## Supplementary material for "TAZ-CAMTA1 and YAP-TFE3 modulate the basal TAZ/YAP transcriptional program by recruiting the ATAC histone acetyltransferase complex": Combined supplemental files

### Figure S1

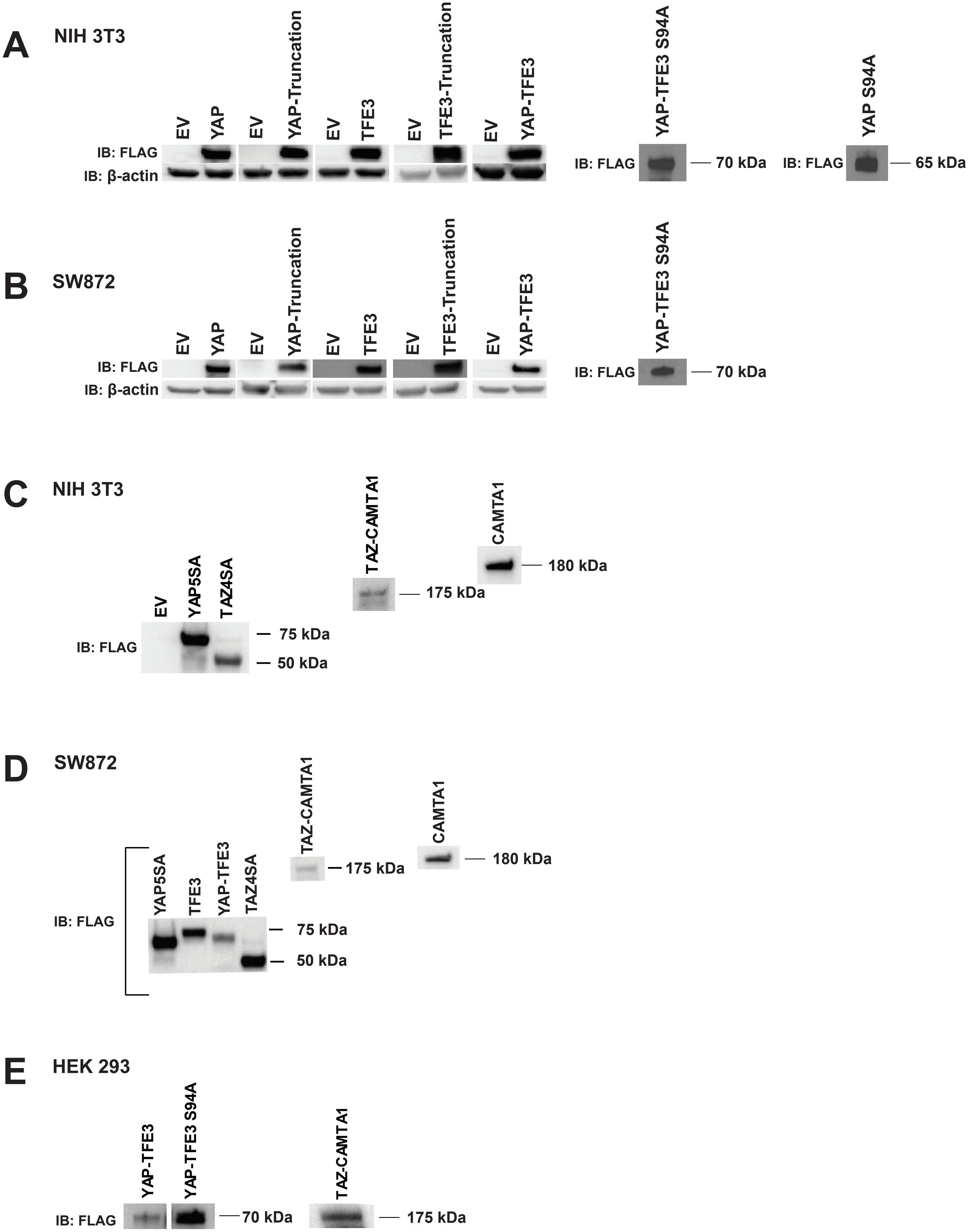
A
NIH 3T3
EV
YAP
EV
YAP-Truncation
EV
TFE3
EV
TFE3-Truncation
EV
YAP-TFE3
IB: FLAG
IB: β-actin
YAP-TFE3 S94A
IB: FLAG
70 kDa
YAP S94A
IB: FLAG
65 kDa
B
SW872
EV
YAP
EV
YAP-Truncation
EV
TFE3
EV
TFE3-Truncation
EV
YAP-TFE3
IB: FLAG
IB: β-actin
YAP-TFE3 S94A
IB: FLAG
70 kDa
C
NIH 3T3
EV
YAP5SA
TAZ4SA
IB: FLAG
75 kDa
50 kDa
TAZ-CAMTA1
175 kDa
CAMTA1
180 kDa
D
SW872
IB: FLAG
YAP5SA
TFE3
YAP-TFE3
TAZ4SA
75 kDa
50 kDa
TAZ-CAMTA1
175 kDa
CAMTA1
180 kDa
E
HEK 293
YAP-TFE3
YAP-TFE3 S94A
IB: FLAG
70 kDa
TAZ-CAMTA1
175 kDa

**Figure S1. YAP-TFE3, TAZ-CAMTA1, and controls are expressed in NIH 3T3 cells, SW 872 cells, and HEK293 cells. (A)** Western blots for Flag tagged YAP, YAP-truncation, TFE3, TFE3 truncation, YAP-TFE3, YAP-TFE3 S94A, and YAP S94A in NIH 3T3 cells. (**B**) Western blots for Flag tagged YAP, YAP-truncation, TFE3, TFE3 truncation, YAP-TFE3, and YAP-TFE3 S94A in SW872 cells. (**C**) Western blots for Flag tagged YAP5SA, TAZ4SA, TAZ-CAMTA1, and CAMTA1 in NIH 3T3 cells. (**D**) Western blots for Flag tagged YAP5SA, TFE3, YAP-TFE3, TAZ4SA, TAZ-CAMTA1, and CAMTA1 in SW872 cells. (**E**) Western blots for Flag tagged YAP-TFE3, YAP-TFE3 S94A, and TAZ-CAMTA1 in HEK293 cells.

### Figure S2

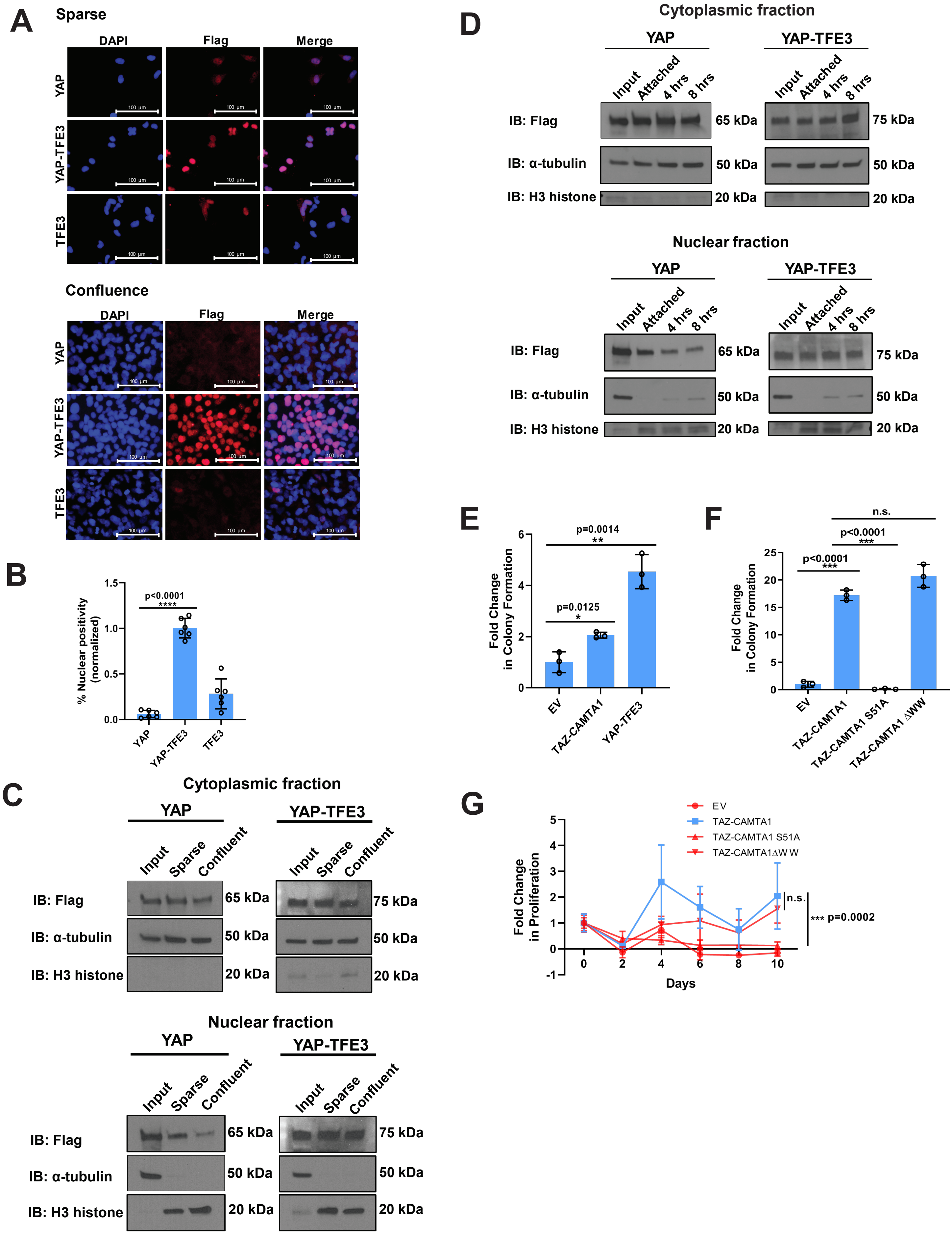
A
Sparse
DAPI
Flag
Merge
YAP
YAP-TFE3
TFE3
Confluence
B
% Nuclear positivity (normalized)
p<0.0001
****
YAP
YAP-TFE3
TFE3
C
Cytoplasmic fraction
Nuclear fraction
Input
Sparse
Confluent
IB: Flag
IB: α-tubulin
IB: H3 histone
65 kDa
75 kDa
50 kDa
20 kDa
D
Attached
4 hrs
8 hrs
E
Fold Change in Colony Formation
p=0.0014
**
p=0.0125
*
EV
TAZ-CAMTA1
YAP-TFE3
F
n.s.
p<0.0001
***
TAZ-CAMTA1 S51A
TAZ-CAMTA1 ΔWW
G
EV
TAZ-CAMTA1
TAZ-CAMTA1 S51A
TAZ-CAMTA1ΔW W
Fold Change in Proliferation
Days
*** p=0.0002

**Figure S2. YAP-TFE3 is not regulated by the Hippo pathway and required binding to TEAD transcription factors to drive its oncogenic transcriptional program.** (**A**) YAP and YAP-TFE3 are localized within the nucleus under spare conditions in SW872 cells. YAP-TFE3 but not YAP is localized within the nucleus under confluent conditions in SW872 cells. (**B**) Graphical representation of results in part (**A**) with % nuclear positivity during cell confluence normalized to % nuclear positivity during sparse conditions. (**C**) NIH 3T3 cells expressing YAP and YAP-TFE3 are plated under sparse and confluent conditions. YAP accumulation in the nucleus is diminished during confluence while YAP-TFE3 levels in the nucleus remain constant. α-tubulin is the cytoplasmic loading control; H3 histone is the nuclear loading control. (**D**) NIH 3T3 cells are detached in a time course. Nuclear:cytoplasmic fractionation show YAP-TFE3 remains in the nucleus during detachment while YAP localization in the nucleus diminishes with time. (**E**) TAZ-CAMTA1 and YAP-TFE3 promote anchorage independent growth in SW872 cells in soft agar colony formation assay. (**F**) The TAZ-CAMTA1 S51A mutant (TEAD binding domain mutant) abrogates colony formation in soft agar in NIH 3T3 cells. Mutating the WW domain in TAZ-CAMTA1 (TAZ-CAMTA1 ΔWW mutant) does not reduce colony formation in soft agar. (**G**) The TAZ-CAMTA1 S51A mutant but not the ΔWW mutant reduces anchorage independent proliferation on poly-HEMA.

### Figure S3

**A** NIH 3T3 cells

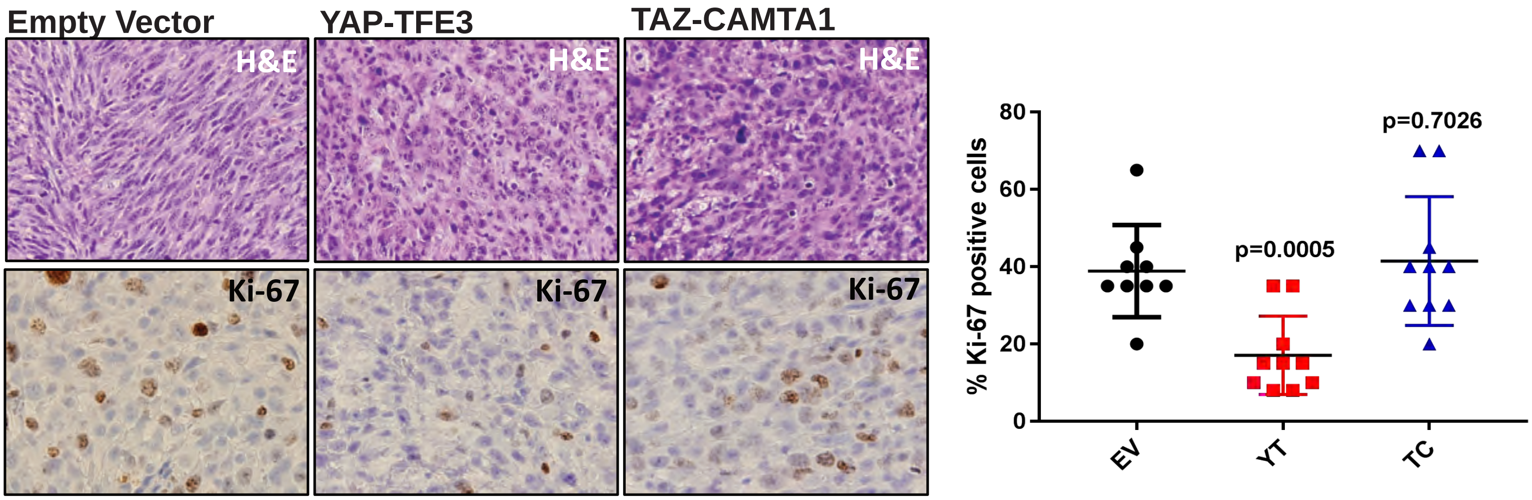


**B** SW 872 cells

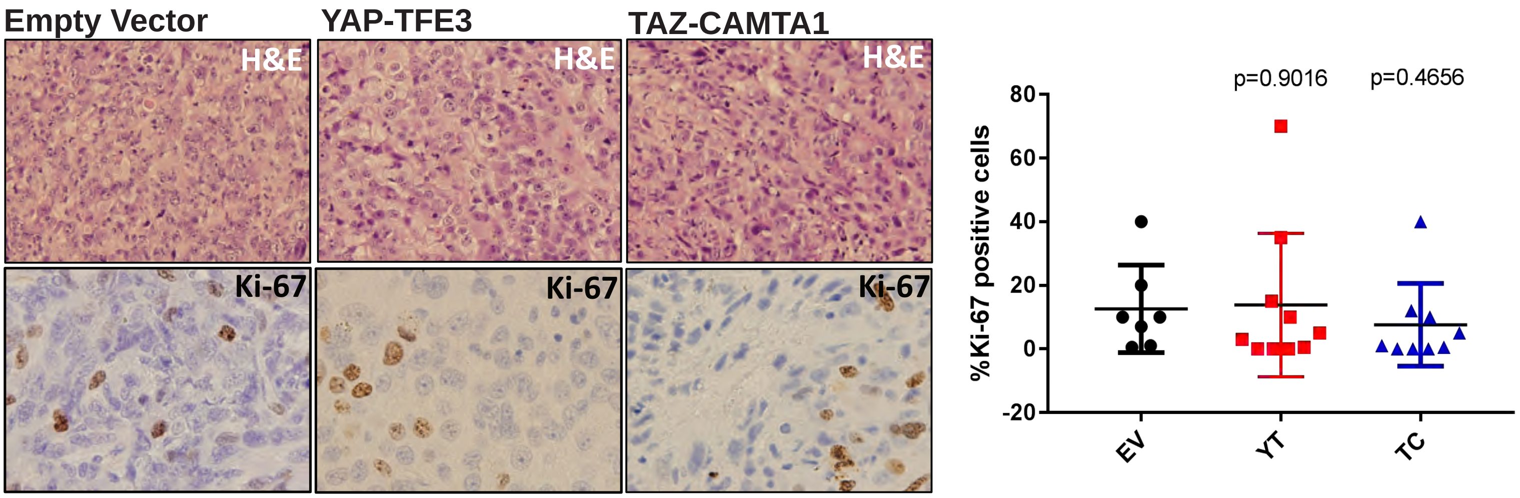

**Figure S3. YAP-TFE3 and TAZ-CAMTA1 driven xenograft in NOD.*Cg-Prkdc$^{scid}$Il2rg$^{tm1Wjl}$/SzJ* (NSG) mice form tumors *in vivo*.** (**A**) H&E sections of NIH 3T3 xenografts transduced with EV, YAP-TFE3 or TAZ-CAMTA1. Tumors expressing YAP-TFE3 or TAZ-CAMTA1 showed a more epithelioid cytomorphology and a greater degree of pleomorphism than EV control. Ki-67 labeling was not increased in YAP-TFE3 or TAZ-CAMTA1 tumors compared to EV. (**B**) H&E sections of SW872 xenografts transduced with EV, YAP-TFE3, or TAZ-CAMTA1. Tumors expressing YAP-TFE3 or TAZ-CAMTA1 showed a greater degree of pleomorphism, however this change was less pronounced than it was in NIH 3T3 cells. Ki-67 labeling was not increased in YAP-TFE3 or TAZ-CAMTA1 tumors compared to EV.

### Figure S4

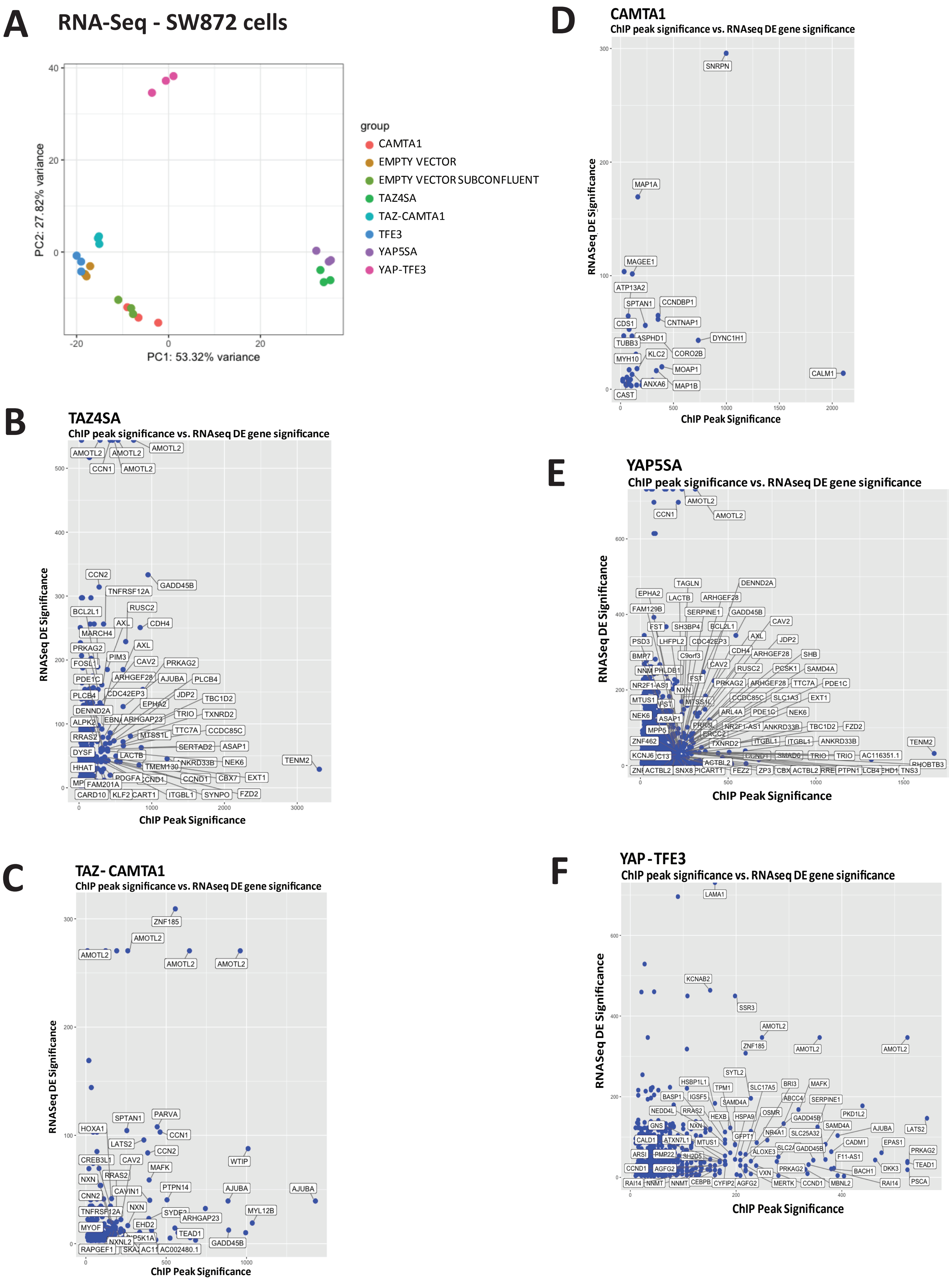
A
RNA-Seq - SW872 cells
PC2: 27.82% variance
PC1: 53.32% variance
group
CAMTA1
EMPTY VECTOR
EMPTY VECTOR SUBCONFLUENT
TAZ4SA
TAZ-CAMTA1
TFE3
YAP5SA
YAP-TFE3
B
TAZ4SA
ChIP peak significance vs. RNAseq DE gene significance
RNAseq DE Significance
ChIP Peak Significance
C
TAZ- CAMTA1
ChIP peak significance vs. RNAseq DE gene significance
RNAseq DE Significance
ChIP Peak Significance
D
CAMTA1
ChIP peak significance vs. RNAseq DE gene significance
RNAseq DE Significance
ChIP Peak Significance
E
YAP5SA
ChIP peak significance vs. RNAseq DE gene significance
RNAseq DE Significance
ChIP Peak Significance
F
YAP - TFE3
ChIP peak significance vs. RNAseq DE gene significance
RNAseq DE Significance
ChIP Peak Significance

**Figure S4. Differential expression of YAP-TFE3 and TAZ-CAMTA1 in SW872 cells.** (**A**) Principal component analysis of RNA expression after variance stabilizing transformation in SW872 cells. (**B-F**) Scatterplot arraying differentially expressed genes in terms of increasing statistical significance along the Y axis, and genes in terms of increasing ChIP peak "score" (i.e. statistical significance) along the X axis. Many TAZ-CAMTA1 (**C**) and YAP-TFE3 (**F**) bound genes are also differentially expressed. TAZ4SA (**B**), CAMTA1 (**D**), and YAP5SA (**E**) controls included.

### Figure S5

A

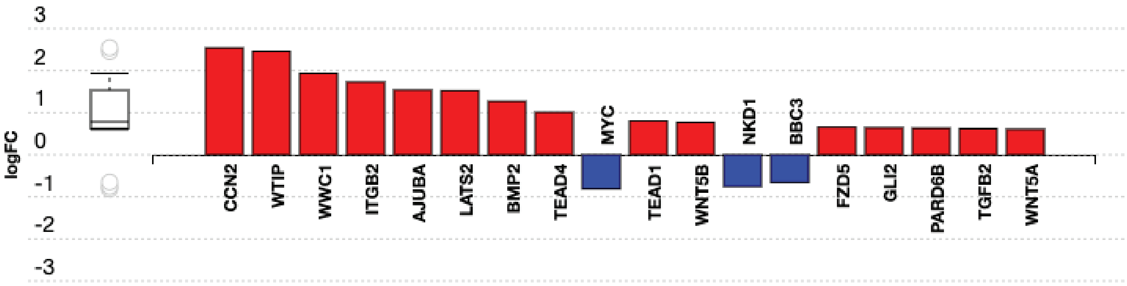


B

TAZ-CAMTA1

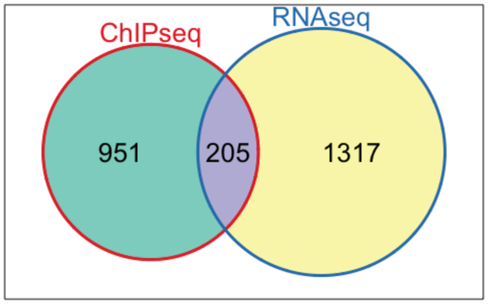


C

YAP-TFE3

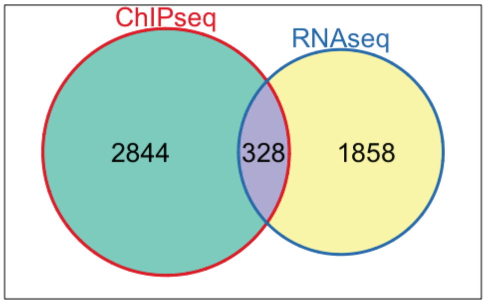


D

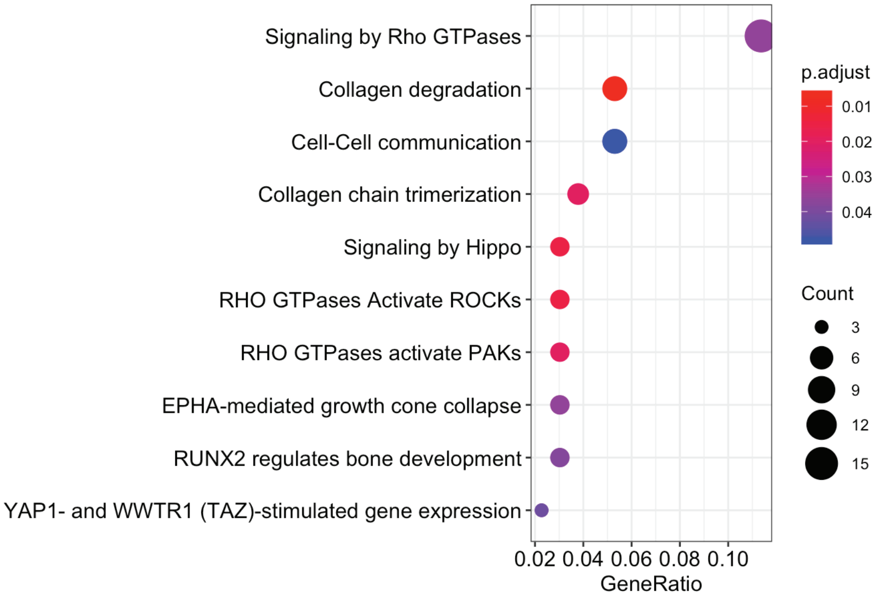

**Figure S5. Pathway analysis of TAZ-CAMTA1 differential expression and DNA binding shows enrichment of Hippo pathway genes as well as other Hippo related pathways.** (**A**) iPathwayGuide analysis showing enrichment of Hippo pathway genes in DE expression analysis in SW872 cells expressing TAZ-CAMTA1. (**B**) Venn diagram showing intersection of DE expressed genes in SW872 TAZ-CAMTA1 with TAZ-CAMTA1 bound genes. (**C**) Venn diagram showing intersection of DE expressed genes in SW872 YAP-TFE3 with YAP-TFE3 bound genes. (**D)** KEGG pathway enrichment analysis of the collection of genes representing the intersection of TAZ-CAMTA1 bound genes (by ChIP-Seq) and DE genes in TAZ-CAMTA1 expressing SW872 cells.

### Figure S6

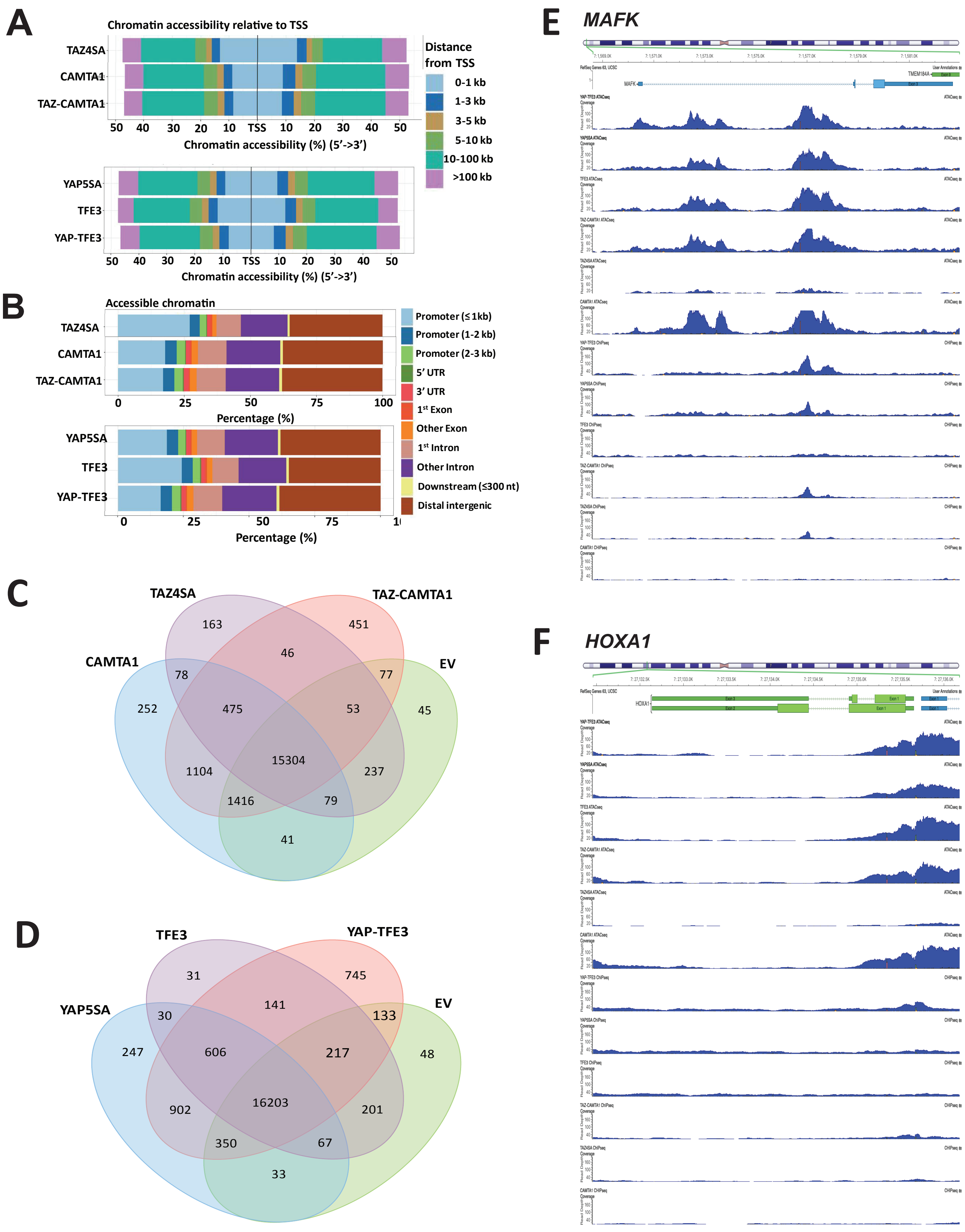
A
Chromatin accessibility relative to TSS
TAZ4SA
CAMTA1
TAZ-CAMTA1
YAP5SA
TFE3
YAP-TFE3
Chromatin accessibility (%) (5'->3')
Distance from TSS
0-1 kb
1-3 kb
3-5 kb
5-10 kb
10-100 kb
>100 kb
B
Accessible chromatin
Percentage (%)
Promoter (≤1kb)
Promoter (1-2 kb)
Promoter (2-3 kb)
5' UTR
3' UTR
1st Exon
Other Exon
1st Intron
Other Intron
Downstream (≤300 nt)
Distal intergenic
C
TAZ4SA
TAZ-CAMTA1
CAMTA1
EV
163
451
46
78
77
252
475
53
45
15304
1104
237
1416
79
41
D
TFE3
YAP-TFE3
YAP5SA
EV
31
745
141
30
133
247
606
217
48
16203
902
201
350
67
33
E
MAFK
F
HOXA1

**Figure S6. TAZ-CAMTA1 and YAP-TFE3 alter the chromatin landscape.** Distribution of transposase accessible chromatin relative to transcriptional start site (**A**) and functional annotation (**B**). (**C**) Venn diagram analysis for TAZ-CAMTA1 after ATAC peaks annotated to their nearest gene feature and including empty vector to demonstrate the baseline level of accessible chromatin. (**D**) Venn diagram analysis for YAP-TFE3 after ATAC peaks annotated to their nearest gene feature and including empty vector to demonstrate the baseline level of accessible chromatin. Combined ATAC-seq and ChIP-Seq signal tracks for *MAFK* (**E**) and *HOX1A* (**F**).

### Figure S7

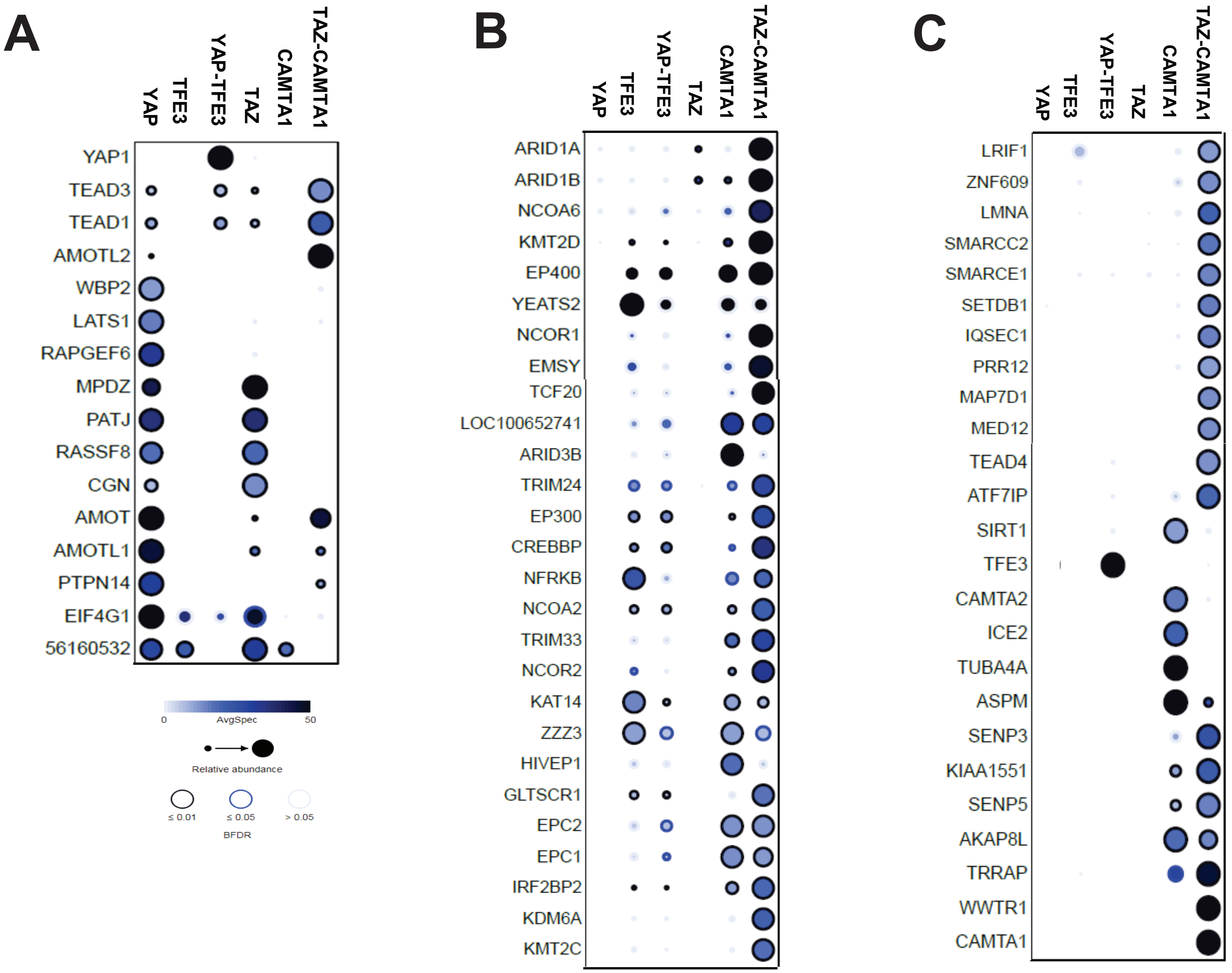
A
YAP
TFE3
YAP-TFE3
TAZ
CAMTA1
TAZ-CAMTA1
YAP1
TEAD3
TEAD1
AMOTL2
WBP2
LATS1
RAPGEF6
MPDZ
PATJ
RASSF8
CGN
AMOT
AMOTL1
PTPN14
EIF4G1
56160532
0
AvgSpec
50
Relative abundance
≤ 0.01
≤ 0.05
> 0.05
BFDR
B
YAP
TFE3
YAP-TFE3
TAZ
CAMTA1
TAZ-CAMTA1
ARID1A
ARID1B
NCOA6
KMT2D
EP400
YEATS2
NCOR1
EMSY
TCF20
LOC100652741
ARID3B
TRIM24
EP300
CREBBP
NFRKB
NCOA2
TRIM33
NCOR2
KAT14
ZZZ3
HIVEP1
GLTSCR1
EPC2
EPC1
IRF2BP2
KDM6A
KMT2C
C
YAP
TFE3
YAP-TFE3
TAZ
CAMTA1
TAZ-CAMTA1
LRIF1
ZNF609
LMNA
SMARCC2
SMARCE1
SETDB1
IQSEC1
PRR12
MAP7D1
MED12
TEAD4
ATF7IP
SIRT1
TFE3
CAMTA2
ICE2
TUBA4A
ASPM
SENP3
KIAA1551
SENP5
AKAP8L
TRRAP
WWTR1
CAMTA1

**Figure S7. BioID Mass Spectrometry shows that YAP-TFE3 and TAZ-CAMTA1 have an altered interactome compared to YAP and TAZ.** Group (**A**) prey proteins often demonstrated a loss of interaction with the fusion proteins. Group (**B**) prey proteins demonstrated enhanced interaction with the fusion proteins relative to TAZ and YAP and generally consisted of epigenetic modifiers and also interacted with CAMTA1 and TFE3. Group (**C**) prey proteins predominantly demonstrated enhanced interactions with TAZ-CAMTA1 relative to TAZ, and predominantly represented epigenetic modifiers. Results demonstrated in dot plot form. Size of dot is proportionate to relative abundance. Darkness of dot is proportionate to the spectral count. Darkness of the ring indicates the Bayesian False Discovery Rate (BFDR); black (FDR $\leq$ 0.01), blue (FDR$\leq$0.05), light blue (FDR> 0.05).

### Figure S8

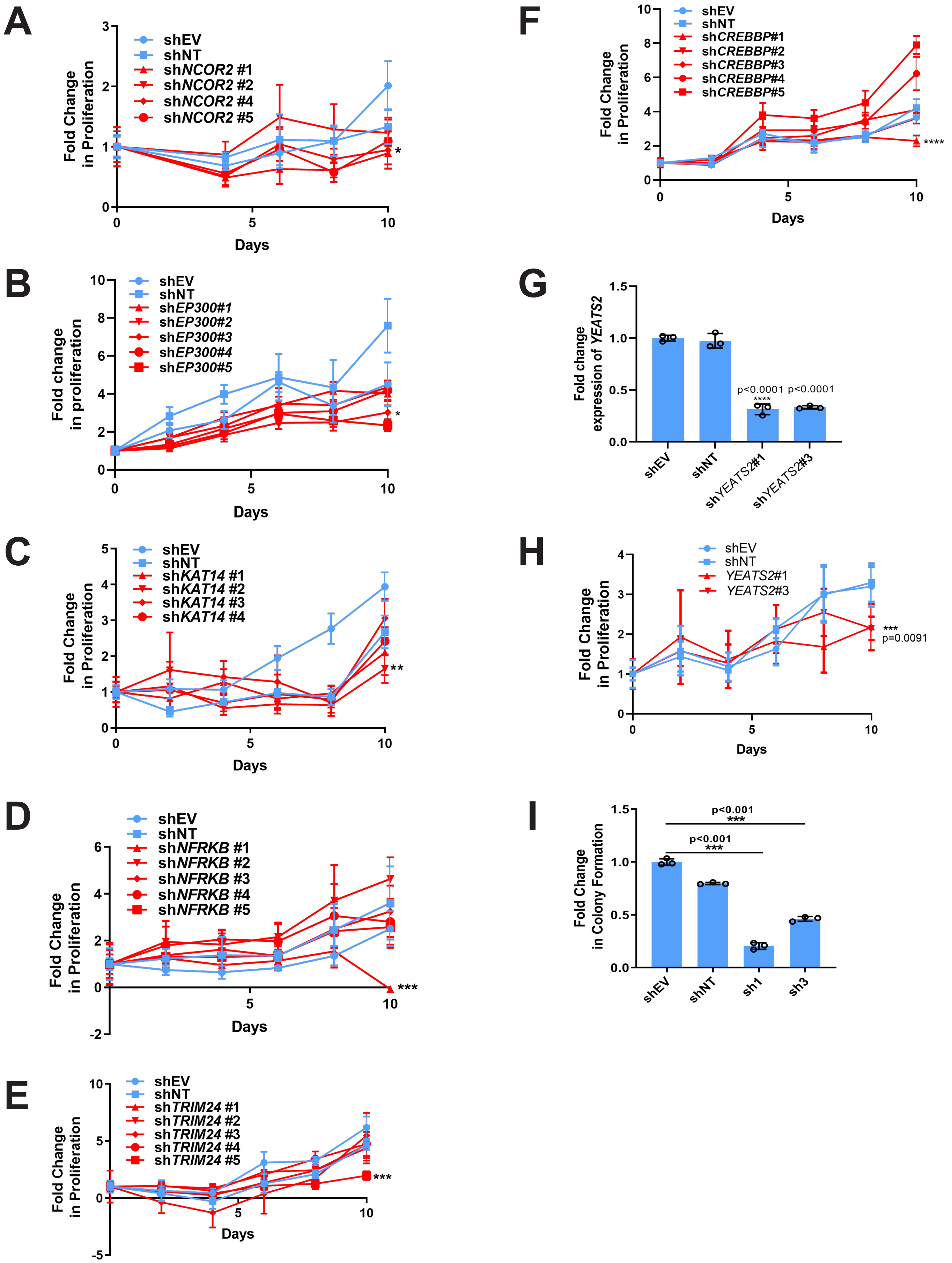
A
Fold Change in Proliferation
Days
shEV
shNT
shNCOR2 #1
shNCOR2 #2
shNCOR2 #4
shNCOR2 #5
*
B
Fold change in proliferation
Days
shEV
shNT
shEP300#1
shEP300#2
shEP300#3
shEP300#4
shEP300#5
*
C
Fold Change in Proliferation
Days
shEV
shNT
shKAT14 #1
shKAT14 #2
shKAT14 #3
shKAT14 #4
**
D
Fold Change in Proliferation
Days
shEV
shNT
shNFRKB #1
shNFRKB #2
shNFRKB #3
shNFRKB #4
shNFRKB #5
***
E
Fold Change in Proliferation
Days
shEV
shNT
shTRIM24 #1
shTRIM24 #2
shTRIM24 #3
shTRIM24 #4
shTRIM24 #5
***
F
Fold Change in Proliferation
Days
shEV
shNT
shCREBBP#1
shCREBBP#2
shCREBBP#3
shCREBBP#4
shCREBBP#5
****
G
Fold change expression of YEATS2
p<0.0001
****
p<0.0001
shEV
shNT
shYEATS2#1
shYEATS2#3
H
Fold Change in Proliferation
Days
shEV
shNT
YEATS2#1
YEATS2#3
***
p=0.0091
I
Fold Change in Colony Formation
p<0.001
***
p<0.001
***
shEV
shNT
sh1
sh3

**Figure S8. RNAi screen identifies epigenetic modifiers important for the activity of TAZ-CAMTA1 and YAP-TFE3.** (**A-F**) Additional epigenetic modifiers with at least 1 shRNA showing a deficit in anchorage independent growth in SW872 TAZ-CAMTA1 cells. (**G**) Knock-down of *YEATS2* in SW872 YAP-TFE3 cells by qRT-PCR. Knock-down of *YEATS2* demonstrated a deficit in anchorage independent growth in SW872 YAP-TFE3 cells when assayed by poly-HEMA assay (**H**) and soft agar (**I**).

| MISSION shRNA name | TRCN# |
|---|---|
| shTead#1 | TRCN0000374164 |
| shTead#2 | TRCN0000085865 |
| shTead#3 | TRCN0000304525 |
| shTead#4 | TRCN0000301799 |
| shTead#5 | TRCN0000085867 |

**Oligo sequence (5'->3')**

CCGGGTGTATTTGAAGTCTCGAATACTCGAGTATTCGAGACTTCAAATACACTTTTTG
CCGGGCTCGCCAATGTGTGAATATACTCGAGTATATTCACACATTGGCGAGCTTTTTG
CCGGTCAAATTCTGGGCGGACTTAACTCGAGTTAAGTCCGCCCAGAATTTGATTTTTG
CCGGGCTTGGTTGAACTATCCTAATCTCGAGATTAGGATAGTTCAACCAAGCTTTTTG
CCGGCAGAAGGAAATCTCGTGATTTCTCGAGAAATCACGAGATTTCCTTCTGTTTTTG

| Gene | baseMean | log2 Fold Change | padj |
|---|---|---|---|
| Upk1b | 1749.009226 | 14.30107985 | 1.39E-25 |
| Slc6a12 | 232.6272342 | 12.33636988 | 6.78E-23 |
| Wnt7a | 83.29495754 | 11.58439297 | 7.61E-20 |
| Wnt7b | 75.66538815 | 11.50930968 | 1.16E-18 |
| Anxa8 | 232.8381859 | 11.32736888 | 3.77E-19 |
| Cdh15 | 124.6645698 | 11.2630599 | 1.06E-18 |
| Vip | 761.4057475 | 11.16688422 | 5.91E-17 |
| Serpinb9g | 65.5698485 | 11.15373451 | 4.43E-19 |
| Il31ra | 119.7831368 | 10.41670498 | 1.72E-16 |
| Robo3 | 32.48397439 | 9.89120176 | 3.56E-12 |
| 4933402E13Rik | 48.25961404 | 9.676572747 | 3.14E-14 |
| Lamb3 | 47.93842017 | 9.675674248 | 1.14E-13 |
| Crct1 | 218.1985879 | 9.639510667 | 3.68E-14 |
| Tmem150b | 157.484637 | 9.611411257 | 2.91E-07 |
| Birc7 | 495.5983212 | 9.469546058 | 2.53E-75 |
| Jph2 | 553.3177943 | 9.026395178 | 1.73E-88 |
| Serpinb1c | 17.20225213 | 9.007033467 | 1.39E-10 |
| Dnah11 | 127.6062225 | 8.93768167 | 2.65E-07 |
| Serpinb9e | 15.75118303 | 8.915192892 | 2.90E-11 |
| Psca | 64.92123435 | 8.879521831 | 1.53E-11 |
| Gm15056 | 33.57462403 | 8.792694124 | 8.52E-03 |
| Ankk1 | 13.33636605 | 8.691275171 | 1.13E-10 |
| Cnfn | 24.09477608 | 8.566008664 | 2.76E-10 |
| Gm1110 | 11.92620283 | 8.560012019 | 7.90E-09 |
| Doc2a | 128.9404018 | 8.539536063 | 1.11E-03 |
| Adamts13 | 189.314739 | 8.521699452 | 2.67E-03 |
| Ikzf4 | 76.17939403 | 8.361250739 | 1.38E-21 |
| Sult4a1 | 8.298499443 | 8.301898961 | 1.77E-08 |
| Dppa4 | 15.36586433 | 8.299635129 | 4.15E-09 |
| Sh3tc1 | 213.0734353 | 8.268226402 | 4.67E-29 |
| Got1l1 | 8.440417381 | 8.231979752 | 1.61E-08 |
| Gm9428 | 14.61896193 | 8.202039492 | 1.09E-09 |
| Vmn1r44 | 18.97883308 | 8.193140413 | 3.44E-10 |
| 1700012B07Rik | 12.47027256 | 8.191048854 | 1.48E-09 |
| NA | 188.127415 | 8.143788227 | 1.81E-40 |
| Pkhd1 | 14.16137139 | 8.124158502 | 6.90E-06 |
| Gm13415 | 175.3337851 | 8.114223188 | 5.18E-40 |
| Tmem171 | 242.2820833 | 8.110703972 | 1.98E-20 |
| Ccl20 | 50.88600856 | 8.109557256 | 2.38E-09 |
| Atg9b | 216.6082704 | 8.103734108 | 4.89E-54 |
| Adam22 | 234.5747723 | 8.074453117 | 5.50E-27 |
| Serpinb1a | 435.1696668 | 8.046094471 | 3.15E-76 |
| Serpinb9f | 8.681433932 | 8.016377479 | 4.00E-07 |
| Gm13418 | 7.051919701 | 7.999986014 | 9.50E-08 |
| Limch1 | 706.4773048 | 7.91836824 | 4.76E-72 |
| Hist1h4n | 13.89900442 | 7.711326345 | 3.79E-08 |

| | | | |
|---|---|---|---|
| Gdf15 | 40.77172247 | 7.690732508 | 1.75E-11 |
| Gm27286 | 31.94524763 | 7.683793549 | 1.23E-07 |
| Pax8 | 75.11368423 | 7.664875431 | 1.50E-22 |
| Serpina5 | 6.801924643 | 7.658077193 | 3.20E-07 |
| Arhgap9 | 58.21154531 | 7.602875599 | 2.89E-19 |
| Prg2 | 8.098456987 | 7.56715957 | 5.18E-07 |
| Ces2e | 38.3662501 | 7.561647216 | 2.25E-11 |
| Ces2f | 8.035604011 | 7.552934179 | 2.12E-07 |
| Gm25043 | 5.794616842 | 7.50332422 | 1.14E-06 |
| Rapgef5 | 8.056496296 | 7.482065318 | 5.84E-07 |
| Ccdc113 | 31.53830894 | 7.47353641 | 8.60E-19 |
| Ctnnd2 | 13.32903101 | 7.468901027 | 3.37E-08 |
| 1520401A03Rik | 8.114182141 | 7.449440248 | 2.04E-06 |
| Parm1 | 50.62742008 | 7.414263366 | 1.27E-08 |
| Krt14 | 42.84356732 | 7.391149385 | 7.95E-19 |
| Serpinb9b | 3482.508606 | 7.327778527 | 1.27E-86 |
| Epn3 | 33.31382918 | 7.326830013 | 3.76E-14 |
| Snca | 7.396202048 | 7.322186564 | 2.10E-05 |
| Gm13284 | 12.99212276 | 7.30839468 | 1.84E-07 |
| Sntg1 | 26.65447637 | 7.274364532 | 3.58E-08 |
| Cyp3a13 | 57.57513375 | 7.268981671 | 3.78E-21 |
| Gm12153 | 16.73241645 | 7.184725709 | 5.37E-08 |
| 1700009N14Rik | 112.4486022 | 7.18175579 | 9.58E-28 |
| Gm43154 | 35.32626595 | 7.167935735 | 1.19E-07 |
| Nkain4 | 34.42273738 | 7.160616157 | 1.40E-12 |
| Snx20 | 7.501681989 | 7.158751649 | 1.26E-05 |
| Upk3bl | 49.55256089 | 7.11848073 | 1.84E-19 |
| Vnn1 | 94.96773462 | 7.109056149 | 8.33E-17 |
| Msln | 8968.235968 | 7.058648991 | 3.70E-34 |
| B230206L02Rik | 12.43408177 | 7.011996104 | 9.44E-07 |
| Cyp24a1 | 5.817295899 | 6.993899268 | 8.45E-06 |
| Cblc | 13.39498369 | 6.984777902 | 1.20E-06 |
| Trav13-2 | 8.537472155 | 6.966726419 | 1.29E-06 |
| Gm21293 | 16.49303806 | 6.948985403 | 3.10E-06 |
| Btn2a2 | 21.67262263 | 6.911921243 | 1.88E-09 |
| NA | 26.06308258 | 6.900284725 | 6.33E-07 |
| Tmem246 | 6.736769451 | 6.899853979 | 1.73E-05 |
| Gm5286 | 7.531885012 | 6.895764877 | 7.57E-07 |
| Gm15389 | 6.567666468 | 6.880965174 | 7.14E-06 |
| Psd4 | 5.219049876 | 6.861377119 | 5.48E-05 |
| Serping1 | 32.77937136 | 6.845654806 | 1.21E-07 |
| Gm12295 | 5.965441193 | 6.844818704 | 2.30E-05 |
| Hbb-bh1 | 14.94205789 | 6.82003518 | 3.82E-08 |
| Tnnt3 | 106.6121583 | 6.809568461 | 1.15E-20 |
| NA | 240.538982 | 6.783701352 | 7.11E-04 |
| Spo11 | 29.75667518 | 6.766346199 | 7.40E-07 |
| Krt7 | 1408.919025 | 6.742540902 | 1.44E-124 |

| | | | |
|---|---|---|---|
| Slc47a1 | 16.32805259 | 6.741404275 | 2.27E-06 |
| Slc2a12 | 153.088269 | 6.733717727 | 1.14E-02 |
| Ptprt | 174.3279632 | 6.717866001 | 1.19E-09 |
| Fras1 | 50.54206584 | 6.706467303 | 1.97E-06 |
| Myom2 | 1179.530624 | 6.668948943 | 6.94E-72 |
| Dsp | 6.380203445 | 6.651173882 | 3.97E-04 |
| Gm5812 | 53.47797956 | 6.627944522 | 6.23E-03 |
| B3gnt3 | 5.759179752 | 6.612211714 | 1.50E-05 |
| Plekhs1 | 102.3016648 | 6.597757169 | 1.33E-36 |
| Dnaic1 | 8.46165829 | 6.561523772 | 6.38E-05 |
| Hepacam | 5.666744147 | 6.456352143 | 1.08E-04 |
| Arg1 | 5.523346439 | 6.434660359 | 1.87E-05 |
| Foxa3 | 6.492805962 | 6.430896609 | 6.56E-06 |
| Vill | 20.01966172 | 6.406891378 | 4.42E-06 |
| Kdr | 75.92150854 | 6.396779655 | 5.53E-22 |
| Far2os1 | 51.84053146 | 6.37524601 | 1.55E-25 |
| Ninj2 | 25.84237259 | 6.33896603 | 2.77E-15 |
| Gm13710 | 14.91984042 | 6.324008686 | 1.76E-10 |
| Ptprq | 158.5957609 | 6.288642217 | 1.57E-25 |
| Hist1h2br | 20.36525225 | 6.272462581 | 8.56E-07 |
| Sgce | 5.719415256 | 6.264311891 | 1.33E-05 |
| 4933427D06Rik | 13.64440714 | 6.254374279 | 4.64E-07 |
| Gm53 | 5.854823975 | 6.245148894 | 3.92E-05 |
| Serpinb6b | 303.4332572 | 6.219750877 | 3.12E-85 |
| Gm10505 | 10.24804569 | 6.210419801 | 7.34E-07 |
| Ttn | 108.5364841 | 6.083779565 | 2.69E-07 |
| Zfp750 | 9.306518556 | 6.075989314 | 5.56E-07 |
| Inpp5d | 43.74884049 | 6.048120752 | 1.60E-10 |
| Slamf9 | 31.67214215 | 6.043191098 | 1.59E-13 |
| Sh3rf2 | 283.6625641 | 6.025221581 | 3.56E-39 |
| Fgfbp1 | 21.39660242 | 6.024870468 | 7.23E-09 |
| Mogat2 | 37.49728946 | 6.023344833 | 1.79E-16 |
| Siglecg | 4363.02185 | 5.909202173 | 3.82E-132 |
| Gpr174 | 13.33517572 | 5.901600577 | 1.76E-06 |
| Csf1r | 229.3071014 | 5.882390181 | 1.03E-51 |
| BC051019 | 6.81730173 | 5.836129596 | 6.33E-05 |
| Fyb2 | 205.4623369 | 5.832322181 | 2.43E-29 |
| Prl3d2 | 7.108386125 | 5.810612513 | 2.84E-05 |
| Lama5 | 488.2888004 | 5.800834131 | 6.75E-26 |
| Klra4 | 79.2878235 | 5.79941945 | 1.27E-19 |
| Fgf21 | 49.48268511 | 5.765967652 | 1.62E-11 |
| Synpr | 13.66684869 | 5.741745414 | 5.01E-05 |
| Ifitm10 | 12.42299472 | 5.725610662 | 1.02E-08 |
| Psme2b | 18.26708444 | 5.720034769 | 1.08E-05 |
| Syk | 89.27270301 | 5.698535033 | 6.78E-18 |
| Tnfsf18 | 94.04928164 | 5.697567296 | 4.60E-21 |
| Upk3b | 18.38391089 | 5.68024178 | 1.04E-06 |

| | | | |
|---|---|---|---|
| Scel | 35.56464911 | 5.64970329 | 2.00E-04 |
| Card10 | 5.87499347 | 5.649678761 | 2.93E-04 |
| Fcgbp | 17.64993208 | 5.628348688 | 3.79E-12 |
| Wdfy4 | 18.09422915 | 5.626376072 | 2.23E-05 |
| 9530053A07Rik | 64.37286542 | 5.611364945 | 4.50E-21 |
| Btn1a1 | 43.15252764 | 5.590590084 | 1.07E-16 |
| Hephl1 | 25.60224789 | 5.562179207 | 1.04E-04 |
| Smarca5-ps | 60.03761091 | 5.560892552 | 1.11E-02 |
| Klk8 | 65.88130734 | 5.553804415 | 3.73E-23 |
| Lamc3 | 81.73661022 | 5.529824173 | 4.07E-25 |
| Ptgs2 | 3952.831909 | 5.52218542 | 6.25E-35 |
| Hspb7 | 111.5849735 | 5.520168119 | 2.01E-33 |
| Slc15a3 | 5.507672821 | 5.495445932 | 4.27E-05 |
| 2610028E06Rik | 10.97570129 | 5.452097305 | 4.99E-05 |
| Myoz2 | 26.20642193 | 5.444794145 | 1.06E-17 |
| AU021092 | 16.8814082 | 5.444774801 | 1.30E-04 |
| Slc30a2 | 5.640362476 | 5.442533107 | 1.32E-04 |
| NA | 43.86976017 | 5.439708404 | 1.25E-07 |
| Emcn | 5.59271389 | 5.424067661 | 6.48E-05 |
| Ttll9 | 5.377351426 | 5.411105306 | 6.05E-04 |
| Gm14341 | 16.93756775 | 5.403602494 | 4.85E-10 |
| Nppb | 446.9261959 | 5.365044283 | 1.97E-44 |
| Tgtp1 | 20.61468666 | 5.341395096 | 1.99E-11 |
| Fst | 208.7049669 | 5.323171467 | 6.85E-37 |
| Fcna | 17.16685909 | 5.319444284 | 2.37E-05 |
| Dpp4 | 5.265900201 | 5.317744204 | 5.73E-04 |
| Cobl | 32.63939068 | 5.264819236 | 2.25E-10 |
| Gm10696 | 6.667621713 | 5.255147184 | 9.73E-04 |
| Lims2 | 179.0157548 | 5.254796926 | 1.39E-40 |
| Sbk3 | 20.10660947 | 5.239943736 | 2.02E-08 |
| Ccdc81 | 6.318104581 | 5.237983249 | 1.59E-04 |
| S100a7a | 436.7014925 | 5.224297668 | 2.36E-81 |
| Ablim3 | 364.5652957 | 5.211781733 | 2.97E-49 |
| Des | 60.75912734 | 5.202673377 | 1.40E-16 |
| Gm26635 | 8.223066354 | 5.180600362 | 3.12E-02 |
| Gadd45g | 1363.863302 | 5.177330238 | 1.44E-90 |
| Ccn1 | 16942.56933 | 5.164325791 | 1.41E-33 |
| Tmeff2 | 105.2957531 | 5.16156083 | 1.63E-27 |
| Als2cl | 101.8281501 | 5.158608672 | 9.31E-10 |
| Cyp2ab1 | 5.554438667 | 5.147187024 | 7.56E-04 |
| Mat1a | 7.834774998 | 5.141796563 | 2.03E-06 |
| Vmn1r-ps128 | 315.8095013 | 5.137501185 | 3.31E-44 |
| Serpine1 | 8069.154667 | 5.097855338 | 1.40E-53 |
| Adgrl4 | 5.885316791 | 5.09477592 | 3.87E-04 |
| Tmem182 | 44.63172361 | 5.084619529 | 8.45E-16 |
| Folr1 | 25.64995036 | 5.075694571 | 1.37E-14 |
| Dppa2 | 32.68223449 | 5.072716019 | 3.51E-16 |

| | | | |
|---|---|---|---|
| Suox | 3135.576887 | 5.047471125 | 8.78E-164 |
| Rnd1 | 586.3380609 | 5.028208196 | 7.61E-58 |
| Gm27224 | 10.41317876 | 5.021031995 | 8.44E-08 |
| Vtn | 30.23869185 | 4.997389038 | 3.00E-08 |
| Csrnp1 | 607.3843703 | 4.988042371 | 3.69E-55 |
| Lman1l | 19.17703041 | 4.986137146 | 3.63E-12 |
| Tcp11x2 | 18.44972698 | 4.962394057 | 4.27E-09 |
| Gprc5a | 2003.573682 | 4.957091951 | 1.76E-70 |
| Slc2a3 | 86.49054597 | 4.951428566 | 6.32E-14 |
| NA | 87.8527127 | 4.934487065 | 1.02E-29 |
| Misp | 6.782132558 | 4.916761262 | 4.84E-04 |
| Pim1 | 2348.868326 | 4.91246664 | 9.86E-51 |
| Tnni2 | 25.52205906 | 4.894451007 | 1.68E-14 |
| Klhl30 | 170.2009204 | 4.893783583 | 1.63E-28 |
| Pmaip1 | 85.15518226 | 4.873505269 | 2.12E-33 |
| Cdc42bpg | 72.4716449 | 4.869888995 | 1.97E-14 |
| Pkp2 | 365.3274075 | 4.855622903 | 1.10E-76 |
| Eps8l2 | 14.97634174 | 4.8460185 | 7.63E-05 |
| Hapln1 | 63.3439414 | 4.837400082 | 5.20E-03 |
| Foxn1 | 11.80917538 | 4.823258566 | 2.32E-08 |
| Syt8 | 16.57992572 | 4.796984654 | 5.96E-07 |
| Xlr3a | 7.716902717 | 4.758071395 | 2.53E-04 |
| Gsta1 | 191.6070784 | 4.756337466 | 9.41E-33 |
| 3425401B19Rik | 10.93216987 | 4.752551415 | 2.42E-03 |
| Sdcbp2 | 61.62660346 | 4.751907515 | 4.02E-26 |
| Gm15854 | 18.23662287 | 4.750352699 | 1.08E-04 |
| Rtn4r | 11.45555949 | 4.746133404 | 1.84E-08 |
| Sorl1 | 6.29132497 | 4.71191586 | 2.39E-03 |
| Eml2 | 184.9655233 | 4.706412266 | 2.31E-51 |
| Gm20521 | 108.1932135 | 4.682497254 | 1.22E-23 |
| Gadd45b | 5683.916854 | 4.675246851 | 6.41E-65 |
| Adrb3 | 11.41728903 | 4.674842275 | 1.60E-08 |
| Itih4 | 19.70594933 | 4.651226954 | 1.41E-12 |
| Sh3bgr | 6.330771203 | 4.644210212 | 5.59E-04 |
| Prss27 | 17.7205746 | 4.639279783 | 1.14E-09 |
| Gm19345 | 11.4622406 | 4.632728263 | 5.80E-08 |
| Gm6088 | 11.0474142 | 4.625013877 | 2.42E-03 |
| Cdh5 | 11.00256545 | 4.611667897 | 1.68E-03 |
| Gm11351 | 79.42538293 | 4.604644379 | 3.35E-29 |
| Otoa | 5.72929926 | 4.603731159 | 3.61E-03 |
| Klra5 | 12.83281442 | 4.600944424 | 4.98E-07 |
| Itga7 | 815.1436197 | 4.591299552 | 4.77E-56 |
| Aldh1l1 | 17.6599573 | 4.583470695 | 3.72E-07 |
| Oprk1 | 32.03179652 | 4.581109886 | 4.30E-04 |
| Tmem184a | 18.15966957 | 4.57450856 | 8.13E-08 |
| Sbsn | 1060.826381 | 4.572049071 | 2.85E-200 |
| Cdkn1a | 10457.59614 | 4.568843911 | 1.76E-73 |

| | | | |
|---|---|---|---|
| Tuba8 | 7.274931265 | 4.564838902 | 9.41E-06 |
| Gm13988 | 25.63523994 | 4.560389146 | 2.84E-10 |
| Serpinb9c | 5.608671978 | 4.519361872 | 8.20E-03 |
| Mamld1 | 9.313569485 | 4.511190842 | 1.49E-05 |
| Gata3 | 165.7026761 | 4.504764728 | 2.84E-36 |
| Cdhr2 | 8.715260545 | 4.49439062 | 8.97E-06 |
| Gm6650 | 24.45313423 | 4.489519275 | 7.80E-08 |
| Tmem40 | 10.09618904 | 4.477471237 | 5.06E-08 |
| Panx3 | 37.5768691 | 4.463686883 | 3.32E-03 |
| Bach2os | 7.936230742 | 4.462702813 | 3.65E-04 |
| Plcxd1 | 187.9861981 | 4.451507357 | 3.91E-69 |
| Cfap70 | 23.3829331 | 4.400517947 | 1.11E-07 |
| Gli1 | 5.477929601 | 4.369149795 | 4.56E-03 |
| Cnksr1 | 26.39635045 | 4.365855036 | 5.16E-09 |
| Cd209c | 391.2517191 | 4.357732098 | 1.33E-30 |
| Clca3b | 16.002642 | 4.356404336 | 3.41E-07 |
| Gm10256 | 43.00942646 | 4.354527727 | 9.24E-11 |
| Cd68 | 539.0889678 | 4.345128046 | 1.27E-33 |
| Mef2b | 78.07998856 | 4.340513799 | 1.54E-02 |
| Slc17a7 | 5.760315762 | 4.331257371 | 2.31E-03 |
| Psg17 | 21.47017957 | 4.328641527 | 3.56E-08 |
| Gm15287 | 92.60236988 | 4.326673953 | 1.21E-37 |
| Ncf2 | 13.41769721 | 4.318381561 | 2.69E-07 |
| Gm10109 | 6.409401559 | 4.296364127 | 1.72E-04 |
| Klk1b27 | 12.04208527 | 4.295783441 | 9.14E-03 |
| Rpl29 | 569.7586007 | 4.294482336 | 5.67E-62 |
| 1110017D15Rik | 13.74255596 | 4.27703936 | 3.34E-07 |
| Drc1 | 7.205444409 | 4.275170368 | 4.73E-05 |
| Wnt11 | 29.78098512 | 4.271320158 | 8.88E-10 |
| 2310002F09Rik | 28.54212603 | 4.262753536 | 9.16E-11 |
| Cyp4f40 | 16.78817656 | 4.261405318 | 4.76E-04 |
| 2900026A02Rik | 18.97069 | 4.258287953 | 1.12E-08 |
| Fgd5 | 6.720168263 | 4.258144054 | 8.54E-04 |
| Best3 | 5.268568397 | 4.244545033 | 2.02E-03 |
| Itga3 | 4169.255291 | 4.235519408 | 8.66E-61 |
| Perp | 26.99007095 | 4.223925103 | 1.06E-12 |
| Trpm5 | 54.98351022 | 4.215033281 | 5.59E-33 |
| Lncenc1 | 53.16397997 | 4.191008102 | 3.47E-33 |
| Gal3st2 | 33.39687285 | 4.186307192 | 7.48E-03 |
| Ccn2 | 7484.582414 | 4.181995924 | 1.01E-19 |
| Chac1 | 2414.321663 | 4.178609028 | 8.87E-71 |
| Rassf7 | 636.3971329 | 4.171043578 | 1.95E-57 |
| Cd80 | 43.14414784 | 4.165878413 | 2.35E-16 |
| Pkp1 | 13.7724583 | 4.150964112 | 2.74E-09 |
| C530050E15Rik | 28.3053703 | 4.148442195 | 9.31E-11 |
| Zbtb8b | 15.33196433 | 4.135877245 | 8.60E-09 |
| Bcam | 6.821030757 | 4.130190602 | 2.94E-03 |

| | | | |
|---|---|---|---|
| Ripor2 | 382.9148722 | 4.128678095 | 1.12E-31 |
| Islr2 | 6.801156223 | 4.12867069 | 2.00E-04 |
| Casp14 | 18.04409302 | 4.126613685 | 1.12E-07 |
| Aif1l | 2280.404424 | 4.123678926 | 8.31E-90 |
| Aldh1a1 | 6563.728029 | 4.118001549 | 6.52E-30 |
| U90926 | 10.05769719 | 4.117648097 | 1.90E-02 |
| Rtn4rl2 | 154.9682333 | 4.114616349 | 9.82E-25 |
| Olfr456 | 23.66145086 | 4.111409109 | 1.09E-07 |
| Acvrl1 | 34.10597704 | 4.110110186 | 2.08E-14 |
| Mypn | 6.086652581 | 4.091970656 | 3.81E-02 |
| Tnn | 22.38362316 | 4.08575513 | 3.14E-09 |
| Prkag3 | 47.08753681 | 4.07616635 | 7.23E-17 |
| Mme | 42.60564629 | 4.075871053 | 3.77E-18 |
| Hist1h4h | 20.58006819 | 4.07109804 | 2.35E-07 |
| Pard6b | 681.0243997 | 4.060326113 | 1.46E-43 |
| NA | 1301.961584 | 4.045268225 | 1.57E-57 |
| Llgl2 | 523.0277484 | 4.044843961 | 1.21E-64 |
| Enkur | 17.5217497 | 4.032130457 | 6.00E-06 |
| Speer3 | 6.085578054 | 4.018407444 | 2.40E-02 |
| Gm26640 | 7.68673924 | 4.017888983 | 1.97E-06 |
| Ddx4 | 6.74894573 | 4.005987879 | 3.69E-02 |
| Muc15 | 5.022882533 | 4.005052815 | 1.13E-03 |
| NA | 141.937738 | 4.004865098 | 1.04E-33 |
| Prrg3 | 9.649158634 | 4.000815398 | 9.31E-04 |
| Hist1h1e | 12.45311968 | 3.991375147 | 1.23E-08 |
| Tjp2 | 3236.987169 | 3.98692811 | 2.76E-83 |
| Hist1h4i | 68.69874854 | 3.970944649 | 2.83E-14 |
| Itgb7 | 197.2536301 | 3.970776018 | 2.26E-26 |
| Dusp27 | 32.80363271 | 3.968038255 | 7.90E-09 |
| Tm4sf4 | 8.031164054 | 3.967517645 | 3.52E-05 |
| Gm20400 | 779.139461 | 3.959220337 | 4.72E-26 |
| Klk14 | 5.470717404 | 3.93975536 | 3.71E-03 |
| Dppa5a | 5.547308591 | 3.926539758 | 1.88E-03 |
| Serpina1c | 24.40367768 | 3.925428867 | 1.56E-03 |
| Exph5 | 13.80776321 | 3.913470202 | 1.49E-03 |
| Nr1d1 | 3745.690596 | 3.910777089 | 1.92E-15 |
| Gm26579 | 5.928312409 | 3.909725218 | 6.35E-05 |
| Adra1d | 14.40500804 | 3.907547122 | 1.51E-05 |
| Fbxw10 | 18.3927969 | 3.894860865 | 3.10E-06 |
| Slc2a5 | 32.51930166 | 3.889822424 | 2.65E-05 |
| BC051077 | 62.17491931 | 3.881965712 | 3.11E-20 |
| Krt80 | 423.3382572 | 3.879307168 | 3.68E-40 |
| Tinagl1 | 7597.988131 | 3.876033789 | 1.10E-61 |
| Ces2g | 117.9433417 | 3.875820477 | 1.38E-23 |
| Dusp13 | 6.94157199 | 3.860406793 | 4.83E-05 |
| Gm12840 | 2928.5719 | 3.853146482 | 2.47E-14 |
| Tgm2 | 2829.880322 | 3.852810047 | 3.81E-102 |

| | | | |
|---|---|---|---|
| Serpinb1b | 80.38277849 | 3.843349775 | 2.01E-22 |
| Rap1gap2 | 970.5392269 | 3.842705365 | 2.78E-51 |
| Itgam | 17.41154262 | 3.828627511 | 3.58E-05 |
| Pdlim1 | 2636.81262 | 3.811454552 | 8.12E-37 |
| Trim7 | 396.6823301 | 3.800166429 | 1.74E-105 |
| Cpa4 | 91.48931586 | 3.793758309 | 6.96E-12 |
| Slc14a2 | 5.198235582 | 3.793453322 | 1.85E-03 |
| Hist1h1c | 1898.756039 | 3.79297089 | 5.00E-29 |
| Inhba | 546.6979894 | 3.789826527 | 1.63E-11 |
| Sulf1 | 1797.810171 | 3.788690589 | 1.18E-38 |
| Crybg2 | 69.94425025 | 3.787979704 | 5.83E-27 |
| Prg4 | 107.0476853 | 3.777471723 | 2.99E-13 |
| Snhg7os | 10.44086061 | 3.775396128 | 4.49E-05 |
| Prelid3a | 26.92713764 | 3.7584967 | 3.79E-11 |
| Itgb4 | 228.6910735 | 3.738403897 | 1.30E-07 |
| 4930590J08Rik | 8.309802627 | 3.727506147 | 3.19E-04 |
| Ripk3 | 33.31369269 | 3.725197247 | 3.57E-12 |
| Sorbs2 | 490.7709526 | 3.723603449 | 4.13E-24 |
| Serpina1a | 229.6910667 | 3.718482191 | 1.29E-23 |
| Acer1 | 15.46558608 | 3.716248073 | 1.31E-09 |
| Btg3 | 59.40803996 | 3.71590407 | 1.44E-03 |
| Amd-ps3 | 45.53365464 | 3.715416971 | 1.66E-05 |
| Sptbn4 | 27.97214524 | 3.711128955 | 2.24E-08 |
| Clca4a | 15.01272408 | 3.709857608 | 5.78E-07 |
| Gm13067 | 13.02922408 | 3.699700676 | 1.59E-07 |
| Erbb3 | 69.46120585 | 3.695299312 | 2.92E-09 |
| 4921536K21Rik | 33.37512953 | 3.693285299 | 3.70E-13 |
| Klf5 | 911.9925518 | 3.690224739 | 9.99E-89 |
| Fmnl1 | 6.076091521 | 3.667646195 | 3.97E-02 |
| Acmsd | 84.38312468 | 3.665722267 | 4.72E-23 |
| Apol7d | 28.86154236 | 3.663174613 | 2.59E-14 |
| Svop | 24.57369793 | 3.660575525 | 3.82E-08 |
| Gm9750 | 83.52959486 | 3.656923048 | 3.40E-22 |
| Rassf6 | 30.46412628 | 3.650364366 | 1.73E-06 |
| Tmem71 | 6.057387095 | 3.649626851 | 7.55E-03 |
| Mill2 | 7.236578422 | 3.635375314 | 1.32E-02 |
| Ptprj | 72.06266337 | 3.630182636 | 1.34E-14 |
| Frat2 | 305.7883249 | 3.630002988 | 6.77E-48 |
| Ano3 | 117.0441969 | 3.629898024 | 6.03E-39 |
| Arhgap45 | 18.56644359 | 3.627112279 | 7.33E-09 |
| Clic3 | 21.76777198 | 3.62415405 | 3.26E-11 |
| Gm16702 | 278.1798623 | 3.6192516 | 1.87E-34 |
| Mmp15 | 11.32755157 | 3.615657919 | 9.77E-06 |
| Slc1a7 | 20.64700538 | 3.611796324 | 3.08E-10 |
| Cntfr | 7.670953597 | 3.611350273 | 1.38E-04 |
| Dll4 | 13.871069 | 3.603914244 | 7.37E-07 |
| Tpd52l1 | 5.150976508 | 3.603881955 | 1.53E-03 |

| | | | |
|---|---|---|---|
| Eda2r | 1123.269013 | 3.595476274 | 7.04E-50 |
| Aloxe3 | 163.7815803 | 3.590013041 | 8.29E-28 |
| Gria3 | 1179.231791 | 3.579867153 | 6.10E-61 |
| Nes | 211.7096441 | 3.578728552 | 1.34E-11 |
| NA | 101.8655657 | 3.574955776 | 2.00E-35 |
| Synpo2l | 127.4131202 | 3.569251341 | 4.73E-13 |
| Dbil5 | 14.31520151 | 3.563661934 | 1.38E-10 |
| NA | 125.6560357 | 3.561363334 | 3.84E-18 |
| Acta1 | 8.400779174 | 3.551599791 | 4.24E-03 |
| Ccl6 | 20.15019644 | 3.55117615 | 4.63E-05 |
| Mreg | 281.1053841 | 3.550776949 | 5.94E-22 |
| Ccdc106 | 72.36994274 | 3.545402367 | 2.08E-33 |
| NA | 324.7999154 | 3.54298006 | 6.39E-25 |
| Traf3ip3 | 5.654108574 | 3.536922824 | 2.98E-02 |
| Ak7 | 5.835582438 | 3.533182673 | 1.37E-03 |
| Mgll | 24.84962995 | 3.533060091 | 3.18E-07 |
| Pla1a | 89.47502873 | 3.521420706 | 1.05E-15 |
| Cavin2 | 13533.10625 | 3.513836109 | 7.32E-27 |
| Cxcl1 | 119.9745271 | 3.509009499 | 1.35E-17 |
| Slc17a9 | 18.00504775 | 3.506829449 | 9.05E-06 |
| Hist1h2bl | 33.18392678 | 3.498142224 | 4.67E-08 |
| Ecscr | 21.55153097 | 3.495543777 | 1.88E-02 |
| Tfap2b | 5.181267629 | 3.490042634 | 1.45E-02 |
| Shank2 | 22.58620902 | 3.488054483 | 1.24E-03 |
| Plb1 | 53.16380267 | 3.484804731 | 5.22E-15 |
| Cystm1 | 169.2870408 | 3.482053213 | 1.08E-21 |
| AU018091 | 5.236497473 | 3.478770384 | 1.27E-02 |
| Gm6133 | 18.62084645 | 3.472863913 | 1.95E-07 |
| Plaur | 1149.588067 | 3.463302878 | 3.09E-39 |
| Stox2 | 10.87126108 | 3.463177061 | 1.60E-06 |
| Cdsn | 1502.659716 | 3.458552919 | 1.69E-50 |
| Scn3a | 16.94562687 | 3.456357632 | 7.19E-05 |
| Pkhd1l1 | 6.114660324 | 3.445134478 | 1.66E-02 |
| Pdzd2 | 406.3513246 | 3.443742688 | 8.13E-08 |
| Vmn1r43 | 19.18440843 | 3.438449854 | 6.19E-05 |
| Car14 | 32.99689159 | 3.43476861 | 4.48E-08 |
| Dnah7a | 5.729662646 | 3.429630455 | 1.78E-02 |
| Efhd1 | 6.477393889 | 3.427762995 | 1.81E-03 |
| Cryab | 13202.45602 | 3.421129239 | 2.06E-53 |
| Gm12037 | 11.05132918 | 3.417692173 | 1.85E-07 |
| Dusp14 | 1099.248265 | 3.412570038 | 1.14E-65 |
| Ppm1h | 131.7107471 | 3.39874403 | 1.89E-16 |
| Cacna1b | 7.340094445 | 3.37343782 | 1.45E-02 |
| Trim72 | 12.45876701 | 3.363146663 | 1.16E-05 |
| Gm12505 | 700.5275066 | 3.360274768 | 1.76E-24 |
| Nrap | 5.646324322 | 3.35664018 | 9.31E-03 |
| Vmn1r45 | 6.489789663 | 3.346145255 | 5.56E-04 |

| | | | |
|---|---|---|---|
| Chmp4c | 13.45073978 | 3.343112518 | 6.71E-07 |
| Gm16998 | 21.06629374 | 3.342078964 | 1.38E-10 |
| Hes7 | 30.3534822 | 3.340676815 | 5.45E-18 |
| Htatip2 | 7.892515834 | 3.34062179 | 8.48E-04 |
| Anxa3 | 7482.75502 | 3.338469542 | 4.85E-52 |
| Gm38393 | 571.0226571 | 3.33721716 | 2.25E-29 |
| Adgrd1 | 8.97434972 | 3.336880367 | 3.16E-03 |
| Tmem144 | 24.43046245 | 3.330314595 | 5.43E-09 |
| Usp17le | 14.67590836 | 3.32978295 | 1.08E-05 |
| Lzts1 | 36.27116963 | 3.329142046 | 2.82E-17 |
| Gm27406 | 15.93084718 | 3.328436134 | 3.55E-05 |
| 2200002D01Rik | 188.5911851 | 3.323780787 | 1.46E-43 |
| Vgf | 80.48610746 | 3.317599804 | 9.18E-25 |
| Hist1h2bc | 898.5619503 | 3.315838605 | 1.22E-15 |
| Igf2os | 450.3914571 | 3.313556856 | 4.85E-16 |
| Gch1 | 62.90561554 | 3.311096574 | 7.66E-19 |
| Lnx1 | 47.50474965 | 3.309616049 | 8.03E-12 |
| Dyrk4 | 47.6636991 | 3.304144736 | 1.83E-15 |
| Dusp8 | 245.7185379 | 3.299323333 | 6.00E-27 |
| NA | 63.62668047 | 3.298782181 | 1.50E-09 |
| Gm23849 | 8.016410947 | 3.290098574 | 9.39E-04 |
| Stxbp2 | 15.11401419 | 3.288183469 | 3.51E-05 |
| Diras2 | 135.873439 | 3.285611188 | 2.07E-18 |
| Pla2g5 | 14.77056158 | 3.276175525 | 2.18E-03 |
| Wnt9a | 916.1471833 | 3.271803539 | 4.84E-47 |
| Mab21l3 | 6.546200284 | 3.270761633 | 2.67E-03 |
| Gm16531 | 8.914604206 | 3.269825594 | 2.52E-02 |
| Lamc2 | 395.7838555 | 3.24719328 | 9.50E-35 |
| Gm12415 | 16.9163411 | 3.240296167 | 5.47E-08 |
| Gpr39 | 133.0862302 | 3.234748578 | 2.11E-16 |
| Gabre | 296.5455035 | 3.233078419 | 3.99E-42 |
| Cited2 | 5203.839702 | 3.225600958 | 4.65E-124 |
| Gm11380 | 56.60624028 | 3.222495159 | 4.74E-16 |
| Pgk1-rs7 | 137.6276523 | 3.222288364 | 3.21E-02 |
| Il3ra | 21.87015892 | 3.221428522 | 2.16E-08 |
| Prrt3 | 27.4990097 | 3.21996353 | 4.55E-12 |
| Gm6166 | 5392.037064 | 3.216835212 | 9.47E-65 |
| Tmem266 | 7.99764906 | 3.2162134 | 2.35E-04 |
| H19 | 1822.874962 | 3.215707387 | 2.51E-05 |
| Prrt2 | 13.99413336 | 3.215618304 | 1.61E-02 |
| Mylpf | 41.0963031 | 3.213650802 | 1.31E-17 |
| Gm11331 | 59.22672314 | 3.199657859 | 1.16E-07 |
| Gm11520 | 9.983968603 | 3.197033761 | 7.99E-07 |
| Ephx1 | 9309.25785 | 3.1951804 | 5.62E-57 |
| Klra1 | 6.79691526 | 3.19154549 | 3.99E-02 |
| Insl6 | 38.19475117 | 3.190504122 | 2.52E-19 |
| Ppp1r32 | 13.45526339 | 3.184847913 | 1.04E-05 |

| | | | |
|---|---|---|---|
| Hoxb6 | 11.04812908 | 3.181621783 | 3.71E-04 |
| Spns3 | 13.83643692 | 3.181469928 | 2.13E-04 |
| Mbp | 219.5097508 | 3.17718839 | 1.08E-31 |
| Fabp5 | 6643.323886 | 3.172041348 | 6.90E-72 |
| Kctd4 | 12.52468892 | 3.158141172 | 1.18E-03 |
| Disp2 | 132.6711548 | 3.157896123 | 6.73E-33 |
| Nkpd1 | 5.186491559 | 3.149019149 | 2.04E-02 |
| Crabp2 | 68.19681442 | 3.14626525 | 1.53E-18 |
| Txk | 7.539374443 | 3.143678133 | 1.96E-03 |
| Klhl40 | 24.70386697 | 3.142862723 | 4.56E-10 |
| Tg | 42.06323385 | 3.137574554 | 2.23E-05 |
| Krt88 | 15.95000092 | 3.133249924 | 4.85E-06 |
| Tmem37 | 109.5363239 | 3.127086332 | 1.76E-27 |
| Ihh | 7.218110808 | 3.1214485 | 3.90E-04 |
| Psrc1 | 2282.570702 | 3.110225771 | 2.17E-41 |
| NA | 15.15424967 | 3.10781879 | 1.08E-05 |
| 1700013F07Rik | 7.109137326 | 3.107108602 | 3.40E-03 |
| Sh3tc2 | 46.01764535 | 3.105640893 | 4.08E-13 |
| Hist1h2ae | 5.952941222 | 3.102720862 | 9.94E-04 |
| Cgn | 236.0254742 | 3.096697515 | 3.08E-18 |
| Smyd1 | 7.587107186 | 3.096118512 | 1.68E-03 |
| Gm11525 | 36.80127275 | 3.096089926 | 6.94E-14 |
| Shisa6 | 8.904196846 | 3.09553637 | 2.31E-03 |
| 4930486L24Rik | 435.5390485 | 3.091059936 | 1.36E-17 |
| Ifit1bl1 | 78.34996578 | 3.086164498 | 2.00E-15 |
| Tmem130 | 8.431316085 | 3.081429124 | 4.20E-05 |
| Chrnb1 | 351.1485445 | 3.079336608 | 6.52E-23 |
| Arrdc4 | 1906.491093 | 3.078539296 | 1.46E-19 |
| Afap1l2 | 7.391967593 | 3.072773016 | 2.59E-02 |
| Plcd3 | 1876.451148 | 3.072230847 | 1.46E-36 |
| Thsd7b | 12.27170345 | 3.0704938 | 2.11E-04 |
| Kank4os | 15.91605683 | 3.05985255 | 2.97E-06 |
| Syt17 | 46.84650688 | 3.055839181 | 1.41E-10 |
| Prrg2 | 490.5911814 | 3.05485047 | 7.53E-40 |
| Nrcam | 42.60013369 | 3.048374444 | 1.56E-11 |
| Gdnf | 878.6590879 | 3.044743997 | 1.47E-36 |
| Cth | 393.5185974 | 3.043678691 | 3.02E-06 |
| Hes1 | 186.8574295 | 3.032542572 | 2.75E-09 |
| 2310061I04Rik | 1207.433837 | 3.019903822 | 1.14E-65 |
| Clcnkb | 11.32601771 | 3.019551274 | 1.09E-02 |
| Snhg15 | 979.4343226 | 3.018003328 | 6.53E-48 |
| Mmel1 | 15.80909654 | 3.016538484 | 2.32E-07 |
| Ngf | 909.5389495 | 3.013326789 | 2.96E-44 |
| Brsk2 | 7.690449615 | 3.002818643 | 6.38E-04 |
| Apob | 19.03152808 | 3.00067297 | 1.04E-03 |
| 4933406C10Rik | 13.8312681 | 2.999745459 | 3.07E-07 |
| Gm14057 | 34.74908516 | 2.988768098 | 1.95E-14 |

| | | | |
|---|---|---|---|
| Gm13713 | 5.072477726 | 2.9845077 | 3.69E-02 |
| Gm26867 | 112.2321674 | 2.980728485 | 3.47E-14 |
| Fgd6 | 331.6053799 | 2.978064153 | 1.63E-28 |
| NA | 5.21045359 | 2.977018493 | 4.86E-03 |
| Trpm1 | 12.59951623 | 2.976760527 | 2.26E-06 |
| Cd96 | 6.045846529 | 2.956154787 | 1.08E-03 |
| Ankrd1 | 4164.102927 | 2.955904399 | 4.18E-20 |
| Slco3a1 | 1020.790683 | 2.941581207 | 1.73E-61 |
| Cbx3-ps7 | 16.14635048 | 2.93918338 | 2.84E-05 |
| Myrfl | 5.472787012 | 2.938878952 | 7.27E-03 |
| Dusp18 | 16.55368329 | 2.937466926 | 7.47E-05 |
| Rhpn2 | 103.2238498 | 2.931996869 | 1.27E-16 |
| Gm6311 | 19.58354765 | 2.929129604 | 3.05E-04 |
| Lmntd2 | 7.734375031 | 2.928476324 | 6.74E-04 |
| Prickle4 | 8.014106294 | 2.925553695 | 1.78E-04 |
| Myh11 | 10.27823204 | 2.922654206 | 9.06E-04 |
| Pde1c | 5.536475538 | 2.921705281 | 9.71E-03 |
| Spdef | 6.009277668 | 2.918456967 | 2.34E-03 |
| Eif2c5 | 9.618557942 | 2.916107368 | 2.85E-02 |
| Efcab8 | 17.7494641 | 2.9123173 | 7.36E-06 |
| Rspo4os | 11.50713663 | 2.911541885 | 1.33E-05 |
| Pvr | 3501.591273 | 2.902064589 | 1.65E-59 |
| Lrrn4cl | 2335.883644 | 2.89716846 | 7.60E-23 |
| 1700028J19Rik | 26.72022137 | 2.888180144 | 2.33E-12 |
| Gm26651 | 25.61219024 | 2.882729054 | 3.51E-06 |
| Cpz | 10.48574976 | 2.875496154 | 3.08E-03 |
| Map3k9 | 11.53418457 | 2.873709618 | 8.03E-04 |
| Gsdmc4 | 5.219483731 | 2.870679957 | 2.11E-03 |
| Arhgef28 | 758.8625099 | 2.869634755 | 4.66E-60 |
| 3830403N18Rik | 23.80216124 | 2.867787894 | 3.44E-10 |
| Snord55 | 6.834009265 | 2.867627842 | 5.30E-04 |
| Fas | 523.473713 | 2.866178684 | 7.20E-38 |
| Vpreb1 | 24.29579557 | 2.858692233 | 2.83E-13 |
| Camk2a | 97.06394336 | 2.858174802 | 1.57E-13 |
| Kcp | 5.256357168 | 2.855996848 | 3.85E-03 |
| Hspb2 | 59.28042065 | 2.85563869 | 4.18E-22 |
| Txlnb | 29.58797101 | 2.851222329 | 5.28E-07 |
| Aaas | 2372.654229 | 2.85084109 | 5.85E-32 |
| B4galt4 | 584.3450299 | 2.849739186 | 4.01E-82 |
| Cyp2c55 | 890.3392128 | 2.849687887 | 1.71E-46 |
| Trap1a | 12.33813831 | 2.844690189 | 2.06E-04 |
| Ppp1r18os | 8.829648244 | 2.844021945 | 4.48E-05 |
| Rapgef3os2 | 8.64117429 | 2.839081604 | 1.08E-03 |
| Eno3 | 744.5122267 | 2.835870535 | 5.28E-22 |
| Stard8 | 6.074829507 | 2.83175855 | 2.30E-02 |
| Catsperd | 26.54257683 | 2.827338203 | 6.11E-08 |
| Hoxb3 | 9.193710774 | 2.826186507 | 1.46E-03 |

| | | | |
|---|---|---|---|
| Pla2g4b | 67.61780263 | 2.824343534 | 2.83E-08 |
| Vmn1r40 | 12.07000502 | 2.816747543 | 1.14E-03 |
| Papss2 | 2566.376957 | 2.815544312 | 4.91E-79 |
| Cgnl1 | 1941.398104 | 2.811015576 | 1.56E-15 |
| Arl4d | 342.751126 | 2.810009656 | 5.19E-18 |
| Igf2bp1 | 2960.658169 | 2.804687681 | 2.14E-181 |
| Lrrc75a | 924.7491795 | 2.804584027 | 4.27E-95 |
| Hist1h2ac | 24.74191821 | 2.802511606 | 2.59E-06 |
| Fam71f2 | 47.87919002 | 2.802077583 | 4.68E-09 |
| Hbegf | 543.5379025 | 2.80196911 | 4.94E-31 |
| Gdf5 | 22.57180371 | 2.80176263 | 7.24E-05 |
| Dglucy | 1002.104884 | 2.801446214 | 2.38E-48 |
| Klra2 | 109.4765134 | 2.799484132 | 2.60E-13 |
| Hoxb3os | 9.42460263 | 2.799134887 | 4.80E-03 |
| E130218I03Rik | 17.73443953 | 2.795775451 | 6.79E-04 |
| Hist1h2bk | 54.97336704 | 2.794786611 | 4.19E-06 |
| Colec11 | 15.15317968 | 2.792619141 | 1.68E-05 |
| Zfp341 | 461.7226419 | 2.789843244 | 7.10E-39 |
| Dennd2d | 8.919399004 | 2.787098153 | 7.85E-03 |
| Igf2 | 11022.73197 | 2.785870184 | 5.72E-06 |
| Olfr1318 | 12.8937525 | 2.783936929 | 6.55E-04 |
| Heph | 11.43738456 | 2.77134803 | 1.80E-03 |
| Cfap57 | 7.191033356 | 2.770331851 | 1.31E-03 |
| Parvg | 9.60610989 | 2.770077754 | 2.39E-03 |
| Ddah1 | 1706.491516 | 2.767189296 | 2.45E-22 |
| Ddias | 367.2417911 | 2.765523762 | 5.68E-13 |
| Stbd1 | 242.832905 | 2.765273647 | 3.58E-27 |
| BC051665 | 8.5578041 | 2.762488902 | 4.50E-04 |
| Stc2 | 605.8941595 | 2.757522078 | 1.11E-22 |
| Gm12185 | 28.38347216 | 2.753955 | 2.18E-05 |
| Gm11944 | 37.06617334 | 2.747071471 | 6.27E-12 |
| Avpi1 | 1533.78618 | 2.746099126 | 1.42E-84 |
| Gm26574 | 14.54837472 | 2.744696092 | 1.27E-05 |
| Bcar3 | 810.1377157 | 2.741472057 | 1.45E-42 |
| Hacd4 | 175.9725215 | 2.740957117 | 9.66E-18 |
| Tnnt2 | 43.75984317 | 2.735295767 | 6.64E-05 |
| Grb14 | 1511.859625 | 2.734673331 | 5.23E-32 |
| Prss36 | 320.8623567 | 2.732097332 | 1.72E-34 |
| Zfp941 | 7.943074269 | 2.728601193 | 6.11E-03 |
| Hist2h2aa2 | 243.0853257 | 2.721732466 | 1.73E-08 |
| Pnpla2 | 1540.394814 | 2.719983578 | 1.19E-41 |
| Susd2 | 15.13366232 | 2.712633354 | 9.54E-03 |
| Procr | 826.0507845 | 2.707209269 | 2.50E-38 |
| Hyal3 | 30.56485389 | 2.705850428 | 5.58E-08 |
| Nfam1 | 6.77097658 | 2.704944973 | 3.48E-03 |
| Gipc2 | 16.09109545 | 2.703392503 | 3.86E-06 |
| Gpr68 | 23.11995535 | 2.70333358 | 6.45E-06 |

| | | | |
|---|---|---|---|
| Platr14 | 62.69361273 | 2.701669359 | 1.65E-15 |
| Lipg | 535.1730908 | 2.700324452 | 2.69E-35 |
| Nectin1 | 1412.399235 | 2.698304712 | 5.21E-56 |
| Apol9b | 84.6420391 | 2.696478328 | 2.95E-11 |
| Flnc | 35973.77426 | 2.6949593 | 1.71E-57 |
| Pim3 | 2043.372491 | 2.694566539 | 1.68E-31 |
| Gm26885 | 5.436629015 | 2.694025597 | 4.99E-03 |
| Gm12138 | 2019.192784 | 2.689086525 | 5.74E-19 |
| NA | 5.722001699 | 2.688037719 | 7.84E-04 |
| Gm17090 | 14.75252409 | 2.687848248 | 6.51E-04 |
| Ifnlr1 | 8.948726448 | 2.686038752 | 2.16E-02 |
| Bmp6 | 5.901513132 | 2.68441513 | 1.01E-02 |
| Tnfrsf10b | 1728.080669 | 2.683818355 | 1.47E-49 |
| Sema6b | 29.71095728 | 2.680703044 | 9.71E-08 |
| Matn3 | 9.122129289 | 2.678368537 | 7.87E-04 |
| Ass1 | 199.8319632 | 2.678250899 | 4.81E-23 |
| Gm6292 | 5.549756566 | 2.677579537 | 5.08E-03 |
| Gm16166 | 9.035890847 | 2.676859499 | 2.26E-03 |
| Tigit | 30.28802111 | 2.673499873 | 3.49E-09 |
| Tbc1d2 | 1016.924895 | 2.672091477 | 2.32E-33 |
| Gm10639 | 17.14337393 | 2.667177178 | 8.42E-05 |
| Cst6 | 60.21863085 | 2.666291589 | 1.37E-09 |
| Gja1 | 1405.954553 | 2.663296786 | 1.79E-15 |
| Btg2 | 1025.665619 | 2.661773075 | 4.53E-20 |
| 1110002E22Rik | 5.899518649 | 2.660336415 | 4.09E-02 |
| Gm11642 | 21.13040859 | 2.654264977 | 4.71E-09 |
| Otud1 | 340.3620018 | 2.652738071 | 5.34E-36 |
| 1700125H20Rik | 6.112302566 | 2.639712694 | 2.16E-03 |
| Bbc3 | 624.0895791 | 2.629067398 | 6.19E-41 |
| Serpina9 | 31.42166562 | 2.627480506 | 3.01E-08 |
| Aqp5 | 257.8230496 | 2.623841597 | 1.21E-17 |
| Muc2 | 8.809115183 | 2.623520037 | 5.27E-04 |
| Cstl1 | 15.00122308 | 2.622966125 | 4.53E-05 |
| Rbmy | 70.47372177 | 2.617422491 | 5.19E-05 |
| S100a11 | 28004.48893 | 2.616165592 | 9.41E-36 |
| Ica1 | 26.09752958 | 2.615393859 | 1.90E-05 |
| Gm26397 | 15.37497149 | 2.612576921 | 8.92E-06 |
| Asns | 8358.728063 | 2.612411373 | 3.13E-32 |
| Erc2 | 648.4103087 | 2.612404533 | 1.92E-53 |
| Gm10499 | 78.66015587 | 2.610527715 | 7.48E-22 |
| Inka2 | 122.816321 | 2.609206613 | 4.49E-10 |
| Atf3 | 486.7080334 | 2.605339197 | 1.43E-34 |
| Dlc1 | 2597.470192 | 2.603403578 | 9.88E-61 |
| 4833427G06Rik | 113.596248 | 2.598754696 | 2.26E-28 |
| Jmjd7 | 318.0362651 | 2.596699869 | 8.23E-36 |
| Akr1b3 | 27184.12226 | 2.593744214 | 2.08E-16 |
| Tmem43 | 6334.664426 | 2.593167216 | 1.24E-41 |

| | | | |
|---|---|---|---|
| Gm10230 | 10.1897906 | 2.590978943 | 3.18E-02 |
| Gm26708 | 7.912831015 | 2.589436434 | 3.28E-03 |
| Atg16l2 | 1186.275372 | 2.589181487 | 6.66E-18 |
| Fam110a | 459.3283501 | 2.584909905 | 1.25E-26 |
| Fam71f1 | 49.35392991 | 2.583719298 | 2.34E-08 |
| Steap1 | 1192.998083 | 2.581074537 | 1.70E-18 |
| Ms4a4d | 289.8013129 | 2.578870403 | 2.14E-09 |
| Gtse1 | 869.2872648 | 2.578510477 | 2.76E-13 |
| 2310043M15Rik | 5.025728522 | 2.575935963 | 3.82E-02 |
| Gm15694 | 10.95927481 | 2.571139571 | 4.32E-05 |
| Plcxd2 | 1602.480094 | 2.569746532 | 7.70E-54 |
| Rin1 | 4778.34829 | 2.567136714 | 1.97E-66 |
| Podnl1 | 3817.279448 | 2.564354488 | 1.35E-57 |
| Cfap43 | 26.34884187 | 2.564240831 | 1.87E-05 |
| Cltb | 2400.270914 | 2.562323333 | 4.60E-39 |
| Dgat2 | 90.86863884 | 2.558511461 | 1.69E-18 |
| Ehd1 | 9278.785003 | 2.558012858 | 1.82E-45 |
| Zfp36 | 1397.592095 | 2.555269675 | 2.98E-15 |
| Gm15264 | 33.15051985 | 2.549479781 | 6.73E-09 |
| Jag1 | 1491.58118 | 2.543331677 | 2.48E-27 |
| NA | 5457.007199 | 2.541729163 | 1.36E-14 |
| Rhox5 | 444.7664169 | 2.540091195 | 4.73E-21 |
| Ttll7 | 695.451499 | 2.539929085 | 3.81E-41 |
| Angptl6 | 251.3102899 | 2.537216054 | 1.03E-24 |
| Gm13285 | 29.53639999 | 2.536900074 | 4.61E-08 |
| Usp2 | 268.4067797 | 2.535999001 | 1.16E-14 |
| Prss23os | 1155.334691 | 2.535663769 | 1.40E-22 |
| Kcnab3 | 152.1664527 | 2.534496565 | 6.68E-21 |
| Ablim1 | 3555.853548 | 2.531653639 | 2.42E-35 |
| 1110046J04Rik | 236.2657601 | 2.531344963 | 2.56E-22 |
| Has1 | 15.99957924 | 2.530864664 | 5.01E-03 |
| Foxf1 | 338.939874 | 2.528495061 | 6.13E-61 |
| Gm12254 | 5.360247685 | 2.519768293 | 3.35E-02 |
| Slco2a1 | 4121.36187 | 2.51863383 | 3.84E-49 |
| Fhl1 | 107.2708123 | 2.518001915 | 6.71E-06 |
| Patl2 | 5.171266621 | 2.514983356 | 1.55E-02 |
| Tslp | 20.98326568 | 2.512592642 | 1.78E-04 |
| Cct2 | 26032.40918 | 2.510756668 | 2.32E-82 |
| Mst1 | 6.564989527 | 2.508227622 | 3.34E-03 |
| Plac8 | 10.13342526 | 2.503814686 | 1.26E-03 |
| Abcb1b | 1465.604472 | 2.503671902 | 9.35E-51 |
| Cep295nl | 32.72658667 | 2.503472552 | 1.33E-10 |
| Maff | 1125.074098 | 2.501182101 | 2.54E-24 |
| Apold1 | 41.24765489 | 2.497586024 | 2.79E-13 |
| Nomo1 | 5376.942128 | 2.49354494 | 1.87E-40 |
| Gm5570 | 6.543427362 | 2.492027301 | 1.30E-02 |
| Adgrg6 | 18.66629187 | 2.491033553 | 1.84E-04 |

| | | | |
|---|---|---|---|
| Egln3 | 507.4247638 | 2.490604485 | 3.92E-40 |
| Rpl12 | 5438.515123 | 2.490271127 | 3.58E-30 |
| Gm13981 | 7.337667877 | 2.488238635 | 2.31E-03 |
| Gm14760 | 24.30930879 | 2.48698164 | 3.45E-04 |
| Gm5106 | 7.079855874 | 2.485781884 | 7.69E-03 |
| Klf10 | 2846.196887 | 2.481942531 | 2.30E-18 |
| Cep72 | 201.8607482 | 2.478794211 | 1.14E-34 |
| Fap | 55.22080039 | 2.478182968 | 5.65E-05 |
| Lrrc31 | 14.73142716 | 2.476523851 | 7.29E-03 |
| Npr3 | 1290.290208 | 2.473911864 | 1.12E-09 |
| Oplah | 158.7571282 | 2.472783313 | 2.23E-20 |
| Kcnu1 | 19.04652871 | 2.47212558 | 1.14E-03 |
| Lrp2bp | 10.02509851 | 2.47079966 | 4.19E-04 |
| Gm4654 | 5.85885103 | 2.464679488 | 3.71E-03 |
| Pfkfb3 | 882.241222 | 2.46372348 | 3.98E-18 |
| Aqp6 | 6.596852005 | 2.463449907 | 7.20E-03 |
| Pram1 | 33.24727736 | 2.462495224 | 1.72E-04 |
| Gm26543 | 9.395393219 | 2.454778686 | 4.01E-03 |
| Higd2a | 2214.016855 | 2.454572645 | 6.65E-72 |
| Prdx6 | 12709.33805 | 2.453489385 | 5.09E-39 |
| Gm16170 | 35.38246777 | 2.453247012 | 2.10E-07 |
| 1700047I17Rik2 | 769.1116899 | 2.449088573 | 6.02E-61 |
| Per1 | 5421.880749 | 2.448601867 | 7.58E-17 |
| Adgrf5 | 8.050799548 | 2.447612957 | 4.08E-02 |
| Plch2 | 6.803676316 | 2.445453628 | 1.96E-02 |
| Piwil2 | 15.37645003 | 2.445215794 | 2.06E-03 |
| Soat2 | 50.10265489 | 2.441890858 | 1.10E-08 |
| Hist1h2bq | 18.51075948 | 2.441253223 | 2.12E-03 |
| Hspb6 | 825.7361147 | 2.440405623 | 1.32E-22 |
| Nlrp4e | 17.54639026 | 2.438575717 | 5.89E-03 |
| Unc5c | 5.83703364 | 2.434651427 | 1.82E-02 |
| Plk2 | 10524.64627 | 2.431687216 | 6.19E-65 |
| Rpl3l | 10.45497779 | 2.431410391 | 4.82E-03 |
| Nhej1 | 239.237029 | 2.423982792 | 1.17E-28 |
| Gm26809 | 35780.51269 | 2.423893409 | 4.99E-15 |
| Rmrp | 8.643832047 | 2.423392275 | 1.38E-02 |
| Mafk | 2125.926029 | 2.419815473 | 5.20E-29 |
| Lyrm9 | 234.5217743 | 2.417930936 | 3.49E-21 |
| Rps2-ps6 | 64.97681542 | 2.417275093 | 1.60E-09 |
| Bst2 | 1204.425443 | 2.414421511 | 2.64E-21 |
| Nectin4 | 8.839627048 | 2.409765896 | 5.19E-03 |
| C130074G19Rik | 6.267890918 | 2.406246677 | 4.48E-03 |
| St18 | 9.877112847 | 2.399138246 | 9.11E-03 |
| Nhlrc2 | 1888.903339 | 2.39853137 | 4.99E-116 |
| Jag2 | 9.330326422 | 2.396485186 | 4.57E-04 |
| Crx | 15.38764971 | 2.39614404 | 3.65E-03 |
| Cnn1 | 11.15129988 | 2.388694867 | 2.94E-04 |

| | | | |
|---|---|---|---|
| Gm15246 | 19.65445391 | 2.387699299 | 5.03E-06 |
| Gm1141 | 15.33114807 | 2.385685618 | 7.79E-05 |
| Abca7 | 2260.675367 | 2.383539643 | 1.96E-49 |
| Gm6139 | 18.06609433 | 2.378293479 | 3.00E-04 |
| Il15ra | 123.6384944 | 2.377650626 | 3.74E-12 |
| Itga6 | 3086.584435 | 2.377426136 | 8.77E-54 |
| Tspan13 | 14.97697967 | 2.376684121 | 1.66E-05 |
| Serpini1 | 53.62556717 | 2.375166432 | 1.44E-13 |
| Hspbap1 | 670.3335746 | 2.371269604 | 2.18E-71 |
| Gm11599 | 19.48675284 | 2.371149701 | 1.83E-05 |
| Lenep | 70.28657422 | 2.370748541 | 3.57E-06 |
| Slc35e4 | 1871.809364 | 2.370273865 | 9.93E-33 |
| Pcdh12 | 5.129888463 | 2.369898534 | 1.55E-02 |
| Hcfc1r1 | 8234.320913 | 2.3697949 | 5.85E-29 |
| Uck2 | 2613.500918 | 2.367657717 | 6.54E-49 |
| Hecw2 | 21.57444205 | 2.358270043 | 1.39E-02 |
| Tlr1 | 42.23168809 | 2.358122174 | 2.29E-09 |
| Hist1h2al | 73.64496331 | 2.357711531 | 5.08E-09 |
| Odc1 | 6015.135819 | 2.35471777 | 5.95E-66 |
| Tuft1 | 1030.058028 | 2.35222253 | 3.87E-27 |
| Golga7b | 16.54206909 | 2.351449308 | 7.13E-05 |
| Grin3a | 5.74326454 | 2.346029832 | 1.88E-02 |
| Glns-ps1 | 14.7088613 | 2.345389983 | 9.92E-05 |
| Dnah6 | 60.00583954 | 2.344138717 | 1.32E-09 |
| Fry | 21.32990924 | 2.343474586 | 4.78E-05 |
| Ltc4s | 33.90047656 | 2.342224709 | 3.49E-05 |
| Ccdc180 | 8.668083621 | 2.340144839 | 3.12E-02 |
| Als2cr12 | 58.85300664 | 2.33366084 | 3.02E-09 |
| Ccng1 | 10250.65217 | 2.33016922 | 2.19E-51 |
| Rras2 | 2707.729271 | 2.330136974 | 3.06E-36 |
| 9130008F23Rik | 333.9612646 | 2.32946099 | 9.03E-29 |
| Ccdc78 | 27.35552252 | 2.328643492 | 3.67E-08 |
| Slc43a3 | 2964.656793 | 2.321488824 | 6.46E-21 |
| Ndufaf6 | 307.0560471 | 2.31948351 | 4.30E-33 |
| Rgs17 | 87.00021087 | 2.317808486 | 1.18E-09 |
| Pnkp | 1016.334908 | 2.317069253 | 1.73E-51 |
| Cgref1 | 204.5506992 | 2.314449888 | 7.17E-21 |
| Adrb2 | 121.6545444 | 2.308772025 | 6.54E-20 |
| Lta | 33.84052544 | 2.307670638 | 4.17E-08 |
| Acox2 | 9.843345696 | 2.30528264 | 9.29E-03 |
| Fstl3 | 1713.818872 | 2.305238686 | 3.70E-29 |
| Pbld2 | 34.86217392 | 2.300516382 | 1.31E-05 |
| Thoc6 | 1074.782664 | 2.296439882 | 3.30E-52 |
| Kif5a | 70.92537186 | 2.294073369 | 1.92E-09 |
| Ikzf2 | 190.7222877 | 2.293682801 | 2.47E-09 |
| Tnnc1 | 109.1518449 | 2.293050858 | 3.80E-19 |
| Ace2 | 6.182647258 | 2.291785789 | 9.09E-03 |

| | | | |
|---|---|---|---|
| Eif3d | 11371.15065 | 2.29092207 | 3.87E-45 |
| Klf4 | 2395.995965 | 2.29026407 | 3.47E-36 |
| Aph1b | 607.0118737 | 2.289994117 | 8.62E-40 |
| Nfkb2 | 5751.40937 | 2.286799447 | 4.62E-43 |
| Esd | 14951.22292 | 2.285081119 | 8.87E-40 |
| Dyrk3 | 209.7965907 | 2.283432503 | 9.99E-28 |
| Usp17la | 30.31757862 | 2.281305469 | 3.11E-07 |
| Mfge8 | 10440.85802 | 2.278450896 | 1.54E-27 |
| Alx4 | 5.88386325 | 2.277267972 | 1.25E-02 |
| D630029K05Rik | 40.79021596 | 2.276292432 | 1.88E-08 |
| Lag3 | 35.71725018 | 2.272730901 | 1.80E-08 |
| Pmm1 | 3567.219961 | 2.272486977 | 5.86E-36 |
| Mslnl | 8.191156947 | 2.271933707 | 6.06E-03 |
| Gm25053 | 5.004406347 | 2.271493418 | 1.52E-02 |
| Tspan9 | 8905.533771 | 2.266189671 | 5.73E-46 |
| Thy1 | 3489.688816 | 2.264901209 | 2.72E-28 |
| Slc9a3r1 | 541.036435 | 2.262680214 | 9.41E-22 |
| Styk1 | 452.3674977 | 2.259849636 | 3.02E-14 |
| Dlk2 | 145.1833932 | 2.259732797 | 4.80E-10 |
| Epha2 | 1944.113419 | 2.259008888 | 2.18E-14 |
| Fitm2 | 1939.475976 | 2.258749172 | 2.14E-41 |
| Phactr2 | 110.9007105 | 2.257767362 | 5.92E-09 |
| Mcub | 378.0966601 | 2.257683921 | 4.18E-28 |
| Gm16062 | 85.00001513 | 2.25704479 | 2.37E-10 |
| Igfbp6 | 10099.86965 | 2.256439479 | 5.17E-33 |
| Slc1a2 | 18.64910711 | 2.256429002 | 2.42E-02 |
| Mycl | 20.21135286 | 2.254826018 | 3.33E-04 |
| Nyap1 | 908.6854763 | 2.252618421 | 6.46E-29 |
| 5-Sep | 7626.58109 | 2.25245095 | 4.51E-41 |
| Optc | 22.31321699 | 2.252256409 | 2.39E-05 |
| Synpo2 | 219.9091425 | 2.25037298 | 3.56E-18 |
| Ap2a2 | 11186.22479 | 2.24952646 | 2.33E-72 |
| AA467197 | 26.56671516 | 2.246513352 | 9.46E-08 |
| Map3k20 | 2221.020669 | 2.245608357 | 4.12E-59 |
| Syngr4 | 10.875277 | 2.245424204 | 1.43E-03 |
| Gm27206 | 7.56169608 | 2.244602802 | 3.64E-03 |
| Clpb | 1889.540012 | 2.2435045 | 2.04E-28 |
| Lgals9 | 3984.469823 | 2.241801787 | 5.17E-39 |
| Slc43a2 | 56.28890673 | 2.24073076 | 1.34E-06 |
| Amotl2 | 15490.23543 | 2.24014888 | 6.93E-36 |
| Tnfrsf21 | 266.5152627 | 2.240009152 | 3.54E-25 |
| Nabp2 | 2302.580906 | 2.238801971 | 9.82E-47 |
| Prr11 | 1150.453785 | 2.234917534 | 6.46E-15 |
| Pdgfb | 7.20281354 | 2.234750278 | 4.80E-02 |
| 4933427I04Rik | 39.54466148 | 2.232521461 | 1.17E-09 |
| Gm15708 | 52.21141378 | 2.231223932 | 8.04E-10 |
| Tmprss6 | 29.40343431 | 2.225586476 | 1.00E-04 |

| | | | |
|---|---|---|---|
| Amy1 | 9.16464884 | 2.223415046 | 4.85E-03 |
| Il11 | 23.34212275 | 2.220974805 | 2.45E-07 |
| Blvrb | 2497.902833 | 2.217931507 | 1.97E-35 |
| Sorbs1 | 833.0862304 | 2.216710194 | 2.40E-35 |
| Rps2-ps10 | 66191.96649 | 2.216239906 | 1.05E-48 |
| Gm10291 | 6.772280103 | 2.21497978 | 1.58E-02 |
| Spire1 | 861.3974626 | 2.214562331 | 2.81E-42 |
| Brsk1 | 879.0230187 | 2.211870575 | 1.62E-39 |
| Ybx3 | 12605.60181 | 2.211146003 | 2.64E-44 |
| Pold4 | 1270.663176 | 2.208252881 | 9.59E-27 |
| Nek2 | 1268.788324 | 2.20633726 | 7.73E-13 |
| Selp | 38.63983632 | 2.204230697 | 1.24E-02 |
| Muc13 | 7.743443714 | 2.200820109 | 8.05E-03 |
| Arl14epl | 50.54681463 | 2.198890238 | 3.03E-10 |
| Alkbh2 | 215.7559227 | 2.197625204 | 9.05E-40 |
| Emx2 | 282.5000379 | 2.196685262 | 5.17E-19 |
| Pgf | 5.848257713 | 2.195808245 | 3.06E-02 |
| Stambpl1 | 1280.981375 | 2.195040707 | 8.14E-36 |
| Rpl10a | 10385.15404 | 2.193175775 | 2.29E-23 |
| Gm6736 | 924.2982575 | 2.190923313 | 4.06E-10 |
| Wwc1 | 1254.514596 | 2.19032379 | 8.29E-20 |
| Taldo1 | 5508.674467 | 2.188543768 | 1.33E-32 |
| Dusp1 | 2323.364801 | 2.188539137 | 2.95E-13 |
| Ulbp1 | 1078.85357 | 2.186529808 | 1.03E-16 |
| Cd74 | 18.46669239 | 2.186234928 | 8.00E-03 |
| Rnf43 | 74.77146117 | 2.185940918 | 1.65E-09 |
| Tuba1b | 52546.48143 | 2.185669591 | 5.49E-16 |
| Hecw1 | 7.383878072 | 2.185574183 | 8.57E-03 |
| Pcsk4 | 254.3779611 | 2.185041506 | 1.64E-37 |
| Grk5 | 918.5251472 | 2.184340775 | 3.41E-19 |
| Wdr45 | 2171.76663 | 2.181393014 | 1.37E-42 |
| Mif | 7576.735311 | 2.179210075 | 3.46E-31 |
| Gm15730 | 7.292559368 | 2.179077357 | 4.64E-03 |
| Gm15541 | 42.64852593 | 2.178494478 | 1.23E-10 |
| Kcnj15 | 706.1352588 | 2.178361445 | 1.20E-09 |
| Snupn | 481.3671527 | 2.177663714 | 1.88E-93 |
| Relt | 217.4731898 | 2.176402229 | 8.02E-26 |
| Stim1 | 2475.820194 | 2.175342413 | 4.01E-47 |
| Fbxo15 | 9.25004082 | 2.172341684 | 7.89E-03 |
| 1700015F17Rik | 5.549174145 | 2.168651644 | 2.27E-02 |
| Sphk1 | 627.9533835 | 2.167618274 | 2.05E-13 |
| Gm2a | 1019.030594 | 2.165625015 | 1.02E-41 |
| Prss23 | 12096.74038 | 2.165508134 | 2.40E-39 |
| Ppfia3 | 146.6688839 | 2.16439577 | 1.10E-21 |
| Eef2 | 390505.9252 | 2.16101469 | 3.16E-25 |
| Fendrr | 16.4166356 | 2.159195128 | 1.10E-03 |
| Gm14659 | 5.051219153 | 2.159055719 | 4.60E-02 |

| | | | |
|---|---|---|---|
| Plekhn1 | 640.7350483 | 2.158411428 | 1.55E-17 |
| Gm10566 | 25.43802156 | 2.154951742 | 3.66E-06 |
| Tubb4a | 66.11470353 | 2.154299164 | 5.38E-14 |
| Fam189b | 38.7043353 | 2.153621324 | 3.36E-07 |
| Lrwd1 | 1708.037956 | 2.150302294 | 4.06E-37 |
| Rab11fip1 | 1416.019755 | 2.149486761 | 5.70E-36 |
| C3ar1 | 28.45683189 | 2.148613571 | 2.18E-03 |
| Gm5620 | 281.5414567 | 2.14765002 | 1.44E-09 |
| Gm10420 | 383.0517429 | 2.145035738 | 3.84E-18 |
| Pfkp | 1611.630881 | 2.143772872 | 4.24E-23 |
| Myo1g | 7.060165232 | 2.139764377 | 4.74E-02 |
| Tmem132b | 17.53167815 | 2.139357049 | 6.70E-03 |
| Gm16982 | 30.26455867 | 2.13932662 | 6.06E-09 |
| Ccdc9b | 3998.146773 | 2.134502518 | 1.15E-66 |
| Inafm2 | 1281.565853 | 2.127238035 | 2.84E-95 |
| NA | 13.11091694 | 2.126482649 | 1.01E-03 |
| Gpank1 | 589.3973124 | 2.125894875 | 5.53E-19 |
| Prps1 | 1657.907258 | 2.124904209 | 1.21E-40 |
| Gm11263 | 10.2203104 | 2.124874973 | 8.07E-04 |
| Mtfr2 | 211.4222988 | 2.123733822 | 4.98E-08 |
| Rras | 2672.576845 | 2.12279315 | 6.15E-27 |
| Ppp1r2-ps4 | 7.020669578 | 2.122644291 | 2.21E-02 |
| Pstpip1 | 90.41301441 | 2.121654466 | 1.16E-09 |
| Tmem245 | 7288.006629 | 2.118063389 | 7.17E-53 |
| Chn2 | 8.322440648 | 2.116169384 | 3.35E-02 |
| D630023F18Rik | 6.989008852 | 2.115227459 | 4.80E-02 |
| Cacna1g | 708.8000386 | 2.110524416 | 4.57E-08 |
| Zfas1 | 5923.00628 | 2.10843349 | 1.98E-20 |
| Rps2-ps13 | 65962.345 | 2.104563567 | 8.44E-44 |
| Txnip | 34225.99991 | 2.103205334 | 9.44E-06 |
| Gm15510 | 5.627902167 | 2.102819574 | 3.55E-02 |
| Mief2 | 738.7242967 | 2.102529954 | 3.76E-30 |
| Nuak2 | 683.3125158 | 2.102521978 | 1.85E-30 |
| NA | 10.87827974 | 2.102298493 | 1.72E-02 |
| Atp8b5 | 13.2202444 | 2.101863657 | 4.04E-03 |
| Hspa12a | 126.8535222 | 2.101367967 | 1.30E-17 |
| Mfap3l | 581.8453732 | 2.101174391 | 1.30E-16 |
| Dcxr | 824.7511236 | 2.100136579 | 1.53E-19 |
| Rusc2 | 4030.034565 | 2.09836723 | 8.96E-20 |
| Gss | 1651.838005 | 2.097614344 | 1.22E-29 |
| Fat1 | 9394.675609 | 2.094005397 | 4.33E-22 |
| Dele1 | 1859.40767 | 2.092270436 | 5.09E-30 |
| Exoc6 | 416.7822632 | 2.091046555 | 6.22E-27 |
| Gm5921 | 644.8903188 | 2.090338255 | 3.60E-25 |
| Ptgis | 3691.451478 | 2.088980752 | 3.53E-20 |
| Ggct | 110.6567063 | 2.088766386 | 3.65E-13 |
| Cd151 | 4413.725542 | 2.088722909 | 2.02E-44 |

| | | | |
|---|---|---|---|
| Hist1h1d | 29.16335522 | 2.083262966 | 8.39E-06 |
| P2ry6 | 8.544628265 | 2.083222573 | 4.80E-02 |
| Slmapos2 | 3326.360358 | 2.081418728 | 2.99E-14 |
| Tspan15 | 23.84249731 | 2.081148978 | 5.04E-03 |
| Glul | 897.3710217 | 2.080766447 | 5.16E-16 |
| Rpl13a | 71987.69167 | 2.080203019 | 4.72E-27 |
| Rps2 | 1054.55721 | 2.075862643 | 8.69E-29 |
| Rasgrp3 | 117.9121061 | 2.073365901 | 3.91E-14 |
| Reep6 | 146.6378485 | 2.07192814 | 2.73E-21 |
| Gm11884 | 8.165736454 | 2.069960971 | 1.38E-02 |
| Ctps | 9423.67962 | 2.068434695 | 1.15E-35 |
| Stard13 | 1677.000687 | 2.065888646 | 1.14E-11 |
| Gm16061 | 14.26029884 | 2.065180091 | 1.21E-02 |
| Sord | 8476.138057 | 2.064782161 | 8.28E-18 |
| Gm6410 | 85.28209271 | 2.064372777 | 4.04E-13 |
| Ndufs2 | 12117.44549 | 2.064231214 | 2.40E-43 |
| Gm7665 | 5.938223775 | 2.062346335 | 2.40E-02 |
| Gm12933 | 53.05099324 | 2.059253526 | 3.70E-06 |
| Mucl3 | 111.7139975 | 2.058622919 | 1.48E-12 |
| Slc6a15 | 33.49232237 | 2.058222095 | 2.79E-04 |
| Hcrtr1 | 11.63728349 | 2.058160405 | 1.03E-03 |
| Lrrc1 | 506.016248 | 2.057552255 | 5.85E-35 |
| Ier3 | 749.7978944 | 2.057422184 | 1.53E-17 |
| Upk1a | 25.37899221 | 2.057350784 | 2.88E-06 |
| Zfp286os | 14.25886093 | 2.056964079 | 1.07E-04 |
| Trac | 20.1751909 | 2.055317031 | 1.03E-04 |
| Itpkc | 1053.756826 | 2.054657394 | 1.13E-22 |
| Cdkn2d | 517.3707037 | 2.053493479 | 3.02E-16 |
| Lama2 | 873.8408923 | 2.053094208 | 2.53E-12 |
| Gm11427 | 43.61658226 | 2.052995942 | 1.83E-07 |
| Cfap100 | 434.4066328 | 2.052150381 | 2.49E-15 |
| Vps36 | 2824.129634 | 2.05199704 | 1.98E-88 |
| Cdkl1 | 21.11717046 | 2.051496977 | 1.72E-04 |
| Fam241a | 67.45370295 | 2.051404211 | 2.30E-15 |
| Dbhos | 23.80023757 | 2.051162588 | 3.57E-05 |
| Gfer | 1246.79568 | 2.050535431 | 2.12E-40 |
| Nlrp4f | 12.24097274 | 2.049056381 | 5.14E-03 |
| Nmnat1 | 220.6102235 | 2.048728346 | 6.15E-45 |
| Nrm | 626.7617768 | 2.045590294 | 2.56E-13 |
| Ezr | 2125.964047 | 2.044551223 | 6.48E-48 |
| Ppp1r15a | 3481.359866 | 2.04315319 | 4.79E-35 |
| Slc25a25 | 821.5438689 | 2.043140197 | 4.07E-17 |
| Cdk18 | 2832.68387 | 2.041441687 | 7.36E-25 |
| Wdr12 | 1978.768705 | 2.041123921 | 1.18E-47 |
| Gm11496 | 7.041601801 | 2.037572827 | 2.47E-02 |
| Pcdh9 | 103.422577 | 2.036302972 | 3.87E-07 |
| Ints12 | 1233.993372 | 2.035023334 | 5.34E-32 |

| | | | |
|---|---|---|---|
| NA | 7.035009564 | 2.030203631 | 1.95E-02 |
| Cdiptos | 63.14051489 | 2.02893466 | 1.75E-09 |
| Ube2c | 1898.382155 | 2.02837565 | 1.05E-10 |
| Cct4 | 19364.54992 | 2.02765583 | 4.22E-67 |
| Unc13d | 41.0409234 | 2.026814448 | 1.90E-08 |
| Pde8b | 714.9937037 | 2.026572728 | 1.47E-20 |
| Elp6 | 429.5943132 | 2.026182405 | 7.55E-29 |
| Arrb2 | 918.8118723 | 2.025731914 | 8.23E-34 |
| Pdlim7 | 2855.17683 | 2.025664325 | 2.20E-38 |
| Tob1 | 562.3538878 | 2.024134438 | 3.77E-22 |
| Map6 | 3232.025626 | 2.023808969 | 8.62E-31 |
| Snhg6 | 1874.929114 | 2.022237175 | 4.80E-20 |
| Wt1 | 33.01804751 | 2.021143385 | 2.80E-03 |
| Dbndd2 | 321.9744503 | 2.021043244 | 2.46E-17 |
| Nop53 | 8144.379275 | 2.020257192 | 5.36E-18 |
| Tfec | 10.99540601 | 2.01816805 | 3.90E-02 |
| Aen | 3318.36924 | 2.0161137 | 8.80E-26 |
| Ttc41 | 109.4536472 | 2.015563053 | 5.41E-11 |
| C1qtnf2 | 17.27844736 | 2.014991584 | 3.30E-04 |
| Cdh17 | 219.8371349 | 2.014986962 | 1.17E-24 |
| Trim12a | 37.33199526 | 2.011701024 | 4.38E-07 |
| AW047730 | 11.98434172 | 2.010250548 | 2.72E-02 |
| Gm15228 | 7.647457266 | 2.010013031 | 1.29E-02 |
| Glrx3 | 6612.60578 | 2.008851266 | 3.89E-47 |
| Gm26788 | 40.96349029 | 2.007982352 | 2.38E-09 |
| Gm27021 | 16.6205354 | 2.006149149 | 3.29E-04 |
| Gm4609 | 20.16889004 | 2.005214938 | 9.28E-04 |
| Slc4a4 | 911.506914 | 2.004653826 | 7.43E-17 |
| Nalcn | 35.68450605 | 2.004093002 | 2.22E-04 |
| Mfsd9 | 752.6085643 | 2.002064687 | 2.07E-39 |
| Aph1c | 281.8841578 | 2.001673568 | 5.58E-17 |
| Pbx4 | 103.3083903 | 1.999203756 | 9.44E-17 |
| Gm10169 | 48.90505126 | 1.998767529 | 2.12E-06 |
| A630095E13Rik | 9.983925072 | 1.997162067 | 1.55E-02 |
| Gm7334 | 682.962771 | 1.996871797 | 2.34E-47 |
| 9930111J21Rik2 | 470.8629972 | 1.996636291 | 1.12E-08 |
| Tpd52 | 282.1749715 | 1.993777631 | 1.29E-14 |
| Noc2l | 3844.883424 | 1.993562338 | 2.16E-55 |
| Gramd1a | 2969.753099 | 1.992180259 | 1.70E-19 |
| Zfp428 | 275.3299011 | 1.991900242 | 7.17E-29 |
| Phactr4 | 1891.975684 | 1.987387622 | 5.49E-31 |
| Faf1 | 5358.729333 | 1.98485019 | 9.07E-66 |
| Spata5l1 | 220.8232095 | 1.984034755 | 3.43E-14 |
| Nmnat3 | 174.6720184 | 1.983701632 | 1.22E-18 |
| Tc2n | 5.42349818 | 1.982963448 | 3.40E-02 |
| Synb | 17.59181136 | 1.982507695 | 2.32E-03 |
| Ckap2 | 2101.539709 | 1.98102666 | 3.82E-11 |

| | | | |
|---|---|---|---|
| Gm10293 | 22.84175474 | 1.980931589 | 2.17E-05 |
| Gm3376 | 148.220178 | 1.979923661 | 9.69E-10 |
| Thpo | 50.84521309 | 1.979850916 | 3.26E-07 |
| Abcc12 | 7.638658383 | 1.978277561 | 1.92E-02 |
| Gjc2 | 6.210392759 | 1.976236857 | 2.97E-02 |
| Gm11518 | 16388.69852 | 1.97621093 | 8.18E-34 |
| Rpl8 | 66813.58999 | 1.975965978 | 3.13E-32 |
| Cnbd2 | 352.579731 | 1.975893649 | 2.26E-15 |
| Chrm1 | 15.25123971 | 1.97501316 | 1.23E-02 |
| Kctd13 | 656.9195456 | 1.9739811 | 6.23E-21 |
| Limk2 | 1846.809037 | 1.971408321 | 7.34E-51 |
| Gm16253 | 14.99152782 | 1.971333039 | 1.09E-04 |
| Zfp809 | 553.769558 | 1.970843513 | 6.72E-29 |
| Pidd1 | 231.8430029 | 1.969253883 | 1.17E-11 |
| Myo1c | 17433.95275 | 1.96901615 | 2.00E-45 |
| Ahr | 292.2985283 | 1.967551971 | 3.06E-17 |
| Sp2 | 1110.544619 | 1.966892962 | 1.02E-21 |
| Poln | 47.95986064 | 1.966682466 | 1.63E-06 |
| Dynlt1a | 769.8087586 | 1.966004963 | 4.78E-26 |
| Arvcf | 1255.272406 | 1.96356535 | 3.99E-42 |
| Psmd4 | 6484.294445 | 1.962605882 | 2.12E-31 |
| Adipor1 | 9075.210439 | 1.961868127 | 1.06E-34 |
| Efna1 | 1322.865678 | 1.96047496 | 1.05E-10 |
| Gcnt3 | 16.25473207 | 1.959676977 | 1.14E-03 |
| Tubb3 | 51.35408728 | 1.958495945 | 1.47E-05 |
| Cyp2s1 | 8.772565514 | 1.95744067 | 4.10E-02 |
| NA | 59602.75301 | 1.955898907 | 1.74E-45 |
| Cmip | 3604.487031 | 1.955217316 | 4.02E-53 |
| Hsf2bp | 28.16029851 | 1.954251071 | 7.95E-05 |
| Gm11687 | 35.35260304 | 1.953231454 | 6.28E-07 |
| Btnl7-ps | 9.554801893 | 1.948905222 | 4.43E-03 |
| Clk3 | 1992.491787 | 1.946242101 | 3.98E-34 |
| Zbtb7b | 2933.12507 | 1.945638603 | 6.43E-27 |
| Htr1b | 5.212664692 | 1.944858489 | 4.35E-02 |
| Rnf128 | 22.02821797 | 1.94187315 | 5.29E-03 |
| Car6 | 110.8106253 | 1.939031755 | 4.53E-03 |
| P2rx3 | 1054.952462 | 1.938016734 | 1.15E-04 |
| 4933438K21Rik | 20.61022562 | 1.937114318 | 1.59E-04 |
| Cd93 | 188.2576324 | 1.936683239 | 9.85E-07 |
| Ctc1 | 2558.238951 | 1.936553715 | 3.83E-15 |
| Klrg2 | 65.68197859 | 1.936408293 | 2.65E-10 |
| Rhbdd1 | 2472.901059 | 1.934271192 | 2.33E-36 |
| Gm12091 | 282.2413441 | 1.933232246 | 1.24E-08 |
| Usp3 | 2309.151463 | 1.932237654 | 1.76E-20 |
| Ttc21a | 7.552495532 | 1.931089116 | 4.20E-02 |
| Fam220a | 353.5374561 | 1.92868874 | 2.30E-05 |
| Arl5c | 11.25248985 | 1.926479757 | 2.46E-03 |

| | | | |
|---|---|---|---|
| Sync | 42.33056854 | 1.922690146 | 4.73E-07 |
| Myl12a | 24114.22838 | 1.922627096 | 4.53E-47 |
| Rab3c | 286.3804103 | 1.919938083 | 1.47E-05 |
| Psmd8 | 7697.479585 | 1.918954124 | 3.84E-25 |
| Atp6v0e2 | 330.9666067 | 1.918415091 | 4.05E-09 |
| Rrm1 | 7755.011656 | 1.917366713 | 7.84E-15 |
| 9230111E07Rik | 13.96764762 | 1.916546168 | 1.08E-03 |
| Stk32c | 7.691422658 | 1.915993343 | 6.12E-03 |
| Ptprf | 4297.628625 | 1.915274353 | 8.90E-99 |
| Pef1 | 4262.675557 | 1.915271906 | 3.11E-27 |
| Zfp143 | 1097.907523 | 1.915248989 | 3.63E-38 |
| Hspa8 | 63221.41105 | 1.914677046 | 1.44E-46 |
| Kcmf1 | 5176.903926 | 1.91387051 | 1.63E-69 |
| Dennd5b | 74.30478023 | 1.912354156 | 1.46E-05 |
| Entpd1 | 10.40574415 | 1.911380362 | 1.30E-02 |
| Fam110c | 419.3831265 | 1.910591182 | 1.57E-19 |
| 2310022A10Rik | 765.7830668 | 1.910247706 | 6.33E-30 |
| Gm28042 | 211.2134342 | 1.9068394 | 3.84E-10 |
| F11r | 12.52591921 | 1.906449267 | 1.47E-02 |
| Chst7 | 291.6271761 | 1.905683638 | 1.12E-15 |
| Mocos | 401.6732574 | 1.905637725 | 2.64E-27 |
| Hagh | 1322.182493 | 1.902308668 | 2.27E-45 |
| Syn3 | 37.65473483 | 1.901434019 | 2.51E-04 |
| Mras | 3862.451972 | 1.900497372 | 4.27E-28 |
| Nppa | 6.942840074 | 1.899834223 | 3.54E-02 |
| Gpat3 | 333.521046 | 1.899295396 | 6.17E-17 |
| 4933417D19Rik | 27.85225599 | 1.899230522 | 1.01E-04 |
| Gm10481 | 14.12705343 | 1.897481259 | 1.60E-03 |
| Syn1 | 1096.842102 | 1.895605823 | 4.37E-38 |
| Smg9 | 787.7115908 | 1.895523838 | 3.55E-28 |
| Utp4 | 3049.021415 | 1.895166537 | 2.22E-42 |
| Gm14471 | 11.34534093 | 1.894823919 | 9.00E-03 |
| Gm23969 | 8.056606807 | 1.894449852 | 2.12E-02 |
| Ifrd1 | 4709.926875 | 1.89308193 | 3.49E-21 |
| Gm14494 | 1272.813194 | 1.892479869 | 4.68E-14 |
| Dlgap4 | 3400.879371 | 1.890806852 | 3.20E-26 |
| Pycr2 | 1798.030111 | 1.888387252 | 3.49E-44 |
| Arhgap19 | 667.7567467 | 1.886337733 | 7.96E-11 |
| Gm9726 | 134.0824814 | 1.885530895 | 1.92E-10 |
| Dusp9 | 278.6759648 | 1.885318038 | 2.79E-10 |
| Nr4a1 | 921.088903 | 1.883444488 | 1.91E-06 |
| Me3 | 835.3947927 | 1.883334131 | 3.01E-30 |
| Adora2b | 955.5375759 | 1.882364052 | 2.76E-25 |
| Pdgfa | 1217.553266 | 1.88126698 | 3.46E-51 |
| Kifc5b | 429.9223858 | 1.880789701 | 1.13E-09 |
| Acot2 | 886.3418002 | 1.879942099 | 1.45E-23 |
| Klhdc8a | 1128.698314 | 1.877921679 | 1.27E-10 |

| | | | |
|---|---|---|---|
| Lysmd1 | 338.1310677 | 1.874406259 | 1.01E-30 |
| Rpl18 | 36466.21172 | 1.87179055 | 5.22E-26 |
| Snrnp48 | 727.3748628 | 1.871182874 | 1.43E-28 |
| Dapk3 | 1412.104759 | 1.870742542 | 4.29E-28 |
| Snord83b | 17.56343378 | 1.869151657 | 1.79E-03 |
| Mgam | 514.7573389 | 1.866382417 | 1.56E-04 |
| Atxn10 | 13554.53568 | 1.864894504 | 1.27E-33 |
| Sgms2 | 1191.807906 | 1.864811841 | 4.91E-38 |
| Adcy1 | 1520.53709 | 1.864302014 | 6.96E-13 |
| Gmds | 1941.410771 | 1.862510835 | 3.28E-41 |
| Pttg1 | 1028.152404 | 1.859834516 | 5.67E-14 |
| Gm26825 | 36665.95223 | 1.857011579 | 2.84E-35 |
| Gm13394 | 70314.65906 | 1.856975899 | 2.87E-29 |
| Arid3a | 862.6284981 | 1.856744848 | 3.65E-19 |
| Gm13832 | 8.699723904 | 1.856041539 | 1.45E-02 |
| Rnf169 | 1270.185398 | 1.854984747 | 1.01E-24 |
| Tspan33 | 25.32257853 | 1.854809414 | 3.24E-04 |
| Rdh9 | 51.98665338 | 1.853650056 | 3.82E-05 |
| Chfr | 2461.953026 | 1.85255615 | 8.28E-29 |
| 2010015M23Rik | 28.65831723 | 1.852191815 | 1.26E-04 |
| Sqor | 2866.171031 | 1.852049468 | 4.05E-52 |
| Ly6i | 6.094536957 | 1.851561362 | 4.55E-02 |
| Mpi | 945.5748437 | 1.851214346 | 1.53E-29 |
| Flt1 | 1329.706088 | 1.851058376 | 4.67E-54 |
| Rasa4 | 4237.672709 | 1.850591043 | 4.40E-29 |
| Loxl4 | 984.284874 | 1.849109314 | 3.29E-13 |
| Tmprss11d | 8.313460733 | 1.848160243 | 1.26E-02 |
| Kars | 8104.296081 | 1.84705987 | 3.14E-41 |
| Palm3 | 21.3394262 | 1.845785246 | 2.53E-02 |
| Ide | 5488.789588 | 1.844280949 | 1.06E-40 |
| Dus3l | 1405.012927 | 1.841768494 | 6.03E-41 |
| As3mt | 32.98295994 | 1.840178462 | 4.45E-04 |
| Csnk1e | 4058.989069 | 1.838654932 | 2.19E-68 |
| Gm16577 | 5.330348081 | 1.837665567 | 4.82E-02 |
| Thap1 | 316.451354 | 1.837559036 | 3.90E-19 |
| Serpinb9 | 1571.525424 | 1.836238228 | 3.55E-31 |
| Mdm2 | 3807.437062 | 1.835353045 | 5.57E-39 |
| Abcb6 | 523.6394462 | 1.834873687 | 2.04E-24 |
| D6Ertd527e | 17.77531328 | 1.83341319 | 5.69E-03 |
| 4921524J17Rik | 641.6105448 | 1.833193033 | 5.74E-47 |
| Gm6316 | 9.358180361 | 1.830780427 | 1.87E-02 |
| 1700061G19Rik | 8.01976113 | 1.830618144 | 8.27E-03 |
| Rpl28-ps1 | 40054.69604 | 1.830615236 | 8.65E-18 |
| Rpsa | 106522.1942 | 1.829280354 | 1.42E-23 |
| Ttc1 | 1980.789827 | 1.828929072 | 7.13E-44 |
| Ipo5 | 14120.82957 | 1.825174329 | 6.70E-46 |
| Pik3cb | 1633.017527 | 1.824567186 | 1.05E-28 |

| | | | |
|---|---|---|---|
| Creb3l3 | 9.152823137 | 1.824530289 | 3.12E-02 |
| Spock2 | 18.69978441 | 1.821859778 | 3.08E-04 |
| Samhd1 | 3737.536878 | 1.821745635 | 9.58E-49 |
| Ppef1 | 316.4882102 | 1.821563573 | 3.41E-20 |
| Eif3g | 4110.815225 | 1.82096826 | 1.85E-39 |
| Rnf114 | 2517.365589 | 1.820784371 | 6.54E-21 |
| Rab3ip | 2085.051427 | 1.820371356 | 3.35E-44 |
| Plek | 6.237955819 | 1.819294019 | 3.66E-02 |
| Col10a1 | 24.89550555 | 1.812158306 | 3.04E-06 |
| Gm10036 | 36.68042067 | 1.812026079 | 8.23E-06 |
| Wdr66 | 88.12706011 | 1.811434103 | 1.86E-09 |
| Rps18-ps3 | 520.1252683 | 1.81109509 | 8.12E-11 |
| Hoxb1 | 35.62416815 | 1.810701761 | 3.99E-02 |
| Trmt10a | 1090.231032 | 1.804704711 | 4.42E-26 |
| Gm10029 | 43.04999321 | 1.804534552 | 1.06E-03 |
| Gm12033 | 19598.80379 | 1.804393231 | 1.28E-22 |
| Cisd1 | 1216.639511 | 1.801899893 | 1.46E-25 |
| Gm15640 | 32.6140749 | 1.801525154 | 4.45E-04 |
| Tas1r2 | 31.23794289 | 1.801251223 | 1.83E-06 |
| Abcc1 | 6563.18358 | 1.7992 | 6.47E-48 |
| Hmcn2 | 50.91192735 | 1.798814335 | 8.19E-03 |
| Rab39b | 192.5716271 | 1.797908744 | 5.65E-07 |
| Rps27l | 3039.048456 | 1.797515041 | 4.31E-33 |
| Gm13068 | 8.713129617 | 1.795704498 | 2.23E-02 |
| Gm24119 | 8.338371078 | 1.795556358 | 4.13E-02 |
| Prdx5 | 5188.911969 | 1.795421166 | 1.92E-37 |
| Ube2t | 295.3875627 | 1.794455024 | 8.30E-08 |
| Selenoh | 777.6593273 | 1.794178906 | 8.14E-21 |
| Trip6 | 4067.669331 | 1.793403542 | 1.13E-25 |
| Tent5c | 31.88437475 | 1.792907521 | 4.86E-04 |
| Slc22a13b-ps | 8.826899059 | 1.792887801 | 3.26E-02 |
| Yjefn3 | 129.0269012 | 1.792591407 | 6.89E-10 |
| Babam2 | 2414.199913 | 1.791412942 | 6.48E-34 |
| Rnh1 | 9811.714619 | 1.789988737 | 5.13E-22 |
| Hist1h2be | 35.43583052 | 1.789539633 | 1.36E-04 |
| E030030I06Rik | 73.11210014 | 1.789522036 | 9.86E-06 |
| Ip6k3 | 40.3684319 | 1.78931595 | 9.59E-04 |
| Ptpn14 | 6806.862281 | 1.787756721 | 1.20E-70 |
| Gm24044 | 25.64572557 | 1.787703365 | 1.22E-04 |
| Raet1e | 152.5635618 | 1.787420013 | 3.71E-14 |
| Irf7 | 51.08821135 | 1.787235087 | 2.81E-06 |
| Nectin2 | 3231.192606 | 1.786262376 | 1.21E-41 |
| Trpt1 | 309.8301647 | 1.786161962 | 6.08E-24 |
| Rasgef1a | 8.735346562 | 1.784037761 | 2.79E-02 |
| Gm3200 | 26.1239579 | 1.783915743 | 2.76E-05 |
| Thap4 | 2190.18433 | 1.783552252 | 4.94E-29 |
| Gars | 14768.92027 | 1.780782264 | 3.68E-35 |

| | | | |
|---|---|---|---|
| Snx15 | 878.8217529 | 1.780731138 | 2.96E-77 |
| Tbc1d7 | 554.0725872 | 1.780231162 | 1.24E-32 |
| Wwox | 869.2521846 | 1.779756374 | 4.92E-29 |
| 9230114K14Rik | 104.7462116 | 1.778008223 | 4.64E-10 |
| 2410006H16Rik | 10631.39287 | 1.777337097 | 1.18E-10 |
| Ripor3 | 41.96255131 | 1.777089436 | 1.69E-03 |
| Rpl10 | 12459.82895 | 1.776986079 | 4.54E-14 |
| Lsm11 | 725.4015622 | 1.776840055 | 4.65E-23 |
| Hspa4l | 3271.537121 | 1.776253565 | 2.28E-13 |
| Mcrip2 | 335.8526933 | 1.776123632 | 2.86E-16 |
| NA | 56.00787468 | 1.77547768 | 3.98E-05 |
| Acot9 | 3567.356035 | 1.774765984 | 2.76E-30 |
| Kctd7 | 323.977193 | 1.77451331 | 1.20E-19 |
| Gm12366 | 374.6077916 | 1.772863414 | 1.16E-09 |
| Adamts4 | 1372.936021 | 1.772145926 | 8.00E-16 |
| Syngr3 | 13.5009954 | 1.771365667 | 4.38E-02 |
| Zfp474 | 8.436150026 | 1.769745109 | 2.61E-02 |
| Ptprd | 25.42487391 | 1.769434483 | 4.02E-02 |
| Nudt15 | 117.3997007 | 1.768845437 | 1.70E-13 |
| Adamtsl1 | 75.62285662 | 1.768192963 | 6.51E-03 |
| Bax | 4107.204276 | 1.767178499 | 1.22E-29 |
| Srp72 | 10229.00723 | 1.765935564 | 2.99E-35 |
| Gm5786 | 119.6563614 | 1.765624033 | 2.98E-13 |
| Il7 | 12.90409676 | 1.765209312 | 2.20E-02 |
| Klhl26 | 2008.492919 | 1.764959527 | 3.20E-45 |
| Flrt3 | 33.14674922 | 1.764055385 | 5.40E-03 |
| Prkd1 | 1467.094425 | 1.764024749 | 3.42E-46 |
| Hdgf | 20795.9107 | 1.762966057 | 6.90E-42 |
| Gm15191 | 9.540595841 | 1.76224139 | 1.28E-02 |
| Gm10602 | 148.9557599 | 1.761660233 | 9.05E-22 |
| Gm11249 | 1092.346685 | 1.760881535 | 1.83E-10 |
| Lmbr1l | 980.757885 | 1.760319593 | 1.35E-17 |
| Dst | 14726.69931 | 1.759155754 | 6.85E-25 |
| Plekha8 | 1302.28539 | 1.758823515 | 2.46E-42 |
| Rpusd3 | 280.6170281 | 1.757767151 | 1.83E-18 |
| Mknk2 | 5451.658763 | 1.756199866 | 2.14E-17 |
| Efnb1 | 347.491773 | 1.755803252 | 2.74E-20 |
| 4-Sep | 46.97005786 | 1.75453958 | 2.94E-05 |
| Lias | 1391.859481 | 1.754376431 | 5.97E-67 |
| Cfap61 | 19.52881633 | 1.754251209 | 6.59E-03 |
| BC022960 | 75.28455684 | 1.753715192 | 5.08E-08 |
| Cavin4 | 290.9811514 | 1.752547328 | 4.37E-12 |
| Rgl3 | 384.4796404 | 1.749498934 | 4.56E-28 |
| Tent5b | 1620.453803 | 1.749154601 | 9.55E-25 |
| Rnu1a1 | 7.850509731 | 1.749138022 | 4.51E-02 |
| Eif3i | 12035.09517 | 1.749112372 | 3.56E-26 |
| Gm11478 | 30.23898364 | 1.748643181 | 3.09E-04 |

| | | | |
|---|---|---|---|
| Atp1a1 | 13088.27316 | 1.748194034 | 5.93E-33 |
| Ppp2r5d | 4897.843244 | 1.747590659 | 1.86E-35 |
| Cct5 | 20882.64327 | 1.74715823 | 4.35E-60 |
| Gm8349 | 64.45013855 | 1.745503872 | 8.35E-13 |
| Mrpl32 | 969.4241825 | 1.743276854 | 5.97E-48 |
| Epb41l4aos | 846.5376575 | 1.742530771 | 2.51E-10 |
| Rpl27-ps1 | 134.8717961 | 1.741440373 | 1.01E-07 |
| Nxf3 | 15.38053015 | 1.741178407 | 2.92E-02 |
| Nosip | 1483.982081 | 1.738696814 | 3.21E-31 |
| Nme2 | 20448.66004 | 1.737165189 | 2.13E-49 |
| Clu | 185.6709907 | 1.736381037 | 3.38E-08 |
| Ccdc170 | 17.32655283 | 1.736320214 | 8.22E-04 |
| Gmip | 1120.355618 | 1.735803489 | 1.52E-22 |
| Junb | 1589.766988 | 1.735630237 | 2.80E-08 |
| Ddit3 | 6642.720834 | 1.735120363 | 2.07E-13 |
| Slc19a2 | 1917.481573 | 1.73426764 | 6.01E-32 |
| Atp6v1g2 | 90.09851119 | 1.734046997 | 1.06E-12 |
| Cnn2 | 23457.26203 | 1.732108699 | 3.45E-49 |
| Cabp4 | 6.743589546 | 1.731925784 | 2.71E-02 |
| Mcu | 1703.920113 | 1.731116107 | 2.71E-56 |
| Bsg | 26626.6997 | 1.731074457 | 1.21E-23 |
| Mrvi1 | 124.3727741 | 1.730895405 | 7.77E-05 |
| Zmat3 | 4898.113607 | 1.730089466 | 1.90E-30 |
| Dusp4 | 2570.483965 | 1.728845958 | 2.51E-09 |
| Disp1 | 2262.130775 | 1.728356398 | 1.47E-61 |
| Slc39a10 | 3524.383162 | 1.727569825 | 1.95E-13 |
| Slc2a9 | 175.1610797 | 1.724486625 | 8.42E-17 |
| Gm4737 | 8568.322965 | 1.724327339 | 3.91E-22 |
| Mrps18b | 1968.321664 | 1.723848311 | 2.52E-38 |
| Tsen54 | 797.3115622 | 1.722797121 | 3.50E-30 |
| Rps2-ps4 | 80.95409934 | 1.720873823 | 7.69E-04 |
| Krt86 | 11.25936999 | 1.720646493 | 3.56E-03 |
| Mybbp1a | 15401.0598 | 1.720519787 | 1.15E-33 |
| Tmc7 | 16.65327218 | 1.72020088 | 6.48E-03 |
| Rpl35a-ps5 | 15.41563685 | 1.720158769 | 1.49E-03 |
| Flt3 | 12.61002332 | 1.719561174 | 6.60E-03 |
| A230020J21Rik | 18.87247229 | 1.718790286 | 1.98E-03 |
| Ufc1 | 3015.992891 | 1.717872821 | 1.93E-26 |
| Mnd1 | 246.5320564 | 1.716621384 | 5.77E-25 |
| Slc12a4 | 4348.57105 | 1.715937981 | 2.29E-56 |
| Car5b | 116.6171212 | 1.714329127 | 5.64E-13 |
| Aim2 | 214.7586752 | 1.714217235 | 6.13E-15 |
| Rabggtb | 5299.556326 | 1.713773218 | 7.57E-48 |
| Tnpo2 | 9752.197038 | 1.71371538 | 6.06E-35 |
| Gm26619 | 12811.53859 | 1.712573759 | 2.71E-12 |
| Retreg1 | 1973.380373 | 1.71144313 | 2.10E-21 |
| Phlda1 | 939.9688801 | 1.709438549 | 3.03E-11 |

| | | | |
|---|---|---|---|
| Fam177a | 1383.292923 | 1.709377282 | 9.64E-30 |
| Fau | 8902.564704 | 1.708993476 | 7.60E-16 |
| Pld2 | 1123.434152 | 1.708861437 | 3.02E-27 |
| NA | 329.9046769 | 1.707335455 | 1.35E-09 |
| Siglece | 8.051011043 | 1.706797831 | 1.38E-02 |
| Rps3 | 56649.04585 | 1.705632085 | 6.04E-19 |
| Gm6433 | 39.0436362 | 1.705507493 | 4.51E-04 |
| Znrf2 | 1311.359839 | 1.704992545 | 7.56E-22 |
| Nudt22 | 470.2166979 | 1.704688204 | 7.86E-37 |
| Fxr2 | 4780.176202 | 1.704040795 | 5.28E-30 |
| Gm5805 | 361.4001265 | 1.703998176 | 3.63E-07 |
| Gm13075 | 19.69455583 | 1.703934069 | 1.22E-03 |
| Chp1 | 6682.072199 | 1.702841 | 5.84E-24 |
| Nuggc | 50.09471182 | 1.702581628 | 3.44E-08 |
| Mrpl17 | 2145.416491 | 1.702205264 | 1.93E-46 |
| Nepro | 974.7880945 | 1.700288628 | 5.84E-33 |
| Gm26917 | 957.8739768 | 1.699893471 | 2.35E-31 |
| Crim1 | 4958.715791 | 1.699303978 | 1.66E-14 |
| Rpl18-ps2 | 29.06764237 | 1.699153492 | 5.37E-05 |
| Fgf18 | 10.05836057 | 1.698516392 | 4.98E-02 |
| Me1 | 3504.254363 | 1.698432095 | 8.11E-38 |
| Rpl27 | 2340.730614 | 1.698137659 | 7.60E-13 |
| Col5a2 | 40728.55901 | 1.697641753 | 1.42E-14 |
| Raet1d | 118.6635537 | 1.696830469 | 4.35E-09 |
| Mrpl45 | 1814.459067 | 1.696685918 | 2.14E-32 |
| Rps6-ps4 | 132832.5808 | 1.696540139 | 1.17E-19 |
| Zfp579 | 654.3509177 | 1.696329652 | 1.84E-17 |
| Gm12070 | 115.8778542 | 1.695904873 | 8.89E-04 |
| Sgsm1 | 11.82230782 | 1.694902406 | 1.98E-02 |
| Ncam1 | 1630.791622 | 1.694551944 | 4.27E-21 |
| Ubl4a | 2372.32782 | 1.694398459 | 4.42E-43 |
| Ifi205 | 92.81123866 | 1.694265941 | 9.19E-03 |
| Tnfaip3 | 769.7438906 | 1.694099282 | 1.24E-11 |
| Bcl2l1 | 4294.000393 | 1.693454179 | 5.50E-27 |
| Tmem170 | 255.3052847 | 1.693305575 | 2.64E-24 |
| Cdipt | 3223.076682 | 1.692998131 | 2.64E-57 |
| Gm5915 | 8.232533777 | 1.692663656 | 3.63E-02 |
| Pqlc3 | 1007.222603 | 1.692644295 | 7.68E-26 |
| Gm3534 | 28.95131471 | 1.69254895 | 1.67E-04 |
| Aurka | 1220.677723 | 1.691423734 | 3.28E-06 |
| Chchd5 | 838.8662133 | 1.690730671 | 1.12E-16 |
| Gm10382 | 31.87078083 | 1.689427697 | 8.48E-03 |
| Il15 | 186.9354469 | 1.68826699 | 2.63E-13 |
| Bambi | 119.379043 | 1.687690552 | 1.21E-08 |
| Rack1 | 120396.584 | 1.687239356 | 3.45E-17 |
| Numbl | 3202.627356 | 1.686993958 | 6.22E-31 |
| Gm10108 | 146.643854 | 1.68672148 | 7.15E-06 |

| | | | |
|---|---|---|---|
| Gm16470 | 10.75496756 | 1.684506138 | 5.00E-03 |
| Drap1 | 6991.132614 | 1.684473311 | 1.22E-42 |
| Gm6682 | 155.9065787 | 1.682756315 | 4.10E-07 |
| Cd24a | 2574.110999 | 1.682056109 | 1.64E-06 |
| Ajuba | 2948.78609 | 1.681119275 | 2.80E-33 |
| Gm2546 | 31.89944019 | 1.681097592 | 1.15E-06 |
| Wbp2 | 5052.649699 | 1.680644261 | 2.39E-14 |
| Orai1 | 454.0094432 | 1.680638037 | 3.94E-16 |
| Fblim1 | 958.4750544 | 1.680587758 | 6.61E-21 |
| Gm10275 | 71607.79352 | 1.680567111 | 1.08E-15 |
| Tmbim1 | 4932.481657 | 1.680320206 | 2.32E-31 |
| Hint3 | 384.6312068 | 1.677846186 | 5.17E-14 |
| Ubxn1 | 7751.977327 | 1.67632858 | 2.51E-49 |
| 3110009E18Rik | 177.7241509 | 1.673538303 | 1.21E-08 |
| Utp14a | 2739.522103 | 1.670984154 | 4.46E-40 |
| Gm11953 | 12.31386492 | 1.670677155 | 1.29E-02 |
| Slc22a4 | 48.52663821 | 1.67066582 | 1.70E-04 |
| Cdc20 | 2852.257054 | 1.669959892 | 1.33E-07 |
| Rnf31 | 992.4717778 | 1.668454393 | 1.67E-18 |
| Thbs1 | 56074.01074 | 1.66722974 | 4.71E-07 |
| Oser1 | 2251.689214 | 1.666217091 | 2.90E-19 |
| Gm42743 | 86.79263655 | 1.665691557 | 5.69E-04 |
| Ercc6l2 | 120.8231153 | 1.66461122 | 1.14E-08 |
| Ncmap | 11.34816572 | 1.664592425 | 2.76E-02 |
| Rhbdf2 | 448.6446709 | 1.663863779 | 2.73E-21 |
| Ftl1-ps1 | 92.97599514 | 1.66268548 | 2.73E-04 |
| Cdk6 | 743.1289463 | 1.66259267 | 4.26E-26 |
| Ddb1 | 78266.76622 | 1.662147754 | 1.68E-45 |
| Rpl13-ps6 | 16.44435408 | 1.661490551 | 1.25E-02 |
| Gm2541 | 306.2248907 | 1.660966188 | 2.35E-06 |
| Mvp | 5662.63892 | 1.660811508 | 5.04E-30 |
| Kif18b | 709.5316689 | 1.659551001 | 8.83E-07 |
| Rps16 | 14597.25608 | 1.65731748 | 2.22E-20 |
| Sms-ps | 1178.87176 | 1.657194519 | 7.06E-19 |
| Gm10653 | 235.1147325 | 1.656132476 | 3.39E-04 |
| Psat1 | 15457.79927 | 1.655199218 | 5.56E-09 |
| Ier5 | 3710.218611 | 1.654033282 | 3.25E-28 |
| Pfkm | 3671.085007 | 1.653725745 | 4.70E-31 |
| Gm12286 | 430.9828818 | 1.653583466 | 4.36E-10 |
| Cdc34 | 2896.485483 | 1.651955708 | 1.81E-23 |
| Stxbp3-ps | 13.88435339 | 1.651566604 | 2.18E-02 |
| Ddit4 | 1013.742835 | 1.650380335 | 1.08E-19 |
| Sgcd | 202.5062698 | 1.649356737 | 5.78E-06 |
| Gm4968 | 86.28465387 | 1.649323284 | 1.53E-07 |
| Wdr93 | 28.35747803 | 1.648955934 | 1.64E-05 |
| Dbp | 721.6468037 | 1.648494638 | 3.32E-07 |
| Gm11185 | 17.00304434 | 1.647960326 | 6.16E-03 |

| | | | |
|---|---|---|---|
| Lin37 | 543.5409492 | 1.647608696 | 4.04E-20 |
| Tmem63b | 2442.58208 | 1.646900379 | 9.89E-53 |
| Supt3 | 1376.160322 | 1.646108127 | 2.46E-21 |
| Prorsd1 | 28.67614291 | 1.643472288 | 5.25E-04 |
| Arl14ep | 4362.339537 | 1.643149981 | 4.38E-21 |
| Cd81 | 15598.87096 | 1.643086529 | 4.00E-21 |
| Vrk2 | 912.3379689 | 1.64305524 | 5.47E-46 |
| Clic4 | 7974.140184 | 1.642433559 | 5.83E-21 |
| Myl6b | 236.3366549 | 1.64046161 | 3.27E-13 |
| Kcna4 | 440.1062529 | 1.639460695 | 1.54E-11 |
| Uhrf1bp1l | 3287.025469 | 1.639310609 | 1.08E-49 |
| Gm13050 | 29.96616328 | 1.637210847 | 4.20E-05 |
| Tubb4b | 9221.441394 | 1.636800891 | 8.05E-10 |
| Plekha7 | 19.49701154 | 1.636373611 | 1.13E-02 |
| Trib3 | 1385.443885 | 1.635906588 | 2.24E-06 |
| Irak3 | 70.04821296 | 1.63577205 | 8.03E-09 |
| Prkag2 | 1252.873195 | 1.634826066 | 1.68E-48 |
| Tpk1 | 262.2730087 | 1.634293277 | 4.85E-21 |
| Mrrf | 1845.383387 | 1.63362915 | 2.04E-33 |
| B3gnt5 | 34.6745235 | 1.630362346 | 1.11E-02 |
| Tap1 | 754.3650937 | 1.630027105 | 1.70E-16 |
| Ccdc107 | 815.763532 | 1.629369084 | 1.44E-37 |
| Gm10243 | 15969.90068 | 1.62868622 | 9.24E-21 |
| Sbno2 | 2509.386322 | 1.628439822 | 1.86E-28 |
| Hsdl2 | 1811.389637 | 1.62637905 | 3.41E-44 |
| Smpd1 | 2430.281075 | 1.626333428 | 4.84E-25 |
| Gm13464 | 317.0423644 | 1.625573716 | 9.06E-11 |
| Tmem209 | 1826.003205 | 1.625112904 | 8.41E-37 |
| Oxsr1 | 2997.297148 | 1.624588601 | 2.55E-40 |
| Farp2 | 1084.046954 | 1.624129864 | 9.03E-37 |
| Hspg2 | 28446.74545 | 1.624037177 | 1.29E-16 |
| Upp1 | 35.47148637 | 1.623773514 | 7.55E-03 |
| Ryr1 | 20.13106927 | 1.623442125 | 4.66E-03 |
| Ntn4 | 184.2709408 | 1.622281989 | 6.33E-10 |
| Sp6 | 12.2715641 | 1.622243496 | 3.23E-02 |
| Gm12176 | 45.17947775 | 1.622139834 | 3.47E-05 |
| Eif3h | 21119.97463 | 1.616310226 | 2.71E-15 |
| Sik3 | 2887.657643 | 1.61623858 | 1.16E-31 |
| Hbs1l | 4908.209871 | 1.616179342 | 1.58E-94 |
| F8a | 147.1483612 | 1.615304533 | 5.08E-10 |
| Por | 4956.615258 | 1.615175651 | 2.10E-18 |
| Ubxn6 | 3031.372245 | 1.614799594 | 2.00E-29 |
| Rps15 | 55482.86243 | 1.614289772 | 2.48E-15 |
| Eif4ebp1 | 7809.692884 | 1.614227354 | 1.39E-34 |
| Faim2 | 94.52067842 | 1.61399565 | 2.23E-08 |
| Gm17383 | 49.24206705 | 1.613784132 | 2.38E-04 |
| Col5a3 | 679.7155384 | 1.612013319 | 1.82E-07 |

| | | | |
|---|---|---|---|
| Gpx4 | 15070.71857 | 1.61100743 | 1.10E-21 |
| C9orf72 | 868.8703599 | 1.609482546 | 1.40E-78 |
| Gm14252 | 6.16975334 | 1.609252175 | 4.62E-02 |
| Fam107b | 3625.918197 | 1.609070932 | 7.76E-21 |
| Cngb1 | 14.26182037 | 1.608466957 | 4.92E-02 |
| Sh2d4a | 65.96527793 | 1.608349986 | 1.14E-12 |
| Kcnj2 | 21.53821937 | 1.60724832 | 2.21E-02 |
| Nrg1 | 1866.437419 | 1.60710598 | 1.71E-21 |
| Awat2 | 33.20162048 | 1.606966024 | 3.21E-05 |
| Dot1l | 2904.513899 | 1.606207719 | 4.13E-25 |
| Dedd2 | 306.1735718 | 1.605951337 | 4.97E-16 |
| Zdhhc2 | 222.3920511 | 1.605034588 | 5.20E-07 |
| Naa38 | 1144.535359 | 1.604764241 | 5.27E-32 |
| C2cd2l | 454.1034844 | 1.604566333 | 5.43E-11 |
| Gm13509 | 20.7459726 | 1.604388952 | 2.21E-03 |
| Gm26850 | 12.53616781 | 1.603705057 | 4.52E-02 |
| Ttc16 | 24.87972731 | 1.60331537 | 1.10E-03 |
| Maoa | 11869.71092 | 1.602390596 | 7.20E-19 |
| Parl | 1832.883324 | 1.602337065 | 2.14E-33 |
| 1810037I17Rik | 1837.466688 | 1.602049297 | 3.50E-35 |
| Rmc1 | 1218.193521 | 1.599684059 | 7.63E-25 |
| Gm26578 | 320.2566482 | 1.599436172 | 3.14E-23 |
| Kat5 | 1999.455412 | 1.59877799 | 1.32E-27 |
| Gm25442 | 9.790656419 | 1.598642172 | 2.97E-02 |
| Tom1l1 | 742.8401817 | 1.59843834 | 8.52E-29 |
| Gm12416 | 46.48077812 | 1.598361888 | 4.48E-04 |
| Grk2 | 3292.244991 | 1.598103931 | 1.21E-44 |
| Rps11 | 38460.82027 | 1.598037858 | 6.87E-18 |
| Specc1 | 3432.279109 | 1.597652039 | 2.10E-19 |
| Prnp | 14550.6052 | 1.597439531 | 1.53E-22 |
| Copg2 | 4967.60205 | 1.596367742 | 2.76E-35 |
| Sbds | 2941.875552 | 1.595059281 | 4.00E-38 |
| Castor1 | 337.8062982 | 1.595047614 | 2.53E-19 |
| Mgmt | 694.135641 | 1.594734136 | 9.46E-22 |
| Gm26909 | 21.61032335 | 1.594352382 | 1.11E-04 |
| Ptx3 | 4202.894405 | 1.594151964 | 3.50E-06 |
| Bloc1s2 | 1188.418543 | 1.593738847 | 1.42E-79 |
| Qsox1 | 10100.16956 | 1.59283343 | 1.37E-21 |
| Trim16 | 1755.34812 | 1.592006634 | 6.75E-21 |
| Lrrc8e | 165.339491 | 1.591360873 | 5.54E-13 |
| Rps6-ps3 | 125.0573789 | 1.590655817 | 4.98E-08 |
| Rasl11b | 155.4030989 | 1.589545657 | 9.82E-08 |
| Gapdh | 2734.936209 | 1.587558761 | 1.51E-03 |
| Tbx6 | 29.79739049 | 1.587319108 | 9.37E-04 |
| NA | 737.3410238 | 1.58731505 | 2.63E-18 |
| Tkt | 7929.85388 | 1.58730058 | 6.40E-29 |
| Ptchd4 | 348.94966 | 1.586497456 | 6.51E-04 |

| | | | |
|---|---|---|---|
| Gm13436 | 72.53817952 | 1.58644799 | 1.40E-05 |
| Bex3 | 8754.027769 | 1.586371969 | 4.55E-42 |
| Aldh18a1 | 7021.690676 | 1.585223333 | 1.30E-19 |
| Tubd1 | 135.3189566 | 1.585105613 | 7.23E-15 |
| Cox5a | 4353.767426 | 1.584881963 | 1.46E-28 |
| Ppp1r3a | 78.06682045 | 1.584234288 | 3.97E-02 |
| Rpl7a-ps5 | 17.20044573 | 1.583301755 | 9.09E-04 |
| Haus8 | 750.9704817 | 1.583123252 | 1.05E-11 |
| Cox6a2 | 24.89091578 | 1.582329177 | 2.64E-03 |
| Rpl3 | 68.93257305 | 1.581550822 | 8.27E-04 |
| Adrm1 | 5007.958963 | 1.58123541 | 9.80E-24 |
| Spon1 | 17.90306055 | 1.580260771 | 1.80E-02 |
| Sh3bp1 | 1826.718969 | 1.579891513 | 3.04E-28 |
| Kat8 | 866.0276859 | 1.579229671 | 1.44E-25 |
| Cav1 | 38978.11609 | 1.57879232 | 9.22E-37 |
| Kif2a | 2279.023051 | 1.577940137 | 5.07E-32 |
| Rpl9 | 44618.973 | 1.577909951 | 4.04E-20 |
| 4932414N04Rik | 12.45760852 | 1.57764856 | 9.09E-03 |
| Dkk2 | 170.6475019 | 1.57717855 | 3.61E-02 |
| Taf10 | 2599.103771 | 1.575713901 | 6.66E-23 |
| Inppl1 | 6885.394772 | 1.574804444 | 1.59E-56 |
| 5430416N02Rik | 1000.533301 | 1.572289603 | 1.42E-39 |
| Ighmbp2 | 888.9972279 | 1.571769667 | 2.60E-40 |
| Fam118b | 1286.976374 | 1.571244954 | 1.84E-33 |
| NA | 1727.994687 | 1.569608477 | 1.36E-03 |
| Gm11517 | 11.31534709 | 1.568144289 | 2.84E-02 |
| Rpl9-ps7 | 28.22713166 | 1.568059373 | 3.84E-04 |
| Gm13226 | 135.786182 | 1.568046141 | 7.61E-08 |
| Vdac1 | 9711.986753 | 1.567909874 | 6.97E-48 |
| Strbp | 1590.73397 | 1.567512389 | 1.56E-23 |
| Fam129b | 15591.52094 | 1.567228327 | 1.54E-21 |
| Gm8355 | 4111.759531 | 1.566701087 | 4.69E-06 |
| Rad51d | 1139.790286 | 1.56590146 | 1.15E-27 |
| Acsl1 | 1302.461648 | 1.565704723 | 1.32E-02 |
| Txnl4a | 1437.889867 | 1.565469635 | 7.24E-18 |
| Wdr36 | 2910.009891 | 1.564465525 | 8.44E-44 |
| Acot13 | 2081.691453 | 1.5639905 | 3.22E-24 |
| Pdcd2l | 631.8121723 | 1.562693924 | 1.13E-18 |
| Nob1 | 1945.869429 | 1.562072663 | 2.83E-40 |
| Ttll2 | 60.65077863 | 1.561866586 | 4.90E-02 |
| Schip1 | 526.8642713 | 1.561428962 | 4.34E-22 |
| Pkmyt1 | 487.067601 | 1.559980166 | 8.67E-08 |
| Tpmt | 384.4346636 | 1.559965627 | 4.67E-32 |
| Gm13864 | 254.081662 | 1.559371013 | 2.26E-09 |
| Dstn | 11056.37856 | 1.559281618 | 4.23E-38 |
| Mta3 | 1941.017603 | 1.559052245 | 4.02E-39 |
| Dis3 | 2028.200728 | 1.558122972 | 2.38E-20 |

| | | | |
|---|---|---|---|
| Slc25a30 | 2177.751789 | 1.558044082 | 2.97E-16 |
| Mmp16 | 66.21102913 | 1.557880927 | 3.89E-04 |
| 1110038B12Rik | 8194.910558 | 1.557878561 | 3.54E-16 |
| Ell2 | 681.4752625 | 1.555886654 | 3.38E-14 |
| Grsf1 | 4328.906575 | 1.555528674 | 4.24E-33 |
| Tmem88 | 67.02043446 | 1.554817079 | 2.84E-05 |
| Sgtb | 669.2050801 | 1.553690409 | 4.48E-26 |
| Bnc1 | 128.6434788 | 1.55368099 | 2.62E-04 |
| Gk | 712.7420813 | 1.553252559 | 2.08E-10 |
| Cav2 | 3663.325958 | 1.552400383 | 7.95E-19 |
| Lrrc57 | 651.3940836 | 1.551143359 | 1.83E-29 |
| Snta1 | 63.91870191 | 1.551006818 | 7.62E-05 |
| Cfl2 | 7309.759639 | 1.549350419 | 5.81E-23 |
| Zfp598 | 2577.338794 | 1.548273141 | 4.34E-41 |
| Fez2 | 3426.090875 | 1.54763578 | 2.41E-19 |
| Txnl1 | 3876.007935 | 1.547193939 | 1.19E-48 |
| Slc25a39 | 8371.758463 | 1.546732117 | 6.48E-22 |
| Gm12922 | 36.52443524 | 1.546319136 | 9.00E-04 |
| Acvr2b | 107.6232016 | 1.546301267 | 2.91E-09 |
| Gm5093 | 17.47709868 | 1.546115104 | 4.71E-03 |
| Gm15489 | 7.30828796 | 1.545962302 | 2.86E-02 |
| Rab11fip5 | 6386.276154 | 1.545670704 | 8.68E-19 |
| Trmt1 | 3494.043104 | 1.545363049 | 1.51E-25 |
| Rnps1 | 1965.837119 | 1.544948312 | 2.33E-21 |
| Gm12915 | 36.12145581 | 1.544881883 | 6.54E-06 |
| Prkrip1 | 783.1122703 | 1.54479935 | 5.92E-24 |
| Gm4335 | 27.54501694 | 1.544634674 | 5.42E-04 |
| Fxyd5 | 177.0089438 | 1.541734476 | 8.69E-05 |
| Gtpbp4 | 3254.645096 | 1.541339753 | 6.57E-48 |
| Whamm | 496.7500502 | 1.540358494 | 2.47E-14 |
| Gm20517 | 50.83540261 | 1.540097342 | 1.43E-07 |
| Mak16 | 2982.450982 | 1.539926395 | 1.05E-32 |
| Fancc | 434.9790087 | 1.539474942 | 9.99E-20 |
| Eif2b5 | 4375.143382 | 1.539025491 | 1.37E-35 |
| Sap25 | 22.93362904 | 1.537780935 | 3.39E-02 |
| Olfr1314 | 22.59357489 | 1.534853856 | 1.27E-03 |
| Prkaa2 | 23.60887112 | 1.534052159 | 1.16E-02 |
| Ints11 | 1652.185538 | 1.530662002 | 5.35E-27 |
| Ctif | 1560.762332 | 1.529717702 | 2.88E-37 |
| Slc35g1 | 93.21178618 | 1.529381716 | 9.78E-07 |
| Mthfd1l | 1964.85493 | 1.529138189 | 4.38E-10 |
| Wdr91 | 824.4786997 | 1.528818241 | 8.11E-17 |
| Lzic | 586.7404707 | 1.528500814 | 1.46E-19 |
| Tagln | 2708.688587 | 1.52691662 | 1.99E-07 |
| Shmt2 | 7386.878222 | 1.526801656 | 3.88E-09 |
| Knstrn | 1791.183893 | 1.526609557 | 1.30E-09 |
| Pik3cd | 389.1741217 | 1.526314455 | 7.96E-15 |

| | | | |
|---|---|---|---|
| Aimp2 | 1282.29294 | 1.523878428 | 4.34E-18 |
| Rpl15 | 5028.833574 | 1.523322225 | 3.31E-21 |
| Zc3hc1 | 1726.952055 | 1.522126322 | 1.98E-36 |
| Pgam5 | 2236.525896 | 1.521954022 | 1.14E-38 |
| Rusc1 | 481.2122263 | 1.521302792 | 1.54E-10 |
| Zfp335 | 1805.266162 | 1.521022805 | 4.01E-18 |
| Map4k1 | 81.71397557 | 1.520695496 | 2.23E-10 |
| Rpl21-ps15 | 1077.507034 | 1.52035084 | 2.56E-05 |
| Rpl3-ps1 | 93867.25035 | 1.519058624 | 1.12E-15 |
| Rpl13 | 43964.4684 | 1.518768638 | 1.94E-13 |
| Creld1 | 74.5768632 | 1.51837056 | 6.09E-05 |
| Acyp2 | 175.9484139 | 1.517430767 | 4.09E-18 |
| Flnb | 18993.59883 | 1.516836897 | 1.19E-26 |
| Ifitm1 | 15.2790241 | 1.515693402 | 4.08E-02 |
| Bpnt1 | 749.3892407 | 1.515516848 | 3.10E-19 |
| Rpl15-ps2 | 26.45417817 | 1.515012486 | 5.38E-03 |
| Pithd1 | 813.3192727 | 1.513505895 | 4.10E-19 |
| Dlgap5 | 995.0483588 | 1.512523287 | 9.71E-08 |
| Asnsd1 | 5010.281596 | 1.511538152 | 6.92E-28 |
| Rb1 | 846.9033777 | 1.510880849 | 8.21E-19 |
| Polr2h | 1088.850475 | 1.509973733 | 2.63E-30 |
| Gm2574 | 455.518203 | 1.509701091 | 1.25E-18 |
| Klc4 | 995.6251541 | 1.50937937 | 5.41E-11 |
| 0610010K14Rik | 1202.24141 | 1.508455709 | 1.16E-18 |
| Gm3222 | 53.99952612 | 1.507299292 | 3.06E-07 |
| Cntnap1 | 62.99681169 | 1.506975914 | 1.69E-07 |
| Mrpl38 | 1679.111537 | 1.506672809 | 6.62E-31 |
| Lpcat4 | 1034.700321 | 1.50660524 | 1.47E-20 |
| BC016548 | 15.36212957 | 1.506141936 | 5.27E-03 |
| Gabpb1 | 1205.660233 | 1.505966473 | 1.25E-32 |
| Zfp622 | 2476.499162 | 1.505779114 | 7.24E-11 |
| Ppp6c | 1863.56707 | 1.505544701 | 2.17E-43 |
| Slc48a1 | 9279.752174 | 1.505030669 | 6.53E-21 |
| Esyt1 | 2873.959478 | 1.50496767 | 3.58E-31 |
| Rps14 | 37307.50526 | 1.504777186 | 5.24E-10 |
| Abhd11 | 803.7195628 | 1.504423571 | 3.00E-22 |
| Rbks | 25.29987498 | 1.503897371 | 6.61E-04 |
| Lats2 | 3075.227714 | 1.503863282 | 1.48E-26 |
| Tor4a | 194.1873465 | 1.503499582 | 1.16E-09 |
| Grik5 | 256.8579891 | 1.503311917 | 2.17E-13 |
| Emd | 2493.987764 | 1.503166052 | 5.54E-23 |
| Ppme1 | 2062.268588 | 1.502866646 | 9.04E-31 |
| Polr2c | 2424.517876 | 1.501274876 | 5.81E-27 |
| Gm15337 | 9.801948551 | 1.501124809 | 2.49E-02 |
| Sars | 17432.90906 | 1.501005755 | 3.65E-34 |
| Eif2b3 | 1233.244349 | 1.500574504 | 1.36E-40 |
| Ttc25 | 33.34011969 | 1.498794766 | 1.62E-02 |

| | | | |
|---|---|---|---|
| Sil1 | 1576.433198 | 1.497448052 | 4.79E-13 |
| Asl | 1505.36333 | 1.49731273 | 3.70E-17 |
| Tubg1 | 1655.844266 | 1.496679577 | 1.25E-11 |
| Phlda3 | 4700.215127 | 1.494678854 | 7.66E-11 |
| Isg20 | 692.6214078 | 1.494560048 | 2.79E-12 |
| Spag4 | 73.35005613 | 1.494004306 | 1.11E-06 |
| Mettl21e | 9.771196451 | 1.493321634 | 2.52E-02 |
| Pwp1 | 2355.646022 | 1.493063573 | 1.13E-36 |
| Slc3a2 | 11311.11817 | 1.492363462 | 7.02E-31 |
| Serbp1 | 24175.27377 | 1.491054275 | 1.52E-41 |
| Hebp2 | 255.3948829 | 1.490403418 | 3.03E-17 |
| Gtf3c1 | 5130.112731 | 1.489429784 | 3.42E-26 |
| Htra1 | 323.732752 | 1.489141755 | 1.79E-20 |
| Usp39 | 3405.710317 | 1.48870427 | 1.03E-19 |
| Gm9083 | 22.23643344 | 1.487477159 | 2.91E-03 |
| Lrp2 | 44.96142077 | 1.486603787 | 3.07E-03 |
| Wnt4 | 10.64991564 | 1.486063383 | 4.45E-02 |
| Sod1 | 7945.096107 | 1.485853286 | 1.54E-30 |
| Man1a | 1574.812904 | 1.485682448 | 2.43E-12 |
| Ghitm | 11802.31619 | 1.485160002 | 1.30E-27 |
| Gm17501 | 28.85545055 | 1.484914139 | 1.27E-02 |
| Phgdh | 14268.86496 | 1.484439617 | 2.24E-12 |
| Fuk | 922.1422652 | 1.483400728 | 1.10E-30 |
| Gm8318 | 56.05561955 | 1.482954698 | 1.27E-05 |
| Vps11 | 2731.764542 | 1.482572354 | 2.46E-12 |
| Pkd1l3 | 40.65792714 | 1.48210762 | 3.20E-05 |
| Fam83h | 457.8759574 | 1.481304285 | 9.22E-29 |
| 6430562O15Rik | 12.17978969 | 1.479685348 | 5.77E-03 |
| Eif2b2 | 1629.982564 | 1.478404933 | 1.76E-22 |
| Vwa1 | 45.31304822 | 1.477980115 | 2.28E-04 |
| Rpl10a-ps1 | 49044.15001 | 1.477588525 | 8.66E-13 |
| Gng3 | 55.78018148 | 1.477552551 | 1.79E-03 |
| Col20a1 | 635.2226194 | 1.476948067 | 8.86E-08 |
| Dcaf15 | 1619.339583 | 1.476435676 | 5.27E-18 |
| Fbxl8 | 383.1510087 | 1.476264076 | 1.04E-12 |
| Rpl24 | 25807.90155 | 1.475346682 | 1.82E-15 |
| Pde5a | 58.33909461 | 1.475000861 | 6.23E-07 |
| Brinp1 | 311.655202 | 1.474956748 | 7.54E-05 |
| Fzd6 | 270.1231641 | 1.474832858 | 3.25E-13 |
| Fuca2 | 3318.401005 | 1.474637518 | 1.17E-19 |
| Fbxl20 | 3801.092862 | 1.474371277 | 4.79E-11 |
| Atxn2l | 6670.119001 | 1.472364082 | 6.50E-37 |
| Arhgef39 | 102.9442882 | 1.471565294 | 1.35E-04 |
| Ftl1 | 156446.3262 | 1.471546307 | 7.12E-15 |
| Camk2b | 29.49686363 | 1.47073463 | 2.95E-02 |
| Gm9013 | 38.9956223 | 1.470384064 | 5.97E-04 |
| Gm13632 | 65.5249491 | 1.470371688 | 9.95E-08 |

| | | | |
|---|---|---|---|
| Nkiras2 | 1742.751952 | 1.469726951 | 1.28E-15 |
| Map1lc3a | 2634.199358 | 1.469515977 | 4.01E-17 |
| Psmd11 | 5834.119852 | 1.469052122 | 1.19E-41 |
| Cit | 699.3070544 | 1.468480816 | 2.72E-05 |
| Bysl | 1467.654049 | 1.467565266 | 7.04E-17 |
| Plpp2 | 931.3288588 | 1.467020528 | 5.14E-21 |
| Il17rc | 2513.616066 | 1.466692345 | 8.40E-65 |
| Wincr1 | 360.4301665 | 1.466624753 | 4.33E-06 |
| Cox18 | 676.0390405 | 1.466344555 | 4.23E-25 |
| NA | 3226.764882 | 1.46612527 | 3.75E-18 |
| Ppa1 | 2005.850286 | 1.466006162 | 3.13E-13 |
| Pgpep1 | 2247.346939 | 1.465012937 | 4.21E-25 |
| Cox7a2l | 10854.98014 | 1.464432663 | 4.15E-12 |
| Pde6g | 13.41208273 | 1.461696695 | 1.83E-02 |
| Gm10080 | 25.45478434 | 1.460056146 | 2.08E-04 |
| Inca1 | 136.1457142 | 1.459371171 | 2.97E-08 |
| Mpp6 | 1620.758067 | 1.459090893 | 1.92E-10 |
| 1810009A15Rik | 1592.766553 | 1.458527758 | 3.81E-28 |
| A530013C23Rik | 96.11709964 | 1.458325915 | 5.84E-07 |
| Qars | 5609.254646 | 1.457157767 | 3.87E-17 |
| Cpxm1 | 32.08295723 | 1.456774752 | 2.32E-04 |
| Ctbs | 628.3728415 | 1.456610055 | 4.89E-37 |
| Pcbp1 | 15320.23063 | 1.45646168 | 1.02E-30 |
| Polr1d | 2742.962488 | 1.455349461 | 7.39E-44 |
| Wwp2 | 3084.461446 | 1.454899383 | 3.25E-19 |
| Mecp2 | 3791.304598 | 1.454051261 | 9.51E-17 |
| Aldh1a7 | 739.2403274 | 1.453827624 | 5.47E-03 |
| S100a3 | 56.32424442 | 1.453135949 | 1.82E-04 |
| Rpl17-ps9 | 231.5005683 | 1.45313375 | 6.78E-10 |
| Rbms2 | 13507.54112 | 1.453071873 | 7.36E-14 |
| Gm4204 | 1280.220195 | 1.4529283 | 5.78E-12 |
| Cib2 | 11.95957914 | 1.452496595 | 1.09E-02 |
| Ssh1 | 2810.464612 | 1.452466704 | 2.86E-38 |
| Dvl1 | 2494.315404 | 1.452393796 | 8.14E-29 |
| Zhx2 | 795.6214179 | 1.452127562 | 2.38E-12 |
| Nfil3 | 1259.058948 | 1.452089238 | 1.07E-12 |
| Gm26767 | 23.92654766 | 1.451501503 | 1.41E-03 |
| Tnfsf9 | 487.3199691 | 1.450693305 | 8.78E-09 |
| Aoc2 | 159.8164997 | 1.450536874 | 9.38E-08 |
| Ube2cbp | 39.28225239 | 1.450350624 | 1.41E-04 |
| Abcg4 | 19.25427227 | 1.450336172 | 8.24E-03 |
| Spindoc | 1533.971125 | 1.449133491 | 2.98E-17 |
| Bscl2 | 225.8361494 | 1.446849719 | 2.87E-08 |
| Polk | 816.6986151 | 1.44652659 | 1.18E-13 |
| Btf3 | 18129.28541 | 1.446344323 | 5.64E-23 |
| Ptprv | 67.14689553 | 1.44601718 | 4.60E-02 |
| Cmtr1 | 4053.649139 | 1.445132406 | 1.86E-37 |

| | | | |
|---|---|---|---|
| Wdr74 | 1689.807573 | 1.444671397 | 1.04E-21 |
| Rps6-ps1 | 64.52353795 | 1.443665935 | 4.10E-05 |
| 1190005I06Rik | 16.86881948 | 1.441767708 | 1.47E-02 |
| Rps5 | 42435.71707 | 1.441593161 | 3.14E-11 |
| Slc7a5 | 7418.657169 | 1.441536565 | 1.66E-05 |
| Usp20 | 1287.726103 | 1.440138133 | 1.09E-23 |
| Stip1 | 10290.17675 | 1.439788088 | 8.91E-17 |
| Bach2 | 553.8509816 | 1.439497126 | 8.92E-06 |
| E2f7 | 788.1276993 | 1.439439381 | 3.53E-05 |
| Limk1 | 1848.013808 | 1.438618156 | 2.30E-16 |
| Rps23 | 23042.03205 | 1.438437641 | 6.20E-10 |
| Gm7536 | 46411.44855 | 1.437798018 | 3.47E-15 |
| Idh3b | 5787.050988 | 1.435708389 | 8.87E-23 |
| Gpi1 | 20361.72384 | 1.435661972 | 3.96E-19 |
| Pam16 | 808.4039737 | 1.435464788 | 8.62E-30 |
| Cobll1 | 2804.002598 | 1.435164794 | 1.67E-28 |
| Cpt1c | 488.9285033 | 1.43503193 | 4.25E-26 |
| Thnsl1 | 299.7345792 | 1.435004756 | 1.41E-19 |
| Eif3l | 10508.6021 | 1.434854956 | 3.12E-26 |
| Fmo3 | 67.82999797 | 1.434697143 | 3.63E-02 |
| Eif4b | 44123.87184 | 1.434458567 | 7.56E-18 |
| Plekhh3 | 1084.0824 | 1.434099926 | 1.49E-18 |
| Rps10 | 48037.18352 | 1.432852648 | 3.37E-13 |
| Rab26 | 6.78843172 | 1.432060824 | 3.61E-02 |
| Rin3 | 39.43852099 | 1.431470647 | 3.56E-04 |
| Impdh1 | 2647.662952 | 1.431427195 | 6.68E-35 |
| Sod3 | 262.7305939 | 1.431239293 | 3.96E-03 |
| Pnpla6 | 1143.850243 | 1.430811563 | 1.83E-40 |
| Tuba4a | 1663.613125 | 1.429748778 | 3.69E-09 |
| Tns2 | 5211.113391 | 1.429634681 | 1.31E-25 |
| Ndufb6 | 1520.582627 | 1.429067931 | 3.46E-31 |
| Rps25 | 29032.20276 | 1.428913402 | 2.97E-15 |
| Wnk4 | 1179.840556 | 1.428522968 | 5.64E-13 |
| Tmem106c | 1500.53999 | 1.428183294 | 3.97E-19 |
| Pabpc1 | 86856.51255 | 1.427929898 | 6.68E-23 |
| Bhlhe41 | 899.5494575 | 1.427707311 | 9.06E-09 |
| Zfp473 | 122.7050172 | 1.426887857 | 2.39E-05 |
| Rbm28 | 3905.034538 | 1.426131026 | 1.54E-22 |
| Rpl26 | 50023.25159 | 1.425908223 | 9.57E-18 |
| Atp7b | 74.21699247 | 1.425886237 | 5.39E-09 |
| Gm11960 | 82.05329476 | 1.425406165 | 4.27E-05 |
| Slc30a6 | 877.1499439 | 1.425327245 | 4.51E-40 |
| Fam228b | 22.34992606 | 1.424927953 | 6.01E-04 |
| Gm6969 | 2684.561492 | 1.424464876 | 4.54E-11 |
| Hgs | 3316.298431 | 1.424206784 | 4.42E-19 |
| Dnm3 | 207.3505454 | 1.423579555 | 1.37E-05 |
| Prrg4 | 14.31062335 | 1.42198485 | 2.75E-02 |

| | | | |
|---|---|---|---|
| Rpl41 | 80605.63953 | 1.421897 | 1.57E-12 |
| Popdc3 | 16.72023235 | 1.4216749 | 3.99E-02 |
| Gamt | 997.1896559 | 1.421477004 | 1.38E-16 |
| Amacr | 720.3402139 | 1.420994485 | 4.56E-18 |
| Emc3 | 5381.192997 | 1.420629114 | 2.87E-26 |
| Serpinb6a | 9297.117704 | 1.419643057 | 1.26E-35 |
| Tbrg1 | 7943.990575 | 1.418645576 | 7.09E-23 |
| Ttc32 | 115.9407862 | 1.417369619 | 2.57E-13 |
| Emc6 | 1268.950995 | 1.417337694 | 1.18E-56 |
| Gm11808 | 30010.49529 | 1.416622103 | 2.87E-16 |
| Calr3 | 72.66829931 | 1.415020575 | 3.08E-09 |
| Rpsa-ps9 | 649.5998619 | 1.41474491 | 8.99E-07 |
| Gm12454 | 22.74329518 | 1.413954513 | 2.42E-02 |
| Rps15a-ps6 | 8.092617238 | 1.413492329 | 4.57E-02 |
| Platr20 | 22.75396567 | 1.412421273 | 3.23E-03 |
| Adra1b | 80.73107556 | 1.412212929 | 1.95E-04 |
| Dcaf4 | 1122.333632 | 1.411861456 | 5.93E-17 |
| Mta1 | 6436.498902 | 1.411451028 | 4.24E-28 |
| Cpeb2 | 370.5002161 | 1.410910617 | 4.29E-13 |
| Cdc42ep5 | 621.0593718 | 1.410683082 | 7.43E-12 |
| Gm8172 | 44.9062134 | 1.410611826 | 2.41E-04 |
| Tpx2 | 3448.017744 | 1.410215279 | 2.58E-06 |
| Adss | 4524.176795 | 1.409918295 | 6.33E-34 |
| Il18r1 | 15.30815582 | 1.409066419 | 2.74E-02 |
| NA | 28.34641966 | 1.408872116 | 3.74E-04 |
| Rpp30 | 619.7082099 | 1.40877807 | 1.92E-18 |
| Uroc1 | 10.6706882 | 1.408552366 | 3.90E-02 |
| Ndufb4 | 577.8670532 | 1.40837605 | 4.94E-13 |
| Rassf1 | 1982.544538 | 1.408362855 | 4.13E-25 |
| Gm5847 | 8.502743325 | 1.407905799 | 4.06E-02 |
| Tll1 | 41.96951859 | 1.40775991 | 1.17E-02 |
| Reep4 | 971.1311621 | 1.407236699 | 2.88E-12 |
| Trmo | 211.4049252 | 1.407138295 | 6.60E-11 |
| Mmaa | 989.4904332 | 1.406441428 | 7.32E-12 |
| Ifi35 | 678.7565277 | 1.406162516 | 1.23E-16 |
| Arl2 | 2679.750191 | 1.405943578 | 1.57E-14 |
| Rabl3 | 368.0500421 | 1.405568085 | 5.69E-15 |
| Gm5526 | 15.93317991 | 1.405291267 | 7.32E-03 |
| Zfand2b | 745.0809418 | 1.405006676 | 2.69E-32 |
| Mcts2 | 402.5475994 | 1.403800588 | 2.67E-22 |
| Rpp40 | 775.7960989 | 1.402735477 | 2.79E-13 |
| E2f6 | 108.6963516 | 1.402161886 | 1.04E-08 |
| Zfp511 | 616.6235673 | 1.402009261 | 3.62E-20 |
| Rpl17 | 47230.04156 | 1.401962269 | 5.87E-12 |
| Gm4691 | 61.07780483 | 1.401621921 | 5.13E-04 |
| Sympk | 8088.670725 | 1.401443068 | 8.31E-21 |
| Gm20636 | 8.32813839 | 1.400697598 | 3.72E-02 |

| | | | |
|---|---|---|---|
| Nmt1 | 7745.587509 | 1.400525782 | 2.56E-25 |
| Zfp593 | 763.2498669 | 1.400093606 | 8.12E-17 |
| Necap1 | 2428.802313 | 1.399262304 | 1.39E-32 |
| Mapre3 | 1426.057277 | 1.398699446 | 3.44E-13 |
| Slc8a3 | 48.28956434 | 1.39802151 | 1.05E-03 |
| Slc29a2 | 224.3345335 | 1.397853545 | 2.78E-14 |
| Ciao2b | 1227.316825 | 1.396691914 | 5.21E-21 |
| Gabarapl2 | 2747.465548 | 1.396265752 | 6.48E-15 |
| Tuba1a | 32146.48033 | 1.396064727 | 3.22E-18 |
| Usp22 | 5532.790043 | 1.395681443 | 6.43E-14 |
| Paxx | 701.5538367 | 1.39556603 | 1.86E-13 |
| Aip | 2860.21654 | 1.394964253 | 2.88E-22 |
| Hras | 1694.369396 | 1.392877843 | 3.31E-15 |
| 2310039H08Rik | 526.4348822 | 1.392828893 | 3.82E-21 |
| Rpl28 | 2509.809573 | 1.392089476 | 5.24E-07 |
| Pif1 | 283.1872027 | 1.391968689 | 3.91E-04 |
| Snrnp200 | 11710.44943 | 1.391917887 | 6.43E-29 |
| Rpl7a | 25955.19859 | 1.391848303 | 3.74E-13 |
| Pdzd11 | 1920.318247 | 1.391805837 | 3.42E-24 |
| Tcirg1 | 2505.669485 | 1.39109615 | 3.37E-17 |
| Rplp0 | 148995.8377 | 1.390573046 | 4.47E-16 |
| Rarg | 5414.781718 | 1.390562895 | 1.03E-22 |
| Ap2a1 | 5510.979936 | 1.389570643 | 1.84E-26 |
| Aup1 | 6846.043004 | 1.38930135 | 6.44E-32 |
| Adk | 3190.013385 | 1.387261779 | 1.07E-29 |
| Sap30 | 554.9416513 | 1.386778626 | 8.13E-11 |
| Ogfod1 | 1856.045924 | 1.385743696 | 5.76E-27 |
| Eif3k | 5580.642354 | 1.385091931 | 5.52E-14 |
| Gm8730 | 13578.63654 | 1.384921385 | 2.90E-12 |
| Zpr1 | 2032.138192 | 1.384552545 | 5.27E-21 |
| Pes1 | 5474.440582 | 1.384408254 | 3.89E-32 |
| C1qbp | 6609.162753 | 1.384372854 | 1.24E-29 |
| Hist3h2a | 838.6480394 | 1.384012032 | 5.97E-11 |
| Ccnk | 2008.508271 | 1.383274013 | 2.78E-12 |
| Mars | 4381.005849 | 1.382810511 | 1.47E-16 |
| Tpst2 | 2312.601895 | 1.382325171 | 6.11E-19 |
| Snap47 | 3433.560277 | 1.382133559 | 1.75E-24 |
| Rps2-ps5 | 10.85470053 | 1.381856164 | 1.25E-02 |
| Pih1d1 | 1075.565427 | 1.380103035 | 1.98E-14 |
| Hoxc4 | 166.5136363 | 1.380011398 | 1.07E-15 |
| Vopp1 | 3424.567913 | 1.379398343 | 9.96E-21 |
| Ppp1r35 | 376.7552798 | 1.378549879 | 1.13E-10 |
| Aldoa | 31312.43539 | 1.378385128 | 3.47E-14 |
| Apbb1 | 1088.659518 | 1.378184883 | 3.10E-25 |
| Immp2l | 226.8841086 | 1.377882493 | 6.09E-12 |
| Notch3 | 14.41931232 | 1.377393108 | 3.51E-02 |
| Gm14438 | 233.5442823 | 1.37721031 | 2.01E-04 |

| | | | |
|---|---|---|---|
| Cs | 17852.67489 | 1.376983825 | 2.13E-22 |
| Gm11627 | 21.59124812 | 1.376882496 | 2.66E-03 |
| Map3k13 | 64.74950077 | 1.376806645 | 1.39E-07 |
| Pla2g12a | 876.4735081 | 1.376184591 | 5.67E-16 |
| Gjc3 | 18.46004011 | 1.375801801 | 4.61E-03 |
| Agpat4 | 1161.530843 | 1.375006252 | 2.00E-17 |
| Rpl38 | 13527.73583 | 1.373070462 | 7.34E-14 |
| Plekhf2 | 391.5956648 | 1.372901359 | 1.39E-17 |
| Lgmn | 8054.730872 | 1.372475702 | 2.95E-12 |
| Gm7488 | 1833.106868 | 1.37226454 | 4.71E-27 |
| Yipf3 | 3447.546377 | 1.371535788 | 5.66E-11 |
| Naa10 | 1512.749773 | 1.370775423 | 1.38E-22 |
| Yif1b | 1695.896623 | 1.370531713 | 1.53E-19 |
| Ak1 | 2929.214187 | 1.369554097 | 3.25E-23 |
| Mettl22 | 320.1298717 | 1.369521273 | 1.77E-18 |
| Eif2d | 2660.698327 | 1.3694699 | 2.06E-18 |
| Lsm12 | 2001.617156 | 1.368972553 | 2.79E-50 |
| Kifc1 | 820.5197881 | 1.367986109 | 2.91E-06 |
| Lrrc28 | 607.671651 | 1.366749656 | 1.77E-30 |
| Map1b | 6864.42189 | 1.366676022 | 1.10E-17 |
| Brf2 | 644.3112501 | 1.366330555 | 4.13E-17 |
| Gm10247 | 32.75537283 | 1.366161129 | 1.49E-03 |
| Mrps31 | 966.2834087 | 1.365103133 | 1.50E-15 |
| Mdh2 | 12364.87538 | 1.364542218 | 4.32E-22 |
| Dusp11 | 7910.266802 | 1.364195695 | 4.04E-19 |
| Zfp703 | 916.2166658 | 1.364170779 | 2.26E-10 |
| Basp1 | 5262.232606 | 1.363382235 | 4.56E-24 |
| Hsd3b7 | 1224.252911 | 1.362323324 | 2.42E-13 |
| Rpl7a-ps11 | 16425.80428 | 1.361970515 | 2.56E-12 |
| Fggy | 872.0594798 | 1.361712767 | 1.92E-17 |
| Rab4a | 910.1117394 | 1.360667394 | 1.62E-25 |
| Mrps16 | 1130.112843 | 1.360596429 | 3.89E-25 |
| Utp23 | 570.5468979 | 1.359631368 | 6.77E-48 |
| Eef1d | 9782.425807 | 1.357909546 | 1.49E-17 |
| Gm9396 | 66.43147602 | 1.356771601 | 3.16E-04 |
| Gnb5 | 631.122664 | 1.355570006 | 6.30E-19 |
| Snx7 | 2260.56405 | 1.354807302 | 5.66E-12 |
| Psmb7 | 5329.244185 | 1.354630698 | 2.04E-34 |
| Serp2 | 10.36553764 | 1.354189488 | 4.24E-02 |
| Rplp1 | 72292.15409 | 1.352383159 | 3.14E-12 |
| Ptrh1 | 1785.441485 | 1.351755353 | 4.23E-16 |
| Mtg1 | 484.3456982 | 1.351493821 | 1.20E-12 |
| Idh3g | 3552.809488 | 1.351200378 | 7.34E-51 |
| Tns4 | 36.47295026 | 1.351196921 | 2.03E-02 |
| Cacnb3 | 4366.285053 | 1.351187793 | 6.04E-11 |
| 2810025M15Rik | 454.5514841 | 1.350689336 | 7.21E-13 |
| Spryd7 | 748.305391 | 1.350269531 | 5.39E-35 |

| | | | |
|---|---|---|---|
| Naa25 | 2845.116492 | 1.350021492 | 5.79E-31 |
| Ercc3 | 2067.814332 | 1.349049507 | 1.55E-27 |
| Rps6ka5 | 321.2408474 | 1.348057827 | 2.85E-12 |
| Pcdh1 | 2459.618889 | 1.34764351 | 2.27E-19 |
| Inpp5k | 2174.199723 | 1.347477562 | 4.96E-17 |
| Actr10 | 3839.071474 | 1.34747742 | 5.05E-40 |
| Trim11 | 1637.383168 | 1.347270277 | 1.33E-26 |
| Psma7 | 5994.916682 | 1.3471359 | 6.48E-23 |
| Gm10327 | 77.92351245 | 1.3470518 | 3.80E-05 |
| Gtpbp2 | 2282.051956 | 1.34618434 | 1.05E-23 |
| Rpl15-ps3 | 37991.77318 | 1.345603964 | 5.19E-11 |
| Slc37a2 | 33.44387964 | 1.345356132 | 8.23E-04 |
| 2310022B05Rik | 12775.90222 | 1.345285815 | 7.62E-11 |
| Ufsp1 | 105.012883 | 1.345155574 | 6.89E-09 |
| Spry1 | 847.9921191 | 1.345083254 | 3.51E-07 |
| Nsmce1 | 1028.989147 | 1.344509783 | 1.65E-21 |
| Rfc4 | 532.2784859 | 1.343957575 | 1.92E-12 |
| Gm15793 | 214.717008 | 1.343807472 | 1.50E-05 |
| Rpl35 | 28922.30866 | 1.343484265 | 2.34E-16 |
| Ccni | 10061.03268 | 1.34311672 | 1.44E-16 |
| Slc20a1 | 2917.806275 | 1.342693012 | 2.31E-14 |
| Gm14150 | 809.4434399 | 1.342682799 | 1.56E-14 |
| 4632427E13Rik | 66.32009799 | 1.342068891 | 6.29E-03 |
| Susd6 | 4237.928786 | 1.341113617 | 2.39E-11 |
| Gm13292 | 28.69762668 | 1.340795596 | 1.88E-02 |
| Rpl19 | 61202.65009 | 1.340434819 | 5.69E-12 |
| Bcorl1 | 1056.977538 | 1.340146609 | 8.98E-13 |
| Nmt2 | 3256.678842 | 1.3400445 | 4.35E-18 |
| Col18a1 | 133.7544371 | 1.33922339 | 2.29E-03 |
| Baz2a | 4071.956571 | 1.338865815 | 6.67E-72 |
| Timmdc1 | 721.404571 | 1.338058829 | 1.22E-22 |
| Stap2 | 79.90043653 | 1.336320681 | 1.21E-06 |
| Coq10b | 1114.821112 | 1.336297211 | 2.09E-13 |
| Tm4sf1 | 4004.71189 | 1.335807647 | 2.76E-13 |
| Pwwp2b | 1031.015521 | 1.335439473 | 1.26E-11 |
| Mea1 | 2622.416807 | 1.334942739 | 4.78E-24 |
| Gm20900 | 83.4022082 | 1.334705856 | 2.59E-04 |
| NA | 20.71352463 | 1.334664805 | 1.41E-02 |
| Ang2 | 58.05561455 | 1.334647417 | 6.94E-04 |
| Rrp12 | 1698.775959 | 1.334090853 | 1.35E-08 |
| Snord37 | 16.51974151 | 1.333989576 | 2.32E-02 |
| Eed | 1239.554501 | 1.333857144 | 4.99E-25 |
| Git2 | 3927.938835 | 1.333028062 | 1.88E-15 |
| Pex16 | 676.2702163 | 1.33296381 | 1.60E-16 |
| Pitpnm1 | 3326.041469 | 1.332902918 | 1.34E-37 |
| Gm10073 | 9.998377766 | 1.332689456 | 3.74E-02 |
| Npepps | 9806.645939 | 1.332078924 | 3.51E-32 |

| | | | |
|---|---|---|---|
| Rpl4 | 100332.3709 | 1.331307729 | 2.81E-14 |
| Dnajc4 | 650.8374999 | 1.330957644 | 4.02E-11 |
| Ddx39 | 3293.012759 | 1.330703936 | 1.70E-09 |
| Thumpd3 | 1520.849078 | 1.330422113 | 3.60E-29 |
| Nploc4 | 4231.422089 | 1.330075895 | 1.17E-16 |
| Lap3 | 4765.227468 | 1.328900791 | 2.58E-31 |
| Mrps30 | 1413.123487 | 1.328292702 | 1.80E-28 |
| Ap2m1 | 16486.59881 | 1.327734849 | 2.86E-28 |
| Irs2 | 2222.67984 | 1.327627357 | 3.67E-20 |
| Dtwd1 | 574.3759161 | 1.327505874 | 9.82E-27 |
| Phb2 | 9596.173477 | 1.32677688 | 1.06E-22 |
| Ptgs2os | 66.7624299 | 1.326507451 | 8.31E-04 |
| Gm17709 | 122.5539367 | 1.325390661 | 6.12E-12 |
| Gm7808 | 50.83150304 | 1.325359736 | 1.61E-04 |
| Gm9843 | 23508.2696 | 1.32535348 | 8.10E-11 |
| NA | 4706.046236 | 1.325296345 | 2.17E-21 |
| Fbxo47 | 20.02647274 | 1.325188622 | 1.14E-02 |
| Psmb6 | 4995.39886 | 1.324196946 | 5.46E-28 |
| Agpat3 | 4907.691748 | 1.32387887 | 5.01E-32 |
| Ndufb4c | 1556.461565 | 1.323818006 | 2.35E-20 |
| Rpl6 | 74724.03916 | 1.323453212 | 1.33E-13 |
| Prc1 | 2935.87793 | 1.323437482 | 2.91E-05 |
| Nadk | 3967.334799 | 1.322096437 | 7.60E-23 |
| Pabpc4 | 13810.91171 | 1.322064686 | 9.11E-23 |
| Psmd12 | 4257.969189 | 1.321524043 | 4.17E-32 |
| Ssr4 | 4717.047134 | 1.32124468 | 1.63E-12 |
| Tox4 | 2329.373657 | 1.321237675 | 3.07E-16 |
| Cnpy4 | 1150.911627 | 1.32101339 | 1.30E-17 |
| Drg2 | 2531.129456 | 1.3209215 | 2.25E-22 |
| Mllt3 | 4797.047418 | 1.320565839 | 7.80E-16 |
| NA | 8103.204573 | 1.320494655 | 1.34E-12 |
| Ptcd2 | 866.1635995 | 1.319949329 | 2.87E-26 |
| Spsb1 | 515.9985486 | 1.319165477 | 6.97E-18 |
| Psmd13 | 2802.297219 | 1.319154538 | 1.31E-13 |
| Ecsit | 800.8495056 | 1.318863513 | 1.07E-16 |
| Ppp1r11 | 1908.709021 | 1.318279667 | 9.80E-12 |
| Rpl11 | 26732.08196 | 1.317927485 | 7.12E-14 |
| Nfu1 | 1848.556139 | 1.317744227 | 2.90E-58 |
| Hyal1 | 467.9810951 | 1.317145726 | 2.38E-06 |
| Zc3h8 | 210.3834763 | 1.31680481 | 1.44E-09 |
| Ddr1 | 5318.493109 | 1.31623834 | 1.34E-19 |
| Prex2 | 704.5135401 | 1.315466624 | 3.63E-06 |
| Gm8116 | 1189.5011 | 1.315225697 | 1.88E-36 |
| Ecm1 | 1410.00657 | 1.314665422 | 7.78E-06 |
| Zswim7 | 240.4272276 | 1.313957519 | 1.63E-18 |
| Tjp1 | 4773.974161 | 1.313875792 | 1.67E-18 |
| Tarsl2 | 774.4453427 | 1.313423596 | 1.25E-29 |

| | | | |
|---|---|---|---|
| Ahcy | 200.6339253 | 1.313363529 | 1.83E-03 |
| Abhd17a | 2211.350324 | 1.313152252 | 9.97E-16 |
| Rwdd2b | 220.2707525 | 1.312674248 | 1.12E-15 |
| Nhp2 | 1896.70157 | 1.311599798 | 1.73E-13 |
| Eno1b | 12787.54532 | 1.311334309 | 5.12E-13 |
| Ssu72 | 1549.612509 | 1.311098274 | 4.05E-27 |
| Dctn2 | 11012.49689 | 1.310822784 | 6.19E-21 |
| Dnpep | 4104.734475 | 1.310789972 | 6.39E-22 |
| Cct6b | 24.23556002 | 1.310071043 | 2.23E-03 |
| Rrn3 | 3178.831608 | 1.309987509 | 9.85E-28 |
| Pmpcb | 3918.866943 | 1.309475916 | 5.53E-28 |
| Uba52-ps | 41.60175151 | 1.309241911 | 6.86E-04 |
| Hid1 | 87.39260748 | 1.309134818 | 5.23E-04 |
| Mrpl21 | 1397.633345 | 1.309078045 | 4.84E-41 |
| Ppp1r13l | 651.9201341 | 1.308592481 | 8.57E-09 |
| Syn2 | 38.51540974 | 1.30716428 | 2.89E-03 |
| Gm26724 | 228.142172 | 1.306403861 | 2.60E-06 |
| Adprm | 564.4792591 | 1.30626538 | 9.04E-19 |
| Txnrd1 | 11050.67123 | 1.306111341 | 2.98E-14 |
| Aldh7a1 | 2850.269756 | 1.305298356 | 8.48E-28 |
| 1190007I07Rik | 76.5727787 | 1.304504617 | 7.44E-10 |
| Gm45837 | 28.69853484 | 1.304443048 | 1.51E-02 |
| Aprt | 6193.680378 | 1.30436225 | 8.19E-17 |
| Plod3 | 6604.231022 | 1.303639931 | 4.30E-18 |
| Rpl36 | 15843.79799 | 1.303057021 | 2.89E-11 |
| Cenpp | 150.4209713 | 1.302209 | 8.42E-08 |
| Ncbp1 | 3020.179347 | 1.301937893 | 1.08E-20 |
| Gm10284 | 47.68678524 | 1.301195839 | 2.06E-03 |
| Tbl3 | 1798.960926 | 1.300898129 | 2.17E-11 |
| Rps7 | 10324.97548 | 1.300474341 | 3.24E-08 |
| Gsto2 | 540.3671304 | 1.300455694 | 1.78E-09 |
| Ube2g2 | 3636.513381 | 1.299298185 | 6.09E-14 |
| Abhd5 | 2340.155512 | 1.299020835 | 1.10E-14 |
| Cyth2 | 2204.307457 | 1.298622443 | 1.79E-16 |
| Gtpbp6 | 1438.376684 | 1.29844934 | 3.23E-18 |
| Paip2b | 3971.739471 | 1.298146607 | 1.24E-16 |
| Gm15050 | 189.9018835 | 1.298077696 | 1.83E-09 |
| Ltbp1 | 3648.116466 | 1.297970626 | 4.00E-10 |
| Nup43 | 725.9634131 | 1.296996243 | 2.09E-13 |
| Eif3c | 17700.40432 | 1.296588975 | 1.25E-33 |
| Gm9855 | 2483.951233 | 1.295602883 | 8.98E-28 |
| Ethe1 | 653.1180357 | 1.294499775 | 1.03E-23 |
| Vezt | 1941.869543 | 1.294156189 | 3.23E-29 |
| Cad | 5610.102911 | 1.293745764 | 5.92E-12 |
| Pold2 | 1477.91555 | 1.292599044 | 5.86E-12 |
| Plac1 | 35.76022485 | 1.292517096 | 3.49E-03 |
| Poc1a | 204.6897645 | 1.291401404 | 1.86E-06 |

| | | | |
|---|---|---|---|
| Rpl18a | 66237.11451 | 1.291351704 | 7.11E-09 |
| Poldip3 | 5976.958649 | 1.291098453 | 2.35E-18 |
| Gm7367 | 116.3501826 | 1.290425919 | 1.55E-02 |
| Ascc1 | 1690.168605 | 1.290346414 | 1.22E-11 |
| Rrp9 | 952.0180092 | 1.289851011 | 1.07E-12 |
| 3110040N11Rik | 458.3785251 | 1.289050169 | 9.73E-29 |
| Rpl14 | 39168.7694 | 1.287892115 | 1.95E-09 |
| E230025N22Rik | 56.16707186 | 1.287849656 | 7.22E-04 |
| Pcgf5 | 1180.442953 | 1.287409158 | 1.01E-19 |
| Rnasek | 2284.435357 | 1.286218762 | 7.65E-11 |
| Rplp2 | 29391.29674 | 1.285822892 | 1.89E-13 |
| Dpm1 | 807.4452685 | 1.285484428 | 4.51E-08 |
| Uqcrfs1 | 2399.105649 | 1.284422816 | 5.18E-29 |
| Gm10076 | 56.07162855 | 1.284311264 | 2.80E-04 |
| Zdhhc9 | 2534.383131 | 1.284233977 | 2.50E-21 |
| Crlf1 | 353.0096371 | 1.284030618 | 1.54E-04 |
| Cops4 | 4067.416077 | 1.283698703 | 2.37E-23 |
| Tdrp | 456.9726997 | 1.283436096 | 7.65E-16 |
| Stk17b | 1172.316199 | 1.283379196 | 9.35E-13 |
| 2610318N02Rik | 78.5731003 | 1.283244344 | 1.70E-03 |
| Usp30 | 1743.102786 | 1.283099581 | 3.35E-14 |
| Pla2g4a | 4766.449097 | 1.283088065 | 1.92E-05 |
| Abca3 | 2846.907726 | 1.282820755 | 9.94E-24 |
| 2-Mar | 2784.399668 | 1.282506133 | 4.48E-17 |
| Rpl37 | 20885.2681 | 1.282120296 | 8.38E-12 |
| Gng2 | 472.9891909 | 1.281787387 | 1.54E-09 |
| Polb | 1158.380286 | 1.281033268 | 7.34E-24 |
| Tjap1 | 2545.037818 | 1.280457279 | 2.89E-26 |
| Actg1 | 96158.53644 | 1.280036763 | 6.56E-19 |
| Sh3gl3 | 191.0386475 | 1.279902109 | 3.85E-10 |
| Gm6472 | 47311.23449 | 1.279844108 | 1.01E-11 |
| Mrps35 | 1483.24933 | 1.279508043 | 1.61E-25 |
| Tnfaip1 | 4512.561739 | 1.279036918 | 7.94E-28 |
| Neil1 | 265.1797796 | 1.277486821 | 5.29E-09 |
| Per3 | 660.4660491 | 1.276569661 | 7.31E-10 |
| Fam214b | 1462.593913 | 1.275926191 | 1.47E-16 |
| Gm26525 | 46.56506484 | 1.275382874 | 1.08E-02 |
| Eno1 | 16599.21767 | 1.275057858 | 9.48E-12 |
| Ppm1g | 5386.882069 | 1.274517632 | 2.33E-16 |
| Ncan | 14.08838223 | 1.274359518 | 4.38E-02 |
| Map2k3os | 158.9798746 | 1.27352315 | 7.52E-05 |
| Vcam1 | 8733.754356 | 1.273193209 | 6.76E-26 |
| Gna13 | 4550.858975 | 1.272629278 | 3.38E-22 |
| Mttp | 478.4641153 | 1.272491536 | 1.49E-25 |
| Zfp972 | 96.72997032 | 1.272164995 | 2.31E-05 |
| Ccdc40 | 33.18276155 | 1.271702454 | 5.74E-03 |
| Carhsp1 | 4864.078369 | 1.271002618 | 2.44E-17 |

| | | | |
|---|---|---|---|
| Ccdc43 | 996.7600218 | 1.270586656 | 7.39E-36 |
| Smap1 | 2848.464316 | 1.270366929 | 2.86E-51 |
| Gm10260 | 27463.1406 | 1.270107424 | 5.37E-11 |
| Rcc1 | 1466.074838 | 1.270095998 | 2.37E-11 |
| Gm17828 | 1219.480847 | 1.270092227 | 5.08E-06 |
| Parp6 | 1359.090638 | 1.270020878 | 1.11E-15 |
| Ptprk | 4819.221325 | 1.269349058 | 4.89E-14 |
| Rpl7a-ps3 | 1301.1174 | 1.268648685 | 1.40E-09 |
| Hibadh | 3736.286012 | 1.268538639 | 2.20E-21 |
| Zc2hc1c | 99.23641284 | 1.268408996 | 1.06E-05 |
| Klf11 | 571.7586071 | 1.268336267 | 1.45E-06 |
| Pkn3 | 159.00837 | 1.267965353 | 2.29E-04 |
| Jup | 2353.842657 | 1.26796015 | 2.12E-08 |
| Prss53 | 73.1378317 | 1.267634385 | 2.16E-04 |
| Msh5 | 95.26155785 | 1.266572668 | 8.55E-04 |
| Slc7a11 | 1232.289792 | 1.26631991 | 4.35E-21 |
| Gm11560 | 44.62081345 | 1.265929484 | 2.16E-03 |
| Tdp2 | 876.6310077 | 1.263775056 | 1.44E-15 |
| Rps12 | 48476.15642 | 1.263341365 | 6.48E-16 |
| Gm13413 | 64.46038927 | 1.26273003 | 1.79E-05 |
| Lgalsl | 3882.880886 | 1.262619902 | 9.52E-19 |
| Cpsf1 | 3715.112604 | 1.261962621 | 3.15E-23 |
| 2610206C17Rik | 15.87470752 | 1.260904452 | 4.39E-02 |
| Epha4 | 1347.621821 | 1.260762584 | 5.31E-20 |
| Tmem161a | 1680.07129 | 1.260125947 | 1.65E-26 |
| Rpl35a | 516.7149947 | 1.260060833 | 1.22E-10 |
| Gm12918 | 11988.44703 | 1.26005287 | 1.50E-04 |
| Hotairm1 | 163.7859472 | 1.25990544 | 1.11E-08 |
| Guca1a | 12.28516956 | 1.25988373 | 3.86E-02 |
| Abca5 | 1770.476615 | 1.259860413 | 3.56E-09 |
| Bag1 | 4120.741985 | 1.259564929 | 1.59E-19 |
| Gps1 | 4800.277109 | 1.259287773 | 4.61E-14 |
| Lonp2 | 4330.10061 | 1.258625214 | 4.26E-26 |
| Srpk1 | 4585.424015 | 1.257423231 | 9.65E-37 |
| Riok3 | 2754.746068 | 1.256433696 | 1.73E-23 |
| Ciao1 | 1904.407259 | 1.256430501 | 3.59E-19 |
| H2-T-ps | 256.5588408 | 1.256313073 | 5.77E-06 |
| Traf4 | 3041.560397 | 1.255707823 | 3.87E-12 |
| Tmem206 | 879.2146497 | 1.255067514 | 6.33E-29 |
| Mark4 | 1158.332822 | 1.254660041 | 2.86E-11 |
| Prmt2 | 1770.370006 | 1.254205221 | 1.93E-35 |
| Sdhb | 3446.628872 | 1.254045853 | 6.11E-26 |
| Cdc42se2 | 2787.71278 | 1.253640765 | 3.75E-49 |
| Crybg1 | 279.3516519 | 1.252981857 | 6.63E-05 |
| Dnph1 | 336.876214 | 1.251996007 | 6.63E-05 |
| Pck2 | 4321.410982 | 1.251360379 | 1.02E-11 |
| Exd2 | 1441.583752 | 1.250806009 | 2.70E-19 |

| | | | |
|---|---|---|---|
| Aga | 1959.623389 | 1.250485678 | 7.85E-19 |
| Ndufaf5 | 624.1161059 | 1.250009135 | 7.22E-15 |
| Gm26686 | 46.31665202 | 1.248995503 | 1.40E-03 |
| Gm13205 | 30.44429336 | 1.248689758 | 1.46E-03 |
| Tsg101 | 2849.982475 | 1.248526698 | 4.62E-18 |
| Eef1b2 | 46128.15298 | 1.248079646 | 1.63E-12 |
| Gm16409 | 267.0783991 | 1.248023296 | 9.62E-05 |
| Dlst | 7252.782551 | 1.247583706 | 9.46E-25 |
| Fam104a | 1760.116884 | 1.247233674 | 8.32E-20 |
| H2-Q7 | 211.7583553 | 1.247192316 | 5.45E-06 |
| Rpl34-ps1 | 26642.99647 | 1.246991289 | 1.03E-11 |
| Depdc7 | 242.1642713 | 1.246863135 | 7.85E-08 |
| Gm5566 | 46.62608169 | 1.246486725 | 2.64E-05 |
| Scamp3 | 3093.025348 | 1.245867237 | 1.12E-18 |
| Cenpw | 161.2582704 | 1.244945547 | 1.09E-05 |
| Gsto1 | 9133.181255 | 1.244483624 | 1.07E-11 |
| Oxa1l | 3698.955215 | 1.243288612 | 2.65E-08 |
| Mpp1 | 3836.686581 | 1.242236905 | 8.79E-19 |
| Htra2 | 5540.271863 | 1.241565383 | 6.92E-25 |
| Smyd3 | 1007.188368 | 1.241443932 | 9.80E-11 |
| Bcl3 | 359.6483038 | 1.241252481 | 4.17E-06 |
| Gstm4 | 203.1581612 | 1.241183455 | 8.79E-13 |
| Rpl9-ps6 | 49.21276014 | 1.240771409 | 4.61E-02 |
| Rpsa-ps11 | 28.09659456 | 1.239704152 | 3.77E-02 |
| Gm15501 | 77795.36435 | 1.238877909 | 1.84E-11 |
| Gcnt2 | 266.7781291 | 1.238760174 | 1.76E-07 |
| CAAA01118383.1 | 707.8337304 | 1.238709475 | 9.96E-16 |
| Rbmx2 | 337.2129035 | 1.237975129 | 2.50E-11 |
| Slc5a6 | 627.4269561 | 1.23787302 | 3.47E-14 |
| Pa2g4 | 8738.523288 | 1.237805351 | 7.68E-14 |
| Esrra | 994.4250692 | 1.237690548 | 2.43E-10 |
| Mrpl22 | 994.9769437 | 1.237606379 | 3.67E-27 |
| Apol9a | 154.7926263 | 1.236332123 | 5.35E-04 |
| Trp53cor1 | 155.8446845 | 1.236138685 | 6.73E-10 |
| Myc | 3282.758284 | 1.235802501 | 2.43E-16 |
| Pak1ip1 | 1946.659596 | 1.235668319 | 2.82E-28 |
| Gpd2 | 3642.520661 | 1.234430532 | 5.05E-40 |
| Mettl1 | 1327.406253 | 1.232961642 | 3.33E-16 |
| Flrt2 | 1425.858476 | 1.232834746 | 3.29E-06 |
| Rpl19-ps12 | 21.89225118 | 1.232374953 | 1.28E-02 |
| Hipk2 | 2809.337586 | 1.232188176 | 3.05E-17 |
| Rps25-ps1 | 24.66073837 | 1.232029058 | 1.85E-02 |
| Ripk2 | 653.0041488 | 1.231560354 | 1.69E-24 |
| Dnajb2 | 1081.510458 | 1.230372504 | 7.38E-19 |
| Ndufa9 | 3674.367459 | 1.229917956 | 1.83E-32 |
| Rad23b | 17055.86365 | 1.229878712 | 2.25E-19 |
| Rn7s1 | 84.50678455 | 1.228963926 | 7.07E-04 |

| | | | |
|---|---|---|---|
| Psmb1 | 9738.740386 | 1.228933114 | 4.97E-18 |
| Zbtb11 | 1086.873096 | 1.228849367 | 1.89E-14 |
| Dazap1 | 4156.04218 | 1.228617231 | 9.81E-38 |
| Acss2 | 638.0747025 | 1.228455585 | 9.34E-12 |
| B3gat3 | 1341.032423 | 1.22841224 | 5.80E-31 |
| Memo1 | 2828.465884 | 1.228226824 | 3.16E-28 |
| Ddx56 | 2294.187769 | 1.227713704 | 8.97E-17 |
| Slc52a2 | 724.533322 | 1.227652696 | 1.94E-16 |
| Pigf | 401.9097768 | 1.227324498 | 1.49E-24 |
| Eif3m | 5716.330193 | 1.227269171 | 7.90E-20 |
| Pdha1 | 9868.078405 | 1.227227688 | 1.63E-48 |
| Catsperg1 | 24.37350102 | 1.22473962 | 1.35E-02 |
| Adprh | 2437.400221 | 1.22406208 | 7.34E-15 |
| Fhdc1 | 1544.683828 | 1.22378741 | 2.99E-19 |
| Traf2 | 1462.06301 | 1.223144537 | 2.69E-10 |
| Eif6 | 3599.494945 | 1.223114147 | 6.08E-16 |
| Mrpl15 | 2833.345375 | 1.22290289 | 1.47E-25 |
| Pdss2 | 496.4537985 | 1.222848598 | 9.71E-12 |
| Gm16178 | 25.22230611 | 1.220666706 | 1.47E-02 |
| Polr2g | 3004.711592 | 1.220632753 | 1.13E-35 |
| Eef1a1 | 672249.7408 | 1.220293992 | 8.37E-10 |
| Spire2 | 545.5638821 | 1.2198726 | 1.32E-19 |
| Cops5 | 4298.493893 | 1.219527311 | 2.18E-49 |
| Col4a5 | 13744.73591 | 1.219459841 | 2.24E-22 |
| Psmd5 | 3176.614808 | 1.219290281 | 8.36E-21 |
| Asap3 | 387.600447 | 1.218850042 | 1.92E-07 |
| Sypl | 3290.631028 | 1.218299095 | 9.43E-31 |
| Cars | 3768.251332 | 1.218111318 | 2.37E-23 |
| Abhd8 | 2737.344855 | 1.21715388 | 3.72E-11 |
| Slc26a11 | 326.5897719 | 1.216255633 | 1.70E-10 |
| Tpra1 | 1917.311456 | 1.216244268 | 2.05E-27 |
| Duox2 | 35.60345511 | 1.215457975 | 1.60E-02 |
| Pcid2 | 1369.909591 | 1.214935546 | 8.76E-15 |
| Rps16-ps2 | 25503.26783 | 1.213993958 | 3.36E-07 |
| Mrpl35 | 1306.855469 | 1.213818407 | 1.35E-36 |
| Dlg4 | 3652.121914 | 1.213502558 | 7.47E-17 |
| Mgat4b | 6635.711925 | 1.212529334 | 1.60E-13 |
| Ruvbl2 | 1919.938162 | 1.212490572 | 6.66E-09 |
| Gm2606 | 23.06282492 | 1.212416689 | 1.74E-02 |
| Phf13 | 1356.38196 | 1.212314427 | 3.75E-08 |
| Gemin2 | 424.6443906 | 1.212303796 | 4.59E-23 |
| Gm27913 | 37.99607744 | 1.211008482 | 1.07E-03 |
| Mrps5 | 2126.38738 | 1.209832366 | 1.27E-36 |
| Prkx | 992.1556184 | 1.209334951 | 1.53E-14 |
| Myzap | 43.56695522 | 1.209118668 | 1.89E-03 |
| Morc4 | 3288.740245 | 1.209109865 | 9.19E-12 |
| Mid1ip1 | 939.9451529 | 1.208819975 | 5.67E-07 |

| | | | |
|---|---|---|---|
| Agpat1 | 3534.213831 | 1.20868751 | 6.41E-23 |
| Ube2j2 | 1199.238568 | 1.208472711 | 1.10E-25 |
| Klc1 | 5592.891025 | 1.20843172 | 5.32E-35 |
| Pttg1ip | 15567.85936 | 1.207871019 | 2.70E-14 |
| Timm17a | 2158.37418 | 1.207821801 | 7.05E-49 |
| Trim24 | 3746.206389 | 1.207593156 | 1.78E-09 |
| Prpf6 | 5480.800313 | 1.206920315 | 8.62E-14 |
| Ifi47 | 69.23413701 | 1.206806016 | 1.38E-06 |
| Ddx27 | 2854.313142 | 1.206760315 | 3.51E-21 |
| Rps12-ps3 | 32513.16414 | 1.206551437 | 1.09E-08 |
| Afmid | 240.573853 | 1.206204182 | 1.53E-06 |
| Sugt1 | 2423.145237 | 1.205972969 | 1.04E-19 |
| Chd5 | 78.15936498 | 1.205592018 | 1.33E-02 |
| Bcl7b | 963.352409 | 1.204180635 | 1.35E-14 |
| Usp14 | 3649.869986 | 1.203132344 | 8.79E-25 |
| Glud1 | 8559.981822 | 1.202868754 | 5.25E-70 |
| Helz | 2920.114213 | 1.201744329 | 7.24E-30 |
| Gnb2 | 15018.84145 | 1.201377827 | 4.83E-19 |
| Dnajb12 | 2755.669575 | 1.201157924 | 3.22E-22 |
| Gm16477 | 34.24256402 | 1.201122878 | 2.44E-04 |
| Gemin5 | 2847.233799 | 1.200101481 | 9.57E-16 |
| Gm10020 | 173.4954606 | 1.199870343 | 1.31E-04 |
| Rala | 2465.88936 | 1.199437782 | 1.84E-33 |
| Gm6436 | 10382.66031 | 1.199199241 | 8.83E-08 |
| Gm26707 | 18.84366641 | 1.199044137 | 1.89E-02 |
| Gm15590 | 184.0391023 | 1.198865485 | 1.06E-04 |
| Eif3f | 13440.69317 | 1.198436201 | 9.32E-07 |
| Cgrrf1 | 795.8021606 | 1.198235569 | 1.81E-19 |
| Gnb1 | 10433.30772 | 1.197731393 | 4.18E-82 |
| Ccnf | 1313.249746 | 1.197495257 | 8.56E-04 |
| Nsa2 | 9495.710085 | 1.196897802 | 1.44E-10 |
| Hint1 | 7364.354729 | 1.196631014 | 9.06E-19 |
| Eif4e2 | 4880.464822 | 1.19648155 | 1.34E-30 |
| Mrpl16 | 1103.742269 | 1.196352273 | 4.55E-16 |
| Myl6 | 22034.81211 | 1.195146394 | 9.90E-57 |
| Rpl23a | 76915.38591 | 1.195141609 | 1.85E-09 |
| Vta1 | 2205.334183 | 1.19478487 | 3.87E-23 |
| Sertad3 | 577.2030855 | 1.194681289 | 1.69E-09 |
| Tssc4 | 969.7799251 | 1.193769914 | 1.84E-12 |
| Sod2 | 2401.870402 | 1.193254707 | 2.24E-21 |
| Tlcd2 | 144.2625361 | 1.192927689 | 3.11E-06 |
| Mtfr1 | 2428.446967 | 1.192447806 | 4.74E-25 |
| Gm773 | 40.14795369 | 1.19244098 | 1.60E-02 |
| Mavs | 1763.108388 | 1.192265633 | 1.18E-27 |
| Trim36 | 26.62347626 | 1.191875263 | 1.98E-02 |
| Med1 | 3533.303566 | 1.191863134 | 2.65E-13 |
| Uimc1 | 1375.567608 | 1.191231366 | 7.66E-18 |

| | | | |
|---|---|---|---|
| Fkbp10 | 8574.958963 | 1.189956762 | 1.35E-13 |
| Rap1gds1 | 3574.090192 | 1.189558771 | 1.24E-35 |
| Sap18b | 2815.50859 | 1.189404144 | 2.32E-20 |
| Opa1 | 3963.090742 | 1.188547789 | 3.32E-51 |
| Gm5121 | 57.77765749 | 1.18847149 | 6.81E-03 |
| Gm3531 | 19.26519257 | 1.187794974 | 3.32E-02 |
| Rps12-ps1 | 89.85863566 | 1.187788314 | 1.40E-04 |
| Gm14414 | 932.8384723 | 1.187697416 | 4.33E-07 |
| Tmem38b | 711.4493136 | 1.187552797 | 2.50E-08 |
| Klc3 | 265.3530538 | 1.187153653 | 1.36E-08 |
| Cops6 | 4398.622067 | 1.186608847 | 1.42E-21 |
| Ptgs2os2 | 206.3760549 | 1.186607866 | 3.58E-03 |
| Gm4735 | 2471.198211 | 1.186431858 | 2.16E-08 |
| Sema3c | 667.7097169 | 1.186115775 | 3.64E-02 |
| Gdpd5 | 258.3174992 | 1.186038171 | 1.76E-10 |
| Ezh1 | 1677.187912 | 1.185302311 | 2.47E-20 |
| Mrpl9 | 1944.256036 | 1.185264125 | 9.08E-35 |
| Cyp4f17 | 372.9455404 | 1.184641939 | 2.35E-16 |
| Iars | 7024.962915 | 1.184005786 | 2.09E-13 |
| Atp10d | 515.9679941 | 1.182701112 | 9.41E-04 |
| 4430402I18Rik | 54.46899431 | 1.1823637 | 2.13E-02 |
| Gm8213 | 418.508554 | 1.182016547 | 1.90E-07 |
| Rnf138 | 732.8314506 | 1.181902149 | 5.06E-13 |
| Zfat | 598.1955488 | 1.181077595 | 1.21E-07 |
| Rps8 | 13664.61444 | 1.180969406 | 3.53E-08 |
| Rps26-ps1 | 19459.09523 | 1.180404004 | 1.01E-08 |
| Gm6286 | 108.3360089 | 1.180153742 | 9.75E-05 |
| Mkrn2 | 1675.962948 | 1.180039001 | 9.28E-13 |
| Nat9 | 476.1271399 | 1.179817196 | 1.35E-07 |
| Atp5c1 | 10686.56103 | 1.178815032 | 1.38E-15 |
| Narfl | 1220.368788 | 1.178290294 | 6.96E-19 |
| 2010204K13Rik | 97.13126604 | 1.177995841 | 2.81E-05 |
| Angel1 | 396.2138089 | 1.177843187 | 7.60E-08 |
| Nipsnap1 | 1460.490521 | 1.177288618 | 3.03E-21 |
| Amn1 | 462.9509753 | 1.177053866 | 4.35E-09 |
| Recql4 | 294.0795684 | 1.176944111 | 2.30E-03 |
| Rtcb | 11773.29027 | 1.176548129 | 6.77E-16 |
| Sh3bp5l | 2141.079408 | 1.175661958 | 7.39E-16 |
| Hikeshi | 958.3911583 | 1.175644697 | 4.88E-25 |
| Gm9625 | 405.7501673 | 1.175499814 | 3.18E-06 |
| Me2 | 1361.748678 | 1.1744103 | 3.73E-14 |
| Gpsm1 | 1158.941864 | 1.174086379 | 7.14E-12 |
| C130046K22Rik | 42.56062004 | 1.173941587 | 2.86E-04 |
| Hsp90ab1 | 120629.8161 | 1.173865811 | 6.19E-36 |
| Clcn2 | 458.1085294 | 1.173831572 | 2.70E-12 |
| Rps19 | 31869.82538 | 1.17191789 | 9.23E-09 |
| Cyp39a1 | 736.9729686 | 1.171913093 | 1.89E-19 |

| | | | |
|---|---|---|---|
| Dnajb6 | 7299.017882 | 1.171767166 | 1.98E-19 |
| Tmem120b | 354.8541929 | 1.17147944 | 1.70E-12 |
| Gm10051 | 181.0733855 | 1.171171762 | 1.07E-02 |
| Kif20a | 2457.420305 | 1.171001448 | 6.59E-04 |
| Pfdn4 | 874.6926454 | 1.170952503 | 4.35E-18 |
| Gm13835 | 2487.733138 | 1.170836688 | 1.00E-20 |
| Nub1 | 6026.984575 | 1.170605486 | 3.94E-10 |
| Coq5 | 1504.412996 | 1.16962317 | 4.27E-28 |
| Drg1 | 4096.902164 | 1.169221958 | 5.20E-20 |
| Gm26668 | 16.88403157 | 1.168989295 | 3.34E-02 |
| Ror1 | 243.8030807 | 1.168715455 | 4.08E-04 |
| Afdn | 6938.094101 | 1.168521184 | 1.93E-14 |
| Gm10040 | 874.6698116 | 1.16732604 | 6.11E-20 |
| Cox17 | 741.8334911 | 1.167312533 | 8.72E-32 |
| Bcat2 | 1292.256022 | 1.16690678 | 8.05E-16 |
| Gm21817 | 26.91196458 | 1.166440637 | 6.18E-03 |
| Farsb | 3179.035491 | 1.166107351 | 1.50E-18 |
| Nol10 | 2056.588005 | 1.166002595 | 8.86E-25 |
| Mri1 | 697.8904914 | 1.165842613 | 1.27E-09 |
| Chka | 2275.514018 | 1.165772547 | 1.90E-12 |
| Gm6451 | 47.40372897 | 1.1655882 | 1.14E-02 |
| Mrps28 | 608.7447475 | 1.165217474 | 1.67E-17 |
| Rpl13-ps3 | 18.70398708 | 1.165191947 | 2.47E-02 |
| Tmem269 | 12.75665785 | 1.16504843 | 2.90E-02 |
| Nufip1 | 993.2100757 | 1.164656305 | 1.41E-15 |
| Get4 | 3291.279844 | 1.164523311 | 6.61E-17 |
| Scnm1 | 789.5838704 | 1.163745454 | 1.15E-09 |
| Prpf40b | 1411.837332 | 1.163421733 | 7.09E-11 |
| Gm20695 | 55.31005275 | 1.163042355 | 3.18E-03 |
| Dgka | 489.6759605 | 1.162880971 | 2.12E-11 |
| Gm9027 | 16.48389127 | 1.162300203 | 3.91E-02 |
| Gab2 | 872.0524057 | 1.162242732 | 1.07E-08 |
| Cct6a | 10672.83687 | 1.162163963 | 8.66E-19 |
| Rhod | 1593.061284 | 1.160218786 | 3.64E-10 |
| Atp5b | 49226.93766 | 1.16002944 | 4.42E-17 |
| Acer3 | 753.5966644 | 1.159916126 | 7.95E-09 |
| Csnk2b | 3640.167179 | 1.159746975 | 1.38E-16 |
| Apaf1 | 2157.688006 | 1.159350489 | 3.04E-15 |
| Usf2 | 2888.615629 | 1.159264786 | 2.64E-19 |
| Atg9a | 2536.843685 | 1.159118694 | 5.69E-20 |
| Akirin2 | 2250.172154 | 1.159110551 | 1.30E-20 |
| Smim26 | 239.5610761 | 1.157760902 | 9.32E-09 |
| Bcl2l13 | 3503.679248 | 1.157580571 | 3.16E-27 |
| Stradb | 1050.361299 | 1.15751468 | 1.93E-11 |
| Zfp330 | 1591.298012 | 1.157298508 | 2.75E-20 |
| Mir6236 | 4368.612436 | 1.157173854 | 1.08E-07 |
| Cbr3 | 934.7984105 | 1.156957851 | 4.18E-17 |

| | | | |
|---|---|---|---|
| Rel | 189.2158242 | 1.156532307 | 3.91E-08 |
| Trappc1 | 1596.35037 | 1.156526384 | 1.95E-14 |
| Cubn | 333.1584532 | 1.156187489 | 1.69E-05 |
| Nprl2 | 675.6085235 | 1.156110752 | 1.85E-14 |
| Trabd | 1764.693389 | 1.155991175 | 4.98E-22 |
| Ldlr | 7632.961331 | 1.155826886 | 5.17E-06 |
| Nrde2 | 1207.833915 | 1.154979978 | 3.76E-12 |
| Rpl3-ps2 | 54.61022222 | 1.154354734 | 5.40E-03 |
| Ube2j1 | 2831.915582 | 1.153643232 | 1.89E-42 |
| Zfpm1 | 851.7480048 | 1.153499152 | 7.59E-11 |
| Serpine2 | 1071.175796 | 1.153185964 | 1.62E-06 |
| Polr3b | 1812.205495 | 1.15286939 | 1.13E-31 |
| Mtor | 2627.083558 | 1.152345288 | 3.22E-24 |
| Rpl17-ps4 | 69.9664268 | 1.151912258 | 2.45E-03 |
| Gm5292 | 16.18044631 | 1.151025061 | 1.41E-02 |
| Med25 | 2722.795482 | 1.150624992 | 8.58E-23 |
| Plk3 | 880.6175897 | 1.149638449 | 2.31E-12 |
| Gm10110 | 45.84062827 | 1.149583864 | 1.86E-02 |
| Tdp1 | 913.0317336 | 1.149477655 | 4.20E-13 |
| Rpl17-ps10 | 39.5528456 | 1.149263484 | 3.56E-02 |
| Tnfaip8l1 | 90.31987106 | 1.148804244 | 1.04E-03 |
| Gm13456 | 238.4475069 | 1.147336267 | 6.09E-05 |
| Ifngr1 | 2047.302005 | 1.147086617 | 1.56E-25 |
| Bmp2k | 1711.020882 | 1.146238568 | 3.56E-16 |
| Mlx | 1355.07942 | 1.146219629 | 7.92E-24 |
| NA | 1282.239263 | 1.145913468 | 3.75E-05 |
| Ppp2r3c | 930.1243978 | 1.145735759 | 2.72E-31 |
| Arpc3 | 6622.626879 | 1.145374661 | 7.02E-21 |
| Klf3 | 3490.931244 | 1.145090503 | 3.76E-18 |
| Lrch4 | 1036.656072 | 1.144832897 | 3.03E-15 |
| E4f1 | 934.5039487 | 1.144795924 | 1.08E-12 |
| 2210016F16Rik | 2097.051085 | 1.144591826 | 2.96E-07 |
| Pfdn1 | 1895.188969 | 1.143973555 | 7.72E-30 |
| Slit3 | 7087.344904 | 1.1431829 | 8.30E-08 |
| Naxe | 3709.451477 | 1.142676905 | 5.50E-13 |
| Sh3glb2 | 1959.605868 | 1.142419695 | 4.67E-26 |
| Gpr157 | 124.3548755 | 1.142064171 | 2.82E-04 |
| Ciapin1 | 1962.754716 | 1.141917655 | 3.97E-09 |
| Vdac3-ps1 | 39.06140609 | 1.141653155 | 8.51E-03 |
| H2-K1 | 1381.165026 | 1.141539015 | 4.77E-10 |
| Prkd2 | 613.8018891 | 1.140399828 | 1.51E-10 |
| Phldb1 | 4483.938021 | 1.140316518 | 2.34E-13 |
| Rpl37a | 29354.97811 | 1.140256047 | 5.96E-07 |
| Gm15441 | 17.41878548 | 1.140043807 | 1.99E-02 |
| Dgkd | 3573.208434 | 1.139010742 | 1.78E-49 |
| Gm7331 | 343.4208214 | 1.138419327 | 3.10E-03 |
| Vim | 152127.4244 | 1.138163575 | 1.03E-15 |

| | | | |
|---|---|---|---|
| Il18rap | 98.60169899 | 1.137878358 | 9.31E-04 |
| Vldlr | 180.1056824 | 1.13772774 | 1.07E-03 |
| Dnaja3 | 3016.129967 | 1.137686542 | 1.24E-22 |
| Mkln1os | 17.31510653 | 1.137262876 | 2.75E-02 |
| H13 | 6024.342285 | 1.136750655 | 2.31E-18 |
| Usp36 | 1623.508219 | 1.136628822 | 2.15E-11 |
| Stk4 | 2349.963146 | 1.136503744 | 2.63E-50 |
| Rps6ka2 | 2190.081064 | 1.136159133 | 1.72E-20 |
| Ccdc163 | 481.2533435 | 1.135500117 | 1.36E-05 |
| Ccnb2 | 359.5961839 | 1.13516841 | 1.61E-04 |
| Parn | 3178.519304 | 1.134876223 | 5.99E-22 |
| Actn4 | 15981.88887 | 1.13373759 | 8.43E-17 |
| Nsun2 | 5120.925682 | 1.131619255 | 4.25E-14 |
| Slc12a9 | 656.8429679 | 1.131480189 | 9.27E-16 |
| Mapt | 26.16495815 | 1.131223019 | 2.76E-02 |
| Ypel3 | 1637.852338 | 1.131139428 | 9.97E-09 |
| Gm6136 | 182.5775809 | 1.131138729 | 6.94E-04 |
| Gm9825 | 3797.027924 | 1.129143742 | 2.09E-13 |
| Lipt1 | 375.2285292 | 1.128923087 | 1.16E-12 |
| Prr5 | 206.0395211 | 1.128548693 | 9.02E-07 |
| Hdgfl2 | 4078.471706 | 1.128527409 | 1.17E-32 |
| Trpc4ap | 8668.387395 | 1.128431803 | 2.00E-14 |
| Sh3glb1 | 7842.113378 | 1.127362067 | 4.04E-24 |
| Uba1 | 17656.13637 | 1.127300537 | 4.79E-17 |
| Gm12712 | 27.7636762 | 1.126997281 | 4.06E-02 |
| Mtln | 167.9560115 | 1.126895872 | 4.46E-09 |
| Fetub | 34.84454848 | 1.125586506 | 2.36E-02 |
| Dixdc1 | 787.0275851 | 1.125530631 | 5.83E-08 |
| Slc25a28 | 779.0647433 | 1.125309745 | 1.03E-11 |
| Slc6a9 | 2482.059137 | 1.123041517 | 4.26E-17 |
| Rpl39l | 5625.606209 | 1.122806888 | 1.41E-12 |
| Reps1 | 2264.486585 | 1.122702933 | 5.41E-25 |
| Gm9493 | 223.7226295 | 1.121816254 | 2.03E-03 |
| AA474408 | 8595.94056 | 1.121797393 | 6.00E-08 |
| Fads3 | 4292.424506 | 1.121419602 | 3.25E-06 |
| Alpk2 | 16.48575008 | 1.121350326 | 2.54E-02 |
| Phf1 | 1872.957205 | 1.121249829 | 7.62E-09 |
| Kcnh2 | 22.92218311 | 1.12113526 | 1.90E-02 |
| Eno4 | 35.79923314 | 1.120843506 | 9.86E-04 |
| Haus4 | 657.3397337 | 1.120804042 | 2.13E-06 |
| Dimt1 | 665.6401902 | 1.120655774 | 2.17E-28 |
| Syne2 | 2077.972178 | 1.11999907 | 5.75E-09 |
| Kansl2 | 5499.25737 | 1.119768188 | 6.60E-18 |
| Alas1 | 1110.822137 | 1.119707583 | 3.99E-09 |
| Lig3 | 2759.69298 | 1.119226131 | 9.78E-28 |
| Ska1 | 241.1235717 | 1.119198647 | 7.87E-03 |
| Mrpl53 | 903.2269651 | 1.119124993 | 3.78E-22 |

| | | | |
|---|---|---|---|
| Zyx | 23861.9633 | 1.119053073 | 8.54E-22 |
| Plk1 | 1159.243141 | 1.117846017 | 4.25E-03 |
| Gm15427 | 13183.07084 | 1.117499984 | 8.24E-04 |
| Clasrp | 883.122175 | 1.117406185 | 1.96E-15 |
| Sin3b | 4130.398059 | 1.116607804 | 1.26E-26 |
| Psmb5 | 2568.243617 | 1.115771194 | 2.59E-15 |
| Nans | 1668.575675 | 1.115153103 | 4.66E-10 |
| Bbs4 | 321.4642689 | 1.114642059 | 1.02E-15 |
| Rpl37rt | 7114.695718 | 1.114443978 | 7.64E-10 |
| Egfr | 1382.648293 | 1.114117365 | 5.26E-06 |
| BC052040 | 653.4512046 | 1.113310834 | 4.73E-25 |
| Gm17638 | 49.71635285 | 1.113101312 | 4.05E-05 |
| Hspa5 | 48769.31287 | 1.113082836 | 8.81E-07 |
| Uba52 | 18378.21562 | 1.112580585 | 6.16E-06 |
| Slc50a1 | 1503.064834 | 1.112550521 | 5.67E-17 |
| Cd55 | 298.5806079 | 1.111210918 | 2.23E-02 |
| Ndufv1 | 4450.278284 | 1.111032357 | 2.94E-19 |
| Akr1b8 | 5004.962685 | 1.110165778 | 8.28E-17 |
| Sdf4 | 4735.660727 | 1.110137292 | 1.14E-14 |
| Def6 | 175.6286037 | 1.109703786 | 2.98E-12 |
| Acaa2 | 2550.415331 | 1.109601675 | 1.32E-09 |
| Nop14 | 3249.947278 | 1.109262983 | 7.81E-51 |
| Lmna | 24900.25804 | 1.109094184 | 3.93E-11 |
| Fip1l1 | 6051.922285 | 1.109062486 | 1.78E-10 |
| Ribc1 | 38.24492566 | 1.108977828 | 5.48E-03 |
| Rps26 | 31879.79427 | 1.108619505 | 3.70E-07 |
| Uqcrq | 2719.276559 | 1.108165167 | 1.02E-13 |
| Ndufs8 | 2449.680964 | 1.108036402 | 1.21E-10 |
| Nfs1 | 1585.844528 | 1.107979858 | 1.11E-21 |
| Ticam1 | 1747.623083 | 1.107976183 | 3.41E-07 |
| Ybx1 | 42624.67727 | 1.107404201 | 1.93E-13 |
| Aptx | 863.7687715 | 1.107273019 | 1.26E-17 |
| Neurl2 | 135.0104615 | 1.107097326 | 1.88E-05 |
| Sec14l1 | 5437.380374 | 1.105609344 | 1.35E-24 |
| Rps17 | 34211.18673 | 1.105008528 | 4.07E-10 |
| Pkn2 | 5004.679856 | 1.104970055 | 2.53E-21 |
| Atp5e | 1767.933432 | 1.1048593 | 1.04E-09 |
| Smim3 | 135.0460714 | 1.104739896 | 4.32E-06 |
| Actr1a | 7248.29814 | 1.104704748 | 1.32E-13 |
| Msn | 18054.98124 | 1.104661706 | 7.33E-09 |
| Slc25a5 | 8607.553285 | 1.103674528 | 7.94E-09 |
| Ptpn23 | 5720.306209 | 1.103499083 | 8.93E-09 |
| Dlx2 | 556.9964188 | 1.103321567 | 8.16E-09 |
| Cyb561d2 | 332.507694 | 1.102962973 | 8.12E-11 |
| Herc3 | 407.8347654 | 1.102813256 | 1.23E-07 |
| Leng1 | 441.7638383 | 1.102353411 | 1.36E-11 |
| Dhps | 1793.496769 | 1.102332657 | 6.06E-15 |

| | | | |
|---|---|---|---|
| Aurkaip1 | 1428.016499 | 1.101995722 | 6.69E-21 |
| Foxp1 | 2437.44245 | 1.101439077 | 1.58E-04 |
| NA | 144.6696948 | 1.101149434 | 4.44E-05 |
| Elac2 | 1667.303953 | 1.100989532 | 5.43E-13 |
| Fam98c | 410.091738 | 1.100912893 | 2.87E-12 |
| Flii | 7693.446933 | 1.100016628 | 1.29E-33 |
| Ndufa7 | 3232.605156 | 1.098869288 | 1.61E-09 |
| Eef1e1 | 830.8756283 | 1.098049992 | 5.24E-11 |
| Lrrc8c | 1417.4025 | 1.097541196 | 9.19E-09 |
| Uqcc1 | 1536.397941 | 1.09751004 | 2.91E-17 |
| Ankrd55 | 177.8979238 | 1.097386983 | 1.85E-04 |
| Osgin1 | 375.3213527 | 1.097249912 | 9.78E-05 |
| Itga1 | 65.1732428 | 1.09691666 | 6.82E-03 |
| Eif2b1 | 1612.710149 | 1.096731562 | 4.57E-20 |
| Plekhg3 | 2284.954782 | 1.096638175 | 1.54E-11 |
| Vps37b | 1439.0067 | 1.096417609 | 2.61E-06 |
| Gm14286 | 25.7290986 | 1.096007092 | 2.13E-02 |
| Wdtc1 | 2442.822428 | 1.095135578 | 3.71E-15 |
| Mms19 | 2815.736031 | 1.094587358 | 1.58E-19 |
| Pgm2 | 1881.276836 | 1.092931177 | 3.85E-19 |
| Eif2b4 | 1155.389 | 1.092750943 | 7.97E-19 |
| Got1 | 2662.479814 | 1.092720362 | 2.62E-14 |
| Tmem184b | 4196.309966 | 1.09272026 | 1.21E-17 |
| Kank2 | 6902.794724 | 1.092118751 | 7.18E-23 |
| Psmc1 | 6304.674307 | 1.092036463 | 7.50E-24 |
| Gtf3a | 2196.178491 | 1.09156041 | 2.54E-11 |
| Isca2 | 752.0861875 | 1.090804263 | 1.31E-21 |
| Rpl22 | 11046.40294 | 1.09078271 | 1.45E-06 |
| Relb | 763.8076857 | 1.090445612 | 1.99E-08 |
| Vdac3 | 5778.986181 | 1.088864469 | 1.87E-25 |
| Gm14149 | 106.2779207 | 1.088630509 | 1.81E-06 |
| Tep1 | 1733.768891 | 1.08859447 | 9.32E-22 |
| 1810032O08Rik | 226.283254 | 1.088585219 | 5.50E-07 |
| Atp5d | 7992.530441 | 1.088461867 | 1.06E-14 |
| Tbc1d22a | 1870.58861 | 1.088381257 | 5.05E-10 |
| Smn1 | 972.1731807 | 1.088079952 | 5.80E-12 |
| Tcf25 | 13497.59425 | 1.08796716 | 5.57E-25 |
| NA | 1944.15319 | 1.087924414 | 1.12E-10 |
| Ubr4 | 11843.35144 | 1.087722472 | 9.62E-14 |
| Psma1 | 3673.371411 | 1.08716917 | 5.23E-32 |
| Phkg1 | 37.75219016 | 1.086140204 | 1.41E-02 |
| Rad9a | 2033.493668 | 1.08557515 | 1.16E-08 |
| Gm1840 | 48.69248908 | 1.085220075 | 1.01E-02 |
| Cdpf1 | 451.3269781 | 1.084867387 | 1.84E-10 |
| Psmb4 | 6707.696983 | 1.084851194 | 6.30E-21 |
| Epb41l3 | 11414.40963 | 1.084245016 | 5.35E-33 |
| Gm10136 | 300.3157053 | 1.084169551 | 1.05E-04 |

| | | | |
|---|---|---|---|
| Epb41 | 92.26991572 | 1.083623441 | 4.92E-04 |
| Prr33 | 24.9195298 | 1.083216153 | 3.72E-02 |
| Chid1 | 1865.137384 | 1.082770718 | 3.09E-09 |
| H2-K2 | 70.25926091 | 1.08252431 | 3.60E-04 |
| Acox1 | 3724.20556 | 1.082102687 | 5.49E-15 |
| Kyat1 | 280.1672108 | 1.081722929 | 5.02E-08 |
| 4930442H23Rik | 17.36876745 | 1.080051117 | 2.59E-02 |
| Gdi1 | 5499.526865 | 1.078897815 | 1.89E-14 |
| Galk1 | 1811.403168 | 1.078828157 | 2.01E-09 |
| Nr1h5 | 425.4240054 | 1.078774322 | 2.99E-02 |
| H60b | 795.9867706 | 1.07778059 | 6.08E-08 |
| Hprt | 2471.179009 | 1.077617042 | 4.57E-12 |
| Alad | 1880.043307 | 1.077399504 | 2.55E-13 |
| Gm14399 | 30.01144222 | 1.077135205 | 1.24E-02 |
| Clptm1 | 4833.878488 | 1.076703095 | 1.11E-13 |
| Cox6a1 | 5935.091383 | 1.076362184 | 1.57E-09 |
| Ganab | 13601.05205 | 1.076186689 | 2.30E-27 |
| Rps23-ps2 | 93.85077079 | 1.076145827 | 1.83E-04 |
| Asna1 | 3940.798485 | 1.076070273 | 6.13E-10 |
| Srfbp1 | 1357.544805 | 1.075719211 | 2.68E-09 |
| Tmem68 | 608.1000298 | 1.075209767 | 2.06E-18 |
| Nedd4l | 2500.331586 | 1.074996854 | 8.95E-10 |
| R3hdm2 | 5526.132124 | 1.074852277 | 3.19E-18 |
| Mapk8ip3 | 4942.510858 | 1.074821159 | 7.95E-12 |
| Actr8 | 1337.352931 | 1.074412045 | 4.02E-20 |
| Mrpl47 | 715.1774425 | 1.074230271 | 2.15E-11 |
| Gtf2f2 | 887.8471566 | 1.074050943 | 4.92E-16 |
| Atad3a | 1366.86159 | 1.073917978 | 1.47E-16 |
| Gca | 78.20706386 | 1.07360298 | 5.67E-03 |
| Psph | 959.6907578 | 1.073395959 | 8.34E-04 |
| Chchd4 | 753.277916 | 1.072409948 | 4.41E-12 |
| Pgk1 | 25876.91073 | 1.072115187 | 3.21E-08 |
| Edn1 | 882.9784013 | 1.071748352 | 9.59E-05 |
| Crlf3 | 1281.734592 | 1.07084909 | 6.22E-14 |
| Ndufb9 | 3591.013312 | 1.070601142 | 9.31E-12 |
| 2700062C07Rik | 561.1969031 | 1.070108166 | 4.79E-16 |
| Ptar1 | 207.0188926 | 1.069951208 | 9.63E-07 |
| Plekhb2 | 3753.770953 | 1.068855389 | 2.05E-12 |
| Gm16425 | 17.03954714 | 1.06816816 | 3.45E-02 |
| Prkcg | 214.7714828 | 1.067810862 | 2.13E-05 |
| Sri | 3410.645404 | 1.067371097 | 5.79E-33 |
| Gtdc1 | 588.2460699 | 1.067297557 | 1.42E-09 |
| Ppia | 832.0567905 | 1.067149514 | 8.77E-14 |
| Rpl35a-ps2 | 2188.965404 | 1.066007272 | 9.41E-04 |
| Cenpm | 389.4200119 | 1.065065845 | 2.13E-03 |
| Ddx54 | 4300.181664 | 1.064963072 | 6.03E-12 |
| Egfl7 | 61.93243651 | 1.064590842 | 1.43E-03 |

| | | | |
|---|---|---|---|
| Smpd3 | 867.4805472 | 1.06434392 | 1.74E-07 |
| Mrps24 | 1143.813471 | 1.0643116 | 5.63E-11 |
| 4930539J05Rik | 104.1542753 | 1.062634605 | 5.68E-05 |
| Ankrd9 | 197.5218888 | 1.062480145 | 3.58E-06 |
| Pear1 | 2676.766374 | 1.06223925 | 1.40E-14 |
| Gm26620 | 24.75500295 | 1.062237795 | 4.92E-02 |
| Pdia5 | 2747.707923 | 1.061840316 | 2.87E-07 |
| C920009B18Rik | 939.532051 | 1.061379419 | 9.63E-05 |
| Mrs2 | 469.060725 | 1.06071996 | 1.08E-11 |
| Ube2s | 4362.208061 | 1.060286994 | 5.74E-07 |
| Rpsa-ps10 | 455.3269898 | 1.060188106 | 2.04E-04 |
| Ell | 1456.306905 | 1.060172716 | 1.00E-10 |
| Epha1 | 75.86205553 | 1.060019428 | 3.10E-04 |
| Zfp524 | 482.7409231 | 1.059747794 | 3.64E-08 |
| Snora34 | 25.72971137 | 1.059745171 | 1.55E-02 |
| Prkcsh | 5914.118506 | 1.059110186 | 2.24E-15 |
| Stard3 | 1188.605779 | 1.059055732 | 8.99E-13 |
| 5730405O15Rik | 32.60095963 | 1.05830083 | 3.21E-03 |
| Uqcrc1 | 5283.028191 | 1.057548026 | 1.67E-15 |
| 2900052L18Rik | 59.70445642 | 1.057389715 | 1.08E-03 |
| Ttll12 | 2912.203449 | 1.057322614 | 3.46E-12 |
| Hdac11 | 2157.610884 | 1.057232989 | 5.68E-06 |
| Gba | 4711.112014 | 1.056985465 | 4.20E-12 |
| Uxt | 499.6757564 | 1.056633279 | 6.56E-10 |
| Ahnak | 67844.71225 | 1.056393963 | 1.83E-14 |
| Supv3l1 | 1487.778323 | 1.055691334 | 1.15E-34 |
| Trmt112-ps2 | 2121.34201 | 1.055601371 | 8.34E-17 |
| Pip4k2c | 5308.496544 | 1.055405906 | 9.33E-10 |
| Sec61g | 587.7553782 | 1.055165831 | 6.74E-17 |
| Zfp575 | 56.61006131 | 1.054960828 | 1.10E-04 |
| Emc4 | 2098.421114 | 1.054311904 | 2.70E-26 |
| Pla2g16 | 886.3125233 | 1.052864016 | 4.10E-06 |
| Cox4i1 | 12755.47037 | 1.052440507 | 3.48E-10 |
| Zfp993 | 76.49799003 | 1.051654852 | 4.44E-03 |
| Mrip-ps | 1177.21672 | 1.05163988 | 8.76E-11 |
| Mfsd4b5 | 246.1474623 | 1.051540088 | 1.06E-04 |
| Plin2 | 9880.879654 | 1.050942329 | 7.51E-09 |
| Wasf1 | 460.3368112 | 1.050715435 | 2.37E-18 |
| Mllt11 | 896.5428316 | 1.050451387 | 2.26E-13 |
| Cyba | 1385.240954 | 1.050379642 | 3.49E-12 |
| Sike1 | 1752.912471 | 1.050269173 | 1.42E-15 |
| Cdh2 | 5735.006876 | 1.049432395 | 1.17E-11 |
| NA | 27.02896418 | 1.048247385 | 9.69E-03 |
| Acsl6 | 253.6968893 | 1.048084076 | 9.44E-08 |
| Arhgef4 | 384.8597554 | 1.048058987 | 3.56E-05 |
| NA | 11703.17523 | 1.047617058 | 2.19E-06 |
| Polr2i | 588.2928749 | 1.047136192 | 1.12E-10 |

| | | | |
|---|---|---|---|
| Phkg2 | 610.2063309 | 1.046983685 | 1.05E-16 |
| Slc25a1 | 5391.834169 | 1.04610247 | 6.66E-11 |
| Atg101 | 738.9112077 | 1.045664307 | 1.59E-13 |
| Grpel1 | 2345.33627 | 1.045267775 | 6.15E-11 |
| Tmem202 | 48.67880422 | 1.045259565 | 1.36E-03 |
| Tbca | 3268.050332 | 1.045249982 | 7.53E-24 |
| Ascc2 | 3721.48564 | 1.045192173 | 7.07E-22 |
| Tagln2 | 5657.746603 | 1.044450488 | 1.05E-06 |
| Sf3a2 | 2291.998884 | 1.044172629 | 5.78E-09 |
| Iqcb1 | 717.9089585 | 1.04399782 | 1.36E-12 |
| Slc8b1 | 1315.172544 | 1.043730897 | 2.93E-09 |
| Ndufb2 | 1099.511065 | 1.042961182 | 2.49E-14 |
| Uchl3 | 918.5000259 | 1.042766527 | 5.67E-16 |
| Oxld1 | 130.1722169 | 1.042650076 | 4.11E-06 |
| Rassf5 | 557.4393745 | 1.042489887 | 1.75E-06 |
| Eif2s1 | 4474.748387 | 1.04240255 | 3.37E-16 |
| Cd109 | 12991.36315 | 1.041973762 | 1.94E-11 |
| Ngdn | 1047.951766 | 1.041420942 | 8.56E-20 |
| Timp1 | 3421.454883 | 1.040808343 | 1.18E-06 |
| Hspe1 | 7725.560891 | 1.040786232 | 3.97E-16 |
| Smu1 | 3414.245524 | 1.040601239 | 4.08E-30 |
| Stamos | 17.11432608 | 1.040278457 | 1.87E-02 |
| Nek3 | 321.6148187 | 1.040254206 | 7.57E-14 |
| Snrpc | 2036.48846 | 1.039728793 | 4.87E-13 |
| Rps6 | 9260.197394 | 1.039475012 | 2.84E-07 |
| Tfb1m | 210.7171039 | 1.039340108 | 6.93E-06 |
| Lsp1 | 852.1553722 | 1.039059357 | 1.14E-07 |
| Nudt14 | 66.89279419 | 1.038902266 | 7.59E-05 |
| Gm11353 | 146.2762761 | 1.038728377 | 2.36E-03 |
| Kif22 | 949.2951466 | 1.038592561 | 1.27E-03 |
| Ttc12 | 284.5390079 | 1.038490659 | 1.95E-05 |
| Lmtk2 | 2086.187159 | 1.038453284 | 2.14E-26 |
| Tspan31 | 6006.152092 | 1.038316445 | 1.97E-11 |
| Dync1h1 | 28582.84367 | 1.038051641 | 2.38E-17 |
| Gdf9 | 43.03072072 | 1.0379136 | 7.36E-03 |
| Mrpl43 | 1580.403356 | 1.037281578 | 8.91E-19 |
| Pdcd5 | 1694.809403 | 1.037119443 | 2.61E-19 |
| Ptpn1 | 2274.268902 | 1.036833748 | 1.59E-12 |
| Psma6 | 5135.561452 | 1.036371415 | 5.02E-32 |
| Zfp446 | 215.0941646 | 1.036263975 | 1.62E-06 |
| Trappc9 | 2452.715589 | 1.035960339 | 4.30E-11 |
| Cbr1 | 1090.694584 | 1.035276299 | 3.97E-08 |
| Gm12953 | 38.853626 | 1.035080268 | 1.43E-02 |
| Foxo3 | 3075.973274 | 1.034899195 | 6.10E-11 |
| H60c | 251.3013379 | 1.034771087 | 6.81E-06 |
| Edf1 | 4065.529406 | 1.034518782 | 3.38E-14 |
| T2 | 37.33308043 | 1.034425625 | 3.07E-03 |

| | | | |
|---|---|---|---|
| Espl1 | 1089.594393 | 1.034309388 | 5.48E-03 |
| C77080 | 1863.179005 | 1.034153082 | 2.51E-14 |
| Gm6313 | 19.41210927 | 1.033912702 | 2.51E-02 |
| Rps20 | 53567.39921 | 1.033506123 | 8.77E-07 |
| Klhdc2 | 4037.018707 | 1.033398522 | 2.57E-19 |
| Gm12696 | 377.0348595 | 1.033361001 | 7.65E-10 |
| Rps28 | 15164.62267 | 1.033351704 | 4.19E-06 |
| Pigyl | 425.4755021 | 1.032633161 | 2.24E-09 |
| Rps15a-ps7 | 44.62255626 | 1.032485214 | 1.17E-02 |
| 4930430F08Rik | 478.4478484 | 1.032254767 | 3.73E-08 |
| Uvrag | 2392.79397 | 1.032049361 | 8.04E-14 |
| Ccnb1 | 1707.028829 | 1.031995256 | 3.88E-03 |
| Med18 | 182.6379288 | 1.030913673 | 1.76E-07 |
| Pvt1 | 2024.36745 | 1.030885661 | 3.67E-13 |
| Anln | 4893.272358 | 1.030053878 | 4.39E-03 |
| Spata6 | 2206.374903 | 1.029739916 | 3.16E-15 |
| Ankrd61 | 29.46407003 | 1.029236401 | 2.46E-02 |
| Tars | 6511.285888 | 1.02907234 | 7.11E-24 |
| Dbn1 | 5523.632008 | 1.028360114 | 1.85E-16 |
| Polr2e | 3801.975991 | 1.0281644 | 1.52E-15 |
| Sec13 | 6633.822026 | 1.027487838 | 1.03E-13 |
| Csnk1g1 | 2131.842807 | 1.027430286 | 3.38E-02 |
| Tmbim4 | 1686.78166 | 1.026996694 | 1.45E-25 |
| Surf1 | 1397.802757 | 1.026848182 | 3.46E-10 |
| Erich1 | 560.5336326 | 1.026636117 | 9.37E-12 |
| Tbc1d8 | 59.18404534 | 1.026464844 | 4.16E-04 |
| Plrg1 | 2765.068295 | 1.025172089 | 7.01E-17 |
| Yif1a | 3116.067448 | 1.023720403 | 8.75E-10 |
| Steap2 | 5089.313269 | 1.023534857 | 3.13E-07 |
| Osbpl3 | 2074.055951 | 1.023441962 | 5.27E-12 |
| Asb3 | 997.0760849 | 1.023250577 | 2.38E-18 |
| Arfip2 | 2214.951548 | 1.023062934 | 8.39E-16 |
| Rps15a | 30497.77941 | 1.022753153 | 1.26E-08 |
| Ehd2 | 12732.25354 | 1.02250757 | 4.78E-18 |
| Ptcd1 | 1214.415313 | 1.02249154 | 1.03E-13 |
| Stx11 | 348.3990913 | 1.022225718 | 6.51E-09 |
| Clic1 | 9855.984042 | 1.022120583 | 7.41E-16 |
| Gm15772 | 85.38857994 | 1.022068771 | 1.29E-02 |
| Gm9701 | 21363.22693 | 1.022028588 | 3.52E-13 |
| Chn1 | 68.77845635 | 1.021912611 | 5.89E-03 |
| Rpl39 | 31120.35881 | 1.021521418 | 4.34E-06 |
| Sh2d3c | 108.5494914 | 1.021354341 | 6.46E-04 |
| Gm5879 | 60.46901209 | 1.021273493 | 4.34E-02 |
| Ssx2ip | 1161.638843 | 1.020884689 | 4.45E-11 |
| Ppm1d | 820.5467079 | 1.020415177 | 3.60E-10 |
| Gm7027 | 42.16724364 | 1.020326431 | 2.35E-03 |
| Ggnbp1 | 248.4931598 | 1.020111068 | 7.60E-05 |

| | | | |
|---|---|---|---|
| Pdcd2 | 1168.649565 | 1.019837732 | 5.00E-16 |
| Gdi2 | 14707.93982 | 1.01865125 | 9.91E-18 |
| Tes | 1801.715462 | 1.018637802 | 2.82E-06 |
| Nop16 | 880.4090839 | 1.018006017 | 5.87E-10 |
| Tom1 | 1367.922974 | 1.016130247 | 5.40E-03 |
| Gga1 | 2714.168085 | 1.015904093 | 1.94E-13 |
| Rdm1 | 418.9154272 | 1.015325391 | 8.05E-04 |
| Aifm2 | 1026.37371 | 1.015100051 | 3.76E-19 |
| Mrps9 | 1479.100855 | 1.014901755 | 9.40E-36 |
| Ppp1r18 | 4671.279419 | 1.014522664 | 1.25E-22 |
| Rps9 | 54843.11362 | 1.013278537 | 3.85E-09 |
| Miip | 631.6917154 | 1.01302021 | 2.09E-06 |
| Klf7 | 2129.134471 | 1.012861288 | 9.88E-10 |
| Eci2 | 2234.646018 | 1.012852763 | 7.41E-17 |
| Scarb1 | 4510.378786 | 1.012831282 | 1.11E-10 |
| Ska3 | 318.64434 | 1.012601074 | 1.06E-03 |
| Csk | 2408.111774 | 1.012442347 | 1.46E-12 |
| Lat2 | 39.69412982 | 1.011508766 | 1.41E-02 |
| Slc25a33 | 464.665477 | 1.011450788 | 1.19E-10 |
| Fyn | 3259.615135 | 1.011443113 | 1.47E-26 |
| Btrc | 1416.886445 | 1.011439766 | 1.52E-10 |
| Cyp4f41-ps | 24.80738338 | 1.011389314 | 8.71E-03 |
| Nfkbil1 | 740.5464751 | 1.011143033 | 1.41E-14 |
| Zcwpw1 | 487.8432153 | 1.011081688 | 8.17E-10 |
| Usp35 | 407.4218091 | 1.010996147 | 9.19E-05 |
| Rps18 | 7591.717016 | 1.010853815 | 4.68E-04 |
| Dynlt1f | 1410.208679 | 1.010341558 | 3.09E-09 |
| Dlgap1 | 42.72816766 | 1.010266746 | 1.20E-02 |
| Rabgef1 | 1307.570608 | 1.010231578 | 5.37E-22 |
| Ptch1 | 475.913733 | 1.009372359 | 3.74E-11 |
| Gm19705 | 81.49688775 | 1.009305852 | 1.16E-03 |
| Polrmt | 1417.394738 | 1.008994082 | 4.45E-11 |
| Rae1 | 2709.734454 | 1.008899493 | 2.59E-12 |
| Sac3d1 | 362.1060275 | 1.008863968 | 4.18E-04 |
| Gadd45a | 2223.304238 | 1.008668248 | 1.44E-04 |
| Jmjd4 | 1066.637866 | 1.008525188 | 1.69E-23 |
| Pdgfc | 1278.888861 | 1.00848869 | 1.92E-09 |
| Trp53i13 | 637.8712156 | 1.008390857 | 5.44E-09 |
| Rpl39-ps | 200.9664968 | 1.008189592 | 1.89E-03 |
| Gmpr2 | 704.4970757 | 1.008049318 | 1.34E-21 |
| Daglb | 227.5009461 | 1.007751267 | 4.54E-07 |
| Mterf4 | 757.3895142 | 1.007704248 | 2.29E-24 |
| Cox7c | 1341.314878 | 1.007616409 | 2.83E-13 |
| Patz1 | 1776.586514 | 1.007451305 | 1.70E-09 |
| Ostf1 | 2423.365697 | 1.007434221 | 9.10E-24 |
| Ctu2 | 965.207299 | 1.006853313 | 5.50E-10 |
| Ttll5 | 1440.65249 | 1.006566653 | 1.31E-02 |

| | | | |
|---|---|---|---|
| Fosl1 | 1007.39669 | 1.00570528 | 1.62E-03 |
| Lrrc42 | 3350.571451 | 1.005563971 | 1.56E-16 |
| Ryk | 8106.711931 | 1.005268013 | 5.24E-22 |
| Gm17315 | 21.55675448 | 1.005149937 | 2.32E-02 |
| Vars2 | 1444.707556 | 1.004927911 | 3.18E-19 |
| Rpl36-ps12 | 66.11856536 | 1.004528172 | 2.57E-03 |
| Pmpca | 4581.083073 | 1.004164583 | 2.09E-39 |
| Tmem253 | 30.43885092 | 1.00408814 | 5.18E-03 |
| Sidt2 | 5631.235272 | 1.003311904 | 1.42E-11 |
| Psmb3 | 3320.528605 | 1.002933561 | 2.60E-10 |
| Gm9762 | 589.6013071 | 1.002821447 | 1.16E-10 |
| Tpm1 | 28225.26008 | 1.001927279 | 6.92E-19 |
| Mbd1 | 1415.633505 | 1.001896181 | 1.85E-06 |
| Msra | 160.0002431 | 1.001332907 | 3.15E-07 |
| Foxk2 | 3737.143576 | 1.001157492 | 5.51E-10 |
| Rps13-ps2 | 21055.30253 | 1.00052991 | 4.71E-08 |
| Galc | 1514.601027 | 0.999238769 | 6.02E-13 |
| Cd2bp2 | 2807.343141 | 0.998919935 | 7.10E-23 |
| Med15 | 3551.143144 | 0.998608197 | 4.36E-10 |
| Coq8b | 489.0724424 | 0.996974315 | 1.85E-08 |
| Edrf1 | 931.4257018 | 0.996957452 | 1.16E-11 |
| Sorbs3 | 4375.993113 | 0.996477478 | 2.57E-12 |
| 9330136K24Rik | 58.8334473 | 0.995465226 | 8.66E-04 |
| Tomm20 | 7274.548066 | 0.994543974 | 1.09E-11 |
| Metrnl | 1886.117273 | 0.994108866 | 3.54E-13 |
| Arhgap29 | 5638.40206 | 0.994003555 | 2.92E-05 |
| Retreg2 | 4599.688868 | 0.993433156 | 5.73E-07 |
| Snrpd2 | 2347.998224 | 0.99295897 | 4.07E-10 |
| Gm10443 | 18934.25035 | 0.992809336 | 1.27E-05 |
| Fbxw8 | 5950.711225 | 0.992808852 | 2.65E-14 |
| Mrpl39 | 1700.080335 | 0.992599687 | 6.36E-28 |
| Cog6 | 1639.903711 | 0.992560847 | 3.02E-15 |
| Rce1 | 1079.36259 | 0.992352154 | 6.19E-24 |
| Saraf | 4956.861244 | 0.991788971 | 8.85E-10 |
| Gpkow | 1945.850323 | 0.990972418 | 1.27E-16 |
| Cct8 | 12262.24507 | 0.990944198 | 2.48E-27 |
| Lancl1 | 2052.092334 | 0.990684052 | 4.34E-13 |
| Snx6 | 4503.412023 | 0.990586902 | 7.98E-54 |
| Sirt2 | 4842.503244 | 0.990405631 | 5.65E-10 |
| Hspb8 | 6743.145759 | 0.990003368 | 1.53E-13 |
| Tsr3 | 1281.295736 | 0.98973932 | 1.41E-15 |
| Ppp2r5b | 1603.372505 | 0.989535452 | 2.74E-15 |
| Edc4 | 3070.827727 | 0.989080091 | 2.22E-10 |
| Cep170b | 2978.572285 | 0.988961656 | 8.30E-14 |
| Gm17029 | 17.53182691 | 0.988165327 | 4.64E-02 |
| Gm7658 | 524.6113651 | 0.98791883 | 1.49E-07 |
| Mical1 | 409.2832872 | 0.987080603 | 3.18E-07 |

| | | | |
|---|---|---|---|
| Etfbkmt | 562.6280982 | 0.987079895 | 1.62E-05 |
| Gm13680 | 112.4082536 | 0.986425989 | 1.59E-03 |
| Rpl7 | 74255.39802 | 0.986211976 | 2.63E-10 |
| Stk35 | 1822.968386 | 0.986182064 | 3.24E-17 |
| Cenpn | 611.6297696 | 0.986051109 | 4.47E-04 |
| Grik2 | 1429.423583 | 0.985592096 | 2.69E-06 |
| Aktip | 1968.524584 | 0.985374749 | 3.12E-19 |
| Farp1 | 7954.256896 | 0.984979487 | 1.02E-19 |
| Gm9755 | 75.68857213 | 0.984909501 | 9.64E-03 |
| Cdca3 | 1209.281701 | 0.984510541 | 1.91E-03 |
| Psmc2 | 6666.244406 | 0.984299168 | 1.66E-19 |
| Fdx1 | 723.8667476 | 0.984292932 | 3.13E-08 |
| Vapb | 4955.166935 | 0.984282397 | 1.50E-16 |
| Gatad2a | 4485.776346 | 0.983954663 | 2.20E-18 |
| Gm13393 | 20.06726507 | 0.983560503 | 4.16E-02 |
| Tmem128 | 899.2173669 | 0.983469548 | 4.78E-15 |
| Ino80e | 1732.624095 | 0.983430182 | 4.07E-10 |
| Sinhcaf | 1136.208845 | 0.98340101 | 8.81E-06 |
| Rdh13 | 1335.03426 | 0.983374046 | 4.08E-14 |
| Cd44 | 4483.253208 | 0.983330639 | 4.04E-07 |
| Csnk2a2 | 2886.982549 | 0.983289105 | 1.28E-20 |
| Trim47 | 1273.010191 | 0.98328099 | 6.03E-08 |
| Axl | 36075.15712 | 0.983036022 | 6.94E-12 |
| Banf1 | 3837.995394 | 0.983019921 | 5.43E-12 |
| Prkab2 | 1120.185202 | 0.982794944 | 6.26E-10 |
| Psmd3 | 7020.635315 | 0.982635891 | 4.89E-14 |
| Peli1 | 768.5211593 | 0.982409087 | 1.21E-12 |
| Umps | 1718.713927 | 0.982381865 | 4.18E-09 |
| Ttc39b | 2306.341585 | 0.982381499 | 2.59E-08 |
| Rpl23a-ps3 | 79.47664741 | 0.982104328 | 2.72E-02 |
| Uevld | 1160.400693 | 0.980892728 | 5.77E-09 |
| Cox6b1 | 4254.104049 | 0.980755558 | 1.17E-13 |
| Rpl34 | 304.9125979 | 0.980420689 | 1.09E-08 |
| Med28 | 1860.448472 | 0.980226812 | 4.42E-15 |
| Scrn2 | 419.7699392 | 0.980207799 | 1.80E-09 |
| Gm5069 | 25.00969687 | 0.979963813 | 7.41E-03 |
| Fhl2 | 3219.077124 | 0.979577465 | 2.65E-10 |
| Mvb12a | 2343.567481 | 0.979460923 | 1.96E-07 |
| Fdxacb1 | 210.4348022 | 0.979077393 | 9.43E-05 |
| Sik1 | 2788.494883 | 0.979036878 | 1.13E-04 |
| Alkbh7 | 365.0892985 | 0.978961454 | 6.60E-14 |
| Tecr | 7238.644524 | 0.978244491 | 5.27E-13 |
| Ndufb8 | 3365.96496 | 0.978059137 | 3.99E-19 |
| Exoc4 | 6670.852414 | 0.977312279 | 2.44E-16 |
| Slc39a6 | 3751.096942 | 0.976795829 | 2.77E-12 |
| Ranbp9 | 3139.587622 | 0.976652835 | 8.60E-07 |
| Abhd2 | 3646.914905 | 0.976332893 | 1.09E-11 |

| | | | |
|---|---|---|---|
| Ddx49 | 1574.01169 | 0.97582722 | 4.08E-13 |
| Ywhae | 20410.46669 | 0.975706926 | 4.44E-24 |
| Pxdc1 | 198.4170374 | 0.975489396 | 2.26E-05 |
| Selenoo | 1274.742205 | 0.974280009 | 1.07E-13 |
| Map3k14 | 972.1145369 | 0.973619936 | 8.74E-08 |
| Tmsb4x | 9263.718803 | 0.973122175 | 3.67E-15 |
| Ei24 | 3209.591063 | 0.972847572 | 3.28E-11 |
| Leprotl1 | 1717.958866 | 0.972556844 | 9.27E-25 |
| Surf2 | 1370.085633 | 0.972460192 | 7.96E-22 |
| Ppid | 3377.358914 | 0.972170353 | 3.61E-08 |
| Nop2 | 4072.658811 | 0.971755513 | 8.50E-10 |
| Palm2 | 231.3428786 | 0.971578264 | 1.33E-03 |
| Pde10a | 470.0570704 | 0.971454194 | 2.47E-12 |
| Polr2l | 617.9798244 | 0.971317918 | 5.30E-06 |
| Pxmp2 | 57.82182366 | 0.970982941 | 9.15E-04 |
| Abhd4 | 7163.071034 | 0.970892217 | 3.93E-06 |
| Brinp3 | 129.3249775 | 0.970445803 | 2.84E-02 |
| Ddx18 | 3710.623133 | 0.970308877 | 2.42E-15 |
| Tmem62 | 479.9034088 | 0.969829183 | 2.25E-08 |
| G6pc3 | 1338.998851 | 0.969645286 | 8.56E-10 |
| Ppp2r2a | 2867.173136 | 0.969417053 | 8.79E-21 |
| Zfp689 | 181.2686209 | 0.969377637 | 1.79E-07 |
| Rhebl1 | 244.5837871 | 0.968854834 | 7.05E-06 |
| Arhgef18 | 3617.647802 | 0.96795311 | 3.21E-12 |
| Nup133 | 2240.282604 | 0.967357396 | 1.35E-09 |
| Nubp1 | 1342.338329 | 0.967289481 | 2.36E-11 |
| Aco2 | 10683.99062 | 0.967096743 | 4.44E-14 |
| Asrgl1 | 595.939998 | 0.966968284 | 1.54E-05 |
| Prr7 | 85.92663974 | 0.966852759 | 4.20E-04 |
| Xpc | 1557.390835 | 0.966585463 | 1.26E-14 |
| Zfp354a | 365.4228856 | 0.96644398 | 4.10E-05 |
| Agrn | 5204.164369 | 0.966336837 | 7.04E-08 |
| Tigar | 293.8160429 | 0.966272749 | 3.67E-05 |
| Tubb5 | 51167.76877 | 0.965512963 | 3.02E-08 |
| Tnfrsf23 | 1998.923511 | 0.965353391 | 1.15E-04 |
| Atic | 6279.229534 | 0.965223621 | 1.34E-11 |
| Imp3 | 1479.412076 | 0.965168613 | 1.58E-10 |
| Trip10 | 3204.625039 | 0.964855517 | 4.08E-10 |
| Rplp0-ps1 | 63.10554626 | 0.964553207 | 1.41E-02 |
| Vgll4 | 2778.211313 | 0.964472463 | 1.39E-10 |
| Abhd17c | 2306.603289 | 0.963905791 | 1.12E-10 |
| Espnl | 43.3323916 | 0.963551819 | 1.93E-02 |
| Ak6 | 673.3172241 | 0.96295897 | 2.62E-14 |
| Zfpl1 | 1580.649371 | 0.96269451 | 3.86E-07 |
| Zbtb20 | 2019.282866 | 0.962568915 | 4.76E-02 |
| Gtf2e1 | 984.1282225 | 0.962444899 | 1.62E-04 |
| Ddx11 | 404.9730421 | 0.962075042 | 1.49E-03 |

| | | | |
|---|---|---|---|
| Psma3 | 4329.011223 | 0.961934766 | 4.68E-09 |
| Pafah1b1 | 8649.10865 | 0.961576844 | 9.87E-22 |
| Gm9830 | 252.6703547 | 0.960670683 | 1.18E-04 |
| Hgsnat | 2033.324024 | 0.959931774 | 1.96E-18 |
| Rps4x | 93555.43077 | 0.959755447 | 3.33E-07 |
| Gsdmd | 431.2673849 | 0.959398152 | 1.22E-15 |
| Rtn3 | 10304.32456 | 0.959218584 | 2.68E-12 |
| Fam219a | 921.6076423 | 0.958566834 | 1.33E-12 |
| Id4 | 162.2104674 | 0.958155877 | 4.98E-03 |
| Rabepk | 865.8826236 | 0.957767999 | 1.42E-08 |
| Mastl | 676.3577733 | 0.957674328 | 7.53E-03 |
| Mtfr1l | 3041.82241 | 0.956267577 | 4.37E-07 |
| Kcnn1 | 233.8205423 | 0.956088241 | 1.72E-06 |
| Slc25a20 | 624.4631335 | 0.95524872 | 1.46E-09 |
| Rpl38-ps2 | 415.2384649 | 0.954834408 | 4.96E-02 |
| Smg8 | 1733.268969 | 0.95461315 | 1.61E-10 |
| Txn1 | 10959.5229 | 0.95445788 | 4.75E-11 |
| Wdr5 | 2950.251821 | 0.954305466 | 2.83E-20 |
| Afg3l1 | 3276.76621 | 0.954277677 | 1.66E-23 |
| Bdnf | 1365.54172 | 0.954110525 | 5.84E-06 |
| Ndufa10 | 5483.308946 | 0.953598541 | 8.83E-09 |
| Fbxo6 | 2022.688948 | 0.952804504 | 9.69E-11 |
| NA | 486.6545116 | 0.952497937 | 1.23E-09 |
| Msh3 | 550.292747 | 0.952406994 | 3.54E-08 |
| Hmgcl | 973.914097 | 0.951916326 | 6.23E-09 |
| Cers2 | 6182.42433 | 0.951800174 | 1.92E-27 |
| Bri3 | 2945.239017 | 0.950901243 | 4.99E-05 |
| Lrp10 | 4039.665181 | 0.949878695 | 1.06E-12 |
| Ralgps1 | 724.2808194 | 0.949635366 | 1.89E-11 |
| Rnf10 | 11588.78172 | 0.949339233 | 8.30E-12 |
| Zwilch | 669.3877584 | 0.948872389 | 2.89E-03 |
| Psmd1 | 10295.53513 | 0.94875753 | 7.52E-16 |
| Cwc15 | 2545.719953 | 0.948538947 | 3.02E-30 |
| Smndc1 | 2122.331944 | 0.948423548 | 3.47E-13 |
| Lzts2 | 5933.576252 | 0.94837695 | 1.80E-12 |
| 9530077C05Rik | 123.6192175 | 0.947891619 | 3.99E-06 |
| Gm26648 | 42.53327553 | 0.947736796 | 6.09E-03 |
| Rnf181 | 3995.128789 | 0.947712373 | 5.91E-10 |
| G430049J08Rik | 23.43563077 | 0.947336419 | 3.13E-02 |
| Rps21 | 17061.14801 | 0.947106884 | 1.55E-06 |
| Prmt7 | 1502.437278 | 0.946759872 | 3.66E-16 |
| 4833439L19Rik | 3511.377629 | 0.946728293 | 3.17E-10 |
| Cand1 | 10381.58173 | 0.946041893 | 2.17E-14 |
| Cherp | 3589.831824 | 0.945556001 | 5.54E-12 |
| Enho | 74.8155069 | 0.945512168 | 4.72E-03 |
| Cwf19l1 | 759.216675 | 0.944723086 | 9.19E-12 |
| Trpm2 | 50.50789183 | 0.944548766 | 3.16E-02 |

| | | | |
|---|---|---|---|
| Gm5931 | 88.3662409 | 0.944430366 | 1.91E-02 |
| Adh5 | 4642.32907 | 0.944264303 | 4.18E-31 |
| Ccdc51 | 211.5969972 | 0.944217044 | 1.53E-05 |
| H2-T23 | 367.1066819 | 0.943998485 | 3.98E-04 |
| Snd1 | 12835.65461 | 0.943794684 | 1.05E-17 |
| Galt | 472.9768465 | 0.943787334 | 1.32E-09 |
| Dnajb1 | 2104.812103 | 0.943525451 | 4.98E-08 |
| Arv1 | 297.5590793 | 0.943204015 | 5.83E-13 |
| Rpl32 | 65196.27315 | 0.943081639 | 1.85E-06 |
| Fbxo25 | 1442.277771 | 0.942868406 | 6.67E-14 |
| Fam216a | 1524.130343 | 0.942413521 | 5.41E-24 |
| Tead1 | 8305.982112 | 0.942239099 | 4.06E-13 |
| Hyou1 | 5671.534428 | 0.942107717 | 3.21E-08 |
| Tmem237 | 787.3719009 | 0.94201037 | 5.68E-20 |
| Ddb2 | 437.666468 | 0.941451598 | 6.56E-07 |
| Siva1 | 1713.39971 | 0.94089941 | 2.78E-06 |
| Fyb | 230.8656686 | 0.940575417 | 1.45E-02 |
| Pcbp4 | 5072.455738 | 0.9404338 | 5.25E-24 |
| Cln3 | 168.8465078 | 0.940042054 | 3.99E-06 |
| Mroh2a | 196.7055094 | 0.939712518 | 1.45E-02 |
| Hsf2 | 1151.428926 | 0.939429362 | 6.10E-12 |
| Akr1e1 | 217.9345246 | 0.939257816 | 3.17E-05 |
| Gm8399 | 62.43835672 | 0.9388499 | 4.44E-03 |
| Tbc1d31 | 1630.86014 | 0.938763471 | 4.14E-05 |
| Gpr107 | 3622.865967 | 0.938594113 | 1.54E-13 |
| Zfp365 | 1958.032679 | 0.93830732 | 2.28E-09 |
| Rfxap | 458.8020367 | 0.9377027 | 1.24E-06 |
| Tor1a | 1466.830404 | 0.937625731 | 1.30E-13 |
| Ints10 | 2126.187577 | 0.937192266 | 1.73E-19 |
| Tmem238 | 87.91425267 | 0.937086154 | 1.26E-02 |
| H2-D1 | 9845.207692 | 0.936264139 | 1.78E-10 |
| Fbxw9 | 1293.75865 | 0.936161352 | 2.29E-13 |
| Zbtb40 | 700.9420987 | 0.936122687 | 3.08E-09 |
| Pde4dip | 2967.356983 | 0.936099683 | 5.73E-35 |
| Cmss1 | 343.7516773 | 0.935901798 | 1.05E-08 |
| Ormdl2 | 671.3612949 | 0.935441594 | 4.52E-08 |
| Rsl1d1 | 5971.149172 | 0.935262549 | 1.11E-16 |
| Pea15a | 11198.47623 | 0.934118623 | 3.08E-14 |
| Mrpl2 | 1684.064819 | 0.933870001 | 1.62E-12 |
| Fez1 | 805.2676056 | 0.93306126 | 9.01E-09 |
| Maea | 6459.924432 | 0.933047447 | 1.72E-16 |
| Pex14 | 1215.974622 | 0.932978138 | 9.71E-10 |
| Ubb | 30944.43698 | 0.932949815 | 2.49E-06 |
| Tagap1 | 1599.643518 | 0.932778993 | 3.18E-11 |
| Gm12751 | 1729.560456 | 0.932465605 | 4.81E-07 |
| Gm340 | 499.4293114 | 0.931686895 | 1.26E-09 |
| Wwc2 | 6641.559661 | 0.930938941 | 1.12E-14 |

| | | | |
|---|---|---|---|
| Zfand3 | 5760.058426 | 0.930514368 | 5.26E-08 |
| E130308A19Rik | 977.5793705 | 0.930475312 | 7.24E-11 |
| Azin2 | 2031.835172 | 0.930472208 | 1.67E-07 |
| Rnpep | 1933.761328 | 0.930262617 | 4.31E-16 |
| Ndufb3 | 1297.060705 | 0.929871364 | 1.26E-22 |
| Atp13a1 | 2374.27894 | 0.929683574 | 1.87E-13 |
| Pmel | 43.3739947 | 0.929225635 | 4.22E-03 |
| Cry1 | 919.8294528 | 0.928996416 | 3.61E-07 |
| Mrpl11 | 1587.25396 | 0.928519077 | 7.16E-16 |
| Stk39 | 1399.447575 | 0.927366649 | 1.01E-08 |
| Gatad2b | 1668.082965 | 0.92731503 | 1.54E-08 |
| Impdh2 | 6001.17937 | 0.927012205 | 6.33E-10 |
| Stk11 | 3742.313692 | 0.926892373 | 5.44E-10 |
| Irgm1 | 1281.382103 | 0.926782702 | 5.69E-13 |
| Sys1 | 1599.216733 | 0.926679392 | 1.64E-08 |
| NA | 77.0601651 | 0.926442638 | 3.16E-03 |
| 4933430I17Rik | 27.15124252 | 0.92638321 | 3.05E-02 |
| Rabl6 | 4751.701307 | 0.926377151 | 9.14E-14 |
| Gm10132 | 193.158572 | 0.926255435 | 1.15E-02 |
| Zfp385a | 1283.547352 | 0.925852744 | 8.34E-09 |
| Tyw1 | 1171.744952 | 0.925750955 | 7.99E-09 |
| Nfkbia | 1595.494175 | 0.92559196 | 2.67E-07 |
| Rad54b | 268.0386838 | 0.925257324 | 2.42E-02 |
| Mrpl14 | 1006.247001 | 0.925118778 | 1.20E-06 |
| Pdia4 | 11618.21839 | 0.924711002 | 1.33E-08 |
| Gnai2 | 16160.10343 | 0.92434861 | 1.92E-11 |
| Xab2 | 2424.782106 | 0.924317843 | 2.17E-09 |
| Utp11 | 3221.327935 | 0.924150357 | 2.59E-20 |
| Atg4d | 558.9833682 | 0.924078601 | 1.77E-11 |
| Nop10 | 1906.592631 | 0.923618876 | 1.20E-12 |
| Malsu1 | 636.9300842 | 0.923398193 | 1.68E-09 |
| Mapk12 | 322.9227326 | 0.923276327 | 2.63E-12 |
| Mapk3 | 6423.195426 | 0.922495965 | 6.33E-08 |
| Nol9 | 1501.271379 | 0.922447997 | 2.09E-11 |
| Mrpl57 | 631.7549906 | 0.922061851 | 2.35E-07 |
| Zfp981 | 48.727574 | 0.922022244 | 4.13E-02 |
| Dhfr | 781.3575348 | 0.921656309 | 5.25E-03 |
| Slc41a3 | 517.5453823 | 0.921188514 | 1.88E-10 |
| Srpr | 5889.393341 | 0.921152129 | 2.05E-10 |
| Acap3 | 1180.426838 | 0.920447209 | 2.59E-10 |
| Sp1 | 6965.188488 | 0.920093 | 4.05E-10 |
| Prpf38a | 1921.609227 | 0.919777791 | 8.98E-22 |
| Wdr89 | 406.7336064 | 0.919130188 | 2.78E-03 |
| Heatr5b | 3231.838328 | 0.918929005 | 4.84E-18 |
| Hspa9 | 32616.301 | 0.918849461 | 7.30E-21 |
| Ccdc80 | 6478.722348 | 0.918740439 | 2.51E-04 |
| Plod2 | 1368.669902 | 0.918669979 | 3.62E-06 |

| | | | |
|---|---|---|---|
| Katna1 | 1574.205449 | 0.91858446 | 2.18E-14 |
| Impdh2-ps | 1489.025213 | 0.918562516 | 1.37E-11 |
| Cyc1 | 3631.315047 | 0.918370841 | 1.90E-11 |
| H3f3b | 19605.31291 | 0.917839944 | 2.62E-15 |
| Chmp2a | 3257.77723 | 0.917746227 | 1.35E-25 |
| Ddx58 | 1237.338292 | 0.917523584 | 1.39E-08 |
| Gm10774 | 279.4015294 | 0.917148186 | 5.30E-07 |
| Ubxn8 | 665.5000472 | 0.916701483 | 4.09E-12 |
| 1110065P20Rik | 1148.910686 | 0.916558517 | 3.30E-10 |
| Slc25a3 | 21829.57629 | 0.916269001 | 1.43E-25 |
| Cpox | 1136.258528 | 0.916229179 | 1.70E-04 |
| Farsa | 2227.240026 | 0.915622712 | 5.10E-16 |
| Nsun5 | 676.7308702 | 0.915517616 | 1.08E-08 |
| Sqstm1 | 31647.65135 | 0.915340712 | 4.54E-07 |
| Abhd14a | 565.1011237 | 0.915201977 | 9.12E-12 |
| Ttc17 | 2517.874017 | 0.914483741 | 2.86E-07 |
| Taf1d | 1247.952012 | 0.914449456 | 6.24E-06 |
| Snhg12 | 1711.852806 | 0.913622076 | 1.12E-05 |
| Cdt1 | 2176.814691 | 0.913554983 | 5.30E-04 |
| Taf12 | 648.1120911 | 0.912627632 | 6.90E-12 |
| NA | 457.4716934 | 0.912592283 | 1.01E-04 |
| Wtip | 1616.112467 | 0.912586933 | 3.30E-09 |
| Fam117a | 85.5208589 | 0.91257026 | 2.83E-02 |
| Med30 | 695.4020104 | 0.912237445 | 6.76E-15 |
| Tcaim | 463.9162715 | 0.911848437 | 6.55E-09 |
| Atg14 | 874.8182583 | 0.911801765 | 2.60E-04 |
| H2-Q5 | 32.8399492 | 0.911753167 | 5.32E-03 |
| Cstf1 | 2074.099098 | 0.911681648 | 2.82E-07 |
| 4930452B06Rik | 303.1615407 | 0.911630605 | 5.38E-06 |
| Vcpkmt | 777.1607539 | 0.911431022 | 1.28E-07 |
| Drosha | 4357.54497 | 0.91124853 | 1.33E-28 |
| 5330438D12Rik | 62.65591807 | 0.911230767 | 1.16E-02 |
| Pkn1 | 3328.989249 | 0.911026015 | 9.62E-15 |
| Mtch2 | 7896.965041 | 0.910841873 | 9.22E-32 |
| Vcl | 15499.81698 | 0.910560672 | 1.01E-13 |
| AU040320 | 3791.60754 | 0.909672106 | 9.95E-17 |
| Kdm8 | 161.4759552 | 0.909653527 | 5.56E-04 |
| Trnau1ap | 1335.035752 | 0.909171915 | 1.49E-15 |
| Atf6 | 5289.469533 | 0.908761129 | 1.80E-09 |
| Uchl5 | 1036.057855 | 0.908755565 | 1.64E-07 |
| Snx18 | 1837.625647 | 0.90838159 | 3.94E-18 |
| Prmt5 | 1974.839417 | 0.908103229 | 7.89E-12 |
| Rnf113a1 | 147.4647827 | 0.907532789 | 7.62E-06 |
| Atp6v1d | 4059.992121 | 0.906980263 | 4.65E-08 |
| Ptpn21 | 3194.640395 | 0.906887934 | 1.94E-08 |
| Adcy6 | 3708.430534 | 0.906660465 | 2.27E-26 |
| Tmed4 | 3139.271629 | 0.906261934 | 5.07E-10 |

| | | | |
|---|---|---|---|
| Smim8 | 236.3226799 | 0.906026342 | 1.62E-10 |
| Gm15500 | 76446.03163 | 0.905910207 | 1.83E-06 |
| Cenpa | 1628.559306 | 0.905797002 | 1.18E-03 |
| Ahi1 | 1541.580211 | 0.905462322 | 1.34E-07 |
| NA | 236.8173459 | 0.905264513 | 1.41E-05 |
| Desi1 | 3182.457457 | 0.905225102 | 2.13E-08 |
| Hdlbp | 31467.4146 | 0.905143616 | 9.15E-15 |
| Dip2a | 3267.183888 | 0.904954419 | 4.89E-19 |
| Lasp1 | 9498.425232 | 0.904229572 | 8.57E-17 |
| Dnm2 | 5735.557998 | 0.90386137 | 3.28E-10 |
| Ccdc159 | 201.3206634 | 0.903161259 | 4.45E-05 |
| Gcn1l1 | 5972.383838 | 0.902826143 | 2.38E-14 |
| Cep57l1 | 538.9271566 | 0.902736524 | 6.38E-08 |
| Snhg5 | 748.1058599 | 0.902423108 | 7.85E-06 |
| Gpr108 | 2920.507615 | 0.902309867 | 8.25E-10 |
| Atxn2 | 4115.093461 | 0.902104066 | 4.60E-12 |
| Cops7a | 6190.887762 | 0.9020707 | 1.42E-19 |
| Myo19 | 568.3216705 | 0.902060293 | 3.61E-04 |
| Med11 | 515.6146003 | 0.901945838 | 6.24E-10 |
| Taf9 | 2514.048924 | 0.90176088 | 2.22E-10 |
| Gstm5 | 2305.479204 | 0.901682923 | 1.38E-10 |
| Ghdc | 1058.772751 | 0.901282296 | 1.22E-06 |
| Aldh9a1 | 4090.448895 | 0.901220935 | 4.49E-16 |
| Arl6ip1 | 275.4071157 | 0.900892852 | 1.44E-08 |
| Pdcd6ip | 8464.98572 | 0.90076395 | 7.32E-27 |
| Neurl1a | 101.6784878 | 0.900663236 | 2.86E-03 |
| Fam160a2 | 1223.39513 | 0.899636665 | 1.85E-16 |
| Slc35d1 | 859.5984113 | 0.899160313 | 4.83E-16 |
| Rela | 3095.57112 | 0.898039779 | 8.42E-16 |
| Gtpbp10 | 161.1492663 | 0.897594789 | 1.36E-03 |
| Prdm15 | 720.4881743 | 0.897139585 | 3.42E-06 |
| C1qtnf12 | 46.38102859 | 0.896954359 | 2.00E-02 |
| Zfp692 | 591.0013863 | 0.896904376 | 5.88E-11 |
| Cdca8 | 1439.881188 | 0.896808943 | 6.95E-03 |
| Prelid3b | 1774.650704 | 0.89677866 | 4.42E-09 |
| Cdc42ep3 | 3187.14258 | 0.896421755 | 1.61E-10 |
| Psmc5 | 3850.307525 | 0.896332942 | 9.74E-10 |
| Abcf3 | 2478.910211 | 0.896318832 | 2.84E-10 |
| Ppp1r2 | 6493.223235 | 0.896171715 | 7.07E-11 |
| Mettl25 | 462.2847605 | 0.896015267 | 1.04E-08 |
| Ahcyl2 | 2542.94842 | 0.895964587 | 5.54E-07 |
| Amz2 | 3080.784919 | 0.895912824 | 3.28E-11 |
| Cstb | 9856.088638 | 0.895601761 | 4.62E-15 |
| Anxa2 | 27933.32394 | 0.895453133 | 1.94E-08 |
| Opn3 | 112.0638704 | 0.894838036 | 2.87E-05 |
| Nolc1 | 4595.577339 | 0.894810747 | 1.09E-07 |
| Cnp | 33.65963087 | 0.89409837 | 1.20E-02 |

| | | | |
|---|---|---|---|
| Dffb | 114.904738 | 0.893168351 | 3.60E-05 |
| Gm10157 | 272.6510177 | 0.893123233 | 4.29E-03 |
| Gm10177 | 3215.751042 | 0.892915894 | 4.19E-15 |
| Supt4a | 1530.662931 | 0.892406754 | 4.02E-09 |
| Gm3362 | 137.0655522 | 0.892352052 | 4.26E-03 |
| Ddx19a | 2503.906907 | 0.892149964 | 6.71E-09 |
| Qrsl1 | 552.2016329 | 0.891867043 | 7.24E-12 |
| Adnp2 | 2115.907237 | 0.891699261 | 1.40E-16 |
| Ptges3-ps | 1167.527342 | 0.891506496 | 1.38E-06 |
| Heatr6 | 2310.476748 | 0.891103906 | 7.88E-12 |
| Utp18 | 1810.570753 | 0.89099119 | 3.81E-10 |
| Setd6 | 561.7236217 | 0.890877833 | 5.49E-14 |
| Saysd1 | 190.5484555 | 0.890749567 | 9.23E-07 |
| Fbxo17 | 722.6181176 | 0.890676927 | 4.45E-12 |
| Lrrc41 | 6294.169907 | 0.890293623 | 1.22E-10 |
| Mprip | 17779.67662 | 0.890233594 | 3.52E-31 |
| Slc39a7 | 10355.74721 | 0.890036839 | 6.19E-09 |
| G6pdx | 4365.833196 | 0.889884181 | 3.95E-08 |
| Spryd3 | 1383.849558 | 0.889047148 | 4.68E-09 |
| Zfp691 | 296.0729184 | 0.888870976 | 2.18E-03 |
| Angpt4 | 781.819534 | 0.888786647 | 3.92E-02 |
| Nvl | 1837.408422 | 0.888697144 | 3.98E-26 |
| Pias4 | 1560.177006 | 0.888679172 | 1.33E-05 |
| Lpar2 | 1343.831789 | 0.888361481 | 1.82E-08 |
| Ogfod3 | 112.2973235 | 0.888144168 | 1.46E-04 |
| Atp11a | 5318.325548 | 0.887572775 | 3.66E-10 |
| Gm12013 | 265.5132074 | 0.887323738 | 4.16E-03 |
| Grb2 | 2639.41625 | 0.887184525 | 4.22E-14 |
| Klhdc10 | 3315.489594 | 0.886988039 | 6.53E-16 |
| Fdft1 | 6536.743541 | 0.886703484 | 5.16E-04 |
| Rxrb | 2724.534752 | 0.886257052 | 1.68E-12 |
| Coq6 | 914.9612862 | 0.885383897 | 1.19E-08 |
| Acp1 | 3181.095991 | 0.885367771 | 3.93E-17 |
| Mrps23 | 1720.567113 | 0.885271295 | 7.26E-09 |
| Aurkb | 1122.511781 | 0.884751565 | 9.99E-03 |
| Srrm1 | 6012.177293 | 0.883834016 | 5.69E-15 |
| Rab3gap2 | 4136.31955 | 0.883818272 | 1.54E-18 |
| Abca2 | 3887.217761 | 0.883723755 | 1.84E-16 |
| Phc1 | 2184.532526 | 0.883495835 | 1.96E-10 |
| Grhl1 | 28.30210692 | 0.883235994 | 2.96E-02 |
| Polg2 | 290.1108251 | 0.882818827 | 1.21E-05 |
| Gas2l3 | 1203.193311 | 0.882505638 | 8.53E-04 |
| Sssca1 | 1262.870255 | 0.882443126 | 5.83E-12 |
| Kpna6 | 7686.854538 | 0.882168048 | 5.64E-17 |
| Slc25a19 | 461.7725604 | 0.881141026 | 3.30E-07 |
| Cdc25b | 1752.307462 | 0.880198776 | 8.00E-05 |
| Anapc15 | 633.2408699 | 0.87913371 | 3.13E-06 |

| | | | |
|---|---|---|---|
| Zxdc | 3218.603639 | 0.878829938 | 6.64E-08 |
| Smdt1 | 1962.312894 | 0.878766976 | 3.82E-12 |
| Zfyve19 | 792.6092984 | 0.878655802 | 5.27E-06 |
| Calcoco1 | 4591.223224 | 0.878326905 | 1.02E-04 |
| Ppm1f | 3507.661585 | 0.87820527 | 1.62E-14 |
| Copa | 20491.01741 | 0.878179172 | 9.40E-12 |
| Rfx2 | 152.4655248 | 0.877924195 | 1.41E-05 |
| Ubxn4 | 6611.923936 | 0.877506799 | 5.91E-27 |
| Actb | 74206.67695 | 0.877366597 | 1.73E-12 |
| Gps2 | 1053.198647 | 0.876992903 | 2.56E-09 |
| Emg1 | 2548.431435 | 0.876905852 | 2.28E-13 |
| 1110012L19Rik | 1013.889739 | 0.876211523 | 6.33E-09 |
| Dgkh | 329.3063989 | 0.875629396 | 3.20E-04 |
| Rpl36al | 5933.505914 | 0.875162039 | 4.00E-19 |
| Faf2 | 3343.387474 | 0.874453667 | 1.48E-20 |
| Pycr1 | 948.3752732 | 0.87433552 | 9.59E-04 |
| Tmem120a | 2131.025476 | 0.873974365 | 2.70E-13 |
| Fbxl4 | 863.6181875 | 0.873734185 | 1.13E-15 |
| Prkag1 | 2253.546472 | 0.873588201 | 6.36E-09 |
| Hpcal1 | 926.3645226 | 0.873339029 | 4.73E-05 |
| Micu1 | 6570.285669 | 0.873182673 | 3.89E-10 |
| Tmpo | 7007.127582 | 0.872822537 | 7.64E-07 |
| Ifitm3 | 3638.162294 | 0.872648479 | 1.15E-04 |
| Abca4 | 66.55109914 | 0.872561469 | 2.02E-02 |
| Prrc1 | 2984.459546 | 0.872189006 | 4.36E-09 |
| Sirt3 | 334.231396 | 0.872029088 | 5.69E-07 |
| Phpt1 | 1260.695996 | 0.871911564 | 1.92E-09 |
| Timm23 | 2565.708129 | 0.871138178 | 8.60E-14 |
| Exoc3l | 208.8741871 | 0.870022021 | 4.53E-07 |
| Ppm1m | 1008.031276 | 0.869356285 | 1.48E-11 |
| Eif1 | 18854.81854 | 0.868958861 | 2.26E-09 |
| Rpl31 | 2778.260788 | 0.868870475 | 1.11E-12 |
| Celf4 | 267.6869814 | 0.868577958 | 2.58E-04 |
| Cdc37 | 6153.441156 | 0.86857296 | 9.04E-11 |
| Dctpp1 | 464.4771838 | 0.868418271 | 3.63E-04 |
| Tmem109 | 3376.214754 | 0.868125497 | 2.38E-06 |
| Apba3 | 2076.974687 | 0.867811611 | 3.90E-12 |
| Ncstn | 4112.498853 | 0.867714753 | 6.43E-16 |
| Nudt9 | 3226.818754 | 0.867673242 | 7.68E-08 |
| Sms | 1954.671987 | 0.867580415 | 7.82E-08 |
| Jmjd6 | 1587.362824 | 0.866609404 | 9.14E-12 |
| Coa4 | 295.6661041 | 0.866414452 | 1.46E-04 |
| Gm6055 | 349.7896906 | 0.866241793 | 1.23E-02 |
| Slc37a4 | 401.0618275 | 0.865866377 | 1.32E-10 |
| Pwp2 | 1822.29441 | 0.865831359 | 1.97E-09 |
| Vps39 | 2002.970673 | 0.86570292 | 1.44E-06 |
| Mad2l2 | 595.8755217 | 0.864942764 | 1.43E-11 |

| | | | |
|---|---|---|---|
| Tatdn3 | 338.0046401 | 0.864736472 | 8.84E-06 |
| Fubp1 | 6171.280554 | 0.864445561 | 6.86E-13 |
| Pla2g6 | 1759.603545 | 0.864276907 | 7.97E-06 |
| Nifk | 2606.036144 | 0.864113239 | 3.60E-11 |
| NA | 5575.684898 | 0.863797418 | 1.18E-11 |
| Trit1 | 873.3104816 | 0.863598844 | 1.83E-10 |
| Ubqln4 | 3372.00673 | 0.863475723 | 3.13E-12 |
| Ergic3 | 4114.58219 | 0.863251406 | 3.71E-07 |
| Caskin2 | 1078.194197 | 0.862961007 | 1.14E-16 |
| Arih2 | 2519.227217 | 0.862863626 | 1.21E-10 |
| Tpm4 | 16366.22209 | 0.862548731 | 2.28E-10 |
| Smtn | 3576.331185 | 0.862491561 | 1.29E-08 |
| Sh3bp4 | 1970.38902 | 0.862301163 | 9.74E-09 |
| Lyrm1 | 97.40410017 | 0.86206966 | 3.64E-04 |
| Rps15a-ps3 | 65.56143529 | 0.862008153 | 7.37E-03 |
| Hsp90aa1 | 28139.47731 | 0.86199213 | 1.97E-03 |
| Ttc27 | 1412.862851 | 0.861151773 | 1.14E-15 |
| Actr1b | 4863.150793 | 0.860657377 | 4.58E-10 |
| Ubc | 68322.4058 | 0.860358483 | 1.57E-05 |
| Dus1l | 1059.078537 | 0.859029812 | 5.57E-11 |
| Fis1 | 3116.094015 | 0.858910052 | 2.51E-10 |
| Timm50 | 1105.524902 | 0.858544274 | 2.69E-06 |
| Sumf2 | 1672.193117 | 0.858345857 | 5.25E-09 |
| Tut1 | 1688.031275 | 0.858197801 | 1.69E-12 |
| Rbm18 | 1817.797828 | 0.857954871 | 1.70E-08 |
| Coq7 | 524.6389928 | 0.857044782 | 5.32E-09 |
| Gpn2 | 393.9806168 | 0.85684275 | 4.93E-07 |
| Idh2 | 11638.97368 | 0.856790464 | 2.47E-09 |
| Napg | 996.6463771 | 0.856641907 | 5.93E-10 |
| Ddx19b | 450.9567018 | 0.856612088 | 2.76E-15 |
| Tuba1c | 4795.987546 | 0.856586102 | 1.39E-03 |
| Cpsf4 | 763.0921259 | 0.856030011 | 2.45E-11 |
| Lamc1 | 22727.02501 | 0.855733999 | 2.21E-09 |
| Ercc8 | 336.7685356 | 0.855625379 | 2.32E-06 |
| Prcp | 2518.163059 | 0.855353476 | 6.64E-07 |
| Map2k3 | 4419.736716 | 0.855300737 | 5.01E-09 |
| Mrpl1 | 717.9429599 | 0.855268283 | 1.41E-04 |
| Thra | 5095.172409 | 0.855090832 | 2.62E-04 |
| Cbwd1 | 629.8398015 | 0.854710111 | 9.85E-07 |
| Gfm2 | 1007.986387 | 0.854637654 | 4.44E-23 |
| Ccdc15 | 119.2153648 | 0.854419057 | 3.43E-04 |
| Gsn | 46199.21147 | 0.853947403 | 4.44E-09 |
| Slc44a2 | 6308.547057 | 0.853771602 | 2.22E-11 |
| Gm1821 | 5869.390963 | 0.853662858 | 2.89E-05 |
| Gm8113 | 678.7557631 | 0.853555607 | 3.03E-03 |
| Nr4a2 | 174.1972914 | 0.853319044 | 7.37E-04 |
| Tedc2 | 465.082066 | 0.852940399 | 3.27E-04 |

| | | | |
|---|---|---|---|
| Scmh1 | 2538.520534 | 0.852485201 | 1.67E-08 |
| Dhx29 | 1698.366442 | 0.852468412 | 5.84E-25 |
| Mthfd1 | 3288.788194 | 0.851744776 | 5.16E-07 |
| Gm26762 | 29.75269382 | 0.851083704 | 3.06E-02 |
| Akap8l | 1442.125993 | 0.850609048 | 4.50E-08 |
| Luc7l | 3039.936213 | 0.850179514 | 2.21E-09 |
| Lrp8 | 1552.953999 | 0.849860628 | 9.85E-03 |
| Dnaja2 | 8057.883399 | 0.849832469 | 7.33E-14 |
| Bub3 | 3338.139578 | 0.849822116 | 1.69E-06 |
| Ankrd54 | 751.29234 | 0.848297712 | 4.88E-06 |
| Map2k2 | 5007.849089 | 0.847577422 | 6.95E-12 |
| Mlkl | 1417.964644 | 0.847541843 | 3.07E-08 |
| Phykpl | 589.4291438 | 0.847444831 | 3.53E-13 |
| Ube2f | 1729.361484 | 0.84725712 | 3.97E-14 |
| Proser3 | 387.0409439 | 0.84658468 | 5.70E-04 |
| Kxd1 | 1435.726318 | 0.846198347 | 7.46E-05 |
| Cdk1 | 3299.072583 | 0.845742581 | 8.95E-03 |
| Trappc5 | 1301.529436 | 0.845669426 | 9.65E-09 |
| Git1 | 4693.589768 | 0.845250739 | 9.37E-10 |
| Twf2 | 817.7616965 | 0.845176812 | 8.85E-09 |
| Mipol1 | 608.7016312 | 0.845121448 | 8.88E-13 |
| Pcbp2 | 19401.57002 | 0.845023931 | 9.77E-11 |
| Cdca5 | 431.2039611 | 0.844859886 | 2.94E-02 |
| Gm2000 | 6602.439031 | 0.843975102 | 6.97E-04 |
| Rpa1 | 7115.964406 | 0.843644743 | 9.83E-05 |
| Pir | 1475.999153 | 0.843539883 | 1.02E-04 |
| Tmbim6 | 15673.1219 | 0.843338868 | 1.25E-11 |
| Ndfip2 | 1306.937112 | 0.843298235 | 1.81E-25 |
| Alg2 | 1510.645998 | 0.842875249 | 1.90E-09 |
| Ahcyl1 | 10647.42186 | 0.842529338 | 9.38E-32 |
| Spc24 | 355.8336016 | 0.842438085 | 1.90E-02 |
| Zfp7 | 302.867491 | 0.842059529 | 9.93E-05 |
| Slc25a40 | 773.2498353 | 0.841660688 | 7.64E-05 |
| Akt1s1 | 3858.360262 | 0.841650034 | 1.97E-06 |
| Shroom3 | 71.88219185 | 0.841085059 | 1.64E-02 |
| Tfe3 | 4096.617787 | 0.840958937 | 2.00E-09 |
| Ubxn7 | 3892.875648 | 0.840391356 | 4.89E-12 |
| Ccdc97 | 1381.339871 | 0.840229814 | 7.93E-10 |
| Fam198b | 10855.77158 | 0.839978853 | 1.96E-04 |
| Fntb | 1105.956478 | 0.839174662 | 1.76E-06 |
| Bicd2 | 3364.546628 | 0.839072124 | 5.40E-13 |
| Creb3l1 | 1785.610876 | 0.838767924 | 2.82E-04 |
| Stx7 | 2576.759255 | 0.838257319 | 5.10E-13 |
| Gm9828 | 55.4899361 | 0.837904918 | 7.90E-03 |
| Lars2 | 26122.06978 | 0.837641828 | 5.05E-05 |
| Psen1 | 3740.441008 | 0.837482901 | 1.36E-12 |
| Txndc17 | 3979.458947 | 0.837035297 | 6.10E-13 |

| | | | |
|---|---|---|---|
| S100a13 | 308.5479394 | 0.836960433 | 1.38E-07 |
| Bloc1s6 | 2040.363911 | 0.836814003 | 7.20E-09 |
| Nap1l1 | 24172.49471 | 0.836564326 | 2.39E-13 |
| Ncoa4 | 4582.091116 | 0.836521198 | 7.10E-20 |
| Ercc5 | 1751.423907 | 0.83612904 | 1.17E-15 |
| Gga3 | 928.5292727 | 0.835663842 | 6.09E-06 |
| Cep104 | 1205.104973 | 0.834688359 | 1.54E-15 |
| 4933434E20Rik | 1868.94148 | 0.834582185 | 6.93E-18 |
| Prkci | 7019.667731 | 0.834514764 | 3.51E-07 |
| Tcaf2 | 513.7387912 | 0.833745303 | 6.46E-04 |
| Trappc4 | 1400.935846 | 0.833730933 | 7.88E-11 |
| Thoc5 | 2117.128019 | 0.833478351 | 5.21E-11 |
| Ube2d3 | 9445.414009 | 0.832914459 | 1.70E-29 |
| Lactb2 | 1674.430397 | 0.832758276 | 5.43E-07 |
| Zrsr1 | 2272.804588 | 0.832393464 | 7.11E-05 |
| Patl1 | 4016.791295 | 0.831902749 | 1.14E-05 |
| Slc12a7 | 1768.921695 | 0.831838558 | 6.90E-11 |
| Pdxdc1 | 8226.574578 | 0.831057846 | 8.23E-09 |
| Hk1 | 5099.447276 | 0.830668501 | 7.87E-07 |
| Wdr75 | 2758.597647 | 0.830561432 | 1.50E-08 |
| Ccdc130 | 585.6354187 | 0.830413545 | 1.06E-05 |
| Mcph1 | 1213.891184 | 0.830132696 | 1.59E-11 |
| Gfm1 | 3165.757619 | 0.82992507 | 3.13E-13 |
| Tex2 | 1768.876038 | 0.82976656 | 4.19E-11 |
| Sugct | 28.28930194 | 0.828879738 | 4.20E-02 |
| Psenen | 1775.204145 | 0.828730767 | 3.50E-08 |
| Atg2a | 2180.865891 | 0.828421309 | 1.62E-09 |
| Gbe1 | 696.8299439 | 0.828290907 | 1.44E-05 |
| Tufm | 2564.006724 | 0.828240315 | 6.81E-09 |
| Mthfd2 | 2675.163557 | 0.828023027 | 1.09E-02 |
| Puf60 | 5332.702283 | 0.827686874 | 7.78E-10 |
| Dctn4 | 3196.598025 | 0.827566428 | 2.63E-13 |
| Ube2i | 9079.566651 | 0.827414016 | 2.38E-15 |
| Rtraf | 4191.009413 | 0.82643713 | 2.00E-05 |
| Bok | 494.9314589 | 0.826408598 | 9.70E-03 |
| Cycs | 1585.90036 | 0.826085343 | 1.37E-05 |
| Nupr1 | 2029.051611 | 0.825514579 | 1.40E-03 |
| Hadh | 4701.504074 | 0.825509492 | 9.63E-11 |
| Ndufa11 | 1512.079023 | 0.825308118 | 2.83E-07 |
| Gstp2 | 131.7087738 | 0.824861737 | 1.10E-02 |
| Kif1b | 4905.83088 | 0.824837447 | 1.34E-16 |
| Stam | 2412.624451 | 0.824814326 | 9.15E-08 |
| Cpped1 | 2173.563252 | 0.824727147 | 4.52E-12 |
| Sf3b2 | 15940.53564 | 0.824562965 | 3.50E-22 |
| Dym | 1835.376692 | 0.824331589 | 3.67E-06 |
| Mcm3ap | 3567.789929 | 0.823966225 | 7.99E-19 |
| Zswim1 | 590.8453736 | 0.823624263 | 4.89E-09 |

| | | | |
|---|---|---|---|
| Gng12 | 10176.62266 | 0.823401306 | 4.85E-09 |
| Prr13 | 2601.997656 | 0.823250509 | 3.08E-08 |
| Exosc10 | 2718.984245 | 0.823208224 | 1.83E-14 |
| Elp2 | 4679.842283 | 0.822981793 | 3.27E-15 |
| Rrp1b | 1636.99729 | 0.822925059 | 4.22E-14 |
| Tial1 | 1884.714745 | 0.822698217 | 4.12E-10 |
| Ephx2 | 595.9836792 | 0.822398567 | 1.49E-05 |
| Ltv1 | 1629.7818 | 0.821798765 | 5.09E-10 |
| Rexo2 | 2426.783116 | 0.821693638 | 2.65E-15 |
| NA | 5024.128592 | 0.820476525 | 9.31E-04 |
| Aimp1 | 2724.724341 | 0.820199504 | 1.38E-08 |
| Rtn4 | 18961.53588 | 0.819932048 | 2.53E-10 |
| Cpsf3 | 2747.549532 | 0.818925326 | 8.71E-12 |
| Mrps36 | 791.1543842 | 0.818578003 | 1.12E-08 |
| Cldnd1 | 2104.132741 | 0.818356079 | 1.74E-10 |
| Nol7 | 912.3460862 | 0.818069484 | 5.19E-07 |
| Tollip | 3274.862957 | 0.81777997 | 1.58E-07 |
| Dysf | 3326.040047 | 0.817622575 | 5.83E-05 |
| Bcas3 | 993.4164983 | 0.817619625 | 4.00E-08 |
| Lrrc47 | 1617.599376 | 0.817425357 | 2.88E-10 |
| Thap3 | 441.7221161 | 0.81635255 | 9.51E-14 |
| Tnfrsf22 | 1190.813432 | 0.816300209 | 2.69E-04 |
| Mpnd | 2802.889083 | 0.816251249 | 1.95E-09 |
| Rogdi | 701.325496 | 0.81621538 | 1.40E-07 |
| Rangap1 | 3843.086733 | 0.816116154 | 5.26E-03 |
| Cds2 | 6656.30948 | 0.815837003 | 6.59E-15 |
| Tnfrsf12a | 4419.471402 | 0.815439698 | 2.56E-04 |
| Sesn2 | 1711.20642 | 0.815210695 | 3.96E-06 |
| Mtmr4 | 3714.192411 | 0.815110733 | 7.43E-09 |
| Nt5dc3 | 2881.841793 | 0.815028134 | 9.26E-05 |
| Dcaf12 | 2233.394066 | 0.814222577 | 4.03E-16 |
| Casp7 | 402.0609059 | 0.813791461 | 1.16E-06 |
| Rars | 6691.937495 | 0.813555593 | 2.26E-10 |
| Arfgef2 | 4840.915022 | 0.8134163 | 5.73E-09 |
| Nadsyn1 | 273.4059839 | 0.813413638 | 7.86E-05 |
| 9830144P21Rik | 73.20824837 | 0.812858396 | 2.00E-02 |
| Fig4 | 2668.683324 | 0.812647909 | 2.33E-07 |
| Ubac2 | 1504.085057 | 0.812395199 | 2.17E-08 |
| Braf | 1890.433423 | 0.811694738 | 7.53E-06 |
| Fbxo38 | 2616.09343 | 0.811592241 | 2.54E-15 |
| Ap4b1 | 1040.56706 | 0.811079241 | 3.07E-16 |
| Sra1 | 1578.75584 | 0.809758341 | 3.89E-07 |
| Zc3hav1 | 5239.317525 | 0.809683176 | 7.09E-09 |
| Nfe2l1 | 17080.71886 | 0.809649239 | 5.69E-10 |
| Pou2f1 | 712.764363 | 0.809505356 | 2.87E-05 |
| Ccl27a | 208.3353949 | 0.809305383 | 4.34E-04 |
| Rab3a | 716.4417644 | 0.807761905 | 4.50E-03 |

| | | | |
|---|---|---|---|
| Crtc2 | 1839.734255 | 0.807661564 | 1.31E-06 |
| Ccdc124 | 1462.20588 | 0.807183495 | 2.60E-06 |
| Syf2 | 1649.935848 | 0.806940842 | 6.47E-11 |
| Snx3 | 7101.723968 | 0.806661145 | 6.26E-07 |
| Aqr | 3503.576024 | 0.806510595 | 6.73E-03 |
| Fbxl18 | 1195.193914 | 0.80453753 | 8.25E-12 |
| Plp2 | 8742.786313 | 0.804252003 | 5.83E-07 |
| Prdx3 | 4040.344397 | 0.804173807 | 4.82E-16 |
| Agap3 | 3243.418147 | 0.80359847 | 1.68E-07 |
| Ddrgk1 | 2594.72518 | 0.803530679 | 4.48E-11 |
| Cmas | 1604.032091 | 0.803297595 | 2.55E-22 |
| Cdk2ap2 | 1795.173273 | 0.802984889 | 2.01E-06 |
| Hspb11 | 1261.435509 | 0.801544614 | 5.48E-06 |
| Grpel2 | 916.2086707 | 0.801523489 | 3.46E-09 |
| Ndufb10 | 2793.144385 | 0.801475727 | 1.32E-06 |
| Igbp1 | 3698.89057 | 0.801194249 | 3.41E-05 |
| Sgta | 5423.891798 | 0.800979789 | 5.41E-11 |
| Rps12-ps19 | 164.2671423 | 0.800805507 | 1.05E-03 |
| Gpr89 | 1644.684102 | 0.800759209 | 3.59E-14 |
| Akt2 | 5161.192976 | 0.800757489 | 1.05E-10 |
| Ube2a | 1452.93935 | 0.800168362 | 2.23E-18 |
| Npm3 | 2710.905344 | 0.79985846 | 1.01E-11 |
| Cbl | 1012.925113 | 0.799788945 | 2.86E-08 |
| Nudt2 | 706.1488302 | 0.799280742 | 9.52E-21 |
| Cmc1 | 656.2945244 | 0.799103736 | 6.16E-07 |
| Gtf3c3 | 1632.751785 | 0.799068439 | 1.43E-20 |
| Rpl34-ps2 | 184.2154543 | 0.798845893 | 1.07E-02 |
| Man1b1 | 5748.065786 | 0.798063888 | 9.06E-15 |
| Srebf1 | 2530.686061 | 0.797804983 | 5.46E-04 |
| Ube2k | 2857.850823 | 0.797505859 | 2.58E-12 |
| Rab40c | 1203.858473 | 0.797112073 | 2.37E-07 |
| Stx1a | 409.9590611 | 0.796641791 | 8.53E-07 |
| Lmf2 | 2890.177535 | 0.796542865 | 1.10E-08 |
| Fcf1 | 1579.920549 | 0.796519944 | 1.73E-24 |
| Eif4a3 | 3389.716389 | 0.796154611 | 8.15E-09 |
| Gm12346 | 480.5896986 | 0.795326024 | 6.47E-03 |
| Haghl | 919.2093131 | 0.795305682 | 1.48E-07 |
| Mre11a | 1333.239055 | 0.794805805 | 4.83E-05 |
| Itfg2 | 542.4298973 | 0.794511261 | 6.20E-10 |
| Txndc9 | 3522.902588 | 0.793941925 | 4.65E-15 |
| Acot1 | 957.7954011 | 0.793603762 | 1.61E-04 |
| Plekho2 | 4026.295225 | 0.793052008 | 3.85E-07 |
| Napb | 476.5876789 | 0.79261837 | 8.78E-07 |
| Map4 | 13272.96546 | 0.791606171 | 1.11E-11 |
| Atp6v0b | 9934.293287 | 0.791413978 | 5.26E-06 |
| Gls2 | 118.3808906 | 0.791241504 | 7.16E-03 |
| Flad1 | 1014.220375 | 0.790701616 | 2.59E-06 |

| | | | |
|---|---|---|---|
| Ddt | 536.6455984 | 0.789705842 | 1.05E-06 |
| Drc3 | 61.559682 | 0.789307048 | 3.66E-03 |
| S100a10 | 14257.32699 | 0.789168939 | 5.56E-09 |
| Mknk1 | 2154.3649 | 0.789021694 | 2.29E-10 |
| Haus2 | 1057.827371 | 0.788953681 | 1.42E-09 |
| Psmb2 | 5869.994809 | 0.788220274 | 2.20E-15 |
| Plekhj1 | 1287.09312 | 0.788143747 | 6.48E-12 |
| Gm8186 | 1195.312773 | 0.788120181 | 1.33E-07 |
| H2-Bl | 144.2958941 | 0.787673675 | 3.39E-04 |
| Ftsj1 | 1016.642197 | 0.786977772 | 4.64E-08 |
| 2700097O09Rik | 262.8757636 | 0.786938898 | 4.31E-05 |
| Suv39h1 | 1282.53505 | 0.786683407 | 3.00E-04 |
| Osbpl9 | 6203.981448 | 0.786104514 | 4.81E-16 |
| Mto1 | 581.4842172 | 0.785618012 | 5.04E-08 |
| Rnf220 | 2987.612839 | 0.785498628 | 3.60E-14 |
| Rsph3b | 1349.967464 | 0.785444551 | 3.31E-06 |
| Kctd5 | 2795.822994 | 0.785376045 | 3.70E-13 |
| Cacnb1 | 471.9894397 | 0.78513557 | 1.43E-05 |
| Klhdc3 | 2766.400916 | 0.784766909 | 3.33E-11 |
| Kmt5a | 3867.372276 | 0.784705678 | 5.21E-09 |
| Ppan | 1552.316091 | 0.784568289 | 6.82E-09 |
| Cyfip1 | 6745.902934 | 0.7843808 | 2.14E-16 |
| Cyb5r3 | 22966.34465 | 0.784139402 | 3.44E-08 |
| Bri3bp | 3340.230155 | 0.784089217 | 3.74E-11 |
| Fh1 | 3616.006475 | 0.783742167 | 2.43E-09 |
| Yaf2 | 830.9713549 | 0.7837002 | 1.54E-06 |
| Exoc7 | 1976.272419 | 0.783419928 | 8.72E-09 |
| Lsm2 | 1237.134903 | 0.782612673 | 2.28E-04 |
| Sdad1 | 2442.35499 | 0.782508697 | 4.59E-15 |
| Fbxo24 | 56.11228112 | 0.782359484 | 8.68E-03 |
| Mettl15 | 267.6661028 | 0.782077156 | 2.65E-04 |
| Gata4 | 849.8210775 | 0.781979777 | 4.07E-05 |
| Ficd | 496.8706782 | 0.781387568 | 1.89E-04 |
| Nars | 9176.2195 | 0.781167503 | 4.51E-07 |
| Fkbp4 | 9091.266618 | 0.781131246 | 2.25E-07 |
| Lamb2 | 7100.083165 | 0.780965359 | 5.27E-04 |
| Gpn3 | 839.2302068 | 0.780781153 | 2.94E-09 |
| Sap30bp | 1774.467389 | 0.780766967 | 2.32E-08 |
| Nae1 | 2185.814268 | 0.780405596 | 3.89E-10 |
| Nit2 | 1118.356382 | 0.780241829 | 3.57E-08 |
| Tbc1d17 | 2049.150822 | 0.780141219 | 6.63E-04 |
| Lemd2 | 2831.27415 | 0.779920788 | 7.52E-08 |
| 1810010H24Rik | 33.02850824 | 0.779799868 | 1.62E-02 |
| Ano10 | 2755.935856 | 0.778767134 | 2.56E-10 |
| Alg14 | 549.7607113 | 0.778446309 | 2.79E-08 |
| Naca | 33543.9046 | 0.778213928 | 3.52E-07 |
| Tmsb10 | 24402.93356 | 0.778067603 | 1.12E-07 |

| | | | |
|---|---|---|---|
| Mrps27 | 872.321991 | 0.778021345 | 1.01E-06 |
| Ephb3 | 4231.935626 | 0.777677037 | 1.14E-07 |
| Diaph1 | 1722.652336 | 0.777432882 | 6.46E-08 |
| Ufl1 | 1661.102463 | 0.777239324 | 3.97E-07 |
| Nudt5 | 1618.254727 | 0.777223247 | 3.31E-06 |
| Yju2 | 424.7330676 | 0.777117245 | 4.36E-07 |
| Ran | 17075.7888 | 0.776863406 | 3.79E-03 |
| Arfgap1 | 3463.669508 | 0.77669236 | 2.37E-09 |
| Rpl31-ps8 | 39422.84122 | 0.776256408 | 2.27E-04 |
| Mir22hg | 1001.832139 | 0.775909463 | 1.33E-03 |
| Eaf1 | 2071.798067 | 0.775645471 | 4.17E-11 |
| Hdhd2 | 1257.142152 | 0.774242911 | 6.18E-12 |
| Hax1 | 2992.579162 | 0.773875315 | 3.94E-15 |
| Paqr3 | 356.3402682 | 0.772874465 | 3.62E-05 |
| Psmd7 | 6631.576451 | 0.772526816 | 2.90E-09 |
| Otud7b | 4504.227944 | 0.77222462 | 9.97E-20 |
| Cdk5rap3 | 2181.115954 | 0.77175072 | 1.85E-06 |
| Mier2 | 787.9011876 | 0.771529552 | 5.30E-08 |
| Eef1g | 80391.92701 | 0.770671831 | 1.41E-06 |
| Clip1 | 4554.958969 | 0.770103352 | 1.61E-07 |
| Ppp1ca | 14230.22317 | 0.769670466 | 1.56E-10 |
| Palm | 5324.7755 | 0.768982815 | 9.26E-08 |
| Anapc1 | 10971.14232 | 0.768522033 | 7.03E-17 |
| Klc2 | 1334.112269 | 0.767817134 | 8.17E-10 |
| Nipal1 | 860.9984183 | 0.766761669 | 1.26E-03 |
| Rhog | 1417.782828 | 0.766617464 | 8.20E-05 |
| Sp140 | 412.3072278 | 0.765989353 | 1.68E-03 |
| Gm9294 | 71.67654231 | 0.765822279 | 3.31E-02 |
| Itgb1bp1 | 1464.062863 | 0.76513443 | 3.23E-08 |
| Gm6030 | 218.7687813 | 0.76457881 | 4.26E-02 |
| Utrn | 8786.265038 | 0.764370393 | 3.00E-08 |
| Herc4 | 2473.94669 | 0.76423197 | 1.13E-10 |
| Cd274 | 79.84677291 | 0.763971431 | 6.32E-03 |
| Hnrnpc | 10522.29268 | 0.763934663 | 2.37E-10 |
| Tle1 | 1922.88961 | 0.761424998 | 1.71E-08 |
| Npdc1 | 6730.563996 | 0.76134333 | 2.55E-05 |
| Thop1 | 1815.914219 | 0.761036282 | 1.27E-08 |
| Nup37 | 843.5753032 | 0.760815095 | 9.82E-05 |
| Preb | 3063.586097 | 0.760778896 | 4.64E-19 |
| Naa16 | 558.3411683 | 0.760624676 | 1.30E-05 |
| Rnf166 | 1504.843752 | 0.759878227 | 1.46E-05 |
| Stoml2 | 1850.321277 | 0.759671948 | 3.63E-12 |
| Gm7102 | 42.86306938 | 0.758884852 | 1.17E-02 |
| Usp5 | 7780.615986 | 0.758789255 | 9.43E-10 |
| Swt1 | 741.4746105 | 0.758571339 | 9.50E-04 |
| Raly | 10314.59865 | 0.758376369 | 1.72E-11 |
| Nudcd2 | 1721.561492 | 0.75786963 | 6.14E-09 |

| | | | |
|---|---|---|---|
| Fkbp1a | 9585.237864 | 0.757456382 | 1.11E-08 |
| Rbm42 | 2022.691058 | 0.756938898 | 2.83E-06 |
| Rnf126 | 1969.762057 | 0.7566578 | 7.76E-07 |
| Ifnar2 | 1462.423568 | 0.75664116 | 3.00E-05 |
| Nfkbib | 688.5445045 | 0.756011544 | 4.11E-08 |
| Scamp5 | 1179.827721 | 0.755537547 | 6.86E-08 |
| Unc119 | 1277.409738 | 0.755432464 | 3.31E-04 |
| Sec22c | 967.296682 | 0.75519515 | 2.60E-05 |
| Stx4a | 2718.125959 | 0.754839827 | 4.23E-10 |
| Scyl1 | 5743.234042 | 0.754790232 | 1.31E-07 |
| Pfdn2 | 1804.340573 | 0.754374022 | 1.53E-11 |
| NA | 184.5233713 | 0.754272026 | 6.92E-03 |
| Selenoi | 2343.480976 | 0.754031519 | 2.03E-08 |
| Snrpa1 | 1239.871439 | 0.75393294 | 2.63E-06 |
| Gnb1l | 365.8711781 | 0.753718391 | 2.31E-03 |
| Fgfrl1 | 2587.189807 | 0.753093796 | 2.83E-05 |
| Etfb | 2261.36003 | 0.752574377 | 1.85E-08 |
| Rbm8a | 2712.709074 | 0.752517289 | 8.47E-11 |
| Jade3 | 903.244704 | 0.752334968 | 7.19E-10 |
| Wdr70 | 1348.139472 | 0.75229483 | 2.98E-11 |
| Usp53 | 923.2688407 | 0.752201199 | 3.67E-04 |
| Uqcc2 | 2774.225035 | 0.751831848 | 5.74E-05 |
| Polr1e | 447.9290646 | 0.751620837 | 2.65E-04 |
| Clta | 6971.978575 | 0.751595984 | 9.74E-05 |
| Cnnm4 | 356.2181148 | 0.751391244 | 6.95E-07 |
| Rhoa | 12392.3443 | 0.751365122 | 3.19E-11 |
| Racgap1 | 3856.467284 | 0.751349208 | 9.12E-03 |
| Tmem38a | 516.3255784 | 0.751302836 | 3.68E-08 |
| Rsph3a | 1682.445855 | 0.750159715 | 4.01E-06 |
| Gm15983 | 68.29334638 | 0.750153576 | 3.46E-03 |
| Kctd9 | 1161.401207 | 0.750132947 | 5.71E-03 |
| Psma2 | 4410.673088 | 0.749359386 | 2.08E-12 |
| Pcdh10 | 140.2969449 | 0.748981726 | 7.13E-03 |
| Urod | 2234.310979 | 0.748872046 | 1.34E-09 |
| Abtb1 | 677.458367 | 0.748454096 | 1.52E-05 |
| Polr1c | 2348.159737 | 0.74841904 | 6.09E-06 |
| Kdm3b | 3364.46103 | 0.748342942 | 3.17E-24 |
| Morn4 | 760.2486884 | 0.748325313 | 3.29E-16 |
| Acy1 | 719.2862101 | 0.748077855 | 1.63E-05 |
| Rps24 | 46619.81007 | 0.747643207 | 8.86E-06 |
| Chtf18 | 539.4430916 | 0.747577533 | 4.85E-02 |
| Arl8a | 4599.940929 | 0.747430054 | 6.37E-06 |
| Msrb1 | 847.4834858 | 0.747024877 | 9.25E-07 |
| Arf4 | 9849.937236 | 0.746920124 | 8.88E-07 |
| Eml5 | 284.730193 | 0.746711576 | 2.64E-02 |
| Polr3d | 1127.105134 | 0.746266195 | 8.60E-10 |
| Smad4 | 3283.698347 | 0.745991039 | 9.70E-07 |

| | | | |
|---|---|---|---|
| Lrrc51 | 118.5855804 | 0.745301129 | 3.16E-04 |
| Unc45a | 1699.276498 | 0.745210176 | 4.78E-08 |
| Mapk6 | 5890.268225 | 0.744945769 | 1.89E-08 |
| Rps3a1 | 109609.6564 | 0.744021272 | 1.95E-06 |
| Use1 | 4339.895075 | 0.743893913 | 1.99E-03 |
| Aldh3a2 | 3225.872424 | 0.743553485 | 2.96E-14 |
| Nit1 | 1001.7084 | 0.743537031 | 5.13E-08 |
| Ola1 | 4769.480168 | 0.743480808 | 5.75E-13 |
| U2af1 | 2772.347274 | 0.743307903 | 8.48E-11 |
| Csnk2a1 | 5927.167477 | 0.743125413 | 3.13E-14 |
| Icmt | 2198.9569 | 0.741997101 | 9.23E-07 |
| Rer1 | 2247.001878 | 0.74190792 | 8.09E-11 |
| Oard1 | 738.8061374 | 0.741679631 | 1.40E-09 |
| Otub1 | 4405.542155 | 0.741643759 | 4.53E-11 |
| Ppp2cb | 3059.960561 | 0.741067803 | 9.52E-12 |
| Dhx34 | 694.6250922 | 0.7405933 | 6.15E-08 |
| Nudt7 | 341.5085116 | 0.740331556 | 1.01E-06 |
| Rcl1 | 1451.162847 | 0.74030914 | 1.29E-07 |
| Scand1 | 571.2306615 | 0.74015409 | 4.06E-03 |
| Tead2 | 4318.690094 | 0.740060186 | 1.46E-04 |
| Efhc1 | 235.6947125 | 0.739774991 | 1.27E-03 |
| Spata5 | 1125.39688 | 0.739698174 | 9.97E-06 |
| Rps27rt | 30142.86293 | 0.738868257 | 1.17E-03 |
| Ddx24 | 5239.950081 | 0.738370918 | 4.49E-06 |
| Gm12112 | 87.35281576 | 0.738185759 | 4.35E-02 |
| Brpf1 | 2256.159578 | 0.738032805 | 4.54E-09 |
| Bcar1 | 3817.148731 | 0.737615651 | 1.17E-10 |
| Mettl23 | 306.5676268 | 0.737457458 | 7.79E-04 |
| Tcea3 | 238.4744052 | 0.737424399 | 3.54E-02 |
| Larp7 | 1751.920278 | 0.737362515 | 2.32E-10 |
| Pex10 | 185.311056 | 0.736302964 | 1.60E-03 |
| Myadm | 15438.02828 | 0.736170212 | 6.20E-16 |
| Coro1b | 12286.48232 | 0.735559151 | 5.26E-08 |
| Atp5h | 4474.721524 | 0.734903749 | 1.55E-10 |
| Sec62 | 6353.816318 | 0.734737284 | 3.19E-06 |
| Sh3bp2 | 1443.810206 | 0.734555587 | 2.57E-04 |
| Tns3 | 8946.526377 | 0.73436334 | 4.01E-07 |
| Stk10 | 1458.098386 | 0.733605393 | 6.13E-08 |
| Rpl27a | 3086.246768 | 0.733550181 | 3.52E-06 |
| Radil | 568.4919136 | 0.733036537 | 2.98E-05 |
| Zc3h10 | 826.6331516 | 0.732649554 | 1.04E-06 |
| Arf1 | 15591.73516 | 0.732418956 | 1.26E-07 |
| Pold1 | 894.4656266 | 0.731955738 | 7.00E-03 |
| Nudcd1 | 1384.714775 | 0.731607658 | 3.36E-06 |
| Patj | 672.2244754 | 0.731544107 | 4.79E-05 |
| Hdac6 | 1485.523679 | 0.731157802 | 5.68E-09 |
| Tbc1d15 | 3591.297068 | 0.731077622 | 1.51E-08 |

| | | | |
|---|---|---|---|
| Lrig3 | 1775.697697 | 0.730683794 | 3.55E-05 |
| Ppie | 1371.92535 | 0.72961298 | 3.26E-09 |
| Hoxa1 | 320.536735 | 0.729439744 | 3.89E-03 |
| Snf8 | 1664.037907 | 0.728506757 | 3.39E-12 |
| Vrk3 | 1213.537914 | 0.728453524 | 6.28E-13 |
| Smad1 | 1578.21778 | 0.728050116 | 1.55E-06 |
| B4galt3 | 1066.306636 | 0.728019266 | 2.51E-10 |
| Wdr45b | 3049.093296 | 0.726102612 | 2.81E-11 |
| Mfap1b | 2130.549542 | 0.726067242 | 4.04E-08 |
| Nop9 | 1165.628715 | 0.725483143 | 1.95E-07 |
| Aasdhppt | 981.0537025 | 0.724954988 | 6.50E-12 |
| Tiprl | 2259.29702 | 0.724810605 | 3.01E-12 |
| Zc3h14 | 5985.988222 | 0.724075991 | 4.24E-13 |
| Gfod1 | 804.2426049 | 0.723811545 | 1.11E-02 |
| Pbx2 | 2268.811586 | 0.723280363 | 5.96E-09 |
| Arnt | 5519.889564 | 0.722830866 | 3.18E-06 |
| Ogfod2 | 485.1672615 | 0.722462064 | 6.17E-06 |
| Mnat1 | 1466.460687 | 0.722402692 | 7.15E-06 |
| Ube2m | 2947.986069 | 0.721723531 | 3.64E-08 |
| Kifc2 | 266.15885 | 0.721557041 | 7.03E-03 |
| Dnajc27 | 615.2318411 | 0.721235273 | 5.24E-07 |
| Ubxn2a | 924.4544045 | 0.720952435 | 3.41E-04 |
| Phb | 4971.402228 | 0.720824261 | 3.15E-05 |
| Prpf8 | 19463.90492 | 0.720397534 | 4.32E-12 |
| Grasp | 391.1139554 | 0.720175717 | 2.92E-07 |
| Rnf141 | 1366.065122 | 0.719722004 | 3.60E-06 |
| Mrpl27 | 1066.397577 | 0.719489116 | 3.12E-10 |
| Snip1 | 1187.676737 | 0.719351237 | 4.90E-08 |
| Fchsd2 | 1458.876734 | 0.719224096 | 2.14E-04 |
| Tmem167b | 3267.309721 | 0.718967438 | 2.76E-05 |
| Car7 | 47.50530047 | 0.718623388 | 2.41E-02 |
| Hmgxb3 | 1878.722133 | 0.718593512 | 1.78E-07 |
| Hoxa4 | 123.1936353 | 0.718572535 | 6.00E-03 |
| Ncapd2 | 2905.234723 | 0.718482508 | 2.42E-02 |
| Ahsa1 | 4009.452714 | 0.718437252 | 1.53E-06 |
| Pdhb | 2648.913108 | 0.718366781 | 4.97E-12 |
| Efl1 | 1294.99223 | 0.718351076 | 1.33E-09 |
| Dnajc9 | 984.8756573 | 0.718137277 | 2.89E-03 |
| Ccdc59 | 1338.787267 | 0.718032465 | 9.55E-15 |
| Usmg5 | 2737.433256 | 0.717451261 | 1.60E-13 |
| Ehmt2 | 10097.78051 | 0.717345925 | 2.68E-09 |
| Pacsin3 | 1779.598782 | 0.717188849 | 2.56E-06 |
| Ets2 | 2152.537196 | 0.716579234 | 6.44E-06 |
| Ncaph2 | 5516.037748 | 0.716543228 | 2.62E-05 |
| Slc38a9 | 414.9045467 | 0.716498299 | 1.56E-05 |
| Dpf2 | 3923.177458 | 0.716295054 | 6.67E-09 |
| Msi2 | 1364.372491 | 0.71591685 | 5.51E-07 |

| | | | |
|---|---|---|---|
| Bard1 | 291.9104034 | 0.715908668 | 3.69E-02 |
| Cage1 | 131.6143912 | 0.715435439 | 1.09E-05 |
| Upb1 | 36.43430221 | 0.715406573 | 3.79E-02 |
| Rdh11 | 2494.14244 | 0.71450851 | 4.46E-06 |
| Acot8 | 248.0830666 | 0.714204643 | 6.62E-05 |
| Anp32b | 8481.918321 | 0.714131006 | 7.82E-09 |
| Araf | 2783.283806 | 0.713926172 | 1.06E-15 |
| A430078G23Rik | 32.30141715 | 0.713877118 | 3.90E-02 |
| Trp53 | 5498.367133 | 0.71341868 | 4.32E-05 |
| Samm50 | 3134.279396 | 0.712827427 | 8.05E-18 |
| Yrdc | 1246.308307 | 0.712088591 | 2.42E-09 |
| Sfr1 | 9916.619043 | 0.712071884 | 1.51E-10 |
| Atg16l1 | 1600.559711 | 0.711965785 | 2.96E-08 |
| Sat1 | 5732.563773 | 0.711790652 | 9.21E-04 |
| Bag6 | 9394.64185 | 0.710906229 | 6.06E-09 |
| Cbfa2t2 | 2370.940178 | 0.710789379 | 3.87E-08 |
| Clip3 | 4191.373142 | 0.710456684 | 1.79E-03 |
| Vapa | 4514.815785 | 0.710450743 | 2.91E-08 |
| Lsm5 | 534.3659574 | 0.709893684 | 2.44E-06 |
| Stau1 | 4739.563788 | 0.709729513 | 3.08E-08 |
| Orc3 | 1568.554497 | 0.709627871 | 2.81E-09 |
| Raph1 | 579.1077271 | 0.708822106 | 3.03E-03 |
| Supt5 | 6937.965658 | 0.708733345 | 6.95E-07 |
| Heatr1 | 2150.07386 | 0.70844316 | 5.77E-06 |
| Rps29 | 19692.01595 | 0.708257685 | 4.13E-06 |
| Tmem198 | 59.85625987 | 0.708219689 | 1.35E-02 |
| Rnf123 | 1593.468219 | 0.708032417 | 2.36E-08 |
| Gm10288 | 2402.863389 | 0.707686078 | 4.77E-02 |
| Eloc | 1343.046711 | 0.707663336 | 2.99E-05 |
| Mrps18a | 1418.794395 | 0.707516524 | 6.59E-06 |
| Atp5f1 | 8259.418685 | 0.707484694 | 1.81E-15 |
| Ndufs3 | 2510.570969 | 0.707287905 | 1.25E-09 |
| Mrpl37 | 2575.873984 | 0.706192572 | 7.05E-10 |
| Gm7008 | 42.62583199 | 0.705597133 | 2.11E-02 |
| Copb2 | 10535.47111 | 0.705328007 | 4.89E-06 |
| Wdr18 | 2059.905686 | 0.704160892 | 1.20E-05 |
| Wdr35 | 1847.48163 | 0.704150725 | 7.70E-07 |
| Pitpnc1 | 829.297569 | 0.704064514 | 5.66E-03 |
| R3hdm4 | 3380.236701 | 0.703918589 | 4.06E-06 |
| Ptbp1 | 15707.70459 | 0.703885838 | 2.34E-07 |
| Xpnpep1 | 4934.26555 | 0.703575291 | 2.26E-10 |
| Snrnp25 | 373.4357615 | 0.70325033 | 2.55E-03 |
| Tef | 3446.398834 | 0.702822701 | 3.06E-04 |
| Hyal2 | 1065.562101 | 0.702658189 | 8.31E-05 |
| Irf2bpl | 2111.470138 | 0.702319971 | 3.46E-07 |
| Il1rl1 | 26338.39734 | 0.702170958 | 2.20E-02 |
| Rita1 | 573.5582389 | 0.701540451 | 2.03E-06 |

| | | | |
|---|---|---|---|
| Fbxo7 | 1341.227542 | 0.701498945 | 1.65E-04 |
| Nampt | 3299.598355 | 0.700959547 | 7.37E-07 |
| Plxnb2 | 19455.25205 | 0.700843219 | 1.04E-08 |
| Ginm1 | 1392.945406 | 0.700781249 | 9.39E-07 |
| Syvn1 | 2152.597269 | 0.700753346 | 8.84E-08 |
| Rpap2 | 1188.573752 | 0.700748326 | 3.57E-08 |
| NA | 859.2090966 | 0.700649384 | 2.38E-04 |
| Mark2 | 2420.298645 | 0.700388851 | 2.66E-15 |
| Cebpzos | 554.3928388 | 0.700308516 | 3.73E-14 |
| Trim27 | 2284.639948 | 0.700208397 | 6.27E-07 |
| Tesk2 | 669.5921419 | 0.700178767 | 6.89E-05 |
| Gm9531 | 2715.543962 | 0.700132867 | 1.86E-06 |
| NA | 1983.288498 | 0.700028516 | 2.45E-06 |
| Gm13841 | 29018.88927 | 0.69990813 | 3.97E-04 |
| Psmd14 | 2462.854411 | 0.699383636 | 1.69E-05 |
| NA | 50.27877924 | 0.699203804 | 2.14E-02 |
| Arih1 | 4317.986198 | 0.698800337 | 2.39E-13 |
| Tram2 | 3451.186882 | 0.698751982 | 5.56E-12 |
| Golga2 | 4785.074064 | 0.698690394 | 2.48E-06 |
| Bsdc1 | 2498.400934 | 0.698032895 | 2.38E-04 |
| Pygo2 | 2490.294348 | 0.697593531 | 2.59E-11 |
| Gmpr | 213.3268615 | 0.697329732 | 1.14E-03 |
| Ppp2r2d | 1434.57114 | 0.696505098 | 3.10E-05 |
| Itm2c | 7431.97345 | 0.696453515 | 6.38E-04 |
| Baiap2l1 | 282.5346504 | 0.696439463 | 1.50E-02 |
| Insig2 | 2774.628956 | 0.696386767 | 6.25E-07 |
| Gnl1 | 3426.643249 | 0.696113529 | 1.46E-05 |
| Coa7 | 609.5024897 | 0.696018567 | 7.95E-12 |
| Pitrm1 | 6900.706977 | 0.695954362 | 1.06E-11 |
| Cript | 1250.406291 | 0.695902352 | 1.08E-10 |
| Bap1 | 2457.617171 | 0.695138806 | 3.08E-09 |
| Creb3l4 | 51.97945249 | 0.694720676 | 3.06E-02 |
| Tmem141 | 232.8755583 | 0.694178776 | 1.21E-05 |
| Secisbp2 | 567.6859357 | 0.692911189 | 1.76E-07 |
| Tbc1d23 | 1888.512151 | 0.69267782 | 1.10E-08 |
| Col12a1 | 2667.384937 | 0.692541284 | 7.40E-07 |
| Ttf2 | 1497.723252 | 0.692258733 | 9.37E-05 |
| Rad23a | 3073.219033 | 0.691748768 | 1.48E-07 |
| Repin1 | 1500.785639 | 0.691665535 | 8.47E-06 |
| Clybl | 692.1619379 | 0.691250818 | 7.95E-05 |
| Rad18 | 808.0924725 | 0.690686429 | 8.93E-03 |
| Tmem183a | 2815.550326 | 0.690547278 | 2.03E-13 |
| Tead3 | 2341.916993 | 0.690308276 | 5.12E-06 |
| Sp4 | 343.7880857 | 0.690174772 | 3.00E-04 |
| Fbxl5 | 2118.341111 | 0.689675996 | 4.78E-14 |
| Eif4h | 13353.88039 | 0.689594373 | 1.07E-08 |
| Glis2 | 2505.628587 | 0.68891467 | 7.75E-07 |

| | | | |
|---|---|---|---|
| Ttc37 | 2099.340995 | 0.688844121 | 2.22E-04 |
| Rtca | 1608.30568 | 0.68852198 | 1.13E-11 |
| Fn3krp | 425.7525463 | 0.68837227 | 1.95E-06 |
| Rbm38 | 607.9930154 | 0.68793361 | 1.19E-02 |
| Parva | 10879.78875 | 0.687702389 | 1.27E-10 |
| Ubap1 | 2119.323972 | 0.687498888 | 2.45E-03 |
| Rpf1 | 1264.591 | 0.687378022 | 1.41E-06 |
| A430005L14Rik | 411.6249794 | 0.686759067 | 1.09E-06 |
| Grina | 10349.31318 | 0.686647211 | 7.24E-04 |
| Washc1 | 1985.195212 | 0.68631613 | 2.02E-05 |
| Frg1 | 2103.851192 | 0.686048227 | 2.63E-08 |
| Mtmr10 | 1541.372356 | 0.685906718 | 1.61E-04 |
| Mien1 | 1410.361947 | 0.685817051 | 1.93E-05 |
| Socs7 | 1466.776444 | 0.685635057 | 1.28E-11 |
| Cul3 | 5673.330745 | 0.685554506 | 2.27E-09 |
| Ccar2 | 1822.781877 | 0.685527222 | 1.94E-08 |
| Mad2l1 | 1774.898828 | 0.685328437 | 2.94E-02 |
| Vars | 7296.234314 | 0.683894242 | 1.23E-04 |
| Pop5 | 528.7413162 | 0.683803112 | 8.97E-05 |
| Zfp346 | 2029.029437 | 0.683672133 | 9.57E-04 |
| Dok1 | 2983.994436 | 0.683387216 | 1.92E-04 |
| Cd99l2 | 3754.262879 | 0.68327912 | 7.97E-09 |
| Gnl3l | 4240.911019 | 0.682985495 | 4.12E-18 |
| Shmt1 | 1266.240147 | 0.682945349 | 9.63E-05 |
| Zfhx3 | 1535.860148 | 0.682413792 | 1.32E-08 |
| Sec24a | 2291.392764 | 0.682068744 | 1.99E-04 |
| Zc3h3 | 967.5588601 | 0.681646194 | 7.32E-04 |
| Fbl | 2993.699857 | 0.681225298 | 1.35E-07 |
| Rfc2 | 1332.762709 | 0.6812002 | 9.38E-04 |
| U2af2 | 7519.96701 | 0.68117059 | 9.51E-07 |
| Dvl3 | 1937.725035 | 0.680620964 | 4.79E-05 |
| Zfp574 | 814.8553804 | 0.680593672 | 9.04E-04 |
| Eif4a2 | 15379.29281 | 0.680410004 | 2.83E-05 |
| Cenpu | 251.9751922 | 0.680094462 | 4.87E-02 |
| Ero1lb | 197.7904632 | 0.679880201 | 4.71E-04 |
| Mcrs1 | 2054.270711 | 0.679541956 | 5.47E-07 |
| Cryz | 2067.245893 | 0.679226204 | 1.94E-03 |
| Kptn | 495.5398175 | 0.679030484 | 3.49E-06 |
| Suds3 | 3742.606582 | 0.678895388 | 3.72E-08 |
| Psmd2 | 21874.64041 | 0.678666722 | 9.64E-08 |
| Ptges3 | 6256.261295 | 0.678536895 | 1.21E-06 |
| Hinfp | 406.8236308 | 0.678303919 | 1.94E-06 |
| Fchsd1 | 780.790929 | 0.677551812 | 1.41E-03 |
| Pex13 | 1616.351738 | 0.67748369 | 7.64E-09 |
| 1190002N15Rik | 866.5027261 | 0.677315736 | 1.68E-03 |
| Mrpl10 | 2215.87896 | 0.67693345 | 1.91E-12 |
| Tln1 | 13133.47837 | 0.676895748 | 3.48E-10 |

| | | | |
|---|---|---|---|
| Chmp5 | 2578.82645 | 0.676241728 | 1.32E-10 |
| Itpr2 | 1991.961663 | 0.676189355 | 2.37E-06 |
| Kmt5c | 716.1049833 | 0.67618059 | 2.44E-04 |
| Cpt2 | 2107.034094 | 0.675738664 | 2.78E-10 |
| Ywhah | 6364.92468 | 0.675655339 | 8.56E-07 |
| Cbx4 | 917.6543838 | 0.675512389 | 5.18E-04 |
| Dis3l | 1248.96123 | 0.675406081 | 1.79E-06 |
| Brat1 | 1070.094288 | 0.675247102 | 2.10E-07 |
| Gm13166 | 77.61614625 | 0.675228146 | 6.43E-03 |
| Pml | 1978.102798 | 0.674676189 | 2.73E-05 |
| Srr | 2490.166765 | 0.674254325 | 3.99E-06 |
| Gyg | 5935.453933 | 0.673771223 | 3.12E-03 |
| Bcdin3d | 199.363341 | 0.673501747 | 4.09E-04 |
| Ngrn | 916.2905254 | 0.672550024 | 6.09E-12 |
| Lsm14a | 3705.854334 | 0.672307789 | 1.41E-07 |
| Acbd4 | 1094.853639 | 0.672282687 | 4.83E-10 |
| 4930451G09Rik | 46.88870039 | 0.671810454 | 2.00E-02 |
| Pomgnt1 | 3559.582041 | 0.671574429 | 1.62E-10 |
| Cox7a2 | 2411.57902 | 0.671350484 | 1.38E-09 |
| Kdm6b | 4030.264971 | 0.671284968 | 1.84E-03 |
| Upf3a | 1578.50849 | 0.671152726 | 4.16E-13 |
| Wrnip1 | 1355.163956 | 0.670942569 | 6.46E-12 |
| Ubl7 | 1626.244088 | 0.670462291 | 2.03E-04 |
| Slc46a1 | 317.5478592 | 0.670418093 | 2.36E-04 |
| Abt1 | 516.2441004 | 0.669873452 | 1.60E-06 |
| Snapc2 | 1266.040637 | 0.669133271 | 4.34E-04 |
| Agl | 3359.570353 | 0.669042139 | 1.34E-20 |
| Ripor1 | 4272.017785 | 0.668836149 | 7.49E-09 |
| Pi4k2b | 1510.332724 | 0.668811851 | 3.04E-07 |
| Dus2 | 436.7762165 | 0.668720846 | 2.86E-05 |
| Pycrl | 980.0945065 | 0.6686528 | 1.03E-04 |
| Dgcr6 | 1533.711583 | 0.66859815 | 4.80E-06 |
| Txn2 | 4732.276052 | 0.668318064 | 1.01E-06 |
| Rpl36a | 23382.0782 | 0.668093851 | 1.77E-03 |
| Rbms3 | 1823.229044 | 0.668049936 | 1.57E-05 |
| 1700030K09Rik | 586.5005332 | 0.667990843 | 3.37E-05 |
| Slu7 | 2633.493033 | 0.667216928 | 2.39E-05 |
| Elp4 | 759.6580291 | 0.667077304 | 1.43E-06 |
| Tspo | 2413.333548 | 0.667062733 | 1.32E-08 |
| Rwdd2a | 82.87345654 | 0.666852667 | 1.69E-02 |
| Sec31a | 15080.55081 | 0.666755515 | 1.29E-05 |
| Gipc1 | 3013.934014 | 0.66665486 | 1.22E-06 |
| Col8a1 | 3683.903915 | 0.666554487 | 1.86E-02 |
| Elp5 | 2617.836782 | 0.666307692 | 2.00E-11 |
| Axin1 | 4077.021414 | 0.6662923 | 4.89E-12 |
| Capzb | 8468.560577 | 0.666165086 | 9.05E-06 |
| Isl2 | 63.99342943 | 0.666113475 | 2.85E-02 |

| | | | |
|---|---|---|---|
| Slc7a7 | 105.7139369 | 0.665862205 | 1.17E-02 |
| Klf6 | 6415.94845 | 0.665785656 | 8.25E-12 |
| Tubb6 | 4737.383742 | 0.665615828 | 6.94E-03 |
| Tsta3 | 1507.503282 | 0.665562957 | 2.24E-09 |
| Bop1 | 2336.321526 | 0.665493933 | 3.31E-04 |
| Ncaph | 1055.329841 | 0.665377023 | 2.32E-02 |
| Fads1 | 13261.24545 | 0.665347812 | 3.36E-08 |
| Gnas | 44076.85637 | 0.665027228 | 2.01E-06 |
| Fbxw4 | 1212.574999 | 0.664772205 | 2.51E-10 |
| Efemp1 | 43132.42209 | 0.664087925 | 4.09E-05 |
| Gm20498 | 557.4901711 | 0.663684507 | 2.89E-06 |
| Rapgef1 | 5967.043459 | 0.663361428 | 3.01E-08 |
| Ilkap | 2207.153689 | 0.66331367 | 3.70E-13 |
| Rnaseh2a | 1209.479254 | 0.662879272 | 8.33E-04 |
| Pecr | 434.5207829 | 0.662041299 | 5.74E-10 |
| Dynlrb1 | 4195.992513 | 0.661935224 | 6.38E-05 |
| Usp19 | 5211.507926 | 0.661653198 | 6.20E-07 |
| Tmem143 | 540.7880561 | 0.661413331 | 1.25E-03 |
| Npc1 | 3965.691488 | 0.661254215 | 1.58E-04 |
| Rabac1 | 2922.165955 | 0.661175608 | 9.28E-04 |
| Ubqln1 | 4261.004222 | 0.661031423 | 7.76E-09 |
| Nicn1 | 1087.416872 | 0.660512887 | 1.04E-03 |
| Wipi2 | 4767.419219 | 0.659992356 | 2.47E-09 |
| Magohb | 625.3616612 | 0.659588667 | 6.39E-03 |
| Ppp2r1a | 11659.34767 | 0.659344588 | 6.27E-06 |
| Crem | 172.0738427 | 0.659161397 | 2.23E-05 |
| Gm19680 | 282.3254426 | 0.658469184 | 2.99E-04 |
| Nip7 | 2066.690499 | 0.6582887 | 1.18E-05 |
| Kpna1 | 2050.904323 | 0.657515242 | 1.33E-02 |
| Spast | 2112.317484 | 0.657372166 | 1.30E-06 |
| Vps16 | 1845.3321 | 0.657316303 | 4.63E-10 |
| Capn2 | 13394.05317 | 0.657274025 | 1.48E-06 |
| Atg2b | 4251.553425 | 0.657095664 | 6.97E-07 |
| Ndufa8 | 2398.782292 | 0.656870351 | 8.27E-07 |
| Fosl2 | 7134.778544 | 0.656818133 | 1.08E-04 |
| Mob2 | 788.0670922 | 0.656449841 | 4.95E-05 |
| Exosc1 | 1110.733076 | 0.656306879 | 5.30E-07 |
| Uso1 | 6535.449643 | 0.655956837 | 1.83E-06 |
| Polh | 607.3972268 | 0.655893648 | 1.43E-03 |
| Kpnb1 | 26279.66806 | 0.655691246 | 4.44E-07 |
| Kmt2d | 3395.264604 | 0.655633955 | 1.39E-03 |
| Hdac7 | 6665.610222 | 0.655253984 | 9.30E-16 |
| Naga | 3447.187179 | 0.655227076 | 2.43E-04 |
| Ankrd13d | 440.9352955 | 0.655169185 | 1.65E-03 |
| Hmces | 1311.48031 | 0.65435603 | 9.04E-11 |
| Snrpb | 5637.203642 | 0.654138328 | 1.06E-05 |
| Xpo4 | 1005.969614 | 0.654123966 | 6.93E-06 |

| | | | |
|---|---|---|---|
| Nol11 | 2131.47981 | 0.653815997 | 4.68E-13 |
| Rnpc3 | 644.1823741 | 0.653653065 | 7.48E-03 |
| Cib1 | 847.3343329 | 0.65342882 | 2.39E-07 |
| Rpl30 | 7654.741905 | 0.653035451 | 1.01E-04 |
| Rnf214 | 1252.647553 | 0.652858848 | 1.23E-11 |
| Hspa4 | 15381.02481 | 0.652828222 | 1.89E-05 |
| Fkbp2 | 1281.795309 | 0.65262369 | 1.02E-06 |
| Ddah2 | 4558.89888 | 0.651445196 | 6.58E-04 |
| Chchd3 | 4298.451419 | 0.651437464 | 1.77E-06 |
| Strn4 | 2571.884023 | 0.651405127 | 2.57E-09 |
| Sumo2 | 2921.158398 | 0.650996647 | 5.20E-05 |
| Cdk11b | 2039.570556 | 0.650539951 | 1.10E-11 |
| Clpp | 2273.411807 | 0.650288859 | 3.52E-05 |
| Creb3 | 1212.471112 | 0.650287136 | 1.31E-05 |
| Ap1b1 | 3907.493539 | 0.649911788 | 4.14E-07 |
| Tbc1d25 | 541.5737425 | 0.649658064 | 1.81E-06 |
| Dohh | 1632.800307 | 0.649476948 | 6.93E-07 |
| Optn | 361.5681594 | 0.649137943 | 1.83E-04 |
| Wdr46 | 1834.741141 | 0.64908338 | 3.35E-05 |
| Gart | 2651.950456 | 0.649083277 | 1.84E-03 |
| Zfp526 | 555.9011632 | 0.648898433 | 2.19E-03 |
| Cnot8 | 2936.25042 | 0.648759226 | 2.49E-08 |
| Pgam1 | 19698.03657 | 0.648503107 | 9.30E-04 |
| Ctsd | 41499.83588 | 0.648379917 | 2.63E-03 |
| Dcbld2 | 3824.810857 | 0.648038876 | 2.59E-04 |
| Comt | 3881.085594 | 0.647800512 | 6.06E-10 |
| Gmps | 4740.881068 | 0.647098438 | 8.67E-10 |
| Dis3l2 | 992.1240902 | 0.646961697 | 1.57E-07 |
| Tube1 | 230.4590009 | 0.646948672 | 1.38E-02 |
| Phf23 | 3785.987225 | 0.646736284 | 5.39E-06 |
| Pnpo | 218.6551061 | 0.646372867 | 1.30E-03 |
| Naa35 | 1827.19018 | 0.646293362 | 2.60E-10 |
| Map3k11 | 1003.039186 | 0.64622625 | 7.94E-09 |
| Btbd1 | 3241.787789 | 0.646054483 | 4.08E-06 |
| Gmeb1 | 1118.296004 | 0.645825515 | 2.10E-08 |
| Mff | 3918.125802 | 0.64579953 | 6.47E-12 |
| Gm10704 | 127.3962194 | 0.645692751 | 1.57E-02 |
| Ruvbl1 | 2816.222171 | 0.645632365 | 1.22E-05 |
| Sirt6 | 1008.851613 | 0.64558227 | 2.03E-07 |
| Irak1bp1 | 511.6876268 | 0.64541518 | 3.29E-06 |
| Sec24c | 4984.872007 | 0.64508125 | 3.28E-07 |
| Zfp219 | 1438.940271 | 0.64501826 | 3.70E-09 |
| Hilpda | 740.6665374 | 0.644980884 | 1.42E-07 |
| Map2k7 | 2944.395202 | 0.64484446 | 5.30E-06 |
| Arrdc1 | 446.1481351 | 0.644723883 | 2.98E-06 |
| Inpp5f | 1261.195768 | 0.644477238 | 2.53E-08 |
| Spsb2 | 835.3781798 | 0.64446567 | 8.77E-08 |

| | | | |
|---|---|---|---|
| Krtcap2 | 3311.574394 | 0.643761308 | 7.75E-06 |
| St6galnac4 | 1396.81264 | 0.643708989 | 2.34E-04 |
| Flot2 | 3914.985823 | 0.643700813 | 1.06E-09 |
| Pdk3 | 1684.471163 | 0.643698928 | 2.13E-07 |
| Tmed3 | 3867.412961 | 0.643342131 | 2.30E-03 |
| Usp46 | 1648.110825 | 0.642473804 | 4.49E-07 |
| Slc41a1 | 1950.284416 | 0.642026123 | 1.22E-05 |
| Slc39a14 | 1634.769816 | 0.641868305 | 4.48E-06 |
| Dirc2 | 1518.295363 | 0.641623025 | 9.72E-06 |
| Dbf4 | 1797.842921 | 0.641420813 | 3.49E-02 |
| Dnajc17 | 243.5826044 | 0.641242018 | 4.30E-05 |
| Naa80 | 610.3779038 | 0.640987641 | 8.01E-04 |
| Yipf6 | 840.4900824 | 0.640939967 | 1.00E-07 |
| Ptpmt1 | 905.1759236 | 0.640837805 | 4.09E-06 |
| Cyp20a1 | 1599.590162 | 0.640356177 | 4.70E-08 |
| Pelo | 681.6239708 | 0.640103685 | 9.87E-06 |
| Nme4 | 944.0496733 | 0.640083022 | 6.86E-04 |
| Atp6v1f | 4383.57506 | 0.639847675 | 4.97E-04 |
| Hsd17b10 | 3298.465071 | 0.639247156 | 2.79E-05 |
| Tmem230 | 1598.27255 | 0.639229302 | 1.15E-06 |
| Setd1a | 2741.234231 | 0.639093 | 9.39E-06 |
| Gtpbp3 | 1278.258545 | 0.638872702 | 4.81E-05 |
| Lsm4 | 1011.076088 | 0.638262086 | 3.60E-06 |
| Abhd1 | 424.5594923 | 0.638223813 | 2.51E-06 |
| Ess2 | 1232.75127 | 0.638197495 | 1.89E-04 |
| Mpst | 1039.896141 | 0.637759945 | 2.42E-03 |
| Ulk3 | 367.2827981 | 0.637214334 | 1.20E-03 |
| Gnpda2 | 1029.052467 | 0.636910215 | 7.90E-05 |
| Exoc3 | 2109.633632 | 0.636689147 | 1.27E-06 |
| Rnf40 | 2469.510568 | 0.636053884 | 7.34E-09 |
| Smurf1 | 4586.089216 | 0.635201742 | 6.45E-06 |
| Yipf1 | 1602.069816 | 0.6350498 | 3.19E-05 |
| Acadl | 4515.648214 | 0.634838316 | 2.80E-11 |
| Wbp1 | 2160.848682 | 0.634405327 | 2.24E-03 |
| Nup155 | 2585.298343 | 0.634222977 | 1.06E-04 |
| Pld3 | 2140.500516 | 0.633668592 | 5.08E-04 |
| Ube2v1 | 6995.372513 | 0.633376671 | 2.03E-05 |
| Mcm7 | 4085.380433 | 0.63254477 | 7.60E-03 |
| Dhdds | 1430.297237 | 0.631872842 | 1.98E-04 |
| Lrrfip2 | 1836.448746 | 0.631643637 | 6.60E-08 |
| Psma5 | 4475.505167 | 0.631494203 | 8.37E-10 |
| Ccdc86 | 1531.939262 | 0.631316431 | 1.41E-05 |
| Mis18a | 305.8991117 | 0.631272983 | 3.73E-03 |
| Smg5 | 4366.585321 | 0.630746764 | 2.80E-04 |
| Cul4a | 6931.867817 | 0.630606706 | 4.35E-32 |
| Lyplal1 | 1068.156704 | 0.630234992 | 9.79E-06 |
| Ltbp2 | 1061.332223 | 0.629149333 | 3.87E-02 |

| | | | |
|---|---|---|---|
| Mkrn1 | 1831.344411 | 0.628539403 | 2.43E-04 |
| Ampd2 | 2558.553788 | 0.627387144 | 5.47E-06 |
| Mtpap | 1104.094798 | 0.627322174 | 5.81E-09 |
| Poc5 | 569.4921546 | 0.627306216 | 3.04E-05 |
| Bcl6 | 1044.315841 | 0.627298787 | 1.70E-03 |
| Slc25a37 | 3381.662168 | 0.6271664 | 5.84E-04 |
| Ehd4 | 2099.212753 | 0.626371691 | 6.36E-05 |
| Tbc1d13 | 2285.124383 | 0.626282665 | 8.30E-06 |
| Tkfc | 1275.455666 | 0.626128507 | 3.44E-05 |
| Plgrkt | 1248.255989 | 0.626097018 | 5.58E-14 |
| Aph1a | 5523.669385 | 0.625784301 | 2.78E-06 |
| Rp9 | 470.7603178 | 0.625311178 | 3.51E-03 |
| Kat7 | 3759.136679 | 0.624840281 | 1.25E-06 |
| Wdr62 | 764.1814903 | 0.624830294 | 2.01E-02 |
| Irx1 | 1133.860938 | 0.624499326 | 4.04E-03 |
| Syngr2 | 2533.946804 | 0.62448902 | 1.96E-07 |
| Emc7 | 2809.673745 | 0.624252916 | 7.85E-06 |
| Fads2 | 5458.473187 | 0.624162354 | 9.34E-07 |
| Cry2 | 1311.129949 | 0.623100234 | 9.24E-07 |
| Tmem147 | 1932.796916 | 0.622712102 | 3.24E-08 |
| Park7 | 2984.256339 | 0.622458347 | 1.35E-06 |
| Cebpb | 1041.170112 | 0.622383628 | 1.27E-02 |
| Yars | 7597.314974 | 0.621720726 | 8.48E-07 |
| Sh3gl1 | 4195.01312 | 0.621637602 | 1.02E-07 |
| Dhx30 | 2802.794432 | 0.621409468 | 1.62E-05 |
| Sfmbt2 | 667.3441797 | 0.621187832 | 3.75E-03 |
| Etfdh | 2166.481833 | 0.620601861 | 3.34E-21 |
| Ndfip1 | 2962.313867 | 0.6198287 | 5.14E-06 |
| Il6st | 23732.48129 | 0.619823193 | 5.44E-04 |
| Echs1 | 2288.2654 | 0.619708536 | 1.30E-06 |
| Dcps | 1314.550216 | 0.619650614 | 3.13E-06 |
| Mad2l1bp | 381.6900226 | 0.61964968 | 1.76E-04 |
| Nsmce2 | 1712.981215 | 0.618972364 | 1.29E-07 |
| Eif3b | 10504.03029 | 0.618940843 | 1.46E-04 |
| Mrps12 | 608.7077497 | 0.617506983 | 2.81E-06 |
| Ubl5 | 2285.251775 | 0.617209742 | 7.84E-05 |
| Ccdc61 | 496.8746763 | 0.617140401 | 6.19E-03 |
| Vps45 | 804.2893788 | 0.616940219 | 5.35E-05 |
| Herc1 | 3285.880765 | 0.616798294 | 2.14E-07 |
| Mrpl28 | 2037.882619 | 0.616540709 | 3.10E-04 |
| Cldn12 | 2413.444114 | 0.616430939 | 6.58E-04 |
| Prpsap1 | 2055.416811 | 0.616416696 | 3.05E-10 |
| Nelfe | 1986.089958 | 0.61600898 | 5.74E-07 |
| Dynlt1b | 1798.335892 | 0.615381311 | 8.77E-08 |
| Ralgapa2 | 995.1711884 | 0.615305567 | 4.79E-04 |
| Brox | 3007.5817 | 0.614833142 | 2.43E-17 |
| Ndufb11 | 3018.005221 | 0.614234076 | 1.74E-04 |

| | | | |
|---|---|---|---|
| Ppp1r12c | 1921.830022 | 0.614194606 | 3.91E-09 |
| Daxx | 1244.430088 | 0.614188253 | 5.07E-08 |
| Rgs19 | 1525.312029 | 0.614174168 | 4.51E-14 |
| Slc9a5 | 964.4551971 | 0.614068748 | 8.66E-03 |
| Ctsz | 6039.909013 | 0.612825742 | 1.20E-05 |
| Akt3 | 3456.107411 | 0.612603438 | 1.03E-04 |
| Helz2 | 1049.268653 | 0.612515867 | 9.23E-07 |
| Gm16576 | 237.6025889 | 0.61246301 | 3.46E-03 |
| Mapkapk2 | 5768.945182 | 0.612446231 | 1.83E-06 |
| Pex7 | 519.9801991 | 0.612390788 | 2.30E-07 |
| Tchp | 576.9511312 | 0.612278704 | 8.52E-04 |
| Npm1 | 45543.89849 | 0.61225641 | 1.88E-04 |
| Pip4p1 | 2929.907263 | 0.612207628 | 1.43E-03 |
| Ccdc85b | 694.6340861 | 0.611779806 | 1.29E-03 |
| Cnppd1 | 3116.994516 | 0.611752761 | 2.37E-04 |
| Atrnl1 | 4479.794656 | 0.611539291 | 1.87E-05 |
| Dzip1l | 1361.683775 | 0.611466797 | 1.05E-07 |
| Vwa5a | 5468.402827 | 0.61103644 | 3.84E-03 |
| Rraga | 5750.079961 | 0.610076251 | 1.43E-07 |
| Rps27a | 38338.23439 | 0.609816393 | 2.38E-03 |
| Sh3yl1 | 217.2738414 | 0.60978526 | 2.64E-04 |
| Srgap2 | 3776.658232 | 0.609131258 | 5.88E-17 |
| Fkbpl | 364.4896429 | 0.609016155 | 4.30E-04 |
| Acads | 1920.723929 | 0.608962573 | 1.53E-05 |
| Ap3b1 | 4777.203822 | 0.6087824 | 1.64E-09 |
| Cbll1 | 784.2011838 | 0.608715139 | 2.95E-04 |
| Gtf2a2 | 1249.80594 | 0.608498473 | 9.91E-06 |
| Lcmt1 | 805.5401037 | 0.608483137 | 5.39E-06 |
| Vps26b | 3604.127261 | 0.60822905 | 2.32E-04 |
| Rsl24d1 | 1367.856925 | 0.608223082 | 9.57E-09 |
| Tmem203 | 416.9914192 | 0.608097139 | 5.22E-04 |
| Prkra | 1707.47307 | 0.607943493 | 3.31E-04 |
| Narf | 1213.76747 | 0.607801682 | 8.65E-04 |
| Pcgf2 | 2343.36249 | 0.607273228 | 1.14E-03 |
| Pan2 | 1537.473845 | 0.606758323 | 7.39E-11 |
| Isyna1 | 639.1786947 | 0.606543801 | 2.02E-07 |
| Rab5c | 6665.436685 | 0.606123291 | 1.32E-06 |
| B4gat1 | 1831.688547 | 0.605840516 | 2.80E-05 |
| Tmem234 | 4147.137472 | 0.605831444 | 2.56E-08 |
| Slc7a6 | 3393.128208 | 0.605558556 | 7.99E-09 |
| Cwc27 | 742.8550555 | 0.605259053 | 9.75E-05 |
| Tmem181a | 524.5608539 | 0.605137928 | 2.47E-09 |
| Gan | 285.4934913 | 0.604756742 | 2.22E-05 |
| Zfp784 | 178.7752622 | 0.604629508 | 1.16E-03 |
| Tnip2 | 759.4249082 | 0.604584705 | 4.09E-05 |
| Tnxb | 839.0427032 | 0.604491048 | 2.24E-04 |
| Mfsd8 | 360.6439686 | 0.604469015 | 1.74E-03 |

| | | | |
|---|---|---|---|
| Ccdc12 | 559.7686421 | 0.604171077 | 5.90E-07 |
| Plin3 | 5523.193935 | 0.603863315 | 5.79E-04 |
| Mcat | 964.4933201 | 0.60312671 | 1.09E-06 |
| Arhgap23 | 3979.574992 | 0.602479915 | 1.45E-06 |
| Dars | 4070.211979 | 0.60238768 | 2.23E-10 |
| Tango6 | 700.5811635 | 0.602039306 | 5.11E-05 |
| Frmd6 | 6537.824608 | 0.602019036 | 5.99E-05 |
| Tmem199 | 785.0747928 | 0.601702361 | 2.85E-08 |
| Pramef8 | 1116.244024 | 0.601571614 | 1.70E-05 |
| Nudt16 | 122.149193 | 0.601307741 | 1.39E-02 |
| Anapc16 | 1878.679111 | 0.601092974 | 4.44E-08 |
| Cdk5r1 | 132.2529447 | 0.600861746 | 4.47E-02 |
| Mtmr14 | 1510.787619 | 0.600698445 | 7.83E-06 |
| Chmp2b | 2474.73103 | 0.599667973 | 5.81E-07 |
| Ddost | 6638.293536 | 0.599264718 | 1.58E-04 |
| Litaf | 5188.580695 | 0.598902609 | 1.79E-03 |
| Odf2 | 3143.041756 | 0.598484626 | 8.79E-07 |
| Snw1 | 3904.766616 | 0.598147567 | 2.52E-07 |
| Il10rb | 939.8976722 | 0.597827081 | 8.54E-09 |
| Nup214 | 3369.149841 | 0.597782001 | 5.00E-04 |
| Rwdd4a | 1263.978367 | 0.597745353 | 9.29E-08 |
| Psmc4 | 3375.438816 | 0.597214007 | 1.03E-04 |
| Tinf2 | 460.587957 | 0.596944902 | 1.51E-03 |
| Rock2 | 6286.389359 | 0.596570369 | 3.55E-02 |
| Guk1 | 1445.832017 | 0.59645724 | 2.18E-05 |
| Hdac5 | 4813.627712 | 0.596204168 | 2.18E-02 |
| 1700021F05Rik | 907.3929152 | 0.596082848 | 1.55E-11 |
| Magoh | 2065.067574 | 0.595094262 | 7.70E-08 |
| Gm10123 | 58159.74363 | 0.595088951 | 3.50E-06 |
| Med17 | 1362.023412 | 0.595031729 | 1.31E-09 |
| Capns1 | 11304.90824 | 0.595023778 | 8.16E-07 |
| Nxt1 | 655.9980238 | 0.595011325 | 7.54E-04 |
| Plod1 | 3492.523358 | 0.59488429 | 5.44E-04 |
| Zfp653 | 251.8025585 | 0.594340336 | 2.18E-03 |
| Eid2 | 551.6096972 | 0.594080032 | 1.20E-07 |
| Lin7c | 1704.717814 | 0.593741415 | 5.20E-03 |
| Arl3 | 1484.978537 | 0.59361755 | 5.67E-08 |
| Gm10163 | 19766.21336 | 0.593471379 | 2.07E-03 |
| Hsdl1 | 1948.417286 | 0.593413886 | 1.42E-05 |
| Ap3m2 | 731.4968151 | 0.593205444 | 3.81E-03 |
| Dennd2c | 573.6539703 | 0.593179302 | 7.65E-08 |
| Scaf1 | 4485.533781 | 0.592861003 | 8.87E-08 |
| Alkbh3 | 1383.005993 | 0.592677886 | 1.96E-09 |
| Dcaf10 | 1139.197036 | 0.592142012 | 3.05E-06 |
| Stambp | 1548.578292 | 0.591872079 | 1.10E-06 |
| Simc1 | 369.8851408 | 0.590583918 | 1.64E-03 |
| Setd3 | 4338.931259 | 0.590506149 | 2.97E-07 |

| | | | |
|---|---|---|---|
| Mrps17 | 1874.149448 | 0.590178094 | 1.19E-06 |
| Atn1 | 9447.780035 | 0.589566027 | 6.14E-06 |
| Slc16a1 | 2748.76285 | 0.589522983 | 9.69E-03 |
| R3hdm1 | 3476.07313 | 0.589496776 | 1.20E-08 |
| Wrap73 | 404.8710006 | 0.589292816 | 2.34E-03 |
| Gm10231 | 6874.757385 | 0.589143004 | 1.27E-04 |
| Eif2ak1 | 2626.861785 | 0.588742403 | 1.23E-05 |
| Tec | 115.0557458 | 0.588697667 | 2.41E-02 |
| Pick1 | 781.9958229 | 0.588557003 | 1.85E-05 |
| Tpm2 | 4623.603642 | 0.587897844 | 1.16E-07 |
| Strada | 827.2796994 | 0.587743856 | 6.90E-07 |
| Palld | 2774.077585 | 0.587171841 | 1.10E-03 |
| Bcat1 | 3993.521955 | 0.587098316 | 4.89E-02 |
| Fam171a2 | 1340.633396 | 0.586666422 | 1.15E-03 |
| Trappc10 | 2918.060848 | 0.586338709 | 2.87E-08 |
| Plaa | 5924.78038 | 0.586217595 | 4.68E-05 |
| Rbfa | 690.2126453 | 0.586198653 | 1.21E-04 |
| Ccndbp1 | 1400.492824 | 0.586088008 | 8.06E-06 |
| Mccc1 | 1236.34865 | 0.58547677 | 4.20E-05 |
| Yipf2 | 568.8044309 | 0.585049025 | 9.32E-04 |
| Ints5 | 1926.240369 | 0.584977815 | 1.78E-05 |
| Smpd2 | 1105.312988 | 0.584930822 | 9.81E-06 |
| Nt5c3 | 901.5486206 | 0.583805505 | 3.55E-06 |
| Cnot10 | 1671.516662 | 0.583554451 | 6.18E-09 |
| P4ha1 | 9161.320401 | 0.583364073 | 2.67E-03 |
| Gm14681 | 1360.044557 | 0.582669572 | 3.00E-03 |
| Rbm33 | 3346.777096 | 0.582652629 | 7.73E-06 |
| Arpc2 | 10253.14542 | 0.582075935 | 1.22E-11 |
| Haus5 | 343.3806765 | 0.582041932 | 4.31E-02 |
| Ermard | 436.0657237 | 0.581501957 | 2.48E-05 |
| Ift46 | 2411.826845 | 0.580965346 | 8.98E-06 |
| Rgs2 | 984.5440039 | 0.580710377 | 2.26E-02 |
| 2310033P09Rik | 321.5540812 | 0.580411018 | 2.44E-05 |
| Ppil3 | 1048.930102 | 0.580308783 | 3.55E-06 |
| Pi4k2a | 2798.58575 | 0.579582353 | 8.67E-06 |
| Tex10 | 1746.463961 | 0.579018585 | 9.32E-08 |
| Nek8 | 529.5072841 | 0.578293958 | 2.09E-07 |
| Ndufa12 | 1429.626157 | 0.578127386 | 5.80E-04 |
| Gm12191 | 13447.05379 | 0.578076624 | 8.11E-04 |
| 0610037L13Rik | 1757.031948 | 0.5780171 | 4.97E-06 |
| Psme3 | 5358.956441 | 0.577391613 | 3.56E-08 |
| Gle1 | 2111.626911 | 0.577273775 | 5.26E-04 |
| Cox8a | 8974.358587 | 0.577066449 | 1.12E-04 |
| Rpusd4 | 523.4815444 | 0.576930108 | 2.14E-07 |
| Mef2d | 4971.28913 | 0.576662129 | 1.37E-05 |
| 1700025G04Rik | 2025.145476 | 0.576196577 | 7.47E-05 |
| Klf16 | 1257.19304 | 0.575241359 | 2.26E-03 |

| | | | |
|---|---|---|---|
| Stam2 | 2719.319093 | 0.575198124 | 3.14E-12 |
| Sphk2 | 1000.143385 | 0.57500904 | 1.14E-05 |
| Herpud2 | 1800.931372 | 0.574956414 | 5.96E-06 |
| Neu1 | 2682.126546 | 0.57493932 | 3.61E-03 |
| Mocs2 | 1122.778625 | 0.574847164 | 1.69E-07 |
| Kif1c | 11260.85304 | 0.574647766 | 8.61E-06 |
| Layn | 1804.865648 | 0.574566395 | 1.64E-03 |
| Snrpg | 1662.606002 | 0.574526704 | 3.95E-04 |
| Rnf187 | 6422.38981 | 0.574273481 | 1.09E-05 |
| Dtx2 | 985.8070884 | 0.573263312 | 1.29E-03 |
| Ccdc9 | 1160.732769 | 0.572102059 | 4.05E-06 |
| Tmem94 | 1964.367392 | 0.571965028 | 2.47E-05 |
| Map1a | 6290.433199 | 0.571812499 | 5.22E-05 |
| Pigv | 317.9398644 | 0.571762175 | 1.16E-02 |
| Sec63 | 6038.823214 | 0.571696526 | 3.21E-05 |
| Cenpo | 452.4141085 | 0.571667207 | 1.49E-03 |
| Smad7 | 439.850284 | 0.571446999 | 3.88E-02 |
| Tsku | 1205.341302 | 0.571426773 | 6.24E-06 |
| Ergic1 | 11142.20075 | 0.571164517 | 1.30E-04 |
| Tbc1d10b | 1987.395714 | 0.570983483 | 1.46E-05 |
| Rwdd1 | 3045.160573 | 0.570941484 | 7.85E-08 |
| Slc22a5 | 722.5646486 | 0.570770623 | 3.01E-04 |
| Anapc2 | 5642.744713 | 0.570659408 | 2.36E-07 |
| Canx | 28016.75408 | 0.570534484 | 3.08E-05 |
| Zfp64 | 1050.308417 | 0.570498463 | 2.71E-04 |
| Cyp4f13 | 464.5427587 | 0.57023415 | 5.62E-03 |
| Cacna1a | 1112.647342 | 0.569696364 | 3.11E-02 |
| Tbcb | 1604.288802 | 0.569667965 | 2.32E-03 |
| Rps3a2 | 220.5441406 | 0.569072958 | 3.11E-02 |
| Trpm4 | 642.7733671 | 0.569035445 | 2.03E-05 |
| Copg1 | 12031.54603 | 0.56902042 | 1.34E-07 |
| Dbr1 | 635.1497289 | 0.568895621 | 2.61E-04 |
| Wasf2 | 5249.227574 | 0.568302765 | 2.60E-05 |
| Fuz | 552.4981124 | 0.568090784 | 9.24E-05 |
| Ophn1 | 1096.550011 | 0.568029454 | 5.24E-05 |
| Bet1l | 959.1702443 | 0.567876009 | 2.32E-03 |
| Rab1b | 4911.964953 | 0.567813392 | 5.18E-05 |
| 9130221H12Rik | 286.8279774 | 0.567424641 | 1.39E-02 |
| Stau2 | 1353.634259 | 0.567015148 | 3.78E-12 |
| Ptprn | 7409.737366 | 0.566943261 | 1.40E-06 |
| Cfap300 | 360.5795128 | 0.566923163 | 4.72E-03 |
| Xpo5 | 4096.279036 | 0.566629918 | 6.81E-05 |
| Foxp4 | 2395.208816 | 0.566290209 | 1.81E-05 |
| Sipa1l3 | 3854.926667 | 0.565977946 | 5.73E-06 |
| Map2k1 | 2663.239994 | 0.565967638 | 2.14E-08 |
| Med8 | 1643.850492 | 0.565580919 | 1.21E-05 |
| Tmem59 | 14616.98512 | 0.565531184 | 1.41E-03 |

| | | | |
|---|---|---|---|
| Cavin1 | 29736.21876 | 0.565101725 | 6.54E-05 |
| Aarsd1 | 1053.296333 | 0.565048158 | 1.92E-03 |
| Zranb1 | 1747.798794 | 0.564362135 | 3.34E-07 |
| Pfn1 | 17269.4251 | 0.564340594 | 5.43E-04 |
| Polr3a | 1153.652272 | 0.563747582 | 2.45E-06 |
| Acp2 | 2896.824916 | 0.563734914 | 2.98E-06 |
| Gdap2 | 944.6345225 | 0.563433279 | 4.91E-09 |
| Cap1 | 19890.28716 | 0.563417194 | 1.17E-09 |
| Polr1a | 4966.441682 | 0.563337701 | 1.19E-05 |
| Ppp2r5c | 4554.30962 | 0.562893151 | 4.16E-11 |
| Ndrg4 | 1002.618499 | 0.562484297 | 2.36E-03 |
| Ldha | 13824.04381 | 0.562480478 | 9.47E-03 |
| Snrnp27 | 1828.287886 | 0.56229409 | 4.72E-06 |
| Cyld | 2898.729581 | 0.562212085 | 2.90E-04 |
| Becn1 | 3487.621834 | 0.56214016 | 1.88E-16 |
| Rab8a | 3357.260812 | 0.562076734 | 1.44E-06 |
| Fam214a | 744.1921239 | 0.561934877 | 2.00E-02 |
| Ddhd1 | 899.9729154 | 0.561687726 | 1.15E-05 |
| Mettl13 | 1248.651196 | 0.561297943 | 2.74E-05 |
| Yipf4 | 2227.605612 | 0.561101844 | 1.54E-04 |
| Hmg20b | 1606.6635 | 0.560288354 | 6.58E-05 |
| Ufsp2 | 1807.769792 | 0.560181103 | 4.12E-08 |
| Rab5if | 2691.810294 | 0.560161426 | 1.80E-03 |
| Fam213a | 757.8558732 | 0.560054221 | 2.47E-02 |
| Rnf167 | 1913.105388 | 0.559215671 | 4.33E-04 |
| Ccdc189 | 104.5542721 | 0.558983682 | 1.84E-02 |
| Focad | 3203.348512 | 0.558594465 | 2.40E-07 |
| Smurf2 | 3989.71208 | 0.558527799 | 9.85E-10 |
| Prr14 | 1622.26639 | 0.557910526 | 2.64E-09 |
| Cdc16 | 4416.767764 | 0.557593183 | 4.85E-07 |
| Baz1a | 2082.890024 | 0.557484075 | 1.68E-03 |
| Metap1 | 2869.366179 | 0.557182556 | 3.86E-12 |
| Bcl10 | 1391.517893 | 0.556801568 | 4.93E-06 |
| Mrpl19 | 1414.367424 | 0.55675745 | 2.77E-03 |
| Gatad1 | 2934.150248 | 0.556498023 | 3.22E-08 |
| Gcsh | 2534.925227 | 0.556038737 | 1.34E-04 |
| Gsk3b | 3602.663052 | 0.555840531 | 5.29E-07 |
| Grin2d | 155.278892 | 0.555516594 | 4.80E-02 |
| Rsu1 | 5433.912262 | 0.555504514 | 6.76E-09 |
| Gm9864 | 1039.948219 | 0.55547846 | 1.04E-05 |
| Sat2 | 217.4762623 | 0.555209662 | 1.06E-02 |
| Cln6 | 1612.759689 | 0.554721214 | 3.33E-03 |
| Ube4b | 4488.5307 | 0.554615425 | 3.29E-05 |
| Ubr3 | 4475.974516 | 0.554578235 | 3.33E-07 |
| Ppp1r9b | 4501.93599 | 0.554054194 | 1.05E-06 |
| Cndp2 | 2438.22089 | 0.553543209 | 3.86E-02 |
| Mgst3 | 998.4397806 | 0.552547171 | 2.60E-03 |

| | | | |
|---|---|---|---|
| Gmeb2 | 1450.329167 | 0.552497153 | 9.15E-05 |
| Abcc5 | 6534.980853 | 0.552110068 | 3.07E-03 |
| Cyb5r4 | 1403.43879 | 0.551869334 | 6.05E-07 |
| Acd | 2131.77091 | 0.551860644 | 1.91E-03 |
| Atg7 | 2072.141009 | 0.55105827 | 2.83E-04 |
| Ndufs4 | 1702.476031 | 0.551018262 | 6.87E-06 |
| Paox | 462.702227 | 0.550214106 | 7.31E-05 |
| H2-T10 | 831.3455959 | 0.550173618 | 2.15E-02 |
| Itpa | 1348.474225 | 0.550091691 | 1.18E-06 |
| Helb | 926.5999354 | 0.549688059 | 1.32E-07 |
| Moap1 | 164.6982918 | 0.549627093 | 1.48E-02 |
| 8430429K09Rik | 165.2112728 | 0.549616642 | 3.78E-03 |
| Spag7 | 2417.33747 | 0.549583408 | 1.88E-10 |
| Mrpl40 | 1051.796623 | 0.549500439 | 2.01E-06 |
| 2510002D24Rik | 276.2069138 | 0.549370123 | 4.42E-04 |
| Tob2 | 2102.527049 | 0.549257697 | 4.34E-04 |
| Dync1i2 | 7167.443248 | 0.54912681 | 2.27E-12 |
| Commd10 | 1041.836611 | 0.549006402 | 1.15E-06 |
| Mlf2 | 9541.962556 | 0.548594172 | 2.84E-05 |
| Nup54 | 1688.162602 | 0.548513976 | 6.00E-05 |
| Slc35c2 | 1816.266464 | 0.548341182 | 5.19E-05 |
| Skap2 | 2394.603323 | 0.547905429 | 6.08E-12 |
| Gm5611 | 403.0955032 | 0.547869004 | 3.31E-03 |
| Ankrd13a | 3982.563234 | 0.547835028 | 4.18E-12 |
| Mrps14 | 1169.475178 | 0.547483323 | 1.93E-06 |
| Rhoc | 7213.832226 | 0.547467438 | 3.01E-04 |
| Mrps7 | 1445.006038 | 0.547320106 | 3.96E-05 |
| Sppl2a | 3377.428579 | 0.546816075 | 1.64E-06 |
| Rac1 | 12684.70984 | 0.545558996 | 1.05E-07 |
| Thap11 | 1739.529214 | 0.545328816 | 6.58E-04 |
| Naip2 | 1161.031535 | 0.544754339 | 2.32E-06 |
| Plpbp | 2115.5974 | 0.54467634 | 4.23E-14 |
| Snrpe | 2802.549934 | 0.544269811 | 1.59E-05 |
| Llph | 1359.624563 | 0.544265458 | 3.56E-06 |
| Dgke | 540.5581551 | 0.543884358 | 2.92E-05 |
| Agpat5 | 1466.315111 | 0.543874319 | 3.88E-05 |
| Hebp1 | 1013.408107 | 0.543793683 | 1.61E-03 |
| Xbp1 | 4888.960651 | 0.543631415 | 9.30E-03 |
| Utp20 | 4892.506841 | 0.543361015 | 7.09E-04 |
| Primpol | 719.4967497 | 0.543107423 | 1.53E-04 |
| Alkbh6 | 589.5487126 | 0.542689787 | 2.29E-03 |
| Mta2 | 6317.531041 | 0.542221152 | 1.16E-08 |
| Cyb5r1 | 3302.743638 | 0.542195568 | 2.50E-02 |
| Arf5 | 4503.439777 | 0.541721358 | 2.24E-05 |
| Wdfy2 | 314.5094219 | 0.541393541 | 6.38E-04 |
| Mrpl30 | 2494.206912 | 0.541223296 | 1.46E-03 |
| Mast2 | 8459.255157 | 0.54071782 | 2.18E-04 |

| | | | |
|---|---|---|---|
| Adck5 | 731.9848639 | 0.540628737 | 2.11E-03 |
| Fiz1 | 1176.565028 | 0.540597361 | 5.11E-06 |
| Txndc12 | 3770.539548 | 0.54028332 | 7.74E-06 |
| Cdc42ep2 | 2880.635182 | 0.540243766 | 1.65E-03 |
| Psma4 | 4648.273589 | 0.539992317 | 2.44E-09 |
| Ubr5 | 12270.45866 | 0.539313019 | 2.10E-09 |
| Zfp688 | 338.1326918 | 0.539238741 | 4.66E-03 |
| Morc3 | 1497.828179 | 0.539060137 | 1.19E-03 |
| Sart1 | 3217.739631 | 0.538473674 | 6.34E-08 |
| Inpp5a | 3109.503164 | 0.538243167 | 9.94E-03 |
| Itga5 | 8198.609151 | 0.538224966 | 1.29E-03 |
| Tmem70 | 867.4598913 | 0.538205838 | 7.18E-06 |
| Mecr | 644.0354512 | 0.537925014 | 1.02E-03 |
| Zdhhc7 | 1702.309656 | 0.53727108 | 2.04E-05 |
| Ppox | 965.157333 | 0.537247645 | 4.54E-03 |
| Akr1b10 | 376.3471069 | 0.536993989 | 3.80E-03 |
| Srp14 | 4319.453531 | 0.536941726 | 1.88E-09 |
| Afg3l2 | 2671.085849 | 0.535992574 | 4.95E-07 |
| Fzd8 | 383.6272413 | 0.534814038 | 1.37E-04 |
| Hmox2 | 2425.930296 | 0.534749386 | 1.29E-04 |
| Hibch | 1051.562632 | 0.534708764 | 7.10E-04 |
| Zfp462 | 2551.882414 | 0.534256992 | 4.78E-03 |
| Afap1 | 3020.859185 | 0.533795567 | 5.41E-05 |
| Unkl | 1023.23768 | 0.5336044 | 2.38E-04 |
| Myo6 | 987.1695516 | 0.533588163 | 8.46E-04 |
| Marf1 | 4341.805183 | 0.533550636 | 3.88E-04 |
| Cdk10 | 1543.456391 | 0.533508431 | 4.83E-09 |
| Mfn2 | 3174.330346 | 0.533055406 | 2.56E-07 |
| Fopnl | 1523.346677 | 0.532639245 | 2.34E-07 |
| Fam120b | 2980.824706 | 0.532634535 | 1.32E-04 |
| Abcf2 | 4906.179787 | 0.532086226 | 1.46E-03 |
| Scap | 4677.143336 | 0.531932086 | 3.30E-05 |
| 1110008L16Rik | 693.3561909 | 0.531497814 | 1.61E-04 |
| Pdhx | 940.8641112 | 0.531424298 | 4.74E-03 |
| AU022252 | 1364.971733 | 0.5314046 | 1.15E-04 |
| Aldh1l2 | 7819.791201 | 0.531001047 | 3.61E-02 |
| Trappc11 | 2425.129797 | 0.530681638 | 8.66E-13 |
| Rprd1a | 1030.14927 | 0.530554808 | 5.16E-04 |
| Klhl18 | 706.0119398 | 0.529637606 | 4.67E-04 |
| Zfr2 | 251.1434612 | 0.528988097 | 4.91E-02 |
| Shb | 889.4084882 | 0.528979753 | 1.47E-04 |
| Ogdh | 6423.124912 | 0.528544614 | 3.08E-04 |
| Mrps15 | 2715.97954 | 0.528184996 | 2.56E-07 |
| Gm11868 | 349.9990098 | 0.528162911 | 3.36E-06 |
| Rgs12 | 1802.3496 | 0.528083941 | 2.13E-03 |
| Stx8 | 1318.595004 | 0.528010895 | 5.51E-04 |
| Mtmr9 | 1803.405493 | 0.52788812 | 5.65E-14 |

| | | | |
|---|---|---|---|
| Xrcc6 | 1228.827517 | 0.527650125 | 1.15E-03 |
| Khnyn | 1116.250918 | 0.527412224 | 1.74E-04 |
| Ino80c | 526.2001063 | 0.526819785 | 2.06E-03 |
| Rnf5 | 945.3833538 | 0.526596671 | 1.12E-05 |
| Slc15a4 | 1139.09426 | 0.526546197 | 1.91E-02 |
| Tmem248 | 2966.009369 | 0.52636553 | 3.79E-04 |
| Zfp369 | 776.9165251 | 0.526155425 | 1.21E-05 |
| Elf4 | 2047.77655 | 0.526002497 | 2.94E-08 |
| Vcp | 24296.04105 | 0.525756229 | 3.86E-06 |
| Tmem231 | 1590.140116 | 0.525616304 | 1.29E-04 |
| Rab3gap1 | 3040.80505 | 0.524899902 | 9.05E-12 |
| Tpd52l2 | 3633.98065 | 0.524427976 | 2.76E-06 |
| Slc25a11 | 3169.733048 | 0.524183658 | 3.05E-06 |
| Rpf2 | 971.3745042 | 0.523910522 | 4.92E-02 |
| Napa | 5584.664807 | 0.523765116 | 4.50E-04 |
| Sema3f | 4201.918047 | 0.52355802 | 9.09E-04 |
| Abi1 | 4282.059818 | 0.523507912 | 2.85E-05 |
| Slc25a4 | 16951.5366 | 0.523438434 | 2.07E-12 |
| Ddx23 | 3775.542555 | 0.523001472 | 5.11E-05 |
| Zfp236 | 2213.260685 | 0.522998581 | 1.10E-04 |
| Serac1 | 662.4093304 | 0.522701673 | 6.11E-04 |
| Sdha | 7204.468236 | 0.5223601 | 4.23E-10 |
| Rsc1a1 | 1091.624438 | 0.522357879 | 2.87E-04 |
| Bag2 | 1009.73514 | 0.522182258 | 2.87E-03 |
| 1700037H04Rik | 1863.740915 | 0.522171487 | 1.03E-04 |
| Hyi | 318.3341667 | 0.522154763 | 3.09E-03 |
| Sptan1 | 24705.76715 | 0.52214167 | 7.89E-05 |
| Atp5a1 | 24534.04847 | 0.522096826 | 2.60E-06 |
| Chmp7 | 1677.684671 | 0.521970142 | 8.24E-05 |
| Mok | 173.5492469 | 0.521464651 | 6.82E-03 |
| Gm49342 | 3671.500329 | 0.521229052 | 3.56E-03 |
| Pank4 | 603.6682192 | 0.521123778 | 1.96E-04 |
| Nckipsd | 805.9801633 | 0.519784931 | 2.76E-04 |
| Plppr2 | 1300.032739 | 0.519015255 | 6.77E-04 |
| Nup35 | 1055.107469 | 0.518972444 | 1.13E-02 |
| Lrrc61 | 428.7985426 | 0.518491426 | 3.92E-02 |
| Fam20a | 1473.199596 | 0.518040562 | 1.85E-02 |
| Sh3rf1 | 1249.064827 | 0.517851172 | 4.85E-04 |
| Tmod3 | 2418.238521 | 0.51771172 | 1.81E-03 |
| Pstk | 460.105305 | 0.517685547 | 2.22E-03 |
| Dolpp1 | 505.6016964 | 0.517570179 | 2.77E-04 |
| Cast | 5248.563907 | 0.517474926 | 4.16E-08 |
| Lrrk1 | 3325.760949 | 0.517472239 | 1.64E-04 |
| Fam136a | 1289.210965 | 0.51694644 | 4.58E-04 |
| Ccnt1 | 2298.363621 | 0.516855528 | 3.10E-03 |
| Zfpm2 | 579.7477361 | 0.516677223 | 6.53E-03 |
| AI837181 | 2095.389026 | 0.516294623 | 3.41E-03 |

| | | | |
|---|---|---|---|
| Adsl | 2404.650685 | 0.516290206 | 7.06E-04 |
| Ccdc93 | 3957.529583 | 0.51620722 | 1.14E-04 |
| P3h4 | 3930.457188 | 0.516025914 | 4.25E-03 |
| Erf | 2001.173073 | 0.515685838 | 1.28E-02 |
| Tmem259 | 4431.668597 | 0.515678546 | 3.08E-11 |
| St13 | 14945.84821 | 0.51551584 | 1.78E-03 |
| Tacc2 | 2141.000492 | 0.515379051 | 4.06E-07 |
| Uqcr10 | 2097.495222 | 0.51499355 | 2.31E-04 |
| Arhgdia | 21848.33896 | 0.514640068 | 8.35E-04 |
| Ifi30 | 2933.154737 | 0.514169133 | 4.60E-02 |
| Ift52 | 1834.55518 | 0.513847275 | 7.42E-07 |
| Nrd1 | 9559.450072 | 0.513303736 | 1.80E-06 |
| Lonp1 | 6941.8007 | 0.513054103 | 6.26E-04 |
| Cct7 | 14582.72543 | 0.512952901 | 2.39E-04 |
| Atg5 | 1278.933549 | 0.512824576 | 4.51E-06 |
| Enc1 | 4373.996949 | 0.512768585 | 2.08E-02 |
| Exosc5 | 597.2657099 | 0.512668416 | 2.22E-03 |
| Prep | 3836.793026 | 0.512571223 | 1.40E-02 |
| Zdhhc12 | 144.6380655 | 0.512347748 | 2.52E-02 |
| Pdcl3 | 2311.344659 | 0.511952589 | 1.83E-07 |
| Polr2b | 4760.275088 | 0.511733566 | 2.41E-04 |
| Necap2 | 1811.57524 | 0.511731102 | 9.07E-08 |
| Prpf19 | 7538.045994 | 0.511627914 | 6.57E-05 |
| 9930012K11Rik | 537.6727168 | 0.511547774 | 2.87E-03 |
| Acot7 | 1795.605865 | 0.511463587 | 1.59E-04 |
| Mfhas1 | 2892.570226 | 0.51100388 | 4.12E-09 |
| Mrps10 | 1521.663651 | 0.510579784 | 8.15E-06 |
| Dhx8 | 2608.475505 | 0.510389786 | 2.58E-05 |
| Gstm2 | 1617.930043 | 0.510059807 | 4.17E-02 |
| Gm49405 | 509.8473078 | 0.509464327 | 1.46E-03 |
| Acyp1 | 248.5914622 | 0.509060826 | 6.68E-03 |
| Asb6 | 1407.281146 | 0.509040632 | 1.03E-03 |
| Mzt2 | 762.5559631 | 0.508223484 | 3.25E-03 |
| Gm15920 | 1814.698209 | 0.508107304 | 8.83E-03 |
| Srbd1 | 1448.629917 | 0.507959674 | 1.12E-05 |
| Pgap2 | 1147.432946 | 0.507903324 | 4.77E-04 |
| Nudt18 | 604.0577606 | 0.50790152 | 4.22E-03 |
| Pxn | 3926.522418 | 0.507195449 | 1.18E-04 |
| Apobec1 | 552.4754437 | 0.506174493 | 1.95E-02 |
| Wdr20 | 732.3281428 | 0.505851485 | 2.92E-09 |
| Poll | 515.957832 | 0.505098099 | 1.16E-03 |
| Hnrnpa1 | 23929.72459 | 0.504693168 | 9.30E-04 |
| Xrcc4 | 301.2973331 | 0.504287039 | 7.49E-03 |
| Otub2 | 372.7939584 | 0.503319587 | 1.45E-02 |
| Rcan1 | 795.1523126 | 0.503250045 | 1.07E-02 |
| Sirt7 | 672.5356625 | 0.502578557 | 8.55E-05 |
| Ppt2 | 1697.89408 | 0.501414709 | 5.66E-03 |

| | | | |
|---|---|---|---|
| Prim2 | 794.0553518 | 0.501054312 | 3.69E-02 |
| mt-Co1 | 763641.7603 | 0.500770184 | 5.52E-04 |
| Anapc7 | 2454.939837 | 0.500668536 | 7.30E-04 |
| Gm28043 | 890.2744121 | 0.500655437 | 7.61E-03 |
| Plcb3 | 3361.405742 | 0.500374638 | 1.43E-10 |
| Tent4a | 1488.434797 | 0.500264117 | 5.63E-04 |
| Gabarapl1 | 11663.63086 | 0.500184962 | 9.53E-03 |
| Fam234a | 4598.43371 | 0.499896125 | 5.12E-03 |
| Frmd8 | 2457.447622 | 0.499019779 | 1.54E-02 |
| Zfp592 | 1895.845173 | 0.498814738 | 3.30E-04 |
| Gng5 | 2852.253698 | 0.498609028 | 1.36E-08 |
| Eif5a | 21555.42109 | 0.498539487 | 1.71E-02 |
| Mrpl12 | 1862.811651 | 0.497908348 | 7.20E-03 |
| Tomm70a | 4143.787542 | 0.497788128 | 1.15E-04 |
| Iqgap1 | 14617.56628 | 0.497659438 | 2.57E-07 |
| Mgea5 | 6513.668935 | 0.496956067 | 2.59E-03 |
| Rab13 | 815.1880269 | 0.49682451 | 1.63E-05 |
| Larp1 | 8675.057462 | 0.49671336 | 6.28E-06 |
| Dap3 | 2531.53342 | 0.496421433 | 3.21E-04 |
| Ppp3cc | 406.7928018 | 0.496196806 | 9.82E-04 |
| Ctsa | 21262.24103 | 0.495975607 | 1.28E-02 |
| Mthfd2l | 443.9979493 | 0.495651693 | 7.28E-05 |
| Zfp956 | 415.6674772 | 0.495145325 | 2.07E-03 |
| Cmtr2 | 516.4360552 | 0.495112051 | 1.25E-05 |
| Lurap1l | 1882.936888 | 0.494329438 | 2.61E-03 |
| Rpl23 | 66491.88099 | 0.494269173 | 5.90E-03 |
| Aifm1 | 2622.394853 | 0.494222801 | 1.93E-05 |
| Endov | 1049.701626 | 0.493897291 | 1.62E-04 |
| Tmem214 | 3151.052134 | 0.493743058 | 2.58E-04 |
| Letmd1 | 1317.473792 | 0.493242005 | 1.15E-07 |
| Arhgap21 | 4091.646736 | 0.493181459 | 2.15E-02 |
| Pde6d | 1448.894996 | 0.492330122 | 1.37E-04 |
| Taok3 | 2501.86846 | 0.491926654 | 2.74E-07 |
| Chchd1 | 748.5531693 | 0.491859029 | 1.70E-03 |
| Tmem159 | 526.5163809 | 0.491601326 | 4.07E-03 |
| Unc13b | 378.855196 | 0.491596285 | 3.86E-02 |
| Ntan1 | 2090.352579 | 0.491370236 | 7.55E-04 |
| Snap29 | 2083.038228 | 0.490111107 | 7.35E-05 |
| Cuta | 2544.799639 | 0.490017095 | 3.78E-03 |
| Stard3nl | 981.4973654 | 0.48948269 | 2.02E-07 |
| Dusp7 | 1971.188281 | 0.488982684 | 2.62E-05 |
| Sdhc | 3946.444783 | 0.488514697 | 6.42E-04 |
| Anxa7 | 4754.841561 | 0.487833076 | 4.76E-07 |
| Dcaf13 | 2171.351908 | 0.487662233 | 3.78E-06 |
| Gm43517 | 631.9198304 | 0.48726816 | 2.72E-02 |
| Phrf1 | 3217.922762 | 0.487201064 | 7.90E-04 |
| Cops3 | 3367.131836 | 0.487033846 | 2.59E-07 |

| | | | |
|---|---|---|---|
| Prdx1 | 26184.93752 | 0.486926156 | 1.21E-06 |
| Cdc40 | 2423.015152 | 0.486898665 | 3.71E-06 |
| Usp10 | 2331.73001 | 0.48646028 | 7.57E-05 |
| Exosc4 | 794.7367626 | 0.486452674 | 2.16E-03 |
| Phf5a | 1104.508101 | 0.48616123 | 5.39E-04 |
| Brd2 | 13357.93295 | 0.485861683 | 2.18E-04 |
| Camsap1 | 3025.579919 | 0.485846355 | 9.39E-06 |
| Supt20 | 1894.238868 | 0.485706522 | 2.10E-03 |
| S100a6 | 44660.9589 | 0.485656937 | 9.85E-03 |
| Madd | 889.1617141 | 0.48548695 | 7.35E-04 |
| Eif1ad | 1505.772972 | 0.485316375 | 1.76E-03 |
| Myh9 | 47052.31137 | 0.48500552 | 1.28E-04 |
| Tnpo1 | 8965.551593 | 0.484536384 | 1.78E-08 |
| Snx14 | 1287.976179 | 0.484427624 | 5.39E-05 |
| D16Ertd472e | 232.6290134 | 0.484288094 | 1.56E-03 |
| Mau2 | 4038.002112 | 0.483671372 | 5.32E-05 |
| Med31 | 305.7132664 | 0.483407313 | 2.86E-04 |
| Coa6 | 428.7493738 | 0.483275943 | 3.26E-03 |
| Mrpl48 | 486.4467665 | 0.483009505 | 2.59E-02 |
| Vps72 | 1872.257713 | 0.482999539 | 2.35E-02 |
| Map2k5 | 1196.801708 | 0.482789995 | 2.58E-07 |
| Xrn2 | 6587.658003 | 0.482610864 | 7.31E-07 |
| Dnajc21 | 953.490279 | 0.482288795 | 1.02E-02 |
| Nfatc2ip | 343.9678999 | 0.482282875 | 2.09E-02 |
| Sestd1 | 2933.03197 | 0.481844425 | 1.51E-05 |
| Dmac1 | 1447.805556 | 0.481648172 | 2.25E-08 |
| Gm10053 | 1698.198741 | 0.481510117 | 4.49E-02 |
| Eif3e | 15575.42035 | 0.481455606 | 5.13E-03 |
| Osbp | 4497.550677 | 0.481408474 | 6.11E-05 |
| Tada3 | 1642.533578 | 0.481378389 | 2.62E-03 |
| Twnk | 1587.171026 | 0.480825556 | 1.55E-04 |
| Riox2 | 304.3405249 | 0.480527688 | 1.84E-02 |
| Acox3 | 1204.483995 | 0.480445965 | 9.29E-04 |
| Nup107 | 1865.084275 | 0.480195122 | 8.42E-03 |
| Utp15 | 1382.972538 | 0.479564513 | 4.16E-04 |
| Elob | 5284.404596 | 0.479301713 | 1.72E-04 |
| Trappc6a | 772.3366347 | 0.479103978 | 9.56E-03 |
| Phf20 | 3151.247539 | 0.479049813 | 1.27E-07 |
| Chmp3 | 6440.19342 | 0.478661507 | 2.55E-05 |
| Ypel5 | 2403.272004 | 0.478528943 | 2.89E-03 |
| Pex5 | 3893.377517 | 0.478509105 | 6.60E-07 |
| Wars | 2773.816283 | 0.478461143 | 3.46E-03 |
| Fbxl12 | 679.757772 | 0.478234568 | 1.05E-03 |
| Vps51 | 1405.641349 | 0.478020038 | 1.14E-04 |
| Zranb2 | 4108.229627 | 0.477978592 | 8.71E-03 |
| Kdelr1 | 5927.242726 | 0.477829971 | 1.04E-02 |
| Pum3 | 4557.771787 | 0.477786025 | 6.57E-06 |

| | | | |
|---|---|---|---|
| Mrpl3 | 1782.428916 | 0.47740031 | 7.50E-07 |
| Tmem134 | 2708.878956 | 0.477212095 | 7.39E-04 |
| Crnde | 141.719853 | 0.477206618 | 4.85E-02 |
| Tead4 | 2019.099038 | 0.47685592 | 1.21E-02 |
| Psmg1 | 640.1736031 | 0.476087164 | 1.23E-06 |
| En1 | 1200.582007 | 0.475934732 | 1.11E-02 |
| Mul1 | 856.8914795 | 0.475529529 | 2.03E-02 |
| Clcn7 | 3441.410259 | 0.475350292 | 4.10E-03 |
| Atp6v1h | 5874.496336 | 0.475320466 | 7.41E-04 |
| Lars | 4924.644824 | 0.474605198 | 2.74E-02 |
| Elovl1 | 5057.052182 | 0.474448535 | 6.39E-04 |
| Tspyl1 | 4410.886126 | 0.474439588 | 3.44E-03 |
| Epn1 | 2966.878104 | 0.474351894 | 2.11E-04 |
| Gsk3a | 3027.222386 | 0.474065898 | 4.19E-05 |
| Phospho2 | 943.8096213 | 0.473931596 | 3.30E-04 |
| Dgkq | 1168.892023 | 0.473643626 | 1.10E-03 |
| Emc9 | 106.7389187 | 0.47338518 | 3.80E-02 |
| Hook2 | 814.6719502 | 0.473352298 | 3.26E-03 |
| Atp6ap1 | 11667.39151 | 0.473342168 | 5.01E-04 |
| Lage3 | 686.8422425 | 0.473169418 | 7.72E-05 |
| Bbs2 | 880.2996349 | 0.473134871 | 2.51E-02 |
| Nxpe3 | 351.1963934 | 0.473088829 | 3.14E-03 |
| Fpgs | 1316.666915 | 0.472898726 | 8.49E-03 |
| Trim46 | 801.3602403 | 0.472792463 | 1.12E-02 |
| Akap1 | 2550.866096 | 0.472783757 | 3.66E-04 |
| Fnta | 2765.044566 | 0.472547979 | 1.29E-07 |
| Klhl21 | 1593.791091 | 0.472244899 | 4.81E-02 |
| Dedd | 1622.876603 | 0.471651976 | 2.60E-10 |
| Ptgr2 | 619.5256586 | 0.471600368 | 2.59E-04 |
| Harbi1 | 406.9577452 | 0.471571316 | 1.38E-03 |
| Tmem18 | 963.5596269 | 0.471494537 | 7.89E-07 |
| Abl2 | 5144.461312 | 0.471025715 | 1.70E-07 |
| Nif3l1 | 915.7202951 | 0.470594027 | 7.72E-03 |
| Frrs1 | 1733.117738 | 0.469770513 | 1.46E-05 |
| Fzr1 | 2045.795168 | 0.469491269 | 1.83E-02 |
| Klf13 | 2027.299638 | 0.469350991 | 2.70E-03 |
| Tdrkh | 618.162234 | 0.469220825 | 6.60E-03 |
| Fam160a1 | 1093.71908 | 0.469066511 | 2.28E-03 |
| Ftsj3 | 4055.065065 | 0.468741053 | 9.29E-04 |
| Bud23 | 1182.391185 | 0.468117127 | 2.90E-04 |
| Akirin1 | 2974.695353 | 0.467763842 | 5.24E-05 |
| Sec23ip | 3395.713208 | 0.467687869 | 1.20E-03 |
| Dhx38 | 2719.106964 | 0.467616677 | 2.31E-05 |
| Pqbp1 | 1870.459313 | 0.467299844 | 2.59E-03 |
| Wdr55 | 862.4348251 | 0.46684394 | 3.48E-04 |
| Zswim6 | 572.7350864 | 0.466557926 | 5.30E-03 |
| Pgm1 | 4305.875486 | 0.46641058 | 3.81E-04 |

| | | | |
|---|---|---|---|
| Surf6 | 1223.042779 | 0.46635652 | 2.03E-05 |
| Gnpat | 3120.524208 | 0.466029527 | 1.34E-08 |
| Crbn | 1780.102223 | 0.465945529 | 5.70E-04 |
| Zfp414 | 942.5399202 | 0.465858728 | 2.23E-04 |
| Trim3 | 2043.175664 | 0.465683724 | 8.65E-05 |
| Ap1s1 | 3404.57732 | 0.465674242 | 6.41E-04 |
| Eif2a | 4013.91275 | 0.46538701 | 2.54E-05 |
| Cd47 | 2998.687264 | 0.465022054 | 6.99E-03 |
| Pakap | 1146.85027 | 0.465009543 | 3.68E-02 |
| Tomm22 | 2970.207624 | 0.464965334 | 7.84E-07 |
| Stx5a | 3441.259391 | 0.464917707 | 6.69E-05 |
| Kdm4a | 6293.193988 | 0.464628262 | 3.31E-07 |
| Mrpl51 | 2225.268468 | 0.464585227 | 3.02E-04 |
| Rpap1 | 1187.497531 | 0.464530375 | 1.26E-05 |
| Tmem41a | 421.3225088 | 0.464141471 | 4.93E-03 |
| Atrip | 465.3775041 | 0.463394082 | 2.53E-02 |
| Ppm1a | 3456.516388 | 0.462495038 | 4.29E-09 |
| Gak | 5311.988674 | 0.461864654 | 2.60E-05 |
| Ccdc181 | 639.3289339 | 0.461628645 | 1.90E-03 |
| Sugp1 | 1683.026125 | 0.460710888 | 2.96E-04 |
| Zfp282 | 1414.152334 | 0.46043199 | 2.15E-04 |
| Kyat3 | 444.3764685 | 0.460389075 | 1.12E-02 |
| Large1 | 4373.999407 | 0.460057451 | 1.55E-06 |
| Ppp2r5a | 2519.975068 | 0.459954726 | 3.12E-04 |
| Mogs | 3717.350253 | 0.459850933 | 4.30E-03 |
| Tsacc | 100.5470491 | 0.458908486 | 4.58E-02 |
| Pxk | 1881.676776 | 0.458754858 | 9.15E-08 |
| Ntpcr | 1088.202802 | 0.4585968 | 9.22E-04 |
| Ccdc120 | 311.5972816 | 0.4584216 | 1.93E-02 |
| Osr1 | 5513.095233 | 0.457992651 | 2.71E-02 |
| Nipsnap3b | 762.8209259 | 0.457974183 | 1.36E-03 |
| Gtpbp1 | 2056.74881 | 0.457064062 | 3.66E-03 |
| Cox5b | 4421.831308 | 0.457050792 | 4.35E-05 |
| Tprgl | 2416.991549 | 0.455909875 | 1.07E-02 |
| Llgl1 | 4105.61428 | 0.455881117 | 6.83E-05 |
| Epn2 | 3510.416008 | 0.455874757 | 9.19E-03 |
| Idh3a | 2254.942093 | 0.455826342 | 2.98E-03 |
| Hsd17b4 | 3036.782046 | 0.455717253 | 1.11E-07 |
| Etv3 | 1203.077921 | 0.455595436 | 1.94E-04 |
| Commd9 | 1041.174624 | 0.454997736 | 2.00E-02 |
| Elk1 | 1611.0863 | 0.45472826 | 1.11E-02 |
| Ndufa13 | 3518.981647 | 0.454708241 | 2.57E-03 |
| Abcg2 | 2509.433036 | 0.454554561 | 5.30E-03 |
| Pafah1b3 | 1426.134526 | 0.454227386 | 1.43E-03 |
| Dusp16 | 2072.5505 | 0.454125401 | 4.74E-03 |
| Dpm3 | 443.9062842 | 0.453921742 | 8.16E-03 |
| 1810013L24Rik | 2909.228309 | 0.453638022 | 5.14E-04 |

| | | | |
|---|---|---|---|
| Gm45713 | 119.8688671 | 0.453610721 | 4.29E-02 |
| Pkm | 49897.4209 | 0.453517698 | 8.83E-03 |
| Uap1 | 2457.583466 | 0.453437356 | 1.02E-07 |
| Cyhr1 | 3502.365889 | 0.453226437 | 7.57E-03 |
| Borcs7 | 429.8216862 | 0.453194161 | 9.21E-04 |
| Ppp1r14b | 5784.837077 | 0.453145131 | 1.41E-02 |
| Kdelr3 | 1636.509425 | 0.452592369 | 2.30E-02 |
| Dctn3 | 1006.383455 | 0.452410705 | 6.37E-07 |
| Pacsin2 | 4361.917334 | 0.452145267 | 1.53E-05 |
| Wiz | 3587.376658 | 0.452061719 | 3.29E-05 |
| Wfs1 | 1732.71301 | 0.451601481 | 1.61E-03 |
| Ndufb5 | 2748.937973 | 0.451299852 | 5.86E-03 |
| Ipo4 | 4162.893495 | 0.451285097 | 9.87E-03 |
| Sec61b | 1615.257574 | 0.451269917 | 2.66E-04 |
| Gm7887 | 398.1521651 | 0.451171935 | 8.34E-04 |
| Fxn | 355.7300904 | 0.450984867 | 9.36E-03 |
| 1110004F10Rik | 2600.912527 | 0.450782137 | 1.98E-05 |
| Psmf1 | 1876.021687 | 0.450262536 | 4.16E-03 |
| Vps9d1 | 897.8907511 | 0.449324495 | 1.75E-05 |
| Nup98 | 2513.759391 | 0.449082274 | 8.99E-04 |
| Zswim3 | 309.3906797 | 0.448652953 | 2.15E-02 |
| Jade1 | 3558.396045 | 0.448636442 | 3.96E-03 |
| Pcmt1 | 2281.828033 | 0.448424155 | 1.91E-04 |
| Shf | 903.0088903 | 0.448217479 | 2.25E-02 |
| Arl4a | 298.3612635 | 0.448069597 | 3.45E-03 |
| Rundc3a | 652.5243746 | 0.447898509 | 4.44E-02 |
| Os9 | 7755.641324 | 0.447608148 | 2.94E-03 |
| NA | 1647.516016 | 0.447227185 | 5.04E-03 |
| Slc17a5 | 1194.706084 | 0.446408623 | 1.00E-02 |
| Zfp787 | 485.2637602 | 0.446154944 | 8.83E-03 |
| Tfcp2 | 1065.086979 | 0.445556057 | 6.81E-03 |
| Rilpl1 | 1857.379582 | 0.444851209 | 1.06E-03 |
| Tbc1d22b | 1063.836405 | 0.444730944 | 1.66E-05 |
| Ctnnbl1 | 1369.037269 | 0.444064682 | 3.37E-04 |
| Gm21988 | 501.1876847 | 0.44403314 | 1.37E-02 |
| Cebpz | 3061.302049 | 0.443562481 | 8.32E-05 |
| Pcif1 | 2628.461221 | 0.44339254 | 5.71E-03 |
| Fuom | 175.3865216 | 0.44337047 | 3.54E-02 |
| Bloc1s3 | 433.2225642 | 0.443265453 | 2.47E-02 |
| Nadk2 | 1271.547494 | 0.443005779 | 2.49E-02 |
| Ppp2ca | 10386.53634 | 0.442771561 | 3.38E-06 |
| Med14 | 2332.440218 | 0.442061884 | 2.29E-04 |
| Als2 | 1499.841059 | 0.441934239 | 1.61E-04 |
| Mcrip1 | 3277.7431 | 0.441436291 | 2.32E-02 |
| Zc2hc1a | 2130.526618 | 0.441311572 | 4.74E-03 |
| Zswim4 | 2006.776445 | 0.441026186 | 4.25E-03 |
| Stat5b | 2117.456746 | 0.439372884 | 5.36E-07 |

| | | | |
|---|---|---|---|
| Washc3 | 1500.94525 | 0.439047244 | 4.10E-04 |
| Nrf1 | 1279.08247 | 0.438814039 | 2.84E-03 |
| Naf1 | 1224.57437 | 0.43879628 | 1.07E-04 |
| BC023719 | 452.5926543 | 0.437907043 | 1.30E-02 |
| Chst12 | 2131.435361 | 0.437737102 | 7.99E-03 |
| Zfp553 | 1197.130045 | 0.437180009 | 3.81E-02 |
| Rfx3 | 232.2533201 | 0.436536982 | 3.45E-02 |
| Pdcd11 | 3527.629205 | 0.436282147 | 2.79E-05 |
| Tpgs1 | 719.3056991 | 0.436114933 | 8.16E-03 |
| Retsat | 1353.471053 | 0.434790292 | 6.70E-03 |
| Irak4 | 619.5218289 | 0.434549721 | 3.85E-04 |
| Cbx2 | 830.0959647 | 0.434480182 | 2.56E-02 |
| Hadha | 6973.56177 | 0.434345377 | 1.60E-03 |
| Gpc1 | 3053.071229 | 0.434176707 | 2.02E-02 |
| Arfgap3 | 2436.475969 | 0.433784562 | 9.95E-03 |
| Copb1 | 7208.232283 | 0.433378967 | 4.62E-08 |
| Tma16 | 794.3992447 | 0.433001202 | 3.42E-03 |
| Slc26a6 | 286.9579948 | 0.432953699 | 1.07E-02 |
| Sptbn1 | 59834.79882 | 0.432574683 | 6.88E-03 |
| Borcs6 | 770.1972836 | 0.432097677 | 2.60E-02 |
| Zfp865 | 1022.925073 | 0.431846611 | 6.71E-05 |
| Wdr33 | 2511.881024 | 0.431552116 | 2.40E-04 |
| Mocs1 | 2925.075305 | 0.431184231 | 3.08E-02 |
| Pot1a | 895.8685969 | 0.430825682 | 4.47E-03 |
| Psmd9 | 1420.823399 | 0.430412903 | 4.72E-03 |
| Rnf215 | 897.5323402 | 0.429683849 | 4.24E-05 |
| Ccdc77 | 463.8315296 | 0.428881257 | 2.65E-02 |
| Igf1r | 2901.97391 | 0.428874236 | 1.53E-04 |
| Klhl42 | 1851.794143 | 0.428817168 | 5.35E-06 |
| Nucb1 | 11251.76517 | 0.427789865 | 2.04E-04 |
| Dtd2 | 514.8764301 | 0.427394607 | 5.35E-04 |
| Rchy1 | 1824.914905 | 0.427347388 | 2.82E-03 |
| Zfp263 | 1718.195765 | 0.426392046 | 5.85E-03 |
| Wwp1 | 1335.549327 | 0.425866434 | 4.21E-02 |
| Med26 | 634.3945367 | 0.425386126 | 2.70E-03 |
| Rap2b | 1894.249099 | 0.42523276 | 2.23E-04 |
| Ube2e2 | 954.284212 | 0.42503862 | 1.48E-03 |
| Eloa | 2936.855797 | 0.424899966 | 1.79E-06 |
| Tpt1 | 131718.7681 | 0.42465021 | 3.56E-02 |
| Ldb1 | 7917.754413 | 0.424200917 | 7.53E-03 |
| 8-Sep | 10704.68529 | 0.424066341 | 3.23E-04 |
| Atp5l | 4692.184 | 0.423982574 | 1.38E-02 |
| Fam185a | 517.0409463 | 0.423875693 | 1.87E-02 |
| Got2 | 9551.023887 | 0.423764824 | 1.41E-02 |
| Fbln5 | 11685.97594 | 0.423647075 | 3.04E-02 |
| Aars | 20365.17343 | 0.423640616 | 2.41E-04 |
| Kmt2b | 2483.737405 | 0.422795399 | 1.31E-02 |

| | | | |
|---|---|---|---|
| Psmg4 | 256.9783017 | 0.422627949 | 1.16E-02 |
| NA | 713.1694764 | 0.422386406 | 1.02E-02 |
| Pgs1 | 1768.015867 | 0.421948516 | 1.52E-03 |
| Ate1 | 1420.82635 | 0.421424426 | 2.15E-03 |
| Brap | 2566.325462 | 0.421127913 | 4.94E-03 |
| Tbpl1 | 1549.163799 | 0.420952316 | 1.16E-04 |
| Plpp1 | 5028.842612 | 0.420904389 | 2.97E-02 |
| Smarca4 | 6267.623213 | 0.42019526 | 4.48E-04 |
| Ddx47 | 2966.178771 | 0.419929235 | 1.01E-04 |
| Fbxo34 | 895.2201394 | 0.4198992 | 9.92E-04 |
| Mex3d | 2980.139648 | 0.419677316 | 4.31E-02 |
| Tsfm | 720.1176681 | 0.419568866 | 1.17E-02 |
| Lamtor5 | 1897.270012 | 0.419516374 | 1.86E-04 |
| Mtif3 | 797.1508615 | 0.419317326 | 1.69E-02 |
| Xpot | 6963.581493 | 0.419213245 | 3.25E-03 |
| Snx29 | 375.6562173 | 0.419066178 | 4.29E-02 |
| Cdc42 | 11259.05128 | 0.418374092 | 5.88E-08 |
| Cep76 | 375.9843548 | 0.418014094 | 2.12E-02 |
| Rfk | 2308.638808 | 0.41795157 | 2.09E-04 |
| Cers4 | 1306.675198 | 0.417666679 | 4.47E-04 |
| Prpf4 | 1801.281111 | 0.417658044 | 2.54E-03 |
| Dguok | 1124.366048 | 0.417595171 | 3.68E-03 |
| Timm13 | 1691.713944 | 0.417333133 | 6.56E-03 |
| Epg5 | 1506.603937 | 0.416882222 | 5.12E-07 |
| Zdhhc18 | 1133.727077 | 0.416833714 | 2.00E-02 |
| Derl1 | 5766.467668 | 0.416517486 | 1.98E-03 |
| Cln5 | 1783.706728 | 0.415955767 | 5.69E-03 |
| Ptpn2 | 736.8581651 | 0.415319361 | 3.15E-03 |
| Fkbp8 | 6857.949609 | 0.414975331 | 2.28E-03 |
| Metap2 | 6994.685675 | 0.414727719 | 1.37E-03 |
| Nudcd3 | 5085.866745 | 0.414342159 | 6.66E-03 |
| Dnajb4 | 1668.421695 | 0.413760998 | 3.88E-03 |
| Cc2d1a | 2511.210551 | 0.413526571 | 3.94E-03 |
| Maf1 | 2954.734566 | 0.413482237 | 1.64E-02 |
| Dnaaf5 | 936.3481781 | 0.413071134 | 2.10E-03 |
| Xpo7 | 3778.843741 | 0.413038396 | 7.58E-06 |
| Foxn3 | 857.0623571 | 0.412856837 | 2.01E-02 |
| Entr1 | 2471.686962 | 0.412722205 | 7.80E-05 |
| Sec23b | 2213.107425 | 0.411721421 | 3.55E-05 |
| Wdr3 | 1937.156413 | 0.411558191 | 3.83E-03 |
| Txnrd3 | 885.4457633 | 0.411156835 | 6.79E-03 |
| Huwe1 | 19835.01404 | 0.410837799 | 3.97E-06 |
| Eci1 | 1615.091732 | 0.410833723 | 1.41E-03 |
| Tbrg4 | 2034.760982 | 0.410666041 | 1.14E-02 |
| Crat | 2486.540887 | 0.410357896 | 8.00E-05 |
| Pias1 | 1417.343596 | 0.410304842 | 6.73E-05 |
| Fbxo22 | 1204.372058 | 0.410146265 | 8.90E-06 |

| | | | |
|---|---|---|---|
| Snrpb2 | 1610.992456 | 0.40991746 | 6.21E-03 |
| Zbtb17 | 993.1594324 | 0.40870984 | 2.99E-02 |
| Pgls | 2664.443408 | 0.408574052 | 3.57E-02 |
| Cpne1 | 5044.131924 | 0.407997293 | 1.92E-07 |
| Habp4 | 1289.065897 | 0.407950526 | 1.66E-08 |
| Angel2 | 1995.898618 | 0.407873 | 5.18E-03 |
| Mapre1 | 6524.825861 | 0.407685004 | 7.05E-05 |
| AU020206 | 146.3601913 | 0.407434859 | 2.92E-02 |
| Retreg3 | 2865.0767 | 0.406707366 | 1.28E-03 |
| Usf3 | 1381.631125 | 0.406694663 | 1.44E-02 |
| Csnk1d | 6941.480387 | 0.406237127 | 9.57E-04 |
| Ubald1 | 2040.828702 | 0.405082002 | 4.30E-02 |
| Gm12338 | 2281.166651 | 0.405022563 | 1.82E-02 |
| Kctd10 | 13098.86895 | 0.404860609 | 9.44E-05 |
| Dpp3 | 3369.61079 | 0.40484569 | 1.19E-02 |
| Uggt1 | 8254.05786 | 0.404754473 | 1.16E-04 |
| Hjurp | 5943.468661 | 0.404693864 | 2.43E-02 |
| Sdhaf1 | 433.54347 | 0.404692072 | 4.73E-02 |
| Fastkd1 | 692.7686316 | 0.404106629 | 3.63E-04 |
| Abcf1 | 5975.295718 | 0.404098625 | 3.85E-05 |
| Cbx7 | 739.4739341 | 0.403864808 | 4.56E-03 |
| Sfxn4 | 238.9702114 | 0.403412762 | 4.61E-02 |
| Ecd | 1522.526592 | 0.402585866 | 2.70E-04 |
| Ggnbp2 | 4298.82299 | 0.402578722 | 1.24E-02 |
| Tpp2 | 4539.424409 | 0.402525993 | 6.59E-05 |
| Dnajc24 | 378.3848799 | 0.401869183 | 2.71E-03 |
| Plcg1 | 7566.406721 | 0.401592912 | 6.52E-03 |
| Nelfb | 2470.739406 | 0.401552179 | 4.47E-04 |
| Dxo | 631.7096353 | 0.401362457 | 7.48E-04 |
| Scpep1 | 18619.66479 | 0.401251677 | 3.58E-03 |
| Eapp | 1456.644053 | 0.400801101 | 4.77E-03 |
| Wipi1 | 5221.186575 | 0.400518561 | 2.27E-04 |
| Wipf2 | 1118.249451 | 0.40051781 | 3.74E-02 |
| Pias3 | 1708.261894 | 0.400304533 | 2.02E-03 |
| Sdhaf2 | 1871.044111 | 0.400012633 | 8.03E-04 |
| Rangrf | 427.5682787 | 0.399920222 | 1.08E-02 |
| Fbxw17 | 660.5170251 | 0.399747474 | 2.39E-03 |
| Dand5 | 170.4514732 | 0.399568939 | 1.64E-02 |
| Gm10698 | 709.1241477 | 0.399233865 | 4.63E-02 |
| Zfp9 | 1864.128904 | 0.399192168 | 1.64E-03 |
| Hmbs | 1209.092363 | 0.398737719 | 1.25E-02 |
| Polr2j | 1633.58678 | 0.397901087 | 7.12E-05 |
| Rcan3 | 1082.07026 | 0.397354557 | 3.49E-03 |
| Ipo11 | 1848.149906 | 0.397285387 | 2.03E-02 |
| Haus7 | 723.0560045 | 0.396762039 | 4.69E-03 |
| Zcrb1 | 1391.467671 | 0.396575538 | 2.70E-05 |
| Immp1l | 814.9343149 | 0.396544727 | 6.50E-03 |

| | | | |
|---|---|---|---|
| Pgm3 | 874.3873228 | 0.396402439 | 2.25E-03 |
| Rrp8 | 1637.732775 | 0.396146027 | 4.62E-02 |
| Papss1 | 3988.897528 | 0.39590515 | 2.22E-03 |
| Atp2a2 | 15822.23775 | 0.395750308 | 1.13E-02 |
| Pter | 781.8123917 | 0.395497457 | 7.76E-04 |
| Stxbp1 | 3426.852098 | 0.395491148 | 1.21E-02 |
| Yod1 | 191.6414735 | 0.395268397 | 2.14E-02 |
| Cdk17 | 1107.511682 | 0.394942011 | 2.78E-04 |
| Zcchc10 | 277.3639513 | 0.394887569 | 3.70E-02 |
| Serinc3 | 12902.53338 | 0.394662312 | 7.89E-04 |
| Kremen1 | 3328.728218 | 0.394378559 | 2.02E-03 |
| Edem2 | 2018.732178 | 0.394238269 | 8.74E-04 |
| Srebf2 | 11304.7876 | 0.394192284 | 1.15E-02 |
| Pdap1 | 5185.52205 | 0.394185265 | 4.28E-04 |
| Rbm22 | 1683.263056 | 0.394003108 | 4.16E-03 |
| Fdx1l | 902.2452556 | 0.393952896 | 1.11E-02 |
| Auh | 961.4463672 | 0.393943855 | 3.33E-02 |
| Gm13443 | 688.7594814 | 0.393530413 | 9.54E-04 |
| Rpusd1 | 979.1902002 | 0.392172499 | 7.08E-03 |
| Rpain | 423.7311747 | 0.391677229 | 3.22E-02 |
| 2310009B15Rik | 380.4070517 | 0.391663245 | 3.92E-02 |
| Atpif1 | 2983.892833 | 0.391460469 | 1.36E-03 |
| Ctdnep1 | 4303.325777 | 0.391328248 | 1.22E-02 |
| Glce | 2652.834273 | 0.391289131 | 3.94E-02 |
| Enoph1 | 817.0058006 | 0.391012615 | 1.29E-02 |
| Fbxo31 | 875.7258759 | 0.3909941 | 3.22E-02 |
| Kdm5a | 4716.918006 | 0.390903717 | 3.21E-03 |
| Trmt6 | 1442.51025 | 0.39081475 | 9.46E-03 |
| Mthfsd | 1470.292497 | 0.390558496 | 7.37E-07 |
| Pop7 | 475.5727012 | 0.390285471 | 1.19E-02 |
| Zfp106 | 11251.11847 | 0.390147233 | 7.79E-09 |
| Calhm5 | 378.443981 | 0.390082347 | 2.96E-02 |
| Kdm5c | 4324.335109 | 0.390015628 | 1.59E-05 |
| Top3a | 748.5691303 | 0.389771908 | 2.66E-02 |
| 11-Sep | 12235.26164 | 0.389685797 | 3.11E-03 |
| Gpt2 | 2629.31278 | 0.389468024 | 3.71E-02 |
| E2f4 | 3205.637373 | 0.389427827 | 3.91E-04 |
| Pigk | 4249.556601 | 0.389289414 | 4.40E-03 |
| Tab1 | 1195.635095 | 0.38907146 | 1.24E-02 |
| Ccdc127 | 2076.615621 | 0.388989664 | 1.97E-05 |
| Irf2bp1 | 1235.539186 | 0.388929455 | 1.25E-02 |
| Trrap | 7929.435386 | 0.388787253 | 4.96E-02 |
| Lmbrd1 | 2417.679745 | 0.388628395 | 2.49E-03 |
| Pgrmc1 | 9425.547064 | 0.388259301 | 3.45E-03 |
| Fam53c | 2241.692611 | 0.388069159 | 9.07E-06 |
| NA | 998.8343986 | 0.387130728 | 1.11E-02 |
| Gpat4 | 4294.080592 | 0.387102098 | 2.32E-03 |

| | | | |
|---|---|---|---|
| Mink1 | 2720.963999 | 0.387083182 | 1.45E-03 |
| Tsen34 | 2980.73204 | 0.386510571 | 3.67E-03 |
| Lsm3 | 1116.592052 | 0.386333593 | 3.17E-02 |
| Ttc21b | 2440.384871 | 0.386298252 | 2.12E-05 |
| Flot1 | 2483.620995 | 0.386172127 | 2.68E-02 |
| Gcc1 | 1481.838615 | 0.386123902 | 1.81E-02 |
| Uck1 | 1754.310657 | 0.385951315 | 1.99E-03 |
| Serf2 | 9414.865504 | 0.385701301 | 6.13E-05 |
| Glb1 | 2857.56341 | 0.385553691 | 7.32E-04 |
| Strap | 7041.803168 | 0.385044911 | 1.83E-04 |
| Mapk1 | 9186.482385 | 0.384596816 | 1.21E-05 |
| Hivep1 | 1746.576588 | 0.384296444 | 2.05E-04 |
| Arl6ip4 | 1389.976539 | 0.383943908 | 1.90E-04 |
| Adamtsl5 | 3366.847999 | 0.383784543 | 2.62E-03 |
| Map1s | 2799.329107 | 0.383202216 | 2.18E-02 |
| Tnip1 | 1654.714596 | 0.383101055 | 2.61E-02 |
| Oma1 | 581.1175536 | 0.382985591 | 1.53E-02 |
| Ap2b1 | 9895.76698 | 0.382816851 | 2.74E-05 |
| Med10 | 1079.70181 | 0.382491411 | 8.79E-04 |
| Il17ra | 1539.540835 | 0.382117088 | 6.07E-04 |
| Sbf2 | 4819.618936 | 0.380961584 | 1.66E-07 |
| Rnf38 | 1227.714967 | 0.380279127 | 4.41E-02 |
| Sec61a1 | 12275.49602 | 0.380028032 | 5.23E-04 |
| Mtfmt | 419.5935587 | 0.380000865 | 4.55E-03 |
| Utp6 | 2172.945915 | 0.379685573 | 1.07E-02 |
| Clns1a | 2100.979167 | 0.379273481 | 2.62E-04 |
| Mbd6 | 2281.456436 | 0.379088713 | 2.28E-02 |
| Ccnd3 | 4542.864049 | 0.378670031 | 3.78E-02 |
| Mrgbp | 899.3957712 | 0.378280082 | 7.81E-05 |
| Paics | 14439.95988 | 0.37737776 | 7.85E-08 |
| Tfpt | 416.8764704 | 0.377364292 | 2.10E-02 |
| Tomm7 | 2106.696315 | 0.377272246 | 8.45E-03 |
| Sdhd | 3094.16066 | 0.376465279 | 1.84E-02 |
| Rxylt1 | 2203.764838 | 0.376439425 | 1.50E-03 |
| Snrpf | 1289.863239 | 0.376262173 | 4.66E-02 |
| Nup50 | 3476.731492 | 0.376236711 | 1.16E-02 |
| Mtrr | 953.9515698 | 0.376004592 | 4.41E-03 |
| Stard5 | 614.9927071 | 0.375972165 | 1.50E-02 |
| Fbxo28 | 1806.420033 | 0.375941937 | 6.46E-04 |
| Eva1b | 2403.016199 | 0.374690028 | 3.05E-02 |
| Mrps34 | 1027.085385 | 0.374266973 | 1.95E-02 |
| Triqk | 268.8786932 | 0.374163122 | 2.11E-02 |
| Bbip1 | 680.608969 | 0.374113337 | 4.44E-02 |
| Ralgapb | 4220.545798 | 0.373850898 | 5.72E-07 |
| Elavl1 | 3244.196393 | 0.372730974 | 8.93E-03 |
| Nkx2-5 | 211.3079351 | 0.372574759 | 4.57E-02 |
| Foxo4 | 1470.022901 | 0.37219785 | 3.52E-03 |

| | | | |
|---|---|---|---|
| Elp3 | 2915.615554 | 0.372125605 | 9.50E-03 |
| Apc2 | 1048.209407 | 0.371697885 | 4.73E-03 |
| Eef2kmt | 674.7200596 | 0.371355167 | 3.23E-03 |
| Cacul1 | 4722.619289 | 0.371315777 | 6.60E-05 |
| Ints14 | 965.0288745 | 0.370917961 | 1.92E-03 |
| Dock6 | 1238.663641 | 0.370765457 | 9.64E-03 |
| Stt3b | 6004.274142 | 0.370526183 | 2.51E-04 |
| Fbxw2 | 2586.135429 | 0.369024715 | 3.64E-07 |
| Ltn1 | 2194.469987 | 0.3686944 | 1.26E-05 |
| Arpc4 | 4891.018247 | 0.367509932 | 8.27E-03 |
| N4bp1 | 3589.986201 | 0.367257145 | 2.49E-06 |
| NA | 256.5573422 | 0.367168911 | 1.31E-02 |
| Stub1 | 5148.132786 | 0.367131562 | 1.49E-02 |
| Slc38a1 | 5119.465128 | 0.366881709 | 3.93E-05 |
| Scarb2 | 3430.447569 | 0.366525068 | 1.20E-02 |
| Trim25 | 2953.14383 | 0.366194794 | 1.92E-03 |
| Rrp36 | 1042.505097 | 0.366055675 | 1.54E-03 |
| Rbm15 | 572.4601641 | 0.366006443 | 3.96E-02 |
| Ap1m1 | 3076.866517 | 0.365493759 | 2.73E-03 |
| Smyd5 | 3133.949759 | 0.36505324 | 1.16E-02 |
| Zfp438 | 226.8653021 | 0.364665452 | 1.69E-02 |
| Fam117b | 1725.697617 | 0.363956489 | 2.07E-03 |
| Zfp830 | 698.123764 | 0.36366638 | 7.31E-04 |
| Dop1b | 1500.10808 | 0.363347508 | 3.33E-02 |
| Banp | 1329.834469 | 0.363344816 | 3.61E-02 |
| Prrc2b | 13572.01384 | 0.363297476 | 7.88E-04 |
| Sik2 | 1615.083702 | 0.362832484 | 1.74E-02 |
| Aspscr1 | 1850.262874 | 0.362799405 | 9.86E-04 |
| Fam207a | 1843.838055 | 0.362680608 | 1.29E-02 |
| Adipor2 | 3974.141308 | 0.362609588 | 1.09E-03 |
| Pcgf1 | 643.0125964 | 0.361858666 | 3.41E-03 |
| Dnajc8 | 3861.33955 | 0.361841749 | 2.33E-05 |
| Ric1 | 2773.699318 | 0.361575991 | 3.86E-03 |
| Tpp1 | 8880.766729 | 0.361142756 | 3.23E-02 |
| Taf1c | 527.9761851 | 0.360601279 | 1.06E-02 |
| Rps6kb2 | 734.5964013 | 0.359671281 | 2.05E-02 |
| Tmem33 | 2683.243717 | 0.359382143 | 7.77E-05 |
| Arfgap2 | 3236.509744 | 0.359042028 | 7.69E-04 |
| 5-Mar | 3832.102311 | 0.358472469 | 1.54E-06 |
| Piga | 562.1998323 | 0.356664594 | 4.18E-02 |
| Armt1 | 1217.678356 | 0.35636703 | 1.46E-03 |
| Lmo4 | 1591.794629 | 0.356036333 | 2.93E-04 |
| Mark3 | 3507.222576 | 0.355709595 | 6.82E-06 |
| Terf2ip | 998.8334094 | 0.355627707 | 3.07E-02 |
| Mapk9 | 2993.864 | 0.355157635 | 1.58E-04 |
| Mtr | 885.9411703 | 0.355041659 | 3.64E-02 |
| Srp19 | 1021.369308 | 0.355035675 | 4.83E-02 |

| | | | |
|---|---|---|---|
| Dapp1 | 542.9780254 | 0.354484834 | 1.74E-02 |
| Dennd5a | 6370.596711 | 0.352725523 | 2.25E-06 |
| Gpr173 | 432.4446883 | 0.35229183 | 1.16E-02 |
| Arpc5l | 2244.355316 | 0.35217748 | 4.40E-03 |
| Nnt | 4601.606286 | 0.35197501 | 2.37E-07 |
| Synj1 | 1193.434325 | 0.351934003 | 2.92E-02 |
| Klhdc4 | 649.4677115 | 0.351123764 | 1.13E-02 |
| Ldlrad4 | 327.7250835 | 0.350992658 | 4.50E-02 |
| Hsf1 | 2362.532022 | 0.350938459 | 4.70E-03 |
| Wnk1 | 21009.8334 | 0.350832322 | 2.78E-02 |
| Immt | 7540.562334 | 0.350678018 | 2.82E-03 |
| Rnf130 | 1604.760353 | 0.350106821 | 1.03E-02 |
| Capza2 | 8096.038169 | 0.349616546 | 1.55E-02 |
| Ndel1 | 3898.669197 | 0.349399348 | 1.96E-02 |
| Tulp3 | 4525.053787 | 0.349273062 | 1.79E-03 |
| Cct3 | 10769.55899 | 0.349055982 | 4.82E-02 |
| Eif4g1 | 31562.38989 | 0.348328455 | 1.70E-03 |
| Sars2 | 394.5444787 | 0.348212844 | 3.33E-02 |
| Mtg2 | 797.9187037 | 0.347135813 | 6.14E-03 |
| Sin3a | 3188.735018 | 0.347092307 | 2.82E-02 |
| Nisch | 16444.12916 | 0.34679464 | 2.01E-03 |
| Rps6kc1 | 1738.429506 | 0.346639675 | 1.17E-02 |
| Trim32 | 3371.688236 | 0.346593277 | 4.95E-03 |
| Ppp5c | 2025.354471 | 0.346501948 | 2.61E-02 |
| Dyrk1a | 2184.20678 | 0.346434389 | 9.31E-03 |
| Tpi1 | 16501.50437 | 0.346315156 | 3.67E-02 |
| Coasy | 1306.789236 | 0.346055356 | 2.35E-03 |
| Psmd6 | 2606.420377 | 0.345858892 | 4.26E-03 |
| Tsc2 | 3734.861746 | 0.345679197 | 1.51E-03 |
| Srprb | 2065.424096 | 0.345070216 | 2.86E-02 |
| Mettl2 | 1593.731798 | 0.34492884 | 1.69E-03 |
| Brk1 | 5138.533732 | 0.344878716 | 4.16E-03 |
| Upf1 | 2598.878259 | 0.34464424 | 1.15E-02 |
| Lypla2 | 1388.392472 | 0.344251185 | 8.18E-03 |
| Tgds | 791.4115282 | 0.344039679 | 3.66E-03 |
| Nfrkb | 1182.921367 | 0.34396696 | 4.50E-03 |
| Stat6 | 5625.010172 | 0.343412314 | 3.19E-05 |
| Rab6a | 3433.18195 | 0.343071534 | 1.29E-03 |
| Acp6 | 792.6797913 | 0.342853195 | 4.63E-03 |
| Atxn7l1 | 1301.845341 | 0.342820743 | 2.13E-02 |
| 4930453N24Rik | 1049.629045 | 0.342814906 | 1.10E-02 |
| Myo9b | 8108.594078 | 0.342283277 | 5.42E-04 |
| Apeh | 1943.694711 | 0.342012988 | 5.98E-03 |
| Stk25 | 4558.968745 | 0.341934879 | 1.73E-05 |
| B3gntl1 | 339.1393466 | 0.341911038 | 9.91E-03 |
| BC002163 | 1895.01447 | 0.341357123 | 3.59E-02 |
| Tarbp2 | 899.8778158 | 0.341334561 | 1.21E-02 |

| | | | |
|---|---|---|---|
| Prdm10 | 358.6740737 | 0.340441457 | 4.35E-02 |
| Sptlc2 | 3897.874693 | 0.340122155 | 1.32E-03 |
| Ik | 3973.641719 | 0.339634461 | 4.12E-05 |
| Samd4 | 3008.136996 | 0.339514959 | 1.16E-02 |
| Fbxl6 | 1647.638682 | 0.339245781 | 2.00E-02 |
| Tada2a | 1216.281985 | 0.339223242 | 2.24E-02 |
| Nup188 | 4987.880274 | 0.338779229 | 1.04E-02 |
| Gaa | 16101.78382 | 0.338567119 | 4.09E-02 |
| Slf2 | 3579.020865 | 0.338136519 | 4.50E-02 |
| Wdr41 | 1253.631657 | 0.337811244 | 9.96E-05 |
| Reep5 | 3384.547633 | 0.337724927 | 4.58E-02 |
| Gm29394 | 1296.624358 | 0.336985196 | 1.26E-02 |
| Rnf7 | 1866.571505 | 0.336440508 | 6.05E-04 |
| Hars | 2334.883029 | 0.33550691 | 3.40E-02 |
| Tdrd7 | 1895.989433 | 0.335404269 | 1.07E-03 |
| Trappc6b | 1441.26128 | 0.334879009 | 5.04E-04 |
| Bcl7a | 1302.545937 | 0.334700522 | 5.44E-03 |
| Gins4 | 1158.512093 | 0.334618899 | 2.15E-02 |
| NA | 1786.724464 | 0.334589744 | 3.06E-02 |
| Elovl5 | 2213.912141 | 0.333803674 | 2.72E-03 |
| Med4 | 736.5183557 | 0.333007564 | 1.05E-02 |
| Pigb | 356.954535 | 0.332860467 | 3.07E-02 |
| Ufd1 | 2067.713197 | 0.332395914 | 3.06E-03 |
| Smim7 | 2421.951949 | 0.332058206 | 7.85E-03 |
| Hspbp1 | 928.0207555 | 0.331982095 | 1.18E-02 |
| Vhl | 1585.185787 | 0.331883942 | 2.89E-02 |
| Hmgn1 | 7690.882493 | 0.331812261 | 3.46E-02 |
| Mpc2 | 1599.611664 | 0.331791864 | 2.56E-02 |
| Atp13a2 | 1312.41158 | 0.331172456 | 6.12E-03 |
| Fermt2 | 3801.900806 | 0.330969805 | 2.03E-02 |
| Tmcc1 | 1566.82676 | 0.329822274 | 1.35E-02 |
| Surf4 | 13492.55385 | 0.329662192 | 1.77E-02 |
| Add1 | 14766.81535 | 0.329604639 | 2.39E-03 |
| Mkln1 | 4393.243402 | 0.329593334 | 4.79E-05 |
| Coq9 | 1669.857785 | 0.329560367 | 3.11E-02 |
| Actr3 | 13135.49264 | 0.328421507 | 3.35E-03 |
| Bud31 | 2133.400063 | 0.328279009 | 3.94E-03 |
| Ndufs7 | 1838.714035 | 0.328060103 | 5.88E-03 |
| Yeats4 | 1397.088105 | 0.328030735 | 1.77E-03 |
| Gpx8 | 2857.904981 | 0.32768844 | 1.15E-04 |
| Tubgcp4 | 2134.913717 | 0.327598842 | 9.91E-04 |
| Slc38a6 | 9317.07918 | 0.327329213 | 1.46E-03 |
| Rmnd5b | 897.9283373 | 0.327023011 | 1.50E-02 |
| Crlf2 | 952.9025188 | 0.326211867 | 3.61E-02 |
| Tcp1 | 15233.79817 | 0.326197003 | 1.35E-04 |
| Fam114a2 | 3449.452351 | 0.326166157 | 2.42E-05 |
| Nras | 3618.572248 | 0.325346101 | 1.06E-04 |

| | | | |
|---|---|---|---|
| Rhot1 | 2347.280075 | 0.325085162 | 1.02E-04 |
| Usp4 | 3655.583413 | 0.324725343 | 1.48E-03 |
| Elp1 | 4443.627689 | 0.32454245 | 1.88E-02 |
| Smg7 | 3227.148695 | 0.323759154 | 3.11E-03 |
| Rbpms | 1460.507555 | 0.323674884 | 4.25E-02 |
| Bicra | 720.7496501 | 0.323396197 | 2.80E-02 |
| Rgp1 | 2788.044564 | 0.323099589 | 2.44E-03 |
| Zfp513 | 1098.824088 | 0.322827376 | 1.59E-04 |
| Noa1 | 980.0937759 | 0.322597863 | 2.60E-02 |
| Chkb | 1184.070151 | 0.322158597 | 4.34E-03 |
| Fbxo4 | 853.3693442 | 0.322054714 | 1.25E-02 |
| Snrpa | 2781.442247 | 0.321631298 | 9.36E-03 |
| Lrsam1 | 1151.868033 | 0.321247269 | 3.04E-02 |
| Lsm14b | 2293.235444 | 0.321165844 | 4.30E-02 |
| Mxi1 | 1578.380318 | 0.319754594 | 4.50E-02 |
| Brd4 | 6239.995293 | 0.319551267 | 1.29E-02 |
| Gys1 | 2048.955579 | 0.31878368 | 1.20E-02 |
| Atp5s | 258.5845938 | 0.318710437 | 2.16E-02 |
| Qtrt1 | 428.9958719 | 0.318620716 | 3.55E-02 |
| Bcl9l | 4861.60902 | 0.318438291 | 2.01E-03 |
| Gorasp2 | 5275.470705 | 0.318310963 | 5.85E-03 |
| Cdk9 | 2985.525803 | 0.317822071 | 7.01E-03 |
| D17Wsu92e | 5640.635753 | 0.317533284 | 5.03E-03 |
| Slc20a2 | 2786.725506 | 0.317296096 | 4.21E-02 |
| Rapgef3 | 465.241526 | 0.31650529 | 3.63E-02 |
| Mtrf1 | 201.4749012 | 0.315820216 | 4.30E-02 |
| Rnf41 | 3468.949675 | 0.315528555 | 2.42E-02 |
| Acad11 | 584.7985704 | 0.315073539 | 2.58E-02 |
| Nudt3 | 2818.544077 | 0.314334803 | 1.18E-02 |
| Praf2 | 2315.534915 | 0.314170819 | 1.04E-02 |
| Ranbp10 | 2194.498125 | 0.314027456 | 7.41E-04 |
| Ptp4a2 | 11669.3148 | 0.31370781 | 1.66E-02 |
| Sf3b6 | 2064.9888 | 0.311373366 | 2.67E-04 |
| Ipp | 573.9750255 | 0.311245838 | 1.30E-02 |
| Ilf2 | 3066.686516 | 0.311128469 | 2.17E-02 |
| Thumpd2 | 335.1465948 | 0.31074927 | 4.92E-03 |
| Pdia6 | 14361.81594 | 0.309638095 | 3.34E-02 |
| Trmt2b | 1368.574183 | 0.309413895 | 1.28E-02 |
| Rbm17 | 4210.243158 | 0.309295439 | 1.92E-03 |
| NA | 387.3489469 | 0.307439028 | 2.60E-02 |
| Zfp277 | 1107.154918 | 0.306947094 | 7.60E-03 |
| Usp21 | 1961.571321 | 0.306758557 | 4.20E-03 |
| Zbtb10 | 572.374976 | 0.306625986 | 4.48E-02 |
| Ccm2 | 2148.139071 | 0.306583327 | 2.41E-02 |
| Map4k4 | 13173.18194 | 0.305833048 | 4.24E-04 |
| App | 18994.67596 | 0.305398752 | 4.02E-02 |
| Rbbp5 | 1649.334514 | 0.30391412 | 2.70E-03 |

| | | | |
|---|---|---|---|
| Atp6v1c1 | 4552.101365 | 0.303772537 | 4.73E-02 |
| Cdk16 | 7880.240212 | 0.303422803 | 6.54E-03 |
| Srsf4 | 2159.848775 | 0.302471576 | 9.89E-03 |
| Sprtn | 812.7061567 | 0.302234876 | 1.83E-02 |
| Tmub2 | 1734.236763 | 0.301998723 | 4.06E-02 |
| Mief1 | 1553.193963 | 0.301727692 | 1.27E-03 |
| Siah1a | 745.6340288 | 0.301649628 | 1.10E-02 |
| Mettl8 | 622.6240002 | 0.3014557 | 2.95E-02 |
| Sap130 | 2107.661239 | 0.301226988 | 1.70E-02 |
| Map3k10 | 1637.445681 | 0.300784466 | 4.75E-02 |
| Ankrd13c | 1592.952556 | 0.300746462 | 9.36E-03 |
| Chmp4b | 6534.757523 | 0.300031056 | 2.18E-02 |
| Khsrp | 4308.732968 | 0.299566641 | 1.63E-03 |
| Clptm1l | 4291.729412 | 0.299372306 | 3.14E-02 |
| Alg1 | 1213.748362 | 0.299301948 | 1.38E-02 |
| 6-Mar | 3809.295394 | 0.298519254 | 1.27E-02 |
| Zrsr2 | 1460.694365 | 0.298025203 | 1.25E-02 |
| Dda1 | 2103.385569 | 0.297841572 | 2.27E-02 |
| Kif3c | 2739.168318 | 0.297183941 | 1.49E-03 |
| Aco1 | 2755.25536 | 0.29669251 | 1.08E-03 |
| Mon2 | 5512.525877 | 0.296442908 | 2.47E-03 |
| Arel1 | 2008.331544 | 0.296326209 | 3.33E-03 |
| Pdcl | 1348.652634 | 0.296084938 | 5.26E-03 |
| Tm9sf4 | 3990.326375 | 0.296000637 | 3.40E-02 |
| Tmem63a | 1994.571198 | 0.29548759 | 1.12E-02 |
| Lpcat3 | 2591.326237 | 0.2953871 | 3.36E-02 |
| Neo1 | 4545.56012 | 0.295054163 | 8.49E-03 |
| D11Wsu47e | 413.81489 | 0.294934528 | 3.73E-02 |
| Aatf | 2596.894846 | 0.294300319 | 1.04E-02 |
| Sec24b | 3696.789891 | 0.294175162 | 4.89E-03 |
| Noct | 8107.901444 | 0.293975441 | 1.06E-02 |
| Emp2 | 4062.72683 | 0.293960471 | 2.26E-03 |
| Mtrex | 2716.987977 | 0.292593987 | 2.07E-03 |
| Ccdc71 | 1481.215184 | 0.292318849 | 2.09E-02 |
| Tex264 | 2795.650085 | 0.291126123 | 2.36E-02 |
| Plxna1 | 20953.16237 | 0.290807529 | 1.02E-04 |
| Ccz1 | 2351.573949 | 0.290493381 | 5.02E-04 |
| Dazap2 | 7672.178723 | 0.289963919 | 4.60E-02 |
| Nup88 | 3246.624486 | 0.28987968 | 2.68E-04 |
| Zfyve27 | 1433.895386 | 0.289681504 | 1.27E-03 |
| Dync1li1 | 2155.249373 | 0.28962949 | 5.39E-03 |
| Impa1 | 1974.053515 | 0.288049306 | 9.66E-03 |
| Hddc2 | 804.2135858 | 0.287113489 | 1.43E-02 |
| Rfx7 | 1726.032941 | 0.286808958 | 5.95E-03 |
| Myd88 | 1070.318763 | 0.286337964 | 2.91E-02 |
| Src | 2935.32324 | 0.285799734 | 1.38E-02 |
| Exoc2 | 1768.947934 | 0.284899542 | 6.29E-03 |

| | | | |
|---|---|---|---|
| Psmg3 | 501.873255 | 0.284844385 | 4.68E-02 |
| Atp5g3 | 7645.161236 | 0.283175326 | 2.85E-02 |
| Vdac2 | 9351.945713 | 0.282470546 | 1.77E-02 |
| Rnf20 | 2420.762924 | 0.281499774 | 3.95E-04 |
| Wdfy1 | 2965.331295 | 0.281270351 | 5.45E-03 |
| Mycbp2 | 4214.136191 | 0.28007727 | 1.16E-02 |
| Slx1b | 369.2781855 | 0.279824106 | 1.34E-02 |
| Cox16 | 493.8339891 | 0.279294811 | 1.27E-02 |
| Senp3 | 3435.215381 | 0.278791014 | 1.28E-03 |
| Atg4b | 2480.845886 | 0.278758623 | 9.73E-03 |
| Rbck1 | 3641.012485 | 0.277509033 | 1.98E-02 |
| Eif4a1 | 28084.96844 | 0.276966415 | 1.30E-02 |
| Agfg1 | 2748.187289 | 0.276310261 | 5.03E-03 |
| Prdx2 | 10872.40196 | 0.276175791 | 4.93E-02 |
| Lhfp | 4843.198567 | 0.275727409 | 4.06E-03 |
| Cpsf7 | 3483.078608 | 0.275363229 | 4.81E-04 |
| Dhx35 | 1030.525777 | 0.274993342 | 2.13E-02 |
| Ccdc6 | 2420.855025 | 0.274303405 | 9.98E-03 |
| Mfsd10 | 2347.017728 | 0.27407548 | 3.51E-02 |
| Hnrnpul1 | 6740.962743 | 0.2734644 | 4.42E-03 |
| Rnmt | 1215.05708 | 0.271070092 | 4.15E-02 |
| Arpc1a | 4104.665188 | 0.269991816 | 4.16E-02 |
| Pank2 | 2024.787923 | 0.269527177 | 4.67E-03 |
| Spg7 | 2088.585806 | 0.26896952 | 4.43E-02 |
| Zbtb12 | 936.8639599 | 0.267651097 | 2.67E-02 |
| Ttc13 | 1929.317458 | 0.267303533 | 3.24E-02 |
| Timm44 | 1806.621639 | 0.265773777 | 1.15E-02 |
| Rnf44 | 2695.652928 | 0.265260961 | 4.54E-02 |
| Fyco1 | 2113.385571 | 0.265097181 | 3.66E-02 |
| Cep112it | 847.0867356 | 0.263711296 | 4.07E-02 |
| Uqcr11 | 2043.452294 | 0.263540901 | 2.90E-02 |
| Setdb1 | 3317.723671 | 0.263417108 | 1.93E-03 |
| Mtmr3 | 4507.121796 | 0.261934113 | 2.15E-02 |
| Ldah | 2630.414737 | 0.260953846 | 2.41E-02 |
| Sf1 | 8718.970602 | 0.260740092 | 3.61E-02 |
| Med19 | 1033.326527 | 0.260319539 | 3.34E-02 |
| Dnajc15 | 497.9328464 | 0.258986186 | 3.65E-02 |
| Atg3 | 1744.113181 | 0.257835206 | 3.61E-02 |
| Map4k3 | 1741.257735 | 0.25775367 | 2.47E-02 |
| Dctn5 | 1807.865925 | 0.257499411 | 4.59E-02 |
| Tsnax | 2533.133708 | 0.256716734 | 4.79E-03 |
| Mboat7 | 2519.861218 | 0.255728878 | 4.49E-02 |
| Spg20 | 6566.427666 | 0.255625565 | 1.97E-02 |
| Zyg11b | 5267.774086 | 0.254400562 | 2.04E-02 |
| Virma | 3718.754435 | 0.25404043 | 5.11E-03 |
| Mrpl46 | 412.9064619 | 0.25384294 | 3.86E-02 |
| Nbas | 2321.441303 | 0.253694402 | 4.34E-02 |

| | | | |
|---|---|---|---|
| Fam98a | 1905.214616 | 0.253535226 | 2.16E-02 |
| Spata7 | 434.5194125 | 0.253510326 | 3.37E-02 |
| Cnot2 | 3155.015605 | 0.251914676 | 7.69E-04 |
| Supt6 | 7336.991304 | 0.251243264 | 3.62E-02 |
| Amfr | 5482.908097 | 0.250823505 | 7.25E-03 |
| Golga7 | 2979.170609 | 0.24956365 | 1.02E-02 |
| Ndufs6 | 1052.213876 | 0.247822703 | 3.34E-02 |
| Tor1b | 2653.994304 | 0.247122973 | 3.04E-02 |
| Arhgap17 | 1804.268498 | 0.246112642 | 1.88E-02 |
| Tm2d3 | 567.3585455 | 0.244979778 | 2.20E-02 |
| Nudc | 3584.828604 | 0.24328125 | 4.22E-02 |
| Zfp384 | 2438.431101 | 0.242174413 | 3.58E-02 |
| Ppp3r1 | 4512.171715 | 0.241518354 | 2.54E-02 |
| Pdcd6 | 2365.713327 | 0.240623595 | 4.78E-03 |
| Osbpl2 | 2282.302263 | 0.239134522 | 7.81E-04 |
| Cwc25 | 1224.52217 | 0.236730523 | 4.03E-02 |
| Rrp1 | 7200.402786 | 0.236679193 | 2.68E-02 |
| Msl3 | 2208.540722 | 0.236108027 | 3.75E-02 |
| Cmpk1 | 3161.859525 | 0.234357762 | 9.47E-03 |
| Itprid2 | 2850.831262 | 0.233583577 | 3.82E-02 |
| Tulp4 | 16880.04521 | 0.233447103 | 3.75E-02 |
| Fbxo3 | 2578.681611 | 0.230649643 | 2.49E-02 |
| Uspl1 | 1969.222052 | 0.230633202 | 2.47E-02 |
| Ppp4c | 2200.376241 | 0.230043038 | 3.42E-02 |
| Mmadhc | 2680.153335 | 0.228618139 | 9.67E-03 |
| Slc7a6os | 1293.861495 | 0.222752441 | 3.05E-02 |
| Ddx59 | 608.5547225 | 0.222185543 | 4.90E-02 |
| Rnf19a | 1760.078004 | 0.220180647 | 1.79E-02 |
| Nsf | 2095.639979 | 0.219474847 | 4.02E-02 |
| Josd1 | 4707.625447 | 0.215749637 | 4.56E-02 |
| Gpd1l | 1232.297381 | 0.21513394 | 2.02E-02 |
| Mrps21 | 1364.811082 | 0.213790043 | 2.90E-02 |
| Ubr2 | 5346.693588 | 0.213764859 | 6.50E-03 |
| Dhx16 | 2218.027029 | 0.212753891 | 4.20E-02 |
| Mcl1 | 6120.725986 | 0.211678366 | 3.52E-02 |
| Etfa | 3192.680966 | 0.209013046 | 4.18E-02 |
| Utp3 | 2514.145431 | 0.208739393 | 1.94E-02 |
| Ube2q1 | 5225.360789 | 0.207149236 | 3.90E-02 |
| Wdr59 | 1368.374064 | 0.203148221 | 3.98E-02 |
| Pak2 | 5721.701201 | 0.195761664 | 1.19E-03 |
| Ythdf1 | 3589.274297 | 0.194083577 | 4.60E-02 |
| Rnf14 | 3214.705751 | 0.181171504 | 1.71E-02 |
| Wwtr1 | 5195.543564 | 0.172003878 | 3.86E-02 |
| Thumpd1 | 2170.674412 | 0.168728133 | 3.92E-02 |
| Kctd3 | 1744.740515 | 0.166214463 | 1.51E-02 |
| Ube2d2a | 3927.040904 | -0.161726796 | 2.64E-02 |
| Paf1 | 1759.690936 | -0.172401028 | 2.85E-02 |

| | | | |
|---|---|---|---|
| Bnip2 | 3652.520686 | -0.182675993 | 4.84E-02 |
| Akap10 | 1520.79788 | -0.183645446 | 2.85E-02 |
| Cers5 | 3272.249933 | -0.188297547 | 8.24E-03 |
| BC005537 | 7337.972699 | -0.190250465 | 3.22E-02 |
| Ptdss1 | 2133.806868 | -0.192595185 | 3.58E-02 |
| Tor1aip2 | 6209.143612 | -0.194645184 | 2.26E-02 |
| Ehmt1 | 2812.096587 | -0.198510857 | 3.11E-02 |
| Fam149b | 889.385841 | -0.203667812 | 2.99E-02 |
| Fam120a | 9571.131555 | -0.20517407 | 2.86E-02 |
| Zscan29 | 790.0709857 | -0.207160875 | 4.82E-02 |
| Rnf145 | 4349.170828 | -0.207311766 | 3.45E-02 |
| Wdr48 | 2346.652087 | -0.209581288 | 1.85E-02 |
| Cebpg | 2322.035649 | -0.210725825 | 4.92E-02 |
| Capza1 | 5169.359605 | -0.212384227 | 1.10E-02 |
| Atxn7l3b | 7440.579257 | -0.212484814 | 2.70E-02 |
| Sar1a | 8919.114577 | -0.219454723 | 3.31E-02 |
| Zmynd8 | 5278.19677 | -0.219893901 | 6.33E-03 |
| Stx12 | 1913.740575 | -0.220258814 | 1.26E-02 |
| 2610301B20Rik | 1235.416847 | -0.220436638 | 4.01E-02 |
| Rexo1 | 2127.055018 | -0.223791404 | 4.15E-02 |
| Rnf115 | 1861.018874 | -0.225996106 | 5.00E-02 |
| Atmin | 1212.579264 | -0.228284346 | 1.39E-02 |
| Cog7 | 1094.294008 | -0.228355562 | 9.92E-03 |
| Armh3 | 1013.355979 | -0.230111434 | 1.26E-02 |
| Ctnna1 | 13605.32906 | -0.231044159 | 4.98E-03 |
| Vasp | 2678.87239 | -0.231264923 | 2.30E-02 |
| Arl2bp | 3849.906749 | -0.236240972 | 2.57E-02 |
| Mat2b | 3057.082385 | -0.23736618 | 1.93E-02 |
| Kbtbd4 | 953.0351392 | -0.237478698 | 5.58E-03 |
| Zfp623 | 763.4393593 | -0.239652386 | 3.77E-02 |
| Trim26 | 1760.62455 | -0.239849138 | 1.79E-02 |
| Slc25a51 | 3150.308466 | -0.241693947 | 8.87E-03 |
| Hars2 | 834.9974646 | -0.242771144 | 1.40E-02 |
| Limd1 | 3109.454773 | -0.243429549 | 3.74E-03 |
| Eps15 | 5455.733955 | -0.244269396 | 1.09E-02 |
| Elmo2 | 2691.4094 | -0.246049143 | 8.42E-03 |
| Cltc | 24544.13461 | -0.247628453 | 5.14E-05 |
| Gskip | 1077.810338 | -0.247819753 | 3.74E-02 |
| Scd2 | 61452.75163 | -0.248084898 | 4.45E-02 |
| Zzef1 | 3506.293149 | -0.248098103 | 1.79E-02 |
| Cdk5 | 1892.485916 | -0.248982356 | 1.50E-02 |
| Cdc26 | 2437.359862 | -0.250071361 | 1.61E-02 |
| Vps26a | 5958.591577 | -0.250083467 | 4.89E-02 |
| Zc3h13 | 2165.313132 | -0.250913113 | 9.54E-03 |
| Rc3h1 | 2093.844484 | -0.250949863 | 2.09E-02 |
| Mmgt1 | 1671.149507 | -0.252577305 | 2.68E-02 |
| 5830417I10Rik | 2116.658395 | -0.253539419 | 1.12E-02 |

| | | | |
|---|---|---|---|
| Zgpat | 1572.13059 | -0.25409362 | 7.96E-03 |
| Hdac2 | 5826.991218 | -0.254098613 | 2.13E-02 |
| Gab1 | 1286.750464 | -0.255124769 | 2.73E-02 |
| Ibtk | 1777.858862 | -0.255142708 | 1.90E-02 |
| Tpbg | 2142.107889 | -0.255726427 | 2.56E-02 |
| Cpne8 | 1083.712569 | -0.255747073 | 2.80E-02 |
| Ppp2r5e | 2577.239633 | -0.256078325 | 2.32E-04 |
| Atf6b | 3107.322539 | -0.257228126 | 4.33E-02 |
| Dpy19l3 | 2100.747006 | -0.258142716 | 4.46E-02 |
| Dctn6 | 1522.17295 | -0.259154111 | 2.05E-02 |
| 4930402H24Rik | 5341.149355 | -0.261019286 | 1.99E-02 |
| Rab12 | 2907.184409 | -0.2611997 | 1.74E-04 |
| Shoc2 | 2739.440178 | -0.266102882 | 1.26E-02 |
| Zdhhc24 | 757.5320873 | -0.26634906 | 4.33E-02 |
| Abcd3 | 3631.045164 | -0.266518568 | 1.83E-04 |
| Fech | 1997.126972 | -0.267106558 | 1.41E-03 |
| Ndufv2 | 2829.125466 | -0.267981634 | 2.43E-02 |
| Tbl1x | 7690.858636 | -0.268077194 | 5.62E-03 |
| Bccip | 1315.490618 | -0.268124848 | 1.89E-02 |
| Fbf1 | 2196.630713 | -0.268162712 | 3.04E-02 |
| Smarcc1 | 5292.980505 | -0.269071383 | 1.01E-02 |
| Ocel1 | 830.5687289 | -0.270905279 | 4.50E-02 |
| Taok2 | 2417.6946 | -0.27350482 | 3.12E-02 |
| Sgpl1 | 13380.67463 | -0.274645826 | 3.40E-02 |
| BC031181 | 1034.180156 | -0.276000469 | 1.51E-02 |
| Gm16286 | 2912.987106 | -0.276169513 | 1.44E-02 |
| Wbp4 | 1679.499856 | -0.276896062 | 2.45E-02 |
| Vps4a | 1813.209297 | -0.277621758 | 1.99E-02 |
| Chmp1a | 5050.613376 | -0.277637301 | 2.95E-02 |
| Nfatc3 | 4109.575676 | -0.277893938 | 3.50E-06 |
| Socs6 | 1293.79635 | -0.279031709 | 2.31E-02 |
| Pigm | 1186.751818 | -0.279306917 | 1.95E-03 |
| Gpatch8 | 3437.946536 | -0.280107133 | 1.86E-03 |
| Commd3 | 3470.393101 | -0.280508958 | 9.12E-03 |
| Scyl3 | 1120.710529 | -0.280560664 | 2.68E-03 |
| Smpd4 | 1683.926286 | -0.28161104 | 1.33E-02 |
| Sppl3 | 3400.317528 | -0.282781179 | 3.51E-03 |
| Fbxw11 | 4239.429717 | -0.283368474 | 2.27E-04 |
| Btbd9 | 833.0426777 | -0.283596095 | 2.84E-02 |
| Prelid1 | 8505.205021 | -0.284304387 | 1.36E-02 |
| Gadd45gip1 | 1198.857908 | -0.285001039 | 1.49E-02 |
| Vegfc | 674.5119938 | -0.285509849 | 4.46E-02 |
| Soat1 | 3693.855559 | -0.285665183 | 5.85E-03 |
| Gripap1 | 1321.857965 | -0.286353953 | 3.60E-02 |
| Smarcd3 | 630.858313 | -0.287196527 | 1.95E-02 |
| Mbtps1 | 9771.008573 | -0.287253116 | 4.99E-03 |
| 2810004N23Rik | 1657.133658 | -0.287571102 | 3.28E-03 |

| | | | |
|---|---|---|---|
| Prkca | 3873.741095 | -0.290056218 | 3.26E-02 |
| Mrps26 | 1220.384571 | -0.290164841 | 2.01E-02 |
| Pprc1 | 3591.050554 | -0.290385849 | 7.73E-03 |
| Kat14 | 1963.154269 | -0.29075766 | 1.76E-02 |
| Rbbp7 | 9676.724973 | -0.290965944 | 3.86E-03 |
| C87436 | 843.8956946 | -0.291528999 | 2.89E-02 |
| Crot | 1867.061663 | -0.294285649 | 9.75E-03 |
| Diexf | 951.2860776 | -0.295221041 | 3.73E-03 |
| D10Jhu81e | 1329.248274 | -0.296455351 | 3.39E-02 |
| Tesk1 | 1587.617809 | -0.296733418 | 7.90E-03 |
| Cipc | 1168.346399 | -0.296778111 | 8.80E-03 |
| Tmem127 | 5979.127084 | -0.296948051 | 4.04E-02 |
| Ipmk | 2418.776262 | -0.298972769 | 1.34E-03 |
| Toe1 | 901.2311272 | -0.300011777 | 6.32E-03 |
| Sucla2 | 2404.682437 | -0.301729703 | 1.25E-02 |
| Pex3 | 966.7864216 | -0.301854236 | 3.52E-02 |
| Fam76a | 1691.346603 | -0.301915813 | 4.35E-02 |
| Sccpdh | 1211.67741 | -0.302236053 | 1.72E-02 |
| Tlk2 | 3022.803034 | -0.302726938 | 4.59E-02 |
| Foxj2 | 2108.968859 | -0.304681719 | 8.77E-05 |
| Kifap3 | 3772.714154 | -0.304807282 | 2.57E-03 |
| Cdc123 | 2172.327021 | -0.305501001 | 1.94E-02 |
| Slc35c1 | 794.2651529 | -0.306346227 | 2.60E-02 |
| Rragc | 10259.96991 | -0.306474565 | 4.93E-02 |
| BC003965 | 804.6201398 | -0.306524401 | 1.80E-02 |
| Nr2f2 | 2990.503183 | -0.307390386 | 2.88E-02 |
| Senp2 | 2448.303089 | -0.307616387 | 1.44E-03 |
| Pcsk7 | 888.8351781 | -0.308060848 | 2.91E-03 |
| Tmem101 | 489.0305594 | -0.30815467 | 2.25E-02 |
| Arhgap1 | 4297.533144 | -0.308367003 | 7.73E-04 |
| Ltbr | 4575.460131 | -0.308497727 | 5.75E-03 |
| Ralbp1 | 3528.08954 | -0.308989981 | 1.43E-04 |
| Cog1 | 1510.656347 | -0.309743597 | 2.21E-02 |
| Jmy | 1698.02112 | -0.309759526 | 9.66E-04 |
| Trmt5 | 584.4873569 | -0.310093321 | 3.06E-02 |
| Tmem41b | 1689.446509 | -0.310478554 | 3.16E-02 |
| Usp28 | 1013.741207 | -0.311124151 | 2.99E-03 |
| Notch2 | 5564.204759 | -0.31275673 | 6.83E-03 |
| Sec16a | 6100.782522 | -0.313693987 | 1.88E-02 |
| Ash2l | 2579.786406 | -0.313895149 | 1.78E-03 |
| Gpn1 | 1307.42094 | -0.314416249 | 3.28E-02 |
| Sec22b | 3178.347462 | -0.315259706 | 6.10E-03 |
| Mpg | 550.6470613 | -0.315489435 | 4.66E-02 |
| Mmgt2 | 671.2994786 | -0.317977966 | 3.78E-02 |
| Egln2 | 2115.922258 | -0.320432193 | 9.84E-03 |
| Rreb1 | 1281.82888 | -0.320474945 | 1.63E-02 |
| Gnao1 | 4073.149345 | -0.321081816 | 5.77E-03 |

| | | | |
|---|---|---|---|
| Dph1 | 914.7849843 | -0.321139767 | 1.57E-02 |
| Zfyve9 | 2465.724035 | -0.321872202 | 2.37E-03 |
| Lrch3 | 1722.549272 | -0.322223903 | 9.79E-06 |
| Mtmr12 | 1044.113782 | -0.322396082 | 3.12E-02 |
| Unc50 | 1307.062746 | -0.322662264 | 1.22E-02 |
| Agtrap | 1417.40242 | -0.323371137 | 1.14E-02 |
| Hnrnpk | 21286.09285 | -0.323387206 | 1.29E-04 |
| Ccser2 | 3364.433109 | -0.323413531 | 5.51E-03 |
| Mypop | 419.0754148 | -0.324094191 | 4.31E-02 |
| Nek9 | 8266.50416 | -0.32479474 | 5.77E-03 |
| Gm13736 | 1587.120877 | -0.325123171 | 7.85E-03 |
| Msantd4 | 1976.487313 | -0.325265188 | 3.94E-05 |
| Bnip3l | 4675.143122 | -0.326535527 | 1.77E-02 |
| Lsm1 | 795.3759526 | -0.326824832 | 3.98E-03 |
| Erlin1 | 2251.338134 | -0.32723124 | 1.46E-02 |
| Ewsr1 | 10998.79773 | -0.329243269 | 1.74E-02 |
| Ttf1 | 1026.73623 | -0.33017915 | 2.18E-02 |
| Cybc1 | 1087.851641 | -0.331706001 | 4.11E-04 |
| Dusp12 | 1078.997366 | -0.332305255 | 3.31E-02 |
| Dmac2 | 437.6982699 | -0.332794338 | 2.15E-02 |
| Gpatch2 | 1041.002905 | -0.33305529 | 4.04E-03 |
| Snap23 | 1353.916862 | -0.333119557 | 9.82E-03 |
| Ppp3ca | 3841.872004 | -0.333134735 | 3.26E-03 |
| Golga5 | 2169.683447 | -0.333880091 | 3.94E-03 |
| Commd2 | 951.5123925 | -0.334257108 | 2.37E-02 |
| Specc1l | 4786.27288 | -0.334321668 | 2.90E-05 |
| Zfp451 | 2269.649158 | -0.33437342 | 2.65E-04 |
| Mcur1 | 737.7989981 | -0.335084062 | 7.17E-03 |
| Tent2 | 1578.662334 | -0.335181739 | 7.12E-03 |
| Cfap20 | 1689.539204 | -0.335616702 | 6.51E-03 |
| Gcdh | 962.2082415 | -0.335881453 | 6.50E-03 |
| Mrpl4 | 1025.618602 | -0.336376144 | 2.51E-02 |
| Mtap | 23447.27157 | -0.337086855 | 4.71E-02 |
| Cop1 | 3510.093014 | -0.337326096 | 9.19E-04 |
| Nbdy | 1309.915387 | -0.337625981 | 9.92E-04 |
| Ccdc32 | 828.4896004 | -0.337677906 | 1.82E-02 |
| G3bp1 | 9404.431586 | -0.338239049 | 8.09E-03 |
| Psme2 | 2209.647031 | -0.338683593 | 3.32E-03 |
| Gna11 | 4652.144884 | -0.339579289 | 3.22E-02 |
| Vti1b | 2649.138363 | -0.340008921 | 2.19E-03 |
| Fam160b2 | 868.648153 | -0.341978407 | 1.73E-03 |
| Ssr3 | 17468.73704 | -0.342091054 | 5.86E-04 |
| Prkce | 764.757114 | -0.342556274 | 2.61E-02 |
| NA | 4537.193566 | -0.342925973 | 3.31E-05 |
| Cep350 | 2934.829578 | -0.34308585 | 2.13E-03 |
| Phf2 | 3036.045331 | -0.343355931 | 2.57E-04 |
| Plpp5 | 772.2669265 | -0.343459069 | 4.65E-02 |

| | | | |
|---|---|---|---|
| Prkdc | 1324.769727 | -0.343717209 | 2.12E-02 |
| Cdc14b | 821.3770431 | -0.343919548 | 2.51E-03 |
| Zbed3 | 1298.091305 | -0.344012805 | 1.40E-06 |
| Smug1 | 791.1906468 | -0.344447863 | 5.53E-04 |
| Mrtfa | 2448.317165 | -0.345456814 | 7.32E-03 |
| Setd1b | 1635.701756 | -0.345694448 | 2.83E-02 |
| Ankib1 | 2741.451452 | -0.346655148 | 4.64E-03 |
| Zfp668 | 533.3538571 | -0.347024404 | 3.88E-02 |
| Jcad | 4643.802879 | -0.347431384 | 3.10E-02 |
| Trim35 | 6233.884212 | -0.348276535 | 9.59E-04 |
| Zfp189 | 296.7393454 | -0.348796791 | 4.52E-02 |
| Snx16 | 596.553418 | -0.349499714 | 3.04E-02 |
| Acvr1b | 825.908788 | -0.350073928 | 1.16E-03 |
| Rab22a | 4367.407689 | -0.350337216 | 9.03E-04 |
| Tprkb | 2647.743254 | -0.351701157 | 1.29E-02 |
| Tmem167 | 2323.710803 | -0.351851199 | 4.39E-03 |
| Gigyf1 | 2070.367659 | -0.352383491 | 5.49E-03 |
| Akna | 1024.927798 | -0.353256273 | 1.00E-02 |
| Gon4l | 2759.220346 | -0.353434989 | 8.61E-04 |
| Rab28 | 2040.054985 | -0.354923043 | 9.65E-05 |
| Rab33b | 906.8254427 | -0.356211631 | 3.32E-03 |
| Zfp870 | 203.7841449 | -0.356241009 | 4.44E-02 |
| Letm1 | 2063.672833 | -0.356461 | 1.08E-02 |
| Asph | 3636.471194 | -0.356728698 | 2.29E-02 |
| Tax1bp3 | 4764.282672 | -0.357256022 | 2.50E-02 |
| Zbtb5 | 524.8705552 | -0.357526595 | 1.33E-02 |
| Rab18 | 4215.54324 | -0.357983604 | 1.32E-04 |
| Clstn1 | 7598.808291 | -0.35811675 | 2.33E-03 |
| Pard3 | 3039.571374 | -0.358138104 | 1.65E-04 |
| Ndn | 1355.557359 | -0.358719971 | 1.45E-02 |
| Sem1 | 4653.426344 | -0.359059636 | 3.05E-02 |
| Hspa1b | 1228.877042 | -0.359275448 | 1.63E-02 |
| Safb2 | 1979.333132 | -0.359304609 | 2.03E-02 |
| Adarb1 | 849.0097309 | -0.359678828 | 2.89E-02 |
| Rai1 | 2652.568346 | -0.359854061 | 3.00E-04 |
| Rbpj-ps3 | 2159.928873 | -0.359996946 | 2.62E-03 |
| Aplf | 766.3187187 | -0.360195612 | 1.04E-02 |
| BC005624 | 2059.921733 | -0.36020335 | 4.41E-06 |
| BC029722 | 1983.763936 | -0.360210266 | 1.68E-02 |
| Josd2 | 899.9900018 | -0.360840266 | 7.66E-03 |
| Cyb5d1 | 1009.24945 | -0.361238413 | 4.86E-04 |
| Rapgef2 | 1760.896602 | -0.361830956 | 4.19E-02 |
| Ankrd11 | 9266.580924 | -0.362108732 | 2.17E-05 |
| Dcun1d5 | 1542.903361 | -0.362109634 | 1.64E-03 |
| Pum1 | 4713.99963 | -0.362113299 | 7.26E-05 |
| Rbmxl1 | 3706.246739 | -0.362856948 | 1.11E-03 |
| Minos1 | 1389.650092 | -0.362864777 | 5.51E-03 |

| | | | |
|---|---|---|---|
| Coa5 | 4878.455822 | -0.363029297 | 5.04E-04 |
| Dld | 3979.457662 | -0.363471018 | 4.27E-02 |
| Ube4a | 2494.407906 | -0.363612173 | 4.83E-04 |
| Arhgap22 | 1622.996624 | -0.365028589 | 1.65E-02 |
| Ppp1r37 | 2197.916903 | -0.365109518 | 8.63E-03 |
| Foxred1 | 683.0766349 | -0.365490051 | 9.16E-03 |
| Trappc13 | 1405.841872 | -0.365555775 | 4.03E-02 |
| Fbxw5 | 1900.946667 | -0.365807796 | 2.62E-05 |
| Uba5 | 2011.905068 | -0.366211812 | 4.08E-03 |
| Hpf1 | 810.3108242 | -0.366396513 | 3.06E-02 |
| Prickle2 | 610.6101831 | -0.366672244 | 1.91E-02 |
| Rnf8 | 695.2111225 | -0.366902809 | 7.08E-04 |
| Secisbp2l | 3505.578734 | -0.367860501 | 3.24E-02 |
| Mtf1 | 1418.940464 | -0.367984299 | 6.94E-04 |
| Ctr9 | 3056.119718 | -0.368429346 | 9.39E-04 |
| Dnlz | 934.93158 | -0.368474513 | 3.15E-03 |
| Pik3r4 | 1596.763516 | -0.369006326 | 1.84E-03 |
| Arpc1b | 12666.70678 | -0.369473752 | 2.79E-03 |
| Nufip2 | 2755.24518 | -0.369501139 | 7.08E-03 |
| Phc2 | 4897.151543 | -0.372076208 | 1.05E-02 |
| Ap4m1 | 644.4259784 | -0.372123704 | 2.97E-02 |
| Swi5 | 5268.153823 | -0.372320559 | 7.55E-06 |
| Tmed7 | 4080.361082 | -0.372459123 | 1.38E-04 |
| Eme2 | 679.3381335 | -0.373050966 | 3.60E-02 |
| Zfp408 | 1020.215457 | -0.373302861 | 8.87E-03 |
| Fkbp15 | 2776.85651 | -0.373415627 | 1.34E-04 |
| Fzd2 | 5450.697507 | -0.373436544 | 1.33E-02 |
| Gbf1 | 6069.662746 | -0.374431748 | 3.65E-03 |
| Ap1g1 | 3420.398712 | -0.375159263 | 2.33E-05 |
| Cd9 | 9138.524752 | -0.375478363 | 2.91E-04 |
| Pigu | 1268.087199 | -0.375542021 | 8.50E-05 |
| Vipas39 | 1504.93256 | -0.376235339 | 4.57E-04 |
| Oaz1 | 9771.830723 | -0.376297279 | 4.81E-03 |
| Zfp287 | 249.7870389 | -0.376905996 | 1.54E-02 |
| Alg9 | 906.9621859 | -0.377062435 | 7.30E-04 |
| Lyrm2 | 460.0390523 | -0.377148942 | 2.59E-02 |
| Ubqln2 | 3270.808587 | -0.377170196 | 7.82E-05 |
| Naa20 | 2437.665542 | -0.377195337 | 6.71E-06 |
| Bnip3 | 1792.535201 | -0.377669205 | 2.31E-02 |
| Lman2l | 996.9305942 | -0.379229087 | 1.44E-03 |
| Prpsap2 | 830.6666767 | -0.379235294 | 9.71E-03 |
| Mfap1a | 3352.589017 | -0.379579 | 4.35E-03 |
| Mcfd2 | 5084.76864 | -0.380121464 | 8.68E-03 |
| Ttpal | 2000.246656 | -0.380592999 | 1.16E-02 |
| Sfswap | 2364.600261 | -0.380632667 | 1.59E-05 |
| Ints3 | 3305.873649 | -0.381539991 | 6.28E-03 |
| Luzp1 | 3837.373776 | -0.381866093 | 4.88E-03 |

| | | | |
|---|---|---|---|
| Cblb | 1328.176995 | -0.38239206 | 9.04E-03 |
| Ccdc92 | 341.0632489 | -0.3825156 | 4.42E-02 |
| Cables2 | 840.8736199 | -0.382607521 | 4.40E-03 |
| Fbxo30 | 1577.825112 | -0.382703506 | 1.85E-02 |
| Map2k4 | 2943.46728 | -0.384593798 | 5.14E-04 |
| Atp5j2 | 3346.217142 | -0.384732694 | 3.28E-03 |
| Siae | 1505.165601 | -0.384939621 | 5.60E-03 |
| Kpna4 | 4074.367138 | -0.385575217 | 4.75E-02 |
| Ift88 | 627.0852278 | -0.38557782 | 1.10E-02 |
| Zfp507 | 2655.215056 | -0.385665389 | 5.13E-05 |
| Brd1 | 2882.283459 | -0.385768904 | 2.70E-08 |
| Thrap3 | 10251.1699 | -0.385934077 | 2.03E-03 |
| Gnl2 | 3384.380656 | -0.386197885 | 4.79E-03 |
| Zfp952 | 434.2315256 | -0.386356813 | 1.16E-03 |
| Usp7 | 5978.042099 | -0.387125061 | 1.78E-04 |
| Myo10 | 6334.157457 | -0.38839606 | 3.20E-02 |
| Emc8 | 1371.50203 | -0.388731855 | 4.44E-02 |
| Ywhaz | 11408.90752 | -0.388924033 | 1.58E-04 |
| Gna12 | 3909.739383 | -0.389695977 | 6.86E-04 |
| Cdk2 | 961.5697005 | -0.389946729 | 1.44E-02 |
| Gm3608 | 2249.543933 | -0.391020228 | 1.02E-04 |
| Sdc2 | 7669.630869 | -0.392159739 | 1.61E-02 |
| Ufm1 | 2432.671543 | -0.392307391 | 2.96E-03 |
| Arfip1 | 2355.251809 | -0.392488553 | 2.18E-05 |
| Vamp8 | 2018.695151 | -0.393094811 | 3.48E-04 |
| Nt5c2 | 1953.179617 | -0.393132267 | 7.16E-06 |
| Gtf2h3 | 971.4168647 | -0.393471346 | 4.98E-03 |
| Sdhaf4 | 475.0879917 | -0.395019911 | 3.57E-03 |
| Slc25a16 | 1028.92458 | -0.395383606 | 2.39E-03 |
| Myh10 | 12571.26723 | -0.395509428 | 6.36E-03 |
| Gm20604 | 399.9364725 | -0.395547124 | 1.65E-03 |
| Dclk2 | 983.6272227 | -0.395571809 | 5.77E-03 |
| Desi2 | 2829.811094 | -0.396600266 | 6.43E-06 |
| Ywhab | 10120.15948 | -0.3968195 | 2.87E-03 |
| Sf3a1 | 4586.280995 | -0.396861597 | 2.13E-02 |
| Tusc3 | 3182.406469 | -0.397367101 | 2.27E-05 |
| Numa1 | 7864.573362 | -0.397574853 | 2.23E-04 |
| Pmm2 | 1576.310651 | -0.397804324 | 2.51E-04 |
| Vps26c | 2458.394981 | -0.397925885 | 4.56E-03 |
| Zfp1 | 503.9292446 | -0.398033431 | 2.20E-03 |
| Iqce | 1194.492392 | -0.398128325 | 9.00E-03 |
| Tmem110 | 707.2892674 | -0.398408791 | 1.33E-03 |
| Gphn | 863.3605628 | -0.398755244 | 1.66E-03 |
| Srrd | 319.5444584 | -0.398759519 | 1.05E-02 |
| Romo1 | 1261.431817 | -0.399442881 | 1.42E-02 |
| Ing5 | 967.3715042 | -0.399508188 | 2.46E-04 |
| Btbd3 | 870.3274116 | -0.399902816 | 6.05E-04 |

| | | | |
|---|---|---|---|
| Itsn1 | 2752.843847 | -0.399935258 | 1.62E-04 |
| Nupl1 | 1449.7265 | -0.400484496 | 7.06E-03 |
| Ints1 | 2784.850217 | -0.400698078 | 9.28E-03 |
| Celf5 | 475.0065591 | -0.400823716 | 4.35E-02 |
| Ubn2 | 1931.029383 | -0.400864178 | 2.18E-02 |
| Slc12a6 | 3549.421791 | -0.401310326 | 3.75E-03 |
| Ccdc102a | 1567.023143 | -0.401625835 | 5.97E-03 |
| NA | 3354.889398 | -0.401885031 | 4.20E-06 |
| Zfp30 | 422.1195397 | -0.402468442 | 2.65E-03 |
| Mafg | 3468.758033 | -0.403320383 | 1.50E-04 |
| Soga1 | 1586.138467 | -0.404117563 | 4.64E-03 |
| Entpd4 | 1607.124128 | -0.404488978 | 4.57E-03 |
| Dnajc16 | 942.4498415 | -0.40462742 | 2.16E-03 |
| Ndufaf1 | 402.0330849 | -0.404758166 | 1.09E-02 |
| Eid2b | 415.8998296 | -0.405249122 | 3.76E-03 |
| Npnt | 449.3441054 | -0.405312922 | 4.74E-02 |
| Mesd | 4178.825069 | -0.405624325 | 1.26E-02 |
| Xpa | 512.5686762 | -0.405883917 | 4.89E-02 |
| Chuk | 2436.929155 | -0.406057677 | 2.52E-02 |
| Cfdp1 | 4640.495616 | -0.406505423 | 1.94E-04 |
| Ccnj | 254.3550901 | -0.40658604 | 4.14E-02 |
| Ttl | 702.6056377 | -0.407046939 | 1.84E-03 |
| Dnmbp | 2254.506423 | -0.407595617 | 1.32E-05 |
| Ube2g1 | 1838.730964 | -0.407719846 | 2.61E-03 |
| Commd8 | 1529.596254 | -0.409064334 | 1.06E-03 |
| Sgsm3 | 1109.231287 | -0.409350315 | 4.26E-03 |
| Slc4a2 | 6711.940086 | -0.409420544 | 6.29E-03 |
| Pkp4 | 2837.958 | -0.409717174 | 1.95E-02 |
| Bckdk | 2049.280461 | -0.410019642 | 6.93E-04 |
| Rptor | 2361.625741 | -0.410190601 | 4.00E-07 |
| Them4 | 258.1979688 | -0.410381455 | 6.72E-03 |
| Eif4ebp2 | 3020.522678 | -0.410732756 | 8.07E-05 |
| Zmiz2 | 2535.331176 | -0.410939887 | 1.98E-04 |
| Fam91a1 | 5484.339588 | -0.411387934 | 5.44E-05 |
| Abcd1 | 1111.639964 | -0.412561224 | 2.03E-02 |
| Sap18 | 518.3172933 | -0.412723057 | 1.39E-02 |
| Pkd1 | 8593.732781 | -0.412731959 | 6.07E-05 |
| 4931414P19Rik | 239.6934697 | -0.412752114 | 1.87E-02 |
| Eif1b | 955.7925393 | -0.413114823 | 1.12E-02 |
| Mynn | 1124.562014 | -0.413681162 | 3.81E-03 |
| Cbx8 | 308.4306254 | -0.41387013 | 3.67E-02 |
| Slc30a5 | 1945.250866 | -0.414340813 | 6.25E-07 |
| 1110038F14Rik | 735.5469736 | -0.414792809 | 1.09E-04 |
| Cstf2t | 2665.498894 | -0.415379539 | 1.31E-04 |
| Fem1a | 1367.887487 | -0.416029224 | 3.40E-04 |
| Irf3 | 1284.181054 | -0.416149417 | 6.97E-05 |
| Slc35b2 | 1827.407569 | -0.416192675 | 8.08E-03 |

| | | | |
|---|---|---|---|
| Zbtb24 | 960.6102902 | -0.416617074 | 2.72E-04 |
| Rbpj | 6189.6897 | -0.416812468 | 3.00E-04 |
| Nek6 | 2770.3449 | -0.417237314 | 2.75E-04 |
| Nr3c1 | 3466.11795 | -0.41796737 | 1.29E-03 |
| Srpk2 | 4173.862921 | -0.418044367 | 7.33E-04 |
| Zfp202 | 333.0936493 | -0.418534201 | 5.88E-03 |
| Ppp1r21 | 1610.304203 | -0.418670094 | 1.97E-08 |
| Bphl | 452.6391813 | -0.418820237 | 9.53E-04 |
| Hoxd9 | 476.4829214 | -0.41890656 | 2.90E-03 |
| Sh3bgrl3 | 2819.46658 | -0.419561277 | 4.30E-02 |
| D5Ertd579e | 4848.886748 | -0.419952181 | 7.15E-03 |
| Erlec1 | 2362.333871 | -0.42028782 | 3.44E-04 |
| Ankrd10 | 1201.376825 | -0.420639914 | 3.04E-06 |
| Dag1 | 10803.36973 | -0.420646798 | 1.27E-04 |
| H6pd | 4608.678006 | -0.421888597 | 4.37E-02 |
| Dr1 | 1586.629601 | -0.422461485 | 3.85E-04 |
| Anxa6 | 6987.123827 | -0.422618064 | 2.21E-02 |
| Zfp398 | 382.5512451 | -0.423349617 | 7.37E-03 |
| Anapc5 | 11211.90137 | -0.423389838 | 7.49E-04 |
| NA | 2939.508412 | -0.423453671 | 4.47E-04 |
| Ehbp1 | 2247.789155 | -0.423658424 | 2.23E-04 |
| Srrm2 | 21325.05914 | -0.423884149 | 2.62E-03 |
| Fcho2 | 1868.138627 | -0.423893018 | 3.72E-02 |
| Scd1 | 8875.775287 | -0.423930128 | 1.86E-03 |
| Txndc5 | 14360.93511 | -0.424330839 | 7.55E-03 |
| Opa3 | 1502.032382 | -0.424338208 | 4.24E-02 |
| Paip1 | 2505.193787 | -0.424453602 | 6.78E-05 |
| Ndst1 | 5649.967415 | -0.425429355 | 1.23E-04 |
| Zdhhc5 | 4681.471334 | -0.425699414 | 1.07E-05 |
| Raver1 | 2661.25353 | -0.42607166 | 3.39E-03 |
| Ano6 | 9272.629877 | -0.426216818 | 5.54E-04 |
| Atxn7l3 | 3554.494769 | -0.42652994 | 2.81E-06 |
| Osgep | 1715.128448 | -0.426808446 | 5.57E-03 |
| Spata24 | 121.247919 | -0.427166431 | 2.80E-02 |
| Ddx42 | 3726.502989 | -0.427384065 | 3.62E-05 |
| Tm7sf3 | 4188.572659 | -0.428194713 | 4.63E-05 |
| Lactb | 625.5953829 | -0.428420242 | 1.33E-02 |
| Abhd16a | 1314.595388 | -0.428958673 | 4.79E-04 |
| Chordc1 | 1486.870167 | -0.429423595 | 3.67E-02 |
| 2410004B18Rik | 470.626286 | -0.429745308 | 4.05E-03 |
| Tut7 | 3327.102852 | -0.430814082 | 1.07E-03 |
| Tmem39b | 450.7819741 | -0.431151582 | 7.86E-03 |
| Nt5m | 402.8418024 | -0.431591477 | 6.84E-03 |
| Srsf9 | 1778.892419 | -0.431670473 | 1.13E-06 |
| Zfp87 | 507.9830536 | -0.431800747 | 4.63E-03 |
| Ano8 | 1962.656392 | -0.432027131 | 2.68E-06 |
| Wdr7 | 1610.402645 | -0.432172387 | 3.84E-04 |

| | | | |
|---|---|---|---|
| Dynll2 | 6105.990932 | -0.432213149 | 9.21E-04 |
| Rasa3 | 6805.378419 | -0.432626275 | 9.77E-03 |
| Srsf7 | 4012.366097 | -0.432783717 | 6.41E-03 |
| Vmac | 1013.466863 | -0.432846711 | 9.79E-03 |
| Ccdc136 | 1423.32675 | -0.432955198 | 3.43E-02 |
| Alpk1 | 418.5502308 | -0.433081731 | 3.02E-02 |
| Zfp326 | 1099.788456 | -0.433492897 | 3.41E-02 |
| Aars2 | 599.8250935 | -0.433526022 | 3.94E-04 |
| Ctnnbip1 | 507.0421115 | -0.434776312 | 1.26E-03 |
| Slc35b3 | 449.8835323 | -0.435383661 | 8.02E-03 |
| Iws1 | 1708.971764 | -0.435514657 | 2.50E-03 |
| Ebag9 | 758.3804734 | -0.435697943 | 5.88E-04 |
| Parvb | 2285.010317 | -0.435702092 | 5.02E-03 |
| Fam32a | 3348.599776 | -0.436860276 | 7.66E-06 |
| Mapk7 | 1811.638448 | -0.437123441 | 2.04E-04 |
| Sh3bp5 | 1876.843321 | -0.437402647 | 3.38E-03 |
| Eid1 | 4134.703214 | -0.437893006 | 2.63E-03 |
| Tomm5 | 1139.643768 | -0.438624557 | 7.74E-03 |
| Hnrnpf | 22991.53822 | -0.438878604 | 3.98E-05 |
| BC017158 | 795.086068 | -0.438883304 | 9.48E-05 |
| Dyrk2 | 1360.699862 | -0.439342656 | 3.41E-04 |
| Sfxn3 | 5162.307059 | -0.439626819 | 2.37E-05 |
| mt-Atp6 | 187651.4583 | -0.439724025 | 5.69E-03 |
| Ppp6r3 | 4632.142651 | -0.439886555 | 6.09E-12 |
| Bcl9 | 1951.774164 | -0.439991871 | 1.58E-03 |
| Fbxo45 | 992.1009613 | -0.440057066 | 8.59E-05 |
| Pomk | 854.3945557 | -0.440384779 | 1.73E-04 |
| Tmem173 | 257.4625801 | -0.440477832 | 4.48E-02 |
| Lyrm4 | 469.9816086 | -0.440509293 | 1.31E-04 |
| Tsen2 | 691.2193844 | -0.441557876 | 3.45E-03 |
| N4bp2l2 | 2115.566049 | -0.441651711 | 6.96E-06 |
| Ctdsp2 | 23274.70546 | -0.441652982 | 1.09E-03 |
| NA | 1773.329601 | -0.441916282 | 2.83E-05 |
| Slc25a14 | 246.6469147 | -0.442141882 | 6.21E-03 |
| Rab34 | 4682.192332 | -0.442295143 | 7.29E-05 |
| Eral1 | 699.6738377 | -0.442435087 | 4.93E-05 |
| Ss18l2 | 298.5950248 | -0.442640139 | 8.24E-03 |
| Kazald1 | 321.4247234 | -0.442873509 | 1.36E-02 |
| Abracl | 1310.993802 | -0.443209187 | 1.05E-02 |
| Ints8 | 1963.027738 | -0.443672582 | 9.02E-03 |
| Arcn1 | 8216.940913 | -0.44371452 | 8.73E-07 |
| Dgat1 | 845.3611873 | -0.444282293 | 6.09E-07 |
| Scaf11 | 6880.726904 | -0.444352182 | 3.80E-05 |
| Mllt10 | 2069.02611 | -0.444580647 | 6.15E-07 |
| 1600012H06Rik | 1609.676472 | -0.445142498 | 4.92E-06 |
| Txndc11 | 1459.372549 | -0.445180195 | 1.54E-05 |
| Yae1d1 | 672.7722005 | -0.446869916 | 1.26E-02 |

| | | | |
|---|---|---|---|
| 6820431F20Rik | 4434.102125 | -0.447489425 | 6.32E-03 |
| Parp16 | 205.5236357 | -0.447577313 | 7.65E-03 |
| Arhgap32 | 2413.87787 | -0.447589908 | 4.80E-04 |
| Dtx3 | 3727.879191 | -0.448000662 | 1.32E-02 |
| Bad | 1258.755194 | -0.448371346 | 3.85E-04 |
| C1galt1 | 851.1254799 | -0.448374788 | 1.80E-02 |
| Snrpd3 | 2992.359328 | -0.449120882 | 4.26E-02 |
| Fbrsl1 | 745.1517284 | -0.449140615 | 3.90E-04 |
| Zfp867 | 524.7941376 | -0.44915267 | 3.54E-02 |
| Dtnbp1 | 2425.756691 | -0.449230406 | 6.63E-03 |
| Tanc1 | 5268.526244 | -0.449851422 | 5.58E-06 |
| Uhrf1bp1 | 1844.893817 | -0.44994017 | 5.71E-04 |
| Commd4 | 942.1458007 | -0.449969761 | 4.96E-04 |
| Rars2 | 673.8057716 | -0.450004893 | 6.31E-03 |
| Taf15 | 4039.925904 | -0.450093623 | 2.30E-02 |
| Map3k2 | 1933.527456 | -0.450580709 | 3.95E-03 |
| Carmil1 | 968.5312716 | -0.450674915 | 2.02E-03 |
| Elof1 | 1123.40013 | -0.450937608 | 1.21E-03 |
| Gm5898 | 599.5779994 | -0.45159584 | 1.33E-02 |
| Zfp772 | 327.9103734 | -0.451737837 | 1.69E-03 |
| Zfp251 | 1144.446552 | -0.452402851 | 1.82E-07 |
| Lrpprc | 5310.296473 | -0.452964622 | 7.44E-04 |
| Fbxo8 | 727.2861706 | -0.453604733 | 1.09E-04 |
| Gprc5b | 5063.518833 | -0.453949723 | 2.30E-02 |
| Rmnd5a | 3357.074221 | -0.45461209 | 3.13E-05 |
| Ppfia1 | 3284.486732 | -0.454621963 | 8.97E-08 |
| Vps29 | 2535.509627 | -0.454694781 | 5.71E-05 |
| A530017D24Rik | 207.4785903 | -0.45517219 | 1.84E-03 |
| Rdh5 | 275.4133273 | -0.455766843 | 9.32E-03 |
| Nuak1 | 2656.800253 | -0.455796012 | 1.06E-02 |
| Zfp869 | 866.2160119 | -0.455883483 | 1.21E-05 |
| Ring1 | 784.773901 | -0.456290329 | 1.69E-02 |
| Slc16a10 | 806.7478132 | -0.456663157 | 4.15E-02 |
| Mbd2 | 1918.447548 | -0.457116501 | 7.48E-07 |
| Zfp110 | 1694.471274 | -0.457260315 | 2.46E-10 |
| Zcchc17 | 1182.648684 | -0.457609037 | 2.72E-03 |
| Fam234b | 435.9697922 | -0.457654544 | 2.81E-03 |
| Gga2 | 4453.49319 | -0.457989732 | 5.58E-03 |
| Cttn | 14805.07831 | -0.45815521 | 2.27E-10 |
| Pde4d | 597.7003963 | -0.458230838 | 1.24E-02 |
| Rexo4 | 1818.23831 | -0.459854056 | 1.17E-04 |
| Gid8 | 1780.261878 | -0.45993904 | 1.57E-05 |
| Atp5j | 2735.573287 | -0.460791166 | 1.10E-06 |
| Ppic | 4106.544221 | -0.461060457 | 3.40E-04 |
| Serp1 | 6384.5106 | -0.461236466 | 2.35E-03 |
| Qdpr | 1935.818044 | -0.461416758 | 4.42E-03 |
| Rab1a | 7539.257883 | -0.461658229 | 7.13E-06 |

| | | | |
|---|---|---|---|
| Zfp444 | 542.1527879 | -0.461689614 | 2.25E-03 |
| Camk2d | 6564.725921 | -0.462350317 | 2.36E-04 |
| Cdk5rap2 | 4844.501444 | -0.462680367 | 6.61E-03 |
| Ndufab1 | 1673.332492 | -0.462810235 | 8.83E-03 |
| Swap70 | 2724.175662 | -0.463203841 | 2.99E-03 |
| Hacd2 | 1385.12878 | -0.463713992 | 1.05E-02 |
| Rab29 | 491.9800228 | -0.464637583 | 3.19E-03 |
| Cradd | 175.9835452 | -0.465958356 | 1.23E-02 |
| Cdk13 | 2416.935395 | -0.466050022 | 6.36E-07 |
| Rfx5 | 647.994861 | -0.467134633 | 1.27E-03 |
| Fah | 863.0818308 | -0.467206324 | 2.58E-04 |
| Acap2 | 6513.117941 | -0.46802573 | 1.55E-04 |
| Irx2 | 1120.625039 | -0.468419383 | 2.13E-02 |
| Pdzd8 | 3340.981836 | -0.468548874 | 3.90E-02 |
| Rev1 | 1466.258566 | -0.468755191 | 8.69E-05 |
| Ubfd1 | 2088.508959 | -0.469422166 | 4.16E-04 |
| Polr3c | 1444.932593 | -0.469448419 | 1.73E-04 |
| Med22 | 2582.347251 | -0.47096762 | 1.15E-03 |
| 4931406P16Rik | 2722.4401 | -0.472191903 | 9.80E-06 |
| Fam204a | 644.7460821 | -0.472939184 | 2.53E-02 |
| Dtymk | 1449.195743 | -0.473005226 | 1.29E-02 |
| Qrich1 | 3515.068439 | -0.473700229 | 1.13E-08 |
| Mgat5 | 813.1866966 | -0.474036656 | 6.50E-04 |
| Qser1 | 2903.660055 | -0.474249824 | 8.04E-06 |
| Cdc42ep4 | 1198.950253 | -0.474251606 | 2.80E-03 |
| Ezh2 | 1645.93811 | -0.474270361 | 2.59E-02 |
| Gtf2e2 | 1104.94076 | -0.474513237 | 1.47E-06 |
| Gemin7 | 920.3277622 | -0.474639296 | 9.75E-05 |
| Eri1 | 1694.265833 | -0.474643128 | 1.84E-03 |
| Spata2 | 1154.432802 | -0.476265447 | 7.19E-05 |
| Mxra7 | 2456.24539 | -0.476481125 | 4.95E-04 |
| Ap3s1 | 6475.604168 | -0.476898398 | 2.62E-02 |
| Orai2 | 1421.004878 | -0.476964494 | 3.07E-04 |
| Clasp1 | 4060.473029 | -0.477270036 | 1.95E-09 |
| Usp38 | 1425.829906 | -0.477610874 | 3.98E-08 |
| Meis3 | 878.4392067 | -0.478077493 | 2.50E-04 |
| Twistnb | 808.5278395 | -0.478205255 | 2.08E-02 |
| Epdr1 | 994.1146596 | -0.478349192 | 1.88E-03 |
| Zfp212 | 832.4312734 | -0.478399721 | 2.24E-04 |
| Usp48 | 2411.170558 | -0.478845323 | 2.16E-04 |
| Ldlrap1 | 698.7591419 | -0.479345246 | 1.31E-02 |
| Rtl8b | 1525.92564 | -0.479809562 | 3.35E-03 |
| Rpl7l1 | 3740.19368 | -0.480507496 | 1.34E-03 |
| Ptbp2 | 996.7176837 | -0.480598358 | 2.08E-02 |
| 1110051M20Rik | 1345.512438 | -0.480697012 | 1.11E-04 |
| Sirt4 | 287.7350307 | -0.481288054 | 2.78E-02 |
| Rmdn1 | 288.7588206 | -0.481486127 | 5.70E-04 |

| | | | |
|---|---|---|---|
| Nkrf | 418.5610682 | -0.481688812 | 5.19E-04 |
| Pcmtd2 | 1549.253239 | -0.482015204 | 1.12E-02 |
| Rara | 1780.058804 | -0.48277357 | 3.57E-04 |
| Pigc | 1164.872634 | -0.483472427 | 1.77E-03 |
| Pla2r1 | 1271.627081 | -0.48352203 | 2.44E-02 |
| 9430038I01Rik | 156.3527109 | -0.483523656 | 5.39E-03 |
| Rab24 | 1438.55139 | -0.483992652 | 4.73E-07 |
| Rnd2 | 944.1092971 | -0.484230347 | 1.98E-02 |
| Alg11 | 1483.530021 | -0.484622071 | 1.50E-03 |
| Ap3m1 | 2675.647402 | -0.485006554 | 1.72E-02 |
| Scarf2 | 3742.419041 | -0.485132686 | 3.46E-04 |
| Stx17 | 1277.852869 | -0.48531765 | 8.82E-04 |
| Prpf38b | 3109.602672 | -0.485479595 | 3.43E-03 |
| Slc35a2 | 1147.598411 | -0.485652476 | 8.09E-04 |
| Mycbp | 444.75304 | -0.486455737 | 4.63E-02 |
| Man2a2 | 5486.323806 | -0.487553473 | 1.46E-05 |
| Senp7 | 1089.772864 | -0.488214782 | 3.99E-03 |
| Ston1 | 4324.364224 | -0.488216554 | 1.79E-03 |
| Zfp746 | 1625.00071 | -0.488397852 | 2.69E-07 |
| Nck1 | 1126.62883 | -0.488573594 | 2.12E-03 |
| Taf2 | 2167.708385 | -0.488753555 | 4.18E-17 |
| Atg10 | 242.0275732 | -0.489116176 | 1.11E-03 |
| Mgrn1 | 4288.647559 | -0.489295558 | 5.13E-11 |
| Pon2 | 996.9557705 | -0.489683452 | 1.61E-02 |
| Ammecr1 | 543.3185001 | -0.490230359 | 2.08E-02 |
| Gorab | 1255.96873 | -0.490257647 | 1.34E-04 |
| Pheta1 | 721.9805754 | -0.490566215 | 1.94E-02 |
| Itch | 3325.939321 | -0.490921168 | 7.26E-06 |
| Tmem87a | 1703.135091 | -0.491544428 | 1.58E-03 |
| Abhd17b | 807.0306078 | -0.491736836 | 3.31E-02 |
| Ormdl1 | 569.7657093 | -0.491815814 | 4.22E-02 |
| Bptf | 6943.697425 | -0.49188522 | 1.00E-04 |
| Rnf135 | 468.7873319 | -0.492088029 | 1.59E-04 |
| Psd | 627.8684929 | -0.492160578 | 3.68E-04 |
| Sptlc1 | 1235.658035 | -0.492584015 | 4.26E-09 |
| Cnot11 | 1342.876554 | -0.492984179 | 6.71E-06 |
| Gkap1 | 1017.367931 | -0.493116503 | 7.69E-04 |
| Zfp316 | 1016.687164 | -0.493139849 | 1.28E-03 |
| Ipo13 | 5588.190074 | -0.493382813 | 7.57E-05 |
| Fam122a | 464.1460504 | -0.494466934 | 1.12E-04 |
| Fra10ac1 | 379.3987442 | -0.49450138 | 1.52E-02 |
| Mum1 | 1919.342188 | -0.494552816 | 7.81E-07 |
| Ebpl | 299.9519528 | -0.496456015 | 1.12E-02 |
| Mrpl36 | 758.7165102 | -0.49692875 | 2.49E-05 |
| Pcdh18 | 4140.879768 | -0.496987494 | 4.52E-04 |
| Cse1l | 4120.440244 | -0.497667607 | 6.39E-03 |
| Bcl7c | 1233.600283 | -0.497709747 | 1.30E-03 |

| | | | |
|---|---|---|---|
| Uhmk1 | 4265.778395 | -0.497714875 | 1.35E-07 |
| Mphosph8 | 1179.310236 | -0.497892205 | 2.83E-02 |
| NA | 479.4478713 | -0.498341668 | 4.86E-04 |
| Hectd1 | 10105.91826 | -0.498450043 | 1.46E-04 |
| Thada | 1328.834289 | -0.498497446 | 2.75E-05 |
| Acadsb | 2549.580084 | -0.498729551 | 3.77E-02 |
| Zkscan5 | 710.4717409 | -0.498762532 | 2.13E-03 |
| Zfp992 | 359.8629747 | -0.498823777 | 7.48E-03 |
| Erlin2 | 2954.042327 | -0.500603003 | 1.17E-04 |
| Phf3 | 5509.061134 | -0.500627559 | 1.55E-05 |
| Pip4k2b | 4426.450694 | -0.50065757 | 7.43E-09 |
| Mlst8 | 977.1244368 | -0.500715522 | 8.55E-05 |
| Washc5 | 9453.018285 | -0.50132075 | 3.64E-08 |
| Man1a2 | 4199.116273 | -0.501426054 | 2.49E-03 |
| Hmgn2 | 1321.386804 | -0.502216698 | 7.02E-03 |
| Pip4p2 | 1878.030152 | -0.502502549 | 1.04E-04 |
| Cwc22 | 1548.880878 | -0.502703139 | 7.88E-03 |
| 2-Mar | 1502.229125 | -0.502946075 | 2.22E-04 |
| Elmod3 | 569.4908347 | -0.503198942 | 1.69E-05 |
| Pcnt | 1725.263666 | -0.504097666 | 1.24E-02 |
| Prpf3 | 1244.119026 | -0.504117961 | 1.87E-03 |
| Ift122 | 1556.937666 | -0.504129502 | 8.39E-05 |
| Ptma | 34544.23717 | -0.504492934 | 5.56E-04 |
| Kpna3 | 2411.979668 | -0.505008101 | 6.44E-03 |
| D17H6S53E | 459.8831001 | -0.505296507 | 2.67E-03 |
| Amer1 | 1350.755714 | -0.505341055 | 3.06E-09 |
| Bckdhb | 830.3338723 | -0.505627821 | 2.71E-03 |
| Wdr19 | 1255.580238 | -0.505972351 | 1.86E-04 |
| Sntb2 | 2942.491811 | -0.506131741 | 6.95E-03 |
| Osbpl8 | 1429.936323 | -0.506153827 | 7.95E-03 |
| Hmgn3 | 770.6502188 | -0.506869293 | 1.42E-02 |
| Trafd1 | 1802.932578 | -0.507194404 | 1.80E-03 |
| Timp2 | 44603.91543 | -0.507210738 | 2.07E-04 |
| Trim56 | 2778.079535 | -0.5082975 | 4.51E-05 |
| Armc9 | 957.2896761 | -0.508689832 | 1.01E-06 |
| Pex2 | 1610.317343 | -0.508712784 | 1.63E-06 |
| Ankrd52 | 3927.4539 | -0.509026886 | 9.84E-04 |
| Pigl | 298.5422387 | -0.509650582 | 1.06E-03 |
| Dtx3l | 451.8181588 | -0.509770994 | 1.13E-04 |
| Trub2 | 1464.184844 | -0.510211715 | 9.63E-07 |
| Mfn1 | 2369.80232 | -0.510717454 | 7.10E-04 |
| Bcor | 1899.950226 | -0.511113188 | 4.58E-05 |
| Pigs | 4158.40818 | -0.511194259 | 5.38E-04 |
| Ttc7 | 1120.005252 | -0.511194331 | 1.68E-03 |
| Eif4g3 | 5017.46316 | -0.511395222 | 5.58E-10 |
| Tada2b | 1040.973773 | -0.511704221 | 5.62E-08 |
| Dbt | 333.5748819 | -0.511726972 | 1.86E-02 |

| | | | |
|---|---|---|---|
| Zfyve16 | 1187.000652 | -0.512278304 | 1.71E-02 |
| Add3 | 3445.316954 | -0.512328307 | 3.75E-04 |
| Eif5 | 9245.504961 | -0.512596912 | 1.36E-05 |
| Rab43 | 2992.193124 | -0.513111547 | 8.46E-05 |
| Pank1 | 534.8184598 | -0.513231625 | 1.67E-02 |
| Vamp5 | 899.8965301 | -0.513456837 | 3.98E-02 |
| Rprd2 | 1838.435115 | -0.513635615 | 7.06E-08 |
| Prr3 | 728.9177642 | -0.51371064 | 1.57E-06 |
| Sec61a2 | 1335.17424 | -0.513927572 | 6.12E-03 |
| Snapc1 | 358.4501178 | -0.514329376 | 1.65E-04 |
| Cope | 2935.825434 | -0.514791347 | 3.31E-05 |
| Ddr2 | 21485.53935 | -0.514877875 | 6.68E-03 |
| Mlycd | 405.9267261 | -0.51509979 | 1.01E-03 |
| Afg1l | 436.7627528 | -0.515574277 | 1.03E-03 |
| Ptpa | 5174.820688 | -0.515818126 | 8.66E-04 |
| Trmt61a | 494.8076108 | -0.516216308 | 4.72E-03 |
| Slc29a3 | 2607.580111 | -0.51673192 | 7.67E-04 |
| Ddx28 | 579.0195724 | -0.516740224 | 3.20E-04 |
| Arhgef12 | 4275.790492 | -0.516776323 | 9.50E-10 |
| Msl2 | 1328.305038 | -0.51700422 | 9.54E-08 |
| Dab2 | 7942.183086 | -0.517235741 | 1.23E-02 |
| Kansl1 | 3445.684526 | -0.517359402 | 4.32E-07 |
| Itgb5 | 5005.644612 | -0.517581756 | 9.18E-03 |
| Znhit3 | 321.3577153 | -0.517720703 | 1.05E-02 |
| R74862 | 140.0729216 | -0.517932965 | 1.62E-02 |
| Chaf1b | 1033.445061 | -0.518006922 | 4.39E-02 |
| Mettl17 | 418.7816212 | -0.518173061 | 1.27E-02 |
| Amd1 | 4601.579232 | -0.518224821 | 1.95E-03 |
| AW554918 | 348.6668057 | -0.518805323 | 1.97E-03 |
| Smim19 | 793.360551 | -0.518909349 | 5.47E-04 |
| AI413582 | 932.4708856 | -0.518991655 | 3.81E-03 |
| Ice1 | 3774.633263 | -0.519511456 | 1.22E-05 |
| Inpp1 | 473.1785364 | -0.519841818 | 1.54E-03 |
| Dars2 | 894.803958 | -0.520648593 | 2.37E-02 |
| Lig4 | 448.5148621 | -0.520840236 | 1.13E-02 |
| Rft1 | 677.9886492 | -0.520919901 | 5.57E-05 |
| Ncdn | 2384.110763 | -0.522150878 | 6.88E-06 |
| Mterf1a | 128.382231 | -0.522714268 | 3.95E-02 |
| Lrig1 | 3532.647732 | -0.523474282 | 2.97E-03 |
| Acadm | 2856.942427 | -0.523490948 | 7.42E-07 |
| Rap1a | 1336.699842 | -0.524414054 | 1.16E-02 |
| Ccnq | 612.2135084 | -0.524951147 | 1.33E-04 |
| Zfp664 | 3680.499792 | -0.525738747 | 5.92E-08 |
| Stk38l | 824.657163 | -0.525992817 | 2.31E-06 |
| Zmym1 | 884.2906158 | -0.526008354 | 7.48E-04 |
| Sc5d | 2351.772826 | -0.526647012 | 2.96E-02 |
| Mfsd14b | 1609.43286 | -0.526881894 | 1.07E-03 |

| | | | |
|---|---|---|---|
| Brms1l | 1453.82211 | -0.527956915 | 3.33E-04 |
| Klhl8 | 432.6864808 | -0.527989542 | 1.29E-04 |
| Atp10a | 1864.778291 | -0.528213479 | 3.77E-02 |
| Ccdc115 | 665.7497048 | -0.528348545 | 1.01E-03 |
| Tmem218 | 321.622946 | -0.528672034 | 3.14E-03 |
| Usp11 | 2202.578828 | -0.529030855 | 4.94E-03 |
| Chd4 | 15088.84888 | -0.529394082 | 1.44E-09 |
| Rtf1 | 2978.284882 | -0.529468869 | 7.79E-06 |
| Cdk20 | 349.3497726 | -0.529728625 | 1.39E-03 |
| Polr3e | 952.8042013 | -0.52986793 | 1.23E-03 |
| Zcchc7 | 971.1907976 | -0.530725585 | 1.31E-02 |
| Fbxl17 | 1422.489761 | -0.530925281 | 8.54E-07 |
| Ehbp1l1 | 2904.929473 | -0.531049183 | 1.55E-05 |
| Mrpl41 | 755.618332 | -0.531051595 | 3.01E-04 |
| Trmt2a | 1583.31861 | -0.531147189 | 4.79E-08 |
| Ints4 | 1468.076545 | -0.53136081 | 9.38E-07 |
| Fam118a | 816.1656045 | -0.531898512 | 5.66E-06 |
| Vps4b | 3274.311852 | -0.531938282 | 3.47E-05 |
| Lca5 | 237.679466 | -0.532067671 | 1.22E-02 |
| Kdm1b | 834.1471258 | -0.532419366 | 3.39E-03 |
| Fam210b | 928.8968634 | -0.532740103 | 8.65E-03 |
| Tirap | 1020.400692 | -0.532958841 | 1.09E-04 |
| Apoo | 571.4592001 | -0.533253669 | 3.42E-04 |
| Zfp27 | 492.8091239 | -0.533608474 | 2.51E-06 |
| Ranbp1 | 4886.197583 | -0.534477787 | 2.76E-02 |
| Ipo8 | 2808.34261 | -0.534525128 | 5.27E-07 |
| Fam110b | 769.9936195 | -0.534672902 | 3.82E-02 |
| 9330104G04Rik | 82.39673676 | -0.535523524 | 2.24E-02 |
| Prdm2 | 2816.518592 | -0.535548073 | 7.58E-09 |
| Hypk | 259.71492 | -0.535991321 | 1.10E-02 |
| Zbtb8a | 353.867421 | -0.536366057 | 4.26E-03 |
| Plekho1 | 1741.480657 | -0.536683743 | 3.33E-02 |
| Fam120c | 578.4235692 | -0.537296125 | 7.80E-03 |
| Dexi | 703.8502771 | -0.537412659 | 1.49E-04 |
| Abtb2 | 1084.725545 | -0.537580616 | 3.52E-03 |
| Pars2 | 317.5111591 | -0.53771539 | 4.14E-02 |
| Depdc5 | 1318.126855 | -0.537979408 | 1.47E-03 |
| Zc3h12a | 425.3628521 | -0.53822177 | 8.00E-05 |
| Rab9 | 2610.758043 | -0.538726572 | 1.54E-02 |
| Coro1c | 5207.156324 | -0.539055664 | 1.34E-04 |
| Trmu | 326.8927546 | -0.539208527 | 1.96E-03 |
| NA | 270.2693905 | -0.540273138 | 2.54E-02 |
| Zmat5 | 369.1131801 | -0.540651013 | 2.22E-04 |
| Parp2 | 627.2086813 | -0.541554916 | 1.99E-03 |
| Srsf11 | 4467.49277 | -0.541682295 | 6.65E-04 |
| Spred3 | 503.1761065 | -0.541729666 | 3.70E-03 |
| Arl5b | 1105.834095 | -0.542061304 | 3.23E-03 |

| | | | |
|---|---|---|---|
| Abl1 | 7240.633084 | -0.542584164 | 4.57E-08 |
| Elf2 | 3398.181671 | -0.542610279 | 3.82E-08 |
| Mdn1 | 3461.176169 | -0.544067276 | 1.68E-02 |
| NA | 2691.384087 | -0.544130095 | 2.06E-04 |
| E130317F20Rik | 111.3208225 | -0.545185538 | 3.34E-02 |
| Cep19 | 695.464579 | -0.546178518 | 8.16E-08 |
| Ank2 | 3380.138198 | -0.546477316 | 1.03E-03 |
| Peli3 | 459.9064544 | -0.547287751 | 2.80E-02 |
| Ampd3 | 3598.275107 | -0.550045807 | 1.66E-04 |
| Snapc3 | 1027.631758 | -0.550689553 | 3.13E-05 |
| Mpdu1 | 1210.812562 | -0.551020729 | 1.64E-05 |
| Usb1 | 1737.627401 | -0.551438837 | 8.14E-06 |
| Kidins220 | 6720.791685 | -0.551681069 | 3.81E-06 |
| Lrrc45 | 624.2525475 | -0.551853525 | 1.09E-03 |
| Selenot | 3177.042245 | -0.551880222 | 1.19E-02 |
| Laptm4a | 29285.87798 | -0.551889687 | 6.80E-04 |
| Eny2 | 1448.613816 | -0.552106675 | 2.71E-06 |
| Pim2 | 183.1195995 | -0.553032782 | 4.04E-02 |
| Pard6g | 691.6014837 | -0.553076613 | 7.41E-05 |
| Pyroxd1 | 888.8072622 | -0.553251389 | 9.75E-03 |
| Creg1 | 2258.352822 | -0.553671463 | 1.03E-02 |
| Zfp617 | 1019.171138 | -0.554128396 | 1.88E-02 |
| Ninl | 1227.198457 | -0.554188371 | 1.37E-03 |
| Ncs1 | 2519.874797 | -0.55532277 | 5.61E-05 |
| Scfd2 | 558.8131694 | -0.555715095 | 2.86E-04 |
| Atat1 | 753.3053063 | -0.555781622 | 1.14E-02 |
| Hp1bp3 | 6438.328602 | -0.556060385 | 1.92E-06 |
| Dpysl2 | 1095.968635 | -0.556146024 | 2.34E-03 |
| Slc39a9 | 2865.46119 | -0.556323016 | 6.17E-10 |
| Zfp871 | 1310.679291 | -0.556507199 | 7.36E-04 |
| Lima1 | 14608.60545 | -0.557227387 | 3.08E-04 |
| Pgghg | 1003.379207 | -0.557246964 | 6.67E-07 |
| Gzf1 | 2233.832383 | -0.557486041 | 6.37E-05 |
| Nphp4 | 152.6202732 | -0.557709415 | 3.83E-03 |
| Nfya | 2749.990945 | -0.557907211 | 2.28E-05 |
| Gas2l1 | 1866.627813 | -0.558476456 | 3.29E-06 |
| Zmym2 | 2205.020865 | -0.558685095 | 2.30E-05 |
| Taf4 | 981.5170063 | -0.55886077 | 3.85E-05 |
| Pole3 | 2616.86679 | -0.559043161 | 8.25E-05 |
| Ep300 | 3594.368947 | -0.559443757 | 6.12E-14 |
| Gm10182 | 1077.283482 | -0.559547385 | 2.70E-05 |
| Irak2 | 771.0028241 | -0.559597726 | 9.19E-04 |
| Hdhd3 | 255.8322952 | -0.560310394 | 2.23E-02 |
| Brd7 | 3005.751299 | -0.56051069 | 1.43E-04 |
| Tmem220 | 63.81390771 | -0.560635327 | 4.64E-02 |
| Snx19 | 1980.259491 | -0.560933607 | 2.77E-10 |
| Sh2b2 | 263.5269605 | -0.56109882 | 8.93E-03 |

| | | | |
|---|---|---|---|
| Zbtb39 | 784.3304255 | -0.561303346 | 1.97E-06 |
| Gm6483 | 1650.189577 | -0.561610678 | 2.95E-03 |
| Eipr1 | 580.7879384 | -0.562983397 | 1.39E-03 |
| Hnrnpab | 12537.02073 | -0.56320045 | 1.28E-03 |
| Zfp112 | 141.6587009 | -0.563227101 | 1.50E-02 |
| Ubn1 | 5569.528947 | -0.563321034 | 6.82E-09 |
| Kdsr | 1932.623425 | -0.563371667 | 9.40E-06 |
| Naip5 | 268.7274309 | -0.563991319 | 8.87E-04 |
| Pgd | 3686.528964 | -0.564519488 | 2.77E-06 |
| Antxr1 | 10062.55027 | -0.564859163 | 3.00E-03 |
| Zkscan17 | 2056.602394 | -0.564965225 | 3.27E-03 |
| Ado | 1738.480241 | -0.565011922 | 5.22E-09 |
| NA | 83.4285938 | -0.565057882 | 2.69E-02 |
| Tmx1 | 1818.606557 | -0.565086721 | 2.58E-05 |
| Pnpla8 | 2288.601537 | -0.565375047 | 1.04E-03 |
| Nol8 | 839.9327086 | -0.56593094 | 4.72E-02 |
| Zfp46 | 933.397489 | -0.565982563 | 1.76E-04 |
| Pptc7 | 1901.222242 | -0.566208026 | 8.12E-07 |
| Mettl5 | 275.6754539 | -0.566691481 | 6.10E-03 |
| Gm10039 | 530.8826632 | -0.566702507 | 2.93E-02 |
| Tbc1d32 | 741.9345639 | -0.567135586 | 5.58E-04 |
| Uba6 | 1719.595584 | -0.56743535 | 2.86E-02 |
| Ptpn11 | 9188.837608 | -0.568001303 | 4.74E-05 |
| Rab20 | 279.5572422 | -0.568425633 | 1.88E-02 |
| Gm20716 | 4001.886924 | -0.568829385 | 3.23E-12 |
| Chmp6 | 1112.882257 | -0.568843465 | 2.84E-04 |
| Cdc25a | 821.8925884 | -0.569064055 | 2.22E-04 |
| Tmem242 | 1643.421866 | -0.569079244 | 4.24E-04 |
| Abhd14b | 229.6613872 | -0.569111061 | 2.65E-02 |
| F2r | 8616.173877 | -0.569577359 | 1.92E-06 |
| Vkorc1l1 | 1525.062969 | -0.56997078 | 4.42E-12 |
| Glipr2 | 2939.975943 | -0.570227687 | 1.66E-04 |
| Rab11fip3 | 2701.591854 | -0.570249243 | 6.45E-10 |
| Hdac1 | 4349.289025 | -0.570444716 | 9.17E-06 |
| Tmem87b | 2010.229776 | -0.571409523 | 6.42E-04 |
| Pawr | 993.2221019 | -0.572052488 | 5.74E-06 |
| Nacc1 | 5060.814783 | -0.572191495 | 9.36E-06 |
| Apool | 575.8688515 | -0.572945725 | 8.32E-05 |
| Nup93 | 2169.44978 | -0.573471828 | 4.39E-03 |
| 4930579G18Rik | 123.347476 | -0.574079397 | 8.18E-03 |
| Tmem123 | 3880.257383 | -0.574098822 | 2.19E-06 |
| Rcbtb1 | 2189.384948 | -0.574105663 | 1.65E-06 |
| Cpeb1 | 899.5519733 | -0.574301795 | 2.75E-03 |
| Iqsec2 | 575.0814242 | -0.574930755 | 1.37E-03 |
| Zfp747 | 399.2398778 | -0.575110785 | 2.82E-05 |
| Zfp207 | 5415.144154 | -0.575119415 | 2.20E-08 |
| Cyb561d1 | 293.9033658 | -0.57546673 | 3.58E-05 |

| | | | |
|---|---|---|---|
| Zfand1 | 845.8720894 | -0.575676418 | 2.34E-04 |
| Oat | 4512.35951 | -0.575822033 | 8.05E-05 |
| 1110059G10Rik | 340.4845622 | -0.576425946 | 1.83E-03 |
| 8030462N17Rik | 494.1981898 | -0.577002273 | 6.53E-04 |
| Zfand5 | 4632.448716 | -0.577322768 | 4.46E-05 |
| NA | 1559.71249 | -0.577433576 | 7.69E-04 |
| Nfatc4 | 1786.320943 | -0.57801284 | 1.17E-03 |
| Xdh | 15174.39706 | -0.578453794 | 1.02E-02 |
| Stt3a | 9093.140975 | -0.57851821 | 1.00E-06 |
| Fem1c | 1546.41211 | -0.578747442 | 6.87E-13 |
| Scp2 | 5871.714806 | -0.579401392 | 5.42E-07 |
| Slc33a1 | 1241.532103 | -0.579407173 | 1.37E-05 |
| Crybg3 | 324.3155032 | -0.57991812 | 5.69E-03 |
| Mppe1 | 337.864424 | -0.580006268 | 2.17E-04 |
| Pafah1b2 | 3932.142107 | -0.580873362 | 2.21E-11 |
| Rgmb | 2135.748027 | -0.581574171 | 1.69E-02 |
| Slc25a46 | 1399.314162 | -0.581622142 | 1.11E-03 |
| H2-Ke6 | 877.4545958 | -0.582021523 | 1.26E-03 |
| Smim11 | 611.446549 | -0.582439663 | 4.84E-08 |
| Cyth3 | 4789.656829 | -0.582632366 | 8.26E-05 |
| Cptp | 462.7882176 | -0.582918388 | 1.07E-03 |
| Tsga10 | 265.0241548 | -0.583185688 | 2.29E-03 |
| Uqcrb | 2131.306192 | -0.58351797 | 1.89E-06 |
| Fancl | 733.7646602 | -0.583932306 | 1.85E-02 |
| Hif1an | 2520.763791 | -0.58411954 | 2.40E-11 |
| Nsdhl | 4567.873814 | -0.584242744 | 4.14E-02 |
| Fmn1 | 2418.368649 | -0.584292461 | 4.80E-03 |
| Mthfr | 3144.198278 | -0.584594221 | 7.94E-06 |
| Rdx | 5160.05572 | -0.58475395 | 1.32E-05 |
| Trak2 | 2626.579049 | -0.584928713 | 1.22E-06 |
| Tefm | 293.5605631 | -0.58603624 | 1.19E-03 |
| Sbk1 | 915.23951 | -0.586137721 | 8.32E-03 |
| Tmem250-ps | 1722.302464 | -0.586444874 | 3.63E-07 |
| Uprt | 713.1006371 | -0.586870571 | 6.60E-04 |
| Dnal4 | 1352.547094 | -0.587157685 | 2.47E-06 |
| Osbpl1a | 1582.537579 | -0.587460633 | 1.77E-05 |
| Sipa1 | 1737.308702 | -0.587676668 | 5.91E-05 |
| Traf3 | 1527.743095 | -0.587908592 | 3.10E-03 |
| Tiam1 | 806.6891846 | -0.588671745 | 9.41E-03 |
| Zfp706 | 4663.743055 | -0.588922522 | 3.96E-06 |
| Ctnnal1 | 2217.420049 | -0.589699312 | 9.71E-04 |
| Slc25a44 | 2191.360797 | -0.590595466 | 1.64E-04 |
| Vegfb | 3825.597118 | -0.590799731 | 1.34E-03 |
| Exoc5 | 1533.417989 | -0.590830694 | 7.77E-03 |
| Zcchc4 | 584.7811622 | -0.59096568 | 6.96E-05 |
| Ang | 240.4623036 | -0.59117521 | 1.14E-02 |
| Ckap5 | 6224.732297 | -0.591223517 | 2.17E-02 |

| | | | |
|---|---|---|---|
| Dhx36 | 3037.663644 | -0.592090523 | 4.27E-03 |
| Sirt1 | 358.785771 | -0.592615225 | 3.03E-02 |
| Dync2li1 | 1037.764832 | -0.592822105 | 3.59E-04 |
| Ift22 | 607.373969 | -0.592835607 | 1.33E-07 |
| Rab23 | 1604.157666 | -0.593299194 | 5.92E-05 |
| Gpatch11 | 1362.790505 | -0.593357891 | 6.47E-07 |
| R3hcc1 | 995.4229444 | -0.593441827 | 2.93E-04 |
| Mocs3 | 161.1274591 | -0.593676594 | 1.43E-03 |
| Setd7 | 5112.78778 | -0.593780204 | 2.24E-06 |
| Arid1a | 3254.783003 | -0.593847177 | 1.74E-07 |
| Zfp932 | 375.9339726 | -0.593924377 | 4.39E-02 |
| Galnt11 | 1119.990764 | -0.593996813 | 2.66E-06 |
| Pced1b | 224.7482532 | -0.594717593 | 2.24E-04 |
| Arhgap39 | 893.335859 | -0.596035789 | 4.74E-04 |
| Pphln1 | 1556.478777 | -0.59625031 | 2.12E-12 |
| Cd34 | 10535.55549 | -0.596286848 | 1.76E-04 |
| Tmem165 | 2684.294746 | -0.59676413 | 8.68E-09 |
| Gm6741 | 320.3714133 | -0.597270738 | 1.56E-02 |
| Cox7b | 3207.21518 | -0.597562624 | 6.57E-04 |
| Zfp961 | 486.3026627 | -0.597761499 | 1.00E-02 |
| Dcp1a | 1156.200016 | -0.598160202 | 6.37E-12 |
| Osbpl7 | 425.4315274 | -0.598495975 | 8.64E-07 |
| Mzt1 | 737.1184684 | -0.598526563 | 2.32E-05 |
| Aggf1 | 1481.03892 | -0.598621287 | 1.36E-06 |
| Gdpgp1 | 236.3563167 | -0.598905001 | 1.33E-03 |
| Apip | 617.6113811 | -0.599062508 | 8.76E-05 |
| Rab8b | 1190.981597 | -0.599593586 | 3.65E-04 |
| Otulin | 961.990477 | -0.599896737 | 4.52E-08 |
| Atp6ap2 | 4358.08888 | -0.600412831 | 2.01E-05 |
| Zc3h4 | 3335.352754 | -0.600980239 | 2.45E-06 |
| Rufy2 | 1450.023558 | -0.601322656 | 2.58E-04 |
| Chpf2 | 2871.583349 | -0.60136685 | 1.82E-07 |
| Tdg | 173.5710662 | -0.601431961 | 1.98E-02 |
| Gm17344 | 712.596112 | -0.601712085 | 2.03E-11 |
| Tsn | 3126.119514 | -0.602306144 | 3.04E-09 |
| Zfp667 | 89.21251769 | -0.602347531 | 3.06E-02 |
| Ccdc91 | 1289.115254 | -0.602744054 | 1.15E-09 |
| Wapl | 2654.610917 | -0.603055924 | 3.61E-03 |
| St3gal2 | 8902.023726 | -0.603251182 | 2.78E-09 |
| Lifr | 229.2415172 | -0.603528036 | 7.50E-03 |
| Thsd7a | 4019.127381 | -0.603716139 | 2.11E-03 |
| A330035P11Rik | 70.16852895 | -0.60423228 | 2.03E-02 |
| Chst14 | 1836.493334 | -0.604274753 | 2.89E-07 |
| Nek1 | 1368.716511 | -0.604386157 | 1.39E-02 |
| Tbck | 763.4982955 | -0.604783576 | 1.51E-06 |
| Anapc13 | 612.8802342 | -0.604785875 | 7.16E-08 |
| Tecpr2 | 835.5171694 | -0.604923956 | 4.56E-06 |

| | | | |
|---|---|---|---|
| Timm9 | 538.7327621 | -0.605402093 | 1.66E-05 |
| Ppp3cb | 2558.783048 | -0.605417089 | 1.85E-08 |
| Nfix | 9711.486408 | -0.6057442 | 1.80E-06 |
| Btd | 882.1449736 | -0.605808158 | 5.81E-05 |
| Decr1 | 1500.980543 | -0.60583932 | 1.70E-09 |
| Dync2h1 | 1413.254867 | -0.605865096 | 3.41E-03 |
| Ammecr1l | 2249.366946 | -0.606014986 | 3.07E-12 |
| Cfb | 68.25731523 | -0.606653305 | 2.94E-02 |
| Mob3a | 1207.127894 | -0.607120908 | 1.25E-04 |
| Slc1a5 | 4845.639286 | -0.607832715 | 1.96E-03 |
| Aldh6a1 | 1590.691443 | -0.608107201 | 2.67E-03 |
| Rwdd3 | 232.2336999 | -0.60822715 | 1.84E-02 |
| Papola | 6677.657305 | -0.608296638 | 4.39E-07 |
| Cdk7 | 2374.916507 | -0.608581977 | 3.54E-07 |
| Ino80 | 2504.264289 | -0.6090234 | 3.06E-08 |
| Tmem60 | 583.2390495 | -0.609149316 | 4.82E-06 |
| Tfb2m | 1122.816783 | -0.609349233 | 3.81E-06 |
| Zfp36l2 | 7867.303204 | -0.609764819 | 6.14E-04 |
| Star | 63.2158291 | -0.610091699 | 4.91E-02 |
| Med6 | 834.6516389 | -0.610284869 | 6.48E-05 |
| NA | 167.7309758 | -0.610348333 | 6.42E-03 |
| Eif2ak2 | 1733.612018 | -0.610405957 | 9.28E-04 |
| Rex1bd | 1738.186848 | -0.610466721 | 2.42E-05 |
| Naglu | 4106.854751 | -0.610561203 | 4.41E-03 |
| Pip4k2a | 1015.444415 | -0.610821085 | 1.87E-07 |
| Plekhg5 | 645.5685303 | -0.610987639 | 3.00E-02 |
| Serf1 | 393.2430891 | -0.611208147 | 9.00E-03 |
| Sf3b1 | 11144.92847 | -0.611344021 | 2.85E-09 |
| Kctd21 | 352.0213232 | -0.611700319 | 2.28E-03 |
| Hacd1 | 603.4796318 | -0.612006891 | 1.43E-03 |
| NA | 161.3501806 | -0.612071467 | 7.55E-04 |
| Chtop | 5837.65029 | -0.612475368 | 2.26E-11 |
| Pde4a | 1127.432746 | -0.612478152 | 5.58E-06 |
| Tfap4 | 806.8709696 | -0.613310004 | 2.16E-03 |
| Carf | 168.4458168 | -0.614717002 | 4.90E-02 |
| Heatr5a | 3537.032457 | -0.614801613 | 4.39E-14 |
| Nnmt | 470.8866788 | -0.615376219 | 3.39E-03 |
| Rnf13 | 3890.78372 | -0.615427184 | 9.06E-05 |
| Zfp12 | 1260.57049 | -0.616314197 | 1.31E-08 |
| Dhx57 | 1977.793762 | -0.616358884 | 3.72E-12 |
| Cnnm3 | 667.6426996 | -0.616577996 | 8.47E-04 |
| Rundc1 | 970.3247728 | -0.616677531 | 8.60E-08 |
| Gas8 | 1271.176052 | -0.61670467 | 3.77E-05 |
| Skil | 1068.678727 | -0.616746192 | 6.78E-03 |
| Rbm26 | 1857.080921 | -0.61709098 | 1.05E-03 |
| Dtwd2 | 147.383029 | -0.617203206 | 5.21E-04 |
| Zfyve21 | 755.2797833 | -0.617319428 | 2.27E-05 |

| | | | |
|---|---|---|---|
| Nmral1 | 513.5951069 | -0.617349244 | 1.44E-03 |
| Zbtb18 | 764.6264989 | -0.617572759 | 1.18E-02 |
| N4bp2l1 | 517.1616349 | -0.617819231 | 4.32E-02 |
| Herc6 | 237.6921114 | -0.619848989 | 3.17E-02 |
| Zfp407 | 895.1626937 | -0.620024595 | 5.26E-06 |
| Esyt2 | 5933.474857 | -0.620135299 | 4.90E-09 |
| Tpst1 | 2809.181828 | -0.621378375 | 1.95E-05 |
| Azi2 | 1115.706096 | -0.621437484 | 4.84E-08 |
| Lonrf1 | 505.0783429 | -0.621845585 | 4.48E-05 |
| Pcyt1a | 4562.392465 | -0.621873277 | 6.56E-12 |
| Ppp4r1l-ps | 1152.662507 | -0.622304191 | 2.16E-05 |
| Fbxl19 | 1033.975476 | -0.622313741 | 5.12E-07 |
| Man2b1 | 3916.818943 | -0.622800731 | 2.21E-04 |
| Tctn3 | 629.4105587 | -0.622892443 | 9.02E-08 |
| Tmem168 | 2136.488341 | -0.623481271 | 1.81E-04 |
| Zfp532 | 1559.243215 | -0.624050349 | 1.49E-04 |
| Gm17131 | 1124.972203 | -0.624120187 | 3.66E-02 |
| Pls3 | 9293.72303 | -0.624176021 | 5.83E-12 |
| Tsc1 | 3398.460772 | -0.624542937 | 8.08E-07 |
| Gm15398 | 164.4360715 | -0.62464018 | 3.00E-02 |
| Abraxas1 | 442.9505407 | -0.624709554 | 1.68E-04 |
| Erh | 4021.046823 | -0.625185088 | 6.82E-04 |
| Ak3 | 4242.137544 | -0.62551894 | 5.81E-09 |
| Dnaja1 | 8039.758531 | -0.625633159 | 2.33E-08 |
| Lxn | 2852.121736 | -0.625803788 | 7.18E-03 |
| Casp3 | 1397.104307 | -0.626137408 | 1.03E-06 |
| Crk | 5752.903002 | -0.626268178 | 1.31E-07 |
| Yme1l1 | 4626.937224 | -0.626287793 | 2.86E-03 |
| Gpx7 | 623.8937088 | -0.626662828 | 4.76E-04 |
| Zbtb34 | 893.6006717 | -0.626951362 | 3.19E-04 |
| Tusc2 | 636.7772395 | -0.627192685 | 2.45E-04 |
| E130307A14Rik | 356.3034166 | -0.62757262 | 7.30E-04 |
| Srrt | 3532.906012 | -0.627645956 | 3.44E-05 |
| Nhlrc3 | 2088.125293 | -0.629029618 | 2.87E-09 |
| Mllt1 | 2904.39071 | -0.629318792 | 8.16E-09 |
| Ankrd40 | 3457.239698 | -0.629335417 | 4.63E-09 |
| Mfsd14a | 1911.459662 | -0.630089439 | 9.92E-08 |
| Slc43a1 | 365.6670623 | -0.630269544 | 2.73E-02 |
| Ddx5 | 36977.81738 | -0.630573259 | 8.73E-08 |
| Gns | 20094.75228 | -0.630596206 | 1.59E-04 |
| Nrbp2 | 771.4583412 | -0.630828951 | 1.96E-02 |
| Tbcc | 642.9037322 | -0.631708796 | 1.12E-04 |
| Svbp | 1041.238098 | -0.632547166 | 1.55E-06 |
| Nelfa | 1367.005053 | -0.632997142 | 2.17E-11 |
| Glyr1 | 5537.263863 | -0.6336083 | 5.45E-12 |
| Garem2 | 297.7203972 | -0.635018025 | 7.84E-04 |
| Mtcl1 | 1053.202858 | -0.635250906 | 7.62E-05 |

| | | | |
|---|---|---|---|
| Dync1li2 | 5420.762549 | -0.637084308 | 2.86E-10 |
| Cul5 | 1666.263736 | -0.637095509 | 5.78E-03 |
| Tmem126a | 982.003366 | -0.637883111 | 5.17E-03 |
| Fbxo11 | 3313.191893 | -0.637900085 | 1.15E-06 |
| Slmap | 3527.376361 | -0.638311426 | 3.67E-05 |
| Tgs1 | 1089.093533 | -0.638321732 | 1.74E-06 |
| Akt1 | 12758.06405 | -0.638878028 | 1.20E-06 |
| Ext1 | 3885.770872 | -0.639950598 | 2.12E-06 |
| Jarid2 | 391.5625149 | -0.640106859 | 9.39E-03 |
| Pon3 | 413.776097 | -0.640244954 | 9.12E-05 |
| Ddx17 | 14422.11125 | -0.640619944 | 2.27E-06 |
| Rnf213 | 3156.486219 | -0.640631916 | 2.13E-06 |
| Txndc16 | 3149.373031 | -0.640633166 | 2.19E-04 |
| 2500002B13Rik | 80.42307177 | -0.64137108 | 3.29E-02 |
| Zfp800 | 699.3697943 | -0.642042852 | 4.37E-03 |
| Arhgef10l | 1765.531792 | -0.642290033 | 4.90E-04 |
| Tmx3 | 1991.990171 | -0.642416804 | 1.58E-02 |
| Atf2 | 2251.338789 | -0.642643316 | 8.08E-06 |
| Cabin1 | 2885.667804 | -0.643002465 | 3.05E-10 |
| Nrip1 | 567.8159237 | -0.643167139 | 1.79E-04 |
| Anxa5 | 17795.31783 | -0.643199352 | 4.77E-07 |
| Tgfbr2 | 6135.16223 | -0.643929522 | 3.14E-04 |
| Garem1 | 770.4422413 | -0.644152364 | 1.32E-06 |
| Klhl11 | 480.4745573 | -0.64418443 | 4.04E-05 |
| Birc3 | 495.0236977 | -0.644858892 | 1.21E-04 |
| Rtel1 | 1078.20905 | -0.644916191 | 3.35E-02 |
| Asxl2 | 1701.60244 | -0.644949553 | 2.72E-07 |
| Slc25a26 | 255.1476367 | -0.645659264 | 1.02E-04 |
| Dctn1 | 6439.938184 | -0.645854433 | 1.01E-09 |
| Ttc39c | 557.9704609 | -0.646382444 | 2.24E-04 |
| A130010J15Rik | 350.9587272 | -0.646465374 | 1.80E-06 |
| Ttc4 | 1740.905347 | -0.646550695 | 2.26E-11 |
| Gigyf2 | 4619.799954 | -0.648166724 | 1.31E-16 |
| Yap1 | 5868.317144 | -0.648353079 | 6.04E-20 |
| Rnf139 | 1837.564298 | -0.648566759 | 1.39E-11 |
| Mospd2 | 1181.516859 | -0.648649029 | 2.74E-02 |
| Zfp639 | 1185.948539 | -0.648758964 | 1.18E-16 |
| Ppp1r15b | 4964.1046 | -0.648887449 | 4.26E-11 |
| Tor1aip1 | 3198.975944 | -0.649231088 | 1.78E-12 |
| Cstf2 | 1462.967735 | -0.649963609 | 1.59E-03 |
| Kiss1r | 85.47449588 | -0.65007943 | 6.99E-03 |
| 1110032A03Rik | 326.5533345 | -0.650162764 | 3.28E-03 |
| Uggt2 | 1534.658459 | -0.650358722 | 5.33E-04 |
| Pros1 | 1181.249787 | -0.650398293 | 6.15E-05 |
| Nktr | 4432.706255 | -0.650804831 | 4.19E-03 |
| Kif3b | 2552.903667 | -0.650953427 | 1.03E-20 |
| Pin1 | 1061.889102 | -0.651102209 | 9.79E-05 |

| | | | |
|---|---|---|---|
| Atg12 | 4017.335377 | -0.651200264 | 2.80E-03 |
| Mia3 | 7047.789822 | -0.651337842 | 2.89E-09 |
| Foxg1 | 4338.937908 | -0.651396631 | 2.32E-05 |
| Gt(ROSA)26Sor | 115.8026385 | -0.651717669 | 2.25E-03 |
| Tbc1d14 | 2671.591197 | -0.652263409 | 4.85E-07 |
| Dnal1 | 1014.638004 | -0.652562177 | 4.24E-04 |
| Fam219b | 604.1462657 | -0.6526688 | 1.50E-05 |
| R3hcc1l | 960.2885558 | -0.652760565 | 1.97E-05 |
| Cirbp | 1092.334366 | -0.652927566 | 1.89E-04 |
| Kdelc1 | 1895.28066 | -0.653218045 | 5.73E-11 |
| Tymp | 200.2920391 | -0.653369971 | 8.04E-03 |
| Pcyt2 | 535.7741907 | -0.654076977 | 1.52E-02 |
| Cdk5rap1 | 416.9799843 | -0.654997848 | 3.28E-07 |
| Nsrp1 | 872.0705176 | -0.655298816 | 8.70E-03 |
| Dstyk | 1943.600084 | -0.655514641 | 7.51E-10 |
| Gm10146 | 505.8695599 | -0.656417105 | 4.68E-03 |
| Lrfn4 | 2023.205322 | -0.656869571 | 8.68E-07 |
| Myo1e | 5416.527455 | -0.65688092 | 3.88E-04 |
| Nedd9 | 2791.535274 | -0.657085233 | 8.18E-03 |
| Kdm5b | 5947.881651 | -0.657102095 | 4.76E-03 |
| Tle3 | 2473.32434 | -0.657141249 | 2.14E-05 |
| Dram2 | 830.4501593 | -0.657537633 | 4.76E-03 |
| Gnb4 | 263.9667297 | -0.657564569 | 1.34E-02 |
| Phyhd1 | 936.9221417 | -0.658874767 | 2.26E-02 |
| Gm16199 | 125.1724322 | -0.659037948 | 3.79E-02 |
| Mettl27 | 1096.641165 | -0.659318305 | 2.23E-04 |
| Gnai3 | 6342.144305 | -0.659402708 | 4.23E-05 |
| Chtf8 | 5434.622375 | -0.659523328 | 1.37E-07 |
| Eif2ak4 | 3546.105363 | -0.659716827 | 3.63E-11 |
| Cep120 | 1497.538669 | -0.659949143 | 1.32E-10 |
| Idh1 | 8516.619482 | -0.659993418 | 1.89E-08 |
| Gmcl1 | 1500.19973 | -0.660045281 | 1.65E-17 |
| Zfp13 | 87.85007492 | -0.660404523 | 1.85E-02 |
| Zbtb14 | 946.9784755 | -0.660504048 | 1.12E-08 |
| Cep44 | 426.9071182 | -0.660615965 | 7.35E-04 |
| Apc | 3191.587842 | -0.66071471 | 3.81E-03 |
| 2410002F23Rik | 2404.042911 | -0.660748381 | 1.47E-07 |
| Zfp874b | 473.1071347 | -0.661539242 | 3.11E-08 |
| Gm16973 | 160.4522238 | -0.66212368 | 5.02E-04 |
| Hsd17b12 | 4957.384465 | -0.662383725 | 2.28E-04 |
| Ap5s1 | 405.3152319 | -0.663055223 | 2.87E-04 |
| Rpp25l | 408.0873489 | -0.663174583 | 1.05E-05 |
| Trnt1 | 2212.121754 | -0.663380689 | 5.14E-04 |
| Lrrc58 | 3830.129758 | -0.663662746 | 3.02E-08 |
| Spout1 | 972.9814925 | -0.664435879 | 5.95E-04 |
| Gm12473 | 155.3930602 | -0.66456478 | 5.41E-04 |
| Tmem14a | 58.42559324 | -0.664730718 | 2.63E-02 |

| | | | |
|---|---|---|---|
| Tmem17 | 204.5718151 | -0.665249967 | 5.03E-05 |
| Sgpp1 | 3316.645421 | -0.665446907 | 3.78E-09 |
| Fryl | 2801.659489 | -0.665981586 | 8.77E-05 |
| 3110082I17Rik | 709.1107669 | -0.666010995 | 3.81E-05 |
| Taf1b | 795.1423822 | -0.666507527 | 7.26E-05 |
| Lrrc8b | 376.9902679 | -0.666667154 | 5.56E-03 |
| Ppwd1 | 530.3046537 | -0.666854209 | 6.58E-04 |
| Gng10 | 2800.643553 | -0.667403224 | 1.68E-13 |
| Ss18 | 2071.284024 | -0.667965014 | 7.05E-07 |
| Oaz2 | 2466.83586 | -0.668026755 | 4.35E-10 |
| Cacng7 | 2011.225826 | -0.668049756 | 2.09E-03 |
| Ubiad1 | 901.4301508 | -0.669665366 | 3.00E-07 |
| Atxn1l | 3689.513877 | -0.669927637 | 2.35E-27 |
| 4930581F22Rik | 94.75752944 | -0.670792315 | 2.53E-02 |
| Scn1b | 147.6156986 | -0.670869563 | 3.54E-02 |
| Twist1 | 1323.592562 | -0.671721231 | 1.22E-08 |
| Tmem126b | 728.0357211 | -0.672125012 | 8.81E-12 |
| Cuedc1 | 1729.792206 | -0.672364075 | 3.39E-07 |
| Ash1l | 4678.45912 | -0.672849857 | 6.49E-07 |
| Trim13 | 272.2413146 | -0.673089136 | 1.04E-05 |
| Uri1 | 1845.880355 | -0.673378186 | 1.05E-25 |
| Tyro3 | 1115.648355 | -0.67344715 | 1.49E-05 |
| Kif1bp | 2098.509956 | -0.673581118 | 2.32E-07 |
| Dnajc25 | 1205.90302 | -0.67366532 | 2.70E-06 |
| Nmi | 245.734847 | -0.67385527 | 5.01E-05 |
| Ppm1e | 886.2907223 | -0.673938077 | 3.13E-03 |
| Btf3l4 | 4495.978449 | -0.67395366 | 6.61E-14 |
| Nom1 | 1882.71559 | -0.674862574 | 7.77E-11 |
| Ppm1l | 2353.623018 | -0.675112512 | 1.27E-09 |
| Ciao2a | 1186.863786 | -0.675532327 | 5.82E-08 |
| Vmn1r65 | 92.26155685 | -0.675648933 | 8.80E-03 |
| Smad3 | 3160.732874 | -0.676196881 | 3.41E-08 |
| Tubb2b | 2259.346938 | -0.676777806 | 1.95E-04 |
| Nfkbie | 307.6811974 | -0.677012244 | 1.66E-03 |
| Cep128 | 695.3413724 | -0.677117372 | 3.79E-02 |
| Psap | 78612.42307 | -0.677125044 | 8.42E-04 |
| Ip6k2 | 1213.466928 | -0.677361011 | 2.99E-05 |
| Ptges2 | 754.8783035 | -0.677478017 | 2.93E-04 |
| Med13 | 3961.167349 | -0.677670565 | 7.01E-05 |
| Hspa1a | 228.6343034 | -0.678554949 | 2.83E-03 |
| Clip2 | 3685.485315 | -0.678738625 | 1.55E-06 |
| Tgif2 | 708.2075399 | -0.67874931 | 1.16E-04 |
| Pccb | 1287.765021 | -0.679278235 | 4.93E-05 |
| Ccdc25 | 818.1589063 | -0.679671874 | 6.13E-05 |
| Rlim | 4055.693503 | -0.679678278 | 1.26E-04 |
| Iglon5 | 4473.234465 | -0.679814448 | 8.68E-04 |
| Acaca | 6220.310423 | -0.680313474 | 5.43E-06 |

| | | | |
|---|---|---|---|
| Peak1 | 6982.986032 | -0.680688015 | 4.70E-11 |
| Guf1 | 1373.90788 | -0.681017708 | 3.47E-03 |
| 3300002I08Rik | 78.85212045 | -0.681515343 | 7.59E-03 |
| Calm1 | 13158.15346 | -0.681655272 | 2.78E-07 |
| Unc13a | 386.4394156 | -0.682805059 | 4.15E-03 |
| Hltf | 1302.480206 | -0.682874743 | 5.54E-03 |
| Rsf1 | 1494.238977 | -0.682878982 | 4.57E-06 |
| Hnrnpu | 20347.87798 | -0.683060395 | 5.34E-07 |
| Atraid | 2965.766926 | -0.683485415 | 4.43E-06 |
| Tspan17 | 1736.884601 | -0.683737588 | 3.60E-08 |
| Klhl29 | 577.8649113 | -0.683743943 | 2.16E-06 |
| Prr14l | 2104.405396 | -0.68421512 | 9.77E-03 |
| Casp9 | 1187.691822 | -0.68459831 | 1.61E-06 |
| Dbnl | 5109.073654 | -0.685052113 | 3.56E-06 |
| Gm14230 | 113.3153658 | -0.685098043 | 2.96E-02 |
| Ccdc47 | 3476.211324 | -0.685476883 | 6.90E-04 |
| Fam8a1 | 2114.147731 | -0.685947573 | 2.65E-04 |
| Mfap3 | 1621.114974 | -0.686324354 | 5.04E-11 |
| Zfyve26 | 2479.501653 | -0.686680654 | 9.42E-10 |
| Camta2 | 1178.937289 | -0.686718869 | 5.83E-12 |
| Arhgef11 | 2254.102821 | -0.686942199 | 3.36E-19 |
| Ptger4 | 492.752494 | -0.687154629 | 5.64E-04 |
| Adamtsl4 | 642.4400238 | -0.687764664 | 1.32E-04 |
| Rpl22l1 | 5120.112213 | -0.687794315 | 1.41E-04 |
| Ap1ar | 1516.331122 | -0.687896352 | 1.51E-07 |
| Scaf8 | 3158.438473 | -0.688095982 | 1.81E-19 |
| Zfp128 | 209.8468391 | -0.688183823 | 9.28E-05 |
| Fkbp5 | 2306.504311 | -0.689138961 | 1.31E-09 |
| 7-Sep | 6788.598028 | -0.689340286 | 5.55E-03 |
| Gm28036 | 1213.811392 | -0.689669298 | 3.69E-03 |
| S100pbp | 2801.716801 | -0.689783772 | 1.23E-08 |
| Phka1 | 969.7610533 | -0.68987308 | 2.33E-07 |
| Enkd1 | 439.8664695 | -0.689974685 | 2.99E-04 |
| Fam129a | 1172.688172 | -0.690135972 | 6.90E-06 |
| Crtap | 4391.875566 | -0.690266924 | 9.74E-05 |
| Tlcd1 | 600.2415488 | -0.690321235 | 8.21E-05 |
| Lrrc14 | 1426.419279 | -0.691166786 | 1.25E-06 |
| Armcx2 | 5543.690073 | -0.691209125 | 1.38E-12 |
| Zbtb4 | 1221.091028 | -0.691311586 | 1.40E-07 |
| G2e3 | 810.0107457 | -0.691423059 | 4.11E-02 |
| Ap5z1 | 1165.206489 | -0.691874631 | 6.93E-09 |
| Ugdh | 6304.035598 | -0.692497432 | 2.38E-12 |
| Cmc4 | 100.5006959 | -0.693118578 | 2.36E-03 |
| Cbx5 | 6654.599166 | -0.693148757 | 2.13E-03 |
| Clock | 1899.267566 | -0.693901371 | 5.01E-03 |
| Prkar2a | 2154.478436 | -0.693914799 | 2.08E-03 |
| Prps2 | 2712.855431 | -0.694308834 | 8.77E-08 |

| | | | |
|---|---|---|---|
| Zdhhc21 | 1121.368188 | -0.694383826 | 1.44E-02 |
| Gins3 | 349.528308 | -0.695386469 | 4.06E-03 |
| Tcn2 | 1543.355567 | -0.695694619 | 3.02E-03 |
| Mif4gd | 450.899296 | -0.695827744 | 1.83E-07 |
| 2-Sep | 11980.90362 | -0.696487541 | 3.98E-05 |
| Gm4419 | 105.9221896 | -0.696723978 | 1.87E-02 |
| Traf3ip2 | 927.1277928 | -0.697210948 | 6.22E-07 |
| Nsun3 | 307.1312189 | -0.697225293 | 7.02E-03 |
| Speg | 1856.726043 | -0.697253363 | 2.36E-06 |
| Lrrc20 | 643.8913043 | -0.699379918 | 1.52E-03 |
| Arpc5 | 5430.297978 | -0.699542654 | 2.95E-18 |
| Zfp275 | 2068.451814 | -0.699860324 | 1.53E-13 |
| Slc36a4 | 1156.576941 | -0.700237079 | 6.05E-19 |
| Rbfox2 | 9966.948387 | -0.700273196 | 6.29E-10 |
| Ankrd24 | 376.7422993 | -0.700422029 | 1.47E-05 |
| Mdc1 | 2605.907214 | -0.700475458 | 3.10E-03 |
| Npr2 | 2586.247944 | -0.701789798 | 2.34E-15 |
| Spcs2 | 3414.660533 | -0.702309222 | 3.32E-12 |
| Gm13977 | 61.77917577 | -0.70284395 | 2.20E-02 |
| H3f3a | 13435.63948 | -0.702855763 | 2.56E-08 |
| Tmem186 | 774.7522416 | -0.703238354 | 7.19E-09 |
| Scaf4 | 1389.036488 | -0.704290046 | 2.13E-11 |
| 1600014C10Rik | 740.8140799 | -0.704350719 | 9.40E-05 |
| Tcp11l2 | 1451.319384 | -0.704417766 | 1.41E-02 |
| Heg1 | 6138.242062 | -0.704521278 | 4.30E-05 |
| Dennd1a | 1904.296513 | -0.704770149 | 9.67E-07 |
| Cldn34c1 | 161.8468038 | -0.704925212 | 3.49E-02 |
| Cdc27 | 4023.936669 | -0.705107426 | 1.19E-04 |
| Chchd7 | 559.246393 | -0.705789559 | 3.02E-07 |
| Gapvd1 | 3649.129331 | -0.706847707 | 3.66E-13 |
| Furin | 4097.260466 | -0.706905139 | 1.35E-07 |
| Yeats2 | 2822.94201 | -0.707730127 | 1.47E-14 |
| Ocrl | 1987.16698 | -0.707814668 | 6.61E-11 |
| Mcm9 | 451.883513 | -0.707874705 | 4.38E-08 |
| Mlec | 15108.74515 | -0.708297023 | 6.99E-12 |
| Cep68 | 2000.679055 | -0.70917773 | 1.74E-18 |
| Acadvl | 2866.060471 | -0.709373616 | 1.12E-07 |
| Ndufc2 | 1245.322101 | -0.709595515 | 1.34E-05 |
| Gm21092 | 10769.17919 | -0.710088249 | 1.16E-04 |
| Vps13d | 2923.790283 | -0.71032033 | 3.59E-05 |
| Trip12 | 11199.36888 | -0.71035468 | 2.97E-15 |
| Nptn | 6929.496085 | -0.710817071 | 3.24E-13 |
| Hira | 2627.73973 | -0.710880155 | 1.53E-17 |
| Gprasp1 | 801.852662 | -0.710973222 | 1.70E-05 |
| Baz1b | 8196.336798 | -0.711188035 | 4.95E-15 |
| Pdzd4 | 1139.545824 | -0.711728056 | 1.67E-08 |
| Mkks | 849.6034523 | -0.712276253 | 3.18E-05 |

| | | | |
|---|---|---|---|
| Cpa6 | 890.522578 | -0.712582973 | 2.02E-04 |
| Wdr90 | 668.2938756 | -0.712585113 | 2.40E-02 |
| Gusb | 2912.311294 | -0.712827604 | 2.55E-06 |
| LTO1 | 699.7176963 | -0.713441384 | 6.20E-03 |
| Borcs8 | 860.7623581 | -0.713513102 | 1.12E-06 |
| Akap11 | 3406.170716 | -0.713515542 | 4.62E-08 |
| Dynlt3 | 2657.833029 | -0.713812573 | 1.53E-02 |
| Tfam | 1401.597515 | -0.714038002 | 5.83E-07 |
| Tspan14 | 1344.749055 | -0.714751966 | 1.49E-10 |
| Trim68 | 280.7631145 | -0.716166782 | 1.30E-05 |
| Zfp825 | 163.4175946 | -0.716167718 | 1.86E-03 |
| Phlpp2 | 1467.445767 | -0.716541554 | 2.16E-15 |
| Coprs | 814.1861832 | -0.716722742 | 8.89E-10 |
| Ythdc1 | 2829.349268 | -0.717611167 | 2.20E-09 |
| Zdhhc8 | 3036.914019 | -0.717657759 | 4.03E-10 |
| Stx18 | 996.9462278 | -0.717758914 | 1.42E-06 |
| NA | 232.7467139 | -0.718141576 | 1.52E-02 |
| Bach1 | 2273.465941 | -0.718187001 | 1.10E-15 |
| Gm15764 | 62.42387083 | -0.718330163 | 4.57E-02 |
| Crip2 | 63.44237168 | -0.718758339 | 2.98E-02 |
| 1700003F12Rik | 81.47917467 | -0.719081948 | 4.52E-03 |
| Zdhhc20 | 2954.604114 | -0.719554806 | 2.70E-04 |
| Gatb | 891.9343914 | -0.719768746 | 1.83E-03 |
| Timp3 | 19304.01546 | -0.720096712 | 1.27E-03 |
| Angpt2 | 316.2272496 | -0.720166079 | 1.44E-02 |
| Wbp1l | 6033.497275 | -0.720419297 | 1.30E-11 |
| Gm17018 | 524.6653148 | -0.720497604 | 3.12E-13 |
| Polr3h | 617.8748839 | -0.720678448 | 3.75E-04 |
| Tmem251 | 678.3297086 | -0.721229748 | 6.86E-07 |
| Pask | 665.4025986 | -0.721759231 | 3.31E-05 |
| Asxl3 | 95.93112386 | -0.722768888 | 3.94E-02 |
| 1700017B05Rik | 1654.177394 | -0.723031066 | 2.91E-05 |
| Fgd3 | 456.6207369 | -0.72312287 | 4.08E-02 |
| Tyw5 | 397.5307478 | -0.723150862 | 2.11E-02 |
| Gtf2i | 8843.674879 | -0.723483709 | 5.93E-09 |
| Adar | 2479.583721 | -0.72415119 | 4.96E-11 |
| Acbd5 | 1379.102261 | -0.725049522 | 4.50E-03 |
| Kirrel | 13677.00951 | -0.725146487 | 3.45E-26 |
| Hnrnpdl | 5096.050461 | -0.725750769 | 4.33E-06 |
| Fam102b | 2904.554024 | -0.725780976 | 2.16E-12 |
| Inha | 254.1528339 | -0.725786629 | 1.06E-05 |
| AI597479 | 1129.332889 | -0.726151853 | 6.13E-22 |
| Ankrd13b | 2771.960592 | -0.726434474 | 2.13E-07 |
| Plekhm1 | 2389.867812 | -0.726461739 | 6.42E-08 |
| Api5 | 5016.463876 | -0.728610354 | 1.02E-13 |
| Gpalpp1 | 1808.305997 | -0.728908034 | 4.08E-14 |
| Ints6 | 1189.813151 | -0.729625963 | 2.70E-04 |

| | | | |
|---|---|---|---|
| Rab32 | 1574.14668 | -0.730004468 | 3.26E-06 |
| Nlgn2 | 3845.598607 | -0.730253216 | 1.21E-07 |
| Naa30 | 708.3590329 | -0.730671427 | 5.56E-11 |
| Ifitm2 | 6827.596898 | -0.730722246 | 2.62E-05 |
| Scrib | 2356.920438 | -0.730896841 | 1.22E-06 |
| Hivep2 | 1729.316059 | -0.731338422 | 2.48E-03 |
| Mdm4 | 1505.114202 | -0.732414117 | 3.92E-04 |
| Ikzf5 | 763.4734331 | -0.73267292 | 1.49E-07 |
| Ttyh2 | 1997.972056 | -0.732766226 | 1.68E-05 |
| Ppp4r1 | 3187.238595 | -0.733548992 | 1.99E-04 |
| Mib1 | 3545.099851 | -0.734790764 | 2.93E-04 |
| Fzd7 | 1499.058727 | -0.735146535 | 6.33E-07 |
| Hdac10 | 484.6780465 | -0.735696417 | 1.76E-06 |
| Fndc4 | 669.6069823 | -0.735769949 | 5.59E-04 |
| Ggcx | 1596.794998 | -0.73609311 | 7.77E-09 |
| Rfc1 | 3100.460772 | -0.736538205 | 5.79E-07 |
| Ermp1 | 2303.615021 | -0.736793422 | 3.26E-07 |
| Zfp319 | 991.9383572 | -0.737056879 | 8.17E-10 |
| A430057M04Rik | 95.38244642 | -0.73708201 | 6.37E-03 |
| Bzw1 | 11535.9414 | -0.737803916 | 5.80E-08 |
| Sacm1l | 2042.468251 | -0.738367472 | 5.47E-04 |
| Dtl | 972.7640328 | -0.738645868 | 4.62E-02 |
| Usp9x | 10545.38415 | -0.738775595 | 4.77E-05 |
| Ephb2 | 1039.48774 | -0.738953976 | 6.49E-05 |
| Hspa2 | 705.0792204 | -0.739236036 | 7.06E-05 |
| Ralgapa1 | 2002.317679 | -0.739490664 | 3.64E-05 |
| Isca1 | 1912.366855 | -0.739925002 | 3.87E-12 |
| Fgfr2 | 2705.677105 | -0.739968019 | 5.74E-04 |
| Parp8 | 1704.661461 | -0.740002073 | 3.41E-08 |
| Igtp | 114.2451841 | -0.740627291 | 8.18E-03 |
| Kif6 | 161.0754494 | -0.740887603 | 4.80E-03 |
| Ldhb | 3916.76833 | -0.741044281 | 1.27E-05 |
| Pkia | 759.5393534 | -0.741058523 | 4.30E-03 |
| Tbk1 | 1987.788939 | -0.741107122 | 6.93E-07 |
| Suco | 2030.134186 | -0.741157168 | 1.73E-03 |
| 2610002M06Rik | 1470.102267 | -0.741354392 | 6.02E-15 |
| Taf1 | 2938.602297 | -0.741476353 | 6.70E-05 |
| Lox | 121572.0046 | -0.741777833 | 3.99E-03 |
| Zfp580 | 250.268861 | -0.742062846 | 8.91E-04 |
| Lrig2 | 1243.676386 | -0.74221067 | 2.72E-09 |
| Fnbp4 | 2349.6636 | -0.742219097 | 1.00E-04 |
| Tor2a | 1252.70039 | -0.742457659 | 6.70E-08 |
| Acat1 | 4212.253723 | -0.742609765 | 1.90E-09 |
| Zfp874a | 257.4111334 | -0.742864248 | 7.94E-04 |
| Dennd4b | 511.3383527 | -0.742925271 | 3.14E-06 |
| Pdxk | 5185.761374 | -0.743345598 | 3.26E-05 |
| Klhl24 | 4261.066099 | -0.743437994 | 3.47E-05 |

| | | | |
|---|---|---|---|
| Arhgap44 | 266.8234364 | -0.743501199 | 9.85E-03 |
| Ints2 | 1439.53712 | -0.743729765 | 1.97E-06 |
| Pum2 | 6337.250185 | -0.743782958 | 1.89E-10 |
| Gm17132 | 459.1111935 | -0.7440455 | 1.87E-02 |
| Dusp6 | 789.6529648 | -0.744205031 | 8.58E-03 |
| Ppfibp1 | 13504.46439 | -0.744253394 | 1.27E-08 |
| Ctsb | 36468.22462 | -0.744658632 | 2.70E-04 |
| Acsf3 | 664.047027 | -0.745275511 | 1.76E-06 |
| Lrp5 | 2822.529842 | -0.745423354 | 5.48E-04 |
| Pik3ca | 3935.265814 | -0.746083331 | 2.48E-26 |
| Ncor1 | 9518.98468 | -0.746246114 | 2.42E-07 |
| Adgra3 | 2363.887936 | -0.746611082 | 6.36E-10 |
| Pex26 | 694.5375104 | -0.74709443 | 3.87E-08 |
| Mitd1 | 486.7458551 | -0.747204144 | 2.28E-05 |
| Ywhaq | 13241.07001 | -0.74773032 | 4.96E-16 |
| Rere | 6426.410439 | -0.747926579 | 4.46E-08 |
| Gm12655 | 31.02432306 | -0.748172626 | 5.00E-02 |
| Fkbp1b | 126.766473 | -0.748542897 | 2.89E-03 |
| Slc6a6 | 8237.226832 | -0.748603983 | 2.64E-05 |
| Zswim9 | 494.6910546 | -0.748630754 | 2.13E-04 |
| Vcpip1 | 2503.113416 | -0.748651638 | 1.13E-10 |
| Rbm43 | 682.9030732 | -0.748753888 | 1.69E-10 |
| Zfp84 | 572.4479467 | -0.749144506 | 1.34E-11 |
| Cdr2l | 1223.894987 | -0.749275844 | 2.77E-16 |
| Lynx1 | 1430.13896 | -0.7493233 | 8.88E-07 |
| Gm26792 | 177.3943853 | -0.749412429 | 5.53E-04 |
| Zfp651 | 2092.260882 | -0.749608818 | 7.98E-08 |
| Clec16a | 2108.067705 | -0.749943351 | 2.30E-22 |
| Uvssa | 1162.7755 | -0.749973812 | 8.53E-04 |
| Evi5l | 1300.320633 | -0.750032287 | 2.94E-12 |
| Gm15387 | 5255.01591 | -0.750043315 | 4.56E-07 |
| Hnrnpa0 | 3624.462535 | -0.750769858 | 2.18E-06 |
| Tfdp2 | 1412.093919 | -0.750859294 | 1.85E-05 |
| Bfar | 1557.571232 | -0.75105535 | 2.15E-13 |
| Gm14117 | 173.7043936 | -0.751119561 | 9.30E-03 |
| Tmem170b | 491.2964213 | -0.751152676 | 5.64E-04 |
| Stk16 | 1996.715038 | -0.752183341 | 4.40E-08 |
| Rnaseh1 | 673.6648435 | -0.752908044 | 8.43E-19 |
| Usp6nl | 1363.084088 | -0.753028621 | 7.50E-12 |
| Rhobtb1 | 610.439932 | -0.753072146 | 8.99E-03 |
| Dgcr2 | 5162.081899 | -0.753533868 | 1.85E-12 |
| A930005H10Rik | 101.6160897 | -0.753548169 | 2.35E-03 |
| Smchd1 | 3941.626738 | -0.753752027 | 4.06E-06 |
| Vmp1 | 3794.103478 | -0.753850884 | 3.53E-04 |
| Snrk | 1370.299256 | -0.754008599 | 4.81E-10 |
| Setx | 3233.215275 | -0.755018536 | 8.54E-03 |
| Gm2824 | 44.1800568 | -0.75529015 | 2.94E-02 |

| | | | |
|---|---|---|---|
| Dnajc10 | 5082.055813 | -0.756085312 | 3.22E-22 |
| Rsbn1 | 753.7959019 | -0.75618207 | 2.99E-06 |
| Acad8 | 707.5740021 | -0.756234518 | 1.88E-09 |
| Cdkn2aip | 683.7261103 | -0.756656313 | 3.49E-08 |
| Wrb | 675.6283793 | -0.756779171 | 5.15E-07 |
| Efna3 | 212.0063188 | -0.756989887 | 1.25E-03 |
| Cercam | 171.2368976 | -0.757096386 | 5.65E-03 |
| Xpnpep3 | 989.0992619 | -0.757616775 | 4.08E-18 |
| Srgap3 | 3597.599338 | -0.758178452 | 3.72E-05 |
| D230017M19Rik | 82.99630944 | -0.758821436 | 1.15E-02 |
| Pus10 | 791.0007265 | -0.759544286 | 1.25E-03 |
| Bmt2 | 1303.453939 | -0.75989655 | 2.75E-04 |
| Evc | 3246.728565 | -0.759949281 | 1.83E-09 |
| Syap1 | 1746.197058 | -0.760201799 | 4.65E-05 |
| Arntl | 722.4816621 | -0.760769135 | 1.15E-07 |
| Ccdc22 | 1040.823011 | -0.760865037 | 3.76E-13 |
| Tet3 | 2987.453673 | -0.760933167 | 3.61E-09 |
| Fam126a | 2910.896476 | -0.761046118 | 2.65E-15 |
| Ncoa7 | 841.4675163 | -0.761089752 | 3.26E-08 |
| Zfp280d | 615.38634 | -0.761092485 | 1.20E-03 |
| Fam53a | 1630.046894 | -0.761969851 | 1.18E-10 |
| Eef1akmt1 | 477.1061299 | -0.762061983 | 2.23E-04 |
| Rhobtb2 | 871.4547682 | -0.76216287 | 3.57E-08 |
| Antxr2 | 3509.943442 | -0.762545278 | 4.98E-06 |
| Rnf146 | 1837.081402 | -0.762837973 | 5.15E-11 |
| Kif13a | 1516.705461 | -0.762947702 | 5.29E-10 |
| Tnc | 3603.199261 | -0.762964294 | 7.31E-04 |
| Gm26610 | 45.93485586 | -0.762980699 | 1.67E-02 |
| D630044L22Rik | 67.84507621 | -0.763181626 | 6.53E-03 |
| Hbp1 | 5566.199986 | -0.76378942 | 3.66E-06 |
| Polr2d | 570.7006928 | -0.764088652 | 9.27E-08 |
| Qtrt2 | 643.2243804 | -0.764121943 | 2.32E-04 |
| 6030458C11Rik | 897.7734304 | -0.76420386 | 1.16E-09 |
| Ubp1 | 2230.908591 | -0.764360718 | 1.31E-07 |
| Taf1a | 403.3021228 | -0.764680641 | 1.21E-06 |
| B4galt2 | 3537.6615 | -0.764780928 | 6.53E-10 |
| Gm14276 | 62.27855161 | -0.764869394 | 3.39E-02 |
| Cic | 8307.861902 | -0.765617986 | 3.34E-09 |
| Rbl1 | 738.9772961 | -0.765689603 | 4.54E-03 |
| Nr2c1 | 630.205208 | -0.766303903 | 2.70E-07 |
| Zbtb8os | 1050.643546 | -0.766382119 | 6.46E-05 |
| Ppt1 | 7244.146237 | -0.766528561 | 8.67E-15 |
| Pgm2l1 | 185.7113534 | -0.766647881 | 3.04E-04 |
| Hspa13 | 1659.899174 | -0.767058918 | 1.24E-09 |
| P4htm | 35.99990357 | -0.767120681 | 1.95E-02 |
| Otud4 | 3740.54233 | -0.7676236 | 1.15E-04 |
| Colgalt1 | 6309.791937 | -0.767686707 | 5.05E-09 |

| | | | |
|---|---|---|---|
| Sft2d1 | 798.2097011 | -0.768120403 | 3.11E-12 |
| Creb3l2 | 4400.318381 | -0.768254587 | 3.17E-08 |
| NA | 459.8289227 | -0.768714334 | 3.41E-12 |
| Pigp | 767.26785 | -0.768861666 | 1.26E-06 |
| Helq | 674.3319847 | -0.769067921 | 4.46E-05 |
| Hoga1 | 295.3476107 | -0.769265078 | 6.34E-04 |
| Rpa3 | 645.0287183 | -0.769270186 | 2.02E-06 |
| Atp13a3 | 9000.749949 | -0.769271987 | 6.06E-06 |
| Kdm2a | 7140.470013 | -0.769554309 | 2.60E-19 |
| Vti1a | 1513.738788 | -0.76968298 | 2.61E-10 |
| Vangl1 | 1595.126302 | -0.769932204 | 4.62E-05 |
| Ube2q2 | 1050.628754 | -0.770252175 | 1.49E-04 |
| Zdhhc1 | 1130.427471 | -0.770675229 | 1.85E-07 |
| Golga1 | 1274.09039 | -0.772303525 | 6.86E-04 |
| Sgk3 | 113.1127876 | -0.772644008 | 4.00E-02 |
| Slc11a2 | 1839.744541 | -0.772834236 | 9.60E-06 |
| Capn15 | 724.6406254 | -0.772914794 | 3.91E-11 |
| Cnot9 | 2635.142109 | -0.773822959 | 1.21E-07 |
| Dnajc30 | 610.0677149 | -0.774024467 | 8.11E-09 |
| Hcfc2 | 1637.57446 | -0.774095621 | 6.05E-07 |
| Cep131 | 587.196652 | -0.774164647 | 3.56E-05 |
| Gm13722 | 178.4238107 | -0.774421891 | 1.55E-03 |
| Spata2l | 99.79410299 | -0.775690526 | 2.79E-04 |
| Smarca5 | 5548.939349 | -0.775833195 | 1.79E-06 |
| Ift172 | 1593.456915 | -0.776388083 | 6.62E-09 |
| Rtl8c | 186.4143832 | -0.776709601 | 8.75E-04 |
| Cmtm4 | 10031.66492 | -0.776933303 | 1.12E-15 |
| Zfp523 | 1205.100763 | -0.776974169 | 1.66E-07 |
| Zc4h2 | 400.4251748 | -0.777876911 | 3.65E-11 |
| Ilk | 4670.521348 | -0.778255433 | 6.15E-09 |
| Glis3 | 1111.322553 | -0.778811775 | 5.74E-05 |
| Taco1 | 417.8883259 | -0.778957501 | 1.05E-08 |
| Gm14584 | 141.3280251 | -0.78020797 | 3.50E-04 |
| Golgb1 | 5482.119432 | -0.782211461 | 3.46E-11 |
| Slc39a11 | 1072.037658 | -0.782613058 | 1.35E-07 |
| Cpq | 2992.352624 | -0.782742828 | 1.80E-07 |
| Mpc1 | 1267.168125 | -0.783276514 | 6.60E-09 |
| Akap8 | 2567.406678 | -0.783520345 | 1.15E-10 |
| Rcor1 | 2598.897465 | -0.783613496 | 4.50E-29 |
| B4galt5 | 4562.137611 | -0.783850041 | 8.51E-04 |
| Manea | 992.7930942 | -0.78440357 | 2.03E-03 |
| Comtd1 | 549.7820272 | -0.784583273 | 5.55E-08 |
| Plxna2 | 3143.852049 | -0.784815628 | 6.36E-06 |
| Ppdpf | 1355.859809 | -0.784947594 | 6.71E-15 |
| Fam193a | 2712.375027 | -0.785008016 | 3.52E-18 |
| Rsbn1l | 600.0395247 | -0.78515554 | 6.57E-04 |
| Zfp951 | 126.5205167 | -0.785429131 | 1.03E-02 |

| | | | |
|---|---|---|---|
| Lmbrd2 | 671.6944806 | -0.787315622 | 7.74E-03 |
| Uckl1 | 1220.761514 | -0.787455626 | 7.54E-11 |
| Slc35a4 | 2217.830955 | -0.788632689 | 6.65E-13 |
| Bcr | 3158.078807 | -0.788854614 | 3.77E-16 |
| Eef2k | 2828.119429 | -0.789293828 | 3.11E-09 |
| Ern1 | 1411.742797 | -0.789372144 | 3.27E-13 |
| Slc25a35 | 132.6975674 | -0.789568292 | 1.57E-03 |
| Rc3h2 | 1945.035031 | -0.790186134 | 4.25E-05 |
| Mios | 1271.525436 | -0.790192801 | 5.01E-08 |
| Tlr3 | 335.217897 | -0.790223363 | 1.65E-04 |
| Jak2 | 1748.376278 | -0.791544585 | 2.21E-07 |
| Psg16 | 1040.230629 | -0.792040599 | 2.60E-06 |
| Srp54c | 1928.786045 | -0.792669335 | 2.92E-05 |
| Zfp605 | 275.4334845 | -0.792905934 | 9.30E-03 |
| Tmem30a | 3454.308382 | -0.79351538 | 7.19E-05 |
| Mpdz | 8375.563949 | -0.793555075 | 1.89E-03 |
| 4931440P22Rik | 33.62408602 | -0.793962838 | 4.72E-02 |
| Supt16 | 3884.352525 | -0.794317988 | 3.78E-06 |
| Gm5514 | 63.12948495 | -0.794617818 | 1.36E-02 |
| Phyh | 208.6455419 | -0.795315869 | 2.75E-03 |
| Top1mt | 841.059034 | -0.795350768 | 1.70E-10 |
| Enah | 2404.280719 | -0.796180647 | 3.69E-10 |
| Zfp626 | 49.10786744 | -0.796580389 | 1.57E-02 |
| Resf1 | 4767.446193 | -0.797281945 | 2.17E-03 |
| Cep295 | 1053.007575 | -0.797785289 | 1.03E-03 |
| Tbcel | 1611.699035 | -0.798059314 | 2.55E-20 |
| Hcn3 | 50.97651329 | -0.798222163 | 2.94E-02 |
| Fancf | 204.1776999 | -0.798442086 | 1.34E-05 |
| Slc25a36 | 2060.018335 | -0.798792606 | 3.07E-05 |
| Zfp28 | 611.0405196 | -0.799794023 | 5.90E-11 |
| Pole2 | 557.9420967 | -0.800431828 | 5.22E-03 |
| Trub1 | 627.2813923 | -0.800589301 | 2.26E-10 |
| Gpatch1 | 528.5209635 | -0.800751487 | 2.84E-13 |
| Rest | 3692.63625 | -0.801020165 | 6.08E-23 |
| Gm6563 | 1607.864764 | -0.801679174 | 2.54E-04 |
| Nrn1 | 2360.363595 | -0.801942429 | 1.07E-04 |
| Anp32a | 2932.882658 | -0.80224265 | 2.44E-13 |
| Ddx46 | 2557.119627 | -0.802875361 | 1.16E-04 |
| Sez6l2 | 293.3386285 | -0.803368426 | 2.22E-03 |
| Tipin | 782.6723888 | -0.803744188 | 4.90E-04 |
| Wdr81 | 3346.664813 | -0.803876084 | 7.67E-07 |
| Tpr | 8918.937942 | -0.804614087 | 6.75E-03 |
| Pin4 | 625.4302369 | -0.805162182 | 3.84E-04 |
| Mcc | 1935.549035 | -0.805176618 | 1.17E-08 |
| Kmt2c | 2822.831742 | -0.805476293 | 3.39E-10 |
| Ino80b | 616.8921082 | -0.806679398 | 2.02E-06 |
| NA | 121.2677958 | -0.807215603 | 6.13E-05 |

| | | | |
|---|---|---|---|
| Kmt5b | 1784.508974 | -0.807415545 | 2.66E-05 |
| Lrrfip1 | 4238.945427 | -0.807449352 | 2.48E-06 |
| Magt1 | 3336.975739 | -0.807553826 | 3.53E-07 |
| 1810030O07Rik | 1755.512131 | -0.80775959 | 4.77E-23 |
| Anp32e | 3696.837051 | -0.807963699 | 2.08E-05 |
| Slc2a1 | 3768.575499 | -0.808246978 | 2.31E-10 |
| Nenf | 2679.17814 | -0.808827897 | 2.25E-08 |
| Nek7 | 4505.140843 | -0.809438595 | 3.97E-06 |
| AI838599 | 78.18413624 | -0.809872168 | 1.47E-02 |
| Shpk | 226.3556193 | -0.811837424 | 2.22E-06 |
| Dnajc7 | 3526.747744 | -0.812185211 | 1.01E-05 |
| Plk4 | 915.3174009 | -0.812223317 | 8.81E-03 |
| Tapbpl | 512.5127428 | -0.812468897 | 5.64E-11 |
| Slc36a1 | 1573.335444 | -0.81259483 | 8.22E-09 |
| Dennd1b | 638.5635192 | -0.812696734 | 7.47E-04 |
| Bmp1 | 4680.338256 | -0.812709107 | 7.50E-15 |
| Foxd2os | 411.5132541 | -0.812984436 | 1.25E-08 |
| Zbtb41 | 1325.200177 | -0.813159488 | 1.36E-03 |
| Camkk2 | 1761.969751 | -0.813552388 | 2.13E-15 |
| 9130019O22Rik | 353.1637933 | -0.814495768 | 3.44E-05 |
| Ddx51 | 1561.419541 | -0.814569361 | 4.64E-10 |
| Ankrd12 | 2220.360824 | -0.814737258 | 2.13E-04 |
| Sash1 | 2475.130362 | -0.814848724 | 5.59E-12 |
| Stk11ip | 1416.486596 | -0.815359109 | 1.96E-09 |
| Tmem175 | 1175.19773 | -0.815501259 | 9.12E-15 |
| Galk2 | 2673.268011 | -0.81567355 | 1.76E-11 |
| Zc3h7a | 2250.277667 | -0.815958642 | 3.15E-04 |
| Ndst2 | 1120.700593 | -0.816207781 | 9.15E-21 |
| Cenpb | 8519.626358 | -0.816311627 | 9.89E-10 |
| Tgfb1 | 1304.531232 | -0.816704038 | 1.65E-03 |
| Ccny | 4186.355601 | -0.816783383 | 1.33E-23 |
| Dse | 4466.695367 | -0.817083613 | 7.56E-06 |
| Uba3 | 3570.617151 | -0.817176698 | 2.74E-03 |
| Gm6211 | 148.7366503 | -0.81763722 | 1.14E-03 |
| Fmo1 | 97.89348174 | -0.817933985 | 6.80E-03 |
| Acin1 | 5414.640843 | -0.818596949 | 1.24E-25 |
| Gm10125 | 80.42503309 | -0.818992611 | 5.85E-03 |
| Magee1 | 2530.570754 | -0.818992721 | 4.95E-12 |
| Rnf149 | 1199.129938 | -0.819224278 | 6.61E-17 |
| Mnt | 3501.854391 | -0.81983527 | 4.58E-05 |
| Itsn2 | 2226.775109 | -0.820003372 | 2.74E-05 |
| Col6a2 | 28456.27254 | -0.820976174 | 1.01E-02 |
| Emp3 | 1776.507322 | -0.821478726 | 1.01E-06 |
| 1700123M08Rik | 104.1186089 | -0.821865519 | 9.78E-03 |
| Myo1h | 414.7993453 | -0.822075278 | 3.88E-05 |
| Ints6l | 343.212195 | -0.822554757 | 2.42E-03 |
| Lrfn3 | 523.8739616 | -0.822945227 | 6.82E-09 |

| | | | |
|---|---|---|---|
| Nckap5l | 1105.351254 | -0.823218433 | 5.74E-07 |
| Rnf157 | 1391.358528 | -0.823486658 | 1.07E-10 |
| Odf2l | 557.1822106 | -0.824704962 | 3.07E-04 |
| Slc22a21 | 399.1708764 | -0.825846785 | 3.11E-06 |
| Fth1 | 88712.82582 | -0.826076598 | 5.97E-05 |
| Gm17201 | 126.9571466 | -0.826276344 | 1.64E-03 |
| Mex3a | 2190.570495 | -0.828300873 | 4.19E-04 |
| Gm7803 | 29.42446638 | -0.82900069 | 4.16E-02 |
| Ice2 | 642.1654413 | -0.829503452 | 3.29E-05 |
| Dcun1d4 | 1164.457837 | -0.829906834 | 1.13E-06 |
| Sarnp | 583.9157138 | -0.829951236 | 7.32E-05 |
| Acta2 | 22267.60312 | -0.830461507 | 1.32E-04 |
| Ensa | 4415.155983 | -0.831337987 | 1.89E-11 |
| Gopc | 1774.957849 | -0.832704293 | 1.16E-09 |
| Galnt2 | 8979.608691 | -0.833170413 | 2.36E-08 |
| Zfp459 | 54.92350494 | -0.834387239 | 6.49E-03 |
| Arf3 | 7135.48518 | -0.835076334 | 1.56E-35 |
| Cetn3 | 1261.039176 | -0.836162748 | 4.07E-04 |
| Rassf8 | 5424.313331 | -0.83623907 | 7.54E-10 |
| Kctd18 | 586.3295036 | -0.836439178 | 7.50E-06 |
| Glcci1 | 283.1218107 | -0.836894369 | 1.44E-02 |
| Otud3 | 277.8243889 | -0.836909153 | 7.91E-08 |
| Zkscan7 | 200.504492 | -0.836960538 | 8.64E-06 |
| 3110053B16Rik | 271.5926797 | -0.837054884 | 2.63E-06 |
| Fuca1 | 1794.844636 | -0.83764219 | 1.53E-05 |
| Pdzrn3 | 1450.783321 | -0.838201247 | 4.01E-06 |
| Eif4g2 | 37082.43035 | -0.838900563 | 3.22E-17 |
| Pik3c2b | 54.85501759 | -0.839033997 | 4.80E-02 |
| Tmem65 | 2586.368853 | -0.840002507 | 1.04E-12 |
| Mia2 | 2335.340629 | -0.840069226 | 1.41E-10 |
| Cntf | 90.70659904 | -0.84018903 | 2.37E-02 |
| Ube3a | 2165.048968 | -0.840234845 | 6.61E-04 |
| Pnkd | 893.3433196 | -0.840464908 | 7.05E-14 |
| Acsl3 | 2747.009398 | -0.840657182 | 4.68E-04 |
| Naa15 | 3151.720685 | -0.840739201 | 1.34E-03 |
| Cklf | 151.6952611 | -0.841137117 | 5.61E-03 |
| Lrp12 | 1151.827332 | -0.841503503 | 1.40E-11 |
| Dram1 | 593.7149253 | -0.841606471 | 1.18E-07 |
| NA | 31.52558462 | -0.842310601 | 4.87E-02 |
| Vegfd | 6147.027 | -0.842606013 | 8.75E-03 |
| Sqle | 9213.229076 | -0.843513567 | 2.02E-04 |
| Mrpl50 | 1080.201533 | -0.843547789 | 5.89E-09 |
| Dock7 | 10072.46313 | -0.843815795 | 6.02E-22 |
| Kat6a | 4809.065282 | -0.844148268 | 2.05E-14 |
| Slc35a5 | 538.059794 | -0.844769052 | 4.44E-08 |
| Tmc5 | 75.57411912 | -0.845050347 | 2.82E-02 |
| Pex11b | 937.9156307 | -0.845409282 | 1.69E-19 |

| | | | |
|---|---|---|---|
| Mark1 | 807.1047026 | -0.845539133 | 2.50E-05 |
| Urgcp | 1854.416943 | -0.845542133 | 1.97E-11 |
| Ybey | 168.7312436 | -0.846835958 | 2.77E-04 |
| Unc119b | 2447.389577 | -0.846908065 | 6.26E-16 |
| Eif3j1 | 236.6541089 | -0.846971536 | 1.89E-02 |
| Vamp1 | 618.7179027 | -0.847553703 | 2.17E-06 |
| Gm16300 | 124.443785 | -0.848124961 | 1.08E-02 |
| Acsl4 | 2428.085415 | -0.848503853 | 5.87E-04 |
| Keap1 | 1638.540529 | -0.848614975 | 6.36E-11 |
| Slc25a24 | 3916.415838 | -0.850149687 | 4.30E-04 |
| Dnajb9 | 908.8568972 | -0.850243936 | 3.24E-07 |
| Trp53rka | 613.6532897 | -0.85062932 | 4.75E-08 |
| Elovl7 | 28.61036178 | -0.852029024 | 4.13E-02 |
| Hdx | 176.7894411 | -0.852561682 | 6.18E-03 |
| Rgl1 | 2957.467567 | -0.853342032 | 3.71E-04 |
| Ptgs1 | 3977.955639 | -0.854207915 | 2.69E-06 |
| Agpat2 | 111.9244998 | -0.854295245 | 1.80E-04 |
| Loxl3 | 4747.787426 | -0.855172594 | 1.01E-05 |
| Sgsm2 | 1683.587835 | -0.855236917 | 3.92E-07 |
| Kif16b | 1057.721431 | -0.855701097 | 4.95E-12 |
| Rhbdd3 | 570.80931 | -0.855783355 | 6.20E-07 |
| Tubb2a | 6574.060184 | -0.856176861 | 1.52E-07 |
| Mapk1ip1 | 378.0234365 | -0.856757287 | 9.06E-07 |
| Cd164 | 10632.62841 | -0.857657943 | 1.75E-24 |
| Gm1818 | 103.6610247 | -0.857915002 | 4.52E-03 |
| Pdzd7 | 210.0352206 | -0.857940657 | 8.51E-04 |
| Kcnab1 | 1245.884438 | -0.858302884 | 2.44E-04 |
| Men1 | 2488.414312 | -0.858509519 | 1.11E-22 |
| Cep170 | 6223.844815 | -0.858954561 | 1.30E-11 |
| Ap3b2 | 72.29580284 | -0.859730048 | 4.65E-03 |
| Upf2 | 1590.124209 | -0.859831923 | 6.41E-04 |
| Kdm5d | 903.6611732 | -0.859929927 | 1.17E-08 |
| Larp4 | 3176.471045 | -0.860068872 | 6.68E-05 |
| Tmem106b | 10840.02422 | -0.860436305 | 8.13E-07 |
| Ptpn9 | 2903.919013 | -0.861336901 | 9.38E-16 |
| Tigd2 | 928.8889991 | -0.861387036 | 5.38E-09 |
| St3gal4 | 2084.028363 | -0.861471958 | 1.24E-04 |
| Btbd7 | 2855.621626 | -0.862423019 | 1.74E-44 |
| Tex261 | 4268.859971 | -0.863026469 | 9.65E-09 |
| Napepld | 430.2690843 | -0.863904348 | 2.69E-08 |
| Matr3 | 6757.454026 | -0.863916441 | 4.92E-04 |
| Slc31a1 | 3001.893987 | -0.864083702 | 4.57E-36 |
| Gosr1 | 1647.177887 | -0.864117676 | 2.41E-05 |
| Myo5a | 10004.30674 | -0.864141999 | 1.06E-07 |
| Mtif2 | 862.6809578 | -0.864595883 | 8.71E-07 |
| Fbxo10 | 1508.820967 | -0.86467226 | 3.94E-09 |
| Fam229b | 171.6780668 | -0.865307038 | 1.06E-04 |

| | | | |
|---|---|---|---|
| Rdh16 | 64.36347481 | -0.865395976 | 3.76E-03 |
| Vps13b | 3442.13905 | -0.865683892 | 1.91E-08 |
| Synj2 | 1645.375646 | -0.866414192 | 3.11E-22 |
| G3bp2 | 3581.96833 | -0.866644829 | 1.04E-15 |
| Zeb2os | 155.3605102 | -0.866683504 | 4.10E-04 |
| Ago4 | 324.5381755 | -0.869524624 | 5.65E-05 |
| Rab3d | 425.5947872 | -0.869688341 | 5.51E-04 |
| Asap1 | 15094.41644 | -0.869875463 | 2.28E-11 |
| Zfp652 | 3085.865232 | -0.870038674 | 2.37E-21 |
| Ttc8 | 1298.530567 | -0.871208994 | 1.93E-21 |
| Rbm6 | 2487.44003 | -0.87137081 | 1.53E-06 |
| Cbx1 | 2142.534803 | -0.871572596 | 1.56E-13 |
| Shroom4 | 831.9222619 | -0.871918804 | 6.81E-07 |
| Mtch1 | 11913.88654 | -0.872260849 | 1.01E-09 |
| Zmym6 | 2149.481954 | -0.872351866 | 1.12E-04 |
| Caprin2 | 403.2197993 | -0.872407881 | 4.35E-06 |
| Tmcc2 | 1387.568865 | -0.872664924 | 7.55E-07 |
| Slc25a47 | 102.6606474 | -0.873310918 | 8.90E-04 |
| Itgav | 6626.652067 | -0.873680366 | 6.53E-10 |
| Zfp362 | 2229.096229 | -0.874018263 | 1.01E-13 |
| Btbd6 | 204.0903082 | -0.874023418 | 8.57E-08 |
| B230217C12Rik | 62.92638536 | -0.874198634 | 3.85E-02 |
| Wdfy3 | 4331.225501 | -0.874343664 | 1.15E-16 |
| Rcor3 | 1322.653627 | -0.875655029 | 7.28E-08 |
| Six1 | 1079.039133 | -0.876174434 | 2.24E-03 |
| Trmt11 | 423.7101061 | -0.876503484 | 4.55E-04 |
| Aar2 | 1658.318704 | -0.876673649 | 1.24E-11 |
| Hoxaas3 | 58.30301202 | -0.876690167 | 2.93E-02 |
| Zhx3 | 1585.705602 | -0.876814084 | 3.99E-08 |
| Camsap2 | 3519.542233 | -0.877298888 | 4.51E-22 |
| Usp37 | 982.1370996 | -0.877365489 | 8.18E-07 |
| Slc2a4rg-ps | 147.8306169 | -0.877691219 | 2.07E-03 |
| Fndc3b | 7112.954594 | -0.878371908 | 3.22E-06 |
| Ing4 | 1691.109084 | -0.878419563 | 1.08E-12 |
| Amigo2 | 624.6922769 | -0.878479971 | 4.50E-09 |
| Mapkapk3 | 612.1223251 | -0.878688327 | 1.79E-04 |
| Kdm6a | 1109.474701 | -0.879073396 | 1.62E-10 |
| Metap1d | 1532.634791 | -0.879341581 | 4.06E-10 |
| Tab3 | 871.1103317 | -0.879537034 | 1.19E-06 |
| Mosmo | 928.7988729 | -0.879840333 | 2.48E-07 |
| Rnf217 | 747.2323047 | -0.880066232 | 1.13E-11 |
| Pcna | 3425.363174 | -0.880198954 | 3.06E-04 |
| Ccpg1 | 2606.364703 | -0.880711257 | 4.50E-07 |
| Gm16153 | 64.45525906 | -0.881073823 | 1.23E-02 |
| Pacs2 | 7227.003698 | -0.881195365 | 6.81E-23 |
| Zfp142 | 2034.722645 | -0.88132091 | 1.67E-12 |
| Tmem164 | 4384.562062 | -0.882308184 | 3.85E-11 |

| | | | |
|---|---|---|---|
| Zfp51 | 342.0893331 | -0.883069895 | 1.40E-02 |
| Nr2c2 | 2635.174019 | -0.883579009 | 8.04E-12 |
| Pura | 1313.30586 | -0.884029471 | 1.26E-04 |
| Chd6 | 3406.923992 | -0.884193673 | 2.69E-02 |
| Ltbp4 | 1251.339727 | -0.885747416 | 5.09E-05 |
| Entpd7 | 479.0742444 | -0.886317669 | 7.21E-09 |
| Alg5 | 1251.071303 | -0.886681701 | 1.89E-08 |
| C630043F03Rik | 71.42610825 | -0.886974915 | 7.44E-03 |
| Tmtc4 | 1184.027016 | -0.888071808 | 6.64E-06 |
| Mtmr2 | 3520.614254 | -0.888970788 | 2.95E-08 |
| Itm2b | 23528.2636 | -0.889566854 | 1.43E-05 |
| Tmem176a | 167.3003744 | -0.88976695 | 6.25E-03 |
| Agfg2 | 1357.380105 | -0.890209821 | 5.36E-10 |
| Ackr3 | 559.0724489 | -0.890424586 | 6.49E-03 |
| Zmym4 | 3703.024196 | -0.890624398 | 3.50E-10 |
| Hectd4 | 2188.400038 | -0.891399809 | 9.35E-10 |
| Gldc | 24.74146371 | -0.891507221 | 3.22E-02 |
| Tbc1d1 | 1872.956571 | -0.89205485 | 4.59E-06 |
| Smim15 | 1057.349609 | -0.89227578 | 5.92E-04 |
| Cyren | 435.9422685 | -0.892506522 | 1.38E-03 |
| Efhd2 | 1565.299968 | -0.892814394 | 2.47E-09 |
| Errfi1 | 2068.090918 | -0.893030823 | 4.46E-05 |
| Rnaset2b | 1643.229936 | -0.893930752 | 1.52E-12 |
| Kcnn4 | 208.2608762 | -0.894087535 | 3.69E-03 |
| Tnrc6c | 2070.107347 | -0.894322248 | 7.71E-20 |
| Zfp558 | 139.6941417 | -0.895016973 | 7.02E-04 |
| Rai14 | 4032.0779 | -0.895135047 | 2.46E-08 |
| Lpgat1 | 5562.591021 | -0.895189359 | 3.71E-13 |
| Whrn | 1183.150778 | -0.895358491 | 1.50E-07 |
| Lpar6 | 212.9759482 | -0.896085506 | 2.59E-03 |
| Hipk4 | 55.60434058 | -0.896510318 | 1.78E-02 |
| Qk | 4353.559499 | -0.896810892 | 8.44E-10 |
| Nfyc | 2165.842063 | -0.896966858 | 5.32E-19 |
| Hnrnpa3 | 10251.47568 | -0.89725868 | 1.27E-03 |
| Tspan3 | 3878.105411 | -0.897331279 | 9.20E-16 |
| Npas3 | 130.7817696 | -0.897512136 | 3.91E-05 |
| Ints7 | 1422.713731 | -0.897797922 | 2.47E-11 |
| Mettl7a1 | 836.3886156 | -0.89796033 | 8.26E-06 |
| Ifi203 | 486.7251377 | -0.898048099 | 7.35E-06 |
| Mob1a | 10016.4113 | -0.898159984 | 6.52E-16 |
| Camk2n2 | 678.9875033 | -0.898231351 | 7.35E-06 |
| Rab11fip2 | 1364.111661 | -0.898299153 | 1.67E-04 |
| Max | 2035.937597 | -0.898388895 | 4.01E-19 |
| Xntrpc | 480.116466 | -0.898804595 | 2.34E-13 |
| Ppp1r12b | 4138.333762 | -0.899139404 | 2.19E-09 |
| Crebzf | 1134.690437 | -0.899149533 | 1.67E-05 |
| Cog3 | 2017.359627 | -0.899564457 | 3.81E-14 |

| | | | |
|---|---|---|---|
| Lmnb1 | 1761.407172 | -0.900420106 | 1.31E-03 |
| Vat1 | 27960.27472 | -0.901959789 | 2.17E-05 |
| Zfp334 | 1151.134813 | -0.902926414 | 8.92E-18 |
| Epc1 | 1094.48254 | -0.903143251 | 3.59E-11 |
| Etv6 | 1583.328039 | -0.903309841 | 4.21E-19 |
| Traf6 | 1586.592483 | -0.903351419 | 1.09E-19 |
| Zfp39 | 324.6592607 | -0.904524306 | 7.84E-11 |
| NA | 125.8365537 | -0.904665429 | 1.21E-02 |
| Mterf3 | 690.8179778 | -0.90474293 | 1.19E-07 |
| Stk36 | 385.152876 | -0.904949258 | 1.86E-06 |
| Calm3 | 9475.102056 | -0.904985203 | 1.24E-10 |
| Phtf1 | 2088.129967 | -0.90500942 | 1.31E-17 |
| Fbln2 | 51001.24849 | -0.905057038 | 9.73E-05 |
| Glmn | 555.0524008 | -0.905126196 | 4.81E-05 |
| Cpd | 4849.178331 | -0.905188051 | 1.41E-11 |
| mt-Nd2 | 99498.30713 | -0.905293857 | 3.12E-17 |
| Slc35a1 | 1244.299512 | -0.906047075 | 2.94E-15 |
| Gpr85 | 486.5449696 | -0.906105407 | 2.00E-05 |
| Aacs | 4095.139985 | -0.906110808 | 3.21E-06 |
| Nedd4 | 84397.06472 | -0.906230673 | 1.86E-17 |
| Brca1 | 832.1662432 | -0.906234139 | 8.15E-03 |
| Rad50 | 1970.461368 | -0.906563747 | 2.65E-08 |
| Fam133b | 677.0444772 | -0.906613476 | 1.30E-07 |
| NA | 1997.531107 | -0.907020659 | 3.34E-09 |
| Irf1 | 1630.516384 | -0.907415723 | 4.61E-08 |
| Lrrcc1 | 1408.285589 | -0.907582124 | 7.59E-03 |
| Emsy | 1493.02595 | -0.909051118 | 7.97E-20 |
| Cdh23 | 67.32634277 | -0.909230564 | 4.45E-02 |
| Selenon | 4253.1088 | -0.909597137 | 4.77E-13 |
| Dffa | 430.3054641 | -0.910255606 | 7.28E-15 |
| Sp3 | 4082.066464 | -0.91049712 | 1.34E-05 |
| Vps50 | 1798.051151 | -0.910991708 | 3.21E-09 |
| Hfe | 52.7239069 | -0.911208601 | 4.08E-02 |
| Dusp5 | 674.9902809 | -0.91166934 | 1.71E-04 |
| Perm1 | 74.82401421 | -0.911847398 | 2.26E-03 |
| 9330151L19Rik | 193.2703544 | -0.911921791 | 1.88E-03 |
| Rtp4 | 57.72055596 | -0.913226636 | 2.80E-03 |
| Tcf12 | 4651.7336 | -0.913315635 | 3.12E-17 |
| Sdc1 | 4334.086121 | -0.913410397 | 6.15E-06 |
| Asb7 | 543.7279623 | -0.914330689 | 5.29E-19 |
| Pik3c2a | 2963.767554 | -0.915156659 | 3.37E-03 |
| Prpf18 | 1264.394982 | -0.915297531 | 4.00E-10 |
| H2afv | 1309.866945 | -0.91554291 | 1.78E-08 |
| NA | 132.3463999 | -0.916225563 | 1.06E-02 |
| Gm10093 | 872.0229362 | -0.917008937 | 3.67E-13 |
| Coq4 | 358.7442056 | -0.91728414 | 2.04E-05 |
| Erc1 | 3022.322006 | -0.917352321 | 3.03E-32 |

| | | | |
|---|---|---|---|
| Wdr34 | 712.2965379 | -0.91784997 | 4.83E-09 |
| Gm10575 | 53.05328232 | -0.917896413 | 1.75E-02 |
| Adgra2 | 9959.089649 | -0.918379822 | 3.35E-13 |
| Gm15867 | 82.20330109 | -0.918940615 | 1.56E-03 |
| Ranbp2 | 8070.714676 | -0.918962205 | 6.63E-04 |
| Dtna | 3290.17433 | -0.919614622 | 1.73E-11 |
| Eif2s3y | 1012.072768 | -0.919785414 | 2.76E-04 |
| Akap9 | 2311.061138 | -0.91986471 | 1.09E-03 |
| Cul7 | 4030.507774 | -0.920507616 | 1.45E-11 |
| Zik1 | 289.836167 | -0.920843031 | 4.27E-08 |
| Tbccd1 | 1403.677915 | -0.922601501 | 1.72E-24 |
| Gm14005 | 151.5950979 | -0.923030387 | 3.27E-02 |
| Pogk | 1686.347408 | -0.923104268 | 2.13E-12 |
| Cisd2 | 773.2577454 | -0.923167655 | 3.84E-06 |
| Zfp266 | 1891.611082 | -0.923935882 | 4.68E-07 |
| Nfkb1 | 4076.642103 | -0.924211775 | 7.27E-16 |
| C8g | 63.07082444 | -0.924929098 | 3.38E-04 |
| Ptbp3 | 3849.275156 | -0.925183947 | 3.04E-08 |
| Shc1 | 9740.164831 | -0.925275838 | 6.08E-10 |
| Ptges3l | 205.1528227 | -0.926462383 | 3.00E-06 |
| Adcy9 | 386.8584794 | -0.926629865 | 7.94E-09 |
| Col28a1 | 133.400327 | -0.927526766 | 2.02E-02 |
| 4930515G01Rik | 25.56917121 | -0.927531066 | 2.04E-02 |
| Exoc3l4 | 133.8271647 | -0.927534368 | 5.93E-04 |
| Ids | 2262.136179 | -0.927546316 | 2.12E-17 |
| Iah1 | 1772.443177 | -0.927746764 | 7.09E-07 |
| Car13 | 2496.327558 | -0.927999302 | 8.16E-15 |
| Xkr6 | 187.590343 | -0.928040353 | 4.51E-06 |
| Chpf | 4455.956934 | -0.928360855 | 4.50E-06 |
| Cars2 | 395.453121 | -0.928637903 | 8.84E-09 |
| Ogfrl1 | 619.3082176 | -0.929231978 | 2.33E-05 |
| Mier1 | 2199.416579 | -0.929344364 | 1.93E-07 |
| Slc23a1 | 34.46118179 | -0.92975073 | 3.51E-02 |
| Tnfsf13 | 53.02449501 | -0.930231071 | 9.29E-03 |
| Fbxo41 | 890.8379166 | -0.930671769 | 1.05E-08 |
| Leprot | 5202.282116 | -0.930965102 | 2.64E-25 |
| Synj2bp | 736.8357298 | -0.932460419 | 2.38E-10 |
| Setbp1 | 1813.691562 | -0.933885682 | 4.31E-06 |
| Gm14325 | 470.7536262 | -0.934406232 | 4.37E-04 |
| Igfbp7 | 2473.484216 | -0.934464872 | 2.15E-04 |
| Acvr1 | 1714.787564 | -0.934942262 | 5.28E-12 |
| Prpf39 | 1718.798443 | -0.935032536 | 1.18E-03 |
| Wsb1 | 4822.029222 | -0.935413104 | 3.26E-03 |
| Pde7a | 1818.082955 | -0.935534212 | 4.68E-20 |
| Gpatch4 | 1008.352246 | -0.935759267 | 6.82E-04 |
| 5730480H06Rik | 325.1206865 | -0.936406632 | 1.75E-10 |
| Taf7 | 271.7514652 | -0.936717884 | 1.76E-05 |

| | | | |
|---|---|---|---|
| Catsper2 | 170.3526591 | -0.936742718 | 7.68E-10 |
| Plekhm3 | 751.7569629 | -0.937949451 | 1.84E-16 |
| 1810014B01Rik | 81.46576989 | -0.938199791 | 2.92E-03 |
| Wdr47 | 680.4647398 | -0.938216317 | 7.22E-05 |
| Mgat2 | 2476.636202 | -0.938389508 | 3.17E-14 |
| Gm28050 | 30.14436521 | -0.938692048 | 3.31E-02 |
| Dock4 | 204.615827 | -0.938831242 | 3.96E-05 |
| Zfp90 | 681.0618636 | -0.94011116 | 2.10E-09 |
| Tns1 | 7416.997207 | -0.940814996 | 4.54E-12 |
| Eml6 | 386.0830764 | -0.940987108 | 7.11E-04 |
| Cab39 | 2954.878358 | -0.941064528 | 2.79E-16 |
| Ppp1r8 | 1171.081422 | -0.941228128 | 7.30E-05 |
| Atp7a | 1656.515308 | -0.941298183 | 4.53E-14 |
| Ncbp2 | 2446.490191 | -0.941327448 | 3.84E-13 |
| Lrrc49 | 603.8737907 | -0.941355707 | 5.21E-13 |
| Zfp235 | 197.9093928 | -0.941390725 | 2.02E-04 |
| Lsm6 | 915.3953535 | -0.941946863 | 1.35E-07 |
| Ulk2 | 3079.236902 | -0.943448906 | 1.89E-31 |
| Wasf3 | 609.9759654 | -0.943609446 | 1.69E-10 |
| Ccp110 | 807.4716263 | -0.944299654 | 5.92E-05 |
| Mindy4 | 685.858107 | -0.944412992 | 8.96E-10 |
| Bet1 | 1021.278895 | -0.944658366 | 2.17E-13 |
| Tra2b | 4384.951138 | -0.944949957 | 1.94E-08 |
| Cr1l | 1416.508505 | -0.945397215 | 3.04E-10 |
| Syncrip | 5515.504038 | -0.946655291 | 6.18E-12 |
| Tra2a | 3145.174792 | -0.947362996 | 7.92E-07 |
| Senp6 | 2994.154322 | -0.947791004 | 4.28E-06 |
| Ccdc167 | 216.2758511 | -0.948338779 | 8.16E-07 |
| Arid5a | 111.8645725 | -0.948818606 | 3.59E-02 |
| Smim13 | 747.323615 | -0.949339512 | 1.25E-12 |
| Stard7 | 2441.522627 | -0.950226723 | 1.02E-13 |
| Spef1 | 267.7432895 | -0.950402466 | 9.09E-05 |
| C130036L24Rik | 41.61183727 | -0.950829202 | 1.28E-02 |
| Zfp488 | 181.8487077 | -0.952339855 | 6.87E-04 |
| Exo5 | 1308.513939 | -0.952825461 | 6.71E-17 |
| Ubtf | 5461.848329 | -0.952837974 | 3.53E-10 |
| Nme7 | 496.5352665 | -0.953185393 | 2.28E-10 |
| Atp2c1 | 2132.770983 | -0.95347661 | 6.25E-09 |
| Map7d3 | 53.69648233 | -0.953697978 | 4.55E-02 |
| 2610001J05Rik | 2297.6783 | -0.95385907 | 1.26E-16 |
| Laptm4b | 5740.524772 | -0.95407251 | 3.26E-07 |
| Mrps2 | 1751.345024 | -0.954268323 | 8.18E-30 |
| 9930021J03Rik | 2322.785857 | -0.955089538 | 3.40E-08 |
| Ngly1 | 876.5239073 | -0.955312971 | 1.28E-09 |
| Fign | 239.6826974 | -0.956744177 | 2.64E-03 |
| Syna | 17.54066673 | -0.956822433 | 3.35E-02 |
| Gm4540 | 805.2713897 | -0.95687481 | 2.54E-08 |

| | | | | |
|---|---|---|---|---|
| | 7-Mar | 2746.3225 | -0.956953675 | 1.66E-04 |
| Cd302 | | 1091.553765 | -0.957236897 | 2.20E-09 |
| AI480526 | | 225.4831329 | -0.957768134 | 7.39E-03 |
| NA | | 1811.90362 | -0.957834412 | 8.90E-16 |
| Gm26599 | | 26.08773149 | -0.958175929 | 2.77E-02 |
| N6amt1 | | 522.967724 | -0.958260136 | 1.73E-11 |
| mt-Co3 | | 264639.126 | -0.958401163 | 3.32E-11 |
| Suclg2 | | 5554.452872 | -0.958584749 | 6.04E-21 |
| Fam57a | | 572.5291046 | -0.958840184 | 2.40E-07 |
| Alg8 | | 450.3239946 | -0.959318059 | 1.92E-07 |
| Pex11g | | 135.442031 | -0.959910338 | 7.98E-04 |
| Magi1 | | 863.00933 | -0.960056072 | 5.11E-12 |
| Pan3 | | 1649.932503 | -0.960354596 | 1.60E-06 |
| Gng8 | | 129.1031785 | -0.960606344 | 5.69E-06 |
| Gm15728 | | 120.8445457 | -0.961221416 | 1.94E-04 |
| Taf11 | | 1033.711611 | -0.961551169 | 1.17E-13 |
| Fam102a | | 1293.326522 | -0.962686203 | 1.43E-05 |
| Tm9sf3 | | 10160.30778 | -0.962837246 | 4.25E-09 |
| Sp100 | | 229.87064 | -0.963012251 | 7.75E-03 |
| Cyb5b | | 7929.007222 | -0.963053165 | 2.56E-12 |
| Ube2v2 | | 1076.557813 | -0.96339852 | 1.21E-02 |
| Pds5a | | 3377.579484 | -0.964233341 | 5.12E-06 |
| Zmym5 | | 1054.81283 | -0.964358769 | 6.84E-09 |
| Iqck | | 94.08072981 | -0.964439767 | 1.71E-05 |
| Il18 | | 182.5136154 | -0.964523659 | 1.12E-03 |
| Slc25a32 | | 612.1776734 | -0.964625641 | 2.47E-08 |
| Slc35b4 | | 3374.352088 | -0.964626958 | 4.39E-09 |
| Tmem263 | | 2118.154995 | -0.966011087 | 6.21E-15 |
| | 8-Mar | 6434.713929 | -0.966151447 | 7.66E-90 |
| Zfp518b | | 797.7422678 | -0.966434699 | 1.12E-09 |
| Ulk4 | | 167.008026 | -0.966925467 | 2.39E-05 |
| Zkscan1 | | 2415.13319 | -0.967850917 | 9.77E-23 |
| Fgf10 | | 1182.615126 | -0.968061452 | 1.68E-05 |
| Cntrl | | 1076.170369 | -0.968168539 | 4.09E-04 |
| Irx5 | | 374.8914831 | -0.968530302 | 2.29E-09 |
| Ddx55 | | 523.2516774 | -0.968775825 | 8.58E-08 |
| Arl13b | | 708.5328787 | -0.969201906 | 7.52E-13 |
| Gm4673 | | 94.57211719 | -0.969222873 | 5.22E-04 |
| Prcc | | 1702.755892 | -0.969405459 | 1.39E-14 |
| Ppp4r2 | | 4872.112859 | -0.969418633 | 1.49E-12 |
| Hlcs | | 514.8003438 | -0.969858524 | 3.17E-11 |
| Nxpe4 | | 417.4256941 | -0.969903095 | 6.03E-15 |
| Arrb1 | | 618.1247361 | -0.96991665 | 4.88E-06 |
| Prmt6 | | 763.8135835 | -0.970109789 | 5.92E-16 |
| Taok1 | | 3582.691983 | -0.970148296 | 7.66E-05 |
| Zfp54 | | 145.6278503 | -0.970532714 | 4.48E-03 |
| Mettl9 | | 2407.464825 | -0.970765981 | 1.74E-24 |

| | | | |
|---|---|---|---|
| Rab31 | 6124.422698 | -0.971140715 | 3.10E-19 |
| Selenos | 1618.134668 | -0.971540365 | 1.14E-08 |
| Ostm1 | 1935.936349 | -0.972408113 | 6.48E-12 |
| Nav1 | 2802.729917 | -0.97342574 | 1.14E-14 |
| Nipa2 | 1056.294581 | -0.973529143 | 1.09E-09 |
| 2810433D01Rik | 81.57471888 | -0.973701191 | 3.31E-03 |
| Rnft1 | 1646.026657 | -0.974178103 | 2.11E-06 |
| Slc10a7 | 1417.196697 | -0.974512098 | 8.60E-09 |
| Gm6565 | 182.3058885 | -0.974883151 | 4.11E-02 |
| Maged1 | 25928.54256 | -0.975000558 | 1.54E-19 |
| Accs | 415.1684912 | -0.975411289 | 6.37E-05 |
| Dhrs3 | 1200.156624 | -0.975645558 | 1.95E-02 |
| Dnajc3 | 2949.642665 | -0.976154172 | 1.01E-07 |
| Ccl7 | 1521.615401 | -0.976790601 | 2.25E-04 |
| Extl3 | 3064.630989 | -0.97756739 | 2.18E-15 |
| Slc4a3 | 999.3973339 | -0.977568229 | 1.43E-10 |
| Ccnl1 | 2321.999919 | -0.977573904 | 2.74E-06 |
| Cryl1 | 233.4306095 | -0.978367865 | 3.15E-07 |
| Zfp868 | 720.900899 | -0.978508788 | 5.60E-09 |
| Rdh10 | 613.1453821 | -0.978862605 | 1.11E-08 |
| Lysmd2 | 420.1955019 | -0.979344874 | 1.47E-07 |
| Fam122b | 417.4900481 | -0.979837353 | 5.18E-13 |
| 4732491K20Rik | 50.26132976 | -0.979859486 | 1.64E-02 |
| Selenof | 8426.348698 | -0.980311417 | 1.47E-19 |
| Adam19 | 3337.629642 | -0.98043749 | 9.35E-13 |
| Sub1 | 3397.985703 | -0.980495834 | 1.63E-14 |
| Lacc1 | 400.6671085 | -0.981075574 | 8.32E-05 |
| BC039966 | 29.60548201 | -0.981106616 | 2.11E-02 |
| Col4a1 | 20658.11282 | -0.981217094 | 4.06E-06 |
| 9330020H09Rik | 49.04901894 | -0.982191118 | 1.32E-02 |
| Ttc5 | 737.7181432 | -0.98232181 | 2.69E-19 |
| Gm21897 | 269.4514575 | -0.983468033 | 5.28E-03 |
| Ssb | 5065.476175 | -0.983827955 | 1.98E-03 |
| U2surp | 2613.824615 | -0.983837828 | 8.78E-05 |
| Nfia | 1183.604419 | -0.984733408 | 2.43E-05 |
| Hmgcr | 4300.914669 | -0.986318216 | 7.80E-04 |
| Tnrc6b | 3087.399165 | -0.986535087 | 8.09E-19 |
| L3hypdh | 996.178476 | -0.986848427 | 1.50E-15 |
| Il4i1 | 184.7277245 | -0.987585671 | 1.24E-05 |
| Enox2 | 311.907405 | -0.988259994 | 5.78E-09 |
| Gpcpd1 | 953.453731 | -0.989334745 | 5.49E-05 |
| Atm | 1655.842703 | -0.989602527 | 2.68E-04 |
| 4833418N02Rik | 87.81324026 | -0.989921673 | 3.14E-03 |
| Flvcr1 | 683.9766244 | -0.990481446 | 2.01E-06 |
| Smo | 9359.591469 | -0.990600383 | 2.47E-25 |
| Raf1 | 5257.897708 | -0.991057051 | 5.63E-30 |
| 2610021A01Rik | 455.860301 | -0.991072987 | 1.41E-03 |

| | | | |
|---|---|---|---|
| Lix1l | 6902.240498 | -0.991490301 | 6.07E-10 |
| Gm14324 | 32.8070965 | -0.991619975 | 3.80E-02 |
| Bcl2l2 | 1753.498301 | -0.991786277 | 6.52E-13 |
| Uhrf1 | 2138.211695 | -0.99202684 | 4.40E-02 |
| Bhlhb9 | 1587.323127 | -0.992178813 | 6.43E-10 |
| Hspa12b | 551.3314607 | -0.992184614 | 1.38E-10 |
| Rnf2 | 2451.487291 | -0.992512852 | 4.60E-13 |
| Slc35e1 | 2802.975292 | -0.992525023 | 3.13E-15 |
| Ctcf | 2881.745882 | -0.992740218 | 6.20E-27 |
| Tnfaip2 | 3658.703536 | -0.99293028 | 2.12E-09 |
| Dhodh | 572.5121825 | -0.993061506 | 1.14E-05 |
| Nipbl | 4012.210935 | -0.993107908 | 3.98E-04 |
| Atp11b | 2214.282937 | -0.993153116 | 6.91E-05 |
| Smarcc2 | 6613.270965 | -0.993208396 | 1.44E-25 |
| Endod1 | 423.568897 | -0.993767299 | 8.11E-06 |
| Zbed4 | 1014.724302 | -0.99423947 | 2.73E-15 |
| B3galnt2 | 1195.272558 | -0.994279246 | 6.19E-08 |
| Ubtd2 | 899.0570191 | -0.995342008 | 4.60E-11 |
| Lss | 5106.435912 | -0.995403886 | 5.70E-03 |
| Clspn | 928.4095926 | -0.995722685 | 6.23E-03 |
| Gtpbp8 | 618.9859008 | -0.99572334 | 1.17E-06 |
| Zfp512b | 2113.08326 | -0.995783134 | 1.48E-10 |
| Dalrd3 | 632.742894 | -0.995796061 | 2.91E-12 |
| Dapk1 | 317.0635772 | -0.996118789 | 1.62E-03 |
| Cflar | 3787.80611 | -0.996376871 | 9.47E-16 |
| Samd1 | 1577.418184 | -0.996523351 | 1.39E-10 |
| Smpd5 | 32.81608022 | -0.996777037 | 9.48E-03 |
| L3mbtl3 | 963.0967322 | -0.996828012 | 1.86E-07 |
| Trim34a | 72.94271336 | -0.997297934 | 3.94E-04 |
| Rmdn3 | 951.2252578 | -0.997352111 | 8.96E-16 |
| Ppp2r1b | 1945.402469 | -0.997399486 | 1.22E-13 |
| Sox12 | 1777.455189 | -0.998010431 | 6.51E-10 |
| Ostc | 2694.914602 | -0.998063192 | 7.15E-15 |
| Abcd2 | 730.8520276 | -0.998265117 | 1.16E-06 |
| Ankrd34a | 289.6306119 | -0.998466547 | 1.18E-07 |
| Atp9b | 2468.969284 | -0.998630428 | 2.87E-26 |
| Fam131b | 2012.653011 | -0.998760475 | 2.29E-10 |
| Zfp120 | 529.9207457 | -0.998806483 | 1.01E-04 |
| Pot1b | 357.1730331 | -0.999172354 | 9.83E-07 |
| Rasl11a | 634.5078439 | -0.999211622 | 1.00E-04 |
| Slk | 5336.649698 | -0.999733924 | 2.97E-07 |
| Serpinb8 | 975.2511855 | -1.000176107 | 1.45E-04 |
| Zfp521 | 2160.446859 | -1.000246916 | 2.29E-07 |
| Fkbp3 | 3751.554593 | -1.000351355 | 2.91E-04 |
| Ccdc85c | 228.5972203 | -1.000538244 | 5.19E-07 |
| Abca1 | 1441.23125 | -1.000883762 | 3.45E-03 |
| Tsc22d1 | 5481.602433 | -1.001104826 | 8.53E-15 |

| | | | |
|---|---|---|---|
| Daam1 | 2029.928172 | -1.002878925 | 5.02E-11 |
| Cemip2 | 1597.684135 | -1.003088125 | 3.85E-19 |
| Gcc2 | 1712.595495 | -1.003217303 | 6.04E-05 |
| Mccc1os | 80.25130271 | -1.003385003 | 5.17E-04 |
| Zfp94 | 133.1127041 | -1.003483615 | 6.49E-06 |
| Morn1 | 87.48653537 | -1.003543131 | 5.10E-05 |
| Nkapd1 | 1223.372902 | -1.004003679 | 1.87E-11 |
| Elmod2 | 768.1670159 | -1.004270863 | 1.70E-06 |
| Tmem69 | 430.4526778 | -1.004666987 | 1.08E-09 |
| Pigw | 191.815086 | -1.005288808 | 4.05E-03 |
| Aaed1 | 890.5933786 | -1.006164533 | 1.22E-11 |
| Nipa1 | 460.541922 | -1.006260422 | 1.07E-09 |
| Ces2b | 20.89041723 | -1.006406772 | 4.94E-02 |
| Caap1 | 304.6149182 | -1.006462331 | 3.63E-06 |
| Adnp | 1561.291486 | -1.006485797 | 3.44E-11 |
| Gm27029 | 107.2998506 | -1.007440947 | 1.99E-02 |
| Fam172a | 1437.904506 | -1.007631684 | 1.35E-16 |
| Nemf | 1753.799341 | -1.007991139 | 3.41E-03 |
| Snx13 | 2199.007322 | -1.008401137 | 1.79E-06 |
| Trim23 | 667.5805986 | -1.008408943 | 2.15E-04 |
| Lancl3 | 159.4956977 | -1.008704821 | 3.28E-03 |
| Pggt1b | 571.5865341 | -1.008839136 | 4.73E-06 |
| Sowahc | 830.3609363 | -1.009130362 | 4.23E-10 |
| Gm26669 | 87.35143234 | -1.009620983 | 4.17E-02 |
| Pfkfb4 | 268.5857426 | -1.009689701 | 1.50E-06 |
| Zfp184 | 226.740299 | -1.009805224 | 3.38E-08 |
| Pank3 | 4644.409009 | -1.010136368 | 1.19E-06 |
| Acvr2a | 745.8765756 | -1.010500775 | 4.16E-05 |
| Plxnb1 | 240.1316655 | -1.010594932 | 4.78E-02 |
| Luc7l2 | 5353.744953 | -1.010847354 | 2.21E-06 |
| Fbxw7 | 724.7639213 | -1.011318875 | 6.75E-10 |
| Tlk1 | 4354.547591 | -1.011392942 | 1.98E-16 |
| Cdkn2c | 2309.987574 | -1.012462422 | 1.37E-13 |
| Fam168b | 9749.745025 | -1.012735635 | 9.71E-21 |
| Fam199x | 378.9540079 | -1.013227538 | 5.21E-03 |
| Paxbp1 | 1368.923884 | -1.013463091 | 1.69E-03 |
| Arl6ip6 | 702.06621 | -1.013485206 | 5.07E-11 |
| Cep192 | 1814.92121 | -1.013564567 | 2.86E-05 |
| Mgme1 | 457.9350659 | -1.013643891 | 2.59E-16 |
| Rp2 | 620.5085526 | -1.013847625 | 7.74E-04 |
| Lsm10 | 1095.464393 | -1.01418926 | 5.29E-19 |
| Rnf34 | 1154.614628 | -1.014917184 | 3.38E-21 |
| Tk1 | 1172.738402 | -1.015578026 | 9.12E-03 |
| Prps1l3 | 896.6416978 | -1.016161934 | 5.03E-09 |
| Gm15270 | 34.01967855 | -1.016210944 | 2.71E-02 |
| Phf6 | 1129.146427 | -1.01769366 | 2.31E-05 |
| Znrf3 | 529.8891783 | -1.018957733 | 4.59E-14 |

| | | | |
|---|---|---|---|
| Rtl10 | 88.75112838 | -1.019037223 | 8.58E-05 |
| Irgm2 | 164.2745168 | -1.019572792 | 3.84E-05 |
| A930035D04Rik | 22.72611546 | -1.019841706 | 2.79E-02 |
| Pi4kb | 3352.487585 | -1.021793992 | 2.07E-14 |
| Zfp799 | 721.4912572 | -1.021937854 | 4.78E-20 |
| Pisd-ps2 | 162.680632 | -1.022938281 | 1.33E-05 |
| Spin4 | 112.9802311 | -1.023002685 | 1.63E-03 |
| Zfp429 | 53.739323 | -1.023028161 | 2.59E-03 |
| Mmp11 | 3113.336865 | -1.023190682 | 2.48E-18 |
| Runx1t1 | 722.8523518 | -1.023577586 | 1.25E-06 |
| Plk-ps1 | 24.66026024 | -1.025004838 | 3.05E-02 |
| Macrod2 | 179.0981956 | -1.025115578 | 1.75E-07 |
| Sorcs2 | 114.3662152 | -1.02562101 | 1.07E-02 |
| Rbbp6 | 3361.835241 | -1.026001245 | 7.19E-11 |
| Htatsf1 | 1879.559997 | -1.026104958 | 3.49E-08 |
| Gm5909 | 37.52129575 | -1.026578288 | 5.29E-03 |
| Ppip5k1 | 941.1084221 | -1.027058591 | 7.16E-16 |
| Dpy19l4 | 1294.644785 | -1.027624399 | 1.26E-08 |
| A930012L18Rik | 23.28516798 | -1.027760883 | 2.08E-02 |
| Fam13b | 1672.285168 | -1.028131184 | 1.37E-08 |
| Gm17477 | 26.30376676 | -1.028189309 | 4.54E-02 |
| Parp9 | 681.8767794 | -1.028296504 | 9.16E-21 |
| Pcca | 1116.587658 | -1.028573395 | 3.12E-13 |
| Thoc1 | 1156.504266 | -1.028590208 | 4.41E-06 |
| Gm10516 | 98.22767106 | -1.029751377 | 4.57E-05 |
| AW146154 | 327.3249277 | -1.03002107 | 3.53E-04 |
| 2410018L13Rik | 753.6620775 | -1.030211663 | 8.60E-16 |
| Pcolce | 25050.1764 | -1.030906647 | 1.98E-14 |
| Ldlrad3 | 1390.98449 | -1.03093963 | 5.62E-11 |
| Adamts1 | 17979.02683 | -1.03100209 | 5.94E-04 |
| Fbxl3 | 1735.020904 | -1.031552754 | 1.91E-10 |
| Psen2 | 389.5631355 | -1.031820035 | 1.71E-10 |
| Hexim2 | 171.0155295 | -1.03235135 | 2.88E-07 |
| Dmxl1 | 1786.98676 | -1.032659524 | 2.56E-04 |
| Dido1 | 3460.394899 | -1.032733595 | 2.73E-14 |
| Gm9530 | 34.82102145 | -1.032919177 | 2.98E-02 |
| Bvht | 149.9887075 | -1.033157067 | 4.19E-05 |
| Tarbp1 | 227.408269 | -1.033681548 | 7.05E-05 |
| Med20 | 1331.253788 | -1.034090379 | 1.92E-12 |
| Klf9 | 3975.253106 | -1.03450207 | 6.40E-11 |
| 1700086O06Rik | 134.763061 | -1.034998868 | 6.77E-05 |
| Coq3 | 453.259282 | -1.035011443 | 8.60E-09 |
| Chek1 | 275.9564489 | -1.035366234 | 1.80E-02 |
| Nprl3 | 1164.930093 | -1.035786271 | 2.84E-17 |
| Sft2d2 | 796.2368986 | -1.035879649 | 3.34E-09 |
| Dmxl2 | 711.1668361 | -1.036020316 | 5.98E-04 |
| Alg10b | 2845.75944 | -1.036032384 | 2.50E-12 |

| | | | |
|---|---|---|---|
| Pign | 3567.705886 | -1.03614849 | 8.12E-17 |
| Gm8615 | 190.0903932 | -1.036329434 | 2.32E-05 |
| Rcc2 | 9407.022186 | -1.036569938 | 6.01E-15 |
| Rfesd | 285.2920985 | -1.037192072 | 3.88E-09 |
| Rnf219 | 689.5253814 | -1.037485651 | 3.61E-15 |
| Msantd2 | 431.847042 | -1.03782747 | 6.72E-13 |
| Lamp2 | 21188.1191 | -1.038094772 | 3.71E-14 |
| Katnb1 | 817.8951214 | -1.038933876 | 2.57E-09 |
| Fktn | 1315.024605 | -1.039059037 | 2.03E-11 |
| C430049B03Rik | 299.2756912 | -1.039252562 | 1.91E-04 |
| Cand2 | 2044.67946 | -1.03954539 | 1.38E-14 |
| Mbtd1 | 1716.420246 | -1.039967926 | 4.20E-10 |
| Hes6 | 612.1037007 | -1.040001944 | 1.44E-09 |
| Gm17066 | 107.9290911 | -1.040351084 | 1.46E-03 |
| Mmachc | 686.9795778 | -1.041006073 | 2.12E-13 |
| Plin4 | 983.3697174 | -1.041537698 | 1.63E-02 |
| Meis1 | 1497.087718 | -1.041607809 | 4.20E-11 |
| Skp2 | 912.3949815 | -1.041867716 | 9.02E-05 |
| Zfp661 | 298.2016297 | -1.044169996 | 3.06E-09 |
| Dip2c | 2363.258568 | -1.045161749 | 2.03E-23 |
| Gm26516 | 33.49088012 | -1.045262353 | 3.69E-02 |
| Crebl2 | 1856.82481 | -1.045265317 | 1.64E-11 |
| Gm9922 | 24.5227298 | -1.047136877 | 2.96E-02 |
| Rcn1 | 7279.957607 | -1.047232605 | 1.61E-21 |
| Morf4l2 | 12523.02325 | -1.047234389 | 5.91E-14 |
| Smc1a | 5445.475379 | -1.04729314 | 8.67E-14 |
| B230219D22Rik | 3240.706529 | -1.047490548 | 3.35E-11 |
| Aida | 2825.873098 | -1.047566792 | 1.12E-16 |
| Asic1 | 2363.956811 | -1.047738048 | 8.33E-08 |
| Crocc | 472.3468662 | -1.047800252 | 2.82E-06 |
| AI429214 | 333.0356269 | -1.048586675 | 3.19E-11 |
| Srek1 | 1768.790581 | -1.049337944 | 6.27E-11 |
| Pctp | 173.5637158 | -1.049620551 | 2.48E-03 |
| Abr | 2974.771084 | -1.049935161 | 2.47E-18 |
| Mvk | 1649.647814 | -1.050340183 | 1.36E-05 |
| Ifit3b | 229.6966565 | -1.050451589 | 9.64E-03 |
| Gm13288 | 22.68868773 | -1.051660135 | 4.09E-02 |
| Rngtt | 654.9001586 | -1.05230393 | 1.25E-08 |
| Cnep1r1 | 1321.404235 | -1.052332593 | 2.52E-13 |
| Fam173a | 718.2912736 | -1.052564048 | 1.11E-16 |
| Idua | 698.7732812 | -1.053172979 | 2.90E-17 |
| Hs2st1 | 3965.428802 | -1.053330784 | 7.81E-21 |
| Slc1a4 | 3460.178396 | -1.054454932 | 9.04E-19 |
| Eif5b | 5660.941743 | -1.054820438 | 1.84E-10 |
| Dennd4c | 5771.928255 | -1.055022205 | 9.98E-20 |
| Gdap10 | 114.4604381 | -1.055222515 | 1.30E-03 |
| Gm20712 | 69.74493002 | -1.055281559 | 1.12E-04 |

| | | | |
|---|---|---|---|
| Gm6548 | 191.9369595 | -1.055412001 | 1.52E-07 |
| Mrpl23-ps1 | 524.2313248 | -1.056017118 | 4.13E-08 |
| Akr1c12 | 25.24508091 | -1.056128685 | 3.42E-02 |
| Gm8991 | 29.80053889 | -1.056142078 | 4.98E-02 |
| Cryzl1 | 961.3287174 | -1.056795663 | 1.10E-11 |
| Pcna-ps2 | 616.8396399 | -1.057292456 | 1.51E-05 |
| 3110056K07Rik | 100.0527371 | -1.05862601 | 1.03E-03 |
| Ssh2 | 685.016075 | -1.058704028 | 7.76E-08 |
| Suv39h2 | 217.1038152 | -1.059701264 | 2.04E-02 |
| Ddx10 | 1380.39871 | -1.059812631 | 2.19E-08 |
| Ncoa1 | 1626.174698 | -1.060779739 | 1.31E-21 |
| Zbtb45 | 334.1316636 | -1.060847108 | 1.13E-10 |
| Dnajc13 | 5378.577471 | -1.060866522 | 1.83E-16 |
| Zfp101 | 297.5685569 | -1.061339314 | 1.72E-04 |
| Lclat1 | 2716.89391 | -1.061895454 | 1.52E-23 |
| Gm3555 | 33.44381624 | -1.062029685 | 1.23E-02 |
| Tardbp | 5588.960402 | -1.062177142 | 2.03E-10 |
| H2-M5 | 55.87435966 | -1.062305276 | 2.07E-03 |
| Brwd3 | 700.4136653 | -1.063379681 | 4.98E-05 |
| Ttc26 | 659.8498206 | -1.063694197 | 4.80E-11 |
| Ncapg2 | 1246.754677 | -1.06389728 | 2.39E-03 |
| Xaf1 | 354.2355811 | -1.065099458 | 7.94E-06 |
| Mmab | 2103.661292 | -1.065101633 | 8.88E-07 |
| Fam135a | 1961.021116 | -1.066336619 | 3.36E-05 |
| Gm15411 | 49.73035422 | -1.066739647 | 1.72E-03 |
| Hykk | 297.2623057 | -1.066996512 | 2.23E-15 |
| Limd2 | 1732.780265 | -1.067009847 | 3.16E-10 |
| Cep70 | 410.6996312 | -1.067279992 | 1.17E-04 |
| Mysm1 | 1703.180241 | -1.067553413 | 2.18E-05 |
| Zeb2 | 3746.786289 | -1.067822974 | 3.78E-24 |
| NA | 152.4716177 | -1.068002984 | 7.16E-06 |
| Fmnl3 | 2696.537561 | -1.068208107 | 1.10E-14 |
| Six5 | 615.5519533 | -1.068376882 | 5.38E-12 |
| Zc3h6 | 614.4255871 | -1.068712782 | 7.48E-09 |
| Jmjd8 | 638.2911338 | -1.068754984 | 3.38E-27 |
| Slc38a7 | 2944.818323 | -1.06896236 | 1.13E-14 |
| S1pr3 | 3918.192315 | -1.069104112 | 4.52E-12 |
| Tmem241 | 182.3624427 | -1.069750087 | 3.96E-05 |
| Katnbl1 | 1110.930944 | -1.070793034 | 3.47E-11 |
| Gemin6 | 230.8022335 | -1.070841373 | 5.81E-04 |
| Zbtb46 | 81.22314712 | -1.072037177 | 1.24E-03 |
| Enpp3 | 439.6794586 | -1.072270324 | 7.90E-05 |
| Dagla | 576.2981415 | -1.072981969 | 3.47E-16 |
| Zfp979 | 556.4311338 | -1.0730857 | 3.56E-03 |
| Gm27042 | 75.90676536 | -1.073252255 | 5.11E-06 |
| Slc22a17 | 205.312658 | -1.073785364 | 1.32E-07 |
| Zfp324 | 561.8524386 | -1.073796386 | 1.36E-18 |

| | | | |
|---|---|---|---|
| Gpbp1 | 2390.321323 | -1.074332363 | 1.98E-06 |
| Wdr11 | 1814.743598 | -1.074699259 | 6.58E-20 |
| Zscan12 | 310.7903595 | -1.074936219 | 1.25E-12 |
| Rpap3 | 708.2044117 | -1.075765976 | 5.78E-15 |
| Ccdc183 | 20.08276762 | -1.076357394 | 3.89E-02 |
| Hsph1 | 6295.873221 | -1.076467862 | 8.49E-19 |
| Leo1 | 880.8601505 | -1.077004937 | 6.90E-12 |
| Zfp367 | 599.2259266 | -1.077818039 | 1.39E-04 |
| Phip | 2280.134654 | -1.07875217 | 1.95E-04 |
| Hexdc | 950.8161435 | -1.079375808 | 2.41E-11 |
| Naip6 | 212.161741 | -1.079400943 | 3.30E-10 |
| Nid1 | 41135.2065 | -1.079426068 | 8.59E-25 |
| 1700066M21Rik | 329.5716986 | -1.080472145 | 8.37E-04 |
| Clk1 | 3445.62983 | -1.080515611 | 8.72E-06 |
| Mcm4 | 3562.673628 | -1.080814732 | 4.05E-07 |
| Crebbp | 2005.517214 | -1.08122121 | 5.45E-22 |
| NA | 48.83985045 | -1.081224898 | 1.72E-02 |
| Lins1 | 515.0195333 | -1.081372251 | 2.41E-06 |
| Sema5a | 3228.140088 | -1.081667897 | 5.33E-05 |
| Ylpm1 | 3107.707483 | -1.081670857 | 3.51E-14 |
| C530005A16Rik | 28.57807176 | -1.081903509 | 1.29E-02 |
| Zdhhc17 | 1143.789916 | -1.082327904 | 3.39E-11 |
| Trim33 | 2609.996824 | -1.082394381 | 1.55E-10 |
| Dhcr24 | 6989.867164 | -1.082791261 | 6.28E-04 |
| Zfp654 | 364.2675256 | -1.083761657 | 1.08E-06 |
| Rbm4b | 986.1917831 | -1.084431018 | 4.09E-12 |
| Mrpl24 | 995.155896 | -1.08448812 | 1.16E-08 |
| Bod1 | 1941.315102 | -1.084548106 | 2.37E-20 |
| Mcm6 | 3739.888641 | -1.084708118 | 3.10E-04 |
| Serinc5 | 1871.588337 | -1.084889839 | 1.29E-23 |
| 4930513N10Rik | 46.10247408 | -1.085415281 | 3.44E-04 |
| Dolk | 812.9649783 | -1.085496731 | 5.29E-20 |
| Aldh3a1 | 155.6027006 | -1.086737869 | 3.99E-02 |
| Gpsm2 | 2322.001977 | -1.087902131 | 1.28E-14 |
| Pskh1 | 1911.4363 | -1.088589511 | 7.80E-27 |
| Mzf1 | 33.13338563 | -1.088989347 | 2.03E-02 |
| Peli2 | 1908.882325 | -1.089901146 | 8.87E-17 |
| Zfp24 | 1016.182409 | -1.090107538 | 3.77E-13 |
| Ccar1 | 3219.749532 | -1.090616565 | 4.00E-07 |
| F3 | 10673.0851 | -1.090751525 | 2.61E-07 |
| Zzz3 | 5136.327403 | -1.090949636 | 9.47E-13 |
| Trmt61b | 146.4603856 | -1.09112062 | 5.43E-06 |
| Ydjc | 121.0858377 | -1.092257539 | 1.80E-04 |
| Gramd1c | 21.27151137 | -1.09239455 | 2.91E-02 |
| Pibf1 | 375.6418153 | -1.092426577 | 7.04E-08 |
| Dicer1 | 2835.407638 | -1.093265885 | 2.66E-26 |
| Gm16183 | 29.51470691 | -1.093592831 | 1.05E-02 |

| | | | |
|---|---|---|---|
| Tceal8 | 3073.445068 | -1.094699855 | 7.75E-29 |
| Homez | 326.2089349 | -1.094811888 | 4.11E-06 |
| Trak1 | 2002.750485 | -1.095187891 | 7.12E-22 |
| Masp1 | 1959.534601 | -1.095671992 | 2.86E-05 |
| Fbxl14 | 1290.48478 | -1.095763538 | 1.11E-18 |
| Tbx15 | 3782.490487 | -1.09660921 | 1.33E-19 |
| Zfp768 | 743.7030957 | -1.096865371 | 1.77E-24 |
| Fen1 | 1040.024734 | -1.096989117 | 1.88E-04 |
| Pbdc1 | 1501.992805 | -1.097319501 | 3.57E-15 |
| Vangl2 | 622.778806 | -1.097668855 | 5.12E-11 |
| Son | 10931.49173 | -1.098270238 | 6.54E-20 |
| Ctdspl2 | 1328.663166 | -1.099003193 | 7.19E-09 |
| Grcc10 | 2167.032303 | -1.099353462 | 1.59E-08 |
| Gm26816 | 34.25874803 | -1.09937198 | 8.66E-03 |
| 1600020E01Rik | 376.3471128 | -1.099992823 | 2.53E-14 |
| Foxd2 | 47.01500774 | -1.100534542 | 4.11E-04 |
| Znhit2 | 471.4218009 | -1.101892183 | 2.18E-09 |
| Snx22 | 20.06032273 | -1.103723511 | 4.52E-02 |
| Cfap97 | 1089.736 | -1.103813579 | 2.25E-09 |
| D10Wsu102e | 2431.206995 | -1.106408049 | 6.31E-22 |
| Zfp839 | 451.8448377 | -1.106472596 | 3.11E-17 |
| Pik3r2 | 5877.98043 | -1.10671228 | 8.27E-17 |
| Gm21992 | 240.7428873 | -1.107205374 | 2.88E-04 |
| NA | 72.61252199 | -1.107391707 | 1.48E-02 |
| Usp27x | 226.8415996 | -1.107800544 | 1.09E-10 |
| Gramd4 | 2354.504038 | -1.108253565 | 1.97E-19 |
| Zbtb33 | 1203.252921 | -1.108920056 | 3.10E-13 |
| Sdcbp | 7570.072267 | -1.109599319 | 1.24E-20 |
| Gm26621 | 27.54801655 | -1.110292844 | 3.07E-02 |
| Ltbp3 | 11015.44366 | -1.111777211 | 7.71E-21 |
| Arsk | 464.5614322 | -1.112077089 | 9.96E-14 |
| Abhd12 | 3305.451109 | -1.114209419 | 6.15E-16 |
| Ccdc90b | 690.3988291 | -1.115023168 | 3.19E-07 |
| Fam76b | 695.9564414 | -1.115258478 | 1.15E-06 |
| Snord49a | 38.68955715 | -1.115478614 | 3.98E-02 |
| Obsl1 | 4116.268852 | -1.115623488 | 9.31E-12 |
| Dnm1l | 3671.652075 | -1.116034019 | 3.57E-05 |
| Tmx4 | 2058.445111 | -1.116162161 | 5.54E-27 |
| Rad54l2 | 1696.94065 | -1.116303432 | 1.17E-33 |
| Acsbg1 | 82.3921383 | -1.116627032 | 1.46E-02 |
| Evl | 2952.632705 | -1.117606835 | 4.11E-08 |
| Tnrc18 | 4357.493149 | -1.117946032 | 1.92E-13 |
| AA465934 | 40.81545396 | -1.118014015 | 2.22E-03 |
| Cd40 | 29.42286657 | -1.118020578 | 6.24E-03 |
| Per2 | 279.5536971 | -1.118987716 | 2.18E-07 |
| Cdc6 | 389.1138025 | -1.119377303 | 1.33E-02 |
| Tmppe | 488.8157654 | -1.119401859 | 2.09E-13 |

| | | | |
|---|---|---|---|
| Ssh3 | 139.3431978 | -1.11970733 | 7.55E-05 |
| Hace1 | 931.2661007 | -1.120188968 | 1.63E-17 |
| Paxip1 | 2358.455763 | -1.120465097 | 2.31E-18 |
| Galnt1 | 3097.996652 | -1.12124934 | 1.05E-16 |
| Mex3b | 174.4803361 | -1.121879336 | 9.92E-05 |
| Ly6c1 | 1095.469361 | -1.122044792 | 1.34E-06 |
| Igf2bp2 | 8644.418417 | -1.122293262 | 1.48E-32 |
| Sema4c | 1046.850603 | -1.122749434 | 6.37E-15 |
| Cyb561 | 80.17326152 | -1.122890968 | 5.74E-03 |
| Kdm3a | 3959.529429 | -1.122967613 | 9.33E-21 |
| C330011M18Rik | 41.66936782 | -1.123030442 | 1.31E-03 |
| Ska2 | 986.6881394 | -1.123070908 | 2.20E-04 |
| Gm9996 | 20.33827661 | -1.123335494 | 3.27E-02 |
| Ppip5k2 | 1909.281978 | -1.124484707 | 5.31E-06 |
| Sfxn2 | 329.2799816 | -1.124818698 | 7.05E-06 |
| Zcchc14 | 3410.077933 | -1.124900437 | 3.81E-41 |
| Armc8 | 1164.412789 | -1.128607642 | 4.52E-21 |
| Nfatc1 | 796.3990061 | -1.129558353 | 1.74E-11 |
| Gm15784 | 32.74440799 | -1.129726124 | 8.43E-03 |
| Dip2b | 2838.90961 | -1.12973392 | 1.54E-11 |
| Ppig | 3219.137977 | -1.129747669 | 1.13E-04 |
| Insig1 | 7189.3603 | -1.13031869 | 1.68E-05 |
| 2410004P03Rik | 25.1661059 | -1.131744536 | 3.05E-02 |
| Gm49322 | 185.7261263 | -1.131818992 | 2.28E-08 |
| Cul1 | 6224.737814 | -1.132677357 | 3.37E-14 |
| Tmem132a | 1717.660449 | -1.132711 | 3.11E-14 |
| Gnpda1 | 2534.474834 | -1.132865035 | 3.92E-12 |
| Foxo1 | 589.3772702 | -1.133111538 | 8.69E-21 |
| Lats1 | 3099.689664 | -1.133602765 | 1.06E-14 |
| B3glct | 917.7545041 | -1.133605149 | 3.60E-24 |
| Atad2b | 628.6636208 | -1.134120558 | 3.18E-09 |
| Pxdn | 12242.77878 | -1.134386549 | 8.16E-12 |
| Gm12258 | 435.4925812 | -1.134809953 | 8.79E-05 |
| Smim27 | 95.77154053 | -1.135497152 | 3.47E-05 |
| Tfrc | 1850.025373 | -1.136040429 | 1.97E-02 |
| Mbip | 596.8817882 | -1.136267409 | 4.71E-10 |
| Lcmt2 | 150.6669994 | -1.136683117 | 3.08E-06 |
| Folr2 | 18.99814039 | -1.137092687 | 3.85E-02 |
| Sult6b1 | 15.676378 | -1.137249106 | 4.24E-02 |
| Aak1 | 1915.969594 | -1.137824996 | 8.80E-25 |
| Rnf6 | 2353.676709 | -1.1380952 | 3.25E-07 |
| Adam17 | 3233.205911 | -1.13874336 | 2.04E-43 |
| Sipa1l2 | 3528.093693 | -1.139564379 | 3.03E-09 |
| Setd2 | 3613.390968 | -1.139777148 | 8.48E-09 |
| 4933421O10Rik | 122.4029757 | -1.139852134 | 4.54E-05 |
| Tbx18 | 1240.526144 | -1.140069231 | 4.46E-31 |
| Slc17a8 | 25.91453632 | -1.140133695 | 3.38E-02 |

| | | | |
|---|---|---|---|
| Paqr4 | 951.0335697 | -1.140270091 | 1.04E-07 |
| Ssr1 | 7136.202432 | -1.140337436 | 1.71E-19 |
| Mn1 | 1568.859744 | -1.140875232 | 1.64E-08 |
| AW549877 | 3473.329196 | -1.142120257 | 3.36E-12 |
| Fli1 | 48.07928747 | -1.142704383 | 2.75E-03 |
| Zfp248 | 467.7710312 | -1.143051988 | 3.25E-07 |
| E130208F15Rik | 44.87845881 | -1.143313469 | 4.52E-03 |
| 4632404H12Rik | 210.9051878 | -1.143408519 | 4.58E-08 |
| Rsrc2 | 3040.044271 | -1.14343434 | 1.93E-06 |
| Pemt | 84.99351402 | -1.143442244 | 1.55E-04 |
| Gm17435 | 160.567384 | -1.143661364 | 4.44E-10 |
| 1700030J22Rik | 46.49523078 | -1.144114652 | 2.47E-04 |
| Rab10os | 762.5082916 | -1.144883451 | 1.28E-09 |
| Zc3h7b | 5316.848278 | -1.14684528 | 2.69E-31 |
| Ireb2 | 2896.385537 | -1.147819346 | 4.91E-10 |
| Prox2 | 90.38021678 | -1.148177364 | 3.69E-05 |
| Tbc1d19 | 941.0508285 | -1.148670322 | 1.82E-17 |
| Zc3h12c | 402.0650147 | -1.148699766 | 3.87E-09 |
| Krt10 | 176.393471 | -1.149201647 | 2.56E-08 |
| Akap6 | 117.141912 | -1.150166728 | 4.32E-04 |
| Pld1 | 1259.68068 | -1.15129647 | 3.40E-05 |
| Aspa | 75.09945744 | -1.151359381 | 1.91E-02 |
| Suz12 | 2260.611263 | -1.152417975 | 5.67E-05 |
| Brip1 | 378.3607621 | -1.154041992 | 4.91E-03 |
| Irx3 | 178.4497727 | -1.155057729 | 9.62E-08 |
| Bola1 | 299.4104525 | -1.155548289 | 8.40E-12 |
| Tnik | 413.6788858 | -1.155553916 | 2.90E-07 |
| Ccdc30 | 89.38024234 | -1.156535445 | 7.51E-05 |
| Alyref2 | 123.0794176 | -1.1565937 | 3.14E-07 |
| Zfp37 | 576.1216051 | -1.157257171 | 1.31E-03 |
| 2310001H17Rik | 317.2797806 | -1.157257631 | 2.53E-07 |
| Phlpp1 | 938.9364218 | -1.157520294 | 1.09E-17 |
| Chrac1 | 757.1424589 | -1.158207294 | 5.80E-20 |
| Plekha5 | 1255.587539 | -1.158882738 | 1.67E-14 |
| Ttc14 | 1834.000703 | -1.159110938 | 8.32E-04 |
| Anxa9 | 109.4151629 | -1.159452872 | 1.55E-06 |
| Mpzl1 | 3792.936615 | -1.160465048 | 2.99E-10 |
| Brdt | 227.9030075 | -1.161234011 | 1.90E-07 |
| Jun | 4890.668098 | -1.16184099 | 3.09E-06 |
| Aspm | 1755.329385 | -1.161893604 | 3.92E-02 |
| 4732440D04Rik | 22.54247112 | -1.162533529 | 2.92E-02 |
| Ncoa5 | 1911.343697 | -1.162560653 | 2.34E-20 |
| Mdm4-ps | 1511.456696 | -1.162913628 | 4.89E-09 |
| Mob1b | 1690.668524 | -1.163355093 | 1.45E-12 |
| Dscc1 | 138.1416843 | -1.163842139 | 5.05E-04 |
| Gm14735 | 40.31413703 | -1.164085749 | 4.11E-03 |
| Gm16211 | 299.0104868 | -1.164547652 | 8.48E-14 |

| | | | |
|---|---|---|---|
| Kif9 | 130.1670536 | -1.164961502 | 1.35E-08 |
| Sft2d3 | 128.2159729 | -1.166479373 | 3.80E-07 |
| Stag2 | 5411.882155 | -1.166708778 | 6.52E-07 |
| Slc2a6 | 121.7876624 | -1.16693127 | 1.05E-04 |
| 0610009E02Rik | 39.17645434 | -1.1669754 | 1.58E-03 |
| Zfp930 | 228.3535006 | -1.167518957 | 2.84E-03 |
| Ikbip | 3735.889795 | -1.167656318 | 1.84E-10 |
| mt-Nd1 | 144168.3645 | -1.16843571 | 6.09E-14 |
| Cdrt4 | 25.61038649 | -1.168470616 | 3.81E-03 |
| Cstf3 | 1249.131363 | -1.168479261 | 4.86E-12 |
| Pip5k1c | 2979.353255 | -1.169246183 | 1.27E-21 |
| mt-Rnr2 | 13741.51398 | -1.169254409 | 5.39E-04 |
| Ofd1 | 934.0707456 | -1.169463436 | 4.10E-11 |
| Gm6356 | 69.88684248 | -1.169538103 | 5.98E-04 |
| Loxl1 | 13086.55011 | -1.169572973 | 1.22E-11 |
| Agap1 | 3485.368846 | -1.169659033 | 4.51E-22 |
| BC024978 | 232.6752538 | -1.169744392 | 6.64E-11 |
| Pole4 | 1287.876454 | -1.169901926 | 9.05E-16 |
| Snapin | 2650.194462 | -1.17096775 | 1.71E-19 |
| Rbm10 | 2075.724467 | -1.170971642 | 4.67E-16 |
| Gm26659 | 43.34765276 | -1.171325716 | 2.68E-03 |
| Tmub1 | 212.8439358 | -1.171349222 | 5.04E-10 |
| Lrr1 | 63.46400517 | -1.171415568 | 4.33E-02 |
| Tm7sf2 | 703.3797047 | -1.17168541 | 6.81E-06 |
| Pkdcc | 437.5342631 | -1.171769624 | 5.02E-08 |
| Nat14 | 49.35207722 | -1.172050956 | 1.06E-03 |
| P4ha3 | 952.268623 | -1.172342084 | 1.62E-05 |
| Npat | 1337.634223 | -1.172480879 | 1.26E-12 |
| Plekha2 | 266.0834896 | -1.172745417 | 7.99E-04 |
| Tfdp1 | 4208.88843 | -1.173148707 | 2.96E-08 |
| 4833417C18Rik | 23.3663168 | -1.173243761 | 4.73E-02 |
| Akap13 | 5008.332383 | -1.173700848 | 2.15E-15 |
| Ccne2 | 288.3521814 | -1.174123743 | 1.48E-02 |
| Tbl1xr1 | 2044.586522 | -1.174263219 | 1.46E-23 |
| Rcor2 | 232.6964551 | -1.174308245 | 5.85E-05 |
| Gm10425 | 67.09687368 | -1.174396464 | 2.72E-04 |
| P3h3 | 5126.717251 | -1.174406442 | 9.33E-24 |
| Maml2 | 1522.735573 | -1.175626769 | 6.97E-11 |
| Zfp141 | 460.0176366 | -1.175822574 | 7.13E-05 |
| Gm12183 | 25.24142721 | -1.175954428 | 3.06E-02 |
| Mrps18c | 677.5568705 | -1.176047642 | 1.34E-13 |
| Gm23201 | 29.8277375 | -1.176172088 | 2.29E-02 |
| Pgam2 | 56.35623073 | -1.176238066 | 1.93E-04 |
| Ssbp2 | 1469.826249 | -1.178468874 | 3.12E-05 |
| Capn10 | 780.8179271 | -1.178764732 | 9.69E-15 |
| Nudt19 | 340.0343303 | -1.178837586 | 2.47E-10 |
| mt-Tl1 | 66.97544178 | -1.178944649 | 6.32E-03 |

| | | | |
|---|---|---|---|
| Adamts10 | 953.871541 | -1.179091571 | 4.01E-20 |
| Fgfbp3 | 84.82346159 | -1.179204313 | 3.63E-04 |
| Tfap2a | 730.9542063 | -1.179311124 | 3.10E-07 |
| Clec11a | 34.24030649 | -1.180103505 | 1.15E-02 |
| A630052C17Rik | 85.93514374 | -1.180150191 | 1.70E-04 |
| Kat2b | 1853.046154 | -1.180532298 | 1.99E-09 |
| Slc30a4 | 4466.816693 | -1.181336083 | 3.49E-06 |
| Emilin2 | 6079.768848 | -1.181437529 | 2.07E-23 |
| Gm14418 | 318.6864151 | -1.181600134 | 5.44E-09 |
| Lmln | 194.6595696 | -1.181673675 | 2.62E-02 |
| Uaca | 2130.933084 | -1.182345783 | 4.97E-17 |
| Stard4 | 2046.999912 | -1.182652542 | 1.74E-04 |
| Fundc1 | 1249.116808 | -1.183226955 | 1.38E-28 |
| Relch | 1655.605876 | -1.183579069 | 7.79E-17 |
| Casc4 | 602.869766 | -1.183737303 | 1.32E-07 |
| Dnajb14 | 1687.272902 | -1.184767912 | 3.43E-23 |
| Nlrc3 | 519.874105 | -1.184833682 | 7.74E-11 |
| Fnip2 | 5464.462773 | -1.185151476 | 2.47E-13 |
| Pcyt1b | 608.1630088 | -1.185211632 | 1.59E-06 |
| Smoc1 | 319.0475636 | -1.185333763 | 1.27E-02 |
| Trim21 | 337.3701587 | -1.185997385 | 2.78E-17 |
| Ap5m1 | 552.9572744 | -1.186854974 | 1.14E-09 |
| Scrn1 | 542.6863496 | -1.187165615 | 1.82E-04 |
| Fn1 | 474467.2828 | -1.187488376 | 1.65E-11 |
| Prrx1 | 19059.58088 | -1.188596777 | 1.12E-23 |
| Gm17396 | 29.03386503 | -1.188793361 | 8.95E-03 |
| Zfp449 | 559.0070296 | -1.189716139 | 1.63E-06 |
| Rnf170 | 766.78241 | -1.190700899 | 3.84E-10 |
| Gm16124 | 23.47259755 | -1.191695583 | 2.61E-02 |
| Nrip2 | 87.48086505 | -1.192483005 | 1.77E-03 |
| Kdelr2 | 5081.551697 | -1.192746968 | 1.46E-19 |
| Npas2 | 67.53822572 | -1.1928425 | 1.66E-02 |
| Gm9913 | 38.08840571 | -1.193035902 | 2.12E-02 |
| Oxct1 | 16865.66235 | -1.193159406 | 4.01E-14 |
| Plat | 4991.615202 | -1.193264852 | 5.10E-08 |
| Srsf5 | 9108.135251 | -1.193598846 | 4.57E-08 |
| Csgalnact2 | 1490.922805 | -1.19372081 | 9.01E-11 |
| Usp34 | 6882.952531 | -1.193963366 | 9.55E-07 |
| Klhdc8b | 78.31204509 | -1.194341786 | 2.21E-05 |
| Lemd3 | 799.9605492 | -1.195537803 | 1.30E-18 |
| Spag1 | 148.3787446 | -1.195918783 | 9.45E-05 |
| Brf1 | 1638.014583 | -1.196296569 | 2.00E-35 |
| Ece1 | 1437.72767 | -1.19681078 | 4.10E-09 |
| Asf1a | 736.1877707 | -1.197050167 | 1.50E-11 |
| Pcdhgc4 | 30257.72995 | -1.197671613 | 6.30E-16 |
| Vsig10 | 686.1056348 | -1.197691986 | 2.33E-20 |
| Srp9 | 340.4882686 | -1.200095795 | 1.90E-16 |

| | | | |
|---|---|---|---|
| Brd8 | 2043.868302 | -1.200297493 | 1.86E-15 |
| Gm17251 | 159.5166582 | -1.200461686 | 5.92E-07 |
| 2810408I11Rik | 99.88867526 | -1.200728037 | 2.10E-02 |
| Rbm12 | 1195.18032 | -1.201667847 | 2.33E-10 |
| Kdm7a | 2156.934782 | -1.202221031 | 3.00E-08 |
| Tmem64 | 1663.412228 | -1.20228854 | 7.64E-18 |
| Hivep3 | 515.0723489 | -1.203814747 | 2.21E-09 |
| Kcnd1 | 74.87306616 | -1.204143283 | 1.78E-03 |
| Dhx33 | 2116.155745 | -1.204768273 | 1.24E-25 |
| Slc26a2 | 908.0177184 | -1.205486895 | 5.95E-22 |
| C1d | 926.7708056 | -1.205862076 | 2.46E-09 |
| Fbxo32 | 1611.242564 | -1.206229663 | 3.86E-07 |
| Stxbp4 | 1774.063882 | -1.206419116 | 2.87E-23 |
| Dpm2 | 1496.657824 | -1.206941393 | 7.36E-16 |
| Ift57 | 787.3549709 | -1.207228065 | 1.30E-19 |
| Ppp1r16a | 1023.013923 | -1.208197821 | 2.62E-21 |
| Fam13a | 488.6827203 | -1.208808278 | 1.45E-08 |
| Zfp317 | 753.4419576 | -1.209524869 | 6.47E-16 |
| Rmdn2 | 355.5715743 | -1.209633349 | 9.34E-09 |
| Dock5 | 2703.985943 | -1.209927941 | 1.13E-15 |
| Bdp1 | 1218.233805 | -1.210168584 | 1.69E-05 |
| Soga3 | 486.8497861 | -1.210267671 | 7.37E-14 |
| Gtf3c5 | 1178.49991 | -1.210807278 | 4.44E-30 |
| Bag4 | 698.9273242 | -1.21083907 | 7.71E-24 |
| Bhmt2 | 20.87048708 | -1.212102244 | 3.34E-02 |
| 3010003L21Rik | 23.41418922 | -1.212248894 | 2.49E-02 |
| Wipf1 | 3029.779313 | -1.212581559 | 6.43E-48 |
| Lrrc56 | 115.614384 | -1.212652493 | 8.37E-06 |
| Hoxd3os1 | 173.6527096 | -1.213316695 | 7.26E-07 |
| Csnk1g3 | 1352.910275 | -1.215283353 | 1.26E-19 |
| Pnrc2 | 1547.386536 | -1.215537171 | 3.72E-16 |
| Gpr155 | 418.0128172 | -1.215604042 | 1.87E-09 |
| NA | 78.76176561 | -1.215878718 | 3.31E-02 |
| Actr6 | 294.6156024 | -1.216194136 | 1.22E-04 |
| Mrfap1 | 12176.20273 | -1.216842131 | 4.42E-26 |
| Zfp318 | 1909.003472 | -1.21747386 | 8.92E-08 |
| mt-Nd4 | 129372.965 | -1.217572376 | 1.52E-16 |
| Sacs | 898.5330918 | -1.217772321 | 2.36E-03 |
| Jmjd1c | 4950.142215 | -1.218863779 | 6.68E-09 |
| Etfrf1 | 953.9923609 | -1.219342109 | 8.27E-16 |
| AC157931.1 | 78.99463552 | -1.219644556 | 8.78E-05 |
| Fus | 8428.016269 | -1.219730642 | 8.43E-09 |
| Sdccag8 | 700.0101104 | -1.220759193 | 1.06E-13 |
| Itpripl1 | 456.6583994 | -1.221227424 | 1.81E-07 |
| Yes1 | 1175.543531 | -1.221989283 | 1.87E-05 |
| Rufy1 | 2070.818133 | -1.223695617 | 2.77E-12 |
| Oscp1 | 498.6865751 | -1.223808595 | 3.32E-10 |

| | | | |
|---|---|---|---|
| Gmfb | 3530.728036 | -1.224346456 | 6.44E-08 |
| Socs4 | 969.7173352 | -1.22442649 | 7.33E-12 |
| Efna4 | 476.6852267 | -1.224938841 | 2.82E-13 |
| Samd15 | 15.18295231 | -1.225166523 | 3.63E-02 |
| Shprh | 1740.163833 | -1.225730596 | 2.54E-08 |
| Smarcd1 | 1668.953612 | -1.22699608 | 1.49E-21 |
| Lbh | 5291.647432 | -1.227079479 | 1.24E-05 |
| Ccdc157 | 793.9460529 | -1.227794777 | 5.45E-22 |
| Dnajc18 | 1352.375434 | -1.22785088 | 1.17E-18 |
| Naalad2 | 1266.955718 | -1.229590795 | 3.23E-22 |
| Natd1 | 2182.198035 | -1.229985913 | 5.89E-13 |
| Cdk19 | 2233.948062 | -1.230462428 | 1.58E-39 |
| Cspp1 | 957.6473057 | -1.231081429 | 5.17E-07 |
| Smim12 | 756.8155337 | -1.232216506 | 4.44E-12 |
| Gm5641 | 10531.33056 | -1.232501105 | 8.81E-09 |
| Ap1s3 | 90.19535064 | -1.233401931 | 3.92E-05 |
| Gm14379 | 22.4080526 | -1.233560554 | 2.60E-02 |
| Tmtc3 | 2089.237018 | -1.23431104 | 2.12E-08 |
| Morn2 | 184.5289294 | -1.235228205 | 1.64E-07 |
| Pou6f1 | 1919.566169 | -1.235326372 | 2.61E-27 |
| Cc2d2a | 1162.851323 | -1.235544688 | 1.01E-11 |
| Gm11847 | 90.45244817 | -1.235819923 | 4.25E-02 |
| Arl6 | 456.3562911 | -1.237145714 | 9.15E-13 |
| Sav1 | 3229.415502 | -1.237659375 | 4.12E-42 |
| Gm3839 | 40.37264345 | -1.237879743 | 2.63E-02 |
| Epb41l1 | 6358.9137 | -1.238066544 | 2.71E-21 |
| Tdrd3 | 952.5802765 | -1.238282633 | 8.07E-23 |
| Tasp1 | 468.6773577 | -1.238297218 | 1.08E-19 |
| Snhg20 | 229.742394 | -1.239013588 | 2.34E-06 |
| Foxn2 | 1053.302421 | -1.239709785 | 3.32E-16 |
| C1qtnf6 | 2407.485434 | -1.239830686 | 1.53E-16 |
| Rarb | 39.25205665 | -1.240737616 | 3.46E-02 |
| Ccnc | 460.2356239 | -1.241182587 | 2.54E-05 |
| Mxra8 | 6216.491192 | -1.241902304 | 7.44E-17 |
| Gm26547 | 74.60976844 | -1.241972639 | 2.06E-03 |
| Man2c1 | 1776.054593 | -1.242192766 | 2.73E-21 |
| Arhgef1 | 2053.505929 | -1.242465522 | 1.59E-22 |
| Meaf6 | 1077.454916 | -1.243122588 | 1.02E-29 |
| Mlh1 | 506.401336 | -1.243146318 | 1.12E-12 |
| Dubr | 27.90810858 | -1.243574105 | 2.82E-03 |
| Recql | 672.013649 | -1.243906455 | 1.22E-09 |
| Adam23 | 4333.929332 | -1.244361547 | 1.03E-09 |
| Fnip1 | 4227.643221 | -1.24444089 | 2.02E-17 |
| St3gal5 | 2651.830812 | -1.244722676 | 5.13E-21 |
| Ebf3 | 1369.264586 | -1.244727543 | 1.16E-11 |
| Klhl36 | 240.6418848 | -1.245645556 | 1.11E-10 |
| Ppil4 | 1478.851273 | -1.245939698 | 2.39E-11 |

| | | | |
|---|---|---|---|
| Dpep2 | 58.14460163 | -1.245950536 | 5.22E-03 |
| Gpr176 | 2045.393277 | -1.246062738 | 1.07E-16 |
| Fgfr3 | 193.996665 | -1.247613247 | 4.20E-08 |
| Tnfrsf18 | 32.81611636 | -1.24893471 | 1.46E-02 |
| Lrp6 | 8327.93429 | -1.249946503 | 1.69E-19 |
| Leng9 | 144.2703895 | -1.250086148 | 2.49E-08 |
| Nphp3 | 459.6841505 | -1.250124354 | 1.36E-13 |
| Zfp955a | 638.3229474 | -1.251228599 | 4.91E-08 |
| Rhoq | 7241.961221 | -1.251251392 | 9.24E-30 |
| Ppp2r3a | 1273.429635 | -1.251302372 | 3.59E-13 |
| Sf3b3 | 10639.11885 | -1.251717127 | 1.04E-24 |
| Zmiz1 | 4690.270148 | -1.251819915 | 4.41E-17 |
| Zfp983 | 210.3850671 | -1.253602581 | 2.21E-14 |
| Dcun1d2 | 872.1359931 | -1.25372875 | 1.35E-29 |
| Wdr82 | 3294.636451 | -1.253761013 | 5.64E-32 |
| Zfp938 | 398.0537243 | -1.254288093 | 1.29E-06 |
| Cops2 | 5354.090409 | -1.254514007 | 2.25E-05 |
| 5730409E04Rik | 1776.754366 | -1.255159577 | 7.69E-33 |
| Cd248 | 6880.534459 | -1.256373788 | 1.98E-09 |
| Ublcp1 | 1024.594223 | -1.256378672 | 3.11E-17 |
| D3Ertd254e | 1068.137171 | -1.256466918 | 6.71E-15 |
| Rock1 | 1927.129653 | -1.256573899 | 2.80E-04 |
| Zfp708 | 70.2331927 | -1.258536254 | 1.83E-03 |
| Mmd | 4325.604263 | -1.258579051 | 8.24E-04 |
| Mex3c | 2408.706296 | -1.259826479 | 2.21E-30 |
| Zfand4 | 198.1185896 | -1.260395493 | 1.45E-05 |
| B9d1 | 529.0966433 | -1.260459939 | 1.22E-12 |
| Zfp68 | 2035.29604 | -1.260915816 | 5.31E-13 |
| E330009J07Rik | 844.1774462 | -1.262285348 | 9.20E-22 |
| Cenpc1 | 1290.091489 | -1.262322119 | 2.18E-11 |
| Cbx3 | 4932.723007 | -1.262504523 | 4.91E-17 |
| Elf1 | 1872.473563 | -1.262898458 | 1.43E-31 |
| Il34 | 440.8805053 | -1.262949862 | 1.89E-07 |
| B130055M24Rik | 163.0194118 | -1.263607172 | 2.59E-06 |
| Plscr4 | 108.3537055 | -1.263772358 | 5.48E-05 |
| Gm26838 | 16.74123391 | -1.263899961 | 1.73E-02 |
| Tfpi | 73.57390232 | -1.264093483 | 1.03E-03 |
| E430018J23Rik | 232.8995405 | -1.264359832 | 8.41E-12 |
| Hmgb1-ps3 | 192.7784252 | -1.265862961 | 3.11E-09 |
| Bend3 | 818.5543217 | -1.265943037 | 2.19E-16 |
| Mtus2 | 261.3824621 | -1.266376716 | 2.33E-04 |
| Hoxa5 | 641.1710565 | -1.266881407 | 4.47E-13 |
| Il20rb | 23.27753996 | -1.267672422 | 4.17E-02 |
| Cpeb4 | 981.9575186 | -1.268122353 | 4.19E-07 |
| Dab2ip | 2202.223697 | -1.268308119 | 3.63E-13 |
| 4632415L05Rik | 318.8005624 | -1.268696633 | 2.10E-13 |
| Fkbp14 | 1386.081438 | -1.269530636 | 1.39E-10 |

| | | | |
|---|---|---|---|
| Dennd2a | 5263.659133 | -1.269558843 | 3.27E-14 |
| Gm26797 | 27.78923782 | -1.272028215 | 1.29E-02 |
| Flicr | 17.9057832 | -1.272347566 | 4.70E-02 |
| Hdhd5 | 498.4192023 | -1.272773569 | 4.12E-17 |
| Sepsecs | 743.2789513 | -1.273153765 | 3.93E-12 |
| Slc5a4b | 28.62253057 | -1.273249753 | 1.86E-03 |
| 2810032G03Rik | 133.6360171 | -1.273355269 | 1.08E-10 |
| Ptk2b | 79.64446995 | -1.275173678 | 2.75E-07 |
| Hipk3 | 5355.807055 | -1.275630632 | 1.20E-14 |
| Rnaseh2c | 449.8484921 | -1.276139234 | 1.05E-11 |
| NA | 22.99628667 | -1.276340739 | 2.30E-02 |
| Has3 | 89.44932683 | -1.276485675 | 3.31E-06 |
| Adgre5 | 1076.317522 | -1.277057195 | 2.02E-11 |
| Cyb5a | 3603.739228 | -1.277067409 | 3.62E-14 |
| Serpinf1 | 25986.98908 | -1.277523924 | 1.38E-08 |
| Zfp229 | 252.3107468 | -1.277991151 | 1.15E-04 |
| Hmgb1 | 6151.969447 | -1.278274531 | 3.02E-12 |
| Marveld1 | 6344.789281 | -1.278722308 | 8.97E-58 |
| Gm15879 | 29.63890879 | -1.281099207 | 3.41E-03 |
| Zfp551 | 180.988907 | -1.281288828 | 1.05E-09 |
| Morc2a | 4216.368248 | -1.281404903 | 2.84E-18 |
| Kmt2e | 4574.680012 | -1.281419189 | 4.32E-25 |
| Pdf | 290.2676097 | -1.281784204 | 9.95E-09 |
| Fam171b | 768.6035689 | -1.282091462 | 1.05E-10 |
| Gm12743 | 64.09499747 | -1.282439907 | 1.25E-04 |
| Arl15 | 412.0136059 | -1.282555353 | 2.10E-15 |
| Nid2 | 6347.243041 | -1.282744111 | 4.93E-03 |
| Svil | 5457.138753 | -1.282746234 | 1.29E-20 |
| Ap3d1 | 7686.894989 | -1.283382774 | 9.72E-62 |
| Jtb | 1085.524652 | -1.283457042 | 8.87E-16 |
| Vdr | 1256.985056 | -1.28355253 | 7.58E-07 |
| Slc30a7 | 2473.28765 | -1.283626389 | 3.12E-54 |
| Itga2b | 570.2372038 | -1.285062539 | 1.93E-11 |
| Cbx6 | 10839.95353 | -1.285120508 | 2.05E-36 |
| Rab2b | 836.6849826 | -1.287299387 | 1.74E-18 |
| 2210408I21Rik | 146.384477 | -1.287309645 | 7.15E-12 |
| Ankrd16 | 929.5128238 | -1.287523798 | 4.22E-14 |
| E230001N04Rik | 20.98671475 | -1.287961119 | 9.06E-03 |
| Dcaf6 | 5277.412717 | -1.288168877 | 2.68E-14 |
| Pradc1 | 384.2185328 | -1.288638972 | 1.12E-10 |
| Aamdc | 197.088835 | -1.288799013 | 3.58E-11 |
| Gm11033 | 338.8365071 | -1.289877732 | 7.48E-09 |
| Srsf6 | 7575.428591 | -1.289945345 | 1.83E-31 |
| Lipt2 | 116.9283302 | -1.290817761 | 1.83E-06 |
| Ift27 | 800.9781334 | -1.290951765 | 2.60E-20 |
| Amot | 5972.569539 | -1.291056944 | 2.91E-23 |
| Clec2d | 2151.763597 | -1.292168341 | 2.27E-06 |

| | | | |
|---|---|---|---|
| Mrc2 | 8828.930084 | -1.292250765 | 3.76E-13 |
| S1pr2 | 3510.869616 | -1.292736497 | 1.76E-25 |
| Npm2 | 23.26991696 | -1.292801305 | 1.94E-02 |
| Mdfic | 2585.20032 | -1.293247334 | 2.42E-56 |
| Ndufb7 | 1320.50736 | -1.293935064 | 1.39E-13 |
| Washc4 | 3491.836837 | -1.293935256 | 1.97E-06 |
| Rmi2 | 98.74410112 | -1.294782649 | 1.45E-03 |
| E030042O20Rik | 33.24426692 | -1.295700513 | 6.17E-04 |
| Ctbp1 | 5894.416865 | -1.295708502 | 3.76E-23 |
| Galm | 284.6776046 | -1.296530868 | 2.58E-10 |
| Cetn2 | 985.5248549 | -1.296650304 | 1.59E-13 |
| Emc2 | 2388.117355 | -1.296877223 | 1.21E-09 |
| Gdf11 | 1582.164993 | -1.297494652 | 6.64E-15 |
| Gm16754 | 42.88730402 | -1.298132411 | 1.57E-03 |
| Tspan11 | 2916.150625 | -1.298658762 | 7.47E-12 |
| Gm21994 | 134.2549258 | -1.298665271 | 2.53E-09 |
| Rbm5 | 4614.006628 | -1.299983896 | 1.80E-37 |
| Ccdc103 | 53.9477953 | -1.300598974 | 7.52E-05 |
| Il13ra1 | 219.0472217 | -1.300912562 | 7.28E-13 |
| Kctd11 | 386.5987816 | -1.30136589 | 7.28E-10 |
| Gtf3c6 | 754.2313491 | -1.301680908 | 1.25E-14 |
| Gng7 | 249.2332224 | -1.302091434 | 1.20E-12 |
| Stx2 | 1865.461332 | -1.302332936 | 3.11E-16 |
| Crebrf | 2283.692067 | -1.302788543 | 6.97E-10 |
| Cst3 | 29679.40866 | -1.303906697 | 1.57E-10 |
| Lyst | 3273.741853 | -1.303958299 | 2.72E-23 |
| Glt8d1 | 1660.943834 | -1.304740835 | 2.00E-18 |
| Pds5b | 3572.252805 | -1.306234407 | 1.61E-15 |
| Fer | 2906.447521 | -1.306274353 | 1.81E-31 |
| Zfp148 | 2485.900454 | -1.30650036 | 6.91E-12 |
| 2810021J22Rik | 299.5547927 | -1.306902495 | 2.25E-12 |
| Lrp4 | 529.2244838 | -1.306970014 | 3.59E-05 |
| Msmo1 | 8301.39548 | -1.307581517 | 2.07E-05 |
| A430110L20Rik | 37.73548911 | -1.307995165 | 1.93E-03 |
| Rap2c | 1453.772771 | -1.308684442 | 9.79E-13 |
| Vamp7 | 964.5436274 | -1.308802852 | 1.31E-09 |
| Zfp358 | 1303.342284 | -1.309514132 | 1.48E-30 |
| Zfp964 | 78.48029832 | -1.310579871 | 1.69E-06 |
| Gm27209 | 17.6173853 | -1.311409589 | 1.72E-02 |
| Gm17300 | 172.1249343 | -1.312270394 | 3.21E-05 |
| Mtrf1l | 255.3618304 | -1.313309099 | 3.01E-08 |
| Zmpste24 | 3169.95583 | -1.313328198 | 9.97E-21 |
| Trp53rkb | 133.0751093 | -1.313676489 | 3.19E-11 |
| 4930539E08Rik | 68.16425014 | -1.313688399 | 5.15E-06 |
| BC065403 | 92.11563179 | -1.314233825 | 3.52E-07 |
| Chd2 | 3693.709782 | -1.314768651 | 2.38E-10 |
| Sumf1 | 4147.413684 | -1.314813366 | 4.08E-26 |

| | | | |
|---|---|---|---|
| Slit2 | 4587.564039 | -1.315176253 | 8.62E-09 |
| Wdr5b | 137.9910893 | -1.315224982 | 1.17E-09 |
| Pcdha11 | 424.0706161 | -1.315451619 | 1.66E-04 |
| Ttc33 | 677.2382922 | -1.32117029 | 7.84E-19 |
| Slc45a3 | 239.2303382 | -1.321295118 | 2.32E-06 |
| Kat6b | 1896.384666 | -1.321774297 | 2.36E-23 |
| Fgfr1 | 26119.72218 | -1.324153269 | 3.51E-16 |
| Thap12 | 2285.134238 | -1.324398345 | 1.07E-31 |
| Mbnl2 | 5029.844149 | -1.325525416 | 5.87E-13 |
| Rybp | 2361.128019 | -1.326084308 | 9.74E-24 |
| Slc23a2 | 2101.09374 | -1.326745243 | 3.23E-14 |
| Mus81 | 696.6969706 | -1.32678613 | 2.95E-38 |
| Msrb2 | 106.301837 | -1.327184527 | 3.86E-06 |
| Rbl2 | 2066.659329 | -1.328232466 | 1.10E-55 |
| Zdhhc6 | 1741.824061 | -1.328287668 | 1.85E-21 |
| Kdm2b | 1932.525567 | -1.328832506 | 1.34E-19 |
| Col4a2 | 13125.40325 | -1.330822605 | 1.70E-11 |
| 2410131K14Rik | 1774.50699 | -1.331065283 | 1.31E-30 |
| Pcf11 | 1771.039127 | -1.331163861 | 6.37E-08 |
| Gm26704 | 17.69182856 | -1.331165754 | 3.42E-02 |
| Dpp8 | 3923.176834 | -1.331474548 | 1.20E-61 |
| Fignl2 | 141.930956 | -1.331768509 | 6.24E-10 |
| Ust | 2007.144156 | -1.33349357 | 1.09E-14 |
| Gm17611 | 18.62857657 | -1.334101494 | 1.24E-02 |
| Gm22642 | 14.1373254 | -1.334206678 | 2.82E-02 |
| 4930480K23Rik | 94.19300099 | -1.335538288 | 1.06E-07 |
| Pbrm1 | 3746.543043 | -1.336542697 | 3.57E-11 |
| Gm26606 | 60.32037353 | -1.33728804 | 1.54E-04 |
| Arntl2 | 104.313339 | -1.337469814 | 8.50E-06 |
| Gm17491 | 91.17287409 | -1.337578163 | 9.04E-05 |
| Nxt2 | 344.3911455 | -1.33788936 | 2.56E-06 |
| Edem3 | 2887.189314 | -1.338743241 | 6.92E-13 |
| BC028528 | 309.7950399 | -1.338758394 | 5.49E-16 |
| Parvaos | 101.9905319 | -1.338939331 | 1.07E-03 |
| Slirp | 1059.190464 | -1.339037916 | 5.32E-20 |
| Nagk | 2436.902089 | -1.339120854 | 7.70E-32 |
| Olfml3 | 3715.895388 | -1.339854117 | 4.03E-16 |
| Zmym3 | 1752.30601 | -1.339912631 | 5.18E-25 |
| Gxylt1 | 1462.867959 | -1.339978024 | 1.72E-10 |
| Hoxd8 | 1212.483579 | -1.340100121 | 7.07E-20 |
| Adamts12 | 1032.731012 | -1.342116342 | 1.63E-09 |
| Exo1 | 451.9536838 | -1.342652216 | 1.26E-03 |
| Gm10814 | 28.90121507 | -1.342986443 | 8.17E-03 |
| Sall2 | 1033.6407 | -1.343662495 | 1.36E-27 |
| Brwd1 | 3301.856348 | -1.343877674 | 3.10E-07 |
| Fer1l6 | 38.79742372 | -1.344251535 | 1.07E-03 |
| Ift140 | 1141.357097 | -1.344288795 | 2.67E-12 |

| | | | |
|---|---|---|---|
| Card19 | 646.5482948 | -1.345337341 | 1.58E-13 |
| NA | 13.22047084 | -1.345475201 | 3.80E-02 |
| Zbtb38 | 1479.838627 | -1.346101841 | 5.20E-11 |
| 3110001I22Rik | 71.57358443 | -1.347705357 | 2.25E-06 |
| Ddx6 | 7985.738967 | -1.349095252 | 2.74E-60 |
| Dcp2 | 1731.728548 | -1.349448944 | 8.87E-19 |
| Ccdc28b | 161.1195197 | -1.35164564 | 2.32E-10 |
| 2700046G09Rik | 44.87274014 | -1.351977207 | 5.38E-04 |
| Platr25 | 127.2381172 | -1.352763957 | 5.07E-04 |
| Pde3a | 117.8255858 | -1.352944157 | 2.55E-07 |
| Nfat5 | 11458.97903 | -1.353831222 | 4.68E-19 |
| Hsd17b1 | 30.82236224 | -1.35436194 | 6.54E-03 |
| Gm11991 | 26.04455271 | -1.35437168 | 3.07E-03 |
| Ivns1abp | 3699.912295 | -1.354805637 | 3.80E-10 |
| 0610010F05Rik | 801.5454022 | -1.355653664 | 1.28E-04 |
| Ankrd28 | 3180.342278 | -1.356171631 | 8.03E-15 |
| Cask | 3506.668855 | -1.356256058 | 5.67E-24 |
| Celf2 | 6245.986694 | -1.356857112 | 6.27E-14 |
| Atl3 | 9377.879914 | -1.357959831 | 7.16E-21 |
| Lrba | 1694.232808 | -1.358165393 | 2.70E-26 |
| Slbp | 1414.315151 | -1.358960274 | 4.27E-11 |
| Zfp395 | 1588.050862 | -1.359268139 | 7.92E-15 |
| Esf1 | 1086.489617 | -1.359425906 | 7.31E-04 |
| Gm9796 | 10.08008836 | -1.359761453 | 4.54E-02 |
| Slc29a1 | 1079.904629 | -1.359819933 | 5.65E-09 |
| 1110019D14Rik | 97.35392094 | -1.359936059 | 3.16E-06 |
| Ripk1 | 1870.091494 | -1.360101531 | 1.17E-32 |
| Akr1c14 | 1643.65333 | -1.360269672 | 3.87E-04 |
| Cacna1c | 477.6875855 | -1.360443567 | 6.41E-10 |
| Zfp866 | 1101.719992 | -1.360783163 | 8.25E-16 |
| Klhl9 | 5226.60583 | -1.361836116 | 9.03E-29 |
| Gm15774 | 16.91903401 | -1.363174968 | 1.54E-02 |
| Mmp19 | 2149.521448 | -1.363354432 | 1.14E-17 |
| Impad1 | 4133.85745 | -1.363851293 | 2.04E-21 |
| Esco1 | 506.7553112 | -1.363989373 | 8.68E-06 |
| Gm9752 | 52.85376842 | -1.364388046 | 5.20E-03 |
| Gm26896 | 25.24819437 | -1.364865604 | 1.66E-02 |
| Tlnrd1 | 1548.830688 | -1.365057324 | 2.47E-19 |
| Gm9978 | 77.15633169 | -1.365058878 | 1.73E-06 |
| Cfap46 | 33.74864826 | -1.365387485 | 2.91E-03 |
| Asap2 | 581.2720119 | -1.366047122 | 5.66E-13 |
| Atpaf1 | 1066.490644 | -1.366076263 | 2.43E-27 |
| Ehhadh | 38.52226287 | -1.367568609 | 1.43E-04 |
| Ccdc8 | 17.37507298 | -1.367789271 | 2.25E-02 |
| Gng11 | 381.5507162 | -1.368025759 | 8.41E-09 |
| Ccdc173 | 120.6844617 | -1.368902138 | 7.56E-04 |
| Bmpr1a | 3988.482522 | -1.369033211 | 2.12E-25 |

| | | | |
|---|---|---|---|
| Tcta | 597.1160012 | -1.369842721 | 3.29E-18 |
| Zfp963 | 272.2670617 | -1.370141306 | 1.06E-10 |
| Txlna | 4884.051793 | -1.370429048 | 5.01E-30 |
| Slc38a2 | 19026.7174 | -1.371171421 | 9.99E-20 |
| Zfp637 | 1645.301751 | -1.371545781 | 3.62E-18 |
| Ift80 | 1433.054562 | -1.372280495 | 4.64E-15 |
| Deaf1 | 690.6344136 | -1.372429893 | 5.21E-21 |
| Zfp60 | 821.8985304 | -1.372675541 | 4.71E-12 |
| Gm20036 | 44.30314038 | -1.373339184 | 1.73E-04 |
| Icam5 | 90.53152646 | -1.373642226 | 7.72E-05 |
| Sfn | 26.79580954 | -1.374635763 | 2.05E-03 |
| Ttc39aos1 | 48.76699706 | -1.374696187 | 3.43E-03 |
| NA | 208.8689783 | -1.377626376 | 2.25E-10 |
| Ly6h | 28.54279703 | -1.377857582 | 4.62E-04 |
| Mast1 | 37.90958504 | -1.377869649 | 6.96E-03 |
| Slc2a10 | 452.5441176 | -1.378131845 | 3.40E-15 |
| 1700082M22Rik | 108.1681906 | -1.37857285 | 5.29E-07 |
| Zfp418 | 129.6532408 | -1.379400547 | 9.90E-12 |
| Tnfsf12 | 1683.302223 | -1.379699335 | 1.06E-23 |
| Mga | 4009.852013 | -1.379772216 | 7.05E-10 |
| Cerk | 1102.588167 | -1.380185267 | 3.76E-27 |
| Epha7 | 250.107713 | -1.380411064 | 3.57E-12 |
| Mgat4a | 25.58784331 | -1.380940914 | 2.90E-02 |
| Zfp827 | 2374.032775 | -1.381744765 | 4.07E-23 |
| Rps6kb1 | 1533.635537 | -1.38248823 | 2.00E-12 |
| Zfx | 1758.364518 | -1.382941306 | 5.12E-10 |
| Chac2 | 165.5154092 | -1.383580724 | 1.94E-06 |
| B130034C11Rik | 28.09028887 | -1.384790271 | 2.40E-03 |
| Atp2b1 | 7973.205069 | -1.385195301 | 3.06E-18 |
| 1600002K03Rik | 136.2918221 | -1.385692145 | 1.73E-11 |
| Ddi2 | 2416.311868 | -1.387207523 | 8.44E-52 |
| Zfp655 | 3110.698296 | -1.389280387 | 1.72E-46 |
| 5031425E22Rik | 290.5449219 | -1.389325847 | 1.26E-18 |
| Casd1 | 1764.77328 | -1.389646275 | 6.21E-20 |
| Arhgap18 | 896.4577738 | -1.389833665 | 1.27E-04 |
| Gm14444 | 29.84845564 | -1.39231671 | 3.70E-02 |
| Gm17484 | 94.6692011 | -1.392338734 | 4.09E-05 |
| Tmed1 | 963.0524274 | -1.392588966 | 2.34E-20 |
| Nck2 | 490.0943103 | -1.392753244 | 2.01E-17 |
| Kctd1 | 652.3447104 | -1.393412527 | 1.38E-17 |
| Cenpj | 331.1364679 | -1.393587853 | 2.86E-09 |
| Nynrin | 2362.227013 | -1.393864023 | 3.18E-26 |
| Eif5a2 | 835.2089142 | -1.394202094 | 1.06E-10 |
| Ikbke | 873.2403649 | -1.394551558 | 4.33E-18 |
| Dhcr7 | 1440.708452 | -1.394647512 | 1.11E-09 |
| Bcap29 | 901.6323732 | -1.395102196 | 4.18E-09 |
| Zpbp | 33.14557248 | -1.395854591 | 4.81E-05 |

| | | | |
|---|---|---|---|
| Gm10130 | 13.49872942 | -1.396292554 | 3.06E-02 |
| Gm26750 | 86.09606895 | -1.396309133 | 2.96E-07 |
| Shcbp1 | 529.1576106 | -1.396408441 | 1.61E-02 |
| B630019K06Rik | 230.2762103 | -1.397535204 | 7.15E-20 |
| Rbm39 | 8622.554061 | -1.397830876 | 6.29E-07 |
| Faap100 | 615.3568584 | -1.39877477 | 2.52E-17 |
| Fan1 | 251.048372 | -1.399126323 | 1.73E-17 |
| Zeb1 | 3675.931373 | -1.399178062 | 1.07E-16 |
| Rnls | 41.93373739 | -1.399189464 | 4.93E-04 |
| mt-Atp8 | 24945.38778 | -1.399804982 | 3.93E-05 |
| Nek4 | 454.1602901 | -1.400616966 | 7.91E-18 |
| Zgrf1 | 386.5955645 | -1.40090142 | 2.21E-07 |
| Mafa | 18.2822908 | -1.401319437 | 2.14E-02 |
| Gm14391 | 930.6181873 | -1.401355697 | 3.48E-09 |
| Plpp3 | 5481.615621 | -1.401625206 | 1.77E-13 |
| 1700109K24Rik | 65.79683356 | -1.401760311 | 7.93E-04 |
| Stk38 | 4121.885645 | -1.401821443 | 6.00E-39 |
| Naa50 | 3422.302758 | -1.402247036 | 1.49E-13 |
| Dock1 | 3924.573139 | -1.403228306 | 3.17E-36 |
| Stag1 | 2203.340069 | -1.40343728 | 1.56E-21 |
| Usp16 | 1373.588607 | -1.403906223 | 1.86E-10 |
| Gm26691 | 125.394815 | -1.404007826 | 2.44E-07 |
| Mir7-1 | 12.19980414 | -1.404101297 | 4.47E-02 |
| Det1 | 414.467086 | -1.404399144 | 1.42E-14 |
| Mbtps2 | 1552.93338 | -1.404836143 | 7.61E-10 |
| Gm4924 | 170.5538508 | -1.404998888 | 8.35E-04 |
| Inpp4b | 39.07537712 | -1.405069564 | 2.39E-02 |
| Ndufaf3 | 55.58015648 | -1.406218362 | 1.09E-06 |
| St7l | 643.9586616 | -1.406271755 | 2.15E-26 |
| Stmn1 | 1582.862217 | -1.407205362 | 2.68E-07 |
| Ccdc88a | 1689.147043 | -1.408061709 | 2.64E-04 |
| Tlr2 | 500.01692 | -1.408327232 | 7.18E-10 |
| C330018D20Rik | 298.9653095 | -1.408477719 | 3.19E-20 |
| Gm2670 | 10.29472409 | -1.408751397 | 4.30E-02 |
| Slc31a2 | 992.1188663 | -1.408791352 | 6.23E-21 |
| Zfp931 | 208.6056449 | -1.409225693 | 4.00E-06 |
| Rab27b | 133.4508688 | -1.410266508 | 4.16E-04 |
| Man2a1 | 13475.00019 | -1.410297587 | 3.10E-51 |
| Hoxd3 | 456.5880826 | -1.410548632 | 6.03E-07 |
| Efnb2 | 1976.379573 | -1.411193289 | 1.40E-15 |
| C330007P06Rik | 1979.096959 | -1.411681949 | 1.26E-16 |
| Eya1 | 701.8287389 | -1.41285473 | 2.77E-30 |
| Ndufa5 | 1256.806341 | -1.413410817 | 1.56E-21 |
| Vps41 | 3917.059342 | -1.413554986 | 1.80E-34 |
| Sh3rf3 | 2120.548004 | -1.413801384 | 3.08E-32 |
| Vcan | 10736.2105 | -1.414181632 | 4.39E-32 |
| NA | 11.74088147 | -1.41484055 | 2.72E-02 |

| | | | |
|---|---|---|---|
| Zfp59 | 117.0891697 | -1.41557424 | 2.03E-11 |
| Sfpq | 8941.106294 | -1.415992924 | 2.51E-20 |
| Gm22270 | 16.49993171 | -1.4162088 | 1.81E-02 |
| Apbb2 | 3282.670896 | -1.417931461 | 1.54E-19 |
| Gm12018 | 45.77515052 | -1.41801914 | 8.69E-04 |
| 1700037C18Rik | 79.88967796 | -1.418337258 | 1.60E-05 |
| Wrn | 1098.5201 | -1.41883213 | 5.27E-17 |
| Frmd4a | 3299.821544 | -1.419002957 | 2.31E-13 |
| Spon2 | 2156.979328 | -1.419011129 | 3.97E-06 |
| Gm9987 | 116.0074177 | -1.419219759 | 1.00E-11 |
| Gm15777 | 83.7950099 | -1.420166996 | 7.01E-05 |
| Pcdh19 | 2133.007913 | -1.420410689 | 1.34E-13 |
| Zfp960 | 443.9527272 | -1.421352537 | 2.45E-06 |
| Rnpepl1 | 315.374151 | -1.421371381 | 2.41E-16 |
| Gtf2ird2 | 552.0214477 | -1.421618889 | 2.01E-16 |
| Gm14634 | 11.82749569 | -1.421773007 | 1.76E-02 |
| Ier5l | 736.6308499 | -1.422639263 | 1.94E-13 |
| Zfp949 | 680.9002348 | -1.422881535 | 2.89E-07 |
| Stk3 | 2814.161093 | -1.422951718 | 4.22E-11 |
| Top1 | 3900.261935 | -1.423340955 | 1.50E-20 |
| Gm26853 | 56.97836267 | -1.423444869 | 3.26E-05 |
| Ptk7 | 1571.76109 | -1.424390002 | 1.16E-14 |
| Id3 | 1711.956127 | -1.427863519 | 8.04E-03 |
| Lnpk | 631.473907 | -1.429077083 | 1.33E-06 |
| Acat3 | 97.28512709 | -1.429088783 | 7.45E-06 |
| Xkr5 | 41.05272347 | -1.429126863 | 1.05E-02 |
| Gm2115 | 916.8914908 | -1.429195152 | 7.51E-08 |
| Nucks1 | 4521.486297 | -1.430913079 | 2.16E-14 |
| Wtap | 2388.347222 | -1.431242784 | 7.90E-30 |
| 9330111N05Rik | 10.97158586 | -1.432313774 | 3.58E-02 |
| Ppil1 | 752.6917392 | -1.432507411 | 9.68E-09 |
| Nphp1 | 936.7893837 | -1.433027143 | 5.71E-20 |
| Cyp2j13 | 20.6584464 | -1.433416888 | 7.96E-03 |
| Fastkd2 | 1129.972847 | -1.433764209 | 3.75E-22 |
| Abca8b | 235.2305154 | -1.434333109 | 9.81E-08 |
| G730003C15Rik | 29.50403608 | -1.434552373 | 3.01E-03 |
| Pde1b | 115.0782532 | -1.434733742 | 1.10E-04 |
| Slc25a10 | 990.5896264 | -1.434959032 | 1.64E-09 |
| 9030407P20Rik | 53.53998062 | -1.435312107 | 8.75E-04 |
| Mbnl1 | 5934.787029 | -1.4364227 | 6.24E-13 |
| Slc25a53 | 238.7168006 | -1.43699877 | 5.20E-08 |
| 5530601H04Rik | 352.289377 | -1.438185419 | 7.31E-11 |
| Rhob | 2399.85739 | -1.438463537 | 4.48E-09 |
| Gm7964 | 24.99744468 | -1.440374568 | 1.90E-03 |
| Kcnk2 | 223.8511014 | -1.440674815 | 3.57E-09 |
| E230016M11Rik | 61.17761226 | -1.441697918 | 4.61E-04 |
| Runx1 | 1094.061865 | -1.441720117 | 2.12E-36 |

| | | | |
|---|---|---|---|
| Scara3 | 1941.279625 | -1.442324311 | 3.96E-19 |
| Trpm3 | 35.24716459 | -1.442792842 | 5.17E-04 |
| Gm14326 | 251.3502862 | -1.44317809 | 6.64E-06 |
| Myo5c | 18.82323414 | -1.443365999 | 1.41E-02 |
| Ankrd6 | 596.1378015 | -1.443417203 | 6.94E-23 |
| Ncbp3 | 1301.175897 | -1.443713876 | 7.99E-24 |
| Kif15 | 556.6763184 | -1.443854126 | 2.40E-03 |
| Zfp322a | 971.5690419 | -1.443879545 | 3.66E-08 |
| Fmnl2 | 2152.240826 | -1.444173555 | 1.71E-27 |
| Acat2 | 6980.262021 | -1.44499633 | 1.04E-07 |
| Adam5 | 19.51957787 | -1.445145324 | 8.89E-03 |
| Mis12 | 1396.433358 | -1.446260695 | 2.98E-10 |
| Mtx3 | 733.5799151 | -1.446360047 | 2.00E-13 |
| Begain | 238.4066239 | -1.446647068 | 2.45E-07 |
| Phf14 | 2376.386027 | -1.447489676 | 1.00E-15 |
| Gm6645 | 54.99621619 | -1.448197419 | 1.33E-04 |
| Baz2b | 2192.623411 | -1.44879208 | 2.00E-21 |
| 0610031O16Rik | 23.83237418 | -1.449665851 | 1.02E-02 |
| Hsd17b11 | 1801.223666 | -1.449710431 | 5.51E-27 |
| Kif13b | 1354.851538 | -1.449710912 | 3.02E-19 |
| E130311K13Rik | 560.9654514 | -1.449776147 | 1.79E-15 |
| Mtcp1 | 143.2347273 | -1.449898399 | 2.81E-05 |
| Ninj1 | 599.4857441 | -1.451861391 | 1.60E-12 |
| Epm2aip1 | 2273.909362 | -1.452160198 | 2.26E-17 |
| 9330159F19Rik | 260.4380525 | -1.453274027 | 1.55E-07 |
| Gm17690 | 111.0720358 | -1.453508939 | 7.21E-08 |
| Gm3604 | 37.39336299 | -1.453782563 | 1.04E-02 |
| Gen1 | 267.1936314 | -1.454185506 | 9.23E-04 |
| Zfp719 | 839.4258072 | -1.454506437 | 1.19E-17 |
| Gm12925 | 25.35113162 | -1.454734866 | 6.50E-03 |
| Ptger1 | 1375.579965 | -1.454790436 | 8.20E-24 |
| Abi2 | 1463.946605 | -1.454915919 | 1.55E-24 |
| Nanos1 | 79.92065785 | -1.455290505 | 2.44E-03 |
| Taz | 1239.733738 | -1.455527384 | 1.18E-23 |
| BC037034 | 465.9142831 | -1.455809652 | 8.96E-05 |
| Alms1 | 2337.478094 | -1.455838112 | 7.50E-16 |
| NA | 41.98412907 | -1.45623696 | 4.54E-04 |
| C030037D09Rik | 117.7660195 | -1.456248547 | 8.16E-09 |
| Lysmd3 | 498.995161 | -1.456424586 | 1.58E-06 |
| Gm14292 | 357.5749687 | -1.456479739 | 3.08E-21 |
| Pnisr | 2217.702301 | -1.456989039 | 5.44E-08 |
| Agbl2 | 25.22031612 | -1.457023973 | 6.06E-04 |
| Zmat2 | 2291.274399 | -1.457162987 | 1.50E-58 |
| Alg3 | 402.4435632 | -1.457496871 | 2.87E-10 |
| Sptbn2 | 22.14600034 | -1.457796667 | 6.16E-03 |
| Rbm4 | 516.6815499 | -1.459209796 | 2.36E-20 |
| Ppp4r3a | 2146.905736 | -1.459980057 | 6.22E-24 |

| | | | |
|---|---|---|---|
| Tbc1d12 | 677.2638307 | -1.461000605 | 2.00E-11 |
| Gm20878 | 18.0253798 | -1.461747574 | 5.32E-03 |
| Tmem138 | 722.3299743 | -1.461861267 | 1.08E-24 |
| NA | 63.84407624 | -1.462521076 | 2.06E-03 |
| Depp1 | 214.6965571 | -1.462540272 | 4.15E-04 |
| Zfp955b | 368.1558793 | -1.462998153 | 3.54E-07 |
| Dlg5 | 3568.439899 | -1.464990255 | 1.22E-60 |
| Pwwp2a | 695.825925 | -1.465138315 | 8.67E-12 |
| Rbm12b2 | 327.5625957 | -1.465596724 | 5.71E-06 |
| Myo9a | 2346.723944 | -1.466039576 | 3.47E-09 |
| Tekt2 | 30.20879387 | -1.467154433 | 1.59E-03 |
| Gm26982 | 51.70163914 | -1.467337196 | 1.08E-05 |
| Rap2a | 3349.270391 | -1.467761857 | 9.34E-17 |
| Gm17057 | 15.87823374 | -1.467845296 | 3.69E-03 |
| Cdrt4os1 | 11.58248016 | -1.468543684 | 3.20E-02 |
| Ptgir | 52.09243532 | -1.469278491 | 6.85E-05 |
| Slx4ip | 322.4792239 | -1.470449033 | 4.60E-16 |
| Rhbdd2 | 1090.819054 | -1.470520174 | 8.54E-24 |
| Smc3 | 4460.02087 | -1.470598902 | 8.36E-09 |
| Ggh | 449.0071855 | -1.472257637 | 2.79E-17 |
| NA | 146.9254783 | -1.473888221 | 1.82E-12 |
| Rnf168 | 1483.705527 | -1.475652273 | 9.64E-14 |
| Gm26858 | 34.62157132 | -1.476722861 | 2.32E-02 |
| Gm20403 | 14.36654804 | -1.477763459 | 1.99E-02 |
| Far1 | 2576.261243 | -1.477892455 | 1.66E-07 |
| Zfp971 | 202.4386453 | -1.478267878 | 1.11E-07 |
| B230369F24Rik | 12.92141657 | -1.478812137 | 3.71E-02 |
| Grk1 | 121.1867528 | -1.479561273 | 4.47E-06 |
| Tmem8 | 134.9929926 | -1.48095075 | 4.99E-14 |
| Gnptab | 4858.629519 | -1.480993611 | 8.05E-33 |
| Mum1l1 | 1107.86875 | -1.480999526 | 1.21E-06 |
| Zfp976 | 153.6269941 | -1.481154183 | 2.60E-03 |
| Cdc14a | 289.2179467 | -1.481210435 | 2.22E-15 |
| Hnrnph2 | 3745.444933 | -1.481827751 | 4.33E-29 |
| Mkx | 371.1273353 | -1.481975708 | 1.20E-08 |
| Bbs7 | 975.2240167 | -1.482353198 | 4.04E-09 |
| Gm12359 | 30.84415271 | -1.483318128 | 2.02E-03 |
| Zfp933 | 326.5717679 | -1.48338585 | 2.55E-08 |
| Kalrn | 1301.719211 | -1.483436342 | 4.40E-22 |
| Nyx | 17.73463253 | -1.483869147 | 2.37E-02 |
| Topbp1 | 2858.394934 | -1.484414183 | 1.89E-07 |
| Gm12648 | 261.1845011 | -1.484833154 | 7.87E-16 |
| Nsun7 | 86.05227179 | -1.485289699 | 2.79E-05 |
| Smarca2 | 2276.346727 | -1.485556778 | 7.38E-18 |
| Pikfyve | 2011.636706 | -1.489818375 | 1.26E-25 |
| Irf5 | 142.3257791 | -1.490012416 | 8.19E-05 |
| Serpina3g | 65.50415713 | -1.49026531 | 6.25E-03 |

| | | | |
|---|---|---|---|
| Paqr7 | 35.8722446 | -1.490309381 | 1.38E-02 |
| Uba7 | 268.7536281 | -1.490704113 | 4.99E-10 |
| Pspc1 | 505.5553659 | -1.492778926 | 1.67E-09 |
| Hip1 | 3266.38303 | -1.492798062 | 1.09E-37 |
| 1700048M11Rik | 10.49803794 | -1.493191585 | 4.94E-02 |
| Shc4 | 137.7357455 | -1.493636984 | 6.01E-08 |
| Haglr | 19.13102496 | -1.494344336 | 1.16E-02 |
| Pdpr | 1055.824546 | -1.494365 | 1.84E-38 |
| Slc10a6 | 188.9386759 | -1.494481508 | 3.19E-03 |
| Mblac2 | 232.0593899 | -1.49456494 | 2.16E-09 |
| Smad6 | 222.526087 | -1.494677084 | 4.57E-06 |
| Cep164 | 1466.016671 | -1.495586304 | 4.68E-28 |
| Lgr4 | 1019.428449 | -1.496677847 | 3.29E-16 |
| Zfp422 | 441.3517438 | -1.496861836 | 7.97E-09 |
| Abcd4 | 634.1555892 | -1.497331183 | 1.96E-20 |
| 2500004C02Rik | 115.4313729 | -1.498881385 | 2.34E-04 |
| Bspry | 18.37848977 | -1.499126096 | 7.06E-03 |
| Lrriq3 | 33.70355744 | -1.499284918 | 2.28E-04 |
| Clcc1 | 1204.501927 | -1.500002892 | 1.22E-18 |
| 4930516B21Rik | 34.01233477 | -1.500420876 | 2.34E-04 |
| Gm10222 | 23166.10098 | -1.500833985 | 1.19E-16 |
| Zfp945 | 839.8369458 | -1.500942873 | 1.47E-10 |
| Pomt2 | 1195.558329 | -1.501100229 | 3.32E-34 |
| Psd3 | 883.8155306 | -1.501302751 | 8.54E-17 |
| Tmem86a | 734.3320314 | -1.501303257 | 5.86E-13 |
| Gm17586 | 152.4386316 | -1.501379795 | 5.79E-14 |
| Gabpa | 2042.440526 | -1.50154431 | 1.17E-15 |
| Gm14027 | 23.21964448 | -1.501741582 | 2.33E-03 |
| Ddx3x | 12841.37381 | -1.501902858 | 7.85E-21 |
| Rgcc | 230.2419975 | -1.503285651 | 3.73E-07 |
| Glis1 | 258.3571055 | -1.50365283 | 4.33E-05 |
| H2afy2 | 1900.864898 | -1.504155809 | 1.34E-19 |
| Rnf125 | 21.54989324 | -1.504222287 | 1.16E-02 |
| mt-Cytb | 212693.2349 | -1.504289794 | 1.27E-19 |
| Ift74 | 2106.799547 | -1.505152376 | 2.96E-08 |
| Zfp11 | 209.7158581 | -1.506081596 | 1.79E-18 |
| Gm20707 | 119.736565 | -1.506104868 | 5.25E-07 |
| Eml4 | 2243.380341 | -1.506326282 | 9.03E-29 |
| Ccdc126 | 260.6403871 | -1.507085449 | 1.40E-16 |
| Mrps25 | 1661.276357 | -1.507119713 | 7.74E-19 |
| Mrgpra4 | 199.1215619 | -1.507198806 | 1.05E-04 |
| Creb1 | 1592.2065 | -1.507366378 | 7.11E-09 |
| Ptchd1 | 55.72263066 | -1.507835227 | 2.28E-02 |
| Entpd5 | 1361.739639 | -1.508448522 | 3.38E-22 |
| Stab2 | 34.5652292 | -1.509815551 | 4.75E-03 |
| Cdc42ep1 | 2564.879225 | -1.510311871 | 2.73E-13 |
| Atxn7 | 639.3804785 | -1.510794336 | 1.54E-13 |

| | | | |
|---|---|---|---|
| Traf5 | 535.9410057 | -1.511179118 | 5.48E-20 |
| Tusc1 | 20.89561595 | -1.511388632 | 1.36E-02 |
| Usp47 | 10646.51981 | -1.511830279 | 8.27E-47 |
| Rtl9 | 47.36193539 | -1.512576366 | 3.02E-05 |
| Pdp1 | 4819.970157 | -1.513054788 | 1.10E-11 |
| Tepp | 46.4196485 | -1.513104957 | 8.15E-06 |
| Ifi27l2a | 63.9925599 | -1.514191888 | 2.16E-05 |
| Gm20482 | 178.0800543 | -1.514413155 | 4.76E-15 |
| Tpcn1 | 2505.938347 | -1.514431557 | 1.59E-31 |
| Mdh1 | 5766.576341 | -1.51508732 | 1.09E-25 |
| Ctdspl | 1253.236411 | -1.51533239 | 6.08E-14 |
| Pcdhb11 | 29.99806765 | -1.515899321 | 1.93E-03 |
| AC132444.6 | 61.30137669 | -1.516134833 | 4.77E-03 |
| Lpin2 | 2438.108658 | -1.516332625 | 1.90E-45 |
| Hus1 | 719.1719753 | -1.516394821 | 1.85E-10 |
| Wdr83os | 1481.78488 | -1.516415454 | 6.13E-22 |
| Gm12276 | 19.03335426 | -1.516797803 | 1.17E-02 |
| Satb2 | 298.8094884 | -1.516890026 | 1.02E-13 |
| Zbtb6 | 597.3816576 | -1.519524617 | 3.77E-15 |
| Hnrnpa2b1 | 19860.98294 | -1.519803659 | 1.41E-12 |
| Adamts2 | 9146.817329 | -1.521290705 | 5.43E-11 |
| Trib2 | 1232.911453 | -1.522088487 | 9.15E-15 |
| Ccdc71l | 320.4996366 | -1.523003193 | 3.14E-18 |
| Nqo2 | 413.4205542 | -1.523434872 | 2.61E-13 |
| Adamts9 | 285.3258477 | -1.523492234 | 6.37E-05 |
| Eldr | 67.99065804 | -1.523703799 | 4.82E-04 |
| Fnbp1l | 2320.759181 | -1.524050217 | 2.54E-08 |
| Adgrb2 | 56.35235799 | -1.525125887 | 6.94E-03 |
| Nbl1 | 1782.408442 | -1.525580304 | 2.34E-08 |
| Pfdn5 | 3955.074207 | -1.525728549 | 3.01E-15 |
| Rasa2 | 836.285905 | -1.527966582 | 1.30E-19 |
| NA | 106.2130085 | -1.528463907 | 6.89E-10 |
| Ttyh3 | 8901.200216 | -1.528670165 | 2.38E-13 |
| Adam10 | 4023.172379 | -1.52879921 | 1.83E-24 |
| Gm14443 | 88.67641707 | -1.529755213 | 1.53E-06 |
| Slc18b1 | 718.7074161 | -1.530071323 | 8.07E-16 |
| Sohlh2 | 9.321089239 | -1.530078438 | 2.96E-02 |
| Ddc | 24.03719735 | -1.530559346 | 1.54E-03 |
| Socs1 | 288.2224024 | -1.534203548 | 2.03E-11 |
| Zfp948 | 823.2408788 | -1.534513768 | 9.68E-06 |
| Rrbp1 | 16274.11852 | -1.535178287 | 1.18E-63 |
| Mgst2 | 162.1947575 | -1.535672563 | 4.01E-09 |
| Gpr137b | 6771.907523 | -1.536176008 | 5.92E-10 |
| Fam173b | 346.8684431 | -1.538576458 | 2.73E-20 |
| Armcx1 | 1688.155142 | -1.539719793 | 1.11E-22 |
| Rbm12b1 | 398.9236447 | -1.54064041 | 9.12E-11 |
| Thoc2 | 2635.484445 | -1.541224687 | 2.42E-05 |

| | | | |
|---|---|---|---|
| Wdr13 | 1853.351441 | -1.541621172 | 1.14E-27 |
| Pcbd2 | 191.883457 | -1.542203995 | 1.25E-10 |
| Kif7 | 398.6317459 | -1.543599356 | 5.24E-22 |
| Nol3 | 15.7892644 | -1.544543229 | 4.72E-02 |
| NA | 17.57245839 | -1.544622012 | 2.01E-03 |
| Dpyd | 147.048122 | -1.545615459 | 8.39E-05 |
| Gm13698 | 41.2442785 | -1.545709648 | 1.22E-04 |
| 4732419C18Rik | 17.65707049 | -1.545892447 | 5.13E-03 |
| Tmed8 | 1367.151775 | -1.546258803 | 3.23E-44 |
| Ptp4a3 | 157.9606979 | -1.546353577 | 4.52E-09 |
| Rslcan18 | 70.301619 | -1.54647796 | 4.98E-05 |
| Cox20 | 380.4983109 | -1.546843662 | 4.20E-19 |
| Zfp472 | 207.0491988 | -1.546928858 | 3.82E-12 |
| Bbs12 | 294.4829504 | -1.546933668 | 1.15E-07 |
| Fgd4 | 592.2398547 | -1.547281192 | 1.03E-13 |
| Slc44a1 | 6489.8044 | -1.547626719 | 1.80E-19 |
| NA | 153.1519126 | -1.548261829 | 6.17E-10 |
| Ctdsp1 | 4964.789485 | -1.549098047 | 1.71E-31 |
| Ccdc62 | 84.71422288 | -1.549505089 | 9.13E-06 |
| Mirg | 1133.798695 | -1.549576004 | 1.72E-11 |
| Cacna2d1 | 1239.235042 | -1.549591417 | 1.46E-19 |
| C920006O11Rik | 12.30282308 | -1.549740929 | 2.28E-02 |
| Srsf10 | 2729.370307 | -1.549749216 | 8.28E-10 |
| Gm10433 | 28.16759338 | -1.550013979 | 4.28E-04 |
| Chst11 | 5163.815561 | -1.551856494 | 7.27E-21 |
| Slc7a3 | 69.61508554 | -1.553502804 | 3.00E-04 |
| 1300002E11Rik | 233.5490192 | -1.554071108 | 6.47E-13 |
| Fancm | 513.5863206 | -1.554150792 | 1.22E-07 |
| A730020E08Rik | 31.93034117 | -1.554187379 | 6.77E-03 |
| Meis2 | 2452.885529 | -1.554307014 | 8.72E-11 |
| Togaram1 | 2240.824097 | -1.554488555 | 2.04E-15 |
| Gm4117 | 584.0060264 | -1.554522997 | 6.13E-07 |
| Gm15834 | 35.5211194 | -1.554580091 | 8.78E-04 |
| Chm | 1244.672927 | -1.554878887 | 1.71E-08 |
| Arhgap5 | 3044.182628 | -1.557452163 | 4.29E-05 |
| Zfp950 | 460.7675227 | -1.557601207 | 5.41E-04 |
| Zfp773 | 15.52759869 | -1.557863222 | 5.63E-03 |
| Amigo3 | 59.57360313 | -1.558848827 | 1.59E-06 |
| Fsip1 | 20.73777009 | -1.560762346 | 5.53E-03 |
| Pgm5 | 58.86253734 | -1.560897774 | 1.74E-02 |
| Tbc1d8b | 1093.135708 | -1.560977045 | 1.75E-08 |
| Dcst1 | 38.85705804 | -1.56102773 | 9.21E-06 |
| 5730596B20Rik | 137.2936872 | -1.561222868 | 5.09E-06 |
| Arid2 | 2802.678595 | -1.56145078 | 1.08E-18 |
| Zkscan3 | 2374.665102 | -1.562350935 | 3.47E-37 |
| Dbi | 3225.641979 | -1.563003614 | 1.08E-41 |
| Gm6712 | 31.73004018 | -1.563423563 | 1.40E-02 |

| | | | |
|---|---|---|---|
| A930002H24Rik | 37.79997615 | -1.564193992 | 7.39E-04 |
| Pdk1 | 1503.806453 | -1.564807657 | 1.45E-10 |
| Ubxn11 | 137.916065 | -1.565329903 | 2.78E-08 |
| Far1os | 17.20304771 | -1.565342983 | 3.46E-02 |
| BC002059 | 127.0815277 | -1.565854189 | 2.59E-04 |
| Bace1 | 2787.184137 | -1.566427033 | 2.59E-33 |
| Ung | 266.879367 | -1.566702373 | 8.18E-11 |
| Zfp658 | 480.0067555 | -1.567962775 | 1.12E-10 |
| Mfsd4b4 | 543.5534433 | -1.568009013 | 1.08E-20 |
| Srd5a1 | 148.0881979 | -1.568537792 | 1.09E-08 |
| Rbbp9 | 1104.425234 | -1.569245553 | 1.38E-26 |
| Mettl26 | 461.2555934 | -1.569631225 | 1.02E-13 |
| Chadl | 33.16524172 | -1.569888354 | 2.42E-04 |
| Gm17354 | 67.70481599 | -1.570280179 | 1.31E-08 |
| C2cd5 | 1214.242853 | -1.570401022 | 5.58E-26 |
| Dzip3 | 1177.649643 | -1.572550342 | 8.70E-05 |
| Aldh3b1 | 101.3746066 | -1.572729586 | 3.67E-09 |
| 2610008E11Rik | 1808.69585 | -1.574332132 | 2.09E-05 |
| Jrkl | 316.2799906 | -1.574374435 | 6.54E-07 |
| Dlx1 | 451.0195392 | -1.574743342 | 1.66E-10 |
| Igf1 | 89.98250005 | -1.575938667 | 3.35E-03 |
| Zfp984 | 306.0112459 | -1.57667809 | 1.43E-06 |
| Cdk14 | 5133.388251 | -1.577572463 | 9.07E-08 |
| Ggact | 310.702016 | -1.578241955 | 1.52E-13 |
| Taf9b | 211.1311733 | -1.578846176 | 9.57E-13 |
| Med16 | 1503.89229 | -1.578858798 | 2.80E-30 |
| B3galnt1 | 1459.923705 | -1.580448033 | 5.05E-34 |
| Megf9 | 1364.252732 | -1.580649505 | 1.40E-14 |
| Armcx3 | 3815.423845 | -1.580752823 | 4.72E-30 |
| Nab1 | 1203.85222 | -1.580845694 | 2.88E-10 |
| Trim65 | 262.1264386 | -1.580883416 | 4.06E-14 |
| Wdpcp | 403.2797746 | -1.581386151 | 4.01E-16 |
| Gm9958 | 29.2944882 | -1.581640992 | 5.17E-03 |
| Bclaf3 | 423.8993827 | -1.582024758 | 4.71E-08 |
| Mpzl2 | 41.79260049 | -1.58272203 | 6.63E-04 |
| Rsad1 | 456.0324608 | -1.582960657 | 1.26E-34 |
| Gm16185 | 101.2788479 | -1.584511646 | 1.39E-03 |
| Tbc1d2b | 5994.726219 | -1.584612157 | 5.94E-08 |
| 5730455P16Rik | 1509.639387 | -1.584923438 | 7.37E-23 |
| Nudt12 | 265.2947616 | -1.587552802 | 1.42E-14 |
| Izumo4 | 98.40570222 | -1.587580277 | 1.35E-11 |
| Spred1 | 1968.043063 | -1.58858243 | 7.79E-39 |
| Csad | 1257.892946 | -1.588740473 | 4.59E-25 |
| St5 | 1526.821015 | -1.590162644 | 1.31E-19 |
| Zfp991 | 54.63737771 | -1.590313333 | 4.72E-03 |
| Gm21989 | 45.534082 | -1.590829689 | 2.24E-04 |
| NA | 21.64261849 | -1.591300089 | 5.10E-03 |

| | | | |
|---|---|---|---|
| Rbm25 | 4138.697711 | -1.59178649 | 1.66E-10 |
| Mfsd7a | 51.5810738 | -1.592040022 | 1.52E-06 |
| Ttbk2 | 1297.767438 | -1.592958511 | 3.02E-16 |
| Zfp715 | 596.4060816 | -1.593259234 | 3.73E-15 |
| Bhlhe40 | 3834.220772 | -1.594235372 | 4.14E-12 |
| Tctn2 | 715.0863891 | -1.595202554 | 1.77E-09 |
| NA | 574.234844 | -1.596082328 | 2.73E-21 |
| Dpagt1 | 948.9650305 | -1.596143774 | 1.65E-31 |
| Kantr | 75.93088182 | -1.596392075 | 1.87E-06 |
| Gm14139 | 207.1634126 | -1.597252279 | 1.31E-14 |
| Uty | 529.0603127 | -1.59830532 | 1.20E-07 |
| Naprt | 66.15425766 | -1.598857062 | 7.39E-09 |
| Arhgef40 | 3841.130444 | -1.600300564 | 3.43E-28 |
| Lvrn | 146.2069254 | -1.600912996 | 1.05E-09 |
| Alkbh8 | 540.3213084 | -1.60161935 | 1.21E-10 |
| Itgb2 | 505.7153736 | -1.602746193 | 2.25E-09 |
| Tnrc6a | 3803.20519 | -1.603098665 | 5.10E-23 |
| Zfp810 | 353.3936938 | -1.603248095 | 2.85E-13 |
| Phospho1 | 692.1845389 | -1.603688668 | 1.08E-07 |
| Gm16249 | 81.29125866 | -1.605460429 | 6.77E-08 |
| Cyp26c1 | 9.040582588 | -1.607181326 | 1.58E-02 |
| Gm15726 | 15.84810327 | -1.607851284 | 3.28E-02 |
| Fam43a | 1435.289342 | -1.608223555 | 2.10E-21 |
| Cep41 | 537.163494 | -1.608259833 | 2.12E-18 |
| Pcx | 1264.131113 | -1.608535689 | 1.65E-07 |
| A930012O16Rik | 25.52054419 | -1.60857753 | 4.88E-04 |
| Zscan22 | 382.3077886 | -1.609818675 | 3.62E-24 |
| Synm | 371.6020985 | -1.610871248 | 8.12E-07 |
| Slc1a3 | 1356.973152 | -1.611135864 | 2.93E-17 |
| Plscr1 | 527.9546792 | -1.611748308 | 4.08E-18 |
| Ifi208 | 45.86600101 | -1.612529477 | 1.12E-04 |
| BC037032 | 28.00622575 | -1.612584295 | 2.54E-03 |
| Purb | 3528.782382 | -1.613266851 | 1.79E-23 |
| Epop | 29.97060165 | -1.613340707 | 1.14E-02 |
| Itih5 | 10.38686915 | -1.614482132 | 4.24E-02 |
| Rbak | 676.8083874 | -1.61453286 | 3.26E-29 |
| Cdyl2 | 218.9938247 | -1.61534457 | 2.28E-19 |
| 4930526L06Rik | 20.81954448 | -1.615586922 | 2.32E-03 |
| A930038B10Rik | 19.0527149 | -1.615594406 | 4.26E-03 |
| Has2os | 14.78403193 | -1.61561065 | 4.78E-03 |
| Hexim1 | 1678.805287 | -1.61586198 | 2.25E-48 |
| Zfp518a | 234.9199991 | -1.616040577 | 1.36E-04 |
| Gm16907 | 51.26309275 | -1.616163775 | 4.33E-04 |
| Myl4 | 52.50706572 | -1.616388602 | 7.67E-07 |
| 4930506C21Rik | 19.3739159 | -1.617553624 | 5.10E-03 |
| Gm6793 | 34.41213098 | -1.61919186 | 2.25E-03 |
| Kansl1l | 717.0289781 | -1.619203818 | 2.04E-19 |

| | | | |
|---|---|---|---|
| Fadd | 424.1906884 | -1.620178152 | 3.28E-25 |
| Rapgefl1 | 180.997305 | -1.620807662 | 2.31E-11 |
| Dop1a | 969.9602123 | -1.620842806 | 6.39E-23 |
| Aqp11 | 57.42947331 | -1.621544319 | 1.17E-04 |
| Fmod | 352.8327308 | -1.621678188 | 3.47E-02 |
| Tnfrsf1b | 2946.17122 | -1.62167844 | 1.02E-34 |
| Gm13230 | 51.25331689 | -1.622379532 | 1.92E-02 |
| Pnn | 2572.950217 | -1.622568185 | 6.42E-13 |
| Spata48 | 29.05752533 | -1.623180093 | 6.58E-04 |
| Sh3d19 | 6514.789156 | -1.623517555 | 9.37E-23 |
| Hmg20a | 1475.554886 | -1.62393831 | 6.40E-58 |
| Gm13615 | 9.512650198 | -1.624544447 | 3.34E-02 |
| Cachd1 | 1495.499593 | -1.626529043 | 3.77E-31 |
| Ythdf3 | 3180.029988 | -1.627018952 | 1.84E-20 |
| Ksr1 | 974.9766751 | -1.627060714 | 7.29E-20 |
| Rmi1 | 653.8504849 | -1.627484924 | 6.47E-19 |
| Sirpa | 3616.513785 | -1.629441353 | 2.19E-27 |
| 2010001A14Rik | 129.6180682 | -1.629612644 | 1.23E-17 |
| Gm12106 | 18.19262209 | -1.629788602 | 9.85E-03 |
| Cavin3 | 2957.834925 | -1.630176069 | 4.43E-20 |
| E2f8 | 355.3960181 | -1.630315308 | 4.74E-06 |
| Gsap | 597.175052 | -1.630580013 | 4.51E-30 |
| Nceh1 | 3203.829638 | -1.631847914 | 1.40E-20 |
| Ppp4r3b | 2886.064442 | -1.63189463 | 4.75E-14 |
| Foxl1 | 95.73223871 | -1.632384842 | 1.77E-08 |
| Mipep | 470.7267755 | -1.632918167 | 1.08E-12 |
| Ranbp6 | 626.1189375 | -1.635065923 | 6.56E-10 |
| Cbr4 | 464.8334136 | -1.63516756 | 1.03E-16 |
| Eefsec | 760.5553838 | -1.635506314 | 4.48E-22 |
| Slc12a2 | 4935.850474 | -1.63560945 | 2.12E-29 |
| Trim30a | 433.047542 | -1.637284896 | 3.39E-09 |
| Ank3 | 2241.447597 | -1.638201969 | 1.19E-16 |
| Sdc4 | 3683.171926 | -1.639282258 | 2.41E-26 |
| Invs | 851.0288157 | -1.639587172 | 3.29E-32 |
| Erg28 | 1761.041711 | -1.639698777 | 2.96E-17 |
| Polr3k | 906.3976112 | -1.641981831 | 4.40E-14 |
| Iqcd | 20.73632352 | -1.642511099 | 1.55E-03 |
| Wdyhv1 | 343.7885209 | -1.642975135 | 1.00E-25 |
| Gm38394 | 4556.884526 | -1.645037701 | 1.72E-11 |
| Bdh2 | 1089.036205 | -1.645291452 | 4.85E-13 |
| NA | 341.1389176 | -1.645937519 | 1.37E-11 |
| Vav3 | 109.5812845 | -1.646840909 | 2.97E-07 |
| Prl8a1 | 197.3899827 | -1.647621687 | 2.91E-11 |
| Hoxa7 | 1586.992664 | -1.649178733 | 7.38E-47 |
| Prkcz | 18.35611058 | -1.650587325 | 1.22E-02 |
| 1010001B22Rik | 11.61181883 | -1.651409067 | 2.73E-02 |
| Zfp81 | 566.5113587 | -1.6525785 | 2.71E-10 |

| | | | |
|---|---|---|---|
| S100a4 | 42457.13509 | -1.653007126 | 1.55E-10 |
| Ptgfr | 515.8706601 | -1.653658028 | 8.58E-09 |
| Plekhg4 | 303.5491276 | -1.655398167 | 9.50E-05 |
| Tgfbr3 | 8535.55638 | -1.65583275 | 1.20E-30 |
| Arid4a | 961.9188335 | -1.656716777 | 6.76E-13 |
| Bahcc1 | 1260.799357 | -1.656944204 | 2.85E-22 |
| 1700029F12Rik | 12.22197706 | -1.657939973 | 2.98E-02 |
| Ptpn13 | 1286.598036 | -1.658301826 | 1.57E-12 |
| Cpe | 17622.21883 | -1.659199518 | 4.40E-24 |
| Zfp944 | 404.0550411 | -1.659878728 | 1.36E-10 |
| Smc2 | 1507.282032 | -1.660060591 | 7.24E-03 |
| Gm15860 | 15.13392669 | -1.660062827 | 1.16E-02 |
| Cdc7 | 594.7862899 | -1.660485942 | 1.01E-06 |
| Bola3 | 767.7054442 | -1.661011879 | 2.52E-20 |
| Gm10501 | 41.08916846 | -1.663734934 | 2.88E-04 |
| Gm26719 | 16.00054035 | -1.663757986 | 9.46E-03 |
| Epm2a | 133.8997319 | -1.664062356 | 7.18E-08 |
| Pkd2l2 | 165.7486455 | -1.664807311 | 2.56E-08 |
| Adhfe1 | 58.16465703 | -1.665401891 | 4.64E-07 |
| Gm26785 | 24.64401246 | -1.665410511 | 2.54E-03 |
| Ccdc191 | 288.9313192 | -1.665590059 | 9.38E-10 |
| Zfc3h1 | 2508.455457 | -1.665898137 | 2.39E-06 |
| Gm10025 | 36.1476908 | -1.666250774 | 1.99E-05 |
| Casp12 | 2453.996963 | -1.667450657 | 1.33E-17 |
| Snora41 | 14.15229559 | -1.668526043 | 2.00E-02 |
| Ppp1r3e | 56.64665345 | -1.669528229 | 9.57E-07 |
| 4930563E22Rik | 46.10621929 | -1.669681931 | 7.54E-05 |
| Xpnpep2 | 463.2230938 | -1.670713778 | 3.88E-09 |
| Nanos3 | 11.84376575 | -1.670727457 | 6.98E-03 |
| Mxd3 | 180.75113 | -1.671661818 | 7.83E-05 |
| Rac3 | 64.09443732 | -1.672047115 | 4.39E-06 |
| Lrrc74b | 21.95606927 | -1.673081336 | 5.45E-03 |
| Ebf1 | 2433.234753 | -1.673313119 | 1.97E-08 |
| Glrb | 356.4512423 | -1.674306314 | 3.14E-13 |
| Armcx4 | 547.9847829 | -1.674732082 | 5.73E-09 |
| Hoxa3 | 715.999191 | -1.675049318 | 2.22E-19 |
| Gm26869 | 331.3716579 | -1.675159874 | 8.11E-07 |
| Npas1 | 10.86969447 | -1.675466261 | 1.41E-02 |
| 4921504A21Rik | 64.24312908 | -1.676725435 | 5.33E-06 |
| Gm17655 | 8.882211556 | -1.677003405 | 2.93E-02 |
| Rnase10 | 47.92102259 | -1.677275416 | 8.64E-06 |
| Fzd3 | 172.7317828 | -1.677832119 | 7.47E-16 |
| Adal | 268.8178673 | -1.677962401 | 8.47E-16 |
| D930016D06Rik | 227.053603 | -1.678716542 | 1.09E-06 |
| Slc23a3 | 79.80737097 | -1.679170842 | 5.64E-04 |
| Echdc3 | 481.9063913 | -1.68128244 | 3.24E-19 |
| Gm13237 | 39.05376099 | -1.683483296 | 1.09E-02 |

| | | | |
|---|---|---|---|
| Capn7 | 1695.101627 | -1.683549292 | 1.93E-19 |
| Foxc1 | 342.3062006 | -1.68377895 | 9.75E-23 |
| Zfp157 | 1050.188746 | -1.683922684 | 3.42E-14 |
| Zfp131 | 1252.290088 | -1.685019382 | 2.52E-09 |
| Zfp563 | 188.7486021 | -1.687808666 | 8.27E-18 |
| Gm23130 | 69.16721741 | -1.688117928 | 9.07E-03 |
| Extl2 | 1343.270838 | -1.688873199 | 1.51E-27 |
| Scfd1 | 1563.777827 | -1.689170132 | 1.68E-12 |
| Npepl1 | 10.70708853 | -1.694856443 | 3.32E-02 |
| Gm5884 | 1399.330987 | -1.695409271 | 3.79E-03 |
| Enpp1 | 302.9825204 | -1.695637489 | 7.49E-12 |
| Mob3c | 513.3066035 | -1.695774558 | 2.43E-18 |
| Gpr161 | 765.4282915 | -1.696317497 | 6.35E-23 |
| 1810064F22Rik | 29.02977189 | -1.696551289 | 8.65E-04 |
| Hspa1l | 63.89465712 | -1.696840382 | 1.14E-07 |
| Tcf4 | 3628.891416 | -1.69839514 | 5.29E-52 |
| Slc37a3 | 2707.708926 | -1.699581698 | 1.33E-80 |
| Gm16121 | 16.22995493 | -1.699891963 | 2.59E-02 |
| Pdik1l | 420.2720688 | -1.700239952 | 2.95E-11 |
| Pclaf | 978.5195751 | -1.70027426 | 6.97E-04 |
| Zfp503 | 2242.630648 | -1.700274358 | 1.73E-30 |
| Zfp780b | 192.5945161 | -1.701031028 | 2.50E-10 |
| Slc22a23 | 1141.901598 | -1.701430135 | 8.91E-23 |
| Bbx | 2721.317942 | -1.702124366 | 2.44E-22 |
| Trdmt1 | 240.7301506 | -1.702160504 | 5.81E-07 |
| Tspyl4 | 789.2797587 | -1.702399136 | 2.42E-11 |
| Ube2d-ps | 793.7075186 | -1.703414606 | 1.94E-29 |
| Cdadc1 | 577.8821487 | -1.705378867 | 7.32E-29 |
| Tssk4 | 22.16839144 | -1.705829942 | 1.98E-03 |
| Hnrnpr | 3057.474168 | -1.706880114 | 9.18E-26 |
| Gm16295 | 26.92563693 | -1.707709199 | 1.40E-04 |
| Gpd1 | 13.62226953 | -1.708231609 | 1.70E-02 |
| Mfsd3 | 248.4748801 | -1.710838652 | 2.85E-16 |
| Dhrs7 | 751.5447118 | -1.711218262 | 6.04E-19 |
| Cyb561a3 | 744.7278893 | -1.711456183 | 7.23E-29 |
| Htr2a | 66.90647818 | -1.711593319 | 8.40E-06 |
| Prss35 | 25.22353741 | -1.713002976 | 5.00E-03 |
| Hdac9 | 113.3884376 | -1.713039735 | 5.95E-04 |
| Sspn | 1024.357293 | -1.713059539 | 1.50E-24 |
| Trim63 | 604.4480012 | -1.714888243 | 9.15E-05 |
| Pcdhb15 | 375.5700413 | -1.72030884 | 1.30E-19 |
| Oaf | 3976.294308 | -1.720368405 | 1.03E-20 |
| Galnt5 | 11.63583862 | -1.72260947 | 3.64E-02 |
| Fndc3a | 4745.827846 | -1.72274369 | 2.43E-75 |
| Mxra8os | 7.148754445 | -1.723648431 | 2.65E-02 |
| 4930556M19Rik | 82.05104413 | -1.724569818 | 7.41E-07 |
| Gla | 4390.097848 | -1.72548352 | 8.46E-18 |

| | | | |
|---|---|---|---|
| Dgcr8 | 1706.26144 | -1.726203252 | 6.68E-10 |
| Podn | 20.13299284 | -1.726461747 | 3.64E-02 |
| Lhfpl2 | 6465.270021 | -1.726494446 | 6.29E-34 |
| Lst1 | 26.82333931 | -1.726839469 | 2.48E-04 |
| Fam227a | 82.93503029 | -1.727048015 | 6.49E-07 |
| Pfkfb2 | 1070.695959 | -1.729091317 | 6.91E-26 |
| Pcm1 | 5197.668132 | -1.729914738 | 5.82E-11 |
| Styx | 48.45081395 | -1.731446553 | 1.17E-04 |
| Cep152 | 366.564991 | -1.73370704 | 5.01E-12 |
| Tpcn2 | 753.9566622 | -1.734002002 | 7.60E-47 |
| A930007I19Rik | 40.04911523 | -1.73424298 | 5.48E-04 |
| Adam12 | 2470.942642 | -1.734975699 | 5.59E-10 |
| Gm15128 | 352.174942 | -1.735502036 | 8.09E-05 |
| Ap5b1 | 295.1850734 | -1.735970238 | 3.12E-26 |
| Irs1 | 2008.224803 | -1.737131781 | 5.34E-19 |
| Gm16861 | 183.4760558 | -1.737687716 | 1.32E-16 |
| Nasp | 2787.702527 | -1.738754154 | 5.30E-10 |
| Degs2 | 79.4466478 | -1.738968721 | 9.82E-09 |
| Arl10 | 239.535473 | -1.739644742 | 8.35E-18 |
| Gm16365 | 67.34365029 | -1.740569051 | 9.67E-07 |
| Erich6 | 9.01126769 | -1.741506614 | 4.24E-02 |
| Adck2 | 829.9544706 | -1.741881454 | 5.32E-41 |
| Pik3ip1 | 2251.596988 | -1.742129083 | 1.52E-12 |
| Grin1 | 7.823242048 | -1.742620299 | 3.40E-02 |
| Gm10561 | 210.1280875 | -1.742803308 | 3.13E-22 |
| Fam89a | 14.33546896 | -1.746437159 | 8.38E-03 |
| N4bp2 | 751.2548306 | -1.74735557 | 4.29E-10 |
| Rif1 | 2369.223093 | -1.748023444 | 1.18E-03 |
| Gm26601 | 11.23064245 | -1.748937216 | 1.00E-02 |
| Cnpy2 | 2958.714011 | -1.749117811 | 8.26E-48 |
| 6-Sep | 1358.601144 | -1.749739165 | 1.67E-23 |
| Snn | 2448.99706 | -1.750566641 | 8.93E-33 |
| Gm12580 | 9.951218649 | -1.750717566 | 2.47E-02 |
| Gm11851 | 45.809336 | -1.750825689 | 1.70E-05 |
| Xiap | 2319.815008 | -1.750842122 | 3.93E-18 |
| Adamts6 | 404.2434059 | -1.751439924 | 6.00E-08 |
| Stoml1 | 563.1153803 | -1.752120583 | 2.05E-19 |
| 1110002J07Rik | 12.818549 | -1.752829529 | 5.61E-03 |
| Nudt6 | 176.1843673 | -1.755117759 | 6.92E-11 |
| Csrp1 | 2952.741716 | -1.75622501 | 5.08E-51 |
| Kif26b | 704.3016406 | -1.756519886 | 2.34E-09 |
| Lmod1 | 164.8111626 | -1.756550297 | 2.16E-05 |
| Mtmr1 | 788.909476 | -1.756650429 | 6.12E-29 |
| Abcc4 | 345.0528919 | -1.756965714 | 4.26E-10 |
| Gm27003 | 15.40465391 | -1.757217675 | 5.40E-03 |
| Tfcp2l1 | 14.22561057 | -1.757287668 | 6.22E-03 |
| Snord45b | 12.13719533 | -1.757957854 | 4.68E-02 |

| | | | |
|---|---|---|---|
| Socs5 | 5784.952356 | -1.758710704 | 1.99E-49 |
| Mmp14 | 40429.12938 | -1.759011659 | 3.73E-26 |
| 3830406C13Rik | 528.8087456 | -1.759137133 | 1.72E-27 |
| Csrp2 | 1603.401949 | -1.759480038 | 2.35E-09 |
| Bmi1 | 3119.393451 | -1.759504176 | 1.87E-22 |
| Slc35a3 | 927.6897423 | -1.760397255 | 3.30E-19 |
| 1700020I14Rik | 3599.969541 | -1.7609857 | 3.63E-18 |
| Gm11223 | 1494.168631 | -1.763087896 | 7.48E-10 |
| Gm9905 | 11.71472939 | -1.764003409 | 1.93E-02 |
| Rftn2 | 1288.815036 | -1.764476523 | 4.53E-21 |
| Spin2c | 57.0212798 | -1.766837197 | 6.88E-08 |
| Col17a1 | 62.0117311 | -1.76719359 | 3.66E-05 |
| Top2b | 4599.091835 | -1.768842133 | 3.61E-08 |
| mt-Nd3 | 16305.7006 | -1.76905434 | 4.00E-21 |
| Gm2093 | 35.49229119 | -1.769217058 | 2.83E-05 |
| Tacc1 | 884.4454082 | -1.771385994 | 7.88E-21 |
| Prcd | 10.9762808 | -1.772739374 | 7.54E-03 |
| Ghr | 7386.390632 | -1.773736073 | 7.23E-22 |
| Zfp788 | 503.9710848 | -1.774380495 | 3.05E-08 |
| Gm17382 | 35.29974084 | -1.77500343 | 4.49E-04 |
| Dnajc28 | 239.7158735 | -1.775096365 | 1.06E-14 |
| mt-Nd5 | 120857.742 | -1.77529338 | 2.05E-42 |
| Rps19-ps3 | 21.03043894 | -1.775914686 | 3.75E-04 |
| Celsr1 | 29.54169175 | -1.77592556 | 2.38E-03 |
| Car11 | 122.9055571 | -1.776438956 | 2.90E-08 |
| Twsg1 | 5761.860578 | -1.776548993 | 1.63E-54 |
| Pola1 | 1268.841336 | -1.777269339 | 1.46E-05 |
| Wnk3 | 29.28258561 | -1.777655523 | 1.31E-02 |
| Smarcad1 | 2090.096887 | -1.778266338 | 1.22E-15 |
| Trim44 | 4685.908872 | -1.778497633 | 4.73E-31 |
| Zfp397 | 1409.011093 | -1.778773352 | 5.72E-11 |
| Slfn2 | 22.2414266 | -1.779288086 | 8.75E-03 |
| C1qtnf1 | 929.0184909 | -1.779480262 | 1.94E-33 |
| Gbp4 | 14.23422782 | -1.779502711 | 9.73E-04 |
| Mtf2 | 1124.744827 | -1.779657425 | 7.66E-15 |
| Fbxo33 | 557.3720924 | -1.78000279 | 3.18E-19 |
| Gli3 | 1922.023409 | -1.781853872 | 9.00E-32 |
| Hk1os | 14.68810408 | -1.781901554 | 1.06E-02 |
| Dclk1 | 6246.021643 | -1.782346964 | 9.75E-36 |
| Sgsh | 942.5070194 | -1.783166366 | 1.86E-20 |
| Galnt15 | 67.07683312 | -1.784501143 | 4.46E-04 |
| Zbtb26 | 467.1449705 | -1.786145325 | 4.32E-08 |
| Gm15443 | 102.0140851 | -1.786478318 | 2.12E-10 |
| Gm21981 | 86.72813471 | -1.786659367 | 8.38E-09 |
| Zmat1 | 277.1067175 | -1.787213465 | 1.50E-09 |
| Ankrd46 | 2144.475571 | -1.787286182 | 4.41E-31 |
| Slc4a11 | 115.9732355 | -1.787534395 | 1.70E-14 |

| | | | |
|---|---|---|---|
| Fam217b | 836.7603683 | -1.78805397 | 7.26E-20 |
| Gm12248 | 9.109659201 | -1.788836267 | 4.47E-02 |
| Gm18360 | 8.20057758 | -1.78912151 | 4.14E-02 |
| Tyw3 | 278.3643857 | -1.789764493 | 2.67E-09 |
| Clcn4 | 1622.727894 | -1.790030965 | 4.36E-38 |
| Dynll1 | 5076.522784 | -1.79061256 | 2.93E-19 |
| Galnt17 | 1070.268328 | -1.790684633 | 1.54E-24 |
| Lancl2 | 747.3264616 | -1.791656416 | 2.81E-17 |
| C1rb | 431.70044 | -1.791731478 | 6.17E-13 |
| Chn1os3 | 57.36693281 | -1.792714552 | 2.42E-05 |
| Nkap | 795.2055193 | -1.793571987 | 6.79E-15 |
| Wdr78 | 241.0414196 | -1.795407965 | 7.80E-13 |
| 4921531C22Rik | 54.37804619 | -1.796413058 | 5.67E-07 |
| Srsf2 | 5846.326793 | -1.797076346 | 1.51E-17 |
| Zfp26 | 803.5414538 | -1.797116921 | 4.93E-11 |
| Carmn | 43.03451096 | -1.798958702 | 4.00E-03 |
| Gm12424 | 18.16555463 | -1.799099451 | 3.04E-03 |
| Slc22a15 | 345.992738 | -1.799656295 | 1.15E-36 |
| Erbin | 6002.117223 | -1.801132479 | 2.81E-11 |
| Rab7b | 1493.162856 | -1.801148991 | 1.93E-44 |
| Arsb | 3780.501608 | -1.801153429 | 3.03E-41 |
| Mtmr7 | 29.58287104 | -1.802898051 | 6.58E-05 |
| Carmil2 | 23.53441272 | -1.803413815 | 3.00E-03 |
| Gm15408 | 18.81798496 | -1.803636097 | 4.90E-03 |
| Vwa8 | 2171.685012 | -1.803872016 | 3.35E-35 |
| Gchfr | 290.9261129 | -1.804815169 | 2.30E-09 |
| Rsrp1 | 7585.578583 | -1.806267145 | 1.63E-12 |
| mt-Nd6 | 36849.03868 | -1.808199199 | 3.42E-38 |
| Tagap | 14.85566499 | -1.808782834 | 9.04E-03 |
| Otulinl | 214.2498603 | -1.808865734 | 5.27E-24 |
| Rida | 433.8767028 | -1.809726606 | 1.02E-16 |
| Pdcd4 | 2480.242851 | -1.810300384 | 3.60E-23 |
| 4930594M22Rik | 28.25743457 | -1.812791554 | 1.43E-03 |
| Tbc1d5 | 2332.869746 | -1.813336585 | 1.79E-66 |
| Slc3a1 | 16.02250516 | -1.81348797 | 1.22E-03 |
| Fmr1 | 3504.187624 | -1.815805042 | 6.13E-10 |
| Ptpre | 155.6299662 | -1.816335596 | 6.99E-10 |
| Jam2 | 337.7129228 | -1.817130778 | 1.06E-11 |
| Cdh26 | 290.3703007 | -1.818172391 | 9.18E-07 |
| Zfp58 | 250.0516868 | -1.818439134 | 1.24E-19 |
| Gm19569 | 24.3437234 | -1.819049689 | 8.16E-04 |
| Reep3 | 3673.165162 | -1.821039678 | 6.19E-99 |
| Pdk4 | 19724.77029 | -1.821953246 | 6.65E-10 |
| Draxin | 16.69298831 | -1.822342255 | 1.27E-03 |
| Scn2a | 379.9991975 | -1.822836954 | 2.23E-07 |
| Cep162 | 817.932987 | -1.825639231 | 8.66E-06 |
| Lzts3 | 586.2326723 | -1.825856553 | 1.95E-21 |

| | | | |
|---|---|---|---|
| Vps8 | 2880.10801 | -1.825948993 | 5.37E-34 |
| Gm16675 | 32.05963952 | -1.826408728 | 2.84E-05 |
| Mir99ahg | 106.1611718 | -1.826480027 | 4.75E-08 |
| Krtcap3 | 56.95226486 | -1.826854961 | 1.06E-07 |
| Arhgap24 | 1837.038701 | -1.827119614 | 2.70E-16 |
| Gm11769 | 12.57310372 | -1.827485236 | 1.27E-02 |
| Kifc3 | 995.0069766 | -1.827649422 | 3.20E-13 |
| Gm10778 | 129.4366747 | -1.82987739 | 7.77E-08 |
| Gm26786 | 83.47726905 | -1.830788299 | 7.37E-10 |
| Mfsd4b3 | 26.36714971 | -1.831486017 | 1.76E-04 |
| Zfp119b | 118.8405696 | -1.831812612 | 3.90E-09 |
| Art3 | 18.30251111 | -1.831825393 | 2.03E-03 |
| Cd276 | 4044.747532 | -1.833127358 | 4.51E-52 |
| Zc3hav1l | 1399.674989 | -1.834217637 | 6.40E-77 |
| A830005F24Rik | 10.58896151 | -1.834722245 | 1.95E-02 |
| Nap1l3 | 28.84721273 | -1.834722484 | 4.83E-05 |
| Cetn4 | 155.0230006 | -1.834999808 | 2.18E-12 |
| Zfp296 | 38.59202607 | -1.83591008 | 2.67E-05 |
| Zfp763 | 273.8263546 | -1.836624391 | 6.69E-14 |
| Hdac8 | 481.6676714 | -1.837505413 | 2.28E-25 |
| Gm13187 | 17.5193156 | -1.838472694 | 6.84E-03 |
| Gm6345 | 21.15217882 | -1.839631677 | 1.40E-03 |
| Zfp712 | 74.56902935 | -1.840811696 | 2.05E-07 |
| Gjc1 | 1240.108962 | -1.841318322 | 4.22E-27 |
| Map3k1 | 164.9162762 | -1.841756747 | 8.10E-07 |
| Cyp46a1 | 102.4882055 | -1.842284009 | 9.23E-09 |
| Peg12 | 309.1835184 | -1.842318005 | 1.01E-18 |
| Nat8f1 | 120.6153721 | -1.842707182 | 1.18E-07 |
| Epha5 | 18.84096967 | -1.843327451 | 1.74E-03 |
| Wnt10b | 29.53377535 | -1.843678839 | 5.15E-04 |
| Pcdhb4 | 29.69585276 | -1.84449603 | 8.18E-06 |
| Bsn | 48.5312342 | -1.845034458 | 5.26E-09 |
| Fhl4 | 42.64920327 | -1.848600561 | 3.72E-06 |
| Riox1 | 464.5125167 | -1.848630937 | 4.97E-13 |
| Hist2h2be | 103.5079661 | -1.851405624 | 3.00E-11 |
| BC005561 | 334.3062812 | -1.851415674 | 4.86E-06 |
| Gm10509 | 7.378122108 | -1.851454556 | 4.33E-02 |
| Zfp764 | 357.7441711 | -1.85162179 | 9.09E-33 |
| Efcab7 | 494.6609103 | -1.852056064 | 5.46E-09 |
| Stat1 | 1423.127776 | -1.852111488 | 5.23E-34 |
| Gm26779 | 11.84712196 | -1.85248637 | 1.65E-02 |
| Srgap1 | 1826.660683 | -1.852644674 | 9.50E-14 |
| Nsg1 | 1730.070035 | -1.854707854 | 3.69E-40 |
| Ttc28 | 3338.3816 | -1.855808311 | 8.45E-31 |
| Arhgef19 | 326.3317744 | -1.856274541 | 5.95E-12 |
| Gm6977 | 21.7987696 | -1.857087937 | 5.55E-03 |
| Ctnnb1 | 26830.96708 | -1.857955504 | 3.80E-99 |

| | | | |
|---|---|---|---|
| Zfp455 | 70.70734888 | -1.858716779 | 2.54E-11 |
| Ifi207 | 781.2702642 | -1.859265992 | 5.10E-23 |
| H2-DMa | 231.9741204 | -1.859946615 | 5.43E-17 |
| Atr | 1202.631887 | -1.860044173 | 3.72E-11 |
| Gm26108 | 19.72662174 | -1.860741689 | 2.91E-03 |
| Igf2bp3 | 4305.408781 | -1.862336416 | 3.93E-48 |
| Pkig | 750.3973578 | -1.862451768 | 3.55E-26 |
| Mblac1 | 76.13773596 | -1.863845582 | 5.58E-10 |
| Eif1a | 6421.978995 | -1.86497633 | 5.84E-22 |
| NA | 76.02244038 | -1.865619365 | 5.51E-04 |
| Cnih2 | 222.9740135 | -1.866212813 | 3.21E-14 |
| Snord53 | 11.38426083 | -1.866792959 | 3.58E-02 |
| Slc26a1 | 15.21426017 | -1.867639756 | 1.09E-02 |
| Gm5784 | 30.3737508 | -1.869461756 | 8.18E-05 |
| C1ra | 10143.28128 | -1.871669095 | 9.22E-17 |
| 3300005D01Rik | 873.8809125 | -1.872088133 | 2.86E-11 |
| Zcchc24 | 8085.911851 | -1.872230808 | 2.91E-58 |
| Cdh18 | 12.16129654 | -1.873014217 | 2.76E-02 |
| Sdsl | 62.01515941 | -1.874321395 | 6.95E-05 |
| Sclt1 | 532.3854206 | -1.874650487 | 2.56E-11 |
| Il4 | 67.4342044 | -1.876063996 | 9.08E-07 |
| NA | 63.66815679 | -1.876100407 | 3.45E-08 |
| Gas2 | 879.7185941 | -1.876818418 | 8.68E-16 |
| Akap5 | 25.59355521 | -1.877381904 | 4.42E-03 |
| 2610044O15Rik8 | 77.27477444 | -1.882010864 | 2.60E-09 |
| Slco1a5 | 23.68395135 | -1.882249721 | 1.07E-02 |
| Zfp442 | 237.9315236 | -1.883604493 | 3.62E-08 |
| Gm12222 | 7.278448278 | -1.883828984 | 4.03E-02 |
| Ifi204 | 1140.215139 | -1.884396279 | 1.12E-17 |
| Armcx6 | 583.5901757 | -1.88556729 | 1.29E-30 |
| Zfp260 | 2209.690586 | -1.886019387 | 1.01E-20 |
| Uckl1os | 18.48553068 | -1.886540247 | 8.24E-04 |
| Golm1 | 1094.89663 | -1.887461153 | 2.27E-19 |
| 6330403L08Rik | 423.3189736 | -1.887991978 | 6.44E-22 |
| Hoxa6 | 269.2072896 | -1.888011652 | 7.58E-15 |
| Tanc2 | 4417.326718 | -1.888110792 | 6.23E-43 |
| Tmem47 | 608.4736339 | -1.889685516 | 1.53E-08 |
| Gm9776 | 112.7083575 | -1.889782705 | 4.66E-12 |
| Arhgef15 | 50.24344551 | -1.889891386 | 8.91E-05 |
| 4933431E20Rik | 881.878442 | -1.889933696 | 7.09E-22 |
| Plcb4 | 1673.625399 | -1.890899176 | 4.56E-14 |
| Cep135 | 362.1027343 | -1.892271139 | 2.58E-10 |
| Gm8494 | 9.59836089 | -1.895195852 | 1.28E-02 |
| Pmepa1 | 1219.457425 | -1.895381795 | 1.36E-25 |
| Rora | 549.7334082 | -1.896198203 | 2.00E-14 |
| Rbm41 | 364.9897645 | -1.897011367 | 2.59E-08 |
| Prkd3 | 3344.207344 | -1.898623285 | 3.42E-41 |

| | | | |
|---|---|---|---|
| Arhgap20 | 61.50283451 | -1.899198493 | 1.76E-03 |
| Atad5 | 543.9945073 | -1.89959532 | 1.82E-06 |
| Tgfbi | 34.33221477 | -1.901699647 | 8.54E-04 |
| Prelp | 2798.880483 | -1.90175727 | 2.52E-05 |
| Gm26690 | 12.97637611 | -1.902466379 | 1.72E-02 |
| Tnfrsf14 | 35.33998753 | -1.903408968 | 3.18E-06 |
| Prune2 | 851.2860509 | -1.90454649 | 5.88E-15 |
| Ypel4 | 290.1410355 | -1.904811376 | 4.90E-18 |
| Ube2d1 | 1311.582683 | -1.90678458 | 4.45E-40 |
| D030068K23Rik | 56.07754723 | -1.907087485 | 9.73E-09 |
| Zfp937 | 394.8604838 | -1.908178379 | 2.27E-11 |
| Gm17399 | 12.21236887 | -1.908264674 | 7.99E-03 |
| Tmem150a | 1636.619374 | -1.909077165 | 1.42E-34 |
| Gm15446 | 215.8839463 | -1.91010036 | 7.01E-11 |
| Lrrc8d | 953.5605398 | -1.910820929 | 1.91E-24 |
| Bmf | 1679.651699 | -1.91138084 | 4.98E-08 |
| Tnfrsf11a | 123.4947407 | -1.91281207 | 1.10E-20 |
| Lama3 | 7.991540785 | -1.912986704 | 4.71E-02 |
| Gm7598 | 27.13875223 | -1.913322428 | 1.63E-05 |
| Arid4b | 1455.105327 | -1.913583043 | 1.25E-12 |
| Meg3 | 12210.99322 | -1.916792769 | 1.37E-10 |
| Gm10762 | 70.63600912 | -1.916798256 | 5.12E-10 |
| NA | 126.927072 | -1.917294107 | 2.13E-11 |
| 1700088E04Rik | 90.25557833 | -1.918050543 | 4.43E-09 |
| Map9 | 417.747882 | -1.919357096 | 5.37E-12 |
| Tmc3 | 56.59669875 | -1.919492562 | 3.70E-06 |
| Gm15163 | 28.34791949 | -1.921406294 | 1.28E-05 |
| Kiz | 936.1381678 | -1.922015342 | 8.22E-21 |
| Pts | 474.3278678 | -1.923373855 | 2.97E-24 |
| Gm10638 | 16.6363986 | -1.923454174 | 3.85E-03 |
| Hmgcs1 | 10146.3423 | -1.923464033 | 3.81E-08 |
| Zfp677 | 74.51178123 | -1.924023051 | 2.49E-05 |
| Pafah2 | 1268.397562 | -1.924402392 | 1.59E-43 |
| Fam120aos | 58.1147555 | -1.92457824 | 2.01E-06 |
| Phkb | 2588.754316 | -1.924802695 | 2.99E-41 |
| C2cd4c | 10.15695358 | -1.925195254 | 1.77E-02 |
| Gpc6 | 435.6774066 | -1.925773746 | 3.82E-12 |
| Fbxl22 | 36.52026256 | -1.925955601 | 2.49E-05 |
| Elavl2 | 196.7850386 | -1.926686003 | 2.48E-09 |
| Hif1a | 7405.238458 | -1.927070162 | 5.35E-27 |
| Gm42688 | 427.2518859 | -1.927689058 | 6.96E-41 |
| Plag1 | 548.7458397 | -1.928233667 | 3.51E-18 |
| Erp27 | 12.20386998 | -1.92848886 | 3.48E-02 |
| Gm26737 | 245.1761763 | -1.928970894 | 2.75E-13 |
| Rsl1 | 57.97589736 | -1.929245665 | 2.03E-08 |
| Gm16070 | 26.39325451 | -1.930009226 | 8.07E-05 |
| Trim2 | 2299.108785 | -1.934370371 | 2.52E-15 |

| | | | |
|---|---|---|---|
| Zhx1 | 1595.963463 | -1.934605657 | 6.65E-18 |
| Snord123 | 14.8425164 | -1.937965101 | 3.20E-03 |
| Idi1 | 4535.218215 | -1.938169587 | 4.16E-04 |
| Gm26873 | 34.36725225 | -1.939120656 | 9.77E-04 |
| Ctns | 2362.555936 | -1.940012545 | 6.69E-43 |
| Rbmx | 972.7927229 | -1.940215879 | 4.49E-25 |
| Gm10702 | 9.176440757 | -1.941710202 | 2.04E-02 |
| Plpp7 | 57.83169076 | -1.943625623 | 3.65E-09 |
| A430018G15Rik | 38.4698772 | -1.944513605 | 3.34E-05 |
| Tcf7l1 | 1795.450597 | -1.94480706 | 3.65E-21 |
| Tut4 | 2806.378558 | -1.9455718 | 3.29E-18 |
| Gm20775 | 22.95694495 | -1.945694539 | 1.91E-03 |
| Decr2 | 584.6527844 | -1.946422927 | 5.71E-29 |
| Pjvk | 9.273777323 | -1.946992493 | 2.85E-02 |
| Gm12205 | 9.93294853 | -1.947100176 | 1.45E-02 |
| Utp14b | 229.3382319 | -1.947210474 | 6.88E-10 |
| Sesn1 | 1627.682062 | -1.947404888 | 1.22E-42 |
| 4930522L14Rik | 312.3836131 | -1.947502015 | 5.26E-08 |
| Scd4 | 53.94463503 | -1.948722707 | 2.37E-08 |
| Etaa1 | 461.0245111 | -1.950316565 | 4.58E-08 |
| Camk4 | 17.90380593 | -1.951108661 | 2.75E-03 |
| Kmo | 16.90059884 | -1.951566322 | 1.48E-03 |
| Gm25791 | 29.41108585 | -1.951826459 | 1.27E-03 |
| Gm4791 | 10.92142209 | -1.95185096 | 8.55E-03 |
| Atf5 | 11430.63398 | -1.951969454 | 2.60E-44 |
| Gm20541 | 28.26502666 | -1.95271956 | 3.03E-03 |
| Tmem106a | 958.7992917 | -1.953222238 | 2.16E-31 |
| Zfp445 | 1254.964781 | -1.953259976 | 1.75E-13 |
| Tent5a | 42.14233375 | -1.954459811 | 4.12E-06 |
| Strip2 | 14.36443478 | -1.954606772 | 1.62E-02 |
| Bnc2 | 1922.400801 | -1.95479706 | 2.70E-39 |
| Socs2 | 153.4955773 | -1.956466892 | 6.31E-07 |
| Sox4 | 7067.281048 | -1.956651828 | 1.19E-16 |
| Zfp292 | 1751.446612 | -1.957415075 | 3.52E-14 |
| Tmem267 | 228.135723 | -1.958766316 | 8.97E-19 |
| Gm12657 | 10.27908881 | -1.961381747 | 2.21E-02 |
| Gm10399 | 11.62803883 | -1.962472422 | 7.82E-03 |
| A730017L22Rik | 398.5430896 | -1.962605197 | 3.37E-18 |
| Pla2g7 | 967.292464 | -1.962793948 | 2.59E-16 |
| NA | 72.73572633 | -1.963380782 | 5.20E-07 |
| Zfp160 | 566.4007685 | -1.96402589 | 5.04E-11 |
| 9130023H24Rik | 247.178609 | -1.964114234 | 3.04E-28 |
| Mtss1l | 4009.15688 | -1.964240314 | 1.04E-30 |
| Smim10l1 | 2712.536032 | -1.964351709 | 4.16E-18 |
| B230354K17Rik | 1037.407643 | -1.964566545 | 1.08E-24 |
| Etv4 | 307.4908993 | -1.965804218 | 4.59E-07 |
| Gm24539 | 27.90993882 | -1.968441433 | 2.30E-03 |

| | | | |
|---|---|---|---|
| Ptgfrn | 7767.767633 | -1.969561662 | 1.27E-44 |
| Tnr | 97.7195546 | -1.969675284 | 3.06E-04 |
| Pdss1 | 204.7238761 | -1.970046204 | 1.38E-07 |
| 1700034H15Rik | 27.77523794 | -1.970141789 | 1.65E-04 |
| Tcea2 | 213.1302564 | -1.970583822 | 5.51E-18 |
| 9030624G23Rik | 33.45556206 | -1.972323738 | 1.97E-06 |
| Rasl12 | 7.779363372 | -1.972352787 | 1.47E-02 |
| Ace | 11.96119416 | -1.972355356 | 2.95E-02 |
| Bicdl1 | 356.0545461 | -1.973091837 | 1.10E-26 |
| Zfp469 | 681.3675061 | -1.973217703 | 5.56E-17 |
| Emid1 | 31.69594959 | -1.974666497 | 1.30E-04 |
| Atp11c | 750.7779488 | -1.97516427 | 1.73E-09 |
| Rtl1 | 52.28000868 | -1.975189597 | 4.42E-07 |
| Gpr162 | 212.3158726 | -1.975764198 | 1.11E-13 |
| Srrm4os | 14.95289305 | -1.977016409 | 1.52E-02 |
| Id2 | 912.7516454 | -1.979811295 | 7.87E-04 |
| Gm14698 | 642.0530449 | -1.980036764 | 3.15E-09 |
| 4933404O12Rik | 648.6074273 | -1.980643772 | 4.60E-16 |
| Thap2 | 667.6400008 | -1.981090088 | 1.70E-26 |
| Anpep | 300.0514348 | -1.981544187 | 6.34E-06 |
| Elobl | 14.0838148 | -1.981754045 | 1.55E-02 |
| Tppp3 | 30.71678209 | -1.982394634 | 1.92E-05 |
| Gm12586 | 24.70570618 | -1.982590451 | 3.17E-04 |
| Snx32 | 113.7111598 | -1.982811734 | 2.26E-10 |
| Atrx | 5044.519523 | -1.984032417 | 2.62E-05 |
| Esyt3 | 17.29674193 | -1.984226755 | 1.79E-04 |
| Hnrnpd | 4309.091954 | -1.984493304 | 3.96E-19 |
| Nfib | 4927.428113 | -1.984987935 | 1.33E-24 |
| Mrgprf | 1953.146531 | -1.98568407 | 1.09E-14 |
| Gm24357 | 23.09912928 | -1.985987721 | 6.52E-04 |
| Gm14951 | 6.722864299 | -1.986444863 | 4.52E-02 |
| 1810021B22Rik | 22.88639876 | -1.986568427 | 7.93E-04 |
| Dzank1 | 68.2702937 | -1.986754973 | 4.63E-04 |
| Snx24 | 596.8955764 | -1.987625918 | 3.64E-32 |
| 6330403K07Rik | 19.3962192 | -1.98869497 | 2.64E-03 |
| Shtn1 | 23.34544253 | -1.989875761 | 1.13E-04 |
| Il1r1 | 1964.698977 | -1.990081692 | 2.30E-22 |
| Cbfb | 3583.177904 | -1.990230352 | 3.68E-88 |
| Gm16759 | 14.23985702 | -1.991900024 | 3.09E-02 |
| NA | 757.4786446 | -1.99215126 | 6.23E-06 |
| Angptl2 | 12262.23054 | -1.994863482 | 3.11E-18 |
| Il17d | 37.22225942 | -1.995460378 | 2.88E-05 |
| Medag | 7575.216717 | -1.995637977 | 1.71E-20 |
| Magi2 | 526.5829282 | -1.996718615 | 2.03E-48 |
| Gm10143 | 35.22900184 | -1.997038395 | 1.64E-04 |
| Gli2 | 250.1196828 | -1.99754331 | 2.30E-16 |
| Arsg | 389.4454428 | -1.997632817 | 5.94E-33 |

| | | | |
|---|---|---|---|
| C230035I16Rik | 12.02907447 | -1.998560441 | 1.93E-02 |
| Dennd4a | 6083.669471 | -1.99884714 | 3.21E-25 |
| Pcdhb17 | 304.6149099 | -2.001308378 | 1.27E-29 |
| Marcks | 14713.78478 | -2.002131012 | 2.17E-19 |
| Mmp23 | 14.79485096 | -2.003100885 | 3.07E-03 |
| Ifi211 | 544.3470546 | -2.00310317 | 5.71E-12 |
| Lztfl1 | 853.8656872 | -2.003560432 | 2.98E-15 |
| Zfp111 | 586.8419565 | -2.005131194 | 5.67E-16 |
| Slc24a5 | 20.57805544 | -2.006826129 | 2.43E-03 |
| Nod1 | 274.7364886 | -2.007698098 | 9.92E-18 |
| Gm14124 | 65.72396806 | -2.008879977 | 6.13E-08 |
| Gm10478 | 10.73763532 | -2.011951057 | 8.07E-03 |
| Reck | 1994.171189 | -2.01424564 | 1.25E-28 |
| Ror2 | 442.1456838 | -2.015466026 | 2.75E-23 |
| Ftx | 591.5860801 | -2.015796949 | 2.20E-17 |
| Aig1 | 239.7079789 | -2.015850945 | 1.79E-16 |
| 9430091E24Rik | 71.37412386 | -2.017000051 | 1.36E-12 |
| Sytl1 | 58.78219199 | -2.017167051 | 1.63E-05 |
| Plxnd1 | 6200.522837 | -2.017708535 | 4.80E-30 |
| Fut10 | 709.5832085 | -2.017811355 | 1.56E-35 |
| Pappa | 4604.374407 | -2.017996328 | 5.65E-16 |
| Glb1l | 1400.038872 | -2.018574409 | 2.04E-33 |
| NA | 667.970585 | -2.019827111 | 7.83E-51 |
| 1700018A04Rik | 23.62567072 | -2.020010667 | 1.08E-02 |
| Gas7 | 13591.79219 | -2.020294069 | 1.64E-20 |
| Ogt | 5848.32168 | -2.020316321 | 5.46E-15 |
| Ugt3a1 | 8.452433688 | -2.020927115 | 1.29E-02 |
| Ccdc87 | 25.43668448 | -2.021190638 | 1.45E-05 |
| Inpp5e | 1283.030225 | -2.021258732 | 4.19E-52 |
| Zfp942 | 555.0205701 | -2.021365263 | 2.42E-07 |
| Mast4 | 3107.692726 | -2.021669874 | 1.86E-43 |
| Polr3f | 941.8123888 | -2.021724358 | 3.39E-24 |
| A330009N23Rik | 54.46191423 | -2.021964266 | 2.97E-09 |
| Zc3h12d | 7.659289797 | -2.023383821 | 1.19E-02 |
| Ypel1 | 252.6287883 | -2.0236464 | 1.04E-12 |
| Msrb3 | 3410.420807 | -2.024372692 | 4.51E-20 |
| Mamdc2 | 140.4200342 | -2.025195116 | 1.44E-06 |
| Ubxn2b | 458.5861195 | -2.025576581 | 1.41E-18 |
| Zfp935 | 205.3922271 | -2.026006198 | 6.17E-10 |
| Cplx2 | 1224.275436 | -2.027448048 | 1.81E-51 |
| D030028A08Rik | 174.8608595 | -2.027817053 | 2.59E-20 |
| 3830408C21Rik | 13.15067126 | -2.031159854 | 3.26E-02 |
| Trpm7 | 6048.847824 | -2.0311892 | 1.91E-13 |
| Adgrg1 | 72.72172502 | -2.033108857 | 7.41E-07 |
| Alms1-ps2 | 88.59135599 | -2.033660929 | 1.41E-09 |
| NA | 216.623365 | -2.034018426 | 2.03E-23 |
| Agbl3 | 142.3879339 | -2.036098659 | 7.28E-16 |

| | | | |
|---|---|---|---|
| Tsr2 | 1165.014558 | -2.036401252 | 2.82E-46 |
| Mb21d2 | 423.4672271 | -2.039252946 | 7.76E-28 |
| 2310009A05Rik | 19.63698033 | -2.039328801 | 2.27E-03 |
| Jam3 | 380.139953 | -2.042178683 | 2.15E-34 |
| Zfp273 | 74.37311127 | -2.042378837 | 7.06E-08 |
| Pcdhb18 | 74.91592395 | -2.043507025 | 1.45E-10 |
| Aff3 | 1563.295887 | -2.04380625 | 9.49E-14 |
| B3galt1 | 227.8123498 | -2.044146707 | 1.11E-24 |
| Zfp946 | 237.5303763 | -2.044936897 | 3.50E-07 |
| Tshz3 | 29.61253025 | -2.046768141 | 3.36E-03 |
| mt-Ta | 78.99941108 | -2.046887314 | 5.61E-05 |
| Itgb3bp | 300.9143278 | -2.047681451 | 4.76E-12 |
| 5033430I15Rik | 56.08870669 | -2.050370674 | 1.83E-09 |
| Tle6 | 416.6767866 | -2.0505246 | 1.60E-31 |
| Gpx3 | 8.412329513 | -2.05059181 | 1.15E-02 |
| Mid1 | 2636.808961 | -2.05290186 | 3.99E-24 |
| Tenm4 | 841.8870128 | -2.053932581 | 1.71E-03 |
| A230056P14Rik | 61.05474717 | -2.054527514 | 1.01E-09 |
| Gm5977 | 48.68085952 | -2.054692559 | 1.71E-07 |
| Dgki | 31.1515082 | -2.055852379 | 8.01E-04 |
| Evi5 | 2624.44236 | -2.057110675 | 4.64E-16 |
| Pnma1 | 34.96893406 | -2.058249353 | 4.27E-05 |
| Pdgfrb | 13123.50877 | -2.060471538 | 4.32E-60 |
| Hdgfl3 | 1749.187573 | -2.060552215 | 1.15E-18 |
| Zfp456 | 23.67828685 | -2.06075138 | 2.50E-04 |
| 2210408F21Rik | 135.6814517 | -2.060973783 | 1.56E-15 |
| Adamts7 | 1045.159352 | -2.060977354 | 3.21E-33 |
| Chchd10 | 24.56151814 | -2.062242213 | 6.62E-06 |
| NA | 6.652014732 | -2.06388408 | 2.49E-02 |
| Gm15413 | 17.33829686 | -2.064551061 | 3.45E-04 |
| Cbarp | 525.0437716 | -2.066379993 | 3.14E-41 |
| Zfp974 | 285.1415955 | -2.066582081 | 1.90E-15 |
| Atg4c | 643.0173726 | -2.067509423 | 4.14E-26 |
| Slc16a11 | 113.1379731 | -2.06927183 | 6.67E-11 |
| Gm20544 | 23.19780166 | -2.070188823 | 9.73E-04 |
| Pcmtd1 | 2794.947783 | -2.073446714 | 3.71E-14 |
| Eea1 | 3828.799504 | -2.073884796 | 2.11E-14 |
| Hsd17b7 | 1863.090416 | -2.074222479 | 4.46E-12 |
| Phf11b | 9.223932221 | -2.074302518 | 1.79E-02 |
| Xxylt1 | 1691.672168 | -2.075152265 | 3.08E-53 |
| Zfp729b | 395.1662651 | -2.077048683 | 5.99E-12 |
| Phactr1 | 1833.467544 | -2.078163787 | 6.30E-37 |
| Nacad | 47.47538368 | -2.079667473 | 1.30E-05 |
| Lrif1 | 631.3650522 | -2.080405297 | 8.88E-11 |
| Gm10548 | 14.20004954 | -2.080594562 | 8.04E-03 |
| Nrp2 | 11248.52068 | -2.080831388 | 4.50E-54 |
| Gbp11 | 15.68366756 | -2.08090226 | 3.06E-02 |

| | | | |
|---|---|---|---|
| Trim12c | 714.1829737 | -2.081385328 | 1.59E-19 |
| Unc93a | 52.636787 | -2.083190086 | 3.00E-02 |
| Gm15298 | 211.5333334 | -2.083349499 | 4.62E-16 |
| NA | 11.55450061 | -2.083960804 | 4.48E-03 |
| Zfp882 | 177.2345233 | -2.086955509 | 3.20E-10 |
| Gm26548 | 14.86370093 | -2.087595859 | 2.43E-02 |
| Cideb | 9.944776261 | -2.089044253 | 1.22E-02 |
| Ankrd44 | 1362.280154 | -2.090845068 | 3.00E-31 |
| Slc5a3 | 5155.170795 | -2.091335408 | 2.56E-23 |
| Znrd1 | 628.2231869 | -2.092697473 | 3.46E-27 |
| Ppp1r3d | 238.405961 | -2.093196955 | 2.82E-19 |
| Zbtb16 | 8.168993781 | -2.094630061 | 3.77E-02 |
| Zfp109 | 105.0394414 | -2.095100913 | 4.08E-13 |
| NA | 154.0539091 | -2.09537338 | 6.49E-18 |
| Ptprm | 1104.193032 | -2.095921297 | 1.58E-49 |
| Eps8 | 11574.16555 | -2.096163968 | 5.84E-55 |
| Hnrnph1 | 8004.785467 | -2.096438658 | 6.14E-16 |
| Padi3 | 10.25855959 | -2.096889089 | 3.10E-02 |
| Ntn1 | 231.1255559 | -2.097360988 | 4.12E-15 |
| Zfp182 | 311.6166338 | -2.102377172 | 3.47E-09 |
| Nova1 | 277.0734488 | -2.102406497 | 4.25E-10 |
| Mir1949 | 13.61835505 | -2.103374577 | 1.38E-02 |
| Cyp1b1 | 3968.844327 | -2.103816347 | 1.73E-29 |
| H2-T24 | 90.28178638 | -2.10567924 | 1.70E-09 |
| Mir8112 | 30.71183278 | -2.105841216 | 6.21E-04 |
| Sppl2b | 851.0028724 | -2.105878892 | 4.88E-24 |
| Pde4c | 11.66519836 | -2.110872261 | 7.96E-03 |
| Zfp943 | 335.8360668 | -2.111908915 | 5.36E-11 |
| Sort1 | 907.944398 | -2.112892384 | 1.94E-13 |
| Faddos | 6.170598786 | -2.115533481 | 2.26E-02 |
| Zfp457 | 8.610118667 | -2.115579742 | 4.62E-02 |
| Zfp959 | 168.894081 | -2.115614525 | 1.70E-19 |
| Ceacam1 | 5.720254876 | -2.117067133 | 4.97E-02 |
| Lhx6 | 339.1144646 | -2.117245288 | 3.20E-21 |
| Cyp26a1 | 12.85867698 | -2.118451806 | 2.47E-03 |
| Mat2a | 11860.39905 | -2.119165039 | 7.10E-29 |
| Slc25a29 | 113.262095 | -2.121544588 | 1.61E-11 |
| Gm13147 | 17.96081472 | -2.12286091 | 1.61E-03 |
| Ccl28 | 69.41710484 | -2.123534724 | 1.84E-12 |
| AI182371 | 13.38423638 | -2.123729666 | 3.16E-03 |
| BC107364 | 7.454424741 | -2.124490198 | 2.64E-02 |
| Zfp74 | 465.250991 | -2.12451765 | 2.51E-16 |
| Hnrnpm | 7454.04858 | -2.125228512 | 5.03E-21 |
| Plch1 | 34.94020001 | -2.125307898 | 7.02E-04 |
| 6720427I07Rik | 137.3231581 | -2.125609394 | 1.06E-06 |
| Pi15 | 156.8925324 | -2.12703909 | 1.02E-07 |
| Nucb2 | 1252.406383 | -2.127212645 | 3.39E-31 |

| | | | |
|---|---|---|---|
| Vwa5b2 | 28.56756157 | -2.127735575 | 2.29E-02 |
| Cdh11 | 8716.03324 | -2.128084171 | 1.01E-24 |
| Slc39a13 | 2542.695428 | -2.129243113 | 1.64E-34 |
| Acnat1 | 19.47805909 | -2.129414239 | 1.45E-03 |
| Zfp97 | 105.3791855 | -2.131689377 | 5.05E-06 |
| Rian | 11881.87867 | -2.132915782 | 3.81E-31 |
| Gm20431 | 201.6035838 | -2.133740704 | 1.22E-02 |
| 9330154J02Rik | 40.21475152 | -2.134352996 | 1.20E-05 |
| Gm3086 | 5.19932287 | -2.134541528 | 2.86E-02 |
| Gulp1 | 1061.560392 | -2.134847396 | 1.01E-27 |
| Gm5518 | 20.12754576 | -2.135289042 | 1.36E-03 |
| Gm5432 | 8.334657074 | -2.13627471 | 1.71E-02 |
| Emc1 | 889.6482432 | -2.136942606 | 1.18E-47 |
| Zfp644 | 1613.466685 | -2.137534057 | 7.96E-20 |
| AI427809 | 6.421281759 | -2.137681141 | 4.85E-02 |
| Cd46 | 40.97987189 | -2.138517703 | 2.88E-04 |
| Ttc30a1 | 153.2911787 | -2.139650947 | 7.84E-23 |
| Cited1 | 16.05689947 | -2.139743987 | 9.30E-04 |
| Ttyh1 | 71.46831245 | -2.141495308 | 2.99E-04 |
| Castor2 | 1817.867489 | -2.14164755 | 6.23E-56 |
| Atp8b4 | 29.3419901 | -2.142946665 | 1.20E-03 |
| Ankle1 | 132.7054838 | -2.143782953 | 3.20E-05 |
| Ccdc116 | 16.45258406 | -2.145395609 | 6.38E-03 |
| C030014I23Rik | 243.4663144 | -2.146259326 | 1.51E-25 |
| mt-Tq | 47.7235201 | -2.146490379 | 5.96E-07 |
| Cnot6 | 3465.79491 | -2.146810943 | 3.74E-26 |
| Ndufa4l2 | 9.031759966 | -2.149402375 | 1.86E-02 |
| Ccnb2-ps | 54.8298888 | -2.151369825 | 1.64E-07 |
| Tmem117 | 124.9016676 | -2.15175585 | 1.45E-10 |
| Slc27a1 | 1298.849242 | -2.15182248 | 2.62E-19 |
| Apobr | 620.5613799 | -2.157140187 | 1.63E-12 |
| Snapc5 | 308.3123639 | -2.157628364 | 2.68E-23 |
| Lrrc73 | 44.40294994 | -2.158631214 | 9.67E-09 |
| Zfp433 | 217.8127299 | -2.162488072 | 2.64E-09 |
| Gm11613 | 86.03133756 | -2.165581857 | 2.61E-12 |
| Gm14416 | 31.20614591 | -2.167343371 | 8.90E-04 |
| Gm9747 | 221.4334643 | -2.16767306 | 2.44E-12 |
| Unc5b | 390.1679829 | -2.168280457 | 6.48E-13 |
| Matn4 | 26.3751028 | -2.168724301 | 4.87E-05 |
| Col14a1 | 21.17008947 | -2.170046994 | 1.00E-02 |
| Larp6 | 54.75592821 | -2.170758198 | 3.90E-10 |
| D130040H23Rik | 50.8991406 | -2.171368281 | 1.00E-06 |
| Zfp638 | 3672.885195 | -2.171870058 | 2.31E-12 |
| Col6a3 | 17819.46317 | -2.173051098 | 1.42E-24 |
| Nkain1 | 814.8730286 | -2.17385041 | 2.44E-37 |
| Gbp2 | 472.493555 | -2.174454224 | 1.12E-23 |
| Pcdhb22 | 261.1195135 | -2.17596843 | 4.02E-30 |

| | | | |
|---|---|---|---|
| Ankrd42 | 231.1595588 | -2.176595735 | 4.84E-29 |
| 3000002C10Rik | 44.84368038 | -2.176653506 | 1.34E-07 |
| Asb4 | 41.09296277 | -2.179055128 | 9.58E-04 |
| Fam208a | 522.2502803 | -2.179619528 | 1.98E-09 |
| B230118H07Rik | 425.1159329 | -2.180465473 | 1.42E-18 |
| Fzd4 | 527.5932981 | -2.180763555 | 6.85E-13 |
| 1700067K01Rik | 19.25634044 | -2.180997585 | 1.28E-04 |
| Nlrp1b | 59.85443185 | -2.181620061 | 5.10E-11 |
| Pmp2 | 16.17721345 | -2.182456509 | 2.50E-02 |
| Golph3l | 941.6517126 | -2.183803633 | 5.27E-65 |
| Usp1 | 2727.71314 | -2.184948291 | 3.71E-11 |
| Dlec1 | 30.45660228 | -2.185112428 | 1.78E-04 |
| Bicc1 | 3603.644497 | -2.186021632 | 5.14E-64 |
| Mir351 | 18.63857239 | -2.186187767 | 1.22E-03 |
| Klf12 | 71.36571759 | -2.187410777 | 4.20E-14 |
| Zfp820 | 151.3426224 | -2.188400661 | 7.21E-06 |
| Casp8ap2 | 835.5596121 | -2.189599915 | 2.93E-09 |
| Gm26688 | 12.27901547 | -2.194779221 | 3.84E-03 |
| Ppargc1a | 542.9043571 | -2.198887598 | 2.27E-10 |
| Zfp493 | 62.7561201 | -2.199707441 | 5.22E-08 |
| Cd3d | 19.4225017 | -2.202729638 | 3.22E-03 |
| Gm26490 | 11.39270309 | -2.203526676 | 6.30E-03 |
| Gm13212 | 790.6580958 | -2.203747279 | 5.50E-13 |
| Gm45351 | 101.0112841 | -2.205528593 | 4.93E-17 |
| Pik3r3 | 1442.838135 | -2.205563449 | 6.33E-34 |
| Pgbd5 | 323.7462955 | -2.208602339 | 4.00E-23 |
| Slfn8 | 18.73354758 | -2.208784249 | 4.53E-03 |
| Raver2 | 8.675742128 | -2.209492528 | 2.23E-02 |
| Hoxa2 | 364.402932 | -2.20990427 | 2.99E-20 |
| Mmp28 | 209.9282617 | -2.210747563 | 6.91E-10 |
| Chic1 | 323.0857059 | -2.210803115 | 3.19E-23 |
| Gm10676 | 25.58480143 | -2.211804645 | 2.92E-05 |
| Gabpb2 | 2957.735198 | -2.212689721 | 3.18E-29 |
| Brca2 | 529.5132143 | -2.214271476 | 4.71E-07 |
| Ddx3y | 1595.987767 | -2.214654067 | 6.73E-11 |
| Atp1b1 | 1462.526931 | -2.215114913 | 5.28E-27 |
| Gpr63 | 64.54603199 | -2.217986965 | 1.45E-11 |
| B430203G13Rik | 111.3993379 | -2.218213667 | 1.24E-12 |
| Macc1 | 17.40048592 | -2.218754827 | 7.41E-03 |
| Npr1 | 26.77724999 | -2.219826702 | 2.85E-04 |
| Klrb1c | 12.08738048 | -2.220302738 | 3.58E-03 |
| Gm16617 | 22.46511473 | -2.222220661 | 2.93E-05 |
| Lrrc17 | 862.6812303 | -2.222809951 | 2.45E-10 |
| Gm15704 | 10.21925034 | -2.225949313 | 1.38E-02 |
| Gm12080 | 10.62390129 | -2.226269736 | 1.15E-02 |
| 4930483K19Rik | 9.197652411 | -2.22644497 | 1.63E-02 |
| Gpr135 | 158.3505966 | -2.226555319 | 3.76E-25 |

| | | | |
|---|---|---|---|
| Mtmr11 | 1713.709962 | -2.228946387 | 4.42E-43 |
| A430033K04Rik | 372.7872773 | -2.229846158 | 4.47E-29 |
| Gm7162 | 14.15814186 | -2.230275941 | 4.58E-03 |
| Zfp760 | 404.3092604 | -2.230704842 | 1.91E-10 |
| AI606473 | 18.78385635 | -2.233187459 | 6.09E-04 |
| Srsf1 | 6220.767209 | -2.233567083 | 7.42E-13 |
| Grin3b | 31.43570586 | -2.233594528 | 8.89E-07 |
| Arglu1 | 1665.246254 | -2.233680851 | 4.30E-19 |
| Ccnb1ip1 | 16.25829992 | -2.234480124 | 3.79E-05 |
| Sptssa | 1180.152327 | -2.234559282 | 3.52E-33 |
| Gstm1 | 8315.734253 | -2.235661785 | 1.03E-18 |
| Fbxo36 | 335.8821186 | -2.23739671 | 1.65E-22 |
| Cdkn2aipnl | 929.6184746 | -2.23780002 | 1.51E-20 |
| Chd9 | 2494.76083 | -2.238274699 | 1.47E-31 |
| Ly96 | 148.4629841 | -2.239838413 | 7.69E-18 |
| Galr2 | 29.98084627 | -2.242064686 | 1.47E-05 |
| Zfp61 | 290.8195837 | -2.242895108 | 1.62E-23 |
| Gm4425 | 83.20614124 | -2.243561941 | 2.18E-13 |
| Kbtbd7 | 200.5646128 | -2.247585527 | 2.79E-14 |
| Thtpa | 224.2179658 | -2.24923343 | 1.29E-35 |
| Gm9968 | 18.46010921 | -2.249541757 | 8.33E-04 |
| Thg1l | 413.4328502 | -2.249717601 | 2.13E-39 |
| Pcdhb8 | 54.15088343 | -2.250193458 | 7.40E-11 |
| Zfp213 | 424.7323897 | -2.251218852 | 1.98E-16 |
| Wdr60 | 933.6140883 | -2.255556344 | 1.69E-31 |
| Zfp595 | 165.6562495 | -2.255890399 | 6.29E-15 |
| Dnase1l2 | 34.76247815 | -2.257114899 | 4.15E-06 |
| Cmtm3 | 7129.083408 | -2.260000885 | 2.95E-45 |
| Frg2f1 | 213.8372542 | -2.26092402 | 2.37E-26 |
| Mbd4 | 461.5537666 | -2.262415612 | 4.88E-27 |
| Tet2 | 657.7186831 | -2.263452938 | 8.52E-31 |
| Homer1 | 244.6755818 | -2.264514176 | 6.60E-12 |
| Zc3h12b | 308.6497987 | -2.265994831 | 4.66E-38 |
| Dlx1as | 29.61315962 | -2.266206296 | 3.25E-04 |
| Rilp | 7.065350126 | -2.270107088 | 3.68E-02 |
| Il17rd | 972.8019314 | -2.27131322 | 1.99E-19 |
| Zfp345 | 250.9356857 | -2.271783783 | 2.41E-15 |
| Rabl2 | 424.7187683 | -2.27305786 | 2.47E-21 |
| Actn2 | 23.22558396 | -2.273812768 | 1.19E-04 |
| Gm6871 | 49.20881328 | -2.275418992 | 1.97E-06 |
| Tbc1d9 | 40.79818821 | -2.281066534 | 1.77E-05 |
| Ajm1 | 101.9957552 | -2.283441911 | 2.71E-04 |
| Trps1 | 1353.888354 | -2.285312169 | 8.48E-10 |
| Zfp354c | 1061.502211 | -2.285464726 | 2.13E-33 |
| Spry2 | 928.0977301 | -2.285871892 | 6.33E-69 |
| Zdhhc14 | 26.56961032 | -2.287034833 | 7.47E-06 |
| Cpne7 | 6.76200543 | -2.288782244 | 2.69E-02 |

| | | | |
|---|---|---|---|
| Pigo | 175.9082829 | -2.28940753 | 4.17E-12 |
| 4930481B07Rik | 25.10703027 | -2.29162279 | 5.34E-06 |
| Etnk1 | 4408.390478 | -2.293847555 | 8.05E-53 |
| Sertad4 | 988.7002916 | -2.294890994 | 1.73E-10 |
| Ism1 | 148.8317432 | -2.295869095 | 3.55E-08 |
| Zmiz1os1 | 10.16799365 | -2.297336911 | 7.40E-03 |
| 6030445D17Rik | 6.833795613 | -2.297847142 | 4.63E-02 |
| Zfp934 | 59.10685606 | -2.301052298 | 9.31E-05 |
| Trem2 | 96.77326875 | -2.301081878 | 1.17E-10 |
| 5430403G16Rik | 257.0774916 | -2.30171885 | 7.44E-10 |
| Il1rap | 2529.832075 | -2.303898475 | 2.12E-40 |
| Zfp386 | 962.4306407 | -2.304975402 | 8.77E-14 |
| Nme3 | 171.0026249 | -2.306296337 | 2.80E-23 |
| Hgf | 466.3773992 | -2.307294901 | 2.87E-14 |
| Ptdss2 | 1024.474847 | -2.308207634 | 6.57E-48 |
| Rev3l | 2594.882945 | -2.309440105 | 3.61E-15 |
| Galnt13 | 425.0375608 | -2.310589464 | 1.82E-32 |
| Itgb3 | 406.4148934 | -2.311655403 | 2.54E-11 |
| Dsel | 941.8407461 | -2.311866743 | 2.54E-15 |
| Rnft2 | 369.3814148 | -2.312956175 | 7.64E-18 |
| NA | 20.46702798 | -2.313083442 | 2.14E-03 |
| Zfp970 | 672.497198 | -2.313295579 | 6.23E-16 |
| Trmt12 | 249.3859157 | -2.315441232 | 4.62E-33 |
| Itgal | 47.97439676 | -2.316022998 | 3.82E-08 |
| Srsf3 | 8264.123244 | -2.316669049 | 4.51E-34 |
| Ell3 | 8.738846931 | -2.316749935 | 2.66E-02 |
| Fxyd1 | 14.72790562 | -2.318549575 | 1.61E-02 |
| AA543186 | 24.50231718 | -2.319378221 | 1.13E-05 |
| Ucp2 | 1083.734914 | -2.320138164 | 9.67E-32 |
| Hic1 | 1990.450822 | -2.320255637 | 5.42E-21 |
| Impact | 6753.387895 | -2.320417238 | 1.65E-49 |
| Phf20l1 | 1651.961763 | -2.321393494 | 5.08E-15 |
| Tmem191c | 138.1462665 | -2.321898274 | 3.72E-15 |
| Rhou | 4021.987846 | -2.323712845 | 2.82E-46 |
| Far2 | 36.7669344 | -2.323734846 | 1.86E-05 |
| Phtf1os | 43.702832 | -2.323759588 | 1.43E-04 |
| Zfp985 | 227.584577 | -2.324932101 | 6.93E-06 |
| Lrguk | 36.65908511 | -2.324940803 | 7.31E-05 |
| Mecom | 1419.704379 | -2.325290772 | 7.07E-46 |
| mt-Ti | 38.83048144 | -2.325854199 | 4.26E-04 |
| C4b | 168.9332119 | -2.326342661 | 4.58E-10 |
| Cacfd1 | 1602.512009 | -2.327968775 | 3.23E-75 |
| Cd1d1 | 192.488829 | -2.328417876 | 3.71E-20 |
| Bclaf1 | 5262.010873 | -2.329079304 | 9.53E-21 |
| Gm14127 | 12.86916418 | -2.329974591 | 3.39E-03 |
| Rgs9bp | 289.0652429 | -2.330567698 | 1.14E-25 |
| Prpf4b | 2196.000112 | -2.33388452 | 1.12E-15 |

| | | | |
|---|---|---|---|
| BC006965 | 6.703558381 | -2.333931816 | 1.12E-02 |
| Gm24299 | 23.96221992 | -2.334050464 | 3.04E-04 |
| B3gnt9 | 1121.027082 | -2.335978389 | 2.77E-46 |
| Tgfb3 | 3717.223368 | -2.338866051 | 1.17E-37 |
| Ccdc151 | 30.69612096 | -2.339868611 | 3.45E-06 |
| Senp8 | 238.1056629 | -2.340229848 | 9.71E-18 |
| Sox5 | 174.3593763 | -2.341425581 | 1.99E-07 |
| Ctso | 1002.433076 | -2.344184256 | 1.81E-54 |
| Ank | 9508.851247 | -2.346741048 | 5.82E-93 |
| Mep1a | 25.58712896 | -2.347101935 | 2.33E-02 |
| Ecm2 | 56.32071213 | -2.347603415 | 1.25E-09 |
| Slf1 | 437.365383 | -2.347964128 | 1.13E-10 |
| Klhl23 | 494.130889 | -2.349476689 | 2.08E-16 |
| Nab2 | 4607.437328 | -2.350393513 | 1.47E-73 |
| Tada1 | 953.2882728 | -2.352186085 | 5.28E-39 |
| Gm12940 | 515.8451704 | -2.353735172 | 2.46E-12 |
| Lpar1 | 5879.931993 | -2.354619933 | 2.94E-60 |
| Sh3bgrl | 3106.270232 | -2.356444019 | 8.32E-11 |
| Zfp947 | 97.00750437 | -2.357697965 | 1.83E-07 |
| Zfp419 | 31.98303988 | -2.358658821 | 2.61E-07 |
| Igsf10 | 1498.548461 | -2.361231998 | 1.04E-15 |
| Gm15758 | 33.86750818 | -2.362270153 | 7.86E-07 |
| Slc15a2 | 73.21034976 | -2.362438798 | 1.43E-05 |
| Gm26733 | 24.26345678 | -2.364249161 | 2.27E-03 |
| Zfp995 | 106.1881916 | -2.364277946 | 2.14E-09 |
| Prox2os | 18.94389743 | -2.364622419 | 6.17E-04 |
| Clip4 | 144.42984 | -2.365973948 | 7.93E-18 |
| Col16a1 | 10534.38836 | -2.367320608 | 9.75E-36 |
| Slc25a45 | 460.588334 | -2.367725138 | 3.73E-42 |
| Ttc30b | 441.1625201 | -2.367874097 | 4.15E-45 |
| Gbp3 | 49.15600115 | -2.369746131 | 2.33E-10 |
| Mab21l1 | 801.0403118 | -2.370079846 | 4.20E-54 |
| Camkk1 | 480.61585 | -2.370904292 | 7.95E-25 |
| Twist2 | 1503.498649 | -2.371655128 | 6.06E-43 |
| Myo1b | 2440.909108 | -2.371875025 | 1.18E-43 |
| Lef1 | 146.4483945 | -2.375331883 | 5.26E-09 |
| Fam208b | 1989.088708 | -2.375355892 | 1.03E-16 |
| Miga2 | 802.4941053 | -2.378132947 | 1.21E-30 |
| Gm13410 | 9.648131191 | -2.380883554 | 1.49E-02 |
| Gm26874 | 6.163567678 | -2.383079212 | 3.80E-02 |
| Gm20457 | 45.64204163 | -2.383483432 | 6.01E-08 |
| Tmod4 | 28.18849812 | -2.383605422 | 4.54E-04 |
| Gm2399 | 12.05274708 | -2.385356851 | 2.69E-03 |
| Neat1 | 13268.97801 | -2.387964268 | 3.21E-18 |
| Lekr1 | 154.8880671 | -2.38801899 | 6.83E-08 |
| Ankrd49 | 493.1015181 | -2.390680392 | 2.90E-18 |
| Luc7l3 | 2630.403889 | -2.390922254 | 8.74E-13 |

| | | | |
|---|---|---|---|
| Ptafr | 9.392685792 | -2.390989145 | 1.97E-02 |
| Gm11008 | 25.36760532 | -2.394349383 | 2.55E-04 |
| Dock11 | 1878.430365 | -2.396680039 | 1.95E-04 |
| 1700055D18Rik | 8.249492801 | -2.39752124 | 5.66E-03 |
| Zbtb1 | 1295.613615 | -2.399560216 | 7.56E-37 |
| A330074K22Rik | 266.6373111 | -2.401793056 | 5.57E-18 |
| Fam71e1 | 11.96666864 | -2.402109517 | 1.52E-03 |
| Tmem260 | 658.9304686 | -2.402295982 | 6.17E-51 |
| Agap2 | 1004.939107 | -2.403896009 | 8.62E-18 |
| Rnf150 | 1744.58427 | -2.40449773 | 1.45E-46 |
| Gm26387 | 13.19810809 | -2.404922185 | 9.29E-04 |
| Il11ra1 | 3333.985703 | -2.405960856 | 1.37E-30 |
| Zfp85 | 115.2997844 | -2.407936335 | 9.11E-12 |
| Cyb5rl | 377.8189539 | -2.409306292 | 1.01E-17 |
| Vax2 | 16.92850451 | -2.409814879 | 1.53E-03 |
| Dlk1 | 29943.381 | -2.411162781 | 4.46E-12 |
| Sh3kbp1 | 3211.059854 | -2.411733279 | 2.62E-91 |
| Gm26730 | 29.49930531 | -2.412459026 | 1.01E-06 |
| AV039307 | 11.6712299 | -2.412644957 | 9.06E-03 |
| Zfp770 | 544.5327502 | -2.414049807 | 5.04E-16 |
| Pla2g4c | 10.85244054 | -2.415594947 | 3.24E-02 |
| B430212C06Rik | 14.01429096 | -2.415676499 | 3.29E-04 |
| Slc16a4 | 67.2259842 | -2.417531164 | 6.04E-07 |
| 4933440N22Rik | 14.45191676 | -2.417695785 | 9.74E-04 |
| Dpy19l1 | 4064.222751 | -2.417942056 | 1.94E-67 |
| Nol4 | 339.5172744 | -2.418388907 | 1.88E-19 |
| Cep290 | 586.5849252 | -2.420656434 | 2.89E-05 |
| Gm16023 | 30.5915826 | -2.420805154 | 1.08E-06 |
| Snai2 | 1626.102104 | -2.422368473 | 2.12E-49 |
| Lmo7 | 1891.856719 | -2.423592718 | 1.73E-59 |
| Gpr153 | 1837.287844 | -2.425842048 | 1.18E-35 |
| Fbxl21 | 8.014840798 | -2.426016531 | 3.80E-02 |
| Anks6 | 655.9616026 | -2.426764247 | 1.48E-21 |
| Fgf14 | 155.1480338 | -2.427222217 | 3.22E-10 |
| 4931422A03Rik | 10.25819116 | -2.429334735 | 6.04E-03 |
| Mgat3 | 1523.241024 | -2.431036467 | 2.64E-51 |
| Mdfi | 37.12431873 | -2.432546486 | 6.15E-09 |
| Hs6st1 | 2054.227925 | -2.434369687 | 5.67E-52 |
| Xrn1 | 931.0024997 | -2.436023231 | 1.96E-24 |
| Lppos | 143.0048581 | -2.436737829 | 2.06E-20 |
| Cr2 | 8.580331134 | -2.439320994 | 7.14E-03 |
| Trpv3 | 7.69065872 | -2.440282925 | 6.76E-03 |
| E230013L22Rik | 57.3366247 | -2.44070808 | 4.90E-14 |
| Tnfrsf25 | 17.04286528 | -2.441773084 | 4.76E-04 |
| Gse1 | 312.4825439 | -2.444306749 | 3.11E-18 |
| Cdh1 | 10.89184837 | -2.44616095 | 3.90E-03 |
| Il1rn | 36.407963 | -2.448556828 | 9.63E-05 |

| | | | |
|---|---|---|---|
| Yy2 | 149.4951159 | -2.449488907 | 1.34E-09 |
| Anks1b | 74.61954556 | -2.450188487 | 4.49E-06 |
| Fstl1 | 39494.87916 | -2.450893735 | 1.06E-47 |
| Samd14 | 1239.574654 | -2.452423805 | 1.27E-57 |
| Ccn4 | 3715.730625 | -2.454599688 | 2.29E-08 |
| Lca5l | 18.56150822 | -2.454962951 | 2.03E-04 |
| Gm22392 | 13.95865186 | -2.459506089 | 3.89E-03 |
| NA | 54.53741336 | -2.46113151 | 4.78E-07 |
| Dhrs9 | 1469.943128 | -2.465865131 | 2.32E-25 |
| Mmp2 | 89388.04831 | -2.468131914 | 3.36E-22 |
| G630016G05Rik | 35.92889756 | -2.468755033 | 2.69E-06 |
| Zfp619 | 187.8261664 | -2.469251586 | 3.38E-16 |
| Pdgfra | 29231.84787 | -2.470237979 | 5.27E-18 |
| Tbc1d4 | 572.5515038 | -2.470958709 | 1.68E-22 |
| Rasa1 | 2097.985419 | -2.472975275 | 1.76E-20 |
| Scg2 | 1103.15175 | -2.474079383 | 9.22E-05 |
| Zfp35 | 601.7816323 | -2.474797163 | 2.98E-43 |
| Spry3 | 464.077843 | -2.474991081 | 1.30E-17 |
| Nin | 2334.99443 | -2.475599372 | 1.53E-36 |
| Zfp953 | 159.3076468 | -2.47673108 | 1.62E-13 |
| Glt8d2 | 963.6849507 | -2.47783317 | 1.62E-25 |
| Gp5 | 7.194843487 | -2.479424918 | 3.22E-03 |
| Apopt1 | 749.4078064 | -2.479875525 | 6.90E-56 |
| NA | 12.28994974 | -2.480494858 | 6.86E-03 |
| Wdr83 | 632.260428 | -2.48077287 | 2.88E-69 |
| Dek | 2477.319996 | -2.480998842 | 4.02E-13 |
| Tmem67 | 422.0083957 | -2.481852155 | 1.87E-22 |
| Zfp458 | 135.3705613 | -2.488849203 | 1.50E-12 |
| Insyn1 | 1056.11518 | -2.489195666 | 1.84E-38 |
| Agk | 886.1543945 | -2.490853305 | 1.57E-50 |
| Gm10643 | 42.40207878 | -2.494035304 | 6.00E-10 |
| Sh3pxd2b | 11222.03918 | -2.494585859 | 3.20E-37 |
| Gm9951 | 8.324679201 | -2.494775439 | 2.37E-02 |
| Gm14403 | 121.8702542 | -2.500914784 | 1.17E-08 |
| Gata2 | 333.1625954 | -2.501540634 | 6.35E-30 |
| Kctd17 | 1532.436301 | -2.502079369 | 7.36E-45 |
| Gm16835 | 21.2767008 | -2.504615697 | 4.83E-05 |
| Spred2 | 1836.996101 | -2.507263847 | 9.48E-92 |
| Eya4 | 341.0867101 | -2.507886571 | 3.61E-33 |
| Rerg | 14.52101119 | -2.513055291 | 1.51E-03 |
| Arxes1 | 501.9970735 | -2.514829421 | 3.35E-10 |
| Tia1 | 2607.982324 | -2.517650322 | 9.77E-19 |
| Adamts14 | 1745.125069 | -2.518895624 | 6.50E-23 |
| Prss30 | 9.310014953 | -2.523483994 | 8.56E-03 |
| NA | 11.55182351 | -2.523748597 | 1.13E-02 |
| Mef2c | 246.1899401 | -2.52728906 | 8.80E-10 |
| Cabyr | 15.12677381 | -2.527540302 | 3.92E-04 |

| | | | |
|---|---|---|---|
| Car15 | 64.28556731 | -2.529964562 | 7.72E-12 |
| Gm5141 | 44.57070115 | -2.530368034 | 2.11E-05 |
| Btnl10 | 6.951584442 | -2.532112203 | 2.72E-02 |
| Mycn | 50.26697572 | -2.533715824 | 1.08E-06 |
| Itga11 | 1425.902456 | -2.536857576 | 4.45E-12 |
| Phxr4 | 8.275227431 | -2.536879255 | 4.82E-03 |
| B4galt6 | 1066.319975 | -2.538540053 | 8.80E-35 |
| Hvcn1 | 604.403497 | -2.540423044 | 7.74E-42 |
| 1700060J05Rik | 10.45083137 | -2.54332525 | 1.61E-03 |
| Epc2 | 1151.623631 | -2.54580932 | 1.59E-37 |
| Gm26546 | 16.82540964 | -2.549568317 | 1.08E-04 |
| Zfp808 | 153.8817309 | -2.550581062 | 2.48E-08 |
| Atxn1 | 1217.972466 | -2.552560944 | 5.74E-33 |
| Krcc1 | 1725.674243 | -2.552615627 | 5.11E-21 |
| Gm16263 | 10.50272982 | -2.553018696 | 1.79E-02 |
| Trio | 8424.77309 | -2.553455832 | 1.16E-75 |
| Zfp62 | 1535.939692 | -2.553612588 | 1.19E-14 |
| Parp14 | 278.3693229 | -2.558757613 | 4.18E-26 |
| Onecut2 | 225.0983773 | -2.559065197 | 2.06E-23 |
| Zfp422-ps | 64.25166758 | -2.560095958 | 4.57E-10 |
| Cntln | 2121.188526 | -2.560508568 | 2.32E-16 |
| Gm2245 | 5.67545899 | -2.562835966 | 1.49E-02 |
| Fam69a | 2022.668361 | -2.563033533 | 3.38E-31 |
| Zfp748 | 408.1503428 | -2.566508076 | 1.63E-14 |
| Gm26771 | 21.68010433 | -2.570089617 | 1.27E-05 |
| 2310015A10Rik | 393.9563693 | -2.571516482 | 5.75E-31 |
| Rnasel | 68.25410949 | -2.572261732 | 8.53E-10 |
| Gdpd2 | 416.9752147 | -2.572485509 | 1.27E-11 |
| Islr | 180.9664839 | -2.574315753 | 8.27E-07 |
| Trim62 | 389.1707065 | -2.575179346 | 1.65E-45 |
| Gm10524 | 163.6017619 | -2.576094905 | 7.86E-21 |
| Slfn9 | 1195.956977 | -2.576848463 | 1.54E-06 |
| Eepd1 | 56.11607845 | -2.58063018 | 2.32E-07 |
| D630033O11Rik | 7.891037492 | -2.580875676 | 5.01E-03 |
| Zfp281 | 2185.820373 | -2.582819827 | 1.22E-48 |
| Osbpl10 | 87.89074291 | -2.583017276 | 2.33E-10 |
| Zfp146 | 1880.563755 | -2.585047247 | 3.47E-32 |
| Frk | 907.1646372 | -2.585443031 | 1.30E-20 |
| Etaa1os | 11.32037963 | -2.586660048 | 1.51E-03 |
| Zfp618 | 1231.162364 | -2.589667144 | 3.98E-18 |
| Gm26540 | 8.589600188 | -2.5914575 | 3.32E-03 |
| Gm10226 | 13.74316307 | -2.591485378 | 3.39E-04 |
| NA | 16.14919928 | -2.591919459 | 3.49E-03 |
| Cyp2j9 | 75.57103561 | -2.592654185 | 3.46E-12 |
| Cotl1 | 536.8515866 | -2.593901642 | 4.47E-25 |
| Fam84b | 9.611565536 | -2.595543152 | 9.68E-03 |
| Amigo1 | 89.49915741 | -2.596422324 | 4.66E-14 |

| | | | |
|---|---|---|---|
| Negr1 | 66.82959444 | -2.596488924 | 3.73E-13 |
| Pofut1 | 2332.656105 | -2.598869127 | 2.41E-34 |
| Gm15283 | 31.78380944 | -2.599233874 | 1.88E-07 |
| Pi16 | 9.391261602 | -2.6024312 | 2.15E-02 |
| Gcnt4 | 773.2398383 | -2.603502321 | 3.60E-37 |
| Plagl2 | 1099.649691 | -2.60738082 | 5.49E-43 |
| Tmem229b | 14.0780733 | -2.60893021 | 7.44E-03 |
| Prr22 | 7.558916078 | -2.612090191 | 7.99E-03 |
| Fam198a | 33.40205875 | -2.612220346 | 3.71E-10 |
| Bcan | 8.867739542 | -2.612424681 | 7.85E-03 |
| Arhgef3 | 59.83266333 | -2.618658124 | 2.11E-10 |
| Rnase4 | 11680.33536 | -2.61923922 | 9.89E-35 |
| Usp40 | 1922.118719 | -2.61996407 | 1.70E-58 |
| A830082K12Rik | 468.2476811 | -2.620724917 | 5.72E-38 |
| 4933406P04Rik | 9.227396209 | -2.626842927 | 4.83E-03 |
| Timm22 | 1003.238842 | -2.632718067 | 2.04E-21 |
| DQ267102 | 5.610198586 | -2.63369796 | 4.93E-02 |
| Tmem42 | 153.7982272 | -2.636049671 | 8.61E-29 |
| Olr1 | 36.00857858 | -2.641766606 | 3.87E-03 |
| Zup1 | 502.8916938 | -2.641928666 | 4.86E-23 |
| Rnase1 | 36.55767924 | -2.642446868 | 9.27E-07 |
| Timm21 | 480.2239543 | -2.643881094 | 3.43E-37 |
| Nat8f4 | 100.8466667 | -2.643913316 | 1.14E-11 |
| Gstm7 | 9.666511752 | -2.645142987 | 5.39E-03 |
| Arhgef33 | 6.08745988 | -2.646749142 | 1.62E-02 |
| Zfp2 | 138.1886809 | -2.647173692 | 2.44E-14 |
| Gm16229 | 22.95610143 | -2.647639298 | 2.65E-05 |
| Btc | 11.83361267 | -2.647960431 | 6.69E-04 |
| Znrd1as | 91.526769 | -2.648011525 | 1.56E-18 |
| Gm17509 | 9.815080105 | -2.648595417 | 4.50E-03 |
| Zfhx2 | 31.19525386 | -2.650162488 | 2.23E-05 |
| D630045J12Rik | 1186.490416 | -2.654999236 | 8.65E-38 |
| Mab21l2 | 405.6572256 | -2.657261443 | 1.09E-27 |
| Gm20518 | 49.03831615 | -2.657997221 | 1.85E-06 |
| AU041133 | 94.90491122 | -2.661562853 | 2.40E-10 |
| Zfp354b | 32.87930447 | -2.662215993 | 5.00E-05 |
| Tmem179 | 20.54769501 | -2.664976832 | 1.58E-03 |
| Mylk | 284.7805888 | -2.66517008 | 3.48E-22 |
| Fam3c | 4044.640668 | -2.66588717 | 2.62E-91 |
| Zfp966 | 23.05339885 | -2.668392289 | 2.04E-03 |
| Map2k6 | 397.271134 | -2.668859244 | 8.36E-36 |
| Arc | 134.080199 | -2.670717542 | 2.15E-18 |
| 1700020L24Rik | 39.09349375 | -2.670810972 | 2.42E-05 |
| Kdelc2 | 3012.784827 | -2.672860946 | 3.10E-54 |
| Gm6288 | 19.23685061 | -2.673035105 | 3.26E-05 |
| Gm17195 | 18.69066063 | -2.673259864 | 1.44E-05 |
| Mgst1 | 1067.813549 | -2.674175545 | 1.27E-36 |

| | | | |
|---|---|---|---|
| Nr2f1 | 1515.258968 | -2.674252121 | 9.64E-58 |
| Cspg4 | 1149.427576 | -2.674570423 | 3.83E-24 |
| 9-Sep | 7495.015017 | -2.674716534 | 5.50E-78 |
| Ggt7 | 21.30559824 | -2.67724921 | 3.37E-06 |
| Gata5 | 7.901781493 | -2.677713878 | 1.24E-02 |
| Gfra1 | 41.83640276 | -2.678060468 | 2.85E-04 |
| Smad9 | 231.529786 | -2.679022142 | 2.99E-13 |
| Gm13470 | 63.86688211 | -2.679560273 | 4.35E-14 |
| Zfp758 | 213.2476453 | -2.682426203 | 1.03E-12 |
| 2210016L21Rik | 958.8047931 | -2.683318446 | 4.82E-57 |
| Gm14295 | 40.67472279 | -2.686332194 | 1.76E-03 |
| Tmem44 | 491.8757937 | -2.688889507 | 3.52E-13 |
| Camk2n1 | 11.24597342 | -2.689645415 | 1.92E-03 |
| Tmem254a | 133.019222 | -2.690386644 | 5.96E-10 |
| mt-Tn | 66.01413775 | -2.694669135 | 4.26E-07 |
| Scly | 161.3156578 | -2.697113176 | 5.92E-22 |
| Ctxn1 | 645.1595188 | -2.702132235 | 2.73E-37 |
| Fcrlb | 50.76172523 | -2.702489828 | 2.43E-08 |
| Pianp | 1185.174268 | -2.703418856 | 1.65E-42 |
| Dio2 | 87.5272909 | -2.703570031 | 2.43E-10 |
| Asb13 | 427.8861044 | -2.707925472 | 6.98E-28 |
| Arhgap42 | 1172.465651 | -2.708135764 | 5.23E-38 |
| Sardh | 902.4568354 | -2.708904703 | 7.81E-28 |
| 2700081O15Rik | 2585.082579 | -2.711327449 | 1.76E-58 |
| Thrb | 322.1208178 | -2.711329261 | 3.09E-19 |
| Kirrel3 | 34.68484594 | -2.711723491 | 7.25E-06 |
| Gsdme | 190.6699199 | -2.714106109 | 5.80E-18 |
| Tlr7 | 12.65124356 | -2.71617238 | 2.61E-02 |
| NA | 141.1068307 | -2.716514756 | 6.07E-19 |
| P3h2 | 17.77383867 | -2.718165302 | 1.80E-04 |
| Ciita | 7.733065526 | -2.718371246 | 8.66E-03 |
| Kng2 | 30.33286104 | -2.719793479 | 3.16E-05 |
| Il1rl2 | 175.7426724 | -2.720435594 | 1.90E-24 |
| Zfp119a | 132.5293073 | -2.722607596 | 3.91E-16 |
| Igdcc4 | 80.19494662 | -2.722731369 | 2.47E-13 |
| Adh7 | 1804.860141 | -2.724810547 | 1.47E-25 |
| Cd82 | 121.0116936 | -2.725410232 | 9.39E-19 |
| Zfp454 | 65.32644374 | -2.726003127 | 1.64E-15 |
| Prrg1 | 395.4345146 | -2.730971212 | 1.80E-17 |
| Gm2415 | 6.484549693 | -2.731690771 | 1.59E-02 |
| Il12rb1 | 41.79475726 | -2.732032893 | 5.07E-08 |
| Tlr4 | 2569.985835 | -2.734040991 | 2.87E-44 |
| Siglec15 | 19.76496087 | -2.734713108 | 2.70E-05 |
| Gm20939 | 156.2065304 | -2.735007692 | 1.85E-09 |
| Mindy2 | 1585.001885 | -2.735212127 | 5.87E-23 |
| P2rx4 | 2480.638167 | -2.738137095 | 2.58E-48 |
| Pabpc4l | 55.49155236 | -2.739748576 | 5.63E-10 |

| | | | |
|---|---|---|---|
| 4930478K11Rik | 10.82729642 | -2.742315854 | 5.32E-04 |
| Platr22 | 32.98595665 | -2.743059595 | 5.70E-08 |
| Zfp629 | 757.4171943 | -2.747362805 | 1.60E-37 |
| Gm13836 | 8.255389852 | -2.749715344 | 2.32E-03 |
| Gm15907 | 6.608722355 | -2.750178679 | 1.63E-02 |
| 2810039B14Rik | 112.6447876 | -2.7524145 | 5.07E-29 |
| Fcor | 26.09012877 | -2.753436363 | 1.79E-06 |
| Mir186 | 9.472613992 | -2.754948115 | 1.31E-03 |
| Emilin1 | 5288.590701 | -2.760039487 | 2.59E-47 |
| 4931428F04Rik | 479.9064024 | -2.760642569 | 1.07E-34 |
| Snap25 | 20.82983854 | -2.760948397 | 2.85E-05 |
| Zfp239 | 614.0938397 | -2.761497975 | 2.97E-35 |
| Sparcl1 | 1663.612457 | -2.764151206 | 6.72E-14 |
| Dhx9 | 8577.897448 | -2.766447556 | 3.75E-37 |
| Eml1 | 3052.707289 | -2.767214876 | 8.58E-80 |
| Zfp738 | 241.9782701 | -2.767890005 | 4.46E-14 |
| Neurl4 | 1596.034851 | -2.769672708 | 7.02E-74 |
| Nap1l2 | 52.4545836 | -2.77409212 | 2.18E-07 |
| Adamts15 | 21.18110807 | -2.774639379 | 3.85E-06 |
| Ankdd1b | 30.83395126 | -2.776615372 | 2.03E-08 |
| Zfp729a | 400.7358496 | -2.777833703 | 1.11E-17 |
| E030013I19Rik | 11.43396328 | -2.780910428 | 4.51E-03 |
| Zfp133-ps | 30.65106094 | -2.781048015 | 1.47E-08 |
| Uqcc3 | 436.9863654 | -2.78142843 | 1.42E-40 |
| 4930577N17Rik | 13.86405934 | -2.782418733 | 4.63E-04 |
| Gfra2 | 25.3859268 | -2.784256757 | 5.44E-06 |
| Zfp65 | 459.1248073 | -2.787144852 | 1.45E-23 |
| Ift81 | 1302.686909 | -2.788891667 | 5.06E-16 |
| Prdm11 | 475.2305451 | -2.793023413 | 2.00E-57 |
| Ccdc162 | 5.798399572 | -2.795624577 | 1.38E-02 |
| Faah | 126.5236943 | -2.796143933 | 2.09E-13 |
| Sfrp2 | 592.0206281 | -2.798540367 | 3.72E-14 |
| Lgals4 | 78.28538866 | -2.800783827 | 3.45E-13 |
| Tril | 35.05868021 | -2.801498886 | 3.30E-04 |
| Zfp994 | 554.6194131 | -2.801548728 | 5.58E-16 |
| Pcdhb2 | 19.19826051 | -2.802485794 | 8.10E-07 |
| Hhipl1 | 257.2349416 | -2.803815322 | 4.10E-20 |
| Rassf2 | 2868.908583 | -2.8039895 | 1.93E-30 |
| Gm17096 | 12.90294207 | -2.80560914 | 4.37E-04 |
| Hif3a | 7.878060011 | -2.805667387 | 8.98E-03 |
| Gm15792 | 8.665228527 | -2.805925858 | 1.95E-02 |
| Pcdhb5 | 354.373418 | -2.808921474 | 4.09E-44 |
| Zfp873 | 157.1558645 | -2.809810379 | 4.34E-17 |
| Snx9 | 6913.287897 | -2.813759022 | 5.08E-93 |
| Gm17084 | 46.48215339 | -2.815540508 | 3.04E-02 |
| Gbp9 | 51.96187374 | -2.816341595 | 7.45E-15 |
| Cdon | 2024.016055 | -2.816640251 | 1.45E-30 |

| | | | |
|---|---|---|---|
| 4930547M16Rik | 12.48702434 | -2.817708927 | 1.57E-03 |
| Sfxn5 | 564.0641619 | -2.822158604 | 2.75E-58 |
| Ccdc38 | 9.527172374 | -2.826223322 | 3.69E-03 |
| Clec4a2 | 19.35931451 | -2.827879232 | 2.53E-05 |
| Cped1 | 8743.703581 | -2.830717816 | 8.20E-46 |
| Gm26945 | 12.42640804 | -2.832518853 | 3.21E-03 |
| Gm13305 | 188.831561 | -2.834068982 | 3.08E-24 |
| Stra6l | 28.27227403 | -2.835852711 | 4.52E-04 |
| Plekha4 | 436.889388 | -2.837398511 | 3.06E-21 |
| Ptpn3 | 659.1680817 | -2.838127991 | 6.44E-56 |
| Zfp606 | 511.9724779 | -2.843223452 | 3.80E-22 |
| Gm16096 | 18.53812567 | -2.84344745 | 4.47E-05 |
| Gm3739 | 55.08943698 | -2.84560226 | 1.61E-10 |
| A930024E05Rik | 70.97068599 | -2.848159176 | 9.47E-16 |
| Nlrp2 | 6.369497996 | -2.852342746 | 1.53E-02 |
| 2700033N17Rik | 66.80711653 | -2.853530319 | 4.39E-15 |
| Ccdc39 | 190.5301171 | -2.855561216 | 4.72E-16 |
| Fzd1 | 3526.570472 | -2.857092473 | 6.07E-49 |
| Zadh2 | 2436.775149 | -2.857172928 | 1.22E-70 |
| Spats1 | 7.028321187 | -2.857646379 | 1.89E-02 |
| Hr | 118.7699506 | -2.858140454 | 3.78E-08 |
| Notch4 | 22.94704906 | -2.859477166 | 4.10E-04 |
| Arhgap28 | 1085.405443 | -2.861913399 | 1.49E-52 |
| Gm11377 | 12.83673501 | -2.862204912 | 4.98E-03 |
| Ict1os | 13.02405551 | -2.868760216 | 1.10E-03 |
| Tvp23a | 784.6639239 | -2.871107247 | 9.18E-39 |
| Matk | 59.77616886 | -2.872311887 | 3.05E-12 |
| 3110039I08Rik | 2289.917775 | -2.87440605 | 2.72E-10 |
| 2810403D21Rik | 60.92745888 | -2.876182568 | 1.33E-11 |
| Igsf3 | 583.6384486 | -2.879228496 | 1.15E-46 |
| 2210418O10Rik | 246.2963268 | -2.879505289 | 5.25E-23 |
| Glra1 | 7.980948167 | -2.881624806 | 3.44E-02 |
| Hmga2 | 10182.13318 | -2.882703657 | 1.33E-29 |
| Pxylp1 | 827.890857 | -2.890675065 | 8.06E-27 |
| A730056A06Rik | 112.1017604 | -2.89225797 | 1.22E-13 |
| Mfap4 | 212.0521077 | -2.893061981 | 5.45E-04 |
| Gm10139 | 66.38850078 | -2.89409298 | 4.66E-07 |
| Dnajc12 | 45.61121461 | -2.894273423 | 1.88E-08 |
| Tmem140 | 492.4092491 | -2.895545714 | 9.51E-22 |
| E130018N17Rik | 5.325017621 | -2.895780326 | 4.52E-02 |
| Pcdhb3 | 9.868502243 | -2.899352613 | 6.37E-04 |
| Synpo | 5453.723669 | -2.905978259 | 1.90E-61 |
| Mest | 25.66506474 | -2.908863345 | 1.84E-07 |
| Ociad2 | 337.1734645 | -2.90984158 | 6.81E-45 |
| 4933413J09Rik | 6.786959589 | -2.910763567 | 1.08E-02 |
| Slc35g2 | 9.457983186 | -2.914559675 | 3.88E-03 |
| Ccbe1 | 1756.029322 | -2.918656417 | 2.60E-35 |

| | | | |
|---|---|---|---|
| Trip11 | 1539.402913 | -2.918723086 | 1.06E-17 |
| mt-Tc | 174.937795 | -2.919141364 | 1.72E-12 |
| Nxph3 | 56.41829472 | -2.92036887 | 9.52E-12 |
| Polm | 3037.083453 | -2.922046501 | 3.19E-98 |
| Gm11537 | 9.445219704 | -2.923493624 | 4.12E-03 |
| NA | 24.57661118 | -2.925382757 | 5.96E-06 |
| Gm16209 | 7.103626664 | -2.926675465 | 1.88E-02 |
| Hddc3 | 237.1640719 | -2.92921656 | 3.64E-28 |
| Ehd3 | 485.4889265 | -2.931797919 | 7.00E-26 |
| Pcdhb16 | 566.0208578 | -2.931928191 | 1.19E-57 |
| Gas6 | 3563.445454 | -2.933077119 | 4.13E-25 |
| Gm20692 | 7.4490656 | -2.94201164 | 7.61E-03 |
| Omd | 64.4193638 | -2.945305027 | 1.63E-06 |
| Gm14410 | 104.8186658 | -2.946822935 | 4.94E-13 |
| Chsy1 | 1508.249314 | -2.946864283 | 8.61E-113 |
| Elovl2 | 19.10085203 | -2.947483572 | 5.57E-04 |
| Adamts16 | 30.27965716 | -2.950984641 | 1.90E-05 |
| Gm3298 | 5.144971386 | -2.952212954 | 4.44E-02 |
| Tnfaip6 | 646.6209697 | -2.954431987 | 1.54E-34 |
| Rspo2 | 7.266487856 | -2.959317091 | 2.41E-03 |
| 9330161L09Rik | 7.055396606 | -2.960269935 | 1.02E-02 |
| Zfp40 | 305.3181922 | -2.96874581 | 5.53E-09 |
| A430035B10Rik | 7.243714529 | -2.971048049 | 2.04E-03 |
| Rhoj | 1534.443716 | -2.971708364 | 6.68E-58 |
| NA | 14.79549638 | -2.972137013 | 5.32E-05 |
| Osr2 | 352.6919725 | -2.975992717 | 2.44E-16 |
| Elovl6 | 3066.198611 | -2.977571038 | 2.61E-09 |
| Gm17055 | 35.10683914 | -2.982641476 | 6.31E-10 |
| Gpr75 | 16.08289918 | -2.985658136 | 2.17E-05 |
| Ttc39a | 153.8251168 | -2.990279813 | 7.86E-15 |
| Rragd | 324.476639 | -2.990524243 | 1.27E-14 |
| Gm14412 | 35.41472427 | -2.991230285 | 1.32E-05 |
| 4930570G19Rik | 12.97709924 | -2.993202216 | 4.21E-03 |
| Gm23455 | 33.12231138 | -2.995076739 | 1.46E-05 |
| Slc23a4 | 14.09889523 | -2.996505156 | 6.38E-04 |
| Pmp22 | 6001.44192 | -2.997023544 | 2.48E-63 |
| Acot11 | 161.2842215 | -2.998012523 | 1.10E-10 |
| Bbs1 | 345.4501395 | -2.998162985 | 4.54E-50 |
| Ankrd35 | 13.53436989 | -2.999214266 | 5.20E-04 |
| Capn6 | 8375.626269 | -3.000023307 | 2.91E-40 |
| Zfp607a | 91.92954207 | -3.002486738 | 1.86E-18 |
| Gm22249 | 9.956659232 | -3.004949599 | 4.61E-03 |
| Gm12219 | 9.137256872 | -3.005306566 | 8.44E-03 |
| Arxes2 | 825.7504101 | -3.006761469 | 2.28E-29 |
| Gm22935 | 8.206150922 | -3.00959463 | 9.91E-03 |
| Man1c1 | 553.3265867 | -3.009870986 | 6.38E-66 |
| Gxylt2 | 1431.643095 | -3.012887448 | 1.74E-29 |

| | | | |
|---|---|---|---|
| 4933431K23Rik | 11.94158742 | -3.015043709 | 4.15E-04 |
| Cdo1 | 5428.454521 | -3.015778068 | 5.12E-21 |
| NA | 48.41532146 | -3.016941938 | 8.36E-10 |
| Zfp975 | 158.7501867 | -3.020333545 | 1.08E-13 |
| Agt | 230.4174338 | -3.021383106 | 3.22E-11 |
| Gm25829 | 10.54213017 | -3.022179374 | 1.74E-02 |
| Tmem98 | 1290.414908 | -3.026916675 | 1.10E-94 |
| Gm11261 | 171.9841549 | -3.035610502 | 2.88E-14 |
| Gm11766 | 15.91984041 | -3.037838834 | 4.86E-05 |
| Adgrl3 | 128.3042568 | -3.040099406 | 1.24E-15 |
| Gm26840 | 19.51410435 | -3.04024947 | 3.75E-05 |
| Smco3 | 15.58925897 | -3.042298517 | 8.42E-04 |
| Slc30a1 | 484.7571357 | -3.043936575 | 8.80E-15 |
| Setmar | 64.67531696 | -3.047408487 | 2.12E-18 |
| Tbxa2r | 12.46534744 | -3.048881827 | 7.19E-03 |
| Gm16046 | 54.5878734 | -3.049895047 | 4.03E-07 |
| Lama4 | 3042.134624 | -3.050491287 | 1.56E-08 |
| Slc41a2 | 949.467359 | -3.05121918 | 3.77E-67 |
| Colec12 | 20930.91238 | -3.051435503 | 1.59E-113 |
| Shroom2 | 8.787730123 | -3.058365894 | 2.06E-03 |
| Mpp4 | 6.753419752 | -3.05890905 | 1.42E-02 |
| Adcy7 | 42.46266469 | -3.059222891 | 5.08E-08 |
| Gm14409 | 23.87494513 | -3.060183927 | 3.73E-03 |
| Tek | 47.78473353 | -3.060565475 | 2.68E-09 |
| Abca9 | 166.1965339 | -3.06349354 | 7.60E-13 |
| Slc12a5 | 95.98187009 | -3.065086831 | 1.50E-11 |
| Paqr8 | 1206.296549 | -3.067767879 | 1.22E-42 |
| Dpep1 | 950.7647622 | -3.067973156 | 9.25E-17 |
| NA | 85.90070398 | -3.068163141 | 6.95E-16 |
| Grem2 | 1174.534392 | -3.070120265 | 1.25E-32 |
| Prr36 | 21.45874571 | -3.073010221 | 5.99E-07 |
| Pik3r1 | 5071.593751 | -3.074505543 | 3.95E-61 |
| Unc93a2 | 67.49710805 | -3.075372157 | 6.79E-05 |
| Dleu2 | 153.9845958 | -3.075806272 | 3.13E-23 |
| Gm10220 | 11.21547464 | -3.077069646 | 8.13E-03 |
| Gm10330 | 164.1859516 | -3.077527211 | 2.66E-18 |
| Pcdhb6 | 173.737983 | -3.079793849 | 5.75E-29 |
| Fgf7 | 4782.992174 | -3.084769716 | 8.90E-38 |
| Gm20493 | 7.930003298 | -3.086280787 | 4.92E-03 |
| Lrrd1 | 7.978361091 | -3.086325591 | 1.46E-02 |
| Dkk3 | 1119.94565 | -3.086654774 | 6.72E-35 |
| Rassf4 | 19.56252476 | -3.089368847 | 1.27E-06 |
| Gm16574 | 9.62552461 | -3.090550056 | 1.65E-02 |
| Dhrs4 | 481.3038845 | -3.092072526 | 3.66E-43 |
| Gm14322 | 342.7270789 | -3.095484278 | 7.70E-42 |
| Htra3 | 82.77514087 | -3.095636575 | 3.74E-14 |
| Prl2c3 | 113.2380744 | -3.096872413 | 1.29E-03 |

| | | | |
|---|---|---|---|
| Ccdc74a | 117.2288527 | -3.098043854 | 4.78E-17 |
| Zfp968 | 6.045798492 | -3.098661729 | 3.67E-02 |
| NA | 21.29929916 | -3.101740359 | 6.10E-05 |
| Gpc3 | 11.30597956 | -3.101998422 | 1.84E-03 |
| Adam11 | 158.3843609 | -3.10209096 | 7.72E-15 |
| 2810408A11Rik | 84.21173571 | -3.103304027 | 6.74E-19 |
| Apba1 | 301.8235088 | -3.103312276 | 1.36E-30 |
| Elac1 | 354.5344417 | -3.109988478 | 3.82E-41 |
| Cldn20 | 7.614908487 | -3.113800185 | 2.99E-03 |
| Slc16a9 | 86.65457978 | -3.114492495 | 2.09E-13 |
| Ccl17 | 8.136382902 | -3.117575101 | 6.20E-03 |
| Pcdhb14 | 233.1268323 | -3.118850621 | 3.40E-33 |
| Gm11816 | 245.2771472 | -3.120972699 | 1.13E-19 |
| Il16 | 43.63478451 | -3.126849209 | 9.80E-09 |
| C1s2 | 1092.038141 | -3.130594336 | 7.84E-24 |
| Tiam2 | 1483.526478 | -3.13096286 | 3.11E-41 |
| Tmem268 | 2601.301169 | -3.133924197 | 1.25E-52 |
| Cdcp1 | 16.20798329 | -3.134498186 | 8.70E-05 |
| Zfp113 | 518.8178997 | -3.140667616 | 2.16E-17 |
| Gm9903 | 6.713650102 | -3.143239912 | 1.66E-02 |
| Itgb8 | 478.0701506 | -3.148441424 | 5.95E-35 |
| Rbm15b | 151.7920532 | -3.149439589 | 4.47E-05 |
| Chrdl1 | 9.252815843 | -3.150537229 | 5.44E-04 |
| Rnf112 | 13.03420123 | -3.150862283 | 3.17E-02 |
| Snord69 | 17.3634094 | -3.154371172 | 2.64E-05 |
| Irf6 | 5.620115735 | -3.158674039 | 2.55E-02 |
| Plp1 | 415.718745 | -3.158684259 | 1.06E-15 |
| Flvcr2 | 5.065362858 | -3.159687392 | 2.27E-02 |
| Ypel2 | 260.1805721 | -3.163614402 | 8.09E-26 |
| NA | 35.3883632 | -3.163690797 | 2.37E-11 |
| Gm26645 | 20.70438726 | -3.165465188 | 1.12E-06 |
| mt-Ty | 79.47743535 | -3.165731965 | 4.30E-11 |
| Gm15998 | 16.83269665 | -3.168964229 | 6.94E-06 |
| Kbtbd8 | 94.49187229 | -3.17168645 | 1.35E-08 |
| Gm10254 | 21.60174155 | -3.172716044 | 1.69E-03 |
| Gm27509 | 7.270988291 | -3.173395339 | 2.60E-02 |
| C1s1 | 15631.97993 | -3.179040346 | 5.44E-31 |
| Sesn3 | 451.770964 | -3.179323705 | 5.65E-51 |
| Lgi4 | 6.261486722 | -3.181271883 | 4.20E-02 |
| Rab27a | 205.8409117 | -3.181564445 | 1.24E-25 |
| 4921534H16Rik | 7.363642845 | -3.182304833 | 1.98E-02 |
| Cstad | 40.98467497 | -3.183734497 | 2.24E-09 |
| AY512931 | 27.67092346 | -3.184149016 | 4.19E-04 |
| Prrx2 | 980.971953 | -3.187635402 | 8.47E-24 |
| Col27a1 | 963.2021591 | -3.189857004 | 1.52E-45 |
| Creg2 | 7.409406499 | -3.191318383 | 1.65E-02 |
| Aoc3 | 1354.327432 | -3.194327464 | 2.97E-28 |

| | | | |
|---|---|---|---|
| Gm12591 | 9.177330515 | -3.197489913 | 3.15E-03 |
| Slc6a7 | 36.75567192 | -3.199659276 | 1.27E-12 |
| Klf2 | 230.8305466 | -3.200208447 | 1.74E-14 |
| Gm6594 | 10.76466459 | -3.20201919 | 1.92E-03 |
| Ccdc57 | 258.3154069 | -3.209551426 | 3.21E-32 |
| Zscan26 | 1306.932827 | -3.209631953 | 8.76E-36 |
| Gm10544 | 12.32905783 | -3.211025753 | 1.10E-04 |
| Fjx1 | 266.2076356 | -3.217094761 | 6.73E-50 |
| D630036H23Rik | 6.049307055 | -3.221343635 | 1.35E-02 |
| NA | 97.40321005 | -3.223601833 | 1.10E-13 |
| Tmem216 | 308.3576184 | -3.224349079 | 2.69E-60 |
| Gm27321 | 8.927264981 | -3.224557961 | 3.33E-03 |
| Gm26829 | 19.23961468 | -3.227117224 | 1.82E-07 |
| Gbp7 | 50.27437464 | -3.232248971 | 2.86E-11 |
| 5830432E09Rik | 23.64403577 | -3.233619568 | 2.87E-07 |
| Gm12963 | 65.53533855 | -3.236142006 | 9.51E-15 |
| Gm43302 | 33.491425 | -3.236928319 | 8.02E-08 |
| Gm15958 | 23.51822755 | -3.23705397 | 1.86E-04 |
| Galnt4 | 1123.766541 | -3.244569875 | 4.14E-54 |
| Il6ra | 1020.18581 | -3.250107441 | 9.39E-110 |
| 1700006J14Rik | 16.77111565 | -3.254184346 | 3.78E-04 |
| Tinag | 23.1948581 | -3.258479047 | 2.42E-03 |
| Pcdh7 | 730.661321 | -3.260558858 | 3.83E-51 |
| Rin2 | 6488.596361 | -3.262002991 | 2.32E-57 |
| Tshz1 | 831.544133 | -3.266536246 | 3.84E-55 |
| Selenop | 9438.313738 | -3.266782452 | 8.42E-28 |
| Rarres2 | 54.65008237 | -3.273934499 | 6.15E-11 |
| NA | 14.20874993 | -3.274966641 | 8.01E-03 |
| Efcab9 | 37.28015627 | -3.279199814 | 1.05E-09 |
| Ralgds | 5291.738528 | -3.282012314 | 1.40E-53 |
| Akr1c18 | 34.14361785 | -3.285088117 | 4.79E-04 |
| AW551984 | 1346.712062 | -3.286453909 | 2.72E-24 |
| Rnf144a | 1573.389355 | -3.28838015 | 1.33E-22 |
| Ephb6 | 218.4738707 | -3.291136177 | 1.75E-16 |
| Cep83os | 409.9112091 | -3.293742543 | 6.42E-47 |
| Gm9869 | 51.84992159 | -3.300590704 | 2.17E-12 |
| Myl9 | 237.5650558 | -3.301398524 | 1.18E-22 |
| Mlana | 173.3382716 | -3.303775908 | 1.23E-08 |
| Myo7a | 1099.229425 | -3.304521672 | 2.95E-58 |
| 2010315B03Rik | 196.2225042 | -3.307115544 | 2.29E-15 |
| Srpx2 | 547.7024361 | -3.30844131 | 7.09E-09 |
| Gm3558 | 41.23278577 | -3.311432766 | 5.78E-06 |
| E230015B07Rik | 16.39336344 | -3.312543027 | 1.45E-05 |
| Pappa2 | 2623.486677 | -3.316114436 | 3.01E-50 |
| Zfp14 | 70.96744208 | -3.316917757 | 4.49E-20 |
| Ckb | 5.205320121 | -3.32361218 | 1.99E-02 |
| Gm27517 | 14.62030774 | -3.328361297 | 6.91E-03 |

| | | | |
|---|---|---|---|
| Tcim | 182.9808592 | -3.329953029 | 1.64E-23 |
| Phf2os1 | 5.264763034 | -3.335817865 | 1.89E-02 |
| Olfml2b | 2018.256671 | -3.336098096 | 2.06E-44 |
| 2510009E07Rik | 671.3141264 | -3.34108973 | 8.02E-55 |
| NA | 10.53704637 | -3.341472197 | 1.27E-02 |
| Il11ra2 | 54.33013918 | -3.342453782 | 3.31E-11 |
| Gm25500 | 6.528691617 | -3.342593972 | 2.67E-02 |
| Clec3b | 429.6188855 | -3.343290549 | 9.44E-07 |
| Clvs1 | 109.3908839 | -3.344440623 | 3.82E-10 |
| Tet1 | 358.3477956 | -3.345684138 | 3.10E-25 |
| Trnp1 | 989.6323959 | -3.348964007 | 6.45E-61 |
| Crocc2 | 99.75445273 | -3.349217513 | 2.12E-09 |
| Flrt1 | 7.038971897 | -3.351637465 | 4.34E-02 |
| Pcdhb19 | 163.5809958 | -3.353143042 | 2.33E-35 |
| Ccl8 | 242.947781 | -3.356023885 | 2.11E-13 |
| 5730507C01Rik | 114.2588328 | -3.361658504 | 1.43E-22 |
| Plxna4 | 92.75828692 | -3.365734383 | 2.80E-08 |
| Pcdhb20 | 156.4655691 | -3.366195543 | 3.00E-33 |
| Ifi202b | 2370.951925 | -3.370598322 | 9.08E-13 |
| Samd9l | 629.3467263 | -3.370923894 | 1.79E-20 |
| Ntrk2 | 6.767507947 | -3.371294496 | 1.31E-02 |
| Fkrp | 819.6783063 | -3.37165086 | 1.04E-49 |
| Ccl11 | 14.11907563 | -3.372530826 | 4.54E-04 |
| Rhov | 20.06302063 | -3.373517022 | 2.38E-05 |
| Col22a1 | 44.64731885 | -3.382429789 | 2.22E-10 |
| Tug1 | 5617.309804 | -3.385409409 | 2.50E-39 |
| Rab3il1 | 19.55437173 | -3.38550759 | 4.88E-05 |
| Aox3 | 125.6643492 | -3.388501285 | 3.43E-07 |
| Pcdhb7 | 281.6238328 | -3.392644467 | 2.19E-46 |
| Mansc4 | 11.73314333 | -3.39465005 | 4.38E-05 |
| Gm4631 | 911.8427902 | -3.395332894 | 1.42E-16 |
| A330023F24Rik | 61.15415772 | -3.395508103 | 3.50E-12 |
| Gm2381 | 276.5731117 | -3.396877861 | 1.17E-13 |
| A230028O05Rik | 24.80296605 | -3.401308528 | 7.13E-07 |
| Zfp125 | 122.523314 | -3.406472864 | 1.65E-23 |
| St6gal1 | 1757.152116 | -3.406816449 | 1.22E-57 |
| Zkscan16 | 70.2059796 | -3.407169663 | 2.85E-15 |
| Ctsk | 939.5622255 | -3.407416366 | 1.73E-54 |
| Camk1d | 90.9566728 | -3.414113375 | 3.94E-10 |
| Tex9 | 290.3539929 | -3.418739295 | 5.79E-29 |
| Ssbp4 | 614.8805247 | -3.421912947 | 1.21E-54 |
| Zcchc18 | 5.88133833 | -3.423245828 | 7.93E-03 |
| Sned1 | 9956.936022 | -3.42660387 | 9.86E-40 |
| Eef1akmt2 | 275.4335541 | -3.43107891 | 2.39E-15 |
| Gm12590 | 87.10131386 | -3.435590385 | 4.48E-17 |
| Gm12480 | 7.167525394 | -3.437687558 | 8.80E-03 |
| Gm20633 | 69.1003007 | -3.441252796 | 1.85E-13 |

| | | | |
|---|---|---|---|
| Mdp1 | 559.3208089 | -3.445130063 | 2.10E-70 |
| Gm14569 | 7.156873918 | -3.447053069 | 6.73E-04 |
| 2700069I18Rik | 246.3421382 | -3.447252701 | 1.60E-20 |
| Cd300lb | 309.9653894 | -3.448827414 | 1.31E-19 |
| Gm11992 | 6.530987242 | -3.452758544 | 7.24E-03 |
| Gm14421 | 160.9356899 | -3.45561066 | 2.40E-19 |
| Xk | 7.137264591 | -3.456969973 | 8.97E-04 |
| Cx3cl1 | 722.2112652 | -3.465339824 | 9.52E-45 |
| Csf2rb2 | 11.60028118 | -3.46686287 | 2.25E-04 |
| Rhpn1 | 32.93132459 | -3.479434212 | 2.30E-12 |
| 1700071M16Rik | 6.45197741 | -3.48026943 | 1.78E-02 |
| Ccdc148 | 58.82159294 | -3.481463098 | 1.47E-07 |
| Gm7361 | 115.296168 | -3.482020063 | 3.08E-13 |
| Wdr63 | 7.18728154 | -3.490319479 | 1.20E-03 |
| Rbp1 | 2005.260463 | -3.492302882 | 1.88E-93 |
| Tnfrsf11b | 121.1051387 | -3.501056673 | 2.03E-11 |
| Etv5 | 692.3569151 | -3.501428564 | 8.68E-39 |
| Jrk | 382.3720798 | -3.5082426 | 2.02E-47 |
| Carmil3 | 108.3115967 | -3.51105167 | 4.93E-18 |
| Aox1 | 478.0590306 | -3.512151383 | 1.52E-52 |
| Bag5 | 1327.489717 | -3.512693358 | 3.93E-57 |
| B3galt4 | 33.62953106 | -3.514016586 | 3.89E-10 |
| Cfap161 | 19.83244907 | -3.523978811 | 7.41E-07 |
| Slc2a13 | 76.15910103 | -3.529460553 | 2.40E-16 |
| Mettl18 | 120.06686 | -3.52975705 | 1.56E-22 |
| Gria1 | 276.9010458 | -3.530205797 | 1.76E-17 |
| Tha1 | 19.03802371 | -3.53031381 | 1.97E-06 |
| Maml3 | 728.8511573 | -3.530549542 | 5.89E-61 |
| Gm9923 | 49.22259011 | -3.531257056 | 6.79E-07 |
| Klhl4 | 12.53402334 | -3.534374809 | 1.62E-03 |
| Itgbl1 | 4703.422305 | -3.53822716 | 1.38E-36 |
| Tnk1 | 13.45238113 | -3.548381081 | 2.89E-02 |
| 9630013D21Rik | 65.25458142 | -3.549089266 | 5.88E-08 |
| Spag8 | 8.074328966 | -3.552519843 | 3.25E-03 |
| Gm16845 | 68.96377463 | -3.554262396 | 1.53E-17 |
| Tspan18 | 194.9774176 | -3.558186375 | 1.30E-21 |
| Ranbp3l | 21.28438069 | -3.571257906 | 4.51E-02 |
| E130102H24Rik | 90.14504892 | -3.57285136 | 1.28E-12 |
| Rtl5 | 11.39763817 | -3.576871698 | 1.16E-02 |
| Gm26775 | 8.002624879 | -3.577721301 | 5.03E-03 |
| Plxnc1 | 5.257759957 | -3.577981793 | 1.50E-02 |
| Lpar4 | 27.03383507 | -3.580637631 | 1.45E-05 |
| Gm14440 | 165.1266027 | -3.580651243 | 2.68E-08 |
| Mylip | 3518.788342 | -3.581376198 | 5.90E-59 |
| 1700024B05Rik | 11.97341044 | -3.586962279 | 1.20E-03 |
| 5930403N24Rik | 408.5526991 | -3.587483266 | 8.33E-32 |
| Zfhx4 | 9921.32866 | -3.591327065 | 1.59E-36 |

| | | | |
|---|---|---|---|
| Klrb1a | 6.733824506 | -3.592689797 | 6.41E-04 |
| Mitf | 1042.401336 | -3.594640742 | 1.48E-52 |
| Pcdhb9 | 216.2214447 | -3.602237442 | 1.59E-39 |
| Gm3173 | 12.70900116 | -3.60280796 | 4.62E-04 |
| Spry4 | 239.4302629 | -3.610480333 | 2.26E-30 |
| Mrgpre | 112.8488705 | -3.617712552 | 3.92E-15 |
| Dnase1 | 109.3277492 | -3.624781053 | 7.25E-20 |
| Rep15 | 19.80358759 | -3.645054572 | 1.64E-07 |
| Plcb1 | 374.5524231 | -3.64930515 | 3.95E-54 |
| Aebp1 | 72201.79584 | -3.649571639 | 3.92E-97 |
| Spaar | 6.692330531 | -3.661694224 | 1.02E-02 |
| Catip | 21.30652682 | -3.6633177 | 1.65E-07 |
| Jade2 | 1489.987394 | -3.671486202 | 6.48E-181 |
| Mir17hg | 152.1074557 | -3.67764529 | 1.55E-10 |
| Mt1 | 6812.419424 | -3.681035708 | 3.80E-23 |
| Traf3ip1 | 465.2906491 | -3.681795288 | 2.98E-47 |
| Cel | 7.378356627 | -3.682483695 | 1.32E-02 |
| Coro2b | 6.45669062 | -3.683177891 | 1.26E-02 |
| Inmt | 199.4518932 | -3.693434508 | 1.52E-10 |
| Klhl41 | 8.79281984 | -3.694665993 | 3.44E-03 |
| Dlgap2 | 221.2029046 | -3.695251057 | 2.80E-37 |
| Gm26682 | 24.22475906 | -3.697070353 | 7.23E-08 |
| Rhbdl3 | 44.7420194 | -3.698075384 | 7.14E-08 |
| Zfp108 | 34.41460638 | -3.703990624 | 2.85E-11 |
| Sycp2 | 67.3916275 | -3.704578212 | 1.27E-08 |
| Zkscan8 | 842.0575177 | -3.709088763 | 3.70E-30 |
| Lmnb2 | 823.7442095 | -3.717092179 | 7.02E-18 |
| Fam13c | 1244.440345 | -3.726735813 | 1.49E-54 |
| Ubd | 22.06665641 | -3.727634855 | 2.61E-05 |
| Agtr2 | 1656.388944 | -3.728244392 | 1.23E-07 |
| Rpgrip1l | 758.612363 | -3.730548617 | 3.83E-44 |
| Gm3667 | 10.55630223 | -3.734882591 | 2.35E-03 |
| Pygo1 | 741.5294784 | -3.737895655 | 2.29E-70 |
| Pigh | 312.2573961 | -3.738985274 | 1.04E-54 |
| Timp4 | 80.55768285 | -3.752615617 | 4.19E-11 |
| Gm4890 | 10.73927302 | -3.754709824 | 3.41E-04 |
| Col15a1 | 10.29721222 | -3.755885535 | 1.06E-02 |
| Abi3 | 24.75257189 | -3.756363572 | 4.39E-03 |
| Mrnip | 63.17783418 | -3.759045175 | 9.38E-15 |
| 8430419K02Rik | 5.106795061 | -3.766834019 | 2.63E-02 |
| Gfap | 7.471407455 | -3.77780547 | 2.45E-02 |
| Gm10822 | 7.318685899 | -3.786829782 | 3.53E-03 |
| Rab36 | 271.0864376 | -3.790136915 | 3.20E-36 |
| Gm16701 | 13.96346115 | -3.793643411 | 1.33E-04 |
| Tmem200a | 180.2586998 | -3.808520352 | 8.83E-30 |
| Sult5a1 | 65.16037106 | -3.810229512 | 6.58E-12 |
| Gm5751 | 5.367922004 | -3.816051708 | 1.08E-02 |

| | | | |
|---|---|---|---|
| Scara5 | 982.0551509 | -3.823472702 | 1.23E-12 |
| Trim30d | 101.5243115 | -3.8255847 | 1.23E-11 |
| Zfp300 | 75.47022026 | -3.82760785 | 1.16E-14 |
| Enpp2 | 759.7675373 | -3.828234993 | 3.30E-40 |
| Prtg | 275.2554638 | -3.83538899 | 1.87E-39 |
| Pcdhb21 | 187.4813216 | -3.837835325 | 6.57E-41 |
| Adh1 | 110.5458724 | -3.842058201 | 3.08E-12 |
| Gm13091 | 17.69307921 | -3.845812977 | 5.95E-06 |
| C530008M17Rik | 222.325705 | -3.850223276 | 2.11E-21 |
| Scarletltr | 7.10325454 | -3.850323806 | 1.57E-02 |
| Pkd2 | 8695.457457 | -3.850446864 | 2.52E-78 |
| Trpm6 | 7.21072813 | -3.851065065 | 5.11E-04 |
| Shc2 | 1586.018871 | -3.862242811 | 8.58E-80 |
| Gm10390 | 11.7170738 | -3.87246734 | 1.30E-02 |
| Sytl2 | 158.0277612 | -3.884776319 | 8.45E-16 |
| Cxxc4 | 5.811367535 | -3.885091076 | 7.80E-03 |
| H1fx | 45.98158594 | -3.888287141 | 2.24E-12 |
| Lyve1 | 9.828957395 | -3.89228198 | 3.38E-04 |
| P2rx6 | 61.83965373 | -3.903027492 | 3.30E-16 |
| Tgfbr1 | 1819.850756 | -3.906981982 | 2.00E-52 |
| Angptl4 | 1038.167366 | -3.912195373 | 4.58E-41 |
| Thbs2 | 76114.02438 | -3.916252512 | 7.13E-48 |
| Adgrl1 | 1938.183396 | -3.922001281 | 1.57E-76 |
| Mettl5os | 7.059956268 | -3.923433993 | 6.12E-03 |
| NA | 5.566068501 | -3.931715441 | 1.15E-02 |
| Zfp709 | 209.956385 | -3.936912202 | 2.44E-31 |
| Tmem252 | 7.498715846 | -3.943949011 | 2.26E-03 |
| Gm3629 | 23.23421655 | -3.955606213 | 5.50E-05 |
| Dpf1 | 35.92731429 | -3.955753387 | 7.57E-09 |
| Gm27845 | 5.357455858 | -3.970115831 | 1.87E-02 |
| Gm8126 | 8.405641104 | -3.979379204 | 4.66E-03 |
| Zfp704 | 652.3489939 | -3.984334616 | 8.27E-47 |
| B930094E09Rik | 8.478385925 | -3.984847631 | 1.35E-03 |
| Gm15911 | 6.812046895 | -3.986402232 | 5.84E-03 |
| Cd200 | 115.5327228 | -3.987443079 | 2.51E-16 |
| Gm10701 | 7.826260597 | -3.993848983 | 1.64E-03 |
| 4930426L09Rik | 16.02716752 | -3.995005667 | 3.40E-05 |
| Gm21847 | 17.84747835 | -3.997225237 | 5.16E-04 |
| Arhgap6 | 33.54524912 | -4.003133962 | 7.40E-06 |
| Arhgap8 | 18.23760369 | -4.003431607 | 5.81E-06 |
| Lrrc4b | 17.38360114 | -4.007570087 | 2.63E-03 |
| Ppl | 22.69188888 | -4.027867161 | 6.04E-06 |
| Hlf | 720.1008505 | -4.037229505 | 1.41E-12 |
| Gm26923 | 29.45211618 | -4.037635957 | 4.70E-06 |
| Prob1 | 109.771206 | -4.043380388 | 2.11E-21 |
| Sec16b | 930.6328352 | -4.047830235 | 6.91E-57 |
| Gm14387 | 74.45587935 | -4.047934558 | 9.36E-13 |

| | | | |
|---|---|---|---|
| Mertk | 609.7724486 | -4.055104378 | 2.87E-69 |
| Gas1 | 20046.86774 | -4.056310271 | 1.44E-102 |
| BC106179 | 76.11099145 | -4.056991335 | 8.93E-20 |
| Pknox2 | 149.7278646 | -4.058018975 | 1.37E-25 |
| Slc40a1 | 380.186994 | -4.059602039 | 4.45E-28 |
| Cd33 | 68.59571646 | -4.060342808 | 1.62E-16 |
| Caps2 | 114.6304859 | -4.06122583 | 1.28E-21 |
| 2610035D17Rik | 96.21859554 | -4.069328878 | 8.58E-16 |
| Dhx58os | 12.54913426 | -4.071121442 | 5.13E-04 |
| 3222401L13Rik | 7.347962629 | -4.079585327 | 1.13E-03 |
| Kctd15 | 882.2490959 | -4.08142942 | 1.35E-49 |
| Ogn | 19778.96094 | -4.091757738 | 1.09E-14 |
| Grid1 | 22.69881994 | -4.093853933 | 6.16E-07 |
| Zfp759 | 126.1544654 | -4.100791908 | 1.18E-20 |
| Lum | 974.5741517 | -4.101129518 | 1.51E-12 |
| 1700125H03Rik | 12.54704266 | -4.111210848 | 5.61E-05 |
| Zscan30 | 5.346686754 | -4.111821244 | 4.79E-03 |
| Chst2 | 135.1732714 | -4.127742597 | 3.03E-24 |
| Ang5 | 6.410110454 | -4.130000801 | 9.44E-03 |
| NA | 14.33436608 | -4.132197478 | 3.49E-06 |
| Ccdc65 | 26.87887487 | -4.132293845 | 4.25E-08 |
| Trim66 | 32.12966506 | -4.13691161 | 5.39E-07 |
| Itga8 | 1363.794526 | -4.14489594 | 8.81E-42 |
| Plagl1 | 737.429685 | -4.148099571 | 3.21E-40 |
| Gm26763 | 7.555053541 | -4.149408776 | 3.79E-03 |
| Scube3 | 14.00198406 | -4.157681307 | 1.96E-04 |
| Gm26680 | 7.197796939 | -4.183752434 | 2.50E-02 |
| Runx2os1 | 7.032015264 | -4.186230041 | 4.08E-03 |
| Sult1a1 | 55.76000516 | -4.191867717 | 3.72E-07 |
| Gm3194 | 17.06507034 | -4.194383924 | 3.48E-05 |
| Serpina3n | 668.2845824 | -4.194760593 | 1.94E-34 |
| Prl2c1 | 5.79024499 | -4.197041642 | 1.12E-02 |
| Aspn | 5188.83664 | -4.203546797 | 1.36E-12 |
| NA | 45.07335818 | -4.220151614 | 5.85E-09 |
| Gpnmb | 3590.839261 | -4.222988908 | 2.42E-43 |
| NA | 9.225849333 | -4.225854306 | 5.11E-03 |
| Rnf152 | 9.581791921 | -4.233094863 | 4.57E-04 |
| Plppr3 | 487.5192114 | -4.23500593 | 1.33E-35 |
| Gm3411 | 95.97843402 | -4.241012072 | 4.54E-11 |
| Galnt9 | 9.989789218 | -4.246834257 | 8.26E-05 |
| Apod | 983.8203818 | -4.252743788 | 8.32E-15 |
| Tmem100 | 6.230469592 | -4.253998597 | 2.38E-03 |
| Etv1 | 303.4475497 | -4.262629904 | 2.71E-25 |
| Matn2 | 13519.17868 | -4.268367598 | 2.12E-83 |
| Csgalnact1 | 1244.091072 | -4.272242501 | 6.86E-93 |
| Tmem119 | 3163.057139 | -4.296258954 | 9.04E-101 |
| Ccdc152 | 31.66243982 | -4.317769513 | 2.18E-08 |

| | | | |
|---|---|---|---|
| Pcdhb12 | 18.19238944 | -4.324276504 | 5.80E-07 |
| Steap3 | 17632.6646 | -4.324498105 | 1.65E-62 |
| Proz | 26.7661844 | -4.341717571 | 1.32E-08 |
| Egr1 | 4443.449226 | -4.356999626 | 1.17E-18 |
| Foxred2 | 88.56860549 | -4.371760917 | 8.63E-16 |
| Dpt | 3000.84147 | -4.375775899 | 1.70E-14 |
| Opcml | 366.9743187 | -4.37656347 | 2.13E-39 |
| Cxcl12 | 14907.90677 | -4.379710401 | 1.75E-47 |
| Gm14401 | 16.81454501 | -4.380582702 | 1.59E-04 |
| Gm22119 | 5.404121612 | -4.384187572 | 2.97E-03 |
| Slc24a1 | 28.67118619 | -4.38572725 | 3.22E-10 |
| Adam1a | 40.12227664 | -4.387086648 | 2.28E-10 |
| Osbpl6 | 120.9530659 | -4.394137145 | 4.52E-20 |
| 3110070M22Rik | 24.64606842 | -4.398974182 | 1.32E-07 |
| Gm14445 | 7.644904696 | -4.41098126 | 1.69E-03 |
| Ugt1a7c | 392.0152248 | -4.418607374 | 7.43E-37 |
| Cadm4 | 7.869765111 | -4.426560475 | 2.14E-03 |
| Fam78b | 275.8695293 | -4.432061205 | 3.42E-45 |
| Zfp965 | 8.037022261 | -4.439269412 | 6.54E-03 |
| Cldn19 | 7.792817537 | -4.442844789 | 7.89E-04 |
| Dok3 | 10.13367908 | -4.479886252 | 1.50E-03 |
| Krt13 | 6.738745077 | -4.484868135 | 3.66E-03 |
| Dyx1c1 | 41.08029909 | -4.490107785 | 1.18E-13 |
| Npff | 17.81950082 | -4.495033562 | 3.59E-06 |
| Csf2rb | 20.90762805 | -4.50090999 | 3.26E-05 |
| Gm2464 | 5.686631402 | -4.532404722 | 2.57E-03 |
| Nacc2 | 1412.305969 | -4.54196434 | 2.92E-127 |
| 10-Mar | 64.82310953 | -4.574851364 | 2.59E-15 |
| Gm12105 | 5.951807365 | -4.578185942 | 2.70E-03 |
| Osbp2 | 32.12984721 | -4.581327839 | 1.51E-06 |
| Gm14411 | 5.065863336 | -4.589589852 | 7.81E-03 |
| Zcchc3 | 416.4233873 | -4.593086377 | 3.45E-43 |
| C130071C03Rik | 75.01846086 | -4.608941831 | 4.75E-18 |
| Epha10 | 7.545852402 | -4.614259913 | 2.59E-04 |
| Gm15401 | 8.9244427 | -4.621739266 | 9.45E-04 |
| Plekhb1 | 22.62480202 | -4.630086806 | 2.45E-08 |
| Emb | 6148.231165 | -4.646572726 | 2.03E-140 |
| Fmr1os | 6.08347099 | -4.654232634 | 9.66E-04 |
| NA | 11.41496147 | -4.656704633 | 9.11E-05 |
| Syngr1 | 40.08116894 | -4.662610822 | 1.75E-07 |
| Gm26650 | 6.680380822 | -4.664617245 | 2.44E-03 |
| NA | 6.161689882 | -4.67729358 | 1.02E-03 |
| Vash1 | 29.13997177 | -4.683235755 | 1.40E-06 |
| Trhde | 38.6088199 | -4.722507319 | 1.28E-06 |
| Prkg2 | 2708.349054 | -4.724067707 | 9.74E-48 |
| Speer4d | 12.14569302 | -4.736891483 | 4.46E-03 |
| Mgp | 2186.4868 | -4.744773428 | 1.48E-49 |

| | | | |
|---|---|---|---|
| Bmper | 999.7225729 | -4.748948419 | 2.22E-49 |
| Cldn22 | 6.464303654 | -4.751944374 | 8.37E-04 |
| Sdc3 | 1488.655788 | -4.762800641 | 2.87E-98 |
| Zfp72 | 103.7844434 | -4.774246671 | 1.38E-15 |
| Ptn | 40530.58054 | -4.785144687 | 2.92E-26 |
| Htra4 | 6.102645879 | -4.801694972 | 2.40E-03 |
| Wfdc17 | 23.56673593 | -4.81277211 | 5.41E-06 |
| Mc2r | 13.56370526 | -4.833378602 | 8.82E-05 |
| Cldn2 | 11.0341259 | -4.834545656 | 3.07E-03 |
| 9430002A10Rik | 10.20621605 | -4.85635506 | 3.61E-04 |
| Maf | 987.6224961 | -4.857676472 | 1.14E-47 |
| Cabp7 | 8.452464321 | -4.871010103 | 7.14E-03 |
| C3 | 1005.759055 | -4.881773699 | 1.29E-33 |
| Epyc | 9.018814475 | -4.884816693 | 5.63E-04 |
| Has2 | 320.7677252 | -4.937774199 | 1.77E-45 |
| Ugt1a6b | 1056.631265 | -4.949004676 | 2.81E-114 |
| Nat8f6 | 9.726106815 | -4.949378045 | 1.04E-04 |
| Edil3 | 1365.668624 | -4.950725368 | 2.57E-55 |
| Sema7a | 84.82804028 | -4.953668592 | 4.92E-19 |
| NA | 20.64834532 | -4.958979636 | 1.40E-05 |
| Mirlet7b | 6.125468596 | -4.959492477 | 5.92E-04 |
| NA | 10.88839713 | -4.964681935 | 2.74E-04 |
| Tecta | 36.05411746 | -4.972031856 | 1.02E-10 |
| Gm4724 | 16.04273291 | -4.97411055 | 7.14E-04 |
| Tmem178b | 75.53419669 | -4.974626922 | 3.07E-11 |
| 4833422C13Rik | 31.02501048 | -4.978491598 | 1.33E-08 |
| Clmp | 2125.373568 | -4.987049489 | 7.12E-208 |
| Egr3 | 140.915746 | -5.009708738 | 1.14E-15 |
| Acpp | 8.017078266 | -5.011008531 | 1.53E-03 |
| Gm14002 | 28.28916881 | -5.023705889 | 3.64E-08 |
| Gm13610 | 5.013164379 | -5.024367651 | 3.54E-03 |
| Gm2366 | 40.57644542 | -5.024452148 | 5.90E-03 |
| Gm2026 | 125.1518551 | -5.029288168 | 7.47E-06 |
| Gm14406 | 5.570444499 | -5.041926114 | 1.28E-02 |
| Ear10 | 5.072886965 | -5.051269113 | 6.20E-03 |
| Ces1a | 829.2986385 | -5.077070346 | 3.54E-21 |
| Zfp977 | 12.83380859 | -5.087090485 | 1.64E-04 |
| Clec1a | 18.53247397 | -5.099273078 | 4.93E-05 |
| Klhl33 | 14.40315675 | -5.103815298 | 1.66E-04 |
| Dmp1 | 15.30594467 | -5.119240569 | 3.45E-05 |
| Gm15996 | 11.99713073 | -5.126511513 | 1.63E-04 |
| 4930519F16Rik | 13.44671808 | -5.129255463 | 3.83E-04 |
| E2f2 | 41.48521494 | -5.144859696 | 2.18E-05 |
| Endou | 5.25502425 | -5.158845641 | 2.77E-04 |
| Ryr3 | 5.624264745 | -5.16107398 | 3.38E-03 |
| Cela1 | 174.5106123 | -5.168313967 | 8.59E-31 |
| Sox9 | 2547.006272 | -5.170282988 | 5.34E-103 |

| | | | |
|---|---|---|---|
| Gm11767 | 9.219788467 | -5.179814258 | 1.18E-04 |
| Dlx4 | 6.030862526 | -5.193099394 | 1.82E-04 |
| Ssc5d | 1608.633546 | -5.19876116 | 7.45E-90 |
| Gpr88 | 11.30675403 | -5.203224854 | 6.25E-04 |
| Ccl2 | 2277.435037 | -5.20646257 | 1.24E-53 |
| Atoh8 | 455.8228685 | -5.224500217 | 2.49E-18 |
| Axin2 | 27.73513628 | -5.226059574 | 1.77E-11 |
| Ripply1 | 17.40893938 | -5.228487875 | 2.59E-04 |
| Nlrp10 | 17.4659925 | -5.234991056 | 3.34E-04 |
| NA | 7.364960641 | -5.235410522 | 1.59E-04 |
| Nmur1 | 13.54843957 | -5.247743725 | 9.17E-07 |
| Gm16596 | 16.02890155 | -5.258229714 | 1.08E-04 |
| 2700012I20Rik | 5.993940404 | -5.267864662 | 2.53E-04 |
| Gm20319 | 6.961059712 | -5.345389789 | 1.19E-04 |
| Fam135b | 222.7442954 | -5.347081363 | 2.49E-21 |
| Foxo6 | 14.26804265 | -5.368492938 | 7.59E-05 |
| Actr3b | 183.7091602 | -5.378045367 | 3.75E-28 |
| Lpcat2 | 612.0918981 | -5.395860773 | 8.10E-40 |
| Ednra | 698.5583548 | -5.415483606 | 5.02E-58 |
| Smoc2 | 5606.559969 | -5.420868771 | 2.67E-51 |
| Prl2c2 | 22898.70079 | -5.432163927 | 6.19E-29 |
| Fgf2 | 34.6754416 | -5.464873698 | 1.88E-05 |
| Gm11841 | 7.340654826 | -5.467785566 | 1.74E-04 |
| Angpt1 | 126.2034669 | -5.467803097 | 3.76E-16 |
| Gper1 | 5.596836191 | -5.478869971 | 2.41E-04 |
| Dcn | 41417.76249 | -5.481906516 | 7.36E-27 |
| Egfl6 | 11.47141831 | -5.505711579 | 1.06E-04 |
| Gm26587 | 44.99508691 | -5.506379327 | 3.76E-08 |
| Serpina3m | 6.914672563 | -5.512134108 | 2.12E-04 |
| Eln | 24.79874943 | -5.531938646 | 4.97E-05 |
| Pnldc1 | 9.570498215 | -5.535618883 | 3.72E-05 |
| Prtn3 | 13.25797432 | -5.538210411 | 7.51E-05 |
| Bcl2 | 622.3489031 | -5.547802729 | 3.55E-51 |
| Postn | 49906.43674 | -5.554972671 | 8.48E-31 |
| Prl2c5 | 68.59226915 | -5.555206742 | 1.06E-12 |
| Nat8f7 | 24.83797719 | -5.557112165 | 2.30E-09 |
| Galnt12 | 136.076518 | -5.559559223 | 2.68E-22 |
| NA | 18.24678081 | -5.559916849 | 1.66E-04 |
| Sfrp1 | 704.6605608 | -5.564757827 | 1.24E-70 |
| Nr1i3 | 116.7264137 | -5.567517938 | 2.75E-04 |
| Slc25a23 | 164.104295 | -5.577988065 | 4.46E-18 |
| Slc12a1 | 5.052709889 | -5.586358868 | 1.74E-04 |
| Samd11 | 155.349818 | -5.602262497 | 9.20E-03 |
| Cytip | 32.07785982 | -5.647038439 | 4.99E-07 |
| Fam180a | 160.3125889 | -5.675397331 | 1.92E-27 |
| Gm21671 | 8.561994867 | -5.702864795 | 3.05E-04 |
| Mmp8 | 5.245523941 | -5.703184952 | 8.51E-05 |

| | | | |
|---|---|---|---|
| Gm12216 | 7.974735615 | -5.70414633 | 3.62E-05 |
| Gm5799 | 11.406214 | -5.745781779 | 2.30E-05 |
| Tle2 | 1136.122372 | -5.798150101 | 2.45E-69 |
| Dnmt3b | 153.2781329 | -5.825079538 | 5.04E-19 |
| Ggt5 | 37.76480545 | -5.838223037 | 2.02E-04 |
| Ugt1a6a | 1314.632066 | -5.891964248 | 2.90E-66 |
| Ddx60 | 6.938250587 | -5.902693418 | 2.34E-04 |
| Kcnk3 | 92.21429487 | -5.912232663 | 1.12E-15 |
| Lrrn3 | 42.94924044 | -5.941050882 | 2.52E-07 |
| Dnm3os | 3686.208543 | -5.946498753 | 2.53E-39 |
| Cldn1 | 6.784948916 | -6.018931711 | 3.59E-05 |
| Capns2 | 41.29573201 | -6.032968827 | 4.82E-10 |
| B930082K07Rik | 10.59285588 | -6.043266119 | 5.26E-06 |
| 4930589L23Rik | 9.662169631 | -6.067325255 | 3.17E-05 |
| Gm11638 | 6.989282699 | -6.073457567 | 8.83E-06 |
| Prokr1 | 338.3992619 | -6.09496537 | 9.54E-44 |
| Gm12263 | 29.33682477 | -6.097923184 | 2.02E-02 |
| Fam169a | 9.893249726 | -6.104183185 | 1.00E-05 |
| Zfa-ps | 45.81363219 | -6.121352007 | 6.84E-03 |
| Ccdc177 | 18.1886864 | -6.125603731 | 4.21E-06 |
| Gm6829 | 6.218569805 | -6.160046064 | 4.34E-03 |
| Spp1 | 33894.67107 | -6.167850454 | 2.84E-71 |
| Fam19a5 | 11.74775917 | -6.196463869 | 7.65E-05 |
| Ccn3 | 191.5912267 | -6.206536417 | 2.80E-22 |
| Mt2 | 2687.292186 | -6.215413568 | 1.85E-37 |
| Apln | 371.7087337 | -6.232913238 | 1.51E-40 |
| Hc | 29.33803059 | -6.260567269 | 5.64E-11 |
| Cp | 1547.646794 | -6.267395113 | 1.01E-74 |
| Nrep | 127.7232309 | -6.274609124 | 2.03E-22 |
| 5530401A14Rik | 6.491227945 | -6.282975141 | 2.08E-03 |
| Mmp9 | 324.7980202 | -6.310313027 | 3.22E-17 |
| Wfdc3 | 92.4419757 | -6.32873563 | 7.15E-13 |
| Gm17308 | 13.38750999 | -6.370614776 | 8.62E-07 |
| Trabd2b | 1980.320248 | -6.389347004 | 4.67E-90 |
| Il33 | 385.9276264 | -6.423021949 | 4.34E-19 |
| Ear1 | 28.60473268 | -6.433897676 | 1.04E-05 |
| Gm1979 | 16.54561795 | -6.460616617 | 8.76E-07 |
| Gm26552 | 14.03270238 | -6.499738902 | 1.62E-06 |
| Zfp41 | 1083.240847 | -6.505796294 | 2.35E-79 |
| Fam19a1 | 75.29991681 | -6.553342596 | 2.60E-16 |
| Gm15422 | 17.50296185 | -6.568808934 | 1.08E-06 |
| mt-Te | 93.59404751 | -6.576388303 | 1.76E-11 |
| Speer4b | 14.69744655 | -6.594421754 | 7.42E-07 |
| Rgs4 | 1205.292047 | -6.606859294 | 1.76E-32 |
| Pla2g2e | 18.47206542 | -6.615508871 | 3.89E-07 |
| Atf1-ps | 23.95725453 | -6.626488654 | 3.88E-02 |
| Nat8f3 | 96.81673939 | -6.627073152 | 1.14E-15 |

| | | | |
|---|---|---|---|
| Mir124a-1hg | 9.198990273 | -6.668214739 | 1.79E-05 |
| Znf41-ps | 70.54421761 | -6.672184636 | 9.21E-09 |
| Gm6158 | 12.97793091 | -6.674852154 | 2.20E-02 |
| 5031410I06Rik | 317.2038905 | -6.768390303 | 3.66E-23 |
| Kctd14 | 13.33077476 | -6.769276982 | 4.05E-06 |
| Kctd12b | 28.18537233 | -6.789691599 | 1.91E-07 |
| Gm27624 | 17.52873322 | -6.811276791 | 7.57E-07 |
| B630019A10Rik | 22.44071648 | -6.876416846 | 1.92E-07 |
| Aspg | 128.0287642 | -6.877574291 | 5.17E-19 |
| Nat8 | 9.423743375 | -7.05992694 | 1.39E-07 |
| Egr2 | 252.8800803 | -7.081634163 | 2.65E-29 |
| Cd200r3 | 139.6286064 | -7.112151788 | 1.06E-03 |
| Gm21698 | 27.37379651 | -7.126090463 | 1.29E-07 |
| Ear2 | 11.40384437 | -7.241308177 | 5.14E-05 |
| Klhdc9 | 17.89947461 | -7.364907546 | 8.79E-04 |
| Rgs5 | 30.59780507 | -7.421135022 | 7.19E-09 |
| Dmrtc1c2 | 1708.160336 | -7.491681913 | 4.40E-28 |
| Gm10401 | 31.09799767 | -7.572568223 | 1.41E-04 |
| Lingo2 | 18.63616195 | -7.682144598 | 4.61E-08 |
| Gm7347 | 27.39865267 | -7.682161267 | 4.02E-09 |
| Cdk15 | 129.3148332 | -7.704559309 | 4.32E-17 |
| Plxdc1 | 183.9040077 | -7.803938261 | 8.06E-20 |
| Gm21083 | 126.1258943 | -7.956181667 | 1.48E-12 |
| Megf10 | 616.628578 | -8.120693498 | 1.67E-41 |
| Ear-ps2 | 25.66046438 | -8.267050242 | 7.57E-07 |
| Scn7a | 62.80044543 | -8.363897768 | 2.43E-10 |
| Tspan10 | 235.8434891 | -8.453517463 | 9.27E-08 |
| Gm10221 | 118.8385297 | -8.605153616 | 2.59E-02 |
| Gm5862 | 499.4662236 | -8.622219457 | 7.92E-31 |
| NA | 68.2755783 | -8.78915436 | 2.45E-11 |
| Gm26649 | 48.11705398 | -8.82055988 | 6.48E-05 |
| Igfbp5 | 819.6479129 | -9.101062843 | 3.44E-12 |
| Gm26751 | 46.39692143 | -9.40366823 | 1.16E-03 |
| Gm14567 | 65.48514594 | -9.612085656 | 5.25E-06 |
| Speer4a | 330.0211079 | -9.924473671 | 3.44E-13 |
| Wfdc18 | 86.62622672 | -9.957394378 | 8.30E-14 |
| Sla2 | 584.6313518 | -10.41141551 | 6.53E-10 |
| Zfp781 | 19.65596644 | -23.30973411 | 2.09E-04 |

| Gene | baseMean | log2 Fold Change | padj |
|---|---|---|---|
| Crct1 | 218.1985879 | 11.7708862 | 3.93E-19 |
| Gm43154 | 35.32626595 | 10.3647144 | 1.04E-13 |
| Lypd5 | 36.53736749 | 10.2789084 | 2.22E-12 |
| Gal3st2 | 33.39687285 | 10.09259813 | 6.35E-12 |
| Ptprt | 174.3279632 | 9.71125709 | 8.79E-18 |
| Fras1 | 50.54206584 | 9.68217334 | 4.27E-11 |
| Rbpms2 | 29.23539116 | 9.161995941 | 2.64E-12 |
| Ankrd29 | 28.1865502 | 9.137301618 | 2.20E-03 |
| Prr32 | 25.91099265 | 8.971513848 | 2.53E-09 |
| Dthd1 | 17.05478183 | 8.956538809 | 1.55E-08 |
| Scel | 35.56464911 | 8.854784745 | 2.03E-08 |
| Lyz2 | 828.2358917 | 8.604986778 | 1.02E-91 |
| 3425401B19Rik | 10.93216987 | 8.556735658 | 4.24E-08 |
| AU021092 | 16.8814082 | 8.278543127 | 1.26E-08 |
| Abcc2 | 27.61579762 | 8.197671492 | 3.29E-14 |
| Mgam | 514.7573389 | 8.002556303 | 6.67E-69 |
| Hephl1 | 25.60224789 | 7.942455136 | 8.11E-08 |
| Gm6083 | 6.947319638 | 7.786887968 | 1.14E-06 |
| Sntg1 | 26.65447637 | 7.767605737 | 3.62E-08 |
| Als2cl | 101.8281501 | 7.760566776 | 1.84E-19 |
| Vill | 20.01966172 | 7.742999512 | 1.70E-07 |
| Cfap52 | 5.900962001 | 7.697504458 | 8.63E-06 |
| Anxa8 | 232.8381859 | 7.629502259 | 3.96E-08 |
| Atp9a | 5.328480303 | 7.524725766 | 2.62E-04 |
| Gm6088 | 11.0474142 | 7.27491265 | 4.71E-06 |
| Cyp4b1 | 7.700139718 | 7.257002333 | 8.06E-05 |
| Cyp4f14 | 11.05060069 | 7.244557552 | 1.41E-06 |
| Tmem150b | 157.484637 | 7.112846048 | 1.17E-03 |
| Ttll2 | 60.65077863 | 7.056646749 | 7.56E-26 |
| Klk1b27 | 12.04208527 | 7.008867263 | 2.84E-05 |
| Serpinb6c | 18.60767378 | 6.993868943 | 1.17E-09 |
| Sh3tc1 | 213.0734353 | 6.989503429 | 1.44E-19 |
| Hist1h4n | 13.89900442 | 6.829781055 | 1.25E-05 |
| Slc27a6 | 6.663998212 | 6.812892175 | 8.00E-05 |
| Lctl | 9.524116933 | 6.772058532 | 1.89E-05 |
| Gpr174 | 13.33517572 | 6.766063442 | 1.91E-07 |
| Slc2a12 | 153.088269 | 6.706721288 | 3.97E-02 |
| Mypn | 6.086652581 | 6.698409937 | 1.61E-03 |
| Rapgef5 | 8.056496296 | 6.692430896 | 7.25E-05 |
| Ecscr | 21.55153097 | 6.668322541 | 9.41E-06 |
| Ppp1r3a | 78.06682045 | 6.605992768 | 1.74E-22 |
| Slc6a12 | 232.6272342 | 6.556617577 | 4.01E-06 |
| Mef2b | 78.07998856 | 6.554322135 | 9.26E-04 |
| Lamb3 | 47.93842017 | 6.383437877 | 1.71E-05 |
| Col4a4 | 11.5650756 | 6.382734248 | 6.24E-05 |
| Gm15127 | 7.210988012 | 6.257661173 | 9.39E-05 |

| | | | |
|---|---|---|---|
| Gbp8 | 10.32154589 | 6.214107608 | 2.33E-05 |
| Vnn1 | 94.96773462 | 6.114932811 | 2.11E-11 |
| Cpne4 | 29.44823632 | 6.072805446 | 1.68E-08 |
| Tph1 | 18.10438353 | 6.029606037 | 2.96E-10 |
| Serpinb1a | 435.1696668 | 6.010830913 | 5.16E-41 |
| Mup5 | 8.555026896 | 6.008885231 | 5.80E-04 |
| Cyp24a1 | 5.817295899 | 5.966658737 | 1.06E-03 |
| Ptprn2 | 18.06401101 | 5.905266112 | 4.39E-07 |
| Havcr2 | 173.0157584 | 5.892108066 | 1.46E-26 |
| Slco2b1 | 11.7499612 | 5.881206378 | 5.58E-04 |
| Spo11 | 29.75667518 | 5.866900145 | 1.59E-04 |
| Gm9428 | 14.61896193 | 5.814865818 | 1.98E-04 |
| Insc | 11.98764112 | 5.705461321 | 2.43E-04 |
| Prg4 | 107.0476853 | 5.676900602 | 2.46E-27 |
| Lhx4 | 18.42303899 | 5.64704524 | 3.95E-07 |
| Elmo1 | 276.9450485 | 5.616164762 | 1.43E-17 |
| Il31ra | 119.7831368 | 5.596666422 | 1.79E-04 |
| Gm16531 | 8.914604206 | 5.59094079 | 2.24E-04 |
| Eif2c5 | 9.618557942 | 5.585657944 | 3.30E-05 |
| NA | 10.54876441 | 5.551317896 | 1.89E-07 |
| Rufy4 | 5.678289834 | 5.537536728 | 9.85E-05 |
| Gm7609 | 148.5996821 | 5.500533692 | 5.61E-37 |
| Limch1 | 706.4773048 | 5.473933773 | 3.70E-33 |
| Serpind1 | 71.4025159 | 5.466851901 | 1.77E-28 |
| Plekha6 | 15.60175078 | 5.465284878 | 2.09E-09 |
| Gm15854 | 18.23662287 | 5.465221305 | 6.46E-05 |
| Arhgap9 | 58.21154531 | 5.384944674 | 6.13E-09 |
| Itln1 | 5.005083366 | 5.363365644 | 9.18E-03 |
| Csprs | 36.23461021 | 5.323667773 | 3.33E-12 |
| Unc93a | 52.636787 | 5.323274143 | 8.01E-18 |
| Lgr6 | 16.14818361 | 5.314340268 | 3.16E-06 |
| Serping1 | 32.77937136 | 5.308370047 | 3.89E-04 |
| Adam22 | 234.5747723 | 5.290544763 | 6.01E-11 |
| Chst1 | 44.51961143 | 5.226471893 | 5.74E-10 |
| Hbb-bh1 | 14.94205789 | 5.071613422 | 4.22E-04 |
| Ranbp3l | 21.28438069 | 5.0662902 | 6.15E-05 |
| Ikzf4 | 76.17939403 | 5.045289483 | 2.11E-07 |
| Ifnlr1 | 8.948726448 | 5.03392073 | 1.52E-05 |
| Myom2 | 1179.530624 | 5.02102967 | 2.16E-39 |
| Ankk1 | 13.33636605 | 5.010741012 | 2.47E-03 |
| Mgll | 24.84962995 | 4.985059782 | 1.79E-12 |
| Cdh5 | 11.00256545 | 4.980032145 | 2.92E-03 |
| Gm25043 | 5.794616842 | 4.966598039 | 9.59E-03 |
| 2610028E06Rik | 10.97570129 | 4.961321988 | 1.45E-03 |
| A730049H05Rik | 20.80637073 | 4.912181919 | 1.56E-05 |
| Ccl6 | 20.15019644 | 4.900620314 | 6.39E-08 |
| Gm15080 | 6.203336083 | 4.896430016 | 1.62E-02 |

| | | | |
|---|---|---|---|
| Enkur | 17.5217497 | 4.873446459 | 2.47E-07 |
| Wdr72 | 6.573811052 | 4.864592252 | 8.40E-05 |
| Sorl1 | 6.29132497 | 4.845142057 | 8.14E-03 |
| Serpinb1c | 17.20225213 | 4.839236691 | 5.92E-03 |
| Hist1h2bg | 15.84317161 | 4.81076889 | 9.42E-04 |
| Psca | 64.92123435 | 4.808049372 | 2.77E-03 |
| Upk1b | 1749.009226 | 4.783887902 | 4.74E-03 |
| Dnaic1 | 8.46165829 | 4.773695142 | 1.98E-02 |
| Dnah7a | 5.729662646 | 4.760310871 | 2.67E-03 |
| Muc15 | 5.022882533 | 4.750594113 | 3.89E-04 |
| NA | 26.06308258 | 4.74747405 | 4.74E-03 |
| Slc2a3 | 86.49054597 | 4.717910456 | 1.76E-11 |
| Rpl3l | 10.45497779 | 4.711582284 | 2.38E-08 |
| Fam186a | 5.780700193 | 4.686660964 | 5.45E-03 |
| Crmp1 | 10.17559597 | 4.658952561 | 1.20E-07 |
| Fcna | 17.16685909 | 4.628702799 | 1.60E-03 |
| Slc30a2 | 5.640362476 | 4.625680106 | 6.46E-03 |
| Klra1 | 6.79691526 | 4.60902189 | 7.06E-03 |
| Ptprj | 72.06266337 | 4.576233064 | 1.09E-21 |
| Nrcam | 42.60013369 | 4.541913365 | 4.86E-24 |
| Calcb | 38.52361346 | 4.491352358 | 8.80E-04 |
| Pdzd2 | 406.3513246 | 4.44715981 | 4.34E-11 |
| Tfec | 10.99540601 | 4.435805048 | 2.21E-06 |
| Gm5286 | 7.531885012 | 4.428783183 | 1.24E-02 |
| Serpinb9b | 3482.508606 | 4.425241586 | 1.67E-30 |
| Wt1 | 33.01804751 | 4.410898737 | 2.10E-11 |
| Unc93a2 | 67.49710805 | 4.403149688 | 2.34E-23 |
| Des | 60.75912734 | 4.389879793 | 8.13E-11 |
| Gm10230 | 10.1897906 | 4.372076553 | 3.94E-04 |
| Lrrc31 | 14.73142716 | 4.356978479 | 4.27E-06 |
| Gm26749 | 17.48551257 | 4.323140899 | 1.20E-02 |
| Gm10256 | 43.00942646 | 4.316240803 | 1.68E-09 |
| Cacnb4 | 40.60664551 | 4.287667924 | 4.94E-18 |
| Gm53 | 5.854823975 | 4.263481111 | 2.69E-02 |
| Rtn4r | 11.45555949 | 4.227591744 | 6.26E-06 |
| Otoa | 5.72929926 | 4.215599041 | 2.63E-02 |
| Fndc1 | 5.820280764 | 4.20893478 | 4.12E-02 |
| Klhl30 | 170.2009204 | 4.143891155 | 2.99E-19 |
| Cngb1 | 14.26182037 | 4.138957784 | 5.02E-08 |
| Gm17067 | 9.657809531 | 4.122352461 | 1.39E-05 |
| Trpc5os | 6.190066466 | 4.122349201 | 6.52E-04 |
| Gm13713 | 5.072477726 | 4.11558564 | 9.16E-03 |
| Loxl2 | 5.43924584 | 4.089952489 | 2.55E-02 |
| Tfeb | 959.7623694 | 4.075622358 | 1.97E-97 |
| Gm26740 | 5.601699982 | 4.068000489 | 3.25E-03 |
| Card10 | 5.87499347 | 4.065372295 | 2.96E-02 |
| Sgce | 5.719415256 | 4.037179081 | 2.87E-02 |

| | | | |
|---|---|---|---|
| Gm13415 | 175.3337851 | 3.984731594 | 5.27E-09 |
| Parm1 | 50.62742008 | 3.981163436 | 1.80E-02 |
| Speer4cos | 8.082481503 | 3.978497998 | 2.23E-06 |
| Ifit1bl1 | 78.34996578 | 3.965833417 | 3.04E-24 |
| Ccdc3 | 14.86370165 | 3.926440119 | 7.77E-07 |
| Tmem71 | 6.057387095 | 3.922795687 | 1.39E-02 |
| Klra5 | 12.83281442 | 3.920790269 | 1.58E-04 |
| Gm2619 | 5.817097055 | 3.906223706 | 1.53E-03 |
| Birc7 | 495.5983212 | 3.905035281 | 5.36E-12 |
| Col6a4 | 50.04165493 | 3.900210204 | 1.60E-16 |
| Hpgds | 7.146207006 | 3.880772323 | 9.92E-04 |
| Slc2a5 | 32.51930166 | 3.878279785 | 1.91E-04 |
| Klra4 | 79.2878235 | 3.875740655 | 3.82E-08 |
| P2ry6 | 8.544628265 | 3.85097144 | 3.44E-04 |
| Eps8l1 | 19.97258385 | 3.830413283 | 2.79E-08 |
| Arxes1 | 501.9970735 | 3.811365944 | 7.41E-28 |
| Ggt5 | 37.76480545 | 3.783519171 | 3.77E-04 |
| Fmo3 | 67.82999797 | 3.78125025 | 8.63E-09 |
| Snta1 | 63.91870191 | 3.778759117 | 9.46E-25 |
| Rnf128 | 22.02821797 | 3.763847494 | 1.63E-08 |
| Prss12 | 16.0826015 | 3.749006489 | 4.69E-08 |
| Slc4a10 | 26.51771149 | 3.74132572 | 6.27E-04 |
| Tmem255a | 15.76381383 | 3.737305579 | 1.18E-07 |
| Gpr35 | 13.25276182 | 3.717892155 | 1.98E-09 |
| Fmo2 | 37.14481697 | 3.700599304 | 1.68E-06 |
| Tgtp1 | 20.61468666 | 3.691175711 | 6.81E-05 |
| Gm12153 | 16.73241645 | 3.688725416 | 3.62E-02 |
| Gm7592 | 64.29282183 | 3.683906056 | 3.13E-15 |
| Hist1h2ac | 24.74191821 | 3.680242303 | 2.16E-09 |
| Rbmy | 70.47372177 | 3.651683419 | 7.14E-08 |
| Olr1 | 36.00857858 | 3.642724007 | 6.38E-10 |
| Gngt2 | 39.73962039 | 3.630146743 | 9.83E-22 |
| Selp | 38.63983632 | 3.611638954 | 1.18E-04 |
| Dusp27 | 32.80363271 | 3.610713455 | 1.80E-06 |
| Kdr | 75.92150854 | 3.599976873 | 1.92E-06 |
| Lama5 | 488.2888004 | 3.581465216 | 2.35E-09 |
| Siglecg | 4363.02185 | 3.524808017 | 1.12E-45 |
| Bmp6 | 5.901513132 | 3.519259136 | 2.15E-03 |
| Mettl7b | 5.575022441 | 3.504839638 | 2.77E-02 |
| Upk3b | 18.38391089 | 3.4758107 | 1.96E-02 |
| Ms4a4d | 289.8013129 | 3.475085995 | 6.12E-15 |
| Adgrd1 | 8.97434972 | 3.46465334 | 8.17E-03 |
| Nxf7 | 10.40513731 | 3.460997776 | 1.69E-02 |
| Lims2 | 179.0157548 | 3.454385055 | 1.60E-16 |
| Mamld1 | 9.313569485 | 3.446994064 | 6.35E-03 |
| Slc9a9 | 20.87683751 | 3.417151065 | 7.29E-07 |
| Sybu | 5.676802523 | 3.371328882 | 3.54E-03 |

| | | | |
|---|---|---|---|
| Gch1 | 62.90561554 | 3.361917186 | 3.26E-18 |
| Jph2 | 553.3177943 | 3.348857538 | 2.77E-11 |
| Nlrp10 | 17.4659925 | 3.340136262 | 4.18E-05 |
| D630023F18Rik | 6.989008852 | 3.322636847 | 2.07E-03 |
| Gsta3 | 16.36459403 | 3.304993936 | 4.28E-05 |
| Ccdc180 | 8.668083621 | 3.276644287 | 6.03E-03 |
| Itga7 | 815.1436197 | 3.275349832 | 2.39E-27 |
| Klra17 | 8.115897874 | 3.27298281 | 4.58E-03 |
| Nlrc5 | 42.39167115 | 3.259087037 | 3.11E-14 |
| Il6 | 22.46548116 | 3.253944826 | 2.06E-05 |
| Phactr2 | 110.9007105 | 3.220537095 | 1.60E-16 |
| Syk | 89.27270301 | 3.214485985 | 2.45E-05 |
| Fn3k | 24.39829699 | 3.202527362 | 3.11E-05 |
| Crb2 | 8.069357588 | 3.186444876 | 1.10E-03 |
| Clcnkb | 11.32601771 | 3.186412663 | 2.79E-02 |
| Folr1 | 25.64995036 | 3.178813035 | 3.17E-05 |
| Gm15433 | 6.517950262 | 3.161438012 | 1.60E-02 |
| Eml2 | 184.9655233 | 3.160533423 | 8.02E-22 |
| P2ry10b | 6.207368498 | 3.148882804 | 2.03E-02 |
| Nxf3 | 15.38053015 | 3.144951967 | 1.32E-04 |
| Cstl1 | 15.00122308 | 3.142761031 | 4.29E-06 |
| Upk3bl | 49.55256089 | 3.136165183 | 1.21E-03 |
| Rhpn2 | 103.2238498 | 3.128433836 | 9.56E-18 |
| Trpv4 | 13.02702406 | 3.127106435 | 6.50E-06 |
| C1qtnf2 | 17.27844736 | 3.115782676 | 3.47E-08 |
| Lrrc24 | 8.09439088 | 3.115623613 | 2.66E-02 |
| Slc6a4 | 6.592371236 | 3.112889847 | 1.15E-02 |
| Kctd4 | 12.52468892 | 3.085266056 | 6.48E-03 |
| Cavin4 | 290.9811514 | 3.085093782 | 1.53E-34 |
| Enho | 74.8155069 | 3.076812265 | 7.25E-24 |
| Tmeff2 | 105.2957531 | 3.04506632 | 3.33E-08 |
| Obscn | 23.8335051 | 3.040803241 | 3.42E-04 |
| Tspan15 | 23.84249731 | 3.020703018 | 9.83E-05 |
| Hist1h4i | 68.69874854 | 3.012213212 | 1.65E-07 |
| Usp17le | 14.67590836 | 2.990493367 | 5.25E-04 |
| Gm12138 | 2019.192784 | 2.982187387 | 1.42E-21 |
| Sp110 | 64.32583629 | 2.977391615 | 1.60E-11 |
| Gfpt2 | 60.13655232 | 2.972712172 | 3.43E-17 |
| Ifitm10 | 12.42299472 | 2.957633445 | 2.18E-02 |
| Cyp3a13 | 57.57513375 | 2.954376542 | 2.21E-03 |
| Mfsd6 | 21.51319353 | 2.940147763 | 1.32E-04 |
| Cd74 | 18.46669239 | 2.924350511 | 1.04E-03 |
| Susd2 | 15.13366232 | 2.921137606 | 1.69E-02 |
| Ptprv | 67.14689553 | 2.92062386 | 1.01E-04 |
| Hes1 | 186.8574295 | 2.916747648 | 1.36E-07 |
| C1qtnf5 | 193.5238496 | 2.914501198 | 6.60E-21 |
| Popdc3 | 16.72023235 | 2.903168842 | 1.94E-05 |

| | | | |
|---|---|---|---|
| Col14a1 | 21.17008947 | 2.896407175 | 2.37E-06 |
| Sod3 | 262.7305939 | 2.886002171 | 1.25E-08 |
| Tmem184a | 18.15966957 | 2.884996339 | 6.18E-03 |
| Akr1b3 | 27184.12226 | 2.87542495 | 1.87E-18 |
| Mamdc2 | 140.4200342 | 2.874081741 | 1.38E-18 |
| Cxcl1 | 119.9745271 | 2.858735337 | 8.94E-11 |
| Hecw2 | 21.57444205 | 2.849520049 | 1.08E-02 |
| Ppfibp2 | 6.683566386 | 2.846585545 | 3.74E-02 |
| Aloxe3 | 163.7815803 | 2.833082288 | 2.28E-16 |
| Icam1 | 7.776917133 | 2.832997622 | 4.89E-03 |
| Tceal5 | 16.32229525 | 2.825568297 | 3.63E-03 |
| Rasl10b | 32.86393239 | 2.818887601 | 3.05E-06 |
| Tnfsf15 | 5.532606882 | 2.811481218 | 2.63E-02 |
| Sh3bgrl2 | 10.59069966 | 2.807114272 | 1.48E-03 |
| Hist1h4h | 20.58006819 | 2.798149261 | 3.29E-03 |
| Ptchd1 | 55.72263066 | 2.780567103 | 2.33E-06 |
| Tinagl1 | 7597.988131 | 2.779732167 | 1.50E-30 |
| Tspan32 | 38.52298728 | 2.765401524 | 5.86E-07 |
| Prss27 | 17.7205746 | 2.765050902 | 3.25E-03 |
| Tspan12 | 74.6818032 | 2.759944093 | 4.27E-11 |
| Has1 | 15.99957924 | 2.751553455 | 8.53E-03 |
| Adamtsl1 | 75.62285662 | 2.750896244 | 7.39E-05 |
| B3gnt5 | 34.6745235 | 2.750212233 | 3.30E-05 |
| Ripk3 | 33.31369269 | 2.736868667 | 5.94E-06 |
| C3ar1 | 28.45683189 | 2.731347445 | 3.18E-04 |
| Gm10642 | 6.906922096 | 2.730848174 | 4.35E-02 |
| Heph | 11.43738456 | 2.71017226 | 8.65E-03 |
| Nes | 211.7096441 | 2.708382795 | 4.80E-06 |
| Slc6a15 | 33.49232237 | 2.707877795 | 4.01E-06 |
| Gm16998 | 21.06629374 | 2.707566518 | 2.88E-06 |
| Hsd11b1 | 26.914393 | 2.705229177 | 5.03E-03 |
| Apob | 19.03152808 | 2.703572984 | 1.31E-02 |
| Eln | 24.79874943 | 2.688622439 | 8.99E-06 |
| Fyb2 | 205.4623369 | 2.666763748 | 7.30E-06 |
| Anpep | 300.0514348 | 2.664845631 | 1.54E-10 |
| Acsbg1 | 82.3921383 | 2.662268457 | 7.17E-13 |
| Ip6k3 | 40.3684319 | 2.653829739 | 1.78E-06 |
| Tns4 | 36.47295026 | 2.647502861 | 9.36E-06 |
| Catsperd | 26.54257683 | 2.634523624 | 4.01E-06 |
| Dpep2 | 58.14460163 | 2.633769783 | 1.11E-12 |
| Htr1b | 5.212664692 | 2.629993122 | 1.37E-02 |
| Sp100 | 229.87064 | 2.626295664 | 5.58E-15 |
| Prrt4 | 6.235099417 | 2.621067947 | 1.62E-02 |
| Plekha2 | 266.0834896 | 2.616018217 | 6.49E-16 |
| Tg | 42.06323385 | 2.607975442 | 2.62E-03 |
| Cd28 | 13.18085716 | 2.599821555 | 8.18E-04 |
| Gm12059 | 5.20445124 | 2.598912607 | 4.62E-02 |

| | | | |
|---|---|---|---|
| Gm3376 | 148.220178 | 2.594059522 | 4.11E-15 |
| Pmaip1 | 85.15518226 | 2.590313147 | 1.25E-08 |
| Clcnka | 6.209978955 | 2.58991038 | 1.41E-02 |
| Serpinb6b | 303.4332572 | 2.587707567 | 2.28E-13 |
| Ankrd1 | 4164.102927 | 2.586711737 | 2.70E-14 |
| Ppef1 | 316.4882102 | 2.586692471 | 3.68E-39 |
| Arxes2 | 825.7504101 | 2.577460233 | 7.56E-26 |
| Ifi205 | 92.81123866 | 2.577155817 | 2.60E-04 |
| NA | 7.9647822 | 2.571469646 | 9.88E-03 |
| As3mt | 32.98295994 | 2.562969355 | 2.21E-06 |
| Hist1h2bl | 33.18392678 | 2.55914403 | 5.86E-04 |
| Dusp9 | 278.6759648 | 2.556457989 | 7.55E-17 |
| Adra1d | 14.40500804 | 2.54482102 | 2.88E-02 |
| Hrc | 7.301964216 | 2.538245205 | 1.05E-02 |
| Gm5570 | 6.543427362 | 2.536403521 | 3.32E-02 |
| Sema3c | 667.7097169 | 2.534626272 | 1.76E-05 |
| Btn1a1 | 43.15252764 | 2.516833053 | 2.67E-03 |
| Nipal3 | 40.66859929 | 2.51171152 | 4.01E-06 |
| Aff2 | 6.182382875 | 2.510980825 | 2.27E-02 |
| Abcc3 | 10.35850329 | 2.50976366 | 1.11E-02 |
| Gm6736 | 924.2982575 | 2.497433004 | 1.56E-11 |
| Gpr179 | 12.91737115 | 2.485336775 | 1.38E-02 |
| Prkg1 | 134.4890151 | 2.480725203 | 3.41E-13 |
| Inmt | 199.4518932 | 2.47001607 | 8.13E-08 |
| Hist1h2bc | 898.5619503 | 2.466197882 | 5.06E-08 |
| Pltp | 1305.024307 | 2.465805634 | 5.99E-21 |
| Mogat2 | 37.49728946 | 2.452862435 | 1.04E-02 |
| Camk2a | 97.06394336 | 2.451953936 | 4.69E-09 |
| Syt17 | 46.84650688 | 2.448983324 | 3.94E-06 |
| Crybg2 | 69.94425025 | 2.443674414 | 4.17E-10 |
| Itga3 | 4169.255291 | 2.429327847 | 2.99E-19 |
| Usp17la | 30.31757862 | 2.420828825 | 3.60E-07 |
| Erc2 | 648.4103087 | 2.42073921 | 1.63E-44 |
| Cth | 393.5185974 | 2.417520956 | 1.48E-03 |
| Dkk2 | 170.6475019 | 2.414537354 | 4.14E-03 |
| Zfp600 | 19.86953804 | 2.41416786 | 2.85E-02 |
| Chn2 | 8.322440648 | 2.402009269 | 3.94E-02 |
| Abca8a | 34.13438541 | 2.401183189 | 1.33E-02 |
| Mreg | 281.1053841 | 2.397474962 | 2.26E-09 |
| Lman1l | 19.17703041 | 2.393942724 | 9.68E-03 |
| Grk5 | 918.5251472 | 2.390791194 | 2.35E-21 |
| Alx4 | 5.88386325 | 2.389194951 | 3.02E-02 |
| Gm12840 | 2928.5719 | 2.379098398 | 3.30E-05 |
| Abcb11 | 9.530869549 | 2.367210646 | 8.12E-04 |
| Cobl | 32.63939068 | 2.364278684 | 3.54E-02 |
| Ica1l | 23.42769138 | 2.344230047 | 7.31E-04 |
| Sorbs2 | 490.7709526 | 2.342485989 | 7.93E-09 |

| | | | |
|---|---|---|---|
| Map3k9 | 11.53418457 | 2.341329092 | 2.77E-02 |
| Gipc2 | 16.09109545 | 2.338561431 | 4.50E-04 |
| Adgrg6 | 18.66629187 | 2.336665879 | 2.15E-03 |
| Plscr2 | 362.8998898 | 2.335588547 | 1.42E-10 |
| Lamc3 | 81.73661022 | 2.322506609 | 2.97E-04 |
| Gm20400 | 779.139461 | 2.307605482 | 2.08E-08 |
| Gm11884 | 8.165736454 | 2.306878458 | 1.56E-02 |
| Gria3 | 1179.231791 | 2.303221388 | 2.94E-24 |
| Oit3 | 92.97206647 | 2.291930496 | 4.60E-05 |
| Golga7b | 16.54206909 | 2.28229052 | 6.37E-04 |
| Cst6 | 60.21863085 | 2.266900728 | 3.10E-06 |
| Slc45a3 | 239.2303382 | 2.261900415 | 2.99E-19 |
| Adra1b | 80.73107556 | 2.256641637 | 2.32E-09 |
| Fry | 21.32990924 | 2.251021283 | 5.11E-04 |
| Tgm2 | 2829.880322 | 2.24942206 | 1.11E-33 |
| C130026I21Rik | 69.24352748 | 2.23610166 | 8.39E-08 |
| Il3ra | 21.87015892 | 2.227868257 | 1.21E-03 |
| Cgref1 | 204.5506992 | 2.227480761 | 3.80E-18 |
| Nhsl2 | 58.44723449 | 2.224119436 | 1.21E-02 |
| Rex2 | 58.76235083 | 2.219505377 | 6.12E-05 |
| Mucl3 | 111.7139975 | 2.201686837 | 3.29E-13 |
| Gm11496 | 7.041601801 | 2.195380744 | 4.20E-02 |
| Acta2 | 22267.60312 | 2.194269766 | 2.94E-24 |
| Cgnl1 | 1941.398104 | 2.185755016 | 1.13E-08 |
| Ephx1 | 9309.25785 | 2.179419864 | 1.69E-25 |
| Fgf18 | 10.05836057 | 2.175569628 | 2.87E-02 |
| Mmp28 | 209.9282617 | 2.17108167 | 2.49E-11 |
| Tent5c | 31.88437475 | 2.167551935 | 8.06E-05 |
| Pkp2 | 365.3274075 | 2.163826796 | 4.87E-14 |
| Paqr7 | 35.8722446 | 2.155204526 | 1.29E-04 |
| Hist1h2bq | 18.51075948 | 2.151169515 | 2.66E-02 |
| Plxna4 | 92.75828692 | 2.133444116 | 5.51E-06 |
| Gm4841 | 16.51594958 | 2.115191855 | 2.88E-03 |
| Ereg | 462.3479018 | 2.109706636 | 1.70E-07 |
| Cd93 | 188.2576324 | 2.101361136 | 9.61E-07 |
| Defb41 | 36.08056729 | 2.099953024 | 1.35E-09 |
| Matn4 | 26.3751028 | 2.095005912 | 2.19E-06 |
| Lmcd1 | 190.9655838 | 2.081171557 | 1.50E-05 |
| Bst2 | 1204.425443 | 2.079182196 | 9.80E-15 |
| Fst | 208.7049669 | 2.076770379 | 2.01E-05 |
| Synpo2l | 127.4131202 | 2.076182821 | 3.31E-04 |
| Herc3 | 407.8347654 | 2.060555192 | 4.54E-23 |
| Fgfbp3 | 84.82346159 | 2.055504858 | 2.13E-15 |
| Cyp3a41a | 110.7883611 | 2.055449071 | 2.69E-03 |
| Olfm1 | 251.9775343 | 2.049075193 | 4.90E-07 |
| Fbxl7 | 12.4386985 | 2.044374714 | 1.97E-03 |
| Gm38393 | 571.0226571 | 2.037695484 | 2.68E-10 |

| | | | |
|---|---|---|---|
| Ptgs2 | 3952.831909 | 2.034685052 | 8.73E-05 |
| Mcam | 18.62577123 | 2.034027488 | 1.52E-02 |
| Sema6b | 29.71095728 | 2.031612452 | 5.07E-04 |
| Gm12505 | 700.5275066 | 2.027544005 | 1.98E-08 |
| Sorbs1 | 833.0862304 | 2.025574186 | 4.10E-28 |
| Dennd5b | 74.30478023 | 2.019094934 | 2.55E-05 |
| Plac8 | 10.13342526 | 2.017694537 | 3.63E-02 |
| Hspb7 | 111.5849735 | 2.014381766 | 2.76E-04 |
| Acvrl1 | 34.10597704 | 2.008734712 | 2.25E-03 |
| Slc22a13b-ps | 8.826899059 | 2.006025286 | 4.37E-02 |
| Veph1 | 84.62075448 | 2.005466427 | 4.59E-04 |
| Tmem37 | 109.5363239 | 2.001819471 | 2.39E-10 |
| Aim2 | 214.7586752 | 1.999006475 | 5.52E-19 |
| Mmp3 | 85.42811866 | 1.992256978 | 5.74E-04 |
| Prrg4 | 14.31062335 | 1.990561858 | 5.21E-03 |
| AA467197 | 26.56671516 | 1.989972394 | 1.88E-05 |
| 1700020L24Rik | 39.09349375 | 1.98473673 | 1.39E-06 |
| Pla1a | 89.47502873 | 1.978796506 | 1.02E-04 |
| Gm10825 | 19.23658637 | 1.97253919 | 5.25E-04 |
| P2rx3 | 1054.952462 | 1.970797808 | 5.46E-04 |
| Csrnp1 | 607.3843703 | 1.970767366 | 2.82E-08 |
| Ifit1bl2 | 54.30370509 | 1.965329798 | 5.34E-08 |
| Gbp11 | 15.68366756 | 1.957445534 | 7.50E-03 |
| Rap1gap2 | 970.5392269 | 1.956756139 | 1.04E-12 |
| Dgat2 | 90.86863884 | 1.954118369 | 5.98E-10 |
| Hist1h1d | 29.16335522 | 1.953982893 | 1.79E-04 |
| A530016L24Rik | 46.43521955 | 1.951305109 | 7.14E-03 |
| Abca4 | 66.55109914 | 1.945203165 | 3.38E-07 |
| Dock8 | 275.2900409 | 1.944037147 | 3.95E-05 |
| Cystm1 | 169.2870408 | 1.942552798 | 2.21E-06 |
| Cdkl1 | 21.11717046 | 1.942472338 | 1.87E-03 |
| Hist1h1c | 1898.756039 | 1.93964937 | 2.57E-07 |
| Slc23a3 | 79.80737097 | 1.925319837 | 8.13E-05 |
| Pdlim1 | 2636.81262 | 1.924780307 | 5.01E-09 |
| Cd59a | 250.5614553 | 1.920780917 | 1.04E-13 |
| Aldh3a1 | 155.6027006 | 1.920633895 | 2.75E-04 |
| Mrgpre | 112.8488705 | 1.916218132 | 3.19E-09 |
| Rassf7 | 636.3971329 | 1.915619179 | 1.68E-11 |
| Pid1 | 12.18015165 | 1.914272486 | 2.97E-03 |
| Igfbp7 | 2473.484216 | 1.909943726 | 5.73E-14 |
| Ntn4 | 184.2709408 | 1.909885659 | 2.25E-12 |
| Hsf2bp | 28.16029851 | 1.905256971 | 5.27E-04 |
| Csdc2 | 64.87675074 | 1.903110159 | 2.85E-06 |
| Kank4os | 15.91605683 | 1.902737858 | 2.37E-02 |
| Lamc2 | 395.7838555 | 1.877626413 | 5.50E-11 |
| Efcab8 | 17.7494641 | 1.877334936 | 2.31E-02 |
| 3830403N18Rik | 23.80216124 | 1.872077308 | 4.50E-04 |

| | | | |
|---|---|---|---|
| Gm26912 | 15.59095208 | 1.871742897 | 6.57E-04 |
| Tshz3 | 29.61253025 | 1.86888251 | 1.38E-03 |
| Adamts9 | 285.3258477 | 1.86413341 | 5.48E-07 |
| Gm26651 | 25.61219024 | 1.858844414 | 1.72E-02 |
| Slc7a2 | 1073.849895 | 1.857813694 | 4.48E-11 |
| Gm28044 | 9.247468107 | 1.85383843 | 4.88E-02 |
| 2310002F09Rik | 28.54212603 | 1.852612349 | 3.93E-02 |
| Fibin | 47.24750194 | 1.846106996 | 2.99E-03 |
| Car9 | 41.71092485 | 1.844870196 | 1.39E-03 |
| Igfbp6 | 10099.86965 | 1.841455576 | 7.06E-21 |
| NA | 188.127415 | 1.835133391 | 3.27E-02 |
| Prrt3 | 27.4990097 | 1.834586372 | 1.20E-03 |
| Brinp3 | 129.3249775 | 1.83345065 | 7.51E-05 |
| Gm15128 | 352.174942 | 1.832411985 | 1.19E-04 |
| Nlrp4f | 12.24097274 | 1.826964597 | 4.20E-02 |
| Sap25 | 22.93362904 | 1.826795759 | 3.00E-02 |
| Raet1d | 118.6635537 | 1.826323407 | 2.01E-09 |
| Cdh19 | 11.64226065 | 1.825812897 | 4.82E-02 |
| Tmc7 | 16.65327218 | 1.824138878 | 9.83E-03 |
| Raet1e | 152.5635618 | 1.821788094 | 1.04E-13 |
| Tjp2 | 3236.987169 | 1.820914318 | 9.00E-17 |
| 4933438K21Rik | 20.61022562 | 1.819830802 | 1.84E-03 |
| Mustn1 | 214.872157 | 1.812248718 | 4.77E-13 |
| Cd53 | 21.19451119 | 1.810853483 | 2.66E-02 |
| Vwf | 71.18653804 | 1.804909 | 1.46E-04 |
| Olfr1314 | 22.59357489 | 1.802216417 | 4.36E-04 |
| Hist1h2be | 35.43583052 | 1.797061131 | 6.07E-04 |
| Pfkfb3 | 882.241222 | 1.79655324 | 5.53E-09 |
| Cryab | 13202.45602 | 1.788215945 | 4.80E-14 |
| Mical1 | 409.2832872 | 1.787398229 | 2.77E-20 |
| Ercc6l2 | 120.8231153 | 1.780398144 | 6.76E-09 |
| Klf4 | 2395.995965 | 1.778330585 | 8.99E-21 |
| Akap6 | 117.141912 | 1.778172218 | 1.72E-09 |
| Myzap | 43.56695522 | 1.777018172 | 1.05E-05 |
| Chrnb1 | 351.1485445 | 1.776999599 | 3.12E-07 |
| Prlr | 9.919996375 | 1.774337432 | 2.32E-02 |
| Suox | 3135.576887 | 1.76632155 | 2.99E-19 |
| Dmtn | 35.78379888 | 1.765193314 | 3.81E-05 |
| Trim63 | 604.4480012 | 1.762781518 | 2.75E-04 |
| Ptgir | 52.09243532 | 1.761652135 | 8.33E-09 |
| Klk8 | 65.88130734 | 1.757152124 | 2.11E-02 |
| Lmod1 | 164.8111626 | 1.750047228 | 2.81E-05 |
| Mndal | 41.59689426 | 1.747069617 | 3.82E-04 |
| Cntf | 90.70659904 | 1.743362901 | 5.41E-07 |
| Cox6a2 | 24.89091578 | 1.735680388 | 3.43E-03 |
| Rflnb | 613.7469556 | 1.726488921 | 1.42E-15 |
| Fuca2 | 3318.401005 | 1.725919993 | 5.41E-25 |

| | | | |
|---|---|---|---|
| Krt80 | 423.3382572 | 1.72302253 | 1.35E-07 |
| Cxxc5 | 218.8335225 | 1.718354713 | 4.07E-13 |
| Prr5 | 206.0395211 | 1.716242688 | 1.43E-13 |
| Nhsl1 | 543.1360852 | 1.703239386 | 1.40E-05 |
| Apol9b | 84.6420391 | 1.69988989 | 3.51E-04 |
| Cldn34c1 | 161.8468038 | 1.698339618 | 7.39E-09 |
| Rnd1 | 586.3380609 | 1.692767419 | 2.84E-06 |
| Zdhhc2 | 222.3920511 | 1.686227367 | 9.22E-07 |
| Prorsd1 | 28.67614291 | 1.685839085 | 1.22E-03 |
| Pfkp | 1611.630881 | 1.683762273 | 2.38E-13 |
| Zfp980 | 23.43098719 | 1.682385495 | 2.18E-02 |
| Thpo | 50.84521309 | 1.679163268 | 1.25E-04 |
| Mfrp | 16.50642772 | 1.677466211 | 7.90E-03 |
| Uroc1 | 10.6706882 | 1.674126494 | 3.56E-02 |
| E230025N22Rik | 56.16707186 | 1.672985096 | 4.55E-05 |
| Gm14444 | 29.84845564 | 1.672200794 | 3.11E-03 |
| Tspan33 | 25.32257853 | 1.671064044 | 4.99E-03 |
| Trim30d | 101.5243115 | 1.670309742 | 2.82E-04 |
| Gdpd5 | 258.3174992 | 1.666104109 | 7.59E-19 |
| Rhox5 | 444.7664169 | 1.665268668 | 1.68E-08 |
| Gpr39 | 133.0862302 | 1.660764386 | 3.75E-04 |
| Svep1 | 11238.12662 | 1.659579413 | 6.38E-10 |
| Arhgap20 | 61.50283451 | 1.659219231 | 4.33E-03 |
| Pi15 | 156.8925324 | 1.652333007 | 1.85E-05 |
| Ccn1 | 16942.56933 | 1.649167301 | 1.28E-03 |
| Dhrs13os | 15.23519987 | 1.646952839 | 1.80E-02 |
| Fgd6 | 331.6053799 | 1.644647709 | 3.23E-08 |
| Tnfsf9 | 487.3199691 | 1.642327671 | 8.69E-10 |
| Fstl3 | 1713.818872 | 1.641244054 | 5.73E-14 |
| Gm1141 | 15.33114807 | 1.639111118 | 3.08E-02 |
| Sphk1 | 627.9533835 | 1.630429751 | 5.32E-07 |
| Zfp981 | 48.727574 | 1.629372452 | 3.88E-04 |
| Chac1 | 2414.321663 | 1.624504642 | 2.50E-10 |
| Tinag | 23.1948581 | 1.621857844 | 2.55E-02 |
| Serpinb1b | 80.38277849 | 1.621030437 | 7.37E-04 |
| Kcnj15 | 706.1352588 | 1.619559661 | 6.66E-05 |
| Mgp | 2186.4868 | 1.616380861 | 1.73E-06 |
| Gm10382 | 31.87078083 | 1.601780524 | 4.22E-02 |
| Pcdh17 | 32.83385484 | 1.598194998 | 3.80E-03 |
| Smim3 | 135.0460714 | 1.597652601 | 6.04E-11 |
| Otud1 | 340.3620018 | 1.594732639 | 2.64E-12 |
| 4430402I18Rik | 54.46899431 | 1.589138244 | 6.22E-03 |
| Slc37a2 | 33.44387964 | 1.584199705 | 2.36E-04 |
| Duox2 | 35.60345511 | 1.583277438 | 5.17E-03 |
| Serpine1 | 8069.154667 | 1.580152463 | 3.30E-05 |
| Dlk2 | 145.1833932 | 1.577149752 | 1.57E-04 |
| Ptgs2os2 | 206.3760549 | 1.575976689 | 4.29E-04 |

| | | | |
|---|---|---|---|
| Nmnat3 | 174.6720184 | 1.57510726 | 5.50E-11 |
| Thbd | 4214.750375 | 1.572861376 | 2.36E-10 |
| Cavin2 | 13533.10625 | 1.568484 | 2.73E-05 |
| Slc16a1 | 2748.76285 | 1.566683665 | 5.70E-12 |
| Gcnt3 | 16.25473207 | 1.557970263 | 3.80E-02 |
| Ube2t | 295.3875627 | 1.557020834 | 3.15E-05 |
| Plscr4 | 108.3537055 | 1.555307018 | 1.72E-08 |
| Fads3 | 4292.424506 | 1.547414916 | 1.16E-09 |
| Fhl1 | 107.2708123 | 1.547403155 | 2.88E-02 |
| Txlnb | 29.58797101 | 1.546907004 | 3.98E-02 |
| Gm12439 | 13.97232501 | 1.542349043 | 9.42E-03 |
| Zfp52 | 703.5987578 | 1.539425446 | 5.84E-04 |
| Tnr | 97.7195546 | 1.527969587 | 1.29E-02 |
| Tmem117 | 124.9016676 | 1.526480131 | 1.84E-07 |
| Tril | 35.05868021 | 1.525384227 | 4.50E-04 |
| Hrct1 | 51.68066241 | 1.52413637 | 1.58E-04 |
| Gm12454 | 22.74329518 | 1.523082761 | 4.18E-02 |
| Samd10 | 92.46193541 | 1.520253121 | 5.32E-07 |
| Rtn4rl2 | 154.9682333 | 1.518939313 | 2.02E-03 |
| Cd55 | 298.5806079 | 1.518726963 | 6.13E-03 |
| Sh3rf2 | 283.6625641 | 1.511409411 | 1.13E-02 |
| Neurl1a | 101.6784878 | 1.506506443 | 1.14E-06 |
| Cd80 | 43.14414784 | 1.50263212 | 2.83E-02 |
| Eda2r | 1123.269013 | 1.497847783 | 2.38E-08 |
| Pstpip1 | 90.41301441 | 1.496565636 | 1.88E-04 |
| Pim1 | 2348.868326 | 1.492817497 | 9.21E-05 |
| Gstt1 | 157.0937604 | 1.49228177 | 7.47E-04 |
| Acy3 | 101.9828741 | 1.486610368 | 5.79E-07 |
| Dnph1 | 336.876214 | 1.48368236 | 1.42E-05 |
| Slco3a1 | 1020.790683 | 1.482195501 | 6.12E-15 |
| Ryr1 | 20.13106927 | 1.480980274 | 3.45E-02 |
| S100a11 | 28004.48893 | 1.475794137 | 7.02E-11 |
| Loxl4 | 984.284874 | 1.473651497 | 1.08E-07 |
| Colq | 10.81989595 | 1.47273582 | 1.86E-02 |
| Zfp40 | 305.3181922 | 1.471338754 | 1.29E-02 |
| Ogn | 19778.96094 | 1.467755355 | 3.18E-02 |
| Mtss1 | 37.26253904 | 1.466647327 | 4.10E-04 |
| Dusp8 | 245.7185379 | 1.465663252 | 4.18E-05 |
| Dmpk | 271.5900462 | 1.464379311 | 7.51E-07 |
| Arrb1 | 618.1247361 | 1.463062876 | 3.77E-12 |
| Inhba | 546.6979894 | 1.4612221 | 4.92E-02 |
| Igfbp4 | 605.6600899 | 1.457019967 | 1.56E-03 |
| 2210016F16Rik | 2097.051085 | 1.444671396 | 9.96E-10 |
| Glt8d2 | 963.6849507 | 1.443274035 | 2.11E-09 |
| Gpat3 | 333.521046 | 1.440645639 | 3.90E-09 |
| Lancl3 | 159.4956977 | 1.439868622 | 2.69E-05 |
| E2f6 | 108.6963516 | 1.439030386 | 2.78E-08 |

| | | | |
|---|---|---|---|
| Fat1 | 9394.675609 | 1.437478772 | 8.78E-10 |
| Fam241a | 67.45370295 | 1.436433039 | 5.00E-07 |
| Pcdh10 | 140.2969449 | 1.432402787 | 3.59E-07 |
| Thy1 | 3489.688816 | 1.428652242 | 1.14E-10 |
| Col7a1 | 345.9172629 | 1.42584676 | 2.06E-04 |
| Sgcd | 202.5062698 | 1.425713758 | 6.18E-04 |
| Nppb | 446.9261959 | 1.424408357 | 2.60E-03 |
| Cltb | 2400.270914 | 1.422912025 | 1.72E-11 |
| Maff | 1125.074098 | 1.421538567 | 1.70E-07 |
| Hist2h2aa2 | 243.0853257 | 1.420665922 | 1.95E-02 |
| Gm17382 | 35.29974084 | 1.419470897 | 3.98E-04 |
| Gimap6 | 24.98875556 | 1.419258868 | 7.81E-04 |
| Gadd45g | 1363.863302 | 1.416459823 | 1.28E-06 |
| Gpsm3 | 22.92218898 | 1.413827036 | 2.41E-03 |
| Optn | 361.5681594 | 1.413155917 | 7.71E-17 |
| NA | 37.82154378 | 1.412538165 | 2.50E-02 |
| Nalcn | 35.68450605 | 1.411124013 | 4.00E-02 |
| Gm16178 | 25.22230611 | 1.408618667 | 1.41E-02 |
| Nabp1 | 861.6662125 | 1.407441205 | 3.43E-08 |
| 2610316D01Rik | 31.67538577 | 1.407377247 | 2.93E-02 |
| Slc9a3r1 | 541.036435 | 1.406135055 | 6.39E-08 |
| Ddah1 | 1706.491516 | 1.400595366 | 1.44E-05 |
| Ngf | 909.5389495 | 1.400082287 | 3.76E-09 |
| Cd274 | 79.84677291 | 1.398522193 | 5.00E-07 |
| Angpt4 | 781.819534 | 1.395421991 | 3.78E-03 |
| Ikzf2 | 190.7222877 | 1.3914236 | 2.43E-03 |
| Serpinb9 | 1571.525424 | 1.391240284 | 5.42E-17 |
| Dnah6 | 60.00583954 | 1.390135629 | 3.03E-03 |
| Pla2g4b | 67.61780263 | 1.388711433 | 3.87E-02 |
| Tmem88 | 67.02043446 | 1.386803513 | 1.07E-03 |
| Pla2g16 | 886.3125233 | 1.386476467 | 1.00E-08 |
| Ablim1 | 3555.853548 | 1.38232877 | 4.52E-10 |
| S100a7a | 436.7014925 | 1.377267433 | 2.04E-05 |
| Nexn | 586.7586746 | 1.376849526 | 5.78E-03 |
| Thnsl2 | 60.98386303 | 1.373515186 | 4.67E-06 |
| Tnxb | 839.0427032 | 1.372298887 | 3.05E-17 |
| C2cd2 | 903.6257456 | 1.367637259 | 4.83E-16 |
| Ptgds | 167.4192768 | 1.366745306 | 7.08E-04 |
| Htr2a | 66.90647818 | 1.364758965 | 1.94E-04 |
| Dlc1 | 2597.470192 | 1.364143124 | 4.90E-16 |
| Cemip | 8576.297188 | 1.362594367 | 1.26E-03 |
| Arhgap18 | 896.4577738 | 1.362369349 | 7.95E-04 |
| Spon2 | 2156.979328 | 1.361307695 | 6.65E-05 |
| Paqr4 | 951.0335697 | 1.359631523 | 8.64E-10 |
| Cbr2 | 50.33587452 | 1.359467836 | 1.37E-02 |
| Trib3 | 1385.443885 | 1.357125255 | 6.48E-04 |
| Thbs1 | 56074.01074 | 1.356638634 | 3.42E-04 |

| | | | |
|---|---|---|---|
| Neu3 | 186.830771 | 1.356013897 | 9.08E-10 |
| Slco2a1 | 4121.36187 | 1.355854378 | 1.14E-13 |
| Gm26809 | 35780.51269 | 1.355250769 | 1.44E-04 |
| Calhm5 | 378.443981 | 1.354727444 | 8.03E-16 |
| Kank1 | 1147.219222 | 1.34373632 | 1.68E-10 |
| Atp10d | 515.9679941 | 1.33847942 | 9.16E-04 |
| Cgn | 236.0254742 | 1.338103898 | 1.99E-03 |
| Hacd4 | 175.9725215 | 1.33747688 | 3.90E-04 |
| Vmn1r-ps128 | 315.8095013 | 1.335624932 | 3.97E-03 |
| Fam117a | 85.5208589 | 1.333420851 | 3.57E-03 |
| Sulf1 | 1797.810171 | 1.332722222 | 8.09E-05 |
| Ptgs1 | 3977.955639 | 1.331347365 | 1.22E-12 |
| Herc6 | 237.6921114 | 1.327479579 | 2.84E-06 |
| Man1a | 1574.812904 | 1.326918297 | 6.15E-09 |
| Mfsd4b5 | 246.1474623 | 1.320029704 | 6.24E-06 |
| 4930458D05Rik | 61.19370456 | 1.318284699 | 4.94E-03 |
| Sync | 42.33056854 | 1.31622153 | 4.17E-03 |
| Sult1a1 | 55.76000516 | 1.309878353 | 3.11E-02 |
| Cyp2j9 | 75.57103561 | 1.307032972 | 3.11E-05 |
| Aig1 | 239.7079789 | 1.30592679 | 3.31E-08 |
| Abhd6 | 166.9699771 | 1.305104789 | 8.67E-09 |
| Rab30 | 327.3676624 | 1.302805457 | 2.10E-03 |
| Tlr1 | 42.23168809 | 1.299151263 | 8.69E-03 |
| Fzd4 | 527.5932981 | 1.296473952 | 7.89E-05 |
| Stac3 | 27.44952208 | 1.29295149 | 9.14E-03 |
| Kcnab3 | 152.1664527 | 1.291780909 | 3.43E-05 |
| Hist2h3c2 | 70.03396305 | 1.290872813 | 1.72E-02 |
| Zc2hc1c | 99.23641284 | 1.288277105 | 4.63E-05 |
| Ccl9 | 880.6190831 | 1.287410945 | 2.98E-03 |
| 4833417C18Rik | 23.3663168 | 1.281694788 | 1.94E-02 |
| Gm16364 | 22.25215599 | 1.281458649 | 1.34E-02 |
| Trpm2 | 50.50789183 | 1.276763184 | 1.26E-02 |
| Ctla2b | 860.0005034 | 1.274775269 | 1.64E-03 |
| Agmo | 28.0253374 | 1.274770082 | 1.91E-03 |
| Atp1a3 | 132.1001762 | 1.272353107 | 4.24E-03 |
| Rin3 | 39.43852099 | 1.270564552 | 7.18E-03 |
| Serpina1a | 229.6910667 | 1.267767109 | 6.49E-03 |
| Papss2 | 2566.376957 | 1.264700534 | 2.23E-15 |
| Dusp1 | 2323.364801 | 1.262593681 | 2.79E-04 |
| Il15 | 186.9354469 | 1.2573783 | 7.82E-07 |
| Fam13a | 488.6827203 | 1.255930961 | 6.18E-09 |
| 4930486L24Rik | 435.5390485 | 1.252394511 | 4.74E-03 |
| Anxa3 | 7482.75502 | 1.248873915 | 3.75E-07 |
| Stc2 | 605.8941595 | 1.248666723 | 1.32E-04 |
| Tmem38b | 711.4493136 | 1.247582054 | 5.18E-08 |
| Dennd3 | 112.4646379 | 1.247561051 | 3.30E-04 |
| Ano3 | 117.0441969 | 1.245874196 | 3.27E-04 |

| | | | |
|---|---|---|---|
| Col11a1 | 3443.82184 | 1.243367551 | 3.46E-02 |
| Rasl11b | 155.4030989 | 1.241634064 | 2.71E-04 |
| Clvs1 | 109.3908839 | 1.239966086 | 1.22E-02 |
| Ttc39b | 2306.341585 | 1.239179114 | 2.02E-11 |
| Malat1 | 24971.98118 | 1.237780023 | 8.92E-04 |
| Ube2l6 | 711.5714298 | 1.237170964 | 9.06E-06 |
| Ripor2 | 382.9148722 | 1.236934375 | 4.95E-03 |
| Rarb | 39.25205665 | 1.232286761 | 2.24E-02 |
| Inava | 109.6608324 | 1.230322739 | 8.74E-07 |
| Foxf1 | 338.939874 | 1.22811635 | 2.20E-13 |
| Stap2 | 79.90043653 | 1.227408342 | 5.57E-05 |
| Sox5 | 174.3593763 | 1.219544351 | 1.69E-02 |
| Wnt9a | 916.1471833 | 1.218007791 | 2.27E-06 |
| Gabarapl2 | 2747.465548 | 1.217220594 | 2.32E-10 |
| Dqx1 | 97.9924442 | 1.214911859 | 6.47E-05 |
| Baiap2l1 | 282.5346504 | 1.213217342 | 5.90E-05 |
| Rab27b | 133.4508688 | 1.211786164 | 3.53E-03 |
| Ahi1 | 1541.580211 | 1.206531032 | 1.88E-11 |
| Hoxa1 | 320.536735 | 1.20549577 | 5.50E-06 |
| Atp2b4 | 128.2057106 | 1.205079281 | 3.25E-02 |
| Steap1 | 1192.998083 | 1.204599727 | 4.89E-04 |
| Cd14 | 165.7940412 | 1.20455909 | 7.44E-05 |
| Shroom3 | 71.88219185 | 1.204091191 | 1.42E-03 |
| Ifit1 | 122.4955534 | 1.201283259 | 2.01E-04 |
| Crlf1 | 353.0096371 | 1.198276276 | 2.35E-03 |
| Dpep1 | 950.7647622 | 1.192429235 | 6.82E-03 |
| Col5a2 | 40728.55901 | 1.192096377 | 1.07E-06 |
| Ccn4 | 3715.730625 | 1.190746332 | 3.41E-02 |
| Stx11 | 348.3990913 | 1.18965033 | 1.01E-10 |
| Celsr3 | 31.26221965 | 1.188853438 | 2.41E-02 |
| Stard13 | 1677.000687 | 1.188402568 | 8.92E-04 |
| Steap2 | 5089.313269 | 1.185437504 | 3.23E-08 |
| Npr3 | 1290.290208 | 1.184827102 | 2.11E-02 |
| Ass1 | 199.8319632 | 1.181139285 | 2.15E-04 |
| Ehd1 | 9278.785003 | 1.176613087 | 2.60E-09 |
| Tln2 | 2751.788901 | 1.174758776 | 7.99E-11 |
| Ppp1r12b | 4138.333762 | 1.174341787 | 3.66E-14 |
| Xylb | 84.70507461 | 1.172646661 | 2.99E-05 |
| Csf1r | 229.3071014 | 1.172157089 | 2.58E-02 |
| Depdc7 | 242.1642713 | 1.171376915 | 4.04E-06 |
| Cd24a | 2574.110999 | 1.170682871 | 5.38E-03 |
| Pard6b | 681.0243997 | 1.170514001 | 9.38E-04 |
| Afmid | 240.573853 | 1.166751665 | 2.56E-05 |
| Spaca6 | 1058.427464 | 1.166721074 | 1.01E-03 |
| Fgd3 | 456.6207369 | 1.163865732 | 2.22E-03 |
| Phactr4 | 1891.975684 | 1.161081028 | 4.67E-10 |
| Sdsl | 62.01515941 | 1.160420494 | 1.77E-02 |

| | | | |
|---|---|---|---|
| Arhgap29 | 5638.40206 | 1.153088353 | 9.32E-06 |
| Gm11427 | 43.61658226 | 1.152486831 | 2.15E-02 |
| Gm20634 | 348.1245484 | 1.15210114 | 1.10E-04 |
| Usp2 | 268.4067797 | 1.145560443 | 4.41E-03 |
| Trpm5 | 54.98351022 | 1.140489073 | 1.90E-02 |
| Diras2 | 135.873439 | 1.136736274 | 1.99E-02 |
| Tob1 | 562.3538878 | 1.133397153 | 1.28E-06 |
| Gm20632 | 184.1790835 | 1.132579776 | 2.26E-05 |
| Morc4 | 3288.740245 | 1.120275728 | 4.33E-09 |
| Slmapos2 | 3326.360358 | 1.116711023 | 4.79E-04 |
| Zfp36 | 1397.592095 | 1.111467803 | 4.90E-03 |
| Gm15287 | 92.60236988 | 1.108348062 | 1.57E-02 |
| Gprc5a | 2003.573682 | 1.103968368 | 1.09E-03 |
| Maoa | 11869.71092 | 1.099257442 | 2.50E-08 |
| Pkib | 43.16851639 | 1.098559982 | 2.14E-02 |
| Apol9a | 154.7926263 | 1.098418489 | 9.71E-03 |
| Asah2 | 1867.355462 | 1.097476161 | 1.06E-08 |
| Rbm47 | 58.66599152 | 1.093368131 | 4.41E-02 |
| Vldlr | 180.1056824 | 1.091922309 | 7.37E-03 |
| Cep295nl | 32.72658667 | 1.091561526 | 4.18E-02 |
| Mbp | 219.5097508 | 1.087686966 | 9.17E-04 |
| Arl6ip1 | 275.4071157 | 1.087097432 | 2.87E-11 |
| Prnp | 14550.6052 | 1.08427077 | 8.97E-10 |
| Slc16a11 | 113.1379731 | 1.084215494 | 1.65E-04 |
| Slc35g1 | 93.21178618 | 1.083887602 | 3.43E-03 |
| Mocos | 401.6732574 | 1.081309649 | 2.55E-08 |
| Jph1 | 69.63530886 | 1.080411097 | 3.66E-03 |
| Fam110c | 419.3831265 | 1.078188684 | 6.02E-06 |
| Lama2 | 873.8408923 | 1.076108204 | 2.14E-03 |
| Hist1h2al | 73.64496331 | 1.072281124 | 4.39E-02 |
| Gpc6 | 435.6774066 | 1.069818632 | 2.02E-04 |
| Stat2 | 1625.247407 | 1.069411526 | 4.69E-08 |
| Nadk2 | 1271.547494 | 1.069354085 | 8.45E-08 |
| Gm16062 | 85.00001513 | 1.069287834 | 1.85E-02 |
| Boc | 364.9113936 | 1.068564329 | 2.40E-03 |
| Tacc1 | 884.4454082 | 1.067501428 | 6.35E-08 |
| Fign | 239.6826974 | 1.06738023 | 1.88E-03 |
| Isoc1 | 868.2467197 | 1.066799872 | 7.64E-06 |
| Alcam | 657.7870578 | 1.062374754 | 1.60E-04 |
| Znrf2 | 1311.359839 | 1.060462033 | 4.70E-08 |
| Adam11 | 158.3843609 | 1.058698668 | 1.63E-02 |
| Pla2g12a | 876.4735081 | 1.058064005 | 9.28E-09 |
| Raph1 | 579.1077271 | 1.057806095 | 4.00E-05 |
| Crip2 | 63.44237168 | 1.055354272 | 3.43E-04 |
| Hbegf | 543.5379025 | 1.055239133 | 2.22E-04 |
| Ahr | 292.2985283 | 1.053984311 | 8.96E-05 |
| Plaur | 1149.588067 | 1.053525468 | 9.01E-04 |

| | | | |
|---|---|---|---|
| Tnfrsf10b | 1728.080669 | 1.05352032 | 1.94E-07 |
| Nr4a1 | 921.088903 | 1.052608838 | 3.78E-02 |
| Polk | 816.6986151 | 1.051892298 | 1.05E-06 |
| Grk4 | 204.3808361 | 1.051067583 | 1.24E-05 |
| Gm21750 | 59.65533783 | 1.041106634 | 4.20E-02 |
| Aif1l | 2280.404424 | 1.038270641 | 1.08E-05 |
| Cited2 | 5203.839702 | 1.032407038 | 2.76E-12 |
| Ism1 | 148.8317432 | 1.030836529 | 3.14E-02 |
| E230016K23Rik | 97.97888687 | 1.02839447 | 2.29E-02 |
| Gm14296 | 55.8943979 | 1.025980207 | 3.56E-02 |
| Dmxl2 | 711.1668361 | 1.025193444 | 2.75E-03 |
| Fcgr4 | 95.65933917 | 1.025164367 | 9.18E-03 |
| Avil | 62.46903402 | 1.019782279 | 3.46E-02 |
| Myom1 | 84.62851477 | 1.017268884 | 6.11E-03 |
| Slc25a25 | 821.5438689 | 1.012066948 | 3.77E-04 |
| Stbd1 | 242.832905 | 1.009756712 | 1.48E-03 |
| Lipg | 535.1730908 | 1.009551608 | 7.18E-05 |
| Serpinb8 | 975.2511855 | 1.008190909 | 6.12E-04 |
| Acot2 | 886.3418002 | 1.00696255 | 1.76E-06 |
| Platr14 | 62.69361273 | 1.006173878 | 2.45E-02 |
| Vgll3 | 2688.695715 | 1.005866558 | 3.03E-03 |
| Ephx2 | 595.9836792 | 1.00365395 | 8.36E-07 |
| Chst7 | 291.6271761 | 1.002735193 | 3.11E-04 |
| Stim1 | 2475.820194 | 0.998731803 | 1.46E-09 |
| Tfpi | 73.57390232 | 0.996384395 | 1.90E-02 |
| Diaph3 | 1284.11 | 0.995029083 | 4.96E-02 |
| Stk17b | 1172.316199 | 0.99219975 | 4.99E-07 |
| Gm5620 | 281.5414567 | 0.988480522 | 3.06E-02 |
| Zfp951 | 126.5205167 | 0.988215718 | 1.98E-03 |
| Hip1 | 3266.38303 | 0.987921144 | 1.06E-16 |
| Abca9 | 166.1965339 | 0.985975906 | 4.47E-02 |
| Rin1 | 4778.34829 | 0.984975101 | 1.69E-09 |
| Nrp1 | 3718.261942 | 0.984475911 | 6.13E-07 |
| Slc43a3 | 2964.656793 | 0.98434115 | 7.65E-04 |
| Prex2 | 704.5135401 | 0.984231806 | 3.43E-03 |
| Atp1a1 | 13088.27316 | 0.98304153 | 5.92E-10 |
| Adora2b | 955.5375759 | 0.980786757 | 1.34E-06 |
| Dhrs13 | 291.7875454 | 0.980299232 | 4.58E-03 |
| Gabre | 296.5455035 | 0.979044689 | 6.39E-04 |
| Slc25a40 | 773.2498353 | 0.978279445 | 2.67E-05 |
| Dusp14 | 1099.248265 | 0.969521828 | 2.83E-05 |
| Mpp5 | 1955.405807 | 0.967704537 | 1.87E-05 |
| Il15ra | 123.6384944 | 0.967547463 | 3.04E-02 |
| Fam111a | 2899.275804 | 0.967425442 | 5.62E-03 |
| Slc39a10 | 3524.383162 | 0.965802924 | 4.13E-04 |
| Col1a1 | 121831.0483 | 0.965248304 | 7.53E-04 |
| Tent5b | 1620.453803 | 0.964554688 | 3.39E-07 |

| | | | |
|---|---|---|---|
| Chst12 | 2131.435361 | 0.963630821 | 1.04E-08 |
| Pgrmc1 | 9425.547064 | 0.963302649 | 4.91E-13 |
| Pde8b | 714.9937037 | 0.963112944 | 1.39E-04 |
| Fbn1 | 5465.355355 | 0.962609709 | 2.38E-03 |
| Slc25a42 | 514.849024 | 0.96260944 | 4.22E-07 |
| Gm11816 | 245.2771472 | 0.961791013 | 1.26E-03 |
| Chrd | 441.3963201 | 0.960975487 | 6.37E-05 |
| Cd248 | 6880.534459 | 0.960797835 | 4.51E-05 |
| Mgst3 | 998.4397806 | 0.959123509 | 5.79E-07 |
| Zfp53 | 773.8964099 | 0.958556976 | 1.29E-03 |
| Bmp2k | 1711.020882 | 0.958024487 | 1.88E-10 |
| Gm15764 | 62.42387083 | 0.95556762 | 9.11E-03 |
| Gm17167 | 121.3447426 | 0.953395519 | 1.27E-02 |
| Serpine2 | 1071.175796 | 0.948345746 | 5.99E-04 |
| Gm2115 | 916.8914908 | 0.947086894 | 1.93E-03 |
| Serpinb6a | 9297.117704 | 0.945662528 | 4.57E-15 |
| Tubb4a | 66.11470353 | 0.943642855 | 1.07E-02 |
| Lyrm9 | 234.5217743 | 0.942847422 | 2.40E-03 |
| Btg2 | 1025.665619 | 0.941179084 | 9.42E-03 |
| Ncam1 | 1630.791622 | 0.940577601 | 3.06E-06 |
| Ptpn21 | 3194.640395 | 0.937787658 | 7.45E-08 |
| Prickle1 | 1065.178979 | 0.93435747 | 8.52E-05 |
| Mindy2 | 1585.001885 | 0.933987666 | 4.74E-03 |
| Ulbp1 | 1078.85357 | 0.933504274 | 3.53E-03 |
| Rras2 | 2707.729271 | 0.932601092 | 9.65E-06 |
| Hacd2 | 1385.12878 | 0.928473336 | 5.79E-07 |
| Myo1c | 17433.95275 | 0.928400959 | 9.56E-10 |
| Cracr2a | 41.73135551 | 0.926232486 | 1.31E-02 |
| Fas | 523.473713 | 0.926159403 | 5.09E-04 |
| Tmsb4x | 9263.718803 | 0.926093961 | 1.55E-12 |
| Phtf2 | 418.8036949 | 0.925396153 | 1.45E-04 |
| Lyn | 252.2098934 | 0.924554372 | 3.77E-03 |
| Col12a1 | 2667.384937 | 0.924550329 | 3.42E-10 |
| Dbndd2 | 321.9744503 | 0.923996648 | 1.22E-03 |
| Kcnk2 | 223.8511014 | 0.923539994 | 3.25E-04 |
| Ddr1 | 5318.493109 | 0.922849978 | 4.98E-09 |
| Ehd3 | 485.4889265 | 0.92282505 | 2.36E-03 |
| Ier3 | 749.7978944 | 0.922071557 | 1.38E-03 |
| Gja1 | 1405.954553 | 0.914215812 | 3.67E-02 |
| Gstm2 | 1617.930043 | 0.91403762 | 7.41E-04 |
| Acvr2a | 745.8765756 | 0.91341481 | 9.67E-04 |
| Styk1 | 452.3674977 | 0.912933733 | 1.56E-02 |
| Pla2g4a | 4766.449097 | 0.912828995 | 1.27E-02 |
| Dtd1 | 634.7652521 | 0.912675875 | 5.51E-06 |
| Pcx | 1264.131113 | 0.909561767 | 1.58E-02 |
| Fgf10 | 1182.615126 | 0.909287333 | 2.94E-04 |
| Atp11a | 5318.325548 | 0.906689673 | 2.09E-09 |

| | | | |
|---|---|---|---|
| Sp140 | 412.3072278 | 0.904604281 | 9.61E-04 |
| C1qtnf1 | 929.0184909 | 0.904521557 | 2.04E-09 |
| Tjp1 | 4773.974161 | 0.903788888 | 3.23E-08 |
| Stambpl1 | 1280.981375 | 0.902570923 | 5.96E-06 |
| Rusc2 | 4030.034565 | 0.902405389 | 9.64E-04 |
| Emx2 | 282.5000379 | 0.901868349 | 2.72E-03 |
| Bag2 | 1009.73514 | 0.900075382 | 9.04E-07 |
| Ltbp1 | 3648.116466 | 0.898362117 | 1.54E-04 |
| Cdk18 | 2832.68387 | 0.897901872 | 8.64E-05 |
| Gm19705 | 81.49688775 | 0.895873214 | 1.71E-02 |
| Tor4a | 194.1873465 | 0.892541622 | 2.61E-03 |
| Sgms1 | 1435.255104 | 0.890795602 | 6.95E-06 |
| Lgals9 | 3984.469823 | 0.890620972 | 4.47E-06 |
| Eml5 | 284.730193 | 0.888885016 | 2.58E-02 |
| Stom | 2151.225051 | 0.884946043 | 3.96E-04 |
| Akap12 | 11002.38031 | 0.882166639 | 8.06E-05 |
| Jag1 | 1491.58118 | 0.881259851 | 1.88E-03 |
| Rasgrp3 | 117.9121061 | 0.88054492 | 1.14E-02 |
| Itga6 | 3086.584435 | 0.879165082 | 3.39E-07 |
| 2500002B13Rik | 80.42307177 | 0.87906422 | 3.03E-03 |
| Lgmn | 8054.730872 | 0.875676576 | 9.67E-05 |
| Sparc | 83467.83876 | 0.875469505 | 1.01E-04 |
| Trim30a | 433.047542 | 0.874563423 | 6.86E-03 |
| Cdkn1a | 10457.59614 | 0.872103339 | 5.27E-03 |
| Cobll1 | 2804.002598 | 0.870430148 | 5.88E-10 |
| Slc46a3 | 77.82817835 | 0.869766629 | 3.32E-02 |
| Cd151 | 4413.725542 | 0.868171534 | 1.77E-07 |
| Efemp1 | 43132.42209 | 0.864999698 | 6.71E-07 |
| Osgin1 | 375.3213527 | 0.864470111 | 1.12E-02 |
| Dynll2 | 6105.990932 | 0.863422778 | 9.25E-11 |
| Hebp2 | 255.3948829 | 0.861546572 | 1.72E-05 |
| Gzf1 | 2233.832383 | 0.859934046 | 2.31E-09 |
| Rrp8 | 1637.732775 | 0.859644394 | 3.24E-05 |
| Tnfaip8 | 770.730871 | 0.858853557 | 2.06E-14 |
| Prss23os | 1155.334691 | 0.856587568 | 8.01E-03 |
| Adamts1 | 17979.02683 | 0.855347489 | 1.98E-02 |
| Col4a5 | 13744.73591 | 0.852488592 | 2.84E-10 |
| Mpp6 | 1620.758067 | 0.849830554 | 1.77E-03 |
| Yif1b | 1695.896623 | 0.849064804 | 4.22E-07 |
| 2210408F21Rik | 135.6814517 | 0.847817222 | 1.58E-04 |
| Ero1l | 4526.176064 | 0.847591361 | 6.18E-03 |
| Gsn | 46199.21147 | 0.846103966 | 7.99E-08 |
| Ppp1r15a | 3481.359866 | 0.845795194 | 6.45E-06 |
| Epb41l5 | 589.5562055 | 0.845190123 | 4.43E-06 |
| Pdgfrl | 4182.667061 | 0.84475714 | 3.40E-03 |
| Tdg | 173.5710662 | 0.842622246 | 2.75E-03 |
| Apobec1 | 552.4754437 | 0.840104515 | 3.31E-04 |

| | | | |
|---|---|---|---|
| Tbc1d2 | 1016.924895 | 0.839800448 | 1.80E-03 |
| Lin54 | 751.046264 | 0.837916693 | 1.06E-03 |
| 1110046J04Rik | 236.2657601 | 0.836296876 | 1.17E-02 |
| Clip1 | 4554.958969 | 0.832439145 | 1.58E-07 |
| Prss23 | 12096.74038 | 0.832410194 | 8.99E-06 |
| Ccng1 | 10250.65217 | 0.831418671 | 1.80E-06 |
| Pdgfc | 1278.888861 | 0.831304945 | 8.24E-06 |
| Rsu1 | 5433.912262 | 0.829883259 | 3.43E-17 |
| Lpin3 | 493.5528399 | 0.829474582 | 2.70E-03 |
| Sord | 8476.138057 | 0.829427446 | 4.66E-03 |
| Psmd8 | 7697.479585 | 0.827466315 | 1.12E-04 |
| 2810433D01Rik | 81.57471888 | 0.826309223 | 2.56E-02 |
| Ackr3 | 559.0724489 | 0.823459281 | 3.60E-02 |
| Hsp90aa1 | 28139.47731 | 0.822683907 | 1.37E-02 |
| Sorbs3 | 4375.993113 | 0.82235449 | 1.20E-07 |
| Efnb1 | 347.491773 | 0.821428781 | 2.32E-04 |
| Tmem45a | 630.8650328 | 0.819543606 | 4.99E-03 |
| Sh3bp5l | 2141.079408 | 0.817447791 | 3.79E-07 |
| Ptprg | 2392.25637 | 0.816690347 | 3.98E-04 |
| Zfp953 | 159.3076468 | 0.814230037 | 2.50E-02 |
| Deptor | 1172.456592 | 0.813766428 | 5.55E-03 |
| Map2k3os | 158.9798746 | 0.812452261 | 4.89E-02 |
| P4ha1 | 9161.320401 | 0.811112764 | 1.40E-04 |
| Stard10 | 184.9670242 | 0.809933261 | 3.86E-02 |
| Sash1 | 2475.130362 | 0.808761343 | 5.51E-11 |
| Tns3 | 8946.526377 | 0.801471216 | 3.30E-07 |
| Wwp1 | 1335.549327 | 0.801155387 | 3.38E-04 |
| Hoxc5 | 124.7217338 | 0.800485386 | 4.66E-02 |
| Col1a2 | 322977.1213 | 0.799777072 | 5.16E-04 |
| Csf1 | 14262.92663 | 0.799330702 | 9.33E-03 |
| Pdzd7 | 210.0352206 | 0.794070595 | 5.92E-03 |
| Fzd7 | 1499.058727 | 0.793408494 | 4.02E-07 |
| Lzts3 | 586.2326723 | 0.793083678 | 1.17E-04 |
| Adamtsl4 | 642.4400238 | 0.792267776 | 2.36E-05 |
| Col5a1 | 27771.21141 | 0.79208498 | 1.31E-03 |
| Nudt16 | 122.149193 | 0.789452664 | 3.67E-03 |
| Pxdc1 | 198.4170374 | 0.78884863 | 3.53E-03 |
| Mtmr10 | 1541.372356 | 0.788435432 | 8.40E-05 |
| Cdh2 | 5735.006876 | 0.786902466 | 4.82E-06 |
| Nectin1 | 1412.399235 | 0.786726698 | 8.20E-05 |
| Lgals3bp | 961.5598894 | 0.786298269 | 9.50E-03 |
| H6pd | 4608.678006 | 0.785854669 | 4.58E-04 |
| Tmem173 | 257.4625801 | 0.784960288 | 5.40E-04 |
| Qsox1 | 10100.16956 | 0.784450965 | 3.86E-05 |
| Tuba1b | 52546.48143 | 0.783726815 | 2.32E-02 |
| Rnf113a1 | 147.4647827 | 0.783665088 | 6.51E-04 |
| Ank2 | 3380.138198 | 0.782014794 | 1.22E-05 |

| | | | |
|---|---|---|---|
| Enpp4 | 380.5708592 | 0.781188519 | 1.45E-04 |
| Pofut1 | 2332.656105 | 0.781031171 | 1.55E-03 |
| Itpr3 | 5436.339997 | 0.779547032 | 5.61E-07 |
| Prdx6 | 12709.33805 | 0.779256771 | 4.36E-04 |
| Ext2 | 5673.335571 | 0.778785891 | 4.14E-05 |
| Myo1d | 3631.060795 | 0.777592405 | 7.06E-06 |
| Grem1 | 116.2492527 | 0.77594912 | 2.58E-02 |
| Slc35e4 | 1871.809364 | 0.775174031 | 1.16E-03 |
| Calhm2 | 211.5215987 | 0.772200086 | 9.48E-05 |
| Tlcd2 | 144.2625361 | 0.771964466 | 1.42E-02 |
| Cgrrf1 | 795.8021606 | 0.765226182 | 1.55E-07 |
| Camk1 | 3159.841714 | 0.764968988 | 1.34E-07 |
| Vopp1 | 3424.567913 | 0.763759104 | 4.10E-06 |
| Plagl1 | 737.429685 | 0.761626719 | 3.79E-02 |
| Tuft1 | 1030.058028 | 0.761568481 | 4.52E-03 |
| Nnmt | 470.8866788 | 0.761264274 | 8.87E-04 |
| Daglb | 227.5009461 | 0.760826693 | 1.02E-03 |
| Lgals4 | 78.28538866 | 0.760219897 | 3.23E-02 |
| Ghitm | 11802.31619 | 0.758943585 | 5.68E-07 |
| Psmb10 | 782.003635 | 0.758295826 | 5.21E-04 |
| Ccnyl1 | 2407.209529 | 0.758269725 | 3.05E-06 |
| Slc41a3 | 517.5453823 | 0.755815554 | 2.02E-06 |
| Cyb5r1 | 3302.743638 | 0.7556956 | 6.13E-03 |
| Tmbim1 | 4932.481657 | 0.75497081 | 3.39E-06 |
| Dnajb4 | 1668.421695 | 0.754464378 | 4.02E-07 |
| Macrod1 | 694.0355666 | 0.749399927 | 2.28E-02 |
| Grik5 | 256.8579891 | 0.747948701 | 2.50E-03 |
| Colca2 | 129.8506087 | 0.747775307 | 3.91E-03 |
| Tns2 | 5211.113391 | 0.745223849 | 1.03E-06 |
| Cpox | 1136.258528 | 0.742660821 | 1.15E-02 |
| A530040E14Rik | 89.26674794 | 0.741758358 | 3.42E-02 |
| Osbpl10 | 87.89074291 | 0.741627064 | 1.91E-02 |
| Grn | 30705.7805 | 0.741092861 | 6.55E-04 |
| Adamts5 | 15137.11031 | 0.740228175 | 9.29E-03 |
| Npc1 | 3965.691488 | 0.736589559 | 1.60E-04 |
| Enpp3 | 439.6794586 | 0.735868735 | 2.43E-02 |
| Uvrag | 2392.79397 | 0.734564801 | 1.62E-06 |
| Zhx3 | 1585.705602 | 0.733973537 | 2.76E-05 |
| Iqsec2 | 575.0814242 | 0.733794018 | 7.83E-05 |
| Tlr4 | 2569.985835 | 0.732176316 | 1.34E-03 |
| Col8a1 | 3683.903915 | 0.732122165 | 3.28E-02 |
| Epn2 | 3510.416008 | 0.729406555 | 1.17E-04 |
| Itpr1 | 2029.82746 | 0.728777551 | 1.58E-05 |
| Atp10a | 1864.778291 | 0.727357478 | 1.27E-02 |
| Radil | 568.4919136 | 0.726696673 | 2.21E-04 |
| Sms-ps | 1178.87176 | 0.726549023 | 1.07E-03 |
| Galnt10 | 2209.826918 | 0.725474219 | 2.25E-04 |

| | | | |
|---|---|---|---|
| Map3k20 | 2221.020669 | 0.724632046 | 4.30E-06 |
| Pcdh1 | 2459.618889 | 0.722868217 | 2.14E-05 |
| Mgat3 | 1523.241024 | 0.722711561 | 4.05E-05 |
| Plod2 | 1368.669902 | 0.721877915 | 1.84E-03 |
| Pcgf5 | 1180.442953 | 0.720253788 | 6.13E-06 |
| Rdh10 | 613.1453821 | 0.719549836 | 1.15E-04 |
| Angptl4 | 1038.167366 | 0.719428224 | 3.11E-02 |
| Dab2 | 7942.183086 | 0.71909158 | 1.91E-03 |
| Gm12346 | 480.5896986 | 0.716620808 | 4.88E-02 |
| Hoxa4 | 123.1936353 | 0.716298622 | 2.00E-02 |
| Akap2 | 11618.68398 | 0.716199966 | 2.81E-05 |
| Cav1 | 38978.11609 | 0.715230969 | 2.40E-07 |
| Zfp809 | 553.769558 | 0.714971944 | 6.97E-04 |
| Adipor1 | 9075.210439 | 0.713124644 | 1.20E-04 |
| Crebl2 | 1856.82481 | 0.712835897 | 3.61E-05 |
| Arvcf | 1255.272406 | 0.712833376 | 1.74E-05 |
| Flnb | 18993.59883 | 0.71155057 | 9.41E-06 |
| Abhd14b | 229.6613872 | 0.710944192 | 1.22E-02 |
| Syne2 | 2077.972178 | 0.709545701 | 1.79E-03 |
| Plekhf2 | 391.5956648 | 0.709180157 | 1.38E-04 |
| Il6st | 23732.48129 | 0.707376082 | 4.43E-04 |
| Man2b1 | 3916.818943 | 0.706134483 | 1.59E-04 |
| Amotl1 | 7811.016398 | 0.705401102 | 5.73E-06 |
| Dock4 | 204.615827 | 0.704646286 | 4.25E-03 |
| Cfl2 | 7309.759639 | 0.702962921 | 1.04E-04 |
| Nqo2 | 413.4205542 | 0.701872959 | 2.21E-03 |
| Spire1 | 861.3974626 | 0.701382572 | 2.71E-04 |
| Sap30 | 554.9416513 | 0.701235011 | 7.37E-03 |
| 2200002D01Rik | 188.5911851 | 0.700603304 | 3.55E-02 |
| Cyb5r3 | 22966.34465 | 0.697091632 | 9.91E-06 |
| Lrrc51 | 118.5855804 | 0.696897504 | 3.45E-03 |
| Asap3 | 387.600447 | 0.696722071 | 1.70E-02 |
| Fez2 | 3426.090875 | 0.696495318 | 5.80E-04 |
| Tango2 | 725.8753531 | 0.695492247 | 3.43E-06 |
| Ifi203 | 486.7251377 | 0.694442992 | 2.31E-03 |
| Tmem245 | 7288.006629 | 0.693602767 | 1.11E-05 |
| Dlg3 | 2321.002913 | 0.693059395 | 3.02E-09 |
| Hipk2 | 2809.337586 | 0.692273347 | 2.88E-05 |
| Fth1 | 88712.82582 | 0.691273203 | 4.48E-03 |
| Pank1 | 534.8184598 | 0.690848106 | 3.43E-03 |
| Prkci | 7019.667731 | 0.690639495 | 2.12E-04 |
| A930004D18Rik | 436.6953306 | 0.690491064 | 8.23E-03 |
| Gm10516 | 98.22767106 | 0.690399841 | 7.58E-03 |
| Mink1 | 2720.963999 | 0.69039169 | 4.72E-08 |
| Ino80c | 526.2001063 | 0.69001875 | 1.91E-04 |
| Utrn | 8786.265038 | 0.689890862 | 6.13E-06 |
| Hotairm1 | 163.7859472 | 0.686839025 | 1.21E-02 |

| | | | |
|---|---|---|---|
| Gsto2 | 540.3671304 | 0.686466166 | 1.00E-02 |
| Abca5 | 1770.476615 | 0.685148099 | 8.83E-03 |
| Neo1 | 4545.56012 | 0.68483463 | 1.60E-09 |
| Nop10 | 1906.592631 | 0.683491718 | 2.03E-06 |
| Endod1 | 423.568897 | 0.683379308 | 7.07E-03 |
| Cfap157 | 270.2462511 | 0.682607743 | 7.20E-04 |
| Lats2 | 3075.227714 | 0.681634276 | 2.25E-05 |
| Pros1 | 1181.249787 | 0.681577465 | 1.28E-04 |
| Dysf | 3326.040047 | 0.679825375 | 4.71E-03 |
| Rab20 | 279.5572422 | 0.679541363 | 1.36E-02 |
| Creb3l1 | 1785.610876 | 0.679532574 | 1.56E-02 |
| Crim1 | 4958.715791 | 0.679234777 | 1.46E-02 |
| Arhgef18 | 3617.647802 | 0.678623667 | 1.34E-05 |
| Usp13 | 226.2858383 | 0.675279598 | 4.71E-02 |
| Osbpl5 | 2420.062007 | 0.674696062 | 1.61E-04 |
| Cda | 445.5243863 | 0.673031229 | 1.03E-02 |
| Hikeshi | 958.3911583 | 0.667674949 | 9.44E-08 |
| Fam110a | 459.3283501 | 0.66696955 | 3.85E-02 |
| Hspb6 | 825.7361147 | 0.666159142 | 4.47E-02 |
| Gstt3 | 236.4840369 | 0.665396234 | 7.66E-03 |
| Ttc12 | 284.5390079 | 0.665088995 | 2.98E-02 |
| Slc1a5 | 4845.639286 | 0.663839757 | 3.32E-03 |
| Prr5l | 589.662235 | 0.663503878 | 4.38E-02 |
| Disp1 | 2262.130775 | 0.662589232 | 9.41E-09 |
| Anxa1 | 40517.67387 | 0.662232976 | 3.15E-05 |
| Pgam1 | 19698.03657 | 0.660918869 | 3.69E-03 |
| Spsb1 | 515.9985486 | 0.66029956 | 2.03E-04 |
| Large1 | 4373.999407 | 0.659570687 | 4.27E-11 |
| Macf1 | 20441.87418 | 0.659078339 | 1.31E-02 |
| Plod1 | 3492.523358 | 0.658542548 | 7.03E-04 |
| Atp1b3 | 3296.651774 | 0.658184453 | 5.50E-04 |
| Enpp5 | 660.2317262 | 0.657774665 | 2.04E-02 |
| Rbms2 | 13507.54112 | 0.657309326 | 5.56E-03 |
| AI429214 | 333.0356269 | 0.656766275 | 4.60E-05 |
| Chfr | 2461.953026 | 0.655022965 | 9.61E-04 |
| Lmbr1l | 980.757885 | 0.652366351 | 1.17E-02 |
| Hspg2 | 28446.74545 | 0.652274085 | 6.98E-03 |
| Gclc | 1433.972265 | 0.650246389 | 3.14E-03 |
| 4930523C07Rik | 776.0317708 | 0.650148989 | 3.09E-02 |
| Orai1 | 454.0094432 | 0.647982645 | 1.27E-02 |
| Adam9 | 10091.06238 | 0.647861068 | 1.48E-03 |
| Naaa | 680.2746552 | 0.647291193 | 2.34E-03 |
| Plpp1 | 5028.842612 | 0.646645043 | 2.77E-03 |
| Mafk | 2125.926029 | 0.646345365 | 2.02E-02 |
| Osgin2 | 294.2721574 | 0.645954995 | 4.12E-03 |
| H2-K1 | 1381.165026 | 0.645603611 | 3.36E-03 |
| Erich1 | 560.5336326 | 0.644753845 | 1.93E-04 |

| | | | | |
|---|---|---|---|---|
| Ap1s2 | | 981.3291552 | 0.644073902 | 1.46E-03 |
| | 3-Mar | 99.19508776 | 0.64277883 | 3.87E-02 |
| Abca3 | | 2846.907726 | 0.640992918 | 9.06E-06 |
| Fbxo32 | | 1611.242564 | 0.638160725 | 3.39E-02 |
| Apcdd1 | | 937.5246247 | 0.637516105 | 1.14E-03 |
| Dnaja4 | | 286.7765266 | 0.636319251 | 3.02E-02 |
| Myl12a | | 24114.22838 | 0.636315062 | 3.33E-05 |
| Slc39a13 | | 2542.695428 | 0.636028098 | 1.84E-03 |
| Pld2 | | 1123.434152 | 0.634485772 | 7.37E-04 |
| Lrrc27 | | 419.7854774 | 0.633965712 | 4.05E-02 |
| Tesk2 | | 669.5921419 | 0.632810594 | 1.84E-03 |
| Dhrs9 | | 1469.943128 | 0.631942475 | 3.55E-02 |
| Apba1 | | 301.8235088 | 0.630959616 | 2.98E-02 |
| Slc19a1 | | 824.1036615 | 0.629180662 | 3.92E-02 |
| Gmpr | | 213.3268615 | 0.628992627 | 1.37E-02 |
| Paics | | 14439.95988 | 0.628798729 | 1.98E-18 |
| Zfp595 | | 165.6562495 | 0.62788886 | 4.87E-02 |
| Bsg | | 26626.6997 | 0.626806678 | 2.72E-03 |
| Slc16a13 | | 668.4799613 | 0.626668764 | 1.99E-03 |
| Pqlc3 | | 1007.222603 | 0.626537955 | 1.16E-03 |
| Prps1 | | 1657.907258 | 0.62440086 | 1.11E-03 |
| Hras | | 1694.369396 | 0.62312053 | 3.65E-03 |
| Fyn | | 3259.615135 | 0.621388665 | 1.53E-09 |
| Tmem62 | | 479.9034088 | 0.620888211 | 2.56E-03 |
| Zfp57 | | 2113.079957 | 0.620469037 | 3.31E-02 |
| Ftl1 | | 156446.3262 | 0.62039849 | 7.81E-03 |
| Haus8 | | 750.9704817 | 0.620219224 | 4.17E-02 |
| Pex13 | | 1616.351738 | 0.620010392 | 1.31E-06 |
| Ptges | | 1888.104658 | 0.619997686 | 1.43E-02 |
| Snap47 | | 3433.560277 | 0.618561124 | 7.25E-05 |
| Esyt1 | | 2873.959478 | 0.617356982 | 3.30E-05 |
| Plpp2 | | 931.3288588 | 0.616713169 | 8.97E-04 |
| Ubl3 | | 1139.357592 | 0.616011178 | 4.03E-06 |
| Rtn3 | | 10304.32456 | 0.615865446 | 8.17E-05 |
| Man2b2 | | 7485.806257 | 0.614765038 | 2.26E-03 |
| Hoxa6 | | 269.2072896 | 0.614111022 | 2.47E-02 |
| Gm26691 | | 125.394815 | 0.613316832 | 4.07E-02 |
| Cyb5a | | 3603.739228 | 0.611768717 | 2.28E-03 |
| Acaa2 | | 2550.415331 | 0.610676804 | 6.02E-03 |
| Tex2 | | 1768.876038 | 0.609302223 | 1.52E-05 |
| Mcub | | 378.0966601 | 0.60876127 | 2.48E-02 |
| Itgb1 | | 37771.62253 | 0.60870336 | 9.40E-09 |
| Ldlrad4 | | 327.7250835 | 0.607907888 | 1.11E-03 |
| Dst | | 14726.69931 | 0.607363302 | 3.50E-03 |
| Nomo1 | | 5376.942128 | 0.605716484 | 1.02E-02 |
| Slc48a1 | | 9279.752174 | 0.604767443 | 1.64E-03 |
| Wdcp | | 317.2671453 | 0.603900039 | 8.65E-03 |

| | | | |
|---|---|---|---|
| Rb1 | 846.9033777 | 0.602680516 | 3.79E-03 |
| Zhx2 | 795.6214179 | 0.601976456 | 2.28E-02 |
| Glce | 2652.834273 | 0.601572233 | 4.74E-03 |
| Ifngr1 | 2047.302005 | 0.600885437 | 9.27E-07 |
| Fbxl2 | 422.3385071 | 0.600854252 | 2.02E-02 |
| Pdzrn3 | 1450.783321 | 0.600603283 | 5.01E-03 |
| Aph1b | 607.0118737 | 0.599281383 | 5.25E-03 |
| Glb1l | 1400.038872 | 0.599180571 | 2.24E-03 |
| Irak2 | 771.0028241 | 0.598205392 | 1.48E-03 |
| Telo2 | 1265.208066 | 0.597546607 | 1.06E-03 |
| Cyp39a1 | 736.9729686 | 0.597186818 | 6.12E-05 |
| Usp12 | 2202.538901 | 0.596035277 | 5.76E-06 |
| Trim7 | 396.6823301 | 0.595658692 | 1.08E-02 |
| Exoc6 | 416.7822632 | 0.595269803 | 1.79E-02 |
| Zfp763 | 273.8263546 | 0.595191163 | 4.06E-02 |
| Mtus1 | 3124.588945 | 0.594003153 | 2.02E-02 |
| Flot1 | 2483.620995 | 0.593734839 | 2.21E-03 |
| Ide | 5488.789588 | 0.593444207 | 2.45E-04 |
| Thsd7a | 4019.127381 | 0.592973805 | 1.07E-02 |
| Ppip5k1 | 941.1084221 | 0.591949704 | 1.52E-05 |
| Ehd4 | 2099.212753 | 0.591937807 | 9.46E-04 |
| Mlh3 | 176.7584739 | 0.591534765 | 4.09E-02 |
| Sec62 | 6353.816318 | 0.590540084 | 1.27E-03 |
| Naga | 3447.187179 | 0.589956261 | 4.99E-03 |
| Cers6 | 4036.257231 | 0.588258537 | 4.22E-04 |
| Heg1 | 6138.242062 | 0.587978373 | 3.61E-03 |
| Acyp2 | 175.9484139 | 0.587963764 | 7.85E-03 |
| Abcb1b | 1465.604472 | 0.587130442 | 4.71E-03 |
| Tef | 3446.398834 | 0.586952869 | 1.26E-02 |
| Sms | 1954.671987 | 0.586241339 | 2.08E-03 |
| Tpm2 | 4623.603642 | 0.58597966 | 1.33E-06 |
| Lrch1 | 792.2566081 | 0.58227873 | 5.45E-05 |
| Dzip1l | 1361.683775 | 0.581502363 | 3.99E-06 |
| Nagpa | 1014.528249 | 0.58003661 | 3.16E-03 |
| Cacna1c | 477.6875855 | 0.579802127 | 2.95E-02 |
| Cuedc1 | 1729.792206 | 0.5796951 | 7.28E-05 |
| Rap2b | 1894.249099 | 0.578696259 | 2.61E-06 |
| Abcg2 | 2509.433036 | 0.578355738 | 1.61E-03 |
| Gm43517 | 631.9198304 | 0.57783053 | 2.78E-02 |
| Lamb2 | 7100.083165 | 0.576064314 | 4.31E-02 |
| Slc35d1 | 859.5984113 | 0.575964537 | 2.84E-06 |
| Cbx5 | 6654.599166 | 0.57593936 | 4.05E-02 |
| Me1 | 3504.254363 | 0.57566681 | 1.97E-04 |
| Cdk6 | 743.1289463 | 0.574958707 | 2.70E-03 |
| Ptpmt1 | 905.1759236 | 0.574523908 | 2.49E-04 |
| Cdc42bpa | 4204.491388 | 0.574244465 | 3.62E-02 |
| Txnrd1 | 11050.67123 | 0.574095263 | 6.45E-03 |

| | | | |
|---|---|---|---|
| Slc38a4 | 4435.554761 | 0.573068404 | 3.16E-02 |
| Nucb1 | 11251.76517 | 0.572501845 | 4.16E-06 |
| Nhlrc2 | 1888.903339 | 0.572268096 | 1.92E-06 |
| Magi3 | 1822.469333 | 0.572259607 | 1.80E-02 |
| Tsc22d2 | 3177.01915 | 0.571685429 | 1.44E-02 |
| Pigl | 298.5422387 | 0.571033324 | 3.52E-04 |
| 1110012L19Rik | 1013.889739 | 0.570841757 | 1.25E-03 |
| Hsd17b10 | 3298.465071 | 0.570778106 | 1.17E-03 |
| Syngr2 | 2533.946804 | 0.569413358 | 1.94E-05 |
| Fam177a | 1383.292923 | 0.568387036 | 1.84E-03 |
| Clic4 | 7974.140184 | 0.56746807 | 9.17E-03 |
| Gpr155 | 418.0128172 | 0.567313538 | 1.76E-02 |
| Vcl | 15499.81698 | 0.566774836 | 4.59E-05 |
| Ifrd1 | 4709.926875 | 0.565766156 | 2.97E-02 |
| Ccn3 | 191.5912267 | 0.565346925 | 4.57E-02 |
| Fut11 | 609.8350578 | 0.565045278 | 4.07E-03 |
| Klc1 | 5592.891025 | 0.564267954 | 2.12E-07 |
| Stip1 | 10290.17675 | 0.563550056 | 8.60E-03 |
| Tep1 | 1733.768891 | 0.561574112 | 1.27E-05 |
| Rpain | 423.7311747 | 0.561485075 | 6.48E-03 |
| Azin2 | 2031.835172 | 0.561212297 | 9.94E-03 |
| F2r | 8616.173877 | 0.561004913 | 2.19E-05 |
| Il10rb | 939.8976722 | 0.55899581 | 6.42E-07 |
| Mrc2 | 8828.930084 | 0.558372982 | 1.14E-02 |
| Napb | 476.5876789 | 0.55757035 | 3.43E-03 |
| Nudt7 | 341.5085116 | 0.556788588 | 1.48E-03 |
| Ezr | 2125.964047 | 0.556465691 | 1.01E-03 |
| Ctbs | 628.3728415 | 0.556283326 | 2.62E-05 |
| Cpeb3 | 346.758479 | 0.554835534 | 1.54E-02 |
| Tmpo | 7007.127582 | 0.5545181 | 1.01E-02 |
| Gpx4 | 15070.71857 | 0.554310869 | 7.97E-03 |
| Tmco3 | 1560.168356 | 0.553647614 | 2.44E-05 |
| Itpripl2 | 5913.84639 | 0.552475063 | 1.28E-04 |
| Rhod | 1593.061284 | 0.551544511 | 1.78E-02 |
| Exoc3l | 208.8741871 | 0.551462781 | 8.44E-03 |
| Samd4 | 3008.136996 | 0.550390353 | 1.45E-04 |
| Prkce | 764.757114 | 0.550147468 | 7.71E-04 |
| Map1b | 6864.42189 | 0.548075006 | 4.94E-03 |
| Ctps | 9423.67962 | 0.547685301 | 8.23E-03 |
| Cd47 | 2998.687264 | 0.546968217 | 5.95E-03 |
| Fut8 | 1050.838888 | 0.545385934 | 6.57E-04 |
| Nudt18 | 604.0577606 | 0.544309521 | 8.65E-03 |
| Scamp1 | 1792.177025 | 0.542607807 | 2.69E-03 |
| Prdx5 | 5188.911969 | 0.542174143 | 1.30E-03 |
| Galm | 284.6776046 | 0.541802835 | 2.55E-02 |
| Galk1 | 1811.403168 | 0.541071433 | 1.63E-02 |
| Adamts10 | 953.871541 | 0.540649763 | 1.01E-04 |

| | | | |
|---|---|---|---|
| Ptch1 | 475.913733 | 0.539008653 | 3.35E-03 |
| Mosmo | 928.7988729 | 0.538587358 | 7.51E-03 |
| Abca7 | 2260.675367 | 0.538571384 | 7.86E-03 |
| Cdipt | 3223.076682 | 0.537211479 | 9.49E-06 |
| Myo6 | 987.1695516 | 0.536990002 | 3.72E-03 |
| Lamc1 | 22727.02501 | 0.536652779 | 1.46E-03 |
| Pde4dip | 2967.356983 | 0.536385586 | 4.85E-11 |
| Tspan9 | 8905.533771 | 0.536310618 | 6.86E-03 |
| H2-DMa | 231.9741204 | 0.5361076 | 3.98E-02 |
| Dusp3 | 2333.354456 | 0.535916148 | 6.17E-03 |
| Sptbn1 | 59834.79882 | 0.532563483 | 3.66E-03 |
| S100a6 | 44660.9589 | 0.531598194 | 1.78E-02 |
| Tmem60 | 583.2390495 | 0.531277118 | 2.16E-04 |
| Ctif | 1560.762332 | 0.531203631 | 1.53E-04 |
| Abi2 | 1463.946605 | 0.529089622 | 1.25E-03 |
| Kremen1 | 3328.728218 | 0.528921522 | 1.63E-04 |
| Wwc2 | 6641.559661 | 0.528867554 | 1.35E-04 |
| 1810037I17Rik | 1837.466688 | 0.527902182 | 5.93E-04 |
| Lgr4 | 1019.428449 | 0.527739794 | 1.88E-02 |
| Sptan1 | 24705.76715 | 0.527734828 | 4.24E-04 |
| Pcdh7 | 730.661321 | 0.526678029 | 4.37E-02 |
| Tnfaip1 | 4512.561739 | 0.526338148 | 1.00E-04 |
| Ube2h | 7839.735982 | 0.525944918 | 3.98E-02 |
| Slc20a2 | 2786.725506 | 0.525581468 | 2.26E-03 |
| Cd81 | 15598.87096 | 0.525137782 | 1.78E-02 |
| Tns1 | 7416.997207 | 0.524728668 | 9.74E-04 |
| Ap4b1 | 1040.56706 | 0.523381694 | 2.01E-06 |
| Ttc7b | 1433.927736 | 0.522868831 | 6.37E-04 |
| Emilin1 | 5288.590701 | 0.522158661 | 3.56E-02 |
| Snx7 | 2260.56405 | 0.52175808 | 4.24E-02 |
| Pml | 1978.102798 | 0.521703371 | 6.71E-03 |
| Emilin2 | 6079.768848 | 0.519392081 | 1.31E-04 |
| Pcdhb22 | 261.1195135 | 0.51886731 | 5.98E-03 |
| Tfdp2 | 1412.093919 | 0.518429164 | 1.43E-02 |
| Ncoa3 | 2075.835614 | 0.517543474 | 4.39E-06 |
| Cd99l2 | 3754.262879 | 0.514687716 | 1.32E-04 |
| Sike1 | 1752.912471 | 0.514453742 | 9.13E-04 |
| Tpk1 | 262.2730087 | 0.514408339 | 2.43E-02 |
| Dlg1 | 7243.816205 | 0.513461124 | 8.53E-03 |
| Ostf1 | 2423.365697 | 0.513066831 | 5.51E-06 |
| Nrip1 | 567.8159237 | 0.512399419 | 9.59E-03 |
| Abhd17c | 2306.603289 | 0.509935489 | 4.79E-03 |
| Smco4 | 219.148333 | 0.509827322 | 3.59E-02 |
| Lonp1 | 6941.8007 | 0.507960094 | 3.63E-03 |
| Coq7 | 524.6389928 | 0.507950278 | 3.84E-03 |
| Net1 | 1218.273579 | 0.507910011 | 1.15E-02 |
| 1700047I17Rik2 | 769.1116899 | 0.504137071 | 7.65E-03 |

| | | | |
|---|---|---|---|
| Cald1 | 39536.21402 | 0.50350282 | 1.75E-02 |
| Cav2 | 3663.325958 | 0.502699989 | 2.62E-02 |
| Lrrk1 | 3325.760949 | 0.502643712 | 1.45E-03 |
| Usp32 | 4236.091678 | 0.501567212 | 2.00E-02 |
| Gng8 | 129.1031785 | 0.500940246 | 3.51E-02 |
| Ccdc181 | 639.3289339 | 0.499378675 | 3.32E-03 |
| Sdhaf2 | 1871.044111 | 0.498967482 | 1.44E-04 |
| Hoxa5 | 641.1710565 | 0.498459513 | 1.53E-02 |
| Clcn5 | 3296.841631 | 0.497146236 | 2.32E-09 |
| Fyco1 | 2113.385571 | 0.496663897 | 2.34E-04 |
| Magi1 | 863.00933 | 0.496587005 | 1.61E-03 |
| Mief2 | 738.7242967 | 0.496552212 | 4.39E-02 |
| Galc | 1514.601027 | 0.49577866 | 2.98E-03 |
| Ebna1bp2 | 3539.682864 | 0.495586489 | 3.48E-02 |
| Mfsd8 | 360.6439686 | 0.493660607 | 3.88E-02 |
| Uxt | 499.6757564 | 0.492398258 | 2.37E-02 |
| Arhgdib | 13907.08035 | 0.491965793 | 1.46E-02 |
| Specc1 | 3432.279109 | 0.490903397 | 3.39E-02 |
| Ahnak | 67844.71225 | 0.49059535 | 3.21E-03 |
| Sec14l1 | 5437.380374 | 0.489653091 | 8.36E-05 |
| Chchd4 | 753.277916 | 0.487763711 | 1.14E-02 |
| Gm11110 | 223.6628417 | 0.486893867 | 3.61E-02 |
| Jmjd4 | 1066.637866 | 0.486537397 | 2.33E-05 |
| Htra1 | 323.732752 | 0.486512925 | 1.89E-02 |
| Cachd1 | 1495.499593 | 0.485730365 | 3.25E-03 |
| Aldoa | 31312.43539 | 0.485100576 | 4.16E-02 |
| Slc19a2 | 1917.481573 | 0.484743225 | 8.76E-03 |
| Fig4 | 2668.683324 | 0.484007061 | 1.24E-02 |
| Limk2 | 1846.809037 | 0.483792749 | 2.71E-03 |
| Tead1 | 8305.982112 | 0.482215752 | 1.84E-03 |
| Npnt | 449.3441054 | 0.48060413 | 4.16E-02 |
| Bnip3 | 1792.535201 | 0.479526219 | 1.25E-02 |
| Vcam1 | 8733.754356 | 0.478599343 | 8.65E-04 |
| Rapgef2 | 1760.896602 | 0.478317977 | 2.10E-02 |
| Slc9a1 | 2227.40331 | 0.476839842 | 1.42E-05 |
| Ankrd40 | 3457.239698 | 0.475927291 | 6.06E-05 |
| Cox5a | 4353.767426 | 0.475918158 | 7.45E-03 |
| Amotl2 | 15490.23543 | 0.475110963 | 4.64E-02 |
| Fdx1 | 723.8667476 | 0.474210556 | 3.88E-02 |
| Znrd1 | 628.2231869 | 0.47265565 | 4.90E-02 |
| Narf | 1213.76747 | 0.472328036 | 3.83E-02 |
| Mvp | 5662.63892 | 0.472042493 | 9.88E-03 |
| Slc22a5 | 722.5646486 | 0.471752933 | 1.28E-02 |
| Inpp5k | 2174.199723 | 0.471261404 | 2.18E-02 |
| Tcf7l2 | 966.9965045 | 0.47073123 | 1.11E-02 |
| Acap3 | 1180.426838 | 0.470372907 | 8.65E-03 |
| S100a10 | 14257.32699 | 0.469829919 | 3.81E-03 |

| | | | |
|---|---|---|---|
| Gng12 | 10176.62266 | 0.469681262 | 5.82E-03 |
| Epb41l4a | 2231.957676 | 0.469126054 | 2.97E-04 |
| Fat4 | 5165.505281 | 0.468997328 | 4.93E-02 |
| Ccdc9b | 3998.146773 | 0.468757868 | 1.86E-03 |
| Slc23a2 | 2101.09374 | 0.468545757 | 3.65E-02 |
| Lactb2 | 1674.430397 | 0.466091492 | 2.64E-02 |
| Tmsb10 | 24402.93356 | 0.465817126 | 9.42E-03 |
| Ddr2 | 21485.53935 | 0.46535193 | 4.96E-02 |
| Nmnat1 | 220.6102235 | 0.465260844 | 1.69E-02 |
| Hoxc4 | 166.5136363 | 0.465109515 | 4.35E-02 |
| Mfsd12 | 933.9556439 | 0.463649949 | 4.28E-02 |
| Ogfod1 | 1856.045924 | 0.463487272 | 3.15E-03 |
| Rab29 | 491.9800228 | 0.462862867 | 1.01E-02 |
| Fn3krp | 425.7525463 | 0.459560858 | 8.35E-03 |
| Atl2 | 1798.458299 | 0.458969892 | 6.02E-04 |
| Pfdn4 | 874.6926454 | 0.458326461 | 5.84E-03 |
| Emp3 | 1776.507322 | 0.45681035 | 3.03E-02 |
| Arhgef5 | 944.6905667 | 0.455365121 | 3.19E-03 |
| Slc8b1 | 1315.172544 | 0.455075417 | 4.79E-02 |
| Smpd1 | 2430.281075 | 0.45493175 | 2.56E-02 |
| 4921524J17Rik | 641.6105448 | 0.45479881 | 4.32E-03 |
| Cers4 | 1306.675198 | 0.454566505 | 6.56E-04 |
| Insig2 | 2774.628956 | 0.454090997 | 7.14E-03 |
| Fkbp10 | 8574.958963 | 0.453020496 | 2.87E-02 |
| Otud7b | 4504.227944 | 0.452553206 | 1.66E-06 |
| Abcc1 | 6563.18358 | 0.452518383 | 2.75E-03 |
| Dtna | 3290.17433 | 0.450005629 | 6.55E-03 |
| Prrx1 | 19059.58088 | 0.449939244 | 1.43E-03 |
| Ctsl | 40956.20104 | 0.449938211 | 1.36E-02 |
| Ampd3 | 3598.275107 | 0.449871272 | 1.01E-02 |
| Acot9 | 3567.356035 | 0.449652475 | 2.55E-02 |
| Rtn4 | 18961.53588 | 0.449057803 | 4.05E-03 |
| Cnnm4 | 356.2181148 | 0.448621943 | 1.66E-02 |
| Kank2 | 6902.794724 | 0.447819435 | 6.18E-04 |
| Ctnnal1 | 2217.420049 | 0.447726162 | 4.65E-02 |
| NA | 1559.71249 | 0.447562069 | 3.46E-02 |
| Map1a | 6290.433199 | 0.446291939 | 8.44E-03 |
| Thnsl1 | 299.7345792 | 0.445408389 | 3.33E-02 |
| Bcl2l1 | 4294.000393 | 0.44467719 | 3.03E-02 |
| Tmem106b | 10840.02422 | 0.444604181 | 4.88E-02 |
| Tpm1 | 28225.26008 | 0.444241421 | 8.78E-04 |
| Etfb | 2261.36003 | 0.443307174 | 6.26E-03 |
| Bag1 | 4120.741985 | 0.441914489 | 1.14E-02 |
| Got1 | 2662.479814 | 0.4415166 | 1.43E-02 |
| Sema3f | 4201.918047 | 0.440462437 | 2.28E-02 |
| Cdk5rap1 | 416.9799843 | 0.44029568 | 1.62E-03 |
| Col4a3bp | 1675.621637 | 0.440090006 | 4.55E-03 |

| | | | |
|---|---|---|---|
| Irgm1 | 1281.382103 | 0.439186481 | 4.95E-03 |
| Ak1 | 2929.214187 | 0.439146135 | 1.14E-02 |
| Hprt | 2471.179009 | 0.438637448 | 2.84E-02 |
| Pear1 | 2676.766374 | 0.437976841 | 1.08E-02 |
| Rbm18 | 1817.797828 | 0.437765277 | 2.26E-02 |
| Nsmaf | 1785.080286 | 0.437712172 | 3.82E-04 |
| Ap2a2 | 11186.22479 | 0.437626782 | 4.68E-03 |
| Add3 | 3445.316954 | 0.436945237 | 1.11E-02 |
| Adarb1 | 849.0097309 | 0.435509885 | 2.34E-02 |
| Necap1 | 2428.802313 | 0.434206109 | 2.40E-03 |
| Arfgap1 | 3463.669508 | 0.433260693 | 6.09E-03 |
| Ganab | 13601.05205 | 0.432467081 | 1.81E-04 |
| Traf6 | 1586.592483 | 0.432147229 | 8.06E-05 |
| Mitd1 | 486.7458551 | 0.431764858 | 4.61E-02 |
| Akap13 | 5008.332383 | 0.430962291 | 2.15E-02 |
| Erlin2 | 2954.042327 | 0.430013631 | 4.65E-03 |
| Itprid2 | 2850.831262 | 0.428899212 | 3.77E-04 |
| Eya1 | 701.8287389 | 0.428860945 | 1.26E-03 |
| Pdia6 | 14361.81594 | 0.428527056 | 1.06E-02 |
| Rab10 | 6118.675221 | 0.428490931 | 2.04E-04 |
| Map2k3 | 4419.736716 | 0.428288927 | 2.00E-02 |
| Tmem80 | 217.7392199 | 0.427632028 | 4.05E-02 |
| Dagla | 576.2981415 | 0.427502589 | 3.34E-03 |
| Gin1 | 216.2621153 | 0.425951845 | 4.86E-02 |
| Lrrc75a | 924.7491795 | 0.424464679 | 1.75E-02 |
| Vwa8 | 2171.685012 | 0.424043258 | 1.94E-02 |
| Dag1 | 10803.36973 | 0.423923008 | 6.57E-04 |
| NA | 387.3489469 | 0.423201644 | 6.10E-03 |
| Mprip | 17779.67662 | 0.423033016 | 7.46E-07 |
| Tmem135 | 1215.614319 | 0.421616494 | 2.13E-02 |
| Mapk14 | 3703.827213 | 0.42111008 | 3.16E-04 |
| Stx16 | 1833.507757 | 0.420711815 | 4.65E-02 |
| Isca2 | 752.0861875 | 0.419232956 | 2.44E-03 |
| 11-Sep | 12235.26164 | 0.419187647 | 6.46E-03 |
| Galnt1 | 3097.996652 | 0.418170841 | 1.18E-02 |
| Adar | 2479.583721 | 0.417853094 | 1.01E-03 |
| Pde10a | 470.0570704 | 0.414899897 | 1.56E-02 |
| P4hb | 39144.28767 | 0.413217457 | 3.48E-02 |
| Piezo1 | 14797.30952 | 0.409177891 | 2.91E-02 |
| Myadm | 15438.02828 | 0.408393314 | 9.11E-05 |
| Edem1 | 4433.87396 | 0.408136505 | 3.76E-04 |
| Ripor1 | 4272.017785 | 0.407205293 | 3.25E-03 |
| Plekhb2 | 3753.770953 | 0.406775104 | 4.02E-02 |
| Eci1 | 1615.091732 | 0.406196241 | 7.14E-03 |
| Cd109 | 12991.36315 | 0.406003521 | 4.61E-02 |
| Mr1 | 1889.33807 | 0.40532054 | 2.47E-02 |
| Sri | 3410.645404 | 0.404159068 | 8.54E-05 |

| | | | |
|---|---|---|---|
| Nat10 | 2589.324134 | 0.403994249 | 2.20E-02 |
| Phka2 | 2086.31599 | 0.40299882 | 1.58E-02 |
| AC149090.1 | 18121.43559 | 0.401887674 | 6.36E-03 |
| Dnajb6 | 7299.017882 | 0.401835648 | 1.43E-02 |
| Cyld | 2898.729581 | 0.401814934 | 4.00E-02 |
| Mthfr | 3144.198278 | 0.401097955 | 1.10E-02 |
| Ptp4a2 | 11669.3148 | 0.399987709 | 8.17E-03 |
| Atl3 | 9377.879914 | 0.398398409 | 3.40E-02 |
| Entpd4 | 1607.124128 | 0.398308703 | 1.86E-02 |
| Wfs1 | 1732.71301 | 0.397444722 | 2.28E-02 |
| Fbxo10 | 1508.820967 | 0.397244064 | 3.22E-02 |
| Ajuba | 2948.78609 | 0.396295469 | 3.04E-02 |
| Snap23 | 1353.916862 | 0.395844033 | 7.10E-03 |
| Gpx1 | 3782.337533 | 0.395843612 | 3.60E-02 |
| Taok3 | 2501.86846 | 0.395427596 | 2.93E-04 |
| Sqor | 2866.171031 | 0.395106142 | 1.05E-02 |
| Arhgap17 | 1804.268498 | 0.394165647 | 5.16E-04 |
| Fuz | 552.4981124 | 0.392324294 | 3.01E-02 |
| Cpd | 4849.178331 | 0.392114831 | 1.95E-02 |
| Kpna6 | 7686.854538 | 0.391815091 | 1.89E-03 |
| Hspb8 | 6743.145759 | 0.391223665 | 2.19E-02 |
| Dcaf17 | 807.4886767 | 0.391051855 | 3.46E-02 |
| Elob | 5284.404596 | 0.390685869 | 1.11E-02 |
| Ehbp1l1 | 2904.929473 | 0.390618097 | 7.37E-03 |
| Uaca | 2130.933084 | 0.390157241 | 2.96E-02 |
| Cnnm2 | 1162.640313 | 0.389981127 | 3.41E-02 |
| E130308A19Rik | 977.5793705 | 0.388642277 | 3.56E-02 |
| Vti1a | 1513.738788 | 0.388457878 | 7.60E-03 |
| Higd2a | 2214.016855 | 0.387410516 | 3.28E-02 |
| Gm10561 | 210.1280875 | 0.387340416 | 4.55E-02 |
| Cdkn2c | 2309.987574 | 0.387114199 | 2.40E-02 |
| Arpc2 | 10253.14542 | 0.38667313 | 7.56E-05 |
| Rad23b | 17055.86365 | 0.384944324 | 2.98E-02 |
| Helz | 2920.114213 | 0.38436686 | 2.79E-03 |
| Iqsec1 | 6250.869472 | 0.383158333 | 3.41E-02 |
| Samd8 | 1859.769831 | 0.381578661 | 1.66E-02 |
| Cnn2 | 23457.26203 | 0.380562804 | 1.01E-02 |
| Tmem38a | 516.3255784 | 0.380359068 | 2.85E-02 |
| Ugp2 | 4229.526194 | 0.380065987 | 7.90E-03 |
| Trmt1l | 1562.328931 | 0.379970519 | 7.89E-04 |
| Nudcd3 | 5085.866745 | 0.379826281 | 4.57E-02 |
| Faf1 | 5358.729333 | 0.379652148 | 9.43E-03 |
| Fam104a | 1760.116884 | 0.379437204 | 3.39E-02 |
| Lmbrd1 | 2417.679745 | 0.378924331 | 1.33E-02 |
| Aifm2 | 1026.37371 | 0.378683825 | 6.95E-03 |
| Creb3 | 1212.471112 | 0.378372737 | 4.87E-02 |
| Cast | 5248.563907 | 0.37805782 | 5.09E-04 |

| | | | |
|---|---|---|---|
| Elf4 | 2047.77655 | 0.377959605 | 5.33E-04 |
| Nectin2 | 3231.192606 | 0.375478324 | 3.02E-02 |
| Wdr44 | 567.3078697 | 0.374539927 | 2.91E-02 |
| Agpat5 | 1466.315111 | 0.374391237 | 2.19E-02 |
| Ptpn12 | 2592.977983 | 0.373702421 | 3.11E-02 |
| Dnm2 | 5735.557998 | 0.373652947 | 4.73E-02 |
| Hibadh | 3736.286012 | 0.3727742 | 3.22E-02 |
| Rdh13 | 1335.03426 | 0.371862497 | 2.65E-02 |
| Tnrc6c | 2070.107347 | 0.371557835 | 9.92E-04 |
| Ptgfrn | 7767.767633 | 0.371453576 | 4.57E-02 |
| Mpp1 | 3836.686581 | 0.370188242 | 4.64E-02 |
| Aph1a | 5523.669385 | 0.370080758 | 2.84E-02 |
| Aak1 | 1915.969594 | 0.368327842 | 5.26E-03 |
| Larp4b | 2101.209701 | 0.367415303 | 8.84E-03 |
| Cpt1a | 5815.364835 | 0.36721066 | 2.77E-03 |
| Slc39a6 | 3751.096942 | 0.366994732 | 4.54E-02 |
| Notch2 | 5564.204759 | 0.365818663 | 6.09E-03 |
| Pip5k1a | 3867.199522 | 0.365771425 | 1.21E-02 |
| Pigg | 682.5190938 | 0.365637992 | 4.66E-02 |
| Xpo5 | 4096.279036 | 0.365532767 | 4.39E-02 |
| Ywhah | 6364.92468 | 0.365438721 | 3.78E-02 |
| Vps53 | 1711.372237 | 0.364664152 | 1.52E-03 |
| Osbpl1a | 1582.537579 | 0.364586439 | 3.24E-02 |
| Vps36 | 2824.129634 | 0.364112393 | 4.58E-03 |
| Trim8 | 1911.483462 | 0.363701966 | 4.08E-03 |
| Zfp91 | 7065.381387 | 0.363399407 | 6.33E-03 |
| Eif3c | 17700.40432 | 0.362631474 | 6.37E-03 |
| Tsku | 1205.341302 | 0.358866936 | 2.29E-02 |
| Ehd2 | 12732.25354 | 0.358014869 | 1.67E-02 |
| Cdc42se2 | 2787.71278 | 0.357489123 | 3.93E-04 |
| Cdc42 | 11259.05128 | 0.354689387 | 4.10E-05 |
| Pygo2 | 2490.294348 | 0.352486175 | 5.66E-03 |
| Rubcn | 3167.973214 | 0.351941834 | 1.62E-02 |
| Rapgef6 | 1995.599441 | 0.351170493 | 3.86E-02 |
| Ptprf | 4297.628625 | 0.351073754 | 1.51E-03 |
| Kif1c | 11260.85304 | 0.34978144 | 3.25E-02 |
| Fam98a | 1905.214616 | 0.348749848 | 5.18E-03 |
| Pacsin2 | 4361.917334 | 0.348603317 | 4.96E-03 |
| Plxnb2 | 19455.25205 | 0.347735664 | 2.54E-02 |
| Tmem128 | 899.2173669 | 0.345684991 | 3.41E-02 |
| Pnpla6 | 1143.850243 | 0.345551383 | 1.18E-02 |
| Zfp39 | 324.6592607 | 0.345028127 | 3.16E-02 |
| Rab7b | 1493.162856 | 0.343905259 | 3.04E-02 |
| Txndc12 | 3770.539548 | 0.343579262 | 2.27E-02 |
| Chmp2b | 2474.73103 | 0.343434673 | 2.23E-02 |
| Cipc | 1168.346399 | 0.342667918 | 7.65E-03 |
| Prkag2 | 1252.873195 | 0.342583711 | 1.76E-02 |

| | | | |
|---|---|---|---|
| Bcl9l | 4861.60902 | 0.341587693 | 4.21E-03 |
| Snap29 | 2083.038228 | 0.341266564 | 2.66E-02 |
| Nfe2l1 | 17080.71886 | 0.341185863 | 4.57E-02 |
| Pdha1 | 9868.078405 | 0.340507505 | 6.37E-04 |
| Parva | 10879.78875 | 0.340178473 | 9.88E-03 |
| Gpd2 | 3642.520661 | 0.339947037 | 2.79E-03 |
| Nsf | 2095.639979 | 0.339806742 | 4.31E-03 |
| Polr2e | 3801.975991 | 0.339497565 | 4.60E-02 |
| Zdhhc3 | 3899.941909 | 0.338591302 | 3.05E-02 |
| 1700021F05Rik | 907.3929152 | 0.335732669 | 1.13E-03 |
| Tor1aip2 | 6209.143612 | 0.334210055 | 2.64E-04 |
| Vapa | 4514.815785 | 0.333756403 | 4.49E-02 |
| Pi4ka | 5631.269742 | 0.330665001 | 4.29E-03 |
| Ddb1 | 78266.76622 | 0.330347131 | 3.18E-02 |
| Ncstn | 4112.498853 | 0.326656689 | 1.60E-02 |
| Anxa7 | 4754.841561 | 0.325815418 | 4.94E-03 |
| Bmp1 | 4680.338256 | 0.324685556 | 1.21E-02 |
| Pea15a | 11198.47623 | 0.321712347 | 4.71E-02 |
| Myl6 | 22034.81211 | 0.320818829 | 2.98E-04 |
| Sh3pxd2a | 15740.33814 | 0.320386538 | 3.42E-04 |
| Aldh3a2 | 3225.872424 | 0.319785251 | 8.03E-03 |
| Ankrd13c | 1592.952556 | 0.316235737 | 2.17E-02 |
| Ascc2 | 3721.48564 | 0.316177181 | 2.37E-02 |
| Man1b1 | 5748.065786 | 0.315102281 | 1.49E-02 |
| Emp2 | 4062.72683 | 0.314230594 | 4.81E-03 |
| Smarcd3 | 630.858313 | 0.312333104 | 2.94E-02 |
| Rapgef1 | 5967.043459 | 0.311721205 | 4.53E-02 |
| Npepps | 9806.645939 | 0.31104835 | 3.59E-02 |
| Utp23 | 570.5468979 | 0.309468424 | 9.14E-03 |
| Slc1a4 | 3460.178396 | 0.308777972 | 4.69E-02 |
| Serinc3 | 12902.53338 | 0.307503021 | 3.65E-02 |
| Osbpl11 | 1140.020702 | 0.30510506 | 1.58E-02 |
| Lmtk2 | 2086.187159 | 0.302220141 | 1.49E-02 |
| Zzef1 | 3506.293149 | 0.300326671 | 1.37E-02 |
| Ptpn14 | 6806.862281 | 0.299179107 | 2.20E-02 |
| Stat6 | 5625.010172 | 0.299153632 | 1.79E-03 |
| Ccser2 | 3364.433109 | 0.297681348 | 3.74E-02 |
| Kif1b | 4905.83088 | 0.297231303 | 1.91E-02 |
| Vrk3 | 1213.537914 | 0.296537283 | 2.15E-02 |
| Zfyve9 | 2465.724035 | 0.293986956 | 2.02E-02 |
| Pomgnt1 | 3559.582041 | 0.292833802 | 2.99E-02 |
| Socs6 | 1293.79635 | 0.290524541 | 4.91E-02 |
| Stat5b | 2117.456746 | 0.288682944 | 6.02E-03 |
| Plgrkt | 1248.255989 | 0.288402821 | 4.15E-03 |
| Tln1 | 13133.47837 | 0.28504189 | 4.24E-02 |
| BC031181 | 1034.180156 | 0.284462296 | 3.49E-02 |
| Adam17 | 3233.205911 | 0.27913145 | 4.51E-03 |

| | | | |
|---|---|---|---|
| Hipk1 | 4679.274894 | 0.277688963 | 2.61E-02 |
| Araf | 2783.283806 | 0.27573647 | 1.35E-02 |
| Pitpnm1 | 3326.041469 | 0.274571592 | 4.89E-02 |
| Minpp1 | 2115.190894 | 0.271772417 | 2.61E-02 |
| Ccdc43 | 996.7600218 | 0.271288768 | 4.72E-02 |
| Epb41l3 | 11414.40963 | 0.270760639 | 2.00E-02 |
| Cyfip1 | 6745.902934 | 0.27060747 | 2.82E-02 |
| Cdk11b | 2039.570556 | 0.270066969 | 2.75E-02 |
| 4931406P16Rik | 2722.4401 | 0.269372691 | 4.79E-02 |
| Adcy6 | 3708.430534 | 0.268779487 | 1.26E-02 |
| Sfxn3 | 5162.307059 | 0.264614226 | 4.63E-02 |
| Utp11 | 3221.327935 | 0.262553697 | 4.77E-02 |
| Man2a1 | 13475.00019 | 0.25941099 | 3.39E-02 |
| Tmbim4 | 1686.78166 | 0.258015216 | 4.91E-02 |
| Fam114a2 | 3449.452351 | 0.256961025 | 4.82E-03 |
| Zfyve27 | 1433.895386 | 0.248889303 | 2.17E-02 |
| Habp4 | 1289.065897 | 0.248810314 | 3.65E-03 |
| Tmem131 | 4629.57427 | 0.246120511 | 1.95E-02 |
| Ubr5 | 12270.45866 | 0.243176112 | 3.62E-02 |
| Gpx8 | 2857.904981 | 0.241608875 | 2.04E-02 |
| Cers2 | 6182.42433 | 0.237681831 | 4.00E-02 |
| Heatr5a | 3537.032457 | 0.230496145 | 2.39E-02 |
| Nras | 3618.572248 | 0.223920464 | 3.24E-02 |
| Eml3 | 1886.203232 | 0.223485611 | 3.75E-02 |
| Gnb1 | 10433.30772 | 0.223119688 | 3.67E-03 |
| Agl | 3359.570353 | 0.213324832 | 1.99E-02 |
| Glud1 | 8559.981822 | 0.211510855 | 1.52E-02 |
| Zfp106 | 11251.11847 | 0.208047842 | 1.27E-02 |
| Kif3b | 2552.903667 | 0.187415104 | 3.28E-02 |
| Uri1 | 1845.880355 | -0.184471572 | 1.73E-02 |
| Ppp2r5e | 2577.239633 | -0.186804377 | 2.85E-02 |
| Yap1 | 5868.317144 | -0.187134381 | 4.27E-02 |
| Kdm1a | 3091.893595 | -0.194058127 | 3.08E-02 |
| Fem1c | 1546.41211 | -0.203771382 | 4.71E-02 |
| Abcd3 | 3631.045164 | -0.203933279 | 1.81E-02 |
| Wwtr1 | 5195.543564 | -0.204087768 | 4.25E-02 |
| Slc36a4 | 1156.576941 | -0.205466213 | 3.42E-02 |
| Plxna1 | 20953.16237 | -0.211877679 | 2.22E-02 |
| Fcf1 | 1579.920549 | -0.213209418 | 4.04E-02 |
| Ddx6 | 7985.738967 | -0.214675073 | 4.87E-02 |
| Qrich1 | 3515.068439 | -0.217060488 | 4.14E-02 |
| Ppfia1 | 3284.486732 | -0.229999191 | 3.28E-02 |
| Armc1 | 1584.697443 | -0.235134773 | 3.50E-02 |
| Men1 | 2488.414312 | -0.235842221 | 3.55E-02 |
| Kirrel | 13677.00951 | -0.23907577 | 4.07E-03 |
| Ccny | 4186.355601 | -0.241403575 | 1.85E-02 |
| Zmat2 | 2291.274399 | -0.241414577 | 3.48E-02 |

| | | | |
|---|---|---|---|
| AI597479 | 1129.332889 | -0.241844915 | 5.66E-03 |
| Prdm2 | 2816.518592 | -0.242736102 | 4.16E-02 |
| Ctbp2 | 1278.494096 | -0.243641455 | 2.79E-02 |
| Ago1 | 6615.470307 | -0.243786557 | 4.07E-03 |
| Swi5 | 5268.153823 | -0.244501041 | 1.67E-02 |
| Cdc5l | 3272.304668 | -0.245550558 | 3.95E-02 |
| Wdr59 | 1368.374064 | -0.247651714 | 3.73E-02 |
| Chd4 | 15088.84888 | -0.249483009 | 2.46E-02 |
| Hdgfl2 | 4078.471706 | -0.252983278 | 4.64E-02 |
| 2610002M06Rik | 1470.102267 | -0.253170509 | 3.48E-02 |
| Ddi2 | 2416.311868 | -0.253460025 | 2.83E-02 |
| Smug1 | 791.1906468 | -0.253949817 | 3.81E-02 |
| Gng10 | 2800.643553 | -0.254894095 | 2.58E-02 |
| 2610301B20Rik | 1235.416847 | -0.258415396 | 4.49E-02 |
| Trip12 | 11199.36888 | -0.258961409 | 2.42E-02 |
| Chtop | 5837.65029 | -0.260253123 | 2.48E-02 |
| Taf2 | 2167.708385 | -0.260347598 | 4.68E-05 |
| Pprc1 | 3591.050554 | -0.266021164 | 4.95E-02 |
| Rpl36al | 5933.505914 | -0.266660114 | 3.83E-02 |
| Cdk5 | 1892.485916 | -0.266921716 | 3.03E-02 |
| Smarce1 | 5139.95045 | -0.267426263 | 1.75E-02 |
| Ythdf2 | 2107.953455 | -0.270268214 | 3.74E-02 |
| Ctcf | 2881.745882 | -0.270977354 | 1.95E-02 |
| 2810004N23Rik | 1657.133658 | -0.271256952 | 2.06E-02 |
| Thumpd1 | 2170.674412 | -0.272777857 | 2.68E-03 |
| Cttn | 14805.07831 | -0.274013605 | 1.23E-03 |
| Armc10 | 1676.034362 | -0.274502051 | 4.82E-02 |
| Mrps9 | 1479.100855 | -0.276120561 | 7.63E-03 |
| Synj2 | 1645.375646 | -0.277246152 | 1.05E-02 |
| Letmd1 | 1317.473792 | -0.277267871 | 1.79E-02 |
| Wdr92 | 843.1441166 | -0.277530481 | 4.66E-02 |
| Slc7a6os | 1293.861495 | -0.278977549 | 2.23E-02 |
| Btf3l4 | 4495.978449 | -0.280831717 | 1.12E-02 |
| Rbl2 | 2066.659329 | -0.281202258 | 4.99E-03 |
| Brd1 | 2882.283459 | -0.281688734 | 3.56E-04 |
| Lclat1 | 2716.89391 | -0.281827704 | 4.03E-02 |
| Vipas39 | 1504.93256 | -0.282051623 | 3.39E-02 |
| Dock7 | 10072.46313 | -0.282330579 | 9.41E-03 |
| Wdr5 | 2950.251821 | -0.282856265 | 3.83E-02 |
| Ppa2 | 1042.259052 | -0.282959541 | 3.99E-02 |
| Gmps | 4740.881068 | -0.283600501 | 3.83E-02 |
| Klhl42 | 1851.794143 | -0.284177036 | 1.46E-02 |
| Trub2 | 1464.184844 | -0.284242954 | 2.89E-02 |
| Ibtk | 1777.858862 | -0.285184718 | 2.83E-02 |
| Pcgf3 | 3689.92062 | -0.286067811 | 4.59E-03 |
| Rmnd5a | 3357.074221 | -0.286117817 | 3.80E-02 |
| Smcr8 | 3771.341043 | -0.286835947 | 4.25E-02 |

| | | | |
|---|---|---|---|
| Clns1a | 2100.979167 | -0.288199768 | 2.57E-02 |
| Csnk1g2 | 5738.741204 | -0.291456499 | 4.88E-02 |
| Rundc1 | 970.3247728 | -0.291634767 | 4.73E-02 |
| Nek6 | 2770.3449 | -0.292709126 | 4.32E-02 |
| Xpnpep3 | 989.0992619 | -0.293275393 | 4.05E-03 |
| Camkk2 | 1761.969751 | -0.293618328 | 2.24E-02 |
| Zc3hav1l | 1399.674989 | -0.293796038 | 1.09E-02 |
| Zfp334 | 1151.134813 | -0.294951696 | 2.42E-02 |
| Tbc1d24 | 2492.496938 | -0.295072814 | 2.80E-02 |
| Elp2 | 4679.842283 | -0.295982431 | 2.86E-02 |
| Rnf115 | 1861.018874 | -0.299330268 | 2.69E-02 |
| Rtf1 | 2978.284882 | -0.299518198 | 4.87E-02 |
| Ank | 9508.851247 | -0.300260501 | 4.87E-02 |
| Timm8b | 906.7143595 | -0.300807133 | 1.21E-02 |
| Mex3c | 2408.706296 | -0.301178054 | 3.20E-02 |
| Tbl1xr1 | 2044.586522 | -0.301664168 | 4.82E-02 |
| BC048403 | 738.7293582 | -0.301943034 | 4.21E-02 |
| Nhlrc3 | 2088.125293 | -0.304270049 | 2.15E-02 |
| Egln2 | 2115.922258 | -0.304628957 | 4.73E-02 |
| Tsta3 | 1507.503282 | -0.304914663 | 3.50E-02 |
| Mrps5 | 2126.38738 | -0.305611807 | 1.24E-02 |
| Max | 2035.937597 | -0.305689664 | 1.36E-02 |
| Jmy | 1698.02112 | -0.307249288 | 4.53E-03 |
| Sco1 | 557.039177 | -0.307584175 | 2.10E-02 |
| Zbtb9 | 615.7391535 | -0.308174711 | 4.59E-02 |
| Sart3 | 2034.691382 | -0.308206892 | 4.16E-02 |
| Usp42 | 1288.769798 | -0.308263172 | 3.79E-03 |
| Zmiz2 | 2535.331176 | -0.309014478 | 2.29E-02 |
| Tmem186 | 774.7522416 | -0.309154443 | 4.48E-02 |
| Zfp358 | 1303.342284 | -0.310048281 | 3.11E-02 |
| Trnau1ap | 1335.035752 | -0.311939062 | 3.83E-02 |
| Fam53a | 1630.046894 | -0.312994646 | 3.96E-02 |
| Tubgcp2 | 1519.63264 | -0.314416249 | 4.72E-02 |
| Atxn7l3 | 3554.494769 | -0.315787478 | 3.25E-03 |
| Zyx | 23861.9633 | -0.315885582 | 3.94E-02 |
| Acap2 | 6513.117941 | -0.316543548 | 4.39E-02 |
| Fastkd1 | 692.7686316 | -0.317306313 | 2.46E-02 |
| Gigyf2 | 4619.799954 | -0.319007152 | 4.16E-04 |
| Nono | 17185.03395 | -0.319128247 | 3.30E-02 |
| Eif4a1 | 28084.96844 | -0.321038514 | 1.48E-02 |
| Zfp799 | 721.4912572 | -0.321091071 | 1.69E-02 |
| Gadd45gip1 | 1198.857908 | -0.321315174 | 2.01E-02 |
| Rab11fip3 | 2701.591854 | -0.321421343 | 3.35E-03 |
| Ado | 1738.480241 | -0.324308046 | 4.63E-03 |
| Rnf220 | 2987.612839 | -0.324813479 | 1.26E-02 |
| Prr3 | 728.9177642 | -0.325461381 | 1.13E-02 |
| Slc37a3 | 2707.708926 | -0.326015145 | 1.81E-03 |

| | | | |
|---|---|---|---|
| Srsf9 | 1778.892419 | -0.326062656 | 1.40E-03 |
| Prmt6 | 763.8135835 | -0.326171435 | 2.85E-02 |
| Vkorc1l1 | 1525.062969 | -0.326582451 | 3.44E-04 |
| Nsmce2 | 1712.981215 | -0.327024956 | 2.91E-02 |
| Cog3 | 2017.359627 | -0.32741426 | 3.06E-02 |
| Washc5 | 9453.018285 | -0.327716892 | 2.30E-03 |
| Sertad2 | 3856.909765 | -0.327888416 | 1.69E-02 |
| Cep170 | 6223.844815 | -0.328003274 | 4.83E-02 |
| Yeats2 | 2822.94201 | -0.329213356 | 2.59E-03 |
| Pip5k1c | 2979.353255 | -0.331784492 | 3.55E-02 |
| Stx6 | 2150.494347 | -0.331896753 | 2.29E-02 |
| Gmcl1 | 1500.19973 | -0.332555964 | 1.17E-04 |
| Polr1a | 4966.441682 | -0.332751981 | 4.44E-02 |
| Dcun1d5 | 1542.903361 | -0.333067316 | 1.50E-02 |
| Pcnx | 4490.427285 | -0.333388016 | 1.33E-02 |
| Vdac2 | 9351.945713 | -0.334101625 | 1.78E-02 |
| Cacfd1 | 1602.512009 | -0.33422471 | 3.49E-02 |
| Smim13 | 747.323615 | -0.334640932 | 4.86E-02 |
| Mrm3 | 359.86988 | -0.334712275 | 4.31E-02 |
| Tet3 | 2987.453673 | -0.334854296 | 4.48E-02 |
| Ccar2 | 1822.781877 | -0.335176417 | 3.34E-02 |
| Hnrnpf | 22991.53822 | -0.335385175 | 9.20E-03 |
| Mrpl9 | 1944.256036 | -0.335712254 | 5.33E-03 |
| Armcx2 | 5543.690073 | -0.335898959 | 4.24E-03 |
| Dnlz | 934.93158 | -0.336270594 | 2.59E-02 |
| Amer1 | 1350.755714 | -0.336287317 | 4.89E-04 |
| Sbf1 | 5212.739294 | -0.336691891 | 3.00E-02 |
| Actn1 | 13390.55678 | -0.336738137 | 1.85E-02 |
| Fbxl19 | 1033.975476 | -0.337789894 | 2.94E-02 |
| Anks3 | 1314.908854 | -0.337805984 | 4.84E-02 |
| Pou6f1 | 1919.566169 | -0.338006539 | 1.80E-02 |
| Znrf1 | 2102.508541 | -0.338635939 | 2.33E-02 |
| Surf6 | 1223.042779 | -0.339197923 | 1.15E-02 |
| Gm7488 | 1833.106868 | -0.339840853 | 4.73E-02 |
| Ccdc22 | 1040.823011 | -0.341665482 | 6.22E-03 |
| Farp1 | 7954.256896 | -0.344994101 | 1.12E-02 |
| Mgat5 | 813.1866966 | -0.3454413 | 4.73E-02 |
| Sall2 | 1033.6407 | -0.345481028 | 2.45E-02 |
| Tcf3 | 4577.602104 | -0.345980624 | 1.37E-02 |
| Pdzd4 | 1139.545824 | -0.346971645 | 2.87E-02 |
| Enah | 2404.280719 | -0.347932213 | 3.17E-02 |
| Carm1 | 4057.020687 | -0.348450735 | 5.84E-03 |
| Ttc21b | 2440.384871 | -0.349732688 | 8.35E-04 |
| Zbtb5 | 524.8705552 | -0.350281769 | 4.69E-02 |
| Nlgn2 | 3845.598607 | -0.351900942 | 4.94E-02 |
| Trak1 | 2002.750485 | -0.351943099 | 1.25E-02 |
| Zbtb12 | 936.8639599 | -0.352207321 | 1.28E-02 |

| | | | |
|---|---|---|---|
| Zmynd19 | 1294.299713 | -0.352680658 | 4.96E-02 |
| Mfsd14a | 1911.459662 | -0.353867245 | 1.46E-02 |
| Commd10 | 1041.836611 | -0.35442891 | 1.10E-02 |
| mt-Nd2 | 99498.30713 | -0.354792545 | 7.31E-03 |
| Ap5z1 | 1165.206489 | -0.354855063 | 1.56E-02 |
| Tada2b | 1040.973773 | -0.355217308 | 8.66E-04 |
| Khdrbs1 | 7784.512588 | -0.355373907 | 1.28E-02 |
| Uhrf2 | 4017.205027 | -0.355781749 | 3.56E-02 |
| Bach1 | 2273.465941 | -0.356217767 | 5.39E-04 |
| Mpnd | 2802.889083 | -0.357210192 | 4.50E-02 |
| Gigyf1 | 2070.367659 | -0.358511276 | 1.81E-02 |
| Polr2h | 1088.850475 | -0.358588283 | 4.47E-02 |
| Bcl10 | 1391.517893 | -0.358768383 | 1.84E-02 |
| Ehmt1 | 2812.096587 | -0.358991059 | 2.75E-04 |
| NA | 5575.684898 | -0.359096114 | 2.85E-02 |
| Tmem14c | 2037.871057 | -0.359492671 | 2.96E-02 |
| Lrpprc | 5310.296473 | -0.35977439 | 3.07E-02 |
| Tomm34 | 1451.330428 | -0.35998079 | 6.70E-03 |
| Eif2ak4 | 3546.105363 | -0.359981961 | 2.30E-03 |
| Llgl1 | 4105.61428 | -0.361370736 | 8.82E-03 |
| Nab2 | 4607.437328 | -0.361894917 | 2.99E-02 |
| Cnbp | 12294.52008 | -0.362048838 | 3.05E-02 |
| Fbxl6 | 1647.638682 | -0.363284825 | 4.27E-02 |
| Zbtb21 | 1062.931372 | -0.363538699 | 1.14E-02 |
| Spast | 2112.317484 | -0.363642015 | 3.70E-02 |
| Set | 10211.37983 | -0.364316005 | 4.82E-02 |
| Timm9 | 538.7327621 | -0.36511688 | 3.48E-02 |
| Adam19 | 3337.629642 | -0.365481254 | 4.02E-02 |
| Cep19 | 695.464579 | -0.365599756 | 1.59E-03 |
| Gpsm2 | 2322.001977 | -0.36585478 | 4.67E-02 |
| Pex5 | 3893.377517 | -0.366161155 | 1.13E-03 |
| Tmx4 | 2058.445111 | -0.366860881 | 2.86E-03 |
| Spata7 | 434.5194125 | -0.366884503 | 7.51E-03 |
| Tsc1 | 3398.460772 | -0.367100084 | 1.96E-02 |
| Wipf1 | 3029.779313 | -0.367186647 | 1.01E-04 |
| Lmo4 | 1591.794629 | -0.368529373 | 1.09E-03 |
| Mrpl36 | 758.7165102 | -0.368757427 | 7.98E-03 |
| H1f0 | 12531.5451 | -0.368923154 | 7.55E-03 |
| Ppm1l | 2353.623018 | -0.369114092 | 5.68E-03 |
| Slc4a2 | 6711.940086 | -0.369202168 | 4.85E-02 |
| Elavl1 | 3244.196393 | -0.369850917 | 3.43E-02 |
| Fads1 | 13261.24545 | -0.372019386 | 1.26E-02 |
| Hectd3 | 2568.610222 | -0.372116572 | 2.06E-02 |
| Prdx2 | 10872.40196 | -0.372394261 | 2.46E-02 |
| Atic | 6279.229534 | -0.372528027 | 4.71E-02 |
| Nob1 | 1945.869429 | -0.373843241 | 1.35E-02 |
| Met | 7204.015306 | -0.374117815 | 1.56E-02 |

| | | | |
|---|---|---|---|
| Tecr | 7238.644524 | -0.374957285 | 3.27E-02 |
| Aatf | 2596.894846 | -0.375066027 | 4.39E-03 |
| Txlna | 4884.051793 | -0.375589845 | 1.26E-02 |
| 2310033P09Rik | 321.5540812 | -0.375703216 | 3.56E-02 |
| Psme2 | 2209.647031 | -0.375760088 | 4.74E-03 |
| Spin1 | 3118.156723 | -0.376024434 | 3.40E-03 |
| Rnf34 | 1154.614628 | -0.376559746 | 2.80E-03 |
| Spry2 | 928.0977301 | -0.377135401 | 1.35E-02 |
| Gas8 | 1271.176052 | -0.377359857 | 4.75E-02 |
| Zfp655 | 3110.698296 | -0.378769273 | 7.47E-04 |
| Deaf1 | 690.6344136 | -0.379195401 | 3.95E-02 |
| Ctnnbl1 | 1369.037269 | -0.380531538 | 1.10E-02 |
| Fadd | 424.1906884 | -0.381003637 | 4.66E-02 |
| Sipa1 | 1737.308702 | -0.381326672 | 3.90E-02 |
| Prepl | 2110.742543 | -0.381468204 | 3.90E-04 |
| Slc38a7 | 2944.818323 | -0.381895191 | 3.12E-02 |
| Noa1 | 980.0937759 | -0.38189587 | 2.88E-02 |
| Ywhaz | 11408.90752 | -0.382888304 | 1.16E-03 |
| Ebp | 1605.346395 | -0.383059193 | 4.79E-02 |
| Plk2 | 10524.64627 | -0.383830053 | 4.48E-02 |
| Ift43 | 828.8161268 | -0.383966371 | 1.63E-03 |
| Mtfmt | 419.5935587 | -0.384560636 | 1.75E-02 |
| Diexf | 951.2860776 | -0.384581316 | 5.75E-04 |
| Dock1 | 3924.573139 | -0.386569399 | 4.28E-03 |
| Ttc27 | 1412.862851 | -0.388132542 | 3.19E-03 |
| Nat2 | 437.4994644 | -0.388220896 | 2.87E-02 |
| Ppp1r10 | 2578.197764 | -0.388384171 | 2.70E-02 |
| Josd2 | 899.9900018 | -0.388419155 | 1.50E-02 |
| Tmem251 | 678.3297086 | -0.38864415 | 3.08E-02 |
| Nrp2 | 11248.52068 | -0.388700076 | 2.53E-02 |
| Midn | 4349.625845 | -0.388720632 | 3.96E-02 |
| Plekha5 | 1255.587539 | -0.390034006 | 4.61E-02 |
| Fam118a | 816.1656045 | -0.390514134 | 4.16E-03 |
| Mrps26 | 1220.384571 | -0.391357062 | 5.78E-03 |
| Rcl1 | 1451.162847 | -0.391857407 | 2.94E-02 |
| Nol6 | 2151.227173 | -0.391953419 | 3.86E-02 |
| Mum1 | 1919.342188 | -0.392495075 | 5.74E-04 |
| Pus7l | 363.0645938 | -0.392992923 | 2.67E-02 |
| Vim | 152127.4244 | -0.3930814 | 3.26E-02 |
| Mrpl52 | 1057.21332 | -0.393479248 | 3.56E-03 |
| Mafg | 3468.758033 | -0.393529443 | 1.27E-03 |
| Nit2 | 1118.356382 | -0.393899583 | 3.07E-02 |
| Echs1 | 2288.2654 | -0.394960197 | 1.24E-02 |
| B4galt2 | 3537.6615 | -0.395670227 | 8.97E-03 |
| Ralb | 3073.453009 | -0.39682609 | 2.36E-02 |
| Cttnbp2nl | 3034.29804 | -0.396887868 | 5.52E-03 |
| Usp47 | 10646.51981 | -0.397466925 | 1.64E-03 |

| | | | |
|---|---|---|---|
| Pdgfrb | 13123.50877 | -0.398043182 | 1.23E-02 |
| Gaa | 16101.78382 | -0.39842416 | 4.83E-02 |
| Elf2 | 3398.181671 | -0.399318791 | 4.09E-04 |
| Fes | 449.9226603 | -0.400100342 | 3.35E-02 |
| Zcchc14 | 3410.077933 | -0.400101678 | 2.10E-05 |
| Dclre1b | 430.4982949 | -0.400233696 | 3.24E-02 |
| Magee1 | 2530.570754 | -0.401184366 | 4.82E-03 |
| Cd34 | 10535.55549 | -0.401277502 | 4.79E-02 |
| Pip4p2 | 1878.030152 | -0.401579385 | 9.42E-03 |
| Slc35e3 | 2219.989806 | -0.401593314 | 1.40E-02 |
| Ctnnd1 | 14049.60831 | -0.401631642 | 2.30E-02 |
| Naa10 | 1512.749773 | -0.402003913 | 2.91E-02 |
| Coro1c | 5207.156324 | -0.402052048 | 2.05E-02 |
| 5031425E22Rik | 290.5449219 | -0.403333542 | 2.92E-02 |
| Ing2 | 511.6968124 | -0.404254656 | 3.95E-02 |
| Zfp827 | 2374.032775 | -0.404974482 | 2.15E-02 |
| Cdkn2aip | 683.7261103 | -0.405042616 | 1.45E-02 |
| Zfp706 | 4663.743055 | -0.405186965 | 8.58E-03 |
| Rpp38 | 445.7698222 | -0.405194117 | 3.46E-02 |
| Myc | 3282.758284 | -0.405279743 | 4.18E-02 |
| Coq6 | 914.9612862 | -0.405462667 | 4.79E-02 |
| Rrp15 | 1350.946807 | -0.405543624 | 3.03E-02 |
| Zbtb45 | 334.1316636 | -0.40659234 | 4.82E-02 |
| Armc9 | 957.2896761 | -0.407147949 | 4.13E-04 |
| Xrcc1 | 1307.691342 | -0.407344477 | 4.43E-02 |
| Washc2 | 10254.84457 | -0.40742721 | 2.30E-02 |
| Mob3a | 1207.127894 | -0.408013094 | 4.06E-02 |
| Phc1 | 2184.532526 | -0.40815623 | 2.08E-02 |
| Hinfp | 406.8236308 | -0.408488267 | 2.62E-02 |
| Gpatch2l | 1358.907756 | -0.40889995 | 1.99E-02 |
| Rbm4 | 516.6815499 | -0.409135194 | 3.45E-02 |
| Bcl7a | 1302.545937 | -0.409165602 | 3.15E-03 |
| Phf23 | 3785.987225 | -0.409212172 | 2.14E-02 |
| D10Wsu102e | 2431.206995 | -0.409640286 | 2.82E-03 |
| Pik3r2 | 5877.98043 | -0.410201177 | 1.36E-02 |
| Acaca | 6220.310423 | -0.410386523 | 2.96E-02 |
| Pcdh18 | 4140.879768 | -0.410459764 | 1.72E-02 |
| Hnrnpa1 | 23929.72459 | -0.411297319 | 2.96E-02 |
| Sdad1 | 2442.35499 | -0.411422355 | 4.64E-04 |
| Pdpr | 1055.824546 | -0.411478885 | 1.59E-03 |
| Pi4kb | 3352.487585 | -0.411666804 | 1.34E-02 |
| Syncrip | 5515.504038 | -0.412487988 | 1.65E-02 |
| Itsn1 | 2752.843847 | -0.412736564 | 5.33E-04 |
| 3110082I17Rik | 709.1107669 | -0.413503514 | 4.20E-02 |
| Lgals1 | 85771.53346 | -0.41391397 | 3.08E-02 |
| Nkrf | 418.5610682 | -0.413974262 | 9.86E-03 |
| Stard5 | 614.9927071 | -0.416505446 | 2.66E-02 |

| | | | |
|---|---|---|---|
| Inf2 | 6535.964029 | -0.41664733 | 2.12E-02 |
| Stat1 | 1423.127776 | -0.416967165 | 3.08E-02 |
| Armcx3 | 3815.423845 | -0.417018175 | 1.71E-02 |
| Irx5 | 374.8914831 | -0.417696817 | 3.86E-02 |
| Pabpc1 | 86856.51255 | -0.41817233 | 2.54E-02 |
| Bola1 | 299.4104525 | -0.418797222 | 4.63E-02 |
| Parp8 | 1704.661461 | -0.419085897 | 1.02E-02 |
| Maz | 5576.002949 | -0.419155112 | 3.25E-03 |
| Coprs | 814.1861832 | -0.419282978 | 1.95E-03 |
| Wdr82 | 3294.636451 | -0.419380296 | 7.59E-04 |
| Irs2 | 2222.67984 | -0.420349006 | 2.48E-02 |
| Pitrm1 | 6900.706977 | -0.420675293 | 4.14E-04 |
| Ogdhl | 4923.875176 | -0.421862779 | 1.18E-02 |
| Bcl7c | 1233.600283 | -0.42319217 | 2.48E-02 |
| Hnrnpa0 | 3624.462535 | -0.424447214 | 3.51E-02 |
| Ppp1r13b | 826.0314193 | -0.424458098 | 1.67E-02 |
| Polr1c | 2348.159737 | -0.424897502 | 4.74E-02 |
| Hbp1 | 5566.199986 | -0.425034778 | 4.55E-02 |
| Tiparp | 2669.572333 | -0.425361449 | 8.18E-03 |
| Faap100 | 615.3568584 | -0.42657202 | 4.16E-02 |
| Usp28 | 1013.741207 | -0.426606 | 1.78E-04 |
| Taf6l | 798.7607961 | -0.426761405 | 4.39E-02 |
| Kat2a | 2700.153633 | -0.426915342 | 1.99E-02 |
| Zfp362 | 2229.096229 | -0.427338745 | 2.08E-03 |
| Hsdl1 | 1948.417286 | -0.427875214 | 1.03E-02 |
| Fam136a | 1289.210965 | -0.429132481 | 1.76E-02 |
| Capn10 | 780.8179271 | -0.429568541 | 2.38E-02 |
| Pip4k2b | 4426.450694 | -0.43000598 | 6.89E-06 |
| Stxbp1 | 3426.852098 | -0.430146661 | 2.35E-02 |
| Sox12 | 1777.455189 | -0.430375288 | 3.79E-02 |
| Tax1bp3 | 4764.282672 | -0.430607869 | 2.30E-02 |
| Impact | 6753.387895 | -0.430621277 | 3.54E-02 |
| Btd | 882.1449736 | -0.430926102 | 1.91E-02 |
| Sema4c | 1046.850603 | -0.431054667 | 1.57E-02 |
| Mrpl2 | 1684.064819 | -0.43398337 | 8.23E-03 |
| Lpar1 | 5879.931993 | -0.435073181 | 1.69E-02 |
| 4930402H24Rik | 5341.149355 | -0.435465035 | 3.27E-04 |
| Mpdu1 | 1210.812562 | -0.435917309 | 3.67E-03 |
| Rbm3 | 13054.23758 | -0.436114337 | 7.41E-03 |
| Gtf3c5 | 1178.49991 | -0.437345883 | 2.60E-04 |
| Edc3 | 1285.870104 | -0.438039818 | 2.14E-03 |
| Ap4m1 | 644.4259784 | -0.438254825 | 3.24E-02 |
| Cic | 8307.861902 | -0.438278652 | 4.97E-03 |
| Mrps33 | 1660.844546 | -0.440014546 | 1.79E-02 |
| Apobec3 | 1537.869773 | -0.440207538 | 3.66E-03 |
| Usb1 | 1737.627401 | -0.441479027 | 2.17E-03 |
| NA | 1773.329601 | -0.442542951 | 1.71E-04 |

| | | | |
|---|---|---|---|
| Rbp1 | 2005.260463 | -0.442995643 | 4.18E-02 |
| Xrcc5 | 2169.701875 | -0.444655921 | 7.78E-04 |
| Slc12a7 | 1768.921695 | -0.444746777 | 4.16E-03 |
| Rpl31 | 2778.260788 | -0.444760691 | 2.49E-03 |
| Nmnat2 | 1973.93081 | -0.444770432 | 3.92E-02 |
| Tsr2 | 1165.014558 | -0.445256007 | 8.24E-03 |
| Fbxo41 | 890.8379166 | -0.445428686 | 3.03E-02 |
| Cbarp | 525.0437716 | -0.445456767 | 1.03E-02 |
| Gtf2ird1 | 1929.055866 | -0.446180787 | 3.97E-04 |
| Bcs1l | 748.8985829 | -0.446825194 | 2.99E-02 |
| Rdh11 | 2494.14244 | -0.447531698 | 2.17E-02 |
| Orai2 | 1421.004878 | -0.449008283 | 3.42E-03 |
| Ick | 1414.685462 | -0.449095245 | 6.55E-04 |
| Zfp319 | 991.9383572 | -0.450234762 | 1.19E-03 |
| Tsen2 | 691.2193844 | -0.450413996 | 1.08E-02 |
| Ip6k2 | 1213.466928 | -0.450484644 | 2.54E-02 |
| Agpat4 | 1161.530843 | -0.450539568 | 3.50E-02 |
| Gm8186 | 1195.312773 | -0.450567123 | 1.62E-02 |
| Fam126a | 2910.896476 | -0.45083789 | 2.99E-05 |
| Samd1 | 1577.418184 | -0.450973946 | 2.02E-02 |
| Tmed1 | 963.0524274 | -0.45175152 | 1.47E-02 |
| Lta4h | 6976.776006 | -0.452251232 | 2.05E-02 |
| Eif3e | 15575.42035 | -0.452835971 | 3.22E-02 |
| Mettl27 | 1096.641165 | -0.453792547 | 4.34E-02 |
| Lrfn4 | 2023.205322 | -0.454203481 | 4.13E-03 |
| Vmac | 1013.466863 | -0.45440839 | 2.38E-02 |
| Atp6v0b | 9934.293287 | -0.454615866 | 4.16E-02 |
| Nrg1 | 1866.437419 | -0.455196856 | 4.26E-02 |
| Zfp202 | 333.0936493 | -0.456187098 | 8.65E-03 |
| Zfp687 | 1806.062977 | -0.4565075 | 1.25E-06 |
| Hdac3 | 3639.465084 | -0.456737202 | 8.99E-04 |
| Rpl35 | 28922.30866 | -0.456778656 | 3.14E-02 |
| Pcdhb17 | 304.6149099 | -0.456793189 | 2.67E-02 |
| Ilf2 | 3066.686516 | -0.457846891 | 2.67E-03 |
| Mccc2 | 227.1006885 | -0.458487637 | 3.57E-02 |
| Ttyh2 | 1997.972056 | -0.459072142 | 3.22E-02 |
| Tubb2a | 6574.060184 | -0.459534422 | 2.58E-02 |
| Ddx54 | 4300.181664 | -0.459538975 | 1.92E-02 |
| Cog4 | 4133.200435 | -0.460024822 | 2.02E-02 |
| Tle1 | 1922.88961 | -0.460055663 | 4.90E-03 |
| Wdr46 | 1834.741141 | -0.461166041 | 1.72E-02 |
| Smad1 | 1578.21778 | -0.462854513 | 1.36E-02 |
| Zfp142 | 2034.722645 | -0.463764239 | 1.53E-03 |
| Ints7 | 1422.713731 | -0.464110573 | 3.66E-03 |
| Soga3 | 486.8497861 | -0.464147982 | 1.86E-02 |
| Anxa6 | 6987.123827 | -0.464202945 | 3.90E-02 |
| Tmcc2 | 1387.568865 | -0.464919138 | 3.87E-02 |

| | | | |
|---|---|---|---|
| Ppp1r14b | 5784.837077 | -0.465339205 | 4.01E-02 |
| Poc1b | 1164.775363 | -0.466748811 | 4.68E-03 |
| Ptpn9 | 2903.919013 | -0.467136277 | 1.32E-04 |
| Nsg1 | 1730.070035 | -0.46731816 | 5.52E-03 |
| Mmp11 | 3113.336865 | -0.467446679 | 6.33E-04 |
| Ddx39b | 10038.84561 | -0.468124184 | 5.15E-03 |
| Top3a | 748.5691303 | -0.46879383 | 2.66E-02 |
| Cfap20 | 1689.539204 | -0.468984189 | 5.64E-04 |
| Usp11 | 2202.578828 | -0.469057339 | 4.56E-02 |
| Cdk5rap2 | 4844.501444 | -0.469077642 | 2.23E-02 |
| Cpne8 | 1083.712569 | -0.469078343 | 1.40E-04 |
| Ptma | 34544.23717 | -0.470052434 | 6.47E-03 |
| Mecom | 1419.704379 | -0.470852774 | 2.08E-02 |
| Ctnnb1 | 26830.96708 | -0.470992242 | 1.97E-06 |
| Noct | 8107.901444 | -0.47153306 | 1.59E-04 |
| Bcl2l12 | 332.2104542 | -0.472235877 | 3.46E-02 |
| Gm29394 | 1296.624358 | -0.47312511 | 1.90E-03 |
| Cblb | 1328.176995 | -0.474000935 | 4.58E-03 |
| Dock5 | 2703.985943 | -0.474001716 | 1.13E-02 |
| Rps29 | 19692.01595 | -0.474476133 | 1.17E-02 |
| Rufy3 | 357.8406021 | -0.475518333 | 1.89E-02 |
| Rpl8 | 66813.58999 | -0.475772799 | 2.89E-02 |
| Arhgap32 | 2413.87787 | -0.47630664 | 1.04E-03 |
| Gm4540 | 805.2713897 | -0.476477557 | 2.69E-02 |
| Fnip2 | 5464.462773 | -0.476739596 | 1.98E-02 |
| Ears2 | 383.1650498 | -0.477236964 | 7.64E-03 |
| Phc2 | 4897.151543 | -0.477997219 | 3.96E-03 |
| Exosc5 | 597.2657099 | -0.478593352 | 2.00E-02 |
| Taf5l | 1587.864323 | -0.479642152 | 3.23E-03 |
| Atp2b1 | 7973.205069 | -0.47997649 | 1.69E-02 |
| Gm10243 | 15969.90068 | -0.480086026 | 3.50E-02 |
| Lsm14b | 2293.235444 | -0.480301169 | 7.99E-03 |
| B3gnt9 | 1121.027082 | -0.480500057 | 1.56E-02 |
| Dph1 | 914.7849843 | -0.480596596 | 1.03E-03 |
| Acy1 | 719.2862101 | -0.48197562 | 2.93E-02 |
| Garem2 | 297.7203972 | -0.482482148 | 3.73E-02 |
| Kdm2b | 1932.525567 | -0.484099468 | 6.86E-03 |
| Amigo2 | 624.6922769 | -0.484133995 | 6.65E-03 |
| Dbnl | 5109.073654 | -0.484174412 | 6.33E-03 |
| Gm42688 | 427.2518859 | -0.484429081 | 2.20E-03 |
| Ksr1 | 974.9766751 | -0.484759462 | 3.22E-02 |
| Acd | 2131.77091 | -0.484910721 | 2.70E-02 |
| Mtss1l | 4009.15688 | -0.485188448 | 2.74E-02 |
| Snrpg | 1662.606002 | -0.485844548 | 1.36E-02 |
| Rps4x | 93555.43077 | -0.4866279 | 4.61E-02 |
| Vash2 | 2507.446433 | -0.48719523 | 1.69E-02 |
| Fbxl14 | 1290.48478 | -0.488839968 | 6.37E-04 |

| | | | |
|---|---|---|---|
| Cdk19 | 2233.948062 | -0.488977457 | 1.86E-06 |
| Qdpr | 1935.818044 | -0.489119753 | 1.05E-02 |
| Grhpr | 457.3816047 | -0.490051275 | 1.26E-02 |
| Pus1 | 1110.89264 | -0.490808902 | 2.07E-03 |
| Abr | 2974.771084 | -0.49095676 | 4.36E-04 |
| Lpar2 | 1343.831789 | -0.491202086 | 1.25E-02 |
| C1qbp | 6609.162753 | -0.491279798 | 7.48E-04 |
| Pgp | 525.9322498 | -0.491602053 | 4.31E-02 |
| Adgra2 | 9959.089649 | -0.492488136 | 8.88E-04 |
| Rufy1 | 2070.818133 | -0.492926596 | 2.71E-02 |
| Psme1 | 1774.506782 | -0.493291329 | 1.80E-03 |
| Parp12 | 537.1403291 | -0.493755728 | 1.51E-03 |
| Fmnl3 | 2696.537561 | -0.494968408 | 2.70E-03 |
| Got2 | 9551.023887 | -0.496186065 | 1.50E-02 |
| Lima1 | 14608.60545 | -0.498848247 | 6.38E-03 |
| Aprt | 6193.680378 | -0.498931785 | 1.08E-02 |
| Smarcc1 | 5292.980505 | -0.499057459 | 5.84E-06 |
| D16Ertd472e | 232.6290134 | -0.499105561 | 6.71E-03 |
| Rps16 | 14597.25608 | -0.499261008 | 3.24E-02 |
| Rps15a | 30497.77941 | -0.500145624 | 2.94E-02 |
| Gar1 | 912.2814622 | -0.5002999 | 1.52E-02 |
| Rpl23 | 66491.88099 | -0.500909722 | 2.06E-02 |
| Map2k6 | 397.271134 | -0.501235731 | 4.61E-02 |
| 5-Sep | 7626.58109 | -0.501525005 | 2.06E-02 |
| Ppp3ca | 3841.872004 | -0.502150245 | 3.95E-05 |
| Zfp821 | 579.2143519 | -0.502407885 | 3.96E-03 |
| Adck5 | 731.9848639 | -0.503431792 | 1.97E-02 |
| NA | 216.623365 | -0.504692351 | 2.67E-02 |
| Urb2 | 1491.82971 | -0.507409932 | 8.96E-05 |
| Acvr1 | 1714.787564 | -0.508160066 | 1.38E-03 |
| Wdr6 | 3416.318234 | -0.509790657 | 3.57E-03 |
| Wdr34 | 712.2965379 | -0.50999974 | 6.67E-03 |
| Tbcc | 642.9037322 | -0.510288844 | 8.44E-03 |
| Tcf4 | 3628.891416 | -0.511034 | 5.15E-05 |
| H2afy2 | 1900.864898 | -0.511772885 | 1.30E-02 |
| Rps3a1 | 109609.6564 | -0.513577789 | 6.37E-03 |
| Jund | 2487.535816 | -0.513650019 | 4.69E-02 |
| Rps12 | 48476.15642 | -0.514005355 | 7.77E-03 |
| Mapk8ip1 | 2586.65763 | -0.514215896 | 2.18E-02 |
| Samd14 | 1239.574654 | -0.514991233 | 4.17E-03 |
| Hyal2 | 1065.562101 | -0.515976281 | 2.00E-02 |
| Ankrd54 | 751.29234 | -0.516196422 | 2.96E-02 |
| Bahcc1 | 1260.799357 | -0.51670623 | 1.46E-02 |
| Aacs | 4095.139985 | -0.516738401 | 3.73E-02 |
| Trib2 | 1232.911453 | -0.517343225 | 4.14E-02 |
| B9d1 | 529.0966433 | -0.51752588 | 1.68E-02 |
| Rnd2 | 944.1092971 | -0.518018738 | 4.07E-02 |

| | | | |
|---|---|---|---|
| Ilf3 | 5444.124218 | -0.518111936 | 2.76E-05 |
| Ttc26 | 659.8498206 | -0.51838073 | 7.12E-03 |
| Nradd | 1116.064392 | -0.519690956 | 2.67E-04 |
| Tmeff1 | 2004.0604 | -0.519944906 | 9.05E-04 |
| Asic1 | 2363.956811 | -0.521177752 | 3.74E-02 |
| Trp53 | 5498.367133 | -0.521930621 | 1.46E-02 |
| Gm15500 | 76446.03163 | -0.522690929 | 2.99E-02 |
| Rita1 | 573.5582389 | -0.523703389 | 3.37E-03 |
| 1500015A07Rik | 235.5015399 | -0.523725559 | 1.39E-02 |
| Entpd6 | 901.5502184 | -0.523946414 | 2.56E-02 |
| NA | 256.5573422 | -0.524866173 | 2.08E-03 |
| Dip2b | 2838.90961 | -0.524973182 | 1.10E-02 |
| Snx8 | 2047.563724 | -0.525077267 | 5.97E-04 |
| Rps3 | 56649.04585 | -0.525517072 | 3.59E-02 |
| Srebf2 | 11304.7876 | -0.525750426 | 2.98E-03 |
| Ccdc115 | 665.7497048 | -0.526834315 | 4.56E-03 |
| Dmac2 | 437.6982699 | -0.526909598 | 9.22E-04 |
| Bri3bp | 3340.230155 | -0.52727948 | 1.06E-04 |
| Ldhb | 3916.76833 | -0.527979594 | 1.03E-02 |
| Nr2c2ap | 954.217837 | -0.528131372 | 1.72E-02 |
| Ppan | 1552.316091 | -0.530294759 | 9.13E-04 |
| Gas1 | 20046.86774 | -0.530683368 | 3.14E-02 |
| 2410002F23Rik | 2404.042911 | -0.531391478 | 1.80E-04 |
| Polr1e | 447.9290646 | -0.531566583 | 4.58E-02 |
| Tdrkh | 618.162234 | -0.53178219 | 9.26E-03 |
| Slc6a6 | 8237.226832 | -0.532086181 | 1.46E-02 |
| Gm9701 | 21363.22693 | -0.532431216 | 1.41E-03 |
| Dpysl2 | 1095.968635 | -0.533510414 | 1.44E-02 |
| Atp1b1 | 1462.526931 | -0.534587422 | 4.61E-02 |
| Smarcd1 | 1668.953612 | -0.534973154 | 2.93E-04 |
| Ppp1cc | 8121.670663 | -0.535247844 | 2.84E-08 |
| Gm10260 | 27463.1406 | -0.535384507 | 3.18E-02 |
| Tmem106a | 958.7992917 | -0.536423048 | 7.61E-03 |
| Eef1b2 | 46128.15298 | -0.536875554 | 1.56E-02 |
| Sf3b3 | 10639.11885 | -0.537772296 | 1.33E-04 |
| H2-Q4 | 205.1152131 | -0.538638822 | 4.08E-02 |
| Micall2 | 1659.531975 | -0.538651947 | 1.93E-02 |
| Impdh2-ps | 1489.025213 | -0.539447127 | 8.12E-04 |
| Nthl1 | 448.1070006 | -0.539686553 | 4.19E-03 |
| Igf2bp2 | 8644.418417 | -0.539889805 | 2.22E-07 |
| Gm15501 | 77795.36435 | -0.540100016 | 2.08E-02 |
| Pdcd2 | 1168.649565 | -0.540848932 | 2.71E-04 |
| Atp6v1g1 | 12568.78181 | -0.541557261 | 9.43E-03 |
| Rps21 | 17061.14801 | -0.543429177 | 2.98E-02 |
| Zfp184 | 226.740299 | -0.543599945 | 1.24E-02 |
| Serinc5 | 1871.588337 | -0.543755923 | 6.02E-06 |
| Ilvbl | 2448.329512 | -0.543850653 | 2.30E-03 |

| | | | |
|---|---|---|---|
| Snn | 2448.99706 | -0.543879608 | 1.84E-03 |
| 4931428F04Rik | 479.9064024 | -0.54426074 | 4.96E-02 |
| Hook2 | 814.6719502 | -0.545597229 | 3.56E-03 |
| Sh3pxd2b | 11222.03918 | -0.546212558 | 3.26E-02 |
| Ddx10 | 1380.39871 | -0.546246093 | 2.07E-02 |
| Suclg2 | 5554.452872 | -0.547248839 | 1.32E-06 |
| Rpl32 | 65196.27315 | -0.548900916 | 2.84E-02 |
| Gamt | 997.1896559 | -0.549817577 | 1.17E-02 |
| Wdr81 | 3346.664813 | -0.551792021 | 4.42E-03 |
| Rpl14 | 39168.7694 | -0.552154868 | 4.95E-02 |
| Mrps6 | 3383.155268 | -0.553275807 | 3.22E-02 |
| Gga2 | 4453.49319 | -0.553363518 | 3.45E-03 |
| Dph2 | 675.7724169 | -0.555320678 | 3.80E-03 |
| Rplp2 | 29391.29674 | -0.555382788 | 1.06E-02 |
| Abhd11 | 803.7195628 | -0.55574023 | 4.05E-03 |
| Akr7a5 | 1052.456729 | -0.556013099 | 2.35E-02 |
| Sbk1 | 915.23951 | -0.556586415 | 4.22E-02 |
| Zfp637 | 1645.301751 | -0.556819779 | 3.21E-03 |
| Tspan14 | 1344.749055 | -0.557178813 | 5.52E-06 |
| 3110053B16Rik | 271.5926797 | -0.558802133 | 7.42E-03 |
| Ltbp4 | 1251.339727 | -0.560788089 | 4.36E-02 |
| Rps6-ps4 | 132832.5808 | -0.561641241 | 1.81E-02 |
| Dctd | 1097.232119 | -0.563778847 | 4.64E-03 |
| Rps19 | 31869.82538 | -0.564138787 | 3.07E-02 |
| Hs1bp3 | 3425.439165 | -0.565316824 | 9.01E-03 |
| Sfxn5 | 564.0641619 | -0.565465933 | 3.45E-03 |
| Zbtb1 | 1295.613615 | -0.566015371 | 1.52E-02 |
| Arhgef40 | 3841.130444 | -0.566165543 | 9.71E-04 |
| Rps10 | 48037.18352 | -0.566231498 | 2.45E-02 |
| Sp4 | 343.7880857 | -0.567207069 | 1.66E-02 |
| Gm17586 | 152.4386316 | -0.569523265 | 7.91E-03 |
| Ntmt1 | 1200.274153 | -0.569788147 | 7.35E-04 |
| Ncs1 | 2519.874797 | -0.571203583 | 2.15E-04 |
| Impdh2 | 6001.17937 | -0.571727985 | 1.23E-03 |
| Gtf2i | 8843.674879 | -0.572270921 | 4.25E-05 |
| Rcc2 | 9407.022186 | -0.572346606 | 1.81E-04 |
| Selenow | 3078.602105 | -0.572561819 | 7.22E-03 |
| Gm6472 | 47311.23449 | -0.573419941 | 1.49E-02 |
| Tbcel | 1611.699035 | -0.575121732 | 2.82E-10 |
| Timp3 | 19304.01546 | -0.576006994 | 3.96E-02 |
| Rps28 | 15164.62267 | -0.57709268 | 4.66E-02 |
| Rpp40 | 775.7960989 | -0.577254092 | 1.78E-02 |
| Cry1 | 919.8294528 | -0.577323514 | 1.07E-02 |
| Chsy1 | 1508.249314 | -0.578884266 | 3.17E-05 |
| Trmt61b | 146.4603856 | -0.579344383 | 4.97E-02 |
| Plxnd1 | 6200.522837 | -0.579538923 | 8.42E-03 |
| Zswim6 | 572.7350864 | -0.579569462 | 2.69E-03 |

| | | | |
|---|---|---|---|
| Foxk1 | 3191.645725 | -0.581614453 | 1.73E-04 |
| Clmp | 2125.373568 | -0.581952161 | 1.78E-04 |
| Rps13-ps2 | 21055.30253 | -0.582047955 | 9.57E-03 |
| Qtrt1 | 428.9958719 | -0.582112862 | 4.98E-04 |
| Sdc3 | 1488.655788 | -0.582869317 | 3.16E-02 |
| Rps9 | 54843.11362 | -0.583515009 | 4.95E-03 |
| Trp53rkb | 133.0751093 | -0.584663704 | 3.93E-03 |
| Garnl3 | 869.1182456 | -0.58536522 | 4.13E-03 |
| Rpl39 | 31120.35881 | -0.585719228 | 3.97E-02 |
| Spred2 | 1836.996101 | -0.586226227 | 9.91E-06 |
| Efna3 | 212.0063188 | -0.586330963 | 4.57E-02 |
| Fam43a | 1435.289342 | -0.587462362 | 3.82E-03 |
| Wasf3 | 609.9759654 | -0.587917437 | 4.36E-04 |
| Arhgap39 | 893.335859 | -0.588617319 | 2.70E-03 |
| Satb2 | 298.8094884 | -0.58898815 | 1.36E-02 |
| Tmem121 | 208.8525126 | -0.589215538 | 9.46E-03 |
| Cebpg | 2322.035649 | -0.589275218 | 2.75E-08 |
| Rapgef3 | 465.241526 | -0.589348565 | 3.78E-04 |
| Slc25a1 | 5391.834169 | -0.589920517 | 2.02E-03 |
| Tead2 | 4318.690094 | -0.592249963 | 1.22E-02 |
| Psmg4 | 256.9783017 | -0.592866762 | 2.24E-03 |
| Slc6a8 | 7871.670639 | -0.594381458 | 1.18E-02 |
| Pcdhb7 | 281.6238328 | -0.59457798 | 1.88E-02 |
| Tmem97 | 1767.275724 | -0.596592403 | 3.78E-02 |
| Atp6v1b2 | 9772.289405 | -0.596665602 | 6.39E-03 |
| Snai2 | 1626.102104 | -0.598061244 | 1.81E-03 |
| Pdcd4 | 2480.242851 | -0.598148738 | 7.67E-03 |
| Lsp1 | 852.1553722 | -0.598709236 | 1.46E-02 |
| Hsf4 | 234.9047654 | -0.598762691 | 9.15E-03 |
| Tes | 1801.715462 | -0.598867471 | 3.05E-02 |
| Cuta | 2544.799639 | -0.599157214 | 1.89E-03 |
| Whrn | 1183.150778 | -0.599989125 | 2.84E-03 |
| Prdm11 | 475.2305451 | -0.600314312 | 1.85E-03 |
| Vat1 | 27960.27472 | -0.600377094 | 2.34E-02 |
| Ctns | 2362.555936 | -0.601366533 | 2.03E-04 |
| Clip2 | 3685.485315 | -0.603946419 | 1.52E-04 |
| Gdpd1 | 1125.495183 | -0.604333116 | 3.00E-02 |
| Rps7 | 10324.97548 | -0.604636422 | 4.90E-02 |
| Pafah1b3 | 1426.134526 | -0.60579648 | 1.26E-04 |
| Trmt61a | 494.8076108 | -0.60584124 | 3.62E-03 |
| Gnpda1 | 2534.474834 | -0.606889145 | 1.61E-03 |
| Gm11175 | 589.7593106 | -0.607018428 | 4.38E-02 |
| 9-Sep | 7495.015017 | -0.607210689 | 2.76E-04 |
| Fam3c | 4044.640668 | -0.607908178 | 4.15E-05 |
| Vegfb | 3825.597118 | -0.608167377 | 4.57E-03 |
| Marcks | 14713.78478 | -0.608255066 | 3.59E-02 |
| Adgrl1 | 1938.183396 | -0.608372487 | 1.64E-02 |

| | | | |
|---|---|---|---|
| Tprkb | 2647.743254 | -0.608839824 | 5.71E-05 |
| 2410018L13Rik | 753.6620775 | -0.609106086 | 1.30E-05 |
| Camkk1 | 480.61585 | -0.609713345 | 2.96E-02 |
| Plscr3 | 3839.393331 | -0.61149092 | 1.30E-04 |
| Ly6c1 | 1095.469361 | -0.611946736 | 3.79E-02 |
| Cask | 3506.668855 | -0.612689569 | 6.19E-05 |
| Eps8 | 11574.16555 | -0.612766018 | 7.58E-05 |
| H2afv | 1309.866945 | -0.613377381 | 1.16E-03 |
| Cln8 | 3141.229382 | -0.614239902 | 1.21E-02 |
| B3galt1 | 227.8123498 | -0.615112239 | 2.26E-03 |
| Ift140 | 1141.357097 | -0.615148622 | 8.40E-03 |
| Sh3bp2 | 1443.810206 | -0.616435145 | 1.13E-02 |
| Ank3 | 2241.447597 | -0.619939513 | 1.14E-02 |
| Sgk1 | 8387.420148 | -0.620545496 | 1.24E-02 |
| Acly | 18875.66242 | -0.621680886 | 5.65E-04 |
| Gm17251 | 159.5166582 | -0.622185991 | 3.35E-02 |
| Ccng2 | 2087.349482 | -0.622459685 | 9.18E-03 |
| Ptges3l | 205.1528227 | -0.622734248 | 6.71E-03 |
| Rgma | 3295.806106 | -0.623336213 | 1.06E-02 |
| Zbed5 | 182.5059509 | -0.62636147 | 9.94E-03 |
| Zfp553 | 1197.130045 | -0.626829921 | 9.88E-03 |
| Limd2 | 1732.780265 | -0.626907153 | 1.68E-03 |
| Cacng7 | 2011.225826 | -0.627310259 | 1.63E-02 |
| Erf | 2001.173073 | -0.627701488 | 9.60E-03 |
| Sarm1 | 300.3287627 | -0.628129846 | 8.12E-03 |
| Rps27rt | 30142.86293 | -0.628574682 | 2.47E-02 |
| Etv6 | 1583.328039 | -0.630381832 | 4.71E-09 |
| Bin1 | 3469.194523 | -0.630839976 | 2.71E-06 |
| Rnf157 | 1391.358528 | -0.635176795 | 3.73E-06 |
| Casp9 | 1187.691822 | -0.635770465 | 6.28E-05 |
| Gdf11 | 1582.164993 | -0.63702259 | 1.06E-03 |
| Ulk4 | 167.008026 | -0.637206168 | 1.93E-02 |
| Fkbp4 | 9091.266618 | -0.637437289 | 2.12E-04 |
| Gm10443 | 18934.25035 | -0.637628988 | 2.54E-02 |
| Lhfpl2 | 6465.270021 | -0.63791672 | 9.88E-05 |
| Nckap5l | 1105.351254 | -0.63841291 | 7.37E-04 |
| Urb1 | 1070.200587 | -0.638653441 | 1.07E-02 |
| Scd2 | 61452.75163 | -0.640890021 | 3.12E-07 |
| Pogk | 1686.347408 | -0.64283929 | 1.00E-05 |
| Pcdhb20 | 156.4655691 | -0.644025347 | 1.94E-02 |
| Rps12-ps19 | 164.2671423 | -0.644311629 | 4.30E-02 |
| Grasp | 391.1139554 | -0.645066071 | 8.52E-05 |
| Cx3cl1 | 722.2112652 | -0.647752099 | 3.35E-02 |
| P2rx4 | 2480.638167 | -0.647796205 | 4.23E-03 |
| Ralgds | 5291.738528 | -0.647942373 | 1.64E-02 |
| Nme3 | 171.0026249 | -0.648823295 | 7.88E-03 |
| Sgip1 | 339.6091924 | -0.650830478 | 2.54E-04 |

| | | | |
|---|---|---|---|
| 1700003F12Rik | 81.47917467 | -0.651945587 | 2.77E-02 |
| Gcat | 1335.897836 | -0.652092286 | 8.87E-04 |
| Slc5a3 | 5155.170795 | -0.652657517 | 1.33E-02 |
| 2410131K14Rik | 1774.50699 | -0.654604642 | 1.94E-07 |
| Magi2 | 526.5829282 | -0.654647236 | 2.21E-06 |
| Tnfrsf11a | 123.4947407 | -0.655746608 | 1.21E-03 |
| Lhx9 | 259.9429502 | -0.656216721 | 1.17E-02 |
| Psd3 | 883.8155306 | -0.658234084 | 1.60E-03 |
| Zfp703 | 916.2166658 | -0.658288753 | 1.55E-02 |
| Cbfb | 3583.177904 | -0.659373189 | 5.83E-10 |
| Nudt16l1 | 1616.991711 | -0.660092708 | 4.14E-04 |
| Ldlrad3 | 1390.98449 | -0.661843146 | 2.21E-04 |
| Zik1 | 289.836167 | -0.661914576 | 3.34E-04 |
| Acsl3 | 2747.009398 | -0.664644462 | 2.50E-02 |
| Tlnrd1 | 1548.830688 | -0.664737954 | 1.18E-04 |
| Sqle | 9213.229076 | -0.665828293 | 1.62E-02 |
| Maml3 | 728.8511573 | -0.66740278 | 5.52E-03 |
| Pdp2 | 461.4955656 | -0.668468395 | 1.48E-03 |
| Ankrd44 | 1362.280154 | -0.670682651 | 1.48E-03 |
| Rpl36 | 15843.79799 | -0.673588761 | 4.53E-03 |
| Emc2 | 2388.117355 | -0.673715992 | 1.00E-02 |
| Otulinl | 214.2498603 | -0.674845288 | 2.55E-04 |
| Rhebl1 | 244.5837871 | -0.674995314 | 1.35E-02 |
| Pbdc1 | 1501.992805 | -0.676040415 | 1.33E-05 |
| Shc1 | 9740.164831 | -0.676503718 | 6.42E-05 |
| Ddx51 | 1561.419541 | -0.676887574 | 2.27E-06 |
| Tanc2 | 4417.326718 | -0.678305503 | 1.12E-05 |
| Gjc1 | 1240.108962 | -0.67840195 | 5.71E-04 |
| Enc1 | 4373.996949 | -0.68006602 | 7.83E-03 |
| Arrdc2 | 510.3397326 | -0.680751138 | 1.22E-02 |
| Fgfr1 | 26119.72218 | -0.683229536 | 2.93E-04 |
| Traf4 | 3041.560397 | -0.685569537 | 1.44E-03 |
| Agfg2 | 1357.380105 | -0.686038262 | 1.58E-05 |
| Podnl1 | 3817.279448 | -0.686239505 | 3.16E-04 |
| Myo1b | 2440.909108 | -0.686722909 | 5.85E-04 |
| Coq8a | 152.0031913 | -0.68761518 | 4.67E-02 |
| Dynll1 | 5076.522784 | -0.690230985 | 4.41E-03 |
| AI413582 | 932.4708856 | -0.691774689 | 5.00E-04 |
| Sc5d | 2351.772826 | -0.695488476 | 1.34E-02 |
| Epha7 | 250.107713 | -0.696330815 | 1.75E-03 |
| Top1mt | 841.059034 | -0.697820489 | 1.96E-07 |
| Tcp11l2 | 1451.319384 | -0.699524072 | 4.87E-02 |
| Gm1673 | 194.6979044 | -0.701828113 | 3.65E-02 |
| Orc5 | 1511.84814 | -0.704431457 | 7.52E-03 |
| Ermp1 | 2303.615021 | -0.704907439 | 8.86E-06 |
| Irs1 | 2008.224803 | -0.708129297 | 2.37E-03 |
| Pcyt2 | 535.7741907 | -0.708414312 | 2.93E-02 |

| | | | |
|---|---|---|---|
| S1pr3 | 3918.192315 | -0.708731707 | 4.94E-05 |
| Mvk | 1649.647814 | -0.70905592 | 1.69E-02 |
| Apex1 | 3288.753952 | -0.709480101 | 2.22E-07 |
| Pcdhgc4 | 30257.72995 | -0.709831318 | 2.31E-05 |
| Fasn | 11598.25822 | -0.711713092 | 7.14E-03 |
| Pcyt1b | 608.1630088 | -0.711785595 | 1.91E-02 |
| 9430091E24Rik | 71.37412386 | -0.712666506 | 1.34E-02 |
| Nol4 | 339.5172744 | -0.713811987 | 3.21E-02 |
| Glrb | 356.4512423 | -0.715554087 | 7.69E-03 |
| Bzw2 | 2826.086381 | -0.715737426 | 5.90E-05 |
| Serf1 | 393.2430891 | -0.718556567 | 7.78E-03 |
| Tfap4 | 806.8709696 | -0.718733432 | 1.48E-03 |
| Cpe | 17622.21883 | -0.721061115 | 1.42E-04 |
| Amot | 5972.569539 | -0.723248926 | 4.72E-07 |
| Fhl3 | 1894.248643 | -0.728542031 | 1.36E-04 |
| Gm16211 | 299.0104868 | -0.729473621 | 1.09E-05 |
| Fmo1 | 97.89348174 | -0.729919571 | 4.66E-02 |
| Slc16a10 | 806.7478132 | -0.731304546 | 3.35E-03 |
| Rnd3 | 3804.865344 | -0.733148904 | 2.25E-06 |
| Efna4 | 476.6852267 | -0.738440803 | 7.81E-05 |
| Lbh | 5291.647432 | -0.744910476 | 3.69E-02 |
| Gpr176 | 2045.393277 | -0.745706706 | 8.49E-06 |
| Ociad2 | 337.1734645 | -0.74631808 | 4.64E-04 |
| Mmp14 | 40429.12938 | -0.746839756 | 9.94E-05 |
| Bhlhe40 | 3834.220772 | -0.747262124 | 8.02E-03 |
| Ypel1 | 252.6287883 | -0.748276316 | 2.93E-02 |
| Mast4 | 3107.692726 | -0.749629742 | 4.32E-06 |
| Rnase4 | 11680.33536 | -0.750521577 | 3.93E-03 |
| Sox4 | 7067.281048 | -0.75654872 | 9.87E-03 |
| Siah2 | 587.9083439 | -0.760066901 | 1.68E-05 |
| Asap2 | 581.2720119 | -0.761060957 | 3.89E-04 |
| Mpzl1 | 3792.936615 | -0.762011639 | 3.30E-04 |
| Rtkn | 1336.449565 | -0.762508511 | 1.26E-04 |
| Synpo | 5453.723669 | -0.765833393 | 1.70E-04 |
| Fam69b | 943.3674919 | -0.768577889 | 1.41E-04 |
| Hivep3 | 515.0723489 | -0.770868622 | 1.33E-04 |
| Dusp6 | 789.6529648 | -0.771074823 | 2.36E-02 |
| Dvl2 | 3542.891552 | -0.77206438 | 2.67E-05 |
| Dpy19l1 | 4064.222751 | -0.777918188 | 3.79E-07 |
| Txnrd3 | 885.4457633 | -0.780922161 | 1.34E-06 |
| Apln | 371.7087337 | -0.781852691 | 4.02E-02 |
| P4htm | 35.99990357 | -0.783999602 | 4.35E-02 |
| Zfp521 | 2160.446859 | -0.78513227 | 3.88E-04 |
| Gm16861 | 183.4760558 | -0.787699808 | 3.32E-04 |
| Mertk | 609.7724486 | -0.791308779 | 1.24E-03 |
| Tmem200a | 180.2586998 | -0.793809996 | 2.37E-02 |
| Gm12013 | 265.5132074 | -0.794329566 | 4.25E-02 |

| | | | |
|---|---|---|---|
| Kdm5b | 5947.881651 | -0.794409993 | 2.82E-03 |
| Tnfaip6 | 646.6209697 | -0.799798903 | 4.80E-03 |
| Atxn1 | 1217.972466 | -0.800241451 | 1.37E-03 |
| Jun | 4890.668098 | -0.800509386 | 7.81E-03 |
| Fam131b | 2012.653011 | -0.801794788 | 3.98E-06 |
| Rpl22l1 | 5120.112213 | -0.805352168 | 5.35E-05 |
| Nln | 2979.601328 | -0.805580409 | 5.28E-13 |
| Shmt1 | 1266.240147 | -0.808240308 | 3.33E-05 |
| 9430038I01Rik | 156.3527109 | -0.808857169 | 7.29E-06 |
| Nkain1 | 814.8730286 | -0.809973477 | 1.26E-05 |
| 2810039B14Rik | 112.6447876 | -0.810346537 | 1.77E-04 |
| Sfrp1 | 704.6605608 | -0.817429028 | 9.67E-03 |
| Olfml2b | 2018.256671 | -0.818314015 | 4.61E-03 |
| Klf16 | 1257.19304 | -0.82114059 | 7.54E-05 |
| Plat | 4991.615202 | -0.822218076 | 1.35E-03 |
| Tgfb2 | 3781.852125 | -0.824814931 | 3.13E-03 |
| Sesn3 | 451.770964 | -0.826288277 | 2.21E-04 |
| Hvcn1 | 604.403497 | -0.830672321 | 4.36E-05 |
| Prps2 | 2712.855431 | -0.830923405 | 1.55E-09 |
| Shroom1 | 179.9661314 | -0.834565436 | 1.03E-02 |
| Armcx6 | 583.5901757 | -0.835910407 | 1.49E-06 |
| Slc29a1 | 1079.904629 | -0.839544632 | 2.21E-03 |
| Gm15427 | 13183.07084 | -0.839558052 | 4.74E-02 |
| Gla | 4390.097848 | -0.840612833 | 3.05E-04 |
| Clip4 | 144.42984 | -0.840996098 | 3.21E-03 |
| Shf | 903.0088903 | -0.846883109 | 5.91E-05 |
| Gm13226 | 135.786182 | -0.848716953 | 3.09E-02 |
| Acot1 | 957.7954011 | -0.849280588 | 3.88E-04 |
| Arhgef19 | 326.3317744 | -0.849945051 | 7.64E-03 |
| Tubb2b | 2259.346938 | -0.851212722 | 1.75E-05 |
| Maf | 987.6224961 | -0.853741391 | 3.99E-02 |
| Txndc16 | 3149.373031 | -0.85956696 | 4.32E-06 |
| Prob1 | 109.771206 | -0.860032995 | 2.70E-02 |
| Dennd2a | 5263.659133 | -0.860126296 | 3.94E-06 |
| Gm12763 | 245.8440299 | -0.86092486 | 3.61E-03 |
| Hmga1 | 10223.72691 | -0.86412738 | 5.07E-03 |
| Polm | 3037.083453 | -0.864714595 | 6.58E-09 |
| A930024E05Rik | 70.97068599 | -0.86483612 | 1.31E-02 |
| Tfap2a | 730.9542063 | -0.866415996 | 1.13E-03 |
| Efnb2 | 1976.379573 | -0.867667477 | 1.09E-05 |
| 1700056E22Rik | 47.30814952 | -0.86840274 | 1.73E-02 |
| Lonrf1 | 505.0783429 | -0.868785462 | 6.59E-08 |
| Acat2 | 6980.262021 | -0.871362728 | 8.47E-03 |
| Dcaf6 | 5277.412717 | -0.876972745 | 3.12E-06 |
| Arhgap24 | 1837.038701 | -0.87864173 | 7.59E-04 |
| Gm9747 | 221.4334643 | -0.879805874 | 1.91E-02 |
| Hexdc | 950.8161435 | -0.887898564 | 4.43E-07 |

| | | | |
|---|---|---|---|
| Gbp2 | 472.493555 | -0.889226612 | 2.73E-04 |
| Hmgcr | 4300.914669 | -0.889923855 | 1.14E-02 |
| Cyp51 | 10822.29261 | -0.890844861 | 6.05E-03 |
| Myo1e | 5416.527455 | -0.893410861 | 8.24E-06 |
| Rnf150 | 1744.58427 | -0.895038674 | 1.29E-06 |
| Ptger4 | 492.752494 | -0.897643404 | 3.48E-05 |
| Inka1 | 187.7790777 | -0.900494294 | 1.21E-03 |
| Foxc2 | 481.5561448 | -0.901544957 | 1.24E-03 |
| Ptk7 | 1571.76109 | -0.902228163 | 1.22E-05 |
| Gcnt4 | 773.2398383 | -0.904256867 | 5.69E-05 |
| Faah | 126.5236943 | -0.904259451 | 2.88E-02 |
| Hcn3 | 50.97651329 | -0.905149838 | 4.05E-02 |
| Ramp3 | 1853.838291 | -0.906368002 | 1.65E-02 |
| Tgif2 | 708.2075399 | -0.907227655 | 1.62E-06 |
| Mmp2 | 89388.04831 | -0.90915986 | 3.30E-03 |
| A330009N23Rik | 54.46191423 | -0.913156025 | 1.49E-02 |
| Rps15a-ps3 | 65.56143529 | -0.913746857 | 2.96E-02 |
| Amdhd2 | 1696.242283 | -0.913771599 | 4.50E-04 |
| Shc2 | 1586.018871 | -0.916335 | 4.41E-05 |
| 4933431E20Rik | 881.878442 | -0.92014438 | 2.61E-05 |
| Evl | 2952.632705 | -0.920709639 | 5.93E-05 |
| Pmvk | 936.202711 | -0.924289504 | 9.51E-04 |
| Nsdhl | 4567.873814 | -0.925202005 | 3.85E-03 |
| Csgalnact1 | 1244.091072 | -0.926363643 | 2.28E-05 |
| Mylip | 3518.788342 | -0.927665364 | 3.07E-04 |
| Dcakd | 2695.360278 | -0.930059964 | 7.11E-08 |
| Gli2 | 250.1196828 | -0.931625366 | 6.75E-04 |
| Aqp5 | 257.8230496 | -0.93199895 | 2.52E-02 |
| Cmtm3 | 7129.083408 | -0.933972471 | 1.26E-07 |
| Hmgcs1 | 10146.3423 | -0.938904339 | 3.69E-02 |
| Slc2a6 | 121.7876624 | -0.939555634 | 7.12E-03 |
| Gas7 | 13591.79219 | -0.941599169 | 1.86E-04 |
| Ucp2 | 1083.734914 | -0.9418467 | 1.97E-05 |
| 2810408A11Rik | 84.21173571 | -0.943524376 | 6.45E-03 |
| NA | 126.927072 | -0.944671068 | 3.30E-03 |
| Plekha4 | 436.889388 | -0.9458532 | 9.06E-03 |
| Ror2 | 442.1456838 | -0.951530346 | 1.58E-05 |
| Noc4l | 1562.446352 | -0.95204491 | 5.46E-09 |
| Sco2 | 266.3938489 | -0.952159987 | 7.06E-03 |
| Prr7 | 85.92663974 | -0.953905511 | 7.56E-03 |
| Mex3d | 2980.139648 | -0.954299297 | 9.43E-06 |
| Fdft1 | 6536.743541 | -0.960873881 | 9.45E-04 |
| Mtmr2 | 3520.614254 | -0.966026802 | 1.98E-08 |
| Spry4 | 239.4302629 | -0.969564099 | 2.87E-03 |
| Prtg | 275.2554638 | -0.972075867 | 6.80E-04 |
| Rhou | 4021.987846 | -0.975036013 | 4.32E-08 |
| Capn6 | 8375.626269 | -0.977014082 | 2.02E-04 |

| | | | |
|---|---|---|---|
| Cd200 | 115.5327228 | -0.977025732 | 4.60E-02 |
| Il34 | 440.8805053 | -0.980369227 | 3.77E-04 |
| Pde7a | 1818.082955 | -0.980609659 | 4.43E-21 |
| Dlk1 | 29943.381 | -0.983965915 | 2.77E-02 |
| Hmga1b | 5808.517452 | -0.99001272 | 1.40E-03 |
| Rtp4 | 57.72055596 | -0.990666828 | 3.44E-03 |
| Egr2 | 252.8800803 | -0.992322772 | 3.38E-02 |
| Fgf14 | 155.1480338 | -0.994713189 | 3.59E-02 |
| Opcml | 366.9743187 | -0.996370449 | 3.61E-03 |
| NA | 167.7309758 | -1.000314674 | 2.88E-05 |
| Runx1 | 1094.061865 | -1.002274236 | 6.85E-18 |
| Slpi | 339.1667458 | -1.006373023 | 1.09E-02 |
| Stard4 | 2046.999912 | -1.008619555 | 7.04E-03 |
| Pik3r3 | 1442.838135 | -1.016078475 | 2.77E-07 |
| Ak5 | 1153.641956 | -1.019643926 | 4.19E-10 |
| Srgap1 | 1826.660683 | -1.022523237 | 3.71E-04 |
| Ccl2 | 2277.435037 | -1.02494865 | 1.28E-02 |
| Erg28 | 1761.041711 | -1.027435422 | 1.62E-06 |
| Cdo1 | 5428.454521 | -1.029317919 | 9.87E-03 |
| Il18 | 182.5136154 | -1.030075167 | 2.14E-03 |
| Cdk15 | 129.3148332 | -1.03076686 | 9.86E-03 |
| Lpin1 | 2970.0158 | -1.032148953 | 7.15E-04 |
| Rab27a | 205.8409117 | -1.035028928 | 4.09E-04 |
| Prkg2 | 2708.349054 | -1.040385805 | 8.75E-03 |
| Chchd10 | 24.56151814 | -1.040781214 | 3.57E-02 |
| Gm5862 | 499.4662236 | -1.044738324 | 2.08E-02 |
| Pcdhb9 | 216.2214447 | -1.044967042 | 7.58E-05 |
| Chst2 | 135.1732714 | -1.04500485 | 5.60E-03 |
| Glis1 | 258.3571055 | -1.048088519 | 1.92E-02 |
| Msmo1 | 8301.39548 | -1.050575413 | 3.77E-03 |
| St6gal1 | 1757.152116 | -1.05265183 | 7.88E-06 |
| Tiam2 | 1483.526478 | -1.053719445 | 5.90E-05 |
| Sh2b2 | 263.5269605 | -1.054236356 | 3.31E-06 |
| Fjx1 | 266.2076356 | -1.054603036 | 8.16E-08 |
| Tgfb3 | 3717.223368 | -1.074797848 | 7.91E-08 |
| Kif26b | 704.3016406 | -1.084488751 | 1.65E-03 |
| Gm4617 | 58.09501644 | -1.085901595 | 3.29E-02 |
| Gm15280 | 482.2429882 | -1.08790814 | 2.21E-06 |
| Igsf3 | 583.6384486 | -1.092295866 | 2.34E-07 |
| Slc27a1 | 1298.849242 | -1.102657681 | 4.68E-05 |
| Mex3a | 2190.570495 | -1.103713352 | 1.52E-05 |
| Nceh1 | 3203.829638 | -1.112965762 | 4.40E-09 |
| Pdk4 | 19724.77029 | -1.118962331 | 1.27E-03 |
| Il17rd | 972.8019314 | -1.124377341 | 8.44E-05 |
| Mtus2 | 261.3824621 | -1.124830613 | 4.81E-03 |
| Trps1 | 1353.888354 | -1.125508575 | 1.44E-02 |
| Unc13a | 386.4394156 | -1.131007206 | 7.76E-06 |

| | | | |
|---|---|---|---|
| Gpr137b | 6771.907523 | -1.138337464 | 4.81E-05 |
| Axin2 | 27.73513628 | -1.139172758 | 3.70E-02 |
| Dusp10 | 1991.634529 | -1.146041367 | 2.36E-03 |
| Insig1 | 7189.3603 | -1.147637798 | 9.23E-05 |
| Pla2g7 | 967.292464 | -1.149362917 | 1.69E-05 |
| Il1rl1 | 26338.39734 | -1.156251765 | 5.51E-04 |
| Kcnab1 | 1245.884438 | -1.157179145 | 4.32E-06 |
| Gm7964 | 24.99744468 | -1.157352227 | 2.94E-02 |
| Nat8f1 | 120.6153721 | -1.158237574 | 4.67E-03 |
| Kctd15 | 882.2490959 | -1.161471869 | 1.45E-04 |
| Csrp2 | 1603.401949 | -1.162566482 | 6.64E-04 |
| Ednra | 698.5583548 | -1.16432512 | 1.09E-03 |
| H2-K2 | 70.25926091 | -1.180642453 | 1.68E-03 |
| Fdps | 10188.03978 | -1.183562598 | 1.24E-03 |
| Tm7sf2 | 703.3797047 | -1.199127005 | 3.11E-05 |
| 5033430I15Rik | 56.08870669 | -1.199740347 | 4.25E-04 |
| St3gal4 | 2084.028363 | -1.202680413 | 5.45E-07 |
| Steap3 | 17632.6646 | -1.21608518 | 4.51E-05 |
| mt-Tq | 47.7235201 | -1.217513779 | 9.78E-03 |
| Gm9392 | 25.55154648 | -1.218673866 | 1.79E-02 |
| Lss | 5106.435912 | -1.231568939 | 2.70E-03 |
| Gm7729 | 37.07732584 | -1.232226251 | 3.30E-02 |
| Dlgap2 | 221.2029046 | -1.245142454 | 7.35E-06 |
| Sorcs2 | 114.3662152 | -1.246735581 | 6.37E-03 |
| Gm5643 | 21.10175863 | -1.250599214 | 1.70E-02 |
| Tnfrsf11b | 121.1051387 | -1.255580909 | 3.96E-02 |
| Mex3b | 174.4803361 | -1.26595167 | 6.45E-05 |
| Depp1 | 214.6965571 | -1.270593549 | 9.92E-03 |
| Mycn | 50.26697572 | -1.276137433 | 2.70E-02 |
| Hsd17b7 | 1863.090416 | -1.276471017 | 2.01E-04 |
| Il1rap | 2529.832075 | -1.276686336 | 3.78E-12 |
| Masp1 | 1959.534601 | -1.287537911 | 6.53E-06 |
| Col28a1 | 133.400327 | -1.293340656 | 4.17E-03 |
| Rgs5 | 30.59780507 | -1.296412508 | 8.02E-03 |
| Zfp41 | 1083.240847 | -1.30652133 | 1.40E-04 |
| Il16 | 43.63478451 | -1.309166286 | 3.28E-03 |
| Smoc1 | 319.0475636 | -1.328692302 | 1.89E-02 |
| 3010003L21Rik | 23.41418922 | -1.331494577 | 3.67E-02 |
| Etv4 | 307.4908993 | -1.333354362 | 3.66E-03 |
| Itgb8 | 478.0701506 | -1.337892902 | 4.71E-07 |
| Rhpn1 | 32.93132459 | -1.339326036 | 1.23E-03 |
| Cela1 | 174.5106123 | -1.339372474 | 6.18E-04 |
| Plxnb1 | 240.1316655 | -1.34639214 | 2.54E-02 |
| Fam135b | 222.7442954 | -1.365607769 | 3.83E-02 |
| Etv5 | 692.3569151 | -1.366343373 | 2.18E-06 |
| Spata48 | 29.05752533 | -1.368098609 | 9.73E-03 |
| Gm7361 | 115.296168 | -1.371528339 | 1.26E-02 |

| | | | |
|---|---|---|---|
| Ctxn1 | 645.1595188 | -1.385052391 | 3.89E-10 |
| Arhgap28 | 1085.405443 | -1.403770796 | 3.41E-13 |
| Angpt1 | 126.2034669 | -1.409052265 | 3.98E-02 |
| Rgs4 | 1205.292047 | -1.415372861 | 4.41E-02 |
| Has2os | 14.78403193 | -1.432694627 | 2.22E-02 |
| Fam19a1 | 75.29991681 | -1.4330406 | 1.22E-02 |
| Apol10b | 37.80520983 | -1.436142927 | 2.24E-03 |
| Sox9 | 2547.006272 | -1.445770034 | 6.89E-09 |
| Edil3 | 1365.668624 | -1.450031586 | 3.01E-05 |
| BC106179 | 76.11099145 | -1.455612089 | 6.64E-05 |
| Pknox2 | 149.7278646 | -1.463935671 | 2.16E-05 |
| Cux2 | 16.35978818 | -1.465964398 | 3.37E-02 |
| Mvd | 2840.656231 | -1.468614713 | 9.29E-06 |
| Casq2 | 10.12003193 | -1.478018252 | 3.85E-02 |
| Gbp9 | 51.96187374 | -1.485477493 | 7.48E-06 |
| Foxl1 | 95.73223871 | -1.486063025 | 1.08E-06 |
| Gfra2 | 25.3859268 | -1.494533496 | 3.26E-02 |
| Ggt7 | 21.30559824 | -1.496738664 | 1.20E-02 |
| 2610035D17Rik | 96.21859554 | -1.507483335 | 2.42E-03 |
| Nrep | 127.7232309 | -1.509978185 | 5.45E-03 |
| Tmem176a | 167.3003744 | -1.512417668 | 1.52E-05 |
| Slc30a1 | 484.7571357 | -1.519852629 | 7.16E-04 |
| Plp1 | 415.718745 | -1.520973366 | 5.36E-04 |
| Gsta1 | 191.6070784 | -1.523332858 | 7.67E-03 |
| Cxcl12 | 14907.90677 | -1.525719599 | 8.84E-06 |
| Nat8 | 9.423743375 | -1.527764856 | 4.47E-02 |
| Nat8f4 | 100.8466667 | -1.535709977 | 2.91E-04 |
| Tmem268 | 2601.301169 | -1.551904408 | 8.26E-13 |
| Paqr8 | 1206.296549 | -1.579129607 | 2.89E-11 |
| Rhov | 20.06302063 | -1.584679193 | 4.69E-02 |
| NA | 13.22047084 | -1.596038328 | 3.62E-02 |
| Tmem176b | 522.89459 | -1.608332601 | 7.97E-08 |
| B630019A10Rik | 22.44071648 | -1.611728389 | 1.02E-02 |
| Tfcp2l1 | 14.22561057 | -1.618148481 | 2.79E-02 |
| Gm14323 | 29.8499962 | -1.632298925 | 2.36E-03 |
| Zan | 76.59000702 | -1.633865403 | 1.21E-02 |
| Lef1 | 146.4483945 | -1.663019203 | 2.13E-04 |
| Spp1 | 33894.67107 | -1.667623369 | 2.53E-05 |
| Pgbd5 | 323.7462955 | -1.673252854 | 1.04E-13 |
| Nat8f3 | 96.81673939 | -1.696747884 | 2.70E-02 |
| Cfap161 | 19.83244907 | -1.703872592 | 4.81E-03 |
| Has2 | 320.7677252 | -1.706778628 | 4.39E-07 |
| mt-Ti | 38.83048144 | -1.721186534 | 2.48E-02 |
| Igf2 | 11022.73197 | -1.726028498 | 2.54E-02 |
| Duox1 | 119.8789528 | -1.750190734 | 2.42E-06 |
| Sema7a | 84.82804028 | -1.754443866 | 4.84E-04 |
| NA | 11.72957923 | -1.772118578 | 2.85E-02 |

| | | | |
|---|---|---|---|
| Plppr3 | 487.5192114 | -1.8277039 | 3.41E-07 |
| Gbp4 | 14.23422782 | -1.835554125 | 1.46E-03 |
| Gpr137c | 36.62518333 | -1.845260248 | 4.68E-04 |
| Kirrel3 | 34.68484594 | -1.851030783 | 3.70E-03 |
| Flrt3 | 33.14674922 | -1.867045102 | 3.00E-02 |
| Adamts15 | 21.18110807 | -1.897662776 | 2.11E-03 |
| Serpina3n | 668.2845824 | -1.938255986 | 1.21E-07 |
| Gpnmb | 3590.839261 | -1.949538272 | 1.39E-09 |
| Dmrtc1c2 | 1708.160336 | -1.956244809 | 1.46E-02 |
| Megf10 | 616.628578 | -1.964979333 | 6.53E-06 |
| Kcnk3 | 92.21429487 | -1.973685611 | 6.45E-05 |
| Prl2c5 | 68.59226915 | -1.974470602 | 2.13E-03 |
| Hc | 29.33803059 | -1.98119287 | 4.93E-05 |
| Zc3h12d | 7.659289797 | -1.991380042 | 2.79E-02 |
| 1700006J14Rik | 16.77111565 | -1.992375312 | 1.59E-02 |
| Rnase1 | 36.55767924 | -1.995710115 | 4.16E-04 |
| Nat8f7 | 24.83797719 | -1.997380837 | 3.65E-03 |
| Tcim | 182.9808592 | -2.020033512 | 1.23E-09 |
| Dpt | 3000.84147 | -2.041600027 | 2.87E-03 |
| Prtn3 | 13.25797432 | -2.063736598 | 1.64E-02 |
| Prl2c2 | 22898.70079 | -2.067944174 | 2.72E-04 |
| Gm14165 | 18.44418343 | -2.092964738 | 2.03E-02 |
| Tecta | 36.05411746 | -2.143319737 | 7.06E-05 |
| Il33 | 385.9276264 | -2.149152588 | 7.37E-03 |
| Tnni1 | 8.579755006 | -2.251664121 | 1.83E-02 |
| Atp8a2 | 5.479564426 | -2.287155283 | 4.32E-02 |
| H2-T3 | 6.159642751 | -2.313741649 | 4.80E-02 |
| Klrb1a | 6.733824506 | -2.338720534 | 2.29E-02 |
| Nmur1 | 13.54843957 | -2.381733009 | 2.44E-03 |
| Tdrd9 | 51.03735793 | -2.429018055 | 3.01E-08 |
| Galnt9 | 9.989789218 | -2.56740209 | 3.58E-03 |
| Gm22935 | 8.206150922 | -2.662823451 | 3.87E-02 |
| Igf2os | 450.3914571 | -2.872503733 | 6.87E-10 |
| Myo5b | 13.71114935 | -2.957526029 | 4.96E-02 |
| Aspg | 128.0287642 | -3.477843411 | 2.16E-11 |
| NA | 9.225849333 | -3.524636248 | 2.36E-02 |
| Tspan10 | 235.8434891 | -3.548190822 | 4.63E-03 |
| Nr1i3 | 116.7264137 | -3.90724801 | 3.35E-02 |
| Homer2 | 11.8001883 | -4.341052358 | 3.33E-02 |
| Cabp7 | 8.452464321 | -5.220610089 | 1.34E-02 |
| Gm2366 | 40.57644542 | -5.542631544 | 9.88E-03 |
| Cd200r3 | 139.6286064 | -5.979969237 | 1.52E-02 |
| Gm15655 | 5.77256052 | -6.212476794 | 4.81E-02 |
| Gm12263 | 29.33682477 | -7.288969043 | 2.14E-02 |
| Klhdc9 | 17.89947461 | -7.353902472 | 4.07E-03 |
| Samd11 | 155.349818 | -8.840541265 | 2.89E-04 |
| Sla2 | 584.6313518 | -9.401121571 | 8.12E-08 |

| | | | |
|---|---|---|---|
| Gm7666 | 51.9416385 | -44.88723913 | 4.45E-30 |

| Gene | baseMean | log2 Fold Change | padj |
|---|---|---|---|
| Adamts13 | 189.314739 | 10.24904711 | 4.32E-03 |
| Plekhh1 | 212.2371776 | 8.05062859 | 3.28E-06 |
| Tmem150b | 157.484637 | 6.958019897 | 4.85E-03 |
| Cyp4f14 | 11.05060069 | 5.264457867 | 3.58E-03 |
| Gm27286 | 31.94524763 | 5.060941799 | 1.09E-02 |
| Tfeb | 959.7623694 | 4.827490541 | 9.31E-139 |
| Abcc2 | 27.61579762 | 4.77783541 | 1.96E-04 |
| Gm26749 | 17.48551257 | 4.46924387 | 2.34E-02 |
| Tspan10 | 235.8434891 | 4.451879005 | 2.49E-04 |
| Gpr174 | 13.33517572 | 4.401606959 | 6.48E-03 |
| Igfbp5 | 819.6479129 | 3.858200243 | 1.04E-11 |
| Mep1a | 25.58712896 | 3.47846926 | 2.40E-04 |
| Kdr | 75.92150854 | 3.468283411 | 2.39E-05 |
| Gpnmb | 3590.839261 | 3.207506524 | 2.38E-25 |
| Mamld1 | 9.313569485 | 3.12599478 | 4.07E-02 |
| Dusp15 | 15.63811429 | 2.73328188 | 1.33E-04 |
| Agt | 230.4174338 | 2.637812988 | 1.35E-09 |
| Rragd | 324.476639 | 2.630012131 | 9.85E-16 |
| Pmel | 43.3739947 | 2.597591535 | 4.42E-18 |
| Mgam | 514.7573389 | 2.538494312 | 3.98E-06 |
| Rnf43 | 74.77146117 | 2.502925012 | 1.66E-10 |
| Gm10256 | 43.00942646 | 2.467505542 | 8.05E-03 |
| Mreg | 281.1053841 | 2.415480436 | 8.37E-09 |
| Ptgds | 167.4192768 | 2.395725597 | 2.64E-10 |
| Celsr1 | 29.54169175 | 2.352994788 | 5.81E-06 |
| St3gal6 | 28.19900837 | 2.301850139 | 2.44E-02 |
| Plp1 | 415.718745 | 2.140060823 | 6.57E-07 |
| Erbb3 | 69.46120585 | 2.1107842 | 1.60E-02 |
| Tspan12 | 74.6818032 | 2.000081748 | 2.62E-05 |
| Kbtbd11 | 17.4369085 | 1.978548326 | 1.09E-02 |
| Rnf128 | 22.02821797 | 1.948622138 | 3.84E-02 |
| Trim63 | 604.4480012 | 1.928119638 | 1.90E-04 |
| Rab27a | 205.8409117 | 1.912785657 | 2.22E-13 |
| Renbp | 533.6090566 | 1.89847119 | 9.32E-22 |
| Hyal1 | 467.9810951 | 1.897811149 | 3.63E-10 |
| Notch4 | 22.94704906 | 1.891249849 | 2.46E-02 |
| Peg3 | 3145.504318 | 1.881239809 | 3.21E-18 |
| Aloxe3 | 163.7815803 | 1.876976114 | 2.20E-06 |
| Dmrtc1c2 | 1708.160336 | 1.87397093 | 4.91E-02 |
| Ccn3 | 191.5912267 | 1.842331091 | 3.08E-16 |
| Pla2g2e | 18.47206542 | 1.740931289 | 1.33E-03 |
| Hhatl | 22.27895666 | 1.723288973 | 1.26E-02 |
| Mlana | 173.3382716 | 1.631517529 | 4.23E-02 |
| Optn | 361.5681594 | 1.617204413 | 1.89E-21 |
| E330034G19Rik | 21.08984885 | 1.589017733 | 4.19E-04 |
| Gm17382 | 35.29974084 | 1.583424965 | 1.84E-04 |

| | | | |
|---|---|---|---|
| Mx2 | 49.68098047 | 1.581890795 | 5.59E-03 |
| Tfe3 | 4096.617787 | 1.565710654 | 2.23E-27 |
| Dmxl2 | 711.1668361 | 1.558362846 | 3.27E-06 |
| Tnfrsf21 | 266.5152627 | 1.557635083 | 1.06E-10 |
| Gm10010 | 87.03730997 | 1.547604738 | 2.61E-05 |
| Garem2 | 297.7203972 | 1.524699891 | 5.21E-17 |
| Kcnk3 | 92.21429487 | 1.512580815 | 6.45E-03 |
| Selenop | 9438.313738 | 1.492363212 | 3.86E-05 |
| Ttc39a | 153.8251168 | 1.476009082 | 6.91E-06 |
| Ccdc40 | 33.18276155 | 1.457472838 | 1.12E-02 |
| Ttc39aos1 | 48.76699706 | 1.455826869 | 1.10E-03 |
| Vgf | 80.48610746 | 1.45434385 | 5.79E-04 |
| Nlrc5 | 42.39167115 | 1.441348443 | 1.50E-02 |
| Carmil3 | 108.3115967 | 1.440296036 | 1.93E-04 |
| Hsd17b1 | 30.82236224 | 1.422956068 | 7.16E-03 |
| Gm16861 | 183.4760558 | 1.421125715 | 9.19E-14 |
| Sytl2 | 158.0277612 | 1.376256702 | 1.30E-02 |
| Col9a3 | 88.80701058 | 1.306501083 | 5.28E-06 |
| Itgb3 | 406.4148934 | 1.304345025 | 2.72E-03 |
| AA467197 | 26.56671516 | 1.29545511 | 3.67E-02 |
| Atf3 | 486.7080334 | 1.294266898 | 1.92E-07 |
| 2610318N02Rik | 78.5731003 | 1.287712528 | 1.75E-02 |
| Adgrg1 | 72.72172502 | 1.25194502 | 3.24E-03 |
| Sap30 | 554.9416513 | 1.235088164 | 4.66E-07 |
| Fbxo32 | 1611.242564 | 1.220760103 | 9.48E-06 |
| Fuca2 | 3318.401005 | 1.20493615 | 1.63E-11 |
| Lyst | 3273.741853 | 1.199078536 | 2.78E-18 |
| 3010001F23Rik | 41.86125242 | 1.188149025 | 1.18E-02 |
| Ramp3 | 1853.838291 | 1.185636308 | 2.38E-03 |
| Ankrd12 | 2220.360824 | 1.151771883 | 3.36E-06 |
| Dusp4 | 2570.483965 | 1.145230363 | 2.27E-03 |
| Bhlhe41 | 899.5494575 | 1.143641578 | 1.54E-04 |
| Dpf1 | 35.92731429 | 1.140451263 | 4.82E-02 |
| Pnpla7 | 844.5630092 | 1.138052179 | 8.37E-09 |
| Adcy1 | 1520.53709 | 1.134676226 | 4.83E-04 |
| C77080 | 1863.179005 | 1.129496658 | 7.05E-15 |
| Smad9 | 231.529786 | 1.128689244 | 1.05E-02 |
| Klrg2 | 65.68197859 | 1.122397751 | 7.23E-03 |
| Lrguk | 36.65908511 | 1.122291417 | 2.67E-02 |
| Hebp2 | 255.3948829 | 1.116572334 | 2.44E-08 |
| 2500002B13Rik | 80.42307177 | 1.112645472 | 2.33E-04 |
| Myo7a | 1099.229425 | 1.112122195 | 9.87E-07 |
| Fnip2 | 5464.462773 | 1.105368061 | 6.55E-10 |
| Creg1 | 2258.352822 | 1.098916556 | 3.84E-06 |
| Slc25a44 | 2191.360797 | 1.098380439 | 2.74E-11 |
| Tcn2 | 1543.355567 | 1.091036905 | 4.88E-05 |
| Gabre | 296.5455035 | 1.089606859 | 3.51E-04 |

| | | | |
|---|---|---|---|
| Cry1 | 919.8294528 | 1.082996245 | 1.45E-07 |
| Myo5a | 10004.30674 | 1.073874958 | 1.87E-09 |
| Washc2 | 10254.84457 | 1.066308408 | 3.08E-12 |
| Mrgpre | 112.8488705 | 1.061462927 | 1.04E-02 |
| Tbc1d2 | 1016.924895 | 1.057896611 | 1.33E-04 |
| Hk2 | 3039.306829 | 1.052887963 | 3.56E-10 |
| Gas2l3 | 1203.193311 | 1.051421341 | 1.23E-03 |
| Cndp2 | 2438.22089 | 1.047482649 | 8.49E-04 |
| Dpp7 | 5371.98911 | 1.046197052 | 2.51E-06 |
| Elovl4 | 159.9698766 | 1.045529783 | 2.34E-03 |
| Gm7008 | 42.62583199 | 1.045226632 | 3.56E-03 |
| Chaf1b | 1033.445061 | 1.03757063 | 4.04E-04 |
| Lrp12 | 1151.827332 | 1.036077366 | 2.82E-16 |
| Mcoln1 | 2770.651326 | 1.020789449 | 3.51E-08 |
| Mfsd1 | 8319.866312 | 1.013708171 | 5.44E-14 |
| C530008M17Rik | 222.325705 | 1.011132702 | 3.80E-02 |
| Pappa2 | 2623.486677 | 1.003692776 | 1.41E-04 |
| 5031439G07Rik | 7244.859997 | 0.990900761 | 1.21E-07 |
| Gm8615 | 190.0903932 | 0.987075773 | 4.60E-04 |
| Naglu | 4106.854751 | 0.982817325 | 7.22E-05 |
| Pxylp1 | 827.890857 | 0.973009607 | 4.30E-03 |
| Tmem268 | 2601.301169 | 0.966105821 | 1.15E-04 |
| Dbp | 721.6468037 | 0.965686098 | 3.92E-02 |
| Akap6 | 117.141912 | 0.953513714 | 1.28E-02 |
| Sirpa | 3616.513785 | 0.952589135 | 1.90E-08 |
| Phlda1 | 939.9688801 | 0.948187031 | 5.84E-03 |
| 1700003F12Rik | 81.47917467 | 0.946148806 | 2.94E-04 |
| Macrod2 | 179.0981956 | 0.946094203 | 1.23E-06 |
| Lrp8 | 1552.953999 | 0.944420905 | 3.76E-02 |
| Ctns | 2362.555936 | 0.940217725 | 1.52E-09 |
| Unc13b | 378.855196 | 0.938059444 | 6.95E-04 |
| Tifa | 255.8989682 | 0.931721972 | 4.30E-03 |
| Apba1 | 301.8235088 | 0.928198526 | 1.27E-03 |
| Cfap157 | 270.2462511 | 0.91892898 | 4.96E-06 |
| Gnpda1 | 2534.474834 | 0.91790926 | 9.87E-07 |
| Sulf1 | 1797.810171 | 0.917773003 | 3.30E-02 |
| Shmt1 | 1266.240147 | 0.914935306 | 4.72E-06 |
| P2ry2 | 104.4183659 | 0.909296723 | 2.34E-02 |
| Mical1 | 409.2832872 | 0.907552548 | 9.00E-05 |
| Impact | 6753.387895 | 0.902229781 | 8.14E-07 |
| Tmem140 | 492.4092491 | 0.899044813 | 3.30E-02 |
| Mfsd11 | 2064.247758 | 0.897660818 | 1.90E-07 |
| Stk17b | 1172.316199 | 0.883494428 | 4.51E-05 |
| Wdr62 | 764.1814903 | 0.88117062 | 1.01E-02 |
| Ezr | 2125.964047 | 0.878985851 | 8.23E-08 |
| Npc1 | 3965.691488 | 0.877423292 | 1.42E-05 |
| Naa80 | 610.3779038 | 0.877422894 | 8.80E-05 |

| | | | |
|---|---|---|---|
| Lamp1 | 56088.28006 | 0.868649787 | 2.52E-05 |
| Slc39a8 | 483.1111622 | 0.868310254 | 6.95E-05 |
| Sord | 8476.138057 | 0.85986362 | 8.79E-03 |
| Gla | 4390.097848 | 0.855492591 | 7.43E-04 |
| Eda2r | 1123.269013 | 0.843660842 | 1.38E-02 |
| Ass1 | 199.8319632 | 0.842015977 | 3.97E-02 |
| Atp6v1c1 | 4552.101365 | 0.841431026 | 1.96E-07 |
| Bhlhe40 | 3834.220772 | 0.833813526 | 6.71E-03 |
| Wdr81 | 3346.664813 | 0.833022576 | 1.11E-05 |
| Etl4 | 562.100196 | 0.831321176 | 2.06E-02 |
| Hacd2 | 1385.12878 | 0.830121161 | 4.60E-05 |
| Ang | 240.4623036 | 0.82752138 | 3.09E-03 |
| Pcdh10 | 140.2969449 | 0.827461657 | 2.50E-02 |
| Grn | 30705.7805 | 0.817804917 | 4.29E-04 |
| Enpp2 | 759.7675373 | 0.817529998 | 3.65E-02 |
| Vash2 | 2507.446433 | 0.81515687 | 2.49E-05 |
| Zfp704 | 652.3489939 | 0.814073142 | 2.10E-02 |
| Neu1 | 2682.126546 | 0.812867871 | 6.00E-04 |
| Zc3h8 | 210.3834763 | 0.809626975 | 4.91E-03 |
| Hspa12a | 126.8535222 | 0.805842031 | 2.57E-02 |
| Gstp1 | 12119.5976 | 0.804957864 | 1.67E-06 |
| Angel1 | 396.2138089 | 0.79985016 | 5.88E-03 |
| Zfand2a | 2124.767632 | 0.797834234 | 4.30E-05 |
| Atp6v1h | 5874.496336 | 0.796473614 | 3.50E-07 |
| Aim2 | 214.7586752 | 0.796083368 | 8.83E-03 |
| Ctsa | 21262.24103 | 0.796012217 | 8.20E-04 |
| Slc39a10 | 3524.383162 | 0.794512307 | 1.48E-02 |
| Rell1 | 1320.85393 | 0.794141051 | 2.61E-06 |
| Fgfbp3 | 84.82346159 | 0.793183426 | 3.39E-02 |
| Abcc4 | 345.0528919 | 0.792660927 | 3.76E-02 |
| Cpeb2 | 370.5002161 | 0.791556452 | 1.66E-03 |
| Opn3 | 112.0638704 | 0.79108794 | 3.41E-03 |
| Znrf2 | 1311.359839 | 0.788522775 | 4.17E-04 |
| Trim37 | 1840.754342 | 0.788360063 | 1.14E-09 |
| Gm4673 | 94.57211719 | 0.78791053 | 2.24E-02 |
| Rufy3 | 357.8406021 | 0.768524868 | 4.07E-05 |
| Ung | 266.879367 | 0.767934501 | 1.05E-02 |
| Atp6v0d1 | 8071.893965 | 0.765115319 | 2.90E-03 |
| Tuba4a | 1663.613125 | 0.762544288 | 2.70E-02 |
| Smyd3 | 1007.188368 | 0.757183782 | 2.35E-03 |
| Dnase2a | 1680.910599 | 0.751770316 | 2.29E-08 |
| Cerk | 1102.588167 | 0.751639841 | 8.61E-08 |
| Pnpo | 218.6551061 | 0.744390879 | 2.89E-03 |
| Clcn7 | 3441.410259 | 0.740822005 | 1.25E-04 |
| Rnase4 | 11680.33536 | 0.735894025 | 1.30E-02 |
| Maoa | 11869.71092 | 0.734494638 | 1.78E-03 |
| Washc5 | 9453.018285 | 0.728593676 | 3.98E-14 |

| | | | |
|---|---|---|---|
| Nudt16l1 | 1616.991711 | 0.726295031 | 2.33E-04 |
| Cutc | 134.6270003 | 0.72263145 | 9.14E-04 |
| Dock4 | 204.615827 | 0.720887579 | 9.22E-03 |
| Fam214a | 744.1921239 | 0.720731662 | 2.46E-02 |
| Snai2 | 1626.102104 | 0.719371192 | 2.98E-04 |
| Bid | 1806.508215 | 0.717355868 | 1.17E-05 |
| Nagpa | 1014.528249 | 0.717038168 | 4.28E-04 |
| Slc36a1 | 1573.335444 | 0.71612409 | 1.11E-05 |
| Hexa | 11840.38452 | 0.715834246 | 3.50E-02 |
| Gdf11 | 1582.164993 | 0.71532583 | 4.94E-04 |
| Vps18 | 2476.558079 | 0.71167032 | 4.30E-03 |
| Camkk1 | 480.61585 | 0.710226641 | 1.88E-02 |
| Zbed5 | 182.5059509 | 0.707529289 | 4.47E-03 |
| Lurap1 | 310.6166841 | 0.707368872 | 1.71E-05 |
| Ppfia3 | 146.6688839 | 0.705883695 | 3.82E-02 |
| Ss18l2 | 298.5950248 | 0.70091505 | 1.07E-04 |
| Slc25a33 | 464.665477 | 0.694861458 | 3.46E-04 |
| Fam171b | 768.6035689 | 0.694103454 | 7.04E-03 |
| Slc39a6 | 3751.096942 | 0.693632768 | 3.73E-05 |
| Mroh1 | 6120.311947 | 0.693372387 | 4.71E-04 |
| Tceal9 | 6722.958843 | 0.685159799 | 1.01E-03 |
| Pwwp2b | 1031.015521 | 0.679452044 | 1.22E-02 |
| Klhl21 | 1593.791091 | 0.676237362 | 3.50E-02 |
| 4632404H12Rik | 210.9051878 | 0.675928649 | 8.86E-03 |
| Ctsb | 36468.22462 | 0.674503741 | 1.36E-02 |
| Hsd3b7 | 1224.252911 | 0.668739595 | 8.01E-03 |
| Acy1 | 719.2862101 | 0.666967997 | 2.59E-03 |
| Hsdl1 | 1948.417286 | 0.665705359 | 3.41E-05 |
| Slc29a3 | 2607.580111 | 0.665617594 | 2.49E-04 |
| Uap1l1 | 10255.8949 | 0.665106867 | 2.11E-02 |
| Gaa | 16101.78382 | 0.663836341 | 5.64E-04 |
| Chrnb2 | 134.4276621 | 0.661873217 | 7.19E-03 |
| Pogk | 1686.347408 | 0.661831028 | 1.59E-05 |
| Atp6v0b | 9934.293287 | 0.659259347 | 3.29E-03 |
| Mapk8ip1 | 2586.65763 | 0.656421362 | 5.11E-03 |
| Gba | 4711.112014 | 0.652412714 | 6.96E-04 |
| Pikfyve | 2011.636706 | 0.652208111 | 1.54E-04 |
| Pip4k2c | 5308.496544 | 0.650751932 | 4.21E-03 |
| Ephx2 | 595.9836792 | 0.650469813 | 1.01E-02 |
| Man2b1 | 3916.818943 | 0.649545584 | 2.17E-03 |
| Hectd3 | 2568.610222 | 0.646383275 | 1.73E-05 |
| Atp6v1a | 10459.35955 | 0.644834495 | 2.16E-03 |
| Gns | 20094.75228 | 0.644548419 | 2.25E-03 |
| Rnf157 | 1391.358528 | 0.644410412 | 7.49E-06 |
| Tsc1 | 3398.460772 | 0.641195278 | 1.34E-05 |
| 6430548M08Rik | 814.8945097 | 0.640560121 | 3.51E-04 |
| Map3k6 | 224.2736214 | 0.640450546 | 1.99E-02 |

| | | | |
|---|---|---|---|
| Vps8 | 2880.10801 | 0.639834321 | 6.96E-04 |
| Galc | 1514.601027 | 0.63472699 | 2.08E-04 |
| St6galnac4 | 1396.81264 | 0.630117118 | 5.25E-03 |
| Col4a3bp | 1675.621637 | 0.62996901 | 4.34E-05 |
| Oxct1 | 16865.66235 | 0.628768357 | 2.20E-03 |
| Ptrh1 | 1785.441485 | 0.628709412 | 4.61E-03 |
| Vegfb | 3825.597118 | 0.628575802 | 8.71E-03 |
| Prkx | 992.1556184 | 0.627507005 | 2.25E-03 |
| Lnx2 | 610.26002 | 0.626819374 | 9.14E-04 |
| Ahcyl2 | 2542.94842 | 0.623350562 | 9.39E-03 |
| Nkain1 | 814.8730286 | 0.623327195 | 3.85E-03 |
| Gclc | 1433.972265 | 0.621060658 | 1.38E-02 |
| Ogdhl | 4923.875176 | 0.620886162 | 1.73E-04 |
| Acap3 | 1180.426838 | 0.618184597 | 7.44E-04 |
| Plbd2 | 8599.309391 | 0.618169903 | 5.14E-03 |
| Slc20a1 | 2917.806275 | 0.614838545 | 1.08E-02 |
| Vps13c | 3424.132169 | 0.613672759 | 1.78E-03 |
| Spns1 | 1815.839164 | 0.613145886 | 2.37E-04 |
| Gga2 | 4453.49319 | 0.611187385 | 2.91E-03 |
| Fbxl2 | 422.3385071 | 0.610835835 | 4.29E-02 |
| Fv1 | 303.0703483 | 0.610561436 | 5.77E-03 |
| Myo6 | 987.1695516 | 0.609301684 | 2.26E-03 |
| Nat10 | 2589.324134 | 0.608808447 | 4.36E-04 |
| Mnt | 3501.854391 | 0.608475211 | 3.15E-02 |
| Specc1 | 3432.279109 | 0.607286375 | 1.38E-02 |
| Ostm1 | 1935.936349 | 0.606279553 | 4.91E-04 |
| Osbpl1a | 1582.537579 | 0.605986163 | 1.90E-04 |
| Ctsd | 41499.83588 | 0.605747897 | 4.85E-02 |
| Gramd3 | 607.1091723 | 0.604762484 | 9.31E-03 |
| Tmem192 | 1385.041556 | 0.604337848 | 8.85E-03 |
| Rragc | 10259.96991 | 0.602751079 | 9.41E-04 |
| Mgst1 | 1067.813549 | 0.601049044 | 4.79E-02 |
| Tmem251 | 678.3297086 | 0.599338904 | 5.01E-04 |
| Adi1 | 3350.226368 | 0.596713257 | 3.55E-03 |
| Gnptab | 4858.629519 | 0.592279644 | 9.03E-05 |
| Mif | 7576.735311 | 0.58911061 | 3.25E-02 |
| Ccdc115 | 665.7497048 | 0.588576628 | 2.89E-03 |
| Atxn3 | 1054.949583 | 0.587959237 | 2.87E-02 |
| Bax | 4107.204276 | 0.582218744 | 6.06E-03 |
| Igf2r | 15440.49464 | 0.581448014 | 3.07E-03 |
| Glmp | 7292.413588 | 0.580814587 | 4.79E-03 |
| Col12a1 | 2667.384937 | 0.58080339 | 8.89E-04 |
| Hook2 | 814.6719502 | 0.580736473 | 4.12E-03 |
| Rmnd1 | 1544.503591 | 0.579308727 | 9.69E-03 |
| Sec11c | 466.8889073 | 0.578891648 | 1.67E-03 |
| Fam160a2 | 1223.39513 | 0.576516647 | 9.57E-06 |
| Arfgef2 | 4840.915022 | 0.575571928 | 1.13E-03 |

| | | | |
|---|---|---|---|
| Ugp2 | 4229.526194 | 0.573025379 | 4.35E-05 |
| Sgpl1 | 13380.67463 | 0.570823913 | 9.92E-05 |
| Magi2 | 526.5829282 | 0.568882264 | 1.32E-04 |
| Atp6v1d | 4059.992121 | 0.56471123 | 1.25E-02 |
| Ogdh | 6423.124912 | 0.5644125 | 2.23E-03 |
| Parvb | 2285.010317 | 0.561916049 | 3.68E-03 |
| Soat1 | 3693.855559 | 0.561630115 | 6.57E-07 |
| Slc17a5 | 1194.706084 | 0.560314557 | 1.25E-02 |
| Fnip1 | 4227.643221 | 0.559794398 | 3.60E-03 |
| Cdc14a | 289.2179467 | 0.558876174 | 1.50E-02 |
| Shpk | 226.3556193 | 0.55720371 | 8.73E-03 |
| Cuta | 2544.799639 | 0.556935574 | 1.18E-02 |
| Pianp | 1185.174268 | 0.556868954 | 4.96E-02 |
| Akap13 | 5008.332383 | 0.556236914 | 4.50E-03 |
| Ap5z1 | 1165.206489 | 0.555609341 | 7.83E-05 |
| Zfyve26 | 2479.501653 | 0.555574557 | 2.52E-05 |
| Dhtkd1 | 505.0186144 | 0.553292988 | 3.23E-03 |
| Tmem199 | 785.0747928 | 0.551933153 | 1.38E-05 |
| Diaph2 | 713.883724 | 0.550201283 | 3.57E-02 |
| Ccdc171 | 404.936003 | 0.548554865 | 1.93E-02 |
| Gtf2h1 | 3210.848105 | 0.546640178 | 3.23E-03 |
| Kif16b | 1057.721431 | 0.545675457 | 2.08E-04 |
| Wdr7 | 1610.402645 | 0.542079242 | 1.33E-04 |
| Tmem150a | 1636.619374 | 0.541467702 | 9.55E-03 |
| Commd4 | 942.1458007 | 0.540663187 | 3.51E-04 |
| Sbno1 | 4742.795818 | 0.540131958 | 3.44E-03 |
| Rai14 | 4032.0779 | 0.537907093 | 1.36E-02 |
| Itpr3 | 5436.339997 | 0.536597275 | 4.26E-03 |
| Mrs2 | 469.060725 | 0.534014687 | 1.28E-02 |
| Plpbp | 2115.5974 | 0.53119035 | 9.61E-12 |
| Mgat4b | 6635.711925 | 0.531073798 | 2.29E-02 |
| Nln | 2979.601328 | 0.530578227 | 2.49E-05 |
| Nr2c2ap | 954.217837 | 0.529310494 | 3.83E-02 |
| Prkdc | 1324.769727 | 0.526050282 | 3.71E-03 |
| Nars2 | 357.7982091 | 0.524996686 | 5.63E-03 |
| Fam50a | 3368.128408 | 0.524883919 | 2.03E-03 |
| Gnl3 | 3183.598349 | 0.522764655 | 2.50E-02 |
| Vps33a | 2892.012132 | 0.522695922 | 4.09E-07 |
| Fndc10 | 367.9322098 | 0.520682622 | 4.76E-02 |
| Dcun1d4 | 1164.457837 | 0.51812118 | 2.88E-02 |
| Tpp1 | 8880.766729 | 0.517481838 | 1.86E-02 |
| Htt | 2803.662578 | 0.514465035 | 3.22E-04 |
| Txnrd3 | 885.4457633 | 0.514436272 | 7.92E-03 |
| Plpp2 | 931.3288588 | 0.514136503 | 2.12E-02 |
| Snx8 | 2047.563724 | 0.511004483 | 2.67E-03 |
| Slc35f6 | 2258.273448 | 0.509732497 | 7.46E-03 |
| Rdh13 | 1335.03426 | 0.508857801 | 2.96E-03 |

| | | | |
|---|---|---|---|
| 8-Mar | 6434.713929 | 0.505102599 | 2.33E-25 |
| Serac1 | 662.4093304 | 0.504526848 | 1.15E-02 |
| AI413582 | 932.4708856 | 0.50447687 | 4.34E-02 |
| Orai2 | 1421.004878 | 0.502362107 | 2.21E-03 |
| mt-Nd6 | 36849.03868 | 0.500378681 | 9.45E-03 |
| Pno1 | 1764.155483 | 0.499157006 | 3.99E-02 |
| Scpep1 | 18619.66479 | 0.495790534 | 4.44E-03 |
| Tmem87b | 2010.229776 | 0.495284302 | 3.30E-02 |
| Armcx6 | 583.5901757 | 0.495082559 | 2.31E-02 |
| Cd3eap | 705.0522757 | 0.493730443 | 4.79E-02 |
| Cln5 | 1783.706728 | 0.4902495 | 1.23E-02 |
| Flt1 | 1329.706088 | 0.490044762 | 2.17E-03 |
| Tmeff1 | 2004.0604 | 0.489167014 | 5.76E-03 |
| Pnpla6 | 1143.850243 | 0.48778462 | 3.65E-04 |
| Snx30 | 5144.862324 | 0.487695802 | 6.37E-03 |
| Ank2 | 3380.138198 | 0.484745329 | 3.76E-02 |
| Slc20a2 | 2786.725506 | 0.484710033 | 1.52E-02 |
| Lrpprc | 5310.296473 | 0.484237323 | 4.69E-03 |
| Scarb2 | 3430.447569 | 0.483704459 | 9.67E-03 |
| Nr6a1 | 421.218285 | 0.48365331 | 1.90E-02 |
| Tbc1d16 | 1866.941337 | 0.482479118 | 9.41E-03 |
| Zfp655 | 3110.698296 | 0.481460868 | 2.52E-05 |
| mt-Nd5 | 120857.742 | 0.47637975 | 7.43E-03 |
| Bcas3 | 993.4164983 | 0.471434638 | 2.45E-02 |
| Lta4h | 6976.776006 | 0.469789634 | 3.62E-02 |
| Foxn2 | 1053.302421 | 0.465703413 | 2.68E-02 |
| Fbxw7 | 724.7639213 | 0.464859907 | 4.57E-02 |
| Rnf115 | 1861.018874 | 0.46381256 | 3.67E-04 |
| Clcn5 | 3296.841631 | 0.462737779 | 1.88E-07 |
| Nepro | 974.7880945 | 0.461890115 | 2.54E-02 |
| Phf20 | 3151.247539 | 0.460129692 | 1.40E-05 |
| Tctn3 | 629.4105587 | 0.459696115 | 9.14E-04 |
| Elf2 | 3398.181671 | 0.459646769 | 9.74E-05 |
| Vac14 | 1852.536674 | 0.457450663 | 3.13E-02 |
| Kmt2c | 2822.831742 | 0.456758498 | 7.26E-03 |
| Vps26a | 5958.591577 | 0.452398487 | 3.13E-03 |
| Tbc1d24 | 2492.496938 | 0.445934892 | 6.76E-04 |
| Atg3 | 1744.113181 | 0.445299796 | 2.59E-03 |
| Nsmce2 | 1712.981215 | 0.443313618 | 3.60E-03 |
| 2410131K14Rik | 1774.50699 | 0.44278778 | 3.03E-03 |
| Atg7 | 2072.141009 | 0.442425515 | 4.05E-02 |
| Abhd12 | 3305.451109 | 0.441358736 | 2.31E-02 |
| Paxip1 | 2358.455763 | 0.440841994 | 1.09E-02 |
| Slc26a2 | 908.0177184 | 0.44048549 | 5.45E-03 |
| Hnrnph2 | 3745.444933 | 0.438333104 | 1.73E-02 |
| Slc25a15 | 757.3726621 | 0.435469521 | 3.03E-03 |
| Coprs | 814.1861832 | 0.434273997 | 3.03E-03 |

| | | | | |
|---|---|---|---|---|
| Nup50 | | 3476.731492 | 0.434150128 | 3.30E-02 |
| Bex3 | | 8754.027769 | 0.433050839 | 6.57E-03 |
| Wdr12 | | 1978.768705 | 0.429597263 | 4.35E-02 |
| Cstb | | 9856.088638 | 0.425593271 | 5.43E-03 |
| Utp18 | | 1810.570753 | 0.424762095 | 4.20E-02 |
| Hivep1 | | 1746.576588 | 0.424619693 | 7.51E-04 |
| Celf1 | | 4225.590777 | 0.424102329 | 3.49E-02 |
| Ccdc93 | | 3957.529583 | 0.423445201 | 2.04E-02 |
| Adamtsl5 | | 3366.847999 | 0.42161615 | 1.15E-02 |
| Jak1 | | 13668.72976 | 0.421331831 | 2.07E-02 |
| Lrp11 | | 3762.571826 | 0.418649356 | 2.70E-02 |
| Degs1 | | 9219.28485 | 0.418455043 | 3.65E-02 |
| Zfand5 | | 4632.448716 | 0.417807191 | 3.74E-02 |
| Als2 | | 1499.841059 | 0.414845672 | 5.98E-03 |
| Irak1bp1 | | 511.6876268 | 0.413417815 | 3.65E-02 |
| Nckap1 | | 11777.13914 | 0.412592115 | 2.31E-02 |
| Ift88 | | 627.0852278 | 0.412085376 | 4.64E-02 |
| Kdm4c | | 2657.626562 | 0.411749681 | 1.72E-02 |
| | 8-Sep | 10704.68529 | 0.410180705 | 7.87E-03 |
| Bcl7a | | 1302.545937 | 0.410066386 | 7.52E-03 |
| Rab43 | | 2992.193124 | 0.408857823 | 2.14E-02 |
| Pcyt1a | | 4562.392465 | 0.408082635 | 2.10E-04 |
| Atp6ap2 | | 4358.08888 | 0.406214524 | 4.56E-02 |
| Mtrr | | 953.9515698 | 0.404071725 | 2.18E-02 |
| Asxl2 | | 1701.60244 | 0.402723868 | 1.63E-02 |
| Nup188 | | 4987.880274 | 0.400557313 | 2.36E-02 |
| Max | | 2035.937597 | 0.400396244 | 1.61E-03 |
| Ube2q1 | | 5225.360789 | 0.399832822 | 6.01E-04 |
| Hps3 | | 1032.072822 | 0.398516702 | 1.37E-02 |
| Tmem181a | | 524.5608539 | 0.397597092 | 2.21E-03 |
| Dolk | | 812.9649783 | 0.396620511 | 9.55E-03 |
| Parp8 | | 1704.661461 | 0.393326283 | 4.16E-02 |
| Pusl1 | | 543.0205238 | 0.391702048 | 3.26E-02 |
| Dnajc13 | | 5378.577471 | 0.385782001 | 4.15E-02 |
| Golgb1 | | 5482.119432 | 0.385620811 | 1.87E-02 |
| Rab3gap2 | | 4136.31955 | 0.385483118 | 3.98E-03 |
| Dzip1 | | 1896.61168 | 0.385041858 | 3.03E-03 |
| Rhoq | | 7241.961221 | 0.383865053 | 1.15E-02 |
| Ppp1r37 | | 2197.916903 | 0.383723516 | 4.82E-02 |
| Lrfn4 | | 2023.205322 | 0.382949445 | 4.82E-02 |
| Tulp4 | | 16880.04521 | 0.379754068 | 6.45E-03 |
| Zfp28 | | 611.0405196 | 0.379682131 | 1.79E-02 |
| mt-Nd2 | | 99498.30713 | 0.379245292 | 9.70E-03 |
| Aak1 | | 1915.969594 | 0.378764936 | 1.06E-02 |
| Agfg1 | | 2748.187289 | 0.377343461 | 1.69E-03 |
| Uggt1 | | 8254.05786 | 0.377078976 | 5.69E-03 |
| Rcbtb1 | | 2189.384948 | 0.376050508 | 2.20E-02 |

| | | | |
|---|---|---|---|
| mt-Co2 | 249817.806 | 0.375800581 | 9.82E-03 |
| Focad | 3203.348512 | 0.373465581 | 1.04E-02 |
| Gid4 | 1936.274946 | 0.370284538 | 7.15E-03 |
| Agtrap | 1417.40242 | 0.364530801 | 3.14E-02 |
| Atp7a | 1656.515308 | 0.362741827 | 4.53E-02 |
| Tceal8 | 3073.445068 | 0.361601601 | 5.10E-03 |
| Akap10 | 1520.79788 | 0.360885835 | 8.90E-05 |
| Mtch2 | 7896.965041 | 0.358530864 | 2.19E-04 |
| Taok3 | 2501.86846 | 0.354875718 | 4.63E-03 |
| Gpr89 | 1644.684102 | 0.354318856 | 1.49E-02 |
| Rsad1 | 456.0324608 | 0.352879152 | 3.23E-02 |
| Usp28 | 1013.741207 | 0.350896152 | 7.43E-03 |
| Tmed8 | 1367.151775 | 0.347289699 | 2.02E-02 |
| Smcr8 | 3771.341043 | 0.344300091 | 2.47E-02 |
| Pi4ka | 5631.269742 | 0.342719255 | 8.06E-03 |
| Mycbp2 | 4214.136191 | 0.342218828 | 1.98E-02 |
| D6Wsu163e | 991.3243772 | 0.341018611 | 1.03E-02 |
| Noct | 8107.901444 | 0.33924421 | 3.02E-02 |
| Nhlrc3 | 2088.125293 | 0.338221638 | 2.04E-02 |
| Pdpr | 1055.824546 | 0.336110391 | 3.30E-02 |
| Timm8b | 906.7143595 | 0.335726904 | 8.85E-03 |
| Fbxl17 | 1422.489761 | 0.334233923 | 2.33E-02 |
| Anapc1 | 10971.14232 | 0.329289484 | 8.71E-03 |
| Kif3b | 2552.903667 | 0.322581823 | 9.69E-05 |
| Cipc | 1168.346399 | 0.322506659 | 3.38E-02 |
| 2610301B20Rik | 1235.416847 | 0.320482351 | 1.89E-02 |
| Nf2 | 4991.381476 | 0.319746102 | 4.83E-02 |
| Spg11 | 3357.707005 | 0.318995128 | 5.16E-03 |
| Bach1 | 2273.465941 | 0.317936206 | 7.10E-03 |
| Pitrm1 | 6900.706977 | 0.314491054 | 3.46E-02 |
| Huwe1 | 19835.01404 | 0.311546504 | 8.73E-03 |
| Ncoa3 | 2075.835614 | 0.311492471 | 3.50E-02 |
| Micall1 | 3562.670056 | 0.310767683 | 4.66E-02 |
| Kif1b | 4905.83088 | 0.306751045 | 3.58E-02 |
| Slc38a1 | 5119.465128 | 0.306561156 | 9.55E-03 |
| Sfxn1 | 2367.25292 | 0.305005585 | 6.24E-03 |
| Prepl | 2110.742543 | 0.302898681 | 1.85E-02 |
| Mafg | 3468.758033 | 0.302765048 | 4.79E-02 |
| Tbcel | 1611.699035 | 0.296459842 | 9.50E-03 |
| Acadl | 4515.648214 | 0.2919735 | 3.50E-02 |
| Dnajc11 | 1545.72422 | 0.291634297 | 2.14E-02 |
| Ago2 | 7856.332648 | 0.289277531 | 4.06E-02 |
| Tmem126b | 728.0357211 | 0.27829379 | 3.66E-02 |
| Pacs2 | 7227.003698 | 0.274974497 | 3.50E-02 |
| Zfp652 | 3085.865232 | 0.269912124 | 4.37E-02 |
| Nop14 | 3249.947278 | 0.254443836 | 1.51E-02 |
| Ankrd13a | 3982.563234 | 0.235365594 | 4.35E-02 |

| | | | |
|---|---|---|---|
| Zfp106 | 11251.11847 | 0.228926374 | 1.30E-02 |
| Btbd7 | 2855.621626 | 0.226548779 | 4.07E-03 |
| Cltc | 24544.13461 | 0.224903691 | 4.30E-03 |
| Tgfbrap1 | 1916.492283 | 0.218408079 | 4.76E-02 |
| Kctd3 | 1744.740515 | -0.191656794 | 4.31E-02 |
| Marveld1 | 6344.789281 | -0.25678584 | 2.35E-02 |
| Wwtr1 | 5195.543564 | -0.256953593 | 1.67E-02 |
| Ugdh | 6304.035598 | -0.292473202 | 4.40E-02 |
| Man2a1 | 13475.00019 | -0.296279666 | 2.97E-02 |
| Cd9 | 9138.524752 | -0.2984316 | 4.42E-02 |
| Wipf1 | 3029.779313 | -0.30355277 | 6.05E-03 |
| Fyn | 3259.615135 | -0.306520784 | 2.62E-02 |
| Zbed3 | 1298.091305 | -0.315102029 | 1.92E-04 |
| Smim11 | 611.446549 | -0.315588725 | 3.25E-02 |
| Dnmbp | 2254.506423 | -0.320186489 | 9.55E-03 |
| Hdac7 | 6665.610222 | -0.326595387 | 2.26E-03 |
| Myl6 | 22034.81211 | -0.328532855 | 6.96E-04 |
| Farp1 | 7954.256896 | -0.329768902 | 4.04E-02 |
| St3gal2 | 8902.023726 | -0.329849304 | 1.98E-02 |
| Comt | 3881.085594 | -0.331313761 | 2.69E-02 |
| Calu | 43534.96136 | -0.347234481 | 4.56E-02 |
| Atf4 | 16684.06758 | -0.351057406 | 3.77E-02 |
| Sh3gl1 | 4195.01312 | -0.352366606 | 3.76E-02 |
| Cpne1 | 5044.131924 | -0.354108684 | 2.27E-04 |
| Tpm1 | 28225.26008 | -0.35477285 | 3.09E-02 |
| Gypc | 1383.765939 | -0.356168469 | 4.61E-02 |
| E2f4 | 3205.637373 | -0.362521602 | 1.37E-02 |
| Emp2 | 4062.72683 | -0.364110476 | 2.55E-03 |
| Tmem63a | 1994.571198 | -0.365964375 | 1.84E-02 |
| Nfix | 9711.486408 | -0.368978729 | 4.64E-02 |
| Parva | 10879.78875 | -0.372858089 | 1.04E-02 |
| Scd2 | 61452.75163 | -0.382106489 | 1.61E-02 |
| Gtf3c5 | 1178.49991 | -0.38309289 | 5.59E-03 |
| Fbxl14 | 1290.48478 | -0.384915627 | 2.84E-02 |
| Synj2 | 1645.375646 | -0.389420623 | 3.51E-04 |
| Rab3ip | 2085.051427 | -0.395620826 | 4.77E-02 |
| Lmo4 | 1591.794629 | -0.396337161 | 1.20E-03 |
| Foxd2os | 411.5132541 | -0.398229901 | 4.68E-02 |
| Gsap | 597.175052 | -0.398900517 | 4.64E-02 |
| Dnajc24 | 378.3848799 | -0.399227382 | 3.57E-02 |
| Adamts10 | 953.871541 | -0.401419687 | 2.13E-02 |
| Ehd2 | 12732.25354 | -0.406721285 | 1.30E-02 |
| Trim8 | 1911.483462 | -0.409593039 | 3.22E-03 |
| Il17rc | 2513.616066 | -0.409671763 | 2.19E-04 |
| Cpne8 | 1083.712569 | -0.416413472 | 3.03E-03 |
| Tnfsf12 | 1683.302223 | -0.417760329 | 3.54E-02 |
| Trio | 8424.77309 | -0.418824353 | 4.31E-02 |

| | | | |
|---|---|---|---|
| Tram2 | 3451.186882 | -0.419106561 | 1.31E-03 |
| Dynlt1b | 1798.335892 | -0.419234754 | 6.45E-03 |
| Rbpj | 6189.6897 | -0.421345452 | 4.30E-03 |
| Ajuba | 2948.78609 | -0.422076249 | 4.68E-02 |
| Rbpj-ps3 | 2159.928873 | -0.422505023 | 5.34E-03 |
| Mab21l1 | 801.0403118 | -0.423194736 | 4.98E-02 |
| Ctso | 1002.433076 | -0.427816164 | 4.71E-02 |
| Tpst1 | 2809.181828 | -0.429450233 | 3.87E-02 |
| Fam3c | 4044.640668 | -0.430007139 | 1.92E-02 |
| Tns2 | 5211.113391 | -0.430212299 | 3.22E-02 |
| Slc6a9 | 2482.059137 | -0.430935863 | 2.57E-02 |
| Tspo | 2413.333548 | -0.435955458 | 5.39E-03 |
| Fuz | 552.4981124 | -0.436854913 | 3.71E-02 |
| Jdp2 | 2313.384115 | -0.437216627 | 2.44E-02 |
| Parp12 | 537.1403291 | -0.438867741 | 1.60E-02 |
| Man1c1 | 553.3265867 | -0.440015394 | 4.68E-02 |
| Pcdh18 | 4140.879768 | -0.440028486 | 2.32E-02 |
| Ext1 | 3885.770872 | -0.440472895 | 1.72E-02 |
| Cplx2 | 1224.275436 | -0.446790146 | 1.22E-02 |
| Cavin1 | 29736.21876 | -0.447754849 | 2.16E-02 |
| Sbno2 | 2509.386322 | -0.447964668 | 4.35E-02 |
| Rgs10 | 422.0667231 | -0.453650329 | 4.29E-02 |
| Ebp | 1605.346395 | -0.458924449 | 3.13E-02 |
| Osbpl3 | 2074.055951 | -0.464939221 | 3.09E-02 |
| Tecr | 7238.644524 | -0.467895404 | 1.23E-02 |
| Acadvl | 2866.060471 | -0.467897756 | 8.85E-03 |
| Anxa2 | 27933.32394 | -0.468794314 | 4.58E-02 |
| Epb41l2 | 9926.271712 | -0.469564294 | 9.14E-04 |
| Efemp1 | 43132.42209 | -0.46992671 | 4.49E-02 |
| Sqor | 2866.171031 | -0.476508831 | 4.15E-03 |
| Tgfbr3 | 8535.55638 | -0.477651139 | 1.89E-02 |
| Peli2 | 1908.882325 | -0.480978808 | 5.59E-03 |
| Acly | 18875.66242 | -0.483311637 | 3.01E-02 |
| Fads2 | 5458.473187 | -0.486325639 | 3.24E-03 |
| 9930012K11Rik | 537.6727168 | -0.488726877 | 4.79E-02 |
| Cask | 3506.668855 | -0.488755493 | 6.89E-03 |
| Mcc | 1935.549035 | -0.490156242 | 9.55E-03 |
| Alkbh7 | 365.0892985 | -0.490959762 | 7.98E-03 |
| Scd1 | 8875.775287 | -0.494441087 | 4.26E-03 |
| Rnf150 | 1744.58427 | -0.495584273 | 4.52E-02 |
| Csad | 1257.892946 | -0.50104285 | 1.86E-02 |
| Rasa4 | 4237.672709 | -0.505573768 | 4.11E-02 |
| Cog4 | 4133.200435 | -0.507496682 | 2.14E-02 |
| Atf5 | 11430.63398 | -0.512186829 | 7.15E-03 |
| Mars | 4381.005849 | -0.517472105 | 3.54E-02 |
| Nfkb2 | 5751.40937 | -0.518522166 | 3.60E-02 |
| Synpo | 5453.723669 | -0.51928859 | 4.92E-02 |

| | | | |
|---|---|---|---|
| Lpar1 | 5879.931993 | -0.519491021 | 7.94E-03 |
| Fez1 | 805.2676056 | -0.520408527 | 2.46E-02 |
| Tbx18 | 1240.526144 | -0.520803509 | 3.36E-06 |
| Fhl2 | 3219.077124 | -0.524402441 | 1.47E-02 |
| Papss2 | 2566.376957 | -0.52785732 | 1.32E-02 |
| Gsdmd | 431.2673849 | -0.528818999 | 1.06E-03 |
| Gpr176 | 2045.393277 | -0.538512036 | 7.61E-03 |
| Il1rap | 2529.832075 | -0.539155748 | 3.16E-02 |
| Lurap1l | 1882.936888 | -0.539592461 | 1.29E-02 |
| Trp53i13 | 637.8712156 | -0.53984021 | 3.30E-02 |
| Rufy1 | 2070.818133 | -0.545329743 | 2.94E-02 |
| Tdrkh | 618.162234 | -0.545913871 | 1.86E-02 |
| Noc4l | 1562.446352 | -0.546029315 | 7.26E-03 |
| Runx1 | 1094.061865 | -0.548619006 | 4.35E-05 |
| Il4ra | 1231.92372 | -0.550110543 | 1.70E-02 |
| Shb | 889.4084882 | -0.550345481 | 2.02E-03 |
| Sertad1 | 765.3883428 | -0.552179309 | 2.70E-02 |
| Nol7 | 912.3460862 | -0.552768016 | 1.37E-02 |
| E430018J23Rik | 232.8995405 | -0.557452186 | 2.04E-02 |
| Mmp11 | 3113.336865 | -0.55989098 | 8.80E-05 |
| Pced1b | 224.7482532 | -0.562026114 | 5.72E-03 |
| Plekha1 | 3449.235513 | -0.569809048 | 4.74E-03 |
| Ccdc9b | 3998.146773 | -0.570497005 | 3.16E-04 |
| Ifi35 | 678.7565277 | -0.574981745 | 1.90E-02 |
| Itga2b | 570.2372038 | -0.575644228 | 3.54E-02 |
| Srebf2 | 11304.7876 | -0.57699348 | 2.78E-03 |
| Tgfb3 | 3717.223368 | -0.584311197 | 2.54E-02 |
| Cdc42ep2 | 2880.635182 | -0.589507335 | 8.65E-03 |
| Foxc1 | 342.3062006 | -0.591263785 | 5.35E-03 |
| Tmem119 | 3163.057139 | -0.594278717 | 3.52E-02 |
| Frmd6 | 6537.824608 | -0.594518977 | 1.67E-03 |
| Tgfbr2 | 6135.16223 | -0.595062161 | 1.23E-02 |
| Cavin3 | 2957.834925 | -0.595262397 | 1.58E-02 |
| Abcd2 | 730.8520276 | -0.595538697 | 4.35E-02 |
| Htra1 | 323.732752 | -0.595579404 | 1.19E-02 |
| Rpl35a | 516.7149947 | -0.597864554 | 4.07E-02 |
| Vangl1 | 1595.126302 | -0.609043604 | 1.70E-02 |
| Pdlim2 | 3693.375744 | -0.613290584 | 3.61E-02 |
| Tmem62 | 479.9034088 | -0.616999536 | 1.04E-02 |
| Spry2 | 928.0977301 | -0.618844848 | 2.54E-05 |
| Sdc4 | 3683.171926 | -0.621420083 | 1.90E-03 |
| Rnaseh2c | 449.8484921 | -0.621687687 | 1.27E-02 |
| Erg28 | 1761.041711 | -0.6227552 | 2.29E-02 |
| Trp53cor1 | 155.8446845 | -0.624144412 | 4.92E-02 |
| Tm4sf1 | 4004.71189 | -0.636452606 | 1.15E-02 |
| Il1r1 | 1964.698977 | -0.638617215 | 2.94E-02 |
| Npnt | 449.3441054 | -0.640351899 | 1.24E-02 |

| | | | |
|---|---|---|---|
| Pygo2 | 2490.294348 | -0.64060094 | 1.21E-07 |
| Irs1 | 2008.224803 | -0.642207981 | 1.88E-02 |
| Medag | 7575.216717 | -0.645536508 | 4.31E-02 |
| Fhl3 | 1894.248643 | -0.646969964 | 3.03E-03 |
| Mettl26 | 461.2555934 | -0.650327475 | 2.36E-02 |
| Plpp1 | 5028.842612 | -0.652796509 | 7.33E-03 |
| Myc | 3282.758284 | -0.652925947 | 7.09E-04 |
| Igfbp6 | 10099.86965 | -0.654821338 | 1.30E-02 |
| Arhgap28 | 1085.405443 | -0.66098008 | 7.15E-03 |
| Lmo7 | 1891.856719 | -0.662574026 | 2.33E-04 |
| Plpp5 | 772.2669265 | -0.662932031 | 1.06E-03 |
| Lhx9 | 259.9429502 | -0.664268859 | 2.70E-02 |
| Adamts4 | 1372.936021 | -0.66876965 | 4.13E-02 |
| Cdc42ep5 | 621.0593718 | -0.668880782 | 2.50E-02 |
| Tspan11 | 2916.150625 | -0.675488199 | 7.92E-03 |
| Tiam2 | 1483.526478 | -0.676381333 | 4.85E-02 |
| Jund | 2487.535816 | -0.67787572 | 1.22E-02 |
| Nuak1 | 2656.800253 | -0.692294915 | 1.31E-03 |
| Enpp1 | 302.9825204 | -0.69237063 | 4.33E-02 |
| Ror2 | 442.1456838 | -0.692584791 | 9.07E-03 |
| Map3k14 | 972.1145369 | -0.69317523 | 4.15E-03 |
| Olfml3 | 3715.895388 | -0.696177098 | 8.38E-04 |
| Gas2 | 879.7185941 | -0.69618256 | 3.76E-02 |
| Kcnk2 | 223.8511014 | -0.700543776 | 3.58E-02 |
| Rgs3 | 1513.56933 | -0.701975296 | 1.70E-02 |
| Plk2 | 10524.64627 | -0.703966626 | 7.32E-05 |
| Hspa1a | 228.6343034 | -0.704845891 | 1.93E-02 |
| Galnt17 | 1070.268328 | -0.710146542 | 1.38E-03 |
| Ablim1 | 3555.853548 | -0.71203505 | 1.31E-02 |
| Fzd2 | 5450.697507 | -0.712767428 | 2.89E-05 |
| Isg20 | 692.6214078 | -0.713010501 | 1.98E-02 |
| Trib2 | 1232.911453 | -0.713957941 | 6.19E-03 |
| Pcyt1b | 608.1630088 | -0.718520547 | 4.31E-02 |
| Timp3 | 19304.01546 | -0.720625155 | 1.58E-02 |
| Hspa1b | 1228.877042 | -0.729915407 | 1.29E-05 |
| Gng2 | 472.9891909 | -0.732304734 | 1.21E-02 |
| Clip4 | 144.42984 | -0.732885226 | 3.19E-02 |
| Mvk | 1649.647814 | -0.733749464 | 3.13E-02 |
| Klc3 | 265.3530538 | -0.737301388 | 1.28E-02 |
| Pcdh19 | 2133.007913 | -0.743232317 | 2.85E-03 |
| Fam43a | 1435.289342 | -0.763996441 | 2.40E-04 |
| Arntl | 722.4816621 | -0.771824544 | 2.61E-06 |
| Lrig1 | 3532.647732 | -0.778341271 | 1.72E-04 |
| Ngf | 909.5389495 | -0.783281975 | 1.24E-02 |
| Gm11816 | 245.2771472 | -0.784898488 | 4.04E-02 |
| Inka1 | 187.7790777 | -0.800197397 | 1.33E-02 |
| Ldlr | 7632.961331 | -0.80269187 | 2.32E-02 |

| | | | |
|---|---|---|---|
| Il13ra1 | 219.0472217 | -0.819217564 | 5.31E-05 |
| Itgb5 | 5005.644612 | -0.820599124 | 4.83E-04 |
| Angptl6 | 251.3102899 | -0.826088465 | 3.99E-02 |
| Hmgcr | 4300.914669 | -0.828816083 | 4.93E-02 |
| Gpr85 | 486.5449696 | -0.831174256 | 1.71E-03 |
| Grem2 | 1174.534392 | -0.83258462 | 1.90E-02 |
| H2-T23 | 367.1066819 | -0.836882695 | 2.46E-02 |
| Amigo2 | 624.6922769 | -0.837275947 | 8.93E-07 |
| Klf4 | 2395.995965 | -0.84262181 | 3.16E-04 |
| Cx3cl1 | 722.2112652 | -0.845047101 | 8.11E-03 |
| Gm45713 | 119.8688671 | -0.845903552 | 3.55E-03 |
| Kcna4 | 440.1062529 | -0.855738986 | 1.28E-02 |
| Mn1 | 1568.859744 | -0.859365496 | 6.31E-04 |
| Cercam | 171.2368976 | -0.860905144 | 1.38E-02 |
| Lbh | 5291.647432 | -0.864108208 | 2.88E-02 |
| Podnl1 | 3817.279448 | -0.865750005 | 9.48E-06 |
| Nfil3 | 1259.058948 | -0.866170452 | 9.56E-04 |
| Hexdc | 950.8161435 | -0.873222588 | 3.28E-06 |
| Cyp51 | 10822.29261 | -0.873581054 | 1.92E-02 |
| Fdft1 | 6536.743541 | -0.874214559 | 9.31E-03 |
| Nsdhl | 4567.873814 | -0.879858385 | 1.76E-02 |
| Itgbl1 | 4703.422305 | -0.885847361 | 2.89E-02 |
| Csrp2 | 1603.401949 | -0.886409719 | 3.80E-02 |
| Gbp2 | 472.493555 | -0.888848932 | 9.14E-04 |
| Gpr63 | 64.54603199 | -0.902433783 | 2.18E-02 |
| S1pr3 | 3918.192315 | -0.90457556 | 3.46E-07 |
| Kank3 | 220.0939253 | -0.905275117 | 2.04E-02 |
| Olfml2b | 2018.256671 | -0.91087866 | 3.75E-03 |
| Lsp1 | 852.1553722 | -0.913238767 | 1.53E-04 |
| Adm | 2177.281661 | -0.935004168 | 1.01E-02 |
| Fosl1 | 1007.39669 | -0.956994162 | 3.30E-02 |
| Gm13056 | 41.42652466 | -0.95905091 | 3.10E-02 |
| Rgs16 | 180.0814537 | -0.967253725 | 4.98E-02 |
| Dhcr24 | 6989.867164 | -0.976916436 | 2.46E-02 |
| Ecm1 | 1410.00657 | -0.989876701 | 1.37E-02 |
| Pmvk | 936.202711 | -1.003532859 | 8.89E-04 |
| Acy3 | 101.9828741 | -1.013282845 | 2.02E-02 |
| NA | 84.09963672 | -1.018126083 | 1.72E-02 |
| Fdps | 10188.03978 | -1.019045076 | 1.93E-02 |
| Ccdc141 | 352.3930217 | -1.021672685 | 6.32E-03 |
| Ccl7 | 1521.615401 | -1.032588261 | 1.85E-03 |
| AW551984 | 1346.712062 | -1.044336961 | 2.00E-02 |
| Fbln5 | 11685.97594 | -1.055643319 | 5.56E-07 |
| Tm7sf2 | 703.3797047 | -1.078092729 | 8.20E-04 |
| Col5a3 | 679.7155384 | -1.082073297 | 1.04E-02 |
| Phex | 174.470391 | -1.090065956 | 2.31E-02 |
| Cdsn | 1502.659716 | -1.090250477 | 3.16E-04 |

| | | | |
|---|---|---|---|
| Neurl1a | 101.6784878 | -1.093857941 | 1.36E-02 |
| Gprc5a | 2003.573682 | -1.101494494 | 5.11E-03 |
| Xpnpep2 | 463.2230938 | -1.101570492 | 2.26E-03 |
| Il34 | 440.8805053 | -1.117058815 | 1.23E-04 |
| Osr2 | 352.6919725 | -1.1426853 | 1.94E-02 |
| Chrnb1 | 351.1485445 | -1.150196279 | 1.16E-02 |
| Ifi27l2a | 63.9925599 | -1.1525191 | 1.08E-02 |
| Clu | 185.6709907 | -1.169933425 | 1.01E-02 |
| Tnfaip8l1 | 90.31987106 | -1.201715973 | 1.90E-02 |
| Hap1 | 31.38598453 | -1.207638731 | 2.32E-02 |
| Gm15280 | 482.2429882 | -1.252912121 | 1.45E-07 |
| H2-K2 | 70.25926091 | -1.288306062 | 2.11E-03 |
| Gm17057 | 15.87823374 | -1.317304381 | 4.58E-02 |
| Ifi47 | 69.23413701 | -1.327841924 | 1.41E-04 |
| Tek | 47.78473353 | -1.369536036 | 4.52E-02 |
| Ccl2 | 2277.435037 | -1.379387364 | 1.02E-03 |
| Fam19a1 | 75.29991681 | -1.41710875 | 3.40E-02 |
| Depp1 | 214.6965571 | -1.446800225 | 7.13E-03 |
| Angptl4 | 1038.167366 | -1.450744751 | 2.61E-06 |
| Cd55 | 298.5806079 | -1.47252534 | 2.61E-02 |
| Gm15710 | 93.55804308 | -1.481850097 | 7.43E-04 |
| Cfap161 | 19.83244907 | -1.540600596 | 2.86E-02 |
| Crabp1 | 530.6832668 | -1.571379348 | 3.84E-06 |
| Ngef | 111.9462157 | -1.590073391 | 4.31E-02 |
| Serpina3g | 65.50415713 | -1.607926506 | 3.13E-02 |
| Krt87 | 33.77427656 | -1.639707128 | 3.76E-02 |
| Gfra2 | 25.3859268 | -1.789329654 | 2.04E-02 |
| Cpa4 | 91.48931586 | -1.968914638 | 2.31E-02 |
| Cytip | 32.07785982 | -1.989218609 | 3.09E-02 |
| Sorcs2 | 114.3662152 | -1.991649695 | 1.11E-05 |
| Nes | 211.7096441 | -2.098119021 | 4.56E-02 |
| Serpina3h | 32.62380238 | -2.148205777 | 1.03E-02 |
| Aqp1 | 1103.311567 | -2.352770256 | 1.41E-04 |
| Msln | 8968.235968 | -2.429449234 | 1.54E-03 |
| Rhbdl2 | 8.373552228 | -2.790221704 | 4.59E-02 |
| Palm3 | 21.3394262 | -3.050875448 | 9.55E-03 |
| Serpina3i | 55.90345178 | -3.136739451 | 6.11E-05 |
| Lamc3 | 81.73661022 | -3.536065052 | 4.41E-02 |
| Nr1i3 | 116.7264137 | -4.197588128 | 4.83E-02 |
| Fgf21 | 49.48268511 | -4.510885479 | 3.84E-02 |
| Sla2 | 584.6313518 | -9.715147328 | 1.69E-07 |
| NA | 19.95430791 | -24.2127121 | 4.30E-05 |

| Gene | baseMean | log2 Fold Change | padj |
|---|---|---|---|
| Vip | 761.4057475 | 14.85041013 | 1.22E-28 |
| Slc6a12 | 232.6272342 | 12.03766372 | 2.09E-21 |
| Il31ra | 119.7831368 | 11.85597865 | 1.32E-20 |
| Upk1b | 1749.009226 | 11.17382424 | 1.16E-15 |
| Anxa8 | 232.8381859 | 11.06192071 | 6.76E-18 |
| Ccl20 | 50.88600856 | 10.74558061 | 2.63E-15 |
| Arhgap36 | 36.01428515 | 10.60666255 | 4.96E-03 |
| Panx3 | 37.5768691 | 10.5179598 | 7.12E-15 |
| Hapln1 | 63.3439414 | 10.45741111 | 2.14E-10 |
| Slc2a12 | 153.088269 | 10.42627461 | 6.97E-05 |
| 4933402E13Rik | 48.25961404 | 10.23212126 | 2.11E-15 |
| Parm1 | 50.62742008 | 9.828138195 | 4.76E-14 |
| Gm15056 | 33.57462403 | 9.812762255 | 4.00E-03 |
| Sntg1 | 26.65447637 | 9.48956457 | 4.90E-13 |
| Cnfn | 24.09477608 | 9.206703463 | 2.10E-11 |
| H2-Aa | 11.95773125 | 8.999747339 | 2.85E-02 |
| Gm5812 | 53.47797956 | 8.755846532 | 2.87E-04 |
| Serping1 | 32.77937136 | 8.674900113 | 2.06E-11 |
| Wnt7a | 83.29495754 | 8.668167466 | 3.22E-11 |
| Vill | 20.01966172 | 8.665622743 | 3.20E-10 |
| Wnt7b | 75.66538815 | 8.432683301 | 3.66E-10 |
| Robo3 | 32.48397439 | 8.363164534 | 1.03E-08 |
| Pkhd1 | 14.16137139 | 8.348519788 | 5.61E-06 |
| Dppa4 | 15.36586433 | 8.345110658 | 6.41E-09 |
| Serpinb9g | 65.5698485 | 8.3199287 | 1.04E-10 |
| Gm27286 | 31.94524763 | 8.267139858 | 2.02E-08 |
| Csmd1 | 7.471778892 | 8.210860573 | 3.07E-07 |
| Limch1 | 706.4773048 | 8.054763004 | 5.26E-74 |
| Cdh15 | 124.6645698 | 7.992958981 | 1.25E-09 |
| Slc47a1 | 16.32805259 | 7.978427328 | 2.54E-08 |
| Oprk1 | 32.03179652 | 7.976430607 | 3.22E-10 |
| Ddx4 | 6.74894573 | 7.953080249 | 6.04E-06 |
| Ces2e | 38.3662501 | 7.836745673 | 8.23E-12 |
| Ptprq | 158.5957609 | 7.82794001 | 1.54E-38 |
| Lamb3 | 47.93842017 | 7.746549151 | 8.82E-09 |
| U90926 | 10.05769719 | 7.739360309 | 2.93E-06 |
| 1700012B07Rik | 12.47027256 | 7.69439449 | 2.85E-08 |
| Gdf15 | 40.77172247 | 7.680422056 | 3.90E-11 |
| Cntn2 | 22.21630626 | 7.559950384 | 1.77E-12 |
| Hist1h4n | 13.89900442 | 7.554959742 | 1.31E-07 |
| Scel | 35.56464911 | 7.485581167 | 7.02E-07 |
| Gm9428 | 14.61896193 | 7.391701873 | 8.81E-08 |
| Sorbs2 | 490.7709526 | 7.338384894 | 3.36E-91 |
| Serpinb1a | 435.1696668 | 7.30237979 | 3.15E-62 |
| AU021092 | 16.8814082 | 7.28488654 | 1.70E-07 |
| Snx20 | 7.501681989 | 7.272761266 | 1.37E-05 |

| | | | |
|---|---|---|---|
| Map3k7cl | 13.82843052 | 7.230406972 | 3.51E-08 |
| Hepacam | 5.666744147 | 7.200436102 | 1.90E-05 |
| Serpinb1c | 17.20225213 | 7.159429234 | 8.73E-07 |
| Serpinb9e | 15.75118303 | 7.147586913 | 2.67E-07 |
| Dsp | 6.380203445 | 7.128990393 | 1.99E-04 |
| Snca | 7.396202048 | 7.091227425 | 5.94E-05 |
| Hephl1 | 25.60224789 | 7.068677052 | 5.59E-07 |
| Gm12153 | 16.73241645 | 7.010154763 | 2.05E-07 |
| 1520401A03Rik | 8.114182141 | 6.994295976 | 1.43E-05 |
| Prg2 | 8.098456987 | 6.957047933 | 7.15E-06 |
| Jph2 | 553.3177943 | 6.946063736 | 5.52E-52 |
| Kdr | 75.92150854 | 6.933515789 | 3.12E-25 |
| Arg1 | 5.523346439 | 6.894415006 | 5.89E-06 |
| Cblc | 13.39498369 | 6.876129947 | 2.99E-06 |
| NA | 188.127415 | 6.834350729 | 3.06E-28 |
| Gvin1 | 17.14007767 | 6.794506768 | 5.70E-05 |
| Speer3 | 6.085578054 | 6.778376289 | 6.26E-05 |
| Sh3tc1 | 213.0734353 | 6.776343969 | 2.12E-19 |
| Cacna1s | 6.269835238 | 6.753932633 | 1.22E-03 |
| Synpr | 13.66684869 | 6.743573524 | 2.13E-06 |
| Tmem171 | 242.2820833 | 6.718491076 | 5.51E-14 |
| Btn2a2 | 21.67262263 | 6.669727149 | 1.34E-08 |
| B230206L02Rik | 12.43408177 | 6.632943613 | 6.19E-06 |
| Birc7 | 495.5983212 | 6.632128131 | 1.15E-36 |
| Cyp3a13 | 57.57513375 | 6.613946984 | 2.97E-17 |
| 1700009N14Rik | 112.4486022 | 6.60853892 | 3.72E-23 |
| Gm21293 | 16.49303806 | 6.581701662 | 1.69E-05 |
| Ces2f | 8.035604011 | 6.578806424 | 1.29E-05 |
| Ikzf4 | 76.17939403 | 6.523012968 | 3.79E-13 |
| Gm4070 | 36.86624693 | 6.521880541 | 5.13E-09 |
| Gm5286 | 7.531885012 | 6.502311971 | 5.59E-06 |
| Des | 60.75912734 | 6.488551813 | 4.91E-25 |
| NA | 26.06308258 | 6.449157403 | 5.76E-06 |
| Plekhs1 | 102.3016648 | 6.429426399 | 3.28E-34 |
| Ctnnd2 | 13.32903101 | 6.39893974 | 4.73E-06 |
| Fgfbp1 | 21.39660242 | 6.394770628 | 1.38E-09 |
| Serpinb6b | 303.4332572 | 6.388571253 | 2.41E-89 |
| Gm6650 | 24.45313423 | 6.376438122 | 1.02E-14 |
| Vmn1r44 | 18.97883308 | 6.33258317 | 3.07E-06 |
| Fcna | 17.16685909 | 6.264176291 | 7.59E-07 |
| Nkain4 | 34.42273738 | 6.235032245 | 1.70E-09 |
| Cyp4f40 | 16.78817656 | 6.198139141 | 2.35E-07 |
| Mogat2 | 37.49728946 | 6.198102597 | 5.07E-17 |
| Ankk1 | 13.33636605 | 6.180210519 | 1.36E-05 |
| Crct1 | 218.1985879 | 6.160110426 | 4.09E-06 |
| Upk3b | 18.38391089 | 6.123510118 | 2.02E-07 |
| Psme2b | 18.26708444 | 6.116378429 | 3.61E-06 |

| | | | |
|---|---|---|---|
| Ube4bos1 | 31.36650529 | 6.102930809 | 7.45E-03 |
| NA | 240.538982 | 6.09547576 | 3.39E-03 |
| 4933427D06Rik | 13.64440714 | 6.088297873 | 1.60E-06 |
| Gm26635 | 8.223066354 | 6.056044855 | 1.26E-02 |
| Arhgap9 | 58.21154531 | 6.026691785 | 4.41E-12 |
| Hist1h2br | 20.36525225 | 6.023141821 | 3.93E-06 |
| Serpinb9f | 8.681433932 | 6.023132804 | 3.10E-04 |
| Fmo2 | 37.14481697 | 5.989270605 | 6.67E-18 |
| Cobl | 32.63939068 | 5.988687209 | 6.50E-13 |
| Sbk3 | 20.10660947 | 5.937110353 | 2.74E-10 |
| Vtn | 30.23869185 | 5.926878381 | 6.80E-11 |
| Inpp5d | 43.74884049 | 5.923905693 | 7.61E-10 |
| Slamf9 | 31.67214215 | 5.902262196 | 1.32E-12 |
| Dpp4 | 5.265900201 | 5.868457594 | 1.83E-04 |
| Tcp11x2 | 18.44972698 | 5.829847888 | 5.61E-12 |
| Serpina5 | 6.801924643 | 5.807124449 | 2.45E-04 |
| Serpinb9c | 5.608671978 | 5.784758474 | 6.57E-04 |
| Serpinb9b | 3482.508606 | 5.776622549 | 1.62E-53 |
| Atg9b | 216.6082704 | 5.742517627 | 5.78E-27 |
| Gm10696 | 6.667621713 | 5.706122361 | 4.14E-04 |
| Doc2a | 128.9404018 | 5.680026858 | 4.51E-02 |
| Upk3bl | 49.55256089 | 5.650732801 | 3.09E-12 |
| Susd2 | 15.13366232 | 5.628922553 | 5.76E-09 |
| Gli1 | 5.477929601 | 5.616339583 | 2.17E-04 |
| Gm15854 | 18.23662287 | 5.606308199 | 6.58E-06 |
| Tmeff2 | 105.2957531 | 5.602664166 | 6.13E-32 |
| Col4a4 | 11.5650756 | 5.589647767 | 1.63E-04 |
| B3gnt3 | 5.759179752 | 5.51781335 | 5.94E-04 |
| Gpc2 | 10.67028678 | 5.508086991 | 2.60E-03 |
| Als2cl | 101.8281501 | 5.478959704 | 1.33E-10 |
| Trav13-2 | 8.537472155 | 5.440252525 | 3.23E-04 |
| Card10 | 5.87499347 | 5.438304747 | 3.02E-04 |
| Sh3rf2 | 283.6625641 | 5.416400675 | 2.55E-31 |
| Ablim3 | 364.5652957 | 5.411342393 | 1.93E-52 |
| Ttll9 | 5.377351426 | 5.409903036 | 8.85E-04 |
| Msln | 8968.235968 | 5.393148926 | 5.76E-20 |
| Wdfy4 | 18.09422915 | 5.370992762 | 1.07E-04 |
| Gal3st2 | 33.39687285 | 5.360002332 | 3.89E-04 |
| Acta1 | 8.400779174 | 5.334985791 | 7.50E-06 |
| Epn3 | 33.31382918 | 5.331638053 | 1.24E-07 |
| Myom2 | 1179.530624 | 5.331141397 | 1.35E-45 |
| Dnaic1 | 8.46165829 | 5.284705495 | 2.19E-03 |
| Pdzd2 | 406.3513246 | 5.271766324 | 1.52E-16 |
| Dppa2 | 32.68223449 | 5.262799093 | 5.07E-17 |
| Got1l1 | 8.440417381 | 5.25805877 | 8.46E-04 |
| Vnn1 | 94.96773462 | 5.248231036 | 3.07E-09 |
| Gm11351 | 79.42538293 | 5.209361675 | 7.56E-37 |

| | | | |
|---|---|---|---|
| Cyp2ab1 | 5.554438667 | 5.200957913 | 6.68E-04 |
| Olfr456 | 23.66145086 | 5.169299159 | 2.34E-11 |
| Lama5 | 488.2888004 | 5.153877104 | 3.13E-20 |
| Foxa3 | 6.492805962 | 5.150195472 | 5.98E-04 |
| Slc2a3 | 86.49054597 | 5.134382383 | 1.37E-14 |
| Psca | 64.92123435 | 5.01464186 | 4.11E-04 |
| Fmnl1 | 6.076091521 | 5.014511616 | 4.57E-03 |
| Adgrl4 | 5.885316791 | 5.012945623 | 6.37E-04 |
| Sgce | 5.719415256 | 4.99273759 | 1.13E-03 |
| Misp | 6.782132558 | 4.981298368 | 5.57E-04 |
| Gm25043 | 5.794616842 | 4.965119135 | 3.03E-03 |
| 2900026A02Rik | 18.97069 | 4.952951628 | 3.67E-11 |
| Klra4 | 79.2878235 | 4.952739564 | 3.61E-14 |
| Inhba | 546.6979894 | 4.945904052 | 1.97E-18 |
| Tnnt3 | 106.6121583 | 4.940814654 | 4.92E-11 |
| Mmp16 | 66.21102913 | 4.938050947 | 1.12E-35 |
| Siglecg | 4363.02185 | 4.937440951 | 1.70E-91 |
| Slc30a2 | 5.640362476 | 4.925930892 | 8.43E-04 |
| Car6 | 110.8106253 | 4.92143544 | 2.38E-14 |
| Pmaip1 | 85.15518226 | 4.918203022 | 1.61E-33 |
| Bcam | 6.821030757 | 4.915467163 | 4.00E-04 |
| Gm10505 | 10.24804569 | 4.905846439 | 1.90E-04 |
| Gm8430 | 288.0183719 | 4.899318452 | 2.59E-02 |
| Mef2b | 78.07998856 | 4.897915568 | 7.42E-03 |
| Rnd1 | 586.3380609 | 4.896860053 | 2.64E-54 |
| Fmo3 | 67.82999797 | 4.886241281 | 1.46E-15 |
| Tll1 | 41.96951859 | 4.884693594 | 4.38E-24 |
| Tnfsf18 | 94.04928164 | 4.865612608 | 3.18E-15 |
| Folr1 | 25.64995036 | 4.861205776 | 4.37E-13 |
| Tmem246 | 6.736769451 | 4.861159403 | 4.53E-03 |
| Ninj2 | 25.84237259 | 4.835801248 | 6.12E-09 |
| BC051019 | 6.81730173 | 4.829833627 | 1.59E-03 |
| Scn3a | 16.94562687 | 4.82749899 | 1.05E-08 |
| Gm43154 | 35.32626595 | 4.804143533 | 1.19E-03 |
| Shank2 | 22.58620902 | 4.78451872 | 8.41E-06 |
| Hist1h4h | 20.58006819 | 4.773387724 | 1.41E-09 |
| Adam22 | 234.5747723 | 4.765821724 | 1.14E-09 |
| Slc17a7 | 5.760315762 | 4.762638379 | 9.44E-04 |
| Wnt11 | 29.78098512 | 4.752258733 | 1.43E-11 |
| Klk14 | 5.470717404 | 4.733853693 | 4.68E-04 |
| Cd74 | 18.46669239 | 4.728924848 | 4.54E-10 |
| Klk8 | 65.88130734 | 4.718445718 | 1.67E-16 |
| Sh3bgr | 6.330771203 | 4.695228742 | 6.75E-04 |
| Fyb2 | 205.4623369 | 4.690017997 | 7.48E-19 |
| Afap1l2 | 7.391967593 | 4.675597948 | 4.22E-04 |
| Lims2 | 179.0157548 | 4.662084763 | 1.62E-31 |
| Syk | 89.27270301 | 4.661282757 | 6.14E-12 |

| | | | |
|---|---|---|---|
| Sorl1 | 6.29132497 | 4.64776196 | 3.61E-03 |
| Prl3d2 | 7.108386125 | 4.640962373 | 1.47E-03 |
| Tslp | 20.98326568 | 4.639861666 | 2.00E-13 |
| Fras1 | 50.54206584 | 4.624454153 | 2.29E-03 |
| Hp | 45.64326259 | 4.612649046 | 1.26E-16 |
| Hist1h4i | 68.69874854 | 4.60917075 | 1.31E-18 |
| Aldh1a1 | 6563.728029 | 4.587863065 | 3.09E-36 |
| Ttn | 108.5364841 | 4.586746719 | 2.00E-04 |
| St3gal6 | 28.19900837 | 4.582623556 | 1.05E-10 |
| C530050E15Rik | 28.3053703 | 4.570910598 | 1.34E-12 |
| Btn1a1 | 43.15252764 | 4.568271364 | 4.29E-11 |
| Gm10256 | 43.00942646 | 4.555569569 | 1.96E-11 |
| Pdgfb | 7.20281354 | 4.553531631 | 1.17E-05 |
| Ripor2 | 382.9148722 | 4.553109181 | 9.38E-38 |
| Ptgs2 | 3952.831909 | 4.522331128 | 2.60E-23 |
| Emcn | 5.59271389 | 4.519456478 | 1.66E-03 |
| Crybg2 | 69.94425025 | 4.498337147 | 3.55E-37 |
| Fgf21 | 49.48268511 | 4.495053941 | 3.87E-07 |
| Zfp750 | 9.306518556 | 4.48716079 | 4.85E-04 |
| Nkpd1 | 5.186491559 | 4.482081875 | 7.15E-04 |
| Vmn1r-ps128 | 315.8095013 | 4.481316988 | 3.07E-33 |
| Best3 | 5.268568397 | 4.476817516 | 1.39E-03 |
| C1qtnf3 | 20.4941898 | 4.476235363 | 1.70E-09 |
| Clca3b | 16.002642 | 4.473144163 | 2.54E-07 |
| Fap | 55.22080039 | 4.45216677 | 1.06E-13 |
| Ifitm10 | 12.42299472 | 4.443365849 | 2.13E-05 |
| Krt14 | 42.84356732 | 4.442833305 | 4.36E-07 |
| Nes | 211.7096441 | 4.434299131 | 5.13E-17 |
| Pkhd1l1 | 6.114660324 | 4.432356285 | 1.89E-03 |
| Ccdc81 | 6.318104581 | 4.431683172 | 2.31E-03 |
| Pkp1 | 13.7724583 | 4.421306123 | 3.52E-10 |
| Tmem71 | 6.057387095 | 4.400321247 | 1.19E-03 |
| Spo11 | 29.75667518 | 4.360927398 | 4.14E-03 |
| Ip6k3 | 40.3684319 | 4.360050134 | 1.21E-18 |
| A730049H05Rik | 20.80637073 | 4.333945267 | 4.89E-05 |
| Gata3 | 165.7026761 | 4.300127431 | 1.41E-32 |
| Cmklr1 | 11.05913899 | 4.298453833 | 1.96E-05 |
| Gpr39 | 133.0862302 | 4.293171667 | 5.83E-28 |
| 4930590J08Rik | 8.309802627 | 4.282526309 | 3.74E-05 |
| Ccl6 | 20.15019644 | 4.264980915 | 8.58E-07 |
| Pkp2 | 365.3274075 | 4.245571422 | 5.80E-58 |
| Islr2 | 6.801156223 | 4.245385843 | 1.74E-04 |
| Chst1 | 44.51961143 | 4.243827852 | 2.41E-07 |
| Fst | 208.7049669 | 4.233539864 | 3.75E-23 |
| Sema3d | 11.73243349 | 4.231593248 | 7.65E-08 |
| Gm53 | 5.854823975 | 4.22868895 | 1.15E-02 |
| Atp2a3 | 10.67440491 | 4.217026752 | 5.94E-05 |

| | | | |
|---|---|---|---|
| Slc39a4 | 5.328099942 | 4.208207159 | 5.67E-04 |
| Svop | 24.57369793 | 4.194271243 | 3.67E-10 |
| Gm6088 | 11.0474142 | 4.18877605 | 8.75E-03 |
| Erbb3 | 69.46120585 | 4.186127885 | 2.41E-11 |
| Gm13713 | 5.072477726 | 4.186012179 | 2.47E-03 |
| Gm27406 | 15.93084718 | 4.151776619 | 2.16E-07 |
| Sdcbp2 | 61.62660346 | 4.150378942 | 1.22E-19 |
| Gm20521 | 108.1932135 | 4.147299845 | 2.59E-18 |
| Stbd1 | 242.832905 | 4.131346964 | 1.36E-60 |
| NA | 87.8527127 | 4.118683199 | 1.94E-20 |
| Prelid3a | 26.92713764 | 4.110266591 | 6.46E-13 |
| 1110017D15Rik | 13.74255596 | 4.10816605 | 1.62E-06 |
| Prss27 | 17.7205746 | 4.104722205 | 1.76E-07 |
| Itgb7 | 197.2536301 | 4.100098624 | 1.31E-27 |
| Serpina1a | 229.6910667 | 4.088794809 | 6.44E-28 |
| Mamld1 | 9.313569485 | 4.083625674 | 1.67E-04 |
| Cdh5 | 11.00256545 | 4.078715234 | 8.33E-03 |
| Sh3tc2 | 46.01764535 | 4.076878212 | 5.82E-22 |
| Gm13284 | 12.99212276 | 4.065770175 | 9.06E-03 |
| Fendrr | 16.4166356 | 4.061630154 | 4.94E-11 |
| Gm10230 | 10.1897906 | 4.06056713 | 3.48E-04 |
| Npr3 | 1290.290208 | 4.055909016 | 1.05E-23 |
| Klra5 | 12.83281442 | 4.04007171 | 1.88E-05 |
| Cyp24a1 | 5.817295899 | 4.039782231 | 2.41E-02 |
| Tgtp1 | 20.61468666 | 4.020427886 | 1.57E-06 |
| 1700013F07Rik | 7.109137326 | 4.013656291 | 1.14E-04 |
| Htatip2 | 7.892515834 | 4.009627515 | 5.51E-05 |
| Atp2b4 | 128.2057106 | 3.994649564 | 1.65E-20 |
| 1110002E22Rik | 5.899518649 | 3.986983589 | 1.87E-03 |
| Hspb7 | 111.5849735 | 3.979243606 | 2.82E-17 |
| Mab21l3 | 6.546200284 | 3.972351784 | 2.33E-04 |
| Gm13415 | 175.3337851 | 3.965771489 | 1.02E-09 |
| Hbb-bh1 | 14.94205789 | 3.963091049 | 3.51E-03 |
| Vwce | 9.513615881 | 3.948712444 | 1.02E-04 |
| Serpina1c | 24.40367768 | 3.943156393 | 1.99E-03 |
| Fcgbp | 17.64993208 | 3.930239797 | 5.45E-06 |
| Fgd5 | 6.720168263 | 3.93006176 | 2.93E-03 |
| Itgam | 17.41154262 | 3.919379207 | 3.36E-05 |
| Ccn2 | 7484.582414 | 3.896862753 | 7.00E-17 |
| Pla2g5 | 14.77056158 | 3.88961121 | 2.84E-04 |
| Itgb4 | 228.6910735 | 3.885250307 | 6.97E-08 |
| Slc1a7 | 20.64700538 | 3.865205354 | 2.49E-11 |
| Lamc3 | 81.73661022 | 3.864366904 | 2.73E-12 |
| Rassf6 | 30.46412628 | 3.864085135 | 6.04E-07 |
| 2310002F09Rik | 28.54212603 | 3.862416534 | 1.06E-08 |
| Serpinb1b | 80.38277849 | 3.851453291 | 3.91E-22 |
| Gm16702 | 278.1798623 | 3.834786877 | 4.66E-38 |

| | | | |
|---|---|---|---|
| Tmem182 | 44.63172361 | 3.825207811 | 4.85E-09 |
| Tuba8 | 7.274931265 | 3.808029754 | 4.19E-04 |
| Gprc5a | 2003.573682 | 3.807250992 | 2.51E-41 |
| Klra1 | 6.79691526 | 3.802636345 | 1.44E-02 |
| Arhgap45 | 18.56644359 | 3.80170149 | 2.06E-09 |
| Cdc42bpg | 72.4716449 | 3.800975122 | 6.65E-09 |
| Tnn | 22.38362316 | 3.797446328 | 8.02E-08 |
| Gm19345 | 11.4622406 | 3.788870391 | 2.10E-05 |
| Gjb5 | 5.452997903 | 3.780839679 | 1.21E-02 |
| Gm17090 | 14.75252409 | 3.76534529 | 1.14E-06 |
| Cth | 393.5185974 | 3.764674454 | 9.86E-09 |
| Slc2a5 | 32.51930166 | 3.754573912 | 1.12E-04 |
| Txlnb | 29.58797101 | 3.741691843 | 3.22E-11 |
| Foxn1 | 11.80917538 | 3.740186176 | 3.79E-05 |
| Tm4sf4 | 8.031164054 | 3.735176716 | 1.57E-04 |
| Slc17a9 | 18.00504775 | 3.730233702 | 2.71E-06 |
| Soat2 | 50.10265489 | 3.7215514 | 3.93E-19 |
| Pim1 | 2348.868326 | 3.718704511 | 5.25E-29 |
| Hoxb3os | 9.42460263 | 3.710382725 | 1.39E-04 |
| Rtn4rl2 | 154.9682333 | 3.709093699 | 7.72E-20 |
| Ptprv | 67.14689553 | 3.680417203 | 4.78E-08 |
| Brsk2 | 7.690449615 | 3.67750264 | 2.97E-05 |
| Card14 | 94.21879325 | 3.674177906 | 7.26E-03 |
| Car14 | 32.99689159 | 3.67128172 | 6.35E-09 |
| Ppp1r9a | 32.14076349 | 3.668668065 | 2.78E-03 |
| Kank4os | 15.91605683 | 3.667680359 | 1.60E-08 |
| Hoxb3 | 9.193710774 | 3.665866623 | 3.05E-05 |
| Hacd4 | 175.9725215 | 3.653708286 | 1.37E-30 |
| Ano3 | 117.0441969 | 3.650446599 | 4.65E-39 |
| Sult4a1 | 8.298499443 | 3.649426398 | 3.89E-02 |
| Tmem144 | 24.43046245 | 3.647370275 | 2.09E-10 |
| Cavin2 | 13533.10625 | 3.643249982 | 2.76E-28 |
| Mycl | 20.21135286 | 3.638913831 | 1.98E-09 |
| Efcab8 | 17.7494641 | 3.637561101 | 1.40E-08 |
| Eda2r | 1123.269013 | 3.634506531 | 2.28E-50 |
| Bach2os | 7.936230742 | 3.630469212 | 6.24E-03 |
| Gm6133 | 18.62084645 | 3.618962228 | 8.83E-08 |
| S100a7a | 436.7014925 | 3.602236284 | 3.56E-38 |
| Gm20400 | 779.139461 | 3.583517951 | 4.29E-21 |
| Dusp18 | 16.55368329 | 3.581712896 | 1.20E-06 |
| Mmp15 | 11.32755157 | 3.580090008 | 1.81E-05 |
| Cd68 | 539.0889678 | 3.567709068 | 1.56E-22 |
| Pde1a | 91.39232492 | 3.560683012 | 2.10E-19 |
| Trap1a | 12.33813831 | 3.559490329 | 2.59E-06 |
| Nfam1 | 6.77097658 | 3.555155669 | 9.48E-05 |
| Rasl10a | 6.669734342 | 3.552382893 | 1.25E-04 |
| Csrnp1 | 607.3843703 | 3.550608021 | 7.22E-28 |

| | | | |
|---|---|---|---|
| Piwil2 | 15.37645003 | 3.547766945 | 4.81E-06 |
| Serpine1 | 8069.154667 | 3.542682851 | 5.95E-26 |
| Fam71f2 | 47.87919002 | 3.542092554 | 1.10E-13 |
| Vmn1r40 | 12.07000502 | 3.537561512 | 3.64E-05 |
| Drc1 | 7.205444409 | 3.536040219 | 1.37E-03 |
| Crx | 15.38764971 | 3.532297531 | 8.41E-06 |
| Adra1d | 14.40500804 | 3.516108275 | 1.74E-04 |
| Cyp4f14 | 11.05060069 | 3.510713698 | 2.67E-02 |
| BC051077 | 62.17491931 | 3.494555977 | 3.87E-16 |
| Gm14341 | 16.93756775 | 3.47913078 | 1.75E-04 |
| Gm15287 | 92.60236988 | 3.469060955 | 8.85E-24 |
| Inka2 | 122.816321 | 3.45461342 | 1.12E-16 |
| Gm12840 | 2928.5719 | 3.452011267 | 2.15E-11 |
| Cd80 | 43.14414784 | 3.450834692 | 4.02E-11 |
| Rbmy | 70.47372177 | 3.449542236 | 7.89E-08 |
| Pcdh9 | 103.422577 | 3.445393211 | 1.40E-18 |
| 9530053A07Rik | 64.37286542 | 3.443925988 | 3.20E-08 |
| Cdkn1a | 10457.59614 | 3.438167913 | 1.93E-41 |
| Dcdc2a | 6.01120644 | 3.437174445 | 2.34E-03 |
| Fam71f1 | 49.35392991 | 3.434835596 | 7.78E-14 |
| Ecscr | 21.55153097 | 3.430210249 | 2.99E-02 |
| Ak7 | 5.835582438 | 3.429939774 | 2.56E-03 |
| Tmem184a | 18.15966957 | 3.426434049 | 1.31E-04 |
| Gadd45b | 5683.916854 | 3.423902607 | 1.03E-34 |
| Parvg | 9.60610989 | 3.410807476 | 1.39E-04 |
| Tpd52l1 | 5.150976508 | 3.406105563 | 3.81E-03 |
| Hes1 | 186.8574295 | 3.401638787 | 4.02E-11 |
| Itih4 | 19.70594933 | 3.392298032 | 8.37E-07 |
| Ces2g | 117.9433417 | 3.390685081 | 7.78E-18 |
| Gadd45g | 1363.863302 | 3.375362439 | 2.85E-38 |
| Ppp1r3a | 78.06682045 | 3.368228092 | 1.32E-06 |
| Hist1h1c | 1898.756039 | 3.364674695 | 1.31E-22 |
| Shisa6 | 8.904196846 | 3.364152508 | 1.28E-03 |
| Plac8 | 10.13342526 | 3.35923191 | 7.18E-06 |
| Acox2 | 9.843345696 | 3.353265172 | 6.40E-05 |
| Aldh1l1 | 17.6599573 | 3.346903645 | 4.33E-04 |
| 4921536K21Rik | 33.37512953 | 3.335622054 | 1.43E-10 |
| Ankrd1 | 4164.102927 | 3.32985581 | 9.87E-25 |
| Efhd1 | 6.477393889 | 3.327067371 | 3.32E-03 |
| Stard8 | 6.074829507 | 3.319792124 | 4.87E-03 |
| Chn2 | 8.322440648 | 3.294221284 | 4.34E-04 |
| Tinagl1 | 7597.988131 | 3.289299301 | 4.26E-44 |
| Gm12505 | 700.5275066 | 3.281759413 | 5.98E-23 |
| Cpz | 10.48574976 | 3.274080025 | 8.18E-04 |
| Rspo4os | 11.50713663 | 3.270861738 | 1.02E-06 |
| Hist1h2bc | 898.5619503 | 3.26945695 | 7.53E-15 |
| Muc2 | 8.809115183 | 3.235011287 | 1.46E-05 |

| | | | |
|---|---|---|---|
| Gm13067 | 13.02922408 | 3.230280749 | 9.71E-06 |
| Mreg | 281.1053841 | 3.226684147 | 6.69E-18 |
| Eps8l2 | 14.97634174 | 3.203328801 | 1.53E-02 |
| Chac1 | 2414.321663 | 3.199044799 | 3.02E-41 |
| Mgll | 24.84962995 | 3.195771434 | 7.46E-06 |
| Slc14a2 | 5.198235582 | 3.195735121 | 1.26E-02 |
| NA | 1301.961584 | 3.194498537 | 1.25E-35 |
| Adgrg6 | 18.66629187 | 3.191249642 | 9.03E-07 |
| Ms4a4c | 8.55212296 | 3.189531202 | 5.45E-03 |
| Hist1h2bq | 18.51075948 | 3.184231756 | 4.97E-05 |
| Catsperd | 26.54257683 | 3.180474334 | 1.05E-09 |
| NA | 101.8655657 | 3.179295375 | 1.35E-27 |
| Myh11 | 10.27823204 | 3.174351305 | 3.79E-04 |
| Dyrk4 | 47.6636991 | 3.172001694 | 6.61E-14 |
| Tnnt2 | 43.75984317 | 3.166790615 | 4.45E-06 |
| Suox | 3135.576887 | 3.163547198 | 8.18E-64 |
| Stxbp2 | 15.11401419 | 3.162291016 | 8.83E-05 |
| Id4 | 162.2104674 | 3.152360736 | 1.65E-23 |
| Hoxb6 | 11.04812908 | 3.14475844 | 6.13E-04 |
| Eml2 | 184.9655233 | 3.129363745 | 2.45E-22 |
| AU018091 | 5.236497473 | 3.12555769 | 3.43E-02 |
| Gdf5 | 22.57180371 | 3.123957445 | 1.17E-05 |
| Ccdc106 | 72.36994274 | 3.104176276 | 5.42E-25 |
| Gpr68 | 23.11995535 | 3.102114547 | 2.99E-07 |
| Gm26579 | 5.928312409 | 3.101556026 | 2.81E-03 |
| Mcam | 18.62577123 | 3.098340533 | 4.91E-06 |
| Chrm1 | 15.25123971 | 3.08568758 | 6.33E-05 |
| Tlr1 | 42.23168809 | 3.085099528 | 1.51E-15 |
| Tceal5 | 16.32229525 | 3.077598436 | 3.08E-04 |
| Tubb4a | 66.11470353 | 3.070902381 | 3.34E-28 |
| Hrc | 7.301964216 | 3.059079459 | 2.74E-04 |
| Adrb3 | 11.41728903 | 3.05785301 | 5.73E-04 |
| Acvrl1 | 34.10597704 | 3.053935242 | 7.09E-08 |
| Crabp2 | 68.19681442 | 3.049960485 | 4.10E-17 |
| Slc24a3 | 7.064572383 | 3.048580629 | 4.26E-02 |
| Hist1h1e | 12.45311968 | 3.047576369 | 3.70E-05 |
| Camk2a | 97.06394336 | 3.044275991 | 7.29E-15 |
| Gm12185 | 28.38347216 | 3.042788375 | 3.88E-06 |
| Fgd6 | 331.6053799 | 3.042346114 | 2.82E-29 |
| Apol9b | 84.6420391 | 3.039810726 | 7.74E-14 |
| Prss23os | 1155.334691 | 3.035969867 | 2.03E-31 |
| Mrvi1 | 124.3727741 | 3.035565477 | 1.65E-12 |
| Taf7l | 8.416501097 | 3.034620926 | 3.67E-03 |
| Cyp2c55 | 890.3392128 | 3.032517273 | 5.27E-52 |
| Fbxw10 | 18.3927969 | 3.031211319 | 5.21E-04 |
| Nr1d1 | 3745.690596 | 3.028763481 | 2.05E-09 |
| Csf1r | 229.3071014 | 3.023782738 | 8.49E-14 |

| | | | |
|---|---|---|---|
| Il3ra | 21.87015892 | 3.021746151 | 3.09E-07 |
| Itga3 | 4169.255291 | 3.019137998 | 7.21E-31 |
| Il7 | 12.90409676 | 3.016978831 | 3.36E-05 |
| NA | 5.21045359 | 3.015192799 | 5.36E-03 |
| Ccdc113 | 31.53830894 | 3.002134923 | 1.70E-03 |
| Cntfr | 7.670953597 | 2.994935624 | 2.66E-03 |
| Gm26708 | 7.912831015 | 2.994243105 | 6.34E-04 |
| Apol7d | 28.86154236 | 2.991413146 | 1.77E-09 |
| Enkur | 17.5217497 | 2.99113341 | 1.77E-03 |
| Lrrc24 | 8.09439088 | 2.983562734 | 1.21E-02 |
| Dusp8 | 245.7185379 | 2.979983516 | 1.17E-21 |
| Gm38393 | 571.0226571 | 2.977726582 | 4.24E-23 |
| Mslnl | 8.191156947 | 2.97497327 | 2.47E-04 |
| Olfr1318 | 12.8937525 | 2.97465713 | 3.17E-04 |
| Zbtb7c | 40.91004377 | 2.966961597 | 4.62E-12 |
| Gm10109 | 6.409401559 | 2.960064843 | 1.74E-02 |
| Has1 | 15.99957924 | 2.957851211 | 1.09E-03 |
| Nlrp4f | 12.24097274 | 2.953696611 | 2.78E-05 |
| Hist1h2ac | 24.74191821 | 2.945245944 | 9.88E-07 |
| Kcnu1 | 19.04652871 | 2.944922749 | 1.12E-04 |
| Rap1gap2 | 970.5392269 | 2.944619055 | 5.94E-30 |
| Ptpru | 9.867573582 | 2.940236555 | 1.91E-03 |
| Hist1h2bg | 15.84317161 | 2.940043312 | 3.59E-02 |
| Casp14 | 18.04409302 | 2.937938483 | 3.74E-04 |
| Myrfl | 5.472787012 | 2.93790169 | 9.06E-03 |
| Rhpn2 | 103.2238498 | 2.937784342 | 2.40E-16 |
| Slc43a3 | 2964.656793 | 2.932487895 | 3.52E-32 |
| Tmem132b | 17.53167815 | 2.92806767 | 1.73E-04 |
| A330033J07Rik | 5.576475669 | 2.927361158 | 1.01E-02 |
| Pgf | 5.848257713 | 2.926306681 | 3.14E-03 |
| NA | 7.9647822 | 2.924537 | 6.15E-04 |
| Pdlim1 | 2636.81262 | 2.91998466 | 1.36E-21 |
| Aif1l | 2280.404424 | 2.9191765 | 1.01E-44 |
| Stox2 | 10.87126108 | 2.915141672 | 1.15E-04 |
| Lman1l | 19.17703041 | 2.910790355 | 1.96E-04 |
| Cxcl1 | 119.9745271 | 2.894275775 | 7.11E-12 |
| Prkaa2 | 23.60887112 | 2.891444198 | 3.81E-07 |
| Ppp1r32 | 13.45526339 | 2.887430922 | 1.13E-04 |
| Tmem37 | 109.5363239 | 2.886185177 | 5.12E-23 |
| Trim7 | 396.6823301 | 2.876770282 | 1.93E-59 |
| Insl6 | 38.19475117 | 2.875963907 | 2.03E-15 |
| Trib3 | 1385.443885 | 2.874049624 | 4.65E-17 |
| Usp17lc | 5.351707464 | 2.873656843 | 3.19E-02 |
| Pla1a | 89.47502873 | 2.870871382 | 2.00E-10 |
| H19 | 1822.874962 | 2.866426105 | 2.78E-04 |
| Disp2 | 132.6711548 | 2.861252262 | 1.32E-26 |
| Grin3a | 5.74326454 | 2.857882158 | 3.97E-03 |

| | | | |
|---|---|---|---|
| Sulf1 | 1797.810171 | 2.855558382 | 6.00E-22 |
| Apob | 19.03152808 | 2.854807002 | 2.47E-03 |
| Plaur | 1149.588067 | 2.844884795 | 2.73E-26 |
| Llgl2 | 523.0277484 | 2.84456997 | 1.16E-31 |
| Txk | 7.539374443 | 2.842635583 | 7.22E-03 |
| Mylpf | 41.0963031 | 2.840467248 | 1.61E-13 |
| A230072E10Rik | 9.529520467 | 2.836438184 | 5.42E-03 |
| Gm23849 | 8.016410947 | 2.832904365 | 6.72E-03 |
| Gm6166 | 5392.037064 | 2.82780097 | 1.23E-49 |
| Nalcn | 35.68450605 | 2.823126298 | 1.26E-07 |
| Brinp1 | 311.655202 | 2.819738619 | 1.10E-14 |
| Cystm1 | 169.2870408 | 2.816608935 | 4.01E-14 |
| Gm3376 | 148.220178 | 2.816144538 | 9.51E-19 |
| Far2os1 | 51.84053146 | 2.810192393 | 2.66E-05 |
| Psg17 | 21.47017957 | 2.810155886 | 9.19E-04 |
| Elavl3 | 5.174486242 | 2.809876048 | 9.42E-03 |
| Calcrl | 135.2839369 | 2.796939609 | 6.07E-08 |
| Psrc1 | 2282.570702 | 2.792776073 | 5.32E-33 |
| Perp | 26.99007095 | 2.787326459 | 9.82E-06 |
| Gm12037 | 11.05132918 | 2.782573684 | 4.95E-05 |
| Adgrd1 | 8.97434972 | 2.781086843 | 1.88E-02 |
| Lag3 | 35.71725018 | 2.779189432 | 4.33E-12 |
| Acer2 | 5.820463227 | 2.778729683 | 9.56E-03 |
| Gm11525 | 36.80127275 | 2.778652857 | 5.31E-11 |
| Mapk4 | 10.79076515 | 2.777325178 | 1.73E-03 |
| NA | 324.7999154 | 2.777177539 | 2.64E-15 |
| Itga2 | 5.07357676 | 2.774272592 | 2.54E-02 |
| 1110035H17Rik | 5.169770735 | 2.761437015 | 1.09E-02 |
| Cited2 | 5203.839702 | 2.760309241 | 3.46E-90 |
| Cdsn | 1502.659716 | 2.75879411 | 6.51E-32 |
| Cgnl1 | 1941.398104 | 2.756335666 | 1.37E-14 |
| Acer1 | 15.46558608 | 2.745947556 | 2.28E-05 |
| Adamts9 | 285.3258477 | 2.743887974 | 8.00E-16 |
| Fabp5 | 6643.323886 | 2.741083281 | 2.50E-53 |
| Ephx1 | 9309.25785 | 2.739383574 | 1.53E-41 |
| Gm13710 | 14.91984042 | 2.736818679 | 1.55E-02 |
| Btg3 | 59.40803996 | 2.736655552 | 2.79E-02 |
| Trpm5 | 54.98351022 | 2.732300956 | 1.81E-13 |
| Rasgef1a | 8.735346562 | 2.728495419 | 4.30E-04 |
| Fabp4 | 102.7972834 | 2.726767238 | 1.63E-14 |
| Rasgrp3 | 117.9121061 | 2.726234358 | 8.70E-24 |
| Klf5 | 911.9925518 | 2.723142682 | 7.52E-48 |
| Mfap3l | 581.8453732 | 2.714102978 | 1.18E-26 |
| Nxf3 | 15.38053015 | 2.713180598 | 4.66E-04 |
| Cspg5 | 5.282491474 | 2.712968855 | 2.39E-02 |
| 3830403N18Rik | 23.80216124 | 2.712480654 | 4.97E-09 |
| Rpl29 | 569.7586007 | 2.70663664 | 1.34E-24 |

| | | | |
|---|---|---|---|
| 4933417D19Rik | 27.85225599 | 2.706169965 | 1.55E-08 |
| Wdr66 | 88.12706011 | 2.700865608 | 5.54E-20 |
| Serpinb6c | 18.60767378 | 2.698495297 | 3.20E-02 |
| D630029K05Rik | 40.79021596 | 2.694932741 | 2.67E-11 |
| Angptl6 | 251.3102899 | 2.689213002 | 2.95E-27 |
| Atp8b3 | 5.397180196 | 2.685926091 | 2.87E-02 |
| Notch3 | 14.41931232 | 2.681034883 | 8.78E-06 |
| Gria3 | 1179.231791 | 2.674189651 | 8.95E-34 |
| Pax8 | 75.11368423 | 2.67397825 | 3.29E-03 |
| Arl9 | 5.951293518 | 2.670100393 | 1.57E-02 |
| Styk1 | 452.3674977 | 2.66918683 | 4.30E-19 |
| Gm27224 | 10.41317876 | 2.665653878 | 1.20E-02 |
| Ifit1bl1 | 78.34996578 | 2.665215158 | 2.49E-11 |
| Tgm2 | 2829.880322 | 2.662121703 | 1.67E-48 |
| Unc5c | 5.83703364 | 2.651074901 | 8.29E-03 |
| Atf3 | 486.7080334 | 2.650574565 | 3.00E-35 |
| Snhg15 | 979.4343226 | 2.647367466 | 1.78E-36 |
| Ccn1 | 16942.56933 | 2.64053098 | 2.57E-09 |
| Plch2 | 6.803676316 | 2.640496155 | 1.49E-02 |
| Adcyap1r1 | 11.14523057 | 2.639550258 | 3.48E-03 |
| Fas | 523.473713 | 2.631970555 | 1.51E-31 |
| Cfap70 | 23.3829331 | 2.62863794 | 3.25E-03 |
| Pard6b | 681.0243997 | 2.626711339 | 2.39E-18 |
| Ddias | 367.2417911 | 2.626543018 | 1.75E-11 |
| Esr2 | 5.598042516 | 2.624365918 | 2.52E-03 |
| Cnksr1 | 26.39635045 | 2.62051622 | 1.18E-03 |
| Hist1h2ae | 5.952941222 | 2.617573503 | 8.53E-03 |
| Pygm | 20.09698139 | 2.615793886 | 6.46E-04 |
| Gm5570 | 6.543427362 | 2.613282988 | 1.01E-02 |
| Clca4a | 15.01272408 | 2.608236631 | 9.78E-04 |
| Dennd2d | 8.919399004 | 2.606521243 | 1.72E-02 |
| Sema6b | 29.71095728 | 2.602949837 | 3.90E-07 |
| Tnni2 | 25.52205906 | 2.601150072 | 1.68E-04 |
| Trim72 | 12.45876701 | 2.599346259 | 1.38E-03 |
| Hes7 | 30.3534822 | 2.587181153 | 1.35E-10 |
| Vmn1r43 | 19.18440843 | 2.583849442 | 4.54E-03 |
| Arl4d | 342.751126 | 2.581875884 | 5.62E-15 |
| Klf15 | 6.447471502 | 2.579923901 | 6.67E-03 |
| Dyrk3 | 209.7965907 | 2.579656573 | 5.39E-35 |
| Dll4 | 13.871069 | 2.578589031 | 8.50E-04 |
| Jag1 | 1491.58118 | 2.576796003 | 1.57E-27 |
| Ncf2 | 13.41769721 | 2.575489283 | 5.13E-03 |
| Ripk3 | 33.31369269 | 2.575348925 | 6.18E-06 |
| Pla2g4b | 67.61780263 | 2.574847987 | 7.78E-07 |
| Gm13988 | 25.63523994 | 2.573949321 | 1.06E-03 |
| Gm13981 | 7.337667877 | 2.57390833 | 1.89E-03 |
| Dlc1 | 2597.470192 | 2.571130986 | 1.13E-58 |

| | | | |
|---|---|---|---|
| Acmsd | 84.38312468 | 2.566865289 | 2.33E-11 |
| Tnfaip3 | 769.7438906 | 2.554543849 | 1.19E-24 |
| NA | 10.87827974 | 2.551555041 | 3.68E-03 |
| Wnt9a | 916.1471833 | 2.550902763 | 2.01E-28 |
| Hist1h2bk | 54.97336704 | 2.547829637 | 4.60E-05 |
| A3galt2 | 6.703661174 | 2.545905595 | 1.07E-02 |
| Foxf1 | 338.939874 | 2.545566872 | 1.28E-61 |
| Tigit | 30.28802111 | 2.543374923 | 3.72E-08 |
| Hist1h2bl | 33.18392678 | 2.537297694 | 1.66E-04 |
| Fam107a | 8.437155477 | 2.536703628 | 3.96E-03 |
| Fgf18 | 10.05836057 | 2.536370314 | 2.18E-03 |
| Gm16166 | 9.035890847 | 2.534531893 | 5.22E-03 |
| Mylk2 | 10.00166894 | 2.534273103 | 4.19E-04 |
| A230050P20Rik | 6.105782914 | 2.528433664 | 5.03E-03 |
| Itga7 | 815.1436197 | 2.524751297 | 3.60E-17 |
| Colec11 | 15.15317968 | 2.521414289 | 1.74E-04 |
| Arhgef28 | 758.8625099 | 2.520060994 | 7.72E-46 |
| Plekha6 | 15.60175078 | 2.519521611 | 9.79E-03 |
| Ifi44 | 10.76225655 | 2.514673286 | 9.06E-04 |
| Itga1 | 65.1732428 | 2.506211664 | 2.10E-11 |
| Lama2 | 873.8408923 | 2.505712445 | 1.95E-17 |
| Pram1 | 33.24727736 | 2.503190362 | 1.88E-04 |
| NA | 141.937738 | 2.50244752 | 3.18E-13 |
| Tg | 42.06323385 | 2.500976662 | 1.29E-03 |
| Vmn1r45 | 6.489789663 | 2.49489508 | 1.72E-02 |
| Slc6a15 | 33.49232237 | 2.491933691 | 7.80E-06 |
| Hist1h2be | 35.43583052 | 2.491455331 | 5.40E-08 |
| Lta | 33.84052544 | 2.491421938 | 4.22E-09 |
| Sema3e | 12.67231211 | 2.487660429 | 4.08E-03 |
| Ttpa | 5.406595632 | 2.486820251 | 4.17E-02 |
| Foxs1 | 115.5421906 | 2.485072043 | 9.76E-14 |
| Fxyd5 | 177.0089438 | 2.483730927 | 8.50E-11 |
| Fat2 | 8.056016086 | 2.481665531 | 3.16E-02 |
| Snhg6 | 1874.929114 | 2.480464776 | 3.77E-29 |
| Ablim1 | 3555.853548 | 2.476035925 | 2.85E-33 |
| Plb1 | 53.16380267 | 2.472928849 | 9.47E-08 |
| 1700015F17Rik | 5.549174145 | 2.472329543 | 9.81E-03 |
| Cfap43 | 26.34884187 | 2.471871573 | 5.09E-05 |
| Nkain3 | 6.736363694 | 2.469807259 | 7.45E-03 |
| E330013P04Rik | 7.302192827 | 2.464425377 | 8.08E-03 |
| P2ry6 | 8.544628265 | 2.462761399 | 1.95E-02 |
| Mirt1 | 5.892982918 | 2.461687488 | 3.06E-02 |
| P2rx3 | 1054.952462 | 2.460860429 | 1.14E-06 |
| Tjp2 | 3236.987169 | 2.457696441 | 1.16E-31 |
| NA | 125.6560357 | 2.457318121 | 7.66E-09 |
| Tent5c | 31.88437475 | 2.454516443 | 1.12E-06 |
| F11r | 12.52591921 | 2.450363981 | 1.52E-03 |

| | | | |
|---|---|---|---|
| Cnn1 | 11.15129988 | 2.444960535 | 2.67E-04 |
| Syt17 | 46.84650688 | 2.442840913 | 8.38E-07 |
| Klhl30 | 170.2009204 | 2.442782863 | 1.78E-07 |
| Gja1 | 1405.954553 | 2.442377244 | 7.86E-13 |
| Dgat2 | 90.86863884 | 2.437653295 | 1.69E-16 |
| Spon1 | 17.90306055 | 2.435065041 | 1.28E-04 |
| Gm6311 | 19.58354765 | 2.434106665 | 4.24E-03 |
| Cd209c | 391.2517191 | 2.432558626 | 6.72E-10 |
| Ppm1h | 131.7107471 | 2.429710284 | 1.28E-08 |
| Hyal3 | 30.56485389 | 2.429211191 | 2.18E-06 |
| Amd-ps3 | 45.53365464 | 2.42235296 | 8.85E-03 |
| Snord55 | 6.834009265 | 2.420504565 | 5.64E-03 |
| Upp1 | 35.47148637 | 2.419827177 | 4.28E-05 |
| Prss23 | 12096.74038 | 2.418154783 | 3.80E-48 |
| Zbtb8b | 15.33196433 | 2.417633996 | 2.06E-03 |
| Spdef | 6.009277668 | 2.416553525 | 1.78E-02 |
| Skap1 | 6.501013336 | 2.415277195 | 9.93E-03 |
| Klra17 | 8.115897874 | 2.414340393 | 2.60E-02 |
| Cdhr2 | 8.715260545 | 2.411167862 | 3.76E-02 |
| Dusp14 | 1099.248265 | 2.410840257 | 1.41E-32 |
| Gm12454 | 22.74329518 | 2.409580392 | 5.04E-05 |
| Hist2h2aa2 | 243.0853257 | 2.407804649 | 1.21E-06 |
| Pappa2 | 2623.486677 | 2.407484323 | 2.62E-29 |
| Nlrp4e | 17.54639026 | 2.405511779 | 8.51E-03 |
| Ppfia4 | 8.420556161 | 2.39991984 | 4.25E-03 |
| Tmc7 | 16.65327218 | 2.398227157 | 5.12E-05 |
| Hbegf | 543.5379025 | 2.391452059 | 2.23E-22 |
| Cd37 | 12.38251855 | 2.389796539 | 3.26E-03 |
| Ikzf2 | 190.7222877 | 2.386506061 | 9.67E-10 |
| Lyrm9 | 234.5217743 | 2.386136252 | 2.02E-20 |
| Popdc3 | 16.72023235 | 2.385279442 | 2.36E-04 |
| D6Ertd527e | 17.77531328 | 2.383210123 | 2.87E-04 |
| Sptbn4 | 27.97214524 | 2.376119184 | 7.63E-04 |
| Hecw1 | 7.383878072 | 2.375156969 | 4.54E-03 |
| Pmp2 | 16.17721345 | 2.373940891 | 1.52E-03 |
| Hcrtr1 | 11.63728349 | 2.372238573 | 1.29E-04 |
| Aqp6 | 6.596852005 | 2.371980539 | 1.26E-02 |
| Flt3 | 12.61002332 | 2.369423253 | 1.44E-04 |
| Sbsn | 1060.826381 | 2.362323298 | 3.33E-52 |
| Klf4 | 2395.995965 | 2.361513557 | 8.09E-38 |
| Pim3 | 2043.372491 | 2.361411873 | 5.73E-24 |
| Rhbdf2 | 448.6446709 | 2.360938168 | 9.87E-42 |
| Arl5c | 11.25248985 | 2.360871487 | 1.38E-04 |
| Zfp941 | 7.943074269 | 2.355541396 | 2.51E-02 |
| D630023F18Rik | 6.989008852 | 2.35454285 | 2.48E-02 |
| Cyp2s1 | 8.772565514 | 2.352446898 | 2.18E-02 |
| 2200002D01Rik | 188.5911851 | 2.35101427 | 3.16E-21 |

| | | | |
|---|---|---|---|
| Rpl12 | 5438.515123 | 2.350271652 | 1.66E-26 |
| Alas2 | 5.704140781 | 2.349710444 | 2.84E-02 |
| Mst1 | 6.564989527 | 2.348595316 | 8.20E-03 |
| Rnf128 | 22.02821797 | 2.348192401 | 5.89E-04 |
| Rassf7 | 636.3971329 | 2.347463552 | 2.77E-18 |
| Proca1 | 10.97401158 | 2.340448247 | 1.01E-03 |
| Trac | 20.1751909 | 2.338755276 | 9.51E-06 |
| Ifitm1 | 15.2790241 | 2.337917802 | 1.00E-03 |
| Gm5106 | 7.079855874 | 2.336862884 | 1.61E-02 |
| Gdap1l1 | 51.61483492 | 2.33655455 | 9.24E-05 |
| Tnnc1 | 109.1518449 | 2.33547505 | 1.59E-19 |
| Gabre | 296.5455035 | 2.335256568 | 8.26E-22 |
| Gtse1 | 869.2872648 | 2.334822452 | 8.93E-11 |
| Bcar3 | 810.1377157 | 2.333978736 | 1.49E-30 |
| Obscn | 23.8335051 | 2.331398282 | 3.45E-03 |
| Gm26809 | 35780.51269 | 2.32939334 | 1.35E-13 |
| Steap1 | 1192.998083 | 2.328328741 | 7.26E-15 |
| Gm16061 | 14.26029884 | 2.320874607 | 5.42E-03 |
| Kcnj14 | 9.643672781 | 2.32038722 | 2.12E-03 |
| Btg2 | 1025.665619 | 2.318991637 | 4.16E-15 |
| 5430416N02Rik | 1000.533301 | 2.3149104 | 7.69E-87 |
| Igfbp4 | 605.6600899 | 2.313042878 | 5.85E-09 |
| Anxa3 | 7482.75502 | 2.312436474 | 4.83E-25 |
| Rdh9 | 51.98665338 | 2.312196373 | 2.68E-07 |
| Pde6g | 13.41208273 | 2.310139657 | 8.17E-05 |
| Prickle4 | 8.014106294 | 2.308712732 | 5.86E-03 |
| Gsdmc4 | 5.219483731 | 2.303622189 | 2.14E-02 |
| Frat2 | 305.7883249 | 2.301344335 | 3.43E-19 |
| Slc22a13b-ps | 8.826899059 | 2.298733574 | 4.95E-03 |
| Avpi1 | 1533.78618 | 2.297451975 | 1.13E-58 |
| Gdnf | 878.6590879 | 2.294575843 | 9.76E-21 |
| Eln | 24.79874943 | 2.291490353 | 6.37E-05 |
| Cldn15 | 6.758990275 | 2.29089286 | 1.56E-02 |
| Lenep | 70.28657422 | 2.281501506 | 1.31E-05 |
| Spns3 | 13.83643692 | 2.274421354 | 1.58E-02 |
| Epha2 | 1944.113419 | 2.274091663 | 3.44E-14 |
| Sema4a | 10.76380038 | 2.271553866 | 6.14E-03 |
| Atg16l2 | 1186.275372 | 2.270615274 | 1.20E-13 |
| Dusp1 | 2323.364801 | 2.261926566 | 1.01E-13 |
| Arl14epl | 50.54681463 | 2.261247891 | 1.48E-10 |
| Ddit4 | 1013.742835 | 2.261028492 | 1.26E-35 |
| Gm1141 | 15.33114807 | 2.259616896 | 2.45E-04 |
| Cxcl10 | 13.09469926 | 2.258214919 | 3.06E-03 |
| Npy1r | 34.61345112 | 2.254773771 | 2.58E-06 |
| Nectin4 | 8.839627048 | 2.25314365 | 1.21E-02 |
| Tspan12 | 74.6818032 | 2.240445104 | 5.31E-08 |
| Rab3c | 286.3804103 | 2.238268992 | 5.74E-07 |

| | | | |
|---|---|---|---|
| Hist2h2aa1 | 50.36297655 | 2.238185015 | 2.33E-02 |
| 9930111J21Rik2 | 470.8629972 | 2.237985597 | 2.46E-10 |
| Spock2 | 18.69978441 | 2.236585588 | 3.97E-06 |
| Ptpn22 | 7.501573766 | 2.235748867 | 4.36E-02 |
| Lmntd2 | 7.734375031 | 2.235560033 | 1.61E-02 |
| Il11 | 23.34212275 | 2.228694851 | 3.26E-07 |
| 4930486L24Rik | 435.5390485 | 2.227798371 | 2.36E-09 |
| Ccl5 | 19.03032931 | 2.22679886 | 4.23E-04 |
| 2410006H16Rik | 10631.39287 | 2.226224354 | 1.10E-15 |
| Heph | 11.43738456 | 2.220530291 | 1.92E-02 |
| Dbhos | 23.80023757 | 2.21606076 | 9.43E-06 |
| Mbp | 219.5097508 | 2.215665522 | 2.11E-15 |
| Aqp5 | 257.8230496 | 2.214407134 | 1.70E-12 |
| Emx2 | 282.5000379 | 2.21201752 | 6.91E-19 |
| Faim2 | 94.52067842 | 2.201844733 | 1.17E-14 |
| Cgn | 236.0254742 | 2.20179331 | 2.28E-09 |
| Grk5 | 918.5251472 | 2.200473864 | 4.74E-19 |
| Cep295nl | 32.72658667 | 2.198117655 | 4.97E-08 |
| Ptafr | 9.392685792 | 2.195572547 | 7.60E-04 |
| Zfp36 | 1397.592095 | 2.195470245 | 3.22E-11 |
| Zfp341 | 461.7226419 | 2.19421321 | 7.22E-24 |
| Poln | 47.95986064 | 2.19157127 | 1.09E-07 |
| Mfsd6 | 21.51319353 | 2.19006617 | 3.25E-03 |
| Flt1 | 1329.706088 | 2.187971808 | 4.00E-75 |
| Ifit1 | 122.4955534 | 2.18672124 | 7.04E-15 |
| Sla | 6.121986309 | 2.181767657 | 4.82E-02 |
| Tubb3 | 51.35408728 | 2.173693892 | 1.79E-06 |
| Optc | 22.31321699 | 2.173397282 | 6.71E-05 |
| Gm11520 | 9.983968603 | 2.170300223 | 1.94E-03 |
| Ptx3 | 4202.894405 | 2.169705493 | 3.07E-10 |
| Ttc25 | 33.34011969 | 2.16666483 | 4.03E-04 |
| Gm15640 | 32.6140749 | 2.161600114 | 2.58E-05 |
| Procr | 826.0507845 | 2.1582683 | 3.15E-24 |
| Syngr3 | 13.5009954 | 2.15625079 | 1.53E-02 |
| Thsd7b | 12.27170345 | 2.151492786 | 1.83E-02 |
| Sema3a | 2160.937361 | 2.146800691 | 8.55E-12 |
| Map2 | 61.11523575 | 2.144397105 | 7.34E-09 |
| AW047730 | 11.98434172 | 2.136215082 | 2.08E-02 |
| Ppp1r2-ps4 | 7.020669578 | 2.136150878 | 2.55E-02 |
| Lrp2 | 44.96142077 | 2.134731196 | 1.84E-05 |
| Noxred1 | 5.257158721 | 2.133312733 | 1.03E-02 |
| Cd96 | 6.045846529 | 2.129092896 | 3.09E-02 |
| Gipc2 | 16.09109545 | 2.126750919 | 6.11E-04 |
| Creb3l3 | 9.152823137 | 2.126409491 | 9.82E-03 |
| Tbx6 | 29.79739049 | 2.123171647 | 7.42E-06 |
| Wwc1 | 1254.514596 | 2.120382883 | 3.23E-18 |
| Bbc3 | 624.0895791 | 2.106890307 | 4.33E-26 |

| | | | |
|---|---|---|---|
| Icam1 | 7.776917133 | 2.101832915 | 2.48E-02 |
| Gm11642 | 21.13040859 | 2.101492984 | 9.40E-06 |
| Mroh6 | 11.76675397 | 2.100571091 | 4.07E-03 |
| Gm11476 | 21.68780768 | 2.100494361 | 2.13E-04 |
| Ckap2 | 2101.539709 | 2.095934289 | 5.22E-12 |
| Ppp1r15a | 3481.359866 | 2.093529702 | 3.91E-36 |
| Ovgp1 | 83.50523948 | 2.092571118 | 4.07E-06 |
| Abat | 7.958698331 | 2.090996293 | 3.04E-02 |
| Jag2 | 9.330326422 | 2.084031801 | 3.69E-03 |
| Gm14057 | 34.74908516 | 2.082394183 | 3.97E-07 |
| Lrrc75a | 924.7491795 | 2.080968329 | 9.51E-52 |
| Slc16a8 | 7.869228903 | 2.080302603 | 1.27E-02 |
| Mcub | 378.0966601 | 2.073849943 | 2.06E-23 |
| Epha1 | 75.86205553 | 2.072552065 | 1.26E-13 |
| Fhl1 | 107.2708123 | 2.066616238 | 3.86E-04 |
| Platr14 | 62.69361273 | 2.065769562 | 4.15E-09 |
| 4933406C10Rik | 13.8312681 | 2.064666292 | 1.08E-03 |
| Usp17le | 14.67590836 | 2.062166932 | 1.47E-02 |
| Gca | 78.20706386 | 2.058352742 | 3.58E-08 |
| Aaas | 2372.654229 | 2.052529949 | 1.06E-16 |
| Pstpip1 | 90.41301441 | 2.05210817 | 7.02E-09 |
| Gpat3 | 333.521046 | 2.051294014 | 2.08E-19 |
| Mpeg1 | 23.16351824 | 2.05064911 | 7.46E-04 |
| Ngf | 909.5389495 | 2.050372539 | 1.63E-20 |
| Gch1 | 62.90561554 | 2.048317432 | 2.57E-07 |
| Ddit3 | 6642.720834 | 2.046547141 | 8.28E-18 |
| Gm12693 | 66.34240934 | 2.044765321 | 1.66E-04 |
| Gm12643 | 5.091002663 | 2.044252134 | 1.95E-02 |
| Sap25 | 22.93362904 | 2.042030182 | 4.05E-03 |
| Pkd1l3 | 40.65792714 | 2.041354735 | 4.80E-09 |
| 2010300F17Rik | 13.13878252 | 2.040294541 | 5.79E-03 |
| Car9 | 41.71092485 | 2.03786192 | 6.38E-05 |
| Nyap1 | 908.6854763 | 2.037778346 | 2.24E-23 |
| Car5b | 116.6171212 | 2.032873995 | 5.88E-18 |
| Hist1h2al | 73.64496331 | 2.032791547 | 1.03E-06 |
| Rasgrp2 | 25.33324843 | 2.032470127 | 3.58E-02 |
| Ripk2 | 653.0041488 | 2.02886886 | 2.65E-66 |
| Thbs1 | 56074.01074 | 2.026743048 | 1.30E-09 |
| Fry | 21.32990924 | 2.025983986 | 7.69E-04 |
| Camk2b | 29.49686363 | 2.025788178 | 2.13E-03 |
| Prkcg | 214.7714828 | 2.024462396 | 8.24E-17 |
| Ccdc18 | 133.6308939 | 2.022019304 | 1.07E-04 |
| Dbndd2 | 321.9744503 | 2.021308029 | 5.66E-17 |
| B4galt4 | 584.3450299 | 2.016743738 | 1.78E-40 |
| Rnu1a1 | 7.850509731 | 2.015932524 | 2.18E-02 |
| Gm16170 | 35.38246777 | 2.013981065 | 4.23E-05 |
| Ptgs2os | 66.7624299 | 2.006520298 | 1.45E-07 |

| | | | |
|---|---|---|---|
| Slc43a2 | 56.28890673 | 2.002529749 | 2.56E-05 |
| Reep6 | 146.6378485 | 2.000341549 | 1.68E-19 |
| Stap2 | 79.90043653 | 1.98933403 | 9.07E-14 |
| Lipg | 535.1730908 | 1.985929596 | 4.87E-19 |
| Gm9967 | 10.06285755 | 1.985363786 | 6.76E-03 |
| Rhof | 50.58345819 | 1.984697198 | 6.33E-07 |
| Synb | 17.59181136 | 1.980215155 | 2.99E-03 |
| Krt7 | 1408.919025 | 1.976191558 | 4.08E-11 |
| Ncam1 | 1630.791622 | 1.972669977 | 8.51E-28 |
| Gm16998 | 21.06629374 | 1.971604659 | 6.01E-04 |
| Arhgap19 | 667.7567467 | 1.970613406 | 2.16E-11 |
| Il2rb | 49.38214337 | 1.968477989 | 5.63E-06 |
| Gm11427 | 43.61658226 | 1.967913539 | 9.46E-07 |
| Gm11425 | 8.581756494 | 1.965404514 | 2.62E-02 |
| Prg4 | 107.0476853 | 1.964724938 | 4.73E-04 |
| Gm9750 | 83.52959486 | 1.96441608 | 1.09E-06 |
| Gm2541 | 306.2248907 | 1.96295325 | 3.20E-08 |
| Mapk15 | 11.12845642 | 1.960477606 | 1.10E-02 |
| Olfr1314 | 22.59357489 | 1.960047561 | 2.39E-05 |
| Tlcd2 | 144.2625361 | 1.959531326 | 4.50E-15 |
| Tob1 | 562.3538878 | 1.955883333 | 2.45E-20 |
| Rpl10a | 10385.15404 | 1.953160737 | 2.37E-18 |
| Zfas1 | 5923.00628 | 1.952980172 | 2.78E-17 |
| Pfkfb3 | 882.241222 | 1.948397341 | 2.07E-11 |
| Gm16253 | 14.99152782 | 1.945731646 | 1.81E-04 |
| 1110046J04Rik | 236.2657601 | 1.942833726 | 3.76E-13 |
| Spire1 | 861.3974626 | 1.941685011 | 3.82E-32 |
| Adra1b | 80.73107556 | 1.94141298 | 1.24E-07 |
| Gjc2 | 6.210392759 | 1.936690582 | 4.05E-02 |
| Abcb1b | 1465.604472 | 1.933388151 | 4.32E-30 |
| Gng3 | 55.78018148 | 1.926582594 | 4.17E-05 |
| Gm42743 | 86.79263655 | 1.92447286 | 8.25E-05 |
| Dglucy | 1002.104884 | 1.922103792 | 1.02E-22 |
| Plcd3 | 1876.451148 | 1.92169115 | 1.41E-14 |
| Myoz2 | 26.20642193 | 1.920479284 | 1.01E-02 |
| Epb41l4aos | 846.5376575 | 1.920465544 | 5.57E-12 |
| Mmel1 | 15.80909654 | 1.920209751 | 2.40E-03 |
| Gm26917 | 957.8739768 | 1.915900652 | 3.13E-39 |
| Cd93 | 188.2576324 | 1.911024814 | 2.25E-06 |
| Ntn4 | 184.2709408 | 1.910143387 | 3.37E-13 |
| Abca7 | 2260.675367 | 1.908342478 | 1.70E-31 |
| D930019O06Rik | 18.49910595 | 1.903856065 | 7.51E-05 |
| Matn3 | 9.122129289 | 1.899322527 | 3.04E-02 |
| Ccng1 | 10250.65217 | 1.899164548 | 6.64E-34 |
| Gm13436 | 72.53817952 | 1.897592252 | 2.22E-07 |
| Ggct | 110.6567063 | 1.897154883 | 1.19E-10 |
| Plcxd2 | 1602.480094 | 1.896752091 | 3.07E-29 |

| | | | |
|---|---|---|---|
| Slc9a3r1 | 541.036435 | 1.895016522 | 3.73E-15 |
| Slmapos2 | 3326.360358 | 1.894837825 | 1.15E-11 |
| Cep72 | 201.8607482 | 1.891069433 | 6.59E-20 |
| Gm11399 | 8.864055192 | 1.890655734 | 1.26E-02 |
| Pkn3 | 159.00837 | 1.890303275 | 2.87E-08 |
| Zdhhc2 | 222.3920511 | 1.888945598 | 3.55E-09 |
| Ddah1 | 1706.491516 | 1.888378959 | 1.14E-10 |
| Als2cr12 | 58.85300664 | 1.885633897 | 3.86E-06 |
| Sgms2 | 1191.807906 | 1.882630833 | 3.17E-38 |
| Nek2 | 1268.788324 | 1.879356048 | 2.52E-09 |
| Mdm2 | 3807.437062 | 1.877888155 | 3.84E-40 |
| 9330171B17Rik | 7.554386795 | 1.877495755 | 7.49E-03 |
| Mucl3 | 111.7139975 | 1.876963167 | 2.61E-10 |
| Gm15730 | 7.292559368 | 1.874961687 | 2.11E-02 |
| Oasl2 | 31.36978808 | 1.874927578 | 8.37E-04 |
| Hpx | 24.31517719 | 1.87286375 | 4.21E-04 |
| Pllp | 6.162697618 | 1.869398926 | 3.81E-02 |
| Pnpla2 | 1540.394814 | 1.867000024 | 1.02E-19 |
| Dbp | 721.6468037 | 1.864287212 | 1.18E-08 |
| Ripor3 | 41.96255131 | 1.864064681 | 1.24E-03 |
| Gm15694 | 10.95927481 | 1.861135971 | 6.14E-03 |
| Slc25a33 | 464.665477 | 1.859554815 | 7.37E-34 |
| Aoc2 | 159.8164997 | 1.859307156 | 6.74E-12 |
| Prr11 | 1150.453785 | 1.858207689 | 2.41E-10 |
| Dusp15 | 15.63811429 | 1.858011586 | 3.72E-03 |
| Gm11517 | 11.31534709 | 1.857243653 | 9.53E-03 |
| Slc9a9 | 20.87683751 | 1.855022664 | 8.97E-03 |
| Kifc5b | 429.9223858 | 1.854773048 | 3.67E-09 |
| Tnfrsf21 | 266.5152627 | 1.85462988 | 4.10E-17 |
| Ass1 | 199.8319632 | 1.851185714 | 4.01E-11 |
| Polk | 816.6986151 | 1.851017679 | 2.41E-21 |
| Slc4a4 | 911.506914 | 1.850521261 | 3.66E-14 |
| 9230114K14Rik | 104.7462116 | 1.849991433 | 1.29E-10 |
| Zbtb7b | 2933.12507 | 1.849914526 | 5.73E-24 |
| Arrdc4 | 1906.491093 | 1.849434391 | 1.65E-07 |
| Lamc2 | 395.7838555 | 1.849100662 | 1.58E-11 |
| Avil | 62.46903402 | 1.846858358 | 1.24E-06 |
| Gm11263 | 10.2203104 | 1.83726112 | 5.69E-03 |
| Mtss1 | 37.26253904 | 1.836253086 | 6.37E-07 |
| Chmp4c | 13.45073978 | 1.833956178 | 1.46E-02 |
| Nhlrc2 | 1888.903339 | 1.833650946 | 3.35E-67 |
| Rhbdd1 | 2472.901059 | 1.833420044 | 3.20E-32 |
| Gm11884 | 8.165736454 | 1.833041788 | 3.83E-02 |
| Mafk | 2125.926029 | 1.832273236 | 1.06E-16 |
| Ttll2 | 60.65077863 | 1.828330402 | 2.03E-02 |
| Zfp143 | 1097.907523 | 1.826773366 | 2.94E-34 |
| Gm11934 | 9.445736696 | 1.826761407 | 5.64E-03 |

| | | | |
|---|---|---|---|
| NA | 13.11091694 | 1.826484051 | 7.17E-03 |
| Rgs17 | 87.00021087 | 1.822872701 | 3.95E-06 |
| Tns4 | 36.47295026 | 1.819313631 | 1.55E-03 |
| Cybrd1 | 13.86455351 | 1.816494394 | 1.66E-03 |
| Gm12138 | 2019.192784 | 1.814601617 | 6.04E-09 |
| Myc | 3282.758284 | 1.813674851 | 2.11E-33 |
| Eef2 | 390505.9252 | 1.81352992 | 1.09E-17 |
| Gm773 | 40.14795369 | 1.812970191 | 1.43E-04 |
| Kif18b | 709.5316689 | 1.812111724 | 1.22E-07 |
| 9230111E07Rik | 13.96764762 | 1.811125726 | 2.78E-03 |
| Bst2 | 1204.425443 | 1.80668793 | 4.63E-12 |
| Slc37a2 | 33.44387964 | 1.80645532 | 2.67E-06 |
| Ube2c | 1898.382155 | 1.806399128 | 1.89E-08 |
| Crybg1 | 279.3516519 | 1.806295975 | 7.32E-09 |
| Cct2 | 26032.40918 | 1.805252529 | 1.91E-42 |
| Gm26788 | 40.96349029 | 1.804793961 | 1.73E-07 |
| Spata5l1 | 220.8232095 | 1.802511725 | 1.35E-11 |
| Klra2 | 109.4765134 | 1.801370244 | 7.34E-06 |
| Slc25a25 | 821.5438689 | 1.797799102 | 3.89E-13 |
| Slc35g1 | 93.21178618 | 1.794285193 | 8.91E-09 |
| 2610318N02Rik | 78.5731003 | 1.793378643 | 7.79E-06 |
| Hspb6 | 825.7361147 | 1.791564429 | 2.55E-12 |
| Gm13050 | 29.96616328 | 1.789719877 | 8.55E-06 |
| NA | 63.62668047 | 1.789094983 | 2.47E-03 |
| Gm13532 | 26.6412113 | 1.787585036 | 1.15E-03 |
| Prkag3 | 47.08753681 | 1.785181722 | 1.33E-03 |
| Col20a1 | 635.2226194 | 1.78425547 | 1.45E-10 |
| Usp2 | 268.4067797 | 1.783703107 | 1.63E-07 |
| 4833427G06Rik | 113.596248 | 1.782108166 | 3.68E-13 |
| Col5a2 | 40728.55901 | 1.781684149 | 1.53E-15 |
| Lgals9 | 3984.469823 | 1.781580327 | 1.52E-24 |
| 1110038B12Rik | 8194.910558 | 1.78082361 | 2.38E-20 |
| Siglece | 8.051011043 | 1.776884598 | 1.19E-02 |
| Hist1h1d | 29.16335522 | 1.77667631 | 2.58E-04 |
| 1700061G19Rik | 8.01976113 | 1.775579161 | 1.32E-02 |
| Ppbp | 8.064294932 | 1.77522212 | 3.65E-02 |
| Pidd1 | 231.8430029 | 1.775031581 | 2.29E-09 |
| Ccdc170 | 17.32655283 | 1.772116995 | 8.11E-04 |
| Gm4841 | 16.51594958 | 1.769336723 | 6.95E-03 |
| Tspan32 | 38.52298728 | 1.767267642 | 1.42E-03 |
| Fam110a | 459.3283501 | 1.766135204 | 1.39E-12 |
| Gm26574 | 14.54837472 | 1.763564817 | 1.04E-02 |
| Tnfrsf10b | 1728.080669 | 1.763020796 | 1.72E-21 |
| Gpank1 | 589.3973124 | 1.759736037 | 5.55E-13 |
| Fam241a | 67.45370295 | 1.753800881 | 3.92E-11 |
| Fam169b | 128.3296111 | 1.751570532 | 1.73E-06 |
| Aim2 | 214.7586752 | 1.750891686 | 3.29E-15 |

| | | | |
|---|---|---|---|
| Stk32c | 7.691422658 | 1.750168715 | 1.66E-02 |
| Wt1 | 33.01804751 | 1.749878369 | 1.49E-02 |
| As3mt | 32.98295994 | 1.748329945 | 1.18E-03 |
| Clic3 | 21.76777198 | 1.748254519 | 3.96E-03 |
| Ahr | 292.2985283 | 1.747296225 | 1.84E-13 |
| Clpb | 1889.540012 | 1.745287249 | 3.22E-17 |
| Fat1 | 9394.675609 | 1.741778937 | 3.06E-15 |
| Krt80 | 423.3382572 | 1.736439305 | 1.66E-08 |
| Upk1a | 25.37899221 | 1.736160794 | 1.53E-04 |
| Mx2 | 49.68098047 | 1.735298531 | 1.15E-04 |
| Blvrb | 2497.902833 | 1.735051762 | 1.24E-21 |
| Rpl15-ps2 | 26.45417817 | 1.733957801 | 1.56E-03 |
| Snhg12 | 1711.852806 | 1.732200208 | 2.76E-17 |
| Vldlr | 180.1056824 | 1.732146619 | 4.01E-07 |
| Irf7 | 51.08821135 | 1.730557603 | 9.15E-06 |
| Spag4 | 73.35005613 | 1.727974069 | 1.77E-08 |
| Arhgef39 | 102.9442882 | 1.727923919 | 8.29E-06 |
| Akr1b3 | 27184.12226 | 1.727538495 | 1.22E-07 |
| Gm11944 | 37.06617334 | 1.726877599 | 5.05E-05 |
| Gm5921 | 644.8903188 | 1.726161164 | 4.41E-17 |
| Serpinb9 | 1571.525424 | 1.725022781 | 3.64E-27 |
| Asns | 8358.728063 | 1.724913794 | 2.64E-14 |
| Myo1a | 26.02622937 | 1.724827374 | 7.03E-04 |
| Per1 | 5421.880749 | 1.724608757 | 1.22E-08 |
| Rpl13a | 71987.69167 | 1.722744053 | 1.72E-18 |
| Gm13832 | 8.699723904 | 1.721666934 | 2.95E-02 |
| Pfkp | 1611.630881 | 1.721019812 | 7.04E-15 |
| Ecm1 | 1410.00657 | 1.718437235 | 5.92E-09 |
| Pcsk4 | 254.3779611 | 1.716397822 | 7.09E-23 |
| Gm45837 | 28.69853484 | 1.716072859 | 1.23E-03 |
| Gm15246 | 19.65445391 | 1.715372304 | 2.20E-03 |
| Vwa1 | 45.31304822 | 1.714475284 | 2.05E-05 |
| Znrf2 | 1311.359839 | 1.714368292 | 1.12E-21 |
| Eci3 | 9.306711395 | 1.713780826 | 1.39E-02 |
| Ybx3 | 12605.60181 | 1.712896882 | 1.58E-26 |
| Alcam | 657.7870578 | 1.712131781 | 6.22E-12 |
| Rpl35a-ps2 | 2188.965404 | 1.712126249 | 8.23E-08 |
| Trpm1 | 12.59951623 | 1.710656873 | 1.02E-02 |
| Extl1 | 31.44522133 | 1.710045653 | 2.94E-02 |
| Bhlhe41 | 899.5494575 | 1.704903925 | 9.67E-12 |
| Cyp26b1 | 1058.188833 | 1.704803885 | 9.68E-06 |
| Gm11380 | 56.60624028 | 1.704794895 | 7.49E-05 |
| Adrb2 | 121.6545444 | 1.70447317 | 7.19E-11 |
| Ell2 | 681.4752625 | 1.702910174 | 1.87E-16 |
| Vgf | 80.48610746 | 1.702389826 | 9.81E-07 |
| Mcrip2 | 335.8526933 | 1.702011221 | 1.15E-14 |
| Il15 | 186.9354469 | 1.70029622 | 3.34E-13 |

| | | | |
|---|---|---|---|
| NA | 27.02896418 | 1.700071369 | 9.54E-06 |
| Fblim1 | 958.4750544 | 1.699728332 | 5.83E-21 |
| Pold4 | 1270.663176 | 1.69500474 | 8.47E-16 |
| Fhdc1 | 1544.683828 | 1.693231601 | 1.74E-35 |
| Crlf1 | 353.0096371 | 1.692180029 | 6.35E-07 |
| Gm11478 | 30.23898364 | 1.691393573 | 6.80E-04 |
| Thpo | 50.84521309 | 1.68930783 | 2.71E-05 |
| Cnbd2 | 352.579731 | 1.687582016 | 3.59E-11 |
| Rps6-ps1 | 64.52353795 | 1.687079425 | 1.74E-06 |
| Gm11127 | 32.61799423 | 1.685956302 | 3.68E-03 |
| 9530026P05Rik | 12.09696974 | 1.685307271 | 2.12E-03 |
| Fam189b | 38.7043353 | 1.684995489 | 1.52E-04 |
| Lrp2bp | 10.02509851 | 1.684838374 | 2.91E-02 |
| Map3k20 | 2221.020669 | 1.684694965 | 4.08E-33 |
| Rps2-ps6 | 64.97681542 | 1.682560123 | 6.63E-05 |
| Fibin | 47.24750194 | 1.682106709 | 2.59E-03 |
| Gm10499 | 78.66015587 | 1.681281729 | 4.54E-09 |
| Gm10653 | 235.1147325 | 1.680928871 | 3.87E-04 |
| Nppb | 446.9261959 | 1.680883705 | 4.78E-05 |
| Rpl21-ps15 | 1077.507034 | 1.680130615 | 4.59E-06 |
| Hmcn2 | 50.91192735 | 1.679139918 | 1.78E-02 |
| Lnx1 | 47.50474965 | 1.675450244 | 1.45E-03 |
| Bscl2 | 225.8361494 | 1.675343133 | 1.64E-10 |
| Gm26810 | 8.924958287 | 1.6752747 | 4.19E-02 |
| Fzd6 | 270.1231641 | 1.675059952 | 1.59E-16 |
| Aldh1a7 | 739.2403274 | 1.67216008 | 1.66E-03 |
| Recql4 | 294.0795684 | 1.67118168 | 1.31E-05 |
| Gm7334 | 682.962771 | 1.667033126 | 1.22E-32 |
| Gm15541 | 42.64852593 | 1.666195139 | 2.60E-06 |
| Gm11331 | 59.22672314 | 1.665182337 | 1.20E-02 |
| Plk2 | 10524.64627 | 1.660657829 | 2.45E-30 |
| Gm5822 | 14.5937471 | 1.660432457 | 1.97E-02 |
| Rpl23a-ps3 | 79.47664741 | 1.659224048 | 1.18E-04 |
| Lrrc71 | 28.98119436 | 1.658932426 | 9.95E-03 |
| Zfp811 | 11.95022398 | 1.65831247 | 6.44E-03 |
| Ces1g | 19.45501609 | 1.658223258 | 2.42E-02 |
| Mtfr2 | 211.4222988 | 1.658083315 | 4.21E-05 |
| Fstl3 | 1713.818872 | 1.656201572 | 3.64E-15 |
| Plekha7 | 19.49701154 | 1.654548619 | 1.23E-02 |
| Kifc1 | 820.5197881 | 1.654449727 | 2.07E-08 |
| Dusp4 | 2570.483965 | 1.651750621 | 2.44E-08 |
| Rras2 | 2707.729271 | 1.651329559 | 2.61E-18 |
| Cgref1 | 204.5506992 | 1.651204648 | 1.08E-10 |
| Gm9806 | 17.38597716 | 1.648445954 | 2.32E-02 |
| Dkk2 | 170.6475019 | 1.645474219 | 3.46E-02 |
| Gm4204 | 1280.220195 | 1.640949824 | 1.30E-14 |
| 2310061I04Rik | 1207.433837 | 1.64010684 | 1.52E-19 |

| | | | |
|---|---|---|---|
| Gm6969 | 2684.561492 | 1.639319479 | 6.08E-14 |
| Unc13d | 41.0409234 | 1.638964301 | 8.88E-06 |
| 1810032O08Rik | 226.283254 | 1.637552406 | 2.19E-14 |
| Snord37 | 16.51974151 | 1.635407899 | 5.07E-03 |
| Slc7a5 | 7418.657169 | 1.63123116 | 1.55E-06 |
| 4933430I17Rik | 27.15124252 | 1.629940934 | 3.97E-05 |
| Cd53 | 21.19451119 | 1.627722705 | 2.21E-02 |
| Gm6736 | 924.2982575 | 1.625470888 | 7.70E-06 |
| Jsrp1 | 11.60423913 | 1.624850676 | 2.58E-02 |
| Fgf11 | 9.959730422 | 1.624565091 | 1.24E-02 |
| Cenpw | 161.2582704 | 1.624104646 | 6.55E-09 |
| Snrnp48 | 727.3748628 | 1.624078934 | 2.41E-21 |
| Rps2 | 1054.55721 | 1.620674554 | 1.59E-17 |
| Gm10029 | 43.04999321 | 1.619924401 | 4.68E-03 |
| Gm10420 | 383.0517429 | 1.615241649 | 2.06E-10 |
| Il15ra | 123.6384944 | 1.614982357 | 6.90E-06 |
| Inafm2 | 1281.565853 | 1.614475715 | 2.77E-54 |
| Slc39a10 | 3524.383162 | 1.613387654 | 1.57E-11 |
| Igf2bp1 | 2960.658169 | 1.611606118 | 3.25E-59 |
| Scarf1 | 6.608561061 | 1.611345376 | 4.77E-02 |
| E2f6 | 108.6963516 | 1.609925857 | 4.30E-11 |
| NA | 10.0590874 | 1.608729748 | 4.21E-02 |
| Itga6 | 3086.584435 | 1.608606255 | 1.12E-24 |
| Ripply1 | 17.40893938 | 1.607983591 | 3.23E-02 |
| Ccdc163 | 481.2533435 | 1.607267659 | 6.37E-10 |
| 2900052L18Rik | 59.70445642 | 1.607129986 | 3.02E-07 |
| Gm13573 | 14.31675529 | 1.60703353 | 2.00E-02 |
| Prdx6 | 12709.33805 | 1.606491431 | 5.77E-17 |
| Rhbdl1 | 40.6676154 | 1.605379477 | 4.08E-04 |
| Kcnj15 | 706.1352588 | 1.605073683 | 1.60E-05 |
| Ttc41 | 109.4536472 | 1.602256346 | 4.77E-07 |
| Pcgf5 | 1180.442953 | 1.602221799 | 1.15E-29 |
| NA | 737.3410238 | 1.6016084 | 3.07E-18 |
| Aen | 3318.36924 | 1.600983233 | 2.81E-16 |
| Plk3 | 880.6175897 | 1.60082711 | 1.22E-22 |
| Chd5 | 78.15936498 | 1.599926713 | 9.51E-04 |
| Gm27913 | 37.99607744 | 1.598006554 | 1.07E-05 |
| Ica1 | 26.09752958 | 1.598003289 | 1.63E-02 |
| Gtpbp2 | 2282.051956 | 1.595297131 | 2.43E-32 |
| Slco2a1 | 4121.36187 | 1.594916991 | 6.89E-20 |
| Ezr | 2125.964047 | 1.594152086 | 5.88E-29 |
| Amotl2 | 15490.23543 | 1.591887672 | 2.95E-18 |
| Fau | 8902.564704 | 1.590693542 | 1.64E-13 |
| Eno3 | 744.5122267 | 1.59034276 | 2.20E-07 |
| Nop53 | 8144.379275 | 1.590269151 | 3.02E-11 |
| Kcnab3 | 152.1664527 | 1.588987666 | 1.98E-08 |
| 4932414N04Rik | 12.45760852 | 1.588887508 | 1.08E-02 |

| | | | |
|---|---|---|---|
| Ptprj | 72.06266337 | 1.587039375 | 3.40E-03 |
| Dbil5 | 14.31520151 | 1.586313904 | 1.38E-02 |
| Apol9a | 154.7926263 | 1.583785052 | 8.57E-06 |
| Dusp9 | 278.6759648 | 1.582633285 | 2.71E-07 |
| Rack1 | 120396.584 | 1.581562005 | 7.49E-15 |
| Pde8b | 714.9937037 | 1.581146344 | 1.45E-12 |
| Me3 | 835.3947927 | 1.58096223 | 3.51E-21 |
| Hspa4l | 3271.537121 | 1.580390494 | 1.64E-10 |
| Cpxm1 | 32.08295723 | 1.579288437 | 6.96E-05 |
| Diras2 | 135.873439 | 1.578163693 | 9.31E-05 |
| Rpsa-ps9 | 649.5998619 | 1.577719562 | 6.44E-08 |
| Nmnat3 | 174.6720184 | 1.576067277 | 1.27E-11 |
| Tmem63c | 11.0959347 | 1.576052831 | 8.10E-03 |
| Gm13083 | 9.705678089 | 1.576030219 | 4.06E-02 |
| Odc1 | 6015.135819 | 1.575527985 | 1.46E-29 |
| Hsd3b3 | 12.47940476 | 1.575489544 | 1.49E-02 |
| Rnf169 | 1270.185398 | 1.575334906 | 1.03E-17 |
| Flrt2 | 1425.858476 | 1.572982557 | 3.58E-09 |
| Apold1 | 41.24765489 | 1.572334237 | 1.72E-05 |
| Gm15559 | 33.97139489 | 1.571339754 | 1.82E-04 |
| Aif1 | 20.87617467 | 1.570789549 | 5.39E-04 |
| Egln3 | 507.4247638 | 1.567920601 | 5.46E-16 |
| Cdkn2d | 517.3707037 | 1.567164943 | 1.34E-09 |
| Prrg2 | 490.5911814 | 1.564812877 | 8.15E-11 |
| Gdpd5 | 258.3174992 | 1.564161821 | 2.29E-17 |
| Gsta1 | 191.6070784 | 1.563694684 | 3.24E-04 |
| Thy1 | 3489.688816 | 1.563559855 | 1.09E-13 |
| Rps27a-ps2 | 26.39735323 | 1.562320084 | 1.17E-02 |
| Pvr | 3501.591273 | 1.56156351 | 1.41E-17 |
| Pttg1 | 1028.152404 | 1.558650504 | 7.71E-10 |
| Selenoh | 777.6593273 | 1.558427655 | 1.41E-15 |
| Plekha2 | 266.0834896 | 1.557885583 | 1.63E-06 |
| Lncenc1 | 53.16397997 | 1.557862928 | 1.10E-04 |
| Gm13226 | 135.786182 | 1.557228948 | 1.55E-07 |
| Gm6139 | 18.06609433 | 1.556264372 | 3.07E-02 |
| Acyp2 | 175.9484139 | 1.554900249 | 8.73E-19 |
| Veph1 | 84.62075448 | 1.553498686 | 3.52E-03 |
| Gm5805 | 361.4001265 | 1.553025586 | 6.29E-06 |
| Txnip | 34225.99991 | 1.552956232 | 1.83E-03 |
| Hid1 | 87.39260748 | 1.552784052 | 4.28E-05 |
| Slc19a2 | 1917.481573 | 1.551952684 | 2.66E-25 |
| Gm15793 | 214.717008 | 1.551554274 | 7.34E-07 |
| Gm16174 | 11.37027811 | 1.551301499 | 6.60E-03 |
| Loxl4 | 984.284874 | 1.550693226 | 2.58E-09 |
| Gm26510 | 11.30750862 | 1.550684537 | 2.97E-02 |
| Tmem169 | 34.23990879 | 1.549378052 | 4.22E-04 |
| Tmcc3 | 374.2348867 | 1.547994531 | 1.49E-09 |

| | | | |
|---|---|---|---|
| Junb | 1589.766988 | 1.547339779 | 1.42E-06 |
| Pmm1 | 3567.219961 | 1.546577918 | 7.62E-17 |
| Ehd1 | 9278.785003 | 1.545433318 | 6.81E-17 |
| Mettl21e | 9.771196451 | 1.545035776 | 2.26E-02 |
| Gm10247 | 32.75537283 | 1.544934874 | 3.42E-04 |
| Hmgcll1 | 8.897186381 | 1.543497189 | 4.92E-02 |
| Tcaf2 | 513.7387912 | 1.543175048 | 1.07E-10 |
| Gstm6 | 8.21388758 | 1.54317214 | 4.71E-02 |
| Rpl8 | 66813.58999 | 1.54101233 | 1.30E-19 |
| Papss2 | 2566.376957 | 1.540791845 | 7.92E-24 |
| Glul | 897.3710217 | 1.540394057 | 5.65E-09 |
| Gm13632 | 65.5249491 | 1.538931872 | 2.68E-08 |
| Uhrf1bp1l | 3287.025469 | 1.538186154 | 2.76E-43 |
| Rb1 | 846.9033777 | 1.536602604 | 4.97E-19 |
| Tpx2 | 3448.017744 | 1.534230563 | 4.81E-07 |
| Dnah6 | 60.00583954 | 1.534111649 | 2.07E-04 |
| Rpl18 | 36466.21172 | 1.531600723 | 2.23E-17 |
| Itpkc | 1053.756826 | 1.531561033 | 1.02E-12 |
| Gm11249 | 1092.346685 | 1.529894527 | 6.61E-08 |
| Lysmd1 | 338.1310677 | 1.529646235 | 2.89E-20 |
| Disp1 | 2262.130775 | 1.528306779 | 1.07E-47 |
| Col11a2 | 157.3560411 | 1.527649568 | 8.24E-05 |
| NA | 1727.994687 | 1.52722881 | 2.50E-03 |
| Rpl28-ps1 | 40054.69604 | 1.526116239 | 2.33E-12 |
| Atf7ip2 | 31.79534999 | 1.525528084 | 3.63E-02 |
| Trim12a | 37.33199526 | 1.525324495 | 2.78E-04 |
| Fbln1 | 65.66334147 | 1.524850539 | 2.41E-03 |
| NA | 2233.700196 | 1.523446057 | 1.24E-12 |
| Gm15708 | 52.21141378 | 1.521542212 | 7.46E-05 |
| Tec | 115.0557458 | 1.521009652 | 2.43E-10 |
| Gm8623 | 21.75656886 | 1.519493707 | 1.22E-02 |
| Stx11 | 348.3990913 | 1.519089723 | 2.14E-18 |
| Acot2 | 886.3418002 | 1.517691515 | 2.55E-15 |
| Abcg4 | 19.25427227 | 1.515382408 | 6.59E-03 |
| Gm11599 | 19.48675284 | 1.515152652 | 1.21E-02 |
| Rpsa-ps11 | 28.09659456 | 1.513527821 | 1.16E-02 |
| Cyp2t4 | 9.343747417 | 1.511660495 | 2.05E-02 |
| Gm10275 | 71607.79352 | 1.510855608 | 1.52E-12 |
| Wdr45 | 2171.76663 | 1.509898214 | 1.78E-20 |
| Oxr1 | 3434.000898 | 1.509439982 | 1.47E-09 |
| Cfap100 | 434.4066328 | 1.509140754 | 1.66E-08 |
| Tspan33 | 25.32257853 | 1.509054084 | 5.19E-03 |
| Gm6169 | 8.709692281 | 1.508681452 | 4.07E-02 |
| Sh2d6 | 55.41535397 | 1.508393203 | 1.17E-04 |
| Esd | 14951.22292 | 1.50651986 | 1.66E-17 |
| Ap2a2 | 11186.22479 | 1.505563856 | 2.01E-32 |
| Rps2-ps10 | 66191.96649 | 1.50523151 | 1.34E-22 |

| | | | |
|---|---|---|---|
| Slc22a4 | 48.52663821 | 1.503261669 | 1.05E-03 |
| Isg15 | 100.3018409 | 1.503025754 | 5.75E-07 |
| Ak1 | 2929.214187 | 1.502593217 | 2.94E-27 |
| Oplah | 158.7571282 | 1.502509016 | 7.98E-08 |
| Klf10 | 2846.196887 | 1.502213167 | 3.53E-07 |
| Rps6-ps4 | 132832.5808 | 1.500778448 | 3.06E-15 |
| Sp2 | 1110.544619 | 1.5001053 | 1.01E-12 |
| NA | 329.9046769 | 1.500076721 | 2.11E-07 |
| A630031M04Rik | 11.42034849 | 1.49958265 | 2.51E-02 |
| Rusc2 | 4030.034565 | 1.498941881 | 2.49E-10 |
| Col28a1 | 133.400327 | 1.497677072 | 7.56E-05 |
| Msh5 | 95.26155785 | 1.495391525 | 8.76E-05 |
| Gm16364 | 22.25215599 | 1.49398672 | 7.44E-04 |
| Acy3 | 101.9828741 | 1.491909314 | 1.03E-07 |
| Stard13 | 1677.000687 | 1.491782904 | 2.23E-06 |
| 9130008F23Rik | 333.9612646 | 1.491582158 | 6.14E-12 |
| Cyp39a1 | 736.9729686 | 1.489778121 | 1.03E-30 |
| 4933427I04Rik | 39.54466148 | 1.488710526 | 1.50E-04 |
| Cav2 | 3663.325958 | 1.487675372 | 5.47E-17 |
| Noc2l | 3844.883424 | 1.48734793 | 9.64E-31 |
| Paqr3 | 356.3402682 | 1.485768899 | 2.85E-16 |
| Dot1l | 2904.513899 | 1.482134556 | 4.17E-21 |
| Arl14ep | 4362.339537 | 1.482070392 | 5.62E-17 |
| Rnf114 | 2517.365589 | 1.481675014 | 7.48E-14 |
| Ercc6l2 | 120.8231153 | 1.481311572 | 7.56E-07 |
| 1700028J19Rik | 26.72022137 | 1.480896374 | 1.27E-03 |
| Gm11824 | 14.12119418 | 1.479069816 | 2.88E-02 |
| Zfp286os | 14.25886093 | 1.47541998 | 1.02E-02 |
| Gm9493 | 223.7226295 | 1.472190222 | 5.17E-05 |
| Rpsa | 106522.1942 | 1.470851724 | 2.98E-15 |
| Wincr1 | 360.4301665 | 1.470833646 | 6.20E-06 |
| Prorsd1 | 28.67614291 | 1.470673543 | 2.52E-03 |
| Rps12-ps1 | 89.85863566 | 1.469812833 | 2.25E-06 |
| Col4a5 | 13744.73591 | 1.469116085 | 1.84E-31 |
| NA | 58.68890451 | 1.46882902 | 3.08E-05 |
| Abcc1 | 6563.18358 | 1.467546889 | 1.14E-31 |
| Hspbap1 | 670.3335746 | 1.466998344 | 4.89E-27 |
| Trmt10a | 1090.231032 | 1.466304473 | 3.63E-17 |
| Rps15 | 55482.86243 | 1.465841641 | 1.74E-12 |
| Usp3 | 2309.151463 | 1.465268839 | 6.61E-12 |
| Slco3a1 | 1020.790683 | 1.464233577 | 1.97E-15 |
| Fcer2a | 17.18996379 | 1.462328595 | 3.48E-03 |
| Gm26686 | 46.31665202 | 1.460938207 | 1.86E-04 |
| Gm19705 | 81.49688775 | 1.45856252 | 1.65E-06 |
| Prelp | 2798.880483 | 1.457545809 | 1.95E-03 |
| Rpl17-ps10 | 39.5528456 | 1.456593994 | 7.64E-03 |
| Gm16409 | 267.0783991 | 1.45567108 | 6.61E-06 |

| | | | | |
|---|---|---|---|---|
| Gm14149 | | 106.2779207 | 1.4552922 | 8.82E-11 |
| Rpl9-ps6 | | 49.21276014 | 1.454341721 | 2.14E-02 |
| Rpsa-ps5 | | 10.03686359 | 1.453533168 | 4.93E-02 |
| Klrg2 | | 65.68197859 | 1.452139689 | 6.27E-06 |
| Zhx2 | | 795.6214179 | 1.451849822 | 5.05E-12 |
| Hsf2bp | | 28.16029851 | 1.449284051 | 6.07E-03 |
| BC022960 | | 75.28455684 | 1.449117108 | 1.37E-05 |
| Clk3 | | 1992.491787 | 1.44893745 | 5.98E-19 |
| Gm9855 | | 2483.951233 | 1.448892878 | 6.00E-34 |
| Eif3d | | 11371.15065 | 1.448517256 | 2.75E-18 |
| Arrb2 | | 918.8118723 | 1.447936949 | 2.61E-17 |
| Zmat3 | | 4898.113607 | 1.447314487 | 3.44E-21 |
| Rgl3 | | 384.4796404 | 1.447228986 | 5.31E-19 |
| Jmjd7 | | 318.0362651 | 1.446215065 | 2.60E-11 |
| Gm16201 | | 31.28094923 | 1.445713438 | 1.79E-03 |
| Rps16 | | 14597.25608 | 1.444866487 | 2.28E-15 |
| Specc1 | | 3432.279109 | 1.44445094 | 1.14E-15 |
| Rpl17-ps4 | | 69.9664268 | 1.442797711 | 1.46E-04 |
| Por | | 4956.615258 | 1.441399749 | 1.69E-14 |
| Lmcd1 | | 190.9655838 | 1.441329396 | 1.69E-03 |
| Rps3 | | 56649.04585 | 1.440050491 | 1.83E-13 |
| Mamstr | | 61.73933429 | 1.439312904 | 2.07E-02 |
| Rps6-ps3 | | 125.0573789 | 1.43854803 | 1.51E-06 |
| Crim1 | | 4958.715791 | 1.437590385 | 2.16E-10 |
| Necab3 | | 62.92270688 | 1.436635258 | 1.64E-03 |
| Yjefn3 | | 129.0269012 | 1.435954991 | 1.79E-06 |
| | 5-Sep | 7626.58109 | 1.434152628 | 6.58E-17 |
| Whamm | | 496.7500502 | 1.433276094 | 3.18E-12 |
| Rhox5 | | 444.7664169 | 1.432538628 | 3.92E-07 |
| Kcmf1 | | 5176.903926 | 1.430727733 | 1.26E-38 |
| Gm10036 | | 36.68042067 | 1.42949499 | 8.05E-04 |
| Ttc16 | | 24.87972731 | 1.429105598 | 5.52E-03 |
| Gm4968 | | 86.28465387 | 1.429072221 | 1.03E-05 |
| Gm7367 | | 116.3501826 | 1.429048736 | 8.70E-03 |
| Gm11518 | | 16388.69852 | 1.428393234 | 8.71E-18 |
| Rdh12 | | 38.3276781 | 1.427634053 | 2.98E-05 |
| Gm26651 | | 25.61219024 | 1.427455468 | 4.52E-02 |
| Ndufaf6 | | 307.0560471 | 1.427398393 | 1.14E-12 |
| Hcfc1r1 | | 8234.320913 | 1.427175162 | 6.76E-11 |
| Gm14494 | | 1272.813194 | 1.426901641 | 3.42E-08 |
| Hagh | | 1322.182493 | 1.426585723 | 2.15E-25 |
| Abca5 | | 1770.476615 | 1.426481575 | 3.86E-11 |
| Mknk2 | | 5451.658763 | 1.425579257 | 1.66E-11 |
| Gm15264 | | 33.15051985 | 1.425442709 | 2.98E-03 |
| Wfikkn1 | | 16.62283162 | 1.425435358 | 4.36E-03 |
| Gm12242 | | 18.90098167 | 1.424735599 | 2.23E-02 |
| Agpat4 | | 1161.530843 | 1.424712292 | 3.10E-18 |

| | | | |
|---|---|---|---|
| Ldb3 | 22.13416126 | 1.422929309 | 3.64E-02 |
| Cxcl5 | 153.6533999 | 1.422638048 | 4.87E-04 |
| Gm15050 | 189.9018835 | 1.422038708 | 5.60E-11 |
| Guca1a | 12.28516956 | 1.421933924 | 2.09E-02 |
| Chfr | 2461.953026 | 1.42086758 | 5.48E-17 |
| Prss53 | 73.1378317 | 1.420852485 | 4.06E-05 |
| Ctc1 | 2558.238951 | 1.42067992 | 2.21E-08 |
| Grip2 | 30.08385853 | 1.420392131 | 3.55E-02 |
| Tmem43 | 6334.664426 | 1.419625073 | 6.82E-13 |
| Pqlc3 | 1007.222603 | 1.419497908 | 4.43E-18 |
| Rpl13 | 43964.4684 | 1.419133742 | 1.53E-11 |
| 2610206C17Rik | 15.87470752 | 1.417168966 | 2.47E-02 |
| Gm10020 | 173.4954606 | 1.416545244 | 7.34E-06 |
| Mllt3 | 4797.047418 | 1.415892183 | 1.21E-17 |
| Strbp | 1590.73397 | 1.414604256 | 5.88E-19 |
| Rps2-ps13 | 65962.345 | 1.412278541 | 6.79E-20 |
| 4933438K21Rik | 20.61022562 | 1.410083816 | 1.05E-02 |
| Cdc25b | 1752.307462 | 1.409954985 | 1.85E-10 |
| Rpl18-ps2 | 29.06764237 | 1.409343448 | 1.33E-03 |
| Awat2 | 33.20162048 | 1.409296852 | 4.27E-04 |
| Ltc4s | 33.90047656 | 1.408457611 | 2.27E-02 |
| Maff | 1125.074098 | 1.408334543 | 3.58E-08 |
| Fermt3 | 34.13683324 | 1.407141691 | 1.20E-04 |
| Cd151 | 4413.725542 | 1.404594379 | 3.24E-20 |
| Cd274 | 79.84677291 | 1.402217132 | 1.03E-07 |
| Flrt3 | 33.14674922 | 1.40209929 | 3.81E-02 |
| Slc35e4 | 1871.809364 | 1.401251043 | 8.62E-12 |
| Aurka | 1220.677723 | 1.401222657 | 2.03E-04 |
| Gm10136 | 300.3157053 | 1.399891061 | 5.72E-07 |
| Rps26 | 31879.79427 | 1.397152143 | 2.00E-10 |
| Hmgn5 | 168.6581029 | 1.394508404 | 2.50E-02 |
| Ifi205 | 92.81123866 | 1.393195535 | 4.39E-02 |
| Otud1 | 340.3620018 | 1.390819282 | 3.67E-10 |
| Gfer | 1246.79568 | 1.390139192 | 1.18E-18 |
| Ccser1 | 471.615598 | 1.389474466 | 1.57E-10 |
| Cltb | 2400.270914 | 1.388771013 | 6.98E-12 |
| Exoc6 | 416.7822632 | 1.387498061 | 5.22E-12 |
| Gm4462 | 7.257038782 | 1.387082472 | 4.72E-02 |
| Gm17315 | 21.55675448 | 1.386719884 | 1.22E-03 |
| Slc25a28 | 779.0647433 | 1.384681043 | 7.13E-17 |
| Faah | 126.5236943 | 1.38456255 | 1.81E-05 |
| Gm26555 | 27.36621759 | 1.383581042 | 9.54E-03 |
| Stambpl1 | 1280.981375 | 1.383544868 | 1.87E-14 |
| Gm9625 | 405.7501673 | 1.38240839 | 5.61E-08 |
| Slc25a30 | 2177.751789 | 1.382298096 | 1.12E-12 |
| Gm12366 | 374.6077916 | 1.380565952 | 4.66E-06 |
| Rpl13-ps3 | 18.70398708 | 1.380375884 | 7.91E-03 |

| | | | |
|---|---|---|---|
| Rpl17-ps9 | 231.5005683 | 1.380179553 | 8.77E-09 |
| Dcxr | 824.7511236 | 1.378341348 | 9.61E-09 |
| Gm8213 | 418.508554 | 1.377681082 | 1.69E-09 |
| Pid1 | 12.18015165 | 1.377407863 | 2.79E-02 |
| Tbc1d31 | 1630.86014 | 1.377272055 | 1.60E-09 |
| Gm13205 | 30.44429336 | 1.376908798 | 4.79E-04 |
| Hrct1 | 51.68066241 | 1.376841176 | 2.48E-04 |
| Nuak2 | 683.3125158 | 1.375909835 | 3.36E-13 |
| Kcna4 | 440.1062529 | 1.375704237 | 3.52E-08 |
| Tap1 | 754.3650937 | 1.37557699 | 1.04E-11 |
| Mapre3 | 1426.057277 | 1.374620691 | 1.94E-12 |
| Phactr2 | 110.9007105 | 1.372870793 | 1.00E-03 |
| Rps12 | 48476.15642 | 1.37275634 | 3.61E-18 |
| Cox7a2l | 10854.98014 | 1.37225877 | 1.84E-10 |
| A230060F14Rik | 15.63411288 | 1.371731187 | 2.16E-02 |
| Gm5828 | 12.07803403 | 1.371723176 | 3.13E-02 |
| Gm9083 | 22.23643344 | 1.371497549 | 8.26E-03 |
| Zwilch | 669.3877584 | 1.367241133 | 1.56E-05 |
| C77080 | 1863.179005 | 1.367203292 | 7.94E-24 |
| Rpsa-ps10 | 455.3269898 | 1.366254745 | 1.83E-06 |
| Gm7808 | 50.83150304 | 1.366136223 | 1.28E-04 |
| Gm12712 | 27.7636762 | 1.365194184 | 1.36E-02 |
| Gm16062 | 85.00001513 | 1.364911714 | 3.30E-04 |
| Acsbg1 | 82.3921383 | 1.364723757 | 3.43E-04 |
| Dennd5b | 74.30478023 | 1.364047196 | 3.21E-03 |
| Cct6b | 24.23556002 | 1.363979305 | 1.67E-03 |
| NA | 10.10771867 | 1.363681507 | 3.45E-02 |
| Usp18 | 22.06187172 | 1.362192972 | 2.38E-03 |
| Rps12-ps24 | 14.81712885 | 1.361975267 | 4.78E-02 |
| Glns-ps1 | 14.7088613 | 1.361857271 | 4.30E-02 |
| Gm7536 | 46411.44855 | 1.3615571 | 2.27E-13 |
| Ccdc78 | 27.35552252 | 1.361535817 | 3.32E-03 |
| Rps12-ps3 | 32513.16414 | 1.361119393 | 1.92E-10 |
| Gm10382 | 31.87078083 | 1.359919595 | 4.75E-02 |
| Tmbim1 | 4932.481657 | 1.357615671 | 2.11E-20 |
| Slc2a9 | 175.1610797 | 1.357464419 | 2.01E-10 |
| Krt86 | 11.25936999 | 1.357272966 | 3.29E-02 |
| Adcy1 | 1520.53709 | 1.356898592 | 4.30E-07 |
| Phlda1 | 939.9688801 | 1.355574519 | 3.18E-07 |
| Gm13413 | 64.46038927 | 1.354977149 | 4.87E-06 |
| Apaf1 | 2157.688006 | 1.353958036 | 5.11E-20 |
| Gm3531 | 19.26519257 | 1.353650962 | 1.57E-02 |
| Castor1 | 337.8062982 | 1.351193867 | 9.26E-14 |
| Nabp2 | 2302.580906 | 1.350706419 | 3.03E-17 |
| Rpl36-ps12 | 66.11856536 | 1.349666025 | 4.13E-05 |
| Kif2a | 2279.023051 | 1.349534544 | 3.16E-23 |
| Gm10108 | 146.643854 | 1.347752935 | 5.83E-04 |

| | | | |
|---|---|---|---|
| 4921524J17Rik | 641.6105448 | 1.346701798 | 4.23E-25 |
| Egfr | 1382.648293 | 1.345840635 | 4.80E-08 |
| Cyp4x1 | 10.86680834 | 1.345606207 | 4.80E-02 |
| Impdh1 | 2647.662952 | 1.345463792 | 1.93E-30 |
| Gm12176 | 45.17947775 | 1.34535908 | 9.96E-04 |
| Gm10243 | 15969.90068 | 1.344852507 | 3.85E-14 |
| Spry1 | 847.9921191 | 1.344664483 | 6.04E-07 |
| Rps10 | 48037.18352 | 1.344640925 | 2.01E-11 |
| Ifi35 | 678.7565277 | 1.344554946 | 6.28E-15 |
| Rps16-ps2 | 25503.26783 | 1.343702634 | 2.60E-08 |
| Rpl27-ps1 | 134.8717961 | 1.343571515 | 7.91E-05 |
| Rabggtb | 5299.556326 | 1.343488284 | 2.79E-29 |
| Hotairm1 | 163.7859472 | 1.343031479 | 1.50E-09 |
| Sord | 8476.138057 | 1.342585839 | 6.62E-08 |
| Phkg1 | 37.75219016 | 1.341971698 | 2.49E-03 |
| Ltbp1 | 3648.116466 | 1.34186714 | 1.93E-10 |
| Cpox | 1136.258528 | 1.340785094 | 3.22E-08 |
| Gm4294 | 216.0372457 | 1.340689107 | 4.35E-03 |
| Rpl4 | 100332.3709 | 1.340466559 | 4.38E-14 |
| Slc3a2 | 11311.11817 | 1.340363523 | 1.23E-24 |
| Col8a1 | 3683.903915 | 1.340041171 | 1.07E-06 |
| Gm20900 | 83.4022082 | 1.339927179 | 3.34E-04 |
| Gimap6 | 24.98875556 | 1.33911975 | 5.08E-04 |
| Col5a3 | 679.7155384 | 1.337939357 | 2.81E-05 |
| Aloxe3 | 163.7815803 | 1.337525272 | 2.63E-04 |
| Rin1 | 4778.34829 | 1.337169783 | 2.33E-18 |
| Eed | 1239.554501 | 1.336950939 | 9.61E-25 |
| Rps14 | 37307.50526 | 1.33513973 | 7.49E-08 |
| Cpne9 | 17.409485 | 1.334881498 | 3.59E-02 |
| Sinhcaf | 1136.208845 | 1.334353484 | 1.70E-09 |
| Sorbs1 | 833.0862304 | 1.333332926 | 5.49E-13 |
| NA | 5457.007199 | 1.33311373 | 1.30E-04 |
| Sema3c | 667.7097169 | 1.331864865 | 2.18E-02 |
| Gm10051 | 181.0733855 | 1.331355435 | 4.36E-03 |
| Uck2 | 2613.500918 | 1.32782057 | 1.08E-15 |
| Herc3 | 407.8347654 | 1.327748862 | 2.45E-10 |
| Cep55 | 681.4845732 | 1.327081215 | 1.97E-02 |
| Prr5 | 206.0395211 | 1.326711648 | 7.99E-09 |
| Rps25 | 29032.20276 | 1.326239784 | 6.20E-13 |
| Wdr12 | 1978.768705 | 1.326199409 | 3.27E-20 |
| Cst6 | 60.21863085 | 1.325316804 | 7.38E-03 |
| Slc29a2 | 224.3345335 | 1.32493274 | 1.31E-12 |
| Sec31b | 230.9272441 | 1.324552992 | 1.99E-06 |
| Rpl23a | 76915.38591 | 1.323956196 | 4.90E-11 |
| Rplp0-ps1 | 63.10554626 | 1.323135519 | 6.58E-04 |
| C1qtnf12 | 46.38102859 | 1.321240744 | 4.08E-04 |
| Gm15427 | 13183.07084 | 1.320212607 | 9.50E-05 |

| | | | |
|---|---|---|---|
| Pck2 | 4321.410982 | 1.319769728 | 1.46E-12 |
| Gm10169 | 48.90505126 | 1.319473831 | 3.43E-03 |
| Gm6433 | 39.0436362 | 1.319400016 | 1.05E-02 |
| Rpl17 | 47230.04156 | 1.318894789 | 2.10E-10 |
| Lrwd1 | 1708.037956 | 1.318336225 | 3.23E-14 |
| C2cd2l | 454.1034844 | 1.317973301 | 1.65E-07 |
| Trim24 | 3746.206389 | 1.317016187 | 9.43E-11 |
| Knstrn | 1791.183893 | 1.31697167 | 3.42E-07 |
| Rpl22 | 11046.40294 | 1.316432934 | 8.48E-09 |
| Limk2 | 1846.809037 | 1.315885097 | 1.21E-22 |
| Tmem63b | 2442.58208 | 1.315369278 | 2.30E-33 |
| Uba52 | 18378.21562 | 1.314693569 | 1.28E-07 |
| Tuft1 | 1030.058028 | 1.314162497 | 6.46E-09 |
| Rras | 2672.576845 | 1.314122313 | 1.10E-10 |
| Prdx5 | 5188.911969 | 1.3139871 | 4.12E-20 |
| Pbx4 | 103.3083903 | 1.313632725 | 2.10E-07 |
| Rps26-ps1 | 19459.09523 | 1.313073939 | 3.16E-10 |
| Rpl9 | 44618.973 | 1.31261737 | 6.66E-14 |
| Mfsd2b | 28.2560697 | 1.312463818 | 2.75E-02 |
| Baz2a | 4071.956571 | 1.311436541 | 1.55E-68 |
| Snhg5 | 748.1058599 | 1.311346079 | 6.75E-11 |
| Tmem245 | 7288.006629 | 1.310797356 | 1.78E-20 |
| Naa25 | 2845.116492 | 1.3098735 | 9.06E-29 |
| Rpl17-ps5 | 73.10362811 | 1.309569152 | 4.98E-02 |
| Cdk18 | 2832.68387 | 1.30948955 | 1.46E-10 |
| A230020J21Rik | 18.87247229 | 1.309412132 | 2.81E-02 |
| Nomo1 | 5376.942128 | 1.308979752 | 1.32E-11 |
| Espnl | 43.3323916 | 1.307782798 | 1.27E-03 |
| Gm13075 | 19.69455583 | 1.307199372 | 2.03E-02 |
| Ager | 26.41507912 | 1.30680921 | 4.71E-03 |
| Ide | 5488.789588 | 1.306645702 | 1.59E-20 |
| Rps2-ps4 | 80.95409934 | 1.306089018 | 1.63E-02 |
| Gm9828 | 55.4899361 | 1.30564609 | 1.03E-05 |
| Dynlt1a | 769.8087586 | 1.304986484 | 1.06E-11 |
| Dlgap5 | 995.0483588 | 1.304762457 | 7.97E-06 |
| Crnde | 141.719853 | 1.303086904 | 4.25E-09 |
| Gm10132 | 193.158572 | 1.302787498 | 3.47E-04 |
| Plekhn1 | 640.7350483 | 1.302407661 | 8.12E-07 |
| Efna1 | 1322.865678 | 1.30236562 | 3.97E-05 |
| Gmip | 1120.355618 | 1.302249508 | 8.72E-13 |
| Nrg1 | 1866.437419 | 1.302134615 | 4.09E-14 |
| Rpl26 | 50023.25159 | 1.302109003 | 1.34E-14 |
| Ifrd1 | 4709.926875 | 1.301767778 | 2.68E-10 |
| Pdgfa | 1217.553266 | 1.300812028 | 2.48E-24 |
| Gm14438 | 233.5442823 | 1.300723059 | 6.53E-04 |
| Prss36 | 320.8623567 | 1.29967529 | 3.15E-08 |
| Lsm11 | 725.4015622 | 1.299644089 | 1.63E-12 |

| | | | |
|---|---|---|---|
| Wdr91 | 824.4786997 | 1.297648711 | 4.57E-12 |
| Rps19-ps6 | 69.44201172 | 1.297413307 | 1.37E-02 |
| Gm26707 | 18.84366641 | 1.297399235 | 1.22E-02 |
| Zfr2 | 251.1434612 | 1.29737907 | 3.12E-07 |
| Yaf2 | 830.9713549 | 1.296678352 | 7.91E-16 |
| Me1 | 3504.254363 | 1.296240565 | 5.31E-22 |
| Eif2b2 | 1629.982564 | 1.294622268 | 4.75E-17 |
| Cct4 | 19364.54992 | 1.293759587 | 1.85E-27 |
| Rpl39-ps | 200.9664968 | 1.292598611 | 6.87E-05 |
| Vezt | 1941.869543 | 1.292479844 | 8.91E-29 |
| Erc2 | 648.4103087 | 1.292216705 | 3.71E-13 |
| Zfp598 | 2577.338794 | 1.291677104 | 2.19E-28 |
| Vps36 | 2824.129634 | 1.291413009 | 8.17E-35 |
| Gm26578 | 320.2566482 | 1.291247914 | 4.55E-15 |
| Tbrg1 | 7943.990575 | 1.291224233 | 1.00E-18 |
| Cd24a | 2574.110999 | 1.289257231 | 4.29E-04 |
| Gm12933 | 53.05099324 | 1.289119246 | 7.18E-03 |
| Rps12-ps19 | 164.2671423 | 1.288921865 | 6.34E-08 |
| Rps27a | 38338.23439 | 1.287624696 | 4.69E-11 |
| Lif | 170.5025553 | 1.286402821 | 7.14E-04 |
| Ankrd61 | 29.46407003 | 1.286278958 | 4.68E-03 |
| Fam117a | 85.5208589 | 1.286031927 | 1.63E-03 |
| Prps1 | 1657.907258 | 1.285469852 | 4.11E-15 |
| Gm11808 | 30010.49529 | 1.284510572 | 3.26E-13 |
| Ifit3b | 229.6966565 | 1.284455831 | 1.22E-03 |
| Pvt1 | 2024.36745 | 1.284069034 | 1.95E-19 |
| Syn1 | 1096.842102 | 1.283107288 | 1.54E-17 |
| Thap4 | 2190.18433 | 1.282215782 | 3.95E-15 |
| Nmnat1 | 220.6102235 | 1.281997103 | 3.25E-17 |
| Rps5 | 42435.71707 | 1.281666765 | 7.90E-09 |
| Milr1 | 11.1371401 | 1.281346939 | 4.48E-02 |
| Col10a1 | 24.89550555 | 1.281306146 | 2.10E-03 |
| Tex26 | 48.26017168 | 1.281136149 | 1.29E-03 |
| Syne2 | 2077.972178 | 1.281126399 | 4.59E-11 |
| Gm5459 | 15.18266592 | 1.279135272 | 4.98E-02 |
| Ppp2r5d | 4897.843244 | 1.279037989 | 4.63E-19 |
| Gpr157 | 124.3548755 | 1.278617622 | 5.77E-05 |
| Alkbh2 | 215.7559227 | 1.278600425 | 2.23E-13 |
| Gm17828 | 1219.480847 | 1.278426326 | 6.92E-06 |
| Gm15772 | 85.38857994 | 1.277873177 | 1.94E-03 |
| Dhx58 | 52.4208171 | 1.277459224 | 2.46E-05 |
| Gm5931 | 88.3662409 | 1.275438578 | 1.39E-03 |
| Zc3h8 | 210.3834763 | 1.27526 | 8.31E-09 |
| Rpl15-ps3 | 37991.77318 | 1.274086596 | 1.09E-09 |
| Ifih1 | 56.97521354 | 1.271263547 | 8.91E-04 |
| Dvl1 | 2494.315404 | 1.269469938 | 8.39E-22 |
| Edn1 | 882.9784013 | 1.269435922 | 4.80E-06 |

| | | | |
|---|---|---|---|
| Sh2d3c | 108.5494914 | 1.269423604 | 2.31E-05 |
| Nova2 | 53.87519582 | 1.269333925 | 4.12E-04 |
| Thap1 | 316.451354 | 1.265790906 | 2.75E-09 |
| Rps18 | 7591.717016 | 1.265714704 | 1.37E-05 |
| Gdf9 | 43.03072072 | 1.264576755 | 1.16E-03 |
| NA | 56.00787468 | 1.263231351 | 5.98E-03 |
| Rplp2 | 29391.29674 | 1.263160759 | 1.15E-12 |
| Rpl24 | 25807.90155 | 1.261572694 | 2.80E-11 |
| Cdh3 | 15.56937876 | 1.261487948 | 1.97E-02 |
| Rpl32-ps | 16.76282452 | 1.261078194 | 1.88E-02 |
| Rab3ip | 2085.051427 | 1.260684647 | 2.96E-21 |
| Gab2 | 872.0524057 | 1.259459875 | 9.61E-10 |
| Tjp1 | 4773.974161 | 1.25927424 | 1.06E-16 |
| H2-Q7 | 211.7583553 | 1.258611151 | 6.86E-06 |
| Gm6030 | 218.7687813 | 1.258512774 | 6.25E-04 |
| Pnkp | 1016.334908 | 1.258348966 | 2.12E-15 |
| Rps15a-ps7 | 44.62255626 | 1.257657981 | 2.07E-03 |
| Gss | 1651.838005 | 1.256685853 | 5.35E-11 |
| Gm12091 | 282.2413441 | 1.254945822 | 4.57E-04 |
| Trim69 | 27.47810504 | 1.254145684 | 3.23E-03 |
| NA | 16.7581873 | 1.25414345 | 4.71E-02 |
| Lpcat4 | 1034.700321 | 1.253807898 | 3.45E-14 |
| 1700001O22Rik | 11.23019865 | 1.252999437 | 3.97E-02 |
| Bpnt1 | 749.3892407 | 1.252765663 | 3.89E-13 |
| Rpl18a | 66237.11451 | 1.250207127 | 4.03E-08 |
| Il18rap | 98.60169899 | 1.249685395 | 3.69E-04 |
| Gm9726 | 134.0824814 | 1.249166127 | 6.11E-05 |
| Mfrp | 16.50642772 | 1.248698472 | 3.44E-02 |
| Gm9396 | 66.43147602 | 1.247873826 | 1.34E-03 |
| Igfbp6 | 10099.86965 | 1.246829336 | 1.53E-10 |
| Paip2b | 3971.739471 | 1.245293404 | 5.40E-15 |
| Rpl28 | 2509.809573 | 1.24507217 | 1.27E-05 |
| Oxsr1 | 2997.297148 | 1.245017119 | 1.16E-23 |
| Ifi47 | 69.23413701 | 1.244673637 | 8.31E-07 |
| Sesn2 | 1711.20642 | 1.24382266 | 1.55E-12 |
| Pgpep1 | 2247.346939 | 1.242733157 | 5.92E-18 |
| Rps7 | 10324.97548 | 1.24218736 | 2.41E-07 |
| Dst | 14726.69931 | 1.242053969 | 1.28E-12 |
| Anln | 4893.272358 | 1.240211582 | 6.99E-04 |
| Gstt1 | 157.0937604 | 1.240128149 | 2.29E-03 |
| Dqx1 | 97.9924442 | 1.239748524 | 9.54E-06 |
| Adipor1 | 9075.210439 | 1.239660733 | 3.81E-14 |
| Kctd13 | 656.9195456 | 1.23888236 | 1.34E-08 |
| Per3 | 660.4660491 | 1.23747516 | 4.62E-09 |
| Arvcf | 1255.272406 | 1.237266167 | 7.27E-17 |
| Rpl35 | 28922.30866 | 1.236894544 | 1.14E-13 |
| Sphk1 | 627.9533835 | 1.236382538 | 6.94E-05 |

| | | | |
|---|---|---|---|
| Rpl10a-ps1 | 49044.15001 | 1.236163837 | 5.31E-09 |
| Rps6ka5 | 321.2408474 | 1.236094774 | 3.33E-10 |
| Gm20683 | 64.00408063 | 1.235881599 | 1.46E-02 |
| Mif | 7576.735311 | 1.235814634 | 1.79E-10 |
| Mme | 42.60564629 | 1.234887972 | 2.49E-02 |
| Serpina9 | 31.42166562 | 1.232999741 | 2.23E-02 |
| Dpm1 | 807.4452685 | 1.232035209 | 2.86E-07 |
| Creb3l4 | 51.97945249 | 1.231542328 | 4.98E-05 |
| Zfp972 | 96.72997032 | 1.229027383 | 6.21E-05 |
| Gm8116 | 1189.5011 | 1.22781163 | 1.68E-31 |
| Ptges | 1888.104658 | 1.227735454 | 2.94E-09 |
| 2310022A10Rik | 765.7830668 | 1.227119875 | 1.47E-12 |
| Ifi213 | 21.28483327 | 1.226941229 | 1.45E-02 |
| Mcu | 1703.920113 | 1.226210233 | 4.05E-28 |
| Gm13456 | 238.4475069 | 1.225674801 | 2.48E-05 |
| Fam220a | 353.5374561 | 1.225357417 | 1.21E-02 |
| Mroh2a | 196.7055094 | 1.225235576 | 1.46E-03 |
| Rps23 | 23042.03205 | 1.224734642 | 2.95E-07 |
| Zfpm1 | 851.7480048 | 1.224645827 | 8.89E-12 |
| Pla2g4a | 4766.449097 | 1.224174849 | 7.13E-05 |
| Rps25-ps1 | 24.66073837 | 1.22368117 | 2.32E-02 |
| Cdc34 | 2896.485483 | 1.223072507 | 5.29E-13 |
| Gm10335 | 326.9928239 | 1.221958369 | 1.66E-03 |
| Schip1 | 526.8642713 | 1.221406172 | 1.56E-13 |
| Isg20 | 692.6214078 | 1.221098513 | 2.76E-08 |
| Glis2 | 2505.628587 | 1.220470254 | 8.27E-19 |
| Rpl27 | 2340.730614 | 1.220453407 | 6.37E-07 |
| Arhgap33 | 330.2406032 | 1.220104907 | 3.00E-07 |
| Mttp | 478.4641153 | 1.219721689 | 3.11E-23 |
| 6430562O15Rik | 12.17978969 | 1.219218866 | 3.21E-02 |
| Cs | 17852.67489 | 1.219072464 | 2.15E-17 |
| Htra1 | 323.732752 | 1.218979795 | 1.29E-13 |
| Lamb2 | 7100.083165 | 1.218668349 | 5.15E-08 |
| Dmtn | 35.78379888 | 1.218473133 | 3.66E-03 |
| Cav1 | 38978.11609 | 1.218399616 | 6.93E-22 |
| Irf2bpl | 2111.470138 | 1.217777399 | 4.12E-19 |
| Cdc20 | 2852.257054 | 1.217680991 | 2.32E-04 |
| Primpol | 719.4967497 | 1.21706279 | 1.24E-18 |
| Srp72 | 10229.00723 | 1.216689238 | 6.45E-17 |
| Nectin1 | 1412.399235 | 1.21646331 | 7.99E-12 |
| Stim1 | 2475.820194 | 1.21588078 | 5.04E-15 |
| Usp17la | 30.31757862 | 1.215354728 | 1.57E-02 |
| Hspa5 | 48769.31287 | 1.214551108 | 1.28E-07 |
| Rplp0 | 148995.8377 | 1.214526778 | 3.48E-12 |
| Gm12922 | 36.52443524 | 1.214501169 | 1.39E-02 |
| Pdlim7 | 2855.17683 | 1.214498314 | 4.15E-14 |
| Spsb1 | 515.9985486 | 1.2140893 | 5.19E-15 |

| | | | |
|---|---|---|---|
| Rpl15 | 5028.833574 | 1.213502254 | 1.66E-13 |
| Fam107b | 3625.918197 | 1.212522956 | 5.72E-12 |
| Sap18b | 2815.50859 | 1.210667134 | 1.23E-20 |
| 2010015M23Rik | 28.65831723 | 1.209361946 | 2.14E-02 |
| Mras | 3862.451972 | 1.208586761 | 1.09E-11 |
| Gm15500 | 76446.03163 | 1.207840561 | 2.38E-10 |
| Ube2cbp | 39.28225239 | 1.207495361 | 2.58E-03 |
| Pik3cb | 1633.017527 | 1.207043936 | 8.53E-13 |
| Cdh2 | 5735.006876 | 1.20592521 | 1.15E-14 |
| Mss51 | 56.32042687 | 1.205575743 | 3.37E-05 |
| Retreg1 | 1973.380373 | 1.205247076 | 7.52E-11 |
| Cit | 699.3070544 | 1.205007844 | 9.19E-04 |
| Ccdc9b | 3998.146773 | 1.204611397 | 1.91E-21 |
| C9orf72 | 868.8703599 | 1.20421622 | 2.76E-43 |
| Gm5526 | 15.93317991 | 1.203871072 | 2.98E-02 |
| Dele1 | 1859.40767 | 1.202673599 | 2.48E-10 |
| Fitm2 | 1939.475976 | 1.202370081 | 3.91E-12 |
| Nr4a1 | 921.088903 | 1.20225219 | 4.25E-03 |
| Sbds | 2941.875552 | 1.201160039 | 1.30E-21 |
| Ctps | 9423.67962 | 1.20072952 | 2.08E-12 |
| Bex3 | 8754.027769 | 1.200295829 | 4.58E-24 |
| Gm2000 | 6602.439031 | 1.198948025 | 1.42E-06 |
| Gm9843 | 23508.2696 | 1.198931091 | 8.93E-09 |
| Ubxn1 | 7751.977327 | 1.197527295 | 3.42E-25 |
| Cox6a2 | 24.89091578 | 1.197343942 | 3.55E-02 |
| Thoc6 | 1074.782664 | 1.19649941 | 1.98E-14 |
| Sqor | 2866.171031 | 1.195129275 | 8.14E-22 |
| Gm28042 | 211.2134342 | 1.194192351 | 2.03E-04 |
| Gm5121 | 57.77765749 | 1.193909313 | 8.23E-03 |
| Mrps31 | 966.2834087 | 1.193284257 | 8.36E-12 |
| Plpp2 | 931.3288588 | 1.192695471 | 7.21E-14 |
| Grb14 | 1511.859625 | 1.192069621 | 1.03E-06 |
| Ier3 | 749.7978944 | 1.191539104 | 2.36E-06 |
| Tpd52 | 282.1749715 | 1.191361107 | 1.12E-05 |
| Serpini1 | 53.62556717 | 1.191056861 | 8.12E-04 |
| Gm10260 | 27463.1406 | 1.190098716 | 1.70E-09 |
| Mthfd2 | 2675.163557 | 1.189462749 | 2.39E-04 |
| Rpl30 | 7654.741905 | 1.189211937 | 7.16E-13 |
| Ska1 | 241.1235717 | 1.188544467 | 5.80E-03 |
| Tmem170 | 255.3052847 | 1.188229557 | 5.11E-12 |
| Gm10040 | 874.6698116 | 1.188117619 | 2.86E-20 |
| Rpl3-ps2 | 54.61022222 | 1.187644563 | 5.19E-03 |
| Gm10602 | 148.9557599 | 1.186976692 | 5.98E-10 |
| Rps19 | 31869.82538 | 1.186969669 | 1.11E-08 |
| Gm13421 | 26.55001828 | 1.186626137 | 1.63E-02 |
| Ccl9 | 880.6190831 | 1.186254436 | 2.16E-03 |
| Flnc | 35973.77426 | 1.186044776 | 1.05E-11 |

| | | | |
|---|---|---|---|
| Wdr36 | 2910.009891 | 1.185829772 | 4.31E-25 |
| 4930458D05Rik | 61.19370456 | 1.185750982 | 4.95E-03 |
| Reep4 | 971.1311621 | 1.185363937 | 9.62E-09 |
| Zfp354a | 365.4228856 | 1.185324923 | 5.38E-07 |
| Gk | 712.7420813 | 1.185161726 | 2.83E-06 |
| Dgka | 489.6759605 | 1.184746748 | 1.70E-11 |
| Rasl11b | 155.4030989 | 1.183021216 | 1.48E-04 |
| Tcirg1 | 2505.669485 | 1.182132141 | 2.25E-12 |
| 2810025M15Rik | 454.5514841 | 1.181206217 | 8.43E-10 |
| Aldh18a1 | 7021.690676 | 1.180331402 | 4.73E-11 |
| Cenpv | 267.4788372 | 1.180310507 | 5.89E-09 |
| Tbc1d2 | 1016.924895 | 1.179861932 | 4.29E-07 |
| Prpf40b | 1411.837332 | 1.179857515 | 7.81E-11 |
| Adarb2 | 76.97265872 | 1.179634417 | 8.20E-03 |
| Rpl37rt | 7114.695718 | 1.179312085 | 1.40E-10 |
| Susd6 | 4237.928786 | 1.179249778 | 9.57E-09 |
| Mpp6 | 1620.758067 | 1.178903226 | 5.99E-07 |
| Gm8730 | 13578.63654 | 1.17889907 | 6.52E-09 |
| Gm7331 | 343.4208214 | 1.178038007 | 2.84E-03 |
| Eef1b2 | 46128.15298 | 1.17705692 | 6.15E-11 |
| Zfpm2 | 579.7477361 | 1.176511817 | 1.11E-10 |
| Rpl32 | 65196.27315 | 1.176097549 | 3.65E-09 |
| Trim36 | 26.62347626 | 1.176063315 | 2.59E-02 |
| Lrrn4cl | 2335.883644 | 1.175958905 | 1.75E-04 |
| Gramd1a | 2969.753099 | 1.175516343 | 2.95E-07 |
| Zfp335 | 1805.266162 | 1.175504051 | 6.09E-11 |
| Rbm38 | 607.9930154 | 1.175153746 | 1.15E-05 |
| Ipo5 | 14120.82957 | 1.175003099 | 3.15E-19 |
| Utp4 | 3049.021415 | 1.174327541 | 1.60E-16 |
| Esyt1 | 2873.959478 | 1.173326776 | 6.33E-19 |
| Mrpl32 | 969.4241825 | 1.173221973 | 1.14E-21 |
| Nr1h5 | 425.4240054 | 1.172802955 | 2.16E-02 |
| Rpl38-ps2 | 415.2384649 | 1.172479506 | 1.75E-02 |
| Bcorl1 | 1056.977538 | 1.172007762 | 9.89E-10 |
| Gm13864 | 254.081662 | 1.171161485 | 1.55E-05 |
| Rpl19 | 61202.65009 | 1.170813662 | 4.12E-09 |
| Ccdc107 | 815.763532 | 1.170282287 | 2.41E-19 |
| Fbxo47 | 20.02647274 | 1.170119134 | 3.02E-02 |
| Pif1 | 283.1872027 | 1.169992508 | 4.37E-03 |
| Ghitm | 11802.31619 | 1.168112945 | 4.16E-17 |
| Tmem240 | 41.68437518 | 1.167525943 | 1.78E-02 |
| Esrra | 994.4250692 | 1.167320569 | 4.94E-09 |
| Rpl36 | 15843.79799 | 1.165547238 | 5.95E-09 |
| Myl6b | 236.3366549 | 1.164890012 | 6.59E-07 |
| Rpl35a | 516.7149947 | 1.164710056 | 5.17E-09 |
| Slc38a9 | 414.9045467 | 1.163838531 | 8.85E-13 |
| Faf1 | 5358.729333 | 1.163778372 | 8.34E-23 |

| | | | |
|---|---|---|---|
| Grsf1 | 4328.906575 | 1.162493965 | 1.53E-18 |
| Zc2hc1c | 99.23641284 | 1.162386007 | 8.67E-05 |
| Cmip | 3604.487031 | 1.161153072 | 5.74E-19 |
| Abcb6 | 523.6394462 | 1.160972157 | 4.76E-10 |
| Cacnb4 | 40.60664551 | 1.159313398 | 4.52E-02 |
| Rpl6 | 74724.03916 | 1.158774449 | 2.19E-10 |
| Rps19-ps2 | 57.10865603 | 1.157371884 | 9.24E-03 |
| Gabpb1 | 1205.660233 | 1.156855562 | 2.83E-19 |
| Gm7027 | 42.16724364 | 1.156591836 | 5.89E-04 |
| Rtp4 | 57.72055596 | 1.15598532 | 1.31E-05 |
| Synpo2l | 127.4131202 | 1.155975878 | 4.42E-02 |
| Tkt | 7929.85388 | 1.155504031 | 1.90E-15 |
| Gm12918 | 11988.44703 | 1.155303612 | 7.67E-04 |
| Prex2 | 704.5135401 | 1.154756352 | 8.16E-05 |
| Dcbld2 | 3824.810857 | 1.153723754 | 4.18E-11 |
| Lat2 | 39.69412982 | 1.152301258 | 5.53E-03 |
| Rpl11 | 26732.08196 | 1.152198886 | 1.48E-10 |
| Snx15 | 878.8217529 | 1.151998452 | 9.49E-32 |
| Chchd5 | 838.8662133 | 1.151507807 | 4.88E-08 |
| Gm6472 | 47311.23449 | 1.151483559 | 2.06E-09 |
| Wdr89 | 406.7336064 | 1.151318343 | 1.95E-04 |
| Nob1 | 1945.869429 | 1.149390953 | 8.89E-22 |
| Etfbkmt | 562.6280982 | 1.149134893 | 6.70E-07 |
| Gm5786 | 119.6563614 | 1.149085415 | 6.38E-06 |
| Fxr2 | 4780.176202 | 1.148564484 | 7.39E-14 |
| Gm13841 | 29018.88927 | 1.148464267 | 4.50E-09 |
| Tm4sf1 | 4004.71189 | 1.148358723 | 8.27E-10 |
| Gm11353 | 146.2762761 | 1.145580983 | 9.63E-04 |
| Rpl3-ps1 | 93867.25035 | 1.145032756 | 4.22E-09 |
| Bcl2l1 | 4294.000393 | 1.144729443 | 1.42E-12 |
| Pde4b | 1859.321615 | 1.143620028 | 2.25E-04 |
| Gm8172 | 44.9062134 | 1.143615731 | 4.54E-03 |
| Lyz2 | 828.2358917 | 1.143281019 | 2.50E-02 |
| Higd2a | 2214.016855 | 1.142983222 | 6.13E-16 |
| Nkiras2 | 1742.751952 | 1.140004075 | 1.55E-09 |
| 4930558J18Rik | 24.7967093 | 1.139709824 | 3.31E-02 |
| Oas1b | 52.61821612 | 1.138885633 | 2.34E-03 |
| Tpk1 | 262.2730087 | 1.138610642 | 2.83E-10 |
| Sik1 | 2788.494883 | 1.137764589 | 9.31E-06 |
| Gm6177 | 33.69468597 | 1.137354019 | 3.20E-02 |
| Ska3 | 318.64434 | 1.13717898 | 2.85E-04 |
| Neurl1a | 101.6784878 | 1.136535029 | 1.53E-04 |
| Pycr2 | 1798.030111 | 1.135981973 | 3.27E-16 |
| Zyx | 23861.9633 | 1.135730876 | 5.86E-22 |
| Gm3362 | 137.0655522 | 1.135012049 | 2.78E-04 |
| Ccdc80 | 6478.722348 | 1.134777406 | 7.25E-06 |
| Grik5 | 256.8579891 | 1.134693962 | 8.81E-08 |

| | | | |
|---|---|---|---|
| Tarsl2 | 774.4453427 | 1.134128719 | 6.52E-22 |
| Eef1a1 | 672249.7408 | 1.133954098 | 2.41E-08 |
| Tnfrsf22 | 1190.813432 | 1.133845839 | 3.92E-07 |
| Rpl14 | 39168.7694 | 1.133715959 | 2.60E-07 |
| NA | 127.6455783 | 1.133537937 | 5.08E-05 |
| Afmid | 240.573853 | 1.133534289 | 9.89E-06 |
| Rpl10 | 12459.82895 | 1.13350992 | 3.74E-06 |
| Adprm | 564.4792591 | 1.133011293 | 4.90E-14 |
| Prc1 | 2935.87793 | 1.131973738 | 5.66E-04 |
| Usp30 | 1743.102786 | 1.131956732 | 5.57E-11 |
| Nupr1 | 2029.051611 | 1.130282083 | 1.22E-05 |
| Gga3 | 928.5292727 | 1.129793362 | 1.01E-09 |
| Glrx3 | 6612.60578 | 1.129707314 | 2.97E-15 |
| Nfkb2 | 5751.40937 | 1.128756177 | 5.34E-11 |
| Gm15542 | 49.28181558 | 1.128740507 | 1.54E-02 |
| Adora2b | 955.5375759 | 1.127883243 | 1.93E-09 |
| Csnk1e | 4058.989069 | 1.127749643 | 7.92E-26 |
| Amacr | 720.3402139 | 1.127584247 | 1.99E-11 |
| Gm11449 | 50.67927013 | 1.127579909 | 4.87E-02 |
| Psat1 | 15457.79927 | 1.127358794 | 1.47E-04 |
| Cntnap1 | 62.99681169 | 1.126721744 | 1.95E-04 |
| Gm6767 | 22.91698675 | 1.126514989 | 3.71E-02 |
| Atxn2l | 6670.119001 | 1.126372962 | 1.34E-21 |
| Cdiptos | 63.14051489 | 1.125090326 | 2.11E-03 |
| Zfat | 598.1955488 | 1.124989332 | 8.52E-07 |
| Klhl26 | 2008.492919 | 1.124678674 | 1.68E-18 |
| Rplp1 | 72292.15409 | 1.123610984 | 1.64E-08 |
| Sgtb | 669.2050801 | 1.122726351 | 1.04E-13 |
| Gm21817 | 26.91196458 | 1.120430069 | 1.06E-02 |
| Gm11687 | 35.35260304 | 1.120236307 | 9.17E-03 |
| Clcn3 | 952.4309388 | 1.119775943 | 1.73E-05 |
| AU020206 | 146.3601913 | 1.118025296 | 2.41E-11 |
| Gm10076 | 56.07162855 | 1.116995243 | 2.36E-03 |
| Lats2 | 3075.227714 | 1.116749119 | 9.85E-15 |
| Dlgap4 | 3400.879371 | 1.116661769 | 1.44E-09 |
| Rpl38 | 13527.73583 | 1.116409571 | 2.98E-09 |
| Baiap2l1 | 282.5346504 | 1.116318856 | 6.60E-05 |
| Gm26694 | 109.4297847 | 1.116113144 | 3.43E-04 |
| Clic4 | 7974.140184 | 1.115712839 | 5.67E-10 |
| Bax | 4107.204276 | 1.114594695 | 4.16E-12 |
| Synpo2 | 219.9091425 | 1.112613093 | 5.70E-05 |
| Cyb5r1 | 3302.743638 | 1.112077802 | 1.98E-06 |
| Gm26909 | 21.61032335 | 1.111806895 | 1.28E-02 |
| Gm12599 | 17.24211812 | 1.110536007 | 4.27E-02 |
| Eif4ebp1 | 7809.692884 | 1.110041586 | 1.53E-16 |
| Klf7 | 2129.134471 | 1.108574056 | 3.65E-11 |
| Fez2 | 3426.090875 | 1.108222755 | 3.76E-10 |

| | | | |
|---|---|---|---|
| Vpreb1 | 24.29579557 | 1.107319327 | 1.50E-02 |
| Napb | 476.5876789 | 1.10678199 | 4.66E-12 |
| Fam110c | 419.3831265 | 1.106691309 | 5.72E-07 |
| Fosl1 | 1007.39669 | 1.106211639 | 6.64E-04 |
| Tmpo | 7007.127582 | 1.106038138 | 4.78E-10 |
| Podnl1 | 3817.279448 | 1.105681347 | 2.98E-11 |
| Cpt1c | 488.9285033 | 1.105592923 | 2.02E-15 |
| Rpl19-ps11 | 96.05830387 | 1.105560012 | 8.73E-03 |
| Tmem209 | 1826.003205 | 1.104819348 | 3.91E-17 |
| Ptcd2 | 866.1635995 | 1.104646977 | 2.66E-18 |
| S100a11 | 28004.48893 | 1.104560702 | 4.84E-07 |
| Rnf31 | 992.4717778 | 1.104242092 | 2.03E-08 |
| Platr20 | 22.75396567 | 1.103743597 | 3.16E-02 |
| Ube2g2 | 3636.513381 | 1.103605984 | 4.59E-10 |
| NA | 8103.204573 | 1.103132427 | 7.55E-09 |
| Sh3bp1 | 1826.718969 | 1.103007483 | 6.25E-14 |
| Gm9013 | 38.9956223 | 1.102917608 | 1.58E-02 |
| Tsen54 | 797.3115622 | 1.10190089 | 1.55E-12 |
| Mak16 | 2982.450982 | 1.099888514 | 8.16E-17 |
| Ddx11 | 404.9730421 | 1.099316009 | 3.43E-04 |
| Traf2 | 1462.06301 | 1.098940952 | 2.94E-08 |
| Rpl41 | 80605.63953 | 1.098773671 | 1.13E-07 |
| Lrrc1 | 506.016248 | 1.097477744 | 2.88E-10 |
| Rps11 | 38460.82027 | 1.096277909 | 9.92E-09 |
| Cdc42ep3 | 3187.14258 | 1.096214373 | 7.98E-15 |
| Gm17638 | 49.71635285 | 1.096203217 | 6.92E-05 |
| Gm20517 | 50.83540261 | 1.095846828 | 4.17E-04 |
| Efnb3 | 30.29227065 | 1.095567755 | 6.98E-03 |
| Rpl34-ps1 | 26642.99647 | 1.095108508 | 5.28E-09 |
| Slc5a5 | 16.03111384 | 1.09470642 | 3.62E-02 |
| Brsk1 | 879.0230187 | 1.093968807 | 4.23E-10 |
| Ccnb1 | 1707.028829 | 1.093955004 | 2.79E-03 |
| Pxdc1 | 198.4170374 | 1.0933414 | 2.53E-06 |
| Psmd8 | 7697.479585 | 1.092963115 | 1.24E-08 |
| Gm16477 | 34.24256402 | 1.091178591 | 1.24E-03 |
| Ube2t | 295.3875627 | 1.090208005 | 2.25E-03 |
| Prdm15 | 720.4881743 | 1.090160473 | 2.07E-08 |
| Klf11 | 571.7586071 | 1.090114396 | 6.05E-05 |
| Rpl7a-ps11 | 16425.80428 | 1.089552381 | 5.11E-08 |
| Mapk8ip3 | 4942.510858 | 1.08908903 | 8.89E-12 |
| Rpp30 | 619.7082099 | 1.088957496 | 4.07E-11 |
| Slc25a39 | 8371.758463 | 1.088697909 | 4.24E-11 |
| Grk2 | 3292.244991 | 1.086614858 | 9.87E-21 |
| Pef1 | 4262.675557 | 1.086474851 | 3.19E-09 |
| NA | 1282.239263 | 1.086333451 | 1.44E-04 |
| Tnpo2 | 9752.197038 | 1.083928711 | 2.89E-14 |
| Lrrc8e | 165.339491 | 1.082347769 | 2.89E-06 |

| | | | |
|---|---|---|---|
| Steap2 | 5089.313269 | 1.082104444 | 1.04E-07 |
| Cxxc5 | 218.8335225 | 1.081934429 | 5.21E-06 |
| AA467197 | 26.56671516 | 1.081243835 | 2.56E-02 |
| Fam198b | 10855.77158 | 1.080280129 | 1.92E-06 |
| Rpl30-ps10 | 596.1498409 | 1.079596791 | 1.53E-02 |
| Pcdh1 | 2459.618889 | 1.079158133 | 1.83E-12 |
| Gm6451 | 47.40372897 | 1.077787171 | 2.50E-02 |
| Rpl31-ps11 | 31.77537998 | 1.076951802 | 1.50E-02 |
| Clcn2 | 458.1085294 | 1.076627033 | 3.34E-10 |
| Sik3 | 2887.657643 | 1.076534412 | 2.88E-14 |
| Rps20 | 53567.39921 | 1.075810096 | 5.05E-07 |
| Pam16 | 808.4039737 | 1.075748766 | 1.04E-16 |
| 3110009E18Rik | 177.7241509 | 1.075231022 | 5.37E-04 |
| Eif3h | 21119.97463 | 1.075103809 | 3.89E-07 |
| Ints12 | 1233.993372 | 1.07469311 | 2.22E-09 |
| Zfp809 | 553.769558 | 1.074477392 | 5.82E-09 |
| Gm14226 | 33.86813149 | 1.073920504 | 2.78E-03 |
| Col11a1 | 3443.82184 | 1.073642792 | 3.43E-02 |
| Gm1840 | 48.69248908 | 1.07363978 | 1.37E-02 |
| Rcc1 | 1466.074838 | 1.073617026 | 3.71E-08 |
| Naa16 | 558.3411683 | 1.073500762 | 6.07E-10 |
| Tbpl1 | 1549.163799 | 1.072791579 | 2.13E-24 |
| 1190002N15Rik | 866.5027261 | 1.072655261 | 4.74E-07 |
| Ptgis | 3691.451478 | 1.071998825 | 6.63E-06 |
| Mybl1 | 202.1011964 | 1.071308067 | 2.04E-02 |
| Hspb11 | 1261.435509 | 1.071166622 | 1.32E-09 |
| Ttc1 | 1980.789827 | 1.070857002 | 2.64E-15 |
| Rpl37 | 20885.2681 | 1.070776909 | 2.68E-08 |
| Troap | 510.2589497 | 1.070313421 | 7.51E-03 |
| Dlg4 | 3652.121914 | 1.069722808 | 5.43E-13 |
| Dusp11 | 7910.266802 | 1.068895368 | 7.75E-12 |
| Gm14633 | 22.79272291 | 1.068480087 | 3.15E-02 |
| Nans | 1668.575675 | 1.067744749 | 4.99E-09 |
| Gm15501 | 77795.36435 | 1.06746263 | 1.61E-08 |
| Lzic | 586.7404707 | 1.067120931 | 9.89E-10 |
| Prkd1 | 1467.094425 | 1.06699689 | 4.58E-17 |
| Tbc1d8 | 59.18404534 | 1.06650795 | 3.03E-04 |
| A330069E16Rik | 17.58090328 | 1.064970949 | 4.27E-02 |
| Pithd1 | 813.3192727 | 1.064601275 | 1.11E-09 |
| 4-Sep | 46.97005786 | 1.064145003 | 2.09E-02 |
| Spag5 | 1260.809925 | 1.064024997 | 3.84E-03 |
| Gm5879 | 60.46901209 | 1.062659587 | 4.21E-02 |
| Kif5a | 70.92537186 | 1.062612236 | 1.21E-02 |
| Nosip | 1483.982081 | 1.06255976 | 6.01E-12 |
| Prkg1 | 134.4890151 | 1.062302831 | 3.00E-03 |
| Pitpnm1 | 3326.041469 | 1.062131051 | 9.22E-24 |
| Mical1 | 409.2832872 | 1.061879667 | 5.59E-08 |

| | | | |
|---|---|---|---|
| Shmt2 | 7386.878222 | 1.060469218 | 8.90E-05 |
| Lin37 | 543.5409492 | 1.059082369 | 1.33E-08 |
| Heca | 1189.435065 | 1.058787003 | 3.03E-11 |
| Map3k13 | 64.74950077 | 1.05807822 | 1.11E-04 |
| Ifit1bl2 | 54.30370509 | 1.057053273 | 4.70E-03 |
| Chp1 | 6682.072199 | 1.056540525 | 1.31E-09 |
| Actn3 | 77.20788642 | 1.056482115 | 4.94E-04 |
| Magohb | 625.3616612 | 1.056211826 | 8.89E-06 |
| Gm16982 | 30.26455867 | 1.055598069 | 1.09E-02 |
| Gm7488 | 1833.106868 | 1.055557992 | 4.66E-16 |
| Adss | 4524.176795 | 1.054472467 | 5.08E-19 |
| Zcwpw2 | 14.80335001 | 1.054254236 | 3.51E-02 |
| Rps8 | 13664.61444 | 1.054169748 | 1.65E-06 |
| Amn1 | 462.9509753 | 1.053973809 | 2.85E-07 |
| NA | 140.031756 | 1.053349095 | 6.34E-08 |
| Rpl7a | 25955.19859 | 1.05319352 | 9.56E-08 |
| Fhl2 | 3219.077124 | 1.052975905 | 2.06E-11 |
| Mnd1 | 246.5320564 | 1.052786952 | 1.46E-09 |
| Grhl1 | 28.30210692 | 1.0520402 | 9.22E-03 |
| Gcnt2 | 266.7781291 | 1.05196818 | 1.71E-05 |
| Ticam1 | 1747.623083 | 1.051740198 | 2.27E-06 |
| Pycr1 | 948.3752732 | 1.051017174 | 8.47E-05 |
| Gm26703 | 14.15106203 | 1.048331626 | 4.24E-02 |
| Gm26619 | 12811.53859 | 1.048045134 | 4.34E-05 |
| Aldh1l2 | 7819.791201 | 1.047411387 | 1.93E-05 |
| Fip1l1 | 6051.922285 | 1.04736051 | 3.54E-09 |
| Elp6 | 429.5943132 | 1.047056125 | 4.12E-08 |
| Gm20628 | 35.31350781 | 1.046793438 | 3.31E-02 |
| Gm6436 | 10382.66031 | 1.046610941 | 5.69E-06 |
| Ptpn14 | 6806.862281 | 1.046504063 | 2.16E-24 |
| Dlgap1 | 42.72816766 | 1.046482507 | 1.13E-02 |
| Gm14427 | 29.03266401 | 1.045381092 | 3.01E-02 |
| Map6 | 3232.025626 | 1.045127713 | 9.91E-09 |
| Lurap1l | 1882.936888 | 1.044867064 | 5.60E-11 |
| Atp1a1 | 13088.27316 | 1.044142242 | 3.96E-12 |
| Eif3i | 12035.09517 | 1.04392276 | 9.61E-10 |
| Rps15a-ps3 | 65.56143529 | 1.043302336 | 1.14E-03 |
| Emd | 2493.987764 | 1.041496634 | 2.87E-11 |
| Gas5 | 17534.6299 | 1.040951462 | 3.76E-06 |
| Usp36 | 1623.508219 | 1.040613794 | 1.93E-09 |
| Pa2g4 | 8738.523288 | 1.039645242 | 8.60E-10 |
| A130014A01Rik | 79.0835431 | 1.039126183 | 1.18E-02 |
| Rpl31-ps16 | 94.33671546 | 1.038926342 | 8.75E-03 |
| Plekhf2 | 391.5956648 | 1.038638292 | 3.48E-10 |
| Efnb1 | 347.491773 | 1.036532944 | 1.96E-07 |
| Gpx4 | 15070.71857 | 1.036167944 | 2.46E-09 |
| Fam214b | 1462.593913 | 1.035959302 | 5.79E-11 |

| | | | |
|---|---|---|---|
| Hbs1l | 4908.209871 | 1.034964043 | 1.47E-38 |
| Rps13-ps5 | 26.70885885 | 1.034895454 | 2.41E-02 |
| Atp5d | 7992.530441 | 1.034785193 | 4.90E-13 |
| Sod1 | 7945.096107 | 1.034511918 | 5.54E-15 |
| Mthfd1l | 1964.85493 | 1.034337697 | 5.29E-05 |
| Mrpl38 | 1679.111537 | 1.033722034 | 1.07E-14 |
| Nr4a2 | 174.1972914 | 1.033131923 | 3.67E-05 |
| Tex30 | 300.7795105 | 1.033054477 | 1.65E-03 |
| Pfkm | 3671.085007 | 1.032463137 | 2.03E-12 |
| Hmga1 | 10223.72691 | 1.032400135 | 1.08E-04 |
| Gm17709 | 122.5539367 | 1.031637901 | 2.40E-07 |
| Serpinb6a | 9297.117704 | 1.030896132 | 7.07E-19 |
| Opn3 | 112.0638704 | 1.030303877 | 1.36E-06 |
| Rps15a | 30497.77941 | 1.029901587 | 1.86E-08 |
| Samhd1 | 3737.536878 | 1.02958158 | 7.21E-16 |
| Afdn | 6938.094101 | 1.029250764 | 3.90E-11 |
| Nsa2 | 9495.710085 | 1.029082499 | 7.68E-08 |
| Zfp428 | 275.3299011 | 1.028577704 | 5.07E-08 |
| Eef1g | 80391.92701 | 1.028524121 | 1.47E-10 |
| Pdgfc | 1278.888861 | 1.02804733 | 1.68E-09 |
| Wdr74 | 1689.807573 | 1.027777354 | 3.40E-11 |
| Gm12191 | 13447.05379 | 1.026706333 | 1.58E-09 |
| Trim47 | 1273.010191 | 1.026333795 | 2.66E-08 |
| Gm6286 | 108.3360089 | 1.026262645 | 1.08E-03 |
| Lrrc8c | 1417.4025 | 1.026232195 | 1.52E-07 |
| Fbxl8 | 383.1510087 | 1.025447596 | 1.97E-06 |
| Ppme1 | 2062.268588 | 1.024990727 | 1.73E-14 |
| Smndc1 | 2122.331944 | 1.024473645 | 7.27E-15 |
| Nfe2l1 | 17080.71886 | 1.022958743 | 7.26E-15 |
| Ighmbp2 | 888.9972279 | 1.022881051 | 4.15E-17 |
| Agpat3 | 4907.691748 | 1.022272327 | 4.05E-19 |
| Ssr4 | 4717.047134 | 1.022078289 | 1.13E-07 |
| Ccdc15 | 119.2153648 | 1.021566081 | 1.77E-05 |
| Gm340 | 499.4293114 | 1.021493103 | 4.02E-11 |
| Mta3 | 1941.017603 | 1.02143056 | 5.92E-17 |
| Rpl39 | 31120.35881 | 1.021317241 | 6.95E-06 |
| Serbp1 | 24175.27377 | 1.020794066 | 1.26E-19 |
| Usp35 | 407.4218091 | 1.02076463 | 1.11E-04 |
| Btf3 | 18129.28541 | 1.020363979 | 1.15E-11 |
| Gm12696 | 377.0348595 | 1.020147865 | 2.15E-09 |
| Cthrc1 | 357.533666 | 1.019998037 | 5.98E-07 |
| Rnps1 | 1965.837119 | 1.018638803 | 1.34E-09 |
| Hsd3b7 | 1224.252911 | 1.018612611 | 1.12E-07 |
| Pml | 1978.102798 | 1.018218425 | 2.02E-10 |
| Gm8113 | 678.7557631 | 1.018071221 | 4.72E-04 |
| Pwwp2b | 1031.015521 | 1.017550069 | 5.92E-07 |
| Mocos | 401.6732574 | 1.017028477 | 3.92E-08 |

| | | | |
|---|---|---|---|
| Eif3g | 4110.815225 | 1.016414261 | 1.11E-12 |
| Eml5 | 284.730193 | 1.016043354 | 2.42E-03 |
| Cars | 3768.251332 | 1.015862896 | 3.46E-16 |
| Mgat4b | 6635.711925 | 1.015353594 | 1.60E-09 |
| Tpst2 | 2312.601895 | 1.014851201 | 2.13E-10 |
| Fads3 | 4292.424506 | 1.014453998 | 4.31E-05 |
| Eif3f | 13440.69317 | 1.014420451 | 5.88E-05 |
| Vegfa | 2252.329387 | 1.014139046 | 4.75E-13 |
| Rps6 | 9260.197394 | 1.013958466 | 9.63E-07 |
| Mark4 | 1158.332822 | 1.013745952 | 1.76E-07 |
| Phactr4 | 1891.975684 | 1.013729603 | 1.41E-08 |
| Rpl6l | 113.5010521 | 1.013497947 | 3.00E-02 |
| Tubg1 | 1655.844266 | 1.012044606 | 1.07E-05 |
| C920021L13Rik | 43.17377465 | 1.010779562 | 5.54E-03 |
| Trim11 | 1637.383168 | 1.009667597 | 5.18E-15 |
| Qars | 5609.254646 | 1.009521665 | 1.59E-08 |
| Cobll1 | 2804.002598 | 1.00898361 | 3.08E-14 |
| Cct5 | 20882.64327 | 1.00896518 | 2.48E-20 |
| Spire2 | 545.5638821 | 1.008584011 | 2.44E-13 |
| Ms4a4d | 289.8013129 | 1.007624028 | 3.78E-02 |
| Ulbp1 | 1078.85357 | 1.007055411 | 3.39E-04 |
| Oser1 | 2251.689214 | 1.006584461 | 1.81E-07 |
| Nfil3 | 1259.058948 | 1.006285035 | 1.98E-06 |
| Kdm6b | 4030.264971 | 1.005725836 | 2.69E-06 |
| Cisd1 | 1216.639511 | 1.005225403 | 2.09E-08 |
| Klc4 | 995.6251541 | 1.004460505 | 2.90E-05 |
| Abca4 | 66.55109914 | 1.004311272 | 9.45E-03 |
| Rps24 | 46619.81007 | 1.003647662 | 2.81E-09 |
| Nrp1 | 3718.261942 | 1.003245638 | 5.46E-08 |
| Nme2 | 20448.66004 | 1.002895842 | 8.26E-17 |
| Sync | 42.33056854 | 1.002820812 | 2.04E-02 |
| Asap3 | 387.600447 | 1.002361046 | 3.56E-05 |
| Rps8-ps4 | 109.3114323 | 1.002008483 | 1.27E-02 |
| Mybl2 | 567.7128471 | 1.001790562 | 7.38E-03 |
| Plekhh3 | 1084.0824 | 1.001179019 | 2.71E-09 |
| Ogfod1 | 1856.045924 | 1.000675941 | 3.37E-14 |
| Sac3d1 | 362.1060275 | 1.000289891 | 6.50E-04 |
| Neil1 | 265.1797796 | 0.99983591 | 1.05E-05 |
| Oxa1l | 3698.955215 | 0.99979609 | 1.50E-05 |
| Ptprf | 4297.628625 | 0.99970749 | 5.53E-27 |
| Ppm1d | 820.5467079 | 0.999422058 | 1.54E-09 |
| D16Ertd472e | 232.6290134 | 0.999312656 | 3.45E-12 |
| Cbx4 | 917.6543838 | 0.999016118 | 2.36E-07 |
| Vps37b | 1439.0067 | 0.998585757 | 3.17E-05 |
| Zfp579 | 654.3509177 | 0.998501595 | 1.72E-06 |
| Carhsp1 | 4864.078369 | 0.997887604 | 8.51E-11 |
| Kmt5a | 3867.372276 | 0.995944173 | 1.68E-13 |

| | | | |
|---|---|---|---|
| Kif20a | 2457.420305 | 0.99591516 | 5.54E-03 |
| Rpp40 | 775.7960989 | 0.995729698 | 5.56E-07 |
| Tubb4b | 9221.441394 | 0.995635954 | 3.93E-04 |
| Ppp1r11 | 1908.709021 | 0.995494898 | 6.43E-07 |
| Gfpt2 | 60.13655232 | 0.995201998 | 1.18E-02 |
| Ndufs2 | 12117.44549 | 0.995155121 | 1.30E-10 |
| 1700047I17Rik2 | 769.1116899 | 0.995068342 | 1.98E-10 |
| Map1a | 6290.433199 | 0.994629148 | 1.09E-12 |
| Aph1b | 607.0118737 | 0.994604876 | 5.26E-08 |
| Sil1 | 1576.433198 | 0.994227178 | 3.91E-06 |
| Cdca8 | 1439.881188 | 0.993582852 | 3.38E-03 |
| Irgm1 | 1281.382103 | 0.99344514 | 2.07E-14 |
| Rbms2 | 13507.54112 | 0.992789848 | 8.08E-07 |
| Gtf3c1 | 5130.112731 | 0.991956482 | 6.80E-12 |
| NA | 28.31675729 | 0.991561587 | 2.45E-02 |
| Smim26 | 239.5610761 | 0.991277048 | 1.70E-06 |
| NA | 468.7515269 | 0.991046643 | 1.44E-02 |
| Rbm28 | 3905.034538 | 0.990985807 | 3.94E-11 |
| Plcxd1 | 187.9861981 | 0.99087546 | 5.95E-04 |
| Sars | 17432.90906 | 0.990856172 | 3.99E-15 |
| Rbks | 25.29987498 | 0.990504985 | 4.08E-02 |
| Lig3 | 2759.69298 | 0.990246975 | 1.58E-21 |
| Gars | 14768.92027 | 0.989704907 | 2.66E-11 |
| Trip6 | 4067.669331 | 0.988922054 | 2.66E-08 |
| Psph | 959.6907578 | 0.988832944 | 2.97E-03 |
| Ptch1 | 475.913733 | 0.988714431 | 1.76E-10 |
| Emc6 | 1268.950995 | 0.988649275 | 2.10E-27 |
| Myo1c | 17433.95275 | 0.98862533 | 6.13E-12 |
| Cdt1 | 2176.814691 | 0.98860171 | 2.35E-04 |
| Slc48a1 | 9279.752174 | 0.98775171 | 2.37E-09 |
| Prkd2 | 613.8018891 | 0.986741923 | 6.62E-08 |
| Cenpn | 611.6297696 | 0.986722731 | 6.10E-04 |
| Rpl31-ps8 | 39422.84122 | 0.986513007 | 3.24E-06 |
| S100a3 | 56.32424442 | 0.986454936 | 1.86E-02 |
| Riok3 | 2754.746068 | 0.986125553 | 1.63E-14 |
| Fam57b | 17.95663098 | 0.986093237 | 4.61E-02 |
| Pan2 | 1537.473845 | 0.985988185 | 8.63E-27 |
| Rassf5 | 557.4393745 | 0.98533826 | 9.95E-06 |
| Polb | 1158.380286 | 0.985159097 | 3.61E-14 |
| 4930451G09Rik | 46.88870039 | 0.984877731 | 3.92E-04 |
| H2-T-ps | 256.5588408 | 0.984691163 | 6.68E-04 |
| Nap1l1 | 24172.49471 | 0.983832124 | 1.26E-17 |
| Rasa4 | 4237.672709 | 0.981506569 | 1.08E-08 |
| Cdc42se2 | 2787.71278 | 0.981139975 | 6.17E-30 |
| Mrpl45 | 1814.459067 | 0.981112159 | 3.46E-11 |
| Ptprk | 4819.221325 | 0.980679679 | 1.51E-08 |
| Cdk6 | 743.1289463 | 0.979008271 | 2.08E-09 |

| | | | |
|---|---|---|---|
| Sms-ps | 1178.87176 | 0.978675532 | 5.03E-07 |
| Parp6 | 1359.090638 | 0.977949508 | 1.91E-09 |
| Espl1 | 1089.594393 | 0.976593015 | 1.17E-02 |
| Vrk2 | 912.3379689 | 0.976106236 | 2.51E-16 |
| Spata6 | 2206.374903 | 0.976072142 | 1.92E-13 |
| Cmtr1 | 4053.649139 | 0.97574475 | 3.04E-17 |
| Txnrd1 | 11050.67123 | 0.975645361 | 3.57E-08 |
| Mfge8 | 10440.85802 | 0.975411418 | 9.71E-06 |
| Mtmr10 | 1541.372356 | 0.975214024 | 7.11E-08 |
| Smg9 | 787.7115908 | 0.973571003 | 6.62E-08 |
| Zfp575 | 56.61006131 | 0.972045011 | 5.34E-04 |
| Gm7910 | 157.1334931 | 0.971955378 | 1.17E-02 |
| Ets2 | 2152.537196 | 0.969991931 | 1.12E-09 |
| Rps17 | 34211.18673 | 0.969812097 | 8.70E-08 |
| Ppp2r5b | 1603.372505 | 0.968717468 | 2.47E-14 |
| Fam131a | 31.76925957 | 0.968692058 | 7.10E-03 |
| Unc5b | 390.1679829 | 0.968631027 | 1.10E-03 |
| Hoxc4 | 166.5136363 | 0.967723073 | 7.15E-08 |
| Klhdc8a | 1128.698314 | 0.967560957 | 1.95E-03 |
| Zfp599 | 148.7800721 | 0.967540574 | 2.09E-04 |
| Atp7b | 74.21699247 | 0.96663183 | 1.78E-04 |
| Nsl1 | 311.0216032 | 0.966144778 | 8.17E-03 |
| Aifm2 | 1026.37371 | 0.965902052 | 3.93E-17 |
| Smpd3 | 867.4805472 | 0.965897439 | 3.87E-06 |
| Gtpbp4 | 3254.645096 | 0.965140127 | 5.79E-19 |
| Dgkd | 3573.208434 | 0.964839739 | 3.50E-35 |
| Map1b | 6864.42189 | 0.964751113 | 4.44E-09 |
| Traf4 | 3041.560397 | 0.963217291 | 2.41E-07 |
| Sidt2 | 5631.235272 | 0.96317513 | 1.92E-10 |
| Phlda3 | 4700.215127 | 0.963158241 | 6.08E-05 |
| Vsir | 66.02772379 | 0.962563231 | 8.73E-03 |
| Slc8b1 | 1315.172544 | 0.962193168 | 8.81E-08 |
| Slc8a3 | 48.28956434 | 0.962064566 | 3.94E-02 |
| Sbno2 | 2509.386322 | 0.961466914 | 2.60E-10 |
| Dixdc1 | 787.0275851 | 0.96069597 | 6.73E-06 |
| Pcdh10 | 140.2969449 | 0.959461538 | 5.33E-04 |
| Hectd2 | 366.6262872 | 0.959391192 | 4.49E-04 |
| Slc6a9 | 2482.059137 | 0.959274963 | 2.03E-12 |
| Pik3cd | 389.1741217 | 0.95925066 | 3.33E-06 |
| Gpi1 | 20361.72384 | 0.95846314 | 7.32E-09 |
| Gm13680 | 112.4082536 | 0.958281549 | 2.84E-03 |
| Eef1d | 9782.425807 | 0.957313702 | 5.37E-09 |
| Rps27rt | 30142.86293 | 0.956443647 | 2.91E-05 |
| Cubn | 333.1584532 | 0.95525626 | 6.32E-04 |
| Sgo1 | 501.443209 | 0.955021607 | 3.41E-02 |
| Cacnb3 | 4366.285053 | 0.954913392 | 8.42E-06 |
| Gm9701 | 21363.22693 | 0.954743358 | 2.57E-11 |

| | | | |
|---|---|---|---|
| Utp14a | 2739.522103 | 0.953941753 | 2.12E-13 |
| Psmd4 | 6484.294445 | 0.953355681 | 5.47E-08 |
| Arhgap4 | 43.72618217 | 0.952976762 | 2.15E-02 |
| Mbd1 | 1415.633505 | 0.952754424 | 9.53E-06 |
| Rps27l | 3039.048456 | 0.95219781 | 9.17E-10 |
| Lias | 1391.859481 | 0.951542165 | 9.70E-20 |
| Txnl1 | 3876.007935 | 0.950997883 | 1.36E-18 |
| Cd81 | 15598.87096 | 0.950647657 | 1.48E-07 |
| Gm5566 | 46.62608169 | 0.950552981 | 2.37E-03 |
| Mrps5 | 2126.38738 | 0.949792434 | 1.56E-22 |
| Gm15710 | 93.55804308 | 0.949351221 | 6.70E-03 |
| C2 | 32.68304212 | 0.948456092 | 1.61E-02 |
| Rps23-ps2 | 93.85077079 | 0.948182794 | 1.44E-03 |
| 1810010H24Rik | 33.02850824 | 0.94792057 | 2.98E-03 |
| Copg2 | 4967.60205 | 0.947321744 | 8.99E-13 |
| Pabpc4 | 13810.91171 | 0.947234153 | 6.77E-12 |
| Gm6410 | 85.28209271 | 0.946985742 | 2.56E-03 |
| Cenpa | 1628.559306 | 0.946717223 | 9.22E-04 |
| Taf5 | 394.9900112 | 0.946016018 | 8.42E-07 |
| Eif3m | 5716.330193 | 0.945909709 | 6.77E-12 |
| Uba52-ps | 41.60175151 | 0.945582428 | 2.21E-02 |
| Tmem161a | 1680.07129 | 0.944946262 | 5.93E-15 |
| NA | 216.7207786 | 0.944507435 | 1.34E-02 |
| Gpt2 | 2629.31278 | 0.943564126 | 1.17E-07 |
| Foxo3 | 3075.973274 | 0.943479848 | 5.37E-09 |
| Rbmx2 | 337.2129035 | 0.943367498 | 9.14E-07 |
| Ccnb2 | 359.5961839 | 0.942514145 | 2.66E-03 |
| Setd6 | 561.7236217 | 0.942415047 | 3.22E-15 |
| Reps1 | 2264.486585 | 0.941401358 | 1.71E-17 |
| Zc3hav1 | 5239.317525 | 0.941276774 | 2.71E-11 |
| Dedd2 | 306.1735718 | 0.940118038 | 6.36E-06 |
| Mpi | 945.5748437 | 0.938725952 | 4.44E-08 |
| Mrpl30 | 2494.206912 | 0.938372916 | 2.04E-08 |
| Trp53cor1 | 155.8446845 | 0.93735547 | 6.90E-06 |
| Asrgl1 | 595.939998 | 0.937119297 | 4.20E-05 |
| Gm26724 | 228.142172 | 0.936985864 | 1.36E-03 |
| Cox18 | 676.0390405 | 0.936810634 | 1.73E-10 |
| Sh3glb1 | 7842.113378 | 0.93666849 | 1.29E-16 |
| Psmd5 | 3176.614808 | 0.936645873 | 2.19E-12 |
| Rrp1b | 1636.99729 | 0.936572922 | 1.22E-17 |
| Snupn | 481.3671527 | 0.936167959 | 1.02E-16 |
| Sp1 | 6965.188488 | 0.935794687 | 3.93E-10 |
| Zfp993 | 76.49799003 | 0.935257484 | 1.65E-02 |
| Polr1d | 2742.962488 | 0.934968942 | 2.91E-18 |
| 1700021F05Rik | 907.3929152 | 0.934783449 | 3.40E-27 |
| Rad23b | 17055.86365 | 0.934640544 | 2.48E-11 |
| Ccdc43 | 996.7600218 | 0.934172739 | 2.34E-19 |

| | | | |
|---|---|---|---|
| Apoe | 75.61519687 | 0.933914456 | 1.23E-03 |
| Dtx2 | 985.8070884 | 0.932714722 | 1.12E-07 |
| Gadd45a | 2223.304238 | 0.932560827 | 6.64E-04 |
| Patl1 | 4016.791295 | 0.931791235 | 1.25E-06 |
| Tmem38b | 711.4493136 | 0.931535527 | 2.46E-05 |
| Tube1 | 230.4590009 | 0.931072521 | 3.24E-04 |
| Asnsd1 | 5010.281596 | 0.930802066 | 6.10E-11 |
| S100a13 | 308.5479394 | 0.929921277 | 5.89E-09 |
| Ppp6c | 1863.56707 | 0.92947747 | 8.92E-17 |
| Mta1 | 6436.498902 | 0.927305316 | 2.08E-12 |
| NA | 59602.75301 | 0.92642757 | 9.32E-11 |
| Usp20 | 1287.726103 | 0.925954484 | 4.31E-10 |
| Dcp1b | 84.1534547 | 0.924864131 | 2.47E-02 |
| Zfp703 | 916.2166658 | 0.924615417 | 3.83E-05 |
| Nhej1 | 239.237029 | 0.924516068 | 9.33E-05 |
| Gfod1 | 804.2426049 | 0.924330826 | 1.26E-03 |
| Nmt2 | 3256.678842 | 0.924158439 | 6.78E-09 |
| Plscr2 | 362.8998898 | 0.923930557 | 1.48E-02 |
| Alas1 | 1110.822137 | 0.923510324 | 2.51E-06 |
| Prx | 427.4098247 | 0.923162057 | 4.71E-05 |
| NA | 184.5233713 | 0.923107214 | 9.55E-04 |
| Itpr2 | 1991.961663 | 0.92305784 | 1.17E-10 |
| Utrn | 8786.265038 | 0.92269043 | 3.34E-11 |
| Prkab2 | 1120.185202 | 0.921923075 | 1.31E-08 |
| Tmem62 | 479.9034088 | 0.921657989 | 1.93E-07 |
| Polr3gl | 623.184602 | 0.921555015 | 1.94E-09 |
| Dimt1 | 665.6401902 | 0.921401338 | 3.76E-19 |
| Gm7658 | 524.6113651 | 0.920863219 | 1.70E-06 |
| Hoxa4 | 123.1936353 | 0.920481146 | 3.66E-04 |
| Gm11560 | 44.62081345 | 0.92038892 | 3.88E-02 |
| Eif2b3 | 1233.244349 | 0.918887468 | 2.21E-15 |
| Mfsd9 | 752.6085643 | 0.915882007 | 1.03E-08 |
| Bloc1s2 | 1188.418543 | 0.915188756 | 6.72E-26 |
| Dclre1a | 1218.627447 | 0.91506792 | 2.18E-10 |
| Rgs2 | 984.5440039 | 0.914897362 | 2.68E-04 |
| 4930430F08Rik | 478.4478484 | 0.913769522 | 2.07E-06 |
| Limk1 | 1848.013808 | 0.912619577 | 5.51E-07 |
| Sap30 | 554.9416513 | 0.912152956 | 4.43E-05 |
| Usp22 | 5532.790043 | 0.911946057 | 2.42E-06 |
| Tor4a | 194.1873465 | 0.911454302 | 5.66E-04 |
| Taf1d | 1247.952012 | 0.911346691 | 1.03E-05 |
| H2-K1 | 1381.165026 | 0.910286752 | 1.51E-06 |
| Thnsl1 | 299.7345792 | 0.909883099 | 3.72E-08 |
| Fdxacb1 | 210.4348022 | 0.909231587 | 4.38E-04 |
| Ciao2b | 1227.316825 | 0.908068712 | 3.36E-09 |
| Tsg101 | 2849.982475 | 0.907878838 | 9.14E-10 |
| Epha4 | 1347.621821 | 0.907842035 | 1.42E-10 |

| | | | |
|---|---|---|---|
| Helz | 2920.114213 | 0.907810814 | 4.04E-17 |
| Eif4b | 44123.87184 | 0.907453107 | 1.49E-07 |
| Hyou1 | 5671.534428 | 0.907452637 | 1.86E-07 |
| Plk1 | 1159.243141 | 0.907347487 | 2.84E-02 |
| Wdr93 | 28.35747803 | 0.907248347 | 3.47E-02 |
| Mir6236 | 4368.612436 | 0.906959257 | 5.97E-05 |
| Ppa1 | 2005.850286 | 0.906408504 | 1.61E-05 |
| Cep76 | 375.9843548 | 0.906391593 | 1.40E-07 |
| Tspan9 | 8905.533771 | 0.905806395 | 5.30E-08 |
| Hspa8 | 63221.41105 | 0.905745321 | 5.70E-11 |
| 9530077C05Rik | 123.6192175 | 0.905417595 | 1.53E-05 |
| Rtcb | 11773.29027 | 0.904428377 | 1.49E-09 |
| Slc26a11 | 326.5897719 | 0.903702086 | 5.14E-06 |
| Arl6ip1 | 275.4071157 | 0.903433843 | 1.88E-08 |
| Rps13-ps2 | 21055.30253 | 0.903140759 | 1.56E-06 |
| Angel1 | 396.2138089 | 0.902892855 | 7.80E-05 |
| Ttll7 | 695.451499 | 0.902625782 | 7.60E-06 |
| Epn2 | 3510.416008 | 0.90241111 | 1.15E-07 |
| Rpl36a | 23382.0782 | 0.902084187 | 2.51E-05 |
| Gatad2b | 1668.082965 | 0.901774857 | 6.98E-08 |
| Rel | 189.2158242 | 0.901215552 | 3.77E-05 |
| Git2 | 3927.938835 | 0.901174355 | 2.10E-07 |
| Fam149a | 2567.250985 | 0.901020831 | 8.61E-14 |
| Dnah17 | 62.31221111 | 0.901001004 | 1.61E-02 |
| Hipk2 | 2809.337586 | 0.900862466 | 1.96E-09 |
| Eif3k | 5580.642354 | 0.900818458 | 2.54E-06 |
| Srpk1 | 4585.424015 | 0.900359083 | 6.18E-19 |
| Helz2 | 1049.268653 | 0.900292465 | 4.05E-13 |
| Tdp2 | 876.6310077 | 0.900123173 | 3.74E-08 |
| Hebp2 | 255.3948829 | 0.899782086 | 1.36E-06 |
| NA | 1944.15319 | 0.898124434 | 2.24E-07 |
| Ero1lb | 197.7904632 | 0.897180749 | 3.12E-06 |
| Pdzd7 | 210.0352206 | 0.89717007 | 3.47E-04 |
| Gm9762 | 589.6013071 | 0.897136811 | 1.60E-08 |
| Adrm1 | 5007.958963 | 0.896269679 | 4.14E-08 |
| Arid3a | 862.6284981 | 0.895979402 | 4.41E-05 |
| Kars | 8104.296081 | 0.895455984 | 3.16E-10 |
| Stk11 | 3742.313692 | 0.894901436 | 4.19E-09 |
| Nfkbil1 | 740.5464751 | 0.894702852 | 2.49E-11 |
| Diaph1 | 1722.652336 | 0.894618553 | 7.53E-10 |
| Rmc1 | 1218.193521 | 0.894519128 | 3.23E-08 |
| Dapk3 | 1412.104759 | 0.893949265 | 5.53E-07 |
| Pdcd2l | 631.8121723 | 0.893527689 | 1.46E-06 |
| Polr2c | 2424.517876 | 0.893377009 | 6.12E-10 |
| Yipf3 | 3447.546377 | 0.891910798 | 4.56E-05 |
| Arhgef4 | 384.8597554 | 0.891546579 | 7.01E-04 |
| Ddx58 | 1237.338292 | 0.891526274 | 6.55E-08 |

| | | | |
|---|---|---|---|
| Depdc7 | 242.1642713 | 0.891417158 | 2.51E-04 |
| Prmt7 | 1502.437278 | 0.890076136 | 5.00E-14 |
| Leprotl1 | 1717.958866 | 0.889098739 | 1.75E-20 |
| Adm | 2177.281661 | 0.88851256 | 1.77E-03 |
| Rnf138 | 732.8314506 | 0.888271 | 1.40E-07 |
| Bcat1 | 3993.521955 | 0.887570608 | 2.62E-03 |
| Gemin2 | 424.6443906 | 0.88688343 | 2.06E-12 |
| Atp6v1g2 | 90.09851119 | 0.886861559 | 8.59E-04 |
| Inca1 | 136.1457142 | 0.886827795 | 1.63E-03 |
| Rrn3 | 3178.831608 | 0.886185451 | 6.41E-13 |
| NA | 144.6696948 | 0.886095178 | 1.67E-03 |
| Snta1 | 63.91870191 | 0.885307064 | 4.55E-02 |
| Gm6136 | 182.5775809 | 0.884681349 | 1.20E-02 |
| Polg2 | 290.1108251 | 0.884521628 | 1.68E-05 |
| Gata4 | 849.8210775 | 0.884054076 | 4.59E-06 |
| Ptges3-ps | 1167.527342 | 0.883879622 | 2.68E-06 |
| Rdh1 | 35.13277364 | 0.88375131 | 1.28E-02 |
| Ubxn2a | 924.4544045 | 0.883690418 | 1.27E-05 |
| Ktn1 | 2814.57127 | 0.883098792 | 3.76E-03 |
| Hspb2 | 59.28042065 | 0.883004598 | 1.09E-02 |
| Iqcb1 | 717.9089585 | 0.881817939 | 4.90E-09 |
| Rad18 | 808.0924725 | 0.881699616 | 8.85E-04 |
| Trmt1 | 3494.043104 | 0.880651283 | 9.36E-09 |
| Bag2 | 1009.73514 | 0.879943716 | 3.04E-07 |
| Pabpc1 | 86856.51255 | 0.879410718 | 4.25E-09 |
| Pcid2 | 1369.909591 | 0.879391544 | 5.44E-08 |
| Mybbp1a | 15401.0598 | 0.879164244 | 2.53E-09 |
| Kpna1 | 2050.904323 | 0.87839536 | 9.79E-04 |
| Pla2g12a | 876.4735081 | 0.877996963 | 7.01E-07 |
| Rpl39l | 5625.606209 | 0.877848321 | 7.35E-08 |
| Spindoc | 1533.971125 | 0.877626548 | 9.04E-07 |
| Pold2 | 1477.91555 | 0.877466274 | 7.13E-06 |
| Arih2 | 2519.227217 | 0.87674761 | 1.18E-10 |
| Kat8 | 866.0276859 | 0.876650542 | 2.61E-08 |
| Gpsm1 | 1158.941864 | 0.876082564 | 7.56E-07 |
| Ccl25 | 167.8819223 | 0.875025556 | 1.78E-05 |
| Zfyve19 | 792.6092984 | 0.874656905 | 9.05E-06 |
| Spn | 32.34573545 | 0.874610946 | 4.40E-02 |
| Xylb | 84.70507461 | 0.873910513 | 1.37E-03 |
| Rarg | 5414.781718 | 0.873404611 | 2.43E-09 |
| Sec24a | 2291.392764 | 0.872814785 | 2.15E-06 |
| Cda | 445.5243863 | 0.872617291 | 8.69E-05 |
| Ranbp9 | 3139.587622 | 0.872401188 | 1.92E-05 |
| Plod2 | 1368.669902 | 0.870803125 | 1.84E-05 |
| Osgin2 | 294.2721574 | 0.869605538 | 9.29E-06 |
| Mecp2 | 3791.304598 | 0.869244858 | 1.78E-06 |
| Man1a | 1574.812904 | 0.868840021 | 9.71E-05 |

| | | | |
|---|---|---|---|
| Rbm33 | 3346.777096 | 0.868259015 | 2.25E-11 |
| Ruvbl2 | 1919.938162 | 0.867752122 | 6.84E-05 |
| Zpr1 | 2032.138192 | 0.867694136 | 1.25E-08 |
| Osgin1 | 375.3213527 | 0.867584697 | 3.34E-03 |
| Ttc32 | 115.9407862 | 0.867438392 | 2.62E-05 |
| Bdnf | 1365.54172 | 0.865485661 | 6.50E-05 |
| Rusc1 | 481.2122263 | 0.864947903 | 6.10E-04 |
| Ifngr1 | 2047.302005 | 0.864330042 | 1.47E-14 |
| Mrps18b | 1968.321664 | 0.863536438 | 4.35E-10 |
| Rabl3 | 368.0500421 | 0.862546033 | 4.79E-06 |
| Plod3 | 6604.231022 | 0.862424012 | 2.88E-08 |
| Map2k3os | 158.9798746 | 0.862157107 | 1.23E-02 |
| Gm16576 | 237.6025889 | 0.861640742 | 3.17E-05 |
| Hint3 | 384.6312068 | 0.861169706 | 2.89E-04 |
| Baz1a | 2082.890024 | 0.860986885 | 9.96E-07 |
| Ghdc | 1058.772751 | 0.860913663 | 6.00E-06 |
| Klf3 | 3490.931244 | 0.860504619 | 2.00E-10 |
| Atp6v0e2 | 330.9666067 | 0.860353819 | 1.63E-02 |
| H2-Bl | 144.2958941 | 0.860345928 | 1.08E-04 |
| Ypel3 | 1637.852338 | 0.860151128 | 2.68E-05 |
| Vopp1 | 3424.567913 | 0.859816826 | 1.89E-08 |
| Pigf | 401.9097768 | 0.859639841 | 3.83E-12 |
| Calr3 | 72.66829931 | 0.859633971 | 8.62E-04 |
| Plekha8 | 1302.28539 | 0.858223235 | 1.50E-10 |
| Eif3l | 10508.6021 | 0.858020604 | 8.51E-10 |
| AA474408 | 8595.94056 | 0.857645377 | 6.67E-05 |
| Lgalsl | 3882.880886 | 0.857520163 | 5.91E-09 |
| Rps18-ps3 | 520.1252683 | 0.857490224 | 4.36E-03 |
| Ercc5 | 1751.423907 | 0.857150918 | 4.88E-16 |
| Aurkb | 1122.511781 | 0.857145857 | 1.62E-02 |
| Tinf2 | 460.587957 | 0.85538387 | 4.41E-06 |
| Slc7a7 | 105.7139369 | 0.855190062 | 1.17E-03 |
| 0610010K14Rik | 1202.24141 | 0.854398675 | 1.78E-06 |
| Ddx27 | 2854.313142 | 0.854262953 | 7.69E-11 |
| Taldo1 | 5508.674467 | 0.854180937 | 1.10E-05 |
| Mars | 4381.005849 | 0.85412853 | 9.35E-07 |
| Gm10157 | 272.6510177 | 0.853919446 | 8.32E-03 |
| Sumo2 | 2921.158398 | 0.853590815 | 1.24E-07 |
| Ribc1 | 38.24492566 | 0.853178937 | 4.63E-02 |
| Acrbp | 73.24599371 | 0.8527978 | 1.68E-03 |
| Rpl23 | 66491.88099 | 0.852693982 | 1.38E-06 |
| Dbf4 | 1797.842921 | 0.852172566 | 5.20E-03 |
| Dus3l | 1405.012927 | 0.851761068 | 2.98E-09 |
| Lrch4 | 1036.656072 | 0.851332159 | 1.31E-08 |
| Racgap1 | 3856.467284 | 0.850513863 | 3.76E-03 |
| Cryab | 13202.45602 | 0.850384993 | 3.86E-04 |
| Agpat1 | 3534.213831 | 0.850126653 | 1.49E-11 |

| | | | |
|---|---|---|---|
| Tma16 | 794.3992447 | 0.848883042 | 2.59E-09 |
| Lyar | 1204.940602 | 0.848687079 | 1.48E-04 |
| Cpsf4 | 763.0921259 | 0.84829427 | 6.87E-11 |
| E2f7 | 788.1276993 | 0.84816842 | 2.45E-02 |
| Nploc4 | 4231.422089 | 0.848152899 | 3.53E-07 |
| Vdac1 | 9711.986753 | 0.848114063 | 2.13E-14 |
| Ccni | 10061.03268 | 0.847802783 | 5.08E-07 |
| 2300009A05Rik | 106.6990656 | 0.847364598 | 2.71E-05 |
| Iars | 7024.962915 | 0.847339777 | 3.69E-07 |
| Ciapin1 | 1962.754716 | 0.846843068 | 2.66E-05 |
| Fyn | 3259.615135 | 0.846806342 | 1.55E-18 |
| Plekhb2 | 3753.770953 | 0.844499552 | 6.65E-08 |
| Alg13 | 429.2832683 | 0.84442222 | 1.74E-04 |
| Dnajc9 | 984.8756573 | 0.842818383 | 5.53E-04 |
| Lrrc57 | 651.3940836 | 0.842336795 | 4.60E-09 |
| Gtpbp6 | 1438.376684 | 0.842139119 | 5.22E-08 |
| Zfp473 | 122.7050172 | 0.840793108 | 2.15E-02 |
| Fbxl20 | 3801.092862 | 0.839930238 | 3.88E-04 |
| Mier2 | 787.9011876 | 0.839839896 | 4.96E-09 |
| Ccnt1 | 2298.363621 | 0.839543358 | 1.14E-06 |
| Kcnn1 | 233.8205423 | 0.839527156 | 4.65E-05 |
| Zfp692 | 591.0013863 | 0.838739576 | 1.89E-09 |
| Pgam5 | 2236.525896 | 0.838160276 | 4.24E-12 |
| Ubl4a | 2372.32782 | 0.837930488 | 5.21E-11 |
| Stc2 | 605.8941595 | 0.837895635 | 6.93E-03 |
| Nprl2 | 675.6085235 | 0.837838269 | 7.89E-08 |
| Mri1 | 697.8904914 | 0.837775838 | 2.87E-05 |
| Ints11 | 1652.185538 | 0.837506566 | 1.52E-08 |
| Sqstm1 | 31647.65135 | 0.837407479 | 6.93E-06 |
| Bcl2l13 | 3503.679248 | 0.837291065 | 2.22E-14 |
| Pwp1 | 2355.646022 | 0.837019374 | 6.80E-12 |
| Hint1 | 7364.354729 | 0.836351703 | 1.93E-09 |
| 2310039H08Rik | 526.4348822 | 0.835602112 | 5.76E-08 |
| Gls2 | 118.3808906 | 0.835266038 | 5.52E-03 |
| Bcl3 | 359.6483038 | 0.834019955 | 3.53E-03 |
| Orai1 | 454.0094432 | 0.833610069 | 1.50E-04 |
| Smad4 | 3283.698347 | 0.833425938 | 6.78E-08 |
| Eif2d | 2660.698327 | 0.833368217 | 2.96E-07 |
| Hspg2 | 28446.74545 | 0.833361264 | 5.51E-05 |
| Akap8l | 1442.125993 | 0.832626335 | 1.54E-07 |
| Socs7 | 1466.776444 | 0.832574844 | 2.29E-16 |
| Gm12013 | 265.5132074 | 0.832296955 | 9.56E-03 |
| Rps21 | 17061.14801 | 0.831616599 | 4.29E-05 |
| Fbxl5 | 2118.341111 | 0.831504243 | 1.37E-19 |
| Ankrd55 | 177.8979238 | 0.831423229 | 7.16E-03 |
| Lin7c | 1704.717814 | 0.830124653 | 8.89E-05 |
| Chka | 2275.514018 | 0.829055912 | 1.34E-06 |

| | | | |
|---|---|---|---|
| Oxld1 | 130.1722169 | 0.828428828 | 4.55E-04 |
| NA | 3226.764882 | 0.827683634 | 2.67E-06 |
| Rps29 | 19692.01595 | 0.827573049 | 1.05E-07 |
| Gm10443 | 18934.25035 | 0.827187939 | 4.60E-04 |
| Morn4 | 760.2486884 | 0.827153161 | 2.17E-19 |
| 1810009A15Rik | 1592.766553 | 0.826756915 | 1.87E-09 |
| Drg2 | 2531.129456 | 0.826149702 | 4.03E-09 |
| Egfl7 | 61.93243651 | 0.82568935 | 2.08E-02 |
| Nat9 | 476.1271399 | 0.824069924 | 4.56E-04 |
| CAAA01118383.1 | 707.8337304 | 0.823603354 | 2.78E-07 |
| Gm9825 | 3797.027924 | 0.822660872 | 2.22E-07 |
| Dcaf15 | 1619.339583 | 0.8217221 | 4.27E-06 |
| Trip10 | 3204.625039 | 0.821367555 | 2.21E-07 |
| Kansl2 | 5499.25737 | 0.821057072 | 7.76E-10 |
| H2-Q5 | 32.8399492 | 0.82095407 | 1.62E-02 |
| Psmd11 | 5834.119852 | 0.820787964 | 2.20E-13 |
| Uimc1 | 1375.567608 | 0.820152194 | 9.89E-09 |
| Ankrd23 | 160.0658246 | 0.819975704 | 1.42E-03 |
| Kctd9 | 1161.401207 | 0.81981566 | 3.11E-03 |
| Ppp1r2 | 6493.223235 | 0.819428473 | 5.37E-09 |
| Plin2 | 9880.879654 | 0.818499532 | 1.37E-05 |
| Rpl7a-ps3 | 1301.1174 | 0.818425631 | 1.99E-04 |
| Zfp7 | 302.867491 | 0.817644035 | 2.28E-04 |
| Kif22 | 949.2951466 | 0.817364613 | 1.64E-02 |
| Rps28 | 15164.62267 | 0.817258172 | 4.77E-04 |
| Agpat5 | 1466.315111 | 0.817232988 | 4.68E-10 |
| Nop2 | 4072.658811 | 0.817055922 | 5.31E-07 |
| Ncaph | 1055.329841 | 0.816994046 | 5.80E-03 |
| Gm14414 | 932.8384723 | 0.815611885 | 9.69E-04 |
| Gm10705 | 193.1483854 | 0.815556347 | 3.98E-02 |
| Wbp2 | 5052.649699 | 0.81539054 | 5.01E-04 |
| Ube2j1 | 2831.915582 | 0.814887437 | 2.63E-21 |
| Rps4x | 93555.43077 | 0.814865698 | 2.69E-05 |
| Apba3 | 2076.974687 | 0.814500445 | 1.61E-10 |
| Mkrn2 | 1675.962948 | 0.814403796 | 2.04E-06 |
| Smim1 | 81.00682783 | 0.813983273 | 3.75E-02 |
| Rab4a | 910.1117394 | 0.813964021 | 1.78E-09 |
| Rasip1 | 66.82205557 | 0.813768376 | 1.43E-03 |
| Cfl2 | 7309.759639 | 0.81367282 | 6.82E-07 |
| Mllt11 | 896.5428316 | 0.812762398 | 3.63E-08 |
| Pzp | 23.6449918 | 0.812039443 | 3.38E-02 |
| Prnp | 14550.6052 | 0.811920491 | 2.04E-06 |
| Desi1 | 3182.457457 | 0.811887153 | 9.73E-07 |
| Gpd2 | 3642.520661 | 0.811871934 | 1.71E-17 |
| Peli1 | 768.5211593 | 0.811739113 | 1.03E-08 |
| Cep57l1 | 538.9271566 | 0.811513702 | 2.11E-06 |
| Irs2 | 2222.67984 | 0.810930838 | 6.16E-08 |

| | | | |
|---|---|---|---|
| Adamts5 | 15137.11031 | 0.810620575 | 9.44E-04 |
| Mid1ip1 | 939.9451529 | 0.810002419 | 1.49E-03 |
| Rrm1 | 7755.011656 | 0.809023141 | 2.31E-03 |
| Mettl22 | 320.1298717 | 0.808669677 | 8.44E-07 |
| Il6st | 23732.48129 | 0.808329164 | 7.14E-06 |
| Gm10163 | 19766.21336 | 0.808192521 | 2.81E-05 |
| Babam2 | 2414.199913 | 0.807825993 | 1.74E-07 |
| Golga4 | 4342.991697 | 0.807798432 | 2.58E-07 |
| Cbr3 | 934.7984105 | 0.807355666 | 1.37E-08 |
| Atxn10 | 13554.53568 | 0.806968243 | 5.96E-07 |
| Dlx2 | 556.9964188 | 0.806522941 | 5.23E-05 |
| Cbx2 | 830.0959647 | 0.806137212 | 1.97E-05 |
| Fdxr | 851.9175355 | 0.80611667 | 4.10E-04 |
| Myl12a | 24114.22838 | 0.805171054 | 6.88E-09 |
| Psmd12 | 4257.969189 | 0.804945095 | 3.02E-12 |
| Immp2l | 226.8841086 | 0.804454375 | 1.53E-04 |
| Nim1k | 75.94060927 | 0.804217355 | 9.58E-04 |
| Relt | 217.4731898 | 0.80417912 | 4.27E-04 |
| Phf13 | 1356.38196 | 0.803963083 | 5.23E-04 |
| Dtwd1 | 574.3759161 | 0.803949302 | 4.69E-10 |
| Tubd1 | 135.3189566 | 0.80387499 | 2.79E-04 |
| Adat2 | 575.114688 | 0.803849012 | 8.64E-05 |
| Acp6 | 792.6797913 | 0.803750274 | 3.45E-12 |
| Hdgf | 20795.9107 | 0.803630955 | 2.69E-09 |
| Prkrip1 | 783.1122703 | 0.803102311 | 5.67E-07 |
| Cdc25c | 279.9314129 | 0.801965377 | 4.43E-02 |
| Zfp365 | 1958.032679 | 0.801903004 | 6.72E-07 |
| Slc25a27 | 344.4537154 | 0.800971537 | 7.09E-04 |
| Tnfrsf23 | 1998.923511 | 0.800322502 | 2.17E-03 |
| Ahcyl1 | 10647.42186 | 0.799669325 | 3.10E-28 |
| Gtpbp10 | 161.1492663 | 0.798857683 | 5.30E-03 |
| Hsf2 | 1151.428926 | 0.797811652 | 1.18E-08 |
| Tsc22d3 | 2297.21333 | 0.797222913 | 2.62E-04 |
| Nepro | 974.7880945 | 0.796929016 | 9.26E-08 |
| Idh3b | 5787.050988 | 0.795733239 | 1.66E-07 |
| Sypl | 3290.631028 | 0.795649166 | 2.22E-13 |
| Gm26819 | 158.0427173 | 0.795606176 | 7.26E-03 |
| Fgfrl1 | 2587.189807 | 0.794825315 | 1.43E-05 |
| Aimp2 | 1282.29294 | 0.794393906 | 1.75E-05 |
| Cops4 | 4067.416077 | 0.794065551 | 2.52E-09 |
| Snrpa1 | 1239.871439 | 0.793933563 | 1.13E-06 |
| Rxrb | 2724.534752 | 0.793170754 | 6.29E-10 |
| Mpp1 | 3836.686581 | 0.792951575 | 4.96E-08 |
| Smtn | 3576.331185 | 0.792895552 | 3.33E-07 |
| Dctn2 | 11012.49689 | 0.791651682 | 4.47E-08 |
| Agrn | 5204.164369 | 0.790868694 | 1.96E-05 |
| Pir | 1475.999153 | 0.789856974 | 4.12E-04 |

| | | | |
|---|---|---|---|
| Apbb1 | 1088.659518 | 0.789376343 | 1.11E-08 |
| Tent5b | 1620.453803 | 0.789296651 | 1.14E-05 |
| Wnt5a | 600.197003 | 0.78926845 | 6.19E-07 |
| Tbc1d15 | 3591.297068 | 0.78914663 | 1.68E-09 |
| Pcbp1 | 15320.23063 | 0.788431404 | 1.70E-09 |
| Rps9 | 54843.11362 | 0.788179066 | 9.41E-06 |
| Tle1 | 1922.88961 | 0.788090098 | 9.19E-09 |
| R3hdm2 | 5526.132124 | 0.787995532 | 5.30E-10 |
| Phb2 | 9596.173477 | 0.787974889 | 1.87E-08 |
| Shroom3 | 71.88219185 | 0.787519235 | 3.22E-02 |
| Prkx | 992.1556184 | 0.787257278 | 1.53E-06 |
| Tmem206 | 879.2146497 | 0.78603451 | 1.41E-11 |
| Rflnb | 613.7469556 | 0.785565227 | 3.97E-04 |
| Nudt22 | 470.2166979 | 0.78551463 | 3.48E-08 |
| Slc4a7 | 1372.608239 | 0.785189074 | 2.47E-05 |
| Ddb1 | 78266.76622 | 0.785017638 | 1.05E-10 |
| Msi2 | 1364.372491 | 0.784812836 | 5.32E-08 |
| Cdkn3 | 244.1485424 | 0.784563241 | 2.75E-02 |
| Bcl2l11 | 1515.764537 | 0.78412667 | 2.23E-02 |
| Taf10 | 2599.103771 | 0.784058142 | 2.91E-06 |
| Mrrf | 1845.383387 | 0.783997997 | 3.20E-08 |
| Deptor | 1172.456592 | 0.783591439 | 2.46E-03 |
| Rock2 | 6286.389359 | 0.783534582 | 5.99E-03 |
| Ube2s | 4362.208061 | 0.783101551 | 4.11E-04 |
| Ryk | 8106.711931 | 0.78253377 | 2.03E-13 |
| Foxk2 | 3737.143576 | 0.782447999 | 2.68E-06 |
| Ankrd9 | 197.5218888 | 0.781883435 | 1.20E-03 |
| Ccnk | 2008.508271 | 0.781602055 | 1.82E-04 |
| Pmpcb | 3918.866943 | 0.781546466 | 2.35E-10 |
| Nectin2 | 3231.192606 | 0.781376084 | 1.50E-08 |
| Eef1e1 | 830.8756283 | 0.780927647 | 6.96E-06 |
| Tmem68 | 608.1000298 | 0.779624995 | 6.97E-10 |
| Acsl6 | 253.6968893 | 0.779284452 | 1.33E-04 |
| Ufl1 | 1661.102463 | 0.779090256 | 6.04E-07 |
| Rn7s1 | 84.50678455 | 0.779035767 | 4.93E-02 |
| Ethe1 | 653.1180357 | 0.778529603 | 6.67E-09 |
| Fam98c | 410.091738 | 0.778190739 | 1.93E-06 |
| Rpl34-ps2 | 184.2154543 | 0.777823833 | 1.64E-02 |
| NA | 11703.17523 | 0.777155545 | 7.90E-04 |
| Gm26825 | 36665.95223 | 0.776300635 | 7.68E-07 |
| Aup1 | 6846.043004 | 0.776233615 | 2.06E-10 |
| Pltp | 1305.024307 | 0.776174992 | 5.05E-03 |
| Bysl | 1467.654049 | 0.775876027 | 2.76E-05 |
| Slc25a37 | 3381.662168 | 0.775720301 | 2.44E-05 |
| Plekhg6 | 45.970297 | 0.775691325 | 2.19E-02 |
| Celf4 | 267.6869814 | 0.773518178 | 1.69E-03 |
| Rpl7 | 74255.39802 | 0.773447536 | 1.57E-06 |

| | | | |
|---|---|---|---|
| Tom1l1 | 742.8401817 | 0.772832599 | 3.04E-07 |
| Cyth2 | 2204.307457 | 0.772612282 | 2.64E-06 |
| Miip | 631.6917154 | 0.772484034 | 5.40E-04 |
| Mlh3 | 176.7584739 | 0.772047789 | 1.16E-03 |
| Nup43 | 725.9634131 | 0.77163802 | 3.19E-05 |
| Polr2h | 1088.850475 | 0.771451364 | 2.23E-08 |
| Rpl37a | 29354.97811 | 0.771355153 | 1.34E-03 |
| Npepps | 9806.645939 | 0.771309361 | 3.35E-11 |
| Tmod3 | 2418.238521 | 0.770909383 | 2.92E-06 |
| Srrm1 | 6012.177293 | 0.770062995 | 2.64E-11 |
| Kdm8 | 161.4759552 | 0.769713867 | 5.67E-03 |
| Adk | 3190.013385 | 0.769126333 | 1.42E-09 |
| Cenpu | 251.9751922 | 0.768600407 | 3.00E-02 |
| Cyp2d22 | 1717.641267 | 0.768549272 | 7.37E-03 |
| Tcaim | 463.9162715 | 0.76842469 | 1.99E-06 |
| Taf4b | 222.6082141 | 0.768326116 | 4.67E-03 |
| Mtln | 167.9560115 | 0.767478685 | 1.49E-04 |
| Rpl31 | 2778.260788 | 0.765900603 | 8.24E-10 |
| Rprd1a | 1030.14927 | 0.765753628 | 4.57E-07 |
| F730043M19Rik | 811.4228063 | 0.765720699 | 8.27E-03 |
| Mastl | 676.3577733 | 0.765190856 | 4.46E-02 |
| Map3k14 | 972.1145369 | 0.765134582 | 5.09E-05 |
| Sirt3 | 334.231396 | 0.764651466 | 1.97E-05 |
| Ncbp1 | 3020.179347 | 0.764472 | 1.36E-07 |
| Tmem253 | 30.43885092 | 0.764257421 | 4.85E-02 |
| 1810037I17Rik | 1837.466688 | 0.763788841 | 1.61E-08 |
| Exd2 | 1441.583752 | 0.76374401 | 1.32E-07 |
| Zfp511 | 616.6235673 | 0.763191784 | 1.87E-06 |
| Myo19 | 568.3216705 | 0.76310924 | 3.79E-03 |
| Ctla2b | 860.0005034 | 0.763023906 | 4.71E-02 |
| Ssu72 | 1549.612509 | 0.76222397 | 1.53E-09 |
| Mtfr1 | 2428.446967 | 0.762156072 | 1.44E-10 |
| Dnajc4 | 650.8374999 | 0.760567008 | 3.70E-04 |
| Abhd8 | 2737.344855 | 0.760077904 | 8.14E-05 |
| Sfmbt2 | 667.3441797 | 0.759291723 | 4.39E-04 |
| Got1 | 2662.479814 | 0.75877367 | 3.24E-07 |
| Lmbr1l | 980.757885 | 0.758665807 | 5.95E-04 |
| Ccdc77 | 463.8315296 | 0.758473471 | 5.17E-05 |
| NA | 77.0601651 | 0.758350789 | 2.22E-02 |
| Nub1 | 6026.984575 | 0.757388203 | 1.11E-04 |
| Mxi1 | 1578.380318 | 0.757312191 | 6.04E-07 |
| Nudcd1 | 1384.714775 | 0.75665598 | 2.33E-06 |
| Sympk | 8088.670725 | 0.75602151 | 1.33E-06 |
| Car7 | 47.50530047 | 0.755167166 | 2.08E-02 |
| Gm14681 | 1360.044557 | 0.755055125 | 1.25E-04 |
| Ctbs | 628.3728415 | 0.7548476 | 3.34E-10 |
| Smim3 | 135.0460714 | 0.753945358 | 3.31E-03 |

| | | | |
|---|---|---|---|
| Ubxn6 | 3031.372245 | 0.753572856 | 5.27E-07 |
| Mad2l1 | 1774.898828 | 0.753354946 | 1.97E-02 |
| Tmem243 | 153.1688877 | 0.753020517 | 4.08E-05 |
| Rabl6 | 4751.701307 | 0.752929299 | 3.51E-09 |
| Acvr2b | 107.6232016 | 0.752663614 | 8.73E-03 |
| Ascc1 | 1690.168605 | 0.752477944 | 1.76E-04 |
| Gm10288 | 2402.863389 | 0.752326692 | 4.20E-02 |
| Morc3 | 1497.828179 | 0.752041034 | 5.74E-06 |
| Acot13 | 2081.691453 | 0.751938185 | 3.33E-06 |
| Raet1e | 152.5635618 | 0.751587694 | 4.34E-03 |
| Rad51c | 374.2213069 | 0.751243605 | 4.74E-05 |
| Nfkbiz | 359.018599 | 0.751181869 | 1.85E-02 |
| Trpt1 | 309.8301647 | 0.750679888 | 8.43E-05 |
| Mgmt | 694.135641 | 0.750212518 | 2.11E-05 |
| Plcb3 | 3361.405742 | 0.749908654 | 4.71E-22 |
| Cep104 | 1205.104973 | 0.749808953 | 2.13E-12 |
| Heatr6 | 2310.476748 | 0.749769031 | 1.98E-08 |
| Snd1 | 12835.65461 | 0.749762996 | 2.98E-11 |
| Acbd6 | 1946.20672 | 0.749688169 | 1.93E-04 |
| Dnm2 | 5735.557998 | 0.749438173 | 4.06E-07 |
| Mapk3 | 6423.195426 | 0.748850321 | 2.16E-05 |
| Maoa | 11869.71092 | 0.748801156 | 8.81E-05 |
| Pes1 | 5474.440582 | 0.748674986 | 7.25E-10 |
| Crtc2 | 1839.734255 | 0.74858284 | 1.25E-05 |
| Gm26602 | 167.2911212 | 0.748470331 | 3.65E-02 |
| Fam83h | 457.8759574 | 0.748324628 | 1.08E-07 |
| Impdh2-ps | 1489.025213 | 0.748300636 | 8.37E-08 |
| Kifc2 | 266.15885 | 0.748283803 | 6.50E-03 |
| Ppfia3 | 146.6688839 | 0.747925231 | 3.05E-03 |
| Nolc1 | 4595.577339 | 0.746244004 | 1.78E-05 |
| Pam | 7016.485341 | 0.744587283 | 1.87E-07 |
| Gna13 | 4550.858975 | 0.744584308 | 4.64E-08 |
| Polrmt | 1417.394738 | 0.744402788 | 2.70E-06 |
| Mrpl15 | 2833.345375 | 0.744127658 | 7.73E-10 |
| Coq5 | 1504.412996 | 0.743927531 | 1.25E-11 |
| Bend6 | 173.0386063 | 0.743703389 | 2.86E-03 |
| Ube2j2 | 1199.238568 | 0.743581198 | 4.69E-10 |
| Cldn12 | 2413.444114 | 0.743510702 | 4.66E-05 |
| Mpst | 1039.896141 | 0.742974592 | 4.92E-04 |
| Thbd | 4214.750375 | 0.742495085 | 2.94E-03 |
| Rpa1 | 7115.964406 | 0.742246143 | 9.43E-04 |
| Ada | 254.4672862 | 0.741936849 | 7.04E-04 |
| Uqcrfs1 | 2399.105649 | 0.741741573 | 4.43E-10 |
| H2-D1 | 9845.207692 | 0.741211869 | 9.73E-07 |
| Chchd4 | 753.277916 | 0.741150928 | 4.22E-06 |
| Nrde2 | 1207.833915 | 0.740866643 | 2.00E-05 |
| Mrpl35 | 1306.855469 | 0.740740395 | 7.83E-14 |

| | | | |
|---|---|---|---|
| Sertad3 | 577.2030855 | 0.740579341 | 4.06E-04 |
| Cdca2 | 627.3753724 | 0.740059358 | 4.83E-02 |
| Sf3a2 | 2291.998884 | 0.739904464 | 7.63E-05 |
| Prpf6 | 5480.800313 | 0.739818889 | 1.17E-05 |
| Edrf1 | 931.4257018 | 0.739233703 | 1.20E-06 |
| H2-T23 | 367.1066819 | 0.739205105 | 8.49E-03 |
| BC052040 | 653.4512046 | 0.739143083 | 3.20E-11 |
| Tjap1 | 2545.037818 | 0.739044274 | 3.60E-09 |
| Rela | 3095.57112 | 0.738412966 | 1.01E-10 |
| Hspa9 | 32616.301 | 0.737892134 | 1.70E-13 |
| Lsm12 | 2001.617156 | 0.737624715 | 7.53E-15 |
| Dtx4 | 6178.699218 | 0.737329978 | 2.26E-03 |
| Mief2 | 738.7242967 | 0.736931477 | 2.06E-04 |
| Cenpp | 150.4209713 | 0.736911371 | 5.04E-03 |
| Rp9 | 470.7603178 | 0.73656986 | 6.72E-04 |
| Plp2 | 8742.786313 | 0.736391515 | 8.30E-06 |
| Drap1 | 6991.132614 | 0.736248533 | 9.52E-09 |
| Akr1b10 | 376.3471069 | 0.736178552 | 6.47E-05 |
| Setd1a | 2741.234231 | 0.735849592 | 4.67E-07 |
| Tcea3 | 238.4744052 | 0.7357491 | 4.38E-02 |
| Nrm | 626.7617768 | 0.735381496 | 1.76E-02 |
| Lemd2 | 2831.27415 | 0.735310836 | 7.26E-07 |
| Ppp2r2a | 2867.173136 | 0.735258394 | 4.33E-12 |
| Psmb1 | 9738.740386 | 0.735141511 | 6.44E-07 |
| Fam216a | 1524.130343 | 0.735110663 | 1.23E-14 |
| Pus7 | 1910.719923 | 0.734943076 | 1.05E-03 |
| Cbl | 1012.925113 | 0.734023511 | 6.50E-07 |
| Mmp28 | 209.9282617 | 0.733423546 | 3.49E-02 |
| Zfp330 | 1591.298012 | 0.733131694 | 1.66E-08 |
| Rassf3 | 475.9706472 | 0.732694415 | 9.06E-06 |
| Csnk2a2 | 2886.982549 | 0.73259792 | 1.34E-11 |
| Cbx7 | 739.4739341 | 0.732464287 | 1.30E-07 |
| Hmces | 1311.48031 | 0.732313792 | 6.14E-13 |
| Zbtb11 | 1086.873096 | 0.732303127 | 1.30E-05 |
| 9330136K24Rik | 58.8334473 | 0.731428745 | 2.17E-02 |
| Ppm1g | 5386.882069 | 0.730887279 | 6.74E-06 |
| Dlg1 | 7243.816205 | 0.730844194 | 9.25E-06 |
| Eaf1 | 2071.798067 | 0.730754265 | 1.05E-09 |
| Atxn2 | 4115.093461 | 0.730392789 | 5.10E-08 |
| Nfu1 | 1848.556139 | 0.7298651 | 5.24E-18 |
| Supt3 | 1376.160322 | 0.729354096 | 7.52E-05 |
| Clasrp | 883.122175 | 0.728914682 | 6.37E-07 |
| Pkn2 | 5004.679856 | 0.728534988 | 1.33E-09 |
| Actg1 | 96158.53644 | 0.728241732 | 1.24E-06 |
| Aprt | 6193.680378 | 0.727396498 | 8.99E-06 |
| Flnb | 18993.59883 | 0.727333562 | 9.60E-07 |
| Ckap2l | 1587.056983 | 0.727017358 | 4.11E-02 |

| | | | |
|---|---|---|---|
| A430046D13Rik | 73.22061336 | 0.726545343 | 2.00E-02 |
| Bag1 | 4120.741985 | 0.726407385 | 5.59E-07 |
| Stradb | 1050.361299 | 0.72632523 | 6.17E-05 |
| Snora7a | 41.98139657 | 0.725840626 | 2.06E-02 |
| Ddx49 | 1574.01169 | 0.725730389 | 1.81E-07 |
| Ergic1 | 11142.20075 | 0.725633324 | 1.35E-06 |
| Nol9 | 1501.271379 | 0.725450748 | 3.17E-07 |
| Kat5 | 1999.455412 | 0.725202736 | 2.66E-06 |
| Gm4737 | 8568.322965 | 0.725153566 | 1.27E-04 |
| Mrpl21 | 1397.633345 | 0.725124113 | 7.57E-13 |
| Abhd2 | 3646.914905 | 0.724710316 | 1.08E-06 |
| Klc1 | 5592.891025 | 0.724304606 | 7.03E-13 |
| Tbkbp1 | 102.9282373 | 0.72388936 | 4.37E-02 |
| Prr7 | 85.92663974 | 0.723859887 | 1.28E-02 |
| Cacnb1 | 471.9894397 | 0.723584465 | 9.96E-05 |
| Lrrc28 | 607.671651 | 0.723205698 | 7.10E-09 |
| Grasp | 391.1139554 | 0.72287473 | 4.11E-07 |
| Tmem11 | 1225.6397 | 0.722705579 | 1.43E-06 |
| Plekhg3 | 2284.954782 | 0.722609613 | 2.03E-05 |
| Aptx | 863.7687715 | 0.721570768 | 7.90E-08 |
| Cyba | 1385.240954 | 0.721280903 | 4.31E-06 |
| Tox4 | 2329.373657 | 0.721032233 | 2.15E-05 |
| Pex16 | 676.2702163 | 0.720805422 | 2.33E-05 |
| Rrp9 | 952.0180092 | 0.720080349 | 1.72E-04 |
| Isl2 | 63.99342943 | 0.719209113 | 2.05E-02 |
| Stip1 | 10290.17675 | 0.718832372 | 8.29E-05 |
| Usp39 | 3405.710317 | 0.718802026 | 3.15E-05 |
| Exoc4 | 6670.852414 | 0.718620121 | 4.69E-09 |
| Inppl1 | 6885.394772 | 0.718467618 | 2.87E-12 |
| Slc20a1 | 2917.806275 | 0.718409853 | 1.09E-04 |
| Jmjd6 | 1587.362824 | 0.718363192 | 3.64E-08 |
| Nav2 | 1894.646959 | 0.718201904 | 4.77E-02 |
| Atad3a | 1366.86159 | 0.718120718 | 1.02E-07 |
| Rps6ka2 | 2190.081064 | 0.718009105 | 1.50E-08 |
| Cds2 | 6656.30948 | 0.717546851 | 1.91E-11 |
| Gm2a | 1019.030594 | 0.717417598 | 2.77E-05 |
| Cpeb2 | 370.5002161 | 0.716307955 | 5.81E-04 |
| Lgmn | 8054.730872 | 0.716298843 | 5.98E-04 |
| Fam177a | 1383.292923 | 0.716040976 | 7.30E-06 |
| Gtf2f2 | 887.8471566 | 0.715939409 | 1.86E-07 |
| Cox5a | 4353.767426 | 0.715924563 | 1.83E-06 |
| Swt1 | 741.4746105 | 0.715907961 | 2.53E-03 |
| Mcts2 | 402.5475994 | 0.71566178 | 3.01E-06 |
| Prkag2 | 1252.873195 | 0.715409149 | 1.12E-09 |
| Gpr146 | 226.6367334 | 0.715053703 | 4.00E-02 |
| Akap6 | 117.141912 | 0.714938042 | 2.20E-02 |
| Gspt2 | 338.4968154 | 0.714409551 | 6.60E-07 |

| | | | |
|---|---|---|---|
| Hibadh | 3736.286012 | 0.713786792 | 2.86E-07 |
| Stom | 2151.225051 | 0.713495971 | 1.97E-03 |
| Rnf141 | 1366.065122 | 0.713453728 | 6.58E-06 |
| Slc39a8 | 483.1111622 | 0.713282818 | 1.82E-04 |
| Cad | 5610.102911 | 0.713169916 | 3.34E-04 |
| Sh3bp4 | 1970.38902 | 0.713087166 | 4.28E-06 |
| Ptar1 | 207.0188926 | 0.712254389 | 2.14E-03 |
| Gmds | 1941.410771 | 0.712127498 | 1.12E-06 |
| Ufc1 | 3015.992891 | 0.711989598 | 3.08E-05 |
| Tef | 3446.398834 | 0.711311338 | 3.68E-04 |
| Wdr5 | 2950.251821 | 0.711284882 | 2.04E-11 |
| Haus8 | 750.9704817 | 0.711100168 | 4.85E-03 |
| Ezh1 | 1677.187912 | 0.710727206 | 9.79E-08 |
| Lars2 | 26122.06978 | 0.710694807 | 9.30E-04 |
| Dnajb1 | 2104.812103 | 0.710556722 | 7.92E-05 |
| Tns2 | 5211.113391 | 0.71018082 | 6.65E-07 |
| Uvrag | 2392.79397 | 0.709754247 | 7.20E-07 |
| Dpf2 | 3923.177458 | 0.709700275 | 1.75E-08 |
| Mtfr1l | 3041.82241 | 0.709664982 | 3.31E-04 |
| Bri3bp | 3340.230155 | 0.709518114 | 4.71E-09 |
| Rala | 2465.88936 | 0.709414329 | 4.83E-12 |
| Kif1b | 4905.83088 | 0.708567376 | 3.33E-12 |
| Aimp1 | 2724.724341 | 0.708315625 | 1.87E-06 |
| Caskin2 | 1078.194197 | 0.706850647 | 3.25E-11 |
| Ggps1 | 715.1528405 | 0.706793351 | 1.31E-07 |
| Ets1 | 2712.616302 | 0.70603825 | 1.76E-05 |
| Drg1 | 4096.902164 | 0.705752301 | 9.92E-08 |
| E4f1 | 934.5039487 | 0.705172992 | 2.86E-05 |
| Rpl27a | 3086.246768 | 0.704800457 | 1.36E-05 |
| Parl | 1832.883324 | 0.704336243 | 4.79E-07 |
| Snrnp200 | 11710.44943 | 0.703909409 | 5.89E-08 |
| Mxd1 | 507.8744242 | 0.702808164 | 2.35E-03 |
| Eif4e2 | 4880.464822 | 0.702749849 | 5.91E-11 |
| Ifitm3 | 3638.162294 | 0.702587887 | 2.97E-03 |
| Tceanc | 180.8782985 | 0.70251479 | 3.60E-04 |
| Tmem106c | 1500.53999 | 0.702426043 | 3.12E-05 |
| Tex2 | 1768.876038 | 0.702233516 | 5.28E-08 |
| Nup37 | 843.5753032 | 0.702039247 | 4.88E-04 |
| AA986860 | 219.9850829 | 0.701994844 | 2.43E-03 |
| Lsm14a | 3705.854334 | 0.701341861 | 6.70E-08 |
| Nadk | 3967.334799 | 0.701236568 | 5.76E-07 |
| Nufip1 | 993.2100757 | 0.701207124 | 4.25E-06 |
| Dus1l | 1059.078537 | 0.700939453 | 1.99E-07 |
| Arv1 | 297.5590793 | 0.700933017 | 2.35E-07 |
| Tuba4a | 1663.613125 | 0.700734218 | 7.51E-03 |
| Smap1 | 2848.464316 | 0.70040114 | 7.53E-16 |
| Nudt15 | 117.3997007 | 0.700354747 | 9.23E-03 |

| | | | | |
|---|---|---|---|---|
| Dnajc21 | | 953.490279 | 0.700154875 | 1.76E-04 |
| Cox6a1 | | 5935.091383 | 0.69996249 | 1.82E-04 |
| Pawr | | 993.2221019 | 0.699426439 | 1.07E-08 |
| Zfp691 | | 296.0729184 | 0.699130041 | 2.31E-02 |
| Mre11a | | 1333.239055 | 0.698772256 | 5.55E-04 |
| Brpf1 | | 2256.159578 | 0.698459655 | 5.57E-08 |
| Xpot | | 6963.581493 | 0.698089877 | 6.71E-07 |
| Rbm18 | | 1817.797828 | 0.697812353 | 8.90E-06 |
| Cwf19l1 | | 759.216675 | 0.697499109 | 1.15E-06 |
| Dgkh | | 329.3063989 | 0.697444325 | 6.34E-03 |
| Mrps30 | | 1413.123487 | 0.697430765 | 2.55E-08 |
| Acss2 | | 638.0747025 | 0.697115309 | 2.64E-04 |
| Tmbim4 | | 1686.78166 | 0.696760983 | 5.52E-12 |
| Pou2f1 | | 712.764363 | 0.696749502 | 4.97E-04 |
| Abca2 | | 3887.217761 | 0.696647828 | 2.48E-10 |
| Crybg3 | | 324.3155032 | 0.696133736 | 6.53E-04 |
| Pnpla6 | | 1143.850243 | 0.69604928 | 6.06E-10 |
| Tcp1 | | 15233.79817 | 0.695799556 | 8.55E-17 |
| Metap2 | | 6994.685675 | 0.695674788 | 5.45E-08 |
| Palm | | 5324.7755 | 0.695640433 | 2.51E-06 |
| Ppm1m | | 1008.031276 | 0.695289247 | 1.64E-07 |
| | 2-Mar | 2784.399668 | 0.695116663 | 1.42E-05 |
| Mms19 | | 2815.736031 | 0.694816467 | 3.03E-08 |
| Akt3 | | 3456.107411 | 0.694395537 | 1.39E-05 |
| Wwp1 | | 1335.549327 | 0.694078321 | 7.27E-04 |
| Arhgap21 | | 4091.646736 | 0.69396505 | 1.19E-03 |
| Ldlrad4 | | 327.7250835 | 0.693950189 | 3.00E-05 |
| Phf1 | | 1872.957205 | 0.693685067 | 7.16E-04 |
| Stk4 | | 2349.963146 | 0.693604768 | 6.79E-19 |
| Txnl4a | | 1437.889867 | 0.693584415 | 3.52E-04 |
| Naa10 | | 1512.749773 | 0.693569903 | 2.46E-06 |
| Zswim7 | | 240.4272276 | 0.692801158 | 1.42E-05 |
| Supv3l1 | | 1487.778323 | 0.692657366 | 4.52E-15 |
| Vps11 | | 2731.764542 | 0.692342254 | 2.30E-03 |
| Tead1 | | 8305.982112 | 0.692238688 | 2.43E-07 |
| Ppan | | 1552.316091 | 0.691997748 | 6.38E-07 |
| Bambi | | 119.379043 | 0.690763224 | 4.03E-02 |
| Rfxap | | 458.8020367 | 0.690612986 | 6.48E-04 |
| Nop14 | | 3249.947278 | 0.690537506 | 7.86E-20 |
| Dcaf4 | | 1122.333632 | 0.689749493 | 1.16E-04 |
| Pld2 | | 1123.434152 | 0.689715512 | 4.29E-05 |
| Chchd1 | | 748.5531693 | 0.689606584 | 9.15E-06 |
| Slc25a19 | | 461.7725604 | 0.689231705 | 1.29E-04 |
| Ubqln1 | | 4261.004222 | 0.68804326 | 3.29E-09 |
| Mtbp | | 724.1374728 | 0.686866731 | 2.64E-02 |
| Sulf2 | | 1429.615205 | 0.686372481 | 1.98E-02 |
| Hspa12a | | 126.8535222 | 0.685230738 | 1.58E-02 |

| | | | |
|---|---|---|---|
| Cgrrf1 | 795.8021606 | 0.685144781 | 8.03E-07 |
| Ccdc186 | 2985.129838 | 0.685076609 | 1.28E-03 |
| Ahi1 | 1541.580211 | 0.684975628 | 1.28E-04 |
| Scnm1 | 789.5838704 | 0.68497406 | 7.27E-04 |
| Pdss2 | 496.4537985 | 0.684828983 | 3.24E-04 |
| Paxx | 701.5538367 | 0.684376632 | 7.36E-04 |
| Impdh2 | 6001.17937 | 0.68436482 | 1.07E-05 |
| Ttll3 | 274.4218314 | 0.683024016 | 1.62E-02 |
| Edf1 | 4065.529406 | 0.682975248 | 1.43E-06 |
| Tpmt | 384.4346636 | 0.682934099 | 1.44E-06 |
| H3f3b | 19605.31291 | 0.682709133 | 1.09E-08 |
| E130308A19Rik | 977.5793705 | 0.682707115 | 3.97E-06 |
| Gtpbp3 | 1278.258545 | 0.682702968 | 1.93E-05 |
| Gm12286 | 430.9828818 | 0.682649253 | 1.99E-02 |
| Akap1 | 2550.866096 | 0.682613228 | 2.42E-07 |
| Sh3glb2 | 1959.605868 | 0.682598695 | 1.12E-09 |
| Cdca3 | 1209.281701 | 0.68210843 | 4.62E-02 |
| 2700062C07Rik | 561.1969031 | 0.681543325 | 7.16E-07 |
| Xbp1 | 4888.960651 | 0.681469959 | 1.23E-03 |
| Amz2 | 3080.784919 | 0.681214894 | 1.05E-06 |
| Hsp90ab1 | 120629.8161 | 0.680521379 | 1.77E-12 |
| Pdia4 | 11618.21839 | 0.680325574 | 5.86E-05 |
| Mrpl1 | 717.9429599 | 0.679813266 | 3.89E-03 |
| Gfm2 | 1007.986387 | 0.679648511 | 1.43E-14 |
| Ostf1 | 2423.365697 | 0.679440177 | 4.74E-11 |
| Nek3 | 321.6148187 | 0.679358034 | 3.17E-06 |
| Gas2l3 | 1203.193311 | 0.679148191 | 1.53E-02 |
| Stard10 | 184.9670242 | 0.678951013 | 4.58E-02 |
| Thap3 | 441.7221161 | 0.678678491 | 1.41E-09 |
| Cnn2 | 23457.26203 | 0.678300702 | 3.28E-08 |
| Elf4 | 2047.77655 | 0.678200375 | 9.67E-13 |
| Exd1 | 114.7818395 | 0.677663838 | 1.50E-02 |
| NA | 4706.046236 | 0.677589516 | 3.56E-06 |
| Bsg | 26626.6997 | 0.677548222 | 2.31E-04 |
| Incenp | 2097.979085 | 0.677436345 | 3.43E-02 |
| Ppef1 | 316.4882102 | 0.677241891 | 1.91E-03 |
| Ubxn7 | 3892.875648 | 0.675242708 | 6.64E-08 |
| Nfs1 | 1585.844528 | 0.674391581 | 2.00E-08 |
| Herpud1 | 2966.583906 | 0.673851614 | 1.86E-05 |
| Sos1 | 2970.646572 | 0.673183969 | 4.59E-06 |
| Dnajb6 | 7299.017882 | 0.672905109 | 6.65E-07 |
| Zfp524 | 482.7409231 | 0.672388817 | 9.67E-04 |
| Ptprm | 1104.193032 | 0.672260986 | 8.12E-07 |
| Fbn1 | 5465.355355 | 0.671698138 | 2.18E-02 |
| Fbxl12 | 679.757772 | 0.67147499 | 3.97E-06 |
| Gm14303 | 4285.914875 | 0.670868521 | 1.91E-02 |
| Pigyl | 425.4755021 | 0.67079911 | 2.27E-04 |

| | | | |
|---|---|---|---|
| Mettl23 | 306.5676268 | 0.670738985 | 3.22E-03 |
| Wdr75 | 2758.597647 | 0.670340297 | 9.68E-06 |
| Ppp2r3c | 930.1243978 | 0.669952163 | 5.49E-11 |
| Aip | 2860.21654 | 0.669931017 | 9.31E-06 |
| Stk35 | 1822.968386 | 0.668648532 | 3.15E-08 |
| Nup107 | 1865.084275 | 0.668311589 | 2.37E-04 |
| Ccl27a | 208.3353949 | 0.668059137 | 5.56E-03 |
| Cald1 | 39536.21402 | 0.667739878 | 1.62E-04 |
| Snx18 | 1837.625647 | 0.667323985 | 5.59E-10 |
| Cebpb | 1041.170112 | 0.667278442 | 9.16E-03 |
| Srfbp1 | 1357.544805 | 0.66711443 | 4.71E-04 |
| Rnf166 | 1504.843752 | 0.666713107 | 2.35E-04 |
| Zfp385a | 1283.547352 | 0.665910997 | 7.08E-05 |
| Dnaja3 | 3016.129967 | 0.665628546 | 3.42E-08 |
| Ccdc59 | 1338.787267 | 0.665367927 | 1.52E-12 |
| Ddx19a | 2503.906907 | 0.66524861 | 3.15E-05 |
| Simc1 | 369.8851408 | 0.665227385 | 4.57E-04 |
| Uevld | 1160.400693 | 0.665161785 | 1.65E-04 |
| Poc1a | 204.6897645 | 0.664632804 | 2.60E-02 |
| Sbf2 | 4819.618936 | 0.664453732 | 2.92E-20 |
| Xpo4 | 1005.969614 | 0.664432798 | 7.12E-06 |
| Ndufb4c | 1556.461565 | 0.664285301 | 1.02E-05 |
| Phkg2 | 610.2063309 | 0.664115841 | 4.55E-07 |
| Nemp1 | 657.820187 | 0.664099993 | 3.20E-05 |
| Herc4 | 2473.94669 | 0.663862801 | 4.55E-08 |
| Rhog | 1417.782828 | 0.663710342 | 1.01E-03 |
| Polh | 607.3972268 | 0.663438892 | 1.64E-03 |
| Plekhj1 | 1287.09312 | 0.663283939 | 1.73E-08 |
| Ero1l | 4526.176064 | 0.662787126 | 1.76E-02 |
| Eif2b5 | 4375.143382 | 0.662661804 | 3.12E-07 |
| Tmem38a | 516.3255784 | 0.66183179 | 2.30E-06 |
| Bcl7b | 963.352409 | 0.661616051 | 6.04E-05 |
| Tnfsf9 | 487.3199691 | 0.661509104 | 1.69E-02 |
| Krtcap2 | 3311.574394 | 0.661130999 | 6.66E-06 |
| Dut | 1071.493419 | 0.660538118 | 2.79E-02 |
| Rab11fip1 | 1416.019755 | 0.660464587 | 3.62E-04 |
| Cand1 | 10381.58173 | 0.660180504 | 2.52E-07 |
| Ubald2 | 692.815041 | 0.660139253 | 9.36E-04 |
| Slc52a2 | 724.533322 | 0.659456244 | 2.86E-05 |
| Dnhd1 | 158.3893005 | 0.659354327 | 2.09E-02 |
| Pfdn4 | 874.6926454 | 0.659128899 | 3.26E-06 |
| Gtdc1 | 588.2460699 | 0.65872633 | 3.94E-04 |
| Ubap1 | 2119.323972 | 0.658457554 | 5.05E-03 |
| Lipe | 395.8530031 | 0.658348091 | 8.24E-04 |
| Tars | 6511.285888 | 0.658034381 | 4.11E-10 |
| Baiap2 | 2265.134962 | 0.658013886 | 9.06E-04 |
| Psmb3 | 3320.528605 | 0.657800237 | 7.40E-05 |

| | | | |
|---|---|---|---|
| Rabgef1 | 1307.570608 | 0.657103298 | 1.33E-09 |
| Mrs2 | 469.060725 | 0.656887634 | 6.40E-05 |
| Rsc1a1 | 1091.624438 | 0.656806357 | 5.60E-06 |
| Trabd | 1764.693389 | 0.656469352 | 1.42E-07 |
| Lsm5 | 534.3659574 | 0.65548588 | 2.05E-05 |
| Smc5 | 2503.855218 | 0.654888748 | 1.52E-02 |
| Coq10b | 1114.821112 | 0.65466598 | 7.49E-04 |
| Smg8 | 1733.268969 | 0.654579748 | 2.58E-05 |
| Ddx56 | 2294.187769 | 0.654075874 | 2.51E-05 |
| Tomm20 | 7274.548066 | 0.65397471 | 1.81E-05 |
| Cdk2ap2 | 1795.173273 | 0.653726694 | 1.94E-04 |
| Msn | 18054.98124 | 0.653586854 | 1.24E-03 |
| Fam104a | 1760.116884 | 0.653398304 | 5.43E-06 |
| Pcbp3 | 1704.285604 | 0.653358882 | 5.60E-05 |
| Snhg3 | 152.1985693 | 0.652767656 | 1.44E-02 |
| Dazap1 | 4156.04218 | 0.652749963 | 4.31E-11 |
| Gdi1 | 5499.526865 | 0.652744031 | 9.03E-06 |
| Slc7a1 | 3571.861693 | 0.652697105 | 3.67E-03 |
| Rpl36al | 5933.505914 | 0.652310295 | 8.37E-11 |
| Naaa | 680.2746552 | 0.652177413 | 5.83E-04 |
| Rps3a1 | 109609.6564 | 0.652086321 | 5.27E-05 |
| Cyp4f17 | 372.9455404 | 0.651545285 | 2.05E-05 |
| Tut1 | 1688.031275 | 0.651443178 | 2.08E-07 |
| Cnnm4 | 356.2181148 | 0.651122088 | 3.01E-05 |
| Ak6 | 673.3172241 | 0.651073284 | 7.19E-07 |
| Trim27 | 2284.639948 | 0.650625703 | 6.27E-06 |
| Hivep1 | 1746.576588 | 0.650327259 | 1.69E-10 |
| Mnat1 | 1466.460687 | 0.649994837 | 8.77E-05 |
| Hgs | 3316.298431 | 0.649871832 | 1.23E-04 |
| Abhd4 | 7163.071034 | 0.649586471 | 3.53E-03 |
| Eif3c | 17700.40432 | 0.649382084 | 5.52E-09 |
| Glud1 | 8559.981822 | 0.648741559 | 1.30E-20 |
| Ccdc130 | 585.6354187 | 0.648592317 | 1.01E-03 |
| Tnfaip1 | 4512.561739 | 0.64827457 | 1.07E-07 |
| Ankrd54 | 751.29234 | 0.648190757 | 8.37E-04 |
| Idh3g | 3552.809488 | 0.647947942 | 3.91E-12 |
| Crlf3 | 1281.734592 | 0.647798594 | 1.44E-05 |
| Psmb6 | 4995.39886 | 0.647573996 | 2.83E-07 |
| Dlst | 7252.782551 | 0.647414578 | 3.18E-07 |
| Saraf | 4956.861244 | 0.646372576 | 1.38E-04 |
| Mapk6 | 5890.268225 | 0.646093177 | 2.11E-06 |
| Ndufb4 | 577.8670532 | 0.645866151 | 2.11E-03 |
| Zc3hc1 | 1726.952055 | 0.644988638 | 3.92E-07 |
| H2-T10 | 831.3455959 | 0.644534126 | 8.07E-03 |
| Nsun2 | 5120.925682 | 0.644368563 | 4.19E-05 |
| Rpl21 | 28626.7454 | 0.644161212 | 9.77E-04 |
| Gin1 | 216.2621153 | 0.6431687 | 2.00E-04 |

| | | | |
|---|---|---|---|
| Klhdc10 | 3315.489594 | 0.643127863 | 1.24E-08 |
| Map4k1 | 81.71397557 | 0.642748882 | 1.77E-02 |
| Slc22a5 | 722.5646486 | 0.642511527 | 5.85E-05 |
| Acbd4 | 1094.853639 | 0.642434166 | 5.78E-09 |
| Gm10557 | 952.751582 | 0.642391869 | 8.22E-05 |
| Parn | 3178.519304 | 0.642130831 | 1.64E-07 |
| Rundc3a | 652.5243746 | 0.641995791 | 3.71E-03 |
| Inpp5f | 1261.195768 | 0.641978054 | 4.92E-08 |
| Pop5 | 528.7413162 | 0.64190312 | 3.48E-04 |
| Gabarapl2 | 2747.465548 | 0.641578297 | 7.99E-04 |
| Cers2 | 6182.42433 | 0.64157742 | 1.03E-12 |
| Rtraf | 4191.009413 | 0.641522665 | 1.55E-03 |
| Kyat1 | 280.1672108 | 0.641216962 | 2.52E-03 |
| F8a | 147.1483612 | 0.641149777 | 2.82E-02 |
| Eif2b4 | 1155.389 | 0.641094068 | 6.37E-07 |
| Asl | 1505.36333 | 0.639916193 | 7.90E-04 |
| Mrpl48-ps | 169.5839347 | 0.639817443 | 1.01E-02 |
| Smpd1 | 2430.281075 | 0.639716696 | 1.37E-04 |
| Gm12751 | 1729.560456 | 0.639512092 | 1.03E-03 |
| Gga1 | 2714.168085 | 0.639468203 | 9.09E-06 |
| Dgke | 540.5581551 | 0.63883143 | 9.85E-07 |
| Def6 | 175.6286037 | 0.638414373 | 1.74E-04 |
| Uchl5 | 1036.057855 | 0.638307746 | 4.51E-04 |
| Ifnar2 | 1462.423568 | 0.637846091 | 6.99E-04 |
| Hsdl2 | 1811.389637 | 0.637743751 | 2.11E-07 |
| Mvp | 5662.63892 | 0.637283779 | 3.79E-05 |
| Stam | 2412.624451 | 0.637134352 | 7.25E-05 |
| Rpl5 | 1094.398523 | 0.636976193 | 5.72E-05 |
| Akr1b8 | 5004.962685 | 0.636069883 | 4.93E-06 |
| Hnrnph3 | 773.6832434 | 0.63593767 | 5.97E-03 |
| Arih1 | 4317.986198 | 0.635349652 | 5.97E-11 |
| Rexo5 | 237.0252487 | 0.634919005 | 2.85E-02 |
| Rad9a | 2033.493668 | 0.634739731 | 1.70E-03 |
| Kpna6 | 7686.854538 | 0.634450368 | 4.94E-09 |
| Mrpl48 | 486.4467665 | 0.633821034 | 3.49E-03 |
| Mipol1 | 608.7016312 | 0.63371206 | 2.03E-07 |
| 4930539J05Rik | 104.1542753 | 0.633711739 | 2.78E-02 |
| Stil | 640.2338351 | 0.633470009 | 4.20E-02 |
| Fuk | 922.1422652 | 0.633042544 | 3.54E-06 |
| Gm10704 | 127.3962194 | 0.632855138 | 2.19E-02 |
| Zfp689 | 181.2686209 | 0.632772128 | 1.37E-03 |
| Pank4 | 603.6682192 | 0.632567554 | 6.63E-06 |
| Prrc1 | 2984.459546 | 0.632439152 | 4.30E-05 |
| Gm10774 | 279.4015294 | 0.632243765 | 1.04E-03 |
| Fosl2 | 7134.778544 | 0.632216284 | 2.90E-04 |
| Mpnd | 2802.889083 | 0.631890728 | 7.11E-06 |
| Fbxo7 | 1341.227542 | 0.631833225 | 1.04E-03 |

| | | | |
|---|---|---|---|
| Jmjd4 | 1066.637866 | 0.631810742 | 1.58E-09 |
| Hpcal1 | 926.3645226 | 0.631682681 | 5.33E-03 |
| Stx7 | 2576.759255 | 0.631665598 | 1.30E-07 |
| Mfsd8 | 360.6439686 | 0.630740342 | 1.33E-03 |
| Tob2 | 2102.527049 | 0.630661064 | 6.69E-05 |
| Vps39 | 2002.970673 | 0.630490995 | 8.14E-04 |
| Agap3 | 3243.418147 | 0.629871955 | 7.83E-05 |
| Abraxas1 | 442.9505407 | 0.629599701 | 9.43E-05 |
| Hyal1 | 467.9810951 | 0.629510236 | 4.16E-02 |
| Hras | 1694.369396 | 0.629108195 | 8.88E-04 |
| Hax1 | 2992.579162 | 0.629022397 | 4.50E-10 |
| Dip2a | 3267.183888 | 0.628364095 | 1.90E-09 |
| Rab11fip5 | 6386.276154 | 0.627953806 | 7.86E-04 |
| Sec63 | 6038.823214 | 0.627942273 | 6.86E-06 |
| Mrpl17 | 2145.416491 | 0.627919192 | 5.84E-07 |
| Farsb | 3179.035491 | 0.627645594 | 6.30E-06 |
| Fam118b | 1286.976374 | 0.627196921 | 5.60E-06 |
| Rrp12 | 1698.775959 | 0.626046246 | 1.46E-02 |
| Pi4k2a | 2798.58575 | 0.625906651 | 2.27E-06 |
| Lrrc47 | 1617.599376 | 0.625339877 | 3.10E-06 |
| Casp7 | 402.0609059 | 0.625321834 | 3.42E-04 |
| Jup | 2353.842657 | 0.624921272 | 1.09E-02 |
| Gchfr | 290.9261129 | 0.624887224 | 4.33E-02 |
| Ndufa6 | 2802.477104 | 0.624468401 | 7.08E-03 |
| H2-T22 | 227.0660543 | 0.624409881 | 2.66E-02 |
| Mavs | 1763.108388 | 0.624330308 | 4.58E-08 |
| Doc2g | 74.53898672 | 0.62426015 | 3.96E-02 |
| Rcl1 | 1451.162847 | 0.623941467 | 1.58E-05 |
| Smyd3 | 1007.188368 | 0.623386473 | 2.46E-03 |
| Pfdn1 | 1895.188969 | 0.62225433 | 2.93E-09 |
| Nol7 | 912.3460862 | 0.621944232 | 2.50E-04 |
| Ppp1r35 | 376.7552798 | 0.621789595 | 7.60E-03 |
| Letm2 | 106.9009209 | 0.621626533 | 8.19E-03 |
| Stard5 | 614.9927071 | 0.621257896 | 3.97E-05 |
| Unc119 | 1277.409738 | 0.621047313 | 4.80E-03 |
| Spryd7 | 748.305391 | 0.62008611 | 8.07E-08 |
| Nipsnap1 | 1460.490521 | 0.619938817 | 2.00E-06 |
| Abhd5 | 2340.155512 | 0.619537303 | 5.41E-04 |
| Zfand2a | 2124.767632 | 0.619368241 | 2.96E-04 |
| Ajuba | 2948.78609 | 0.619185099 | 3.10E-05 |
| Cry2 | 1311.129949 | 0.618571622 | 1.81E-06 |
| Atg9a | 2536.843685 | 0.618427476 | 3.12E-06 |
| Nedd9 | 2791.535274 | 0.618104565 | 1.65E-02 |
| Ier5 | 3710.218611 | 0.617814972 | 1.13E-04 |
| Poldip3 | 5976.958649 | 0.617744042 | 7.59E-05 |
| Tfeb | 959.7623694 | 0.617524854 | 4.25E-03 |
| Uso1 | 6535.449643 | 0.616330574 | 1.23E-05 |

| | | | |
|---|---|---|---|
| Mgea5 | 6513.668935 | 0.615836799 | 2.15E-04 |
| Fbxo31 | 875.7258759 | 0.615556474 | 6.15E-04 |
| Trmo | 211.4049252 | 0.614914962 | 9.62E-03 |
| Gm10177 | 3215.751042 | 0.614629628 | 1.75E-07 |
| Polr2e | 3801.975991 | 0.614504819 | 4.96E-06 |
| Banf1 | 3837.995394 | 0.614409922 | 3.82E-05 |
| Prim1 | 735.9249099 | 0.614143283 | 3.81E-02 |
| N4bp1 | 3589.986201 | 0.61388114 | 1.77E-15 |
| Dis3 | 2028.200728 | 0.613705777 | 6.94E-04 |
| Ttc39b | 2306.341585 | 0.613362458 | 9.81E-04 |
| Fmn1 | 2418.368649 | 0.612971343 | 3.68E-03 |
| Mrps16 | 1130.112843 | 0.61243396 | 1.04E-05 |
| Dnpep | 4104.734475 | 0.61237335 | 1.98E-05 |
| Creb3l1 | 1785.610876 | 0.612236905 | 1.25E-02 |
| Eya1 | 701.8287389 | 0.612045997 | 1.44E-07 |
| Hsp90b1 | 54806.66579 | 0.611994077 | 1.98E-03 |
| 1700029J07Rik | 68.72205517 | 0.611688268 | 3.69E-02 |
| Tent4b | 1416.599917 | 0.611224672 | 2.34E-03 |
| Atf6 | 5289.469533 | 0.611188214 | 1.10E-04 |
| Bmp2k | 1711.020882 | 0.610600473 | 3.72E-05 |
| mt-Rnr1 | 9050.912142 | 0.610429137 | 2.05E-02 |
| Nfatc2ip | 343.9678999 | 0.609402263 | 3.65E-03 |
| Psmc2 | 6666.244406 | 0.609173821 | 6.85E-08 |
| Dnmt3a | 1779.411715 | 0.608750431 | 5.98E-07 |
| Ate1 | 1420.82635 | 0.608715983 | 7.80E-06 |
| Zbtb40 | 700.9420987 | 0.608710393 | 2.50E-04 |
| Sod2 | 2401.870402 | 0.608684594 | 3.89E-06 |
| Tpt1 | 131718.7681 | 0.606828409 | 2.57E-03 |
| Ciao1 | 1904.407259 | 0.606484067 | 4.35E-05 |
| Memo1 | 2828.465884 | 0.606404862 | 1.95E-07 |
| Syvn1 | 2152.597269 | 0.606167192 | 6.94E-06 |
| Tbc1d23 | 1888.512151 | 0.605912541 | 1.12E-06 |
| NA | 457.4716934 | 0.605328996 | 1.61E-02 |
| Ttc17 | 2517.874017 | 0.605124753 | 1.27E-03 |
| Lyrm1 | 97.40410017 | 0.605115245 | 1.99E-02 |
| Tor1a | 1466.830404 | 0.60460119 | 4.65E-06 |
| Opa1 | 3963.090742 | 0.60431547 | 1.24E-13 |
| Zfp622 | 2476.499162 | 0.604213983 | 1.74E-02 |
| Ppid | 3377.358914 | 0.603918335 | 1.21E-03 |
| H13 | 6024.342285 | 0.603847108 | 9.35E-06 |
| Rabepk | 865.8826236 | 0.603636113 | 7.23E-04 |
| Mcph1 | 1213.891184 | 0.603480482 | 2.22E-06 |
| Ywhae | 20410.46669 | 0.603269292 | 1.34E-09 |
| Galc | 1514.601027 | 0.602363383 | 3.51E-05 |
| Med1 | 3533.303566 | 0.601813938 | 5.00E-04 |
| Map3k11 | 1003.039186 | 0.601430758 | 1.50E-07 |
| Cmss1 | 343.7516773 | 0.601369484 | 5.05E-04 |

| | | | |
|---|---|---|---|
| Vapb | 4955.166935 | 0.601060843 | 1.27E-06 |
| Ppp1r18 | 4671.279419 | 0.601017218 | 2.21E-08 |
| Slc39a14 | 1634.769816 | 0.600997009 | 2.80E-05 |
| Dffb | 114.904738 | 0.600705791 | 9.14E-03 |
| Naip2 | 1161.031535 | 0.600657131 | 2.62E-07 |
| Rwdd2a | 82.87345654 | 0.600393958 | 4.03E-02 |
| Sugt1 | 2423.145237 | 0.600290572 | 1.73E-05 |
| Egf | 72.21418597 | 0.600223803 | 3.97E-02 |
| Abca3 | 2846.907726 | 0.599478233 | 8.30E-06 |
| Mvb12a | 2343.567481 | 0.599392701 | 2.72E-03 |
| Nsd2 | 4012.714233 | 0.59930169 | 5.55E-04 |
| Arfip2 | 2214.951548 | 0.59911232 | 6.41E-06 |
| Mef2d | 4971.28913 | 0.59871808 | 9.49E-06 |
| Sat2 | 217.4762623 | 0.598706035 | 7.00E-03 |
| Lcat | 54.65890367 | 0.598532545 | 3.59E-02 |
| Rpusd3 | 280.6170281 | 0.598502935 | 7.18E-03 |
| Trim46 | 801.3602403 | 0.598376397 | 1.40E-03 |
| Gm10698 | 709.1241477 | 0.598258219 | 2.48E-03 |
| Tagap1 | 1599.643518 | 0.597530004 | 4.84E-05 |
| Aasdhppt | 981.0537025 | 0.597438833 | 3.29E-08 |
| Ptges3 | 6256.261295 | 0.597412691 | 3.36E-05 |
| Timm23 | 2565.708129 | 0.597405053 | 7.98E-07 |
| Yipf4 | 2227.605612 | 0.597201017 | 7.60E-05 |
| Ripor1 | 4272.017785 | 0.597165376 | 4.90E-07 |
| Irf9 | 319.9910044 | 0.596930119 | 8.02E-04 |
| Emc3 | 5381.192997 | 0.596230736 | 2.54E-05 |
| Rtn3 | 10304.32456 | 0.595987578 | 3.23E-05 |
| Atg4d | 558.9833682 | 0.595719169 | 3.64E-05 |
| Ube2h | 7839.735982 | 0.595631763 | 5.01E-03 |
| Fam160a2 | 1223.39513 | 0.595610107 | 1.49E-07 |
| Napg | 996.6463771 | 0.595147694 | 3.73E-05 |
| Rnf123 | 1593.468219 | 0.594674741 | 5.33E-06 |
| Exoc3l | 208.8741871 | 0.594662448 | 1.14E-03 |
| Imp3 | 1479.412076 | 0.594326154 | 1.81E-04 |
| Ssh1 | 2810.464612 | 0.594132952 | 4.73E-07 |
| Mrps35 | 1483.24933 | 0.59409467 | 4.20E-06 |
| Gnb2 | 15018.84145 | 0.593947468 | 2.80E-05 |
| Helb | 926.5999354 | 0.593886164 | 1.66E-08 |
| Mettl1 | 1327.406253 | 0.593504706 | 2.20E-04 |
| Rassf1 | 1982.544538 | 0.593419017 | 3.91E-05 |
| Gm45713 | 119.8688671 | 0.59315306 | 7.32E-03 |
| Kctd7 | 323.977193 | 0.592956789 | 6.31E-03 |
| Etv3 | 1203.077921 | 0.592659398 | 1.27E-06 |
| Sycp1 | 75.62340135 | 0.592524364 | 4.32E-02 |
| Dnajc27 | 615.2318411 | 0.592408094 | 6.65E-05 |
| Dstn | 11056.37856 | 0.59229732 | 3.26E-06 |
| Usf2 | 2888.615629 | 0.591649843 | 1.29E-05 |

| | | | |
|---|---|---|---|
| Nr1d2 | 1181.241596 | 0.591438438 | 1.63E-02 |
| Gpbp1l1 | 2823.12087 | 0.591162922 | 9.02E-04 |
| Trpc4ap | 8668.387395 | 0.590862423 | 1.48E-04 |
| Timmdc1 | 721.404571 | 0.590852341 | 4.91E-05 |
| Cep170b | 2978.572285 | 0.590817097 | 2.03E-05 |
| Fggy | 872.0594798 | 0.59059618 | 5.96E-04 |
| Manf | 3144.303668 | 0.590104531 | 5.31E-03 |
| Ak5 | 1153.641956 | 0.58964855 | 1.91E-04 |
| Gk5 | 482.1202121 | 0.58919459 | 6.78E-04 |
| Ell | 1456.306905 | 0.588702249 | 7.18E-04 |
| Tfe3 | 4096.617787 | 0.588654242 | 5.68E-05 |
| Srf | 2338.457348 | 0.588486947 | 1.62E-03 |
| Cox7c | 1341.314878 | 0.58824854 | 4.99E-05 |
| Wrnip1 | 1355.163956 | 0.588202132 | 3.73E-09 |
| Spata5 | 1125.39688 | 0.588041175 | 7.59E-04 |
| Nbeal2 | 273.9990957 | 0.588037341 | 4.01E-02 |
| Rfc4 | 532.2784859 | 0.58791218 | 4.58E-03 |
| Eif2b1 | 1612.710149 | 0.587550056 | 2.73E-06 |
| Phykpl | 589.4291438 | 0.587193668 | 1.23E-06 |
| Katna1 | 1574.205449 | 0.587154885 | 2.73E-06 |
| Mknk1 | 2154.3649 | 0.587113516 | 5.18E-06 |
| Nudcd2 | 1721.561492 | 0.586872365 | 1.36E-05 |
| Zfp593 | 763.2498669 | 0.586775212 | 1.20E-03 |
| Csnk2a1 | 5927.167477 | 0.58671258 | 5.18E-09 |
| Gclc | 1433.972265 | 0.586506915 | 2.89E-03 |
| Prkcd | 2207.110446 | 0.586411032 | 4.43E-06 |
| Gm9830 | 252.6703547 | 0.585903966 | 3.04E-02 |
| Gstm4 | 203.1581612 | 0.585903033 | 2.08E-03 |
| Mdh2 | 12364.87538 | 0.585749112 | 9.00E-05 |
| Grpel1 | 2345.33627 | 0.58573193 | 5.41E-04 |
| Canx | 28016.75408 | 0.585473859 | 2.81E-05 |
| Fbxw9 | 1293.75865 | 0.585385287 | 1.15E-05 |
| Ppp2r2d | 1434.57114 | 0.585291227 | 7.46E-04 |
| Mettl25 | 462.2847605 | 0.584891722 | 3.99E-04 |
| Tns3 | 8946.526377 | 0.584891444 | 1.00E-04 |
| Mettl15 | 267.6661028 | 0.584810366 | 1.00E-02 |
| Cdk17 | 1107.511682 | 0.583776635 | 5.48E-08 |
| Slc12a4 | 4348.57105 | 0.583685135 | 3.26E-07 |
| Bsdc1 | 2498.400934 | 0.583381907 | 3.25E-03 |
| Cpsf1 | 3715.112604 | 0.582989858 | 1.33E-05 |
| Mrpl53 | 903.2269651 | 0.582974273 | 1.64E-06 |
| Ufsp1 | 105.012883 | 0.582929234 | 2.66E-02 |
| Ddx19b | 450.9567018 | 0.582462545 | 1.78E-07 |
| Phtf2 | 418.8036949 | 0.582297297 | 1.32E-02 |
| Rps3a2 | 220.5441406 | 0.582264514 | 3.28E-02 |
| Pdzd11 | 1920.318247 | 0.581497131 | 6.61E-05 |
| 3110040N11Rik | 458.3785251 | 0.580998045 | 2.73E-06 |

| | | | |
|---|---|---|---|
| Prpsap1 | 2055.416811 | 0.580806261 | 6.02E-09 |
| Sdf4 | 4735.660727 | 0.580602447 | 1.31E-04 |
| Gnl3l | 4240.911019 | 0.580550335 | 4.70E-13 |
| Rars | 6691.937495 | 0.58045053 | 1.32E-05 |
| Wdr45b | 3049.093296 | 0.580134493 | 2.41E-07 |
| Pfdn2 | 1804.340573 | 0.579654974 | 5.17E-07 |
| Rad51d | 1139.790286 | 0.579482487 | 1.69E-04 |
| Eif6 | 3599.494945 | 0.578946365 | 3.17E-04 |
| Trmt13 | 347.1095397 | 0.578857825 | 2.48E-02 |
| Metrnl | 1886.117273 | 0.578711353 | 5.59E-05 |
| Dym | 1835.376692 | 0.57858983 | 2.04E-03 |
| Igbp1 | 3698.89057 | 0.577518031 | 4.64E-03 |
| Jade3 | 903.244704 | 0.576984641 | 4.94E-06 |
| Ccdc117 | 1297.355029 | 0.576893607 | 1.35E-05 |
| Nsun5 | 676.7308702 | 0.576390804 | 6.77E-04 |
| Psmb7 | 5329.244185 | 0.575924793 | 7.21E-07 |
| Eif1 | 18854.81854 | 0.575789715 | 1.56E-04 |
| Nkx2-5 | 211.3079351 | 0.574818336 | 1.55E-03 |
| Thap11 | 1739.529214 | 0.574423752 | 4.45E-04 |
| Igf1r | 2901.97391 | 0.574115504 | 4.15E-07 |
| Atl2 | 1798.458299 | 0.574051271 | 1.38E-06 |
| Arl2 | 2679.750191 | 0.573923714 | 3.75E-03 |
| Rnf126 | 1969.762057 | 0.573796767 | 3.30E-04 |
| Prcp | 2518.163059 | 0.573698766 | 1.57E-03 |
| Sh3rf1 | 1249.064827 | 0.573542959 | 1.41E-04 |
| Ahnak | 67844.71225 | 0.573060225 | 7.81E-05 |
| Btbd1 | 3241.787789 | 0.572983535 | 7.37E-05 |
| Alg14 | 549.7607113 | 0.572887383 | 8.76E-05 |
| Saysd1 | 190.5484555 | 0.57218598 | 3.12E-03 |
| Csk | 2408.111774 | 0.571488282 | 1.52E-04 |
| Mad2l2 | 595.8755217 | 0.571265078 | 2.06E-05 |
| Qsox1 | 10100.16956 | 0.571225878 | 1.49E-03 |
| Wwc2 | 6641.559661 | 0.570218889 | 5.69E-06 |
| Wnk1 | 21009.8334 | 0.570117385 | 2.83E-04 |
| Gm9531 | 2715.543962 | 0.56996851 | 1.83E-04 |
| Ganab | 13601.05205 | 0.569915365 | 3.34E-08 |
| Dnajb12 | 2755.669575 | 0.569825044 | 1.24E-05 |
| Otub2 | 372.7939584 | 0.569770346 | 6.53E-03 |
| Stk39 | 1399.447575 | 0.569605232 | 8.76E-04 |
| Erf | 2001.173073 | 0.569250835 | 7.21E-03 |
| Fbxo38 | 2616.09343 | 0.568991098 | 8.00E-08 |
| Osr1 | 5513.095233 | 0.568928527 | 6.60E-03 |
| Zbtb18 | 764.6264989 | 0.568797999 | 2.44E-02 |
| Mlx | 1355.07942 | 0.568506341 | 2.06E-06 |
| Lonp1 | 6941.8007 | 0.568214517 | 1.97E-04 |
| Cct6a | 10672.83687 | 0.568152716 | 4.03E-05 |
| Ccdc85b | 694.6340861 | 0.56810955 | 3.87E-03 |

| | | | |
|---|---|---|---|
| Stx1a | 409.9590611 | 0.567922684 | 8.43E-04 |
| Usf3 | 1381.631125 | 0.567779109 | 5.35E-04 |
| Med15 | 3551.143144 | 0.567761593 | 8.16E-04 |
| Rsl1d1 | 5971.149172 | 0.567623226 | 1.33E-06 |
| Usp19 | 5211.507926 | 0.566451479 | 3.56E-05 |
| Ybx1 | 42624.67727 | 0.566291604 | 3.90E-04 |
| Eci2 | 2234.646018 | 0.56536536 | 8.85E-06 |
| Actr8 | 1337.352931 | 0.56531498 | 4.15E-06 |
| Numbl | 3202.627356 | 0.56474184 | 3.09E-04 |
| Nop16 | 880.4090839 | 0.564497769 | 1.29E-03 |
| Srpr | 5889.393341 | 0.564414949 | 2.12E-04 |
| Tmem70 | 867.4598913 | 0.564395248 | 3.48E-06 |
| Slc39a7 | 10355.74721 | 0.564241866 | 4.67E-04 |
| Mrps24 | 1143.813471 | 0.564235852 | 1.12E-03 |
| Snx14 | 1287.976179 | 0.564173511 | 3.10E-06 |
| Tmem120b | 354.8541929 | 0.564058239 | 1.63E-03 |
| Thumpd3 | 1520.849078 | 0.563370499 | 7.37E-06 |
| Actn4 | 15981.88887 | 0.563238494 | 8.91E-05 |
| Aktip | 1968.524584 | 0.563205885 | 9.11E-07 |
| Herpud2 | 1800.931372 | 0.562916699 | 1.43E-05 |
| Bbs4 | 321.4642689 | 0.562848415 | 1.54E-04 |
| Dnaja2 | 8057.883399 | 0.562640624 | 1.83E-06 |
| Luc7l | 3039.936213 | 0.562554729 | 1.56E-04 |
| Kctd6 | 240.6580014 | 0.561485833 | 1.13E-04 |
| Ints5 | 1926.240369 | 0.561096707 | 5.97E-05 |
| Rnf38 | 1227.714967 | 0.560899332 | 2.72E-03 |
| Zfp526 | 555.9011632 | 0.559960113 | 1.16E-02 |
| Daxx | 1244.430088 | 0.558857183 | 1.38E-06 |
| Clip1 | 4554.958969 | 0.558463552 | 2.77E-04 |
| Prelid3b | 1774.650704 | 0.558147874 | 5.41E-04 |
| Tle4 | 360.9712993 | 0.558033998 | 1.21E-03 |
| Dennd2c | 573.6539703 | 0.557644113 | 7.16E-07 |
| Gpn3 | 839.2302068 | 0.557562527 | 4.83E-05 |
| Adamtsl4 | 642.4400238 | 0.557411752 | 1.86E-03 |
| Ankrd13d | 440.9352955 | 0.556459139 | 1.08E-02 |
| Uba1 | 17656.13637 | 0.556427585 | 8.71E-05 |
| Adamts4 | 1372.936021 | 0.555146112 | 2.36E-02 |
| Polr1c | 2348.159737 | 0.555073312 | 1.38E-03 |
| Cdpf1 | 451.3269781 | 0.554743541 | 2.49E-03 |
| Radil | 568.4919136 | 0.554242903 | 2.70E-03 |
| Atp5c1 | 10686.56103 | 0.553829673 | 4.18E-04 |
| Pak1ip1 | 1946.659596 | 0.553723122 | 2.72E-06 |
| Fastkd5 | 450.8853233 | 0.553636973 | 5.77E-03 |
| Ermard | 436.0657237 | 0.553100552 | 8.94E-05 |
| Arhgef18 | 3617.647802 | 0.553000683 | 1.59E-04 |
| Utp23 | 570.5468979 | 0.552908775 | 3.23E-08 |
| Tspan31 | 6006.152092 | 0.552878903 | 7.69E-04 |

| | | | |
|---|---|---|---|
| Mrps23 | 1720.567113 | 0.552723653 | 6.24E-04 |
| Actr1a | 7248.29814 | 0.552661418 | 4.88E-04 |
| 1110002L01Rik | 207.9137928 | 0.552413738 | 1.12E-02 |
| Sys1 | 1599.216733 | 0.552365336 | 1.52E-03 |
| Gm1821 | 5869.390963 | 0.552208742 | 1.14E-02 |
| Ppp1r12c | 1921.830022 | 0.552193868 | 2.46E-07 |
| Ltv1 | 1629.7818 | 0.552169618 | 6.42E-05 |
| Ubr2 | 5346.693588 | 0.552162678 | 2.15E-13 |
| Ubb | 30944.43698 | 0.552056448 | 9.36E-03 |
| Surf2 | 1370.085633 | 0.551897732 | 1.79E-07 |
| Tollip | 3274.862957 | 0.551730357 | 7.70E-04 |
| Hnrnpc | 10522.29268 | 0.551549165 | 1.03E-05 |
| Rita1 | 573.5582389 | 0.551536903 | 3.41E-04 |
| Cox6b1 | 4254.104049 | 0.550752979 | 7.45E-05 |
| Mrpl14 | 1006.247001 | 0.550423614 | 6.99E-03 |
| Clptm1 | 4833.878488 | 0.550239967 | 3.45E-04 |
| Scyl1 | 5743.234042 | 0.550225157 | 2.30E-04 |
| Ftl1 | 156446.3262 | 0.550064382 | 7.59E-03 |
| Bcl6 | 1044.315841 | 0.550043328 | 8.35E-03 |
| Akirin2 | 2250.172154 | 0.549320013 | 2.92E-05 |
| Sos2 | 1289.168662 | 0.549157046 | 6.33E-05 |
| Tubb6 | 4737.383742 | 0.54865802 | 3.57E-02 |
| Casc3 | 2226.789352 | 0.548597452 | 1.13E-07 |
| Tmem183a | 2815.550326 | 0.548452367 | 1.31E-08 |
| Cox17 | 741.8334911 | 0.548133701 | 1.68E-07 |
| Gnb1 | 10433.30772 | 0.54719396 | 1.65E-17 |
| Arfgap3 | 2436.475969 | 0.546807764 | 1.26E-03 |
| Dhx29 | 1698.366442 | 0.546757334 | 1.46E-10 |
| Kcnab1 | 1245.884438 | 0.546723069 | 2.88E-02 |
| Prmt5 | 1974.839417 | 0.546602988 | 8.94E-05 |
| Actr10 | 3839.071474 | 0.546595251 | 3.15E-07 |
| Supt5 | 6937.965658 | 0.546546782 | 2.38E-04 |
| Zfand2b | 745.0809418 | 0.546373437 | 1.74E-05 |
| Bicd2 | 3364.546628 | 0.546171235 | 6.51E-06 |
| Pcbp2 | 19401.57002 | 0.546005966 | 6.37E-05 |
| Rhebl1 | 244.5837871 | 0.545900764 | 2.00E-02 |
| Naca | 33543.9046 | 0.545766256 | 6.63E-04 |
| Gatad2a | 4485.776346 | 0.545633177 | 3.50E-06 |
| Diaph2 | 713.883724 | 0.545261396 | 5.33E-03 |
| Cyc1 | 3631.315047 | 0.545184053 | 1.52E-04 |
| Sdf2l1 | 506.6027358 | 0.545145481 | 2.11E-02 |
| Nmt1 | 7745.587509 | 0.545122125 | 1.43E-04 |
| Fastkd1 | 692.7686316 | 0.545116019 | 1.26E-06 |
| Timm17a | 2158.37418 | 0.545083108 | 2.15E-10 |
| Slc41a1 | 1950.284416 | 0.545013801 | 3.40E-04 |
| Rab40c | 1203.858473 | 0.544873302 | 7.97E-04 |
| Lap3 | 4765.227468 | 0.544353852 | 6.17E-06 |

| | | | |
|---|---|---|---|
| Gmeb1 | 1118.296004 | 0.544025353 | 4.40E-06 |
| Sirt2 | 4842.503244 | 0.543886959 | 1.37E-03 |
| Zfand3 | 5760.058426 | 0.543851272 | 2.81E-03 |
| Fam129b | 15591.52094 | 0.543715318 | 2.22E-03 |
| Hprt | 2471.179009 | 0.543575822 | 1.07E-03 |
| Cox4i1 | 12755.47037 | 0.542671928 | 2.47E-03 |
| Pbx3 | 618.9527378 | 0.542525853 | 7.38E-04 |
| Gdi2 | 14707.93982 | 0.542337288 | 1.32E-05 |
| Cage1 | 131.6143912 | 0.542227666 | 1.44E-03 |
| Magi3 | 1822.469333 | 0.542100051 | 9.58E-03 |
| Zc3h10 | 826.6331516 | 0.542055876 | 5.63E-04 |
| Txn1 | 10959.5229 | 0.541574765 | 4.13E-04 |
| Ehbp1l1 | 2904.929473 | 0.541470348 | 1.23E-05 |
| Hikeshi | 958.3911583 | 0.541179434 | 7.10E-06 |
| Snora43 | 461.7845665 | 0.541060724 | 2.76E-02 |
| Sssca1 | 1262.870255 | 0.540897031 | 5.61E-05 |
| Ormdl2 | 671.3612949 | 0.540564903 | 3.09E-03 |
| Mplkip | 339.1711707 | 0.540405838 | 8.09E-03 |
| U2af1 | 2772.347274 | 0.539823455 | 5.42E-06 |
| Lars | 4924.644824 | 0.539823301 | 1.42E-02 |
| Zrsr1 | 2272.804588 | 0.539753287 | 1.62E-02 |
| Ddx50 | 4596.223589 | 0.539392668 | 9.87E-04 |
| Eif1ad | 1505.772972 | 0.539086398 | 6.31E-04 |
| Map2k1 | 2663.239994 | 0.539006554 | 1.77E-07 |
| Gtf3a | 2196.178491 | 0.538904721 | 2.11E-03 |
| Atg14 | 874.8182583 | 0.538700928 | 4.82E-02 |
| Gtpbp1 | 2056.74881 | 0.538602544 | 7.22E-04 |
| Npm1 | 45543.89849 | 0.538361659 | 1.55E-03 |
| Hdgfl2 | 4078.471706 | 0.537925647 | 5.56E-08 |
| Zc3h14 | 5985.988222 | 0.537820604 | 1.81E-07 |
| Prkci | 7019.667731 | 0.537790158 | 1.91E-03 |
| Zdhhc9 | 2534.383131 | 0.53778957 | 1.94E-04 |
| Ndufa7 | 3232.605156 | 0.536891403 | 6.28E-03 |
| Nhp2 | 1896.70157 | 0.53679506 | 5.48E-03 |
| Hspb8 | 6743.145759 | 0.536582162 | 1.48E-04 |
| Mterf4 | 757.3895142 | 0.536426583 | 2.51E-07 |
| Tchp | 576.9511312 | 0.536089678 | 5.04E-03 |
| Rpain | 423.7311747 | 0.535706464 | 3.27E-03 |
| Ncoa3 | 2075.835614 | 0.5355192 | 3.32E-07 |
| Sec61g | 587.7553782 | 0.534945805 | 6.97E-05 |
| Atp5b | 49226.93766 | 0.534718485 | 2.65E-04 |
| Ubald1 | 2040.828702 | 0.534605056 | 7.86E-03 |
| Tbca | 3268.050332 | 0.534468298 | 8.42E-07 |
| Use1 | 4339.895075 | 0.534330473 | 3.84E-02 |
| Dph7 | 114.5500558 | 0.534125152 | 2.18E-02 |
| Grb2 | 2639.41625 | 0.5339264 | 1.37E-05 |
| Ngdn | 1047.951766 | 0.533715337 | 9.47E-06 |

| | | | |
|---|---|---|---|
| Gnb5 | 631.122664 | 0.53297885 | 1.27E-03 |
| Ahctf1 | 4874.929657 | 0.531895699 | 4.48E-04 |
| Umps | 1718.713927 | 0.531769022 | 2.93E-03 |
| Pde4dip | 2967.356983 | 0.531640454 | 1.18E-11 |
| Mtif3 | 797.1508615 | 0.531444643 | 2.61E-03 |
| Eif4enif1 | 3055.847692 | 0.531423613 | 1.32E-06 |
| Camsap1 | 3025.579919 | 0.531409318 | 1.81E-06 |
| Akr1e1 | 217.9345246 | 0.531369979 | 3.12E-02 |
| Rce1 | 1079.36259 | 0.531369559 | 2.55E-07 |
| Kmt5c | 716.1049833 | 0.530836967 | 6.11E-03 |
| Hmg20b | 1606.6635 | 0.530781116 | 2.34E-04 |
| Mrps27 | 872.321991 | 0.530407797 | 1.58E-03 |
| Csnk2b | 3640.167179 | 0.530042028 | 3.90E-04 |
| Zxdc | 3218.603639 | 0.529221622 | 2.20E-03 |
| R3hdm1 | 3476.07313 | 0.529161506 | 6.05E-07 |
| Smyd4 | 635.3620075 | 0.528736082 | 2.23E-03 |
| Tagln2 | 5657.746603 | 0.528715282 | 2.36E-02 |
| Edc4 | 3070.827727 | 0.52834915 | 1.47E-03 |
| Surf1 | 1397.802757 | 0.528161611 | 2.57E-03 |
| Adat1 | 343.821926 | 0.528086027 | 5.60E-03 |
| NA | 1983.288498 | 0.52743277 | 6.80E-04 |
| Mlkl | 1417.964644 | 0.527394218 | 1.13E-03 |
| Psma7 | 5994.916682 | 0.527353669 | 3.02E-04 |
| Pwp2 | 1822.29441 | 0.527182851 | 5.44E-04 |
| Utp18 | 1810.570753 | 0.527124614 | 4.55E-04 |
| Slc46a1 | 317.5478592 | 0.527056804 | 6.31E-03 |
| Zcwpw1 | 487.8432153 | 0.527022204 | 2.93E-03 |
| Ccdc97 | 1381.339871 | 0.526899221 | 2.53E-04 |
| Hspe1 | 7725.560891 | 0.526834471 | 9.45E-05 |
| Praf2 | 2315.534915 | 0.526489331 | 1.18E-05 |
| Nifk | 2606.036144 | 0.526148331 | 1.26E-04 |
| Stk17b | 1172.316199 | 0.5259276 | 7.30E-03 |
| Ndfip2 | 1306.937112 | 0.524847909 | 3.27E-10 |
| Zbtb10 | 572.374976 | 0.524783404 | 3.93E-04 |
| Ndc1 | 2175.202521 | 0.524719939 | 1.88E-02 |
| Srebf1 | 2530.686061 | 0.524621769 | 3.52E-02 |
| Snap47 | 3433.560277 | 0.524607401 | 2.90E-04 |
| Ccdc14 | 270.2503542 | 0.524373204 | 2.72E-02 |
| Cherp | 3589.831824 | 0.52412186 | 3.04E-04 |
| Pgm2 | 1881.276836 | 0.524049911 | 5.06E-05 |
| Zc2hc1a | 2130.526618 | 0.524027432 | 9.14E-04 |
| Pdha1 | 9868.078405 | 0.523998046 | 1.94E-09 |
| Kmt2b | 2483.737405 | 0.523871139 | 2.36E-03 |
| Gmpr2 | 704.4970757 | 0.523676478 | 2.59E-06 |
| Tmem167b | 3267.309721 | 0.523480889 | 3.75E-03 |
| Zxda | 240.0594274 | 0.523468919 | 1.17E-02 |
| Map4k3 | 1741.257735 | 0.523149607 | 2.18E-06 |

| | | | |
|---|---|---|---|
| Gm28043 | 890.2744121 | 0.522084886 | 6.70E-03 |
| Smu1 | 3414.245524 | 0.52186128 | 4.23E-08 |
| Alg2 | 1510.645998 | 0.521501024 | 4.27E-04 |
| Glce | 2652.834273 | 0.521157751 | 6.21E-03 |
| mt-Co1 | 763641.7603 | 0.520901935 | 4.48E-04 |
| Cd44 | 4483.253208 | 0.520702303 | 1.32E-02 |
| Lurap1 | 310.6166841 | 0.520634783 | 4.48E-04 |
| Sptbn1 | 59834.79882 | 0.520462328 | 1.32E-03 |
| Ophn1 | 1096.550011 | 0.520400358 | 3.36E-04 |
| Mrpl16 | 1103.742269 | 0.520283423 | 1.01E-03 |
| Nol10 | 2056.588005 | 0.5200308 | 1.43E-05 |
| Ubr4 | 11843.35144 | 0.519900226 | 8.43E-04 |
| Syf2 | 1649.935848 | 0.518970915 | 5.99E-05 |
| Braf | 1890.433423 | 0.518784202 | 7.16E-03 |
| Snrpc | 2036.48846 | 0.518123874 | 7.22E-04 |
| Eprs | 14576.67394 | 0.517710117 | 1.99E-03 |
| Prkag1 | 2253.546472 | 0.517316258 | 1.18E-03 |
| Bcar1 | 3817.148731 | 0.517312813 | 1.38E-05 |
| Slc30a6 | 877.1499439 | 0.517035641 | 7.32E-06 |
| Ecsit | 800.8495056 | 0.516987489 | 2.76E-03 |
| Ndufaf5 | 624.1161059 | 0.516611053 | 3.09E-03 |
| Slc15a4 | 1139.09426 | 0.516563972 | 2.71E-02 |
| Sec22c | 967.296682 | 0.516484042 | 6.72E-03 |
| Ptrh1 | 1785.441485 | 0.51625491 | 4.26E-03 |
| Ndufb6 | 1520.582627 | 0.516095154 | 8.83E-05 |
| Gsdmd | 431.2673849 | 0.515882197 | 5.29E-05 |
| Slc25a40 | 773.2498353 | 0.51568533 | 2.50E-02 |
| Nadsyn1 | 273.4059839 | 0.515558942 | 2.12E-02 |
| Tceanc2 | 1099.383763 | 0.515545251 | 4.88E-02 |
| Xpc | 1557.390835 | 0.515467545 | 9.95E-05 |
| Eif2s1 | 4474.748387 | 0.515050099 | 1.39E-04 |
| Ndufb9 | 3591.013312 | 0.51501989 | 2.22E-03 |
| NA | 236.8173459 | 0.514979691 | 2.34E-02 |
| Fchsd1 | 780.790929 | 0.514754918 | 2.26E-02 |
| Zfp462 | 2551.882414 | 0.514732218 | 8.72E-03 |
| Nsmce1 | 1028.989147 | 0.514349194 | 7.32E-04 |
| Dhx38 | 2719.106964 | 0.514131471 | 4.52E-06 |
| Gm15417 | 73.13027253 | 0.513739271 | 4.96E-02 |
| Tiprl | 2259.29702 | 0.513422379 | 1.85E-06 |
| Rbm15 | 572.4601641 | 0.512474162 | 3.69E-03 |
| Aurkaip1 | 1428.016499 | 0.512426137 | 3.89E-05 |
| Rubcn | 3167.973214 | 0.512326849 | 2.54E-05 |
| Mto1 | 581.4842172 | 0.512248939 | 7.57E-04 |
| Cenpo | 452.4141085 | 0.51215373 | 6.24E-03 |
| Ttc37 | 2099.340995 | 0.511951672 | 9.46E-03 |
| Acot9 | 3567.356035 | 0.511909449 | 2.45E-03 |
| Cul3 | 5673.330745 | 0.51140194 | 1.72E-05 |

| | | | |
|---|---|---|---|
| Usp10 | 2331.73001 | 0.511263007 | 4.34E-05 |
| Gpkow | 1945.850323 | 0.511026998 | 5.28E-05 |
| Akap7 | 767.851978 | 0.510701474 | 8.12E-04 |
| Polr2i | 588.2928749 | 0.510387672 | 3.59E-03 |
| Wwtr1 | 5195.543564 | 0.510149266 | 6.75E-11 |
| Cdkn1b | 3842.681411 | 0.510148971 | 2.26E-03 |
| Tmem214 | 3151.052134 | 0.509993779 | 2.23E-04 |
| C1qbp | 6609.162753 | 0.509676394 | 9.16E-05 |
| Gm9864 | 1039.948219 | 0.509270268 | 8.36E-05 |
| Palld | 2774.077585 | 0.508509292 | 6.81E-03 |
| Ncoa4 | 4582.091116 | 0.50844923 | 9.01E-08 |
| Mis18a | 305.8991117 | 0.507660828 | 2.75E-02 |
| Ythdf1 | 3589.274297 | 0.507621232 | 3.43E-08 |
| Icmt | 2198.9569 | 0.506987231 | 1.46E-03 |
| Epb41l3 | 11414.40963 | 0.50678968 | 8.26E-08 |
| Calhm2 | 211.5215987 | 0.506604118 | 8.44E-03 |
| Spata7 | 434.5194125 | 0.506399203 | 7.22E-06 |
| Ugcg | 3159.293624 | 0.50632045 | 2.05E-03 |
| Stau2 | 1353.634259 | 0.505982843 | 1.37E-09 |
| Leng1 | 441.7638383 | 0.505842962 | 4.30E-03 |
| Tmem189 | 1348.449585 | 0.505826933 | 4.21E-06 |
| Cry1 | 919.8294528 | 0.505495831 | 1.04E-02 |
| Elmo3 | 103.5252001 | 0.505413826 | 3.54E-02 |
| Vta1 | 2205.334183 | 0.505170701 | 8.16E-05 |
| Rpl10-ps3 | 54386.59665 | 0.505159536 | 1.53E-02 |
| Wdr44 | 567.3078697 | 0.50503636 | 3.76E-04 |
| Mta2 | 6317.531041 | 0.504930103 | 2.07E-07 |
| Edem1 | 4433.87396 | 0.504816233 | 8.08E-07 |
| Tcf25 | 13497.59425 | 0.504448117 | 5.13E-06 |
| Npdc1 | 6730.563996 | 0.504221554 | 8.83E-03 |
| Zcchc10 | 277.3639513 | 0.503702458 | 7.52E-03 |
| Zfp628 | 484.0723769 | 0.503683973 | 1.46E-03 |
| Cep85l | 193.1259737 | 0.503561106 | 3.36E-02 |
| Mcm3ap | 3567.789929 | 0.503025218 | 1.93E-07 |
| Prdx3 | 4040.344397 | 0.50275703 | 1.07E-06 |
| Fam131b | 2012.653011 | 0.502186708 | 2.57E-03 |
| Adcy6 | 3708.430534 | 0.501724169 | 1.54E-08 |
| Ephx2 | 595.9836792 | 0.501612098 | 1.42E-02 |
| Sh3gl3 | 191.0386475 | 0.501335244 | 2.97E-02 |
| Zfp280b | 1049.133878 | 0.501289977 | 4.83E-06 |
| Polr2l | 617.9798244 | 0.50048416 | 3.26E-02 |
| Afg3l2 | 2671.085849 | 0.500254782 | 4.62E-06 |
| Ndrg4 | 1002.618499 | 0.500240431 | 9.59E-03 |
| Stat5b | 2117.456746 | 0.500158141 | 1.67E-08 |
| Abhd6 | 166.9699771 | 0.500130194 | 4.00E-02 |
| Herc1 | 3285.880765 | 0.500107608 | 4.89E-05 |
| Cox8a | 8974.358587 | 0.49963124 | 1.27E-03 |

| | | | |
|---|---|---|---|
| Tead3 | 2341.916993 | 0.499137132 | 1.70E-03 |
| Nvl | 1837.408422 | 0.498995142 | 1.15E-08 |
| Ptpn1 | 2274.268902 | 0.498709678 | 1.47E-03 |
| Pmpca | 4581.083073 | 0.498473582 | 3.40E-10 |
| Slc16a1 | 2748.76285 | 0.498181056 | 3.88E-02 |
| Lhx9 | 259.9429502 | 0.498118725 | 2.99E-02 |
| Tbl3 | 1798.960926 | 0.498094067 | 2.04E-02 |
| Eci1 | 1615.091732 | 0.49775194 | 1.25E-04 |
| Zyg11b | 5267.774086 | 0.497544706 | 2.91E-06 |
| Med26 | 634.3945367 | 0.497378955 | 5.17E-04 |
| Lancl1 | 2052.092334 | 0.497265757 | 6.38E-04 |
| Ei24 | 3209.591063 | 0.497117875 | 1.49E-03 |
| Mrpl47 | 715.1774425 | 0.496960404 | 4.24E-03 |
| Lsm2 | 1237.134903 | 0.496953933 | 3.02E-02 |
| Tmem109 | 3376.214754 | 0.496951211 | 1.22E-02 |
| Pdcd2 | 1168.649565 | 0.496849338 | 2.04E-04 |
| Naf1 | 1224.57437 | 0.496836564 | 1.36E-05 |
| Sdhb | 3446.628872 | 0.495646144 | 9.09E-05 |
| Psmc1 | 6304.674307 | 0.495565727 | 1.42E-05 |
| Ubc | 68322.4058 | 0.495559318 | 2.15E-02 |
| Rhoc | 7213.832226 | 0.49555181 | 1.58E-03 |
| Cbwd1 | 629.8398015 | 0.495544543 | 8.31E-03 |
| Crbn | 1780.102223 | 0.494887079 | 3.33E-04 |
| Abhd17c | 2306.603289 | 0.494584949 | 1.97E-03 |
| Polr2b | 4760.275088 | 0.4939491 | 5.71E-04 |
| Stard3 | 1188.605779 | 0.493890483 | 1.94E-03 |
| Nit1 | 1001.7084 | 0.493886885 | 5.80E-04 |
| Ccdc120 | 311.5972816 | 0.493877026 | 1.47E-02 |
| Acot8 | 248.0830666 | 0.493701229 | 9.74E-03 |
| Ttll12 | 2912.203449 | 0.493081229 | 2.52E-03 |
| Rsph3b | 1349.967464 | 0.493005497 | 6.19E-03 |
| Mtg1 | 484.3456982 | 0.492832131 | 2.01E-02 |
| Gm13394 | 70314.65906 | 0.492585499 | 6.61E-03 |
| Mrpl57 | 631.7549906 | 0.492461355 | 1.08E-02 |
| Kmt2d | 3395.264604 | 0.492164754 | 2.41E-02 |
| Piga | 562.1998323 | 0.492018886 | 4.65E-03 |
| Klc2 | 1334.112269 | 0.49160754 | 1.86E-04 |
| Otud7b | 4504.227944 | 0.491284558 | 2.21E-08 |
| Sp4 | 343.7880857 | 0.491271734 | 1.57E-02 |
| Ino80c | 526.2001063 | 0.491032553 | 5.24E-03 |
| Pear1 | 2676.766374 | 0.49042475 | 8.87E-04 |
| Ndufa10 | 5483.308946 | 0.49042235 | 5.97E-03 |
| Slc44a2 | 6308.547057 | 0.489966096 | 2.74E-04 |
| Ube2i | 9079.566651 | 0.489660098 | 7.13E-06 |
| 2610005L07Rik | 4557.828866 | 0.489458558 | 2.55E-04 |
| Kpnb1 | 26279.66806 | 0.48925111 | 3.07E-04 |
| Zswim6 | 572.7350864 | 0.488869023 | 4.31E-03 |

| | | | |
|---|---|---|---|
| Crcp | 993.0124417 | 0.488821744 | 6.65E-08 |
| Atp13a1 | 2374.27894 | 0.488810816 | 2.61E-04 |
| Tmsb4x | 9263.718803 | 0.488702083 | 1.90E-04 |
| A430005L14Rik | 411.6249794 | 0.48827347 | 9.97E-04 |
| Rpl34 | 304.9125979 | 0.488203684 | 8.96E-03 |
| Snhg17 | 1429.438413 | 0.488060151 | 1.58E-03 |
| Ubqln4 | 3372.00673 | 0.48761261 | 1.90E-04 |
| Slc5a6 | 627.4269561 | 0.487511079 | 6.47E-03 |
| Sap30bp | 1774.467389 | 0.487455417 | 9.53E-04 |
| Tatdn3 | 338.0046401 | 0.487380869 | 2.17E-02 |
| Rnf10 | 11588.78172 | 0.486910159 | 9.90E-04 |
| S100a10 | 14257.32699 | 0.486736396 | 6.57E-04 |
| Rae1 | 2709.734454 | 0.486446732 | 1.62E-03 |
| Selenoi | 2343.480976 | 0.486393022 | 5.85E-04 |
| Sms | 1954.671987 | 0.486288185 | 4.95E-03 |
| Gdpd1 | 1125.495183 | 0.485595814 | 4.48E-02 |
| Yif1a | 3116.067448 | 0.484888442 | 7.27E-03 |
| Ralgapa2 | 995.1711884 | 0.484540916 | 8.77E-03 |
| Dcaf12 | 2233.394066 | 0.48438843 | 3.55E-06 |
| Git1 | 4693.589768 | 0.484127313 | 9.48E-04 |
| Zbtb44 | 864.3557499 | 0.484123235 | 4.03E-05 |
| Unc45a | 1699.276498 | 0.484112359 | 7.65E-04 |
| Scmh1 | 2538.520534 | 0.484048178 | 2.64E-03 |
| Slc25a3 | 21829.57629 | 0.483498095 | 1.15E-07 |
| Acer3 | 753.5966644 | 0.483304173 | 2.98E-02 |
| Cycs | 1585.90036 | 0.483287771 | 1.86E-02 |
| Cog6 | 1639.903711 | 0.483248182 | 3.07E-04 |
| Nae1 | 2185.814268 | 0.483088895 | 2.31E-04 |
| Ccnd2 | 10678.38031 | 0.4824409 | 6.98E-06 |
| Polr1e | 447.9290646 | 0.482299505 | 3.05E-02 |
| Tmem248 | 2966.009369 | 0.481593847 | 1.67E-03 |
| Phgdh | 14268.86496 | 0.481279778 | 4.25E-02 |
| Comt | 3881.085594 | 0.481117677 | 9.19E-06 |
| Ric1 | 2773.699318 | 0.48109159 | 1.24E-04 |
| Arnt | 5519.889564 | 0.480691592 | 3.42E-03 |
| Smurf2 | 3989.71208 | 0.480569436 | 2.93E-07 |
| Exosc1 | 1110.733076 | 0.480565657 | 4.44E-04 |
| Pias4 | 1560.177006 | 0.479914731 | 3.12E-02 |
| Ergic3 | 4114.58219 | 0.479452822 | 8.68E-03 |
| Wwox | 869.2521846 | 0.479302787 | 6.52E-03 |
| Hspa4 | 15381.02481 | 0.479271537 | 2.81E-03 |
| Med25 | 2722.795482 | 0.47887058 | 1.22E-04 |
| Zfp688 | 338.1326918 | 0.478378743 | 1.62E-02 |
| Usp15 | 2743.233149 | 0.478165927 | 2.72E-02 |
| Tdp1 | 913.0317336 | 0.478089424 | 5.61E-03 |
| Aga | 1959.623389 | 0.47754275 | 1.73E-03 |
| Rnf113a1 | 147.4647827 | 0.477110441 | 3.41E-02 |

| | | | |
|---|---|---|---|
| Gypc | 1383.765939 | 0.476516724 | 1.49E-04 |
| Mapk12 | 322.9227326 | 0.476324155 | 7.93E-04 |
| Tep1 | 1733.768891 | 0.476289096 | 8.12E-05 |
| Tmem259 | 4431.668597 | 0.476181711 | 1.81E-09 |
| Ankrd13c | 1592.952556 | 0.476059277 | 2.80E-05 |
| Dnajb2 | 1081.510458 | 0.475398126 | 1.55E-03 |
| Elac2 | 1667.303953 | 0.475366199 | 3.99E-03 |
| Dagla | 576.2981415 | 0.475153649 | 2.15E-04 |
| Dusp16 | 2072.5505 | 0.475047004 | 3.97E-03 |
| Nol11 | 2131.47981 | 0.474952444 | 3.62E-07 |
| Cfap300 | 360.5795128 | 0.474952405 | 2.45E-02 |
| Setd3 | 4338.931259 | 0.474441276 | 7.04E-05 |
| Rab5if | 2691.810294 | 0.47436569 | 1.17E-02 |
| Tspo | 2413.333548 | 0.473804488 | 1.10E-04 |
| Naxe | 3709.451477 | 0.472987073 | 5.92E-03 |
| Trit1 | 873.3104816 | 0.472866715 | 1.04E-03 |
| Ndufa9 | 3674.367459 | 0.472863677 | 1.66E-05 |
| Uqcc1 | 1536.397941 | 0.472556639 | 6.91E-04 |
| Relb | 763.8076857 | 0.4725432 | 2.78E-02 |
| Taf12 | 648.1120911 | 0.472533615 | 8.98E-04 |
| Fam219a | 921.6076423 | 0.47236294 | 1.10E-03 |
| B4galt3 | 1066.306636 | 0.472308411 | 9.11E-05 |
| Pdhb | 2648.913108 | 0.472140791 | 1.32E-05 |
| Trp53i13 | 637.8712156 | 0.472037391 | 1.25E-02 |
| Lmtk2 | 2086.187159 | 0.471913083 | 4.66E-06 |
| Plekho2 | 4026.295225 | 0.471843588 | 4.65E-03 |
| Gng2 | 472.9891909 | 0.471273106 | 4.95E-02 |
| Cebpz | 3061.302049 | 0.471060544 | 4.03E-05 |
| Psma1 | 3673.371411 | 0.470607829 | 1.25E-06 |
| Yrdc | 1246.308307 | 0.470579977 | 1.66E-04 |
| Smg5 | 4366.585321 | 0.470519248 | 1.04E-02 |
| Ginm1 | 1392.945406 | 0.470423069 | 1.83E-03 |
| Gtf3c3 | 1632.751785 | 0.47004295 | 1.49E-07 |
| Rad52 | 832.8818623 | 0.469967386 | 1.20E-02 |
| Lsm4 | 1011.076088 | 0.46992335 | 1.17E-03 |
| Sec31a | 15080.55081 | 0.469807228 | 3.58E-03 |
| Cdipt | 3223.076682 | 0.469545798 | 3.63E-05 |
| Nsd3 | 4509.284761 | 0.469400379 | 1.21E-03 |
| Golga2 | 4785.074064 | 0.469203215 | 2.78E-03 |
| Ino80e | 1732.624095 | 0.469182296 | 5.75E-03 |
| Atp2a2 | 15822.23775 | 0.469154435 | 3.08E-03 |
| Pttg1ip | 15567.85936 | 0.469146089 | 6.53E-03 |
| Zc3h3 | 967.5588601 | 0.468986046 | 3.05E-02 |
| Pi4k2b | 1510.332724 | 0.468733224 | 6.24E-04 |
| Klf16 | 1257.19304 | 0.468463034 | 1.85E-02 |
| Ndufb11 | 3018.005221 | 0.468399232 | 6.55E-03 |
| Slc38a1 | 5119.465128 | 0.467906786 | 1.79E-07 |

| | | | |
|---|---|---|---|
| Crem | 172.0738427 | 0.467899284 | 4.28E-03 |
| Copa | 20491.01741 | 0.46787186 | 6.18E-04 |
| Rer1 | 2247.001878 | 0.467806515 | 9.28E-05 |
| Gcn1l1 | 5972.383838 | 0.467628762 | 1.87E-04 |
| Ccdc9 | 1160.732769 | 0.467601543 | 2.91E-04 |
| Kdm5a | 4716.918006 | 0.467582062 | 4.86E-04 |
| Grpel2 | 916.2086707 | 0.467114366 | 1.19E-03 |
| Pdcd5 | 1694.809403 | 0.46701583 | 1.44E-04 |
| B3gat3 | 1341.032423 | 0.466292633 | 3.99E-05 |
| Heatr1 | 2150.07386 | 0.466279445 | 4.94E-03 |
| Txn2 | 4732.276052 | 0.466231804 | 1.18E-03 |
| Osbpl3 | 2074.055951 | 0.466145963 | 3.56E-03 |
| Cdk11b | 2039.570556 | 0.466035192 | 2.66E-06 |
| Tyw1 | 1171.744952 | 0.465865565 | 7.20E-03 |
| Ago3 | 1945.986236 | 0.465816084 | 4.16E-02 |
| Klhdc3 | 2766.400916 | 0.4650711 | 1.91E-04 |
| Trnau1ap | 1335.035752 | 0.464635836 | 1.21E-04 |
| 1700037H04Rik | 1863.740915 | 0.464540584 | 8.38E-04 |
| Yif1b | 1695.896623 | 0.464481993 | 5.07E-03 |
| Snip1 | 1187.676737 | 0.464158855 | 8.43E-04 |
| Ift46 | 2411.826845 | 0.464154772 | 6.68E-04 |
| Yars | 7597.314974 | 0.463708352 | 4.41E-04 |
| Rabac1 | 2922.165955 | 0.463680096 | 3.03E-02 |
| Xrcc6 | 1228.827517 | 0.463604555 | 6.06E-03 |
| Ndufa12 | 1429.626157 | 0.463563341 | 8.69E-03 |
| Abhd14a | 565.1011237 | 0.46341849 | 1.30E-03 |
| Spryd3 | 1383.849558 | 0.462856575 | 4.61E-03 |
| Haus2 | 1057.827371 | 0.462679858 | 7.93E-04 |
| Akirin1 | 2974.695353 | 0.462411036 | 9.43E-05 |
| Nxpe3 | 351.1963934 | 0.461511334 | 5.04E-03 |
| Hibch | 1051.562632 | 0.461401233 | 5.04E-03 |
| Sf1 | 8718.970602 | 0.46085274 | 1.48E-04 |
| Ercc3 | 2067.814332 | 0.460698162 | 5.73E-04 |
| Lonp2 | 4330.10061 | 0.460558907 | 3.03E-04 |
| 8430429K09Rik | 165.2112728 | 0.460485371 | 2.14E-02 |
| Map1lc3a | 2634.199358 | 0.460401889 | 1.75E-02 |
| Emc7 | 2809.673745 | 0.460261847 | 1.68E-03 |
| Necap1 | 2428.802313 | 0.460217287 | 2.78E-04 |
| Capn2 | 13394.05317 | 0.459898842 | 1.36E-03 |
| Zranb1 | 1747.798794 | 0.458788395 | 5.76E-05 |
| Lamc1 | 22727.02501 | 0.458497248 | 2.69E-03 |
| Lrrc41 | 6294.169907 | 0.458166918 | 1.93E-03 |
| Fkbp2 | 1281.795309 | 0.458052244 | 1.10E-03 |
| Psmd3 | 7020.635315 | 0.457932157 | 1.01E-03 |
| Wdr33 | 2511.881024 | 0.457778823 | 1.29E-04 |
| Tnip2 | 759.4249082 | 0.457537275 | 3.11E-03 |
| Cpt1a | 5815.364835 | 0.457323584 | 1.84E-05 |

| | | | |
|---|---|---|---|
| Raly | 10314.59865 | 0.457203152 | 1.12E-04 |
| Nars | 9176.2195 | 0.457032749 | 5.74E-03 |
| Fam120b | 2980.824706 | 0.456759823 | 1.61E-03 |
| Trim25 | 2953.14383 | 0.456120239 | 1.24E-04 |
| Dync1h1 | 28582.84367 | 0.456094482 | 4.77E-04 |
| Gemin5 | 2847.233799 | 0.455582764 | 5.05E-03 |
| Ascc2 | 3721.48564 | 0.455490322 | 7.79E-05 |
| Mrpl43 | 1580.403356 | 0.455466408 | 2.75E-04 |
| Ddhd1 | 899.9729154 | 0.455277084 | 6.30E-04 |
| Acox1 | 3724.20556 | 0.454965928 | 2.28E-03 |
| Prkra | 1707.47307 | 0.454821934 | 1.12E-02 |
| Zfp219 | 1438.940271 | 0.454784178 | 6.80E-05 |
| Ercc8 | 336.7685356 | 0.453399281 | 2.22E-02 |
| Gng12 | 10176.62266 | 0.45321186 | 2.50E-03 |
| Npm3 | 2710.905344 | 0.452885546 | 2.65E-04 |
| Hilpda | 740.6665374 | 0.452829436 | 4.24E-04 |
| Dbr1 | 635.1497289 | 0.452381331 | 5.60E-03 |
| Mtor | 2627.083558 | 0.452123074 | 1.88E-04 |
| Abcg2 | 2509.433036 | 0.452050095 | 7.16E-03 |
| Snrnp27 | 1828.287886 | 0.451965921 | 4.02E-04 |
| Cbll1 | 784.2011838 | 0.451837952 | 1.10E-02 |
| Vcam1 | 8733.754356 | 0.451679312 | 4.99E-04 |
| Mon2 | 5512.525877 | 0.451569985 | 3.31E-06 |
| Tial1 | 1884.714745 | 0.451087041 | 1.29E-03 |
| Pbx2 | 2268.811586 | 0.450822172 | 5.83E-04 |
| Klf6 | 6415.94845 | 0.45067187 | 8.59E-06 |
| Faf2 | 3343.387474 | 0.450246575 | 4.90E-06 |
| Fchsd2 | 1458.876734 | 0.450030833 | 3.23E-02 |
| Plekha1 | 3449.235513 | 0.449371309 | 6.19E-03 |
| Abcf3 | 2478.910211 | 0.449235658 | 3.23E-03 |
| Nup214 | 3369.149841 | 0.448941268 | 1.36E-02 |
| Atpif1 | 2983.892833 | 0.448880587 | 2.94E-04 |
| Gm13835 | 2487.733138 | 0.448818809 | 8.75E-04 |
| Fam185a | 517.0409463 | 0.448627191 | 1.54E-02 |
| Strn4 | 2571.884023 | 0.448297587 | 8.84E-05 |
| Srgap2 | 3776.658232 | 0.448128914 | 2.13E-09 |
| Ahcyl2 | 2542.94842 | 0.448126588 | 2.17E-02 |
| Tkfc | 1275.455666 | 0.447624576 | 5.06E-03 |
| Pafah1b1 | 8649.10865 | 0.447505501 | 2.28E-05 |
| Gclm | 1768.792317 | 0.447479371 | 3.97E-03 |
| Abt1 | 516.2441004 | 0.447213645 | 2.49E-03 |
| Ogfod2 | 485.1672615 | 0.447202023 | 9.09E-03 |
| Tpra1 | 1917.311456 | 0.446787564 | 2.03E-04 |
| Tmem184b | 4196.309966 | 0.446715662 | 1.13E-03 |
| Rab8a | 3357.260812 | 0.446526099 | 2.34E-04 |
| Utp11 | 3221.327935 | 0.446471174 | 2.31E-05 |
| Mex3d | 2980.139648 | 0.445832835 | 3.77E-02 |

| | | | |
|---|---|---|---|
| Timm50 | 1105.524902 | 0.445744779 | 2.59E-02 |
| Madd | 889.1617141 | 0.444507094 | 2.91E-03 |
| Cdc37 | 6153.441156 | 0.443401715 | 1.95E-03 |
| Cdk5rap3 | 2181.115954 | 0.44325962 | 1.10E-02 |
| Rfx3 | 232.2533201 | 0.442896239 | 3.71E-02 |
| Tmem129 | 1354.313334 | 0.442877431 | 8.38E-03 |
| Sugp1 | 1683.026125 | 0.442762186 | 7.24E-04 |
| Acaa2 | 2550.415331 | 0.442407786 | 2.91E-02 |
| Gm21811 | 6988.957672 | 0.442336357 | 8.75E-04 |
| Ubr5 | 12270.45866 | 0.441974042 | 1.90E-06 |
| Selenoo | 1274.742205 | 0.441892939 | 1.73E-03 |
| Lactb2 | 1674.430397 | 0.441767002 | 1.42E-02 |
| Clic1 | 9855.984042 | 0.440567294 | 1.18E-03 |
| Shox2 | 1446.207875 | 0.440371262 | 1.70E-03 |
| Slu7 | 2633.493033 | 0.440296057 | 8.85E-03 |
| Dus4l | 339.6680114 | 0.440129406 | 4.24E-02 |
| Pex14 | 1215.974622 | 0.440036067 | 7.89E-03 |
| Mob2 | 788.0670922 | 0.440024068 | 1.08E-02 |
| Wdtc1 | 2442.822428 | 0.439501846 | 3.55E-03 |
| Farp1 | 7954.256896 | 0.439146169 | 1.33E-04 |
| Bivm | 1320.752866 | 0.438913947 | 8.44E-04 |
| Mr1 | 1889.33807 | 0.438157873 | 4.07E-03 |
| Plekha3 | 1730.437148 | 0.437910675 | 1.42E-03 |
| Emg1 | 2548.431435 | 0.437804266 | 5.84E-04 |
| Epb41l5 | 589.5562055 | 0.437708021 | 1.74E-02 |
| Gtf2a2 | 1249.80594 | 0.437377825 | 2.52E-03 |
| Tatdn2 | 1825.377767 | 0.436706935 | 2.58E-03 |
| Cep112it | 847.0867356 | 0.43620992 | 5.07E-04 |
| Kat7 | 3759.136679 | 0.436087639 | 1.30E-03 |
| Yy1 | 2360.24052 | 0.435895803 | 5.57E-06 |
| Inpp5a | 3109.503164 | 0.435684737 | 5.00E-02 |
| Get4 | 3291.279844 | 0.434688759 | 4.07E-03 |
| 4833439L19Rik | 3511.377629 | 0.434304884 | 7.75E-03 |
| Vcpkmt | 777.1607539 | 0.433809544 | 2.19E-02 |
| Plaa | 5924.78038 | 0.43379227 | 4.21E-03 |
| Fam98a | 1905.214616 | 0.433547106 | 5.81E-05 |
| Vars2 | 1444.707556 | 0.433518408 | 3.03E-04 |
| Pstk | 460.105305 | 0.433313212 | 1.53E-02 |
| Mrps28 | 608.7447475 | 0.433170514 | 3.91E-03 |
| Gpr108 | 2920.507615 | 0.43302262 | 6.37E-03 |
| 2700049A03Rik | 653.9146772 | 0.432472322 | 8.92E-03 |
| Fbxl4 | 863.6181875 | 0.432023167 | 2.00E-04 |
| Pelo | 681.6239708 | 0.43152604 | 4.99E-03 |
| Stoml2 | 1850.321277 | 0.431487004 | 1.84E-04 |
| Taok3 | 2501.86846 | 0.431322 | 1.16E-05 |
| Afg3l1 | 3276.76621 | 0.431221121 | 1.90E-05 |
| Atg2a | 2180.865891 | 0.431152012 | 3.41E-03 |

| | | | |
|---|---|---|---|
| Ddx31 | 694.0201295 | 0.431138919 | 2.53E-03 |
| Rxylt1 | 2203.764838 | 0.430999067 | 3.33E-04 |
| Zfp106 | 11251.11847 | 0.43038838 | 3.08E-10 |
| Mrip-ps | 1177.21672 | 0.430303803 | 1.59E-02 |
| Stau1 | 4739.563788 | 0.429949281 | 1.55E-03 |
| Cct8 | 12262.24507 | 0.429803197 | 8.06E-06 |
| Prpf8 | 19463.90492 | 0.429773889 | 8.21E-05 |
| Malsu1 | 636.9300842 | 0.429645196 | 1.01E-02 |
| Tab2 | 5843.044862 | 0.429149785 | 1.19E-06 |
| Mrps9 | 1479.100855 | 0.428081722 | 6.50E-07 |
| Tsr3 | 1281.295736 | 0.428061931 | 1.34E-03 |
| Eif2ak1 | 2626.861785 | 0.428054036 | 2.50E-03 |
| Scamp3 | 3093.025348 | 0.427865715 | 5.52E-03 |
| Efemp1 | 43132.42209 | 0.427768714 | 1.36E-02 |
| Ptpn2 | 736.8581651 | 0.427760945 | 2.91E-03 |
| Hdhd2 | 1257.142152 | 0.427717562 | 3.30E-04 |
| Fam136a | 1289.210965 | 0.427543549 | 5.57E-03 |
| Alkbh1 | 1016.532387 | 0.427391404 | 2.19E-02 |
| Anapc1 | 10971.14232 | 0.427306145 | 9.18E-06 |
| Fkbp10 | 8574.958963 | 0.427257012 | 1.58E-02 |
| Rnaseh2c | 449.8484921 | 0.427057695 | 3.19E-02 |
| Kank2 | 6902.794724 | 0.426630479 | 3.17E-04 |
| Trp53 | 5498.367133 | 0.426580376 | 2.37E-02 |
| Atg101 | 738.9112077 | 0.426448958 | 5.81E-03 |
| Plpbp | 2115.5974 | 0.426437346 | 7.88E-09 |
| Mettl14 | 970.764186 | 0.425975401 | 1.61E-03 |
| Casp2 | 1562.016278 | 0.425903927 | 1.14E-02 |
| Rpf1 | 1264.591 | 0.425889175 | 5.04E-03 |
| Zswim1 | 590.8453736 | 0.425322934 | 5.19E-03 |
| Nudt18 | 604.0577606 | 0.4252189 | 2.30E-02 |
| Hipk1 | 4679.274894 | 0.425072357 | 3.09E-05 |
| Polr3b | 1812.205495 | 0.424298773 | 5.53E-05 |
| Eif4a2 | 15379.29281 | 0.423638617 | 1.52E-02 |
| Zfp865 | 1022.925073 | 0.423530566 | 1.35E-04 |
| Khnyn | 1116.250918 | 0.42324398 | 4.00E-03 |
| Slc37a4 | 401.0618275 | 0.422764473 | 3.82E-03 |
| Ipp | 573.9750255 | 0.422648325 | 6.61E-04 |
| Dus2 | 436.7762165 | 0.422386656 | 1.42E-02 |
| Nup54 | 1688.162602 | 0.422382603 | 3.19E-03 |
| Cops6 | 4398.622067 | 0.422136143 | 1.67E-03 |
| Hdlbp | 31467.4146 | 0.421979926 | 6.98E-04 |
| Bloc1s6 | 2040.363911 | 0.421831476 | 6.85E-03 |
| Ap2a1 | 5510.979936 | 0.421643028 | 2.99E-03 |
| Ptcd1 | 1214.415313 | 0.421085125 | 4.85E-03 |
| Ndufb2 | 1099.511065 | 0.421084709 | 4.67E-03 |
| Blmh | 1538.113957 | 0.420980955 | 1.60E-03 |
| Psmd13 | 2802.297219 | 0.420607677 | 3.51E-02 |

| | | | |
|---|---|---|---|
| AU040320 | 3791.60754 | 0.420312004 | 3.11E-04 |
| Snrpg | 1662.606002 | 0.420213255 | 1.46E-02 |
| Txndc17 | 3979.458947 | 0.4199571 | 6.89E-04 |
| Usp46 | 1648.110825 | 0.419945889 | 1.79E-03 |
| Sphk2 | 1000.143385 | 0.419630878 | 2.34E-03 |
| Tmed4 | 3139.271629 | 0.419209409 | 7.98E-03 |
| Rilpl1 | 1857.379582 | 0.419133752 | 2.83E-03 |
| Rnf11 | 4607.610523 | 0.419057862 | 7.35E-05 |
| Ppm1a | 3456.516388 | 0.41898243 | 1.98E-07 |
| Dpm3 | 443.9062842 | 0.418966137 | 1.91E-02 |
| Sin3b | 4130.398059 | 0.418843001 | 1.75E-04 |
| Them4 | 258.1979688 | 0.418745513 | 3.72E-03 |
| Uchl3 | 918.5000259 | 0.418721037 | 2.75E-03 |
| Wdr70 | 1348.139472 | 0.418387609 | 4.90E-04 |
| Asb3 | 997.0760849 | 0.418310653 | 9.20E-04 |
| Polr2g | 3004.711592 | 0.4179901 | 6.38E-05 |
| Foxo4 | 1470.022901 | 0.417782433 | 1.27E-03 |
| Twnk | 1587.171026 | 0.417715204 | 1.53E-03 |
| Crat | 2486.540887 | 0.417414812 | 8.65E-05 |
| Psen1 | 3740.441008 | 0.417308806 | 9.19E-04 |
| Nars2 | 357.7982091 | 0.416992882 | 7.10E-03 |
| Vdac3 | 5778.986181 | 0.416790298 | 1.82E-04 |
| Ccdc86 | 1531.939262 | 0.416403162 | 7.08E-03 |
| Ampd2 | 2558.553788 | 0.416243606 | 4.47E-03 |
| Pdcd6ip | 8464.98572 | 0.416114378 | 2.30E-06 |
| Sprtn | 812.7061567 | 0.416045965 | 1.09E-03 |
| Dnajb4 | 1668.421695 | 0.416039915 | 4.72E-03 |
| Adprh | 2437.400221 | 0.415676495 | 1.69E-02 |
| Kdelr3 | 1636.509425 | 0.415133167 | 4.74E-02 |
| Eloc | 1343.046711 | 0.414829901 | 2.38E-02 |
| Htra2 | 5540.271863 | 0.414650022 | 1.45E-03 |
| Atg5 | 1278.933549 | 0.414618318 | 3.55E-04 |
| Ap2m1 | 16486.59881 | 0.414455835 | 1.45E-03 |
| Psma3 | 4329.011223 | 0.414385993 | 2.17E-02 |
| Kdelr1 | 5927.242726 | 0.414376124 | 3.51E-02 |
| Slc38a6 | 9317.07918 | 0.413896626 | 6.39E-05 |
| Arf1 | 15591.73516 | 0.413604106 | 5.29E-03 |
| Nop10 | 1906.592631 | 0.413548606 | 3.18E-03 |
| Cmas | 1604.032091 | 0.412837673 | 1.98E-06 |
| Tlr6 | 142.0509724 | 0.412461527 | 4.96E-02 |
| Phf20 | 3151.247539 | 0.412090827 | 9.80E-06 |
| 2310057M21Rik | 364.6686249 | 0.411788072 | 6.59E-03 |
| Proser1 | 898.7818854 | 0.411574088 | 1.47E-02 |
| Ctu2 | 965.207299 | 0.411060326 | 2.18E-02 |
| Larp1 | 8675.057462 | 0.4104265 | 3.25E-04 |
| Fbxw17 | 660.5170251 | 0.410381929 | 2.37E-03 |
| Mtrr | 953.9515698 | 0.41014231 | 2.28E-03 |

| | | | |
|---|---|---|---|
| Slc39a6 | 3751.096942 | 0.410051794 | 6.90E-03 |
| Gm20498 | 557.4901711 | 0.40992406 | 6.93E-03 |
| Frrs1 | 1733.117738 | 0.409384788 | 2.53E-04 |
| Pter | 781.8123917 | 0.409305159 | 6.34E-04 |
| Ssx2ip | 1161.638843 | 0.408937746 | 1.66E-02 |
| Usp14 | 3649.869986 | 0.408561882 | 1.23E-03 |
| Mff | 3918.125802 | 0.408379637 | 3.31E-05 |
| Tpm4 | 16366.22209 | 0.408362347 | 5.36E-03 |
| Mfap1b | 2130.549542 | 0.407950035 | 3.92E-03 |
| Ggnbp2 | 4298.82299 | 0.407274258 | 1.44E-02 |
| Cwc15 | 2545.719953 | 0.40711353 | 3.34E-06 |
| Sec24d | 7002.722589 | 0.407069382 | 8.59E-03 |
| Psme3 | 5358.956441 | 0.406869576 | 2.02E-04 |
| Morc4 | 3288.740245 | 0.406692203 | 4.08E-02 |
| Zfp217 | 1499.500526 | 0.406328069 | 4.76E-04 |
| Rnf220 | 2987.612839 | 0.40576335 | 2.21E-04 |
| Dop1b | 1500.10808 | 0.405617188 | 2.04E-02 |
| Aff4 | 11956.73403 | 0.405097479 | 3.00E-02 |
| Cdk8 | 1690.559402 | 0.404626496 | 2.10E-03 |
| Dand5 | 170.4514732 | 0.404102511 | 1.82E-02 |
| Med30 | 695.4020104 | 0.403908493 | 1.39E-03 |
| Rbpj-ps3 | 2159.928873 | 0.403782253 | 7.74E-04 |
| Mthfd2l | 443.9979493 | 0.40377846 | 1.86E-03 |
| Suds3 | 3742.606582 | 0.403778278 | 2.07E-03 |
| Ppp1r9b | 4501.93599 | 0.403654093 | 6.85E-04 |
| Pdxdc1 | 8226.574578 | 0.403477329 | 9.81E-03 |
| Aarsd1 | 1053.296333 | 0.403444015 | 3.91E-02 |
| Wipf2 | 1118.249451 | 0.403425112 | 4.41E-02 |
| Erich1 | 560.5336326 | 0.403400307 | 1.57E-02 |
| Uqcc2 | 2774.225035 | 0.402972715 | 4.97E-02 |
| Cd2bp2 | 2807.343141 | 0.40239261 | 2.02E-04 |
| Ndel1 | 3898.669197 | 0.401446513 | 8.54E-03 |
| Vcl | 15499.81698 | 0.401362742 | 2.27E-03 |
| Eloa | 2936.855797 | 0.401302551 | 1.05E-05 |
| Scrn2 | 419.7699392 | 0.401092308 | 2.73E-02 |
| Ptbp1 | 15707.70459 | 0.400911585 | 5.94E-03 |
| Ddx18 | 3710.623133 | 0.400830697 | 2.42E-03 |
| Zfp784 | 178.7752622 | 0.400675951 | 4.74E-02 |
| Med14 | 2332.440218 | 0.400532607 | 1.21E-03 |
| Klf13 | 2027.299638 | 0.400316186 | 1.48E-02 |
| 2310009B15Rik | 380.4070517 | 0.399668468 | 4.21E-02 |
| Cstf1 | 2074.099098 | 0.399572388 | 4.25E-02 |
| Pdcl3 | 2311.344659 | 0.39950589 | 8.76E-05 |
| 1700025G04Rik | 2025.145476 | 0.398715478 | 9.83E-03 |
| Rnf44 | 2695.652928 | 0.39845108 | 2.39E-03 |
| Mark2 | 2420.298645 | 0.398145058 | 1.73E-05 |
| Clybl | 692.1619379 | 0.398131089 | 3.72E-02 |

| | | | |
|---|---|---|---|
| Tmem128 | 899.2173669 | 0.397893925 | 3.60E-03 |
| Prmt2 | 1770.370006 | 0.397860626 | 2.61E-04 |
| Dnajc19 | 884.2635241 | 0.397858616 | 4.11E-02 |
| Magoh | 2065.067574 | 0.397516597 | 6.38E-04 |
| Zfp710 | 555.8720173 | 0.397482595 | 1.03E-03 |
| Lhfp | 4843.198567 | 0.397073737 | 3.20E-05 |
| Hccs | 884.9279035 | 0.397008456 | 1.86E-02 |
| Gm2574 | 455.518203 | 0.396428818 | 4.44E-02 |
| Nup98 | 2513.759391 | 0.396399476 | 4.83E-03 |
| Ddx23 | 3775.542555 | 0.396198238 | 3.50E-03 |
| Inpp5k | 2174.199723 | 0.396084063 | 2.76E-02 |
| Tln1 | 13133.47837 | 0.395980194 | 5.14E-04 |
| Flot2 | 3914.985823 | 0.395785119 | 3.74E-04 |
| Gm8186 | 1195.312773 | 0.395379432 | 1.50E-02 |
| Gle1 | 2111.626911 | 0.395127617 | 2.70E-02 |
| Prpf4 | 1801.281111 | 0.394252817 | 5.91E-03 |
| Naa35 | 1827.19018 | 0.393930492 | 2.54E-04 |
| Me2 | 1361.748678 | 0.393744267 | 2.27E-02 |
| Zcchc8 | 1549.897255 | 0.393638182 | 4.27E-04 |
| Etfdh | 2166.481833 | 0.393548343 | 6.82E-09 |
| Fn3krp | 425.7525463 | 0.393307599 | 1.20E-02 |
| Arl8a | 4599.940929 | 0.393058232 | 2.98E-02 |
| Rab3gap2 | 4136.31955 | 0.392769749 | 2.44E-04 |
| Tbc1d7 | 554.0725872 | 0.391694802 | 2.21E-02 |
| Wars | 2773.816283 | 0.391558019 | 2.35E-02 |
| Fiz1 | 1176.565028 | 0.390840485 | 1.71E-03 |
| Tnpo1 | 8965.551593 | 0.390297291 | 1.09E-05 |
| Vapa | 4514.815785 | 0.389826762 | 4.47E-03 |
| Exosc4 | 794.7367626 | 0.389823099 | 2.00E-02 |
| Ubl7 | 1626.244088 | 0.389669113 | 4.82E-02 |
| Lnx2 | 610.26002 | 0.389545298 | 1.88E-02 |
| Gm11868 | 349.9990098 | 0.389039069 | 1.03E-03 |
| Galt | 472.9768465 | 0.388666409 | 2.49E-02 |
| Sec14l1 | 5437.380374 | 0.388177387 | 8.41E-04 |
| Osbp | 4497.550677 | 0.387847482 | 1.98E-03 |
| Unkl | 1023.23768 | 0.387519041 | 1.19E-02 |
| Il10rb | 939.8976722 | 0.387458665 | 3.98E-04 |
| Cldnd1 | 2104.132741 | 0.387187095 | 5.17E-03 |
| Mrpl22 | 994.9769437 | 0.387109592 | 2.06E-03 |
| Rsph3a | 1682.445855 | 0.386269715 | 3.01E-02 |
| Mrpl2 | 1684.064819 | 0.385673481 | 7.42E-03 |
| Anapc16 | 1878.679111 | 0.385519801 | 8.75E-04 |
| Thop1 | 1815.914219 | 0.385484927 | 7.66E-03 |
| Ints10 | 2126.187577 | 0.385186061 | 5.42E-04 |
| Acad11 | 584.7985704 | 0.384221197 | 6.93E-03 |
| Rbm8a | 2712.709074 | 0.383429473 | 1.98E-03 |
| Ndufb10 | 2793.144385 | 0.383248298 | 3.58E-02 |

| | | | |
|---|---|---|---|
| Zbtb7a | 2825.843584 | 0.383200204 | 2.61E-03 |
| Fam117b | 1725.697617 | 0.38275626 | 1.52E-03 |
| Fbl | 2993.699857 | 0.381980787 | 5.82E-03 |
| Ftsj3 | 4055.065065 | 0.381866871 | 1.03E-02 |
| Fuca2 | 3318.401005 | 0.381316829 | 3.80E-02 |
| Prdm10 | 358.6740737 | 0.381228942 | 2.66E-02 |
| Rap1gds1 | 3574.090192 | 0.380923611 | 2.02E-04 |
| Mapk9 | 2993.864 | 0.380607416 | 6.57E-05 |
| Plppr2 | 1300.032739 | 0.379540865 | 1.96E-02 |
| 1810013L24Rik | 2909.228309 | 0.379514865 | 5.39E-03 |
| Dolpp1 | 505.6016964 | 0.378871132 | 1.22E-02 |
| Nip7 | 2066.690499 | 0.378829661 | 1.98E-02 |
| Eif4h | 13353.88039 | 0.378434045 | 3.31E-03 |
| Farsa | 2227.240026 | 0.377873526 | 1.92E-03 |
| Rexo1 | 2127.055018 | 0.377780003 | 3.89E-04 |
| Ppm1k | 350.4821301 | 0.377701198 | 3.87E-02 |
| Lrrc40 | 1157.772138 | 0.376955883 | 2.75E-02 |
| Hk1 | 5099.447276 | 0.376802711 | 4.30E-02 |
| Il17rc | 2513.616066 | 0.376725207 | 5.12E-05 |
| Poli | 459.5423416 | 0.376062107 | 5.92E-03 |
| Cpt2 | 2107.034094 | 0.375837683 | 9.58E-04 |
| Dtd2 | 514.8764301 | 0.375622961 | 3.31E-03 |
| Surf6 | 1223.042779 | 0.375440177 | 9.91E-04 |
| Ptp4a2 | 11669.3148 | 0.375373695 | 4.70E-03 |
| Abhd17a | 2211.350324 | 0.375050794 | 4.24E-02 |
| Vps37a | 1652.988048 | 0.374988392 | 4.21E-02 |
| Actb | 74206.67695 | 0.374958333 | 5.31E-03 |
| Ubxn4 | 6611.923936 | 0.374566807 | 1.36E-05 |
| Rell1 | 1320.85393 | 0.374398671 | 1.92E-02 |
| Ptpn21 | 3194.640395 | 0.373709334 | 3.72E-02 |
| Prkcsh | 5914.118506 | 0.373262105 | 1.09E-02 |
| Ppp1r12a | 3585.345434 | 0.373087231 | 4.44E-02 |
| Chchd3 | 4298.451419 | 0.372571914 | 1.12E-02 |
| Ola1 | 4769.480168 | 0.371968808 | 7.04E-04 |
| Unk | 1102.176227 | 0.371782563 | 1.10E-02 |
| Cmc1 | 656.2945244 | 0.371746138 | 3.60E-02 |
| Rraga | 5750.079961 | 0.371328704 | 2.56E-03 |
| Mrpl9 | 1944.256036 | 0.371193822 | 3.66E-04 |
| Tarbp2 | 899.8778158 | 0.371079617 | 7.72E-03 |
| Ubr3 | 4475.974516 | 0.371075171 | 1.17E-03 |
| Dvl3 | 1937.725035 | 0.371043266 | 4.32E-02 |
| Abcf1 | 5975.295718 | 0.370795963 | 2.50E-04 |
| Arfgef2 | 4840.915022 | 0.370624991 | 1.51E-02 |
| Rnpep | 1933.761328 | 0.370166092 | 2.85E-03 |
| Pacsin2 | 4361.917334 | 0.369982445 | 6.68E-04 |
| Cdc16 | 4416.767764 | 0.369809522 | 1.56E-03 |
| Galns | 1404.178413 | 0.369416192 | 2.27E-02 |

| | | | |
|---|---|---|---|
| Smn1 | 972.1731807 | 0.369106418 | 3.77E-02 |
| Foxk1 | 3191.645725 | 0.368998536 | 1.20E-02 |
| Ppat | 1263.284928 | 0.368687955 | 3.70E-02 |
| Gatad1 | 2934.150248 | 0.368630288 | 4.93E-04 |
| Plekhh2 | 2098.280773 | 0.368278635 | 4.06E-02 |
| Rpusd2 | 266.1180622 | 0.368173102 | 2.37E-02 |
| Map2k2 | 5007.849089 | 0.367521648 | 6.04E-03 |
| Ube2k | 2857.850823 | 0.367464774 | 2.69E-03 |
| Tmem230 | 1598.27255 | 0.367457387 | 9.34E-03 |
| Zfp956 | 415.6674772 | 0.367400231 | 3.25E-02 |
| Sppl2a | 3377.428579 | 0.367003457 | 2.30E-03 |
| Mprip | 17779.67662 | 0.366822791 | 5.45E-06 |
| Qtrt1 | 428.9958719 | 0.366464684 | 1.73E-02 |
| Kansl3 | 3444.874046 | 0.366167483 | 1.35E-05 |
| Ube2d3 | 9445.414009 | 0.366153541 | 2.34E-06 |
| Harbi1 | 406.9577452 | 0.366085256 | 1.84E-02 |
| Dock6 | 1238.663641 | 0.36584081 | 1.36E-02 |
| D230025D16Rik | 1404.011213 | 0.364912663 | 3.42E-02 |
| Pde6d | 1448.894996 | 0.364789005 | 7.42E-03 |
| Abhd11 | 803.7195628 | 0.364699264 | 3.93E-02 |
| Fermt2 | 3801.900806 | 0.364694076 | 1.25E-02 |
| Ubl3 | 1139.357592 | 0.364671857 | 5.34E-03 |
| Brd2 | 13357.93295 | 0.363544051 | 8.82E-03 |
| Lrrc42 | 3350.571451 | 0.363539748 | 6.25E-03 |
| Mast3 | 1601.42766 | 0.363457386 | 2.70E-02 |
| Tsta3 | 1507.503282 | 0.363126626 | 2.27E-03 |
| Rpap2 | 1188.573752 | 0.363072263 | 8.22E-03 |
| Cfap298 | 524.7661239 | 0.362925825 | 1.42E-02 |
| Mtpap | 1104.094798 | 0.362735216 | 1.54E-03 |
| Snrpd2 | 2347.998224 | 0.362171376 | 4.16E-02 |
| Kmt2a | 4699.546315 | 0.361835027 | 1.89E-02 |
| Tbc1d10b | 1987.395714 | 0.361541731 | 1.03E-02 |
| Gps2 | 1053.198647 | 0.361355153 | 2.67E-02 |
| Wdr43 | 3511.800235 | 0.36101119 | 2.49E-02 |
| Pcmt1 | 2281.828033 | 0.360187327 | 4.14E-03 |
| Hmgxb3 | 1878.722133 | 0.35925338 | 1.65E-02 |
| Hmgxb4 | 1190.644908 | 0.358774813 | 1.44E-02 |
| NA | 486.6545116 | 0.357983549 | 4.26E-02 |
| Sec23ip | 3395.713208 | 0.357944679 | 1.95E-02 |
| Pprc1 | 3591.050554 | 0.357805536 | 1.06E-03 |
| Ddx54 | 4300.181664 | 0.357724284 | 3.91E-02 |
| Ncstn | 4112.498853 | 0.35735639 | 2.01E-03 |
| Zfp236 | 2213.260685 | 0.357324951 | 1.32E-02 |
| Nedd1 | 1214.112669 | 0.357283231 | 4.90E-02 |
| Cpne1 | 5044.131924 | 0.357186295 | 9.51E-06 |
| Thoc5 | 2117.128019 | 0.356630853 | 1.00E-02 |
| Kdm3b | 3364.46103 | 0.356517457 | 4.23E-06 |

| | | | |
|---|---|---|---|
| Mettl13 | 1248.651196 | 0.356298572 | 1.30E-02 |
| Armt1 | 1217.678356 | 0.356101756 | 1.94E-03 |
| Uap1 | 2457.583466 | 0.355581758 | 5.60E-05 |
| Dhx30 | 2802.794432 | 0.355342526 | 2.30E-02 |
| Ttc27 | 1412.862851 | 0.355032251 | 2.27E-03 |
| Ttc21b | 2440.384871 | 0.354862923 | 1.45E-04 |
| B3gntl1 | 339.1393466 | 0.354862279 | 8.72E-03 |
| Gm7887 | 398.1521651 | 0.35453695 | 1.25E-02 |
| Smad5 | 2597.963491 | 0.354519058 | 2.23E-03 |
| Gmeb2 | 1450.329167 | 0.354440571 | 1.95E-02 |
| Ctif | 1560.762332 | 0.354394331 | 7.99E-03 |
| Gak | 5311.988674 | 0.354040453 | 2.09E-03 |
| Dctn4 | 3196.598025 | 0.353785113 | 3.82E-03 |
| Ppp2r5a | 2519.975068 | 0.353584252 | 8.55E-03 |
| Rbm22 | 1683.263056 | 0.353420009 | 1.38E-02 |
| Arl3 | 1484.978537 | 0.352938783 | 2.42E-03 |
| Srprb | 2065.424096 | 0.352781927 | 3.08E-02 |
| Uqcrc1 | 5283.028191 | 0.352243494 | 1.64E-02 |
| Isyna1 | 639.1786947 | 0.352215287 | 4.94E-03 |
| Klhdc2 | 4037.018707 | 0.352185402 | 4.97E-03 |
| Cct7 | 14582.72543 | 0.3513832 | 1.86E-02 |
| Tbrg4 | 2034.760982 | 0.351323951 | 4.08E-02 |
| Cox7a2 | 2411.57902 | 0.351127397 | 3.09E-03 |
| Tapt1 | 1487.131827 | 0.350956262 | 9.08E-05 |
| Pgap2 | 1147.432946 | 0.349695576 | 2.48E-02 |
| Akt2 | 5161.192976 | 0.348789474 | 9.74E-03 |
| Ndufv1 | 4450.278284 | 0.34815178 | 1.07E-02 |
| Llph | 1359.624563 | 0.347572949 | 5.37E-03 |
| Bcl7a | 1302.545937 | 0.347258331 | 4.91E-03 |
| Gng5 | 2852.253698 | 0.347247348 | 1.53E-04 |
| Elovl5 | 2213.912141 | 0.347086499 | 2.32E-03 |
| Zfp91 | 7065.381387 | 0.346895642 | 3.06E-03 |
| Atg2b | 4251.553425 | 0.346480338 | 1.59E-02 |
| Rbpj | 6189.6897 | 0.346461464 | 3.82E-03 |
| Gid4 | 1936.274946 | 0.346246309 | 1.58E-03 |
| Tctex1d2 | 890.6787767 | 0.346178797 | 2.50E-02 |
| Nus1 | 4625.394918 | 0.346114995 | 1.37E-02 |
| Emp2 | 4062.72683 | 0.345788119 | 3.88E-04 |
| Lrrfip2 | 1836.448746 | 0.345394467 | 5.93E-03 |
| Tex10 | 1746.463961 | 0.345020001 | 2.82E-03 |
| Fbxo28 | 1806.420033 | 0.343565548 | 2.56E-03 |
| Actr1b | 4863.150793 | 0.343499142 | 2.43E-02 |
| Map2k7 | 2944.395202 | 0.34311501 | 2.64E-02 |
| Dzip1l | 1361.683775 | 0.342828774 | 5.54E-03 |
| Bcat2 | 1292.256022 | 0.342555841 | 3.62E-02 |
| Tmem134 | 2708.878956 | 0.341476256 | 2.37E-02 |
| Med28 | 1860.448472 | 0.341377204 | 1.33E-02 |

| | | | |
|---|---|---|---|
| Map2k5 | 1196.801708 | 0.340894677 | 5.20E-04 |
| Slc12a7 | 1768.921695 | 0.340454363 | 1.51E-02 |
| Ifi27 | 1560.472021 | 0.340441588 | 2.06E-02 |
| Siah1a | 745.6340288 | 0.339489898 | 4.73E-03 |
| Brox | 3007.5817 | 0.339132489 | 8.18E-06 |
| Axin1 | 4077.021414 | 0.338811599 | 9.86E-04 |
| Atic | 6279.229534 | 0.338698317 | 3.32E-02 |
| Jak1 | 13668.72976 | 0.337883934 | 1.77E-02 |
| Poc5 | 569.4921546 | 0.337090784 | 4.13E-02 |
| Hadh | 4701.504074 | 0.336984454 | 1.60E-02 |
| Araf | 2783.283806 | 0.336698387 | 3.88E-04 |
| Wdr3 | 1937.156413 | 0.336628033 | 2.52E-02 |
| Pdcd11 | 3527.629205 | 0.336533184 | 2.02E-03 |
| Mrpl11 | 1587.25396 | 0.335309527 | 7.93E-03 |
| Hdac7 | 6665.610222 | 0.334590954 | 1.03E-04 |
| Dnajc1 | 2143.380387 | 0.334524805 | 2.87E-03 |
| Marf1 | 4341.805183 | 0.334070222 | 4.05E-02 |
| 0610030E20Rik | 1672.283913 | 0.333985347 | 1.29E-02 |
| Golph3 | 3356.80977 | 0.333613 | 2.25E-03 |
| Acbd3 | 3022.146409 | 0.332812383 | 1.40E-02 |
| Tcerg1 | 2443.339398 | 0.332095544 | 1.03E-02 |
| Ccdc6 | 2420.855025 | 0.331384928 | 2.08E-03 |
| Ntan1 | 2090.352579 | 0.33112368 | 3.51E-02 |
| Asb6 | 1407.281146 | 0.330973008 | 4.91E-02 |
| Mark3 | 3507.222576 | 0.330663776 | 4.58E-05 |
| Derl1 | 5766.467668 | 0.330612619 | 2.03E-02 |
| Trpm4 | 642.7733671 | 0.330249459 | 2.27E-02 |
| Fkbp1a | 9585.237864 | 0.330203079 | 2.35E-02 |
| Nrf1 | 1279.08247 | 0.330019147 | 3.51E-02 |
| Tomm70a | 4143.787542 | 0.329861473 | 1.70E-02 |
| Phc1 | 2184.532526 | 0.329717312 | 3.30E-02 |
| Rhoa | 12392.3443 | 0.329538378 | 7.24E-03 |
| Tmem63a | 1994.571198 | 0.329519186 | 5.58E-03 |
| Rsl24d1 | 1367.856925 | 0.328973683 | 3.80E-03 |
| Prpsap2 | 830.6666767 | 0.328843234 | 2.98E-02 |
| Cmtr2 | 516.4360552 | 0.327886777 | 6.45E-03 |
| Micu1 | 6570.285669 | 0.327321295 | 3.51E-02 |
| Gnai2 | 16160.10343 | 0.327210439 | 3.29E-02 |
| Snw1 | 3904.766616 | 0.327038347 | 8.83E-03 |
| Wipi2 | 4767.419219 | 0.326944348 | 6.12E-03 |
| Oard1 | 738.8061374 | 0.326094202 | 1.54E-02 |
| Rsrc1 | 3007.886735 | 0.325082815 | 1.30E-02 |
| 4930453N24Rik | 1049.629045 | 0.324832917 | 2.04E-02 |
| Mau2 | 4038.002112 | 0.324758874 | 1.09E-02 |
| Hdac4 | 1038.543867 | 0.324598566 | 4.67E-02 |
| Dbn1 | 5523.632008 | 0.324583349 | 1.92E-02 |
| Dnajc8 | 3861.33955 | 0.324079421 | 2.38E-04 |

| | | | |
|---|---|---|---|
| Ptprn | 7409.737366 | 0.323310896 | 1.06E-02 |
| Stx5a | 3441.259391 | 0.322606908 | 9.17E-03 |
| Ndfip1 | 2962.313867 | 0.322528975 | 3.00E-02 |
| Phf23 | 3785.987225 | 0.322485903 | 3.93E-02 |
| Narfl | 1220.368788 | 0.322008613 | 3.17E-02 |
| Utp3 | 2514.145431 | 0.321852044 | 2.54E-04 |
| Sgta | 5423.891798 | 0.321562615 | 1.65E-02 |
| Pdcl | 1348.652634 | 0.321341984 | 2.99E-03 |
| Capza2 | 8096.038169 | 0.321205889 | 3.40E-02 |
| Vps9d1 | 897.8907511 | 0.320051524 | 4.03E-03 |
| Nop9 | 1165.628715 | 0.320031782 | 3.86E-02 |
| Rad23a | 3073.219033 | 0.319637631 | 2.74E-02 |
| Ralgps1 | 724.2808194 | 0.319267547 | 4.60E-02 |
| Ppp1r10 | 2578.197764 | 0.318827399 | 3.69E-02 |
| Zcchc9 | 831.0395613 | 0.318381642 | 4.58E-02 |
| Tpm1 | 28225.26008 | 0.318130314 | 1.03E-02 |
| Fbxo4 | 853.3693442 | 0.317643802 | 1.72E-02 |
| Aldh3a2 | 3225.872424 | 0.317040303 | 2.66E-03 |
| Golga7 | 2979.170609 | 0.316236101 | 1.20E-03 |
| Cops7a | 6190.887762 | 0.316226895 | 3.47E-03 |
| Cnot10 | 1671.516662 | 0.315787557 | 3.31E-03 |
| Itprid2 | 2850.831262 | 0.315723436 | 5.07E-03 |
| Prpf38a | 1921.609227 | 0.315327463 | 2.52E-03 |
| Yipf6 | 840.4900824 | 0.314907553 | 1.53E-02 |
| Atg16l1 | 1600.559711 | 0.314376527 | 2.65E-02 |
| Sdad1 | 2442.35499 | 0.314357565 | 3.68E-03 |
| Eif4a3 | 3389.716389 | 0.314100911 | 4.14E-02 |
| Lcmt1 | 805.5401037 | 0.313013693 | 3.31E-02 |
| Spg20 | 6566.427666 | 0.312767195 | 4.79E-03 |
| Ago2 | 7856.332648 | 0.312648309 | 2.43E-03 |
| Sec24b | 3696.789891 | 0.312553947 | 3.49E-03 |
| Gmps | 4740.881068 | 0.312150096 | 6.11E-03 |
| Arpc5l | 2244.355316 | 0.311954051 | 1.58E-02 |
| Fam160b1 | 2368.974209 | 0.311592327 | 8.72E-03 |
| Ube2f | 1729.361484 | 0.3115485 | 1.15E-02 |
| Ralgapb | 4220.545798 | 0.311408658 | 5.46E-05 |
| Lrrc8a | 1684.686033 | 0.310659878 | 1.54E-02 |
| Anp32b | 8481.918321 | 0.310658443 | 2.23E-02 |
| Stam2 | 2719.319093 | 0.308965107 | 4.07E-04 |
| Mtfmt | 419.5935587 | 0.306585961 | 2.98E-02 |
| Tmem199 | 785.0747928 | 0.304830659 | 9.65E-03 |
| Gpr89 | 1644.684102 | 0.304611261 | 8.47E-03 |
| Huwe1 | 19835.01404 | 0.30448326 | 1.10E-03 |
| Gpr173 | 432.4446883 | 0.304321167 | 3.87E-02 |
| Ttc19 | 1089.857695 | 0.304099884 | 2.45E-02 |
| Sfr1 | 9916.619043 | 0.304025502 | 1.22E-02 |
| Psmg1 | 640.1736031 | 0.30366401 | 3.55E-03 |

| | | | |
|---|---|---|---|
| Trmt112-ps2 | 2121.34201 | 0.303562304 | 3.33E-02 |
| Wasf2 | 5249.227574 | 0.303521655 | 4.04E-02 |
| Hnrnpul2 | 7133.827205 | 0.303479068 | 1.63E-03 |
| Plrg1 | 2765.068295 | 0.303365629 | 2.73E-02 |
| Cnot2 | 3155.015605 | 0.303089309 | 5.69E-05 |
| Irak4 | 619.5218289 | 0.302416166 | 2.07E-02 |
| Frg1 | 2103.851192 | 0.301595377 | 2.64E-02 |
| Zfp560 | 338.4447528 | 0.301519704 | 4.68E-02 |
| Map4k4 | 13173.18194 | 0.300918648 | 7.39E-04 |
| 4933434E20Rik | 1868.94148 | 0.300828519 | 4.41E-03 |
| Exosc10 | 2718.984245 | 0.299174978 | 1.13E-02 |
| Cops5 | 4298.493893 | 0.298636761 | 8.83E-04 |
| Pxn | 3926.522418 | 0.29862813 | 3.73E-02 |
| Afap1 | 3020.859185 | 0.298463557 | 3.88E-02 |
| Cpped1 | 2173.563252 | 0.297875572 | 2.44E-02 |
| Dda1 | 2103.385569 | 0.297867266 | 2.82E-02 |
| Ehmt2 | 10097.78051 | 0.297313932 | 2.53E-02 |
| Ehd2 | 12732.25354 | 0.297122614 | 2.40E-02 |
| Paics | 14439.95988 | 0.29696529 | 4.54E-05 |
| 5-Mar | 3832.102311 | 0.29634008 | 1.23E-04 |
| Sec23a | 4948.848425 | 0.295866443 | 4.85E-02 |
| Usmg5 | 2737.433256 | 0.295575572 | 5.06E-03 |
| Rap2b | 1894.249099 | 0.29518175 | 1.62E-02 |
| Epb41l4a | 2231.957676 | 0.295099119 | 1.72E-02 |
| Epn1 | 2966.878104 | 0.294476439 | 3.36E-02 |
| Dynlt1b | 1798.335892 | 0.294004973 | 1.94E-02 |
| Cpsf6 | 3087.474864 | 0.293980764 | 4.34E-02 |
| Atp5f1 | 8259.418685 | 0.293394037 | 2.19E-03 |
| Brat1 | 1070.094288 | 0.2931222 | 4.31E-02 |
| Zbtb25 | 464.9360401 | 0.292680242 | 1.47E-02 |
| Mapkapk5 | 1443.221454 | 0.292561258 | 1.01E-02 |
| Ppp2r5c | 4554.30962 | 0.29169411 | 1.34E-03 |
| Rtca | 1608.30568 | 0.291480685 | 8.46E-03 |
| Arpc2 | 10253.14542 | 0.290920753 | 1.51E-03 |
| Eif2s3x | 10181.23195 | 0.2906475 | 1.05E-02 |
| Iqgap1 | 14617.56628 | 0.290522991 | 4.85E-03 |
| Fubp1 | 6171.280554 | 0.290404113 | 3.04E-02 |
| Ppie | 1371.92535 | 0.289630625 | 3.50E-02 |
| Gsk3a | 3027.222386 | 0.289375853 | 2.04E-02 |
| Srsf4 | 2159.848775 | 0.289255384 | 1.77E-02 |
| Med17 | 1362.023412 | 0.28924899 | 6.39E-03 |
| Aco2 | 10683.99062 | 0.288478587 | 4.58E-02 |
| Cul4a | 6931.867817 | 0.288153371 | 2.66E-07 |
| Atp8b2 | 6346.170488 | 0.28808382 | 1.48E-02 |
| Gpr107 | 3622.865967 | 0.287926682 | 4.42E-02 |
| Mtrex | 2716.987977 | 0.287451371 | 3.26E-03 |
| Clint1 | 4999.687017 | 0.28633887 | 3.35E-03 |

| | | | |
|---|---|---|---|
| Psmd1 | 10295.53513 | 0.284981246 | 3.03E-02 |
| Orc3 | 1568.554497 | 0.284920149 | 3.18E-02 |
| Polr1a | 4966.441682 | 0.284901653 | 4.43E-02 |
| Mtdh | 6058.46066 | 0.284147548 | 2.90E-02 |
| Flii | 7693.446933 | 0.283800897 | 4.44E-03 |
| Cast | 5248.563907 | 0.283593274 | 5.04E-03 |
| Hgsnat | 2033.324024 | 0.283414262 | 2.06E-02 |
| Rwdd4a | 1263.978367 | 0.281932427 | 2.11E-02 |
| Gtf2ird1 | 1929.055866 | 0.280660708 | 1.83E-02 |
| Mrpl58 | 767.8759832 | 0.279215255 | 2.59E-02 |
| Lasp1 | 9498.425232 | 0.278245637 | 2.12E-02 |
| Atf4 | 16684.06758 | 0.278041294 | 3.05E-02 |
| Cramp1l | 1364.654941 | 0.277794068 | 4.27E-02 |
| Fbf1 | 2196.630713 | 0.277733344 | 2.90E-02 |
| 1110004F10Rik | 2600.912527 | 0.277439546 | 1.45E-02 |
| Supt6 | 7336.991304 | 0.27681472 | 2.48E-02 |
| Kctd3 | 1744.740515 | 0.276609957 | 3.31E-05 |
| Chmp5 | 2578.82645 | 0.276429832 | 1.69E-02 |
| Snrpe | 2802.549934 | 0.276230597 | 4.68E-02 |
| Pcbp4 | 5072.455738 | 0.276171885 | 6.89E-03 |
| Ncor2 | 6492.036827 | 0.27485592 | 4.64E-02 |
| Anapc2 | 5642.744713 | 0.274809965 | 2.30E-02 |
| Ufd1 | 2067.713197 | 0.274101991 | 2.05E-02 |
| Lmbrd1 | 2417.679745 | 0.273589311 | 4.80E-02 |
| Med13l | 5103.80171 | 0.272922828 | 3.74E-02 |
| Dlat | 2761.146179 | 0.272857852 | 7.79E-03 |
| Gfm1 | 3165.757619 | 0.271265096 | 3.33E-02 |
| Dennd5a | 6370.596711 | 0.269572421 | 5.32E-04 |
| Tmed2 | 11159.26364 | 0.268625747 | 1.95E-02 |
| Prr14 | 1622.26639 | 0.268246558 | 8.37E-03 |
| Sart1 | 3217.739631 | 0.268072503 | 1.32E-02 |
| Preb | 3063.586097 | 0.266450047 | 4.08E-03 |
| Acadl | 4515.648214 | 0.266423367 | 1.05E-02 |
| Mrpl10 | 2215.87896 | 0.266165799 | 1.16E-02 |
| Cebpzos | 554.3928388 | 0.26586755 | 9.31E-03 |
| Med21 | 976.9462806 | 0.265628785 | 1.42E-02 |
| Sf3b2 | 15940.53564 | 0.26545703 | 4.18E-03 |
| Fam53c | 2241.692611 | 0.265179443 | 4.14E-03 |
| Rapgef1 | 5967.043459 | 0.264623013 | 4.77E-02 |
| Khdc4 | 2525.121927 | 0.263338303 | 2.11E-02 |
| Txndc9 | 3522.902588 | 0.261893068 | 1.98E-02 |
| Mkln1 | 4393.243402 | 0.261806508 | 1.95E-03 |
| Ube2a | 1452.93935 | 0.261401844 | 9.71E-03 |
| Rfx7 | 1726.032941 | 0.26130519 | 1.58E-02 |
| Ambra1 | 2446.423628 | 0.260725712 | 3.64E-02 |
| Emc4 | 2098.421114 | 0.260424552 | 1.96E-02 |
| Vezf1 | 3105.724991 | 0.259651486 | 3.57E-02 |

| | | | |
|---|---|---|---|
| Rbm17 | 4210.243158 | 0.259138208 | 1.33E-02 |
| Aars | 20365.17343 | 0.258365404 | 3.92E-02 |
| Scaf1 | 4485.533781 | 0.257492711 | 3.59E-02 |
| Vps16 | 1845.3321 | 0.257067311 | 2.82E-02 |
| D17Wsu92e | 5640.635753 | 0.256660178 | 3.25E-02 |
| Gnaq | 3219.879092 | 0.256370582 | 1.13E-02 |
| Nf2 | 4991.381476 | 0.255635815 | 3.57E-02 |
| Zdhhc16 | 1645.02251 | 0.255270485 | 1.30E-02 |
| Sco1 | 557.039177 | 0.253674387 | 2.87E-02 |
| Elp2 | 4679.842283 | 0.252920574 | 3.04E-02 |
| Nelfb | 2470.739406 | 0.252504276 | 4.19E-02 |
| Rnf24 | 2352.640376 | 0.25244531 | 2.75E-02 |
| Acp1 | 3181.095991 | 0.25209486 | 3.31E-02 |
| Tpm2 | 4623.603642 | 0.250982125 | 4.18E-02 |
| Entr1 | 2471.686962 | 0.250366073 | 2.67E-02 |
| Mmadhc | 2680.153335 | 0.250334694 | 5.46E-03 |
| Metap1 | 2869.366179 | 0.250055283 | 3.89E-03 |
| Dhx35 | 1030.525777 | 0.24903585 | 4.71E-02 |
| Chmp3 | 6440.19342 | 0.247554926 | 4.79E-02 |
| Nup88 | 3246.624486 | 0.247488953 | 2.75E-03 |
| Eif2a | 4013.91275 | 0.247219849 | 4.14E-02 |
| Setd5 | 9969.903168 | 0.246229025 | 3.56E-02 |
| Letmd1 | 1317.473792 | 0.24511904 | 1.57E-02 |
| Rab6a | 3433.18195 | 0.244603952 | 3.20E-02 |
| Ppp3r1 | 4512.171715 | 0.244442571 | 2.91E-02 |
| Rbbp5 | 1649.334514 | 0.243292303 | 2.29E-02 |
| Necap2 | 1811.57524 | 0.241917235 | 2.11E-02 |
| Pdap1 | 5185.52205 | 0.241872157 | 4.70E-02 |
| Fbxo3 | 2578.681611 | 0.241170412 | 2.27E-02 |
| Noct | 8107.901444 | 0.240598534 | 4.91E-02 |
| Amfr | 5482.908097 | 0.239065571 | 1.38E-02 |
| Zfp384 | 2438.431101 | 0.238944604 | 4.70E-02 |
| Kif3c | 2739.168318 | 0.238597835 | 1.56E-02 |
| Tnks2 | 9380.128971 | 0.238178979 | 4.28E-02 |
| Smg7 | 3227.148695 | 0.238152039 | 4.23E-02 |
| Heatr5b | 3231.838328 | 0.237332371 | 4.92E-02 |
| Gpd1l | 1232.297381 | 0.237102001 | 1.19E-02 |
| Arl5a | 2208.885859 | 0.235607218 | 3.29E-02 |
| Rgs19 | 1525.312029 | 0.234802578 | 8.66E-03 |
| Tbc1d22b | 1063.836405 | 0.23300687 | 4.01E-02 |
| Smarcd2 | 2024.989409 | 0.231774884 | 3.20E-02 |
| Tmem131 | 4629.57427 | 0.230041889 | 1.18E-02 |
| Elp5 | 2617.836782 | 0.229868379 | 3.91E-02 |
| Xpo7 | 3778.843741 | 0.227195411 | 2.34E-02 |
| Zcrb1 | 1391.467671 | 0.226344547 | 2.72E-02 |
| Agfg1 | 2748.187289 | 0.222083031 | 3.33E-02 |
| Ppp2ca | 10386.53634 | 0.220540186 | 3.51E-02 |

| | | | |
|---|---|---|---|
| Stat6 | 5625.010172 | 0.21997935 | 1.28E-02 |
| Khsrp | 4308.732968 | 0.219628146 | 3.05E-02 |
| Tmem18 | 963.5596269 | 0.219328246 | 3.78E-02 |
| Cacul1 | 4722.619289 | 0.2189244 | 3.00E-02 |
| Ncoa6 | 4067.91543 | 0.218346116 | 4.13E-02 |
| Spag7 | 2417.33747 | 0.218335752 | 2.20E-02 |
| Ndufb3 | 1297.060705 | 0.216704008 | 4.69E-02 |
| Fbxw2 | 2586.135429 | 0.214871448 | 5.57E-03 |
| Mrgbp | 899.3957712 | 0.214349613 | 4.06E-02 |
| Hnrnpul1 | 6740.962743 | 0.213226853 | 3.71E-02 |
| Rnf111 | 1614.977381 | 0.21273619 | 3.11E-02 |
| Pxk | 1881.676776 | 0.212275854 | 2.45E-02 |
| Cdc23 | 2628.540815 | 0.210622458 | 9.27E-03 |
| Rhot1 | 2347.280075 | 0.207486039 | 2.11E-02 |
| Copb1 | 7208.232283 | 0.207325199 | 1.64E-02 |
| Adh5 | 4642.32907 | 0.20501111 | 2.55E-02 |
| Sri | 3410.645404 | 0.20232305 | 4.84E-02 |
| Rab3gap1 | 3040.80505 | 0.199857053 | 1.86E-02 |
| Setdb1 | 3317.723671 | 0.199792786 | 2.69E-02 |
| Ccdc127 | 2076.615621 | 0.199123303 | 4.61E-02 |
| Rnf20 | 2420.762924 | 0.196937452 | 2.02E-02 |
| Mapk1 | 9186.482385 | 0.195445938 | 4.32E-02 |
| Ankrd13a | 3982.563234 | 0.194564586 | 2.69E-02 |
| Mtch2 | 7896.965041 | 0.190383917 | 3.00E-02 |
| Habp4 | 1289.065897 | 0.189920169 | 1.63E-02 |
| Nfatc3 | 4109.575676 | 0.174624692 | 5.35E-03 |
| Pak2 | 5721.701201 | 0.173443714 | 5.67E-03 |
| Ankle2 | 2199.59166 | 0.167363117 | 4.76E-02 |
| Paf1 | 1759.690936 | 0.164742335 | 4.32E-02 |
| Becn1 | 3487.621834 | 0.154972177 | 4.55E-02 |
| Usp8 | 4956.37385 | -0.144022453 | 4.54E-02 |
| Ppp2r5e | 2577.239633 | -0.158662304 | 3.34E-02 |
| Rab12 | 2907.184409 | -0.172252454 | 1.98E-02 |
| Herc2 | 5066.067327 | -0.178896581 | 4.80E-02 |
| Kbtbd4 | 953.0351392 | -0.186029128 | 3.94E-02 |
| Gigyf2 | 4619.799954 | -0.189793164 | 2.91E-02 |
| Ep300 | 3594.368947 | -0.192282797 | 1.82E-02 |
| Nras | 3618.572248 | -0.19419725 | 3.36E-02 |
| Fam20b | 3653.87584 | -0.19612301 | 3.80E-02 |
| Mapkap1 | 2840.876994 | -0.197952548 | 4.56E-02 |
| Kdm2a | 7140.470013 | -0.199434793 | 3.78E-02 |
| Cat | 3139.699228 | -0.200711457 | 4.70E-02 |
| Ndufs1 | 4503.206639 | -0.204337122 | 3.53E-02 |
| Pafah1b2 | 3932.142107 | -0.206277027 | 3.12E-02 |
| Eps15l1 | 2232.088824 | -0.208163706 | 3.47E-02 |
| Mtpn | 11061.19823 | -0.210089296 | 2.26E-02 |
| Slx4 | 1785.115886 | -0.210530155 | 2.99E-02 |

| | | | |
|---|---|---|---|
| Ube2q1 | 5225.360789 | -0.212467217 | 4.21E-02 |
| Ube2d2a | 3927.040904 | -0.21357824 | 3.40E-03 |
| Gm17344 | 712.596112 | -0.217851042 | 2.16E-02 |
| Zfp251 | 1144.446552 | -0.219325783 | 1.85E-02 |
| Limd1 | 3109.454773 | -0.222392126 | 1.08E-02 |
| Zdhhc5 | 4681.471334 | -0.222903857 | 3.46E-02 |
| Vps53 | 1711.372237 | -0.225109286 | 4.11E-02 |
| Ppp6r3 | 4632.142651 | -0.225886512 | 8.05E-04 |
| Nipsnap2 | 2457.606527 | -0.227233386 | 2.97E-02 |
| Capza1 | 5169.359605 | -0.227401784 | 7.87E-03 |
| Paip1 | 2505.193787 | -0.228206112 | 4.98E-02 |
| Prepl | 2110.742543 | -0.229434612 | 2.58E-02 |
| Zfp275 | 2068.451814 | -0.231682947 | 2.56E-02 |
| Usp4 | 3655.583413 | -0.232247036 | 3.46E-02 |
| Zc3h13 | 2165.313132 | -0.232269853 | 2.12E-02 |
| Dzip1 | 1896.61168 | -0.235204122 | 3.59E-02 |
| NA | 459.8289227 | -0.236827287 | 4.13E-02 |
| Dync1li2 | 5420.762549 | -0.237131738 | 3.40E-02 |
| Xpnpep3 | 989.0992619 | -0.237331588 | 1.16E-02 |
| Fech | 1997.126972 | -0.238189982 | 5.96E-03 |
| Ppil2 | 3213.172897 | -0.239227295 | 4.17E-02 |
| Mbd2 | 1918.447548 | -0.239352738 | 1.58E-02 |
| Gng10 | 2800.643553 | -0.23950117 | 1.50E-02 |
| Pced1a | 1754.726717 | -0.239951263 | 3.11E-02 |
| Otulin | 961.990477 | -0.240604891 | 4.51E-02 |
| Rptor | 2361.625741 | -0.240821259 | 4.98E-03 |
| Irf3 | 1284.181054 | -0.242104724 | 3.17E-02 |
| Ino80 | 2504.264289 | -0.243410638 | 4.54E-02 |
| Slc36a4 | 1156.576941 | -0.24398444 | 2.88E-03 |
| Itfg1 | 5550.990102 | -0.246178169 | 3.50E-02 |
| Pdcd6 | 2365.713327 | -0.24723706 | 5.12E-03 |
| Zfp12 | 1260.57049 | -0.247397848 | 3.79E-02 |
| Gapvd1 | 3649.129331 | -0.247817318 | 2.03E-02 |
| Pex2 | 1610.317343 | -0.247896238 | 3.14E-02 |
| Itgb1 | 37771.62253 | -0.248485318 | 2.21E-02 |
| Ilf3 | 5444.124218 | -0.249007069 | 4.25E-02 |
| Sfswap | 2364.600261 | -0.250382378 | 7.26E-03 |
| Elf2 | 3398.181671 | -0.250552303 | 1.96E-02 |
| Atxn7l3 | 3554.494769 | -0.251818782 | 9.69E-03 |
| Fam160b2 | 868.648153 | -0.253938905 | 2.79E-02 |
| Fam91a1 | 5484.339588 | -0.254681668 | 2.00E-02 |
| M6pr | 6868.970325 | -0.254774088 | 2.78E-02 |
| Alg9 | 906.9621859 | -0.256806057 | 3.13E-02 |
| Qrich1 | 3515.068439 | -0.257207961 | 3.52E-03 |
| Acvr1b | 825.908788 | -0.257780574 | 2.35E-02 |
| Thada | 1328.834289 | -0.259857518 | 4.46E-02 |
| Spcs1 | 3229.514527 | -0.259953754 | 2.76E-02 |

| | | | |
|---|---|---|---|
| Pptc7 | 1901.222242 | -0.260164896 | 3.85E-02 |
| Anapc4 | 3891.529376 | -0.260449236 | 2.16E-02 |
| Zfp507 | 2655.215056 | -0.260553789 | 9.79E-03 |
| D6Wsu163e | 991.3243772 | -0.260645146 | 1.78E-02 |
| Usp6nl | 1363.084088 | -0.261206782 | 2.92E-02 |
| Cab39 | 2954.878358 | -0.261460708 | 4.12E-02 |
| Lman2l | 996.9305942 | -0.261703755 | 3.93E-02 |
| Suclg1 | 3399.964263 | -0.262134403 | 9.22E-03 |
| Commd3 | 3470.393101 | -0.264283301 | 1.82E-02 |
| Tbc1d9b | 5415.2979 | -0.264343971 | 1.14E-02 |
| Srsf9 | 1778.892419 | -0.265084232 | 4.67E-03 |
| Baz1b | 8196.336798 | -0.266915001 | 6.71E-03 |
| Arl6ip5 | 5111.98551 | -0.267613507 | 4.73E-02 |
| Rc3h1 | 2093.844484 | -0.268071019 | 1.59E-02 |
| Ankrd10 | 1201.376825 | -0.268618942 | 4.54E-03 |
| Msl2 | 1328.305038 | -0.268623739 | 8.97E-03 |
| Trappc3 | 2759.576289 | -0.268809797 | 4.10E-03 |
| Fbrsl1 | 745.1517284 | -0.268979875 | 4.92E-02 |
| Rnf8 | 695.2111225 | -0.27019411 | 1.67E-02 |
| Wdr61 | 1665.389284 | -0.270658841 | 3.55E-02 |
| Acin1 | 5414.640843 | -0.27102352 | 1.17E-03 |
| Zbtb39 | 784.3304255 | -0.27241937 | 3.26E-02 |
| Rabif | 689.5941583 | -0.272558133 | 1.20E-02 |
| Aars2 | 599.8250935 | -0.274497313 | 3.58E-02 |
| Pip4k2b | 4426.450694 | -0.274575116 | 2.86E-03 |
| Stx12 | 1913.740575 | -0.274750081 | 2.00E-03 |
| Cnot11 | 1342.876554 | -0.274845602 | 1.90E-02 |
| Slc31a1 | 3001.893987 | -0.275296018 | 1.23E-04 |
| Tmed10 | 10506.46475 | -0.276253896 | 9.84E-03 |
| Txndc11 | 1459.372549 | -0.276520101 | 1.07E-02 |
| Papola | 6677.657305 | -0.276661865 | 3.69E-02 |
| Prcc | 1702.755892 | -0.278464726 | 4.71E-02 |
| Rab1a | 7539.257883 | -0.278917863 | 1.13E-02 |
| Poglut1 | 2986.878341 | -0.279146102 | 1.57E-02 |
| Crk | 5752.903002 | -0.280372153 | 3.14E-02 |
| Psmd6 | 2606.420377 | -0.282706042 | 2.80E-02 |
| Chd4 | 15088.84888 | -0.283331018 | 2.38E-03 |
| Fem1a | 1367.887487 | -0.283381077 | 2.18E-02 |
| Vps52 | 2360.232529 | -0.283763978 | 1.23E-02 |
| Rab7 | 17268.35184 | -0.284483379 | 7.47E-03 |
| Tfb2m | 1122.816783 | -0.284899518 | 4.83E-02 |
| Wbp4 | 1679.499856 | -0.284949332 | 2.50E-02 |
| Btbd3 | 870.3274116 | -0.285019073 | 2.08E-02 |
| Pelp1 | 1937.94971 | -0.28506333 | 4.43E-02 |
| Tmcc1 | 1566.82676 | -0.285291813 | 4.32E-02 |
| Dcp1a | 1156.200016 | -0.285336447 | 1.73E-03 |
| Rtf2 | 2919.948112 | -0.28600573 | 2.31E-02 |

| | | | |
|---|---|---|---|
| Fbxo11 | 3313.191893 | -0.286271707 | 4.83E-02 |
| 2410002F23Rik | 2404.042911 | -0.28671612 | 3.83E-02 |
| Ubfd1 | 2088.508959 | -0.287392497 | 4.61E-02 |
| Adgra3 | 2363.887936 | -0.288090197 | 3.01E-02 |
| Gas2l1 | 1866.627813 | -0.288207381 | 2.71E-02 |
| Sgcb | 2110.307292 | -0.28873328 | 1.59E-02 |
| Cog7 | 1094.294008 | -0.290023651 | 1.12E-03 |
| Txndc15 | 1324.69371 | -0.290096216 | 3.33E-02 |
| Mex3c | 2408.706296 | -0.290281653 | 1.55E-02 |
| Tmem123 | 3880.257383 | -0.291133095 | 2.74E-02 |
| Asb8 | 1164.021748 | -0.29150518 | 1.70E-02 |
| Hnrnpk | 21286.09285 | -0.291581372 | 8.39E-04 |
| Mrtfb | 2231.817183 | -0.292254411 | 1.77E-02 |
| Snx2 | 3747.672178 | -0.29232315 | 3.09E-02 |
| Smad3 | 3160.732874 | -0.292507506 | 2.96E-02 |
| Mfsd14a | 1911.459662 | -0.29263317 | 2.27E-02 |
| Kif13a | 1516.705461 | -0.292696817 | 2.96E-02 |
| Supt7l | 842.860683 | -0.292922245 | 5.35E-03 |
| Scap | 4677.143336 | -0.293406827 | 3.66E-02 |
| Chd3 | 13653.34876 | -0.293486021 | 3.56E-02 |
| Cyb5d1 | 1009.24945 | -0.29353787 | 6.35E-03 |
| Dnlz | 934.93158 | -0.293614958 | 2.56E-02 |
| Mfsd10 | 2347.017728 | -0.294067691 | 2.91E-02 |
| Tmem60 | 583.2390495 | -0.294216371 | 4.08E-02 |
| Zfp84 | 572.4479467 | -0.294331865 | 1.17E-02 |
| Zfp869 | 866.2160119 | -0.295425451 | 6.75E-03 |
| Ifngr2 | 2171.576009 | -0.296056227 | 4.68E-02 |
| Tmem127 | 5979.127084 | -0.296910199 | 4.93E-02 |
| Gigyf1 | 2070.367659 | -0.297079696 | 2.62E-02 |
| Fam173a | 718.2912736 | -0.297135617 | 3.04E-02 |
| Specc1l | 4786.27288 | -0.298551122 | 2.93E-04 |
| Nptn | 6929.496085 | -0.29880298 | 4.48E-03 |
| Tada2b | 1040.973773 | -0.299866193 | 2.30E-03 |
| Aco1 | 2755.25536 | -0.299946591 | 1.41E-03 |
| S100pbp | 2801.716801 | -0.300799906 | 2.29E-02 |
| Vcpip1 | 2503.113416 | -0.301053891 | 1.71E-02 |
| Nnt | 4601.606286 | -0.301238441 | 2.11E-05 |
| Psme4 | 6579.725646 | -0.301494447 | 2.61E-02 |
| Anapc5 | 11211.90137 | -0.301727885 | 2.45E-02 |
| Atmin | 1212.579264 | -0.301768138 | 1.17E-03 |
| Magi1 | 863.00933 | -0.301912776 | 4.74E-02 |
| 9530082P21Rik | 1031.08362 | -0.301929515 | 4.84E-02 |
| Ctnna1 | 13605.32906 | -0.301951036 | 2.60E-04 |
| Klhl22 | 5015.7865 | -0.303618054 | 1.36E-02 |
| Cox5b | 4421.831308 | -0.304044377 | 1.11E-02 |
| Myl12b | 5643.908536 | -0.304189095 | 1.83E-02 |
| Dpy19l3 | 2100.747006 | -0.30422772 | 2.02E-02 |

| | | | |
|---|---|---|---|
| Zbtb21 | 1062.931372 | -0.304925935 | 1.71E-02 |
| Crebbp | 2005.517214 | -0.305410518 | 1.21E-02 |
| Arhgef25 | 5617.048691 | -0.305585305 | 1.74E-03 |
| Cep350 | 2934.829578 | -0.305761008 | 8.47E-03 |
| Ccs | 1597.308004 | -0.306088525 | 1.90E-02 |
| Zfp639 | 1185.948539 | -0.306274225 | 1.40E-04 |
| Foxred1 | 683.0766349 | -0.306361797 | 3.83E-02 |
| Bptf | 6943.697425 | -0.306864538 | 2.42E-02 |
| Ado | 1738.480241 | -0.309683921 | 2.39E-03 |
| Rnf145 | 4349.170828 | -0.309943486 | 1.43E-03 |
| Snx25 | 1932.640755 | -0.310329201 | 2.06E-02 |
| Ccnh | 791.1746844 | -0.311927275 | 1.63E-03 |
| Zfp212 | 832.4312734 | -0.312001944 | 2.40E-02 |
| F2r | 8616.173877 | -0.312044755 | 1.57E-02 |
| Tma7 | 1031.886558 | -0.312212671 | 1.29E-02 |
| Pgrmc2 | 4295.144483 | -0.312317429 | 2.03E-02 |
| Mettl9 | 2407.464825 | -0.312403856 | 1.96E-03 |
| Foxj2 | 2108.968859 | -0.313215879 | 7.57E-05 |
| Sgpp1 | 3316.645421 | -0.313301019 | 1.01E-02 |
| Fam192a | 1935.108348 | -0.313473439 | 3.18E-02 |
| Dhx57 | 1977.793762 | -0.313693641 | 7.63E-04 |
| Rexo4 | 1818.23831 | -0.314046375 | 1.34E-02 |
| Parp9 | 681.8767794 | -0.314377137 | 5.78E-03 |
| Rft1 | 677.9886492 | -0.314724786 | 2.20E-02 |
| Apobec3 | 1537.869773 | -0.315090595 | 2.38E-02 |
| Tmem258 | 1971.731613 | -0.315131219 | 3.05E-02 |
| Rtf1 | 2978.284882 | -0.315383044 | 1.29E-02 |
| Zfp202 | 333.0936493 | -0.315511954 | 4.86E-02 |
| Micu2 | 2127.341309 | -0.3170873 | 1.11E-03 |
| Zfp746 | 1625.00071 | -0.317747881 | 1.40E-03 |
| Plekhm3 | 751.7569629 | -0.318317834 | 7.35E-03 |
| Aplf | 766.3187187 | -0.318348886 | 3.04E-02 |
| Dhx33 | 2116.155745 | -0.318677877 | 1.06E-02 |
| Nrbp1 | 3395.974321 | -0.320258494 | 4.02E-04 |
| Alg1 | 1213.748362 | -0.320610485 | 1.11E-02 |
| Lsm1 | 795.3759526 | -0.32126766 | 5.69E-03 |
| Bag4 | 698.9273242 | -0.322194907 | 1.01E-02 |
| Soat1 | 3693.855559 | -0.322228848 | 2.24E-03 |
| Rfx1 | 784.7024519 | -0.322637118 | 3.23E-02 |
| Fbxw5 | 1900.946667 | -0.323304872 | 3.17E-04 |
| Ssr2 | 4724.151705 | -0.323781854 | 2.02E-02 |
| Ralb | 3073.453009 | -0.323849854 | 3.51E-02 |
| Cers6 | 4036.257231 | -0.323891983 | 4.56E-02 |
| Pik3ca | 3935.265814 | -0.324024081 | 8.84E-06 |
| Slc27a4 | 1532.925228 | -0.324214609 | 7.17E-03 |
| Mfn1 | 2369.80232 | -0.324222469 | 4.69E-02 |
| Dexi | 703.8502771 | -0.32439988 | 3.26E-02 |

| | | | |
|---|---|---|---|
| Pigt | 4933.604732 | -0.324956766 | 1.88E-02 |
| Nr3c1 | 3466.11795 | -0.325460665 | 1.77E-02 |
| NA | 3354.889398 | -0.326015627 | 3.07E-04 |
| Ndufa4 | 6517.052841 | -0.326106044 | 1.34E-04 |
| Fads2 | 5458.473187 | -0.327292476 | 1.83E-02 |
| Vkorc1l1 | 1525.062969 | -0.327355498 | 9.81E-05 |
| Amer1 | 1350.755714 | -0.328126149 | 2.00E-04 |
| Trappc12 | 1816.216958 | -0.328709365 | 2.53E-02 |
| Rps6ka4 | 3774.441517 | -0.328980583 | 4.14E-02 |
| Tmem256 | 1681.651426 | -0.329255768 | 3.94E-02 |
| Serhl | 1090.949015 | -0.33036219 | 1.24E-02 |
| Unc50 | 1307.062746 | -0.330713892 | 1.25E-02 |
| Gtf2f1 | 3305.095847 | -0.330824957 | 2.09E-03 |
| Hnrnpf | 22991.53822 | -0.33103298 | 3.15E-03 |
| Mus81 | 696.6969706 | -0.331206619 | 1.22E-03 |
| Swap70 | 2724.175662 | -0.331982399 | 4.74E-02 |
| Csrnp2 | 676.1118606 | -0.332082687 | 3.29E-02 |
| Asxl2 | 1701.60244 | -0.332285179 | 1.30E-02 |
| Dido1 | 3460.394899 | -0.332865229 | 2.64E-02 |
| Camta2 | 1178.937289 | -0.333003243 | 1.47E-03 |
| Evc2 | 3217.80212 | -0.333243132 | 5.14E-04 |
| Mif4gd | 450.899296 | -0.333433183 | 1.90E-02 |
| Ap1ar | 1516.331122 | -0.333984646 | 1.83E-02 |
| Kidins220 | 6720.791685 | -0.334276509 | 8.77E-03 |
| Asah1 | 13422.84013 | -0.335226943 | 1.40E-02 |
| NA | 1997.531107 | -0.335453287 | 4.92E-02 |
| Rab18 | 4215.54324 | -0.33552802 | 4.92E-04 |
| Zer1 | 1546.7305 | -0.33562023 | 2.67E-02 |
| 4931406C07Rik | 1602.982884 | -0.335724713 | 1.38E-02 |
| Trim28 | 8583.476259 | -0.337782811 | 3.14E-02 |
| Ubp1 | 2230.908591 | -0.338128947 | 3.34E-02 |
| Maged2 | 4335.856591 | -0.3384807 | 4.51E-02 |
| Tti1 | 1037.172499 | -0.338935475 | 3.71E-02 |
| Cfl1 | 29084.9649 | -0.339478128 | 2.83E-02 |
| Usp28 | 1013.741207 | -0.339492473 | 1.43E-03 |
| Acsf3 | 664.047027 | -0.339913532 | 4.59E-02 |
| Ubqln2 | 3270.808587 | -0.340095243 | 5.52E-04 |
| Orai3 | 1110.900108 | -0.340843136 | 2.62E-02 |
| Gm16211 | 299.0104868 | -0.340885663 | 3.92E-02 |
| Msantd4 | 1976.487313 | -0.341180931 | 2.10E-05 |
| Prr3 | 728.9177642 | -0.341540206 | 2.33E-03 |
| Dpp8 | 3923.176834 | -0.341936849 | 4.17E-05 |
| Xrcc1 | 1307.691342 | -0.342180939 | 4.99E-02 |
| Spsb2 | 835.3781798 | -0.342347589 | 9.93E-03 |
| Gnao1 | 4073.149345 | -0.343342476 | 3.91E-03 |
| Cep120 | 1497.538669 | -0.343793123 | 1.48E-03 |
| Gtf2h5 | 1581.79946 | -0.343972026 | 4.83E-03 |

| | | | |
|---|---|---|---|
| Map3k2 | 1933.527456 | -0.343995048 | 3.87E-02 |
| Rsf1 | 1494.238977 | -0.345071811 | 3.32E-02 |
| Commd5 | 445.54609 | -0.345701442 | 4.61E-02 |
| Cope | 2935.825434 | -0.346250082 | 8.51E-03 |
| Akap8 | 2567.406678 | -0.346625947 | 8.06E-03 |
| Rbbp6 | 3361.835241 | -0.346742344 | 4.89E-02 |
| Mfsd5 | 2190.404327 | -0.346784483 | 1.54E-02 |
| Mesd | 4178.825069 | -0.346840755 | 4.38E-02 |
| Peak1 | 6982.986032 | -0.34705618 | 1.61E-03 |
| Spns1 | 1815.839164 | -0.34723513 | 2.11E-02 |
| Blcap | 1673.554479 | -0.347341918 | 1.33E-03 |
| Minpp1 | 2115.190894 | -0.34769198 | 7.15E-04 |
| Slc35f5 | 3506.333334 | -0.348017212 | 9.48E-03 |
| Kdelc1 | 1895.28066 | -0.348311644 | 8.90E-04 |
| Gnai3 | 6342.144305 | -0.348645794 | 4.86E-02 |
| Srd5a3 | 981.177044 | -0.348999922 | 1.04E-03 |
| Kdm6a | 1109.474701 | -0.349019138 | 1.88E-02 |
| Eif4g3 | 5017.46316 | -0.34911713 | 4.50E-05 |
| Ppp4r2 | 4872.112859 | -0.349176019 | 2.04E-02 |
| Dram1 | 593.7149253 | -0.349425345 | 4.43E-02 |
| Tns1 | 7416.997207 | -0.349694293 | 1.93E-02 |
| Anp32a | 2932.882658 | -0.35022988 | 2.74E-03 |
| Btbd10 | 734.5276884 | -0.350820865 | 5.46E-03 |
| Nt5m | 402.8418024 | -0.351488382 | 3.61E-02 |
| Capn15 | 724.6406254 | -0.351562214 | 4.39E-03 |
| Vps35 | 10913.33651 | -0.352181994 | 4.15E-03 |
| C87436 | 843.8956946 | -0.352404403 | 8.29E-03 |
| D17H6S53E | 459.8831001 | -0.352655329 | 4.98E-02 |
| Ncdn | 2384.110763 | -0.352846218 | 3.94E-03 |
| Zdhhc8 | 3036.914019 | -0.353915464 | 3.90E-03 |
| Cep68 | 2000.679055 | -0.35487888 | 1.95E-05 |
| Rcor1 | 2598.897465 | -0.355564311 | 7.56E-07 |
| Rmdn3 | 951.2252578 | -0.355604147 | 7.06E-03 |
| Nfya | 2749.990945 | -0.355872801 | 1.13E-02 |
| Trak2 | 2626.579049 | -0.356081356 | 5.38E-03 |
| Slc35c1 | 794.2651529 | -0.356082118 | 1.10E-02 |
| Gopc | 1774.957849 | -0.356231408 | 1.63E-02 |
| Rnf139 | 1837.564298 | -0.356327016 | 3.74E-04 |
| Abcb10 | 1746.478949 | -0.356496303 | 1.06E-02 |
| Slc35b3 | 449.8835323 | -0.356551507 | 3.90E-02 |
| Lrpap1 | 3320.63054 | -0.356833262 | 3.80E-02 |
| Hps3 | 1032.072822 | -0.35686194 | 5.78E-03 |
| Rai14 | 4032.0779 | -0.357205235 | 4.50E-02 |
| Gpr137 | 943.9495011 | -0.357274663 | 1.16E-02 |
| Ift20 | 885.2811192 | -0.358095841 | 3.87E-02 |
| Clasp1 | 4060.473029 | -0.358440644 | 1.25E-05 |
| Zbtb24 | 960.6102902 | -0.358532026 | 2.43E-03 |

| | | | |
|---|---|---|---|
| Rad54l2 | 1696.94065 | -0.358717121 | 1.76E-04 |
| Osgep | 1715.128448 | -0.359004022 | 2.68E-02 |
| Ash2l | 2579.786406 | -0.359129173 | 4.25E-04 |
| Mfap3 | 1621.114974 | -0.359210336 | 1.07E-03 |
| Ptprs | 10843.05157 | -0.35941107 | 9.59E-05 |
| Btbd9 | 833.0426777 | -0.360130031 | 5.74E-03 |
| Letm1 | 2063.672833 | -0.360957491 | 1.24E-02 |
| Dnaja1 | 8039.758531 | -0.361561652 | 2.36E-03 |
| Rragc | 10259.96991 | -0.361735975 | 2.30E-02 |
| Fkbp15 | 2776.85651 | -0.361786732 | 3.10E-04 |
| Plpp5 | 772.2669265 | -0.362083583 | 4.22E-02 |
| Nedd8 | 2187.083819 | -0.362087431 | 6.68E-04 |
| Zmynd8 | 5278.19677 | -0.362515598 | 4.85E-06 |
| G3bp2 | 3581.96833 | -0.362537165 | 1.63E-03 |
| Zswim8 | 5853.559868 | -0.362553676 | 3.13E-02 |
| Mypop | 419.0754148 | -0.362776411 | 2.63E-02 |
| Atp7a | 1656.515308 | -0.362813718 | 6.95E-03 |
| Ube2z | 5727.588705 | -0.363574908 | 2.04E-03 |
| Nemp2 | 323.1133591 | -0.364071409 | 3.57E-02 |
| Stx2 | 1865.461332 | -0.364189255 | 3.97E-02 |
| Ptms | 12716.67832 | -0.364348532 | 9.49E-03 |
| Sertad2 | 3856.909765 | -0.364724987 | 1.79E-03 |
| Zfp1 | 503.9292446 | -0.365120441 | 6.38E-03 |
| Yeats2 | 2822.94201 | -0.365407749 | 1.50E-04 |
| Snx16 | 596.553418 | -0.365520019 | 2.77E-02 |
| 2610301B20Rik | 1235.416847 | -0.365865877 | 5.21E-04 |
| Tpst1 | 2809.181828 | -0.366214468 | 1.94E-02 |
| Rnf185 | 2243.235099 | -0.366301754 | 1.57E-02 |
| Intu | 504.6476843 | -0.366413227 | 4.32E-02 |
| Jak2 | 1748.376278 | -0.366755325 | 2.77E-02 |
| Tmem165 | 2684.294746 | -0.366965596 | 7.46E-04 |
| Cnpy3 | 2372.834644 | -0.367275281 | 2.49E-02 |
| Tmem168 | 2136.488341 | -0.367675776 | 4.16E-02 |
| Arpc1b | 12666.70678 | -0.367892398 | 3.87E-03 |
| Rpa3 | 645.0287183 | -0.36797531 | 3.54E-02 |
| Anxa4 | 8612.757354 | -0.367997745 | 4.04E-04 |
| Scp2 | 5871.714806 | -0.368158711 | 2.61E-03 |
| Cables2 | 840.8736199 | -0.368333583 | 7.73E-03 |
| 1110038F14Rik | 735.5469736 | -0.368736897 | 8.06E-04 |
| Vps33b | 466.8240164 | -0.369594003 | 3.21E-03 |
| Rad50 | 1970.461368 | -0.369832462 | 3.97E-02 |
| Orai2 | 1421.004878 | -0.369917117 | 7.64E-03 |
| Api5 | 5016.463876 | -0.370556783 | 3.32E-04 |
| Erp29 | 5236.891141 | -0.371308607 | 5.92E-03 |
| Hnrnpab | 12537.02073 | -0.371628786 | 4.96E-02 |
| Ss18 | 2071.284024 | -0.371701696 | 9.89E-03 |
| Abcc10 | 532.3965489 | -0.371826284 | 1.05E-03 |

| | | | |
|---|---|---|---|
| Gab1 | 1286.750464 | -0.372516071 | 1.19E-03 |
| Manbal | 1400.09717 | -0.372817625 | 2.65E-02 |
| Rbmxl1 | 3706.246739 | -0.373045215 | 1.06E-03 |
| Mob1a | 10016.4113 | -0.373765323 | 1.70E-03 |
| Stt3a | 9093.140975 | -0.373944613 | 2.80E-03 |
| 3110082I17Rik | 709.1107669 | -0.374549352 | 3.15E-02 |
| Rbm19 | 1250.338151 | -0.375040838 | 1.02E-02 |
| Fkbp9 | 22454.91561 | -0.375240188 | 1.12E-02 |
| Amd1 | 4601.579232 | -0.375449582 | 3.58E-02 |
| Tgs1 | 1089.093533 | -0.375483527 | 8.15E-03 |
| Atxn1l | 3689.513877 | -0.376025761 | 2.62E-09 |
| Ptpn11 | 9188.837608 | -0.376521981 | 1.14E-02 |
| Map2k4 | 2943.46728 | -0.376718235 | 9.21E-04 |
| Iffo2 | 1434.049209 | -0.376736548 | 3.70E-03 |
| Sfxn3 | 5162.307059 | -0.376852746 | 4.65E-04 |
| Klhl8 | 432.6864808 | -0.377173375 | 8.74E-03 |
| AC149090.1 | 18121.43559 | -0.377202951 | 3.68E-03 |
| Pgap3 | 296.0703753 | -0.377556861 | 4.01E-02 |
| Mthfsd | 1470.292497 | -0.377595391 | 4.99E-06 |
| Smcr8 | 3771.341043 | -0.377616065 | 1.08E-03 |
| Zmiz1 | 4690.270148 | -0.377662141 | 2.18E-02 |
| Ints8 | 1963.027738 | -0.378009889 | 3.50E-02 |
| Zdhhc4 | 1522.705857 | -0.378372042 | 9.16E-03 |
| Cc2d1b | 3391.120348 | -0.378572213 | 3.51E-03 |
| Trub1 | 627.2813923 | -0.379118669 | 4.17E-03 |
| Arl13b | 708.5328787 | -0.379518606 | 7.62E-03 |
| Uhmk1 | 4265.778395 | -0.380513832 | 1.00E-04 |
| Tbk1 | 1987.788939 | -0.380642737 | 1.83E-02 |
| Abl1 | 7240.633084 | -0.380876583 | 2.38E-04 |
| Gm10123 | 58159.74363 | -0.381129347 | 5.20E-03 |
| Lpcat1 | 1186.570369 | -0.381304594 | 2.27E-03 |
| Recql5 | 745.1329941 | -0.381502188 | 5.81E-03 |
| Sema4c | 1046.850603 | -0.381916358 | 1.46E-02 |
| Ndufa1 | 1200.192316 | -0.382099113 | 4.91E-02 |
| Pigu | 1268.087199 | -0.383390994 | 7.88E-05 |
| Chchd6 | 733.0271583 | -0.383679465 | 4.79E-02 |
| Dusp19 | 368.3047924 | -0.3837698 | 6.20E-03 |
| Atp13a3 | 9000.749949 | -0.384734354 | 3.94E-02 |
| Rnf2 | 2451.487291 | -0.384816689 | 9.47E-03 |
| Cabin1 | 2885.667804 | -0.385066127 | 3.22E-04 |
| Zmat5 | 369.1131801 | -0.385150674 | 1.20E-02 |
| Wdsub1 | 1434.266269 | -0.385237001 | 1.10E-02 |
| Pgm1 | 4305.875486 | -0.386448313 | 5.04E-03 |
| Msantd2 | 431.847042 | -0.386942354 | 1.03E-02 |
| Stx18 | 996.9462278 | -0.387202286 | 1.51E-02 |
| Inip | 1433.22946 | -0.387338409 | 2.66E-03 |
| Swi5 | 5268.153823 | -0.387523875 | 4.66E-06 |

| | | | |
|---|---|---|---|
| Pced1b | 224.7482532 | -0.387727121 | 2.19E-02 |
| Mat2b | 3057.082385 | -0.388067988 | 1.03E-04 |
| Pdrg1 | 1875.615487 | -0.388301761 | 3.17E-03 |
| Vti1a | 1513.738788 | -0.38852999 | 2.55E-03 |
| S1pr3 | 3918.192315 | -0.388719426 | 2.23E-02 |
| Ap2s1 | 2093.820475 | -0.388927107 | 9.21E-04 |
| Mtf1 | 1418.940464 | -0.389227675 | 4.00E-04 |
| Lyrm4 | 469.9816086 | -0.389795015 | 8.84E-04 |
| Abhd16a | 1314.595388 | -0.389982977 | 2.15E-03 |
| Tceal9 | 6722.958843 | -0.390036325 | 3.31E-02 |
| Lrp11 | 3762.571826 | -0.390165794 | 6.84E-03 |
| Xrcc5 | 2169.701875 | -0.390415268 | 1.23E-03 |
| Kansl1 | 3445.684526 | -0.39096115 | 2.40E-04 |
| Taf2 | 2167.708385 | -0.391397639 | 2.38E-11 |
| Nckipsd | 805.9801633 | -0.391650034 | 1.08E-02 |
| Crlf2 | 952.9025188 | -0.39170047 | 1.42E-02 |
| Hif1an | 2520.763791 | -0.391764789 | 1.44E-05 |
| Hsbp1 | 4851.142607 | -0.392180106 | 2.11E-04 |
| Skiv2l | 2653.900037 | -0.392322384 | 1.71E-04 |
| Gm20716 | 4001.886924 | -0.392355054 | 3.19E-06 |
| Psmd10 | 850.2875447 | -0.392697227 | 3.79E-05 |
| Psme2 | 2209.647031 | -0.393719911 | 7.65E-04 |
| Zfp874b | 473.1071347 | -0.393860185 | 1.35E-03 |
| Ctnnd1 | 14049.60831 | -0.394191866 | 9.11E-03 |
| Iqsec2 | 575.0814242 | -0.394304752 | 3.46E-02 |
| Xntrpc | 480.116466 | -0.394354335 | 1.92E-03 |
| Fto | 4606.764482 | -0.394541888 | 2.70E-02 |
| Tgif2 | 708.2075399 | -0.394665832 | 3.83E-02 |
| Kat14 | 1963.154269 | -0.394716303 | 1.29E-03 |
| Ptrhd1 | 267.3701113 | -0.394748491 | 4.71E-02 |
| Mcur1 | 737.7989981 | -0.395546929 | 1.67E-03 |
| Tmed7 | 4080.361082 | -0.395812671 | 6.68E-05 |
| Sowahc | 830.3609363 | -0.395898055 | 2.40E-02 |
| Col12a1 | 2667.384937 | -0.396010279 | 8.98E-03 |
| Ccnyl1 | 2407.209529 | -0.396011538 | 1.43E-02 |
| Map4k5 | 3127.866476 | -0.396108225 | 9.66E-04 |
| Irf1 | 1630.516384 | -0.396110835 | 2.96E-02 |
| Stard7 | 2441.522627 | -0.396142215 | 3.75E-03 |
| Dusp3 | 2333.354456 | -0.396255366 | 2.68E-02 |
| Fbxl15 | 233.2235007 | -0.396436517 | 4.69E-02 |
| Zfp668 | 533.3538571 | -0.397026085 | 2.04E-02 |
| Zfp128 | 209.8468391 | -0.397491629 | 3.06E-02 |
| Atox1 | 1778.466527 | -0.397517196 | 1.81E-02 |
| Oaz2 | 2466.83586 | -0.397598239 | 3.93E-04 |
| Zfp27 | 492.8091239 | -0.397786738 | 5.98E-04 |
| Slc25a38 | 454.986801 | -0.397917432 | 8.47E-03 |
| Ephb3 | 4231.935626 | -0.39799661 | 1.25E-02 |

| | | | |
|---|---|---|---|
| Pphln1 | 1556.478777 | -0.398023335 | 4.56E-06 |
| Chchd7 | 559.246393 | -0.39824826 | 5.91E-03 |
| Elmod3 | 569.4908347 | -0.399106658 | 9.08E-04 |
| Coa5 | 4878.455822 | -0.399504961 | 1.68E-04 |
| Pgrmc1 | 9425.547064 | -0.399789716 | 3.43E-03 |
| Eif1b | 955.7925393 | -0.399826153 | 1.78E-02 |
| Zfp87 | 507.9830536 | -0.399906211 | 1.00E-02 |
| Acly | 18875.66242 | -0.400043448 | 1.89E-02 |
| Mppe1 | 337.864424 | -0.400853365 | 1.46E-02 |
| Fam50a | 3368.128408 | -0.400898332 | 4.84E-03 |
| Nfatc1 | 796.3990061 | -0.401368472 | 2.75E-02 |
| Mbd3 | 3436.50102 | -0.401375501 | 2.32E-02 |
| Mterf3 | 690.8179778 | -0.401379072 | 2.99E-02 |
| Prpf3 | 1244.119026 | -0.401470488 | 1.84E-02 |
| Nr2c2 | 2635.174019 | -0.401581978 | 3.65E-03 |
| Yeats4 | 1397.088105 | -0.40193789 | 1.86E-04 |
| Gemin7 | 920.3277622 | -0.402889405 | 1.35E-03 |
| Lman2 | 5637.996807 | -0.403051204 | 4.68E-03 |
| Ciao2a | 1186.863786 | -0.403163157 | 2.02E-03 |
| Atp5g1 | 2775.394835 | -0.403612898 | 1.61E-02 |
| Pgd | 3686.528964 | -0.403768056 | 1.37E-03 |
| Eif5 | 9245.504961 | -0.403843394 | 1.00E-03 |
| Smim11 | 611.446549 | -0.403985598 | 2.18E-04 |
| Atp5k | 1792.535023 | -0.404114739 | 3.67E-04 |
| Atf2 | 2251.338789 | -0.404713906 | 8.16E-03 |
| Ltbp3 | 11015.44366 | -0.404860643 | 1.49E-03 |
| Ppp1r7 | 1580.060918 | -0.404999775 | 4.17E-03 |
| Zmym2 | 2205.020865 | -0.405038053 | 3.42E-03 |
| Unc119b | 2447.389577 | -0.40552607 | 2.32E-04 |
| C2cd3 | 1122.973297 | -0.405933095 | 7.32E-03 |
| Ube4a | 2494.407906 | -0.405998417 | 1.20E-04 |
| Fryl | 2801.659489 | -0.406283435 | 2.66E-02 |
| Rars2 | 673.8057716 | -0.406293172 | 1.71E-02 |
| Sucla2 | 2404.682437 | -0.406496537 | 7.87E-04 |
| Dgat1 | 845.3611873 | -0.406568792 | 7.04E-06 |
| Cltc | 24544.13461 | -0.40674695 | 2.10E-11 |
| Ddx5 | 36977.81738 | -0.407029354 | 1.05E-03 |
| Pex11b | 937.9156307 | -0.407086028 | 1.99E-05 |
| Son | 10931.49173 | -0.407271572 | 1.61E-03 |
| Pbxip1 | 1531.014964 | -0.407737512 | 2.59E-02 |
| Mlec | 15108.74515 | -0.407862487 | 1.73E-04 |
| Gm45902 | 1976.714974 | -0.408034531 | 2.06E-02 |
| Fam122a | 464.1460504 | -0.408079192 | 1.92E-03 |
| Tnfrsf1b | 2946.17122 | -0.408429032 | 3.94E-03 |
| Slc38a10 | 6334.272706 | -0.40944528 | 1.91E-03 |
| Cep19 | 695.464579 | -0.409915426 | 8.24E-05 |
| Rcbtb2 | 2432.790428 | -0.410112684 | 5.83E-04 |

| | | | |
|---|---|---|---|
| Heg1 | 6138.242062 | -0.410350353 | 2.77E-02 |
| Vps4a | 1813.209297 | -0.410665826 | 5.32E-04 |
| Tigd2 | 928.8889991 | -0.4108957 | 8.92E-03 |
| Gorab | 1255.96873 | -0.41171976 | 1.94E-03 |
| Dnajc3 | 2949.642665 | -0.411832607 | 4.21E-02 |
| Zfp46 | 933.397489 | -0.412217257 | 9.68E-03 |
| Sdcbp | 7570.072267 | -0.412249645 | 1.23E-03 |
| Sec61a2 | 1335.17424 | -0.412282618 | 3.85E-02 |
| Prdx4 | 3691.055708 | -0.413185451 | 1.58E-02 |
| Gas8 | 1271.176052 | -0.413420379 | 9.06E-03 |
| Zfp747 | 399.2398778 | -0.413433563 | 3.68E-03 |
| Htt | 2803.662578 | -0.413470029 | 8.30E-04 |
| Eif4e | 4590.336867 | -0.413500066 | 2.33E-05 |
| Fam76a | 1691.346603 | -0.413576014 | 5.69E-03 |
| Dynll2 | 6105.990932 | -0.413581688 | 2.11E-03 |
| Rnf149 | 1199.129938 | -0.413794461 | 4.12E-05 |
| AI597479 | 1129.332889 | -0.414257667 | 4.41E-08 |
| Esyt2 | 5933.474857 | -0.414450086 | 1.82E-04 |
| Tdrd3 | 952.5802765 | -0.414586338 | 1.60E-03 |
| Hars2 | 834.9974646 | -0.414673233 | 2.02E-05 |
| Pum2 | 6337.250185 | -0.414837005 | 7.65E-04 |
| Lemd3 | 799.9605492 | -0.415208442 | 3.47E-03 |
| Noc4l | 1562.446352 | -0.415364179 | 1.18E-02 |
| Fam193a | 2712.375027 | -0.415458026 | 9.01E-06 |
| Apoo | 571.4592001 | -0.415483928 | 7.06E-03 |
| Arhgap31 | 1599.694509 | -0.415521493 | 1.90E-02 |
| Hectd1 | 10105.91826 | -0.415615734 | 2.36E-03 |
| Stk16 | 1996.715038 | -0.415692323 | 4.48E-03 |
| Mfsd11 | 2064.247758 | -0.415893678 | 1.21E-02 |
| Dpysl2 | 1095.968635 | -0.416064881 | 3.23E-02 |
| Pdcd10 | 1292.535562 | -0.417465722 | 5.84E-04 |
| Ppp4r3a | 2146.905736 | -0.417586036 | 7.43E-03 |
| 2810021J22Rik | 299.5547927 | -0.41776396 | 3.57E-02 |
| Ddr2 | 21485.53935 | -0.418580976 | 3.77E-02 |
| Hace1 | 931.2661007 | -0.418878381 | 2.14E-03 |
| Dstyk | 1943.600084 | -0.419122665 | 1.57E-04 |
| A130010J15Rik | 350.9587272 | -0.41974531 | 2.45E-03 |
| Kdelr2 | 5081.551697 | -0.419960147 | 3.08E-03 |
| Glyr1 | 5537.263863 | -0.420427874 | 1.03E-05 |
| Ppp6r2 | 1188.00831 | -0.420718933 | 4.72E-09 |
| Mocs3 | 161.1274591 | -0.420795902 | 2.93E-02 |
| Ppwd1 | 530.3046537 | -0.421130755 | 4.51E-02 |
| Abtb2 | 1084.725545 | -0.421545676 | 3.08E-02 |
| Gon4l | 2759.220346 | -0.42169983 | 8.40E-05 |
| Aaed1 | 890.5933786 | -0.421730788 | 7.82E-03 |
| Vipas39 | 1504.93256 | -0.421783295 | 1.07E-04 |
| Sap30l | 703.9109609 | -0.421798021 | 1.82E-03 |

| | | | |
|---|---|---|---|
| Smarcb1 | 2669.132999 | -0.421837305 | 2.33E-03 |
| Ugdh | 6304.035598 | -0.422229649 | 4.17E-05 |
| Efcab14 | 3236.247204 | -0.422305338 | 1.87E-08 |
| Wdr7 | 1610.402645 | -0.422706688 | 7.06E-04 |
| Degs1 | 9219.28485 | -0.422768966 | 4.32E-03 |
| Invs | 851.0288157 | -0.422787465 | 2.57E-03 |
| Alg10b | 2845.75944 | -0.423457699 | 8.02E-03 |
| Lrrc58 | 3830.129758 | -0.423617333 | 7.60E-04 |
| Nom1 | 1882.71559 | -0.423730487 | 8.31E-05 |
| Tyro3 | 1115.648355 | -0.423838492 | 1.04E-02 |
| Carmil1 | 968.5312716 | -0.423838679 | 5.00E-03 |
| Ap3m1 | 2675.647402 | -0.423880073 | 4.85E-02 |
| Rnf135 | 468.7873319 | -0.424072733 | 1.52E-03 |
| Scaf8 | 3158.438473 | -0.424449435 | 5.58E-08 |
| Fam162a | 1420.891024 | -0.424538175 | 1.26E-02 |
| Hint2 | 338.0065764 | -0.424792936 | 3.18E-02 |
| Brd8 | 2043.868302 | -0.424859789 | 9.37E-03 |
| Gpatch1 | 528.5209635 | -0.425799392 | 1.30E-04 |
| Snx30 | 5144.862324 | -0.425957084 | 3.02E-03 |
| Adpgk | 1682.900064 | -0.426154929 | 4.87E-03 |
| BC028528 | 309.7950399 | -0.426568871 | 1.18E-02 |
| Traf7 | 6372.750636 | -0.426831259 | 5.35E-05 |
| Wdr73 | 690.8516135 | -0.426962382 | 1.95E-02 |
| Tomm6 | 2647.365377 | -0.427227762 | 2.64E-03 |
| 2-Mar | 1502.229125 | -0.427469684 | 2.50E-03 |
| 2310011J03Rik | 1357.96746 | -0.427601387 | 2.94E-03 |
| Wdfy1 | 2965.331295 | -0.427741074 | 2.35E-05 |
| Cstf2t | 2665.498894 | -0.427878018 | 1.13E-04 |
| Raver1 | 2661.25353 | -0.427963482 | 4.24E-03 |
| Cpne8 | 1083.712569 | -0.427995282 | 1.86E-04 |
| Wdr83os | 1481.78488 | -0.428286556 | 1.21E-02 |
| Snx33 | 2482.042903 | -0.428499694 | 3.84E-03 |
| Atp6ap1 | 11667.39151 | -0.428958552 | 2.38E-03 |
| Hexb | 4185.185652 | -0.429070696 | 2.39E-02 |
| Cchcr1 | 826.9712325 | -0.429092728 | 4.15E-02 |
| Wdr11 | 1814.743598 | -0.429241086 | 5.21E-04 |
| Rpap3 | 708.2044117 | -0.429454512 | 2.82E-03 |
| BC017158 | 795.086068 | -0.429980519 | 1.72E-04 |
| Tmcc2 | 1387.568865 | -0.430260378 | 2.49E-02 |
| Adcy9 | 386.8584794 | -0.430280638 | 1.09E-02 |
| Mgme1 | 457.9350659 | -0.430627219 | 6.04E-04 |
| Numa1 | 7864.573362 | -0.431035695 | 8.49E-05 |
| Borcs8 | 860.7623581 | -0.431257341 | 5.39E-03 |
| Spcs2 | 3414.660533 | -0.431607213 | 3.90E-05 |
| Ikzf5 | 763.4734331 | -0.431683557 | 2.97E-03 |
| Mrps2 | 1751.345024 | -0.431704391 | 4.49E-07 |
| Ints3 | 3305.873649 | -0.431738892 | 2.39E-03 |

| | | | |
|---|---|---|---|
| Fam126a | 2910.896476 | -0.431893395 | 1.57E-05 |
| Rab22a | 4367.407689 | -0.431948396 | 4.73E-05 |
| Tmem164 | 4384.562062 | -0.432010041 | 2.42E-03 |
| Slc35b2 | 1827.407569 | -0.43213814 | 7.45E-03 |
| Rpl7l1 | 3740.19368 | -0.432559333 | 5.42E-03 |
| Zfp39 | 324.6592607 | -0.432748933 | 2.16E-03 |
| Depdc5 | 1318.126855 | -0.433173274 | 1.47E-02 |
| Baz2b | 2192.623411 | -0.433736813 | 8.90E-03 |
| Mtif2 | 862.6809578 | -0.434053892 | 2.20E-02 |
| Oat | 4512.35951 | -0.43410396 | 4.66E-03 |
| Rgma | 3295.806106 | -0.434947693 | 4.94E-02 |
| Mrps18c | 677.5568705 | -0.435258039 | 9.66E-03 |
| Zfp871 | 1310.679291 | -0.435948397 | 1.20E-02 |
| Foxd2os | 411.5132541 | -0.436207473 | 3.07E-03 |
| Nipsnap3b | 762.8209259 | -0.436956389 | 3.65E-03 |
| Swsap1 | 242.7743012 | -0.4369653 | 7.04E-03 |
| Rtkn | 1336.449565 | -0.437024175 | 2.28E-02 |
| Rab8b | 1190.981597 | -0.43722864 | 1.39E-02 |
| 1110051M20Rik | 1345.512438 | -0.43724022 | 6.44E-04 |
| Atp6v0c-ps2 | 12789.63813 | -0.437536612 | 1.76E-02 |
| Kmt2c | 2822.831742 | -0.438448977 | 1.21E-03 |
| Pnpla8 | 2288.601537 | -0.439701279 | 1.56E-02 |
| Wdr6 | 3416.318234 | -0.440018857 | 5.04E-03 |
| Fem1c | 1546.41211 | -0.440204092 | 7.61E-08 |
| Ipo8 | 2808.34261 | -0.440246233 | 6.13E-05 |
| Zfp652 | 3085.865232 | -0.440324581 | 3.56E-06 |
| Mrps11 | 708.4432458 | -0.440735428 | 2.24E-03 |
| Acadm | 2856.942427 | -0.441306566 | 5.10E-05 |
| Taf1 | 2938.602297 | -0.441532993 | 2.79E-02 |
| Otud3 | 277.8243889 | -0.441663336 | 6.19E-03 |
| Gfod2 | 607.8982402 | -0.442083491 | 3.91E-02 |
| 1810022K09Rik | 286.9124674 | -0.442094258 | 1.24E-02 |
| Tle3 | 2473.32434 | -0.442263076 | 6.90E-03 |
| Mosmo | 928.7988729 | -0.442815205 | 1.54E-02 |
| Dpy19l4 | 1294.644785 | -0.443337211 | 2.35E-02 |
| Ice2 | 642.1654413 | -0.44413512 | 4.00E-02 |
| Dtnb | 1357.44367 | -0.445265663 | 4.02E-05 |
| Cipc | 1168.346399 | -0.445351578 | 7.59E-05 |
| Ttc4 | 1740.905347 | -0.445402135 | 7.77E-06 |
| Eme2 | 679.3381335 | -0.445402909 | 1.29E-02 |
| 9930021J03Rik | 2322.785857 | -0.445942775 | 1.76E-02 |
| Tamm41 | 731.5477176 | -0.445985329 | 1.16E-02 |
| Ift80 | 1433.054562 | -0.44615118 | 1.95E-02 |
| Smim13 | 747.323615 | -0.446254507 | 1.36E-03 |
| Bet1 | 1021.278895 | -0.446289778 | 8.88E-04 |
| Ip6k2 | 1213.466928 | -0.446290102 | 9.55E-03 |
| Tm9sf1 | 2720.226018 | -0.446380107 | 4.81E-05 |

| | | | |
|---|---|---|---|
| Slc2a10 | 452.5441176 | -0.446667895 | 1.57E-02 |
| Ccser2 | 3364.433109 | -0.447395893 | 1.25E-04 |
| Pot1b | 357.1730331 | -0.447405849 | 4.23E-02 |
| Vps13b | 3442.13905 | -0.447669661 | 6.73E-03 |
| Ctps2 | 3181.451717 | -0.448141826 | 4.55E-04 |
| Zfand5 | 4632.448716 | -0.448163514 | 2.43E-03 |
| Pla2g15 | 2388.719268 | -0.448338197 | 1.03E-02 |
| Znrf1 | 2102.508541 | -0.448601569 | 3.13E-04 |
| Ikbip | 3735.889795 | -0.450002763 | 2.56E-02 |
| Vps26a | 5958.591577 | -0.450176112 | 2.76E-04 |
| Cflar | 3787.80611 | -0.451202444 | 5.80E-04 |
| Keap1 | 1638.540529 | -0.451323563 | 9.66E-04 |
| Ifi207 | 781.2702642 | -0.452733977 | 2.57E-02 |
| Ncoa1 | 1626.174698 | -0.452854046 | 8.59E-05 |
| Ppil4 | 1478.851273 | -0.453004949 | 2.65E-02 |
| Clstn1 | 7598.808291 | -0.45305268 | 1.32E-04 |
| Mia3 | 7047.789822 | -0.453754752 | 6.98E-05 |
| Rnf217 | 747.2323047 | -0.453815738 | 7.40E-04 |
| Trim68 | 280.7631145 | -0.453842927 | 7.86E-03 |
| Aamp | 5473.874888 | -0.45391941 | 1.74E-05 |
| Arhgap1 | 4297.533144 | -0.454048601 | 6.95E-07 |
| Sft2d2 | 796.2368986 | -0.454581431 | 1.60E-02 |
| Msto1 | 1393.541493 | -0.455040582 | 6.79E-04 |
| Nfkb1 | 4076.642103 | -0.455109267 | 1.60E-04 |
| Nrn1 | 2360.363595 | -0.455152418 | 4.32E-02 |
| Vps33a | 2892.012132 | -0.455198582 | 1.26E-06 |
| Higd1a | 580.7001197 | -0.455269722 | 4.04E-02 |
| Gpx8 | 2857.904981 | -0.455270162 | 1.23E-07 |
| Rcc2 | 9407.022186 | -0.455583919 | 1.33E-03 |
| Zfp326 | 1099.788456 | -0.456218392 | 3.11E-02 |
| Elof1 | 1123.40013 | -0.456632145 | 1.37E-03 |
| Akap11 | 3406.170716 | -0.457184637 | 8.61E-04 |
| Slc25a44 | 2191.360797 | -0.457204442 | 5.37E-03 |
| Tacc1 | 884.4454082 | -0.457280374 | 2.50E-02 |
| D5Ertd579e | 4848.886748 | -0.457532061 | 4.13E-03 |
| Prmt6 | 763.8135835 | -0.458013854 | 2.09E-04 |
| Pinx1 | 467.0674322 | -0.458147743 | 1.13E-02 |
| Rfc1 | 3100.460772 | -0.458431919 | 3.27E-03 |
| Chtop | 5837.65029 | -0.458682532 | 1.16E-06 |
| Pign | 3567.705886 | -0.459055966 | 4.86E-04 |
| Tor1aip1 | 3198.975944 | -0.459194596 | 1.17E-06 |
| Eif4g2 | 37082.43035 | -0.459460744 | 9.82E-06 |
| Trim35 | 6233.884212 | -0.45951786 | 1.42E-05 |
| Anapc11 | 1259.126626 | -0.460407739 | 1.01E-02 |
| Ptdss1 | 2133.806868 | -0.460553803 | 1.64E-07 |
| Tbc1d32 | 741.9345639 | -0.460678775 | 7.11E-03 |
| Tsen2 | 691.2193844 | -0.46074812 | 2.91E-03 |

| | | | |
|---|---|---|---|
| Fah | 863.0818308 | -0.461453571 | 4.15E-04 |
| Abcd3 | 3631.045164 | -0.461632351 | 5.58E-11 |
| Hsf4 | 234.9047654 | -0.462028813 | 2.57E-02 |
| Ccpg1 | 2606.364703 | -0.462131146 | 1.42E-02 |
| Gt(ROSA)26Sor | 115.8026385 | -0.462223513 | 3.51E-02 |
| Six5 | 615.5519533 | -0.462560838 | 4.71E-03 |
| Atrn | 3824.366302 | -0.462734267 | 2.73E-08 |
| Tln2 | 2751.788901 | -0.462785101 | 1.33E-02 |
| Pik3r2 | 5877.98043 | -0.462820149 | 1.11E-03 |
| Sars2 | 394.5444787 | -0.46296598 | 6.10E-03 |
| Nav1 | 2802.729917 | -0.46344942 | 4.93E-04 |
| Slc35e1 | 2802.975292 | -0.463481681 | 4.90E-04 |
| Vti1b | 2649.138363 | -0.463580826 | 2.95E-05 |
| Acad9 | 2304.741806 | -0.463761816 | 1.67E-04 |
| Sppl3 | 3400.317528 | -0.464007784 | 1.24E-06 |
| Npr2 | 2586.247944 | -0.46428702 | 3.29E-07 |
| Aldh6a1 | 1590.691443 | -0.46432909 | 3.09E-02 |
| Mlst8 | 977.1244368 | -0.464394307 | 3.87E-04 |
| Cfdp1 | 4640.495616 | -0.464598698 | 2.63E-05 |
| Pank3 | 4644.409009 | -0.464758494 | 4.29E-02 |
| Prps2 | 2712.855431 | -0.464767503 | 6.29E-04 |
| Pogk | 1686.347408 | -0.464773009 | 7.80E-04 |
| Hlx | 144.0303873 | -0.46495494 | 2.96E-02 |
| Tanc1 | 5268.526244 | -0.465028057 | 4.07E-06 |
| Cyb5b | 7929.007222 | -0.465783479 | 1.48E-03 |
| Elmo2 | 2691.4094 | -0.466003346 | 3.23E-07 |
| Hip1r | 805.6628022 | -0.466204246 | 1.61E-02 |
| Hs1bp3 | 3425.439165 | -0.466419301 | 1.55E-02 |
| Dnajc10 | 5082.055813 | -0.466536719 | 5.73E-09 |
| Creb3l2 | 4400.318381 | -0.466786482 | 1.45E-03 |
| Qk | 4353.559499 | -0.466916657 | 2.70E-03 |
| B3galt6 | 196.4103202 | -0.467574317 | 9.79E-03 |
| Ap1s2 | 981.3291552 | -0.46773533 | 1.39E-02 |
| Fat4 | 5165.505281 | -0.468031633 | 1.95E-02 |
| Cd164 | 10632.62841 | -0.468166018 | 7.11E-08 |
| Polr2d | 570.7006928 | -0.468512831 | 1.63E-03 |
| Slc39a9 | 2865.46119 | -0.469137665 | 3.42E-07 |
| Rngtt | 654.9001586 | -0.469293119 | 1.81E-02 |
| 1600012H06Rik | 1609.676472 | -0.469405683 | 1.90E-06 |
| Ccdc28b | 161.1195197 | -0.469407973 | 3.64E-02 |
| Pcdh18 | 4140.879768 | -0.469621578 | 1.31E-03 |
| Rest | 3692.63625 | -0.469812485 | 1.75E-08 |
| Ctdspl2 | 1328.663166 | -0.470131841 | 2.29E-02 |
| Ccdc25 | 818.1589063 | -0.470316439 | 8.42E-03 |
| Rad9b | 181.9457967 | -0.470732758 | 1.11E-02 |
| Dtymk | 1449.195743 | -0.470862932 | 1.68E-02 |
| Tyk2 | 1273.344657 | -0.471140325 | 3.19E-04 |

| | | | |
|---|---|---|---|
| Fbxw7 | 724.7639213 | -0.471217487 | 6.59E-03 |
| Vps4b | 3274.311852 | -0.471409625 | 3.74E-04 |
| Calhm5 | 378.443981 | -0.471665946 | 1.23E-02 |
| Ptpn9 | 2903.919013 | -0.472116659 | 2.35E-05 |
| Dnm1 | 9239.958421 | -0.472132673 | 2.98E-02 |
| Anxa1 | 40517.67387 | -0.472306242 | 1.71E-03 |
| Magt1 | 3336.975739 | -0.472367498 | 5.07E-03 |
| Ostc | 2694.914602 | -0.472451306 | 4.79E-04 |
| Scaper | 920.9526199 | -0.472881428 | 2.32E-02 |
| Tm9sf3 | 10160.30778 | -0.472953363 | 7.42E-03 |
| Ifi203 | 486.7251377 | -0.473105223 | 2.91E-02 |
| Ptges2 | 754.8783035 | -0.473202776 | 1.72E-02 |
| BC005624 | 2059.921733 | -0.473803615 | 1.79E-09 |
| Bbs5 | 411.9729032 | -0.473878392 | 9.09E-03 |
| Slmap | 3527.376361 | -0.474251539 | 3.49E-03 |
| Atf6b | 3107.322539 | -0.474482828 | 1.30E-04 |
| Acaca | 6220.310423 | -0.474514112 | 2.58E-03 |
| Ttc5 | 737.7181432 | -0.474598421 | 2.05E-05 |
| Tmem250-ps | 1722.302464 | -0.474871972 | 6.46E-05 |
| Foxn2 | 1053.302421 | -0.475247528 | 3.03E-03 |
| Ubxn8 | 665.5000472 | -0.4753058 | 1.10E-03 |
| Smarcd3 | 630.858313 | -0.475584793 | 1.10E-04 |
| Zfp90 | 681.0618636 | -0.476993719 | 3.95E-03 |
| Zfp523 | 1205.100763 | -0.477294795 | 2.25E-03 |
| Ak3 | 4242.137544 | -0.477712776 | 1.66E-05 |
| Egln2 | 2115.922258 | -0.477877908 | 1.08E-04 |
| Arhgef11 | 2254.102821 | -0.477913797 | 9.61E-10 |
| Ptma | 34544.23717 | -0.478271574 | 1.52E-03 |
| Tspan5 | 5806.925104 | -0.478751411 | 1.08E-02 |
| Rbm5 | 4614.006628 | -0.479053885 | 6.23E-06 |
| Rab10os | 762.5082916 | -0.479095534 | 1.94E-02 |
| Tprkb | 2647.743254 | -0.479768928 | 7.03E-04 |
| Firre | 651.0119016 | -0.4798291 | 3.71E-02 |
| D10Jhu81e | 1329.248274 | -0.48009232 | 4.93E-04 |
| Zcchc17 | 1182.648684 | -0.480151508 | 2.13E-03 |
| Slc4a2 | 6711.940086 | -0.48050615 | 1.59E-03 |
| Eif5b | 5660.941743 | -0.481074462 | 7.06E-03 |
| Crtap | 4391.875566 | -0.481137709 | 1.05E-02 |
| Polr2f | 1393.352791 | -0.481633485 | 4.45E-03 |
| Aagab | 1352.172232 | -0.481849594 | 4.20E-06 |
| Stx17 | 1277.852869 | -0.482164257 | 1.18E-03 |
| Sdc1 | 4334.086121 | -0.482624267 | 2.84E-02 |
| Trmt5 | 584.4873569 | -0.483390296 | 6.50E-04 |
| Sgpl1 | 13380.67463 | -0.483423549 | 1.36E-04 |
| Pten | 6875.916686 | -0.483588254 | 5.78E-03 |
| Ywhaz | 11408.90752 | -0.483788401 | 3.09E-06 |
| Abcd1 | 1111.639964 | -0.483882645 | 7.47E-03 |

| | | | |
|---|---|---|---|
| Ireb2 | 2896.385537 | -0.484273995 | 1.60E-02 |
| Mbtps1 | 9771.008573 | -0.484646281 | 1.56E-06 |
| Nkapd1 | 1223.372902 | -0.484698051 | 2.17E-03 |
| Tmem19 | 1982.052025 | -0.484773729 | 3.73E-06 |
| 4931406P16Rik | 2722.4401 | -0.485393507 | 8.17E-06 |
| Cblb | 1328.176995 | -0.485393589 | 9.96E-04 |
| Zbed3 | 1298.091305 | -0.485501632 | 1.09E-11 |
| Hp1bp3 | 6438.328602 | -0.48551587 | 5.49E-05 |
| Ptpa | 5174.820688 | -0.485742732 | 2.41E-03 |
| Erlec1 | 2362.333871 | -0.486104411 | 4.28E-05 |
| Eif2ak2 | 1733.612018 | -0.486117014 | 1.21E-02 |
| Ubn1 | 5569.528947 | -0.486758761 | 1.06E-06 |
| Pqlc2 | 321.7284343 | -0.486943367 | 4.08E-03 |
| Zfp992 | 359.8629747 | -0.48721188 | 1.08E-02 |
| Setd2 | 3613.390968 | -0.487274733 | 2.49E-02 |
| 2310001H17Rik | 317.2797806 | -0.487583779 | 4.29E-02 |
| Sephs2 | 1513.013476 | -0.487770903 | 5.68E-03 |
| Iba57 | 239.3181474 | -0.488752593 | 3.60E-02 |
| Bphl | 452.6391813 | -0.488846595 | 1.33E-04 |
| Fam32a | 3348.599776 | -0.490242238 | 7.23E-07 |
| Eif2ak4 | 3546.105363 | -0.490487459 | 1.80E-06 |
| Ebpl | 299.9519528 | -0.490529082 | 1.51E-02 |
| Timm10 | 561.3568094 | -0.490731049 | 5.88E-03 |
| Fut10 | 709.5832085 | -0.491114977 | 2.59E-03 |
| 9130019O22Rik | 353.1637933 | -0.492483204 | 1.81E-02 |
| Pyroxd1 | 888.8072622 | -0.49340014 | 2.78E-02 |
| Fam122b | 417.4900481 | -0.49395036 | 3.54E-04 |
| Jcad | 4643.802879 | -0.49422345 | 2.08E-03 |
| Wsb2 | 4678.99171 | -0.494378132 | 6.86E-03 |
| Zfp316 | 1016.687164 | -0.494382491 | 1.66E-03 |
| Sbk1 | 915.23951 | -0.495488121 | 3.46E-02 |
| Cops9 | 1741.656329 | -0.495773996 | 7.63E-03 |
| Rmdn1 | 288.7588206 | -0.495871651 | 4.17E-04 |
| Dffa | 430.3054641 | -0.496015847 | 2.82E-05 |
| Evi5l | 1300.320633 | -0.496053688 | 6.80E-06 |
| Nckap1 | 11777.13914 | -0.496185338 | 2.50E-04 |
| Mum1 | 1919.342188 | -0.496265712 | 1.13E-06 |
| Ampd3 | 3598.275107 | -0.49649358 | 1.00E-03 |
| Sertad1 | 765.3883428 | -0.496555029 | 9.22E-03 |
| Ctsa | 21262.24103 | -0.496701422 | 1.62E-02 |
| Brpf3 | 1602.387227 | -0.496975451 | 3.03E-03 |
| Acad8 | 707.5740021 | -0.497796943 | 1.29E-04 |
| Camkk2 | 1761.969751 | -0.498061992 | 2.43E-06 |
| Clcn5 | 3296.841631 | -0.499876524 | 5.69E-10 |
| Mios | 1271.525436 | -0.499951324 | 9.84E-04 |
| Glmn | 555.0524008 | -0.500033785 | 3.76E-02 |
| Pkdcc | 437.5342631 | -0.500363563 | 3.16E-02 |

| | | | |
|---|---|---|---|
| NA | 152.4716177 | -0.500664689 | 4.47E-02 |
| Ccdc85c | 228.5972203 | -0.500667126 | 1.66E-02 |
| Polr3e | 952.8042013 | -0.5009727 | 3.04E-03 |
| Mrpl41 | 755.618332 | -0.501149353 | 8.85E-04 |
| Tomm5 | 1139.643768 | -0.501546473 | 2.72E-03 |
| Cradd | 175.9835452 | -0.50157785 | 8.06E-03 |
| Trak1 | 2002.750485 | -0.501928115 | 2.30E-05 |
| Lamtor3 | 1439.284321 | -0.502881042 | 2.60E-04 |
| Armc1 | 1584.697443 | -0.502920533 | 2.27E-08 |
| Isca1 | 1912.366855 | -0.5029603 | 4.62E-06 |
| Med24 | 2175.951609 | -0.503113092 | 3.10E-04 |
| Ndufv3 | 2611.286604 | -0.503540089 | 9.05E-03 |
| Dph1 | 914.7849843 | -0.503670758 | 1.30E-04 |
| Dnmbp | 2254.506423 | -0.504225954 | 8.93E-08 |
| Lrig2 | 1243.676386 | -0.504387346 | 9.39E-05 |
| Nhlrc3 | 2088.125293 | -0.5049301 | 3.47E-06 |
| Maml2 | 1522.735573 | -0.505154386 | 9.31E-03 |
| Lyrm2 | 460.0390523 | -0.505177429 | 2.83E-03 |
| Tmem263 | 2118.154995 | -0.505754977 | 8.90E-05 |
| Ift57 | 787.3549709 | -0.505821362 | 2.10E-04 |
| Ndst1 | 5649.967415 | -0.505845076 | 6.31E-06 |
| Aggf1 | 1481.03892 | -0.505992762 | 7.24E-05 |
| Ltbr | 4575.460131 | -0.506293428 | 4.49E-06 |
| Chst14 | 1836.493334 | -0.506438058 | 2.93E-05 |
| Dera | 752.132981 | -0.506721713 | 6.07E-07 |
| Pcyox1 | 9858.97276 | -0.506959431 | 2.91E-03 |
| Sgsm2 | 1683.587835 | -0.506988345 | 4.67E-03 |
| Porcn | 567.2858887 | -0.507119376 | 1.57E-02 |
| Parvb | 2285.010317 | -0.507651246 | 1.29E-03 |
| Gprasp1 | 801.852662 | -0.508176819 | 3.24E-03 |
| Lonrf1 | 505.0783429 | -0.508340799 | 1.24E-03 |
| Plekha5 | 1255.587539 | -0.508642572 | 1.45E-03 |
| Cdk5rap2 | 4844.501444 | -0.50973384 | 3.37E-03 |
| Vps26c | 2458.394981 | -0.509841092 | 3.09E-04 |
| Ddi2 | 2416.311868 | -0.510383976 | 4.42E-08 |
| Camsap2 | 3519.542233 | -0.510687529 | 4.45E-08 |
| Tpm3 | 11984.02528 | -0.510738414 | 1.47E-04 |
| Trim33 | 2609.996824 | -0.510755371 | 4.82E-03 |
| Eny2 | 1448.613816 | -0.510912889 | 2.11E-05 |
| Itm2b | 23528.2636 | -0.5116182 | 2.10E-02 |
| Eid2b | 415.8998296 | -0.511816587 | 2.68E-04 |
| Sntb2 | 2942.491811 | -0.512300352 | 8.06E-03 |
| Supt16 | 3884.352525 | -0.512452311 | 4.90E-03 |
| Wbp1l | 6033.497275 | -0.512616534 | 3.23E-06 |
| Zfp148 | 2485.900454 | -0.513000445 | 1.33E-02 |
| Rbbp7 | 9676.724973 | -0.513147382 | 2.22E-07 |
| Gpsm2 | 2322.001977 | -0.513242026 | 5.76E-04 |

| | | | |
|---|---|---|---|
| Tmem14c | 2037.871057 | -0.513431454 | 1.56E-04 |
| Katnbl1 | 1110.930944 | -0.513519804 | 2.65E-03 |
| Slc6a6 | 8237.226832 | -0.5137223 | 6.47E-03 |
| Psg16 | 1040.230629 | -0.514236468 | 3.70E-03 |
| Tnfaip8 | 770.730871 | -0.514296353 | 1.42E-05 |
| Cdk5 | 1892.485916 | -0.514386496 | 2.43E-07 |
| BC031181 | 1034.180156 | -0.514437634 | 3.77E-06 |
| Adck1 | 756.8498528 | -0.514984549 | 9.06E-06 |
| AI464131 | 913.7164078 | -0.515016608 | 2.91E-03 |
| Pde4d | 597.7003963 | -0.515085163 | 5.80E-03 |
| Scd1 | 8875.775287 | -0.5153749 | 1.82E-04 |
| Mgat2 | 2476.636202 | -0.516480731 | 6.03E-05 |
| Acat1 | 4212.253723 | -0.517429826 | 5.59E-05 |
| Mettl21a | 1118.409393 | -0.517781703 | 4.13E-05 |
| Dennd6b | 107.576249 | -0.517893535 | 1.39E-02 |
| Ipmk | 2418.776262 | -0.517988338 | 1.77E-08 |
| Zfp362 | 2229.096229 | -0.518018638 | 2.16E-05 |
| Zeb2 | 3746.786289 | -0.518036667 | 2.14E-06 |
| Wdfy3 | 4331.225501 | -0.518411581 | 2.16E-06 |
| Nbdy | 1309.915387 | -0.519011849 | 3.59E-07 |
| P3h1 | 3970.874625 | -0.519238895 | 1.56E-03 |
| Syde1 | 3457.474687 | -0.519654483 | 6.17E-05 |
| Cmtm4 | 10031.66492 | -0.519774815 | 2.11E-07 |
| Kctd18 | 586.3295036 | -0.519792133 | 8.51E-03 |
| Ppp1r21 | 1610.304203 | -0.519958916 | 4.17E-12 |
| Kat6a | 4809.065282 | -0.520127643 | 5.48E-06 |
| Psmc3ip | 403.3394449 | -0.520205098 | 1.32E-03 |
| Gtf2h3 | 971.4168647 | -0.520289621 | 2.15E-04 |
| Mtcl1 | 1053.202858 | -0.520698126 | 1.76E-03 |
| Gmcl1 | 1500.19973 | -0.521283063 | 2.67E-11 |
| Rab4b | 1144.9524 | -0.521294739 | 3.50E-05 |
| Arhgap22 | 1622.996624 | -0.521430999 | 5.97E-04 |
| Bcs1l | 748.8985829 | -0.521836759 | 2.52E-03 |
| Pfkl | 3875.900593 | -0.521926577 | 7.70E-03 |
| Hdhd5 | 498.4192023 | -0.522473047 | 7.80E-04 |
| Fancf | 204.1776999 | -0.523056826 | 5.26E-03 |
| Ercc4 | 754.5038931 | -0.523626932 | 7.78E-05 |
| Ipo13 | 5588.190074 | -0.523829878 | 3.74E-05 |
| Ltbp4 | 1251.339727 | -0.524136567 | 2.65E-02 |
| Mlycd | 405.9267261 | -0.524147339 | 1.05E-03 |
| Zfp398 | 382.5512451 | -0.524805135 | 8.83E-04 |
| Pard3 | 3039.571374 | -0.524862902 | 3.03E-08 |
| Slc22a17 | 205.312658 | -0.525487435 | 1.36E-02 |
| Fam102a | 1293.326522 | -0.525500381 | 2.90E-02 |
| Gskip | 1077.810338 | -0.526048294 | 4.86E-06 |
| Slc35a1 | 1244.299512 | -0.526216 | 8.03E-06 |
| Wdr1 | 9550.788112 | -0.526304727 | 1.59E-07 |

| | | | |
|---|---|---|---|
| Agtrap | 1417.40242 | -0.526427159 | 2.46E-05 |
| Mphosph9 | 656.4240869 | -0.526507467 | 4.26E-02 |
| Spg21 | 3580.650246 | -0.526520312 | 1.87E-03 |
| Fam13b | 1672.285168 | -0.527256691 | 6.43E-03 |
| Zfyve16 | 1187.000652 | -0.527271742 | 1.74E-02 |
| Hmgn3 | 770.6502188 | -0.527503012 | 1.31E-02 |
| Trnt1 | 2212.121754 | -0.528366574 | 8.38E-03 |
| Ppp4r1 | 3187.238595 | -0.528472633 | 1.13E-02 |
| Brms1l | 1453.82211 | -0.528656395 | 4.48E-04 |
| Npat | 1337.634223 | -0.528674884 | 2.51E-03 |
| Fndc3b | 7112.954594 | -0.528968769 | 8.76E-03 |
| Ctdsp2 | 23274.70546 | -0.529074652 | 1.10E-04 |
| Siae | 1505.165601 | -0.52931618 | 1.40E-04 |
| Lactb | 625.5953829 | -0.529386503 | 2.43E-03 |
| Eri3 | 5009.343592 | -0.530139384 | 1.30E-04 |
| Mllt1 | 2904.39071 | -0.530218581 | 2.27E-06 |
| Kif7 | 398.6317459 | -0.531150891 | 1.07E-03 |
| Ccdc32 | 828.4896004 | -0.531201214 | 1.74E-04 |
| Cavin1 | 29736.21876 | -0.531574873 | 2.67E-04 |
| Meis1 | 1497.087718 | -0.531705201 | 1.40E-03 |
| Rnf157 | 1391.358528 | -0.531878825 | 4.16E-05 |
| Gm20482 | 178.0800543 | -0.532006114 | 5.77E-03 |
| Mgst3 | 998.4397806 | -0.532343738 | 5.66E-03 |
| 7-Sep | 6788.598028 | -0.532388189 | 4.48E-02 |
| Zmym5 | 1054.81283 | -0.532440116 | 2.41E-03 |
| 1110059G10Rik | 340.4845622 | -0.532448269 | 5.39E-03 |
| Tmem170b | 491.2964213 | -0.532540051 | 2.09E-02 |
| Cpe | 17622.21883 | -0.532620411 | 2.67E-03 |
| Thyn1 | 917.6132514 | -0.532741165 | 1.32E-02 |
| Sft2d1 | 798.2097011 | -0.533430842 | 2.22E-06 |
| Cfap20 | 1689.539204 | -0.533511285 | 1.23E-05 |
| Nubp2 | 1267.351179 | -0.533852352 | 2.16E-03 |
| Usp34 | 6882.952531 | -0.534329813 | 4.79E-02 |
| Exoc1 | 1809.747397 | -0.534560592 | 4.69E-02 |
| Isoc2a | 444.5362702 | -0.534835466 | 6.03E-03 |
| Etfrf1 | 953.9923609 | -0.534896615 | 6.93E-04 |
| Itpripl1 | 456.6583994 | -0.535069834 | 3.51E-02 |
| Cdc123 | 2172.327021 | -0.53623337 | 2.80E-05 |
| Eid1 | 4134.703214 | -0.536243947 | 2.65E-04 |
| Camk2d | 6564.725921 | -0.537044693 | 2.47E-05 |
| Mob1b | 1690.668524 | -0.537105475 | 1.95E-03 |
| Sepsecs | 743.2789513 | -0.537362834 | 5.11E-03 |
| Grn | 30705.7805 | -0.537522927 | 7.88E-03 |
| Tardbp | 5588.960402 | -0.53784784 | 2.51E-03 |
| Dctd | 1097.232119 | -0.538162164 | 2.36E-03 |
| Arhgdib | 13907.08035 | -0.53854595 | 1.78E-03 |
| Rex1bd | 1738.186848 | -0.538595693 | 3.04E-04 |

| | | | |
|---|---|---|---|
| Ubiad1 | 901.4301508 | -0.539778144 | 5.71E-05 |
| Mib1 | 3545.099851 | -0.539855975 | 1.19E-02 |
| Rundc1 | 970.3247728 | -0.539954096 | 4.48E-06 |
| Ifit2 | 2698.970925 | -0.540044324 | 1.05E-02 |
| Cnnm2 | 1162.640313 | -0.540233281 | 4.09E-04 |
| Tpbg | 2142.107889 | -0.540912623 | 1.06E-06 |
| Tbl1xr1 | 2044.586522 | -0.541216932 | 8.50E-06 |
| Nacc1 | 5060.814783 | -0.5412356 | 4.39E-05 |
| Usp9x | 10545.38415 | -0.541458461 | 4.66E-03 |
| Mmp19 | 2149.521448 | -0.541772643 | 1.40E-03 |
| Crot | 1867.061663 | -0.542106636 | 1.18E-06 |
| Ptprg | 2392.25637 | -0.542181107 | 1.33E-02 |
| Med22 | 2582.347251 | -0.542222067 | 2.20E-04 |
| Ammecr1l | 2249.366946 | -0.542692492 | 7.80E-10 |
| Rbm45 | 1231.891643 | -0.542801946 | 5.75E-04 |
| Bend3 | 818.5543217 | -0.54287565 | 7.25E-04 |
| Prpf31 | 1632.334095 | -0.542885845 | 1.26E-03 |
| Card19 | 646.5482948 | -0.543027342 | 4.56E-03 |
| Vamp8 | 2018.695151 | -0.543215574 | 7.99E-07 |
| Hdac3 | 3639.465084 | -0.543500279 | 8.45E-06 |
| Tle6 | 416.6767866 | -0.543535978 | 1.94E-03 |
| Myh10 | 12571.26723 | -0.543662482 | 1.79E-04 |
| Zkscan7 | 200.504492 | -0.543806771 | 4.83E-03 |
| Rnf146 | 1837.081402 | -0.544389576 | 5.16E-06 |
| Furin | 4097.260466 | -0.54452378 | 9.05E-05 |
| Zfp407 | 895.1626937 | -0.544600867 | 9.62E-05 |
| Spout1 | 972.9814925 | -0.544727345 | 7.08E-03 |
| Ksr1 | 974.9766751 | -0.544734094 | 4.07E-03 |
| Gm49322 | 185.7261263 | -0.544855213 | 8.96E-03 |
| Ppdpf | 1355.859809 | -0.544934091 | 1.25E-07 |
| Rhobtb2 | 871.4547682 | -0.545031617 | 1.38E-04 |
| Caprin2 | 403.2197993 | -0.54510667 | 6.31E-03 |
| Smpd4 | 1683.926286 | -0.545399216 | 9.11E-07 |
| Rcn1 | 7279.957607 | -0.545471405 | 1.84E-06 |
| Tspan3 | 3878.105411 | -0.546068321 | 2.35E-06 |
| Prdm2 | 2816.518592 | -0.54618885 | 6.60E-09 |
| Dicer1 | 2835.407638 | -0.546215867 | 2.60E-07 |
| Flna | 59929.71668 | -0.546305815 | 1.65E-04 |
| Hoxa5 | 641.1710565 | -0.546423756 | 2.53E-03 |
| Cacng7 | 2011.225826 | -0.546610875 | 1.68E-02 |
| Nek9 | 8266.50416 | -0.546753726 | 2.43E-06 |
| 3110053B16Rik | 271.5926797 | -0.547128133 | 3.50E-03 |
| Lrp12 | 1151.827332 | -0.547143323 | 1.91E-05 |
| Ino80b | 616.8921082 | -0.547652183 | 2.07E-03 |
| Mterf1a | 128.382231 | -0.548051224 | 3.43E-02 |
| Tmem126b | 728.0357211 | -0.54810218 | 2.79E-08 |
| Cox7b | 3207.21518 | -0.548366048 | 2.51E-03 |

| | | | |
|---|---|---|---|
| Cryzl1 | 961.3287174 | -0.548987968 | 8.14E-04 |
| Snx11 | 910.8597265 | -0.549158357 | 5.72E-05 |
| Atp6v0a2 | 2069.637557 | -0.549335898 | 1.32E-05 |
| Tmem17 | 204.5718151 | -0.550163381 | 9.78E-04 |
| Notch2 | 5564.204759 | -0.55087342 | 1.26E-06 |
| Rdh10 | 613.1453821 | -0.550980266 | 2.07E-03 |
| Phlpp2 | 1467.445767 | -0.551110725 | 1.92E-09 |
| Hapln4 | 436.7943246 | -0.55122507 | 7.06E-04 |
| Scd2 | 61452.75163 | -0.551230857 | 3.25E-06 |
| Rbm6 | 2487.44003 | -0.551495082 | 4.00E-03 |
| 6030458C11Rik | 897.7734304 | -0.551814075 | 1.82E-05 |
| Ppp1r12b | 4138.333762 | -0.552121067 | 4.69E-04 |
| Pomp | 2998.288686 | -0.552928097 | 2.55E-06 |
| Ndufaf7 | 1011.202297 | -0.552984803 | 2.44E-06 |
| Wls | 19235.93451 | -0.553482437 | 1.17E-06 |
| Tlcd1 | 600.2415488 | -0.554063318 | 2.27E-03 |
| Selenof | 8426.348698 | -0.554399043 | 8.37E-07 |
| Rab24 | 1438.55139 | -0.554418074 | 1.11E-08 |
| Meis3 | 878.4392067 | -0.55502305 | 2.87E-05 |
| Ust | 2007.144156 | -0.555468797 | 2.58E-03 |
| Srsf7 | 4012.366097 | -0.555489221 | 5.07E-04 |
| Smc1a | 5445.475379 | -0.555497601 | 1.70E-04 |
| Jmjd1c | 4950.142215 | -0.55564968 | 1.52E-02 |
| Atp6v1b2 | 9772.289405 | -0.555704268 | 3.97E-03 |
| Rnf13 | 3890.78372 | -0.555728002 | 6.16E-04 |
| Ncoa5 | 1911.343697 | -0.55590759 | 2.05E-05 |
| Asph | 3636.471194 | -0.55603483 | 3.40E-04 |
| Washc2 | 10254.84457 | -0.557012056 | 1.75E-04 |
| Socs4 | 969.7173352 | -0.557334972 | 3.18E-03 |
| Gcc2 | 1712.595495 | -0.557421697 | 4.07E-02 |
| Cd63 | 40987.52582 | -0.557662384 | 2.68E-04 |
| Ccar1 | 3219.749532 | -0.558429802 | 1.65E-02 |
| Commd8 | 1529.596254 | -0.558636002 | 7.91E-06 |
| Akr1a1 | 21668.11073 | -0.558871878 | 1.40E-04 |
| Smug1 | 791.1906468 | -0.55934641 | 1.72E-08 |
| Timp2 | 44603.91543 | -0.559753669 | 5.66E-05 |
| Dennd4c | 5771.928255 | -0.55984821 | 3.56E-06 |
| Clip2 | 3685.485315 | -0.560199068 | 1.28E-04 |
| Tecpr2 | 835.5171694 | -0.560343327 | 3.28E-05 |
| Mga | 4009.852013 | -0.561480959 | 2.23E-02 |
| Lyst | 3273.741853 | -0.562308766 | 4.17E-05 |
| Minos1 | 1389.650092 | -0.562807174 | 1.40E-05 |
| Atp5j | 2735.573287 | -0.563077558 | 3.49E-09 |
| Chmp1a | 5050.613376 | -0.563097882 | 5.17E-06 |
| Cd82 | 121.0116936 | -0.563331184 | 4.48E-02 |
| NA | 121.2677958 | -0.563670785 | 5.79E-03 |
| Kif3b | 2552.903667 | -0.563761849 | 1.16E-15 |

| | | | |
|---|---|---|---|
| Trim45 | 432.7742059 | -0.563785394 | 3.82E-04 |
| Srsf5 | 9108.135251 | -0.563900388 | 1.78E-02 |
| Gmppb | 553.2926187 | -0.56402276 | 1.11E-03 |
| Kifap3 | 3772.714154 | -0.564069189 | 1.36E-08 |
| Bcl9 | 1951.774164 | -0.56421517 | 5.80E-05 |
| Mapk1ip1 | 378.0234365 | -0.564311893 | 1.83E-03 |
| Pigc | 1164.872634 | -0.564771817 | 3.15E-04 |
| Syap1 | 1746.197058 | -0.564980817 | 3.90E-03 |
| Nckap5l | 1105.351254 | -0.565083311 | 1.04E-03 |
| Ext1 | 3885.770872 | -0.56556935 | 4.69E-05 |
| Tspan14 | 1344.749055 | -0.566014386 | 7.31E-07 |
| Ing5 | 967.3715042 | -0.56609418 | 2.06E-07 |
| Zzz3 | 5136.327403 | -0.566202844 | 4.58E-04 |
| Zfp971 | 202.4386453 | -0.56635729 | 4.70E-02 |
| Ttc26 | 659.8498206 | -0.566547397 | 7.61E-04 |
| Twist1 | 1323.592562 | -0.567090752 | 2.57E-06 |
| Gid8 | 1780.261878 | -0.567294891 | 1.24E-07 |
| Max | 2035.937597 | -0.568298209 | 3.31E-08 |
| Rbm43 | 682.9030732 | -0.568840398 | 1.76E-06 |
| Rcbtb1 | 2189.384948 | -0.568984351 | 3.08E-06 |
| Plscr3 | 3839.393331 | -0.569011114 | 1.05E-04 |
| Cdk7 | 2374.916507 | -0.569078666 | 3.19E-06 |
| Pnkd | 893.3433196 | -0.569338941 | 6.73E-07 |
| Vmac | 1013.466863 | -0.569595946 | 7.18E-04 |
| Urm1 | 1412.734785 | -0.569624742 | 7.91E-04 |
| Lamtor4 | 1098.802993 | -0.56990318 | 1.39E-03 |
| Slc35f6 | 2258.273448 | -0.569922075 | 1.44E-04 |
| Shisa4 | 1281.213874 | -0.569941426 | 1.47E-05 |
| Stag2 | 5411.882155 | -0.569969578 | 2.62E-02 |
| Tmx4 | 2058.445111 | -0.569974964 | 8.09E-08 |
| Zfp518b | 797.7422678 | -0.570581604 | 5.43E-04 |
| Gm4540 | 805.2713897 | -0.57090848 | 1.58E-03 |
| Zfp532 | 1559.243215 | -0.571017579 | 7.65E-04 |
| Zcchc14 | 3410.077933 | -0.571369092 | 2.31E-11 |
| Gm5900 | 339.4007408 | -0.571457773 | 7.52E-03 |
| Tfdp1 | 4208.88843 | -0.571897397 | 1.26E-02 |
| Tmem160 | 502.2737753 | -0.572412283 | 1.26E-02 |
| Mcc | 1935.549035 | -0.572855607 | 9.24E-05 |
| Tor2a | 1252.70039 | -0.572873065 | 5.60E-05 |
| Tm7sf3 | 4188.572659 | -0.572990702 | 5.70E-08 |
| Nlgn2 | 3845.598607 | -0.573077297 | 6.04E-05 |
| Zbtb14 | 946.9784755 | -0.573375636 | 1.08E-06 |
| Rbm14 | 1809.83463 | -0.573668121 | 1.83E-03 |
| 4930402H24Rik | 5341.149355 | -0.573846211 | 1.12E-07 |
| Btbd7 | 2855.621626 | -0.574074534 | 1.27E-20 |
| Emc10 | 2968.610409 | -0.575006134 | 1.93E-04 |
| Dcaf11 | 750.4487475 | -0.575211999 | 1.94E-03 |

| | | | |
|---|---|---|---|
| Gng7 | 249.2332224 | -0.575255913 | 1.51E-03 |
| Bin1 | 3469.194523 | -0.575815526 | 4.82E-06 |
| Dnal4 | 1352.547094 | -0.576178533 | 5.99E-06 |
| Pyurf | 964.7439251 | -0.576432174 | 9.37E-07 |
| Cyp4f16 | 592.6168284 | -0.576497794 | 4.22E-04 |
| Zcchc2 | 3565.226416 | -0.576520545 | 1.53E-05 |
| Cep152 | 366.564991 | -0.576597608 | 3.09E-02 |
| Gramd4 | 2354.504038 | -0.576636961 | 6.25E-06 |
| Fdft1 | 6536.743541 | -0.57678872 | 3.67E-02 |
| Taf6l | 798.7607961 | -0.577192579 | 8.87E-04 |
| Inha | 254.1528339 | -0.577239616 | 6.25E-04 |
| Asb7 | 543.7279623 | -0.577245973 | 1.45E-08 |
| Rab11fip3 | 2701.591854 | -0.577662833 | 6.85E-10 |
| Pik3r4 | 1596.763516 | -0.577737776 | 9.21E-07 |
| Efemp2 | 7439.007368 | -0.578106553 | 4.29E-05 |
| Slc35a5 | 538.059794 | -0.578275866 | 3.08E-04 |
| Cep85 | 1088.695654 | -0.578362246 | 9.30E-03 |
| Osbpl7 | 425.4315274 | -0.578661231 | 2.77E-06 |
| Washc5 | 9453.018285 | -0.579192377 | 3.10E-10 |
| Gm13563 | 86.15815411 | -0.579665095 | 3.31E-02 |
| 3300002I08Rik | 78.85212045 | -0.580075139 | 2.69E-02 |
| Cfap97 | 1089.736 | -0.580726736 | 2.99E-03 |
| Fbxl19 | 1033.975476 | -0.580736638 | 4.51E-06 |
| 1700034P13Rik | 389.2191072 | -0.58099539 | 4.34E-03 |
| Pnn | 2572.950217 | -0.581300126 | 1.87E-02 |
| Ppfia1 | 3284.486732 | -0.581945262 | 1.05E-11 |
| Hectd3 | 2568.610222 | -0.58258407 | 1.10E-05 |
| 5730455P16Rik | 1509.639387 | -0.582912659 | 5.53E-04 |
| Ormdl1 | 569.7657093 | -0.583148454 | 1.76E-02 |
| Mrfap1 | 12176.20273 | -0.583339252 | 1.20E-06 |
| Tgfbr2 | 6135.16223 | -0.584104494 | 1.59E-03 |
| Megf8 | 7974.050836 | -0.584155783 | 3.28E-06 |
| Pip5k1c | 2979.353255 | -0.584182206 | 4.24E-06 |
| Zfp983 | 210.3850671 | -0.584408836 | 2.34E-04 |
| Traf5 | 535.9410057 | -0.584695505 | 5.61E-04 |
| Srp54c | 1928.786045 | -0.585237401 | 3.26E-03 |
| Gpatch3 | 203.8086641 | -0.586177716 | 4.66E-03 |
| Bzw1 | 11535.9414 | -0.586225068 | 3.10E-05 |
| Erlin2 | 2954.042327 | -0.586551381 | 8.04E-06 |
| Spg11 | 3357.707005 | -0.586737425 | 3.25E-11 |
| Hnrnpa0 | 3624.462535 | -0.587031173 | 3.70E-04 |
| Dock7 | 10072.46313 | -0.587095203 | 6.37E-11 |
| Lrrc14 | 1426.419279 | -0.587182774 | 6.24E-05 |
| Cdk5rap1 | 416.9799843 | -0.587349679 | 6.86E-06 |
| Pi4ka | 5631.269742 | -0.587489671 | 2.51E-09 |
| Zmym4 | 3703.024196 | -0.588439523 | 6.85E-05 |
| Fam53a | 1630.046894 | -0.588735909 | 1.26E-06 |

| | | | |
|---|---|---|---|
| Ulk2 | 3079.236902 | -0.588753815 | 6.54E-13 |
| Zfp28 | 611.0405196 | -0.589231883 | 1.97E-06 |
| Tnrc6c | 2070.107347 | -0.589380878 | 4.31E-09 |
| Ints2 | 1439.53712 | -0.589453195 | 2.68E-04 |
| Ptgr1 | 2247.24335 | -0.589634803 | 1.37E-05 |
| Bin3 | 1086.867767 | -0.589764509 | 6.05E-07 |
| Sf3b1 | 11144.92847 | -0.590270881 | 1.86E-08 |
| Syncrip | 5515.504038 | -0.590300822 | 3.87E-05 |
| Srd5a1 | 148.0881979 | -0.590724029 | 3.94E-02 |
| Ppip5k1 | 941.1084221 | -0.590926376 | 5.83E-06 |
| Ogdhl | 4923.875176 | -0.591120888 | 2.75E-05 |
| Pqlc1 | 490.6899246 | -0.591308924 | 3.67E-02 |
| Mgat1 | 4378.074304 | -0.591450092 | 1.33E-06 |
| Plbd2 | 8599.309391 | -0.591565785 | 8.06E-04 |
| Atp6v0d1 | 8071.893965 | -0.591821762 | 5.28E-03 |
| Hexdc | 950.8161435 | -0.591896974 | 4.94E-04 |
| Gm10039 | 530.8826632 | -0.591926595 | 2.75E-02 |
| Lrrfip1 | 4238.945427 | -0.592501703 | 9.59E-04 |
| Gm12648 | 261.1845011 | -0.592630755 | 1.20E-03 |
| Ppp1r8 | 1171.081422 | -0.593278734 | 1.95E-02 |
| Tbx15 | 3782.490487 | -0.593471839 | 2.34E-06 |
| Btf3l4 | 4495.978449 | -0.593494694 | 8.49E-11 |
| Fbxo32 | 1611.242564 | -0.59382685 | 2.14E-02 |
| Fam133b | 677.0444772 | -0.594270375 | 8.64E-04 |
| Nsd1 | 1011.722466 | -0.594628787 | 3.17E-02 |
| Capn1 | 3476.697024 | -0.596329183 | 7.79E-07 |
| Zfp395 | 1588.050862 | -0.597043817 | 1.30E-03 |
| Ankrd24 | 376.7422993 | -0.597303524 | 2.50E-04 |
| Nipa2 | 1056.294581 | -0.597681274 | 3.11E-04 |
| Catsper2 | 170.3526591 | -0.597702859 | 3.24E-05 |
| Hlcs | 514.8003438 | -0.597921685 | 6.58E-05 |
| Cep170 | 6223.844815 | -0.598284553 | 5.32E-06 |
| Klhl36 | 240.6418848 | -0.598364897 | 2.62E-03 |
| Adnp | 1561.291486 | -0.598797523 | 1.50E-04 |
| Snx8 | 2047.563724 | -0.600640659 | 1.14E-05 |
| Kif13b | 1354.851538 | -0.601268152 | 4.02E-04 |
| Bid | 1806.508215 | -0.602323203 | 3.69E-05 |
| Zfp324 | 561.8524386 | -0.602365375 | 8.60E-07 |
| Tmem186 | 774.7522416 | -0.602547376 | 1.13E-06 |
| Trim13 | 272.2413146 | -0.602617147 | 9.10E-05 |
| Nfib | 4927.428113 | -0.602624818 | 4.05E-03 |
| Erc1 | 3022.322006 | -0.602825275 | 1.64E-14 |
| Bod1 | 1941.315102 | -0.603221241 | 5.85E-07 |
| Atg12 | 4017.335377 | -0.603506373 | 7.65E-03 |
| Kctd11 | 386.5987816 | -0.604464439 | 6.60E-03 |
| Ccdc91 | 1289.115254 | -0.604569794 | 1.68E-09 |
| Cdk4 | 23550.22214 | -0.604643298 | 1.09E-05 |

| | | | |
|---|---|---|---|
| Ddx51 | 1561.419541 | -0.604788951 | 6.81E-06 |
| Zfp654 | 364.2675256 | -0.604815937 | 9.48E-03 |
| Bcl2l12 | 332.2104542 | -0.604963068 | 1.15E-03 |
| Ttbk2 | 1297.767438 | -0.605188485 | 3.72E-03 |
| Zik1 | 289.836167 | -0.605675782 | 4.00E-04 |
| Atp6v1a | 10459.35955 | -0.605984607 | 4.04E-04 |
| Mkks | 849.6034523 | -0.606467645 | 5.73E-04 |
| B230219D22Rik | 3240.706529 | -0.606531274 | 2.55E-04 |
| Tax1bp3 | 4764.282672 | -0.606952332 | 1.05E-04 |
| Gtf2ird2 | 552.0214477 | -0.607540806 | 6.78E-04 |
| Tbcel | 1611.699035 | -0.6076157 | 3.99E-12 |
| Smc3 | 4460.02087 | -0.607784749 | 3.09E-02 |
| Pask | 665.4025986 | -0.608549061 | 7.25E-04 |
| Akt1 | 12758.06405 | -0.608701706 | 6.30E-06 |
| Gcdh | 962.2082415 | -0.608726566 | 5.00E-07 |
| NA | 1811.90362 | -0.609203742 | 6.49E-07 |
| Gpx7 | 623.8937088 | -0.60925216 | 9.08E-04 |
| Trip12 | 11199.36888 | -0.609311623 | 3.25E-11 |
| Bola1 | 299.4104525 | -0.609528937 | 4.08E-04 |
| Tbx18 | 1240.526144 | -0.609591847 | 6.83E-10 |
| Ttyh2 | 1997.972056 | -0.609717809 | 5.53E-04 |
| Ddx3x | 12841.37381 | -0.61001549 | 3.52E-04 |
| AW146154 | 327.3249277 | -0.610442247 | 4.93E-02 |
| Sdccag8 | 700.0101104 | -0.610612528 | 3.58E-04 |
| Kdm5b | 5947.881651 | -0.610901028 | 1.17E-02 |
| Rbm10 | 2075.724467 | -0.611338322 | 4.82E-05 |
| Naip6 | 212.161741 | -0.611390128 | 3.90E-04 |
| Nr2c1 | 630.205208 | -0.612124491 | 6.15E-05 |
| Anapc13 | 612.8802342 | -0.612579467 | 6.20E-08 |
| Med6 | 834.6516389 | -0.61291652 | 8.37E-05 |
| Rhobtb1 | 610.439932 | -0.613007101 | 4.46E-02 |
| Mccc2 | 227.1006885 | -0.613443352 | 8.39E-04 |
| Rbm25 | 4138.697711 | -0.61418022 | 2.52E-02 |
| Emsy | 1493.02595 | -0.614283486 | 1.31E-09 |
| Bcap29 | 901.6323732 | -0.614352519 | 1.65E-02 |
| Lrfn3 | 523.8739616 | -0.614958518 | 2.34E-05 |
| Atp6ap2 | 4358.08888 | -0.615174267 | 1.82E-05 |
| Cog4 | 4133.200435 | -0.615255603 | 1.91E-04 |
| Det1 | 414.467086 | -0.61537353 | 1.04E-03 |
| Nipa1 | 460.541922 | -0.615766777 | 2.78E-04 |
| Zcchc24 | 8085.911851 | -0.61623188 | 3.69E-07 |
| Dyrk1b | 695.0887966 | -0.616361367 | 4.74E-03 |
| Tbc1d1 | 1872.956571 | -0.61639288 | 2.59E-03 |
| Birc3 | 495.0236977 | -0.616463339 | 3.42E-04 |
| Apool | 575.8688515 | -0.616656484 | 2.83E-05 |
| Lynx1 | 1430.13896 | -0.616756634 | 8.75E-05 |
| Shq1 | 500.8612755 | -0.61675964 | 2.97E-04 |

| | | | |
|---|---|---|---|
| Naa20 | 2437.665542 | -0.617024386 | 1.41E-13 |
| Npnt | 449.3441054 | -0.617146408 | 2.36E-03 |
| H2-DMa | 231.9741204 | -0.618382861 | 6.06E-03 |
| NA | 167.7309758 | -0.619011632 | 6.89E-03 |
| Bclaf3 | 423.8993827 | -0.619123628 | 4.96E-02 |
| Hspa13 | 1659.899174 | -0.619300661 | 1.69E-06 |
| Lsm10 | 1095.464393 | -0.619999313 | 8.52E-08 |
| Lysmd2 | 420.1955019 | -0.620511687 | 1.33E-03 |
| Spata2l | 99.79410299 | -0.621097679 | 4.11E-03 |
| Htatsf1 | 1879.559997 | -0.621437831 | 1.53E-03 |
| Dnase2a | 1680.910599 | -0.621772411 | 6.43E-07 |
| Prelid1 | 8505.205021 | -0.622156009 | 2.44E-08 |
| Epha7 | 250.107713 | -0.622356882 | 1.97E-03 |
| Mdh1 | 5766.576341 | -0.622364503 | 4.34E-05 |
| 5031439G07Rik | 7244.859997 | -0.622866513 | 3.26E-04 |
| Rbfox2 | 9966.948387 | -0.623038517 | 7.75E-08 |
| Rab9 | 2610.758043 | -0.623246848 | 5.91E-03 |
| Ppp2r3a | 1273.429635 | -0.623400258 | 5.57E-04 |
| Abcc4 | 345.0528919 | -0.623486542 | 3.69E-02 |
| Ocrl | 1987.16698 | -0.623846952 | 1.66E-08 |
| Zfp825 | 163.4175946 | -0.624087393 | 7.71E-03 |
| Tbc1d19 | 941.0508285 | -0.624163819 | 6.03E-06 |
| Mn1 | 1568.859744 | -0.624217039 | 3.70E-03 |
| Fam229b | 171.6780668 | -0.624708426 | 6.29E-03 |
| Rbm12 | 1195.18032 | -0.624861266 | 1.88E-03 |
| Prickle1 | 1065.178979 | -0.62493838 | 6.26E-03 |
| Adar | 2479.583721 | -0.625040263 | 2.63E-08 |
| Tusc2 | 636.7772395 | -0.625296809 | 3.52E-04 |
| Hspa2 | 705.0792204 | -0.625301308 | 1.17E-03 |
| Rrp7a | 2906.308174 | -0.625341585 | 1.11E-02 |
| Rab34 | 4682.192332 | -0.625895587 | 2.03E-08 |
| Ilk | 4670.521348 | -0.626012808 | 5.74E-06 |
| Rtl8c | 186.4143832 | -0.626619222 | 9.73E-03 |
| Gsto2 | 540.3671304 | -0.627554084 | 8.92E-03 |
| Tbc1d14 | 2671.591197 | -0.627567321 | 2.13E-06 |
| Ermp1 | 2303.615021 | -0.627755743 | 2.31E-05 |
| Jmjd8 | 638.2911338 | -0.628275603 | 1.32E-10 |
| Kmt2e | 4574.680012 | -0.628562427 | 9.70E-07 |
| Urgcp | 1854.416943 | -0.628572888 | 1.24E-06 |
| Cog3 | 2017.359627 | -0.628733558 | 2.60E-07 |
| Cachd1 | 1495.499593 | -0.628863527 | 1.47E-05 |
| Cnot9 | 2635.142109 | -0.629283305 | 3.02E-05 |
| BC024978 | 232.6752538 | -0.629644929 | 6.66E-04 |
| Topbp1 | 2858.394934 | -0.629745941 | 4.60E-02 |
| Iqck | 94.08072981 | -0.629754224 | 5.55E-03 |
| Hoxd8 | 1212.483579 | -0.629793916 | 3.34E-05 |
| Tspan17 | 1736.884601 | -0.629913884 | 6.64E-07 |

| | | | |
|---|---|---|---|
| Zfp938 | 398.0537243 | -0.630546543 | 2.23E-02 |
| Zfp142 | 2034.722645 | -0.630593298 | 9.08E-07 |
| Parp8 | 1704.661461 | -0.630945408 | 4.51E-06 |
| Pros1 | 1181.249787 | -0.631131778 | 1.44E-04 |
| Frmd4a | 3299.821544 | -0.631334168 | 2.27E-03 |
| Aven | 437.0444546 | -0.631535308 | 5.12E-04 |
| Cnn3 | 23459.34671 | -0.631564208 | 3.84E-06 |
| Ing4 | 1691.109084 | -0.632132659 | 6.23E-07 |
| Zbed4 | 1014.724302 | -0.632236737 | 9.27E-07 |
| NA | 2939.508412 | -0.633800708 | 1.34E-07 |
| Mrpl50 | 1080.201533 | -0.633953893 | 2.07E-05 |
| Kdm5d | 903.6611732 | -0.634014335 | 4.46E-05 |
| Tnrc6b | 3087.399165 | -0.634048758 | 2.96E-08 |
| Osmr | 2002.323368 | -0.634187973 | 4.03E-03 |
| Bcor | 1899.950226 | -0.635475649 | 4.89E-07 |
| Gpalpp1 | 1808.305997 | -0.635635048 | 8.00E-11 |
| Pwwp2a | 695.825925 | -0.636048944 | 5.17E-03 |
| Ap5m1 | 552.9572744 | -0.636068607 | 1.79E-03 |
| Ifitm2 | 6827.596898 | -0.636146856 | 4.04E-04 |
| Wrb | 675.6283793 | -0.636341995 | 3.79E-05 |
| Armcx2 | 5543.690073 | -0.636947665 | 1.43E-10 |
| Coq3 | 453.259282 | -0.636971518 | 5.50E-04 |
| Golgb1 | 5482.119432 | -0.637118071 | 1.43E-07 |
| Iglon5 | 4473.234465 | -0.637354647 | 2.53E-03 |
| Gdpgp1 | 236.3563167 | -0.637933581 | 7.60E-04 |
| Polr3f | 941.8123888 | -0.638069241 | 2.11E-03 |
| Rap2c | 1453.772771 | -0.638822238 | 9.43E-04 |
| Sptlc1 | 1235.658035 | -0.639451301 | 3.33E-14 |
| Zdhhc6 | 1741.824061 | -0.639547885 | 8.99E-06 |
| Cpq | 2992.352624 | -0.639554636 | 3.63E-05 |
| Mmp11 | 3113.336865 | -0.639641599 | 1.22E-07 |
| Pdlim2 | 3693.375744 | -0.639699223 | 3.02E-03 |
| Mcm9 | 451.883513 | -0.640040498 | 9.75E-07 |
| Selenon | 4253.1088 | -0.640145794 | 8.27E-07 |
| Gm4419 | 105.9221896 | -0.64014736 | 3.66E-02 |
| Ppp4r1l-ps | 1152.662507 | -0.640289127 | 1.42E-05 |
| Gm14325 | 470.7536262 | -0.640708823 | 2.11E-02 |
| Mmachc | 686.9795778 | -0.640716278 | 8.84E-06 |
| Macrod2 | 179.0981956 | -0.64083205 | 1.02E-03 |
| Hpf1 | 810.3108242 | -0.640854756 | 1.14E-04 |
| Rnaset2b | 1643.229936 | -0.640950961 | 7.72E-07 |
| Naa50 | 3422.302758 | -0.641149251 | 1.50E-03 |
| Fen1 | 1040.024734 | -0.641171762 | 4.42E-02 |
| Lrpprc | 5310.296473 | -0.641608566 | 1.75E-06 |
| Pskh1 | 1911.4363 | -0.641807529 | 5.47E-10 |
| Zdhhc24 | 757.5320873 | -0.641901685 | 4.27E-07 |
| Phip | 2280.134654 | -0.642125399 | 4.12E-02 |

| | | | |
|---|---|---|---|
| Taz | 1239.733738 | -0.64261697 | 1.69E-05 |
| Iqsec1 | 6250.869472 | -0.643022693 | 1.06E-05 |
| Usp11 | 2202.578828 | -0.643204911 | 7.27E-04 |
| Cyp4v3 | 587.8034895 | -0.643552626 | 3.70E-02 |
| Ptdss2 | 1024.474847 | -0.643556707 | 6.00E-05 |
| Cdc42ep1 | 2564.879225 | -0.644198744 | 3.68E-03 |
| Dnajc16 | 942.4498415 | -0.644460836 | 8.38E-07 |
| Apex1 | 3288.753952 | -0.644484614 | 6.55E-07 |
| Nfix | 9711.486408 | -0.644767945 | 5.91E-07 |
| Adam17 | 3233.205911 | -0.645503834 | 9.22E-15 |
| Zfp81 | 566.5113587 | -0.645774769 | 2.23E-02 |
| Sav1 | 3229.415502 | -0.646049506 | 2.85E-12 |
| Metap1d | 1532.634791 | -0.646060151 | 7.83E-06 |
| Colgalt1 | 6309.791937 | -0.646316498 | 1.70E-06 |
| Rnaseh1 | 673.6648435 | -0.646607964 | 2.47E-14 |
| Gm21992 | 240.7428873 | -0.64706203 | 4.80E-02 |
| Morf4l2 | 12523.02325 | -0.647088938 | 8.30E-06 |
| Kirrel | 13677.00951 | -0.647252707 | 1.01E-20 |
| Usp16 | 1373.588607 | -0.647316789 | 6.18E-03 |
| Zc4h2 | 400.4251748 | -0.64766611 | 3.56E-08 |
| Kat2b | 1853.046154 | -0.64845442 | 1.87E-03 |
| Phlpp1 | 938.9364218 | -0.648595139 | 2.58E-06 |
| Shpk | 226.3556193 | -0.648632345 | 2.07E-04 |
| Trim2 | 2299.108785 | -0.648778478 | 1.50E-02 |
| Fuca1 | 1794.844636 | -0.649298885 | 1.31E-03 |
| Cnep1r1 | 1321.404235 | -0.649493009 | 1.17E-05 |
| Ldlr | 7632.961331 | -0.649519245 | 1.81E-02 |
| Zfp207 | 5415.144154 | -0.649720231 | 4.04E-10 |
| Romo1 | 1261.431817 | -0.650435019 | 5.31E-05 |
| Zfp235 | 197.9093928 | -0.650444809 | 1.41E-02 |
| R3hcc1 | 995.4229444 | -0.651017315 | 9.38E-05 |
| Clk1 | 3445.62983 | -0.651395436 | 1.24E-02 |
| Abcd4 | 634.1555892 | -0.651425782 | 7.82E-05 |
| Fbxo10 | 1508.820967 | -0.651940089 | 1.71E-05 |
| Tbc1d24 | 2492.496938 | -0.65209301 | 1.46E-09 |
| Ints7 | 1422.713731 | -0.652275705 | 2.36E-06 |
| Zfp60 | 821.8985304 | -0.652387673 | 1.83E-03 |
| Tceal1 | 290.1467987 | -0.652534331 | 7.45E-04 |
| Selenow | 3078.602105 | -0.652685829 | 3.93E-04 |
| Mipep | 470.7267755 | -0.652702042 | 6.88E-03 |
| Ncs1 | 2519.874797 | -0.652778389 | 2.86E-06 |
| Cuedc2 | 3606.221772 | -0.652860957 | 2.99E-08 |
| Gm21994 | 134.2549258 | -0.652887042 | 2.11E-03 |
| Tsn | 3126.119514 | -0.653072742 | 2.12E-10 |
| Ttc33 | 677.2382922 | -0.653211736 | 1.78E-05 |
| Uap1l1 | 10255.8949 | -0.653231177 | 2.98E-03 |
| Irs1 | 2008.224803 | -0.653476476 | 1.73E-03 |

| | | | |
|---|---|---|---|
| Csgalnact2 | 1490.922805 | -0.653791222 | 7.16E-04 |
| Pbrm1 | 3746.543043 | -0.65390364 | 2.39E-03 |
| Rpp25l | 408.0873489 | -0.65399782 | 1.92E-05 |
| Epb41l1 | 6358.9137 | -0.65423808 | 1.44E-06 |
| Srek1 | 1768.790581 | -0.654767254 | 8.77E-05 |
| Lgr4 | 1019.428449 | -0.656220871 | 6.07E-04 |
| Cic | 8307.861902 | -0.656236367 | 8.09E-07 |
| Gga2 | 4453.49319 | -0.656624579 | 6.77E-05 |
| Sfpq | 8941.106294 | -0.656665503 | 4.58E-05 |
| D030068K23Rik | 56.07754723 | -0.656894651 | 3.12E-02 |
| Antxr2 | 3509.943442 | -0.657000055 | 1.39E-04 |
| Slbp | 1414.315151 | -0.657402709 | 2.60E-03 |
| Trmt61a | 494.8076108 | -0.657503205 | 3.52E-04 |
| Zfp317 | 753.4419576 | -0.657727779 | 1.76E-05 |
| Phka2 | 2086.31599 | -0.658436488 | 2.44E-06 |
| 2610002M06Rik | 1470.102267 | -0.659094987 | 6.44E-12 |
| Inava | 109.6608324 | -0.659351962 | 1.57E-02 |
| Cutc | 134.6270003 | -0.65948041 | 1.35E-03 |
| Irx3 | 178.4497727 | -0.659654249 | 2.98E-03 |
| Nmnat2 | 1973.93081 | -0.659758055 | 1.63E-04 |
| Vegfb | 3825.597118 | -0.661168488 | 4.20E-04 |
| Rnpepl1 | 315.374151 | -0.661281907 | 1.16E-04 |
| Cstf3 | 1249.131363 | -0.661349114 | 1.76E-04 |
| Sipa1 | 1737.308702 | -0.661757106 | 8.21E-06 |
| Mrpl36 | 758.7165102 | -0.66218163 | 2.21E-08 |
| Cyth3 | 4789.656829 | -0.662522958 | 9.98E-06 |
| Actn1 | 13390.55678 | -0.662707505 | 1.18E-08 |
| C330018D20Rik | 298.9653095 | -0.663531683 | 1.04E-05 |
| Klhl24 | 4261.066099 | -0.663748389 | 3.45E-04 |
| Ift43 | 828.8161268 | -0.664074178 | 5.59E-10 |
| Gstm1 | 8315.734253 | -0.664193063 | 1.77E-02 |
| Selenos | 1618.134668 | -0.664240343 | 1.76E-04 |
| Gabpa | 2042.440526 | -0.665033853 | 8.07E-04 |
| Sox12 | 1777.455189 | -0.665372786 | 7.45E-05 |
| 6330418K02Rik | 74.5174767 | -0.666021216 | 3.53E-02 |
| Erbb2 | 2761.808853 | -0.666255475 | 3.45E-08 |
| Sc5d | 2351.772826 | -0.666840841 | 6.32E-03 |
| Rab7b | 1493.162856 | -0.66703948 | 2.61E-07 |
| Arsk | 464.5614322 | -0.667174679 | 9.11E-06 |
| Cbx3 | 4932.723007 | -0.667411588 | 2.16E-05 |
| Cdk19 | 2233.948062 | -0.667507322 | 1.82E-12 |
| Zfp963 | 272.2670617 | -0.667553619 | 2.20E-03 |
| Zdhhc1 | 1130.427471 | -0.667588477 | 1.07E-05 |
| Pgam2 | 56.35623073 | -0.667688379 | 4.27E-02 |
| Ppm1l | 2353.623018 | -0.667943307 | 3.36E-09 |
| Mpc1 | 1267.168125 | -0.667967779 | 1.30E-06 |
| Ddx46 | 2557.119627 | -0.668242288 | 2.04E-03 |

| | | | |
|---|---|---|---|
| Evc | 3246.728565 | -0.668316661 | 2.48E-07 |
| Zfp768 | 743.7030957 | -0.668376918 | 4.70E-10 |
| Chrac1 | 757.1424589 | -0.670042552 | 1.69E-07 |
| Armc5 | 822.7150533 | -0.670240792 | 7.25E-06 |
| Hexim2 | 171.0155295 | -0.670780441 | 9.18E-04 |
| Ccny | 4186.355601 | -0.670978271 | 4.49E-16 |
| Tmem132a | 1717.660449 | -0.671128864 | 1.43E-05 |
| Setd7 | 5112.78778 | -0.671159817 | 1.32E-07 |
| Ppfibp1 | 13504.46439 | -0.671736203 | 5.50E-07 |
| Traf3 | 1527.743095 | -0.6731886 | 8.32E-04 |
| Unc13a | 386.4394156 | -0.673216775 | 6.00E-03 |
| Ebf3 | 1369.264586 | -0.673654802 | 4.60E-04 |
| Pars2 | 317.5111591 | -0.67369176 | 1.15E-02 |
| Snx19 | 1980.259491 | -0.673762701 | 5.42E-14 |
| Enah | 2404.280719 | -0.673887969 | 2.23E-07 |
| Slc35a4 | 2217.830955 | -0.674183715 | 1.61E-09 |
| Fam120c | 578.4235692 | -0.674669191 | 9.19E-04 |
| Zfp422 | 441.3517438 | -0.675099772 | 1.48E-02 |
| Bcl7c | 1233.600283 | -0.675177912 | 1.30E-05 |
| Fbxo41 | 890.8379166 | -0.675378537 | 6.03E-05 |
| Mtf2 | 1124.744827 | -0.675504602 | 5.87E-03 |
| Zfp651 | 2092.260882 | -0.675725584 | 2.33E-06 |
| Pcsk7 | 888.8351781 | -0.675735876 | 3.24E-11 |
| Hdac10 | 484.6780465 | -0.676025111 | 1.90E-05 |
| Trp53rkb | 133.0751093 | -0.676527439 | 3.53E-04 |
| Ccdc102a | 1567.023143 | -0.67751312 | 2.63E-06 |
| Atl1 | 306.3142026 | -0.677574217 | 6.02E-04 |
| Gm10146 | 505.8695599 | -0.677840129 | 4.38E-03 |
| 2610001J05Rik | 2297.6783 | -0.67802612 | 8.33E-09 |
| Slc25a14 | 246.6469147 | -0.678176855 | 2.22E-05 |
| D330023K18Rik | 75.36005953 | -0.678304469 | 4.19E-02 |
| Pgghg | 1003.379207 | -0.678484995 | 2.06E-09 |
| Msra | 160.0002431 | -0.678488761 | 2.61E-03 |
| Dpagt1 | 948.9650305 | -0.678493818 | 7.85E-07 |
| Pepd | 1464.655481 | -0.678550533 | 1.58E-10 |
| Rbbp9 | 1104.425234 | -0.68004862 | 6.17E-06 |
| Slc6a8 | 7871.670639 | -0.680137172 | 7.42E-04 |
| Elavl2 | 196.7850386 | -0.680334138 | 4.25E-02 |
| Fam168b | 9749.745025 | -0.680886376 | 1.00E-09 |
| Bfar | 1557.571232 | -0.681526547 | 5.37E-11 |
| Ly6a | 4142.015759 | -0.681751542 | 1.13E-02 |
| Ufm1 | 2432.671543 | -0.681861208 | 1.56E-07 |
| Gm17300 | 172.1249343 | -0.681935258 | 3.96E-02 |
| E430018J23Rik | 232.8995405 | -0.682685396 | 2.36E-04 |
| Wasf3 | 609.9759654 | -0.682863618 | 6.20E-06 |
| Pdpn | 5637.459948 | -0.683650597 | 1.83E-04 |
| Hbp1 | 5566.199986 | -0.683893839 | 5.68E-05 |

| | | | |
|---|---|---|---|
| Zfp418 | 129.6532408 | -0.684079353 | 3.98E-04 |
| Gm14326 | 251.3502862 | -0.68438844 | 4.11E-02 |
| Rybp | 2361.128019 | -0.684436098 | 5.01E-07 |
| Slc35a3 | 927.6897423 | -0.685038111 | 7.60E-04 |
| Mex3a | 2190.570495 | -0.685119906 | 5.23E-03 |
| Hdac1 | 4349.289025 | -0.685493309 | 1.31E-07 |
| Ddx10 | 1380.39871 | -0.686695216 | 5.29E-04 |
| Arpc5 | 5430.297978 | -0.686749564 | 2.70E-17 |
| Zfp59 | 117.0891697 | -0.687008794 | 6.25E-04 |
| Lgals4 | 78.28538866 | -0.687148581 | 3.77E-02 |
| Tmem218 | 321.622946 | -0.687159739 | 1.32E-04 |
| Chpf2 | 2871.583349 | -0.687180754 | 3.79E-09 |
| Tspan6 | 853.1154017 | -0.687336439 | 3.59E-08 |
| Ssh2 | 685.016075 | -0.688123464 | 8.17E-04 |
| Crocc | 472.3468662 | -0.68821646 | 3.14E-03 |
| Agpat2 | 111.9244998 | -0.688847979 | 3.12E-03 |
| Gtf3c5 | 1178.49991 | -0.688951051 | 1.24E-10 |
| Fnbp1l | 2320.759181 | -0.68941907 | 2.10E-02 |
| Mpg | 550.6470613 | -0.689528522 | 6.83E-06 |
| Slc10a7 | 1417.196697 | -0.689601419 | 8.45E-05 |
| Zeb1 | 3675.931373 | -0.689643344 | 1.01E-04 |
| 5031425E22Rik | 290.5449219 | -0.689784877 | 7.86E-06 |
| Smim19 | 793.360551 | -0.689894905 | 4.67E-06 |
| Phtf1 | 2088.129967 | -0.689938129 | 1.50E-10 |
| Atp2c1 | 2132.770983 | -0.690456304 | 4.76E-05 |
| Cd9 | 9138.524752 | -0.690532766 | 1.45E-11 |
| Snapin | 2650.194462 | -0.690717776 | 2.20E-07 |
| Dcun1d2 | 872.1359931 | -0.690812276 | 4.15E-10 |
| Dag1 | 10803.36973 | -0.692106512 | 2.10E-10 |
| Dctn1 | 6439.938184 | -0.692332147 | 1.10E-10 |
| Ccng2 | 2087.349482 | -0.692545468 | 7.70E-04 |
| Bach1 | 2273.465941 | -0.693055614 | 1.82E-14 |
| Rab11b | 4006.527392 | -0.693301998 | 3.76E-05 |
| Zfp111 | 586.8419565 | -0.693372057 | 6.85E-03 |
| Decr1 | 1500.980543 | -0.694230853 | 7.56E-12 |
| Gm15867 | 82.20330109 | -0.694466312 | 2.02E-02 |
| Ift172 | 1593.456915 | -0.696589805 | 3.53E-07 |
| Camkmt | 146.0385414 | -0.69679996 | 7.42E-04 |
| Mks1 | 91.58251318 | -0.696895758 | 7.89E-03 |
| Mnt | 3501.854391 | -0.697335337 | 8.34E-04 |
| Zfp626 | 49.10786744 | -0.697510013 | 3.81E-02 |
| H3f3a | 13435.63948 | -0.697822266 | 5.86E-08 |
| Tasp1 | 468.6773577 | -0.69863731 | 2.86E-07 |
| Lfng | 220.6117941 | -0.698681168 | 2.11E-02 |
| Ccdc115 | 665.7497048 | -0.698955574 | 1.50E-05 |
| Zfp960 | 443.9527272 | -0.699142101 | 3.18E-02 |
| Zc3hav1l | 1399.674989 | -0.699268025 | 3.36E-13 |

| | | | |
|---|---|---|---|
| Plpp7 | 57.83169076 | -0.699447042 | 1.98E-02 |
| Nqo2 | 413.4205542 | -0.699653237 | 9.61E-04 |
| Ece1 | 1437.72767 | -0.699735088 | 1.11E-03 |
| Gm20707 | 119.736565 | -0.699749309 | 2.05E-02 |
| Comtd1 | 549.7820272 | -0.700213349 | 2.05E-06 |
| Tepp | 46.4196485 | -0.701077901 | 3.82E-02 |
| Tmem41b | 1689.446509 | -0.701238133 | 4.57E-07 |
| Marveld1 | 6344.789281 | -0.70130859 | 4.49E-18 |
| Ccdc157 | 793.9460529 | -0.701526645 | 5.58E-08 |
| Csnk1g3 | 1352.910275 | -0.70157858 | 3.12E-07 |
| Zfp358 | 1303.342284 | -0.702672302 | 1.20E-09 |
| Arid2 | 2802.678595 | -0.703479496 | 1.56E-04 |
| Ylpm1 | 3107.707483 | -0.703529606 | 1.83E-06 |
| Stag1 | 2203.340069 | -0.703779904 | 3.86E-06 |
| Zmym3 | 1752.30601 | -0.703849945 | 1.27E-07 |
| Gm10425 | 67.09687368 | -0.703951235 | 3.09E-02 |
| Slc25a26 | 255.1476367 | -0.703970865 | 2.84E-05 |
| Cyb561d1 | 293.9033658 | -0.704034619 | 5.71E-07 |
| Inf2 | 6535.964029 | -0.70406051 | 1.90E-06 |
| Zgrf1 | 386.5955645 | -0.704589137 | 1.39E-02 |
| Agfg2 | 1357.380105 | -0.704820105 | 1.67E-06 |
| Itsn1 | 2752.843847 | -0.705554785 | 1.66E-11 |
| Arl10 | 239.535473 | -0.706788176 | 4.82E-04 |
| Yap1 | 5868.317144 | -0.70762738 | 3.55E-23 |
| Pnpla7 | 844.5630092 | -0.708074802 | 1.86E-04 |
| 1110008P14Rik | 539.8675936 | -0.708109574 | 9.71E-05 |
| 1600014C10Rik | 740.8140799 | -0.708756858 | 1.21E-04 |
| Whrn | 1183.150778 | -0.708812088 | 5.60E-05 |
| Arhgap32 | 2413.87787 | -0.709010062 | 2.53E-08 |
| AW549877 | 3473.329196 | -0.70908436 | 3.31E-05 |
| Srrt | 3532.906012 | -0.709986414 | 3.83E-06 |
| Gm17484 | 94.6692011 | -0.710335294 | 4.66E-02 |
| Edem3 | 2887.189314 | -0.710538918 | 2.93E-04 |
| 1810058I24Rik | 4262.602694 | -0.710743815 | 1.45E-03 |
| Chtf8 | 5434.622375 | -0.711002246 | 2.24E-08 |
| NA | 84.09963672 | -0.711984321 | 3.82E-02 |
| Paxip1 | 2358.455763 | -0.712977028 | 6.17E-08 |
| Zbtb6 | 597.3816576 | -0.71333287 | 3.53E-04 |
| Trim23 | 667.5805986 | -0.713346356 | 1.33E-02 |
| Zscan22 | 382.3077886 | -0.714455731 | 4.67E-06 |
| Grcc10 | 2167.032303 | -0.714631496 | 4.63E-04 |
| Ccdc167 | 216.2758511 | -0.714649127 | 2.22E-04 |
| Klhdc8b | 78.31204509 | -0.714951521 | 1.25E-02 |
| Zbtb8a | 353.867421 | -0.715189364 | 1.52E-04 |
| Echdc3 | 481.9063913 | -0.715332823 | 1.95E-04 |
| Nek6 | 2770.3449 | -0.715410276 | 3.18E-10 |
| Tex261 | 4268.859971 | -0.715694482 | 3.78E-06 |

| | | | |
|---|---|---|---|
| Nagpa | 1014.528249 | -0.715714813 | 3.99E-05 |
| Atg10 | 242.0275732 | -0.715736563 | 1.73E-06 |
| Lrp6 | 8327.93429 | -0.715798052 | 6.21E-07 |
| Dop1a | 969.9602123 | -0.715924852 | 2.13E-05 |
| Mcm4 | 3562.673628 | -0.71666514 | 1.41E-03 |
| Ddx55 | 523.2516774 | -0.716750644 | 1.14E-04 |
| Mis12 | 1396.433358 | -0.716998844 | 3.31E-03 |
| Zfp101 | 297.5685569 | -0.717008074 | 1.20E-02 |
| Cr1l | 1416.508505 | -0.717106099 | 3.26E-06 |
| Ndst2 | 1120.700593 | -0.717331413 | 2.93E-16 |
| Usb1 | 1737.627401 | -0.717545752 | 7.91E-09 |
| Uty | 529.0603127 | -0.717793569 | 2.80E-02 |
| Cdk20 | 349.3497726 | -0.717925937 | 1.64E-05 |
| Tmem251 | 678.3297086 | -0.718236119 | 1.08E-06 |
| Iqce | 1194.492392 | -0.718279217 | 1.61E-06 |
| Asic1 | 2363.956811 | -0.718383209 | 4.45E-04 |
| Faap100 | 615.3568584 | -0.718669036 | 2.28E-05 |
| Rnf6 | 2353.676709 | -0.719178191 | 2.25E-03 |
| Mgat5 | 813.1866966 | -0.719448703 | 2.16E-07 |
| Col4a1 | 20658.11282 | -0.719771218 | 1.27E-03 |
| Ogfrl1 | 619.3082176 | -0.719797949 | 1.58E-03 |
| Plau | 1516.461126 | -0.720419883 | 4.07E-02 |
| Npas3 | 130.7817696 | -0.72247504 | 1.11E-03 |
| Tmem241 | 182.3624427 | -0.722563559 | 6.14E-03 |
| Gm10125 | 80.42503309 | -0.723459292 | 1.54E-02 |
| Prps1l3 | 896.6416978 | -0.723670579 | 5.40E-05 |
| Pex11g | 135.442031 | -0.723850356 | 1.64E-02 |
| Fez1 | 805.2676056 | -0.724149657 | 2.46E-05 |
| Ilvbl | 2448.329512 | -0.724263767 | 3.10E-06 |
| Ofd1 | 934.0707456 | -0.725391593 | 7.75E-05 |
| 9930012K11Rik | 537.6727168 | -0.725481843 | 3.79E-05 |
| Dip2c | 2363.258568 | -0.725785906 | 9.65E-12 |
| Tra2a | 3145.174792 | -0.726043497 | 2.75E-04 |
| Tmem106b | 10840.02422 | -0.726080932 | 5.51E-05 |
| Rnf168 | 1483.705527 | -0.726655958 | 4.77E-04 |
| Pdik1l | 420.2720688 | -0.726701939 | 7.13E-03 |
| Pigp | 767.26785 | -0.726978212 | 6.62E-06 |
| Atp6v1g1 | 12568.78181 | -0.727079808 | 3.86E-05 |
| Rnf219 | 689.5253814 | -0.72739123 | 4.91E-08 |
| Tsc1 | 3398.460772 | -0.727854238 | 1.30E-08 |
| Zfp964 | 78.48029832 | -0.728331877 | 6.43E-03 |
| Ublcp1 | 1024.594223 | -0.72884548 | 1.80E-06 |
| Pdzd4 | 1139.545824 | -0.728982044 | 1.26E-08 |
| Nenf | 2679.17814 | -0.72969536 | 8.37E-07 |
| Taf11 | 1033.711611 | -0.729950906 | 3.41E-08 |
| Nrp2 | 11248.52068 | -0.729973487 | 1.91E-07 |
| Zfp319 | 991.9383572 | -0.730222898 | 2.08E-09 |

| | | | |
|---|---|---|---|
| Arhgap39 | 893.335859 | -0.730281 | 2.22E-05 |
| Arhgef10 | 6143.142145 | -0.731419473 | 1.90E-11 |
| Zfp329 | 528.9877162 | -0.731976426 | 4.43E-02 |
| Sergef | 239.6963491 | -0.732257168 | 8.32E-05 |
| Dcp2 | 1731.728548 | -0.732893181 | 3.28E-06 |
| St7l | 643.9586616 | -0.733054338 | 2.67E-08 |
| Zfp266 | 1891.611082 | -0.734659722 | 1.08E-04 |
| Foxd2 | 47.01500774 | -0.734761891 | 1.82E-02 |
| Runx1 | 1094.061865 | -0.735181879 | 1.56E-10 |
| Wdpcp | 403.2797746 | -0.735237624 | 2.29E-04 |
| D3Ertd254e | 1068.137171 | -0.735394548 | 9.42E-06 |
| Epc1 | 1094.48254 | -0.735394623 | 1.27E-07 |
| Pin4 | 625.4302369 | -0.735431957 | 1.63E-03 |
| Trim21 | 337.3701587 | -0.735543913 | 1.19E-07 |
| Gbp3 | 49.15600115 | -0.735552275 | 2.96E-02 |
| Hnrnpa3 | 10251.47568 | -0.736146193 | 1.19E-02 |
| Rac3 | 64.09443732 | -0.736329143 | 4.30E-02 |
| 9430038I01Rik | 156.3527109 | -0.73702589 | 1.65E-05 |
| Arf3 | 7135.48518 | -0.737420884 | 1.26E-27 |
| Scd4 | 53.94463503 | -0.737812199 | 2.50E-02 |
| Grk1 | 121.1867528 | -0.73786034 | 2.86E-02 |
| Plcb4 | 1673.625399 | -0.738151923 | 6.29E-03 |
| Agbl3 | 142.3879339 | -0.738259049 | 7.15E-04 |
| Lzts3 | 586.2326723 | -0.738929863 | 1.47E-04 |
| Atp6v0c | 247.5894249 | -0.739958958 | 5.17E-03 |
| Fignl2 | 141.930956 | -0.740163205 | 5.50E-04 |
| Impad1 | 4133.85745 | -0.740320631 | 6.29E-07 |
| Gtf2i | 8843.674879 | -0.740657309 | 4.76E-09 |
| Ghr | 7386.390632 | -0.741213842 | 1.45E-04 |
| Lrig1 | 3532.647732 | -0.741348564 | 2.52E-05 |
| Ints1 | 2784.850217 | -0.741790109 | 8.95E-07 |
| Ensa | 4415.155983 | -0.742090016 | 4.36E-09 |
| Myo1d | 3631.060795 | -0.742321737 | 4.53E-06 |
| Bnip3 | 1792.535201 | -0.74267245 | 4.40E-06 |
| Ube2d1 | 1311.582683 | -0.742754205 | 3.53E-07 |
| Ctcf | 2881.745882 | -0.743026441 | 1.97E-15 |
| Slc25a24 | 3916.415838 | -0.743434146 | 3.04E-03 |
| Ephb2 | 1039.48774 | -0.744094334 | 7.94E-05 |
| Armc8 | 1164.412789 | -0.74424155 | 8.14E-10 |
| Tnrc18 | 4357.493149 | -0.744586734 | 2.22E-06 |
| Zfp24 | 1016.182409 | -0.744597311 | 1.19E-06 |
| Tut4 | 2806.378558 | -0.745092168 | 1.83E-03 |
| Laptm4a | 29285.87798 | -0.745141349 | 4.72E-06 |
| E030042O20Rik | 33.24426692 | -0.745186135 | 4.46E-02 |
| Glt8d1 | 1660.943834 | -0.746093276 | 1.16E-06 |
| Usp27x | 226.8415996 | -0.746241998 | 1.30E-05 |
| Wdr90 | 668.2938756 | -0.746903446 | 2.11E-02 |

| | | | |
|---|---|---|---|
| Morc2a | 4216.368248 | -0.74720033 | 9.19E-07 |
| Zbtb33 | 1203.252921 | -0.747603312 | 1.62E-06 |
| Samd1 | 1577.418184 | -0.747708813 | 2.88E-06 |
| Cdk5r1 | 132.2529447 | -0.747860885 | 2.25E-02 |
| Bloc1s1 | 2005.739788 | -0.748041963 | 7.68E-11 |
| Wdr19 | 1255.580238 | -0.748330589 | 3.20E-08 |
| 4933404O12Rik | 648.6074273 | -0.748656181 | 3.42E-03 |
| Zfp551 | 180.988907 | -0.749061328 | 3.92E-04 |
| Casp9 | 1187.691822 | -0.74906484 | 2.30E-07 |
| Pomt2 | 1195.558329 | -0.749660682 | 1.51E-09 |
| Lrba | 1694.232808 | -0.750263843 | 8.76E-09 |
| Mdn1 | 3461.176169 | -0.750819093 | 9.65E-04 |
| Clba1 | 2846.204258 | -0.750870837 | 4.25E-03 |
| Fut11 | 609.8350578 | -0.750927325 | 1.80E-05 |
| Txlna | 4884.051793 | -0.751621358 | 1.30E-09 |
| Pibf1 | 375.6418153 | -0.751719212 | 3.11E-04 |
| Ift122 | 1556.937666 | -0.753282033 | 4.25E-09 |
| Thoc1 | 1156.504266 | -0.753333712 | 1.27E-03 |
| Cercam | 171.2368976 | -0.753400821 | 6.38E-03 |
| Lamtor1 | 3855.900239 | -0.753407006 | 1.73E-06 |
| Enpp1 | 302.9825204 | -0.753745094 | 2.80E-03 |
| Nudt16l1 | 1616.991711 | -0.753873709 | 7.10E-06 |
| Zfp229 | 252.3107468 | -0.754106037 | 2.70E-02 |
| Pik3ip1 | 2251.596988 | -0.754107094 | 4.40E-03 |
| Dnajb5 | 479.406288 | -0.754526745 | 1.17E-04 |
| Psmb10 | 782.003635 | -0.754597471 | 1.76E-04 |
| Hoxa2 | 364.402932 | -0.754817034 | 1.70E-03 |
| Hykk | 297.2623057 | -0.755061105 | 8.91E-09 |
| Zfp712 | 74.56902935 | -0.755514518 | 3.20E-02 |
| Mob3a | 1207.127894 | -0.755583379 | 2.21E-06 |
| Synj2bp | 736.8357298 | -0.755670482 | 3.87E-07 |
| Traf6 | 1586.592483 | -0.75637252 | 4.27E-14 |
| Cenpb | 8519.626358 | -0.757220907 | 2.90E-08 |
| Ap5s1 | 405.3152319 | -0.757631049 | 4.57E-05 |
| Capn7 | 1695.101627 | -0.758413407 | 1.01E-04 |
| Slc38a2 | 19026.7174 | -0.758562602 | 1.37E-06 |
| Rbx1 | 2644.786189 | -0.758630832 | 2.26E-10 |
| Golga1 | 1274.09039 | -0.758881703 | 1.16E-03 |
| Ldhb | 3916.76833 | -0.759463948 | 1.15E-05 |
| Dab2 | 7942.183086 | -0.759625337 | 2.21E-04 |
| Borcs5 | 355.7722684 | -0.759771814 | 1.03E-04 |
| Vps29 | 2535.509627 | -0.760165601 | 1.31E-11 |
| Syne3 | 1388.172839 | -0.760166752 | 4.38E-05 |
| A830082K12Rik | 468.2476811 | -0.760443872 | 1.11E-04 |
| Gpr63 | 64.54603199 | -0.760867694 | 2.15E-02 |
| Slc25a53 | 238.7168006 | -0.761041015 | 3.68E-03 |
| Zfp948 | 823.2408788 | -0.761326453 | 4.46E-02 |

| | | | |
|---|---|---|---|
| F8 | 92.05127566 | -0.76136534 | 3.15E-02 |
| BC029722 | 1983.763936 | -0.761671957 | 1.94E-07 |
| Smarcc2 | 6613.270965 | -0.762377793 | 3.18E-15 |
| Pdpr | 1055.824546 | -0.762833838 | 1.96E-11 |
| Chd9 | 2494.76083 | -0.763609871 | 1.46E-04 |
| Slc35a2 | 1147.598411 | -0.763620553 | 1.24E-07 |
| 1600020E01Rik | 376.3471128 | -0.764292606 | 7.58E-08 |
| Psen2 | 389.5631355 | -0.764348897 | 3.23E-06 |
| Hmg20a | 1475.554886 | -0.764448856 | 2.69E-14 |
| Gm14418 | 318.6864151 | -0.765084372 | 1.86E-04 |
| Erh | 4021.046823 | -0.765296943 | 3.83E-05 |
| Slc30a7 | 2473.28765 | -0.765639129 | 2.03E-20 |
| Celf2 | 6245.986694 | -0.765782372 | 5.28E-05 |
| Sdhaf4 | 475.0879917 | -0.76619699 | 1.06E-08 |
| Obsl1 | 4116.268852 | -0.7663064 | 6.23E-06 |
| Ppp4r3b | 2886.064442 | -0.767420979 | 8.06E-04 |
| H2-T24 | 90.28178638 | -0.76778102 | 2.12E-02 |
| Mlh1 | 506.401336 | -0.76813041 | 1.74E-05 |
| Pex26 | 694.5375104 | -0.768207891 | 2.16E-08 |
| Pomk | 854.3945557 | -0.76833606 | 4.70E-11 |
| Mettl7a1 | 836.3886156 | -0.768361818 | 2.15E-04 |
| Fut8 | 1050.838888 | -0.768551524 | 1.09E-07 |
| Ptp4a3 | 157.9606979 | -0.769189027 | 4.24E-03 |
| Tmem64 | 1663.412228 | -0.769193844 | 7.39E-08 |
| Wdyhv1 | 343.7885209 | -0.769372193 | 3.53E-07 |
| Sh2b1 | 1763.552917 | -0.769426762 | 7.82E-14 |
| Plekhm1 | 2389.867812 | -0.769806151 | 1.66E-08 |
| Fbxl17 | 1422.489761 | -0.769951027 | 1.00E-12 |
| Gm20712 | 69.74493002 | -0.770516193 | 4.49E-03 |
| Zfp868 | 720.900899 | -0.770719906 | 7.16E-06 |
| Zkscan1 | 2415.13319 | -0.771136809 | 1.14E-14 |
| Rhbdd3 | 570.80931 | -0.771578898 | 1.14E-05 |
| Ppic | 4106.544221 | -0.771597923 | 1.46E-09 |
| Ldlrad3 | 1390.98449 | -0.771774909 | 1.82E-06 |
| Nfyc | 2165.842063 | -0.772167581 | 3.83E-14 |
| Serpinf1 | 25986.98908 | -0.77246562 | 1.18E-03 |
| Ppa2 | 1042.259052 | -0.772883842 | 2.08E-12 |
| Anxa6 | 6987.123827 | -0.773104916 | 1.81E-05 |
| Agk | 886.1543945 | -0.773182267 | 2.71E-06 |
| Spred2 | 1836.996101 | -0.773295047 | 2.01E-10 |
| Arhgap24 | 1837.038701 | -0.773474263 | 1.15E-03 |
| Pla2g7 | 967.292464 | -0.773799897 | 2.37E-03 |
| Vamp7 | 964.5436274 | -0.774198687 | 5.97E-04 |
| Rnf170 | 766.78241 | -0.774232397 | 6.90E-05 |
| Dennd4b | 511.3383527 | -0.77439645 | 1.97E-06 |
| Fbxl14 | 1290.48478 | -0.774625999 | 8.80E-10 |
| Zfyve26 | 2479.501653 | -0.774706054 | 8.89E-12 |

| | | | |
|---|---|---|---|
| Slx4ip | 322.4792239 | -0.775264949 | 1.60E-05 |
| Snx13 | 2199.007322 | -0.775298784 | 4.15E-04 |
| Ggn | 64.83403107 | -0.77786603 | 1.37E-02 |
| Ids | 2262.136179 | -0.778016523 | 2.29E-12 |
| Maged1 | 25928.54256 | -0.778031065 | 1.63E-12 |
| Hexa | 11840.38452 | -0.778875193 | 1.81E-03 |
| Zfp955a | 638.3229474 | -0.779015779 | 1.14E-03 |
| Ctsl | 40956.20104 | -0.779223306 | 2.28E-07 |
| Pfdn5 | 3955.074207 | -0.779294811 | 1.29E-04 |
| Fam102b | 2904.554024 | -0.779595679 | 8.37E-14 |
| Rbm3 | 13054.23758 | -0.779859633 | 1.02E-08 |
| Fbln5 | 11685.97594 | -0.780153712 | 4.35E-05 |
| Tsc22d1 | 5481.602433 | -0.780858164 | 3.43E-09 |
| Gm10093 | 872.0229362 | -0.781188818 | 9.65E-10 |
| Vac14 | 1852.536674 | -0.781246369 | 3.52E-07 |
| Zfp493 | 62.7561201 | -0.78255135 | 1.34E-02 |
| Gm26792 | 177.3943853 | -0.782814842 | 3.78E-04 |
| Mccc1os | 80.25130271 | -0.783005721 | 7.16E-03 |
| Cirbp | 1092.334366 | -0.783148685 | 9.21E-06 |
| Fndc4 | 669.6069823 | -0.78359048 | 3.09E-04 |
| Vegfd | 6147.027 | -0.783855237 | 1.95E-02 |
| Pi4kb | 3352.487585 | -0.784150796 | 1.06E-08 |
| Osbpl1a | 1582.537579 | -0.784442298 | 1.20E-08 |
| 5730480H06Rik | 325.1206865 | -0.784495341 | 9.85E-08 |
| Rmdn2 | 355.5715743 | -0.784504478 | 2.78E-04 |
| Gm17586 | 152.4386316 | -0.784721136 | 3.57E-05 |
| Paqr4 | 951.0335697 | -0.785143281 | 4.26E-04 |
| Bbx | 2721.317942 | -0.78527671 | 1.71E-05 |
| Fundc1 | 1249.116808 | -0.785490167 | 2.04E-13 |
| Cog1 | 1510.656347 | -0.785536047 | 1.80E-09 |
| Scfd2 | 558.8131694 | -0.785632084 | 3.05E-07 |
| Arl15 | 412.0136059 | -0.785958976 | 1.21E-06 |
| Fam219b | 604.1462657 | -0.786270566 | 2.44E-07 |
| Snx24 | 596.8955764 | -0.786499471 | 2.58E-06 |
| Bhlhb9 | 1587.323127 | -0.786612434 | 1.79E-06 |
| Alyref2 | 123.0794176 | -0.786612743 | 5.34E-04 |
| Lins1 | 515.0195333 | -0.786632335 | 8.70E-04 |
| Dnal1 | 1014.638004 | -0.787054006 | 2.31E-05 |
| Fam227a | 82.93503029 | -0.787363931 | 2.80E-02 |
| Fes | 449.9226603 | -0.78771098 | 2.95E-07 |
| Gdf11 | 1582.164993 | -0.787887724 | 4.72E-06 |
| Extl3 | 3064.630989 | -0.788483607 | 3.73E-10 |
| Nek4 | 454.1602901 | -0.78863007 | 1.39E-06 |
| Cd302 | 1091.553765 | -0.789112555 | 1.45E-06 |
| Efna5 | 479.7293286 | -0.789692185 | 7.62E-05 |
| Dab2ip | 2202.223697 | -0.789699644 | 1.39E-05 |
| Glcci1 | 283.1218107 | -0.789963968 | 2.50E-02 |

| | | | |
|---|---|---|---|
| C2cd5 | 1214.242853 | -0.790176749 | 1.94E-07 |
| Mfsd1 | 8319.866312 | -0.790311466 | 8.29E-10 |
| Cttn | 14805.07831 | -0.790430514 | 4.35E-28 |
| Zscan12 | 310.7903595 | -0.79047902 | 1.51E-07 |
| Tmem47 | 608.4736339 | -0.791716188 | 2.83E-02 |
| Zbtb38 | 1479.838627 | -0.791881692 | 2.31E-04 |
| Epdr1 | 994.1146596 | -0.791989143 | 2.11E-07 |
| Atxn7 | 639.3804785 | -0.792398945 | 1.79E-04 |
| Fbln2 | 51001.24849 | -0.792459878 | 9.95E-04 |
| Cisd2 | 773.2577454 | -0.792565519 | 1.16E-04 |
| Ctf1 | 130.0064015 | -0.792583071 | 6.39E-04 |
| Gm14391 | 930.6181873 | -0.792639458 | 1.45E-03 |
| Bckdhb | 830.3338723 | -0.793156206 | 2.23E-06 |
| Gm1976 | 219.0049381 | -0.793156919 | 3.31E-03 |
| Pcca | 1116.587658 | -0.793443794 | 3.87E-08 |
| Mgrn1 | 4288.647559 | -0.793533383 | 1.83E-26 |
| Ydjc | 121.0858377 | -0.793878067 | 8.62E-03 |
| Atat1 | 753.3053063 | -0.794419775 | 3.00E-04 |
| Slc25a35 | 132.6975674 | -0.794651445 | 1.74E-03 |
| Anxa9 | 109.4151629 | -0.794845993 | 1.34E-03 |
| Suclg2 | 5554.452872 | -0.795041126 | 1.89E-14 |
| Dnajc7 | 3526.747744 | -0.795627438 | 2.42E-05 |
| Lima1 | 14608.60545 | -0.796128686 | 2.50E-07 |
| Lmo7 | 1891.856719 | -0.796234803 | 1.37E-07 |
| Il4i1 | 184.7277245 | -0.796662418 | 5.53E-04 |
| Atl3 | 9377.879914 | -0.796719474 | 1.07E-07 |
| Ywhab | 10120.15948 | -0.797390906 | 8.85E-10 |
| Alkbh8 | 540.3213084 | -0.797489877 | 2.13E-03 |
| mt-Nd1 | 144168.3645 | -0.797581236 | 7.56E-07 |
| Pcyt1b | 608.1630088 | -0.797610261 | 2.07E-03 |
| Mthfsl | 129.1169754 | -0.798605359 | 1.86E-04 |
| mt-Cytb | 212693.2349 | -0.799028448 | 4.12E-06 |
| Mdfic | 2585.20032 | -0.79937092 | 9.85E-23 |
| mt-Nd2 | 99498.30713 | -0.79984621 | 2.42E-13 |
| Pigs | 4158.40818 | -0.799918261 | 5.15E-08 |
| Brdt | 227.9030075 | -0.800006624 | 4.63E-04 |
| Rab20 | 279.5572422 | -0.800619601 | 9.52E-04 |
| Nucks1 | 4521.486297 | -0.800958185 | 4.29E-05 |
| Nedd4 | 84397.06472 | -0.801329903 | 1.57E-13 |
| Hnrnpr | 3057.474168 | -0.80138563 | 2.02E-06 |
| Nln | 2979.601328 | -0.802712718 | 9.34E-14 |
| Fign | 239.6826974 | -0.80282397 | 1.45E-02 |
| Ramp3 | 1853.838291 | -0.803090101 | 1.50E-02 |
| Zfp839 | 451.8448377 | -0.803100557 | 8.81E-10 |
| Tmem173 | 257.4625801 | -0.803100726 | 2.02E-04 |
| Tbcc | 642.9037322 | -0.803140356 | 1.09E-06 |
| Srgap3 | 3597.599338 | -0.803140826 | 1.73E-05 |

| | | | |
|---|---|---|---|
| Clec16a | 2108.067705 | -0.803223898 | 2.02E-25 |
| Shroom4 | 831.9222619 | -0.803628079 | 7.47E-06 |
| Efcab2 | 42.83091983 | -0.804817906 | 4.12E-02 |
| Brf1 | 1638.014583 | -0.805247155 | 8.53E-17 |
| Vps50 | 1798.051151 | -0.805435405 | 2.98E-07 |
| Calm3 | 9475.102056 | -0.805640101 | 2.15E-08 |
| 1810014B01Rik | 81.46576989 | -0.805924944 | 1.21E-02 |
| Ndufa5 | 1256.806341 | -0.806090426 | 9.26E-08 |
| Zfp449 | 559.0070296 | -0.80656287 | 1.86E-03 |
| Rnf34 | 1154.614628 | -0.806683316 | 9.33E-14 |
| Tomm6os | 40.86290194 | -0.807240451 | 1.93E-02 |
| Gpatch11 | 1362.790505 | -0.807537723 | 1.62E-11 |
| Dync2li1 | 1037.764832 | -0.808330211 | 1.23E-06 |
| 1700082M22Rik | 108.1681906 | -0.808678132 | 2.91E-03 |
| Rhoq | 7241.961221 | -0.808841724 | 7.86E-13 |
| Zfp706 | 4663.743055 | -0.809795189 | 2.52E-10 |
| Pou6f1 | 1919.566169 | -0.809912513 | 2.96E-12 |
| Arid4a | 961.9188335 | -0.809963144 | 8.04E-04 |
| Cdkn2aipnl | 929.6184746 | -0.810101652 | 1.32E-03 |
| Lipt2 | 116.9283302 | -0.810567736 | 2.91E-03 |
| Scly | 161.3156578 | -0.810898311 | 2.02E-03 |
| Soga3 | 486.8497861 | -0.811641885 | 7.90E-07 |
| Mrc2 | 8828.930084 | -0.811965609 | 1.16E-05 |
| Dnajc30 | 610.0677149 | -0.812047411 | 2.27E-09 |
| Elf1 | 1872.473563 | -0.812336417 | 9.35E-14 |
| Naglu | 4106.854751 | -0.813122171 | 1.58E-04 |
| Txndc16 | 3149.373031 | -0.814196137 | 3.06E-06 |
| Garem1 | 770.4422413 | -0.8143595 | 1.34E-09 |
| Apip | 617.6113811 | -0.814501097 | 1.11E-07 |
| Dgcr2 | 5162.081899 | -0.814872061 | 4.98E-14 |
| Wdr82 | 3294.636451 | -0.81541794 | 5.03E-14 |
| Zfp799 | 721.4912572 | -0.815525321 | 2.61E-13 |
| Gm1818 | 103.6610247 | -0.815537105 | 8.31E-03 |
| Cd248 | 6880.534459 | -0.815985266 | 1.99E-04 |
| Ocel1 | 830.5687289 | -0.816172012 | 1.65E-10 |
| Zc3h12c | 402.0650147 | -0.816212003 | 3.96E-05 |
| Tusc3 | 3182.406469 | -0.816792096 | 1.18E-18 |
| C1d | 926.7708056 | -0.818011954 | 9.23E-05 |
| Fasn | 11598.25822 | -0.819292536 | 3.30E-04 |
| Sclt1 | 532.3854206 | -0.819592791 | 5.53E-03 |
| Dnm1l | 3671.652075 | -0.819986712 | 3.88E-03 |
| Tpcn1 | 2505.938347 | -0.820087079 | 6.59E-10 |
| Casd1 | 1764.77328 | -0.820341947 | 1.46E-07 |
| Pds5b | 3572.252805 | -0.820533929 | 1.33E-06 |
| Rbl2 | 2066.659329 | -0.821433337 | 2.07E-22 |
| Purb | 3528.782382 | -0.821797821 | 9.15E-07 |
| Cenpc1 | 1290.091489 | -0.822321386 | 2.49E-05 |

| | | | |
|---|---|---|---|
| Thap12 | 2285.134238 | -0.822697452 | 6.53E-13 |
| Bmpr1a | 3988.482522 | -0.823858818 | 9.21E-10 |
| Shc1 | 9740.164831 | -0.824266028 | 7.26E-08 |
| Ccdc22 | 1040.823011 | -0.825056901 | 6.23E-15 |
| Mthfs | 115.8052317 | -0.825059633 | 1.42E-03 |
| Zfand4 | 198.1185896 | -0.825111943 | 5.77E-03 |
| LTO1 | 699.7176963 | -0.825126782 | 1.74E-03 |
| Itgb3bp | 300.9143278 | -0.825505417 | 6.81E-03 |
| Abi2 | 1463.946605 | -0.825712058 | 1.18E-08 |
| Exo5 | 1308.513939 | -0.825972723 | 6.54E-13 |
| Zfp827 | 2374.032775 | -0.826277138 | 7.51E-09 |
| Znhit2 | 471.4218009 | -0.82634483 | 1.12E-05 |
| Adamts6 | 404.2434059 | -0.826398424 | 1.70E-02 |
| Gxylt1 | 1462.867959 | -0.826430494 | 1.55E-04 |
| Ripk1 | 1870.091494 | -0.826775147 | 8.33E-13 |
| 4930515G01Rik | 25.56917121 | -0.827222375 | 4.10E-02 |
| H2afy2 | 1900.864898 | -0.827784567 | 1.43E-06 |
| Zhx3 | 1585.705602 | -0.827932957 | 3.43E-07 |
| Ift27 | 800.9781334 | -0.828217169 | 4.82E-09 |
| NA | 153.1519126 | -0.828239316 | 9.23E-04 |
| Zfp419 | 31.98303988 | -0.828305829 | 3.44E-02 |
| Osbpl5 | 2420.062007 | -0.830069947 | 3.10E-07 |
| Emilin2 | 6079.768848 | -0.831170427 | 6.77E-12 |
| Zfp866 | 1101.719992 | -0.831363035 | 1.71E-06 |
| Gsap | 597.175052 | -0.831573552 | 4.05E-09 |
| Myo9a | 2346.723944 | -0.83164056 | 1.56E-03 |
| Jtb | 1085.524652 | -0.831779927 | 3.51E-07 |
| Gpatch4 | 1008.352246 | -0.832073611 | 3.59E-03 |
| Aida | 2825.873098 | -0.83269083 | 9.96E-11 |
| Zfp970 | 672.497198 | -0.832883911 | 5.34E-03 |
| 1300002E11Rik | 233.5490192 | -0.833064257 | 1.27E-04 |
| Bace1 | 2787.184137 | -0.833279367 | 3.72E-10 |
| Nxpe4 | 417.4256941 | -0.833516959 | 1.57E-11 |
| Pcna | 3425.363174 | -0.834324547 | 8.95E-04 |
| Mapk11 | 597.2809478 | -0.834430604 | 2.02E-04 |
| Prkd3 | 3344.207344 | -0.835051625 | 8.12E-09 |
| Acvr1 | 1714.787564 | -0.835666694 | 1.40E-09 |
| Tmem65 | 2586.368853 | -0.836682986 | 2.41E-12 |
| Spon2 | 2156.979328 | -0.836716072 | 1.10E-02 |
| 1700030J22Rik | 46.49523078 | -0.837049909 | 6.97E-03 |
| Ap5b1 | 295.1850734 | -0.837337047 | 9.27E-08 |
| Tmem267 | 228.135723 | -0.837466981 | 9.74E-05 |
| 2210011C24Rik | 74.38268018 | -0.838792437 | 1.71E-02 |
| Plcd1 | 819.3767576 | -0.839008436 | 5.23E-09 |
| Cstf2 | 1462.967735 | -0.840280112 | 4.83E-05 |
| Mrps6 | 3383.155268 | -0.841000119 | 6.20E-05 |
| Adi1 | 3350.226368 | -0.841007134 | 1.77E-07 |

| | | | |
|---|---|---|---|
| Zfp109 | 105.0394414 | -0.841034656 | 1.92E-03 |
| Acsl4 | 2428.085415 | -0.841348283 | 9.03E-04 |
| Stk11ip | 1416.486596 | -0.842001942 | 1.04E-09 |
| Dubr | 27.90810858 | -0.84243253 | 4.07E-02 |
| Agbl2 | 25.22031612 | -0.842765921 | 4.68E-02 |
| Atm | 1655.842703 | -0.842824984 | 2.83E-03 |
| Ift88 | 627.0852278 | -0.843766043 | 1.41E-08 |
| Phf6 | 1129.146427 | -0.84408173 | 6.63E-04 |
| Ikbke | 873.2403649 | -0.844113639 | 3.15E-07 |
| Pde7a | 1818.082955 | -0.844274469 | 2.59E-16 |
| Glmp | 7292.413588 | -0.844311733 | 1.31E-07 |
| Vcan | 10736.2105 | -0.844525771 | 6.64E-12 |
| Fan1 | 251.048372 | -0.845559001 | 9.69E-08 |
| Gm15387 | 5255.01591 | -0.845927994 | 1.95E-08 |
| Setd4 | 125.7559305 | -0.845944975 | 1.81E-02 |
| Nfat5 | 11458.97903 | -0.847066839 | 6.61E-08 |
| Raf1 | 5257.897708 | -0.847500437 | 6.51E-22 |
| Arsa | 6700.14466 | -0.847540787 | 6.33E-09 |
| Itga2b | 570.2372038 | -0.84795643 | 1.79E-05 |
| Adamts12 | 1032.731012 | -0.849445318 | 2.54E-04 |
| Ptpn13 | 1286.598036 | -0.849711413 | 5.72E-04 |
| Tcta | 597.1160012 | -0.85050415 | 8.98E-08 |
| Pnisr | 2217.702301 | -0.851150521 | 2.73E-03 |
| Eml4 | 2243.380341 | -0.851174099 | 7.28E-10 |
| Zeb2os | 155.3605102 | -0.851557707 | 6.57E-04 |
| Cc2d2a | 1162.851323 | -0.851619337 | 4.98E-06 |
| Nprl3 | 1164.930093 | -0.851835324 | 8.38E-12 |
| Hipk3 | 5355.807055 | -0.852156988 | 6.18E-07 |
| Chpf | 4455.956934 | -0.852243831 | 4.21E-05 |
| Loxl3 | 4747.787426 | -0.85280763 | 1.66E-05 |
| Lrfn4 | 2023.205322 | -0.852820612 | 2.26E-10 |
| Rtl10 | 88.75112838 | -0.853251871 | 1.15E-03 |
| 5530601H04Rik | 352.289377 | -0.853412092 | 1.41E-04 |
| 1700123M08Rik | 104.1186089 | -0.853470464 | 8.68E-03 |
| Pcyt1a | 4562.392465 | -0.853506429 | 5.41E-21 |
| Rgmb | 2135.748027 | -0.85374072 | 4.24E-04 |
| Slc39a13 | 2542.695428 | -0.853760265 | 1.90E-06 |
| Arhgef10l | 1765.531792 | -0.854197552 | 3.96E-06 |
| Fmnl3 | 2696.537561 | -0.854787939 | 1.46E-09 |
| 1700020I14Rik | 3599.969541 | -0.855195946 | 5.35E-05 |
| Cetn3 | 1261.039176 | -0.856343202 | 4.00E-04 |
| Cln8 | 3141.229382 | -0.856873512 | 3.06E-05 |
| Lcmt2 | 150.6669994 | -0.856927769 | 5.21E-04 |
| Atp11c | 750.7779488 | -0.857407772 | 1.53E-02 |
| Ubtf | 5461.848329 | -0.85775371 | 3.35E-08 |
| Ythdf3 | 3180.029988 | -0.858256887 | 2.43E-06 |
| Phf14 | 2376.386027 | -0.858335474 | 4.44E-06 |

| | | | |
|---|---|---|---|
| Stk38 | 4121.885645 | -0.858706148 | 3.86E-15 |
| Med20 | 1331.253788 | -0.859694099 | 8.59E-09 |
| Arid4b | 1455.105327 | -0.860056615 | 2.68E-03 |
| Gm3650 | 185.2176749 | -0.860066606 | 3.07E-06 |
| Top1 | 3900.261935 | -0.860150253 | 5.14E-08 |
| Zfp955b | 368.1558793 | -0.860156956 | 4.28E-03 |
| Hhat | 581.9096131 | -0.860165398 | 6.29E-08 |
| Sf3a1 | 4586.280995 | -0.860662121 | 2.39E-07 |
| Gm12018 | 45.77515052 | -0.861067325 | 4.60E-02 |
| Gm2115 | 916.8914908 | -0.861702875 | 1.98E-03 |
| Deaf1 | 690.6344136 | -0.862274289 | 4.83E-09 |
| Abcb8 | 1416.156753 | -0.862550974 | 5.46E-09 |
| Gm26786 | 83.47726905 | -0.863297984 | 1.87E-03 |
| Hoxa6 | 269.2072896 | -0.86329978 | 2.94E-04 |
| Rtl8b | 1525.92564 | -0.863353382 | 8.52E-08 |
| Shroom1 | 179.9661314 | -0.863799917 | 2.39E-03 |
| Clcc1 | 1204.501927 | -0.86404367 | 7.04E-07 |
| Rap2a | 3349.270391 | -0.864265837 | 2.39E-06 |
| Plxnd1 | 6200.522837 | -0.864815438 | 3.07E-06 |
| Galnt2 | 8979.608691 | -0.864849094 | 1.21E-08 |
| Accs | 415.1684912 | -0.865139088 | 5.75E-04 |
| Foxo1 | 589.3772702 | -0.865453037 | 7.67E-13 |
| Hs2st1 | 3965.428802 | -0.865570189 | 3.62E-14 |
| Myo5a | 10004.30674 | -0.866090764 | 1.72E-07 |
| Nmi | 245.734847 | -0.866401287 | 2.45E-07 |
| Asap1 | 15094.41644 | -0.86651267 | 5.64E-11 |
| Kdm2b | 1932.525567 | -0.866625572 | 8.94E-09 |
| Togaram1 | 2240.824097 | -0.867653766 | 2.06E-05 |
| Rbm4 | 516.6815499 | -0.867930374 | 3.59E-08 |
| NA | 126.927072 | -0.868445896 | 2.61E-03 |
| Elmod2 | 768.1670159 | -0.869143764 | 5.25E-05 |
| Gm27042 | 75.90676536 | -0.869991465 | 1.67E-04 |
| Csrp1 | 2952.741716 | -0.870293808 | 1.91E-13 |
| Arhgef1 | 2053.505929 | -0.870709597 | 1.77E-11 |
| Cpeb1 | 899.5519733 | -0.87100333 | 5.32E-06 |
| Gpc6 | 435.6774066 | -0.871033401 | 1.61E-03 |
| Rab31 | 6124.422698 | -0.871089938 | 2.38E-15 |
| Eea1 | 3828.799504 | -0.871098529 | 2.75E-03 |
| Ctnnal1 | 2217.420049 | -0.872114387 | 1.00E-06 |
| Magee1 | 2530.570754 | -0.8724186 | 3.60E-13 |
| Far1 | 2576.261243 | -0.872469458 | 3.59E-03 |
| Kalrn | 1301.719211 | -0.873736149 | 2.74E-08 |
| Cenpj | 331.1364679 | -0.87495904 | 2.81E-04 |
| Mat2a | 11860.39905 | -0.875224975 | 1.16E-05 |
| Daam1 | 2029.928172 | -0.87523532 | 1.98E-08 |
| Pole4 | 1287.876454 | -0.876081331 | 3.08E-09 |
| Six1 | 1079.039133 | -0.876517537 | 2.96E-03 |

| | | | |
|---|---|---|---|
| Zfp442 | 237.9315236 | -0.876613016 | 1.45E-02 |
| Mrps25 | 1661.276357 | -0.876788067 | 5.41E-07 |
| Armc9 | 957.2896761 | -0.877642396 | 2.51E-17 |
| Cpeb4 | 981.9575186 | -0.877664844 | 7.68E-04 |
| Cul1 | 6224.737814 | -0.878307831 | 9.95E-09 |
| Ttc8 | 1298.530567 | -0.878328273 | 1.27E-21 |
| Cacna2d1 | 1239.235042 | -0.879058249 | 4.77E-07 |
| Zfx | 1758.364518 | -0.879128537 | 1.49E-04 |
| Wrn | 1098.5201 | -0.879874024 | 3.75E-07 |
| Tmem86a | 734.3320314 | -0.879878065 | 4.62E-05 |
| Srp9 | 340.4882686 | -0.880235754 | 5.44E-10 |
| Car15 | 64.28556731 | -0.880510757 | 6.82E-03 |
| Nudt1 | 309.718058 | -0.880525862 | 3.67E-05 |
| Tmppe | 488.8157654 | -0.880574675 | 9.87E-09 |
| Amigo2 | 624.6922769 | -0.881974813 | 6.33E-09 |
| 2900089D17Rik | 112.3122989 | -0.882420741 | 1.60E-02 |
| Adam23 | 4333.929332 | -0.882892485 | 2.99E-05 |
| Hmgb1-ps3 | 192.7784252 | -0.882903163 | 3.29E-05 |
| Hs6st1 | 2054.227925 | -0.883177012 | 6.96E-08 |
| Krtcap3 | 56.95226486 | -0.883178312 | 4.90E-03 |
| Gm14292 | 357.5749687 | -0.883198557 | 2.76E-09 |
| Fus | 8428.016269 | -0.883623353 | 6.01E-05 |
| Zfp933 | 326.5717679 | -0.884063323 | 1.14E-03 |
| Zfp810 | 353.3936938 | -0.884260377 | 7.62E-05 |
| Rbm4b | 986.1917831 | -0.884703949 | 2.64E-08 |
| Gm11991 | 26.04455271 | -0.885032998 | 4.91E-02 |
| Cacfd1 | 1602.512009 | -0.885204222 | 2.04E-12 |
| Plag1 | 548.7458397 | -0.885770761 | 1.00E-04 |
| Zc3h7b | 5316.848278 | -0.887022669 | 7.08E-19 |
| Myo1e | 5416.527455 | -0.887270446 | 1.76E-06 |
| Tspyl4 | 789.2797587 | -0.887309976 | 8.84E-04 |
| Ubtd2 | 899.0570191 | -0.887480402 | 7.88E-09 |
| Sdc4 | 3683.171926 | -0.887817633 | 2.35E-08 |
| Etv6 | 1583.328039 | -0.888214217 | 2.53E-18 |
| Dnajc13 | 5378.577471 | -0.888429896 | 1.37E-11 |
| Hspa1l | 63.89465712 | -0.888708527 | 3.55E-03 |
| Stxbp4 | 1774.063882 | -0.889003375 | 3.78E-13 |
| Fnip1 | 4227.643221 | -0.889132865 | 3.26E-09 |
| Atraid | 2965.766926 | -0.890209408 | 2.81E-09 |
| Gm12758 | 62.41482331 | -0.890228903 | 2.53E-02 |
| 4930522L14Rik | 312.3836131 | -0.890248636 | 1.76E-02 |
| Zfp521 | 2160.446859 | -0.890504912 | 7.24E-06 |
| Cldn34c1 | 161.8468038 | -0.89149382 | 6.31E-03 |
| Sfxn2 | 329.2799816 | -0.891715999 | 5.53E-04 |
| Dse | 4466.695367 | -0.892344898 | 1.49E-06 |
| Idh1 | 8516.619482 | -0.892672825 | 3.74E-14 |
| Ints4 | 1468.076545 | -0.895457822 | 1.35E-16 |

| | | | |
|---|---|---|---|
| Pdk4 | 19724.77029 | -0.895528524 | 4.76E-03 |
| Pcm1 | 5197.668132 | -0.896848879 | 1.40E-03 |
| Ano6 | 9272.629877 | -0.896933386 | 1.19E-13 |
| Adal | 268.8178673 | -0.897043455 | 2.13E-05 |
| Pcbd2 | 191.883457 | -0.897068246 | 1.74E-04 |
| Dlg5 | 3568.439899 | -0.897877912 | 1.13E-23 |
| Dmxl2 | 711.1668361 | -0.898344748 | 4.04E-03 |
| Mxra8 | 6216.491192 | -0.900150033 | 4.00E-09 |
| Kif16b | 1057.721431 | -0.900823997 | 6.06E-13 |
| Entpd6 | 901.5502184 | -0.900879319 | 2.68E-06 |
| Tpgs2 | 676.6281657 | -0.901032725 | 3.85E-08 |
| Dhrs7 | 751.5447118 | -0.901130811 | 4.47E-06 |
| Atpaf1 | 1066.490644 | -0.901242933 | 5.51E-13 |
| Cul7 | 4030.507774 | -0.901463544 | 7.99E-11 |
| Mcm6 | 3739.888641 | -0.902098008 | 4.05E-03 |
| NA | 574.234844 | -0.902225043 | 1.07E-07 |
| Riox1 | 464.5125167 | -0.902320057 | 6.59E-04 |
| Aacs | 4095.139985 | -0.902323874 | 5.67E-06 |
| Kank1 | 1147.219222 | -0.902635408 | 1.68E-05 |
| Man2a1 | 13475.00019 | -0.903332986 | 2.23E-21 |
| Sh3bgrl3 | 2819.46658 | -0.903368236 | 5.99E-06 |
| Ywhaq | 13241.07001 | -0.903430049 | 2.07E-22 |
| Col4a2 | 13125.40325 | -0.9034405 | 1.10E-05 |
| Dhrs9 | 1469.943128 | -0.903591138 | 2.50E-04 |
| Slc5a4b | 28.62253057 | -0.903747958 | 2.57E-02 |
| Svil | 5457.138753 | -0.904119195 | 1.54E-10 |
| Zfp780b | 192.5945161 | -0.904686971 | 8.82E-04 |
| Atp9b | 2468.969284 | -0.905053978 | 1.54E-21 |
| Eif5a2 | 835.2089142 | -0.90520707 | 4.94E-05 |
| Pdxk | 5185.761374 | -0.905629197 | 5.38E-07 |
| Lpin2 | 2438.108658 | -0.905635763 | 4.97E-17 |
| Map2k6 | 397.271134 | -0.9057767 | 7.73E-06 |
| Hnrnph1 | 8004.785467 | -0.905985539 | 1.05E-03 |
| Zfp984 | 306.0112459 | -0.906026284 | 8.03E-03 |
| mt-Nd3 | 16305.7006 | -0.906674843 | 3.74E-06 |
| B3galt1 | 227.8123498 | -0.90704758 | 5.50E-07 |
| Sh3rf3 | 2120.548004 | -0.911870703 | 5.18E-14 |
| Zfp318 | 1909.003472 | -0.912427651 | 1.12E-04 |
| Zfp354c | 1061.502211 | -0.912553452 | 2.44E-06 |
| Aar2 | 1658.318704 | -0.912971802 | 3.24E-12 |
| Lgals1 | 85771.53346 | -0.913632434 | 1.93E-09 |
| Ankrd12 | 2220.360824 | -0.913684765 | 4.20E-05 |
| Top2b | 4599.091835 | -0.914201778 | 8.15E-03 |
| Tceal8 | 3073.445068 | -0.914644938 | 2.49E-20 |
| Lrif1 | 631.3650522 | -0.915274248 | 6.87E-03 |
| Zbtb45 | 334.1316636 | -0.915493049 | 3.58E-08 |
| Lrrc56 | 115.614384 | -0.917068613 | 1.03E-03 |

| | | | |
|---|---|---|---|
| Arhgap26 | 119.8297351 | -0.917272302 | 1.32E-03 |
| Ctsb | 36468.22462 | -0.917739003 | 8.54E-06 |
| B230354K17Rik | 1037.407643 | -0.918237214 | 2.78E-06 |
| Mvd | 2840.656231 | -0.919306092 | 3.92E-03 |
| Ccdc191 | 288.9313192 | -0.91930763 | 1.22E-03 |
| Camkk1 | 480.61585 | -0.919408309 | 6.72E-05 |
| Stoml1 | 563.1153803 | -0.919761158 | 3.30E-06 |
| Bmp1 | 4680.338256 | -0.92006332 | 2.69E-18 |
| Mid1 | 2636.808961 | -0.920088004 | 1.35E-05 |
| NA | 154.0539091 | -0.920235136 | 2.59E-05 |
| Ankrd16 | 929.5128238 | -0.920660905 | 9.67E-08 |
| Fbxl3 | 1735.020904 | -0.921529284 | 2.47E-08 |
| Ly6h | 28.54279703 | -0.922011412 | 1.95E-02 |
| Morn1 | 87.48653537 | -0.923374129 | 2.88E-04 |
| 1600002K03Rik | 136.2918221 | -0.923590872 | 3.80E-06 |
| Gm10282 | 76.85288788 | -0.923629663 | 7.96E-03 |
| Nasp | 2787.702527 | -0.923707262 | 1.90E-03 |
| 4921531C22Rik | 54.37804619 | -0.923866555 | 3.97E-03 |
| Gmfb | 3530.728036 | -0.923867934 | 8.48E-05 |
| Shld1 | 158.5939244 | -0.923988227 | 1.15E-05 |
| Gm27029 | 107.2998506 | -0.924189357 | 4.11E-02 |
| 4632404H12Rik | 210.9051878 | -0.924371699 | 1.17E-05 |
| Azin1 | 5848.456593 | -0.924819551 | 2.99E-09 |
| Gm12743 | 64.09499747 | -0.92508304 | 5.52E-03 |
| Etaa1 | 461.0245111 | -0.925279035 | 1.50E-02 |
| Leo1 | 880.8601505 | -0.927093501 | 6.34E-09 |
| Ssr1 | 7136.202432 | -0.927139453 | 5.84E-13 |
| Tbc1d8b | 1093.135708 | -0.928097313 | 1.43E-03 |
| Sft2d3 | 128.2159729 | -0.928435138 | 5.11E-05 |
| Leprot | 5202.282116 | -0.929061853 | 8.42E-25 |
| 2210016F16Rik | 2097.051085 | -0.929477999 | 6.97E-05 |
| Scn1b | 147.6156986 | -0.930325656 | 3.80E-03 |
| Igf2bp2 | 8644.418417 | -0.930344474 | 2.42E-22 |
| Akap13 | 5008.332383 | -0.930911927 | 7.95E-10 |
| Prickle2 | 610.6101831 | -0.93171728 | 8.61E-10 |
| Zfp770 | 544.5327502 | -0.932416933 | 2.66E-03 |
| Gm17251 | 159.5166582 | -0.933168377 | 1.32E-04 |
| Crebl2 | 1856.82481 | -0.933479329 | 3.48E-09 |
| Ppp2r1b | 1945.402469 | -0.933717185 | 8.39E-12 |
| Trim65 | 262.1264386 | -0.933742152 | 6.67E-06 |
| Gm21981 | 86.72813471 | -0.934245957 | 2.30E-03 |
| Myo10 | 6334.157457 | -0.934453765 | 7.12E-08 |
| Smo | 9359.591469 | -0.934669326 | 2.92E-22 |
| Marcksl1 | 7037.230072 | -0.935509835 | 8.23E-06 |
| Ung | 266.879367 | -0.936337893 | 1.08E-04 |
| Idua | 698.7732812 | -0.936445231 | 8.37E-14 |
| Nxt2 | 344.3911455 | -0.936689396 | 1.35E-03 |

| | | | |
|---|---|---|---|
| Limd2 | 1732.780265 | -0.936875774 | 6.50E-08 |
| Brwd1 | 3301.856348 | -0.937256088 | 6.50E-04 |
| A930012L18Rik | 23.28516798 | -0.937638988 | 3.66E-02 |
| Msrb2 | 106.301837 | -0.937822491 | 1.23E-03 |
| Lclat1 | 2716.89391 | -0.937862537 | 2.29E-18 |
| Ncbp3 | 1301.175897 | -0.938806664 | 1.20E-10 |
| Capn10 | 780.8179271 | -0.939784511 | 1.19E-09 |
| Wdr78 | 241.0414196 | -0.941637845 | 1.31E-04 |
| Usp13 | 226.2858383 | -0.941853585 | 1.06E-03 |
| Pccb | 1287.765021 | -0.94196353 | 2.14E-08 |
| Trim30a | 433.047542 | -0.942252384 | 1.03E-03 |
| Mapk8ip1 | 2586.65763 | -0.942307958 | 2.43E-07 |
| NA | 31.52558462 | -0.942674095 | 2.87E-02 |
| Nod1 | 274.7364886 | -0.94285148 | 5.20E-05 |
| Car13 | 2496.327558 | -0.942988035 | 5.97E-15 |
| A430057M04Rik | 95.38244642 | -0.943396419 | 6.07E-04 |
| Klhl9 | 5226.60583 | -0.943427374 | 3.88E-14 |
| Ptger1 | 1375.579965 | -0.944255126 | 1.29E-10 |
| Slc22a23 | 1141.901598 | -0.944417736 | 8.67E-08 |
| Pradc1 | 384.2185328 | -0.945039327 | 3.53E-06 |
| Mpzl1 | 3792.936615 | -0.945331333 | 6.04E-07 |
| Cyp46a1 | 102.4882055 | -0.946132337 | 2.91E-03 |
| Jam2 | 337.7129228 | -0.94628441 | 4.38E-04 |
| Calm1 | 13158.15346 | -0.946556632 | 1.13E-12 |
| Glrb | 356.4512423 | -0.947081658 | 4.41E-05 |
| Sall2 | 1033.6407 | -0.947352614 | 2.32E-14 |
| Slc36a1 | 1573.335444 | -0.949041336 | 2.81E-11 |
| L3hypdh | 996.178476 | -0.94904846 | 3.04E-14 |
| Btd | 882.1449736 | -0.949560081 | 2.76E-10 |
| Plxna2 | 3143.852049 | -0.950201842 | 6.21E-08 |
| Efhd2 | 1565.299968 | -0.951368937 | 3.69E-10 |
| Recql | 672.013649 | -0.951427535 | 4.95E-06 |
| Dtnbp1 | 2425.756691 | -0.951760256 | 3.57E-09 |
| Zfp119b | 118.8405696 | -0.952099291 | 1.77E-03 |
| Mindy4 | 685.858107 | -0.952228931 | 1.11E-09 |
| Tbccd1 | 1403.677915 | -0.952478882 | 8.29E-26 |
| Gm17018 | 524.6653148 | -0.95255994 | 1.69E-21 |
| Krt10 | 176.393471 | -0.953015031 | 4.26E-06 |
| Nfatc4 | 1786.320943 | -0.953569716 | 6.79E-08 |
| Fzd3 | 172.7317828 | -0.953616969 | 1.67E-06 |
| Cdh17 | 219.8371349 | -0.953947785 | 3.52E-05 |
| Rrbp1 | 16274.11852 | -0.953978424 | 6.42E-25 |
| Lrriq3 | 33.70355744 | -0.95442596 | 1.56E-02 |
| 2810433D01Rik | 81.57471888 | -0.95520402 | 5.28E-03 |
| Fam172a | 1437.904506 | -0.955408318 | 8.55E-15 |
| Atg4c | 643.0173726 | -0.955526283 | 1.19E-06 |
| Pikfyve | 2011.636706 | -0.955746698 | 4.13E-11 |

| | | | |
|---|---|---|---|
| Eef2k | 2828.119429 | -0.955784354 | 1.15E-12 |
| 8-Mar | 6434.713929 | -0.956203479 | 2.41E-89 |
| Ggh | 449.0071855 | -0.956935392 | 3.75E-08 |
| Lgals3 | 26908.25829 | -0.957924584 | 1.95E-04 |
| Hmgb1 | 6151.969447 | -0.95807878 | 3.86E-07 |
| Ddx6 | 7985.738967 | -0.95836185 | 8.49E-31 |
| Serinc5 | 1871.588337 | -0.958467012 | 1.93E-18 |
| Mtmr11 | 1713.709962 | -0.959995916 | 5.18E-09 |
| Nphp3 | 459.6841505 | -0.960641194 | 1.66E-08 |
| Gm12940 | 515.8451704 | -0.96080781 | 7.05E-03 |
| Mblac2 | 232.0593899 | -0.961590931 | 1.39E-04 |
| Gm10561 | 210.1280875 | -0.961864758 | 8.17E-09 |
| St5 | 1526.821015 | -0.961910221 | 9.43E-08 |
| 9430091E24Rik | 71.37412386 | -0.962019755 | 1.60E-04 |
| Tmem163 | 21.88372852 | -0.963443361 | 2.33E-02 |
| Vsig10 | 686.1056348 | -0.963697733 | 9.41E-14 |
| Decr2 | 584.6527844 | -0.963960605 | 3.40E-08 |
| Epm2aip1 | 2273.909362 | -0.964158019 | 4.12E-08 |
| Emp3 | 1776.507322 | -0.964262209 | 1.42E-08 |
| Gm15163 | 28.34791949 | -0.964824794 | 1.95E-02 |
| Ccdc126 | 260.6403871 | -0.964908219 | 5.29E-08 |
| Zfyve21 | 755.2797833 | -0.965187564 | 3.84E-11 |
| Anxa5 | 17795.31783 | -0.966468588 | 3.74E-14 |
| Pisd-ps2 | 162.680632 | -0.96756495 | 4.80E-05 |
| 4930556M19Rik | 82.05104413 | -0.967870204 | 4.51E-03 |
| Slirp | 1059.190464 | -0.968233339 | 6.88E-11 |
| Ostm1 | 1935.936349 | -0.969082584 | 1.46E-11 |
| Atp6v0a1 | 3975.629212 | -0.969100193 | 4.57E-16 |
| Apbb2 | 3282.670896 | -0.969146146 | 1.53E-09 |
| Pkd2l2 | 165.7486455 | -0.971252743 | 1.34E-03 |
| Gm14698 | 642.0530449 | -0.971276565 | 6.16E-03 |
| Zfp763 | 273.8263546 | -0.971362954 | 8.86E-05 |
| Prr36 | 21.45874571 | -0.971837626 | 3.69E-02 |
| Mdrl | 28.27028879 | -0.971858691 | 1.80E-02 |
| Fam173b | 346.8684431 | -0.972026762 | 1.03E-08 |
| Camk2n2 | 678.9875033 | -0.972287961 | 1.75E-06 |
| Usp1 | 2727.71314 | -0.972499791 | 6.15E-03 |
| Hexim1 | 1678.805287 | -0.972644568 | 1.25E-18 |
| Reck | 1994.171189 | -0.972742169 | 1.91E-07 |
| L3mbtl3 | 963.0967322 | -0.972931208 | 6.01E-07 |
| Ctdspl | 1253.236411 | -0.973116937 | 2.78E-06 |
| Lamb1 | 12422.21913 | -0.973743065 | 1.74E-03 |
| Fzd2 | 5450.697507 | -0.973783181 | 1.77E-11 |
| Svbp | 1041.238098 | -0.974039591 | 1.64E-13 |
| Gm26750 | 86.09606895 | -0.974307096 | 2.09E-04 |
| Srsf6 | 7575.428591 | -0.974414603 | 4.01E-18 |
| Mtmr2 | 3520.614254 | -0.975032241 | 1.99E-09 |

| | | | |
|---|---|---|---|
| Adam10 | 4023.172379 | -0.97532649 | 1.99E-10 |
| Relch | 1655.605876 | -0.975731006 | 1.33E-11 |
| 5730409E04Rik | 1776.754366 | -0.975944517 | 2.81E-20 |
| Pacs2 | 7227.003698 | -0.97720501 | 2.09E-27 |
| Sco2 | 266.3938489 | -0.977248292 | 1.61E-03 |
| Adgra2 | 9959.089649 | -0.977566029 | 1.98E-14 |
| 6330403L08Rik | 423.3189736 | -0.978270324 | 3.85E-07 |
| Nudt6 | 176.1843673 | -0.979005243 | 3.93E-04 |
| Ift22 | 607.373969 | -0.979529831 | 7.07E-18 |
| Prrx1 | 19059.58088 | -0.980431799 | 3.94E-16 |
| Ifi204 | 1140.215139 | -0.981332879 | 1.48E-05 |
| Zfp503 | 2242.630648 | -0.981450564 | 8.03E-11 |
| Coq4 | 358.7442056 | -0.982038957 | 6.05E-06 |
| H2afv | 1309.866945 | -0.982145766 | 2.57E-09 |
| Jun | 4890.668098 | -0.982236766 | 1.38E-04 |
| Gpr161 | 765.4282915 | -0.982749405 | 1.61E-08 |
| Amigo3 | 59.57360313 | -0.983092407 | 1.99E-03 |
| Stmn1 | 1582.862217 | -0.983761549 | 5.76E-04 |
| Pbx1 | 3313.477385 | -0.98406103 | 1.51E-05 |
| Gm5514 | 63.12948495 | -0.984389457 | 2.49E-03 |
| Rsl1 | 57.97589736 | -0.98477822 | 1.85E-03 |
| Mbnl2 | 5029.844149 | -0.985591096 | 1.99E-07 |
| Gdpd2 | 416.9752147 | -0.986153371 | 1.54E-02 |
| Bbs7 | 975.2240167 | -0.98692144 | 1.62E-04 |
| Gm17435 | 160.567384 | -0.987832561 | 5.74E-08 |
| E130311K13Rik | 560.9654514 | -0.988249524 | 8.08E-08 |
| Pole2 | 557.9420967 | -0.988276293 | 6.42E-04 |
| Cdadc1 | 577.8821487 | -0.98994759 | 4.60E-11 |
| Rasa2 | 836.285905 | -0.991718851 | 6.33E-09 |
| Wipf1 | 3029.779313 | -0.99189471 | 1.93E-32 |
| Tnrc6a | 3803.20519 | -0.99262232 | 2.56E-09 |
| D10Wsu102e | 2431.206995 | -0.992663246 | 1.31E-17 |
| Stk36 | 385.152876 | -0.992910767 | 2.61E-07 |
| Nnmt | 470.8866788 | -0.993555963 | 2.08E-06 |
| Hivep2 | 1729.316059 | -0.993641859 | 4.11E-05 |
| Eef1akmt1 | 477.1061299 | -0.99369483 | 1.75E-06 |
| 1810030O07Rik | 1755.512131 | -0.993950857 | 1.14E-33 |
| Trim44 | 4685.908872 | -0.994796058 | 2.69E-10 |
| Tnfsf12 | 1683.302223 | -0.994952247 | 1.02E-12 |
| Wdr83 | 632.260428 | -0.99512566 | 2.09E-14 |
| Zfp937 | 394.8604838 | -0.996443729 | 7.24E-04 |
| Zfp397 | 1409.011093 | -0.996553905 | 4.75E-04 |
| Zfp26 | 803.5414538 | -0.996681172 | 4.90E-04 |
| St3gal2 | 8902.023726 | -0.996911439 | 7.10E-23 |
| Fancl | 733.7646602 | -0.997982747 | 4.15E-05 |
| Gm7935 | 28.89187514 | -0.998092636 | 4.52E-02 |
| Kdm3a | 3959.529429 | -0.999848382 | 2.23E-16 |

| | | | |
|---|---|---|---|
| Dolk | 812.9649783 | -0.999909318 | 4.29E-17 |
| Ppp1r16a | 1023.013923 | -1.000401977 | 6.12E-15 |
| Matk | 59.77616886 | -1.001250971 | 4.35E-03 |
| Mettl27 | 1096.641165 | -1.001352523 | 1.81E-08 |
| Pcolce | 25050.1764 | -1.002743748 | 2.33E-13 |
| Laptm4b | 5740.524772 | -1.002787713 | 1.31E-07 |
| Slc16a13 | 668.4799613 | -1.003421727 | 3.78E-08 |
| Xiap | 2319.815008 | -1.003464642 | 1.48E-06 |
| St3gal4 | 2084.028363 | -1.00361667 | 9.99E-06 |
| 1700088E04Rik | 90.25557833 | -1.004200962 | 1.84E-03 |
| Ogt | 5848.32168 | -1.00517887 | 2.31E-04 |
| 2410018L13Rik | 753.6620775 | -1.005241837 | 6.02E-15 |
| Hist2h2be | 103.5079661 | -1.005617437 | 1.91E-04 |
| Ccnb2-ps | 54.8298888 | -1.005675506 | 1.36E-02 |
| Tmem138 | 722.3299743 | -1.00583337 | 1.46E-12 |
| Zfp322a | 971.5690419 | -1.005977799 | 2.16E-04 |
| Alms1 | 2337.478094 | -1.006544509 | 5.84E-08 |
| Esyt3 | 17.29674193 | -1.006629901 | 4.34E-02 |
| Agap1 | 3485.368846 | -1.009263483 | 2.00E-16 |
| Prl8a1 | 197.3899827 | -1.009286948 | 3.80E-05 |
| Kdsr | 1932.623425 | -1.010222131 | 1.34E-15 |
| Lsm6 | 915.3953535 | -1.010832402 | 2.21E-08 |
| Dnajc12 | 45.61121461 | -1.011190086 | 1.67E-02 |
| Tubb2b | 2259.346938 | -1.011363403 | 2.44E-08 |
| Spred1 | 1968.043063 | -1.011943275 | 1.22E-16 |
| Nphp4 | 152.6202732 | -1.012034313 | 2.33E-07 |
| Cdk14 | 5133.388251 | -1.012055901 | 1.15E-03 |
| Zfp719 | 839.4258072 | -1.012061938 | 3.89E-09 |
| Wdr34 | 712.2965379 | -1.012064988 | 1.96E-10 |
| Cep164 | 1466.016671 | -1.01209214 | 1.69E-13 |
| Nme7 | 496.5352665 | -1.012191581 | 2.39E-11 |
| Cfap46 | 33.74864826 | -1.012215359 | 3.02E-02 |
| Zfp455 | 70.70734888 | -1.012513626 | 3.51E-05 |
| Zfp422-ps | 64.25166758 | -1.01280774 | 8.15E-03 |
| D130040H23Rik | 50.8991406 | -1.01295224 | 1.45E-02 |
| Smim27 | 95.77154053 | -1.013031949 | 2.34E-04 |
| F3 | 10673.0851 | -1.014050874 | 3.00E-06 |
| Zfp595 | 165.6562495 | -1.015491652 | 2.09E-04 |
| Tfap2a | 730.9542063 | -1.015720655 | 1.73E-05 |
| E2f8 | 355.3960181 | -1.015948547 | 6.55E-03 |
| Gsta4 | 6626.313589 | -1.015954725 | 6.33E-07 |
| Parp14 | 278.3693229 | -1.016203643 | 1.56E-05 |
| Fam69a | 2022.668361 | -1.016916481 | 8.69E-06 |
| Zfp974 | 285.1415955 | -1.016983415 | 1.22E-04 |
| Arl6 | 456.3562911 | -1.018411678 | 4.80E-09 |
| Foxc1 | 342.3062006 | -1.01864574 | 1.23E-09 |
| Atr | 1202.631887 | -1.019167822 | 5.53E-04 |

| | | | |
|---|---|---|---|
| 6430548M08Rik | 814.8945097 | -1.019230308 | 2.27E-11 |
| Tmed8 | 1367.151775 | -1.019237903 | 2.07E-20 |
| Kat6b | 1896.384666 | -1.019260905 | 3.86E-14 |
| Wtap | 2388.347222 | -1.019794044 | 1.31E-15 |
| Chm | 1244.672927 | -1.019810648 | 4.05E-04 |
| Zfp658 | 480.0067555 | -1.019983199 | 4.43E-05 |
| 4632415L05Rik | 318.8005624 | -1.020475152 | 4.05E-09 |
| Zfp637 | 1645.301751 | -1.020487513 | 2.32E-10 |
| Plscr1 | 527.9546792 | -1.020554555 | 5.07E-08 |
| Uaca | 2130.933084 | -1.020805971 | 1.02E-12 |
| Btbd6 | 204.0903082 | -1.020807669 | 6.78E-10 |
| Gm2093 | 35.49229119 | -1.020816172 | 9.20E-03 |
| Brca2 | 529.5132143 | -1.020876257 | 3.18E-02 |
| 2810430I11Rik | 151.6916456 | -1.022424439 | 2.60E-04 |
| Gm14139 | 207.1634126 | -1.022788543 | 4.27E-07 |
| Smim12 | 756.8155337 | -1.023308316 | 1.61E-08 |
| S1pr2 | 3510.869616 | -1.023679985 | 4.00E-16 |
| Hsd17b11 | 1801.223666 | -1.024907002 | 6.17E-14 |
| Pcna-ps2 | 616.8396399 | -1.025100158 | 3.93E-05 |
| Gm45351 | 101.0112841 | -1.026125618 | 6.62E-06 |
| Aak1 | 1915.969594 | -1.026216861 | 3.85E-20 |
| Zfp58 | 250.0516868 | -1.026216961 | 1.16E-07 |
| B630019K06Rik | 230.2762103 | -1.027530878 | 1.75E-12 |
| Casp8ap2 | 835.5596121 | -1.027564274 | 9.24E-03 |
| Kiz | 936.1381678 | -1.027991248 | 8.66E-07 |
| Dcaf6 | 5277.412717 | -1.028139213 | 3.02E-09 |
| Ddx3y | 1595.987767 | -1.028382273 | 4.59E-03 |
| Srsf1 | 6220.767209 | -1.028638741 | 1.99E-03 |
| Peli2 | 1908.882325 | -1.029362123 | 8.55E-15 |
| Dek | 2477.319996 | -1.029768342 | 5.07E-03 |
| Oxct1 | 16865.66235 | -1.030201463 | 1.63E-10 |
| Tet2 | 657.7186831 | -1.030881448 | 1.26E-07 |
| Cerk | 1102.588167 | -1.031098476 | 9.31E-16 |
| Mfsd4b4 | 543.5534433 | -1.032750051 | 8.49E-10 |
| Socs2 | 153.4955773 | -1.032966167 | 1.21E-02 |
| Lix1l | 6902.240498 | -1.033685562 | 2.08E-10 |
| 5730596B20Rik | 137.2936872 | -1.034530455 | 3.11E-03 |
| Zfp558 | 139.6941417 | -1.03465525 | 8.94E-05 |
| Ptk2b | 79.64446995 | -1.034903666 | 3.95E-05 |
| Lamp2 | 21188.1191 | -1.035289643 | 9.99E-14 |
| Zfp563 | 188.7486021 | -1.035704427 | 3.44E-08 |
| Gm21989 | 45.534082 | -1.036173867 | 1.71E-02 |
| Atrx | 5044.519523 | -1.036940035 | 4.53E-02 |
| Pls3 | 9293.72303 | -1.037763983 | 1.87E-30 |
| Sf3b3 | 10639.11885 | -1.037851874 | 5.49E-17 |
| Gns | 20094.75228 | -1.03792047 | 3.68E-10 |
| Vangl2 | 622.778806 | -1.038252033 | 5.39E-10 |

| | | | |
|---|---|---|---|
| Igf2bp3 | 4305.408781 | -1.038425865 | 1.27E-15 |
| Slf1 | 437.365383 | -1.039501356 | 6.65E-03 |
| Erbin | 6002.117223 | -1.039900161 | 2.60E-04 |
| Tmem175 | 1175.19773 | -1.040009303 | 1.21E-22 |
| C330007P06Rik | 1979.096959 | -1.040129577 | 2.34E-09 |
| Ndufb7 | 1320.50736 | -1.040135742 | 5.52E-09 |
| Luc7l3 | 2630.403889 | -1.040407541 | 3.68E-03 |
| 6720427I07Rik | 137.3231581 | -1.040559925 | 2.12E-02 |
| Cemip2 | 1597.684135 | -1.041302922 | 3.45E-20 |
| Mob3c | 513.3066035 | -1.04147827 | 1.11E-07 |
| Ggact | 310.702016 | -1.041554205 | 1.13E-06 |
| Smim10l1 | 2712.536032 | -1.042760317 | 9.46E-06 |
| Rassf8 | 5424.313331 | -1.04298928 | 2.60E-14 |
| AI413582 | 932.4708856 | -1.043823536 | 3.23E-09 |
| Vasn | 7393.562647 | -1.043846422 | 3.04E-06 |
| Gulp1 | 1061.560392 | -1.044260692 | 1.67E-07 |
| Arhgef40 | 3841.130444 | -1.044654222 | 2.13E-12 |
| Shtn1 | 23.34544253 | -1.045235122 | 2.27E-02 |
| Dpp7 | 5371.98911 | -1.045442625 | 9.10E-08 |
| Fbxo33 | 557.3720924 | -1.045658041 | 1.63E-07 |
| Qdpr | 1935.818044 | -1.045992605 | 3.50E-11 |
| 2010001A14Rik | 129.6180682 | -1.046313905 | 3.41E-09 |
| Gm14488 | 37.31048439 | -1.046422381 | 3.51E-02 |
| Slc26a2 | 908.0177184 | -1.048580987 | 4.21E-17 |
| Cryl1 | 233.4306095 | -1.049906955 | 5.91E-08 |
| Evi5 | 2624.44236 | -1.05022759 | 7.53E-05 |
| Il34 | 440.8805053 | -1.051969923 | 2.35E-05 |
| Slc1a3 | 1356.973152 | -1.05197588 | 7.51E-08 |
| Fmr1 | 3504.187624 | -1.052254972 | 6.74E-04 |
| Atf5 | 11430.63398 | -1.053539681 | 2.01E-13 |
| Adgrg1 | 72.72172502 | -1.054974978 | 7.06E-03 |
| 4930539E08Rik | 68.16425014 | -1.056484184 | 2.48E-04 |
| Pxdn | 12242.77878 | -1.056580332 | 4.23E-10 |
| AI429214 | 333.0356269 | -1.058416509 | 2.26E-11 |
| Mtap | 23447.27157 | -1.058469454 | 3.36E-11 |
| Zfp260 | 2209.690586 | -1.059931162 | 3.57E-07 |
| Hoxa3 | 715.999191 | -1.060057078 | 1.50E-08 |
| Rubcnl | 44.07152091 | -1.06011656 | 2.12E-02 |
| Nphp1 | 936.7893837 | -1.060180272 | 2.23E-11 |
| Efna4 | 476.6852267 | -1.060211357 | 4.38E-10 |
| Napepld | 430.2690843 | -1.06060332 | 1.46E-11 |
| Tmem254c | 143.0849402 | -1.06076169 | 6.74E-03 |
| Sox4 | 7067.281048 | -1.061374426 | 1.74E-05 |
| Zfp160 | 566.4007685 | -1.061582155 | 6.24E-04 |
| Naalad2 | 1266.955718 | -1.06203225 | 7.61E-17 |
| Srsf2 | 5846.326793 | -1.062196319 | 1.19E-06 |
| Tctn3 | 629.4105587 | -1.062418882 | 1.61E-19 |

| | | | |
|---|---|---|---|
| Trpm7 | 6048.847824 | -1.062419195 | 2.66E-04 |
| Scarf2 | 3742.419041 | -1.062521029 | 1.38E-15 |
| Dusp10 | 1991.634529 | -1.06343439 | 1.60E-03 |
| Alg5 | 1251.071303 | -1.063452003 | 2.42E-11 |
| C8g | 63.07082444 | -1.064253385 | 4.43E-05 |
| Il13ra1 | 219.0472217 | -1.064353175 | 2.61E-09 |
| Pdss1 | 204.7238761 | -1.065843701 | 5.65E-03 |
| Bahcc1 | 1260.799357 | -1.066277253 | 9.28E-10 |
| Ap5z1 | 1165.206489 | -1.06661703 | 7.46E-19 |
| Socs1 | 288.2224024 | -1.066641384 | 3.95E-06 |
| Rbak | 676.8083874 | -1.066673763 | 7.47E-14 |
| Smarca2 | 2276.346727 | -1.066703777 | 1.59E-09 |
| Adgre5 | 1076.317522 | -1.066803063 | 4.02E-08 |
| Nudt19 | 340.0343303 | -1.067650037 | 1.35E-08 |
| 3000002C10Rik | 44.84368038 | -1.067726714 | 3.97E-03 |
| Mpdu1 | 1210.812562 | -1.068195044 | 5.46E-17 |
| Ubxn2b | 458.5861195 | -1.068437316 | 3.61E-06 |
| C1qtnf6 | 2407.485434 | -1.06863627 | 2.75E-12 |
| Hoxa7 | 1586.992664 | -1.069909745 | 6.21E-21 |
| Fgd4 | 592.2398547 | -1.070039781 | 4.27E-07 |
| Pclaf | 978.5195751 | -1.070598896 | 4.80E-02 |
| Hhipl1 | 257.2349416 | -1.07147022 | 3.96E-04 |
| NA | 63.66815679 | -1.071623648 | 9.06E-04 |
| Nap1l2 | 52.4545836 | -1.072079143 | 1.95E-02 |
| Pcyt2 | 535.7741907 | -1.072276119 | 5.84E-05 |
| Zfp445 | 1254.964781 | -1.072287129 | 1.03E-04 |
| Gm16249 | 81.29125866 | -1.072917176 | 1.95E-04 |
| Slc16a4 | 67.2259842 | -1.073454414 | 2.09E-02 |
| Rnls | 41.93373739 | -1.0740271 | 5.76E-03 |
| 1110019D14Rik | 97.35392094 | -1.074231841 | 2.12E-04 |
| Pip4k2a | 1015.444415 | -1.074682021 | 9.00E-20 |
| Gusb | 2912.311294 | -1.07514039 | 1.30E-12 |
| Aspa | 75.09945744 | -1.075485963 | 2.81E-02 |
| Pkig | 750.3973578 | -1.076036831 | 1.31E-09 |
| Sesn1 | 1627.682062 | -1.076280016 | 5.45E-14 |
| Meaf6 | 1077.454916 | -1.07746353 | 1.34E-22 |
| Ucp2 | 1083.734914 | -1.077844643 | 9.92E-08 |
| Hnrnpa2b1 | 19860.98294 | -1.077890209 | 1.20E-06 |
| Vsig10l | 521.0060425 | -1.078443699 | 5.15E-06 |
| Tekt2 | 30.20879387 | -1.079317965 | 1.47E-02 |
| B4galt2 | 3537.6615 | -1.079789049 | 3.26E-18 |
| Zfp760 | 404.3092604 | -1.080394209 | 3.18E-03 |
| 4930563E22Rik | 46.10621929 | -1.080518231 | 9.72E-03 |
| Cyb561a3 | 744.7278893 | -1.080539978 | 1.71E-12 |
| Medag | 7575.216717 | -1.081036334 | 1.37E-06 |
| 3300005D01Rik | 873.8809125 | -1.081666765 | 2.34E-04 |
| Cep162 | 817.932987 | -1.081919063 | 1.36E-02 |

| | | | |
|---|---|---|---|
| Ubxn11 | 137.916065 | -1.082233385 | 1.32E-04 |
| Ap3d1 | 7686.894989 | -1.082407261 | 5.64E-44 |
| Hip1 | 3266.38303 | -1.082674722 | 2.60E-20 |
| Gm4673 | 94.57211719 | -1.082773349 | 1.34E-04 |
| Zkscan3 | 2374.665102 | -1.083627154 | 1.68E-18 |
| Zfp655 | 3110.698296 | -1.083665766 | 6.48E-29 |
| Zfp433 | 217.8127299 | -1.083919782 | 3.78E-03 |
| Fam208b | 1989.088708 | -1.084202252 | 3.21E-04 |
| Ppp1r3d | 238.405961 | -1.08444766 | 1.57E-06 |
| Tmem8 | 134.9929926 | -1.085951506 | 9.38E-09 |
| Arglu1 | 1665.246254 | -1.086614316 | 2.89E-05 |
| Vps41 | 3917.059342 | -1.088628467 | 1.13E-20 |
| Pip4p2 | 1878.030152 | -1.089006478 | 1.45E-17 |
| Dalrd3 | 632.742894 | -1.089318457 | 4.29E-14 |
| Zfp239 | 614.0938397 | -1.089388888 | 6.02E-07 |
| Tgfbr3 | 8535.55638 | -1.090410233 | 1.25E-13 |
| Gm13212 | 790.6580958 | -1.091274259 | 5.87E-04 |
| Zfp74 | 465.250991 | -1.092424817 | 3.31E-05 |
| Man1c1 | 553.3265867 | -1.092597859 | 2.27E-12 |
| Il12rb1 | 41.79475726 | -1.092710207 | 1.79E-02 |
| Gse1 | 312.4825439 | -1.092907686 | 7.87E-05 |
| Ppm1e | 886.2907223 | -1.093946936 | 1.35E-06 |
| Tom1 | 1367.922974 | -1.09419879 | 3.58E-03 |
| Ap4s1 | 235.0127263 | -1.094270588 | 5.75E-08 |
| Evl | 2952.632705 | -1.094277988 | 1.41E-07 |
| Smpd5 | 32.81608022 | -1.094326795 | 6.75E-03 |
| Zfp157 | 1050.188746 | -1.094422814 | 1.62E-06 |
| Hmgn2 | 1321.386804 | -1.09490231 | 1.83E-09 |
| Speg | 1856.726043 | -1.095698423 | 1.13E-13 |
| Grem1 | 116.2492527 | -1.096796802 | 8.60E-04 |
| Cdc7 | 594.7862899 | -1.097640447 | 2.01E-03 |
| Kifc3 | 995.0069766 | -1.097770485 | 2.34E-05 |
| Extl2 | 1343.270838 | -1.098177637 | 2.84E-12 |
| 2210016L21Rik | 958.8047931 | -1.098379931 | 1.15E-11 |
| Tbc1d5 | 2332.869746 | -1.098515421 | 9.03E-26 |
| Ncbp2 | 2446.490191 | -1.098580045 | 4.07E-17 |
| Galk2 | 2673.268011 | -1.099860115 | 1.80E-19 |
| Zfp944 | 404.0550411 | -1.100198544 | 3.02E-05 |
| mt-Nd4 | 129372.965 | -1.100252196 | 2.34E-13 |
| Ngly1 | 876.5239073 | -1.100824168 | 4.45E-12 |
| N6amt1 | 522.967724 | -1.100960198 | 1.88E-14 |
| Psap | 78612.42307 | -1.101166593 | 4.26E-08 |
| Cacna1c | 477.6875855 | -1.101176223 | 7.89E-07 |
| Bbs12 | 294.4829504 | -1.101224814 | 2.27E-04 |
| Fancm | 513.5863206 | -1.101725794 | 2.83E-04 |
| Cst3 | 29679.40866 | -1.101802418 | 1.40E-07 |
| Commd4 | 942.1458007 | -1.103017001 | 6.95E-18 |

| | | | |
|---|---|---|---|
| Itgb5 | 5005.644612 | -1.105452772 | 1.01E-08 |
| Art3 | 18.30251111 | -1.10647574 | 4.65E-02 |
| 3830406C13Rik | 528.8087456 | -1.10911722 | 1.99E-12 |
| Plagl2 | 1099.649691 | -1.109374997 | 4.80E-09 |
| NA | 106.2130085 | -1.110027295 | 2.45E-06 |
| Gm26880 | 2822.471388 | -1.111045894 | 9.54E-06 |
| Med16 | 1503.89229 | -1.112944188 | 1.38E-15 |
| Gbp9 | 51.96187374 | -1.113557861 | 3.83E-04 |
| Izumo4 | 98.40570222 | -1.115162697 | 9.39E-07 |
| Emc1 | 889.6482432 | -1.11552 | 1.03E-14 |
| Dtna | 3290.17433 | -1.115768975 | 5.41E-16 |
| Zfp820 | 151.3426224 | -1.116508019 | 2.87E-02 |
| Krcc1 | 1725.674243 | -1.116625866 | 8.00E-05 |
| Dbnl | 5109.073654 | -1.11690943 | 3.33E-14 |
| Ankrd28 | 3180.342278 | -1.117091541 | 3.68E-10 |
| Gm15774 | 16.91903401 | -1.117580753 | 4.77E-02 |
| Prkca | 3873.741095 | -1.117680288 | 2.07E-18 |
| Caps2 | 114.6304859 | -1.117976503 | 1.18E-03 |
| Top1mt | 841.059034 | -1.118523544 | 5.44E-19 |
| Naprt | 66.15425766 | -1.118619442 | 2.48E-05 |
| Ccne2 | 288.3521814 | -1.119364574 | 2.58E-02 |
| Wdr13 | 1853.351441 | -1.119390586 | 6.23E-15 |
| Cbx6 | 10839.95353 | -1.119715858 | 1.85E-27 |
| Dennd2a | 5263.659133 | -1.11978895 | 5.32E-11 |
| Lrp4 | 529.2244838 | -1.121010989 | 6.06E-04 |
| Lpgat1 | 5562.591021 | -1.12146689 | 1.24E-19 |
| Usp40 | 1922.118719 | -1.121479919 | 5.31E-12 |
| Tas1r2 | 31.23794289 | -1.121618809 | 2.03E-02 |
| Slc44a1 | 6489.8044 | -1.124841748 | 1.52E-10 |
| AU041133 | 94.90491122 | -1.125514904 | 4.23E-03 |
| A630052C17Rik | 85.93514374 | -1.125807454 | 4.10E-04 |
| Fam43a | 1435.289342 | -1.126265865 | 6.60E-11 |
| Ier5l | 736.6308499 | -1.126278822 | 9.58E-09 |
| Zfp935 | 205.3922271 | -1.127516804 | 6.49E-04 |
| Mtx3 | 733.5799151 | -1.128117015 | 1.67E-08 |
| Kansl1l | 717.0289781 | -1.128304989 | 4.65E-10 |
| Ctdsp1 | 4964.789485 | -1.128574022 | 6.09E-17 |
| Gm15443 | 102.0140851 | -1.130131885 | 1.90E-05 |
| Lmln | 194.6595696 | -1.130482953 | 4.25E-02 |
| Zmat1 | 277.1067175 | -1.131979461 | 1.72E-04 |
| Hes6 | 612.1037007 | -1.132726602 | 7.81E-11 |
| Pafah2 | 1268.397562 | -1.132775167 | 4.59E-16 |
| Zfp146 | 1880.563755 | -1.133363471 | 4.98E-07 |
| 2410131K14Rik | 1774.50699 | -1.135107992 | 2.07E-22 |
| 4921504A21Rik | 64.24312908 | -1.135480789 | 1.78E-03 |
| Gm12925 | 25.35113162 | -1.136499407 | 2.51E-02 |
| Gm19426 | 43.53584447 | -1.1365514 | 4.73E-03 |

| | | | |
|---|---|---|---|
| Atp2b1 | 7973.205069 | -1.136868465 | 2.46E-12 |
| Zfp248 | 467.7710312 | -1.136882419 | 5.50E-07 |
| Zfp995 | 106.1881916 | -1.137068267 | 2.47E-03 |
| Ppil1 | 752.6917392 | -1.137171195 | 9.24E-06 |
| Hoxd3 | 456.5880826 | -1.137498734 | 8.97E-05 |
| Gm10575 | 53.05328232 | -1.137514981 | 3.19E-03 |
| Hsd17b12 | 4957.384465 | -1.138092701 | 1.59E-10 |
| Zfp273 | 74.37311127 | -1.139014301 | 1.65E-03 |
| Usp47 | 10646.51981 | -1.140212089 | 9.84E-27 |
| Platr25 | 127.2381172 | -1.141555889 | 3.18E-03 |
| Ttc39c | 557.9704609 | -1.141802898 | 5.77E-11 |
| Cd276 | 4044.747532 | -1.141850191 | 8.62E-21 |
| Prpf4b | 2196.000112 | -1.142418717 | 1.88E-04 |
| Mmd | 4325.604263 | -1.142725312 | 3.41E-03 |
| Gm20457 | 45.64204163 | -1.143219295 | 4.14E-03 |
| Prr5l | 589.662235 | -1.14442748 | 1.98E-05 |
| Fam208a | 522.2502803 | -1.144442627 | 2.70E-03 |
| Hnrnph2 | 3745.444933 | -1.145252527 | 1.32E-17 |
| Spry2 | 928.0977301 | -1.145684741 | 7.80E-20 |
| Marcks | 14713.78478 | -1.146569449 | 7.10E-07 |
| Cdh26 | 290.3703007 | -1.146652284 | 2.63E-03 |
| Slc37a3 | 2707.708926 | -1.14667765 | 4.56E-38 |
| Leng9 | 144.2703895 | -1.146858755 | 3.73E-07 |
| Fer | 2906.447521 | -1.1474076 | 2.60E-24 |
| Chn1os3 | 57.36693281 | -1.147466781 | 6.00E-03 |
| Zfp472 | 207.0491988 | -1.147703443 | 2.10E-07 |
| Ahnak2 | 23819.93924 | -1.147788564 | 3.54E-11 |
| Kitl | 3273.736457 | -1.148014841 | 6.18E-07 |
| A230056P14Rik | 61.05474717 | -1.148709467 | 1.49E-04 |
| Clec2d | 2151.763597 | -1.150169742 | 4.35E-05 |
| Smarcd1 | 1668.953612 | -1.150470542 | 8.69E-19 |
| Gtf3c6 | 754.2313491 | -1.150570982 | 1.50E-11 |
| Agap2 | 1004.939107 | -1.151670289 | 7.89E-05 |
| Ssbp2 | 1469.826249 | -1.151962173 | 6.78E-05 |
| Coprs | 814.1861832 | -1.152158167 | 1.81E-22 |
| NA | 41.98412907 | -1.152293437 | 5.21E-03 |
| Rnase10 | 47.92102259 | -1.152791378 | 1.79E-03 |
| Wdr5b | 137.9910893 | -1.153408209 | 7.26E-08 |
| mt-Nd6 | 36849.03868 | -1.154692665 | 7.01E-16 |
| Tgfb2 | 3781.852125 | -1.155297946 | 1.55E-06 |
| Mtch1 | 11913.88654 | -1.155364096 | 8.33E-16 |
| Stat1 | 1423.127776 | -1.155415732 | 5.18E-14 |
| Zfp715 | 596.4060816 | -1.15548288 | 1.87E-08 |
| Kif9 | 130.1670536 | -1.155667191 | 1.97E-08 |
| Aoc3 | 1354.327432 | -1.15593067 | 1.23E-04 |
| Gpr176 | 2045.393277 | -1.156297056 | 3.17E-14 |
| Lca5l | 18.56150822 | -1.156610976 | 4.41E-02 |

| | | | |
|---|---|---|---|
| Scara3 | 1941.279625 | -1.158141545 | 1.72E-12 |
| Mfsd4b5 | 246.1474623 | -1.158946051 | 8.14E-05 |
| Golm1 | 1094.89663 | -1.159123238 | 6.18E-08 |
| Ifi211 | 544.3470546 | -1.159538824 | 1.03E-04 |
| Ccdc103 | 53.9477953 | -1.160533413 | 2.54E-04 |
| Gm16675 | 32.05963952 | -1.161144652 | 3.86E-03 |
| NA | 146.9254783 | -1.162236643 | 1.15E-08 |
| Zpbp | 33.14557248 | -1.162490664 | 4.29E-04 |
| E530011L22Rik | 207.3667489 | -1.163024581 | 7.74E-06 |
| Cetn4 | 155.0230006 | -1.163182136 | 2.65E-06 |
| Ccl28 | 69.41710484 | -1.164020646 | 1.62E-05 |
| Slc35b4 | 3374.352088 | -1.165173455 | 2.14E-12 |
| Hivep3 | 515.0723489 | -1.16574895 | 4.54E-10 |
| Lrrc49 | 603.8737907 | -1.166538017 | 7.27E-19 |
| Zfp296 | 38.59202607 | -1.167291341 | 5.51E-03 |
| Ftx | 591.5860801 | -1.168900882 | 1.20E-06 |
| Sp100 | 229.87064 | -1.169395098 | 1.26E-03 |
| 5430403G16Rik | 257.0774916 | -1.170080311 | 2.38E-03 |
| H2-M5 | 55.87435966 | -1.170584032 | 1.00E-03 |
| Pcmtd1 | 2794.947783 | -1.170766562 | 4.16E-05 |
| Zfp68 | 2035.29604 | -1.172526015 | 3.92E-11 |
| Tapbpl | 512.5127428 | -1.173071391 | 1.27E-20 |
| Epha5 | 18.84096967 | -1.173869286 | 4.33E-02 |
| Zfp788 | 503.9710848 | -1.174072409 | 3.69E-04 |
| Zfp292 | 1751.446612 | -1.174451452 | 1.15E-05 |
| Alms1-ps2 | 88.59135599 | -1.174931168 | 2.49E-04 |
| Tshz1 | 831.544133 | -1.175337797 | 2.55E-09 |
| Zfp946 | 237.5303763 | -1.175396551 | 4.37E-03 |
| Smarcad1 | 2090.096887 | -1.175695952 | 2.78E-07 |
| Tsr2 | 1165.014558 | -1.176176842 | 6.60E-17 |
| Morn2 | 184.5289294 | -1.177150697 | 6.96E-07 |
| Hcn3 | 50.97651329 | -1.177562963 | 1.36E-03 |
| Ankrd6 | 596.1378015 | -1.177628713 | 8.92E-16 |
| Itga11 | 1425.902456 | -1.178961844 | 2.49E-03 |
| Gm10676 | 25.58480143 | -1.179070342 | 1.03E-02 |
| Hspa12b | 551.3314607 | -1.179490376 | 5.15E-14 |
| Kctd17 | 1532.436301 | -1.179599534 | 4.55E-11 |
| Plxnb1 | 240.1316655 | -1.181791256 | 2.34E-02 |
| Slc39a11 | 1072.037658 | -1.18223003 | 2.40E-15 |
| Zfp976 | 153.6269941 | -1.182900461 | 1.98E-02 |
| Adamts2 | 9146.817329 | -1.183102097 | 7.53E-07 |
| G730003C15Rik | 29.50403608 | -1.183344015 | 1.37E-02 |
| Tmed1 | 963.0524274 | -1.183371348 | 7.28E-15 |
| 4930432B10Rik | 19.72505041 | -1.183939689 | 3.76E-02 |
| Gm6793 | 34.41213098 | -1.184714297 | 2.52E-02 |
| Draxin | 16.69298831 | -1.185295824 | 2.85E-02 |
| Amdhd2 | 1696.242283 | -1.186334738 | 3.07E-07 |

| | | | |
|---|---|---|---|
| Ift140 | 1141.357097 | -1.186458222 | 1.24E-09 |
| Zmat2 | 2291.274399 | -1.18676102 | 1.04E-39 |
| Hic1 | 1990.450822 | -1.186881405 | 3.58E-06 |
| Foxc2 | 481.5561448 | -1.187199118 | 1.59E-06 |
| Gm14443 | 88.67641707 | -1.187234853 | 1.10E-04 |
| Ggt7 | 21.30559824 | -1.187859749 | 2.52E-02 |
| Ttc39a | 153.8251168 | -1.188298292 | 2.96E-04 |
| mt-Nd5 | 120857.742 | -1.188425089 | 3.35E-19 |
| Emilin1 | 5288.590701 | -1.190342062 | 1.41E-09 |
| Clvs1 | 109.3908839 | -1.191909227 | 1.69E-02 |
| Mab21l2 | 405.6572256 | -1.192249807 | 6.50E-07 |
| NA | 208.8689783 | -1.193256267 | 3.98E-08 |
| Ncoa2 | 3430.512152 | -1.193275323 | 3.32E-02 |
| Ntn1 | 231.1255559 | -1.193657714 | 7.71E-06 |
| Ptpre | 155.6299662 | -1.193862326 | 4.63E-05 |
| Bclaf1 | 5262.010873 | -1.194420564 | 4.26E-06 |
| Zfp644 | 1613.466685 | -1.195628077 | 6.94E-07 |
| Adamts7 | 1045.159352 | -1.196377801 | 3.96E-12 |
| Tmem44 | 491.8757937 | -1.197580794 | 2.10E-03 |
| Runx1t1 | 722.8523518 | -1.197737963 | 1.80E-08 |
| Fndc3a | 4745.827846 | -1.198277354 | 2.81E-37 |
| Mfsd3 | 248.4748801 | -1.198301497 | 7.86E-09 |
| Pbdc1 | 1501.992805 | -1.199376211 | 1.53E-17 |
| Ankrd46 | 2144.475571 | -1.199540097 | 1.66E-14 |
| Gm11008 | 25.36760532 | -1.201465003 | 3.26E-02 |
| Mum1l1 | 1107.86875 | -1.202305799 | 1.40E-04 |
| Npas2 | 67.53822572 | -1.202695321 | 1.91E-02 |
| Hnrnpd | 4309.091954 | -1.20313155 | 1.55E-07 |
| Chd6 | 3406.923992 | -1.203331069 | 2.59E-03 |
| Dpm2 | 1496.657824 | -1.20334805 | 1.93E-15 |
| Zfp133-ps | 30.65106094 | -1.204013369 | 5.72E-03 |
| B3glct | 917.7545041 | -1.204105879 | 5.33E-27 |
| Zfp607b | 114.4188307 | -1.204500703 | 3.97E-03 |
| Abr | 2974.771084 | -1.204750879 | 2.89E-23 |
| Samd15 | 15.18295231 | -1.20478956 | 2.96E-02 |
| Rsad1 | 456.0324608 | -1.205062333 | 1.54E-21 |
| Armcx3 | 3815.423845 | -1.205075307 | 1.04E-17 |
| Zfp943 | 335.8360668 | -1.206567968 | 2.44E-04 |
| Armcx6 | 583.5901757 | -1.206723554 | 9.41E-14 |
| Gabpb2 | 2957.735198 | -1.207062856 | 2.37E-09 |
| Lvrn | 146.2069254 | -1.207910448 | 4.40E-06 |
| Gm26869 | 331.3716579 | -1.208041442 | 5.56E-04 |
| Cox20 | 380.4983109 | -1.208307238 | 8.13E-13 |
| Iah1 | 1772.443177 | -1.209671774 | 1.34E-10 |
| Zfp959 | 168.894081 | -1.209993937 | 1.89E-08 |
| Wdr60 | 933.6140883 | -1.210047434 | 4.23E-10 |
| Scfd1 | 1563.777827 | -1.210146685 | 8.80E-07 |

| | | | |
|---|---|---|---|
| Csad | 1257.892946 | -1.212328068 | 5.96E-15 |
| Sub1 | 3397.985703 | -1.213459291 | 3.52E-21 |
| Rnd2 | 944.1092971 | -1.214631752 | 1.41E-09 |
| Cbr4 | 464.8334136 | -1.214709411 | 7.50E-10 |
| Gm20939 | 156.2065304 | -1.214916728 | 8.24E-03 |
| P2rx5 | 276.0859069 | -1.215878716 | 1.19E-05 |
| Zfp931 | 208.6056449 | -1.216281211 | 8.25E-05 |
| Gm9392 | 25.55154648 | -1.21787933 | 9.51E-03 |
| Adam19 | 3337.629642 | -1.218542579 | 1.21E-18 |
| Twist2 | 1503.498649 | -1.219677441 | 2.11E-12 |
| Antxr1 | 10062.55027 | -1.219938988 | 4.60E-11 |
| Pts | 474.3278678 | -1.220039011 | 8.82E-11 |
| Fam69b | 943.3674919 | -1.220374212 | 1.55E-11 |
| Abhd12 | 3305.451109 | -1.221595752 | 1.70E-18 |
| Sh3bgrl | 3106.270232 | -1.221772065 | 1.52E-03 |
| 4930513N10Rik | 46.10247408 | -1.224024222 | 4.39E-05 |
| Gjc1 | 1240.108962 | -1.224537732 | 1.43E-12 |
| Gm5641 | 10531.33056 | -1.225854481 | 1.98E-08 |
| Dnajb14 | 1687.272902 | -1.22641015 | 1.84E-24 |
| Natd1 | 2182.198035 | -1.226467713 | 1.49E-12 |
| Zfp2 | 138.1886809 | -1.226713464 | 2.24E-04 |
| Pdf | 290.2676097 | -1.227985295 | 6.28E-08 |
| Ctbp1 | 5894.416865 | -1.229110824 | 1.53E-20 |
| Tmem67 | 422.0083957 | -1.229487445 | 1.22E-06 |
| Slc31a2 | 992.1188663 | -1.230824113 | 3.53E-16 |
| C030014I23Rik | 243.4663144 | -1.230992467 | 2.23E-10 |
| Eipr1 | 580.7879384 | -1.232876306 | 1.57E-12 |
| NA | 216.623365 | -1.234579218 | 1.20E-10 |
| Matn4 | 26.3751028 | -1.235383552 | 2.07E-02 |
| Samd14 | 1239.574654 | -1.235929586 | 3.99E-16 |
| Gm10699 | 19.8947923 | -1.236637552 | 2.20E-02 |
| Mfap5 | 6144.004538 | -1.236982808 | 1.83E-04 |
| Ranbp6 | 626.1189375 | -1.237224098 | 5.01E-06 |
| Golph3l | 941.6517126 | -1.237847646 | 8.73E-24 |
| Ccdc57 | 258.3154069 | -1.238185387 | 9.42E-07 |
| N4bp2 | 751.2548306 | -1.238418672 | 1.58E-05 |
| Lxn | 2852.121736 | -1.238843274 | 5.08E-08 |
| Fam110b | 769.9936195 | -1.239367465 | 6.31E-07 |
| Tubb2a | 6574.060184 | -1.23977752 | 3.25E-14 |
| Rtl9 | 47.36193539 | -1.240849848 | 6.10E-04 |
| Cyp51 | 10822.29261 | -1.241648715 | 7.07E-06 |
| Xrn1 | 931.0024997 | -1.243211695 | 2.90E-07 |
| Dennd4a | 6083.669471 | -1.243218523 | 3.35E-10 |
| Stk3 | 2814.161093 | -1.243447449 | 1.72E-08 |
| Gm15406 | 36.87485789 | -1.244508803 | 8.65E-04 |
| Dynll1 | 5076.522784 | -1.244668015 | 1.26E-09 |
| Gm9987 | 116.0074177 | -1.244840124 | 1.54E-09 |

| | | | |
|---|---|---|---|
| Bhlhe40 | 3834.220772 | -1.244928895 | 1.38E-07 |
| Gpc1 | 3053.071229 | -1.246322908 | 3.61E-12 |
| Gm38394 | 4556.884526 | -1.24709871 | 7.49E-07 |
| Nynrin | 2362.227013 | -1.247247349 | 6.54E-21 |
| Slc27a1 | 1298.849242 | -1.248431457 | 4.34E-07 |
| Taf9b | 211.1311733 | -1.25015878 | 7.05E-09 |
| Slc25a10 | 990.5896264 | -1.251361089 | 2.70E-07 |
| Dhodh | 572.5121825 | -1.252036327 | 4.20E-08 |
| Bmi1 | 3119.393451 | -1.252393655 | 9.97E-12 |
| Gm12424 | 18.16555463 | -1.253122864 | 3.02E-02 |
| 6-Sep | 1358.601144 | -1.253441017 | 1.63E-12 |
| Spry3 | 464.077843 | -1.253951344 | 1.80E-05 |
| Oit3 | 92.97206647 | -1.254207762 | 3.14E-02 |
| Phf20l1 | 1651.961763 | -1.254628827 | 4.78E-05 |
| Adamts15 | 21.18110807 | -1.25571292 | 2.49E-02 |
| Rgcc | 230.2419975 | -1.25609093 | 3.34E-05 |
| Tril | 35.05868021 | -1.256403882 | 2.79E-02 |
| Zfp456 | 23.67828685 | -1.256751644 | 2.00E-02 |
| Klhl23 | 494.130889 | -1.257809158 | 1.45E-05 |
| Srrm4 | 674.1551676 | -1.258189082 | 3.45E-08 |
| Cd63-ps | 20.0657463 | -1.258255286 | 1.96E-02 |
| D030028A08Rik | 174.8608595 | -1.25873835 | 1.49E-09 |
| Sipa1l2 | 3528.093693 | -1.259321192 | 9.92E-11 |
| Tha1 | 19.03802371 | -1.259435977 | 2.00E-02 |
| Arhgap42 | 1172.465651 | -1.260494452 | 2.02E-09 |
| Dgcr8 | 1706.26144 | -1.260588183 | 1.27E-05 |
| Gm15128 | 352.174942 | -1.260934441 | 6.07E-03 |
| Fadd | 424.1906884 | -1.261217877 | 2.65E-16 |
| Zfp975 | 158.7501867 | -1.261529764 | 9.80E-04 |
| Zfp947 | 97.00750437 | -1.262022282 | 4.49E-03 |
| Gm7598 | 27.13875223 | -1.262453493 | 2.11E-03 |
| Ccdc71l | 320.4996366 | -1.262978551 | 2.33E-13 |
| Sspn | 1024.357293 | -1.265868345 | 6.51E-14 |
| Rftn2 | 1288.815036 | -1.265961981 | 2.66E-11 |
| Zfp61 | 290.8195837 | -1.266294966 | 2.22E-09 |
| Etnk1 | 4408.390478 | -1.26668452 | 7.54E-17 |
| Lppos | 143.0048581 | -1.266801782 | 1.65E-07 |
| Clcn4 | 1622.727894 | -1.266927221 | 8.64E-20 |
| Cars2 | 395.453121 | -1.267220429 | 1.17E-14 |
| Gnb4 | 263.9667297 | -1.267445486 | 1.07E-06 |
| Cnih2 | 222.9740135 | -1.268182754 | 2.37E-07 |
| Zfp638 | 3672.885195 | -1.26926463 | 9.00E-05 |
| Gm14634 | 11.82749569 | -1.270550985 | 3.33E-02 |
| A730056A06Rik | 112.1017604 | -1.270978047 | 7.32E-04 |
| Ppt1 | 7244.146237 | -1.273004599 | 5.55E-38 |
| Entpd5 | 1361.739639 | -1.273038902 | 5.21E-16 |
| Nlrc3 | 519.874105 | -1.273128991 | 5.09E-12 |

| | | | |
|---|---|---|---|
| Zfp182 | 311.6166338 | -1.273161191 | 4.89E-04 |
| Zcchc3 | 416.4233873 | -1.273351007 | 8.19E-06 |
| Gm16295 | 26.92563693 | -1.273868409 | 2.95E-03 |
| Gm9776 | 112.7083575 | -1.27450396 | 1.13E-06 |
| Mmab | 2103.661292 | -1.274940779 | 5.75E-09 |
| Yy2 | 149.4951159 | -1.274984027 | 1.12E-03 |
| Chst11 | 5163.815561 | -1.275219365 | 3.96E-14 |
| Zfp345 | 250.9356857 | -1.275348054 | 6.57E-06 |
| Zfp942 | 555.0205701 | -1.275612845 | 1.83E-03 |
| 9330159F19Rik | 260.4380525 | -1.276882808 | 5.88E-06 |
| Slc16a11 | 113.1379731 | -1.277980412 | 1.13E-05 |
| Bok | 494.9314589 | -1.278238161 | 8.14E-05 |
| Kbtbd7 | 200.5646128 | -1.278896807 | 1.07E-05 |
| Rora | 549.7334082 | -1.279397013 | 2.83E-07 |
| Pmepa1 | 1219.457425 | -1.279605587 | 2.01E-12 |
| Gm15411 | 49.73035422 | -1.279807737 | 2.29E-04 |
| Shc4 | 137.7357455 | -1.279980221 | 3.65E-06 |
| Tmem140 | 492.4092491 | -1.281384074 | 3.17E-05 |
| Ndufaf3 | 55.58015648 | -1.281888029 | 7.74E-06 |
| Fkbp14 | 1386.081438 | -1.282977905 | 1.63E-10 |
| Pcdhb15 | 375.5700413 | -1.283708578 | 9.56E-12 |
| Gnpda1 | 2534.474834 | -1.283939082 | 8.54E-15 |
| Gm16140 | 20.7670719 | -1.285150133 | 1.76E-02 |
| Slc4a11 | 115.9732355 | -1.285380065 | 6.92E-09 |
| Angptl2 | 12262.23054 | -1.286230651 | 5.59E-08 |
| Gm22270 | 16.49993171 | -1.28865235 | 3.06E-02 |
| 1700056E22Rik | 47.30814952 | -1.289031235 | 6.85E-05 |
| Utp14b | 229.3382319 | -1.289435784 | 5.52E-05 |
| C430049B03Rik | 299.2756912 | -1.289812202 | 4.72E-06 |
| Wdr81 | 3346.664813 | -1.290296747 | 1.93E-15 |
| B430203G13Rik | 111.3993379 | -1.291085979 | 1.15E-05 |
| Arhgap18 | 896.4577738 | -1.292850246 | 5.23E-04 |
| Ccl7 | 1521.615401 | -1.294386052 | 1.14E-06 |
| Gli3 | 1922.023409 | -1.294829582 | 2.39E-17 |
| Pigo | 175.9082829 | -1.29562139 | 6.39E-05 |
| 9130023H24Rik | 247.178609 | -1.297977739 | 7.53E-15 |
| Ift81 | 1302.686909 | -1.298388918 | 3.15E-04 |
| Timm22 | 1003.238842 | -1.299331061 | 4.94E-06 |
| C1qtnf1 | 929.0184909 | -1.299563721 | 9.98E-19 |
| Trim12c | 714.1829737 | -1.300257358 | 2.61E-08 |
| Gm10182 | 1077.283482 | -1.300395725 | 1.34E-22 |
| Sirpa | 3616.513785 | -1.300399736 | 1.40E-17 |
| Cask | 3506.668855 | -1.301002156 | 8.87E-22 |
| Oscp1 | 498.6865751 | -1.301289647 | 4.93E-11 |
| B9d1 | 529.0966433 | -1.301498668 | 4.37E-13 |
| Zfp808 | 153.8817309 | -1.301888831 | 4.19E-03 |
| Apopt1 | 749.4078064 | -1.302478009 | 6.02E-18 |

| | | | |
|---|---|---|---|
| Gm10222 | 23166.10098 | -1.303486916 | 1.70E-12 |
| NA | 68.2755783 | -1.3048758 | 2.08E-02 |
| Arhgef19 | 326.3317744 | -1.305582622 | 1.59E-06 |
| Kirrel3 | 34.68484594 | -1.305861382 | 2.32E-02 |
| Gpr162 | 212.3158726 | -1.30745501 | 8.23E-07 |
| Neurl4 | 1596.034851 | -1.307605702 | 2.21E-18 |
| Vps8 | 2880.10801 | -1.308010029 | 9.07E-18 |
| Mex3b | 174.4803361 | -1.308267971 | 7.78E-06 |
| Gm26737 | 245.1761763 | -1.310560228 | 4.52E-07 |
| Rtl1 | 52.28000868 | -1.310711803 | 4.50E-04 |
| Cand2 | 2044.67946 | -1.31072098 | 5.95E-22 |
| Gm14403 | 121.8702542 | -1.311154414 | 2.19E-03 |
| Casp12 | 2453.996963 | -1.311155964 | 4.46E-11 |
| Slfn9 | 1195.956977 | -1.311271216 | 2.39E-02 |
| Ecm2 | 56.32071213 | -1.31149299 | 1.96E-04 |
| Man2b1 | 3916.818943 | -1.311532086 | 2.54E-15 |
| Tshz3 | 29.61253025 | -1.311542553 | 4.84E-02 |
| Hif1a | 7405.238458 | -1.312148561 | 7.87E-13 |
| Sqle | 9213.229076 | -1.312815374 | 6.08E-09 |
| Pcdhb18 | 74.91592395 | -1.313044645 | 7.60E-06 |
| Zhx1 | 1595.963463 | -1.313870952 | 9.76E-09 |
| Lztfl1 | 853.8656872 | -1.314299038 | 3.92E-07 |
| Otulinl | 214.2498603 | -1.315139895 | 1.72E-14 |
| Omd | 64.4193638 | -1.316659768 | 1.67E-02 |
| Lhx6 | 339.1144646 | -1.317796978 | 2.74E-09 |
| Gm11223 | 1494.168631 | -1.318674925 | 7.90E-06 |
| Ankrd42 | 231.1595588 | -1.319062253 | 1.90E-13 |
| Zfp213 | 424.7323897 | -1.319260926 | 1.86E-06 |
| Pdgfra | 29231.84787 | -1.319277244 | 1.02E-05 |
| Gm6710 | 114.0969952 | -1.320129961 | 1.26E-02 |
| Mvk | 1649.647814 | -1.320736675 | 5.88E-08 |
| Mettl18 | 120.06686 | -1.32086962 | 2.02E-06 |
| Cdrt4 | 25.61038649 | -1.320908422 | 1.20E-03 |
| 1700037C18Rik | 79.88967796 | -1.321838677 | 8.23E-05 |
| Twsg1 | 5761.860578 | -1.321892169 | 1.76E-30 |
| Zfp40 | 305.3181922 | -1.321899535 | 1.25E-02 |
| Adam1a | 40.12227664 | -1.322333028 | 1.11E-03 |
| Il11ra1 | 3333.985703 | -1.322367347 | 7.91E-10 |
| Dpf1 | 35.92731429 | -1.322792333 | 1.71E-02 |
| Dock1 | 3924.573139 | -1.322877406 | 6.26E-32 |
| Eif1a | 6421.978995 | -1.324872958 | 2.34E-11 |
| Tagln | 2708.688587 | -1.325794229 | 1.29E-05 |
| Tnfrsf11a | 123.4947407 | -1.327281232 | 2.86E-12 |
| Hnrnpm | 7454.04858 | -1.327667852 | 1.20E-08 |
| Tlr2 | 500.01692 | -1.327693778 | 1.05E-08 |
| Cstad | 40.98467497 | -1.327973471 | 3.34E-03 |
| Vwa8 | 2171.685012 | -1.328899786 | 1.24E-19 |

| | | | |
|---|---|---|---|
| Tent5a | 42.14233375 | -1.329135171 | 1.04E-03 |
| Gpr85 | 486.5449696 | -1.329750871 | 4.14E-10 |
| Onecut2 | 225.0983773 | -1.329930263 | 2.41E-08 |
| Cep83os | 409.9112091 | -1.331069912 | 1.33E-10 |
| Lrguk | 36.65908511 | -1.331289633 | 4.95E-03 |
| Timm21 | 480.2239543 | -1.331365567 | 1.74E-11 |
| Cbarp | 525.0437716 | -1.33156836 | 1.22E-19 |
| Map9 | 417.747882 | -1.331895425 | 2.51E-06 |
| Cnot6 | 3465.79491 | -1.332069338 | 1.43E-10 |
| Cdyl2 | 218.9938247 | -1.332273708 | 2.30E-14 |
| Nanos1 | 79.92065785 | -1.336560883 | 6.26E-03 |
| B130055M24Rik | 163.0194118 | -1.33663694 | 8.08E-07 |
| Tlr4 | 2569.985835 | -1.336904027 | 1.54E-11 |
| Rmi1 | 653.8504849 | -1.33717711 | 3.49E-13 |
| Gm16835 | 21.2767008 | -1.337385686 | 1.21E-02 |
| Gpr135 | 158.3505966 | -1.339283603 | 9.05E-12 |
| Scn2a | 379.9991975 | -1.339641242 | 2.15E-04 |
| Ift74 | 2106.799547 | -1.340524585 | 1.46E-06 |
| Bicdl1 | 356.0545461 | -1.342496455 | 1.25E-13 |
| Wnt10b | 29.53377535 | -1.342972691 | 1.01E-02 |
| NA | 54.53741336 | -1.343186922 | 2.83E-03 |
| AI606473 | 18.78385635 | -1.344077791 | 3.14E-02 |
| Socs5 | 5784.952356 | -1.344571612 | 4.18E-29 |
| Dhcr7 | 1440.708452 | -1.344861758 | 8.18E-09 |
| Tcf4 | 3628.891416 | -1.345130827 | 3.33E-33 |
| Ndn | 1355.557359 | -1.345778755 | 1.17E-21 |
| A330023F24Rik | 61.15415772 | -1.34627575 | 2.15E-03 |
| Peg12 | 309.1835184 | -1.346577151 | 4.33E-11 |
| Snapc5 | 308.3123639 | -1.346780441 | 9.80E-11 |
| Mest | 25.66506474 | -1.347306745 | 1.48E-02 |
| B4galt5 | 4562.137611 | -1.348165796 | 6.50E-09 |
| Zfp606 | 511.9724779 | -1.348567651 | 4.88E-06 |
| Bola3 | 767.7054442 | -1.348773346 | 4.82E-14 |
| Ly96 | 148.4629841 | -1.349866446 | 1.97E-08 |
| Gm42688 | 427.2518859 | -1.350721422 | 2.66E-22 |
| Phactr1 | 1833.467544 | -1.350879898 | 2.39E-16 |
| Fgfbp3 | 84.82346159 | -1.350882906 | 4.83E-05 |
| Pcdha11 | 424.0706161 | -1.351869777 | 1.63E-04 |
| Adck2 | 829.9544706 | -1.352347001 | 1.10E-25 |
| mt-Ty | 79.47743535 | -1.352526864 | 2.30E-03 |
| Jam3 | 380.139953 | -1.353943441 | 2.99E-17 |
| 4933431E20Rik | 881.878442 | -1.354729918 | 9.18E-12 |
| Osr2 | 352.6919725 | -1.354799029 | 2.46E-04 |
| Asap2 | 581.2720119 | -1.354900322 | 1.60E-12 |
| P3h3 | 5126.717251 | -1.355876824 | 9.74E-31 |
| Mbd4 | 461.5537666 | -1.356277058 | 7.41E-11 |
| Tyw3 | 278.3643857 | -1.357134633 | 9.33E-06 |

| | | | |
|---|---|---|---|
| Fastkd2 | 1129.972847 | -1.358898669 | 6.83E-20 |
| Stard4 | 2046.999912 | -1.359652019 | 2.05E-05 |
| Gbp7 | 50.27437464 | -1.359723179 | 1.71E-03 |
| Brinp3 | 129.3249775 | -1.359912177 | 4.31E-03 |
| Reep3 | 3673.165162 | -1.360673541 | 4.56E-57 |
| Rcan2 | 964.4870689 | -1.360804815 | 2.67E-04 |
| Megf9 | 1364.252732 | -1.362354335 | 6.54E-11 |
| NA | 24.57661118 | -1.363036231 | 1.27E-02 |
| 2210408F21Rik | 135.6814517 | -1.363744358 | 7.03E-09 |
| Katnb1 | 817.8951214 | -1.363940486 | 1.05E-14 |
| Tcea2 | 213.1302564 | -1.36414704 | 1.30E-09 |
| Zfp748 | 408.1503428 | -1.364813461 | 5.32E-05 |
| Bbs1 | 345.4501395 | -1.367263746 | 2.46E-15 |
| A930012O16Rik | 25.52054419 | -1.369216351 | 2.16E-03 |
| Etv4 | 307.4908993 | -1.369269058 | 7.54E-04 |
| D630045J12Rik | 1186.490416 | -1.369321169 | 4.60E-11 |
| Rassf2 | 2868.908583 | -1.369934879 | 5.51E-08 |
| Gm21596 | 17.61082874 | -1.371876139 | 4.26E-02 |
| Tmem14a | 58.42559324 | -1.373600762 | 7.52E-06 |
| Gpr153 | 1837.287844 | -1.373635854 | 4.00E-12 |
| 9030624G23Rik | 33.45556206 | -1.37444546 | 3.55E-04 |
| Gm16183 | 29.51470691 | -1.374994295 | 1.73E-03 |
| Inpp5e | 1283.030225 | -1.375499492 | 2.62E-25 |
| Cyb5a | 3603.739228 | -1.378020441 | 6.52E-16 |
| Fgf7 | 4782.992174 | -1.378129034 | 2.46E-08 |
| Xxylt1 | 1691.672168 | -1.378878075 | 1.42E-24 |
| Pgbd5 | 323.7462955 | -1.379966298 | 3.13E-10 |
| Vamp5 | 899.8965301 | -1.380657378 | 7.45E-09 |
| Gm26797 | 27.78923782 | -1.380754757 | 7.96E-03 |
| Sgsh | 942.5070194 | -1.382007736 | 1.48E-12 |
| Myl9 | 237.5650558 | -1.383016702 | 1.88E-06 |
| Glt8d2 | 963.6849507 | -1.383018197 | 4.71E-09 |
| 3110039I08Rik | 2289.917775 | -1.383445863 | 4.53E-03 |
| Kbtbd8 | 94.49187229 | -1.384057842 | 1.18E-02 |
| Gm9747 | 221.4334643 | -1.38409824 | 1.01E-05 |
| E230016M11Rik | 61.17761226 | -1.384807467 | 3.55E-04 |
| Pdcd4 | 2480.242851 | -1.385320667 | 8.80E-14 |
| Egr1 | 4443.449226 | -1.38603999 | 1.01E-02 |
| Dsel | 941.8407461 | -1.386796464 | 3.86E-06 |
| Nck2 | 490.0943103 | -1.388337858 | 4.37E-17 |
| Dscc1 | 138.1416843 | -1.389358792 | 4.01E-05 |
| Armcx1 | 1688.155142 | -1.389708839 | 1.99E-18 |
| Apobr | 620.5613799 | -1.389908469 | 8.42E-06 |
| Zfp950 | 460.7675227 | -1.390265541 | 2.64E-03 |
| Msrb3 | 3410.420807 | -1.391890752 | 7.25E-10 |
| B3galnt1 | 1459.923705 | -1.393863193 | 9.28E-27 |
| Sgk1 | 8387.420148 | -1.394775481 | 6.03E-12 |

| | | | |
|---|---|---|---|
| Srsf3 | 8264.123244 | -1.39480952 | 8.50E-13 |
| Rev3l | 2594.882945 | -1.395019505 | 4.68E-06 |
| Dock11 | 1878.430365 | -1.396608473 | 4.62E-02 |
| Cep41 | 537.163494 | -1.396863991 | 1.63E-14 |
| Fam217b | 836.7603683 | -1.397228318 | 1.67E-12 |
| Pcdhgc4 | 30257.72995 | -1.397714152 | 7.99E-21 |
| A730017L22Rik | 398.5430896 | -1.39820323 | 4.12E-10 |
| BC039966 | 29.60548201 | -1.399019552 | 1.52E-03 |
| Slc2a6 | 121.7876624 | -1.400240262 | 4.58E-06 |
| P4htm | 35.99990357 | -1.400730821 | 4.53E-05 |
| Rgs9bp | 289.0652429 | -1.40091287 | 4.12E-11 |
| Zfp35 | 601.7816323 | -1.402416816 | 4.16E-16 |
| Camk1d | 90.9566728 | -1.403372731 | 6.57E-04 |
| Sec16b | 930.6328352 | -1.403534188 | 3.05E-09 |
| Ppp2r2c | 14.44398726 | -1.404467414 | 1.34E-02 |
| Rab27b | 133.4508688 | -1.404614515 | 5.39E-04 |
| Gm11033 | 338.8365071 | -1.405401192 | 5.47E-10 |
| NA | 11.74088147 | -1.40570801 | 3.05E-02 |
| Zfp758 | 213.2476453 | -1.405988716 | 1.58E-04 |
| Cetn2 | 985.5248549 | -1.406521324 | 2.41E-15 |
| Ulk4 | 167.008026 | -1.407100169 | 1.68E-09 |
| Pmp22 | 6001.44192 | -1.40711944 | 9.74E-15 |
| Dip2b | 2838.90961 | -1.408530778 | 6.40E-17 |
| Emc2 | 2388.117355 | -1.409430203 | 7.02E-11 |
| Zfp882 | 177.2345233 | -1.412361631 | 1.96E-05 |
| Pdk1 | 1503.806453 | -1.412902696 | 1.19E-08 |
| Galm | 284.6776046 | -1.41407227 | 7.78E-12 |
| Arsg | 389.4454428 | -1.415876549 | 3.09E-18 |
| 2610044O15Rik8 | 77.27477444 | -1.41593892 | 3.16E-06 |
| Cnpy2 | 2958.714011 | -1.41638344 | 1.33E-31 |
| Neat1 | 13268.97801 | -1.41684667 | 6.61E-07 |
| Mgst1 | 1067.813549 | -1.417679972 | 1.97E-11 |
| Mamdc2 | 140.4200342 | -1.417897696 | 3.66E-04 |
| Zfp764 | 357.7441711 | -1.418823319 | 2.28E-21 |
| Spata48 | 29.05752533 | -1.42180657 | 2.80E-03 |
| Zfp953 | 159.3076468 | -1.422189555 | 7.15E-06 |
| Mdfi | 37.12431873 | -1.422732829 | 3.86E-04 |
| Gm10762 | 70.63600912 | -1.423642373 | 1.03E-06 |
| Dnase1l2 | 34.76247815 | -1.424672362 | 2.42E-03 |
| Prcd | 10.9762808 | -1.424882497 | 2.34E-02 |
| BC037032 | 28.00622575 | -1.425637791 | 5.08E-03 |
| Gm6871 | 49.20881328 | -1.426290698 | 1.85E-03 |
| Cbfb | 3583.177904 | -1.426310123 | 1.82E-46 |
| Nin | 2334.99443 | -1.426471893 | 7.43E-13 |
| Gm17055 | 35.10683914 | -1.426920153 | 8.64E-05 |
| Arsb | 3780.501608 | -1.427947183 | 3.77E-26 |
| A430033K04Rik | 372.7872773 | -1.428406175 | 9.63E-14 |

| | | | |
|---|---|---|---|
| Xdh | 15174.39706 | -1.428639134 | 4.26E-11 |
| Pi15 | 156.8925324 | -1.430079946 | 2.54E-04 |
| Epc2 | 1151.623631 | -1.431830209 | 7.98E-13 |
| Tymp | 200.2920391 | -1.433573109 | 7.59E-09 |
| Degs2 | 79.4466478 | -1.434211966 | 1.76E-06 |
| Zfp386 | 962.4306407 | -1.43504547 | 6.34E-06 |
| Gm20403 | 14.36654804 | -1.435783879 | 2.74E-02 |
| B3gnt9 | 1121.027082 | -1.436513297 | 1.01E-18 |
| Nr2f1 | 1515.258968 | -1.437539797 | 1.73E-18 |
| Ttyh3 | 8901.200216 | -1.438657156 | 1.28E-11 |
| Rhbdd2 | 1090.819054 | -1.439139896 | 8.34E-23 |
| Slc25a29 | 113.262095 | -1.440057246 | 3.98E-06 |
| Bicc1 | 3603.644497 | -1.442571736 | 1.01E-28 |
| Fam151b | 10.37822406 | -1.443906229 | 3.51E-02 |
| 2810408A11Rik | 84.21173571 | -1.445120125 | 2.56E-06 |
| Sema5a | 3228.140088 | -1.445292472 | 7.66E-08 |
| Igdcc4 | 80.19494662 | -1.448350284 | 2.28E-05 |
| Gm10814 | 28.90121507 | -1.449129625 | 5.15E-03 |
| Gm13615 | 9.512650198 | -1.449992852 | 4.95E-02 |
| Rabl2 | 424.7187683 | -1.450453921 | 1.40E-09 |
| Gm27003 | 15.40465391 | -1.452586008 | 2.03E-02 |
| 9030407P20Rik | 53.53998062 | -1.453836364 | 8.77E-04 |
| Ralgds | 5291.738528 | -1.454428876 | 2.92E-11 |
| Fnip2 | 5464.462773 | -1.458206864 | 3.80E-19 |
| Plxna4 | 92.75828692 | -1.458535148 | 3.93E-03 |
| Gatb | 891.9343914 | -1.459049471 | 1.08E-10 |
| Gm10767 | 19.72172038 | -1.460240989 | 8.80E-03 |
| Zfp729b | 395.1662651 | -1.460542152 | 1.87E-06 |
| Trmt12 | 249.3859157 | -1.460660103 | 7.04E-17 |
| Vat1 | 27960.27472 | -1.461025857 | 4.55E-12 |
| Coro1a | 13.10308147 | -1.461078285 | 4.97E-02 |
| Vmn2r-ps54 | 19.44055338 | -1.461244546 | 1.95E-02 |
| Gm3558 | 41.23278577 | -1.461276922 | 1.89E-02 |
| Fmnl2 | 2152.240826 | -1.461869058 | 7.74E-28 |
| Ssbp4 | 614.8805247 | -1.462432127 | 4.76E-13 |
| Snx32 | 113.7111598 | -1.463393715 | 2.00E-06 |
| Znrd1 | 628.2231869 | -1.463403273 | 2.35E-14 |
| Ypel1 | 252.6287883 | -1.464017691 | 2.26E-07 |
| Pabpc4l | 55.49155236 | -1.464791668 | 1.57E-04 |
| Aamdc | 197.088835 | -1.466718542 | 1.41E-13 |
| Trib2 | 1232.911453 | -1.468857656 | 1.59E-13 |
| Gm15413 | 17.33829686 | -1.469778151 | 5.63E-03 |
| Cplx2 | 1224.275436 | -1.470291024 | 2.06E-28 |
| Cracr2a | 41.73135551 | -1.471202774 | 5.18E-04 |
| Magi2 | 526.5829282 | -1.473212886 | 3.24E-29 |
| Gnptab | 4858.629519 | -1.473922804 | 4.34E-32 |
| Il1r1 | 1964.698977 | -1.475009848 | 1.35E-12 |

| | | | |
|---|---|---|---|
| Vangl1 | 1595.126302 | -1.475319306 | 3.09E-15 |
| Prune2 | 851.2860509 | -1.475593689 | 2.38E-09 |
| A930002H24Rik | 37.79997615 | -1.475713356 | 1.61E-03 |
| Begain | 238.4066239 | -1.477669164 | 2.13E-07 |
| Has2os | 14.78403193 | -1.479304779 | 9.02E-03 |
| Mxd3 | 180.75113 | -1.482778089 | 6.05E-04 |
| Zc3h12b | 308.6497987 | -1.484379402 | 3.80E-22 |
| Pappa | 4604.374407 | -1.484650752 | 6.65E-09 |
| Fhl4 | 42.64920327 | -1.485680174 | 9.49E-05 |
| Znrd1as | 91.526769 | -1.486149575 | 2.69E-09 |
| Glipr2 | 2939.975943 | -1.489125917 | 1.03E-23 |
| Il27ra | 37.22449657 | -1.489173662 | 2.52E-04 |
| Nid1 | 41135.2065 | -1.489795621 | 1.55E-45 |
| Cntln | 2121.188526 | -1.490833699 | 4.00E-06 |
| Plekhg4 | 303.5491276 | -1.492504219 | 6.31E-04 |
| Il18 | 182.5136154 | -1.492789516 | 5.06E-07 |
| Itih2 | 173.4232574 | -1.494383678 | 9.99E-05 |
| Arc | 134.080199 | -1.494407509 | 2.99E-08 |
| Ogn | 19778.96094 | -1.495981493 | 9.62E-03 |
| Pfkfb2 | 1070.695959 | -1.498293523 | 9.07E-20 |
| Il1rl2 | 175.7426724 | -1.49918562 | 1.39E-09 |
| Arhgef15 | 50.24344551 | -1.499284928 | 1.17E-03 |
| Sumf1 | 4147.413684 | -1.500484517 | 4.08E-33 |
| Thtpa | 224.2179658 | -1.501426999 | 5.48E-20 |
| Tbc1d4 | 572.5515038 | -1.501670597 | 2.72E-09 |
| Slc12a2 | 4935.850474 | -1.501769337 | 1.37E-24 |
| Kctd1 | 652.3447104 | -1.501956134 | 5.54E-20 |
| Aff3 | 1563.295887 | -1.502187274 | 9.44E-08 |
| Myo5c | 18.82323414 | -1.503194953 | 1.25E-02 |
| NA | 667.970585 | -1.503442484 | 1.32E-30 |
| Gm5784 | 30.3737508 | -1.503495999 | 9.69E-04 |
| Aspn | 5188.83664 | -1.503540604 | 2.13E-02 |
| Gm26490 | 11.39270309 | -1.503639339 | 4.38E-02 |
| Rab2b | 836.6849826 | -1.503879237 | 3.87E-24 |
| Gm16229 | 22.95610143 | -1.50682226 | 5.79E-03 |
| Tmem150a | 1636.619374 | -1.506838052 | 7.01E-22 |
| Cyp2j9 | 75.57103561 | -1.50830909 | 5.01E-06 |
| Fn1 | 474467.2828 | -1.510114085 | 1.66E-17 |
| Mtss1l | 4009.15688 | -1.510350455 | 2.62E-18 |
| Ttc30a1 | 153.2911787 | -1.510639175 | 1.46E-14 |
| Cdh11 | 8716.03324 | -1.512888239 | 9.65E-13 |
| 1700034H15Rik | 27.77523794 | -1.514789953 | 3.25E-03 |
| Chic1 | 323.0857059 | -1.515595299 | 2.80E-12 |
| 4930533I22Rik | 11.42858759 | -1.521644572 | 3.61E-02 |
| Sppl2b | 851.0028724 | -1.522087385 | 4.09E-13 |
| Castor2 | 1817.867489 | -1.523539132 | 2.76E-29 |
| Zfp281 | 2185.820373 | -1.524692677 | 7.54E-18 |

| | | | |
|---|---|---|---|
| Lhfpl2 | 6465.270021 | -1.525646295 | 2.51E-26 |
| Tada1 | 953.2882728 | -1.528492463 | 1.09E-17 |
| A830005F24Rik | 10.58896151 | -1.530703542 | 4.47E-02 |
| Asb13 | 427.8861044 | -1.534098289 | 2.03E-10 |
| Zfp773 | 15.52759869 | -1.534792446 | 6.72E-03 |
| Ube2d-ps | 793.7075186 | -1.534831065 | 2.65E-24 |
| Ankdd1b | 30.83395126 | -1.535715267 | 1.48E-04 |
| Tex9 | 290.3539929 | -1.535826198 | 1.92E-08 |
| Bnc2 | 1922.400801 | -1.536528131 | 9.47E-25 |
| Hdgfl3 | 1749.187573 | -1.536535689 | 1.05E-10 |
| Loxl1 | 13086.55011 | -1.538463671 | 7.02E-19 |
| Mblac1 | 76.13773596 | -1.539531412 | 1.17E-07 |
| Pofut1 | 2332.656105 | -1.541329429 | 5.83E-13 |
| Elovl2 | 19.10085203 | -1.541873657 | 3.41E-02 |
| Gm15996 | 11.99713073 | -1.54224622 | 3.13E-02 |
| Anks1b | 74.61954556 | -1.542624202 | 3.48E-03 |
| Pdgfrb | 13123.50877 | -1.544032228 | 8.86E-34 |
| Mb21d2 | 423.4672271 | -1.54451518 | 2.03E-17 |
| Gm5977 | 48.68085952 | -1.547586839 | 2.94E-05 |
| Prrx2 | 980.971953 | -1.549553752 | 1.57E-06 |
| Prrg1 | 395.4345146 | -1.552653849 | 1.47E-06 |
| B230118H07Rik | 425.1159329 | -1.552795556 | 3.38E-10 |
| Hmgcr | 4300.914669 | -1.554204256 | 9.86E-08 |
| Olfml2b | 2018.256671 | -1.554245593 | 1.75E-10 |
| B4galt6 | 1066.319975 | -1.554456938 | 4.24E-14 |
| Dock5 | 2703.985943 | -1.556843135 | 1.32E-24 |
| Slc22a15 | 345.992738 | -1.558172541 | 4.84E-30 |
| Cd1d1 | 192.488829 | -1.558630355 | 3.06E-10 |
| Nacad | 47.47538368 | -1.558738953 | 1.32E-03 |
| Fbxo36 | 335.8821186 | -1.558782686 | 5.76E-12 |
| Gm15283 | 31.78380944 | -1.559055698 | 2.44E-04 |
| Gm12963 | 65.53533855 | -1.561102786 | 8.61E-06 |
| Tcf7l1 | 1795.450597 | -1.562322918 | 8.64E-14 |
| 4930481B07Rik | 25.10703027 | -1.562724534 | 4.16E-04 |
| Gm14401 | 16.81454501 | -1.563440822 | 8.84E-03 |
| Gm16596 | 16.02890155 | -1.564073808 | 2.67E-02 |
| Ptchd1 | 55.72263066 | -1.565383536 | 2.68E-02 |
| Rufy1 | 2070.818133 | -1.566739643 | 6.18E-19 |
| Amot | 5972.569539 | -1.567242386 | 4.06E-33 |
| Slc41a2 | 949.467359 | -1.567337424 | 7.15E-21 |
| Nagk | 2436.902089 | -1.568486383 | 2.22E-42 |
| Rarres2 | 54.65008237 | -1.568561354 | 5.60E-04 |
| Snn | 2448.99706 | -1.568876104 | 3.03E-26 |
| Gm6288 | 19.23685061 | -1.570692739 | 3.13E-03 |
| Mab21l1 | 801.0403118 | -1.570872933 | 6.61E-26 |
| Lpar1 | 5879.931993 | -1.573467837 | 2.18E-27 |
| mt-Tn | 66.01413775 | -1.573636486 | 2.70E-03 |

| | | | |
|---|---|---|---|
| Gm15998 | 16.83269665 | -1.574823289 | 8.17E-03 |
| Selenop | 9438.313738 | -1.576017457 | 4.02E-07 |
| Gm16845 | 68.96377463 | -1.577535167 | 2.25E-06 |
| Ppp1r3e | 56.64665345 | -1.579946913 | 2.25E-06 |
| 9330154J02Rik | 40.21475152 | -1.580567302 | 4.77E-04 |
| Uckl1os | 18.48553068 | -1.580991727 | 3.26E-03 |
| A130051J06Rik | 19.20010434 | -1.581950339 | 4.23E-03 |
| Epm2a | 133.8997319 | -1.582903956 | 3.52E-07 |
| Ptgfrn | 7767.767633 | -1.583474011 | 7.24E-29 |
| Slpi | 339.1667458 | -1.583657839 | 2.02E-06 |
| Efnb2 | 1976.379573 | -1.585167049 | 6.34E-19 |
| Tdrd9 | 51.03735793 | -1.585355129 | 9.09E-05 |
| Gm43302 | 33.491425 | -1.585971913 | 3.61E-03 |
| Gng11 | 381.5507162 | -1.590187785 | 3.95E-11 |
| Zfp619 | 187.8261664 | -1.591108856 | 5.20E-08 |
| Adam12 | 2470.942642 | -1.591462865 | 2.42E-08 |
| Rbmx | 972.7927229 | -1.593674594 | 2.52E-17 |
| Gm14410 | 104.8186658 | -1.595240974 | 7.54E-06 |
| Il6ra | 1020.18581 | -1.595736689 | 1.20E-32 |
| Lama4 | 3042.134624 | -1.596296673 | 5.79E-03 |
| 2310015A10Rik | 393.9563693 | -1.596600277 | 1.37E-13 |
| Tlnrd1 | 1548.830688 | -1.596691478 | 1.81E-25 |
| Garem2 | 297.7203972 | -1.598142454 | 1.50E-16 |
| Endod1 | 423.568897 | -1.599027782 | 1.93E-12 |
| Zfp72 | 103.7844434 | -1.600361126 | 1.30E-04 |
| Gm22392 | 13.95865186 | -1.603270779 | 4.33E-02 |
| Zbtb1 | 1295.613615 | -1.604267306 | 2.71E-17 |
| Plat | 4991.615202 | -1.604483349 | 3.16E-13 |
| Adgrl1 | 1938.183396 | -1.606099136 | 5.34E-15 |
| Ctxn1 | 645.1595188 | -1.607047095 | 2.42E-14 |
| Tnfrsf25 | 17.04286528 | -1.607253136 | 1.04E-02 |
| Nap1l3 | 28.84721273 | -1.608491763 | 2.12E-04 |
| Ttc30b | 441.1625201 | -1.609365275 | 3.05E-24 |
| Lancl2 | 747.3264616 | -1.610613258 | 5.02E-14 |
| 0610031O16Rik | 23.83237418 | -1.610946392 | 2.93E-03 |
| Adgrb2 | 56.35235799 | -1.611396274 | 6.10E-03 |
| Cyb5rl | 377.8189539 | -1.613494482 | 6.58E-09 |
| Rab32 | 1574.14668 | -1.61374074 | 5.87E-25 |
| Nlrp1b | 59.85443185 | -1.614093233 | 2.21E-07 |
| Sfxn5 | 564.0641619 | -1.614356269 | 4.60E-22 |
| Slc12a5 | 95.98187009 | -1.615187325 | 2.27E-04 |
| Zfp469 | 681.3675061 | -1.615735198 | 1.25E-11 |
| Samd9l | 629.3467263 | -1.616777071 | 1.10E-05 |
| Col6a3 | 17819.46317 | -1.619279907 | 8.49E-14 |
| NA | 21.64261849 | -1.61950405 | 4.90E-03 |
| Polr3k | 906.3976112 | -1.620166716 | 1.34E-13 |
| Duoxa1 | 78.00167486 | -1.621141966 | 3.08E-05 |

| | | | |
|---|---|---|---|
| Gm10524 | 163.6017619 | -1.622871251 | 9.68E-11 |
| 2210408I21Rik | 146.384477 | -1.622991123 | 3.21E-17 |
| Lrrc17 | 862.6812303 | -1.623582695 | 7.25E-06 |
| Zadh2 | 2436.775149 | -1.624322627 | 2.38E-24 |
| Lss | 5106.435912 | -1.624513638 | 4.95E-06 |
| Xkr5 | 41.05272347 | -1.624920216 | 4.28E-03 |
| Psd3 | 883.8155306 | -1.625099654 | 2.91E-19 |
| Mirg | 1133.798695 | -1.625516592 | 3.33E-12 |
| Ctns | 2362.555936 | -1.62744007 | 1.84E-30 |
| Zfp65 | 459.1248073 | -1.627756793 | 1.64E-09 |
| Fstl1 | 39494.87916 | -1.628850976 | 2.87E-21 |
| Srgap1 | 1826.660683 | -1.630023086 | 1.21E-10 |
| Ccdc62 | 84.71422288 | -1.632957408 | 1.46E-06 |
| Fdps | 10188.03978 | -1.634052684 | 3.36E-07 |
| Lancl3 | 159.4956977 | -1.637162485 | 2.75E-06 |
| Zfp618 | 1231.162364 | -1.638273346 | 9.18E-08 |
| Tmem42 | 153.7982272 | -1.638472098 | 2.96E-15 |
| Olr1 | 36.00857858 | -1.638767302 | 4.82E-02 |
| Ankrd49 | 493.1015181 | -1.638783079 | 2.40E-09 |
| Gm3667 | 10.55630223 | -1.640593843 | 3.81E-02 |
| Zfp62 | 1535.939692 | -1.641915745 | 1.52E-06 |
| Cx3cl1 | 722.2112652 | -1.643585483 | 1.22E-11 |
| Zfp759 | 126.1544654 | -1.6438346 | 1.10E-06 |
| Camk2n1 | 11.24597342 | -1.643891399 | 3.95E-02 |
| Spin2c | 57.0212798 | -1.644108813 | 3.29E-07 |
| Impg2 | 34.0966657 | -1.647582166 | 1.28E-02 |
| Cavin3 | 2957.834925 | -1.648543261 | 4.19E-20 |
| Zfp652os | 10.1798594 | -1.649915896 | 3.80E-02 |
| Pdp1 | 4819.970157 | -1.650187998 | 2.51E-13 |
| Trio | 8424.77309 | -1.650763619 | 4.88E-32 |
| Itgb3 | 406.4148934 | -1.652242692 | 2.74E-06 |
| Homer1 | 244.6755818 | -1.65452258 | 2.90E-07 |
| 4930577N17Rik | 13.86405934 | -1.656169844 | 1.31E-02 |
| Cdo1 | 5428.454521 | -1.658265075 | 6.47E-07 |
| Phkb | 2588.754316 | -1.658784984 | 1.08E-30 |
| Gm11766 | 15.91984041 | -1.659853352 | 7.53E-03 |
| Mgat3 | 1523.241024 | -1.660342443 | 2.72E-25 |
| Tenm4 | 841.8870128 | -1.661775096 | 1.61E-02 |
| Sdsl | 62.01515941 | -1.661824288 | 4.03E-04 |
| NA | 132.3463999 | -1.662004673 | 3.56E-06 |
| Trim30d | 101.5243115 | -1.662620672 | 3.21E-04 |
| Mmp14 | 40429.12938 | -1.663719287 | 4.73E-23 |
| B3galt4 | 33.62953106 | -1.66544797 | 3.13E-05 |
| Col16a1 | 10534.38836 | -1.667299453 | 5.90E-18 |
| Frk | 907.1646372 | -1.668286243 | 2.74E-09 |
| Sema3b | 1221.942745 | -1.668713873 | 1.27E-09 |
| 2700081O15Rik | 2585.082579 | -1.668910501 | 4.54E-23 |

| | | | |
|---|---|---|---|
| Tanc2 | 4417.326718 | -1.669387191 | 1.45E-33 |
| Gm12205 | 9.93294853 | -1.670136919 | 2.46E-02 |
| Scube3 | 14.00198406 | -1.673041987 | 4.94E-02 |
| Snai2 | 1626.102104 | -1.673137802 | 1.42E-24 |
| Fyb | 230.8656686 | -1.674338636 | 2.01E-05 |
| Ly6c1 | 1095.469361 | -1.67464271 | 7.15E-13 |
| Prob1 | 109.771206 | -1.67577861 | 3.53E-07 |
| Wnk3 | 29.28258561 | -1.676274311 | 1.62E-02 |
| Senp8 | 238.1056629 | -1.676595146 | 2.14E-10 |
| Trim62 | 389.1707065 | -1.677008428 | 1.37E-22 |
| Nceh1 | 3203.829638 | -1.677327165 | 2.92E-21 |
| Arxes1 | 501.9970735 | -1.677576847 | 1.43E-05 |
| Glis1 | 258.3571055 | -1.67852094 | 6.67E-06 |
| NA | 14.79549638 | -1.680705558 | 5.05E-03 |
| Ank3 | 2241.447597 | -1.681545489 | 4.10E-17 |
| Galnt17 | 1070.268328 | -1.681734334 | 1.30E-21 |
| Cabyr | 15.12677381 | -1.682354384 | 5.69E-03 |
| Trps1 | 1353.888354 | -1.682507378 | 1.18E-05 |
| 2210418O10Rik | 246.2963268 | -1.683369566 | 1.95E-10 |
| Olfml3 | 3715.895388 | -1.684442542 | 2.95E-24 |
| Gm14379 | 22.4080526 | -1.684725071 | 3.83E-03 |
| Nsun7 | 86.05227179 | -1.685608293 | 2.45E-06 |
| 1700101I11Rik | 24.33272945 | -1.686272928 | 1.60E-03 |
| Gm14387 | 74.45587935 | -1.686668899 | 1.09E-05 |
| Atp8b4 | 29.3419901 | -1.687390828 | 1.14E-02 |
| Zfp994 | 554.6194131 | -1.688900203 | 1.46E-06 |
| Gm26816 | 34.25874803 | -1.689485929 | 1.08E-04 |
| Hdac9 | 113.3884376 | -1.690061951 | 8.67E-04 |
| Impact | 6753.387895 | -1.690505599 | 1.46E-26 |
| 4930506C21Rik | 19.3739159 | -1.692176114 | 2.45E-03 |
| Ifi208 | 45.86600101 | -1.694589548 | 5.86E-05 |
| Traf3ip1 | 465.2906491 | -1.694787874 | 2.19E-13 |
| NA | 21.29929916 | -1.696030335 | 1.57E-02 |
| Ang | 240.4623036 | -1.698665853 | 3.07E-13 |
| Cotl1 | 536.8515866 | -1.698753714 | 1.22E-11 |
| Thg1l | 413.4328502 | -1.700417753 | 6.83E-25 |
| Mfsd4b3 | 26.36714971 | -1.702464133 | 3.98E-04 |
| Dnmt3b | 153.2781329 | -1.704585394 | 7.50E-05 |
| Tug1 | 5617.309804 | -1.707211613 | 1.15E-10 |
| Pcdh19 | 2133.007913 | -1.707295813 | 1.16E-18 |
| Slc40a1 | 380.186994 | -1.70758563 | 1.37E-06 |
| Acat3 | 97.28512709 | -1.707877005 | 7.09E-08 |
| 9330111N05Rik | 10.97158586 | -1.70847467 | 1.63E-02 |
| Rerg | 14.52101119 | -1.709283006 | 1.14E-02 |
| Upp2 | 42.13980025 | -1.70943988 | 5.66E-06 |
| Cspg4 | 1149.427576 | -1.709990102 | 1.35E-10 |
| Gm10501 | 41.08916846 | -1.710085089 | 2.34E-04 |

| | | | |
|---|---|---|---|
| Zfp119a | 132.5293073 | -1.710694553 | 3.10E-08 |
| Tbc1d2b | 5994.726219 | -1.710742538 | 8.07E-09 |
| Bdh2 | 1089.036205 | -1.711863552 | 1.11E-13 |
| Gli2 | 250.1196828 | -1.71188782 | 3.00E-12 |
| Meg3 | 12210.99322 | -1.713460115 | 2.04E-08 |
| Tspan11 | 2916.150625 | -1.713602632 | 2.26E-19 |
| Mindy2 | 1585.001885 | -1.715513285 | 1.13E-09 |
| Il4 | 67.4342044 | -1.7160089 | 6.57E-06 |
| Snap25 | 20.82983854 | -1.720468162 | 3.58E-03 |
| Ap3m1-ps | 10.19075366 | -1.722415428 | 3.62E-02 |
| Zc3h12d | 7.659289797 | -1.724425039 | 2.78E-02 |
| Dpy19l1 | 4064.222751 | -1.727173457 | 4.70E-35 |
| Adamts14 | 1745.125069 | -1.728566532 | 3.03E-11 |
| Chadl | 33.16524172 | -1.728715098 | 6.60E-05 |
| Gm13091 | 17.69307921 | -1.730513209 | 1.19E-03 |
| Gm5518 | 20.12754576 | -1.731260878 | 7.70E-03 |
| Mdp1 | 559.3208089 | -1.731700889 | 7.45E-24 |
| Mpzl2 | 41.79260049 | -1.733139417 | 2.40E-04 |
| Zfp458 | 135.3705613 | -1.734212725 | 2.58E-07 |
| Smim17 | 13.1914216 | -1.734619338 | 3.45E-02 |
| Myo1b | 2440.909108 | -1.734896005 | 9.70E-24 |
| Dhx9 | 8577.897448 | -1.735057235 | 5.19E-15 |
| Tmem179 | 20.54769501 | -1.735566013 | 2.53E-02 |
| Lekr1 | 154.8880671 | -1.736098954 | 8.19E-05 |
| St3gal5 | 2651.830812 | -1.738151429 | 7.60E-39 |
| Gm10139 | 66.38850078 | -1.743008593 | 1.91E-03 |
| Tmem260 | 658.9304686 | -1.743953534 | 1.34E-30 |
| Elac1 | 354.5344417 | -1.745849917 | 1.29E-16 |
| Zfp729a | 400.7358496 | -1.746533018 | 6.52E-08 |
| Lyve1 | 9.828957395 | -1.748280159 | 2.78E-02 |
| Mir186 | 9.472613992 | -1.753505701 | 1.65E-02 |
| NA | 20.64834532 | -1.754553959 | 7.70E-04 |
| Rnf152 | 9.581791921 | -1.754692497 | 1.43E-02 |
| Ptpn3 | 659.1680817 | -1.755860809 | 5.33E-24 |
| Ctnnb1 | 26830.96708 | -1.756935465 | 2.45E-88 |
| Sptssa | 1180.152327 | -1.757513615 | 3.81E-21 |
| Mast4 | 3107.692726 | -1.757880292 | 6.94E-33 |
| Tiam2 | 1483.526478 | -1.758197849 | 6.93E-14 |
| Pcdhb17 | 304.6149099 | -1.758883735 | 4.54E-24 |
| Tmem117 | 124.9016676 | -1.759083539 | 4.42E-08 |
| Adhfe1 | 58.16465703 | -1.759152794 | 7.55E-08 |
| Gm14127 | 12.86916418 | -1.759673543 | 1.86E-02 |
| Gas2 | 879.7185941 | -1.76266057 | 9.35E-14 |
| Hvcn1 | 604.403497 | -1.763504633 | 1.06E-21 |
| Gm10544 | 12.32905783 | -1.765665086 | 5.96E-03 |
| Tnfrsf14 | 35.33998753 | -1.765692472 | 5.97E-06 |
| Tmem191c | 138.1462665 | -1.768346305 | 7.07E-10 |

| | | | |
|---|---|---|---|
| Pcdhb22 | 261.1195135 | -1.770278085 | 1.08E-22 |
| Gm9796 | 10.08008836 | -1.770728378 | 1.27E-02 |
| Gm26923 | 29.45211618 | -1.77123435 | 2.33E-03 |
| Insig1 | 7189.3603 | -1.771372315 | 1.31E-11 |
| Plagl1 | 737.429685 | -1.771380949 | 2.03E-09 |
| Sfrp1 | 704.6605608 | -1.772771511 | 1.47E-11 |
| Faddos | 6.170598786 | -1.776042291 | 4.57E-02 |
| Miga2 | 802.4941053 | -1.779044951 | 7.21E-18 |
| Gm8615 | 190.0903932 | -1.779575023 | 1.57E-12 |
| Gm12580 | 9.951218649 | -1.77988307 | 2.24E-02 |
| Gm10548 | 14.20004954 | -1.780422145 | 2.22E-02 |
| 1110002J07Rik | 12.818549 | -1.780423935 | 4.89E-03 |
| Il11ra2 | 54.33013918 | -1.782719283 | 5.46E-05 |
| Dbi | 3225.641979 | -1.785286962 | 2.50E-53 |
| Myo1h | 414.7993453 | -1.785762559 | 4.95E-18 |
| Rsrp1 | 7585.578583 | -1.788303406 | 5.82E-12 |
| Tmem216 | 308.3576184 | -1.78912638 | 4.35E-29 |
| Nab2 | 4607.437328 | -1.792763554 | 2.01E-43 |
| Ttc28 | 3338.3816 | -1.792889288 | 2.45E-28 |
| Erg28 | 1761.041711 | -1.793801804 | 6.22E-20 |
| Ctso | 1002.433076 | -1.797724399 | 1.48E-33 |
| Tctn2 | 715.0863891 | -1.800190365 | 1.59E-11 |
| Sh3pxd2b | 11222.03918 | -1.800773854 | 1.49E-19 |
| Slc5a3 | 5155.170795 | -1.801003257 | 3.05E-17 |
| Ccdc39 | 190.5301171 | -1.801362222 | 1.92E-07 |
| Ror2 | 442.1456838 | -1.802313472 | 7.29E-19 |
| Gm26771 | 21.68010433 | -1.803260435 | 5.70E-04 |
| Fer1l6 | 38.79742372 | -1.806903356 | 3.86E-05 |
| Atoh8 | 455.8228685 | -1.80763507 | 2.35E-03 |
| Slc9a4 | 10.64877529 | -1.80796371 | 1.86E-02 |
| Zscan26 | 1306.932827 | -1.808980768 | 2.14E-12 |
| Gm4631 | 911.8427902 | -1.809320746 | 1.78E-05 |
| Far1os | 17.20304771 | -1.812664774 | 1.98E-02 |
| 2010315B03Rik | 196.2225042 | -1.8131282 | 6.19E-06 |
| E130102H24Rik | 90.14504892 | -1.814792401 | 4.64E-05 |
| Trip11 | 1539.402913 | -1.815256228 | 1.83E-07 |
| Gfra1 | 41.83640276 | -1.815738218 | 5.39E-03 |
| Dleu2 | 153.9845958 | -1.816967094 | 1.05E-11 |
| Eml1 | 3052.707289 | -1.818074307 | 1.12E-35 |
| Gm3739 | 55.08943698 | -1.822411788 | 2.50E-06 |
| Pcdhb8 | 54.15088343 | -1.822879011 | 2.07E-08 |
| Anks6 | 655.9616026 | -1.822958221 | 1.23E-12 |
| Zfp454 | 65.32644374 | -1.824753016 | 5.40E-10 |
| Gm14440 | 165.1266027 | -1.826289562 | 2.86E-03 |
| Nkain1 | 814.8730286 | -1.829272034 | 3.95E-27 |
| Csf2rb2 | 11.60028118 | -1.831783893 | 1.12E-02 |
| Zfp354b | 32.87930447 | -1.833295541 | 6.17E-04 |

| | | | |
|---|---|---|---|
| Myl4 | 52.50706572 | -1.837945066 | 4.32E-08 |
| Atxn1 | 1217.972466 | -1.838797787 | 1.12E-17 |
| Fkrp | 819.6783063 | -1.839129952 | 4.75E-17 |
| Gm7162 | 14.15814186 | -1.840698321 | 1.31E-02 |
| Kif26b | 704.3016406 | -1.841939959 | 6.93E-10 |
| Bsn | 48.5312342 | -1.844220151 | 2.86E-08 |
| Gm16759 | 14.23985702 | -1.846587219 | 4.96E-02 |
| Sertad4 | 988.7002916 | -1.847446828 | 5.16E-07 |
| 2500004C02Rik | 115.4313729 | -1.847794441 | 9.64E-06 |
| Pcdhb4 | 29.69585276 | -1.851019052 | 6.68E-06 |
| Prdm11 | 475.2305451 | -1.851092953 | 1.75E-28 |
| Gm10638 | 16.6363986 | -1.853329705 | 4.95E-03 |
| Adam5 | 19.51957787 | -1.854134684 | 1.34E-03 |
| Nme3 | 171.0026249 | -1.854391193 | 2.19E-17 |
| Rasa1 | 2097.985419 | -1.854894554 | 9.07E-12 |
| Hsd17b7 | 1863.090416 | -1.855258901 | 1.30E-09 |
| Zfp629 | 757.4171943 | -1.855396551 | 3.15E-18 |
| Lrrn3 | 42.94924044 | -1.856118233 | 4.27E-04 |
| Cd200 | 115.5327228 | -1.858557553 | 2.46E-06 |
| 9-Sep | 7495.015017 | -1.858883771 | 5.35E-38 |
| Gm26690 | 12.97637611 | -1.859451795 | 2.03E-02 |
| Osbpl10 | 87.89074291 | -1.860898324 | 1.12E-08 |
| Cped1 | 8743.703581 | -1.862068477 | 3.68E-20 |
| Tmem119 | 3163.057139 | -1.863891399 | 1.59E-21 |
| Klrb1c | 12.08738048 | -1.863926433 | 1.12E-02 |
| Rab36 | 271.0864376 | -1.86622363 | 3.62E-12 |
| Gm26645 | 20.70438726 | -1.867446458 | 2.22E-04 |
| Ypel2 | 260.1805721 | -1.868732627 | 9.13E-11 |
| Dhcr24 | 6989.867164 | -1.869011103 | 2.36E-09 |
| Pitx2 | 8.309447683 | -1.869226393 | 3.83E-02 |
| Ifi202b | 2370.951925 | -1.870990654 | 1.54E-04 |
| Arxes2 | 825.7504101 | -1.87767149 | 2.14E-13 |
| Gm16861 | 183.4760558 | -1.879414262 | 9.96E-19 |
| Gas1 | 20046.86774 | -1.879652249 | 1.13E-22 |
| Tia1 | 2607.982324 | -1.879744366 | 9.30E-11 |
| Synpo | 5453.723669 | -1.879799172 | 3.62E-26 |
| Zfp14 | 70.96744208 | -1.88116923 | 1.40E-11 |
| Gm26688 | 12.27901547 | -1.881172951 | 1.17E-02 |
| Scg2 | 1103.15175 | -1.883452902 | 4.54E-03 |
| Zfp113 | 518.8178997 | -1.883912398 | 2.75E-07 |
| Apoh | 10.70072802 | -1.886551387 | 1.07E-02 |
| mt-Tc | 174.937795 | -1.887861435 | 4.06E-06 |
| Fgfr1 | 26119.72218 | -1.888512099 | 3.50E-31 |
| Gm26829 | 19.23961468 | -1.890917971 | 4.02E-05 |
| Gm14421 | 160.9356899 | -1.893197967 | 7.55E-09 |
| Sult1a1 | 55.76000516 | -1.894356492 | 1.03E-03 |
| Msmo1 | 8301.39548 | -1.894646133 | 6.90E-10 |

| | | | |
|---|---|---|---|
| Klf12 | 71.36571759 | -1.896765186 | 2.70E-12 |
| Gpr75 | 16.08289918 | -1.902059853 | 8.18E-04 |
| Bag5 | 1327.489717 | -1.905544723 | 1.29E-18 |
| Rab3il1 | 19.55437173 | -1.906273092 | 1.14E-02 |
| Scrn1 | 542.6863496 | -1.908705329 | 1.95E-09 |
| Gata2 | 333.1625954 | -1.909503793 | 2.85E-19 |
| Snx9 | 6913.287897 | -1.909832322 | 1.69E-43 |
| Fgf2 | 34.6754416 | -1.910836715 | 1.35E-02 |
| Ispd | 67.05220787 | -1.911673592 | 2.79E-07 |
| C130071C03Rik | 75.01846086 | -1.914315439 | 8.52E-10 |
| Zfp966 | 23.05339885 | -1.915551562 | 2.21E-02 |
| Foxl1 | 95.73223871 | -1.915926828 | 1.08E-10 |
| Lrrc8d | 953.5605398 | -1.916000549 | 7.33E-24 |
| Pkd2 | 8695.457457 | -1.91873669 | 3.04E-20 |
| Etaa1os | 11.32037963 | -1.919215168 | 6.96E-03 |
| Lifr | 229.2415172 | -1.919722061 | 9.34E-17 |
| Glb1l | 1400.038872 | -1.919935215 | 3.35E-30 |
| Ang2 | 58.05561455 | -1.924849779 | 2.13E-05 |
| Smco3 | 15.58925897 | -1.925665116 | 4.25E-03 |
| Fcrlb | 50.76172523 | -1.925968125 | 4.15E-05 |
| Nat8f1 | 120.6153721 | -1.933980177 | 9.39E-08 |
| Gm13305 | 188.831561 | -1.9350406 | 2.74E-13 |
| Nat8f6 | 9.726106815 | -1.935417992 | 1.66E-02 |
| Gm20319 | 6.961059712 | -1.937136548 | 2.14E-02 |
| NA | 10.88839713 | -1.937430652 | 1.16E-02 |
| Ankrd44 | 1362.280154 | -1.939553336 | 9.78E-27 |
| Mycn | 50.26697572 | -1.940384676 | 9.21E-05 |
| Zfp873 | 157.1558645 | -1.943278684 | 8.67E-10 |
| Ttyh1 | 71.46831245 | -1.944734059 | 1.38E-03 |
| Tmem254a | 133.019222 | -1.945381869 | 6.46E-06 |
| NA | 35.3883632 | -1.945481766 | 5.53E-07 |
| Tmem98 | 1290.414908 | -1.94658769 | 1.11E-43 |
| Gm11816 | 245.2771472 | -1.946980947 | 2.49E-11 |
| Gm26540 | 8.589600188 | -1.947777913 | 1.18E-02 |
| Jade2 | 1489.987394 | -1.94794066 | 8.74E-67 |
| Gp5 | 7.194843487 | -1.948275627 | 1.21E-02 |
| Gm2897 | 18.49746913 | -1.949175148 | 3.18E-02 |
| Gm5432 | 8.334657074 | -1.949384821 | 2.56E-02 |
| Setmar | 64.67531696 | -1.94961108 | 2.10E-12 |
| Lmnb2 | 823.7442095 | -1.949801323 | 1.01E-05 |
| Angpt4 | 781.819534 | -1.951226914 | 3.31E-06 |
| Hmga2 | 10182.13318 | -1.956606822 | 5.97E-14 |
| Zfp300 | 75.47022026 | -1.956990125 | 7.71E-07 |
| Nsg1 | 1730.070035 | -1.957410947 | 5.11E-44 |
| Fhod3 | 6.890644467 | -1.959764004 | 4.72E-02 |
| Sparcl1 | 1663.612457 | -1.960546511 | 2.35E-07 |
| Etv1 | 303.4475497 | -1.96133158 | 1.66E-07 |

| | | | |
|---|---|---|---|
| Zfhx4 | 9921.32866 | -1.963024128 | 1.83E-11 |
| Hddc3 | 237.1640719 | -1.963406343 | 3.28E-15 |
| Tpcn2 | 753.9566622 | -1.96452916 | 2.31E-58 |
| Rhoj | 1534.443716 | -1.964923093 | 1.17E-26 |
| BC065403 | 92.11563179 | -1.96599061 | 8.26E-13 |
| Rhou | 4021.987846 | -1.967076578 | 4.08E-33 |
| Unc93a2 | 67.49710805 | -1.96803293 | 1.71E-03 |
| Gsdme | 190.6699199 | -1.968476095 | 4.07E-11 |
| Dlk1 | 29943.381 | -1.96988813 | 3.64E-08 |
| Mecom | 1419.704379 | -1.971485858 | 2.89E-33 |
| NA | 45.07335818 | -1.973931699 | 2.05E-04 |
| Tm7sf2 | 703.3797047 | -1.975301817 | 4.49E-14 |
| 1700006J14Rik | 16.77111565 | -1.977538103 | 8.94E-03 |
| Gm14124 | 65.72396806 | -1.979883047 | 6.46E-08 |
| Gm11767 | 9.219788467 | -1.980903675 | 5.02E-03 |
| Mgst2 | 162.1947575 | -1.981200988 | 5.00E-14 |
| Pik3r3 | 1442.838135 | -1.981242621 | 1.77E-27 |
| Dclk1 | 6246.021643 | -1.98410815 | 2.43E-43 |
| 4933440N22Rik | 14.45191676 | -1.989587076 | 3.96E-03 |
| A430108G06Rik | 8.933382318 | -1.99129946 | 1.14E-02 |
| Mrgpre | 112.8488705 | -1.992128829 | 4.67E-08 |
| Rgs16 | 180.0814537 | -1.992364528 | 2.01E-08 |
| Pde3a | 117.8255858 | -1.997401106 | 4.55E-13 |
| Gm20717 | 8.660994201 | -1.997827769 | 2.64E-02 |
| 4930547M16Rik | 12.48702434 | -1.99803933 | 1.04E-02 |
| 2810403D21Rik | 60.92745888 | -2.004997159 | 8.02E-08 |
| Itih5 | 10.38686915 | -2.005886082 | 1.71E-02 |
| 2510009E07Rik | 671.3141264 | -2.007470979 | 7.00E-23 |
| Mrgprf | 1953.146531 | -2.008696542 | 1.25E-14 |
| Mxra8os | 7.148754445 | -2.010513029 | 1.37E-02 |
| Larp6 | 54.75592821 | -2.014489412 | 2.62E-09 |
| Plcb1 | 374.5524231 | -2.014863958 | 4.29E-22 |
| 9430002A10Rik | 10.20621605 | -2.015239486 | 9.59E-03 |
| Xpnpep2 | 463.2230938 | -2.016067074 | 3.42E-12 |
| Frg2f1 | 213.8372542 | -2.016100088 | 1.21E-22 |
| Chsy1 | 1508.249314 | -2.016193465 | 6.86E-58 |
| Rragd | 324.476639 | -2.016988282 | 2.34E-09 |
| Kcnk2 | 223.8511014 | -2.020837802 | 1.32E-15 |
| Slc25a45 | 460.588334 | -2.022453999 | 7.89E-32 |
| Rnf150 | 1744.58427 | -2.025776358 | 2.74E-33 |
| Mef2c | 246.1899401 | -2.030661233 | 1.16E-06 |
| Nsdhl | 4567.873814 | -2.030943304 | 4.41E-14 |
| Gm14322 | 342.7270789 | -2.031653368 | 9.13E-24 |
| Vav3 | 109.5812845 | -2.040543715 | 1.32E-10 |
| Dnase1 | 109.3277492 | -2.040890348 | 1.83E-13 |
| Gm4425 | 83.20614124 | -2.046097859 | 3.66E-12 |
| Gm24365 | 7.040799217 | -2.047708872 | 2.96E-02 |

| | | | |
|---|---|---|---|
| 1700001K19Rik | 6.929533922 | -2.049748029 | 4.41E-02 |
| Fcor | 26.09012877 | -2.049984571 | 4.09E-05 |
| Spry4 | 239.4302629 | -2.051104785 | 7.55E-13 |
| Akr1c14 | 1643.65333 | -2.051640123 | 8.21E-08 |
| Ptgfr | 515.8706601 | -2.052760655 | 1.71E-12 |
| Hgf | 466.3773992 | -2.053465327 | 2.01E-11 |
| Gm15758 | 33.86750818 | -2.056643564 | 9.61E-06 |
| Gm26587 | 44.99508691 | -2.059081128 | 3.81E-04 |
| 4833422C13Rik | 31.02501048 | -2.061994135 | 1.38E-04 |
| S100a4 | 42457.13509 | -2.063543529 | 2.14E-15 |
| Galnt4 | 1123.766541 | -2.070384817 | 4.02E-24 |
| Gm14412 | 35.41472427 | -2.071259937 | 6.81E-04 |
| Tvp23a | 784.6639239 | -2.071354729 | 1.75E-21 |
| Fgfr2 | 2705.677105 | -2.073982087 | 3.68E-23 |
| Zfp977 | 12.83380859 | -2.074325383 | 4.89E-03 |
| Zfp607a | 91.92954207 | -2.076553071 | 2.97E-12 |
| C330011M18Rik | 41.66936782 | -2.076803729 | 7.04E-08 |
| Il16 | 43.63478451 | -2.080103305 | 8.01E-07 |
| Ccdc74a | 117.2288527 | -2.081819398 | 9.04E-11 |
| Stab2 | 34.5652292 | -2.082025435 | 2.54E-05 |
| NA | 7.740859217 | -2.083555108 | 2.81E-02 |
| Ppargc1a | 542.9043571 | -2.085787679 | 3.07E-09 |
| Itgb2 | 505.7153736 | -2.086812745 | 1.40E-14 |
| D630036H23Rik | 6.049307055 | -2.086875055 | 4.15E-02 |
| Oaf | 3976.294308 | -2.089482781 | 2.05E-29 |
| Bmper | 999.7225729 | -2.089828381 | 1.47E-11 |
| Klhl41 | 8.79281984 | -2.091385214 | 2.05E-02 |
| Gm10330 | 164.1859516 | -2.094764146 | 2.07E-10 |
| Dact1 | 476.0212076 | -2.096030065 | 5.96E-10 |
| 2700033N17Rik | 66.80711653 | -2.101227732 | 6.65E-11 |
| Gm14002 | 28.28916881 | -2.101953048 | 5.37E-07 |
| A930024E05Rik | 70.97068599 | -2.104816505 | 5.07E-11 |
| 3110070M22Rik | 24.64606842 | -2.105123948 | 1.99E-05 |
| Pianp | 1185.174268 | -2.108288345 | 3.54E-27 |
| Zscan30 | 5.346686754 | -2.113593715 | 4.79E-02 |
| Trpv3 | 7.69065872 | -2.114509348 | 1.28E-02 |
| NA | 11.41496147 | -2.114831701 | 2.22E-03 |
| Id3 | 1711.956127 | -2.115888245 | 7.96E-05 |
| Dmp1 | 15.30594467 | -2.116315703 | 3.68E-03 |
| Gm26945 | 12.42640804 | -2.117474642 | 9.93E-03 |
| Bcas3os1 | 7.263128961 | -2.123204034 | 2.03E-02 |
| Gm21847 | 17.84747835 | -2.123237722 | 1.78E-02 |
| Sycp2 | 67.3916275 | -2.125581551 | 8.99E-05 |
| Zfp704 | 652.3489939 | -2.126437316 | 2.88E-16 |
| Tgfb3 | 3717.223368 | -2.12764104 | 6.19E-31 |
| Gpr137b | 6771.907523 | -2.130102761 | 1.15E-17 |
| Il1rap | 2529.832075 | -2.130754467 | 2.20E-34 |

| | | | |
|---|---|---|---|
| Gm2381 | 276.5731117 | -2.130789388 | 2.14E-06 |
| Ces1a | 829.2986385 | -2.136967147 | 1.12E-04 |
| 5031410I06Rik | 317.2038905 | -2.142621567 | 4.59E-06 |
| Lef1 | 146.4483945 | -2.151836336 | 1.79E-07 |
| Matn2 | 13519.17868 | -2.153307551 | 7.70E-22 |
| Rian | 11881.87867 | -2.15461399 | 3.07E-31 |
| Inpp4b | 39.07537712 | -2.155958897 | 3.86E-04 |
| Mir351 | 18.63857239 | -2.15825577 | 1.47E-03 |
| Pcdhb16 | 566.0208578 | -2.158435408 | 1.28E-35 |
| Insyn1 | 1056.11518 | -2.160527485 | 2.35E-29 |
| Gm26763 | 7.555053541 | -2.164048576 | 3.52E-02 |
| Itga8 | 1363.794526 | -2.165367584 | 8.04E-13 |
| Adh7 | 1804.860141 | -2.165586775 | 2.14E-16 |
| Ptk7 | 1571.76109 | -2.166578965 | 2.21E-31 |
| Gm17057 | 15.87823374 | -2.16945073 | 9.10E-05 |
| Id2 | 912.7516454 | -2.169604125 | 3.07E-04 |
| 4930570G19Rik | 12.97709924 | -2.17229013 | 1.98E-02 |
| Eps8 | 11574.16555 | -2.172328254 | 2.02E-58 |
| Gm10643 | 42.40207878 | -2.172690115 | 7.94E-09 |
| Zfp738 | 241.9782701 | -2.175122536 | 2.10E-10 |
| Slc12a1 | 5.052709889 | -2.178341367 | 4.73E-02 |
| Mmp2 | 89388.04831 | -2.178672009 | 3.79E-17 |
| Gm20633 | 69.1003007 | -2.181218816 | 1.66E-08 |
| Gm3629 | 23.23421655 | -2.186728463 | 2.35E-02 |
| 2610035D17Rik | 96.21859554 | -2.194741457 | 6.59E-07 |
| Ccdc151 | 30.69612096 | -2.195421619 | 8.61E-06 |
| Gm27030 | 6.892780479 | -2.197127163 | 2.50E-02 |
| Cd33 | 68.59571646 | -2.197342825 | 5.58E-08 |
| Nanos3 | 11.84376575 | -2.203266251 | 9.82E-04 |
| Tmem106a | 958.7992917 | -2.205937001 | 2.45E-39 |
| Ccdc87 | 25.43668448 | -2.2064397 | 3.30E-06 |
| 5930403N24Rik | 408.5526991 | -2.207811393 | 1.41E-13 |
| Gm26840 | 19.51410435 | -2.20860391 | 3.06E-04 |
| Maml3 | 728.8511573 | -2.210888382 | 1.41E-27 |
| 2810039B14Rik | 112.6447876 | -2.214352655 | 2.57E-24 |
| Gm10220 | 11.21547464 | -2.214949805 | 3.02E-02 |
| Gm1979 | 16.54561795 | -2.215546194 | 8.03E-04 |
| Pcdhb14 | 233.1268323 | -2.216131685 | 6.71E-21 |
| Nxph3 | 56.41829472 | -2.216356359 | 3.38E-08 |
| Arhgef3 | 59.83266333 | -2.219661049 | 9.06E-09 |
| Fam3c | 4044.640668 | -2.224184182 | 1.46E-64 |
| Ptgds | 167.4192768 | -2.224760462 | 4.64E-08 |
| C2cd4c | 10.15695358 | -2.22899008 | 8.73E-03 |
| Mrnip | 63.17783418 | -2.234016732 | 4.89E-10 |
| Gxylt2 | 1431.643095 | -2.237561257 | 9.90E-17 |
| C1ra | 10143.28128 | -2.239746045 | 5.13E-23 |
| A230028O05Rik | 24.80296605 | -2.240527879 | 1.72E-04 |

| | | | |
|---|---|---|---|
| Tet1 | 358.3477956 | -2.242580708 | 1.03E-14 |
| Pcdhb3 | 9.868502243 | -2.243718588 | 2.48E-03 |
| Tbc1d9 | 40.79818821 | -2.247600929 | 4.82E-06 |
| Gm10604 | 15.18683589 | -2.247887282 | 8.65E-04 |
| A330009N23Rik | 54.46191423 | -2.2487151 | 1.18E-10 |
| Pik3r1 | 5071.593751 | -2.249577198 | 3.07E-33 |
| Gm4724 | 16.04273291 | -2.250305759 | 2.24E-02 |
| Aox3 | 125.6643492 | -2.250998606 | 5.28E-04 |
| Tmem178b | 75.53419669 | -2.252094726 | 4.78E-06 |
| NA | 141.1068307 | -2.254133862 | 2.45E-14 |
| Jrk | 382.3720798 | -2.254165706 | 4.27E-26 |
| Gm15298 | 211.5333334 | -2.256725498 | 3.70E-18 |
| Ank | 9508.851247 | -2.259013597 | 5.93E-86 |
| Col27a1 | 963.2021591 | -2.260590845 | 1.41E-24 |
| Zfp709 | 209.956385 | -2.263961361 | 3.83E-16 |
| C530008M17Rik | 222.325705 | -2.266644059 | 2.62E-09 |
| Myo7a | 1099.229425 | -2.272813559 | 1.55E-29 |
| Gm7638 | 5.959151382 | -2.27423964 | 4.35E-02 |
| Carmil3 | 108.3115967 | -2.275939685 | 2.67E-10 |
| Sox9 | 2547.006272 | -2.279157243 | 1.21E-22 |
| Zfp125 | 122.523314 | -2.280092375 | 4.06E-16 |
| Gm16046 | 54.5878734 | -2.282004901 | 9.44E-05 |
| Negr1 | 66.82959444 | -2.28256848 | 1.18E-11 |
| 2700069I18Rik | 246.3421382 | -2.282746832 | 1.70E-10 |
| Eef1akmt2 | 275.4335541 | -2.285052669 | 3.95E-08 |
| Mylip | 3518.788342 | -2.285714389 | 1.17E-24 |
| Gm26682 | 24.22475906 | -2.286802559 | 1.24E-05 |
| Hmgcs1 | 10146.3423 | -2.286803576 | 9.73E-11 |
| Prokr1 | 338.3992619 | -2.287085041 | 6.04E-13 |
| Apod | 983.8203818 | -2.288791634 | 5.89E-05 |
| 5033430I15Rik | 56.08870669 | -2.28908759 | 6.24E-11 |
| Trnp1 | 989.6323959 | -2.290386047 | 1.53E-31 |
| Thrb | 322.1208178 | -2.292020384 | 6.66E-15 |
| Zkscan8 | 842.0575177 | -2.29254616 | 3.82E-13 |
| Gbp2 | 472.493555 | -2.299328031 | 6.66E-26 |
| Sh3kbp1 | 3211.059854 | -2.300924022 | 1.02E-83 |
| Nat8f4 | 100.8466667 | -2.302070635 | 2.74E-09 |
| Aox1 | 478.0590306 | -2.303693039 | 9.75E-29 |
| Sardh | 902.4568354 | -2.303901926 | 4.46E-21 |
| Fam198a | 33.40205875 | -2.303958238 | 3.85E-09 |
| Ccbe1 | 1756.029322 | -2.305616383 | 1.66E-22 |
| Uqcc3 | 436.9863654 | -2.305655999 | 1.93E-30 |
| Idi1 | 4535.218215 | -2.307522283 | 3.27E-05 |
| Acat2 | 6980.262021 | -2.316879879 | 1.26E-17 |
| Plp1 | 415.718745 | -2.321791236 | 3.98E-09 |
| Ebf1 | 2433.234753 | -2.324358436 | 6.40E-15 |
| Trhde | 38.6088199 | -2.329404628 | 1.54E-05 |

| | | | |
|---|---|---|---|
| Atp1b1 | 1462.526931 | -2.335641304 | 2.09E-29 |
| Plekha4 | 436.889388 | -2.336373132 | 8.92E-15 |
| Tmem268 | 2601.301169 | -2.338625644 | 9.63E-30 |
| Cmtm3 | 7129.083408 | -2.342412582 | 6.82E-48 |
| 5730507C01Rik | 114.2588328 | -2.347470614 | 5.38E-16 |
| Prkg2 | 2708.349054 | -2.348164212 | 7.67E-13 |
| P2rx4 | 2480.638167 | -2.349771098 | 1.09E-35 |
| Duox1 | 119.8789528 | -2.361786058 | 3.05E-11 |
| Sdc3 | 1488.655788 | -2.36483346 | 1.26E-28 |
| Col17a1 | 62.0117311 | -2.364911287 | 9.12E-08 |
| Gm11851 | 45.809336 | -2.365539924 | 3.77E-08 |
| Grem2 | 1174.534392 | -2.369066018 | 4.29E-20 |
| Fxyd1 | 14.72790562 | -2.374356901 | 1.45E-02 |
| Gm11638 | 6.989282699 | -2.375551226 | 4.09E-03 |
| Fam169a | 9.893249726 | -2.381649687 | 5.91E-03 |
| Catip | 21.30652682 | -2.385777946 | 6.97E-05 |
| Tnr | 97.7195546 | -2.388878612 | 2.01E-05 |
| Rtl5 | 11.39763817 | -2.394597366 | 3.52E-02 |
| Pgm5 | 58.86253734 | -2.397563015 | 3.66E-04 |
| Rab27a | 205.8409117 | -2.403280614 | 4.24E-17 |
| Rbm15b | 151.7920532 | -2.403692606 | 2.58E-03 |
| Gm13502 | 199.0616539 | -2.407006853 | 3.47E-02 |
| Gas7 | 13591.79219 | -2.407390016 | 3.87E-28 |
| Abca9 | 166.1965339 | -2.413943076 | 5.96E-09 |
| Vash1 | 29.13997177 | -2.419537576 | 3.56E-05 |
| Dlx4 | 6.030862526 | -2.4199864 | 1.33E-02 |
| Ccnb1ip1 | 16.25829992 | -2.422510179 | 9.87E-06 |
| Tgfbr1 | 1819.850756 | -2.422593261 | 1.10E-21 |
| Flvcr2 | 5.065362858 | -2.422895976 | 4.70E-02 |
| Rnf144a | 1573.389355 | -2.435639048 | 9.82E-13 |
| Gla | 4390.097848 | -2.436930506 | 1.21E-33 |
| Casq2 | 10.12003193 | -2.438796395 | 8.95E-05 |
| Mc2r | 13.56370526 | -2.439745888 | 1.68E-03 |
| Kdelc2 | 3012.784827 | -2.440159576 | 3.21E-45 |
| Ociad2 | 337.1734645 | -2.440692609 | 6.26E-35 |
| Galnt13 | 425.0375608 | -2.441560657 | 9.34E-37 |
| E230013L22Rik | 57.3366247 | -2.441944633 | 2.84E-14 |
| Igsf3 | 583.6384486 | -2.442085796 | 1.03E-34 |
| Rin2 | 6488.596361 | -2.442521415 | 2.33E-32 |
| C1s2 | 1092.038141 | -2.447459212 | 6.91E-15 |
| Dlec1 | 30.45660228 | -2.450828622 | 3.69E-05 |
| Fam13c | 1244.440345 | -2.454284117 | 2.38E-25 |
| Ccdc152 | 31.66243982 | -2.457462791 | 1.57E-06 |
| 5830432E09Rik | 23.64403577 | -2.458983572 | 3.85E-06 |
| Pcdhb5 | 354.373418 | -2.461748984 | 3.60E-36 |
| Islr | 180.9664839 | -2.463303923 | 3.76E-06 |
| 4933431K23Rik | 11.94158742 | -2.463722156 | 1.78E-03 |

| | | | |
|---|---|---|---|
| Gm13470 | 63.86688211 | -2.463785465 | 2.91E-13 |
| Gm3194 | 17.06507034 | -2.464040799 | 3.76E-04 |
| Nacc2 | 1412.305969 | -2.466402821 | 1.47E-54 |
| Gm13836 | 8.255389852 | -2.471576515 | 3.72E-03 |
| Plekhb1 | 22.62480202 | -2.475371536 | 6.83E-04 |
| Gm21698 | 27.37379651 | -2.478532049 | 4.45E-04 |
| Lpar4 | 27.03383507 | -2.480232131 | 5.37E-04 |
| 4931428F04Rik | 479.9064024 | -2.492161209 | 5.58E-29 |
| Trem2 | 96.77326875 | -2.50227161 | 1.35E-12 |
| Pigh | 312.2573961 | -2.503251815 | 4.80E-34 |
| Mitf | 1042.401336 | -2.510092376 | 2.02E-27 |
| NA | 9.46447353 | -2.513353626 | 5.39E-03 |
| Foxred2 | 88.56860549 | -2.517547644 | 9.08E-08 |
| Serpinb8 | 975.2511855 | -2.518978861 | 7.22E-22 |
| Ccdc148 | 58.82159294 | -2.520949199 | 2.85E-05 |
| Tnfaip6 | 646.6209697 | -2.521595542 | 7.91E-26 |
| Sort1 | 907.944398 | -2.525819523 | 3.18E-18 |
| Cdh1 | 10.89184837 | -2.526350414 | 8.34E-03 |
| C1rb | 431.70044 | -2.535877973 | 4.11E-23 |
| Ptger4 | 492.752494 | -2.540928625 | 1.26E-35 |
| Rep15 | 19.80358759 | -2.541236063 | 1.13E-06 |
| Colec12 | 20930.91238 | -2.552329496 | 1.76E-79 |
| Dpt | 3000.84147 | -2.56185877 | 1.65E-05 |
| Nat8f7 | 24.83797719 | -2.56735191 | 1.72E-04 |
| 1700024B05Rik | 11.97341044 | -2.572204809 | 5.10E-03 |
| Adgrl3 | 128.3042568 | -2.575097746 | 3.34E-11 |
| Osbpl6 | 120.9530659 | -2.577792059 | 4.22E-10 |
| Gm18360 | 8.20057758 | -2.579127831 | 6.43E-03 |
| Il17rd | 972.8019314 | -2.585344337 | 3.15E-24 |
| Dkk3 | 1119.94565 | -2.589102821 | 5.58E-25 |
| Pcdhb20 | 156.4655691 | -2.590766697 | 4.78E-25 |
| C1qtnf2 | 17.27844736 | -2.593026491 | 1.90E-02 |
| Gm22935 | 8.206150922 | -2.596924813 | 2.06E-02 |
| Elovl6 | 3066.198611 | -2.598828353 | 4.01E-07 |
| 9630013D21Rik | 65.25458142 | -2.600756367 | 1.07E-05 |
| Gm11407 | 5.515355237 | -2.609174153 | 1.94E-02 |
| Dcn | 41417.76249 | -2.60937747 | 1.02E-06 |
| Gm7361 | 115.296168 | -2.609713023 | 2.54E-08 |
| Cfap161 | 19.83244907 | -2.611325042 | 6.95E-04 |
| Zan | 76.59000702 | -2.617910434 | 7.03E-06 |
| C1s1 | 15631.97993 | -2.619620949 | 5.50E-21 |
| Pcdhb2 | 19.19826051 | -2.632412069 | 1.26E-06 |
| A430035B10Rik | 7.243714529 | -2.648317255 | 2.84E-03 |
| Acot11 | 161.2842215 | -2.653570082 | 1.18E-08 |
| Prtg | 275.2554638 | -2.659341747 | 1.74E-23 |
| Rnft2 | 369.3814148 | -2.660923961 | 5.75E-22 |
| Dio2 | 87.5272909 | -2.664395095 | 4.13E-10 |

| | | | |
|---|---|---|---|
| Zkscan16 | 70.2059796 | -2.664972412 | 1.11E-11 |
| Bcl2 | 622.3489031 | -2.666523175 | 1.03E-16 |
| Apba1 | 301.8235088 | -2.666808445 | 4.32E-25 |
| Il1rn | 36.407963 | -2.673753975 | 4.08E-04 |
| 3222401L13Rik | 7.347962629 | -2.691126914 | 2.79E-03 |
| Aig1 | 239.7079789 | -2.692061375 | 3.54E-27 |
| Fam19a1 | 75.29991681 | -2.693528518 | 5.95E-08 |
| Gm17509 | 9.815080105 | -2.693778837 | 3.64E-03 |
| Slc2a13 | 76.15910103 | -2.700883682 | 3.70E-13 |
| Gm3173 | 12.70900116 | -2.701894679 | 8.53E-04 |
| Grid1 | 22.69881994 | -2.703702913 | 1.59E-05 |
| Gm3411 | 95.97843402 | -2.70816822 | 2.89E-06 |
| Slc24a1 | 28.67118619 | -2.709636923 | 1.38E-07 |
| Arhgap8 | 18.23760369 | -2.716164208 | 2.02E-03 |
| Sesn3 | 451.770964 | -2.718512082 | 7.84E-40 |
| Gm16096 | 18.53812567 | -2.722855274 | 3.92E-05 |
| Efcab9 | 37.28015627 | -2.724946223 | 7.58E-09 |
| Pcdhb21 | 187.4813216 | -2.725613784 | 1.51E-31 |
| Csrp2 | 1603.401949 | -2.738192267 | 1.99E-20 |
| Kctd12b | 28.18537233 | -2.741127982 | 8.58E-07 |
| Pcdhb19 | 163.5809958 | -2.743524905 | 5.88E-30 |
| Gm24576 | 5.441975694 | -2.748866601 | 2.65E-02 |
| 4930426L09Rik | 16.02716752 | -2.756659756 | 5.60E-05 |
| Pcdhb12 | 18.19238944 | -2.761249541 | 5.05E-06 |
| Lmx1b | 7.22827015 | -2.762348169 | 3.13E-02 |
| Ugt1a6b | 1056.631265 | -2.770277475 | 8.76E-54 |
| Htr2a | 66.90647818 | -2.776404529 | 1.34E-10 |
| Etv5 | 692.3569151 | -2.789192104 | 9.96E-26 |
| Grpr | 52.83871958 | -2.794774513 | 6.36E-06 |
| Xk | 7.137264591 | -2.798285204 | 1.13E-03 |
| Snord69 | 17.3634094 | -2.801363665 | 6.53E-05 |
| Fjx1 | 266.2076356 | -2.806035565 | 6.15E-43 |
| Emb | 6148.231165 | -2.807768269 | 5.39E-54 |
| Rpgrip1l | 758.612363 | -2.810767689 | 9.57E-29 |
| Rhpn1 | 32.93132459 | -2.822823794 | 2.68E-10 |
| Prtn3 | 13.25797432 | -2.835608387 | 7.25E-04 |
| Fgf14 | 155.1480338 | -2.837319652 | 4.66E-13 |
| Ckb | 5.205320121 | -2.841119201 | 3.12E-02 |
| Gpr88 | 11.30675403 | -2.851519675 | 3.12E-02 |
| Ctsk | 939.5622255 | -2.854495018 | 2.77E-40 |
| Pcdhb7 | 281.6238328 | -2.855031138 | 2.27E-37 |
| Itgbl1 | 4703.422305 | -2.860713321 | 5.58E-24 |
| Ephb6 | 218.4738707 | -2.861234808 | 6.30E-14 |
| Gm25829 | 10.54213017 | -2.862950805 | 2.71E-02 |
| Egfl6 | 11.47141831 | -2.866340424 | 1.42E-03 |
| Pcdhb6 | 173.737983 | -2.875538758 | 1.12E-26 |
| Serpina3h | 32.62380238 | -2.876660852 | 3.75E-05 |

| | | | |
|---|---|---|---|
| 4933406P04Rik | 9.227396209 | -2.887818359 | 2.82E-03 |
| Gm10254 | 21.60174155 | -2.90040983 | 3.58E-03 |
| Slc30a1 | 484.7571357 | -2.903636125 | 2.31E-13 |
| P2rx6 | 61.83965373 | -2.907874637 | 1.63E-13 |
| Ptn | 40530.58054 | -2.90891563 | 3.99E-10 |
| Gm17195 | 18.69066063 | -2.910474375 | 3.90E-06 |
| Tgfbi | 34.33221477 | -2.913186983 | 7.46E-06 |
| Gm2464 | 5.686631402 | -2.922571945 | 1.43E-02 |
| Polm | 3037.083453 | -2.926698423 | 1.11E-98 |
| Dhrs4 | 481.3038845 | -2.927615902 | 9.11E-40 |
| Rspo2 | 7.266487856 | -2.929426204 | 1.93E-03 |
| Znf41-ps | 70.54421761 | -2.930141316 | 1.94E-07 |
| 4930483K19Rik | 9.197652411 | -2.935073748 | 5.22E-03 |
| Cldn20 | 7.614908487 | -2.9373522 | 3.30E-03 |
| Platr22 | 32.98595665 | -2.940741034 | 1.06E-08 |
| Mt1 | 6812.419424 | -2.941057321 | 7.49E-15 |
| Itgb8 | 478.0701506 | -2.946417151 | 3.67E-31 |
| Adam11 | 158.3843609 | -2.946702017 | 3.03E-14 |
| D630033O11Rik | 7.891037492 | -2.971188296 | 2.42E-03 |
| Ugt1a6a | 1314.632066 | -2.987425036 | 6.70E-22 |
| Tcim | 182.9808592 | -2.988559439 | 6.08E-20 |
| Chst2 | 135.1732714 | -2.999122073 | 1.29E-17 |
| Sohlh2 | 9.321089239 | -3.000653517 | 3.02E-04 |
| Hlf | 720.1008505 | -3.004271964 | 2.52E-07 |
| Sned1 | 9956.936022 | -3.005401025 | 1.46E-30 |
| Sfrp2 | 592.0206281 | -3.006402389 | 9.35E-16 |
| Cdon | 2024.016055 | -3.015317298 | 2.35E-34 |
| Gm15907 | 6.608722355 | -3.023530455 | 1.11E-02 |
| Arhgap28 | 1085.405443 | -3.024089771 | 5.80E-58 |
| Mtus2 | 261.3824621 | -3.028591158 | 1.05E-17 |
| Rhbdl3 | 44.7420194 | -3.039785262 | 3.05E-06 |
| Actr3b | 183.7091602 | -3.052720325 | 2.39E-16 |
| Shc2 | 1586.018871 | -3.056205486 | 2.21E-52 |
| Ssc5d | 1608.633546 | -3.062634362 | 9.17E-38 |
| Gcnt4 | 773.2398383 | -3.075376957 | 1.67E-49 |
| Speer4a | 330.0211079 | -3.076464638 | 2.22E-06 |
| Tmem252 | 7.498715846 | -3.076599089 | 3.70E-03 |
| Gm21671 | 8.561994867 | -3.076707247 | 2.09E-02 |
| Slc16a9 | 86.65457978 | -3.087204671 | 5.13E-13 |
| Emid1 | 31.69594959 | -3.090813042 | 1.16E-06 |
| Cela1 | 174.5106123 | -3.09352874 | 1.02E-17 |
| Adh1 | 110.5458724 | -3.097923399 | 6.11E-09 |
| Epha10 | 7.545852402 | -3.098406557 | 8.43E-04 |
| Rnase4 | 11680.33536 | -3.099681544 | 1.79E-47 |
| Gm15958 | 23.51822755 | -3.102094901 | 3.09E-05 |
| Irf6 | 5.620115735 | -3.109379179 | 2.99E-02 |
| Speer4b | 14.69744655 | -3.110049199 | 3.92E-05 |

| | | | |
|---|---|---|---|
| 1700071M16Rik | 6.45197741 | -3.124278976 | 1.50E-02 |
| Crocc2 | 99.75445273 | -3.129214716 | 2.06E-08 |
| Ednra | 698.5583548 | -3.140212452 | 3.21E-24 |
| Gria1 | 276.9010458 | -3.14087357 | 3.08E-15 |
| St6gal1 | 1757.152116 | -3.14753474 | 1.90E-49 |
| Prl2c3 | 113.2380744 | -3.155694555 | 1.32E-03 |
| E030013I19Rik | 11.43396328 | -3.159527587 | 2.66E-04 |
| Tdo2 | 5.007406738 | -3.191911601 | 2.17E-02 |
| Nbl1 | 1782.408442 | -3.193750841 | 1.18E-31 |
| Pla2g2e | 18.47206542 | -3.200550845 | 3.76E-05 |
| Aebp1 | 72201.79584 | -3.209025276 | 8.71E-75 |
| NA | 12.28994974 | -3.21048564 | 1.15E-03 |
| Capn6 | 8375.626269 | -3.217381104 | 1.83E-45 |
| Slc6a7 | 36.75567192 | -3.219306672 | 5.37E-13 |
| Pgam1-ps2 | 32.29934769 | -3.223933278 | 5.04E-04 |
| Postn | 49906.43674 | -3.225722238 | 7.89E-11 |
| Gm7347 | 27.39865267 | -3.22944118 | 2.52E-07 |
| Sema6d | 5.168120737 | -3.23260475 | 4.78E-02 |
| Gm14445 | 7.644904696 | -3.246447597 | 4.16E-03 |
| Plppr3 | 487.5192114 | -3.254381379 | 2.63E-22 |
| Gm17308 | 13.38750999 | -3.258751978 | 5.29E-05 |
| Sox5 | 174.3593763 | -3.261638334 | 1.96E-11 |
| NA | 48.41532146 | -3.262674808 | 1.08E-10 |
| Klhl4 | 12.53402334 | -3.277442514 | 2.10E-03 |
| Mgp | 2186.4868 | -3.29717144 | 3.77E-26 |
| Zfp108 | 34.41460638 | -3.298454019 | 2.16E-11 |
| Lpcat2 | 612.0918981 | -3.310180883 | 4.26E-17 |
| Enpp2 | 759.7675373 | -3.319750972 | 5.48E-32 |
| Steap3 | 17632.6646 | -3.334931192 | 3.58E-37 |
| Siglec15 | 19.76496087 | -3.341967276 | 3.84E-06 |
| Mmp8 | 5.245523941 | -3.355216438 | 5.64E-03 |
| Pygo1 | 741.5294784 | -3.371921835 | 2.84E-61 |
| Edil3 | 1365.668624 | -3.376146443 | 1.60E-27 |
| Pxylp1 | 827.890857 | -3.391493859 | 1.01E-35 |
| Slc25a23 | 164.104295 | -3.394151239 | 2.54E-11 |
| mt-Te | 93.59404751 | -3.394816073 | 2.65E-10 |
| Gm9869 | 51.84992159 | -3.406697071 | 3.66E-13 |
| Gm21083 | 126.1258943 | -3.412125074 | 8.93E-15 |
| Kng2 | 30.33286104 | -3.421533339 | 6.26E-07 |
| Ccl2 | 2277.435037 | -3.429529955 | 7.94E-25 |
| Pcdhb9 | 216.2214447 | -3.431965322 | 6.19E-38 |
| Mdga1 | 7.954351591 | -3.432942335 | 1.86E-02 |
| Gm10409 | 5.644235183 | -3.443047241 | 3.88E-02 |
| Tle2 | 1136.122372 | -3.452130371 | 4.81E-35 |
| Cxcl12 | 14907.90677 | -3.469365789 | 9.53E-30 |
| Ccl8 | 242.947781 | -3.475212827 | 5.28E-14 |
| Egr3 | 140.915746 | -3.479926124 | 1.59E-11 |

| | | | |
|---|---|---|---|
| Cd300lb | 309.9653894 | -3.502609607 | 3.07E-20 |
| Gm11261 | 171.9841549 | -3.514592115 | 8.71E-18 |
| Slfn2 | 22.2414266 | -3.526044948 | 8.25E-06 |
| Mcf2l | 69.89545145 | -3.53007031 | 3.19E-13 |
| Ugt1a7c | 392.0152248 | -3.535484961 | 3.42E-27 |
| Sema7a | 84.82804028 | -3.537474878 | 1.40E-13 |
| H1fx | 45.98158594 | -3.541447302 | 2.70E-12 |
| Thbs2 | 76114.02438 | -3.544994002 | 8.59E-39 |
| Osbp2 | 32.12984721 | -3.553466881 | 1.58E-05 |
| BC106179 | 76.11099145 | -3.559991312 | 8.43E-19 |
| Rbp1 | 2005.260463 | -3.566453608 | 7.04E-97 |
| B930094E09Rik | 8.478385925 | -3.56683092 | 1.17E-03 |
| Rnasel | 68.25410949 | -3.577255009 | 4.53E-18 |
| Igsf10 | 1498.548461 | -3.588494829 | 3.52E-33 |
| Gm5862 | 499.4662236 | -3.594105791 | 4.37E-22 |
| Kctd14 | 13.33077476 | -3.604963521 | 7.38E-04 |
| Serpina3i | 55.90345178 | -3.605781328 | 2.36E-07 |
| Gm2026 | 125.1518551 | -3.633772885 | 1.17E-03 |
| NA | 85.90070398 | -3.642857473 | 4.36E-19 |
| Il33 | 385.9276264 | -3.679491136 | 5.67E-08 |
| Col22a1 | 44.64731885 | -3.705452049 | 2.48E-12 |
| Syngr1 | 40.08116894 | -3.708141416 | 6.92E-06 |
| Sytl2 | 158.0277612 | -3.712549632 | 3.83E-15 |
| Kctd15 | 882.2490959 | -3.73168812 | 2.99E-42 |
| Capns2 | 41.29573201 | -3.731742659 | 1.22E-10 |
| Galnt12 | 136.076518 | -3.733281427 | 5.87E-23 |
| Fzd1 | 3526.570472 | -3.740829819 | 1.47E-80 |
| Nrep | 127.7232309 | -3.747215922 | 1.08E-14 |
| Med12l | 34.90194708 | -3.750787778 | 1.64E-03 |
| NA | 6.161689882 | -3.756327037 | 1.23E-02 |
| Dyx1c1 | 41.08029909 | -3.77615182 | 7.49E-15 |
| Gm12576 | 5.138505381 | -3.78704125 | 5.81E-03 |
| Egr2 | 252.8800803 | -3.80541484 | 8.30E-20 |
| Clmp | 2125.373568 | -3.808619713 | 4.96E-153 |
| Map3k5 | 17.22064227 | -3.818554812 | 1.77E-03 |
| Angpt1 | 126.2034669 | -3.825923168 | 3.38E-11 |
| Gm26552 | 14.03270238 | -3.83817342 | 8.47E-05 |
| Gm16701 | 13.96346115 | -3.858254407 | 1.52E-04 |
| Gpnmb | 3590.839261 | -3.858994983 | 1.30E-36 |
| Tek | 47.78473353 | -3.863335201 | 3.70E-12 |
| Apln | 371.7087337 | -3.873349084 | 3.54E-32 |
| Tspan18 | 194.9774176 | -3.882861678 | 3.15E-24 |
| Ccn3 | 191.5912267 | -3.903457039 | 5.39E-32 |
| Mpp4 | 6.753419752 | -3.905411487 | 2.52E-03 |
| 1700125H03Rik | 12.54704266 | -3.914404423 | 3.68E-05 |
| NA | 97.40321005 | -3.940866186 | 3.51E-17 |
| Lgi4 | 6.261486722 | -3.946977513 | 9.89E-03 |

| | | | |
|---|---|---|---|
| Has2 | 320.7677252 | -3.961389068 | 5.83E-34 |
| Timp4 | 80.55768285 | -3.968270598 | 8.37E-12 |
| Mmp9 | 324.7980202 | -3.971072734 | 1.24E-08 |
| Kcne4 | 7.230896568 | -3.973388653 | 3.04E-03 |
| Mlana | 173.3382716 | -4.017737384 | 2.39E-11 |
| Pcdh7 | 730.661321 | -4.023238103 | 9.11E-70 |
| Csgalnact1 | 1244.091072 | -4.032346004 | 4.59E-87 |
| Serpina3n | 668.2845824 | -4.032374308 | 8.63E-32 |
| Tecta | 36.05411746 | -4.049997152 | 3.91E-12 |
| Tmem200a | 180.2586998 | -4.053941105 | 1.82E-32 |
| Cabp7 | 8.452464321 | -4.063181063 | 2.20E-02 |
| Gfra2 | 25.3859268 | -4.104247432 | 1.00E-08 |
| Ubd | 22.06665641 | -4.119144451 | 7.15E-06 |
| Dnm3os | 3686.208543 | -4.128304386 | 8.12E-20 |
| Ccdc65 | 26.87887487 | -4.137619708 | 2.04E-08 |
| Rnase1 | 36.55767924 | -4.153294599 | 1.34E-10 |
| 10-Mar | 64.82310953 | -4.171824474 | 4.24E-14 |
| Mertk | 609.7724486 | -4.198991781 | 4.04E-74 |
| Akr1c18 | 34.14361785 | -4.245338949 | 3.06E-05 |
| Ang5 | 6.410110454 | -4.268444161 | 8.40E-03 |
| Smoc2 | 5606.559969 | -4.282447788 | 2.27E-32 |
| Gm12590 | 87.10131386 | -4.286823051 | 8.93E-21 |
| Serpina3g | 65.50415713 | -4.294178263 | 2.17E-10 |
| B930082K07Rik | 10.59285588 | -4.316877535 | 4.19E-04 |
| Dlgap2 | 221.2029046 | -4.322963456 | 7.58E-43 |
| Gm27517 | 14.62030774 | -4.340174762 | 2.32E-03 |
| Paqr8 | 1206.296549 | -4.349886818 | 4.72E-79 |
| Mt2 | 2687.292186 | -4.406964827 | 8.27E-20 |
| Opcml | 366.9743187 | -4.416866747 | 6.07E-43 |
| Dhx58os | 12.54913426 | -4.43999338 | 3.88E-04 |
| Fam180a | 160.3125889 | -4.443195391 | 6.84E-27 |
| Igfbp5 | 819.6479129 | -4.447604299 | 6.93E-14 |
| Zfp41 | 1083.240847 | -4.472860125 | 4.57E-48 |
| Maf | 987.6224961 | -4.545069796 | 8.80E-43 |
| Pknox2 | 149.7278646 | -4.630725211 | 3.24E-29 |
| Csf2rb | 20.90762805 | -4.658524068 | 2.59E-05 |
| Plxnc1 | 5.257759957 | -4.690205491 | 1.52E-03 |
| Gm10701 | 7.826260597 | -4.700930893 | 1.07E-03 |
| Nat8f3 | 96.81673939 | -4.717034457 | 1.63E-12 |
| Gm27509 | 7.270988291 | -4.755625363 | 4.53E-03 |
| Wfdc3 | 92.4419757 | -4.81821485 | 4.05E-12 |
| Acpp | 8.017078266 | -4.821446875 | 4.54E-03 |
| Prl2c5 | 68.59226915 | -4.823770724 | 2.67E-11 |
| Gm5799 | 11.406214 | -4.868924675 | 5.45E-04 |
| Wdr63 | 7.18728154 | -4.879650413 | 1.22E-05 |
| AW551984 | 1346.712062 | -4.880958885 | 1.22E-47 |
| Arhgap20 | 61.50283451 | -4.904326951 | 8.44E-08 |

| | | | |
|---|---|---|---|
| Spp1 | 33894.67107 | -4.909268874 | 4.21E-45 |
| Cdk15 | 129.3148332 | -4.911544423 | 2.76E-29 |
| Gm2237 | 6.67013034 | -4.962424126 | 3.38E-02 |
| Gm2366 | 40.57644542 | -5.140188567 | 6.11E-03 |
| Cp | 1547.646794 | -5.148102241 | 8.31E-65 |
| Cd200r3 | 139.6286064 | -5.149953341 | 1.62E-02 |
| Rgs5 | 30.59780507 | -5.173615763 | 8.90E-14 |
| 4930589L23Rik | 9.662169631 | -5.222179792 | 5.52E-04 |
| Tmem100 | 6.230469592 | -5.25656298 | 7.14E-04 |
| Clec1a | 18.53247397 | -5.288690695 | 4.57E-05 |
| NA | 9.225849333 | -5.301221367 | 1.06E-03 |
| Tnfrsf11b | 121.1051387 | -5.303268244 | 8.54E-17 |
| Fam78b | 275.8695293 | -5.312100141 | 3.59E-59 |
| Scn7a | 62.80044543 | -5.334533875 | 1.69E-14 |
| Clip4 | 144.42984 | -5.353513198 | 3.19E-28 |
| Kcnk3 | 92.21429487 | -5.472463189 | 6.62E-17 |
| Ccdc177 | 18.1886864 | -5.516091098 | 1.92E-06 |
| Trabd2b | 1980.320248 | -5.521761776 | 2.58E-74 |
| Hc | 29.33803059 | -5.530592504 | 8.89E-11 |
| Axin2 | 27.73513628 | -5.571863317 | 1.66E-06 |
| Nr1i3 | 116.7264137 | -5.614779936 | 3.55E-04 |
| Serpina3m | 6.914672563 | -5.63889454 | 2.14E-04 |
| Plxdc1 | 183.9040077 | -5.657982478 | 1.03E-19 |
| Mir124a-1hg | 9.198990273 | -5.714219791 | 3.07E-04 |
| Gm15422 | 17.50296185 | -5.849222357 | 1.64E-06 |
| Tinag | 23.1948581 | -5.871842686 | 2.69E-05 |
| Galnt9 | 9.989789218 | -5.880912541 | 3.59E-05 |
| Klhdc9 | 17.89947461 | -6.012010736 | 1.09E-02 |
| Nat8 | 9.423743375 | -6.226214177 | 6.30E-06 |
| Megf10 | 616.628578 | -6.242837793 | 4.50E-45 |
| Fam19a5 | 11.74775917 | -6.323224285 | 7.92E-05 |
| Myo5b | 13.71114935 | -6.339015868 | 9.32E-05 |
| Rgs4 | 1205.292047 | -6.409715916 | 9.55E-31 |
| Prl2c2 | 22898.70079 | -6.461967864 | 1.06E-39 |
| Trpm6 | 7.21072813 | -6.483195659 | 6.26E-06 |
| Cytip | 32.07785982 | -6.590697375 | 1.49E-07 |
| Klrb1a | 6.733824506 | -6.690121693 | 4.88E-06 |
| Samd11 | 155.349818 | -6.732006489 | 2.16E-03 |
| Nmur1 | 13.54843957 | -6.783296023 | 1.34E-06 |
| Aspg | 128.0287642 | -6.930045834 | 2.11E-19 |
| B630019A10Rik | 22.44071648 | -7.022529167 | 1.81E-07 |
| NA | 18.24678081 | -7.624448727 | 3.98E-06 |
| Dmrtc1c2 | 1708.160336 | -7.632915506 | 1.20E-28 |
| Gm10401 | 31.09799767 | -7.699328553 | 1.57E-04 |
| Gm9803 | 36.28989778 | -7.770535888 | 3.00E-02 |
| Lingo2 | 18.63616195 | -7.808905035 | 4.97E-08 |
| Fam135b | 222.7442954 | -8.221417766 | 1.25E-22 |

| | | | |
|---|---|---|---|
| Tspan10 | 235.8434891 | -8.580277869 | 1.05E-07 |
| Gm10221 | 118.8385297 | -8.720361037 | 2.97E-02 |
| Gm26649 | 48.11705398 | -8.936003195 | 7.61E-05 |
| Sla2 | 584.6313518 | -9.539845097 | 1.00E-08 |
| Gm15801 | 42.34653278 | -21.72354016 | 1.31E-12 |
| Zfp781 | 19.65596644 | -22.78722088 | 4.20E-04 |

| Gene | baseMean | log2 Fold Change | padj |
|---|---|---|---|
| Slc2a12 | 153.088269 | 10.45556904 | 2.15E-04 |
| Lamb3 | 47.93842017 | 9.840430458 | 5.24E-13 |
| Psca | 64.92123435 | 9.817207773 | 8.56E-13 |
| Hephl1 | 25.60224789 | 9.354470658 | 5.91E-11 |
| Spo11 | 29.75667518 | 9.114167736 | 1.10E-10 |
| Gm2163 | 20.51038887 | 8.960754739 | 2.79E-02 |
| Gm5812 | 53.47797956 | 8.930462959 | 6.21E-04 |
| Klk1b27 | 12.04208527 | 8.391618486 | 1.38E-07 |
| Vnn1 | 94.96773462 | 8.265320432 | 3.50E-21 |
| Mup5 | 8.555026896 | 8.203730704 | 3.09E-07 |
| Slc47a1 | 16.32805259 | 8.16304669 | 4.97E-08 |
| Ecscr | 21.55153097 | 7.950187799 | 3.39E-08 |
| Ptprt | 174.3279632 | 7.749208496 | 1.50E-11 |
| Sh3tc1 | 213.0734353 | 7.672791635 | 6.90E-24 |
| Dthd1 | 17.05478183 | 7.559335639 | 2.45E-06 |
| AU021092 | 16.8814082 | 7.462845494 | 3.20E-07 |
| Cdh5 | 11.00256545 | 7.320903548 | 1.00E-06 |
| Itln1 | 5.005083366 | 7.316327135 | 6.51E-05 |
| Serping1 | 32.77937136 | 7.200989591 | 1.67E-07 |
| Cyp4b1 | 7.700139718 | 7.13885153 | 7.85E-05 |
| Slc2a5 | 32.51930166 | 7.099740542 | 7.19E-15 |
| Chst1 | 44.51961143 | 7.025678633 | 2.03E-18 |
| Anxa8 | 232.8381859 | 6.992510354 | 4.83E-07 |
| Gm13303 | 7.269012401 | 6.96913055 | 7.46E-04 |
| Gm21293 | 16.49303806 | 6.648255053 | 4.72E-05 |
| Gm9428 | 14.61896193 | 6.510921585 | 1.20E-05 |
| Gm1110 | 11.92620283 | 6.35566665 | 1.42E-04 |
| Gm13284 | 12.99212276 | 6.294846558 | 4.82E-05 |
| Insc | 11.98764112 | 6.270860712 | 2.43E-05 |
| Upk3b | 18.38391089 | 6.265717371 | 4.06E-07 |
| Nes | 211.7096441 | 6.184347283 | 1.15E-31 |
| Tmem150b | 157.484637 | 6.09720879 | 5.34E-03 |
| Gm5286 | 7.531885012 | 6.089200695 | 7.93E-05 |
| 2610028E06Rik | 10.97570129 | 6.024605647 | 3.38E-05 |
| Gm13415 | 175.3337851 | 6.018055463 | 3.67E-21 |
| Gm27286 | 31.94524763 | 6.017865259 | 2.25E-04 |
| Lhx4 | 18.42303899 | 5.980472999 | 3.90E-08 |
| B3gnt3 | 5.759179752 | 5.976573598 | 4.54E-04 |
| Serpinb9b | 3482.508606 | 5.974525383 | 2.69E-56 |
| Tmeff2 | 105.2957531 | 5.973028198 | 5.28E-35 |
| Pdzd2 | 406.3513246 | 5.889814151 | 1.13E-19 |
| Wnt7a | 83.29495754 | 5.87373004 | 5.30E-05 |
| Gm14373 | 26.27208476 | 5.733959944 | 1.41E-02 |
| Lyz2 | 828.2358917 | 5.710741974 | 8.48E-40 |
| A730049H05Rik | 20.80637073 | 5.682746505 | 1.78E-07 |
| Als2cl | 101.8281501 | 5.666017914 | 1.35E-10 |

| | | | |
|---|---|---|---|
| Card10 | 5.87499347 | 5.657735398 | 4.64E-04 |
| Gdf15 | 40.77172247 | 5.567048853 | 1.11E-05 |
| Gm6083 | 6.947319638 | 5.519055949 | 1.09E-03 |
| Adam22 | 234.5747723 | 5.477887094 | 6.31E-12 |
| Erbb3 | 69.46120585 | 5.456278132 | 4.03E-18 |
| Mill2 | 7.236578422 | 5.423342338 | 4.00E-04 |
| Slc6a12 | 232.6272342 | 5.42266826 | 2.04E-04 |
| Gm12153 | 16.73241645 | 5.41109538 | 2.90E-04 |
| Calcb | 38.52361346 | 5.403831744 | 1.98E-05 |
| Gm15452 | 18.49115526 | 5.37282138 | 2.00E-03 |
| Cyp2ab1 | 5.554438667 | 5.327911594 | 1.36E-03 |
| Cobl | 32.63939068 | 5.263096526 | 1.87E-09 |
| Ces2f | 8.035604011 | 5.20549898 | 2.22E-03 |
| Gm10256 | 43.00942646 | 5.187433573 | 5.02E-14 |
| Synpo2l | 127.4131202 | 5.082531805 | 2.50E-24 |
| Ptprj | 72.06266337 | 5.075282579 | 4.01E-27 |
| Crct1 | 218.1985879 | 5.045405324 | 6.56E-04 |
| Cdhr1 | 7.429102719 | 5.03529637 | 4.47E-05 |
| Gm10696 | 6.667621713 | 5.021787461 | 5.38E-03 |
| Limch1 | 706.4773048 | 4.991671774 | 1.05E-27 |
| Gm16531 | 8.914604206 | 4.923688645 | 1.17E-03 |
| Tspan15 | 23.84249731 | 4.911495972 | 1.76E-12 |
| Gm43154 | 35.32626595 | 4.89947655 | 2.08E-03 |
| Fyb2 | 205.4623369 | 4.887356021 | 1.07E-19 |
| Cyp3a41a | 110.7883611 | 4.863252019 | 7.21E-17 |
| Gm13418 | 7.051919701 | 4.854006858 | 6.86E-03 |
| Hist1h4n | 13.89900442 | 4.83886808 | 3.74E-03 |
| Cyp4f40 | 16.78817656 | 4.783537122 | 3.03E-04 |
| Prkag3 | 47.08753681 | 4.776626386 | 6.78E-22 |
| Gal3st2 | 33.39687285 | 4.776020052 | 4.59E-03 |
| Atg9b | 216.6082704 | 4.770279854 | 4.97E-18 |
| Robo3 | 32.48397439 | 4.756639711 | 6.03E-03 |
| Cyp24a1 | 5.817295899 | 4.703859433 | 1.28E-02 |
| NA | 26.06308258 | 4.663417808 | 4.15E-03 |
| Serpinb1c | 17.20225213 | 4.624733017 | 7.04E-03 |
| Elfn1 | 11.57763753 | 4.608900118 | 1.05E-06 |
| Arhgap9 | 58.21154531 | 4.552582991 | 1.32E-06 |
| Ikzf4 | 76.17939403 | 4.512002073 | 3.78E-06 |
| Psg17 | 21.47017957 | 4.509075876 | 6.06E-08 |
| Slc27a6 | 6.663998212 | 4.506540659 | 1.52E-02 |
| Synpr | 13.66684869 | 4.499602696 | 6.49E-03 |
| Hs3st1 | 5.546121186 | 4.476675347 | 3.93E-03 |
| Pkp2 | 365.3274075 | 4.46856739 | 1.12E-63 |
| Ces2e | 38.3662501 | 4.466408217 | 6.42E-04 |
| Sorl1 | 6.29132497 | 4.450891282 | 1.36E-02 |
| Tmem171 | 242.2820833 | 4.441697527 | 4.99E-06 |
| Rnd1 | 586.3380609 | 4.438311752 | 9.68E-44 |

| | | | |
|---|---|---|---|
| Kctd4 | 12.52468892 | 4.425070992 | 1.12E-05 |
| BC051019 | 6.81730173 | 4.423611352 | 1.04E-02 |
| Rapgef5 | 8.056496296 | 4.410554239 | 1.69E-02 |
| Prr32 | 25.91099265 | 4.407999645 | 9.90E-03 |
| Cth | 393.5185974 | 4.398553874 | 7.37E-11 |
| Serpind1 | 71.4025159 | 4.391841593 | 1.56E-18 |
| Fras1 | 50.54206584 | 4.386499989 | 9.20E-03 |
| AU018091 | 5.236497473 | 4.369136197 | 3.67E-03 |
| Ankk1 | 13.33636605 | 4.368805562 | 9.03E-03 |
| Serpinb9c | 5.608671978 | 4.342026891 | 3.01E-02 |
| Cgn | 236.0254742 | 4.30198856 | 4.35E-33 |
| Enkur | 17.5217497 | 4.276182257 | 7.09E-06 |
| Serpina5 | 6.801924643 | 4.272936142 | 2.26E-02 |
| Rhpn2 | 103.2238498 | 4.261941909 | 7.00E-34 |
| Cdhr2 | 8.715260545 | 4.256823053 | 1.33E-04 |
| Pax8 | 75.11368423 | 4.246285956 | 1.24E-06 |
| Gpr179 | 12.91737115 | 4.238132149 | 7.86E-07 |
| Slc30a2 | 5.640362476 | 4.227995626 | 1.21E-02 |
| Lamc3 | 81.73661022 | 4.213803824 | 7.62E-14 |
| Ifit1bl1 | 78.34996578 | 4.186762205 | 2.10E-27 |
| Rbmy | 70.47372177 | 4.178472383 | 1.91E-10 |
| Tacr2 | 9.773124393 | 4.165525214 | 4.09E-07 |
| Cxcl1 | 119.9745271 | 4.163815869 | 1.71E-23 |
| Ctnnd2 | 13.32903101 | 4.145277958 | 1.19E-02 |
| Serpinb9g | 65.5698485 | 4.112065702 | 9.78E-03 |
| NA | 6.543162396 | 4.081555202 | 1.32E-05 |
| Gm53 | 5.854823975 | 4.056173523 | 3.28E-02 |
| Ccl6 | 20.15019644 | 4.052592417 | 1.17E-05 |
| Best3 | 5.268568397 | 4.030932642 | 1.02E-02 |
| Gm17167 | 121.3447426 | 4.015842248 | 1.22E-42 |
| Adgrf5 | 8.050799548 | 3.99467713 | 1.19E-03 |
| Tmem71 | 6.057387095 | 3.983918571 | 8.64E-03 |
| Susd2 | 15.13366232 | 3.957033055 | 2.45E-04 |
| Apob | 19.03152808 | 3.956587052 | 2.97E-05 |
| Dppa5a | 5.547308591 | 3.94837571 | 5.37E-03 |
| Lman1l | 19.17703041 | 3.936242212 | 4.95E-07 |
| Adcyap1r1 | 11.14523057 | 3.932834973 | 1.19E-05 |
| Itga3 | 4169.255291 | 3.908377089 | 2.01E-50 |
| Nlrc5 | 42.39167115 | 3.904636663 | 5.80E-21 |
| Klk14 | 5.470717404 | 3.904562488 | 1.12E-02 |
| Lypd5 | 36.53736749 | 3.892282937 | 3.21E-02 |
| Car6 | 110.8106253 | 3.891730148 | 1.21E-08 |
| Ms4a4d | 289.8013129 | 3.857237088 | 1.12E-18 |
| Sult4a1 | 8.298499443 | 3.856823206 | 4.51E-02 |
| Cd209c | 391.2517191 | 3.837280432 | 1.01E-22 |
| Wt1 | 33.01804751 | 3.829158639 | 8.09E-09 |
| Dusp8 | 245.7185379 | 3.825596768 | 9.63E-35 |

| | | | |
|---|---|---|---|
| Cyp4f14 | 11.05060069 | 3.792257644 | 2.80E-02 |
| Fbxw10 | 18.3927969 | 3.768348377 | 3.32E-05 |
| Btn1a1 | 43.15252764 | 3.757403354 | 3.95E-07 |
| Scel | 35.56464911 | 3.739794639 | 4.67E-02 |
| Oxtr | 5.267159996 | 3.709531682 | 4.40E-02 |
| Myom2 | 1179.530624 | 3.706418924 | 1.43E-21 |
| Gm12505 | 700.5275066 | 3.703800187 | 3.06E-28 |
| Gm13988 | 25.63523994 | 3.692099874 | 2.99E-06 |
| Col6a4 | 50.04165493 | 3.676067168 | 6.70E-15 |
| Birc7 | 495.5983212 | 3.665192466 | 8.77E-11 |
| Atp2a3 | 10.67440491 | 3.657614951 | 1.65E-03 |
| Des | 60.75912734 | 3.62368319 | 1.40E-07 |
| Sod3 | 262.7305939 | 3.607993569 | 1.37E-13 |
| Serpinb1a | 435.1696668 | 3.604572826 | 1.64E-14 |
| Syk | 89.27270301 | 3.603495025 | 8.77E-07 |
| Gm20400 | 779.139461 | 3.592545869 | 1.85E-20 |
| Gpr174 | 13.33517572 | 3.589454551 | 1.92E-02 |
| Gm26870 | 5.060075859 | 3.587433908 | 4.72E-02 |
| Fcna | 17.16685909 | 3.579042592 | 1.81E-02 |
| Adra1b | 80.73107556 | 3.564721827 | 2.50E-24 |
| Plb1 | 53.16380267 | 3.561796491 | 3.50E-14 |
| Fgf21 | 49.48268511 | 3.549940552 | 2.69E-04 |
| Sh3bgrl2 | 10.59069966 | 3.549318693 | 1.27E-05 |
| Ccdc180 | 8.668083621 | 3.549090952 | 1.58E-03 |
| Ms4a4c | 8.55212296 | 3.540742836 | 4.21E-03 |
| Plscr2 | 362.8998898 | 3.536962067 | 1.77E-24 |
| Zfp750 | 9.306518556 | 3.506265892 | 1.86E-02 |
| Fmo2 | 37.14481697 | 3.504030959 | 5.47E-06 |
| Matn3 | 9.122129289 | 3.503893235 | 2.33E-05 |
| Gm27224 | 10.41317876 | 3.503470975 | 1.19E-03 |
| Ereg | 462.3479018 | 3.495577955 | 1.59E-20 |
| Car14 | 32.99689159 | 3.491804683 | 1.25E-07 |
| Itih4 | 19.70594933 | 3.472535736 | 1.45E-06 |
| Apol9b | 84.6420391 | 3.442660109 | 5.06E-17 |
| Il15ra | 123.6384944 | 3.435449835 | 1.13E-23 |
| Slc39a4 | 5.328099942 | 3.423592432 | 1.42E-02 |
| Bcam | 6.821030757 | 3.407248265 | 4.54E-02 |
| Htatip2 | 7.892515834 | 3.401996568 | 2.10E-03 |
| Pla2g5 | 14.77056158 | 3.395112132 | 4.39E-03 |
| Fbln1 | 65.66334147 | 3.395105294 | 1.66E-12 |
| Anpep | 300.0514348 | 3.393246922 | 2.67E-17 |
| Far2os1 | 51.84053146 | 3.386198459 | 6.92E-07 |
| Ica1l | 23.42769138 | 3.37781561 | 7.48E-08 |
| Adamtsl1 | 75.62285662 | 3.375163653 | 2.88E-07 |
| Prss12 | 16.0826015 | 3.3617946 | 1.09E-06 |
| Gm10230 | 10.1897906 | 3.351700536 | 9.10E-03 |
| Gm11884 | 8.165736454 | 3.334847688 | 5.03E-05 |

| | | | |
|---|---|---|---|
| Rassf6 | 30.46412628 | 3.321859665 | 7.59E-05 |
| Arxes1 | 501.9970735 | 3.316195443 | 2.49E-21 |
| Havcr2 | 173.0157584 | 3.301489043 | 1.53E-08 |
| Rassf7 | 636.3971329 | 3.30053995 | 8.58E-35 |
| Gdap1l1 | 51.61483492 | 3.273598937 | 5.36E-08 |
| Vwce | 9.513615881 | 3.269145411 | 4.17E-03 |
| Ly6i | 6.094536957 | 3.264351561 | 2.46E-04 |
| Eps8l2 | 14.97634174 | 3.261963255 | 2.80E-02 |
| Gm3376 | 148.220178 | 3.255482792 | 2.57E-24 |
| Ostn | 5.88146593 | 3.241433588 | 2.21E-02 |
| Gja1 | 1405.954553 | 3.229496653 | 4.98E-21 |
| Phactr2 | 110.9007105 | 3.224814082 | 8.25E-17 |
| Rnf165 | 18.2686871 | 3.212386115 | 1.33E-05 |
| Aloxe3 | 163.7815803 | 3.207861165 | 2.69E-21 |
| Il3ra | 21.87015892 | 3.19549919 | 1.56E-07 |
| Gm12840 | 2928.5719 | 3.194083049 | 3.26E-09 |
| Flrt1 | 7.038971897 | 3.181183798 | 4.26E-03 |
| A530016L24Rik | 46.43521955 | 3.169973661 | 5.72E-07 |
| Ifnlr1 | 8.948726448 | 3.158080532 | 1.34E-02 |
| Hbegf | 543.5379025 | 3.152455184 | 9.37E-38 |
| Gpat3 | 333.521046 | 3.1471234 | 6.48E-45 |
| Snhg7os | 10.44086061 | 3.138620224 | 3.18E-03 |
| S100a7a | 436.7014925 | 3.13690751 | 2.62E-28 |
| Camta1 | 704.1684855 | 3.117612331 | 2.96E-70 |
| Siglecg | 4363.02185 | 3.110453616 | 1.17E-35 |
| Slfn5 | 11.24247403 | 3.099852163 | 1.12E-02 |
| Btn2a2 | 21.67262263 | 3.091026108 | 3.60E-02 |
| Ttn | 108.5364841 | 3.090935609 | 3.31E-02 |
| C3ar1 | 28.45683189 | 3.081679669 | 1.96E-05 |
| Gsta3 | 16.36459403 | 3.030386651 | 1.70E-04 |
| NA | 15.15424967 | 3.026259636 | 8.61E-05 |
| Tfec | 10.99540601 | 3.026177248 | 2.47E-03 |
| Fmo3 | 67.82999797 | 3.013040565 | 7.60E-06 |
| Tmem37 | 109.5363239 | 2.995551134 | 2.57E-24 |
| Tspan32 | 38.52298728 | 2.989616222 | 2.71E-08 |
| Abi3 | 24.75257189 | 2.984442991 | 2.19E-03 |
| Dlk2 | 145.1833932 | 2.972168622 | 8.21E-16 |
| Inpp5d | 43.74884049 | 2.960515534 | 9.55E-03 |
| Gm26651 | 25.61219024 | 2.955372242 | 9.90E-06 |
| Mamld1 | 9.313569485 | 2.951809129 | 2.03E-02 |
| Gm11331 | 59.22672314 | 2.947533201 | 7.05E-06 |
| Rnf112 | 13.03420123 | 2.941754234 | 5.73E-04 |
| Ggt5 | 37.76480545 | 2.939909482 | 6.97E-03 |
| Aldh3a1 | 155.6027006 | 2.933717592 | 1.25E-09 |
| Sybu | 5.676802523 | 2.931906433 | 1.10E-02 |
| Fbxo15 | 9.25004082 | 2.923837438 | 6.29E-04 |
| Tinag | 23.1948581 | 2.913959905 | 1.72E-06 |

| | | | |
|---|---|---|---|
| Ptchd1 | 55.72263066 | 2.913831187 | 4.06E-07 |
| 2200002D01Rik | 188.5911851 | 2.912827236 | 1.75E-32 |
| Stbd1 | 242.832905 | 2.912820845 | 3.29E-29 |
| Upk3bl | 49.55256089 | 2.910882444 | 2.37E-03 |
| Taf7l | 8.416501097 | 2.910486589 | 1.21E-02 |
| Gprc5a | 2003.573682 | 2.910386198 | 1.13E-23 |
| Prrt4 | 6.235099417 | 2.906149166 | 3.86E-03 |
| Clvs1 | 109.3908839 | 2.901181053 | 1.77E-12 |
| Cdsn | 1502.659716 | 2.881135423 | 6.68E-34 |
| Ccdc3 | 14.86370165 | 2.872446408 | 5.40E-04 |
| Loxl4 | 984.284874 | 2.865753138 | 5.09E-29 |
| Casq1 | 13.57339567 | 2.860009982 | 8.95E-04 |
| Acsbg1 | 82.3921383 | 2.857008949 | 4.27E-15 |
| Syt17 | 46.84650688 | 2.85077387 | 1.80E-08 |
| Ptgs2 | 3952.831909 | 2.843767718 | 4.20E-09 |
| Gm11380 | 56.60624028 | 2.835495094 | 1.16E-11 |
| Plekha6 | 15.60175078 | 2.832848174 | 6.24E-03 |
| Hoxb3 | 9.193710774 | 2.82798036 | 4.51E-03 |
| Ptprv | 67.14689553 | 2.81984294 | 1.38E-04 |
| Prss27 | 17.7205746 | 2.817218524 | 1.72E-03 |
| Jph2 | 553.3177943 | 2.815904631 | 3.74E-08 |
| Cox6a2 | 24.89091578 | 2.810632714 | 4.48E-08 |
| Itga1 | 65.1732428 | 2.809233875 | 1.29E-13 |
| Lims2 | 179.0157548 | 2.797766588 | 6.12E-11 |
| Hspa12a | 126.8535222 | 2.791435159 | 6.09E-30 |
| Heph | 11.43738456 | 2.790314597 | 4.48E-03 |
| Ccdc184 | 7.875763359 | 2.787876806 | 1.74E-03 |
| Prg4 | 107.0476853 | 2.764871303 | 1.08E-06 |
| Ulbp1 | 1078.85357 | 2.761060831 | 1.04E-24 |
| 2310002F09Rik | 28.54212603 | 2.742755891 | 3.05E-04 |
| Chac1 | 2414.321663 | 2.742711617 | 1.17E-29 |
| Grk5 | 918.5251472 | 2.742010242 | 2.13E-28 |
| Ccnd1 | 5838.880133 | 2.736578117 | 2.60E-07 |
| Cst6 | 60.21863085 | 2.716059273 | 3.92E-09 |
| Arl9 | 5.951293518 | 2.70576058 | 2.73E-02 |
| Hoxb6 | 11.04812908 | 2.694759236 | 9.11E-03 |
| Tmem88 | 67.02043446 | 2.692321097 | 9.60E-14 |
| Ces2g | 117.9433417 | 2.692305019 | 8.01E-11 |
| Tspan12 | 74.6818032 | 2.691913325 | 8.01E-11 |
| Catsperd | 26.54257683 | 2.663560355 | 1.88E-06 |
| Atp6v0a4 | 40.07859287 | 2.656317523 | 1.57E-07 |
| Epha2 | 1944.113419 | 2.654014078 | 2.77E-18 |
| Gm9750 | 83.52959486 | 2.650816937 | 4.64E-11 |
| Lgr6 | 16.14818361 | 2.650051834 | 4.51E-02 |
| Ninj2 | 25.84237259 | 2.646382311 | 8.05E-03 |
| Erc2 | 648.4103087 | 2.644050365 | 1.08E-53 |
| Hoxb3os | 9.42460263 | 2.636567542 | 2.07E-02 |

| | | | |
|---|---|---|---|
| Thpo | 50.84521309 | 2.623262386 | 2.46E-11 |
| BC051077 | 62.17491931 | 2.622708909 | 1.00E-08 |
| Tspan33 | 25.32257853 | 2.618348197 | 4.36E-07 |
| Plaur | 1149.588067 | 2.616981551 | 1.38E-21 |
| Cd28 | 13.18085716 | 2.61692808 | 5.17E-04 |
| Gm5570 | 6.543427362 | 2.616594138 | 1.89E-02 |
| Sema6b | 29.71095728 | 2.593903407 | 1.38E-06 |
| P2rx3 | 1054.952462 | 2.593888728 | 1.10E-06 |
| Rasl10b | 32.86393239 | 2.592220865 | 1.73E-05 |
| Apol9a | 154.7926263 | 2.577175352 | 2.63E-13 |
| Tinagl1 | 7597.988131 | 2.562738189 | 3.04E-26 |
| Clu | 185.6709907 | 2.555163966 | 1.33E-15 |
| Prr5 | 206.0395211 | 2.551688165 | 5.20E-31 |
| Hist1h4h | 20.58006819 | 2.551364093 | 6.50E-03 |
| Fat2 | 8.056016086 | 2.550148897 | 4.74E-02 |
| Cilp | 7.772029667 | 2.545281741 | 3.40E-02 |
| Gm5106 | 7.079855874 | 2.543076088 | 1.47E-02 |
| Cyp3a13 | 57.57513375 | 2.539481214 | 9.16E-03 |
| Cstl1 | 15.00122308 | 2.534287142 | 3.10E-04 |
| Inmt | 199.4518932 | 2.533699391 | 2.18E-08 |
| Pard6b | 681.0243997 | 2.531566406 | 2.15E-16 |
| Cldn9 | 6.563795951 | 2.520890652 | 5.81E-03 |
| Has1 | 15.99957924 | 2.515390619 | 1.42E-02 |
| Gm17173 | 7.49841357 | 2.50996912 | 5.10E-03 |
| Gm26548 | 14.86370093 | 2.501061729 | 7.23E-05 |
| Ifi205 | 92.81123866 | 2.493768519 | 3.16E-04 |
| Sync | 42.33056854 | 2.480552429 | 1.43E-10 |
| Ccn1 | 16942.56933 | 2.466446156 | 1.32E-07 |
| Cacna1a | 1112.647342 | 2.452520281 | 9.08E-23 |
| Ip6k3 | 40.3684319 | 2.44735032 | 1.01E-05 |
| Scn3a | 16.94562687 | 2.44168899 | 2.38E-02 |
| Gadd45g | 1363.863302 | 2.440773213 | 1.29E-19 |
| Slc9a3r1 | 541.036435 | 2.436071568 | 6.67E-24 |
| Cxcl5 | 153.6533999 | 2.433539215 | 1.90E-09 |
| Mirt1 | 5.892982918 | 2.424550874 | 4.89E-02 |
| Tjp2 | 3236.987169 | 2.423731015 | 5.90E-30 |
| Gm26809 | 35780.51269 | 2.421902133 | 6.98E-14 |
| Perp | 26.99007095 | 2.419570776 | 4.33E-04 |
| Cdkl1 | 21.11717046 | 2.419538671 | 2.63E-05 |
| Gm26867 | 112.2321674 | 2.417584593 | 9.67E-09 |
| Sult1a1 | 55.76000516 | 2.417517072 | 2.25E-06 |
| Olr1 | 36.00857858 | 2.415917115 | 1.14E-04 |
| Igfbp5 | 819.6479129 | 2.407820866 | 3.12E-05 |
| Kank4os | 15.91605683 | 2.38592553 | 1.34E-03 |
| Gm17501 | 28.85545055 | 2.374744353 | 1.03E-04 |
| Sgcd | 202.5062698 | 2.37143632 | 2.53E-10 |
| Kdr | 75.92150854 | 2.368014818 | 4.22E-03 |

| | | | |
|---|---|---|---|
| Camk2a | 97.06394336 | 2.366845798 | 1.16E-08 |
| Nppb | 446.9261959 | 2.365243449 | 1.73E-08 |
| Rnf128 | 22.02821797 | 2.361498931 | 1.14E-03 |
| Ifit1bl2 | 54.30370509 | 2.344068664 | 1.29E-11 |
| Itgb7 | 197.2536301 | 2.338207558 | 8.98E-09 |
| Gngt2 | 39.73962039 | 2.331957992 | 5.92E-09 |
| Bst2 | 1204.425443 | 2.326011454 | 1.04E-18 |
| 4933427I04Rik | 39.54466148 | 2.320021158 | 1.19E-09 |
| Krt14 | 42.84356732 | 2.319253819 | 3.38E-02 |
| Hoxb1 | 35.62416815 | 2.31584146 | 1.86E-02 |
| Entpd2 | 16.74233881 | 2.314198308 | 1.17E-03 |
| Stox2 | 10.87126108 | 2.309736016 | 7.20E-03 |
| Pstpip1 | 90.41301441 | 2.295293304 | 2.14E-10 |
| Mucl3 | 111.7139975 | 2.293240319 | 1.23E-14 |
| Gm11816 | 245.2771472 | 2.292988607 | 3.93E-19 |
| Mogat2 | 37.49728946 | 2.292761684 | 1.42E-02 |
| Ces1g | 19.45501609 | 2.284422554 | 2.38E-03 |
| Eda2r | 1123.269013 | 2.276574248 | 2.08E-19 |
| Itga7 | 815.1436197 | 2.268598703 | 2.63E-13 |
| Spns3 | 13.83643692 | 2.263981747 | 2.83E-02 |
| Abat | 7.958698331 | 2.262393082 | 2.33E-02 |
| Chrnb1 | 351.1485445 | 2.257462682 | 9.78E-12 |
| Shroom3 | 71.88219185 | 2.246717972 | 5.13E-12 |
| Muc13 | 7.743443714 | 2.245981357 | 1.66E-02 |
| Trap1a | 12.33813831 | 2.243199411 | 1.20E-02 |
| Tgtp1 | 20.61468666 | 2.240935154 | 2.99E-02 |
| Slco3a1 | 1020.790683 | 2.240402043 | 1.18E-34 |
| Cep55 | 681.4845732 | 2.236083599 | 1.10E-04 |
| Tceal5 | 16.32229525 | 2.2355686 | 2.49E-02 |
| Gm26543 | 9.395393219 | 2.231048595 | 2.24E-02 |
| Cldn15 | 6.758990275 | 2.230565128 | 3.59E-02 |
| Crip2 | 63.44237168 | 2.220168436 | 6.34E-19 |
| Klf4 | 2395.995965 | 2.200240025 | 4.11E-32 |
| Eln | 24.79874943 | 2.196364112 | 3.74E-04 |
| Pla1a | 89.47502873 | 2.196133047 | 6.93E-06 |
| 4930458D05Rik | 61.19370456 | 2.194715972 | 8.45E-08 |
| Ntn4 | 184.2709408 | 2.188528289 | 1.31E-16 |
| Igfbp6 | 10099.86965 | 2.185615003 | 1.17E-29 |
| Soat2 | 50.10265489 | 2.183504019 | 2.15E-06 |
| NA | 7.035009564 | 2.176566883 | 2.63E-02 |
| Cd80 | 43.14414784 | 2.174573062 | 2.24E-04 |
| Ddah1 | 1706.491516 | 2.173652477 | 4.14E-13 |
| Anln | 4893.272358 | 2.172755435 | 3.11E-09 |
| Elmo1 | 276.9450485 | 2.167154302 | 3.69E-03 |
| Ncf2 | 13.41769721 | 2.162843229 | 4.20E-02 |
| Tmem130 | 8.431316085 | 2.162597443 | 1.61E-02 |
| Serpine1 | 8069.154667 | 2.156711009 | 1.67E-09 |

| | | | |
|---|---|---|---|
| Gm26640 | 7.68673924 | 2.152698613 | 4.50E-02 |
| Diaph3 | 1284.11 | 2.152503762 | 2.24E-07 |
| Aif1l | 2280.404424 | 2.147373947 | 9.13E-24 |
| Cdc42bpg | 72.4716449 | 2.143353949 | 5.05E-03 |
| Mamdc2 | 140.4200342 | 2.136566242 | 1.91E-10 |
| Pygm | 20.09698139 | 2.133296124 | 1.54E-02 |
| Dnah6 | 60.00583954 | 2.132947341 | 2.46E-07 |
| Csprs | 36.23461021 | 2.130547583 | 1.78E-02 |
| Cd74 | 18.46669239 | 2.121745162 | 2.42E-02 |
| NA | 5457.007199 | 2.116702155 | 1.82E-09 |
| Klhdc8a | 1128.698314 | 2.11063029 | 4.49E-12 |
| Tbxa2r | 12.46534744 | 2.11060199 | 6.22E-03 |
| Ly6a | 4142.015759 | 2.109934492 | 7.78E-17 |
| As3mt | 32.98295994 | 2.106620818 | 1.46E-04 |
| Prx | 427.4098247 | 2.097209252 | 9.11E-22 |
| Gm1141 | 15.33114807 | 2.093284568 | 1.84E-03 |
| Cfap70 | 23.3829331 | 2.09252154 | 4.54E-02 |
| Gm7241 | 14.92296646 | 2.091623591 | 3.86E-03 |
| Slc22a13b-ps | 8.826899059 | 2.089499539 | 2.29E-02 |
| Aqp1 | 1103.311567 | 2.085917868 | 2.05E-04 |
| Igfbp4 | 605.6600899 | 2.084725937 | 7.77E-07 |
| Suox | 3135.576887 | 2.084557055 | 3.66E-27 |
| Efcab8 | 17.7494641 | 2.084537777 | 6.11E-03 |
| Dusp9 | 278.6759648 | 2.082138749 | 2.22E-11 |
| Gm23935 | 1035.186958 | 2.078273241 | 8.07E-03 |
| Ptchd4 | 348.94966 | 2.076675605 | 3.04E-05 |
| Fads3 | 4292.424506 | 2.07257171 | 1.86E-17 |
| Gm16998 | 21.06629374 | 2.068697896 | 6.32E-04 |
| Scara5 | 982.0551509 | 2.066291948 | 5.60E-04 |
| Pdlim1 | 2636.81262 | 2.061872504 | 1.60E-10 |
| Ttll2 | 60.65077863 | 2.057718398 | 1.40E-02 |
| Olfr1318 | 12.8937525 | 2.055985073 | 3.62E-02 |
| Gm16702 | 278.1798623 | 2.044956913 | 1.43E-10 |
| Gm14296 | 55.8943979 | 2.041226079 | 3.15E-07 |
| Ablim3 | 364.5652957 | 2.037087653 | 2.48E-07 |
| Gm16178 | 25.22230611 | 2.036858947 | 5.06E-05 |
| Sema3c | 667.7097169 | 2.018236796 | 7.87E-04 |
| Trpv4 | 13.02702406 | 2.01686605 | 7.82E-03 |
| Pof1b | 9.373576499 | 2.016470497 | 4.31E-02 |
| Usp17le | 14.67590836 | 2.014751961 | 3.18E-02 |
| Ica1 | 26.09752958 | 2.014156695 | 3.63E-03 |
| Ankrd1 | 4164.102927 | 2.013626297 | 5.92E-09 |
| Steap1 | 1192.998083 | 2.008988058 | 1.32E-10 |
| Ablim1 | 3555.853548 | 1.995596925 | 3.79E-21 |
| Eml2 | 184.9655233 | 1.991664755 | 9.98E-09 |
| 9530053A07Rik | 64.37286542 | 1.987898997 | 6.34E-03 |
| Cpox | 1136.258528 | 1.986700056 | 3.01E-16 |

| | | | |
|---|---|---|---|
| C1qtnf2 | 17.27844736 | 1.979629196 | 1.32E-03 |
| Klk8 | 65.88130734 | 1.976322775 | 4.58E-03 |
| Slmapos2 | 3326.360358 | 1.974136068 | 7.28E-12 |
| Trib3 | 1385.443885 | 1.973798034 | 7.01E-08 |
| Srpx2 | 547.7024361 | 1.972825637 | 1.74E-03 |
| 4-Sep | 46.97005786 | 1.970650012 | 9.94E-06 |
| Neurl1a | 101.6784878 | 1.964631377 | 1.51E-11 |
| Pi15 | 156.8925324 | 1.959439862 | 1.08E-07 |
| Als2cr12 | 58.85300664 | 1.949060222 | 5.32E-06 |
| Sorbs2 | 490.7709526 | 1.939055772 | 2.45E-06 |
| Cdh19 | 11.64226065 | 1.919367533 | 2.54E-02 |
| Vmn2r96 | 15.73884589 | 1.917198833 | 9.10E-05 |
| Rflnb | 613.7469556 | 1.911143293 | 2.23E-19 |
| Spon1 | 17.90306055 | 1.898387746 | 8.65E-03 |
| Cryab | 13202.45602 | 1.885770306 | 7.32E-16 |
| Psg19 | 9.355351805 | 1.884539802 | 3.67E-02 |
| Gbp11 | 15.68366756 | 1.884020309 | 7.78E-03 |
| Lipg | 535.1730908 | 1.878660068 | 1.86E-16 |
| Plekha2 | 266.0834896 | 1.87796134 | 1.87E-08 |
| Chn1 | 68.77845635 | 1.876006767 | 4.29E-07 |
| Ect2 | 1479.659604 | 1.872957795 | 3.29E-04 |
| Ppef1 | 316.4882102 | 1.872615127 | 1.85E-20 |
| Gnrh1 | 12.96179263 | 1.872572324 | 4.12E-03 |
| Gm7609 | 148.5996821 | 1.872392123 | 1.45E-04 |
| Abcb11 | 9.530869549 | 1.871223633 | 9.98E-03 |
| Fgd6 | 331.6053799 | 1.870470744 | 9.02E-11 |
| Clcn3 | 952.4309388 | 1.869732747 | 7.02E-13 |
| S100a11 | 28004.48893 | 1.864237744 | 1.57E-17 |
| Zdhhc2 | 222.3920511 | 1.862781034 | 2.24E-08 |
| 1810019D21Rik | 9.516214438 | 1.850402225 | 2.76E-02 |
| Gm15337 | 9.801948551 | 1.849467853 | 1.04E-02 |
| Trim71 | 10.11243533 | 1.847184949 | 3.15E-02 |
| 2610316D01Rik | 31.67538577 | 1.845364857 | 1.29E-03 |
| Smim3 | 135.0460714 | 1.840600863 | 6.51E-15 |
| Ephx1 | 9309.25785 | 1.840152872 | 1.83E-18 |
| Arhgap45 | 18.56644359 | 1.835287154 | 1.98E-02 |
| Slc45a3 | 239.2303382 | 1.830737256 | 8.45E-13 |
| 9330185C12Rik | 10.38647242 | 1.825356477 | 2.74E-02 |
| Mgam | 514.7573389 | 1.824874349 | 7.64E-04 |
| Igf2bp1 | 2960.658169 | 1.82259186 | 4.38E-75 |
| Rap1gap2 | 970.5392269 | 1.821330656 | 3.06E-11 |
| Prrt3 | 27.4990097 | 1.820092905 | 8.72E-04 |
| Hist1h2ac | 24.74191821 | 1.81696976 | 1.04E-02 |
| Clec3b | 429.6188855 | 1.8147427 | 2.37E-02 |
| Dnph1 | 336.876214 | 1.810322868 | 2.69E-08 |
| Phactr4 | 1891.975684 | 1.80995678 | 1.45E-24 |
| Mcam | 18.62577123 | 1.806703966 | 2.67E-02 |

| | | | |
|---|---|---|---|
| Platr15 | 27.55767206 | 1.804708201 | 2.38E-04 |
| Pcdh10 | 140.2969449 | 1.799857598 | 1.73E-11 |
| Cavin2 | 13533.10625 | 1.798972228 | 5.61E-07 |
| Rab30 | 327.3676624 | 1.796478951 | 3.76E-06 |
| Inka2 | 122.816321 | 1.79476299 | 1.34E-04 |
| S100a3 | 56.32424442 | 1.794294461 | 1.14E-05 |
| Ptgs2os | 66.7624299 | 1.789688799 | 1.01E-05 |
| Hoxa1 | 320.536735 | 1.78837464 | 4.45E-13 |
| Adam33 | 32.31827042 | 1.783163921 | 6.01E-05 |
| Mmp28 | 209.9282617 | 1.781617729 | 6.78E-08 |
| Nr1h5 | 425.4240054 | 1.780812257 | 7.35E-04 |
| Itgb4 | 228.6910735 | 1.779666193 | 4.23E-02 |
| Gm26525 | 46.56506484 | 1.777124369 | 8.68E-04 |
| E2f7 | 788.1276993 | 1.771761062 | 1.72E-06 |
| A430106G13Rik | 6.653140069 | 1.771739617 | 4.77E-02 |
| Fosl1 | 1007.39669 | 1.771350798 | 7.10E-08 |
| Serpinb1b | 80.38277849 | 1.771231214 | 1.07E-04 |
| Sh3tc2 | 46.01764535 | 1.770219855 | 3.74E-04 |
| Msln | 8968.235968 | 1.768793096 | 1.22E-02 |
| Serpina9 | 31.42166562 | 1.767633293 | 1.17E-03 |
| Eqtn | 8.292335483 | 1.75950755 | 3.49E-02 |
| Gm15731 | 11.98380975 | 1.756297365 | 2.35E-02 |
| Kcnab3 | 152.1664527 | 1.756247899 | 1.43E-09 |
| Dpep2 | 58.14460163 | 1.754301476 | 6.86E-06 |
| Klhl30 | 170.2009204 | 1.751710949 | 7.97E-04 |
| Fendrr | 16.4166356 | 1.748236701 | 2.45E-02 |
| Cd59a | 250.5614553 | 1.74652754 | 1.49E-11 |
| Prune2 | 851.2860509 | 1.744256083 | 1.45E-12 |
| Slc7a5 | 7418.657169 | 1.741455924 | 1.08E-06 |
| 4931428L18Rik | 8.34340106 | 1.741083065 | 3.90E-02 |
| Rhox5 | 444.7664169 | 1.740782498 | 1.82E-09 |
| Phex | 174.470391 | 1.733817514 | 1.12E-06 |
| Lama5 | 488.2888004 | 1.732251804 | 9.71E-03 |
| Tmem204 | 18.79207328 | 1.731332869 | 1.39E-03 |
| Gm5380 | 24.31303982 | 1.72590977 | 2.39E-03 |
| Nrcam | 42.60013369 | 1.724307591 | 1.21E-03 |
| Acy3 | 101.9828741 | 1.723186715 | 1.57E-09 |
| Prelp | 2798.880483 | 1.719429797 | 6.14E-04 |
| Sap25 | 22.93362904 | 1.717014873 | 3.41E-02 |
| Arhgap18 | 896.4577738 | 1.712131813 | 7.00E-06 |
| Krt80 | 423.3382572 | 1.708599708 | 1.15E-07 |
| Ddias | 367.2417911 | 1.706900104 | 7.33E-05 |
| Fstl3 | 1713.818872 | 1.703775229 | 2.77E-15 |
| Vgf | 80.48610746 | 1.702876116 | 2.73E-06 |
| E2f2 | 41.48521494 | 1.700320495 | 1.37E-02 |
| Tmcc3 | 374.2348867 | 1.699334998 | 1.17E-10 |
| Ryr1 | 20.13106927 | 1.691966505 | 7.94E-03 |

| | | | |
|---|---|---|---|
| Igf1 | 89.98250005 | 1.689573407 | 3.51E-04 |
| Ercc6l2 | 120.8231153 | 1.689188568 | 3.15E-08 |
| Crybg2 | 69.94425025 | 1.688183197 | 7.28E-05 |
| Nol4l | 22.67327197 | 1.687044303 | 4.51E-02 |
| Atf3 | 486.7080334 | 1.68677177 | 6.03E-14 |
| Gpr39 | 133.0862302 | 1.686194 | 1.94E-04 |
| Svep1 | 11238.12662 | 1.684205841 | 2.03E-10 |
| Pi16 | 9.391261602 | 1.683791144 | 2.74E-02 |
| Gm20521 | 108.1932135 | 1.682518847 | 2.84E-03 |
| Spc24 | 355.8336016 | 1.680614905 | 4.45E-06 |
| Ccl9 | 880.6190831 | 1.678619581 | 2.79E-05 |
| Ogn | 19778.96094 | 1.67486476 | 7.91E-03 |
| Mbnl3 | 134.2834975 | 1.673992542 | 7.29E-08 |
| Gm22620 | 10.99034939 | 1.673163683 | 4.20E-02 |
| Slc6a15 | 33.49232237 | 1.673065601 | 1.00E-02 |
| Ehd1 | 9278.785003 | 1.66548478 | 1.01E-18 |
| Gm11525 | 36.80127275 | 1.657841255 | 6.56E-04 |
| 2210016F16Rik | 2097.051085 | 1.656628624 | 6.08E-13 |
| Zfp52 | 703.5987578 | 1.654445055 | 1.24E-04 |
| Ctps | 9423.67962 | 1.647369565 | 9.11E-22 |
| Cldn34c1 | 161.8468038 | 1.647316626 | 1.52E-08 |
| Gm12138 | 2019.192784 | 1.645370209 | 6.62E-07 |
| Gm10382 | 31.87078083 | 1.643605026 | 2.61E-02 |
| Arhgap29 | 5638.40206 | 1.642786578 | 1.75E-11 |
| Ccl5 | 19.03032931 | 1.641375752 | 2.54E-02 |
| Atp2b4 | 128.2057106 | 1.638537826 | 1.02E-03 |
| Tfrc | 1850.025373 | 1.638496088 | 1.81E-03 |
| Gm2115 | 916.8914908 | 1.637331502 | 1.81E-09 |
| Gm4876 | 18.04393838 | 1.63523283 | 2.73E-02 |
| Gm13285 | 29.53639999 | 1.630728666 | 2.52E-03 |
| Tmem62 | 479.9034088 | 1.628186927 | 5.49E-21 |
| Csdc2 | 64.87675074 | 1.626818063 | 7.43E-05 |
| Emx2 | 282.5000379 | 1.624094171 | 7.31E-10 |
| Hfe | 52.7239069 | 1.623935466 | 5.29E-05 |
| Plxna4 | 92.75828692 | 1.623470756 | 7.86E-04 |
| Ripk3 | 33.31369269 | 1.623128858 | 1.65E-02 |
| Gm4841 | 16.51594958 | 1.618877185 | 2.79E-02 |
| Gm12128 | 11.44534595 | 1.615712866 | 3.13E-02 |
| Gm19705 | 81.49688775 | 1.615576253 | 3.23E-07 |
| P4ha3 | 952.268623 | 1.615410057 | 7.23E-09 |
| Adarb2 | 76.97265872 | 1.615106093 | 4.47E-04 |
| Csrnp1 | 607.3843703 | 1.614956573 | 7.60E-06 |
| Gtse1 | 869.2872648 | 1.614043789 | 4.26E-05 |
| Camsap3 | 10.74646449 | 1.610108624 | 2.85E-02 |
| NA | 125.6560357 | 1.609792587 | 7.34E-04 |
| Pla2g4b | 67.61780263 | 1.609565679 | 8.09E-03 |
| Bglap3 | 128.1990344 | 1.607631334 | 1.64E-14 |

| | | | |
|---|---|---|---|
| Cd53 | 21.19451119 | 1.606267257 | 4.40E-02 |
| Acer1 | 15.46558608 | 1.605531383 | 4.42E-02 |
| Pfkp | 1611.630881 | 1.604161004 | 2.39E-12 |
| Cystm1 | 169.2870408 | 1.603032999 | 1.25E-04 |
| Lyn | 252.2098934 | 1.593595143 | 1.61E-08 |
| Golga7b | 16.54206909 | 1.592260687 | 2.58E-02 |
| Vldlr | 180.1056824 | 1.590018258 | 1.34E-05 |
| Gm5620 | 281.5414567 | 1.588995732 | 5.58E-05 |
| Zc2hc1c | 99.23641284 | 1.586041545 | 1.05E-07 |
| Psat1 | 15457.79927 | 1.585950381 | 1.97E-07 |
| Cd14 | 165.7940412 | 1.585674684 | 2.42E-08 |
| Gm14444 | 29.84845564 | 1.585146592 | 4.00E-03 |
| Hist1h4i | 68.69874854 | 1.585033001 | 1.49E-02 |
| Nek2 | 1268.788324 | 1.58217768 | 2.71E-06 |
| Steap2 | 5089.313269 | 1.578717642 | 1.33E-14 |
| B3gnt5 | 34.6745235 | 1.57782764 | 3.14E-02 |
| Gm6410 | 85.28209271 | 1.576732862 | 3.03E-07 |
| Rrm2 | 2219.879428 | 1.572306232 | 3.29E-03 |
| Dusp1 | 2323.364801 | 1.570209404 | 1.69E-06 |
| Nup210l | 38.33852126 | 1.56906497 | 3.15E-06 |
| Iqgap3 | 1736.914357 | 1.566066326 | 4.02E-05 |
| Sh3rf2 | 283.6625641 | 1.566027838 | 5.40E-03 |
| Mybl2 | 567.7128471 | 1.549508815 | 5.20E-05 |
| Gsta1 | 191.6070784 | 1.545213673 | 1.02E-03 |
| Odc1 | 6015.135819 | 1.543613889 | 1.33E-27 |
| Rnf225 | 18.6574331 | 1.541270254 | 2.31E-02 |
| Ptgds | 167.4192768 | 1.536544323 | 6.50E-05 |
| Gcnt3 | 16.25473207 | 1.532634924 | 3.13E-02 |
| Sh2d6 | 55.41535397 | 1.53138873 | 2.40E-04 |
| Arl5c | 11.25248985 | 1.529881133 | 4.16E-02 |
| Akr1b3 | 27184.12226 | 1.525573339 | 1.35E-05 |
| Serp2 | 10.36553764 | 1.521990535 | 4.01E-02 |
| Angptl6 | 251.3102899 | 1.515499033 | 1.71E-08 |
| Ism1 | 148.8317432 | 1.511151674 | 3.19E-04 |
| 1700020L24Rik | 39.09349375 | 1.501999345 | 4.16E-04 |
| Arhgap20 | 61.50283451 | 1.49330679 | 9.44E-03 |
| Il11 | 23.34212275 | 1.492209605 | 2.94E-03 |
| Tent5b | 1620.453803 | 1.488511971 | 4.26E-17 |
| Rin3 | 39.43852099 | 1.487956277 | 6.76E-04 |
| Boc | 364.9113936 | 1.486543074 | 3.90E-06 |
| Prkg1 | 134.4890151 | 1.485615048 | 4.71E-05 |
| Tnr | 97.7195546 | 1.485403745 | 1.20E-02 |
| Cd59b | 100.6980068 | 1.484244187 | 2.16E-03 |
| Plac1 | 35.76022485 | 1.48419836 | 2.00E-03 |
| Hrct1 | 51.68066241 | 1.483223456 | 1.77E-04 |
| Calhm5 | 378.443981 | 1.481600883 | 2.90E-19 |
| Atp10a | 1864.778291 | 1.481218917 | 2.62E-09 |

| | | | |
|---|---|---|---|
| Sorbs1 | 833.0862304 | 1.479501369 | 3.46E-15 |
| Smpd3 | 867.4805472 | 1.471760582 | 2.39E-12 |
| Ifit1 | 122.4955534 | 1.471512552 | 1.30E-06 |
| Lfng | 220.6117941 | 1.469825141 | 1.87E-07 |
| Ajuba | 2948.78609 | 1.468208941 | 2.33E-24 |
| Raph1 | 579.1077271 | 1.467712639 | 1.05E-09 |
| Dqx1 | 97.9924442 | 1.466207129 | 3.32E-07 |
| Glt8d2 | 963.6849507 | 1.464222431 | 7.30E-10 |
| Asf1b | 669.9867269 | 1.464210178 | 4.64E-04 |
| Gm16364 | 22.25215599 | 1.463985882 | 2.11E-03 |
| Ppm1h | 131.7107471 | 1.459684323 | 2.39E-03 |
| Gpc6 | 435.6774066 | 1.459499402 | 5.30E-08 |
| Ly6e | 6968.053691 | 1.459494922 | 3.29E-15 |
| Foxm1 | 2886.173548 | 1.458783784 | 2.57E-04 |
| Hells | 1567.737808 | 1.45674147 | 4.55E-02 |
| Gm23130 | 69.16721741 | 1.449408388 | 3.02E-02 |
| Itga6 | 3086.584435 | 1.448605902 | 1.90E-19 |
| Creld1 | 74.5768632 | 1.447554767 | 5.45E-04 |
| Kctd9 | 1161.401207 | 1.44580734 | 1.93E-07 |
| Nqo1 | 596.4674168 | 1.444536986 | 1.51E-05 |
| Tnxb | 839.0427032 | 1.443305286 | 2.25E-19 |
| NA | 49.426301 | 1.437341546 | 7.97E-03 |
| Aunip | 99.4352662 | 1.43723597 | 1.31E-02 |
| Tns4 | 36.47295026 | 1.436642775 | 3.08E-02 |
| Sorbs3 | 4375.993113 | 1.433100869 | 4.42E-23 |
| Cd55 | 298.5806079 | 1.424436207 | 8.41E-03 |
| Neo1 | 4545.56012 | 1.423371306 | 2.71E-41 |
| Clic3 | 21.76777198 | 1.422678696 | 4.43E-02 |
| 5031410I06Rik | 317.2038905 | 1.421071534 | 6.23E-03 |
| Rad54b | 268.0386838 | 1.420004923 | 1.22E-03 |
| Nexn | 586.7586746 | 1.418357537 | 2.98E-03 |
| 4930558J18Rik | 24.7967093 | 1.414768692 | 1.12E-02 |
| Sapcd2 | 288.9189524 | 1.41367249 | 6.26E-03 |
| Fam117a | 85.5208589 | 1.412314654 | 1.22E-03 |
| Lrr1 | 63.46400517 | 1.411115349 | 1.35E-02 |
| Igfbp7 | 2473.484216 | 1.410049642 | 6.43E-08 |
| Eml5 | 284.730193 | 1.408890845 | 4.71E-05 |
| Slc25a25 | 821.5438689 | 1.4076155 | 8.92E-08 |
| Tmpo | 7007.127582 | 1.406989709 | 6.23E-15 |
| Defb41 | 36.08056729 | 1.406144996 | 1.44E-04 |
| Acot2 | 886.3418002 | 1.40613361 | 1.32E-12 |
| Gria3 | 1179.231791 | 1.402743017 | 3.12E-09 |
| Naaladl1 | 22.98190992 | 1.399971233 | 4.89E-02 |
| Gm38393 | 571.0226571 | 1.398903007 | 3.43E-05 |
| Colq | 10.81989595 | 1.397719909 | 2.05E-02 |
| Ckap2 | 2101.539709 | 1.395734901 | 2.66E-05 |
| Cd93 | 188.2576324 | 1.392144903 | 2.13E-03 |

| | | | |
|---|---|---|---|
| Fgd3 | 456.6207369 | 1.391075903 | 9.62E-05 |
| Abca4 | 66.55109914 | 1.387360862 | 5.47E-04 |
| Galnt15 | 67.07683312 | 1.387045688 | 3.64E-03 |
| Prc1 | 2935.87793 | 1.386494081 | 6.01E-05 |
| Mfsd4b5 | 246.1474623 | 1.381646727 | 1.24E-06 |
| Crlf1 | 353.0096371 | 1.372650609 | 2.19E-04 |
| Cip2a | 1058.148088 | 1.372463874 | 1.98E-02 |
| Mtfr2 | 211.4222988 | 1.372073357 | 2.25E-03 |
| Tk1 | 1172.738402 | 1.369127887 | 1.07E-03 |
| Lmcd1 | 190.9655838 | 1.368534625 | 7.19E-03 |
| Ube2c | 1898.382155 | 1.363651982 | 1.04E-04 |
| Ska1 | 241.1235717 | 1.362592539 | 3.29E-03 |
| Dbf4 | 1797.842921 | 1.361640829 | 1.21E-05 |
| Maff | 1125.074098 | 1.360849236 | 4.42E-07 |
| Kif18b | 709.5316689 | 1.360704652 | 3.09E-04 |
| Sp100 | 229.87064 | 1.360096317 | 1.94E-04 |
| Gm10660 | 29.5555364 | 1.359316717 | 3.32E-02 |
| Pmaip1 | 85.15518226 | 1.357471266 | 9.58E-03 |
| NA | 87.8527127 | 1.353770243 | 1.58E-02 |
| Hipk4 | 55.60434058 | 1.353371051 | 1.05E-04 |
| Dennd5b | 74.30478023 | 1.348040548 | 7.97E-03 |
| Gm26672 | 17.76847359 | 1.345964418 | 9.91E-03 |
| Ccna2 | 3121.371366 | 1.341690769 | 1.55E-03 |
| Hipk2 | 2809.337586 | 1.340131171 | 3.93E-19 |
| Sms | 1954.671987 | 1.33604339 | 4.59E-16 |
| NA | 37.82154378 | 1.334534289 | 2.79E-02 |
| Fanca | 670.1156277 | 1.329567018 | 7.71E-04 |
| Fam111a | 2899.275804 | 1.326803015 | 3.17E-05 |
| Cavin4 | 290.9811514 | 1.325980012 | 1.46E-06 |
| Gm20465 | 16.41696527 | 1.324318284 | 4.48E-02 |
| Dmpk | 271.5900462 | 1.323712762 | 7.88E-06 |
| Nmnat3 | 174.6720184 | 1.323065334 | 5.51E-08 |
| Gm10825 | 19.23658637 | 1.318867344 | 3.12E-02 |
| Agt | 230.4174338 | 1.318047575 | 4.24E-03 |
| Rras2 | 2707.729271 | 1.316567902 | 2.91E-11 |
| Eno3 | 744.5122267 | 1.314383287 | 7.93E-05 |
| Mybl1 | 202.1011964 | 1.313343191 | 8.16E-03 |
| Cdk1 | 3299.072583 | 1.312762871 | 1.37E-04 |
| Wdhd1 | 811.8103482 | 1.311731202 | 3.51E-02 |
| Cdc6 | 389.1138025 | 1.311249737 | 8.05E-03 |
| Col12a1 | 2667.384937 | 1.308926783 | 1.22E-20 |
| Gm5526 | 15.93317991 | 1.304139835 | 2.85E-02 |
| Cd151 | 4413.725542 | 1.303407641 | 8.28E-17 |
| Recql4 | 294.0795684 | 1.298543724 | 2.42E-03 |
| Hectd2 | 366.6262872 | 1.294351438 | 3.90E-06 |
| Gata3 | 165.7026761 | 1.293446565 | 3.25E-03 |
| Pla2g16 | 886.3125233 | 1.291895303 | 8.57E-08 |

| | | | |
|---|---|---|---|
| Frat2 | 305.7883249 | 1.290981539 | 4.99E-06 |
| Cd84 | 73.90841491 | 1.290071316 | 1.28E-07 |
| Gm14427 | 29.03266401 | 1.28794744 | 1.10E-02 |
| Stambpl1 | 1280.981375 | 1.287928783 | 5.51E-12 |
| Tmem238 | 87.91425267 | 1.286507448 | 1.32E-03 |
| Olfm1 | 251.9775343 | 1.28508796 | 3.08E-03 |
| Spsb1 | 515.9985486 | 1.283867467 | 3.24E-16 |
| Plk1 | 1159.243141 | 1.283550714 | 3.18E-03 |
| Sgms2 | 1191.807906 | 1.279229695 | 2.49E-17 |
| Mtss1 | 37.26253904 | 1.27820062 | 2.15E-03 |
| Wwc1 | 1254.514596 | 1.276616198 | 1.37E-06 |
| Raet1e | 152.5635618 | 1.276564876 | 6.23E-07 |
| Col4a5 | 13744.73591 | 1.273863935 | 4.50E-23 |
| Polq | 283.6487716 | 1.264476263 | 1.56E-02 |
| Fzd4 | 527.5932981 | 1.263641459 | 9.02E-05 |
| M1ap | 53.57928597 | 1.261916564 | 1.27E-02 |
| Vstm5 | 49.36700862 | 1.261536113 | 4.52E-05 |
| Xpnpep2 | 463.2230938 | 1.260660209 | 3.28E-05 |
| Lmod1 | 164.8111626 | 1.259700483 | 3.75E-03 |
| Tspan9 | 8905.533771 | 1.258364694 | 7.95E-14 |
| D630029K05Rik | 40.79021596 | 1.25705613 | 1.04E-02 |
| Sgo1 | 501.443209 | 1.256125254 | 8.83E-03 |
| Tnfrsf23 | 1998.923511 | 1.255201249 | 2.37E-06 |
| Kifc1 | 820.5197881 | 1.255007366 | 1.00E-04 |
| Troap | 510.2589497 | 1.254310364 | 3.59E-03 |
| 1700034P13Rik | 389.2191072 | 1.25140169 | 3.51E-11 |
| Knstrn | 1791.183893 | 1.250642789 | 5.36E-06 |
| Mthfd2 | 2675.163557 | 1.248065954 | 3.35E-04 |
| Znrf2 | 1311.359839 | 1.246553879 | 3.32E-11 |
| Dhdh | 34.0646641 | 1.245919077 | 1.75E-03 |
| Adamtsl4 | 642.4400238 | 1.244990966 | 5.64E-13 |
| Cdc25c | 279.9314129 | 1.244775785 | 2.39E-03 |
| Chek1 | 275.9564489 | 1.244636139 | 8.59E-03 |
| Ppp1r12b | 4138.333762 | 1.243095273 | 4.05E-16 |
| Fgfbp3 | 84.82346159 | 1.242206544 | 7.34E-06 |
| Hspg2 | 28446.74545 | 1.240942023 | 3.47E-09 |
| Tpx2 | 3448.017744 | 1.239023253 | 2.01E-04 |
| Ggct | 110.6567063 | 1.236042416 | 1.46E-04 |
| Gm6736 | 924.2982575 | 1.235436387 | 2.37E-03 |
| Kif23 | 1634.120811 | 1.234679647 | 1.34E-03 |
| Lrrc1 | 506.016248 | 1.234255164 | 3.70E-12 |
| Trim16 | 1755.34812 | 1.23329466 | 6.20E-12 |
| Gfod1 | 804.2426049 | 1.230561233 | 3.76E-05 |
| 2810408I11Rik | 99.88867526 | 1.229783702 | 2.91E-02 |
| Styk1 | 452.3674977 | 1.227799135 | 2.93E-04 |
| Dpep1 | 950.7647622 | 1.227606653 | 3.57E-03 |
| Lyrm9 | 234.5217743 | 1.227417348 | 1.91E-05 |

| | | | |
|---|---|---|---|
| Stac3 | 27.44952208 | 1.22700578 | 1.11E-02 |
| Wnt9a | 916.1471833 | 1.226669828 | 1.24E-06 |
| Aurka | 1220.677723 | 1.225846832 | 3.40E-03 |
| Ezr | 2125.964047 | 1.225822774 | 9.75E-17 |
| 1700028J19Rik | 26.72022137 | 1.224298884 | 1.86E-02 |
| Hspb7 | 111.5849735 | 1.222738355 | 4.55E-02 |
| Mad2l1 | 1774.898828 | 1.219687123 | 2.23E-04 |
| Arhgap19 | 667.7567467 | 1.216716843 | 2.00E-04 |
| Chd5 | 78.15936498 | 1.21576964 | 3.17E-02 |
| Hist1h2be | 35.43583052 | 1.214748942 | 3.08E-02 |
| Vmn1r-ps128 | 315.8095013 | 1.213144706 | 7.93E-03 |
| Chst7 | 291.6271761 | 1.213013307 | 3.71E-06 |
| Osgin1 | 375.3213527 | 1.212887949 | 7.38E-05 |
| Unc13d | 41.0409234 | 1.208839537 | 4.51E-03 |
| Tuba1b | 52546.48143 | 1.207657297 | 6.77E-05 |
| Ngf | 909.5389495 | 1.205886552 | 4.71E-07 |
| Map1b | 6864.42189 | 1.205273515 | 6.53E-13 |
| Spaca6 | 1058.427464 | 1.204651818 | 4.42E-04 |
| Syt12 | 32.59906048 | 1.203821538 | 2.68E-02 |
| Prdx6 | 12709.33805 | 1.203372278 | 3.03E-09 |
| Parpbp | 379.4779051 | 1.202705218 | 9.55E-03 |
| Cnn2 | 23457.26203 | 1.202173112 | 5.29E-23 |
| Gm5844 | 37.75653119 | 1.199568444 | 3.74E-02 |
| Hmmr | 1694.008987 | 1.199138804 | 7.20E-03 |
| Slc35g1 | 93.21178618 | 1.198606993 | 5.89E-04 |
| Cdc20 | 2852.257054 | 1.197652747 | 8.56E-04 |
| 2700099C18Rik | 244.6458142 | 1.197469792 | 1.07E-02 |
| Samhd1 | 3737.536878 | 1.193783623 | 2.18E-20 |
| Abcb1b | 1465.604472 | 1.192484244 | 2.97E-11 |
| Slc5a5 | 16.03111384 | 1.192270996 | 3.56E-02 |
| Pitpnc1 | 829.297569 | 1.190238136 | 6.68E-06 |
| Tigit | 30.28802111 | 1.186709681 | 3.95E-02 |
| Mcm5 | 1985.10944 | 1.182636997 | 3.36E-03 |
| Endod1 | 423.568897 | 1.181103926 | 9.91E-08 |
| Nsl1 | 311.0216032 | 1.176772097 | 2.51E-03 |
| Mpp6 | 1620.758067 | 1.17509792 | 2.50E-06 |
| Cenpv | 267.4788372 | 1.174702523 | 2.50E-08 |
| Gstt1 | 157.0937604 | 1.174362304 | 9.31E-03 |
| Nuak2 | 683.3125158 | 1.174325485 | 3.21E-09 |
| Sgms1 | 1435.255104 | 1.174020339 | 3.89E-10 |
| Shmt2 | 7386.878222 | 1.172610736 | 4.49E-05 |
| Stard13 | 1677.000687 | 1.172573384 | 7.64E-04 |
| Map2k3os | 158.9798746 | 1.169408956 | 1.11E-03 |
| Cltb | 2400.270914 | 1.167201781 | 5.00E-08 |
| Orc1 | 478.3133488 | 1.16387541 | 5.32E-04 |
| G430049J08Rik | 23.43563077 | 1.1623295 | 1.41E-02 |
| Ly6c1 | 1095.469361 | 1.162068603 | 2.37E-06 |

| | | | |
|---|---|---|---|
| Gm15541 | 42.64852593 | 1.160372576 | 3.97E-03 |
| Tdg | 173.5710662 | 1.160278211 | 5.63E-06 |
| Dusp14 | 1099.248265 | 1.160139997 | 1.50E-07 |
| Mcu | 1703.920113 | 1.158925202 | 1.66E-24 |
| Col8a1 | 3683.903915 | 1.156491843 | 1.04E-04 |
| Psph | 959.6907578 | 1.156283489 | 1.17E-03 |
| Arvcf | 1255.272406 | 1.156130883 | 3.61E-14 |
| Klf2 | 230.8305466 | 1.155705605 | 5.09E-03 |
| Nuf2 | 1145.648729 | 1.155412562 | 9.58E-03 |
| Raet1d | 118.6635537 | 1.154247936 | 3.92E-04 |
| Nectin2 | 3231.192606 | 1.152064826 | 8.05E-17 |
| NA | 127.6455783 | 1.150720438 | 1.09E-04 |
| Paqr4 | 951.0335697 | 1.149940775 | 2.73E-07 |
| Ifngr1 | 2047.302005 | 1.149555743 | 1.66E-24 |
| Apoe | 75.61519687 | 1.148382264 | 1.10E-04 |
| Cdca8 | 1439.881188 | 1.14793184 | 1.60E-03 |
| Zwilch | 669.3877584 | 1.145797704 | 9.99E-04 |
| Slc39a10 | 3524.383162 | 1.145785911 | 1.01E-05 |
| Rad51 | 623.6209141 | 1.145450496 | 2.94E-03 |
| Cmah | 754.9686317 | 1.135805463 | 8.80E-03 |
| Mndal | 41.59689426 | 1.135773394 | 3.15E-02 |
| Ifrd1 | 4709.926875 | 1.135617292 | 1.97E-07 |
| Stc2 | 605.8941595 | 1.134875403 | 4.64E-04 |
| Ptpn21 | 3194.640395 | 1.132212709 | 1.63E-11 |
| Fancb | 219.634365 | 1.131938196 | 2.44E-02 |
| Birc5 | 1048.907529 | 1.13151229 | 1.23E-02 |
| Gmnn | 367.7893567 | 1.129202721 | 9.45E-04 |
| Chrd | 441.3963201 | 1.128266673 | 8.64E-07 |
| Kcnh2 | 22.92218311 | 1.125092667 | 2.54E-02 |
| Cit | 699.3070544 | 1.124932644 | 5.19E-03 |
| Cenpn | 611.6297696 | 1.124560807 | 2.37E-04 |
| Pif1 | 283.1872027 | 1.123505185 | 1.40E-02 |
| Hsp90aa1 | 28139.47731 | 1.119376761 | 2.08E-04 |
| Hspb2 | 59.28042065 | 1.11855303 | 1.69E-03 |
| Lats2 | 3075.227714 | 1.118157928 | 4.31E-14 |
| Kif2c | 1636.606177 | 1.114497624 | 9.51E-03 |
| Gm17709 | 122.5539367 | 1.111998755 | 4.95E-08 |
| Rrm1 | 7755.011656 | 1.110943805 | 5.87E-05 |
| Npnt | 449.3441054 | 1.110874985 | 5.10E-09 |
| Nav3 | 643.5698791 | 1.109718078 | 2.15E-02 |
| Ttc39b | 2306.341585 | 1.109185509 | 2.06E-09 |
| Kifc5b | 429.9223858 | 1.108882199 | 1.89E-03 |
| Abca9 | 166.1965339 | 1.108475106 | 1.41E-02 |
| Otud1 | 340.3620018 | 1.106394267 | 3.10E-06 |
| Cdkn1a | 10457.59614 | 1.106354407 | 1.30E-04 |
| Macf1 | 20441.87418 | 1.103931433 | 2.08E-06 |
| Tbc1d8 | 59.18404534 | 1.100511912 | 4.23E-04 |

| | | | |
|---|---|---|---|
| Exo1 | 451.9536838 | 1.09661927 | 2.13E-02 |
| Top2a | 7403.168567 | 1.094673967 | 4.37E-02 |
| Csf1 | 14262.92663 | 1.093205162 | 8.82E-05 |
| Ier3 | 749.7978944 | 1.090268294 | 5.96E-05 |
| Tril | 35.05868021 | 1.090050415 | 1.78E-02 |
| 2610318N02Rik | 78.5731003 | 1.086372584 | 2.22E-02 |
| Cenpq | 506.4884526 | 1.08591867 | 1.79E-03 |
| 6430590A07Rik | 23.85473417 | 1.08405286 | 4.22E-02 |
| Tjp1 | 4773.974161 | 1.082368159 | 7.29E-12 |
| Gm26880 | 2822.471388 | 1.079963601 | 5.45E-05 |
| Racgap1 | 3856.467284 | 1.078058116 | 5.16E-04 |
| E030030I06Rik | 73.11210014 | 1.078004772 | 2.79E-02 |
| Radil | 568.4919136 | 1.074687536 | 2.81E-09 |
| Pld2 | 1123.434152 | 1.074353385 | 2.07E-10 |
| Lcat | 54.65890367 | 1.07353109 | 8.39E-05 |
| Txnrd1 | 11050.67123 | 1.073402138 | 5.13E-09 |
| Spc25 | 518.5776184 | 1.072161221 | 1.59E-02 |
| Cdc42ep5 | 621.0593718 | 1.071050039 | 1.83E-06 |
| Gcnt2 | 266.7781291 | 1.070915401 | 3.72E-05 |
| Ncapg | 1081.828361 | 1.070478305 | 2.19E-02 |
| Ccnb1 | 1707.028829 | 1.069707679 | 8.36E-03 |
| Chtf18 | 539.4430916 | 1.068545343 | 1.03E-02 |
| Kif20a | 2457.420305 | 1.067872929 | 6.61E-03 |
| Gsta4 | 6626.313589 | 1.065100815 | 6.34E-07 |
| Rasal2 | 873.60698 | 1.064484566 | 7.95E-05 |
| Col7a1 | 345.9172629 | 1.062962005 | 7.37E-03 |
| Espl1 | 1089.594393 | 1.060719781 | 1.26E-02 |
| Nop58 | 4575.776706 | 1.060306915 | 1.13E-03 |
| Ecm1 | 1410.00657 | 1.058745509 | 1.47E-03 |
| Pxdc1 | 198.4170374 | 1.058234232 | 1.75E-05 |
| Pkmyt1 | 487.067601 | 1.057628784 | 1.49E-03 |
| Fat1 | 9394.675609 | 1.056366457 | 1.21E-05 |
| Gsto1 | 9133.181255 | 1.054399522 | 8.71E-08 |
| Tm4sf1 | 4004.71189 | 1.054004525 | 8.94E-08 |
| Plpp2 | 931.3288588 | 1.053194322 | 2.30E-10 |
| AU020206 | 146.3601913 | 1.052271699 | 9.72E-10 |
| Cks2 | 825.4966081 | 1.051783976 | 3.85E-03 |
| Slc7a1 | 3571.861693 | 1.049886666 | 4.42E-06 |
| Hist1h2bc | 898.5619503 | 1.047586378 | 4.47E-02 |
| Gm10499 | 78.66015587 | 1.044641989 | 1.32E-03 |
| Eme1 | 193.7007537 | 1.042769129 | 1.12E-02 |
| Sms-ps | 1178.87176 | 1.041330929 | 3.16E-07 |
| Arf2 | 1611.683155 | 1.040606897 | 4.73E-07 |
| Abcg2 | 2509.433036 | 1.036643379 | 2.07E-10 |
| 4921524J17Rik | 641.6105448 | 1.03455248 | 1.61E-14 |
| Mtbp | 724.1374728 | 1.034347899 | 1.23E-03 |
| Gm26705 | 51.19502999 | 1.032824723 | 1.03E-02 |

| | | | |
|---|---|---|---|
| Efemp1 | 43132.42209 | 1.031921765 | 7.59E-10 |
| Amotl2 | 15490.23543 | 1.030870744 | 1.45E-07 |
| Cdk6 | 743.1289463 | 1.03057304 | 9.73E-10 |
| Cd274 | 79.84677291 | 1.029461575 | 3.90E-04 |
| Pim1 | 2348.868326 | 1.029048201 | 1.01E-02 |
| Tob1 | 562.3538878 | 1.028461962 | 1.06E-05 |
| Ptgs1 | 3977.955639 | 1.025999201 | 8.35E-08 |
| Slc35e4 | 1871.809364 | 1.02501742 | 3.65E-06 |
| Prkcg | 214.7714828 | 1.023228618 | 1.97E-04 |
| Nckap5 | 208.3120577 | 1.023101124 | 4.40E-02 |
| Gm16365 | 67.34365029 | 1.021270517 | 7.78E-04 |
| Prr11 | 1150.453785 | 1.021030169 | 2.30E-03 |
| Gm26909 | 21.61032335 | 1.020637059 | 4.00E-02 |
| Sparc | 83467.83876 | 1.017046905 | 2.36E-06 |
| Stim1 | 2475.820194 | 1.01554178 | 4.31E-10 |
| Zfp57 | 2113.079957 | 1.015213994 | 5.33E-05 |
| Mcm10 | 600.8357867 | 1.014837151 | 3.13E-02 |
| Skida1 | 68.19523049 | 1.014629148 | 1.69E-03 |
| Aurkb | 1122.511781 | 1.012662879 | 8.77E-03 |
| Gdnf | 878.6590879 | 1.012583257 | 2.73E-04 |
| Ndc80 | 681.3837944 | 1.012123998 | 3.13E-02 |
| Cxxc5 | 218.8335225 | 1.011605053 | 6.74E-05 |
| Ttk | 376.1005494 | 1.011103273 | 2.64E-02 |
| Ero1l | 4526.176064 | 1.011074193 | 4.54E-04 |
| Myl9 | 237.5650558 | 1.007782966 | 6.38E-04 |
| Kpna2 | 566.1143964 | 1.006705622 | 2.40E-03 |
| Arhgap26 | 119.8297351 | 1.005065889 | 2.79E-04 |
| Slc1a5 | 4845.639286 | 1.004534723 | 8.64E-07 |
| Cdca3 | 1209.281701 | 1.004292648 | 5.01E-03 |
| Wtip | 1616.112467 | 1.002726231 | 6.29E-10 |
| Gm26788 | 40.96349029 | 1.00215021 | 1.48E-02 |
| Dhfr | 781.3575348 | 1.002006087 | 7.09E-03 |
| Gm11427 | 43.61658226 | 0.995409913 | 4.42E-02 |
| Ppa1 | 2005.850286 | 0.994649902 | 7.15E-06 |
| Neu3 | 186.830771 | 0.993133686 | 1.49E-05 |
| Asns | 8358.728063 | 0.99308474 | 7.44E-05 |
| Wdr66 | 88.12706011 | 0.991293037 | 5.37E-03 |
| Plcxd2 | 1602.480094 | 0.990557014 | 5.78E-08 |
| Phlda1 | 939.9688801 | 0.990155663 | 7.64E-04 |
| Kif22 | 949.2951466 | 0.989469827 | 7.05E-03 |
| Gm21750 | 59.65533783 | 0.989070316 | 4.32E-02 |
| Mafk | 2125.926029 | 0.986862447 | 5.56E-05 |
| Kif4 | 977.9406374 | 0.985788429 | 1.74E-02 |
| Clspn | 928.4095926 | 0.984549038 | 1.73E-02 |
| Bend6 | 173.0386063 | 0.983081783 | 1.23E-04 |
| Nuggc | 50.09471182 | 0.982752848 | 7.13E-03 |
| S100a10 | 14257.32699 | 0.982446037 | 3.04E-12 |

| | | | |
|---|---|---|---|
| Tnfrsf22 | 1190.813432 | 0.981630416 | 4.72E-05 |
| Gm10184 | 6253.77365 | 0.98059322 | 5.19E-03 |
| Tcte2 | 253.6465231 | 0.98038611 | 2.24E-08 |
| Ston1 | 4324.364224 | 0.979961476 | 5.52E-10 |
| Thbd | 4214.750375 | 0.979098536 | 1.89E-04 |
| Tdrp | 456.9726997 | 0.978484037 | 9.05E-09 |
| Cyb5r1 | 3302.743638 | 0.977642757 | 1.15E-04 |
| Magi1 | 863.00933 | 0.977313177 | 1.39E-12 |
| Trip13 | 438.5730302 | 0.976833749 | 1.49E-02 |
| Atad2 | 2781.337186 | 0.973985386 | 7.76E-03 |
| H2-K1 | 1381.165026 | 0.97201921 | 9.86E-07 |
| Fam83d | 571.3319939 | 0.971652615 | 3.40E-02 |
| Tacc3 | 1930.82365 | 0.970055829 | 1.67E-02 |
| Slc30a4 | 4466.816693 | 0.969787945 | 6.67E-04 |
| Efcab11 | 48.08988877 | 0.968023068 | 3.31E-02 |
| Cenpm | 389.4200119 | 0.967882528 | 1.57E-02 |
| Rassf5 | 557.4393745 | 0.965390325 | 5.06E-05 |
| Prps1 | 1657.907258 | 0.963557219 | 2.89E-08 |
| Gm16344 | 100.0569507 | 0.963081358 | 5.32E-05 |
| Fignl1 | 577.6069439 | 0.962971884 | 1.51E-02 |
| Pdgfrl | 4182.667061 | 0.96191688 | 4.04E-04 |
| Gm12346 | 480.5896986 | 0.961736362 | 2.81E-03 |
| Plagl1 | 737.429685 | 0.961647808 | 3.59E-03 |
| Pdgfa | 1217.553266 | 0.961165281 | 5.33E-13 |
| Tead1 | 8305.982112 | 0.961025762 | 1.55E-12 |
| Lyar | 1204.940602 | 0.958364852 | 5.09E-05 |
| Flnb | 18993.59883 | 0.958152419 | 2.73E-10 |
| Pcdh19 | 2133.007913 | 0.958025912 | 3.75E-06 |
| Spdl1 | 447.2875193 | 0.957383858 | 2.15E-02 |
| Ror1 | 243.8030807 | 0.955911705 | 1.27E-02 |
| Emp1 | 29485.48831 | 0.955519818 | 4.86E-06 |
| Kank1 | 1147.219222 | 0.954176486 | 1.18E-05 |
| Gm10080 | 25.45478434 | 0.951343851 | 4.63E-02 |
| Ercc6l | 592.8696419 | 0.950573081 | 2.89E-02 |
| Lmnb1 | 1761.407172 | 0.949450556 | 2.23E-03 |
| Hacd4 | 175.9725215 | 0.946450088 | 1.68E-02 |
| Spag5 | 1260.809925 | 0.9464315 | 2.36E-02 |
| Cdca5 | 431.2039611 | 0.945237688 | 3.41E-02 |
| Them6 | 440.6205857 | 0.945133755 | 2.39E-04 |
| Atp1a1 | 13088.27316 | 0.943015823 | 2.17E-09 |
| Mpi | 945.5748437 | 0.942431324 | 1.56E-07 |
| Nhsl1 | 543.1360852 | 0.941971365 | 2.87E-02 |
| Myo1c | 17433.95275 | 0.941868924 | 3.13E-10 |
| Magohb | 625.3616612 | 0.941175236 | 2.63E-04 |
| Tbc1d31 | 1630.86014 | 0.938419652 | 1.98E-04 |
| Foxf1 | 338.939874 | 0.938381121 | 4.92E-08 |
| Cdipt | 3223.076682 | 0.936925039 | 4.02E-17 |

| | | | |
|---|---|---|---|
| Chaf1a | 1268.21855 | 0.934871236 | 9.95E-03 |
| Cavin1 | 29736.21876 | 0.934775248 | 1.42E-10 |
| Acta2 | 22267.60312 | 0.933798953 | 7.70E-05 |
| Timeless | 737.6886152 | 0.930845613 | 5.72E-03 |
| Ltbp2 | 1061.332223 | 0.929152075 | 5.10E-03 |
| Col5a2 | 40728.55901 | 0.928643787 | 2.03E-04 |
| Ddr2 | 21485.53935 | 0.928637561 | 2.39E-06 |
| Adam9 | 10091.06238 | 0.928472825 | 6.72E-07 |
| Rnf113a1 | 147.4647827 | 0.926808728 | 1.76E-05 |
| Fign | 239.6826974 | 0.924859369 | 7.05E-03 |
| Micu1 | 6570.285669 | 0.923896358 | 3.25E-10 |
| Wdr45 | 2171.76663 | 0.921624769 | 1.42E-07 |
| Ehd4 | 2099.212753 | 0.919893172 | 1.60E-08 |
| Acot9 | 3567.356035 | 0.919358196 | 5.33E-08 |
| Cenpu | 251.9751922 | 0.918396341 | 1.64E-02 |
| Gchfr | 290.9261129 | 0.916221942 | 4.37E-03 |
| Incenp | 2097.979085 | 0.915910959 | 7.09E-03 |
| Spire2 | 545.5638821 | 0.915464467 | 1.41E-10 |
| Dyrk3 | 209.7965907 | 0.913733226 | 1.77E-04 |
| Dlgap5 | 995.0483588 | 0.913430497 | 5.94E-03 |
| 2810429I04Rik | 100.9140103 | 0.913104797 | 1.77E-02 |
| Rad54l | 518.4103323 | 0.911273323 | 1.93E-02 |
| Llgl2 | 523.0277484 | 0.911145353 | 1.44E-03 |
| Me3 | 835.3947927 | 0.91104152 | 4.58E-07 |
| Tesk2 | 669.5921419 | 0.91025723 | 8.71E-07 |
| Sh3rf1 | 1249.064827 | 0.90956345 | 1.84E-09 |
| Plk4 | 915.3174009 | 0.907574378 | 8.47E-03 |
| Ccdc9b | 3998.146773 | 0.904630138 | 8.99E-12 |
| Gm15270 | 34.01967855 | 0.903590383 | 4.30E-02 |
| Tex30 | 300.7795105 | 0.903570897 | 1.47E-02 |
| Ptch1 | 475.913733 | 0.903152085 | 2.44E-08 |
| Pcgf5 | 1180.442953 | 0.903089566 | 2.49E-09 |
| Slc16a1 | 2748.76285 | 0.902948723 | 2.01E-04 |
| Traip | 238.25573 | 0.901043284 | 1.48E-02 |
| Melk | 527.1280361 | 0.899611657 | 4.57E-02 |
| Snx7 | 2260.56405 | 0.898706348 | 4.06E-05 |
| Tent4b | 1416.599917 | 0.897990997 | 1.35E-05 |
| Cenpi | 568.2024036 | 0.897814722 | 2.80E-02 |
| Prkci | 7019.667731 | 0.897759281 | 2.95E-07 |
| Car5b | 116.6171212 | 0.89700569 | 1.09E-03 |
| Arhgap44 | 266.8234364 | 0.896045196 | 4.14E-03 |
| Calr3 | 72.66829931 | 0.895591513 | 1.04E-03 |
| Eno1b | 12787.54532 | 0.894655748 | 7.83E-06 |
| Cdh17 | 219.8371349 | 0.894158006 | 6.45E-05 |
| B4galt4 | 584.3450299 | 0.893886856 | 7.14E-08 |
| Anxa3 | 7482.75502 | 0.893624445 | 4.51E-04 |
| Magi3 | 1822.469333 | 0.892771079 | 2.70E-05 |

| | | | |
|---|---|---|---|
| Irak3 | 70.04821296 | 0.892584193 | 7.97E-03 |
| Loxl3 | 4747.787426 | 0.892545961 | 2.12E-05 |
| Myl12a | 24114.22838 | 0.891867871 | 5.46E-10 |
| Haus8 | 750.9704817 | 0.888655231 | 8.74E-04 |
| Dst | 14726.69931 | 0.887107228 | 2.57E-06 |
| Pkn3 | 159.00837 | 0.887065309 | 3.26E-02 |
| Adss | 4524.176795 | 0.883301172 | 6.53E-13 |
| Ddr1 | 5318.493109 | 0.882598555 | 1.78E-08 |
| Tbc1d2 | 1016.924895 | 0.882034111 | 6.32E-04 |
| Ddx39 | 3293.012759 | 0.881355555 | 4.19E-04 |
| Cenpw | 161.2582704 | 0.878501835 | 6.96E-03 |
| Hotairm1 | 163.7859472 | 0.878150295 | 3.57E-04 |
| Bora | 321.195499 | 0.877991225 | 1.53E-02 |
| Il15 | 186.9354469 | 0.877665675 | 1.03E-03 |
| Myadm | 15438.02828 | 0.877252829 | 5.64E-21 |
| Dlc1 | 2597.470192 | 0.875370654 | 6.45E-07 |
| Ptprf | 4297.628625 | 0.874244989 | 3.84E-20 |
| Arhgef18 | 3617.647802 | 0.872184424 | 3.60E-09 |
| Gm19426 | 43.53584447 | 0.870965845 | 1.98E-02 |
| Cbx5 | 6654.599166 | 0.87059706 | 3.77E-04 |
| Man1a | 1574.812904 | 0.869029051 | 2.91E-04 |
| CAAA01147332.1 | 997.9501685 | 0.86865409 | 1.63E-02 |
| 4833427G06Rik | 113.596248 | 0.867213011 | 2.46E-03 |
| Rock2 | 6286.389359 | 0.867113111 | 5.22E-03 |
| Eno1 | 16599.21767 | 0.866312212 | 3.10E-05 |
| Gas6 | 3563.445454 | 0.86477088 | 1.08E-02 |
| Ets1 | 2712.616302 | 0.864016534 | 4.16E-07 |
| Mns1 | 88.61665514 | 0.863862603 | 4.25E-02 |
| Dhrs9 | 1469.943128 | 0.862920032 | 1.22E-03 |
| Ttc41 | 109.4536472 | 0.862734812 | 2.33E-02 |
| Rad51ap1 | 471.7413004 | 0.862238195 | 3.13E-02 |
| Sema3f | 4201.918047 | 0.860925999 | 1.49E-07 |
| Herc3 | 407.8347654 | 0.860510039 | 2.35E-04 |
| Grk4 | 204.3808361 | 0.860283572 | 4.25E-04 |
| Lrp4 | 529.2244838 | 0.860234264 | 1.99E-02 |
| Arl6ip1 | 275.4071157 | 0.859481667 | 2.68E-07 |
| Fbn1 | 5465.355355 | 0.859442488 | 6.08E-03 |
| A930004D18Rik | 436.6953306 | 0.859348178 | 3.54E-04 |
| Cdk2ap2 | 1795.173273 | 0.858674462 | 2.24E-06 |
| Prkar2b | 2610.181424 | 0.85828239 | 2.07E-06 |
| Plpp1 | 5028.842612 | 0.857718866 | 1.69E-05 |
| Arhgap11a | 2033.144585 | 0.857565132 | 3.09E-02 |
| Bcl2l1 | 4294.000393 | 0.856919712 | 7.48E-07 |
| Cdkn3 | 244.1485424 | 0.856044288 | 2.92E-02 |
| Dtl | 972.7640328 | 0.855616801 | 4.37E-02 |
| Ppat | 1263.284928 | 0.855315364 | 5.15E-07 |
| 2210011C24Rik | 74.38268018 | 0.855110124 | 1.73E-02 |

| | | | |
|---|---|---|---|
| Cenpa | 1628.559306 | 0.85484233 | 7.33E-03 |
| Dnajc9 | 984.8756573 | 0.854651733 | 1.22E-03 |
| Disp1 | 2262.130775 | 0.854022041 | 1.15E-14 |
| Nup160 | 1549.140682 | 0.850444453 | 3.96E-03 |
| Slc18a2 | 77.34349994 | 0.849638444 | 4.72E-02 |
| Tiam1 | 806.6891846 | 0.849126708 | 4.04E-04 |
| Pola2 | 1593.727292 | 0.847740631 | 2.60E-03 |
| Slc48a1 | 9279.752174 | 0.845258205 | 1.69E-06 |
| Lonp1 | 6941.8007 | 0.844527591 | 5.71E-08 |
| Spin4 | 112.9802311 | 0.843970337 | 1.35E-02 |
| Ybx3 | 12605.60181 | 0.843899705 | 1.54E-06 |
| Tns3 | 8946.526377 | 0.842765422 | 4.10E-08 |
| Klrg2 | 65.68197859 | 0.842611985 | 2.80E-02 |
| Yif1b | 1695.896623 | 0.842172027 | 3.54E-07 |
| Haus6 | 1556.803127 | 0.841624597 | 1.14E-02 |
| Nrm | 626.7617768 | 0.841573439 | 1.26E-02 |
| Tagln2 | 5657.746603 | 0.841407448 | 4.55E-04 |
| Deptor | 1172.456592 | 0.841161753 | 2.74E-03 |
| Aqp5 | 257.8230496 | 0.841113109 | 2.84E-02 |
| Rpp30 | 619.7082099 | 0.839274777 | 2.00E-06 |
| Slc2a1 | 3768.575499 | 0.839063922 | 2.73E-10 |
| Zfp53 | 773.8964099 | 0.837899918 | 4.73E-03 |
| Optn | 361.5681594 | 0.834923065 | 4.00E-06 |
| Morc4 | 3288.740245 | 0.833490355 | 2.20E-05 |
| Ptgis | 3691.451478 | 0.833090095 | 1.64E-03 |
| Gsdmd | 431.2673849 | 0.832577428 | 2.97E-11 |
| Kdm8 | 161.4759552 | 0.830834906 | 5.27E-03 |
| Efnb1 | 347.491773 | 0.830141963 | 1.30E-04 |
| Zfp809 | 553.769558 | 0.830124845 | 3.30E-05 |
| Ktn1 | 2814.57127 | 0.829247868 | 1.53E-02 |
| Nomo1 | 5376.942128 | 0.828884548 | 1.04E-04 |
| Osbpl3 | 2074.055951 | 0.826671222 | 2.32E-07 |
| Tmem245 | 7288.006629 | 0.825443466 | 4.95E-08 |
| Fas | 523.473713 | 0.825174153 | 1.85E-03 |
| Asxl3 | 95.93112386 | 0.823827322 | 2.75E-02 |
| Ipo11 | 1848.149906 | 0.823617697 | 2.35E-06 |
| Cercam | 171.2368976 | 0.822598547 | 4.45E-03 |
| Dut | 1071.493419 | 0.822349982 | 1.10E-02 |
| Rpl29 | 569.7586007 | 0.822075908 | 1.03E-02 |
| Gm6645 | 54.99621619 | 0.821377389 | 1.97E-02 |
| Ube2t | 295.3875627 | 0.820313872 | 4.93E-02 |
| E2f6 | 108.6963516 | 0.819048703 | 4.12E-03 |
| NA | 806.7719172 | 0.81889895 | 4.41E-05 |
| Epb41l4a | 2231.957676 | 0.81844146 | 2.64E-12 |
| Stil | 640.2338351 | 0.818067054 | 1.43E-02 |
| Gk | 712.7420813 | 0.817434819 | 4.36E-03 |
| Rbms2 | 13507.54112 | 0.816233723 | 2.05E-04 |

| | | | |
|---|---|---|---|
| Abca5 | 1770.476615 | 0.815844123 | 7.78E-04 |
| Slc38a4 | 4435.554761 | 0.813845189 | 5.55E-04 |
| Ccnb2 | 359.5961839 | 0.813144349 | 2.25E-02 |
| Adk | 3190.013385 | 0.812308802 | 6.53E-10 |
| Ptpn14 | 6806.862281 | 0.812299552 | 2.48E-14 |
| Pak1 | 1454.477288 | 0.811208906 | 3.66E-06 |
| Zfp36 | 1397.592095 | 0.80966011 | 4.72E-02 |
| Tor4a | 194.1873465 | 0.809482669 | 5.51E-03 |
| Fam110a | 459.3283501 | 0.808176218 | 5.54E-03 |
| Aoc2 | 159.8164997 | 0.807907068 | 1.27E-02 |
| Uba7 | 268.7536281 | 0.804331428 | 1.66E-03 |
| Junos | 162.0037706 | 0.804039537 | 8.25E-03 |
| Lamc2 | 395.7838555 | 0.803200739 | 1.43E-02 |
| Pdzd8 | 3340.981836 | 0.803004369 | 8.26E-04 |
| Erich1 | 560.5336326 | 0.800851771 | 8.77E-07 |
| Efcab2 | 42.83091983 | 0.799425004 | 3.36E-02 |
| Pdgfc | 1278.888861 | 0.799424726 | 1.42E-05 |
| Tubb4a | 66.11470353 | 0.797890124 | 3.04E-02 |
| Wwc2 | 6641.559661 | 0.796279022 | 4.84E-10 |
| Exoc6 | 416.7822632 | 0.795833566 | 4.17E-04 |
| Slc39a13 | 2542.695428 | 0.795703226 | 3.00E-05 |
| Lrp8 | 1552.953999 | 0.795190556 | 4.00E-02 |
| Acsl5 | 3339.880617 | 0.794471276 | 1.81E-03 |
| Cd44 | 4483.253208 | 0.793380987 | 2.51E-04 |
| Ifit2 | 2698.970925 | 0.792313665 | 2.74E-04 |
| Cyld | 2898.729581 | 0.792178106 | 1.25E-06 |
| 9130008F23Rik | 333.9612646 | 0.790524341 | 1.37E-03 |
| Abcc1 | 6563.18358 | 0.790423549 | 3.70E-09 |
| Bbc3 | 624.0895791 | 0.788956937 | 5.95E-04 |
| S100a6 | 44660.9589 | 0.788401533 | 7.49E-05 |
| Dkc1 | 2009.176878 | 0.783425289 | 2.31E-02 |
| Cfap69 | 254.5171885 | 0.78302098 | 1.54E-02 |
| Flvcr1 | 683.9766244 | 0.782966268 | 5.43E-04 |
| Ccnyl1 | 2407.209529 | 0.781656879 | 8.84E-07 |
| Asap3 | 387.600447 | 0.78054204 | 4.07E-03 |
| Adamts14 | 1745.125069 | 0.779216763 | 1.01E-02 |
| Dnajb4 | 1668.421695 | 0.778905365 | 9.36E-08 |
| Rasl11a | 634.5078439 | 0.777163911 | 7.47E-03 |
| Fuca2 | 3318.401005 | 0.774984812 | 1.95E-05 |
| Myof | 24621.41901 | 0.773248916 | 1.06E-06 |
| Slc25a40 | 773.2498353 | 0.771846857 | 1.16E-03 |
| Xylb | 84.70507461 | 0.771779369 | 1.04E-02 |
| Apcdd1 | 937.5246247 | 0.771057365 | 2.87E-05 |
| Cdc42bpa | 4204.491388 | 0.769973446 | 1.62E-03 |
| Stip1 | 10290.17675 | 0.768558409 | 7.95E-05 |
| Rrp1b | 1636.99729 | 0.76814046 | 1.38E-11 |
| Gm13092 | 594.121747 | 0.767634681 | 3.17E-05 |

| | | | |
|---|---|---|---|
| Mlkl | 1417.964644 | 0.767011851 | 3.77E-06 |
| Gm13632 | 65.5249491 | 0.766729116 | 2.17E-02 |
| Pxk | 1881.676776 | 0.766221229 | 5.11E-19 |
| Uck2 | 2613.500918 | 0.765442141 | 2.70E-05 |
| Bicd2 | 3364.546628 | 0.76510138 | 5.10E-10 |
| Reep4 | 971.1311621 | 0.763850877 | 9.69E-04 |
| Spats2l | 58.43551475 | 0.763067383 | 3.92E-02 |
| Stap2 | 79.90043653 | 0.760580293 | 2.22E-02 |
| 9230114K14Rik | 104.7462116 | 0.759671989 | 3.26E-02 |
| Hoxc5 | 124.7217338 | 0.758924167 | 4.66E-02 |
| Hdgf | 20795.9107 | 0.758302578 | 9.70E-08 |
| Filip1l | 2075.750886 | 0.756835294 | 2.33E-03 |
| Ccng1 | 10250.65217 | 0.756300986 | 1.34E-05 |
| Ddx11 | 404.9730421 | 0.755947506 | 3.58E-02 |
| Slc9a5 | 964.4551971 | 0.755433919 | 3.45E-03 |
| Lamc1 | 22727.02501 | 0.754138515 | 1.15E-06 |
| Secisbp2l | 3505.578734 | 0.753993246 | 1.89E-05 |
| Cacybp | 3685.283473 | 0.75192066 | 2.35E-02 |
| Avpi1 | 1533.78618 | 0.751293597 | 2.24E-06 |
| NA | 1559.71249 | 0.750988085 | 3.61E-05 |
| Zfp568 | 651.051108 | 0.749237389 | 1.20E-02 |
| Smtn | 3576.331185 | 0.747551903 | 6.29E-06 |
| Prdx4 | 3691.055708 | 0.746861122 | 1.31E-05 |
| Prim1 | 735.9249099 | 0.746105329 | 2.03E-02 |
| Clip1 | 4554.958969 | 0.745605066 | 2.70E-06 |
| Tnfaip1 | 4512.561739 | 0.745510246 | 3.26E-09 |
| Rbpj-ps3 | 2159.928873 | 0.743401594 | 4.17E-10 |
| Etl4 | 562.100196 | 0.742627065 | 1.60E-02 |
| Gm4735 | 2471.198211 | 0.742623469 | 2.39E-03 |
| Vcam1 | 8733.754356 | 0.742579363 | 1.37E-08 |
| Serpinh1 | 36919.64396 | 0.742435263 | 3.42E-05 |
| Glud1 | 8559.981822 | 0.741069918 | 5.90E-26 |
| Vgll3 | 2688.695715 | 0.740636311 | 3.41E-02 |
| Cdkn2c | 2309.987574 | 0.740340275 | 3.42E-07 |
| Il10rb | 939.8976722 | 0.738719141 | 3.76E-12 |
| Thy1 | 3489.688816 | 0.738128571 | 2.26E-03 |
| Dagla | 576.2981415 | 0.73673131 | 8.52E-09 |
| Mdm2 | 3807.437062 | 0.73575338 | 2.46E-06 |
| Ube2l6 | 711.5714298 | 0.733952112 | 1.48E-02 |
| Selenoi | 2343.480976 | 0.732737119 | 3.38E-07 |
| Bcar3 | 810.1377157 | 0.732507478 | 2.16E-03 |
| Samd4 | 3008.136996 | 0.729767233 | 7.88E-08 |
| Atp11a | 5318.325548 | 0.727904159 | 2.27E-06 |
| Cdc45 | 743.4999014 | 0.727158191 | 4.98E-02 |
| Rapgef2 | 1760.896602 | 0.725984936 | 6.87E-05 |
| Pidd1 | 231.8430029 | 0.725938809 | 4.77E-02 |
| Rad18 | 808.0924725 | 0.725347318 | 1.60E-02 |

| | | | |
|---|---|---|---|
| Ncaph | 1055.329841 | 0.723899259 | 3.22E-02 |
| Cfap36 | 1389.217841 | 0.723575348 | 1.15E-02 |
| Pacsin2 | 4361.917334 | 0.72326523 | 1.51E-11 |
| Rbpj | 6189.6897 | 0.721963352 | 8.71E-10 |
| Afdn | 6938.094101 | 0.721143107 | 2.13E-05 |
| Slco2a1 | 4121.36187 | 0.72052092 | 2.56E-04 |
| Htra1 | 323.732752 | 0.720001783 | 8.81E-05 |
| Syne2 | 2077.972178 | 0.719636136 | 1.05E-03 |
| Gabre | 296.5455035 | 0.718101732 | 1.59E-02 |
| A530040E14Rik | 89.26674794 | 0.71704406 | 3.19E-02 |
| Esd | 14951.22292 | 0.716683775 | 3.24E-04 |
| Syde1 | 3457.474687 | 0.716062023 | 5.90E-08 |
| Cav1 | 38978.11609 | 0.714980045 | 1.60E-07 |
| Tpst2 | 2312.601895 | 0.714889383 | 4.17E-05 |
| Cep85 | 1088.695654 | 0.714393086 | 2.52E-03 |
| Gm7102 | 42.86306938 | 0.713563961 | 3.53E-02 |
| Ubr1 | 1990.706341 | 0.713395132 | 1.56E-03 |
| Nedd9 | 2791.535274 | 0.712691452 | 1.11E-02 |
| Ostf1 | 2423.365697 | 0.711584716 | 2.12E-11 |
| Wdr62 | 764.1814903 | 0.710784814 | 2.02E-02 |
| Creb3 | 1212.471112 | 0.709991405 | 9.80E-06 |
| Utp11 | 3221.327935 | 0.709283715 | 1.93E-11 |
| Crim1 | 4958.715791 | 0.70801642 | 7.29E-03 |
| Tmsb10 | 24402.93356 | 0.707605358 | 9.88E-06 |
| Apbb1ip | 2739.651596 | 0.704996068 | 6.26E-04 |
| Bscl2 | 225.8361494 | 0.702656763 | 2.75E-02 |
| Polk | 816.6986151 | 0.702209327 | 1.96E-03 |
| Eif3c | 17700.40432 | 0.698375032 | 1.48E-09 |
| Plcxd1 | 187.9861981 | 0.697772933 | 3.84E-02 |
| Prkag2 | 1252.873195 | 0.697154751 | 1.06E-08 |
| Paqr3 | 356.3402682 | 0.695992317 | 8.26E-04 |
| Lrrc8c | 1417.4025 | 0.694827306 | 1.47E-03 |
| Wars | 2773.816283 | 0.693896017 | 6.86E-05 |
| Tuba1c | 4795.987546 | 0.69263506 | 2.90E-02 |
| Afmid | 240.573853 | 0.691503365 | 2.16E-02 |
| Slc7a2 | 1073.849895 | 0.69118395 | 3.60E-02 |
| Dnmt1 | 4454.586651 | 0.690491899 | 4.40E-03 |
| Stx11 | 348.3990913 | 0.689565999 | 4.99E-04 |
| Csgalnact2 | 1490.922805 | 0.689452929 | 8.26E-04 |
| Adora2b | 955.5375759 | 0.688747945 | 1.12E-03 |
| Procr | 826.0507845 | 0.687026228 | 6.68E-03 |
| Tgm2 | 2829.880322 | 0.686220317 | 1.27E-03 |
| Ghitm | 11802.31619 | 0.68445897 | 6.46E-06 |
| Zbtb18 | 764.6264989 | 0.683966167 | 1.23E-02 |
| Ephb2 | 1039.48774 | 0.683945993 | 7.11E-04 |
| Itpr1 | 2029.82746 | 0.683388788 | 4.49E-05 |
| Fbln5 | 11685.97594 | 0.682094028 | 1.13E-03 |

| | | | |
|---|---|---|---|
| Lpar6 | 212.9759482 | 0.681402268 | 4.48E-02 |
| Gstm2 | 1617.930043 | 0.681332914 | 1.48E-02 |
| Pltp | 1305.024307 | 0.680154911 | 3.15E-02 |
| Lig1 | 1985.265355 | 0.679248123 | 2.10E-02 |
| Pa2g4 | 8738.523288 | 0.676094104 | 3.28E-04 |
| Enpp4 | 380.5708592 | 0.674910325 | 1.06E-03 |
| Ifrd2 | 723.7027877 | 0.674025072 | 3.83E-02 |
| Nol7 | 912.3460862 | 0.673497563 | 1.93E-04 |
| Rapgef3 | 465.241526 | 0.67282184 | 9.27E-06 |
| Col1a1 | 121831.0483 | 0.669560206 | 2.58E-02 |
| Arhgap33 | 330.2406032 | 0.66947807 | 1.69E-02 |
| Fzd6 | 270.1231641 | 0.66871479 | 5.40E-03 |
| Ldlrap1 | 698.7591419 | 0.668562841 | 1.12E-03 |
| Ahr | 292.2985283 | 0.668084619 | 2.04E-02 |
| Calhm2 | 211.5215987 | 0.666844372 | 7.76E-04 |
| Dab2 | 7942.183086 | 0.666581982 | 3.42E-03 |
| Tmem128 | 899.2173669 | 0.666114162 | 1.07E-06 |
| Sptan1 | 24705.76715 | 0.665906435 | 2.28E-06 |
| 9930012K11Rik | 537.6727168 | 0.665727586 | 3.05E-04 |
| Dnajb1 | 2104.812103 | 0.664248995 | 7.01E-04 |
| Ctnnal1 | 2217.420049 | 0.664227949 | 6.80E-04 |
| Smc5 | 2503.855218 | 0.662816384 | 2.83E-02 |
| Hmga1 | 10223.72691 | 0.661050096 | 3.57E-02 |
| Smco4 | 219.148333 | 0.660792379 | 2.27E-03 |
| Slc12a4 | 4348.57105 | 0.660670691 | 2.48E-08 |
| Pcolce | 25050.1764 | 0.660618966 | 8.81E-06 |
| Kpna1 | 2050.904323 | 0.660334381 | 3.28E-02 |
| Banf1 | 3837.995394 | 0.659779363 | 3.11E-05 |
| P4ha1 | 9161.320401 | 0.658942733 | 2.27E-03 |
| Alg13 | 429.2832683 | 0.65868148 | 9.71E-03 |
| Lrrc75a | 924.7491795 | 0.658434767 | 2.54E-05 |
| Gdpd5 | 258.3174992 | 0.657055965 | 2.31E-03 |
| Cfl2 | 7309.759639 | 0.65619017 | 2.52E-04 |
| Myl6 | 22034.81211 | 0.654963708 | 1.13E-16 |
| Fkbp10 | 8574.958963 | 0.654804593 | 3.40E-04 |
| Gm20404 | 84.38756408 | 0.654695573 | 3.88E-02 |
| Dnaja4 | 286.7765266 | 0.65377808 | 1.79E-02 |
| Anxa2 | 27933.32394 | 0.653276559 | 2.63E-04 |
| Esyt1 | 2873.959478 | 0.652538917 | 6.36E-06 |
| Nrp1 | 3718.261942 | 0.652345916 | 1.68E-03 |
| Cdk18 | 2832.68387 | 0.651181836 | 6.03E-03 |
| Lamb2 | 7100.083165 | 0.650603144 | 1.30E-02 |
| Gm13977 | 61.77917577 | 0.649304105 | 3.28E-02 |
| Gsr | 1627.238933 | 0.649186803 | 7.86E-04 |
| Exoc3l4 | 133.8271647 | 0.648511807 | 2.70E-02 |
| Orc2 | 1454.761569 | 0.647715418 | 8.72E-03 |
| Sbno2 | 2509.386322 | 0.647452932 | 1.10E-04 |

| | | | |
|---|---|---|---|
| Usp32 | 4236.091678 | 0.647235969 | 9.13E-04 |
| Sema4b | 529.3265577 | 0.647121054 | 3.05E-06 |
| Pkp4 | 2837.958 | 0.646768676 | 5.24E-04 |
| Cavin3 | 2957.834925 | 0.646674621 | 1.54E-03 |
| Plod2 | 1368.669902 | 0.64541564 | 4.90E-03 |
| Aplp2 | 11967.92074 | 0.645270319 | 5.06E-05 |
| Egln1 | 2310.538171 | 0.645129378 | 6.11E-06 |
| Parva | 10879.78875 | 0.643262999 | 1.68E-08 |
| Strbp | 1590.73397 | 0.642307232 | 3.52E-04 |
| Uchl5 | 1036.057855 | 0.64228719 | 1.10E-03 |
| Gclc | 1433.972265 | 0.641001217 | 2.61E-03 |
| 9530077C05Rik | 123.6192175 | 0.640611989 | 6.85E-03 |
| Elf4 | 2047.77655 | 0.639948127 | 7.73E-11 |
| Ldlrad4 | 327.7250835 | 0.639940394 | 3.43E-04 |
| Rin1 | 4778.34829 | 0.637649889 | 2.09E-04 |
| Gclm | 1768.792317 | 0.636607967 | 7.28E-05 |
| Rars | 6691.937495 | 0.63563884 | 6.06E-06 |
| Pea15a | 11198.47623 | 0.635377573 | 2.44E-06 |
| Fth1 | 88712.82582 | 0.634403583 | 7.90E-03 |
| Tcf20 | 4460.867637 | 0.634339018 | 2.24E-08 |
| Msn | 18054.98124 | 0.631434739 | 4.68E-03 |
| Lrrk1 | 3325.760949 | 0.631316968 | 1.89E-05 |
| Myo19 | 568.3216705 | 0.629438264 | 3.83E-02 |
| Hspa5 | 48769.31287 | 0.627588328 | 2.13E-02 |
| Mthfd1l | 1964.85493 | 0.627318691 | 3.92E-02 |
| Ino80c | 526.2001063 | 0.625111848 | 6.63E-04 |
| Slc19a2 | 1917.481573 | 0.623645442 | 2.29E-04 |
| Sec14l1 | 5437.380374 | 0.623214168 | 1.15E-07 |
| Itgb1 | 37771.62253 | 0.622513776 | 2.53E-09 |
| Man1a2 | 4199.116273 | 0.621832533 | 5.56E-04 |
| Man2b2 | 7485.806257 | 0.621212666 | 1.38E-03 |
| Cd248 | 6880.534459 | 0.621115098 | 1.29E-02 |
| Agps | 3230.015004 | 0.619559187 | 7.72E-04 |
| Prnp | 14550.6052 | 0.618569454 | 1.12E-03 |
| Relt | 217.4731898 | 0.618511469 | 1.71E-02 |
| Rbpms | 1460.507555 | 0.617801374 | 1.82E-04 |
| Ide | 5488.789588 | 0.617547946 | 8.35E-05 |
| Utrn | 8786.265038 | 0.616922737 | 5.24E-05 |
| Ttll7 | 695.451499 | 0.615431593 | 7.47E-03 |
| Mustn1 | 214.872157 | 0.615186099 | 4.12E-02 |
| Ndc1 | 2175.202521 | 0.614792267 | 1.11E-02 |
| Tle4 | 360.9712993 | 0.614571603 | 7.90E-04 |
| Colca2 | 129.8506087 | 0.614541895 | 1.86E-02 |
| Snrpf | 1289.863239 | 0.614132501 | 2.35E-03 |
| Nup43 | 725.9634131 | 0.613926012 | 2.97E-03 |
| Dzip1l | 1361.683775 | 0.613709261 | 5.63E-07 |
| Nabp2 | 2302.580906 | 0.612114855 | 7.76E-04 |

| | | | |
|---|---|---|---|
| Ppid | 3377.358914 | 0.611749895 | 2.67E-03 |
| Jdp2 | 2313.384115 | 0.611165357 | 5.48E-05 |
| Rpf2 | 971.3745042 | 0.610072081 | 4.68E-02 |
| Iqsec1 | 6250.869472 | 0.609874849 | 1.00E-04 |
| Ier5 | 3710.218611 | 0.609616056 | 4.18E-04 |
| H2-Bl | 144.2958941 | 0.60958262 | 1.69E-02 |
| Akap12 | 11002.38031 | 0.608578302 | 9.38E-03 |
| Arxes2 | 825.7504101 | 0.608200852 | 4.37E-02 |
| Sh3bp5 | 1876.843321 | 0.608134314 | 1.23E-04 |
| Cav2 | 3663.325958 | 0.607956206 | 3.21E-03 |
| Map6 | 3232.025626 | 0.607723434 | 3.52E-03 |
| Sqor | 2866.171031 | 0.607591561 | 9.33E-06 |
| Cobll1 | 2804.002598 | 0.606575439 | 3.28E-05 |
| Smurf2 | 3989.71208 | 0.606497575 | 2.34E-10 |
| Nudt15 | 117.3997007 | 0.60580206 | 4.78E-02 |
| Myl12b | 5643.908536 | 0.605429013 | 1.99E-06 |
| Wwp1 | 1335.549327 | 0.605241406 | 8.45E-03 |
| Vps54 | 1685.981771 | 0.604893754 | 4.54E-02 |
| Clic4 | 7974.140184 | 0.603736254 | 3.43E-03 |
| Glce | 2652.834273 | 0.60259952 | 3.36E-03 |
| H2afx | 1067.491448 | 0.602286914 | 3.96E-02 |
| Zfp428 | 275.3299011 | 0.601780539 | 5.42E-03 |
| Napb | 476.5876789 | 0.601487017 | 8.80E-04 |
| Mcrip2 | 335.8526933 | 0.601083571 | 2.62E-02 |
| Tmsb4x | 9263.718803 | 0.600752694 | 1.17E-05 |
| Aldh1l2 | 7819.791201 | 0.600731443 | 4.02E-02 |
| Ttll4 | 1807.441143 | 0.59889544 | 7.01E-03 |
| Ruvbl2 | 1919.938162 | 0.598819279 | 1.75E-02 |
| Klc1 | 5592.891025 | 0.597253592 | 2.02E-08 |
| Nup37 | 843.5753032 | 0.596933062 | 8.12E-03 |
| Orai1 | 454.0094432 | 0.596768434 | 1.84E-02 |
| Rgs3 | 1513.56933 | 0.596629352 | 1.92E-02 |
| Gm340 | 499.4293114 | 0.595866574 | 5.98E-04 |
| Slc3a2 | 11311.11817 | 0.595664465 | 4.44E-05 |
| Gsto2 | 540.3671304 | 0.595113032 | 2.41E-02 |
| C1rb | 431.70044 | 0.594247977 | 4.65E-02 |
| Sbsn | 1060.826381 | 0.59245044 | 1.30E-03 |
| Mid1ip1 | 939.9451529 | 0.592429429 | 4.76E-02 |
| Naaa | 680.2746552 | 0.590002183 | 4.85E-03 |
| Eprs | 14576.67394 | 0.58978309 | 1.03E-03 |
| Aen | 3318.36924 | 0.589222915 | 1.14E-02 |
| Prkce | 764.757114 | 0.588655384 | 1.83E-04 |
| Mthfd1 | 3288.788194 | 0.587732761 | 2.56E-03 |
| Pml | 1978.102798 | 0.586467171 | 1.17E-03 |
| Faf1 | 5358.729333 | 0.586203143 | 6.55E-06 |
| Slc35d1 | 859.5984113 | 0.585568706 | 1.17E-06 |
| C1ra | 10143.28128 | 0.585316516 | 3.67E-02 |

| | | | |
|---|---|---|---|
| Tmem43 | 6334.664426 | 0.58509009 | 1.25E-02 |
| Usp24 | 5368.656531 | 0.584704328 | 4.43E-03 |
| Gm5921 | 644.8903188 | 0.584237557 | 1.93E-02 |
| Ahnak | 67844.71225 | 0.583237189 | 1.86E-04 |
| Rad23b | 17055.86365 | 0.582672683 | 1.76E-04 |
| Rassf3 | 475.9706472 | 0.582206964 | 1.38E-03 |
| Mbtps2 | 1552.93338 | 0.58196388 | 3.62E-02 |
| Btaf1 | 2399.825233 | 0.581550577 | 2.91E-03 |
| Chd1l | 216.3429542 | 0.580397549 | 3.18E-02 |
| Mrps15 | 2715.97954 | 0.579131428 | 9.58E-08 |
| Msantd3 | 455.0885317 | 0.579021999 | 1.00E-02 |
| Lin54 | 751.046264 | 0.578698601 | 3.12E-02 |
| NA | 4706.046236 | 0.578304194 | 2.86E-04 |
| Syde2 | 1207.49208 | 0.578044779 | 6.75E-03 |
| Slc25a33 | 464.665477 | 0.577263409 | 1.33E-03 |
| Chst12 | 2131.435361 | 0.577257196 | 1.31E-03 |
| Dlx2 | 556.9964188 | 0.576454993 | 1.13E-02 |
| Lgr4 | 1019.428449 | 0.576050603 | 6.10E-03 |
| Cks1b | 1676.270287 | 0.576027936 | 3.73E-02 |
| Gm7334 | 682.962771 | 0.575994939 | 3.55E-04 |
| Slc22a23 | 1141.901598 | 0.5754441 | 3.36E-03 |
| Abcb7 | 679.7533069 | 0.57379571 | 1.49E-02 |
| Slc20a1 | 2917.806275 | 0.57265681 | 6.08E-03 |
| Suv39h1 | 1282.53505 | 0.572016675 | 2.74E-02 |
| Pear1 | 2676.766374 | 0.571338866 | 2.63E-04 |
| Plec | 14666.51798 | 0.570719697 | 2.15E-02 |
| Uxs1 | 2643.985863 | 0.568966109 | 2.23E-04 |
| Ppp1r15a | 3481.359866 | 0.568394293 | 3.94E-03 |
| Nnmt | 470.8866788 | 0.566919453 | 1.63E-02 |
| 6330403L08Rik | 423.3189736 | 0.565678654 | 7.57E-03 |
| Cdc34 | 2896.485483 | 0.564879902 | 4.00E-03 |
| Mars | 4381.005849 | 0.564509728 | 4.34E-03 |
| Col5a1 | 27771.21141 | 0.564322658 | 2.79E-02 |
| Cited2 | 5203.839702 | 0.56428694 | 3.88E-04 |
| Xpot | 6963.581493 | 0.5634423 | 2.46E-04 |
| Ptges3-ps | 1167.527342 | 0.563198318 | 9.31E-03 |
| Tnfaip8 | 770.730871 | 0.561875361 | 1.77E-06 |
| Mlh3 | 176.7584739 | 0.561856757 | 4.23E-02 |
| Gtpbp2 | 2282.051956 | 0.561534609 | 2.59E-04 |
| Itpripl2 | 5913.84639 | 0.561373636 | 6.59E-05 |
| Myc | 3282.758284 | 0.561150767 | 1.31E-03 |
| Zfp608 | 707.2103349 | 0.560949954 | 9.80E-03 |
| Txndc12 | 3770.539548 | 0.56050241 | 1.90E-05 |
| Ap4s1 | 235.0127263 | 0.560105656 | 8.80E-03 |
| Asph | 3636.471194 | 0.559306872 | 8.26E-04 |
| Rasa4 | 4237.672709 | 0.559247117 | 4.52E-03 |
| Pnpla2 | 1540.394814 | 0.558965873 | 2.68E-02 |

| | | | |
|---|---|---|---|
| Ptp4a2 | 11669.3148 | 0.557550311 | 4.71E-05 |
| Nadk2 | 1271.547494 | 0.55577631 | 1.15E-02 |
| Mrip-ps | 1177.21672 | 0.554900063 | 3.36E-03 |
| Zfpm1 | 851.7480048 | 0.554343082 | 8.52E-03 |
| Itpr3 | 5436.339997 | 0.55406965 | 6.16E-04 |
| Snhg15 | 979.4343226 | 0.553647778 | 3.63E-02 |
| Sigmar1 | 2775.964074 | 0.553253109 | 3.86E-03 |
| Prim2 | 794.0553518 | 0.553012125 | 4.65E-02 |
| Rps6ka3 | 2296.409602 | 0.551829953 | 1.71E-02 |
| Tns2 | 5211.113391 | 0.550544786 | 4.64E-04 |
| Taf15 | 4039.925904 | 0.5497734 | 1.34E-02 |
| Cdh2 | 5735.006876 | 0.548949617 | 2.23E-03 |
| Antxr2 | 3509.943442 | 0.548367459 | 4.21E-03 |
| Dusp3 | 2333.354456 | 0.546980904 | 3.54E-03 |
| Rassf1 | 1982.544538 | 0.546519281 | 4.89E-04 |
| Sos1 | 2970.646572 | 0.546060893 | 7.34E-04 |
| Lsm2 | 1237.134903 | 0.545465175 | 3.18E-02 |
| Tnfrsf12a | 4419.471402 | 0.543644827 | 4.44E-02 |
| Gata4 | 849.8210775 | 0.543346801 | 1.55E-02 |
| Spire1 | 861.3974626 | 0.542893544 | 5.85E-03 |
| Ube2s | 4362.208061 | 0.542548605 | 3.69E-02 |
| Capza2 | 8096.038169 | 0.542541651 | 4.37E-04 |
| Crybg3 | 324.3155032 | 0.542243313 | 1.97E-02 |
| Gm7879 | 315.5063862 | 0.541909163 | 4.54E-02 |
| Pawr | 993.2221019 | 0.54014385 | 4.48E-05 |
| Tinf2 | 460.587957 | 0.539917766 | 1.21E-02 |
| Pgrmc1 | 9425.547064 | 0.539451391 | 1.56E-04 |
| Mgat3 | 1523.241024 | 0.538594449 | 3.01E-03 |
| Pfdn4 | 874.6926454 | 0.537536953 | 5.17E-04 |
| Kif24 | 253.3085184 | 0.537347006 | 4.59E-02 |
| Map3k20 | 2221.020669 | 0.535694156 | 1.01E-03 |
| Fam177a | 1383.292923 | 0.535434085 | 2.71E-03 |
| Gm4204 | 1280.220195 | 0.535336412 | 4.22E-02 |
| Rtn4 | 18961.53588 | 0.535284125 | 2.50E-04 |
| Tjap1 | 2545.037818 | 0.535165654 | 9.38E-05 |
| Golim4 | 4417.201827 | 0.533804428 | 1.05E-02 |
| Fhod1 | 1074.977248 | 0.532736541 | 1.32E-04 |
| Bmp2k | 1711.020882 | 0.531389735 | 1.04E-03 |
| Hsp90b1 | 54806.66579 | 0.53077869 | 1.79E-02 |
| Kcmf1 | 5176.903926 | 0.529996523 | 1.57E-05 |
| Strn3 | 4349.359229 | 0.529248585 | 3.33E-02 |
| Rab23 | 1604.157666 | 0.528523892 | 1.17E-03 |
| Snrpd1 | 1590.092186 | 0.528215806 | 7.96E-03 |
| Lmbr1l | 980.757885 | 0.526186178 | 4.28E-02 |
| Smarcd3 | 630.858313 | 0.52581156 | 1.73E-05 |
| Nup107 | 1865.084275 | 0.52519136 | 1.08E-02 |
| NA | 140.031756 | 0.524687832 | 2.42E-02 |

| | | | |
|---|---|---|---|
| Arhgdib | 13907.08035 | 0.524335764 | 5.83E-03 |
| Oxsr1 | 2997.297148 | 0.523379649 | 1.89E-04 |
| Nrip1 | 567.8159237 | 0.523233876 | 5.49E-03 |
| Phgdh | 14268.86496 | 0.522806465 | 4.88E-02 |
| Gm10040 | 874.6698116 | 0.522458866 | 3.51E-04 |
| Actn4 | 15981.88887 | 0.522290708 | 8.74E-04 |
| Trappc2 | 271.7114901 | 0.521250173 | 4.80E-02 |
| Senp1 | 2049.743705 | 0.521108328 | 3.07E-04 |
| Ybey | 168.7312436 | 0.520567267 | 4.33E-02 |
| Acot7 | 1795.605865 | 0.520095756 | 4.93E-04 |
| Il6st | 23732.48129 | 0.519999887 | 1.25E-02 |
| Pcdh1 | 2459.618889 | 0.519648963 | 3.25E-03 |
| Gspt2 | 338.4968154 | 0.518894881 | 1.06E-03 |
| Erlin1 | 2251.338134 | 0.518832065 | 2.35E-04 |
| Rfx2 | 152.4655248 | 0.518017083 | 3.38E-02 |
| Ric1 | 2773.699318 | 0.517846481 | 1.04E-04 |
| Ppm1g | 5386.882069 | 0.516716482 | 4.97E-03 |
| Sh3bp4 | 1970.38902 | 0.516194957 | 3.06E-03 |
| Gm26578 | 320.2566482 | 0.515503138 | 8.43E-03 |
| Tacc1 | 884.4454082 | 0.515411156 | 1.98E-02 |
| Plxnb2 | 19455.25205 | 0.514603473 | 1.77E-04 |
| Fkbp2 | 1281.795309 | 0.514511396 | 5.82E-04 |
| Rcc1 | 1466.074838 | 0.513932364 | 2.81E-02 |
| NA | 1944.15319 | 0.513817277 | 1.07E-02 |
| Jmjd4 | 1066.637866 | 0.513572387 | 4.19E-06 |
| Maml1 | 3478.645256 | 0.512581805 | 2.20E-07 |
| Enoph1 | 817.0058006 | 0.512470565 | 2.75E-03 |
| Vcl | 15499.81698 | 0.512371968 | 2.17E-04 |
| Otud7b | 4504.227944 | 0.511107518 | 2.17E-08 |
| Tmem259 | 4431.668597 | 0.511098924 | 3.88E-10 |
| Gars | 14768.92027 | 0.51066399 | 2.68E-03 |
| Wasf1 | 460.3368112 | 0.510051239 | 1.84E-04 |
| Bsg | 26626.6997 | 0.509231224 | 1.58E-02 |
| Eif1ad | 1505.772972 | 0.509178203 | 3.31E-03 |
| Ptprg | 2392.25637 | 0.507095553 | 4.07E-02 |
| Ctif | 1560.762332 | 0.506487477 | 2.46E-04 |
| Tmbim1 | 4932.481657 | 0.505282006 | 3.11E-03 |
| Ankrd13c | 1592.952556 | 0.50526554 | 2.54E-05 |
| Mpp5 | 1955.405807 | 0.504742877 | 4.47E-02 |
| Ehd2 | 12732.25354 | 0.504658484 | 1.66E-04 |
| Scmh1 | 2538.520534 | 0.503672369 | 4.21E-03 |
| Ncoa3 | 2075.835614 | 0.502981409 | 6.12E-06 |
| Agpat3 | 4907.691748 | 0.501797029 | 8.07E-05 |
| Slc6a9 | 2482.059137 | 0.50089586 | 1.20E-03 |
| Gsn | 46199.21147 | 0.500543693 | 3.06E-03 |
| Gm45713 | 119.8688671 | 0.500231293 | 4.73E-02 |
| Stk10 | 1458.098386 | 0.499663367 | 1.17E-03 |

| | | | |
|---|---|---|---|
| Sik3 | 2887.657643 | 0.498658896 | 2.15E-03 |
| Lrch1 | 792.2566081 | 0.4976596 | 5.94E-04 |
| Wdr74 | 1689.807573 | 0.496379969 | 5.88E-03 |
| Ttc32 | 115.9407862 | 0.496376284 | 4.27E-02 |
| Lrig3 | 1775.697697 | 0.496117298 | 1.77E-02 |
| Me1 | 3504.254363 | 0.495762847 | 1.36E-03 |
| Reck | 1994.171189 | 0.494901268 | 2.48E-02 |
| Dcaf17 | 807.4886767 | 0.494819617 | 3.06E-03 |
| Paics | 14439.95988 | 0.494775016 | 1.17E-11 |
| Tln1 | 13133.47837 | 0.494007686 | 3.61E-05 |
| Ubl3 | 1139.357592 | 0.493922905 | 2.79E-04 |
| Col1a2 | 322977.1213 | 0.491548985 | 4.72E-02 |
| Nus1 | 4625.394918 | 0.491010577 | 7.92E-04 |
| Dlg3 | 2321.002913 | 0.490653633 | 5.21E-05 |
| Acyp2 | 175.9484139 | 0.490616976 | 2.64E-02 |
| Ddb2 | 437.666468 | 0.490509125 | 3.31E-02 |
| Ddb1 | 78266.76622 | 0.489922262 | 2.87E-04 |
| Gzf1 | 2233.832383 | 0.489872619 | 1.55E-03 |
| H6pd | 4608.678006 | 0.489587597 | 4.20E-02 |
| Piga | 562.1998323 | 0.489342542 | 1.05E-02 |
| Sestd1 | 2933.03197 | 0.488299438 | 5.53E-05 |
| Rap2b | 1894.249099 | 0.487267791 | 8.84E-05 |
| Klf13 | 2027.299638 | 0.486105256 | 5.87E-03 |
| 2310057M21Rik | 364.6686249 | 0.485718929 | 2.39E-03 |
| Vopp1 | 3424.567913 | 0.484785951 | 5.93E-03 |
| Mast2 | 8459.255157 | 0.484365406 | 3.54E-03 |
| 1810037I17Rik | 1837.466688 | 0.483740223 | 1.43E-03 |
| Lzic | 586.7404707 | 0.483702366 | 2.03E-02 |
| Emilin2 | 6079.768848 | 0.482470811 | 3.29E-04 |
| Epn2 | 3510.416008 | 0.481830605 | 1.61E-02 |
| Cd47 | 2998.687264 | 0.481702232 | 1.42E-02 |
| Gm9855 | 2483.951233 | 0.481361572 | 4.18E-04 |
| Cars | 3768.251332 | 0.481257363 | 6.57E-04 |
| Eif2b2 | 1629.982564 | 0.481132984 | 8.59E-03 |
| Pef1 | 4262.675557 | 0.481083779 | 2.96E-02 |
| Heg1 | 6138.242062 | 0.480875494 | 1.80E-02 |
| Card19 | 646.5482948 | 0.480058948 | 2.41E-02 |
| Slc7a11 | 1232.289792 | 0.477959964 | 2.46E-03 |
| Trip6 | 4067.669331 | 0.477441256 | 2.45E-02 |
| Bcl2l13 | 3503.679248 | 0.476918964 | 8.53E-05 |
| Psmd5 | 3176.614808 | 0.475936558 | 1.76E-03 |
| Coq10b | 1114.821112 | 0.474471511 | 3.60E-02 |
| Trim8 | 1911.483462 | 0.473257732 | 4.89E-05 |
| Ywhae | 20410.46669 | 0.473212909 | 1.04E-05 |
| Lsm14a | 3705.854334 | 0.471295432 | 1.18E-03 |
| Rnf126 | 1969.762057 | 0.470290462 | 8.96E-03 |
| Plekhg2 | 1751.520827 | 0.469368662 | 1.73E-03 |

| | | | |
|---|---|---|---|
| Col4a6 | 6304.756814 | 0.469322418 | 9.45E-03 |
| Noc2l | 3844.883424 | 0.469168605 | 1.79E-03 |
| Adipor1 | 9075.210439 | 0.467967109 | 1.71E-02 |
| Chchd1 | 748.5531693 | 0.467399527 | 8.47E-03 |
| Myo18a | 2886.634282 | 0.466996973 | 1.08E-02 |
| Lmtk2 | 2086.187159 | 0.466966045 | 1.98E-05 |
| Hivep3 | 515.0723489 | 0.466961385 | 2.79E-02 |
| Pold2 | 1477.91555 | 0.466860313 | 4.76E-02 |
| Larp7 | 1751.920278 | 0.466498988 | 3.80E-04 |
| Aldh7a1 | 2850.269756 | 0.4655088 | 7.76E-04 |
| Dse | 4466.695367 | 0.464543375 | 3.59E-02 |
| Hsf2 | 1151.428926 | 0.464063432 | 3.62E-03 |
| Usp14 | 3649.869986 | 0.463881189 | 5.92E-04 |
| Plekhf2 | 391.5956648 | 0.46283569 | 1.92E-02 |
| Lipt1 | 375.2285292 | 0.462627112 | 1.61E-02 |
| Apaf1 | 2157.688006 | 0.462527615 | 8.40E-03 |
| Cpd | 4849.178331 | 0.46190704 | 2.83E-03 |
| Pvr | 3501.591273 | 0.461115752 | 4.25E-02 |
| Atp1b3 | 3296.651774 | 0.460673466 | 2.07E-02 |
| Utp20 | 4892.506841 | 0.46033333 | 1.35E-02 |
| Crtc3 | 1345.590025 | 0.460212806 | 1.95E-02 |
| Usp12 | 2202.538901 | 0.459721236 | 6.36E-04 |
| Rubcn | 3167.973214 | 0.457140539 | 5.73E-04 |
| Arhgap17 | 1804.268498 | 0.456617092 | 2.30E-05 |
| Nap1l1 | 24172.49471 | 0.455698934 | 4.61E-04 |
| Slc39a14 | 1634.769816 | 0.455505694 | 4.71E-03 |
| Tmem63a | 1994.571198 | 0.453943626 | 2.48E-04 |
| Zcchc10 | 277.3639513 | 0.453927936 | 3.21E-02 |
| Htra2 | 5540.271863 | 0.453676602 | 1.21E-03 |
| Pitpnm1 | 3326.041469 | 0.453353196 | 1.40E-04 |
| Hoxc4 | 166.5136363 | 0.453009251 | 3.74E-02 |
| Mnat1 | 1466.460687 | 0.452346531 | 1.80E-02 |
| Hikeshi | 958.3911583 | 0.452331574 | 5.60E-04 |
| Ppic | 4106.544221 | 0.452199721 | 1.58E-03 |
| E2f4 | 3205.637373 | 0.451370704 | 1.56E-04 |
| Hspa1b | 1228.877042 | 0.450549007 | 5.87E-03 |
| Map2k3 | 4419.736716 | 0.450241511 | 9.49E-03 |
| Serpinb6a | 9297.117704 | 0.449659263 | 6.64E-04 |
| Fez2 | 3426.090875 | 0.449178302 | 3.72E-02 |
| Ganab | 13601.05205 | 0.449149345 | 6.26E-05 |
| Tfdp2 | 1412.093919 | 0.448642328 | 3.18E-02 |
| Cuedc1 | 1729.792206 | 0.448245857 | 2.70E-03 |
| Dnmbp | 2254.506423 | 0.446671642 | 6.21E-06 |
| Flnc | 35973.77426 | 0.446436324 | 3.60E-02 |
| Canx | 28016.75408 | 0.446179571 | 4.57E-03 |
| Morc3 | 1497.828179 | 0.445129219 | 2.23E-02 |
| Cdc42ep2 | 2880.635182 | 0.444871925 | 2.82E-02 |

| | | | | |
|---|---|---|---|---|
| B3gnt2 | | 1006.377242 | 0.44360209 | 2.35E-02 |
| Cdk2 | | 961.5697005 | 0.441568489 | 1.25E-02 |
| Dcbld2 | | 3824.810857 | 0.441521501 | 3.92E-02 |
| Rap1b | | 4653.914511 | 0.44000223 | 9.91E-07 |
| U2af1 | | 2772.347274 | 0.439780257 | 7.55E-04 |
| Tln2 | | 2751.788901 | 0.438815809 | 3.76E-02 |
| Ap1s2 | | 981.3291552 | 0.438577187 | 3.93E-02 |
| Asah2 | | 1867.355462 | 0.438049747 | 4.97E-02 |
| Chka | | 2275.514018 | 0.43801238 | 3.27E-02 |
| Wdr12 | | 1978.768705 | 0.437757069 | 1.11E-02 |
| Inafm2 | | 1281.565853 | 0.437440092 | 2.85E-04 |
| | 11-Sep | 12235.26164 | 0.436348677 | 2.99E-03 |
| Spen | | 2670.265912 | 0.436326859 | 1.09E-03 |
| Zfhx3 | | 1535.860148 | 0.436132679 | 1.47E-03 |
| Kmt5a | | 3867.372276 | 0.436000535 | 5.64E-03 |
| Schip1 | | 526.8642713 | 0.435919652 | 3.18E-02 |
| Abhd17c | | 2306.603289 | 0.435751763 | 1.56E-02 |
| Xpo5 | | 4096.279036 | 0.435269236 | 8.33E-03 |
| Cdk7 | | 2374.916507 | 0.434500567 | 1.18E-03 |
| Fam45a | | 1463.362685 | 0.433958252 | 1.51E-02 |
| Galt | | 472.9768465 | 0.432910974 | 2.22E-02 |
| Bid | | 1806.508215 | 0.43280526 | 8.41E-03 |
| Ubxn7 | | 3892.875648 | 0.432524389 | 2.18E-03 |
| NA | | 3226.764882 | 0.431812775 | 4.20E-02 |
| Necap1 | | 2428.802313 | 0.431073158 | 1.85E-03 |
| Fut11 | | 609.8350578 | 0.430632578 | 3.23E-02 |
| Rhbdf1 | | 3389.183734 | 0.430544844 | 7.76E-03 |
| Il17rc | | 2513.616066 | 0.430381046 | 9.94E-06 |
| Aph1a | | 5523.669385 | 0.428248759 | 5.75E-03 |
| Btbd19 | | 938.1618624 | 0.427716782 | 3.89E-02 |
| Caskin2 | | 1078.194197 | 0.426744044 | 3.15E-04 |
| Alg6 | | 639.3946694 | 0.426435026 | 4.16E-02 |
| Cast | | 5248.563907 | 0.425155108 | 4.36E-05 |
| Arl4a | | 298.3612635 | 0.424094077 | 1.21E-02 |
| Iars | | 7024.962915 | 0.423863359 | 3.41E-02 |
| Scamp5 | | 1179.827721 | 0.421086452 | 1.12E-02 |
| Isca2 | | 752.0861875 | 0.419542706 | 1.66E-03 |
| Nup153 | | 3184.816415 | 0.41839135 | 6.07E-03 |
| Abca3 | | 2846.907726 | 0.417657699 | 6.28E-03 |
| Hdac7 | | 6665.610222 | 0.415394966 | 3.65E-06 |
| Pinx1 | | 467.0674322 | 0.415021281 | 3.83E-02 |
| Dnm2 | | 5735.557998 | 0.413479701 | 1.73E-02 |
| Pgpep1 | | 2247.346939 | 0.41339076 | 1.74E-02 |
| Flot1 | | 2483.620995 | 0.412929264 | 4.17E-02 |
| Prdx5 | | 5188.911969 | 0.412344287 | 1.71E-02 |
| Fn3krp | | 425.7525463 | 0.411936221 | 1.60E-02 |
| Rabggtb | | 5299.556326 | 0.411728982 | 3.39E-03 |

| | | | |
|---|---|---|---|
| Mprip | 17779.67662 | 0.411161663 | 1.15E-06 |
| Podnl1 | 3817.279448 | 0.410744237 | 4.46E-02 |
| Sec63 | 6038.823214 | 0.409962572 | 1.07E-02 |
| Thnsl1 | 299.7345792 | 0.409152066 | 4.28E-02 |
| Cept1 | 1366.960895 | 0.408400343 | 2.43E-02 |
| Dstn | 11056.37856 | 0.407882859 | 4.71E-03 |
| Apmap | 1337.145835 | 0.407495206 | 4.86E-03 |
| Nars | 9176.2195 | 0.40726675 | 3.09E-02 |
| Pigl | 298.5422387 | 0.407019872 | 1.46E-02 |
| Gli3 | 1922.023409 | 0.405204468 | 2.70E-02 |
| Uhrf1bp1l | 3287.025469 | 0.404505857 | 2.00E-03 |
| Chchd4 | 753.277916 | 0.403693728 | 3.58E-02 |
| 1810055G02Rik | 1562.499636 | 0.403552204 | 2.32E-02 |
| Fzd7 | 1499.058727 | 0.403296159 | 2.04E-02 |
| Piezo1 | 14797.30952 | 0.403039464 | 2.40E-02 |
| Syngr2 | 2533.946804 | 0.402829049 | 3.70E-03 |
| Cwc27 | 742.8550555 | 0.402195358 | 3.00E-02 |
| Cers2 | 6182.42433 | 0.400354412 | 5.21E-05 |
| Slc7a6 | 3393.128208 | 0.400195469 | 7.84E-04 |
| Gm9531 | 2715.543962 | 0.399247801 | 2.40E-02 |
| Ltv1 | 1629.7818 | 0.39896327 | 1.13E-02 |
| Tmem80 | 217.7392199 | 0.398659313 | 4.58E-02 |
| Abca2 | 3887.217761 | 0.398490077 | 1.35E-03 |
| Nfyb | 844.2811569 | 0.397700253 | 1.35E-02 |
| Ptpn2 | 736.8581651 | 0.397371625 | 1.27E-02 |
| Mfhas1 | 2892.570226 | 0.397244417 | 3.31E-05 |
| Tbpl1 | 1549.163799 | 0.396853407 | 1.05E-03 |
| Tpm4 | 16366.22209 | 0.396785967 | 1.55E-02 |
| Txnl1 | 3876.007935 | 0.396577403 | 1.37E-03 |
| Frmd6 | 6537.824608 | 0.395615259 | 2.80E-02 |
| Plod3 | 6604.231022 | 0.394713499 | 3.55E-02 |
| Anxa1 | 40517.67387 | 0.393288343 | 2.22E-02 |
| Axl | 36075.15712 | 0.393129728 | 2.54E-02 |
| Cdc42se2 | 2787.71278 | 0.392671102 | 4.93E-05 |
| Notch2 | 5564.204759 | 0.39241486 | 1.94E-03 |
| Peli1 | 768.5211593 | 0.391983312 | 1.99E-02 |
| Ppp2r5a | 2519.975068 | 0.391679004 | 7.48E-03 |
| Dync1h1 | 28582.84367 | 0.391585283 | 7.49E-03 |
| Atn1 | 9447.780035 | 0.390937146 | 1.07E-02 |
| Fam83h | 457.8759574 | 0.390398985 | 1.87E-02 |
| Cgrrf1 | 795.8021606 | 0.389253537 | 1.64E-02 |
| Zc3hav1 | 5239.317525 | 0.388809144 | 2.19E-02 |
| Ginm1 | 1392.945406 | 0.388115554 | 2.44E-02 |
| Map1a | 6290.433199 | 0.388075738 | 2.06E-02 |
| Kansl2 | 5499.25737 | 0.3877427 | 1.39E-02 |
| Med25 | 2722.795482 | 0.387422008 | 5.51E-03 |
| Celf1 | 4225.590777 | 0.386253823 | 2.19E-02 |

| | | | |
|---|---|---|---|
| Zfp217 | 1499.500526 | 0.385590054 | 2.43E-03 |
| Cdc40 | 2423.015152 | 0.383674288 | 1.21E-03 |
| Tdrd7 | 1895.989433 | 0.383492123 | 6.13E-04 |
| Txn1 | 10959.5229 | 0.383328081 | 3.22E-02 |
| H2-D1 | 9845.207692 | 0.382637229 | 3.50E-02 |
| Smap1 | 2848.464316 | 0.380421082 | 7.70E-05 |
| Metap2 | 6994.685675 | 0.379973313 | 1.07E-02 |
| Cfap298 | 524.7661239 | 0.379629643 | 1.95E-02 |
| Crlf3 | 1281.734592 | 0.37922404 | 3.21E-02 |
| Nck1 | 1126.62883 | 0.378663504 | 4.30E-02 |
| Vps36 | 2824.129634 | 0.376591861 | 2.19E-03 |
| Zfp91 | 7065.381387 | 0.375330893 | 3.18E-03 |
| Nsmaf | 1785.080286 | 0.374331928 | 2.40E-03 |
| Nid1 | 41135.2065 | 0.373297373 | 2.44E-03 |
| Mknk1 | 2154.3649 | 0.372790667 | 1.22E-02 |
| Qk | 4353.559499 | 0.372726485 | 3.79E-02 |
| Stag1 | 2203.340069 | 0.372524393 | 3.95E-02 |
| Ube2j1 | 2831.915582 | 0.372423042 | 1.04E-04 |
| Usp10 | 2331.73001 | 0.371031213 | 9.19E-03 |
| Tnk2 | 5977.180731 | 0.370482157 | 1.31E-02 |
| Rad23a | 3073.219033 | 0.370377969 | 1.94E-02 |
| Lsm4 | 1011.076088 | 0.369948594 | 2.54E-02 |
| Eaf1 | 2071.798067 | 0.369711656 | 8.00E-03 |
| Erap1 | 966.344512 | 0.36948975 | 9.32E-03 |
| Sipa1l3 | 3854.926667 | 0.369402936 | 1.19E-02 |
| Ppp1r12c | 1921.830022 | 0.368350453 | 2.25E-03 |
| Lrp11 | 3762.571826 | 0.367996859 | 2.31E-02 |
| Hmgn1 | 7690.882493 | 0.367485845 | 4.37E-02 |
| Trak2 | 2626.579049 | 0.364816973 | 9.24E-03 |
| Cpne3 | 3705.747662 | 0.363420737 | 3.41E-02 |
| Ptpn11 | 9188.837608 | 0.363140427 | 3.04E-02 |
| Tab2 | 5843.044862 | 0.362128324 | 1.65E-04 |
| Gltp | 1692.539984 | 0.360553152 | 4.75E-02 |
| Mrpl21 | 1397.633345 | 0.360536965 | 1.78E-03 |
| Uvrag | 2392.79397 | 0.360230119 | 3.61E-02 |
| Mthfsd | 1470.292497 | 0.360229082 | 2.08E-05 |
| Pdia6 | 14361.81594 | 0.360180244 | 3.10E-02 |
| Tmem201 | 1116.37731 | 0.360024079 | 2.97E-03 |
| Cyfip1 | 6745.902934 | 0.359468887 | 1.12E-03 |
| Itgav | 6626.652067 | 0.358118709 | 4.04E-02 |
| Gpsm2 | 2322.001977 | 0.357756484 | 3.95E-02 |
| Phka2 | 2086.31599 | 0.357685356 | 2.88E-02 |
| Dolpp1 | 505.6016964 | 0.356837842 | 3.58E-02 |
| Gng12 | 10176.62266 | 0.356337951 | 4.07E-02 |
| Fuz | 552.4981124 | 0.356219856 | 4.18E-02 |
| Prrc2c | 11572.89267 | 0.354228059 | 2.37E-02 |
| Higd2a | 2214.016855 | 0.354129518 | 4.36E-02 |

| | | | |
|---|---|---|---|
| Grk2 | 3292.244991 | 0.353834752 | 1.09E-02 |
| Rtn3 | 10304.32456 | 0.351590815 | 3.95E-02 |
| Osbpl9 | 6203.981448 | 0.35126696 | 1.80E-03 |
| Tmem131 | 4629.57427 | 0.350538311 | 1.89E-04 |
| Zrsr2 | 1460.694365 | 0.350271896 | 8.58E-03 |
| Mrps28 | 608.7447475 | 0.349256332 | 4.32E-02 |
| Ccdc43 | 996.7600218 | 0.348262733 | 4.19E-03 |
| Vars2 | 1444.707556 | 0.347099781 | 1.03E-02 |
| Prmt5 | 1974.839417 | 0.346913929 | 3.50E-02 |
| Ankib1 | 2741.451452 | 0.346329759 | 1.25E-02 |
| Nucb1 | 11251.76517 | 0.345266238 | 9.71E-03 |
| Stat6 | 5625.010172 | 0.343219423 | 1.53E-04 |
| Socs6 | 1293.79635 | 0.341995847 | 1.07E-02 |
| Hipk1 | 4679.274894 | 0.340747794 | 2.69E-03 |
| Atad3a | 1366.86159 | 0.340373764 | 3.62E-02 |
| Mgat2 | 2476.636202 | 0.340054617 | 2.21E-02 |
| Camk1 | 3159.841714 | 0.339968103 | 4.01E-02 |
| Tubgcp3 | 2045.209214 | 0.339490728 | 6.86E-03 |
| Atl2 | 1798.458299 | 0.338752909 | 1.43E-02 |
| Gde1 | 1219.065937 | 0.338129994 | 4.44E-03 |
| Cactin | 1721.152786 | 0.337095292 | 6.49E-03 |
| Snx15 | 878.8217529 | 0.335927037 | 4.45E-03 |
| Uso1 | 6535.449643 | 0.335503343 | 4.79E-02 |
| Ptpmt1 | 905.1759236 | 0.335150393 | 4.97E-02 |
| Dvl1 | 2494.315404 | 0.334983676 | 4.28E-02 |
| Gemin2 | 424.6443906 | 0.331923771 | 3.13E-02 |
| Wdr44 | 567.3078697 | 0.330505523 | 4.79E-02 |
| Pgd | 3686.528964 | 0.330041352 | 2.11E-02 |
| Yrdc | 1246.308307 | 0.327269783 | 2.40E-02 |
| Ipo5 | 14120.82957 | 0.3268437 | 4.53E-02 |
| Fbl | 2993.699857 | 0.325247563 | 4.13E-02 |
| Tsku | 1205.341302 | 0.324722092 | 3.42E-02 |
| Tial1 | 1884.714745 | 0.3245024 | 4.76E-02 |
| St7 | 1024.711178 | 0.3230117 | 4.24E-02 |
| 5430416N02Rik | 1000.533301 | 0.32205149 | 3.04E-02 |
| Cpsf4 | 763.0921259 | 0.320479281 | 4.55E-02 |
| Pfdn1 | 1895.188969 | 0.318354395 | 9.14E-03 |
| Emc6 | 1268.950995 | 0.318337099 | 2.82E-03 |
| Gypc | 1383.765939 | 0.318011895 | 3.04E-02 |
| Psma1 | 3673.371411 | 0.317756745 | 3.76E-03 |
| Hspa9 | 32616.301 | 0.317586408 | 6.73E-03 |
| Ctbs | 628.3728415 | 0.31750126 | 2.85E-02 |
| Pnpla6 | 1143.850243 | 0.317133662 | 1.76E-02 |
| Zzef1 | 3506.293149 | 0.316556529 | 6.02E-03 |
| Utp23 | 570.5468979 | 0.316124578 | 5.46E-03 |
| Paf1 | 1759.690936 | 0.314426899 | 8.84E-05 |
| Gm13835 | 2487.733138 | 0.314217704 | 4.75E-02 |

| | | | |
|---|---|---|---|
| Sash1 | 2475.130362 | 0.313822175 | 2.81E-02 |
| Rnf145 | 4349.170828 | 0.313637363 | 2.96E-03 |
| Ube2j2 | 1199.238568 | 0.313404211 | 3.05E-02 |
| Pdcd6ip | 8464.98572 | 0.31272485 | 1.39E-03 |
| Drap1 | 6991.132614 | 0.31042907 | 4.78E-02 |
| Axin1 | 4077.021414 | 0.306996829 | 7.35E-03 |
| Ercc4 | 754.5038931 | 0.305534991 | 4.62E-02 |
| Zfyve9 | 2465.724035 | 0.304569717 | 1.08E-02 |
| Gm8116 | 1189.5011 | 0.303704981 | 1.86E-02 |
| Magoh | 2065.067574 | 0.303463068 | 2.32E-02 |
| Phykpl | 589.4291438 | 0.30195967 | 3.81E-02 |
| Spop | 6108.041441 | 0.300933823 | 4.44E-03 |
| Pdcl | 1348.652634 | 0.299645669 | 1.26E-02 |
| Zfp598 | 2577.338794 | 0.298186657 | 4.22E-02 |
| Dimt1 | 665.6401902 | 0.298136049 | 1.68E-02 |
| Cdk11b | 2039.570556 | 0.297260991 | 8.90E-03 |
| Cmtm6 | 929.6733678 | 0.296002848 | 3.21E-02 |
| Psmd1 | 10295.53513 | 0.295755689 | 4.55E-02 |
| Ptk2 | 4021.590193 | 0.295738665 | 1.51E-02 |
| Exoc8 | 988.8659388 | 0.29454709 | 3.55E-02 |
| Ccdc12 | 559.7686421 | 0.294453487 | 4.79E-02 |
| Mbd2 | 1918.447548 | 0.29332463 | 5.03E-03 |
| Tars | 6511.285888 | 0.293240317 | 1.95E-02 |
| Tacc2 | 2141.000492 | 0.292852754 | 1.56E-02 |
| Tep1 | 1733.768891 | 0.291419778 | 4.22E-02 |
| Rwdd1 | 3045.160573 | 0.290072982 | 2.42E-02 |
| Gnb1 | 10433.30772 | 0.287917035 | 5.00E-05 |
| Ube2f | 1729.361484 | 0.287631644 | 3.95E-02 |
| Calm2 | 10854.30123 | 0.286369271 | 2.06E-02 |
| Fyn | 3259.615135 | 0.285138325 | 1.36E-02 |
| Rab8a | 3357.260812 | 0.284573959 | 4.79E-02 |
| Tpbg | 2142.107889 | 0.284415338 | 2.86E-02 |
| Otub1 | 4405.542155 | 0.283639517 | 4.34E-02 |
| Cdk17 | 1107.511682 | 0.282300447 | 2.76E-02 |
| Usp6nl | 1363.084088 | 0.281789529 | 3.13E-02 |
| Dag1 | 10803.36973 | 0.281508683 | 3.25E-02 |
| Sec23b | 2213.107425 | 0.281460685 | 1.61E-02 |
| Wwtr1 | 5195.543564 | 0.278024841 | 1.72E-03 |
| Fam102b | 2904.554024 | 0.276223337 | 2.71E-02 |
| Itprid2 | 2850.831262 | 0.273018334 | 3.40E-02 |
| Rnf19a | 1760.078004 | 0.27285321 | 7.80E-03 |
| Comt | 3881.085594 | 0.272272643 | 3.51E-02 |
| N4bp1 | 3589.986201 | 0.271977598 | 2.16E-03 |
| Srpk1 | 4585.424015 | 0.270092114 | 2.96E-02 |
| Clcn5 | 3296.841631 | 0.265650006 | 3.43E-03 |
| Pdha1 | 9868.078405 | 0.263092601 | 9.90E-03 |
| Arhgef17 | 6255.352482 | 0.262654731 | 3.66E-02 |

| | | | |
|---|---|---|---|
| Cwc15 | 2545.719953 | 0.262267389 | 8.89E-03 |
| Zdhhc5 | 4681.471334 | 0.261939618 | 2.26E-02 |
| Socs7 | 1466.776444 | 0.259298688 | 3.92E-02 |
| Nvl | 1837.408422 | 0.255870612 | 1.22E-02 |
| Dync1li1 | 2155.249373 | 0.25562119 | 3.62E-02 |
| Gpx8 | 2857.904981 | 0.254299182 | 9.52E-03 |
| Slc9a1 | 2227.40331 | 0.254078135 | 3.66E-02 |
| Cpne1 | 5044.131924 | 0.253065007 | 5.57E-03 |
| Heatr5a | 3537.032457 | 0.250140155 | 8.65E-03 |
| Elk3 | 4455.882184 | 0.2496236 | 1.70E-02 |
| Zdhhc16 | 1645.02251 | 0.248696607 | 3.08E-02 |
| Ccdc59 | 1338.787267 | 0.246229667 | 3.08E-02 |
| Sri | 3410.645404 | 0.24591999 | 2.66E-02 |
| Pan2 | 1537.473845 | 0.244897247 | 3.26E-02 |
| Ahcyl1 | 10647.42186 | 0.244840689 | 4.18E-03 |
| Plgrkt | 1248.255989 | 0.241622917 | 1.62E-02 |
| Supv3l1 | 1487.778323 | 0.241072591 | 2.52E-02 |
| Morn4 | 760.2486884 | 0.239023743 | 3.65E-02 |
| Slc38a1 | 5119.465128 | 0.228383692 | 3.38E-02 |
| Timm17a | 2158.37418 | 0.218141299 | 3.66E-02 |
| Tram1 | 2081.460987 | 0.216168377 | 4.65E-02 |
| Zfp106 | 11251.11847 | 0.19646884 | 1.51E-02 |
| Appbp2 | 3439.901311 | 0.194046177 | 1.92E-02 |
| Cnot2 | 3155.015605 | 0.19240562 | 2.85E-02 |
| Ube3b | 6295.625369 | 0.191472085 | 4.65E-02 |
| Nfatc3 | 4109.575676 | 0.16107486 | 2.19E-02 |
| Uri1 | 1845.880355 | -0.160333321 | 3.45E-02 |
| Zfp110 | 1694.471274 | -0.179323902 | 3.86E-02 |
| Mgrn1 | 4288.647559 | -0.181199985 | 4.98E-02 |
| Brd1 | 2882.283459 | -0.183078916 | 2.81E-02 |
| Tgfbrap1 | 1916.492283 | -0.187986528 | 4.39E-02 |
| Kansl3 | 3444.874046 | -0.201356423 | 4.71E-02 |
| Stx12 | 1913.740575 | -0.203200519 | 4.79E-02 |
| Akap10 | 1520.79788 | -0.204329804 | 3.04E-02 |
| Msl1 | 3847.048481 | -0.205029346 | 1.70E-02 |
| Cog7 | 1094.294008 | -0.205695141 | 4.49E-02 |
| Fem1c | 1546.41211 | -0.209415517 | 2.92E-02 |
| Adh5 | 4642.32907 | -0.211643407 | 4.02E-02 |
| Cttn | 14805.07831 | -0.21531247 | 1.25E-02 |
| Abcd3 | 3631.045164 | -0.216321588 | 7.77E-03 |
| Pknox1 | 1152.190822 | -0.216490828 | 4.75E-02 |
| Ppp1cc | 8121.670663 | -0.22009805 | 4.78E-02 |
| Atxn7l3b | 7440.579257 | -0.220843533 | 4.88E-02 |
| Fam193a | 2712.375027 | -0.222851654 | 4.59E-02 |
| Msantd4 | 1976.487313 | -0.224188659 | 1.41E-02 |
| Slc30a7 | 2473.28765 | -0.225836873 | 2.22E-02 |
| Ppp6r2 | 1188.00831 | -0.226816372 | 4.51E-03 |

| | | | |
|---|---|---|---|
| Sdha | 7204.468236 | -0.22762634 | 2.64E-02 |
| Fbrs | 1611.001734 | -0.23142397 | 4.30E-02 |
| Psmd10 | 850.2875447 | -0.233455389 | 3.37E-02 |
| Dhx57 | 1977.793762 | -0.234317559 | 2.85E-02 |
| Erc1 | 3022.322006 | -0.235768185 | 9.90E-03 |
| Xpo7 | 3778.843741 | -0.236815319 | 3.50E-02 |
| Glyr1 | 5537.263863 | -0.238389053 | 3.51E-02 |
| Atp5f1 | 8259.418685 | -0.238481154 | 3.08E-02 |
| Sppl3 | 3400.317528 | -0.241034208 | 3.40E-02 |
| Dmac1 | 1447.805556 | -0.242822003 | 2.00E-02 |
| Gdap2 | 944.6345225 | -0.243101055 | 4.32E-02 |
| Ccdc91 | 1289.115254 | -0.243927398 | 4.23E-02 |
| Trappc11 | 2425.129797 | -0.244607807 | 5.48E-03 |
| Sde2 | 2069.820891 | -0.245281404 | 4.01E-02 |
| Stk25 | 4558.968745 | -0.24649864 | 7.88E-03 |
| Phf2 | 3036.045331 | -0.246701737 | 2.64E-02 |
| Arl2bp | 3849.906749 | -0.248608258 | 4.32E-02 |
| 1600012H06Rik | 1609.676472 | -0.249273043 | 3.08E-02 |
| Rab3gap2 | 4136.31955 | -0.251895799 | 4.79E-02 |
| Ppt1 | 7244.146237 | -0.252442346 | 3.99E-02 |
| Rab11fip3 | 2701.591854 | -0.253103572 | 2.25E-02 |
| Gpalpp1 | 1808.305997 | -0.254105198 | 2.91E-02 |
| Aagab | 1352.172232 | -0.254250632 | 4.00E-02 |
| Fam53c | 2241.692611 | -0.255913887 | 1.34E-02 |
| Rab24 | 1438.55139 | -0.257496411 | 2.38E-02 |
| Taf2 | 2167.708385 | -0.257923547 | 3.75E-05 |
| Fam114a2 | 3449.452351 | -0.259978201 | 3.25E-03 |
| Slc31a1 | 3001.893987 | -0.260073907 | 7.69E-04 |
| Yap1 | 5868.317144 | -0.260227619 | 1.39E-03 |
| Fam149b | 889.385841 | -0.260891873 | 1.10E-02 |
| Dlg5 | 3568.439899 | -0.261068745 | 1.42E-02 |
| Ipo8 | 2808.34261 | -0.263994095 | 4.23E-02 |
| Chd4 | 15088.84888 | -0.26446794 | 1.10E-02 |
| Map3k12 | 890.7020701 | -0.264610831 | 3.48E-02 |
| Ap3d1 | 7686.894989 | -0.265510123 | 3.50E-03 |
| Carm1 | 4057.020687 | -0.265582043 | 3.94E-02 |
| Tmem9b | 1153.311785 | -0.265802463 | 3.51E-02 |
| Impa1 | 1974.053515 | -0.266709792 | 4.22E-02 |
| Zfp869 | 866.2160119 | -0.266721588 | 2.88E-02 |
| Tbx18 | 1240.526144 | -0.266755558 | 2.09E-02 |
| Tmed8 | 1367.151775 | -0.268562833 | 4.46E-02 |
| Slc25a12 | 2589.919914 | -0.269219648 | 4.61E-02 |
| Cep68 | 2000.679055 | -0.269530309 | 3.34E-03 |
| Ulk2 | 3079.236902 | -0.270130045 | 3.85E-03 |
| 5730409E04Rik | 1776.754366 | -0.270315456 | 3.49E-02 |
| Dctn1 | 6439.938184 | -0.270387292 | 3.83E-02 |
| Ap2s1 | 2093.820475 | -0.270413343 | 4.92E-02 |

| | | | |
|---|---|---|---|
| Cript | 1250.406291 | -0.270762963 | 4.49E-02 |
| Cep19 | 695.464579 | -0.271257927 | 2.19E-02 |
| Mafg | 3468.758033 | -0.271293844 | 3.37E-02 |
| Zfp746 | 1625.00071 | -0.271572825 | 1.48E-02 |
| Pcsk7 | 888.8351781 | -0.271855861 | 2.18E-02 |
| Ppp3ca | 3841.872004 | -0.271899345 | 4.32E-02 |
| Prdm5 | 1471.436735 | -0.272042876 | 3.83E-02 |
| Atp9b | 2468.969284 | -0.27212054 | 1.58E-02 |
| Mus81 | 696.6969706 | -0.273205407 | 1.61E-02 |
| Ehmt1 | 2812.096587 | -0.273579538 | 6.73E-03 |
| Mtmr3 | 4507.121796 | -0.273581906 | 3.95E-02 |
| Hadhb | 3032.965333 | -0.275349682 | 5.70E-03 |
| Pip4k2b | 4426.450694 | -0.275433316 | 6.50E-03 |
| Stk38l | 824.657163 | -0.275769613 | 3.50E-02 |
| Synj2 | 1645.375646 | -0.27589879 | 7.88E-03 |
| Umad1 | 626.3534589 | -0.276303547 | 1.14E-02 |
| Clec16a | 2108.067705 | -0.278667197 | 1.31E-03 |
| Nedd4 | 84397.06472 | -0.278920228 | 3.59E-02 |
| Adar | 2479.583721 | -0.280409564 | 3.75E-02 |
| Kif3c | 2739.168318 | -0.280929688 | 8.77E-03 |
| Mllt1 | 2904.39071 | -0.281031489 | 3.51E-02 |
| Ndufaf7 | 1011.202297 | -0.281490022 | 4.21E-02 |
| Anxa4 | 8612.757354 | -0.281603636 | 1.80E-02 |
| Fam118a | 816.1656045 | -0.282061714 | 4.40E-02 |
| Ift52 | 1834.55518 | -0.282739184 | 2.46E-02 |
| Cmpk1 | 3161.859525 | -0.283361931 | 4.62E-03 |
| Jmy | 1698.02112 | -0.283794145 | 7.32E-03 |
| Plxna1 | 20953.16237 | -0.285309017 | 6.05E-04 |
| Zbtb21 | 1062.931372 | -0.285343147 | 4.81E-02 |
| Ppil2 | 3213.172897 | -0.2856138 | 2.59E-02 |
| Actr3 | 13135.49264 | -0.28566499 | 3.04E-02 |
| Kif3b | 2552.903667 | -0.285711425 | 2.23E-04 |
| Gsk3b | 3602.663052 | -0.286864452 | 3.37E-02 |
| Elmod3 | 569.4908347 | -0.286969827 | 3.72E-02 |
| Decr1 | 1500.980543 | -0.287067975 | 1.52E-02 |
| Zeb2 | 3746.786289 | -0.287405427 | 2.56E-02 |
| Egln2 | 2115.922258 | -0.287446811 | 4.98E-02 |
| Zfp513 | 1098.824088 | -0.287859215 | 3.18E-03 |
| Man2a1 | 13475.00019 | -0.288196228 | 1.10E-02 |
| Zfp410 | 1325.253078 | -0.289297842 | 1.08E-02 |
| Usp28 | 1013.741207 | -0.289362933 | 1.49E-02 |
| Wdr59 | 1368.374064 | -0.28961062 | 7.72E-03 |
| Gid8 | 1780.261878 | -0.289838052 | 2.11E-02 |
| Fbxl17 | 1422.489761 | -0.290169056 | 2.32E-02 |
| Poldip2 | 3963.395315 | -0.291679639 | 3.57E-02 |
| Tmem237 | 787.3719009 | -0.2923491 | 2.38E-02 |
| Fam32a | 3348.599776 | -0.292714614 | 1.01E-02 |

| | | | |
|---|---|---|---|
| Ldah | 2630.414737 | -0.29304815 | 2.76E-02 |
| Aldh9a1 | 4090.448895 | -0.293259006 | 3.41E-02 |
| Zc4h2 | 400.4251748 | -0.293600307 | 2.90E-02 |
| Cbx6 | 10839.95353 | -0.29369296 | 1.89E-02 |
| Zfp358 | 1303.342284 | -0.293932578 | 3.29E-02 |
| Sft2d1 | 798.2097011 | -0.294016926 | 2.36E-02 |
| Lamtor5 | 1897.270012 | -0.294216563 | 2.87E-02 |
| Nudt2 | 706.1488302 | -0.294227742 | 4.66E-03 |
| Dcaf7 | 6677.160184 | -0.295674252 | 1.83E-02 |
| Kifap3 | 3772.714154 | -0.296200225 | 1.00E-02 |
| Plekhm3 | 751.7569629 | -0.296862424 | 2.52E-02 |
| Itsn1 | 2752.843847 | -0.296941099 | 1.63E-02 |
| Ist1 | 3319.949775 | -0.29720652 | 1.90E-02 |
| Capn1 | 3476.697024 | -0.298875717 | 3.88E-02 |
| Cox5b | 4421.831308 | -0.299894463 | 2.54E-02 |
| Zfp251 | 1144.446552 | -0.300000794 | 1.96E-03 |
| Abcc10 | 532.3965489 | -0.300344673 | 1.69E-02 |
| Hmces | 1311.48031 | -0.301033119 | 1.41E-02 |
| Max | 2035.937597 | -0.301194753 | 1.12E-02 |
| Chkb | 1184.070151 | -0.301552939 | 2.17E-02 |
| Fbxw5 | 1900.946667 | -0.302048339 | 1.97E-03 |
| Bloc1s1 | 2005.739788 | -0.302053216 | 2.80E-02 |
| Dolk | 812.9649783 | -0.302145136 | 3.16E-02 |
| Rabgap1 | 4533.421953 | -0.302583894 | 2.25E-02 |
| Gcdh | 962.2082415 | -0.302987433 | 3.40E-02 |
| Ccny | 4186.355601 | -0.304508763 | 1.10E-03 |
| Unc119b | 2447.389577 | -0.304686151 | 1.51E-02 |
| Pik3r4 | 1596.763516 | -0.304789111 | 2.80E-02 |
| Dgat1 | 845.3611873 | -0.304965738 | 2.06E-03 |
| Wipi1 | 5221.186575 | -0.305132294 | 1.68E-02 |
| Hp1bp3 | 6438.328602 | -0.30547283 | 3.10E-02 |
| Bin1 | 3469.194523 | -0.306317659 | 4.22E-02 |
| Trim35 | 6233.884212 | -0.306530619 | 1.17E-02 |
| Fastkd1 | 692.7686316 | -0.306872542 | 2.26E-02 |
| Eci2 | 2234.646018 | -0.307302389 | 4.55E-02 |
| Cdc42se1 | 4459.289015 | -0.30740417 | 1.10E-02 |
| Pls3 | 9293.72303 | -0.308089083 | 3.54E-03 |
| Ddx6 | 7985.738967 | -0.308114822 | 1.22E-03 |
| Sgpl1 | 13380.67463 | -0.308251382 | 3.97E-02 |
| Foxo1 | 589.3772702 | -0.308398774 | 2.87E-02 |
| Ccdc22 | 1040.823011 | -0.308791575 | 1.14E-02 |
| Zfp672 | 709.0883327 | -0.31026158 | 3.43E-02 |
| Zfp706 | 4663.743055 | -0.312454149 | 4.58E-02 |
| Stambp | 1548.578292 | -0.312508507 | 3.61E-02 |
| Atf7 | 5470.285064 | -0.312521156 | 3.99E-02 |
| Blvra | 783.0242707 | -0.312624233 | 3.81E-02 |
| Vti1b | 2649.138363 | -0.312630995 | 1.40E-02 |

| | | | |
|---|---|---|---|
| Parp11 | 900.9732298 | -0.312779512 | 3.64E-02 |
| Ugp2 | 4229.526194 | -0.313701318 | 2.91E-02 |
| Spats2 | 2069.741681 | -0.314330546 | 1.81E-02 |
| Rbl2 | 2066.659329 | -0.314506377 | 8.82E-04 |
| Rnf157 | 1391.358528 | -0.314543252 | 3.96E-02 |
| Commd3 | 3470.393101 | -0.314611457 | 9.31E-03 |
| Arpc5 | 5430.297978 | -0.314626375 | 5.65E-04 |
| Cmtm4 | 10031.66492 | -0.314927781 | 6.02E-03 |
| Zcchc14 | 3410.077933 | -0.31499139 | 9.99E-04 |
| Cnot8 | 2936.25042 | -0.315111057 | 2.70E-02 |
| Pex2 | 1610.317343 | -0.315504491 | 1.03E-02 |
| Prdm2 | 2816.518592 | -0.316077741 | 3.00E-03 |
| Vamp8 | 2018.695151 | -0.316145085 | 1.27E-02 |
| Mov10 | 1852.686349 | -0.316411637 | 3.60E-02 |
| Gm20716 | 4001.886924 | -0.316635935 | 5.89E-04 |
| Chd3 | 13653.34876 | -0.316732389 | 4.25E-02 |
| Wdr41 | 1253.631657 | -0.316865019 | 1.16E-03 |
| Cox16 | 493.8339891 | -0.316966652 | 1.07E-02 |
| Smug1 | 791.1906468 | -0.317131524 | 4.08E-03 |
| Exoc2 | 1768.947934 | -0.31762193 | 6.86E-03 |
| Zfp9 | 1864.128904 | -0.318150941 | 3.52E-02 |
| Atg4b | 2480.845886 | -0.318574603 | 8.71E-03 |
| Ift43 | 828.8161268 | -0.318799788 | 8.86E-03 |
| Zmat2 | 2291.274399 | -0.319421493 | 1.79E-03 |
| Rit1 | 1586.676908 | -0.320176333 | 1.09E-02 |
| Ppp1r16a | 1023.013923 | -0.320664376 | 3.77E-02 |
| Spg11 | 3357.707005 | -0.320864164 | 1.20E-03 |
| Ttl | 702.6056377 | -0.320937412 | 3.52E-02 |
| Ltbr | 4575.460131 | -0.321312466 | 1.14E-02 |
| Ubn1 | 5569.528947 | -0.321368918 | 4.43E-03 |
| Llgl1 | 4105.61428 | -0.321531953 | 1.77E-02 |
| P3h3 | 5126.717251 | -0.321756031 | 2.52E-02 |
| Cpped1 | 2173.563252 | -0.321896811 | 2.91E-02 |
| Prkd1 | 1467.094425 | -0.322228591 | 4.36E-02 |
| Naa35 | 1827.19018 | -0.32251007 | 8.08E-03 |
| Fbxo25 | 1442.277771 | -0.323371919 | 4.13E-02 |
| Commd8 | 1529.596254 | -0.323890622 | 2.70E-02 |
| Armcx2 | 5543.690073 | -0.323918417 | 4.51E-03 |
| Atp5s | 258.5845938 | -0.324043461 | 4.55E-02 |
| B4galnt1 | 1418.878612 | -0.325202355 | 3.74E-02 |
| Iqcc | 667.0780684 | -0.326549895 | 4.93E-02 |
| Atp5g3 | 7645.161236 | -0.326931968 | 2.74E-02 |
| Fndc3a | 4745.827846 | -0.326958162 | 2.66E-03 |
| Rcc2 | 9407.022186 | -0.327786464 | 4.93E-02 |
| Tstd3 | 514.7463397 | -0.328646981 | 4.28E-02 |
| Trnau1ap | 1335.035752 | -0.328733991 | 1.95E-02 |
| Slc35e3 | 2219.989806 | -0.329241588 | 4.35E-02 |

| | | | |
|---|---|---|---|
| Rab31 | 6124.422698 | -0.329374394 | 1.12E-02 |
| Dnttip1 | 1338.58901 | -0.329567769 | 1.52E-02 |
| Uspl1 | 1969.222052 | -0.330616864 | 3.20E-03 |
| Zkscan3 | 2374.665102 | -0.331188212 | 2.62E-02 |
| Ift22 | 607.373969 | -0.331870451 | 9.51E-03 |
| Rtf1 | 2978.284882 | -0.332307636 | 1.77E-02 |
| Dyrk2 | 1360.699862 | -0.332385007 | 2.03E-02 |
| Vps29 | 2535.509627 | -0.3329866 | 1.11E-02 |
| Zgpat | 1572.13059 | -0.333505876 | 1.30E-03 |
| Gm42688 | 427.2518859 | -0.333508504 | 4.15E-02 |
| Naxd | 2049.602672 | -0.334033662 | 2.61E-02 |
| Scpep1 | 18619.66479 | -0.334291027 | 4.17E-02 |
| Soat1 | 3693.855559 | -0.335310173 | 3.46E-03 |
| Zfp334 | 1151.134813 | -0.336100571 | 5.41E-03 |
| Gigyf1 | 2070.367659 | -0.336148785 | 2.19E-02 |
| Nbdy | 1309.915387 | -0.336175137 | 3.15E-03 |
| Yeats2 | 2822.94201 | -0.337062833 | 1.32E-03 |
| Prkaca | 4495.591608 | -0.337634862 | 4.33E-02 |
| NA | 2939.508412 | -0.337877307 | 1.62E-02 |
| Fbxl6 | 1647.638682 | -0.338167907 | 4.89E-02 |
| Tmem181a | 524.5608539 | -0.338295463 | 5.33E-03 |
| Prepl | 2110.742543 | -0.339382248 | 1.45E-03 |
| Ago1 | 6615.470307 | -0.340085015 | 1.14E-05 |
| Pdrg1 | 1875.615487 | -0.340221912 | 2.20E-02 |
| Ndufb6 | 1520.582627 | -0.340357702 | 2.82E-02 |
| Zc3h12b | 308.6497987 | -0.340548591 | 4.86E-02 |
| 1700123O20Rik | 1002.793485 | -0.340646018 | 1.22E-02 |
| Clp1 | 553.9320773 | -0.340903202 | 9.49E-03 |
| Pja1 | 1983.521891 | -0.341169839 | 6.82E-03 |
| Tmem98 | 1290.414908 | -0.3417078 | 4.55E-02 |
| Rnf185 | 2243.235099 | -0.341778248 | 4.75E-02 |
| Tsr2 | 1165.014558 | -0.342069006 | 4.53E-02 |
| Sgcb | 2110.307292 | -0.342518819 | 7.94E-03 |
| Cyb5d1 | 1009.24945 | -0.342894574 | 2.76E-03 |
| Smyd2 | 3058.476522 | -0.343164255 | 2.07E-02 |
| Ccnq | 612.2135084 | -0.344171832 | 3.33E-02 |
| Parn | 3178.519304 | -0.344361129 | 1.75E-02 |
| Zbed3 | 1298.091305 | -0.344545969 | 4.14E-06 |
| Tomm6 | 2647.365377 | -0.344715809 | 3.51E-02 |
| Slc35a1 | 1244.299512 | -0.345163373 | 9.95E-03 |
| Hps3 | 1032.072822 | -0.345755173 | 1.60E-02 |
| Dip2c | 2363.258568 | -0.345938524 | 4.56E-03 |
| Dguok | 1124.366048 | -0.346099783 | 4.44E-02 |
| Ptms | 12716.67832 | -0.346370681 | 2.91E-02 |
| Sbf1 | 5212.739294 | -0.346517984 | 1.78E-02 |
| Ehbp1 | 2247.789155 | -0.347191978 | 8.14E-03 |
| Spry2 | 928.0977301 | -0.347311579 | 1.92E-02 |

| | | | |
|---|---|---|---|
| Rchy1 | 1824.914905 | -0.347475648 | 4.22E-02 |
| Jkamp | 1284.292578 | -0.347748061 | 8.79E-03 |
| Timm29 | 1875.151399 | -0.348004795 | 4.74E-02 |
| Ankmy2 | 1264.514491 | -0.34856778 | 4.32E-02 |
| AI597479 | 1129.332889 | -0.348634198 | 1.08E-05 |
| Slc26a2 | 908.0177184 | -0.348787086 | 1.65E-02 |
| L3hypdh | 996.178476 | -0.348814925 | 1.69E-02 |
| Mrps11 | 708.4432458 | -0.34911703 | 3.37E-02 |
| Dnase2a | 1680.910599 | -0.349363337 | 1.60E-02 |
| Eapp | 1456.644053 | -0.349830998 | 3.74E-02 |
| Ap3b1 | 4777.203822 | -0.349913749 | 2.90E-03 |
| Trim21 | 337.3701587 | -0.349933279 | 2.85E-02 |
| Anks3 | 1314.908854 | -0.350001782 | 2.87E-02 |
| Hmg20a | 1475.554886 | -0.350198167 | 1.98E-03 |
| Fam168b | 9749.745025 | -0.350572127 | 6.57E-03 |
| Npr2 | 2586.247944 | -0.351754694 | 4.04E-04 |
| Colec12 | 20930.91238 | -0.352017735 | 3.88E-02 |
| Ttc21b | 2440.384871 | -0.352129136 | 5.26E-04 |
| Zfp28 | 611.0405196 | -0.352627283 | 1.16E-02 |
| Commd10 | 1041.836611 | -0.353062198 | 8.06E-03 |
| Pitrm1 | 6900.706977 | -0.353312868 | 3.19E-03 |
| Slc36a4 | 1156.576941 | -0.353713358 | 2.17E-05 |
| Akap11 | 3406.170716 | -0.354186203 | 2.47E-02 |
| Dis3l2 | 992.1240902 | -0.354317976 | 1.76E-02 |
| Avl9 | 2443.33882 | -0.35439407 | 4.47E-02 |
| Zfp874b | 473.1071347 | -0.355015596 | 7.92E-03 |
| Inf2 | 6535.964029 | -0.355054887 | 4.65E-02 |
| Twsg1 | 5761.860578 | -0.355400023 | 9.55E-03 |
| Retreg3 | 2865.0767 | -0.355621759 | 1.56E-02 |
| Nit2 | 1118.356382 | -0.355805053 | 4.38E-02 |
| Aamp | 5473.874888 | -0.35616003 | 2.45E-03 |
| Zscan12 | 310.7903595 | -0.356813312 | 4.13E-02 |
| Nipsnap2 | 2457.606527 | -0.357131816 | 8.68E-04 |
| Mpc2 | 1599.611664 | -0.357785738 | 3.79E-02 |
| Cul7 | 4030.507774 | -0.357832742 | 3.28E-02 |
| Mrps26 | 1220.384571 | -0.358376709 | 9.62E-03 |
| Hdac3 | 3639.465084 | -0.358558571 | 1.04E-02 |
| Taz | 1239.733738 | -0.358573684 | 4.28E-02 |
| Uhrf2 | 4017.205027 | -0.358995816 | 2.51E-02 |
| Akna | 1024.927798 | -0.359059823 | 2.22E-02 |
| Rcan3 | 1082.07026 | -0.359756119 | 2.32E-02 |
| Prdx2 | 10872.40196 | -0.360085595 | 2.32E-02 |
| Dera | 752.132981 | -0.360156004 | 1.07E-03 |
| Stard9 | 2695.896783 | -0.36054885 | 2.67E-02 |
| Dhps | 1793.496769 | -0.36058279 | 4.32E-02 |
| Slc37a3 | 2707.708926 | -0.360857045 | 2.83E-04 |
| 2610001J05Rik | 2297.6783 | -0.360871258 | 7.66E-03 |

| | | | |
|---|---|---|---|
| Phkb | 2588.754316 | -0.360947024 | 4.25E-02 |
| Coq10a | 716.0566011 | -0.360973149 | 3.28E-02 |
| Ntan1 | 2090.352579 | -0.360977922 | 3.99E-02 |
| Ywhaq | 13241.07001 | -0.361845449 | 5.94E-04 |
| Tusc3 | 3182.406469 | -0.362058262 | 4.77E-04 |
| Rhbdd2 | 1090.819054 | -0.362076713 | 4.20E-02 |
| NA | 1773.329601 | -0.362601946 | 2.24E-03 |
| Fam160b2 | 868.648153 | -0.363262611 | 2.45E-03 |
| Tsc22d1 | 5481.602433 | -0.363387633 | 2.05E-02 |
| Garem1 | 770.4422413 | -0.364035089 | 2.00E-02 |
| Ankrd6 | 596.1378015 | -0.364242034 | 3.54E-02 |
| Rcbtb2 | 2432.790428 | -0.364434691 | 5.73E-03 |
| Atmin | 1212.579264 | -0.364456385 | 1.82E-04 |
| Pisd | 1832.974643 | -0.364462878 | 4.39E-02 |
| Mpnd | 2802.889083 | -0.365014018 | 2.91E-02 |
| Rnf135 | 468.7873319 | -0.365020281 | 1.36E-02 |
| Coq9 | 1669.857785 | -0.365036922 | 4.01E-02 |
| Zfp512 | 1349.954288 | -0.365264436 | 1.12E-03 |
| Rassf8 | 5424.313331 | -0.365938707 | 2.74E-02 |
| Nsmf | 2651.040508 | -0.36605437 | 1.63E-02 |
| Psme2 | 2209.647031 | -0.366735143 | 4.41E-03 |
| Ctsl | 40956.20104 | -0.366821322 | 4.45E-02 |
| Cuedc2 | 3606.221772 | -0.366946531 | 6.68E-03 |
| Tmx4 | 2058.445111 | -0.366952399 | 2.00E-03 |
| Phf23 | 3785.987225 | -0.367296247 | 3.41E-02 |
| Pi4ka | 5631.269742 | -0.367615878 | 8.26E-04 |
| Lonp2 | 4330.10061 | -0.367621044 | 1.11E-02 |
| Jmjd8 | 638.2911338 | -0.368317858 | 5.05E-04 |
| Rab43 | 2992.193124 | -0.368963951 | 1.58E-02 |
| Zkscan1 | 2415.13319 | -0.369332999 | 9.70E-04 |
| 4930402H24Rik | 5341.149355 | -0.370478513 | 2.32E-03 |
| Zmynd8 | 5278.19677 | -0.370583121 | 1.01E-05 |
| Rbfox2 | 9966.948387 | -0.370734181 | 5.22E-03 |
| Zfand5 | 4632.448716 | -0.371442802 | 2.82E-02 |
| Apex1 | 3288.753952 | -0.371607905 | 1.34E-02 |
| Foxp4 | 2395.208816 | -0.371918351 | 1.81E-02 |
| Rell1 | 1320.85393 | -0.371955261 | 4.01E-02 |
| Slc39a11 | 1072.037658 | -0.372133842 | 3.93E-02 |
| Ilf3 | 5444.124218 | -0.37310778 | 3.55E-03 |
| Rmdn1 | 288.7588206 | -0.373854969 | 1.61E-02 |
| Mccc1 | 1236.34865 | -0.374309686 | 3.07E-02 |
| Washc5 | 9453.018285 | -0.37434783 | 2.33E-04 |
| Capzb | 8468.560577 | -0.37457124 | 4.17E-02 |
| D10Jhu81e | 1329.248274 | -0.375313594 | 1.63E-02 |
| Spast | 2112.317484 | -0.375406923 | 2.20E-02 |
| Tbc1d9b | 5415.2979 | -0.376813131 | 5.35E-04 |
| Edc3 | 1285.870104 | -0.376904807 | 7.93E-03 |

| | | | |
|---|---|---|---|
| D10Wsu102e | 2431.206995 | -0.376994266 | 5.08E-03 |
| Josd2 | 899.9900018 | -0.377062191 | 1.37E-02 |
| Soga1 | 1586.138467 | -0.377458508 | 2.22E-02 |
| Tmed1 | 963.0524274 | -0.377497971 | 4.01E-02 |
| Kansl1 | 3445.684526 | -0.377551292 | 1.10E-03 |
| Pigp | 767.26785 | -0.377955597 | 4.89E-02 |
| Rbm43 | 682.9030732 | -0.379820095 | 4.17E-03 |
| Idua | 698.7732812 | -0.38122728 | 7.93E-03 |
| Acox1 | 3724.20556 | -0.381455104 | 2.59E-02 |
| Cox7c | 1341.314878 | -0.381495868 | 2.59E-02 |
| Acap2 | 6513.117941 | -0.381915805 | 7.36E-03 |
| Mindy1 | 1987.66538 | -0.38206413 | 4.37E-03 |
| Zfp799 | 721.4912572 | -0.382245505 | 2.01E-03 |
| Noct | 8107.901444 | -0.38265845 | 2.48E-03 |
| Ndst2 | 1120.700593 | -0.383021523 | 4.95E-05 |
| Armc9 | 957.2896761 | -0.383733315 | 7.07E-04 |
| Cfap20 | 1689.539204 | -0.38401435 | 5.08E-03 |
| Mtch1 | 11913.88654 | -0.384038237 | 2.80E-02 |
| Pskh1 | 1911.4363 | -0.384213297 | 8.29E-04 |
| Ogdh | 6423.124912 | -0.384330577 | 2.80E-02 |
| Gm9701 | 21363.22693 | -0.384498043 | 2.62E-02 |
| Eri3 | 5009.343592 | -0.384740172 | 1.55E-02 |
| Rab2b | 836.6849826 | -0.384800095 | 3.02E-02 |
| Acad11 | 584.7985704 | -0.384957729 | 1.63E-02 |
| Echs1 | 2288.2654 | -0.38624508 | 1.08E-02 |
| Fbxo46 | 415.7050116 | -0.386504402 | 4.81E-02 |
| Zmym3 | 1752.30601 | -0.386711922 | 1.20E-02 |
| Shkbp1 | 798.6678406 | -0.387038448 | 4.33E-02 |
| Pcgf1 | 643.0125964 | -0.387053895 | 5.96E-03 |
| Wdr83 | 632.260428 | -0.387089889 | 9.31E-03 |
| Psmd2 | 21874.64041 | -0.387222378 | 1.02E-02 |
| Fam234b | 435.9697922 | -0.387555762 | 2.84E-02 |
| Zfp316 | 1016.687164 | -0.387648895 | 3.15E-02 |
| Fbxl19 | 1033.975476 | -0.387708204 | 6.56E-03 |
| Gusb | 2912.311294 | -0.387739866 | 3.50E-02 |
| Gm16211 | 299.0104868 | -0.388012481 | 3.09E-02 |
| Apool | 575.8688515 | -0.388280841 | 2.10E-02 |
| Lclat1 | 2716.89391 | -0.388585898 | 1.39E-03 |
| Hk2 | 3039.306829 | -0.388825347 | 4.05E-02 |
| Slc41a1 | 1950.284416 | -0.389155806 | 2.85E-02 |
| Fam53a | 1630.046894 | -0.389159248 | 4.55E-03 |
| Ing4 | 1691.109084 | -0.390256452 | 6.99E-03 |
| Sdf2 | 2087.194487 | -0.390665591 | 4.75E-02 |
| S100pbp | 2801.716801 | -0.390999755 | 5.42E-03 |
| Tlk1 | 4354.547591 | -0.391352759 | 7.24E-03 |
| Slc25a11 | 3169.733048 | -0.391509256 | 2.38E-03 |
| Slc35b3 | 449.8835323 | -0.391573585 | 3.94E-02 |

| | | | |
|---|---|---|---|
| Ptpdc1 | 1192.393303 | -0.391678971 | 1.51E-03 |
| Lrsam1 | 1151.868033 | -0.39196837 | 2.01E-02 |
| Bloc1s5 | 627.5788997 | -0.39197416 | 2.72E-02 |
| Rita1 | 573.5582389 | -0.39206183 | 3.11E-02 |
| Zfyve26 | 2479.501653 | -0.392165555 | 2.31E-03 |
| Urgcp | 1854.416943 | -0.392544491 | 8.05E-03 |
| Wdr82 | 3294.636451 | -0.393044975 | 1.32E-03 |
| 4833420G17Rik | 1254.475335 | -0.393317008 | 1.31E-02 |
| Capn10 | 780.8179271 | -0.394160551 | 3.21E-02 |
| Scfd2 | 558.8131694 | -0.394375937 | 2.64E-02 |
| Pced1b | 224.7482532 | -0.394681053 | 3.36E-02 |
| Atp6v1h | 5874.496336 | -0.394772733 | 1.65E-02 |
| Zfand1 | 845.8720894 | -0.394893948 | 3.31E-02 |
| Spidr | 1784.161671 | -0.395417868 | 6.26E-03 |
| Eif1b | 955.7925393 | -0.395572852 | 3.64E-02 |
| Gm12481 | 10409.92692 | -0.39557635 | 2.46E-02 |
| 8-Mar | 6434.713929 | -0.395955639 | 5.25E-16 |
| Parp16 | 205.5236357 | -0.396225306 | 3.67E-02 |
| Ube2d-ps | 793.7075186 | -0.396509161 | 2.66E-02 |
| Ywhaz | 11408.90752 | -0.396535993 | 4.87E-04 |
| Zdhhc1 | 1130.427471 | -0.396605098 | 2.54E-02 |
| Hsdl1 | 1948.417286 | -0.396701009 | 1.45E-02 |
| Zfp503 | 2242.630648 | -0.396937501 | 2.84E-02 |
| Trappc2l | 1296.296787 | -0.39740024 | 3.54E-02 |
| Shq1 | 500.8612755 | -0.397464872 | 4.55E-02 |
| Tmem19 | 1982.052025 | -0.397885499 | 4.85E-04 |
| NA | 486.6545116 | -0.398581548 | 4.49E-02 |
| Dzip1 | 1896.61168 | -0.398787792 | 4.49E-04 |
| Smcr8 | 3771.341043 | -0.398841061 | 1.38E-03 |
| Arpc4 | 4891.018247 | -0.398958643 | 1.17E-02 |
| Rilpl2 | 842.7504969 | -0.399925545 | 1.99E-02 |
| Wdr7 | 1610.402645 | -0.400019854 | 3.40E-03 |
| Tulp3 | 4525.053787 | -0.400448003 | 1.20E-03 |
| Cog2 | 1721.709194 | -0.401172902 | 7.16E-03 |
| Rangrf | 427.5682787 | -0.401183982 | 2.96E-02 |
| Atp6v0a1 | 3975.629212 | -0.402121721 | 3.45E-03 |
| Tmem186 | 774.7522416 | -0.402425719 | 3.43E-03 |
| Wls | 19235.93451 | -0.40315339 | 1.49E-03 |
| Cox19 | 726.4729738 | -0.40334304 | 4.81E-02 |
| Cyba | 1385.240954 | -0.403423814 | 3.19E-02 |
| Kif7 | 398.6317459 | -0.403667366 | 2.85E-02 |
| Anapc7 | 2454.939837 | -0.403689082 | 2.09E-02 |
| 2610002M06Rik | 1470.102267 | -0.403712348 | 1.04E-04 |
| Hapln4 | 436.7943246 | -0.40431072 | 2.92E-02 |
| Evi5l | 1300.320633 | -0.404349783 | 7.23E-04 |
| Slc25a14 | 246.6469147 | -0.404437191 | 2.36E-02 |
| Hmbs | 1209.092363 | -0.40497055 | 2.96E-02 |

| | | | |
|---|---|---|---|
| Etfrf1 | 953.9923609 | -0.405052818 | 2.46E-02 |
| Abhd11 | 803.7195628 | -0.405518593 | 4.07E-02 |
| Pafah2 | 1268.397562 | -0.405693421 | 1.27E-02 |
| Rnd3 | 3804.865344 | -0.406166718 | 1.62E-02 |
| Igf2bp2 | 8644.418417 | -0.406229064 | 1.49E-04 |
| Cdk19 | 2233.948062 | -0.406258473 | 8.62E-05 |
| Pik3r2 | 5877.98043 | -0.406935553 | 1.06E-02 |
| Vps33a | 2892.012132 | -0.407081316 | 5.21E-05 |
| Ndufb11 | 3018.005221 | -0.407191708 | 4.01E-02 |
| Cep120 | 1497.538669 | -0.407329915 | 3.61E-04 |
| Tmem14c | 2037.871057 | -0.407558187 | 7.54E-03 |
| Papss2 | 2566.376957 | -0.40764532 | 3.28E-02 |
| Dbi | 3225.641979 | -0.407682667 | 2.35E-03 |
| Atp6v1d | 4059.992121 | -0.408141946 | 4.82E-02 |
| A130010J15Rik | 350.9587272 | -0.408321848 | 6.41E-03 |
| Atp6ap2 | 4358.08888 | -0.408369092 | 1.35E-02 |
| R3hdm4 | 3380.236701 | -0.408415807 | 2.74E-02 |
| Slc25a17 | 2861.439304 | -0.408580884 | 2.18E-03 |
| Csad | 1257.892946 | -0.408630038 | 2.85E-02 |
| Gna12 | 3909.739383 | -0.40872064 | 1.32E-03 |
| Ptdss2 | 1024.474847 | -0.40957507 | 2.81E-02 |
| Maged1 | 25928.54256 | -0.409603391 | 1.01E-03 |
| Wasf3 | 609.9759654 | -0.410465845 | 1.79E-02 |
| Trim32 | 3371.688236 | -0.41058599 | 2.74E-03 |
| Zfp983 | 210.3850671 | -0.410602465 | 1.91E-02 |
| Chmp1a | 5050.613376 | -0.410816022 | 3.04E-03 |
| Atp7a | 1656.515308 | -0.41111319 | 4.44E-03 |
| Cdk4 | 23550.22214 | -0.411349661 | 9.01E-03 |
| Csrp1 | 2952.741716 | -0.411738635 | 2.23E-03 |
| Shisa5 | 4347.519976 | -0.412139307 | 4.49E-02 |
| Zfp142 | 2034.722645 | -0.412851731 | 4.43E-03 |
| Camk2d | 6564.725921 | -0.412996755 | 3.79E-03 |
| Pou6f1 | 1919.566169 | -0.413509187 | 1.56E-03 |
| Tle1 | 1922.88961 | -0.413809707 | 1.01E-02 |
| Castor2 | 1817.867489 | -0.414231702 | 9.31E-03 |
| Rnf8 | 695.2111225 | -0.414362924 | 2.74E-04 |
| Fance | 1141.489143 | -0.414457463 | 2.77E-02 |
| H3f3a | 13435.63948 | -0.414841025 | 4.83E-03 |
| Foxc1 | 342.3062006 | -0.41503779 | 3.32E-02 |
| Rpl26 | 50023.25159 | -0.415624335 | 4.74E-02 |
| Harbi1 | 406.9577452 | -0.415683823 | 1.56E-02 |
| Tomm34 | 1451.330428 | -0.415701635 | 8.10E-04 |
| Acot13 | 2081.691453 | -0.415926137 | 3.16E-02 |
| Hdac10 | 484.6780465 | -0.417170097 | 2.05E-02 |
| Thtpa | 224.2179658 | -0.417284126 | 1.93E-02 |
| Dcaf5 | 2929.156897 | -0.417402646 | 2.38E-03 |
| Ankrd46 | 2144.475571 | -0.418070955 | 2.64E-02 |

| | | | |
|---|---|---|---|
| Grasp | 391.1139554 | -0.418081243 | 1.42E-02 |
| Commd7 | 1506.669976 | -0.418248915 | 2.20E-03 |
| Pik3c3 | 1269.108305 | -0.418374556 | 9.92E-03 |
| B3gnt9 | 1121.027082 | -0.418431318 | 3.23E-02 |
| Clpp | 2273.411807 | -0.419189478 | 2.69E-02 |
| Dusp22 | 270.0580327 | -0.41962733 | 1.79E-02 |
| Smim14 | 4036.84733 | -0.419629651 | 2.37E-02 |
| Bcl7a | 1302.545937 | -0.420244593 | 1.54E-03 |
| Tpcn1 | 2505.938347 | -0.4205196 | 5.95E-03 |
| Tbcel | 1611.699035 | -0.420857716 | 6.48E-06 |
| Stx8 | 1318.595004 | -0.421228931 | 1.97E-02 |
| Slc25a44 | 2191.360797 | -0.421400389 | 2.29E-02 |
| Bzw2 | 2826.086381 | -0.42152216 | 2.85E-02 |
| Epha4 | 1347.621821 | -0.42153506 | 1.23E-02 |
| Trak1 | 2002.750485 | -0.422177478 | 1.17E-03 |
| Lsm10 | 1095.464393 | -0.422234606 | 9.31E-04 |
| Ndufb2 | 1099.511065 | -0.422319285 | 1.06E-02 |
| Slc29a3 | 2607.580111 | -0.42232161 | 1.86E-02 |
| Crot | 1867.061663 | -0.42262229 | 5.23E-04 |
| Ifngr2 | 2171.576009 | -0.42263603 | 6.96E-03 |
| Trappc12 | 1816.216958 | -0.422829399 | 7.05E-03 |
| Plscr3 | 3839.393331 | -0.422936814 | 1.13E-02 |
| Tmem208 | 927.097134 | -0.423039861 | 3.40E-03 |
| Ppard | 1643.695207 | -0.423420437 | 5.54E-03 |
| Gmppb | 553.2926187 | -0.423736719 | 3.16E-02 |
| Dusp19 | 368.3047924 | -0.423929845 | 4.36E-03 |
| Pigx | 1227.555917 | -0.423981227 | 2.11E-03 |
| Pdzd4 | 1139.545824 | -0.424304468 | 3.25E-03 |
| Atraid | 2965.766926 | -0.425456203 | 1.58E-02 |
| Anapc11 | 1259.126626 | -0.425600822 | 3.59E-02 |
| Nipsnap3b | 762.8209259 | -0.425752943 | 1.01E-02 |
| Fundc1 | 1249.116808 | -0.425768532 | 2.80E-04 |
| Cacfd1 | 1602.512009 | -0.426122387 | 2.84E-03 |
| Atp5mpl | 1875.045829 | -0.426139239 | 6.56E-03 |
| Nudt9 | 3226.818754 | -0.426308293 | 3.12E-02 |
| Akr1a1 | 21668.11073 | -0.426718942 | 1.05E-02 |
| Tmem231 | 1590.140116 | -0.426981511 | 7.35E-03 |
| Trip12 | 11199.36888 | -0.427242439 | 1.84E-05 |
| Serhl | 1090.949015 | -0.427408499 | 2.19E-03 |
| Acaca | 6220.310423 | -0.427875089 | 1.58E-02 |
| Nenf | 2679.17814 | -0.428119593 | 1.25E-02 |
| 1810030O07Rik | 1755.512131 | -0.428203018 | 8.69E-07 |
| Lamp2 | 21188.1191 | -0.428405716 | 8.84E-03 |
| Adck1 | 756.8498528 | -0.428462393 | 6.34E-04 |
| Vps41 | 3917.059342 | -0.428985186 | 1.25E-03 |
| Slc35a2 | 1147.598411 | -0.428992863 | 9.63E-03 |
| Stx1a | 409.9590611 | -0.429696864 | 3.23E-02 |

| | | | |
|---|---|---|---|
| Ahnak2 | 23819.93924 | -0.429881239 | 4.32E-02 |
| Tcaf1 | 6725.43509 | -0.430132396 | 8.33E-05 |
| Ppa2 | 1042.259052 | -0.430612377 | 3.35E-04 |
| Prmt9 | 1287.77869 | -0.431085063 | 2.82E-06 |
| Ints4 | 1468.076545 | -0.431255473 | 3.09E-04 |
| Stk38 | 4121.885645 | -0.431274175 | 4.19E-04 |
| NA | 256.5573422 | -0.431412369 | 1.06E-02 |
| Sdhaf4 | 475.0879917 | -0.431695529 | 3.50E-03 |
| Tom1l2 | 3543.79026 | -0.431733586 | 1.23E-02 |
| Lpar2 | 1343.831789 | -0.431798171 | 2.56E-02 |
| Polb | 1158.380286 | -0.431809279 | 4.93E-03 |
| Nthl1 | 448.1070006 | -0.43252551 | 2.23E-02 |
| 1810022K09Rik | 286.9124674 | -0.432856691 | 2.60E-02 |
| Dnajc16 | 942.4498415 | -0.433246324 | 2.97E-03 |
| Tceal9 | 6722.958843 | -0.43325107 | 3.28E-02 |
| Plcg1 | 7566.406721 | -0.43327886 | 9.78E-03 |
| Ift27 | 800.9781334 | -0.433338215 | 7.25E-03 |
| Atp6v1c1 | 4552.101365 | -0.43343996 | 1.05E-02 |
| Taf6l | 798.7607961 | -0.433600466 | 2.94E-02 |
| Got2 | 9551.023887 | -0.433992799 | 3.08E-02 |
| Nudt19 | 340.0343303 | -0.434133861 | 4.95E-02 |
| Tmem17 | 204.5718151 | -0.434693523 | 1.81E-02 |
| C1rl | 598.4219494 | -0.434785149 | 4.88E-02 |
| Lpar1 | 5879.931993 | -0.434874937 | 1.24E-02 |
| Taf1c | 527.9761851 | -0.434888733 | 6.24E-03 |
| Pcnx | 4490.427285 | -0.434960973 | 3.85E-04 |
| Slc35f6 | 2258.273448 | -0.435920633 | 1.02E-02 |
| Mindy4 | 685.858107 | -0.435950737 | 1.62E-02 |
| Zxdc | 3218.603639 | -0.436840925 | 2.81E-02 |
| Slc44a1 | 6489.8044 | -0.437047713 | 4.18E-02 |
| Ndufc1 | 833.0367167 | -0.437375196 | 1.17E-02 |
| Guk1 | 1445.832017 | -0.437980515 | 7.68E-03 |
| Thg1l | 413.4328502 | -0.438615065 | 1.77E-02 |
| Vcpkmt | 777.1607539 | -0.438636326 | 4.11E-02 |
| Pafah1b3 | 1426.134526 | -0.438774657 | 7.16E-03 |
| Stxbp4 | 1774.063882 | -0.439297206 | 1.38E-03 |
| Slc35c2 | 1816.266464 | -0.439648123 | 4.96E-03 |
| Dpysl2 | 1095.968635 | -0.440195989 | 4.26E-02 |
| Kdm2b | 1932.525567 | -0.440253862 | 1.21E-02 |
| Gm10243 | 15969.90068 | -0.440277144 | 4.53E-02 |
| Inpp5e | 1283.030225 | -0.440414459 | 3.48E-03 |
| Faap20 | 598.4536698 | -0.440628071 | 2.13E-03 |
| Pex5 | 3893.377517 | -0.440685952 | 3.20E-05 |
| Pfkfb2 | 1070.695959 | -0.44087323 | 2.51E-02 |
| Zfp444 | 542.1527879 | -0.440986745 | 9.38E-03 |
| Rps17 | 34211.18673 | -0.44100698 | 4.54E-02 |
| Gskip | 1077.810338 | -0.441436895 | 3.90E-04 |

| | | | |
|---|---|---|---|
| Naip5 | 268.7274309 | -0.441757481 | 2.23E-02 |
| Vipas39 | 1504.93256 | -0.442166055 | 1.33E-04 |
| Pgap2 | 1147.432946 | -0.442604701 | 8.46E-03 |
| Smarcc1 | 5292.980505 | -0.443228512 | 5.63E-05 |
| Gcat | 1335.897836 | -0.443600865 | 3.08E-02 |
| Efhd2 | 1565.299968 | -0.443634633 | 1.22E-02 |
| Ophn1 | 1096.550011 | -0.443714362 | 6.28E-03 |
| Erp29 | 5236.891141 | -0.444153449 | 2.13E-03 |
| Cmc1 | 656.2945244 | -0.444679998 | 2.27E-02 |
| Dcun1d4 | 1164.457837 | -0.445038931 | 2.93E-02 |
| Phf14 | 2376.386027 | -0.445846779 | 4.75E-02 |
| Ccdc32 | 828.4896004 | -0.445898089 | 4.34E-03 |
| Atxn7l3 | 3554.494769 | -0.446115255 | 5.24E-06 |
| Tbccd1 | 1403.677915 | -0.44757692 | 3.42E-06 |
| Slc44a2 | 6308.547057 | -0.447777576 | 2.60E-03 |
| Ctso | 1002.433076 | -0.44814226 | 9.65E-03 |
| Tfcp2 | 1065.086979 | -0.448760572 | 1.85E-02 |
| Pogk | 1686.347408 | -0.449019992 | 3.00E-03 |
| 2010001A14Rik | 129.6180682 | -0.449312988 | 1.90E-02 |
| Wdr73 | 690.8516135 | -0.449842388 | 2.56E-02 |
| Cdyl2 | 218.9938247 | -0.449855023 | 1.92E-02 |
| Slc41a2 | 949.467359 | -0.449884034 | 2.34E-02 |
| Ntmt1 | 1200.274153 | -0.44989803 | 8.63E-03 |
| Pebp1 | 16359.7524 | -0.450407645 | 2.06E-02 |
| Ogfr | 3906.693324 | -0.451324782 | 5.19E-03 |
| Kdm5d | 903.6611732 | -0.451425912 | 1.02E-02 |
| Mcrip1 | 3277.7431 | -0.452155412 | 4.75E-02 |
| Uqcrh | 8383.033201 | -0.452261441 | 2.44E-02 |
| Borcs8 | 860.7623581 | -0.452932805 | 7.58E-03 |
| Tmem260 | 658.9304686 | -0.45294094 | 8.03E-03 |
| Ppdpf | 1355.859809 | -0.453328122 | 3.97E-05 |
| Ankrd34a | 289.6306119 | -0.454156213 | 4.04E-02 |
| Arl8b | 6917.709562 | -0.454416255 | 8.66E-03 |
| Hexb | 4185.185652 | -0.454658485 | 3.18E-02 |
| Zik1 | 289.836167 | -0.454830669 | 1.71E-02 |
| Wdpcp | 403.2797746 | -0.454869211 | 4.92E-02 |
| Cdk5rap1 | 416.9799843 | -0.455083585 | 1.17E-03 |
| Armcx1 | 1688.155142 | -0.455089885 | 1.56E-02 |
| Itga5 | 8198.609151 | -0.455406555 | 2.01E-02 |
| E330009J07Rik | 844.1774462 | -0.457404684 | 1.92E-03 |
| Psme4 | 6579.725646 | -0.457672346 | 1.12E-03 |
| Elf2 | 3398.181671 | -0.458181283 | 2.14E-05 |
| Gm3650 | 185.2176749 | -0.4584131 | 2.64E-02 |
| Pnpla7 | 844.5630092 | -0.458856839 | 3.76E-02 |
| Nlrc3 | 519.874105 | -0.459227807 | 3.59E-02 |
| Usf1 | 2269.965477 | -0.459277265 | 4.29E-03 |
| Kif9 | 130.1670536 | -0.459427876 | 4.95E-02 |

| | | | |
|---|---|---|---|
| Cuta | 2544.799639 | -0.460363332 | 1.93E-02 |
| Atg10 | 242.0275732 | -0.460381466 | 4.45E-03 |
| Tmub2 | 1734.236763 | -0.460465928 | 4.16E-03 |
| Cdc42ep4 | 1198.950253 | -0.46071321 | 1.07E-02 |
| Cd276 | 4044.747532 | -0.461440158 | 8.72E-04 |
| Myo5a | 10004.30674 | -0.462134524 | 1.77E-02 |
| Dock1 | 3924.573139 | -0.462573359 | 2.56E-04 |
| Stard5 | 614.9927071 | -0.46290637 | 7.91E-03 |
| Fam89b | 2256.63054 | -0.463010268 | 3.08E-02 |
| Cdk20 | 349.3497726 | -0.463116381 | 1.39E-02 |
| Efna5 | 479.7293286 | -0.463250521 | 4.87E-02 |
| Lrfn4 | 2023.205322 | -0.463896313 | 2.27E-03 |
| Tspyl2 | 2557.62995 | -0.463896924 | 3.61E-03 |
| Kansl1l | 717.0289781 | -0.464169599 | 3.05E-02 |
| Smox | 1088.138552 | -0.464173595 | 2.64E-02 |
| Kif3a | 3165.630292 | -0.464217298 | 1.39E-02 |
| Dtnbp1 | 2425.756691 | -0.465303066 | 1.37E-02 |
| Rps13-ps2 | 21055.30253 | -0.4657268 | 3.95E-02 |
| A830082K12Rik | 468.2476811 | -0.465779049 | 4.13E-02 |
| Ptov1 | 4258.284015 | -0.466127647 | 2.26E-02 |
| NA | 667.970585 | -0.466176856 | 1.09E-03 |
| Mical3 | 1070.981969 | -0.466268865 | 4.20E-02 |
| Rpl37 | 20885.2681 | -0.466614092 | 4.72E-02 |
| Eef1b2 | 46128.15298 | -0.466633039 | 3.29E-02 |
| Mcoln1 | 2770.651326 | -0.467367988 | 1.89E-02 |
| Btd | 882.1449736 | -0.467483719 | 6.64E-03 |
| Ap5s1 | 405.3152319 | -0.467976627 | 2.98E-02 |
| Bloc1s3 | 433.2225642 | -0.468320833 | 4.32E-02 |
| Zbtb1 | 1295.613615 | -0.468447101 | 4.36E-02 |
| Glrx | 1441.157528 | -0.468548361 | 3.04E-02 |
| Alms1 | 2337.478094 | -0.468749062 | 3.54E-02 |
| Tyk2 | 1273.344657 | -0.468772674 | 8.86E-04 |
| Pomk | 854.3945557 | -0.469571725 | 2.14E-04 |
| Ttc30b | 441.1625201 | -0.469802553 | 7.75E-03 |
| Hspa2 | 705.0792204 | -0.469901969 | 3.43E-02 |
| Grb10 | 24850.6108 | -0.470253606 | 1.00E-06 |
| Gm20482 | 178.0800543 | -0.47061274 | 2.81E-02 |
| Cutc | 134.6270003 | -0.470802939 | 3.81E-02 |
| Slc36a1 | 1573.335444 | -0.471214856 | 3.60E-03 |
| Rsad1 | 456.0324608 | -0.472251885 | 3.20E-04 |
| Mgat5 | 813.1866966 | -0.472532456 | 2.16E-03 |
| Zmat5 | 369.1131801 | -0.473309037 | 3.47E-03 |
| Gm14322 | 342.7270789 | -0.473699804 | 4.01E-02 |
| Snx21 | 1540.371223 | -0.473873535 | 3.21E-03 |
| Cdk5 | 1892.485916 | -0.474120177 | 6.95E-06 |
| Atg7 | 2072.141009 | -0.474372111 | 6.61E-03 |
| Tldc1 | 738.7605049 | -0.474376868 | 2.83E-02 |

| | | | |
|---|---|---|---|
| Coq8b | 489.0724424 | -0.474387821 | 3.18E-02 |
| Pard6g | 691.6014837 | -0.47542049 | 2.20E-03 |
| Twist2 | 1503.498649 | -0.475436374 | 2.14E-02 |
| Rad9b | 181.9457967 | -0.47577135 | 1.79E-02 |
| Kif16b | 1057.721431 | -0.4759408 | 5.56E-04 |
| Zswim4 | 2006.776445 | -0.475981452 | 6.41E-03 |
| Tbl1xr1 | 2044.586522 | -0.476597825 | 2.93E-04 |
| Pepd | 1464.655481 | -0.477374958 | 2.85E-05 |
| Nme4 | 944.0496733 | -0.477586628 | 3.52E-02 |
| Tle3 | 2473.32434 | -0.47763821 | 7.64E-03 |
| Tk2 | 1286.754723 | -0.477921873 | 3.94E-04 |
| Borcs5 | 355.7722684 | -0.478903124 | 3.41E-02 |
| Nbr1 | 8450.78952 | -0.479244752 | 3.69E-03 |
| Ino80b | 616.8921082 | -0.4793315 | 1.63E-02 |
| Arhgap32 | 2413.87787 | -0.47935065 | 6.62E-04 |
| Rps9 | 54843.11362 | -0.47999919 | 2.15E-02 |
| Kdm1a | 3091.893595 | -0.480359463 | 8.01E-11 |
| Rpl34-ps1 | 26642.99647 | -0.481446333 | 3.39E-02 |
| Farp1 | 7954.256896 | -0.482445248 | 8.34E-05 |
| Dph1 | 914.7849843 | -0.482762548 | 6.63E-04 |
| Foxk1 | 3191.645725 | -0.483111882 | 1.97E-03 |
| Vash2 | 2507.446433 | -0.48361106 | 1.30E-02 |
| Wdsub1 | 1434.266269 | -0.483626681 | 2.68E-03 |
| Arap1 | 2783.487022 | -0.484092649 | 4.00E-02 |
| Nfya | 2749.990945 | -0.48446301 | 1.02E-03 |
| Fnip1 | 4227.643221 | -0.484766733 | 4.90E-03 |
| Dynlt1b | 1798.335892 | -0.484896168 | 1.77E-04 |
| Pheta1 | 721.9805754 | -0.484920285 | 4.80E-02 |
| Zfp362 | 2229.096229 | -0.485995352 | 2.17E-04 |
| Agpat4 | 1161.530843 | -0.486221854 | 1.46E-02 |
| Alg1 | 1213.748362 | -0.486231996 | 1.82E-04 |
| Tax1bp3 | 4764.282672 | -0.486299453 | 5.61E-03 |
| Abr | 2974.771084 | -0.486325065 | 3.58E-04 |
| S1pr1 | 1398.259521 | -0.486873879 | 2.94E-03 |
| D030028A08Rik | 174.8608595 | -0.486940323 | 4.18E-02 |
| Abcd1 | 1111.639964 | -0.487121412 | 1.51E-02 |
| Agtrap | 1417.40242 | -0.487517151 | 2.66E-04 |
| Eml1 | 3052.707289 | -0.487904811 | 4.08E-03 |
| Cog3 | 2017.359627 | -0.487912068 | 2.46E-04 |
| Dnm1 | 9239.958421 | -0.488289664 | 4.57E-02 |
| Rps21 | 17061.14801 | -0.488380428 | 4.44E-02 |
| Zfp68 | 2035.29604 | -0.490165713 | 2.00E-02 |
| Nhlrc3 | 2088.125293 | -0.490367213 | 2.24E-05 |
| Stat3 | 5566.252116 | -0.490531116 | 1.96E-02 |
| Gng10 | 2800.643553 | -0.490561232 | 4.25E-07 |
| Rps3a1 | 109609.6564 | -0.490658386 | 7.32E-03 |
| Rragc | 10259.96991 | -0.490699017 | 3.55E-03 |

| | | | |
|---|---|---|---|
| Atp6v0b | 9934.293287 | -0.491430504 | 1.78E-02 |
| Rab34 | 4682.192332 | -0.49205944 | 4.68E-05 |
| Tmem251 | 678.3297086 | -0.492383206 | 2.45E-03 |
| Arf5 | 4503.439777 | -0.492874005 | 5.61E-04 |
| Igf2bp3 | 4305.408781 | -0.492887383 | 7.70E-04 |
| Vps26c | 2458.394981 | -0.492932668 | 1.31E-03 |
| Stk11ip | 1416.486596 | -0.492961143 | 1.41E-03 |
| Pnkd | 893.3433196 | -0.493143373 | 5.50E-05 |
| Acat1 | 4212.253723 | -0.493154771 | 3.86E-04 |
| Vmac | 1013.466863 | -0.49326197 | 8.63E-03 |
| Mdp1 | 559.3208089 | -0.493410312 | 1.25E-02 |
| Usp47 | 10646.51981 | -0.49355571 | 2.99E-05 |
| Acd | 2131.77091 | -0.493557912 | 1.72E-02 |
| Atp6v0c-ps2 | 12789.63813 | -0.493693877 | 1.46E-02 |
| Zfp719 | 839.4258072 | -0.493752207 | 1.32E-02 |
| Sgsh | 942.5070194 | -0.493846581 | 3.53E-02 |
| Rpl23a | 76915.38591 | -0.49565807 | 4.52E-02 |
| Pkig | 750.3973578 | -0.495744192 | 1.66E-02 |
| Romo1 | 1261.431817 | -0.496024309 | 5.96E-03 |
| Fes | 449.9226603 | -0.496297679 | 3.60E-03 |
| Ift122 | 1556.937666 | -0.496792327 | 4.24E-04 |
| Spin1 | 3118.156723 | -0.496993024 | 2.68E-05 |
| Adck5 | 731.9848639 | -0.498246001 | 1.55E-02 |
| Abhd12 | 3305.451109 | -0.498507164 | 1.72E-03 |
| Rdh5 | 275.4133273 | -0.498559444 | 1.08E-02 |
| Commd4 | 942.1458007 | -0.498822865 | 3.74E-04 |
| Cacnb1 | 471.9894397 | -0.499567752 | 2.26E-02 |
| 2510009E07Rik | 671.3141264 | -0.500889549 | 4.18E-02 |
| Ndufv3 | 2611.286604 | -0.50104379 | 2.01E-02 |
| Casp6 | 849.5980722 | -0.501524682 | 4.40E-03 |
| Nynrin | 2362.227013 | -0.50191509 | 8.33E-04 |
| Ulk1 | 2950.077175 | -0.502013901 | 2.30E-02 |
| Pgrmc2 | 4295.144483 | -0.502057107 | 2.70E-04 |
| Kalrn | 1301.719211 | -0.502679191 | 4.89E-03 |
| Fam76a | 1691.346603 | -0.502805063 | 1.57E-03 |
| Prkra | 1707.47307 | -0.50282436 | 1.07E-02 |
| Pfdn5 | 3955.074207 | -0.503132611 | 3.59E-02 |
| Galnt18 | 1135.099224 | -0.503916938 | 4.09E-02 |
| Akt3 | 3456.107411 | -0.504285631 | 5.41E-03 |
| Map2k6 | 397.271134 | -0.504456509 | 3.29E-02 |
| Lrrc8d | 953.5605398 | -0.505342252 | 2.52E-02 |
| Noc4l | 1562.446352 | -0.505702056 | 4.33E-03 |
| Tmem119 | 3163.057139 | -0.505808745 | 3.51E-02 |
| Pyroxd1 | 888.8072622 | -0.5062421 | 4.38E-02 |
| Eif2ak4 | 3546.105363 | -0.50628594 | 2.84E-06 |
| Rfxank | 604.2452941 | -0.506338047 | 2.46E-03 |
| Selenok | 2017.394881 | -0.50653674 | 3.41E-03 |

| | | | |
|---|---|---|---|
| Dhrs4 | 481.3038845 | -0.50691805 | 4.75E-02 |
| Zfp266 | 1891.611082 | -0.508253331 | 2.04E-02 |
| Eef2k | 2828.119429 | -0.508464423 | 7.40E-04 |
| Tbcb | 1604.288802 | -0.509134749 | 1.99E-02 |
| Leng9 | 144.2703895 | -0.509245387 | 4.87E-02 |
| Laptm4a | 29285.87798 | -0.509340892 | 5.99E-03 |
| Pfkl | 3875.900593 | -0.509374333 | 2.02E-02 |
| Arpc1a | 4104.665188 | -0.509628856 | 2.45E-04 |
| Nprl3 | 1164.930093 | -0.509636425 | 1.97E-04 |
| Gm14139 | 207.1634126 | -0.510285664 | 2.59E-02 |
| Gata6 | 365.1840322 | -0.510652938 | 8.76E-03 |
| Bnc2 | 1922.400801 | -0.510751437 | 3.05E-03 |
| Nr2f1 | 1515.258968 | -0.512033825 | 7.19E-03 |
| Usp49 | 749.9339687 | -0.512064919 | 2.82E-02 |
| Gpatch2l | 1358.907756 | -0.512089952 | 1.36E-03 |
| Rida | 433.8767028 | -0.512252093 | 4.83E-02 |
| Psme1 | 1774.506782 | -0.512255319 | 7.46E-04 |
| Psap | 78612.42307 | -0.512595603 | 3.43E-02 |
| H1f0 | 12531.5451 | -0.513022817 | 4.25E-05 |
| Gmcl1 | 1500.19973 | -0.513039181 | 1.27E-10 |
| Evc | 3246.728565 | -0.514010797 | 2.87E-04 |
| Ntpcr | 1088.202802 | -0.51505634 | 7.98E-04 |
| Etv6 | 1583.328039 | -0.515961272 | 2.12E-06 |
| Rnf34 | 1154.614628 | -0.516447003 | 8.09E-06 |
| Cep83os | 409.9112091 | -0.516551844 | 3.51E-02 |
| Ctdspl | 1253.236411 | -0.516794047 | 3.66E-02 |
| Gaa | 16101.78382 | -0.517335274 | 4.08E-03 |
| Rps20 | 53567.39921 | -0.517882757 | 4.60E-02 |
| Acads | 1920.723929 | -0.517961399 | 1.17E-03 |
| Naglu | 4106.854751 | -0.5181337 | 4.20E-02 |
| Stat1 | 1423.127776 | -0.518151191 | 3.09E-03 |
| Inpp1 | 473.1785364 | -0.518168275 | 4.38E-03 |
| Tspan14 | 1344.749055 | -0.518176462 | 2.03E-05 |
| Tmem192 | 1385.041556 | -0.518251345 | 1.15E-02 |
| Parvb | 2285.010317 | -0.519268145 | 2.46E-03 |
| Scd2 | 61452.75163 | -0.519540229 | 4.48E-05 |
| Itgb5 | 5005.644612 | -0.51977078 | 2.38E-02 |
| Ifi207 | 781.2702642 | -0.5204786 | 1.88E-02 |
| Catsper2 | 170.3526591 | -0.521274607 | 5.88E-04 |
| Traf5 | 535.9410057 | -0.521406786 | 5.15E-03 |
| Tsc1 | 3398.460772 | -0.521443258 | 2.05E-04 |
| Cand2 | 2044.67946 | -0.521525989 | 6.63E-04 |
| Bphl | 452.6391813 | -0.521869582 | 9.87E-05 |
| Pdxk | 5185.761374 | -0.522430018 | 1.28E-02 |
| Zfp827 | 2374.032775 | -0.522728491 | 1.05E-03 |
| Ap4m1 | 644.4259784 | -0.522970552 | 5.15E-03 |
| Bckdha | 1514.42681 | -0.523349721 | 1.99E-02 |

| | | | |
|---|---|---|---|
| Ttc5 | 737.7181432 | -0.523670882 | 6.37E-06 |
| Flad1 | 1014.220375 | -0.524248729 | 8.23E-03 |
| Naga | 3447.187179 | -0.525117592 | 1.16E-02 |
| Mfsd1 | 8319.866312 | -0.526053747 | 2.14E-04 |
| Chrnb2 | 134.4276621 | -0.52620847 | 2.57E-02 |
| Anxa5 | 17795.31783 | -0.526397183 | 2.19E-04 |
| Churc1 | 720.4306281 | -0.526482171 | 6.63E-03 |
| Rnd2 | 944.1092971 | -0.526914785 | 2.72E-02 |
| Pianp | 1185.174268 | -0.527086398 | 2.42E-02 |
| Frg2f1 | 213.8372542 | -0.52719456 | 2.40E-02 |
| Wbp1 | 2160.848682 | -0.527628347 | 3.21E-02 |
| Enc1 | 4373.996949 | -0.528674024 | 4.17E-02 |
| Cryl1 | 233.4306095 | -0.528676898 | 1.55E-02 |
| Tubb2a | 6574.060184 | -0.528769731 | 5.41E-03 |
| Intu | 504.6476843 | -0.529089011 | 4.93E-03 |
| Dennd4a | 6083.669471 | -0.529345423 | 2.61E-02 |
| Zadh2 | 2436.775149 | -0.529499721 | 4.24E-03 |
| Hilpda | 740.6665374 | -0.52956196 | 1.35E-04 |
| Zfyve1 | 3094.323627 | -0.529728011 | 1.64E-02 |
| Aprt | 6193.680378 | -0.530071443 | 4.24E-03 |
| Fstl1 | 39494.87916 | -0.530323145 | 9.62E-03 |
| Cic | 8307.861902 | -0.530438889 | 2.66E-04 |
| Trpm4 | 642.7733671 | -0.530852012 | 4.40E-04 |
| Tmem8 | 134.9929926 | -0.53131519 | 9.15E-03 |
| NA | 574.234844 | -0.53163598 | 5.49E-03 |
| Kdm7a | 2156.934782 | -0.532615375 | 4.69E-02 |
| Atxn7l1 | 1301.845341 | -0.53473888 | 8.56E-04 |
| Mtmr11 | 1713.709962 | -0.535781422 | 4.21E-03 |
| Slc25a1 | 5391.834169 | -0.536137759 | 4.42E-03 |
| 2310015A10Rik | 393.9563693 | -0.536161697 | 3.60E-02 |
| Spata24 | 121.247919 | -0.536905415 | 9.48E-03 |
| 2610301B20Rik | 1235.416847 | -0.537240114 | 5.29E-07 |
| 1110008P14Rik | 539.8675936 | -0.537749546 | 8.12E-03 |
| Nr2c2ap | 954.217837 | -0.538306953 | 1.05E-02 |
| Slc2a8 | 590.8319803 | -0.53847356 | 3.93E-02 |
| Bcl9 | 1951.774164 | -0.539348348 | 3.69E-04 |
| Ten1 | 499.6524531 | -0.540392316 | 4.36E-02 |
| Abcd2 | 730.8520276 | -0.540421391 | 2.76E-02 |
| Timm17b | 948.1288908 | -0.540504285 | 1.35E-02 |
| Ak5 | 1153.641956 | -0.541644365 | 2.00E-03 |
| Slc12a7 | 1768.921695 | -0.541864303 | 1.83E-04 |
| Plin3 | 5523.193935 | -0.542290719 | 7.12E-03 |
| Tmem267 | 228.135723 | -0.542561379 | 2.73E-02 |
| Sspn | 1024.357293 | -0.543138101 | 5.01E-03 |
| Zfp319 | 991.9383572 | -0.543337331 | 3.29E-05 |
| Gdpd1 | 1125.495183 | -0.543480673 | 4.43E-02 |
| Tmem60 | 583.2390495 | -0.545032578 | 1.48E-04 |

| | | | |
|---|---|---|---|
| Ccn3 | 191.5912267 | -0.545458829 | 4.96E-02 |
| Zcchc2 | 3565.226416 | -0.545533559 | 1.47E-04 |
| H60b | 795.9867706 | -0.545573779 | 2.53E-02 |
| Nlgn2 | 3845.598607 | -0.54722522 | 4.02E-04 |
| Cyp2j6 | 1735.618558 | -0.547287241 | 1.30E-04 |
| Krba1 | 685.1161659 | -0.547312977 | 2.09E-02 |
| Pip4p2 | 1878.030152 | -0.547746654 | 9.51E-05 |
| Mrps33 | 1660.844546 | -0.548282712 | 1.23E-03 |
| Arsg | 389.4454428 | -0.548559382 | 1.90E-03 |
| Tmem203 | 416.9914192 | -0.548812202 | 7.05E-03 |
| Hes6 | 612.1037007 | -0.549176305 | 5.48E-03 |
| Mllt3 | 4797.047418 | -0.549210785 | 4.59E-03 |
| Capg | 4579.700126 | -0.549232706 | 1.25E-02 |
| Ndufa6 | 2802.477104 | -0.549288907 | 3.93E-02 |
| Cnpy2 | 2958.714011 | -0.549421296 | 4.06E-05 |
| Amacr | 720.3402139 | -0.549728554 | 5.66E-03 |
| Rpl36a | 23382.0782 | -0.549803131 | 2.97E-02 |
| Car13 | 2496.327558 | -0.549983624 | 2.88E-05 |
| Fer | 2906.447521 | -0.550264718 | 7.27E-06 |
| Sgip1 | 339.6091924 | -0.550594523 | 1.94E-03 |
| Il6ra | 1020.18581 | -0.551515822 | 1.81E-04 |
| Rpl7a-ps3 | 1301.1174 | -0.551692173 | 3.37E-02 |
| Spg21 | 3580.650246 | -0.551994789 | 2.72E-03 |
| Mrpl36 | 758.7165102 | -0.552356512 | 9.83E-06 |
| Vcan | 10736.2105 | -0.552432294 | 3.98E-05 |
| Cask | 3506.668855 | -0.552470885 | 2.85E-04 |
| Ctsb | 36468.22462 | -0.552585026 | 2.25E-02 |
| Rbm4b | 986.1917831 | -0.55263109 | 1.76E-03 |
| Mecom | 1419.704379 | -0.553923414 | 3.19E-03 |
| Mlycd | 405.9267261 | -0.554641492 | 1.14E-03 |
| Ift140 | 1141.357097 | -0.555478958 | 1.52E-02 |
| Rhebl1 | 244.5837871 | -0.556140471 | 3.92E-02 |
| Ddx51 | 1561.419541 | -0.556193968 | 1.21E-04 |
| Irx3 | 178.4497727 | -0.556487834 | 2.46E-02 |
| Commd6 | 577.2059038 | -0.556845544 | 1.36E-04 |
| Atp6v1a | 10459.35955 | -0.557949016 | 3.16E-03 |
| Gm17586 | 152.4386316 | -0.5581287 | 6.49E-03 |
| Vdac2 | 9351.945713 | -0.558274721 | 5.63E-06 |
| Gtf2i | 8843.674879 | -0.558373329 | 4.95E-05 |
| Bola1 | 299.4104525 | -0.558819266 | 2.74E-03 |
| Wdr81 | 3346.664813 | -0.559304275 | 2.71E-03 |
| Lyplal1 | 1068.156704 | -0.559605308 | 4.89E-04 |
| Pakap | 1146.85027 | -0.559732769 | 2.85E-02 |
| Rpl10a-ps1 | 49044.15001 | -0.559930088 | 2.79E-02 |
| Rtl8b | 1525.92564 | -0.560193219 | 1.91E-03 |
| Impact | 6753.387895 | -0.561751426 | 2.21E-03 |
| Vps8 | 2880.10801 | -0.561958305 | 1.13E-03 |

| | | | |
|---|---|---|---|
| Nab2 | 4607.437328 | -0.562553298 | 1.09E-04 |
| Slc4a11 | 115.9732355 | -0.562562271 | 2.21E-02 |
| Ctps2 | 3181.451717 | -0.562599613 | 2.57E-05 |
| Ap5z1 | 1165.206489 | -0.562800902 | 1.37E-05 |
| Armcx3 | 3815.423845 | -0.564373864 | 3.41E-04 |
| Pbx1 | 3313.477385 | -0.564456521 | 3.67E-02 |
| Gtf2ird1 | 1929.055866 | -0.565305539 | 1.84E-06 |
| Adgra2 | 9959.089649 | -0.565324991 | 5.96E-05 |
| Trim45 | 432.7742059 | -0.565358159 | 8.60E-04 |
| Sema4c | 1046.850603 | -0.565715601 | 4.67E-04 |
| Clybl | 692.1619379 | -0.565778994 | 5.43E-03 |
| Fam78b | 275.8695293 | -0.566051913 | 4.22E-02 |
| Rgmb | 2135.748027 | -0.56837932 | 4.69E-02 |
| Pip4p1 | 2929.907263 | -0.568431472 | 1.01E-02 |
| Nemp2 | 323.1133591 | -0.568478577 | 1.34E-03 |
| St3gal5 | 2651.830812 | -0.569319344 | 1.21E-04 |
| Il1rl2 | 175.7426724 | -0.56966054 | 4.96E-02 |
| Klhl36 | 240.6418848 | -0.569702744 | 8.83E-03 |
| Mn1 | 1568.859744 | -0.569903671 | 1.86E-02 |
| Stk36 | 385.152876 | -0.570372574 | 9.34E-03 |
| Lima1 | 14608.60545 | -0.571101901 | 8.70E-04 |
| Rps4x | 93555.43077 | -0.571114572 | 1.02E-02 |
| Triqk | 268.8786932 | -0.571144008 | 1.07E-03 |
| Pbdc1 | 1501.992805 | -0.571332016 | 2.50E-04 |
| Atat1 | 753.3053063 | -0.571545676 | 2.42E-02 |
| Ralb | 3073.453009 | -0.572352316 | 2.34E-04 |
| Inpp4a | 1118.819903 | -0.573305881 | 4.05E-04 |
| Ilvbl | 2448.329512 | -0.573631739 | 7.88E-04 |
| Smarcd1 | 1668.953612 | -0.573925831 | 5.58E-05 |
| Hlcs | 514.8003438 | -0.573945635 | 3.34E-04 |
| Zfp775 | 415.5759076 | -0.57395659 | 1.03E-02 |
| Mgst1 | 1067.813549 | -0.574047232 | 2.21E-02 |
| Atp6v1e1 | 7121.436934 | -0.574127421 | 4.36E-03 |
| Fgfr1 | 26119.72218 | -0.575699031 | 2.39E-03 |
| Tbcc | 642.9037322 | -0.575813868 | 1.49E-03 |
| Med22 | 2582.347251 | -0.576362958 | 2.38E-04 |
| Sdccag8 | 700.0101104 | -0.577700899 | 1.91E-03 |
| Pdlim5 | 5330.342559 | -0.578139911 | 3.03E-05 |
| Vegfb | 3825.597118 | -0.578791733 | 5.55E-03 |
| Oxct1 | 16865.66235 | -0.57917728 | 1.48E-03 |
| Washc2 | 10254.84457 | -0.579608812 | 2.77E-04 |
| Clip2 | 3685.485315 | -0.580181755 | 2.17E-04 |
| Chst11 | 5163.815561 | -0.580820145 | 2.71E-03 |
| Lamtor1 | 3855.900239 | -0.580977498 | 8.33E-04 |
| Arhgap39 | 893.335859 | -0.581107535 | 2.22E-03 |
| Jade1 | 3558.396045 | -0.581324071 | 6.26E-04 |
| Vps13c | 3424.132169 | -0.581348195 | 8.79E-04 |

| | | | |
|---|---|---|---|
| Acvr1 | 1714.787564 | -0.581504622 | 1.13E-04 |
| Rtl8c | 186.4143832 | -0.581673772 | 3.14E-02 |
| Rcbtb1 | 2189.384948 | -0.581913373 | 6.06E-06 |
| Tmem150a | 1636.619374 | -0.581938107 | 1.03E-03 |
| Ip6k2 | 1213.466928 | -0.583275901 | 1.32E-03 |
| Tmem175 | 1175.19773 | -0.583637677 | 1.59E-07 |
| Rpl37a | 29354.97811 | -0.583979165 | 3.71E-02 |
| Tgfb2 | 3781.852125 | -0.584147665 | 4.38E-02 |
| Ttc39c | 557.9704609 | -0.584196559 | 2.77E-03 |
| Meis2 | 2452.885529 | -0.585844247 | 4.96E-02 |
| Rap2a | 3349.270391 | -0.586132611 | 4.74E-03 |
| Bach1 | 2273.465941 | -0.586342511 | 3.76E-10 |
| Cped1 | 8743.703581 | -0.587796112 | 1.58E-02 |
| Cd1d1 | 192.488829 | -0.588021932 | 4.14E-02 |
| 1190002N15Rik | 866.5027261 | -0.588086504 | 2.02E-02 |
| Slc4a8 | 427.6101811 | -0.588954939 | 4.75E-02 |
| Gm17435 | 160.567384 | -0.589242186 | 2.39E-03 |
| Gstm5 | 2305.479204 | -0.589370533 | 2.13E-04 |
| Lppos | 143.0048581 | -0.589860298 | 3.26E-02 |
| Pi4kb | 3352.487585 | -0.590086936 | 7.69E-05 |
| Zfp821 | 579.2143519 | -0.590337746 | 3.03E-04 |
| Galk2 | 2673.268011 | -0.590376217 | 8.14E-06 |
| Pdgfrb | 13123.50877 | -0.590916241 | 3.17E-05 |
| Coprs | 814.1861832 | -0.591406744 | 1.97E-06 |
| Slc25a30 | 2177.751789 | -0.592181961 | 1.02E-02 |
| Usp11 | 2202.578828 | -0.592744528 | 4.88E-03 |
| Nln | 2979.601328 | -0.592800979 | 2.09E-07 |
| Sall2 | 1033.6407 | -0.593108384 | 7.97E-06 |
| Lamtor4 | 1098.802993 | -0.593390843 | 2.13E-03 |
| Pon3 | 413.776097 | -0.594810225 | 7.67E-04 |
| Zfp647 | 141.5050878 | -0.596014798 | 7.25E-03 |
| Hook2 | 814.6719502 | -0.596315456 | 7.84E-04 |
| Nphp3 | 459.6841505 | -0.596840485 | 1.41E-03 |
| Aldh6a1 | 1590.691443 | -0.597193899 | 9.58E-03 |
| Ksr1 | 974.9766751 | -0.597376944 | 3.62E-03 |
| Abhd18 | 258.364168 | -0.598907956 | 1.71E-02 |
| Rufy1 | 2070.818133 | -0.599023371 | 3.23E-03 |
| Gm10443 | 18934.25035 | -0.599063769 | 2.92E-02 |
| Eef1akmt1 | 477.1061299 | -0.599469865 | 1.13E-02 |
| Snrnp25 | 373.4357615 | -0.600959468 | 3.13E-02 |
| Adi1 | 3350.226368 | -0.601015991 | 7.53E-04 |
| Cacng7 | 2011.225826 | -0.601324453 | 1.68E-02 |
| Gpr176 | 2045.393277 | -0.60135549 | 4.00E-04 |
| Slc38a7 | 2944.818323 | -0.601366708 | 9.75E-05 |
| Aldh5a1 | 498.0285701 | -0.601561702 | 1.82E-03 |
| Snx10 | 1086.178709 | -0.601837783 | 1.13E-04 |
| Sarm1 | 300.3287627 | -0.602203077 | 8.65E-03 |

| | | | |
|---|---|---|---|
| 9430091E24Rik | 71.37412386 | -0.602814234 | 3.20E-02 |
| Sdcbp | 7570.072267 | -0.603452417 | 4.16E-06 |
| Lyst | 3273.741853 | -0.603595197 | 3.37E-05 |
| Rps28 | 15164.62267 | -0.603756306 | 2.61E-02 |
| NA | 146.9254783 | -0.603997424 | 5.74E-03 |
| 3110053B16Rik | 271.5926797 | -0.60526273 | 2.18E-03 |
| Mien1 | 1410.361947 | -0.605351272 | 8.29E-04 |
| Fam50a | 3368.128408 | -0.606110682 | 3.48E-05 |
| Traf4 | 3041.560397 | -0.606381365 | 4.47E-03 |
| Hacl1 | 528.7901018 | -0.606476092 | 1.19E-02 |
| Mpg | 550.6470613 | -0.607055921 | 2.22E-04 |
| Cog4 | 4133.200435 | -0.60761601 | 6.72E-04 |
| Mmp14 | 40429.12938 | -0.607824477 | 1.77E-03 |
| Cbfb | 3583.177904 | -0.60809181 | 9.67E-09 |
| Tpt1 | 131718.7681 | -0.609299251 | 6.03E-03 |
| Dhrs11 | 512.9711271 | -0.609493122 | 1.63E-05 |
| Kdm6b | 4030.264971 | -0.609938616 | 1.47E-02 |
| Zfp58 | 250.0516868 | -0.610264757 | 4.16E-03 |
| Atp6v1f | 4383.57506 | -0.610301785 | 3.37E-03 |
| Asap2 | 581.2720119 | -0.610821278 | 4.87E-03 |
| Gm5900 | 339.4007408 | -0.610885589 | 8.49E-03 |
| Chchd6 | 733.0271583 | -0.610995377 | 2.22E-03 |
| Mex3d | 2980.139648 | -0.611785503 | 7.35E-03 |
| Hsd17b11 | 1801.223666 | -0.612489771 | 3.93E-05 |
| Wdr6 | 3416.318234 | -0.612546439 | 1.83E-04 |
| Chsy1 | 1508.249314 | -0.613444094 | 5.73E-06 |
| Wdyhv1 | 343.7885209 | -0.613936121 | 1.28E-04 |
| Fam229b | 171.6780668 | -0.614459191 | 1.35E-02 |
| Gulp1 | 1061.560392 | -0.616228151 | 6.86E-03 |
| Gm26619 | 12811.53859 | -0.616322846 | 4.43E-02 |
| Timm8b | 906.7143595 | -0.616629264 | 2.25E-09 |
| Atpif1 | 2983.892833 | -0.616664478 | 1.66E-06 |
| Adam19 | 3337.629642 | -0.618315184 | 5.06E-05 |
| 9430038I01Rik | 156.3527109 | -0.61834204 | 5.82E-04 |
| Gamt | 997.1896559 | -0.618344025 | 2.39E-03 |
| Klc4 | 995.6251541 | -0.620251051 | 2.98E-02 |
| Zeb2os | 155.3605102 | -0.620688718 | 2.83E-02 |
| Flcn | 3271.906196 | -0.620839355 | 3.65E-03 |
| Tes | 1801.715462 | -0.620913348 | 1.71E-02 |
| Gm4262 | 265.4630085 | -0.622358763 | 8.47E-03 |
| Ank3 | 2241.447597 | -0.623720772 | 7.83E-03 |
| Mrm1 | 929.4432346 | -0.624219909 | 1.55E-02 |
| Pcdhb16 | 566.0208578 | -0.624437011 | 8.60E-04 |
| Tnfaip6 | 646.6209697 | -0.624490791 | 3.01E-02 |
| Inpp5a | 3109.503164 | -0.624698874 | 7.92E-03 |
| Socs5 | 5784.952356 | -0.625371902 | 1.72E-06 |
| Plcb4 | 1673.625399 | -0.625705741 | 4.46E-02 |

| | | | |
|---|---|---|---|
| Pcdhgc4 | 30257.72995 | -0.62813028 | 1.83E-04 |
| Aldh4a1 | 564.9211389 | -0.628389637 | 5.52E-03 |
| Hbp1 | 5566.199986 | -0.628516358 | 6.89E-04 |
| Kdsr | 1932.623425 | -0.628557019 | 3.78E-06 |
| Tmem218 | 321.622946 | -0.629005948 | 1.14E-03 |
| Gm6436 | 10382.66031 | -0.629389336 | 2.00E-02 |
| Rpl39 | 31120.35881 | -0.629503104 | 1.76E-02 |
| Gatb | 891.9343914 | -0.630152259 | 1.79E-02 |
| Akr7a5 | 1052.456729 | -0.630326925 | 5.52E-03 |
| Rps27rt | 30142.86293 | -0.630572561 | 1.76E-02 |
| Usb1 | 1737.627401 | -0.63079862 | 1.69E-06 |
| Grcc10 | 2167.032303 | -0.632044462 | 5.31E-03 |
| Sema4g | 286.9531932 | -0.632492926 | 1.11E-02 |
| Grhpr | 457.3816047 | -0.632621349 | 4.34E-04 |
| Fbln2 | 51001.24849 | -0.633296661 | 2.18E-02 |
| Faap100 | 615.3568584 | -0.633728855 | 5.41E-04 |
| Samd14 | 1239.574654 | -0.634201543 | 1.51E-04 |
| Hddc3 | 237.1640719 | -0.634623704 | 2.59E-02 |
| Pcdhb5 | 354.373418 | -0.634989822 | 2.54E-03 |
| Pygo1 | 741.5294784 | -0.635072382 | 4.37E-03 |
| Cblb | 1328.176995 | -0.635802867 | 3.48E-05 |
| Cbarp | 525.0437716 | -0.636222779 | 4.72E-05 |
| Kdm3a | 3959.529429 | -0.636925789 | 1.15E-06 |
| Pecr | 434.5207829 | -0.636981778 | 1.03E-07 |
| Pcdhb21 | 187.4813216 | -0.637174501 | 5.64E-03 |
| Kctd1 | 652.3447104 | -0.638250073 | 4.10E-04 |
| 2410131K14Rik | 1774.50699 | -0.639047883 | 2.69E-07 |
| Tmem106a | 958.7992917 | -0.639675593 | 6.29E-04 |
| Glmp | 7292.413588 | -0.640865444 | 2.62E-04 |
| Ptpn3 | 659.1680817 | -0.641075702 | 6.37E-04 |
| Parp8 | 1704.661461 | -0.641349498 | 1.05E-05 |
| Pde7a | 1818.082955 | -0.642225996 | 2.04E-09 |
| Orai3 | 1110.900108 | -0.642739746 | 2.76E-05 |
| Mast4 | 3107.692726 | -0.642795803 | 8.97E-05 |
| Fdft1 | 6536.743541 | -0.643055437 | 3.59E-02 |
| Vps18 | 2476.558079 | -0.643145392 | 3.64E-03 |
| Alpk1 | 418.5502308 | -0.6432836 | 2.62E-03 |
| Ap3m2 | 731.4968151 | -0.643366269 | 5.63E-03 |
| Nfatc4 | 1786.320943 | -0.644279576 | 1.05E-03 |
| Pcdhb19 | 163.5809958 | -0.644733005 | 7.20E-03 |
| Gnptab | 4858.629519 | -0.645567943 | 2.10E-06 |
| Slc4a2 | 6711.940086 | -0.646032007 | 4.70E-05 |
| Aebp1 | 72201.79584 | -0.646427165 | 1.68E-03 |
| 4632404H12Rik | 210.9051878 | -0.647704862 | 5.10E-03 |
| Ap3s1 | 6475.604168 | -0.648715916 | 6.12E-03 |
| Porcn | 567.2858887 | -0.649559195 | 3.56E-03 |
| Xrcc5 | 2169.701875 | -0.649872156 | 9.53E-08 |

| | | | |
|---|---|---|---|
| Tom1l1 | 742.8401817 | -0.650713395 | 8.56E-05 |
| Ccdc57 | 258.3154069 | -0.652165179 | 2.57E-02 |
| Arrdc2 | 510.3397326 | -0.652486274 | 1.26E-02 |
| Pdcd4 | 2480.242851 | -0.652862287 | 2.07E-03 |
| Eif3e | 15575.42035 | -0.653011183 | 4.58E-04 |
| Ccng2 | 2087.349482 | -0.653341618 | 4.00E-03 |
| Dvl2 | 3542.891552 | -0.653614187 | 4.07E-04 |
| Aar2 | 1658.318704 | -0.653733726 | 3.05E-06 |
| Homer3 | 2341.705151 | -0.653931507 | 5.29E-04 |
| Klf9 | 3975.253106 | -0.654106222 | 2.37E-04 |
| Efna4 | 476.6852267 | -0.654506032 | 4.24E-04 |
| Rin2 | 6488.596361 | -0.655142186 | 7.50E-03 |
| Tbc1d16 | 1866.941337 | -0.655531723 | 1.67E-05 |
| Mob3a | 1207.127894 | -0.657332583 | 1.35E-04 |
| Bcs1l | 748.8985829 | -0.658417489 | 2.86E-04 |
| Zfp655 | 3110.698296 | -0.658892146 | 7.73E-11 |
| Sh3bp2 | 1443.810206 | -0.659260186 | 4.13E-03 |
| Tmem50a | 3373.80242 | -0.659653156 | 5.32E-06 |
| Tbc1d24 | 2492.496938 | -0.660407435 | 3.35E-09 |
| 9330104G04Rik | 82.39673676 | -0.660598713 | 8.72E-03 |
| Acy1 | 719.2862101 | -0.661038262 | 7.97E-04 |
| Rtkn | 1336.449565 | -0.661915485 | 8.70E-04 |
| NA | 126.927072 | -0.66295145 | 4.48E-02 |
| Dyrk1b | 695.0887966 | -0.662981061 | 5.22E-03 |
| Natd1 | 2182.198035 | -0.663930813 | 6.21E-04 |
| Ttyh3 | 8901.200216 | -0.664085895 | 7.39E-03 |
| Soga3 | 486.8497861 | -0.665402962 | 1.61E-04 |
| Clba1 | 2846.204258 | -0.666320713 | 2.57E-02 |
| Alcam | 657.7870578 | -0.666358904 | 3.10E-02 |
| Zbed5 | 182.5059509 | -0.666793103 | 3.69E-03 |
| Klhl24 | 4261.066099 | -0.667352513 | 8.90E-04 |
| Tpmt | 384.4346636 | -0.668829081 | 2.29E-05 |
| Orai2 | 1421.004878 | -0.669006484 | 1.51E-06 |
| Pgap3 | 296.0703753 | -0.66973767 | 3.17E-04 |
| Txnrd3 | 885.4457633 | -0.670773748 | 3.40E-05 |
| Anxa6 | 6987.123827 | -0.672147602 | 6.64E-04 |
| Naip6 | 212.161741 | -0.672743734 | 1.83E-04 |
| 4732487G21Rik | 167.6550051 | -0.673217602 | 8.65E-03 |
| Fhdc1 | 1544.683828 | -0.673297962 | 1.28E-05 |
| Nagk | 2436.902089 | -0.673608695 | 3.41E-08 |
| Dennd2a | 5263.659133 | -0.675352434 | 3.80E-04 |
| Qdpr | 1935.818044 | -0.6754226 | 9.31E-05 |
| Pts | 474.3278678 | -0.676150384 | 1.14E-03 |
| Maoa | 11869.71092 | -0.676769378 | 1.25E-03 |
| Pcdhb8 | 54.15088343 | -0.67680897 | 4.99E-02 |
| Inava | 109.6608324 | -0.677888439 | 2.22E-02 |
| Mpdu1 | 1210.812562 | -0.678376647 | 5.34E-07 |

| | | | |
|---|---|---|---|
| Acot1 | 957.7954011 | -0.678853523 | 5.06E-03 |
| Hectd3 | 2568.610222 | -0.679600572 | 8.69E-07 |
| Gabarap | 19416.47096 | -0.67971473 | 1.51E-02 |
| 5-Sep | 7626.58109 | -0.680858199 | 4.64E-04 |
| Tpcn2 | 753.9566622 | -0.681845319 | 2.87E-08 |
| Hs1bp3 | 3425.439165 | -0.682804122 | 6.56E-04 |
| Gyg | 5935.453933 | -0.683633127 | 8.49E-03 |
| Fam210b | 928.8968634 | -0.683675077 | 2.09E-03 |
| Ccdc189 | 104.5542721 | -0.684602382 | 1.42E-02 |
| Nmnat2 | 1973.93081 | -0.685538787 | 2.58E-04 |
| Plekhm1 | 2389.867812 | -0.6860923 | 2.15E-06 |
| Iah1 | 1772.443177 | -0.686559159 | 1.17E-03 |
| Gpr153 | 1837.287844 | -0.686560356 | 2.33E-03 |
| Fbxo36 | 335.8821186 | -0.687239122 | 7.43E-03 |
| Mob3b | 109.8020313 | -0.688887261 | 4.17E-02 |
| Hexdc | 950.8161435 | -0.688977852 | 1.22E-04 |
| Ydjc | 121.0858377 | -0.689635094 | 4.35E-02 |
| Hebp1 | 1013.408107 | -0.690423669 | 2.52E-04 |
| Runx1 | 1094.061865 | -0.69231563 | 5.64E-09 |
| Zfp637 | 1645.301751 | -0.692441003 | 8.21E-05 |
| Eno2 | 4035.358671 | -0.692689061 | 2.81E-02 |
| Rhoq | 7241.961221 | -0.692897649 | 4.90E-09 |
| Bahcc1 | 1260.799357 | -0.693071927 | 2.93E-04 |
| Dclk1 | 6246.021643 | -0.693859806 | 1.33E-05 |
| Ccl28 | 69.41710484 | -0.694601423 | 1.74E-02 |
| Tgfb3 | 3717.223368 | -0.694619931 | 9.84E-04 |
| Irx1 | 1133.860938 | -0.695570126 | 4.44E-03 |
| Mtss1l | 4009.15688 | -0.695987436 | 3.47E-04 |
| Emc2 | 2388.117355 | -0.696406293 | 5.19E-03 |
| Mroh1 | 6120.311947 | -0.696408105 | 9.25E-05 |
| Dcakd | 2695.360278 | -0.699705406 | 7.78E-05 |
| Tmem256 | 1681.651426 | -0.700999527 | 8.30E-06 |
| Cfap100 | 434.4066328 | -0.701386556 | 3.42E-02 |
| Klc3 | 265.3530538 | -0.702556212 | 5.34E-03 |
| Gm15280 | 482.2429882 | -0.702749556 | 3.48E-03 |
| Tprkb | 2647.743254 | -0.703519999 | 1.23E-06 |
| Arntl2 | 104.313339 | -0.704332037 | 3.93E-02 |
| Hsf4 | 234.9047654 | -0.704807666 | 9.50E-04 |
| Plekha5 | 1255.587539 | -0.706084496 | 1.96E-05 |
| Bsn | 48.5312342 | -0.706389776 | 3.81E-02 |
| Stxbp1 | 3426.852098 | -0.70826129 | 1.79E-05 |
| Sh3kbp1 | 3211.059854 | -0.708559899 | 1.81E-08 |
| Shc1 | 9740.164831 | -0.709021028 | 1.65E-05 |
| 2810032G03Rik | 133.6360171 | -0.709026679 | 3.96E-04 |
| Igtp | 114.2451841 | -0.709502404 | 2.56E-02 |
| H2afv | 1309.866945 | -0.709646463 | 7.38E-05 |
| Pdk1 | 1503.806453 | -0.710138979 | 1.42E-02 |

| | | | |
|---|---|---|---|
| Tmem164 | 4384.562062 | -0.711919544 | 8.21E-07 |
| Adck2 | 829.9544706 | -0.712305102 | 1.37E-07 |
| Sptssa | 1180.152327 | -0.712371397 | 6.25E-04 |
| Bnip3 | 1792.535201 | -0.712712443 | 3.63E-05 |
| Magee1 | 2530.570754 | -0.7129112 | 1.46E-08 |
| Rab36 | 271.0864376 | -0.713868758 | 1.87E-02 |
| Efnb2 | 1976.379573 | -0.714401269 | 3.42E-04 |
| Cyp1b1 | 3968.844327 | -0.71478193 | 8.55E-04 |
| Decr2 | 584.6527844 | -0.714898457 | 1.52E-04 |
| Mmp2 | 89388.04831 | -0.715369982 | 2.27E-02 |
| Selenow | 3078.602105 | -0.715578884 | 2.75E-04 |
| Sox4 | 7067.281048 | -0.715608152 | 1.18E-02 |
| Abhd1 | 424.5594923 | -0.716581378 | 2.00E-06 |
| Serf1 | 393.2430891 | -0.716637093 | 5.60E-03 |
| Rnf19b | 2914.707296 | -0.717238791 | 8.60E-04 |
| Fam213a | 757.8558732 | -0.718214205 | 1.03E-02 |
| Sbk1 | 915.23951 | -0.718694846 | 3.41E-03 |
| Lgals3 | 26908.25829 | -0.719048192 | 1.46E-02 |
| Tifa | 255.8989682 | -0.72030019 | 1.95E-02 |
| Cops9 | 1741.656329 | -0.720865086 | 1.86E-04 |
| Marcks | 14713.78478 | -0.721763985 | 6.43E-03 |
| 1700017B05Rik | 1654.177394 | -0.721972661 | 1.39E-04 |
| Mtmr2 | 3520.614254 | -0.722156619 | 4.47E-05 |
| Arhgef40 | 3841.130444 | -0.722590018 | 6.83E-06 |
| Vangl2 | 622.778806 | -0.722668439 | 5.64E-05 |
| Rpl10-ps6 | 116.8439331 | -0.72277227 | 2.97E-02 |
| 5930403N24Rik | 408.5526991 | -0.722957549 | 4.51E-02 |
| Csgalnact1 | 1244.091072 | -0.724124439 | 1.16E-03 |
| Sh3bgrl3 | 2819.46658 | -0.725856053 | 9.75E-04 |
| Pex11a | 343.174823 | -0.726371081 | 3.27E-04 |
| Vat1 | 27960.27472 | -0.727033003 | 2.68E-03 |
| Gm14303 | 4285.914875 | -0.727096088 | 2.20E-02 |
| Rpl22l1 | 5120.112213 | -0.727327958 | 2.46E-04 |
| Fmn1 | 2418.368649 | -0.728026225 | 1.30E-03 |
| Fam3c | 4044.640668 | -0.728487889 | 2.70E-07 |
| Npc2 | 24224.6119 | -0.729210528 | 6.17E-03 |
| Map1lc3b | 9584.386169 | -0.731324392 | 1.19E-03 |
| Fbxo32 | 1611.242564 | -0.731620191 | 8.49E-03 |
| Nckap5l | 1105.351254 | -0.732640805 | 4.72E-05 |
| Zfp184 | 226.740299 | -0.733454824 | 2.01E-04 |
| Mettl27 | 1096.641165 | -0.734815251 | 1.48E-04 |
| Rhoj | 1534.443716 | -0.737028848 | 3.43E-04 |
| NA | 154.0539091 | -0.737054595 | 1.45E-03 |
| Fmnl3 | 2696.537561 | -0.738523983 | 8.20E-07 |
| Acsf2 | 1577.993251 | -0.738693428 | 1.23E-04 |
| Prickle2 | 610.6101831 | -0.740068869 | 3.80E-06 |
| Dpp7 | 5371.98911 | -0.742096761 | 6.21E-04 |

| | | | |
|---|---|---|---|
| B3galt1 | 227.8123498 | -0.745060461 | 7.44E-05 |
| Aph1b | 607.0118737 | -0.745174455 | 2.78E-04 |
| Wdr34 | 712.2965379 | -0.746346404 | 1.11E-05 |
| Hist2h2be | 103.5079661 | -0.749283818 | 1.13E-02 |
| Mitf | 1042.401336 | -0.751690692 | 4.91E-03 |
| Emb | 6148.231165 | -0.752822509 | 2.59E-04 |
| Orc5 | 1511.84814 | -0.753227634 | 2.58E-03 |
| Magi2 | 526.5829282 | -0.753393267 | 1.68E-08 |
| Fbxo41 | 890.8379166 | -0.753674088 | 2.15E-05 |
| Spin2c | 57.0212798 | -0.753829245 | 2.24E-02 |
| Myo10 | 6334.157457 | -0.754064877 | 6.12E-05 |
| Amigo2 | 624.6922769 | -0.754359285 | 2.38E-06 |
| Etv5 | 692.3569151 | -0.754552127 | 1.61E-02 |
| Nme3 | 171.0026249 | -0.754595895 | 9.26E-04 |
| Rwdd2a | 82.87345654 | -0.754607314 | 2.42E-02 |
| Pde4d | 597.7003963 | -0.755049634 | 7.87E-05 |
| Mnt | 3501.854391 | -0.755537017 | 7.64E-04 |
| Tead2 | 4318.690094 | -0.755893588 | 4.64E-04 |
| P4htm | 35.99990357 | -0.757966565 | 3.73E-02 |
| Trim62 | 389.1707065 | -0.758585215 | 2.63E-05 |
| Lhfpl2 | 6465.270021 | -0.759839848 | 1.15E-06 |
| Efna3 | 212.0063188 | -0.760149409 | 3.62E-03 |
| Fgfr2 | 2705.677105 | -0.760158454 | 1.49E-03 |
| Ssbp2 | 1469.826249 | -0.760385428 | 2.41E-02 |
| Prdm11 | 475.2305451 | -0.760571689 | 2.23E-05 |
| Nbl1 | 1782.408442 | -0.761483474 | 2.08E-02 |
| Galnt13 | 425.0375608 | -0.763635628 | 9.67E-05 |
| Comtd1 | 549.7820272 | -0.764981958 | 5.47E-07 |
| Whrn | 1183.150778 | -0.76648769 | 3.96E-05 |
| Pappa | 4604.374407 | -0.767116342 | 1.02E-02 |
| Arhgap42 | 1172.465651 | -0.767959519 | 1.04E-03 |
| Pigh | 312.2573961 | -0.768946794 | 2.64E-04 |
| Vdr | 1256.985056 | -0.769640001 | 1.17E-02 |
| Atg12 | 4017.335377 | -0.771016339 | 1.31E-03 |
| Shpk | 226.3556193 | -0.771367521 | 2.06E-05 |
| Selenop | 9438.313738 | -0.773000972 | 3.92E-02 |
| Pcdhb17 | 304.6149099 | -0.773162423 | 1.59E-05 |
| Evl | 2952.632705 | -0.77364738 | 7.99E-04 |
| Arhgef19 | 326.3317744 | -0.773887577 | 1.30E-02 |
| Timp3 | 19304.01546 | -0.77392028 | 1.89E-03 |
| Cst3 | 29679.40866 | -0.774499027 | 8.82E-04 |
| Atl1 | 306.3142026 | -0.774882874 | 1.92E-04 |
| Sesn3 | 451.770964 | -0.777283825 | 4.16E-04 |
| Mex3a | 2190.570495 | -0.777367923 | 3.40E-03 |
| Mertk | 609.7724486 | -0.778810107 | 1.09E-03 |
| Col5a3 | 679.7155384 | -0.780160289 | 4.27E-02 |
| 2410018L13Rik | 753.6620775 | -0.783682705 | 3.32E-09 |

| | | | |
|---|---|---|---|
| Rnft2 | 369.3814148 | -0.783929765 | 1.15E-02 |
| Pqlc1 | 490.6899246 | -0.785780724 | 9.20E-03 |
| Ttyh2 | 1997.972056 | -0.7859086 | 1.99E-05 |
| Dbnl | 5109.073654 | -0.786524919 | 6.10E-07 |
| Flrt2 | 1425.858476 | -0.787235665 | 1.24E-02 |
| Lrrc56 | 115.614384 | -0.788289362 | 8.83E-03 |
| Aig1 | 239.7079789 | -0.78901386 | 2.52E-03 |
| Fjx1 | 266.2076356 | -0.790589368 | 7.23E-05 |
| Litaf | 5188.580695 | -0.793062337 | 1.29E-04 |
| Trappc6a | 772.3366347 | -0.793316888 | 5.56E-05 |
| Iglon5 | 4473.234465 | -0.793920465 | 3.79E-04 |
| Myo1e | 5416.527455 | -0.794600471 | 7.38E-05 |
| Rps18-ps3 | 520.1252683 | -0.796562676 | 2.03E-02 |
| Vsig10l | 521.0060425 | -0.801253969 | 2.24E-03 |
| Hdac5 | 4813.627712 | -0.806247427 | 4.93E-03 |
| H2-DMb2 | 96.6450081 | -0.806750904 | 8.34E-03 |
| Fam43a | 1435.289342 | -0.806778617 | 1.46E-05 |
| Mapk8ip1 | 2586.65763 | -0.807698985 | 4.03E-05 |
| A930024E05Rik | 70.97068599 | -0.810940018 | 1.57E-02 |
| Cpe | 17622.21883 | -0.811327567 | 8.30E-06 |
| Dact1 | 476.0212076 | -0.81178233 | 4.65E-02 |
| Cspg4 | 1149.427576 | -0.812221197 | 8.31E-03 |
| Met | 7204.015306 | -0.813446402 | 4.82E-10 |
| Slc27a1 | 1298.849242 | -0.813841401 | 3.56E-03 |
| Cnih2 | 222.9740135 | -0.813998912 | 2.55E-03 |
| Nat2 | 437.4994644 | -0.815360578 | 5.69E-08 |
| Nudt16l1 | 1616.991711 | -0.816077922 | 3.78E-06 |
| E230013L22Rik | 57.3366247 | -0.816847782 | 5.81E-03 |
| Ccdc159 | 201.3206634 | -0.817195162 | 1.61E-03 |
| Gm1673 | 194.6979044 | -0.81826577 | 7.54E-03 |
| Tanc2 | 4417.326718 | -0.818636368 | 3.08E-08 |
| Erg28 | 1761.041711 | -0.819503417 | 1.70E-04 |
| Clmp | 2125.373568 | -0.823021709 | 1.25E-08 |
| Trib2 | 1232.911453 | -0.823310574 | 1.77E-04 |
| Setmar | 64.67531696 | -0.823637345 | 2.23E-03 |
| Slc7a7 | 105.7139369 | -0.824488931 | 7.64E-03 |
| Ctnnb1 | 26830.96708 | -0.825117873 | 2.11E-19 |
| Pex11g | 135.442031 | -0.825436666 | 1.07E-02 |
| Nradd | 1116.064392 | -0.826165467 | 2.79E-10 |
| Gnpda1 | 2534.474834 | -0.826729596 | 3.33E-06 |
| Dip2b | 2838.90961 | -0.827491186 | 6.20E-06 |
| Sfxn5 | 564.0641619 | -0.829640082 | 2.32E-06 |
| Agap2 | 1004.939107 | -0.829640794 | 1.27E-02 |
| Gm15298 | 211.5333334 | -0.831352646 | 2.91E-03 |
| Nr3c2 | 281.6346757 | -0.83151501 | 1.04E-02 |
| Pcdhb20 | 156.4655691 | -0.831930875 | 8.87E-04 |
| Fam110b | 769.9936195 | -0.832147418 | 2.84E-03 |

| | | | |
|---|---|---|---|
| Tfap2a | 730.9542063 | -0.832503924 | 1.34E-03 |
| NA | 132.3463999 | -0.832515501 | 4.65E-02 |
| Msrb2 | 106.301837 | -0.832543832 | 7.86E-03 |
| Ulk4 | 167.008026 | -0.833064008 | 7.64E-04 |
| Fam20c | 1116.228718 | -0.833104975 | 2.53E-05 |
| Unc13a | 386.4394156 | -0.834064582 | 1.33E-03 |
| Adhfe1 | 58.16465703 | -0.834473299 | 1.49E-02 |
| Rnf144a | 1573.389355 | -0.835479654 | 4.61E-02 |
| Ptgr1 | 2247.24335 | -0.835728916 | 1.43E-09 |
| Xdh | 15174.39706 | -0.839022645 | 5.43E-04 |
| Tmem200a | 180.2586998 | -0.839947074 | 1.07E-02 |
| Ctns | 2362.555936 | -0.841231821 | 2.42E-08 |
| Gga2 | 4453.49319 | -0.841670462 | 8.48E-07 |
| Hgf | 466.3773992 | -0.843494761 | 1.83E-02 |
| Snn | 2448.99706 | -0.843718205 | 9.58E-08 |
| Arhgef28 | 758.8625099 | -0.844651601 | 5.54E-05 |
| Sardh | 902.4568354 | -0.845439891 | 2.29E-03 |
| Ptk2b | 79.64446995 | -0.846247675 | 1.52E-03 |
| Nsg1 | 1730.070035 | -0.847681496 | 1.06E-08 |
| Gbp3 | 49.15600115 | -0.848203673 | 2.08E-02 |
| Hhipl1 | 257.2349416 | -0.849069361 | 1.23E-02 |
| Gm10288 | 2402.863389 | -0.849327883 | 3.89E-02 |
| Dpy19l1 | 4064.222751 | -0.852590162 | 1.07E-08 |
| Gpr85 | 486.5449696 | -0.852623127 | 2.32E-04 |
| Tmem205 | 457.4021942 | -0.853673705 | 1.31E-03 |
| Cotl1 | 536.8515866 | -0.854681963 | 2.44E-03 |
| Atp6v0d1 | 8071.893965 | -0.854710261 | 1.04E-04 |
| Gm9869 | 51.84992159 | -0.856109669 | 3.86E-02 |
| Nceh1 | 3203.829638 | -0.856249845 | 1.01E-05 |
| Gm14414 | 932.8384723 | -0.857529986 | 1.49E-03 |
| Pik3r5 | 1315.666089 | -0.862567714 | 3.79E-03 |
| Pcdhb7 | 281.6238328 | -0.865165487 | 1.29E-04 |
| H2-DMb1 | 135.8915411 | -0.865209057 | 1.51E-02 |
| Prps2 | 2712.855431 | -0.865274862 | 1.55E-10 |
| Camkk1 | 480.61585 | -0.867633671 | 4.85E-04 |
| Amdhd2 | 1696.242283 | -0.871747494 | 6.56E-04 |
| Cd300lb | 309.9653894 | -0.874011619 | 3.63E-02 |
| Tnik | 413.6788858 | -0.874166252 | 3.72E-04 |
| B4galt5 | 4562.137611 | -0.874621789 | 7.37E-04 |
| Ccdc103 | 53.9477953 | -0.874711595 | 1.04E-02 |
| AW551984 | 1346.712062 | -0.877193093 | 2.50E-02 |
| Angptl2 | 12262.23054 | -0.879374661 | 8.63E-04 |
| Elovl4 | 159.9698766 | -0.881869808 | 6.68E-03 |
| Plxnd1 | 6200.522837 | -0.884061459 | 6.83E-06 |
| Gm21981 | 86.72813471 | -0.884101828 | 7.72E-03 |
| Pmepa1 | 1219.457425 | -0.885924177 | 5.66E-06 |
| Ogdhl | 4923.875176 | -0.88592599 | 5.78E-10 |

| | | | |
|---|---|---|---|
| Pde3a | 117.8255858 | -0.888034755 | 1.53E-03 |
| Scn1b | 147.6156986 | -0.888688466 | 1.07E-02 |
| Nsdhl | 4567.873814 | -0.88903841 | 4.33E-03 |
| 1700003F12Rik | 81.47917467 | -0.889079716 | 7.09E-04 |
| Rragd | 324.476639 | -0.891145733 | 1.97E-02 |
| Atxn1 | 1217.972466 | -0.892173744 | 1.83E-04 |
| Fam69b | 943.3674919 | -0.895780653 | 3.44E-06 |
| Chst2 | 135.1732714 | -0.898662336 | 1.58E-02 |
| Tcp11l2 | 1451.319384 | -0.900077603 | 4.57E-03 |
| Tubb2b | 2259.346938 | -0.900171888 | 3.08E-06 |
| Pcdhb6 | 173.737983 | -0.900884554 | 9.72E-04 |
| AI413582 | 932.4708856 | -0.901324704 | 1.30E-06 |
| Suclg2 | 5554.452872 | -0.903537472 | 1.02E-17 |
| Serinc5 | 1871.588337 | -0.905196427 | 5.01E-16 |
| Hvcn1 | 604.403497 | -0.906006033 | 4.03E-06 |
| Limd2 | 1732.780265 | -0.906360035 | 6.96E-07 |
| Caps2 | 114.6304859 | -0.906748802 | 1.86E-02 |
| Slc1a3 | 1356.973152 | -0.907513634 | 1.48E-05 |
| Casp9 | 1187.691822 | -0.908874971 | 9.82E-10 |
| Gm14002 | 28.28916881 | -0.911628966 | 2.10E-02 |
| Dhcr7 | 1440.708452 | -0.91457739 | 3.74E-04 |
| Dcst1 | 38.85705804 | -0.918891616 | 1.04E-02 |
| Ermp1 | 2303.615021 | -0.919341573 | 1.02E-09 |
| H1fx | 45.98158594 | -0.922763394 | 3.40E-02 |
| Ankrd44 | 1362.280154 | -0.926868371 | 1.95E-06 |
| Ctla2b | 860.0005034 | -0.927786428 | 2.82E-02 |
| Sh2d3c | 108.5494914 | -0.927953552 | 1.14E-02 |
| Adgrl2 | 6387.888046 | -0.92889991 | 1.04E-15 |
| Ghr | 7386.390632 | -0.929593295 | 4.87E-06 |
| Gm13470 | 63.86688211 | -0.931116493 | 4.99E-03 |
| Cmtm3 | 7129.083408 | -0.93116428 | 9.27E-08 |
| Mcf2l | 69.89545145 | -0.93265381 | 2.65E-02 |
| Nxph3 | 56.41829472 | -0.933413297 | 3.25E-02 |
| Cyp2j9 | 75.57103561 | -0.934468193 | 8.07E-03 |
| P2rx4 | 2480.638167 | -0.93470869 | 5.62E-06 |
| St3gal4 | 2084.028363 | -0.934799516 | 1.39E-04 |
| Atp6v1g1 | 12568.78181 | -0.935254041 | 3.13E-07 |
| Rnf122 | 450.2380668 | -0.936317259 | 1.58E-02 |
| Grb14 | 1511.859625 | -0.939375186 | 5.42E-04 |
| Top1mt | 841.059034 | -0.939975281 | 2.34E-13 |
| 9330159F19Rik | 260.4380525 | -0.941922647 | 2.35E-03 |
| Dyx1c1 | 41.08029909 | -0.942635069 | 1.62E-02 |
| 2810408A11Rik | 84.21173571 | -0.943152318 | 4.56E-03 |
| Myo1b | 2440.909108 | -0.944805814 | 3.54E-07 |
| Tiam2 | 1483.526478 | -0.945404425 | 2.93E-04 |
| Marcksl1 | 7037.230072 | -0.948799568 | 2.21E-05 |
| Slc6a8 | 7871.670639 | -0.951012104 | 5.02E-06 |

| | | | |
|---|---|---|---|
| Xpr1 | 2311.363627 | -0.954443366 | 3.50E-02 |
| NA | 85.90070398 | -0.955005138 | 8.74E-03 |
| Gm9294 | 71.67654231 | -0.955210311 | 2.52E-02 |
| Gm10010 | 87.03730997 | -0.955877724 | 1.64E-02 |
| Tmem116 | 74.18436638 | -0.956212414 | 3.92E-02 |
| Ypel1 | 252.6287883 | -0.956858702 | 2.02E-03 |
| Tbc1d4 | 572.5515038 | -0.957255261 | 5.56E-04 |
| Tmem140 | 492.4092491 | -0.964012633 | 5.26E-03 |
| Txndc16 | 3149.373031 | -0.965510997 | 9.58E-08 |
| Cracr2a | 41.73135551 | -0.966186958 | 2.38E-02 |
| Fam110c | 419.3831265 | -0.972115171 | 9.40E-05 |
| Adssl1 | 59.72197817 | -0.972715178 | 3.95E-02 |
| NA | 35.3883632 | -0.98109484 | 1.47E-02 |
| Bdh2 | 1089.036205 | -0.983995401 | 1.00E-04 |
| Sdc3 | 1488.655788 | -0.989676668 | 2.25E-05 |
| Prtg | 275.2554638 | -0.990435637 | 3.56E-04 |
| Itgbl1 | 4703.422305 | -0.991369844 | 2.53E-03 |
| Acat2 | 6980.262021 | -0.991686784 | 1.36E-03 |
| Kdm5b | 5947.881651 | -0.992493471 | 6.10E-05 |
| Gpr137b | 6771.907523 | -0.99300113 | 4.00E-04 |
| Bmper | 999.7225729 | -0.993005609 | 5.26E-03 |
| Fgf14 | 155.1480338 | -0.993240005 | 2.71E-02 |
| Acacb | 256.1837198 | -0.993402877 | 9.80E-03 |
| F730043M19Rik | 811.4228063 | -0.993640658 | 1.30E-03 |
| Lonrf1 | 505.0783429 | -0.996262073 | 1.76E-10 |
| Capn6 | 8375.626269 | -1.001514785 | 9.07E-05 |
| Srl | 383.3789868 | -1.002018626 | 6.64E-04 |
| 2700033N17Rik | 66.80711653 | -1.002349934 | 1.54E-03 |
| Tmem163 | 21.88372852 | -1.003366138 | 2.56E-02 |
| Tmeff1 | 2004.0604 | -1.003453199 | 6.96E-13 |
| Nkain1 | 814.8730286 | -1.004589492 | 1.40E-08 |
| Gng2 | 472.9891909 | -1.005772515 | 1.95E-05 |
| Garem2 | 297.7203972 | -1.006184619 | 2.73E-07 |
| Ccdc136 | 1423.32675 | -1.00629014 | 1.17E-06 |
| Parvaos | 101.9905319 | -1.006639386 | 3.48E-02 |
| Hsd17b7 | 1863.090416 | -1.009800626 | 3.84E-03 |
| Rgs17 | 87.00021087 | -1.010359443 | 4.08E-02 |
| Gm8615 | 190.0903932 | -1.012158301 | 1.20E-04 |
| Tmem86a | 734.3320314 | -1.012346875 | 7.88E-06 |
| Cpne8 | 1083.712569 | -1.012877665 | 6.46E-20 |
| Mgst2 | 162.1947575 | -1.012960975 | 2.00E-04 |
| Arhgap24 | 1837.038701 | -1.01375118 | 4.48E-05 |
| Aph1c | 281.8841578 | -1.015135325 | 3.59E-04 |
| Zfp454 | 65.32644374 | -1.019412276 | 6.70E-04 |
| Khk | 219.7090183 | -1.022719792 | 6.98E-03 |
| A930002H24Rik | 37.79997615 | -1.022926469 | 4.59E-02 |
| Grem2 | 1174.534392 | -1.026590857 | 3.51E-04 |

| | | | |
|---|---|---|---|
| 9-Sep | 7495.015017 | -1.027182793 | 1.34E-11 |
| Sytl2 | 158.0277612 | -1.028358522 | 4.49E-02 |
| Ralgds | 5291.738528 | -1.029564814 | 1.44E-05 |
| Ldhb | 3916.76833 | -1.031422515 | 6.11E-09 |
| Mapkapk3 | 612.1223251 | -1.032647007 | 4.48E-05 |
| Sco2 | 266.3938489 | -1.033156712 | 2.00E-03 |
| 2610307P16Rik | 188.2318779 | -1.034852615 | 5.54E-03 |
| Gm16861 | 183.4760558 | -1.035760389 | 5.91E-07 |
| 1810058I24Rik | 4262.602694 | -1.038078219 | 6.29E-06 |
| Mpzl1 | 3792.936615 | -1.039710983 | 1.45E-07 |
| Pcdh18 | 4140.879768 | -1.039854794 | 3.66E-13 |
| Gm17484 | 94.6692011 | -1.040452059 | 5.26E-03 |
| Fdps | 10188.03978 | -1.040982365 | 4.31E-03 |
| NA | 167.7309758 | -1.046096611 | 6.83E-06 |
| Mylip | 3518.788342 | -1.04612227 | 2.14E-05 |
| Armcx6 | 583.5901757 | -1.047435161 | 3.22E-10 |
| Igsf8 | 3977.096197 | -1.048031449 | 4.93E-07 |
| Fam198b | 10855.77158 | -1.048559191 | 1.54E-05 |
| Tgif2 | 708.2075399 | -1.053387608 | 8.45E-09 |
| Fnip2 | 5464.462773 | -1.054299156 | 8.05E-10 |
| Col17a1 | 62.0117311 | -1.055944976 | 2.97E-02 |
| Htr2a | 66.90647818 | -1.061220816 | 9.32E-03 |
| Pcdhb9 | 216.2214447 | -1.064987617 | 3.54E-05 |
| Dlk1 | 29943.381 | -1.065693461 | 1.05E-02 |
| Tnc | 3603.199261 | -1.065880813 | 9.05E-06 |
| Amot | 5972.569539 | -1.069186412 | 2.91E-15 |
| Il27ra | 37.22449657 | -1.070257464 | 1.42E-02 |
| Axin2 | 27.73513628 | -1.078845084 | 3.42E-02 |
| Nap1l2 | 52.4545836 | -1.080544866 | 2.96E-02 |
| Kctd15 | 882.2490959 | -1.082792567 | 3.39E-04 |
| Dusp10 | 1991.634529 | -1.091596099 | 3.01E-03 |
| Bok | 494.9314589 | -1.096201591 | 2.02E-03 |
| Cd82 | 121.0116936 | -1.10005321 | 1.02E-04 |
| Lpin1 | 2970.0158 | -1.106302298 | 1.67E-04 |
| Snai2 | 1626.102104 | -1.107870429 | 8.01E-11 |
| Carmil3 | 108.3115967 | -1.109613542 | 3.64E-03 |
| Sh3pxd2b | 11222.03918 | -1.112201647 | 1.99E-07 |
| Sparcl1 | 1663.612457 | -1.11221155 | 1.14E-02 |
| Kcna4 | 440.1062529 | -1.11348225 | 7.52E-05 |
| Fam180a | 160.3125889 | -1.114775702 | 4.36E-03 |
| Synpo | 5453.723669 | -1.116380041 | 3.37E-09 |
| Myo7a | 1099.229425 | -1.117891554 | 1.59E-07 |
| 4933431E20Rik | 881.878442 | -1.120386143 | 8.45E-08 |
| Atp6v1b2 | 9772.289405 | -1.123290939 | 3.89E-09 |
| Gm11261 | 171.9841549 | -1.123715446 | 1.00E-02 |
| Mef2c | 246.1899401 | -1.125389302 | 1.95E-02 |
| Foxl1 | 95.73223871 | -1.127893685 | 2.01E-04 |

| | | | |
|---|---|---|---|
| Adcy1 | 1520.53709 | -1.130764736 | 1.26E-04 |
| Irs1 | 2008.224803 | -1.130848824 | 7.24E-08 |
| Glis1 | 258.3571055 | -1.131727575 | 6.97E-03 |
| Bhlhe41 | 899.5494575 | -1.133072783 | 4.79E-05 |
| Trim63 | 604.4480012 | -1.136631384 | 2.88E-02 |
| S1pr3 | 3918.192315 | -1.137554653 | 1.71E-12 |
| Lvrn | 146.2069254 | -1.142460556 | 3.56E-05 |
| Pgam2 | 56.35623073 | -1.144691802 | 7.00E-04 |
| 2810039B14Rik | 112.6447876 | -1.146011279 | 2.13E-08 |
| Rnf150 | 1744.58427 | -1.146927311 | 8.12E-11 |
| Tspan11 | 2916.150625 | -1.155236756 | 9.60E-09 |
| Trps1 | 1353.888354 | -1.156877274 | 8.12E-03 |
| Ociad2 | 337.1734645 | -1.15976352 | 3.78E-09 |
| Pabpc4l | 55.49155236 | -1.160572025 | 4.74E-03 |
| Srgap1 | 1826.660683 | -1.160813884 | 2.37E-05 |
| 5033430I15Rik | 56.08870669 | -1.16197655 | 4.10E-04 |
| Unc13b | 378.855196 | -1.162438492 | 2.85E-06 |
| Dcaf6 | 5277.412717 | -1.163452905 | 7.37E-11 |
| Pik3r3 | 1442.838135 | -1.169002626 | 1.05E-09 |
| Serpinb8 | 975.2511855 | -1.170138433 | 3.47E-05 |
| Olfml2b | 2018.256671 | -1.171624117 | 8.08E-06 |
| Sh2b2 | 263.5269605 | -1.174667335 | 8.58E-08 |
| Tspan18 | 194.9774176 | -1.175381345 | 2.22E-03 |
| Otulinl | 214.2498603 | -1.175726028 | 6.27E-12 |
| Mgarp | 47.39826116 | -1.176336094 | 1.10E-03 |
| Galnt17 | 1070.268328 | -1.197405785 | 6.36E-11 |
| Igsf3 | 583.6384486 | -1.21288167 | 3.60E-09 |
| Foxs1 | 115.5421906 | -1.213907733 | 4.73E-03 |
| Rhpn1 | 32.93132459 | -1.215430943 | 2.46E-03 |
| Mvd | 2840.656231 | -1.221680776 | 2.62E-04 |
| Mex3b | 174.4803361 | -1.22257871 | 8.17E-05 |
| Rbp1 | 2005.260463 | -1.224335594 | 2.59E-12 |
| Cp | 1547.646794 | -1.227226049 | 2.16E-04 |
| Plekha4 | 436.889388 | -1.227715193 | 2.17E-04 |
| Mt1 | 6812.419424 | -1.227944818 | 5.34E-03 |
| Rhou | 4021.987846 | -1.228603434 | 7.22E-13 |
| Maml3 | 728.8511573 | -1.229799015 | 6.29E-09 |
| Ifi47 | 69.23413701 | -1.232217 | 8.45E-05 |
| Pxylp1 | 827.890857 | -1.237296592 | 1.65E-05 |
| Dram1 | 593.7149253 | -1.23890231 | 2.30E-14 |
| Slc24a1 | 28.67118619 | -1.241247535 | 1.61E-02 |
| Steap3 | 17632.6646 | -1.249613661 | 1.76E-05 |
| Il18 | 182.5136154 | -1.249869151 | 7.40E-05 |
| Timp4 | 80.55768285 | -1.253877784 | 4.12E-02 |
| Il16 | 43.63478451 | -1.254143841 | 3.21E-03 |
| Tgfb1 | 1304.531232 | -1.260621856 | 3.88E-06 |
| Cxcl12 | 14907.90677 | -1.262965975 | 2.77E-04 |

| | | | |
|---|---|---|---|
| Lpcat2 | 612.0918981 | -1.263123499 | 5.37E-03 |
| Pde1b | 115.0782532 | -1.270933552 | 1.75E-03 |
| Fcrlb | 50.76172523 | -1.27171659 | 1.38E-02 |
| Ucp2 | 1083.734914 | -1.271884381 | 1.02E-09 |
| C530008M17Rik | 222.325705 | -1.275166901 | 1.13E-03 |
| Zfp41 | 1083.240847 | -1.27678302 | 1.49E-04 |
| Il13ra1 | 219.0472217 | -1.282689958 | 8.53E-13 |
| Id3 | 1711.956127 | -1.28389705 | 4.40E-02 |
| Shroom1 | 179.9661314 | -1.286724983 | 1.05E-05 |
| Nol4 | 339.5172744 | -1.287083885 | 6.34E-06 |
| Slc6a7 | 36.75567192 | -1.291267835 | 3.66E-04 |
| Gas7 | 13591.79219 | -1.292212908 | 3.93E-08 |
| Emid1 | 31.69594959 | -1.297082903 | 1.97E-02 |
| Gm16201 | 31.28094923 | -1.302556227 | 2.77E-02 |
| Tm7sf2 | 703.3797047 | -1.30895694 | 2.67E-06 |
| Slit2 | 4587.564039 | -1.309842038 | 8.05E-08 |
| Trem2 | 96.77326875 | -1.310763229 | 4.14E-04 |
| Klf12 | 71.36571759 | -1.311269403 | 1.93E-07 |
| A330009N23Rik | 54.46191423 | -1.314387236 | 1.23E-04 |
| Pkia | 759.5393534 | -1.319272335 | 9.29E-07 |
| Nat8f1 | 120.6153721 | -1.323175023 | 5.71E-04 |
| Polm | 3037.083453 | -1.329598251 | 5.34E-21 |
| Pdk4 | 19724.77029 | -1.338598436 | 4.40E-05 |
| Plp1 | 415.718745 | -1.343303671 | 2.09E-03 |
| Cdk15 | 129.3148332 | -1.348661999 | 2.46E-04 |
| Il1rap | 2529.832075 | -1.349101538 | 8.91E-14 |
| Ifi27l2a | 63.9925599 | -1.349793522 | 3.35E-04 |
| Dlgap2 | 221.2029046 | -1.350541097 | 5.44E-07 |
| Msi1 | 105.8772489 | -1.36574635 | 1.80E-05 |
| NA | 97.40321005 | -1.368209546 | 1.54E-03 |
| Lrrc46 | 44.82983873 | -1.375749349 | 2.09E-04 |
| Ang2 | 58.05561455 | -1.385200632 | 3.50E-03 |
| Glrb | 356.4512423 | -1.387881182 | 3.65E-09 |
| Spata48 | 29.05752533 | -1.390158401 | 5.95E-03 |
| Anks1b | 74.61954556 | -1.390446073 | 1.76E-02 |
| Lxn | 2852.121736 | -1.396819414 | 2.92E-09 |
| Hlf | 720.1008505 | -1.401149757 | 4.66E-02 |
| Col28a1 | 133.400327 | -1.418627709 | 9.21E-04 |
| Tcim | 182.9808592 | -1.418784866 | 2.86E-05 |
| Tnfrsf11a | 123.4947407 | -1.423880054 | 3.73E-14 |
| Fam169b | 128.3296111 | -1.42984086 | 6.91E-04 |
| Epha7 | 250.107713 | -1.440869761 | 3.71E-13 |
| Gm15422 | 17.50296185 | -1.446803776 | 3.83E-02 |
| Gm9747 | 221.4334643 | -1.455716187 | 1.02E-05 |
| Cyp26c1 | 9.040582588 | -1.4580937 | 4.44E-02 |
| Sytl1 | 58.78219199 | -1.464295535 | 3.15E-03 |
| Shc2 | 1586.018871 | -1.466074994 | 1.57E-12 |

| | | | |
|---|---|---|---|
| Cdo1 | 5428.454521 | -1.466933365 | 4.48E-05 |
| Il12rb1 | 41.79475726 | -1.467323957 | 2.61E-03 |
| Gm12590 | 87.10131386 | -1.469376922 | 7.44E-05 |
| NA | 13.22047084 | -1.471243237 | 4.13E-02 |
| Gm11767 | 9.219788467 | -1.471571489 | 4.13E-02 |
| Csrp2 | 1603.401949 | -1.481706724 | 3.78E-06 |
| Gcnt4 | 773.2398383 | -1.484390479 | 8.57E-13 |
| Sox9 | 2547.006272 | -1.50189575 | 9.37E-10 |
| Ednra | 698.5583548 | -1.511745016 | 5.59E-06 |
| Gm13187 | 17.5193156 | -1.516333988 | 3.26E-02 |
| Zfpm2 | 579.7477361 | -1.518397927 | 3.42E-15 |
| Eya4 | 341.0867101 | -1.521050845 | 3.98E-14 |
| Gbp2 | 472.493555 | -1.524098163 | 8.99E-12 |
| Ptger4 | 492.752494 | -1.534963171 | 2.43E-14 |
| Gm5514 | 63.12948495 | -1.537417993 | 5.12E-06 |
| Egr2 | 252.8800803 | -1.540009035 | 1.70E-04 |
| Smoc1 | 319.0475636 | -1.544458547 | 3.19E-03 |
| Gm6166 | 5392.037064 | -1.55241685 | 2.16E-14 |
| Ctxn1 | 645.1595188 | -1.566328572 | 4.14E-13 |
| Rtp4 | 57.72055596 | -1.567985195 | 1.01E-06 |
| Ramp3 | 1853.838291 | -1.583126621 | 1.43E-06 |
| Mmp16 | 66.21102913 | -1.585047527 | 1.24E-02 |
| Gm10155 | 26.37968544 | -1.586280676 | 1.76E-02 |
| Prcd | 10.9762808 | -1.587096726 | 1.62E-02 |
| Gramd1c | 21.27151137 | -1.587334854 | 3.08E-03 |
| Bhlhe40 | 3834.220772 | -1.587870811 | 5.26E-11 |
| Gria1 | 276.9010458 | -1.589872672 | 1.97E-04 |
| Edil3 | 1365.668624 | -1.590752134 | 2.24E-06 |
| Angpt1 | 126.2034669 | -1.595399606 | 1.15E-02 |
| 2610035D17Rik | 96.21859554 | -1.610460521 | 6.89E-04 |
| Gm26552 | 14.03270238 | -1.611273059 | 3.76E-02 |
| Pla2g7 | 967.292464 | -1.612566836 | 1.48E-10 |
| Hr | 118.7699506 | -1.618814216 | 4.52E-03 |
| Hk1os | 14.68810408 | -1.624804859 | 2.54E-02 |
| Megf10 | 616.628578 | -1.631172474 | 2.13E-04 |
| Fam135b | 222.7442954 | -1.642985509 | 6.16E-03 |
| Gla | 4390.097848 | -1.649614892 | 2.64E-15 |
| Rab27a | 205.8409117 | -1.656009734 | 1.35E-09 |
| Nat8f4 | 100.8466667 | -1.656336918 | 5.03E-05 |
| Gm3194 | 17.06507034 | -1.657184428 | 1.46E-02 |
| Rpl27-ps3 | 6890.101037 | -1.658050702 | 1.68E-02 |
| Igf2os | 450.3914571 | -1.675123776 | 4.50E-04 |
| Nrep | 127.7232309 | -1.676842415 | 1.10E-03 |
| Rgs5 | 30.59780507 | -1.680666431 | 2.01E-04 |
| 10-Mar | 64.82310953 | -1.68652545 | 9.74E-04 |
| Pla2g2e | 18.47206542 | -1.686984179 | 4.30E-03 |
| St6gal1 | 1757.152116 | -1.690878575 | 1.23E-14 |

| | | | |
|---|---|---|---|
| Paqr8 | 1206.296549 | -1.69279605 | 3.92E-13 |
| Pde4b | 1859.321615 | -1.700499054 | 9.44E-08 |
| Slc2a6 | 121.7876624 | -1.701562033 | 7.37E-08 |
| Asb4 | 41.09296277 | -1.703348683 | 2.19E-02 |
| C330018A13Rik | 26.57426401 | -1.711408997 | 3.69E-03 |
| Il1rn | 36.407963 | -1.713844085 | 1.04E-02 |
| Tgtp2 | 74.54802873 | -1.71821481 | 3.39E-03 |
| Mtus2 | 261.3824621 | -1.718561658 | 2.07E-06 |
| Foxred2 | 88.56860549 | -1.721490534 | 5.79E-04 |
| Kif26b | 704.3016406 | -1.748290464 | 2.12E-08 |
| Plppr3 | 487.5192114 | -1.759749255 | 6.79E-07 |
| Masp1 | 1959.534601 | -1.764778757 | 6.47E-11 |
| Fabp5 | 6643.323886 | -1.769748941 | 2.75E-21 |
| Tmem268 | 2601.301169 | -1.783579559 | 4.13E-17 |
| Galnt9 | 9.989789218 | -1.801600648 | 2.66E-02 |
| Cd33 | 68.59571646 | -1.80893947 | 1.42E-05 |
| Prl2c2 | 22898.70079 | -1.8205421 | 1.30E-03 |
| Sohlh2 | 9.321089239 | -1.830762065 | 1.83E-02 |
| Serpina3n | 668.2845824 | -1.84093997 | 4.07E-07 |
| Rnase1 | 36.55767924 | -1.847891219 | 8.01E-04 |
| Dmrtc1c2 | 1708.160336 | -1.865254012 | 1.60E-02 |
| Slc9a4 | 10.64877529 | -1.871282313 | 2.17E-02 |
| Cd200 | 115.5327228 | -1.872698873 | 6.60E-06 |
| Lef1 | 146.4483945 | -1.875874724 | 1.58E-05 |
| Gjb3 | 262.9797057 | -1.890138448 | 1.01E-03 |
| Cela1 | 174.5106123 | -1.89202929 | 2.20E-07 |
| Cytip | 32.07785982 | -1.896243171 | 1.51E-02 |
| Has2 | 320.7677252 | -1.905089211 | 6.66E-09 |
| Scn7a | 62.80044543 | -1.907771598 | 1.17E-03 |
| Gjb4 | 54.22542267 | -1.909860957 | 2.61E-03 |
| Tecta | 36.05411746 | -1.910875663 | 3.42E-04 |
| Lenep | 70.28657422 | -1.91523418 | 3.23E-03 |
| Kcnab1 | 1245.884438 | -1.927357382 | 2.29E-16 |
| Mlana | 173.3382716 | -1.977771498 | 2.06E-03 |
| Gbp9 | 51.96187374 | -1.987919557 | 9.15E-10 |
| Rgs4 | 1205.292047 | -2.001333362 | 1.20E-03 |
| BC106179 | 76.11099145 | -2.025306178 | 7.96E-09 |
| Gm11992 | 6.530987242 | -2.04990858 | 4.40E-02 |
| Plekhb1 | 22.62480202 | -2.068235738 | 7.94E-03 |
| Osbpl6 | 120.9530659 | -2.068731687 | 8.07E-07 |
| Cfap161 | 19.83244907 | -2.071984753 | 2.40E-04 |
| Lingo2 | 18.63616195 | -2.081852034 | 1.62E-02 |
| Fabp4 | 102.7972834 | -2.081979135 | 7.60E-06 |
| Wfdc18 | 86.62622672 | -2.102451093 | 1.00E-03 |
| Slc30a1 | 484.7571357 | -2.107834057 | 4.39E-07 |
| Wfdc17 | 23.56673593 | -2.170915014 | 1.97E-02 |
| Tnfrsf11b | 121.1051387 | -2.189591416 | 3.61E-05 |

| | | | |
|---|---|---|---|
| Gbp4 | 14.23422782 | -2.223499674 | 8.81E-05 |
| Gm15640 | 32.6140749 | -2.235726781 | 1.10E-03 |
| Gm3629 | 23.23421655 | -2.236195151 | 2.19E-02 |
| Nat8 | 9.423743375 | -2.254701752 | 1.35E-03 |
| Epha10 | 7.545852402 | -2.276914206 | 1.30E-02 |
| Wdr63 | 7.18728154 | -2.289001959 | 1.04E-02 |
| Prtn3 | 13.25797432 | -2.307561388 | 5.81E-03 |
| Slco1a5 | 23.68395135 | -2.343886858 | 3.54E-03 |
| Flrt3 | 33.14674922 | -2.402030997 | 3.22E-03 |
| Has2os | 14.78403193 | -2.40291477 | 4.89E-05 |
| 4930589L23Rik | 9.662169631 | -2.482226181 | 2.14E-02 |
| Kcnk3 | 92.21429487 | -2.500916019 | 1.50E-07 |
| Tspan10 | 235.8434891 | -2.503434752 | 4.71E-02 |
| Ubd | 22.06665641 | -2.570031309 | 5.02E-03 |
| B630019A10Rik | 22.44071648 | -2.585527469 | 1.43E-05 |
| Gm11351 | 79.42538293 | -2.727084514 | 2.91E-03 |
| 1700071M16Rik | 6.45197741 | -2.752421347 | 4.88E-02 |
| Fbxl21 | 8.014840798 | -2.778490116 | 2.80E-02 |
| Fam19a1 | 75.29991681 | -2.823795849 | 2.10E-08 |
| Nmur1 | 13.54843957 | -2.833915352 | 1.92E-04 |
| Slc12a1 | 5.052709889 | -2.922248512 | 1.90E-02 |
| Gm22935 | 8.206150922 | -3.02144295 | 1.34E-02 |
| Klrb1a | 6.733824506 | -3.035509673 | 3.29E-03 |
| Mmp8 | 5.245523941 | -3.052710517 | 9.12E-03 |
| Tmem100 | 6.230469592 | -3.066775703 | 1.03E-02 |
| Pmp2 | 16.17721345 | -3.13823872 | 5.81E-03 |
| St6galnac2 | 30.52906661 | -3.551840969 | 1.66E-02 |
| 4931431C16Rik | 8.871351992 | -4.026756682 | 3.67E-02 |
| Sema3d | 11.73243349 | -4.465062325 | 9.16E-03 |
| Gm2366 | 40.57644542 | -5.439031973 | 8.08E-03 |
| Cd200r3 | 139.6286064 | -6.671983477 | 4.88E-03 |
| Samd11 | 155.349818 | -8.892620545 | 1.86E-04 |
| Sla2 | 584.6313518 | -9.115649297 | 1.22E-07 |
| Doc2a | 128.9404018 | -16.67154531 | 2.81E-08 |
| Gm15801 | 42.34653278 | -22.14891552 | 2.39E-12 |

| Gene | baseMean | log2 Fold Change | padj |
|---|---|---|---|
| S100b | 13.49662888 | 6.503580837 | 1.55E-08 |
| Nr1i3 | 116.7264137 | 6.189488688 | 7.08E-04 |
| Gm21293 | 16.49303806 | 5.817590587 | 3.30E-03 |
| Vwa5b2 | 28.56756157 | 4.92549064 | 6.11E-10 |
| NA | 26.06308258 | 4.747235538 | 3.80E-08 |
| Slc6a12 | 232.6272342 | 4.234934088 | 3.62E-02 |
| Gm38393 | 571.0226571 | 3.913480392 | 5.32E-41 |
| Lgr6 | 16.14818361 | 3.797623059 | 1.67E-03 |
| NA | 188.127415 | 3.51768053 | 1.11E-06 |
| Fmo3 | 67.82999797 | 3.452521569 | 1.49E-03 |
| Anxa8 | 232.8381859 | 2.839371007 | 2.44E-04 |
| Calcb | 38.52361346 | 2.714968434 | 8.56E-03 |
| Scn1b | 147.6156986 | 2.687818424 | 3.86E-21 |
| Tubb3 | 51.35408728 | 2.681935404 | 2.83E-09 |
| Peg3 | 3145.504318 | 2.609754001 | 5.86E-36 |
| Gpr149 | 14.67585469 | 2.449397492 | 2.05E-02 |
| Macc1 | 17.40048592 | 2.406559298 | 3.20E-02 |
| Dock8 | 275.2900409 | 2.405106229 | 1.83E-07 |
| Ripk3 | 33.31369269 | 2.378409845 | 5.26E-06 |
| Prtn3 | 13.25797432 | 2.361953846 | 3.76E-02 |
| Ces1g | 19.45501609 | 2.335761105 | 2.73E-03 |
| Enkur | 17.5217497 | 2.311759083 | 3.11E-02 |
| Klra4 | 79.2878235 | 2.308798228 | 1.12E-03 |
| Nr4a1 | 921.088903 | 2.296688594 | 5.30E-07 |
| Cacnb4 | 40.60664551 | 2.283695885 | 5.01E-04 |
| Ptprq | 158.5957609 | 2.207051318 | 2.08E-02 |
| Adgrg6 | 18.66629187 | 2.20676006 | 2.17E-04 |
| Klk8 | 65.88130734 | 2.195959074 | 4.33E-05 |
| Rarb | 39.25205665 | 2.176300361 | 9.44E-07 |
| Hes1 | 186.8574295 | 2.151051402 | 9.54E-04 |
| Nr4a3 | 32.35265595 | 2.072152091 | 5.57E-03 |
| Hmcn1 | 25.31932809 | 2.051003872 | 2.72E-02 |
| Camta1 | 704.1684855 | 2.01147948 | 2.47E-28 |
| Ica1l | 23.42769138 | 2.005672828 | 9.40E-03 |
| Nap1l2 | 52.4545836 | 1.988204963 | 9.15E-06 |
| Inmt | 199.4518932 | 1.976530434 | 2.22E-04 |
| Vnn1 | 94.96773462 | 1.957527799 | 8.57E-03 |
| Slc4a4 | 911.506914 | 1.95734803 | 3.24E-14 |
| Pmaip1 | 85.15518226 | 1.943020184 | 2.95E-04 |
| Mfsd6 | 21.51319353 | 1.941113648 | 2.62E-02 |
| 3830403N18Rik | 23.80216124 | 1.91364461 | 2.55E-03 |
| 2310002F09Rik | 28.54212603 | 1.908095552 | 7.24E-03 |
| Btn1a1 | 43.15252764 | 1.896456951 | 3.69E-02 |
| Ptgs2 | 3952.831909 | 1.889797344 | 1.44E-03 |
| Haglr | 19.13102496 | 1.852820954 | 3.06E-03 |
| Atg9b | 216.6082704 | 1.831805062 | 8.56E-03 |

| | | | |
|---|---|---|---|
| Klhl30 | 170.2009204 | 1.826457807 | 1.93E-03 |
| Dusp27 | 32.80363271 | 1.760041602 | 3.06E-02 |
| Ahi1 | 1541.580211 | 1.733112722 | 3.05E-23 |
| Rnf128 | 22.02821797 | 1.732107891 | 7.91E-03 |
| Thbs1 | 56074.01074 | 1.697239598 | 2.21E-05 |
| Gm6736 | 924.2982575 | 1.684970409 | 1.15E-04 |
| Ang2 | 58.05561455 | 1.677557173 | 7.86E-03 |
| Disp2 | 132.6711548 | 1.598962052 | 6.57E-08 |
| Akr1b3 | 27184.12226 | 1.59513333 | 4.85E-05 |
| Trim36 | 26.62347626 | 1.589099794 | 1.24E-02 |
| Npr3 | 1290.290208 | 1.584702376 | 3.12E-03 |
| Dhrs3 | 1200.156624 | 1.581044287 | 2.08E-03 |
| Angpt1 | 126.2034669 | 1.577806222 | 3.94E-02 |
| Ica1 | 26.09752958 | 1.546895896 | 2.40E-02 |
| Gm12840 | 2928.5719 | 1.532425004 | 3.82E-02 |
| Cst6 | 60.21863085 | 1.528263066 | 1.59E-03 |
| Gm12138 | 2019.192784 | 1.524823495 | 4.67E-05 |
| Fam57b | 17.95663098 | 1.516585851 | 2.23E-02 |
| Serpinb9b | 3482.508606 | 1.505633132 | 2.50E-03 |
| NA | 87.8527127 | 1.505516793 | 1.16E-02 |
| Mgat4a | 25.58784331 | 1.476568726 | 4.47E-02 |
| Mamdc2 | 140.4200342 | 1.459512653 | 1.79E-04 |
| Lims2 | 179.0157548 | 1.452414002 | 4.43E-03 |
| Slc6a15 | 33.49232237 | 1.436972552 | 9.22E-03 |
| Nes | 211.7096441 | 1.428568391 | 3.98E-02 |
| Cgnl1 | 1941.398104 | 1.41319176 | 2.68E-03 |
| Lynx1 | 1430.13896 | 1.400135518 | 1.05E-19 |
| Omd | 64.4193638 | 1.372935787 | 4.25E-02 |
| Adamts9 | 285.3258477 | 1.365509685 | 1.87E-03 |
| Aldh1a1 | 6563.728029 | 1.361655733 | 6.24E-03 |
| Rab27b | 133.4508688 | 1.356680718 | 2.55E-03 |
| Plscr2 | 362.8998898 | 1.342121814 | 3.28E-03 |
| Cavin2 | 13533.10625 | 1.34081656 | 2.24E-03 |
| Klf4 | 2395.995965 | 1.324909014 | 1.00E-10 |
| Camk2a | 97.06394336 | 1.316467462 | 2.05E-02 |
| Gm773 | 40.14795369 | 1.312799593 | 3.69E-02 |
| Plxna4 | 92.75828692 | 1.289389808 | 3.38E-02 |
| Magee1 | 2530.570754 | 1.271990732 | 3.69E-26 |
| Prorsd1 | 28.67614291 | 1.266920634 | 2.64E-02 |
| Tmeff2 | 105.2957531 | 1.26663485 | 3.76E-02 |
| Phospho1 | 692.1845389 | 1.251121581 | 9.43E-04 |
| Krt80 | 423.3382572 | 1.207372861 | 1.75E-03 |
| Gm20400 | 779.139461 | 1.202828332 | 2.40E-02 |
| Nexn | 586.7586746 | 1.190660897 | 4.80E-02 |
| Vmn1r-ps128 | 315.8095013 | 1.186893306 | 2.37E-02 |
| Gm12505 | 700.5275066 | 1.185445147 | 8.16E-03 |
| Gm26809 | 35780.51269 | 1.184791739 | 4.60E-03 |

| | | | |
|---|---|---|---|
| Adcy7 | 42.46266469 | 1.177316894 | 2.40E-02 |
| Tnfaip3 | 769.7438906 | 1.173921528 | 2.42E-04 |
| Gm17024 | 26.18378437 | 1.168974159 | 4.56E-02 |
| 4930486L24Rik | 435.5390485 | 1.166641713 | 2.60E-02 |
| Itgb7 | 197.2536301 | 1.166546466 | 1.99E-02 |
| Mpzl2 | 41.79260049 | 1.162338592 | 3.10E-02 |
| Slc30a1 | 484.7571357 | 1.15835384 | 3.87E-02 |
| Ifi47 | 69.23413701 | 1.14597771 | 3.43E-03 |
| Map1a | 6290.433199 | 1.130792558 | 1.97E-14 |
| Pcdh9 | 103.422577 | 1.130739002 | 3.71E-02 |
| Fetub | 34.84454848 | 1.123254734 | 3.67E-02 |
| Dusp4 | 2570.483965 | 1.121328638 | 3.78E-03 |
| Phlda1 | 939.9688801 | 1.120805485 | 6.75E-04 |
| Eml2 | 184.9655233 | 1.119855947 | 4.60E-03 |
| Grem1 | 116.2492527 | 1.10410049 | 7.08E-04 |
| Rin3 | 39.43852099 | 1.095352349 | 1.47E-02 |
| Styk1 | 452.3674977 | 1.08913071 | 5.51E-03 |
| Gm14137 | 67.19258941 | 1.088142526 | 7.44E-04 |
| Rasgrp3 | 117.9121061 | 1.087218114 | 2.90E-03 |
| Olfm1 | 251.9775343 | 1.08459836 | 4.29E-02 |
| Cystm1 | 169.2870408 | 1.064539279 | 4.47E-02 |
| Lamc2 | 395.7838555 | 1.060554799 | 2.59E-03 |
| Endod1 | 423.568897 | 1.059316981 | 2.03E-05 |
| 9930111J21Rik2 | 470.8629972 | 1.059235476 | 3.75E-02 |
| Crim1 | 4958.715791 | 1.049782367 | 1.91E-04 |
| Snta1 | 63.91870191 | 1.049417786 | 1.80E-02 |
| Gm19705 | 81.49688775 | 1.041242805 | 1.30E-02 |
| Xylb | 84.70507461 | 1.040383762 | 2.36E-04 |
| Slmapos2 | 3326.360358 | 1.029672773 | 5.51E-03 |
| Snhg11 | 55.41700975 | 1.029337044 | 4.96E-02 |
| Tril | 35.05868021 | 1.027040607 | 1.32E-02 |
| Sulf1 | 1797.810171 | 1.025625921 | 1.18E-02 |
| Gpat3 | 333.521046 | 1.025127147 | 2.95E-04 |
| Ly6a | 4142.015759 | 1.024382051 | 1.50E-03 |
| Slc40a1 | 380.186994 | 1.024250622 | 3.82E-02 |
| 9830144P21Rik | 73.20824837 | 1.018565128 | 3.25E-02 |
| Malat1 | 24971.98118 | 1.011455679 | 2.65E-02 |
| Gprc5a | 2003.573682 | 1.00338741 | 9.41E-03 |
| Clu | 185.6709907 | 1.00052181 | 2.81E-02 |
| Atp1b1 | 1462.526931 | 0.998610162 | 1.00E-04 |
| Ly6c1 | 1095.469361 | 0.99719783 | 6.34E-04 |
| Tgm2 | 2829.880322 | 0.993121151 | 5.18E-06 |
| Ppargc1a | 542.9043571 | 0.992563775 | 4.25E-02 |
| Thrb | 322.1208178 | 0.989483726 | 8.09E-03 |
| Spry4 | 239.4302629 | 0.983892011 | 7.20E-03 |
| Tnxb | 839.0427032 | 0.976819546 | 1.12E-07 |
| Gchfr | 290.9261129 | 0.973863941 | 8.11E-03 |

| | | | |
|---|---|---|---|
| Loxl4 | 984.284874 | 0.966247269 | 3.92E-03 |
| Crabp1 | 530.6832668 | 0.958942594 | 1.25E-02 |
| Nqo1 | 596.4674168 | 0.955968936 | 2.39E-02 |
| Gm14410 | 104.8186658 | 0.951658241 | 4.05E-02 |
| Etl4 | 562.100196 | 0.945074163 | 5.80E-03 |
| Fam198b | 10855.77158 | 0.935121092 | 1.05E-03 |
| Slc2a6 | 121.7876624 | 0.933233776 | 1.13E-02 |
| Dpysl3 | 7094.565781 | 0.931762817 | 4.41E-05 |
| Dusp1 | 2323.364801 | 0.929582717 | 3.29E-02 |
| Pitpnc1 | 829.297569 | 0.919755695 | 4.65E-03 |
| Hspa4l | 3271.537121 | 0.915309417 | 5.13E-03 |
| Dock4 | 204.615827 | 0.911722599 | 5.49E-04 |
| Ano3 | 117.0441969 | 0.908944552 | 1.01E-02 |
| Ercc6l2 | 120.8231153 | 0.908769108 | 9.45E-03 |
| Krt7 | 1408.919025 | 0.895210423 | 4.83E-02 |
| Plk3 | 880.6175897 | 0.875875106 | 1.05E-05 |
| Dmxl2 | 711.1668361 | 0.873656145 | 3.76E-02 |
| Clip4 | 144.42984 | 0.872149228 | 3.78E-03 |
| Sema5a | 3228.140088 | 0.870016747 | 1.81E-02 |
| Mir22hg | 1001.832139 | 0.865335566 | 6.37E-03 |
| Epha2 | 1944.113419 | 0.861772616 | 4.79E-02 |
| Specc1 | 3432.279109 | 0.861409023 | 1.24E-04 |
| Tinagl1 | 7597.988131 | 0.853785958 | 8.57E-03 |
| Stap2 | 79.90043653 | 0.846983597 | 1.33E-02 |
| Irak2 | 771.0028241 | 0.844894427 | 1.18E-05 |
| NA | 101.8655657 | 0.841739535 | 4.05E-02 |
| F8 | 92.05127566 | 0.841694956 | 4.78E-02 |
| Cpeb4 | 981.9575186 | 0.833101821 | 1.46E-02 |
| Slc7a2 | 1073.849895 | 0.827170497 | 3.14E-02 |
| Skil | 1068.678727 | 0.825645282 | 3.92E-03 |
| Filip1l | 2075.750886 | 0.820688202 | 4.19E-03 |
| Aox1 | 478.0590306 | 0.812211347 | 9.59E-04 |
| Col12a1 | 2667.384937 | 0.80889277 | 6.14E-07 |
| Cpeb3 | 346.758479 | 0.808221176 | 5.51E-04 |
| Mfap3l | 581.8453732 | 0.803371336 | 2.75E-02 |
| Mtmr10 | 1541.372356 | 0.79569639 | 3.87E-04 |
| Rhox5 | 444.7664169 | 0.794319735 | 4.49E-02 |
| Cpeb2 | 370.5002161 | 0.793095661 | 2.57E-03 |
| Herc3 | 407.8347654 | 0.789698949 | 2.73E-03 |
| Slc20a1 | 2917.806275 | 0.784922727 | 5.51E-04 |
| Ppfia3 | 146.6688839 | 0.782649072 | 2.33E-02 |
| Trim16 | 1755.34812 | 0.778454302 | 4.11E-04 |
| Prkcg | 214.7714828 | 0.773397003 | 2.09E-02 |
| Maoa | 11869.71092 | 0.771022729 | 1.14E-03 |
| Aim2 | 214.7586752 | 0.768407788 | 1.24E-02 |
| Tcaf2 | 513.7387912 | 0.766615134 | 2.08E-02 |
| Sorbs1 | 833.0862304 | 0.765270715 | 1.05E-03 |

| | | | |
|---|---|---|---|
| Pde8b | 714.9937037 | 0.765259995 | 1.01E-02 |
| Zfp53 | 773.8964099 | 0.7634673 | 3.83E-02 |
| Errfi1 | 2068.090918 | 0.762213202 | 8.56E-03 |
| Pqlc3 | 1007.222603 | 0.762196849 | 1.91E-04 |
| Prepl | 2110.742543 | 0.755638212 | 1.38E-14 |
| Cndp2 | 2438.22089 | 0.755565326 | 3.76E-02 |
| Gm10125 | 80.42503309 | 0.754366408 | 4.05E-02 |
| Tspyl4 | 789.2797587 | 0.744192505 | 3.92E-02 |
| Bdnf | 1365.54172 | 0.744091121 | 8.29E-03 |
| Ahr | 292.2985283 | 0.741438715 | 2.39E-02 |
| Nr4a2 | 174.1972914 | 0.740173875 | 2.73E-02 |
| Cthrc1 | 357.533666 | 0.735140942 | 3.84E-03 |
| Cacna1c | 477.6875855 | 0.730045591 | 8.73E-03 |
| Limk1 | 1848.013808 | 0.722703686 | 1.90E-03 |
| Dbndd2 | 321.9744503 | 0.705251234 | 4.29E-02 |
| Zrsr1 | 2272.804588 | 0.704388637 | 1.37E-02 |
| Ier3 | 749.7978944 | 0.701418185 | 4.96E-02 |
| NA | 1944.15319 | 0.699374312 | 1.50E-03 |
| Srsf5 | 9108.135251 | 0.693018434 | 2.51E-02 |
| Gsto2 | 540.3671304 | 0.691941521 | 2.58E-02 |
| Prr5 | 206.0395211 | 0.691256339 | 2.39E-02 |
| Acsl4 | 2428.085415 | 0.690541642 | 4.92E-02 |
| Kank1 | 1147.219222 | 0.689248862 | 1.18E-02 |
| Dst | 14726.69931 | 0.688945318 | 2.65E-03 |
| Fblim1 | 958.4750544 | 0.686703863 | 4.66E-03 |
| Prss23 | 12096.74038 | 0.685602468 | 1.93E-03 |
| Coq10b | 1114.821112 | 0.6855818 | 5.67E-03 |
| Gbe1 | 696.8299439 | 0.684735222 | 5.51E-03 |
| Hoxa4 | 123.1936353 | 0.683803318 | 2.49E-02 |
| Slco3a1 | 1020.790683 | 0.672203625 | 5.17E-03 |
| Map1b | 6864.42189 | 0.666967563 | 1.50E-03 |
| Slc22a23 | 1141.901598 | 0.664577789 | 2.99E-03 |
| Tecr | 7238.644524 | 0.662193478 | 1.00E-04 |
| Ptgs1 | 3977.955639 | 0.660528647 | 5.95E-03 |
| Anxa3 | 7482.75502 | 0.653436315 | 4.70E-02 |
| Dgke | 540.5581551 | 0.653140874 | 2.23E-05 |
| Nktr | 4432.706255 | 0.652589117 | 3.87E-02 |
| Hotairm1 | 163.7859472 | 0.650501716 | 3.71E-02 |
| Vps13d | 2923.790283 | 0.649926245 | 3.56E-03 |
| Zfp36l2 | 7867.303204 | 0.648001425 | 5.12E-03 |
| Haghl | 919.2093131 | 0.644405439 | 9.59E-04 |
| Ifit2 | 2698.970925 | 0.642706901 | 1.80E-02 |
| Man1a | 1574.812904 | 0.641725378 | 3.82E-02 |
| Usp53 | 923.2688407 | 0.637905326 | 3.26E-02 |
| Tsc22d2 | 3177.01915 | 0.637310728 | 1.43E-02 |
| NA | 83.4285938 | 0.636445473 | 3.06E-02 |
| Itga6 | 3086.584435 | 0.635115833 | 2.15E-03 |

| | | | |
|---|---|---|---|
| Abca5 | 1770.476615 | 0.62751859 | 4.33E-02 |
| Chuk | 2436.929155 | 0.617965809 | 7.20E-03 |
| Klf7 | 2129.134471 | 0.615162383 | 5.69E-03 |
| Lin7c | 1704.717814 | 0.609570009 | 3.84E-02 |
| Chfr | 2461.953026 | 0.603695715 | 8.98E-03 |
| Plod2 | 1368.669902 | 0.601427234 | 3.44E-02 |
| Akap12 | 11002.38031 | 0.600956963 | 3.65E-02 |
| Mtus1 | 3124.588945 | 0.600451345 | 4.26E-02 |
| Esyt1 | 2873.959478 | 0.596112279 | 3.71E-04 |
| Fuca2 | 3318.401005 | 0.594410321 | 8.29E-03 |
| Swap70 | 2724.175662 | 0.592597602 | 2.57E-03 |
| Net1 | 1218.273579 | 0.591565766 | 6.46E-03 |
| Efna5 | 479.7293286 | 0.590129356 | 2.95E-02 |
| Gdpd5 | 258.3174992 | 0.577501268 | 3.84E-02 |
| Irgm1 | 1281.382103 | 0.576327792 | 5.21E-04 |
| Ggact | 310.702016 | 0.574985489 | 4.44E-02 |
| Snapc5 | 308.3123639 | 0.568603934 | 4.42E-02 |
| Lrp1 | 57751.77543 | 0.565689826 | 1.42E-02 |
| Gsto1 | 9133.181255 | 0.565449938 | 3.37E-02 |
| Irf2bpl | 2111.470138 | 0.563462916 | 1.50E-03 |
| Abca3 | 2846.907726 | 0.561795279 | 7.45E-04 |
| Fam13b | 1672.285168 | 0.561109749 | 2.83E-02 |
| Lpp | 7312.736329 | 0.56107917 | 5.33E-03 |
| mt-Atp6 | 187651.4583 | 0.559456914 | 6.54E-03 |
| Anxa1 | 40517.67387 | 0.557069327 | 2.96E-03 |
| Sgms2 | 1191.807906 | 0.555720963 | 4.44E-03 |
| Fig4 | 2668.683324 | 0.54628047 | 1.06E-02 |
| Anxa6 | 6987.123827 | 0.543666625 | 2.94E-02 |
| Klc2 | 1334.112269 | 0.542918008 | 7.08E-04 |
| Abracl | 1310.993802 | 0.539983256 | 1.72E-02 |
| Osbpl8 | 1429.936323 | 0.539260677 | 4.00E-02 |
| Ppp1r12b | 4138.333762 | 0.536531491 | 8.12E-03 |
| Vcl | 15499.81698 | 0.531812541 | 8.06E-04 |
| Lactb2 | 1674.430397 | 0.529101256 | 2.39E-02 |
| Myo6 | 987.1695516 | 0.528318675 | 1.31E-02 |
| Nfat5 | 11458.97903 | 0.527196974 | 1.27E-02 |
| Flot1 | 2483.620995 | 0.524725814 | 2.40E-02 |
| Atp6v0a1 | 3975.629212 | 0.522019084 | 5.51E-04 |
| Impact | 6753.387895 | 0.520920949 | 2.03E-02 |
| Hoxa5 | 641.1710565 | 0.520326755 | 2.24E-02 |
| Txnrd1 | 11050.67123 | 0.519185977 | 3.92E-02 |
| Tmsb4x | 9263.718803 | 0.513274642 | 1.61E-03 |
| Ttc39b | 2306.341585 | 0.513076685 | 4.47E-02 |
| Fhl2 | 3219.077124 | 0.510111798 | 1.94E-02 |
| Nipa1 | 460.541922 | 0.510032704 | 2.11E-02 |
| Bcl2l1 | 4294.000393 | 0.499950645 | 2.97E-02 |
| Optn | 361.5681594 | 0.498256981 | 3.76E-02 |

| | | | |
|---|---|---|---|
| Etfrf1 | 953.9923609 | 0.497341234 | 1.51E-02 |
| Tjp1 | 4773.974161 | 0.496973549 | 1.98E-02 |
| Myo5a | 10004.30674 | 0.49085186 | 3.69E-02 |
| Tle4 | 360.9712993 | 0.489938217 | 3.70E-02 |
| Hipk2 | 2809.337586 | 0.488962321 | 1.75E-02 |
| Cldnd1 | 2104.132741 | 0.487368397 | 4.44E-03 |
| mt-Nd1 | 144168.3645 | 0.48704274 | 3.11E-02 |
| Cpt1a | 5815.364835 | 0.479620966 | 2.06E-04 |
| Ermp1 | 2303.615021 | 0.478618197 | 1.61E-02 |
| Acyp2 | 175.9484139 | 0.477440868 | 4.47E-02 |
| Myh10 | 12571.26723 | 0.471895437 | 1.37E-02 |
| Vegfa | 2252.329387 | 0.471454486 | 1.51E-02 |
| F2r | 8616.173877 | 0.471257001 | 2.52E-03 |
| Golga4 | 4342.991697 | 0.470179538 | 3.38E-02 |
| Tmod3 | 2418.238521 | 0.466616703 | 4.61E-02 |
| Sptan1 | 24705.76715 | 0.466202354 | 8.29E-03 |
| Birc6 | 8702.308943 | 0.465705253 | 3.71E-02 |
| Slmap | 3527.376361 | 0.465322733 | 3.35E-02 |
| Ahnak | 67844.71225 | 0.463943065 | 1.72E-02 |
| Armcx1 | 1688.155142 | 0.460766904 | 4.37E-02 |
| Tmf1 | 2643.356274 | 0.46066226 | 4.07E-02 |
| Hk2 | 3039.306829 | 0.459398273 | 3.94E-02 |
| Avpi1 | 1533.78618 | 0.459243404 | 2.33E-02 |
| Mef2a | 1661.812283 | 0.454987327 | 1.28E-02 |
| Flnb | 18993.59883 | 0.451455155 | 2.95E-02 |
| Afdn | 6938.094101 | 0.445322639 | 4.87E-02 |
| Pkd1 | 8593.732781 | 0.444524588 | 5.51E-04 |
| 5031425E22Rik | 290.5449219 | 0.443829339 | 2.90E-02 |
| Synj1 | 1193.434325 | 0.439461568 | 4.58E-02 |
| Pikfyve | 2011.636706 | 0.435672002 | 3.44E-02 |
| Csrp1 | 2952.741716 | 0.435054971 | 5.17E-03 |
| Cnn2 | 23457.26203 | 0.430091559 | 8.56E-03 |
| Lats2 | 3075.227714 | 0.429197922 | 4.05E-02 |
| Stx3 | 1465.82188 | 0.428910506 | 2.65E-02 |
| Tbk1 | 1987.788939 | 0.427410687 | 4.54E-02 |
| Cited2 | 5203.839702 | 0.420397404 | 3.76E-02 |
| Ehbp1l1 | 2904.929473 | 0.419071598 | 9.91E-03 |
| Ostm1 | 1935.936349 | 0.418950513 | 4.10E-02 |
| Prkdc | 1324.769727 | 0.416983759 | 3.87E-02 |
| Wwc2 | 6641.559661 | 0.411413913 | 1.47E-02 |
| Calm1 | 13158.15346 | 0.409230689 | 3.06E-02 |
| Eya1 | 701.8287389 | 0.406536864 | 1.07E-02 |
| mt-Co2 | 249817.806 | 0.406245563 | 4.76E-03 |
| Myh9 | 47052.31137 | 0.405066045 | 2.05E-02 |
| Pls3 | 9293.72303 | 0.401399771 | 5.51E-04 |
| Cast | 5248.563907 | 0.398800762 | 9.70E-04 |
| Runx1 | 1094.061865 | 0.397394074 | 1.01E-02 |

| | | | |
|---|---|---|---|
| Kif1b | 4905.83088 | 0.394955062 | 3.07E-03 |
| Yipf6 | 840.4900824 | 0.389695686 | 1.52E-02 |
| Pgd | 3686.528964 | 0.389675198 | 1.80E-02 |
| Wdr7 | 1610.402645 | 0.388420884 | 1.81E-02 |
| Ttc19 | 1089.857695 | 0.37830574 | 2.97E-02 |
| Abi1 | 4282.059818 | 0.376359647 | 3.49E-02 |
| Tpm1 | 28225.26008 | 0.372350217 | 2.09E-02 |
| Slc25a16 | 1028.92458 | 0.368819706 | 3.78E-02 |
| Golgb1 | 5482.119432 | 0.367260049 | 3.02E-02 |
| Picalm | 6753.449164 | 0.366149518 | 4.77E-02 |
| Lrrc58 | 3830.129758 | 0.365890934 | 3.24E-02 |
| Pdlim5 | 5330.342559 | 0.365746008 | 4.49E-02 |
| Osbpl9 | 6203.981448 | 0.356516911 | 7.20E-03 |
| Klf6 | 6415.94845 | 0.351055602 | 8.56E-03 |
| Slc26a2 | 908.0177184 | 0.350737811 | 4.66E-02 |
| Mus81 | 696.6969706 | 0.348537625 | 4.32E-03 |
| Pxk | 1881.676776 | 0.346891842 | 1.81E-03 |
| Epb41l3 | 11414.40963 | 0.346347892 | 5.17E-03 |
| Ehbp1 | 2247.789155 | 0.336643946 | 3.71E-02 |
| Stk38l | 824.657163 | 0.336586499 | 2.31E-02 |
| Als2 | 1499.841059 | 0.335009795 | 4.23E-02 |
| Mia3 | 7047.789822 | 0.332289413 | 3.63E-02 |
| Tulp4 | 16880.04521 | 0.327093978 | 2.95E-02 |
| Iqgap1 | 14617.56628 | 0.32217877 | 1.60E-02 |
| Nt5c2 | 1953.179617 | 0.319065494 | 4.53E-03 |
| Itprid2 | 2850.831262 | 0.317014952 | 3.77E-02 |
| Cdr2l | 1223.894987 | 0.314484888 | 8.11E-03 |
| Calm2 | 10854.30123 | 0.312725187 | 3.43E-02 |
| Gnaq | 3219.879092 | 0.311964283 | 1.52E-02 |
| Cdk11b | 2039.570556 | 0.310246189 | 2.18E-02 |
| Gpd2 | 3642.520661 | 0.309277283 | 2.05E-02 |
| Slc38a1 | 5119.465128 | 0.28840246 | 1.86E-02 |
| Ndufa4 | 6517.052841 | 0.282940989 | 1.07E-02 |
| Herc2 | 5066.067327 | 0.257451908 | 2.40E-02 |
| Reep3 | 3673.165162 | 0.253586147 | 4.15E-02 |
| Arf3 | 7135.48518 | 0.224435063 | 1.75E-02 |
| Cltc | 24544.13461 | 0.184388642 | 3.29E-02 |
| Kdm1a | 3091.893595 | -0.208496991 | 4.25E-02 |
| Zfp639 | 1185.948539 | -0.227643049 | 3.76E-02 |
| Dgkd | 3573.208434 | -0.228727512 | 4.96E-02 |
| Gmcl1 | 1500.19973 | -0.229642261 | 3.82E-02 |
| Sun1 | 4858.801632 | -0.232055317 | 3.73E-02 |
| Fam114a2 | 3449.452351 | -0.238018024 | 2.95E-02 |
| N4bp1 | 3589.986201 | -0.243985672 | 2.72E-02 |
| Zfp687 | 1806.062977 | -0.246550776 | 4.91E-02 |
| AI597479 | 1129.332889 | -0.258027566 | 8.98E-03 |
| Cdc23 | 2628.540815 | -0.260467477 | 1.05E-02 |

| | | | |
|---|---|---|---|
| Gng10 | 2800.643553 | -0.263714665 | 4.44E-02 |
| Zfp110 | 1694.471274 | -0.267103175 | 4.44E-03 |
| Dnmbp | 2254.506423 | -0.269470718 | 4.14E-02 |
| Msl1 | 3847.048481 | -0.272745208 | 4.40E-03 |
| Cdk10 | 1543.456391 | -0.276619967 | 3.76E-02 |
| Rnf145 | 4349.170828 | -0.289843531 | 2.64E-02 |
| Mbtps1 | 9771.008573 | -0.297762587 | 3.65E-02 |
| Srsf9 | 1778.892419 | -0.298830239 | 1.15E-02 |
| Zfp768 | 743.7030957 | -0.302688278 | 3.93E-02 |
| Pip4k2b | 4426.450694 | -0.303790704 | 9.74E-03 |
| Smo | 9359.591469 | -0.305696016 | 2.63E-02 |
| Cep68 | 2000.679055 | -0.305916128 | 3.60E-03 |
| Rbl2 | 2066.659329 | -0.309109674 | 6.70E-03 |
| Usp28 | 1013.741207 | -0.311111404 | 2.94E-02 |
| Carm1 | 4057.020687 | -0.312052799 | 3.91E-02 |
| Nbdy | 1309.915387 | -0.31903677 | 2.14E-02 |
| Wipf1 | 3029.779313 | -0.321542249 | 3.78E-03 |
| Dgcr2 | 5162.081899 | -0.322809703 | 3.82E-02 |
| Pcnx | 4490.427285 | -0.322876925 | 4.14E-02 |
| Evc2 | 3217.80212 | -0.326181076 | 8.56E-03 |
| Emp2 | 4062.72683 | -0.326343145 | 9.14E-03 |
| Exo5 | 1308.513939 | -0.328380874 | 4.44E-02 |
| Amer1 | 1350.755714 | -0.32841508 | 3.00E-03 |
| Rcbtb2 | 2432.790428 | -0.331598358 | 4.21E-02 |
| 9530068E07Rik | 9957.757091 | -0.334748621 | 4.24E-02 |
| Poglut1 | 2986.878341 | -0.33536079 | 2.60E-02 |
| Rsad1 | 456.0324608 | -0.336743878 | 4.36E-02 |
| St3gal2 | 8902.023726 | -0.337760764 | 1.73E-02 |
| Mrpl9 | 1944.256036 | -0.338077981 | 1.37E-02 |
| Vars2 | 1444.707556 | -0.340176642 | 3.94E-02 |
| Tmem201 | 1116.37731 | -0.340409786 | 2.05E-02 |
| Anp32a | 2932.882658 | -0.341100151 | 2.94E-02 |
| Mettl2 | 1593.731798 | -0.345546907 | 2.05E-02 |
| Cdk19 | 2233.948062 | -0.346754618 | 6.69E-03 |
| Tmem206 | 879.2146497 | -0.347116127 | 4.42E-02 |
| Zc4h2 | 400.4251748 | -0.34858226 | 3.11E-02 |
| Kdelc1 | 1895.28066 | -0.349331632 | 9.40E-03 |
| Zfp251 | 1144.446552 | -0.349336624 | 1.61E-03 |
| Adipor2 | 3974.141308 | -0.352324231 | 2.05E-02 |
| Tbcel | 1611.699035 | -0.352718792 | 2.15E-03 |
| Xpnpep3 | 989.0992619 | -0.35332511 | 1.34E-03 |
| Klhdc3 | 2766.400916 | -0.354670325 | 4.14E-02 |
| Zfp790 | 587.5727377 | -0.356947829 | 1.97E-02 |
| Ick | 1414.685462 | -0.356960898 | 2.95E-02 |
| Sf3b3 | 10639.11885 | -0.357245788 | 4.88E-02 |
| Gypc | 1383.765939 | -0.358797256 | 3.76E-02 |
| Capn15 | 724.6406254 | -0.360281248 | 2.72E-02 |

| | | | |
|---|---|---|---|
| Akt2 | 5161.192976 | -0.360958597 | 4.85E-02 |
| Usb1 | 1737.627401 | -0.364436204 | 4.16E-02 |
| Gtf3c4 | 3125.86937 | -0.365624823 | 3.82E-02 |
| Erbb2 | 2761.808853 | -0.36917219 | 2.90E-02 |
| Mllt1 | 2904.39071 | -0.36952733 | 1.37E-02 |
| Rpl31 | 2778.260788 | -0.371264434 | 3.76E-02 |
| Zfp12 | 1260.57049 | -0.372208845 | 1.13E-02 |
| Rpusd4 | 523.4815444 | -0.376126834 | 1.48E-02 |
| Med31 | 305.7132664 | -0.379204969 | 4.62E-02 |
| Gm29394 | 1296.624358 | -0.380347785 | 4.17E-02 |
| Fbxl19 | 1033.975476 | -0.381009277 | 2.83E-02 |
| Maz | 5576.002949 | -0.383247605 | 2.30E-02 |
| S1pr2 | 3510.869616 | -0.383371971 | 3.47E-02 |
| Zfp30 | 422.1195397 | -0.3835842 | 3.91E-02 |
| Chtf8 | 5434.622375 | -0.383958857 | 3.18E-02 |
| Diablo | 1748.100172 | -0.389124979 | 4.84E-03 |
| Txnl4b | 388.3308839 | -0.38922832 | 2.84E-02 |
| Usp21 | 1961.571321 | -0.389340422 | 4.60E-03 |
| Gid4 | 1936.274946 | -0.389526064 | 5.19E-03 |
| Thumpd2 | 335.1465948 | -0.390730375 | 8.11E-03 |
| Spire2 | 545.5638821 | -0.395109234 | 4.81E-02 |
| Klhl22 | 5015.7865 | -0.396068659 | 1.05E-02 |
| Rara | 1780.058804 | -0.397722341 | 3.69E-02 |
| Selenon | 4253.1088 | -0.399057129 | 2.72E-02 |
| Fam53c | 2241.692611 | -0.403046496 | 2.42E-04 |
| Lrrc42 | 3350.571451 | -0.40543601 | 2.03E-02 |
| Fam53a | 1630.046894 | -0.408364899 | 1.23E-02 |
| Pycr2 | 1798.030111 | -0.408923595 | 4.45E-02 |
| Cebpg | 2322.035649 | -0.409002588 | 1.48E-03 |
| Rpia | 1112.425502 | -0.411266565 | 2.11E-02 |
| Zfp93 | 364.0180944 | -0.412862067 | 1.95E-02 |
| Cyp39a1 | 736.9729686 | -0.413168985 | 2.40E-02 |
| Arhgef2 | 10247.78531 | -0.413713179 | 4.15E-02 |
| Parp12 | 537.1403291 | -0.414865272 | 2.93E-02 |
| Dtx3l | 451.8181588 | -0.41610539 | 1.90E-02 |
| Klhl8 | 432.6864808 | -0.418034139 | 3.01E-02 |
| Ankrd6 | 596.1378015 | -0.422825182 | 4.62E-02 |
| Edc3 | 1285.870104 | -0.424027109 | 1.01E-02 |
| Josd1 | 4707.625447 | -0.42531597 | 1.05E-03 |
| Tap2 | 1573.668119 | -0.429321498 | 4.44E-02 |
| Irf2 | 1439.051402 | -0.429345444 | 1.40E-02 |
| AC149090.1 | 18121.43559 | -0.429897771 | 9.58E-03 |
| Pdzd4 | 1139.545824 | -0.43138028 | 1.16E-02 |
| Rnf135 | 468.7873319 | -0.435853049 | 1.05E-02 |
| Zfp316 | 1016.687164 | -0.436835798 | 4.29E-02 |
| Lpcat1 | 1186.570369 | -0.436991608 | 4.72E-03 |
| Speg | 1856.726043 | -0.437592166 | 3.85E-02 |

| | | | |
|---|---|---|---|
| Tango6 | 700.5811635 | -0.438908772 | 3.80E-02 |
| Thop1 | 1815.914219 | -0.439138588 | 1.92E-02 |
| Fam219b | 604.1462657 | -0.440073732 | 3.69E-02 |
| Oard1 | 738.8061374 | -0.44065814 | 8.57E-03 |
| 2510002D24Rik | 276.2069138 | -0.440755788 | 4.05E-02 |
| Vegfc | 674.5119938 | -0.44337532 | 1.59E-02 |
| Tmem39b | 450.7819741 | -0.444041285 | 4.05E-02 |
| Clip2 | 3685.485315 | -0.444644704 | 2.59E-02 |
| Manbal | 1400.09717 | -0.44712901 | 4.42E-02 |
| Parp16 | 205.5236357 | -0.448708156 | 3.87E-02 |
| Zfp706 | 4663.743055 | -0.451021214 | 8.56E-03 |
| Zfyve21 | 755.2797833 | -0.451669157 | 2.58E-02 |
| Nt5m | 402.8418024 | -0.457481728 | 3.22E-02 |
| Ncs1 | 2519.874797 | -0.458538276 | 1.37E-02 |
| 4933427D14Rik | 1047.149216 | -0.458812885 | 2.94E-02 |
| Poli | 459.5423416 | -0.460117908 | 8.06E-03 |
| Mta1 | 6436.498902 | -0.462018532 | 9.91E-03 |
| Iqcc | 667.0780684 | -0.464899084 | 1.07E-02 |
| Ccdc117 | 1297.355029 | -0.465851311 | 8.25E-03 |
| Zfp874b | 473.1071347 | -0.469396947 | 2.59E-03 |
| H2-M3 | 439.5141547 | -0.4709925 | 4.23E-02 |
| Osgep | 1715.128448 | -0.472009111 | 2.24E-02 |
| Ddx39b | 10038.84561 | -0.478870589 | 1.22E-02 |
| Rcc2 | 9407.022186 | -0.47892696 | 8.56E-03 |
| Rxrb | 2724.534752 | -0.480483753 | 4.61E-03 |
| Serpinh1 | 36919.64396 | -0.481185862 | 4.15E-02 |
| Ttc7 | 1120.005252 | -0.481786036 | 2.93E-02 |
| Prdx4 | 3691.055708 | -0.482419682 | 3.22E-02 |
| Zfp324 | 561.8524386 | -0.483216515 | 1.34E-03 |
| Dguok | 1124.366048 | -0.483443033 | 1.20E-02 |
| Nfya | 2749.990945 | -0.48371685 | 5.17E-03 |
| Ppic | 4106.544221 | -0.48437534 | 3.66E-03 |
| Mxi1 | 1578.380318 | -0.485687071 | 2.08E-02 |
| Olfml3 | 3715.895388 | -0.486448454 | 4.36E-02 |
| Exosc5 | 597.2657099 | -0.48922328 | 3.77E-02 |
| Prkca | 3873.741095 | -0.49021458 | 3.40E-03 |
| Pex26 | 694.5375104 | -0.492908469 | 6.55E-03 |
| Lclat1 | 2716.89391 | -0.493475909 | 2.25E-04 |
| Efhd2 | 1565.299968 | -0.498112516 | 1.61E-02 |
| C1qtnf6 | 2407.485434 | -0.498361752 | 1.92E-02 |
| Dlg4 | 3652.121914 | -0.500893346 | 1.45E-02 |
| Scfd2 | 558.8131694 | -0.503338405 | 1.31E-02 |
| Bcl7c | 1233.600283 | -0.503891326 | 1.43E-02 |
| Sox12 | 1777.455189 | -0.505273717 | 2.75E-02 |
| Hs6st1 | 2054.227925 | -0.505429533 | 2.68E-02 |
| Maf1 | 2954.734566 | -0.506483854 | 3.11E-02 |
| Adamtsl4 | 642.4400238 | -0.508257138 | 3.76E-02 |

| | | | |
|---|---|---|---|
| Ptprn | 7409.737366 | -0.51096788 | 6.52E-04 |
| Khnyn | 1116.250918 | -0.511370922 | 5.80E-03 |
| Galnt13 | 425.0375608 | -0.514549652 | 4.30E-02 |
| Noc4l | 1562.446352 | -0.514759079 | 1.46E-02 |
| Zc3h10 | 826.6331516 | -0.517810318 | 1.26E-02 |
| Med22 | 2582.347251 | -0.519419498 | 5.57E-03 |
| Exog | 327.807065 | -0.5208163 | 2.97E-02 |
| Sema4c | 1046.850603 | -0.520899947 | 8.06E-03 |
| Tmem218 | 321.622946 | -0.520950368 | 3.36E-02 |
| Sbsn | 1060.826381 | -0.523961767 | 2.03E-02 |
| Narf | 1213.76747 | -0.528088058 | 4.10E-02 |
| Slx1b | 369.2781855 | -0.528165787 | 7.17E-05 |
| Ttc23 | 768.0262767 | -0.528366957 | 4.73E-02 |
| Rps13-ps2 | 21055.30253 | -0.529096502 | 4.87E-02 |
| Rps15a | 30497.77941 | -0.529941465 | 4.31E-02 |
| Bcl7a | 1302.545937 | -0.532660419 | 2.82E-04 |
| Coq8b | 489.0724424 | -0.532699212 | 4.29E-02 |
| Hmgn2 | 1321.386804 | -0.532778104 | 3.65E-02 |
| Zscan22 | 382.3077886 | -0.533364545 | 8.05E-03 |
| Plxnd1 | 6200.522837 | -0.535743894 | 4.10E-02 |
| Mmp19 | 2149.521448 | -0.536498205 | 1.61E-02 |
| Micall2 | 1659.531975 | -0.537173449 | 4.42E-02 |
| Gm11518 | 16388.69852 | -0.542903777 | 2.05E-02 |
| Tdrkh | 618.162234 | -0.553398801 | 1.46E-02 |
| Loxl1 | 13086.55011 | -0.555130391 | 2.42E-02 |
| Prdm15 | 720.4881743 | -0.555472961 | 4.87E-02 |
| Naip5 | 268.7274309 | -0.556224179 | 1.50E-02 |
| Zfp688 | 338.1326918 | -0.557500386 | 3.94E-02 |
| Cbx2 | 830.0959647 | -0.558965665 | 3.61E-02 |
| Alkbh6 | 589.5487126 | -0.563694435 | 2.05E-02 |
| Tubb2b | 2259.346938 | -0.565166469 | 2.66E-02 |
| Zfp647 | 141.5050878 | -0.568999786 | 4.31E-02 |
| Atf5 | 11430.63398 | -0.569362405 | 2.45E-03 |
| Itpk1 | 518.1719181 | -0.570191641 | 2.95E-02 |
| Zfp395 | 1588.050862 | -0.571782409 | 2.05E-02 |
| Impdh2-ps | 1489.025213 | -0.573665347 | 1.12E-03 |
| Ormdl3 | 2615.743301 | -0.574851377 | 1.45E-02 |
| Pcsk4 | 254.3779611 | -0.575410046 | 2.47E-02 |
| Mab21l1 | 801.0403118 | -0.578479993 | 3.00E-03 |
| Samd14 | 1239.574654 | -0.578908928 | 3.71E-03 |
| Gm4737 | 8568.322965 | -0.580152473 | 2.40E-02 |
| Rtkn | 1336.449565 | -0.580451107 | 1.84E-02 |
| Harbi1 | 406.9577452 | -0.580963436 | 2.27E-03 |
| Impdh2 | 6001.17937 | -0.583129743 | 3.49E-03 |
| Cdk18 | 2832.68387 | -0.585818534 | 4.83E-02 |
| Sh3bgrl3 | 2819.46658 | -0.588647968 | 3.55E-02 |
| Adam19 | 3337.629642 | -0.590542166 | 9.54E-04 |

| | | | |
|---|---|---|---|
| Pdcd4 | 2480.242851 | -0.59475437 | 2.35E-02 |
| Ccng2 | 2087.349482 | -0.595352611 | 3.69E-02 |
| Gm10182 | 1077.283482 | -0.599191367 | 1.75E-04 |
| Mepce | 1758.836327 | -0.602045043 | 9.40E-03 |
| Fam171a2 | 1340.633396 | -0.602328407 | 1.32E-02 |
| Zfp174 | 256.8040551 | -0.616207776 | 3.76E-02 |
| Rgs3 | 1513.56933 | -0.617801907 | 4.56E-02 |
| Zbtb5 | 524.8705552 | -0.620603464 | 3.25E-04 |
| Acot1 | 957.7954011 | -0.620678738 | 3.89E-02 |
| Agpat2 | 111.9244998 | -0.621068088 | 4.54E-02 |
| Cmss1 | 343.7516773 | -0.622983795 | 3.96E-03 |
| Tcf7l1 | 1795.450597 | -0.623991287 | 3.82E-02 |
| Jcad | 4643.802879 | -0.625746035 | 1.50E-03 |
| Zfp775 | 415.5759076 | -0.629132083 | 1.84E-02 |
| Nfkbie | 307.6811974 | -0.629615449 | 3.52E-02 |
| Aplf | 766.3187187 | -0.630607329 | 1.79E-04 |
| Hivep3 | 515.0723489 | -0.631990068 | 8.56E-03 |
| Lhx6 | 339.1144646 | -0.633920081 | 3.87E-02 |
| Igsf8 | 3977.096197 | -0.634984084 | 1.99E-02 |
| Trim45 | 432.7742059 | -0.641203875 | 7.72E-04 |
| Ucp2 | 1083.734914 | -0.645777782 | 1.95E-02 |
| Lxn | 2852.121736 | -0.649154534 | 4.56E-02 |
| Cd34 | 10535.55549 | -0.649415961 | 1.34E-03 |
| Ociad2 | 337.1734645 | -0.650810702 | 1.22E-02 |
| Irx1 | 1133.860938 | -0.653215895 | 3.08E-02 |
| Sh3bp2 | 1443.810206 | -0.661300144 | 1.81E-02 |
| Il13ra1 | 219.0472217 | -0.663522237 | 1.59E-03 |
| Lsp1 | 852.1553722 | -0.6642603 | 1.69E-02 |
| Efna3 | 212.0063188 | -0.666967904 | 4.60E-02 |
| Rgs20 | 444.9216063 | -0.667194517 | 4.44E-03 |
| Otub2 | 372.7939584 | -0.675927047 | 1.16E-02 |
| Timp1 | 3421.454883 | -0.679039453 | 2.46E-02 |
| Pck2 | 4321.410982 | -0.684950231 | 5.81E-03 |
| Fam131b | 2012.653011 | -0.687455171 | 6.65E-04 |
| Bgn | 97843.4361 | -0.690847791 | 1.29E-02 |
| Pla2g15 | 2388.719268 | -0.691602121 | 9.43E-04 |
| Cnih2 | 222.9740135 | -0.696065421 | 3.82E-02 |
| Dennd2a | 5263.659133 | -0.698671115 | 1.50E-03 |
| Il11ra1 | 3333.985703 | -0.702955547 | 1.80E-02 |
| Trim46 | 801.3602403 | -0.703996252 | 2.59E-03 |
| Sipa1l2 | 3528.093693 | -0.704389702 | 6.63E-03 |
| Proser3 | 387.0409439 | -0.708110156 | 3.98E-02 |
| Ralgds | 5291.738528 | -0.708719058 | 2.03E-02 |
| Zfat | 598.1955488 | -0.719739925 | 2.39E-02 |
| Fbxo41 | 890.8379166 | -0.721379286 | 6.94E-04 |
| Slc29a1 | 1079.904629 | -0.723004515 | 2.83E-02 |
| Smpd3 | 867.4805472 | -0.72360777 | 8.56E-03 |

| | | | |
|---|---|---|---|
| Emb | 6148.231165 | -0.731411372 | 2.57E-03 |
| Tnfaip6 | 646.6209697 | -0.732977987 | 3.42E-02 |
| Gli2 | 250.1196828 | -0.744434133 | 3.11E-02 |
| Zfas1 | 5923.00628 | -0.749056662 | 2.19E-02 |
| Cdon | 2024.016055 | -0.750877465 | 3.65E-02 |
| Lysmd1 | 338.1310677 | -0.75691653 | 7.01E-04 |
| Il1r1 | 1964.698977 | -0.757231112 | 6.30E-03 |
| Zfp128 | 209.8468391 | -0.757769714 | 2.25E-04 |
| Scyl2 | 2384.37133 | -0.758847616 | 4.64E-02 |
| Tfap4 | 806.8709696 | -0.761519042 | 2.59E-03 |
| Tmem45a | 630.8650328 | -0.761694084 | 2.97E-02 |
| Cln8 | 3141.229382 | -0.762271766 | 3.89E-03 |
| Ccl7 | 1521.615401 | -0.764272211 | 4.22E-02 |
| Mtss1l | 4009.15688 | -0.766027842 | 5.51E-04 |
| Socs1 | 288.2224024 | -0.77535644 | 9.14E-03 |
| Kctd21 | 352.0213232 | -0.783102713 | 2.65E-03 |
| Fam13c | 1244.440345 | -0.785062831 | 1.64E-02 |
| Sdc3 | 1488.655788 | -0.791097995 | 5.52E-03 |
| Rhoj | 1534.443716 | -0.791864949 | 7.76E-04 |
| Ahdc1 | 1254.188811 | -0.793406994 | 5.51E-04 |
| Pcdhb9 | 216.2214447 | -0.796845338 | 1.63E-02 |
| Irgm2 | 164.2745168 | -0.799937619 | 2.05E-02 |
| Col6a1 | 53797.49195 | -0.801421616 | 2.35E-02 |
| Xaf1 | 354.2355811 | -0.819627984 | 1.20E-02 |
| 2410006H16Rik | 10631.39287 | -0.821097576 | 4.22E-02 |
| Tfap2a | 730.9542063 | -0.821168418 | 8.03E-03 |
| Pvt1 | 2024.36745 | -0.827795193 | 9.13E-07 |
| Gas7 | 13591.79219 | -0.828004585 | 5.13E-03 |
| H2-Q4 | 205.1152131 | -0.82810535 | 3.33E-03 |
| Lbh | 5291.647432 | -0.832323343 | 3.87E-02 |
| Ptk7 | 1571.76109 | -0.848384663 | 3.15E-04 |
| Myo7a | 1099.229425 | -0.851232383 | 1.40E-03 |
| Snhg12 | 1711.852806 | -0.851850971 | 1.50E-03 |
| Cacnb1 | 471.9894397 | -0.8543559 | 2.31E-04 |
| Adamts2 | 9146.817329 | -0.862229385 | 5.96E-03 |
| Gbp2 | 472.493555 | -0.896821433 | 1.36E-03 |
| Sulf2 | 1429.615205 | -0.906544955 | 1.62E-02 |
| Begain | 238.4066239 | -0.907028947 | 1.81E-02 |
| Lrp4 | 529.2244838 | -0.907151161 | 4.63E-02 |
| Tlr2 | 500.01692 | -0.92490658 | 1.50E-03 |
| Rbp1 | 2005.260463 | -0.925326846 | 4.52E-06 |
| Parp14 | 278.3693229 | -0.925389915 | 3.92E-03 |
| Mex3a | 2190.570495 | -0.927375552 | 2.13E-03 |
| Adgrg1 | 72.72172502 | -0.930387604 | 4.44E-02 |
| Igtp | 114.2451841 | -0.931856012 | 1.52E-02 |
| Enox1 | 399.1181144 | -0.945038502 | 6.12E-04 |
| H2-T24 | 90.28178638 | -0.946924773 | 2.60E-02 |

| | | | |
|---|---|---|---|
| Efna1 | 1322.865678 | -0.948646878 | 3.18E-02 |
| Plpp7 | 57.83169076 | -0.962646914 | 1.31E-02 |
| A330074K22Rik | 266.6373111 | -0.965046304 | 7.00E-03 |
| Rnase10 | 47.92102259 | -0.967828041 | 4.42E-02 |
| Adamts14 | 1745.125069 | -0.973130052 | 4.44E-03 |
| Rdh12 | 38.3276781 | -0.981374495 | 3.11E-02 |
| Il18 | 182.5136154 | -1.006647704 | 1.01E-02 |
| P4ha3 | 952.268623 | -1.007228604 | 4.26E-03 |
| Dlgap2 | 221.2029046 | -1.010949268 | 2.68E-03 |
| Cercam | 171.2368976 | -1.011859892 | 4.32E-03 |
| Igsf10 | 1498.548461 | -1.019373872 | 1.24E-02 |
| Ifi27l2a | 63.9925599 | -1.022311786 | 3.78E-02 |
| H2-DMb2 | 96.6450081 | -1.028705807 | 2.88E-03 |
| Gbp9 | 51.96187374 | -1.0515168 | 3.06E-02 |
| Cd82 | 121.0116936 | -1.067537634 | 8.01E-04 |
| Zfp41 | 1083.240847 | -1.070522087 | 9.91E-03 |
| Nat8f1 | 120.6153721 | -1.079797891 | 3.87E-02 |
| H2-Q7 | 211.7583553 | -1.086982732 | 2.74E-03 |
| Sardh | 902.4568354 | -1.087215636 | 3.72E-04 |
| Csrp2 | 1603.401949 | -1.088557084 | 5.98E-03 |
| Gdpd2 | 416.9752147 | -1.08994436 | 4.81E-02 |
| Nuggc | 50.09471182 | -1.103307906 | 4.10E-02 |
| Boc | 364.9113936 | -1.106999166 | 9.50E-03 |
| Ltbp4 | 1251.339727 | -1.11157052 | 1.89E-05 |
| Gm15280 | 482.2429882 | -1.132126387 | 4.39E-06 |
| Sh2d3c | 108.5494914 | -1.139847107 | 2.55E-03 |
| Atp1a3 | 132.1001762 | -1.156735853 | 4.03E-02 |
| Cd200 | 115.5327228 | -1.18131646 | 3.11E-02 |
| Col6a2 | 28456.27254 | -1.197492376 | 2.87E-03 |
| Il17rd | 972.8019314 | -1.200612844 | 1.91E-04 |
| Has2 | 320.7677252 | -1.273083073 | 3.00E-03 |
| Lrrc17 | 862.6812303 | -1.279102363 | 6.81E-03 |
| Glis1 | 258.3571055 | -1.280725813 | 7.98E-03 |
| Clec2d | 2151.763597 | -1.298378816 | 1.30E-04 |
| Cdsn | 1502.659716 | -1.347029493 | 1.18E-06 |
| Adam12 | 2470.942642 | -1.360839039 | 9.29E-05 |
| Fcrlb | 50.76172523 | -1.370177679 | 3.69E-02 |
| Smoc1 | 319.0475636 | -1.371918734 | 3.48E-02 |
| NA | 35.3883632 | -1.399594989 | 1.95E-02 |
| Col5a3 | 679.7155384 | -1.418225531 | 3.25E-04 |
| Sfrp2 | 592.0206281 | -1.426107347 | 3.60E-03 |
| Nxph3 | 56.41829472 | -1.448089904 | 2.88E-03 |
| Mapkapk3 | 612.1223251 | -1.475300166 | 1.96E-08 |
| H2-K2 | 70.25926091 | -1.477459472 | 4.51E-04 |
| Fam135b | 222.7442954 | -1.493149776 | 4.66E-02 |
| Mdfi | 37.12431873 | -1.508000136 | 3.55E-03 |
| Prokr1 | 338.3992619 | -1.51347897 | 1.13E-04 |

| | | | |
|---|---|---|---|
| Timp4 | 80.55768285 | -1.519952165 | 4.80E-02 |
| Itga11 | 1425.902456 | -1.540776717 | 1.14E-03 |
| Trabd2b | 1980.320248 | -1.547363024 | 1.21E-05 |
| Scara5 | 982.0551509 | -1.551252743 | 4.81E-02 |
| Plekha4 | 436.889388 | -1.554304117 | 2.64E-05 |
| Gm7361 | 115.296168 | -1.570906002 | 1.41E-02 |
| Rgcc | 230.2419975 | -1.573104471 | 1.73E-05 |
| Ednra | 698.5583548 | -1.573569375 | 2.64E-05 |
| Txnip | 34225.99991 | -1.598577709 | 1.40E-02 |
| Ccn5 | 3412.349025 | -1.612798952 | 4.54E-02 |
| Oaz3 | 21.83637891 | -1.626765247 | 1.14E-02 |
| Gria1 | 276.9010458 | -1.629557629 | 1.08E-03 |
| Nlrp1b | 59.85443185 | -1.637236991 | 5.58E-06 |
| Aspg | 128.0287642 | -1.66162218 | 2.16E-02 |
| Depp1 | 214.6965571 | -1.739234117 | 1.59E-03 |
| Hr | 118.7699506 | -1.790322633 | 1.26E-02 |
| Dlk1 | 29943.381 | -1.822310583 | 1.87E-05 |
| Scn7a | 62.80044543 | -1.850566742 | 1.05E-02 |
| Cacna1g | 708.8000386 | -1.883425237 | 2.22E-04 |
| Trem2 | 96.77326875 | -1.910767479 | 1.21E-06 |
| Serpina3n | 668.2845824 | -1.942557669 | 3.01E-06 |
| Rgs4 | 1205.292047 | -1.95567626 | 8.08E-03 |
| Asb4 | 41.09296277 | -2.015590172 | 2.05E-02 |
| Notch1 | 117.642874 | -2.058630448 | 2.22E-04 |
| Scg2 | 1103.15175 | -2.077200093 | 2.28E-02 |
| Gfra2 | 25.3859268 | -2.11136736 | 8.06E-03 |
| Postn | 49906.43674 | -2.115868437 | 8.01E-04 |
| Lingo2 | 18.63616195 | -2.193835237 | 4.36E-02 |
| Tmem176b | 522.89459 | -2.224774847 | 6.86E-13 |
| Tmprss6 | 29.40343431 | -2.256185568 | 1.62E-02 |
| C3 | 1005.759055 | -2.263729938 | 3.80E-06 |
| Rarres2 | 54.65008237 | -2.288947317 | 3.60E-03 |
| Tmem176a | 167.3003744 | -2.407122204 | 4.74E-11 |
| H19 | 1822.874962 | -2.533732551 | 2.30E-02 |
| Islr | 180.9664839 | -2.546163033 | 1.30E-04 |
| Nat8f3 | 96.81673939 | -2.566093202 | 1.01E-02 |
| Hhatl | 22.27895666 | -2.59132803 | 1.02E-02 |
| Lhx8 | 17.10623604 | -3.673059398 | 1.33E-02 |
| Mmp9 | 324.7980202 | -3.999111299 | 7.19E-07 |
| Gbp11 | 15.68366756 | -4.174773683 | 3.76E-02 |
| Serpina3h | 32.62380238 | -4.532221307 | 9.13E-07 |
| AI606473 | 18.78385635 | -4.657829406 | 3.30E-04 |
| Gpr26 | 12.44590916 | -4.887231053 | 1.61E-02 |
| Serpina3g | 65.50415713 | -5.069140553 | 2.84E-06 |
| Card14 | 94.21879325 | -6.240865448 | 1.00E-04 |
| Smarca5-ps | 60.03761091 | -6.367283867 | 4.42E-02 |
| Abcb1a | 30.36830819 | -6.373965402 | 2.59E-03 |

| | | | |
|---|---|---|---|
| Rpgrip1 | 375.287743 | -6.665365473 | 1.20E-02 |
| Serpina3i | 55.90345178 | -6.75648003 | 3.25E-06 |
| Gm20796 | 51.42200161 | -7.470863887 | 7.20E-03 |
| Samd11 | 155.349818 | -8.322643694 | 2.88E-03 |
| Gm7666 | 51.9416385 | -17.15462386 | 6.27E-04 |

| Gene | baseMean | log2 Fold Change | padj |
|---|---|---|---|
| AP003419.1 | 46.01432342 | 19.94684904 | 4.32E-10 |
| CXCR5 | 38.38259048 | 19.89685544 | 7.21E-06 |
| NPPB | 2240.650956 | 11.49253916 | 6.33E-85 |
| AC089983.1 | 88.31094755 | 10.69956979 | 6.35E-17 |
| SLC27A2 | 45.4861244 | 10.3902619 | 5.59E-14 |
| SFTA1P | 288.2208761 | 10.00916747 | 8.46E-28 |
| SLC18A3 | 36.75075062 | 9.631689912 | 1.45E-13 |
| RGPD4 | 132.856446 | 8.939636977 | 2.46E-02 |
| MYO7B | 109.193404 | 8.673655066 | 9.63E-22 |
| XIRP1 | 24.70751638 | 8.59486843 | 5.11E-09 |
| MARCO | 27.67416851 | 8.343381727 | 1.09E-10 |
| PSG3 | 19.01407792 | 8.268621729 | 3.60E-10 |
| TNNI1 | 87.9705331 | 8.159720515 | 8.68E-23 |
| KRT34 | 47.26339011 | 8.082639668 | 9.89E-13 |
| IMPDH1P10 | 18.85870357 | 8.049931459 | 1.46E-04 |
| KRT81 | 3638.578358 | 8.03170974 | 3.53E-234 |
| INHBB | 89.82945103 | 8.010420611 | 3.28E-18 |
| RFPL4AL1 | 48.58179161 | 7.99822836 | 7.11E-10 |
| CEACAMP10 | 596.3616348 | 7.979908333 | 4.76E-100 |
| AC010323.1 | 29.99683061 | 7.897357899 | 2.46E-02 |
| LINC02331 | 15.15784713 | 7.759494028 | 2.15E-08 |
| CHRND | 24.58768675 | 7.734988577 | 5.87E-05 |
| PRAMEF1 | 8.708668192 | 7.703952677 | 2.72E-08 |
| KRT80 | 1730.679913 | 7.592655617 | 1.93E-267 |
| LCN2 | 20.89964453 | 7.530755499 | 2.24E-07 |
| KISS1 | 240.3909755 | 7.528133741 | 1.73E-39 |
| ANKRD1 | 2371.13122 | 7.508110522 | 9.13E-69 |
| CXCR4 | 15.34564949 | 7.339905096 | 3.95E-07 |
| AC115522.1 | 25.34233385 | 7.255101144 | 9.16E-08 |
| KRT4 | 36.4381841 | 7.223122813 | 3.94E-17 |
| PSG7 | 92.83788603 | 7.216441657 | 7.51E-29 |
| CAVIN2 | 121.5099942 | 7.175347267 | 9.06E-35 |
| LINC01186 | 8.428108961 | 7.081777268 | 4.29E-07 |
| MC5R | 25.26232457 | 7.024230158 | 2.11E-07 |
| PSG4 | 1067.986903 | 6.995659739 | 2.35E-185 |
| GGTLC2 | 5.282843147 | 6.983651214 | 2.58E-06 |
| LURAP1L-AS1 | 37.65792059 | 6.964013217 | 2.13E-13 |
| TINAGL1 | 244.1329187 | 6.962719931 | 8.04E-92 |
| TNNT2 | 85.46035183 | 6.914288091 | 5.89E-12 |
| AP003498.1 | 6.759281315 | 6.862882225 | 7.97E-05 |
| C17orf102 | 57.63053537 | 6.778432833 | 4.25E-19 |
| WFDC21P | 140.545174 | 6.777575139 | 1.52E-59 |
| LINC00592 | 5.244624385 | 6.758460135 | 5.33E-06 |
| THAP12P8 | 7.763317663 | 6.748549239 | 4.73E-06 |
| RFPL4A | 89.139459 | 6.736773745 | 3.22E-18 |
| AC114488.1 | 8.478510308 | 6.679069134 | 2.14E-06 |

| | | | |
|---|---|---|---|
| SPX | 445.5857189 | 6.651407824 | 2.02E-66 |
| HBG1 | 1243.158158 | 6.647752194 | 2.56E-38 |
| HCN4 | 60.95695921 | 6.575644439 | 5.48E-39 |
| KRT17 | 109.932617 | 6.526630465 | 5.15E-31 |
| KRTAP1-5 | 5.600439711 | 6.487739539 | 2.71E-05 |
| AC245041.1 | 18.51833017 | 6.451786676 | 3.91E-08 |
| KRTAP3-1 | 7.798810147 | 6.447684719 | 9.73E-06 |
| PSG5 | 429.1534162 | 6.418044094 | 4.91E-92 |
| AC026316.2 | 27.22988646 | 6.40374094 | 1.78E-11 |
| PSG2 | 45.18656473 | 6.374938637 | 2.48E-11 |
| NXPH2 | 6.697336569 | 6.361276288 | 8.22E-06 |
| C4BPB | 51.0915616 | 6.296158266 | 4.06E-31 |
| ADCY8 | 7.411685296 | 6.287519751 | 1.62E-05 |
| AC062015.1 | 5.368918224 | 6.269063979 | 4.35E-05 |
| LNCOG | 57.59267757 | 6.264260736 | 1.35E-19 |
| CPA4 | 8124.801267 | 6.169402228 | 0.00E+00 |
| GDF6 | 21.29645127 | 6.163274685 | 5.78E-15 |
| AC135977.1 | 22.06214483 | 6.110820093 | 7.84E-03 |
| FGF19 | 32.84708541 | 6.091869102 | 3.79E-06 |
| AP1M2 | 110.3798198 | 6.075586798 | 1.08E-26 |
| CEACAMP8 | 13.78743094 | 6.046254534 | 9.95E-06 |
| AC100800.1 | 63.02116095 | 5.998650384 | 2.36E-29 |
| DIRC3-AS1 | 14.94687545 | 5.988818522 | 2.41E-07 |
| KCND3 | 287.4144475 | 5.971779987 | 8.20E-39 |
| AL159166.1 | 10.36203043 | 5.948845339 | 2.47E-05 |
| SUSD4 | 14.80441536 | 5.921521619 | 3.30E-08 |
| FGF21 | 23.9633406 | 5.900134231 | 1.07E-05 |
| TGM4 | 11.45184573 | 5.894288627 | 5.02E-05 |
| CHAT | 14.31282578 | 5.868572076 | 7.56E-06 |
| PSG6 | 42.7531939 | 5.833577013 | 9.02E-09 |
| TRPM1 | 8.303459732 | 5.814591699 | 4.91E-06 |
| APCS | 10.98996835 | 5.805623665 | 9.68E-05 |
| NA | 9.834076694 | 5.785662978 | 3.02E-03 |
| PDGFB | 265.4831051 | 5.777653451 | 4.32E-60 |
| MYL2 | 7.674597299 | 5.777485604 | 3.35E-05 |
| AL159987.1 | 10.21878798 | 5.769756266 | 7.79E-08 |
| RPL23AP47 | 7.779613667 | 5.757978152 | 6.48E-03 |
| AL365184.1 | 7.788440408 | 5.700935415 | 1.38E-04 |
| PSG11 | 67.8097526 | 5.699109288 | 1.53E-22 |
| PSG9 | 110.5829147 | 5.690273251 | 8.61E-29 |
| KRTAP2-3 | 13.1496624 | 5.657482386 | 1.51E-04 |
| IGFN1 | 141.2038353 | 5.644557655 | 6.76E-27 |
| AC067773.1 | 17.64783138 | 5.587847577 | 1.53E-08 |
| AL136452.1 | 5.435183264 | 5.587787204 | 3.21E-04 |
| OLR1 | 33.65778704 | 5.58422227 | 3.81E-13 |
| LY6K | 276.1851787 | 5.536047257 | 1.40E-75 |
| FPR3 | 132.3685572 | 5.522766966 | 1.75E-70 |

| | | | |
|---|---|---|---|
| LINC01293 | 57.56332832 | 5.522146032 | 2.18E-12 |
| ADGRG4 | 32.72646161 | 5.463924947 | 6.75E-07 |
| TNFSF18 | 594.8574692 | 5.462185192 | 1.86E-20 |
| AC087393.2 | 6.395911565 | 5.447653585 | 1.67E-02 |
| CSTA | 29.96038685 | 5.417034943 | 2.24E-08 |
| SPNS3 | 24.62107375 | 5.385024135 | 1.35E-06 |
| AL353743.4 | 7.124459301 | 5.381802219 | 3.94E-04 |
| MSLN | 108.5106732 | 5.355271091 | 1.87E-51 |
| KRT87P | 36.57107821 | 5.355002579 | 2.45E-21 |
| MYH2 | 23.18872912 | 5.342873782 | 1.58E-04 |
| OLFML1 | 58.53919204 | 5.3326063 | 5.52E-28 |
| PSG1 | 87.14686265 | 5.298181267 | 4.02E-32 |
| IL7R | 8419.815253 | 5.291867475 | 2.78E-69 |
| OR10Y1P | 5.655546436 | 5.285473774 | 8.81E-04 |
| ALCAM | 37.52071971 | 5.267789821 | 5.97E-15 |
| MYHAS | 12.21043099 | 5.264336067 | 6.30E-05 |
| STXBP6 | 354.2817869 | 5.257976853 | 5.80E-121 |
| FOXB1 | 5.497409793 | 5.196826079 | 1.34E-03 |
| NGF | 85.12586789 | 5.15143005 | 1.21E-23 |
| KRT86 | 789.1722789 | 5.14302061 | 2.71E-160 |
| GPRIN2 | 25.95356115 | 5.122758474 | 6.29E-14 |
| CST6 | 11.6176009 | 5.116679417 | 1.06E-08 |
| CAPSL | 15.12718097 | 5.110586296 | 2.60E-07 |
| LAD1 | 18.12413109 | 5.099625468 | 3.03E-08 |
| ZNF536 | 5.604134159 | 5.073572254 | 8.04E-04 |
| GPR1 | 155.1271788 | 5.043839343 | 3.07E-13 |
| MYOD1 | 190.9762848 | 5.042852443 | 3.59E-14 |
| GPA33 | 14.52145809 | 5.026503454 | 3.29E-08 |
| CLDN9 | 13.60129441 | 5.013965119 | 3.20E-06 |
| AC132068.1 | 5.382205542 | 4.98791066 | 1.61E-03 |
| PCDHGB8P | 11.70899875 | 4.983931549 | 2.96E-02 |
| NA | 6.666296487 | 4.979721228 | 9.59E-03 |
| NPIPA2 | 22.55804979 | 4.949410647 | 1.76E-02 |
| AL009178.2 | 8.641259053 | 4.946749515 | 9.98E-05 |
| CHRNA4 | 6.453752451 | 4.927474572 | 1.42E-04 |
| AC107871.1 | 14.44185302 | 4.925141102 | 4.23E-04 |
| PAPPA2 | 59.55621791 | 4.921552773 | 2.42E-17 |
| TMEM200C | 23.71648983 | 4.90735289 | 5.05E-08 |
| ITIH3 | 47.92121289 | 4.886632418 | 5.37E-15 |
| PSG8 | 13.98429761 | 4.871837953 | 4.39E-07 |
| DRGX | 5.390330915 | 4.863337164 | 1.93E-03 |
| LINC01291 | 43.47043514 | 4.856922505 | 6.36E-11 |
| LGALS9C | 9.780642832 | 4.845529798 | 3.43E-05 |
| CD24 | 529.1194415 | 4.820518813 | 3.01E-127 |
| TCEA1P3 | 6.557677423 | 4.818240007 | 2.46E-04 |
| LINC02535 | 49.64853035 | 4.805061301 | 1.93E-20 |
| AC114296.1 | 5.972914867 | 4.791164956 | 1.68E-03 |

| | | | |
|---|---|---|---|
| IL1RAPL1 | 13.93100269 | 4.780669785 | 3.80E-08 |
| AC018781.1 | 5.21339508 | 4.774231062 | 2.34E-03 |
| KCNE1B | 47.6023731 | 4.77128285 | 3.13E-12 |
| VSTM1 | 56.60755979 | 4.76070116 | 3.06E-10 |
| TMEM52B | 10.55974503 | 4.760641122 | 2.97E-05 |
| AC005832.4 | 89.28362442 | 4.755095562 | 3.79E-05 |
| MFAP5 | 48.10545696 | 4.75228498 | 3.60E-19 |
| KLHDC7B | 28.70749508 | 4.728899407 | 1.95E-03 |
| AC098613.1 | 9.120879227 | 4.72691479 | 1.30E-06 |
| STX11 | 126.2269358 | 4.715652261 | 1.79E-62 |
| AL021153.1 | 5.180709888 | 4.71065763 | 4.71E-04 |
| C6orf132 | 13.19535717 | 4.704226832 | 2.69E-08 |
| CEACAMP7 | 24.51020739 | 4.699155224 | 5.60E-13 |
| MS4A4A | 31.22211546 | 4.692084414 | 4.62E-21 |
| MT3 | 7.356063884 | 4.652648757 | 6.12E-04 |
| KRT8 | 14283.57687 | 4.642117003 | 0.00E+00 |
| ASS1 | 1006.358665 | 4.616953192 | 1.82E-29 |
| LINC01693 | 25.8599621 | 4.613973394 | 3.05E-15 |
| CCND2 | 40.55711956 | 4.607867767 | 8.40E-10 |
| PLAC8 | 321.3230234 | 4.598928116 | 2.71E-69 |
| KCNQ3 | 69.36936743 | 4.597663535 | 1.18E-23 |
| CDH4 | 706.3312431 | 4.582050274 | 4.35E-98 |
| CCN2 | 8122.707037 | 4.572167702 | 6.21E-123 |
| CTNND2 | 5.418341109 | 4.556599254 | 6.70E-04 |
| VLDLR-AS1 | 123.3112072 | 4.543594085 | 1.14E-21 |
| MEI4 | 8.544885276 | 4.539574507 | 6.52E-05 |
| LAMP3 | 296.1481714 | 4.53217281 | 6.80E-61 |
| AC016717.2 | 5.261098293 | 4.513769291 | 1.42E-03 |
| TMEM132E | 67.2505779 | 4.513266139 | 5.62E-19 |
| ITGB2 | 124.990794 | 4.512183942 | 1.05E-43 |
| AL137129.1 | 25.35779872 | 4.509043611 | 8.77E-06 |
| TMEM130 | 81.1847839 | 4.483625617 | 1.16E-34 |
| ANKRD2 | 7.839182386 | 4.452964021 | 7.83E-05 |
| AC092132.1 | 5.105028167 | 4.442578322 | 1.82E-03 |
| RIMS2 | 41.07355111 | 4.439200628 | 3.55E-13 |
| ANXA8L1 | 163.5609482 | 4.427073957 | 4.40E-39 |
| AOX1 | 235.0610148 | 4.422674359 | 6.26E-34 |
| MYH1 | 5.335709278 | 4.415841943 | 3.05E-02 |
| AC245041.2 | 17.47365683 | 4.407005583 | 3.64E-10 |
| CSDC2 | 8.659595836 | 4.387803585 | 9.16E-06 |
| NA | 5.318823847 | 4.385551841 | 4.94E-02 |
| DNAJC6 | 116.1338755 | 4.372433484 | 1.18E-37 |
| GUCY1A2 | 7.369270985 | 4.363549264 | 6.88E-03 |
| AC114550.1 | 11.42977472 | 4.361453554 | 4.26E-06 |
| LTF | 52.07108289 | 4.358267232 | 1.15E-21 |
| HBBP1 | 30.01402287 | 4.356605625 | 7.34E-10 |
| SYNPO2L | 54.0993148 | 4.342816284 | 1.06E-20 |

| | | | |
|---|---|---|---|
| SNX18P24 | 5.717837345 | 4.314470829 | 2.11E-02 |
| FOXG1-AS1 | 5.642949624 | 4.309025441 | 1.26E-04 |
| RPL13AP20 | 27.56620949 | 4.304121141 | 4.30E-11 |
| AC090505.1 | 11.22101694 | 4.286462179 | 1.92E-08 |
| NA | 16.51338953 | 4.257501595 | 6.74E-03 |
| LCE2A | 7.928852411 | 4.251202235 | 1.09E-02 |
| TNNC1 | 56.70589737 | 4.242217362 | 2.40E-24 |
| CLDN1 | 777.3721858 | 4.233543295 | 2.14E-49 |
| HS3ST5 | 7.638299293 | 4.228903842 | 1.27E-04 |
| AL078612.2 | 11.28350549 | 4.221276376 | 9.76E-07 |
| SLC7A3 | 41.66827949 | 4.215734284 | 3.09E-19 |
| GSX2 | 19.00376094 | 4.181740541 | 5.65E-09 |
| 4-Mar | 1178.181843 | 4.179765613 | 3.81E-62 |
| CCN1 | 10596.59591 | 4.163417313 | 2.09E-303 |
| TAGLN | 4862.193472 | 4.154828998 | 3.49E-74 |
| AL451069.1 | 7.150260511 | 4.136280794 | 1.75E-03 |
| MEGF6 | 249.4909584 | 4.129181406 | 3.88E-29 |
| MB | 5.190958042 | 4.128701746 | 3.91E-03 |
| ADAM18 | 22.12303189 | 4.121837309 | 3.72E-12 |
| EMX1 | 5.979855963 | 4.119956352 | 2.01E-05 |
| ALPP | 37.81297781 | 4.117735478 | 1.84E-23 |
| CD177 | 28.34911627 | 4.108572348 | 1.30E-09 |
| SLAMF7 | 176.448223 | 4.106564443 | 2.46E-15 |
| C7orf65 | 7.884882127 | 4.103583178 | 1.54E-04 |
| GFRA1 | 5.639523961 | 4.094628562 | 2.47E-04 |
| CPNE4 | 26.75781287 | 4.093699492 | 4.63E-10 |
| GJB3 | 26.34134941 | 4.085225136 | 2.97E-12 |
| AL590004.3 | 1289.954072 | 4.082227126 | 5.96E-88 |
| PRELP | 480.5999178 | 4.067842539 | 1.61E-21 |
| NYAP2 | 8.4881526 | 4.065071493 | 5.40E-05 |
| AC095350.1 | 25.6081823 | 4.060862708 | 1.97E-10 |
| NTNG1 | 9.82965435 | 4.053238407 | 1.12E-03 |
| TAGAP | 49.23096721 | 4.048079554 | 7.58E-03 |
| KRT7 | 53.45128578 | 4.046290966 | 1.87E-17 |
| HBG2 | 280.0908711 | 4.040063094 | 1.40E-38 |
| MYOSLID | 13.69503477 | 4.010751434 | 5.20E-06 |
| PPP1R14C | 7.441369772 | 4.008872144 | 3.70E-03 |
| LINC02154 | 47.14458306 | 4.006643539 | 9.03E-23 |
| TRIM29 | 531.2890621 | 4.005509121 | 6.06E-63 |
| USP41 | 16.73443121 | 4.004059659 | 4.51E-05 |
| AC079054.1 | 7.797618888 | 4.002647404 | 1.01E-04 |
| MME | 72.18384025 | 3.991209193 | 3.23E-11 |
| TMEM178B | 177.5366143 | 3.98524677 | 1.82E-29 |
| AC112656.1 | 16.72979437 | 3.974768732 | 4.26E-06 |
| XAGE2 | 41.30684287 | 3.968421112 | 7.03E-26 |
| PADI1 | 49.5811494 | 3.962982254 | 1.13E-16 |
| AL161431.1 | 253.7190024 | 3.957343133 | 3.79E-94 |

| | | | |
|---|---|---|---|
| C5orf67 | 8.957053987 | 3.944338369 | 2.29E-06 |
| AC034206.1 | 5.479961284 | 3.932406277 | 9.50E-04 |
| ELFN2 | 490.7239094 | 3.930639324 | 8.90E-76 |
| NA | 6.039292419 | 3.927979826 | 9.07E-04 |
| LRRC19 | 16.48424518 | 3.917685774 | 3.12E-05 |
| NPY4R | 21.74673346 | 3.915815686 | 3.62E-10 |
| COL12A1 | 7146.720884 | 3.914485629 | 2.64E-196 |
| AMOTL2 | 5194.214834 | 3.908633168 | 0.00E+00 |
| RASGRP4 | 13.34237912 | 3.884328867 | 3.36E-07 |
| LYPD5 | 426.1463919 | 3.881794891 | 6.55E-34 |
| SERPINA3 | 18.99021726 | 3.876130215 | 5.57E-03 |
| AC018464.1 | 5.840792081 | 3.872901658 | 3.71E-03 |
| SERPINB7 | 182.2948166 | 3.870968585 | 4.54E-23 |
| LINC01511 | 8.69732256 | 3.868153583 | 8.00E-05 |
| MOV10L1 | 49.20024885 | 3.867316788 | 4.91E-20 |
| AC022028.2 | 14.49200066 | 3.864326144 | 1.60E-06 |
| LINC01638 | 242.896981 | 3.831816555 | 1.75E-65 |
| FRG2C | 27.16775073 | 3.82845407 | 2.74E-09 |
| C7orf69 | 6.788843734 | 3.806163813 | 2.63E-04 |
| AC011510.1 | 10.81268386 | 3.803905594 | 6.15E-06 |
| HBD | 5.800011754 | 3.800575018 | 9.64E-04 |
| HNRNPA1P33 | 8.089797574 | 3.788754589 | 5.26E-04 |
| ASB18 | 18.35133093 | 3.784922085 | 8.37E-04 |
| TMPRSS3 | 63.85416896 | 3.76535917 | 3.60E-29 |
| GPR65 | 19.00070169 | 3.764698313 | 1.88E-06 |
| AP002478.1 | 8.764480814 | 3.760678479 | 7.36E-04 |
| SHOC1 | 8.469872835 | 3.748565501 | 4.18E-04 |
| LIMS2 | 478.4740095 | 3.741328975 | 6.07E-98 |
| GOLGA8IP | 12.87111558 | 3.737688363 | 8.10E-05 |
| TNFSF4 | 327.5208002 | 3.735706856 | 2.97E-48 |
| AC011591.1 | 33.09399697 | 3.726793331 | 2.74E-14 |
| KLHDC7A | 24.91322927 | 3.718760724 | 5.86E-06 |
| BICDL1 | 37.542591 | 3.718293734 | 1.48E-14 |
| AL590644.1 | 26.03942664 | 3.692767168 | 7.07E-13 |
| AC015909.1 | 13.01442352 | 3.692011948 | 6.19E-05 |
| KCNU1 | 6.149298403 | 3.688246302 | 3.77E-02 |
| GPC4 | 272.3054783 | 3.679468892 | 2.10E-71 |
| NA | 13.87474242 | 3.674286642 | 1.58E-07 |
| FRG2B | 5.367969376 | 3.673032733 | 2.42E-02 |
| TENM2 | 351.6327637 | 3.661479987 | 1.49E-14 |
| GRIP1 | 17.48063495 | 3.647837574 | 1.41E-10 |
| CRLF2 | 21.35257814 | 3.645005424 | 1.33E-06 |
| HSPB3 | 33.71635809 | 3.633839936 | 5.51E-12 |
| AP003064.2 | 24.94135603 | 3.629905597 | 1.86E-07 |
| FRMPD2 | 7.454959441 | 3.628619708 | 3.59E-02 |
| RBP7 | 234.865592 | 3.602796307 | 3.92E-101 |
| MYL3 | 32.80788903 | 3.600004601 | 8.02E-18 |

| | | | |
|---|---|---|---|
| LINC01133 | 24.06459905 | 3.593962323 | 2.85E-07 |
| SAA1 | 78.53236772 | 3.57953343 | 2.88E-23 |
| MYO5B | 212.8051185 | 3.577781452 | 9.34E-33 |
| LHX5 | 8.739437811 | 3.57722123 | 4.28E-04 |
| FAM201A | 45.64172295 | 3.564677829 | 2.31E-13 |
| ARHGAP40 | 27.14455982 | 3.563678629 | 1.11E-13 |
| TM6SF1 | 340.5809889 | 3.549306683 | 7.61E-25 |
| NPY4R2 | 11.66202251 | 3.545510356 | 2.02E-03 |
| FKBP9P1 | 272.6623261 | 3.544707836 | 3.55E-32 |
| SEMA3B | 1213.282702 | 3.544449289 | 1.25E-71 |
| TMEM92-AS1 | 16.11057805 | 3.517926499 | 3.85E-06 |
| ITGB2-AS1 | 20.00398093 | 3.492398478 | 4.88E-09 |
| AC005863.1 | 18.09095241 | 3.486612603 | 1.56E-06 |
| AL049839.2 | 69.57473853 | 3.48624442 | 7.24E-05 |
| WDFY4 | 28.36478348 | 3.483573435 | 3.81E-07 |
| AC010247.2 | 13.42529976 | 3.474968405 | 1.73E-06 |
| FST | 410.4366751 | 3.474443967 | 7.98E-38 |
| CSF2RA | 877.8505258 | 3.465909627 | 5.59E-82 |
| GDAP1L1 | 48.33841871 | 3.457024586 | 1.69E-11 |
| RPL12P14 | 37.54375201 | 3.454122018 | 3.28E-15 |
| GDA | 761.696661 | 3.446546149 | 1.84E-72 |
| SYTL5 | 146.9965863 | 3.446257586 | 4.47E-24 |
| SLC12A3 | 6.886708526 | 3.445152066 | 9.80E-03 |
| LINC01198 | 6.335604086 | 3.436551922 | 4.36E-03 |
| AL645608.7 | 11.78460553 | 3.433786834 | 2.93E-04 |
| AC121757.1 | 28.94935304 | 3.428536017 | 5.41E-11 |
| ATP6V0A4 | 28.12777374 | 3.428148729 | 6.82E-11 |
| ILDR2 | 71.76148004 | 3.428102054 | 3.33E-11 |
| GAS2L2 | 7.131076997 | 3.425249523 | 1.31E-02 |
| WNT7A | 74.42957687 | 3.422957096 | 1.51E-16 |
| INHBE | 431.5658161 | 3.42196789 | 3.09E-08 |
| NA | 53.49267065 | 3.409326253 | 3.69E-20 |
| CHRNA1 | 65.20886781 | 3.399481533 | 1.44E-04 |
| ANXA3 | 816.3716511 | 3.398650038 | 7.32E-72 |
| RPL29P19 | 46.34515324 | 3.397825934 | 9.07E-18 |
| AC009238.1 | 10.03320931 | 3.381129571 | 4.26E-05 |
| TSPAN2 | 207.194681 | 3.378359156 | 7.61E-42 |
| UNC13D | 102.386188 | 3.37262408 | 4.17E-24 |
| LINC00707 | 90.45365985 | 3.369521265 | 3.78E-27 |
| TEX37 | 13.34227782 | 3.365450846 | 9.12E-07 |
| AC245100.3 | 41.78719059 | 3.35262677 | 7.99E-09 |
| PSG8-AS1 | 10.13715312 | 3.348603365 | 1.24E-05 |
| LINC02057 | 19.34808816 | 3.345398584 | 5.43E-09 |
| AC092384.2 | 8.422921767 | 3.342959756 | 8.06E-04 |
| TNFRSF12A | 3033.22403 | 3.33949236 | 1.75E-109 |
| BMP4 | 3231.109292 | 3.336868898 | 5.48E-102 |
| PRR20G | 7.206833657 | 3.332295377 | 2.87E-03 |

| | | | |
|---|---|---|---|
| ALOXE3 | 48.23452377 | 3.324294242 | 3.18E-06 |
| ANKRD34C | 8.916883708 | 3.311663127 | 3.79E-04 |
| PADI3 | 60.49052887 | 3.303531637 | 3.54E-11 |
| BRS3 | 15.9865702 | 3.303381332 | 5.02E-07 |
| IL32 | 397.8318467 | 3.297383703 | 1.92E-33 |
| NANOS1 | 479.3315027 | 3.291411568 | 6.62E-93 |
| AWAT2 | 6.560235983 | 3.291031076 | 4.72E-03 |
| THSD7B | 5.906740791 | 3.28922367 | 1.03E-03 |
| HRH2 | 76.37524624 | 3.27961758 | 9.43E-20 |
| ELF3 | 19.19275641 | 3.268045053 | 3.18E-10 |
| HYDIN2 | 17.04818201 | 3.261932585 | 3.06E-03 |
| BDNF | 222.3395398 | 3.259647327 | 2.15E-77 |
| AC084783.1 | 9.848268834 | 3.258930729 | 6.85E-05 |
| AC005150.1 | 7.249293129 | 3.24983138 | 2.49E-04 |
| LUARIS | 40.9259168 | 3.246140375 | 1.09E-16 |
| SMPD4P1 | 9.902617318 | 3.242868811 | 2.38E-04 |
| FAM3B | 10.079902 | 3.240950713 | 3.78E-06 |
| SBF2-AS1 | 216.8026323 | 3.239571956 | 1.30E-57 |
| LINC02243 | 8.297230953 | 3.231787787 | 2.87E-04 |
| UCA1 | 6683.968826 | 3.226631236 | 4.09E-54 |
| INHBC | 11.53838661 | 3.224265135 | 1.47E-04 |
| ZP1 | 57.88032852 | 3.222334389 | 2.32E-08 |
| SERPINB2 | 585.2962819 | 3.220566156 | 1.47E-25 |
| LINC01605 | 45.79967183 | 3.220046003 | 6.22E-12 |
| TMEFF2 | 21.7217407 | 3.210748955 | 8.37E-05 |
| NEK11 | 62.08406486 | 3.208064804 | 3.48E-14 |
| AC105411.1 | 36.104077 | 3.205428651 | 3.27E-10 |
| AXL | 11912.61544 | 3.198676255 | 1.04E-80 |
| HDAC2-AS2 | 45.6529736 | 3.198036406 | 5.16E-16 |
| ITGBL1 | 61.58578191 | 3.195801831 | 8.07E-17 |
| SNAPC1 | 2436.405541 | 3.187567239 | 6.36E-57 |
| AC010457.1 | 11.60862398 | 3.186676018 | 2.16E-04 |
| PIWIL2 | 12.98599122 | 3.18474016 | 1.86E-05 |
| GALNT18 | 150.014102 | 3.17981229 | 6.02E-42 |
| PLCE1-AS1 | 11.35185947 | 3.179527289 | 1.95E-02 |
| AC105105.1 | 257.2624826 | 3.177837788 | 8.74E-11 |
| KRT18 | 26282.33084 | 3.175387246 | 3.95E-171 |
| AP004607.3 | 9.874376473 | 3.174408702 | 3.52E-05 |
| TGFB2-AS1 | 227.6156955 | 3.169188773 | 5.43E-59 |
| PI3 | 17.94074282 | 3.168057468 | 1.07E-06 |
| EFEMP1 | 741.1103555 | 3.16581704 | 7.62E-62 |
| CXCL14 | 171.4861655 | 3.159327204 | 7.93E-23 |
| DUSP13 | 11.52726261 | 3.150079745 | 3.22E-03 |
| FPGT-TNNI3K | 29.63087152 | 3.147636833 | 1.08E-02 |
| CAPN6 | 166.404887 | 3.144683522 | 1.19E-42 |
| CDH1 | 23.87917036 | 3.136967277 | 1.17E-05 |
| OSCAR | 347.1562098 | 3.132953631 | 3.41E-33 |

| | | | |
|---|---|---|---|
| TPO | 119.1349183 | 3.132268239 | 2.89E-16 |
| SUN3 | 13.02687629 | 3.122708834 | 1.49E-04 |
| CES5A | 10.8197907 | 3.122420106 | 1.26E-02 |
| NXF2B | 60.15929052 | 3.12053605 | 2.94E-02 |
| LINC00525 | 14.59269898 | 3.120319358 | 2.92E-06 |
| AC245100.6 | 52.1791593 | 3.111041545 | 8.96E-09 |
| VCX3A | 15.59637735 | 3.107700571 | 2.99E-06 |
| CDC42EP3 | 1854.372981 | 3.105214755 | 1.04E-44 |
| VSTM2L | 409.4949088 | 3.096817337 | 8.25E-66 |
| FBLN5 | 552.2943098 | 3.095879262 | 1.50E-44 |
| ACKR3 | 110.6345116 | 3.093570269 | 6.10E-43 |
| NA | 26.57063328 | 3.085293428 | 4.83E-06 |
| CA9 | 143.2752701 | 3.084619289 | 4.74E-04 |
| AL590483.1 | 14.41730836 | 3.082997395 | 1.43E-07 |
| AC018529.2 | 5.13035025 | 3.082826417 | 3.23E-02 |
| LOX | 1600.914318 | 3.069363489 | 1.77E-61 |
| SLC16A12 | 9.185038545 | 3.069242193 | 8.24E-05 |
| TMEM178A | 50.3489753 | 3.06641657 | 5.24E-17 |
| FGF14 | 12.63602815 | 3.065656971 | 7.37E-04 |
| AC103740.2 | 6.922522524 | 3.061834027 | 3.16E-02 |
| ALPK2 | 2595.841625 | 3.05642035 | 5.31E-43 |
| NA | 43.80094162 | 3.047264888 | 2.74E-09 |
| AC010207.1 | 29.98635144 | 3.045406432 | 3.46E-11 |
| IL21R | 123.0008977 | 3.036878488 | 2.58E-13 |
| AC121758.1 | 17.51648739 | 3.035870143 | 1.18E-02 |
| PHF24 | 57.9882298 | 3.031873556 | 1.02E-04 |
| IGSF23 | 35.73858368 | 3.030178483 | 4.17E-16 |
| NLRP5 | 272.5607272 | 3.018487618 | 3.54E-46 |
| CA3 | 9.544616765 | 3.017399002 | 1.95E-04 |
| PTH1R | 160.1294675 | 3.012364337 | 1.21E-50 |
| PLEKHA7 | 2443.226225 | 3.011971116 | 3.68E-86 |
| FBP1 | 21.31209753 | 3.011071849 | 3.01E-06 |
| AC004233.3 | 52.94805197 | 3.010762903 | 5.65E-19 |
| CNN1 | 297.6933427 | 3.007205427 | 2.92E-21 |
| NCF2 | 11.76390963 | 3.000862165 | 1.43E-04 |
| MYL7 | 7.43415832 | 3.000614722 | 3.76E-04 |
| SIGLEC6 | 13.02459145 | 2.999763985 | 8.30E-06 |
| HRASLS2 | 6.319019622 | 2.997162822 | 1.23E-03 |
| KRT73-AS1 | 27.90722466 | 2.986687965 | 7.06E-07 |
| WWC1 | 602.1446754 | 2.983745505 | 6.80E-83 |
| RBM24 | 126.6531076 | 2.980024832 | 4.11E-22 |
| AJUBA | 9629.715395 | 2.98002213 | 1.52E-69 |
| AC117500.3 | 30.09938499 | 2.976716046 | 1.10E-07 |
| HNF4A | 25.93293679 | 2.976090895 | 3.18E-03 |
| DAW1 | 15.73967927 | 2.970581267 | 3.09E-05 |
| WNT7B | 200.4097672 | 2.967162111 | 1.78E-16 |
| BMP2 | 67.08671781 | 2.96183078 | 3.66E-19 |

| | | | |
|---|---|---|---|
| MYH15 | 58.51415002 | 2.955327876 | 1.52E-05 |
| CHMP4C | 55.12357223 | 2.951622202 | 1.55E-18 |
| SLPI | 49.88731299 | 2.950267041 | 1.13E-17 |
| GADD45B | 977.0968945 | 2.948432547 | 2.81E-150 |
| CLIC6 | 238.7103255 | 2.947044553 | 2.58E-68 |
| MGAM | 11.91207921 | 2.946967731 | 4.01E-03 |
| CPZ | 462.7221784 | 2.940897776 | 2.61E-42 |
| IL21-AS1 | 5.394513016 | 2.940045712 | 2.07E-02 |
| DUX4L45 | 69.43942931 | 2.938257048 | 1.75E-22 |
| USH1C | 6.338202099 | 2.928914084 | 2.44E-02 |
| AC110048.1 | 5.151843255 | 2.92566208 | 8.32E-03 |
| FLNC-AS1 | 13.89514009 | 2.922551837 | 6.29E-05 |
| AL391422.1 | 7.102066668 | 2.919280454 | 1.67E-03 |
| ACOXL | 37.18565397 | 2.918080593 | 9.76E-14 |
| FO681492.1 | 191.8394703 | 2.912317642 | 5.88E-28 |
| LINC01968 | 5.10520881 | 2.910798195 | 3.51E-02 |
| ZNF185 | 2431.652507 | 2.905150958 | 3.22E-164 |
| RASGRF1 | 134.7074287 | 2.893702412 | 2.93E-22 |
| EPS8L2 | 349.987695 | 2.881798069 | 1.24E-42 |
| SERPINE1 | 2089.772652 | 2.879923768 | 1.30E-69 |
| AC018816.1 | 5.312448188 | 2.868125843 | 1.47E-02 |
| GREM1 | 25.98707971 | 2.858354579 | 6.00E-08 |
| C1orf158 | 8.469554404 | 2.85790343 | 8.21E-03 |
| HSD17B2 | 349.7292265 | 2.856530107 | 1.88E-33 |
| UPB1 | 37.15425623 | 2.855349083 | 2.46E-09 |
| RNF144B | 69.84375385 | 2.849492881 | 3.24E-16 |
| IGFL2-AS1 | 125.576691 | 2.849162121 | 1.37E-28 |
| AL683807.1 | 11.74887014 | 2.847151988 | 1.93E-04 |
| TTR | 6.644641824 | 2.845099006 | 7.87E-03 |
| ADAMTS16 | 718.0575453 | 2.844076366 | 3.62E-35 |
| SLC9A4 | 21.29564053 | 2.843056145 | 8.97E-05 |
| AC009292.1 | 5.752936398 | 2.839973732 | 2.97E-03 |
| CHST4 | 31.95928319 | 2.827326499 | 1.21E-06 |
| SAA2 | 14.69633019 | 2.826653276 | 4.95E-03 |
| DPP6 | 11.99597905 | 2.825873077 | 7.33E-04 |
| PROSER2-AS1 | 38.08475189 | 2.823696278 | 6.62E-10 |
| AC008429.1 | 55.50190177 | 2.818945252 | 1.53E-11 |
| BASP1 | 2291.351755 | 2.817583098 | 3.99E-80 |
| LINC00973 | 7.727607361 | 2.816904805 | 4.68E-03 |
| EDIL3 | 8.97240499 | 2.812816245 | 2.07E-05 |
| LINC01836 | 94.12756194 | 2.811936325 | 3.81E-42 |
| C2orf78 | 9.859463982 | 2.811385806 | 4.03E-03 |
| AC087667.1 | 30.95840747 | 2.810397901 | 1.38E-10 |
| DMKN | 409.4101885 | 2.807223888 | 6.69E-79 |
| ELMOD1 | 26.26171552 | 2.806084534 | 3.09E-06 |
| KRTAP5-AS1 | 7.911436712 | 2.802218637 | 2.56E-02 |
| UGT3A1 | 24.54671501 | 2.802084291 | 9.23E-04 |

| | | | |
|---|---|---|---|
| MIR503 | 28.37757972 | 2.799857824 | 1.41E-06 |
| ACTR3C | 65.52093073 | 2.798976965 | 5.93E-11 |
| ROBO4 | 76.28270049 | 2.789894187 | 9.76E-08 |
| C6orf223 | 35.05250985 | 2.788420001 | 1.38E-07 |
| PSORS1C2 | 8.824272258 | 2.786863374 | 3.01E-04 |
| NKD2 | 30.11768859 | 2.78419036 | 1.18E-07 |
| LINC01203 | 6.584875129 | 2.775569246 | 6.01E-04 |
| TMEM45A | 771.370834 | 2.7734829 | 1.17E-49 |
| FUNDC2P2 | 18.74886774 | 2.758823322 | 1.86E-07 |
| SLC4A9 | 9.435170445 | 2.755623714 | 9.45E-04 |
| GAS6-DT | 47.94899173 | 2.755172615 | 5.51E-16 |
| ALDH2 | 6.145122017 | 2.752681174 | 1.74E-02 |
| C9orf131 | 10.69519957 | 2.751874302 | 2.00E-02 |
| PIM1 | 1122.782319 | 2.747930724 | 5.91E-44 |
| FOSL1 | 1816.844697 | 2.747001263 | 6.31E-28 |
| CDA | 83.82678768 | 2.745939553 | 2.17E-22 |
| AC022167.1 | 33.04569225 | 2.745432191 | 3.72E-02 |
| SAMD7 | 13.89098362 | 2.737341382 | 7.49E-07 |
| GRIN2B | 12.85074978 | 2.73718604 | 1.84E-05 |
| TCL1B | 142.1677943 | 2.73336573 | 8.73E-34 |
| MMP23B | 44.95772942 | 2.731589774 | 4.77E-10 |
| SLC45A2 | 191.8192019 | 2.73055948 | 2.06E-37 |
| GSG1 | 11.75434517 | 2.730476933 | 7.97E-05 |
| NMRK2 | 40.23946307 | 2.722532359 | 4.70E-03 |
| AC097059.2 | 16.24885375 | 2.71608171 | 7.88E-07 |
| MMP24 | 326.4706903 | 2.709293728 | 1.31E-52 |
| SERINC2 | 3346.317931 | 2.708692019 | 3.73E-50 |
| AC130304.1 | 149.9867397 | 2.707135 | 9.37E-03 |
| NTN1 | 74.09737759 | 2.700040274 | 2.24E-14 |
| ASNSP1 | 400.9698674 | 2.696084903 | 1.35E-16 |
| COL4A3 | 9.192934283 | 2.691776397 | 1.30E-02 |
| CRYAB | 8154.737916 | 2.691391253 | 2.40E-30 |
| MAGEB2 | 14.51957996 | 2.690053145 | 3.72E-06 |
| CEACAM22P | 11.65861573 | 2.68257236 | 4.61E-04 |
| CCK | 19.10415853 | 2.681985151 | 1.19E-07 |
| RUNDC3B | 6.946831316 | 2.678098141 | 6.81E-03 |
| TGFB2 | 3830.689573 | 2.674734338 | 6.14E-150 |
| CYP2B7P | 34.19759031 | 2.669105606 | 1.31E-06 |
| MYL4 | 57.40459191 | 2.666703377 | 3.89E-14 |
| NOG | 208.1326203 | 2.661632609 | 7.78E-18 |
| ASNS | 14006.90675 | 2.65706432 | 2.32E-28 |
| KCNE1 | 13.69174082 | 2.654292412 | 1.03E-05 |
| AL023881.1 | 23.0706953 | 2.654118478 | 2.20E-05 |
| C19orf33 | 9.271186735 | 2.653393533 | 4.79E-05 |
| IGF2-AS | 6.867922769 | 2.652140574 | 1.07E-03 |
| UNC5CL | 15.07775381 | 2.649908942 | 3.67E-05 |
| THBS1 | 3896.636739 | 2.649153785 | 2.25E-125 |

| | | | |
|---|---|---|---|
| SPANXB1 | 1009.08562 | 2.642333279 | 6.46E-96 |
| AC020934.1 | 20.74450596 | 2.640450774 | 1.90E-07 |
| ADGRE4P | 80.48512716 | 2.637262845 | 7.47E-14 |
| SLC7A10 | 92.31430533 | 2.63689344 | 3.78E-35 |
| OCLN | 724.0860483 | 2.636221107 | 9.49E-34 |
| AC117500.5 | 24.72259737 | 2.636133817 | 9.62E-06 |
| AC008514.1 | 9.822789484 | 2.634270809 | 4.28E-04 |
| STC2 | 5840.24853 | 2.633851825 | 6.22E-19 |
| JDP2 | 1542.857247 | 2.631558239 | 2.52E-37 |
| HHLA1 | 19.15672691 | 2.629999899 | 1.47E-07 |
| HRASLS | 31.71422465 | 2.621385847 | 1.05E-08 |
| KIAA1324 | 83.13749574 | 2.61861783 | 7.45E-18 |
| SLC12A8 | 358.0297075 | 2.616712025 | 3.60E-38 |
| PLAC1 | 82.08555886 | 2.601999582 | 1.62E-24 |
| AP001025.1 | 5.743856054 | 2.601858545 | 8.70E-03 |
| JPH2 | 828.9286868 | 2.601186191 | 9.99E-52 |
| PSCA | 27.29284018 | 2.599935796 | 1.31E-07 |
| EMP1 | 5374.264184 | 2.594419653 | 1.13E-48 |
| TXNRD1 | 33083.72171 | 2.593612209 | 2.98E-47 |
| AP001783.1 | 27.85078654 | 2.593067266 | 3.57E-09 |
| AC008897.3 | 8.561143206 | 2.591207595 | 3.95E-03 |
| C21orf59-TCP10 | 27.03478783 | 2.590142849 | 1.18E-02 |
| NA | 17.66581051 | 2.589718928 | 1.25E-07 |
| UBE2QL1 | 493.8624657 | 2.586052573 | 1.49E-51 |
| HEPACAM | 11.78854112 | 2.583912193 | 2.14E-02 |
| AL162727.1 | 20.90424433 | 2.582598279 | 1.58E-07 |
| GDF15 | 313.7269975 | 2.578408927 | 4.88E-06 |
| FGFR2 | 5.649823958 | 2.576179127 | 3.20E-02 |
| ADM2 | 805.2354835 | 2.575386774 | 5.60E-11 |
| MYCL | 127.9613386 | 2.573036378 | 7.53E-22 |
| LRRC15 | 26.29205623 | 2.559204653 | 4.81E-04 |
| CDH2 | 355.8677844 | 2.557880493 | 2.81E-37 |
| AC022784.1 | 65.50596088 | 2.55467244 | 1.74E-16 |
| AC017002.1 | 16.76283255 | 2.552801915 | 1.38E-05 |
| BX640514.2 | 77.90288435 | 2.552022836 | 2.78E-12 |
| IRX1 | 12.32971566 | 2.551808741 | 1.46E-03 |
| LAMC2 | 82.01010143 | 2.550860728 | 7.73E-19 |
| F3 | 631.683175 | 2.549787214 | 4.60E-39 |
| GNAT1 | 22.34118719 | 2.54914688 | 8.51E-03 |
| KCNK13 | 67.77061597 | 2.548761138 | 1.39E-25 |
| AC016705.2 | 94.42185277 | 2.548118729 | 2.39E-13 |
| HKDC1 | 536.4580666 | 2.547112529 | 6.57E-28 |
| OLIG2 | 29.16821453 | 2.54302645 | 4.23E-05 |
| IGFBPL1 | 580.1775613 | 2.542888495 | 8.04E-92 |
| LINC01615 | 22.14922554 | 2.538009003 | 1.83E-06 |
| FOLR1 | 159.0086188 | 2.537184275 | 1.38E-17 |
| VN2R19P | 6.404195999 | 2.53416899 | 1.02E-02 |

| | | | |
|---|---|---|---|
| AP000446.1 | 357.0586464 | 2.534099235 | 2.47E-55 |
| NDRG1 | 8255.208812 | 2.532771352 | 4.77E-38 |
| LCMT1-AS2 | 33.15623234 | 2.532662353 | 6.79E-08 |
| MAP3K15 | 28.1466474 | 2.532611624 | 7.44E-07 |
| FAM166B | 10.42880559 | 2.530197677 | 7.80E-04 |
| AC002401.4 | 17.60670996 | 2.524638908 | 7.98E-06 |
| MUC3A | 74.07583469 | 2.522494354 | 2.06E-07 |
| AL157896.1 | 13.64846009 | 2.509912907 | 9.98E-03 |
| KLF17 | 5.565692992 | 2.508002829 | 3.58E-02 |
| AC022239.1 | 18.69611026 | 2.507547282 | 1.85E-06 |
| AC044810.2 | 6.129593592 | 2.506800905 | 5.02E-03 |
| SFRP4 | 13.22019335 | 2.505429009 | 2.02E-04 |
| SMIM34B | 6.97659786 | 2.504605826 | 1.88E-02 |
| PPL | 285.5073873 | 2.503084775 | 1.85E-30 |
| TMEM139 | 162.5650754 | 2.498310641 | 6.50E-48 |
| GPRC5D-AS1 | 5.850032767 | 2.491268639 | 1.61E-02 |
| NA | 5.308877443 | 2.484546432 | 6.98E-03 |
| ARC | 110.2728391 | 2.48056709 | 3.82E-13 |
| SLC7A5 | 18550.72411 | 2.477901853 | 1.97E-14 |
| NOC2LP1 | 35.01371521 | 2.476660072 | 5.75E-09 |
| AC009716.2 | 5.056753745 | 2.469634445 | 2.35E-02 |
| FMN2 | 287.7331264 | 2.468913033 | 1.15E-22 |
| KCNH1 | 22.72830459 | 2.467773549 | 5.48E-06 |
| TUFT1 | 886.6362793 | 2.464820911 | 3.44E-97 |
| SULF1 | 13.73868172 | 2.464067491 | 1.78E-02 |
| CEBPA | 131.3984501 | 2.463147695 | 6.61E-28 |
| PLAUR | 813.9719978 | 2.460239011 | 1.83E-47 |
| HES4 | 110.1398204 | 2.458825655 | 8.24E-19 |
| SLCO1B3 | 13.29089053 | 2.458427023 | 1.65E-03 |
| CST7 | 33.33388902 | 2.45740346 | 1.86E-08 |
| LINC00689 | 9.95143427 | 2.455196583 | 2.10E-03 |
| ADAMTS9-AS2 | 15.97353599 | 2.453973682 | 2.12E-04 |
| DUSP1 | 581.9843181 | 2.453163756 | 1.83E-33 |
| MMP3 | 9.60446711 | 2.448068072 | 2.52E-02 |
| AL590399.4 | 12.31094571 | 2.447613887 | 2.33E-04 |
| AL355916.1 | 143.2120041 | 2.445557406 | 2.38E-32 |
| AC138150.2 | 38.54319126 | 2.443911511 | 5.60E-11 |
| AL136131.3 | 28.31490858 | 2.442424395 | 6.12E-05 |
| BPIFA2 | 22.6494482 | 2.441294656 | 2.65E-07 |
| CRIM1 | 9816.058979 | 2.438982844 | 7.79E-106 |
| LAYN | 609.0277932 | 2.436930352 | 8.29E-53 |
| AL844175.1 | 5.243046801 | 2.436671274 | 2.29E-02 |
| AC093616.1 | 454.260492 | 2.433300306 | 1.56E-51 |
| CSF2RB | 37.96421996 | 2.431978602 | 1.56E-07 |
| FAM131B | 590.29546 | 2.423088065 | 1.41E-70 |
| LINC01583 | 42.75130781 | 2.419721545 | 1.99E-09 |
| RPS6P12 | 74.88269617 | 2.418817107 | 5.91E-22 |

| | | | |
|---|---|---|---|
| AC073610.3 | 16.21540822 | 2.418083676 | 5.60E-04 |
| AC117422.1 | 6.67553884 | 2.413707161 | 2.75E-03 |
| HMGA1 | 9467.267639 | 2.413551812 | 1.59E-77 |
| TMEM155 | 10.51064918 | 2.411539656 | 5.31E-03 |
| KIRREL3 | 364.1978972 | 2.409752411 | 9.49E-34 |
| MYRIP | 5.796576788 | 2.408693028 | 3.68E-02 |
| FBN2 | 312.3030735 | 2.406540197 | 1.31E-14 |
| TMEM184A | 186.8034924 | 2.404754102 | 2.30E-16 |
| RIMS4 | 97.05276827 | 2.401177595 | 1.27E-15 |
| CEACAMP11 | 8.440503374 | 2.400624647 | 1.08E-02 |
| AC090409.1 | 76.02384453 | 2.397869131 | 5.51E-21 |
| IGFL1P1 | 18.4142963 | 2.397287658 | 1.05E-05 |
| SERPINB8 | 817.2619824 | 2.395116178 | 4.70E-29 |
| CYP2A6 | 5.287110338 | 2.392598688 | 3.60E-02 |
| ATF3 | 836.543072 | 2.38906888 | 1.42E-18 |
| CHAC1 | 753.1216115 | 2.388385127 | 7.83E-08 |
| SLAMF9 | 7.095931523 | 2.387672868 | 1.46E-02 |
| WDR66 | 107.1767467 | 2.385939353 | 5.73E-13 |
| AC004877.1 | 19.22436289 | 2.381811556 | 1.10E-04 |
| SLC7A11 | 1423.958723 | 2.379026911 | 8.78E-06 |
| ESRG | 287.4514818 | 2.377232954 | 9.04E-33 |
| LINC01764 | 8.559247651 | 2.375605852 | 6.30E-03 |
| AC090152.1 | 15.60249365 | 2.375373551 | 1.46E-04 |
| LIN28A | 640.0340004 | 2.374330869 | 7.25E-70 |
| INHBA | 62.28707009 | 2.369917868 | 1.28E-18 |
| TNFRSF9 | 304.3101605 | 2.369764129 | 1.05E-19 |
| TMEM225B | 15.79911079 | 2.366975671 | 3.27E-06 |
| MIR503HG | 2154.711243 | 2.366899379 | 5.86E-26 |
| LINC00857 | 12.31235457 | 2.364954911 | 1.17E-03 |
| ERBB3 | 99.31327608 | 2.362743931 | 1.75E-14 |
| CRYBG2 | 93.98396619 | 2.36207028 | 6.60E-08 |
| ITIH5 | 73.04147724 | 2.355100052 | 1.32E-09 |
| LINC01436 | 88.90708551 | 2.353524697 | 3.06E-26 |
| TTC9 | 226.5344015 | 2.353025726 | 1.51E-34 |
| Z99943.1 | 6.154884971 | 2.350760388 | 1.39E-02 |
| OR7E14P | 8.152750421 | 2.346373898 | 1.93E-03 |
| ARNT2 | 978.2147149 | 2.345531993 | 1.77E-55 |
| IL6 | 31.65351887 | 2.342234274 | 4.66E-05 |
| DLGAP1-AS2 | 423.8719857 | 2.338926715 | 1.46E-23 |
| WNT5B | 481.1590035 | 2.338572567 | 2.38E-35 |
| AC122710.2 | 8.911190889 | 2.335796736 | 5.74E-03 |
| AC243972.1 | 13.61856719 | 2.333107078 | 3.57E-04 |
| IL20RB | 107.3800024 | 2.331697194 | 6.48E-09 |
| KCNQ1 | 14.72470286 | 2.331533635 | 8.59E-03 |
| CYFIP2 | 654.8815999 | 2.330770875 | 3.38E-22 |
| NA | 26.04020648 | 2.329641373 | 1.20E-08 |
| SPATA8 | 7.77623881 | 2.328870598 | 6.48E-03 |

| | | | |
|---|---|---|---|
| S100A2 | 104.4095725 | 2.3253185 | 3.62E-12 |
| LINC02518 | 7.233358566 | 2.32443442 | 8.50E-03 |
| FOXE1 | 349.40179 | 2.32085975 | 1.18E-37 |
| EDN1 | 1541.884744 | 2.319594134 | 1.86E-27 |
| EXPH5 | 144.9258139 | 2.31929744 | 4.46E-20 |
| DPP4 | 73.62921846 | 2.318370759 | 2.47E-19 |
| S100A3 | 298.3816144 | 2.317897786 | 5.72E-46 |
| EHD4 | 2189.688796 | 2.312936158 | 1.55E-113 |
| CYTOR | 2035.603011 | 2.31117162 | 1.47E-109 |
| WTIP | 955.4419514 | 2.308478611 | 5.49E-35 |
| RPSAP52 | 35.61562351 | 2.306758656 | 1.46E-11 |
| PICART1 | 40.09760807 | 2.306534101 | 1.26E-10 |
| TDRG1 | 14.32537586 | 2.303608036 | 1.71E-04 |
| HBEGF | 861.6443607 | 2.302650393 | 1.34E-72 |
| ANKFN1 | 98.2235767 | 2.300280132 | 1.48E-16 |
| C1orf198 | 2749.169919 | 2.300005042 | 8.94E-194 |
| AC126177.6 | 5.309390916 | 2.298993027 | 4.16E-02 |
| FSTL3 | 2216.418323 | 2.297957136 | 8.13E-39 |
| PLEKHS1 | 100.4445389 | 2.297693265 | 8.85E-17 |
| DUX4L31 | 29.25806601 | 2.292301085 | 1.90E-07 |
| TNIP3 | 7.611684205 | 2.285139379 | 1.80E-02 |
| GPRC5A | 3645.517891 | 2.284443962 | 3.59E-60 |
| BX322557.1 | 5.468117233 | 2.283459063 | 2.88E-02 |
| AC005670.2 | 15.0788706 | 2.281912253 | 1.57E-03 |
| CARD11 | 487.3641494 | 2.279190138 | 1.64E-28 |
| DIO2 | 72.37432467 | 2.277976494 | 4.96E-07 |
| OGFRL1 | 1421.080525 | 2.27778932 | 2.09E-41 |
| PPP1R15A | 3154.007942 | 2.274656178 | 1.18E-28 |
| HIF1A-AS1 | 59.94623766 | 2.26845617 | 4.22E-13 |
| TFPI2 | 2830.248721 | 2.266060685 | 3.64E-70 |
| AC006273.1 | 24.67120854 | 2.265676963 | 2.41E-07 |
| AC026250.1 | 102.7984567 | 2.265063254 | 1.46E-21 |
| DHRS3 | 27.89732788 | 2.258768891 | 3.43E-06 |
| CLEC4M | 17.52321515 | 2.256682444 | 7.87E-05 |
| NA | 8.854667232 | 2.25310731 | 2.38E-03 |
| AC004817.3 | 15.82350215 | 2.252260089 | 7.01E-04 |
| TMEM56 | 329.4539423 | 2.249924083 | 1.84E-26 |
| LINC00601 | 16.03766609 | 2.248763434 | 9.11E-05 |
| NOX1 | 21.84778441 | 2.248327771 | 3.82E-04 |
| BEX2 | 181.7672279 | 2.24783014 | 1.31E-09 |
| FADS3 | 3833.388893 | 2.243039279 | 9.32E-117 |
| ERICH2 | 16.08752266 | 2.242724761 | 9.91E-04 |
| PTPRR | 11.79398755 | 2.241511903 | 4.48E-03 |
| FAM163B | 39.18434864 | 2.240805387 | 1.51E-06 |
| ACTBL2 | 62.37567461 | 2.235655734 | 2.75E-10 |
| FABP3 | 256.5391819 | 2.23214161 | 1.17E-16 |
| ECEL1P2 | 5.851314691 | 2.230888476 | 2.90E-02 |

| | | | |
|---|---|---|---|
| CX3CR1 | 5.956254789 | 2.230298824 | 4.71E-02 |
| AP003717.1 | 9.464883929 | 2.230222773 | 2.89E-02 |
| C7orf57 | 122.2458911 | 2.227433463 | 4.09E-13 |
| FAM19A5 | 5.778785156 | 2.224565556 | 3.53E-02 |
| CALB1 | 16.64156536 | 2.222356539 | 5.10E-03 |
| NUAK1 | 3145.049458 | 2.221828755 | 8.47E-89 |
| UGCG | 4030.044103 | 2.221241813 | 1.43E-80 |
| AL121890.4 | 11.68394512 | 2.220941867 | 6.64E-04 |
| SIK1B | 50.93325372 | 2.216027953 | 5.55E-07 |
| AC011944.1 | 11.91468553 | 2.215882443 | 7.54E-03 |
| TTBK1 | 13.02622851 | 2.214555868 | 2.97E-04 |
| CAV1 | 23107.43986 | 2.211744154 | 4.56E-73 |
| NEGR1 | 205.2653914 | 2.2115463 | 1.16E-10 |
| HBE1 | 415.2750207 | 2.21135496 | 1.66E-19 |
| INSYN2B | 326.6455031 | 2.211171563 | 1.50E-19 |
| SMPD3 | 199.6240386 | 2.211130482 | 1.88E-10 |
| AC073610.1 | 10.58281692 | 2.210344348 | 6.11E-03 |
| EEF1A2 | 5981.383771 | 2.210131093 | 1.27E-77 |
| ANXA1 | 9614.08568 | 2.208865295 | 9.61E-27 |
| NA | 40.08367415 | 2.202430585 | 9.42E-11 |
| REN | 21.64424951 | 2.196362257 | 3.38E-03 |
| NANOS3 | 11.09929873 | 2.19266347 | 2.99E-04 |
| ALS2CL | 852.7311576 | 2.189093429 | 1.60E-36 |
| PSAT1 | 16232.43473 | 2.188384656 | 3.48E-25 |
| TMEM40 | 73.22422857 | 2.18762262 | 2.32E-10 |
| OLAH | 10.38694469 | 2.187492471 | 2.97E-03 |
| TRIM58 | 392.4915009 | 2.187051263 | 1.36E-54 |
| OTUD7A | 19.82865708 | 2.184524785 | 2.48E-02 |
| XAGE3 | 258.2371969 | 2.18325452 | 2.73E-52 |
| AC092645.1 | 38.54954026 | 2.179844928 | 1.73E-05 |
| AC091806.1 | 113.7234827 | 2.176113516 | 9.02E-13 |
| AC148476.1 | 31.43670773 | 2.174484207 | 7.02E-08 |
| AP001189.5 | 7.632947862 | 2.173940209 | 4.61E-02 |
| RBPJL | 20.92174682 | 2.172857572 | 8.35E-04 |
| CES1P2 | 28.23427915 | 2.172538714 | 1.92E-05 |
| AP000894.2 | 11.2305448 | 2.171911577 | 8.57E-03 |
| MYOZ1 | 20.50391645 | 2.171252289 | 4.03E-05 |
| AL357033.1 | 14.66257818 | 2.169815957 | 3.01E-05 |
| CD55 | 1726.450139 | 2.169645121 | 8.05E-113 |
| CLDN3 | 137.1034934 | 2.168188079 | 3.75E-20 |
| AC106760.1 | 323.397699 | 2.165342157 | 2.78E-43 |
| PNCK | 57.10837434 | 2.163906411 | 7.96E-06 |
| GLI1 | 189.0118592 | 2.162909798 | 8.98E-11 |
| FGF18 | 29.63811764 | 2.162906603 | 2.75E-05 |
| VTN | 154.8509553 | 2.161181164 | 2.31E-12 |
| FMOD | 1627.644591 | 2.156187508 | 7.24E-35 |
| PVR | 6076.548739 | 2.155561132 | 3.39E-40 |

| | | | |
|---|---|---|---|
| RPSAP71 | 7.432948907 | 2.155134039 | 1.53E-02 |
| AL035458.2 | 6.031236525 | 2.15449705 | 2.68E-02 |
| LRRC38 | 9.873393408 | 2.151535842 | 1.95E-02 |
| ARFGEF3 | 55.42513659 | 2.144774147 | 3.52E-08 |
| SLC5A4 | 15.92355371 | 2.142962157 | 1.56E-04 |
| CTH | 928.9606088 | 2.140116271 | 2.31E-12 |
| AL021578.1 | 5.965190378 | 2.137562193 | 4.43E-02 |
| SUSD2 | 605.236627 | 2.137551043 | 1.31E-34 |
| RAB32 | 1770.456458 | 2.13394434 | 7.67E-67 |
| AL391422.4 | 577.9328061 | 2.131876021 | 2.40E-23 |
| PRKAG2 | 1255.564186 | 2.129482436 | 3.13E-57 |
| SH3GL3 | 111.9951372 | 2.128728704 | 2.12E-13 |
| LINC00882 | 70.53774856 | 2.127931919 | 1.24E-13 |
| AC060766.6 | 9.190089575 | 2.12748081 | 2.58E-03 |
| ZNF534 | 9.308285298 | 2.126326143 | 1.04E-02 |
| UPP1 | 3542.442731 | 2.125364727 | 2.08E-26 |
| RUBCNL | 165.9995 | 2.124265511 | 6.81E-12 |
| MAP6 | 112.0278279 | 2.123252562 | 1.09E-13 |
| NA | 21.84005856 | 2.120878079 | 2.29E-06 |
| LINC00862 | 29.77997774 | 2.118219789 | 5.61E-05 |
| SLC35F3 | 102.7993497 | 2.115801001 | 7.27E-18 |
| COL8A1 | 2277.352965 | 2.114337203 | 3.24E-27 |
| AC108463.2 | 15.81970056 | 2.113689345 | 1.43E-04 |
| AC116351.1 | 130.5665582 | 2.11330432 | 5.12E-16 |
| AC009264.1 | 24.17787094 | 2.110533458 | 1.15E-05 |
| MYOZ2 | 10.39892218 | 2.109151025 | 4.78E-03 |
| KIF21B | 1486.646113 | 2.108243731 | 1.19E-26 |
| AC114488.2 | 14.72393248 | 2.108107473 | 1.28E-03 |
| AC015987.1 | 29.65507999 | 2.105545038 | 3.76E-06 |
| CLGN | 74.76072684 | 2.103762172 | 3.11E-04 |
| NA | 32.99010685 | 2.102735437 | 8.36E-08 |
| LINC00958 | 7.133378578 | 2.100700058 | 1.26E-02 |
| LIPH | 201.1100145 | 2.100127309 | 2.44E-25 |
| AC025419.1 | 7.159029254 | 2.099645777 | 3.00E-02 |
| AL359510.2 | 5.030386633 | 2.099164646 | 4.71E-02 |
| RBFOX3 | 23.02501928 | 2.09766222 | 3.15E-03 |
| HOXC13-AS | 178.5640571 | 2.097439622 | 6.12E-20 |
| ADARB1 | 2219.524268 | 2.095467691 | 3.69E-20 |
| SPTB | 236.3609432 | 2.094023903 | 2.68E-14 |
| SDC2 | 2063.499858 | 2.093091985 | 1.41E-70 |
| NA | 1365.072079 | 2.085926174 | 7.76E-63 |
| SDR42E1 | 246.9387746 | 2.085926158 | 9.95E-23 |
| LINC00239 | 13.37656744 | 2.083065703 | 1.46E-02 |
| RPS6KA2 | 1707.929685 | 2.077459675 | 1.13E-38 |
| OXTR | 109.0439804 | 2.077123913 | 6.44E-10 |
| KCNIP1 | 288.8772349 | 2.07453034 | 8.46E-23 |
| SH2D5 | 415.6853727 | 2.071146959 | 1.96E-58 |

| | | | |
|---|---|---|---|
| MATN4 | 29.85064027 | 2.07051546 | 1.85E-05 |
| ERVV-2 | 60.76003593 | 2.0705071 | 9.66E-07 |
| AC129492.2 | 18.58770482 | 2.068609819 | 1.72E-05 |
| PCK2 | 6290.645737 | 2.067372452 | 1.89E-16 |
| PRSS8 | 17.4747297 | 2.062348289 | 7.72E-04 |
| AC010255.1 | 15.32312268 | 2.061313778 | 3.39E-03 |
| MAEL | 7.774685191 | 2.053639178 | 2.46E-02 |
| AL365181.3 | 171.3149321 | 2.052085474 | 3.17E-14 |
| GABARAPL1 | 3444.554127 | 2.049393018 | 1.65E-117 |
| AL139352.1 | 15.41388576 | 2.048955489 | 2.44E-04 |
| RXFP3 | 26.79648996 | 2.048776051 | 6.88E-04 |
| AC005736.1 | 74.59682991 | 2.046006028 | 7.51E-15 |
| LINC00964 | 5.892597204 | 2.043213679 | 4.29E-02 |
| MAFF | 607.0319197 | 2.042904178 | 1.07E-28 |
| ENO2 | 1006.576172 | 2.041606002 | 5.01E-21 |
| INKA2-AS1 | 103.6961057 | 2.040346468 | 5.43E-09 |
| AL590666.2 | 45.63170054 | 2.039877137 | 1.06E-09 |
| TMEM156 | 66.58521868 | 2.039348654 | 1.43E-05 |
| SPINT1 | 91.07398261 | 2.039191563 | 1.15E-11 |
| LINC01018 | 72.32292642 | 2.034089968 | 6.77E-17 |
| MIR4435-2HG | 2097.286563 | 2.032182702 | 2.77E-93 |
| FBLN7 | 185.287182 | 2.029579213 | 1.07E-24 |
| IL11 | 145.4784115 | 2.028709633 | 3.20E-16 |
| CATSPER1 | 133.9849911 | 2.027305434 | 4.76E-20 |
| TNFSF15 | 89.03391984 | 2.024287728 | 2.20E-11 |
| KIAA1211L | 13.14754564 | 2.023785949 | 3.83E-03 |
| INHBA-AS1 | 14.99610622 | 2.022758244 | 1.12E-03 |
| AC078905.1 | 7.967882544 | 2.02160268 | 8.83E-03 |
| INMT | 744.6966048 | 2.021037942 | 2.93E-58 |
| DUXA | 17.7868599 | 2.020187061 | 1.18E-02 |
| HR | 283.1205138 | 2.019210996 | 2.97E-09 |
| CALB2 | 75.59255705 | 2.018110535 | 2.36E-10 |
| AP000892.3 | 436.2274301 | 2.014708529 | 1.77E-26 |
| PDLIM1 | 4528.312637 | 2.013559734 | 6.93E-72 |
| MYPN | 979.7692297 | 2.012772751 | 2.19E-27 |
| NOTUM | 82.32665438 | 2.01275183 | 4.04E-14 |
| GOLT1A | 20.85450575 | 2.008480513 | 5.27E-05 |
| UNC5A | 9.614455664 | 2.008341409 | 2.04E-02 |
| LYPD1 | 36.76512797 | 2.00535628 | 2.81E-07 |
| LINC01121 | 12.19044645 | 2.004798381 | 1.56E-02 |
| PTCH1 | 1620.679928 | 2.003098833 | 1.34E-34 |
| S100A16 | 1005.645382 | 2.001644274 | 2.70E-55 |
| RBMS3-AS3 | 6.144421648 | 1.998422101 | 2.34E-02 |
| ARSI | 573.0458329 | 1.998310394 | 5.90E-23 |
| AC124276.1 | 10.9518038 | 1.997864378 | 1.03E-02 |
| LINC01151 | 6.752733732 | 1.997490723 | 1.40E-02 |
| AP003174.1 | 5.735948833 | 1.995460678 | 2.69E-02 |

| | | | |
|---|---|---|---|
| LOXL4 | 806.0990284 | 1.995263152 | 1.90E-55 |
| SLC6A1 | 26.45725069 | 1.9945848 | 8.54E-03 |
| IRF8 | 69.17208856 | 1.994360099 | 5.30E-08 |
| CAV2 | 2124.493216 | 1.994195648 | 3.31E-61 |
| L1CAM | 10255.39231 | 1.991736824 | 2.03E-36 |
| ANXA8 | 20.84443961 | 1.990955602 | 3.26E-05 |
| SLC3A2 | 10651.30897 | 1.990620725 | 6.19E-17 |
| FAM129A | 2074.598291 | 1.989495831 | 7.39E-15 |
| APLN | 612.3809223 | 1.987565964 | 2.28E-14 |
| MAP1LC3C | 6.634733519 | 1.987437628 | 3.14E-02 |
| CLSTN2 | 328.2908032 | 1.981984637 | 2.85E-16 |
| CALCB | 23.00979005 | 1.980749033 | 1.60E-02 |
| TRIB3 | 8474.977623 | 1.98052006 | 1.40E-08 |
| TOX | 45.34735463 | 1.980508022 | 1.19E-09 |
| EPPK1 | 92.84336005 | 1.979089547 | 1.93E-17 |
| AC091182.2 | 17.00484763 | 1.976045262 | 2.56E-04 |
| FLJ45513 | 22.04428444 | 1.974876834 | 1.69E-04 |
| CCL26 | 75.15112479 | 1.974597012 | 3.88E-10 |
| PTGFR | 15.23427547 | 1.972975249 | 1.11E-04 |
| AC005586.2 | 14.13409443 | 1.971278056 | 1.72E-04 |
| CLIC3 | 15.9697621 | 1.97035346 | 4.08E-03 |
| UBASH3B | 966.8217515 | 1.968217676 | 3.14E-23 |
| AL713998.1 | 83.75568185 | 1.966527141 | 1.14E-06 |
| HNF1B | 9.019723547 | 1.966009438 | 4.19E-02 |
| NA | 43.02971145 | 1.965401305 | 5.09E-05 |
| AC092162.3 | 14.46723367 | 1.964496687 | 7.62E-04 |
| FGFR3 | 194.2592122 | 1.963993087 | 3.99E-10 |
| MMP19 | 140.8610266 | 1.962606868 | 9.24E-13 |
| SIK1 | 154.214165 | 1.960935299 | 7.83E-26 |
| MPP4 | 50.02754041 | 1.956910135 | 1.05E-05 |
| PROSER2 | 428.4427461 | 1.956249725 | 1.90E-25 |
| AC104024.2 | 12.74438091 | 1.956008487 | 4.61E-03 |
| TFCP2L1 | 94.35599976 | 1.955599866 | 3.81E-08 |
| FOXO6 | 98.30035387 | 1.952344917 | 2.30E-18 |
| AC099518.6 | 18.71020804 | 1.951518362 | 1.17E-04 |
| NMNAT2 | 86.63767565 | 1.951050109 | 8.40E-10 |
| AC142381.3 | 18.77409383 | 1.950802208 | 3.87E-03 |
| AL355312.4 | 60.37212687 | 1.94795663 | 6.17E-12 |
| NA | 18.72396924 | 1.947314291 | 5.69E-04 |
| PXDNL | 8.079637129 | 1.947160294 | 2.64E-02 |
| IGF2 | 2046.485533 | 1.94672612 | 2.64E-26 |
| CRIM1-DT | 61.0346116 | 1.946046478 | 4.97E-13 |
| NA | 225.5584139 | 1.944717474 | 3.00E-27 |
| LINP1 | 60.87986528 | 1.939509994 | 3.30E-12 |
| DUSP8 | 315.6478431 | 1.939358542 | 5.36E-15 |
| VWF | 121.5698336 | 1.938980156 | 1.70E-09 |
| AP001057.1 | 8.751761603 | 1.936855803 | 4.95E-03 |

| | | | |
|---|---|---|---|
| AC136628.1 | 42.07864024 | 1.936809077 | 5.20E-05 |
| SPANXA1 | 416.0491261 | 1.935402847 | 3.09E-21 |
| NXN | 1791.096427 | 1.934863089 | 1.05E-59 |
| ITGA3 | 22276.61878 | 1.931975816 | 1.10E-80 |
| PLEKHA6 | 883.8866704 | 1.931581274 | 1.07E-19 |
| ACE2 | 59.32408581 | 1.930207337 | 7.75E-08 |
| BEST1 | 300.9674433 | 1.926694889 | 9.51E-15 |
| PLAC9P1 | 16.69223394 | 1.921495474 | 1.70E-04 |
| ABL2 | 2982.838415 | 1.921297173 | 1.29E-72 |
| CREB5 | 770.2728268 | 1.921079673 | 9.98E-13 |
| PKP2 | 1275.453997 | 1.919424835 | 4.12E-73 |
| LINC00869 | 134.1130625 | 1.912907004 | 2.34E-06 |
| ARHGEF28 | 3310.07867 | 1.910476554 | 1.52E-34 |
| AC017002.3 | 10.79083619 | 1.909437241 | 1.28E-02 |
| TMEM71 | 20.65123729 | 1.909332565 | 6.33E-03 |
| SHOX | 186.8715178 | 1.907200749 | 2.08E-22 |
| GRIK5 | 70.41443968 | 1.906376257 | 3.87E-04 |
| TMEM92 | 202.8482872 | 1.90479807 | 1.39E-25 |
| HSD11B1-AS1 | 25.91185308 | 1.904618933 | 9.02E-05 |
| GAD1 | 53.22551783 | 1.903368727 | 4.42E-11 |
| PEAR1 | 1120.147306 | 1.902616643 | 1.68E-48 |
| GDNF | 9.644136723 | 1.901666177 | 4.59E-02 |
| CFAP58 | 18.54124951 | 1.899007168 | 2.17E-04 |
| FLNB | 29152.77127 | 1.894065626 | 1.20E-47 |
| USP43 | 231.4230637 | 1.889042835 | 8.08E-28 |
| ABCA13 | 16.30721454 | 1.884010075 | 3.53E-02 |
| LINC00900 | 27.39226582 | 1.883359434 | 1.53E-03 |
| AC124947.1 | 10.90404445 | 1.880665176 | 4.71E-02 |
| VAV1 | 107.6228808 | 1.880284064 | 7.58E-12 |
| AKR1C7P | 9.921288973 | 1.876614612 | 3.69E-02 |
| HPSE | 457.3034666 | 1.873620426 | 1.57E-50 |
| KHDRBS3 | 71.81950796 | 1.873038859 | 2.16E-13 |
| SIGLEC14 | 107.3074879 | 1.863991344 | 4.92E-11 |
| CES1P1 | 26.62878007 | 1.860695332 | 2.75E-02 |
| LINC01356 | 69.31803959 | 1.859984689 | 6.42E-10 |
| ACAN | 13.49155482 | 1.856809474 | 3.42E-02 |
| AC243964.3 | 235.8240355 | 1.856139157 | 2.84E-29 |
| HRK | 25.95120166 | 1.855575478 | 1.35E-02 |
| PKD1L2 | 885.8875173 | 1.850317752 | 1.28E-10 |
| TMEM154 | 315.1802205 | 1.847606592 | 3.04E-09 |
| SFN | 28.73189058 | 1.847424646 | 1.02E-05 |
| SLC1A4 | 2242.247256 | 1.847287078 | 6.08E-16 |
| RAB11FIP1 | 3684.571645 | 1.847270938 | 2.04E-85 |
| CD1D | 791.2080238 | 1.847231228 | 8.97E-38 |
| ITPK1 | 2503.186321 | 1.845654726 | 1.76E-61 |
| BNC1 | 1562.452238 | 1.845566869 | 2.10E-19 |
| FOXE3 | 14.39157214 | 1.844145213 | 3.05E-03 |

| | | | |
|---|---|---|---|
| PEA15 | 12565.60558 | 1.843423406 | 3.69E-45 |
| ANO4 | 74.03198246 | 1.841659815 | 2.51E-10 |
| PARP8 | 150.9692421 | 1.840902623 | 2.79E-14 |
| ME3 | 73.97688032 | 1.839271683 | 4.64E-08 |
| RGS7 | 47.66134455 | 1.838334875 | 2.15E-09 |
| QRFPR | 70.06962025 | 1.837231271 | 3.38E-08 |
| LRRC20 | 1074.126572 | 1.836504023 | 2.33E-32 |
| NRK | 14.45792918 | 1.835580603 | 2.01E-03 |
| IL31RA | 161.1209566 | 1.835083293 | 9.91E-12 |
| IL18 | 12.86748055 | 1.835011474 | 5.15E-03 |
| PHGDH | 8589.264013 | 1.83424532 | 6.72E-35 |
| LNCOC1 | 35.33999473 | 1.832978959 | 6.58E-05 |
| KSR2 | 287.877213 | 1.831667039 | 2.19E-14 |
| OTOF | 23.81127999 | 1.830739952 | 4.09E-03 |
| SLC52A3 | 59.35415736 | 1.829540127 | 1.63E-07 |
| PRKG1-AS1 | 7.922083969 | 1.82787273 | 2.14E-02 |
| SZT2-AS1 | 13.31741173 | 1.827607391 | 1.92E-02 |
| PRKCE | 1501.609574 | 1.827231423 | 1.06E-18 |
| LPO | 43.94056569 | 1.825822087 | 6.02E-04 |
| PDE1C | 3542.718138 | 1.825624635 | 5.98E-22 |
| AC147067.1 | 13.41097365 | 1.824272209 | 1.68E-03 |
| CGN | 277.5497533 | 1.823808642 | 7.10E-16 |
| AC005291.2 | 13.64347463 | 1.822884974 | 4.59E-02 |
| TSKS | 10.55565234 | 1.822745981 | 1.37E-02 |
| BCL2L1 | 5620.751235 | 1.818596542 | 3.80E-108 |
| GBX2 | 105.7843387 | 1.815286439 | 2.26E-15 |
| XDH | 47.86677611 | 1.814323298 | 3.29E-05 |
| AL035252.4 | 21.61470413 | 1.812863936 | 4.46E-02 |
| AL390195.2 | 16.80462659 | 1.811404362 | 9.14E-04 |
| LINC00623 | 78.41347884 | 1.810152246 | 9.45E-07 |
| ATP1A2 | 143.772855 | 1.808805074 | 4.11E-05 |
| LINC00513 | 40.82278957 | 1.808784013 | 4.79E-07 |
| FLNC | 24286.4346 | 1.807807803 | 2.26E-46 |
| TPM1 | 23789.68 | 1.806728813 | 7.03E-65 |
| AL355103.1 | 9.811611285 | 1.806100158 | 1.80E-02 |
| AREG | 9.792709533 | 1.804999886 | 3.10E-02 |
| IGFBP5 | 6426.850593 | 1.804660381 | 2.99E-05 |
| GPR17 | 17.22731363 | 1.802804761 | 1.63E-03 |
| PPP2R2C | 56.81038469 | 1.802754168 | 1.33E-04 |
| P3H2 | 6277.970193 | 1.799828178 | 2.52E-44 |
| AC005481.1 | 58.82653227 | 1.799348527 | 4.80E-11 |
| FILIP1 | 45.8162629 | 1.797486879 | 2.56E-10 |
| FO393418.1 | 6.878802365 | 1.794558485 | 4.32E-02 |
| PSORS1C1 | 40.71724625 | 1.793579813 | 5.37E-05 |
| SCGB3A2 | 67.32651415 | 1.791351596 | 2.35E-07 |
| LINC01503 | 111.7516947 | 1.79089812 | 6.29E-22 |
| MET | 4328.732849 | 1.787908175 | 1.59E-33 |

| | | | |
|---|---|---|---|
| CHST3 | 1381.804227 | 1.787194253 | 1.25E-66 |
| VXN | 179.481724 | 1.786199157 | 1.14E-16 |
| SEMA7A | 526.884678 | 1.781970248 | 2.73E-23 |
| ADRB2 | 262.2256592 | 1.781820223 | 3.10E-34 |
| SLC7A1 | 3757.962617 | 1.779165617 | 9.92E-21 |
| MAP3K20 | 5636.626502 | 1.777415002 | 7.35E-35 |
| NMU | 16.59498685 | 1.777005895 | 2.47E-03 |
| PPM1H | 121.3528802 | 1.776191869 | 2.64E-08 |
| TGM1 | 90.89196127 | 1.775348263 | 6.26E-10 |
| MATN3 | 531.0146682 | 1.772885735 | 8.01E-35 |
| NMNAT3 | 18.81273681 | 1.770625157 | 1.47E-03 |
| GAL3ST3 | 7.300615404 | 1.77034801 | 4.93E-02 |
| PXDC1 | 1898.914831 | 1.769978249 | 6.49E-87 |
| WWC2 | 2643.317265 | 1.768927945 | 5.90E-24 |
| IL24 | 21.39327091 | 1.76844601 | 2.13E-02 |
| AL121820.1 | 8.32274803 | 1.764768086 | 3.85E-02 |
| SHMT2 | 14299.48947 | 1.764321271 | 2.14E-18 |
| TCAF2P1 | 28.17114646 | 1.763870209 | 1.34E-03 |
| CACNA1I | 43.84665629 | 1.763456632 | 3.46E-06 |
| SLC22A15 | 397.3289944 | 1.759921752 | 1.54E-07 |
| AJAP1 | 312.8253769 | 1.759721108 | 4.58E-16 |
| HMGA2-AS1 | 18.60959412 | 1.758861513 | 8.63E-04 |
| PTPRN | 673.3985235 | 1.758390478 | 3.09E-19 |
| ARL4D | 533.0645091 | 1.758226214 | 4.45E-25 |
| ZFP42 | 83.95063084 | 1.755665021 | 1.88E-11 |
| NA | 310.8966264 | 1.754260607 | 1.22E-26 |
| AC073130.2 | 9.878200911 | 1.752400382 | 2.17E-02 |
| AC080023.1 | 19.56690321 | 1.750955621 | 5.38E-05 |
| CDH7 | 14.05780657 | 1.750262994 | 1.57E-02 |
| RPLP0P2 | 306.6935964 | 1.7472566 | 4.46E-15 |
| AC004917.1 | 9.77783445 | 1.746342594 | 1.30E-02 |
| AC012464.1 | 11.57719136 | 1.745383326 | 1.87E-02 |
| GGT5 | 898.8201285 | 1.745296008 | 1.96E-18 |
| VLDLR | 1352.477514 | 1.744784445 | 1.05E-74 |
| GCSAM | 14.9224264 | 1.743735471 | 4.87E-03 |
| AL121772.1 | 44.2696685 | 1.740206179 | 1.52E-06 |
| GATA3 | 89.10281909 | 1.740072784 | 2.58E-05 |
| LINC00211 | 14.16910301 | 1.738747723 | 1.10E-03 |
| SPINT2 | 13.58599961 | 1.737418541 | 2.00E-02 |
| AC009108.3 | 8.624638713 | 1.735576471 | 2.08E-02 |
| APBA1 | 534.1102272 | 1.734820497 | 2.93E-22 |
| PLS3-AS1 | 36.45111194 | 1.733933212 | 1.70E-06 |
| ERFE | 143.5214673 | 1.73274908 | 2.48E-11 |
| VEGFA | 2322.712089 | 1.73154419 | 2.40E-10 |
| AC027117.1 | 17.37176545 | 1.730628009 | 2.59E-04 |
| ERICH5 | 54.53299792 | 1.727517593 | 4.93E-10 |
| VWA5B2 | 10.94573865 | 1.726196114 | 7.10E-03 |

| | | | |
|---|---|---|---|
| CAMK2A | 27.28062693 | 1.72442689 | 1.82E-03 |
| STPG4 | 10.24940873 | 1.72412146 | 3.01E-02 |
| IP6K3 | 192.4547338 | 1.723464094 | 3.49E-15 |
| P3H2-AS1 | 15.1095985 | 1.722761641 | 2.02E-03 |
| SDC4 | 4176.418327 | 1.721108012 | 3.32E-34 |
| SH2D3C | 192.6025404 | 1.719343297 | 4.52E-11 |
| GPR78 | 42.20307489 | 1.71792947 | 2.79E-07 |
| CU639417.5 | 17.68752859 | 1.716077701 | 1.47E-02 |
| CYTH3 | 8882.569858 | 1.714966959 | 2.69E-114 |
| KLC3 | 240.3054713 | 1.714659072 | 8.59E-21 |
| AC005381.1 | 14.56912461 | 1.714590553 | 1.78E-02 |
| PDE6A | 35.90315679 | 1.714067358 | 5.84E-07 |
| AC107956.1 | 22.9915758 | 1.713723904 | 4.19E-04 |
| NA | 440.3647721 | 1.712955828 | 1.68E-26 |
| BCL11A | 26.1444669 | 1.712361257 | 1.52E-03 |
| NMT2 | 1496.883168 | 1.711800539 | 1.45E-95 |
| MYLK3 | 310.1611264 | 1.711528806 | 2.27E-24 |
| GAS6 | 1171.454117 | 1.710016442 | 1.17E-16 |
| MAGEA4 | 9.336679457 | 1.708599966 | 2.80E-02 |
| SH3BP4 | 4262.928303 | 1.704774805 | 2.86E-65 |
| PODXL | 5988.65413 | 1.704036223 | 1.49E-32 |
| LINC01600 | 10.46319634 | 1.702555158 | 2.27E-02 |
| ANKRD33B | 166.908834 | 1.699949145 | 1.49E-17 |
| AC114947.2 | 30.29873374 | 1.699441545 | 2.81E-03 |
| SOWAHB | 9.765481541 | 1.698411371 | 1.50E-02 |
| MYH9 | 39701.66481 | 1.695973263 | 6.55E-63 |
| MTHFD2 | 8169.227928 | 1.69498952 | 1.42E-17 |
| GRIN1 | 70.83324153 | 1.694564811 | 6.27E-09 |
| DUBR | 562.7047495 | 1.694388384 | 4.31E-30 |
| RCAN1 | 3868.497247 | 1.691601081 | 3.12E-38 |
| CAVIN1 | 22717.39463 | 1.689564489 | 2.24E-38 |
| AC100803.2 | 22.40861951 | 1.68822236 | 1.29E-03 |
| AC092437.1 | 11.49377109 | 1.686470885 | 6.22E-03 |
| C4orf19 | 21.16468249 | 1.685862421 | 5.97E-04 |
| LINC01647 | 23.74194246 | 1.684965291 | 5.36E-04 |
| GRIK4 | 72.0864243 | 1.684822954 | 3.15E-03 |
| ZIC5 | 122.3265531 | 1.682132721 | 3.48E-13 |
| AP002956.1 | 14354.37341 | 1.681809588 | 1.10E-38 |
| CITED4 | 281.9765894 | 1.680391142 | 2.04E-16 |
| ADAP2 | 273.0761693 | 1.679241797 | 1.48E-18 |
| JUN | 2696.774838 | 1.677887543 | 8.72E-18 |
| LINC01405 | 15.01222138 | 1.673643459 | 3.33E-03 |
| ARID3B | 403.876423 | 1.672333187 | 9.01E-33 |
| TMED10P2 | 29.80088923 | 1.671871075 | 1.26E-05 |
| TEX19 | 43.55508042 | 1.671691817 | 3.38E-06 |
| NFE2L3 | 4016.428492 | 1.670967494 | 2.86E-52 |
| APBB1IP | 649.9227515 | 1.67081059 | 3.73E-29 |

| | | | |
|---|---|---|---|
| WFDC3 | 25.00327681 | 1.669573345 | 7.45E-04 |
| AC099568.2 | 26.25689458 | 1.66799197 | 2.17E-03 |
| FENDRR | 1374.189237 | 1.667919283 | 7.45E-22 |
| PSD2 | 18.55882236 | 1.667916377 | 8.06E-04 |
| OCA2 | 53.42871929 | 1.666333283 | 5.53E-06 |
| SLC13A5 | 229.8750143 | 1.665965454 | 9.67E-12 |
| COX6B2 | 38.77772173 | 1.664103588 | 2.66E-04 |
| AC145124.1 | 226.6834811 | 1.663166627 | 4.17E-34 |
| AC144831.1 | 11.12021588 | 1.662500622 | 2.03E-02 |
| AC020656.2 | 34.51942435 | 1.660325476 | 1.82E-05 |
| CDC42EP1 | 3906.296252 | 1.66022256 | 5.55E-20 |
| MATK | 61.01281891 | 1.659421966 | 9.77E-07 |
| GLS2 | 26.64856311 | 1.659333099 | 2.01E-03 |
| HMX2 | 8.63397014 | 1.658800472 | 2.91E-02 |
| MIR646HG | 21.72389345 | 1.658577963 | 1.44E-03 |
| KCNK6 | 269.045259 | 1.658060783 | 9.75E-36 |
| NEDD9 | 1801.09962 | 1.65624283 | 9.60E-49 |
| CSMD2 | 15.3385253 | 1.654957443 | 5.08E-03 |
| AL049650.1 | 34.41115523 | 1.654381597 | 3.52E-04 |
| OPN3 | 842.1645973 | 1.653899377 | 5.43E-46 |
| PTPRF | 14909.99649 | 1.652860401 | 1.04E-43 |
| METRNL | 538.6256251 | 1.649327975 | 6.20E-27 |
| HSPB2 | 71.61158158 | 1.649214855 | 1.49E-03 |
| BACE2 | 2559.985201 | 1.649200546 | 8.98E-40 |
| MIR3936 | 18.15089301 | 1.648343664 | 1.99E-04 |
| AP003469.4 | 15.34615549 | 1.648295832 | 1.37E-03 |
| GRIN3B | 49.69900749 | 1.648009223 | 2.96E-08 |
| MGLL | 4291.086206 | 1.647423337 | 3.52E-38 |
| LMO7 | 2495.326444 | 1.646907239 | 2.92E-20 |
| AIF1L | 1935.226753 | 1.646496248 | 2.03E-36 |
| HOXC12 | 131.3367996 | 1.645747082 | 6.91E-19 |
| NME1-NME2 | 345.9268082 | 1.644608015 | 1.26E-12 |
| RCSD1 | 77.4899084 | 1.644180303 | 7.82E-06 |
| NNMT | 7823.226213 | 1.644060024 | 1.03E-17 |
| ACTN4 | 24964.65553 | 1.64355339 | 1.02E-57 |
| CLU | 7930.293639 | 1.642905719 | 1.24E-41 |
| PNMA8B | 17.9584709 | 1.642141568 | 1.96E-02 |
| NA | 13.14292939 | 1.641913785 | 3.85E-02 |
| RTN4R | 369.6459485 | 1.641380213 | 2.41E-21 |
| TBC1D2 | 462.5151675 | 1.639217343 | 1.29E-26 |
| TRPM3 | 33.5290193 | 1.638016244 | 5.72E-06 |
| TSHZ3 | 74.67057501 | 1.63601811 | 3.48E-07 |
| PPP1R14A | 175.0821301 | 1.63573409 | 2.74E-16 |
| KIAA1210 | 25.70861216 | 1.635029337 | 4.87E-04 |
| NA | 7.980453732 | 1.633576062 | 3.20E-02 |
| PLEK2 | 599.7952445 | 1.630954511 | 1.38E-28 |
| EPHA2 | 5020.238168 | 1.629833058 | 5.31E-43 |

| | | | |
|---|---|---|---|
| AC108463.1 | 17.95471562 | 1.6293751 | 1.04E-03 |
| PDCD6IPP1 | 11.2216562 | 1.628974226 | 4.32E-03 |
| HYI | 1853.666591 | 1.624499058 | 6.43E-37 |
| AC009055.2 | 28.24339746 | 1.624219926 | 4.48E-03 |
| EZR | 9411.26178 | 1.623575388 | 5.50E-83 |
| TUBB6 | 6630.385277 | 1.622740739 | 5.13E-24 |
| SLC1A7 | 15.21491097 | 1.621882739 | 2.08E-03 |
| NPR3 | 210.5822672 | 1.617776122 | 1.61E-10 |
| ESM1 | 570.3932995 | 1.615305923 | 2.27E-06 |
| AC022034.3 | 9.574877301 | 1.615118595 | 3.26E-02 |
| HAVCR2 | 347.7987367 | 1.613578775 | 4.41E-13 |
| MPP7 | 42.74133974 | 1.613094424 | 1.97E-05 |
| DEPP1 | 122.3980038 | 1.612335271 | 4.23E-06 |
| MAFK | 1886.636662 | 1.611584072 | 2.45E-48 |
| CABP7 | 22.87835713 | 1.611315425 | 3.26E-03 |
| RPSAP41 | 17.36452087 | 1.609241616 | 4.34E-02 |
| MAP7D3 | 5669.030078 | 1.60811882 | 8.74E-39 |
| TCP10L | 15.48542202 | 1.606565528 | 3.75E-03 |
| CFAP57 | 85.99377228 | 1.606463743 | 9.84E-05 |
| AC124067.2 | 159.4351296 | 1.601700806 | 1.68E-14 |
| AC092490.1 | 224.8316177 | 1.599257701 | 9.96E-25 |
| ANG | 121.0230573 | 1.598174828 | 3.26E-11 |
| DDIT3 | 1960.790249 | 1.593875031 | 2.38E-04 |
| SYT15 | 456.2697294 | 1.592885688 | 4.71E-27 |
| SOWAHD | 24.47470161 | 1.58998812 | 4.86E-05 |
| AC011453.1 | 8.512167247 | 1.586693977 | 2.26E-02 |
| CEBPG | 1838.694519 | 1.585821947 | 2.03E-09 |
| MTHFD1L | 2628.193976 | 1.585016978 | 9.43E-25 |
| EBI3 | 130.7138912 | 1.579093281 | 3.22E-10 |
| MEF2B | 30.05939738 | 1.578672041 | 4.88E-04 |
| HES2 | 24.47606085 | 1.578555262 | 3.14E-03 |
| SSX4 | 18.60498698 | 1.576365791 | 4.41E-03 |
| LINC02582 | 131.1487649 | 1.575998819 | 1.24E-10 |
| C22orf34 | 161.5042601 | 1.574401302 | 1.70E-17 |
| AC139530.1 | 15.05487964 | 1.574181714 | 2.04E-02 |
| ADRB1 | 31.51575746 | 1.571499897 | 8.08E-05 |
| SAMD12 | 473.3971852 | 1.570383948 | 1.48E-10 |
| BST1 | 122.7096193 | 1.569908495 | 3.09E-09 |
| LINC01777 | 21.41202174 | 1.569391666 | 7.16E-04 |
| AC092687.3 | 14.56324383 | 1.567759151 | 3.28E-03 |
| MYOF | 15695.67332 | 1.567003655 | 6.04E-36 |
| FLNA | 161615.4789 | 1.566609371 | 2.04E-71 |
| AC106744.1 | 14.94496473 | 1.564556567 | 1.54E-02 |
| LINC00662 | 641.6290959 | 1.564255662 | 1.26E-07 |
| STAC | 1317.904973 | 1.563062674 | 3.54E-20 |
| HEG1 | 10780.54503 | 1.561382908 | 1.49E-99 |
| MOCOS | 902.9342014 | 1.559886194 | 1.44E-10 |

| | | | |
|---|---|---|---|
| NFASC | 750.1182267 | 1.559328223 | 4.37E-52 |
| CD99P1 | 133.5769418 | 1.559165624 | 1.29E-13 |
| LINC02469 | 25.02079487 | 1.558166771 | 1.00E-03 |
| NA | 24.85123195 | 1.557463859 | 1.32E-03 |
| SAMD11 | 775.4203297 | 1.553933551 | 5.64E-04 |
| CYP2J2 | 18.03377978 | 1.553668135 | 1.90E-02 |
| HCLS1 | 91.68229821 | 1.547915911 | 4.69E-07 |
| LAMA1 | 1777.372355 | 1.546850966 | 1.94E-17 |
| RAB3IP | 248.084127 | 1.54596467 | 1.07E-10 |
| COL9A2 | 63.23009237 | 1.545957543 | 3.31E-05 |
| UCHL1 | 173.5698622 | 1.542903936 | 4.24E-10 |
| GPR160 | 34.99439473 | 1.539941172 | 1.17E-04 |
| AC245595.1 | 457.8250349 | 1.537293693 | 1.70E-27 |
| CFD | 50.63983391 | 1.536297074 | 7.54E-06 |
| SHB | 1465.471817 | 1.535533423 | 7.73E-26 |
| PLAU | 3831.106497 | 1.534922116 | 4.16E-23 |
| MICAL2 | 7480.93452 | 1.532830488 | 3.87E-31 |
| ACSM5 | 100.294989 | 1.532649693 | 5.93E-09 |
| KLHL5 | 3640.730673 | 1.53182843 | 6.16E-26 |
| CES1 | 60.91092756 | 1.531096968 | 4.33E-04 |
| RNASE4 | 101.2496272 | 1.530582798 | 2.12E-05 |
| LATS2 | 922.0682349 | 1.529782224 | 1.93E-42 |
| LINC01301 | 23.18791021 | 1.528093195 | 9.76E-04 |
| MYLK | 2007.992478 | 1.524406289 | 8.10E-23 |
| AC099066.2 | 26.07679972 | 1.523533847 | 9.20E-04 |
| GOLGA8S | 17.94810252 | 1.522355973 | 1.99E-03 |
| TIMP3 | 33487.39915 | 1.520354797 | 2.52E-26 |
| ADM | 794.9914859 | 1.517893145 | 4.55E-18 |
| AL691426.1 | 14.79326627 | 1.517075409 | 7.26E-03 |
| PPP1R1C | 18.81541995 | 1.514988943 | 1.99E-03 |
| C3orf52 | 168.1696811 | 1.513447092 | 1.75E-14 |
| AC116021.1 | 33.88852242 | 1.511595719 | 1.52E-04 |
| PAQR5 | 477.0127961 | 1.509437339 | 5.96E-14 |
| MSRB3 | 1780.19995 | 1.505181526 | 4.87E-39 |
| AJ011932.1 | 30.9980947 | 1.50505611 | 9.55E-05 |
| FAM189A2 | 29.59519028 | 1.504519328 | 2.30E-03 |
| POU2F2 | 515.4171588 | 1.503783798 | 1.01E-10 |
| CACNG4 | 946.3481827 | 1.503253182 | 4.89E-16 |
| LINC02381 | 568.2933948 | 1.501982408 | 6.08E-37 |
| RGCC | 105.6600784 | 1.500755491 | 1.75E-05 |
| AL096865.1 | 40.03954789 | 1.500108321 | 1.13E-05 |
| SLC16A3 | 3928.098209 | 1.497796337 | 2.48E-38 |
| YAP1 | 7292.138549 | 1.496556748 | 1.76E-73 |
| AC226118.1 | 19.52819123 | 1.496170482 | 4.62E-03 |
| COL16A1 | 5086.360967 | 1.495791317 | 1.03E-29 |
| PLCB4 | 1544.325878 | 1.495357616 | 4.27E-14 |
| AP001363.2 | 10.81595872 | 1.494932262 | 2.58E-02 |

| | | | |
|---|---|---|---|
| CCDC85C | 2884.103042 | 1.49178026 | 7.03E-33 |
| DSP | 1365.602877 | 1.490606762 | 1.13E-36 |
| AC069209.1 | 17.56227689 | 1.489782332 | 7.82E-03 |
| DENND2A | 3519.216737 | 1.488625991 | 1.28E-31 |
| TENM3-AS1 | 240.739707 | 1.488467167 | 3.55E-21 |
| SUSD3 | 356.0271866 | 1.485394922 | 2.04E-17 |
| LINC02454 | 19.94867742 | 1.485131478 | 1.43E-03 |
| BRSK2 | 130.1529311 | 1.484075615 | 4.23E-06 |
| AC005336.2 | 28.78799943 | 1.481041431 | 6.96E-04 |
| RIMKLA | 73.27344799 | 1.480295467 | 3.10E-04 |
| SCN8A | 851.2309317 | 1.479978028 | 2.66E-17 |
| FAM83G | 1496.74242 | 1.479935455 | 1.96E-30 |
| ARL4A | 624.3512308 | 1.479566343 | 4.87E-25 |
| MICB | 1091.298993 | 1.477582927 | 7.86E-15 |
| FCMR | 73.10846455 | 1.474654469 | 6.83E-06 |
| LINC01119 | 98.2556091 | 1.473124067 | 3.98E-08 |
| CYP2B6 | 16.15268785 | 1.472626438 | 1.84E-02 |
| SLC8A1-AS1 | 28.21217156 | 1.470112423 | 5.58E-03 |
| LINC01357 | 79.30204463 | 1.469786005 | 8.93E-10 |
| RGS9 | 137.3520688 | 1.467223846 | 9.56E-08 |
| GALNT9 | 229.1882238 | 1.466602925 | 5.93E-09 |
| OXCT2 | 18.46815646 | 1.463721189 | 1.90E-02 |
| MYRF | 3061.445988 | 1.46330314 | 6.55E-47 |
| POLRMTP1 | 27.84699559 | 1.463268399 | 5.89E-04 |
| CTIF | 3415.140153 | 1.462939533 | 2.47E-32 |
| CCDC9B | 1980.078368 | 1.461675819 | 1.04E-22 |
| AC116407.1 | 18.5476928 | 1.459836544 | 1.39E-02 |
| LINC01219 | 18.91677116 | 1.457904333 | 2.75E-03 |
| VAT1 | 16525.20215 | 1.457615585 | 6.62E-30 |
| PRSS12 | 21.17623043 | 1.456909328 | 1.43E-02 |
| TDRKH-AS1 | 27.67236504 | 1.456618504 | 8.88E-03 |
| AL356515.1 | 15.89611218 | 1.456238285 | 2.78E-02 |
| AC104435.2 | 31.56962772 | 1.455940395 | 1.90E-04 |
| PROC | 45.91681908 | 1.45582821 | 1.31E-04 |
| CRNN | 18.66305069 | 1.45490053 | 3.83E-03 |
| ARHGEF16 | 104.7080639 | 1.45462076 | 1.69E-09 |
| FOXC2 | 196.881966 | 1.451474838 | 2.72E-10 |
| AC105460.1 | 37.50005812 | 1.450271669 | 2.07E-05 |
| RIPOR2 | 1126.46556 | 1.450188523 | 2.90E-12 |
| EBF2 | 15.16971605 | 1.449570628 | 1.32E-02 |
| S100A10 | 12016.01031 | 1.448456552 | 4.10E-39 |
| TBILA | 15.82723053 | 1.446875799 | 8.92E-03 |
| NPTX2 | 1549.746687 | 1.446468962 | 3.94E-17 |
| AC129492.4 | 81.89087006 | 1.445813094 | 1.78E-03 |
| COL1A1 | 951.6101759 | 1.445112198 | 3.52E-09 |
| DES | 57.98814614 | 1.443885358 | 1.84E-05 |
| LLGL2 | 1015.83685 | 1.443722946 | 7.88E-45 |

| | | | |
|---|---|---|---|
| FAM131C | 149.5905496 | 1.442336819 | 2.13E-10 |
| MIR3936HG | 98.17743369 | 1.442200948 | 3.44E-11 |
| TYRP1 | 289.0563429 | 1.440715485 | 5.23E-28 |
| NSG1 | 71.44836128 | 1.438881267 | 7.35E-04 |
| AL603750.1 | 23.85849569 | 1.438141653 | 9.04E-04 |
| AXIN2 | 442.7244193 | 1.4339107 | 1.05E-13 |
| AC093673.1 | 597.9308383 | 1.432424286 | 2.45E-30 |
| DTX4 | 442.4496529 | 1.432262347 | 6.94E-12 |
| SPOCD1 | 189.9108213 | 1.432164814 | 4.42E-05 |
| SOHLH1 | 91.92949427 | 1.431118755 | 6.62E-10 |
| RDM1P5 | 32.0965191 | 1.431072431 | 3.51E-03 |
| MEGF11 | 58.02669947 | 1.429686184 | 6.65E-04 |
| ALMS1-IT1 | 18.70546363 | 1.428354625 | 7.14E-03 |
| MAP3K21 | 160.6828438 | 1.427562324 | 3.56E-10 |
| CD9 | 3271.154132 | 1.427501451 | 2.01E-20 |
| ARL4C | 1081.703425 | 1.426287135 | 9.91E-25 |
| SAMD4A | 2581.581522 | 1.425541248 | 1.11E-29 |
| CEACAM19 | 174.7498355 | 1.424562745 | 1.99E-12 |
| EIF4EBP1 | 6024.259061 | 1.423747157 | 4.07E-19 |
| AC007040.2 | 54.2009864 | 1.423394297 | 2.50E-05 |
| ANXA2P2 | 38.68847783 | 1.423292483 | 2.92E-04 |
| BIK | 12.27743953 | 1.422658656 | 3.06E-02 |
| CFLAR-AS1 | 19.01424679 | 1.419539005 | 6.96E-03 |
| MAL2 | 92.76503004 | 1.417712605 | 4.65E-04 |
| LOXL2 | 22646.42599 | 1.416674064 | 1.62E-23 |
| LRRC61 | 24.33443373 | 1.416282635 | 4.24E-03 |
| PDCD1LG2 | 137.9408949 | 1.414650157 | 1.12E-07 |
| GLRB | 86.56225061 | 1.413672455 | 4.42E-10 |
| LINC01730 | 59.12212474 | 1.412676158 | 1.62E-04 |
| BARX1 | 13.63832694 | 1.411839449 | 3.49E-02 |
| SLC6A11 | 59.07492838 | 1.410876648 | 7.90E-05 |
| CACNA2D3 | 94.70357472 | 1.409532939 | 8.50E-07 |
| PRR5L | 1135.682346 | 1.408426387 | 1.53E-37 |
| LMOD1 | 281.9537545 | 1.408038818 | 7.75E-08 |
| ODF3L2 | 13.19073808 | 1.405951059 | 3.82E-02 |
| TLN2 | 3537.806415 | 1.405664068 | 5.69E-17 |
| DBNDD2 | 498.5138506 | 1.405552591 | 9.40E-22 |
| COL4A4 | 16.52605434 | 1.405424994 | 2.10E-02 |
| SSR3 | 5937.31788 | 1.404651432 | 7.75E-79 |
| RUSC2 | 2608.465768 | 1.40352794 | 2.30E-65 |
| AL031728.1 | 20.12150877 | 1.401994825 | 3.80E-03 |
| LINC02487 | 24.11198403 | 1.401167081 | 2.29E-02 |
| PTPRB | 576.9107832 | 1.400475981 | 6.59E-15 |
| MESP1 | 46.744598 | 1.399691709 | 4.42E-05 |
| SMOC2 | 58.39393359 | 1.399444861 | 4.05E-03 |
| NPAS1 | 1188.241383 | 1.398577543 | 9.29E-40 |
| PIPOX | 53.95477712 | 1.397915836 | 2.09E-06 |

| | | | |
|---|---|---|---|
| PM20D2 | 1660.566496 | 1.396762365 | 3.36E-25 |
| AC025580.2 | 19.20515102 | 1.394486749 | 3.10E-02 |
| CYS1 | 92.6595099 | 1.394413594 | 2.23E-09 |
| CELF2 | 578.2401139 | 1.393876661 | 1.21E-14 |
| AL512785.2 | 18.65984878 | 1.393785359 | 1.36E-02 |
| NLRP12 | 80.46937671 | 1.392911323 | 2.44E-04 |
| MTMR10 | 1017.928693 | 1.392452869 | 8.15E-38 |
| NLRP3 | 205.3152859 | 1.392349059 | 1.53E-07 |
| DNAH10 | 134.5372647 | 1.39095192 | 9.55E-04 |
| WARS | 18975.32743 | 1.390064676 | 1.78E-12 |
| AC073569.2 | 42.59422281 | 1.389148319 | 3.45E-03 |
| SORBS2 | 136.3448756 | 1.388961895 | 7.47E-06 |
| TRNP1 | 1601.234225 | 1.388374179 | 2.40E-22 |
| KAZN | 2090.457273 | 1.387914469 | 2.25E-51 |
| NR2F2-AS1 | 143.3345313 | 1.387774453 | 1.54E-10 |
| AC010327.4 | 19.75050693 | 1.386944676 | 4.93E-02 |
| AC148477.4 | 20.92675249 | 1.386906024 | 1.24E-02 |
| CNN2 | 8063.875845 | 1.385288766 | 5.99E-55 |
| GFI1 | 39.2406586 | 1.383768742 | 4.07E-03 |
| PAPPA | 2557.838757 | 1.383093194 | 2.51E-07 |
| SMPDL3B | 107.507826 | 1.382884601 | 4.66E-07 |
| AC092667.1 | 12.34521846 | 1.382872806 | 3.88E-02 |
| LPAR1 | 468.0407392 | 1.382694069 | 2.04E-17 |
| TPM1-AS | 112.9196416 | 1.381865227 | 7.26E-04 |
| WNT2B | 151.2588281 | 1.381587981 | 2.56E-10 |
| ATP6V1B1-AS1 | 56.91216974 | 1.380355749 | 4.27E-05 |
| GABRE | 2505.179407 | 1.3794731 | 1.84E-13 |
| PPP1R32 | 33.1700174 | 1.378391809 | 5.29E-04 |
| HABP4 | 1578.110636 | 1.377484872 | 1.70E-65 |
| BAIAP2L1 | 1835.617301 | 1.374843398 | 9.68E-48 |
| AC025857.2 | 32.64328677 | 1.373485463 | 1.07E-03 |
| DENND1C | 82.52985998 | 1.373401999 | 2.48E-07 |
| TRIML2 | 231.9553741 | 1.372817028 | 1.01E-07 |
| SARS | 9946.604605 | 1.372236042 | 2.66E-21 |
| LINC02298 | 64.28724493 | 1.371933344 | 4.07E-07 |
| SLC8A1 | 552.6757728 | 1.371815382 | 1.53E-11 |
| C1QTNF1 | 954.1115381 | 1.371550418 | 5.38E-21 |
| CARS | 10244.17979 | 1.371130813 | 4.02E-54 |
| CEACAM5 | 17.62300151 | 1.367799712 | 8.84E-03 |
| UGT3A2 | 1561.296932 | 1.367524269 | 8.40E-12 |
| LINC01666 | 39.06634075 | 1.366859974 | 2.34E-05 |
| C1QTNF1-AS1 | 25.06605869 | 1.366046329 | 3.03E-03 |
| HS6ST1 | 1299.442214 | 1.365340025 | 1.19E-16 |
| GNB1L | 130.7836641 | 1.365174629 | 9.89E-11 |
| TCHH | 60.14085417 | 1.364748755 | 7.62E-04 |
| GPC1 | 4941.093134 | 1.363451512 | 2.01E-33 |
| AL121721.1 | 38.85487816 | 1.362551191 | 1.14E-05 |

| | | | |
|---|---|---|---|
| PCSK9 | 829.7416089 | 1.362226638 | 1.21E-12 |
| RPS6KL1 | 347.9932618 | 1.362125916 | 4.24E-14 |
| ELOVL7 | 65.47929235 | 1.361542689 | 2.56E-03 |
| DRD1 | 26.45660106 | 1.361070962 | 7.17E-03 |
| HSPG2 | 6936.109191 | 1.361032335 | 4.09E-21 |
| TMEM151A | 32.37522276 | 1.359424032 | 5.48E-03 |
| NUAK2 | 210.063746 | 1.357489241 | 6.41E-17 |
| AC124067.4 | 53.45322662 | 1.357432293 | 1.10E-04 |
| AL844908.1 | 27.56971702 | 1.357380223 | 3.25E-04 |
| LINC01094 | 30.00020827 | 1.356627195 | 4.60E-04 |
| MID1 | 1581.448112 | 1.35576264 | 8.51E-32 |
| NA | 39.12265666 | 1.355439317 | 2.07E-03 |
| TREML2 | 26.08468359 | 1.355379685 | 2.36E-03 |
| TRIM7 | 166.7515713 | 1.354490579 | 3.46E-16 |
| PVALB | 24.08534313 | 1.352472553 | 2.34E-02 |
| AC010247.1 | 21.51955622 | 1.351669655 | 2.99E-02 |
| CLCNKB | 120.2619914 | 1.350148947 | 9.10E-09 |
| LINC00632 | 86.52816748 | 1.348295974 | 1.23E-06 |
| COL13A1 | 1295.802804 | 1.348127702 | 8.41E-20 |
| GAL3ST1 | 22.23042295 | 1.348035284 | 1.55E-02 |
| RTKN2 | 192.5215251 | 1.347675109 | 6.31E-07 |
| CALM2 | 26395.33632 | 1.345765093 | 2.53E-49 |
| AP001453.2 | 57.55253509 | 1.345445824 | 1.86E-05 |
| ZNF469 | 430.1866375 | 1.345121251 | 9.16E-12 |
| AC011558.1 | 173.9603832 | 1.344918324 | 3.23E-07 |
| FAM167A | 682.8611599 | 1.344040931 | 2.96E-09 |
| NA | 175.0159474 | 1.343318986 | 5.03E-21 |
| HCFC1R1 | 2331.026297 | 1.343158585 | 6.98E-15 |
| LAMB4 | 116.1014266 | 1.341597788 | 2.27E-07 |
| AC239859.6 | 53.28698724 | 1.3412873 | 3.83E-07 |
| WWC3 | 2266.235865 | 1.338935039 | 1.22E-26 |
| DKK1 | 937.75661 | 1.338568162 | 6.99E-13 |
| EYA2 | 133.8226664 | 1.337671165 | 2.38E-11 |
| NIPAL4 | 209.0449799 | 1.334461563 | 7.54E-06 |
| NR4A1 | 297.1782047 | 1.33324808 | 7.25E-18 |
| FBXL16 | 218.7308951 | 1.331804243 | 7.44E-05 |
| LINC00678 | 17.96233196 | 1.331787497 | 1.18E-02 |
| CREB3L1 | 742.9290274 | 1.331661607 | 4.26E-20 |
| COLGALT2 | 105.74784 | 1.331631134 | 5.19E-09 |
| SYNPO | 2426.228123 | 1.330966963 | 5.06E-31 |
| STX3 | 2995.088942 | 1.329972863 | 2.26E-49 |
| OR2W3 | 22.54171169 | 1.329022759 | 2.08E-03 |
| PKP3 | 138.0497832 | 1.328798655 | 4.40E-05 |
| PAGE1 | 358.4447149 | 1.32783261 | 6.23E-17 |
| MELTF | 431.330045 | 1.325430285 | 1.62E-12 |
| AL161729.1 | 72.37469199 | 1.325282333 | 7.87E-08 |
| INAFM2 | 266.0135533 | 1.32417907 | 9.87E-22 |

| | | | |
|---|---|---|---|
| PARD3B | 69.05787518 | 1.323877954 | 2.81E-03 |
| CEMIP2 | 4440.133686 | 1.323333099 | 1.33E-36 |
| SLC1A3 | 1565.100729 | 1.32149346 | 5.58E-25 |
| IZUMO1 | 20.46040962 | 1.320132633 | 3.64E-02 |
| AC099489.1 | 53.33698487 | 1.319940726 | 2.84E-03 |
| ETHE1 | 1360.286633 | 1.318528446 | 1.92E-36 |
| LMCD1 | 1289.053248 | 1.318462605 | 3.02E-08 |
| WNT3 | 343.0286232 | 1.318012053 | 2.29E-18 |
| FGFR4 | 381.567178 | 1.317642795 | 5.20E-08 |
| EPSTI1 | 1053.012588 | 1.316394798 | 2.19E-28 |
| FHL2 | 2309.54896 | 1.313434832 | 4.59E-32 |
| XK | 72.49002047 | 1.313350579 | 8.92E-07 |
| PFKP | 5727.138889 | 1.312400707 | 9.16E-37 |
| AHNAK | 122265.4934 | 1.312324699 | 7.24E-55 |
| PRSS50 | 13.04396709 | 1.312201667 | 3.52E-02 |
| AC068987.4 | 22.16306705 | 1.311686728 | 4.00E-03 |
| CITED2 | 3299.680565 | 1.311105144 | 3.54E-17 |
| MPP5 | 2731.596394 | 1.310710006 | 1.68E-19 |
| GARS | 17460.14145 | 1.310517244 | 2.78E-14 |
| ADRA2C | 237.9240199 | 1.310406346 | 5.11E-14 |
| BMP6 | 286.6928456 | 1.309262445 | 4.55E-07 |
| CES4A | 56.32910085 | 1.309126464 | 1.04E-05 |
| KRT18P41 | 21.57738418 | 1.308988685 | 1.99E-03 |
| CRISPLD2 | 701.9003059 | 1.306710013 | 1.91E-09 |
| TRIP6 | 1891.929213 | 1.306659749 | 3.65E-23 |
| TCF15 | 164.4727977 | 1.305260448 | 2.89E-09 |
| SV2B | 97.51491438 | 1.303055326 | 4.68E-02 |
| PTPN6 | 55.81649397 | 1.302908996 | 1.62E-04 |
| SYTL2 | 1943.873549 | 1.302652747 | 4.77E-11 |
| AC011603.2 | 323.6017866 | 1.301884133 | 4.97E-02 |
| LURAP1L | 175.7105056 | 1.30137071 | 8.65E-06 |
| AC010547.4 | 58.16470255 | 1.299618346 | 1.67E-02 |
| RNF165 | 34.72656878 | 1.299049839 | 1.14E-02 |
| CHRFAM7A | 55.5674447 | 1.298847437 | 2.75E-04 |
| HSBP1L1 | 284.4549299 | 1.298451394 | 2.24E-14 |
| LACTB | 732.0504195 | 1.298021657 | 1.48E-23 |
| DOK7 | 12.18678831 | 1.297899554 | 3.72E-02 |
| LINC01287 | 37.77036241 | 1.297813324 | 1.83E-02 |
| AC098487.1 | 25.41401803 | 1.297666131 | 1.16E-02 |
| HLA-DQA1 | 47.19583332 | 1.296692793 | 3.51E-03 |
| GABRR2 | 34.6200028 | 1.29536037 | 2.10E-02 |
| CCRL2 | 23.25096366 | 1.294461496 | 1.51E-02 |
| GAS7 | 2811.772974 | 1.294286869 | 5.15E-26 |
| HMGA2 | 5164.377752 | 1.294155138 | 7.88E-20 |
| OR51B5 | 232.9156724 | 1.293765904 | 1.82E-18 |
| NEURL1 | 376.2572735 | 1.293736576 | 2.44E-20 |
| ARLNC1 | 37.47117289 | 1.293165159 | 2.29E-03 |

| | | | |
|---|---|---|---|
| SYDE2 | 235.870281 | 1.291897068 | 1.36E-07 |
| SCG5 | 39.15796545 | 1.291280901 | 4.10E-03 |
| FLNB-AS1 | 98.05334563 | 1.289112157 | 4.77E-08 |
| CCDC80 | 5060.118978 | 1.28714956 | 2.16E-29 |
| KCNJ4 | 60.10916576 | 1.286801008 | 2.29E-05 |
| KCNG1 | 1345.743577 | 1.284995224 | 2.15E-08 |
| AP000523.1 | 53.76492508 | 1.284505321 | 5.18E-06 |
| AC125807.2 | 800.7685966 | 1.284246181 | 3.09E-29 |
| ADAMTS1 | 240.3419719 | 1.281686797 | 9.68E-06 |
| CHURC1-FNTB | 112.0451196 | 1.281580161 | 4.96E-03 |
| NOXA1 | 57.91642516 | 1.281314786 | 2.03E-04 |
| CADM3 | 1193.863173 | 1.281304769 | 1.05E-15 |
| AL358473.1 | 37.43308023 | 1.28030172 | 6.72E-04 |
| SLC19A3 | 219.3959215 | 1.27919607 | 1.67E-19 |
| CCM2L | 68.80571366 | 1.27880572 | 5.06E-04 |
| PGRMC2 | 2270.365819 | 1.27869519 | 1.11E-93 |
| AC135178.5 | 14.78197746 | 1.278164636 | 8.83E-03 |
| AC016687.3 | 39.0356115 | 1.278075022 | 2.70E-03 |
| AMOT | 168.6287222 | 1.274125584 | 2.61E-05 |
| RPSAP46 | 14.07468125 | 1.272662255 | 3.99E-02 |
| RAB3B | 2500.739363 | 1.272301068 | 1.39E-33 |
| KRT8P33 | 61.62685347 | 1.271980725 | 2.36E-06 |
| CHST13 | 38.55414762 | 1.270980865 | 5.66E-04 |
| FSTL1 | 34023.92307 | 1.270946624 | 1.75E-33 |
| AC090617.9 | 15.7887937 | 1.270270846 | 1.98E-02 |
| MYADM | 3260.603661 | 1.269940585 | 1.20E-28 |
| GDPD5 | 1564.667833 | 1.268944829 | 5.07E-21 |
| ZNF560 | 384.619609 | 1.266021855 | 1.95E-15 |
| KBTBD3 | 101.4109539 | 1.265988332 | 1.01E-04 |
| MYL12B | 10882.15315 | 1.265432765 | 2.55E-95 |
| ACKR1 | 26.76636376 | 1.263314671 | 2.04E-03 |
| MOXD1 | 100.2989232 | 1.262306137 | 2.94E-07 |
| PTTG1IP | 16185.65895 | 1.262106492 | 1.36E-34 |
| PAWR | 1395.931334 | 1.261930488 | 5.31E-10 |
| CASC18 | 23.63218306 | 1.261348346 | 1.02E-02 |
| AC083964.1 | 15.50949792 | 1.261212155 | 3.84E-02 |
| SYNGR3 | 539.8025067 | 1.260031499 | 4.00E-13 |
| PIK3CD | 607.3206334 | 1.259313758 | 4.82E-14 |
| PDGFA | 1396.475027 | 1.258517676 | 1.32E-18 |
| NA | 21.75015542 | 1.257338493 | 8.75E-03 |
| TSEN15 | 2116.407831 | 1.255615573 | 5.97E-21 |
| AP002957.1 | 37.61139018 | 1.255098894 | 1.76E-03 |
| DPY19L2P5 | 27.43561117 | 1.254389952 | 3.37E-03 |
| EVA1C | 367.5986656 | 1.253755823 | 8.12E-12 |
| ADAMTSL5 | 346.9843244 | 1.25361478 | 2.90E-12 |
| VSIG1 | 26.36195593 | 1.2526346 | 4.72E-02 |
| RAB33A | 27.18663417 | 1.251799452 | 1.18E-02 |

| | | | |
|---|---|---|---|
| CTPS1 | 4664.975413 | 1.250348184 | 4.91E-30 |
| MSMO1 | 2899.21411 | 1.24966103 | 1.50E-07 |
| TEAD4 | 1502.038523 | 1.248695024 | 2.44E-35 |
| RBPMS | 1720.796055 | 1.247437839 | 3.26E-36 |
| IARS | 24186.3385 | 1.245720759 | 2.96E-21 |
| LINC02575 | 49.33029379 | 1.243307123 | 8.31E-04 |
| CD274 | 323.6243598 | 1.24326696 | 1.51E-09 |
| LIF | 154.155482 | 1.243210766 | 3.53E-05 |
| AARS | 23738.39566 | 1.242585799 | 1.93E-14 |
| SLC8B1 | 1452.09912 | 1.241568317 | 3.98E-23 |
| CPA2 | 11.47748765 | 1.240361326 | 4.61E-02 |
| VSIR | 1076.142038 | 1.239144648 | 6.50E-15 |
| DIRC3 | 25.51275974 | 1.238929169 | 9.30E-03 |
| PCED1B | 73.68965833 | 1.23775866 | 4.82E-05 |
| TMC5 | 49.88234155 | 1.237089825 | 4.03E-02 |
| SMIM10L2B | 102.9373407 | 1.235630811 | 4.21E-05 |
| GAGE12J | 31.75778717 | 1.233885685 | 3.77E-02 |
| FALEC | 20.55045138 | 1.233024933 | 1.86E-02 |
| GALNT15 | 66.94304396 | 1.232856373 | 2.90E-03 |
| NRN1 | 401.8029362 | 1.23271329 | 8.92E-03 |
| DLGAP1-AS5 | 26.94921455 | 1.231517316 | 2.52E-02 |
| CTNNAL1 | 7787.2537 | 1.231201924 | 8.07E-19 |
| SUGT1P1 | 47.60383308 | 1.231120738 | 4.16E-04 |
| MYO1C | 10838.86687 | 1.230741327 | 1.01E-26 |
| SLC6A20 | 25.26900586 | 1.228933849 | 4.67E-02 |
| JPH3 | 76.67241217 | 1.228821337 | 3.00E-07 |
| AL139020.1 | 290.5916199 | 1.227467575 | 6.92E-14 |
| RN7SL1 | 545.8515391 | 1.227112771 | 1.35E-08 |
| ACTN1 | 17677.66597 | 1.226525263 | 4.05E-50 |
| NEDD4L | 4687.21832 | 1.226517732 | 1.89E-36 |
| AC097652.1 | 33.6880894 | 1.224418517 | 5.57E-04 |
| SLC12A4 | 2661.801366 | 1.224363279 | 1.41E-17 |
| BLK | 325.3412728 | 1.223563511 | 3.33E-09 |
| ACOT9 | 3014.294838 | 1.219992912 | 5.26E-46 |
| JPH1 | 95.12718083 | 1.219631418 | 2.18E-06 |
| SEMA3F | 497.6585612 | 1.219425003 | 3.82E-20 |
| SPANXC | 1478.430067 | 1.21846018 | 1.64E-09 |
| PIM3 | 1569.131357 | 1.218220906 | 1.41E-44 |
| PRDX6 | 12969.97135 | 1.21804597 | 1.03E-23 |
| XBP1 | 4964.531383 | 1.218015243 | 2.40E-35 |
| GPT2 | 3740.348619 | 1.217078523 | 4.81E-08 |
| SERP2 | 73.98911883 | 1.216649264 | 2.44E-06 |
| KIF1A | 33.08770637 | 1.215781403 | 3.02E-02 |
| CPNE7 | 438.1070185 | 1.214474139 | 2.13E-13 |
| IL12A | 40.27153141 | 1.21375847 | 4.49E-04 |
| ERCC2 | 3374.31351 | 1.213072723 | 1.05E-26 |
| C1orf116 | 44.68469099 | 1.209380866 | 3.17E-04 |

| | | | |
|---|---|---|---|
| EXT1 | 6550.50112 | 1.207486276 | 9.86E-22 |
| FAM155B | 469.1044739 | 1.206678987 | 7.50E-24 |
| PPARGC1A | 48.48171747 | 1.206575246 | 3.92E-02 |
| MB21D2 | 492.152103 | 1.204761574 | 6.54E-29 |
| ERC2 | 65.38801751 | 1.204561715 | 3.38E-02 |
| DDAH1 | 3716.229868 | 1.204092801 | 5.88E-18 |
| DHRS2 | 1627.919122 | 1.201198531 | 9.06E-06 |
| RRAS2 | 2136.436366 | 1.200576442 | 1.87E-35 |
| AL138478.1 | 119.427306 | 1.200319287 | 3.79E-06 |
| LINC01835 | 24.62093076 | 1.198898308 | 1.78E-02 |
| ITGB4 | 449.0655626 | 1.196811857 | 8.15E-09 |
| TUBA1C | 18892.83266 | 1.196564305 | 1.96E-13 |
| CCDC169-SOHLH | 38.91818841 | 1.196517908 | 2.27E-02 |
| DLX4 | 530.3310774 | 1.195886914 | 4.81E-23 |
| C9orf3 | 1301.867845 | 1.195648926 | 1.72E-33 |
| TM4SF19 | 104.74363 | 1.195599854 | 3.41E-04 |
| RPS20P33 | 11.01529335 | 1.194638144 | 4.91E-02 |
| FEZ1 | 659.6364878 | 1.194195146 | 1.15E-15 |
| AC073111.1 | 24.22844751 | 1.193365903 | 8.30E-03 |
| PHLDA3 | 421.1813707 | 1.192694515 | 4.52E-11 |
| RHCE | 20.12885806 | 1.191405015 | 3.94E-02 |
| RBMS2 | 4001.840397 | 1.190766621 | 9.29E-45 |
| AC138811.2 | 44.94366895 | 1.190312983 | 1.76E-02 |
| ADORA2B | 805.8301713 | 1.190196052 | 1.13E-22 |
| NUP210 | 2474.763351 | 1.189551549 | 3.12E-26 |
| DIAPH3 | 4667.695024 | 1.187855396 | 5.75E-13 |
| ADAM33 | 189.1758325 | 1.187718394 | 1.05E-12 |
| EPB41L3 | 354.3759169 | 1.1874449 | 9.85E-10 |
| SCN4A | 135.5757544 | 1.187103447 | 1.50E-03 |
| MBP | 3639.442215 | 1.186288168 | 8.77E-12 |
| GRID1 | 67.5443194 | 1.183934955 | 1.91E-05 |
| MMP17 | 411.9505481 | 1.182892363 | 8.09E-09 |
| PATJ | 706.5965863 | 1.180882378 | 8.94E-10 |
| SPANXD | 336.2169453 | 1.180527616 | 5.30E-07 |
| LMX1B | 158.1278197 | 1.179515459 | 5.42E-07 |
| YBX3 | 3730.363678 | 1.179167268 | 2.69E-66 |
| PLCE1 | 1326.090032 | 1.177600021 | 3.52E-25 |
| NBEAP3 | 75.26489319 | 1.176346665 | 1.54E-06 |
| SOCS3 | 897.8181479 | 1.175959172 | 2.23E-05 |
| FIBCD1 | 129.1126746 | 1.174746988 | 1.79E-05 |
| ARID3A | 498.0095121 | 1.173336632 | 2.86E-12 |
| SMAD6 | 1299.992546 | 1.17263253 | 1.02E-14 |
| CCND1 | 9113.412653 | 1.172440106 | 1.55E-15 |
| CRYM-AS1 | 32.07064711 | 1.170723891 | 1.08E-02 |
| HCRTR1 | 28.97834538 | 1.170590106 | 5.43E-03 |
| ZBED6CL | 47.85750858 | 1.170266777 | 2.39E-03 |
| CACNA1B | 179.5369723 | 1.170161767 | 2.25E-06 |

| | | | |
|---|---|---|---|
| FOSL2 | 5985.00852 | 1.169947569 | 2.27E-27 |
| NXPH4 | 627.4375109 | 1.169456914 | 4.39E-13 |
| RN7SL2 | 1281.448062 | 1.169179919 | 6.20E-05 |
| TCEAL2 | 28.40002448 | 1.168846218 | 2.24E-02 |
| MAMLD1 | 1047.30731 | 1.166719637 | 3.81E-12 |
| AP000759.1 | 69.88374719 | 1.166713222 | 3.92E-05 |
| ANKRD18A | 120.6515545 | 1.166021722 | 3.64E-04 |
| ATCAY | 1432.949226 | 1.164362947 | 2.97E-09 |
| SLC1A5 | 12852.03691 | 1.163934048 | 1.04E-12 |
| UPK1A | 68.1821967 | 1.162550867 | 4.09E-03 |
| SOWAHA | 80.37691478 | 1.16241802 | 2.89E-06 |
| FAM43A | 796.7891411 | 1.161503025 | 2.92E-24 |
| ZMAT1 | 38.3723145 | 1.161501096 | 8.41E-03 |
| FOXF2 | 394.4896013 | 1.160568959 | 2.01E-23 |
| EPDR1 | 1988.545424 | 1.159099725 | 5.47E-23 |
| SLC2A1 | 3250.842943 | 1.159066506 | 2.91E-18 |
| LYPD6 | 673.2989076 | 1.155477179 | 1.08E-12 |
| FDFT1 | 15026.82734 | 1.153369666 | 2.97E-12 |
| F7 | 60.99527613 | 1.152350738 | 2.86E-04 |
| AC027682.6 | 85.49644904 | 1.151731923 | 8.72E-05 |
| AL132780.2 | 91.80791985 | 1.151481819 | 1.36E-04 |
| SHROOM3 | 3719.952372 | 1.151313416 | 3.16E-29 |
| THEG | 27.75216297 | 1.150287899 | 1.30E-02 |
| EHD1 | 3523.992995 | 1.149829284 | 2.59E-14 |
| SLC46A3 | 306.615023 | 1.148170383 | 1.04E-12 |
| NT5M | 308.3312684 | 1.145936616 | 2.13E-12 |
| LARP1B | 467.2784037 | 1.145056764 | 2.70E-10 |
| LDLR | 5914.413995 | 1.144349939 | 7.13E-17 |
| PLCD3 | 2293.886567 | 1.143611022 | 1.92E-36 |
| PPP1R13L | 292.5531074 | 1.14322047 | 2.26E-10 |
| IL6R | 802.3266812 | 1.143153276 | 1.25E-12 |
| FGGY | 156.5456048 | 1.142824543 | 2.33E-05 |
| AL117378.1 | 78.33490075 | 1.141056785 | 2.09E-04 |
| SCARF2 | 1756.210107 | 1.140464668 | 5.85E-47 |
| ANOS1 | 67.41388281 | 1.14038096 | 1.08E-04 |
| N4BP3 | 111.414007 | 1.139383688 | 1.98E-06 |
| ANKRD55 | 43.24059336 | 1.138823993 | 2.10E-03 |
| BLVRB | 2573.362732 | 1.138733092 | 1.42E-24 |
| SPECC1L-ADORA | 222.3063778 | 1.138417699 | 1.31E-04 |
| CABLES1 | 2714.737389 | 1.137171338 | 4.25E-14 |
| FOXF1 | 786.624986 | 1.136392832 | 5.09E-11 |
| CHRM4 | 157.7202575 | 1.136055742 | 6.45E-07 |
| C10orf55 | 41.45330799 | 1.135685585 | 6.76E-03 |
| CRY1 | 1344.096812 | 1.135342776 | 1.51E-19 |
| AL596223.1 | 39.32068767 | 1.135055207 | 3.85E-02 |
| GSDME | 2169.264017 | 1.134118458 | 8.53E-21 |
| DENND3 | 1469.517193 | 1.133550485 | 1.75E-30 |

| | | | |
|---|---|---|---|
| ANXA2 | 106890.7943 | 1.133497065 | 2.60E-44 |
| UPK1A-AS1 | 54.35396382 | 1.133310521 | 2.06E-02 |
| GOT1 | 2517.089131 | 1.131609373 | 7.94E-13 |
| DDIT4 | 5087.59468 | 1.130582751 | 4.10E-04 |
| AL132656.4 | 18.46287216 | 1.130545928 | 3.59E-02 |
| CEBPD | 169.0188464 | 1.130490801 | 3.83E-07 |
| CLIP4 | 1736.450377 | 1.130461205 | 2.18E-08 |
| FAM85B | 19.23518435 | 1.129808746 | 1.19E-02 |
| SYNE3 | 2045.696794 | 1.12980098 | 1.72E-22 |
| BOK | 3337.704686 | 1.129503414 | 1.15E-13 |
| AC092802.1 | 33.0940247 | 1.129444719 | 1.01E-02 |
| SUPT3H | 446.3907523 | 1.129094004 | 1.77E-22 |
| RNF41 | 3274.28356 | 1.127984306 | 9.81E-11 |
| C16orf95 | 250.6123435 | 1.127807938 | 8.58E-10 |
| F2RL1 | 1117.544653 | 1.127462104 | 1.68E-17 |
| HK2 | 2102.950957 | 1.126684343 | 1.31E-15 |
| BCAT1 | 1936.78461 | 1.126287096 | 4.42E-09 |
| GABRP | 97.88871789 | 1.125874115 | 3.12E-06 |
| AL354718.1 | 27.67496463 | 1.125656737 | 5.75E-03 |
| KCNF1 | 83.96198227 | 1.125194255 | 1.08E-03 |
| MTSS1L | 4140.209056 | 1.124432269 | 3.26E-26 |
| AL451042.2 | 26.7035443 | 1.123562415 | 7.70E-03 |
| AC027237.3 | 72.9435641 | 1.12354605 | 4.32E-03 |
| ZYX | 10900.51147 | 1.123034498 | 2.74E-20 |
| SMC2-AS1 | 104.4127482 | 1.12297533 | 1.03E-04 |
| FKBP9 | 12757.91606 | 1.122308259 | 4.37E-40 |
| ARHGAP23 | 5404.484757 | 1.122243191 | 5.86E-37 |
| ULBP1 | 1094.423407 | 1.121568808 | 1.86E-08 |
| ATF4 | 18601.19462 | 1.120338172 | 1.40E-29 |
| UAP1 | 2438.450767 | 1.119706843 | 3.43E-20 |
| AC006252.1 | 66.20634047 | 1.118906741 | 7.97E-04 |
| NA | 39.88031917 | 1.118899681 | 1.74E-03 |
| ARHGEF18 | 3169.607921 | 1.118725851 | 1.51E-26 |
| C4orf54 | 44.10009235 | 1.117770392 | 1.12E-03 |
| ALDH1L2 | 216.1567516 | 1.117465674 | 4.92E-05 |
| AC125611.3 | 1591.780644 | 1.117171624 | 6.42E-11 |
| SYDE1 | 3531.235541 | 1.114608601 | 7.12E-20 |
| PCID2 | 2143.240142 | 1.114333099 | 3.66E-46 |
| COMT | 3313.736764 | 1.114225147 | 8.03E-21 |
| AC135506.1 | 22.3710912 | 1.113861132 | 1.15E-02 |
| AL160191.1 | 17.79750141 | 1.112525635 | 4.73E-02 |
| LONP1 | 5588.880683 | 1.112190549 | 7.18E-29 |
| EFR3B | 66.27859465 | 1.111830888 | 9.20E-04 |
| PITPNM3 | 292.5927778 | 1.109261877 | 7.83E-15 |
| ARSG | 201.1469179 | 1.10780676 | 5.19E-10 |
| NXNL2 | 101.7112097 | 1.106723649 | 8.29E-08 |
| CALHM6 | 28.56829687 | 1.106439642 | 1.80E-02 |

| | | | |
|---|---|---|---|
| IGF2BP2 | 5316.699877 | 1.105520476 | 8.05E-41 |
| KCNS1 | 313.8676778 | 1.105414981 | 2.35E-08 |
| SPANXA2-OT1 | 50.20779197 | 1.10360411 | 5.67E-03 |
| TTC7A | 1548.824814 | 1.099712786 | 2.23E-29 |
| C1QTNF2 | 299.2471779 | 1.099162537 | 1.15E-04 |
| KIFC3 | 5432.509691 | 1.098203534 | 2.95E-28 |
| FPR1 | 28.60611966 | 1.09792637 | 6.54E-03 |
| NFIL3 | 1500.966468 | 1.09727195 | 2.47E-08 |
| AC084759.3 | 23.81259529 | 1.09644659 | 3.09E-02 |
| CARD10 | 1474.574126 | 1.095791657 | 3.79E-12 |
| FAM167B | 176.2258737 | 1.095789717 | 5.63E-06 |
| PRAG1 | 3099.318571 | 1.095525322 | 1.10E-12 |
| IL1B | 73.1764465 | 1.095227869 | 1.55E-02 |
| FOXA2 | 107.3111875 | 1.094613206 | 5.20E-05 |
| CAP2 | 1928.403831 | 1.093087145 | 1.39E-28 |
| SMCO4 | 160.0160317 | 1.091254659 | 4.36E-04 |
| AC244669.1 | 91.29498098 | 1.091025304 | 1.43E-06 |
| NABP1 | 2078.385631 | 1.090173207 | 3.26E-09 |
| FZD2 | 1127.496047 | 1.089366535 | 1.05E-16 |
| PYGB | 11837.43242 | 1.089084385 | 7.60E-41 |
| ERN1 | 835.8781716 | 1.08895747 | 2.28E-04 |
| CXADR | 430.9583909 | 1.088675926 | 1.43E-08 |
| NOCT | 193.3207984 | 1.088564832 | 9.04E-09 |
| TMEM158 | 357.808961 | 1.088396553 | 1.24E-07 |
| TEX41 | 51.499136 | 1.088032995 | 1.20E-03 |
| NA | 98.28641465 | 1.086682104 | 2.68E-04 |
| CEBPB | 1453.556869 | 1.085974875 | 1.17E-05 |
| AC073365.1 | 60.71790693 | 1.084676559 | 1.31E-05 |
| PTAFR | 29.50072821 | 1.084555919 | 9.65E-03 |
| CYP24A1 | 230.7349006 | 1.084260869 | 1.24E-06 |
| ZSWIM4 | 566.8401579 | 1.081453598 | 3.18E-18 |
| MARS | 14346.5054 | 1.080737923 | 6.14E-51 |
| RHOBTB3 | 58008.22119 | 1.079446649 | 2.11E-15 |
| OTUD1 | 108.5394167 | 1.079256219 | 2.52E-06 |
| SERTAD2 | 1650.89911 | 1.079206742 | 5.94E-24 |
| LSP1P4 | 1758.041799 | 1.078863491 | 6.29E-33 |
| CHRD | 56.17107808 | 1.077425686 | 9.34E-03 |
| NA | 23.5931674 | 1.077120439 | 4.66E-02 |
| ADAP1 | 466.5208821 | 1.076966855 | 2.35E-15 |
| MAL | 65.62425394 | 1.076340961 | 1.12E-03 |
| MYL12A | 9438.49279 | 1.075116123 | 2.05E-37 |
| KLK1 | 68.0157263 | 1.074745271 | 1.84E-05 |
| TMTC2 | 236.964812 | 1.074682223 | 8.88E-11 |
| WNT9A | 123.5863717 | 1.074029096 | 9.28E-06 |
| AC008957.3 | 81.64226081 | 1.072809481 | 6.44E-05 |
| WNT5A | 1223.951769 | 1.071193715 | 1.99E-13 |
| PNMA2 | 3733.387097 | 1.069976495 | 1.64E-19 |

| | | | |
|---|---|---|---|
| SPAAR | 28.07979697 | 1.069205389 | 8.15E-03 |
| AC073585.1 | 61.66117725 | 1.068979741 | 5.49E-04 |
| CGNL1 | 895.5418779 | 1.068789398 | 3.38E-13 |
| HSPB8 | 1436.3874 | 1.06856467 | 4.63E-17 |
| PIK3R6 | 45.75679108 | 1.067170092 | 1.56E-03 |
| NFKB2 | 732.7160331 | 1.06661556 | 5.06E-11 |
| RTBDN | 28.10301601 | 1.06474795 | 1.24E-02 |
| DICER1-AS1 | 124.6884539 | 1.063387455 | 5.32E-09 |
| KLF2 | 119.5906069 | 1.062776554 | 1.23E-05 |
| ANGPT4 | 46.20427387 | 1.060892334 | 1.64E-03 |
| YARS | 12806.03362 | 1.060827715 | 1.71E-15 |
| CTSB | 51746.63918 | 1.060123044 | 2.30E-16 |
| IGFBP4 | 22445.28338 | 1.057829396 | 2.49E-16 |
| ITPR3 | 5904.785391 | 1.057675544 | 3.01E-28 |
| AC005307.1 | 131.4519514 | 1.057055126 | 5.89E-03 |
| SELP | 136.3642521 | 1.05693139 | 3.48E-03 |
| TAX1BP3 | 4806.333429 | 1.056596244 | 7.03E-18 |
| C12orf75 | 2759.517718 | 1.056364505 | 7.55E-29 |
| LNP1 | 176.8767432 | 1.055791645 | 8.54E-11 |
| AL355601.1 | 25.61358171 | 1.054882492 | 2.03E-02 |
| ITGA7 | 128.2115376 | 1.054548866 | 1.85E-03 |
| ADAMTSL4 | 843.0356382 | 1.054531604 | 3.38E-13 |
| S100A11 | 9916.271027 | 1.053351646 | 5.39E-27 |
| CLCN4 | 1217.164316 | 1.052131246 | 4.27E-10 |
| ADRA1B | 405.8511243 | 1.05197292 | 2.62E-03 |
| PHLDB1 | 8195.744789 | 1.051561288 | 3.37E-22 |
| ZNF850 | 295.9530147 | 1.051255352 | 3.49E-04 |
| SDK1 | 1217.636645 | 1.050973017 | 1.21E-14 |
| NUPR1 | 3841.855504 | 1.050428819 | 4.15E-03 |
| INPP4B | 1588.237089 | 1.049844951 | 5.30E-15 |
| PCLO | 431.3526201 | 1.048868186 | 9.03E-05 |
| FTSJ1 | 3401.260365 | 1.047925738 | 8.68E-43 |
| GAS5 | 7842.947098 | 1.047660843 | 3.02E-11 |
| FAM129B | 17789.76161 | 1.047618375 | 1.54E-27 |
| AL162231.2 | 108.8299862 | 1.047421361 | 3.96E-06 |
| GYG2 | 1137.477831 | 1.046567535 | 1.64E-15 |
| KCNK9 | 51.84500957 | 1.045800447 | 2.44E-03 |
| CLTB | 1764.884731 | 1.045293661 | 3.08E-27 |
| PRICKLE1 | 152.583996 | 1.045176606 | 2.71E-05 |
| TJP2 | 4658.735463 | 1.044984668 | 2.33E-28 |
| AC020916.1 | 458.5870598 | 1.044561835 | 3.29E-15 |
| CD44 | 26250.0004 | 1.043074392 | 7.92E-20 |
| SSR4P1 | 122.1326671 | 1.042271934 | 1.76E-06 |
| AL356056.2 | 88.00872493 | 1.041204018 | 3.02E-07 |
| UBE2S | 5368.618216 | 1.040850567 | 1.17E-19 |
| PLCG2 | 1243.876311 | 1.040492155 | 2.04E-11 |
| IL2RG | 101.3761071 | 1.040379885 | 5.01E-04 |

| | | | |
|---|---|---|---|
| FOXL1 | 378.206564 | 1.038767588 | 2.33E-12 |
| AL662907.1 | 56.14748766 | 1.03816477 | 1.04E-04 |
| AC005224.4 | 114.230809 | 1.038051413 | 5.26E-06 |
| LINC01547 | 217.7179449 | 1.037970706 | 1.26E-09 |
| RASSF7 | 937.3718095 | 1.035940664 | 3.05E-08 |
| TP53I11 | 1725.487319 | 1.034106534 | 6.51E-21 |
| BCAM | 3742.931072 | 1.034076763 | 7.56E-16 |
| SLC31A1 | 4679.004963 | 1.033808182 | 4.02E-38 |
| LZTS3 | 1797.583311 | 1.03224503 | 2.60E-37 |
| ICAM5 | 825.1103563 | 1.031215365 | 1.61E-09 |
| MYO1E | 3124.761668 | 1.029089332 | 2.47E-19 |
| KCMF1 | 2322.440073 | 1.028938718 | 1.92E-14 |
| AC092338.1 | 1800.678811 | 1.028118407 | 2.79E-22 |
| AKR1E2 | 18.94289099 | 1.027500197 | 4.56E-02 |
| ABCA4 | 54.74305486 | 1.026348836 | 2.48E-02 |
| CKS1B | 3700.240863 | 1.023929633 | 3.89E-27 |
| FRMD5 | 262.6352491 | 1.023533777 | 5.84E-09 |
| PRSS53 | 192.0090555 | 1.02351511 | 2.18E-02 |
| MRPL23-AS1 | 189.6877581 | 1.022260837 | 5.09E-10 |
| DOCK2 | 1547.341543 | 1.022215297 | 5.65E-11 |
| RRAD | 54.82459311 | 1.022017396 | 7.89E-03 |
| RASD2 | 211.4641263 | 1.021223084 | 6.84E-12 |
| QSOX1 | 8378.691381 | 1.021174596 | 4.77E-23 |
| KLF5 | 608.4056549 | 1.021128451 | 4.17E-05 |
| SESN2 | 1594.171601 | 1.020842057 | 4.11E-04 |
| ALDOC | 1821.65617 | 1.020761026 | 4.54E-02 |
| ASPA | 207.1723575 | 1.020246753 | 9.11E-11 |
| ACSBG1 | 82.41581756 | 1.019058404 | 7.42E-03 |
| HDAC9 | 214.4988075 | 1.018339157 | 2.11E-06 |
| PTGIS | 1269.187612 | 1.017929457 | 4.21E-16 |
| SPTAN1 | 27548.29185 | 1.017738029 | 1.13E-46 |
| SH3D21 | 340.8390984 | 1.017365202 | 3.20E-05 |
| MYLK-AS1 | 37.7056433 | 1.017235327 | 1.24E-02 |
| FOXN3-AS1 | 91.62970335 | 1.016697612 | 4.21E-05 |
| TLN1 | 34235.64152 | 1.015503598 | 2.72E-21 |
| CA5B | 553.48089 | 1.013607212 | 2.50E-11 |
| SERTAD4-AS1 | 147.6901043 | 1.010799615 | 1.10E-07 |
| CD200 | 73.55103506 | 1.009617827 | 1.17E-02 |
| CERNA1 | 67.95049957 | 1.009278453 | 2.49E-04 |
| DUSP14 | 1404.725443 | 1.008550973 | 9.34E-33 |
| SMS | 6603.49377 | 1.008452442 | 2.24E-20 |
| MRAS | 1665.819629 | 1.008421302 | 3.46E-20 |
| HAX1 | 4016.217241 | 1.006706605 | 7.35E-19 |
| GCK | 86.15722206 | 1.00628283 | 2.36E-04 |
| F11R | 1874.633537 | 1.00620146 | 9.57E-10 |
| AC005336.3 | 43.40246572 | 1.005033275 | 1.54E-02 |
| BX322562.1 | 38.96148071 | 1.004521617 | 5.41E-03 |

| | | | |
|---|---|---|---|
| ASAP1 | 7407.398524 | 1.004462889 | 1.99E-23 |
| DUOXA1 | 48.41294367 | 1.003992236 | 1.77E-02 |
| KIF6 | 184.9785171 | 1.003639038 | 2.70E-03 |
| IL17D | 149.0420822 | 1.003282185 | 3.28E-05 |
| MYO1D | 1277.296482 | 1.00269822 | 1.71E-22 |
| LINC02615 | 53.04671976 | 1.002078253 | 2.06E-02 |
| AL365181.2 | 32.29086124 | 1.001660999 | 1.55E-02 |
| PYCR1 | 8366.040574 | 1.001325447 | 4.57E-17 |
| IFITM2 | 1317.633411 | 1.00040085 | 3.63E-06 |
| TCAF2 | 445.3765845 | 0.999998002 | 6.44E-04 |
| GIT1 | 3699.953202 | 0.999548486 | 2.17E-35 |
| C2orf27A | 102.3823596 | 0.999160231 | 4.71E-06 |
| AC006262.1 | 30.28041748 | 0.99870528 | 1.27E-02 |
| AP003119.2 | 21.05916175 | 0.997778964 | 2.42E-02 |
| NFKBIE | 433.0629905 | 0.996842115 | 2.10E-07 |
| LIN54 | 911.5084985 | 0.99482742 | 5.55E-09 |
| SRPX2 | 1641.567739 | 0.99451387 | 3.31E-08 |
| ZFAS1 | 5020.048184 | 0.993429805 | 1.24E-08 |
| CRYBA2 | 26.68788722 | 0.992830319 | 2.10E-02 |
| AVPI1 | 366.8041458 | 0.991259613 | 3.20E-10 |
| GNL3LP1 | 96.30867823 | 0.991213676 | 2.92E-05 |
| ADAMTS12 | 10377.03832 | 0.99066368 | 1.61E-22 |
| NTMT1 | 2667.508817 | 0.990649476 | 4.95E-27 |
| GUCA1B | 26.55959895 | 0.989533346 | 1.03E-02 |
| RHPN2 | 791.741258 | 0.988497584 | 4.44E-17 |
| SPTBN5 | 182.573012 | 0.988476401 | 1.88E-04 |
| SIGLEC5 | 138.3515049 | 0.988329038 | 6.13E-07 |
| CCNB1IP1 | 4135.158345 | 0.988120389 | 8.70E-20 |
| SH3TC1 | 1122.413012 | 0.987996924 | 2.44E-05 |
| TRABD2A | 897.350399 | 0.987950091 | 2.10E-20 |
| AL139289.2 | 61.65032174 | 0.98705681 | 1.52E-03 |
| KLHL29 | 1265.210865 | 0.984978247 | 1.28E-10 |
| CSRNP1 | 686.1282168 | 0.983893673 | 1.03E-19 |
| PARVA | 12811.79845 | 0.983629944 | 3.10E-84 |
| KTN1-AS1 | 455.5717954 | 0.983610913 | 3.37E-09 |
| NUP50-DT | 172.5240029 | 0.983397536 | 2.10E-05 |
| USP32P1 | 78.77943041 | 0.983209748 | 7.42E-03 |
| C6orf48 | 7131.05733 | 0.982991034 | 2.00E-27 |
| ASB4 | 79.74807 | 0.982424912 | 1.47E-03 |
| GSTM4 | 1524.811665 | 0.981670725 | 2.64E-27 |
| ARHGEF10 | 3349.754618 | 0.981261274 | 1.48E-20 |
| RARRES2 | 5285.601134 | 0.981174885 | 1.76E-10 |
| TENM3 | 2755.606116 | 0.98056156 | 1.79E-22 |
| AC083899.1 | 108.9198831 | 0.979849249 | 4.43E-06 |
| PHLDB2 | 5271.381741 | 0.979027565 | 4.40E-09 |
| TNNI3 | 204.9804876 | 0.978308357 | 2.46E-07 |
| CBX7 | 937.6131425 | 0.978174434 | 1.42E-10 |

| | | | |
|---|---|---|---|
| MIR210HG | 249.23102 | 0.977904953 | 2.22E-03 |
| GRIP2 | 288.8625639 | 0.977168536 | 2.13E-03 |
| TMEM255A | 638.2751286 | 0.976992912 | 1.24E-06 |
| SKP2 | 5169.048372 | 0.97690974 | 6.36E-06 |
| RFTN1 | 2508.131971 | 0.976858737 | 4.01E-18 |
| NLRP4 | 140.5209597 | 0.9753925 | 2.97E-02 |
| SQOR | 301.2361767 | 0.975347877 | 5.28E-09 |
| GNGT2 | 60.85698961 | 0.974914257 | 3.30E-03 |
| IGFBP6 | 2175.730591 | 0.974249264 | 3.84E-12 |
| SEMA4A | 163.4203738 | 0.974101632 | 2.25E-04 |
| MIR22HG | 876.9771294 | 0.971412219 | 2.78E-28 |
| TXNRD2 | 1698.664478 | 0.969891323 | 2.00E-21 |
| BEND7 | 460.5163483 | 0.969823497 | 1.46E-13 |
| FHL1 | 18557.55669 | 0.968986433 | 3.33E-41 |
| MGAT3 | 1251.709054 | 0.968942726 | 4.77E-08 |
| MSANTD3 | 2120.168871 | 0.967916059 | 4.86E-25 |
| IGDCC4 | 762.010444 | 0.966768126 | 7.12E-11 |
| SLC39A14 | 6805.43364 | 0.965620554 | 1.36E-17 |
| SHANK1 | 153.5495197 | 0.965573864 | 3.58E-04 |
| HSPA2 | 2639.01931 | 0.965428328 | 7.12E-05 |
| LRRC2 | 129.9918758 | 0.9653049 | 3.13E-06 |
| ASPHD2 | 456.4862353 | 0.96477011 | 7.38E-18 |
| APLP2 | 27182.95191 | 0.96423729 | 7.68E-21 |
| PPP1R26 | 2889.950209 | 0.963999034 | 1.34E-22 |
| CMSS1 | 1886.108128 | 0.963554597 | 9.68E-15 |
| DLC1 | 8727.154963 | 0.962947777 | 4.65E-54 |
| BEX5 | 103.9003807 | 0.962729384 | 1.74E-04 |
| CCN5 | 26.51268906 | 0.96213933 | 2.13E-02 |
| LASP1 | 19014.37754 | 0.961777694 | 4.62E-21 |
| KLF15 | 216.4359999 | 0.961703221 | 1.05E-03 |
| LYRM1 | 775.5876976 | 0.961526137 | 4.50E-09 |
| LY6E-DT | 189.8540731 | 0.96110214 | 1.77E-07 |
| CLMN | 1499.986064 | 0.960337395 | 7.78E-19 |
| HEBP1 | 1059.86254 | 0.959206345 | 5.25E-18 |
| AC020904.2 | 128.7665194 | 0.958914865 | 7.83E-07 |
| BHLHE41 | 332.2020079 | 0.958710264 | 4.57E-03 |
| BMPER | 162.7326693 | 0.95837238 | 2.13E-02 |
| AL162171.1 | 1232.53051 | 0.957847939 | 2.03E-22 |
| KYAT1 | 621.6280245 | 0.95759041 | 4.97E-20 |
| AC005586.1 | 92.41806109 | 0.957185225 | 6.20E-06 |
| KLRG2 | 100.5493213 | 0.956685933 | 7.28E-04 |
| PTTG1 | 6678.880036 | 0.956215797 | 5.56E-21 |
| LINC00664 | 24.11874421 | 0.955700648 | 4.57E-02 |
| HOXC13 | 1296.224136 | 0.955539546 | 1.35E-13 |
| LINC01224 | 480.4551308 | 0.95545912 | 1.42E-05 |
| SLC6A15 | 394.4907478 | 0.954213186 | 4.04E-04 |
| DAG1 | 8402.798769 | 0.953859743 | 3.36E-18 |

| | | | |
|---|---|---|---|
| MAP3K14 | 719.8879108 | 0.953836437 | 5.43E-09 |
| SOCS2-AS1 | 244.3080781 | 0.953458297 | 5.15E-05 |
| DUSP9 | 394.385755 | 0.953361736 | 6.80E-07 |
| AC139491.2 | 236.3875248 | 0.953328751 | 8.07E-04 |
| MVD | 2075.489569 | 0.953162538 | 5.74E-08 |
| NDUFA4L2 | 114.6081155 | 0.9529626 | 1.33E-04 |
| P4HA2 | 4210.481934 | 0.951781107 | 5.20E-21 |
| ALG1L6P | 33.26554236 | 0.951659302 | 5.96E-03 |
| UHRF1BP1 | 4704.980666 | 0.951421402 | 2.71E-08 |
| CAPN2 | 11550.35271 | 0.951319994 | 1.20E-15 |
| AC105206.2 | 53.39233751 | 0.951059174 | 2.12E-03 |
| WHAMM | 724.8447879 | 0.950625558 | 3.26E-12 |
| NUDCD1 | 1948.346597 | 0.95039954 | 1.02E-09 |
| LOXL1 | 102.2955771 | 0.950363499 | 4.84E-04 |
| AL161729.4 | 34.7473894 | 0.949985283 | 1.35E-02 |
| RALGPS2 | 1148.025275 | 0.949885582 | 2.94E-11 |
| FOXN3 | 4410.147751 | 0.949081936 | 3.56E-30 |
| KCNK15 | 31.2411331 | 0.948114273 | 2.67E-02 |
| AKAP2 | 11074.45609 | 0.947825136 | 5.20E-27 |
| MORC4 | 2027.296062 | 0.947570583 | 2.15E-30 |
| PGM5P2 | 53.12602511 | 0.946407345 | 1.02E-03 |
| EBNA1BP2 | 5155.502085 | 0.946285368 | 1.41E-22 |
| DAPK1 | 5388.257129 | 0.946048752 | 1.83E-39 |
| HIST1H2BJ | 99.37829003 | 0.945255325 | 4.38E-05 |
| SPATA6L | 55.35177118 | 0.945192392 | 2.25E-02 |
| PANX2 | 146.5880724 | 0.945123641 | 5.47E-05 |
| AC004816.1 | 101.0139062 | 0.945117042 | 1.66E-05 |
| ACO1 | 6073.137214 | 0.944763296 | 1.15E-25 |
| PYGL | 5642.226534 | 0.944683476 | 3.07E-60 |
| DNAH10OS | 45.64178324 | 0.944286427 | 1.80E-02 |
| REXO2 | 5050.771619 | 0.944257895 | 4.33E-36 |
| FLT4 | 134.8507834 | 0.944023311 | 2.93E-02 |
| PPIAP46 | 150.622229 | 0.943861873 | 8.51E-09 |
| FAM83H | 345.1934365 | 0.943285535 | 1.03E-12 |
| LINC02515 | 71.41042735 | 0.94304454 | 1.44E-03 |
| EVA1A | 932.7295082 | 0.942850051 | 1.83E-25 |
| LINC01747 | 30.54129894 | 0.942745163 | 3.97E-02 |
| LINC00668 | 106.385289 | 0.942715475 | 5.97E-05 |
| WDR1 | 12088.49163 | 0.942525452 | 3.76E-20 |
| IRX4 | 106.6038776 | 0.941567694 | 4.36E-04 |
| TTC39B | 448.9058529 | 0.941045213 | 1.13E-08 |
| SBDS | 6786.193804 | 0.940408144 | 5.48E-19 |
| PGK1 | 28199.85894 | 0.937203787 | 1.24E-08 |
| SGMS1 | 501.840403 | 0.93665762 | 2.63E-07 |
| GNG11 | 5111.312028 | 0.936478497 | 9.99E-15 |
| PLCXD2 | 49.07100378 | 0.934851621 | 1.17E-02 |
| MS4A15 | 168.0965125 | 0.934643973 | 7.56E-08 |

| | | | |
|---|---|---|---|
| AL122035.1 | 44.99986555 | 0.934340582 | 3.63E-03 |
| CNN3 | 12063.72067 | 0.932921945 | 2.40E-30 |
| WNK4 | 1114.048769 | 0.932439337 | 1.40E-11 |
| AC126773.2 | 82.99795842 | 0.931680624 | 6.10E-04 |
| CDHR3 | 288.3718067 | 0.931377046 | 3.16E-06 |
| AL357033.3 | 34.96904904 | 0.930323018 | 2.06E-02 |
| CT45A1 | 1229.578091 | 0.929391128 | 8.01E-20 |
| MARCKS | 3627.182142 | 0.92920756 | 4.57E-23 |
| MOSPD1 | 1494.579896 | 0.927544259 | 8.74E-09 |
| LINC02593 | 656.2390751 | 0.927240813 | 3.94E-03 |
| COL26A1 | 362.579038 | 0.926516967 | 3.46E-09 |
| NF2 | 3616.898035 | 0.926428222 | 2.81E-18 |
| ARHGAP45 | 1747.783185 | 0.92589199 | 9.24E-14 |
| LINC02600 | 73.12744437 | 0.925594764 | 1.97E-04 |
| HS3ST3A1 | 154.9435874 | 0.925225427 | 8.62E-06 |
| SLC25A34-AS1 | 55.99924143 | 0.925097397 | 1.89E-03 |
| C7orf61 | 25.21213737 | 0.92465972 | 4.89E-02 |
| PIP5K1A | 3765.167022 | 0.923842394 | 7.89E-22 |
| AC156455.1 | 100.1937583 | 0.923825766 | 7.02E-05 |
| BNIP3 | 2020.176455 | 0.923659441 | 2.72E-08 |
| AC244153.1 | 322.5237666 | 0.923604601 | 7.00E-14 |
| LRRN4 | 75.66148512 | 0.923267271 | 6.82E-03 |
| PLEKHN1 | 170.6434411 | 0.923121342 | 3.85E-06 |
| EIF3D | 11749.83261 | 0.923004987 | 3.82E-13 |
| NPC1 | 3213.854759 | 0.922659618 | 2.65E-23 |
| AC010618.4 | 100.5779133 | 0.922292791 | 2.41E-03 |
| AHNAK2 | 1587.711431 | 0.921934009 | 1.89E-07 |
| SPAG1 | 418.6973724 | 0.921734035 | 1.18E-09 |
| CASZ1 | 165.5766828 | 0.921312469 | 2.26E-05 |
| AC009404.1 | 51.36748406 | 0.921089328 | 3.36E-03 |
| ID4 | 43.82615756 | 0.920993682 | 1.03E-02 |
| SLC6A9 | 461.5566515 | 0.920959472 | 2.26E-03 |
| NDUFAF2 | 2164.104687 | 0.920916395 | 4.01E-22 |
| GLT8D2 | 304.745755 | 0.920567169 | 1.03E-11 |
| PDXK | 5385.678385 | 0.920425562 | 1.40E-39 |
| AC099548.2 | 253.6681035 | 0.920402783 | 1.54E-03 |
| RAB43P1 | 75.2544523 | 0.919085471 | 9.67E-03 |
| DRAP1 | 3957.910986 | 0.918688685 | 6.66E-21 |
| NRTN | 83.69516378 | 0.918404351 | 2.69E-04 |
| TNS3 | 16971.90437 | 0.918192476 | 3.17E-15 |
| VCL | 13667.92026 | 0.917784924 | 8.01E-09 |
| MEIOB | 1375.792081 | 0.917527883 | 3.53E-11 |
| RAPGEF1 | 4833.32604 | 0.917301875 | 2.78E-23 |
| HCCS | 1607.400186 | 0.917247192 | 4.25E-08 |
| OLFM2 | 487.2499097 | 0.91540744 | 4.03E-06 |
| AC018413.1 | 63.11440807 | 0.914271428 | 6.74E-03 |
| COL8A2 | 659.7014155 | 0.913614384 | 1.09E-06 |

| | | | |
|---|---|---|---|
| ASMTL | 1137.639438 | 0.913541403 | 6.82E-17 |
| AURKB | 2643.519517 | 0.913198114 | 6.34E-13 |
| HPD | 28.55597483 | 0.913173464 | 3.64E-02 |
| GRB10 | 4252.746946 | 0.913078533 | 3.44E-08 |
| SYNC | 139.3471639 | 0.912407245 | 4.08E-07 |
| NEK6 | 8959.611567 | 0.911296503 | 3.24E-18 |
| CMIP | 2119.779756 | 0.910911237 | 1.72E-17 |
| NECTIN2 | 5365.872754 | 0.909319603 | 7.48E-16 |
| ECHDC3 | 285.3637131 | 0.908904161 | 6.14E-13 |
| LMCD1-AS1 | 48.74290335 | 0.908296527 | 4.66E-03 |
| IFRD1 | 3961.506108 | 0.90810324 | 5.63E-04 |
| GJC2 | 45.70083719 | 0.90797513 | 5.14E-03 |
| CDH3 | 1193.523897 | 0.906880617 | 1.79E-12 |
| ABI3 | 137.2170805 | 0.906364019 | 1.03E-03 |
| HOXC8 | 845.3701195 | 0.905686168 | 4.55E-18 |
| RUNX2 | 681.1290161 | 0.9044351 | 4.10E-09 |
| HIST1H2BN | 83.96504118 | 0.904258966 | 2.50E-03 |
| RASGRF2 | 415.7962532 | 0.903270927 | 3.29E-03 |
| AC018755.2 | 97.94631105 | 0.902951041 | 2.52E-02 |
| CLCN1 | 55.22163434 | 0.902711091 | 2.08E-03 |
| IQGAP3 | 3343.91172 | 0.901428578 | 1.21E-18 |
| GREB1L | 343.0554809 | 0.90117278 | 7.52E-05 |
| TSPAN13 | 149.3123275 | 0.901150721 | 5.17E-03 |
| ETS1 | 13708.86452 | 0.899227179 | 6.01E-33 |
| GLI2 | 1186.161085 | 0.899172303 | 6.97E-07 |
| OAZ1 | 19674.06586 | 0.897511438 | 6.98E-14 |
| GSTM3 | 4602.829408 | 0.897243611 | 4.91E-19 |
| CES3 | 106.6239909 | 0.893364117 | 2.20E-03 |
| TUBA1B | 81573.23168 | 0.892567425 | 1.42E-06 |
| AC013394.1 | 193.2051003 | 0.892376837 | 2.94E-03 |
| NSUN7 | 276.9157398 | 0.892116515 | 5.86E-06 |
| ALPL | 124.5448661 | 0.892053345 | 5.02E-04 |
| SEC14L5 | 39.34326117 | 0.891681885 | 4.77E-03 |
| KCTD20 | 6930.129421 | 0.891670343 | 1.46E-77 |
| PCAT6 | 92.64316638 | 0.891663197 | 5.71E-04 |
| HSPA9 | 36794.30816 | 0.890168281 | 1.72E-07 |
| C2orf72 | 79.40764904 | 0.889915796 | 3.68E-02 |
| PPP1R3C | 1863.591223 | 0.889915365 | 5.88E-19 |
| OVAAL | 29.55126187 | 0.888742321 | 2.48E-02 |
| CEBPA-DT | 56.88123024 | 0.888561858 | 3.30E-03 |
| ITGB8 | 92.24891433 | 0.887762897 | 3.31E-04 |
| PDPN | 1362.464471 | 0.88749453 | 2.34E-06 |
| DNAJC15 | 2292.244546 | 0.886829268 | 5.34E-20 |
| AL928654.4 | 120.2726713 | 0.886565343 | 1.42E-05 |
| ZP3 | 653.8483888 | 0.885842955 | 7.03E-09 |
| C1GALT1 | 513.9821536 | 0.885046769 | 3.47E-05 |
| PCDHGA3 | 35.30244861 | 0.885023113 | 1.32E-02 |

| | | | |
|---|---|---|---|
| CACNG6 | 238.4323998 | 0.884880085 | 6.13E-05 |
| AC016596.1 | 36.07615823 | 0.883837297 | 1.64E-02 |
| FAM27C | 79.29343533 | 0.883016855 | 2.09E-04 |
| RSPO4 | 543.3235221 | 0.882067813 | 1.30E-12 |
| CCDC3 | 1071.774426 | 0.881816383 | 1.65E-10 |
| AP002761.3 | 81.18544798 | 0.881499696 | 1.34E-04 |
| TGFBR3L | 82.45148481 | 0.881474437 | 5.72E-04 |
| MEAK7 | 2844.519966 | 0.881368943 | 1.07E-13 |
| AC023024.1 | 138.7335527 | 0.879737432 | 6.25E-04 |
| FUT1 | 131.4538281 | 0.878687147 | 1.90E-02 |
| SLC2A3 | 3270.065993 | 0.878659474 | 2.30E-04 |
| C4orf36 | 228.6572123 | 0.87786187 | 8.41E-07 |
| AL118516.1 | 162.6771387 | 0.875737437 | 5.13E-07 |
| TMCO4 | 1573.630919 | 0.874906788 | 1.83E-12 |
| ALDH3B1 | 1300.36037 | 0.874665701 | 1.49E-17 |
| LINC01679 | 144.8361926 | 0.874099214 | 1.02E-04 |
| KCNQ4 | 29.93609954 | 0.8739181 | 2.50E-02 |
| NKX6-1 | 64.55961741 | 0.872972658 | 1.28E-03 |
| COL5A1 | 25458.72823 | 0.872894659 | 5.90E-19 |
| AC010776.2 | 44.52266423 | 0.872678564 | 6.23E-03 |
| PVT1 | 340.4968136 | 0.872603845 | 9.44E-05 |
| ASTN1 | 2110.214754 | 0.871936349 | 1.52E-11 |
| SUGCT | 319.5476151 | 0.871727421 | 4.40E-06 |
| TENT5B | 961.9812951 | 0.871454251 | 2.10E-07 |
| AZGP1 | 71.03585013 | 0.871324494 | 1.78E-02 |
| MGAT5B | 1657.991925 | 0.870785569 | 6.96E-11 |
| MATN2 | 4934.640224 | 0.869321564 | 1.73E-11 |
| AL390719.1 | 186.2336545 | 0.869277204 | 9.93E-06 |
| TIMM17B | 1538.405681 | 0.869247162 | 7.09E-17 |
| EIF1 | 14837.75257 | 0.86716154 | 6.73E-21 |
| DPYSL4 | 506.4894992 | 0.866657454 | 2.19E-05 |
| ACOT4 | 121.1949792 | 0.865921817 | 1.07E-05 |
| RBM47 | 230.5145896 | 0.865240434 | 3.83E-07 |
| DYNLT3 | 834.4229311 | 0.8649976 | 5.03E-06 |
| EPAS1 | 9233.833509 | 0.864745175 | 9.52E-19 |
| GTPBP2 | 4056.078583 | 0.864682169 | 2.86E-06 |
| SLC22A31 | 78.82971781 | 0.863960836 | 5.63E-04 |
| LCAT | 227.905104 | 0.863307843 | 2.25E-06 |
| NA | 141.3556843 | 0.863146626 | 2.42E-02 |
| TRIM6 | 699.1372443 | 0.861037399 | 1.37E-11 |
| LITAF | 5168.421173 | 0.860757088 | 1.02E-19 |
| LRRC23 | 334.2435134 | 0.860296331 | 3.82E-08 |
| PPM1J | 142.1954387 | 0.86007559 | 2.07E-04 |
| DHCR24 | 12822.6828 | 0.859720869 | 5.72E-05 |
| CHRNA7 | 46.35821401 | 0.859025035 | 4.14E-02 |
| NOS1AP | 114.7690526 | 0.858600666 | 4.10E-04 |
| CBS | 6183.303293 | 0.858589673 | 1.72E-12 |

| | | | |
|---|---|---|---|
| KIF1C | 10083.17981 | 0.858381379 | 2.74E-22 |
| ZIC2 | 467.9089163 | 0.858336609 | 7.28E-06 |
| SPN | 423.4724567 | 0.858179103 | 1.38E-05 |
| SORT1 | 7230.535163 | 0.857655454 | 3.84E-19 |
| AC105942.1 | 246.655278 | 0.857650372 | 2.76E-08 |
| NA | 437.0559334 | 0.856783758 | 5.54E-07 |
| AC018630.2 | 44.03435817 | 0.856636228 | 1.05E-02 |
| ADGRB2 | 2041.469019 | 0.856400422 | 2.70E-11 |
| AL357033.4 | 129.7848764 | 0.855345433 | 5.25E-05 |
| ANLN | 11414.65258 | 0.855314117 | 1.02E-04 |
| EIF3CL | 2513.103972 | 0.854919573 | 2.10E-02 |
| HMGCS1 | 11282.66558 | 0.854723198 | 1.82E-03 |
| CCBE1 | 1908.550007 | 0.854651477 | 3.85E-11 |
| WSCD1 | 732.1810093 | 0.853666509 | 5.83E-08 |
| NAT8L | 892.8349717 | 0.852742376 | 5.20E-14 |
| FDPS | 5927.418536 | 0.852684852 | 6.81E-07 |
| PFKFB4 | 995.5007413 | 0.852287586 | 3.71E-03 |
| AC012531.2 | 53.48214011 | 0.85220937 | 3.83E-02 |
| AL590617.2 | 125.1792277 | 0.852131642 | 7.96E-05 |
| KCTD19 | 164.1371016 | 0.852071617 | 2.70E-04 |
| TRIM31 | 32.74844046 | 0.851342264 | 1.92E-02 |
| ZBTB25 | 608.6663508 | 0.850916181 | 2.20E-14 |
| DNAH11 | 135.1776425 | 0.85028535 | 2.00E-04 |
| TVP23C-CDRT4 | 503.1404065 | 0.850018771 | 7.04E-05 |
| OLMALINC | 520.9366639 | 0.849246579 | 2.37E-13 |
| NT5E | 5643.186994 | 0.848971581 | 4.86E-07 |
| SLC22A5 | 1353.716616 | 0.848112162 | 5.87E-15 |
| DGKA | 1692.88712 | 0.848015194 | 1.54E-10 |
| AC105052.2 | 44.08002568 | 0.847674791 | 1.72E-02 |
| BRI3 | 3521.077973 | 0.847671186 | 9.95E-10 |
| CADM4 | 889.7866067 | 0.847663294 | 4.63E-06 |
| TPST2 | 341.6889748 | 0.84689245 | 2.00E-09 |
| TSC22D2 | 1341.340198 | 0.846821372 | 1.41E-09 |
| DTD1 | 3987.586041 | 0.846640502 | 7.10E-20 |
| HERPUD1 | 5168.49769 | 0.84608487 | 3.54E-10 |
| THEM6 | 1224.94723 | 0.845985639 | 1.39E-05 |
| MXRA7 | 5655.55128 | 0.845571522 | 5.45E-14 |
| ADAMTS15 | 608.3231517 | 0.845501582 | 7.59E-11 |
| ANKRD18B | 178.8942262 | 0.844264292 | 2.33E-03 |
| IL4R | 1223.73997 | 0.844238293 | 3.29E-09 |
| ZNF710-AS1 | 199.9959931 | 0.844106049 | 6.30E-04 |
| AC142086.6 | 63.49481042 | 0.843628596 | 9.34E-04 |
| TMCO3 | 2286.119453 | 0.843574553 | 9.23E-20 |
| TCL6 | 482.6066254 | 0.843466211 | 7.36E-06 |
| PRDM11 | 1207.530965 | 0.84326995 | 2.99E-09 |
| RAB31 | 4038.601522 | 0.843188373 | 5.83E-25 |
| RFK | 851.8053873 | 0.842902488 | 4.49E-06 |

| | | | |
|---|---|---|---|
| FOXO1 | 731.2693503 | 0.84220588 | 1.69E-15 |
| DNAJB6 | 11954.55005 | 0.84205073 | 7.17E-36 |
| STXBP2 | 1158.876155 | 0.842021978 | 1.80E-13 |
| SORCS2 | 333.4430111 | 0.841938027 | 2.09E-05 |
| C10orf25 | 177.3571878 | 0.841924274 | 8.86E-09 |
| CACNA1H | 127.7563133 | 0.841602796 | 1.42E-02 |
| DAPK3 | 2248.152994 | 0.841546381 | 8.80E-11 |
| ERCC6 | 565.1973192 | 0.841342429 | 4.18E-05 |
| BMP8B | 199.0507767 | 0.841170504 | 2.35E-04 |
| AC017104.1 | 30.59044047 | 0.839770765 | 3.56E-02 |
| MAP4 | 17613.87065 | 0.839402251 | 8.73E-32 |
| B3GNT5 | 183.3353908 | 0.839153154 | 6.47E-04 |
| HOXA1 | 545.3612809 | 0.839147808 | 2.14E-13 |
| AC117402.1 | 30.025143 | 0.838464389 | 2.84E-02 |
| ULBP2 | 214.0881506 | 0.838084835 | 9.32E-05 |
| DOCK9 | 3479.797232 | 0.837873577 | 1.18E-12 |
| DYSF | 2529.150232 | 0.837814203 | 8.09E-09 |
| SNX8 | 3274.668815 | 0.837603861 | 2.73E-09 |
| AL450124.1 | 315.5916599 | 0.837404117 | 1.12E-09 |
| DAAM2 | 3400.637135 | 0.836536974 | 7.19E-07 |
| POC1A | 1050.629485 | 0.836473768 | 2.91E-14 |
| GFPT2 | 1047.076629 | 0.83638613 | 2.37E-07 |
| ATL1 | 51.40407205 | 0.836244671 | 4.93E-02 |
| NAV2 | 2387.667239 | 0.836171488 | 1.46E-11 |
| GPR176 | 2600.86289 | 0.836149881 | 1.39E-08 |
| MRPS24 | 3240.035124 | 0.835512921 | 3.31E-23 |
| IGSF8 | 967.3449524 | 0.835119123 | 1.54E-14 |
| AC040977.1 | 44.63630535 | 0.834854602 | 8.14E-03 |
| ZNF350-AS1 | 41.06051394 | 0.833313086 | 3.20E-02 |
| FLII | 12054.38531 | 0.832675768 | 6.23E-16 |
| SPECC1 | 1835.882639 | 0.832647818 | 6.48E-16 |
| ME1 | 734.9613871 | 0.831604814 | 8.26E-09 |
| CTNNA1 | 21483.2315 | 0.830594351 | 5.49E-37 |
| CAVIN3 | 2094.464026 | 0.830231468 | 1.09E-09 |
| LMNA | 16821.17365 | 0.828498611 | 2.37E-15 |
| SOHLH2 | 153.5629023 | 0.828472287 | 2.35E-04 |
| GADD45A | 1177.960422 | 0.827562841 | 4.19E-04 |
| FAM24B | 116.4415182 | 0.827237351 | 1.58E-05 |
| ADCY2 | 133.2167903 | 0.826718752 | 4.27E-03 |
| FGFRL1 | 1762.380023 | 0.826264658 | 1.03E-14 |
| TNFSF9 | 133.0035316 | 0.826078111 | 2.72E-03 |
| PLPPR3 | 186.1600222 | 0.825634882 | 3.70E-06 |
| AC005330.1 | 96.12826441 | 0.825504589 | 2.45E-04 |
| CCDC155 | 74.62847622 | 0.82494149 | 3.96E-02 |
| AC006504.1 | 78.5852465 | 0.824547837 | 1.13E-02 |
| ZBED1 | 2312.89623 | 0.823710537 | 1.75E-13 |
| KCNN4 | 521.3339695 | 0.823695193 | 3.63E-06 |

| | | | |
|---|---|---|---|
| LIMD1 | 1660.134219 | 0.823620651 | 2.43E-08 |
| PTPRU | 1753.900827 | 0.823606809 | 3.97E-09 |
| NFIX | 4322.548575 | 0.823560494 | 2.65E-17 |
| TLE4 | 2216.44133 | 0.82328563 | 3.19E-05 |
| NSD2 | 9039.317095 | 0.823231693 | 1.40E-26 |
| TPBG | 2572.499946 | 0.821926161 | 4.01E-28 |
| SCARA3 | 3354.9439 | 0.82158943 | 1.32E-07 |
| FAM107B | 1775.665814 | 0.821327434 | 1.39E-06 |
| PMP22 | 6670.421425 | 0.821297285 | 5.53E-12 |
| LINC00941 | 153.5825482 | 0.821188254 | 5.50E-05 |
| AC245297.3 | 91.66191408 | 0.821020898 | 2.83E-04 |
| CD151 | 16569.59205 | 0.819927528 | 2.66E-10 |
| MYH14 | 1638.03928 | 0.819746564 | 3.03E-15 |
| ACSM3 | 214.051937 | 0.81934101 | 1.39E-04 |
| FBXO25 | 1326.467273 | 0.819336507 | 2.98E-22 |
| CCDC71L | 1297.79424 | 0.818889506 | 3.08E-10 |
| ADAMTS2 | 1245.830925 | 0.818184066 | 6.15E-07 |
| LCP1 | 1000.928624 | 0.818026546 | 8.84E-03 |
| ASPH | 8470.783867 | 0.817195546 | 7.36E-14 |
| PSPH | 3489.931624 | 0.816676573 | 2.64E-04 |
| SCRN1 | 19372.32534 | 0.816428257 | 1.92E-30 |
| RAP2B | 924.2648138 | 0.814198843 | 1.74E-07 |
| GUCY2C | 48.51206301 | 0.813843954 | 7.24E-03 |
| EIF4E3 | 382.3841659 | 0.813728948 | 1.17E-08 |
| SLC45A3 | 519.9116178 | 0.813329028 | 3.17E-15 |
| PGM5 | 66.64291581 | 0.813232148 | 3.91E-02 |
| SLC41A1 | 2600.516694 | 0.813079297 | 1.22E-17 |
| SPHK1 | 935.0801792 | 0.812338935 | 2.36E-12 |
| RHEB | 6449.634271 | 0.810435783 | 1.48E-44 |
| SIM2 | 439.4469742 | 0.810435565 | 4.10E-08 |
| RPL39L | 913.2693129 | 0.81034196 | 1.20E-19 |
| SRRM3 | 406.1814457 | 0.808692874 | 4.88E-08 |
| TTC7B | 1528.002513 | 0.808062447 | 3.13E-12 |
| PRRG1 | 823.3572282 | 0.807861007 | 1.82E-10 |
| SHF | 228.0871797 | 0.807722343 | 1.23E-04 |
| RND3 | 1363.687625 | 0.807384578 | 3.33E-02 |
| TCEA1 | 7064.069542 | 0.807335896 | 3.98E-04 |
| GRAMD2B | 728.6384239 | 0.8068512 | 1.29E-05 |
| CLTCL1 | 1082.315327 | 0.806637699 | 1.20E-09 |
| XPOT | 9329.971461 | 0.80636204 | 5.99E-06 |
| NEMP2 | 430.5920123 | 0.805988658 | 9.78E-05 |
| CARMIL2 | 174.2976759 | 0.804662 | 1.89E-02 |
| PRSS27 | 41.26702256 | 0.802611926 | 4.24E-02 |
| CDC20 | 4779.118958 | 0.80114022 | 1.83E-09 |
| RNF39 | 58.19866188 | 0.801088552 | 4.54E-03 |
| INKA2 | 917.345134 | 0.801064808 | 6.70E-05 |
| CRADD | 389.3041389 | 0.800483045 | 5.77E-12 |

| | | | |
|---|---|---|---|
| RHOBTB1 | 1185.631432 | 0.800306364 | 4.73E-16 |
| PPM1E | 322.4396099 | 0.800292463 | 8.66E-03 |
| DHCR7 | 5667.85858 | 0.799409336 | 2.38E-07 |
| PDZK1 | 37.14948163 | 0.798590439 | 2.66E-02 |
| ADAMTS17 | 64.94372239 | 0.798480832 | 3.98E-02 |
| LRRC42 | 2265.315389 | 0.798399438 | 1.17E-21 |
| GCLM | 1084.260776 | 0.798283113 | 2.90E-06 |
| REPIN1 | 8966.942661 | 0.7976291 | 5.92E-28 |
| YBX2 | 1436.654908 | 0.797625465 | 5.89E-07 |
| EHD2 | 10312.92724 | 0.797585316 | 7.83E-13 |
| AL442125.1 | 36.23637267 | 0.797233326 | 4.12E-02 |
| NBEAL2 | 1699.565215 | 0.796098226 | 8.95E-09 |
| ADAMTSL1 | 298.7360119 | 0.795366534 | 1.26E-03 |
| ADCK2 | 1160.502158 | 0.794850836 | 1.96E-10 |
| ATP8B3 | 1210.261948 | 0.794389375 | 1.30E-12 |
| TPK1 | 102.279142 | 0.794144054 | 3.05E-03 |
| LGALS8 | 2152.324426 | 0.793412752 | 2.27E-15 |
| CREB3L2 | 4451.672148 | 0.793088811 | 8.12E-23 |
| WDYHV1 | 642.0397183 | 0.792747572 | 7.61E-10 |
| DPY19L2P3 | 201.8146384 | 0.792722292 | 2.30E-03 |
| AC022034.1 | 171.0128156 | 0.791852275 | 5.18E-05 |
| CNKSR3 | 589.3799822 | 0.791488875 | 9.99E-12 |
| ZMAT5 | 512.1863495 | 0.791362859 | 3.01E-10 |
| PMAIP1 | 884.7431058 | 0.790887182 | 6.70E-03 |
| MIR4527HG | 177.5777164 | 0.790486573 | 2.89E-06 |
| FIGN | 438.4513192 | 0.790058348 | 5.89E-07 |
| HCG4 | 34.57144931 | 0.789485317 | 4.57E-02 |
| ENO1 | 85505.67072 | 0.789447698 | 6.16E-09 |
| AL136162.1 | 32.3372411 | 0.788743469 | 4.27E-02 |
| AP003119.3 | 53.08071754 | 0.788256666 | 2.42E-03 |
| LRRC59 | 6683.308593 | 0.78784379 | 3.48E-25 |
| PCDHGA4 | 133.1061788 | 0.787399776 | 6.53E-04 |
| ATXN2L | 9348.969127 | 0.786412488 | 1.10E-10 |
| KIF9-AS1 | 229.0448139 | 0.786313493 | 1.55E-03 |
| AURKA | 3414.550899 | 0.786169917 | 4.90E-12 |
| NACC2 | 1511.713598 | 0.786060453 | 1.12E-16 |
| FBXO27 | 124.0869824 | 0.785704651 | 2.63E-02 |
| PHLPP1 | 1810.927214 | 0.785471557 | 8.72E-10 |
| TSPAN33 | 331.3822405 | 0.785466042 | 2.10E-06 |
| SNX24 | 1531.40493 | 0.785184449 | 1.99E-11 |
| RALGPS1 | 346.6724631 | 0.785153821 | 9.48E-04 |
| AL133352.1 | 66.18805983 | 0.785040512 | 2.59E-02 |
| RBCK1 | 7551.367995 | 0.784889997 | 5.28E-09 |
| PHYHIP | 101.0500315 | 0.78484161 | 2.20E-03 |
| GRHL1 | 65.06406828 | 0.784253593 | 3.35E-02 |
| CDC42EP5 | 204.9522045 | 0.783699965 | 3.59E-04 |
| LIMD1-AS1 | 173.1250136 | 0.783302095 | 4.11E-04 |

| | | | |
|---|---|---|---|
| SLCO3A1 | 123.9865616 | 0.783265841 | 1.06E-02 |
| HOXB9 | 2074.166601 | 0.782980263 | 1.22E-07 |
| ANXA5 | 24506.2297 | 0.781290471 | 1.04E-13 |
| TANGO6 | 1779.528087 | 0.780668159 | 2.88E-15 |
| METTL22 | 1657.307153 | 0.780567351 | 8.32E-19 |
| CNKSR1 | 91.16836387 | 0.780198341 | 8.97E-03 |
| PLCB2 | 233.3908541 | 0.779568513 | 3.68E-03 |
| KIAA1211 | 73.81802387 | 0.77909104 | 2.20E-02 |
| CDH13 | 11715.24575 | 0.779044434 | 5.43E-11 |
| CASKIN2 | 1291.448053 | 0.778883026 | 2.69E-08 |
| RELB | 233.1853472 | 0.778860884 | 1.49E-02 |
| DUSP3 | 4615.69139 | 0.778175634 | 2.32E-18 |
| FLT1 | 2091.054392 | 0.777968871 | 1.35E-11 |
| DLGAP4-AS1 | 149.9801256 | 0.777645624 | 8.07E-05 |
| AL645568.1 | 62.75099522 | 0.77667505 | 1.00E-02 |
| CT45A3 | 529.6423585 | 0.775911478 | 5.87E-10 |
| LEPR | 47.47876556 | 0.774381308 | 1.73E-02 |
| ELF4 | 1789.071958 | 0.77382836 | 3.87E-11 |
| C16orf74 | 376.5939505 | 0.773330705 | 1.48E-08 |
| FAM207A | 755.1657164 | 0.772833908 | 2.52E-12 |
| FARP1 | 10323.71837 | 0.772407065 | 1.97E-23 |
| SERPING1 | 1163.529777 | 0.772365919 | 2.55E-05 |
| SCAT2 | 31.87320801 | 0.772274504 | 4.28E-02 |
| SLC1A1 | 229.0137763 | 0.772169906 | 1.84E-03 |
| UBR4 | 15901.60597 | 0.771680083 | 4.76E-20 |
| FHDC1 | 144.9143232 | 0.771089813 | 1.24E-03 |
| LYN | 763.4264982 | 0.770653202 | 2.95E-08 |
| AC006547.1 | 93.48753224 | 0.770111716 | 3.61E-04 |
| BEX1 | 1622.382863 | 0.769286792 | 2.85E-10 |
| TBC1D9 | 1044.573317 | 0.769038635 | 7.18E-13 |
| APBB2 | 2409.951598 | 0.768676197 | 2.93E-11 |
| SLC35G2 | 48.39117825 | 0.768444574 | 4.63E-02 |
| SPSB1 | 702.3432713 | 0.767828066 | 2.09E-11 |
| ISL2 | 61.72259261 | 0.766695158 | 8.58E-03 |
| SDCBP2 | 87.31079611 | 0.766564231 | 1.79E-03 |
| RAB34 | 4222.806935 | 0.765740341 | 6.81E-13 |
| F11-AS1 | 217.3634243 | 0.765455208 | 1.05E-02 |
| SDC1 | 3446.919541 | 0.764842365 | 3.39E-08 |
| AC021106.1 | 31.7070498 | 0.764715224 | 3.21E-02 |
| ACSL1 | 1243.219884 | 0.764615563 | 5.80E-18 |
| FAM72D | 245.0786245 | 0.763761451 | 1.89E-03 |
| MAOA | 392.4790427 | 0.762201854 | 1.63E-03 |
| DOCK3 | 240.1180089 | 0.761800839 | 4.43E-06 |
| LRP5 | 4117.859614 | 0.760848971 | 6.74E-19 |
| LOXL3 | 3055.781083 | 0.760803912 | 3.99E-08 |
| AC005253.1 | 114.5949675 | 0.760571433 | 1.79E-04 |
| OGDH | 7751.108466 | 0.75911593 | 2.70E-11 |

| | | | |
|---|---|---|---|
| AP002884.1 | 752.7599133 | 0.758089527 | 2.08E-07 |
| PKN3 | 1552.250126 | 0.757849185 | 9.33E-16 |
| SMYD3 | 591.4497563 | 0.757011763 | 8.75E-08 |
| IDH2 | 4025.990048 | 0.756739898 | 1.46E-06 |
| CCDC84 | 692.6060325 | 0.756538282 | 8.15E-21 |
| APOPT1 | 868.812937 | 0.756150204 | 9.91E-11 |
| HILPDA | 944.6319639 | 0.755848804 | 7.62E-04 |
| PTPRM | 1734.337607 | 0.754060155 | 2.46E-06 |
| SNORD17 | 72.81252209 | 0.753406693 | 1.97E-02 |
| EFHD1 | 599.7387341 | 0.752832221 | 7.17E-07 |
| MYDGF | 2881.007494 | 0.75283128 | 1.99E-13 |
| MED8 | 2065.090107 | 0.751949512 | 2.17E-12 |
| JADE1 | 1188.702036 | 0.751534371 | 3.48E-05 |
| PTPN14 | 5617.28597 | 0.751215291 | 2.26E-25 |
| BCAS4 | 1194.623206 | 0.751132225 | 7.13E-08 |
| FAM160A1 | 139.3017091 | 0.750953412 | 4.65E-02 |
| MRPL58 | 996.859681 | 0.750461496 | 6.32E-08 |
| TMEM206 | 691.1648412 | 0.750442061 | 8.42E-11 |
| TMEM200B | 55.71415746 | 0.750407845 | 1.10E-02 |
| SBDSP1 | 833.1214146 | 0.750325639 | 5.07E-04 |
| PINK1 | 5036.849718 | 0.750177596 | 2.74E-08 |
| C5AR1 | 91.91066328 | 0.749414647 | 9.71E-03 |
| FGF7P6 | 157.2628648 | 0.749413373 | 6.44E-03 |
| CCNI | 9677.528827 | 0.749034585 | 3.43E-23 |
| IER5 | 1734.988743 | 0.748777722 | 3.21E-10 |
| FAM72C | 235.6365345 | 0.74691983 | 4.96E-02 |
| RASL11A | 579.9379851 | 0.746795211 | 3.06E-03 |
| FAM210A | 907.4822041 | 0.746711763 | 3.12E-07 |
| ABRACL | 559.2061279 | 0.746654852 | 1.19E-07 |
| MAP1B | 20810.43855 | 0.746524055 | 2.33E-04 |
| LINC00189 | 232.8947623 | 0.746006148 | 2.34E-04 |
| TRIO | 14689.77273 | 0.745652083 | 3.10E-16 |
| AIFM2 | 829.2813547 | 0.745123907 | 2.50E-08 |
| MTFP1 | 587.7366352 | 0.74330476 | 2.21E-05 |
| CTF1 | 182.4779826 | 0.743118065 | 1.07E-05 |
| GTF2IRD1 | 2131.857364 | 0.742953693 | 6.04E-08 |
| SERPINB1 | 1965.599407 | 0.742190867 | 3.25E-17 |
| CYB5A | 1340.256396 | 0.742101328 | 1.12E-11 |
| RMC1 | 875.8919123 | 0.742021767 | 2.19E-11 |
| HSD17B6 | 125.5224641 | 0.741781196 | 1.09E-03 |
| HSF2BP | 157.1567785 | 0.741129528 | 2.78E-05 |
| ANKRD13A | 3316.621453 | 0.740632762 | 1.04E-11 |
| DAZAP1 | 4240.34988 | 0.740268938 | 9.54E-21 |
| MIR621 | 113.5093563 | 0.740146937 | 1.83E-04 |
| FMNL1 | 1214.864249 | 0.73942309 | 3.37E-10 |
| PACS1 | 5469.333388 | 0.739178707 | 5.12E-16 |
| EDNRB | 780.2042551 | 0.738363229 | 4.28E-07 |

| | | | |
|---|---|---|---|
| KLF16 | 371.8280178 | 0.738097543 | 5.78E-05 |
| RAG1 | 104.3953946 | 0.737343457 | 9.20E-03 |
| LMO4 | 1106.20488 | 0.737171753 | 8.62E-06 |
| AC092692.1 | 32.94360854 | 0.737113758 | 4.33E-02 |
| MKNK2 | 4825.496095 | 0.736745636 | 7.90E-09 |
| FRAS1 | 577.0625188 | 0.736302272 | 8.50E-05 |
| C22orf23 | 284.8102428 | 0.736189543 | 8.89E-06 |
| SERPINE2 | 8188.667421 | 0.73617035 | 6.14E-11 |
| ARHGEF2 | 9221.872532 | 0.735793555 | 3.72E-04 |
| CASC10 | 644.5185033 | 0.735496562 | 1.17E-11 |
| MAGEA5 | 44.96564495 | 0.734506162 | 4.03E-02 |
| AL591845.1 | 91.06227821 | 0.733882309 | 6.11E-03 |
| GASK1A | 355.4300528 | 0.733654361 | 1.09E-09 |
| RIPK2 | 724.4734466 | 0.733141074 | 8.85E-08 |
| AL031587.5 | 395.7116765 | 0.732579001 | 2.86E-06 |
| SYT5 | 170.0209336 | 0.732514123 | 1.18E-05 |
| SMAD7 | 443.250496 | 0.732494727 | 1.20E-03 |
| LDLRAD4 | 335.4509987 | 0.732274001 | 9.70E-06 |
| LARGE2 | 219.8105223 | 0.731808765 | 1.16E-03 |
| SEC61G | 3651.15804 | 0.731454175 | 3.62E-11 |
| CDYL2 | 2093.83975 | 0.731313847 | 2.13E-09 |
| SCML1 | 645.1475496 | 0.7312017 | 9.67E-06 |
| SNRPB | 16462.47914 | 0.730751557 | 1.76E-11 |
| MICAL3 | 4913.970363 | 0.73026331 | 1.55E-21 |
| MAP3K9 | 1540.316129 | 0.730200144 | 1.60E-09 |
| PGM2 | 2123.780405 | 0.729366536 | 2.06E-14 |
| PNPLA4 | 1147.358362 | 0.729053648 | 2.04E-12 |
| WFS1 | 1309.328652 | 0.727641082 | 1.49E-06 |
| AC099850.3 | 182.1025745 | 0.727161113 | 1.08E-04 |
| PKIG | 2587.098172 | 0.727044228 | 1.17E-06 |
| PTGR1 | 3609.649913 | 0.726916628 | 3.98E-11 |
| AL671277.2 | 54.94515208 | 0.726086318 | 2.65E-02 |
| BATF3 | 241.6407464 | 0.725759002 | 5.50E-05 |
| CHCHD6 | 358.4714213 | 0.725392345 | 1.04E-10 |
| TENT5C | 137.8763368 | 0.725354772 | 1.03E-03 |
| MINAR1 | 148.9115714 | 0.724710664 | 7.52E-05 |
| TNFRSF10D | 1337.770238 | 0.724435371 | 6.48E-05 |
| SPRYD3 | 1098.046721 | 0.723457917 | 5.46E-09 |
| PDCD2L | 340.1145273 | 0.72215176 | 1.44E-08 |
| SKA1 | 744.58815 | 0.721710458 | 3.14E-04 |
| FAM241B | 259.3376186 | 0.720326124 | 6.12E-04 |
| HRCT1 | 93.63600929 | 0.719387798 | 5.06E-03 |
| MAGEA10-MAGI | 194.1530519 | 0.718715314 | 4.67E-05 |
| PHLDA2 | 934.0650038 | 0.718256796 | 4.30E-05 |
| RTN4 | 17887.31892 | 0.718113442 | 2.97E-12 |
| ANKRD11 | 12651.38724 | 0.717926766 | 2.43E-09 |
| MAD2L2 | 2128.707886 | 0.717910895 | 6.36E-16 |

| | | | |
|---|---|---|---|
| CYB5R3 | 9520.782062 | 0.717359319 | 2.03E-08 |
| ACTG1 | 90431.48433 | 0.716138979 | 5.20E-04 |
| SLC16A5 | 221.2180602 | 0.715991613 | 4.33E-05 |
| NATD1 | 1387.904722 | 0.715719843 | 6.87E-07 |
| ANKZF1 | 2063.874003 | 0.715592898 | 1.06E-13 |
| TMCC2 | 214.3735261 | 0.7155172 | 2.96E-04 |
| NT5C2 | 2575.304732 | 0.71538872 | 2.98E-10 |
| KLF4 | 330.1654842 | 0.714703742 | 2.95E-02 |
| PORCN | 684.9381108 | 0.714530295 | 5.25E-07 |
| SREBF2 | 12774.93631 | 0.713980375 | 1.67E-06 |
| ADAM15 | 5546.526107 | 0.713767095 | 1.11E-09 |
| SINHCAF | 2560.657428 | 0.712090285 | 1.14E-20 |
| AC006538.1 | 49.8630077 | 0.711988252 | 8.90E-03 |
| LIMA1 | 10004.88249 | 0.711921182 | 3.32E-10 |
| PRC1 | 7878.190155 | 0.71174961 | 2.71E-07 |
| SAV1 | 1549.090313 | 0.711056739 | 1.92E-10 |
| GLP2R | 415.3640839 | 0.710876138 | 3.85E-02 |
| SLC25A24 | 1865.58441 | 0.710869304 | 1.05E-04 |
| UBE2C | 4006.039724 | 0.710818979 | 1.15E-09 |
| CKAP4 | 11320.18536 | 0.71061586 | 1.96E-08 |
| CENPA | 1024.489308 | 0.709656116 | 4.99E-04 |
| ROBO3 | 1232.003209 | 0.709435867 | 4.02E-08 |
| ATP1A3 | 357.0233239 | 0.709420492 | 1.13E-02 |
| ID1 | 4464.244433 | 0.709364024 | 2.57E-03 |
| ZNRF2P2 | 85.72672436 | 0.709117459 | 5.21E-03 |
| TPTEP1 | 414.1805845 | 0.708093187 | 2.00E-05 |
| NUDT18 | 423.799999 | 0.708060366 | 1.33E-04 |
| RAB11FIP5 | 2099.786999 | 0.707865361 | 4.46E-07 |
| LAMB2 | 8298.833374 | 0.706764071 | 3.39E-08 |
| NA | 52.26903518 | 0.705977228 | 4.13E-02 |
| UBE2T | 1400.850171 | 0.705189061 | 9.98E-04 |
| KLC2 | 1774.580244 | 0.705150863 | 8.45E-10 |
| GBE1 | 1827.158317 | 0.705000557 | 9.86E-06 |
| SH2D4A | 2100.933704 | 0.704993664 | 3.09E-22 |
| KLHL36 | 2749.072336 | 0.704406566 | 2.20E-14 |
| GRPEL1 | 1981.801444 | 0.704246063 | 2.82E-16 |
| PRKCA | 7132.858038 | 0.704056699 | 1.58E-17 |
| TMSB10 | 36922.00469 | 0.703944831 | 4.56E-07 |
| KIF17 | 286.1148535 | 0.703867006 | 6.73E-04 |
| PRKCH | 883.6995121 | 0.703860761 | 6.26E-10 |
| CCDC50 | 3220.397041 | 0.702756442 | 1.08E-09 |
| HLA-L | 166.7642701 | 0.701687683 | 2.01E-02 |
| THBS4 | 204.431369 | 0.701362105 | 8.91E-06 |
| C8orf58 | 1572.615118 | 0.701178051 | 1.38E-05 |
| RRAS | 2730.533569 | 0.700994899 | 1.66E-06 |
| DNAH3 | 118.1200945 | 0.700898601 | 5.26E-03 |
| CASP4 | 3032.86953 | 0.700666171 | 3.06E-18 |

| | | | |
|---|---|---|---|
| ETS2 | 1570.480883 | 0.700301688 | 7.82E-12 |
| QPCT | 409.1069256 | 0.699977528 | 7.59E-06 |
| VDAC2 | 3904.428237 | 0.699604563 | 8.41E-17 |
| TMEM80 | 695.471718 | 0.699116586 | 1.27E-10 |
| SH3GL2 | 285.0147928 | 0.698900781 | 3.93E-04 |
| FSCN1 | 13820.42552 | 0.698810655 | 3.53E-07 |
| RNF20 | 5561.7265 | 0.698567575 | 2.32E-10 |
| SMIM10 | 1046.732466 | 0.698510406 | 7.91E-11 |
| AGRN | 19104.86146 | 0.697670315 | 4.52E-08 |
| GNAI1 | 819.6351182 | 0.696846567 | 1.65E-07 |
| PRR11 | 1991.797462 | 0.696619386 | 1.08E-08 |
| ZNF219 | 899.6886662 | 0.69597147 | 2.65E-08 |
| DOP1B | 559.0709286 | 0.695930259 | 3.52E-03 |
| RDM1 | 181.5823242 | 0.695796532 | 2.96E-05 |
| SEL1L3 | 3269.858349 | 0.695591737 | 4.45E-07 |
| SND1 | 22442.27026 | 0.695458133 | 2.33E-11 |
| SCN5A | 570.9436693 | 0.695141655 | 1.05E-03 |
| MYBL1 | 683.1035354 | 0.694900657 | 4.47E-03 |
| DVL1 | 1980.054946 | 0.694814441 | 3.99E-13 |
| AC008957.1 | 76.43432309 | 0.694793391 | 5.43E-03 |
| AL589843.1 | 84.43969269 | 0.694624951 | 4.17E-03 |
| CENPH | 2549.82782 | 0.693813155 | 5.31E-07 |
| TIMM44 | 2236.04911 | 0.693697771 | 2.61E-07 |
| IL1RAP | 613.1327742 | 0.693336483 | 6.39E-05 |
| ATP6V1C2 | 111.0224753 | 0.693255573 | 2.57E-02 |
| PLXNB2 | 10147.39702 | 0.693129203 | 4.58E-12 |
| NBR2 | 171.9762602 | 0.693029196 | 1.64E-03 |
| TRIM8 | 1510.617956 | 0.692956737 | 9.70E-06 |
| CSRP2 | 540.994005 | 0.692633799 | 7.06E-07 |
| TRAM2 | 9655.068286 | 0.692228729 | 3.50E-25 |
| TXNRD3 | 920.3708362 | 0.691829753 | 8.78E-12 |
| FXYD5 | 4701.536881 | 0.691464828 | 8.74E-11 |
| KCTD11 | 1277.421524 | 0.691127389 | 9.27E-07 |
| OSBPL3 | 2070.819801 | 0.690941687 | 1.01E-07 |
| DLGAP4 | 3901.025781 | 0.69090093 | 9.61E-27 |
| SDHB | 2974.624177 | 0.690672588 | 1.65E-19 |
| ECI1 | 2757.268471 | 0.690330176 | 4.36E-16 |
| JAG2 | 524.7348244 | 0.6902408 | 2.53E-03 |
| AOC2 | 89.02258926 | 0.689451467 | 4.02E-03 |
| TBX1 | 62.73316178 | 0.688923288 | 1.93E-02 |
| LRRC75A | 138.5402757 | 0.688871647 | 1.13E-03 |
| MAP4K3-DT | 286.7965894 | 0.688477523 | 6.48E-08 |
| LINC00346 | 364.435918 | 0.688051475 | 2.20E-09 |
| PDGFC | 565.39758 | 0.687668686 | 3.34E-05 |
| TBC1D30 | 168.1408535 | 0.687332446 | 1.61E-03 |
| INF2 | 4687.789987 | 0.687061386 | 2.26E-12 |
| EPB41L4B | 494.6600528 | 0.68648704 | 1.34E-04 |

| | | | |
|---|---|---|---|
| CARD16 | 266.3336459 | 0.686185969 | 4.36E-05 |
| STMN1 | 17626.57597 | 0.685300807 | 1.12E-11 |
| CYSTM1 | 1163.563966 | 0.68513273 | 2.18E-07 |
| LINC00476 | 276.6276502 | 0.684984209 | 1.86E-06 |
| SRF | 2490.360946 | 0.684714849 | 2.94E-16 |
| DOCK5 | 2093.893287 | 0.68403964 | 2.63E-05 |
| ARF4 | 9639.726706 | 0.68383598 | 1.45E-09 |
| CORO1C | 8847.320057 | 0.68351959 | 7.24E-13 |
| PINX1 | 749.064652 | 0.683492703 | 2.66E-15 |
| S100A6 | 9517.564517 | 0.681915452 | 3.99E-07 |
| C11orf45 | 327.0867099 | 0.681718568 | 4.95E-05 |
| TIMM9 | 1153.293094 | 0.68142168 | 1.08E-08 |
| NDUFAF3 | 1905.99529 | 0.679687455 | 6.37E-10 |
| FADD | 1921.919656 | 0.679454737 | 1.02E-10 |
| KIF12 | 74.36197482 | 0.678595181 | 2.16E-02 |
| P2RX7 | 1431.080718 | 0.678170827 | 2.45E-05 |
| DGKD | 3157.094476 | 0.67771577 | 1.89E-07 |
| PLS3 | 10420.30682 | 0.677505457 | 1.79E-13 |
| EIF3J | 2363.847504 | 0.676820205 | 1.83E-06 |
| IFNGR2 | 3672.123779 | 0.67656923 | 7.10E-10 |
| NBPF3 | 1660.670122 | 0.67624491 | 1.76E-08 |
| FAM189A1 | 83.82781644 | 0.676232206 | 4.51E-02 |
| AL354892.2 | 44.61264788 | 0.675976534 | 2.83E-02 |
| P2RX4 | 1389.149782 | 0.675084885 | 5.66E-10 |
| TUBB | 73676.804 | 0.673074235 | 6.99E-05 |
| PLA2G16 | 1773.072902 | 0.672006685 | 3.75E-06 |
| CLCF1 | 243.9135885 | 0.67119739 | 1.69E-05 |
| ST3GAL4 | 2762.325065 | 0.670924956 | 3.46E-08 |
| CADM1 | 234.8479271 | 0.670511035 | 4.03E-02 |
| SWAP70 | 3816.151683 | 0.670327923 | 3.69E-15 |
| MTCH1 | 8829.71424 | 0.669843869 | 5.89E-06 |
| CT45A5 | 774.9288964 | 0.669828171 | 2.01E-13 |
| TSPO | 3048.132338 | 0.669750859 | 5.24E-07 |
| HOGA1 | 115.4192823 | 0.669490173 | 2.48E-02 |
| MND1 | 324.1099673 | 0.669420278 | 1.74E-04 |
| E2F5 | 507.7955568 | 0.668939707 | 1.86E-03 |
| SOX7 | 73.75907218 | 0.668806667 | 2.48E-02 |
| TBC1D22B | 1030.373319 | 0.668043011 | 7.24E-13 |
| AL691420.1 | 230.2834602 | 0.667905003 | 1.70E-06 |
| DCBLD1 | 748.5491359 | 0.667710722 | 1.89E-06 |
| AL138756.1 | 98.1393658 | 0.666937909 | 2.61E-03 |
| AL121753.1 | 48.03673125 | 0.666496938 | 3.83E-02 |
| CCDC59 | 724.4599412 | 0.66505293 | 4.38E-07 |
| GPC6 | 245.0733971 | 0.664346792 | 1.99E-02 |
| OR51B4 | 93.62497888 | 0.664108957 | 1.65E-02 |
| WBP2 | 3411.363076 | 0.664026767 | 1.56E-09 |
| MITF | 317.5617059 | 0.66388595 | 4.27E-04 |

| | | | |
|---|---|---|---|
| ARHGAP10 | 734.6739707 | 0.663394859 | 9.24E-14 |
| DGKH | 590.9709515 | 0.663158486 | 5.51E-04 |
| PGF | 1251.66419 | 0.662683357 | 1.54E-04 |
| MRPL14 | 1572.149765 | 0.662170453 | 1.31E-09 |
| BICD2 | 3147.686758 | 0.661748485 | 2.17E-11 |
| CALU | 26908.66603 | 0.661687604 | 3.57E-07 |
| CDKN1A | 368.8877978 | 0.661533823 | 8.76E-06 |
| STARD5 | 260.8407944 | 0.66094157 | 1.88E-05 |
| C17orf51 | 1818.558345 | 0.659918036 | 2.58E-05 |
| PIK3R5 | 80.61375241 | 0.659382067 | 1.65E-02 |
| KPNA2 | 10845.5826 | 0.659332597 | 1.15E-04 |
| LRRC1 | 426.2692342 | 0.659157309 | 2.99E-05 |
| DENND1A | 2021.521882 | 0.65897305 | 1.48E-06 |
| ZNF793-AS1 | 53.5064077 | 0.658528272 | 2.61E-02 |
| TTC28-AS1 | 350.7466498 | 0.658484241 | 6.02E-06 |
| KIF2C | 4458.272751 | 0.657650666 | 3.29E-10 |
| PCDH1 | 2577.072522 | 0.657623151 | 2.31E-03 |
| PHLDB3 | 358.9219198 | 0.657547028 | 3.69E-05 |
| ZPR1 | 5866.201251 | 0.657295592 | 1.66E-06 |
| MAP3K7CL | 150.299139 | 0.657266586 | 1.56E-03 |
| SNHG17 | 1649.128822 | 0.657034712 | 2.48E-03 |
| JOSD1 | 3246.967769 | 0.656907948 | 1.97E-15 |
| CFAP20 | 1402.706065 | 0.656814828 | 2.61E-14 |
| FYCO1 | 2501.029106 | 0.656725622 | 1.56E-15 |
| RFLNB | 1500.315546 | 0.65607592 | 3.96E-09 |
| RASAL2-AS1 | 121.4954753 | 0.655508647 | 3.01E-03 |
| AP001505.1 | 63.01778494 | 0.655043688 | 1.57E-02 |
| FKBP1B | 606.1349468 | 0.654877068 | 6.01E-05 |
| FPR2 | 56.2775836 | 0.654798028 | 3.96E-02 |
| SCARB1 | 3539.739284 | 0.654611345 | 5.27E-12 |
| BID | 1337.216014 | 0.654289524 | 1.08E-04 |
| CNNM1 | 541.0648735 | 0.654167577 | 8.17E-08 |
| SBNO2 | 2829.160965 | 0.653866211 | 2.56E-13 |
| SIAH2 | 1088.269525 | 0.651473155 | 1.80E-09 |
| DIS3L | 2900.587099 | 0.651462805 | 4.33E-11 |
| RANGAP1 | 7733.942925 | 0.651366095 | 4.83E-16 |
| LSP1 | 262.4820471 | 0.651251589 | 2.17E-02 |
| KLHL3 | 639.2424971 | 0.651159682 | 2.78E-02 |
| HDAC11 | 769.1429549 | 0.65114748 | 2.25E-05 |
| LENG1 | 325.7448315 | 0.65113745 | 2.13E-04 |
| AC012645.1 | 59.53374631 | 0.650573362 | 3.98E-02 |
| NOTCH1 | 1196.391895 | 0.650483662 | 9.07E-07 |
| ULK2 | 882.4735602 | 0.650391046 | 7.17E-07 |
| ACAT2 | 4456.954676 | 0.64992014 | 1.23E-03 |
| ARF1 | 11486.82838 | 0.649536335 | 1.35E-14 |
| HAGH | 1298.627079 | 0.648320413 | 2.10E-07 |
| AC092112.1 | 85.72490216 | 0.647943962 | 2.15E-03 |

| | | | |
|---|---|---|---|
| EML2 | 1514.807018 | 0.647561618 | 1.57E-07 |
| NARS | 10148.14967 | 0.647374704 | 1.80E-05 |
| CHPT1 | 1064.870081 | 0.647362174 | 8.52E-10 |
| OOEP | 184.8376679 | 0.646814711 | 7.80E-04 |
| KIRREL1 | 10728.67019 | 0.646225205 | 1.64E-11 |
| TK1 | 5146.104526 | 0.645971198 | 8.28E-04 |
| IRS2 | 208.4093286 | 0.645859457 | 3.60E-04 |
| CCDC169 | 311.1901616 | 0.645424849 | 2.24E-03 |
| ELP4 | 1116.525473 | 0.645216276 | 1.16E-11 |
| RP9P | 225.4968509 | 0.645145177 | 1.43E-05 |
| MAMDC2 | 4343.615439 | 0.64363381 | 2.65E-06 |
| H2AFZ | 12211.53725 | 0.643515536 | 1.80E-12 |
| CCND3 | 1929.467615 | 0.643436685 | 8.62E-07 |
| IER3 | 2184.936652 | 0.643325579 | 2.09E-06 |
| CLUL1 | 116.4611332 | 0.642167942 | 1.64E-02 |
| CEP85 | 1897.607299 | 0.642130415 | 1.91E-09 |
| PDZD7 | 149.1959024 | 0.641730551 | 2.70E-03 |
| PLXNA2 | 1668.333265 | 0.641582632 | 5.83E-05 |
| AC016026.1 | 401.4028746 | 0.641395732 | 2.51E-04 |
| RILPL1 | 800.4589162 | 0.640900332 | 3.68E-05 |
| GPX8 | 5676.176415 | 0.640834267 | 3.51E-07 |
| LINC01694 | 305.0858403 | 0.640590995 | 1.26E-02 |
| TMEM81 | 102.0745934 | 0.64052667 | 3.25E-03 |
| OR2A1-AS1 | 214.8177946 | 0.640358904 | 1.16E-03 |
| ZC3H7B | 8780.439133 | 0.640249262 | 1.99E-16 |
| PRDM13 | 87.34034449 | 0.640237807 | 1.26E-02 |
| RADIL | 174.7008569 | 0.639616464 | 1.82E-03 |
| TMEM205 | 1463.331865 | 0.638734309 | 2.72E-06 |
| PCBD1 | 805.7687183 | 0.637869967 | 1.34E-09 |
| HTR7P1 | 368.6181894 | 0.637742874 | 8.62E-06 |
| PCNX4 | 3959.147168 | 0.637665622 | 1.52E-05 |
| CPM | 499.6059264 | 0.637353259 | 5.21E-03 |
| DUX4L26 | 66.00978395 | 0.637322668 | 3.24E-02 |
| BACH1 | 2018.700707 | 0.637138201 | 1.99E-03 |
| PTP4A2 | 6807.446394 | 0.636684016 | 7.68E-07 |
| NEURL1B | 558.7832072 | 0.634913459 | 1.51E-03 |
| ZCCHC4 | 500.6188419 | 0.634844768 | 4.32E-06 |
| ABCC1 | 5112.111329 | 0.634460685 | 1.74E-09 |
| GPKOW | 1631.232436 | 0.634396822 | 6.54E-10 |
| BHLHE40 | 1231.608194 | 0.634121231 | 3.33E-03 |
| METTL21A | 721.6296818 | 0.633994885 | 6.17E-07 |
| RPS27L | 1580.599612 | 0.633806976 | 4.70E-10 |
| SLC13A3 | 117.8644963 | 0.6335489 | 3.13E-03 |
| KIAA1549L | 2687.335253 | 0.633163244 | 2.30E-08 |
| XAGE1A | 480.1985385 | 0.633049273 | 1.38E-09 |
| SLC22A23 | 249.0103972 | 0.632975868 | 3.38E-03 |
| STX8 | 1628.006817 | 0.632738501 | 3.79E-11 |

| | | | |
|---|---|---|---|
| CYP2S1 | 1720.583561 | 0.632612178 | 1.57E-03 |
| URGCP | 3502.054904 | 0.632415521 | 2.08E-09 |
| ATXN10 | 5946.981823 | 0.632411445 | 2.22E-11 |
| TLNRD1 | 501.8768901 | 0.632405663 | 9.79E-04 |
| AC090204.1 | 619.9733506 | 0.632015261 | 3.59E-10 |
| RNASEH1 | 1425.965528 | 0.631921816 | 1.79E-19 |
| RPL23AP7 | 323.4192596 | 0.631843485 | 1.11E-08 |
| SNHG5 | 2995.960911 | 0.630920536 | 1.27E-07 |
| OR2A20P | 429.4941753 | 0.630414395 | 1.69E-02 |
| COPS4 | 2229.661381 | 0.630249779 | 1.15E-23 |
| ATP6V1D | 2512.368254 | 0.629922219 | 2.22E-06 |
| AHCYL1 | 7022.281892 | 0.629601404 | 2.91E-29 |
| IMP4 | 3203.583644 | 0.629547132 | 1.35E-11 |
| IGFBP7 | 8049.810997 | 0.62885929 | 1.21E-05 |
| FAM219A | 2447.604118 | 0.628664392 | 7.71E-06 |
| PKMYT1 | 1390.690032 | 0.628468772 | 3.63E-03 |
| HERC3 | 716.0548112 | 0.62834276 | 4.91E-02 |
| ALAS1 | 2028.759906 | 0.62830202 | 1.42E-08 |
| FGF2 | 458.0900956 | 0.628276956 | 1.67E-03 |
| ESYT2 | 7763.823023 | 0.628097683 | 9.75E-17 |
| PLEKHB2 | 5020.256771 | 0.627947143 | 1.98E-15 |
| SLC25A19 | 457.3255449 | 0.627874702 | 1.85E-06 |
| KIAA1522 | 1610.958184 | 0.627855234 | 3.60E-07 |
| ADK | 1521.24063 | 0.627615397 | 3.91E-12 |
| TUBB4B | 23483.62734 | 0.627476985 | 1.59E-04 |
| ASPHD1 | 246.0080628 | 0.627378535 | 2.99E-06 |
| LEMD2 | 2746.266522 | 0.627355959 | 4.70E-18 |
| GPN3 | 1009.88344 | 0.627199994 | 1.16E-06 |
| YIF1B | 1626.377802 | 0.627054109 | 1.41E-10 |
| CDCP1 | 1149.145664 | 0.626447436 | 5.53E-05 |
| YIPF1 | 1067.199585 | 0.624862166 | 6.02E-09 |
| STK3 | 890.5682343 | 0.624250771 | 1.43E-08 |
| AMZ2P1 | 165.7323703 | 0.624155307 | 5.17E-04 |
| AGAP1 | 2415.072612 | 0.6240417 | 7.69E-08 |
| CEP250 | 4914.904169 | 0.624033685 | 7.88E-15 |
| CPPED1 | 2575.427025 | 0.623259494 | 4.51E-12 |
| ARL8A | 771.8332599 | 0.6222161 | 1.16E-04 |
| ACTB | 127290.7296 | 0.621729905 | 5.29E-04 |
| TFE3 | 7339.395033 | 0.621548856 | 3.01E-11 |
| MPRIP | 18080.17353 | 0.621307541 | 4.07E-11 |
| P2RX5 | 672.8200669 | 0.62126021 | 5.23E-06 |
| BTG3 | 724.4575775 | 0.620763689 | 3.36E-06 |
| MAD1L1 | 2125.818007 | 0.620671877 | 2.65E-10 |
| COPG1 | 7879.341187 | 0.620290095 | 1.15E-09 |
| FTL | 97215.3597 | 0.620031724 | 1.38E-05 |
| PPDPF | 3817.555591 | 0.619916764 | 1.30E-07 |
| CSNK2A2 | 5047.769308 | 0.619870637 | 6.79E-23 |

| | | | |
|---|---|---|---|
| HEATR5A | 1798.727542 | 0.619749272 | 1.26E-07 |
| INSIG2 | 868.8112722 | 0.619520379 | 4.57E-10 |
| PFDN2 | 1584.135587 | 0.619262892 | 4.26E-07 |
| ATP13A3 | 4319.212019 | 0.619037548 | 8.78E-04 |
| PLGRKT | 793.4817684 | 0.618627883 | 4.57E-10 |
| ING1 | 361.767476 | 0.618468038 | 1.45E-05 |
| HSPD1P11 | 293.0157327 | 0.618263741 | 2.70E-02 |
| FAM86B3P | 213.3333834 | 0.618201244 | 3.24E-03 |
| APCDD1L-DT | 254.384421 | 0.617680589 | 2.33E-02 |
| STEAP3 | 2478.209655 | 0.617266814 | 3.14E-07 |
| SLC7A7 | 165.6952727 | 0.617028109 | 1.53E-02 |
| SPSB4 | 329.8186146 | 0.616923074 | 3.00E-02 |
| TNFAIP1 | 4428.834655 | 0.61589198 | 1.05E-08 |
| MMAB | 1653.207457 | 0.615382904 | 7.55E-06 |
| EVC2 | 505.3089137 | 0.615006545 | 5.96E-06 |
| PELO | 2799.47239 | 0.614980176 | 7.47E-15 |
| HINT1 | 21385.80112 | 0.614493406 | 5.92E-11 |
| C1QL1 | 789.3243264 | 0.614276353 | 4.03E-04 |
| RPS6KA1 | 1985.693064 | 0.613857526 | 5.08E-07 |
| SIRT5 | 725.0325281 | 0.613443276 | 2.71E-06 |
| CDC14B | 2116.738376 | 0.613095321 | 2.33E-11 |
| SNTA1 | 2042.451858 | 0.61291227 | 3.96E-06 |
| IRS1 | 1437.527567 | 0.612279392 | 5.30E-04 |
| BHMT2 | 687.4043628 | 0.612213202 | 8.05E-04 |
| SBF1 | 7935.53834 | 0.61195101 | 5.22E-13 |
| AC125232.1 | 781.9054721 | 0.611938598 | 1.09E-05 |
| MRPL2 | 1431.242601 | 0.611654088 | 6.94E-07 |
| FJX1 | 664.7311372 | 0.611335602 | 2.32E-02 |
| SMURF2 | 2056.949958 | 0.611295872 | 2.55E-08 |
| ATG4D | 562.7709515 | 0.611282354 | 1.33E-06 |
| HIP1R | 1309.841327 | 0.611023277 | 1.71E-05 |
| ARHGEF10L | 688.723664 | 0.610986832 | 2.91E-06 |
| TMEM98 | 1509.247942 | 0.610146915 | 1.78E-05 |
| SH3RF2 | 238.2640979 | 0.609961569 | 6.29E-03 |
| TRIM25 | 7151.293211 | 0.609865694 | 4.37E-07 |
| GAB2 | 601.0589633 | 0.609388694 | 3.56E-03 |
| TFAP2A | 2148.867207 | 0.609317872 | 5.89E-07 |
| CCNB1 | 16691.04164 | 0.609167091 | 2.55E-12 |
| SLC12A7 | 1143.958255 | 0.608642646 | 9.48E-04 |
| RNF126 | 894.0071533 | 0.608305404 | 8.95E-09 |
| ACOT7 | 3467.030846 | 0.608167949 | 2.25E-09 |
| HHAT | 280.182391 | 0.607778746 | 3.06E-05 |
| CLIP1 | 2506.614825 | 0.607637172 | 3.37E-03 |
| SNHG7 | 855.8523069 | 0.607260798 | 1.93E-09 |
| EIF5A2 | 297.4165173 | 0.607083229 | 6.50E-06 |
| LCORL | 400.3564959 | 0.606975543 | 4.57E-03 |
| EXOC6B | 3172.372247 | 0.606561495 | 1.47E-18 |

| | | | |
|---|---|---|---|
| FHAD1 | 349.9471983 | 0.606337834 | 9.70E-05 |
| MED10 | 2839.473397 | 0.606127856 | 5.45E-06 |
| RAB20 | 447.2426815 | 0.60608303 | 8.84E-05 |
| YDJC | 1016.804538 | 0.606009407 | 3.09E-08 |
| PPA1 | 2590.012982 | 0.605877396 | 3.55E-09 |
| SLC20A1 | 3792.236403 | 0.60508113 | 8.38E-05 |
| AFAP1L1 | 1973.109642 | 0.604969137 | 1.30E-08 |
| AXIN1 | 1792.469903 | 0.604465239 | 2.78E-06 |
| SFXN3 | 2024.371708 | 0.604061122 | 4.72E-06 |
| SNHG15 | 1441.968187 | 0.60394602 | 3.75E-03 |
| RBPMS2 | 349.7867648 | 0.603708083 | 3.62E-07 |
| ZNF488 | 398.2968029 | 0.603555711 | 1.64E-02 |
| CHKA | 2021.673855 | 0.603350205 | 2.03E-10 |
| MAP1LC3B | 6359.449737 | 0.603280013 | 1.30E-08 |
| NARS2 | 1356.554678 | 0.602662117 | 5.03E-08 |
| SPIRE1 | 1623.898975 | 0.602374225 | 1.15E-04 |
| ZNF331 | 767.9253616 | 0.602311968 | 1.31E-03 |
| FAXC | 492.2908407 | 0.602245331 | 4.55E-09 |
| PDP2 | 1653.370186 | 0.601947145 | 1.28E-10 |
| SH2D3A | 372.6600886 | 0.60186386 | 1.39E-05 |
| RIPOR1 | 6265.287925 | 0.601822514 | 8.56E-10 |
| CYTL1 | 1870.470186 | 0.601589747 | 6.69E-06 |
| PNPLA2 | 3504.906473 | 0.601463609 | 5.97E-05 |
| EXD3 | 412.4310246 | 0.601449401 | 5.44E-07 |
| THAP9-AS1 | 1053.717623 | 0.601131744 | 2.32E-02 |
| TUBE1 | 792.6081694 | 0.60089195 | 2.03E-02 |
| YIF1A | 3045.656821 | 0.600787034 | 2.05E-08 |
| SEH1L | 1842.288795 | 0.600723788 | 7.06E-10 |
| TYRO3 | 2229.388693 | 0.600612707 | 1.59E-06 |
| TAGLN2 | 18351.75881 | 0.599908829 | 3.66E-04 |
| LRP11 | 2422.615292 | 0.599459122 | 1.98E-06 |
| FGD1 | 1766.826493 | 0.599183449 | 4.67E-08 |
| ARMC4 | 198.3905927 | 0.598954232 | 1.81E-03 |
| RTN4IP1 | 449.1760696 | 0.598561481 | 9.86E-06 |
| HOXB-AS1 | 138.8490023 | 0.598487108 | 6.96E-04 |
| FUOM | 567.2907635 | 0.598344847 | 4.11E-04 |
| AL445363.3 | 241.367527 | 0.598292416 | 1.21E-03 |
| PKN1 | 5938.711978 | 0.598201738 | 2.93E-09 |
| TRMT44 | 441.8763599 | 0.598115549 | 4.29E-05 |
| AC092042.3 | 124.0100688 | 0.596837382 | 1.69E-02 |
| LINC01145 | 651.6649971 | 0.5968138 | 4.47E-05 |
| ATP7B | 692.4149392 | 0.596566266 | 4.44E-04 |
| CDK7 | 2722.817861 | 0.596466737 | 5.55E-19 |
| USP32 | 3325.072913 | 0.596345009 | 3.54E-14 |
| RNPEPL1 | 2127.531071 | 0.596190761 | 6.64E-06 |
| ELFN1 | 408.2898101 | 0.596046158 | 2.70E-05 |
| OSER1 | 2909.229889 | 0.595781932 | 7.42E-14 |

| | | | |
|---|---|---|---|
| NKILA | 183.3136004 | 0.59567556 | 4.50E-03 |
| PRELID2 | 136.0433833 | 0.59563106 | 4.41E-03 |
| AC020763.4 | 153.2361127 | 0.595335832 | 2.19E-02 |
| MMP15 | 1787.760392 | 0.594790761 | 6.22E-04 |
| TUBG1 | 3738.175157 | 0.593773074 | 1.69E-07 |
| TNFRSF8 | 76.01422714 | 0.593760697 | 1.78E-02 |
| PAM16 | 284.7754864 | 0.593259553 | 3.24E-04 |
| RNF181 | 2072.155686 | 0.593073986 | 5.25E-07 |
| ZNF721 | 490.7716122 | 0.593017466 | 3.42E-07 |
| PFDN4 | 1460.258671 | 0.592724394 | 2.93E-08 |
| GOT2 | 7515.082647 | 0.592232616 | 6.78E-11 |
| RPSA | 54318.93969 | 0.591802021 | 1.53E-06 |
| PLS1 | 372.3599433 | 0.590328202 | 2.76E-03 |
| WDSUB1 | 397.7929314 | 0.589526217 | 7.08E-06 |
| EIF2S2 | 12053.63184 | 0.589385956 | 7.36E-05 |
| SLC25A30 | 977.3527418 | 0.588735715 | 7.97E-04 |
| DHDDS | 1753.60419 | 0.588692609 | 6.75E-08 |
| GOLGA2 | 4842.364652 | 0.588628272 | 3.94E-21 |
| TRIP10 | 3342.101072 | 0.588537466 | 8.29E-22 |
| C11orf68 | 2384.928605 | 0.588326477 | 1.71E-05 |
| PPME1 | 11264.96145 | 0.588323127 | 2.28E-07 |
| SKA3 | 713.8002196 | 0.587711545 | 1.93E-04 |
| CDCA8 | 3450.166152 | 0.587083487 | 5.86E-08 |
| TNS1 | 3945.928745 | 0.585847295 | 1.05E-04 |
| TMEM54 | 1494.946011 | 0.585479831 | 3.25E-07 |
| ATP11A | 1843.254605 | 0.585341477 | 1.32E-05 |
| NFKBIB | 708.9637072 | 0.585309355 | 1.16E-03 |
| ALG9 | 1298.005813 | 0.585184106 | 9.75E-09 |
| HDAC6 | 2892.832686 | 0.58505622 | 9.12E-07 |
| BUB1 | 3783.438651 | 0.584907595 | 1.34E-04 |
| SMIM13 | 1014.459578 | 0.584686004 | 1.67E-04 |
| LINC00899 | 351.447866 | 0.584551312 | 7.24E-04 |
| TRAF4 | 2091.976798 | 0.584140877 | 6.86E-09 |
| SPINDOC | 2425.02415 | 0.583872802 | 1.02E-10 |
| TRAP1 | 2737.631956 | 0.583166378 | 1.26E-11 |
| WIPI1 | 1713.174141 | 0.583095881 | 3.67E-07 |
| PSMB6 | 4576.834031 | 0.582886357 | 1.62E-13 |
| SPG21 | 4901.16491 | 0.582607873 | 5.90E-12 |
| ZC3H12A | 238.2279361 | 0.58250174 | 1.45E-02 |
| RRP9 | 1400.270047 | 0.582481302 | 3.10E-09 |
| EIF1AD | 1509.337044 | 0.582396244 | 5.09E-10 |
| GYPC | 5168.208005 | 0.582140664 | 8.07E-06 |
| COPA | 14477.07068 | 0.581249341 | 1.69E-16 |
| AL353751.1 | 231.7121403 | 0.580932462 | 4.10E-02 |
| CHIC2 | 379.1742009 | 0.580825747 | 7.05E-04 |
| AC122710.1 | 176.9889668 | 0.580646294 | 1.11E-03 |
| WDR45B | 3377.208925 | 0.580312167 | 6.91E-19 |

| | | | |
|---|---|---|---|
| MED15 | 3949.807379 | 0.580186947 | 5.55E-10 |
| AFAP1 | 8267.209501 | 0.579882137 | 6.87E-10 |
| C9orf40 | 891.5125514 | 0.579691057 | 2.71E-05 |
| PIK3AP1 | 191.3237924 | 0.579445362 | 9.63E-03 |
| NDST1 | 8021.325827 | 0.579092672 | 6.95E-09 |
| HSPA5 | 56281.23694 | 0.578877373 | 4.73E-06 |
| RAMP1 | 108.3421471 | 0.578642972 | 4.98E-02 |
| ALDH1B1 | 2870.46866 | 0.577563213 | 9.28E-10 |
| HOXC4 | 316.5324271 | 0.57745184 | 1.67E-05 |
| NR2F2 | 5643.974102 | 0.577229389 | 1.27E-06 |
| DNAH14 | 338.8482397 | 0.577155554 | 7.60E-03 |
| TSPAN14 | 2543.409951 | 0.576768321 | 1.41E-05 |
| SLC25A43 | 1114.800884 | 0.576120164 | 2.35E-09 |
| DDX47 | 4767.009017 | 0.574514896 | 2.34E-15 |
| LPP | 1835.54307 | 0.574182321 | 1.14E-02 |
| ZNF706 | 2431.618789 | 0.574180647 | 9.16E-20 |
| EFTUD2 | 8567.38897 | 0.572603634 | 5.72E-06 |
| TACC1 | 7595.804216 | 0.572517882 | 5.51E-07 |
| CCDC61 | 301.5063114 | 0.571606286 | 5.99E-06 |
| CD3EAP | 786.1555899 | 0.571446595 | 1.50E-03 |
| AL645933.2 | 91.62206735 | 0.571386138 | 4.11E-02 |
| FAM114A1 | 1772.247523 | 0.571304936 | 1.09E-04 |
| LINC01234 | 437.4067572 | 0.571112125 | 4.38E-06 |
| SLC6A6 | 5652.703338 | 0.570864029 | 2.67E-07 |
| SRXN1 | 4583.879997 | 0.570834258 | 3.74E-04 |
| TRHDE-AS1 | 181.9325313 | 0.570038275 | 2.27E-02 |
| AGPAT3 | 4177.175525 | 0.570005767 | 4.45E-18 |
| SLC38A1 | 10325.87367 | 0.569009428 | 2.06E-03 |
| RHOB | 1671.093319 | 0.568825159 | 3.25E-03 |
| PNO1 | 1549.618819 | 0.568616546 | 2.78E-11 |
| SNRPA1 | 1845.883866 | 0.568337612 | 1.54E-11 |
| DOCK6 | 1355.620577 | 0.568184694 | 1.12E-04 |
| PTBP3 | 2504.459584 | 0.568183835 | 1.59E-05 |
| URAHP | 165.1413034 | 0.567560032 | 4.50E-02 |
| DUSP10 | 564.3339546 | 0.56727408 | 6.32E-03 |
| CDKN1C | 326.2645282 | 0.567032191 | 1.18E-02 |
| MN1 | 809.3964248 | 0.566619331 | 1.66E-03 |
| EEF2 | 104137.7217 | 0.566553772 | 2.25E-06 |
| OXNAD1 | 604.3305565 | 0.566453314 | 3.99E-05 |
| NELFE | 4523.328843 | 0.566419663 | 1.62E-06 |
| TCP11L1 | 1101.58559 | 0.566197195 | 2.05E-15 |
| PACSIN3 | 1919.961239 | 0.566175331 | 5.70E-10 |
| CGB7 | 208.1396265 | 0.566128572 | 2.65E-02 |
| LRP1 | 11435.56865 | 0.565852581 | 3.03E-05 |
| GPR19 | 145.3282073 | 0.565574349 | 3.76E-02 |
| POTEF | 330.2910566 | 0.565418576 | 4.12E-03 |
| SPTLC3 | 200.1830274 | 0.565343328 | 7.55E-03 |

| | | | |
|---|---|---|---|
| RNF19B | 646.416233 | 0.565244945 | 8.63E-06 |
| LONRF3 | 116.1789548 | 0.565102362 | 2.30E-02 |
| AKT1 | 9635.270702 | 0.564971356 | 1.37E-10 |
| UQCRC1 | 6211.217267 | 0.564548815 | 5.61E-09 |
| TMC3-AS1 | 109.7734588 | 0.5640654 | 4.38E-03 |
| HACD2 | 1942.141445 | 0.563906793 | 1.49E-05 |
| PAFAH1B1 | 7948.894797 | 0.563441708 | 4.88E-17 |
| STARD13 | 1275.002019 | 0.563331652 | 2.96E-04 |
| FAM171A1 | 2363.984497 | 0.563219862 | 2.23E-06 |
| ATP6V1E1 | 4139.268406 | 0.563003797 | 1.29E-16 |
| VAV2 | 2652.404696 | 0.562960328 | 2.27E-09 |
| RAP1GAP2 | 2036.056692 | 0.562702115 | 8.06E-06 |
| MAFG | 2434.517742 | 0.562657398 | 1.78E-09 |
| GEM | 363.9867785 | 0.56223331 | 1.64E-02 |
| LARP6 | 2466.487265 | 0.561702819 | 5.46E-05 |
| DTNB | 563.0925909 | 0.561561531 | 2.47E-07 |
| STPG1 | 457.4197334 | 0.561416331 | 1.13E-04 |
| TIMP2 | 22643.24137 | 0.561238916 | 6.59E-05 |
| HACL1 | 1377.386205 | 0.560946153 | 1.44E-06 |
| SUN2 | 5461.842658 | 0.560368254 | 1.57E-09 |
| DNAJA3 | 2240.392464 | 0.560216464 | 1.07E-05 |
| TEC | 198.6931738 | 0.560103094 | 2.76E-02 |
| RHBDF1 | 1735.657285 | 0.559783851 | 2.08E-06 |
| AP5B1 | 930.0444665 | 0.559744468 | 8.37E-06 |
| H1F0 | 16936.24213 | 0.559367532 | 2.49E-03 |
| CUL4B | 4541.326137 | 0.559290344 | 5.86E-06 |
| LGALS1 | 23005.81072 | 0.559211657 | 4.73E-05 |
| AC092111.1 | 85.97713132 | 0.559046198 | 3.06E-02 |
| TCEAL3 | 1081.559372 | 0.557210114 | 7.78E-05 |
| UBE2D2 | 2275.060115 | 0.557134523 | 6.46E-10 |
| DLL3 | 453.7860412 | 0.556723869 | 2.29E-03 |
| TMEM39A | 1461.701079 | 0.556059575 | 8.35E-07 |
| RIC8A | 8523.894523 | 0.55548648 | 4.08E-06 |
| GOLGA8N | 204.8886715 | 0.555463323 | 1.53E-03 |
| LHX6 | 463.3941 | 0.555255156 | 9.93E-04 |
| MAGOHB | 603.1906548 | 0.555017388 | 2.37E-07 |
| PFKFB2 | 1199.134629 | 0.554626782 | 2.17E-05 |
| KATNB1 | 1890.166344 | 0.554538627 | 1.91E-06 |
| SNX33 | 3457.403843 | 0.554419046 | 5.97E-05 |
| MRPS15 | 2177.285808 | 0.553978191 | 3.82E-11 |
| IL15RA | 273.3091975 | 0.553569311 | 7.15E-03 |
| IQGAP1 | 16665.12339 | 0.553503957 | 1.43E-07 |
| CENPN | 2368.501306 | 0.553452105 | 3.08E-04 |
| TLCD2 | 1283.167735 | 0.55335764 | 7.18E-05 |
| MIF4GD | 956.5208067 | 0.552998496 | 1.33E-05 |
| ACTR1B | 4339.501904 | 0.55286264 | 3.14E-06 |
| WDR25 | 582.4176668 | 0.552318329 | 2.05E-03 |

| | | | |
|---|---|---|---|
| SEC31A | 8459.893365 | 0.551037706 | 6.05E-10 |
| ALKBH3 | 1188.819224 | 0.550567801 | 4.02E-08 |
| STK40 | 4424.176598 | 0.550295094 | 1.08E-13 |
| TBL3 | 2640.040612 | 0.549794717 | 2.62E-05 |
| SSH3 | 1977.929526 | 0.549617913 | 1.22E-04 |
| CALCOCO1 | 3982.64577 | 0.54961422 | 6.78E-05 |
| DAGLB | 1643.438416 | 0.549570069 | 1.16E-09 |
| ANO6 | 6166.89251 | 0.549329643 | 4.36E-08 |
| MAP3K10 | 677.0189669 | 0.548488272 | 1.71E-03 |
| ABLIM1 | 3580.187643 | 0.548218182 | 1.07E-05 |
| PLEKHG3 | 2080.800905 | 0.548116208 | 3.55E-07 |
| SERPINB6 | 5673.984218 | 0.548062436 | 5.64E-08 |
| TDRKH | 1007.401832 | 0.547707271 | 2.70E-08 |
| MAPKAPK2 | 3962.376668 | 0.547607 | 2.73E-10 |
| FERMT3 | 650.0854354 | 0.547091734 | 5.16E-05 |
| LONRF1 | 546.4227996 | 0.546713901 | 7.17E-04 |
| PLEKHH1 | 203.0087702 | 0.546496194 | 4.81E-02 |
| MYZAP | 230.2645852 | 0.545945604 | 5.89E-03 |
| MRPL4 | 2091.730207 | 0.54573313 | 5.55E-08 |
| EIF3C | 53785.43486 | 0.545234938 | 6.86E-06 |
| DCTN2 | 6303.198494 | 0.544925608 | 1.04E-07 |
| LRRC8D | 3345.512279 | 0.544626099 | 5.99E-06 |
| TMEM79 | 312.6854783 | 0.544275326 | 4.21E-03 |
| ERGIC1 | 7955.280628 | 0.544197584 | 7.38E-08 |
| DHRSX | 467.6353937 | 0.544156123 | 9.48E-04 |
| CHCHD7 | 1011.119218 | 0.544035477 | 1.15E-05 |
| MACROD1 | 1467.887466 | 0.543902591 | 5.14E-05 |
| NA | 123.1612894 | 0.54381748 | 4.64E-02 |
| RGS10 | 602.2226751 | 0.543174966 | 2.09E-05 |
| ARHGDIB | 757.048678 | 0.543128744 | 1.94E-02 |
| PRXL2C | 968.9850378 | 0.543110733 | 3.55E-05 |
| SLC17A5 | 973.1245904 | 0.542953893 | 1.08E-03 |
| NPBWR1 | 390.8449915 | 0.542926513 | 1.24E-02 |
| BTBD3 | 2563.138022 | 0.542907734 | 6.27E-12 |
| LRRC17 | 2568.192514 | 0.542454693 | 4.69E-03 |
| LACTB2 | 789.8368156 | 0.542305219 | 2.23E-05 |
| PROB1 | 216.9344506 | 0.542256988 | 1.26E-02 |
| DNASE2 | 2894.136866 | 0.542043201 | 6.17E-04 |
| TJP1 | 3258.84199 | 0.541709301 | 1.12E-06 |
| ERO1A | 3104.479503 | 0.541550713 | 8.24E-04 |
| GNG12 | 8298.18734 | 0.541443182 | 2.71E-07 |
| PSMD8 | 6091.348911 | 0.541361083 | 1.65E-08 |
| TGIF1 | 1706.310532 | 0.541337217 | 6.85E-03 |
| FAM216A | 1300.115299 | 0.541300312 | 1.80E-05 |
| STIP1 | 14572.5874 | 0.541293316 | 9.62E-12 |
| ARSJ | 838.5835605 | 0.540980579 | 6.01E-04 |
| SLC25A3 | 15226.79426 | 0.540806758 | 3.18E-06 |

| | | | |
|---|---|---|---|
| CDK14 | 3306.003124 | 0.540644783 | 2.14E-07 |
| ABT1 | 1107.478359 | 0.540272216 | 1.78E-06 |
| HAUS4 | 2423.93088 | 0.540249051 | 6.29E-06 |
| YWHAZ | 33323.3696 | 0.539900308 | 1.44E-09 |
| DUS2 | 833.3492684 | 0.539582267 | 6.10E-07 |
| USP25 | 1363.579468 | 0.539496835 | 3.57E-03 |
| 8-Sep | 9809.378699 | 0.539273899 | 1.38E-12 |
| PPP2CB | 3807.900051 | 0.539270921 | 2.67E-25 |
| MAGOH | 973.9237622 | 0.539065041 | 2.76E-08 |
| PDLIM4 | 2965.529172 | 0.538477427 | 2.23E-06 |
| AP002387.2 | 105.8432616 | 0.53798479 | 1.74E-02 |
| ALDOA | 42268.47046 | 0.537814681 | 6.63E-05 |
| SDHAF4 | 156.402571 | 0.537619969 | 2.44E-03 |
| CDC45 | 1432.239354 | 0.537224171 | 1.56E-02 |
| RPL12 | 40443.63764 | 0.53717309 | 4.86E-04 |
| TMBIM1 | 4490.409115 | 0.536865011 | 3.59E-06 |
| VOPP1 | 7094.186809 | 0.536667888 | 7.18E-05 |
| OXSR1 | 3175.687546 | 0.536486623 | 1.23E-10 |
| SAE1 | 11296.4598 | 0.535864296 | 1.77E-11 |
| AC027307.2 | 280.7966597 | 0.53577109 | 3.13E-04 |
| SPRYD7 | 559.7320342 | 0.535763553 | 3.16E-05 |
| MCRIP2 | 724.7230538 | 0.535503313 | 3.40E-05 |
| ATF6 | 3656.141178 | 0.53484612 | 1.44E-09 |
| THOC5 | 3377.448188 | 0.534600286 | 1.60E-08 |
| HUNK | 113.2340518 | 0.534446994 | 3.57E-02 |
| SAP30BP | 2342.287681 | 0.534439931 | 1.03E-09 |
| TROAP | 1436.648234 | 0.534436698 | 8.86E-04 |
| EXOSC7 | 1415.908928 | 0.534401631 | 1.82E-06 |
| NOTCH2NLA | 1200.993235 | 0.534389032 | 6.95E-08 |
| CWF19L1 | 851.5140741 | 0.53340992 | 8.60E-06 |
| GID4 | 1136.933439 | 0.53334883 | 3.10E-07 |
| NAMPT | 5630.167846 | 0.533232318 | 2.95E-02 |
| KCTD5 | 3210.37524 | 0.532583676 | 3.85E-08 |
| FXR2 | 2852.945115 | 0.532243576 | 2.26E-11 |
| BYSL | 1631.402346 | 0.532029623 | 2.09E-05 |
| RPS6KA4 | 2634.070272 | 0.531630343 | 7.01E-08 |
| CPT1C | 384.7895272 | 0.531382029 | 4.04E-03 |
| BUD23 | 4063.891293 | 0.531341903 | 3.45E-18 |
| TEAD3 | 1178.905986 | 0.531221632 | 2.24E-06 |
| ZDHHC9 | 2390.142154 | 0.531002113 | 1.29E-09 |
| ABCA3 | 301.3116936 | 0.530996179 | 4.92E-02 |
| UBE2J1 | 3574.235135 | 0.530961735 | 1.45E-11 |
| RNH1 | 8558.943629 | 0.530588528 | 1.61E-08 |
| SLC8A3 | 826.2647581 | 0.530510984 | 4.49E-03 |
| WDR4 | 1435.970238 | 0.530505058 | 6.65E-06 |
| KIF20A | 9603.437791 | 0.530237685 | 1.36E-07 |
| NDUFC2 | 3541.592762 | 0.53001974 | 9.42E-13 |

| | | | |
|---|---|---|---|
| ZNF853 | 558.4164668 | 0.529840436 | 2.47E-03 |
| STK24 | 5540.39702 | 0.52971784 | 1.53E-09 |
| PLAGL1 | 554.3723733 | 0.529715099 | 1.66E-03 |
| MAP6D1 | 244.6323378 | 0.529255356 | 1.08E-03 |
| XPC | 3166.78906 | 0.528629836 | 6.49E-14 |
| PRMT1 | 7046.38091 | 0.52822602 | 4.79E-07 |
| TMEM9 | 4026.650806 | 0.527967967 | 5.72E-05 |
| KLF7 | 1229.558498 | 0.527933111 | 2.04E-04 |
| PLEC | 19438.93034 | 0.527868266 | 5.68E-07 |
| MICA | 1140.36943 | 0.52785816 | 7.11E-03 |
| ASPSCR1 | 867.8872405 | 0.52779844 | 8.95E-07 |
| RNF187 | 6542.358528 | 0.527560068 | 1.42E-06 |
| GHITM | 5118.84873 | 0.52745698 | 1.33E-17 |
| 2-Mar | 481.9723773 | 0.527194513 | 5.68E-03 |
| PA2G4 | 8850.309135 | 0.526882587 | 8.76E-15 |
| THAP4 | 1823.13702 | 0.526712498 | 8.95E-06 |
| SETD3 | 2342.695916 | 0.526673541 | 1.07E-10 |
| PSMA1 | 6765.463374 | 0.526654745 | 6.73E-14 |
| HAUS2 | 847.5349321 | 0.526633386 | 5.27E-04 |
| TBC1D1 | 3064.045464 | 0.526408192 | 3.53E-07 |
| MRTFA | 1872.178254 | 0.526124579 | 1.30E-05 |
| GAPDH | 113084.8488 | 0.526103859 | 4.51E-04 |
| RABEPK | 1331.448188 | 0.526017951 | 3.06E-05 |
| IKBIP | 1150.002844 | 0.525514478 | 3.27E-03 |
| DDR2 | 12220.85219 | 0.525380447 | 4.51E-03 |
| RTN3 | 7686.045019 | 0.525360703 | 7.95E-09 |
| FAM221A | 241.7934268 | 0.525205904 | 5.41E-03 |
| PRSS23 | 21726.22584 | 0.524920452 | 8.21E-05 |
| CIDECP1 | 295.2909922 | 0.52464167 | 7.55E-03 |
| AC021054.1 | 291.9708505 | 0.524567178 | 1.30E-03 |
| UCK2 | 2195.040756 | 0.523482926 | 4.71E-10 |
| LAMA3 | 690.6140048 | 0.523036163 | 7.15E-03 |
| CGAS | 435.9810156 | 0.522982463 | 1.03E-03 |
| LIPC | 496.2256483 | 0.52290644 | 1.25E-03 |
| BOD1 | 2630.333907 | 0.522815705 | 4.07E-06 |
| TRIM62 | 875.1026027 | 0.5224506 | 5.78E-05 |
| PATL1 | 4544.71706 | 0.522244782 | 1.17E-18 |
| HTATIP2 | 3669.802027 | 0.522213579 | 2.07E-10 |
| TRAPPC3 | 2005.937891 | 0.522103508 | 1.87E-09 |
| NLN | 3580.468692 | 0.521941188 | 1.93E-04 |
| NIPAL3 | 1794.482202 | 0.521831834 | 1.29E-09 |
| MRPS9 | 1646.296068 | 0.521012674 | 2.70E-09 |
| TPM2 | 11840.72667 | 0.520807512 | 7.25E-05 |
| ZFAND2A | 542.0624813 | 0.520705521 | 1.24E-02 |
| PSMD2 | 12961.65287 | 0.520647525 | 8.17E-07 |
| ACLY | 17193.67914 | 0.520322161 | 6.24E-06 |
| CNTNAP3 | 149.2453626 | 0.519572062 | 3.62E-02 |

| | | | |
|---|---|---|---|
| LINC02210 | 631.473022 | 0.519437052 | 2.05E-04 |
| LCMT1 | 1635.262159 | 0.519129675 | 1.60E-08 |
| ABHD4 | 4610.109295 | 0.519024066 | 8.36E-05 |
| ZFAND1 | 995.2253224 | 0.518960528 | 3.46E-02 |
| RAD23A | 3349.438841 | 0.518814788 | 1.01E-05 |
| ZG16B | 226.8742984 | 0.518549863 | 9.79E-03 |
| ZEB2 | 1231.237831 | 0.518401432 | 1.93E-02 |
| RUVBL2 | 6983.216087 | 0.51836871 | 1.12E-09 |
| WASF2 | 8001.080965 | 0.517877279 | 2.16E-09 |
| TSEN54 | 1095.294017 | 0.517862972 | 4.20E-05 |
| RAB6B | 890.8632009 | 0.517573895 | 9.98E-04 |
| C1orf226 | 396.125664 | 0.516768141 | 1.39E-03 |
| ACAP1 | 278.1116252 | 0.516570713 | 3.14E-02 |
| CALML4 | 214.1852994 | 0.516375483 | 8.18E-03 |
| DNAJC25 | 601.7296739 | 0.516375212 | 2.16E-04 |
| SNHG3 | 1840.165392 | 0.516194925 | 1.21E-02 |
| GSTO1 | 3403.432352 | 0.516155768 | 1.08E-05 |
| GRK6 | 1476.105821 | 0.515713412 | 3.46E-06 |
| LMNB2 | 11542.53977 | 0.515534353 | 2.23E-05 |
| TBCA | 6087.859397 | 0.515436784 | 5.82E-11 |
| AC105052.3 | 838.6690067 | 0.515277531 | 3.67E-05 |
| RFNG | 1336.381185 | 0.515217376 | 3.54E-04 |
| LRIF1 | 1383.86094 | 0.515214865 | 1.13E-04 |
| REEP4 | 1433.911046 | 0.515087956 | 2.25E-08 |
| ADCY9 | 1485.946218 | 0.515013944 | 7.53E-06 |
| CALHM5 | 354.1155895 | 0.514681564 | 1.32E-02 |
| TIMM50 | 2568.673392 | 0.514676615 | 5.36E-11 |
| AC112178.1 | 129.5750328 | 0.514214047 | 5.80E-03 |
| PFN2 | 10427.95587 | 0.514205275 | 9.97E-11 |
| LDLRAP1 | 1727.553505 | 0.514176202 | 4.23E-07 |
| NDEL1 | 2052.460807 | 0.51409601 | 3.25E-17 |
| DDX19B | 2093.58396 | 0.514031317 | 8.35E-03 |
| GLIS2 | 746.7602133 | 0.513781854 | 1.07E-04 |
| ACP6 | 1453.343694 | 0.513265867 | 1.68E-06 |
| MSX1 | 641.8880923 | 0.512860375 | 7.50E-04 |
| HIST1H1C | 590.5061429 | 0.512677504 | 2.47E-02 |
| MED13L | 4915.063541 | 0.51234321 | 1.03E-05 |
| SIRPA | 3488.218336 | 0.512098551 | 2.46E-04 |
| ELMSAN1 | 2538.049382 | 0.511697944 | 1.86E-08 |
| NRARP | 134.6099929 | 0.511554296 | 2.80E-02 |
| KDELR3 | 2158.686813 | 0.511387279 | 2.55E-03 |
| CMTM7 | 1057.832695 | 0.511017547 | 3.47E-05 |
| BANF1 | 7623.196999 | 0.510937412 | 4.28E-07 |
| MIEF2 | 859.0616354 | 0.510460521 | 5.18E-04 |
| BICC1 | 1248.921737 | 0.51033758 | 2.67E-05 |
| C12orf45 | 368.0857885 | 0.510331501 | 7.59E-04 |
| TMCC1 | 728.6019316 | 0.510303161 | 1.33E-05 |

| | | | |
|---|---|---|---|
| BIRC5 | 4253.014258 | 0.510033603 | 2.34E-05 |
| RPL9 | 21258.71788 | 0.509923172 | 7.12E-06 |
| DDX59 | 684.8136277 | 0.509582821 | 1.08E-04 |
| SORBS3 | 1260.08882 | 0.509069481 | 1.86E-03 |
| PICALM | 8631.628285 | 0.508904921 | 9.70E-06 |
| SLC41A3 | 2189.823322 | 0.508440898 | 3.13E-05 |
| PTGES3 | 10958.19736 | 0.508347493 | 2.46E-10 |
| VSIG10L | 276.6267659 | 0.508155144 | 2.63E-03 |
| ST3GAL3 | 868.2489115 | 0.507866547 | 6.67E-09 |
| CIAO2B | 2234.297862 | 0.507819798 | 3.36E-05 |
| PSMD11 | 5804.341134 | 0.507777408 | 1.32E-15 |
| HOXC11 | 245.5107089 | 0.507687185 | 1.87E-03 |
| AC006329.1 | 118.9663596 | 0.507484531 | 1.46E-02 |
| AZIN2 | 391.0001786 | 0.50744707 | 2.49E-02 |
| TTBK2 | 1066.932728 | 0.507418514 | 2.94E-02 |
| AC008403.1 | 202.196078 | 0.506537427 | 4.12E-03 |
| NSUN5 | 2016.221066 | 0.506083625 | 3.23E-08 |
| EDEM2 | 2837.570935 | 0.505923215 | 1.67E-04 |
| FEZ2 | 2245.374732 | 0.50579209 | 1.38E-08 |
| ARHGEF1 | 4120.570488 | 0.505633819 | 3.05E-06 |
| DLG1 | 3174.228063 | 0.505552659 | 1.73E-03 |
| PROSER1 | 1189.283984 | 0.505514521 | 7.23E-03 |
| HRH1 | 582.8781532 | 0.505394205 | 5.25E-03 |
| KNSTRN | 2096.989694 | 0.505259896 | 7.95E-06 |
| TAZ | 1646.042878 | 0.504963011 | 3.36E-06 |
| RASSF1 | 1191.274444 | 0.504942985 | 3.36E-04 |
| MYBL2 | 6940.146619 | 0.504893234 | 5.65E-03 |
| CDKN3 | 1240.158803 | 0.504795964 | 5.12E-03 |
| HMGB2 | 6919.321379 | 0.504649743 | 4.81E-03 |
| EIF2AK3-DT | 109.8054073 | 0.504051348 | 1.94E-02 |
| AC004477.1 | 149.757773 | 0.503752042 | 4.98E-02 |
| RFESD | 397.7901993 | 0.503611876 | 1.64E-02 |
| NOC2L | 7230.373689 | 0.503410935 | 4.96E-12 |
| MPST | 1990.715552 | 0.503399551 | 1.56E-05 |
| DUSP23 | 365.8499249 | 0.503256532 | 1.18E-02 |
| RCN1 | 15497.19177 | 0.503226823 | 5.31E-14 |
| LPCAT2 | 1115.439456 | 0.502987385 | 6.79E-05 |
| ZNF503-AS1 | 255.9762914 | 0.502565726 | 2.05E-02 |
| TMEM167B | 1344.383655 | 0.501769466 | 1.47E-05 |
| C2orf49 | 1087.85998 | 0.501426498 | 3.54E-07 |
| ANAPC10 | 365.3440448 | 0.501300416 | 3.59E-04 |
| FPGS | 3260.463423 | 0.501261235 | 2.48E-06 |
| MAP2K3 | 1746.737502 | 0.50123414 | 1.32E-02 |
| SORL1 | 4958.874962 | 0.501110345 | 2.54E-04 |
| CPQ | 874.2487683 | 0.501037744 | 1.06E-03 |
| ACBD7 | 133.8693676 | 0.500726795 | 2.38E-02 |
| PKP4 | 3485.234126 | 0.500713642 | 3.38E-05 |

| | | | |
|---|---|---|---|
| ATG7 | 1515.301373 | 0.500654942 | 9.21E-06 |
| NA | 295.3263647 | 0.500630206 | 1.65E-02 |
| ERCC1 | 2607.469316 | 0.500544307 | 1.06E-05 |
| LRRC3 | 598.7888325 | 0.500192201 | 1.94E-02 |
| PIEZO1 | 9000.487395 | 0.499870765 | 4.45E-06 |
| TARS | 20704.77187 | 0.49978625 | 2.33E-04 |
| NIPSNAP1 | 5969.896051 | 0.499443116 | 9.48E-05 |
| PSMD3 | 5874.943555 | 0.499286749 | 4.15E-09 |
| FAM72B | 387.5188204 | 0.499190899 | 2.34E-02 |
| ITSN1 | 4401.632807 | 0.499085337 | 1.38E-15 |
| FASN | 14089.9427 | 0.498897072 | 2.44E-03 |
| AP2B1 | 20918.25531 | 0.498523813 | 2.36E-09 |
| MYO18A | 7255.058026 | 0.498465825 | 1.54E-04 |
| KCTD10 | 3551.12457 | 0.498216661 | 4.23E-05 |
| EGFL7 | 1775.9885 | 0.497877271 | 2.81E-03 |
| PPIE | 2964.366664 | 0.49772979 | 1.39E-06 |
| MIER2 | 921.5115839 | 0.497582231 | 1.06E-04 |
| MBNL2 | 5478.880166 | 0.497481869 | 3.49E-02 |
| EIF4H | 25471.60151 | 0.497292647 | 7.08E-14 |
| GNB1 | 13180.59273 | 0.497168864 | 1.15E-14 |
| FAF1 | 2836.781069 | 0.496971179 | 1.89E-09 |
| GART | 10888.60204 | 0.496923862 | 9.17E-07 |
| IAH1 | 2107.417709 | 0.496843898 | 1.44E-13 |
| MRPS18B | 4057.860062 | 0.496835189 | 1.74E-07 |
| CDCA7 | 2462.20736 | 0.496540485 | 4.60E-03 |
| ERP29 | 5731.028148 | 0.496356401 | 1.74E-07 |
| SERPINH1 | 45395.88893 | 0.496255281 | 1.63E-08 |
| IFT27 | 1262.90455 | 0.495610567 | 1.62E-04 |
| PABPC1 | 53608.13181 | 0.495211903 | 2.35E-07 |
| IRAK1 | 19076.01415 | 0.494831996 | 4.63E-06 |
| CCNA2 | 2526.037041 | 0.494441322 | 5.29E-03 |
| TPD52L2 | 13159.04135 | 0.494375071 | 1.02E-05 |
| PUSL1 | 454.6324044 | 0.494108442 | 6.70E-06 |
| PSMD7 | 7737.490292 | 0.493948123 | 1.77E-20 |
| INPP5K | 1873.425128 | 0.493557961 | 5.77E-05 |
| KMT2E-AS1 | 139.9163431 | 0.493192493 | 5.94E-03 |
| KIAA0930 | 2835.093006 | 0.492801024 | 1.84E-05 |
| AK4 | 1671.133532 | 0.492729033 | 1.70E-02 |
| TNIP1 | 3697.169382 | 0.492370039 | 7.30E-04 |
| AK2 | 8587.731938 | 0.49224141 | 1.51E-07 |
| FAM200B | 455.2856727 | 0.492021145 | 5.50E-05 |
| TPM4 | 21265.75795 | 0.491893225 | 1.97E-07 |
| CIT | 2750.995169 | 0.491451803 | 1.03E-03 |
| TKT | 23860.21316 | 0.491058148 | 8.07E-06 |
| ASRGL1 | 682.0832095 | 0.490760037 | 3.90E-03 |
| H1FX | 4099.276197 | 0.490403822 | 5.47E-07 |
| CT45A10 | 1249.05887 | 0.490285347 | 7.16E-04 |

| | | | |
|---|---|---|---|
| NSMCE3 | 513.061359 | 0.490039835 | 7.81E-06 |
| GAS2L1 | 793.1223109 | 0.489201741 | 2.02E-03 |
| KLHL17 | 698.8288926 | 0.4889923 | 1.82E-04 |
| CCT4 | 15099.11166 | 0.488145272 | 2.02E-16 |
| MICALL1 | 3271.141042 | 0.487982668 | 7.52E-08 |
| GTSF1 | 1663.890902 | 0.487643741 | 3.56E-10 |
| FBL | 5586.954134 | 0.487624806 | 3.05E-06 |
| RPL9P9 | 16918.13946 | 0.487570475 | 4.32E-06 |
| COG5 | 3073.024123 | 0.487429904 | 1.24E-05 |
| SNX25 | 862.6549565 | 0.487166002 | 1.14E-05 |
| SGPL1 | 1970.346108 | 0.486825126 | 2.55E-03 |
| CNRIP1 | 1462.563083 | 0.486478264 | 2.33E-05 |
| PLRG1 | 3630.724933 | 0.486281782 | 1.90E-10 |
| PIK3CB | 1228.520412 | 0.486055343 | 4.98E-06 |
| PGAM1 | 12691.24699 | 0.486035758 | 5.27E-03 |
| EBP | 1987.716815 | 0.485972268 | 5.95E-03 |
| ILVBL | 2135.803995 | 0.48579646 | 1.32E-06 |
| NXT1 | 1469.60535 | 0.485664842 | 4.89E-07 |
| MED30 | 472.5295775 | 0.485628576 | 5.20E-06 |
| EPG5 | 3554.316738 | 0.485306537 | 6.04E-09 |
| EBF4 | 818.8381144 | 0.48511702 | 7.18E-03 |
| ERBB2 | 4218.931161 | 0.484514263 | 3.08E-05 |
| NQO2 | 3009.345582 | 0.484222571 | 7.68E-07 |
| CARS2 | 1179.357325 | 0.484020636 | 2.06E-05 |
| ZNF106 | 3656.174963 | 0.48389058 | 2.55E-08 |
| CHCHD3 | 3891.440959 | 0.483872186 | 5.32E-06 |
| AK3 | 2865.631536 | 0.483501433 | 4.24E-13 |
| BRD4 | 3830.430528 | 0.483416238 | 9.49E-04 |
| RHOA | 15149.00168 | 0.483112393 | 9.19E-06 |
| MAP2K1 | 3084.114411 | 0.48305634 | 5.69E-09 |
| LINC01006 | 968.1904904 | 0.48294442 | 5.75E-05 |
| RNF26 | 5061.4502 | 0.482922826 | 8.16E-05 |
| PRICKLE3 | 449.079314 | 0.482777567 | 1.23E-04 |
| AMZ2 | 2889.472315 | 0.482712548 | 4.83E-18 |
| GOLT1B | 1313.786465 | 0.482610675 | 3.21E-04 |
| RABAC1 | 2206.457957 | 0.482323264 | 1.77E-03 |
| AC097374.1 | 1567.411536 | 0.482237262 | 1.24E-03 |
| TNNT1 | 3015.335765 | 0.481535917 | 2.68E-04 |
| MCU | 1122.807796 | 0.481346198 | 2.75E-08 |
| FAM229B | 637.9454571 | 0.48106584 | 3.20E-05 |
| RTL8A | 3282.957021 | 0.480894965 | 2.18E-05 |
| SNHG1 | 3764.755585 | 0.480287395 | 3.65E-03 |
| AK5 | 674.3521065 | 0.480232905 | 1.45E-02 |
| SCD | 39207.06667 | 0.480025118 | 1.11E-03 |
| KCTD13 | 506.4129761 | 0.478889052 | 5.99E-03 |
| ZNF702P | 841.7746449 | 0.478775657 | 1.89E-02 |
| SYNGR2 | 1829.502754 | 0.478715544 | 1.65E-04 |

| | | | |
|---|---|---|---|
| RPL15 | 46122.89874 | 0.478603852 | 1.99E-04 |
| TMEM43 | 7214.222135 | 0.478393825 | 2.99E-11 |
| PUDP | 1113.873636 | 0.478374028 | 5.00E-05 |
| BNIP1 | 476.0777862 | 0.478146088 | 1.77E-05 |
| FAM84B | 2458.415805 | 0.477639483 | 1.70E-02 |
| SLC44A2 | 6112.518187 | 0.477477796 | 5.29E-07 |
| DNAJC19 | 835.3569696 | 0.477344842 | 3.26E-05 |
| CRYBG1 | 722.0676876 | 0.477315162 | 1.30E-02 |
| GTF3A | 2319.291999 | 0.477304114 | 7.75E-12 |
| SOCS7 | 1710.926291 | 0.476925311 | 8.99E-09 |
| RIN1 | 1365.463973 | 0.476558657 | 2.36E-03 |
| AC008443.1 | 461.6091636 | 0.476087478 | 7.86E-04 |
| CYP2R1 | 801.0685079 | 0.475975329 | 1.29E-02 |
| AHCY | 11764.42684 | 0.475807699 | 4.41E-06 |
| PLEKHG4 | 1389.84851 | 0.475409747 | 4.33E-04 |
| PTPN2 | 1323.929657 | 0.475274259 | 9.80E-07 |
| MFHAS1 | 1322.185779 | 0.47525793 | 2.64E-06 |
| RNPEP | 3309.936809 | 0.475234358 | 7.55E-04 |
| WDR41 | 4995.132278 | 0.47488477 | 1.08E-09 |
| UBC | 59339.49899 | 0.474249903 | 1.75E-05 |
| NFKB1 | 2198.220305 | 0.474203884 | 1.94E-08 |
| ASGR1 | 173.8182471 | 0.474052387 | 1.96E-02 |
| MCRIP1 | 1829.068626 | 0.474019147 | 1.70E-03 |
| DPP9 | 3238.309079 | 0.473794231 | 3.58E-07 |
| ATP9A | 5110.679593 | 0.473766156 | 3.94E-08 |
| PPP3CC | 566.011236 | 0.4735964 | 7.71E-03 |
| NKX2-4 | 192.9784856 | 0.473329328 | 1.26E-02 |
| GRAMD1A | 1599.035645 | 0.473188353 | 1.45E-05 |
| GMPPA | 1577.592379 | 0.473094033 | 1.44E-05 |
| PFKL | 6261.863339 | 0.47299796 | 2.96E-04 |
| SLC25A26 | 1028.574356 | 0.472631716 | 6.70E-05 |
| AEBP1 | 10182.06945 | 0.472579534 | 1.34E-04 |
| SHC1 | 17857.68695 | 0.471747848 | 3.67E-05 |
| SYNGR1 | 3473.006517 | 0.471731904 | 6.22E-04 |
| ZNF280A | 323.1856629 | 0.471476543 | 9.80E-03 |
| S100A4 | 620.5299465 | 0.471269618 | 5.81E-04 |
| HGS | 3477.552588 | 0.471256114 | 1.89E-11 |
| POLR3D | 3632.780335 | 0.470692243 | 1.45E-06 |
| EXOC2 | 2251.914411 | 0.470447024 | 6.67E-09 |
| LARS | 10091.49174 | 0.470417323 | 1.66E-03 |
| LINC00963 | 2907.811582 | 0.470267326 | 9.23E-05 |
| TPPP | 427.9957226 | 0.47021422 | 3.24E-02 |
| TNFRSF10B | 5148.799909 | 0.470199708 | 1.95E-04 |
| SHKBP1 | 2696.583798 | 0.470088644 | 1.63E-05 |
| EEF1AKMT2 | 304.8747012 | 0.470014676 | 1.50E-03 |
| ETNK2 | 2028.091551 | 0.469915302 | 4.08E-05 |
| C7orf50 | 3262.29235 | 0.469799619 | 1.97E-05 |

| | | | |
|---|---|---|---|
| TMEM241 | 503.1300017 | 0.469792285 | 2.39E-03 |
| ANP32B | 12976.31036 | 0.469751128 | 1.30E-06 |
| NLE1 | 1547.763898 | 0.469633353 | 9.30E-05 |
| YOD1 | 331.6001383 | 0.469575813 | 1.37E-02 |
| LIG3 | 2382.176734 | 0.46932026 | 2.90E-06 |
| PSRC1 | 2696.465216 | 0.469155301 | 6.69E-07 |
| ZHX2 | 545.5590592 | 0.469003532 | 4.76E-04 |
| ABTB2 | 1119.031511 | 0.468986712 | 2.69E-08 |
| KSR1 | 1136.231107 | 0.468528309 | 7.16E-03 |
| FAHD2A | 1741.478709 | 0.46843984 | 9.05E-08 |
| TOM1L1 | 731.0211202 | 0.468186995 | 3.17E-03 |
| HOXA11-AS | 322.0818879 | 0.468059825 | 2.10E-03 |
| EIF3M | 9035.068701 | 0.467644202 | 1.23E-14 |
| FBXO22 | 2360.772059 | 0.467630002 | 5.57E-08 |
| ESX1 | 1145.550705 | 0.467394453 | 1.96E-02 |
| MFSD13A | 221.5244945 | 0.467282877 | 1.68E-02 |
| EIF3F | 10611.45856 | 0.467076993 | 4.52E-04 |
| HMGCL | 1754.137431 | 0.46701974 | 4.51E-04 |
| FOXK2 | 3237.994456 | 0.466987112 | 3.21E-04 |
| PCBP2 | 21473.6083 | 0.466981079 | 2.87E-12 |
| C12orf66 | 315.7310445 | 0.46669662 | 4.11E-03 |
| IER5L | 924.1752821 | 0.46661864 | 1.78E-02 |
| AC131212.3 | 157.8229129 | 0.46623833 | 5.70E-03 |
| ACBD6 | 1499.904214 | 0.465818471 | 1.43E-05 |
| RIC1 | 2976.271407 | 0.465589922 | 2.72E-15 |
| VRK2 | 750.8131496 | 0.465541277 | 1.75E-02 |
| POLB | 1462.502926 | 0.465297887 | 2.71E-06 |
| MCMBP | 2516.238275 | 0.464949244 | 7.03E-05 |
| TXNDC11 | 2341.326044 | 0.464756579 | 1.62E-06 |
| TXN | 6436.015601 | 0.46422449 | 1.93E-09 |
| SLC45A4 | 507.6338217 | 0.464188356 | 8.36E-04 |
| PIP4K2C | 2211.016611 | 0.464001523 | 2.89E-06 |
| ZNF365 | 213.2583861 | 0.463513881 | 1.83E-02 |
| GALC | 2588.669822 | 0.463267949 | 4.09E-07 |
| CXorf49 | 337.5002354 | 0.462724767 | 7.42E-03 |
| AKT3 | 2703.948896 | 0.462685679 | 1.73E-04 |
| RBMS3 | 446.5159869 | 0.462597524 | 9.83E-03 |
| RASA3 | 1805.923465 | 0.462363101 | 1.21E-03 |
| MYEF2 | 342.730579 | 0.462220273 | 4.98E-02 |
| PACSIN2 | 4740.17459 | 0.462200211 | 4.34E-07 |
| IDI1 | 1984.814912 | 0.461835245 | 4.39E-02 |
| MORF4L2 | 11614.8889 | 0.46182364 | 1.01E-17 |
| PDXDC1 | 6002.09432 | 0.461815732 | 1.34E-07 |
| SLC37A2 | 644.4016214 | 0.461800844 | 1.01E-02 |
| PPP1R35 | 1131.012291 | 0.461791481 | 2.26E-05 |
| FAM89A | 912.360877 | 0.461349001 | 9.70E-06 |
| KLF9 | 1741.781689 | 0.461335251 | 8.07E-05 |

| | | | |
|---|---|---|---|
| ZNRF1 | 621.6109898 | 0.460715989 | 4.77E-03 |
| SEC24D | 3034.081143 | 0.460480585 | 1.84E-04 |
| PSMC3 | 7429.183721 | 0.460426211 | 5.69E-09 |
| SGMS2 | 606.7809837 | 0.459907818 | 3.68E-02 |
| AARSD1 | 1040.493228 | 0.45981098 | 7.39E-11 |
| ELOVL1 | 2900.615077 | 0.458776245 | 4.01E-04 |
| ZNRF3 | 595.9671143 | 0.458449623 | 4.38E-05 |
| XAB2 | 2830.758882 | 0.457738947 | 1.44E-03 |
| NUDC | 5364.378525 | 0.457651025 | 1.85E-04 |
| NSDHL | 2495.579896 | 0.457336632 | 4.45E-03 |
| ZNF703 | 2016.919739 | 0.457197591 | 1.14E-04 |
| GOLGA2P5 | 171.0467903 | 0.457032811 | 1.03E-02 |
| DRAM2 | 825.8159279 | 0.456903315 | 1.09E-03 |
| SLC25A4 | 1052.454079 | 0.456521641 | 1.02E-03 |
| POLG | 4710.124758 | 0.456429888 | 8.34E-05 |
| C20orf194 | 2675.497658 | 0.456379364 | 7.20E-08 |
| B4GALT1 | 11172.21506 | 0.456320118 | 7.56E-10 |
| HS3ST3B1 | 480.0958608 | 0.456207615 | 2.21E-02 |
| CALCOCO2 | 4846.363505 | 0.456149089 | 1.06E-09 |
| ZNF419 | 808.3470474 | 0.455725725 | 1.56E-03 |
| AC019080.1 | 129.0291347 | 0.455382044 | 2.33E-02 |
| RBM42 | 2701.957043 | 0.455099756 | 4.80E-06 |
| TOM1L2 | 4289.413806 | 0.454992127 | 4.91E-04 |
| ANXA7 | 2952.787176 | 0.454394491 | 3.07E-09 |
| GMPR2 | 4514.04944 | 0.454359303 | 5.91E-08 |
| COX7A2L | 3464.715754 | 0.454233212 | 1.68E-05 |
| METTL26 | 1644.085805 | 0.453891272 | 1.07E-04 |
| PTPN1 | 5738.112123 | 0.453820411 | 4.29E-10 |
| CLPTM1L | 15633.00909 | 0.453293676 | 1.74E-07 |
| TOR4A | 755.6938358 | 0.45307252 | 1.32E-02 |
| ACSS2 | 5287.328464 | 0.452993938 | 3.40E-03 |
| MSL3P1 | 461.1279657 | 0.452456911 | 1.93E-02 |
| POLR3E | 2515.286688 | 0.452361602 | 7.80E-04 |
| SLC25A28 | 399.421377 | 0.452089051 | 8.09E-04 |
| TWF2 | 2812.418961 | 0.452048829 | 1.27E-05 |
| PCSK6 | 1048.029103 | 0.451303499 | 7.45E-07 |
| ZFPL1 | 1177.567137 | 0.451230122 | 2.47E-04 |
| STAU2 | 1901.995673 | 0.451000644 | 4.08E-05 |
| PFDN1 | 2381.99521 | 0.450329274 | 1.77E-06 |
| MRPS18C | 1122.775971 | 0.449601109 | 1.67E-06 |
| PPP1R12B | 1810.477078 | 0.449403283 | 3.32E-07 |
| CHCHD4 | 918.971077 | 0.449161471 | 9.48E-04 |
| EPS15L1 | 1774.911684 | 0.448849169 | 2.62E-08 |
| ATL3 | 3470.507866 | 0.448785717 | 3.10E-10 |
| GTF2E2 | 2020.494839 | 0.448776912 | 6.08E-09 |
| PRRC2B | 23100.42542 | 0.448536548 | 1.71E-10 |
| YRDC | 837.8103711 | 0.448510668 | 5.31E-06 |

| | | | |
|---|---|---|---|
| RAB4B | 443.5158268 | 0.448369203 | 7.73E-04 |
| PREX1 | 4692.440271 | 0.448252634 | 3.20E-06 |
| WDR74 | 2352.357433 | 0.448112077 | 1.93E-04 |
| NT5DC3 | 443.4467148 | 0.447950628 | 3.64E-02 |
| CRYBB2P1 | 487.1810476 | 0.447849598 | 2.01E-03 |
| SAP30 | 507.7406263 | 0.447673142 | 1.01E-05 |
| RPS17 | 42235.26551 | 0.447497511 | 1.19E-06 |
| C16orf58 | 6656.991018 | 0.447451751 | 3.06E-06 |
| CMC1 | 560.2089086 | 0.447447526 | 7.32E-05 |
| HHLA3 | 421.4929855 | 0.44732904 | 8.11E-03 |
| LDHA | 37610.75825 | 0.447321446 | 2.76E-02 |
| OGFOD2 | 630.4163552 | 0.447266705 | 7.68E-04 |
| TTLL4 | 2183.833793 | 0.447171473 | 4.05E-04 |
| GLRX3 | 1925.851693 | 0.446808652 | 8.47E-08 |
| HMGB3 | 3653.261657 | 0.446660869 | 1.77E-04 |
| NCAPD2 | 11336.63226 | 0.446618356 | 1.43E-07 |
| CLK3 | 3618.729538 | 0.446544156 | 1.11E-05 |
| COL4A2 | 5735.041245 | 0.44639437 | 1.21E-04 |
| DGKQ | 1201.710352 | 0.446392594 | 7.91E-03 |
| ZNF330 | 1429.442326 | 0.446362905 | 1.22E-04 |
| APOL6 | 2761.451997 | 0.446309405 | 6.04E-03 |
| ICAM1 | 10019.94945 | 0.445839127 | 8.76E-03 |
| ZDHHC7 | 4188.484347 | 0.445552604 | 2.48E-03 |
| FLOT1 | 6068.185566 | 0.445435204 | 5.98E-04 |
| SEC13 | 4951.057522 | 0.444979722 | 2.33E-02 |
| TNXB | 1293.362643 | 0.444529805 | 2.33E-04 |
| IGF2BP1 | 4568.368856 | 0.444338346 | 2.89E-10 |
| RFT1 | 1944.490929 | 0.444005381 | 4.77E-10 |
| ARHGAP29 | 4982.779967 | 0.443426328 | 4.01E-02 |
| GRK3 | 836.2779387 | 0.443397906 | 5.40E-03 |
| UBE2M | 2402.768163 | 0.44291114 | 1.09E-06 |
| APBA2 | 1434.218644 | 0.442818925 | 7.71E-04 |
| TMEM147 | 1948.56684 | 0.442537748 | 2.32E-05 |
| RAC2 | 3122.912202 | 0.442097044 | 9.43E-04 |
| ELL2 | 3965.592371 | 0.442058443 | 1.70E-03 |
| BTF3L4P2 | 254.3305076 | 0.441857096 | 4.18E-02 |
| TMEM42 | 1109.272036 | 0.441685109 | 3.22E-03 |
| TBRG4 | 6211.250692 | 0.441675035 | 1.06E-05 |
| HAUS7 | 1980.774412 | 0.441668703 | 1.00E-04 |
| TTLL11 | 281.7170078 | 0.441617236 | 3.68E-03 |
| MFGE8 | 7060.251313 | 0.441595404 | 4.43E-04 |
| BTF3L4 | 1946.302754 | 0.441509977 | 4.56E-03 |
| BAG3 | 1430.928166 | 0.441304854 | 2.86E-04 |
| B4GALT4 | 1171.392361 | 0.441226296 | 4.28E-04 |
| MRPS5 | 2726.626492 | 0.441033053 | 1.07E-15 |
| 11-Sep | 10565.38008 | 0.4406449 | 4.41E-03 |
| AC083798.2 | 648.5242041 | 0.440569425 | 3.38E-02 |

| | | | |
|---|---|---|---|
| ATG101 | 1922.096598 | 0.440434825 | 2.43E-04 |
| CTSZ | 14220.45977 | 0.440257123 | 5.52E-05 |
| GIHCG | 338.5023988 | 0.440122453 | 9.12E-04 |
| NA | 220.007574 | 0.439898862 | 2.26E-02 |
| LRCH4 | 2232.681622 | 0.439865886 | 6.41E-04 |
| EMC6 | 677.2712484 | 0.439408072 | 6.66E-04 |
| DCUN1D2 | 458.6557648 | 0.439166016 | 1.01E-02 |
| ESRRA | 1871.490756 | 0.438748287 | 1.66E-05 |
| RSL24D1 | 4258.03113 | 0.438534811 | 1.94E-03 |
| VKORC1L1 | 3787.780139 | 0.438521467 | 1.50E-04 |
| EXOSC1 | 656.5367159 | 0.438500191 | 1.40E-05 |
| PMM2 | 2270.392662 | 0.438479929 | 5.44E-05 |
| SNHG16 | 2900.523027 | 0.4379018 | 9.48E-09 |
| FHL3 | 1027.877847 | 0.437867662 | 5.83E-03 |
| LSM6 | 664.6222914 | 0.437865389 | 1.19E-05 |
| ADIPOR1 | 4348.452011 | 0.437739856 | 1.13E-04 |
| AC068473.5 | 236.2554259 | 0.437629143 | 6.85E-04 |
| KCTD17 | 1231.905598 | 0.437456357 | 6.40E-05 |
| RNASET2 | 534.0947676 | 0.437012463 | 4.48E-03 |
| SQSTM1 | 11897.91029 | 0.437011398 | 2.04E-03 |
| SNAP23 | 1810.39237 | 0.436959234 | 3.78E-06 |
| FLYWCH1 | 2934.327451 | 0.436943926 | 4.93E-06 |
| ELL | 737.5986994 | 0.436879096 | 5.24E-06 |
| TMEM131L | 1289.718128 | 0.436865534 | 5.85E-04 |
| RPUSD3 | 1681.748031 | 0.436783666 | 5.07E-06 |
| PLXNA1 | 4699.089689 | 0.436772047 | 6.49E-05 |
| MAP2K2 | 4092.995995 | 0.436743812 | 1.91E-05 |
| TTF2 | 2656.857746 | 0.436339072 | 8.50E-04 |
| PARVB | 2787.345432 | 0.436196379 | 2.54E-04 |
| B3GLCT | 630.8665745 | 0.436187294 | 6.04E-03 |
| MMS19 | 2806.241532 | 0.435960373 | 2.41E-05 |
| ARIH2 | 4845.810584 | 0.435807386 | 4.13E-05 |
| BEX4 | 216.329814 | 0.435523517 | 2.28E-02 |
| LAMC3 | 1143.755153 | 0.435493608 | 6.88E-03 |
| GORASP2 | 3964.523769 | 0.435448859 | 2.21E-09 |
| MECR | 1202.982923 | 0.435252868 | 1.39E-06 |
| ARHGAP35 | 10670.21885 | 0.434877454 | 2.70E-05 |
| PIGBOS1 | 339.9707877 | 0.43487013 | 1.67E-02 |
| DUSP15 | 247.1476797 | 0.434195224 | 1.89E-02 |
| COX7A2 | 5358.559036 | 0.43401839 | 4.24E-03 |
| SPATC1L | 320.1765012 | 0.433753148 | 5.43E-03 |
| SPA17 | 608.872348 | 0.433305985 | 1.44E-03 |
| CDC25C | 1731.072231 | 0.433255079 | 3.80E-05 |
| SUCLG1 | 3552.834348 | 0.433214455 | 6.80E-08 |
| APOLD1 | 451.9599406 | 0.433022705 | 3.75E-03 |
| PIGU | 1597.231878 | 0.433017477 | 8.85E-06 |
| PLK1 | 5435.480583 | 0.432807481 | 2.76E-03 |

| | | | |
|---|---|---|---|
| RPS3A | 65602.08282 | 0.432713948 | 3.89E-04 |
| TBL1X | 3544.334191 | 0.432707022 | 1.40E-04 |
| CENPW | 732.2832315 | 0.432519653 | 6.27E-03 |
| THOC3 | 3199.426235 | 0.432210549 | 2.28E-03 |
| PLP2 | 4682.094735 | 0.432121773 | 1.60E-04 |
| ZFAND3 | 4293.104697 | 0.432070225 | 5.01E-03 |
| IMMP2L | 342.1216284 | 0.43204261 | 2.34E-02 |
| MRPS23 | 2563.61453 | 0.432014944 | 1.40E-07 |
| DYRK4 | 644.2354557 | 0.431491923 | 7.71E-06 |
| COL18A1 | 7768.279912 | 0.431458875 | 7.23E-04 |
| NPLOC4 | 6856.1453 | 0.431287919 | 1.76E-04 |
| CYCS | 4213.819034 | 0.431246984 | 1.93E-04 |
| NLRP1 | 4175.84274 | 0.431242228 | 3.31E-05 |
| PHB | 7327.068528 | 0.431073637 | 5.72E-05 |
| GMPS | 4155.036423 | 0.430726941 | 1.37E-07 |
| TOMM34 | 5597.477897 | 0.430237682 | 3.58E-06 |
| C6orf106 | 6662.810659 | 0.429911898 | 3.78E-05 |
| CYP51A1 | 5038.630174 | 0.429796127 | 5.81E-04 |
| CARM1 | 4180.03102 | 0.429714058 | 1.66E-06 |
| CDCA3 | 1454.557961 | 0.429267395 | 5.12E-03 |
| HYOU1 | 23967.33716 | 0.429122845 | 4.64E-05 |
| RIOK3 | 2038.172139 | 0.42904334 | 3.69E-03 |
| FOXP4 | 1453.076162 | 0.428949562 | 4.24E-04 |
| AC011481.1 | 2788.474976 | 0.428581107 | 5.85E-05 |
| DNPEP | 2813.70088 | 0.428262244 | 3.45E-05 |
| RPL8 | 63923.6203 | 0.428248523 | 2.32E-04 |
| POLD2 | 6626.569676 | 0.428064504 | 1.05E-07 |
| UBE2H | 13961.67353 | 0.427879274 | 1.22E-13 |
| VAPA | 4644.133299 | 0.427638685 | 1.75E-07 |
| RP9 | 827.2961611 | 0.427504234 | 1.05E-03 |
| DESI2 | 2337.936691 | 0.427459377 | 9.34E-03 |
| XPO6 | 8367.253338 | 0.427344552 | 6.15E-08 |
| COX19 | 685.3613752 | 0.42711957 | 3.55E-02 |
| PCDHGB5 | 192.1740464 | 0.426743429 | 1.03E-02 |
| ADAM9 | 9482.944296 | 0.426628452 | 1.72E-02 |
| ZSWIM6 | 1861.149076 | 0.426135224 | 2.30E-06 |
| MROH1 | 1333.300415 | 0.42554813 | 6.72E-05 |
| ARNTL2 | 885.1172776 | 0.425524705 | 4.07E-03 |
| OSER1-DT | 334.8931331 | 0.425466188 | 1.17E-02 |
| LAMP2 | 7955.899773 | 0.425387271 | 2.44E-07 |
| RPS10 | 23450.91986 | 0.425100802 | 5.51E-04 |
| CCNK | 2190.032846 | 0.425033456 | 2.62E-07 |
| EIF6 | 9045.030297 | 0.424842497 | 1.79E-04 |
| GIPC1 | 4074.727252 | 0.424670255 | 9.95E-06 |
| PLPBP | 2627.182497 | 0.424540002 | 3.20E-04 |
| MFN2 | 7446.363905 | 0.424476755 | 3.82E-07 |
| CDK5RAP1 | 2109.566458 | 0.423540228 | 6.30E-07 |

| | | | |
|---|---|---|---|
| MRPL33 | 1615.604336 | 0.423470827 | 4.84E-03 |
| STX5 | 2654.151479 | 0.423356893 | 2.73E-03 |
| NA | 264.1330256 | 0.42332517 | 4.08E-02 |
| FOXM1 | 5920.981055 | 0.422964718 | 2.68E-04 |
| PXN | 10227.78138 | 0.422918339 | 3.12E-04 |
| IPO5 | 38071.98562 | 0.422883061 | 7.50E-13 |
| TBC1D16 | 6642.917329 | 0.422765953 | 1.49E-05 |
| CLYBL | 500.0273141 | 0.422516529 | 4.05E-03 |
| PEF1 | 2353.697775 | 0.422263952 | 2.36E-03 |
| LRP8 | 1177.83127 | 0.422176851 | 8.90E-03 |
| L3HYPDH | 792.4822493 | 0.421955816 | 9.78E-05 |
| MPV17L2 | 689.7563318 | 0.421584048 | 5.95E-04 |
| SAP30L | 1605.846888 | 0.421526361 | 7.88E-04 |
| CFL2 | 3651.721601 | 0.421394843 | 2.42E-09 |
| DAP3 | 4392.491215 | 0.420958868 | 1.34E-12 |
| ITPKC | 871.852918 | 0.41947967 | 8.71E-05 |
| CHCHD2 | 15154.40285 | 0.419424394 | 1.32E-04 |
| NA | 1744.479115 | 0.419357684 | 1.43E-04 |
| DARS2 | 3464.006142 | 0.419084815 | 6.81E-08 |
| ZNF674-AS1 | 251.204287 | 0.418947595 | 1.44E-03 |
| COX10 | 905.009525 | 0.418938782 | 1.65E-04 |
| STK26 | 523.7241541 | 0.418771168 | 3.23E-02 |
| RNF220 | 2750.734764 | 0.418759365 | 2.28E-04 |
| ANKRD37 | 182.4951309 | 0.418600729 | 9.02E-03 |
| MRPL9 | 1883.802852 | 0.418445251 | 6.38E-03 |
| AASS | 1629.736665 | 0.418365267 | 1.50E-03 |
| USP39 | 3397.710093 | 0.418220895 | 1.82E-05 |
| SLC30A3 | 187.831053 | 0.418205456 | 4.31E-02 |
| PGM3 | 2990.489355 | 0.418100221 | 4.42E-03 |
| TMEM170B | 265.4732646 | 0.417755437 | 3.08E-02 |
| CARD19 | 976.3250092 | 0.417569666 | 9.70E-04 |
| HEIH | 828.3928894 | 0.417473305 | 1.16E-02 |
| P4HTM | 672.6986684 | 0.417322548 | 8.65E-03 |
| JTB | 2513.992003 | 0.417009065 | 5.35E-05 |
| 10-Sep | 3710.919446 | 0.416913184 | 3.89E-03 |
| SLC39A13 | 2800.97386 | 0.416830331 | 9.77E-03 |
| TMEM25 | 1351.60782 | 0.416789614 | 2.06E-02 |
| MRPL37 | 3539.378493 | 0.416489855 | 2.20E-05 |
| NMT1 | 6026.448678 | 0.416213629 | 4.29E-03 |
| RNMT | 3107.790568 | 0.416203605 | 8.90E-03 |
| PDE9A | 550.0523679 | 0.41613068 | 2.37E-03 |
| FAM234B | 967.1611581 | 0.415979202 | 8.85E-05 |
| RFC4 | 1463.728889 | 0.41578508 | 4.88E-03 |
| MSH5 | 896.9720349 | 0.415768915 | 1.67E-03 |
| PSMD9 | 1835.168441 | 0.41573477 | 5.06E-08 |
| TBL2 | 3247.426371 | 0.415588926 | 3.08E-06 |
| IKBKG | 2243.828522 | 0.415475375 | 4.79E-07 |

| | | | |
|---|---|---|---|
| XPNPEP1 | 2583.231566 | 0.415359524 | 1.99E-04 |
| SLC37A4 | 2303.912945 | 0.415309159 | 8.61E-04 |
| RAB3GAP1 | 4031.730799 | 0.41522252 | 3.23E-05 |
| RAI14 | 6837.00494 | 0.414674154 | 3.15E-02 |
| NUP37 | 1711.26298 | 0.414384175 | 2.92E-03 |
| PSMB3 | 4278.148867 | 0.414191185 | 2.05E-03 |
| ADRM1 | 5929.584761 | 0.414191097 | 1.49E-05 |
| NDUFA7 | 788.1011822 | 0.414140344 | 4.33E-02 |
| RPL4 | 113049.4838 | 0.413857759 | 4.84E-04 |
| SIMC1 | 762.4064736 | 0.413761734 | 8.35E-08 |
| EFEMP2 | 5133.916752 | 0.413658379 | 6.37E-03 |
| VDAC1 | 14777.17519 | 0.413626989 | 3.17E-04 |
| DSCR8 | 287.3163435 | 0.413590917 | 2.18E-02 |
| OSBPL10 | 1180.998379 | 0.413148702 | 1.75E-03 |
| TMEM245 | 5277.538846 | 0.413037809 | 4.25E-08 |
| SPAG5 | 4370.625311 | 0.413023457 | 9.20E-03 |
| PLOD2 | 2680.245608 | 0.412971035 | 6.83E-03 |
| GALE | 2706.092626 | 0.412648024 | 2.23E-04 |
| TRAF2 | 1387.830748 | 0.412629048 | 3.61E-03 |
| TMEM184B | 4398.019052 | 0.412590985 | 5.70E-04 |
| TTC9C | 1503.409275 | 0.412406649 | 1.35E-04 |
| RPLP0 | 75599.32004 | 0.412292452 | 5.73E-04 |
| GJC1 | 1901.807077 | 0.412286279 | 5.07E-04 |
| NUP93 | 5090.165098 | 0.412150592 | 4.54E-07 |
| MCUR1 | 1934.263528 | 0.411782512 | 9.32E-07 |
| 5-Sep | 1245.849617 | 0.411773488 | 2.50E-04 |
| NOD1 | 846.3666035 | 0.411740406 | 4.89E-05 |
| DLST | 6606.867678 | 0.411441686 | 3.70E-06 |
| CXorf40A | 833.72996 | 0.411286545 | 2.87E-04 |
| MDH2 | 11598.52765 | 0.411061281 | 1.52E-05 |
| ZNF480 | 608.6021465 | 0.410964799 | 1.28E-03 |
| EXOSC5 | 1246.924041 | 0.410879877 | 5.79E-03 |
| MRPL52 | 1805.217802 | 0.410841323 | 1.09E-06 |
| UQCC2 | 1854.703094 | 0.410812408 | 1.80E-04 |
| FITM2 | 2198.051981 | 0.410657767 | 6.95E-09 |
| ILF2 | 9055.317368 | 0.410451032 | 1.38E-08 |
| AC144652.1 | 365.0300178 | 0.410370419 | 8.46E-03 |
| NSFL1C | 5125.345501 | 0.410283165 | 2.99E-07 |
| AC141586.1 | 467.7885949 | 0.410155887 | 1.31E-02 |
| KLF10 | 1561.85262 | 0.410039997 | 1.16E-03 |
| MTX1 | 2583.38152 | 0.409962937 | 2.80E-03 |
| CDK2AP2 | 1734.372532 | 0.40955593 | 2.92E-04 |
| INSR | 813.980436 | 0.408602117 | 3.85E-02 |
| PRPS1 | 3124.775206 | 0.40849117 | 1.20E-06 |
| POLE4 | 2168.142337 | 0.40845876 | 6.86E-04 |
| CAPN15 | 2350.329042 | 0.408255629 | 2.06E-05 |
| BCAR1 | 5994.364749 | 0.407616191 | 3.93E-04 |

| | | | |
|---|---|---|---|
| RPS19 | 28332.18881 | 0.407544542 | 2.15E-04 |
| GAGE13 | 2562.719045 | 0.407383168 | 7.32E-06 |
| RNF5 | 1615.342707 | 0.407371253 | 1.45E-02 |
| CTSL | 7027.922032 | 0.407144694 | 2.49E-13 |
| ZNF775 | 895.107378 | 0.407126088 | 6.82E-04 |
| HIVEP1 | 1633.6484 | 0.407090776 | 4.90E-03 |
| WASH8P | 904.7545586 | 0.407009128 | 7.33E-03 |
| PQBP1 | 2394.096602 | 0.40664029 | 2.44E-04 |
| LAMA5 | 8544.457214 | 0.406491593 | 7.77E-03 |
| GSTP1 | 12693.12152 | 0.406210233 | 1.72E-03 |
| TSSC2 | 633.1595099 | 0.406120824 | 3.79E-03 |
| P4HA1 | 2000.926193 | 0.405813722 | 5.04E-04 |
| COQ2 | 766.2534857 | 0.405794626 | 9.92E-05 |
| DAP | 15213.24624 | 0.405783245 | 4.18E-04 |
| NUDT21 | 7108.295312 | 0.40561559 | 2.55E-05 |
| GRHPR | 4130.066864 | 0.405585323 | 1.08E-05 |
| LINC01137 | 148.1694956 | 0.405347777 | 1.89E-02 |
| KIAA1191 | 5326.813769 | 0.405248426 | 3.15E-05 |
| ATP6V1E2 | 503.4465659 | 0.405031485 | 3.30E-03 |
| NUTF2 | 4592.139175 | 0.405007194 | 7.72E-03 |
| AC004556.1 | 758.4552135 | 0.40495417 | 2.54E-02 |
| FAM222B | 821.712738 | 0.40489526 | 9.53E-06 |
| RASAL2 | 1538.28984 | 0.404835505 | 2.96E-04 |
| DENND5A | 8765.225991 | 0.404742687 | 2.25E-05 |
| GALK1 | 1388.050362 | 0.40455002 | 1.87E-02 |
| CELSR2 | 2661.06557 | 0.404140955 | 1.27E-03 |
| SF3A2 | 2191.908115 | 0.404129243 | 1.41E-03 |
| DNAH5 | 297.4360865 | 0.404079956 | 2.36E-02 |
| PPID | 1757.95365 | 0.40393555 | 6.42E-04 |
| HNRNPC | 25372.56747 | 0.4039027 | 1.70E-10 |
| TMEM106C | 4149.946741 | 0.403845244 | 4.79E-07 |
| TMEM109 | 5624.877036 | 0.403686525 | 1.53E-03 |
| CCT5 | 38883.96642 | 0.403680845 | 4.60E-08 |
| G6PD | 13437.0078 | 0.40338311 | 3.75E-04 |
| ALDH18A1 | 2818.954037 | 0.403206623 | 3.03E-03 |
| MAP3K2-DT | 170.5922295 | 0.40319925 | 3.02E-02 |
| SAMD4B | 3994.490781 | 0.403195286 | 3.80E-07 |
| RPS2 | 74595.74233 | 0.403143021 | 1.19E-03 |
| DNTTIP1 | 2150.378831 | 0.401882944 | 1.94E-03 |
| BOLA3 | 668.8711214 | 0.401817378 | 2.12E-04 |
| C9orf64 | 1570.97536 | 0.401812409 | 1.39E-06 |
| SLC39A11 | 639.9957675 | 0.401760915 | 1.59E-03 |
| CDC42BPB | 7684.040986 | 0.401748552 | 1.17E-16 |
| AP2M1 | 19896.15906 | 0.401710431 | 9.11E-04 |
| TSPAN17 | 2270.832624 | 0.401636575 | 4.61E-04 |
| POC1B | 724.6914982 | 0.401547829 | 3.51E-03 |
| CYP11A1 | 221.6021817 | 0.401474727 | 3.84E-02 |

| | | | |
|---|---|---|---|
| STK32B | 1361.46195 | 0.401437858 | 4.66E-03 |
| AKAP8L | 2144.844662 | 0.401374681 | 1.84E-06 |
| CCDC9 | 559.97443 | 0.401177565 | 3.28E-03 |
| ZNF773 | 544.1549742 | 0.400968417 | 1.00E-02 |
| CLPP | 1685.642619 | 0.400448746 | 3.31E-04 |
| PSMD13 | 8806.558774 | 0.40040887 | 2.03E-05 |
| SEPHS1 | 2733.449846 | 0.400185762 | 1.90E-09 |
| MLPH | 3090.876894 | 0.40004039 | 1.57E-02 |
| MPP6 | 815.7351252 | 0.399957865 | 6.02E-03 |
| TAF6 | 4881.790056 | 0.399917253 | 1.85E-06 |
| PRKRA | 1493.271741 | 0.399900563 | 9.66E-07 |
| GDAP2 | 761.0711289 | 0.399827681 | 9.90E-04 |
| SLC35F2 | 1705.247956 | 0.399793101 | 9.60E-04 |
| GAGE2A | 3777.159662 | 0.399780796 | 5.99E-05 |
| ANO2 | 401.3700442 | 0.399768513 | 3.42E-02 |
| MTLN | 469.6545991 | 0.399730674 | 6.88E-03 |
| FKBP4 | 6633.202591 | 0.399554796 | 2.88E-04 |
| PDPR | 5125.12976 | 0.399484211 | 1.26E-07 |
| RPS3 | 78001.32135 | 0.399279065 | 1.09E-04 |
| CAD | 8898.770146 | 0.399030426 | 1.10E-06 |
| FDXR | 399.2808891 | 0.398963645 | 1.12E-03 |
| FAM20C | 2024.588472 | 0.398933172 | 4.38E-02 |
| AC245060.4 | 370.64184 | 0.398622497 | 3.94E-04 |
| SEC22B | 4612.525216 | 0.398501777 | 5.03E-07 |
| CEP170B | 2094.005391 | 0.39847825 | 1.07E-06 |
| JMJD6 | 1235.975837 | 0.398372571 | 2.64E-02 |
| TXNL4A | 1733.43983 | 0.398363848 | 8.81E-06 |
| OSBPL6 | 864.281503 | 0.398251441 | 1.01E-02 |
| TPX2 | 13710.93008 | 0.39824804 | 8.53E-05 |
| PPFIA1 | 3610.881528 | 0.398243332 | 1.03E-04 |
| PIGCP1 | 268.8142606 | 0.398179583 | 1.45E-02 |
| KCNAB2 | 1627.146031 | 0.398112997 | 2.82E-02 |
| AIG1 | 902.4128692 | 0.398031943 | 3.70E-04 |
| KARS | 13794.60913 | 0.39802674 | 6.09E-07 |
| CDYL | 1827.428318 | 0.398023778 | 5.52E-04 |
| ZBTB47 | 508.2560449 | 0.398014124 | 4.51E-02 |
| LRRK1 | 2208.045532 | 0.397728791 | 1.01E-04 |
| PFKFB3 | 2049.457462 | 0.397517281 | 1.36E-03 |
| PHB2 | 8408.780695 | 0.397340494 | 1.43E-05 |
| QPCTL | 1074.333906 | 0.397183074 | 4.85E-04 |
| PAM | 16151.86896 | 0.39714218 | 2.05E-04 |
| DPH5 | 1379.944129 | 0.397025951 | 1.58E-03 |
| PRR14 | 1269.197719 | 0.39686645 | 1.35E-03 |
| CCDC86 | 3009.142193 | 0.396820998 | 9.91E-04 |
| DNAJC1 | 790.5283639 | 0.396667879 | 8.83E-04 |
| HDGF | 21317.82589 | 0.396592002 | 7.97E-05 |
| STXBP1 | 3551.05391 | 0.396381094 | 4.60E-04 |

| | | | |
|---|---|---|---|
| CTU2 | 1166.626706 | 0.396013968 | 1.92E-04 |
| INPP5A | 820.5608163 | 0.395938269 | 1.21E-02 |
| CCNY | 2484.643454 | 0.395681988 | 1.16E-07 |
| RNF113A | 757.8717493 | 0.395063907 | 1.03E-03 |
| CCT7 | 18359.8704 | 0.394951157 | 2.58E-07 |
| AC138028.4 | 241.3457325 | 0.394937999 | 3.53E-02 |
| COMMD7 | 2781.927981 | 0.394589533 | 4.33E-04 |
| DNAJB1 | 5494.200964 | 0.394292662 | 5.51E-03 |
| KDELR2 | 11593.04683 | 0.394250216 | 8.25E-04 |
| NME6 | 613.6357016 | 0.394225366 | 5.71E-04 |
| CCDC113 | 1000.208797 | 0.393962294 | 6.84E-03 |
| CDADC1 | 324.1905473 | 0.393955673 | 4.37E-03 |
| FAM83D | 3176.229694 | 0.393632488 | 7.17E-04 |
| EDEM1 | 2254.97478 | 0.393340681 | 4.65E-04 |
| DUSP22 | 1490.329574 | 0.39292398 | 8.58E-07 |
| WDR62 | 1308.836061 | 0.392865897 | 1.92E-02 |
| ODC1 | 5781.739414 | 0.392811902 | 1.98E-02 |
| MBOAT7 | 4718.094911 | 0.392741067 | 3.31E-03 |
| PWP1 | 3685.791824 | 0.392692475 | 1.03E-09 |
| SETD7 | 2957.166962 | 0.392485473 | 6.55E-06 |
| ARHGAP30 | 309.6936613 | 0.3923134 | 4.96E-02 |
| DIPK1B | 1341.473757 | 0.392294267 | 1.41E-02 |
| PSMB2 | 4850.469396 | 0.392247547 | 4.24E-04 |
| RPL27 | 24671.76414 | 0.391831849 | 1.06E-03 |
| OSBPL2 | 1833.379553 | 0.391745688 | 5.95E-04 |
| COTL1 | 13773.83675 | 0.391695774 | 4.45E-04 |
| NANP | 1026.86019 | 0.391360889 | 9.13E-03 |
| PTGFRN | 3794.796387 | 0.391283048 | 8.87E-03 |
| PDCD5 | 1993.549848 | 0.391157702 | 7.31E-07 |
| NNT-AS1 | 922.7219592 | 0.39086036 | 3.93E-02 |
| MRPL21 | 1743.40577 | 0.390755312 | 6.03E-05 |
| DNHD1 | 2181.427692 | 0.390734866 | 7.64E-04 |
| SNAPC2 | 1028.266326 | 0.390715103 | 8.64E-03 |
| PNP | 1512.768907 | 0.390434557 | 5.89E-04 |
| STYXL1 | 1931.959515 | 0.39033709 | 3.18E-03 |
| NSMCE2 | 1140.479055 | 0.390030903 | 4.57E-06 |
| RPS20 | 35647.71181 | 0.389963856 | 1.50E-03 |
| MAPKAPK5-AS1 | 536.4434689 | 0.389937218 | 1.25E-03 |
| TGFBI | 26371.45713 | 0.389657595 | 2.85E-03 |
| RNASEH2A | 2104.874415 | 0.389614352 | 1.08E-02 |
| FBXL12 | 1173.753934 | 0.389134691 | 1.94E-03 |
| DNAJC25-GNG1( | 425.9037544 | 0.389101628 | 5.59E-03 |
| ARHGDIA | 10479.86553 | 0.388957758 | 5.37E-03 |
| ZNF529-AS1 | 151.9124009 | 0.388672631 | 2.30E-02 |
| VANGL2 | 2032.007805 | 0.388495506 | 1.74E-04 |
| SSSCA1 | 1070.342479 | 0.388456596 | 1.80E-03 |
| MNAT1 | 1215.706917 | 0.38836128 | 2.08E-02 |

| | | | |
|---|---|---|---|
| LSM4 | 3280.823502 | 0.38820584 | 1.28E-05 |
| RPS6KB2 | 2424.493208 | 0.388114322 | 2.51E-04 |
| ABCF2 | 6738.907087 | 0.388086812 | 1.35E-07 |
| MAPK9 | 1850.100491 | 0.387963878 | 4.95E-04 |
| ANO10 | 1558.862414 | 0.387893796 | 5.94E-03 |
| CTDSP2 | 12475.24743 | 0.387778399 | 1.37E-07 |
| FAM27E3 | 808.5898466 | 0.387511579 | 1.91E-02 |
| GSPT1 | 9602.68302 | 0.387449023 | 3.38E-11 |
| PPP1R14B | 3533.962955 | 0.387351863 | 1.10E-03 |
| GALNT10 | 5382.628084 | 0.386925517 | 2.41E-04 |
| EEF1D | 14301.58075 | 0.386787617 | 2.00E-04 |
| CNIH4 | 1503.44764 | 0.386762843 | 4.10E-06 |
| TRPT1 | 1109.105546 | 0.386489402 | 2.05E-02 |
| KANK2 | 5584.704178 | 0.386286693 | 1.24E-02 |
| APTX | 2086.750278 | 0.386095777 | 3.78E-06 |
| INPP5B | 1336.420141 | 0.386049973 | 1.43E-02 |
| SLC25A5 | 11977.73219 | 0.385693452 | 2.95E-02 |
| DUSP12 | 1147.714159 | 0.385001203 | 1.42E-02 |
| ATP5F1B | 32311.30569 | 0.384886859 | 2.98E-05 |
| CBARP | 284.9060585 | 0.384885638 | 4.16E-02 |
| HEMK1 | 1344.047739 | 0.384743194 | 3.21E-04 |
| PRMT5 | 6007.645453 | 0.384624402 | 2.54E-05 |
| INPPL1 | 9644.907944 | 0.384566662 | 9.62E-05 |
| ATP6V1B2 | 7378.659165 | 0.384529519 | 3.09E-05 |
| MRTO4 | 2839.374245 | 0.384475248 | 1.33E-04 |
| POLRMT | 1932.426877 | 0.384423104 | 1.87E-03 |
| ZNF788P | 519.4212818 | 0.38441516 | 7.07E-03 |
| NAT9 | 1754.47402 | 0.384288124 | 4.06E-04 |
| DYNLRB1 | 5076.376126 | 0.383807466 | 6.58E-03 |
| MRPL55 | 606.6771076 | 0.383726481 | 7.52E-04 |
| TPI1 | 33065.67208 | 0.38346638 | 4.52E-02 |
| IFRD2 | 2613.653594 | 0.383027762 | 1.55E-04 |
| MTFR2 | 416.4778021 | 0.382911281 | 3.43E-02 |
| AC022211.2 | 275.9392876 | 0.382883134 | 1.08E-02 |
| NBPF10 | 2314.64115 | 0.382865366 | 3.90E-04 |
| SET | 25045.76652 | 0.382769296 | 4.12E-03 |
| XRCC3 | 1695.724094 | 0.382709742 | 4.29E-07 |
| RIN3 | 1523.104933 | 0.38259326 | 7.68E-04 |
| PFAS | 3969.518652 | 0.382471568 | 2.16E-05 |
| IQSEC1 | 2478.420337 | 0.382229125 | 6.27E-06 |
| EPRS | 10315.70588 | 0.382091763 | 3.69E-02 |
| TMEM18 | 1213.800047 | 0.381965486 | 4.90E-03 |
| ECSIT | 1263.236298 | 0.381794298 | 1.05E-03 |
| CDK16 | 7022.735633 | 0.381682137 | 4.91E-04 |
| APPL2 | 2709.579325 | 0.381371619 | 6.58E-07 |
| PRELID3B | 2493.846453 | 0.381340598 | 1.37E-02 |
| KLF6 | 4088.683155 | 0.381205354 | 1.65E-04 |

| | | | |
|---|---|---|---|
| ZNF319 | 661.4451797 | 0.381128639 | 1.78E-02 |
| EEF1B2 | 14457.85453 | 0.380748598 | 4.50E-04 |
| TRADD | 922.3822232 | 0.38059019 | 4.62E-03 |
| THEMIS2 | 327.7141705 | 0.380539171 | 3.69E-02 |
| LYPLAL1 | 519.1981249 | 0.379997624 | 1.13E-02 |
| RELT | 599.7605953 | 0.379910904 | 2.47E-03 |
| HLA-B | 7949.216215 | 0.37961885 | 1.05E-03 |
| STAG3L5P-PVRIG | 1727.688963 | 0.379618108 | 6.87E-03 |
| VAC14 | 3528.466868 | 0.379584972 | 8.28E-04 |
| ZNF281 | 1503.230845 | 0.379125817 | 6.70E-05 |
| COASY | 5076.517411 | 0.378959223 | 7.80E-04 |
| FAU | 10785.67298 | 0.378656846 | 3.49E-03 |
| BRD1 | 1179.651156 | 0.378584941 | 5.44E-05 |
| EIF2AK4 | 3997.221069 | 0.377552257 | 1.39E-10 |
| DOT1L | 1539.184202 | 0.377108949 | 1.53E-03 |
| RPL35 | 27266.11572 | 0.377069266 | 1.60E-03 |
| SCML2 | 373.2409383 | 0.376500258 | 3.83E-02 |
| YIPF6 | 1932.021176 | 0.376147759 | 5.28E-04 |
| BRSK1 | 458.4730848 | 0.376099734 | 2.47E-02 |
| RPL5 | 35286.27166 | 0.375889403 | 3.48E-05 |
| PDCD11 | 5588.612431 | 0.375766799 | 5.27E-03 |
| HTT | 5698.107226 | 0.37568871 | 2.47E-02 |
| RPS15A | 28078.05717 | 0.375628011 | 3.23E-04 |
| MCTS1 | 2241.850458 | 0.375389279 | 9.17E-07 |
| MRPS35 | 1793.927643 | 0.375164619 | 1.78E-04 |
| UBE2I | 4881.546543 | 0.375116204 | 8.79E-06 |
| SLC35B1 | 1668.984745 | 0.374754743 | 2.71E-06 |
| SLC25A25 | 1315.684058 | 0.374604317 | 9.33E-07 |
| JUND | 1908.596123 | 0.374578318 | 3.85E-02 |
| GALNT14 | 1213.5074 | 0.374286946 | 2.75E-03 |
| SZRD1 | 8977.152041 | 0.374257213 | 6.74E-04 |
| UBE2F | 1396.145029 | 0.373960869 | 1.40E-03 |
| BCKDHA | 1382.500501 | 0.373923286 | 4.42E-02 |
| HSP90AB1 | 76450.15073 | 0.373590082 | 5.21E-05 |
| SLC43A3 | 6224.28028 | 0.373063178 | 2.78E-06 |
| MAP3K1 | 6834.516553 | 0.372893754 | 1.45E-03 |
| UBTD1 | 536.0145794 | 0.372658865 | 1.80E-02 |
| BOLA2 | 1180.562019 | 0.372648634 | 4.01E-03 |
| ESPL1 | 2910.925144 | 0.372617523 | 1.54E-02 |
| ZXDC | 1312.034482 | 0.372575405 | 8.24E-05 |
| GNB5 | 2160.045094 | 0.372558433 | 8.77E-07 |
| ACP1 | 2949.308581 | 0.372209733 | 9.43E-06 |
| DDHD2 | 3026.27877 | 0.371894437 | 3.98E-05 |
| FAHD2B | 1473.812251 | 0.371705976 | 6.15E-06 |
| HOXC6 | 874.2215878 | 0.3715965 | 9.70E-06 |
| YARS2 | 1071.129074 | 0.37148805 | 2.44E-03 |
| SLC25A6 | 13376.01443 | 0.371261279 | 6.10E-03 |

| | | | |
|---|---|---|---|
| SAP18 | 2287.019192 | 0.371254123 | 2.37E-04 |
| NCAPH | 2363.792997 | 0.37112767 | 4.17E-02 |
| ZNF71 | 490.1173141 | 0.370951786 | 2.26E-02 |
| SELENOF | 4529.787408 | 0.37074525 | 1.39E-07 |
| VAMP2 | 1712.31311 | 0.37056003 | 6.83E-04 |
| C17orf75 | 860.7018602 | 0.370090424 | 9.48E-04 |
| PHACTR1 | 383.5787134 | 0.37006845 | 1.74E-02 |
| SMO | 2973.87738 | 0.369958558 | 8.80E-03 |
| FARSB | 2957.701085 | 0.369908423 | 2.44E-07 |
| MRPS30 | 2531.712345 | 0.369638498 | 3.44E-08 |
| FAT1 | 14154.47872 | 0.369605735 | 1.95E-03 |
| CCNB2 | 4442.079969 | 0.369473318 | 1.75E-04 |
| BLVRA | 2172.864212 | 0.369375456 | 1.26E-04 |
| ERI3 | 2151.816879 | 0.369332569 | 9.91E-04 |
| TAF7 | 3538.289254 | 0.369260338 | 3.72E-04 |
| NID2 | 2791.988157 | 0.368745997 | 1.47E-02 |
| TBC1D14 | 3382.105884 | 0.368652263 | 9.78E-06 |
| FZD5 | 364.7941564 | 0.368625597 | 1.23E-02 |
| ELMOD3 | 825.5341406 | 0.368245237 | 1.09E-02 |
| PUS1 | 1053.427984 | 0.367608974 | 8.11E-03 |
| SMAD3 | 7634.379311 | 0.367479333 | 5.13E-05 |
| C8orf33 | 4642.341906 | 0.367174818 | 9.38E-05 |
| NKIRAS2 | 2781.961389 | 0.366854444 | 1.35E-05 |
| DTYMK | 2052.910667 | 0.366588738 | 2.76E-04 |
| SH3RF1 | 1138.990913 | 0.366447448 | 1.17E-02 |
| ZSCAN32 | 573.7474721 | 0.36606502 | 2.53E-02 |
| SART1 | 4923.107164 | 0.365803352 | 4.26E-05 |
| DDX54 | 3050.744023 | 0.365770256 | 4.16E-06 |
| PPIA | 74455.26732 | 0.365669267 | 2.00E-02 |
| RAP1A | 1573.438963 | 0.36555383 | 4.19E-06 |
| RAB2A | 3018.845323 | 0.36486684 | 1.07E-09 |
| PSMC1 | 6088.680726 | 0.364172736 | 2.70E-06 |
| MIIP | 828.9580369 | 0.364120475 | 2.88E-03 |
| TFG | 4603.610223 | 0.36411311 | 4.61E-08 |
| GTF2H4 | 781.8735298 | 0.363809776 | 6.46E-03 |
| STX18 | 1205.829051 | 0.363767288 | 1.05E-03 |
| DHX35 | 1575.892245 | 0.363642637 | 3.07E-03 |
| EIF1B | 1124.244198 | 0.363350653 | 1.98E-03 |
| PDLIM7 | 2642.119675 | 0.363304481 | 5.32E-03 |
| APEH | 7239.845918 | 0.363291311 | 1.06E-02 |
| PERP | 2168.913583 | 0.363229737 | 1.34E-05 |
| MTO1 | 1439.627308 | 0.363173002 | 2.34E-02 |
| RPL3 | 72185.34694 | 0.363139565 | 5.26E-03 |
| PRELID1 | 4349.040238 | 0.362791126 | 2.39E-04 |
| TRMT61A | 1134.142873 | 0.362757135 | 2.58E-02 |
| CCDC28A | 816.7254079 | 0.362653001 | 8.97E-04 |
| GNAO1 | 1149.018389 | 0.362179989 | 3.00E-02 |

| | | | |
|---|---|---|---|
| ADSL | 5458.002757 | 0.362169827 | 2.28E-04 |
| SURF4 | 17447.89564 | 0.362008416 | 2.26E-03 |
| PLEKHM2 | 4411.871107 | 0.361847442 | 3.70E-05 |
| NA | 504.5175673 | 0.361761709 | 8.16E-03 |
| SMAD2 | 2978.677024 | 0.361324983 | 2.01E-05 |
| AL133467.2 | 401.6717852 | 0.361306604 | 5.19E-03 |
| REXO1 | 986.0877504 | 0.361051941 | 1.73E-05 |
| NOA1 | 1315.983886 | 0.360953085 | 3.58E-03 |
| RTCB | 5427.681484 | 0.360595145 | 3.01E-05 |
| TCOF1 | 9862.217299 | 0.360307371 | 1.59E-06 |
| CACNB1 | 433.310513 | 0.360157018 | 9.64E-03 |
| LAS1L | 2829.020455 | 0.36011197 | 4.92E-05 |
| USB1 | 2856.737564 | 0.359703704 | 2.69E-04 |
| NCOR2 | 5055.339838 | 0.359340854 | 4.11E-03 |
| URB1 | 5309.208866 | 0.359153967 | 1.88E-05 |
| ELOVL5 | 4586.484995 | 0.359064394 | 1.91E-07 |
| SOGA1 | 7215.020647 | 0.358908394 | 7.67E-04 |
| ATP8B1 | 1504.624557 | 0.358792752 | 1.23E-02 |
| CENPV | 1153.791311 | 0.358742605 | 1.32E-03 |
| PSMA7 | 11075.96575 | 0.358624643 | 7.61E-06 |
| CCL28 | 830.9079579 | 0.358551724 | 3.93E-03 |
| ARCN1 | 17474.68119 | 0.358274051 | 1.60E-06 |
| PXK | 1077.365537 | 0.358255949 | 3.83E-02 |
| EVI5L | 1512.548342 | 0.358211914 | 5.95E-04 |
| MAP2K5 | 1236.431012 | 0.358186293 | 1.43E-03 |
| ZNRD1 | 1174.003622 | 0.35818168 | 9.56E-03 |
| TP53INP2 | 2226.784429 | 0.358130679 | 2.90E-02 |
| MZT2B | 1646.035276 | 0.357945875 | 3.65E-02 |
| EIF2D | 2568.611508 | 0.357914188 | 1.07E-04 |
| SEC61B | 3153.007258 | 0.357844911 | 2.16E-05 |
| ZNF593 | 490.6340416 | 0.357828909 | 2.97E-02 |
| DTNBP1 | 753.5644288 | 0.357444071 | 6.35E-03 |
| EIF4G1 | 29771.85812 | 0.357407977 | 4.57E-04 |
| RPL38 | 12269.32601 | 0.357266159 | 1.32E-02 |
| CRYL1 | 352.6870594 | 0.356401119 | 4.87E-02 |
| LMTK2 | 2508.032902 | 0.356324595 | 4.03E-06 |
| MTCL1 | 1412.753168 | 0.356069084 | 1.12E-03 |
| PLPP2 | 1427.512506 | 0.35581848 | 2.90E-02 |
| USP31 | 1052.765356 | 0.355782473 | 1.72E-02 |
| CUTA | 2762.537115 | 0.355773704 | 3.31E-02 |
| KLHDC10 | 3849.172421 | 0.355763386 | 2.65E-05 |
| PNMA6A | 795.9455409 | 0.355376528 | 3.25E-02 |
| DDX10 | 2148.269443 | 0.35499162 | 2.97E-03 |
| DRG1 | 3037.512976 | 0.3548254 | 3.01E-04 |
| UQCRFS1 | 3568.156548 | 0.354361738 | 5.72E-04 |
| RNF123 | 2422.551839 | 0.353847334 | 6.82E-04 |
| PPFIBP1 | 7522.570197 | 0.353696113 | 1.98E-06 |

| | | | |
|---|---|---|---|
| BCAR3 | 3028.458632 | 0.353489729 | 7.72E-03 |
| AP3D1 | 8143.720888 | 0.35326352 | 5.56E-07 |
| MRPS18A | 1552.70699 | 0.353113913 | 8.91E-04 |
| NUTM2A-AS1 | 303.0746165 | 0.35300167 | 2.65E-02 |
| AC026401.3 | 298.6534259 | 0.352889178 | 3.06E-02 |
| MYO9B | 6638.196893 | 0.35278008 | 5.85E-03 |
| KIAA2013 | 2549.440012 | 0.352515402 | 6.05E-03 |
| BUD31 | 4153.64177 | 0.352136527 | 1.35E-03 |
| FOPNL | 1315.094864 | 0.352111471 | 3.65E-02 |
| FBXW8 | 1751.709567 | 0.351951071 | 2.04E-04 |
| GOLGA3 | 4858.345823 | 0.35178916 | 5.14E-04 |
| TSIX | 369.066555 | 0.351560776 | 2.43E-02 |
| PSMG4 | 858.7223993 | 0.351284817 | 1.08E-03 |
| NDUFS6 | 3119.638103 | 0.351260706 | 2.69E-03 |
| SH3GLB2 | 3523.835095 | 0.351175525 | 6.03E-05 |
| RPL32 | 31619.84787 | 0.350947449 | 2.49E-03 |
| MRPL38 | 1864.760936 | 0.350802122 | 5.66E-04 |
| XRCC6 | 20895.91892 | 0.350617112 | 4.92E-05 |
| PET117 | 794.4960644 | 0.350520044 | 7.75E-05 |
| SETBP1 | 900.8322632 | 0.350454551 | 2.72E-03 |
| ADGRA2 | 6548.879872 | 0.350126644 | 2.40E-04 |
| FAM220A | 1262.311827 | 0.349750192 | 1.74E-02 |
| CSNK2B | 5612.548763 | 0.349745804 | 5.62E-04 |
| PRRC2A | 20762.84586 | 0.349619444 | 2.98E-07 |
| BTBD11 | 910.1491859 | 0.34947759 | 1.33E-03 |
| GOLGA2P7 | 5165.611373 | 0.349337003 | 3.26E-05 |
| NAPA | 3117.244375 | 0.349302762 | 3.25E-03 |
| PPP3CA | 1295.712804 | 0.349256005 | 1.40E-02 |
| NBPF9 | 4785.546614 | 0.349241511 | 2.73E-06 |
| SNRPN | 5362.88291 | 0.3490204 | 5.03E-04 |
| ANAPC16 | 1880.704335 | 0.348919094 | 7.01E-05 |
| DENND4B | 2722.961269 | 0.348869192 | 1.44E-04 |
| SLC25A15 | 687.4465156 | 0.348850478 | 5.20E-04 |
| MICU1 | 2085.264622 | 0.348827358 | 2.01E-03 |
| YWHAG | 14945.9692 | 0.34856001 | 9.94E-09 |
| MARS2 | 475.8319689 | 0.348548144 | 5.06E-03 |
| FBXL5 | 1766.997965 | 0.348279463 | 2.84E-06 |
| TCF3 | 4011.452498 | 0.348219743 | 3.30E-04 |
| EIF5A | 21651.00118 | 0.348195355 | 7.91E-05 |
| MMGT1 | 2362.006228 | 0.347937218 | 1.44E-06 |
| ZNF433-AS1 | 402.7071823 | 0.347860247 | 3.37E-02 |
| STAG3 | 3593.012809 | 0.34761719 | 8.05E-05 |
| CPNE1 | 12635.24221 | 0.34758779 | 3.31E-03 |
| AFG1L | 293.8883154 | 0.347367992 | 4.56E-02 |
| BRF2 | 893.0764052 | 0.347054737 | 4.30E-02 |
| RPL23A | 15036.58917 | 0.346948474 | 1.54E-02 |
| CMTM4 | 3789.945746 | 0.346887844 | 6.00E-07 |

| | | | |
|---|---|---|---|
| SNRPC | 5495.320625 | 0.346884089 | 2.39E-04 |
| RAC1 | 16484.60893 | 0.346691957 | 3.67E-06 |
| TRAPPC1 | 4848.874062 | 0.346528965 | 2.54E-02 |
| PSMB7 | 8583.989746 | 0.346359457 | 5.85E-05 |
| OLA1 | 3800.835144 | 0.346180443 | 5.08E-08 |
| DUSP7 | 822.9598436 | 0.345445161 | 3.23E-03 |
| BSDC1 | 3231.619132 | 0.345380341 | 2.59E-03 |
| TMED3 | 5421.603628 | 0.345365723 | 1.62E-02 |
| ACO2 | 4130.560406 | 0.345343625 | 1.43E-02 |
| COX17 | 554.2075333 | 0.345289274 | 3.44E-02 |
| BTF3 | 37705.71318 | 0.344813874 | 4.57E-05 |
| CHCHD10 | 1240.010778 | 0.344765296 | 8.61E-03 |
| DIP2C | 1132.351831 | 0.344470157 | 1.76E-02 |
| PPT2 | 1542.529366 | 0.344446157 | 3.78E-02 |
| NOP53 | 6803.000535 | 0.344297525 | 3.16E-02 |
| DHX37 | 2003.689058 | 0.34398289 | 5.47E-04 |
| CBL | 5340.886292 | 0.343725234 | 3.02E-06 |
| PTBP1 | 14317.09673 | 0.343686763 | 9.48E-04 |
| RPS7 | 42243.97316 | 0.343161372 | 2.02E-05 |
| OPTN | 3436.814398 | 0.343130726 | 1.19E-03 |
| PLXND1 | 13090.28614 | 0.343019483 | 2.48E-02 |
| TRABD | 1154.065959 | 0.34298077 | 5.78E-05 |
| NAA80 | 540.0798404 | 0.342931629 | 4.71E-02 |
| MAF1 | 4739.284522 | 0.342833446 | 2.29E-03 |
| CTDSPL | 1864.785257 | 0.342797878 | 3.91E-03 |
| CEP104 | 1693.378755 | 0.342770163 | 5.70E-03 |
| UBE2R2 | 4405.95647 | 0.342614911 | 3.80E-05 |
| GALNT2 | 7296.725498 | 0.342587907 | 1.52E-02 |
| RPL21 | 8227.922657 | 0.34247285 | 1.20E-03 |
| NCS1 | 3791.736683 | 0.342412313 | 2.07E-04 |
| GPI | 9672.754168 | 0.342258875 | 4.61E-03 |
| PPTC7 | 1103.285373 | 0.342101431 | 7.87E-03 |
| RPS16 | 19557.72628 | 0.342074846 | 8.72E-03 |
| VARS | 6546.556051 | 0.341995207 | 4.95E-04 |
| PTDSS1 | 5001.392315 | 0.341839408 | 1.02E-03 |
| DDB2 | 1400.451522 | 0.341763007 | 7.03E-04 |
| NAP1L1 | 25109.18802 | 0.34154144 | 5.36E-08 |
| CRYZL1 | 678.2921722 | 0.341431196 | 1.29E-02 |
| CDC42EP2 | 1460.609892 | 0.341404901 | 6.91E-03 |
| DENND5B | 1132.684712 | 0.341265462 | 1.78E-03 |
| ZNF410 | 2800.845383 | 0.341225744 | 2.24E-03 |
| BOLA2-SMG1P6 | 948.6690809 | 0.341110602 | 3.03E-02 |
| GAGE1 | 7974.436024 | 0.341110242 | 1.43E-04 |
| DBF4B | 1125.920736 | 0.341045771 | 5.30E-03 |
| PKM | 127003.2279 | 0.340869648 | 1.70E-02 |
| AGFG1 | 2601.944699 | 0.340793797 | 1.01E-02 |
| GAGE12F | 4858.93278 | 0.340732211 | 4.00E-02 |

| | | | |
|---|---|---|---|
| TPM3 | 16136.67089 | 0.340614671 | 1.41E-03 |
| BCL2L2 | 2667.728938 | 0.340559046 | 9.36E-05 |
| RPL26L1 | 1232.856647 | 0.340326625 | 2.42E-04 |
| TXNDC5 | 23972.78375 | 0.340209662 | 2.03E-03 |
| SCYL1 | 4860.578339 | 0.340099609 | 1.37E-02 |
| IMPDH1 | 3474.006773 | 0.340057928 | 1.96E-04 |
| CDC37 | 9701.163091 | 0.339675083 | 3.87E-05 |
| MORN4 | 623.2349339 | 0.339658905 | 2.25E-03 |
| PEMT | 720.8887767 | 0.339362404 | 1.51E-02 |
| SIX1 | 639.7445453 | 0.339046307 | 3.42E-03 |
| SNX9 | 2199.738733 | 0.33904279 | 2.04E-06 |
| SYNJ2 | 2575.00764 | 0.338964492 | 9.15E-03 |
| C7orf31 | 557.6287 | 0.338708433 | 1.62E-02 |
| EXOC4 | 8628.260536 | 0.338672137 | 6.85E-06 |
| SOX13 | 1139.603012 | 0.338443221 | 3.03E-03 |
| KATNAL1 | 513.5647486 | 0.338351508 | 1.53E-02 |
| PRPF6 | 7435.468691 | 0.338343852 | 2.69E-02 |
| RAB11FIP3 | 2920.906787 | 0.338209913 | 5.65E-04 |
| RPL10 | 91368.98784 | 0.338058878 | 6.47E-03 |
| IFT52 | 2221.042975 | 0.338037198 | 3.72E-05 |
| ARRDC1 | 1211.294091 | 0.337687979 | 2.06E-02 |
| ACVR2B | 614.6329055 | 0.337670689 | 1.53E-02 |
| PAXX | 1435.946098 | 0.337668314 | 3.93E-04 |
| ASXL1 | 11653.72846 | 0.337633548 | 4.27E-07 |
| JPT1 | 2871.360134 | 0.337504755 | 2.54E-04 |
| PITRM1 | 3121.362636 | 0.337498446 | 5.86E-05 |
| FBXO42 | 1571.145269 | 0.337097952 | 4.41E-02 |
| SLC6A8 | 4800.871325 | 0.337004454 | 3.05E-02 |
| POLR2E | 5539.893555 | 0.336979361 | 2.70E-03 |
| KXD1 | 2873.498203 | 0.336900073 | 1.40E-03 |
| GLUD1 | 5125.474283 | 0.336801601 | 1.29E-06 |
| AC010618.1 | 1835.153133 | 0.336625099 | 1.24E-02 |
| MAP1LC3A | 1651.699059 | 0.336528016 | 2.26E-02 |
| FOXD1 | 1076.126703 | 0.336518724 | 5.96E-03 |
| KDSR | 2931.962615 | 0.336443514 | 2.07E-04 |
| POLR1E | 2520.906887 | 0.336346559 | 1.18E-04 |
| SRRD | 1362.097613 | 0.336340994 | 2.56E-03 |
| ATP1A1 | 28999.11334 | 0.336285159 | 7.64E-04 |
| LSM10 | 1191.931926 | 0.335879788 | 3.50E-03 |
| BET1 | 1110.402084 | 0.335792086 | 1.65E-02 |
| UBXN1 | 5831.619773 | 0.335692808 | 3.07E-03 |
| ATP5MC2 | 5676.214694 | 0.335672765 | 1.27E-03 |
| MED27 | 1626.357315 | 0.335672099 | 6.32E-03 |
| RPL18A | 25731.2536 | 0.335559549 | 1.74E-02 |
| HLA-E | 8048.516065 | 0.335523964 | 2.70E-02 |
| SPATA5L1 | 413.1567765 | 0.335203467 | 6.09E-03 |
| STT3A | 12540.12889 | 0.334916186 | 8.60E-05 |

| | | | |
|---|---|---|---|
| CCT2 | 12555.94788 | 0.334790693 | 1.73E-07 |
| RANGRF | 1588.216267 | 0.33471203 | 1.89E-03 |
| TIAL1 | 2251.871338 | 0.334614697 | 3.81E-05 |
| AC079949.2 | 366.5235527 | 0.334380918 | 3.48E-02 |
| PSMB1 | 6788.226364 | 0.334323236 | 2.49E-04 |
| CDC34 | 2227.878526 | 0.334105399 | 6.97E-04 |
| STK39 | 1510.725526 | 0.333836533 | 2.74E-03 |
| VMP1 | 6417.676705 | 0.333750315 | 2.72E-03 |
| TBRG1 | 2566.845992 | 0.333712625 | 1.39E-03 |
| AATF | 3385.796557 | 0.333679547 | 2.11E-04 |
| HACD1 | 437.3528397 | 0.333678018 | 1.91E-03 |
| RPL37A | 41742.41886 | 0.333677999 | 1.53E-02 |
| THAP3 | 667.2347235 | 0.333651964 | 3.84E-02 |
| PHF2 | 2669.663996 | 0.333642729 | 1.29E-07 |
| COQ5 | 1997.429849 | 0.333627501 | 9.16E-06 |
| ATP5F1C | 4379.356069 | 0.333369526 | 1.04E-03 |
| SURF1 | 3085.300649 | 0.333287522 | 4.52E-03 |
| DDA1 | 1367.310378 | 0.333264251 | 1.39E-02 |
| RPP21 | 594.9113622 | 0.333148719 | 1.52E-02 |
| NOP9 | 3695.923527 | 0.332673299 | 2.10E-04 |
| PEX10 | 1005.175471 | 0.332370752 | 3.64E-03 |
| 5-Mar | 1118.604787 | 0.332315429 | 2.85E-02 |
| RHOT1 | 1697.536288 | 0.332137036 | 3.57E-03 |
| PITPNM1 | 3508.712923 | 0.332109537 | 8.62E-03 |
| SUDS3 | 2194.800008 | 0.332099866 | 8.34E-03 |
| ZBTB18 | 811.3954707 | 0.331777957 | 5.23E-03 |
| IGF2BP3 | 3445.067623 | 0.331742981 | 1.83E-03 |
| TRAF3 | 2982.287153 | 0.331653583 | 4.24E-04 |
| PAQR7 | 1043.956206 | 0.331553663 | 8.97E-03 |
| TRIP4 | 1456.396546 | 0.331403774 | 4.03E-04 |
| GCDH | 1021.008364 | 0.331297276 | 7.07E-03 |
| IDH3A | 2872.949552 | 0.331210019 | 3.29E-02 |
| TDP1 | 1043.60661 | 0.331192245 | 1.41E-03 |
| DGUOK | 3302.395692 | 0.331142771 | 1.56E-04 |
| SH3BGRL3 | 7583.347441 | 0.331097946 | 2.24E-02 |
| ZNF385A | 632.1157056 | 0.331079766 | 2.02E-02 |
| JMJD7 | 296.4011285 | 0.330935974 | 2.80E-02 |
| LINC00910 | 264.0356664 | 0.330861678 | 2.93E-02 |
| CARHSP1 | 4813.82067 | 0.330824193 | 3.42E-02 |
| RBFA | 1166.399006 | 0.3308094 | 1.35E-06 |
| TRRAP | 9853.594491 | 0.330746569 | 6.99E-07 |
| CUL4A | 3078.197308 | 0.330661297 | 3.11E-03 |
| TSG101 | 4160.63018 | 0.330551802 | 4.49E-04 |
| TAMM41 | 651.7368094 | 0.329927585 | 1.11E-03 |
| SF3B2 | 19110.27107 | 0.329793333 | 1.18E-04 |
| MAPKAPK5 | 1373.001255 | 0.329678331 | 6.51E-05 |
| DDX41 | 2155.136552 | 0.329231651 | 9.91E-03 |

| | | | |
|---|---|---|---|
| UBN1 | 2592.183221 | 0.329220427 | 7.42E-04 |
| SUN1 | 8621.36509 | 0.328940711 | 8.13E-03 |
| CAP1 | 16233.00955 | 0.328609408 | 8.75E-04 |
| SIVA1 | 2030.777572 | 0.328583883 | 4.02E-03 |
| PDZD2 | 5704.21073 | 0.328400238 | 3.11E-02 |
| MRPL46 | 1238.821764 | 0.328111223 | 9.83E-05 |
| IFT20 | 1159.347537 | 0.328096676 | 1.39E-05 |
| HPS5 | 2635.985793 | 0.327776396 | 3.49E-05 |
| GEMIN7 | 848.4478189 | 0.327736778 | 2.14E-02 |
| SNRPA | 3244.309368 | 0.327612285 | 2.51E-02 |
| PSMD12 | 2523.757316 | 0.327548833 | 1.85E-02 |
| TMEM14C | 4616.94698 | 0.327268543 | 7.30E-04 |
| NDUFA9 | 2952.013751 | 0.327191225 | 1.99E-02 |
| RAN | 15303.38853 | 0.327161411 | 1.03E-03 |
| RPS2P5 | 8753.092583 | 0.327006485 | 1.06E-02 |
| ITCH | 4592.164107 | 0.326966382 | 1.22E-02 |
| MRPL11 | 2079.489287 | 0.326810815 | 4.61E-04 |
| HMG20B | 2798.243651 | 0.326597576 | 6.46E-03 |
| BPNT1 | 1216.423968 | 0.326502182 | 1.07E-02 |
| SERPINB9 | 1503.778653 | 0.326495302 | 1.91E-04 |
| ATP5PO | 6020.894828 | 0.326340836 | 3.51E-04 |
| PPP6R1 | 4169.616317 | 0.326221286 | 3.32E-04 |
| SELENON | 6949.973863 | 0.326158626 | 8.07E-03 |
| TTC27 | 1234.216433 | 0.326056991 | 9.74E-04 |
| PDIA5 | 1145.144501 | 0.325768685 | 3.01E-02 |
| HDDC3 | 769.9143343 | 0.325203226 | 1.10E-02 |
| HJURP | 3243.06767 | 0.325095599 | 1.94E-02 |
| RPL36 | 12121.41414 | 0.324953142 | 1.48E-02 |
| SRP68 | 5045.523619 | 0.324932678 | 2.63E-07 |
| RUBCN | 1148.934787 | 0.324727775 | 1.19E-03 |
| SLC7A6 | 3175.630792 | 0.324688219 | 5.82E-03 |
| COMMD2 | 1374.963537 | 0.324383079 | 1.92E-04 |
| ESS2 | 772.4988779 | 0.324126278 | 1.09E-02 |
| RAB5C | 6053.456357 | 0.32411392 | 2.32E-04 |
| NANS | 3649.193973 | 0.3239733 | 8.17E-06 |
| PEX14 | 1078.492463 | 0.323970794 | 9.31E-03 |
| AC068888.1 | 445.9172072 | 0.323886501 | 1.78E-02 |
| RPS13 | 31513.02511 | 0.323875675 | 9.92E-03 |
| BCL9L | 9610.645557 | 0.323326848 | 1.05E-02 |
| DIP2B | 4273.333063 | 0.323062432 | 6.61E-06 |
| FAH | 2365.929671 | 0.322859461 | 3.83E-04 |
| PTK2 | 6063.525587 | 0.322381986 | 1.06E-03 |
| CC2D1A | 2123.997049 | 0.322283671 | 7.31E-03 |
| MZT2A | 1203.994707 | 0.322008827 | 3.24E-02 |
| TICAM1 | 498.5809167 | 0.321704721 | 4.00E-02 |
| ATP5PD | 3820.956853 | 0.321689826 | 1.92E-03 |
| GOLGA2P10 | 667.6520873 | 0.321494622 | 1.16E-02 |

| | | | |
|---|---|---|---|
| RGS3 | 2410.104891 | 0.321411029 | 2.26E-02 |
| TPRA1 | 1560.994177 | 0.321409526 | 2.99E-03 |
| LYAR | 1412.981459 | 0.321317089 | 3.60E-02 |
| ME2 | 1256.073843 | 0.321183947 | 4.76E-02 |
| PARS2 | 429.7542574 | 0.321128196 | 2.14E-02 |
| DYNC1H1 | 30557.26086 | 0.320835836 | 8.86E-04 |
| TECPR1 | 2202.322973 | 0.320833924 | 4.71E-03 |
| ZBED4 | 1607.762094 | 0.320738066 | 4.22E-05 |
| PHF10 | 1866.438536 | 0.32073538 | 4.51E-04 |
| AKR1A1 | 2291.915055 | 0.320479633 | 3.82E-02 |
| DDX51 | 1618.888558 | 0.320052627 | 1.22E-02 |
| VPS33A | 1967.463228 | 0.319849651 | 6.16E-04 |
| TUFM | 15387.79824 | 0.319801864 | 1.55E-03 |
| KRTCAP2 | 1559.895768 | 0.319755559 | 5.59E-03 |
| LRRC28 | 807.0602025 | 0.319712797 | 3.80E-04 |
| STK32C | 480.8698462 | 0.319608911 | 1.67E-02 |
| CHMP1B | 1645.111841 | 0.3195133 | 7.09E-05 |
| RRNAD1 | 1100.690946 | 0.319466253 | 7.15E-03 |
| CHTF18 | 1995.240987 | 0.319202036 | 4.99E-02 |
| EIF3B | 14629.46064 | 0.319000147 | 2.33E-02 |
| ZCCHC9 | 2215.726008 | 0.31895326 | 6.66E-03 |
| EME1 | 841.7569482 | 0.31893685 | 2.96E-02 |
| GPALPP1 | 1200.665299 | 0.318476173 | 2.17E-03 |
| COX6A1 | 5248.247842 | 0.318475343 | 1.69E-03 |
| MLF1 | 1080.164123 | 0.31845569 | 6.24E-04 |
| TIPARP | 910.2975068 | 0.318183667 | 4.73E-02 |
| ZNF263 | 2605.294114 | 0.318052033 | 6.42E-04 |
| MCL1 | 7487.483715 | 0.318032287 | 4.60E-05 |
| CELSR1 | 3462.62622 | 0.317220198 | 2.99E-03 |
| RPAP1 | 1524.937256 | 0.316708516 | 1.20E-02 |
| NOM1 | 2171.555554 | 0.316562231 | 1.65E-03 |
| AMIGO2 | 914.0654676 | 0.316549142 | 4.96E-02 |
| DLAT | 2983.280395 | 0.316471957 | 1.51E-03 |
| GORASP1 | 2536.009094 | 0.316351733 | 8.44E-03 |
| CD33 | 476.0162093 | 0.316268774 | 3.97E-02 |
| MRPL22 | 1050.652787 | 0.316191043 | 1.34E-03 |
| RPL18 | 22761.13749 | 0.3158611 | 2.20E-02 |
| CSNK1E | 5056.524669 | 0.315612167 | 4.93E-06 |
| PIGL | 581.9689132 | 0.315395577 | 9.99E-03 |
| TRIM35 | 1131.368175 | 0.315077972 | 6.88E-03 |
| EP400 | 4945.291168 | 0.315037476 | 2.63E-07 |
| NDUFAF5 | 604.7519493 | 0.314792197 | 9.68E-03 |
| EIF4A1 | 39066.0312 | 0.314737184 | 8.46E-07 |
| ZSCAN18 | 2383.204961 | 0.314552708 | 1.41E-02 |
| GPR107 | 9096.320411 | 0.31446282 | 5.57E-05 |
| TMEM41B | 992.340041 | 0.314068585 | 3.80E-02 |
| SNX5 | 7442.90652 | 0.314058688 | 8.37E-04 |

| | | | |
|---|---|---|---|
| AKAP1 | 1854.077959 | 0.314013006 | 1.33E-02 |
| RPL13 | 48811.62292 | 0.313893525 | 7.63E-03 |
| DNMT1 | 10550.99223 | 0.313866618 | 3.37E-02 |
| SFXN4 | 585.4507196 | 0.313673646 | 4.53E-03 |
| TMEM14B | 2820.957355 | 0.313603339 | 3.78E-04 |
| UBE2Q1 | 3304.876586 | 0.313225852 | 2.62E-02 |
| CAB39 | 2508.723948 | 0.313097519 | 2.20E-02 |
| TMEM120B | 1090.727739 | 0.312940678 | 8.31E-03 |
| DHX32 | 1575.566691 | 0.312844261 | 4.99E-04 |
| FAM174A | 424.3305524 | 0.312842367 | 2.81E-02 |
| LINC01578 | 1662.10461 | 0.312827619 | 2.17E-02 |
| NCLN | 3912.468455 | 0.312505818 | 4.89E-03 |
| MPC2 | 1051.78884 | 0.312453926 | 1.49E-03 |
| ATP6V0E2-AS1 | 473.5699297 | 0.312326354 | 2.42E-02 |
| RNF216 | 6139.992269 | 0.312206186 | 4.19E-06 |
| TAF3 | 497.5951653 | 0.312061075 | 1.20E-02 |
| SLC25A37 | 6361.631733 | 0.311729596 | 1.94E-07 |
| GSK3B | 3261.575123 | 0.311467294 | 9.58E-04 |
| ZNF544 | 1742.095957 | 0.311060523 | 1.09E-02 |
| NME2 | 13404.07001 | 0.311045587 | 1.73E-03 |
| SSH1 | 4488.68356 | 0.310847598 | 5.09E-05 |
| ARL2 | 3980.279198 | 0.310812629 | 6.42E-03 |
| NRDC | 8374.393388 | 0.310786528 | 5.92E-04 |
| SGF29 | 499.749353 | 0.310784363 | 3.83E-02 |
| TUBGCP4 | 1347.698251 | 0.310609654 | 5.26E-03 |
| STRN4 | 4766.955253 | 0.310380161 | 1.33E-03 |
| GPR108 | 2585.88666 | 0.310219556 | 2.26E-02 |
| EXOSC2 | 2807.209963 | 0.310170513 | 3.86E-07 |
| SELENOS | 1716.629402 | 0.310153106 | 1.16E-04 |
| RNF114 | 4494.094987 | 0.309828929 | 2.66E-04 |
| NOP10 | 2112.578788 | 0.309820739 | 3.58E-03 |
| EEF1A1 | 626427.2418 | 0.309743028 | 6.68E-04 |
| LYRM4 | 1174.273544 | 0.309693327 | 1.64E-02 |
| PJA1 | 2541.974321 | 0.309689048 | 1.26E-04 |
| RRP7A | 1939.453049 | 0.309562425 | 1.35E-03 |
| GAGE12C | 12322.95692 | 0.309512973 | 2.24E-04 |
| ADD2 | 5103.394758 | 0.309460196 | 3.68E-04 |
| DNAJA4 | 883.5233635 | 0.309411507 | 1.06E-02 |
| INO80E | 2856.807769 | 0.30888166 | 1.37E-02 |
| SNAP29 | 1759.719219 | 0.308831725 | 1.04E-04 |
| CLCN6 | 2361.816732 | 0.308801313 | 1.08E-04 |
| CAMKK2 | 2052.006897 | 0.308800679 | 2.43E-04 |
| STAG3L5P | 572.7572891 | 0.308510399 | 1.55E-02 |
| CKAP5 | 15304.98664 | 0.308487423 | 2.77E-02 |
| AFDN | 4425.730501 | 0.308311053 | 3.13E-02 |
| CKB | 5181.807168 | 0.307482102 | 5.57E-03 |
| TUBGCP3 | 960.1919668 | 0.307351863 | 2.29E-02 |

| | | | |
|---|---|---|---|
| MFF | 2603.156878 | 0.307249905 | 4.46E-07 |
| MRPS12 | 1199.187636 | 0.307215567 | 1.02E-02 |
| TNFAIP8L1 | 574.1983135 | 0.307200334 | 2.53E-02 |
| CYC1 | 5704.306008 | 0.307151045 | 1.65E-03 |
| DKK3 | 22309.34202 | 0.307103942 | 2.83E-02 |
| FAM20B | 2759.207908 | 0.307101281 | 2.18E-02 |
| TMA16 | 607.3034725 | 0.307099615 | 3.43E-02 |
| SBF2 | 3829.910888 | 0.306651837 | 5.44E-03 |
| MCCC1 | 1805.188409 | 0.306570988 | 1.03E-02 |
| KIAA1958 | 673.3497986 | 0.306253245 | 2.96E-02 |
| CLEC16A | 2148.484018 | 0.305764486 | 2.84E-04 |
| RPL22L1 | 2206.738069 | 0.305701355 | 2.11E-02 |
| XYLT2 | 1979.267738 | 0.30561388 | 8.06E-03 |
| FRS3 | 243.2988226 | 0.305446769 | 3.55E-02 |
| ZNF277 | 1041.934133 | 0.305315827 | 6.40E-03 |
| SLC39A3 | 931.2260034 | 0.304953957 | 7.53E-03 |
| AC145212.1 | 558.4339615 | 0.304897591 | 2.93E-02 |
| CR381653.1 | 418.8328096 | 0.304168139 | 3.70E-02 |
| PCCB | 2314.132373 | 0.304059482 | 1.55E-04 |
| SLC39A1 | 6731.336194 | 0.30402251 | 3.39E-02 |
| SUPV3L1 | 1062.588736 | 0.303942761 | 3.24E-02 |
| MKRN2 | 1621.236604 | 0.303690751 | 7.46E-06 |
| NACC1 | 4181.963558 | 0.30368953 | 2.46E-02 |
| ZFYVE27 | 817.0816198 | 0.30368501 | 2.77E-03 |
| PPP1R18 | 5146.357623 | 0.303252121 | 2.96E-02 |
| PAIP1 | 7799.808941 | 0.303247834 | 5.24E-04 |
| SCNM1 | 804.5286563 | 0.303193657 | 3.93E-02 |
| RSU1 | 1873.166407 | 0.303088097 | 1.22E-03 |
| PI4K2A | 1765.032642 | 0.302744235 | 3.93E-02 |
| RUVBL1 | 5432.212495 | 0.302411346 | 1.88E-03 |
| UXT | 1749.111555 | 0.302405844 | 2.76E-02 |
| RPL23AP42 | 8204.968219 | 0.30229914 | 2.12E-02 |
| NBAS | 3584.572694 | 0.302095387 | 8.54E-03 |
| PSMC4 | 5691.909785 | 0.301990447 | 7.14E-04 |
| AC025171.1 | 642.5172103 | 0.301957348 | 1.42E-02 |
| ADCY3 | 3045.328296 | 0.301473314 | 1.04E-02 |
| MRS2 | 1525.675607 | 0.300875535 | 5.44E-04 |
| NELFB | 5735.829947 | 0.299848072 | 5.30E-03 |
| TOX4 | 5267.628927 | 0.299730534 | 3.47E-03 |
| AC093525.7 | 393.9993354 | 0.299718977 | 3.32E-02 |
| ZNF598 | 2854.685967 | 0.299635212 | 5.31E-04 |
| NIT2 | 1435.897603 | 0.299582318 | 4.69E-03 |
| HEATR6 | 1746.01362 | 0.299522739 | 8.07E-04 |
| RPS8 | 38197.62831 | 0.299397371 | 1.51E-03 |
| ZBTB17 | 1035.308393 | 0.299350648 | 5.19E-04 |
| CHID1 | 5044.44065 | 0.299009581 | 7.80E-03 |
| SERP1 | 3386.382468 | 0.298977993 | 7.16E-03 |

| | | | |
|---|---|---|---|
| ZRANB1 | 1147.558633 | 0.298959664 | 4.63E-02 |
| FBXW4 | 791.3351442 | 0.298925919 | 3.17E-02 |
| DAZAP2 | 5104.342531 | 0.298728226 | 4.46E-02 |
| CMTR1 | 5231.338082 | 0.29871075 | 8.23E-04 |
| ZNF710 | 1212.046584 | 0.298688747 | 5.43E-03 |
| PEX19 | 4514.041774 | 0.298537773 | 2.33E-03 |
| KDM7A-DT | 360.7802649 | 0.298490728 | 3.97E-02 |
| CAMTA1 | 1421.581369 | 0.297394627 | 1.38E-02 |
| LZTS1 | 3910.639426 | 0.297065491 | 2.13E-02 |
| CDCA7L | 2435.713917 | 0.296923664 | 8.05E-03 |
| KHNYN | 5658.646905 | 0.296922932 | 7.36E-06 |
| IQCB1 | 903.2856941 | 0.296773587 | 4.89E-02 |
| BUD13 | 1709.776183 | 0.296364772 | 1.63E-04 |
| RTL6 | 1982.757286 | 0.296289063 | 8.67E-03 |
| YKT6 | 13205.20478 | 0.296284321 | 3.83E-04 |
| ARHGEF39 | 1059.646998 | 0.296197032 | 4.55E-02 |
| QSOX2 | 2784.490846 | 0.296084296 | 2.96E-05 |
| TRIM65 | 1666.174936 | 0.296058561 | 7.34E-03 |
| UFD1 | 2667.109898 | 0.295913075 | 7.48E-03 |
| LTB4R2 | 430.8196029 | 0.295906958 | 2.28E-02 |
| GPX4 | 7821.112851 | 0.295609546 | 2.10E-02 |
| PIN1 | 1762.228934 | 0.295570489 | 1.54E-02 |
| GNB2 | 11143.86627 | 0.295267845 | 2.09E-02 |
| PRPF3 | 2440.32239 | 0.295160775 | 9.80E-07 |
| C8orf76 | 1193.820138 | 0.295088273 | 9.65E-03 |
| DCAF4 | 983.4014373 | 0.29498189 | 6.02E-03 |
| POLR1D | 2138.160546 | 0.29487624 | 4.28E-02 |
| MRPS34 | 2965.552352 | 0.294475443 | 2.24E-03 |
| RPS23 | 74273.86759 | 0.29413516 | 9.12E-03 |
| CHCHD1 | 767.6253033 | 0.294064179 | 3.22E-02 |
| FADS1 | 13379.58282 | 0.293792412 | 5.76E-03 |
| SMC1A | 8629.188695 | 0.293630058 | 2.04E-02 |
| HUWE1 | 25381.83121 | 0.29358103 | 6.49E-05 |
| PSMG3 | 1824.579345 | 0.293455423 | 5.81E-04 |
| MOAP1 | 1235.398058 | 0.293391907 | 2.01E-03 |
| ALS2 | 2565.168825 | 0.293279511 | 2.59E-02 |
| NOTCH2 | 14889.49296 | 0.293221338 | 2.22E-04 |
| PPM1M | 1735.27223 | 0.29304134 | 3.17E-02 |
| SNAPC3 | 1889.170263 | 0.292961408 | 9.31E-03 |
| COPS6 | 7614.260616 | 0.292759884 | 3.53E-04 |
| VPS72 | 2156.237537 | 0.292732302 | 3.34E-02 |
| LUZP1 | 6874.52116 | 0.292084965 | 1.78E-05 |
| MRPS27 | 6656.267008 | 0.291790867 | 1.46E-02 |
| PIMREG | 1661.753605 | 0.291744759 | 3.61E-02 |
| SRR | 938.275198 | 0.291346059 | 1.92E-02 |
| PDHB | 3611.84747 | 0.291202974 | 1.81E-03 |
| SEC24C | 6118.579717 | 0.290954623 | 1.38E-02 |

| | | | |
|---|---|---|---|
| KRT10 | 834.2809818 | 0.290850892 | 1.03E-02 |
| MTMR3 | 2790.3142 | 0.290849247 | 8.62E-04 |
| PDIA6 | 20872.47759 | 0.290725044 | 3.76E-04 |
| TMEM41A | 1259.205691 | 0.290494426 | 2.50E-02 |
| CTNNBL1 | 3608.026851 | 0.290392977 | 8.21E-06 |
| SRA1 | 1042.19628 | 0.28975339 | 3.59E-02 |
| R3HDM4 | 1971.816613 | 0.289707991 | 1.33E-02 |
| NBPF14 | 4716.123144 | 0.2893899 | 6.99E-03 |
| TUBG2 | 1807.888365 | 0.28922696 | 2.15E-03 |
| LIN52 | 827.3337377 | 0.289024973 | 1.78E-02 |
| PDHA1 | 4227.095497 | 0.288916663 | 8.24E-04 |
| EIF2B5 | 3033.847097 | 0.288842959 | 2.34E-03 |
| SLC7A6OS | 1998.111305 | 0.288759874 | 7.36E-04 |
| ARPC3 | 7377.140275 | 0.288387633 | 4.22E-03 |
| POLR3A | 1936.558238 | 0.288387627 | 4.80E-03 |
| SNW1 | 4347.853979 | 0.288030247 | 3.41E-02 |
| LARP1 | 12477.89571 | 0.287990168 | 5.98E-06 |
| FH | 3105.884307 | 0.287805779 | 1.48E-05 |
| RPS12 | 23474.3027 | 0.287623683 | 1.39E-03 |
| RPLP2 | 26725.70048 | 0.287506542 | 3.64E-02 |
| MAPKBP1 | 1283.789127 | 0.287474282 | 2.94E-02 |
| SPATS2L | 4827.762082 | 0.287446302 | 3.05E-02 |
| NOL7 | 2262.371265 | 0.287378841 | 1.09E-03 |
| RNF44 | 1765.055586 | 0.287261831 | 4.74E-02 |
| RAB5IF | 3288.76951 | 0.287253265 | 1.48E-02 |
| NAPG | 960.7955834 | 0.287218736 | 1.03E-02 |
| SEC23B | 5037.762538 | 0.286768982 | 8.83E-05 |
| TSPAN4 | 4504.292425 | 0.286714485 | 2.20E-02 |
| SLC22A3 | 402.2075182 | 0.28667741 | 4.70E-02 |
| STIM1 | 3076.092754 | 0.286464958 | 7.50E-03 |
| YIPF2 | 2105.78297 | 0.286388162 | 2.29E-02 |
| NR2C2AP | 655.2007936 | 0.286137678 | 1.43E-02 |
| PXMP2 | 902.2577882 | 0.286039823 | 4.76E-02 |
| DDR1 | 3398.200686 | 0.286001906 | 3.78E-02 |
| TALDO1 | 5576.922782 | 0.285968166 | 7.21E-03 |
| DIAPH1 | 8016.050664 | 0.285909281 | 1.35E-02 |
| RRP12 | 2494.959313 | 0.285557283 | 2.58E-02 |
| ZFYVE19 | 888.1861836 | 0.285433675 | 3.26E-02 |
| TIMM10 | 1509.425945 | 0.28524795 | 1.51E-02 |
| PES1 | 4082.165987 | 0.285003349 | 2.10E-03 |
| INTS4 | 2113.040451 | 0.284877963 | 5.14E-05 |
| ARID1A | 5257.33245 | 0.284867089 | 3.73E-04 |
| POGK | 3075.627708 | 0.284836236 | 4.46E-04 |
| PSMA5 | 4318.210261 | 0.284267876 | 3.73E-03 |
| DSTN | 12861.70921 | 0.284040762 | 2.55E-02 |
| COPZ1 | 6355.496849 | 0.283473199 | 1.07E-02 |
| FBXO41 | 1662.033975 | 0.28342684 | 2.95E-02 |

| | | | |
|---|---|---|---|
| ARFGAP3 | 2468.560438 | 0.283421488 | 4.77E-05 |
| STAT2 | 3251.987653 | 0.283279998 | 5.49E-03 |
| AKIP1 | 922.1010123 | 0.283238444 | 3.36E-02 |
| METTL23 | 915.3090469 | 0.282359787 | 4.14E-03 |
| SLC4A1AP | 2028.376907 | 0.282104164 | 2.75E-03 |
| KLHDC4 | 1557.359036 | 0.282077595 | 3.42E-02 |
| PRDX4 | 3675.105295 | 0.281612725 | 1.65E-02 |
| TMLHE | 1940.538019 | 0.281449486 | 2.22E-03 |
| P4HB | 29172.44412 | 0.281286644 | 9.10E-03 |
| CCDC92 | 1103.160319 | 0.281227914 | 1.20E-02 |
| NAIF1 | 858.6424912 | 0.281220169 | 1.85E-02 |
| TRAF7 | 3862.880105 | 0.280830817 | 1.92E-02 |
| ACAD9 | 2006.68078 | 0.280788295 | 3.75E-03 |
| SMNDC1 | 892.9026407 | 0.280668315 | 7.11E-03 |
| COX4I1 | 10109.85953 | 0.280595183 | 4.27E-02 |
| AHSA1 | 8364.353404 | 0.280593812 | 7.48E-05 |
| MAP7D1 | 4939.854588 | 0.280563591 | 4.95E-03 |
| STK11 | 2145.068827 | 0.280482175 | 8.87E-03 |
| PTPMT1 | 1669.947516 | 0.280251145 | 3.46E-03 |
| METAP1 | 2773.955335 | 0.279853961 | 1.48E-03 |
| CLIC1 | 17383.67349 | 0.279608686 | 1.88E-02 |
| UBE2V1 | 9433.955559 | 0.279511468 | 1.40E-02 |
| NDUFV1 | 7369.650581 | 0.279356116 | 1.12E-03 |
| CCDC130 | 833.3427174 | 0.279277276 | 1.94E-02 |
| ISCA2 | 1204.879052 | 0.279186288 | 2.06E-03 |
| GPN1 | 3418.273067 | 0.279110484 | 6.00E-04 |
| R3HCC1 | 1870.466866 | 0.278789442 | 5.27E-03 |
| CMAS | 1197.058059 | 0.278414583 | 2.77E-02 |
| BAZ2A | 7258.618219 | 0.278204791 | 4.53E-05 |
| SLC9A5 | 766.9386846 | 0.278109678 | 2.97E-02 |
| TMED4 | 5466.26898 | 0.277980026 | 9.12E-03 |
| NOB1 | 4105.266417 | 0.27756468 | 2.06E-02 |
| POLR2J | 3629.819299 | 0.277465978 | 4.97E-03 |
| STOML2 | 9108.069344 | 0.277299079 | 1.14E-03 |
| LAPTM4A | 7258.494821 | 0.277155442 | 9.50E-05 |
| TAP2 | 3870.967011 | 0.276837611 | 1.00E-02 |
| KPNA1 | 3329.771721 | 0.276616503 | 4.33E-04 |
| CPSF4 | 2678.248412 | 0.276407904 | 4.52E-04 |
| ZNF496 | 1806.436781 | 0.276398255 | 1.23E-03 |
| C1orf109 | 1140.631876 | 0.276319443 | 3.85E-03 |
| NRF1 | 837.4999078 | 0.275933035 | 2.97E-03 |
| SNRNP27 | 1096.904373 | 0.275710091 | 3.53E-03 |
| METTL5 | 1304.863506 | 0.275458429 | 2.51E-02 |
| ZFYVE21 | 1166.025972 | 0.275208939 | 3.88E-02 |
| C14orf93 | 695.6090895 | 0.274920797 | 3.37E-02 |
| BLCAP | 3692.352387 | 0.27485549 | 1.42E-02 |
| SMARCD3 | 1854.421528 | 0.274818213 | 3.64E-02 |

| | | | |
|---|---|---|---|
| COPB2 | 9595.875253 | 0.274641962 | 1.45E-03 |
| SRGN | 1547.782683 | 0.274626629 | 2.94E-04 |
| NDUFA8 | 3018.998062 | 0.274469447 | 1.88E-02 |
| DTX2 | 890.7965669 | 0.274404667 | 3.95E-02 |
| DDB1 | 22375.96724 | 0.274391284 | 2.24E-02 |
| NRBF2 | 506.3490214 | 0.27437359 | 1.84E-02 |
| SLC35E1 | 3256.555031 | 0.274338902 | 4.49E-03 |
| PLOD3 | 10152.46719 | 0.274263776 | 5.41E-03 |
| C1orf43 | 7614.203548 | 0.273956961 | 7.10E-03 |
| CSTF2 | 2441.068491 | 0.273758961 | 1.11E-04 |
| TMEM44 | 939.3230402 | 0.27358918 | 1.99E-02 |
| PCBP4 | 2249.447405 | 0.273437007 | 3.52E-02 |
| RANBP1 | 4498.47852 | 0.273359719 | 3.31E-02 |
| CINP | 1999.859516 | 0.273309357 | 2.12E-02 |
| RNF7 | 2168.713989 | 0.273263737 | 1.74E-03 |
| ECPAS | 7118.119316 | 0.273179914 | 3.08E-04 |
| SLC35D2 | 1250.038697 | 0.273096443 | 1.93E-02 |
| AHRR | 2060.829402 | 0.273029293 | 1.37E-02 |
| TFIP11 | 2496.731842 | 0.27262489 | 2.01E-02 |
| COX5A | 3745.18832 | 0.272450605 | 1.57E-02 |
| ELOC | 2405.537577 | 0.272333986 | 8.20E-04 |
| EIF2S1 | 4923.226392 | 0.272056434 | 9.57E-03 |
| PXDN | 21810.67112 | 0.271883229 | 2.14E-03 |
| ANAPC1 | 5455.149671 | 0.27151922 | 2.29E-03 |
| CREG1 | 2653.32401 | 0.271310817 | 5.84E-03 |
| PAICS | 10941.13924 | 0.271274053 | 2.85E-02 |
| GOSR2 | 3360.184991 | 0.271127283 | 3.26E-02 |
| HNRNPUL2 | 6953.721425 | 0.270839495 | 1.88E-04 |
| ALG3 | 1585.689512 | 0.270828401 | 3.59E-02 |
| SERINC3 | 7719.314089 | 0.270732311 | 2.09E-05 |
| SNX17 | 6018.72471 | 0.270693632 | 3.78E-02 |
| DHPS | 2252.553779 | 0.270248612 | 6.89E-05 |
| ACACA | 9223.618154 | 0.270166568 | 6.64E-04 |
| ATG3 | 1629.288593 | 0.270017616 | 3.96E-03 |
| TP53I13 | 980.1633656 | 0.269334369 | 2.29E-02 |
| WIPF2 | 2519.523246 | 0.269257908 | 3.42E-03 |
| GCN1 | 15271.39176 | 0.269186105 | 1.12E-02 |
| RNF212 | 2579.106999 | 0.268988444 | 7.82E-05 |
| EIF2B2 | 2814.549157 | 0.268800788 | 3.86E-02 |
| PPP1R7 | 3054.404302 | 0.268656506 | 8.15E-04 |
| CSNK2A1 | 11719.97268 | 0.268588745 | 2.20E-05 |
| MRGBP | 2498.724127 | 0.2683478 | 1.16E-03 |
| RTRAF | 4394.6709 | 0.267625983 | 2.31E-02 |
| MRPS10 | 1948.921082 | 0.267612884 | 1.30E-02 |
| GFM2 | 5520.51197 | 0.267491291 | 9.51E-04 |
| ILKAP | 1385.386995 | 0.267465398 | 7.99E-05 |
| SSRP1 | 19588.32243 | 0.267240121 | 4.01E-04 |

| | | | |
|---|---|---|---|
| HDAC7 | 5153.464871 | 0.267199119 | 1.96E-03 |
| CYTH2 | 1564.392588 | 0.266921466 | 2.99E-02 |
| MELK | 3619.29311 | 0.266797154 | 3.38E-02 |
| TPT1-AS1 | 345.5680486 | 0.266525092 | 4.99E-02 |
| EIF4A3 | 3461.150558 | 0.266419052 | 1.83E-02 |
| PDCD6IP | 6347.384567 | 0.266417229 | 1.73E-03 |
| RFC5 | 1626.892685 | 0.266372001 | 3.16E-02 |
| COA1 | 3731.114246 | 0.266152555 | 2.90E-02 |
| CEP131 | 811.4527466 | 0.265986484 | 3.59E-02 |
| AFG3L2 | 4562.506854 | 0.265748793 | 4.46E-06 |
| DNM2 | 7737.278026 | 0.265657899 | 3.40E-04 |
| OCRL | 3249.787062 | 0.265627569 | 1.88E-04 |
| BBS2 | 3444.019886 | 0.26552131 | 2.81E-03 |
| FBXO34 | 1559.55816 | 0.265505697 | 1.24E-02 |
| NGRN | 7742.66646 | 0.265265256 | 1.68E-03 |
| ZNF274 | 1260.893165 | 0.265026371 | 2.19E-02 |
| RAB8A | 1921.112499 | 0.264972527 | 2.76E-02 |
| AC026464.4 | 1259.810063 | 0.264797047 | 1.96E-02 |
| NOC4L | 875.5322397 | 0.264626425 | 7.10E-03 |
| FCF1 | 2201.561553 | 0.264613793 | 2.20E-02 |
| H3F3B | 13249.7928 | 0.264559693 | 2.93E-04 |
| EXOSC8 | 1008.870093 | 0.264432453 | 2.18E-02 |
| DIS3L2 | 1432.177132 | 0.263941839 | 1.34E-02 |
| USP7 | 5820.658002 | 0.263884611 | 7.84E-06 |
| PSME1 | 7705.982722 | 0.263863337 | 3.95E-02 |
| NDUFB9 | 4292.418128 | 0.263801099 | 4.01E-02 |
| RPL35A | 14501.75282 | 0.263737702 | 4.58E-02 |
| SAPCD2 | 1918.011644 | 0.263679389 | 3.42E-02 |
| RPL36A | 17414.29202 | 0.263650678 | 3.28E-03 |
| PASD1 | 4060.815916 | 0.263547765 | 1.59E-03 |
| LARP4B | 1886.896081 | 0.263399318 | 2.83E-03 |
| CNDP2 | 7468.433963 | 0.262375347 | 1.23E-02 |
| NDUFS7 | 1677.92942 | 0.261899853 | 7.24E-03 |
| PREB | 2909.820778 | 0.261793743 | 1.34E-02 |
| AP3S1 | 3641.825183 | 0.261701205 | 1.85E-03 |
| ENOPH1 | 1391.236757 | 0.261628412 | 1.14E-02 |
| DCUN1D5 | 2156.25617 | 0.261429932 | 1.77E-03 |
| ST7 | 1282.821539 | 0.261303221 | 2.99E-02 |
| SLC33A1 | 1177.423913 | 0.261245902 | 3.58E-02 |
| ATIC | 7866.447087 | 0.261161953 | 1.27E-02 |
| ZDHHC24 | 651.9338299 | 0.261142489 | 4.99E-02 |
| LONP2 | 5795.25613 | 0.261031184 | 2.80E-04 |
| ABHD11 | 864.5830385 | 0.260802564 | 3.67E-02 |
| KIF22 | 5070.961106 | 0.260743307 | 2.54E-02 |
| MRPL18 | 1831.355952 | 0.260702639 | 3.74E-04 |
| COG2 | 1648.39515 | 0.260680641 | 3.58E-03 |
| MYO19 | 2959.134123 | 0.260481098 | 2.25E-02 |

| | | | |
|---|---|---|---|
| PDGFRB | 11707.28027 | 0.260472251 | 4.07E-02 |
| EMC8 | 1576.06358 | 0.26033159 | 1.89E-02 |
| BAIAP2 | 994.5039872 | 0.26016939 | 4.62E-03 |
| CRTAP | 9964.199093 | 0.259846486 | 3.72E-02 |
| FAM104A | 1453.554848 | 0.259779704 | 4.38E-03 |
| PILRB | 1131.747705 | 0.258914776 | 1.89E-02 |
| CLIC4 | 20156.46442 | 0.258808712 | 4.02E-02 |
| MACF1 | 17521.76745 | 0.257643334 | 3.02E-04 |
| RUFY1 | 1541.020346 | 0.257606365 | 2.29E-02 |
| ANKRD27 | 1949.923053 | 0.257362002 | 3.11E-03 |
| CKS2 | 3853.770361 | 0.257028227 | 1.08E-02 |
| FAM50A | 5466.553151 | 0.256952028 | 7.78E-03 |
| YBX1 | 25603.98182 | 0.256937312 | 2.32E-03 |
| GDI1 | 9466.865383 | 0.256755062 | 1.76E-03 |
| KPNA6 | 4279.182556 | 0.256745719 | 2.64E-02 |
| RACK1 | 52295.32079 | 0.256644232 | 3.47E-02 |
| KIF13B | 1664.961045 | 0.256257949 | 1.33E-02 |
| NUBP2 | 2876.219677 | 0.255555725 | 2.89E-02 |
| LINC01029 | 2160.242943 | 0.255382127 | 2.59E-02 |
| KHSRP | 10535.99363 | 0.255318151 | 1.48E-03 |
| TMEM126A | 956.1421872 | 0.254658947 | 2.25E-02 |
| PPCS | 1325.8896 | 0.254556125 | 1.32E-02 |
| SMARCB1 | 4122.10918 | 0.2545002 | 9.66E-03 |
| DIABLO | 2319.996166 | 0.254010198 | 1.23E-02 |
| HADHA | 12749.68248 | 0.253896145 | 4.54E-03 |
| DNAJC8 | 6062.299342 | 0.253587418 | 5.36E-03 |
| NADSYN1 | 1687.908599 | 0.253494187 | 2.32E-02 |
| SEC11C | 605.6753875 | 0.253425218 | 3.88E-02 |
| ZNF787 | 811.1459574 | 0.25333568 | 1.93E-02 |
| DGKZ | 4303.453503 | 0.253322216 | 5.09E-03 |
| PLBD2 | 5126.821147 | 0.252996108 | 3.69E-02 |
| GCC1 | 2344.373322 | 0.252269038 | 1.90E-02 |
| CDKN1B | 2344.932369 | 0.252225346 | 1.38E-02 |
| SMIM26 | 1687.576725 | 0.252151832 | 4.53E-02 |
| RPIA | 1240.066785 | 0.252095437 | 2.33E-03 |
| ZMIZ2 | 2157.698902 | 0.251931021 | 4.51E-04 |
| RPL14 | 17200.18652 | 0.251765099 | 1.27E-02 |
| SMYD5 | 1789.083633 | 0.251634737 | 3.32E-02 |
| SLC25A39 | 4759.181151 | 0.250829143 | 1.80E-02 |
| NNT | 4515.752462 | 0.250686028 | 3.28E-03 |
| SSBP1 | 6663.456012 | 0.250519651 | 5.62E-05 |
| ATXN1L | 3477.835901 | 0.250399844 | 8.83E-03 |
| PPP2R5A | 1252.576734 | 0.250398526 | 4.40E-02 |
| CDKN2AIPNL | 855.1231238 | 0.250118037 | 4.83E-02 |
| SFT2D1 | 1397.633585 | 0.250106335 | 4.38E-04 |
| TRMT2B | 1049.45023 | 0.249738227 | 8.97E-03 |
| GNG5 | 2669.344981 | 0.24955824 | 2.21E-02 |

| | | | |
|---|---|---|---|
| NTPCR | 2144.04092 | 0.249483646 | 4.63E-02 |
| RPN1 | 10565.1925 | 0.249363315 | 3.52E-02 |
| NUP98 | 10252.36378 | 0.249123807 | 1.35E-03 |
| ZCRB1 | 2337.235148 | 0.249111528 | 3.12E-03 |
| INTS11 | 4008.1811 | 0.249088154 | 9.60E-03 |
| ZNF622 | 2500.069831 | 0.248840865 | 1.21E-03 |
| UBE3B | 2910.506242 | 0.248620467 | 3.09E-02 |
| SPTLC1 | 3390.99703 | 0.248333316 | 8.17E-03 |
| TRIT1 | 1099.34828 | 0.248319661 | 4.74E-02 |
| DCAF7 | 7641.624779 | 0.247833553 | 2.95E-02 |
| EEFSEC | 1107.058862 | 0.247636361 | 1.84E-02 |
| NDUFS1 | 6300.475034 | 0.24758603 | 1.68E-05 |
| CERS5 | 1746.370704 | 0.2475595 | 4.53E-02 |
| AL589743.1 | 1078.609246 | 0.247438753 | 1.38E-02 |
| MRPL45 | 2482.560395 | 0.247328102 | 4.82E-03 |
| RPL30 | 35678.43918 | 0.247286653 | 2.94E-02 |
| PNMA3 | 556.7056313 | 0.247161613 | 4.92E-02 |
| THOC6 | 2050.440282 | 0.247120814 | 2.62E-02 |
| ANKRD52 | 4478.484332 | 0.247112747 | 7.45E-03 |
| STARD3NL | 2013.879759 | 0.247033009 | 5.77E-03 |
| HINT2 | 1283.351019 | 0.246914577 | 1.50E-02 |
| DNAJA2 | 4555.427379 | 0.246891624 | 1.64E-05 |
| SRPRA | 13337.9025 | 0.246569849 | 6.76E-03 |
| AKAP13 | 5307.00127 | 0.246303952 | 2.03E-02 |
| TTL | 3635.258305 | 0.246107218 | 2.22E-02 |
| VPS16 | 3276.175891 | 0.245977833 | 2.20E-02 |
| METTL2B | 1796.131861 | 0.245965504 | 3.73E-03 |
| RABL6 | 5448.593957 | 0.245744675 | 8.00E-03 |
| PGS1 | 1688.179157 | 0.245237018 | 2.26E-02 |
| HDHD5 | 2611.229972 | 0.245207381 | 3.99E-02 |
| PRDM4 | 1802.069444 | 0.24507524 | 1.71E-03 |
| RPL23AP82 | 694.2770221 | 0.244806909 | 3.24E-02 |
| ASAP3 | 2641.095712 | 0.244539558 | 1.80E-02 |
| CS | 9835.018622 | 0.244485924 | 4.74E-03 |
| RPTOR | 2416.020726 | 0.243781956 | 2.95E-02 |
| THOC7 | 1848.437883 | 0.243534805 | 5.46E-04 |
| TMED2 | 8167.232608 | 0.243493875 | 7.28E-04 |
| MRPL10 | 2784.242699 | 0.243012792 | 2.79E-02 |
| TENT4A | 2616.581346 | 0.24292151 | 3.69E-02 |
| CSNK1A1 | 6157.901358 | 0.242356057 | 2.06E-02 |
| IDS | 4286.442473 | 0.241930591 | 1.98E-02 |
| FAM45A | 1390.912981 | 0.241877371 | 2.18E-02 |
| STRADB | 1807.717494 | 0.241719472 | 1.27E-02 |
| RAB29 | 1618.049604 | 0.24143927 | 2.49E-02 |
| PDE6D | 1308.982486 | 0.240238864 | 1.39E-02 |
| CEP41 | 1707.755942 | 0.239685783 | 3.95E-02 |
| ARPC2 | 12473.43948 | 0.239502024 | 5.71E-03 |

| | | | |
|---|---|---|---|
| YPEL5 | 2746.761684 | 0.239198054 | 3.44E-02 |
| EIF2AK2 | 3388.221179 | 0.239154853 | 2.99E-02 |
| PDHX | 1539.431444 | 0.238675445 | 2.92E-02 |
| EMC4 | 1681.225534 | 0.238663664 | 1.81E-02 |
| AKIRIN1 | 2134.676385 | 0.238338869 | 2.42E-02 |
| TRIM14 | 2948.397171 | 0.238239858 | 1.13E-02 |
| RNF4 | 3198.021208 | 0.238225876 | 1.79E-03 |
| SEPHS2 | 6477.342138 | 0.238082994 | 1.77E-03 |
| AIMP2 | 1830.027727 | 0.238073769 | 6.33E-03 |
| HSPD1 | 27693.40273 | 0.237582204 | 1.97E-02 |
| TMEM11 | 884.3596645 | 0.237530137 | 1.25E-02 |
| PPP2CA | 10438.57862 | 0.237489647 | 3.21E-02 |
| DCTN1 | 12802.93387 | 0.23747609 | 2.21E-02 |
| RPS6 | 79859.13227 | 0.237467712 | 1.22E-02 |
| CAMSAP1 | 3654.8622 | 0.237467684 | 2.58E-02 |
| FAM118A | 895.3172424 | 0.236638849 | 4.89E-03 |
| NDE1 | 2540.9428 | 0.236339966 | 2.97E-02 |
| IL13RA1 | 5499.46665 | 0.235866796 | 3.17E-03 |
| VPS26C | 2551.853355 | 0.235722545 | 7.31E-04 |
| CTDP1 | 1130.124063 | 0.235683391 | 3.89E-02 |
| DERL1 | 3417.012654 | 0.235483425 | 1.81E-02 |
| GLTP | 1149.725905 | 0.234758926 | 3.58E-02 |
| POFUT1 | 11405.7374 | 0.234679299 | 2.09E-02 |
| DHX16 | 3120.149289 | 0.234627959 | 1.26E-02 |
| CDK5RAP3 | 3684.848713 | 0.234625952 | 8.94E-04 |
| SEC16A | 7783.876163 | 0.234553818 | 7.52E-03 |
| TTC1 | 6219.880764 | 0.234193982 | 5.44E-04 |
| C19orf48 | 2555.273303 | 0.23387378 | 1.33E-02 |
| NT5C3B | 3653.141291 | 0.233708855 | 1.45E-02 |
| REPS1 | 1371.046726 | 0.233692961 | 2.99E-03 |
| PHF5A | 2143.181465 | 0.233539164 | 7.20E-03 |
| AAGAB | 3925.30352 | 0.233467571 | 2.41E-03 |
| RPL17 | 27377.12479 | 0.23334945 | 1.30E-02 |
| ESD | 3673.991792 | 0.233304531 | 4.39E-03 |
| UQCC1 | 3926.027674 | 0.233286874 | 7.54E-03 |
| IWS1 | 2889.136466 | 0.233118746 | 4.79E-02 |
| HIRIP3 | 1467.028443 | 0.233083116 | 6.75E-03 |
| NXPE3 | 1902.101291 | 0.233064345 | 4.00E-02 |
| NA | 1855.274218 | 0.232513838 | 1.47E-02 |
| RRP1 | 1635.850816 | 0.232393068 | 4.44E-02 |
| EIPR1 | 1284.282965 | 0.232390525 | 4.87E-02 |
| MRPS31 | 962.120588 | 0.232283534 | 4.63E-02 |
| CXorf56 | 1124.240623 | 0.232154157 | 2.96E-02 |
| PPP6C | 3935.07532 | 0.232103903 | 1.58E-03 |
| FARSA | 5326.766618 | 0.2320344 | 4.98E-02 |
| PTPRJ | 3731.301994 | 0.231639994 | 5.21E-03 |
| GATAD2A | 4101.766037 | 0.23136672 | 3.19E-02 |

| | | | |
|---|---|---|---|
| SNF8 | 3726.744613 | 0.2312968 | 6.92E-03 |
| CBLL1 | 1559.789929 | 0.231192876 | 2.02E-02 |
| RRP1B | 3504.70681 | 0.230925476 | 1.33E-04 |
| WWP2 | 2287.978635 | 0.230875706 | 2.39E-02 |
| IPO9 | 8938.980323 | 0.230256372 | 3.14E-02 |
| TCP1 | 10078.89212 | 0.230236924 | 6.62E-04 |
| RNF185 | 2804.861805 | 0.230042154 | 1.82E-02 |
| SNU13 | 7127.161613 | 0.229772096 | 2.89E-02 |
| NUDCD2 | 2588.159888 | 0.228541499 | 2.87E-02 |
| NOL6 | 8157.384097 | 0.228468064 | 2.56E-02 |
| TOMM20 | 6224.484129 | 0.228180053 | 1.29E-02 |
| KIAA1147 | 3149.274543 | 0.227946606 | 2.12E-03 |
| CTPS2 | 1411.08626 | 0.227569595 | 4.20E-02 |
| EIF2S3 | 9547.502372 | 0.227486228 | 4.47E-05 |
| PDPK1 | 4241.916576 | 0.227111902 | 3.29E-02 |
| NOP14 | 2656.143157 | 0.226871906 | 2.65E-05 |
| FCHSD2 | 1863.422345 | 0.226729654 | 9.85E-03 |
| PSMA6 | 8369.254575 | 0.226681092 | 1.77E-02 |
| AC073869.1 | 886.8040614 | 0.226069639 | 1.38E-02 |
| USP10 | 5912.579759 | 0.225957477 | 1.64E-05 |
| TMEM161A | 1198.284385 | 0.225915391 | 2.60E-02 |
| POLDIP2 | 7076.671149 | 0.225605004 | 4.84E-02 |
| WRAP73 | 1404.388509 | 0.22553887 | 1.10E-02 |
| MANBAL | 3030.257961 | 0.225307623 | 2.89E-02 |
| WDR46 | 5956.684672 | 0.22499927 | 1.11E-02 |
| SMG5 | 7376.902738 | 0.22495837 | 9.64E-03 |
| FAM192A | 4054.220044 | 0.224896823 | 2.13E-04 |
| MIB2 | 1313.54863 | 0.224711841 | 3.60E-02 |
| AP000525.1 | 2001.229695 | 0.224625069 | 1.98E-02 |
| MAVS | 12155.02289 | 0.224561533 | 1.43E-02 |
| ARID5B | 1968.657859 | 0.22446497 | 1.88E-02 |
| PARP12 | 3009.496546 | 0.22443021 | 6.98E-03 |
| PDAP1 | 9323.361523 | 0.223781944 | 4.81E-03 |
| DCAF8 | 5161.578887 | 0.223213756 | 4.31E-04 |
| CERS2 | 7121.847108 | 0.223136938 | 2.82E-02 |
| GANAB | 50670.80836 | 0.222620263 | 3.84E-02 |
| USP28 | 3710.643919 | 0.222618094 | 2.24E-02 |
| NFE2L1 | 19179.84364 | 0.222571919 | 4.85E-02 |
| BIN3 | 1269.359925 | 0.222436766 | 3.64E-02 |
| CAPN10 | 1051.764605 | 0.222421517 | 2.51E-02 |
| H2AFV | 7403.940441 | 0.222058224 | 1.19E-02 |
| TBC1D20 | 4180.645879 | 0.22194865 | 3.60E-03 |
| GTPBP1 | 3068.954214 | 0.221666264 | 4.14E-02 |
| PGRMC1 | 3576.645105 | 0.221141635 | 9.42E-03 |
| UQCC3 | 1096.155949 | 0.220992664 | 4.36E-02 |
| CSDE1 | 17721.67776 | 0.22023206 | 9.80E-03 |
| RER1 | 5729.27554 | 0.219891411 | 2.58E-02 |

| | | | |
|---|---|---|---|
| SNRNP40 | 2625.67186 | 0.219725623 | 2.49E-02 |
| METTL2A | 1073.794768 | 0.219638457 | 3.61E-02 |
| RBSN | 2154.092521 | 0.219573254 | 2.47E-02 |
| PHF8 | 2614.998685 | 0.219474518 | 2.10E-02 |
| UBE2D3 | 7342.501166 | 0.21936481 | 9.30E-03 |
| STX10 | 2145.396318 | 0.219347149 | 4.82E-02 |
| MOCS1 | 1229.149912 | 0.219072048 | 1.85E-02 |
| IKBKB | 2090.625545 | 0.218468371 | 2.80E-02 |
| ABCB8 | 1540.220527 | 0.218449137 | 2.05E-02 |
| CCDC124 | 1599.856452 | 0.218191641 | 1.49E-02 |
| CMC2 | 1496.242886 | 0.218174354 | 2.53E-02 |
| BTBD2 | 6344.109791 | 0.217658636 | 4.28E-02 |
| EIF2AK1 | 12276.79952 | 0.217632194 | 3.17E-02 |
| PSMC2 | 5871.671733 | 0.217399968 | 1.87E-02 |
| EDC3 | 4250.423147 | 0.216952171 | 1.60E-02 |
| PHKA1 | 1733.661679 | 0.216653201 | 1.68E-03 |
| RNF10 | 5735.766909 | 0.215518182 | 3.15E-02 |
| UTP6 | 3198.610634 | 0.214679073 | 1.34E-02 |
| VPS37C | 1181.127109 | 0.214290876 | 3.30E-02 |
| ELP2 | 3950.602411 | 0.213454306 | 4.76E-02 |
| GLO1 | 5862.37989 | 0.213392467 | 2.19E-02 |
| RHOQ | 3625.302826 | 0.2133861 | 7.30E-03 |
| XPO7 | 6219.276016 | 0.213235584 | 9.65E-03 |
| SMIM20 | 810.4386894 | 0.213137835 | 4.89E-02 |
| AP1B1 | 6559.260492 | 0.212040665 | 2.74E-02 |
| GRSF1 | 4306.976919 | 0.211970814 | 4.46E-02 |
| PPP2R2A | 3979.054954 | 0.211750762 | 5.56E-03 |
| NDUFA10 | 3977.783713 | 0.211579411 | 3.80E-02 |
| RTF2 | 6751.588276 | 0.211439353 | 2.74E-02 |
| ATP6V1G1 | 2695.790776 | 0.211269081 | 7.79E-03 |
| UPF1 | 5584.674034 | 0.21126022 | 1.56E-02 |
| GUSB | 2894.849897 | 0.21104959 | 1.91E-02 |
| NEK9 | 9405.283839 | 0.210883878 | 2.51E-02 |
| PSMA2 | 6852.048468 | 0.209545962 | 2.96E-02 |
| DESI1 | 2612.386155 | 0.208192264 | 1.23E-02 |
| TIMM23 | 2132.272328 | 0.207942056 | 3.24E-02 |
| ATP2A2 | 12810.82753 | 0.207361932 | 1.82E-02 |
| TRIM11 | 1271.475392 | 0.207195893 | 1.47E-02 |
| POMP | 2654.174036 | 0.207028436 | 5.10E-03 |
| HIBADH | 5149.036504 | 0.206759139 | 6.55E-04 |
| SGCD | 2815.580324 | 0.206702667 | 2.33E-03 |
| NALCN | 1270.957975 | 0.20667458 | 2.76E-02 |
| PPP4R1 | 4147.702113 | 0.206333832 | 2.76E-02 |
| SUCLG2 | 2395.918286 | 0.205687141 | 2.96E-02 |
| RDH13 | 730.3832492 | 0.204509324 | 3.55E-02 |
| VPS29 | 1793.056619 | 0.204373459 | 8.09E-03 |
| DPH7 | 1905.818143 | 0.204294722 | 4.00E-02 |

| | | | |
|---|---|---|---|
| MRFAP1 | 10526.41433 | 0.203327332 | 2.97E-03 |
| OSBP | 5786.050547 | 0.203244243 | 2.95E-02 |
| MCFD2 | 14679.18902 | 0.200755791 | 2.18E-03 |
| SNRNP70 | 6284.377571 | 0.200579464 | 4.43E-02 |
| PHACTR4 | 2601.10134 | 0.200137755 | 1.33E-02 |
| TBC1D2B | 6479.875257 | 0.199769435 | 4.25E-02 |
| ETFA | 5630.621912 | 0.198981853 | 1.03E-03 |
| UTP11 | 1951.039154 | 0.198279784 | 4.34E-02 |
| ATF6B | 3233.231276 | 0.197846226 | 2.04E-02 |
| TSN | 3501.252314 | 0.197179483 | 1.56E-02 |
| SRP54 | 3509.000559 | 0.195888012 | 6.75E-03 |
| ECE1 | 7636.537465 | 0.195870486 | 4.70E-02 |
| ARHGEF7 | 1898.615677 | 0.19479166 | 4.08E-03 |
| RCCD1 | 1438.649141 | 0.194500993 | 4.09E-02 |
| NUP155 | 7161.690823 | 0.194254117 | 3.36E-02 |
| TMX2 | 4622.100163 | 0.193556388 | 1.78E-02 |
| TMEM70 | 998.0146355 | 0.193172551 | 4.48E-02 |
| MAGEA10 | 7039.294074 | 0.192100021 | 2.42E-02 |
| HNRNPAB | 5558.229272 | 0.192087304 | 2.06E-02 |
| AGK | 2711.497476 | 0.191961864 | 3.64E-02 |
| DUS1L | 2081.113592 | 0.190382009 | 1.33E-02 |
| RPS27A | 29292.20617 | 0.190216275 | 4.02E-02 |
| CSAG1 | 2662.965097 | 0.189978881 | 3.97E-02 |
| HDLBP | 32128.14887 | 0.189169747 | 4.33E-02 |
| KANK1 | 4383.01843 | 0.188996789 | 2.66E-02 |
| UBE2J2 | 2039.609144 | 0.188246557 | 2.20E-02 |
| DFFA | 3664.020143 | 0.188159827 | 1.71E-02 |
| MAPK14 | 3125.554692 | 0.187185504 | 1.17E-02 |
| JAK1 | 6969.182078 | 0.185779875 | 1.27E-02 |
| PSMD1 | 7577.741112 | 0.18566322 | 2.80E-02 |
| RNF11 | 2230.717256 | 0.183236749 | 2.35E-02 |
| MAPKAP1 | 7259.855184 | 0.182866869 | 4.91E-02 |
| STK35 | 2874.044657 | 0.182807343 | 2.71E-03 |
| EIF4G2 | 45968.60351 | 0.182649485 | 2.02E-02 |
| LSM12 | 2512.723727 | 0.182532727 | 1.46E-02 |
| MKKS | 2227.120336 | 0.181590819 | 1.05E-02 |
| DIDO1 | 6933.614041 | 0.180846594 | 5.25E-04 |
| PSME3 | 7880.901979 | 0.180390158 | 4.18E-02 |
| UNC119B | 3862.281918 | 0.179941991 | 3.13E-02 |
| EIF3H | 12961.00924 | 0.179193376 | 3.50E-02 |
| MRPL15 | 2929.771809 | 0.179186492 | 2.95E-02 |
| HIGD1A | 2206.17762 | 0.178816529 | 2.58E-02 |
| YWHAE | 30929.50022 | 0.178037941 | 4.15E-02 |
| UBAP2 | 3251.690654 | 0.177546997 | 4.43E-02 |
| YWHAB | 18910.22592 | 0.175727609 | 4.43E-03 |
| MRPL3 | 4222.736049 | 0.175289814 | 2.85E-02 |
| ELMO2 | 4319.093273 | 0.174986244 | 4.98E-02 |

| | | | |
|---|---|---|---|
| NDUFV3 | 1383.346286 | 0.174607561 | 1.96E-02 |
| EPN2 | 2916.468412 | 0.171110043 | 3.33E-02 |
| FAM168B | 7463.704228 | 0.169118057 | 1.06E-02 |
| SIN3B | 3202.374621 | 0.168081812 | 7.38E-03 |
| YEATS2 | 5124.563984 | 0.167644254 | 1.47E-02 |
| UBQLN1 | 7209.186043 | 0.167545412 | 2.33E-02 |
| ZNF275 | 1849.137419 | 0.16733028 | 4.24E-02 |
| TCF20 | 3764.351559 | 0.164202068 | 4.71E-02 |
| GEMIN5 | 2211.92909 | 0.16154529 | 3.16E-02 |
| NAT10 | 6953.557033 | 0.154584034 | 2.64E-02 |
| NUP188 | 7233.696582 | 0.154451531 | 2.88E-02 |
| NBR1 | 8980.711034 | 0.149339962 | 1.93E-02 |
| SPECC1L | 5523.200268 | 0.148918383 | 4.06E-02 |
| SGCB | 2046.737694 | 0.144735085 | 3.46E-02 |
| FBXW11 | 3631.29144 | 0.142137703 | 4.73E-02 |
| ERCC3 | 3716.889141 | 0.133196442 | 3.66E-02 |
| TSR1 | 7944.401475 | 0.132068469 | 4.34E-02 |
| FOXO3 | 3030.424993 | -0.117975848 | 4.03E-02 |
| PSMD5 | 3563.031178 | -0.134863696 | 2.20E-02 |
| SSR1 | 4631.397543 | -0.136354809 | 2.72E-02 |
| WDR26 | 5431.769766 | -0.139611997 | 3.06E-02 |
| FAM136A | 2926.438027 | -0.147702329 | 6.78E-03 |
| HNRNPF | 6684.601055 | -0.152441964 | 1.38E-02 |
| ERC1 | 4832.654921 | -0.157825942 | 3.84E-02 |
| SMG7 | 5444.799526 | -0.161981131 | 1.45E-03 |
| STX12 | 1717.797299 | -0.169573008 | 3.84E-02 |
| ELF1 | 1523.077728 | -0.180130411 | 4.15E-02 |
| GIGYF2 | 4244.851325 | -0.180400476 | 4.27E-02 |
| SOD2 | 3376.369198 | -0.180727981 | 2.57E-02 |
| DYNC1I2 | 4866.851288 | -0.181607932 | 3.71E-02 |
| TXNL1 | 2788.791101 | -0.181963452 | 1.61E-02 |
| UBE4A | 5703.241683 | -0.183192138 | 4.50E-02 |
| SUPT6H | 8733.333625 | -0.183652802 | 9.42E-03 |
| CLN6 | 2149.43622 | -0.183657743 | 3.57E-02 |
| GGA1 | 3194.607398 | -0.184581031 | 4.24E-02 |
| TAF11 | 1504.934528 | -0.184613624 | 4.54E-02 |
| NUMA1 | 17142.57348 | -0.185174276 | 3.32E-02 |
| NDFIP1 | 3015.796015 | -0.186369356 | 2.86E-02 |
| KAT6A | 4640.681421 | -0.186385064 | 3.79E-02 |
| PNMA8A | 2550.232377 | -0.188178521 | 2.47E-03 |
| RABGEF1 | 2589.467863 | -0.188625002 | 2.98E-02 |
| SYVN1 | 2776.979789 | -0.189618356 | 3.37E-02 |
| TSPAN3 | 7268.419862 | -0.189706717 | 4.64E-02 |
| LSM14B | 3273.518921 | -0.190770438 | 1.34E-02 |
| RYK | 3166.723865 | -0.190793615 | 2.53E-02 |
| CENPT | 2443.523131 | -0.191152399 | 2.62E-02 |
| PBX2 | 4801.39461 | -0.19167812 | 4.28E-02 |

| | | | |
|---|---|---|---|
| DDRGK1 | 1823.048264 | -0.192197225 | 2.23E-02 |
| HDDC2 | 1891.097272 | -0.192334296 | 1.01E-02 |
| RPL7 | 41042.59646 | -0.194633395 | 2.76E-02 |
| PTRH2 | 1318.331553 | -0.194734724 | 2.97E-02 |
| MTERF4 | 1692.32588 | -0.195861811 | 2.24E-02 |
| PPIL2 | 3565.270233 | -0.196268192 | 3.39E-02 |
| DIP2A | 2335.099636 | -0.196960577 | 4.11E-02 |
| PWWP3A | 2910.551855 | -0.197067128 | 9.69E-03 |
| FOXP1 | 826.7685641 | -0.198240458 | 4.92E-02 |
| KDM5B | 4458.455445 | -0.198854638 | 4.56E-02 |
| YTHDF2 | 3194.209555 | -0.199003692 | 1.44E-02 |
| GATAD2B | 1645.344527 | -0.19924511 | 1.81E-02 |
| EEF1AKNMT | 2434.565317 | -0.20027099 | 4.91E-02 |
| ZNF335 | 1385.685245 | -0.200729708 | 3.33E-02 |
| MTG2 | 2081.396989 | -0.201134812 | 4.33E-02 |
| DERA | 1351.703334 | -0.201352693 | 2.19E-02 |
| ISG20L2 | 2175.728689 | -0.201471787 | 4.62E-02 |
| CCDC97 | 2026.245674 | -0.201910331 | 2.34E-02 |
| HNRNPA3 | 17215.90036 | -0.202636876 | 1.65E-02 |
| SIPA1L1 | 3181.582534 | -0.202807739 | 1.48E-02 |
| CLTC | 25848.11783 | -0.204792695 | 5.13E-03 |
| MTHFSD | 1180.958953 | -0.205706015 | 1.19E-02 |
| ERCC5 | 3744.110558 | -0.207034824 | 2.18E-02 |
| SYPL1 | 4385.788923 | -0.207086789 | 5.77E-03 |
| PHC1 | 2766.398805 | -0.207356673 | 1.24E-02 |
| NKAPD1 | 3040.424315 | -0.207906195 | 4.74E-02 |
| BAZ1B | 12077.77564 | -0.208385713 | 1.54E-02 |
| INTS14 | 3408.072776 | -0.210899324 | 8.92E-04 |
| RAB13 | 2951.583054 | -0.211270326 | 3.40E-02 |
| DDX25 | 1351.778353 | -0.21348286 | 2.34E-02 |
| C9orf78 | 3511.313378 | -0.214149019 | 2.10E-04 |
| PCED1A | 3718.966878 | -0.214872424 | 7.03E-03 |
| ASL | 1647.09413 | -0.215029657 | 5.00E-02 |
| VPS41 | 3381.596704 | -0.215947397 | 1.27E-02 |
| CRAMP1 | 1819.133989 | -0.217261857 | 2.74E-02 |
| ATF7IP | 4087.275155 | -0.217907536 | 3.94E-02 |
| AC026412.1 | 1148.852394 | -0.217937269 | 4.41E-02 |
| ANKS1A | 3891.174726 | -0.218291617 | 4.83E-02 |
| KIF13A | 2692.151826 | -0.218925389 | 3.00E-02 |
| TIMM8A | 916.3763145 | -0.218936534 | 2.90E-02 |
| AP1G1 | 6782.94064 | -0.21973202 | 6.77E-03 |
| CAST | 16712.96545 | -0.219948045 | 2.49E-04 |
| MIS12 | 1596.761517 | -0.219972572 | 2.80E-02 |
| MTHFS | 1303.668712 | -0.220546239 | 1.25E-02 |
| MLLT1 | 5500.18598 | -0.220658567 | 1.32E-02 |
| UBQLN2 | 1858.80665 | -0.221148968 | 1.68E-03 |
| RBL2 | 4576.824447 | -0.221565419 | 2.56E-02 |

| | | | |
|---|---|---|---|
| ZNF276 | 1389.475469 | -0.221810548 | 3.34E-02 |
| NEU3 | 1544.659068 | -0.222762127 | 6.12E-03 |
| SF3B3 | 19040.42579 | -0.223822694 | 3.06E-02 |
| UTP25 | 1455.765387 | -0.224032896 | 4.03E-03 |
| ERAP1 | 6025.539569 | -0.224729106 | 4.42E-02 |
| PUM1 | 5310.868167 | -0.224945538 | 7.52E-04 |
| MAPRE2 | 4609.080376 | -0.225297035 | 1.71E-03 |
| VPS4B | 1727.894412 | -0.225522358 | 1.16E-02 |
| FBRS | 3452.531189 | -0.225665773 | 4.49E-02 |
| MSL3 | 1482.073196 | -0.225762458 | 9.83E-03 |
| PRRC2C | 9875.15191 | -0.226647169 | 2.40E-02 |
| THADA | 2754.68274 | -0.22677932 | 1.09E-02 |
| PWWP2A | 1799.661611 | -0.226820279 | 3.43E-02 |
| RERE | 5375.12534 | -0.228059066 | 5.31E-03 |
| IVD | 2392.483779 | -0.229184181 | 3.77E-02 |
| MTDH | 5816.793673 | -0.229236017 | 4.43E-02 |
| STX6 | 1556.556609 | -0.229366518 | 9.91E-04 |
| ATP6V1A | 3354.193021 | -0.229434488 | 2.61E-02 |
| RBM6 | 3881.850496 | -0.229450344 | 3.16E-02 |
| MAML1 | 2593.422735 | -0.229765667 | 4.22E-03 |
| TIGD1 | 735.47436 | -0.230187013 | 3.42E-02 |
| CBX6 | 5530.008558 | -0.23022371 | 3.95E-02 |
| KIAA0319L | 6294.756255 | -0.23057703 | 3.07E-02 |
| PCNX3 | 5371.421113 | -0.230915438 | 1.14E-02 |
| SPCS2 | 3572.822167 | -0.231496512 | 2.22E-02 |
| USP42 | 1231.999039 | -0.231835412 | 9.61E-03 |
| TM9SF1 | 3915.379062 | -0.232165327 | 4.66E-02 |
| CYP20A1 | 839.5415502 | -0.233058634 | 4.74E-02 |
| HNRNPA1 | 38484.91909 | -0.233090422 | 4.99E-03 |
| CASK | 2190.921557 | -0.233311055 | 3.89E-02 |
| SARAF | 7191.114511 | -0.23368088 | 1.03E-03 |
| SP140L | 1768.433108 | -0.233879307 | 6.92E-03 |
| TOR1B | 3507.737833 | -0.234030146 | 4.83E-03 |
| PHYH | 939.2769998 | -0.235362025 | 1.86E-02 |
| BRD3 | 2610.24856 | -0.236641578 | 1.16E-02 |
| PABPN1 | 6487.789511 | -0.236702858 | 4.95E-03 |
| GATD1 | 2190.233517 | -0.236711353 | 3.46E-02 |
| TMEM39B | 1129.7264 | -0.23753853 | 3.43E-02 |
| DCTN4 | 3405.87974 | -0.237623991 | 2.04E-02 |
| MAPK1IP1L | 3707.931357 | -0.237773403 | 1.93E-02 |
| ZCCHC14 | 2165.42474 | -0.237783455 | 7.04E-03 |
| SH3PXD2B | 1746.191447 | -0.238091694 | 3.30E-02 |
| LMBR1 | 4188.33573 | -0.2383224 | 3.70E-02 |
| CBX5 | 9249.567894 | -0.238376996 | 4.01E-04 |
| RCE1 | 673.4495861 | -0.238653331 | 2.27E-02 |
| RABL2B | 1249.900016 | -0.23871722 | 1.20E-02 |
| LIX1L | 2287.312237 | -0.238735576 | 1.14E-02 |

| | | | |
|---|---|---|---|
| STOM | 8123.48438 | -0.238974626 | 4.00E-03 |
| C11orf49 | 1568.56911 | -0.239353653 | 2.43E-02 |
| ATG14 | 977.1180919 | -0.239564085 | 4.62E-03 |
| KDM2A | 7320.875344 | -0.239834022 | 1.90E-02 |
| PRKACA | 4284.940824 | -0.239976152 | 2.51E-02 |
| HNRNPLL | 2217.846786 | -0.240157416 | 7.86E-03 |
| MAGEA11 | 992.8345221 | -0.240484941 | 1.42E-02 |
| UTP3 | 1873.104973 | -0.240964171 | 1.39E-02 |
| TNPO2 | 4295.059421 | -0.241473602 | 1.07E-02 |
| NDRG3 | 3229.681963 | -0.241479557 | 2.79E-02 |
| ATF5 | 1389.524915 | -0.241702507 | 2.50E-02 |
| DLD | 4716.307916 | -0.241800376 | 4.99E-02 |
| C2orf68 | 2322.099413 | -0.241976887 | 1.69E-05 |
| OSGEP | 996.6215002 | -0.242274404 | 3.97E-02 |
| CHERP | 3014.334714 | -0.242556868 | 1.45E-02 |
| FAM210B | 3172.578834 | -0.242794493 | 8.50E-04 |
| CAPRIN1 | 19543.43926 | -0.243603717 | 2.15E-07 |
| EZH1 | 1512.234473 | -0.244530779 | 5.44E-03 |
| RBMXL1 | 1282.120288 | -0.244601456 | 4.77E-02 |
| ABI1 | 1233.020187 | -0.245138236 | 2.32E-03 |
| EYA3 | 2181.355695 | -0.245611243 | 6.61E-03 |
| FGD5-AS1 | 5228.2829 | -0.24597422 | 1.22E-02 |
| SPPL2A | 1140.966053 | -0.246153629 | 4.03E-02 |
| BTBD1 | 3723.853847 | -0.246192556 | 4.34E-02 |
| BCDIN3D | 426.0239269 | -0.246340349 | 4.79E-02 |
| CLCC1 | 1194.714376 | -0.247041437 | 2.27E-02 |
| ACAT1 | 4651.28969 | -0.248023927 | 5.95E-03 |
| COQ9 | 2727.947584 | -0.24840426 | 1.16E-02 |
| THUMPD3 | 1703.170376 | -0.248544064 | 2.75E-02 |
| CPT2 | 1235.342638 | -0.248616976 | 3.83E-02 |
| STMP1 | 2351.565466 | -0.24916718 | 2.42E-02 |
| SUPT5H | 6827.373794 | -0.249663145 | 1.18E-02 |
| DNAJA1 | 17897.86696 | -0.249747759 | 1.20E-02 |
| RIPK1 | 2531.004431 | -0.250477491 | 3.28E-02 |
| MAP3K7 | 3642.665461 | -0.250539339 | 4.96E-02 |
| SLC50A1 | 2499.022849 | -0.250623288 | 4.09E-02 |
| ZFP36L2 | 1694.899588 | -0.250930154 | 2.02E-02 |
| SUB1 | 7877.607387 | -0.251156347 | 1.43E-02 |
| ANXA4 | 2729.466506 | -0.251550234 | 4.21E-03 |
| TSC22D4 | 4809.816065 | -0.252015167 | 2.24E-02 |
| SPPL2B | 1636.596338 | -0.252346223 | 1.30E-02 |
| GPAA1 | 6075.963465 | -0.25305215 | 3.48E-02 |
| CCNT1 | 2674.828968 | -0.253204665 | 3.70E-02 |
| TMEM59 | 6590.074609 | -0.253553518 | 4.30E-03 |
| GON4L | 2736.602273 | -0.254290711 | 2.18E-03 |
| ARL17A | 1881.979378 | -0.254688032 | 1.14E-02 |
| PPM1A | 1396.887455 | -0.25491193 | 3.11E-02 |

| | | | |
|---|---|---|---|
| SLC23A2 | 3962.862943 | -0.255132873 | 2.54E-04 |
| MTPN | 15377.19465 | -0.255468033 | 1.29E-03 |
| TXNDC17 | 2660.628943 | -0.255854312 | 2.00E-02 |
| KIF24 | 939.5837656 | -0.256016641 | 3.01E-02 |
| DCAF6 | 2454.745029 | -0.256028335 | 4.40E-02 |
| EPHB2 | 2544.09388 | -0.256161908 | 2.52E-02 |
| STEAP1B | 969.6948838 | -0.256413123 | 2.08E-02 |
| VIRMA | 5473.931037 | -0.25654689 | 2.09E-03 |
| SMAD5 | 6225.784563 | -0.257319041 | 2.38E-02 |
| STAG3L3 | 1698.480032 | -0.2573614 | 1.26E-02 |
| CELF1 | 6881.807579 | -0.257886969 | 1.58E-02 |
| GSS | 4871.01293 | -0.258328312 | 2.47E-02 |
| DCAF16 | 2142.847711 | -0.258472182 | 2.10E-02 |
| ZNF316 | 2117.898613 | -0.258550548 | 1.26E-02 |
| TEP1 | 3548.594867 | -0.258558108 | 4.98E-02 |
| DHX9 | 14124.10549 | -0.259907035 | 1.97E-03 |
| PHF21A | 2545.783277 | -0.26059893 | 5.59E-03 |
| WASH2P | 725.33649 | -0.261355664 | 3.28E-02 |
| RALGAPA2 | 2680.8069 | -0.26184015 | 2.77E-02 |
| INTS7 | 1984.401887 | -0.261921682 | 3.15E-02 |
| ANKMY2 | 1123.608775 | -0.262013995 | 1.22E-02 |
| STAG3L2 | 795.2782283 | -0.26234498 | 1.06E-02 |
| KAT14 | 1554.577656 | -0.262678796 | 1.94E-03 |
| RPS6KC1 | 2016.543957 | -0.263285318 | 1.54E-03 |
| ZNF200 | 1367.270175 | -0.263675127 | 1.79E-03 |
| MSH3 | 1602.672514 | -0.263720255 | 1.81E-03 |
| CASP3 | 1463.757118 | -0.263859277 | 1.47E-02 |
| WASHC5 | 4004.657705 | -0.263972466 | 5.42E-03 |
| LIMK2 | 1150.8064 | -0.264257634 | 3.77E-02 |
| XXYLT1 | 1532.304719 | -0.264792978 | 2.18E-02 |
| ZNF623 | 2360.212587 | -0.264814218 | 2.12E-02 |
| NPR2 | 2835.322934 | -0.265646376 | 2.48E-02 |
| BICRAL | 968.5111093 | -0.265752612 | 3.15E-02 |
| METTL9 | 4952.663648 | -0.265801473 | 2.23E-04 |
| C6orf62 | 6945.794868 | -0.266564326 | 3.38E-03 |
| RNF103 | 872.356071 | -0.266706072 | 1.16E-02 |
| SOX12 | 4342.362057 | -0.268085631 | 1.20E-02 |
| PSEN1 | 3588.470478 | -0.268250604 | 1.33E-03 |
| ZNF398 | 1158.971923 | -0.268369286 | 1.49E-03 |
| ATP5MD | 1693.401647 | -0.268416149 | 9.80E-03 |
| CNNM3 | 926.6017936 | -0.268542284 | 1.63E-02 |
| NRSN2 | 5123.912099 | -0.269558471 | 2.47E-02 |
| UBTF | 5711.417987 | -0.269626155 | 4.24E-04 |
| RBPJ | 2693.98748 | -0.27022024 | 1.49E-02 |
| ZC3H7A | 3187.794361 | -0.270611505 | 2.25E-02 |
| SNAPC4 | 1363.319501 | -0.270998762 | 9.11E-03 |
| PIGX | 1133.959941 | -0.271015818 | 2.34E-02 |

| | | | |
|---|---|---|---|
| MESD | 4307.097829 | -0.272060207 | 2.44E-03 |
| MAPK8IP3 | 3358.91059 | -0.272319945 | 1.01E-02 |
| G3BP2 | 7950.497687 | -0.273123963 | 1.84E-03 |
| QDPR | 2166.566869 | -0.273174374 | 2.13E-02 |
| SNHG20 | 472.8352944 | -0.273623414 | 1.58E-02 |
| USP33 | 3250.439962 | -0.274115692 | 3.88E-02 |
| ZSCAN22 | 377.5474583 | -0.274159084 | 2.63E-02 |
| AFF4 | 7606.913083 | -0.274379779 | 3.11E-02 |
| ARMCX2 | 878.4276883 | -0.275030779 | 1.35E-02 |
| POGLUT1 | 1622.036049 | -0.275101971 | 3.80E-03 |
| ZNF542P | 1000.738491 | -0.275634977 | 1.67E-02 |
| SSH2 | 1653.105872 | -0.27616707 | 2.12E-02 |
| GEMIN2 | 718.8377862 | -0.276953345 | 3.16E-02 |
| DERL2 | 1918.396919 | -0.277274831 | 4.98E-04 |
| C1GALT1C1 | 890.1808052 | -0.277318763 | 1.01E-02 |
| KDM5A | 4501.713184 | -0.277407327 | 4.84E-04 |
| ZNF7 | 1564.127802 | -0.277419015 | 1.67E-04 |
| RPRD2 | 2318.718644 | -0.277880668 | 1.09E-05 |
| SARNP | 2010.285177 | -0.278250454 | 2.41E-03 |
| TMC6 | 4347.104689 | -0.278531111 | 3.17E-03 |
| ZNF789 | 838.5068154 | -0.278697118 | 1.48E-03 |
| TEX2 | 2687.825769 | -0.278701551 | 2.03E-04 |
| AC074117.1 | 550.5267816 | -0.278859977 | 3.66E-02 |
| ANKRD13D | 1126.817695 | -0.278861218 | 2.86E-02 |
| AC110079.1 | 522.6979601 | -0.279036925 | 3.94E-02 |
| ADAT1 | 2374.903336 | -0.279846274 | 5.91E-04 |
| UGDH | 1488.604689 | -0.280552505 | 3.05E-02 |
| TK2 | 1898.599849 | -0.281088255 | 1.46E-02 |
| MRPL32 | 2091.698119 | -0.281130969 | 1.43E-03 |
| ZBTB1 | 1574.403671 | -0.281850785 | 4.12E-02 |
| C1RL | 2143.771862 | -0.282177743 | 3.79E-02 |
| STAT6 | 5677.333921 | -0.282596117 | 3.13E-02 |
| CHD8 | 7116.522774 | -0.282677999 | 3.16E-03 |
| ZNF445 | 2350.177662 | -0.28279453 | 5.42E-04 |
| MRI1 | 1491.264152 | -0.28350159 | 1.10E-02 |
| SOCS6 | 907.8110569 | -0.284139267 | 1.00E-02 |
| AC006001.3 | 380.0272958 | -0.284169068 | 4.92E-02 |
| CCDC125 | 1183.595849 | -0.284399784 | 1.62E-02 |
| TMEM184C | 1244.756484 | -0.284503472 | 2.32E-02 |
| ZNF217 | 4777.839968 | -0.284547502 | 3.10E-03 |
| PDZD4 | 1438.460956 | -0.28462586 | 2.38E-02 |
| PNMA1 | 2633.572079 | -0.284652519 | 1.36E-04 |
| PIP4K2A | 1913.589145 | -0.284916124 | 1.54E-02 |
| AC026471.1 | 476.0324479 | -0.284963872 | 9.90E-03 |
| CRIP2 | 4276.416024 | -0.285414202 | 4.43E-02 |
| CD302 | 1041.566279 | -0.285803258 | 8.37E-03 |
| ANKRD42 | 682.3122319 | -0.285902085 | 2.00E-02 |

| | | | |
|---|---|---|---|
| BBS9 | 1146.378736 | -0.285967213 | 2.15E-03 |
| NF1 | 6160.278563 | -0.286287338 | 2.01E-02 |
| MAP3K3 | 2379.952131 | -0.286360888 | 2.30E-02 |
| CCDC120 | 662.5532422 | -0.286399804 | 1.08E-02 |
| PARP1 | 12306.40327 | -0.286410375 | 3.03E-02 |
| OXCT1 | 3184.337989 | -0.286957545 | 1.88E-02 |
| SERF1B | 1933.088998 | -0.287757212 | 1.84E-04 |
| COPS5 | 3660.217827 | -0.287788721 | 2.76E-05 |
| SLX4 | 472.5174403 | -0.287805581 | 3.93E-02 |
| KRBOX1 | 1196.380718 | -0.287865531 | 1.89E-02 |
| COA7 | 1675.441611 | -0.28839227 | 4.26E-02 |
| SV2A | 2278.471485 | -0.288869386 | 3.71E-03 |
| HERC1 | 6565.165456 | -0.290020053 | 2.99E-02 |
| RNF38 | 1806.265725 | -0.290691232 | 1.09E-03 |
| LRRC14 | 2220.494375 | -0.290787528 | 1.18E-02 |
| SLC39A6 | 2723.388463 | -0.290804961 | 1.65E-02 |
| TRIM26 | 1999.723817 | -0.291303696 | 1.53E-02 |
| DYNLL2 | 4327.120161 | -0.291432017 | 9.12E-04 |
| CDK13 | 3103.381094 | -0.291792599 | 4.94E-03 |
| BBS4 | 1662.859635 | -0.292422806 | 1.35E-02 |
| ARFGAP2 | 4523.910026 | -0.292803063 | 1.33E-02 |
| MTMR2 | 2934.886144 | -0.292951282 | 1.12E-05 |
| FBXO11 | 2388.76673 | -0.292958867 | 1.74E-02 |
| TPCN1 | 4974.758491 | -0.293262868 | 2.02E-03 |
| OSGEPL1 | 614.0173471 | -0.293484728 | 2.95E-02 |
| PTPRS | 6222.712973 | -0.29376236 | 1.03E-02 |
| NID1 | 12845.3558 | -0.294138542 | 3.28E-04 |
| MAML2 | 2592.591056 | -0.294391061 | 1.81E-02 |
| DROSHA | 5892.566378 | -0.294455051 | 3.45E-04 |
| DNAJC3 | 5598.113646 | -0.294469936 | 3.97E-03 |
| TMEM181 | 2526.062735 | -0.294814026 | 1.92E-02 |
| YBEY | 319.8980182 | -0.294872616 | 2.81E-02 |
| NA | 707.1082302 | -0.29572507 | 8.00E-03 |
| PAPOLG | 812.3893065 | -0.296417868 | 4.64E-02 |
| ALDH3A2 | 1982.806079 | -0.296465348 | 1.89E-03 |
| HELQ | 538.5349663 | -0.296765684 | 3.43E-02 |
| ICE2 | 2369.142308 | -0.296932511 | 4.20E-02 |
| PRMT2 | 3752.41591 | -0.297050111 | 1.84E-02 |
| RAB18 | 819.3678411 | -0.297082977 | 1.34E-02 |
| UTP20 | 2908.602101 | -0.297326791 | 1.95E-02 |
| ZKSCAN1 | 6233.227887 | -0.297919115 | 1.37E-02 |
| RHNO1 | 1266.859032 | -0.297932052 | 1.14E-02 |
| FAM122C | 495.1261315 | -0.297932181 | 4.69E-02 |
| SCAF8 | 2710.998518 | -0.298602717 | 2.95E-06 |
| ARMC9 | 2187.710154 | -0.298870728 | 3.80E-03 |
| RETREG3 | 2208.190505 | -0.298890896 | 3.93E-02 |
| CCNL2 | 6146.069744 | -0.298897736 | 9.06E-08 |

| | | | |
|---|---|---|---|
| IFFO1 | 1580.185706 | -0.299047888 | 1.88E-02 |
| VCPIP1 | 1187.72626 | -0.2993169 | 3.53E-02 |
| PIDD1 | 1290.180358 | -0.299865527 | 6.43E-03 |
| CAAP1 | 1071.16947 | -0.299934197 | 3.84E-02 |
| LYRM2 | 2767.812549 | -0.299935047 | 3.29E-03 |
| SON | 15233.18633 | -0.299958073 | 6.32E-03 |
| MINDY3 | 690.3990073 | -0.300298756 | 2.44E-02 |
| TTC31 | 2117.998023 | -0.300792844 | 1.30E-03 |
| PCNT | 3658.598164 | -0.300796474 | 6.17E-03 |
| EDA | 1795.900129 | -0.301542002 | 2.99E-02 |
| TAPBP | 6637.985953 | -0.301550577 | 2.45E-02 |
| RCC1L | 4660.194879 | -0.301889909 | 2.06E-03 |
| ARID1B | 3578.147371 | -0.302278691 | 7.11E-08 |
| RBM4 | 5211.12219 | -0.302476964 | 4.44E-03 |
| ATP2B4 | 5211.832201 | -0.302496597 | 4.95E-03 |
| THG1L | 1892.442873 | -0.303217559 | 4.40E-03 |
| ZNF880 | 1055.217913 | -0.303395498 | 1.48E-02 |
| PHKG2 | 1614.940655 | -0.303397091 | 1.94E-02 |
| SETDB1 | 2138.132053 | -0.303576142 | 5.89E-03 |
| WDR75 | 2797.494185 | -0.303803132 | 9.33E-03 |
| CDC40 | 1575.43891 | -0.304334094 | 1.08E-02 |
| ZNF554 | 237.5549339 | -0.304684046 | 4.16E-02 |
| LINC01116 | 674.8071079 | -0.304919225 | 3.04E-02 |
| ZSCAN30 | 924.1394521 | -0.305537389 | 2.57E-02 |
| CREBBP | 3069.964363 | -0.306589734 | 3.20E-02 |
| ZNF512B | 4994.652742 | -0.306840626 | 4.78E-04 |
| SCAMP1-AS1 | 798.8270424 | -0.306921686 | 1.90E-03 |
| AC026470.1 | 2189.407246 | -0.307021336 | 2.43E-02 |
| SP110 | 1100.571643 | -0.307242176 | 6.36E-04 |
| TFB2M | 842.842429 | -0.30744774 | 3.36E-02 |
| DCAKD | 1106.363567 | -0.307465107 | 1.22E-02 |
| ADAMTS7 | 1523.351905 | -0.307521774 | 2.24E-02 |
| VEZT | 1666.310259 | -0.307523383 | 1.43E-02 |
| MOB3C | 776.5128063 | -0.307679459 | 4.84E-02 |
| MTMR4 | 2791.918713 | -0.307943413 | 2.32E-03 |
| ECD | 1457.84952 | -0.308366513 | 6.61E-05 |
| PPP1R10 | 3168.508356 | -0.309302922 | 2.30E-02 |
| NAA40 | 3343.755419 | -0.309434475 | 1.96E-07 |
| NIFK-AS1 | 296.8162705 | -0.309588705 | 4.40E-02 |
| SAP130 | 2350.290221 | -0.309723953 | 6.45E-04 |
| COMMD4 | 2781.957671 | -0.309740294 | 2.50E-02 |
| DHX57 | 2397.342818 | -0.309835032 | 1.88E-04 |
| LRCH1 | 1532.93364 | -0.309852978 | 1.18E-02 |
| NSL1 | 1006.639425 | -0.309861736 | 1.02E-02 |
| MRFAP1L1 | 2334.33261 | -0.309986773 | 3.83E-03 |
| HCFC1 | 8795.74916 | -0.310016055 | 1.14E-02 |
| C8orf82 | 970.646144 | -0.31018047 | 3.35E-02 |

| | | | |
|---|---|---|---|
| ALX4 | 946.9188903 | -0.310314873 | 2.74E-02 |
| COQ4 | 1733.716167 | -0.310316915 | 1.33E-03 |
| HARS2 | 2149.232641 | -0.31039131 | 2.33E-06 |
| LATS1 | 1070.227587 | -0.310817352 | 3.60E-03 |
| TRUB2 | 3441.465998 | -0.31133141 | 2.59E-02 |
| NAGK | 1852.588991 | -0.311508581 | 2.34E-02 |
| TMBIM4 | 1236.621915 | -0.311591927 | 3.83E-03 |
| ELOVL2 | 1985.730308 | -0.311621927 | 1.37E-02 |
| DCAF5 | 3256.108903 | -0.311734405 | 5.77E-04 |
| PHF12 | 1757.024728 | -0.311796959 | 2.72E-02 |
| PTOV1-AS1 | 556.2368947 | -0.312389562 | 2.78E-02 |
| FDX1 | 969.3216053 | -0.31307114 | 3.18E-03 |
| SARDH | 781.3490579 | -0.313157891 | 4.29E-02 |
| NIP7 | 3651.687745 | -0.313463814 | 2.46E-03 |
| LRCH3 | 2423.263043 | -0.31421008 | 2.76E-02 |
| AC018647.2 | 763.6559716 | -0.314507473 | 2.01E-03 |
| CNOT4 | 1183.783448 | -0.314566681 | 9.65E-05 |
| CNPY3 | 3245.507656 | -0.315083411 | 3.38E-02 |
| TMED7 | 4176.355682 | -0.31533661 | 1.37E-02 |
| RETREG2 | 4730.599494 | -0.315801773 | 1.56E-02 |
| JHY | 621.7022963 | -0.31585589 | 9.12E-03 |
| NOP14-AS1 | 749.5561467 | -0.316178546 | 4.81E-02 |
| BACE1 | 6341.742065 | -0.316438877 | 8.88E-04 |
| SELENOI | 2088.637171 | -0.316663156 | 2.00E-02 |
| NSF | 4379.616314 | -0.316829279 | 1.89E-06 |
| CDK12 | 4237.691112 | -0.316883012 | 1.35E-05 |
| NUDT16L1 | 643.9989348 | -0.316992559 | 4.74E-02 |
| AC046185.3 | 1008.471374 | -0.317637291 | 6.69E-04 |
| ZNF780B | 879.49183 | -0.31798083 | 4.92E-02 |
| WDR48 | 2718.702698 | -0.318088306 | 3.56E-04 |
| MTRF1 | 418.9538556 | -0.318490108 | 3.01E-02 |
| INIP | 1067.234987 | -0.318696313 | 2.59E-03 |
| SLC30A6 | 1921.760547 | -0.318717112 | 5.21E-03 |
| BIVM | 3286.082284 | -0.318834386 | 4.06E-02 |
| RABEP1 | 5406.597359 | -0.318871702 | 2.37E-02 |
| ARAP3 | 3262.492922 | -0.318895446 | 7.34E-03 |
| SPOUT1 | 3020.906239 | -0.318944347 | 1.86E-02 |
| FANCL | 998.7763732 | -0.319476978 | 7.13E-03 |
| KIAA1549 | 2899.370619 | -0.319773893 | 2.23E-05 |
| PPP4R1L | 870.1846927 | -0.320346061 | 2.22E-03 |
| POLG2 | 631.5605938 | -0.320359167 | 3.13E-02 |
| EHBP1L1 | 5588.830947 | -0.320378272 | 3.47E-02 |
| SEC14L2 | 627.0691967 | -0.320445084 | 8.53E-03 |
| RNF14 | 1986.791575 | -0.320538744 | 2.82E-03 |
| ATMIN | 3490.625369 | -0.321416762 | 3.57E-04 |
| HSDL1 | 1696.988844 | -0.321597481 | 6.56E-03 |
| TOMM40L | 1091.925039 | -0.321857082 | 3.27E-02 |

| | | | |
|---|---|---|---|
| PRCP | 7324.04715 | -0.322092776 | 1.69E-02 |
| SIX5 | 1314.827357 | -0.322590607 | 2.08E-02 |
| GEMIN6 | 1396.151903 | -0.322894433 | 1.94E-04 |
| CRK | 4662.264636 | -0.323304846 | 4.90E-04 |
| GTF3C4 | 5054.525674 | -0.323453935 | 1.66E-06 |
| PLEKHO2 | 2012.556267 | -0.323535142 | 3.53E-02 |
| WTAP | 2928.312467 | -0.323581809 | 1.21E-05 |
| NKRF | 953.9954924 | -0.32373381 | 4.01E-03 |
| GGPS1 | 1335.938822 | -0.323895854 | 5.05E-03 |
| DYRK1A | 2154.694965 | -0.323941561 | 5.67E-03 |
| TRIAP1 | 1088.600773 | -0.324038661 | 4.18E-03 |
| KAT2A | 3508.866278 | -0.324153367 | 1.59E-04 |
| ZMIZ1 | 3413.461136 | -0.324211931 | 2.92E-04 |
| GOLIM4 | 1524.415909 | -0.324322341 | 4.07E-02 |
| KIFC2 | 1596.31756 | -0.324446571 | 3.15E-02 |
| SPG7 | 3626.18426 | -0.324608871 | 4.63E-03 |
| MCCC2 | 8099.333828 | -0.32465086 | 1.91E-08 |
| CMPK1 | 2535.089272 | -0.324657999 | 2.42E-02 |
| TRAF5 | 901.765231 | -0.325475974 | 1.89E-02 |
| AIDA | 4089.78514 | -0.32561808 | 4.02E-03 |
| THTPA | 987.8526175 | -0.325809686 | 4.94E-02 |
| CROCCP2 | 636.4083808 | -0.325831436 | 4.40E-03 |
| HOXD4 | 312.6919681 | -0.325850045 | 2.26E-02 |
| PLEKHG5 | 1159.440594 | -0.326689638 | 1.26E-02 |
| NHEJ1 | 722.382986 | -0.326996974 | 3.73E-02 |
| MAPK7 | 1119.071792 | -0.327127989 | 3.31E-02 |
| CDC73 | 1412.75039 | -0.327211229 | 3.91E-02 |
| ARSA | 1347.401683 | -0.327279839 | 1.67E-02 |
| DCAF12 | 4233.100473 | -0.32742849 | 1.55E-05 |
| SUZ12P1 | 346.4032018 | -0.327520589 | 2.37E-02 |
| USP20 | 2227.906697 | -0.327605866 | 3.91E-03 |
| SLC38A9 | 1524.406775 | -0.327726088 | 5.87E-03 |
| PPP1R16A | 1407.164711 | -0.328072554 | 2.38E-02 |
| CDK5 | 1685.224274 | -0.329083353 | 1.95E-02 |
| C6orf47 | 1110.798322 | -0.329177219 | 4.04E-03 |
| TP53RK | 1247.742406 | -0.329547883 | 9.30E-03 |
| NCOA5 | 2667.247924 | -0.329559253 | 9.69E-04 |
| UBR3 | 2018.764591 | -0.329844445 | 3.20E-02 |
| PLPP5 | 1498.63886 | -0.330079326 | 2.08E-04 |
| HCN2 | 777.201268 | -0.330195973 | 1.76E-02 |
| TMEM234 | 454.68688 | -0.330618389 | 1.14E-02 |
| SLC35E2A | 570.8039971 | -0.330672338 | 4.85E-03 |
| ZNF619 | 453.2585032 | -0.3313859 | 1.47E-02 |
| SETD1A | 2873.426182 | -0.331420839 | 5.36E-03 |
| PKNOX1 | 1143.533081 | -0.331451121 | 6.79E-03 |
| KDELC1 | 2977.176079 | -0.332088438 | 2.61E-03 |
| ANAPC13 | 1919.232172 | -0.332108585 | 9.44E-04 |

| | | | |
|---|---|---|---|
| FXN | 656.6777015 | -0.332520708 | 3.38E-02 |
| RNF139 | 1481.985634 | -0.332619539 | 2.41E-04 |
| ABCC5 | 1812.770361 | -0.33280707 | 2.85E-05 |
| CTCF | 3014.64559 | -0.333008519 | 8.11E-07 |
| TNFRSF10A | 429.7567669 | -0.333295862 | 4.94E-02 |
| DHFR2 | 351.7586684 | -0.333315901 | 4.55E-02 |
| CHST15 | 1086.666004 | -0.333340324 | 1.27E-02 |
| PRPS2 | 2404.952892 | -0.333554917 | 1.98E-02 |
| UHMK1 | 2866.460124 | -0.33401869 | 2.95E-02 |
| WASHC2C | 1179.143455 | -0.334030275 | 1.33E-03 |
| VGLL4 | 1747.13902 | -0.334142349 | 1.69E-02 |
| TM2D3 | 1237.924129 | -0.33418312 | 6.32E-04 |
| STARD9 | 896.3127905 | -0.334404934 | 2.49E-02 |
| EIF3E | 11249.5084 | -0.334494261 | 1.13E-02 |
| NA | 432.2714357 | -0.334877061 | 3.97E-03 |
| GABBR1 | 1499.290205 | -0.33495325 | 3.12E-04 |
| ARHGEF25 | 2075.177639 | -0.334990896 | 2.14E-02 |
| PRUNE1 | 1101.991682 | -0.336914981 | 2.63E-03 |
| IPO4 | 3946.715919 | -0.336955125 | 9.12E-04 |
| CDKN2AIP | 624.1619595 | -0.337007182 | 1.92E-02 |
| RAMP2 | 1068.842961 | -0.337173562 | 4.16E-02 |
| C12orf76 | 572.4469284 | -0.337454588 | 4.06E-03 |
| RMDN1 | 2329.178846 | -0.337581627 | 2.55E-04 |
| MLYCD | 694.324879 | -0.337666479 | 1.78E-02 |
| SCAMP1 | 2340.239497 | -0.337843762 | 6.66E-03 |
| BAG4 | 1113.912056 | -0.338022775 | 9.55E-04 |
| CSTB | 2993.214322 | -0.338184332 | 1.90E-02 |
| INTS8 | 2842.185022 | -0.338405788 | 3.13E-02 |
| ZNF324B | 402.6473517 | -0.338569573 | 1.26E-02 |
| BTG1 | 2988.2491 | -0.339054295 | 1.30E-02 |
| ASB8 | 972.3934283 | -0.339184714 | 3.64E-04 |
| CSTF2T | 1225.829139 | -0.339322963 | 1.48E-03 |
| GNAI3 | 4325.565096 | -0.339376903 | 1.24E-02 |
| WASF1 | 1399.641949 | -0.339561541 | 7.09E-03 |
| CHKB | 998.5152636 | -0.339836063 | 1.68E-02 |
| TRERF1 | 1637.405227 | -0.33990277 | 2.40E-02 |
| HOMER3 | 2148.505031 | -0.339971031 | 1.79E-02 |
| MICOS10 | 1821.045151 | -0.340620468 | 1.56E-04 |
| BAIAP2-DT | 1190.829781 | -0.341166812 | 1.60E-02 |
| DONSON | 1377.01341 | -0.341288947 | 3.92E-03 |
| SRPK2 | 3839.713398 | -0.341504592 | 9.65E-06 |
| QTRT1 | 1398.971228 | -0.342151877 | 2.77E-03 |
| FAM102A | 1821.808637 | -0.342211098 | 2.60E-02 |
| KIAA0040 | 1749.222652 | -0.342464405 | 1.82E-02 |
| RABGAP1L | 809.043815 | -0.342833291 | 4.62E-02 |
| ARHGAP39 | 505.0589739 | -0.342891047 | 3.88E-03 |
| ZNF79 | 367.4670001 | -0.34396792 | 3.43E-02 |

| | | | |
|---|---|---|---|
| SRRM2 | 25535.81099 | -0.344009611 | 5.44E-04 |
| VAT1L | 15308.2066 | -0.34407068 | 6.94E-03 |
| NUP50 | 3007.034293 | -0.344087648 | 5.75E-03 |
| SENP8 | 218.9915801 | -0.344387677 | 4.50E-02 |
| CDK2AP1 | 4145.313976 | -0.344433498 | 4.33E-06 |
| SRP14 | 6129.603975 | -0.34495154 | 1.75E-03 |
| MEX3A | 1174.644861 | -0.345438741 | 4.10E-02 |
| ZFP90 | 2128.91407 | -0.345515878 | 7.00E-03 |
| FKBP10 | 18208.3748 | -0.345827387 | 5.24E-03 |
| ELOA | 3907.96247 | -0.345873497 | 3.97E-09 |
| APP | 42894.66391 | -0.346300011 | 1.01E-04 |
| L3MBTL2 | 1878.053994 | -0.346814954 | 6.99E-04 |
| NPHP3 | 735.6109425 | -0.347160021 | 4.86E-02 |
| RBM23 | 4441.381134 | -0.347197857 | 4.33E-04 |
| AL513550.1 | 323.2667061 | -0.347440007 | 1.28E-02 |
| RCOR1 | 3100.077146 | -0.347501287 | 9.11E-03 |
| ANKEF1 | 484.1331775 | -0.347599496 | 1.40E-02 |
| RFX1 | 522.128581 | -0.347646588 | 1.76E-02 |
| RBBP4 | 6433.30477 | -0.347835113 | 3.14E-03 |
| PPP4R3B | 3051.126779 | -0.348002271 | 4.65E-02 |
| GTPBP8 | 382.465942 | -0.3480272 | 2.10E-02 |
| ZMPSTE24 | 1890.659318 | -0.34812842 | 1.39E-03 |
| NELFCD | 4856.196635 | -0.348348693 | 7.70E-03 |
| RAD9A | 799.115388 | -0.348362205 | 8.28E-03 |
| KCTD3 | 2461.517131 | -0.349378924 | 2.07E-02 |
| PDP1 | 2884.757114 | -0.349703513 | 1.49E-03 |
| AC009163.6 | 182.0607473 | -0.349760873 | 3.51E-02 |
| ANAPC4 | 975.3825386 | -0.349909982 | 4.32E-02 |
| RBBP6 | 2157.766372 | -0.350759871 | 4.18E-02 |
| CDC42EP4 | 3656.079881 | -0.350813834 | 3.51E-04 |
| NASP | 6492.894079 | -0.350867829 | 2.15E-02 |
| MAPK12 | 3334.757022 | -0.351405598 | 1.06E-03 |
| MICAL1 | 1878.449662 | -0.3515303 | 7.31E-04 |
| L3MBTL1 | 722.4876846 | -0.351779936 | 7.76E-03 |
| CRIPT | 678.109701 | -0.351833484 | 9.70E-04 |
| SLC25A44 | 2071.496953 | -0.351975689 | 1.46E-02 |
| FLVCR1-DT | 243.0547659 | -0.352064924 | 4.79E-02 |
| SLC25A22 | 2360.771682 | -0.352516154 | 2.16E-04 |
| MYCBP | 1295.260831 | -0.352674395 | 1.26E-02 |
| KCTD18 | 614.5891598 | -0.352823702 | 4.38E-03 |
| SMN2 | 2795.058623 | -0.352890971 | 6.82E-04 |
| ENKD1 | 782.8162869 | -0.35300003 | 3.77E-03 |
| ZNF428 | 892.0978553 | -0.353090815 | 9.86E-03 |
| AFMID | 694.8669109 | -0.353142795 | 1.19E-02 |
| C5orf15 | 4632.545109 | -0.353342134 | 1.32E-03 |
| MANBA | 1597.878041 | -0.353579064 | 5.20E-03 |
| TTC30A | 485.3279646 | -0.353647679 | 1.57E-02 |

| | | | |
|---|---|---|---|
| ORC5 | 1547.191739 | -0.353894088 | 1.81E-02 |
| SLX4IP | 516.7187441 | -0.353922563 | 1.35E-02 |
| DHX58 | 287.5296303 | -0.35423278 | 4.45E-02 |
| UBTD2 | 1371.08928 | -0.354662869 | 1.12E-07 |
| HOMER2 | 738.0435219 | -0.355608442 | 1.19E-02 |
| SEMA4C | 2019.090106 | -0.35567965 | 5.03E-04 |
| MAST3 | 693.4806052 | -0.355708628 | 5.49E-04 |
| PRAF2 | 2410.163933 | -0.355968783 | 4.49E-02 |
| TARBP2 | 1194.823027 | -0.356000374 | 2.30E-03 |
| HSF4 | 604.4274697 | -0.356384571 | 1.59E-02 |
| IKZF4 | 437.2775442 | -0.356490842 | 6.63E-03 |
| C11orf95 | 1680.071571 | -0.356552277 | 2.19E-02 |
| NRDE2 | 1127.583602 | -0.357030086 | 4.09E-03 |
| PHAX | 2311.510306 | -0.357045797 | 2.14E-02 |
| PEX13 | 727.2580527 | -0.357074479 | 3.82E-03 |
| HNRNPM | 10158.43398 | -0.357234375 | 3.70E-08 |
| RAVER2 | 998.6587245 | -0.357252058 | 7.16E-04 |
| TFRC | 9364.404424 | -0.357276853 | 1.73E-03 |
| ICK | 878.5792377 | -0.357784223 | 8.90E-03 |
| STMN3 | 8539.398862 | -0.357799792 | 6.35E-03 |
| ZNF891 | 383.1294511 | -0.358378573 | 3.37E-02 |
| ZFYVE1 | 1790.155802 | -0.359175367 | 2.37E-02 |
| ZNF266 | 2147.525112 | -0.359548182 | 4.97E-05 |
| OSBPL1A | 2509.624306 | -0.359549342 | 9.80E-07 |
| FAM92A | 1091.539897 | -0.359663173 | 3.31E-03 |
| TIMM21 | 780.7950079 | -0.359749735 | 4.90E-02 |
| ISCA1 | 1370.129141 | -0.359785534 | 7.79E-04 |
| MAN1A2 | 2382.046761 | -0.359789343 | 2.83E-02 |
| TCTEX1D2 | 229.6803877 | -0.359874452 | 2.94E-02 |
| RTF1 | 2306.221523 | -0.36001673 | 3.31E-03 |
| PARP10 | 2590.466427 | -0.360270502 | 4.49E-03 |
| SCP2 | 2504.721614 | -0.360661069 | 1.06E-02 |
| KLHDC2 | 2270.566987 | -0.36112535 | 2.27E-05 |
| MRPS21 | 2857.515873 | -0.361194787 | 9.20E-04 |
| C1R | 3793.027515 | -0.361513471 | 4.43E-02 |
| DDX6 | 6778.175847 | -0.361591849 | 7.33E-08 |
| CFAP36 | 1119.165321 | -0.361758598 | 2.52E-04 |
| AC010643.1 | 279.3700195 | -0.362450974 | 8.91E-03 |
| CREB3L4 | 569.1732261 | -0.362814169 | 3.58E-03 |
| FGFR1 | 4324.242565 | -0.362876667 | 2.03E-04 |
| FBN1 | 3258.584951 | -0.363334274 | 6.32E-03 |
| RSPRY1 | 1915.819271 | -0.363600619 | 8.59E-03 |
| HOXB7 | 1350.143893 | -0.363638803 | 1.48E-03 |
| ORAI2 | 6379.575777 | -0.363858229 | 3.29E-03 |
| TEX10 | 3490.266542 | -0.36393643 | 2.95E-11 |
| RLIM | 3157.917378 | -0.364251878 | 4.62E-02 |
| FAM133B | 963.4062515 | -0.364513108 | 2.31E-02 |

| | | | |
|---|---|---|---|
| FBXO46 | 574.6112734 | -0.364729229 | 1.01E-02 |
| MXD1 | 447.3520804 | -0.364813451 | 1.55E-02 |
| FUBP1 | 4556.3948 | -0.364991148 | 7.00E-04 |
| DNAJC16 | 1058.461335 | -0.365018661 | 8.17E-03 |
| PI4KAP1 | 543.6085889 | -0.365116452 | 2.22E-03 |
| PGBD1 | 568.7505738 | -0.365136741 | 3.53E-03 |
| MEGF9 | 1097.538604 | -0.365182751 | 1.73E-03 |
| RGL2 | 4766.753353 | -0.36552181 | 6.32E-03 |
| COX11 | 1314.57339 | -0.365578676 | 5.30E-03 |
| SLC35G1 | 581.9719221 | -0.36600541 | 2.10E-03 |
| PIGB | 459.0139734 | -0.366069137 | 1.21E-02 |
| GLT8D1 | 3116.387313 | -0.366427142 | 2.06E-03 |
| CHTF8 | 7617.256486 | -0.366551341 | 8.91E-04 |
| DDI2 | 1498.305752 | -0.36658153 | 2.50E-02 |
| SRCAP | 10671.7796 | -0.366614787 | 3.75E-10 |
| ZMYM4 | 4083.321672 | -0.366642885 | 2.04E-02 |
| CECR7 | 1422.204956 | -0.366792355 | 1.62E-03 |
| C5orf63 | 487.5369067 | -0.366853972 | 1.93E-02 |
| MED7 | 1176.800016 | -0.3669784 | 2.25E-02 |
| AC021092.1 | 246.9309242 | -0.367010551 | 1.27E-02 |
| TRMO | 540.1436586 | -0.367034618 | 1.31E-02 |
| FAM160A2 | 1211.307237 | -0.367036363 | 1.03E-03 |
| ATP6V0D1 | 4652.196853 | -0.367353265 | 1.43E-02 |
| ZFP1 | 767.8083205 | -0.367371906 | 7.30E-03 |
| ZSWIM7 | 523.3338612 | -0.367445552 | 2.10E-03 |
| IFT46 | 2480.303894 | -0.367504357 | 1.93E-05 |
| ZNF652 | 812.4173567 | -0.367534019 | 4.47E-04 |
| FAM98B | 1262.849122 | -0.367877626 | 1.87E-02 |
| TBC1D24 | 1209.420253 | -0.368320797 | 3.26E-03 |
| TMEM254 | 598.0963259 | -0.368477876 | 3.03E-03 |
| BOLA3-AS1 | 290.6439025 | -0.368656292 | 1.19E-02 |
| CSNK1G1 | 2578.916105 | -0.368805354 | 1.14E-06 |
| LRRC27 | 266.4274175 | -0.369140583 | 4.16E-03 |
| ACAA1 | 1616.455569 | -0.36921148 | 8.05E-04 |
| CDC26 | 1403.047321 | -0.36921646 | 5.07E-05 |
| SEMA4G | 559.1740225 | -0.369321805 | 2.64E-02 |
| ITPRIPL1 | 839.3701575 | -0.369425422 | 6.80E-03 |
| NA | 202.7798493 | -0.369659885 | 4.44E-02 |
| CLASP1 | 3823.442414 | -0.369915592 | 6.61E-05 |
| AL136295.7 | 229.180298 | -0.370128521 | 8.89E-03 |
| IL17RC | 1364.10407 | -0.37039876 | 2.27E-02 |
| ITGB3BP | 726.7298882 | -0.370857553 | 3.24E-02 |
| SYNGAP1 | 1172.130205 | -0.370922493 | 1.66E-03 |
| CCDC40 | 197.5519158 | -0.370958087 | 4.92E-02 |
| PAXIP1 | 1709.812529 | -0.371021759 | 2.95E-03 |
| JMJD7-PLA2G4B | 577.9573265 | -0.371060094 | 2.32E-02 |
| DBR1 | 685.1075807 | -0.37107495 | 5.85E-04 |

| | | | |
|---|---|---|---|
| LINC01106 | 1890.349103 | -0.371086024 | 8.55E-05 |
| ZNF512 | 2797.925537 | -0.371287384 | 2.13E-04 |
| CCDC102A | 865.6513979 | -0.371371302 | 3.63E-02 |
| GOLM1 | 4855.950469 | -0.371373766 | 3.81E-07 |
| B3GALNT1 | 1058.702994 | -0.371403001 | 4.52E-03 |
| JARID2 | 1429.283919 | -0.37157448 | 5.15E-06 |
| ENC1 | 10344.19834 | -0.371752109 | 1.25E-04 |
| STK17A | 2263.186549 | -0.371912172 | 2.53E-03 |
| NICN1 | 592.7048162 | -0.372054772 | 1.49E-02 |
| G6PC3 | 3386.25251 | -0.372881624 | 1.15E-03 |
| ZNF689 | 997.135433 | -0.373061725 | 1.61E-02 |
| SIRT1 | 733.7383319 | -0.37312654 | 2.32E-02 |
| STRN | 1774.042078 | -0.373339956 | 1.09E-02 |
| MAT2A | 17360.67703 | -0.373590362 | 3.54E-06 |
| PPIP5K1 | 1251.016395 | -0.373826763 | 1.69E-02 |
| ZNF597 | 311.6737611 | -0.373898736 | 1.15E-02 |
| ARHGAP42 | 723.8663377 | -0.373950795 | 4.80E-02 |
| HSD17B4 | 7591.053364 | -0.374126112 | 9.32E-07 |
| ZKSCAN7 | 396.0174361 | -0.374468668 | 1.44E-02 |
| CHST11 | 1000.865333 | -0.374969725 | 2.47E-02 |
| ZNF280D | 1224.036056 | -0.375005978 | 8.51E-03 |
| CALHM2 | 962.5218492 | -0.375234979 | 1.47E-03 |
| PGGT1B | 904.8288717 | -0.375576983 | 1.93E-02 |
| DNAAF2 | 775.7030037 | -0.375599149 | 5.82E-04 |
| FAM13A | 1552.306742 | -0.375610201 | 8.58E-03 |
| C9orf16 | 2033.48365 | -0.375682114 | 2.79E-02 |
| POPDC3 | 714.293481 | -0.375778577 | 2.72E-03 |
| SSBP4 | 1404.437664 | -0.376007728 | 1.28E-02 |
| VARS2 | 1563.733782 | -0.376207418 | 8.32E-05 |
| IPO13 | 2334.714692 | -0.376414954 | 2.28E-02 |
| NEO1 | 7967.414653 | -0.376546044 | 5.71E-03 |
| TRMT1L | 1145.38981 | -0.376572161 | 8.97E-03 |
| ZNF251 | 1133.94733 | -0.376947341 | 2.09E-06 |
| KIZ | 1237.572865 | -0.377025844 | 5.10E-03 |
| TRIM52 | 1020.521846 | -0.377599047 | 1.71E-02 |
| SLC37A3 | 2567.358471 | -0.377703959 | 5.38E-06 |
| ANKLE1 | 635.5348284 | -0.377979304 | 2.49E-03 |
| MAZ | 11428.16553 | -0.378031381 | 1.45E-03 |
| FBXL20 | 1050.705018 | -0.378215924 | 2.00E-03 |
| FUT10 | 580.7966807 | -0.378228343 | 3.04E-02 |
| NPHP4 | 695.872442 | -0.378266104 | 7.07E-03 |
| PUS3 | 1045.525188 | -0.378319604 | 4.85E-03 |
| IP6K1 | 2965.032039 | -0.378470371 | 4.85E-04 |
| C1orf131 | 519.7394715 | -0.378551571 | 8.69E-04 |
| NIPSNAP3A | 968.4216957 | -0.378949456 | 5.05E-05 |
| MUC20-OT1 | 762.1769723 | -0.379090013 | 1.51E-02 |
| MSL1 | 3890.623421 | -0.379501794 | 1.24E-05 |

| | | | |
|---|---|---|---|
| ABLIM3 | 5868.459542 | -0.379660822 | 2.18E-02 |
| CCDC24 | 384.6898969 | -0.379860545 | 2.96E-02 |
| TAF4 | 859.8748059 | -0.38007754 | 6.46E-03 |
| SLC25A53 | 297.7436183 | -0.380355269 | 1.37E-02 |
| TFCP2 | 1699.603032 | -0.380436264 | 1.08E-07 |
| MRNIP | 1068.786205 | -0.380500489 | 9.49E-05 |
| SMG6 | 4055.32213 | -0.380619062 | 3.32E-04 |
| UBXN2B | 1528.527194 | -0.380764439 | 4.08E-02 |
| FBXL19 | 2478.860355 | -0.381176858 | 4.34E-03 |
| RTEL1-TNFRSF6E | 821.9011062 | -0.381450254 | 3.08E-02 |
| TRAFD1 | 2732.918766 | -0.38153489 | 2.75E-03 |
| C4orf3 | 2769.442182 | -0.381549192 | 2.41E-02 |
| KIAA0753 | 1607.123434 | -0.381624694 | 9.48E-04 |
| EPHX2 | 966.502786 | -0.381684094 | 2.12E-03 |
| PRPF38B | 1919.634009 | -0.381787747 | 4.27E-02 |
| ASB16-AS1 | 304.3514996 | -0.382466047 | 1.78E-02 |
| SS18L2 | 592.0948868 | -0.382688621 | 7.10E-04 |
| CCDC77 | 870.8727388 | -0.382898665 | 1.50E-02 |
| CPT1A | 5309.283983 | -0.38297973 | 6.95E-04 |
| CARMIL1 | 2603.175438 | -0.38305363 | 1.05E-08 |
| RCOR3 | 966.5714162 | -0.383092483 | 1.36E-03 |
| CYB561A3 | 2743.50724 | -0.383208811 | 3.01E-03 |
| LIMD2 | 2211.717988 | -0.383502483 | 1.15E-03 |
| SH3YL1 | 402.1516766 | -0.383917503 | 3.36E-02 |
| HDAC8 | 1003.791273 | -0.383925104 | 2.45E-04 |
| WAC-AS1 | 606.8259948 | -0.384014273 | 1.31E-04 |
| BBS12 | 273.4155551 | -0.385006887 | 1.58E-02 |
| ZNF800 | 909.8293431 | -0.385225401 | 3.55E-02 |
| PAN3 | 841.5834156 | -0.385509423 | 8.70E-03 |
| CBX2 | 1976.013708 | -0.385575093 | 3.42E-03 |
| SPATA20 | 5233.898565 | -0.386681381 | 2.68E-03 |
| GABPB1-AS1 | 2069.678936 | -0.38672558 | 1.13E-02 |
| BLOC1S3 | 480.2032581 | -0.387067402 | 3.83E-02 |
| THAP8 | 365.5154966 | -0.387384762 | 4.45E-02 |
| LAP3 | 2191.36426 | -0.387498569 | 6.14E-05 |
| UBP1 | 4324.363057 | -0.387510534 | 6.41E-07 |
| CYB561 | 2861.485413 | -0.387664045 | 9.32E-05 |
| AC009113.1 | 261.0991221 | -0.387668519 | 5.42E-03 |
| REV1 | 1476.613051 | -0.388212075 | 1.33E-02 |
| TYK2 | 3847.834705 | -0.388339453 | 3.44E-04 |
| XPO1 | 12353.51509 | -0.388384803 | 3.99E-02 |
| ZNF584 | 705.5673573 | -0.388708707 | 3.18E-03 |
| DPY19L2 | 311.8639347 | -0.389094279 | 2.53E-02 |
| USP6NL | 606.8158013 | -0.389485761 | 8.91E-03 |
| UBE2Q2 | 1264.376967 | -0.389676273 | 1.92E-03 |
| FAM104B | 415.0212845 | -0.389850552 | 1.20E-03 |
| SNX29 | 2506.115096 | -0.390199523 | 8.87E-05 |

| | | | |
|---|---|---|---|
| CD276 | 8333.957018 | -0.390224697 | 3.40E-02 |
| ZNF564 | 315.0327144 | -0.390686656 | 9.74E-03 |
| BAGE2 | 711.3235011 | -0.390772453 | 4.34E-02 |
| PLEKHM1P1 | 493.0380594 | -0.39079296 | 3.71E-04 |
| 4-Sep | 536.272959 | -0.390916846 | 3.25E-02 |
| SH2B1 | 2927.777628 | -0.391038252 | 1.74E-04 |
| MDK | 24505.83644 | -0.391110947 | 3.50E-02 |
| TUG1 | 9645.076116 | -0.391173361 | 1.21E-04 |
| LUC7L2 | 3487.963961 | -0.391338043 | 9.14E-04 |
| CAPN5 | 3165.900756 | -0.391466404 | 5.96E-04 |
| LEO1 | 1631.362175 | -0.391659786 | 1.29E-04 |
| PLSCR1 | 1248.540368 | -0.391757305 | 4.40E-04 |
| MMP25-AS1 | 640.8204245 | -0.391986817 | 3.76E-02 |
| MTMR9 | 1043.343317 | -0.392077811 | 5.28E-04 |
| DCBLD2 | 4378.874141 | -0.392619734 | 1.25E-04 |
| HARBI1 | 273.0406226 | -0.392793315 | 1.72E-02 |
| ITGAV | 3581.278314 | -0.39283606 | 4.65E-02 |
| PBX3 | 961.2103878 | -0.392989858 | 6.59E-05 |
| IRF2BPL | 3296.790079 | -0.393025394 | 3.87E-04 |
| ZNF503 | 297.2788085 | -0.393268188 | 4.40E-03 |
| SLX1B-SULT1A4 | 282.9680088 | -0.393393286 | 2.33E-02 |
| NECTIN1 | 3123.160443 | -0.393502595 | 4.76E-02 |
| LINC00265 | 516.0461094 | -0.393566491 | 7.45E-03 |
| AC012467.2 | 176.2267427 | -0.393910018 | 3.74E-02 |
| GPD1L | 1340.603692 | -0.394042325 | 1.11E-03 |
| TIRAP | 621.8397527 | -0.394248468 | 1.29E-04 |
| SOS2 | 1401.00858 | -0.394762896 | 6.06E-03 |
| FUT8 | 3274.135694 | -0.395391278 | 5.04E-05 |
| NME3 | 701.7911262 | -0.39571701 | 3.57E-02 |
| SBNO1 | 2551.156243 | -0.395950465 | 8.78E-03 |
| PNRC1 | 2333.746429 | -0.39595568 | 3.90E-03 |
| CPVL | 3426.438074 | -0.396011616 | 6.88E-03 |
| MTMR9LP | 289.1436766 | -0.39608529 | 2.46E-02 |
| SRSF10 | 4121.068345 | -0.396099723 | 2.47E-03 |
| DNAJC4 | 1151.179245 | -0.396144629 | 1.81E-02 |
| NOVA2 | 439.1966609 | -0.396190042 | 2.59E-02 |
| DTX3 | 1694.078695 | -0.396528573 | 7.75E-05 |
| TRIM3 | 1257.094969 | -0.397275781 | 5.86E-04 |
| POTEC | 621.7288682 | -0.397305797 | 1.82E-04 |
| FAM156A | 1547.074356 | -0.397768038 | 6.91E-03 |
| SEMA4B | 2489.245754 | -0.39778827 | 5.99E-03 |
| EFCAB2 | 314.3175812 | -0.397931606 | 3.92E-02 |
| CBFB | 1922.228719 | -0.398095854 | 3.61E-03 |
| LTK | 604.491803 | -0.398465348 | 7.73E-03 |
| FAM53B | 1491.038315 | -0.398720954 | 1.78E-04 |
| KCTD1 | 748.5229894 | -0.398859826 | 2.46E-03 |
| SLC36A4 | 873.4201689 | -0.3990119 | 2.96E-02 |

| | | | |
|---|---|---|---|
| CORO2A | 1356.442854 | -0.399021698 | 2.18E-06 |
| KDM4D | 220.213436 | -0.399399179 | 3.60E-02 |
| ATP13A1 | 3127.033339 | -0.399541752 | 6.07E-05 |
| PLEKHH3 | 964.0051629 | -0.399606678 | 5.32E-04 |
| NA | 202.4228237 | -0.400005779 | 2.05E-02 |
| HSD17B11 | 1994.692616 | -0.400141458 | 4.68E-05 |
| PDE8A | 1845.673011 | -0.400181126 | 1.56E-04 |
| PRR7 | 203.9707106 | -0.400291348 | 4.10E-02 |
| LRP4 | 1518.677841 | -0.400933424 | 1.48E-02 |
| PATZ1 | 2235.275444 | -0.40111177 | 1.55E-04 |
| FZD3 | 812.4350586 | -0.401232274 | 2.13E-02 |
| POP5 | 1281.270367 | -0.40147389 | 1.44E-06 |
| B3GNT2 | 692.6789905 | -0.401516109 | 7.36E-04 |
| EMP2 | 4797.302819 | -0.401566673 | 8.48E-04 |
| STRN3 | 913.2813553 | -0.401889302 | 9.21E-03 |
| HIPK2 | 6086.845547 | -0.402399459 | 2.89E-09 |
| MUTYH | 512.5623734 | -0.403033913 | 2.98E-02 |
| DNAJC24 | 615.7328119 | -0.403096785 | 3.83E-02 |
| ZNF778 | 856.5401387 | -0.403320861 | 4.07E-02 |
| USP24 | 5365.564556 | -0.403354176 | 2.30E-06 |
| PIP4P2 | 720.376732 | -0.403401124 | 1.28E-05 |
| KLHL9 | 2682.692212 | -0.403415142 | 1.05E-05 |
| FEM1B | 2626.446844 | -0.403445774 | 5.41E-04 |
| C22orf46 | 2615.757336 | -0.403641383 | 8.15E-08 |
| MARVELD1 | 3273.994686 | -0.403920709 | 1.43E-02 |
| RABL3 | 632.95653 | -0.40393332 | 2.18E-03 |
| BRD3OS | 2561.136172 | -0.403992921 | 4.79E-04 |
| TMCO1 | 3626.680009 | -0.404012326 | 4.65E-06 |
| TMEM135 | 1235.723969 | -0.404257975 | 4.74E-02 |
| BMP2K | 1364.424567 | -0.404553003 | 6.00E-03 |
| ZNF337-AS1 | 269.830778 | -0.404577535 | 9.53E-03 |
| ZNF529 | 751.8925068 | -0.404739464 | 1.51E-02 |
| HOXA7 | 998.7093727 | -0.405027575 | 2.40E-05 |
| YJEFN3 | 352.2178593 | -0.405614721 | 3.00E-02 |
| CMTM6 | 3907.919114 | -0.405622768 | 2.88E-06 |
| CAMSAP2 | 2288.391164 | -0.405824408 | 4.14E-02 |
| SHANK3 | 593.1335192 | -0.406062524 | 2.13E-03 |
| WDR83 | 712.4492011 | -0.406205756 | 2.35E-03 |
| WDR73 | 1944.452553 | -0.406349136 | 5.45E-06 |
| TCEAL8 | 1127.435267 | -0.406469496 | 9.00E-03 |
| USP49 | 781.7022822 | -0.406744796 | 1.10E-03 |
| FBXO3 | 1527.174634 | -0.40676326 | 1.09E-03 |
| GOLPH3L | 1038.121567 | -0.406850407 | 1.26E-04 |
| RNF34 | 1430.120546 | -0.406898792 | 5.42E-04 |
| MTATP6P1 | 11442.5503 | -0.406976346 | 3.75E-02 |
| CETN2 | 2499.92649 | -0.407185867 | 1.15E-03 |
| RAB3IL1 | 1180.86828 | -0.40726312 | 1.01E-02 |

| | | | |
|---|---|---|---|
| CIPC | 1125.986789 | -0.407352042 | 2.13E-04 |
| SLC2A4RG | 3962.621191 | -0.407374022 | 1.65E-03 |
| RAB30-AS1 | 321.317745 | -0.407476399 | 1.10E-03 |
| WDFY3 | 2746.737282 | -0.407797328 | 9.77E-04 |
| LMBRD1 | 897.240277 | -0.407939757 | 2.34E-04 |
| ENO3 | 574.3900042 | -0.408008784 | 1.29E-03 |
| NDUFB3 | 1402.880037 | -0.408444464 | 1.82E-03 |
| ACIN1 | 10252.36296 | -0.408527724 | 1.70E-22 |
| ZNF286A | 1332.71699 | -0.408566797 | 1.39E-03 |
| RLF | 1290.363741 | -0.408713484 | 1.12E-02 |
| CORO2B | 318.2158163 | -0.409168381 | 2.96E-02 |
| NKX2-6 | 293.1226131 | -0.409231347 | 2.56E-02 |
| CTSF | 5110.938984 | -0.409352837 | 2.20E-02 |
| ERMARD | 674.76412 | -0.409456597 | 1.25E-03 |
| PBRM1 | 2780.16346 | -0.409512115 | 1.22E-03 |
| DNM1L | 3372.045611 | -0.409896932 | 1.45E-02 |
| TXNDC9 | 936.8841297 | -0.410289572 | 1.29E-02 |
| CCDC57 | 916.7750703 | -0.410389471 | 3.18E-03 |
| NA | 213.9369198 | -0.410486916 | 2.18E-02 |
| ZNF691 | 543.1359512 | -0.410507641 | 1.51E-04 |
| PIK3R2 | 4034.204486 | -0.411326507 | 3.67E-04 |
| HAT1 | 2459.082348 | -0.411494578 | 2.99E-02 |
| CARD8 | 1192.652439 | -0.411511984 | 8.50E-07 |
| ACAP3 | 1532.688697 | -0.411686092 | 7.23E-03 |
| SELENOH | 3534.246655 | -0.411905699 | 1.21E-02 |
| ARHGEF34P | 257.9797001 | -0.412086661 | 2.27E-02 |
| NA | 2018.632065 | -0.412236569 | 9.45E-04 |
| SPDYE3 | 437.3266365 | -0.412578968 | 9.61E-04 |
| RXRA | 6547.240196 | -0.412632166 | 4.57E-04 |
| HEXD | 641.0207043 | -0.412818591 | 4.19E-04 |
| ZNF347 | 425.5866937 | -0.412953027 | 3.68E-02 |
| ITFG2 | 735.0446452 | -0.413065226 | 3.62E-03 |
| LPAR2 | 318.5182653 | -0.413592712 | 3.05E-02 |
| DNASE1 | 1176.911667 | -0.41369369 | 5.72E-05 |
| ZIK1 | 840.9093282 | -0.413696059 | 7.91E-04 |
| AHDC1 | 1790.089992 | -0.413919115 | 8.88E-03 |
| PIAS2 | 969.8538545 | -0.414269257 | 9.27E-04 |
| CLDN12 | 1756.686322 | -0.414335214 | 7.87E-03 |
| HDHD2 | 1699.767881 | -0.414405098 | 1.76E-03 |
| BMS1P1 | 299.021387 | -0.414543606 | 1.01E-02 |
| ASF1A | 781.5517459 | -0.414679295 | 3.72E-02 |
| SDR39U1 | 2016.387592 | -0.41471147 | 2.13E-04 |
| AASDH | 769.6253677 | -0.414815048 | 1.98E-02 |
| KIAA1841 | 487.9104014 | -0.415011816 | 2.19E-02 |
| DTX3L | 2777.600576 | -0.415298198 | 1.02E-03 |
| HFE | 563.7342057 | -0.415397753 | 1.04E-03 |
| SYNJ2BP-COX16 | 287.8347004 | -0.415660455 | 3.90E-02 |

| | | | |
|---|---|---|---|
| AMBRA1 | 3207.388082 | -0.415670638 | 4.97E-03 |
| INTS5 | 1999.085661 | -0.416033604 | 3.19E-03 |
| ANP32E | 1408.613357 | -0.416273754 | 2.99E-02 |
| ST6GALNAC2 | 968.4348124 | -0.416324462 | 8.68E-04 |
| RILPL2 | 1239.886752 | -0.416675321 | 7.05E-04 |
| HECTD1 | 6357.213805 | -0.41723064 | 1.06E-02 |
| TXN2 | 3663.452541 | -0.417986537 | 8.73E-04 |
| TNFRSF25 | 230.3667618 | -0.41823791 | 2.18E-02 |
| SNX21 | 1691.955302 | -0.418378704 | 2.83E-02 |
| FIP1L1 | 2275.696884 | -0.418438402 | 5.08E-04 |
| COX20 | 1221.240216 | -0.41882356 | 4.05E-07 |
| MYCBP2 | 9079.698849 | -0.418878281 | 4.45E-03 |
| FNBP4 | 2786.790921 | -0.419211281 | 2.68E-02 |
| BCKDHB | 817.3871201 | -0.419680194 | 1.53E-03 |
| NEURL2 | 189.9105612 | -0.419843915 | 3.48E-02 |
| FBXL4 | 893.34174 | -0.419912959 | 4.63E-04 |
| PCSK7 | 5369.388781 | -0.419958286 | 7.49E-07 |
| AC009120.2 | 213.2434982 | -0.42008548 | 3.96E-02 |
| TDG | 846.6086933 | -0.420106444 | 5.21E-03 |
| GTDC1 | 589.4500305 | -0.420185492 | 1.12E-03 |
| TNKS2 | 1792.852115 | -0.420661288 | 1.82E-04 |
| PARP14 | 3812.609189 | -0.420687235 | 1.34E-02 |
| DIPK1A | 488.9177531 | -0.421047601 | 2.80E-02 |
| SLC30A7 | 941.1097818 | -0.421203948 | 3.08E-04 |
| AGPAT5 | 2414.072757 | -0.42133695 | 4.65E-05 |
| WRN | 1840.206464 | -0.421480447 | 2.41E-02 |
| ZNF625-ZNF20 | 174.7981346 | -0.421639695 | 2.88E-02 |
| ZBTB44 | 2561.825986 | -0.421737998 | 1.98E-03 |
| TRAPPC8 | 1683.150941 | -0.422039788 | 9.51E-04 |
| BACH2 | 445.9839007 | -0.422281714 | 1.93E-02 |
| RNF149 | 1382.487106 | -0.422653903 | 1.23E-08 |
| PCNX1 | 5764.565374 | -0.422794845 | 7.32E-05 |
| CCDC159 | 150.0120162 | -0.422798683 | 3.79E-02 |
| DNA2 | 356.2619562 | -0.422888282 | 2.79E-02 |
| GALNS | 1902.302002 | -0.42291596 | 1.05E-03 |
| MAPKAPK3 | 1992.429819 | -0.423000116 | 4.70E-03 |
| AC008124.1 | 360.4932966 | -0.423143775 | 1.16E-02 |
| MAGI3 | 464.1381935 | -0.423290307 | 1.04E-02 |
| APOL3 | 1109.386461 | -0.423355459 | 1.16E-05 |
| UCP2 | 2547.826664 | -0.423440818 | 2.51E-04 |
| ZNF549 | 563.6641424 | -0.424002188 | 5.34E-05 |
| MED11 | 514.0417921 | -0.424417984 | 3.14E-02 |
| EPHX1 | 3232.5919 | -0.424539889 | 4.97E-04 |
| PDSS1 | 287.872663 | -0.425005845 | 2.06E-02 |
| MMACHC | 272.7799796 | -0.425553962 | 4.44E-02 |
| NR1D1 | 572.8036485 | -0.425912844 | 1.88E-03 |
| KPTN | 754.5468126 | -0.426487173 | 1.08E-02 |

| | | | |
|---|---|---|---|
| AP001372.2 | 231.832704 | -0.426520791 | 2.97E-02 |
| TMEM53 | 424.5627033 | -0.426595376 | 1.72E-02 |
| FBXL17 | 1252.937548 | -0.426601733 | 1.03E-03 |
| CASP10 | 374.7130969 | -0.426874588 | 1.73E-02 |
| HIP1 | 8387.051766 | -0.427041385 | 9.30E-07 |
| C1orf216 | 1881.654585 | -0.427153881 | 8.34E-03 |
| SLC25A42 | 745.7633681 | -0.427406537 | 1.70E-03 |
| ZNF559-ZNF177 | 287.9463382 | -0.42767346 | 1.15E-02 |
| ABRAXAS1 | 475.8088465 | -0.428244076 | 3.33E-02 |
| RAPGEF6 | 3147.194901 | -0.428639178 | 8.62E-03 |
| MSH2 | 2826.08355 | -0.428656429 | 4.63E-03 |
| TSPAN6 | 4046.676907 | -0.428917018 | 8.92E-05 |
| SDCBP | 7542.179046 | -0.429077344 | 1.38E-07 |
| LINC00863 | 382.1564718 | -0.429743954 | 5.84E-03 |
| ATF2 | 1408.171597 | -0.429827775 | 3.97E-02 |
| ZNF346 | 892.9662731 | -0.42998902 | 2.46E-04 |
| ZNF322 | 1703.612534 | -0.430053225 | 1.16E-02 |
| TCF7L2 | 706.3416085 | -0.430119189 | 1.53E-03 |
| KDM1B | 941.9864652 | -0.430254801 | 3.17E-05 |
| LIN7C | 545.565844 | -0.43049608 | 4.59E-02 |
| PFN1P2 | 183.2416403 | -0.430539212 | 3.90E-02 |
| PALB2 | 1355.274179 | -0.430904909 | 9.60E-03 |
| PRRT3-AS1 | 120.3373431 | -0.430925752 | 2.93E-02 |
| CLUAP1 | 987.7006588 | -0.431230426 | 8.32E-03 |
| WDR6 | 10006.27012 | -0.431231514 | 1.22E-02 |
| AC021087.3 | 146.4656657 | -0.431417025 | 3.72E-02 |
| CHST10 | 1531.826562 | -0.431809006 | 7.53E-03 |
| ZNF354C | 587.1740906 | -0.432024993 | 2.51E-02 |
| C3orf14 | 394.3980219 | -0.432213556 | 6.30E-03 |
| EFNA4 | 501.216771 | -0.432382068 | 1.90E-02 |
| EP300 | 4842.321 | -0.432842759 | 4.03E-23 |
| SERINC1 | 4518.27721 | -0.433256681 | 7.55E-04 |
| ZNF75A | 1038.680233 | -0.433560581 | 1.83E-04 |
| SENP1 | 1760.284492 | -0.433624257 | 3.07E-04 |
| SLC12A9 | 3367.234538 | -0.433778788 | 5.90E-04 |
| SFXN5 | 1044.332545 | -0.433780808 | 1.91E-03 |
| PTMS | 13732.99283 | -0.433973828 | 1.36E-03 |
| SREBF2-AS1 | 139.3932799 | -0.434162095 | 2.90E-02 |
| TLE1 | 2878.688334 | -0.434228303 | 2.69E-03 |
| ZC3HAV1 | 3837.855423 | -0.434344695 | 5.19E-03 |
| GORAB | 305.0897635 | -0.434451693 | 4.10E-02 |
| CRABP2 | 3933.139023 | -0.434552303 | 1.65E-02 |
| NUMB | 4674.727443 | -0.434723297 | 6.99E-13 |
| SUFU | 1234.590452 | -0.435019979 | 1.16E-02 |
| NCBP2 | 4485.947445 | -0.435160602 | 9.63E-07 |
| MARF1 | 3255.495896 | -0.435431936 | 1.40E-06 |
| NDUFA5 | 1338.229747 | -0.435436297 | 6.62E-03 |

| | | | |
|---|---|---|---|
| PTPN12 | 6077.371603 | -0.435501461 | 2.69E-03 |
| PCNX2 | 1178.527075 | -0.435615663 | 1.63E-05 |
| NCDN | 2694.254182 | -0.435777621 | 2.53E-05 |
| TERF1 | 1319.671707 | -0.435946238 | 1.84E-02 |
| PIKFYVE | 2742.920494 | -0.436592738 | 1.66E-02 |
| PRTG | 674.9816943 | -0.436628073 | 3.56E-03 |
| CSPP1 | 914.3606557 | -0.436879989 | 1.86E-02 |
| FAM171A2 | 602.8195332 | -0.436886745 | 6.28E-03 |
| TAPBPL | 874.5015255 | -0.436954635 | 7.30E-03 |
| RBM26 | 3241.279658 | -0.437040044 | 7.13E-04 |
| LMLN | 677.6147435 | -0.437093847 | 9.77E-03 |
| RTN4RL2 | 207.8606213 | -0.437372596 | 4.96E-02 |
| APOBEC3C | 3158.950832 | -0.43753788 | 2.26E-02 |
| PGAP2 | 928.1459418 | -0.438070743 | 2.91E-05 |
| FAM114A2 | 967.3843151 | -0.438076577 | 4.70E-03 |
| FIGNL1 | 1385.407933 | -0.43815608 | 2.42E-02 |
| AJM1 | 297.7773619 | -0.438487866 | 4.13E-02 |
| FBXW9 | 659.5921597 | -0.438506251 | 3.97E-03 |
| TYW3 | 1219.576547 | -0.439017643 | 2.16E-04 |
| RNF146 | 744.1983505 | -0.439156361 | 4.35E-02 |
| LMBR1L | 1267.801377 | -0.439385676 | 3.62E-04 |
| RBM5 | 5755.397797 | -0.439532184 | 5.23E-12 |
| TMEM86A | 534.2173541 | -0.440303908 | 4.93E-02 |
| LINC01089 | 367.5833516 | -0.440367961 | 7.91E-04 |
| SORBS1 | 666.9068481 | -0.440644209 | 4.01E-04 |
| PPP1R13B | 1276.138984 | -0.441010741 | 3.49E-03 |
| TUBB3 | 3262.462153 | -0.441552672 | 5.05E-03 |
| CLIP3 | 2375.243907 | -0.441828498 | 1.78E-03 |
| ARHGAP1 | 3525.834389 | -0.441996827 | 7.39E-05 |
| AC125257.1 | 163.7034956 | -0.44239801 | 2.59E-02 |
| FAM161B | 403.8397917 | -0.442529465 | 6.64E-04 |
| VAMP1 | 983.9967304 | -0.442921483 | 1.11E-03 |
| FGFR1OP | 667.7562339 | -0.443410167 | 1.12E-02 |
| AL358472.2 | 89.91092447 | -0.443421941 | 4.88E-02 |
| MYORG | 1497.758198 | -0.443710208 | 1.27E-02 |
| MED21 | 921.6848111 | -0.443788181 | 3.22E-04 |
| KIDINS220 | 3754.609944 | -0.443848713 | 4.14E-03 |
| SLC25A46 | 2798.417499 | -0.443869406 | 5.77E-03 |
| MREG | 275.7797383 | -0.44390177 | 8.25E-04 |
| PCDHGB2 | 256.1679818 | -0.44459754 | 1.76E-02 |
| MGAT2 | 2529.203773 | -0.44495523 | 3.91E-08 |
| NBPF4 | 231.6908026 | -0.44504012 | 1.48E-02 |
| SREBF1 | 4307.962205 | -0.445104368 | 1.01E-04 |
| DHX29 | 3842.598765 | -0.445277935 | 1.93E-02 |
| RALB | 5036.240045 | -0.445279534 | 1.08E-07 |
| ELF2 | 1626.106279 | -0.445286828 | 7.59E-10 |
| GPRASP2 | 355.0135181 | -0.445392879 | 9.40E-04 |

| | | | |
|---|---|---|---|
| MGST2 | 872.3258944 | -0.44551931 | 6.54E-03 |
| ZADH2 | 2400.126845 | -0.44569809 | 3.27E-11 |
| SPTBN1 | 22759.67806 | -0.445719781 | 4.34E-15 |
| ZBED6 | 1064.580852 | -0.446232016 | 2.31E-03 |
| GAA | 1637.702727 | -0.446296245 | 6.63E-03 |
| SKAP2 | 3278.228177 | -0.446406229 | 1.28E-05 |
| ISYNA1 | 5628.667813 | -0.4467396 | 3.36E-02 |
| GCNT2 | 168.7839907 | -0.447034132 | 3.63E-02 |
| NUDT15 | 1373.191692 | -0.447215462 | 1.49E-04 |
| DENND4A | 1554.563487 | -0.447264797 | 3.00E-02 |
| TOB2 | 2449.947897 | -0.447334047 | 7.03E-04 |
| ZNF160 | 881.0002724 | -0.447370579 | 5.36E-03 |
| MFN1 | 2306.740901 | -0.447477111 | 2.02E-03 |
| POLM | 811.6532932 | -0.447502341 | 9.62E-04 |
| INTS13 | 1642.98614 | -0.447728432 | 9.88E-03 |
| RAD50 | 5501.443131 | -0.447836748 | 4.68E-02 |
| PI4KA | 4563.049434 | -0.447908322 | 6.76E-09 |
| AKT1S1 | 2743.622308 | -0.448002772 | 2.76E-04 |
| NECTIN3 | 569.8866024 | -0.448305867 | 8.27E-03 |
| VSIG10 | 1369.992613 | -0.448358279 | 1.04E-04 |
| HOXB13 | 1427.82097 | -0.448742225 | 5.00E-03 |
| GTF2IP4 | 7182.880039 | -0.448929735 | 3.00E-05 |
| NAPEPLD | 824.1140459 | -0.449008924 | 1.43E-02 |
| ARHGEF5 | 1397.221325 | -0.449298775 | 5.10E-05 |
| CDC27 | 3449.786474 | -0.449437766 | 3.88E-04 |
| AC055811.4 | 242.529967 | -0.449593342 | 7.27E-03 |
| AP5M1 | 1165.533304 | -0.449649822 | 1.76E-04 |
| PSMB8-AS1 | 238.0784475 | -0.449807702 | 4.04E-02 |
| PTPRE | 298.5658975 | -0.450488805 | 4.83E-02 |
| HTR1D | 270.4186298 | -0.450938192 | 4.70E-02 |
| 7-Mar | 2762.733203 | -0.451015862 | 1.14E-02 |
| PJA2 | 7418.704723 | -0.451322577 | 1.35E-04 |
| CSKMT | 343.0414308 | -0.451748009 | 6.95E-04 |
| PRXL2A | 2340.360732 | -0.451775018 | 6.00E-09 |
| PDCL | 1337.598657 | -0.451894095 | 1.61E-08 |
| SPESP1 | 370.4501273 | -0.451931652 | 1.04E-02 |
| DISC1 | 789.4212616 | -0.452065831 | 2.59E-05 |
| USF1 | 2130.590853 | -0.452253836 | 3.54E-05 |
| DISP1 | 267.7010967 | -0.452349136 | 3.36E-02 |
| HIST2H2BF | 235.8078308 | -0.45241309 | 2.40E-02 |
| EFNB1 | 5451.422566 | -0.452485923 | 9.87E-04 |
| PCLAF | 1284.567792 | -0.452765021 | 3.27E-02 |
| DUSP18 | 224.9387926 | -0.453078418 | 2.06E-02 |
| LINC02035 | 194.8129886 | -0.45316239 | 1.41E-02 |
| ZNF324 | 851.7318929 | -0.453318374 | 3.22E-04 |
| HPDL | 404.0767023 | -0.453353093 | 2.50E-03 |
| ZC3H13 | 3640.434787 | -0.453810937 | 7.87E-03 |

| | | | |
|---|---|---|---|
| ULK3 | 1952.350994 | -0.453886202 | 3.95E-06 |
| CYBA | 3231.264754 | -0.45392209 | 5.89E-03 |
| CACFD1 | 1097.142452 | -0.453969336 | 8.46E-03 |
| TRIM33 | 1174.386367 | -0.454490699 | 3.08E-02 |
| TBC1D8 | 2589.114755 | -0.454573307 | 2.87E-09 |
| C2CD6 | 329.5712679 | -0.454650366 | 9.98E-03 |
| KIF11 | 2007.052058 | -0.455022014 | 3.96E-02 |
| AL121832.3 | 132.6188102 | -0.455716468 | 3.97E-02 |
| EID1 | 3347.970584 | -0.455724247 | 5.01E-04 |
| SETD6 | 1072.763408 | -0.455788404 | 1.89E-04 |
| ATAT1 | 537.2356922 | -0.455947492 | 2.62E-06 |
| CHSY1 | 3084.922925 | -0.45602812 | 5.55E-09 |
| ACTR6 | 773.9997215 | -0.456201841 | 4.05E-03 |
| OST4 | 4570.163147 | -0.456462079 | 7.74E-03 |
| POLR2L | 4348.331672 | -0.456496521 | 2.56E-03 |
| PRPF38A | 2292.914314 | -0.4565438 | 1.68E-06 |
| ZNF605 | 777.2780323 | -0.456606137 | 1.26E-02 |
| RUSC1 | 898.0028328 | -0.457310025 | 7.64E-06 |
| PLK2 | 8444.080224 | -0.457675495 | 1.00E-04 |
| MYD88 | 2389.693906 | -0.457733603 | 1.21E-03 |
| GSTZ1 | 1069.563483 | -0.458195936 | 3.56E-04 |
| KCTD21 | 734.3641873 | -0.458434853 | 7.46E-03 |
| CCSAP | 674.7629138 | -0.459005355 | 2.16E-03 |
| ABCB10 | 750.3101236 | -0.459181016 | 1.57E-03 |
| GIPC3 | 892.0876321 | -0.459298653 | 9.42E-03 |
| ZNF799 | 165.882558 | -0.459364359 | 3.04E-02 |
| TMEM131 | 2558.958861 | -0.459477646 | 1.70E-08 |
| IDNK | 151.7021404 | -0.459605604 | 4.24E-02 |
| PEX12 | 557.9871048 | -0.459900309 | 8.84E-05 |
| TRIM9 | 249.1688506 | -0.460338653 | 3.44E-02 |
| PLA2R1 | 868.557516 | -0.460521252 | 3.60E-03 |
| AC239868.1 | 580.739753 | -0.460561886 | 6.77E-03 |
| WDR89 | 465.2162936 | -0.460564076 | 3.13E-02 |
| CDKL5 | 533.1492249 | -0.46067378 | 3.22E-02 |
| SH3BP5 | 1459.805366 | -0.460733557 | 1.10E-03 |
| CNOT6 | 1530.549631 | -0.4607473 | 3.69E-03 |
| SPTY2D1 | 1895.298232 | -0.460776851 | 9.43E-06 |
| FMC1 | 495.7004804 | -0.461269263 | 2.69E-04 |
| C16orf86 | 157.9991481 | -0.461569196 | 3.34E-02 |
| GCSH | 1242.2651 | -0.461683123 | 1.45E-05 |
| OSTM1 | 1362.323596 | -0.462104575 | 4.38E-03 |
| FBXO10 | 1900.060545 | -0.4621674 | 5.02E-03 |
| SNAI2 | 1114.417032 | -0.462784446 | 3.31E-03 |
| UPF3A | 938.2065818 | -0.463284503 | 5.59E-03 |
| OTULINL | 778.6536385 | -0.463676748 | 1.76E-02 |
| ZNF639 | 844.3296518 | -0.463727086 | 3.35E-03 |
| MSRB1 | 1234.914881 | -0.463763379 | 3.93E-04 |

| | | | |
|---|---|---|---|
| NMB | 413.3741812 | -0.464070988 | 6.44E-03 |
| STAG1 | 1643.158292 | -0.46417753 | 2.40E-02 |
| RRAGB | 446.9069824 | -0.464222261 | 1.06E-02 |
| THRB | 889.0566991 | -0.464254054 | 2.27E-04 |
| LRRC37A4P | 1314.268066 | -0.46440088 | 1.47E-05 |
| FRAT1 | 148.3546136 | -0.464409381 | 2.83E-02 |
| POLR2A | 10759.83302 | -0.464671631 | 3.17E-08 |
| MBIP | 613.297196 | -0.464974314 | 2.97E-03 |
| CCDC134 | 408.6200617 | -0.465019027 | 2.32E-05 |
| MT-ATP8 | 15345.88142 | -0.465123199 | 3.08E-02 |
| XIAP | 1292.126033 | -0.465777504 | 3.37E-02 |
| GBP1 | 1438.088516 | -0.465969597 | 1.04E-04 |
| TIMM8B | 2524.292083 | -0.465992825 | 2.67E-03 |
| ZBTB34 | 895.6646402 | -0.466082246 | 3.97E-06 |
| AL390879.1 | 139.316736 | -0.466222636 | 1.44E-02 |
| TAOK1 | 4239.749078 | -0.466602653 | 2.39E-02 |
| HIPK3 | 1459.481618 | -0.466644867 | 4.54E-04 |
| TULP4 | 870.3654159 | -0.466799346 | 1.14E-02 |
| ADGRA3 | 1740.112729 | -0.467486491 | 1.07E-04 |
| ICAM3 | 189.399634 | -0.467542891 | 3.71E-02 |
| BTBD7 | 2004.166963 | -0.467694023 | 1.93E-09 |
| ANKH | 6322.166104 | -0.467820617 | 1.54E-11 |
| COL4A3BP | 2975.02318 | -0.467848962 | 5.12E-07 |
| PRKAR2B | 977.5424306 | -0.468125808 | 7.60E-05 |
| TMEM200A | 540.674388 | -0.468184249 | 7.43E-03 |
| TMEM59L | 339.1337644 | -0.468285777 | 1.45E-02 |
| ZNF26 | 696.8708955 | -0.46860372 | 3.01E-02 |
| AC240565.2 | 150.4166937 | -0.468886468 | 1.86E-02 |
| SCAND1 | 1159.837333 | -0.468955112 | 3.46E-03 |
| HERC2P5 | 595.3123382 | -0.469016782 | 2.85E-03 |
| ZNF736 | 672.2343097 | -0.469017282 | 6.49E-03 |
| SLIT2 | 273.2187921 | -0.46931167 | 1.59E-02 |
| FBXO16 | 183.9879856 | -0.469462124 | 1.46E-02 |
| YTHDC2 | 2799.831602 | -0.469946784 | 3.25E-02 |
| STS | 1112.178348 | -0.469951803 | 9.33E-07 |
| TBCK | 1057.417717 | -0.470161557 | 5.14E-06 |
| LRRC37A17P | 728.9897898 | -0.470333026 | 2.23E-04 |
| CTSS | 708.5670472 | -0.470531836 | 1.55E-02 |
| AGMAT | 377.0349647 | -0.471123863 | 4.95E-04 |
| CMTM3 | 5438.986239 | -0.471557652 | 2.02E-04 |
| WDR90 | 1545.088941 | -0.471560816 | 5.09E-03 |
| ANKDD1A | 276.2525082 | -0.471721684 | 2.90E-03 |
| CLCN3 | 2983.363341 | -0.471853684 | 1.80E-04 |
| TNRC6A | 4408.836177 | -0.471882334 | 1.73E-05 |
| APOBEC3D | 638.3823992 | -0.472194705 | 8.60E-03 |
| ASAH1 | 3432.713782 | -0.472210167 | 1.63E-05 |
| NOL8 | 2499.308708 | -0.473352721 | 2.33E-02 |

| | | | |
|---|---|---|---|
| ZMYND8 | 5516.740769 | -0.473603966 | 3.95E-06 |
| THAP5 | 611.0550646 | -0.473700164 | 2.08E-02 |
| SCRN3 | 689.0235599 | -0.473751798 | 7.19E-03 |
| CBR3 | 396.1745623 | -0.474334611 | 4.83E-02 |
| MBD5 | 732.2921705 | -0.474741301 | 2.16E-03 |
| ZNF440 | 480.4141961 | -0.475193775 | 5.75E-04 |
| BCORL1 | 1237.78771 | -0.475203654 | 4.78E-04 |
| BRD8 | 7516.993316 | -0.475728505 | 1.47E-13 |
| CAPN7 | 1738.504672 | -0.475955022 | 9.86E-05 |
| NSUN5P1 | 2362.073103 | -0.476177935 | 9.31E-07 |
| GALNT1 | 2272.25254 | -0.476256105 | 5.81E-03 |
| HECW1 | 564.3576593 | -0.476382524 | 7.60E-03 |
| SLC35D1 | 878.4067731 | -0.476388592 | 1.31E-07 |
| ZNF33A | 507.7272835 | -0.476708455 | 3.62E-02 |
| TDRD1 | 1309.35831 | -0.476885569 | 3.12E-05 |
| PYCR3 | 1124.038153 | -0.476991709 | 2.78E-04 |
| EMSY | 1435.066954 | -0.477167265 | 1.11E-02 |
| CYHR1 | 3198.186788 | -0.477341322 | 3.44E-04 |
| RBBP9 | 1634.768975 | -0.477547486 | 4.31E-06 |
| ASH1L | 3747.874399 | -0.477604484 | 2.05E-05 |
| ZRANB2 | 2901.416846 | -0.477613649 | 9.32E-03 |
| APTR | 399.1142399 | -0.477833974 | 3.98E-04 |
| GOLGA8A | 1520.217999 | -0.477930215 | 6.53E-03 |
| NUP35 | 774.1365033 | -0.478140296 | 3.02E-03 |
| SLC25A12 | 2043.058397 | -0.478222606 | 3.14E-07 |
| FERMT1 | 589.4759035 | -0.478394673 | 4.29E-03 |
| TEFM | 462.4051509 | -0.478851897 | 7.97E-04 |
| H3F3A | 10452.75397 | -0.479018868 | 3.40E-05 |
| MLLT3 | 633.9740889 | -0.479099624 | 3.15E-03 |
| AC008755.1 | 1945.393163 | -0.479114675 | 1.50E-08 |
| ATP1B3 | 8061.625662 | -0.480050128 | 2.20E-07 |
| R3HCC1L | 545.7380229 | -0.480161163 | 5.67E-03 |
| THUMPD3-AS1 | 493.6368638 | -0.480215759 | 3.29E-02 |
| FRRS1L | 848.3939876 | -0.480279794 | 6.24E-05 |
| FAM76A | 274.522027 | -0.480501176 | 2.34E-02 |
| FAM81A | 243.0128007 | -0.480812843 | 2.72E-02 |
| SIN3A | 5272.090568 | -0.480930136 | 1.54E-15 |
| MT-ND2 | 53315.87307 | -0.481251845 | 4.88E-02 |
| CCDC183 | 146.8971672 | -0.481527574 | 1.31E-02 |
| ARMCX1 | 1313.386076 | -0.481598133 | 8.14E-04 |
| BAHCC1 | 1925.241566 | -0.481660967 | 2.39E-03 |
| BTBD9 | 821.1860443 | -0.48175292 | 5.44E-04 |
| MED4 | 1518.399422 | -0.481858914 | 8.03E-05 |
| ZNF235 | 385.1857377 | -0.481922074 | 6.07E-03 |
| GALNT6 | 1395.585243 | -0.482464722 | 9.01E-03 |
| TUT4 | 1942.183595 | -0.482575205 | 1.87E-02 |
| TSPAN9 | 2172.099223 | -0.482772027 | 8.54E-04 |

| | | | |
|---|---|---|---|
| JAG1 | 2378.045133 | -0.483047948 | 4.20E-03 |
| CUL7 | 4517.472227 | -0.483281761 | 6.35E-03 |
| ARMC8 | 1212.528709 | -0.483525713 | 2.49E-03 |
| RAB9B | 263.064607 | -0.48355726 | 1.36E-03 |
| ING4 | 1112.173601 | -0.483619248 | 8.09E-04 |
| ZNF704 | 2053.150087 | -0.484145329 | 4.03E-02 |
| FAM168A | 5995.039326 | -0.484862589 | 2.61E-04 |
| IQCE | 1179.997058 | -0.484984818 | 4.94E-04 |
| AC026362.1 | 273.9551125 | -0.48537606 | 3.22E-02 |
| RBAK | 798.7510512 | -0.485414749 | 4.40E-02 |
| DHRS11 | 725.2445982 | -0.486166245 | 9.21E-06 |
| KTN1 | 7762.162086 | -0.486555529 | 4.90E-02 |
| AC011815.1 | 181.4747312 | -0.486659904 | 1.16E-02 |
| PIGW | 1124.188094 | -0.486714959 | 5.07E-05 |
| PABPC1L | 2421.736173 | -0.486905763 | 1.04E-02 |
| ZNF211 | 653.8157566 | -0.487207796 | 1.39E-04 |
| LAPTM4B | 16786.53935 | -0.487253331 | 1.33E-04 |
| RABL2A | 460.6065084 | -0.487274731 | 1.85E-04 |
| TRAF3IP2 | 955.3632333 | -0.487381145 | 3.23E-04 |
| ARHGAP27 | 474.1617742 | -0.487731324 | 2.06E-02 |
| PLBD1 | 1735.301632 | -0.487733677 | 5.12E-05 |
| RNF141 | 2039.07365 | -0.487864575 | 1.81E-07 |
| DCUN1D4 | 1235.219247 | -0.487971055 | 5.29E-03 |
| ARID2 | 1422.741654 | -0.488134501 | 1.39E-03 |
| TRIM34 | 459.8810898 | -0.488209133 | 4.84E-03 |
| C7orf43 | 1455.089757 | -0.488368637 | 2.76E-06 |
| TTN-AS1 | 253.0060172 | -0.488685567 | 3.83E-03 |
| ZNF384 | 2497.210853 | -0.488755964 | 6.04E-07 |
| SEPSECS | 589.7060353 | -0.488756536 | 3.42E-02 |
| IQCG | 313.8111129 | -0.489093381 | 5.27E-03 |
| ZNF532 | 4172.318559 | -0.489225849 | 1.30E-05 |
| LSM11 | 1292.011328 | -0.489226606 | 1.32E-05 |
| LHX4 | 142.5888116 | -0.489275993 | 2.50E-02 |
| FAM200A | 562.1375119 | -0.489794725 | 3.17E-04 |
| GPATCH2 | 545.3141153 | -0.489810633 | 3.33E-03 |
| IRF9 | 1259.63647 | -0.490087042 | 4.75E-04 |
| PPARG | 1847.312041 | -0.490727995 | 1.34E-03 |
| DENND4C | 2174.261154 | -0.490865072 | 2.95E-05 |
| ZNF45 | 1552.966555 | -0.490950224 | 6.38E-08 |
| ARL15 | 968.3590657 | -0.491258858 | 3.01E-04 |
| ZNF407 | 755.8998402 | -0.491509508 | 8.71E-07 |
| 3-Mar | 427.8141635 | -0.491914159 | 7.26E-04 |
| AL691432.2 | 135.8271995 | -0.492070694 | 1.72E-02 |
| RBL1 | 1227.749753 | -0.492084344 | 3.35E-02 |
| GTF2IP20 | 291.374783 | -0.492127621 | 1.59E-02 |
| MRPS25 | 2424.720312 | -0.492149048 | 1.37E-10 |
| RGS12 | 1044.633314 | -0.49255633 | 6.34E-04 |

| | | | |
|---|---|---|---|
| SARS2 | 871.2070724 | -0.493160923 | 9.16E-04 |
| CTNND1 | 17473.49328 | -0.493167374 | 1.58E-10 |
| SF3B1 | 12797.31495 | -0.493260205 | 3.41E-04 |
| LRRC29 | 210.1297555 | -0.493715677 | 4.31E-03 |
| TARBP1 | 1916.655118 | -0.493875153 | 3.97E-03 |
| NA | 1596.217383 | -0.49388883 | 1.17E-06 |
| SCD5 | 399.6806143 | -0.494015681 | 2.00E-03 |
| MNT | 1455.531257 | -0.494107675 | 1.92E-04 |
| HMBOX1 | 1002.827149 | -0.494387111 | 6.82E-04 |
| TCTN1 | 1339.541659 | -0.494510583 | 1.06E-08 |
| USP38 | 1006.367779 | -0.495351581 | 4.33E-04 |
| HAPLN3 | 305.6184811 | -0.495513379 | 1.19E-02 |
| PRIMPOL | 515.2385594 | -0.495613039 | 7.87E-03 |
| AL590560.2 | 111.1315513 | -0.496047695 | 3.30E-02 |
| KBTBD6 | 801.5697179 | -0.496293946 | 2.19E-04 |
| ZSCAN12 | 304.742191 | -0.497236292 | 3.53E-04 |
| NFATC3 | 2721.717658 | -0.497355019 | 2.75E-13 |
| VASH1-AS1 | 366.4480593 | -0.497437292 | 1.47E-02 |
| AL732372.2 | 291.0174159 | -0.497815853 | 2.13E-02 |
| CNTLN | 1036.894075 | -0.497897455 | 8.59E-03 |
| SPATA13 | 828.620463 | -0.497977949 | 1.27E-02 |
| PANK1 | 348.7559744 | -0.498386036 | 3.47E-03 |
| GTF2IP1 | 5488.638803 | -0.49844943 | 1.09E-06 |
| APLP1 | 955.4105536 | -0.499068034 | 1.49E-03 |
| KALRN | 216.1013953 | -0.499611947 | 1.42E-02 |
| ADAT2 | 484.6253686 | -0.499672725 | 4.32E-02 |
| SUZ12 | 1749.24164 | -0.499678526 | 6.67E-03 |
| AC024896.1 | 158.5161316 | -0.500523756 | 1.14E-02 |
| CACNB3 | 1731.645184 | -0.500709098 | 1.34E-04 |
| MGA | 1832.066296 | -0.500932119 | 7.01E-04 |
| MED23 | 1471.504237 | -0.500999481 | 1.04E-04 |
| OPRL1 | 280.5460774 | -0.50102831 | 3.52E-02 |
| MZF1 | 921.5328365 | -0.501308932 | 1.44E-06 |
| LTN1 | 2205.287303 | -0.501520773 | 4.29E-02 |
| ADNP | 9338.900182 | -0.502167507 | 4.88E-19 |
| CLBA1 | 602.1476952 | -0.502299465 | 1.66E-04 |
| AC007383.2 | 119.3644019 | -0.502456876 | 4.06E-02 |
| PLXNA3 | 2210.81147 | -0.502631569 | 4.25E-03 |
| MRPL1 | 701.2691386 | -0.502650659 | 7.33E-03 |
| CKAP2 | 2416.929446 | -0.503035097 | 5.20E-03 |
| OAS1 | 861.9539716 | -0.503160123 | 2.90E-02 |
| NA | 4877.191813 | -0.503356058 | 9.59E-23 |
| ZNF471 | 354.7592985 | -0.50340921 | 1.48E-02 |
| KLHL15 | 574.0645139 | -0.503530028 | 9.70E-04 |
| HLTF | 3216.029954 | -0.503691828 | 2.36E-03 |
| NLRC5 | 1288.003349 | -0.503852971 | 1.47E-03 |
| LIN9 | 409.9594675 | -0.503877287 | 1.49E-02 |

| | | | |
|---|---|---|---|
| MON1A | 421.933622 | -0.503884125 | 3.69E-05 |
| LUC7L | 2156.329626 | -0.504017985 | 4.66E-11 |
| MPHOSPH10 | 1233.545703 | -0.50402403 | 1.54E-03 |
| LRRC37A | 215.1036013 | -0.50414339 | 2.30E-03 |
| ZNF493 | 182.3837113 | -0.504182773 | 8.36E-03 |
| ZNF354B | 411.7844575 | -0.504204818 | 4.36E-02 |
| ZNF182 | 399.0536131 | -0.50433516 | 2.84E-02 |
| YES1 | 1836.174064 | -0.504351712 | 1.74E-02 |
| ZNF843 | 97.59420435 | -0.504355431 | 3.72E-02 |
| SYNRG | 2863.187215 | -0.504764784 | 1.14E-08 |
| RBM43 | 426.6895759 | -0.50493261 | 3.49E-04 |
| CCDC74B | 422.4592504 | -0.505208523 | 1.36E-03 |
| CHD7 | 2809.574019 | -0.505331835 | 4.68E-03 |
| S1PR2 | 135.2910706 | -0.505804964 | 2.27E-02 |
| SDSL | 476.1971922 | -0.506053507 | 2.79E-02 |
| CILP2 | 791.2075924 | -0.506069248 | 5.30E-03 |
| PKN2 | 2294.834694 | -0.50720505 | 1.51E-02 |
| ENTPD7 | 448.2118662 | -0.507611293 | 2.20E-05 |
| NR2C1 | 844.9690927 | -0.507617063 | 5.26E-03 |
| TMX1 | 2060.293302 | -0.507665093 | 2.98E-03 |
| ABCA7 | 1472.770357 | -0.507779052 | 7.82E-04 |
| AL157392.3 | 363.3674805 | -0.508173345 | 1.28E-03 |
| AC106782.2 | 196.6371558 | -0.508419445 | 1.72E-02 |
| AC138932.1 | 1791.102047 | -0.508580366 | 3.72E-02 |
| ATR | 1999.15314 | -0.509107476 | 1.23E-02 |
| ZNF530 | 486.0653137 | -0.509117658 | 6.70E-04 |
| ZNF883 | 505.0140234 | -0.509395944 | 2.69E-03 |
| ACOX1 | 1603.338045 | -0.509494446 | 2.11E-05 |
| TRMT10C | 913.7961037 | -0.509530274 | 2.32E-04 |
| ZNF76 | 1282.837094 | -0.509743938 | 4.98E-06 |
| FP671120.2 | 584.4415846 | -0.510194816 | 1.47E-06 |
| A1BG-AS1 | 468.9123604 | -0.510206436 | 1.28E-04 |
| FNDC3B | 3823.40014 | -0.510424873 | 2.18E-02 |
| AC020915.3 | 223.8893518 | -0.510852927 | 9.44E-04 |
| AC118549.1 | 1932.132923 | -0.510890473 | 1.56E-02 |
| NCOA1 | 2266.228525 | -0.510933243 | 3.88E-05 |
| CCDC144B | 1018.828952 | -0.511315652 | 1.69E-02 |
| ZNF121 | 633.2258292 | -0.511886407 | 6.70E-03 |
| REST | 1531.355795 | -0.512100129 | 2.69E-04 |
| TTC33 | 553.6499434 | -0.512409852 | 6.62E-03 |
| SRSF11 | 4244.378978 | -0.512424996 | 7.63E-04 |
| BLOC1S6 | 1357.049245 | -0.512818998 | 1.43E-03 |
| AC008966.1 | 495.8421598 | -0.512843529 | 3.37E-05 |
| NA | 1377.415725 | -0.512895192 | 1.63E-04 |
| TSPAN15 | 535.6987614 | -0.513069034 | 4.13E-03 |
| PTGIR | 176.9157293 | -0.513187899 | 2.12E-02 |
| AC090114.2 | 835.2073606 | -0.513229009 | 6.90E-05 |

| | | | |
|---|---|---|---|
| ERGIC2 | 921.606192 | -0.513350056 | 2.14E-02 |
| TRPV4 | 204.8151447 | -0.513551584 | 2.17E-02 |
| TMEM204 | 546.6914666 | -0.513688562 | 2.75E-04 |
| SNX7 | 1214.597973 | -0.514256374 | 2.33E-06 |
| PYGO1 | 602.9014045 | -0.514401513 | 1.31E-03 |
| ENTPD5 | 1104.144112 | -0.514909313 | 1.15E-04 |
| USP46 | 1234.24571 | -0.514944568 | 1.67E-03 |
| DCTN6 | 1034.325814 | -0.514994029 | 6.52E-06 |
| DDT | 2299.728872 | -0.515040147 | 7.81E-06 |
| PIAS3 | 2689.601268 | -0.515169952 | 5.16E-05 |
| RAB42 | 462.7888802 | -0.515540832 | 5.05E-04 |
| PC | 2346.183981 | -0.515675147 | 7.10E-04 |
| MEIS3P1 | 287.1832009 | -0.515743332 | 2.58E-03 |
| TRBV20-1 | 248.8926942 | -0.516075938 | 7.74E-03 |
| SLC22A4 | 550.2859545 | -0.516108968 | 4.79E-05 |
| ABCB9 | 492.6059606 | -0.516200033 | 3.44E-03 |
| PROCR | 2832.564645 | -0.517496925 | 5.15E-05 |
| SPHK2 | 601.2242349 | -0.517644112 | 6.45E-04 |
| CENPS | 482.2845624 | -0.517989158 | 2.30E-05 |
| AC007066.2 | 118.9970978 | -0.518047672 | 2.05E-02 |
| EIF4E | 1732.254152 | -0.518192121 | 3.86E-05 |
| AGGF1 | 3409.272125 | -0.518250749 | 8.62E-06 |
| NA | 205.8975033 | -0.51830646 | 2.44E-03 |
| CROCC | 613.0046235 | -0.518366654 | 1.81E-04 |
| NA | 887.2209421 | -0.518524223 | 1.82E-04 |
| IPO8 | 3229.342953 | -0.51900476 | 1.20E-08 |
| SLC25A13 | 3952.548398 | -0.519042331 | 2.25E-11 |
| ANGEL2 | 967.7784368 | -0.519192269 | 1.23E-06 |
| SETMAR | 386.4169727 | -0.519390826 | 9.32E-06 |
| EXOC3-AS1 | 613.1024133 | -0.519404211 | 9.74E-07 |
| PLAG1 | 287.0523866 | -0.519551713 | 6.56E-03 |
| ZNF37BP | 389.2242676 | -0.51958191 | 3.19E-02 |
| BRCC3 | 1658.547271 | -0.519596383 | 3.85E-04 |
| THSD1 | 104.9408088 | -0.519669461 | 9.34E-03 |
| TTPAL | 3969.055166 | -0.520023987 | 1.67E-15 |
| SUMF1 | 1209.276204 | -0.520495486 | 2.29E-04 |
| HMGN3-AS1 | 78.76644849 | -0.520533708 | 4.74E-02 |
| AL669831.1 | 130.1423115 | -0.520696763 | 1.77E-02 |
| AC002467.1 | 95.26917571 | -0.520811031 | 1.50E-02 |
| AKAP12 | 6631.627566 | -0.520943209 | 1.51E-04 |
| PAXBP1 | 1660.356288 | -0.521219862 | 2.32E-02 |
| MKLN1-AS | 277.6009155 | -0.521446602 | 2.44E-02 |
| RASA1 | 3077.667782 | -0.521688282 | 1.19E-02 |
| RSRC1 | 823.9064721 | -0.521941804 | 4.70E-04 |
| BX004987.1 | 445.7885911 | -0.522434522 | 1.34E-03 |
| ZNF284 | 227.9168793 | -0.522566417 | 3.05E-04 |
| FAM8A1 | 1246.636344 | -0.52259988 | 1.41E-05 |

| | | | |
|---|---|---|---|
| KIF3B | 5246.530593 | -0.522742683 | 1.70E-15 |
| MFSD11 | 952.6038169 | -0.522945381 | 1.24E-06 |
| RAPGEFL1 | 411.4915503 | -0.523177347 | 7.07E-04 |
| U91328.1 | 298.4707967 | -0.523331733 | 3.68E-04 |
| TFR2 | 211.3813313 | -0.523582191 | 2.02E-02 |
| MC1R | 850.7944033 | -0.523707468 | 7.14E-07 |
| ZNF561-AS1 | 275.8020633 | -0.524123166 | 5.56E-03 |
| FAHD1 | 990.6586558 | -0.524208631 | 4.29E-05 |
| AL928654.1 | 326.5701445 | -0.524356871 | 8.17E-04 |
| U2SURP | 3767.063815 | -0.524367197 | 1.58E-02 |
| WDHD1 | 1399.6444 | -0.524481423 | 3.93E-02 |
| ZNF662 | 318.1287865 | -0.524521203 | 3.58E-02 |
| ERVK9-11 | 199.3391714 | -0.524725529 | 7.13E-03 |
| ZNF792 | 217.2366505 | -0.525104723 | 1.02E-02 |
| PHF23 | 4034.620571 | -0.525114175 | 3.23E-05 |
| RAB43 | 506.4836653 | -0.525258254 | 1.43E-02 |
| GNPTAB | 2239.063169 | -0.525480949 | 5.40E-06 |
| TASOR2 | 3918.209476 | -0.525649661 | 1.66E-02 |
| MIER1 | 835.1737643 | -0.525764885 | 3.11E-03 |
| SMIM30 | 1342.204583 | -0.526109895 | 1.33E-04 |
| UBA6 | 1703.960493 | -0.526129663 | 3.35E-02 |
| AP000873.2 | 168.6041013 | -0.526382254 | 4.04E-03 |
| RFX5 | 4160.592771 | -0.526543375 | 1.80E-15 |
| MFSD6 | 998.7542204 | -0.526761535 | 2.95E-07 |
| STRIP2 | 1045.859077 | -0.52691733 | 4.55E-04 |
| WIPF1 | 1839.234446 | -0.52697383 | 3.29E-10 |
| BIRC6 | 6959.61733 | -0.527081764 | 6.02E-05 |
| SGSM2 | 4223.108752 | -0.527294479 | 8.85E-06 |
| RBM45 | 475.9013446 | -0.527430439 | 4.80E-04 |
| ZBTB3 | 245.4977007 | -0.52754503 | 1.24E-02 |
| C1orf112 | 1218.550267 | -0.527683567 | 8.59E-05 |
| FNIP1 | 3476.825664 | -0.527845781 | 3.44E-04 |
| MYO1B | 8268.616834 | -0.528149406 | 5.44E-06 |
| TBCCD1 | 605.9990156 | -0.528216535 | 2.07E-06 |
| EXOSC4 | 1304.84018 | -0.528363847 | 1.78E-05 |
| MIB1 | 2496.134306 | -0.528463429 | 5.59E-03 |
| VILL | 408.5323477 | -0.528474368 | 4.67E-05 |
| CASTOR3 | 877.9701155 | -0.528587126 | 1.36E-04 |
| HAGHL | 852.8669522 | -0.529154019 | 1.73E-03 |
| CCP110 | 1115.965857 | -0.529332117 | 1.26E-02 |
| RAPGEF3 | 1404.252954 | -0.529698791 | 2.64E-03 |
| PGBD4 | 228.0935864 | -0.530026102 | 2.26E-03 |
| PRRX2 | 217.4448498 | -0.530222217 | 3.47E-02 |
| PIGZ | 330.2327841 | -0.530267208 | 1.03E-02 |
| TNC | 30916.41729 | -0.530968527 | 1.12E-02 |
| AC003665.1 | 447.9924771 | -0.531436774 | 2.17E-02 |
| CORO1A | 260.7712798 | -0.531469029 | 4.74E-03 |

| | | | |
|---|---|---|---|
| PBX1 | 1185.042455 | -0.531492817 | 1.33E-07 |
| PYM1 | 593.1831054 | -0.531892939 | 2.36E-05 |
| ZNF771 | 204.8286513 | -0.531898232 | 7.29E-03 |
| B3GALT6 | 999.9083851 | -0.531972646 | 1.43E-04 |
| ITGB5 | 9170.149928 | -0.532078806 | 3.15E-03 |
| AC026271.3 | 154.9889749 | -0.532170608 | 3.21E-02 |
| SMARCA1 | 5733.867665 | -0.532262326 | 7.72E-11 |
| WBP1L | 2089.230149 | -0.532267557 | 9.33E-04 |
| SHROOM1 | 1653.721303 | -0.532428679 | 6.55E-06 |
| ZNHIT2 | 588.7238783 | -0.532976122 | 1.28E-03 |
| MAPK11 | 1201.739867 | -0.533817997 | 7.35E-09 |
| AL590428.1 | 120.8243549 | -0.533961591 | 9.64E-03 |
| ZNF785 | 729.5996148 | -0.534351794 | 6.29E-06 |
| SYT7 | 257.2568568 | -0.534444139 | 1.11E-03 |
| CENPK | 922.9012945 | -0.534700226 | 3.90E-02 |
| GTF2IRD2B | 908.8548731 | -0.534750752 | 2.74E-03 |
| PHF20L1 | 2256.175305 | -0.534771774 | 9.83E-03 |
| KBTBD4 | 1247.543525 | -0.535029223 | 1.66E-03 |
| MATR3 | 12010.77523 | -0.535231676 | 2.34E-04 |
| TREX1 | 689.4221273 | -0.535720967 | 3.26E-02 |
| NCAM1 | 1326.577166 | -0.535726185 | 3.64E-07 |
| NQO1 | 36091.71617 | -0.535761154 | 1.83E-03 |
| HOXA-AS2 | 648.9155533 | -0.535877068 | 2.39E-04 |
| MYNN | 726.2999099 | -0.536359074 | 8.02E-04 |
| ZNF362 | 968.369252 | -0.536968173 | 1.26E-02 |
| FAM102B | 808.5164647 | -0.537059442 | 9.12E-04 |
| WBP1 | 3743.929191 | -0.53724209 | 9.77E-04 |
| FOXRED2 | 5609.229125 | -0.537256123 | 5.02E-05 |
| JKAMP | 1240.07522 | -0.537344985 | 1.21E-04 |
| GLRX | 2880.011914 | -0.53745767 | 3.28E-08 |
| DPYD | 377.6234819 | -0.53748086 | 2.06E-04 |
| PXYLP1 | 564.4135188 | -0.537702453 | 3.41E-04 |
| AMACR | 1358.410533 | -0.537966315 | 5.39E-19 |
| SNX1 | 8206.680958 | -0.538008413 | 2.01E-08 |
| ZNF75D | 1009.175721 | -0.538042288 | 2.66E-05 |
| PTRHD1 | 611.6881417 | -0.538281047 | 2.33E-02 |
| LNPEP | 4087.045638 | -0.538305564 | 4.24E-06 |
| FAS | 239.2567132 | -0.538337439 | 3.85E-02 |
| CHN1 | 893.6230543 | -0.538538369 | 2.18E-07 |
| MAPK8IP1 | 1166.539598 | -0.538581282 | 1.04E-04 |
| PRPF40A | 3930.555096 | -0.538718114 | 7.24E-03 |
| SCYL3 | 584.6494419 | -0.538745452 | 6.93E-05 |
| SNRNP25 | 1685.632207 | -0.538874013 | 1.72E-03 |
| AL845472.2 | 297.2034584 | -0.538986268 | 8.93E-04 |
| STT3B | 6050.763375 | -0.539063737 | 1.58E-09 |
| AC124864.2 | 105.2312385 | -0.539284544 | 4.07E-02 |
| AC093726.1 | 112.1297998 | -0.539817947 | 3.11E-02 |

| | | | |
|---|---|---|---|
| SNPH | 1189.696174 | -0.540030393 | 2.26E-05 |
| NA | 115.5666921 | -0.540110389 | 2.92E-02 |
| ZBED5 | 1813.585829 | -0.540136131 | 1.79E-03 |
| DGLUCY | 2153.555328 | -0.540319836 | 1.90E-07 |
| ZNF630 | 219.9959955 | -0.540512724 | 1.64E-03 |
| TP53BP1 | 4052.421138 | -0.540560594 | 2.41E-13 |
| CTXN1 | 271.998645 | -0.540658709 | 8.06E-04 |
| AC004803.1 | 262.714056 | -0.540675699 | 2.76E-02 |
| CRELD2 | 1562.4412 | -0.540697669 | 9.75E-05 |
| TRIM66 | 654.3240807 | -0.540936636 | 3.63E-02 |
| ABHD14A | 756.7593186 | -0.541058043 | 6.67E-04 |
| UFC1 | 2361.765324 | -0.541527308 | 1.44E-06 |
| NUDT11 | 434.1893591 | -0.541694328 | 8.33E-07 |
| SSBP3 | 3859.235303 | -0.542267744 | 6.46E-06 |
| IL27RA | 1682.285814 | -0.542274082 | 8.78E-05 |
| CNTNAP1 | 3759.390506 | -0.542863002 | 5.33E-07 |
| PIGF | 619.1242501 | -0.543071717 | 7.23E-06 |
| PPIL3 | 802.2648949 | -0.543203083 | 5.44E-05 |
| PIK3CA | 1376.042323 | -0.543231887 | 1.41E-02 |
| ZNF134 | 1247.108248 | -0.543264549 | 2.11E-05 |
| LRBA | 3125.68714 | -0.54341202 | 7.84E-06 |
| CUL5 | 1325.527547 | -0.543900876 | 3.33E-02 |
| ATRIP | 655.0171284 | -0.544246621 | 8.24E-04 |
| ZNF436-AS1 | 269.1957982 | -0.544383952 | 1.86E-02 |
| FBXL21P | 1580.257631 | -0.544421845 | 3.66E-02 |
| ING5 | 1464.070134 | -0.544614048 | 1.43E-08 |
| ZNF614 | 555.4187751 | -0.54484541 | 2.76E-06 |
| LIMS1 | 1656.651267 | -0.545215901 | 1.70E-04 |
| KLHL11 | 178.0910136 | -0.545351017 | 7.73E-03 |
| MRTFB | 2701.886717 | -0.545367586 | 3.43E-03 |
| RBM4B | 1235.024212 | -0.545517469 | 1.17E-03 |
| DCAF17 | 861.6994227 | -0.545580439 | 1.61E-02 |
| SYT14 | 285.8834422 | -0.545594996 | 1.00E-02 |
| TASOR | 4044.453702 | -0.545996995 | 3.13E-03 |
| FAM47E | 174.1350881 | -0.546089899 | 7.74E-03 |
| MT-ND1 | 34633.62567 | -0.546846782 | 1.30E-02 |
| GTF2A1 | 961.5536787 | -0.546917983 | 9.83E-03 |
| DIS3 | 4813.73347 | -0.54714266 | 2.15E-03 |
| DCP2 | 2102.574905 | -0.54733792 | 4.97E-05 |
| FOXO4 | 737.7610709 | -0.547345643 | 3.25E-05 |
| TTC37 | 7042.255014 | -0.547620983 | 3.88E-02 |
| CCAR1 | 2806.344201 | -0.548021434 | 2.67E-05 |
| RNF219 | 1318.110589 | -0.548101345 | 7.05E-03 |
| WDR81 | 3131.426754 | -0.54813202 | 5.63E-05 |
| ZNF585B | 695.3659731 | -0.548186201 | 7.38E-05 |
| GTF2IRD2 | 351.0317861 | -0.548408204 | 4.32E-02 |
| GALNT7 | 2610.119088 | -0.548491881 | 3.11E-05 |

| | | | |
|---|---|---|---|
| FAM133A | 452.9836405 | -0.548514647 | 9.79E-04 |
| MT-CYB | 107218.2105 | -0.54877988 | 5.62E-03 |
| UBR5 | 9785.128236 | -0.548784871 | 3.51E-16 |
| AP1AR | 523.019085 | -0.548909157 | 3.83E-02 |
| MYO9A | 2981.552426 | -0.549127118 | 8.81E-04 |
| CNPY4 | 2550.890741 | -0.549987502 | 1.92E-05 |
| BTN3A1 | 1442.820939 | -0.550015333 | 1.14E-05 |
| CCDC82 | 785.0585224 | -0.550599249 | 4.83E-02 |
| SLC29A2 | 765.8787347 | -0.550708011 | 2.52E-03 |
| AL031282.2 | 397.8015291 | -0.550971935 | 6.53E-05 |
| NACAD | 1177.507431 | -0.551278972 | 3.38E-03 |
| HIBCH | 1071.09971 | -0.551293369 | 3.56E-04 |
| GLI3 | 3529.66754 | -0.551298388 | 1.49E-06 |
| AC092919.2 | 104.0774614 | -0.552749126 | 3.43E-02 |
| AC243964.2 | 208.6143951 | -0.553266057 | 6.20E-03 |
| BSCL2 | 1044.211796 | -0.553670558 | 4.03E-03 |
| HELZ | 2055.418843 | -0.553797551 | 1.57E-07 |
| THBS3 | 1463.203303 | -0.554091388 | 1.68E-06 |
| AL117332.1 | 115.9506942 | -0.554275516 | 2.90E-02 |
| SRSF6 | 9770.517034 | -0.554300932 | 1.47E-07 |
| BX664615.1 | 109.2441282 | -0.554309878 | 1.18E-02 |
| CDK17 | 880.674751 | -0.554465754 | 4.65E-03 |
| ANKIB1 | 5079.40403 | -0.554512509 | 2.73E-06 |
| SETD9 | 410.3844377 | -0.554715296 | 4.51E-04 |
| SRPX | 4619.232277 | -0.554725984 | 2.68E-06 |
| SPOPL | 436.6793134 | -0.554825309 | 2.46E-02 |
| TNFAIP2 | 440.9777675 | -0.555398477 | 4.06E-03 |
| LEAP2 | 96.45515125 | -0.555462893 | 2.57E-02 |
| TCF12 | 5252.519598 | -0.555759573 | 1.53E-02 |
| ARSD | 1843.274745 | -0.55582583 | 2.54E-07 |
| TNFSF13 | 541.1227978 | -0.555989563 | 3.68E-03 |
| HCAR1 | 765.884658 | -0.556353083 | 2.11E-05 |
| NRG1 | 723.2771418 | -0.556733919 | 3.37E-04 |
| SPOCK1 | 3684.499858 | -0.556776947 | 9.18E-08 |
| ETAA1 | 484.0096387 | -0.556935048 | 3.62E-02 |
| SLC2A6 | 1907.900978 | -0.55755456 | 6.47E-04 |
| FMC1-LUC7L2 | 1583.580303 | -0.557842483 | 8.79E-08 |
| FAM89B | 1379.792366 | -0.558132824 | 4.52E-05 |
| PAXIP1-AS2 | 428.289815 | -0.558695639 | 5.75E-04 |
| FAM234A | 7638.339532 | -0.55892309 | 1.74E-06 |
| ZNF252P | 1467.129611 | -0.559411097 | 5.06E-03 |
| SCFD1 | 2000.414519 | -0.560033128 | 4.84E-03 |
| SLC35A1 | 1520.866646 | -0.560035735 | 2.34E-04 |
| STXBP5 | 998.2124252 | -0.560213449 | 4.05E-04 |
| PKD2 | 2184.663901 | -0.560509177 | 3.59E-08 |
| GPATCH11 | 944.2607256 | -0.560536756 | 3.80E-04 |
| ZEB1 | 1261.88444 | -0.560614346 | 4.01E-02 |

| | | | |
|---|---|---|---|
| NA | 365.8632067 | -0.560767098 | 3.95E-03 |
| SLC38A7 | 1481.333423 | -0.560800139 | 2.97E-06 |
| ERMP1 | 4149.788729 | -0.56086509 | 2.72E-10 |
| ERAP2 | 8236.42801 | -0.561068518 | 1.52E-15 |
| SLC44A1 | 3595.730604 | -0.561105635 | 1.41E-13 |
| SEC62 | 2051.27731 | -0.561887802 | 4.65E-03 |
| RAP1GAP | 474.2442089 | -0.562020128 | 3.04E-03 |
| HLA-DMA | 431.2072074 | -0.562295079 | 4.68E-02 |
| APPBP2 | 2193.313553 | -0.56233738 | 2.52E-04 |
| AP4S1 | 353.5657067 | -0.562596377 | 1.62E-02 |
| RASSF2 | 380.6043208 | -0.562604949 | 2.66E-04 |
| SETD1B | 1514.715378 | -0.563131636 | 6.75E-08 |
| C2orf76 | 103.0839047 | -0.563857641 | 1.70E-02 |
| SCO2 | 668.0786893 | -0.563936264 | 4.88E-04 |
| CAPS | 230.0045912 | -0.564354948 | 3.82E-03 |
| KRCC1 | 655.4614552 | -0.564470154 | 3.01E-02 |
| SEZ6L2 | 3889.516264 | -0.56457083 | 1.08E-04 |
| BPHL | 824.4779233 | -0.564805791 | 6.64E-04 |
| LRRC37B | 470.6949516 | -0.5648557 | 5.33E-05 |
| ZDHHC20 | 1478.023608 | -0.565313244 | 1.12E-03 |
| RBM26-AS1 | 95.22955443 | -0.56555237 | 3.44E-02 |
| HOXB5 | 349.4882309 | -0.566305377 | 1.02E-04 |
| STX1A | 710.3622965 | -0.566327969 | 5.42E-06 |
| PER3 | 895.4161079 | -0.566404498 | 4.86E-04 |
| CABP4 | 464.2960878 | -0.566492738 | 5.92E-05 |
| AL354822.1 | 208.7467612 | -0.566849656 | 3.36E-02 |
| DNTTIP2 | 1683.644075 | -0.56706707 | 1.23E-02 |
| BORCS8 | 545.5502618 | -0.567155647 | 1.37E-04 |
| LINC00652 | 176.6260923 | -0.567329903 | 8.02E-04 |
| INTS3 | 5212.447762 | -0.567358781 | 2.55E-10 |
| CERS6 | 1849.566839 | -0.568215363 | 1.48E-06 |
| HSBP1 | 10345.67201 | -0.568510777 | 1.38E-06 |
| MT-ND6 | 16357.53968 | -0.568685154 | 1.02E-02 |
| PHF3 | 3186.201128 | -0.568745332 | 1.13E-02 |
| TMEM14A | 661.6508551 | -0.56883637 | 7.06E-06 |
| STK38L | 684.3303227 | -0.56884887 | 6.75E-03 |
| AC138035.1 | 323.6251374 | -0.568988946 | 1.17E-03 |
| BFSP1 | 448.8421094 | -0.56928916 | 1.47E-05 |
| PHF6 | 2873.346761 | -0.569320295 | 3.10E-04 |
| IRAK3 | 959.1155894 | -0.57091824 | 8.00E-05 |
| PHLDA1 | 5771.939055 | -0.571282357 | 7.23E-04 |
| TGFBR1 | 4359.922317 | -0.571371817 | 2.90E-04 |
| APH1B | 999.3437867 | -0.571457464 | 4.90E-05 |
| FNDC3A | 2717.186824 | -0.571461876 | 9.80E-05 |
| PANK3 | 2532.270152 | -0.57150806 | 6.18E-03 |
| OBSCN-AS1 | 87.8409625 | -0.571575721 | 2.34E-02 |
| ATE1 | 938.3486093 | -0.571812759 | 3.50E-03 |

| | | | |
|---|---|---|---|
| PCYOX1 | 5760.612629 | -0.571832726 | 2.61E-07 |
| SPATA33 | 423.1964117 | -0.571982917 | 3.87E-04 |
| MT-ND5 | 72234.1821 | -0.572059742 | 3.15E-03 |
| ALG10 | 159.280928 | -0.572187661 | 3.63E-02 |
| MLKL | 890.2288898 | -0.572971397 | 2.76E-07 |
| MEPCE | 5159.706351 | -0.573131182 | 6.99E-12 |
| GNAQ | 3447.817147 | -0.573697929 | 5.10E-06 |
| NA | 104.9217022 | -0.573822421 | 1.23E-02 |
| LGALSL | 1263.152514 | -0.573946586 | 3.84E-06 |
| TMEM132D-AS1 | 225.5041171 | -0.573990874 | 4.14E-02 |
| ZNF436 | 1465.456013 | -0.574035257 | 1.10E-09 |
| TRIM38 | 1546.239524 | -0.57425415 | 3.39E-11 |
| MDM1 | 410.0232745 | -0.575063411 | 1.38E-02 |
| C11orf71 | 513.7332104 | -0.575328271 | 7.79E-06 |
| ZBTB37 | 507.5107261 | -0.575565304 | 3.33E-03 |
| FRMD8 | 2039.255936 | -0.575755797 | 8.07E-06 |
| PRRT1 | 226.6096598 | -0.575925899 | 4.00E-04 |
| AC010132.3 | 1353.721702 | -0.575957689 | 3.77E-05 |
| FBXL3 | 1749.761037 | -0.576308599 | 1.45E-07 |
| SRCIN1 | 313.1286063 | -0.576857553 | 5.67E-04 |
| CCDC136 | 1824.450033 | -0.577037201 | 1.97E-07 |
| ENTPD1 | 1129.684048 | -0.577616325 | 4.67E-07 |
| AFG3L1P | 627.678639 | -0.577884124 | 1.54E-05 |
| PAPSS2 | 187.5959033 | -0.578091827 | 1.09E-02 |
| LIPG | 311.095507 | -0.578198939 | 1.84E-03 |
| TSPYL1 | 2490.20289 | -0.578607576 | 3.09E-07 |
| CTR9 | 4964.492339 | -0.578847949 | 5.23E-07 |
| TTC28 | 2641.399182 | -0.57898085 | 5.15E-04 |
| NFATC4 | 3878.575819 | -0.579003154 | 7.87E-05 |
| LANCL2 | 1253.792335 | -0.579266794 | 1.72E-04 |
| STK36 | 1396.326232 | -0.579442365 | 6.33E-08 |
| COL6A2 | 15718.66709 | -0.579724376 | 1.02E-07 |
| DNM1 | 2280.331942 | -0.580026827 | 8.32E-09 |
| GUSBP1 | 518.3979596 | -0.580532905 | 1.66E-05 |
| ZNF586 | 548.3422355 | -0.580702864 | 9.34E-06 |
| PCMTD1 | 851.989544 | -0.58074108 | 1.43E-03 |
| ZNF229 | 1229.958577 | -0.581037969 | 4.79E-04 |
| TCTA | 1330.44691 | -0.581497064 | 8.26E-07 |
| RCAN3 | 1155.641347 | -0.581873539 | 4.81E-07 |
| BCAP29 | 2042.475783 | -0.582134988 | 1.16E-05 |
| ZNF230 | 247.6155874 | -0.582140845 | 6.24E-03 |
| ZNF189 | 953.5826802 | -0.582160456 | 1.82E-04 |
| ALX1 | 332.6739069 | -0.582163135 | 4.64E-05 |
| CLUHP3 | 309.9195558 | -0.58252875 | 2.06E-03 |
| AFAP1L2 | 2158.646464 | -0.58282046 | 4.94E-06 |
| COL5A3 | 625.5339231 | -0.583078072 | 9.22E-03 |
| WDR5B | 519.3509633 | -0.583608003 | 6.23E-06 |

| | | | |
|---|---|---|---|
| ZNF449 | 394.9404058 | -0.5841491 | 3.91E-03 |
| SLC25A20 | 850.5040378 | -0.584809877 | 1.27E-03 |
| AHCTF1 | 2181.127846 | -0.584944933 | 2.08E-02 |
| ECM1 | 273.6904325 | -0.585311758 | 1.88E-02 |
| LAMTOR3 | 1013.807565 | -0.585772888 | 9.68E-05 |
| ACACB | 1090.006831 | -0.58656548 | 2.02E-07 |
| ARHGEF40 | 6406.672053 | -0.58672964 | 6.48E-05 |
| KRIT1 | 1969.030056 | -0.587389782 | 1.20E-03 |
| MXRA5 | 2340.291441 | -0.587457355 | 6.02E-11 |
| 7-Sep | 8237.371082 | -0.587655631 | 4.86E-03 |
| CD58 | 338.3302354 | -0.587955774 | 1.70E-04 |
| SPRY2 | 1269.954807 | -0.588032876 | 9.34E-07 |
| POMK | 182.4422372 | -0.588043912 | 1.34E-04 |
| FES | 826.5363836 | -0.588641536 | 1.02E-04 |
| MYO5A | 3183.631342 | -0.588642124 | 2.32E-05 |
| HSPB6 | 899.0889698 | -0.589080323 | 1.33E-03 |
| ZNF184 | 394.1955242 | -0.589084105 | 2.52E-04 |
| RNFT1 | 531.588277 | -0.589707592 | 2.53E-04 |
| GTF3C6 | 1875.255331 | -0.589822244 | 8.10E-08 |
| ZNF195 | 778.8841383 | -0.589963866 | 2.78E-03 |
| CPT1B | 640.0772637 | -0.590118635 | 1.75E-05 |
| MON2 | 1932.049245 | -0.590507251 | 1.91E-03 |
| C2CD5 | 1692.653386 | -0.590660192 | 1.47E-03 |
| HGSNAT | 3374.663359 | -0.59066661 | 7.74E-06 |
| CDC37L1 | 623.5580681 | -0.591371421 | 1.59E-03 |
| IFT57 | 1167.083796 | -0.591559899 | 4.20E-05 |
| TIMP1 | 7816.765979 | -0.591631696 | 1.34E-03 |
| NAP1L5 | 362.4858112 | -0.59176242 | 6.14E-04 |
| SAMD10 | 302.1944671 | -0.591853077 | 6.27E-03 |
| CROT | 779.6871208 | -0.593067155 | 8.00E-05 |
| AC234582.1 | 90.25612707 | -0.593242952 | 2.01E-02 |
| DUSP5 | 787.2594332 | -0.593307411 | 1.10E-02 |
| RASA2 | 422.6520801 | -0.593670364 | 3.92E-02 |
| ARMCX4 | 791.492689 | -0.593730922 | 2.67E-05 |
| MAML3 | 1424.798106 | -0.594158929 | 5.70E-06 |
| DTNA | 635.6860956 | -0.594594443 | 1.43E-04 |
| TOGARAM1 | 1182.064204 | -0.594659869 | 3.01E-03 |
| AC002310.6 | 87.54832053 | -0.594781497 | 4.59E-02 |
| HOMER1 | 828.6769183 | -0.595001834 | 1.81E-04 |
| GON7 | 437.6675684 | -0.595066504 | 6.10E-07 |
| OFD1 | 1073.083644 | -0.595198731 | 2.27E-05 |
| RENBP | 435.6778357 | -0.595566318 | 2.98E-02 |
| STX17 | 1044.005854 | -0.59575831 | 6.00E-04 |
| AC060234.2 | 135.2052896 | -0.595888369 | 1.06E-03 |
| ENOX2 | 1065.289968 | -0.59624449 | 2.14E-10 |
| DZIP3 | 1549.200322 | -0.596327573 | 1.19E-02 |
| ZNF608 | 2639.516423 | -0.59642762 | 1.62E-09 |

| | | | |
|---|---|---|---|
| THRA | 4147.420593 | -0.596736634 | 2.89E-08 |
| GLI4 | 718.4885791 | -0.59679081 | 3.37E-05 |
| SVBP | 540.4729975 | -0.596858016 | 2.05E-05 |
| SAFB2 | 1849.688607 | -0.598075602 | 1.00E-06 |
| FOXD2 | 276.8095472 | -0.59811741 | 1.98E-05 |
| ANGPTL2 | 1290.993455 | -0.598207701 | 1.03E-04 |
| PUS7L | 453.9353949 | -0.598300834 | 4.54E-02 |
| GPR137 | 1729.845174 | -0.598319069 | 6.06E-06 |
| OXR1 | 901.5233758 | -0.598421295 | 4.39E-05 |
| ZSWIM5 | 539.9072653 | -0.598937596 | 5.68E-05 |
| XRN1 | 1398.276899 | -0.598963533 | 4.28E-02 |
| LBHD1 | 835.1787337 | -0.599006449 | 1.38E-05 |
| FAM217B | 1604.216906 | -0.600238533 | 3.01E-03 |
| TOPORS | 1768.495239 | -0.600642476 | 1.51E-06 |
| CAND1 | 6229.939359 | -0.600844243 | 5.87E-09 |
| PLA2G6 | 356.9025369 | -0.600926575 | 2.32E-03 |
| HEXIM2 | 229.1603445 | -0.600946441 | 1.88E-04 |
| REL | 362.8332223 | -0.600984798 | 2.61E-03 |
| SASH1 | 1551.444222 | -0.601600717 | 2.11E-13 |
| PHF14 | 2956.198094 | -0.601738468 | 2.53E-05 |
| RIOK2 | 1734.119616 | -0.601877202 | 1.59E-02 |
| GLYCTK | 459.0239151 | -0.601949111 | 3.75E-04 |
| P2RX6 | 798.6419577 | -0.601977628 | 1.74E-06 |
| FBXW7 | 632.8639789 | -0.60215294 | 1.03E-05 |
| SLC35E3 | 1051.924473 | -0.602374973 | 1.92E-05 |
| AP003352.1 | 107.9554894 | -0.602796495 | 2.68E-02 |
| TSPOAP1 | 131.5695784 | -0.602829278 | 3.83E-02 |
| IFT80 | 1020.407039 | -0.603008008 | 4.32E-02 |
| C6orf120 | 1436.962007 | -0.603136913 | 9.24E-07 |
| AL023806.3 | 69.85019706 | -0.603181104 | 3.77E-02 |
| RHPN1 | 347.3131214 | -0.603238674 | 1.39E-03 |
| B4GALNT4 | 1110.705317 | -0.603243977 | 2.09E-03 |
| QSER1 | 3135.860873 | -0.603328045 | 1.83E-03 |
| SAYSD1 | 553.4734968 | -0.604374183 | 3.62E-07 |
| ELK4 | 1399.151224 | -0.604818026 | 2.67E-04 |
| LUC7L3 | 3823.723063 | -0.604917772 | 1.92E-02 |
| TOR3A | 1769.121677 | -0.604942357 | 5.32E-09 |
| ACOT11 | 447.5869448 | -0.605463669 | 5.32E-05 |
| ELAC1 | 322.3283585 | -0.605480486 | 3.00E-05 |
| ZNF557 | 384.8904144 | -0.605626794 | 1.61E-08 |
| SNX4 | 778.4362423 | -0.606005108 | 1.54E-03 |
| RNF168 | 803.0081132 | -0.606057438 | 2.75E-11 |
| VPS45 | 2279.513238 | -0.60630518 | 9.59E-13 |
| RAD51-AS1 | 223.6582899 | -0.606683871 | 1.39E-04 |
| WASHC4 | 2069.96772 | -0.607022175 | 1.88E-02 |
| UBE2D1 | 283.642357 | -0.607293604 | 1.27E-03 |
| FUS | 18454.28601 | -0.607318773 | 9.42E-06 |

| | | | |
|---|---|---|---|
| ARHGAP20 | 387.949941 | -0.608342709 | 9.46E-03 |
| LRRC37A2 | 983.0797025 | -0.608547707 | 5.11E-14 |
| EPHB4 | 7703.29599 | -0.608563425 | 4.77E-06 |
| CTSA | 8271.579524 | -0.608600644 | 6.11E-06 |
| DMTF1 | 2324.734966 | -0.608613614 | 3.39E-04 |
| MSI2 | 3793.636912 | -0.60881878 | 2.18E-14 |
| SLCO2A1 | 1257.747719 | -0.608898378 | 1.76E-08 |
| CENPJ | 546.3815123 | -0.608946349 | 8.39E-03 |
| FAM126A | 3391.101266 | -0.608961511 | 2.47E-04 |
| ZSCAN23 | 119.0486742 | -0.609332847 | 1.18E-02 |
| AC245452.1 | 293.78634 | -0.609386434 | 1.03E-04 |
| RNF138 | 715.7112521 | -0.609492932 | 9.72E-04 |
| GOLGA8B | 542.4888917 | -0.609897507 | 1.99E-04 |
| CPE | 2379.926947 | -0.609934077 | 8.88E-06 |
| ZNF135 | 520.20737 | -0.610806692 | 5.87E-07 |
| FAM106A | 885.1586983 | -0.610842842 | 5.25E-03 |
| CLN5 | 1590.052287 | -0.61087584 | 6.13E-06 |
| AC007382.1 | 86.90946211 | -0.611768129 | 4.46E-02 |
| DBI | 4740.869507 | -0.611796084 | 7.84E-08 |
| NA | 153.5758231 | -0.611810906 | 1.55E-03 |
| LINC01001 | 877.8552253 | -0.61198541 | 1.97E-03 |
| ACSL4 | 4513.913578 | -0.612121993 | 8.11E-07 |
| ELK3 | 2668.256893 | -0.612195589 | 1.02E-08 |
| TET1 | 199.1206885 | -0.612328918 | 3.58E-03 |
| NIPSNAP3B | 109.7404881 | -0.612385896 | 1.56E-02 |
| PRPH | 172.1192953 | -0.612895955 | 2.53E-02 |
| ROGDI | 517.5564016 | -0.613650714 | 5.62E-05 |
| ZNF181 | 409.5785145 | -0.613823594 | 8.79E-03 |
| FAM172A | 1449.935888 | -0.613895157 | 2.44E-09 |
| SMARCA2 | 4976.288903 | -0.614460732 | 8.89E-11 |
| ANKRD17 | 5883.493827 | -0.614570516 | 3.71E-17 |
| MMP14 | 14551.73684 | -0.614923413 | 4.71E-06 |
| AHI1 | 978.9390272 | -0.615980325 | 8.39E-03 |
| DNASE1L2 | 80.57937059 | -0.616015895 | 2.86E-02 |
| PPIP5K2 | 3322.167985 | -0.616062212 | 1.35E-03 |
| ETFRF1 | 176.6651362 | -0.616145556 | 1.31E-02 |
| METRN | 2250.581151 | -0.616217864 | 7.50E-05 |
| IPP | 567.557776 | -0.616357029 | 3.17E-07 |
| NSUN5P2 | 311.8154007 | -0.616662392 | 8.84E-04 |
| LINC00680 | 465.4090567 | -0.616702684 | 2.68E-07 |
| GCC2 | 1356.216985 | -0.616959271 | 2.05E-02 |
| ANGPT1 | 607.9227096 | -0.617202611 | 1.07E-03 |
| AL390728.4 | 395.6292296 | -0.617364333 | 4.00E-03 |
| SLC2A11 | 910.2474593 | -0.61750648 | 3.24E-07 |
| CRLF1 | 1052.296416 | -0.617747423 | 1.78E-05 |
| TKFC | 1546.53106 | -0.618110079 | 3.28E-15 |
| HENMT1 | 1022.022023 | -0.618172955 | 1.38E-07 |

| | | | |
|---|---|---|---|
| TUSC3 | 4463.581761 | -0.618301012 | 1.51E-24 |
| DNM1P41 | 127.8426165 | -0.618377538 | 3.70E-03 |
| ZNF418 | 240.3467327 | -0.618450332 | 4.90E-03 |
| GSAP | 508.953102 | -0.618680283 | 2.26E-05 |
| SRGAP3 | 587.6223722 | -0.618769866 | 7.49E-06 |
| PCM1 | 4712.755582 | -0.619215816 | 1.05E-02 |
| TMSB15A | 178.0663652 | -0.619238019 | 1.55E-03 |
| PHF2P2 | 183.8502381 | -0.619598567 | 2.22E-02 |
| SNRK | 1621.869494 | -0.619856481 | 1.06E-08 |
| VTA1 | 2431.669244 | -0.620190741 | 1.13E-08 |
| TIGD7 | 862.8036905 | -0.620727038 | 1.07E-03 |
| SLC9A3R1 | 2447.527257 | -0.62094383 | 7.49E-07 |
| FRK | 159.893411 | -0.621189624 | 1.27E-02 |
| TGFBR2 | 6562.551738 | -0.621212664 | 1.83E-09 |
| IDUA | 285.6584141 | -0.621293241 | 1.68E-02 |
| PDE7A | 1118.518144 | -0.622118038 | 1.45E-06 |
| NA | 105.2038393 | -0.622234151 | 2.28E-03 |
| HNRNPH2 | 2907.243069 | -0.622857937 | 1.66E-08 |
| AC009126.1 | 120.3707254 | -0.62289183 | 1.23E-02 |
| ORC3 | 1863.49752 | -0.623076948 | 1.35E-06 |
| BPTF | 4182.268897 | -0.623709475 | 5.34E-05 |
| S1PR1 | 611.9125336 | -0.623823312 | 2.34E-04 |
| VPS50 | 1142.578612 | -0.624243966 | 5.96E-03 |
| LTBP3 | 6500.71372 | -0.624296133 | 1.33E-06 |
| ZNF239 | 420.1033766 | -0.624514509 | 3.87E-03 |
| DENND6B | 132.705302 | -0.624911049 | 1.93E-02 |
| FNIP2 | 1389.398079 | -0.625273139 | 1.71E-08 |
| FTX | 118.6607101 | -0.625702174 | 7.87E-03 |
| MAP4K4 | 13426.54364 | -0.626051304 | 3.36E-24 |
| HIST2H3C | 92.93385555 | -0.626143735 | 3.86E-02 |
| AC009053.1 | 341.7337856 | -0.626548056 | 4.38E-07 |
| TPD52L1 | 2117.343253 | -0.62738887 | 2.77E-04 |
| ZNF513 | 1599.06592 | -0.627501408 | 1.69E-08 |
| ZNF485 | 223.0510035 | -0.627986691 | 4.27E-03 |
| CEP350 | 2166.233297 | -0.628226954 | 1.04E-02 |
| SPARC | 56594.83605 | -0.628332195 | 1.75E-03 |
| STON2 | 213.0074989 | -0.629041749 | 8.21E-03 |
| CENPC | 439.1903264 | -0.629078588 | 3.42E-02 |
| ZNF345 | 182.6037155 | -0.629081678 | 6.21E-04 |
| C3orf62 | 371.8425366 | -0.629125183 | 1.09E-02 |
| MRPL50 | 1319.50401 | -0.629650198 | 1.34E-08 |
| C11orf96 | 147.5207521 | -0.629921572 | 6.50E-03 |
| UFL1 | 1471.591244 | -0.630057026 | 1.92E-02 |
| YAE1 | 670.6135456 | -0.630138845 | 4.57E-06 |
| GTF2I | 19621.1412 | -0.630247002 | 3.04E-12 |
| TTC34 | 117.7523898 | -0.630338512 | 7.23E-03 |
| ADGRE5 | 2806.008902 | -0.630529796 | 8.76E-08 |

| | | | |
|---|---|---|---|
| CRYZ | 1531.82456 | -0.630532856 | 2.19E-05 |
| GSTA4 | 526.6998491 | -0.630821637 | 1.68E-06 |
| TNRC6B | 2313.664522 | -0.6310603 | 2.19E-06 |
| ZNF470 | 625.7405749 | -0.631114177 | 1.64E-05 |
| AC073857.1 | 101.7834547 | -0.631361591 | 3.81E-02 |
| BCL9 | 2022.851338 | -0.631473395 | 1.80E-10 |
| AQP3 | 243.1798854 | -0.631546089 | 4.22E-03 |
| ZNF429 | 118.5701516 | -0.632814812 | 5.88E-03 |
| JADE2 | 2540.950106 | -0.632817597 | 5.92E-09 |
| HOXB8 | 188.1046317 | -0.633179335 | 9.81E-04 |
| CCDC138 | 483.0823704 | -0.633701399 | 6.88E-04 |
| TEAD2 | 2552.04389 | -0.633793874 | 4.53E-06 |
| TSBP1-AS1 | 166.0090922 | -0.633861604 | 3.69E-03 |
| AC114811.2 | 115.9227166 | -0.634873252 | 1.69E-02 |
| LMNTD2 | 174.1655401 | -0.635021244 | 7.28E-03 |
| KIAA0586 | 1386.15036 | -0.63506367 | 3.94E-03 |
| TM7SF2 | 2521.278961 | -0.63507601 | 5.61E-03 |
| BRIP1 | 600.5206492 | -0.635147712 | 1.36E-02 |
| ROBO1 | 184.8300666 | -0.635490478 | 4.96E-02 |
| PDIK1L | 258.2112201 | -0.635587849 | 8.28E-04 |
| AC004854.2 | 59.9329362 | -0.635757458 | 4.25E-02 |
| DACT1 | 395.7859807 | -0.635768458 | 2.43E-03 |
| SYT3 | 154.5266104 | -0.635849003 | 2.44E-02 |
| IKZF5 | 323.6334647 | -0.636212608 | 1.49E-02 |
| AMMECR1 | 735.4118195 | -0.636376639 | 6.66E-08 |
| PSMG3-AS1 | 1134.425137 | -0.636747091 | 1.33E-04 |
| MAP9 | 380.48986 | -0.637082952 | 3.58E-02 |
| CADPS2 | 265.7565386 | -0.637297003 | 9.04E-03 |
| PSTPIP2 | 476.2849638 | -0.637623083 | 3.87E-06 |
| POLK | 1122.115659 | -0.637909244 | 2.72E-02 |
| HTR7 | 323.7397031 | -0.638184733 | 1.37E-05 |
| DNMT3B | 895.4449263 | -0.638370609 | 5.72E-07 |
| SLF2 | 1287.386998 | -0.638402486 | 4.23E-05 |
| LMBRD2 | 905.4643708 | -0.638417982 | 1.97E-02 |
| CC2D2A | 1425.087652 | -0.638566131 | 5.44E-09 |
| IP6K2 | 3247.021157 | -0.638831612 | 4.17E-07 |
| COPS2 | 1367.152129 | -0.638847834 | 1.95E-02 |
| U47924.2 | 96.97950533 | -0.639276101 | 9.06E-03 |
| AL137802.2 | 105.9809884 | -0.639530186 | 1.08E-02 |
| CCDC74A | 423.2740133 | -0.639793926 | 4.87E-06 |
| SRBD1 | 1161.090299 | -0.639859349 | 7.14E-04 |
| KIAA2026 | 1257.987222 | -0.639987106 | 1.48E-05 |
| ARMCX5 | 434.7492399 | -0.640256611 | 6.35E-05 |
| RECQL | 2422.766935 | -0.640790242 | 4.72E-03 |
| HBQ1 | 45.14721855 | -0.640810438 | 4.62E-02 |
| CD27-AS1 | 940.6269923 | -0.640854496 | 4.83E-11 |
| CLK4 | 476.8723821 | -0.641046103 | 3.81E-03 |

| | | | |
|---|---|---|---|
| HAUS3 | 535.9848163 | -0.64147246 | 2.16E-03 |
| DPF1 | 212.3406063 | -0.641537872 | 4.00E-03 |
| ACVR2A | 430.0205324 | -0.641655485 | 4.48E-06 |
| AL365203.2 | 100.0752658 | -0.641951857 | 4.12E-03 |
| ZBTB11-AS1 | 138.5096279 | -0.642050336 | 7.96E-04 |
| SOCS1 | 103.8450996 | -0.642143473 | 7.43E-03 |
| APOC1 | 1307.212753 | -0.642159216 | 1.14E-04 |
| CMTR2 | 1344.960866 | -0.642163308 | 2.14E-04 |
| HOXB-AS3 | 267.2395337 | -0.642336009 | 1.84E-04 |
| FAM135A | 586.9742666 | -0.642568549 | 1.26E-02 |
| FIRRE | 111.7449268 | -0.642888743 | 4.05E-03 |
| CASP6 | 603.1914311 | -0.643075514 | 6.57E-06 |
| TMSB15B | 331.5306934 | -0.643092263 | 4.78E-05 |
| BTN2A3P | 353.6383477 | -0.643151551 | 7.19E-08 |
| ADAM10 | 7600.927323 | -0.6431847 | 2.12E-08 |
| PTPN13 | 4334.427018 | -0.643915093 | 6.32E-03 |
| IRX3 | 770.2032991 | -0.643973917 | 2.48E-03 |
| ZNF451 | 1376.809275 | -0.644245983 | 3.18E-04 |
| SLC24A1 | 908.7448531 | -0.644332226 | 8.57E-05 |
| ZNF680 | 151.8603691 | -0.644343882 | 1.68E-02 |
| AC107068.1 | 89.06929586 | -0.645111015 | 2.27E-02 |
| ARHGEF35 | 220.0362306 | -0.646131613 | 1.12E-03 |
| VAMP8 | 385.8872751 | -0.646524686 | 1.26E-02 |
| CYB5D2 | 726.5046726 | -0.64673518 | 1.19E-08 |
| AC004846.1 | 97.33826787 | -0.646796109 | 4.39E-02 |
| ZNF776 | 951.6721861 | -0.647104801 | 4.72E-08 |
| TRA2A | 2138.299833 | -0.647952 | 8.07E-09 |
| SMARCAD1 | 2144.28931 | -0.648017516 | 1.46E-04 |
| AHSA2P | 2124.375938 | -0.648597363 | 2.05E-06 |
| AC009690.2 | 70.56743767 | -0.648697701 | 2.73E-02 |
| COL4A5 | 2752.108464 | -0.648888315 | 2.80E-07 |
| USP15 | 1224.130539 | -0.649240477 | 5.46E-03 |
| ACY3 | 92.98786841 | -0.649273477 | 3.40E-02 |
| APOBEC3G | 2433.261157 | -0.650291641 | 1.41E-07 |
| MEIS2 | 693.3792644 | -0.65030332 | 1.52E-06 |
| SLC26A1 | 198.7225818 | -0.650537813 | 5.99E-03 |
| CCNG2 | 1339.046455 | -0.650722146 | 1.55E-06 |
| ITPRID2 | 7029.408836 | -0.651167177 | 1.44E-06 |
| GSPT2 | 941.9840154 | -0.651226519 | 1.29E-12 |
| KLHL24 | 976.6612335 | -0.651605301 | 7.29E-06 |
| COQ8A | 1679.746005 | -0.651853233 | 5.79E-06 |
| ZNF714 | 190.8902153 | -0.652008382 | 2.55E-03 |
| RSF1 | 2839.203396 | -0.652126497 | 2.28E-03 |
| ADPRHL1 | 67.14277499 | -0.652279174 | 1.27E-02 |
| ZFP69 | 265.9010029 | -0.652439028 | 3.57E-03 |
| HOXA-AS3 | 169.3705619 | -0.652827805 | 4.14E-03 |
| AC135178.2 | 138.3094543 | -0.653217159 | 2.52E-03 |

| | | | |
|---|---|---|---|
| C4orf33 | 225.6424876 | -0.653393479 | 2.42E-03 |
| IFT88 | 335.7827756 | -0.653777945 | 3.12E-03 |
| TCAF1P1 | 529.6204086 | -0.65381677 | 6.15E-05 |
| IQCK | 379.5866682 | -0.654153658 | 8.42E-07 |
| ZNF519 | 152.1448506 | -0.654203741 | 4.96E-02 |
| LINC01667 | 1112.755145 | -0.654317615 | 1.88E-12 |
| ZNF300 | 709.6145832 | -0.654525109 | 8.19E-04 |
| ROCK1 | 2028.443087 | -0.654929554 | 1.28E-02 |
| THY1 | 1129.874395 | -0.65530794 | 1.82E-02 |
| ZNF320 | 1074.743161 | -0.655362598 | 1.38E-07 |
| EPC2 | 1225.634859 | -0.655873311 | 3.16E-05 |
| MINDY4 | 770.5683414 | -0.655953281 | 4.08E-10 |
| IFI16 | 7219.505499 | -0.656017067 | 3.15E-06 |
| ABHD3 | 445.9021462 | -0.656096927 | 2.22E-04 |
| P3H4 | 2876.333403 | -0.656147598 | 1.54E-04 |
| NA | 213.3305447 | -0.656632263 | 3.19E-04 |
| CDK3 | 153.5450677 | -0.656891012 | 1.02E-02 |
| DNAJC13 | 3775.927102 | -0.657368473 | 1.00E-04 |
| KIAA1109 | 3318.943946 | -0.657434635 | 1.31E-02 |
| AC055713.1 | 109.6029545 | -0.657563018 | 1.47E-03 |
| AC000068.1 | 80.57010928 | -0.657644582 | 1.20E-02 |
| AP000529.1 | 73.28206619 | -0.657826918 | 4.66E-02 |
| HERC2P3 | 1250.815945 | -0.658054274 | 1.48E-08 |
| CACNA1G | 138.963321 | -0.65837855 | 1.67E-03 |
| KMT2C | 4465.661044 | -0.658560315 | 8.39E-10 |
| AL627309.6 | 103.0867455 | -0.658579865 | 2.60E-02 |
| GPAT3 | 499.5762651 | -0.659519359 | 5.30E-03 |
| C8orf44 | 173.4833532 | -0.659886934 | 1.66E-03 |
| ZFP30 | 444.4963992 | -0.659957843 | 7.15E-05 |
| MBNL1 | 2385.858077 | -0.660123449 | 5.28E-05 |
| ZNF117 | 141.3690325 | -0.660284339 | 8.00E-03 |
| BRCA1 | 2010.069614 | -0.660287391 | 6.91E-04 |
| DLEU2 | 124.6313412 | -0.660530617 | 2.12E-02 |
| CD68 | 753.459441 | -0.660610132 | 1.81E-03 |
| HERC6 | 1046.33378 | -0.660686486 | 6.52E-09 |
| SH3BGRL | 2014.089491 | -0.660696887 | 6.03E-09 |
| INTS6L | 853.156252 | -0.660838637 | 2.24E-05 |
| PLEKHG2 | 2307.085248 | -0.660856613 | 5.31E-07 |
| FAM161A | 386.9162848 | -0.660895819 | 1.94E-02 |
| DUSP5P1 | 154.5629042 | -0.661271946 | 7.14E-04 |
| CIP2A | 1223.598363 | -0.661706318 | 1.51E-02 |
| HIST4H4 | 75.16508419 | -0.661806669 | 4.19E-02 |
| ZWILCH | 3027.212332 | -0.661912242 | 1.11E-04 |
| DYRK2 | 1234.067166 | -0.661947346 | 5.88E-07 |
| GLCE | 3567.17982 | -0.662757937 | 1.89E-09 |
| CAPZA2 | 3179.046091 | -0.662790942 | 7.88E-05 |
| TAF9B | 1307.227021 | -0.663048231 | 4.31E-04 |

| | | | |
|---|---|---|---|
| NA | 83.64413233 | -0.663480831 | 9.65E-03 |
| UBAC2-AS1 | 265.6527688 | -0.663513035 | 2.42E-04 |
| TMEM50B | 1177.523805 | -0.663594793 | 2.55E-06 |
| GLB1L2 | 252.2022969 | -0.664182247 | 1.95E-03 |
| ACER2 | 79.7112544 | -0.664359553 | 2.81E-02 |
| 6-Mar | 13289.27483 | -0.664396832 | 1.74E-17 |
| RSRP1 | 1513.468845 | -0.664752374 | 3.66E-04 |
| ARL6 | 147.678687 | -0.665635428 | 4.24E-03 |
| AC015799.1 | 53.59283895 | -0.665682924 | 3.61E-02 |
| TTC30B | 553.0113901 | -0.665696916 | 4.74E-10 |
| EFNB3 | 2025.53239 | -0.665825133 | 1.91E-04 |
| TRIM41 | 1896.095923 | -0.666164291 | 6.46E-14 |
| IQSEC2 | 1053.000885 | -0.666174255 | 4.74E-06 |
| HBP1 | 3602.158838 | -0.666428431 | 4.07E-18 |
| TTLL3 | 1264.197568 | -0.666464198 | 1.11E-08 |
| BMI1 | 922.8108579 | -0.667029062 | 2.02E-02 |
| POLN | 112.3528714 | -0.667442948 | 2.91E-02 |
| AP006284.1 | 314.2504097 | -0.667609774 | 1.88E-02 |
| DCHS1 | 6163.576123 | -0.667942953 | 1.50E-05 |
| PRKAR1B | 1954.365488 | -0.668049798 | 6.11E-06 |
| MACROD2 | 76.02264208 | -0.668063508 | 4.19E-02 |
| HIST2H2BE | 613.9657798 | -0.668738814 | 2.28E-02 |
| CACNG8 | 671.2693113 | -0.669182141 | 2.92E-04 |
| RARRES3 | 1750.500845 | -0.669308941 | 5.73E-04 |
| ERI2 | 682.0807066 | -0.669680005 | 2.77E-04 |
| Z97192.2 | 257.6198424 | -0.66968715 | 3.06E-05 |
| AL929601.1 | 134.5228738 | -0.669766104 | 6.74E-04 |
| MROH6 | 143.2816375 | -0.670030335 | 5.14E-03 |
| BX088651.4 | 150.1509367 | -0.670069553 | 7.63E-04 |
| DDTL | 284.9097104 | -0.67007853 | 1.09E-07 |
| HELLS | 1113.450738 | -0.671439938 | 2.93E-03 |
| FAM241A | 959.3567144 | -0.671728026 | 8.56E-07 |
| ST3GAL6 | 110.4910764 | -0.672184147 | 1.60E-02 |
| TNFRSF21 | 5207.604204 | -0.672809882 | 1.86E-16 |
| ZIM2 | 133.2839755 | -0.673160293 | 4.97E-03 |
| AP000808.1 | 86.69107461 | -0.674027809 | 1.95E-02 |
| ZNF638 | 3723.556478 | -0.674393122 | 2.47E-03 |
| VASH1 | 4125.770126 | -0.674413004 | 4.55E-06 |
| TTC8 | 1064.914823 | -0.674649094 | 1.08E-07 |
| IL12RB1 | 128.2359463 | -0.674680237 | 4.09E-03 |
| AL031282.1 | 233.3734107 | -0.674777405 | 1.39E-05 |
| GCNT1 | 951.6921527 | -0.674838905 | 1.44E-03 |
| IFIT3 | 1284.613263 | -0.674869797 | 2.65E-07 |
| ANKRD6 | 555.8492612 | -0.675223721 | 3.63E-04 |
| THOC2 | 2817.232692 | -0.675618358 | 5.16E-03 |
| AC007485.2 | 149.7281123 | -0.676152442 | 4.12E-04 |
| NAPA-AS1 | 80.1956042 | -0.676206885 | 4.54E-02 |

| | | | |
|---|---|---|---|
| ZNF14 | 181.160133 | -0.67640791 | 6.46E-03 |
| AC005046.1 | 173.6432524 | -0.6766158 | 2.41E-02 |
| TOP2B | 4796.333484 | -0.677038697 | 7.64E-06 |
| AL136295.5 | 119.402497 | -0.677412263 | 1.50E-02 |
| PCMTD2 | 3440.324389 | -0.677621802 | 5.89E-08 |
| RAMP2-AS1 | 693.2912316 | -0.677628177 | 4.93E-08 |
| C12orf4 | 972.3065059 | -0.677671533 | 1.96E-04 |
| TPR | 4477.96525 | -0.678421267 | 6.08E-03 |
| FNBP1P1 | 63.53773888 | -0.678429929 | 4.69E-02 |
| ZNF136 | 332.4460914 | -0.67857222 | 5.89E-04 |
| XKR5 | 178.029926 | -0.679003837 | 1.65E-04 |
| NIPAL2 | 1054.596041 | -0.679026093 | 3.97E-04 |
| FIZ1 | 677.6044093 | -0.679278217 | 1.59E-06 |
| AC034236.2 | 124.9206715 | -0.679479789 | 3.43E-03 |
| STYX | 677.5133932 | -0.679660891 | 1.45E-05 |
| ERVW-1 | 167.7450715 | -0.679706316 | 7.68E-04 |
| EML6 | 415.0599144 | -0.680138835 | 2.35E-02 |
| KRBA2 | 422.1868317 | -0.681167665 | 2.67E-09 |
| CLDN4 | 208.9669803 | -0.681196339 | 7.89E-04 |
| ZNF654 | 394.779115 | -0.681220413 | 1.67E-04 |
| POLR2M | 3888.908804 | -0.681860611 | 5.96E-12 |
| AC004148.2 | 154.6999536 | -0.682082262 | 5.39E-04 |
| ARMC5 | 983.2956384 | -0.682195978 | 4.47E-08 |
| ZNF414 | 209.8551727 | -0.682548316 | 1.68E-03 |
| PCDHB15 | 54.5526868 | -0.683130085 | 4.18E-02 |
| CIITA | 215.5088178 | -0.683606058 | 2.97E-02 |
| VAX2 | 144.6886263 | -0.683641124 | 3.11E-03 |
| TBL1XR1 | 2738.467211 | -0.683680794 | 1.20E-04 |
| ANKRD50 | 1173.25523 | -0.684020456 | 1.70E-06 |
| FMR1 | 2502.389221 | -0.68406244 | 4.09E-06 |
| ZNF561 | 1131.693688 | -0.684081617 | 1.22E-13 |
| SIAE | 1123.263764 | -0.684522778 | 4.06E-10 |
| BTN3A3 | 848.1382446 | -0.684678882 | 2.94E-04 |
| NA | 138.6279895 | -0.685346437 | 2.02E-03 |
| DPY19L4 | 1724.934286 | -0.685470962 | 2.11E-05 |
| PTGER4 | 362.933959 | -0.685882544 | 4.46E-02 |
| LPL | 5043.224616 | -0.686157201 | 2.75E-06 |
| ZNF43 | 401.036937 | -0.686999908 | 8.03E-07 |
| ZNF829 | 399.8671455 | -0.6870559 | 4.82E-04 |
| DNAJB14 | 680.7839592 | -0.687441457 | 1.42E-02 |
| G2E3 | 721.7049318 | -0.687622993 | 1.83E-03 |
| CRTC1 | 1021.148261 | -0.688442501 | 3.14E-10 |
| HNRNPDL | 6883.710354 | -0.689151475 | 3.72E-09 |
| CCDC171 | 280.7352548 | -0.689180477 | 4.49E-03 |
| NA | 389.3797951 | -0.689347688 | 1.88E-04 |
| NDUFA6-DT | 233.1582508 | -0.689428809 | 2.36E-05 |
| TSC22D1 | 5776.662557 | -0.689438201 | 2.10E-14 |

| | | | |
|---|---|---|---|
| MTERF2 | 526.1795391 | -0.689850833 | 3.72E-06 |
| HACD4 | 437.4754155 | -0.6899748 | 2.12E-08 |
| AC138866.1 | 312.6038432 | -0.690206808 | 2.06E-04 |
| SNTB2 | 3318.817684 | -0.690728738 | 7.31E-12 |
| CHPF | 6703.093104 | -0.690937346 | 2.71E-06 |
| PRDM8 | 163.9781801 | -0.69147973 | 8.11E-03 |
| IRAK2 | 464.7437238 | -0.69171499 | 6.08E-06 |
| SP3 | 2625.874883 | -0.691740545 | 4.56E-05 |
| SLC26A11 | 414.2417322 | -0.69180622 | 3.76E-04 |
| ZBTB33 | 1269.78301 | -0.691820132 | 3.12E-04 |
| BBOF1 | 106.4689227 | -0.692088242 | 2.64E-02 |
| CCDC66 | 335.7898333 | -0.69210005 | 1.13E-02 |
| ARL9 | 148.4194161 | -0.693186971 | 1.67E-03 |
| PIK3C2B | 986.2967161 | -0.693407785 | 8.26E-06 |
| ESCO1 | 556.9806551 | -0.693593653 | 5.56E-03 |
| AC024075.2 | 79.91306537 | -0.694336065 | 1.14E-02 |
| C6orf226 | 100.1026753 | -0.694340971 | 6.95E-03 |
| PCAT19 | 298.9928344 | -0.6943411 | 9.68E-03 |
| EMP3 | 3670.20766 | -0.694584426 | 1.41E-04 |
| AC009275.1 | 248.9884021 | -0.694639211 | 1.93E-04 |
| AC010894.3 | 225.0721163 | -0.694963264 | 5.29E-05 |
| LPAL2 | 159.1006028 | -0.69501899 | 5.26E-04 |
| ZNF397 | 629.7096529 | -0.695031235 | 2.14E-05 |
| AARD | 73.18662623 | -0.69503254 | 2.03E-02 |
| ZNF10 | 240.692249 | -0.695212003 | 1.87E-02 |
| USP27X-AS1 | 60.68377957 | -0.695360017 | 4.63E-02 |
| NA | 523.1918312 | -0.695406625 | 1.35E-09 |
| D2HGDH | 937.3314087 | -0.695475806 | 1.28E-06 |
| FAM126B | 379.8792403 | -0.695514919 | 5.84E-03 |
| MST1 | 313.337359 | -0.695549817 | 1.94E-04 |
| TSPY26P | 1120.670006 | -0.695581554 | 3.37E-05 |
| STAC2 | 256.3708383 | -0.696517937 | 2.34E-05 |
| PRPF4B | 3417.369769 | -0.696524158 | 2.11E-05 |
| NAA16 | 439.3530062 | -0.697131797 | 1.84E-04 |
| GPX7 | 858.5942305 | -0.697241669 | 4.59E-06 |
| MMRN2 | 292.7979478 | -0.697252473 | 4.77E-03 |
| AC005519.1 | 52.47188024 | -0.697330652 | 1.43E-02 |
| BX005266.2 | 176.8294464 | -0.697590851 | 3.20E-02 |
| FAM215B | 156.6873607 | -0.698347693 | 1.82E-03 |
| CASP8AP2 | 923.2155655 | -0.698357367 | 9.86E-03 |
| CUBN | 240.1036853 | -0.698494694 | 2.07E-04 |
| TMCO1-AS1 | 41.50881023 | -0.699199818 | 3.98E-02 |
| DCUN1D1 | 609.5664724 | -0.699279368 | 1.56E-04 |
| STXBP4 | 406.927206 | -0.699817732 | 4.14E-03 |
| NFYA | 1791.337203 | -0.699826471 | 5.32E-15 |
| CATSPER2 | 113.6358509 | -0.700197385 | 2.32E-02 |
| CD164 | 6736.973925 | -0.700369462 | 2.59E-11 |

| | | | |
|---|---|---|---|
| SETDB2 | 563.5613098 | -0.700415617 | 1.56E-02 |
| PCDHGB1 | 109.8113584 | -0.700515006 | 1.06E-03 |
| NKTR | 2044.428997 | -0.700617108 | 1.06E-02 |
| NAB1 | 2310.147904 | -0.701010369 | 2.56E-07 |
| ST3GAL1 | 3868.982245 | -0.701917076 | 1.11E-08 |
| SCFD2 | 1398.46596 | -0.702062866 | 1.73E-11 |
| MCC | 3773.783562 | -0.702071939 | 3.04E-15 |
| AC007191.1 | 91.39415374 | -0.702350023 | 4.02E-03 |
| AC012640.2 | 113.968136 | -0.702530367 | 2.17E-03 |
| CCNE1 | 424.1412281 | -0.702565915 | 2.09E-06 |
| ATAD2B | 555.2963023 | -0.702870611 | 2.08E-03 |
| VWA1 | 1340.086787 | -0.703767122 | 4.15E-04 |
| KLHL13 | 144.3735064 | -0.703803598 | 6.23E-03 |
| CSAD | 412.1667877 | -0.703833304 | 1.07E-06 |
| ZNF254 | 243.3316025 | -0.704021891 | 1.35E-03 |
| IFT81 | 713.7989242 | -0.704327772 | 2.41E-04 |
| RALGAPA1 | 1361.744049 | -0.704657483 | 2.30E-04 |
| CSRP1 | 17874.18646 | -0.704830182 | 9.20E-08 |
| TOR1AIP1 | 2235.13473 | -0.705000721 | 4.80E-10 |
| GYG1 | 2317.222666 | -0.705336992 | 1.09E-07 |
| ZNF148 | 1250.193518 | -0.705730603 | 1.23E-04 |
| APOBEC3F | 829.4756475 | -0.705832742 | 9.36E-07 |
| SFR1 | 105.61381 | -0.705852821 | 1.27E-02 |
| ZNF790 | 195.3086188 | -0.707001374 | 7.96E-04 |
| AC100861.1 | 179.1552444 | -0.707100569 | 1.86E-03 |
| RAB36 | 529.2506332 | -0.707496075 | 1.68E-08 |
| GLMP | 3840.216257 | -0.707922546 | 2.37E-06 |
| MT1X | 376.4307243 | -0.708003245 | 2.93E-02 |
| UBN2 | 1275.398508 | -0.708253383 | 1.41E-03 |
| LINC00467 | 293.758979 | -0.708263819 | 2.99E-04 |
| DSCC1 | 574.0425517 | -0.708447786 | 1.94E-06 |
| SPAG8 | 62.51808455 | -0.708513875 | 2.94E-02 |
| AP003392.4 | 253.0696273 | -0.7086689 | 1.81E-05 |
| ZBTB8B | 90.42870502 | -0.708725089 | 2.72E-03 |
| CYB5D1 | 624.0858821 | -0.708766399 | 1.52E-06 |
| TWIST1 | 111.7034611 | -0.708886197 | 4.12E-03 |
| NIPBL | 5744.833362 | -0.708998368 | 1.20E-02 |
| ANKRD36C | 322.2887014 | -0.709015884 | 5.75E-04 |
| BRMS1L | 598.3185116 | -0.709286264 | 1.66E-06 |
| CCDC183-AS1 | 300.2302564 | -0.709846992 | 6.12E-05 |
| C2orf69 | 301.3506584 | -0.709851105 | 1.19E-03 |
| C5orf24 | 1607.435982 | -0.710233879 | 7.29E-10 |
| INO80D | 402.4125441 | -0.710735505 | 7.67E-03 |
| EPHB3 | 918.3399535 | -0.711419928 | 5.03E-07 |
| NA | 85.77156651 | -0.711422949 | 1.00E-02 |
| ALG6 | 407.5561029 | -0.71144668 | 6.60E-05 |
| ZFX | 1304.403397 | -0.711851356 | 1.25E-07 |

| | | | |
|---|---|---|---|
| SECISBP2L | 1068.391519 | -0.711856324 | 4.59E-04 |
| ZNF552 | 298.9944392 | -0.711867186 | 4.60E-06 |
| ADAT3 | 105.9149466 | -0.712153725 | 2.94E-02 |
| JUP | 7928.88656 | -0.712190818 | 4.87E-06 |
| TET3 | 1452.642177 | -0.712272374 | 4.79E-11 |
| AC005840.2 | 106.5115936 | -0.71264009 | 4.41E-03 |
| USP32P2 | 439.998467 | -0.712660476 | 6.01E-03 |
| ZNF688 | 448.3177419 | -0.712791832 | 1.43E-05 |
| KPNA5 | 218.1163122 | -0.713178874 | 6.79E-03 |
| SLC25A27 | 441.4577795 | -0.713588379 | 3.29E-04 |
| PRKRA-AS1 | 232.7012941 | -0.713977977 | 3.65E-04 |
| PANO1 | 51.604583 | -0.714304439 | 2.51E-02 |
| DPYSL3 | 1641.152646 | -0.714434447 | 1.12E-07 |
| SOX9 | 2034.137963 | -0.714625467 | 4.09E-08 |
| AC084125.2 | 187.0679811 | -0.714748505 | 8.20E-04 |
| NA | 45.78185892 | -0.715127438 | 3.61E-02 |
| SAMD14 | 319.8574801 | -0.715145474 | 5.12E-03 |
| HMGN3 | 1577.110129 | -0.715890449 | 1.62E-14 |
| PLEKHH2 | 523.7251953 | -0.71669069 | 8.73E-03 |
| AC135776.4 | 95.35057604 | -0.717013743 | 2.85E-02 |
| EOGT | 588.9739614 | -0.717265318 | 5.34E-07 |
| H1FX-AS1 | 94.09648599 | -0.718504266 | 4.27E-04 |
| ZNF724 | 94.80845695 | -0.718651076 | 2.14E-02 |
| SHLD2 | 793.2501601 | -0.719260164 | 4.96E-05 |
| SGSH | 1798.745288 | -0.719574824 | 7.03E-07 |
| AC124312.3 | 178.0884694 | -0.719647205 | 9.91E-04 |
| EEF1A1P19 | 59.86761148 | -0.719967952 | 3.11E-02 |
| AC008736.2 | 70.52549849 | -0.720435988 | 9.39E-03 |
| BVES | 1349.384071 | -0.720457866 | 1.22E-17 |
| AC091132.3 | 70.90349035 | -0.721036943 | 2.04E-02 |
| MR1 | 1383.354792 | -0.721669145 | 6.95E-07 |
| HOXB6 | 896.7173835 | -0.721684749 | 3.53E-09 |
| EBF3 | 94.37900685 | -0.72173498 | 2.18E-02 |
| ACCS | 1275.123585 | -0.72221401 | 1.98E-11 |
| LTBP1 | 2228.493436 | -0.722979524 | 8.31E-05 |
| NA | 96.70339755 | -0.72304176 | 1.74E-03 |
| FRS2 | 817.1134476 | -0.723262385 | 2.24E-05 |
| THNSL1 | 293.0053922 | -0.723280628 | 1.04E-03 |
| HERC2P2 | 2782.210698 | -0.723610326 | 2.00E-08 |
| WNK3 | 384.5832355 | -0.724386809 | 1.11E-06 |
| AL356599.1 | 142.1623234 | -0.724755613 | 8.88E-03 |
| BOC | 1274.836958 | -0.725000132 | 1.09E-07 |
| CU638689.5 | 79.83533743 | -0.725070774 | 5.36E-03 |
| AP5S1 | 887.952493 | -0.725268676 | 1.01E-07 |
| HAPLN4 | 233.8119336 | -0.726004735 | 5.30E-03 |
| STEAP1 | 213.0817292 | -0.726006826 | 8.64E-04 |
| HNMT | 837.2644328 | -0.726050839 | 4.23E-07 |

| | | | |
|---|---|---|---|
| DPF3 | 481.2364414 | -0.726174595 | 2.18E-04 |
| EEPD1 | 580.6320043 | -0.72640529 | 1.67E-04 |
| AC008972.2 | 38.5054591 | -0.726473876 | 4.16E-02 |
| BCLAF1 | 5637.954325 | -0.726852575 | 6.20E-05 |
| AC100786.1 | 73.86546686 | -0.727385742 | 2.10E-02 |
| CU634019.6 | 561.8181909 | -0.727396513 | 3.60E-05 |
| SCN4B | 2747.623872 | -0.727450462 | 1.10E-03 |
| EPC1 | 776.583326 | -0.727620782 | 1.31E-06 |
| GLMN | 563.3816818 | -0.72765793 | 2.69E-05 |
| FAM230B | 183.8312062 | -0.727817579 | 4.23E-04 |
| TRBC2 | 122.21731 | -0.728242184 | 2.68E-02 |
| RBM12B-AS1 | 53.27711546 | -0.728270953 | 4.04E-02 |
| CWC22 | 1209.979763 | -0.728499458 | 5.69E-03 |
| ALG10B | 377.4116927 | -0.728837618 | 1.40E-03 |
| C8orf48 | 96.08804137 | -0.728970639 | 4.74E-03 |
| AC018521.5 | 171.255319 | -0.729675004 | 1.05E-03 |
| AL589843.2 | 59.16284685 | -0.72984196 | 8.09E-03 |
| NECAB1 | 84.7509274 | -0.730716516 | 2.85E-02 |
| TLR2 | 80.40836449 | -0.731073608 | 4.95E-02 |
| C1QTNF3 | 44.26961409 | -0.731305094 | 3.75E-02 |
| ST8SIA6 | 68.51548135 | -0.731443589 | 2.24E-02 |
| RASSF8-AS1 | 287.0778321 | -0.73211794 | 1.41E-06 |
| AC073508.3 | 191.8299393 | -0.732247484 | 3.76E-06 |
| CCDC15 | 552.3203374 | -0.733213992 | 3.19E-03 |
| PLK4 | 998.2687593 | -0.733635752 | 1.25E-02 |
| PPWD1 | 2051.396356 | -0.734972899 | 7.52E-05 |
| UGT8 | 101.0890748 | -0.735140761 | 1.82E-03 |
| LRRC39 | 48.17217601 | -0.735191086 | 4.74E-02 |
| RRN3P2 | 93.46092709 | -0.735483876 | 4.72E-03 |
| RNF207 | 629.8990604 | -0.736269149 | 1.08E-05 |
| AKAP9 | 1494.681228 | -0.736388376 | 1.07E-02 |
| LHX1-DT | 215.5440843 | -0.736540128 | 2.33E-05 |
| MPP1 | 2061.247563 | -0.737356761 | 2.70E-11 |
| MLLT11 | 927.8788135 | -0.73761233 | 1.42E-09 |
| C3orf38 | 678.6615537 | -0.737675681 | 4.32E-12 |
| CCT6B | 86.79091986 | -0.738096155 | 6.55E-03 |
| ACP5 | 1625.175419 | -0.738177564 | 5.44E-04 |
| AC007686.3 | 43.47349693 | -0.738235887 | 2.20E-02 |
| ASIC1 | 1465.170741 | -0.738387043 | 5.10E-07 |
| ZNF793 | 305.6492898 | -0.738533802 | 1.64E-05 |
| TMEM106A | 336.5545532 | -0.738614303 | 9.85E-07 |
| TMX4 | 4179.591097 | -0.739129304 | 1.43E-10 |
| ZNF462 | 2159.207217 | -0.739220654 | 2.15E-16 |
| ZNF107 | 246.4897489 | -0.739936634 | 1.99E-03 |
| TNFRSF14 | 898.3680434 | -0.740034848 | 5.75E-07 |
| COL27A1 | 4166.309454 | -0.740578715 | 5.36E-11 |
| AC140479.2 | 142.8946411 | -0.741225994 | 9.88E-04 |

| | | | |
|---|---|---|---|
| FAM187A | 233.2630473 | -0.741629777 | 4.06E-06 |
| PPP4R2 | 1886.522598 | -0.741865426 | 5.55E-06 |
| ALMS1 | 1742.721093 | -0.742387265 | 2.56E-04 |
| METTL25 | 105.2190927 | -0.742544994 | 3.23E-03 |
| CCDC18-AS1 | 304.8541376 | -0.742551947 | 6.89E-06 |
| RNF150 | 107.7773362 | -0.742695489 | 1.08E-02 |
| SLC12A5-AS1 | 153.3221957 | -0.743503107 | 1.50E-04 |
| IFT74 | 706.580519 | -0.743535751 | 7.03E-03 |
| SCAF11 | 4087.643525 | -0.744018922 | 1.49E-05 |
| PLIN2 | 5843.809196 | -0.744178374 | 1.03E-07 |
| ZNF100 | 192.1707816 | -0.744283494 | 1.97E-02 |
| PARP9 | 1481.054975 | -0.744317484 | 8.60E-07 |
| ANK1 | 275.01387 | -0.745391875 | 2.90E-02 |
| ADGRG6 | 2889.841449 | -0.745836324 | 2.54E-05 |
| DNAJC28 | 72.00157702 | -0.746214723 | 2.37E-02 |
| AC007406.5 | 157.0453592 | -0.746256355 | 5.07E-04 |
| ZNF527 | 289.327272 | -0.746424266 | 2.01E-05 |
| ZNF682 | 145.7408049 | -0.746447177 | 6.40E-04 |
| MDFIC | 378.7965505 | -0.746449795 | 1.21E-03 |
| AC003102.1 | 62.22034967 | -0.746772001 | 1.34E-02 |
| EXTL2 | 2261.873093 | -0.746853635 | 7.00E-11 |
| HAUS1 | 1379.441748 | -0.746946218 | 1.90E-16 |
| AL359643.3 | 126.3212182 | -0.746974175 | 2.05E-04 |
| ZGRF1 | 425.2006383 | -0.748026941 | 1.54E-02 |
| CYP2T1P | 71.51101725 | -0.748823266 | 7.52E-03 |
| CENPQ | 301.7951478 | -0.74994467 | 7.50E-05 |
| SALL2 | 1175.392959 | -0.750514682 | 1.22E-04 |
| FAM186B | 70.38773791 | -0.750556903 | 1.51E-02 |
| RASSF8 | 897.5966724 | -0.751048807 | 1.17E-03 |
| AL592435.1 | 78.35341732 | -0.752129565 | 2.66E-02 |
| ZBTB45P2 | 717.7323167 | -0.752153257 | 3.98E-08 |
| PKDCC | 957.8413157 | -0.752161429 | 3.55E-08 |
| CEBPZ | 2916.110111 | -0.752380706 | 4.55E-04 |
| NEK7 | 933.9272831 | -0.752653976 | 1.20E-04 |
| TMEM87A | 1484.561536 | -0.752937377 | 2.43E-15 |
| GPR75 | 82.90617554 | -0.752986186 | 3.30E-03 |
| ZNF334 | 302.7358372 | -0.753132713 | 1.47E-04 |
| AL162311.3 | 39.90486061 | -0.75388548 | 3.42E-02 |
| NR3C1 | 4920.832451 | -0.754132456 | 2.62E-12 |
| RUFY2 | 371.2303911 | -0.754254903 | 1.56E-04 |
| SKIDA1 | 152.151256 | -0.754819232 | 1.59E-04 |
| LRIG1 | 1253.713048 | -0.75501387 | 3.03E-14 |
| ZNF624 | 132.3153627 | -0.755466913 | 2.02E-02 |
| AL589666.1 | 81.42536243 | -0.755848322 | 9.03E-03 |
| SHLD1 | 405.9876561 | -0.756336997 | 4.31E-07 |
| C1orf21 | 2338.356182 | -0.756496662 | 1.46E-08 |
| ZFP3 | 574.7828544 | -0.757213223 | 1.19E-12 |

| | | | |
|---|---|---|---|
| PMS2P2 | 63.2047173 | -0.758007154 | 1.91E-02 |
| KCND1 | 722.1343561 | -0.758503563 | 1.51E-05 |
| AC104667.2 | 71.44130672 | -0.758795905 | 1.05E-02 |
| TNRC6C | 889.66465 | -0.759387775 | 2.62E-06 |
| SH3GL1P1 | 57.18052606 | -0.759676786 | 4.12E-02 |
| TRIM23 | 971.8216866 | -0.759761697 | 3.44E-03 |
| SSC5D | 282.0855275 | -0.759870605 | 7.44E-04 |
| AC007390.2 | 108.6961985 | -0.759918235 | 6.70E-03 |
| RGS2 | 189.9077921 | -0.760318193 | 7.41E-04 |
| AL935212.1 | 53.38040857 | -0.760350323 | 3.32E-02 |
| AL512310.11 | 40.74662864 | -0.760505539 | 2.90E-02 |
| SRD5A3 | 467.3307845 | -0.760510673 | 1.07E-07 |
| PRMT6 | 1118.533194 | -0.7606864 | 3.02E-15 |
| ZNF490 | 133.7375409 | -0.760785424 | 8.29E-03 |
| SPATA6 | 471.5236482 | -0.76081158 | 2.67E-08 |
| DERL3 | 144.1694928 | -0.761832836 | 8.20E-03 |
| JAK2 | 767.597303 | -0.762141248 | 4.66E-04 |
| ALDH16A1 | 1223.646242 | -0.762242016 | 2.97E-08 |
| FAM78A | 383.5158475 | -0.762362665 | 8.66E-09 |
| ABCA5 | 556.3366742 | -0.762594787 | 4.65E-03 |
| EEA1 | 1040.930789 | -0.76329852 | 2.52E-02 |
| FAM153B | 429.3182156 | -0.763544791 | 5.13E-04 |
| TTK | 1450.743421 | -0.763845387 | 1.57E-02 |
| SLC7A8 | 1004.591585 | -0.763899133 | 1.48E-06 |
| USP46-AS1 | 131.0591345 | -0.76396704 | 2.41E-04 |
| TGFBR3 | 1252.634429 | -0.764091099 | 1.27E-10 |
| TCAF1 | 6221.734152 | -0.764776978 | 4.42E-06 |
| TEX9 | 137.6996453 | -0.765806215 | 7.01E-04 |
| GSKIP | 690.2447286 | -0.765966954 | 5.95E-04 |
| NBN | 2080.013296 | -0.766585425 | 5.70E-04 |
| AC022400.5 | 62.08055453 | -0.767021085 | 1.64E-02 |
| ZNF30 | 247.4851225 | -0.767078709 | 4.23E-04 |
| ETV5 | 3045.552529 | -0.767311695 | 2.90E-07 |
| SHPRH | 739.154152 | -0.767322597 | 3.05E-05 |
| ASNSD1 | 1743.763404 | -0.767556862 | 3.66E-04 |
| CFB | 187.4850491 | -0.767653098 | 1.74E-03 |
| AP002495.2 | 126.5569363 | -0.767919958 | 9.60E-04 |
| AIFM3 | 126.0632533 | -0.768239742 | 1.83E-03 |
| ARG2 | 2408.765317 | -0.768324006 | 4.02E-10 |
| AC006333.2 | 155.503166 | -0.768383774 | 5.96E-05 |
| ZBTB6 | 524.4678034 | -0.768482682 | 2.13E-04 |
| IGLON5 | 232.1725575 | -0.768490139 | 2.41E-03 |
| COL9A3 | 4082.534637 | -0.769084839 | 2.54E-06 |
| RIDA | 596.8398994 | -0.769457333 | 7.48E-07 |
| MAN1A1 | 698.1988176 | -0.769881141 | 2.12E-08 |
| DPY19L2P2 | 694.3173376 | -0.770307631 | 8.07E-06 |
| ANTXR1 | 11230.02048 | -0.770718931 | 4.96E-16 |

| | | | |
|---|---|---|---|
| AL512625.1 | 409.7626639 | -0.770819665 | 2.87E-02 |
| ZMYM2 | 1599.798682 | -0.771659832 | 2.08E-08 |
| ZNF432 | 386.5222351 | -0.772072154 | 3.09E-09 |
| IREB2 | 3969.02169 | -0.772274706 | 1.37E-10 |
| LINC00504 | 145.4894275 | -0.772413254 | 3.79E-02 |
| THUMPD1 | 2204.022143 | -0.772785426 | 8.63E-08 |
| ETFBKMT | 71.93548066 | -0.772789561 | 7.92E-03 |
| HMGN2P15 | 98.57012837 | -0.772813983 | 3.22E-03 |
| CMTM8 | 58.12048249 | -0.773248073 | 6.03E-03 |
| PHTF1 | 1158.451581 | -0.773604811 | 7.70E-15 |
| AL022328.4 | 56.17580024 | -0.773783949 | 3.17E-02 |
| AC027419.2 | 33.09885901 | -0.774487961 | 4.99E-02 |
| LINC01366 | 70.30869446 | -0.774796473 | 3.82E-03 |
| PELI2 | 304.1972169 | -0.774966055 | 5.12E-04 |
| CPNE8 | 535.1743269 | -0.775174178 | 7.67E-04 |
| CLK1 | 1374.728739 | -0.775406891 | 3.51E-04 |
| LIFR-AS1 | 357.2237635 | -0.775598225 | 8.92E-08 |
| LRRC8C | 1363.062089 | -0.776043585 | 9.32E-12 |
| GPR68 | 67.37087682 | -0.776992552 | 4.26E-02 |
| TMEM67 | 702.5993906 | -0.777432885 | 1.19E-03 |
| AC112702.1 | 125.4890082 | -0.777722138 | 7.60E-03 |
| ANKRD26 | 244.6223901 | -0.777931004 | 1.19E-02 |
| TGFA | 403.9619259 | -0.777937605 | 2.61E-03 |
| AC009061.2 | 39.94211966 | -0.77794982 | 3.86E-02 |
| EXOC5 | 2309.050245 | -0.77821552 | 7.95E-04 |
| MTMR11 | 725.5845595 | -0.778785081 | 3.25E-07 |
| MT-TY | 275.5489081 | -0.779782886 | 3.84E-02 |
| BRWD3 | 947.0454821 | -0.780245739 | 9.34E-05 |
| FBXO6 | 243.709265 | -0.780461078 | 9.02E-05 |
| STARD4 | 2398.647591 | -0.780807854 | 7.04E-08 |
| LRRC49 | 476.3752078 | -0.780842354 | 3.30E-02 |
| APOE | 6958.007481 | -0.781056292 | 2.53E-04 |
| PDE5A | 1577.849555 | -0.781463805 | 8.68E-05 |
| EPM2AIP1 | 1443.908971 | -0.781518298 | 8.09E-04 |
| ZNF606 | 571.9029524 | -0.782256919 | 1.79E-07 |
| SLC5A3 | 5415.192645 | -0.782443205 | 1.59E-07 |
| SLC27A3 | 368.6739809 | -0.782575204 | 3.40E-04 |
| KMT2E | 3440.83109 | -0.783020224 | 8.93E-13 |
| LINC01297 | 1123.725903 | -0.783087177 | 8.30E-03 |
| AL033529.1 | 69.57657961 | -0.783090211 | 3.70E-02 |
| BEAN1 | 243.2446155 | -0.783188441 | 3.57E-05 |
| NA | 99.97772378 | -0.783277567 | 1.92E-02 |
| ORAI3 | 3085.151218 | -0.783604537 | 1.30E-04 |
| RBP5 | 155.6224082 | -0.783688871 | 7.96E-07 |
| PROS1 | 404.9411068 | -0.783806965 | 4.00E-05 |
| CSGALNACT1 | 578.8533399 | -0.784335815 | 1.64E-07 |
| PNISR | 4426.738011 | -0.784984775 | 2.14E-04 |

| | | | |
|---|---|---|---|
| HDX | 246.3347493 | -0.784999415 | 3.30E-04 |
| SLC18B1 | 776.1425478 | -0.785796465 | 1.97E-14 |
| E2F8 | 533.6095413 | -0.786048423 | 4.33E-06 |
| OBSCN | 2244.375511 | -0.787030106 | 1.26E-08 |
| ZDHHC17 | 996.3512409 | -0.78707379 | 7.53E-06 |
| CLSTN3 | 6610.167495 | -0.787391345 | 1.42E-05 |
| ARHGAP5-AS1 | 297.6510154 | -0.787960198 | 8.08E-07 |
| TMEM232 | 61.95177943 | -0.788433286 | 1.17E-02 |
| DRC3 | 166.3603861 | -0.788809799 | 1.37E-02 |
| POLI | 891.7794694 | -0.789264111 | 2.21E-05 |
| ZNF708 | 103.7787774 | -0.789366353 | 6.54E-03 |
| NCALD | 398.8456302 | -0.789485452 | 2.00E-02 |
| ERMAP | 476.2680804 | -0.789842442 | 1.45E-12 |
| AL450326.1 | 41.15195871 | -0.790292566 | 1.67E-02 |
| NA | 221.0300842 | -0.790546367 | 8.15E-04 |
| AL358334.2 | 74.69140229 | -0.791118651 | 2.52E-03 |
| RFX7 | 2054.624167 | -0.791207401 | 2.28E-12 |
| AC040162.1 | 268.8290852 | -0.791518758 | 2.75E-03 |
| TRIM22 | 535.1244149 | -0.791602283 | 1.61E-05 |
| CYP26A1 | 160.8035835 | -0.791665124 | 2.79E-04 |
| DLX2 | 377.9123392 | -0.792234399 | 5.51E-04 |
| DBP | 393.6402923 | -0.792391139 | 7.47E-04 |
| MANEAL | 1265.66944 | -0.795199323 | 5.78E-08 |
| NF1P10 | 67.28745981 | -0.795265238 | 2.20E-02 |
| AC112229.3 | 62.10993256 | -0.795677186 | 1.18E-02 |
| GCFC2 | 532.1667663 | -0.795698287 | 6.70E-05 |
| QRICH2 | 106.3556761 | -0.796547478 | 3.02E-03 |
| MIER3 | 2154.762199 | -0.796713522 | 1.65E-04 |
| AC080038.1 | 8743.391557 | -0.796915355 | 2.17E-08 |
| RAB26 | 425.0303471 | -0.796967648 | 2.07E-04 |
| FAM153A | 178.9188772 | -0.797078972 | 1.45E-03 |
| GBAP1 | 243.0011944 | -0.797153336 | 6.15E-06 |
| MPHOSPH9 | 1122.599799 | -0.797175221 | 4.14E-04 |
| PCBD2 | 158.4686784 | -0.797261313 | 3.96E-04 |
| ZNF700 | 334.5364929 | -0.797932503 | 3.43E-04 |
| TAL1 | 117.5851983 | -0.798622841 | 3.22E-03 |
| C2orf16 | 308.8560384 | -0.798855166 | 7.49E-09 |
| ABHD14A-ACY1 | 52.86248385 | -0.799252899 | 3.93E-02 |
| AJ003147.3 | 35.719397 | -0.799929083 | 4.51E-02 |
| CICP27 | 149.4400659 | -0.79999199 | 4.81E-06 |
| SCAMP5 | 945.5999557 | -0.80091698 | 7.53E-06 |
| AF254983.1 | 196.670656 | -0.801196032 | 1.73E-03 |
| DDAH2 | 1845.955686 | -0.801197196 | 1.39E-10 |
| LINC01123 | 472.9229212 | -0.801391167 | 4.47E-07 |
| PSPC1 | 1501.329787 | -0.801496812 | 1.15E-12 |
| AC027228.2 | 92.37529988 | -0.801508412 | 1.43E-02 |
| CMAHP | 193.4739888 | -0.801806408 | 5.54E-04 |

| | | | |
|---|---|---|---|
| PHC2 | 12014.71889 | -0.802135917 | 9.37E-11 |
| C17orf97 | 179.3364957 | -0.80229478 | 6.99E-05 |
| LRRCC1 | 377.3329368 | -0.802420365 | 2.28E-02 |
| SUCO | 1308.492429 | -0.80252712 | 7.76E-05 |
| AC018653.3 | 136.7528489 | -0.803238574 | 2.79E-05 |
| AKAP11 | 1962.018282 | -0.80344362 | 4.96E-04 |
| TBK1 | 788.5180789 | -0.803940223 | 1.28E-03 |
| SCNN1D | 251.7293197 | -0.804267905 | 2.77E-04 |
| SALL1 | 1727.630831 | -0.80432642 | 7.87E-08 |
| ERICH6-AS1 | 44.62730875 | -0.805707432 | 1.55E-02 |
| AC006435.2 | 48.96497455 | -0.805726013 | 1.47E-02 |
| PSMA3-AS1 | 956.8211118 | -0.80599563 | 2.08E-06 |
| ZNF772 | 795.6098817 | -0.806814735 | 1.58E-12 |
| HECTD2 | 302.8748965 | -0.807043074 | 8.84E-04 |
| CCDC186 | 315.7125108 | -0.807696128 | 1.48E-02 |
| MYLIP | 1308.827892 | -0.808503778 | 1.37E-09 |
| CCDC34 | 1220.437521 | -0.808753896 | 2.54E-09 |
| ZNF292 | 1527.571376 | -0.808887988 | 1.90E-03 |
| AL732372.3 | 267.4658968 | -0.809478691 | 5.09E-08 |
| AC008013.2 | 40.82931398 | -0.810769607 | 3.94E-02 |
| ABAT | 360.9278879 | -0.811351399 | 6.02E-07 |
| MTERF1 | 599.7245431 | -0.811606284 | 2.26E-05 |
| LRP6 | 2588.5102 | -0.812213603 | 7.70E-11 |
| FAM230E | 99.91948562 | -0.812826567 | 1.29E-03 |
| SPIN2A | 48.63188211 | -0.812993729 | 1.65E-02 |
| MTF1 | 1054.800846 | -0.813161985 | 5.85E-07 |
| AC009283.1 | 58.64454459 | -0.813531735 | 3.51E-03 |
| ATXN7L2 | 355.4580206 | -0.813798081 | 1.65E-08 |
| AC135048.1 | 219.3732508 | -0.813861019 | 1.34E-03 |
| LRRC8C-DT | 59.89078642 | -0.813917425 | 8.36E-03 |
| PDS5B | 1981.632488 | -0.814298048 | 4.00E-08 |
| PTBP2 | 289.7813242 | -0.814520803 | 2.16E-03 |
| NEMF | 1174.676923 | -0.814949865 | 1.18E-02 |
| LRRC58 | 1475.387555 | -0.815348854 | 3.47E-04 |
| RARB | 1624.994019 | -0.81558313 | 5.75E-09 |
| SUOX | 1358.723531 | -0.815880018 | 1.44E-06 |
| APOD | 125.6884614 | -0.815959658 | 9.30E-03 |
| CCDC142 | 726.9646701 | -0.816014389 | 9.19E-13 |
| MZF1-AS1 | 123.7415297 | -0.816245965 | 3.92E-05 |
| IGFBP2 | 14621.13362 | -0.816353948 | 3.85E-07 |
| C11orf1 | 496.8051139 | -0.816688165 | 2.25E-11 |
| BMF | 718.4286987 | -0.816872759 | 1.77E-06 |
| RHBDL1 | 79.67093928 | -0.817078013 | 7.50E-03 |
| ATM | 4856.471657 | -0.817122303 | 8.31E-04 |
| TUFMP1 | 318.1004999 | -0.817400222 | 2.08E-03 |
| BLZF1 | 801.6579964 | -0.817637506 | 3.10E-04 |
| MINDY2 | 1015.699395 | -0.817667972 | 9.90E-06 |

| | | | |
|---|---|---|---|
| ANTXR2 | 1357.097896 | -0.818071541 | 3.63E-11 |
| KAZALD1 | 613.8092983 | -0.818629934 | 4.41E-06 |
| SASS6 | 296.0806781 | -0.818821757 | 4.81E-03 |
| FRG1-DT | 78.2155875 | -0.818877803 | 4.26E-03 |
| CRISPLD1 | 185.2859406 | -0.819053214 | 2.63E-03 |
| ADAMTS14 | 70.35092211 | -0.819662191 | 4.08E-02 |
| HOTTIP | 31.11813955 | -0.819671932 | 3.78E-02 |
| TTC3 | 9113.083029 | -0.821432762 | 1.76E-10 |
| ZNF567 | 254.7474616 | -0.822011259 | 1.64E-03 |
| TMEM198B | 935.5913628 | -0.82232785 | 3.66E-19 |
| AC012313.10 | 41.82089725 | -0.823632455 | 2.34E-02 |
| AL162258.2 | 44.64770892 | -0.824003612 | 1.20E-02 |
| SHLD3 | 185.7210417 | -0.824367868 | 1.03E-06 |
| USP16 | 1293.924721 | -0.824941261 | 1.62E-04 |
| ZFPM2-AS1 | 556.3674849 | -0.824980569 | 3.70E-04 |
| BDH2 | 851.1838392 | -0.825487995 | 8.31E-15 |
| ARID4B | 1086.545711 | -0.825687382 | 5.01E-04 |
| PGM2L1 | 488.073069 | -0.825863377 | 2.82E-06 |
| AL121594.1 | 307.8997471 | -0.826341171 | 6.56E-04 |
| CHD1 | 2713.459699 | -0.826769384 | 3.83E-04 |
| AC106820.5 | 64.56184719 | -0.827379402 | 4.12E-03 |
| AC233723.2 | 80.15059256 | -0.827391709 | 5.42E-04 |
| HOXA5 | 769.5073558 | -0.827536549 | 1.43E-13 |
| USP34 | 5304.481039 | -0.828120341 | 9.49E-05 |
| CLHC1 | 102.9765619 | -0.828150515 | 1.20E-02 |
| ZNF302 | 1153.441241 | -0.828742141 | 1.99E-05 |
| SPRY3 | 329.3369305 | -0.82875272 | 1.16E-11 |
| NAIP | 402.0713656 | -0.829196645 | 1.35E-08 |
| CRYBG3 | 1054.976464 | -0.82935588 | 3.61E-03 |
| SETX | 6083.649708 | -0.830134329 | 1.18E-09 |
| SPIN3 | 350.5108314 | -0.830316337 | 3.02E-06 |
| DKKL1 | 62.86708431 | -0.830849502 | 7.42E-03 |
| SYNJ1 | 938.8449281 | -0.831371895 | 2.33E-07 |
| AC109587.1 | 51.32532979 | -0.831551913 | 1.68E-02 |
| ANKRD12 | 1010.776964 | -0.832603028 | 3.83E-03 |
| AC004951.1 | 210.411599 | -0.832774102 | 9.04E-08 |
| ZNF326 | 1281.750808 | -0.83303539 | 1.10E-06 |
| PCDHGA7 | 577.8186976 | -0.833219121 | 4.85E-08 |
| MEIS3P2 | 55.25686464 | -0.833602725 | 2.90E-02 |
| AC211433.1 | 34.35776253 | -0.83397559 | 4.60E-02 |
| BORCS7 | 376.7341782 | -0.834854828 | 1.86E-07 |
| RTKN | 1095.937803 | -0.835846219 | 2.02E-11 |
| C20orf144 | 31.27395043 | -0.837196345 | 4.27E-02 |
| NA | 130.3090386 | -0.837223096 | 1.40E-04 |
| SIRPB1 | 258.9285303 | -0.837286971 | 6.09E-03 |
| IQGAP2 | 1477.33195 | -0.837432862 | 3.60E-05 |
| SOX18 | 172.617791 | -0.837589323 | 6.50E-04 |

| | | | |
|---|---|---|---|
| FKBP5 | 2560.809448 | -0.838159256 | 7.85E-30 |
| TIGD3 | 41.09771545 | -0.838303127 | 4.48E-02 |
| SP6 | 120.9574354 | -0.840151577 | 2.49E-03 |
| FAM227A | 248.8095749 | -0.84020462 | 5.66E-04 |
| NPAT | 1282.054313 | -0.840364728 | 1.06E-05 |
| GAL3ST4 | 1047.330578 | -0.840398381 | 6.19E-08 |
| ASIC3 | 49.17937142 | -0.840521807 | 1.94E-02 |
| TRO | 3618.311323 | -0.84063952 | 6.02E-10 |
| AC137767.1 | 53.94270911 | -0.840641735 | 2.63E-03 |
| CASP7 | 1127.428638 | -0.841213808 | 1.14E-10 |
| KAT2B | 1031.112119 | -0.841449478 | 1.40E-13 |
| ZNF879 | 144.5956642 | -0.841549631 | 9.07E-03 |
| AC007952.6 | 69.57289609 | -0.841708787 | 4.72E-03 |
| L3MBTL4-AS1 | 61.20790155 | -0.841906019 | 1.48E-02 |
| SREK1 | 3158.431512 | -0.843252599 | 1.21E-03 |
| KBTBD7 | 448.7049977 | -0.844043669 | 4.60E-08 |
| HLA-DMB | 409.4052964 | -0.845055719 | 3.05E-02 |
| IQCH-AS1 | 120.9287616 | -0.84530513 | 2.93E-02 |
| COLEC11 | 893.9691598 | -0.845524103 | 1.69E-06 |
| PQLC2L | 60.00408119 | -0.845963354 | 6.32E-03 |
| RABEP2 | 542.7569312 | -0.846044815 | 3.70E-09 |
| HIST2H2AA4 | 253.4964895 | -0.846209374 | 1.22E-03 |
| ZNF41 | 598.0445322 | -0.84664428 | 1.40E-07 |
| SLC25A5-AS1 | 91.85803581 | -0.84690076 | 2.22E-04 |
| NUDT12 | 724.1957269 | -0.847035453 | 1.42E-09 |
| LINC01087 | 87.97856487 | -0.847142517 | 1.49E-03 |
| ZNF137P | 75.91940324 | -0.84720265 | 3.34E-02 |
| CHM | 1261.466858 | -0.847392344 | 2.40E-04 |
| EXOG | 670.6987006 | -0.847659277 | 6.70E-18 |
| NXT2 | 357.2870437 | -0.847701209 | 7.61E-08 |
| SYNE2 | 2558.962612 | -0.847757919 | 2.37E-09 |
| AL096870.2 | 67.05680501 | -0.848447687 | 2.14E-03 |
| AC027796.3 | 301.6582265 | -0.848955019 | 1.66E-06 |
| AC007996.1 | 132.0789388 | -0.849667205 | 2.35E-04 |
| PDE3B | 420.4228213 | -0.849844232 | 6.23E-06 |
| FAM49A | 399.0533098 | -0.850246579 | 2.50E-08 |
| ZNF566 | 241.2730474 | -0.850914501 | 5.70E-03 |
| TMEM220 | 89.7203473 | -0.850972289 | 1.76E-03 |
| PIGM | 786.4326629 | -0.850988028 | 1.07E-24 |
| TBC1D19 | 280.9976935 | -0.852200296 | 7.28E-06 |
| AL512791.2 | 40.08376199 | -0.85238209 | 4.08E-02 |
| LAT | 302.3278347 | -0.853124946 | 3.58E-07 |
| STC1 | 1128.86879 | -0.853362612 | 1.60E-03 |
| RMDN2 | 97.31966742 | -0.853948688 | 9.91E-04 |
| LPIN2 | 5759.958478 | -0.854873999 | 4.23E-16 |
| TTYH2 | 540.5647135 | -0.855098483 | 1.11E-04 |
| ZNF224 | 508.5609643 | -0.855223857 | 1.62E-06 |

| | | | |
|---|---|---|---|
| AC093627.4 | 67.41982005 | -0.855625109 | 7.75E-03 |
| LINC01238 | 72.75569433 | -0.856126244 | 1.11E-03 |
| TRIM2 | 1676.098241 | -0.856207763 | 1.49E-03 |
| ODR4 | 586.5015614 | -0.856590172 | 5.81E-04 |
| MBTD1 | 632.4450253 | -0.856603016 | 1.35E-06 |
| GABPA | 1270.757625 | -0.85682785 | 9.50E-07 |
| SNX13 | 1939.795473 | -0.857407288 | 4.46E-06 |
| FPGT | 345.5712041 | -0.857496287 | 7.65E-05 |
| TEK | 960.4093854 | -0.858158333 | 3.81E-04 |
| TMEM150A | 656.5324793 | -0.858936413 | 5.37E-10 |
| NA | 130.178278 | -0.86120427 | 2.94E-04 |
| ZSCAN31 | 211.2724114 | -0.861652307 | 7.81E-05 |
| RAPGEF4 | 1768.711438 | -0.861659202 | 8.30E-12 |
| AC009244.1 | 77.55731589 | -0.862010012 | 2.59E-03 |
| SIPA1L2 | 2095.321232 | -0.862082228 | 6.75E-13 |
| FDXACB1 | 160.5580444 | -0.862552176 | 3.49E-04 |
| AC005703.6 | 31.16690174 | -0.862592286 | 3.17E-02 |
| KCP | 42.91701193 | -0.862726129 | 4.70E-02 |
| TRIP11 | 1200.792687 | -0.86288252 | 1.76E-05 |
| ZNF569 | 229.4674729 | -0.862897781 | 4.10E-04 |
| TPTE | 162.9454147 | -0.863246136 | 5.86E-04 |
| RBM25 | 4151.031687 | -0.863281805 | 6.87E-11 |
| ZC3H12C | 987.4220162 | -0.863355914 | 1.37E-13 |
| CYP4F12 | 335.7560523 | -0.863665798 | 2.77E-06 |
| RC3H1 | 637.8260933 | -0.86409756 | 1.04E-07 |
| NCR1 | 49.57956449 | -0.864917591 | 2.75E-02 |
| TRMT10A | 167.6673084 | -0.865318905 | 8.91E-03 |
| COLEC10 | 58.18578616 | -0.865911431 | 1.19E-02 |
| SLC4A7 | 1002.091398 | -0.866993135 | 1.54E-04 |
| POLR2J4 | 326.6944127 | -0.868224834 | 1.73E-11 |
| PLEKHA3 | 389.0818929 | -0.868542592 | 2.56E-05 |
| PGBD5 | 220.9593528 | -0.868883697 | 7.30E-04 |
| AC006449.5 | 65.52035688 | -0.869258643 | 2.13E-03 |
| NA | 92.09315874 | -0.869487698 | 6.42E-04 |
| MINPP1 | 1359.435503 | -0.869928561 | 1.64E-13 |
| TPGS1 | 169.8498911 | -0.87011636 | 4.73E-05 |
| AC008622.2 | 52.92455998 | -0.870298354 | 4.02E-03 |
| CHST2 | 604.9230275 | -0.871288261 | 6.46E-21 |
| MED15P9 | 38.96548172 | -0.871419464 | 1.02E-02 |
| RUNDC3A-AS1 | 128.8383308 | -0.871485864 | 1.11E-04 |
| LINC02482 | 133.6517581 | -0.871526346 | 3.26E-06 |
| NCOA2 | 1947.675613 | -0.871715378 | 2.55E-12 |
| ZNF644 | 1095.23322 | -0.871928782 | 1.01E-05 |
| TTLL7 | 129.1930967 | -0.872168868 | 2.42E-02 |
| SPRED2 | 2021.337216 | -0.872390227 | 1.94E-15 |
| CCDC180 | 210.7067393 | -0.872546063 | 6.67E-04 |
| DLX1 | 957.5198547 | -0.872867702 | 3.30E-06 |

| | | | |
|---|---|---|---|
| ATP7A | 1068.071532 | -0.872953736 | 8.20E-10 |
| PBDC1 | 1042.742786 | -0.873917311 | 6.43E-24 |
| CFAP70 | 58.14805916 | -0.873996532 | 3.78E-02 |
| UPF2 | 1004.08589 | -0.875168227 | 4.21E-05 |
| ZNF260 | 617.7063357 | -0.875266576 | 5.83E-04 |
| AF127577.4 | 90.7668684 | -0.875294096 | 1.92E-02 |
| FNDC10 | 630.1756976 | -0.875662201 | 3.75E-08 |
| CNFN | 49.70271958 | -0.875936756 | 6.71E-03 |
| ZFYVE16 | 2545.697203 | -0.876368115 | 1.19E-02 |
| TRMT13 | 719.993185 | -0.876805962 | 6.12E-04 |
| CEP295 | 1241.620742 | -0.877376369 | 9.34E-04 |
| PPIG | 1943.519505 | -0.877750174 | 3.54E-05 |
| CCDC146 | 94.61730271 | -0.879144473 | 1.01E-02 |
| SLC9A3-AS1 | 632.0318409 | -0.879734819 | 6.94E-07 |
| MYO7A | 246.39633 | -0.882477911 | 5.23E-04 |
| KANSL1L | 442.1730578 | -0.883257061 | 1.04E-07 |
| GUSBP2 | 71.99712807 | -0.884676514 | 4.28E-03 |
| AC105339.2 | 180.435808 | -0.884692244 | 5.99E-03 |
| RAB22A | 2724.769956 | -0.885241032 | 3.87E-25 |
| AC024060.1 | 46.15207851 | -0.886550037 | 4.64E-02 |
| DISP2 | 460.1966062 | -0.88660638 | 1.17E-03 |
| AC087292.1 | 79.73504335 | -0.886852583 | 4.35E-02 |
| AC010761.1 | 252.8642833 | -0.887372651 | 3.43E-06 |
| ISPD | 52.78470494 | -0.887706223 | 1.25E-02 |
| MCTP1 | 299.6263461 | -0.888787442 | 2.35E-02 |
| NFAT5 | 5313.455934 | -0.890284345 | 1.12E-06 |
| C7orf25 | 138.5021058 | -0.890643233 | 6.53E-04 |
| NLRC4 | 40.9423328 | -0.890980234 | 2.46E-02 |
| ROR1 | 803.2210795 | -0.89125324 | 1.72E-09 |
| KCNH3 | 436.3653327 | -0.893164119 | 1.26E-06 |
| ARHGAP12 | 906.8280344 | -0.893486377 | 2.11E-05 |
| LAMB3 | 537.2880298 | -0.894207961 | 1.99E-05 |
| SCIMP | 42.69225703 | -0.894814276 | 3.57E-02 |
| AC069061.2 | 47.29811414 | -0.895018671 | 3.55E-03 |
| SAP30L-AS1 | 31.89079869 | -0.895119171 | 3.32E-02 |
| DENND1B | 124.7418185 | -0.896085642 | 9.05E-03 |
| AC098820.1 | 29.61244796 | -0.896626676 | 3.49E-02 |
| TXNDC16 | 735.2004753 | -0.897055497 | 1.49E-14 |
| AP001528.2 | 69.98146508 | -0.897096328 | 1.16E-02 |
| MTUS1 | 3968.754988 | -0.89726615 | 1.54E-20 |
| TUBD1 | 294.8775476 | -0.897395781 | 4.51E-08 |
| AL928970.1 | 299.1134828 | -0.897751787 | 8.09E-10 |
| CYP27C1 | 309.7320071 | -0.897948167 | 2.00E-07 |
| AC068279.1 | 52.80865783 | -0.898190796 | 5.56E-03 |
| SPDYE1 | 47.23973887 | -0.898362838 | 2.44E-03 |
| AC012321.1 | 316.3933628 | -0.898797044 | 6.13E-04 |
| MPZL3 | 518.1093127 | -0.899945418 | 2.33E-09 |

| | | | |
|---|---|---|---|
| AL355388.2 | 43.64300837 | -0.9001739 | 3.37E-03 |
| EPN3 | 37.87810074 | -0.900473782 | 2.37E-02 |
| CHIC1 | 312.476867 | -0.900835331 | 4.96E-06 |
| GMFB | 1903.598644 | -0.900927285 | 2.13E-04 |
| AL031123.3 | 37.00347853 | -0.903073778 | 2.77E-02 |
| SESN1 | 517.2971486 | -0.903099094 | 4.15E-08 |
| RF00017 | 34.86901515 | -0.903428352 | 2.04E-02 |
| TSPYL2 | 2040.348022 | -0.903536349 | 7.56E-08 |
| AGBL2 | 104.4122325 | -0.904332428 | 2.54E-03 |
| ATF7 | 63.46670973 | -0.904367291 | 4.31E-04 |
| SYT13 | 311.0869747 | -0.904546612 | 6.60E-03 |
| AC068338.2 | 87.31231627 | -0.90468239 | 2.94E-05 |
| KDM8 | 224.6049505 | -0.906422391 | 1.19E-07 |
| CACNB2 | 46.1342047 | -0.906787946 | 3.63E-02 |
| BTBD19 | 501.2512357 | -0.906864194 | 4.76E-12 |
| AC015802.1 | 42.72525233 | -0.906991756 | 1.61E-02 |
| SUGT1P4-STRA6 | 208.5434078 | -0.907077387 | 1.23E-06 |
| HS2ST1 | 2383.667486 | -0.908009868 | 3.90E-10 |
| ZNF641 | 822.7066855 | -0.909708452 | 1.98E-07 |
| SWT1 | 219.6789585 | -0.909953002 | 1.82E-04 |
| SSBP2 | 2717.946164 | -0.910204075 | 2.89E-15 |
| LINC00511 | 271.0163228 | -0.910502525 | 1.12E-07 |
| ZNF837 | 70.46561219 | -0.910952995 | 8.33E-04 |
| NA | 921.6526783 | -0.911081883 | 8.01E-07 |
| LMO2 | 1198.776963 | -0.911317543 | 3.79E-14 |
| ZNF214 | 149.0621941 | -0.911391592 | 2.02E-03 |
| TBXA2R | 175.3616382 | -0.911623059 | 1.32E-04 |
| CASC15 | 47.77630025 | -0.911802546 | 1.90E-02 |
| CA11 | 828.0280318 | -0.912535686 | 4.72E-06 |
| DDX60 | 693.0435674 | -0.912781474 | 1.21E-03 |
| PLTP | 6996.727233 | -0.912972386 | 2.29E-10 |
| GJC3 | 27.20107655 | -0.913589603 | 4.08E-02 |
| RSBN1L | 1605.52993 | -0.914023123 | 5.55E-23 |
| HSPA6 | 28.92150367 | -0.914349415 | 4.24E-02 |
| PRAP1 | 135.3358466 | -0.914924378 | 2.47E-02 |
| FLI1 | 3174.945059 | -0.915436911 | 4.50E-24 |
| PDCD4 | 2431.171143 | -0.915456883 | 1.51E-11 |
| RAB40B | 390.349998 | -0.915673804 | 2.18E-08 |
| NRG2 | 272.570081 | -0.915728301 | 7.45E-08 |
| MXD4 | 4041.346101 | -0.916249264 | 8.18E-05 |
| ZFP14 | 308.8122709 | -0.916369787 | 7.68E-10 |
| MASP2 | 36.75894355 | -0.916464099 | 3.18E-02 |
| NPW | 37.18962742 | -0.917258836 | 2.87E-02 |
| RIF1 | 2064.822231 | -0.917777171 | 2.68E-02 |
| HERC2P9 | 445.6592513 | -0.918085434 | 4.09E-10 |
| LINC01347 | 72.27855982 | -0.918167329 | 9.07E-04 |
| ATPSCKMT | 793.0329605 | -0.918365416 | 3.68E-16 |

| | | | |
|---|---|---|---|
| NUTM2A | 89.88279148 | -0.919118264 | 9.34E-05 |
| CRYGS | 33.5931757 | -0.91984394 | 2.31E-02 |
| AL450384.2 | 50.7999228 | -0.920210953 | 2.46E-02 |
| ATP2A3 | 1211.316063 | -0.920266726 | 1.10E-13 |
| RTP4 | 188.0812098 | -0.920317397 | 2.06E-06 |
| DOK3 | 101.2399448 | -0.920785872 | 3.63E-04 |
| DNAJC12 | 164.2504545 | -0.921034104 | 2.07E-06 |
| NEIL1 | 519.4268325 | -0.921287763 | 2.71E-07 |
| C21orf91 | 509.9850167 | -0.921683946 | 3.81E-06 |
| ERVK13-1 | 1300.392176 | -0.921725779 | 2.50E-26 |
| SNX22 | 156.8170892 | -0.922005884 | 3.53E-05 |
| EFNB2 | 435.0140339 | -0.922314632 | 2.31E-15 |
| FXYD6 | 2385.941432 | -0.922585752 | 3.30E-09 |
| CAT | 4283.077306 | -0.92294258 | 4.50E-24 |
| ZNF484 | 334.2493911 | -0.92390282 | 5.39E-05 |
| SPOCK2 | 70.07807607 | -0.924291022 | 1.87E-02 |
| PRR34 | 41.40607072 | -0.924582594 | 3.42E-02 |
| CCDC28B | 704.535287 | -0.924853389 | 9.29E-08 |
| ADAMTS4 | 395.6671777 | -0.925118017 | 1.75E-04 |
| TOX2 | 3322.582474 | -0.925159355 | 1.52E-06 |
| SHOC2 | 769.3246051 | -0.925172978 | 6.42E-04 |
| ZNF180 | 312.6482421 | -0.925893194 | 9.67E-05 |
| CCDC18 | 362.2515424 | -0.926639577 | 1.03E-02 |
| GIN1 | 179.1267096 | -0.928009771 | 1.90E-02 |
| RAD9B | 41.45801765 | -0.928302524 | 2.80E-02 |
| AC247036.1 | 655.3655526 | -0.928800823 | 3.52E-06 |
| NMI | 638.8842373 | -0.928890648 | 1.47E-10 |
| NYNRIN | 5607.528608 | -0.929294639 | 7.34E-05 |
| FAM129C | 53.32007866 | -0.929894099 | 4.49E-03 |
| SLC4A11 | 468.4280727 | -0.929985866 | 1.18E-07 |
| SMPD1 | 2976.700773 | -0.930423311 | 3.89E-11 |
| ZNF271P | 785.0492098 | -0.930794755 | 2.10E-07 |
| LRRC56 | 715.9976465 | -0.930816453 | 1.49E-09 |
| TRIM24 | 2116.562968 | -0.931209365 | 5.60E-34 |
| EP400P1 | 318.6927657 | -0.932921281 | 3.42E-06 |
| MYL5 | 176.9455721 | -0.933422752 | 7.23E-06 |
| ZBTB10 | 1109.922168 | -0.933646221 | 1.51E-12 |
| ADAMTS13 | 316.9800591 | -0.93382294 | 2.88E-09 |
| 6-Sep | 2821.75174 | -0.933982268 | 3.19E-56 |
| AC006254.1 | 73.41288277 | -0.934868062 | 1.89E-03 |
| GDPGP1 | 127.8574598 | -0.93560451 | 2.56E-05 |
| AC008103.1 | 69.65190709 | -0.93570876 | 1.58E-03 |
| G2E3-AS1 | 31.7803958 | -0.936424458 | 4.84E-02 |
| NA | 168.0298735 | -0.936452328 | 1.75E-06 |
| DNAAF4 | 158.9747061 | -0.937098017 | 2.03E-05 |
| CREBRF | 445.0915497 | -0.937340981 | 4.01E-05 |
| AC109322.1 | 83.15132005 | -0.937768229 | 1.50E-04 |

| | | | |
|---|---|---|---|
| CYP4V2 | 348.4362526 | -0.938577956 | 1.72E-04 |
| CYP3A5 | 68.36392406 | -0.940337134 | 6.67E-04 |
| TMEM173 | 1611.363547 | -0.94057045 | 1.20E-14 |
| AC138409.2 | 185.7386091 | -0.940922915 | 1.48E-06 |
| LINC01977 | 90.72125105 | -0.941477061 | 2.79E-05 |
| SCUBE2 | 118.6778719 | -0.941636588 | 1.95E-02 |
| AL365205.1 | 676.4159332 | -0.941949269 | 3.07E-08 |
| PNRC2 | 3356.074831 | -0.94212356 | 1.99E-16 |
| MAPK8IP2 | 998.1611941 | -0.942379809 | 2.69E-11 |
| TACR2 | 92.22274491 | -0.942493437 | 1.34E-03 |
| ANPEP | 1081.191916 | -0.94333389 | 1.51E-15 |
| AL133355.1 | 44.43468705 | -0.943409194 | 7.16E-03 |
| NA | 53.08899995 | -0.943642037 | 1.32E-02 |
| AC090739.1 | 23.09489251 | -0.944505287 | 3.83E-02 |
| ANXA9 | 166.5640661 | -0.944965284 | 1.02E-03 |
| MTND1P23 | 155.9045096 | -0.945026273 | 9.39E-04 |
| SULT1C2 | 407.8494656 | -0.945235413 | 3.63E-03 |
| TTC14 | 1432.076423 | -0.945398844 | 2.33E-06 |
| MIS18BP1 | 607.5874957 | -0.945504078 | 7.61E-05 |
| AC009506.1 | 50.51966715 | -0.945630375 | 4.57E-03 |
| PTGER2 | 910.6436438 | -0.946000243 | 5.65E-07 |
| CCDC189 | 255.0331238 | -0.946245783 | 3.17E-07 |
| CCNT2 | 1129.848901 | -0.947625506 | 1.71E-05 |
| ATXN1 | 3041.292894 | -0.947733958 | 2.04E-22 |
| CCDC191 | 86.06487863 | -0.948858682 | 2.37E-03 |
| FHOD3 | 292.6071871 | -0.948952637 | 5.12E-08 |
| PAQR8 | 949.7430723 | -0.949923509 | 8.22E-15 |
| RHBDL3 | 329.7370229 | -0.950009349 | 4.35E-06 |
| C1GALT1C1L | 229.7544466 | -0.950993126 | 1.01E-06 |
| RN7SL832P | 28.57002741 | -0.951945457 | 3.84E-02 |
| PCOLCE | 18291.75263 | -0.952752049 | 1.36E-11 |
| MICE | 40.47422612 | -0.953523656 | 1.36E-02 |
| SRSF8 | 1740.258175 | -0.954193536 | 1.55E-20 |
| PRKAR1A | 9864.643983 | -0.954244599 | 3.32E-18 |
| CEP19 | 554.178459 | -0.954975746 | 3.36E-15 |
| MT1F | 112.7334355 | -0.955351391 | 1.31E-02 |
| GUSBP3 | 41.01851898 | -0.956227737 | 1.21E-02 |
| PIK3CG | 245.0376781 | -0.957083009 | 7.94E-11 |
| STPG3-AS1 | 34.17829675 | -0.959928222 | 1.43E-02 |
| ARHGAP5 | 3812.275453 | -0.96157935 | 1.13E-04 |
| SNTB1 | 288.127424 | -0.961619954 | 5.77E-03 |
| CCDC121 | 148.176966 | -0.962256125 | 8.45E-05 |
| NPIPA7 | 80.15957517 | -0.962297167 | 6.75E-03 |
| TAF1A-AS1 | 52.66931189 | -0.962499968 | 1.59E-03 |
| AC021504.1 | 106.4338101 | -0.963175491 | 2.52E-04 |
| TOB1 | 570.1577494 | -0.963416645 | 2.11E-08 |
| ADGRE2 | 203.9491662 | -0.964366808 | 1.36E-03 |

| | | | |
|---|---|---|---|
| PAK1 | 4174.330043 | -0.964946701 | 3.62E-27 |
| TEN1-CDK3 | 280.7781373 | -0.965249589 | 2.74E-07 |
| ARHGAP28 | 1096.61935 | -0.965266967 | 2.69E-06 |
| PRR22 | 99.43339654 | -0.965489731 | 2.38E-04 |
| KIAA1324L | 213.9158876 | -0.965716545 | 1.25E-03 |
| CPED1 | 752.439028 | -0.967301064 | 1.01E-11 |
| WFDC1 | 322.5959827 | -0.967486745 | 4.93E-10 |
| LINC02557 | 375.7815063 | -0.968630264 | 4.90E-07 |
| GLUD1P3 | 36.56028664 | -0.968800871 | 1.44E-02 |
| TUBA1A | 19721.63494 | -0.969581563 | 1.07E-05 |
| ZFP2 | 31.7747584 | -0.970177786 | 2.34E-02 |
| ADNP-AS1 | 43.48973831 | -0.971462152 | 4.19E-03 |
| KRT15 | 92.32944513 | -0.971590062 | 3.44E-02 |
| MMP2 | 12012.51999 | -0.971653195 | 9.13E-12 |
| KCNC3 | 168.2532864 | -0.971662181 | 1.83E-04 |
| PGAP3 | 538.2201203 | -0.971713442 | 3.08E-08 |
| CR381670.1 | 68.18514066 | -0.971839688 | 4.67E-03 |
| RGS11 | 225.5788872 | -0.971855474 | 1.85E-05 |
| WDR35 | 1138.627068 | -0.971868894 | 1.90E-05 |
| LRG1 | 182.2253611 | -0.971912272 | 3.74E-03 |
| ZNF658 | 263.7837688 | -0.972204421 | 1.77E-06 |
| LGR4 | 3291.633279 | -0.973562827 | 2.31E-15 |
| GMPR | 1897.304968 | -0.974123253 | 4.38E-10 |
| EPCAM | 29.64192716 | -0.974500669 | 3.37E-02 |
| PTP4A3 | 372.1260805 | -0.974858065 | 1.18E-06 |
| SCN10A | 31.43251154 | -0.975007073 | 2.23E-02 |
| CEP85L | 295.6479056 | -0.975434364 | 4.92E-05 |
| TTC41P | 38.05365793 | -0.975603217 | 1.53E-02 |
| SLC6A16 | 260.3475046 | -0.975661691 | 1.13E-06 |
| TIAM1 | 2164.633176 | -0.976468787 | 2.08E-24 |
| LHX1 | 640.0009217 | -0.976798209 | 2.71E-09 |
| IGSF9 | 81.60414804 | -0.978317055 | 1.25E-03 |
| VSIG2 | 41.62470534 | -0.979567209 | 1.16E-02 |
| NA | 39.83765738 | -0.979866829 | 3.37E-03 |
| FAM117B | 322.1446807 | -0.980041726 | 6.15E-10 |
| AL353622.1 | 109.5816489 | -0.980274708 | 1.39E-05 |
| STPG3 | 31.27329893 | -0.980669054 | 8.59E-03 |
| TNRC6C-AS1 | 1271.337069 | -0.98074883 | 1.62E-12 |
| AC087741.1 | 188.2125206 | -0.980811508 | 2.30E-05 |
| NWD1 | 21.24870833 | -0.980952228 | 4.54E-02 |
| STAG2 | 4891.459207 | -0.981221281 | 3.15E-03 |
| MAP3K12 | 1271.312178 | -0.981281978 | 2.14E-08 |
| SHISA3 | 161.7304846 | -0.981505847 | 6.52E-03 |
| AC010595.1 | 65.32033256 | -0.981553983 | 1.45E-03 |
| AC010733.2 | 127.1405434 | -0.981793781 | 3.18E-02 |
| MFSD1 | 1429.551435 | -0.982024451 | 7.50E-17 |
| AC084018.2 | 65.15693012 | -0.982678271 | 1.14E-03 |

| | | | |
|---|---|---|---|
| AL162171.2 | 35.06777543 | -0.983415343 | 1.08E-02 |
| AL137003.1 | 206.3088742 | -0.984367938 | 6.31E-07 |
| AC025171.2 | 287.6890618 | -0.984526266 | 9.17E-09 |
| TEX15 | 71.56116553 | -0.986507189 | 2.43E-02 |
| EMID1 | 737.0210262 | -0.987205758 | 7.40E-09 |
| BCRP3 | 45.78330543 | -0.987208349 | 2.96E-03 |
| MPZL2 | 259.3641946 | -0.987634962 | 2.51E-06 |
| ACP2 | 3153.085252 | -0.987775361 | 8.75E-20 |
| STXBP3 | 829.6540019 | -0.987879757 | 5.39E-05 |
| TMTC3 | 861.6792463 | -0.988064616 | 4.91E-04 |
| SPRN | 304.4446147 | -0.988155413 | 2.14E-08 |
| CTSK | 134.8465847 | -0.989080884 | 1.91E-03 |
| AC118754.1 | 83.43697291 | -0.989342887 | 3.57E-04 |
| TANC2 | 3184.769522 | -0.990256778 | 8.77E-12 |
| ZDHHC11 | 107.5151782 | -0.991016553 | 9.20E-04 |
| TMSB15B-AS1 | 41.70051277 | -0.991029875 | 3.59E-02 |
| NLGN3 | 48.82620059 | -0.991411108 | 2.41E-03 |
| ING3 | 694.3881223 | -0.991728183 | 8.30E-07 |
| PID1 | 439.8178603 | -0.992459491 | 2.63E-11 |
| MDFI | 1896.304313 | -0.99307581 | 2.35E-13 |
| RICTOR | 2048.459973 | -0.99468004 | 1.77E-04 |
| NA | 75.43671028 | -0.995500101 | 9.14E-05 |
| AC140134.1 | 277.8328783 | -0.996423373 | 2.27E-05 |
| EDEM3 | 1391.633374 | -0.996921124 | 2.01E-08 |
| CACHD1 | 561.4740815 | -0.997162341 | 8.34E-15 |
| PTX3 | 4981.988845 | -0.997632406 | 5.19E-07 |
| ZNF492 | 38.86841734 | -0.997905824 | 1.66E-02 |
| HIST1H4H | 35.41351939 | -0.998122126 | 5.94E-03 |
| ATP11C | 2323.681458 | -0.99830314 | 2.56E-08 |
| ZNF491 | 42.3394643 | -0.998401748 | 2.39E-02 |
| CNTF | 43.98519678 | -0.998413762 | 4.35E-03 |
| VPS13B | 2695.859008 | -0.998497722 | 1.27E-15 |
| PCDHB13 | 90.25920142 | -1.00093419 | 2.06E-04 |
| AC068870.2 | 40.06505962 | -1.001613218 | 9.66E-03 |
| ZNF124 | 122.9671591 | -1.002332845 | 1.59E-05 |
| BMT2 | 620.0680209 | -1.002716801 | 6.69E-13 |
| SNCG | 667.1494932 | -1.003213387 | 1.30E-06 |
| EFCAB12 | 77.01796851 | -1.003315772 | 2.02E-03 |
| ZNF660 | 284.1719379 | -1.003401169 | 1.33E-04 |
| UBXN8 | 546.4351699 | -1.00449718 | 8.72E-15 |
| CD74 | 214.5084595 | -1.004782366 | 5.67E-03 |
| ZFP37 | 238.528989 | -1.005243254 | 2.13E-05 |
| LINC01126 | 42.41739339 | -1.005642039 | 3.16E-03 |
| CCDC188 | 246.0132551 | -1.007519063 | 8.17E-08 |
| PHIP | 1703.51194 | -1.007539732 | 5.04E-04 |
| AC092279.1 | 230.0617854 | -1.007858613 | 2.92E-13 |
| ZNF443 | 150.5584489 | -1.007893167 | 2.16E-06 |

| | | | |
|---|---|---|---|
| SLC25A48 | 75.47198056 | -1.00801223 | 1.31E-03 |
| NEAT1 | 4927.823974 | -1.00875949 | 2.24E-05 |
| KCNMB2-AS1 | 30.03868167 | -1.009914125 | 4.04E-02 |
| AP003774.3 | 29.72754542 | -1.010138213 | 4.04E-02 |
| SLK | 1599.19667 | -1.010799469 | 2.04E-05 |
| NR4A3 | 62.52509383 | -1.011326336 | 1.10E-03 |
| PIK3IP1 | 311.7503973 | -1.011566361 | 7.45E-05 |
| FBXL22 | 54.18573168 | -1.012046864 | 1.26E-03 |
| LARP7 | 955.0836205 | -1.012374822 | 1.07E-04 |
| AC048341.1 | 119.2109249 | -1.013074189 | 2.68E-05 |
| DPY19L2P1 | 178.5518984 | -1.013468233 | 6.39E-04 |
| CDC7 | 1042.852377 | -1.013545796 | 1.46E-09 |
| CCDC150 | 150.6043612 | -1.015141451 | 1.02E-03 |
| APOBR | 38.09673852 | -1.015267624 | 1.79E-02 |
| MAP7 | 29.05664401 | -1.015306221 | 3.38E-02 |
| ECM2 | 30.17136374 | -1.01567362 | 4.88E-02 |
| BATF2 | 455.2453915 | -1.015890671 | 1.09E-07 |
| SLF1 | 1156.522204 | -1.016374261 | 1.27E-05 |
| LCP2 | 54.88249637 | -1.019906825 | 9.61E-03 |
| SERPINA5 | 1276.950546 | -1.019926009 | 1.73E-08 |
| AL662795.2 | 323.8813937 | -1.020100928 | 8.00E-06 |
| AMDHD2 | 1252.631935 | -1.020372256 | 4.13E-21 |
| AGL | 980.3410774 | -1.020477152 | 2.42E-07 |
| SULT1B1 | 66.07862671 | -1.020519374 | 1.19E-03 |
| GIMAP2 | 643.0757087 | -1.020778138 | 6.77E-20 |
| PON2 | 4494.206146 | -1.021766324 | 4.43E-40 |
| COL6A3 | 4751.934988 | -1.023217403 | 2.55E-05 |
| RDX | 9248.951892 | -1.024058381 | 1.96E-15 |
| ZNF625 | 116.0250302 | -1.024183095 | 1.74E-03 |
| PER2 | 398.8658407 | -1.02436635 | 7.33E-09 |
| EREG | 68.09509223 | -1.024440686 | 3.42E-02 |
| CXCL8 | 99.7196793 | -1.028171803 | 2.91E-02 |
| AC093110.1 | 87.1898104 | -1.028276504 | 8.93E-05 |
| AL513327.1 | 97.34746054 | -1.028822744 | 1.97E-05 |
| ELL3 | 67.9189708 | -1.029707087 | 2.11E-04 |
| PTOV1-AS2 | 99.07661411 | -1.030099001 | 1.42E-05 |
| SEMA6C | 475.9537505 | -1.030485655 | 3.99E-07 |
| CFAP300 | 138.3243478 | -1.030916047 | 7.84E-06 |
| DCLK1 | 439.8908667 | -1.031549547 | 8.38E-04 |
| FCGR2A | 305.3492327 | -1.032100479 | 1.47E-09 |
| PRICKLE4 | 26.23740993 | -1.033305421 | 4.32E-02 |
| BAIAP3 | 385.5975178 | -1.035367511 | 1.77E-05 |
| ATRX | 2948.433124 | -1.03620867 | 3.31E-03 |
| NPIPA5 | 155.7742575 | -1.036973685 | 7.88E-03 |
| CTSO | 316.7800897 | -1.037209016 | 8.99E-06 |
| ZBTB41 | 298.4376423 | -1.037382591 | 7.17E-03 |
| ALPK3 | 123.8681661 | -1.038335046 | 2.14E-02 |

| | | | |
|---|---|---|---|
| CEP162 | 363.5001121 | -1.039272255 | 1.42E-03 |
| MMEL1 | 32.26212035 | -1.039501389 | 4.58E-02 |
| ITGA11 | 5052.132686 | -1.039570796 | 1.58E-12 |
| FSD1L | 138.4189189 | -1.039651304 | 6.01E-03 |
| AC010226.1 | 53.78791204 | -1.03977473 | 2.67E-03 |
| PSD3 | 2830.348069 | -1.040706938 | 3.32E-20 |
| STRA6LP | 197.9736375 | -1.041880628 | 7.79E-06 |
| PPM1L | 96.48854693 | -1.042103895 | 1.27E-03 |
| NA | 94.5777297 | -1.043997873 | 6.20E-07 |
| AC012441.1 | 24.68799952 | -1.044226856 | 3.42E-02 |
| GDAP1 | 1189.14057 | -1.044890414 | 3.54E-13 |
| SPRED1 | 1716.526315 | -1.045201205 | 4.41E-22 |
| FMO4 | 51.19801347 | -1.045472631 | 4.14E-02 |
| VPS13A | 2553.723416 | -1.045486715 | 6.33E-03 |
| SH2B3 | 6559.800718 | -1.046072818 | 5.94E-21 |
| XAF1 | 1197.67208 | -1.046359446 | 2.20E-10 |
| LDHD | 69.55842111 | -1.046682652 | 1.35E-03 |
| AC104447.1 | 94.69098291 | -1.046750508 | 1.91E-05 |
| AL132639.2 | 45.55583347 | -1.047135863 | 1.76E-03 |
| AC104667.1 | 34.0317262 | -1.047450045 | 7.33E-03 |
| KIAA0895L | 893.3242262 | -1.048123927 | 7.04E-20 |
| AC126768.3 | 60.52388494 | -1.04867429 | 1.05E-02 |
| OAS2 | 1649.202108 | -1.049250347 | 3.04E-07 |
| MAJIN | 69.31755048 | -1.049473023 | 3.63E-03 |
| C8orf31 | 287.7631837 | -1.049506788 | 1.59E-09 |
| AC018553.1 | 92.95374125 | -1.049910248 | 4.85E-07 |
| MT-TL2 | 78.84596201 | -1.050031371 | 2.94E-02 |
| LINC00565 | 61.82331782 | -1.050212358 | 9.14E-04 |
| SULF2 | 7414.451633 | -1.050481498 | 1.34E-36 |
| FAM111B | 742.562208 | -1.051008214 | 3.56E-02 |
| AL136368.1 | 23.15736852 | -1.051497351 | 1.87E-02 |
| NF1P7 | 19.5947773 | -1.053705289 | 3.30E-02 |
| LRRIQ3 | 42.99257129 | -1.054631107 | 1.92E-03 |
| DDX58 | 932.5016077 | -1.055252554 | 1.06E-25 |
| AC132812.1 | 80.9606679 | -1.055457078 | 1.19E-04 |
| CD207 | 40.47290139 | -1.055751252 | 1.51E-03 |
| AC112484.3 | 34.69759026 | -1.056111885 | 7.49E-03 |
| FAM222A | 185.6045506 | -1.05703175 | 1.96E-07 |
| CYB5B | 13391.33376 | -1.057499141 | 2.11E-14 |
| TBXAS1 | 153.5800672 | -1.057737197 | 9.12E-07 |
| ADAM22 | 711.1995431 | -1.058035647 | 1.01E-08 |
| STOX1 | 103.1043832 | -1.058319929 | 1.42E-05 |
| AC131392.1 | 146.7752183 | -1.058405676 | 4.55E-05 |
| ABCC9 | 49.86244202 | -1.05848761 | 2.96E-02 |
| CD163L1 | 1044.256401 | -1.061401734 | 8.17E-10 |
| AC009902.2 | 45.96325468 | -1.061417721 | 1.25E-03 |
| RGPD6 | 2488.190399 | -1.061595935 | 8.07E-09 |

| | | | |
|---|---|---|---|
| A2M-AS1 | 30.00575528 | -1.061746895 | 1.46E-02 |
| AC009120.3 | 29.06908059 | -1.062305837 | 7.45E-03 |
| HTR2B | 48.42776478 | -1.062480907 | 5.76E-03 |
| EFCAB7 | 208.1890505 | -1.062584102 | 5.26E-04 |
| GRAPL | 65.18748568 | -1.065355841 | 1.25E-04 |
| JCAD | 5110.161346 | -1.065387113 | 3.81E-30 |
| 2-Mar | 358.2724243 | -1.066429876 | 1.23E-06 |
| ZNF84 | 831.139286 | -1.066877361 | 1.38E-09 |
| DOCK11 | 2817.853034 | -1.067582497 | 1.69E-05 |
| DNM1P47 | 297.1681704 | -1.067984408 | 2.74E-07 |
| SENP7 | 434.7236845 | -1.06828271 | 1.30E-04 |
| AC044802.1 | 26.95094169 | -1.06970122 | 2.47E-02 |
| AL133215.2 | 51.61394003 | -1.071812475 | 5.17E-04 |
| S100B | 57.72904892 | -1.071836887 | 6.29E-03 |
| UST | 680.7892115 | -1.072053989 | 1.49E-09 |
| AC011379.1 | 25.8671817 | -1.075468485 | 9.29E-03 |
| LINC01963 | 343.5003131 | -1.075928107 | 4.60E-16 |
| ABCG1 | 981.4096477 | -1.076479101 | 2.59E-07 |
| Z83843.1 | 106.7766771 | -1.076727453 | 2.95E-04 |
| SCUBE3 | 863.7147381 | -1.077196612 | 9.32E-13 |
| AC008610.1 | 16.74355191 | -1.079441726 | 4.76E-02 |
| AC012513.3 | 165.7443447 | -1.079779833 | 4.58E-10 |
| ATRNL1 | 133.6598312 | -1.080670213 | 8.04E-06 |
| AL590399.1 | 405.0080006 | -1.082262641 | 1.01E-04 |
| MINDY1 | 1011.666293 | -1.082623016 | 2.92E-10 |
| PDCD4-AS1 | 327.1842865 | -1.082925782 | 3.98E-09 |
| AL590867.1 | 49.35383082 | -1.083212286 | 1.58E-02 |
| AC026356.1 | 50.63709256 | -1.083407867 | 5.36E-04 |
| SPDYE5 | 63.22792987 | -1.084117584 | 1.28E-04 |
| GOLGB1 | 3111.727492 | -1.084795632 | 1.44E-03 |
| CCDC192 | 19.63658269 | -1.085495363 | 4.88E-02 |
| NBL1 | 2346.529476 | -1.085615361 | 2.17E-15 |
| CAMKV | 31.31748181 | -1.086324241 | 9.73E-03 |
| FAM13B | 5560.921553 | -1.087149409 | 7.54E-22 |
| IZUMO4 | 100.2450413 | -1.087553816 | 3.77E-06 |
| AUTS2 | 1479.629524 | -1.089612099 | 2.40E-15 |
| AC240565.1 | 50.67336097 | -1.089734631 | 1.05E-02 |
| AC148477.3 | 81.05410233 | -1.090581878 | 6.17E-05 |
| AC018665.1 | 157.0923195 | -1.091308078 | 1.64E-10 |
| KIAA1586 | 323.4141024 | -1.09173574 | 5.01E-07 |
| APOL4 | 164.0383577 | -1.09179035 | 8.06E-08 |
| SHC3 | 593.2962704 | -1.092896568 | 1.35E-20 |
| TIA1 | 2320.581695 | -1.093243653 | 1.67E-08 |
| MED15P6 | 29.92975537 | -1.093243796 | 2.73E-02 |
| CLDND2 | 28.29589719 | -1.093488452 | 2.45E-02 |
| GPT | 53.48594233 | -1.093567056 | 9.98E-04 |
| C4A-AS1 | 29.21201907 | -1.094931297 | 4.76E-02 |

| | | | |
|---|---|---|---|
| AL137003.2 | 405.2240052 | -1.096390018 | 1.41E-09 |
| CAMK1 | 633.1255623 | -1.096577488 | 3.10E-09 |
| AC107982.3 | 18.97466292 | -1.096707829 | 4.40E-02 |
| DCLK2 | 707.8872453 | -1.098517583 | 4.57E-08 |
| FMNL2 | 4188.966288 | -1.098814995 | 6.88E-24 |
| LINC01431 | 20.83443025 | -1.100339604 | 2.48E-02 |
| FOXR2 | 367.8161971 | -1.100346235 | 7.70E-11 |
| NA | 209.1836675 | -1.100397669 | 9.23E-09 |
| CX3CL1 | 347.8028956 | -1.100407035 | 3.55E-06 |
| ARHGEF3 | 994.1096122 | -1.100622784 | 3.70E-20 |
| MAF | 556.0220594 | -1.102240023 | 8.53E-15 |
| TSHZ1 | 2579.087137 | -1.102357624 | 2.45E-14 |
| AL606760.1 | 46.42269404 | -1.102408657 | 2.64E-03 |
| CD4 | 153.7742743 | -1.103679455 | 4.09E-05 |
| SOS1-IT1 | 41.19267983 | -1.103829001 | 3.98E-02 |
| ZNF518A | 469.4591773 | -1.104361144 | 9.52E-05 |
| SLC44A5 | 50.24975973 | -1.104912107 | 1.06E-03 |
| RNF32 | 194.6641487 | -1.105709263 | 3.83E-07 |
| SPEF1 | 25.56375867 | -1.105879489 | 2.42E-02 |
| RGL1 | 1815.256907 | -1.107588893 | 1.23E-17 |
| ZDHHC8P1 | 115.4879533 | -1.110360725 | 2.26E-06 |
| AP000254.1 | 23.68196743 | -1.111243464 | 2.96E-02 |
| TRANK1 | 2119.181798 | -1.112093439 | 2.26E-10 |
| DIXDC1 | 1830.006981 | -1.112812215 | 1.06E-13 |
| AC010422.6 | 130.2044948 | -1.114340176 | 3.78E-08 |
| DMXL2 | 3319.662109 | -1.114653111 | 3.91E-06 |
| AC006330.1 | 20.83963745 | -1.115787933 | 3.09E-02 |
| AL353593.1 | 40.48596538 | -1.117467302 | 2.54E-03 |
| AC026691.1 | 44.42698353 | -1.118673291 | 1.00E-03 |
| LGI4 | 122.8848522 | -1.118939274 | 7.03E-06 |
| AC008937.3 | 41.52574998 | -1.11908836 | 4.09E-03 |
| EME2 | 888.6364107 | -1.119242501 | 1.70E-11 |
| ABCC6 | 487.0257148 | -1.119273301 | 2.58E-10 |
| CHST6 | 150.7951797 | -1.120687641 | 1.33E-04 |
| BSN | 168.5919291 | -1.121536823 | 8.16E-08 |
| PBLD | 150.0356914 | -1.122140923 | 1.62E-06 |
| TCEANC | 77.99028319 | -1.123300898 | 1.00E-04 |
| DYNC2LI1 | 454.8326072 | -1.125830216 | 7.74E-09 |
| P4HA3 | 312.285872 | -1.130420138 | 4.09E-12 |
| CCDC88A | 3210.87298 | -1.131238401 | 2.57E-05 |
| ST3GAL5 | 3369.022929 | -1.131885254 | 2.66E-25 |
| AP001992.1 | 25.90380369 | -1.13253965 | 1.88E-02 |
| LINC00893 | 98.4657195 | -1.132617943 | 1.06E-04 |
| FOS | 91.33837697 | -1.13286231 | 2.09E-02 |
| ARHGAP9 | 264.0659115 | -1.133724208 | 5.28E-09 |
| FOXC1 | 860.7224109 | -1.133989187 | 1.46E-16 |
| TET2 | 1173.805508 | -1.134198149 | 2.82E-16 |

| | | | |
|---|---|---|---|
| GABRG2 | 26.96306728 | -1.134571235 | 2.12E-02 |
| RAET1G | 88.40783674 | -1.134871552 | 2.24E-03 |
| SSPO | 225.5519511 | -1.135450502 | 4.12E-03 |
| SLC26A10 | 106.1172386 | -1.135765514 | 2.72E-05 |
| C15orf65 | 39.22759693 | -1.135930745 | 2.25E-02 |
| AC016526.3 | 48.75006189 | -1.1360838 | 1.87E-04 |
| TNR | 24.03847016 | -1.136455604 | 3.58E-02 |
| TBC1D10C | 46.80751802 | -1.13659127 | 7.61E-03 |
| NDRG2 | 3068.619395 | -1.136632693 | 2.51E-10 |
| LPP-AS2 | 110.5716903 | -1.136759031 | 6.92E-08 |
| NMUR1 | 23.97944588 | -1.13689281 | 1.46E-02 |
| COL11A2 | 524.1032316 | -1.137754493 | 1.75E-06 |
| RUNDC3A | 72.75626673 | -1.137812436 | 8.26E-04 |
| CHN2 | 294.5890476 | -1.139020612 | 3.40E-16 |
| STIM2 | 1398.883632 | -1.139572085 | 9.38E-22 |
| AL162171.4 | 23.20073892 | -1.14010548 | 2.04E-02 |
| CYP2D6 | 86.52775803 | -1.140797598 | 9.78E-06 |
| ZMAT4 | 504.9050422 | -1.143468666 | 1.81E-10 |
| IRF6 | 110.6818255 | -1.144490349 | 3.72E-03 |
| AC008494.3 | 51.81223161 | -1.145382903 | 1.18E-03 |
| AC011450.1 | 89.0077873 | -1.14635088 | 6.79E-06 |
| JMJD1C | 1264.147845 | -1.147539346 | 8.55E-06 |
| HIST1H3H | 87.21353459 | -1.1476925 | 4.19E-04 |
| SPIN2B | 409.744545 | -1.148021827 | 1.15E-26 |
| ZNF658B | 53.47303321 | -1.148255098 | 8.17E-03 |
| AC027031.2 | 487.0588179 | -1.148541849 | 4.03E-21 |
| FAM71F2 | 50.614311 | -1.149602723 | 1.40E-03 |
| CARNS1 | 82.0911703 | -1.150133463 | 3.33E-04 |
| FZD9 | 356.4608133 | -1.150386979 | 7.10E-10 |
| AC004943.1 | 20.64228712 | -1.150948379 | 2.94E-02 |
| AL353150.1 | 115.1320375 | -1.151698989 | 6.05E-08 |
| PAX9 | 66.37338436 | -1.151985404 | 3.40E-04 |
| AL121672.2 | 17.12607123 | -1.152634343 | 3.77E-02 |
| SATB1 | 25.96066018 | -1.153474514 | 3.28E-02 |
| DACT3 | 383.0030205 | -1.153773162 | 3.06E-10 |
| ARHGAP24 | 86.39537728 | -1.154196022 | 7.41E-06 |
| CPEB3 | 39.77725826 | -1.154415341 | 1.38E-03 |
| RASSF3 | 2924.247126 | -1.154706761 | 4.55E-41 |
| REV3L | 1912.9575 | -1.158061763 | 1.40E-08 |
| AC022007.1 | 130.4691102 | -1.161575788 | 1.83E-08 |
| NDRG4 | 400.4921828 | -1.161629776 | 2.58E-13 |
| DYRK1B | 543.7450149 | -1.161812087 | 1.12E-11 |
| CCDC78 | 384.8868074 | -1.162987221 | 1.45E-13 |
| TSPAN7 | 39.81758527 | -1.163167988 | 1.12E-03 |
| CD52 | 26.23631844 | -1.163651284 | 3.57E-02 |
| AL391069.2 | 56.01725847 | -1.163757865 | 1.18E-04 |
| NA | 137.0107692 | -1.165205716 | 7.92E-06 |

| | | | |
|---|---|---|---|
| FAM84A | 44.9306386 | -1.165313447 | 2.69E-03 |
| AC002116.1 | 48.13561349 | -1.165391562 | 3.70E-02 |
| ARHGAP18 | 3596.763254 | -1.165558169 | 7.64E-44 |
| ADD3 | 3597.694114 | -1.165651706 | 2.66E-37 |
| PHOSPHO2 | 116.2748943 | -1.171022754 | 7.85E-06 |
| AC005041.1 | 156.4021284 | -1.172220079 | 2.24E-07 |
| RIMBP3B | 55.67544132 | -1.172877125 | 1.66E-02 |
| JAK3 | 626.4733155 | -1.173377368 | 1.51E-07 |
| AL353803.2 | 202.4477704 | -1.173834193 | 2.91E-07 |
| GPR137C | 168.6258987 | -1.175915113 | 3.08E-09 |
| AC006058.4 | 114.8916737 | -1.176752699 | 3.50E-06 |
| RASSF5 | 662.6694676 | -1.177525657 | 3.12E-18 |
| TMEM145 | 253.2058034 | -1.178339109 | 1.20E-08 |
| TSC22D3 | 3742.290724 | -1.17938016 | 2.67E-14 |
| AC022107.1 | 577.3918954 | -1.179390536 | 2.98E-34 |
| TMEM150C | 53.30532729 | -1.180051169 | 2.29E-02 |
| AC097376.2 | 104.004178 | -1.181922633 | 2.39E-06 |
| AC004540.1 | 39.99496744 | -1.182559774 | 2.30E-03 |
| AC008555.2 | 21.86004834 | -1.183317302 | 1.26E-02 |
| BICD1 | 1218.161353 | -1.184171496 | 8.78E-37 |
| LRRC24 | 60.46001221 | -1.184733771 | 3.56E-03 |
| GRAMD1B | 1146.120968 | -1.18579881 | 4.64E-17 |
| EPS8 | 4496.746593 | -1.186763003 | 5.23E-18 |
| IL1R1 | 1260.323681 | -1.187184376 | 3.27E-20 |
| AKAP5 | 222.2346906 | -1.187998507 | 2.00E-09 |
| KCNK2 | 141.6576935 | -1.188257999 | 8.39E-05 |
| SGSM1 | 82.61987736 | -1.188987793 | 1.50E-03 |
| ARID4A | 694.162518 | -1.189428707 | 5.18E-07 |
| AL356489.2 | 24.55033681 | -1.189577338 | 3.72E-02 |
| EHD3 | 6392.771573 | -1.189668697 | 2.04E-16 |
| VMO1 | 25.53474 | -1.192813635 | 1.91E-02 |
| PCDHGC3 | 8838.789475 | -1.193137547 | 5.90E-18 |
| ROM1 | 228.2174532 | -1.194162405 | 2.06E-07 |
| GPCPD1 | 2138.032665 | -1.195336303 | 3.40E-08 |
| PSMD6-AS2 | 40.60371709 | -1.201253419 | 6.77E-03 |
| C1QTNF6 | 3661.584941 | -1.201320489 | 1.35E-08 |
| ZNF253 | 186.1191461 | -1.202910071 | 4.41E-05 |
| RANBP2 | 6080.340109 | -1.203198855 | 4.71E-07 |
| TAS1R1 | 28.52816343 | -1.203524641 | 1.65E-02 |
| CITED1 | 34.09631409 | -1.203810983 | 5.22E-03 |
| AP001107.9 | 109.1125027 | -1.20440192 | 3.54E-07 |
| DOCK4 | 1431.176833 | -1.205722395 | 2.53E-21 |
| RAB11FIP1P1 | 56.13576336 | -1.206444567 | 9.66E-03 |
| WDR72 | 57.53109666 | -1.207463378 | 5.22E-03 |
| TMEM249 | 28.67557228 | -1.208205902 | 3.97E-02 |
| IL6ST | 22565.49008 | -1.208308474 | 1.20E-04 |
| ASB9 | 574.9228447 | -1.208790344 | 1.97E-12 |

| | | | |
|---|---|---|---|
| PES1P1 | 21.79278102 | -1.209212252 | 2.63E-02 |
| NA | 14.84169264 | -1.210226648 | 4.37E-02 |
| USP51 | 104.9721947 | -1.211640767 | 3.02E-05 |
| NPAS3 | 33.85604416 | -1.213055533 | 8.31E-03 |
| AC009902.3 | 19.06579071 | -1.213082819 | 2.82E-02 |
| CPXM1 | 3098.915536 | -1.213952766 | 3.06E-16 |
| AC127024.5 | 44.36894571 | -1.215834315 | 2.87E-04 |
| AC012511.1 | 21.46889958 | -1.216363581 | 6.08E-03 |
| SYN1 | 187.3211338 | -1.216475414 | 2.68E-10 |
| ELOVL3 | 25.0446582 | -1.218671121 | 1.93E-02 |
| AL161891.1 | 53.9731438 | -1.22267573 | 8.89E-03 |
| AC008750.8 | 19.93012182 | -1.222698323 | 2.45E-02 |
| DDN | 37.56274831 | -1.223041722 | 1.90E-02 |
| SLC22A17 | 525.7143164 | -1.22415273 | 8.05E-07 |
| SPDYE2 | 89.92486882 | -1.224665202 | 1.00E-05 |
| CPXM2 | 397.4177067 | -1.225113871 | 2.03E-08 |
| IFIT1 | 1401.914147 | -1.225755897 | 3.16E-09 |
| PEG3 | 8882.770975 | -1.226430768 | 1.18E-18 |
| ATP2B1 | 2269.200867 | -1.227070328 | 1.46E-09 |
| ZNF233 | 106.6235961 | -1.2273692 | 2.93E-10 |
| MT-TQ | 46.69971454 | -1.229396137 | 2.02E-02 |
| HESX1 | 24.15406492 | -1.229479082 | 1.07E-02 |
| NA | 120.1022931 | -1.230901642 | 4.84E-04 |
| WNT6 | 159.8552245 | -1.231666125 | 1.05E-05 |
| KCNN3 | 360.2014472 | -1.232186567 | 7.18E-10 |
| AL353763.1 | 33.38763978 | -1.233579987 | 5.34E-03 |
| CD37 | 311.4740181 | -1.234305897 | 3.66E-13 |
| SH3BP5-AS1 | 201.3828996 | -1.236875671 | 1.92E-07 |
| MAOB | 49.54773042 | -1.238667743 | 7.30E-04 |
| AC096537.1 | 58.48260446 | -1.238966499 | 3.42E-04 |
| CMBL | 2580.311176 | -1.238990383 | 1.20E-17 |
| NPAS2 | 2199.95089 | -1.23990344 | 3.85E-20 |
| PDGFRL | 578.6213394 | -1.240880244 | 7.11E-10 |
| PIEZO2 | 3798.179918 | -1.241257333 | 7.72E-49 |
| FNBP1L | 2268.026796 | -1.241615343 | 4.80E-45 |
| FANCM | 393.9957169 | -1.243088536 | 4.68E-05 |
| CACTIN-AS1 | 46.52365665 | -1.243461748 | 2.34E-04 |
| TMEM198 | 334.0886835 | -1.243644387 | 2.10E-14 |
| CASD1 | 796.3896898 | -1.243658982 | 1.57E-14 |
| CHST1 | 142.5358579 | -1.24388673 | 2.17E-09 |
| HLA-DPA1 | 429.1936339 | -1.244006291 | 2.55E-06 |
| SERINC5 | 1988.56082 | -1.244726313 | 9.07E-35 |
| DOP1A | 788.5080826 | -1.245548883 | 4.25E-08 |
| C3orf18 | 20.94029741 | -1.245614366 | 3.05E-02 |
| MBNL1-AS1 | 33.10521091 | -1.246109848 | 2.01E-02 |
| TP53INP1 | 249.9843985 | -1.246505343 | 8.32E-12 |
| KITLG | 103.264813 | -1.246946041 | 3.55E-04 |

| | | | |
|---|---|---|---|
| GBP4 | 532.7411693 | -1.24750015 | 7.72E-08 |
| AC067930.4 | 32.3071449 | -1.247530679 | 6.76E-03 |
| NA | 17.82293091 | -1.249547112 | 1.98E-02 |
| AC103760.1 | 21.44567501 | -1.249815649 | 2.40E-02 |
| TCTEX1D1 | 67.45108166 | -1.250355995 | 4.64E-04 |
| TP53TG3B | 23.77117958 | -1.25329175 | 3.36E-02 |
| KCNAB3 | 147.848877 | -1.255144322 | 6.54E-10 |
| AP003170.1 | 15.44750304 | -1.257102873 | 4.76E-02 |
| CHD9 | 2148.656633 | -1.257208278 | 3.85E-08 |
| SELENBP1 | 2651.618252 | -1.257237483 | 1.67E-10 |
| SOBP | 699.2504882 | -1.257416587 | 1.09E-27 |
| SUSD5 | 41.31474875 | -1.257948341 | 2.13E-03 |
| AK3P3 | 17.21847133 | -1.26209365 | 4.36E-02 |
| C6orf163 | 32.03509104 | -1.26283886 | 2.80E-03 |
| PARD6G-AS1 | 83.02067989 | -1.263811917 | 1.09E-05 |
| LINC02166 | 38.8899247 | -1.263914104 | 6.16E-04 |
| SEMA6A | 182.063587 | -1.264164702 | 4.17E-07 |
| PLAC9 | 94.03662567 | -1.264838499 | 2.00E-05 |
| PCDHB14 | 50.50684348 | -1.265593401 | 3.16E-04 |
| EML4 | 8450.371969 | -1.266649032 | 6.79E-37 |
| ITPKB | 985.5116325 | -1.268915801 | 3.24E-14 |
| PCDH18 | 57.24297472 | -1.271417692 | 7.87E-03 |
| KIF26A | 548.4895354 | -1.271752122 | 1.31E-16 |
| PRRT3 | 215.8516639 | -1.271938361 | 1.01E-14 |
| LRRC46 | 47.82199529 | -1.27224283 | 8.45E-04 |
| SLC22A20P | 38.07364341 | -1.272837386 | 1.01E-03 |
| RNASEL | 490.163533 | -1.273049777 | 4.20E-26 |
| AC093677.2 | 17.96716077 | -1.27348123 | 2.59E-02 |
| AL513548.1 | 37.02258029 | -1.274243587 | 4.15E-03 |
| PNPLA7 | 124.432657 | -1.275037613 | 4.35E-08 |
| RNF112 | 75.02755394 | -1.276765777 | 1.86E-02 |
| KCNH2 | 218.6381721 | -1.28177549 | 1.97E-07 |
| AC092375.2 | 32.97172105 | -1.283163017 | 1.35E-03 |
| NA | 22.00261677 | -1.283593335 | 4.72E-03 |
| SVIL | 3315.265826 | -1.283631412 | 7.47E-32 |
| NA | 13.41999917 | -1.283719632 | 4.22E-02 |
| AC021087.1 | 21.45280627 | -1.284354138 | 1.27E-02 |
| RIBC1 | 84.71474259 | -1.284755517 | 1.21E-07 |
| TNIK | 2274.191439 | -1.285656452 | 1.49E-23 |
| PPP1R12A-AS1 | 32.66543125 | -1.287492309 | 4.39E-03 |
| AC139491.3 | 70.41335232 | -1.287636157 | 1.88E-04 |
| IL4I1 | 44.42488749 | -1.288070118 | 9.64E-04 |
| ZNF467 | 538.7262357 | -1.288274425 | 1.05E-09 |
| GSN-AS1 | 75.19028754 | -1.291999848 | 4.38E-08 |
| GPR162 | 140.5579805 | -1.292249184 | 5.37E-07 |
| AC000123.2 | 14.95406326 | -1.292363916 | 4.68E-02 |
| AL139274.2 | 45.84750634 | -1.292367164 | 1.12E-03 |

| | | | |
|---|---|---|---|
| LINC00920 | 110.360093 | -1.29290422 | 5.14E-08 |
| C3orf20 | 24.41494674 | -1.293193614 | 4.14E-02 |
| INSRR | 12.9101483 | -1.294093194 | 4.72E-02 |
| CDH12P3 | 37.29249809 | -1.294457568 | 9.03E-03 |
| Z92544.2 | 182.6149303 | -1.29487626 | 1.42E-07 |
| PTCH2 | 132.6977029 | -1.295808342 | 1.72E-05 |
| LAMA4 | 12155.3392 | -1.295834758 | 2.03E-84 |
| ZC3H12D | 48.81604572 | -1.296133427 | 7.24E-03 |
| HTRA1 | 853.162337 | -1.296412117 | 1.57E-14 |
| FOXD4L5 | 21.56223852 | -1.297186796 | 3.07E-02 |
| LINC02503 | 75.87403053 | -1.297892183 | 1.05E-03 |
| NADK2 | 3676.638597 | -1.297937041 | 9.04E-27 |
| KPNA2P3 | 51.10949066 | -1.298510488 | 1.92E-02 |
| AF131216.3 | 48.22332352 | -1.299615777 | 5.98E-04 |
| FZD1 | 1665.798152 | -1.299956766 | 3.19E-30 |
| AC012254.3 | 17.6987559 | -1.300238066 | 1.12E-02 |
| ARRB1 | 3818.529655 | -1.303048385 | 2.06E-22 |
| AL590552.1 | 21.11172157 | -1.303530393 | 2.26E-02 |
| SHE | 294.9360923 | -1.306145799 | 1.52E-11 |
| ETV4 | 2074.53517 | -1.30621195 | 3.43E-09 |
| NAAA | 534.8626594 | -1.306396485 | 5.72E-34 |
| CDH11 | 43.29998427 | -1.306865967 | 6.68E-03 |
| BCAS1 | 91.26170515 | -1.30690497 | 1.42E-04 |
| IGHV3-19 | 30.14242899 | -1.307832717 | 6.45E-04 |
| ST6GAL1 | 1260.199539 | -1.307862309 | 1.48E-14 |
| AC092143.3 | 19.80251712 | -1.309681319 | 1.37E-02 |
| SLC6A17 | 261.5610214 | -1.310444108 | 7.19E-08 |
| B3GNT3 | 176.5313853 | -1.311934725 | 1.85E-06 |
| VPS13C | 3551.851662 | -1.31292723 | 4.98E-05 |
| ZHX1 | 2058.307391 | -1.318055628 | 2.14E-13 |
| USP53 | 1004.853788 | -1.319119779 | 1.49E-07 |
| AL031673.1 | 19.97831105 | -1.319424307 | 1.64E-02 |
| NA | 51.51323304 | -1.320263197 | 1.30E-04 |
| AC005837.1 | 17.76815366 | -1.320494575 | 2.33E-02 |
| RNF125 | 78.72295167 | -1.320795632 | 6.28E-04 |
| LAMP5 | 323.5776342 | -1.322787968 | 4.41E-12 |
| GMFG | 271.7051033 | -1.323427843 | 1.29E-09 |
| AL031716.1 | 24.43679082 | -1.324684598 | 1.05E-02 |
| SLC25A45 | 87.16993623 | -1.327457132 | 5.30E-04 |
| AC015849.5 | 41.31593725 | -1.327474261 | 3.36E-02 |
| SULT1A4 | 358.2553373 | -1.32920184 | 5.35E-03 |
| NARF-IT1 | 9.897559897 | -1.330138042 | 3.80E-02 |
| C6orf141 | 203.3132315 | -1.332163079 | 1.10E-09 |
| SGK1 | 5483.663424 | -1.332816224 | 3.04E-21 |
| CCDC13 | 15.62344012 | -1.333153432 | 1.86E-02 |
| MIR3142HG | 184.9959843 | -1.33534161 | 1.09E-11 |
| ICA1L | 195.1054069 | -1.336550245 | 3.85E-13 |

| | | | |
|---|---|---|---|
| METTL7B | 30.52122426 | -1.337018547 | 7.25E-03 |
| RARRES1 | 136.06074 | -1.337990969 | 1.26E-05 |
| CNIH2 | 324.591191 | -1.33937442 | 4.06E-07 |
| MEIS1 | 435.9633765 | -1.339840423 | 8.25E-11 |
| ENDOD1 | 5289.674656 | -1.341222554 | 1.26E-11 |
| AC008467.1 | 33.37212616 | -1.341516187 | 2.56E-04 |
| ZBTB46 | 1939.837957 | -1.341813046 | 9.00E-21 |
| CKMT1A | 221.78047 | -1.342835776 | 1.25E-06 |
| AC140125.2 | 68.76072382 | -1.343251978 | 4.61E-05 |
| ACPP | 27.55663385 | -1.343922101 | 3.10E-03 |
| AC026740.1 | 48.90043332 | -1.344393373 | 3.79E-05 |
| KAT6B | 1431.533217 | -1.344773939 | 2.27E-29 |
| APC | 2832.53114 | -1.345885818 | 3.99E-08 |
| TDRD10 | 80.80343204 | -1.346041644 | 2.03E-07 |
| ROPN1L | 83.08733187 | -1.346195211 | 1.15E-08 |
| 14-Sep | 29.05544493 | -1.347526463 | 2.88E-03 |
| BCL2L11 | 920.1726165 | -1.34761943 | 3.58E-25 |
| AP005131.6 | 30.74446118 | -1.349252711 | 6.64E-04 |
| PLCD4 | 1142.359525 | -1.349455157 | 2.18E-12 |
| RB1 | 3726.742893 | -1.351719994 | 3.83E-20 |
| PLPP3 | 2030.88115 | -1.353076293 | 3.21E-31 |
| AC009159.3 | 20.44347584 | -1.35707111 | 3.46E-03 |
| EPHX4 | 58.81641382 | -1.361093788 | 1.39E-05 |
| AL022316.1 | 18.85500911 | -1.361293648 | 2.66E-02 |
| AC025766.1 | 17.13948499 | -1.363082109 | 2.60E-02 |
| PRRT2 | 400.9286235 | -1.363978788 | 9.60E-16 |
| IFI30 | 2930.300089 | -1.36547832 | 3.94E-19 |
| LHFPL2 | 4882.279716 | -1.36727052 | 2.14E-97 |
| BX571846.1 | 39.76887745 | -1.369258569 | 3.48E-04 |
| CCDC87 | 25.69209508 | -1.369380955 | 4.81E-03 |
| AC021321.1 | 29.50129298 | -1.370452364 | 2.10E-03 |
| NRP2 | 3214.243923 | -1.370865531 | 2.05E-61 |
| GNG4 | 685.0541399 | -1.372394483 | 2.84E-17 |
| CEACAM21 | 17.26810419 | -1.372509932 | 1.95E-02 |
| FAXDC2 | 500.5299023 | -1.375018915 | 9.58E-16 |
| AC003973.2 | 21.04102182 | -1.376544728 | 3.21E-02 |
| AC021242.3 | 17.8700325 | -1.377913185 | 4.62E-02 |
| GVQW2 | 19.57657147 | -1.378975138 | 9.45E-03 |
| AC254562.2 | 30.44230704 | -1.379406752 | 4.68E-03 |
| BAMBI | 4010.746087 | -1.379603073 | 1.13E-15 |
| LINC02289 | 34.71716412 | -1.380327746 | 5.85E-03 |
| LINC02073 | 22.06696947 | -1.380755352 | 1.21E-02 |
| ZNF92 | 182.2272354 | -1.383021746 | 2.33E-05 |
| C5orf49 | 13.73202357 | -1.383367838 | 3.69E-02 |
| HTR2C | 14.49740132 | -1.383751349 | 2.30E-02 |
| ABCC6P2 | 34.63096341 | -1.384157591 | 2.92E-03 |
| RHOJ | 2033.908093 | -1.384324131 | 1.36E-21 |

| | | | |
|---|---|---|---|
| ABCA8 | 289.3288945 | -1.385086182 | 4.77E-05 |
| PRDM1 | 260.727632 | -1.388628727 | 1.59E-06 |
| ARHGEF6 | 2573.200329 | -1.391258665 | 2.92E-77 |
| AC137894.1 | 39.95266791 | -1.391612755 | 1.57E-04 |
| MSC | 323.6871895 | -1.392344727 | 1.38E-03 |
| LINGO1 | 53.96417116 | -1.392393881 | 2.17E-03 |
| EFCAB5 | 26.20293695 | -1.394230005 | 1.76E-02 |
| AC091563.1 | 20.66601571 | -1.395096785 | 1.52E-02 |
| TSPOAP1-AS1 | 18.80222949 | -1.395376063 | 1.31E-02 |
| EMILIN3 | 78.86613811 | -1.395989217 | 1.94E-04 |
| KIAA0825 | 59.54724542 | -1.396806436 | 3.57E-04 |
| AC006111.2 | 16.47899019 | -1.397727837 | 1.81E-02 |
| MIMT1 | 17.01502868 | -1.403360214 | 2.26E-02 |
| GVINP1 | 117.3805028 | -1.404309898 | 4.44E-08 |
| OASL | 87.89182137 | -1.404647023 | 8.07E-05 |
| A2M | 8519.613877 | -1.405849721 | 4.68E-09 |
| AF233439.1 | 16.16706051 | -1.405975649 | 3.84E-02 |
| FAM78B | 308.7966026 | -1.406804167 | 1.78E-12 |
| AKR1C3 | 83.72430564 | -1.407214567 | 1.24E-03 |
| PECAM1 | 169.5651196 | -1.407909786 | 9.56E-10 |
| ZCCHC24 | 4931.736738 | -1.408642583 | 2.49E-27 |
| C3orf70 | 235.4032375 | -1.40943064 | 2.14E-20 |
| MDH1B | 51.25257486 | -1.412455732 | 1.90E-05 |
| GAB1 | 870.6201106 | -1.412724029 | 3.07E-27 |
| SELL | 65.28807749 | -1.413788304 | 3.95E-04 |
| TBC1D32 | 228.4429051 | -1.41502694 | 1.86E-07 |
| CICP4 | 27.71170368 | -1.4151201 | 6.05E-03 |
| AC084083.1 | 13.67494269 | -1.415650125 | 3.65E-02 |
| IL2RB | 1447.939605 | -1.416011141 | 8.39E-27 |
| SNAP25 | 73.26660247 | -1.41725031 | 1.40E-03 |
| NR1H3 | 1127.557574 | -1.41816155 | 1.30E-17 |
| AC006058.1 | 319.7637519 | -1.418320954 | 3.28E-15 |
| COL6A1 | 17941.64959 | -1.418438933 | 3.85E-51 |
| LINC02076 | 54.9376968 | -1.418714254 | 1.10E-06 |
| FBXO40 | 30.39114751 | -1.419915577 | 9.12E-04 |
| AL606760.3 | 15.33031838 | -1.42052651 | 2.60E-02 |
| PARP15 | 35.78849082 | -1.421170752 | 9.84E-04 |
| AC019254.1 | 10.78643097 | -1.421563651 | 4.99E-02 |
| MUC13 | 39.74750968 | -1.422624601 | 3.27E-04 |
| MARCKSL1 | 6894.580256 | -1.423301949 | 2.84E-23 |
| LINC00461 | 138.3576983 | -1.423501537 | 4.62E-06 |
| CLEC4F | 53.65404115 | -1.424186287 | 3.98E-06 |
| EXOSC10-AS1 | 25.29981927 | -1.42443178 | 6.70E-03 |
| HLA-DRB1 | 80.24218844 | -1.42510824 | 1.19E-03 |
| CFI | 1692.65687 | -1.426094091 | 6.10E-05 |
| AC007000.1 | 15.9064266 | -1.426262389 | 4.30E-02 |
| FN1 | 116782.2439 | -1.428510827 | 3.29E-09 |

| | | | |
|---|---|---|---|
| AL596202.1 | 165.9496982 | -1.428953018 | 5.20E-13 |
| SYPL2 | 83.161266 | -1.429375343 | 6.27E-06 |
| NTAN1P2 | 10.97870799 | -1.43006312 | 4.92E-02 |
| HSD11B2 | 26.99250525 | -1.430884459 | 2.33E-03 |
| GALNT5 | 69.69342666 | -1.430995545 | 8.38E-05 |
| AP000787.1 | 90.97820889 | -1.431241525 | 3.91E-06 |
| RTCA-AS1 | 15.71405731 | -1.43219424 | 2.34E-02 |
| LINC02260 | 53.3414811 | -1.434154298 | 5.78E-04 |
| BAZ2B | 879.1717393 | -1.434436497 | 1.75E-08 |
| PCDHB5 | 79.84968752 | -1.43542315 | 1.46E-06 |
| AL358852.1 | 15.31578304 | -1.435595869 | 1.75E-02 |
| AC078880.3 | 15.99001634 | -1.436420212 | 1.29E-02 |
| GBP2 | 578.7100366 | -1.436767921 | 1.08E-14 |
| SYP | 149.7220486 | -1.437381009 | 4.82E-10 |
| NINJ2 | 86.32428491 | -1.437565664 | 1.24E-06 |
| AL008729.2 | 15.84724234 | -1.438166393 | 1.26E-02 |
| GALNT4 | 422.8417552 | -1.442092692 | 1.38E-15 |
| C2CD4B | 29.23482065 | -1.44259025 | 1.83E-03 |
| KCNRG | 21.37247774 | -1.443583478 | 1.37E-02 |
| TLR7 | 44.58411541 | -1.444512512 | 1.72E-03 |
| TMEM121B | 53.36772833 | -1.445117926 | 2.17E-05 |
| TSNAXIP1 | 79.41687888 | -1.447550721 | 1.32E-06 |
| C4B | 1756.412125 | -1.449505006 | 3.28E-02 |
| PGAP1 | 1273.05056 | -1.450759786 | 1.56E-13 |
| NA | 27.53951476 | -1.452043234 | 9.27E-04 |
| TLR3 | 366.6552858 | -1.452511707 | 1.86E-08 |
| LINC01684 | 16.86587379 | -1.453720104 | 2.42E-02 |
| BBOX1-AS1 | 25.44159368 | -1.454712888 | 5.86E-03 |
| SERPINF1 | 435.8532755 | -1.455416043 | 4.74E-17 |
| LINC02657 | 19.24585261 | -1.455635283 | 1.08E-02 |
| NFATC2 | 610.572034 | -1.45682295 | 4.91E-12 |
| AL359555.1 | 15.27189961 | -1.457040569 | 3.00E-02 |
| NRIP1 | 1058.701831 | -1.457919585 | 6.06E-09 |
| LRRC10B | 64.95964827 | -1.461897623 | 7.05E-04 |
| NA | 14.66958513 | -1.464315331 | 3.31E-02 |
| KCNMB2 | 23.37913114 | -1.46573804 | 1.56E-02 |
| NOX5 | 244.0562303 | -1.466729645 | 2.75E-11 |
| MGAT4C | 13.57337256 | -1.466815765 | 4.27E-02 |
| LNX1 | 162.9220059 | -1.466998174 | 1.50E-14 |
| MARVELD3 | 105.4161867 | -1.47190961 | 1.15E-08 |
| AL391987.4 | 23.88118373 | -1.474849355 | 1.25E-03 |
| AC090627.1 | 17.38586849 | -1.475925065 | 7.34E-03 |
| CD247 | 25.01308638 | -1.476007334 | 2.51E-03 |
| AC002128.1 | 38.25855138 | -1.477004455 | 3.58E-04 |
| SULT2A1 | 70.46590007 | -1.479184277 | 4.67E-04 |
| BTG2 | 1018.081876 | -1.479332239 | 1.09E-20 |
| PCDHGC5 | 14.16082268 | -1.479434102 | 2.79E-02 |

| | | | |
|---|---|---|---|
| C1orf54 | 51.82230516 | -1.484013901 | 3.05E-04 |
| CACNA1E | 31.3844744 | -1.486320741 | 1.06E-02 |
| NA | 32.71385346 | -1.486375529 | 6.46E-03 |
| KIAA1217 | 1220.659916 | -1.486391647 | 2.98E-35 |
| AC106820.3 | 46.58828501 | -1.486421875 | 5.55E-05 |
| WWTR1 | 6889.014365 | -1.486807348 | 1.30E-92 |
| GDF11 | 3916.115371 | -1.48851655 | 1.83E-25 |
| FZD8 | 29.0787521 | -1.488965515 | 1.18E-03 |
| PDE7B | 23.34403892 | -1.490224774 | 1.15E-02 |
| ZNF521 | 1676.551674 | -1.492876788 | 3.54E-27 |
| COL2A1 | 456.0622899 | -1.492984966 | 5.44E-09 |
| DIPK2B | 15.15863809 | -1.496093011 | 1.14E-02 |
| LINC01686 | 57.01245041 | -1.496263146 | 5.49E-06 |
| ABCA9 | 44.27440312 | -1.496622364 | 5.91E-04 |
| CKMT1B | 286.8831445 | -1.496820862 | 6.29E-14 |
| RN7SL465P | 10.37104083 | -1.49764753 | 3.51E-02 |
| TCF4 | 3210.623798 | -1.503474402 | 6.82E-05 |
| NGFR | 2528.28695 | -1.504779621 | 3.56E-14 |
| AC083843.3 | 226.7278591 | -1.505050159 | 3.58E-07 |
| HGFAC | 24.78828744 | -1.507389039 | 1.38E-03 |
| AL356585.1 | 10.47909926 | -1.50882349 | 4.58E-02 |
| CAMK1G | 592.4143727 | -1.51038584 | 8.83E-12 |
| CCNE2 | 275.1172858 | -1.511859586 | 1.16E-09 |
| VASH2 | 115.4401106 | -1.512939694 | 1.13E-05 |
| CLEC18B | 62.85794889 | -1.513457911 | 4.81E-06 |
| NA | 32.79874571 | -1.513595816 | 5.45E-03 |
| ITM2B | 18797.51183 | -1.51384672 | 3.00E-23 |
| NAP1L3 | 138.9603458 | -1.514690219 | 4.84E-07 |
| TRPS1 | 933.1259396 | -1.51479809 | 1.99E-14 |
| ACTL8 | 26.23675356 | -1.515068784 | 5.86E-03 |
| PCOTH | 24.61420542 | -1.516264314 | 1.24E-02 |
| PCDHA1 | 13.97894304 | -1.51688381 | 3.67E-02 |
| AC023421.2 | 9.598318077 | -1.520189511 | 3.83E-02 |
| AL445931.1 | 15.68498476 | -1.520717877 | 1.30E-02 |
| TMEM37 | 476.6411504 | -1.522314064 | 5.41E-12 |
| LRRC18 | 14.9253248 | -1.523034402 | 2.15E-02 |
| MIR1-1HG-AS1 | 31.01433589 | -1.524096166 | 5.30E-03 |
| S1PR3 | 4057.598699 | -1.524319931 | 8.70E-32 |
| MDGA1 | 555.6681225 | -1.525167667 | 4.27E-06 |
| PKNOX2 | 537.5619923 | -1.525324899 | 9.88E-21 |
| AC011290.2 | 43.40247706 | -1.525822712 | 1.40E-03 |
| RAB17 | 2197.833679 | -1.525990398 | 2.51E-22 |
| AC015818.2 | 21.48308687 | -1.527229964 | 2.25E-03 |
| RN7SL268P | 15.40962059 | -1.529384711 | 8.67E-03 |
| AL121601.1 | 13.81718734 | -1.52950609 | 1.97E-02 |
| TAS2R4 | 13.53052874 | -1.52956666 | 3.40E-02 |
| AC009812.4 | 17.00709006 | -1.534079725 | 1.29E-02 |

| | | | |
|---|---|---|---|
| AL513534.2 | 18.4908913 | -1.534551114 | 3.47E-03 |
| AC016738.1 | 27.13772455 | -1.53524764 | 1.14E-02 |
| ANKRD18EP | 104.7435619 | -1.536616258 | 1.89E-09 |
| ELMO1 | 314.1325128 | -1.53850431 | 1.35E-10 |
| ADGRF1 | 58.70965185 | -1.540814323 | 5.18E-07 |
| LANCL1-AS1 | 14.20954401 | -1.541528307 | 2.47E-02 |
| SMOC1 | 69.53858863 | -1.542568205 | 5.50E-06 |
| ST3GAL5-AS1 | 58.4679868 | -1.543030054 | 2.03E-08 |
| C16orf89 | 40.33127915 | -1.543805158 | 9.30E-03 |
| FZD4 | 3295.196726 | -1.544040338 | 4.45E-17 |
| SEMA5B | 29.336469 | -1.544073735 | 2.44E-02 |
| AC073130.3 | 40.02148935 | -1.544458345 | 2.45E-05 |
| DLEC1 | 18.46210474 | -1.544737461 | 4.48E-02 |
| AC092143.2 | 43.03820752 | -1.544956655 | 1.89E-02 |
| TLE2 | 901.3412673 | -1.545169867 | 1.96E-15 |
| PAXBP1-AS1 | 94.88010745 | -1.548529311 | 4.34E-12 |
| EPX | 19.60526602 | -1.548820464 | 2.36E-03 |
| CEND1 | 59.0622389 | -1.553620672 | 4.07E-06 |
| WT1 | 29.36100941 | -1.5544649 | 2.07E-03 |
| TXLNB | 42.38748809 | -1.554773772 | 4.61E-04 |
| RCBTB2 | 954.637502 | -1.554784297 | 9.77E-36 |
| DNMBP-AS1 | 22.86594608 | -1.557468922 | 1.39E-03 |
| SCN11A | 27.86262684 | -1.55985579 | 5.73E-04 |
| TMC8 | 293.0250489 | -1.56204127 | 1.17E-10 |
| DSTNP1 | 29.83426616 | -1.562256225 | 1.00E-03 |
| C1QTNF3-AMAC | 63.15032994 | -1.563019867 | 3.54E-03 |
| MTSS1 | 1117.759609 | -1.56354208 | 4.38E-39 |
| AL589993.1 | 9.839506058 | -1.563816517 | 4.59E-02 |
| TPTE2P6 | 16.97085291 | -1.564409504 | 1.05E-02 |
| SPRY4 | 1960.932788 | -1.565086237 | 1.31E-33 |
| CRIP3 | 31.00547865 | -1.566695393 | 7.63E-04 |
| MAP3K5 | 1322.675559 | -1.568910016 | 1.22E-73 |
| CMPK2 | 34.3634101 | -1.569169102 | 1.52E-03 |
| RAPGEF5 | 299.1615067 | -1.56976216 | 3.63E-13 |
| TMC7 | 119.7353915 | -1.569778856 | 9.66E-15 |
| MAPK10 | 51.86224909 | -1.571199034 | 3.29E-05 |
| MCRIP2P1 | 9.424742252 | -1.571395302 | 4.87E-02 |
| NA | 13.3543947 | -1.57143582 | 3.88E-02 |
| FCGR2C | 56.33897174 | -1.571595462 | 1.40E-07 |
| AL354993.1 | 10.52663601 | -1.572419319 | 3.46E-02 |
| PDZRN3 | 211.3588518 | -1.572584604 | 8.53E-15 |
| LAG3 | 53.74991194 | -1.57444424 | 2.41E-06 |
| GFRA3 | 106.5806063 | -1.576831568 | 2.03E-08 |
| C2CD4A | 10.978227 | -1.577315419 | 2.00E-02 |
| AC080038.2 | 35.49383439 | -1.578611772 | 9.18E-06 |
| CFAP206 | 10.33592942 | -1.582071694 | 4.88E-02 |
| MIS18A-AS1 | 12.05924686 | -1.582077613 | 2.85E-02 |

| | | | |
|---|---|---|---|
| LINC00342 | 173.0711505 | -1.585769396 | 2.44E-10 |
| CR383656.4 | 19.28138109 | -1.589033145 | 1.04E-02 |
| AP000842.3 | 13.45252172 | -1.593666692 | 1.13E-02 |
| BTN1A1 | 14.31306556 | -1.594085924 | 1.75E-02 |
| DYNC2H1 | 1286.378422 | -1.594375456 | 9.04E-08 |
| KIAA1755 | 395.0065777 | -1.595157802 | 1.29E-03 |
| HLA-DPB1 | 47.646873 | -1.596085342 | 1.16E-03 |
| STRA6 | 466.7762795 | -1.596373508 | 1.13E-14 |
| AL592424.1 | 19.3597784 | -1.598397094 | 7.62E-04 |
| TCN2 | 1055.694199 | -1.600900249 | 2.50E-17 |
| FAM110B | 546.9806472 | -1.601703578 | 3.77E-08 |
| GAS1 | 64.47266057 | -1.603437316 | 8.42E-08 |
| C4A | 1312.82826 | -1.60425696 | 3.81E-03 |
| HMCN1 | 50.16564326 | -1.604707599 | 4.77E-02 |
| MAGEE2 | 17.34802969 | -1.606209377 | 3.17E-02 |
| TMEM191C | 53.90717508 | -1.606550788 | 9.75E-04 |
| SLC43A2 | 1323.412679 | -1.606818324 | 1.01E-25 |
| CNIH3 | 994.9039675 | -1.608012874 | 6.86E-25 |
| NR2F1-AS1 | 377.042445 | -1.6087087 | 3.32E-22 |
| GPER1 | 115.2248845 | -1.610157121 | 9.12E-07 |
| AL031666.1 | 32.03510505 | -1.615146067 | 2.19E-05 |
| EYA1 | 61.04274887 | -1.615267741 | 1.17E-06 |
| TMEM119 | 206.1753641 | -1.615932303 | 1.65E-10 |
| AC112694.2 | 15.03655311 | -1.616212118 | 1.14E-02 |
| MLANA | 25.42606915 | -1.617689609 | 9.93E-04 |
| CYP39A1 | 219.4277118 | -1.618146019 | 2.06E-15 |
| NRXN2 | 691.5951493 | -1.619106215 | 4.77E-27 |
| CD101 | 54.93496383 | -1.620367185 | 7.38E-04 |
| FRMD4A | 2369.142518 | -1.620589534 | 2.97E-66 |
| GALNT12 | 1294.735295 | -1.621449528 | 1.39E-15 |
| IMPG2 | 21.46058389 | -1.622137207 | 1.51E-03 |
| MEF2C | 193.607049 | -1.62249318 | 1.06E-12 |
| KCNE4 | 2965.933508 | -1.626756701 | 7.20E-35 |
| LINC00327 | 20.45128611 | -1.629482313 | 7.86E-03 |
| TMEM163 | 27.6127971 | -1.630222553 | 9.43E-04 |
| MAP2K6 | 286.8384036 | -1.632300037 | 5.63E-22 |
| AC012508.2 | 9.930179044 | -1.633825783 | 1.54E-02 |
| NFIA | 845.235121 | -1.635271269 | 3.41E-19 |
| TNFRSF19 | 517.7426435 | -1.639701332 | 4.88E-19 |
| RYR3 | 63.81883754 | -1.640881251 | 5.95E-03 |
| CFAP44 | 349.6453516 | -1.650567846 | 2.28E-12 |
| NA | 9.251941923 | -1.653311134 | 4.20E-02 |
| CACNA1D | 40.85940222 | -1.654952079 | 7.58E-03 |
| PRKCB | 1065.831152 | -1.655506666 | 3.34E-20 |
| LCN15 | 13.40241657 | -1.659216618 | 1.73E-02 |
| AC010273.2 | 19.41594626 | -1.66067133 | 4.80E-03 |
| SELE | 16.48521074 | -1.660934965 | 4.11E-03 |

| | | | |
|---|---|---|---|
| GIMAP4 | 15.60675452 | -1.66104095 | 1.53E-02 |
| TRAF3IP3 | 99.96308918 | -1.662410387 | 2.33E-10 |
| LTB | 51.92894334 | -1.663752498 | 2.34E-04 |
| AL031710.2 | 12.472377 | -1.664034215 | 3.33E-02 |
| FAM171B | 279.1691248 | -1.665623116 | 4.06E-07 |
| TRPM2-AS | 87.9797349 | -1.667043807 | 5.91E-10 |
| AC087632.1 | 30.62216116 | -1.667251379 | 1.08E-03 |
| ITGA2 | 6584.396695 | -1.669482216 | 5.54E-36 |
| ITGB1BP2 | 15.71813818 | -1.670018079 | 1.68E-02 |
| AC012501.2 | 74.6821662 | -1.670967848 | 2.03E-07 |
| PTHLH | 279.5986255 | -1.671276786 | 2.78E-11 |
| VCAM1 | 2482.479407 | -1.672463614 | 1.96E-06 |
| FAP | 92.15692023 | -1.672700916 | 3.77E-04 |
| PCDH19 | 17.98510115 | -1.675481618 | 1.76E-03 |
| NA | 11.84966354 | -1.680310123 | 1.51E-02 |
| CLEC1A | 9.206066054 | -1.682960857 | 3.24E-02 |
| H19 | 25168.32442 | -1.683433113 | 1.82E-04 |
| LINC01063 | 10.14136766 | -1.683572898 | 1.63E-02 |
| PYGM | 80.99656343 | -1.684040662 | 1.07E-05 |
| AL132989.2 | 17.86313348 | -1.687616076 | 8.65E-03 |
| AL139393.2 | 31.78021253 | -1.690047028 | 3.80E-04 |
| NEK2P2 | 32.61469177 | -1.69041013 | 1.35E-03 |
| AC066613.1 | 23.66354476 | -1.691110044 | 2.99E-03 |
| AL513365.2 | 8.494539678 | -1.69207181 | 3.89E-02 |
| AC004973.1 | 15.21382652 | -1.693388107 | 5.95E-03 |
| FAM214A | 916.7426868 | -1.694406538 | 6.73E-23 |
| AC008555.4 | 16.62721988 | -1.695416919 | 8.44E-03 |
| DNALI1 | 156.4162185 | -1.696551667 | 2.35E-10 |
| TTC23L | 18.95128468 | -1.69684027 | 1.34E-02 |
| DRD2 | 181.3447405 | -1.699514989 | 1.07E-12 |
| AC012501.1 | 23.96649049 | -1.699699823 | 5.19E-03 |
| C11orf21 | 13.42042057 | -1.699827604 | 2.30E-02 |
| ANK3 | 152.7292105 | -1.70451962 | 4.19E-06 |
| SAMD5 | 74.58486099 | -1.70547135 | 5.69E-06 |
| AP001972.5 | 1533.814559 | -1.707750769 | 3.52E-19 |
| MBL1P | 30.16663922 | -1.709635832 | 1.72E-03 |
| SLC14A2 | 38.02249123 | -1.709996199 | 5.25E-05 |
| LONRF2 | 240.7121002 | -1.713047134 | 3.05E-14 |
| LRRC74B | 22.31592487 | -1.713255722 | 9.30E-04 |
| ARHGEF15 | 18.36256496 | -1.713818991 | 1.43E-02 |
| PLAT | 9788.326261 | -1.714947402 | 1.06E-18 |
| DUSP6 | 3860.577203 | -1.716081564 | 4.92E-36 |
| ENPP2 | 849.7703046 | -1.716576927 | 1.35E-13 |
| AC096921.2 | 33.69750771 | -1.71764194 | 4.00E-05 |
| PRKXP1 | 23.97776507 | -1.719770904 | 1.12E-04 |
| TBX3 | 4851.962125 | -1.721652532 | 4.71E-53 |
| IFIT2 | 1178.033649 | -1.721907145 | 1.79E-35 |

| | | | |
|---|---|---|---|
| RGS14 | 562.7905907 | -1.723336603 | 4.52E-32 |
| NOSTRIN | 322.9129418 | -1.729112812 | 1.50E-17 |
| CASC8 | 11.89497762 | -1.730283221 | 6.34E-03 |
| NRP1 | 5641.832865 | -1.730431722 | 3.05E-41 |
| GTF2IP7 | 37.94287805 | -1.731783896 | 1.36E-04 |
| NA | 45.0838267 | -1.732808313 | 2.13E-07 |
| SAMD9L | 932.6396089 | -1.734730171 | 1.14E-08 |
| LINC00271 | 18.92309629 | -1.73801273 | 5.28E-03 |
| EPHB6 | 1812.119747 | -1.738484287 | 7.21E-20 |
| DACT3-AS1 | 8.567044404 | -1.738692503 | 3.77E-02 |
| KCNJ6 | 165.3882011 | -1.739285246 | 6.49E-14 |
| MYB | 40.76901378 | -1.740723311 | 6.13E-05 |
| AC092683.2 | 24.27190235 | -1.741909342 | 1.96E-03 |
| UBXN10 | 251.9791355 | -1.742460408 | 1.20E-22 |
| PAK3 | 94.81753912 | -1.7435124 | 4.18E-05 |
| HIC1 | 996.4257033 | -1.743680536 | 8.69E-15 |
| IQCN | 27.27984108 | -1.743960618 | 6.25E-03 |
| CLEC7A | 23.75435793 | -1.748248217 | 6.66E-03 |
| CD93 | 23.58250152 | -1.748847225 | 1.68E-03 |
| TNN | 27.80903769 | -1.748968623 | 6.82E-04 |
| LIFR | 13533.60101 | -1.751473269 | 9.88E-18 |
| LRRC73 | 66.17494783 | -1.753832026 | 3.28E-08 |
| AC024909.1 | 67.99772793 | -1.755450527 | 7.79E-07 |
| RFX3-AS1 | 11.79853105 | -1.755951027 | 2.87E-02 |
| RAB37 | 569.4378075 | -1.75897833 | 4.27E-24 |
| CCDC69 | 1021.222201 | -1.760009205 | 1.53E-25 |
| MBNL3 | 313.5751306 | -1.761623626 | 6.84E-07 |
| AP002784.1 | 15.36806646 | -1.76479076 | 4.07E-03 |
| TPI1P2 | 29.76508584 | -1.770938053 | 4.56E-05 |
| ATP8A1 | 225.1808029 | -1.771388208 | 1.48E-03 |
| IFI44L | 45.59889335 | -1.77808178 | 2.33E-02 |
| CHRNG | 12.01570675 | -1.78046881 | 6.72E-03 |
| NR2E3 | 27.75191453 | -1.781785375 | 3.31E-04 |
| SNED1 | 2396.810234 | -1.783518733 | 5.47E-31 |
| GRIN2A | 673.7685502 | -1.790878046 | 1.17E-18 |
| SCG3 | 17.83135647 | -1.792985633 | 5.19E-03 |
| U82695.1 | 9.740562148 | -1.793830605 | 1.83E-02 |
| SAMD9 | 759.2126716 | -1.796078226 | 1.45E-10 |
| AC010809.2 | 18.6237117 | -1.797312758 | 1.62E-03 |
| NA | 8.643307677 | -1.797969396 | 3.22E-02 |
| AC138123.1 | 17.09661336 | -1.799069504 | 4.01E-03 |
| CYP4F24P | 11.30408815 | -1.799407453 | 2.74E-02 |
| TESC | 28.26307814 | -1.799666518 | 7.63E-05 |
| OR7E126P | 11.34187201 | -1.804295431 | 1.53E-02 |
| MMD | 3317.81114 | -1.8052227 | 1.26E-64 |
| GOLGA8VP | 17.19740158 | -1.80688248 | 1.41E-03 |
| IFI44 | 187.1328943 | -1.807244731 | 2.53E-10 |

| | | | |
|---|---|---|---|
| AC010245.2 | 16.07305884 | -1.809225214 | 3.31E-03 |
| NFAM1 | 176.4578415 | -1.81050882 | 6.20E-10 |
| FOXD4 | 8.7431396 | -1.811973738 | 3.99E-02 |
| ESAM | 145.5043028 | -1.812598697 | 5.94E-10 |
| AC010642.1 | 7.978678307 | -1.813755263 | 4.48E-02 |
| CCN3 | 283.7792535 | -1.813927743 | 7.76E-21 |
| LDB2 | 2892.946269 | -1.816372941 | 2.76E-56 |
| AC004801.2 | 21.85766275 | -1.819994726 | 7.45E-03 |
| RFTN2 | 36.09268729 | -1.82364573 | 4.97E-03 |
| SOX4 | 406.2559629 | -1.82387082 | 9.52E-20 |
| AC245884.8 | 11.17547272 | -1.826381001 | 1.26E-02 |
| AP003068.4 | 67.36195346 | -1.827262911 | 1.17E-05 |
| AC004067.1 | 10.2570936 | -1.83087645 | 4.99E-02 |
| UNC5D | 11.94778688 | -1.831109819 | 3.38E-02 |
| LGI2 | 17.67800082 | -1.833303864 | 2.69E-02 |
| SLC6A13 | 22.01920613 | -1.834954046 | 6.75E-03 |
| ITGA8 | 483.5758338 | -1.835696617 | 5.36E-18 |
| AC004846.2 | 7.837252615 | -1.83785399 | 4.34E-02 |
| LINC01060 | 80.61008245 | -1.838864574 | 1.11E-11 |
| SLC7A4 | 76.41360638 | -1.842869543 | 2.49E-06 |
| TSPAN1 | 25.18067891 | -1.843216266 | 4.44E-03 |
| NHSL2 | 362.3466133 | -1.844490834 | 1.81E-24 |
| EMILIN1 | 5275.584214 | -1.845511474 | 1.19E-25 |
| INKA1 | 307.9654041 | -1.846166223 | 4.21E-21 |
| NA | 24.42026338 | -1.850322331 | 1.29E-04 |
| SLC5A9 | 31.88326152 | -1.853031268 | 1.49E-02 |
| CCL5 | 161.9253664 | -1.854463731 | 8.40E-09 |
| AC018638.7 | 8.397737919 | -1.855874251 | 2.15E-02 |
| RTL5 | 2173.925386 | -1.860238051 | 6.16E-86 |
| SCN2A | 55.63473643 | -1.862899285 | 8.23E-05 |
| HSPA7 | 29.38649302 | -1.86468449 | 1.30E-05 |
| ADAMTS3 | 506.1523499 | -1.866537952 | 4.03E-24 |
| AC254562.1 | 10.04046911 | -1.868308602 | 2.35E-02 |
| LINC02163 | 10.9415487 | -1.869049303 | 2.63E-02 |
| CAMP | 9.532341454 | -1.870119869 | 3.84E-02 |
| VAV3 | 21.65778097 | -1.872906936 | 1.56E-02 |
| ADGRF3 | 53.77481944 | -1.872923181 | 3.64E-05 |
| AC023300.2 | 9.952432568 | -1.877945661 | 1.34E-02 |
| GLP1R | 28.92280827 | -1.880954066 | 1.63E-05 |
| HAS2-AS1 | 12.5539903 | -1.881360553 | 7.09E-03 |
| FAM185BP | 15.85875594 | -1.882287065 | 9.49E-03 |
| HLA-DRB5 | 31.70324631 | -1.882996935 | 2.64E-03 |
| EXTL3-AS1 | 56.09264869 | -1.883180165 | 1.39E-06 |
| AC026355.2 | 13.91176656 | -1.883447654 | 3.06E-03 |
| CARD6 | 1819.649875 | -1.885044833 | 4.60E-49 |
| SEC14L6 | 14.29596388 | -1.886279727 | 6.70E-04 |
| ISM1 | 7.500361104 | -1.890671368 | 4.64E-02 |

| | | | |
|---|---|---|---|
| AC108673.2 | 18.07552634 | -1.891360072 | 2.66E-03 |
| RF02271 | 10.29439901 | -1.892493495 | 2.08E-02 |
| AC120057.4 | 8.014390505 | -1.893192791 | 2.04E-02 |
| BX119917.1 | 35.28777911 | -1.894466642 | 1.44E-06 |
| TNFAIP8L3 | 111.5799797 | -1.896339262 | 2.74E-10 |
| HEPH | 29.34641488 | -1.897843802 | 2.98E-03 |
| AC025171.4 | 21.72926794 | -1.905779629 | 2.91E-04 |
| ADAMTS9 | 372.2011459 | -1.908756862 | 1.12E-16 |
| MPZ | 270.8214232 | -1.909833028 | 1.79E-11 |
| LINC02562 | 16.41945962 | -1.914179813 | 4.79E-03 |
| TUBA8 | 48.32664225 | -1.91579545 | 7.46E-03 |
| AC007950.3 | 9.0963026 | -1.918896038 | 3.89E-02 |
| PPP1R16B | 527.7404052 | -1.919088859 | 6.46E-21 |
| GJA3 | 323.6162197 | -1.919173467 | 5.18E-33 |
| GALR1 | 16.93168639 | -1.920086837 | 2.20E-02 |
| AMBP | 14.23918813 | -1.921500968 | 9.71E-03 |
| PLEKHB1 | 266.0658083 | -1.925676303 | 7.29E-12 |
| AC073621.1 | 9.543323116 | -1.92652929 | 2.00E-02 |
| KCNC1 | 90.05506287 | -1.927107137 | 6.52E-11 |
| AC093620.1 | 7.115466543 | -1.929115021 | 3.23E-02 |
| AL356488.3 | 34.82499263 | -1.929712017 | 1.39E-07 |
| NR2F1 | 1908.419877 | -1.933926718 | 1.12E-23 |
| EID3 | 84.86515522 | -1.9345391 | 2.68E-13 |
| CCDC194 | 12.17004459 | -1.93778904 | 8.74E-03 |
| NAP1L2 | 60.86264806 | -1.937853352 | 6.30E-04 |
| NA | 10.75678427 | -1.944907912 | 1.94E-02 |
| CHRNB4 | 22.88941574 | -1.946397199 | 2.10E-04 |
| UBOX5-AS1 | 6.508770489 | -1.947200057 | 3.73E-02 |
| CLEC3B | 312.9420554 | -1.949215387 | 1.17E-08 |
| PDE4D | 2698.714731 | -1.949685301 | 4.98E-24 |
| CHRM1 | 20.60036107 | -1.952644992 | 2.87E-04 |
| PINLYP | 28.72624574 | -1.952863997 | 1.22E-04 |
| SAMD15 | 178.1043318 | -1.957405433 | 3.76E-17 |
| AC002451.1 | 7.972422639 | -1.958654162 | 2.54E-02 |
| AC092683.1 | 166.0784964 | -1.959612473 | 1.22E-12 |
| CAMK4 | 96.91924048 | -1.961123882 | 9.44E-09 |
| F2RL3 | 45.08989024 | -1.961847653 | 2.26E-05 |
| SYNPO2 | 528.338954 | -1.964306402 | 1.18E-36 |
| MIR3681HG | 18.96520959 | -1.965549089 | 1.61E-03 |
| HPS3 | 6068.142851 | -1.965679975 | 8.73E-52 |
| FAM110D | 49.12210533 | -1.966694115 | 2.49E-06 |
| CLMP | 3028.617727 | -1.970741033 | 3.16E-38 |
| RGS5 | 1408.439858 | -1.976083829 | 7.81E-08 |
| AC079465.1 | 12.33074335 | -1.977478832 | 1.04E-02 |
| PIK3R1 | 4595.722417 | -1.977630202 | 5.60E-69 |
| HEY2 | 65.51168791 | -1.981138137 | 4.95E-09 |
| CNTFR | 700.6352166 | -1.984016901 | 1.29E-19 |

| | | | |
|---|---|---|---|
| ASAH2 | 71.01401298 | -1.988877938 | 7.01E-04 |
| CD38 | 109.1153045 | -1.991984207 | 2.03E-07 |
| PLEKHG1 | 194.9176734 | -1.995757942 | 9.95E-10 |
| AL121574.1 | 8.127876777 | -2.000636115 | 3.23E-02 |
| AC007038.2 | 8.778476666 | -2.003175711 | 2.73E-02 |
| LRFN2 | 11.69458686 | -2.003900697 | 7.58E-03 |
| IRS4 | 15.16197689 | -2.007393118 | 2.82E-02 |
| ETV1 | 1912.447401 | -2.012710628 | 2.31E-64 |
| ZNF663P | 25.43069399 | -2.012790754 | 1.80E-04 |
| NA | 33.09930813 | -2.023298428 | 5.13E-06 |
| HLA-DRA | 272.0727889 | -2.028790383 | 3.61E-14 |
| METTL7A | 666.3257155 | -2.030364452 | 2.19E-14 |
| CA12 | 123.9598698 | -2.031736785 | 5.21E-09 |
| TACR1 | 14.86252692 | -2.037752205 | 3.86E-02 |
| RASEF | 196.4761225 | -2.044126736 | 1.14E-27 |
| PCYT1B | 54.97020519 | -2.045409132 | 3.13E-11 |
| SPAG17 | 14.86134536 | -2.047297594 | 2.60E-03 |
| STARD4-AS1 | 775.5088185 | -2.053811798 | 1.69E-15 |
| LGALS12 | 389.1815947 | -2.053937228 | 7.07E-13 |
| CEACAM1 | 714.1367716 | -2.058696852 | 8.25E-22 |
| AC006213.5 | 7.711689508 | -2.064870862 | 2.80E-02 |
| AC104051.2 | 48.84536052 | -2.071437636 | 1.29E-04 |
| CCT8P1 | 14.76212583 | -2.072180162 | 1.54E-03 |
| KRT75 | 23.29573459 | -2.073507313 | 1.02E-03 |
| AC021945.1 | 17.7759937 | -2.074182519 | 9.06E-04 |
| C16orf96 | 8.604072096 | -2.074898629 | 2.17E-02 |
| JMJD1C-AS1 | 18.31117869 | -2.075355449 | 3.54E-04 |
| NA | 791.4238412 | -2.080175979 | 6.74E-37 |
| AC103691.1 | 22.88767344 | -2.084995637 | 1.08E-03 |
| TLL2 | 230.6792461 | -2.085017238 | 1.33E-24 |
| ENHO | 164.5618616 | -2.086447818 | 9.52E-18 |
| AC137932.2 | 28.57861597 | -2.091755639 | 1.69E-04 |
| IL17RD | 1697.446375 | -2.095366609 | 7.56E-30 |
| CLDN5 | 29.88470087 | -2.102763213 | 1.41E-03 |
| AJ011931.1 | 14.10779318 | -2.106909827 | 4.02E-03 |
| LINC01237 | 22.92773012 | -2.11264384 | 1.34E-04 |
| SGPP2 | 21.98751951 | -2.113950131 | 2.75E-03 |
| NHLRC4 | 70.47882681 | -2.118458514 | 7.69E-10 |
| DRC7 | 9.706805279 | -2.119512709 | 1.44E-02 |
| ADCY10P1 | 106.8816125 | -2.120802912 | 3.56E-09 |
| LINC01132 | 37.93370493 | -2.121943637 | 1.19E-08 |
| NRROS | 203.4804337 | -2.123690908 | 2.96E-21 |
| LINC01967 | 8.849505287 | -2.125972011 | 3.92E-02 |
| CYP19A1 | 37.64401046 | -2.133582949 | 1.17E-03 |
| VWA5A | 131.4514081 | -2.140912511 | 9.05E-11 |
| LRRC37A11P | 9.6427187 | -2.144290949 | 3.11E-02 |
| NA | 224.2178336 | -2.145250925 | 4.07E-18 |

| | | | |
|---|---|---|---|
| IL9R | 20.15512317 | -2.146024226 | 6.72E-03 |
| SEMA3A | 14.92527264 | -2.15020846 | 4.28E-03 |
| MTUS2 | 74.85023664 | -2.152926851 | 4.75E-10 |
| AC253536.5 | 9.237527182 | -2.153867352 | 1.17E-02 |
| PLA2G7 | 21.73822793 | -2.157684863 | 4.53E-05 |
| MANSC1 | 461.6010775 | -2.158484356 | 9.01E-26 |
| ABCC11 | 26.48989666 | -2.162656406 | 9.00E-05 |
| ENO1-AS1 | 7.014030703 | -2.16414951 | 1.93E-02 |
| PLCL2 | 39.70894612 | -2.165368435 | 1.32E-05 |
| CRHBP | 46.71371443 | -2.165738276 | 7.09E-05 |
| TNFSF10 | 463.9076168 | -2.166613806 | 1.42E-10 |
| BFSP2-AS1 | 17.70852641 | -2.170385193 | 6.21E-04 |
| SLC18A1 | 15.55119098 | -2.170418528 | 7.80E-04 |
| RYR2 | 420.6593994 | -2.172801775 | 8.14E-04 |
| PEAK3 | 21.50540919 | -2.175295929 | 3.86E-04 |
| ITGA10 | 182.6699131 | -2.175508851 | 3.79E-18 |
| CP | 65.43405638 | -2.181634186 | 3.49E-09 |
| SLC9A9 | 84.18489021 | -2.190499384 | 1.11E-06 |
| GPRIN3 | 52.6600565 | -2.191882727 | 9.98E-06 |
| ACVR1C | 5.461098509 | -2.199083851 | 3.25E-02 |
| AC025164.1 | 16.69760917 | -2.199267498 | 1.69E-02 |
| UNC13A | 437.1254544 | -2.20356668 | 2.41E-27 |
| GALNT17 | 25.55343124 | -2.203883004 | 4.73E-06 |
| IL1R2 | 12.0580723 | -2.205566462 | 2.99E-03 |
| AL137847.1 | 17.62640037 | -2.207180089 | 5.88E-04 |
| CHGB | 280.9084651 | -2.211392994 | 2.23E-39 |
| AC020663.2 | 9.161786122 | -2.215763051 | 5.75E-03 |
| AL627171.1 | 8.008995733 | -2.217035825 | 1.34E-02 |
| DPT | 7.418061856 | -2.22137113 | 3.48E-02 |
| PCDH12 | 238.254452 | -2.223583041 | 1.03E-16 |
| CYP4F8 | 24.3394758 | -2.228942334 | 3.37E-04 |
| RSAD2 | 15.70848725 | -2.234190302 | 2.81E-02 |
| RSPH4A | 9.050537237 | -2.236622935 | 9.80E-03 |
| DRAXIN | 93.60471832 | -2.237114056 | 9.23E-14 |
| GPRC5C | 1051.716387 | -2.238156342 | 1.24E-25 |
| AC008555.1 | 13.9843821 | -2.241341675 | 5.05E-04 |
| MARK1 | 12.82743026 | -2.246268939 | 1.17E-02 |
| GRM2 | 7.113752725 | -2.246570732 | 1.20E-02 |
| LRRTM2 | 15.22254316 | -2.25060731 | 4.45E-03 |
| RTN1 | 9.232349879 | -2.2507008 | 3.38E-03 |
| HSD11B1 | 40.05443864 | -2.252478967 | 7.87E-08 |
| ZMYND15 | 10.71038702 | -2.252780031 | 1.32E-02 |
| PLXDC1 | 87.55184205 | -2.25546664 | 2.15E-07 |
| POU3F2 | 6.773130487 | -2.25959676 | 2.84E-02 |
| AC093788.1 | 6.901960667 | -2.26216887 | 4.96E-02 |
| AL135818.1 | 13.52423364 | -2.265885911 | 3.94E-04 |
| KRT8P30 | 7.699045847 | -2.27256818 | 1.29E-02 |

| | | | |
|---|---|---|---|
| TNFRSF11B | 3103.669719 | -2.272947496 | 5.81E-70 |
| LGALS9 | 267.2477864 | -2.276834012 | 5.00E-10 |
| AQP6 | 26.44147426 | -2.28890812 | 4.93E-05 |
| LINC01262 | 74.40238564 | -2.300905122 | 6.64E-15 |
| NADK2-AS1 | 55.41232106 | -2.301634613 | 1.14E-10 |
| AC010547.1 | 28.59682538 | -2.304984877 | 7.07E-03 |
| DUSP27 | 48.58308619 | -2.306985461 | 3.38E-06 |
| BPI | 155.9343946 | -2.310327048 | 1.02E-16 |
| AC243773.1 | 15.24207664 | -2.312274671 | 8.08E-03 |
| NREP | 12921.79755 | -2.31527136 | 5.50E-26 |
| CHRNB2 | 21.60480173 | -2.31567849 | 1.07E-05 |
| GIMAP8 | 25.83136293 | -2.316813448 | 1.62E-02 |
| GTF2IRD1P1 | 11.75515321 | -2.322457748 | 8.10E-03 |
| TRIB2 | 5941.218021 | -2.324117918 | 4.29E-73 |
| SIRPG-AS1 | 14.55610769 | -2.324390899 | 6.53E-03 |
| TMC1 | 14.90855238 | -2.324723512 | 1.10E-03 |
| TNFSF13B | 34.70978961 | -2.32522636 | 7.23E-06 |
| AC022558.2 | 12.2776348 | -2.329785039 | 1.45E-02 |
| RRS1-AS1 | 20.2277811 | -2.333219057 | 3.45E-05 |
| HNRNPA3P9 | 8.69930565 | -2.334446072 | 4.17E-02 |
| SPP1 | 3358.989557 | -2.334938526 | 4.42E-22 |
| PCDHB12 | 24.55439884 | -2.335073387 | 1.59E-06 |
| AC092070.3 | 6.920013472 | -2.343151898 | 1.36E-02 |
| HLX | 777.8677108 | -2.349644885 | 5.37E-27 |
| GNG13 | 36.65081425 | -2.354558594 | 6.28E-09 |
| DEPTOR | 574.1833815 | -2.360249869 | 1.11E-29 |
| PDE11A | 1409.744978 | -2.360572484 | 2.97E-21 |
| AC007906.2 | 35.53760287 | -2.366860795 | 1.38E-09 |
| AC005790.1 | 5.466737288 | -2.368764819 | 3.72E-02 |
| CTNNA2 | 10.07638971 | -2.384173773 | 2.28E-02 |
| LINC01197 | 9.765364234 | -2.38731698 | 2.03E-02 |
| LINC02370 | 8.606796761 | -2.398600415 | 4.53E-02 |
| GJA5 | 604.0411684 | -2.400013125 | 3.59E-09 |
| AL109945.1 | 5.389499524 | -2.404103366 | 3.22E-02 |
| SLCO6A1 | 11.45394403 | -2.407670731 | 9.65E-03 |
| BUD13P1 | 14.41791251 | -2.410942156 | 1.51E-03 |
| NOX3 | 28.38747355 | -2.411124226 | 1.34E-06 |
| DOCK10 | 51.60308168 | -2.412313202 | 3.00E-05 |
| RAB27B | 854.5006837 | -2.41478165 | 1.38E-25 |
| PPP1R1B | 37.78421622 | -2.421785964 | 2.73E-07 |
| AL122023.1 | 13.6568424 | -2.422619348 | 9.38E-04 |
| RTN4RL1 | 146.9324584 | -2.430160317 | 1.72E-23 |
| TMTC1 | 1452.335955 | -2.430532992 | 1.49E-41 |
| PRSS35 | 16.35980289 | -2.433352565 | 2.64E-03 |
| SLC44A4 | 43.19448386 | -2.434199274 | 5.47E-07 |
| NA | 106.8658323 | -2.434212301 | 2.98E-09 |
| C1orf115 | 406.0582373 | -2.443319631 | 1.14E-14 |

| | | | |
|---|---|---|---|
| PDGFRA | 1613.900805 | -2.45461394 | 6.19E-59 |
| UBXN10-AS1 | 15.03613687 | -2.465502937 | 4.49E-05 |
| BMP7 | 617.2385659 | -2.465790858 | 3.97E-24 |
| RESF1 | 1525.676642 | -2.470672427 | 3.89E-20 |
| AC023490.1 | 5.774690302 | -2.47381415 | 2.94E-02 |
| PHBP6 | 8.166162406 | -2.475356494 | 1.79E-02 |
| SIGLEC17P | 28.70313536 | -2.480472983 | 2.27E-06 |
| AL365204.2 | 5.640036712 | -2.486955062 | 4.87E-02 |
| AC093525.10 | 14.64149699 | -2.495903689 | 4.78E-02 |
| AL109930.1 | 42.7093915 | -2.49846499 | 3.57E-02 |
| TGFB3 | 607.8050464 | -2.502060361 | 4.86E-38 |
| AL008726.1 | 10.79443075 | -2.509697311 | 5.94E-03 |
| LPAR6 | 141.6397686 | -2.510001564 | 9.60E-12 |
| PIK3R3 | 2884.989418 | -2.518241655 | 2.66E-63 |
| LINC00589 | 13.02961938 | -2.523965874 | 1.28E-03 |
| ACOX2 | 57.51644016 | -2.524687708 | 1.16E-08 |
| GABRA2 | 53.37695158 | -2.547974915 | 8.51E-06 |
| GZMA | 20.76270662 | -2.549956245 | 8.63E-04 |
| AC159540.2 | 46.64017187 | -2.558937435 | 1.35E-05 |
| AL357060.1 | 57.41758522 | -2.57143045 | 6.04E-14 |
| SPRY4-AS1 | 9.432133691 | -2.573160585 | 4.16E-03 |
| TSPAN11 | 2754.87485 | -2.573437827 | 1.15E-30 |
| PEG10 | 103399.184 | -2.584186165 | 5.49E-37 |
| AC048344.4 | 10.38222054 | -2.587377828 | 8.70E-03 |
| BFSP2 | 15.52825355 | -2.594319284 | 1.38E-02 |
| PTENP1 | 23.71083276 | -2.598214384 | 1.31E-06 |
| AANAT | 8.096579261 | -2.604359329 | 1.63E-02 |
| CXCL10 | 5.229519643 | -2.61308565 | 4.78E-02 |
| AC010809.1 | 20.89926275 | -2.614622634 | 5.90E-05 |
| TMEM100 | 132.9671535 | -2.617003233 | 1.70E-26 |
| AC117453.1 | 8.645326187 | -2.628726481 | 2.18E-02 |
| SOD3 | 336.2709669 | -2.630106459 | 1.59E-22 |
| ANGPT2 | 85.43122701 | -2.635816037 | 1.09E-14 |
| OLFML2B | 1524.891652 | -2.63745821 | 1.83E-27 |
| LINC00578 | 8.053509764 | -2.63823571 | 9.54E-03 |
| TMOD1 | 1497.162289 | -2.640222775 | 7.98E-38 |
| AC021517.1 | 5.424603503 | -2.64842481 | 2.18E-02 |
| AC091153.4 | 6.999199867 | -2.648432351 | 1.70E-02 |
| TMEM273 | 14.01436715 | -2.649661079 | 1.75E-02 |
| MUC6 | 14.65399839 | -2.649823053 | 1.78E-02 |
| TRGV5 | 22.90767887 | -2.650294161 | 8.20E-06 |
| CNTFR-AS1 | 13.20071498 | -2.650623197 | 1.22E-03 |
| AP002768.1 | 6.270874198 | -2.653017535 | 3.73E-02 |
| LINC02387 | 53.29999358 | -2.656879493 | 3.58E-09 |
| SLC26A4 | 67.63469275 | -2.666774695 | 2.45E-09 |
| ZBTB7C | 327.535564 | -2.66754657 | 2.33E-24 |
| AC009486.1 | 13.83506336 | -2.67135034 | 8.43E-04 |

| | | | |
|---|---|---|---|
| LINC02620 | 7.019322023 | -2.673844633 | 9.11E-03 |
| AC013472.1 | 6.085497053 | -2.681348653 | 4.58E-02 |
| GNRH2 | 11.35355812 | -2.681383144 | 4.78E-04 |
| FAIM2 | 317.161598 | -2.686602932 | 2.16E-23 |
| TSHZ2 | 2202.117157 | -2.688413724 | 9.41E-86 |
| AL135818.2 | 5.597184778 | -2.694059841 | 1.09E-02 |
| AKR1B10 | 66.94033728 | -2.694149592 | 6.54E-04 |
| AL136084.3 | 71.27286851 | -2.694424139 | 1.90E-07 |
| SLC5A12 | 7.134164637 | -2.695009405 | 1.90E-02 |
| NA | 12.81908503 | -2.698576567 | 1.43E-03 |
| NRAP | 18.38544263 | -2.698960918 | 2.00E-05 |
| TTC3P1 | 64.45909731 | -2.701018597 | 9.52E-12 |
| STEAP4 | 35.06140897 | -2.706330838 | 8.66E-07 |
| FYB2 | 6.811654143 | -2.711581784 | 3.12E-03 |
| AC087045.2 | 9.78579654 | -2.712268009 | 9.48E-04 |
| LMOD2 | 18.51326571 | -2.712769693 | 3.63E-06 |
| PDK4 | 122.2626061 | -2.716256072 | 1.70E-18 |
| GRP | 8.013379437 | -2.718155321 | 2.65E-03 |
| VSTM4 | 321.1915059 | -2.720516931 | 6.28E-29 |
| LINC01929 | 20.14398567 | -2.72256673 | 4.13E-04 |
| GRM6 | 10.24229563 | -2.724034201 | 2.99E-02 |
| NT5C1A | 27.98056727 | -2.725132092 | 1.08E-06 |
| A3GALT2 | 14.22687366 | -2.72608349 | 1.25E-03 |
| COL15A1 | 15563.86287 | -2.728375656 | 1.59E-30 |
| SERPINE3 | 16.91027922 | -2.738882658 | 1.48E-05 |
| TDRD6 | 10.17681565 | -2.755114197 | 1.34E-02 |
| SLC12A2 | 5479.546749 | -2.759743314 | 6.12E-95 |
| NTNG2 | 150.2818736 | -2.765355777 | 2.26E-13 |
| CHI3L1 | 17.46316668 | -2.768602301 | 6.11E-04 |
| EBF1 | 927.3327661 | -2.773559922 | 3.53E-38 |
| FAM124B | 12.99155367 | -2.781343017 | 1.36E-03 |
| SESN3 | 604.4270253 | -2.781692168 | 6.80E-70 |
| MT-TH | 24.1473077 | -2.783703681 | 1.39E-03 |
| EWSAT1 | 144.2678212 | -2.790666875 | 1.02E-14 |
| SLC49A3 | 8.681313307 | -2.800269013 | 3.12E-03 |
| ATP2B2 | 75.43341306 | -2.80336833 | 2.68E-09 |
| COL14A1 | 56.31998566 | -2.805289272 | 2.31E-09 |
| DEPDC1-AS1 | 8.111181073 | -2.808289234 | 2.16E-02 |
| C3AR1 | 62.64796635 | -2.813343144 | 1.43E-11 |
| SCG2 | 992.0410165 | -2.823011497 | 1.57E-29 |
| RGS4 | 778.1227395 | -2.82628235 | 5.90E-23 |
| GNG2 | 1893.590023 | -2.846649957 | 3.72E-70 |
| ITGA9 | 2422.723682 | -2.851264342 | 8.45E-35 |
| ABCA6 | 33.80518392 | -2.852633911 | 1.09E-06 |
| SPP2 | 9.031139875 | -2.860575056 | 3.73E-02 |
| CCDC152 | 5.512540049 | -2.881028676 | 1.88E-02 |
| CDH5 | 1043.252914 | -2.889834081 | 3.57E-33 |

| | | | |
|---|---|---|---|
| AC022182.1 | 5.906331977 | -2.891708631 | 2.83E-02 |
| PLXNA4 | 515.4222672 | -2.898214396 | 4.74E-33 |
| IL33 | 11.32575042 | -2.900287438 | 1.03E-02 |
| CREG2 | 57.31823733 | -2.912328547 | 9.07E-13 |
| C3orf80 | 32.31383395 | -2.923947981 | 5.37E-07 |
| VNN2 | 32.72060292 | -2.932569697 | 8.45E-07 |
| JPH4 | 834.6588439 | -2.9369663 | 4.74E-33 |
| ISLR | 87.15946199 | -2.947592304 | 2.92E-13 |
| LINC01352 | 18.42901211 | -2.950692806 | 1.33E-05 |
| GNA14 | 109.087556 | -2.952194286 | 7.38E-22 |
| SCARA5 | 29.23635387 | -2.953056641 | 1.72E-06 |
| KCNK17 | 7.93099649 | -2.959321636 | 1.50E-03 |
| RXFP4 | 11.38414106 | -2.959701471 | 4.63E-04 |
| SYNGR4 | 26.26163114 | -2.970298542 | 1.07E-05 |
| PCDHB8 | 25.38385958 | -2.987372573 | 9.64E-06 |
| MMP1 | 74.36681813 | -2.988620756 | 1.47E-08 |
| AC104809.2 | 6.881124712 | -2.992911808 | 3.45E-03 |
| HABP2 | 5.536422472 | -3.011757338 | 4.35E-02 |
| DNER | 848.2215533 | -3.013913331 | 5.83E-84 |
| VCAN | 908.1972434 | -3.015758968 | 5.95E-35 |
| NPTX1 | 3190.575678 | -3.020337091 | 1.29E-60 |
| MMP9 | 160.911846 | -3.023629624 | 3.61E-19 |
| LINC00877 | 15.57358359 | -3.024029234 | 1.07E-03 |
| CCL15-CCL14 | 48.89464838 | -3.037540282 | 1.31E-04 |
| RCAN2 | 2097.152447 | -3.04265791 | 1.13E-58 |
| MAP1A | 1216.104392 | -3.043752783 | 1.51E-77 |
| AC013275.1 | 5.193056198 | -3.056619989 | 4.66E-02 |
| SPRY1 | 121.128746 | -3.089485541 | 8.86E-12 |
| AL499616.1 | 13.79995614 | -3.095168595 | 5.08E-04 |
| FXYD2 | 41.98422852 | -3.095841976 | 4.71E-11 |
| PRODH | 16.67697053 | -3.101629486 | 5.16E-03 |
| PCSK1 | 1402.5926 | -3.123620584 | 4.26E-53 |
| MAFB | 2006.013682 | -3.125302218 | 2.03E-62 |
| CYSLTR2 | 151.0712334 | -3.12674863 | 9.29E-21 |
| TM4SF18 | 64.41946996 | -3.133625417 | 2.40E-10 |
| MEP1A | 8.233482334 | -3.136643479 | 2.65E-02 |
| UNC5C | 16.04686619 | -3.146222652 | 1.52E-04 |
| TRPA1 | 331.5429719 | -3.147069572 | 8.63E-26 |
| SLC14A1 | 545.9822054 | -3.14767186 | 3.51E-06 |
| GPM6B | 424.3810447 | -3.152917445 | 2.32E-43 |
| NBPF22P | 6.433062377 | -3.156517878 | 6.23E-03 |
| TMEM72 | 5.552220368 | -3.164347868 | 3.25E-02 |
| EVX1 | 13.13102745 | -3.174635325 | 1.68E-04 |
| AP003559.1 | 5.73232179 | -3.185926061 | 2.83E-02 |
| PLVAP | 42.61214939 | -3.196234619 | 7.35E-09 |
| AC025262.3 | 41.05260034 | -3.197379644 | 1.26E-06 |
| TMEM233 | 90.45126575 | -3.19930858 | 5.98E-08 |

| | | | |
|---|---|---|---|
| CXCL12 | 179.5598405 | -3.206801253 | 7.09E-21 |
| AC103808.2 | 11.94070214 | -3.220692958 | 7.54E-06 |
| AC243960.1 | 7.082069548 | -3.226084645 | 1.76E-02 |
| BAALC | 163.1251946 | -3.238505924 | 4.35E-21 |
| AC005280.1 | 7.066605926 | -3.247150527 | 1.67E-02 |
| AL354824.2 | 10.13187429 | -3.249366637 | 1.64E-02 |
| AC093423.2 | 5.941516339 | -3.261975047 | 2.52E-02 |
| NA | 32.00394573 | -3.261994478 | 3.05E-06 |
| CFP | 142.794171 | -3.292436932 | 1.33E-26 |
| GAB3 | 903.944943 | -3.302940183 | 7.31E-68 |
| SLC30A10 | 9.108997572 | -3.303275795 | 9.20E-04 |
| CACNA2D1 | 5.59929796 | -3.323590622 | 4.45E-02 |
| AL445423.1 | 7.941162631 | -3.328171502 | 7.12E-03 |
| PIP5K1B | 6.120998755 | -3.331237009 | 2.74E-02 |
| C7 | 11.38273905 | -3.350265204 | 1.41E-03 |
| C1QTNF4 | 9.09059817 | -3.366634302 | 1.21E-03 |
| AC134043.2 | 93.25974137 | -3.383286402 | 1.31E-17 |
| HAS2 | 104.2657699 | -3.390147698 | 2.04E-16 |
| RGS16 | 706.038458 | -3.399208053 | 1.98E-27 |
| NAV2-AS4 | 43.2774696 | -3.419199718 | 1.90E-11 |
| DLK1 | 278.9725961 | -3.433925479 | 4.29E-17 |
| F13A1 | 43.43632321 | -3.43872648 | 1.21E-08 |
| LYZ | 14.04044372 | -3.444032538 | 7.61E-05 |
| HEY1 | 1560.216718 | -3.45631366 | 1.26E-71 |
| EDN3 | 6.521416936 | -3.45697427 | 1.24E-02 |
| ZAP70 | 104.1860247 | -3.478833978 | 7.96E-22 |
| AC016957.2 | 7.752886535 | -3.480265298 | 9.83E-03 |
| AL031665.2 | 7.040762134 | -3.483847305 | 9.80E-03 |
| ANKRD44 | 619.7998958 | -3.498423241 | 1.61E-35 |
| LINC02198 | 13.44948038 | -3.503401677 | 1.28E-03 |
| FAM20A | 268.4477144 | -3.522420536 | 1.85E-27 |
| SMTNL2 | 14.55934715 | -3.523467863 | 3.64E-05 |
| STK32A | 19.69460038 | -3.524310328 | 2.61E-06 |
| FNDC5 | 839.7235065 | -3.539503023 | 3.59E-14 |
| GPR88 | 5.493524169 | -3.550975096 | 1.88E-02 |
| SIGLEC15 | 122.8361965 | -3.557281941 | 2.00E-29 |
| POSTN | 840.5191803 | -3.568347197 | 7.39E-21 |
| MAGEL2 | 23.89919631 | -3.588237592 | 2.52E-02 |
| AC012499.1 | 8.245841383 | -3.589529475 | 7.76E-03 |
| TF | 518.8802447 | -3.609069432 | 1.89E-45 |
| TRAC | 26.18792344 | -3.613791638 | 6.12E-06 |
| PRRX1 | 15.5569343 | -3.626312939 | 8.30E-03 |
| MGP | 1888.148666 | -3.627461676 | 2.84E-17 |
| MRVI1 | 377.0708102 | -3.630347766 | 7.90E-49 |
| AC112206.2 | 10.72136123 | -3.658386524 | 5.23E-03 |
| ZP2 | 35.43078166 | -3.68043692 | 2.93E-09 |
| INPP5D | 8.203517566 | -3.684921696 | 3.57E-04 |

| | | | |
|---|---|---|---|
| LINC02202 | 74.77782157 | -3.685029653 | 1.45E-18 |
| ERG | 136.5897879 | -3.695645612 | 1.66E-21 |
| 1-Mar | 48.32321599 | -3.697002775 | 7.33E-07 |
| CADPS | 45.90525098 | -3.700071669 | 3.15E-08 |
| AC083843.4 | 12.60092653 | -3.71805565 | 5.17E-04 |
| MRAP2 | 60.19755106 | -3.782544597 | 4.02E-12 |
| ANKRD30B | 6.197607501 | -3.828157163 | 1.04E-02 |
| SPARCL1 | 344.9499686 | -3.83691241 | 6.45E-26 |
| AC104365.2 | 5.073819365 | -3.857837471 | 1.27E-02 |
| LINC01091 | 34.16317966 | -3.874173877 | 5.74E-15 |
| MIR3125 | 7.117429883 | -3.876098581 | 6.03E-03 |
| OVCH1 | 6.504097974 | -3.88292596 | 2.15E-02 |
| PI15 | 14.13753266 | -3.895370432 | 1.41E-03 |
| IL1RAPL2 | 7.265012822 | -3.900784663 | 5.21E-03 |
| RB1-DT | 5.811625749 | -3.917300587 | 3.35E-02 |
| REEP1 | 645.6708003 | -3.917493165 | 8.23E-70 |
| SHISA2 | 384.9045469 | -3.925486689 | 6.06E-83 |
| AC009063.2 | 45.99358757 | -3.973133702 | 2.12E-15 |
| AC055854.1 | 5.136884525 | -3.983229558 | 1.55E-02 |
| GIMAP6 | 27.79027384 | -3.99784004 | 2.00E-07 |
| KDR | 808.088877 | -3.999588672 | 6.60E-51 |
| AP000787.2 | 6.966646499 | -4.017118766 | 2.76E-03 |
| SFRP5 | 9.335463047 | -4.084798865 | 6.66E-03 |
| PPEF2 | 14.46247732 | -4.123568961 | 1.35E-03 |
| RANBP3L | 670.2879169 | -4.124477325 | 4.42E-39 |
| ADGRF5 | 151.418413 | -4.157853804 | 6.19E-33 |
| AC064875.1 | 87.04053269 | -4.158935061 | 1.04E-19 |
| UGT1A9 | 6.632546904 | -4.215865435 | 2.58E-02 |
| TBC1D3I | 5.769888259 | -4.315247371 | 2.63E-02 |
| HSPB2-C11orf52 | 15.99196793 | -4.333442115 | 2.12E-02 |
| HFM1 | 6.484954432 | -4.384516438 | 6.97E-03 |
| AC026369.2 | 7.990228208 | -4.425245574 | 5.10E-03 |
| CMKLR1 | 1380.27909 | -4.443636164 | 4.59E-58 |
| LYVE1 | 7.222638394 | -4.475694638 | 3.83E-03 |
| SELENOP | 510.7451053 | -4.491310732 | 6.45E-20 |
| TMEM176A | 7.449611994 | -4.528223598 | 2.31E-03 |
| COX6CP13 | 5.294205534 | -4.532613612 | 2.95E-03 |
| AL355987.2 | 12.14815253 | -4.538887504 | 4.15E-03 |
| HLX-AS1 | 5.864391477 | -4.561445287 | 4.15E-03 |
| KRT127P | 5.310953075 | -4.622364519 | 3.32E-03 |
| CYP4X1 | 528.1658392 | -4.630600643 | 6.78E-69 |
| CYP4F23P | 18.20499733 | -4.678019542 | 3.19E-05 |
| TM4SF1 | 371.1106316 | -4.737191003 | 8.14E-20 |
| PCDHGA11 | 39.1975755 | -4.778202874 | 4.67E-04 |
| CPHL1P | 12.06318751 | -4.900114791 | 5.55E-04 |
| OR6W1P | 7.088915303 | -4.918571558 | 1.06E-03 |
| LINC02126 | 16.96698924 | -4.969630853 | 1.10E-06 |

| | | | |
|---|---|---|---|
| CCL14 | 36.66059148 | -4.976816376 | 2.06E-03 |
| BMX | 147.3418536 | -5.147896496 | 4.71E-32 |
| H2AFB3 | 5.067498031 | -5.165865697 | 2.19E-03 |
| ZC3H11B | 16.96159532 | -5.170682608 | 1.34E-05 |
| CD180 | 81.62185089 | -5.177747191 | 7.87E-17 |
| PRND | 26.16392127 | -5.17998937 | 1.74E-07 |
| SPINK4 | 6.515444977 | -5.21256478 | 2.87E-04 |
| AC093392.2 | 6.424742308 | -5.286519186 | 4.80E-04 |
| AC010420.1 | 5.023018429 | -5.321269663 | 3.41E-04 |
| AP005263.1 | 8.043627279 | -5.335215146 | 1.26E-02 |
| SRP9P1 | 27.7738132 | -5.435434196 | 2.59E-02 |
| RGS8 | 9.840947122 | -5.46795742 | 1.93E-04 |
| CLEC10A | 5.718442853 | -5.581530577 | 3.64E-04 |
| LINC02458 | 11.69928983 | -5.693622387 | 4.68E-05 |
| APLNR | 28.83062591 | -6.063161822 | 4.51E-07 |
| ANKRD35 | 7.132370749 | -6.122783623 | 1.48E-05 |
| GABRR1 | 17.11856491 | -6.198503331 | 7.29E-06 |
| AC078777.1 | 8.915935471 | -6.388982137 | 2.66E-03 |
| SPDYE11 | 8.84405202 | -6.782949601 | 1.62E-03 |
| LINC00892 | 26.51675302 | -6.925994711 | 1.18E-06 |
| PRSS2 | 39.46445423 | -7.175382017 | 1.36E-07 |
| AC087164.2 | 292.3801368 | -14.21766926 | 2.39E-04 |
| NA | 19.37100855 | -21.915399 | 1.13E-03 |
| AC093668.2 | 22.02253282 | -29.10417566 | 7.77E-06 |

| Gene | baseMean | log2 Fold Change | padj |
|---|---|---|---|
| AP003419.1 | 46.01432342 | 21.78167805 | 5.03E-12 |
| CXCR5 | 38.38259048 | 19.08061235 | 1.53E-05 |
| TAGAP | 49.23096721 | 9.530110951 | 1.06E-12 |
| DHRS9 | 55.24635714 | 9.256800999 | 1.88E-17 |
| RFPL4AL1 | 48.58179161 | 9.088479184 | 1.12E-12 |
| LCN2 | 20.89964453 | 8.978360448 | 2.22E-10 |
| MAPK4 | 12.45556935 | 8.715415761 | 7.29E-03 |
| CRYAB | 8154.737916 | 8.475317447 | 4.34E-303 |
| CST2 | 34.37746149 | 8.356914788 | 1.07E-02 |
| TNFAIP6 | 16.9490992 | 8.262074528 | 1.31E-04 |
| KLHDC7B | 28.70749508 | 8.222256229 | 4.89E-09 |
| SFTA1P | 288.2208761 | 8.123813863 | 1.54E-18 |
| MYH2 | 23.18872912 | 8.107552808 | 1.16E-09 |
| AC091951.1 | 6.7835681 | 8.004925611 | 2.65E-05 |
| SLAMF7 | 176.448223 | 7.95762747 | 1.00E-58 |
| TRPC7 | 102.66675 | 7.692732198 | 7.40E-46 |
| NA | 12.04352763 | 7.659927709 | 1.64E-08 |
| RFPL4A | 89.139459 | 7.639019398 | 2.17E-23 |
| FGF19 | 32.84708541 | 7.608210369 | 2.42E-09 |
| LCE2A | 7.928852411 | 7.477030894 | 2.27E-07 |
| AL353743.4 | 7.124459301 | 7.449354286 | 1.33E-07 |
| GPR1 | 155.1271788 | 7.443891523 | 2.00E-28 |
| FGF21 | 23.9633406 | 7.417138264 | 1.02E-08 |
| MYH1 | 5.335709278 | 7.40656881 | 3.88E-05 |
| NMRK2 | 40.23946307 | 7.368248279 | 2.84E-19 |
| NA | 5.318823847 | 7.289490754 | 2.06E-04 |
| IGKV3-11 | 7.975026233 | 7.268221938 | 1.35E-02 |
| AC135977.1 | 22.06214483 | 7.235231603 | 1.01E-03 |
| SPINK1 | 11.81860843 | 7.225198474 | 5.29E-09 |
| NA | 9.834076694 | 7.184025434 | 1.04E-04 |
| NR1H4 | 131.3960236 | 7.171662402 | 2.53E-32 |
| LCP1 | 1000.928624 | 7.100958539 | 2.30E-180 |
| PHF24 | 57.9882298 | 7.055414464 | 1.05E-23 |
| NA | 16.51338953 | 6.981485406 | 1.17E-06 |
| MRO | 19.70905316 | 6.932232474 | 1.39E-08 |
| IGFL1 | 6.419057012 | 6.930624459 | 1.42E-05 |
| GDF15 | 313.7269975 | 6.835286185 | 2.00E-39 |
| NA | 6.666296487 | 6.697848569 | 1.64E-04 |
| SV2B | 97.51491438 | 6.653116603 | 1.05E-45 |
| GALP | 19.60835474 | 6.636232184 | 4.85E-09 |
| AP1M2 | 110.3798198 | 6.628590287 | 7.22E-32 |
| CSTA | 29.96038685 | 6.624089123 | 1.83E-12 |
| AC006435.4 | 5.208514891 | 6.577826986 | 5.78E-03 |
| TMEM144 | 32.41508643 | 6.55268695 | 9.10E-07 |
| LUM | 18.69450239 | 6.429069311 | 9.77E-10 |
| MAT1A | 731.6526786 | 6.412502691 | 1.70E-195 |

| | | | |
|---|---|---|---|
| FBLN5 | 552.2943098 | 6.407357982 | 6.79E-204 |
| LAMA1 | 1777.372355 | 6.348613798 | 0.00E+00 |
| GFI1B | 5.107360867 | 6.343342968 | 9.48E-06 |
| LINC01443 | 13.66143475 | 6.254347393 | 3.14E-09 |
| MARCO | 27.67416851 | 6.235173665 | 2.37E-06 |
| UGT3A2 | 1561.296932 | 6.220019709 | 2.57E-251 |
| SEMA3E | 5.00787004 | 6.173062719 | 3.22E-04 |
| LILRA2 | 15.14502316 | 6.144008663 | 4.62E-11 |
| LINC01293 | 57.56332832 | 6.108804904 | 3.20E-15 |
| NPIPA2 | 22.55804979 | 6.076584564 | 2.08E-03 |
| NPHS1 | 7.592583401 | 6.072570943 | 1.36E-05 |
| AC089983.1 | 88.31094755 | 6.05758382 | 7.26E-06 |
| ADCY8 | 7.411685296 | 6.048792502 | 3.39E-05 |
| KLHDC7A | 24.91322927 | 6.035597587 | 2.05E-15 |
| LINC01186 | 8.428108961 | 6.022018939 | 2.11E-05 |
| NMRAL2P | 113.1144229 | 6.015437142 | 3.00E-17 |
| AL159166.1 | 10.36203043 | 5.974446368 | 1.74E-05 |
| RPL23AP47 | 7.779613667 | 5.841928827 | 4.79E-03 |
| LINC01033 | 13.57754568 | 5.822229076 | 8.37E-06 |
| ALOXE3 | 48.23452377 | 5.808692824 | 3.32E-18 |
| CCL3 | 7.465544844 | 5.790918508 | 1.27E-05 |
| WFDC21P | 140.545174 | 5.729446053 | 1.60E-42 |
| DRGX | 5.390330915 | 5.694816844 | 1.08E-04 |
| PPP1R14C | 7.441369772 | 5.692942119 | 8.18E-06 |
| AC005307.1 | 131.4519514 | 5.687065449 | 2.26E-83 |
| AC114296.1 | 5.972914867 | 5.682190953 | 8.98E-05 |
| TRPM1 | 8.303459732 | 5.656639718 | 5.57E-06 |
| ZNF536 | 5.604134159 | 5.588347938 | 1.14E-04 |
| AC003092.1 | 11.66842353 | 5.585316582 | 2.14E-10 |
| LRRC55 | 12.44895496 | 5.558350731 | 5.15E-08 |
| UGT3A1 | 24.54671501 | 5.54682554 | 1.16E-13 |
| LINC00520 | 22.88133806 | 5.527180133 | 9.53E-15 |
| TRIML2 | 231.9553741 | 5.484747265 | 1.49E-148 |
| LRRC15 | 26.29205623 | 5.483668787 | 4.40E-18 |
| MCTP1 | 299.6263461 | 5.427966896 | 1.46E-97 |
| LINC01291 | 43.47043514 | 5.423838518 | 1.03E-13 |
| GAS2L2 | 7.131076997 | 5.388227737 | 1.22E-05 |
| FRMD7 | 28.169423 | 5.369148694 | 7.31E-19 |
| HLA-DMB | 409.4052964 | 5.333199328 | 3.84E-84 |
| GASK1B | 12.09932004 | 5.321164194 | 1.82E-06 |
| HNF4A | 25.93293679 | 5.296613385 | 6.08E-09 |
| IGSF5 | 89.72243329 | 5.277069009 | 1.40E-49 |
| DUSP26 | 18.65841433 | 5.205869214 | 4.08E-13 |
| AC026785.3 | 6.536643418 | 5.198115841 | 3.90E-06 |
| HNF4G | 14.34417802 | 5.195894429 | 6.39E-06 |
| MYHAS | 12.21043099 | 5.169017449 | 7.15E-05 |
| AC005832.4 | 89.28362442 | 5.150233413 | 5.27E-06 |

| | | | |
|---|---|---|---|
| HYAL4 | 7.589053687 | 5.141344095 | 1.26E-05 |
| CES5A | 10.8197907 | 5.119189269 | 5.68E-06 |
| POM121L2 | 11.56421178 | 5.086258171 | 2.48E-04 |
| AL157896.1 | 13.64846009 | 5.075260957 | 8.95E-10 |
| PKD1L2 | 885.8875173 | 5.072434636 | 9.51E-78 |
| AL136962.1 | 29.14557855 | 5.064048466 | 3.80E-21 |
| FPGT-TNNI3K | 29.63087152 | 5.060929726 | 5.34E-06 |
| C1S | 1174.241073 | 5.048314786 | 4.94E-88 |
| SLC27A2 | 45.4861244 | 5.01883424 | 9.94E-04 |
| OR51E2 | 9.551715234 | 5.016440535 | 2.51E-08 |
| AC079760.2 | 14.67783629 | 4.998853572 | 1.62E-08 |
| SPTA1 | 18.6919457 | 4.993659543 | 1.72E-04 |
| CLDN1 | 777.3721858 | 4.985662281 | 1.20E-68 |
| RRAGD | 701.8675929 | 4.921531208 | 2.00E-170 |
| ELMOD1 | 26.26171552 | 4.879206461 | 9.08E-19 |
| INHBE | 431.5658161 | 4.876147214 | 3.98E-16 |
| BCL2A1 | 18.62870038 | 4.873372056 | 5.06E-15 |
| SULT1C2 | 407.8494656 | 4.871523599 | 2.60E-86 |
| PRKG1 | 7.249378884 | 4.868561959 | 1.74E-05 |
| C17orf102 | 57.63053537 | 4.856800861 | 3.98E-10 |
| CLEC3A | 27.21407875 | 4.850407839 | 3.00E-15 |
| MYH15 | 58.51415002 | 4.836043813 | 4.98E-14 |
| IGHE | 6.838546937 | 4.831202341 | 2.91E-03 |
| LURAP1L-AS1 | 37.65792059 | 4.816332231 | 8.82E-07 |
| IP6K3 | 192.4547338 | 4.799670876 | 4.79E-135 |
| AC103740.2 | 6.922522524 | 4.78908632 | 1.28E-04 |
| AC132068.1 | 5.382205542 | 4.788143278 | 2.13E-03 |
| MYH8 | 12.93766983 | 4.757058325 | 5.54E-11 |
| NAT16 | 35.39645748 | 4.728909276 | 6.41E-20 |
| ESRP1 | 58.58254683 | 4.716914296 | 6.79E-16 |
| FOXB1 | 5.497409793 | 4.711454634 | 3.32E-03 |
| PCP4 | 5.451390876 | 4.687243221 | 4.22E-04 |
| NRXN3 | 33.47344038 | 4.666887498 | 1.70E-04 |
| SPNS3 | 24.62107375 | 4.659344979 | 3.45E-05 |
| AL356234.1 | 11.88596717 | 4.64826181 | 1.76E-07 |
| CD5L | 11.64689063 | 4.645439605 | 1.30E-06 |
| ERVV-2 | 60.76003593 | 4.623944987 | 3.44E-34 |
| STXBP6 | 354.2817869 | 4.617658238 | 1.53E-93 |
| CALCB | 23.00979005 | 4.61563526 | 1.80E-11 |
| LNCOG | 57.59267757 | 4.600524671 | 8.48E-11 |
| AC104257.1 | 7.057852466 | 4.585350151 | 4.52E-06 |
| IFI27 | 399.8962451 | 4.583733786 | 9.63E-40 |
| CASP17P | 13.40716843 | 4.565946842 | 3.40E-09 |
| CSF2RA | 877.8505258 | 4.52852455 | 4.89E-142 |
| PSG3 | 19.01407792 | 4.527969737 | 1.68E-03 |
| COL4A3 | 9.192934283 | 4.505529818 | 1.02E-06 |
| KRT34 | 47.26339011 | 4.49958522 | 1.86E-04 |

| | | | |
|---|---|---|---|
| KCNAB2 | 1627.146031 | 4.484308938 | 3.22E-202 |
| TCHH | 60.14085417 | 4.470135267 | 8.26E-41 |
| DUXA | 17.7868599 | 4.468680961 | 3.71E-11 |
| AC112656.1 | 16.72979437 | 4.459147261 | 9.52E-08 |
| LMOD1 | 281.9537545 | 4.455886156 | 8.76E-79 |
| MMP3 | 9.60446711 | 4.442737351 | 2.06E-06 |
| AC011503.1 | 12.20112246 | 4.435808784 | 1.02E-08 |
| ESR2 | 84.85778021 | 4.423410333 | 8.46E-39 |
| TRIM63 | 9.40242175 | 4.405348338 | 7.81E-08 |
| AP002478.1 | 8.764480814 | 4.391522418 | 3.39E-05 |
| HKDC1 | 536.4580666 | 4.382509337 | 1.31E-83 |
| TGM4 | 11.45184573 | 4.382350087 | 4.32E-03 |
| TMEM255A | 638.2751286 | 4.352890951 | 3.96E-131 |
| RPLP0P2 | 306.6935964 | 4.345622975 | 8.27E-99 |
| SLC6A12 | 262.088674 | 4.34243923 | 1.60E-47 |
| XIRP1 | 24.70751638 | 4.316762785 | 1.00E-02 |
| DSG2 | 34.95009439 | 4.302023616 | 5.55E-16 |
| CHST4 | 31.95928319 | 4.297286649 | 2.56E-15 |
| AL645608.7 | 11.78460553 | 4.2904691 | 1.57E-06 |
| ENPP5 | 15.13011806 | 4.283060956 | 7.70E-09 |
| PPARGC1A | 48.48171747 | 4.278363666 | 5.97E-20 |
| USH1C | 6.338202099 | 4.277403579 | 2.08E-04 |
| LOX | 1600.914318 | 4.273093179 | 6.40E-120 |
| CEACAMP10 | 596.3616348 | 4.266814475 | 2.17E-28 |
| TSPAN13 | 149.3123275 | 4.261797998 | 3.01E-60 |
| DOK5 | 5.434187448 | 4.260304717 | 3.37E-05 |
| GUCY1A2 | 7.369270985 | 4.253058552 | 4.46E-03 |
| VLDLR-AS1 | 123.3112072 | 4.252803384 | 4.21E-19 |
| SH3GL3 | 111.9951372 | 4.248383922 | 1.14E-56 |
| KCNQ1 | 14.72470286 | 4.232103264 | 4.59E-08 |
| AP003498.1 | 6.759281315 | 4.230523047 | 2.57E-02 |
| RASGRP3 | 66.3907259 | 4.226998726 | 9.62E-12 |
| GPR65 | 19.00070169 | 4.1675565 | 4.84E-08 |
| DAPL1 | 8.678892955 | 4.160798547 | 1.92E-06 |
| LINC00571 | 5.734873955 | 4.142632442 | 1.47E-04 |
| CELF2 | 578.2401139 | 4.141190779 | 8.99E-135 |
| RHCG | 42.36352371 | 4.115912232 | 2.51E-25 |
| LAMP3 | 296.1481714 | 4.081082479 | 1.91E-49 |
| PINCR | 8.621945023 | 4.073262615 | 2.30E-04 |
| BTK | 23.77544446 | 4.073102489 | 9.67E-10 |
| FAM90A12P | 21.08359169 | 4.057712544 | 3.94E-05 |
| BTNL8 | 8.487747057 | 4.05604775 | 1.52E-05 |
| TUBA3E | 22.9586516 | 4.043686966 | 7.48E-19 |
| MEGF6 | 249.4909584 | 4.038285121 | 4.78E-28 |
| CLGN | 74.76072684 | 4.035520363 | 2.63E-14 |
| UPP1 | 3542.442731 | 4.034437348 | 1.27E-94 |
| AC126768.2 | 6.376656943 | 4.033661971 | 8.56E-05 |

| | | | |
|---|---|---|---|
| AC107871.1 | 14.44185302 | 4.021667578 | 4.72E-03 |
| TSLP | 16.83474388 | 4.015507056 | 1.13E-08 |
| AC069120.1 | 17.9285258 | 4.013323241 | 2.22E-13 |
| SUN3 | 13.02687629 | 4.009425194 | 2.20E-07 |
| EPB41L4A | 28.59384182 | 4.004431283 | 5.39E-08 |
| GPNMB | 7636.758085 | 3.994657364 | 6.58E-139 |
| ASS1 | 1006.358665 | 3.987550769 | 3.67E-22 |
| DYNC1I1 | 48.1708623 | 3.980935082 | 1.52E-25 |
| FAM163B | 39.18434864 | 3.980628674 | 7.50E-21 |
| AC018529.2 | 5.13035025 | 3.959775832 | 2.53E-03 |
| HSPB2-C11orf5 | 15.99196793 | 3.948479465 | 8.42E-04 |
| CCL15 | 16.12851781 | 3.944376772 | 1.03E-08 |
| LINC00862 | 29.77997774 | 3.937434881 | 5.04E-17 |
| BIRC7 | 6.408022844 | 3.924210368 | 3.11E-05 |
| AC079298.3 | 24.94196526 | 3.919408574 | 3.59E-17 |
| AL392023.1 | 7.410504344 | 3.888675676 | 4.80E-02 |
| KLK8 | 6.266111381 | 3.887969548 | 1.27E-04 |
| GCNT3 | 25.18319113 | 3.885305537 | 5.50E-06 |
| CPNE4 | 26.75781287 | 3.874275939 | 3.50E-09 |
| LINC01665 | 8.039503424 | 3.861126336 | 9.27E-04 |
| TMEM191B | 32.2656479 | 3.854833619 | 5.20E-14 |
| IL32 | 397.8318467 | 3.849874002 | 8.14E-46 |
| CDX2 | 10.13813606 | 3.842777157 | 1.54E-04 |
| AL117378.1 | 78.33490075 | 3.833874931 | 8.52E-55 |
| ITGB3 | 1822.991296 | 3.831582414 | 2.48E-200 |
| LINC01638 | 242.896981 | 3.824542701 | 6.52E-66 |
| LINC02412 | 11.24665306 | 3.817625398 | 3.65E-06 |
| IL6 | 31.65351887 | 3.812851049 | 5.80E-13 |
| SLC9A4 | 21.29564053 | 3.802246111 | 1.52E-08 |
| AC018781.1 | 5.21339508 | 3.794824329 | 1.91E-02 |
| EFEMP1 | 741.1103555 | 3.790163487 | 1.78E-89 |
| DIRC3-AS1 | 14.94687545 | 3.785359546 | 2.14E-03 |
| KLK10 | 11.57887164 | 3.770359107 | 2.66E-05 |
| GRIK5 | 70.41443968 | 3.768061386 | 7.77E-15 |
| IGFN1 | 141.2038353 | 3.758026535 | 2.81E-12 |
| GPC4 | 272.3054783 | 3.754261228 | 3.14E-75 |
| MUC4 | 25.83873891 | 3.751981508 | 4.93E-05 |
| C7orf57 | 122.2458911 | 3.743733659 | 1.80E-38 |
| LINC02273 | 6.807350032 | 3.74191312 | 9.10E-06 |
| MME | 72.18384025 | 3.736114147 | 5.17E-10 |
| AC097359.2 | 179.0408846 | 3.726404792 | 6.34E-68 |
| GREB1 | 310.3314579 | 3.700774296 | 1.23E-69 |
| UGT2B7 | 35.22609493 | 3.700280951 | 1.31E-18 |
| MAFF | 607.0319197 | 3.699491821 | 1.47E-96 |
| GJB3 | 26.34134941 | 3.685524183 | 3.10E-10 |
| SYTL2 | 1943.873549 | 3.684805743 | 8.27E-86 |
| IL21-AS1 | 5.394513016 | 3.679778357 | 1.82E-03 |

| | | | |
|---|---|---|---|
| PPFIBP2 | 404.1679279 | 3.677707634 | 5.80E-44 |
| STX11 | 126.2269358 | 3.645657524 | 4.59E-37 |
| MSC | 323.6871895 | 3.644809289 | 1.01E-22 |
| SLC45A2 | 191.8192019 | 3.637798243 | 4.47E-69 |
| SLC6A15 | 394.4907478 | 3.635518369 | 1.12E-52 |
| DEGS2 | 11.73401342 | 3.629887916 | 2.49E-05 |
| FREM1 | 20.55740731 | 3.628382764 | 2.27E-09 |
| PIWIL2 | 12.98599122 | 3.622726658 | 1.66E-07 |
| AL035425.1 | 9.156911154 | 3.621605851 | 5.57E-07 |
| GIMAP8 | 25.83136293 | 3.595276445 | 1.49E-09 |
| KRT17 | 109.932617 | 3.588177946 | 9.62E-10 |
| TEX29 | 11.59831696 | 3.580267353 | 5.85E-07 |
| LINC01198 | 6.335604086 | 3.552676313 | 2.27E-03 |
| AC007406.3 | 8.753190183 | 3.551544035 | 2.29E-04 |
| MYO7B | 109.193404 | 3.549030924 | 3.50E-04 |
| TMEM63C | 137.7208928 | 3.545688946 | 2.54E-29 |
| ARFGEF3 | 55.42513659 | 3.544831138 | 7.68E-23 |
| SLC34A2 | 6.877009725 | 3.542259155 | 1.12E-02 |
| SUSD4 | 14.80441536 | 3.541499409 | 2.46E-03 |
| FGF9 | 5.680541362 | 3.537581144 | 2.36E-02 |
| RTN4R | 369.6459485 | 3.535921806 | 1.89E-105 |
| HSPB3 | 33.71635809 | 3.531406642 | 1.36E-11 |
| KIF6 | 184.9785171 | 3.528673486 | 2.28E-34 |
| TMEM200C | 23.71648983 | 3.52840388 | 1.42E-04 |
| PPM1H | 121.3528802 | 3.525358825 | 2.57E-32 |
| AC005336.1 | 13.39810125 | 3.52520426 | 7.20E-08 |
| AZGP1 | 71.03585013 | 3.524478123 | 1.41E-33 |
| TMEM233 | 90.45126575 | 3.520547161 | 8.33E-23 |
| SLCO5A1 | 8.834629614 | 3.514040155 | 4.69E-07 |
| SPX | 445.5857189 | 3.512095426 | 1.14E-18 |
| LIMS2 | 478.4740095 | 3.512004826 | 6.61E-87 |
| GPX5 | 5.593014953 | 3.508678196 | 1.24E-03 |
| LINC00592 | 5.244624385 | 3.49617106 | 4.22E-02 |
| PRAMEF1 | 8.708668192 | 3.491888038 | 3.58E-02 |
| GDNF | 9.644136723 | 3.489788284 | 6.59E-06 |
| AC124947.1 | 10.90404445 | 3.483984111 | 1.17E-05 |
| TM6SF1 | 340.5809889 | 3.476419824 | 5.82E-24 |
| STC2 | 5840.24853 | 3.47611433 | 1.43E-32 |
| TMEM132E | 67.2505779 | 3.472527277 | 1.53E-11 |
| PKD1L3 | 9.867057583 | 3.471486328 | 4.66E-07 |
| INSYN1-AS1 | 10.62605999 | 3.466326196 | 1.19E-04 |
| BLVRB | 2573.362732 | 3.453048232 | 1.74E-230 |
| FLNC-AS1 | 13.89514009 | 3.448849389 | 6.02E-07 |
| AL023881.1 | 23.0706953 | 3.447115897 | 4.54E-09 |
| AL136018.1 | 8.411072915 | 3.436821884 | 3.56E-06 |
| CTH | 928.9606088 | 3.427246377 | 6.10E-31 |
| GDAP1L1 | 48.33841871 | 3.426782423 | 1.71E-11 |

| | | | |
|---|---|---|---|
| SIRLNT | 31.65264106 | 3.417118928 | 4.36E-12 |
| DDIT3 | 1960.790249 | 3.413352651 | 2.68E-17 |
| CNTNAP4 | 6.299186061 | 3.401669695 | 4.31E-03 |
| LINC02535 | 49.64853035 | 3.376440937 | 2.61E-10 |
| ESR1 | 33.66109608 | 3.370837389 | 3.22E-05 |
| DEFB1 | 27.27250216 | 3.363508234 | 1.32E-15 |
| AC090192.2 | 8.548042765 | 3.359888465 | 2.40E-06 |
| AC092132.1 | 5.105028167 | 3.35887481 | 2.36E-02 |
| ELOVL7 | 65.47929235 | 3.333074834 | 3.91E-17 |
| RIMS4 | 97.05276827 | 3.331603372 | 7.23E-31 |
| IL34 | 16.76840544 | 3.318851171 | 5.32E-06 |
| SLC7A11 | 1423.958723 | 3.317685551 | 1.27E-10 |
| GIMAP5 | 56.13836726 | 3.31729362 | 1.44E-22 |
| KCNF1 | 83.96198227 | 3.316969373 | 6.56E-30 |
| CHRM4 | 157.7202575 | 3.315318901 | 5.85E-62 |
| OSCAR | 347.1562098 | 3.313626957 | 1.50E-37 |
| KLF15 | 216.4359999 | 3.313102484 | 3.19E-38 |
| AC079209.1 | 7.975488055 | 3.306977483 | 2.34E-05 |
| LINC01133 | 24.06459905 | 3.2980569 | 2.30E-06 |
| SLC5A1 | 28.6409774 | 3.297629993 | 5.26E-16 |
| TNFSF9 | 133.0035316 | 3.294851963 | 1.01E-47 |
| CTNND2 | 5.418341109 | 3.293271343 | 1.94E-02 |
| CD209 | 25.41772667 | 3.28213628 | 4.63E-10 |
| CYP4F11 | 312.2638214 | 3.281963221 | 4.34E-90 |
| OSGIN1 | 933.2239717 | 3.278517884 | 1.74E-42 |
| TMEM151A | 32.37522276 | 3.27781144 | 7.84E-16 |
| NEBL | 269.678414 | 3.274145263 | 3.79E-24 |
| AL035425.3 | 8.587364089 | 3.273937701 | 3.66E-05 |
| BEX2 | 181.7672279 | 3.26573326 | 5.58E-20 |
| INMT | 744.6966048 | 3.2640808 | 8.26E-160 |
| PAPPA2 | 59.55621791 | 3.263328646 | 4.76E-08 |
| OTOF | 23.81127999 | 3.258194027 | 9.28E-09 |
| AKR1C3 | 83.72430564 | 3.249799149 | 1.41E-26 |
| TNMD | 57.18971116 | 3.24933275 | 6.43E-12 |
| POF1B | 5.720769074 | 3.240371796 | 7.67E-05 |
| AC034213.1 | 7.875937771 | 3.232062378 | 1.66E-03 |
| AL596223.1 | 39.32068767 | 3.230880872 | 4.22E-13 |
| NUPR1 | 3841.855504 | 3.22884443 | 9.59E-23 |
| AC068234.2 | 7.244235066 | 3.223779337 | 2.72E-04 |
| TM4SF19 | 104.74363 | 3.218527065 | 1.88E-28 |
| TNFSF15 | 89.03391984 | 3.207426941 | 1.33E-29 |
| ADM2 | 805.2354835 | 3.202211879 | 1.01E-16 |
| EFNA5 | 8.792344761 | 3.186897396 | 3.67E-04 |
| GRM1 | 7.557448603 | 3.186312347 | 9.51E-04 |
| CX3CL1 | 347.8028956 | 3.184573221 | 3.35E-63 |
| CLIC6 | 238.7103255 | 3.182825817 | 6.96E-82 |
| CPA4 | 8124.801267 | 3.172005188 | 3.78E-86 |

| | | | |
|---|---|---|---|
| AL590004.3 | 1289.954072 | 3.171819966 | 4.09E-53 |
| AC116351.1 | 130.5665582 | 3.170881367 | 4.07E-37 |
| SLCO2B1 | 28.30708115 | 3.168706497 | 2.89E-03 |
| RASAL1 | 6.47576965 | 3.161032739 | 6.31E-03 |
| CHMP4C | 55.12357223 | 3.156350721 | 9.77E-22 |
| OLIG2 | 29.16821453 | 3.153979434 | 6.76E-08 |
| FAM129A | 2074.598291 | 3.152909525 | 1.42E-36 |
| ANK2 | 166.3824866 | 3.152869287 | 6.95E-16 |
| KRT73-AS1 | 27.90722466 | 3.152333916 | 4.81E-08 |
| DLGAP1-AS2 | 423.8719857 | 3.15221673 | 4.42E-43 |
| PLCH1 | 39.15457025 | 3.15147341 | 6.71E-18 |
| NFASC | 750.1182267 | 3.150547627 | 1.14E-229 |
| LINC00677 | 5.476099145 | 3.149538824 | 2.36E-03 |
| SLC6A4 | 13.84678406 | 3.145341732 | 3.28E-05 |
| AL353608.3 | 5.077825569 | 3.145221064 | 7.49E-03 |
| NAP1L3 | 138.9603458 | 3.129002259 | 5.12E-51 |
| USP41 | 16.73443121 | 3.128040343 | 6.14E-04 |
| FLRT2 | 25.58439069 | 3.125250114 | 3.61E-04 |
| CCN2 | 8122.707037 | 3.123332247 | 1.80E-57 |
| FOLR1 | 159.0086188 | 3.110754027 | 1.32E-26 |
| FAM87A | 25.24779033 | 3.105364062 | 9.57E-06 |
| NOX4 | 6.12413806 | 3.09711734 | 8.30E-04 |
| PPP1R15A | 3154.007942 | 3.096872625 | 7.37E-53 |
| NTN1 | 74.09737759 | 3.094023363 | 2.81E-19 |
| SNTB1 | 288.127424 | 3.093844886 | 1.30E-26 |
| CCR2 | 6.629681133 | 3.093464511 | 8.79E-04 |
| CLEC4M | 17.52321515 | 3.089103511 | 3.48E-09 |
| NA | 10.28867259 | 3.085462322 | 1.50E-07 |
| RPL12P14 | 37.54375201 | 3.08437198 | 1.93E-12 |
| KRT80 | 1730.679913 | 3.075015596 | 1.85E-42 |
| GSG1L | 10.27277008 | 3.066925498 | 1.43E-03 |
| FAM201A | 45.64172295 | 3.064587928 | 3.53E-10 |
| NPAS1 | 1188.241383 | 3.063313838 | 1.33E-200 |
| COBLL1 | 11.23169443 | 3.062042528 | 1.23E-03 |
| AC026369.2 | 7.990228208 | 3.060653789 | 6.70E-05 |
| AP001189.5 | 7.632947862 | 3.051508312 | 1.53E-03 |
| PSG4 | 1067.986903 | 3.051106279 | 3.91E-34 |
| AL138828.1 | 7.572296246 | 3.047149561 | 2.85E-03 |
| MSC-AS1 | 127.9931706 | 3.046807536 | 5.86E-16 |
| ACTN2 | 42.19199169 | 3.046392199 | 2.40E-12 |
| FUT9 | 6.495356224 | 3.043643565 | 2.95E-04 |
| AL356515.1 | 15.89611218 | 3.041637546 | 2.16E-08 |
| BHLHE41 | 332.2020079 | 3.038421497 | 1.22E-24 |
| AL157871.2 | 14.49669009 | 3.037048605 | 8.70E-07 |
| PSG7 | 92.83788603 | 3.036469514 | 2.23E-05 |
| VSX1 | 30.17747751 | 3.034933178 | 5.21E-18 |
| MB | 5.190958042 | 3.025789133 | 4.48E-02 |

| | | | |
|---|---|---|---|
| RASGRF2 | 415.7962532 | 3.025265076 | 5.30E-29 |
| AKR1C7P | 9.921288973 | 3.021377373 | 8.67E-05 |
| INSYN1 | 368.9945437 | 3.020586943 | 3.69E-80 |
| AC026316.2 | 27.22988646 | 3.011995927 | 4.64E-03 |
| AL162582.1 | 8.238777489 | 3.011370996 | 2.72E-04 |
| AC016717.2 | 5.261098293 | 3.006306465 | 4.83E-02 |
| ZNF114 | 507.1171622 | 3.001826097 | 5.27E-47 |
| CADM1 | 234.8479271 | 2.999556428 | 2.17E-28 |
| SLC22A15 | 397.3289944 | 2.998605974 | 7.32E-21 |
| P2RX7 | 1431.080718 | 2.998001482 | 5.37E-95 |
| CALB2 | 75.59255705 | 2.997981701 | 7.90E-25 |
| FAM198B-AS1 | 6.610869286 | 2.995222154 | 3.55E-04 |
| HES4 | 110.1398204 | 2.993455908 | 8.45E-29 |
| PLIN4 | 68.6546174 | 2.991895554 | 1.91E-35 |
| AKR1C1 | 12.1303419 | 2.983204271 | 8.95E-07 |
| HSBP1L1 | 284.4549299 | 2.970641489 | 1.98E-80 |
| AC093752.2 | 6.831945174 | 2.970418089 | 2.40E-02 |
| AC245100.3 | 41.78719059 | 2.965028996 | 3.44E-07 |
| LUARIS | 40.9259168 | 2.963826139 | 3.05E-14 |
| SDS | 76.30977003 | 2.962450922 | 7.68E-25 |
| AC010457.1 | 11.60862398 | 2.956705876 | 5.05E-04 |
| PRDM16 | 35.20819683 | 2.947937805 | 1.34E-07 |
| PNPLA1 | 6.649082666 | 2.947808367 | 1.70E-03 |
| TRIM29 | 531.2890621 | 2.944215728 | 4.44E-34 |
| CTSS | 708.5670472 | 2.944057549 | 1.02E-79 |
| FAM166B | 10.42880559 | 2.943924844 | 2.69E-05 |
| AL136131.3 | 28.31490858 | 2.941839018 | 4.11E-07 |
| HS3ST5 | 7.638299293 | 2.939881197 | 1.16E-02 |
| AGT | 33.3491476 | 2.937513313 | 1.36E-16 |
| ERICH2 | 16.08752266 | 2.936855812 | 3.28E-06 |
| UCA1 | 6683.968826 | 2.936424416 | 6.74E-45 |
| TNFRSF12A | 3033.22403 | 2.936404904 | 7.07E-85 |
| CYP2A6 | 5.287110338 | 2.934011427 | 4.66E-03 |
| FAM110C | 6.641058996 | 2.933485617 | 2.09E-03 |
| AC015987.1 | 29.65507999 | 2.932864388 | 3.07E-12 |
| CRLF2 | 21.35257814 | 2.926463527 | 1.34E-04 |
| LINC00973 | 7.727607361 | 2.925701402 | 2.25E-03 |
| LINC01914 | 16.02441718 | 2.924980145 | 4.42E-06 |
| ERC2 | 65.38801751 | 2.924512944 | 1.06E-09 |
| ANKRD1 | 2371.13122 | 2.922114295 | 6.95E-11 |
| AL354872.2 | 5.409528032 | 2.920319235 | 1.59E-02 |
| AL360181.1 | 9.884059553 | 2.917656017 | 1.60E-05 |
| DLL3 | 453.7860412 | 2.912495012 | 1.11E-79 |
| PTGDS | 58.43516836 | 2.910230064 | 9.33E-13 |
| AC245100.6 | 52.1791593 | 2.904033407 | 6.91E-08 |
| ATP1A4 | 7.598800226 | 2.901411023 | 1.59E-03 |
| CELF2-AS1 | 7.070457865 | 2.89841346 | 3.99E-04 |

| | | | |
|---|---|---|---|
| BARX1 | 13.63832694 | 2.897133048 | 1.86E-07 |
| AL162731.1 | 8.490562927 | 2.895888897 | 3.16E-04 |
| ASNSP1 | 400.9698674 | 2.895296318 | 3.61E-19 |
| AC005291.2 | 13.64347463 | 2.88840445 | 3.70E-04 |
| AL035681.1 | 15.38520111 | 2.886546615 | 2.01E-05 |
| PTPRR | 11.79398755 | 2.88515671 | 7.18E-05 |
| PSG2 | 45.18656473 | 2.884684867 | 7.22E-03 |
| UROC1 | 5.986472261 | 2.879595961 | 2.55E-03 |
| BANF1P2 | 14.75365124 | 2.878140498 | 3.87E-07 |
| AC018816.1 | 5.312448188 | 2.87056644 | 1.13E-02 |
| RENBP | 435.6778357 | 2.862825998 | 1.04E-38 |
| C10orf90 | 19.78044098 | 2.862210066 | 8.26E-06 |
| PPP4R4 | 23.02234291 | 2.856884288 | 3.75E-07 |
| AL445470.1 | 24.97949879 | 2.856444038 | 8.43E-11 |
| AC016559.2 | 5.415044847 | 2.856156165 | 1.25E-02 |
| JAML | 5.844980626 | 2.847128975 | 1.99E-02 |
| NA | 53.49267065 | 2.846159067 | 2.57E-14 |
| CASP12 | 10.77797636 | 2.842637971 | 4.72E-06 |
| CEACAM1 | 714.1367716 | 2.838757135 | 2.17E-51 |
| CLIC2 | 109.9732088 | 2.83641542 | 8.98E-37 |
| AC100802.1 | 90.99252145 | 2.834595786 | 7.60E-06 |
| FABP3 | 256.5391819 | 2.830982348 | 6.76E-27 |
| AC093330.2 | 11.4322902 | 2.826488126 | 8.56E-06 |
| AL009178.2 | 8.641259053 | 2.826277381 | 4.55E-02 |
| UNC5A | 9.614455664 | 2.824698934 | 2.52E-04 |
| AL355312.1 | 21.51973065 | 2.822449563 | 2.44E-11 |
| FAM90A24P | 23.25397222 | 2.821239996 | 1.03E-06 |
| LINC01730 | 59.12212474 | 2.821202203 | 1.78E-17 |
| SLC27A6 | 5.313018257 | 2.815571445 | 1.92E-03 |
| IL6R | 802.3266812 | 2.81509853 | 4.41E-77 |
| CNN1 | 297.6933427 | 2.813508193 | 8.92E-19 |
| SLC47A1 | 75.64051756 | 2.813227329 | 2.86E-24 |
| NMNAT3 | 18.81273681 | 2.812837539 | 1.46E-08 |
| CES1P1 | 26.62878007 | 2.812605233 | 2.14E-04 |
| TMPRSS3 | 63.85416896 | 2.81082422 | 1.99E-16 |
| NAT2 | 41.51491537 | 2.810284072 | 9.85E-18 |
| ARMH4 | 65.3878362 | 2.802926587 | 6.29E-08 |
| AL451069.1 | 7.150260511 | 2.802758784 | 4.86E-02 |
| PAX7 | 19.03668264 | 2.801240481 | 4.77E-09 |
| AL035665.1 | 22.94008987 | 2.800956047 | 1.58E-09 |
| NA | 26.57063328 | 2.799260241 | 3.17E-05 |
| AC136628.1 | 42.07864024 | 2.790777008 | 4.18E-10 |
| FEZ1 | 659.6364878 | 2.780388047 | 1.24E-86 |
| SERPINA1 | 129.2485732 | 2.778209999 | 7.73E-30 |
| LINC00239 | 13.37656744 | 2.777328466 | 4.04E-04 |
| AC084783.1 | 9.848268834 | 2.776096363 | 7.43E-04 |
| LHX5 | 8.739437811 | 2.775359845 | 7.84E-03 |

| | | | |
|---|---|---|---|
| IL31RA | 161.1209566 | 2.772430053 | 3.42E-27 |
| HSPB8 | 1436.3874 | 2.770295007 | 7.56E-116 |
| LINC00632 | 86.52816748 | 2.76982743 | 1.60E-29 |
| PSCA | 27.29284018 | 2.767445721 | 6.70E-09 |
| CST7 | 33.33388902 | 2.76322342 | 4.59E-11 |
| AC095350.1 | 25.6081823 | 2.762597289 | 3.43E-05 |
| IRS4 | 15.16197689 | 2.759027178 | 1.32E-07 |
| CLIP4 | 1736.450377 | 2.75868926 | 2.01E-47 |
| NUDT10 | 44.68841082 | 2.756312133 | 2.48E-17 |
| SLC40A1 | 15.47794911 | 2.754284461 | 1.89E-06 |
| C6orf223 | 35.05250985 | 2.744286279 | 1.26E-07 |
| AP003064.2 | 24.94135603 | 2.744092032 | 1.19E-04 |
| BICDL1 | 37.542591 | 2.742854188 | 3.52E-08 |
| MBP | 3639.442215 | 2.742375306 | 6.54E-61 |
| JAKMIP3 | 45.86104186 | 2.741398333 | 1.59E-18 |
| PRR20G | 7.206833657 | 2.740557504 | 1.60E-02 |
| FHIT | 33.51067215 | 2.735273505 | 2.14E-04 |
| HSPB2 | 71.61158158 | 2.728869677 | 1.30E-08 |
| SHISA3 | 161.7304846 | 2.725952372 | 4.55E-21 |
| TNIP3 | 7.611684205 | 2.725113696 | 2.30E-03 |
| TMEM92-AS1 | 16.11057805 | 2.724296991 | 4.66E-04 |
| AC091806.1 | 113.7234827 | 2.723596465 | 2.24E-20 |
| COL2A1 | 456.0622899 | 2.719588282 | 2.16E-33 |
| ALCAM | 37.52071971 | 2.706980707 | 2.14E-04 |
| LINC00689 | 9.95143427 | 2.706146099 | 3.66E-04 |
| AC023813.3 | 9.998479629 | 2.705729401 | 1.22E-05 |
| OLFM2 | 487.2499097 | 2.705441729 | 1.87E-50 |
| PIFO | 13.24295233 | 2.698170281 | 3.95E-03 |
| ANO4 | 74.03198246 | 2.695108308 | 4.20E-23 |
| RETREG1 | 298.0442197 | 2.692692839 | 2.13E-28 |
| KCNK10 | 7.695928486 | 2.69043035 | 4.56E-04 |
| COL4A4 | 16.52605434 | 2.682901595 | 1.92E-07 |
| ADAMTS14 | 70.35092211 | 2.681302436 | 9.15E-21 |
| HRH2 | 76.37524624 | 2.67984405 | 1.85E-13 |
| APOL1 | 4140.760867 | 2.676340025 | 3.16E-103 |
| AC090241.2 | 9.365140553 | 2.675044093 | 4.24E-03 |
| CXADR | 430.9583909 | 2.673380869 | 1.93E-52 |
| MMP19 | 140.8610266 | 2.672551909 | 4.89E-24 |
| INHBC | 11.53838661 | 2.672220595 | 1.88E-03 |
| TRPM8 | 19.63283509 | 2.666933655 | 9.44E-07 |
| SMIM34B | 6.97659786 | 2.665694935 | 9.07E-03 |
| NEK11 | 62.08406486 | 2.664859222 | 4.77E-10 |
| MRGPRX3 | 70.91891523 | 2.664402312 | 1.07E-13 |
| ICAM1 | 10019.94945 | 2.660203763 | 1.44E-72 |
| AC004817.3 | 15.82350215 | 2.659943742 | 2.08E-05 |
| HRASLS2 | 6.319019622 | 2.655697656 | 4.06E-03 |
| ATF3 | 836.543072 | 2.65263947 | 7.49E-23 |

| | | | |
|---|---|---|---|
| ILDR2 | 71.76148004 | 2.648422897 | 5.11E-07 |
| PMEL | 53.48470474 | 2.646928136 | 3.27E-16 |
| CES1 | 60.91092756 | 2.646927384 | 1.03E-11 |
| SMPDL3B | 107.507826 | 2.64500802 | 8.60E-26 |
| TUNAR | 7.534935806 | 2.644544934 | 3.39E-03 |
| LINC01822 | 12.26246863 | 2.644377908 | 1.28E-05 |
| AL353807.3 | 42.662876 | 2.64358772 | 1.60E-19 |
| GLI1 | 189.0118592 | 2.63813259 | 3.86E-16 |
| AC069209.1 | 17.56227689 | 2.625042033 | 5.64E-08 |
| ZNF185 | 2431.652507 | 2.623454688 | 1.95E-134 |
| LINC01693 | 25.8599621 | 2.621426528 | 2.92E-05 |
| AMOTL2 | 5194.214834 | 2.617020907 | 2.42E-151 |
| COL12A1 | 7146.720884 | 2.615659848 | 1.55E-87 |
| AL713998.1 | 83.75568185 | 2.610135596 | 1.26E-11 |
| CHAC1 | 753.1216115 | 2.609993735 | 2.56E-09 |
| CPEB1 | 414.4696126 | 2.60996399 | 2.51E-43 |
| C1orf158 | 8.469554404 | 2.600736322 | 1.25E-02 |
| AKR1B10 | 66.94033728 | 2.598743076 | 8.14E-06 |
| CXCL16 | 487.4807416 | 2.59749043 | 1.10E-60 |
| SH2D4B | 5.657365801 | 2.596616621 | 1.04E-02 |
| BTF3P14 | 5.48923557 | 2.590855589 | 4.61E-03 |
| NA | 5.058469588 | 2.590561186 | 1.24E-02 |
| EXPH5 | 144.9258139 | 2.587222536 | 1.54E-25 |
| AIPL1 | 5.956943928 | 2.58431461 | 7.51E-03 |
| TP63 | 52.16142317 | 2.583910111 | 1.50E-16 |
| AC091114.1 | 7.893051909 | 2.583670964 | 3.23E-03 |
| SPP1 | 3358.989557 | 2.582306721 | 5.87E-28 |
| TEX37 | 13.34227782 | 2.58218728 | 2.33E-04 |
| STX7 | 2097.799501 | 2.581016305 | 9.20E-98 |
| AKR1C2 | 5.273570741 | 2.578010889 | 2.79E-02 |
| TMEM71 | 20.65123729 | 2.575711915 | 6.09E-05 |
| CAMK2A | 27.28062693 | 2.57363953 | 4.52E-07 |
| AC007405.1 | 12.08922671 | 2.570674629 | 1.99E-03 |
| GALNT18 | 150.014102 | 2.57057763 | 1.38E-27 |
| LUCAT1 | 124.7362473 | 2.569189365 | 3.03E-09 |
| PARD3B | 69.05787518 | 2.569097427 | 1.06E-10 |
| SULT1C2P1 | 6.30677781 | 2.568688977 | 1.26E-02 |
| IFRD1 | 3961.506108 | 2.567191132 | 1.48E-26 |
| WARS | 18975.32743 | 2.564686955 | 4.23E-41 |
| ACSM5 | 100.294989 | 2.558512926 | 1.45E-25 |
| SERPINE1 | 2089.772652 | 2.557298491 | 5.61E-55 |
| CLSTN2 | 328.2908032 | 2.552127133 | 3.33E-27 |
| SHANK2 | 32.47462154 | 2.551190101 | 1.21E-02 |
| AC139530.1 | 15.05487964 | 2.54910661 | 1.62E-05 |
| CD55 | 1726.450139 | 2.549036792 | 6.42E-159 |
| KLF4 | 330.1654842 | 2.548196564 | 5.31E-20 |
| KCNJ11 | 20.34534791 | 2.547089093 | 2.56E-04 |

| | | | |
|---|---|---|---|
| ASB18 | 18.35133093 | 2.543597489 | 3.21E-02 |
| DUOX2 | 14.32215838 | 2.543505412 | 4.97E-02 |
| SNORD63 | 6.649018311 | 2.54096716 | 7.30E-03 |
| CNKSR2 | 12.93025897 | 2.536560186 | 4.20E-04 |
| SAA2 | 14.69633019 | 2.534171673 | 1.17E-02 |
| MMP28 | 50.44223442 | 2.533013055 | 2.94E-12 |
| FAM180A | 81.88165982 | 2.531951624 | 1.46E-10 |
| FKBP9P1 | 272.6623261 | 2.531183109 | 1.13E-16 |
| OCLN | 724.0860483 | 2.52656351 | 3.30E-31 |
| AC126177.6 | 5.309390916 | 2.524811262 | 1.79E-02 |
| MYOZ2 | 10.39892218 | 2.521880843 | 2.52E-04 |
| RBPJL | 20.92174682 | 2.519440163 | 4.82E-05 |
| USH2A | 6.134410686 | 2.51872505 | 5.14E-03 |
| LINC02105 | 10.17191937 | 2.516429642 | 1.71E-04 |
| RUNDC3B | 6.946831316 | 2.511037039 | 8.19E-03 |
| MGAM | 11.91207921 | 2.50808559 | 1.31E-02 |
| AL133445.2 | 5.029610343 | 2.505100923 | 5.61E-03 |
| TNNI1 | 87.9705331 | 2.504322895 | 1.38E-02 |
| SEL1L3 | 3269.858349 | 2.499455494 | 4.22E-85 |
| BMP2 | 67.08671781 | 2.489345876 | 7.62E-14 |
| LINC02234 | 6.166080428 | 2.485825649 | 3.24E-02 |
| LRRC2 | 129.9918758 | 2.48416035 | 2.16E-44 |
| AC007611.1 | 51.12879833 | 2.476157348 | 8.72E-15 |
| ITGAM | 10.17882247 | 2.473648355 | 2.42E-02 |
| CCND2 | 40.55711956 | 2.468713162 | 2.38E-03 |
| KCNJ4 | 60.10916576 | 2.468002521 | 5.20E-20 |
| SLC38A8 | 6.684010108 | 2.465587997 | 5.57E-03 |
| AC087667.1 | 30.95840747 | 2.464423567 | 1.66E-08 |
| NLRP3 | 205.3152859 | 2.46296033 | 2.39E-23 |
| DCDC2 | 6.263599993 | 2.462077623 | 1.09E-03 |
| NA | 43.80094162 | 2.459681369 | 2.07E-06 |
| RASGRF1 | 134.7074287 | 2.459396605 | 1.99E-16 |
| IL21R | 123.0008977 | 2.457262611 | 4.82E-09 |
| IL20RB | 107.3800024 | 2.456706343 | 5.26E-10 |
| SMIM2-AS1 | 22.93913187 | 2.454250858 | 9.59E-08 |
| SLC16A4 | 951.4176711 | 2.454037311 | 6.31E-104 |
| OCA2 | 53.42871929 | 2.452776159 | 6.85E-13 |
| LYPD6B | 296.0477993 | 2.452659541 | 8.27E-23 |
| DUOXA1 | 48.41294367 | 2.451376671 | 9.23E-12 |
| RNF165 | 34.72656878 | 2.44905597 | 4.49E-08 |
| TNFRSF9 | 304.3101605 | 2.445239293 | 3.11E-21 |
| TPRG1LP1 | 5.182973913 | 2.442463365 | 1.31E-02 |
| GOLGA8IP | 12.87111558 | 2.442203771 | 1.48E-02 |
| LINC00900 | 27.39226582 | 2.43778509 | 1.30E-05 |
| IFITM10 | 41.85794267 | 2.436137777 | 4.63E-10 |
| ZSCAN4 | 7.350727045 | 2.432405132 | 6.11E-04 |
| TMEM40 | 73.22422857 | 2.425828372 | 6.73E-13 |

| | | | |
|---|---|---|---|
| ERVFRD-1 | 10.99069373 | 2.424858023 | 4.72E-05 |
| SLC39A8 | 375.9403401 | 2.420275534 | 6.93E-47 |
| IFI6 | 1717.788303 | 2.419964794 | 3.86E-43 |
| AL162430.1 | 7.260013261 | 2.416591987 | 5.44E-03 |
| HPN | 16.52860829 | 2.414953701 | 1.88E-04 |
| NKD2 | 30.11768859 | 2.414462717 | 4.71E-06 |
| CAVIN2 | 121.5099942 | 2.41419518 | 2.85E-04 |
| AJUBA | 9629.715395 | 2.41294946 | 1.04E-45 |
| XAGE2 | 41.30684287 | 2.411045343 | 1.06E-09 |
| AL669918.1 | 106.7115265 | 2.4110393 | 1.28E-02 |
| AC104958.2 | 10.58228854 | 2.406893037 | 1.60E-04 |
| MORC4 | 2027.296062 | 2.406047315 | 1.44E-205 |
| LINC02487 | 24.11198403 | 2.403942039 | 5.76E-06 |
| NYAP2 | 8.4881526 | 2.401991824 | 2.95E-02 |
| LGALS12 | 389.1815947 | 2.40038278 | 1.08E-20 |
| MYCL | 127.9613386 | 2.398106994 | 2.80E-19 |
| AC237221.1 | 6.092602119 | 2.39365758 | 3.79E-03 |
| CDH7 | 14.05780657 | 2.391290589 | 2.00E-04 |
| TFCP2L1 | 94.35599976 | 2.387235782 | 2.86E-12 |
| AP001025.1 | 5.743856054 | 2.383892611 | 1.46E-02 |
| SULT2A1 | 70.46590007 | 2.383713512 | 1.10E-14 |
| SLC7A11-AS1 | 23.40959561 | 2.383075648 | 3.33E-02 |
| ANGPTL1 | 9.294027893 | 2.381197961 | 4.89E-04 |
| HLA-F-AS1 | 194.5623124 | 2.378757641 | 1.64E-34 |
| HRASLS | 31.71422465 | 2.374467029 | 1.69E-07 |
| JAKMIP2 | 22.77357986 | 2.369853628 | 7.56E-05 |
| PRRG3 | 58.4442266 | 2.367748229 | 7.63E-05 |
| AC069528.2 | 12.19333953 | 2.367723997 | 8.12E-05 |
| ENPP1 | 221.0228781 | 2.367060381 | 1.09E-50 |
| KIAA1211 | 73.81802387 | 2.360085368 | 1.22E-17 |
| LINC01659 | 10.88984238 | 2.356704863 | 2.09E-04 |
| MOV10L1 | 49.20024885 | 2.354426091 | 8.16E-08 |
| TENM2 | 351.6327637 | 2.353174058 | 1.66E-06 |
| PCMTD1P2 | 10.24315486 | 2.350372752 | 2.56E-04 |
| RPSAP70 | 9.182282981 | 2.347886871 | 1.23E-03 |
| AC148477.4 | 20.92675249 | 2.346842406 | 1.16E-06 |
| SLC16A14 | 23.96117964 | 2.341554396 | 6.98E-06 |
| ABCG2 | 617.2687721 | 2.338867394 | 2.02E-27 |
| PRKCB | 1065.831152 | 2.338768189 | 3.79E-45 |
| SBDS | 6786.193804 | 2.338371682 | 3.97E-116 |
| DTWD2 | 108.1399521 | 2.337869619 | 1.30E-14 |
| SRPX2 | 1641.567739 | 2.336378812 | 5.43E-43 |
| KITLG | 103.264813 | 2.335696972 | 3.51E-17 |
| NFKBIB | 708.9637072 | 2.333386597 | 1.96E-49 |
| OASL | 87.89182137 | 2.329913958 | 3.00E-17 |
| KCNB1 | 28.10504894 | 2.329888023 | 2.92E-10 |
| AL583722.1 | 7.092551311 | 2.329473284 | 7.26E-03 |

| | | | |
|---|---|---|---|
| NEFL | 99.35758727 | 2.329455349 | 3.74E-09 |
| SLC25A30 | 977.3527418 | 2.328987786 | 9.68E-51 |
| AC130371.2 | 6.428091829 | 2.328750107 | 1.98E-02 |
| FAM149A | 27.54054559 | 2.328236541 | 5.28E-08 |
| SLIT3 | 447.7973117 | 2.327048846 | 2.68E-46 |
| CCDC170 | 11.11378406 | 2.326413947 | 1.19E-03 |
| BHMT2 | 687.4043628 | 2.323315504 | 9.28E-47 |
| NGF | 85.12586789 | 2.322823416 | 2.95E-05 |
| NA | 98.28641465 | 2.321866812 | 1.94E-18 |
| LHX8 | 280.7864281 | 2.321637232 | 5.17E-23 |
| DRP2 | 72.51218544 | 2.321617515 | 2.78E-23 |
| FAM19A2 | 35.64483847 | 2.316962464 | 2.02E-10 |
| CREB5 | 770.2728268 | 2.315893592 | 1.94E-18 |
| AC009185.1 | 7.801160267 | 2.314906848 | 1.14E-02 |
| FBN2 | 312.3030735 | 2.314417032 | 1.13E-13 |
| ABCB4 | 14.89418082 | 2.313432247 | 9.42E-03 |
| RIMS3 | 590.9822661 | 2.31191863 | 1.87E-13 |
| AC079160.1 | 5.187502164 | 2.311066057 | 1.51E-02 |
| DNAH10 | 134.5372647 | 2.308018938 | 4.07E-09 |
| ZFP91-CNTF | 25.65313705 | 2.305530009 | 3.64E-02 |
| TRIB3 | 8474.977623 | 2.304368623 | 1.87E-11 |
| PLA2G4C | 169.8016121 | 2.303926451 | 3.43E-17 |
| LBH | 542.265206 | 2.300676761 | 1.59E-20 |
| NCF2 | 11.76390963 | 2.298454838 | 4.78E-03 |
| POU3F3 | 5.207919673 | 2.297897495 | 2.45E-02 |
| ADGRE4P | 80.48512716 | 2.297608142 | 7.58E-11 |
| ANKFN1 | 98.2235767 | 2.297071853 | 7.62E-17 |
| BDKRB2 | 161.3889826 | 2.294366536 | 5.46E-23 |
| CEACAMP11 | 8.440503374 | 2.294259255 | 1.28E-02 |
| AC104051.2 | 48.84536052 | 2.292999401 | 7.86E-09 |
| RAET1E | 123.7618426 | 2.292615683 | 6.90E-34 |
| AC107884.1 | 35.00949873 | 2.290820452 | 7.65E-06 |
| BASP1 | 2291.351755 | 2.289317224 | 5.22E-53 |
| GUCY1A1 | 12.01694402 | 2.288946876 | 2.96E-03 |
| TUBA8 | 48.32664225 | 2.285145511 | 2.29E-04 |
| EBI3 | 130.7138912 | 2.285110565 | 8.83E-22 |
| ATP1A3 | 357.0233239 | 2.283969671 | 1.21E-22 |
| MAGI2-AS3 | 9.256636258 | 2.283509882 | 9.07E-03 |
| SLC7A3 | 41.66827949 | 2.279929048 | 5.06E-06 |
| NANOGNB | 5.163597743 | 2.278640656 | 2.93E-02 |
| GADD45G | 16.86319634 | 2.276384755 | 5.22E-05 |
| AC107956.1 | 22.9915758 | 2.273634844 | 3.13E-07 |
| AC025754.2 | 7.417419864 | 2.273019197 | 4.66E-03 |
| LINC00623 | 78.41347884 | 2.268344496 | 1.42E-10 |
| LINC01473 | 10.67875329 | 2.267842388 | 8.26E-04 |
| HLA-DRB5 | 31.70324631 | 2.267226275 | 1.05E-07 |
| AC011944.1 | 11.91468553 | 2.26552731 | 4.55E-03 |

| | | | |
|---|---|---|---|
| LPO | 43.94056569 | 2.263375151 | 6.86E-06 |
| CRYZL2P-SEC16 | 94.80652887 | 2.263318869 | 1.40E-10 |
| GBP1P1 | 7.21517128 | 2.260617057 | 2.84E-03 |
| FAP | 92.15692023 | 2.257331084 | 2.22E-12 |
| CASP4 | 3032.86953 | 2.256333783 | 2.03E-193 |
| LINC01511 | 8.69732256 | 2.252538337 | 3.64E-02 |
| RPS6KA2 | 1707.929685 | 2.252062695 | 1.24E-45 |
| AC073610.3 | 16.21540822 | 2.251053313 | 1.12E-03 |
| SPRYD7 | 559.7320342 | 2.24973093 | 4.09E-89 |
| MPO | 8.830366871 | 2.249434888 | 1.39E-02 |
| PDCD1LG2 | 137.9408949 | 2.244504139 | 1.67E-19 |
| CARD11 | 487.3641494 | 2.240315199 | 7.68E-28 |
| WTIP | 955.4419514 | 2.230720722 | 5.28E-33 |
| AC010247.2 | 13.42529976 | 2.229640967 | 3.91E-03 |
| APOE | 6958.007481 | 2.227132628 | 4.49E-30 |
| RAB32 | 1770.456458 | 2.226300613 | 3.19E-73 |
| CACNA1B | 179.5369723 | 2.225614846 | 3.61E-22 |
| VEGFA | 2322.712089 | 2.220392974 | 1.08E-16 |
| TRBV11-2 | 5.028597876 | 2.220013271 | 3.05E-02 |
| SULT1A2 | 161.2956704 | 2.215103427 | 5.03E-26 |
| AC104129.1 | 41.03407315 | 2.214622288 | 3.88E-07 |
| MYOSLID | 13.69503477 | 2.20775759 | 2.46E-02 |
| AC090673.1 | 7.812118437 | 2.207718751 | 6.82E-03 |
| AC148476.1 | 31.43670773 | 2.206853713 | 1.77E-08 |
| MALRD1 | 7.12536122 | 2.20664071 | 4.57E-03 |
| LCNL1 | 5.08364685 | 2.204951807 | 4.64E-02 |
| TSPAN33 | 331.3822405 | 2.204114322 | 7.34E-50 |
| RND3 | 1363.687625 | 2.203251229 | 3.16E-11 |
| SLC9A3 | 209.2046598 | 2.200221394 | 1.50E-23 |
| PARP8 | 150.9692421 | 2.198961658 | 6.59E-22 |
| SSR3 | 5937.31788 | 2.196613441 | 4.24E-196 |
| LINC01220 | 13.87085948 | 2.196446389 | 1.18E-02 |
| PLXDC2 | 20.00721933 | 2.194025857 | 1.93E-05 |
| SLC17A5 | 973.1245904 | 2.192337972 | 2.18E-50 |
| VENTX | 11.24059506 | 2.184504325 | 4.81E-04 |
| LINC01477 | 7.235124661 | 2.18345036 | 6.48E-03 |
| AP000446.1 | 357.0586464 | 2.183118236 | 1.87E-41 |
| CDH12 | 46.78834468 | 2.178259265 | 2.02E-10 |
| SLC2A3 | 3270.065993 | 2.168291612 | 7.52E-23 |
| PTCHD1 | 9.559718963 | 2.165687068 | 7.26E-03 |
| SLC4A9 | 9.435170445 | 2.158833395 | 9.66E-03 |
| PLA2G16 | 1773.072902 | 2.1586325 | 1.25E-58 |
| OTUD1 | 108.5394167 | 2.157290703 | 1.65E-26 |
| DHRS3 | 27.89732788 | 2.153932611 | 5.41E-06 |
| CAVIN4 | 73.99844034 | 2.152343792 | 1.42E-19 |
| AL138902.1 | 7.240013858 | 2.151730392 | 1.69E-02 |
| ITGAD | 14.89201776 | 2.151358954 | 6.43E-03 |

| | | | |
|---|---|---|---|
| AC016924.1 | 5.24745008 | 2.146752474 | 2.94E-02 |
| KCNJ10 | 12.23022389 | 2.145920024 | 3.00E-03 |
| TMEM56 | 329.4539423 | 2.143748158 | 1.65E-24 |
| LINC01764 | 8.559247651 | 2.143695473 | 1.24E-02 |
| AC078905.1 | 7.967882544 | 2.142976697 | 3.40E-03 |
| AC099786.2 | 59.76050552 | 2.141743665 | 4.55E-15 |
| TMEM176B | 11.60900438 | 2.14139694 | 6.07E-05 |
| ALK | 6.847829701 | 2.138394192 | 1.89E-02 |
| SOX8 | 41.97552585 | 2.137839459 | 1.49E-14 |
| CX3CR1 | 5.956254789 | 2.136253546 | 4.70E-02 |
| C1QTNF5 | 673.4987851 | 2.13591705 | 2.35E-33 |
| AC011511.2 | 6.936147743 | 2.133624369 | 1.31E-02 |
| AC243972.1 | 13.61856719 | 2.130622257 | 9.32E-04 |
| ACP5 | 1625.175419 | 2.128946614 | 1.04E-28 |
| PRSS16 | 79.4703776 | 2.128080067 | 1.67E-14 |
| NAMPT | 5630.167846 | 2.127902395 | 6.02E-24 |
| RNF180 | 25.27695137 | 2.123725471 | 3.28E-05 |
| AC009238.1 | 10.03320931 | 2.123677937 | 1.67E-02 |
| AL138955.1 | 8.756164881 | 2.121886707 | 1.16E-03 |
| SNCA | 34.72054321 | 2.121077016 | 2.26E-09 |
| PLIN5 | 41.59867721 | 2.120524428 | 5.97E-08 |
| EGF | 82.7844327 | 2.120077661 | 1.45E-10 |
| OOEP | 184.8376679 | 2.119210087 | 5.43E-39 |
| IL33 | 11.32575042 | 2.117094863 | 3.40E-03 |
| AC105105.1 | 257.2624826 | 2.115453364 | 2.81E-05 |
| ITIH5 | 73.04147724 | 2.112780451 | 1.62E-08 |
| PADI3 | 60.49052887 | 2.112012505 | 5.24E-05 |
| TEX15 | 71.56116553 | 2.111641868 | 1.06E-11 |
| LINC01546 | 5.257423978 | 2.106892988 | 3.08E-02 |
| NA | 11.87824001 | 2.106061002 | 1.32E-03 |
| LINC02663 | 5.132443263 | 2.105599706 | 1.64E-02 |
| AC008663.2 | 6.853971202 | 2.103049132 | 8.44E-03 |
| ULBP1 | 1094.423407 | 2.102861946 | 1.91E-28 |
| CSMD2 | 15.3385253 | 2.099809461 | 1.15E-04 |
| AC092384.2 | 8.422921767 | 2.097782088 | 4.45E-02 |
| SAMD5 | 74.58486099 | 2.096679324 | 5.17E-15 |
| PALMD | 192.6518655 | 2.093965972 | 2.84E-24 |
| HINT3 | 485.57052 | 2.093436479 | 1.42E-20 |
| SEC16B | 132.8719069 | 2.09319736 | 6.13E-11 |
| RSAD2 | 15.70848725 | 2.092930286 | 8.13E-03 |
| IL17D | 149.0420822 | 2.091334835 | 6.49E-22 |
| SLC13A5 | 229.8750143 | 2.08986452 | 1.77E-18 |
| GLB1L | 3888.373573 | 2.087345621 | 5.60E-63 |
| AC009041.2 | 47.19130822 | 2.085799467 | 1.02E-10 |
| ARC | 110.2728391 | 2.085021113 | 1.15E-09 |
| HRK | 25.95120166 | 2.084567655 | 4.06E-03 |
| CYP1A1 | 206.1942386 | 2.083676927 | 1.03E-24 |

| | | | |
|---|---|---|---|
| FMO4 | 51.19801347 | 2.082751826 | 4.90E-08 |
| EDNRB | 780.2042551 | 2.082298048 | 9.89E-55 |
| SBDSP1 | 833.1214146 | 2.081716157 | 1.52E-26 |
| AC007406.4 | 5.499413409 | 2.079972534 | 1.50E-02 |
| AC078845.1 | 11.09755694 | 2.079957627 | 1.57E-03 |
| MAP3K15 | 28.1466474 | 2.079187189 | 5.04E-05 |
| LINC02613 | 5.090352264 | 2.077292017 | 2.79E-02 |
| NA | 23.5931674 | 2.075716912 | 5.09E-06 |
| STON1-GTF2A1 | 83.30213425 | 2.074142345 | 1.95E-10 |
| CACNG3 | 6.716815847 | 2.074114034 | 4.68E-02 |
| COBL | 25.11154967 | 2.073462469 | 9.90E-07 |
| SHC4 | 92.74697178 | 2.073413639 | 2.56E-14 |
| BMP6 | 286.6928456 | 2.07076116 | 3.62E-17 |
| GHR | 39.67772673 | 2.069449836 | 1.58E-08 |
| CSNK2A3 | 12.69435714 | 2.069371384 | 1.34E-02 |
| MYH13 | 7.057222428 | 2.068794627 | 7.52E-03 |
| AC010327.4 | 19.75050693 | 2.068226069 | 1.07E-03 |
| WDFY4 | 28.36478348 | 2.067507305 | 4.70E-03 |
| KCNE3 | 74.4885636 | 2.066742628 | 3.08E-06 |
| SLC16A12 | 9.185038545 | 2.066347804 | 1.56E-02 |
| AC073072.1 | 5.524705053 | 2.066043608 | 4.07E-02 |
| OTOA | 11.52031922 | 2.062370225 | 1.19E-02 |
| AC090358.1 | 11.99193219 | 2.061965768 | 1.23E-03 |
| AL591926.5 | 6.905487616 | 2.060762335 | 1.64E-02 |
| ASNS | 14006.90675 | 2.059281944 | 2.47E-17 |
| TMEM273 | 14.01436715 | 2.057211032 | 1.29E-03 |
| C8orf34 | 8.266408055 | 2.055851635 | 1.07E-03 |
| DPP6 | 11.99597905 | 2.051472766 | 1.70E-02 |
| CES1P2 | 28.23427915 | 2.051326851 | 3.84E-05 |
| PDGFA | 1396.475027 | 2.048728128 | 2.93E-49 |
| PLP1 | 431.1818229 | 2.047489778 | 8.24E-20 |
| ZC3H12B | 78.85435396 | 2.047129266 | 3.72E-11 |
| AL132780.2 | 91.80791985 | 2.046599767 | 8.76E-14 |
| MYL4 | 57.40459191 | 2.041285453 | 1.35E-08 |
| PCOTH | 24.61420542 | 2.040715733 | 7.70E-08 |
| LINC01474 | 31.74958539 | 2.039753248 | 2.24E-08 |
| AC010255.1 | 15.32312268 | 2.039147033 | 2.69E-03 |
| MYPN | 979.7692297 | 2.036662175 | 2.95E-28 |
| NAMPTP1 | 291.5978854 | 2.035767013 | 1.32E-16 |
| GALNT9 | 229.1882238 | 2.033695065 | 4.79E-17 |
| AC126696.3 | 5.391050442 | 2.033528375 | 2.54E-02 |
| HLA-DRB1 | 80.24218844 | 2.033124328 | 1.81E-08 |
| PHOSPHO1 | 77.15191877 | 2.031464195 | 2.25E-11 |
| CPVL | 3426.438074 | 2.030084602 | 1.21E-57 |
| AC007405.3 | 63.73903115 | 2.02772393 | 5.34E-10 |
| SETDB2 | 563.5613098 | 2.026870467 | 3.45E-16 |
| MYOF | 15695.67332 | 2.025635732 | 7.56E-60 |

| | | | |
|---|---|---|---|
| ADGRD1 | 51.50381324 | 2.025443028 | 1.92E-05 |
| LY96 | 154.5175083 | 2.02297991 | 1.51E-17 |
| ANKRD34C | 8.916883708 | 2.022390759 | 4.73E-02 |
| AL121772.1 | 44.2696685 | 2.018848233 | 4.20E-09 |
| AC116407.1 | 18.5476928 | 2.01783981 | 1.63E-04 |
| LEMD1 | 11.03924572 | 2.017735831 | 7.04E-04 |
| FSTL3 | 2216.418323 | 2.015892525 | 4.67E-30 |
| MATN4 | 29.85064027 | 2.015718173 | 2.00E-05 |
| SNCB | 14.57954609 | 2.012025912 | 1.29E-03 |
| CASC15 | 47.77630025 | 2.011485375 | 2.56E-14 |
| GFI1 | 39.2406586 | 2.011458343 | 4.68E-06 |
| MATN3 | 531.0146682 | 2.011154986 | 9.00E-46 |
| CD68 | 753.459441 | 2.010545614 | 1.51E-27 |
| OR1F1 | 8.998703734 | 2.007329183 | 1.21E-03 |
| BEST1 | 300.9674433 | 2.006855578 | 3.22E-16 |
| SLC16A10 | 46.30609065 | 2.004457942 | 3.90E-08 |
| RFTN2 | 36.09268729 | 2.002038251 | 3.29E-04 |
| CYP7B1 | 12.66439045 | 2.001068818 | 1.14E-04 |
| CHRD | 56.17107808 | 2.000711044 | 4.24E-08 |
| TXNRD1 | 33083.72171 | 2.000025042 | 2.58E-28 |
| LINC02159 | 20.49523315 | 1.998988305 | 5.02E-05 |
| FBXO27 | 124.0869824 | 1.997740671 | 4.29E-11 |
| ERN1 | 835.8781716 | 1.997471093 | 4.79E-13 |
| SLC12A7 | 1143.958255 | 1.996559206 | 1.19E-33 |
| LINC02050 | 9.730089576 | 1.99503138 | 3.29E-03 |
| NLRP5 | 272.5607272 | 1.991016661 | 3.17E-20 |
| AL391422.1 | 7.102066668 | 1.987648275 | 4.43E-02 |
| ANGPTL4 | 106.2736684 | 1.986833672 | 3.28E-04 |
| DTNA | 635.6860956 | 1.98663363 | 3.20E-49 |
| OLFML1 | 58.53919204 | 1.986484029 | 2.21E-04 |
| PSG9 | 110.5829147 | 1.98324166 | 6.17E-04 |
| AC008514.1 | 9.822789484 | 1.980500211 | 8.80E-03 |
| PSG11 | 67.8097526 | 1.980215932 | 3.54E-03 |
| MAFK | 1886.636662 | 1.979224901 | 9.59E-74 |
| CD177 | 28.34911627 | 1.979015496 | 8.84E-03 |
| NR4A1 | 297.1782047 | 1.978560639 | 6.93E-41 |
| SLC15A1 | 6.827630458 | 1.978340137 | 2.29E-02 |
| AC006970.2 | 5.939617258 | 1.978279683 | 2.84E-02 |
| EGR1 | 398.2324942 | 1.976263194 | 1.08E-03 |
| ADGRG5 | 6.237538367 | 1.976182054 | 4.72E-02 |
| CCRL2 | 23.25096366 | 1.974284486 | 2.91E-05 |
| FUOM | 567.2907635 | 1.973850493 | 2.13E-39 |
| AC010247.1 | 21.51955622 | 1.973447952 | 3.77E-04 |
| MIR616 | 10.47098309 | 1.971962389 | 6.50E-03 |
| GABARAPL1 | 3444.554127 | 1.971099719 | 2.08E-109 |
| NAALAD2 | 21.61366241 | 1.970840726 | 7.05E-05 |
| LINC01933 | 13.42810455 | 1.970816645 | 2.40E-02 |

| | | | |
|---|---|---|---|
| NOCT | 193.3207984 | 1.970583186 | 4.36E-30 |
| GBA | 4636.234166 | 1.96953089 | 1.12E-49 |
| RRAD | 54.82459311 | 1.969219865 | 3.13E-09 |
| NOS1 | 28.20981896 | 1.9678388 | 1.62E-06 |
| ELFN1 | 408.2898101 | 1.967459786 | 1.34E-56 |
| SLIT1 | 10.50331622 | 1.967271787 | 3.89E-02 |
| AC022509.3 | 15.6517351 | 1.966985644 | 1.19E-04 |
| CDR1 | 15.96781407 | 1.963151349 | 2.28E-02 |
| SNX10 | 285.7625478 | 1.959706214 | 1.80E-15 |
| LINC02057 | 19.34808816 | 1.95663075 | 1.51E-03 |
| GPR17 | 17.22731363 | 1.956455778 | 2.54E-04 |
| PANX2 | 146.5880724 | 1.956276514 | 8.42E-21 |
| AC067852.1 | 11.12127708 | 1.954485112 | 4.98E-02 |
| LINC01085 | 12.75865266 | 1.952760967 | 2.49E-04 |
| MAGEB6 | 5.782625089 | 1.951983117 | 3.85E-02 |
| LSR | 324.5816345 | 1.950283303 | 1.49E-26 |
| AC005670.2 | 15.0788706 | 1.94999622 | 7.08E-03 |
| FCRLA | 42.33965956 | 1.949440006 | 6.25E-12 |
| GRIP1 | 17.48063495 | 1.949430785 | 1.66E-03 |
| AL158055.1 | 6.236570408 | 1.947491789 | 2.84E-02 |
| PRLR | 20.23412662 | 1.946758477 | 2.12E-03 |
| DUX4L45 | 69.43942931 | 1.944679578 | 4.28E-10 |
| MYCT1 | 39.3580197 | 1.943393428 | 4.08E-05 |
| NTF4 | 74.5460595 | 1.943200618 | 4.50E-17 |
| AC084880.1 | 9.559536845 | 1.939824672 | 5.56E-03 |
| LINC01356 | 69.31803959 | 1.937407813 | 4.34E-11 |
| MED12L | 66.54139389 | 1.936917736 | 5.82E-07 |
| CYP2J2 | 18.03377978 | 1.936353289 | 1.51E-03 |
| C1orf127 | 6.669118029 | 1.934468902 | 2.33E-02 |
| LINC02453 | 7.064652985 | 1.93138767 | 1.45E-02 |
| AC002480.1 | 40.60269188 | 1.929100963 | 1.93E-09 |
| USH1G | 10.05305436 | 1.928896122 | 1.36E-03 |
| FGFR4 | 381.567178 | 1.928582215 | 1.08E-16 |
| FBXL7 | 181.7020041 | 1.927306273 | 5.35E-39 |
| CCDC169 | 311.1901616 | 1.925003749 | 7.99E-26 |
| NAP1L6 | 10.72768114 | 1.923206328 | 1.86E-03 |
| SLC30A2 | 14.29975304 | 1.923199312 | 2.78E-04 |
| ENPEP | 9.959916459 | 1.922051367 | 5.32E-03 |
| SLC25A30-AS1 | 7.296996916 | 1.92178949 | 1.12E-02 |
| AC120036.1 | 20.426193 | 1.920115816 | 2.47E-06 |
| MIR503 | 28.37757972 | 1.918348184 | 1.48E-03 |
| MTHFD2 | 8169.227928 | 1.917973751 | 2.29E-22 |
| SLC38A6 | 490.2878983 | 1.917966732 | 4.40E-59 |
| RCAN1 | 3868.497247 | 1.916557243 | 3.42E-49 |
| HDAC9 | 214.4988075 | 1.914264857 | 3.17E-22 |
| DNAJC6 | 116.1338755 | 1.913605529 | 1.66E-07 |
| FAM49A | 399.0533098 | 1.913201171 | 4.01E-51 |

| | | | |
|---|---|---|---|
| AC108515.1 | 38.85659606 | 1.912154881 | 2.00E-08 |
| IGFL2-AS1 | 125.576691 | 1.90989863 | 4.25E-13 |
| PCDH1 | 2577.072522 | 1.90886714 | 1.17E-22 |
| CCL15-CCL14 | 48.89464838 | 1.903410042 | 6.39E-03 |
| CLCN5 | 680.0817195 | 1.90276419 | 1.17E-31 |
| LINC02561 | 10.77705008 | 1.901326272 | 5.30E-03 |
| AC004877.1 | 19.22436289 | 1.898430343 | 2.38E-03 |
| MBL1P | 30.16663922 | 1.898358364 | 1.25E-06 |
| RAB33A | 27.18663417 | 1.895234745 | 1.56E-05 |
| AC131011.1 | 9.333704356 | 1.894829271 | 2.41E-03 |
| NALCN-AS1 | 10.77935959 | 1.8939947 | 3.14E-04 |
| MOGAT3 | 27.3877624 | 1.888064623 | 2.93E-08 |
| SAT1 | 2227.264793 | 1.884207402 | 3.91E-66 |
| SGTB | 1244.16621 | 1.8819074 | 9.07E-26 |
| FIGN | 438.4513192 | 1.881609734 | 1.67E-39 |
| CXCL2 | 13.90457934 | 1.881274845 | 6.70E-03 |
| LINC01470 | 8.989339598 | 1.880240245 | 4.70E-02 |
| CYFIP2 | 654.8815999 | 1.87876886 | 1.01E-14 |
| FSTL4 | 52.65497819 | 1.878388797 | 1.19E-07 |
| AL117335.1 | 26.96280158 | 1.876745495 | 1.75E-05 |
| AC017002.1 | 16.76283255 | 1.8764399 | 1.93E-03 |
| AC226118.1 | 19.52819123 | 1.875753371 | 9.91E-05 |
| DNAJB5 | 1661.627669 | 1.873730768 | 1.01E-39 |
| NIPAL4 | 209.0449799 | 1.873353116 | 3.15E-11 |
| PVALB | 24.08534313 | 1.86782804 | 5.48E-04 |
| ALMS1-IT1 | 18.70546363 | 1.867339168 | 8.26E-05 |
| LINC01602 | 20.38071161 | 1.866802659 | 9.64E-05 |
| MASP1 | 35.16124202 | 1.862763048 | 2.02E-03 |
| EEF1A2 | 5981.383771 | 1.860175507 | 4.04E-55 |
| KCNJ5 | 31.40138865 | 1.858128089 | 1.83E-06 |
| PLS3 | 10420.30682 | 1.856724848 | 1.60E-98 |
| LATS2 | 922.0682349 | 1.856232447 | 2.44E-64 |
| AC005736.1 | 74.59682991 | 1.856028046 | 9.73E-13 |
| GPR160 | 34.99439473 | 1.853171249 | 6.22E-07 |
| SULT1A1 | 2597.775991 | 1.852493619 | 5.26E-24 |
| RGS9 | 137.3520688 | 1.851845947 | 1.91E-12 |
| AC005291.1 | 6.33143469 | 1.851731676 | 1.83E-02 |
| AC003975.1 | 9.906849579 | 1.848678078 | 1.19E-03 |
| SSPN | 336.9878138 | 1.847252546 | 3.69E-17 |
| PPP1R1C | 18.81541995 | 1.847139736 | 5.32E-05 |
| LINC00662 | 641.6290959 | 1.846094211 | 1.74E-10 |
| AL590235.1 | 24.40579455 | 1.845873054 | 5.11E-07 |
| DMRTA1 | 131.8968401 | 1.845543926 | 9.58E-15 |
| LRCH2 | 46.29558708 | 1.844356364 | 4.25E-08 |
| PLAC1 | 82.08555886 | 1.842281053 | 1.21E-12 |
| CSF3R | 14.09332699 | 1.840288448 | 1.18E-03 |
| CACNG2 | 5.500643226 | 1.840244492 | 3.75E-02 |

| | | | |
|---|---|---|---|
| KRT8P33 | 61.62685347 | 1.839438744 | 6.91E-14 |
| PI3 | 17.94074282 | 1.837921538 | 8.92E-03 |
| AC110373.1 | 6.172230091 | 1.837587985 | 2.00E-02 |
| AL135905.1 | 19.51739144 | 1.836839494 | 1.69E-05 |
| SFRP5 | 9.335463047 | 1.836471607 | 3.67E-03 |
| FAT2 | 52.76982091 | 1.835430219 | 1.10E-08 |
| RAB3D | 372.3493686 | 1.834017181 | 3.47E-26 |
| HMGB1P31 | 7.564357389 | 1.83106916 | 1.51E-02 |
| ADAMTS9-AS2 | 15.97353599 | 1.831006042 | 6.60E-03 |
| MFSD2A | 204.18906 | 1.83011551 | 4.46E-32 |
| DTX4 | 442.4496529 | 1.828895597 | 2.84E-19 |
| SLC3A2 | 10651.30897 | 1.828220318 | 1.89E-14 |
| MGAT4A | 18.11281546 | 1.82566774 | 4.23E-02 |
| GPRIN2 | 25.95356115 | 1.824786421 | 2.44E-02 |
| VGLL3 | 29.00567059 | 1.824417818 | 5.46E-05 |
| TRAM1L1 | 19.91426417 | 1.824195242 | 1.17E-04 |
| GOLGA4 | 1736.588028 | 1.8207949 | 1.28E-07 |
| IL6R-AS1 | 5.967976111 | 1.82056505 | 1.46E-02 |
| PSAT1 | 16232.43473 | 1.819631831 | 1.18E-17 |
| KAZN | 2090.457273 | 1.819562339 | 6.02E-90 |
| SLCO1B3 | 13.29089053 | 1.819349427 | 2.51E-02 |
| MICE | 40.47422612 | 1.818982425 | 2.64E-12 |
| MEIS3P2 | 55.25686464 | 1.817659707 | 4.73E-11 |
| MAGEL2 | 23.89919631 | 1.813959072 | 2.52E-02 |
| LINC02577 | 30.04782846 | 1.811556491 | 6.97E-06 |
| ITPR1-DT | 36.37859191 | 1.810942107 | 4.71E-08 |
| NANOS3 | 11.09929873 | 1.80344626 | 2.81E-03 |
| TAT | 5.783451986 | 1.80076792 | 2.67E-02 |
| ASAH1 | 3432.713782 | 1.800690764 | 2.90E-74 |
| RPL23AP49 | 10.95563735 | 1.797796802 | 6.79E-03 |
| AC099786.1 | 15.87473704 | 1.79766025 | 1.99E-04 |
| VLDLR | 1352.477514 | 1.796795412 | 1.38E-80 |
| LAIR1 | 55.86959733 | 1.792523652 | 6.54E-04 |
| GADD45A | 1177.960422 | 1.791544304 | 1.30E-16 |
| CCPG1 | 2140.271698 | 1.791470609 | 1.08E-06 |
| GPR137B | 473.0936551 | 1.790895341 | 4.94E-50 |
| METRNL | 538.6256251 | 1.789221388 | 3.04E-32 |
| IRF8 | 69.17208856 | 1.788054546 | 9.44E-07 |
| AL139002.1 | 119.7836048 | 1.787660973 | 6.31E-22 |
| MYH14 | 1638.03928 | 1.78574904 | 8.98E-73 |
| AJ003147.2 | 8.565431462 | 1.78476153 | 1.35E-02 |
| MYO5B | 212.8051185 | 1.784029748 | 1.25E-08 |
| FPR2 | 56.2775836 | 1.782071124 | 3.00E-12 |
| AREG | 9.792709533 | 1.781890057 | 2.73E-02 |
| B3GALT5 | 20.11888974 | 1.781456782 | 7.71E-03 |
| NLRP6 | 7.283248337 | 1.781157695 | 1.10E-02 |
| UCHL1 | 173.5698622 | 1.780710618 | 1.30E-13 |

| | | | |
|---|---|---|---|
| LINC01436 | 88.90708551 | 1.775839025 | 2.63E-15 |
| ENPP4 | 104.0774545 | 1.775791408 | 4.91E-09 |
| KCNK15 | 31.2411331 | 1.774072553 | 7.88E-07 |
| PIPOX | 53.95477712 | 1.773956829 | 9.04E-11 |
| WWC1 | 602.1446754 | 1.772144904 | 6.46E-29 |
| L3HYPDH | 792.4822493 | 1.771462585 | 3.86E-80 |
| SPHK1 | 935.0801792 | 1.771326359 | 1.22E-59 |
| SLFN14 | 5.902439543 | 1.770693128 | 4.05E-02 |
| PIP5K1B | 6.120998755 | 1.770161162 | 4.80E-02 |
| CARD16 | 266.3336459 | 1.769629456 | 1.13E-33 |
| SGK3 | 372.4773493 | 1.76955282 | 1.79E-24 |
| GFPT1 | 3629.248179 | 1.769347611 | 1.89E-26 |
| BACE2 | 2559.985201 | 1.768323196 | 5.58E-46 |
| BEX5 | 103.9003807 | 1.764554565 | 5.46E-15 |
| GAREM1 | 159.1268217 | 1.763741004 | 1.21E-21 |
| HTRA1 | 853.162337 | 1.763589782 | 1.18E-29 |
| PVR | 6076.548739 | 1.763573746 | 3.97E-27 |
| SEMA3B | 1213.282702 | 1.7631064 | 5.08E-18 |
| KCNH1 | 22.72830459 | 1.7605755 | 1.77E-03 |
| HSD17B14 | 7.846998504 | 1.75717281 | 3.49E-02 |
| NA | 26.04020648 | 1.754343373 | 2.47E-05 |
| FRG2C | 27.16775073 | 1.754230891 | 1.74E-02 |
| NA | 7.609373479 | 1.753552226 | 2.00E-02 |
| RGS5 | 1408.439858 | 1.750948713 | 1.43E-06 |
| PCK2 | 6290.645737 | 1.748554 | 4.94E-12 |
| AC068580.4 | 32.74960803 | 1.74704407 | 3.02E-03 |
| LGALS8-AS1 | 15.24455223 | 1.746792101 | 5.95E-04 |
| FLNC | 24286.4346 | 1.746121803 | 2.51E-43 |
| ICAM2 | 134.2705805 | 1.744255181 | 2.73E-17 |
| CCN1 | 10596.59591 | 1.740805097 | 1.11E-52 |
| C1QTNF1-AS1 | 25.06605869 | 1.739740772 | 4.04E-05 |
| CD82 | 1265.274499 | 1.739215562 | 1.78E-20 |
| CAP2 | 1928.403831 | 1.736274759 | 2.48E-73 |
| KANK3 | 90.57746237 | 1.736213252 | 1.84E-11 |
| SCN4A | 135.5757544 | 1.735570169 | 7.44E-07 |
| FAM160A1 | 139.3017091 | 1.734244075 | 1.04E-07 |
| HLA-F | 512.0742704 | 1.733852171 | 3.15E-22 |
| SMPDL3A | 94.57710573 | 1.733570224 | 8.65E-18 |
| L3MBTL4 | 51.64615564 | 1.732006663 | 2.69E-05 |
| CACNA1I | 43.84665629 | 1.731642194 | 2.79E-06 |
| NA | 16.9656287 | 1.730963785 | 2.82E-03 |
| FGD4 | 160.9786872 | 1.730356326 | 4.10E-13 |
| TMEM205 | 1463.331865 | 1.729598916 | 6.81E-43 |
| AL109809.1 | 7.549586939 | 1.729261167 | 1.85E-02 |
| AC018755.2 | 97.94631105 | 1.728260499 | 1.35E-06 |
| ARAP2 | 8.314747089 | 1.728119738 | 2.25E-02 |
| LINC01303 | 56.42147542 | 1.728093675 | 9.06E-04 |

| | | | |
|---|---|---|---|
| IFIT2 | 1178.033649 | 1.727838554 | 4.89E-42 |
| TMEM191A | 42.51659581 | 1.721818308 | 2.58E-08 |
| TRIM16L | 2552.382183 | 1.721302012 | 6.19E-14 |
| TGIF1 | 1706.310532 | 1.720728263 | 3.18E-22 |
| AC090409.1 | 76.02384453 | 1.718747697 | 4.00E-11 |
| LINC01606 | 42.6193548 | 1.718516996 | 2.28E-04 |
| USP18 | 291.189923 | 1.718363619 | 5.08E-28 |
| MSANTD3-TME | 272.4334585 | 1.717203869 | 2.03E-08 |
| HOMER3-AS1 | 17.99111596 | 1.716600865 | 4.08E-04 |
| AC009549.1 | 45.61235665 | 1.716381768 | 1.50E-10 |
| ARHGEF16 | 104.7080639 | 1.714992173 | 1.07E-13 |
| SNORD63B | 18.0784338 | 1.713726826 | 1.55E-03 |
| HDAC2-AS2 | 45.6529736 | 1.712697189 | 4.56E-05 |
| ARSI | 573.0458329 | 1.712420018 | 3.78E-17 |
| PRKAA2 | 150.5467352 | 1.711743004 | 1.23E-13 |
| BRI3 | 3521.077973 | 1.71096102 | 4.50E-38 |
| FUNDC2P2 | 18.74886774 | 1.70933881 | 2.22E-03 |
| TMEM117 | 26.69301115 | 1.709028875 | 2.09E-04 |
| REN | 21.64424951 | 1.708295239 | 2.60E-02 |
| CASP1 | 701.6894892 | 1.708226336 | 1.90E-23 |
| ZFPM2-AS1 | 556.3674849 | 1.707928224 | 1.35E-16 |
| SHMT2 | 14299.48947 | 1.704322134 | 2.93E-17 |
| CDH1 | 23.87917036 | 1.703672002 | 2.89E-02 |
| KCNK9 | 51.84500957 | 1.703260404 | 2.75E-08 |
| AC016705.2 | 94.42185277 | 1.703016093 | 2.03E-06 |
| CTSD | 13511.99258 | 1.701627168 | 9.72E-44 |
| GAPLINC | 47.0292831 | 1.697791806 | 7.92E-09 |
| SLC7A5 | 18550.72411 | 1.697583334 | 3.05E-07 |
| IRX6 | 9.061079984 | 1.696454006 | 6.90E-03 |
| CCDC169-SOHL | 38.91818841 | 1.693707828 | 3.97E-04 |
| SHANK1 | 153.5495197 | 1.692392978 | 5.35E-12 |
| F11-AS1 | 217.3634243 | 1.691148918 | 1.43E-10 |
| GPRIN3 | 52.6600565 | 1.688713413 | 1.28E-05 |
| FAM184B | 43.04346961 | 1.688193435 | 1.43E-08 |
| PCDHA10 | 9.781250887 | 1.68642782 | 2.29E-02 |
| GCNT2 | 168.7839907 | 1.686376175 | 4.68E-25 |
| SEMA6B | 655.8863531 | 1.684696171 | 2.54E-19 |
| MIR22HG | 876.9771294 | 1.684552682 | 1.07E-90 |
| HECW2 | 136.979685 | 1.684529376 | 6.65E-05 |
| PPM1K | 858.4480811 | 1.683392864 | 4.20E-36 |
| DRD1 | 26.45660106 | 1.683372035 | 2.73E-04 |
| DNAH6 | 35.71945557 | 1.682823974 | 6.60E-04 |
| CTSV | 629.8242174 | 1.682433867 | 2.56E-37 |
| ME1 | 734.9613871 | 1.681628997 | 1.47E-35 |
| OSGIN2 | 858.4617756 | 1.680914359 | 3.36E-31 |
| AL355338.1 | 755.4215629 | 1.680818666 | 1.90E-11 |
| TENT5B | 961.9812951 | 1.679180932 | 3.31E-26 |

| | | | |
|---|---|---|---|
| ITGA1 | 11661.05432 | 1.677980974 | 4.70E-42 |
| CYP4V2 | 348.4362526 | 1.675151612 | 1.82E-14 |
| KIRREL2 | 62.22891329 | 1.672350299 | 1.66E-08 |
| APOL2 | 2137.586548 | 1.669563382 | 1.38E-116 |
| LINC02525 | 15.86356257 | 1.668882466 | 1.56E-03 |
| LMOD3 | 34.88929649 | 1.668816917 | 2.65E-05 |
| WNT5B | 481.1590035 | 1.668509358 | 2.86E-18 |
| ERVH-1 | 8.801856477 | 1.667105996 | 4.62E-02 |
| WIPI1 | 1713.174141 | 1.666973425 | 1.74E-55 |
| AC144831.1 | 11.12021588 | 1.666587158 | 1.53E-02 |
| CTNS | 1997.987538 | 1.666233052 | 2.18E-44 |
| P2RX4 | 1389.149782 | 1.665705363 | 6.85E-60 |
| EAF2 | 18.1112927 | 1.665180468 | 1.79E-03 |
| TINCR | 80.46563483 | 1.664982996 | 1.38E-07 |
| DMKN | 409.4101885 | 1.664261036 | 1.54E-27 |
| AC002401.4 | 17.60670996 | 1.662889794 | 5.31E-03 |
| EREG | 68.09509223 | 1.662674574 | 8.18E-06 |
| ABL2 | 2982.838415 | 1.661872514 | 1.20E-54 |
| AL359237.1 | 12.93591957 | 1.661505844 | 3.95E-03 |
| AC098487.1 | 25.41401803 | 1.660160126 | 4.31E-04 |
| ARL4A | 624.3512308 | 1.658663951 | 4.31E-32 |
| IFIT3 | 1284.613263 | 1.656164495 | 1.48E-44 |
| SRSF12 | 22.635608 | 1.655967048 | 1.41E-02 |
| CD70 | 205.6557704 | 1.6551421 | 1.61E-22 |
| CFAP61 | 17.9428359 | 1.654987647 | 5.58E-03 |
| DPP4 | 73.62921846 | 1.654898579 | 2.77E-10 |
| AL121987.2 | 26.93484969 | 1.652450665 | 2.20E-05 |
| SQOR | 301.2361767 | 1.652168252 | 3.16E-26 |
| KREMEN2 | 296.5025655 | 1.651971719 | 7.75E-18 |
| AC022730.4 | 11.60389552 | 1.650613465 | 5.41E-03 |
| BTNL9 | 31.82631364 | 1.649493473 | 9.49E-05 |
| AL139288.1 | 8.236759422 | 1.647454158 | 1.79E-02 |
| AC010207.1 | 29.98635144 | 1.646331703 | 9.65E-04 |
| MELTF | 431.330045 | 1.644830228 | 2.22E-19 |
| IGSF23 | 35.73858368 | 1.644353837 | 4.25E-05 |
| NEU1 | 2876.213876 | 1.644262906 | 6.95E-25 |
| ACKR3 | 110.6345116 | 1.643267568 | 5.22E-12 |
| PKIB | 7.995446444 | 1.642976044 | 3.00E-02 |
| LINC00243 | 39.0991338 | 1.642747898 | 1.55E-05 |
| GSTO1 | 3403.432352 | 1.642634003 | 3.95E-52 |
| MUC16 | 88.7544573 | 1.640849273 | 4.92E-03 |
| GABRR2 | 34.6200028 | 1.640199087 | 1.63E-03 |
| DMGDH | 358.5659548 | 1.63886567 | 1.05E-11 |
| AL078590.3 | 45.52013897 | 1.638709305 | 2.67E-07 |
| AC024022.1 | 9.387587135 | 1.637536648 | 7.68E-03 |
| COL28A1 | 18.51399564 | 1.637334573 | 1.02E-03 |
| CEBPG | 1838.694519 | 1.635044377 | 4.45E-10 |

| | | | |
|---|---|---|---|
| AL035530.2 | 7.570460569 | 1.634334445 | 2.97E-02 |
| FLNB | 29152.77127 | 1.634231923 | 1.33E-35 |
| CDKN2B | 1954.875598 | 1.633431743 | 5.73E-14 |
| GJA3 | 323.6162197 | 1.632804844 | 1.09E-43 |
| AC005479.1 | 7.73917013 | 1.630237318 | 2.90E-02 |
| SLC2A9 | 168.7358437 | 1.6278492 | 1.25E-23 |
| NUP50-DT | 172.5240029 | 1.627166893 | 3.32E-14 |
| TBX15 | 27.29113919 | 1.62615701 | 2.84E-03 |
| AC010547.4 | 58.16470255 | 1.624556059 | 1.31E-03 |
| MYADM | 3260.603661 | 1.622775922 | 7.29E-47 |
| AL353751.1 | 231.7121403 | 1.620374116 | 2.45E-11 |
| ZMYND12 | 26.09480414 | 1.620360596 | 2.15E-03 |
| GNAL | 206.461877 | 1.620308088 | 2.31E-24 |
| SPINT1 | 91.07398261 | 1.61975006 | 9.84E-08 |
| IL3RA | 225.9906258 | 1.619686904 | 8.11E-23 |
| IER3 | 2184.936652 | 1.618783256 | 1.32E-37 |
| PNMA8B | 17.9584709 | 1.618662884 | 2.03E-02 |
| GABRA2 | 53.37695158 | 1.617815241 | 2.77E-05 |
| WWC2-AS2 | 13.98774191 | 1.617468807 | 3.53E-03 |
| ZP1 | 57.88032852 | 1.615366998 | 1.01E-02 |
| TMED10P2 | 29.80088923 | 1.614640584 | 1.23E-05 |
| AP002884.1 | 752.7599133 | 1.613424152 | 2.38E-32 |
| AC068580.1 | 7.569025765 | 1.613052475 | 2.12E-02 |
| NLRP12 | 80.46937671 | 1.612761351 | 9.75E-06 |
| SIGLEC5 | 138.3515049 | 1.612317623 | 7.23E-19 |
| SVEP1 | 143.2123067 | 1.611916112 | 1.89E-07 |
| CEACAMP7 | 24.51020739 | 1.611863485 | 4.06E-02 |
| SLFN5 | 2817.627108 | 1.611233011 | 1.88E-21 |
| LINC00869 | 134.1130625 | 1.60950914 | 7.07E-05 |
| ATP6V1C1 | 2317.987931 | 1.608665454 | 1.84E-28 |
| TSPAN1 | 25.18067891 | 1.608513311 | 1.14E-04 |
| MTFR2 | 416.4778021 | 1.607964067 | 1.19E-27 |
| ADAMTS1 | 240.3419719 | 1.607883782 | 8.20E-09 |
| TMEM156 | 66.58521868 | 1.607876848 | 7.64E-04 |
| TSKS | 10.55565234 | 1.606943623 | 2.84E-02 |
| IFI44L | 45.59889335 | 1.606197561 | 2.22E-02 |
| STX3 | 2995.088942 | 1.606128097 | 1.05E-72 |
| HSD11B1-AS1 | 25.91185308 | 1.605908765 | 9.40E-04 |
| IDH1 | 9760.191328 | 1.605594877 | 6.94E-41 |
| ITGB1-DT | 24.64010817 | 1.605591294 | 1.99E-04 |
| AC005540.1 | 17.4025111 | 1.604675089 | 1.59E-02 |
| OPTN | 3436.814398 | 1.602589947 | 1.48E-65 |
| PRSS50 | 13.04396709 | 1.602466987 | 4.07E-03 |
| THBS1 | 3896.636739 | 1.602463656 | 5.64E-46 |
| FAM215A | 50.51698922 | 1.601128966 | 5.35E-05 |
| MYL3 | 32.80788903 | 1.600762983 | 5.73E-04 |
| HLA-DPA1 | 429.1936339 | 1.600720374 | 4.63E-11 |

| | | | |
|---|---|---|---|
| SLC16A11 | 75.82061411 | 1.599483369 | 1.09E-12 |
| SYDE2 | 235.870281 | 1.598244045 | 1.15E-11 |
| RNF6 | 714.398433 | 1.592821487 | 1.43E-07 |
| TMEM179 | 21.2981889 | 1.592534586 | 1.57E-03 |
| CLIP1 | 2506.614825 | 1.592436122 | 1.75E-17 |
| ACTN3 | 14.19951352 | 1.591480154 | 6.30E-03 |
| IGF2R | 12286.58132 | 1.590054364 | 6.69E-45 |
| ARHGEF10 | 3349.754618 | 1.589110071 | 1.15E-53 |
| ALPK2 | 2595.841625 | 1.585857684 | 4.62E-12 |
| MRAS | 1665.819629 | 1.583329445 | 6.56E-50 |
| ELFN2 | 490.7239094 | 1.582635489 | 1.70E-12 |
| AC121757.1 | 28.94935304 | 1.582316016 | 6.79E-03 |
| ST6GALNAC3 | 127.2482074 | 1.58207844 | 3.77E-06 |
| TSHZ3 | 74.67057501 | 1.581762356 | 4.55E-07 |
| AP001596.1 | 20.39713961 | 1.581733458 | 4.90E-04 |
| SPANXA1 | 416.0491261 | 1.580829551 | 1.67E-14 |
| NUDT16P1 | 19.55888412 | 1.579690394 | 4.62E-04 |
| MAGEA4 | 9.336679457 | 1.579065914 | 3.63E-02 |
| AC025580.2 | 19.20515102 | 1.577037357 | 8.76E-03 |
| LONRF1 | 546.4227996 | 1.576818807 | 2.93E-28 |
| SLC19A3 | 219.3959215 | 1.5763944 | 1.35E-31 |
| LRRC37A7P | 23.79231287 | 1.576024753 | 3.66E-05 |
| SAMD4A | 2581.581522 | 1.573953635 | 2.48E-36 |
| IL2RG | 101.3761071 | 1.57390293 | 9.66E-09 |
| AC007068.1 | 23.29256962 | 1.572305266 | 6.08E-04 |
| AL513165.2 | 33.63872056 | 1.572243855 | 4.48E-03 |
| AC012158.1 | 8.167687206 | 1.571693107 | 3.51E-02 |
| SAT2 | 1759.277794 | 1.57164847 | 1.91E-34 |
| FAM241B | 259.3376186 | 1.571438933 | 1.14E-16 |
| MAPK13 | 160.0474767 | 1.570920325 | 8.68E-18 |
| FAM27B | 30.52650418 | 1.570730707 | 2.93E-06 |
| RNF122 | 196.3554192 | 1.57036563 | 3.87E-19 |
| SMCO4 | 160.0160317 | 1.570036184 | 6.55E-08 |
| NOC2LP1 | 35.01371521 | 1.567555836 | 4.28E-04 |
| HEPH | 29.34641488 | 1.567460288 | 1.87E-03 |
| AC243830.2 | 13.75084541 | 1.565670073 | 1.04E-02 |
| LINC01354 | 15.60851471 | 1.565668995 | 5.01E-03 |
| AC083862.2 | 85.25145897 | 1.565040889 | 4.83E-14 |
| SLC39A14 | 6805.43364 | 1.564906722 | 1.55E-45 |
| AC105411.1 | 36.104077 | 1.563808942 | 6.66E-03 |
| POSTN | 840.5191803 | 1.563161271 | 3.77E-05 |
| EPM2A | 231.7732836 | 1.562466772 | 2.45E-33 |
| FAM155B | 469.1044739 | 1.562373504 | 1.29E-41 |
| FMN2 | 287.7331264 | 1.561539776 | 4.06E-09 |
| RTN4 | 17887.31892 | 1.560851145 | 6.80E-56 |
| VXN | 179.481724 | 1.56034171 | 3.32E-13 |
| CTSL | 7027.922032 | 1.559928366 | 4.24E-196 |

| | | | |
|---|---|---|---|
| MSRB3 | 1780.19995 | 1.5565635 | 3.11E-42 |
| FTL | 97215.3597 | 1.55601543 | 2.00E-31 |
| AC023906.5 | 19.01930279 | 1.555426683 | 1.66E-03 |
| CCL5 | 161.9253664 | 1.554054274 | 4.18E-08 |
| AL590666.2 | 45.63170054 | 1.553869045 | 4.01E-06 |
| RFTN1 | 2508.131971 | 1.553417001 | 1.40E-45 |
| PEA15 | 12565.60558 | 1.552644688 | 3.29E-32 |
| TPK1 | 102.279142 | 1.552061353 | 4.64E-11 |
| CLCNKB | 120.2619914 | 1.551735427 | 7.10E-12 |
| TCAF2P1 | 28.17114646 | 1.551712874 | 4.53E-03 |
| AC104435.2 | 31.56962772 | 1.551549137 | 2.81E-05 |
| ITGAX | 188.9758 | 1.551293357 | 6.95E-17 |
| IL23R | 9.827229111 | 1.547794616 | 2.09E-02 |
| KRT8P12 | 423.0876632 | 1.547661546 | 2.84E-47 |
| CEACAM21 | 17.26810419 | 1.54746194 | 1.51E-03 |
| ASPA | 207.1723575 | 1.546859986 | 1.54E-25 |
| EDEM1 | 2254.97478 | 1.545551536 | 1.16E-53 |
| PKP1 | 27.47331584 | 1.545019732 | 1.15E-04 |
| NA | 13.14292939 | 1.542299458 | 4.67E-02 |
| AC139149.1 | 11.92405406 | 1.540344234 | 3.09E-02 |
| CYSTM1 | 1163.563966 | 1.539650782 | 1.33E-35 |
| MTX1P1 | 58.64938482 | 1.538861405 | 1.16E-07 |
| LYST | 1092.085017 | 1.536558073 | 1.81E-14 |
| SORL1 | 4958.874962 | 1.536327103 | 1.22E-34 |
| RND1 | 94.31001901 | 1.535397128 | 3.76E-08 |
| FOXD3 | 10.61840442 | 1.535390922 | 9.09E-03 |
| C4orf54 | 44.10009235 | 1.534391036 | 8.93E-07 |
| CDH15 | 192.4055587 | 1.534355794 | 9.73E-12 |
| XPOT | 9329.971461 | 1.533517966 | 1.09E-19 |
| SUGT1P1 | 47.60383308 | 1.532286751 | 1.94E-06 |
| NXPH4 | 627.4375109 | 1.532114476 | 1.63E-22 |
| CES4A | 56.32910085 | 1.531643537 | 4.22E-08 |
| MPP7 | 42.74133974 | 1.531478677 | 3.28E-05 |
| MIR181A2HG | 24.1636788 | 1.530872184 | 1.88E-04 |
| AC012512.1 | 28.87379805 | 1.52970503 | 9.01E-05 |
| NXN | 1791.096427 | 1.527285754 | 2.00E-37 |
| KIF1A | 33.08770637 | 1.526885036 | 3.37E-03 |
| TUFT1 | 886.6362793 | 1.526812017 | 3.45E-37 |
| AL160191.3 | 10.7348622 | 1.526054645 | 1.75E-02 |
| PLXNC1 | 412.2176614 | 1.525985264 | 6.20E-16 |
| MCOLN2 | 40.13451401 | 1.52499706 | 6.99E-04 |
| CLTB | 1764.884731 | 1.524664989 | 9.33E-59 |
| MIXL1 | 18.82158452 | 1.523588268 | 1.42E-03 |
| PROSER2-AS1 | 38.08475189 | 1.52186198 | 2.45E-03 |
| FOS | 91.33837697 | 1.521643413 | 3.49E-04 |
| BMP7 | 617.2385659 | 1.520959265 | 7.39E-11 |
| ALS2CL | 852.7311576 | 1.519575589 | 6.58E-18 |

| | | | |
|---|---|---|---|
| GLP2R | 415.3640839 | 1.518894154 | 6.00E-07 |
| NPTX2 | 1549.746687 | 1.518790569 | 5.79E-19 |
| TFE3 | 7339.395033 | 1.516929617 | 9.93E-65 |
| AL161908.1 | 16.68269058 | 1.515403664 | 1.45E-02 |
| TMEM140 | 1535.959175 | 1.515056563 | 4.24E-47 |
| WHAMM | 724.8447879 | 1.514378994 | 2.75E-31 |
| LAMP2 | 7955.899773 | 1.512402662 | 9.55E-87 |
| RHCE | 20.12885806 | 1.512040886 | 3.36E-03 |
| JPH2 | 828.9286868 | 1.511775115 | 9.16E-18 |
| SESN2 | 1594.171601 | 1.509662466 | 3.81E-08 |
| TDRKH-AS1 | 27.67236504 | 1.508771818 | 4.85E-03 |
| UNC5B | 2009.275815 | 1.505878767 | 9.87E-06 |
| OSTM1 | 1362.323596 | 1.505472629 | 7.49E-27 |
| GOLT1A | 20.85450575 | 1.504707878 | 3.40E-03 |
| FTH1P8 | 14.91430424 | 1.503321435 | 2.64E-02 |
| MIR4482 | 11.97326268 | 1.502136733 | 5.25E-03 |
| HSPA9 | 36794.30816 | 1.501686722 | 4.27E-20 |
| GPR135 | 106.0111647 | 1.500826237 | 5.56E-11 |
| FOXE1 | 349.40179 | 1.499902651 | 4.77E-16 |
| IMMP2L | 342.1216284 | 1.499318114 | 1.03E-20 |
| SQSTM1 | 11897.91029 | 1.498475105 | 6.28E-32 |
| FICD | 254.4358081 | 1.498439642 | 4.45E-23 |
| KCNQ3 | 69.36936743 | 1.498130026 | 5.00E-03 |
| SLC46A3 | 306.615023 | 1.498063527 | 2.94E-22 |
| ZDHHC11B | 68.95118631 | 1.497215334 | 3.57E-07 |
| LINC00513 | 40.82278957 | 1.49707213 | 2.83E-05 |
| PIM1 | 1122.782319 | 1.496184939 | 1.92E-13 |
| NOXA1 | 57.91642516 | 1.495404803 | 4.16E-06 |
| ARHGAP42 | 723.8663377 | 1.49485001 | 5.67E-22 |
| CCSER2 | 913.7991343 | 1.494557646 | 8.30E-18 |
| GLRX2 | 325.8434628 | 1.494356635 | 2.70E-18 |
| GRHL1 | 65.06406828 | 1.493755454 | 2.82E-06 |
| AC009055.2 | 28.24339746 | 1.491590609 | 8.22E-03 |
| ADGRG6 | 2889.841449 | 1.491048654 | 2.02E-19 |
| PRKCE | 1501.609574 | 1.487310913 | 1.08E-12 |
| PRRG4 | 102.5023206 | 1.486069203 | 7.66E-10 |
| NKX2-2 | 22.54412958 | 1.485032815 | 5.79E-04 |
| FTCDNL1 | 96.17166783 | 1.483941115 | 1.01E-10 |
| RIPK2 | 724.4734466 | 1.482595228 | 3.13E-31 |
| ATP6V1B1-AS1 | 56.91216974 | 1.481963123 | 4.19E-06 |
| EPHX1 | 3232.5919 | 1.481031684 | 5.34E-42 |
| IL23A | 13.00103384 | 1.479027127 | 4.92E-03 |
| FAS | 239.2567132 | 1.478887562 | 1.72E-12 |
| AL590644.1 | 26.03942664 | 1.477084884 | 1.35E-02 |
| HLA-DMA | 431.2072074 | 1.475955815 | 7.40E-10 |
| GAL3ST2 | 8.135584626 | 1.474978287 | 4.99E-02 |
| CERS1 | 358.8475784 | 1.474877396 | 4.63E-06 |

| | | | |
|---|---|---|---|
| SUSD5 | 41.31474875 | 1.470835457 | 5.16E-07 |
| AL365181.3 | 171.3149321 | 1.469675503 | 9.72E-08 |
| KRT81 | 3638.578358 | 1.46905715 | 4.68E-08 |
| MRAP2 | 60.19755106 | 1.467039418 | 1.47E-05 |
| SLC35F6 | 2445.04949 | 1.466724367 | 8.71E-15 |
| AC138356.1 | 14.46702467 | 1.466346379 | 8.28E-04 |
| TNFSF4 | 327.5208002 | 1.466072313 | 9.15E-08 |
| BCL6 | 700.8646573 | 1.465380127 | 1.23E-45 |
| SCN1B | 334.4603313 | 1.464489679 | 1.08E-13 |
| ADGRD1-AS1 | 12.59987802 | 1.46402177 | 3.66E-03 |
| PXDC1 | 1898.914831 | 1.464004693 | 5.78E-60 |
| AL139260.1 | 63.32928104 | 1.46311365 | 4.30E-08 |
| CATSPERG | 35.31769266 | 1.462349022 | 1.77E-04 |
| FMOD | 1627.644591 | 1.462178176 | 2.12E-16 |
| UNC13D | 102.386188 | 1.462042648 | 4.77E-05 |
| AC096637.2 | 78.23099629 | 1.459608831 | 3.22E-12 |
| NEDD4L | 4687.21832 | 1.458057464 | 1.11E-51 |
| AC025857.2 | 32.64328677 | 1.457116757 | 2.79E-04 |
| LSMEM1 | 38.37769753 | 1.456925203 | 2.37E-03 |
| CACNA1H | 127.7563133 | 1.456403573 | 2.37E-06 |
| AOX1 | 235.0610148 | 1.456201093 | 2.79E-04 |
| SLC52A1 | 10.20398591 | 1.456075084 | 1.52E-02 |
| PTCH1 | 1620.679928 | 1.455093934 | 1.33E-18 |
| AL049650.1 | 34.41115523 | 1.454857267 | 1.49E-03 |
| NA | 52.26903518 | 1.4540127 | 3.92E-07 |
| SEMA4A | 163.4203738 | 1.45352004 | 3.33E-09 |
| AC087651.1 | 9.612092168 | 1.451256256 | 4.10E-02 |
| C1orf61 | 30.00348649 | 1.450500658 | 6.97E-05 |
| STOM | 8123.48438 | 1.450155304 | 5.04E-90 |
| NSUN7 | 276.9157398 | 1.45006461 | 2.56E-15 |
| TMEM192 | 1449.13137 | 1.448308286 | 9.76E-31 |
| SLPI | 49.88731299 | 1.448101545 | 1.04E-04 |
| EIF2S2 | 12053.63184 | 1.447878385 | 1.10E-25 |
| HEYL | 117.7585554 | 1.446060259 | 9.34E-05 |
| HHLA1 | 19.15672691 | 1.445122942 | 8.25E-03 |
| AL136084.3 | 71.27286851 | 1.444860747 | 8.93E-04 |
| RNF41 | 3274.28356 | 1.444728516 | 2.65E-17 |
| IL7R | 8419.815253 | 1.444019125 | 7.51E-06 |
| MAP7 | 29.05664401 | 1.443580091 | 2.56E-05 |
| ACE2 | 59.32408581 | 1.443077 | 7.90E-05 |
| SLC25A18 | 63.98096952 | 1.440713507 | 2.01E-06 |
| C4BPB | 51.0915616 | 1.440290748 | 3.49E-02 |
| ITPR1 | 1284.633357 | 1.440068965 | 4.61E-16 |
| AC147067.1 | 13.41097365 | 1.439160552 | 1.44E-02 |
| CDC37P1 | 25.58415421 | 1.437927574 | 5.81E-04 |
| AC093616.1 | 454.260492 | 1.43446561 | 4.33E-18 |
| PRELID3A | 164.0322339 | 1.433175403 | 2.40E-10 |

| | | | |
|---|---|---|---|
| KIF5C | 131.4518488 | 1.432943029 | 6.43E-06 |
| ARLNC1 | 37.47117289 | 1.432889625 | 4.55E-04 |
| SDK2 | 192.6470232 | 1.430542248 | 5.52E-05 |
| ANO6 | 6166.89251 | 1.430246488 | 1.03E-51 |
| FAM189A2 | 29.59519028 | 1.429187543 | 3.00E-03 |
| SPART | 2174.26844 | 1.428717342 | 2.66E-54 |
| WWC2 | 2643.317265 | 1.428628515 | 6.27E-16 |
| SRRM4 | 9.317143724 | 1.428321604 | 1.30E-02 |
| PNCK | 57.10837434 | 1.427863382 | 4.80E-03 |
| CSF2RB | 37.96421996 | 1.427852787 | 3.59E-03 |
| TMEM191C | 53.90717508 | 1.424086708 | 7.78E-04 |
| KDSR | 2931.962615 | 1.423279213 | 3.62E-69 |
| UFM1 | 957.9219923 | 1.422295763 | 1.28E-18 |
| SLC4A10 | 10.05535113 | 1.422029681 | 3.87E-02 |
| SLC52A3 | 59.35415736 | 1.421900945 | 5.35E-05 |
| SLC5A5 | 18.76023996 | 1.421610292 | 1.84E-02 |
| DCSTAMP | 13.89413438 | 1.421465375 | 5.00E-02 |
| LINC01088 | 28.81277826 | 1.419758822 | 1.09E-04 |
| AL354740.1 | 98.70817287 | 1.419466887 | 2.87E-04 |
| RPS6P8 | 16.72394368 | 1.418271091 | 8.31E-03 |
| FCMR | 73.10846455 | 1.41781856 | 9.59E-06 |
| AC104123.1 | 9.22245432 | 1.417590409 | 4.06E-02 |
| AC010307.2 | 7.62732822 | 1.416886295 | 3.16E-02 |
| SYBU | 311.9660073 | 1.416099076 | 1.87E-07 |
| LHFPL6 | 422.6305817 | 1.41585898 | 1.91E-22 |
| CHN2 | 294.5890476 | 1.415731724 | 1.05E-32 |
| CAMSAP3 | 101.4902695 | 1.414490822 | 3.20E-06 |
| RBM24 | 126.6531076 | 1.414082649 | 1.74E-05 |
| AC092287.1 | 8.384605128 | 1.413149215 | 2.95E-02 |
| PAGE4 | 12.69744612 | 1.412149455 | 1.49E-02 |
| LONP1 | 5588.880683 | 1.41211946 | 1.66E-46 |
| LINC02582 | 131.1487649 | 1.410182151 | 5.76E-09 |
| LAYN | 609.0277932 | 1.409029496 | 6.31E-18 |
| LINC02551 | 15.80481159 | 1.408915034 | 2.38E-02 |
| SYNPO | 2426.228123 | 1.408445403 | 5.63E-35 |
| RAB3IP | 248.084127 | 1.406615089 | 3.30E-09 |
| VAT1 | 16525.20215 | 1.4065107 | 6.15E-28 |
| ZNF385D-AS1 | 17.75479594 | 1.405942735 | 3.56E-03 |
| MIR646HG | 21.72389345 | 1.405250584 | 6.39E-03 |
| AC245100.1 | 8.802468238 | 1.405043453 | 3.87E-02 |
| OR2A1-AS1 | 214.8177946 | 1.404068744 | 3.28E-16 |
| LMCD1-AS1 | 48.74290335 | 1.403810382 | 1.21E-06 |
| ABRACL | 559.2061279 | 1.402255649 | 9.38E-27 |
| GARS | 17460.14145 | 1.401852686 | 2.56E-16 |
| DNAH14 | 338.8482397 | 1.400039772 | 1.51E-13 |
| AC108463.2 | 15.81970056 | 1.399958592 | 1.67E-02 |
| MCF2L | 207.9155546 | 1.399574278 | 3.08E-05 |

| | | | |
|---|---|---|---|
| MNX1-AS1 | 32.26355736 | 1.399187086 | 1.16E-04 |
| SERPINB8 | 817.2619824 | 1.398013549 | 2.45E-10 |
| SLC1A6 | 42.11919478 | 1.396198954 | 1.43E-04 |
| PLEKHF2 | 603.1114698 | 1.39582989 | 1.57E-17 |
| AL445490.1 | 16.22433025 | 1.394647665 | 6.03E-03 |
| CSMD1 | 132.1473999 | 1.393847565 | 7.59E-03 |
| MYT1 | 13.7927867 | 1.393650189 | 2.90E-02 |
| NECAB2 | 101.487437 | 1.391877624 | 1.21E-06 |
| CHST3 | 1381.804227 | 1.391873789 | 9.42E-41 |
| AL450345.1 | 18.11126184 | 1.390213604 | 7.37E-04 |
| FTH1 | 76509.70998 | 1.388208292 | 4.67E-30 |
| F2RL2 | 2246.079531 | 1.387590624 | 4.10E-33 |
| TPSB2 | 9.626241925 | 1.386197264 | 3.24E-02 |
| PLA2R1 | 868.557516 | 1.385332543 | 4.50E-24 |
| PRAP1 | 135.3358466 | 1.385031224 | 1.03E-04 |
| LYN | 763.4264982 | 1.383921042 | 4.87E-26 |
| AL391422.4 | 577.9328061 | 1.383894381 | 2.73E-10 |
| HEXB | 16325.57173 | 1.383755226 | 4.72E-54 |
| NA | 98.56486979 | 1.383280664 | 9.31E-13 |
| PYY | 26.72723626 | 1.383116284 | 4.20E-04 |
| AC010719.1 | 42.8172365 | 1.382277817 | 5.28E-05 |
| CDS1 | 52.31633794 | 1.38221477 | 3.02E-03 |
| NA | 141.3556843 | 1.38181287 | 6.62E-05 |
| CYTOR | 2035.603011 | 1.381393728 | 3.13E-39 |
| LY6E-DT | 189.8540731 | 1.381054605 | 8.98E-16 |
| HLA-V | 43.93514239 | 1.380709261 | 2.90E-03 |
| EYA4 | 172.560277 | 1.379778862 | 2.92E-05 |
| GALNT16 | 52.81985852 | 1.379176476 | 1.45E-04 |
| TMEM138 | 2071.515528 | 1.378549263 | 7.28E-61 |
| APOBR | 38.09673852 | 1.375797752 | 5.25E-06 |
| AL162171.1 | 1232.53051 | 1.375748917 | 3.73E-47 |
| SPOCK1 | 3684.499858 | 1.375563351 | 6.58E-46 |
| STAC | 1317.904973 | 1.374191155 | 7.49E-16 |
| LGALS8 | 2152.324426 | 1.373912432 | 4.99E-46 |
| ADGRG3 | 68.61698108 | 1.372259481 | 2.21E-08 |
| PQLC2L | 60.00408119 | 1.370684116 | 4.56E-09 |
| ADH5 | 8305.898574 | 1.369424174 | 1.81E-71 |
| SLC8A3 | 826.2647581 | 1.369404924 | 2.12E-16 |
| BACH1-IT1 | 12.98698502 | 1.369223485 | 2.32E-02 |
| MIR503HG | 2154.711243 | 1.369199385 | 3.33E-09 |
| AC078925.4 | 12.18734213 | 1.369141099 | 1.51E-02 |
| CHST15 | 1086.666004 | 1.368420889 | 5.32E-34 |
| TMEM86A | 534.2173541 | 1.367938549 | 2.96E-13 |
| GPD1 | 24.67354325 | 1.36732767 | 1.10E-03 |
| C4orf19 | 21.16468249 | 1.366483746 | 5.68E-03 |
| AL645939.2 | 14.40127914 | 1.364395639 | 2.69E-02 |
| MAN1C1 | 266.1644314 | 1.362229109 | 8.61E-10 |

| | | | |
|---|---|---|---|
| GATM | 9.70851274 | 1.361607006 | 2.25E-02 |
| MAOA | 392.4790427 | 1.361554964 | 1.12E-09 |
| PFKFB2 | 1199.134629 | 1.361500304 | 6.03E-30 |
| LINC02298 | 64.28724493 | 1.361270394 | 1.99E-07 |
| ITGBL1 | 61.58578191 | 1.360256698 | 1.45E-03 |
| AVPI1 | 366.8041458 | 1.359429399 | 1.72E-19 |
| ARHGAP20 | 387.949941 | 1.359251104 | 1.79E-11 |
| TGFB2-AS1 | 227.6156955 | 1.359174043 | 7.96E-11 |
| DDR2 | 12220.85219 | 1.358985702 | 4.43E-16 |
| SMTNL1 | 45.52353378 | 1.35858244 | 5.53E-05 |
| EPOR | 858.3410396 | 1.358448452 | 1.68E-26 |
| NFE2L1 | 19179.84364 | 1.354680369 | 1.10E-46 |
| SMOC2 | 58.39393359 | 1.352505799 | 4.57E-03 |
| IFIH1 | 381.1688632 | 1.352466209 | 1.37E-12 |
| MXRA7 | 5655.55128 | 1.351782142 | 4.78E-35 |
| AL031058.1 | 21.8809268 | 1.351525235 | 5.87E-03 |
| B4GALNT3 | 353.3939525 | 1.350398709 | 2.73E-22 |
| MAP1LC3B | 6359.449737 | 1.350289585 | 6.13E-41 |
| GCLM | 1084.260776 | 1.349588441 | 6.12E-17 |
| MIR4435-2HG | 2097.286563 | 1.349087559 | 2.65E-41 |
| AL121904.1 | 28.52617405 | 1.347308698 | 2.49E-03 |
| AC026436.1 | 24.5464379 | 1.346817529 | 3.00E-03 |
| DHRS11 | 725.2445982 | 1.346422179 | 1.08E-46 |
| ITGB2 | 124.990794 | 1.34529355 | 2.51E-04 |
| CTNNA1 | 21483.2315 | 1.345031922 | 7.32E-97 |
| AC011591.1 | 33.09399697 | 1.344928539 | 1.84E-02 |
| AC097059.2 | 16.24885375 | 1.344838761 | 3.04E-02 |
| ACAP1 | 278.1116252 | 1.344330385 | 8.03E-11 |
| NA | 158.6725789 | 1.343833657 | 6.30E-18 |
| AC068580.3 | 46.49094972 | 1.342119429 | 2.55E-05 |
| NXF3 | 122.3733355 | 1.340400403 | 4.97E-05 |
| PAG1 | 564.5162007 | 1.338610427 | 5.51E-11 |
| ABCC4 | 5268.937555 | 1.337365986 | 1.33E-58 |
| MICB | 1091.298993 | 1.337318082 | 2.17E-12 |
| TPM1 | 23789.68 | 1.33657541 | 1.10E-35 |
| EIF2S2P4 | 29.80356116 | 1.336391483 | 1.61E-02 |
| SLC2A4 | 550.3986875 | 1.336319128 | 4.16E-17 |
| MARVELD3 | 105.4161867 | 1.33581724 | 2.94E-10 |
| CYREN | 3033.595272 | 1.333068503 | 4.40E-18 |
| WBP2 | 3411.363076 | 1.332792414 | 5.64E-37 |
| FAM45A | 1390.912981 | 1.332053819 | 1.15E-52 |
| KIAA1210 | 25.70861216 | 1.331442866 | 4.43E-03 |
| SLC6A20 | 25.26900586 | 1.329070097 | 1.91E-02 |
| AC011511.5 | 30.79983502 | 1.328573417 | 1.81E-04 |
| IRAK4 | 600.6550032 | 1.326874197 | 1.42E-22 |
| AC112220.2 | 53.09066079 | 1.326407201 | 3.08E-07 |
| NA | 149.0104143 | 1.325208868 | 1.88E-10 |

| | | | |
|---|---|---|---|
| ATP1A2 | 143.772855 | 1.324074193 | 3.57E-03 |
| NEXN | 357.8514529 | 1.32365974 | 1.37E-13 |
| HTATSF1P2 | 155.7800433 | 1.322615536 | 7.04E-12 |
| THNSL2 | 53.591121 | 1.322535112 | 2.51E-04 |
| KLF8 | 39.97652723 | 1.322187832 | 1.82E-05 |
| FUT1 | 131.4538281 | 1.321349795 | 1.32E-04 |
| C3orf20 | 24.41494674 | 1.321006963 | 1.27E-02 |
| PTP4A1 | 2535.03776 | 1.320487085 | 1.50E-06 |
| EFCAB5 | 26.20293695 | 1.320422048 | 5.59E-03 |
| GALNT5 | 69.69342666 | 1.320170517 | 2.12E-05 |
| AL158825.2 | 9.612895418 | 1.318550699 | 4.08E-02 |
| DSEL | 19.97265315 | 1.318367831 | 4.64E-03 |
| ATP6V1E1 | 4139.268406 | 1.317515399 | 1.78E-91 |
| MUC3A | 74.07583469 | 1.317446064 | 1.14E-02 |
| CEBPB | 1453.556869 | 1.317084198 | 4.56E-08 |
| WDSUB1 | 397.7929314 | 1.316709789 | 4.06E-29 |
| SORT1 | 7230.535163 | 1.316206531 | 1.27E-44 |
| ABCC9 | 49.86244202 | 1.316096725 | 1.10E-03 |
| ARHGDIG | 19.83265057 | 1.315686985 | 1.27E-02 |
| KCNQ4 | 29.93609954 | 1.315302402 | 9.82E-05 |
| MORN3 | 30.72541857 | 1.314982997 | 2.84E-05 |
| NATD1 | 1387.904722 | 1.314884665 | 5.64E-22 |
| AF196972.1 | 16.53544774 | 1.314788391 | 6.83E-03 |
| CERNA1 | 67.95049957 | 1.314136846 | 2.23E-07 |
| NR1D2 | 2187.670759 | 1.313873873 | 1.21E-12 |
| TRIM37 | 2440.200354 | 1.312874502 | 3.76E-39 |
| HS6ST2 | 758.5218787 | 1.310468212 | 8.65E-18 |
| LACTB2 | 789.8368156 | 1.309778268 | 1.72E-29 |
| INAVA | 330.5912767 | 1.308607516 | 3.57E-12 |
| CEBPD | 169.0188464 | 1.308077288 | 6.92E-10 |
| KLHL29 | 1265.210865 | 1.307548046 | 1.73E-18 |
| COL15A1 | 15563.86287 | 1.307425347 | 1.15E-07 |
| AL161772.1 | 38.97171819 | 1.306206419 | 9.59E-05 |
| ANXA1 | 9614.08568 | 1.304616588 | 7.62E-10 |
| PIGY | 36.65229495 | 1.304429052 | 2.67E-02 |
| AC010186.2 | 280.3512769 | 1.303627091 | 8.24E-15 |
| SLC30A1 | 650.6866827 | 1.303538152 | 8.95E-10 |
| DUSP5 | 787.2594332 | 1.302938139 | 2.80E-10 |
| TLR1 | 15.8981197 | 1.302623469 | 3.56E-02 |
| CHMP2B | 883.1991909 | 1.302029021 | 3.32E-31 |
| CHP2 | 31.8069951 | 1.301649089 | 6.48E-05 |
| HIST1H2BN | 83.96504118 | 1.301563599 | 1.31E-06 |
| C1R | 3793.027515 | 1.301355904 | 2.01E-17 |
| AC010601.1 | 55.14621582 | 1.301267865 | 1.07E-06 |
| P4HA2 | 4210.481934 | 1.300788204 | 4.18E-39 |
| MSANTD3 | 2120.168871 | 1.299513871 | 9.94E-46 |
| ZBTB47 | 508.2560449 | 1.298740457 | 8.68E-15 |

| | | | |
|---|---|---|---|
| KCNE4 | 2965.933508 | 1.297645115 | 1.24E-23 |
| PLAU | 3831.106497 | 1.294299599 | 1.06E-16 |
| SPANXB1 | 1009.08562 | 1.29421555 | 4.90E-23 |
| USP54 | 1239.206021 | 1.294172493 | 1.02E-43 |
| LYPLAL1 | 519.1981249 | 1.294074531 | 3.82E-24 |
| ARSG | 201.1469179 | 1.293489823 | 3.24E-14 |
| P4HA1 | 2000.926193 | 1.292957351 | 6.30E-35 |
| GAD1 | 53.22551783 | 1.292779793 | 1.31E-05 |
| HLA-L | 166.7642701 | 1.292189066 | 1.46E-06 |
| CDC42EP1 | 3906.296252 | 1.29127043 | 2.02E-12 |
| AARS | 23738.39566 | 1.290960952 | 1.32E-15 |
| C20orf202 | 13.62698784 | 1.290902168 | 2.25E-02 |
| FLJ45513 | 22.04428444 | 1.290335976 | 2.00E-02 |
| METTL21A | 721.6296818 | 1.287782433 | 4.80E-28 |
| RSPO4 | 543.3235221 | 1.287753535 | 1.30E-27 |
| PLEKHF1 | 202.8274138 | 1.286891439 | 6.01E-05 |
| KIAA1324 | 83.13749574 | 1.286576241 | 7.71E-05 |
| ZNF469 | 430.1866375 | 1.286291157 | 4.49E-11 |
| RRAS2 | 2136.436366 | 1.285937127 | 3.00E-41 |
| RAI14 | 6837.00494 | 1.284856627 | 1.90E-14 |
| PCNX4 | 3959.147168 | 1.283803288 | 2.00E-20 |
| ECHDC1 | 1506.612531 | 1.283670179 | 1.37E-13 |
| UBE2J1 | 3574.235135 | 1.283348436 | 1.14E-66 |
| GPM6A | 20.45239369 | 1.282484355 | 2.60E-02 |
| ZMAT1 | 38.3723145 | 1.282302248 | 1.69E-03 |
| C16orf71 | 39.22617546 | 1.282159014 | 1.92E-04 |
| ADAMTS16 | 718.0575453 | 1.281765673 | 1.09E-07 |
| AC006252.1 | 66.20634047 | 1.281638737 | 5.24E-05 |
| AGAP11 | 44.86727608 | 1.279946353 | 1.76E-04 |
| FOXD3-AS1 | 24.57546857 | 1.27937188 | 3.53E-03 |
| SERPINH1 | 45395.88893 | 1.278681967 | 4.90E-53 |
| TMEM74B | 138.1700874 | 1.278448399 | 1.11E-04 |
| P2RY6 | 13.91387081 | 1.278071525 | 3.32E-02 |
| TUBE1 | 792.6081694 | 1.277268504 | 3.44E-08 |
| HSPA5 | 56281.23694 | 1.277252433 | 9.43E-27 |
| NMRK1 | 236.4018946 | 1.276238236 | 5.01E-17 |
| CCL26 | 75.15112479 | 1.276226923 | 9.37E-05 |
| ECM2 | 30.17136374 | 1.275498421 | 2.32E-03 |
| TMEFF1 | 177.2918341 | 1.27445004 | 3.14E-04 |
| PLCG2 | 1243.876311 | 1.272575304 | 5.45E-17 |
| AL139352.1 | 15.41388576 | 1.272038709 | 3.33E-02 |
| CCDC9B | 1980.078368 | 1.270712268 | 2.13E-17 |
| IQANK1 | 28.11485551 | 1.270653952 | 2.83E-02 |
| GRN | 14788.05412 | 1.26989912 | 1.45E-17 |
| SH3RF2 | 238.2640979 | 1.269275011 | 1.16E-10 |
| FTH1P2 | 16.42404809 | 1.268347561 | 3.64E-02 |
| PRSS3 | 32.73514005 | 1.267967279 | 1.18E-03 |

| | | | |
|---|---|---|---|
| NOTCH3 | 3538.212932 | 1.267816555 | 8.45E-14 |
| LTF | 52.07108289 | 1.267729005 | 1.98E-02 |
| RBM20 | 491.8489347 | 1.267303372 | 1.89E-13 |
| ITGA3 | 22276.61878 | 1.26640558 | 6.48E-35 |
| SNAPC1 | 2436.405541 | 1.2662474 | 1.55E-09 |
| PSD2 | 18.55882236 | 1.266241491 | 1.24E-02 |
| NPC1 | 3213.854759 | 1.266075649 | 4.87E-44 |
| CNST | 892.6135292 | 1.265369222 | 6.77E-18 |
| FRRS1 | 170.6658154 | 1.264755468 | 2.54E-05 |
| C12orf66 | 315.7310445 | 1.26473512 | 8.78E-20 |
| DUSP3 | 4615.69139 | 1.264621862 | 3.90E-48 |
| LDLRAD4 | 335.4509987 | 1.264558659 | 9.96E-16 |
| NA | 12.0417174 | 1.264334052 | 4.61E-02 |
| C2 | 638.2838083 | 1.263851795 | 1.44E-06 |
| EXOC3L4 | 86.63631761 | 1.261960233 | 1.02E-10 |
| ZEB2 | 1231.237831 | 1.261631265 | 1.23E-10 |
| LINC01666 | 39.06634075 | 1.261585782 | 3.15E-05 |
| UPK1A | 68.1821967 | 1.260550445 | 1.22E-03 |
| TNFRSF10B | 5148.799909 | 1.259905365 | 1.85E-27 |
| CU633904.1 | 124.7368645 | 1.259350701 | 1.03E-02 |
| TMEM178B | 177.5366143 | 1.258395052 | 1.52E-03 |
| EFR3B | 66.27859465 | 1.258280721 | 7.96E-05 |
| PAQR5 | 477.0127961 | 1.255642238 | 5.21E-10 |
| FMN1 | 253.4874342 | 1.255583246 | 1.16E-05 |
| SDC1 | 3446.919541 | 1.254517015 | 4.24E-21 |
| ENO2 | 1006.576172 | 1.254354702 | 1.95E-08 |
| ENPP7 | 27.35216795 | 1.253716492 | 1.03E-03 |
| GEM | 363.9867785 | 1.253682023 | 1.24E-09 |
| ZFAND3 | 4293.104697 | 1.251075025 | 1.20E-19 |
| LINC00261 | 28.26260737 | 1.250414048 | 5.23E-03 |
| ZNF711 | 258.9397943 | 1.25040494 | 3.17E-05 |
| AC092645.1 | 38.54954026 | 1.24811817 | 2.22E-02 |
| CLEC11A | 1129.023465 | 1.248005845 | 1.60E-14 |
| RELB | 233.1853472 | 1.246873852 | 2.01E-05 |
| AL450124.1 | 315.5916599 | 1.246322946 | 3.46E-22 |
| TEAD4 | 1502.038523 | 1.245989113 | 1.02E-35 |
| DNASE2 | 2894.136866 | 1.245699596 | 1.19E-17 |
| RAG1 | 104.3953946 | 1.245522049 | 6.79E-07 |
| CDC37P2 | 28.65264598 | 1.244972067 | 9.39E-03 |
| CPEB1-AS1 | 15.47801072 | 1.244024474 | 4.23E-02 |
| LINC02289 | 34.71716412 | 1.243762207 | 2.81E-03 |
| FAM83H | 345.1934365 | 1.243599013 | 6.16E-23 |
| SBF2-AS1 | 216.8026323 | 1.242718318 | 8.56E-09 |
| HOXA1 | 545.3612809 | 1.242660095 | 1.85E-30 |
| EDNRA | 37.02980821 | 1.242576987 | 3.70E-02 |
| AC010618.4 | 100.5779133 | 1.240893349 | 1.22E-05 |
| FGF13 | 17.17901773 | 1.239827048 | 1.06E-02 |

| | | | |
|---|---|---|---|
| NA | 32.99010685 | 1.239474411 | 2.91E-03 |
| MGST1 | 2201.138041 | 1.238838177 | 1.55E-37 |
| CES3 | 106.6239909 | 1.23879375 | 4.06E-06 |
| SMIM3 | 518.8367427 | 1.238482266 | 1.57E-20 |
| OSMR | 15770.68321 | 1.237806056 | 1.92E-25 |
| AC022784.1 | 65.50596088 | 1.236905072 | 1.99E-04 |
| SLC7A7 | 165.6952727 | 1.235550525 | 2.81E-08 |
| LINC02438 | 24.09064818 | 1.234369404 | 3.06E-03 |
| FAM27C | 79.29343533 | 1.233624119 | 1.20E-08 |
| AC020934.1 | 20.74450596 | 1.23205419 | 3.23E-02 |
| TP53I11 | 1725.487319 | 1.231446317 | 7.95E-30 |
| MBNL2 | 5478.880166 | 1.231430013 | 3.16E-09 |
| LINC01611 | 16.09218297 | 1.231393516 | 1.87E-02 |
| IPMK | 190.0956108 | 1.231233626 | 3.71E-09 |
| IL1RAP | 613.1327742 | 1.230353466 | 2.59E-14 |
| SOHLH2 | 153.5629023 | 1.229220451 | 3.28E-09 |
| ESRG | 287.4514818 | 1.228625671 | 3.65E-09 |
| VTN | 154.8509553 | 1.227999277 | 1.47E-04 |
| CALHM6 | 28.56829687 | 1.22764696 | 4.40E-03 |
| FAM171B | 279.1691248 | 1.227309383 | 9.23E-05 |
| BNC1 | 1562.452238 | 1.226491598 | 4.97E-09 |
| TDH | 10.90157754 | 1.226257031 | 2.38E-02 |
| ZNF280A | 323.1856629 | 1.226228407 | 6.76E-15 |
| CCL28 | 830.9079579 | 1.226186152 | 4.95E-30 |
| FTSJ1 | 3401.260365 | 1.225756341 | 3.23E-59 |
| ASTN1 | 2110.214754 | 1.225754582 | 2.04E-22 |
| IARS | 24186.3385 | 1.225731677 | 1.23E-20 |
| SLC1A4 | 2242.247256 | 1.224764205 | 1.71E-07 |
| NCK1-DT | 221.6192102 | 1.224044795 | 1.01E-21 |
| NA | 19.13286646 | 1.223198588 | 1.00E-02 |
| LINC01278 | 1055.169286 | 1.22185263 | 3.64E-54 |
| MIA2 | 1072.719923 | 1.221562 | 1.43E-10 |
| TLR2 | 80.40836449 | 1.220654201 | 4.26E-05 |
| DNAJC3 | 5598.113646 | 1.220415143 | 3.41E-42 |
| ADAMTSL3 | 34.09242262 | 1.219955383 | 6.92E-03 |
| FAM43A | 796.7891411 | 1.21922194 | 2.02E-27 |
| APBA1 | 534.1102272 | 1.218822718 | 2.06E-11 |
| IGFBPL1 | 580.1775613 | 1.218458601 | 6.53E-21 |
| CTBS | 682.3989996 | 1.217600177 | 7.44E-36 |
| HK2 | 2102.950957 | 1.217043785 | 2.90E-18 |
| CACNG6 | 238.4323998 | 1.216284021 | 4.89E-09 |
| COQ10B | 657.958327 | 1.214329082 | 1.72E-25 |
| RNY3P8 | 37.84131551 | 1.214321363 | 3.30E-04 |
| PLCE1 | 1326.090032 | 1.213470989 | 4.38E-27 |
| CCT6B | 86.79091986 | 1.212467106 | 8.90E-09 |
| ONECUT2 | 116.8377289 | 1.212439841 | 3.07E-02 |
| SH2D5 | 415.6853727 | 1.212366663 | 4.52E-20 |

| | | | |
|---|---|---|---|
| TRBV20-1 | 248.8926942 | 1.212313091 | 3.05E-14 |
| RHOQ | 3625.302826 | 1.212189992 | 2.18E-71 |
| AC068987.4 | 22.16306705 | 1.212175493 | 6.38E-03 |
| LRG1 | 182.2253611 | 1.211223876 | 6.91E-05 |
| MYCBP2 | 9079.698849 | 1.210110336 | 5.03E-20 |
| RB1CC1 | 3116.397545 | 1.210066531 | 5.99E-06 |
| PIK3CG | 245.0376781 | 1.209840163 | 1.72E-25 |
| SYNE1 | 3339.578059 | 1.207964486 | 6.87E-10 |
| KCNH5 | 165.5275543 | 1.207694451 | 1.97E-07 |
| AC009502.1 | 25.57399259 | 1.206912526 | 4.13E-02 |
| HAR1A | 46.38065491 | 1.206054828 | 1.11E-03 |
| FADS3 | 3833.388893 | 1.205900817 | 6.32E-34 |
| C11orf74 | 508.0284312 | 1.205866212 | 1.18E-12 |
| SORBS2 | 136.3448756 | 1.205253548 | 1.14E-04 |
| NA | 43.53099848 | 1.204930992 | 9.49E-04 |
| IGSF1 | 60.08272131 | 1.204659208 | 2.28E-05 |
| STEAP1 | 213.0817292 | 1.204442477 | 4.61E-11 |
| KLHL5 | 3640.730673 | 1.204372528 | 2.32E-16 |
| CTSB | 51746.63918 | 1.20430749 | 5.76E-21 |
| CPEB2 | 156.4561522 | 1.203614876 | 3.14E-04 |
| CD79B | 132.1895311 | 1.203545355 | 1.69E-09 |
| SARS | 9946.604605 | 1.203345742 | 1.34E-16 |
| RNF187 | 6542.358528 | 1.203160869 | 1.55E-31 |
| CYS1 | 92.6595099 | 1.202780276 | 1.74E-07 |
| PGM3 | 2990.489355 | 1.201297974 | 6.25E-20 |
| SVIP | 315.9252992 | 1.200279499 | 1.42E-11 |
| UPK1A-AS1 | 54.35396382 | 1.200172739 | 1.09E-02 |
| RBM47 | 230.5145896 | 1.200135429 | 4.46E-14 |
| MOCOS | 902.9342014 | 1.199705631 | 1.16E-06 |
| SLC2A13 | 99.18506297 | 1.199624755 | 4.12E-08 |
| THAP9-AS1 | 1053.717623 | 1.199621976 | 4.56E-07 |
| ISPD | 52.78470494 | 1.197637075 | 5.79E-06 |
| TMEM266 | 66.73934989 | 1.196976453 | 1.56E-06 |
| NA | 29.32595176 | 1.196686646 | 1.54E-02 |
| FAM91A1 | 3131.372127 | 1.195662201 | 8.68E-11 |
| PRDM5 | 571.6075791 | 1.195628181 | 3.21E-19 |
| BEX4 | 216.329814 | 1.195070828 | 9.19E-14 |
| PHYH | 939.2769998 | 1.194670021 | 2.53E-51 |
| LINC02256 | 15.471134 | 1.19340006 | 4.46E-02 |
| NUDT18 | 423.799999 | 1.193277192 | 4.16E-12 |
| LINC01275 | 38.03555998 | 1.193097564 | 5.99E-04 |
| ARHGEF26 | 403.7307648 | 1.192238257 | 3.05E-14 |
| CD72 | 28.97096591 | 1.191912107 | 1.52E-02 |
| TACC1 | 7595.804216 | 1.190369067 | 4.05E-28 |
| CTSC | 4818.804818 | 1.190076601 | 3.17E-22 |
| AC008686.1 | 60.91805457 | 1.189943921 | 2.31E-05 |
| AC012501.2 | 74.6821662 | 1.189669037 | 8.44E-08 |

| | | | |
|---|---|---|---|
| STON1 | 979.0446458 | 1.189566136 | 3.29E-29 |
| LINC01301 | 23.18791021 | 1.189004666 | 1.07E-02 |
| TENT5C | 137.8763368 | 1.187607329 | 1.87E-09 |
| PNMA6A | 795.9455409 | 1.187380245 | 3.99E-17 |
| TEAD1 | 10939.96256 | 1.187366447 | 1.64E-16 |
| RN7SL689P | 27.47548498 | 1.186027438 | 3.20E-03 |
| AHNAK2 | 1587.711431 | 1.184558013 | 4.69E-12 |
| ADAP1 | 466.5208821 | 1.18415622 | 4.52E-19 |
| PAM | 16151.86896 | 1.183564214 | 1.67E-33 |
| REEP3 | 450.3834737 | 1.1832298 | 4.62E-17 |
| DLGAP1-AS1 | 142.860099 | 1.182975185 | 1.65E-09 |
| NNT-AS1 | 922.7219592 | 1.182513281 | 3.56E-13 |
| SEL1L | 4823.460727 | 1.181186057 | 5.70E-17 |
| ZFYVE28 | 231.0358165 | 1.181035586 | 3.26E-14 |
| AL355601.1 | 25.61358171 | 1.18029744 | 5.10E-03 |
| UHRF1BP1 | 4704.980666 | 1.180034585 | 1.75E-12 |
| SLC31A1 | 4679.004963 | 1.179942487 | 5.24E-50 |
| RRN3P3 | 316.7446903 | 1.179848927 | 1.36E-06 |
| PRKAG2 | 1255.564186 | 1.178963824 | 7.48E-18 |
| MICA | 1140.36943 | 1.178546319 | 2.59E-11 |
| PPIC | 3752.098519 | 1.178472357 | 1.88E-51 |
| FOXN3-AS1 | 91.62970335 | 1.176844204 | 5.19E-07 |
| CIPC | 1125.986789 | 1.176763062 | 7.58E-35 |
| RAB17 | 2197.833679 | 1.175923963 | 3.48E-14 |
| AC083805.2 | 26.76387639 | 1.175209432 | 1.32E-03 |
| GFRA2 | 79.95301758 | 1.175122626 | 9.23E-05 |
| AL160191.1 | 17.79750141 | 1.173700725 | 2.53E-02 |
| C2orf88 | 65.43652657 | 1.173117479 | 1.34E-06 |
| DAG1 | 8402.798769 | 1.172653743 | 2.71E-27 |
| KIF26B | 25.08496738 | 1.172521272 | 7.73E-03 |
| ADGRB1 | 36.18287436 | 1.172438148 | 2.02E-02 |
| INSYN2B | 326.6455031 | 1.172267644 | 4.87E-06 |
| NFKBIE | 433.0629905 | 1.171383388 | 3.08E-10 |
| BACH1 | 2018.700707 | 1.171362318 | 8.09E-10 |
| ABCA1 | 1561.312943 | 1.170975453 | 1.34E-06 |
| AL356056.1 | 31.62686919 | 1.169226606 | 5.21E-03 |
| MCOLN3 | 35.09865512 | 1.168436932 | 3.17E-03 |
| MAPT | 583.6495694 | 1.167887455 | 7.72E-05 |
| CEBPA | 131.3984501 | 1.167821202 | 1.07E-06 |
| LGI3 | 21.16732851 | 1.16764587 | 4.14E-02 |
| USP2 | 233.5515925 | 1.167332262 | 4.36E-12 |
| FBXO32 | 739.3971269 | 1.167208827 | 8.04E-11 |
| XBP1 | 4964.531383 | 1.166284248 | 1.33E-32 |
| HLA-B | 7949.216215 | 1.165440422 | 1.24E-28 |
| BCL2 | 308.996463 | 1.164816339 | 7.95E-08 |
| DCAF8 | 5161.578887 | 1.164709129 | 7.91E-96 |
| AC243964.3 | 235.8240355 | 1.164608956 | 6.38E-12 |

| | | | |
|---|---|---|---|
| SLC1A5 | 12852.03691 | 1.164118498 | 8.54E-13 |
| CLN8 | 972.5212827 | 1.163616803 | 2.78E-23 |
| AC007389.1 | 30.05570423 | 1.162662287 | 7.93E-04 |
| SIRPA | 3488.218336 | 1.16116181 | 2.45E-19 |
| FOSL1 | 1816.844697 | 1.161091262 | 1.23E-05 |
| FBXO10 | 1900.060545 | 1.16066654 | 2.98E-15 |
| TDRD5 | 15.65920173 | 1.160584854 | 4.43E-02 |
| TCEA1 | 7064.069542 | 1.160553932 | 9.25E-08 |
| AL022323.1 | 13.31647754 | 1.159166106 | 3.78E-02 |
| PAIP1 | 7799.808941 | 1.157719062 | 6.03E-49 |
| AL603750.1 | 23.85849569 | 1.157143125 | 7.49E-03 |
| ZNF165 | 17.84093696 | 1.15701425 | 1.26E-02 |
| TSC22D2 | 1341.340198 | 1.15663176 | 1.32E-17 |
| ST8SIA2 | 194.4998905 | 1.15518774 | 4.23E-06 |
| NA | 18.99427585 | 1.153945621 | 2.19E-02 |
| FAM114A1 | 1772.247523 | 1.153173264 | 4.41E-17 |
| FENDRR | 1374.189237 | 1.152454163 | 7.23E-11 |
| SCOC | 836.9758959 | 1.152377674 | 2.67E-06 |
| JPH1 | 95.12718083 | 1.152354023 | 4.70E-06 |
| SHISA9 | 161.7235904 | 1.152157036 | 1.46E-04 |
| AL645568.1 | 62.75099522 | 1.151727767 | 1.88E-05 |
| JDP2 | 1542.857247 | 1.151712568 | 9.61E-08 |
| OPA1 | 2577.858291 | 1.150731173 | 7.99E-08 |
| ARHGAP31 | 2564.11328 | 1.150210051 | 5.20E-20 |
| RPP25 | 1596.165225 | 1.150034973 | 1.69E-46 |
| ZNF391 | 378.830913 | 1.149324261 | 2.35E-06 |
| PSG5 | 429.1534162 | 1.149090117 | 3.00E-03 |
| ZIC5 | 122.3265531 | 1.149000076 | 1.16E-06 |
| PCYT2 | 2161.944403 | 1.148969905 | 5.73E-14 |
| ACSL1 | 1243.219884 | 1.148815537 | 9.92E-42 |
| NA | 29.05715039 | 1.148471688 | 9.10E-04 |
| MEIOC | 288.8670266 | 1.14806498 | 9.14E-04 |
| SIL1 | 1229.856605 | 1.148004861 | 7.79E-45 |
| CNTNAP3 | 149.2453626 | 1.14715122 | 6.79E-08 |
| DNAJB9 | 824.6897093 | 1.146568712 | 7.47E-09 |
| SYPL2 | 83.161266 | 1.146192973 | 1.26E-05 |
| SLC19A2 | 517.6445248 | 1.145931724 | 3.23E-18 |
| CALU | 26908.66603 | 1.145849032 | 3.47E-20 |
| ARL4D | 533.0645091 | 1.144622518 | 4.43E-11 |
| ABHD4 | 4610.109295 | 1.143863046 | 1.36E-20 |
| FBLN7 | 185.287182 | 1.143823029 | 2.54E-08 |
| CAMSAP2 | 2288.391164 | 1.143580026 | 2.70E-11 |
| FAM167A | 682.8611599 | 1.142150369 | 5.16E-07 |
| NIPAL3 | 1794.482202 | 1.141754345 | 3.45E-45 |
| PLTP | 6996.727233 | 1.141428872 | 4.55E-16 |
| AGFG2 | 3050.773205 | 1.140481596 | 1.80E-13 |
| KSR2 | 287.877213 | 1.140103684 | 4.97E-06 |

| | | | |
|---|---|---|---|
| SLC25A32 | 1191.517078 | 1.139828819 | 6.48E-20 |
| CREB3L2 | 4451.672148 | 1.139240187 | 3.50E-47 |
| ARRDC4 | 850.4171308 | 1.139032436 | 2.36E-10 |
| CCM2L | 68.80571366 | 1.138419062 | 1.81E-03 |
| SPTB | 236.3609432 | 1.138217476 | 8.40E-05 |
| NOTCH2 | 14889.49296 | 1.138101292 | 3.77E-56 |
| CCN3 | 283.7792535 | 1.137310125 | 1.18E-12 |
| ZMIZ1-AS1 | 20.98118937 | 1.137077389 | 2.23E-02 |
| BHLHE40 | 1231.608194 | 1.136977961 | 1.20E-08 |
| ABHD6 | 420.4684283 | 1.13610492 | 5.60E-22 |
| CDK18 | 1261.470114 | 1.13588089 | 8.12E-12 |
| ANKRD18A | 120.6515545 | 1.135565499 | 4.12E-04 |
| NBR2 | 171.9762602 | 1.134697538 | 1.28E-08 |
| INSR | 813.980436 | 1.134566416 | 2.64E-11 |
| FOXN3 | 4410.147751 | 1.134024322 | 2.51E-43 |
| ZSWIM4 | 566.8401579 | 1.133928185 | 1.38E-20 |
| AKAP2 | 11074.45609 | 1.133898792 | 1.26E-38 |
| FAM53A | 165.8827813 | 1.132872195 | 3.22E-05 |
| ADAM8 | 44.72445104 | 1.132605145 | 2.09E-03 |
| PLEKHB2 | 5020.256771 | 1.13244586 | 1.07E-49 |
| DOCK3 | 240.1180089 | 1.131961921 | 1.97E-13 |
| HCG4 | 34.57144931 | 1.131585934 | 1.15E-03 |
| CALD1 | 17077.04623 | 1.130978105 | 4.02E-18 |
| H2AFY2 | 115.8074385 | 1.130760438 | 4.04E-09 |
| SPTAN1 | 27548.29185 | 1.129892226 | 1.62E-57 |
| GLMP | 3840.216257 | 1.12962492 | 2.30E-15 |
| IRF5 | 86.67911774 | 1.129408199 | 6.28E-05 |
| SAMD12 | 473.3971852 | 1.129370258 | 6.57E-06 |
| EIF5 | 9765.14699 | 1.129117204 | 2.36E-10 |
| TMEM251 | 542.6480132 | 1.128027968 | 1.11E-26 |
| AC096733.2 | 28.16482893 | 1.127737933 | 1.62E-02 |
| TG | 75.66413324 | 1.127379268 | 2.04E-02 |
| CACNA2D2 | 61.41632791 | 1.127288457 | 5.08E-05 |
| KCP | 42.91701193 | 1.126159506 | 7.14E-04 |
| PTPDC1 | 927.0040926 | 1.125933647 | 7.09E-10 |
| CDH23 | 14.58999267 | 1.12576731 | 3.54E-02 |
| TMEM106C | 4149.946741 | 1.125400583 | 1.77E-51 |
| PARVA | 12811.79845 | 1.125315857 | 3.16E-111 |
| AC090159.1 | 39.8118475 | 1.125202064 | 8.53E-04 |
| S100A3 | 298.3816144 | 1.124937458 | 4.80E-11 |
| TBC1D9 | 1044.573317 | 1.124912361 | 7.46E-28 |
| SLC6A9 | 461.5566515 | 1.123794024 | 1.04E-04 |
| CHPT1 | 1064.870081 | 1.122944952 | 1.67E-29 |
| FAM221A | 241.7934268 | 1.122926569 | 8.16E-12 |
| PTK2B | 244.6017206 | 1.122408158 | 8.46E-06 |
| MAL2 | 92.76503004 | 1.122216707 | 6.41E-03 |
| NAP1L2 | 60.86264806 | 1.121590019 | 6.92E-03 |

| | | | |
|---|---|---|---|
| BPIFA2 | 22.6494482 | 1.121215865 | 3.69E-02 |
| ATXN7L1 | 780.669098 | 1.121204671 | 3.10E-18 |
| AC009086.2 | 455.0561247 | 1.121172078 | 2.68E-03 |
| TTLL6 | 61.41262209 | 1.120400322 | 4.23E-03 |
| TOM1L1 | 731.0211202 | 1.11883526 | 2.79E-15 |
| TM4SF19-AS1 | 41.31412618 | 1.118245369 | 2.86E-02 |
| AL357054.4 | 21.85839292 | 1.118240205 | 1.17E-02 |
| ATP6V1B2 | 7378.659165 | 1.118088751 | 2.28E-39 |
| SP100 | 2292.363697 | 1.118044242 | 5.17E-47 |
| EPDR1 | 1988.545424 | 1.116919268 | 1.38E-21 |
| SFXN3 | 2024.371708 | 1.116693836 | 3.21E-19 |
| CHURC1-FNTB | 112.0451196 | 1.115517936 | 1.48E-02 |
| CMTM8 | 58.12048249 | 1.11535658 | 7.81E-08 |
| CPQ | 874.2487683 | 1.113436826 | 1.22E-15 |
| PACSIN2 | 4740.17459 | 1.113042633 | 2.11E-38 |
| NNMT | 7823.226213 | 1.112634451 | 1.43E-08 |
| ID4 | 43.82615756 | 1.110962116 | 7.14E-04 |
| SLC7A1 | 3757.962617 | 1.110683135 | 1.41E-08 |
| SOCS3 | 897.8181479 | 1.11022357 | 5.69E-05 |
| ITGA5 | 9206.177774 | 1.110086219 | 6.88E-14 |
| CMC2 | 1496.242886 | 1.109853366 | 3.62E-43 |
| GPR3 | 101.6903154 | 1.10896582 | 5.00E-07 |
| SLCO3A1 | 123.9865616 | 1.108700714 | 7.31E-05 |
| AC004889.1 | 128.105198 | 1.107263425 | 2.96E-04 |
| STEAP2 | 51.61545693 | 1.106761583 | 1.82E-05 |
| AL121758.1 | 72.32962357 | 1.106608137 | 2.02E-02 |
| TBC1D20 | 4180.645879 | 1.106590474 | 1.66E-62 |
| KCNJ12 | 307.7566808 | 1.106535362 | 1.21E-06 |
| TVP23B | 715.0767378 | 1.106395442 | 7.22E-11 |
| DSE | 960.8994602 | 1.106040552 | 9.37E-34 |
| ANKRD6 | 555.8492612 | 1.10541664 | 7.83E-11 |
| B4GALT4 | 1171.392361 | 1.104559457 | 1.88E-22 |
| SLC12A8 | 358.0297075 | 1.104186814 | 3.37E-07 |
| ULBP2 | 214.0881506 | 1.103985733 | 4.72E-08 |
| FOXF1 | 786.624986 | 1.103500738 | 1.23E-10 |
| KCNK6 | 269.045259 | 1.102740035 | 3.30E-16 |
| MMP17 | 411.9505481 | 1.101944254 | 6.02E-08 |
| SCARB1 | 3539.739284 | 1.100869459 | 2.78E-33 |
| F2RL1 | 1117.544653 | 1.100347667 | 6.12E-17 |
| ANKRD7 | 117.3084134 | 1.100232718 | 4.19E-05 |
| PHEX | 35.22895379 | 1.10008651 | 4.27E-02 |
| RELL2 | 675.3055991 | 1.099440863 | 4.55E-10 |
| KHDRBS3 | 71.81950796 | 1.098906086 | 5.30E-05 |
| ZFAND1 | 995.2253224 | 1.097967274 | 4.69E-07 |
| AC020656.2 | 34.51942435 | 1.097407779 | 6.49E-03 |
| NA | 39.12265666 | 1.097303992 | 1.29E-02 |
| ZNF204P | 105.068296 | 1.097272714 | 1.01E-03 |

| | | | |
|---|---|---|---|
| AC092338.1 | 1800.678811 | 1.097147067 | 1.03E-25 |
| LAMC2 | 82.01010143 | 1.096741604 | 5.70E-04 |
| LRRC32 | 365.4997774 | 1.096715926 | 2.19E-06 |
| RORA | 241.2367859 | 1.096400748 | 3.55E-08 |
| OR2A20P | 429.4941753 | 1.095294826 | 4.88E-06 |
| DKK3 | 22309.34202 | 1.093975211 | 2.03E-19 |
| RNASE4 | 101.2496272 | 1.093855006 | 3.03E-03 |
| SPTLC3 | 200.1830274 | 1.093488339 | 7.80E-09 |
| AF117829.1 | 217.2515211 | 1.093150945 | 1.54E-05 |
| TCL1B | 142.1677943 | 1.093127299 | 8.90E-06 |
| BCAT1 | 1936.78461 | 1.092950497 | 1.01E-08 |
| AC012501.1 | 23.96649049 | 1.092922131 | 1.27E-02 |
| GNS | 11274.3795 | 1.092439841 | 6.90E-24 |
| NUDCD1 | 1948.346597 | 1.092193429 | 8.80E-13 |
| ASPH | 8470.783867 | 1.091884516 | 2.28E-24 |
| IGSF22 | 26.45185137 | 1.091762826 | 7.07E-03 |
| AC024475.4 | 21.46977055 | 1.091737043 | 2.44E-02 |
| LINC01881 | 83.25249054 | 1.090851266 | 3.53E-03 |
| RAB6A | 4718.383046 | 1.090390962 | 2.70E-30 |
| PHACTR1 | 383.5787134 | 1.090375096 | 1.18E-16 |
| CADPS2 | 265.7565386 | 1.089297654 | 2.64E-07 |
| PTH1R | 160.1294675 | 1.087857646 | 7.83E-07 |
| CRYM | 46.70972607 | 1.087439686 | 1.28E-03 |
| LINC01139 | 24.08748545 | 1.087345568 | 4.86E-03 |
| NFIL3 | 1500.966468 | 1.085474399 | 2.68E-08 |
| CFAP54 | 62.85937517 | 1.085031302 | 3.20E-03 |
| MNX1 | 35.38930747 | 1.085026094 | 4.38E-02 |
| RAB11FIP1 | 3684.571645 | 1.084769507 | 1.08E-29 |
| SMIM10L2A | 80.11043027 | 1.084439276 | 6.84E-03 |
| NQO1 | 36091.71617 | 1.0842033 | 8.96E-12 |
| NAGLU | 2962.678407 | 1.084129389 | 6.25E-20 |
| MFAP3L | 92.71318696 | 1.083620859 | 3.50E-04 |
| ALPK1 | 381.0090307 | 1.083447584 | 4.75E-08 |
| MID1 | 1581.448112 | 1.083407041 | 1.28E-20 |
| HMOX1 | 1720.384188 | 1.08289026 | 4.73E-14 |
| SPATA3-AS1 | 16.67560027 | 1.082641083 | 4.89E-02 |
| TMEM154 | 315.1802205 | 1.082398241 | 9.09E-04 |
| SLC38A7 | 1481.333423 | 1.082164835 | 7.80E-23 |
| VPS37A | 1221.264074 | 1.08164404 | 4.33E-13 |
| ZNF69 | 124.6082643 | 1.081483112 | 3.31E-07 |
| PIR | 1579.361561 | 1.081416542 | 2.93E-18 |
| WFDC2 | 34.33119959 | 1.080297193 | 8.28E-03 |
| TMEM39A | 1461.701079 | 1.079541603 | 2.53E-24 |
| AP000560.1 | 18.54898105 | 1.078993311 | 4.80E-02 |
| C11orf96 | 147.5207521 | 1.078930128 | 5.42E-09 |
| ZBED3-AS1 | 237.4057946 | 1.078634433 | 3.76E-06 |
| AEBP1 | 10182.06945 | 1.078263544 | 7.27E-21 |

| | | | |
|---|---|---|---|
| CCND1 | 9113.412653 | 1.078201034 | 2.55E-13 |
| SLC30A3 | 187.831053 | 1.076804404 | 4.40E-10 |
| FSTL5 | 120.0386041 | 1.076362301 | 1.77E-05 |
| NACC2 | 1511.713598 | 1.075927652 | 1.10E-31 |
| HTATIP2 | 3669.802027 | 1.075800784 | 3.60E-43 |
| USP48 | 3298.933719 | 1.075492148 | 7.36E-24 |
| LINC00471 | 85.39286009 | 1.074032774 | 5.26E-05 |
| ROBO1 | 184.8300666 | 1.073469167 | 8.82E-05 |
| MIEF2 | 859.0616354 | 1.072617513 | 1.70E-15 |
| ZCCHC8 | 1037.586952 | 1.072587891 | 4.19E-08 |
| AC007114.1 | 34.79464803 | 1.071705492 | 1.04E-03 |
| SCN3B | 55.20198548 | 1.071615229 | 2.38E-03 |
| LINC01239 | 109.0223971 | 1.069974711 | 4.06E-07 |
| ADAM9 | 9482.944296 | 1.069424991 | 1.96E-11 |
| ADAL | 456.3891718 | 1.068891931 | 7.22E-11 |
| CYBRD1 | 6049.057482 | 1.067102002 | 2.60E-44 |
| HERPUD1 | 5168.49769 | 1.064802539 | 7.38E-16 |
| EPHA2 | 5020.238168 | 1.064095 | 1.06E-18 |
| GADD45B | 977.0968945 | 1.064075522 | 3.12E-19 |
| MROH1 | 1333.300415 | 1.064000945 | 6.80E-28 |
| CPM | 499.6059264 | 1.063543652 | 3.93E-07 |
| EIF4EBP1 | 6024.259061 | 1.063121779 | 4.89E-11 |
| MCUR1 | 1934.263528 | 1.062583603 | 2.17E-43 |
| GRAP2 | 97.99441749 | 1.062465667 | 6.50E-05 |
| AHR | 1112.314882 | 1.06242016 | 2.96E-08 |
| KLHL21 | 1989.331136 | 1.061953799 | 1.32E-24 |
| LINC02609 | 162.2086078 | 1.061538479 | 2.19E-11 |
| MATN1-AS1 | 99.83044349 | 1.061127665 | 1.27E-04 |
| NAT8L | 892.8349717 | 1.060933311 | 4.79E-22 |
| SMC3 | 2270.366911 | 1.060573016 | 6.39E-05 |
| TKTL1 | 81.13948348 | 1.059884194 | 3.05E-05 |
| HUNK | 113.2340518 | 1.059730268 | 1.23E-06 |
| SLC13A4 | 36.08991941 | 1.059583986 | 4.23E-02 |
| NCR3LG1 | 1883.189571 | 1.059426145 | 3.11E-61 |
| GPD2 | 2495.005008 | 1.059236702 | 1.78E-11 |
| VASH2 | 115.4401106 | 1.05911903 | 8.42E-04 |
| AC125807.2 | 800.7685966 | 1.058104688 | 2.37E-20 |
| KRT75 | 23.29573459 | 1.055707941 | 1.36E-02 |
| GTPBP2 | 4056.078583 | 1.055639785 | 4.47E-09 |
| CLCN4 | 1217.164316 | 1.054741061 | 2.58E-10 |
| PDE6B | 188.8286447 | 1.054148728 | 1.24E-08 |
| GNE | 4850.27859 | 1.053562542 | 1.66E-25 |
| ARHGEF2 | 9221.872532 | 1.052537313 | 9.40E-08 |
| SLC16A7 | 237.0406151 | 1.052371958 | 1.18E-05 |
| SPSB1 | 702.3432713 | 1.052350768 | 1.01E-21 |
| MERTK | 532.0190772 | 1.052085424 | 6.79E-09 |
| WWC3 | 2266.235865 | 1.051255 | 8.25E-17 |

| | | | |
|---|---|---|---|
| HOMER3 | 2148.505031 | 1.051107821 | 2.81E-17 |
| LINC01252 | 31.49057911 | 1.051042204 | 9.78E-03 |
| LNCOC1 | 35.33999473 | 1.050859351 | 3.43E-02 |
| HES1 | 556.0751233 | 1.050078277 | 8.14E-09 |
| AC005481.1 | 58.82653227 | 1.049326667 | 2.70E-04 |
| PARM1 | 19.48175827 | 1.049279865 | 4.00E-02 |
| CYP39A1 | 219.4277118 | 1.04893197 | 1.31E-09 |
| NAV3 | 102.7180682 | 1.048112458 | 2.29E-02 |
| IL15RA | 273.3091975 | 1.047928847 | 1.23E-08 |
| ACBD3 | 1599.890682 | 1.047654071 | 2.87E-23 |
| SH3BGR | 93.78411106 | 1.045797477 | 1.05E-03 |
| DEPP1 | 122.3980038 | 1.045777535 | 4.28E-03 |
| PECR | 523.2439058 | 1.045658916 | 2.04E-11 |
| AC105942.1 | 246.655278 | 1.045549166 | 1.00E-12 |
| EPAS1 | 9233.833509 | 1.044855878 | 3.23E-27 |
| EIF4E3 | 382.3841659 | 1.044821816 | 1.07E-14 |
| RHBDF1 | 1735.657285 | 1.044026739 | 5.26E-21 |
| LINC01257 | 134.8471246 | 1.043919602 | 2.62E-04 |
| MOSPD1 | 1494.579896 | 1.043681864 | 4.01E-11 |
| AC012618.3 | 97.74069137 | 1.043672575 | 2.25E-10 |
| QPCT | 409.1069256 | 1.042550681 | 1.22E-12 |
| SH3GLB1 | 2854.355959 | 1.042545551 | 1.05E-18 |
| AC093010.3 | 72.5179379 | 1.041807881 | 1.05E-03 |
| EVI5 | 724.956478 | 1.041626057 | 8.69E-11 |
| TMEM45A | 771.370834 | 1.041588595 | 1.48E-07 |
| PPP1R32 | 33.1700174 | 1.041458918 | 1.05E-02 |
| ATF6 | 3656.141178 | 1.04143286 | 4.00E-35 |
| ALS2 | 2565.168825 | 1.041239844 | 4.55E-20 |
| RALA | 4010.82315 | 1.041236209 | 1.00E-10 |
| SAA1 | 78.53236772 | 1.041142666 | 1.89E-02 |
| LINC02393 | 82.53397108 | 1.03964471 | 2.64E-03 |
| AL121944.1 | 25.22065618 | 1.039592986 | 4.09E-03 |
| PAPLN | 1234.774959 | 1.039135886 | 1.59E-18 |
| TREML2 | 26.08468359 | 1.038983515 | 2.17E-02 |
| NSMCE1-DT | 52.0761297 | 1.038910423 | 2.65E-02 |
| DNAJC12 | 164.2504545 | 1.037663055 | 1.17E-10 |
| SMOX | 2279.030744 | 1.036663145 | 1.06E-04 |
| TMCC3 | 498.972307 | 1.035501021 | 1.62E-10 |
| ZNF763 | 55.32918658 | 1.035423115 | 3.48E-03 |
| SERP1 | 3386.382468 | 1.034046653 | 3.95E-26 |
| NA | 30.12605748 | 1.034036885 | 6.08E-03 |
| PFKFB3 | 2049.457462 | 1.03377559 | 3.49E-20 |
| AP000769.1 | 648.2871949 | 1.03370411 | 1.32E-23 |
| SMIM14 | 1099.159565 | 1.033661034 | 1.23E-20 |
| APOL6 | 2761.451997 | 1.032874739 | 2.04E-12 |
| TRIM16 | 4045.094715 | 1.032757538 | 1.19E-07 |
| TAX1BP3 | 4806.333429 | 1.032618893 | 3.28E-17 |

| | | | |
|---|---|---|---|
| EIF2AK3 | 1083.717756 | 1.03244026 | 3.70E-13 |
| SPAG1 | 418.6973724 | 1.031367182 | 2.01E-12 |
| GSR | 4194.454514 | 1.031248118 | 1.15E-35 |
| SERAC1 | 461.5293554 | 1.031130509 | 3.61E-08 |
| RRAGC | 1986.326998 | 1.030642017 | 2.87E-29 |
| SLC38A1 | 10325.87367 | 1.0304113 | 2.01E-09 |
| RAD51D | 1062.319911 | 1.029420133 | 6.44E-27 |
| ATXN2L | 9348.969127 | 1.029409113 | 6.11E-18 |
| C1QTNF2 | 299.2471779 | 1.029191367 | 2.70E-04 |
| NNT | 4515.752462 | 1.028028427 | 8.21E-43 |
| M6PR | 8085.993348 | 1.027859911 | 1.27E-51 |
| LONRF3 | 116.1789548 | 1.027524014 | 1.77E-06 |
| PMP22 | 6670.421425 | 1.027034086 | 1.72E-18 |
| KDM7A | 570.1778443 | 1.026702567 | 2.22E-03 |
| GPAT3 | 499.5762651 | 1.026543653 | 1.58E-06 |
| TEX19 | 43.55508042 | 1.026348168 | 6.82E-03 |
| FERMT2 | 5028.125652 | 1.02627683 | 5.72E-27 |
| PCED1B | 73.68965833 | 1.026029969 | 8.23E-04 |
| AC141586.1 | 467.7885949 | 1.025932857 | 1.06E-12 |
| KCNAB1 | 18.66573606 | 1.025284276 | 4.02E-02 |
| TMEM64 | 564.8664716 | 1.025135631 | 5.95E-13 |
| NA | 225.5584139 | 1.024473271 | 5.09E-08 |
| NAXE | 2649.379321 | 1.024443702 | 1.36E-18 |
| DDIT4 | 5087.59468 | 1.02425479 | 1.38E-03 |
| TRAF5 | 901.765231 | 1.022796808 | 1.59E-18 |
| AMIGO2 | 914.0654676 | 1.022598107 | 5.89E-14 |
| CTSH | 1717.103531 | 1.021341481 | 1.57E-12 |
| FILIP1 | 45.8162629 | 1.020757796 | 6.44E-04 |
| COX19 | 685.3613752 | 1.02047547 | 9.08E-09 |
| PROS1 | 404.9411068 | 1.020434385 | 3.11E-09 |
| NDRG1 | 8255.208812 | 1.020338107 | 7.91E-07 |
| KANK1 | 4383.01843 | 1.019399234 | 1.04E-45 |
| GCHFR | 321.9882202 | 1.019171721 | 1.26E-08 |
| MTUS1 | 3968.754988 | 1.019009773 | 3.29E-27 |
| OSER1-DT | 334.8931331 | 1.018728451 | 3.39E-12 |
| AC027237.3 | 72.9435641 | 1.018549781 | 9.04E-03 |
| MAP3K21 | 160.6828438 | 1.017755023 | 1.09E-05 |
| PLD1 | 942.7710749 | 1.017036102 | 1.23E-08 |
| RWDD2A | 631.5284039 | 1.016294654 | 4.33E-27 |
| LINC00189 | 232.8947623 | 1.016137815 | 8.34E-08 |
| NAPB | 804.640324 | 1.016064062 | 2.42E-12 |
| SLC7A2 | 2030.212812 | 1.015230464 | 5.59E-05 |
| RFK | 851.8053873 | 1.014500019 | 1.19E-08 |
| SGMS1 | 501.840403 | 1.014209487 | 9.98E-09 |
| PCSK9 | 829.7416089 | 1.012632985 | 2.00E-07 |
| KRT86 | 789.1722789 | 1.011367361 | 2.63E-06 |
| ADHFE1 | 33.12983558 | 1.011129046 | 3.63E-02 |

| | | | |
|---|---|---|---|
| SULT1A3 | 577.5328229 | 1.010747454 | 3.17E-04 |
| AC008556.1 | 76.87959207 | 1.010695649 | 9.60E-06 |
| NAB2 | 1038.514666 | 1.009385508 | 4.05E-15 |
| CRYL1 | 352.6870594 | 1.009294631 | 2.88E-11 |
| MAST4 | 393.8249765 | 1.00741388 | 3.61E-06 |
| CPNE5 | 74.59636606 | 1.007287498 | 1.16E-03 |
| KCNG1 | 1345.743577 | 1.006903897 | 1.51E-05 |
| LINC02125 | 15.97238253 | 1.006591923 | 3.20E-02 |
| NQO2 | 3009.345582 | 1.005538885 | 7.44E-28 |
| TYRO3 | 2229.388693 | 1.004770285 | 3.20E-17 |
| BNIP2 | 2528.105442 | 1.004405784 | 1.78E-12 |
| ZDBF2 | 858.605472 | 1.003867659 | 5.96E-06 |
| PATL1 | 4544.71706 | 1.003690776 | 1.23E-69 |
| ZHX2 | 545.5590592 | 1.003618203 | 9.67E-17 |
| C4orf46 | 611.5640314 | 1.003348813 | 4.01E-09 |
| AC243965.2 | 18.02824951 | 1.003166651 | 2.68E-02 |
| HOGA1 | 115.4192823 | 1.002766552 | 2.05E-04 |
| CD63 | 15208.4414 | 1.002557067 | 8.37E-14 |
| GSDME | 2169.264017 | 1.002551948 | 1.55E-16 |
| DUX4L31 | 29.25806601 | 1.002431322 | 4.73E-02 |
| GRB10 | 4252.746946 | 1.002294802 | 7.78E-10 |
| LIF | 154.155482 | 1.002146009 | 9.57E-04 |
| PTPRH | 506.6079635 | 1.002125059 | 7.08E-04 |
| NA | 44.41616088 | 1.001916316 | 1.80E-02 |
| KIF21A | 532.2354078 | 1.001831699 | 2.01E-04 |
| CPEB4 | 593.1823739 | 0.999073266 | 5.65E-07 |
| CD59 | 30184.01042 | 0.998576761 | 2.96E-12 |
| RAB9A | 1303.88799 | 0.99847409 | 8.92E-26 |
| EHD2 | 10312.92724 | 0.997428537 | 8.44E-20 |
| RCHY1 | 708.5633167 | 0.996722126 | 1.19E-07 |
| CPNE8 | 535.1743269 | 0.995457306 | 2.56E-06 |
| TCEAL4 | 3073.74654 | 0.994847144 | 5.85E-38 |
| MFGE8 | 7060.251313 | 0.993631002 | 1.19E-17 |
| PNO1 | 1549.618819 | 0.993311667 | 9.28E-35 |
| SLC39A10 | 1544.495775 | 0.99287357 | 1.16E-09 |
| KRT8P46 | 15.12617742 | 0.992683993 | 4.87E-02 |
| NT5M | 308.3312684 | 0.992341978 | 9.56E-10 |
| FERMT3 | 650.0854354 | 0.992068415 | 1.58E-15 |
| ANXA8L1 | 163.5609482 | 0.99185164 | 1.39E-02 |
| AL158152.1 | 88.50634893 | 0.990798464 | 3.73E-03 |
| SLC41A2 | 385.2660685 | 0.990695092 | 1.06E-12 |
| ARSB | 2222.90203 | 0.99062011 | 2.59E-14 |
| EIF1 | 14837.75257 | 0.989804557 | 3.95E-27 |
| RNASEK | 3391.534042 | 0.989718751 | 1.44E-12 |
| PLGRKT | 793.4817684 | 0.989482644 | 2.70E-26 |
| GOLGA5 | 2293.257672 | 0.988843574 | 3.05E-36 |
| RNF145 | 10583.21953 | 0.988666389 | 2.18E-103 |

| | | | |
|---|---|---|---|
| ACHE | 203.4309423 | 0.988190365 | 1.24E-04 |
| TLCD2 | 1283.167735 | 0.988045258 | 3.59E-14 |
| SLC25A19 | 457.3255449 | 0.98801776 | 9.09E-16 |
| NA | 35.16822783 | 0.987111314 | 3.00E-02 |
| LIN54 | 911.5084985 | 0.987083701 | 4.86E-09 |
| CCDC71L | 1297.79424 | 0.986541545 | 7.41E-15 |
| RPS6P12 | 74.88269617 | 0.98654061 | 3.90E-04 |
| SDC4 | 4176.418327 | 0.98644986 | 1.01E-11 |
| PDP2 | 1653.370186 | 0.986076208 | 2.45E-28 |
| NUDT4P2 | 286.9845609 | 0.985792942 | 7.40E-09 |
| TEKT3 | 27.13747258 | 0.98575575 | 1.38E-02 |
| MAFG | 2434.517742 | 0.985156215 | 2.57E-28 |
| PRDM11 | 1207.530965 | 0.984670404 | 1.43E-12 |
| NA | 32.95623974 | 0.983743528 | 7.04E-03 |
| SYT14 | 285.8834422 | 0.983257285 | 4.36E-08 |
| GYPC | 5168.208005 | 0.983093098 | 2.29E-15 |
| SLC43A3 | 6224.28028 | 0.982705424 | 3.75E-40 |
| STARD4 | 2398.647591 | 0.981823078 | 1.72E-12 |
| LINC02610 | 37.27312853 | 0.98165118 | 1.97E-02 |
| 3-Sep | 358.3963111 | 0.981108155 | 6.97E-07 |
| CPZ | 462.7221784 | 0.980915183 | 2.45E-05 |
| PTGR2 | 520.8082814 | 0.979490605 | 8.98E-08 |
| BEND7 | 460.5163483 | 0.97838945 | 2.82E-14 |
| NLRP4 | 140.5209597 | 0.977619748 | 2.59E-02 |
| ZNF330 | 1429.442326 | 0.977494251 | 4.93E-20 |
| ALDH1L1 | 633.3709527 | 0.9770841 | 8.93E-08 |
| VGF | 785.8211023 | 0.976446286 | 2.83E-06 |
| TPST2 | 341.6889748 | 0.976130658 | 7.44E-13 |
| NKIRAS1 | 379.5405499 | 0.97588714 | 8.17E-15 |
| DAB2 | 9628.493215 | 0.975317237 | 8.59E-32 |
| CCDC50 | 3220.397041 | 0.975108345 | 2.86E-18 |
| NECTIN2 | 5365.872754 | 0.975104532 | 3.04E-18 |
| ANG | 121.0230573 | 0.974721103 | 1.07E-04 |
| AC092111.1 | 85.97713132 | 0.97416506 | 1.21E-05 |
| KIF27 | 205.5454771 | 0.973264 | 8.56E-06 |
| MCCC1 | 1805.188409 | 0.971056535 | 1.31E-20 |
| KIAA0930 | 2835.093006 | 0.970143748 | 2.78E-19 |
| MAP1B | 20810.43855 | 0.970053492 | 6.79E-07 |
| DDX43 | 474.1818904 | 0.969134353 | 3.27E-15 |
| CIDECP1 | 295.2909922 | 0.96892267 | 3.79E-08 |
| ARHGAP29 | 4982.779967 | 0.96831431 | 4.47E-07 |
| ZDHHC22 | 23.72903958 | 0.967956276 | 1.26E-02 |
| UBR4 | 15901.60597 | 0.967127578 | 2.93E-31 |
| LPCAT2 | 1115.439456 | 0.966322167 | 1.08E-16 |
| PSAP | 37195.08887 | 0.964670832 | 3.21E-16 |
| SLC16A6 | 45.46440852 | 0.963060887 | 4.71E-02 |
| LINC00882 | 70.53774856 | 0.961948522 | 2.02E-03 |

| | | | |
|---|---|---|---|
| FKBP14 | 2076.395674 | 0.961546133 | 6.07E-17 |
| LY6K | 276.1851787 | 0.961066772 | 9.81E-03 |
| PTP4A2 | 6807.446394 | 0.960931307 | 8.71E-15 |
| SOD2 | 3376.369198 | 0.960596221 | 3.52E-46 |
| USP32 | 3325.072913 | 0.960285286 | 1.44E-36 |
| FLNB-AS1 | 98.05334563 | 0.95990866 | 5.65E-05 |
| GAPDHP14 | 68.65157365 | 0.95909683 | 1.84E-04 |
| ARF4 | 9639.726706 | 0.959069039 | 2.85E-18 |
| TMBIM1 | 4490.409115 | 0.958692621 | 2.83E-18 |
| AC096887.2 | 43.01662171 | 0.958491396 | 3.47E-03 |
| PIGBOS1 | 339.9707877 | 0.958217183 | 1.45E-09 |
| SYN3 | 87.05323327 | 0.958190204 | 4.01E-04 |
| ANGPTL2 | 1290.993455 | 0.958040529 | 1.93E-11 |
| B3GALNT1 | 1058.702994 | 0.957888084 | 3.26E-17 |
| AP001528.1 | 51.90560115 | 0.957455598 | 6.22E-03 |
| SIGLEC14 | 107.3074879 | 0.95602141 | 1.55E-03 |
| HSPA13 | 1936.915409 | 0.955685003 | 6.28E-08 |
| PATJ | 706.5965863 | 0.955408684 | 8.92E-07 |
| STK26 | 523.7241541 | 0.955154329 | 2.03E-08 |
| NARS | 10148.14967 | 0.954428437 | 4.30E-11 |
| ADRB1 | 31.51575746 | 0.953903428 | 2.42E-02 |
| LCMT1-AS1 | 36.14641386 | 0.953575908 | 3.64E-03 |
| AP000866.4 | 43.72058498 | 0.95355291 | 5.34E-04 |
| GOT1 | 2517.089131 | 0.953421305 | 2.03E-09 |
| SLC17A7 | 243.5815998 | 0.952945108 | 2.22E-04 |
| COPG1 | 7879.341187 | 0.952752614 | 4.84E-22 |
| FNDC3B | 3823.40014 | 0.95136457 | 2.36E-06 |
| SPANXA2-OT1 | 50.20779197 | 0.95055614 | 1.65E-02 |
| SH3RF3 | 1202.633035 | 0.950311203 | 6.26E-05 |
| RADX | 47.45559083 | 0.950252578 | 2.15E-03 |
| TSTD2 | 2047.132262 | 0.95009214 | 1.01E-21 |
| PLEKHM1 | 1647.368416 | 0.94990246 | 5.69E-18 |
| AC040977.1 | 44.63630535 | 0.949553562 | 1.10E-03 |
| FBXL13 | 69.01456167 | 0.948617807 | 2.81E-03 |
| Z97634.1 | 66.90428637 | 0.948606147 | 2.16E-04 |
| PHGDH | 8589.264013 | 0.948517006 | 6.96E-10 |
| LAMP1 | 5646.809261 | 0.948416768 | 2.89E-20 |
| EIF1B | 1124.244198 | 0.948194284 | 1.37E-19 |
| RNH1 | 8558.943629 | 0.947547449 | 6.49E-26 |
| GLDN | 160.1301569 | 0.947324804 | 9.72E-05 |
| BCDIN3D-AS1 | 120.9273887 | 0.947277333 | 9.42E-03 |
| IRF1 | 2729.050431 | 0.946032451 | 8.85E-10 |
| SSTR2 | 106.5947595 | 0.94587347 | 2.31E-04 |
| THEGL | 17.28143118 | 0.945670639 | 4.01E-02 |
| AGAP2 | 138.4898913 | 0.945014776 | 7.90E-06 |
| AL138478.1 | 119.427306 | 0.944888609 | 2.88E-04 |
| TPO | 119.1349183 | 0.943931357 | 3.54E-02 |

| | | | |
|---|---|---|---|
| SETD7 | 2957.166962 | 0.943220319 | 1.24E-31 |
| TVP23C-CDRT4 | 503.1404065 | 0.942963008 | 5.62E-06 |
| AL031320.2 | 23.78461717 | 0.942532549 | 3.97E-02 |
| MSI1 | 50.33745093 | 0.942275534 | 1.72E-02 |
| ZFAND2A | 542.0624813 | 0.941469514 | 5.89E-07 |
| MT2A | 3108.118129 | 0.941349043 | 1.63E-04 |
| NFKB2 | 732.7160331 | 0.94127116 | 6.33E-09 |
| LMO2 | 1198.776963 | 0.941218502 | 1.83E-16 |
| TBX2 | 83.70495432 | 0.940687855 | 1.33E-03 |
| NA | 80.29581489 | 0.940630752 | 8.80E-06 |
| C21orf91 | 509.9850167 | 0.940619152 | 3.29E-07 |
| SRP19 | 2464.037603 | 0.94025015 | 1.54E-18 |
| AP000459.2 | 350.6615644 | 0.939514538 | 1.25E-04 |
| ADAM23 | 710.607505 | 0.939083084 | 2.06E-13 |
| ST14 | 50.55600932 | 0.939058818 | 3.80E-03 |
| PLSCR1 | 1248.540368 | 0.937863061 | 2.13E-21 |
| PIP4P2 | 720.376732 | 0.93766809 | 1.58E-34 |
| RGS3 | 2410.104891 | 0.937326478 | 2.56E-14 |
| PITPNM3 | 292.5927778 | 0.937311596 | 3.23E-11 |
| MPZ | 270.8214232 | 0.936859121 | 7.49E-04 |
| SPAG4 | 960.0394197 | 0.936592014 | 1.21E-08 |
| RBBP8 | 1615.696352 | 0.935545725 | 4.19E-06 |
| CYLD | 1888.42743 | 0.935021102 | 3.20E-18 |
| ECM1 | 273.6904325 | 0.934811384 | 1.32E-05 |
| FAM89A | 912.360877 | 0.934740182 | 1.83E-22 |
| JAG2 | 524.7348244 | 0.934311894 | 1.55E-05 |
| LINC01285 | 46.63433688 | 0.933799127 | 5.29E-03 |
| CD164 | 6736.973925 | 0.933252407 | 4.93E-20 |
| AC087203.3 | 32.47593761 | 0.933106741 | 4.90E-02 |
| HIF1A | 14396.45639 | 0.933078898 | 3.27E-07 |
| HLA-E | 8048.516065 | 0.932996066 | 2.55E-12 |
| AC013394.1 | 193.2051003 | 0.932466022 | 1.38E-03 |
| C9orf50 | 19.73529643 | 0.932214174 | 2.26E-02 |
| RAB38 | 37.37841539 | 0.931865589 | 3.33E-02 |
| MPP5 | 2731.596394 | 0.930781848 | 2.93E-10 |
| LRRC1 | 426.2692342 | 0.930277684 | 3.64E-10 |
| CYP27A1 | 893.2282248 | 0.93023651 | 7.41E-14 |
| ADAM11 | 409.5004617 | 0.930041795 | 2.27E-12 |
| PDHX | 1539.431444 | 0.929947989 | 6.82E-24 |
| TNFAIP2 | 440.9777675 | 0.929740781 | 5.08E-08 |
| TVP23C | 392.3235822 | 0.928417214 | 4.18E-13 |
| FAM27E3 | 808.5898466 | 0.928167054 | 1.73E-10 |
| ABCG1 | 981.4096477 | 0.927687023 | 6.13E-06 |
| SNX2 | 4505.561683 | 0.927104365 | 3.91E-11 |
| NCOA3 | 5143.412747 | 0.926917014 | 4.14E-13 |
| GOLGA2 | 4842.364652 | 0.926847288 | 9.78E-53 |
| TNFAIP1 | 4428.834655 | 0.926747107 | 4.96E-19 |

| | | | |
|---|---|---|---|
| FGF1 | 246.2227221 | 0.926659644 | 4.46E-04 |
| AC134312.5 | 157.3323084 | 0.926419369 | 4.65E-04 |
| AC008957.3 | 81.64226081 | 0.926252106 | 4.81E-04 |
| SEC22B | 4612.525216 | 0.926214911 | 1.01E-35 |
| PHACTR4 | 2601.10134 | 0.926066882 | 1.30E-41 |
| PEPD | 2743.434718 | 0.924601696 | 1.24E-16 |
| ADPRM | 325.3197862 | 0.924273426 | 4.03E-08 |
| PLXNA1 | 4699.089689 | 0.9239536 | 1.34E-19 |
| TGFB2 | 3830.689573 | 0.923820555 | 4.02E-18 |
| THEMIS2 | 327.7141705 | 0.923754579 | 2.77E-09 |
| LYNX1 | 851.7560327 | 0.92350112 | 3.37E-07 |
| ATP6V1H | 2279.078079 | 0.923309568 | 1.53E-29 |
| AC021054.1 | 291.9708505 | 0.92316361 | 3.33E-10 |
| NFKB1 | 2198.220305 | 0.923145922 | 3.13E-31 |
| BOD1 | 2630.333907 | 0.922223402 | 8.30E-18 |
| PIK3C2A | 3270.990908 | 0.92146789 | 5.60E-03 |
| AC097515.1 | 23.6821678 | 0.920702069 | 3.18E-02 |
| NRBF2 | 506.3490214 | 0.919958806 | 5.74E-22 |
| UHRF1BP1L | 1014.008429 | 0.919762581 | 2.81E-07 |
| IGFBP7 | 8049.810997 | 0.919546819 | 2.61E-11 |
| TPPP | 427.9957226 | 0.919283624 | 2.19E-06 |
| CCDC113 | 1000.208797 | 0.91903385 | 1.68E-12 |
| PELO | 2799.47239 | 0.918646951 | 3.66E-33 |
| NT5E | 5643.186994 | 0.918573943 | 3.20E-08 |
| MAP3K2 | 1478.666948 | 0.917594441 | 4.71E-04 |
| SLC25A25 | 1315.684058 | 0.917131104 | 6.48E-40 |
| FCRLB | 100.9744833 | 0.916837247 | 1.85E-04 |
| NA | 123.1612894 | 0.91566695 | 1.22E-04 |
| AL121721.1 | 38.85487816 | 0.915544662 | 4.21E-03 |
| SRXN1 | 4583.879997 | 0.915388277 | 1.49E-09 |
| AC008109.1 | 67.38834464 | 0.915348681 | 3.24E-04 |
| YARS | 12806.03362 | 0.915291476 | 8.63E-12 |
| GNPDA1 | 5171.192375 | 0.913490147 | 9.41E-18 |
| LIPC | 496.2256483 | 0.913338756 | 8.87E-10 |
| EDARADD | 47.05528653 | 0.912237613 | 7.86E-04 |
| MYZAP | 230.2645852 | 0.912192101 | 3.10E-07 |
| GABRE | 2505.179407 | 0.9118048 | 2.18E-06 |
| TMEM33 | 2111.21072 | 0.911465139 | 1.80E-20 |
| TAS2R14 | 54.60065095 | 0.911347826 | 3.41E-04 |
| ZNF25 | 75.88499036 | 0.91093089 | 6.95E-03 |
| NPR3 | 210.5822672 | 0.910544432 | 6.26E-04 |
| GLB1 | 4696.962364 | 0.909728617 | 1.97E-12 |
| HLA-C | 3125.12319 | 0.907852499 | 3.60E-11 |
| SLC25A29 | 1574.473808 | 0.90716792 | 3.96E-17 |
| AC020916.1 | 458.5870598 | 0.906499058 | 6.16E-12 |
| RIPOR1 | 6265.287925 | 0.905462341 | 1.63E-21 |
| ZNF563 | 145.358265 | 0.904788494 | 1.13E-04 |

| | | | |
|---|---|---|---|
| AL109976.1 | 46.47939616 | 0.904022219 | 2.30E-03 |
| KRT10 | 834.2809818 | 0.903929806 | 1.33E-20 |
| SLC13A3 | 117.8644963 | 0.903518506 | 3.43E-06 |
| ALAS1 | 2028.759906 | 0.903176653 | 2.56E-17 |
| IL15 | 64.48444539 | 0.902228706 | 1.98E-03 |
| USP25 | 1363.579468 | 0.902039545 | 1.46E-07 |
| GTPBP10 | 966.5168096 | 0.901939278 | 8.02E-04 |
| DNM1P46 | 46.31398542 | 0.901819511 | 1.56E-02 |
| AL671277.2 | 54.94515208 | 0.9006288 | 2.50E-03 |
| PRRT4 | 63.24072006 | 0.90056126 | 1.19E-02 |
| LINC02387 | 53.29999358 | 0.899913064 | 9.79E-03 |
| FLCN | 2181.401598 | 0.898602631 | 1.32E-19 |
| CISD2 | 931.2943319 | 0.898538046 | 3.33E-09 |
| TYMP | 1033.445286 | 0.898391765 | 9.87E-09 |
| TPP1 | 10994.53836 | 0.898183531 | 1.31E-12 |
| C11orf45 | 327.0867099 | 0.898128957 | 1.44E-08 |
| KCMF1 | 2322.440073 | 0.897956696 | 2.66E-11 |
| PCAT6 | 92.64316638 | 0.897705749 | 3.14E-04 |
| NA | 100.9126736 | 0.896371254 | 6.76E-07 |
| AL133352.1 | 66.18805983 | 0.896230819 | 6.83E-03 |
| CANX | 22952.90753 | 0.896120394 | 6.46E-19 |
| FAM177A1 | 1022.512052 | 0.895453896 | 4.87E-26 |
| DUSP15 | 247.1476797 | 0.895141206 | 2.29E-08 |
| UST | 680.7892115 | 0.895087421 | 1.93E-07 |
| AC096992.2 | 55.87612262 | 0.893525088 | 3.14E-04 |
| DENND5B | 1132.684712 | 0.891125356 | 1.49E-20 |
| DENND2C | 97.63736781 | 0.890755322 | 2.21E-04 |
| USP27X-AS1 | 60.68377957 | 0.890283117 | 1.43E-03 |
| AF127577.4 | 90.7668684 | 0.890272224 | 8.30E-03 |
| AP003119.2 | 21.05916175 | 0.889570186 | 3.78E-02 |
| PORCN | 684.9381108 | 0.889338121 | 8.36E-11 |
| FRK | 159.893411 | 0.888472202 | 2.68E-05 |
| TMCC2 | 214.3735261 | 0.888282184 | 1.94E-06 |
| DNAJB4 | 526.3585122 | 0.888154987 | 3.74E-05 |
| AL078587.1 | 63.64593881 | 0.887589542 | 8.54E-05 |
| HOMER1 | 828.6769183 | 0.887373331 | 1.14E-09 |
| LPP | 1835.54307 | 0.887367297 | 2.52E-05 |
| SIRT5 | 725.0325281 | 0.887059281 | 7.41E-13 |
| ZNHIT6 | 1164.820065 | 0.886817953 | 6.71E-08 |
| STAC3 | 65.85717021 | 0.886768867 | 2.95E-02 |
| ST3GAL6 | 110.4910764 | 0.88604136 | 2.45E-04 |
| C11orf52 | 106.2142975 | 0.885068483 | 1.30E-03 |
| OSER1 | 2909.229889 | 0.884913024 | 1.64E-30 |
| CCL2 | 1059.694641 | 0.884567966 | 7.41E-04 |
| SLC39A7 | 7440.550849 | 0.883805191 | 5.71E-16 |
| ABCA5 | 556.3366742 | 0.883661815 | 4.81E-04 |
| RAB43P1 | 75.2544523 | 0.883538102 | 1.07E-02 |

| | | | |
|---|---|---|---|
| PEAR1 | 1120.147306 | 0.882494083 | 6.74E-11 |
| UBLCP1 | 2106.228726 | 0.88240736 | 5.66E-05 |
| GCAT | 807.8466605 | 0.88200593 | 4.09E-14 |
| GOLT1B | 1313.786465 | 0.881977816 | 1.20E-12 |
| AC099850.1 | 27.33321147 | 0.881554819 | 1.93E-02 |
| PQLC3 | 634.4680855 | 0.881134988 | 4.37E-15 |
| COX20 | 1221.240216 | 0.881078688 | 4.73E-34 |
| LARP4 | 2417.405961 | 0.880446075 | 1.75E-05 |
| BET1 | 1110.402084 | 0.880078794 | 5.89E-13 |
| DYNC2H1 | 1286.378422 | 0.879864071 | 4.61E-03 |
| ZNF433 | 225.4216414 | 0.879856973 | 7.53E-10 |
| COPA | 14477.07068 | 0.87923445 | 3.22E-37 |
| GRIN2C | 50.62769367 | 0.878864039 | 6.18E-04 |
| C22orf34 | 161.5042601 | 0.878754061 | 5.39E-06 |
| SYTL4 | 710.6228113 | 0.878424886 | 8.30E-07 |
| CARS | 10244.17979 | 0.878173189 | 1.60E-22 |
| RAD51B | 406.0748261 | 0.876636044 | 7.81E-08 |
| TPD52L1 | 2117.343253 | 0.876522938 | 7.98E-08 |
| MYO1C | 10838.86687 | 0.875834545 | 6.54E-14 |
| RMC1 | 875.8919123 | 0.87551483 | 4.41E-16 |
| LURAP1L | 175.7105056 | 0.875285642 | 3.84E-03 |
| PCSK5 | 671.7166073 | 0.874764099 | 2.16E-04 |
| AC008957.1 | 76.43432309 | 0.874243606 | 1.14E-04 |
| B3GAT3 | 2749.384402 | 0.873705617 | 6.36E-13 |
| CAV2 | 2124.493216 | 0.873415117 | 3.42E-12 |
| WDR13 | 3160.87114 | 0.873356778 | 7.49E-11 |
| SDC2 | 2063.499858 | 0.873158562 | 9.81E-13 |
| COPB2 | 9595.875253 | 0.872511323 | 3.08E-29 |
| CYGB | 71.69235275 | 0.872494062 | 1.21E-03 |
| SLC33A1 | 1177.423913 | 0.872375735 | 1.36E-16 |
| ANKRD18B | 178.8942262 | 0.87208613 | 1.13E-03 |
| C7orf31 | 557.6287 | 0.871651902 | 6.75E-13 |
| ANO2 | 401.3700442 | 0.8715427 | 1.16E-07 |
| SDK1 | 1217.636645 | 0.871332518 | 2.04E-10 |
| PRRG1 | 823.3572282 | 0.871330124 | 1.86E-12 |
| CASQ1 | 33.16502397 | 0.871245266 | 1.04E-02 |
| CCNH | 1624.743121 | 0.871143235 | 1.30E-08 |
| SMPD3 | 199.6240386 | 0.870947101 | 2.49E-02 |
| TUBG2 | 1807.888365 | 0.870800321 | 1.93E-25 |
| KIAA1324L | 213.9158876 | 0.870574481 | 2.38E-03 |
| PIK3AP1 | 191.3237924 | 0.869979989 | 2.06E-05 |
| INTS12 | 1102.674669 | 0.869663078 | 1.23E-17 |
| AP002026.1 | 54.44646705 | 0.86958411 | 1.93E-02 |
| AC012254.2 | 129.1850703 | 0.869480062 | 1.98E-04 |
| MFSD12 | 3145.648508 | 0.869436436 | 7.43E-26 |
| ISG15 | 575.9943197 | 0.86879941 | 5.71E-10 |
| CAVIN1 | 22717.39463 | 0.86863101 | 1.08E-10 |

| | | | |
|---|---|---|---|
| CRIM1 | 9816.058979 | 0.868135503 | 6.69E-14 |
| ZFP69B | 336.1088211 | 0.867846742 | 7.23E-04 |
| ZMAT3 | 865.7743738 | 0.867356969 | 4.36E-11 |
| MFAP2 | 231.7639533 | 0.867344989 | 9.49E-04 |
| RNF19B | 646.416233 | 0.8668904 | 3.13E-13 |
| AC245297.3 | 91.66191408 | 0.866324465 | 4.53E-05 |
| PDXK | 5385.678385 | 0.866134397 | 1.97E-35 |
| SMO | 2973.87738 | 0.865995614 | 8.96E-12 |
| C9orf3 | 1301.867845 | 0.865959534 | 4.79E-18 |
| DTNB | 563.0925909 | 0.865945234 | 1.22E-17 |
| DLG2 | 44.97812953 | 0.865742929 | 6.26E-03 |
| TMEM79 | 312.6854783 | 0.865214831 | 6.79E-07 |
| BX640514.2 | 77.90288435 | 0.865053096 | 4.26E-02 |
| SGMS2 | 606.7809837 | 0.864893344 | 1.03E-05 |
| LGMN | 4006.816281 | 0.864271169 | 1.72E-21 |
| TUB | 3581.772259 | 0.864230436 | 2.91E-21 |
| AL139289.1 | 27.90948198 | 0.864135433 | 3.01E-02 |
| PPARG | 1847.312041 | 0.863654897 | 7.97E-10 |
| CT45A1 | 1229.578091 | 0.863417982 | 1.50E-17 |
| AC016831.1 | 169.9544083 | 0.863328211 | 4.02E-03 |
| C5orf22 | 1986.879617 | 0.862993791 | 1.56E-14 |
| UROD | 4803.217769 | 0.862880892 | 1.67E-15 |
| IGF2BP3 | 3445.067623 | 0.862540598 | 3.31E-19 |
| AC022167.2 | 249.2204537 | 0.862434469 | 1.83E-04 |
| PDXDC1 | 6002.09432 | 0.861889583 | 4.38E-25 |
| ZNF697 | 1155.792288 | 0.861394429 | 3.85E-12 |
| CREBL2 | 2141.478236 | 0.86038925 | 7.35E-23 |
| NOMO2 | 9476.424631 | 0.859577292 | 1.97E-16 |
| PPP1R12B | 1810.477078 | 0.859399183 | 1.54E-25 |
| B2M | 24324.29942 | 0.857311333 | 8.83E-31 |
| MMP11 | 1058.85883 | 0.857265646 | 8.54E-07 |
| LARGE1 | 1127.72581 | 0.856830628 | 1.62E-11 |
| CD40 | 1553.021143 | 0.855865051 | 3.96E-18 |
| CYB5RL | 602.6005154 | 0.855301937 | 3.66E-09 |
| HIP1R | 1309.841327 | 0.854840172 | 3.12E-10 |
| SLC38A10 | 4247.429646 | 0.854709817 | 1.22E-14 |
| ARMCX2 | 878.4276883 | 0.854524481 | 2.87E-20 |
| SEPHS2 | 6477.342138 | 0.854263734 | 3.01E-36 |
| METTL22 | 1657.307153 | 0.854176179 | 8.26E-23 |
| NOTCH2NLA | 1200.993235 | 0.854078925 | 7.65E-20 |
| HYOU1 | 23967.33716 | 0.852981585 | 7.36E-18 |
| ELOVL5 | 4586.484995 | 0.851801652 | 5.03E-40 |
| AL031985.3 | 202.0950318 | 0.851688226 | 5.25E-05 |
| A4GALT | 406.9695032 | 0.850902784 | 1.11E-08 |
| C1QTNF12 | 23.96526483 | 0.850869126 | 4.21E-02 |
| FUT10 | 580.7966807 | 0.850571032 | 1.22E-08 |
| ZFP42 | 83.95063084 | 0.850123566 | 2.65E-03 |

| | | | |
|---|---|---|---|
| ZFAND5 | 6320.016712 | 0.85005514 | 1.62E-18 |
| PSPH | 3489.931624 | 0.849498015 | 1.14E-04 |
| MPHOSPH8 | 1367.059281 | 0.849166913 | 4.05E-11 |
| S100A16 | 1005.645382 | 0.848847321 | 2.17E-10 |
| ZFP36 | 459.2400111 | 0.847649408 | 1.53E-09 |
| CAMK2N1 | 1039.323332 | 0.847160064 | 1.06E-12 |
| RBPMS | 1720.796055 | 0.846543322 | 4.04E-17 |
| HEIH | 828.3928894 | 0.846467778 | 1.15E-08 |
| CHMP1B2P | 118.943088 | 0.846274515 | 1.76E-02 |
| PIP5K1A | 3765.167022 | 0.846135253 | 1.50E-18 |
| ARHGEF6 | 2573.200329 | 0.845813755 | 1.63E-33 |
| NDUFA4L2 | 114.6081155 | 0.845554099 | 5.35E-04 |
| SPRED1 | 1716.526315 | 0.845084793 | 4.42E-16 |
| PICART1 | 40.09760807 | 0.844841745 | 4.38E-02 |
| SELP | 136.3642521 | 0.844719548 | 2.12E-02 |
| CRY1 | 1344.096812 | 0.844712656 | 2.96E-11 |
| SLC16A1-AS1 | 314.3438713 | 0.844545834 | 3.15E-03 |
| CHRNB1 | 684.9731031 | 0.844108064 | 7.22E-07 |
| HMGA1 | 9467.267639 | 0.843506591 | 4.71E-10 |
| TTC39B | 448.9058529 | 0.843225735 | 2.31E-07 |
| CDC42BPA | 3574.299221 | 0.842776845 | 1.85E-06 |
| MFHAS1 | 1322.185779 | 0.842776773 | 6.27E-19 |
| LIPH | 201.1100145 | 0.841949835 | 1.13E-04 |
| PNPLA3 | 392.3513662 | 0.841246494 | 4.25E-09 |
| LPL | 5043.224616 | 0.840836164 | 3.02E-09 |
| SHC2 | 1490.065423 | 0.840603012 | 9.85E-11 |
| GCLC | 1634.099911 | 0.840392759 | 9.62E-12 |
| RAB18 | 819.3678411 | 0.840026935 | 3.58E-17 |
| KTN1-AS1 | 455.5717954 | 0.839746718 | 4.06E-07 |
| HAX1 | 4016.217241 | 0.839006418 | 2.14E-13 |
| SH3KBP1 | 6473.2779 | 0.839002217 | 1.02E-20 |
| GLS | 3403.138827 | 0.838804808 | 2.34E-05 |
| LCORL | 400.3564959 | 0.838749773 | 2.58E-05 |
| CDYL2 | 2093.83975 | 0.8385624 | 2.40E-12 |
| PAEP | 37.82752161 | 0.837843633 | 1.89E-02 |
| AIFM2 | 829.2813547 | 0.837230919 | 1.25E-10 |
| DUSP8 | 315.6478431 | 0.835746743 | 1.89E-03 |
| INPP4B | 1588.237089 | 0.835046533 | 6.58E-10 |
| GRINA | 7128.431244 | 0.834923843 | 9.64E-13 |
| CBR4 | 778.8245348 | 0.83391133 | 1.89E-10 |
| HMGCL | 1754.137431 | 0.833761761 | 1.73E-11 |
| COL4A3BP | 2975.02318 | 0.833529876 | 7.51E-22 |
| SOS1 | 2503.778196 | 0.83338018 | 3.33E-05 |
| LMO4 | 1106.20488 | 0.832845532 | 2.50E-07 |
| VPS18 | 1223.589116 | 0.832521052 | 4.30E-08 |
| AC137894.1 | 39.95266791 | 0.83235878 | 4.51E-03 |
| NUDT4 | 2455.67536 | 0.832018022 | 7.31E-09 |

| | | | |
|---|---|---|---|
| MAGEH1 | 643.9168594 | 0.831984255 | 3.15E-06 |
| ZNF419 | 808.3470474 | 0.831720204 | 3.09E-10 |
| FUCA2 | 6047.718625 | 0.831374157 | 5.34E-37 |
| LMX1B | 158.1278197 | 0.831232777 | 5.55E-04 |
| DNAH10OS | 45.64178324 | 0.831201575 | 3.50E-02 |
| AC008014.1 | 134.2352891 | 0.831183062 | 1.39E-02 |
| GTF2IRD1 | 2131.857364 | 0.830953933 | 6.22E-10 |
| SCYL2 | 1874.750772 | 0.830841089 | 1.34E-04 |
| NAPA | 3117.244375 | 0.830593372 | 1.08E-14 |
| KCTD11 | 1277.421524 | 0.830246735 | 1.33E-09 |
| DRAM1 | 1731.290051 | 0.829382646 | 1.83E-25 |
| PRSS27 | 41.26702256 | 0.829087822 | 2.73E-02 |
| WDR11 | 2263.770168 | 0.828427715 | 6.92E-10 |
| FNIP1 | 3476.825664 | 0.82829651 | 2.01E-09 |
| TCL6 | 482.6066254 | 0.828193263 | 7.72E-06 |
| CCDC59 | 724.4599412 | 0.827871558 | 5.16E-11 |
| NA | 53.39350063 | 0.82756585 | 1.33E-03 |
| MYO1E | 3124.761668 | 0.827434426 | 7.61E-13 |
| MLLT10P1 | 40.66822991 | 0.827190833 | 6.93E-03 |
| ADRA1B | 405.8511243 | 0.826908519 | 2.09E-02 |
| NA | 243.5273505 | 0.825449285 | 2.29E-11 |
| EPB41 | 2483.745836 | 0.825365837 | 1.35E-10 |
| YDJC | 1016.804538 | 0.824935503 | 2.87E-15 |
| PROSER1 | 1189.283984 | 0.824468973 | 2.12E-06 |
| CSF1 | 3189.285983 | 0.824466135 | 2.69E-15 |
| AL161431.1 | 253.7190024 | 0.824043376 | 2.82E-04 |
| RAB5A | 1392.42105 | 0.823143952 | 2.58E-11 |
| CDC37L1 | 623.5580681 | 0.822712256 | 1.65E-06 |
| AC124798.1 | 317.8971155 | 0.822315985 | 6.74E-12 |
| CERS6 | 1849.566839 | 0.822050276 | 1.13E-13 |
| CD47 | 3412.049834 | 0.821872037 | 2.56E-29 |
| MTMR8 | 121.7720386 | 0.821605588 | 1.08E-03 |
| FAM229B | 637.9454571 | 0.82062037 | 1.43E-14 |
| PRXL2A | 2340.360732 | 0.819867317 | 3.40E-30 |
| EEA1 | 1040.930789 | 0.819823908 | 1.25E-02 |
| USP12 | 609.8723212 | 0.819807737 | 1.37E-06 |
| RBM34 | 1316.827311 | 0.819343643 | 2.28E-08 |
| LRRC8B | 317.6187643 | 0.819120229 | 1.59E-05 |
| PRR5L | 1135.682346 | 0.818923793 | 3.98E-13 |
| ABCC6P1 | 93.64657159 | 0.81885163 | 1.93E-03 |
| PLA2G5 | 62.8058783 | 0.818736242 | 2.50E-02 |
| NFE2L2 | 4317.516972 | 0.818324346 | 1.38E-06 |
| RTN4IP1 | 449.1760696 | 0.818307141 | 1.22E-10 |
| USP43 | 231.4230637 | 0.818305989 | 9.62E-06 |
| POFUT2 | 4196.210503 | 0.818021369 | 6.70E-08 |
| FIG4 | 421.5997914 | 0.817794127 | 6.09E-08 |
| KIF21B | 1486.646113 | 0.817534526 | 1.09E-04 |

| | | | |
|---|---|---|---|
| LBX2-AS1 | 690.5999235 | 0.816996246 | 1.81E-08 |
| PLPPR2 | 1979.931161 | 0.816985899 | 4.37E-13 |
| GVINP1 | 117.3805028 | 0.816900547 | 1.98E-04 |
| TCEAL9 | 1197.992242 | 0.815965685 | 5.75E-09 |
| AC010894.2 | 35.73780666 | 0.815873188 | 1.78E-02 |
| CHRFAM7A | 55.5674447 | 0.815842446 | 2.57E-02 |
| B3GNT5 | 183.3353908 | 0.815824883 | 6.01E-04 |
| GPC1 | 4941.093134 | 0.81527819 | 2.00E-12 |
| SLC9A1 | 1343.188308 | 0.814765297 | 8.44E-09 |
| RBMS3 | 446.5159869 | 0.813791835 | 4.94E-07 |
| FAM118B | 1981.856079 | 0.813312305 | 1.41E-19 |
| NSMAF | 2623.86319 | 0.813095964 | 1.79E-15 |
| ZNF720 | 478.7604336 | 0.812276814 | 7.64E-08 |
| VSTM2L | 409.4949088 | 0.811385207 | 5.20E-05 |
| CCNG1 | 13038.2605 | 0.809862706 | 1.11E-27 |
| AL392172.1 | 99.14790272 | 0.809245004 | 2.10E-03 |
| TMEM126B | 1389.611773 | 0.808507452 | 9.57E-11 |
| SNX8 | 3274.668815 | 0.807964888 | 8.24E-09 |
| TTC33 | 553.6499434 | 0.807899387 | 1.59E-06 |
| CHST1 | 142.5358579 | 0.807854202 | 4.65E-06 |
| PPIAP46 | 150.622229 | 0.807828338 | 5.16E-07 |
| MS4A15 | 168.0965125 | 0.80745025 | 2.49E-06 |
| KBTBD3 | 101.4109539 | 0.807353467 | 1.69E-02 |
| ZNF570 | 266.3864458 | 0.807351825 | 5.48E-09 |
| PTDSS1 | 5001.392315 | 0.806729264 | 2.47E-17 |
| FLYWCH1 | 2934.327451 | 0.806526789 | 3.73E-19 |
| DAGLB | 1643.438416 | 0.805290511 | 1.49E-20 |
| PDE2A | 399.4921856 | 0.805270805 | 3.72E-06 |
| ZNF468 | 605.9100128 | 0.804604822 | 5.59E-05 |
| SEC24D | 3034.081143 | 0.804593017 | 3.13E-12 |
| AL021392.1 | 31.55875096 | 0.803547926 | 2.39E-02 |
| LAMB2 | 8298.833374 | 0.803534871 | 1.66E-10 |
| PTGFRN | 3794.796387 | 0.803242359 | 3.33E-09 |
| KLHDC1 | 74.0780579 | 0.803124519 | 3.12E-03 |
| NEMP2 | 430.5920123 | 0.803110144 | 7.13E-05 |
| CHST13 | 38.55414762 | 0.801374088 | 3.86E-02 |
| HOXC13-AS | 178.5640571 | 0.800999671 | 1.49E-03 |
| CYB5R1 | 1271.066427 | 0.799146075 | 1.09E-12 |
| TAF13 | 605.4475085 | 0.798942747 | 3.03E-09 |
| KBTBD11 | 254.4879915 | 0.798836787 | 1.93E-11 |
| ENAH | 5040.728136 | 0.798228733 | 3.07E-05 |
| ADSSL1 | 339.1096646 | 0.797654436 | 5.29E-05 |
| USP32P1 | 78.77943041 | 0.797440481 | 2.97E-02 |
| MMAA | 134.8384962 | 0.797329459 | 2.50E-03 |
| FNIP2 | 1389.398079 | 0.79728149 | 1.73E-14 |
| CCDC43 | 1158.071977 | 0.797036696 | 4.69E-15 |
| RABAC1 | 2206.457957 | 0.796764027 | 3.20E-08 |

| | | | |
|---|---|---|---|
| AIMP1 | 1565.903194 | 0.796197002 | 1.50E-07 |
| SRGAP2C | 904.1371559 | 0.796159364 | 9.65E-07 |
| LAPTM4A | 7258.494821 | 0.795267186 | 2.31E-34 |
| CERS4 | 990.9291724 | 0.795148657 | 2.49E-07 |
| QKI | 4899.422462 | 0.794820888 | 5.34E-09 |
| ARL8B | 2283.464947 | 0.79455715 | 7.52E-16 |
| HSD11B1 | 40.05443864 | 0.794359646 | 1.30E-02 |
| ADAM17 | 2891.299778 | 0.794010619 | 7.06E-17 |
| AC092119.2 | 53.10539414 | 0.793845673 | 1.64E-02 |
| RBP7 | 234.865592 | 0.793703148 | 4.43E-05 |
| C17orf58 | 574.9988208 | 0.793525098 | 8.87E-19 |
| UTP3 | 1873.104973 | 0.792976229 | 1.80E-21 |
| FAM102A | 1821.808637 | 0.792673423 | 4.28E-09 |
| GSTO2 | 130.9767255 | 0.792610126 | 2.86E-05 |
| TMEM167B | 1344.383655 | 0.792573517 | 3.78E-13 |
| MPC1 | 774.0013907 | 0.792524628 | 7.17E-30 |
| C9orf116 | 163.5122909 | 0.792469417 | 6.75E-06 |
| RTN3 | 7686.045019 | 0.792350824 | 2.19E-19 |
| ELOVL6 | 1116.973458 | 0.792249717 | 4.01E-06 |
| KRT8 | 14283.57687 | 0.791706812 | 1.40E-11 |
| HSPA4 | 19032.1917 | 0.791595984 | 1.97E-32 |
| AC068491.4 | 36.13138188 | 0.791555187 | 3.06E-02 |
| KDELR2 | 11593.04683 | 0.791126746 | 4.81E-13 |
| SDAD1 | 3020.402864 | 0.790108315 | 8.81E-11 |
| CTSA | 8271.579524 | 0.790010301 | 1.19E-09 |
| RIOK3 | 2038.172139 | 0.788731718 | 6.70E-09 |
| RAPGEF2 | 1880.163938 | 0.788708812 | 8.12E-08 |
| TAX1BP1 | 10757.76244 | 0.78853496 | 2.31E-09 |
| GATAD1 | 3115.269313 | 0.788256344 | 8.40E-30 |
| CALHM5 | 354.1155895 | 0.787732848 | 3.39E-05 |
| CBLB | 355.9339537 | 0.787362153 | 4.39E-05 |
| NPC2 | 9358.589391 | 0.787291879 | 1.48E-06 |
| PPM1L | 96.48854693 | 0.78693961 | 6.67E-03 |
| MANBA | 1597.878041 | 0.786541188 | 2.66E-12 |
| C5AR2 | 63.75946987 | 0.786197872 | 2.82E-02 |
| FER1L6 | 34.70666307 | 0.785941227 | 4.10E-02 |
| AL139020.1 | 290.5916199 | 0.785370442 | 3.03E-06 |
| CBWD5 | 852.034212 | 0.784765904 | 1.38E-06 |
| GGACT | 293.6186656 | 0.784335178 | 8.06E-08 |
| DIAPH2 | 574.5007948 | 0.784096465 | 3.01E-10 |
| DRAM2 | 825.8159279 | 0.783094468 | 1.16E-09 |
| ARL8A | 771.8332599 | 0.78302398 | 4.10E-07 |
| RAB6B | 890.8632009 | 0.782947668 | 1.09E-07 |
| AKTIP | 780.4366473 | 0.782503702 | 4.60E-11 |
| PRR5 | 714.3942261 | 0.782280049 | 9.40E-07 |
| SPECC1 | 1835.882639 | 0.782071663 | 2.48E-14 |
| ATP11A | 1843.254605 | 0.781898725 | 1.37E-09 |

| | | | |
|---|---|---|---|
| FAM174A | 424.3305524 | 0.781799158 | 9.88E-11 |
| GBP3 | 349.9001315 | 0.781485829 | 8.30E-03 |
| ZNF626 | 98.32174551 | 0.781391321 | 3.80E-05 |
| USP46-AS1 | 131.0591345 | 0.780620236 | 5.02E-06 |
| OSBPL1A | 2509.624306 | 0.780117519 | 1.15E-31 |
| MAGEA5 | 44.96564495 | 0.779912302 | 2.08E-02 |
| LGALSL | 1263.152514 | 0.779538668 | 1.78E-11 |
| TMEM126A | 956.1421872 | 0.778803641 | 1.72E-16 |
| AL731577.2 | 75.59622226 | 0.778605608 | 1.54E-02 |
| PERP | 2168.913583 | 0.778593525 | 4.89E-24 |
| NHLRC3 | 495.9969832 | 0.77859236 | 2.99E-13 |
| AC005330.1 | 96.12826441 | 0.778394122 | 3.51E-04 |
| ZSCAN31 | 211.2724114 | 0.778002154 | 1.15E-04 |
| RNF113A | 757.8717493 | 0.777896712 | 1.03E-12 |
| CREG1 | 2653.32401 | 0.777220307 | 4.74E-19 |
| NKILA | 183.3136004 | 0.776363686 | 6.80E-05 |
| AC091435.2 | 63.24787029 | 0.775669522 | 2.18E-02 |
| AC015922.2 | 73.21311534 | 0.774881188 | 4.16E-04 |
| TSKU | 1641.844363 | 0.774691227 | 3.69E-06 |
| MCC | 3773.783562 | 0.773671409 | 3.10E-19 |
| MSANTD4 | 958.8036689 | 0.773468311 | 6.94E-07 |
| SLC22A18 | 622.2696081 | 0.773339325 | 6.41E-11 |
| PPP2R5A | 1252.576734 | 0.773250456 | 2.11E-13 |
| BNC2 | 1174.348305 | 0.772513714 | 8.22E-07 |
| AKNA | 2135.46623 | 0.772058423 | 3.76E-02 |
| MGLL | 4291.086206 | 0.77202081 | 5.81E-09 |
| CYP2R1 | 801.0685079 | 0.772007227 | 1.09E-05 |
| NEO1 | 7967.414653 | 0.771809334 | 5.15E-10 |
| SCOC-AS1 | 40.66257964 | 0.771636164 | 2.57E-02 |
| HYAL1 | 251.009159 | 0.771525513 | 4.04E-03 |
| TLR3 | 366.6552858 | 0.77137972 | 2.81E-03 |
| SLC35A2 | 1819.685814 | 0.770951749 | 1.77E-10 |
| L1CAM | 10255.39231 | 0.770917086 | 3.96E-06 |
| EAF1 | 1414.178432 | 0.769215085 | 1.40E-09 |
| LRP3 | 178.7844076 | 0.768174358 | 1.21E-02 |
| LMAN1 | 5640.517466 | 0.7681276 | 1.06E-06 |
| GLRX | 2880.011914 | 0.767103017 | 9.60E-17 |
| YIPF6 | 1932.021176 | 0.767044525 | 1.23E-14 |
| CDK5R2 | 57.6406424 | 0.766886681 | 3.45E-02 |
| YPEL5 | 2746.761684 | 0.766436204 | 3.20E-15 |
| LINC02055 | 118.9367343 | 0.766357982 | 2.15E-04 |
| RILPL1 | 800.4589162 | 0.766083578 | 3.11E-07 |
| PTPN11 | 8170.647612 | 0.765472709 | 4.46E-09 |
| PNPLA6 | 3135.262182 | 0.765008607 | 6.88E-09 |
| PNMA3 | 556.7056313 | 0.76484077 | 8.46E-14 |
| ARHGEF10L | 688.723664 | 0.764013372 | 1.04E-09 |
| DUX4L9 | 36.41798153 | 0.763628257 | 3.19E-02 |

| | | | |
|---|---|---|---|
| SCN10A | 31.43251154 | 0.763537405 | 3.35E-02 |
| MILR1 | 125.3341487 | 0.763123823 | 3.41E-06 |
| PTPRK | 2144.776039 | 0.762664406 | 2.51E-10 |
| RAB11FIP2 | 445.2120675 | 0.762339783 | 5.61E-03 |
| GPR4 | 237.5154985 | 0.762117131 | 3.59E-03 |
| ALDH3B1 | 1300.36037 | 0.761985608 | 9.60E-14 |
| NOC3L | 1736.520792 | 0.760590516 | 2.82E-04 |
| RERG | 405.3434287 | 0.760423424 | 4.37E-08 |
| ERO1B | 267.8671326 | 0.760358491 | 2.62E-04 |
| PLEKHH2 | 523.7251953 | 0.760032247 | 3.64E-03 |
| VASN | 2614.474492 | 0.759654541 | 2.19E-09 |
| CPD | 5957.815368 | 0.759466839 | 1.96E-12 |
| PFKP | 5727.138889 | 0.758960728 | 8.95E-13 |
| TRPA1 | 331.5429719 | 0.758552408 | 4.62E-03 |
| NA | 72.2974065 | 0.758321088 | 2.11E-03 |
| AC007040.2 | 54.2009864 | 0.758012989 | 3.77E-02 |
| C1GALT1 | 513.9821536 | 0.757948453 | 3.98E-04 |
| ARL1 | 2047.579279 | 0.757141662 | 4.61E-16 |
| GAB2 | 601.0589633 | 0.756744133 | 1.46E-04 |
| SEC62 | 2051.27731 | 0.756480332 | 5.14E-05 |
| SH3BGRL2 | 622.823889 | 0.756083643 | 1.89E-06 |
| ABHD2 | 5282.318574 | 0.755977263 | 2.01E-09 |
| FUCA1 | 1775.850881 | 0.755976792 | 5.92E-06 |
| YIPF4 | 1869.380808 | 0.755908124 | 1.02E-07 |
| AC022916.1 | 53.82228074 | 0.755851585 | 2.77E-02 |
| LSM14A | 3482.093964 | 0.755551304 | 4.46E-08 |
| CASP8 | 2110.41681 | 0.754891331 | 7.04E-12 |
| TM2D1 | 516.8569344 | 0.75439956 | 2.15E-11 |
| SRPRA | 13337.9025 | 0.754263599 | 1.41E-20 |
| SCAI | 404.6599689 | 0.754083688 | 1.73E-03 |
| AFAP1 | 8267.209501 | 0.754050466 | 2.06E-16 |
| ZNF397 | 629.7096529 | 0.753932025 | 9.39E-07 |
| PTPRU | 1753.900827 | 0.753905695 | 6.56E-08 |
| APIP | 1029.168976 | 0.753576027 | 2.84E-13 |
| ATP6V0A1 | 1849.279857 | 0.753519868 | 3.03E-13 |
| IRS2 | 208.4093286 | 0.753419535 | 9.01E-06 |
| ADAMTSL4 | 843.0356382 | 0.753327809 | 3.23E-07 |
| PFDN2 | 1584.135587 | 0.753122918 | 2.17E-10 |
| PDLIM5 | 3144.819155 | 0.752558852 | 5.42E-09 |
| DTNBP1 | 753.5644288 | 0.752450446 | 9.89E-11 |
| AL021068.1 | 38.23277759 | 0.752205541 | 2.14E-02 |
| HLA-H | 207.193018 | 0.752054437 | 1.59E-06 |
| ZBED3 | 1837.70315 | 0.752038389 | 6.50E-17 |
| CBWD6 | 305.3151335 | 0.751882615 | 3.57E-03 |
| ZNF18 | 496.1851024 | 0.751811296 | 2.90E-07 |
| SOAT1 | 2199.622234 | 0.751778474 | 7.55E-15 |
| SLC15A3 | 873.7281996 | 0.751732912 | 3.13E-10 |

| | | | |
|---|---|---|---|
| CSGALNACT2 | 595.8894276 | 0.75163838 | 2.72E-08 |
| SLC35A3 | 470.8080076 | 0.75156992 | 1.38E-03 |
| NLN | 3580.468692 | 0.751562722 | 1.72E-08 |
| SLCO4A1 | 967.7629917 | 0.751439449 | 3.61E-06 |
| SCD | 39207.06667 | 0.751059631 | 6.55E-08 |
| COG5 | 3073.024123 | 0.750610545 | 1.58E-12 |
| TMEM38B | 819.965342 | 0.748982703 | 9.43E-06 |
| NAPG | 960.7955834 | 0.748838845 | 1.01E-14 |
| PRKCA | 7132.858038 | 0.748664989 | 7.09E-20 |
| CLCN7 | 2382.387063 | 0.748234792 | 6.11E-06 |
| DUX4L46 | 65.62997736 | 0.747841577 | 3.13E-03 |
| CCDC149 | 285.9557911 | 0.747736928 | 5.27E-04 |
| DUSP4 | 6532.80606 | 0.747398958 | 2.16E-05 |
| G0S2 | 2016.718559 | 0.747071675 | 1.19E-02 |
| AC244669.1 | 91.29498098 | 0.747004327 | 1.18E-03 |
| CYB5R2 | 274.8195746 | 0.746937247 | 1.64E-09 |
| CBWD3 | 927.2573785 | 0.74637673 | 1.43E-13 |
| MID2 | 540.4827019 | 0.746357533 | 5.08E-09 |
| ZBTB7B | 765.8803121 | 0.745946108 | 1.75E-07 |
| GPN3 | 1009.88344 | 0.745230854 | 2.08E-09 |
| TBC1D8B | 395.7373092 | 0.745102381 | 9.25E-04 |
| PMM2 | 2270.392662 | 0.74489133 | 3.18E-13 |
| PAWR | 1395.931334 | 0.744885261 | 4.54E-04 |
| WASHC3 | 1013.095231 | 0.744400522 | 1.16E-17 |
| HAP1 | 265.2190354 | 0.744386828 | 4.64E-03 |
| LINC01503 | 111.7516947 | 0.744257253 | 2.49E-04 |
| AC004967.1 | 110.4628644 | 0.744237243 | 1.29E-03 |
| TRIP10 | 3342.101072 | 0.744088165 | 2.13E-35 |
| TMEM139 | 162.5650754 | 0.743402342 | 1.29E-04 |
| NCKAP1 | 6666.369709 | 0.742974261 | 1.49E-06 |
| TTC9 | 226.5344015 | 0.742858049 | 4.72E-04 |
| ACTA2 | 163.5945454 | 0.742575714 | 1.34E-03 |
| RAB2A | 3018.845323 | 0.74225844 | 2.52E-40 |
| PHF7 | 290.484141 | 0.742095208 | 7.08E-07 |
| S1PR2 | 135.2910706 | 0.741945337 | 6.09E-05 |
| RCN3 | 1540.546493 | 0.741640479 | 6.79E-05 |
| ZNF438 | 178.9399402 | 0.741206009 | 4.90E-06 |
| NA | 310.8966264 | 0.740648687 | 2.49E-05 |
| SFMBT1 | 939.1757533 | 0.740515918 | 5.47E-11 |
| ACOT4 | 121.1949792 | 0.740514138 | 1.20E-04 |
| CYFIP1 | 4682.357595 | 0.740387722 | 6.40E-16 |
| LPIN3 | 151.6288702 | 0.739868617 | 8.22E-03 |
| SLC16A1 | 4643.305031 | 0.739449429 | 3.10E-10 |
| ARHGAP10 | 734.6739707 | 0.739271942 | 5.81E-18 |
| SEC63 | 3196.314316 | 0.739249754 | 4.97E-05 |
| COG6 | 1004.635117 | 0.738697512 | 1.33E-06 |
| ADAMTS2 | 1245.830925 | 0.738483943 | 6.49E-06 |

| | | | |
|---|---|---|---|
| C3 | 235.9605219 | 0.73836716 | 1.75E-02 |
| PDPN | 1362.464471 | 0.738122334 | 9.68E-05 |
| CYP2U1 | 1376.719402 | 0.738121992 | 9.48E-27 |
| NEK6 | 8959.611567 | 0.738104794 | 2.86E-12 |
| OAS2 | 1649.202108 | 0.738043135 | 3.82E-04 |
| MAGEA10-MA( | 194.1530519 | 0.737949685 | 1.36E-05 |
| SUPV3L1 | 1062.588736 | 0.737931237 | 2.39E-09 |
| TIMMDC1 | 3831.349576 | 0.737874812 | 6.43E-24 |
| CDC14B | 2116.738376 | 0.737176968 | 1.65E-16 |
| RAP2B | 924.2648138 | 0.73675736 | 1.95E-06 |
| ADGRG1 | 794.4778913 | 0.736564585 | 1.03E-02 |
| MYL12B | 10882.15315 | 0.735810141 | 1.47E-32 |
| CUTC | 391.8384636 | 0.734716524 | 3.62E-07 |
| SHB | 1465.471817 | 0.734618475 | 1.58E-06 |
| PFKM | 5080.728656 | 0.734577468 | 2.39E-18 |
| AFDN | 4425.730501 | 0.734398329 | 6.45E-09 |
| LDLR | 5914.413995 | 0.733517411 | 1.92E-07 |
| ZNF396 | 54.79317222 | 0.7333355 | 1.36E-02 |
| MED8 | 2065.090107 | 0.732922783 | 5.34E-12 |
| FLNA | 161615.4789 | 0.73285839 | 4.16E-16 |
| AC091982.3 | 64.41156226 | 0.732586793 | 5.45E-03 |
| SRPX | 4619.232277 | 0.732535236 | 1.10E-10 |
| GPR107 | 9096.320411 | 0.73253083 | 5.69E-24 |
| 11-Sep | 10565.38008 | 0.732487179 | 3.78E-07 |
| BAG2 | 930.8421993 | 0.732453373 | 3.82E-05 |
| COX7A2L | 3464.715754 | 0.731956073 | 2.73E-13 |
| CDYL | 1827.428318 | 0.731568757 | 7.10E-12 |
| LGALS3 | 3865.647832 | 0.731440504 | 3.84E-14 |
| SEMA3F | 497.6585612 | 0.730798004 | 8.81E-08 |
| GALNT14 | 1213.5074 | 0.730440481 | 1.35E-10 |
| KCNN1 | 105.2077303 | 0.730240763 | 4.92E-02 |
| PRMT3 | 1495.28773 | 0.729144895 | 4.01E-05 |
| ARNT2 | 978.2147149 | 0.729126604 | 5.72E-06 |
| ZDHHC9 | 2390.142154 | 0.729052335 | 6.42E-18 |
| AC209154.1 | 130.7499052 | 0.728444412 | 7.86E-05 |
| SLC37A1 | 253.5659223 | 0.727959352 | 1.74E-05 |
| POPDC3 | 714.293481 | 0.727866395 | 2.39E-11 |
| RASEF | 196.4761225 | 0.727841516 | 3.02E-07 |
| ZNF503-AS1 | 255.9762914 | 0.727724277 | 2.37E-04 |
| PIEZO1 | 9000.487395 | 0.727675125 | 3.61E-12 |
| RBM18 | 1709.971679 | 0.727505952 | 1.19E-14 |
| TXN | 6436.015601 | 0.727054157 | 1.71E-22 |
| AL691420.1 | 230.2834602 | 0.726998173 | 2.95E-08 |
| YES1 | 1836.174064 | 0.726559838 | 2.13E-04 |
| CACNA2D3 | 94.70357472 | 0.726548558 | 1.94E-02 |
| TEX41 | 51.499136 | 0.726290757 | 3.86E-02 |
| CLCF1 | 243.9135885 | 0.726066186 | 9.69E-07 |

| | | | |
|---|---|---|---|
| PLCB1 | 346.4078545 | 0.726044585 | 3.12E-05 |
| AP002956.1 | 14354.37341 | 0.725516736 | 7.95E-08 |
| MTIF2 | 1455.220719 | 0.725180911 | 4.53E-04 |
| TNIK | 2274.191439 | 0.725161638 | 1.98E-08 |
| GPT2 | 3740.348619 | 0.724977269 | 1.93E-03 |
| LINC01357 | 79.30204463 | 0.724285539 | 5.12E-03 |
| RNF24 | 6145.979505 | 0.723346253 | 1.63E-15 |
| TCN2 | 1055.694199 | 0.723089826 | 2.09E-04 |
| ATCAY | 1432.949226 | 0.722933997 | 3.99E-04 |
| ENTPD1 | 1129.684048 | 0.722916832 | 1.39E-11 |
| AGO4 | 998.3810896 | 0.722286478 | 2.03E-07 |
| VHL | 2076.084677 | 0.722055802 | 8.00E-10 |
| PIGK | 2062.992312 | 0.721781206 | 7.76E-09 |
| EYA2 | 133.8226664 | 0.72168064 | 6.68E-04 |
| LRMDA | 117.0090295 | 0.72166383 | 8.06E-03 |
| PTPRJ | 3731.301994 | 0.721452363 | 7.12E-23 |
| AMN1 | 251.6585952 | 0.721226635 | 1.25E-04 |
| GAS7 | 2811.772974 | 0.721043326 | 1.27E-08 |
| KLK1 | 68.0157263 | 0.720745582 | 5.01E-03 |
| GFM1 | 2359.802822 | 0.7207324 | 4.15E-06 |
| AC090204.1 | 619.9733506 | 0.720369827 | 8.72E-14 |
| APLF | 162.0757454 | 0.720345853 | 2.28E-03 |
| LINC01145 | 651.6649971 | 0.719849241 | 2.77E-07 |
| AL158212.3 | 189.4677657 | 0.719825342 | 6.49E-06 |
| KLC3 | 240.3054713 | 0.719548621 | 2.78E-04 |
| LRRC8D | 3345.512279 | 0.719081843 | 5.73E-10 |
| SAR1B | 1511.101139 | 0.718915496 | 1.49E-08 |
| CERS2 | 7121.847108 | 0.718858847 | 3.69E-16 |
| LINC01679 | 144.8361926 | 0.718620643 | 1.36E-03 |
| NA | 220.007574 | 0.718424498 | 2.84E-05 |
| DUX4L26 | 66.00978395 | 0.718063637 | 8.90E-03 |
| BCL2L13 | 4891.044754 | 0.717850485 | 2.87E-16 |
| GBE1 | 1827.158317 | 0.717605521 | 4.99E-06 |
| SMAD9 | 316.1031622 | 0.716986902 | 9.87E-03 |
| SNX9 | 2199.738733 | 0.716170803 | 1.74E-27 |
| TRAM1 | 4228.399092 | 0.715575011 | 4.04E-17 |
| AMZ2P1 | 165.7323703 | 0.715419801 | 2.15E-05 |
| FOXN2 | 499.502951 | 0.715343332 | 5.98E-03 |
| PTPRM | 1734.337607 | 0.715332823 | 7.10E-06 |
| TLR4 | 1003.801474 | 0.715287703 | 3.65E-09 |
| NFE2L3 | 4016.428492 | 0.714475201 | 4.08E-10 |
| PPARA | 1593.036496 | 0.712859926 | 1.41E-11 |
| LINC00265 | 516.0461094 | 0.712791224 | 2.90E-08 |
| MBTPS2 | 1935.13835 | 0.712542328 | 1.49E-07 |
| TOB1 | 570.1577494 | 0.711750527 | 1.73E-05 |
| AC016590.3 | 93.95251053 | 0.711408987 | 7.80E-05 |
| TANGO2 | 883.7555233 | 0.711066572 | 6.12E-04 |

| | | | |
|---|---|---|---|
| ARFGEF2 | 6624.906484 | 0.710949833 | 2.16E-13 |
| RFESD | 397.7901993 | 0.710827195 | 2.40E-04 |
| EID2 | 658.2350262 | 0.710730809 | 2.61E-15 |
| BAALC-AS1 | 90.98072801 | 0.710171532 | 1.45E-03 |
| CT45A10 | 1249.05887 | 0.710122425 | 2.00E-07 |
| SPTY2D1OS | 81.55678222 | 0.709816676 | 2.48E-03 |
| PASD1 | 4060.815916 | 0.709502502 | 4.43E-21 |
| NOD2 | 182.4262089 | 0.709237376 | 4.74E-05 |
| MAN2A2 | 3877.249022 | 0.709216049 | 2.08E-11 |
| COL4A5 | 2752.108464 | 0.709168059 | 7.40E-09 |
| WWP1 | 1340.319215 | 0.708962248 | 5.06E-06 |
| SCRN1 | 19372.32534 | 0.708709038 | 3.74E-23 |
| C12orf4 | 972.3065059 | 0.70862845 | 4.89E-05 |
| GRPEL2 | 798.7788215 | 0.708251134 | 3.31E-03 |
| TRNP1 | 1601.234225 | 0.708163428 | 2.02E-06 |
| EHD4 | 2189.688796 | 0.707567453 | 5.10E-11 |
| AL353625.1 | 89.32845993 | 0.707447359 | 1.61E-02 |
| ACBD5 | 839.2854957 | 0.707296747 | 3.38E-12 |
| ACOT9 | 3014.294838 | 0.707021069 | 6.15E-16 |
| STAM2 | 1135.964921 | 0.705738053 | 1.52E-06 |
| HIVEP2 | 987.86558 | 0.705427773 | 3.01E-05 |
| KATNAL2 | 328.0324392 | 0.705378716 | 4.78E-04 |
| NR4A3 | 62.52509383 | 0.705164324 | 6.32E-03 |
| ERBIN | 5871.196358 | 0.704526587 | 2.03E-03 |
| RASSF8 | 897.5966724 | 0.704400285 | 1.76E-03 |
| AL139384.1 | 35.08370117 | 0.704008088 | 4.64E-02 |
| NA | 80.11913439 | 0.703936605 | 3.86E-02 |
| LINC00508 | 78.87403728 | 0.703907076 | 1.67E-02 |
| E2F5 | 507.7955568 | 0.703681442 | 7.71E-04 |
| IGFBP3 | 141.1190452 | 0.70361366 | 1.71E-02 |
| PGAM2 | 238.1250556 | 0.703472887 | 1.16E-02 |
| YIF1A | 3045.656821 | 0.703389944 | 1.87E-11 |
| AC091390.4 | 55.41285315 | 0.702723839 | 1.48E-02 |
| ZRANB1 | 1147.558633 | 0.702592419 | 7.14E-08 |
| RAB7A | 10166.49167 | 0.702581393 | 1.01E-12 |
| FAM49B | 2948.405696 | 0.702473199 | 4.51E-17 |
| MAX | 2278.562027 | 0.702415797 | 3.26E-13 |
| GOLGA1 | 2042.492574 | 0.702168705 | 1.40E-20 |
| IGHV4-34 | 109.8038037 | 0.701327912 | 1.07E-04 |
| IL13RA1 | 5499.46665 | 0.701209657 | 9.82E-23 |
| GCSAML | 54.05539749 | 0.701115454 | 1.64E-02 |
| KRCC1 | 655.4614552 | 0.700285458 | 3.88E-03 |
| AHCYL1 | 7022.281892 | 0.700168988 | 1.42E-36 |
| KCNIP1 | 288.8772349 | 0.699810039 | 2.71E-03 |
| PTHLH | 279.5986255 | 0.699714671 | 5.31E-03 |
| PRPS1 | 3124.775206 | 0.699688114 | 1.46E-18 |
| OSBPL9 | 5854.524737 | 0.699267786 | 5.69E-12 |

| | | | |
|---|---|---|---|
| IBTK | 2116.937579 | 0.699150465 | 1.61E-04 |
| AC073578.1 | 60.53561471 | 0.698941547 | 1.51E-02 |
| ALOX12-AS1 | 192.4843423 | 0.698382911 | 4.69E-06 |
| KATNB1 | 1890.166344 | 0.698123815 | 5.36E-10 |
| SEC24A | 1917.776468 | 0.698075188 | 4.81E-10 |
| DDX19A | 6044.285641 | 0.69795363 | 3.66E-25 |
| SLC16A3 | 3928.098209 | 0.697701816 | 6.70E-09 |
| ZNF571-AS1 | 328.9492408 | 0.69731956 | 4.47E-04 |
| GDAP2 | 761.0711289 | 0.697043229 | 2.67E-10 |
| AC116666.1 | 88.96662613 | 0.696010992 | 5.99E-04 |
| UMAD1 | 477.9267454 | 0.695927134 | 3.67E-05 |
| IWS1 | 2889.136466 | 0.694053806 | 6.46E-12 |
| PNMA8A | 2550.232377 | 0.693763236 | 3.02E-39 |
| SIRPB1 | 258.9285303 | 0.693516209 | 1.83E-02 |
| AGTPBP1 | 952.3310009 | 0.693421721 | 4.52E-05 |
| ITSN2 | 1449.752516 | 0.693394879 | 1.12E-04 |
| LONRF2 | 240.7121002 | 0.693338061 | 7.19E-04 |
| MRPL58 | 996.859681 | 0.693306495 | 4.68E-07 |
| ETV5 | 3045.552529 | 0.693267419 | 2.80E-06 |
| DNAH3 | 118.1200945 | 0.692650599 | 4.96E-03 |
| PMAIP1 | 884.7431058 | 0.692650471 | 1.83E-02 |
| MT-TN | 372.0712389 | 0.692325526 | 1.86E-02 |
| ATP6V1D | 2512.368254 | 0.692131054 | 1.11E-07 |
| TIPARP | 910.2975068 | 0.692020579 | 7.05E-07 |
| NAGPA | 640.8452558 | 0.691952386 | 6.78E-08 |
| TNFRSF14-AS1 | 83.41506238 | 0.691948946 | 3.44E-02 |
| TMC6 | 4347.104689 | 0.691854365 | 2.79E-16 |
| BATF3 | 241.6407464 | 0.691781723 | 7.79E-05 |
| FLT1 | 2091.054392 | 0.691590168 | 1.77E-09 |
| GBP4 | 532.7411693 | 0.691517095 | 3.42E-03 |
| TBK1 | 788.5180789 | 0.691208993 | 4.90E-03 |
| IFI16 | 7219.505499 | 0.690490723 | 5.80E-07 |
| TMEM38A | 657.6662256 | 0.689835638 | 3.48E-06 |
| CHIC2 | 379.1742009 | 0.689552058 | 2.35E-05 |
| ITCH | 4592.164107 | 0.689466798 | 4.74E-09 |
| PICALM | 8631.628285 | 0.688994958 | 5.29E-10 |
| TRIP6 | 1891.929213 | 0.68884245 | 4.94E-07 |
| ZNF702P | 841.7746449 | 0.688656187 | 2.72E-04 |
| CHCHD3 | 3891.440959 | 0.688591863 | 1.43E-11 |
| PLEKHA8 | 1554.564933 | 0.688256336 | 3.85E-08 |
| SLMAP | 1654.45592 | 0.68801837 | 2.55E-05 |
| ARCN1 | 17474.68119 | 0.687990686 | 2.78E-22 |
| LARP6 | 2466.487265 | 0.687843391 | 3.39E-07 |
| ACADM | 1314.411771 | 0.68673684 | 6.41E-06 |
| AC015922.3 | 166.9699555 | 0.685574695 | 5.52E-06 |
| NBPF19 | 1631.618351 | 0.685509471 | 3.16E-12 |
| ODF3B | 128.5541532 | 0.685259985 | 8.11E-03 |

| | | | |
|---|---|---|---|
| BLCAP | 3692.352387 | 0.685101747 | 6.12E-12 |
| PSMD8 | 6091.348911 | 0.684635037 | 2.47E-13 |
| IFI27L2 | 606.4278873 | 0.684282457 | 1.71E-05 |
| MAN2A1 | 1946.42913 | 0.683975397 | 1.13E-07 |
| TSEN15 | 2116.407831 | 0.683805255 | 8.40E-07 |
| ICAM5 | 825.1103563 | 0.683213036 | 1.01E-04 |
| TMEM206 | 691.1648412 | 0.683168151 | 1.94E-09 |
| AC021078.1 | 620.0647068 | 0.682859156 | 2.60E-04 |
| AC008771.1 | 302.7548959 | 0.682814701 | 5.34E-09 |
| NKAP | 996.2150323 | 0.682434324 | 5.22E-08 |
| BTG3 | 724.4575775 | 0.682393683 | 1.26E-07 |
| COPB1 | 10943.70151 | 0.682261112 | 4.18E-15 |
| SLC6A8 | 4800.871325 | 0.681563579 | 1.07E-06 |
| GHITM | 5118.84873 | 0.681228838 | 1.60E-29 |
| RAPH1 | 2044.357541 | 0.680949637 | 7.82E-09 |
| RSC1A1 | 237.9286545 | 0.680867159 | 4.69E-04 |
| CDCA7L | 2435.713917 | 0.680776402 | 1.18E-11 |
| RPS6KL1 | 347.9932618 | 0.680129777 | 3.69E-04 |
| GUCY2C | 48.51206301 | 0.679896961 | 2.22E-02 |
| SMIM29 | 831.6872953 | 0.679721125 | 4.89E-06 |
| IFI35 | 1324.673963 | 0.6795617 | 1.97E-07 |
| PGRMC2 | 2270.365819 | 0.679383409 | 8.34E-27 |
| CCDC183-AS1 | 300.2302564 | 0.679161314 | 2.68E-05 |
| LMF1 | 437.1314473 | 0.67898693 | 5.29E-04 |
| CPXM2 | 397.4177067 | 0.678065632 | 1.96E-03 |
| ETV7 | 52.83399387 | 0.677327562 | 3.67E-02 |
| GUF1 | 1364.463103 | 0.676985471 | 1.16E-07 |
| ECI1 | 2757.268471 | 0.676529926 | 9.59E-16 |
| ALDH1L2 | 216.1567516 | 0.676421073 | 2.03E-02 |
| LRRC20 | 1074.126572 | 0.67600939 | 4.92E-05 |
| IQGAP2 | 1477.33195 | 0.675856352 | 8.49E-04 |
| ZNF770 | 860.8796294 | 0.675707854 | 3.37E-03 |
| GORASP2 | 3964.523769 | 0.675114448 | 4.75E-22 |
| ROCK1P1 | 82.49286083 | 0.675113373 | 3.78E-02 |
| SHISAL1 | 218.6309467 | 0.674966642 | 3.89E-03 |
| NFKBIZ | 242.4207178 | 0.674846909 | 1.42E-04 |
| YPEL4 | 225.6446491 | 0.674665703 | 1.88E-03 |
| LRRC58 | 1475.387555 | 0.674400461 | 3.02E-03 |
| VWA5A | 131.4514081 | 0.673768737 | 3.15E-02 |
| AC012645.1 | 59.53374631 | 0.673635796 | 2.40E-02 |
| HS3ST3A1 | 154.9435874 | 0.673621117 | 1.37E-03 |
| GSTM3 | 4602.829408 | 0.673467495 | 3.99E-11 |
| CLCN3 | 2983.363341 | 0.673424048 | 1.42E-08 |
| TRAF6 | 743.104133 | 0.673155037 | 5.89E-10 |
| STT3A | 12540.12889 | 0.67306172 | 3.34E-17 |
| CT45A3 | 529.6423585 | 0.671873345 | 5.83E-08 |
| AC008267.5 | 109.8802808 | 0.671731535 | 4.86E-03 |

| | | | |
|---|---|---|---|
| METTL18 | 221.5883232 | 0.671556576 | 6.87E-03 |
| SLC9A8 | 1815.924839 | 0.671033872 | 7.67E-10 |
| IL10RB | 1854.424156 | 0.670484521 | 3.03E-08 |
| PRDM4 | 1802.069444 | 0.670326823 | 2.73E-22 |
| LRCH1 | 1532.93364 | 0.670103582 | 5.91E-10 |
| FAM98A | 2562.067213 | 0.670081392 | 7.53E-05 |
| EMP1 | 5374.264184 | 0.669833605 | 5.42E-04 |
| HCFC1R1 | 2331.026297 | 0.669093113 | 2.42E-04 |
| GOLGA3 | 4858.345823 | 0.668199835 | 1.25E-12 |
| ZFAND4 | 114.6766914 | 0.668148013 | 2.98E-02 |
| B4GALNT1 | 1140.587409 | 0.66705712 | 1.98E-06 |
| OPN1SW | 132.2734607 | 0.666643378 | 1.82E-02 |
| MYDGF | 2881.007494 | 0.66636972 | 7.89E-11 |
| DIS3L | 2900.587099 | 0.666339242 | 9.22E-12 |
| RAB27A | 829.7545444 | 0.66601734 | 6.68E-05 |
| RNMT | 3107.790568 | 0.665983197 | 6.38E-06 |
| 2-Mar | 481.9723773 | 0.66571708 | 2.17E-04 |
| ARMCX3 | 2118.590937 | 0.665659665 | 2.65E-14 |
| KDELR3 | 2158.686813 | 0.665551031 | 3.82E-05 |
| IFT27 | 1262.90455 | 0.665085007 | 9.70E-08 |
| UBE2L6 | 5006.393617 | 0.664999179 | 4.73E-07 |
| IMPA1 | 565.8611817 | 0.664766219 | 3.48E-06 |
| CNDP2 | 7468.433963 | 0.664741544 | 1.14E-12 |
| SLC7A8 | 1004.591585 | 0.664418114 | 1.71E-05 |
| MT-TC | 333.2264633 | 0.664321134 | 3.63E-02 |
| MX2 | 8210.402761 | 0.664300071 | 4.26E-26 |
| TSPAN4 | 4504.292425 | 0.663520611 | 2.57E-09 |
| SNX24 | 1531.40493 | 0.663509752 | 1.55E-08 |
| ZCRB1 | 2337.235148 | 0.663104407 | 8.61E-19 |
| RFFL | 1279.067536 | 0.662965207 | 5.70E-05 |
| BRD4 | 3830.430528 | 0.662482027 | 2.00E-06 |
| AC112178.1 | 129.5750328 | 0.661796722 | 7.50E-05 |
| TM7SF3 | 2482.855936 | 0.661739158 | 6.30E-09 |
| WLS | 5603.222117 | 0.661397217 | 1.75E-11 |
| CTIF | 3415.140153 | 0.661026889 | 3.22E-07 |
| GALE | 2706.092626 | 0.660995152 | 3.37E-10 |
| NBEAL1 | 420.3062366 | 0.660760387 | 4.62E-02 |
| KLHDC10 | 3849.172421 | 0.660620653 | 8.39E-17 |
| NA | 1744.479115 | 0.660466764 | 1.84E-10 |
| TMEM70 | 998.0146355 | 0.660264628 | 6.07E-17 |
| TXNDC11 | 2341.326044 | 0.660189294 | 1.07E-12 |
| ZBTB24 | 622.6159114 | 0.659997452 | 2.18E-08 |
| DVL3 | 4546.708344 | 0.659928889 | 4.89E-12 |
| GSKIP | 690.2447286 | 0.659452658 | 2.56E-03 |
| C18orf32 | 615.4090517 | 0.658767893 | 1.12E-07 |
| SGSH | 1798.745288 | 0.658437879 | 3.79E-06 |
| RAD23A | 3349.438841 | 0.658260732 | 6.91E-09 |

| | | | |
|---|---|---|---|
| SLC39A13 | 2800.97386 | 0.65821199 | 1.10E-05 |
| BAIAP2L2 | 100.9679441 | 0.658143784 | 4.48E-02 |
| PRELID3B | 2493.846453 | 0.658075296 | 3.38E-06 |
| C2orf74 | 329.7861284 | 0.65772409 | 1.79E-03 |
| PTPRG | 3213.930849 | 0.657645192 | 1.02E-07 |
| ALKBH8 | 742.8315573 | 0.657607479 | 1.25E-02 |
| PSMC2 | 5871.671733 | 0.657346554 | 4.09E-16 |
| SRGAP2B | 1652.292853 | 0.657241567 | 1.61E-03 |
| WDR41 | 4995.132278 | 0.656991832 | 3.13E-18 |
| MAPK6 | 2936.015372 | 0.656848608 | 4.49E-03 |
| HID1 | 83.06625357 | 0.656808877 | 2.37E-02 |
| SYNE3 | 2045.696794 | 0.656772006 | 3.76E-08 |
| LSM12P1 | 211.9783025 | 0.656423261 | 3.70E-02 |
| ELOA-AS1 | 286.7072976 | 0.65573262 | 6.90E-04 |
| CDH3 | 1193.523897 | 0.655416682 | 5.20E-07 |
| C1orf43 | 7614.203548 | 0.654545408 | 9.62E-13 |
| CEP57 | 2270.226192 | 0.653947983 | 2.42E-04 |
| TRIM5 | 2409.462025 | 0.65374866 | 3.18E-16 |
| CREB3L1 | 742.9290274 | 0.653605091 | 1.86E-05 |
| RCN1 | 15497.19177 | 0.652855453 | 2.82E-23 |
| RRBP1 | 17700.10449 | 0.651298841 | 2.53E-17 |
| DNAJC1 | 790.5283639 | 0.650745158 | 2.97E-09 |
| GALC | 2588.669822 | 0.650562157 | 1.19E-13 |
| DHDDS | 1753.60419 | 0.650176399 | 1.08E-09 |
| PDGFRB | 11707.28027 | 0.650139468 | 6.03E-09 |
| ACY3 | 92.98786841 | 0.649817921 | 1.70E-02 |
| SIRT4 | 114.5330614 | 0.649219922 | 3.66E-03 |
| ZC3H12C | 987.4220162 | 0.649077268 | 4.97E-09 |
| ZFYVE9 | 2196.825218 | 0.64905487 | 1.23E-13 |
| ZC3H15 | 2388.071453 | 0.649041904 | 2.19E-02 |
| DPM1 | 2222.431242 | 0.648987942 | 1.80E-04 |
| SEC31A | 8459.893365 | 0.648817457 | 1.16E-13 |
| LRRC23 | 334.2435134 | 0.648493709 | 3.66E-05 |
| C17orf51 | 1818.558345 | 0.647923187 | 2.99E-05 |
| DYRK4 | 644.2354557 | 0.647854256 | 3.67E-13 |
| U2AF1L4 | 153.7995225 | 0.647764996 | 1.76E-03 |
| SPCS3 | 2551.177372 | 0.647578059 | 7.29E-09 |
| ABCC1 | 5112.111329 | 0.647417634 | 5.62E-10 |
| GAS2L3 | 828.3362503 | 0.64731422 | 2.04E-04 |
| TFB2M | 842.842429 | 0.646646965 | 1.98E-07 |
| ZG16B | 226.8742984 | 0.646625659 | 5.56E-04 |
| AC019080.1 | 129.0291347 | 0.646584043 | 3.49E-04 |
| SPECC1L-ADOR | 222.3063778 | 0.64627126 | 4.35E-02 |
| MED15 | 3949.807379 | 0.646000441 | 2.17E-12 |
| UTP15 | 1493.468396 | 0.645500914 | 3.35E-04 |
| CHCHD6 | 358.4714213 | 0.64526583 | 3.94E-09 |
| EIF3J | 2363.847504 | 0.645248141 | 4.59E-06 |

| | | | |
|---|---|---|---|
| AC022034.1 | 171.0128156 | 0.645221175 | 9.04E-04 |
| ENPP2 | 849.7703046 | 0.644792105 | 8.85E-03 |
| AC008687.4 | 62.22366419 | 0.644670521 | 9.05E-03 |
| XYLT1 | 2293.659344 | 0.644372084 | 5.62E-16 |
| MRI1 | 1491.264152 | 0.644368837 | 4.38E-11 |
| PYCR1 | 8366.040574 | 0.644021311 | 1.45E-07 |
| DZIP1 | 7215.684019 | 0.643777121 | 2.68E-13 |
| HPSE | 457.3034666 | 0.642897393 | 2.18E-06 |
| LRP5 | 4117.859614 | 0.642727074 | 8.24E-14 |
| TMEM267 | 464.8387547 | 0.64261404 | 5.45E-03 |
| NUFIP2 | 3983.671083 | 0.642404335 | 1.27E-04 |
| LRP12 | 1039.779628 | 0.64215439 | 4.91E-06 |
| NRIP1 | 1058.701831 | 0.641866292 | 1.67E-02 |
| OSBPL10 | 1180.998379 | 0.641827111 | 1.63E-07 |
| CHM | 1261.466858 | 0.641780592 | 5.85E-03 |
| ADCY6 | 3447.816817 | 0.64150825 | 2.47E-08 |
| AFG1L | 293.8883154 | 0.641504139 | 1.59E-05 |
| ZXDA | 133.579191 | 0.641479126 | 2.64E-04 |
| F11R | 1874.633537 | 0.641474409 | 1.64E-04 |
| AC132825.4 | 118.8953263 | 0.640868286 | 4.96E-03 |
| NTMT1 | 2667.508817 | 0.640767941 | 8.42E-12 |
| CXXC5 | 1901.801502 | 0.640422231 | 9.93E-13 |
| PFN2 | 10427.95587 | 0.639163811 | 2.23E-16 |
| DDR1 | 3398.200686 | 0.639137608 | 1.52E-07 |
| RAB29 | 1618.049604 | 0.63785587 | 8.35E-12 |
| ZNF234 | 700.7598395 | 0.637178022 | 1.25E-03 |
| TTC39C | 430.9709289 | 0.636310517 | 5.46E-10 |
| FAM83G | 1496.74242 | 0.636057989 | 2.95E-06 |
| FGGY | 156.5456048 | 0.636046116 | 2.53E-02 |
| C1QTNF1 | 954.1115381 | 0.635994091 | 3.81E-05 |
| SRP14-AS1 | 113.3393621 | 0.635725587 | 2.11E-02 |
| ZNF773 | 544.1549742 | 0.635463971 | 7.62E-06 |
| DYRK3 | 586.7166291 | 0.635121772 | 1.27E-10 |
| GINM1 | 1586.931872 | 0.634427267 | 1.82E-17 |
| DDHD1 | 1115.601227 | 0.634195586 | 2.26E-03 |
| EPHX4 | 58.81641382 | 0.634073972 | 1.25E-02 |
| ZNF564 | 315.0327144 | 0.633905563 | 8.04E-07 |
| SMG7-AS1 | 61.16649876 | 0.633790025 | 3.01E-02 |
| EVC2 | 505.3089137 | 0.633323245 | 1.55E-06 |
| CEBPA-DT | 56.88123024 | 0.633113492 | 4.05E-02 |
| SERPINB9 | 1503.778653 | 0.632274401 | 2.62E-15 |
| CSRNP1 | 686.1282168 | 0.632266044 | 1.01E-08 |
| SMCR8 | 1335.19602 | 0.631638763 | 4.24E-18 |
| CEACAM19 | 174.7498355 | 0.631378522 | 4.14E-03 |
| BIRC3 | 138.5705745 | 0.631285216 | 9.83E-03 |
| VDR | 890.1110362 | 0.631097465 | 2.90E-03 |
| FBXO25 | 1326.467273 | 0.63047525 | 9.34E-14 |

| | | | |
|---|---|---|---|
| ZNF674 | 224.7291535 | 0.630109415 | 3.00E-02 |
| KCTD5 | 3210.37524 | 0.629963864 | 2.71E-11 |
| ACTN1 | 17677.66597 | 0.629752812 | 1.07E-13 |
| RNF11 | 2230.717256 | 0.62966589 | 5.00E-20 |
| FLII | 12054.38531 | 0.629469931 | 1.69E-09 |
| NRARP | 134.6099929 | 0.629324153 | 3.20E-03 |
| PTPN14 | 5617.28597 | 0.62897559 | 3.68E-18 |
| ZNF410 | 2800.845383 | 0.628893506 | 1.00E-09 |
| MYOM2 | 221.8497511 | 0.628405259 | 1.63E-04 |
| TMEM184B | 4398.019052 | 0.627788533 | 2.92E-08 |
| DERL1 | 3417.012654 | 0.627708998 | 7.57E-13 |
| AC005586.1 | 92.41806109 | 0.627588828 | 3.73E-03 |
| ZNF280B | 290.2742717 | 0.62751808 | 4.63E-03 |
| JAK2 | 767.597303 | 0.627266179 | 3.59E-03 |
| OXNAD1 | 604.3305565 | 0.627220715 | 2.26E-06 |
| SUMF2 | 9316.853468 | 0.627218909 | 3.81E-17 |
| AC097532.2 | 288.9884563 | 0.626936128 | 1.34E-05 |
| GNL3LP1 | 96.30867823 | 0.6267511 | 1.09E-02 |
| AC092112.1 | 85.72490216 | 0.626623688 | 1.78E-03 |
| GFPT2 | 1047.076629 | 0.626441255 | 1.37E-04 |
| PMS2CL | 760.247039 | 0.625871045 | 2.71E-04 |
| PSMD14 | 3695.568545 | 0.625580244 | 1.64E-09 |
| VAPA | 4644.133299 | 0.625389735 | 1.69E-15 |
| CLU | 7930.293639 | 0.625342694 | 1.08E-06 |
| DYM | 2560.880205 | 0.625177157 | 8.12E-12 |
| ATF4 | 18601.19462 | 0.625072866 | 9.44E-10 |
| JPH3 | 76.67241217 | 0.625062701 | 1.59E-02 |
| ARNT | 2275.330859 | 0.624964782 | 8.35E-10 |
| UBE2W | 670.5578627 | 0.624809597 | 1.30E-08 |
| DNAJB6 | 11954.55005 | 0.624722973 | 4.30E-20 |
| CLMN | 1499.986064 | 0.623855056 | 1.80E-08 |
| ZBTB8A | 300.8609357 | 0.623802947 | 7.75E-05 |
| PXK | 1077.365537 | 0.623088247 | 6.19E-05 |
| SCPEP1 | 3442.188977 | 0.622898238 | 1.21E-08 |
| ASAH2B | 169.8476485 | 0.622576326 | 3.01E-02 |
| MRPL45P2 | 92.66883192 | 0.622234256 | 6.03E-03 |
| ARL13B | 374.1019177 | 0.62215216 | 1.87E-02 |
| SLC25A46 | 2798.417499 | 0.622061787 | 3.55E-05 |
| ATP13A3 | 4319.212019 | 0.622044802 | 7.01E-04 |
| WWOX | 588.6199029 | 0.62136324 | 8.52E-05 |
| RARRES3 | 1750.500845 | 0.621338763 | 1.20E-03 |
| CDH4 | 706.3312431 | 0.619929518 | 2.00E-02 |
| TAF1A | 434.588444 | 0.619657026 | 2.58E-03 |
| WDR19 | 1924.718523 | 0.619615024 | 3.66E-05 |
| EDEM2 | 2837.570935 | 0.619569377 | 1.79E-06 |
| DR1 | 2134.418743 | 0.619556237 | 6.16E-06 |
| STAG3L4 | 1119.960811 | 0.619369397 | 2.20E-06 |

| | | | |
|---|---|---|---|
| PSEN2 | 718.9357932 | 0.618958351 | 1.08E-05 |
| SH3TC1 | 1122.413012 | 0.618204968 | 1.21E-02 |
| AC004803.1 | 262.714056 | 0.618091641 | 5.93E-03 |
| UBE2QL1 | 493.8624657 | 0.618074508 | 1.33E-03 |
| RNF217 | 448.4124261 | 0.618058393 | 6.55E-03 |
| LINC01138 | 170.844276 | 0.617921331 | 3.93E-03 |
| LAMB4 | 116.1014266 | 0.617368071 | 2.84E-02 |
| UBC | 59339.49899 | 0.617239432 | 7.80E-09 |
| TMED2 | 8167.232608 | 0.617164724 | 6.21E-21 |
| RALGPS2 | 1148.025275 | 0.617150228 | 2.54E-05 |
| DNAJB11 | 2408.536378 | 0.616855538 | 2.85E-10 |
| DLG3 | 615.1176337 | 0.616589962 | 1.77E-02 |
| ZC3H6 | 458.3552978 | 0.61648766 | 2.97E-02 |
| FBXW4 | 791.3351442 | 0.616387017 | 4.60E-07 |
| SRA1 | 1042.19628 | 0.61631054 | 3.78E-07 |
| AC106791.1 | 300.1116305 | 0.616273264 | 1.75E-05 |
| IDNK | 151.7021404 | 0.616161768 | 1.29E-03 |
| RARG | 3374.636552 | 0.616159008 | 1.35E-08 |
| LITAF | 5168.421173 | 0.615854986 | 1.58E-10 |
| FLVCR2 | 215.8963486 | 0.615759421 | 1.41E-02 |
| ZNF354A | 498.5568838 | 0.615634467 | 6.11E-03 |
| FBXO42 | 1571.145269 | 0.615482323 | 4.11E-05 |
| CSAG2 | 96.63433714 | 0.615373086 | 4.83E-03 |
| SPNS1 | 2026.014241 | 0.615107643 | 5.43E-09 |
| KIF16B | 1154.574181 | 0.614167508 | 4.34E-04 |
| WHAMMP3 | 172.3458216 | 0.61384909 | 2.99E-02 |
| EXOSC8 | 1008.870093 | 0.6131955 | 1.00E-09 |
| SASH1 | 1551.444222 | 0.61262497 | 1.41E-15 |
| SWI5 | 1113.086776 | 0.612135243 | 4.05E-15 |
| TMEM51 | 617.4009433 | 0.611943592 | 5.29E-10 |
| SOD3 | 336.2709669 | 0.611523982 | 2.37E-02 |
| PDCD6IP | 6347.384567 | 0.611517249 | 3.22E-15 |
| TMEM221 | 72.69405852 | 0.611493081 | 2.80E-02 |
| AL356599.1 | 142.1623234 | 0.61092858 | 1.73E-02 |
| AC126755.2 | 57.24534748 | 0.610790128 | 3.76E-02 |
| RBPMS2 | 349.7867648 | 0.610733634 | 8.08E-08 |
| CT45A5 | 774.9288964 | 0.609938651 | 8.88E-12 |
| PSMD11 | 5804.341134 | 0.60986071 | 1.30E-22 |
| TMF1 | 1309.052105 | 0.609820194 | 1.10E-02 |
| EPS15L1 | 1774.911684 | 0.609703086 | 2.84E-15 |
| PANX1 | 1303.080535 | 0.6096016 | 3.64E-15 |
| F8 | 532.5279691 | 0.609489579 | 6.22E-07 |
| ALG2 | 2031.964824 | 0.609419899 | 4.87E-22 |
| MGST3 | 1187.077861 | 0.608722677 | 7.91E-07 |
| TRMT11 | 543.8145435 | 0.608522627 | 6.56E-03 |
| SWAP70 | 3816.151683 | 0.608301629 | 8.66E-13 |
| MLXIPL | 379.4299946 | 0.608292618 | 1.52E-02 |

| | | | |
|---|---|---|---|
| SNX25 | 862.6549565 | 0.608219284 | 8.59E-09 |
| MTDH | 5816.793673 | 0.60813287 | 7.53E-10 |
| PCDHGA4 | 133.1061788 | 0.607552448 | 9.04E-03 |
| GRSF1 | 4306.976919 | 0.607113796 | 2.51E-11 |
| LRRC59 | 6683.308593 | 0.607013195 | 2.41E-15 |
| RREB1 | 1630.997352 | 0.607000165 | 1.07E-12 |
| RALGPS1 | 346.6724631 | 0.6067733 | 1.23E-02 |
| AKIRIN2 | 1775.583602 | 0.60642877 | 5.24E-06 |
| ST3GAL1 | 3868.982245 | 0.606295561 | 6.70E-07 |
| STAMBPL1 | 514.9312811 | 0.606192046 | 2.78E-03 |
| SPSB2 | 325.8821285 | 0.605966963 | 2.82E-06 |
| XAF1 | 1197.67208 | 0.605759293 | 2.91E-04 |
| APOLD1 | 451.9599406 | 0.605655413 | 9.85E-06 |
| MRPL39 | 1037.840497 | 0.605529709 | 1.33E-11 |
| PYGB | 11837.43242 | 0.605465154 | 4.09E-13 |
| C19orf71 | 213.6471134 | 0.604963986 | 1.85E-04 |
| HSPG2 | 6936.109191 | 0.604106323 | 7.77E-05 |
| BTF3L4P2 | 254.3305076 | 0.604084484 | 2.28E-03 |
| HTRA3 | 2052.944216 | 0.604073145 | 1.45E-04 |
| CYP2S1 | 1720.583561 | 0.603993058 | 2.35E-03 |
| LARP1B | 467.2784037 | 0.603871192 | 1.64E-03 |
| NUCB2 | 1933.069367 | 0.603664779 | 6.46E-06 |
| GCH1 | 403.235998 | 0.603529161 | 9.06E-04 |
| MACROD1 | 1467.887466 | 0.602525233 | 4.04E-06 |
| GNG12 | 8298.18734 | 0.602383436 | 5.60E-09 |
| ERLEC1 | 3042.316058 | 0.602232745 | 1.13E-12 |
| APOPT1 | 868.812937 | 0.601777399 | 2.96E-07 |
| TGDS | 1360.459436 | 0.601538169 | 1.43E-04 |
| MAP4K3-DT | 286.7965894 | 0.60120914 | 1.68E-06 |
| AC009303.4 | 97.14289571 | 0.599499736 | 3.37E-03 |
| DECR2 | 1338.4698 | 0.599434457 | 2.10E-09 |
| FAM219A | 2447.604118 | 0.599041977 | 1.74E-05 |
| RBMXL1 | 1282.120288 | 0.599025869 | 1.22E-08 |
| NANS | 3649.193973 | 0.598514984 | 1.52E-18 |
| SLC49A4 | 686.2493069 | 0.598484814 | 1.63E-09 |
| NA | 210.4474725 | 0.59847254 | 7.60E-03 |
| PLEKHA3 | 389.0818929 | 0.598381871 | 3.60E-03 |
| ITFG1 | 3584.265264 | 0.597567906 | 2.94E-08 |
| TBCA | 6087.859397 | 0.597271765 | 1.08E-14 |
| SLC26A11 | 414.2417322 | 0.597042292 | 1.45E-03 |
| MEX3C | 1765.034227 | 0.596945708 | 9.14E-06 |
| CCDC90B | 1970.233988 | 0.596509886 | 1.22E-07 |
| STX4 | 2129.364156 | 0.596364217 | 2.14E-08 |
| AC009093.1 | 685.8585947 | 0.59632526 | 2.14E-03 |
| RNF2 | 712.6284226 | 0.595773762 | 1.88E-05 |
| STXBP2 | 1158.876155 | 0.595632298 | 2.93E-07 |
| ATG101 | 1922.096598 | 0.595526371 | 1.89E-07 |

| | | | |
|---|---|---|---|
| SNX33 | 3457.403843 | 0.595251323 | 1.11E-05 |
| CLN3 | 1559.612366 | 0.594961221 | 2.60E-05 |
| ACSF2 | 2502.660311 | 0.594950624 | 1.89E-07 |
| JHY | 621.7022963 | 0.59473511 | 1.04E-08 |
| STAG3L5P | 572.7572891 | 0.594669811 | 1.19E-07 |
| P2RX5 | 672.8200669 | 0.594518212 | 9.03E-06 |
| RPS6KA3 | 4793.968676 | 0.594338936 | 4.72E-07 |
| TATDN3 | 402.676729 | 0.594323742 | 4.50E-04 |
| ADAT2 | 484.6253686 | 0.593734232 | 9.74E-03 |
| C22orf23 | 284.8102428 | 0.593658954 | 3.11E-04 |
| CNTFR | 700.6352166 | 0.593527775 | 1.05E-02 |
| IDH3A | 2872.949552 | 0.5935181 | 2.36E-05 |
| C5orf51 | 3030.487835 | 0.593176887 | 8.90E-03 |
| EIF2AK2 | 3388.221179 | 0.593047701 | 8.17E-10 |
| LGALS3BP | 26830.57345 | 0.593019566 | 4.41E-06 |
| KLF3 | 1692.97105 | 0.592625012 | 1.96E-09 |
| ZNF23 | 548.6402448 | 0.592616056 | 2.10E-02 |
| ARHGEF28 | 3310.07867 | 0.592278964 | 4.74E-04 |
| KPNA4 | 3635.1137 | 0.592042833 | 2.07E-15 |
| YRDC | 837.8103711 | 0.591619472 | 1.94E-10 |
| CDC42EP5 | 204.9522045 | 0.591464889 | 8.06E-03 |
| STAT1 | 5986.588145 | 0.590530781 | 3.41E-10 |
| PDZD7 | 149.1959024 | 0.590498833 | 4.40E-03 |
| OR7E38P | 675.7756901 | 0.590397117 | 1.06E-04 |
| CFAP97 | 1560.868825 | 0.589856952 | 2.29E-02 |
| PDZD8 | 1073.044129 | 0.58962522 | 6.28E-04 |
| CARD19 | 976.3250092 | 0.589571145 | 6.58E-07 |
| CHURC1 | 1669.86386 | 0.589325856 | 4.17E-11 |
| FKBP15 | 3792.11178 | 0.589268091 | 8.13E-22 |
| CLUHP3 | 309.9195558 | 0.588901866 | 8.08E-04 |
| IFNGR1 | 1233.623812 | 0.58875085 | 3.07E-08 |
| GRTP1 | 152.5704356 | 0.588316081 | 1.42E-03 |
| TP53INP2 | 2226.784429 | 0.588158192 | 8.75E-05 |
| SNX16 | 332.872172 | 0.587859223 | 5.68E-04 |
| PMS2 | 1548.094037 | 0.587389008 | 3.98E-06 |
| ZNF275 | 1849.137419 | 0.587361684 | 1.56E-17 |
| EIF5A2 | 297.4165173 | 0.587308622 | 2.05E-06 |
| UGGT2 | 2899.149004 | 0.586769047 | 1.31E-02 |
| HLCS | 2418.065507 | 0.586603465 | 3.63E-05 |
| RTRAF | 4394.6709 | 0.586553331 | 2.38E-08 |
| MYO1D | 1277.296482 | 0.586411571 | 3.03E-08 |
| NBPF10 | 2314.64115 | 0.586323992 | 7.74E-09 |
| FER1L4 | 915.777995 | 0.586259161 | 4.90E-02 |
| SNN | 1753.573762 | 0.585223018 | 2.29E-03 |
| TP53TG1 | 318.3974273 | 0.584839915 | 9.47E-05 |
| CYP11A1 | 221.6021817 | 0.584559593 | 7.70E-04 |
| YIPF2 | 2105.78297 | 0.584411473 | 2.07E-07 |

| | | | |
|---|---|---|---|
| PHLDA3 | 421.1813707 | 0.584219943 | 2.51E-03 |
| ALDOA | 42268.47046 | 0.584183833 | 1.01E-05 |
| OGT | 16811.32788 | 0.584122121 | 2.35E-04 |
| GNG12-AS1 | 76.90898532 | 0.583856707 | 2.10E-02 |
| MAPRE3 | 973.794316 | 0.583838103 | 7.36E-08 |
| CAMTA2 | 2017.593104 | 0.583786747 | 2.66E-03 |
| SRFBP1 | 747.7201859 | 0.583594631 | 1.98E-02 |
| AGFG1 | 2601.944699 | 0.583539561 | 1.56E-06 |
| CACHD1 | 561.4740815 | 0.583391845 | 1.52E-06 |
| GOSR2 | 3360.184991 | 0.583383161 | 2.26E-07 |
| SLC44A1 | 3595.730604 | 0.583123098 | 1.74E-15 |
| ZNF222 | 185.4861938 | 0.582095365 | 5.42E-03 |
| PLPP1 | 1972.58768 | 0.581931197 | 1.59E-03 |
| USO1 | 3733.170279 | 0.581759562 | 2.91E-04 |
| COMMD2 | 1374.963537 | 0.581719023 | 2.62E-13 |
| SIRT2 | 2025.60668 | 0.580959484 | 5.31E-05 |
| RMDN1 | 2329.178846 | 0.580911802 | 6.12E-12 |
| RAPGEF1 | 4833.32604 | 0.580885697 | 7.56E-10 |
| FBXO48 | 85.98279878 | 0.580649837 | 2.40E-02 |
| FAM149B1 | 1431.963422 | 0.580537732 | 2.16E-16 |
| EXD3 | 412.4310246 | 0.580261861 | 5.33E-07 |
| RNF181 | 2072.155686 | 0.580222496 | 6.88E-07 |
| TLN2 | 3537.806415 | 0.578837899 | 1.39E-03 |
| BNIP3L | 5784.038634 | 0.578718072 | 5.46E-09 |
| PDPK2P | 424.9377736 | 0.578222199 | 3.49E-05 |
| ANXA5 | 24506.2297 | 0.578142328 | 6.37E-08 |
| OCIAD2 | 557.958916 | 0.577942178 | 3.74E-05 |
| AC007620.2 | 357.518442 | 0.577941219 | 6.09E-04 |
| SH3GL2 | 285.0147928 | 0.577560791 | 3.44E-03 |
| APOOL | 959.5388291 | 0.577550305 | 8.02E-06 |
| ANKRD11 | 12651.38724 | 0.577234411 | 2.10E-06 |
| TIMM9 | 1153.293094 | 0.577130672 | 1.22E-06 |
| ZNF252P | 1467.129611 | 0.576806213 | 2.90E-03 |
| STK3 | 890.5682343 | 0.576683775 | 1.10E-07 |
| ITGB1 | 24729.84288 | 0.576523615 | 2.76E-11 |
| AL356056.2 | 88.00872493 | 0.576371413 | 7.00E-03 |
| ORC4 | 1160.988213 | 0.576340603 | 5.15E-04 |
| SIAH1 | 978.7728873 | 0.576204519 | 3.56E-10 |
| PIP4K2C | 2211.016611 | 0.576166751 | 1.75E-09 |
| SCARB2 | 7661.922881 | 0.576032706 | 2.39E-08 |
| ERO1A | 3104.479503 | 0.575727902 | 2.83E-04 |
| YIPF1 | 1067.199585 | 0.575205601 | 6.08E-08 |
| GPR137C | 168.6258987 | 0.575016589 | 1.07E-03 |
| HSF2 | 913.681019 | 0.574496102 | 1.28E-06 |
| PTGS1 | 2837.117892 | 0.573945832 | 5.72E-03 |
| COMT | 3313.736764 | 0.573708129 | 3.89E-06 |
| UGP2 | 2885.330738 | 0.573361629 | 1.12E-04 |

| | | | |
|---|---|---|---|
| ARHGEF37 | 360.1984528 | 0.573319148 | 3.32E-03 |
| GPR157 | 512.3955584 | 0.572974181 | 4.98E-03 |
| NCOA7 | 2232.279058 | 0.572730074 | 4.15E-04 |
| RAB21 | 835.0701283 | 0.572519904 | 4.74E-09 |
| ATXN2 | 1989.701599 | 0.57174314 | 2.94E-10 |
| L3MBTL3 | 432.0412705 | 0.571407641 | 2.58E-03 |
| C15orf39 | 1454.321721 | 0.571321136 | 2.14E-05 |
| AL390719.1 | 186.2336545 | 0.570556487 | 4.77E-03 |
| KCTD18 | 614.5891598 | 0.570360213 | 1.18E-07 |
| DNAJC10 | 4213.902468 | 0.569982804 | 1.32E-03 |
| SHC1 | 17857.68695 | 0.569674391 | 2.96E-07 |
| SYNM | 2381.484167 | 0.569474817 | 2.80E-05 |
| CCDC183 | 146.8971672 | 0.569422841 | 7.54E-04 |
| AC093726.1 | 112.1297998 | 0.569186182 | 8.20E-03 |
| SCARF2 | 1756.210107 | 0.569075994 | 2.86E-12 |
| ME2 | 1256.073843 | 0.569025268 | 8.97E-05 |
| SPPL2A | 1140.966053 | 0.568972385 | 2.75E-08 |
| NUTM2B-AS1 | 315.9997057 | 0.568758203 | 1.53E-03 |
| TMED7 | 4176.355682 | 0.56865195 | 9.88E-07 |
| RARRES2 | 5285.601134 | 0.568502355 | 4.05E-04 |
| AC006001.4 | 124.4658008 | 0.568343979 | 1.36E-02 |
| CDKN2B-AS1 | 171.1849375 | 0.568196807 | 4.59E-03 |
| ZCCHC4 | 500.6188419 | 0.568165037 | 3.03E-05 |
| JADE1 | 1188.702036 | 0.568034608 | 2.19E-03 |
| IER3IP1 | 854.4095808 | 0.567928019 | 3.30E-05 |
| CTTN | 9646.555605 | 0.567764704 | 2.95E-18 |
| PRPF18 | 424.9952789 | 0.567701219 | 2.47E-03 |
| DCAF10 | 1959.108398 | 0.567614345 | 5.06E-07 |
| FAM204A | 645.4275913 | 0.567477876 | 1.13E-04 |
| CDC42EP3 | 1854.372981 | 0.567388911 | 2.79E-02 |
| INPP5F | 810.7696978 | 0.567235372 | 4.09E-04 |
| CGN | 277.5497533 | 0.567220665 | 2.82E-02 |
| UNC50 | 1554.82129 | 0.567131217 | 2.69E-15 |
| TIGD2 | 358.8860689 | 0.566561446 | 9.63E-03 |
| PCAT19 | 298.9928344 | 0.566284778 | 3.16E-02 |
| AC138028.4 | 241.3457325 | 0.566106752 | 8.21E-04 |
| MED10 | 2839.473397 | 0.566089457 | 1.97E-05 |
| C12orf29 | 687.7136283 | 0.565604517 | 2.32E-04 |
| FAM114A2 | 967.3843151 | 0.565080573 | 8.91E-05 |
| MIF4GD | 956.5208067 | 0.565041824 | 5.28E-06 |
| AIF1L | 1935.226753 | 0.564648008 | 5.62E-05 |
| EXOC2 | 2251.914411 | 0.564518515 | 6.76E-13 |
| SDHB | 2974.624177 | 0.564400971 | 1.84E-13 |
| RBCK1 | 7551.367995 | 0.564269658 | 4.04E-05 |
| ZNF589 | 962.260922 | 0.564234267 | 8.95E-10 |
| SPG21 | 4901.16491 | 0.563978283 | 1.99E-11 |
| WASHC2A | 1913.235673 | 0.563911604 | 2.72E-08 |

| | | | |
|---|---|---|---|
| PSMG1 | 2121.809787 | 0.563610194 | 2.43E-16 |
| PVT1 | 340.4968136 | 0.563404641 | 1.58E-02 |
| EFR3A | 3105.551569 | 0.563344015 | 7.61E-07 |
| LINC00467 | 293.758979 | 0.563243475 | 2.59E-03 |
| HOXC6 | 874.2215878 | 0.563147276 | 4.05E-13 |
| E2F7 | 1312.39899 | 0.56308364 | 3.63E-04 |
| NLRP1 | 4175.84274 | 0.562926946 | 1.80E-08 |
| PALM | 1592.918682 | 0.562923298 | 5.08E-04 |
| PQLC2 | 640.4156606 | 0.562678703 | 3.34E-05 |
| SEC61A1 | 19518.79438 | 0.562581732 | 1.08E-11 |
| POLD4 | 1469.888723 | 0.562390582 | 3.73E-06 |
| VPS8 | 2121.563019 | 0.562243851 | 1.98E-06 |
| LOXL2 | 22646.42599 | 0.562098146 | 2.07E-04 |
| OGDHL | 863.9834681 | 0.562040129 | 7.36E-05 |
| MRPS10 | 1948.921082 | 0.561753522 | 5.28E-09 |
| HDAC11 | 769.1429549 | 0.561495603 | 2.52E-04 |
| GID4 | 1136.933439 | 0.561460753 | 2.84E-08 |
| TMEM214 | 5234.987819 | 0.561302768 | 7.84E-05 |
| TMEM8A | 4587.609126 | 0.561221963 | 4.51E-06 |
| SEC61G | 3651.15804 | 0.560654965 | 5.45E-07 |
| SMIM4 | 523.0536447 | 0.560520152 | 2.15E-04 |
| MTX1 | 2583.38152 | 0.559962697 | 1.62E-05 |
| PCGF1 | 527.1915392 | 0.559881495 | 1.71E-05 |
| RIMKLB | 1383.799397 | 0.559871057 | 2.19E-03 |
| ZRANB2 | 2901.416846 | 0.559849195 | 1.44E-03 |
| R3HCC1L | 545.7380229 | 0.559769838 | 4.66E-04 |
| DNAL1 | 607.5280497 | 0.559158704 | 3.39E-05 |
| ANKRD13A | 3316.621453 | 0.558631109 | 4.22E-07 |
| SYDE1 | 3531.235541 | 0.558399649 | 1.29E-05 |
| OSTF1 | 1331.429526 | 0.558375024 | 1.14E-06 |
| PNPLA4 | 1147.358362 | 0.55816501 | 8.44E-08 |
| NA | 1427.962526 | 0.558141914 | 9.69E-15 |
| PPCS | 1325.8896 | 0.55777669 | 7.53E-10 |
| AC005520.2 | 60.31597595 | 0.557655835 | 3.69E-02 |
| ABTB2 | 1119.031511 | 0.557623876 | 4.88E-12 |
| ZNF542P | 1000.738491 | 0.557587196 | 2.72E-08 |
| SERTAD1 | 500.6192425 | 0.557504687 | 5.19E-04 |
| PLEKHA6 | 883.8866704 | 0.557480838 | 2.14E-02 |
| OSBP2 | 391.2814485 | 0.557434447 | 1.45E-03 |
| GALK2 | 1013.100879 | 0.556528204 | 2.38E-15 |
| DNAJC27 | 207.4879329 | 0.556380255 | 7.33E-03 |
| ARF1 | 11486.82838 | 0.556344708 | 5.35E-11 |
| ASAP2 | 1882.799983 | 0.556284587 | 4.19E-03 |
| GGT1 | 231.5159883 | 0.556276785 | 5.31E-04 |
| FAH | 2365.929671 | 0.556216469 | 4.29E-11 |
| INTS6 | 1176.900312 | 0.556192095 | 4.62E-05 |
| ABI1 | 1233.020187 | 0.55530601 | 8.47E-16 |

| | | | |
|---|---|---|---|
| MNAT1 | 1215.706917 | 0.55515661 | 3.68E-04 |
| RFLNB | 1500.315546 | 0.555135157 | 6.41E-07 |
| SUSD1 | 1885.99721 | 0.555069968 | 2.45E-07 |
| CADM4 | 889.7866067 | 0.554929273 | 3.92E-03 |
| PTPN12 | 6077.371603 | 0.554616294 | 6.19E-05 |
| HACE1 | 409.0077023 | 0.554527301 | 4.40E-03 |
| CTSO | 316.7800897 | 0.554410125 | 1.96E-02 |
| IQSEC1 | 2478.420337 | 0.554356886 | 5.88E-12 |
| TIMM44 | 2236.04911 | 0.554212495 | 4.73E-05 |
| FBXW8 | 1751.709567 | 0.553944282 | 4.17E-10 |
| CASP10 | 374.7130969 | 0.553446534 | 5.85E-04 |
| NANP | 1026.86019 | 0.553305977 | 7.50E-05 |
| RHBDD3 | 1223.115071 | 0.553088593 | 1.38E-10 |
| ABLIM3 | 5868.459542 | 0.552709367 | 3.28E-04 |
| GNB4 | 1581.570179 | 0.552264905 | 1.30E-05 |
| ARPC1B | 9222.847879 | 0.55199912 | 8.93E-07 |
| MAN1A2 | 2382.046761 | 0.551893062 | 2.35E-04 |
| CLEC2D | 144.1037452 | 0.551447832 | 2.31E-02 |
| GMPR2 | 4514.04944 | 0.551341978 | 1.47E-11 |
| PRRC1 | 3640.671727 | 0.55110129 | 5.63E-05 |
| ADK | 1521.24063 | 0.550956061 | 8.22E-10 |
| UBE2D3-AS1 | 70.66496968 | 0.5503469 | 3.65E-02 |
| AARSD1 | 1040.493228 | 0.550157981 | 2.99E-16 |
| DHRS7 | 1710.897479 | 0.54994154 | 1.51E-06 |
| CREM | 385.7283766 | 0.549452207 | 1.98E-07 |
| QRSL1 | 944.9892823 | 0.549401144 | 2.25E-05 |
| AKAP6 | 411.3853393 | 0.549370039 | 4.75E-02 |
| TMEM98 | 1509.247942 | 0.549154102 | 1.08E-04 |
| C14orf132 | 1060.585725 | 0.5486913 | 5.69E-03 |
| SLC35B3 | 642.8102402 | 0.548434652 | 1.22E-05 |
| MKNK2 | 4825.496095 | 0.54686915 | 2.59E-05 |
| MGME1 | 2856.925219 | 0.546463847 | 9.79E-14 |
| CD302 | 1041.566279 | 0.546124271 | 7.71E-09 |
| ZNF226 | 1041.694412 | 0.545886052 | 4.29E-05 |
| TFPI | 400.6441754 | 0.545514208 | 6.67E-04 |
| HS2ST1 | 2383.667486 | 0.545335493 | 2.44E-04 |
| BTF3L4 | 1946.302754 | 0.545326581 | 2.44E-04 |
| EMC7 | 1604.537184 | 0.544960868 | 1.78E-06 |
| LIPA | 3892.554295 | 0.544915233 | 3.05E-05 |
| HHEX | 480.0358831 | 0.544518558 | 1.88E-03 |
| RANBP9 | 2089.439573 | 0.544361998 | 3.80E-13 |
| TRUB1 | 555.5771618 | 0.544329193 | 9.41E-05 |
| QPRT | 11694.36959 | 0.544182563 | 7.94E-03 |
| ZNFX1 | 5548.017493 | 0.544117757 | 2.49E-05 |
| DESI2 | 2337.936691 | 0.544011671 | 4.94E-04 |
| MORF4L2 | 11614.8889 | 0.543886502 | 1.31E-24 |
| ATP6V0E1 | 3554.564393 | 0.543760091 | 2.86E-05 |

| | | | |
|---|---|---|---|
| SPIRE1 | 1623.898975 | 0.543727663 | 4.86E-04 |
| RAB1A | 4088.452228 | 0.543480846 | 4.30E-14 |
| MTIF3 | 463.7794568 | 0.543137165 | 1.37E-04 |
| AC016582.3 | 142.7343043 | 0.542996332 | 5.57E-03 |
| RASL10B | 1608.736611 | 0.542774195 | 1.75E-05 |
| TKT | 23860.21316 | 0.542205402 | 5.16E-07 |
| CAMLG | 1170.821212 | 0.542140382 | 3.09E-07 |
| SMIM13 | 1014.459578 | 0.541816176 | 4.15E-04 |
| RHOT1 | 1697.536288 | 0.541753092 | 2.63E-07 |
| IQSEC2 | 1053.000885 | 0.541679494 | 1.60E-04 |
| HOXD8 | 615.7860238 | 0.541401983 | 1.89E-02 |
| TRAPPC3 | 2005.937891 | 0.541108079 | 2.20E-10 |
| SLC43A1 | 1313.017204 | 0.540948175 | 2.92E-03 |
| ANAPC10 | 365.3440448 | 0.540918373 | 4.98E-05 |
| CDC73 | 1412.75039 | 0.540703882 | 1.43E-04 |
| JMY | 1626.96102 | 0.540680315 | 3.84E-02 |
| JTB | 2513.992003 | 0.54048924 | 4.83E-08 |
| ATF1 | 622.7838902 | 0.540445401 | 2.21E-02 |
| LIFR | 13533.60101 | 0.540251919 | 1.82E-02 |
| VPS26C | 2551.853355 | 0.540112498 | 9.44E-18 |
| AC156455.1 | 100.1937583 | 0.539570381 | 2.72E-02 |
| NA | 202.7798493 | 0.53860774 | 4.78E-04 |
| SECISBP2L | 1068.391519 | 0.538360063 | 8.57E-03 |
| DPP7 | 4715.09999 | 0.538316656 | 2.20E-05 |
| PPM1M | 1735.27223 | 0.538282056 | 1.15E-05 |
| XIAP | 1292.126033 | 0.538216946 | 9.83E-03 |
| REST | 1531.355795 | 0.538155315 | 6.71E-05 |
| ITGB4 | 449.0655626 | 0.537910374 | 1.77E-02 |
| STAG3L5P-PVR | 1727.688963 | 0.537340819 | 4.48E-05 |
| TRAPPC6B | 1174.086258 | 0.536798783 | 1.53E-02 |
| SELENOS | 1716.629402 | 0.536792964 | 5.50E-13 |
| NRTN | 83.69516378 | 0.536766987 | 4.38E-02 |
| AFG3L2 | 4562.506854 | 0.536751093 | 2.70E-23 |
| SUPT7L | 3056.145985 | 0.53614417 | 2.81E-17 |
| PRKAG2-AS1 | 54.82964751 | 0.536073052 | 4.35E-02 |
| MAP1LC3A | 1651.699059 | 0.53585389 | 7.54E-05 |
| TBL2 | 3247.426371 | 0.5358488 | 4.55E-10 |
| DDX19B | 2093.58396 | 0.535698353 | 4.92E-03 |
| MYH9 | 39701.66481 | 0.535607416 | 6.02E-07 |
| LINC01271 | 81.39153448 | 0.53556332 | 3.34E-02 |
| TMEM41B | 992.340041 | 0.535374136 | 7.70E-05 |
| CAPRIN2 | 846.4307427 | 0.534761969 | 3.62E-02 |
| DNAJA2 | 4555.427379 | 0.534754216 | 5.48E-24 |
| NA | 1871.574947 | 0.534451192 | 1.24E-10 |
| KCTD20 | 6930.129421 | 0.534212851 | 2.45E-28 |
| TBRG1 | 2566.845992 | 0.534118611 | 4.18E-08 |
| TK2 | 1898.599849 | 0.534045502 | 2.02E-07 |

| | | | |
|---|---|---|---|
| SLC4A1AP | 2028.376907 | 0.533998348 | 4.90E-10 |
| TIGAR | 630.8242222 | 0.533880517 | 2.18E-04 |
| PARP14 | 3812.609189 | 0.533776321 | 9.15E-04 |
| TTYH3 | 13891.95316 | 0.533754472 | 1.23E-06 |
| CORO1C | 8847.320057 | 0.533592488 | 3.16E-08 |
| EFNA1 | 986.7455075 | 0.533451168 | 6.98E-03 |
| NUMBL | 1600.540194 | 0.533192747 | 2.03E-04 |
| EGFL7 | 1775.9885 | 0.533111097 | 1.03E-03 |
| SRRM3 | 406.1814457 | 0.532856724 | 4.50E-04 |
| TLCD1 | 494.9847647 | 0.532439435 | 2.93E-04 |
| RAP1GAP | 474.2442089 | 0.532175912 | 3.18E-03 |
| DNAJC18 | 922.7044842 | 0.53213905 | 1.16E-07 |
| PITX2 | 89.53508605 | 0.532011157 | 2.78E-02 |
| NR1H2 | 1443.06312 | 0.531575905 | 9.11E-06 |
| ZCCHC9 | 2215.726008 | 0.531380014 | 9.00E-07 |
| FAT1 | 14154.47872 | 0.53135041 | 2.63E-06 |
| SEC61A2 | 352.1459018 | 0.531083047 | 4.04E-04 |
| SPARC | 56594.83605 | 0.530712896 | 9.08E-03 |
| TIMM23B | 344.5474395 | 0.530640987 | 1.05E-04 |
| DAPK3 | 2248.152994 | 0.530600586 | 7.46E-05 |
| MIA3 | 2860.699507 | 0.530180717 | 7.51E-04 |
| CLINT1 | 9716.206222 | 0.529830927 | 3.86E-05 |
| SRP54 | 3509.000559 | 0.529226576 | 1.08E-16 |
| PALB2 | 1355.274179 | 0.529100067 | 7.18E-04 |
| PLPP5 | 1498.63886 | 0.528665135 | 4.70E-11 |
| LRP5L | 159.6593645 | 0.528480931 | 1.04E-02 |
| MED13L | 4915.063541 | 0.528375769 | 3.90E-06 |
| C6orf48 | 7131.05733 | 0.528294562 | 1.69E-08 |
| TJP1 | 3258.84199 | 0.528058107 | 1.62E-06 |
| RALGAPA2 | 2680.8069 | 0.527917325 | 5.93E-07 |
| ZNF721 | 490.7716122 | 0.527738033 | 1.80E-06 |
| TAOK3 | 1225.38155 | 0.527470892 | 1.84E-06 |
| ZNF460 | 99.66333522 | 0.527434666 | 1.61E-02 |
| SLC37A3 | 2567.358471 | 0.527385037 | 1.19E-11 |
| REXO2 | 5050.771619 | 0.527343491 | 8.51E-12 |
| ANO8 | 402.3322437 | 0.527261449 | 2.61E-06 |
| TRPT1 | 1109.105546 | 0.527072517 | 7.12E-04 |
| CHCHD4 | 918.971077 | 0.526937554 | 5.14E-05 |
| SAGE1 | 622.5687556 | 0.526512217 | 6.21E-05 |
| ALKBH3 | 1188.819224 | 0.526409379 | 8.50E-08 |
| AC244153.1 | 322.5237666 | 0.526400809 | 3.28E-05 |
| SRPK2 | 3839.713398 | 0.526287957 | 3.66E-13 |
| NOL6 | 8157.384097 | 0.526273237 | 7.27E-09 |
| GTF2F2 | 842.4991189 | 0.52619167 | 4.67E-04 |
| GTF2H3 | 1819.584187 | 0.526185686 | 1.05E-03 |
| TYW5 | 484.9749854 | 0.525811813 | 3.58E-03 |
| UTRN | 4972.912532 | 0.525563649 | 7.25E-04 |

| | | | |
|---|---|---|---|
| PTDSS2 | 1997.212239 | 0.525341373 | 1.29E-05 |
| NA | 108.8565664 | 0.525150544 | 3.54E-02 |
| BBS10 | 716.8808489 | 0.525015273 | 9.57E-03 |
| PPFIA1 | 3610.881528 | 0.524912177 | 9.59E-08 |
| CALCOCO2 | 4846.363505 | 0.524646709 | 7.61E-13 |
| STK40 | 4424.176598 | 0.52404304 | 1.15E-12 |
| GOPC | 1413.898633 | 0.523904953 | 8.82E-03 |
| ASAP1 | 7407.398524 | 0.523850195 | 5.67E-07 |
| ALG11 | 446.8904915 | 0.523782099 | 1.73E-04 |
| LSM12 | 2512.723727 | 0.523702746 | 5.25E-16 |
| TMEM108 | 147.75432 | 0.52356322 | 2.70E-02 |
| CRTAP | 9964.199093 | 0.523551133 | 2.75E-06 |
| KCTD3 | 2461.517131 | 0.523483887 | 1.64E-04 |
| RARS | 5457.586302 | 0.523477555 | 1.89E-16 |
| PRKAB2 | 1352.821842 | 0.523009276 | 4.60E-04 |
| RAB3GAP2 | 2766.260583 | 0.52285978 | 5.90E-03 |
| NT5DC1 | 1286.054086 | 0.522805843 | 2.31E-07 |
| ACAD11 | 1034.096395 | 0.522792156 | 3.92E-03 |
| ACOT7 | 3467.030846 | 0.522647972 | 2.90E-07 |
| WBP1L | 2089.230149 | 0.522605901 | 9.17E-04 |
| SERPINB6 | 5673.984218 | 0.52231135 | 1.92E-07 |
| NOMO1 | 11370.47813 | 0.522137409 | 1.53E-06 |
| CHKA | 2021.673855 | 0.521649173 | 3.86E-08 |
| CWF19L1 | 851.5140741 | 0.52162888 | 8.56E-06 |
| AEN | 1754.595128 | 0.520992465 | 1.35E-02 |
| EYA1 | 61.04274887 | 0.520778585 | 4.71E-02 |
| SND1 | 22442.27026 | 0.5204623 | 8.76E-07 |
| CGRRF1 | 421.589538 | 0.52002522 | 7.09E-03 |
| LRRFIP2 | 1680.533681 | 0.519972255 | 2.63E-04 |
| PPP1R15B | 2476.940228 | 0.519574476 | 1.36E-14 |
| POC1B | 724.6914982 | 0.519448406 | 5.22E-05 |
| MROH8 | 164.8533653 | 0.519004528 | 3.74E-02 |
| CRYBB2P1 | 487.1810476 | 0.518841045 | 1.61E-04 |
| THAP6 | 409.1386829 | 0.518771644 | 2.93E-02 |
| C14orf28 | 151.003708 | 0.517887539 | 8.33E-03 |
| STX8 | 1628.006817 | 0.517539037 | 6.96E-08 |
| STAT2 | 3251.987653 | 0.517250554 | 3.32E-08 |
| OXCT1 | 3184.337989 | 0.517188117 | 2.62E-06 |
| NUDT7 | 199.4224604 | 0.517058009 | 3.11E-03 |
| ZNF449 | 394.9404058 | 0.517056229 | 7.17E-03 |
| SAMD8 | 962.9109496 | 0.515903957 | 6.05E-05 |
| TCP11L2 | 360.9121766 | 0.515748449 | 1.49E-02 |
| CR381653.1 | 418.8328096 | 0.515536808 | 5.56E-05 |
| SPON2 | 97.66510103 | 0.515356102 | 2.16E-02 |
| ACAT1 | 4651.28969 | 0.514737157 | 2.34E-10 |
| MEF2A | 1827.254252 | 0.514174276 | 2.40E-05 |
| TMEM159 | 936.2046296 | 0.513962773 | 1.61E-05 |

| | | | |
|---|---|---|---|
| ZNF281 | 1503.230845 | 0.513884778 | 1.13E-08 |
| ATG2B | 2630.035045 | 0.513597722 | 6.12E-03 |
| TRIM25 | 7151.293211 | 0.512959276 | 2.46E-05 |
| WASL | 2325.29594 | 0.51261967 | 1.83E-05 |
| CHIC1 | 312.476867 | 0.512448845 | 7.70E-03 |
| WFS1 | 1309.328652 | 0.512201252 | 9.62E-04 |
| PCBP3 | 259.4287125 | 0.512160019 | 4.10E-02 |
| DUSP12 | 1147.714159 | 0.512058269 | 5.04E-04 |
| SLC39A11 | 639.9957675 | 0.512041868 | 2.07E-05 |
| GPR180 | 1990.300594 | 0.511916936 | 5.96E-04 |
| PTPRF | 14909.99649 | 0.511779254 | 6.46E-05 |
| SHTN1 | 750.5620927 | 0.511613688 | 6.37E-05 |
| RHBDF2 | 1009.728671 | 0.511174165 | 5.10E-06 |
| GTPBP4 | 3698.449712 | 0.511098048 | 3.23E-05 |
| SYT7 | 257.2568568 | 0.510993552 | 7.38E-04 |
| GK | 504.2900034 | 0.510914934 | 2.42E-05 |
| PAXIP1-AS2 | 428.289815 | 0.510570707 | 7.95E-04 |
| TATDN1 | 866.9645407 | 0.510547356 | 3.92E-02 |
| GNAO1 | 1149.018389 | 0.510161354 | 1.01E-03 |
| SYF2 | 1559.790301 | 0.510065463 | 1.46E-07 |
| RTCA | 1024.867536 | 0.510054189 | 1.21E-02 |
| PKP4 | 3485.234126 | 0.509815905 | 1.80E-05 |
| PON3 | 552.0018198 | 0.509387306 | 3.64E-05 |
| ELOVL4 | 381.9483083 | 0.509358787 | 1.86E-04 |
| FYCO1 | 2501.029106 | 0.509162227 | 7.68E-10 |
| NDEL1 | 2052.460807 | 0.509142979 | 1.49E-17 |
| LDLRAP1 | 1727.553505 | 0.509124334 | 3.58E-07 |
| COPZ2 | 778.6428004 | 0.509093279 | 1.13E-03 |
| ASF1A | 781.5517459 | 0.508803254 | 5.90E-03 |
| TBC1D23 | 1030.21793 | 0.508660601 | 3.39E-03 |
| DNAJA3 | 2240.392464 | 0.508659792 | 6.03E-05 |
| PLEKHA8P1 | 283.532566 | 0.50844355 | 2.11E-03 |
| GPCPD1 | 2138.032665 | 0.508442404 | 3.16E-02 |
| NDUFAF4 | 634.1814009 | 0.507966325 | 2.44E-03 |
| AC099778.1 | 87.65766901 | 0.507264152 | 2.96E-02 |
| PRPSAP2 | 2086.22234 | 0.506664391 | 3.96E-06 |
| ATP6V0E2-AS1 | 473.5699297 | 0.506073476 | 4.22E-05 |
| HBEGF | 861.6443607 | 0.505897709 | 3.79E-04 |
| FAM228B | 292.6928435 | 0.505785224 | 3.26E-03 |
| SLC39A1 | 6731.336194 | 0.505596365 | 1.13E-04 |
| MARS | 14346.5054 | 0.505452993 | 1.11E-11 |
| TNFRSF8 | 76.01422714 | 0.505195015 | 3.76E-02 |
| TRIM22 | 535.1244149 | 0.505156641 | 6.36E-03 |
| CGB7 | 208.1396265 | 0.504977121 | 4.48E-02 |
| ERBB2 | 4218.931161 | 0.504871738 | 1.01E-05 |
| SFT2D1 | 1397.633585 | 0.504833198 | 2.08E-15 |
| TMEM268 | 1773.573704 | 0.504766862 | 1.37E-02 |

| | | | |
|---|---|---|---|
| WNT5A | 1223.951769 | 0.504581973 | 1.17E-03 |
| ASAP3 | 2641.095712 | 0.504323676 | 4.87E-08 |
| NOMO3 | 6166.030026 | 0.504182647 | 5.75E-06 |
| MTO1 | 1439.627308 | 0.503972209 | 7.23E-04 |
| DSP | 1365.602877 | 0.503894779 | 7.04E-05 |
| ATP6V1F | 3728.914197 | 0.503809192 | 2.54E-06 |
| UEVLD | 829.6436829 | 0.503634924 | 6.05E-04 |
| SLC20A1 | 3792.236403 | 0.50326232 | 1.20E-03 |
| SLC2A10 | 1667.107831 | 0.503176225 | 1.29E-07 |
| AS3MT | 520.8533791 | 0.502984178 | 8.39E-04 |
| ZNF518B | 1323.76808 | 0.502938686 | 5.83E-05 |
| LRPPRC | 6676.377957 | 0.502852838 | 8.77E-04 |
| IMPACT | 1408.132623 | 0.502069173 | 4.44E-08 |
| MCEE | 233.2220779 | 0.50205156 | 5.68E-03 |
| SERTAD2 | 1650.89911 | 0.501998148 | 7.82E-06 |
| MET | 4328.732849 | 0.5018248 | 2.14E-03 |
| AXL | 11912.61544 | 0.501586982 | 8.81E-03 |
| ZNF317 | 2152.413878 | 0.501426254 | 1.10E-07 |
| ZNF630 | 219.9959955 | 0.501243268 | 1.09E-03 |
| EPOP | 1196.081005 | 0.501108063 | 2.04E-03 |
| TTC7B | 1528.002513 | 0.501107757 | 2.75E-05 |
| RAB10 | 4799.435422 | 0.501033675 | 2.21E-07 |
| GLIS2 | 746.7602133 | 0.501000365 | 1.13E-04 |
| LRIF1 | 1383.86094 | 0.499419225 | 1.40E-04 |
| SMAD2 | 2978.677024 | 0.499363197 | 6.11E-10 |
| MICAL3 | 4913.970363 | 0.499115875 | 1.44E-10 |
| EPRS | 10315.70588 | 0.499110097 | 3.84E-03 |
| DYRK1A | 2154.694965 | 0.499101417 | 3.63E-06 |
| SLC37A2 | 644.4016214 | 0.498388202 | 3.91E-03 |
| PLK3 | 284.7440062 | 0.498376285 | 1.04E-03 |
| HLA-A | 14384.32314 | 0.498343084 | 5.19E-05 |
| CCDC47 | 3711.747249 | 0.497692722 | 2.62E-06 |
| SPAG9 | 5033.827121 | 0.497417037 | 7.36E-07 |
| AC137932.3 | 111.9368522 | 0.497386321 | 8.45E-03 |
| PAIP2B | 247.9293627 | 0.497055883 | 4.54E-03 |
| TRAM2 | 9655.068286 | 0.496972345 | 2.11E-13 |
| TMEM208 | 1264.667512 | 0.496594337 | 1.38E-05 |
| MMADHC | 4024.231194 | 0.496318656 | 1.75E-15 |
| PNPLA7 | 124.432657 | 0.496256382 | 2.63E-02 |
| FAM111A-DT | 181.9856441 | 0.496177546 | 9.20E-04 |
| MOAP1 | 1235.398058 | 0.496135936 | 1.11E-08 |
| AC010442.1 | 720.7233888 | 0.496057459 | 6.78E-06 |
| PIK3CA | 1376.042323 | 0.495881756 | 2.34E-02 |
| SSH1 | 4488.68356 | 0.495688791 | 8.05E-12 |
| DOCK6 | 1355.620577 | 0.495664682 | 7.59E-04 |
| RTN4RL2 | 207.8606213 | 0.495375811 | 1.32E-02 |
| SENP5 | 1734.480153 | 0.495123418 | 4.12E-04 |

| | | | |
|---|---|---|---|
| BBIP1 | 337.4951943 | 0.49498758 | 1.04E-04 |
| RIT1 | 1170.596501 | 0.494891531 | 2.47E-06 |
| NCK1 | 1081.918685 | 0.494258037 | 1.42E-04 |
| DGKG | 127.0190231 | 0.493982989 | 2.91E-02 |
| RPF2 | 1752.066944 | 0.493926078 | 2.36E-02 |
| HSPBAP1 | 442.5596767 | 0.493729948 | 3.83E-06 |
| LSM8 | 1401.246654 | 0.493291976 | 1.18E-06 |
| BUD31 | 4153.64177 | 0.493242704 | 2.18E-06 |
| AC007325.4 | 431.1826631 | 0.49316856 | 5.96E-03 |
| GTF2B | 1132.640177 | 0.493125382 | 1.97E-07 |
| GARS-DT | 1058.115265 | 0.492797093 | 5.28E-06 |
| SMIM20 | 810.4386894 | 0.492556726 | 9.20E-08 |
| GNB5 | 2160.045094 | 0.492545214 | 1.03E-11 |
| ZFAND6 | 1726.062275 | 0.492517684 | 4.53E-07 |
| CDCP1 | 1149.145664 | 0.492345712 | 1.76E-03 |
| RPF1 | 1397.979862 | 0.492174623 | 5.51E-05 |
| VAC14 | 3528.466868 | 0.492112345 | 5.87E-06 |
| SLC25A37 | 6361.631733 | 0.492019341 | 7.14E-18 |
| CEP120 | 1616.91796 | 0.49198188 | 2.65E-02 |
| ATP6V1G1 | 2695.790776 | 0.491745874 | 3.10E-12 |
| DHRS7B | 459.8924902 | 0.491724694 | 8.03E-05 |
| SELENOM | 467.2787855 | 0.491079823 | 3.59E-03 |
| PRPS1P2 | 96.21341954 | 0.490995424 | 3.01E-02 |
| ZNF257 | 172.0790604 | 0.490838683 | 1.22E-02 |
| DLX4 | 530.3310774 | 0.490661231 | 1.64E-04 |
| SOCS6 | 907.8110569 | 0.490652048 | 3.45E-07 |
| TBC1D15 | 1376.672471 | 0.490536732 | 6.75E-03 |
| PTGES | 469.4516679 | 0.490231537 | 2.32E-02 |
| RNF13 | 1302.496039 | 0.490091952 | 3.51E-03 |
| TMEM92 | 202.8482872 | 0.489933543 | 2.15E-02 |
| IFNGR2 | 3672.123779 | 0.489913556 | 1.19E-05 |
| LAMTOR3 | 1013.807565 | 0.489783619 | 8.44E-04 |
| AHNAK | 122265.4934 | 0.489747831 | 2.83E-08 |
| GTF2E2 | 2020.494839 | 0.489727747 | 6.91E-11 |
| PPP1R3C | 1863.591223 | 0.4896986 | 2.35E-06 |
| ACOT2 | 1475.773493 | 0.48935149 | 1.60E-04 |
| IL1R1 | 1260.323681 | 0.4887745 | 1.90E-04 |
| NLRP11 | 220.6543597 | 0.488751843 | 7.63E-03 |
| PARN | 3180.356521 | 0.488161226 | 5.42E-12 |
| CATSPER2P1 | 101.1056114 | 0.488137036 | 3.55E-02 |
| PITPNA-AS1 | 362.7373036 | 0.487936788 | 8.48E-03 |
| USF3 | 829.8752061 | 0.487856116 | 5.67E-03 |
| CLIC4 | 20156.46442 | 0.487720216 | 1.77E-05 |
| MDN1 | 7460.91208 | 0.48761635 | 5.19E-03 |
| EGLN1 | 1845.305623 | 0.487519266 | 1.48E-04 |
| INPP5K | 1873.425128 | 0.487509722 | 5.58E-05 |
| CCDC174 | 895.7703077 | 0.486782979 | 2.10E-04 |

| | | | |
|---|---|---|---|
| REL | 362.8332223 | 0.48641543 | 1.12E-02 |
| OGFRL1 | 1421.080525 | 0.486162952 | 1.18E-02 |
| UBN1 | 2592.183221 | 0.485657525 | 1.29E-07 |
| CYB5R4 | 759.2449732 | 0.485465987 | 2.00E-02 |
| SPRYD3 | 1098.046721 | 0.485133629 | 1.36E-04 |
| GOLM1 | 4855.950469 | 0.485022155 | 3.69E-12 |
| ANGPT1 | 607.9227096 | 0.484991743 | 9.08E-03 |
| CYCS | 4213.819034 | 0.484968502 | 1.63E-05 |
| CLDN12 | 1756.686322 | 0.484798442 | 1.09E-03 |
| YTHDF3 | 3985.073013 | 0.484798033 | 5.18E-10 |
| LRP1 | 11435.56865 | 0.484758716 | 3.87E-04 |
| CREB3 | 2795.420509 | 0.484645373 | 1.10E-04 |
| FZD5 | 364.7941564 | 0.484576786 | 3.18E-04 |
| CECR2 | 599.8277745 | 0.484474139 | 6.77E-03 |
| GSK3B | 3261.575123 | 0.484060887 | 4.17E-08 |
| TMED3 | 5421.603628 | 0.483898953 | 3.27E-04 |
| SNRPN | 5362.88291 | 0.483047001 | 4.16E-07 |
| R3HDM2 | 1816.887832 | 0.48276397 | 4.95E-04 |
| BET1L | 4729.619093 | 0.482714622 | 9.41E-06 |
| PDIA5 | 1145.144501 | 0.482681342 | 4.77E-04 |
| PSMA3 | 2944.602149 | 0.482420646 | 1.79E-05 |
| PHLPP2 | 1636.290208 | 0.482334968 | 1.14E-09 |
| DYNC1LI1 | 1867.123395 | 0.482198223 | 4.01E-04 |
| DOCK1 | 3315.886736 | 0.481919063 | 6.15E-11 |
| GATA4 | 1468.75688 | 0.481888284 | 7.49E-03 |
| ELMSAN1 | 2538.049382 | 0.481018938 | 9.10E-08 |
| FPGS | 3260.463423 | 0.480468324 | 5.25E-06 |
| SNAPC5 | 1007.21619 | 0.480420667 | 3.52E-08 |
| TECPR1 | 2202.322973 | 0.48024595 | 6.10E-06 |
| ZNF853 | 558.4164668 | 0.480230935 | 5.71E-03 |
| SMAD3 | 7634.379311 | 0.480194318 | 3.98E-08 |
| PLSCR4 | 241.6913682 | 0.480109779 | 1.31E-02 |
| GRAMD1B | 1146.120968 | 0.480076538 | 1.07E-03 |
| MORN4 | 623.2349339 | 0.479838414 | 3.46E-06 |
| NA | 646.9206097 | 0.479472941 | 3.74E-05 |
| ANKRD28 | 4485.042749 | 0.479314504 | 5.58E-03 |
| USP38 | 1006.367779 | 0.478846398 | 3.79E-04 |
| RETSAT | 2730.549009 | 0.478824352 | 4.20E-05 |
| FAM76A | 274.522027 | 0.478809596 | 1.46E-02 |
| PON2 | 4494.206146 | 0.47856915 | 6.60E-10 |
| MYO5A | 3183.631342 | 0.478459908 | 5.96E-04 |
| ZHX3 | 3668.787153 | 0.478392082 | 2.06E-05 |
| FRG1JP | 122.945195 | 0.478251748 | 1.27E-02 |
| AVL9 | 3433.850467 | 0.477881035 | 4.30E-04 |
| ATP6V0C | 2785.293244 | 0.477533056 | 2.43E-04 |
| ABCB6 | 2468.804131 | 0.477389941 | 2.56E-04 |
| MITD1 | 822.8792502 | 0.477131269 | 1.28E-05 |

| | | | |
|---|---|---|---|
| UBE2Q1 | 3304.876586 | 0.476512663 | 2.49E-04 |
| SPTLC1 | 3390.99703 | 0.476401548 | 2.25E-08 |
| ENTPD4 | 3597.043861 | 0.476393222 | 1.67E-05 |
| LARP1 | 12477.89571 | 0.476264985 | 3.52E-15 |
| CES2 | 4372.401235 | 0.476018909 | 7.78E-09 |
| DYNLL2 | 4327.120161 | 0.47585022 | 5.19E-09 |
| SYNE2 | 2558.962612 | 0.475447521 | 1.26E-03 |
| RNASEH1 | 1425.965528 | 0.47538847 | 1.01E-11 |
| AC009237.3 | 226.5611666 | 0.47523283 | 2.70E-02 |
| PINX1 | 749.064652 | 0.475177241 | 4.56E-08 |
| SMIM19 | 1014.322732 | 0.475126768 | 1.82E-08 |
| LINC00847 | 682.5496375 | 0.475055717 | 4.22E-05 |
| HEATR5A | 1798.727542 | 0.474924243 | 6.03E-05 |
| MBD4 | 1471.174463 | 0.474813393 | 2.92E-05 |
| PTPMT1 | 1669.947516 | 0.47458872 | 6.48E-08 |
| TNKS1BP1 | 7024.243343 | 0.474491401 | 2.21E-08 |
| ENTPD1-AS1 | 196.7310503 | 0.473930914 | 1.85E-02 |
| CCDC107 | 476.26538 | 0.473879112 | 1.90E-02 |
| RBSN | 2154.092521 | 0.473374102 | 4.57E-08 |
| PPT1 | 10242.991 | 0.473247002 | 2.64E-06 |
| ARL14EP | 914.3959616 | 0.473188861 | 4.56E-02 |
| UBL3 | 632.1579712 | 0.473155824 | 9.50E-04 |
| PEX3 | 815.4626573 | 0.473139946 | 5.42E-09 |
| ANKIB1 | 5079.40403 | 0.473063087 | 6.03E-05 |
| ZMYM5 | 236.1654716 | 0.47269562 | 4.86E-02 |
| PPARGC1B | 475.6106913 | 0.47265753 | 7.07E-03 |
| ZNF738 | 275.9543995 | 0.472427541 | 3.73E-03 |
| TUBB6 | 6630.385277 | 0.471997898 | 8.34E-03 |
| MRPS18C | 1122.775971 | 0.471855152 | 1.42E-07 |
| PBDC1 | 1042.742786 | 0.471371517 | 5.19E-09 |
| MEIS3P1 | 287.1832009 | 0.471268777 | 2.95E-03 |
| SEC11C | 605.6753875 | 0.471198136 | 8.50E-06 |
| XRRA1 | 800.521416 | 0.471185115 | 1.10E-02 |
| RNF114 | 4494.094987 | 0.470990217 | 3.98E-09 |
| SAR1A | 3123.489316 | 0.470733612 | 2.11E-05 |
| PANK3 | 2532.270152 | 0.47062716 | 2.59E-02 |
| ZNF12 | 1848.754705 | 0.470537678 | 2.07E-03 |
| FBLIM1 | 6080.812389 | 0.47043224 | 2.82E-03 |
| ARHGEF12 | 14026.13928 | 0.470352118 | 2.08E-06 |
| UBQLN1 | 7209.186043 | 0.470181499 | 3.36E-13 |
| CAD | 8898.770146 | 0.469838201 | 4.15E-09 |
| ERP44 | 4292.313309 | 0.469616112 | 2.16E-13 |
| TMEM184C | 1244.756484 | 0.46950615 | 2.56E-05 |
| PPP1R2 | 1195.383891 | 0.469078833 | 1.23E-02 |
| PSMC1 | 6088.680726 | 0.469027267 | 3.91E-10 |
| HEXA | 7635.630594 | 0.468975513 | 7.96E-05 |
| C5orf56 | 283.159307 | 0.468895238 | 1.20E-02 |

| | | | |
|---|---|---|---|
| CLOCK | 1168.238529 | 0.468768879 | 2.15E-02 |
| ATP6AP1 | 9199.459948 | 0.468763193 | 1.03E-04 |
| OSBPL6 | 864.281503 | 0.468727695 | 1.44E-03 |
| TNRC6C-AS1 | 1271.337069 | 0.468697771 | 1.10E-03 |
| SAE1 | 11296.4598 | 0.46856077 | 4.56E-09 |
| PPM1B | 1180.743235 | 0.468304638 | 8.06E-03 |
| AGA | 441.1289954 | 0.468127092 | 5.21E-04 |
| BLOC1S6 | 1357.049245 | 0.468020323 | 2.98E-03 |
| PIK3CB | 1228.520412 | 0.46771711 | 7.49E-06 |
| FBXL17 | 1252.937548 | 0.467378957 | 1.59E-04 |
| NAP1L5 | 362.4858112 | 0.467252576 | 4.36E-03 |
| FER | 2018.000342 | 0.466980054 | 4.70E-02 |
| SDE2 | 656.1897337 | 0.466501206 | 5.97E-04 |
| PANK2 | 3143.512483 | 0.466217359 | 3.21E-17 |
| YIPF5 | 2324.500251 | 0.466187659 | 8.40E-05 |
| MAP3K13 | 776.1410187 | 0.466032507 | 1.18E-06 |
| SLC23A2 | 3962.862943 | 0.465940542 | 3.47E-13 |
| AGO2 | 1010.035818 | 0.465692892 | 2.45E-03 |
| WDR7 | 1333.184344 | 0.465669036 | 9.23E-05 |
| UTP18 | 2043.71906 | 0.465577871 | 2.32E-11 |
| FAM234B | 967.1611581 | 0.465547828 | 5.08E-06 |
| AXIN1 | 1792.469903 | 0.465501759 | 3.74E-04 |
| MTSS1L | 4140.209056 | 0.465045494 | 3.64E-05 |
| TRAF4 | 2091.976798 | 0.464789194 | 4.67E-06 |
| CBLL1 | 1559.789929 | 0.464752655 | 1.23E-07 |
| NEDD9 | 1801.09962 | 0.464686406 | 1.56E-04 |
| MTHFD1L | 2628.193976 | 0.464619436 | 6.87E-03 |
| HAUS2 | 847.5349321 | 0.464411769 | 2.06E-03 |
| CNNM2 | 636.4353897 | 0.464409565 | 2.58E-05 |
| AKAP5 | 222.2346906 | 0.464370011 | 1.37E-02 |
| SH3BP4 | 4262.928303 | 0.464034197 | 1.66E-05 |
| SLC15A4 | 708.4678334 | 0.463888415 | 9.60E-09 |
| TMEM241 | 503.1300017 | 0.463587309 | 2.15E-03 |
| GJC1 | 1901.807077 | 0.463474356 | 5.52E-05 |
| SCAMP2 | 3244.65783 | 0.463432481 | 4.84E-04 |
| JUP | 7928.88656 | 0.46328958 | 4.23E-03 |
| ZDHHC2 | 859.2562326 | 0.462746491 | 6.12E-04 |
| LINC00963 | 2907.811582 | 0.462742533 | 9.91E-05 |
| C1orf52 | 736.1965473 | 0.462697069 | 1.68E-06 |
| RP9 | 827.2961611 | 0.462645648 | 2.37E-04 |
| IL17RA | 1078.041984 | 0.462566885 | 8.02E-03 |
| POTEF | 330.2910566 | 0.462408642 | 1.98E-02 |
| FKBP9 | 12757.91606 | 0.461806179 | 1.93E-07 |
| KIF1C | 10083.17981 | 0.461606845 | 4.93E-07 |
| WDR45B | 3377.208925 | 0.461083791 | 2.07E-12 |
| USP9X | 8356.864961 | 0.460750281 | 2.77E-04 |
| DHX36 | 2895.374078 | 0.460635868 | 4.28E-03 |

| | | | |
|---|---|---|---|
| DYNC1H1 | 30557.26086 | 0.46055774 | 5.23E-07 |
| RALGAPB | 7349.93362 | 0.460549441 | 8.61E-08 |
| NECTIN3 | 569.8866024 | 0.460120669 | 4.00E-03 |
| PNPLA8 | 1719.226756 | 0.460077932 | 1.92E-02 |
| FMNL1 | 1214.864249 | 0.459887089 | 1.57E-04 |
| NA | 138.6279895 | 0.459830362 | 2.84E-02 |
| RNF130 | 2620.853033 | 0.459661561 | 5.02E-07 |
| NDUFA4 | 4585.705816 | 0.459619888 | 2.46E-02 |
| CBWD1 | 901.7599615 | 0.459312873 | 1.38E-03 |
| C2orf49 | 1087.85998 | 0.45930104 | 2.04E-06 |
| LINC00839 | 428.792773 | 0.459270799 | 1.23E-05 |
| AP003486.1 | 241.7248542 | 0.459186178 | 1.89E-02 |
| ZNF587 | 1833.970438 | 0.45888416 | 5.27E-06 |
| PIP4P1 | 1266.099527 | 0.458579609 | 1.43E-03 |
| TMCC1 | 728.6019316 | 0.458273834 | 5.53E-05 |
| RABGGTB | 1841.804917 | 0.458223502 | 2.32E-02 |
| COG3 | 1626.367913 | 0.458196304 | 2.27E-04 |
| DDI2 | 1498.305752 | 0.457800376 | 2.84E-03 |
| RFT1 | 1944.490929 | 0.457540909 | 5.52E-11 |
| ERGIC2 | 921.606192 | 0.457382597 | 3.80E-02 |
| AGPAT3 | 4177.175525 | 0.457048725 | 4.39E-12 |
| PDE4D | 2698.714731 | 0.456965357 | 3.70E-02 |
| ZNF706 | 2431.618789 | 0.45658175 | 3.70E-13 |
| TENT2 | 2303.57107 | 0.456295064 | 5.32E-03 |
| MTFR1 | 1478.258109 | 0.456149986 | 6.40E-05 |
| SLC31A2 | 408.2149728 | 0.45583851 | 1.82E-02 |
| MAPRE2 | 4609.080376 | 0.455654359 | 2.87E-12 |
| FGF2 | 458.0900956 | 0.455308297 | 2.70E-02 |
| ATP7A | 1068.071532 | 0.454999517 | 1.69E-03 |
| CDH13 | 11715.24575 | 0.454735379 | 2.43E-04 |
| ZNF486 | 252.3443474 | 0.454091789 | 1.28E-02 |
| IFITM1 | 3198.919102 | 0.454050292 | 2.56E-02 |
| PET117 | 794.4960644 | 0.454025805 | 3.83E-08 |
| TRNT1 | 945.3644513 | 0.453696623 | 3.08E-03 |
| AK6 | 844.4994681 | 0.453658528 | 4.95E-03 |
| AL133467.2 | 401.6717852 | 0.453461039 | 1.51E-04 |
| RNF14 | 1986.791575 | 0.453380223 | 5.24E-06 |
| LRP8 | 1177.83127 | 0.453012441 | 3.71E-03 |
| APBB2 | 2409.951598 | 0.452968496 | 1.64E-04 |
| PTTG1IP | 16185.65895 | 0.452561403 | 3.75E-05 |
| PBLD | 150.0356914 | 0.452540984 | 4.78E-02 |
| HOXC8 | 845.3701195 | 0.452517054 | 3.55E-05 |
| NCKAP5L | 3265.097525 | 0.452440754 | 3.56E-05 |
| IMPAD1 | 4303.641024 | 0.452403128 | 4.34E-06 |
| FOXK2 | 3237.994456 | 0.452340093 | 4.25E-04 |
| KLHL2 | 687.0025985 | 0.452313443 | 2.89E-02 |
| MAGOH | 973.9237622 | 0.452273168 | 2.37E-06 |

| | | | |
|---|---|---|---|
| SH3D19 | 1811.616778 | 0.451707948 | 5.25E-04 |
| AZIN2 | 391.0001786 | 0.451278749 | 4.61E-02 |
| PAM16 | 284.7754864 | 0.451137978 | 6.49E-03 |
| NDUFAF5 | 604.7519493 | 0.450886365 | 4.77E-05 |
| APOL4 | 164.0383577 | 0.450369275 | 2.11E-02 |
| VPS13D | 3534.148094 | 0.450340515 | 8.43E-06 |
| ERP29 | 5731.028148 | 0.450315475 | 2.02E-06 |
| PGPEP1 | 1359.067798 | 0.450108367 | 2.24E-02 |
| ULBP3 | 139.2906982 | 0.449996251 | 3.83E-02 |
| HOXB9 | 2074.166601 | 0.449978018 | 3.90E-03 |
| ZMYM6 | 440.9584887 | 0.449668639 | 7.42E-03 |
| AC012313.1 | 785.7816603 | 0.449288453 | 7.11E-04 |
| MEGF9 | 1097.538604 | 0.449245163 | 3.21E-05 |
| CBX4 | 1634.077251 | 0.449173354 | 1.03E-03 |
| TRIM56 | 1938.37613 | 0.449142652 | 1.40E-06 |
| TP53BP2 | 2437.809138 | 0.448800592 | 1.22E-04 |
| P4HB | 29172.44412 | 0.448672721 | 7.72E-06 |
| NIT2 | 1435.897603 | 0.448569161 | 4.81E-06 |
| RAP1A | 1573.438963 | 0.448281225 | 3.33E-09 |
| EXT2 | 10034.73328 | 0.4478153 | 4.95E-04 |
| ADIRF-AS1 | 624.3818623 | 0.447402688 | 5.91E-04 |
| POLR3C | 1690.913297 | 0.44725913 | 3.94E-03 |
| HDHD3 | 862.3746448 | 0.447101009 | 4.57E-03 |
| ZMYM4 | 4083.321672 | 0.446796425 | 2.96E-03 |
| RRAGB | 446.9069824 | 0.446560749 | 9.09E-03 |
| CREB1 | 1266.207202 | 0.446134516 | 4.14E-02 |
| TMLHE | 1940.538019 | 0.446055714 | 1.57E-07 |
| LSM10 | 1191.931926 | 0.445572863 | 3.79E-05 |
| NMD3 | 1663.788087 | 0.445158409 | 1.33E-02 |
| SUCLG2 | 2395.918286 | 0.445005132 | 9.07E-08 |
| KRTCAP2 | 1559.895768 | 0.444865545 | 3.78E-05 |
| MAPK8IP2 | 998.1611941 | 0.444855399 | 2.18E-03 |
| HSPB6 | 899.0889698 | 0.444683856 | 1.64E-02 |
| NBPF14 | 4716.123144 | 0.444581096 | 8.64E-06 |
| FAM220A | 1262.311827 | 0.444520283 | 1.34E-03 |
| C11orf98 | 2552.043132 | 0.44441477 | 2.78E-06 |
| ANXA7 | 2952.787176 | 0.44428225 | 4.10E-09 |
| SOX13 | 1139.603012 | 0.444187343 | 3.43E-05 |
| WSB2 | 2525.982471 | 0.444170762 | 1.09E-06 |
| GNA13 | 2404.762405 | 0.443870636 | 6.20E-04 |
| ZFPL1 | 1177.567137 | 0.443837668 | 2.36E-04 |
| UBXN4 | 4202.091355 | 0.443796193 | 2.83E-03 |
| SAP30 | 507.7406263 | 0.443758845 | 4.14E-06 |
| DDX60L | 1220.764135 | 0.443683504 | 1.35E-02 |
| NSMCE3 | 513.061359 | 0.443586638 | 3.29E-05 |
| SUPT3H | 446.3907523 | 0.443407189 | 3.71E-04 |
| SNX4 | 778.4362423 | 0.443211548 | 2.13E-02 |

| | | | |
|---|---|---|---|
| TJP2 | 4658.735463 | 0.442866888 | 9.25E-06 |
| SNAP29 | 1759.719219 | 0.442759969 | 2.35E-09 |
| BTAF1 | 2298.431969 | 0.442671989 | 1.03E-02 |
| B4GALT6 | 552.4881887 | 0.442390264 | 2.10E-02 |
| JMJD4 | 1074.760631 | 0.442341861 | 4.83E-04 |
| CGREF1 | 748.9593742 | 0.442183273 | 5.75E-03 |
| ACBD4 | 227.6965106 | 0.442033882 | 4.22E-03 |
| SNAP23 | 1810.39237 | 0.441945824 | 1.85E-06 |
| HNRNPF | 6684.601055 | 0.44176275 | 2.76E-16 |
| GCC1 | 2344.373322 | 0.441715579 | 5.84E-06 |
| FAM50A | 5466.553151 | 0.44168905 | 6.80E-07 |
| TMEM128 | 489.3146157 | 0.441640562 | 3.38E-03 |
| MVP | 3819.471907 | 0.441296922 | 9.44E-03 |
| PPP1CB | 4116.18422 | 0.44128652 | 4.02E-02 |
| FAHD2A | 1741.478709 | 0.441160354 | 3.37E-07 |
| KCNN3 | 360.2014472 | 0.441112084 | 3.52E-02 |
| PCGF2 | 2220.258047 | 0.440761805 | 1.23E-04 |
| ZBTB2 | 1131.647043 | 0.440672261 | 2.39E-02 |
| CDC40 | 1575.43891 | 0.440500992 | 5.42E-05 |
| CFL2 | 3651.721601 | 0.440403435 | 1.88E-10 |
| TRIB1 | 1261.800018 | 0.440295169 | 1.65E-02 |
| ZIC4 | 194.7867583 | 0.440119188 | 2.59E-02 |
| C20orf194 | 2675.497658 | 0.440109722 | 1.43E-07 |
| BDH1 | 398.2595237 | 0.440059768 | 4.77E-04 |
| CAMTA1 | 1421.581369 | 0.439988813 | 7.07E-05 |
| MKLN1 | 5424.076468 | 0.439633192 | 4.03E-07 |
| TMEM44-AS1 | 169.8116783 | 0.439403696 | 3.38E-03 |
| GTF2A2 | 1816.018377 | 0.439207997 | 9.92E-07 |
| IFITM3 | 20144.00674 | 0.439132905 | 1.11E-02 |
| DENND4A | 1554.563487 | 0.43901439 | 2.95E-02 |
| MYORG | 1497.758198 | 0.438683798 | 1.15E-02 |
| FOXF2 | 394.4896013 | 0.438597999 | 4.87E-04 |
| ZNF805 | 126.9341375 | 0.43853161 | 4.71E-02 |
| SMAP1 | 1417.633339 | 0.437732847 | 1.60E-06 |
| CEMIP2 | 4440.133686 | 0.437606989 | 1.02E-04 |
| RHEB | 6449.634271 | 0.437177896 | 1.65E-13 |
| CAMK2D | 1210.58187 | 0.437025885 | 5.73E-03 |
| RAB4B | 443.5158268 | 0.437006306 | 7.01E-04 |
| ITPK1 | 2503.186321 | 0.436799314 | 3.66E-04 |
| MTRR | 3180.935578 | 0.436521524 | 2.49E-03 |
| DBT | 1105.356408 | 0.435956516 | 1.30E-02 |
| NF2 | 3616.898035 | 0.435106343 | 1.08E-04 |
| BRAP | 1210.93741 | 0.43502021 | 1.21E-08 |
| RPS6KA4 | 2634.070272 | 0.434819268 | 1.12E-05 |
| ADIPOR1 | 4348.452011 | 0.434657866 | 1.03E-04 |
| TFG | 4603.610223 | 0.434534942 | 1.87E-11 |
| MTERF2 | 526.1795391 | 0.434278308 | 3.01E-03 |

| | | | |
|---|---|---|---|
| GPX8 | 5676.176415 | 0.433929555 | 8.32E-04 |
| HIBADH | 5149.036504 | 0.433737447 | 5.85E-15 |
| LRP10 | 8748.08372 | 0.432963908 | 1.01E-03 |
| NSMCE2 | 1140.479055 | 0.432696722 | 1.14E-07 |
| CBS | 6183.303293 | 0.432477408 | 7.95E-04 |
| MAGED1 | 16745.61713 | 0.432318607 | 3.87E-02 |
| C11orf24 | 1879.979204 | 0.432253334 | 1.48E-04 |
| RYBP | 2158.376547 | 0.432231821 | 1.08E-02 |
| GPALPP1 | 1200.665299 | 0.432195282 | 8.69E-06 |
| PHF10 | 1866.438536 | 0.432178761 | 5.36E-07 |
| IGF2BP2 | 5316.699877 | 0.432172134 | 6.73E-07 |
| DNAJC19 | 835.3569696 | 0.431732088 | 1.31E-04 |
| WDR26 | 5431.769766 | 0.431727022 | 5.22E-15 |
| IFIT5 | 791.850336 | 0.431726991 | 1.08E-03 |
| ZP3 | 653.8483888 | 0.431650802 | 8.70E-03 |
| C16orf58 | 6656.991018 | 0.431593334 | 5.70E-06 |
| GANAB | 50670.80836 | 0.430875386 | 8.41E-06 |
| LIMS1 | 1656.651267 | 0.430695943 | 3.04E-03 |
| C16orf74 | 376.5939505 | 0.430464916 | 2.59E-03 |
| DTD1 | 3987.586041 | 0.43018413 | 9.14E-06 |
| FEZ2 | 2245.374732 | 0.430124152 | 1.30E-06 |
| CDK5RAP1 | 2109.566458 | 0.429991651 | 2.39E-07 |
| ZC2HC1A | 281.9343025 | 0.429649892 | 4.91E-02 |
| CPSF2 | 4290.404317 | 0.429578711 | 4.52E-02 |
| NR2F1-AS1 | 377.042445 | 0.429442523 | 8.33E-03 |
| FAF1 | 2836.781069 | 0.429055226 | 1.98E-07 |
| AP5Z1 | 1954.951571 | 0.429026091 | 2.68E-03 |
| GMPPA | 1577.592379 | 0.428940462 | 7.38E-05 |
| PSMA5 | 4318.210261 | 0.428632982 | 2.94E-06 |
| EMC3 | 1790.290029 | 0.428583335 | 1.32E-04 |
| BBS5 | 481.4587906 | 0.428341344 | 5.97E-05 |
| CNOT10 | 1611.329825 | 0.428250396 | 4.14E-03 |
| LHFPL2 | 4882.279716 | 0.428225906 | 4.78E-11 |
| ZNF215 | 464.4303147 | 0.428121057 | 4.12E-03 |
| GTF3C3 | 2564.112253 | 0.427965871 | 1.81E-03 |
| RPP38 | 393.6215106 | 0.427538011 | 6.79E-05 |
| UBE2E2 | 803.3808045 | 0.427367235 | 3.77E-03 |
| KIF1B | 4029.593743 | 0.427251233 | 5.32E-08 |
| PRR13 | 3653.230678 | 0.427187623 | 3.28E-04 |
| WDR36 | 3975.291948 | 0.426862808 | 1.15E-03 |
| INPPL1 | 9644.907944 | 0.426860019 | 9.39E-06 |
| FXR1 | 4988.513076 | 0.426582158 | 2.93E-03 |
| MTMR10 | 1017.928693 | 0.42653971 | 3.08E-04 |
| USP31 | 1052.765356 | 0.42630651 | 2.61E-03 |
| SERINC1 | 4518.27721 | 0.426083538 | 7.41E-04 |
| NDUFAF7 | 981.6976472 | 0.426035796 | 2.19E-04 |
| HIST2H2BF | 235.8078308 | 0.425941675 | 2.19E-02 |

| | | | |
|---|---|---|---|
| NSRP1 | 670.5981166 | 0.425353877 | 4.96E-02 |
| RAB24 | 1488.677025 | 0.425111302 | 7.51E-05 |
| WHAMMP2 | 146.3450519 | 0.425000682 | 4.10E-02 |
| AK2 | 8587.731938 | 0.424973265 | 6.23E-06 |
| ETF1 | 16211.09724 | 0.424801023 | 4.25E-05 |
| PTER | 667.7124145 | 0.424579948 | 2.47E-02 |
| RABGAP1L | 809.043815 | 0.424548412 | 7.41E-03 |
| SLC25A33 | 579.6992284 | 0.424474115 | 1.49E-04 |
| OTUD7B | 1444.623169 | 0.424375939 | 1.49E-05 |
| FAM13A | 1552.306742 | 0.424168673 | 1.79E-03 |
| RXYLT1 | 1054.450903 | 0.424073133 | 2.67E-02 |
| SH3BP5 | 1459.805366 | 0.424061006 | 2.16E-03 |
| DDX52 | 2152.99997 | 0.424014481 | 7.03E-06 |
| RAD23B | 8479.786235 | 0.423950791 | 2.08E-04 |
| ITPR2 | 1727.732014 | 0.423871851 | 2.08E-02 |
| GPAM | 1106.907471 | 0.423845615 | 2.91E-04 |
| MRPS12 | 1199.187636 | 0.4229052 | 1.46E-04 |
| FBXO6 | 243.709265 | 0.422815733 | 3.53E-02 |
| HYKK | 262.855178 | 0.422401141 | 7.35E-03 |
| TPST1 | 2070.732166 | 0.422216628 | 1.69E-04 |
| GTF2H1 | 3538.505438 | 0.422069753 | 1.68E-04 |
| PPP1R26 | 2889.950209 | 0.421928843 | 5.23E-05 |
| WDFY1 | 2048.290992 | 0.421863351 | 7.45E-04 |
| GTF2H2B | 334.5807758 | 0.421788452 | 4.67E-03 |
| PAPOLA | 6682.579099 | 0.421649386 | 1.46E-05 |
| VPS26A | 1738.082387 | 0.421589835 | 1.83E-02 |
| BCL2L1 | 5620.751235 | 0.42146886 | 1.82E-06 |
| PKIG | 2587.098172 | 0.42142066 | 7.65E-03 |
| LAPTM5 | 330.5302971 | 0.421334976 | 2.75E-02 |
| RAB37 | 569.4378075 | 0.420985468 | 2.18E-02 |
| AC011481.4 | 113.9890397 | 0.420525969 | 2.92E-02 |
| ZBTB43 | 985.9247111 | 0.420137198 | 4.59E-02 |
| COL5A1 | 25458.72823 | 0.420028181 | 4.88E-05 |
| ADAMTS10 | 829.7042696 | 0.419741586 | 1.01E-02 |
| GPSM1 | 4074.013977 | 0.419546681 | 1.69E-04 |
| ZNF622 | 2500.069831 | 0.419535007 | 3.37E-09 |
| GIHCG | 338.5023988 | 0.419532022 | 1.02E-03 |
| SUPT16H | 16245.5832 | 0.419089085 | 3.83E-08 |
| SUZ12 | 1749.24164 | 0.418315028 | 2.35E-02 |
| CSNK2A2 | 5047.769308 | 0.41777016 | 5.89E-11 |
| BROX | 895.3496209 | 0.41775418 | 1.69E-02 |
| GRK5 | 1308.605948 | 0.417713406 | 1.12E-05 |
| MCPH1 | 1085.253236 | 0.417617281 | 3.65E-03 |
| SYNGR2 | 1829.502754 | 0.417576374 | 1.02E-03 |
| FCGRT | 3493.295156 | 0.417539714 | 3.39E-02 |
| SET | 25045.76652 | 0.417035324 | 1.36E-03 |
| XYLT2 | 1979.267738 | 0.416471342 | 1.19E-04 |

| | | | |
|---|---|---|---|
| ZCCHC2 | 526.4422095 | 0.416217302 | 2.40E-02 |
| ATXN1L | 3477.835901 | 0.416125924 | 2.23E-06 |
| GALNT1 | 2272.25254 | 0.415966975 | 1.58E-02 |
| BPGM | 883.7041193 | 0.415883577 | 1.07E-06 |
| BCOR | 2863.827776 | 0.415574149 | 7.54E-06 |
| TRIT1 | 1099.34828 | 0.415463128 | 2.10E-04 |
| AC026412.1 | 1148.852394 | 0.415407507 | 1.01E-05 |
| ANKRD27 | 1949.923053 | 0.415313975 | 2.16E-07 |
| AAK1 | 2456.29884 | 0.415221344 | 8.07E-03 |
| TMC7 | 119.7353915 | 0.415155533 | 1.79E-02 |
| SLC41A1 | 2600.516694 | 0.413801174 | 3.22E-05 |
| SDCCAG8 | 630.3575857 | 0.413091673 | 4.37E-02 |
| SPTBN1 | 22759.67806 | 0.412756953 | 3.02E-13 |
| UBE2E1 | 2623.630516 | 0.412703293 | 1.07E-04 |
| VTI1A | 776.1752698 | 0.412671664 | 1.88E-03 |
| FBXO30 | 835.150853 | 0.412323957 | 3.01E-03 |
| IL18BP | 506.5528303 | 0.412111815 | 1.39E-02 |
| TTC17 | 4956.933534 | 0.412098388 | 8.74E-06 |
| SLC30A9 | 2686.38585 | 0.412067139 | 3.30E-02 |
| CSDE1 | 17721.67776 | 0.411975438 | 1.27E-07 |
| TNKS | 4426.097578 | 0.4118647 | 4.92E-05 |
| ZNF44 | 400.8301765 | 0.411856579 | 4.02E-02 |
| NFAT5 | 5313.455934 | 0.411777399 | 3.98E-02 |
| SLC38A3 | 241.9542686 | 0.411334487 | 4.68E-02 |
| MICALL1 | 3271.141042 | 0.410439264 | 6.38E-06 |
| AC027307.2 | 280.7966597 | 0.410406445 | 5.85E-03 |
| UAP1L1 | 2532.817372 | 0.410051351 | 2.59E-02 |
| SLC37A4 | 2303.912945 | 0.409424929 | 8.59E-04 |
| PDIA6 | 20872.47759 | 0.409150781 | 1.55E-07 |
| HLA-DQB1 | 303.1547478 | 0.409117735 | 8.12E-03 |
| ARFGAP1 | 6666.931935 | 0.408973304 | 1.03E-04 |
| ATG4A | 490.5422648 | 0.408961369 | 1.50E-02 |
| RHBDL2 | 184.1327411 | 0.408814189 | 1.82E-02 |
| ST13 | 13724.51909 | 0.408689051 | 5.73E-07 |
| ELF4 | 1789.071958 | 0.408201275 | 9.43E-04 |
| LRP6 | 2588.5102 | 0.408010551 | 1.76E-03 |
| RBM28 | 3312.820675 | 0.407611113 | 2.41E-04 |
| SGPP1 | 535.6014895 | 0.407336567 | 4.73E-03 |
| EEF1E1 | 820.3777301 | 0.407303834 | 4.38E-02 |
| PLRG1 | 3630.724933 | 0.407071268 | 1.00E-07 |
| PSMD7 | 7737.490292 | 0.406971807 | 2.47E-14 |
| CCDC28A | 816.7254079 | 0.406865079 | 9.24E-05 |
| NA | 110.1530595 | 0.406705661 | 4.11E-02 |
| MAP3K2-DT | 170.5922295 | 0.406637745 | 2.11E-02 |
| TMEM30A | 6283.824916 | 0.405512621 | 6.88E-04 |
| C9orf72 | 587.3200381 | 0.405397221 | 1.62E-02 |
| LINC00910 | 264.0356664 | 0.405229432 | 3.38E-03 |

| | | | |
|---|---|---|---|
| NCSTN | 7147.975551 | 0.405080737 | 4.81E-04 |
| PPP3CC | 566.011236 | 0.404505533 | 2.25E-02 |
| ALX1 | 332.6739069 | 0.404374107 | 2.28E-03 |
| NUDT14 | 692.5963255 | 0.403862587 | 4.72E-03 |
| NRIP3 | 1681.226296 | 0.403697561 | 2.05E-06 |
| CLN6 | 2149.43622 | 0.403681229 | 8.75E-08 |
| SIAH2 | 1088.269525 | 0.403590326 | 3.05E-04 |
| STYXL1 | 1931.959515 | 0.402954842 | 1.83E-03 |
| LRRC28 | 807.0602025 | 0.402527332 | 1.53E-06 |
| ZNF131 | 2746.314322 | 0.402393959 | 2.72E-03 |
| LINC00667 | 1026.945253 | 0.402217074 | 7.26E-04 |
| SGCE | 1697.348195 | 0.401500018 | 6.45E-05 |
| MAP2K2 | 4092.995995 | 0.401405354 | 7.89E-05 |
| MITF | 317.5617059 | 0.401358712 | 4.35E-02 |
| IMP4 | 3203.583644 | 0.400889044 | 2.84E-05 |
| CLTC | 25848.11783 | 0.400449193 | 2.35E-09 |
| SHISA4 | 760.8688191 | 0.400448812 | 8.00E-03 |
| SMG1P3 | 308.3634136 | 0.400242928 | 2.37E-02 |
| AP001372.2 | 231.832704 | 0.400094338 | 2.56E-02 |
| PWP1 | 3685.791824 | 0.400056607 | 2.30E-10 |
| HDGFL3 | 1750.855512 | 0.399799237 | 4.53E-02 |
| SPSB3 | 1598.016998 | 0.399637281 | 6.73E-04 |
| KMT5B | 1956.088618 | 0.399470372 | 1.62E-03 |
| MRPS22 | 2251.012462 | 0.399251706 | 3.42E-06 |
| WDR25 | 582.4176668 | 0.398524603 | 3.14E-02 |
| MTMR6 | 792.3833963 | 0.39851383 | 3.75E-03 |
| PREPL | 3920.905607 | 0.398511404 | 2.19E-02 |
| TIMP2 | 22643.24137 | 0.398251803 | 6.17E-03 |
| EIF3M | 9035.068701 | 0.398159768 | 5.71E-11 |
| B3GNT2 | 692.6789905 | 0.398129213 | 3.41E-04 |
| SERINC3 | 7719.314089 | 0.398005141 | 5.21E-11 |
| CCNB1IP1 | 4135.158345 | 0.397756505 | 6.45E-04 |
| HPS1 | 1129.99209 | 0.397494092 | 2.40E-03 |
| TMEM150A | 656.5324793 | 0.397462563 | 4.93E-03 |
| ARPC2 | 12473.43948 | 0.3971909 | 7.89E-07 |
| NUP58 | 1380.731867 | 0.397015892 | 1.93E-02 |
| EXOSC1 | 656.5367159 | 0.396894832 | 5.52E-05 |
| HHLA3 | 421.4929855 | 0.39682303 | 1.80E-02 |
| NUAK1 | 3145.049458 | 0.396689076 | 1.41E-03 |
| DGKQ | 1201.710352 | 0.396379505 | 1.84E-02 |
| POLR3G | 623.9924706 | 0.39510961 | 4.13E-02 |
| AAGAB | 3925.30352 | 0.39495192 | 2.82E-08 |
| RFXANK | 1757.375361 | 0.394671164 | 2.15E-03 |
| CMC1 | 560.2089086 | 0.394626912 | 3.28E-04 |
| LY6E | 19754.43352 | 0.394185219 | 2.00E-02 |
| KIN | 403.2926376 | 0.394128012 | 3.55E-02 |
| LINC00630 | 179.5448628 | 0.393927169 | 2.87E-02 |

| | | | |
|---|---|---|---|
| CNOT6L | 810.8697715 | 0.393854136 | 3.98E-02 |
| EDA | 1795.900129 | 0.393442555 | 2.37E-03 |
| LINC01234 | 437.4067572 | 0.393423976 | 1.90E-03 |
| MBD2 | 1768.177903 | 0.393260556 | 2.86E-02 |
| NXPE3 | 1902.101291 | 0.393147557 | 1.23E-04 |
| CHCHD10 | 1240.010778 | 0.392798833 | 1.78E-03 |
| YKT6 | 13205.20478 | 0.392721315 | 9.03E-07 |
| GLG1 | 15942.58437 | 0.392697043 | 5.62E-05 |
| SGPL1 | 1970.346108 | 0.392450956 | 1.70E-02 |
| ETS1 | 13708.86452 | 0.392404279 | 6.43E-07 |
| RHNO1 | 1266.859032 | 0.392001881 | 3.20E-04 |
| CDK8 | 669.859065 | 0.391955569 | 1.36E-03 |
| NPTN | 4805.499864 | 0.391679299 | 2.59E-06 |
| CXorf38 | 1402.157958 | 0.391551862 | 1.05E-05 |
| HK1 | 11421.58405 | 0.391161372 | 9.87E-06 |
| NME4 | 10104.26584 | 0.391096616 | 4.53E-03 |
| PNRC1 | 2333.746429 | 0.390827206 | 3.55E-03 |
| ARHGAP30 | 309.6936613 | 0.39062892 | 4.35E-02 |
| SMNDC1 | 892.9026407 | 0.390592781 | 4.74E-05 |
| GZF1 | 2171.95145 | 0.390486669 | 6.79E-04 |
| LRRC42 | 2265.315389 | 0.390425182 | 7.68E-06 |
| DYNLRB1 | 5076.376126 | 0.390178475 | 4.91E-03 |
| ZNF672 | 729.8899097 | 0.389860524 | 8.91E-04 |
| TXNL4B | 951.0131225 | 0.389853256 | 3.02E-02 |
| UBE2D3 | 7342.501166 | 0.38981734 | 4.61E-07 |
| CCBE1 | 1908.550007 | 0.389607089 | 5.25E-03 |
| HSPH1 | 5633.831136 | 0.389590963 | 7.38E-03 |
| LENG1 | 325.7448315 | 0.388978984 | 3.64E-02 |
| CEPT1 | 951.8770677 | 0.388846371 | 2.79E-03 |
| DMXL1 | 2178.059926 | 0.388763116 | 1.66E-02 |
| TRIP4 | 1456.396546 | 0.387976541 | 1.32E-05 |
| DUSP14 | 1404.725443 | 0.387724117 | 1.60E-05 |
| SRGN | 1547.782683 | 0.387492513 | 3.76E-08 |
| APP | 42894.66391 | 0.387233278 | 8.53E-06 |
| TIGD6 | 286.9580293 | 0.386810618 | 2.16E-03 |
| PDIA3 | 18821.93843 | 0.386797581 | 2.16E-04 |
| C10orf88 | 309.8950589 | 0.386671857 | 2.37E-02 |
| CABLES1 | 2714.737389 | 0.38655092 | 2.20E-02 |
| ACSS3 | 176.7728467 | 0.386550454 | 4.95E-02 |
| TMX1 | 2060.293302 | 0.386508539 | 2.65E-02 |
| PPP2CB | 3807.900051 | 0.386496556 | 1.21E-13 |
| 5-Mar | 1118.604787 | 0.386486613 | 7.30E-03 |
| MINAR1 | 148.9115714 | 0.386429532 | 4.84E-02 |
| ZNF585A | 616.617988 | 0.386385399 | 2.29E-04 |
| C1orf109 | 1140.631876 | 0.386329409 | 1.34E-05 |
| SCLT1 | 582.6754598 | 0.386326301 | 2.56E-02 |
| TMEM182 | 297.9939013 | 0.386105917 | 4.71E-02 |

| | | | |
|---|---|---|---|
| SNRPB2 | 3684.199833 | 0.386079876 | 5.67E-05 |
| ZBTB25 | 608.6663508 | 0.385828946 | 9.86E-04 |
| ZNF426 | 509.8565456 | 0.385766556 | 3.12E-03 |
| C8orf76 | 1193.820138 | 0.385753389 | 2.91E-04 |
| BECN1 | 2776.217333 | 0.385405008 | 8.56E-10 |
| SERPINE2 | 8188.667421 | 0.385116388 | 1.22E-03 |
| HADHA | 12749.68248 | 0.384898183 | 4.49E-06 |
| NR1D1 | 572.8036485 | 0.384867413 | 3.08E-03 |
| ALG9 | 1298.005813 | 0.384730563 | 2.26E-04 |
| HCN2 | 777.201268 | 0.38413301 | 2.93E-03 |
| UIMC1 | 1163.043945 | 0.383982374 | 1.01E-02 |
| PHKB | 4756.907039 | 0.383962282 | 2.49E-09 |
| PCDHGB5 | 192.1740464 | 0.383822503 | 1.68E-02 |
| BCL9L | 9610.645557 | 0.383742609 | 1.56E-03 |
| YIF1B | 1626.377802 | 0.383678923 | 1.46E-04 |
| TMEM42 | 1109.272036 | 0.383572879 | 1.07E-02 |
| TMEM68 | 502.3024622 | 0.383504386 | 2.70E-04 |
| HARBI1 | 273.0406226 | 0.383338296 | 1.07E-02 |
| NORAD | 14921.27391 | 0.383167829 | 1.86E-04 |
| WDR86-AS1 | 427.2589428 | 0.383130843 | 1.05E-02 |
| NDUFAF2 | 2164.104687 | 0.383035382 | 1.57E-04 |
| RBM3 | 4706.629631 | 0.382938359 | 1.07E-02 |
| RUFY3 | 638.5464759 | 0.382659823 | 2.51E-03 |
| REPIN1 | 8966.942661 | 0.382498718 | 4.60E-07 |
| HDLBP | 32128.14887 | 0.382498261 | 4.84E-06 |
| VMA21 | 3061.229425 | 0.382490538 | 3.52E-04 |
| GORASP1 | 2536.009094 | 0.382193524 | 8.67E-04 |
| GTF2H2C | 2923.736029 | 0.381777044 | 3.37E-03 |
| EXOC1 | 2427.226678 | 0.381708065 | 1.36E-02 |
| KSR1 | 1136.231107 | 0.381420162 | 3.14E-02 |
| PLBD2 | 5126.821147 | 0.38136925 | 6.49E-04 |
| CFLAR | 2074.937118 | 0.380766823 | 1.61E-10 |
| COX7A2 | 5358.559036 | 0.380226957 | 1.28E-02 |
| RIOX2 | 1333.224624 | 0.379876806 | 2.32E-02 |
| UVSSA | 368.1860719 | 0.379876172 | 8.99E-03 |
| MIPEP | 509.2321409 | 0.379721514 | 6.88E-03 |
| MTMR3 | 2790.3142 | 0.379674054 | 4.75E-06 |
| BBS7 | 793.6486539 | 0.379470555 | 2.08E-02 |
| FAM3C2 | 399.8232784 | 0.379173863 | 1.60E-02 |
| USP40 | 3112.399187 | 0.378944102 | 2.50E-04 |
| AC145124.1 | 226.6834811 | 0.378250694 | 1.92E-02 |
| MAP4 | 17613.87065 | 0.378200823 | 4.35E-07 |
| CC2D1B | 2873.320909 | 0.378137602 | 1.71E-02 |
| FOSL2 | 5985.00852 | 0.377731809 | 1.35E-03 |
| TBPL1 | 690.9482082 | 0.377604636 | 4.78E-02 |
| METTL6 | 983.6936275 | 0.377405906 | 5.49E-04 |
| GAS5 | 7842.947098 | 0.377314437 | 3.28E-02 |

| | | | |
|---|---|---|---|
| TACO1 | 1177.894129 | 0.377194895 | 4.11E-04 |
| TGFBR2 | 6562.551738 | 0.377173615 | 4.13E-04 |
| DDX3X | 15192.34195 | 0.377106974 | 5.40E-05 |
| AC093525.7 | 393.9993354 | 0.37630219 | 3.67E-03 |
| RBP1 | 684.6441289 | 0.375990325 | 4.23E-02 |
| LTO1 | 862.7388776 | 0.375861512 | 2.32E-03 |
| MAGT1 | 4499.864434 | 0.375855155 | 6.32E-04 |
| ARIH2 | 4845.810584 | 0.375809492 | 4.32E-04 |
| DENND3 | 1469.517193 | 0.375767281 | 4.53E-04 |
| PKD2 | 2184.663901 | 0.37535119 | 2.32E-04 |
| LEMD2 | 2746.266522 | 0.375330577 | 4.44E-07 |
| TAF1D | 3595.679805 | 0.37523369 | 1.35E-02 |
| AFAP1L2 | 2158.646464 | 0.375193339 | 4.25E-03 |
| CUEDC1 | 2613.806841 | 0.375085199 | 5.79E-03 |
| LMTK2 | 2508.032902 | 0.37498461 | 6.36E-07 |
| C6orf106 | 6662.810659 | 0.374980247 | 3.40E-04 |
| PARP16 | 1070.397486 | 0.374941718 | 5.07E-03 |
| ULK2 | 882.4735602 | 0.374785276 | 6.61E-03 |
| SUPT20H | 1593.510877 | 0.374127526 | 2.28E-03 |
| TRAK2 | 3071.599689 | 0.373906304 | 9.35E-09 |
| TXNDC5 | 23972.78375 | 0.373836289 | 5.10E-04 |
| NAA10 | 4808.115404 | 0.373333225 | 1.68E-03 |
| STPG1 | 457.4197334 | 0.373244403 | 1.34E-02 |
| C18orf21 | 443.4218284 | 0.373188171 | 1.25E-03 |
| SLC36A1 | 875.5995958 | 0.373079048 | 3.39E-03 |
| MMP14 | 14551.73684 | 0.373076229 | 8.27E-03 |
| FOXO4 | 737.7610709 | 0.372666145 | 4.57E-03 |
| CAPN7 | 1738.504672 | 0.372649607 | 2.18E-03 |
| KLF10 | 1561.85262 | 0.372562521 | 3.00E-03 |
| PAFAH1B1 | 7948.894797 | 0.372367932 | 5.17E-08 |
| TMBIM6 | 24594.40474 | 0.372083662 | 1.06E-05 |
| HOXB-AS1 | 138.8490023 | 0.371939873 | 4.10E-02 |
| LINC-PINT | 542.6879352 | 0.371823487 | 3.99E-02 |
| CRELD1 | 1238.502371 | 0.371064594 | 3.43E-02 |
| PPID | 1757.95365 | 0.37056911 | 1.58E-03 |
| RAB23 | 599.4729796 | 0.370314585 | 2.46E-02 |
| LINC01578 | 1662.10461 | 0.37016483 | 4.29E-03 |
| AC245297.1 | 206.1312181 | 0.370119789 | 4.17E-02 |
| UHRF2 | 2746.586196 | 0.370072593 | 1.18E-02 |
| PIGH | 917.6961508 | 0.369975917 | 5.36E-06 |
| PARVB | 2787.345432 | 0.369750849 | 2.09E-03 |
| RAB3GAP1 | 4031.730799 | 0.369711568 | 2.12E-04 |
| LINC02210 | 631.473022 | 0.369318858 | 1.01E-02 |
| AC005229.4 | 186.0188007 | 0.368825324 | 1.49E-02 |
| TMED10 | 13060.58854 | 0.368302635 | 2.62E-08 |
| TNFRSF1A | 4509.026064 | 0.367365411 | 2.14E-02 |
| MED13 | 2899.319374 | 0.367348297 | 1.22E-02 |

| | | | |
|---|---|---|---|
| EIF3C | 53785.43486 | 0.367274331 | 3.53E-03 |
| METTL15 | 531.7385375 | 0.367127321 | 2.06E-02 |
| DENND1A | 2021.521882 | 0.367005925 | 1.18E-02 |
| PDE9A | 550.0523679 | 0.366784435 | 6.85E-03 |
| PRDX6 | 12969.97135 | 0.36674896 | 6.40E-03 |
| VEZF1 | 2545.055002 | 0.366112789 | 2.46E-04 |
| EXTL2 | 2261.873093 | 0.366089516 | 2.28E-03 |
| MAPK9 | 1850.100491 | 0.365979295 | 8.85E-04 |
| RPL39L | 913.2693129 | 0.365502864 | 1.02E-04 |
| RGMB | 1091.065118 | 0.36548926 | 1.00E-05 |
| PRICKLE3 | 449.079314 | 0.365379706 | 3.61E-03 |
| LTBR | 4379.978864 | 0.365321727 | 2.21E-04 |
| VPS11 | 3275.966273 | 0.365270201 | 1.45E-03 |
| AC105052.3 | 838.6690067 | 0.364981447 | 4.26E-03 |
| KANK2 | 5584.704178 | 0.364311447 | 1.77E-02 |
| TTC1 | 6219.880764 | 0.364273515 | 1.01E-08 |
| PTPN23 | 3372.476872 | 0.364171127 | 1.08E-03 |
| AMZ2 | 2889.472315 | 0.364168127 | 7.34E-11 |
| PPTC7 | 1103.285373 | 0.363977512 | 3.37E-03 |
| RBM33 | 2975.818229 | 0.363892763 | 2.85E-03 |
| ATP1A1 | 28999.11334 | 0.363846435 | 1.99E-04 |
| TAB2 | 4301.645528 | 0.363499484 | 3.05E-03 |
| C8orf58 | 1572.615118 | 0.363019533 | 3.82E-02 |
| RPN1 | 10565.1925 | 0.362985104 | 9.62E-04 |
| OPA3 | 1728.535822 | 0.362687043 | 5.99E-03 |
| TESK1 | 1077.548336 | 0.362245013 | 3.42E-03 |
| PIGN | 1278.798581 | 0.362102951 | 3.66E-03 |
| ZBTB21 | 1045.562051 | 0.361859986 | 2.39E-02 |
| RFNG | 1336.381185 | 0.361594346 | 1.54E-02 |
| CNNM1 | 541.0648735 | 0.361371228 | 4.73E-03 |
| SNAPC3 | 1889.170263 | 0.360680884 | 7.35E-04 |
| CA13 | 181.8979236 | 0.360285963 | 3.98E-02 |
| MORF4L1 | 15830.39905 | 0.359961334 | 9.53E-06 |
| TMEM263 | 1624.078972 | 0.359647784 | 3.75E-03 |
| ARHGAP35 | 10670.21885 | 0.359634481 | 5.93E-04 |
| SCP2 | 2504.721614 | 0.359436726 | 8.96E-03 |
| GATA6 | 439.6014834 | 0.359357806 | 2.70E-02 |
| ARMCX6 | 1690.845751 | 0.359306591 | 2.42E-03 |
| 10-Sep | 3710.919446 | 0.359224136 | 1.36E-02 |
| KPNA3 | 2751.19483 | 0.359158559 | 1.83E-04 |
| ARMCX5-GPRA | 226.1098284 | 0.358781203 | 3.12E-02 |
| CGNL1 | 895.5418779 | 0.358474111 | 3.04E-02 |
| CD9 | 3271.154132 | 0.358463413 | 4.45E-02 |
| BUD23 | 4063.891293 | 0.357839376 | 8.09E-09 |
| YWHAG | 14945.9692 | 0.357811738 | 2.62E-09 |
| C11orf58 | 5322.868975 | 0.35776153 | 2.94E-03 |
| SPPL3 | 1822.428239 | 0.357609884 | 1.44E-02 |

| | | | |
|---|---|---|---|
| OGFOD2 | 630.4163552 | 0.357603236 | 7.75E-03 |
| RHBDD1 | 1068.658761 | 0.357411121 | 3.75E-02 |
| PRDX5 | 11669.25722 | 0.357172896 | 7.81E-03 |
| SLAIN2 | 1317.445147 | 0.357058957 | 7.17E-03 |
| PDLIM3 | 622.703077 | 0.357048094 | 1.10E-03 |
| TARS | 20704.77187 | 0.356763331 | 1.12E-02 |
| JOSD1 | 3246.967769 | 0.356621052 | 3.50E-05 |
| SNX3 | 5109.333516 | 0.356478013 | 3.68E-05 |
| PPP1R3D | 379.5106735 | 0.356293923 | 6.03E-03 |
| CGAS | 435.9810156 | 0.355858806 | 3.11E-02 |
| BBS4 | 1662.859635 | 0.355848093 | 1.35E-03 |
| ADORA2B | 805.8301713 | 0.355743285 | 8.87E-03 |
| BLOC1S2 | 510.6569008 | 0.355741575 | 9.00E-03 |
| NUDT17 | 302.2081899 | 0.355740417 | 2.16E-02 |
| PEX10 | 1005.175471 | 0.355573301 | 1.22E-03 |
| CAAP1 | 1071.16947 | 0.355519884 | 8.60E-03 |
| PEX13 | 727.2580527 | 0.355104038 | 2.17E-03 |
| CEP104 | 1693.378755 | 0.354951513 | 3.29E-03 |
| TLE3 | 1335.834862 | 0.35488294 | 4.57E-02 |
| ADGRE5 | 2806.008902 | 0.354735938 | 3.86E-03 |
| NDUFB6 | 2354.12987 | 0.354668409 | 9.52E-03 |
| COX6C | 2864.57091 | 0.354476249 | 3.60E-04 |
| DHFR2 | 351.7586684 | 0.354407409 | 1.84E-02 |
| RAF1 | 4631.727471 | 0.354394401 | 3.52E-07 |
| HABP4 | 1578.110636 | 0.353684997 | 5.68E-05 |
| IMP3 | 2504.421106 | 0.353661119 | 8.31E-03 |
| HADHB | 5881.409386 | 0.353398252 | 3.05E-04 |
| MTHFR | 2157.163497 | 0.353262541 | 2.63E-02 |
| VAMP7 | 2563.18571 | 0.353030102 | 4.86E-03 |
| HSPA14 | 1041.680726 | 0.352917715 | 8.02E-04 |
| ATP5PD | 3820.956853 | 0.35289177 | 4.59E-04 |
| ZCCHC10 | 662.269274 | 0.352754779 | 1.48E-03 |
| AC011462.1 | 3328.314682 | 0.351888255 | 6.54E-04 |
| UBAP1 | 3759.769201 | 0.351793158 | 1.35E-03 |
| UHMK1 | 2866.460124 | 0.35157874 | 1.78E-02 |
| FADS1 | 13379.58282 | 0.351452902 | 5.96E-04 |
| HDDC2 | 1891.097272 | 0.351133424 | 9.81E-08 |
| ZNF24 | 2810.113033 | 0.351117588 | 3.27E-02 |
| OIP5-AS1 | 1986.522582 | 0.350910409 | 7.01E-06 |
| MKRN1 | 5568.431711 | 0.35084449 | 4.94E-04 |
| KIAA0232 | 1363.339717 | 0.350782614 | 3.27E-03 |
| RNF216 | 6139.992269 | 0.350698348 | 1.14E-07 |
| YME1L1 | 5387.912874 | 0.350610238 | 4.74E-03 |
| POGLUT1 | 1622.036049 | 0.350609784 | 6.45E-05 |
| DRAP1 | 3957.910986 | 0.350397434 | 9.18E-04 |
| PDE8A | 1845.673011 | 0.350360466 | 7.28E-04 |
| FOXD1 | 1076.126703 | 0.350230941 | 3.09E-03 |

| | | | |
|---|---|---|---|
| PIGP | 575.1401178 | 0.350135698 | 8.28E-04 |
| GMPPB | 1612.893144 | 0.350069789 | 1.39E-03 |
| TMEM99 | 358.1260625 | 0.349867001 | 7.11E-03 |
| SLC35F5 | 1967.09848 | 0.349296843 | 2.71E-02 |
| IFT20 | 1159.347537 | 0.349295448 | 1.32E-06 |
| OSTC | 3548.714486 | 0.349285934 | 1.76E-05 |
| HS1BP3 | 1553.181874 | 0.349199851 | 1.09E-02 |
| ZNF841 | 569.68734 | 0.349119136 | 8.69E-03 |
| ETV1 | 1912.447401 | 0.349051327 | 5.47E-03 |
| CD151 | 16569.59205 | 0.348940019 | 1.40E-02 |
| AP1S2 | 2272.975722 | 0.348073439 | 3.40E-02 |
| GPX4 | 7821.112851 | 0.348041754 | 4.57E-03 |
| ATP6AP2 | 6279.90048 | 0.348005649 | 1.21E-10 |
| COA7 | 1675.441611 | 0.347864172 | 9.04E-03 |
| TEX2 | 2687.825769 | 0.347645649 | 7.93E-07 |
| TMEM127 | 3467.423247 | 0.347559643 | 2.69E-03 |
| ATP8B3 | 1210.261948 | 0.347456185 | 3.99E-03 |
| STIMATE | 1402.581338 | 0.347203007 | 6.29E-03 |
| MAN2B2 | 2532.968815 | 0.347048856 | 1.11E-03 |
| ITGA2 | 6584.396695 | 0.347005457 | 2.17E-02 |
| GTF2H2 | 360.6277818 | 0.346858134 | 4.60E-02 |
| ZNF433-AS1 | 402.7071823 | 0.346844926 | 2.73E-02 |
| CSAG1 | 2662.965097 | 0.346812402 | 2.66E-05 |
| SLC11A2 | 1980.419384 | 0.346799307 | 1.88E-05 |
| FBN1 | 3258.584951 | 0.346755461 | 8.14E-03 |
| RAB11FIP3 | 2920.906787 | 0.34671166 | 3.05E-04 |
| NBPF8 | 1942.365623 | 0.346381264 | 1.02E-04 |
| CSAG3 | 2154.5927 | 0.346137578 | 1.32E-03 |
| BTBD1 | 3723.853847 | 0.345809899 | 2.13E-03 |
| OTUB2 | 248.5069851 | 0.345419632 | 3.80E-02 |
| TIPRL | 2490.142079 | 0.345358207 | 9.13E-05 |
| FAR2 | 1062.330359 | 0.345334496 | 1.02E-03 |
| NUBPL | 852.1598089 | 0.344421194 | 1.56E-04 |
| PARD3 | 3322.902457 | 0.344372802 | 5.86E-03 |
| PLPBP | 2627.182497 | 0.344337685 | 3.89E-03 |
| NUPL2 | 1012.659613 | 0.343947023 | 6.59E-03 |
| CPT1A | 5309.283983 | 0.343622828 | 2.24E-03 |
| OSBPL2 | 1833.379553 | 0.343395802 | 2.57E-03 |
| YEATS4 | 615.0240725 | 0.343196067 | 1.91E-02 |
| DEGS1 | 2886.061069 | 0.343006966 | 2.20E-02 |
| VDAC2 | 3904.428237 | 0.34294353 | 1.06E-04 |
| ELOC | 2405.537577 | 0.342738152 | 8.81E-06 |
| AC093525.6 | 218.5605161 | 0.342715087 | 2.88E-02 |
| CAV1 | 23107.43986 | 0.342347308 | 1.49E-02 |
| UQCRFS1 | 3568.156548 | 0.341852142 | 7.66E-04 |
| ACO1 | 6073.137214 | 0.341845394 | 4.34E-04 |
| TMEM134 | 704.4764652 | 0.341835375 | 2.17E-02 |

| | | | |
|---|---|---|---|
| SETD4 | 917.8819247 | 0.3415683 | 8.30E-03 |
| NA | 284.9394827 | 0.341558751 | 4.91E-02 |
| PSMA4 | 7281.516475 | 0.341349335 | 1.27E-05 |
| MSRB2 | 725.0240156 | 0.341342941 | 1.48E-02 |
| TOX4 | 5267.628927 | 0.341216817 | 5.81E-04 |
| LZTS2 | 2628.988224 | 0.341174758 | 7.73E-03 |
| USF2 | 2621.385513 | 0.341086386 | 1.88E-04 |
| DLD | 4716.307916 | 0.340839853 | 2.77E-03 |
| TIMP3 | 33487.39915 | 0.340794813 | 4.01E-02 |
| RHBDD2 | 4031.725876 | 0.340732424 | 1.62E-03 |
| CPE | 2379.926947 | 0.34045344 | 1.90E-02 |
| ISCU | 1888.172076 | 0.339821679 | 2.20E-07 |
| ZNF865 | 651.0894696 | 0.339037277 | 3.25E-03 |
| C1orf122 | 1607.075983 | 0.338877181 | 2.39E-02 |
| USE1 | 928.9630378 | 0.338546655 | 3.29E-02 |
| GOSR1 | 2158.58845 | 0.338129643 | 6.36E-04 |
| C16orf72 | 3259.203774 | 0.337817684 | 1.45E-02 |
| MCOLN1 | 614.275788 | 0.337724127 | 2.30E-03 |
| DHX35 | 1575.892245 | 0.337717257 | 5.46E-03 |
| SNX11 | 807.4830096 | 0.33759486 | 3.49E-04 |
| ORMDL1 | 1730.741365 | 0.337405054 | 5.11E-04 |
| AIMP2 | 1830.027727 | 0.337374036 | 2.99E-05 |
| TANGO6 | 1779.528087 | 0.337305952 | 1.47E-03 |
| FEM1B | 2626.446844 | 0.337266988 | 3.70E-03 |
| PLOD3 | 10152.46719 | 0.337072209 | 3.61E-04 |
| ALG13 | 1081.099549 | 0.337050751 | 4.64E-02 |
| PLPP2 | 1427.512506 | 0.337022141 | 3.67E-02 |
| MSX1 | 641.8880923 | 0.336711099 | 3.39E-02 |
| SPRY4 | 1960.932788 | 0.336382486 | 1.87E-02 |
| CLPX | 2147.540169 | 0.336366021 | 1.39E-02 |
| PUM2 | 4470.30099 | 0.336273222 | 9.53E-03 |
| STIP1 | 14572.5874 | 0.336071711 | 4.34E-05 |
| SLC35B1 | 1668.984745 | 0.335841829 | 1.95E-05 |
| CAST | 16712.96545 | 0.335821816 | 3.12E-09 |
| KLHL7 | 2045.471299 | 0.33560354 | 1.57E-04 |
| WDR31 | 314.5976903 | 0.335516639 | 4.24E-02 |
| COQ5 | 1997.429849 | 0.335447736 | 4.72E-06 |
| TRIOBP | 4479.917163 | 0.335346898 | 4.91E-03 |
| RASAL2 | 1538.28984 | 0.335284117 | 2.83E-03 |
| RPN2 | 26174.83075 | 0.335020426 | 6.35E-13 |
| CKAP4 | 11320.18536 | 0.335009068 | 1.46E-02 |
| ASB7 | 1029.321998 | 0.334718601 | 2.29E-02 |
| TPM3P9 | 730.5224163 | 0.334711516 | 1.44E-03 |
| VAMP1 | 983.9967304 | 0.33461742 | 1.37E-02 |
| HFE | 563.7342057 | 0.334332596 | 5.96E-03 |
| STN1 | 566.7971959 | 0.334319979 | 2.73E-04 |
| ANKMY2 | 1123.608775 | 0.334236155 | 4.31E-04 |

| | | | |
|---|---|---|---|
| TM9SF2 | 21150.58403 | 0.334059069 | 4.54E-03 |
| RPS27L | 1580.599612 | 0.333905853 | 1.85E-03 |
| IFNAR1 | 2453.459114 | 0.333808139 | 4.79E-02 |
| TES | 6310.989074 | 0.333477369 | 2.00E-04 |
| PPP1R3B | 1558.68049 | 0.333250927 | 1.40E-05 |
| RAB28 | 518.5789756 | 0.333129015 | 2.39E-02 |
| DHX16 | 3120.149289 | 0.333006024 | 1.46E-04 |
| RPS6KC1 | 2016.543957 | 0.332994302 | 1.62E-05 |
| MT-RNR2 | 147786.8412 | 0.332736154 | 2.07E-02 |
| ZNF717 | 470.4797863 | 0.332463681 | 1.59E-03 |
| SNRNP27 | 1096.904373 | 0.332348749 | 1.86E-04 |
| C12orf49 | 2455.796683 | 0.331881901 | 1.35E-04 |
| VEGFB | 3022.252006 | 0.331586881 | 4.20E-02 |
| TNFRSF14 | 898.3680434 | 0.331502739 | 3.61E-02 |
| COA4 | 2229.22841 | 0.331432654 | 2.84E-03 |
| ZNF529-AS1 | 151.9124009 | 0.330847879 | 4.58E-02 |
| SLC30A4 | 349.638387 | 0.330720907 | 2.56E-02 |
| MARK3 | 3930.182116 | 0.330557752 | 4.92E-04 |
| NPLOC4 | 6856.1453 | 0.330254785 | 4.94E-03 |
| PPFIBP1 | 7522.570197 | 0.330182407 | 7.86E-06 |
| PUSL1 | 454.6324044 | 0.329990186 | 2.96E-03 |
| C12orf73 | 289.4162533 | 0.329615601 | 1.02E-02 |
| WASH8P | 904.7545586 | 0.329586034 | 3.24E-02 |
| CBFA2T2 | 1814.649614 | 0.329457092 | 1.16E-03 |
| EBAG9 | 824.6448612 | 0.329384981 | 1.31E-02 |
| TBC1D7 | 760.8227745 | 0.329344575 | 9.07E-03 |
| ROBO3 | 1232.003209 | 0.328621084 | 1.91E-02 |
| TMEM147 | 1948.56684 | 0.328348271 | 2.08E-03 |
| KRT18 | 26282.33084 | 0.328343827 | 1.25E-02 |
| ZDHHC24 | 651.9338299 | 0.328112683 | 7.56E-03 |
| NDST1 | 8021.325827 | 0.328097094 | 1.82E-03 |
| DTD2 | 575.1748008 | 0.327877576 | 1.06E-02 |
| TRAPPC2B | 319.9620755 | 0.327845995 | 2.34E-02 |
| NSUN4 | 1220.122466 | 0.327667327 | 1.13E-03 |
| KTI12 | 644.5004716 | 0.326807348 | 3.44E-03 |
| SLC30A7 | 941.1097818 | 0.326597494 | 4.07E-03 |
| NUFIP1 | 765.1757443 | 0.326547662 | 1.68E-04 |
| ZNF217 | 4777.839968 | 0.326259199 | 4.12E-04 |
| TAF2 | 2850.725467 | 0.325851712 | 1.28E-02 |
| ZNF277 | 1041.934133 | 0.32546508 | 2.47E-03 |
| DPYSL2 | 5563.378319 | 0.325215503 | 6.32E-05 |
| OCRL | 3249.787062 | 0.325137191 | 1.72E-06 |
| TMEM167A | 4949.646247 | 0.325068842 | 1.60E-04 |
| ZFYVE26 | 2573.202655 | 0.325034137 | 1.90E-03 |
| IRF9 | 1259.63647 | 0.323207552 | 2.55E-02 |
| ZFP82 | 392.497152 | 0.322979211 | 2.73E-02 |
| C1D | 518.6044639 | 0.322911456 | 8.33E-03 |

| | | | |
|---|---|---|---|
| DNAJB12 | 1293.375467 | 0.322713325 | 3.22E-03 |
| ELF1 | 1523.077728 | 0.322629293 | 2.49E-05 |
| PMS2P1 | 319.8335432 | 0.322153226 | 3.30E-02 |
| LAMB1 | 49693.47009 | 0.321388103 | 8.01E-03 |
| EML2 | 1514.807018 | 0.32126055 | 1.58E-02 |
| CEP250 | 4914.904169 | 0.321233407 | 1.40E-04 |
| ELP3 | 3058.743108 | 0.321103769 | 1.19E-06 |
| MYL12A | 9438.49279 | 0.32101776 | 4.38E-04 |
| C16orf45 | 734.6864757 | 0.320999329 | 4.47E-03 |
| ATG3 | 1629.288593 | 0.320144053 | 3.09E-04 |
| ZNF655 | 3980.994069 | 0.320097622 | 1.93E-02 |
| CASC4 | 1140.995251 | 0.320063134 | 4.45E-02 |
| SMG1P1 | 784.4459553 | 0.31984488 | 2.57E-02 |
| VPS29 | 1793.056619 | 0.319717708 | 5.17E-06 |
| ZDHHC4 | 3625.181031 | 0.318601645 | 6.60E-04 |
| FH | 3105.884307 | 0.318020332 | 7.37E-07 |
| BBS9 | 1146.378736 | 0.317621484 | 2.43E-04 |
| WAC | 3497.059522 | 0.317574706 | 2.00E-02 |
| FEM1C | 1014.330276 | 0.317555868 | 7.95E-03 |
| TRIM38 | 1546.239524 | 0.317320006 | 2.51E-04 |
| TEX30 | 1442.847285 | 0.317151618 | 3.71E-02 |
| TNKS2 | 1792.852115 | 0.316877662 | 4.77E-03 |
| LIG3 | 2382.176734 | 0.316757974 | 2.24E-03 |
| POMP | 2654.174036 | 0.316651707 | 3.25E-06 |
| HOTAIRM1 | 1189.090175 | 0.316640373 | 3.68E-03 |
| MAP7D3 | 5669.030078 | 0.31605778 | 2.68E-02 |
| PDCD5 | 1993.549848 | 0.315290917 | 6.22E-05 |
| AC079949.2 | 366.5235527 | 0.31527468 | 4.03E-02 |
| TMEM120B | 1090.727739 | 0.315065086 | 6.41E-03 |
| SLC35E4 | 401.9891485 | 0.314663766 | 2.37E-02 |
| BCL7A | 490.2871555 | 0.314567346 | 4.06E-02 |
| CD46 | 4929.697759 | 0.314550274 | 2.66E-02 |
| SEC22A | 837.2172181 | 0.314491818 | 1.40E-03 |
| XPA | 555.6036263 | 0.314440984 | 2.16E-02 |
| RABEP1 | 5406.597359 | 0.314411117 | 2.33E-02 |
| FBXO8 | 615.4023594 | 0.314297901 | 2.89E-03 |
| TBKBP1 | 1994.792743 | 0.314202638 | 1.53E-03 |
| ATXN7 | 1109.239865 | 0.313804231 | 1.37E-02 |
| PRDX4 | 3675.105295 | 0.313629327 | 5.60E-03 |
| DUSP22 | 1490.329574 | 0.313531367 | 8.26E-05 |
| PSMA1 | 6765.463374 | 0.313385575 | 1.62E-05 |
| ATG2A | 2615.448837 | 0.312643172 | 1.08E-02 |
| LARS | 10091.49174 | 0.312572771 | 4.78E-02 |
| CTDP1 | 1130.124063 | 0.31246 | 3.04E-03 |
| FBXO31 | 2020.634313 | 0.312289115 | 1.42E-02 |
| ZDHHC5 | 5504.125348 | 0.312143013 | 1.79E-04 |
| DCPS | 1817.087287 | 0.312002916 | 1.25E-03 |

| | | | |
|---|---|---|---|
| SIK3 | 2445.245228 | 0.311986927 | 3.10E-04 |
| NDUFS1 | 6300.475034 | 0.311696365 | 1.65E-08 |
| MKNK1 | 1050.384741 | 0.311544894 | 3.42E-03 |
| ZNF480 | 608.6021465 | 0.311217014 | 1.40E-02 |
| AC008443.1 | 461.6091636 | 0.311012784 | 3.46E-02 |
| RDH14 | 549.1452708 | 0.311006796 | 1.51E-02 |
| FMNL2 | 4188.966288 | 0.310978828 | 9.41E-03 |
| WNK1 | 11229.30839 | 0.310485726 | 1.48E-03 |
| POLR3GL | 875.5551453 | 0.31024635 | 2.03E-03 |
| EFCAB14 | 3751.538257 | 0.310172794 | 1.02E-02 |
| FOXJ3 | 2806.709873 | 0.310138776 | 2.50E-04 |
| FP671120.2 | 584.4415846 | 0.310025582 | 2.72E-03 |
| MED30 | 472.5295775 | 0.309876839 | 4.35E-03 |
| KDM7A-DT | 360.7802649 | 0.309801819 | 2.45E-02 |
| OARD1 | 789.8360287 | 0.309724121 | 4.14E-02 |
| AASS | 1629.736665 | 0.309715345 | 2.25E-02 |
| FCF1 | 2201.561553 | 0.30958901 | 4.87E-03 |
| ADD2 | 5103.394758 | 0.309555232 | 2.88E-04 |
| UBE2Q2 | 1264.376967 | 0.309077048 | 1.30E-02 |
| NUP210 | 2474.763351 | 0.308939908 | 1.50E-02 |
| ARFGAP3 | 2468.560438 | 0.308590066 | 4.38E-06 |
| ZFP91 | 6060.027481 | 0.308534561 | 2.44E-06 |
| MRS2 | 1525.675607 | 0.308243081 | 2.49E-04 |
| SSH2 | 1653.105872 | 0.308198354 | 6.62E-03 |
| RUSC2 | 2608.465768 | 0.30815797 | 7.07E-04 |
| FAM129B | 17789.76161 | 0.307956908 | 3.78E-03 |
| PCCA | 4759.862482 | 0.307636483 | 4.60E-03 |
| MARCKS | 3627.182142 | 0.307347275 | 2.83E-03 |
| EMC1 | 6779.200104 | 0.306915714 | 3.43E-03 |
| PLEKHA1 | 499.4409068 | 0.306898163 | 3.36E-02 |
| RP2 | 1027.688176 | 0.306774885 | 4.26E-02 |
| GIPC1 | 4074.727252 | 0.3066711 | 1.85E-03 |
| SMIM30 | 1342.204583 | 0.3066019 | 3.29E-02 |
| RABGEF1 | 2589.467863 | 0.306497279 | 8.34E-05 |
| HPS4 | 1781.078037 | 0.306330664 | 8.16E-04 |
| LARP4B | 1886.896081 | 0.306243708 | 2.83E-04 |
| TRIM44 | 11804.00258 | 0.306207355 | 3.49E-02 |
| ZNF250 | 430.0699983 | 0.306176188 | 1.78E-02 |
| MICU1 | 2085.264622 | 0.305900097 | 6.74E-03 |
| CELSR2 | 2661.06557 | 0.305885894 | 1.76E-02 |
| TNPO2 | 4295.059421 | 0.305561418 | 6.20E-04 |
| VPS53 | 2334.714473 | 0.305420668 | 2.83E-02 |
| APEH | 7239.845918 | 0.304967069 | 3.47E-02 |
| HIPK3 | 1459.481618 | 0.304755495 | 2.59E-02 |
| PRXL2C | 968.9850378 | 0.304635984 | 2.95E-02 |
| GYG2 | 1137.477831 | 0.304525139 | 4.33E-02 |
| EMILIN2 | 884.5796368 | 0.304506878 | 4.84E-02 |

| | | | |
|---|---|---|---|
| ATG4D | 562.7709515 | 0.30445639 | 2.51E-02 |
| DLC1 | 8727.154963 | 0.304403338 | 4.27E-06 |
| STS | 1112.178348 | 0.304119483 | 1.18E-03 |
| ADPGK | 4912.460783 | 0.304106666 | 2.31E-03 |
| EIF4G3 | 7933.865896 | 0.304010708 | 4.29E-03 |
| SURF4 | 17447.89564 | 0.303687698 | 1.15E-02 |
| NDUFC1 | 1027.759632 | 0.303053439 | 3.40E-03 |
| UBE2F | 1396.145029 | 0.302763005 | 1.03E-02 |
| MRPS17 | 1038.638066 | 0.302710345 | 3.82E-04 |
| SSBP1 | 6663.456012 | 0.302529446 | 4.48E-07 |
| MRPS15 | 2177.285808 | 0.302290427 | 5.54E-04 |
| TOMM34 | 5597.477897 | 0.302213955 | 1.54E-03 |
| FBXO22 | 2360.772059 | 0.302185278 | 6.36E-04 |
| TGOLN2 | 11684.65617 | 0.301665372 | 6.33E-04 |
| CFDP1 | 2477.404977 | 0.301432956 | 1.34E-04 |
| HEMK1 | 1344.047739 | 0.301235709 | 5.16E-03 |
| GPR108 | 2585.88666 | 0.300878952 | 2.50E-02 |
| MECP2 | 6385.898364 | 0.30012771 | 2.03E-03 |
| IKBKG | 2243.828522 | 0.30003261 | 3.41E-04 |
| PILRB | 1131.747705 | 0.299542763 | 4.06E-03 |
| MAGI1 | 1754.890765 | 0.299468788 | 8.61E-03 |
| 8-Mar | 1120.594271 | 0.299439851 | 1.34E-04 |
| MINK1 | 4742.302078 | 0.299038689 | 1.52E-02 |
| TRIP12 | 14270.81744 | 0.298947105 | 3.03E-03 |
| TOM1 | 1752.806836 | 0.298475425 | 1.18E-02 |
| AKIP1 | 922.1010123 | 0.29842167 | 2.03E-02 |
| SLC48A1 | 1063.484791 | 0.298184627 | 2.03E-02 |
| METTL5 | 1304.863506 | 0.297472106 | 1.16E-02 |
| DAD1 | 4266.170628 | 0.297452358 | 1.18E-02 |
| XXYLT1 | 1532.304719 | 0.296723743 | 6.63E-03 |
| ITPKC | 871.852918 | 0.296569943 | 6.56E-03 |
| SLC1A3 | 1565.100729 | 0.296568388 | 4.81E-02 |
| ARSA | 1347.401683 | 0.296145396 | 2.74E-02 |
| PPP1R37 | 912.1806825 | 0.296123024 | 1.36E-02 |
| PTK2 | 6063.525587 | 0.295897345 | 2.58E-03 |
| TMEM219 | 2731.416102 | 0.295733306 | 2.51E-02 |
| GGCX | 2529.463059 | 0.295570529 | 7.24E-03 |
| CRTC2 | 1579.016731 | 0.295514221 | 6.45E-03 |
| EVI5L | 1512.548342 | 0.295103442 | 4.87E-03 |
| MXD1 | 447.3520804 | 0.294886118 | 4.10E-02 |
| ATP6V0B | 2681.18054 | 0.294404332 | 2.74E-02 |
| CARD9 | 538.1160319 | 0.294316274 | 4.20E-02 |
| MCFD2 | 14679.18902 | 0.294164554 | 1.94E-06 |
| GGH | 2304.859882 | 0.294097618 | 4.73E-02 |
| RAP2C | 1055.919875 | 0.293978012 | 3.48E-02 |
| NSMCE1 | 1244.15062 | 0.293670161 | 5.03E-03 |
| RNF10 | 5735.766909 | 0.293278136 | 1.78E-03 |

| | | | |
|---|---|---|---|
| WASF3 | 984.7548171 | 0.293142891 | 5.71E-05 |
| MCU | 1122.807796 | 0.292659141 | 1.07E-03 |
| AFF4 | 7606.913083 | 0.292484131 | 1.80E-02 |
| PIGL | 581.9689132 | 0.291650695 | 1.45E-02 |
| PPP2R3A | 791.7218731 | 0.291549306 | 4.31E-02 |
| MIIP | 828.9580369 | 0.291364537 | 1.81E-02 |
| TMEM242 | 779.789896 | 0.291100689 | 2.37E-03 |
| ZBTB8OS | 542.9329176 | 0.290995173 | 2.34E-02 |
| TMEM60 | 883.7742257 | 0.290825317 | 3.38E-02 |
| NBDY | 1029.916909 | 0.290740111 | 2.70E-02 |
| UBL7-AS1 | 231.1872658 | 0.290610238 | 3.68E-02 |
| CCNYL1 | 1291.923322 | 0.290531215 | 4.97E-03 |
| MFSD6 | 998.7542204 | 0.290465156 | 4.85E-03 |
| TOP3A | 3683.398939 | 0.290414421 | 1.10E-04 |
| BTN2A2 | 1021.308363 | 0.290355195 | 7.44E-03 |
| MFSD4B | 323.1202287 | 0.289897893 | 3.13E-02 |
| MAN1B1 | 7196.532018 | 0.289482667 | 2.44E-02 |
| OSBP | 5786.050547 | 0.289450427 | 8.51E-04 |
| ABRAXAS2 | 726.1411251 | 0.288903521 | 3.40E-02 |
| SEM1 | 3715.928863 | 0.288626707 | 6.15E-04 |
| CLCN6 | 2361.816732 | 0.288110176 | 2.48E-04 |
| MYO10 | 14327.81064 | 0.287619691 | 6.70E-03 |
| CIAO2B | 2234.297862 | 0.287517619 | 2.80E-02 |
| MPRIP | 18080.17353 | 0.287444795 | 4.52E-03 |
| PIGO | 2514.073758 | 0.287293967 | 2.24E-03 |
| PGRMC1 | 3576.645105 | 0.286836308 | 3.47E-04 |
| MFF | 2603.156878 | 0.286602037 | 1.36E-06 |
| SNW1 | 4347.853979 | 0.286587906 | 3.21E-02 |
| TM9SF4 | 6205.588196 | 0.286481995 | 1.91E-02 |
| SMARCD3 | 1854.421528 | 0.286331593 | 2.48E-02 |
| PDRG1 | 1660.637 | 0.285974113 | 2.75E-02 |
| PLXNB2 | 10147.39702 | 0.285429547 | 9.02E-03 |
| SCAF4 | 1560.497146 | 0.285282878 | 1.96E-03 |
| PPP1R11 | 2646.361088 | 0.28514769 | 1.24E-03 |
| DDX59 | 684.8136277 | 0.285022809 | 4.24E-02 |
| MKKS | 2227.120336 | 0.284671188 | 1.15E-05 |
| GSTA4 | 526.6998491 | 0.284396731 | 3.64E-02 |
| LMBRD1 | 897.240277 | 0.284315307 | 9.60E-03 |
| WDR53 | 375.6925069 | 0.284187364 | 3.14E-02 |
| FAM192A | 4054.220044 | 0.284140788 | 8.82E-07 |
| SRRD | 1362.097613 | 0.284017926 | 1.12E-02 |
| GTPBP1 | 3068.954214 | 0.283949802 | 5.49E-03 |
| ARPC3 | 7377.140275 | 0.283734225 | 4.35E-03 |
| HSD17B12 | 2891.077653 | 0.283607436 | 3.23E-02 |
| FAM122A | 825.2903686 | 0.283566046 | 2.14E-03 |
| TIMM17A | 2036.133331 | 0.283323845 | 9.09E-03 |
| TGFBR3 | 1252.634429 | 0.283316964 | 2.64E-02 |

| | | | |
|---|---|---|---|
| NCOA1 | 2266.228525 | 0.283068892 | 3.13E-02 |
| KLHL42 | 1254.378234 | 0.283064045 | 8.70E-04 |
| SURF1 | 3085.300649 | 0.282985644 | 1.70E-02 |
| BCKDHB | 817.3871201 | 0.282935509 | 3.49E-02 |
| CCDC9 | 559.97443 | 0.282764139 | 4.43E-02 |
| E2F6 | 1460.523896 | 0.282733238 | 1.48E-03 |
| KLC2 | 1774.580244 | 0.282673734 | 2.62E-02 |
| ASMTL | 1137.639438 | 0.282329726 | 2.22E-02 |
| LINC00960 | 736.0030382 | 0.282025865 | 1.30E-03 |
| CUL1 | 4998.254058 | 0.281711611 | 1.97E-06 |
| AC245060.4 | 370.64184 | 0.281678085 | 1.23E-02 |
| TFDP2 | 1297.077178 | 0.281519602 | 4.13E-02 |
| RNF111 | 1914.181521 | 0.28149625 | 2.81E-05 |
| CHMP1B | 1645.111841 | 0.281395391 | 3.91E-04 |
| YAP1 | 7292.138549 | 0.281271917 | 2.24E-03 |
| SEC61B | 3153.007258 | 0.280939943 | 9.65E-04 |
| CRCP | 2261.256039 | 0.280371672 | 1.76E-02 |
| NKX3-1 | 1379.272934 | 0.279590584 | 1.39E-02 |
| TMEM54 | 1494.946011 | 0.279485508 | 2.46E-02 |
| EXOSC7 | 1415.908928 | 0.279254069 | 2.00E-02 |
| KIAA0391 | 2161.833236 | 0.279233152 | 3.30E-02 |
| PCNP | 3681.071871 | 0.279128349 | 6.16E-03 |
| CACNB1 | 433.310513 | 0.278837122 | 4.72E-02 |
| SDHAF3 | 554.5680846 | 0.278429879 | 1.99E-02 |
| PCBP2 | 21473.6083 | 0.278250291 | 6.11E-05 |
| GOLGA7 | 2655.781304 | 0.277853164 | 3.70E-04 |
| MEF2D | 1707.54484 | 0.277661132 | 2.20E-02 |
| STK24 | 5540.39702 | 0.277550336 | 2.83E-03 |
| DCUN1D5 | 2156.25617 | 0.277434503 | 5.89E-04 |
| INSIG2 | 868.8112722 | 0.27722317 | 9.05E-03 |
| GSS | 4871.01293 | 0.277137062 | 1.28E-02 |
| SCAMP1-AS1 | 798.8270424 | 0.277110631 | 3.07E-03 |
| SEC23A | 5558.829986 | 0.276835866 | 8.46E-03 |
| PXDN | 21810.67112 | 0.276764719 | 1.51E-03 |
| SNX19 | 5696.146326 | 0.276699954 | 1.10E-03 |
| CELSR3 | 2048.93203 | 0.276609105 | 1.88E-03 |
| EPG5 | 3554.316738 | 0.276576557 | 1.48E-03 |
| PDLIM1 | 4528.312637 | 0.276464672 | 3.71E-02 |
| PSMD1 | 7577.741112 | 0.275869737 | 4.13E-04 |
| TMEM106B | 1864.784951 | 0.275807242 | 4.41E-02 |
| NDUFA8 | 3018.998062 | 0.275802641 | 1.59E-02 |
| DCP1A | 1566.772076 | 0.275694531 | 7.00E-03 |
| CSNK1A1 | 6157.901358 | 0.275382846 | 6.25E-03 |
| NBR1 | 8980.711034 | 0.274998972 | 1.94E-06 |
| KIF13B | 1664.961045 | 0.274840372 | 5.71E-03 |
| FAHD2B | 1473.812251 | 0.274441007 | 9.44E-04 |
| BAG5 | 1783.062211 | 0.274348313 | 2.22E-02 |

| | | | |
|---|---|---|---|
| ECD | 1457.84952 | 0.274144567 | 1.77E-04 |
| NOP10 | 2112.578788 | 0.273975064 | 9.84E-03 |
| ZXDB | 459.6379814 | 0.273867683 | 4.85E-02 |
| RAB31 | 4038.601522 | 0.273777478 | 2.19E-03 |
| NFU1 | 1357.271741 | 0.273596381 | 2.17E-02 |
| LRP11 | 2422.615292 | 0.272795649 | 4.97E-02 |
| MRPS35 | 1793.927643 | 0.272768565 | 7.55E-03 |
| FXR2 | 2852.945115 | 0.272750964 | 1.16E-03 |
| UCK2 | 2195.040756 | 0.272275029 | 2.18E-03 |
| B4GALT1 | 11172.21506 | 0.272134417 | 4.25E-04 |
| NEK9 | 9405.283839 | 0.272062703 | 2.24E-03 |
| PLAA | 2566.524153 | 0.271993775 | 2.17E-02 |
| SNHG16 | 2900.523027 | 0.271456434 | 5.72E-04 |
| SLC30A6 | 1921.760547 | 0.271245701 | 1.65E-02 |
| STAU2 | 1901.995673 | 0.270490515 | 1.98E-02 |
| TMEM63B | 1468.220273 | 0.270316016 | 3.15E-02 |
| TXNL4A | 1733.43983 | 0.270163429 | 3.34E-03 |
| MAP2K4 | 1534.37162 | 0.269584521 | 6.77E-03 |
| GOLPH3 | 5308.918063 | 0.269172199 | 5.97E-07 |
| ANAPC13 | 1919.232172 | 0.269100823 | 6.86E-03 |
| TPT1-AS1 | 345.5680486 | 0.268168293 | 3.90E-02 |
| PHLDB1 | 8195.744789 | 0.268161602 | 3.10E-02 |
| SLC7A6OS | 1998.111305 | 0.267609796 | 1.55E-03 |
| IFT52 | 2221.042975 | 0.266750324 | 1.24E-03 |
| BRAF | 1683.390518 | 0.266525004 | 4.13E-02 |
| RNF103 | 872.356071 | 0.266459828 | 6.97E-03 |
| ACP1 | 2949.308581 | 0.266431296 | 1.91E-03 |
| WASHC2C | 1179.143455 | 0.266242372 | 9.49E-03 |
| MRPS9 | 1646.296068 | 0.26567677 | 4.13E-03 |
| MRPL19 | 1851.910839 | 0.265569455 | 1.65E-02 |
| EXOC4 | 8628.260536 | 0.265536213 | 5.06E-04 |
| SLC50A1 | 2499.022849 | 0.265116558 | 2.49E-02 |
| MTA3 | 2140.289836 | 0.265042886 | 6.65E-04 |
| TRAK1 | 3100.499793 | 0.264828433 | 3.73E-04 |
| SLC7A6 | 3175.630792 | 0.264752317 | 2.73E-02 |
| PPM1A | 1396.887455 | 0.26471662 | 1.88E-02 |
| AFTPH | 1389.954349 | 0.264630412 | 2.62E-02 |
| SYAP1 | 2470.590071 | 0.264350057 | 9.70E-03 |
| LRPAP1 | 5556.645268 | 0.264338896 | 1.30E-02 |
| CDC42SE2 | 2790.469273 | 0.263381845 | 6.44E-03 |
| WDR44 | 860.5443349 | 0.262610471 | 1.96E-02 |
| P3H1 | 4737.492285 | 0.262434326 | 4.17E-02 |
| COA1 | 3731.114246 | 0.261727841 | 2.91E-02 |
| KIAA1671 | 2181.6701 | 0.261421873 | 1.64E-03 |
| MRPS30 | 2531.712345 | 0.261344374 | 1.11E-04 |
| ARMC10 | 2257.733891 | 0.26111971 | 5.32E-03 |
| CNN3 | 12063.72067 | 0.260802381 | 3.81E-03 |

| | | | |
|---|---|---|---|
| PCCB | 2314.132373 | 0.260165456 | 1.16E-03 |
| LSG1 | 3989.680754 | 0.259714674 | 6.62E-03 |
| QDPR | 2166.566869 | 0.259314268 | 2.56E-02 |
| CHMP4A | 1281.138968 | 0.259224169 | 6.63E-03 |
| RNF126 | 894.0071533 | 0.258992798 | 2.48E-02 |
| CACUL1 | 1176.624673 | 0.258992288 | 2.94E-02 |
| ENG | 22586.22234 | 0.25802363 | 2.10E-02 |
| HUS1 | 841.6123964 | 0.258011066 | 4.18E-02 |
| AK3 | 2865.631536 | 0.258010086 | 2.05E-04 |
| MTX2 | 1898.537794 | 0.257984952 | 1.32E-02 |
| LPIN2 | 5759.958478 | 0.25794537 | 2.88E-02 |
| NBAS | 3584.572694 | 0.257869624 | 2.60E-02 |
| XAGE1A | 480.1985385 | 0.25654459 | 2.43E-02 |
| C3orf38 | 678.6615537 | 0.25644692 | 1.96E-02 |
| IFT43 | 833.1068617 | 0.255673707 | 4.49E-02 |
| AP3B1 | 6576.449798 | 0.255542159 | 2.36E-06 |
| ACSL5 | 2072.118794 | 0.255462149 | 3.62E-02 |
| NDUFS4 | 2202.636562 | 0.255056634 | 4.40E-03 |
| F2R | 9742.24398 | 0.254917589 | 4.40E-04 |
| SAV1 | 1549.090313 | 0.254864663 | 4.24E-02 |
| GTSF1 | 1663.890902 | 0.254444756 | 1.81E-03 |
| SELENOF | 4529.787408 | 0.254430324 | 4.13E-04 |
| CTNNBL1 | 3608.026851 | 0.25442304 | 8.30E-05 |
| AKIRIN1 | 2134.676385 | 0.253704652 | 1.30E-02 |
| FAF2 | 4718.721789 | 0.253459818 | 9.31E-04 |
| RCBTB1 | 1382.029973 | 0.253057491 | 9.83E-03 |
| CMIP | 2119.779756 | 0.252990538 | 3.94E-02 |
| OXSM | 545.8298956 | 0.2527258 | 2.26E-02 |
| ERAP1 | 6025.539569 | 0.252704726 | 1.83E-02 |
| ADCY7 | 1794.366675 | 0.252618881 | 2.78E-02 |
| SIN3B | 3202.374621 | 0.252591727 | 1.29E-05 |
| CYTH3 | 8882.569858 | 0.252543447 | 2.99E-03 |
| MAT2B | 4086.391294 | 0.252350124 | 2.43E-04 |
| AP3S1 | 3641.825183 | 0.251558631 | 2.38E-03 |
| PLEKHM2 | 4411.871107 | 0.251166952 | 5.54E-03 |
| PSMD6 | 3876.685776 | 0.250813191 | 4.41E-03 |
| VTI1B | 3334.280894 | 0.250540929 | 1.28E-03 |
| VPS41 | 3381.596704 | 0.25052429 | 2.24E-03 |
| RER1 | 5729.27554 | 0.250098412 | 8.34E-03 |
| SLC25A43 | 1114.800884 | 0.249153908 | 1.73E-02 |
| SUN2 | 5461.842658 | 0.248812723 | 1.39E-02 |
| CHMP5 | 3982.975966 | 0.248160302 | 4.54E-04 |
| G6PD | 13437.0078 | 0.247795012 | 4.02E-02 |
| TTC27 | 1234.216433 | 0.24775197 | 1.33E-02 |
| CLEC16A | 2148.484018 | 0.247444 | 3.47E-03 |
| PFDN4 | 1460.258671 | 0.247373815 | 3.57E-02 |
| NSFL1C | 5125.345501 | 0.246757578 | 3.23E-03 |

| | | | |
|---|---|---|---|
| TMEM59 | 6590.074609 | 0.246720692 | 4.70E-03 |
| ETHE1 | 1360.286633 | 0.246673091 | 4.56E-02 |
| TMED4 | 5466.26898 | 0.246524119 | 2.12E-02 |
| NGLY1 | 1647.620889 | 0.246424054 | 4.38E-03 |
| AP3D1 | 8143.720888 | 0.246332081 | 6.69E-04 |
| SUCLG1 | 3552.834348 | 0.245712341 | 3.57E-03 |
| C9orf64 | 1570.97536 | 0.244923933 | 4.53E-03 |
| ERCC5 | 3744.110558 | 0.244857335 | 4.12E-03 |
| ANAPC16 | 1880.704335 | 0.244734031 | 6.45E-03 |
| NA | 283.3964559 | 0.244558149 | 4.69E-02 |
| ACSL3 | 7289.522609 | 0.243851999 | 1.98E-02 |
| UBXN2A | 1343.339787 | 0.243753277 | 3.95E-02 |
| CTHRC1 | 861.6752021 | 0.243701263 | 3.51E-02 |
| RUBCN | 1148.934787 | 0.243208049 | 1.70E-02 |
| MAPK1 | 5682.840336 | 0.242965864 | 9.92E-04 |
| CECR7 | 1422.204956 | 0.242892843 | 4.29E-02 |
| ATP5PO | 6020.894828 | 0.242506753 | 9.64E-03 |
| MCL1 | 7487.483715 | 0.242197174 | 2.34E-03 |
| SHKBP1 | 2696.583798 | 0.241988602 | 4.09E-02 |
| ARHGEF18 | 3169.607921 | 0.241553045 | 4.87E-02 |
| NFATC1 | 833.7809692 | 0.241221324 | 3.25E-02 |
| RBM17 | 1897.252331 | 0.24104068 | 6.63E-03 |
| POLB | 1462.502926 | 0.240993906 | 2.30E-02 |
| TDRKH | 1007.401832 | 0.240593312 | 2.50E-02 |
| EIF2AK4 | 3997.221069 | 0.240368359 | 6.53E-05 |
| MAP2K1 | 3084.114411 | 0.239869838 | 6.74E-03 |
| RAP1B | 7934.008772 | 0.239801421 | 2.02E-02 |
| INTS10 | 3396.001548 | 0.239710508 | 3.13E-03 |
| GNPTG | 2089.161259 | 0.239178287 | 4.28E-02 |
| JAM3 | 6397.462548 | 0.239005863 | 3.04E-02 |
| GLOD4 | 2925.096266 | 0.238483531 | 2.91E-02 |
| CCDC25 | 5562.181212 | 0.238325493 | 3.84E-02 |
| STEAP1B | 969.6948838 | 0.238212314 | 2.49E-02 |
| GRIPAP1 | 2385.540867 | 0.237508406 | 6.11E-03 |
| UBXN1 | 5831.619773 | 0.237489733 | 4.49E-02 |
| REXO4 | 2624.892417 | 0.237050736 | 2.88E-02 |
| EIF4G2 | 45968.60351 | 0.236977009 | 1.47E-03 |
| PTPN2 | 1323.929657 | 0.236918478 | 2.30E-02 |
| MAD1L1 | 2125.818007 | 0.235882057 | 3.07E-02 |
| PEX11A | 650.2384399 | 0.235560746 | 3.43E-02 |
| ARHGEF9 | 1519.411048 | 0.234680689 | 3.75E-02 |
| LAMC1 | 24069.24177 | 0.234452775 | 1.02E-02 |
| POLR3D | 3632.780335 | 0.234383749 | 2.69E-02 |
| CHCHD2 | 15154.40285 | 0.234380158 | 4.78E-02 |
| ECE1 | 7636.537465 | 0.233982485 | 1.27E-02 |
| PRKRA | 1493.271741 | 0.23351354 | 5.95E-03 |
| DECR1 | 1827.820885 | 0.233463443 | 2.08E-02 |

| | | | |
|---|---|---|---|
| ALDH5A1 | 1180.765843 | 0.233417871 | 5.00E-02 |
| CEP41 | 1707.755942 | 0.232778012 | 4.18E-02 |
| TOMM7 | 3903.636684 | 0.232006438 | 4.18E-02 |
| HERPUD2 | 1607.403671 | 0.231364749 | 3.09E-03 |
| DCTN2 | 6303.198494 | 0.230800329 | 4.22E-02 |
| METTL23 | 915.3090469 | 0.230424607 | 1.89E-02 |
| ZNF106 | 3656.174963 | 0.230267079 | 1.40E-02 |
| LSM1 | 1371.961436 | 0.230102206 | 5.49E-03 |
| PCBP1 | 10328.69218 | 0.229607886 | 1.30E-02 |
| FBXO9 | 2299.674237 | 0.229053734 | 1.82E-02 |
| NDUFV3 | 1383.346286 | 0.228753959 | 8.25E-04 |
| RAB12 | 994.6987341 | 0.228549466 | 2.64E-02 |
| SV2A | 2278.471485 | 0.228272019 | 2.26E-02 |
| GLA | 1702.524625 | 0.228229309 | 4.46E-02 |
| TIMM8A | 916.3763145 | 0.227814821 | 1.38E-02 |
| NDUFA6 | 2209.094568 | 0.227683355 | 4.05E-02 |
| NUDCD2 | 2588.159888 | 0.226683977 | 2.65E-02 |
| RNF185 | 2804.861805 | 0.226542926 | 1.78E-02 |
| HMGXB3 | 4166.29827 | 0.226470277 | 1.72E-02 |
| APTX | 2086.750278 | 0.226375056 | 9.70E-03 |
| SSU72 | 3414.035404 | 0.225972794 | 2.82E-02 |
| OCIAD1 | 4290.362214 | 0.225209657 | 7.03E-04 |
| FRG1HP | 786.3343413 | 0.224938293 | 3.98E-02 |
| PRRC2C | 9875.15191 | 0.224707127 | 2.29E-02 |
| SSR1 | 4631.397543 | 0.224508087 | 4.69E-05 |
| DCTN4 | 3405.87974 | 0.224136613 | 2.61E-02 |
| TMEM245 | 5277.538846 | 0.224133114 | 4.92E-03 |
| SELENOT | 2207.40553 | 0.224109227 | 2.80E-02 |
| RPUSD3 | 1681.748031 | 0.223984258 | 2.97E-02 |
| COPS7A | 2415.779054 | 0.223860429 | 1.94E-02 |
| DDX17 | 25267.52818 | 0.223039245 | 2.61E-04 |
| MMGT1 | 2362.006228 | 0.222992661 | 2.67E-03 |
| PRKAG1 | 3564.972411 | 0.221648508 | 1.74E-04 |
| DYNC1LI2 | 8557.972053 | 0.22092976 | 3.01E-02 |
| ZXDC | 1312.034482 | 0.220581246 | 2.68E-02 |
| WRNIP1 | 3155.759016 | 0.219896787 | 3.54E-03 |
| ZNF263 | 2605.294114 | 0.219487159 | 2.30E-02 |
| AKAP8L | 2144.844662 | 0.219318306 | 1.39E-02 |
| PSME4 | 4808.399483 | 0.21901312 | 4.18E-02 |
| CD27-AS1 | 940.6269923 | 0.218024125 | 3.65E-02 |
| ZNF621 | 1590.824832 | 0.217864132 | 2.40E-02 |
| AIFM1 | 3633.466305 | 0.216756389 | 1.97E-02 |
| SAP30BP | 2342.287681 | 0.215308849 | 2.57E-02 |
| ESYT2 | 7763.823023 | 0.214597798 | 1.03E-02 |
| ATP9A | 5110.679593 | 0.213332171 | 2.35E-02 |
| SRP68 | 5045.523619 | 0.213136168 | 1.04E-03 |
| BFAR | 4729.283267 | 0.212864752 | 8.67E-04 |

| | | | |
|---|---|---|---|
| CNPY2 | 2428.666619 | 0.212379541 | 1.45E-02 |
| PJA1 | 2541.974321 | 0.212246995 | 1.08E-02 |
| NCBP2 | 4485.947445 | 0.211033696 | 2.71E-02 |
| NFX1 | 3764.769076 | 0.209927755 | 5.07E-04 |
| CEP170B | 2094.005391 | 0.209074896 | 1.61E-02 |
| FAM120AOS | 2297.168803 | 0.208939461 | 2.82E-02 |
| VPS35 | 9627.418689 | 0.208892965 | 1.23E-02 |
| ADRM1 | 5929.584761 | 0.208053161 | 4.62E-02 |
| SLC35E1 | 3256.555031 | 0.207922051 | 3.60E-02 |
| CBL | 5340.886292 | 0.207786768 | 7.04E-03 |
| TRIM27 | 4409.412563 | 0.207105722 | 3.70E-02 |
| MTCH2 | 5501.939807 | 0.206826087 | 2.52E-02 |
| UBE2R2 | 4405.95647 | 0.206281675 | 1.88E-02 |
| UQCRC2 | 8408.55013 | 0.205768266 | 1.13E-03 |
| RNF4 | 3198.021208 | 0.2056745 | 6.94E-03 |
| TSN | 3501.252314 | 0.203906566 | 9.98E-03 |
| ANKHD1 | 3607.744618 | 0.203610712 | 3.37E-02 |
| GSPT1 | 9602.68302 | 0.203339598 | 9.45E-04 |
| UNC13B | 4255.834304 | 0.202532776 | 1.56E-02 |
| STT3B | 6050.763375 | 0.201980877 | 4.18E-02 |
| TTPAL | 3969.055166 | 0.201970624 | 3.54E-03 |
| MMUT | 1660.940296 | 0.201918954 | 4.90E-02 |
| HIKESHI | 2232.270826 | 0.200771893 | 1.58E-02 |
| PHKA1 | 1733.661679 | 0.200302052 | 2.82E-03 |
| SLC39A9 | 4446.358022 | 0.200156184 | 1.54E-02 |
| NA | 1855.274218 | 0.20003997 | 3.56E-02 |
| MRPS36 | 1255.991792 | 0.199729569 | 4.77E-02 |
| SLC25A51 | 716.101912 | 0.199100445 | 4.62E-02 |
| BMS1 | 3236.803498 | 0.198544939 | 3.79E-02 |
| TSG101 | 4160.63018 | 0.198487601 | 4.79E-02 |
| EPS15 | 2833.132169 | 0.198117953 | 2.32E-02 |
| OGA | 5560.027732 | 0.196696873 | 4.13E-02 |
| USP36 | 1986.048346 | 0.195239859 | 5.00E-02 |
| SLC9A6 | 2218.455759 | 0.194859236 | 4.74E-02 |
| ADAR | 16246.27729 | 0.192459023 | 4.05E-03 |
| TTC19 | 2471.205667 | 0.190192578 | 9.31E-03 |
| ETFA | 5630.621912 | 0.188057993 | 1.64E-03 |
| FAM104A | 1453.554848 | 0.185218434 | 4.80E-02 |
| MFN2 | 7446.363905 | 0.185014544 | 4.55E-02 |
| SYPL1 | 4385.788923 | 0.184123464 | 1.29E-02 |
| SDCBP | 7542.179046 | 0.184013586 | 3.96E-02 |
| TPBG | 2572.499946 | 0.183854743 | 3.29E-02 |
| PSMB6 | 4576.834031 | 0.183662299 | 4.07E-02 |
| FBXO7 | 6186.671808 | 0.183076528 | 4.21E-02 |
| SMG5 | 7376.902738 | 0.181617194 | 4.06E-02 |
| SETD5 | 6418.971384 | 0.180510681 | 3.76E-02 |
| UBQLN2 | 1858.80665 | 0.179464932 | 8.80E-03 |

| | | | |
|---|---|---|---|
| MCTS1 | 2241.850458 | 0.177603368 | 3.20E-02 |
| CCNI | 9677.528827 | 0.175784398 | 4.46E-02 |
| SNF8 | 3726.744613 | 0.174592492 | 4.75E-02 |
| RYK | 3166.723865 | 0.174087474 | 3.81E-02 |
| DIDO1 | 6933.614041 | 0.173934856 | 6.67E-04 |
| SYVN1 | 2776.979789 | 0.173859401 | 4.79E-02 |
| IAH1 | 2107.417709 | 0.171447368 | 2.13E-02 |
| PPHLN1 | 2345.279835 | 0.170780683 | 3.23E-02 |
| UBE2K | 2831.245226 | 0.170720401 | 2.33E-03 |
| ACVR1 | 1442.854223 | 0.170613818 | 4.63E-02 |
| USP14 | 4055.224992 | 0.169667798 | 5.89E-03 |
| SARAF | 7191.114511 | 0.164244832 | 2.53E-02 |
| FITM2 | 2198.051981 | 0.163237074 | 3.71E-02 |
| FBXL5 | 1766.997965 | 0.160975318 | 4.52E-02 |
| SP1 | 5444.375441 | 0.160791595 | 4.89E-03 |
| XPC | 3166.78906 | 0.16039608 | 4.60E-02 |
| KLHL12 | 3560.651364 | 0.15632956 | 3.33E-02 |
| UBE2H | 13961.67353 | 0.154776665 | 1.54E-02 |
| SGCB | 2046.737694 | 0.153997038 | 1.74E-02 |
| TXNL1 | 2788.791101 | 0.153637991 | 3.97E-02 |
| FNBP1 | 5457.462751 | 0.139211903 | 2.89E-02 |
| ERCC3 | 3716.889141 | -0.137027315 | 2.68E-02 |
| TCP1 | 10078.89212 | -0.147295208 | 3.96E-02 |
| PUM1 | 5310.868167 | -0.149601109 | 3.11E-02 |
| AP3M1 | 2106.545134 | -0.153644485 | 4.85E-02 |
| STRIP1 | 2498.412763 | -0.160038156 | 4.76E-02 |
| TRIM4 | 2817.363487 | -0.160229261 | 4.48E-02 |
| DFFA | 3664.020143 | -0.161908673 | 4.24E-02 |
| DCAF1 | 2282.453564 | -0.163559259 | 1.08E-02 |
| INPP4A | 1802.976542 | -0.163908577 | 4.64E-02 |
| PLAGL2 | 2515.730917 | -0.170606283 | 2.63E-02 |
| CTNND1 | 17473.49328 | -0.17306957 | 4.63E-02 |
| MED20 | 1921.206544 | -0.173157436 | 1.72E-02 |
| NUP88 | 3423.063769 | -0.173342595 | 4.48E-02 |
| DUXAP8 | 8045.737965 | -0.173836154 | 2.46E-03 |
| CAPRIN1 | 19543.43926 | -0.175509176 | 2.44E-04 |
| RPL36AL | 2425.183161 | -0.175582285 | 4.97E-02 |
| TMEM69 | 1083.687159 | -0.175804921 | 4.51E-02 |
| ZNF7 | 1564.127802 | -0.1764293 | 1.85E-02 |
| WDR46 | 5956.684672 | -0.178463752 | 4.97E-02 |
| RALBP1 | 3901.009253 | -0.178664343 | 1.95E-02 |
| SERF1B | 1933.088998 | -0.178905644 | 2.46E-02 |
| STK35 | 2874.044657 | -0.179365356 | 2.77E-03 |
| STK38 | 3124.942503 | -0.179453015 | 3.42E-02 |
| NUP155 | 7161.690823 | -0.182042961 | 4.58E-02 |
| TRRAP | 9853.594491 | -0.183683274 | 9.57E-03 |
| NELFA | 1190.639151 | -0.18525932 | 4.98E-02 |

| | | | |
|---|---|---|---|
| PCSK7 | 5369.388781 | -0.185741091 | 4.72E-02 |
| FANCF | 773.4633916 | -0.186012193 | 3.88E-02 |
| PPP1R12C | 2655.67742 | -0.188594197 | 1.37E-02 |
| CHMP7 | 4210.63395 | -0.188827758 | 3.06E-02 |
| CHTOP | 3372.68113 | -0.189021598 | 1.79E-02 |
| CSTF2 | 2441.068491 | -0.189059998 | 9.95E-03 |
| ANKRD52 | 4478.484332 | -0.189336341 | 4.70E-02 |
| CHMP3 | 5334.293861 | -0.189463177 | 4.22E-02 |
| VAPB | 5890.824968 | -0.189918177 | 4.02E-02 |
| GON4L | 2736.602273 | -0.190118169 | 2.50E-02 |
| BLMH | 2902.148101 | -0.190483658 | 1.02E-02 |
| ANKH | 6322.166104 | -0.191566696 | 1.07E-02 |
| PDE4DIP | 4512.478732 | -0.191734623 | 3.75E-02 |
| FAM53C | 7850.025975 | -0.191994919 | 1.67E-02 |
| TSR1 | 7944.401475 | -0.19211146 | 1.48E-03 |
| SART3 | 3428.365544 | -0.19281259 | 6.30E-04 |
| ATG10 | 1168.58865 | -0.193463757 | 3.85E-02 |
| NAA40 | 3343.755419 | -0.193680059 | 1.44E-03 |
| PCNX3 | 5371.421113 | -0.193996764 | 3.54E-02 |
| GAGE1 | 7974.436024 | -0.194106491 | 4.44E-02 |
| GTF3C4 | 5054.525674 | -0.194835607 | 5.44E-03 |
| YBX1 | 25603.98182 | -0.196315497 | 2.40E-02 |
| NT5C3B | 3653.141291 | -0.196545532 | 4.30E-02 |
| ANP32A | 7068.971783 | -0.196641931 | 3.18E-02 |
| PTGES3 | 10958.19736 | -0.196995732 | 2.74E-02 |
| LSM4 | 3280.823502 | -0.197845123 | 4.13E-02 |
| NA | 8125.450766 | -0.199452244 | 2.52E-02 |
| RABL6 | 5448.593957 | -0.199587232 | 3.49E-02 |
| RPRD2 | 2318.718644 | -0.199818793 | 1.52E-03 |
| UBE2N | 4070.400118 | -0.200386994 | 2.83E-02 |
| PRRC2B | 23100.42542 | -0.200662993 | 8.41E-03 |
| USP42 | 1231.999039 | -0.200894432 | 2.20E-02 |
| HNRNPU | 19845.21396 | -0.200898523 | 1.31E-02 |
| TOMM5 | 5140.633679 | -0.201361114 | 3.59E-02 |
| FAM120A | 8203.42013 | -0.201444971 | 3.51E-04 |
| DUXAP9 | 5141.360363 | -0.202070051 | 5.22E-03 |
| TOMM70 | 5225.000315 | -0.202078505 | 1.71E-02 |
| UQCC1 | 3926.027674 | -0.202116461 | 2.19E-02 |
| GPN1 | 3418.273067 | -0.202229272 | 1.62E-02 |
| XRCC6 | 20895.91892 | -0.2027912 | 2.81E-02 |
| SPINDOC | 2425.02415 | -0.203496867 | 4.74E-02 |
| KAT14 | 1554.577656 | -0.204133764 | 1.64E-02 |
| YTHDF2 | 3194.209555 | -0.204213347 | 9.54E-03 |
| WAC-AS1 | 606.8259948 | -0.205068832 | 4.67E-02 |
| KMT2D | 6520.858359 | -0.205322392 | 1.27E-02 |
| SEC11A | 6740.047607 | -0.205589464 | 3.75E-02 |
| TSFM | 1745.174459 | -0.206110833 | 3.89E-02 |

| | | | |
|---|---|---|---|
| ACAD9 | 2006.68078 | -0.206557931 | 4.00E-02 |
| RCL1 | 1212.460695 | -0.206764519 | 3.75E-02 |
| HP1BP3 | 7941.351933 | -0.207385284 | 4.45E-03 |
| COPS6 | 7614.260616 | -0.207478895 | 1.47E-02 |
| PAFAH1B2 | 5201.458627 | -0.20777766 | 9.25E-03 |
| UQCRH | 6986.14891 | -0.208469781 | 4.35E-02 |
| HKR1 | 1861.870123 | -0.208471529 | 6.11E-03 |
| WTAP | 2928.312467 | -0.209181825 | 5.79E-03 |
| SIPA1L1 | 3181.582534 | -0.209198321 | 9.46E-03 |
| NUP214 | 5589.627266 | -0.210470414 | 3.01E-02 |
| SRSF3 | 10893.79417 | -0.210723733 | 6.58E-03 |
| RAB13 | 2951.583054 | -0.211078588 | 3.00E-02 |
| APOL3 | 1109.386461 | -0.211639754 | 3.94E-02 |
| TSPAN3 | 7268.419862 | -0.212043307 | 2.07E-02 |
| SMARCA2 | 4976.288903 | -0.212244153 | 4.68E-02 |
| AHSA1 | 8364.353404 | -0.212409638 | 3.42E-03 |
| MAT2A | 17360.67703 | -0.2130052 | 1.28E-02 |
| SCMH1 | 2564.152182 | -0.21329311 | 3.16E-02 |
| FAM219B | 2418.25522 | -0.213645444 | 5.64E-03 |
| RIN2 | 3881.998031 | -0.214229017 | 2.52E-02 |
| TTL | 3635.258305 | -0.214983447 | 4.78E-02 |
| TERF2 | 2077.136829 | -0.215570138 | 1.83E-02 |
| DHRS4 | 1323.074625 | -0.215753182 | 4.52E-02 |
| ZNF232 | 692.3280867 | -0.215843586 | 4.55E-02 |
| ITSN1 | 4401.632807 | -0.21618437 | 1.32E-03 |
| COPS5 | 3660.217827 | -0.216569058 | 1.75E-03 |
| DPH7 | 1905.818143 | -0.216658473 | 2.52E-02 |
| SCAP | 4183.080889 | -0.21697252 | 4.15E-02 |
| MTERF4 | 1692.32588 | -0.217524616 | 7.28E-03 |
| PARK7 | 7399.089819 | -0.217546936 | 3.90E-02 |
| MTG2 | 2081.396989 | -0.217998456 | 2.23E-02 |
| BTBD7 | 2004.166963 | -0.218311656 | 8.00E-03 |
| MRPL20 | 2256.101299 | -0.21837196 | 2.10E-02 |
| PRIM2 | 1382.020413 | -0.218371973 | 2.81E-02 |
| RSU1 | 1873.166407 | -0.218619526 | 2.45E-02 |
| WDR82 | 7613.00644 | -0.219428322 | 6.63E-03 |
| MIEF1 | 4448.973513 | -0.220371429 | 9.52E-03 |
| GLO1 | 5862.37989 | -0.22115279 | 1.52E-02 |
| PGM2 | 2123.780405 | -0.221433568 | 4.34E-02 |
| EIF2B5 | 3033.847097 | -0.221753579 | 2.34E-02 |
| IK | 5694.729371 | -0.221928887 | 3.96E-03 |
| WDR48 | 2718.702698 | -0.222557293 | 1.49E-02 |
| SEH1L | 1842.288795 | -0.223116432 | 4.26E-02 |
| SIGMAR1 | 5799.589283 | -0.223455886 | 3.20E-02 |
| RXRB | 3345.042761 | -0.223885747 | 3.77E-02 |
| NFIX | 4322.548575 | -0.224279377 | 4.59E-02 |
| CDC26 | 1403.047321 | -0.224952265 | 1.64E-02 |

| | | | |
|---|---|---|---|
| RNF168 | 803.0081132 | -0.225022064 | 1.73E-02 |
| POGK | 3075.627708 | -0.22527445 | 6.45E-03 |
| SON | 15233.18633 | -0.225404973 | 4.74E-02 |
| TMED8 | 2389.896747 | -0.226058153 | 1.61E-02 |
| ASB8 | 972.3934283 | -0.226137493 | 1.95E-02 |
| ELP5 | 2602.09496 | -0.226534558 | 1.86E-02 |
| OXSR1 | 3175.687546 | -0.226596437 | 1.32E-02 |
| NUP205 | 10693.41834 | -0.226726226 | 1.01E-02 |
| GNA12 | 6774.681606 | -0.226853014 | 2.65E-02 |
| ZNF45 | 1552.966555 | -0.226981151 | 1.85E-02 |
| BAG4 | 1113.912056 | -0.227671568 | 2.91E-02 |
| SYNRG | 2863.187215 | -0.227945701 | 1.66E-02 |
| R3HCC1 | 1870.466866 | -0.228042675 | 2.51E-02 |
| ADNP | 9338.900182 | -0.228076138 | 1.13E-04 |
| MRPS25 | 2424.720312 | -0.228629626 | 4.63E-03 |
| SLC25A15 | 687.4465156 | -0.228996298 | 3.09E-02 |
| KHDRBS1 | 9211.008352 | -0.228998902 | 2.65E-04 |
| MRPL4 | 2091.730207 | -0.230054653 | 3.96E-02 |
| GNPAT | 2481.947473 | -0.230167264 | 1.90E-03 |
| POM121C | 7805.729312 | -0.230753758 | 3.99E-02 |
| TMEM183A | 2850.845206 | -0.230885358 | 8.99E-03 |
| WARS2 | 469.0013481 | -0.23101891 | 3.34E-02 |
| MPLKIP | 1609.920512 | -0.23110222 | 3.71E-03 |
| SCAF1 | 3012.004697 | -0.23133957 | 1.67E-02 |
| MAPRE1 | 10884.81933 | -0.231897695 | 1.26E-02 |
| PREX1 | 4692.440271 | -0.231951225 | 2.60E-02 |
| MRPL57 | 750.8230746 | -0.232498164 | 1.09E-02 |
| RIC1 | 2976.271407 | -0.232582331 | 1.89E-04 |
| NRF1 | 837.4999078 | -0.232633045 | 1.43E-02 |
| THRAP3 | 8007.967293 | -0.232654375 | 3.81E-03 |
| GUSB | 2894.849897 | -0.232662641 | 7.50E-03 |
| CRAT | 2562.999731 | -0.232734997 | 4.89E-02 |
| RHOT2 | 5883.720344 | -0.232994675 | 1.80E-02 |
| BAP1 | 5048.804008 | -0.233034888 | 1.95E-02 |
| PDHB | 3611.84747 | -0.233204161 | 1.45E-02 |
| CCT5 | 38883.96642 | -0.23348907 | 2.69E-03 |
| CKLF | 1221.047736 | -0.233835734 | 4.41E-03 |
| YWHAE | 30929.50022 | -0.234125649 | 4.42E-03 |
| C7orf26 | 1655.471349 | -0.234250207 | 1.61E-02 |
| PHB2 | 8408.780695 | -0.234281634 | 1.61E-02 |
| MGAT5 | 6876.268674 | -0.234820939 | 3.57E-02 |
| C2orf68 | 2322.099413 | -0.235855801 | 1.62E-05 |
| UBFD1 | 7481.738186 | -0.23587992 | 8.22E-03 |
| NFATC3 | 2721.717658 | -0.236255535 | 8.30E-04 |
| EIF3H | 12961.00924 | -0.236694007 | 3.13E-03 |
| UTP14A | 3064.894564 | -0.237362843 | 2.63E-02 |
| DNAJC15 | 2292.244546 | -0.237585159 | 3.30E-02 |

| | | | |
|---|---|---|---|
| GABBR1 | 1499.290205 | -0.238078102 | 1.13E-02 |
| KDM3A | 2614.767864 | -0.238182398 | 4.38E-02 |
| TTC7A | 1548.824814 | -0.238374186 | 3.83E-02 |
| NSF | 4379.616314 | -0.239096742 | 3.54E-04 |
| KAT5 | 2011.410731 | -0.239649403 | 4.85E-02 |
| SMARCC2 | 5581.460016 | -0.239723082 | 1.41E-02 |
| NSMF | 5938.632953 | -0.240101904 | 2.65E-02 |
| NAP1L4 | 7720.284071 | -0.240445066 | 1.22E-02 |
| USP22 | 19092.274 | -0.240752513 | 2.36E-03 |
| C5orf15 | 4632.545109 | -0.240771686 | 3.62E-02 |
| ZNF789 | 838.5068154 | -0.241246158 | 3.78E-03 |
| RPIA | 1240.066785 | -0.241276034 | 3.24E-03 |
| BTN2A3P | 353.6383477 | -0.241311686 | 4.95E-02 |
| ZMYND8 | 5516.740769 | -0.241560508 | 2.90E-02 |
| BAIAP2L1 | 1835.617301 | -0.241685919 | 3.09E-02 |
| LASP1 | 19014.37754 | -0.241772739 | 4.02E-02 |
| ATF7 | 1907.724653 | -0.241872075 | 2.38E-03 |
| ZMIZ2 | 2157.698902 | -0.242057707 | 6.67E-04 |
| MLH1 | 2396.676964 | -0.242581472 | 8.61E-03 |
| VPS4A | 5291.241345 | -0.243021189 | 7.19E-03 |
| FBXO21 | 2781.616339 | -0.243294186 | 4.14E-02 |
| KIAA1147 | 3149.274543 | -0.244123773 | 7.52E-04 |
| TBC1D8 | 2589.114755 | -0.244213498 | 2.10E-03 |
| NDC1 | 2827.639761 | -0.244432759 | 1.72E-02 |
| PDPR | 5125.12976 | -0.245018226 | 1.96E-03 |
| ILKAP | 1385.386995 | -0.245267481 | 2.79E-04 |
| TMEM63A | 2402.01206 | -0.245356489 | 2.24E-03 |
| PFDN6 | 2605.546708 | -0.245867129 | 2.27E-02 |
| G6PC3 | 3386.25251 | -0.246581745 | 4.04E-02 |
| MRPL38 | 1864.760936 | -0.246606368 | 2.01E-02 |
| ANAPC7 | 3020.887082 | -0.247522881 | 2.49E-02 |
| PBX2 | 4801.39461 | -0.247568889 | 5.34E-03 |
| SPIN2B | 409.744545 | -0.247851256 | 2.06E-02 |
| RAI1 | 4292.083718 | -0.248241045 | 9.04E-03 |
| FBXW2 | 5336.381978 | -0.248388615 | 2.25E-06 |
| WASH4P | 1272.645148 | -0.248445833 | 1.30E-02 |
| VPS52 | 3114.345053 | -0.249148847 | 2.54E-03 |
| NOC4L | 875.5322397 | -0.249152836 | 1.05E-02 |
| SEMA4C | 2019.090106 | -0.249230079 | 1.75E-02 |
| KCTD17 | 1231.905598 | -0.249583957 | 3.40E-02 |
| PITRM1 | 3121.362636 | -0.24966712 | 3.81E-03 |
| PPP4R1 | 4147.702113 | -0.250393628 | 5.03E-03 |
| DIS3L2 | 1432.177132 | -0.250501419 | 1.82E-02 |
| BRD2 | 10327.68486 | -0.250539004 | 4.25E-03 |
| PPP1CC | 7270.810122 | -0.250730772 | 7.47E-03 |
| ADI1 | 4301.525489 | -0.250788003 | 4.35E-02 |
| C11orf49 | 1568.56911 | -0.250798824 | 1.45E-02 |

| | | | |
|---|---|---|---|
| RNF139 | 1481.985634 | -0.250892433 | 5.57E-03 |
| SMARCC1 | 8451.95218 | -0.251518846 | 5.48E-07 |
| ZNF598 | 2854.685967 | -0.251797902 | 3.93E-03 |
| CCDC51 | 1006.545882 | -0.251928627 | 2.89E-02 |
| ZSCAN22 | 377.5474583 | -0.251946863 | 3.16E-02 |
| WBP11 | 4842.758116 | -0.252100244 | 1.41E-02 |
| TUFM | 15387.79824 | -0.252250377 | 1.48E-02 |
| SNX7 | 1214.597973 | -0.252250869 | 2.91E-02 |
| SARNP | 2010.285177 | -0.252706196 | 5.13E-03 |
| TRAPPC2L | 2075.314082 | -0.253171861 | 4.59E-02 |
| CSNK1D | 7358.64663 | -0.253731442 | 1.66E-02 |
| PHF13 | 1800.433566 | -0.253740956 | 4.77E-02 |
| ACACA | 9223.618154 | -0.253852986 | 1.33E-03 |
| SYNGAP1 | 1172.130205 | -0.254715516 | 3.58E-02 |
| ABL1 | 5711.530818 | -0.254982908 | 2.68E-02 |
| CDC42EP4 | 3656.079881 | -0.255260567 | 1.12E-02 |
| ARPC4 | 5112.83738 | -0.25528182 | 2.83E-02 |
| SERBP1 | 15173.87957 | -0.255322551 | 1.37E-02 |
| AZIN1 | 7430.088457 | -0.255678301 | 5.01E-04 |
| NUMB | 4674.727443 | -0.25572051 | 3.64E-05 |
| MAPK1IP1L | 3707.931357 | -0.255806681 | 9.30E-03 |
| KLHL18 | 1508.869788 | -0.256368174 | 1.70E-02 |
| HMGN1 | 7832.934936 | -0.256511708 | 2.76E-05 |
| HOMEZ | 941.8807832 | -0.256840395 | 1.25E-02 |
| NLK | 1029.221641 | -0.257243072 | 5.41E-03 |
| CCS | 1502.729385 | -0.257530323 | 1.48E-02 |
| CCDC144NL-AS | 1631.969598 | -0.257972216 | 3.13E-02 |
| PPP3CB | 1084.96136 | -0.258954348 | 8.69E-03 |
| EYA3 | 2181.355695 | -0.25920572 | 3.03E-03 |
| POLR2A | 10759.83302 | -0.259574728 | 3.35E-03 |
| ZNF768 | 2739.634064 | -0.259960152 | 1.39E-02 |
| THBS3 | 1463.203303 | -0.260159749 | 3.72E-02 |
| TOMM6 | 3943.972343 | -0.260239633 | 3.81E-02 |
| LINC00680 | 465.4090567 | -0.260869295 | 3.69E-02 |
| CPSF4 | 2678.248412 | -0.2610403 | 8.82E-04 |
| AHCY | 11764.42684 | -0.261180111 | 1.88E-02 |
| TBC1D5 | 6004.939625 | -0.261318731 | 1.30E-04 |
| XRN2 | 14878.00599 | -0.261389226 | 1.98E-03 |
| STX10 | 2145.396318 | -0.261511029 | 1.36E-02 |
| LYRM4 | 1174.273544 | -0.26156957 | 4.66E-02 |
| ZC3H7B | 8780.439133 | -0.261594838 | 1.90E-03 |
| IPO9 | 8938.980323 | -0.26165112 | 1.12E-02 |
| ANAPC11 | 1440.404166 | -0.261990887 | 4.24E-02 |
| ZNF746 | 1278.679077 | -0.26312402 | 1.63E-03 |
| PPP1R9B | 4228.42343 | -0.263376805 | 4.31E-03 |
| DHX38 | 5131.050193 | -0.263503712 | 4.84E-04 |
| PIH1D1 | 2491.157385 | -0.263789319 | 4.77E-02 |

| | | | |
|---|---|---|---|
| RELT | 599.7605953 | -0.26441423 | 4.59E-02 |
| KDM2A | 7320.875344 | -0.264862208 | 7.36E-03 |
| EIF4B | 22106.45278 | -0.264926135 | 3.27E-02 |
| UTP4 | 5907.120824 | -0.264950753 | 3.24E-03 |
| C1orf50 | 316.5943599 | -0.265064098 | 4.77E-02 |
| R3HDM1 | 2482.454151 | -0.26621913 | 2.75E-02 |
| ANKFY1 | 7580.467361 | -0.266319719 | 8.32E-03 |
| TAP2 | 3870.967011 | -0.266664978 | 1.24E-02 |
| PKN3 | 1552.250126 | -0.266745693 | 1.13E-02 |
| DUS1L | 2081.113592 | -0.266755714 | 2.14E-04 |
| IRF2BPL | 3296.790079 | -0.266961847 | 2.01E-02 |
| ZCCHC17 | 2099.925354 | -0.267064065 | 4.77E-03 |
| GATC | 2817.492369 | -0.267344667 | 1.29E-03 |
| POLR2F | 1917.43816 | -0.267964227 | 2.39E-02 |
| LENG8 | 3791.016704 | -0.268002613 | 6.81E-03 |
| HTATSF1 | 6072.239422 | -0.268133314 | 2.96E-02 |
| IRAK1 | 19076.01415 | -0.268137101 | 2.09E-02 |
| PDCL3 | 827.3346447 | -0.26817675 | 8.99E-03 |
| MTAP | 3392.856448 | -0.268268364 | 1.70E-02 |
| MRTFA | 1872.178254 | -0.268370585 | 4.18E-02 |
| SF1 | 10578.12563 | -0.268380439 | 1.70E-04 |
| FAU | 10785.67298 | -0.268868832 | 4.74E-02 |
| URB1 | 5309.208866 | -0.268931825 | 1.78E-03 |
| DYNC1I2 | 4866.851288 | -0.269053799 | 7.95E-04 |
| RNPS1 | 7816.659833 | -0.269390875 | 6.84E-03 |
| SH3PXD2B | 1746.191447 | -0.269542971 | 1.13E-02 |
| AGAP3 | 4495.009324 | -0.269571525 | 1.28E-03 |
| MSL3 | 1482.073196 | -0.270302283 | 1.00E-03 |
| RCC2 | 8669.224488 | -0.27064525 | 2.06E-03 |
| MTHFD1 | 8224.678989 | -0.270906501 | 5.57E-04 |
| EEF2KMT | 406.8081592 | -0.271087802 | 3.62E-02 |
| KIF3B | 5246.530593 | -0.271441354 | 6.43E-05 |
| ASB3 | 947.7723291 | -0.271985255 | 3.63E-04 |
| RNF8 | 1200.25097 | -0.272300821 | 7.39E-03 |
| PGAP2 | 928.1459418 | -0.272403249 | 1.10E-02 |
| GNB2 | 11143.86627 | -0.272556602 | 3.30E-02 |
| PIGW | 1124.188094 | -0.272584908 | 3.01E-02 |
| CARD8 | 1192.652439 | -0.272655371 | 1.01E-03 |
| ERVK13-1 | 1300.392176 | -0.273649112 | 2.63E-03 |
| FBXO3 | 1527.174634 | -0.273930916 | 3.38E-02 |
| KDM4A | 4070.725124 | -0.274334748 | 1.91E-02 |
| BOLA2 | 1180.562019 | -0.274388738 | 4.18E-02 |
| TMEM43 | 7214.222135 | -0.274513912 | 2.69E-04 |
| PIGX | 1133.959941 | -0.274601612 | 1.69E-02 |
| EIF2A | 3488.390567 | -0.275240731 | 3.69E-02 |
| MRPS34 | 2965.552352 | -0.275276325 | 4.16E-03 |
| KMT5A | 1885.209063 | -0.275687458 | 4.13E-02 |

| | | | |
|---|---|---|---|
| RNF212 | 2579.106999 | -0.275905909 | 3.86E-05 |
| ESYT1 | 9950.813896 | -0.275955129 | 1.78E-02 |
| MRPL32 | 2091.698119 | -0.27616311 | 1.31E-03 |
| KPNB1 | 23901.24342 | -0.276185224 | 6.60E-03 |
| GBA2 | 3875.278018 | -0.27625548 | 1.20E-03 |
| GPATCH1 | 715.8843927 | -0.276331492 | 2.26E-02 |
| LRCH4 | 2232.681622 | -0.27649607 | 4.39E-02 |
| SNHG7 | 855.8523069 | -0.276558391 | 1.25E-02 |
| CSTF2T | 1225.829139 | -0.276661965 | 9.24E-03 |
| RECQL5 | 1712.668547 | -0.27696405 | 2.56E-02 |
| ATP5F1E | 10326.63516 | -0.277009535 | 4.41E-02 |
| RPL21 | 8227.922657 | -0.277244297 | 1.01E-02 |
| AC073869.1 | 886.8040614 | -0.277526016 | 1.52E-03 |
| CLP1 | 1304.941902 | -0.27754422 | 1.27E-04 |
| DDX51 | 1618.888558 | -0.277730179 | 3.13E-02 |
| RNF166 | 1068.399366 | -0.277816403 | 3.89E-03 |
| PELP1 | 3679.224611 | -0.27789556 | 3.13E-03 |
| ACACB | 1090.006831 | -0.2781013 | 2.01E-02 |
| MOK | 1296.653583 | -0.278115823 | 1.36E-02 |
| C17orf100 | 413.4607647 | -0.278320924 | 3.33E-02 |
| SIMC1 | 762.4064736 | -0.278394086 | 5.53E-04 |
| TMA7 | 3015.831578 | -0.278402903 | 3.14E-02 |
| SPPL2B | 1636.596338 | -0.278533361 | 3.97E-03 |
| DCAF16 | 2142.847711 | -0.278564998 | 9.82E-03 |
| ZNF687 | 1910.367372 | -0.278622696 | 1.09E-02 |
| DHX15 | 7490.4444 | -0.278732812 | 4.15E-03 |
| SPTSSA | 1145.077999 | -0.278830076 | 4.38E-02 |
| ARG2 | 2408.765317 | -0.278980925 | 4.12E-02 |
| ATPSCKMT | 793.0329605 | -0.27899872 | 1.97E-02 |
| SIX5 | 1314.827357 | -0.279036302 | 4.52E-02 |
| KDELC1 | 2977.176079 | -0.279409528 | 1.19E-02 |
| ANGEL2 | 967.7784368 | -0.279540267 | 1.16E-02 |
| KHDC1 | 323.2992763 | -0.280127211 | 4.17E-02 |
| LSM14B | 3273.518921 | -0.280330347 | 8.57E-05 |
| RAD21 | 11043.70758 | -0.281617726 | 3.96E-02 |
| FAM104B | 415.0212845 | -0.281694499 | 1.70E-02 |
| ZNF74 | 870.2220145 | -0.281864216 | 5.19E-03 |
| JPT1 | 2871.360134 | -0.282164206 | 2.52E-03 |
| ENGASE | 1590.466553 | -0.282252589 | 2.37E-02 |
| UBA1 | 23907.21668 | -0.282389473 | 7.44E-03 |
| LUC7L | 2156.329626 | -0.282390201 | 2.93E-04 |
| HYLS1 | 1297.849303 | -0.282567573 | 7.19E-03 |
| EIF3F | 10611.45856 | -0.282648093 | 4.72E-02 |
| ARPC1A | 8304.036239 | -0.283329233 | 2.79E-02 |
| LINC02381 | 568.2933948 | -0.283571012 | 4.98E-02 |
| HNRNPH3 | 5091.643026 | -0.283586343 | 2.95E-02 |
| NFIC | 3080.234408 | -0.283647787 | 1.17E-03 |

| | | | |
|---|---|---|---|
| RPL30 | 35678.43918 | -0.283670055 | 9.49E-03 |
| CARM1 | 4180.03102 | -0.283854912 | 2.37E-03 |
| TXNDC17 | 2660.628943 | -0.284272454 | 7.10E-03 |
| TBC1D1 | 3064.045464 | -0.284426681 | 1.02E-02 |
| NADK | 3529.607159 | -0.285057328 | 9.14E-04 |
| ZNF775 | 895.107378 | -0.285205545 | 2.29E-02 |
| 6-Mar | 13289.27483 | -0.285259851 | 6.01E-04 |
| SNX29 | 2506.115096 | -0.28529789 | 4.89E-03 |
| AGO1 | 2852.696688 | -0.285497859 | 1.55E-04 |
| MINPP1 | 1359.435503 | -0.285711967 | 2.67E-02 |
| MRPL51 | 4519.757422 | -0.285839995 | 1.55E-03 |
| VDAC1 | 14777.17519 | -0.286061277 | 1.69E-02 |
| MOV10 | 4645.557548 | -0.286118848 | 6.92E-03 |
| CRAMP1 | 1819.133989 | -0.286487038 | 1.82E-03 |
| NIPSNAP1 | 5969.896051 | -0.286598496 | 3.71E-02 |
| RPL11 | 36795.27254 | -0.286901492 | 6.78E-04 |
| UBALD1 | 463.8856587 | -0.287541207 | 4.41E-02 |
| ADPRHL2 | 1566.834803 | -0.287765414 | 3.98E-02 |
| TPCN1 | 4974.758491 | -0.287958001 | 2.10E-03 |
| QSOX2 | 2784.490846 | -0.288082799 | 4.19E-05 |
| DARS2 | 3464.006142 | -0.288158119 | 3.22E-04 |
| COG4 | 5080.752884 | -0.288184366 | 8.79E-03 |
| RGL2 | 4766.753353 | -0.288262828 | 3.56E-02 |
| TDP2 | 1202.826721 | -0.288385833 | 3.59E-02 |
| FKBP4 | 6633.202591 | -0.288923016 | 1.13E-02 |
| LEO1 | 1631.362175 | -0.289327643 | 5.03E-03 |
| CSKMT | 343.0414308 | -0.289563544 | 3.00E-02 |
| BZW1 | 9563.85045 | -0.289895084 | 1.16E-02 |
| LRRC41 | 4882.189823 | -0.290334415 | 1.06E-02 |
| METTL16 | 2336.129699 | -0.290608407 | 3.12E-04 |
| NARF | 2110.426255 | -0.290750654 | 1.54E-02 |
| RPL6 | 45697.77222 | -0.291014081 | 4.30E-05 |
| NOL12 | 814.8016187 | -0.291062232 | 4.02E-02 |
| MRPL12 | 3247.097346 | -0.291303009 | 4.15E-03 |
| CCNQ | 1123.60927 | -0.291524853 | 1.89E-02 |
| TMSB4X | 28768.79103 | -0.291914848 | 3.03E-02 |
| ISOC2 | 1214.042127 | -0.292241911 | 4.20E-02 |
| PIGM | 786.4326629 | -0.292270595 | 3.68E-04 |
| DAGLA | 1136.539145 | -0.292326991 | 1.91E-02 |
| PPP1CA | 7704.076957 | -0.292419645 | 2.22E-02 |
| FARP1 | 10323.71837 | -0.292700477 | 4.54E-04 |
| POMT1 | 2123.905987 | -0.2927328 | 2.37E-02 |
| C12orf10 | 2096.887451 | -0.293037915 | 3.46E-02 |
| BTF3 | 37705.71318 | -0.293120163 | 5.91E-04 |
| ITPA | 3951.47197 | -0.293250228 | 1.15E-02 |
| UBE2S | 5368.618216 | -0.293462232 | 2.46E-02 |
| KLHDC4 | 1557.359036 | -0.293728411 | 2.44E-02 |

| | | | |
|---|---|---|---|
| RNF182 | 646.7070756 | -0.293801814 | 4.55E-02 |
| SLFN11 | 2201.403726 | -0.294108811 | 4.52E-02 |
| RPL19 | 41793.06361 | -0.294127331 | 5.27E-03 |
| SMAD5 | 6225.784563 | -0.294176873 | 7.10E-03 |
| POLE3 | 5076.864435 | -0.294298785 | 5.83E-03 |
| FOXRED1 | 3760.919058 | -0.294509231 | 4.59E-04 |
| AL133367.1 | 255.4937845 | -0.294682822 | 3.64E-02 |
| ZNF76 | 1282.837094 | -0.294743105 | 1.13E-02 |
| SIK2 | 3153.93286 | -0.294930978 | 4.78E-02 |
| KCTD2 | 2597.186571 | -0.295082019 | 1.82E-02 |
| GALNS | 1902.302002 | -0.295153018 | 2.71E-02 |
| DGKZ | 4303.453503 | -0.295804138 | 7.15E-04 |
| RNGTT | 1700.91481 | -0.295919154 | 4.87E-02 |
| FBXL19-AS1 | 285.6288051 | -0.296467508 | 1.64E-02 |
| CAPN10 | 1051.764605 | -0.296659799 | 1.48E-03 |
| ZNF335 | 1385.685245 | -0.296753216 | 5.50E-04 |
| MAP2K7 | 1772.492535 | -0.296840221 | 4.55E-02 |
| ZNF646 | 2349.21993 | -0.296966903 | 1.35E-02 |
| METTL1 | 692.8469278 | -0.297026712 | 1.47E-02 |
| MSI2 | 3793.636912 | -0.297076353 | 3.45E-04 |
| SH3BP5L | 1408.159396 | -0.297557044 | 1.08E-02 |
| NOP9 | 3695.923527 | -0.297907093 | 9.40E-04 |
| ANKS6 | 4072.304987 | -0.29794268 | 6.60E-03 |
| NDUFB2 | 3858.029035 | -0.298315542 | 2.26E-02 |
| SLC2A8 | 513.2020919 | -0.298483753 | 2.98E-02 |
| NAP1L1 | 25109.18802 | -0.298862207 | 2.14E-06 |
| GAS8 | 1544.346104 | -0.298982197 | 3.66E-03 |
| PMPCA | 4348.934324 | -0.299140181 | 4.10E-02 |
| SPEN | 5159.913258 | -0.29931811 | 4.95E-09 |
| RPS6KB2 | 2424.493208 | -0.299366676 | 5.85E-03 |
| GIGYF1 | 4330.870716 | -0.29990828 | 5.80E-06 |
| TMEM164 | 2976.713852 | -0.300032665 | 9.43E-03 |
| UNC45A | 5748.358077 | -0.300159971 | 2.22E-02 |
| PDCD6 | 8304.407416 | -0.300270135 | 9.73E-05 |
| HOXA7 | 998.7093727 | -0.300402518 | 1.69E-03 |
| MLXIP | 3771.357326 | -0.300577941 | 1.51E-05 |
| HOXD9 | 382.8018641 | -0.300824303 | 4.59E-02 |
| FAM136A | 2926.438027 | -0.300967209 | 6.30E-10 |
| ZFAND2B | 806.4967146 | -0.30102634 | 7.88E-03 |
| ADD1 | 10092.77545 | -0.301543 | 5.69E-03 |
| ARL15 | 968.3590657 | -0.302181908 | 3.27E-02 |
| ECI2 | 3140.262434 | -0.302575805 | 7.42E-03 |
| DBI | 4740.869507 | -0.302641646 | 1.31E-02 |
| MED24 | 2857.549009 | -0.302789646 | 3.20E-02 |
| IGSF3 | 4190.625472 | -0.303343179 | 1.44E-02 |
| FAM173A | 371.9964491 | -0.303796709 | 4.02E-02 |
| TSPEAR-AS2 | 381.5583813 | -0.303926488 | 3.71E-02 |

| | | | |
|---|---|---|---|
| MYRF | 3061.445988 | -0.304195246 | 8.74E-03 |
| PGP | 1252.965826 | -0.304196742 | 4.65E-02 |
| FUT4 | 525.2112305 | -0.304718576 | 3.92E-02 |
| SNRNP200 | 26799.02674 | -0.30477092 | 3.43E-04 |
| ERF | 3013.278853 | -0.304884167 | 1.64E-03 |
| NIFK-AS1 | 296.8162705 | -0.305494322 | 3.58E-02 |
| SIN3A | 5272.090568 | -0.305571229 | 6.12E-07 |
| FRS3 | 243.2988226 | -0.305718687 | 3.20E-02 |
| RITA1 | 1304.280185 | -0.305893512 | 1.64E-03 |
| USP11 | 6955.716365 | -0.306151283 | 2.59E-02 |
| CEP85 | 1897.607299 | -0.306377119 | 8.23E-03 |
| STK16 | 1384.410778 | -0.306526993 | 3.20E-03 |
| POLDIP3 | 5683.207276 | -0.30682006 | 3.81E-03 |
| PMPCB | 5559.104433 | -0.306908413 | 2.19E-02 |
| DMWD | 1492.062478 | -0.306972055 | 2.36E-03 |
| PABPC4 | 6179.496463 | -0.307185582 | 5.06E-04 |
| CISD3 | 1447.725078 | -0.307188121 | 4.35E-02 |
| PI4K2B | 830.2728568 | -0.307301047 | 1.23E-02 |
| LSM7 | 1436.691062 | -0.307353646 | 4.77E-02 |
| THSD4 | 3166.704389 | -0.307453291 | 2.27E-02 |
| EEF1AKNMT | 2434.565317 | -0.30785126 | 9.12E-04 |
| METTL26 | 1644.085805 | -0.307901186 | 1.19E-02 |
| UBE2G2 | 6206.458586 | -0.308070842 | 1.20E-05 |
| KDM5C | 7269.373489 | -0.308511982 | 1.93E-05 |
| FXYD5 | 4701.536881 | -0.308657106 | 7.64E-03 |
| TBCCD1 | 605.9990156 | -0.308721976 | 5.88E-03 |
| CRADD | 389.3041389 | -0.308854666 | 1.93E-02 |
| MLX | 2711.999625 | -0.309231674 | 5.01E-03 |
| EIF4ENIF1 | 1435.10004 | -0.309298841 | 3.87E-02 |
| ARL6IP1 | 9774.351742 | -0.309384925 | 4.45E-04 |
| TBC1D9B | 7238.498145 | -0.309446218 | 3.13E-03 |
| ZNF444 | 935.445066 | -0.309538218 | 2.44E-03 |
| GEMIN5 | 2211.92909 | -0.309577938 | 4.78E-06 |
| CDC25B | 13099.33858 | -0.309644843 | 2.71E-04 |
| SUPT4H1 | 3356.805422 | -0.309688699 | 3.20E-02 |
| DNAJC14 | 2859.106033 | -0.309779148 | 2.33E-04 |
| KANSL2 | 1885.157961 | -0.309825521 | 7.27E-07 |
| 8-Sep | 9809.378699 | -0.30999184 | 9.69E-05 |
| CCDC36 | 528.8254423 | -0.310069652 | 2.28E-02 |
| COL4A2 | 5735.041245 | -0.310627386 | 1.01E-02 |
| WDR75 | 2797.494185 | -0.310909171 | 6.29E-03 |
| NCBP2-AS2 | 619.0231114 | -0.311379193 | 2.15E-02 |
| ZNF432 | 386.5222351 | -0.311862128 | 2.13E-02 |
| GLI4 | 718.4885791 | -0.311867678 | 4.04E-02 |
| PCBP4 | 2249.447405 | -0.311892361 | 1.27E-02 |
| GPATCH4 | 2639.640989 | -0.31228269 | 4.97E-03 |
| XRCC3 | 1695.724094 | -0.312941116 | 4.89E-05 |

| | | | |
|---|---|---|---|
| GALT | 1702.165667 | -0.313012883 | 1.39E-02 |
| STX6 | 1556.556609 | -0.313158947 | 1.23E-06 |
| EPN1 | 6033.942424 | -0.313161726 | 4.26E-03 |
| ZNF174 | 375.442338 | -0.313214398 | 3.29E-02 |
| SNRPA | 3244.309368 | -0.313329908 | 3.13E-02 |
| CS | 9835.018622 | -0.31335848 | 1.53E-04 |
| ARHGDIA | 10479.86553 | -0.31340046 | 2.84E-02 |
| SHANK3 | 593.1335192 | -0.31352421 | 1.72E-02 |
| TMEM203 | 2294.626469 | -0.313821438 | 6.20E-03 |
| HNRNPAB | 5558.229272 | -0.314272764 | 3.92E-05 |
| DDX24 | 7689.582107 | -0.314424514 | 1.25E-03 |
| EIPR1 | 1284.282965 | -0.314512956 | 4.36E-03 |
| GRK6 | 1476.105821 | -0.31477017 | 7.41E-03 |
| CCAR2 | 13079.8938 | -0.314906705 | 2.96E-03 |
| ELK1 | 2389.169204 | -0.314967006 | 3.67E-03 |
| HEXIM1 | 2270.508308 | -0.315052366 | 1.08E-02 |
| GADD45GIP1 | 1768.229665 | -0.315155513 | 1.14E-02 |
| ILF2 | 9055.317368 | -0.315331267 | 1.83E-05 |
| PITPNM1 | 3508.712923 | -0.315333102 | 1.22E-02 |
| BFSP1 | 448.8421094 | -0.315433543 | 1.94E-02 |
| MLLT1 | 5500.18598 | -0.315615944 | 1.47E-04 |
| ZKSCAN4 | 386.1799195 | -0.315699225 | 4.80E-02 |
| GRAMD1A | 1599.035645 | -0.315830359 | 5.57E-03 |
| AC005523.2 | 1554.025878 | -0.315939585 | 6.00E-03 |
| IMPDH2 | 17822.23971 | -0.316032572 | 1.50E-03 |
| STRADA | 1484.97176 | -0.316383602 | 9.09E-05 |
| BORCS8 | 545.5502618 | -0.316394548 | 4.20E-02 |
| FAM120B | 2619.454493 | -0.31646155 | 2.07E-02 |
| MAP3K11 | 4440.429376 | -0.316508 | 9.14E-03 |
| CKS2 | 3853.770361 | -0.316607546 | 1.02E-03 |
| EARS2 | 2767.420521 | -0.316776388 | 5.60E-03 |
| PGBD1 | 568.7505738 | -0.316861477 | 9.23E-03 |
| G3BP1 | 7461.652877 | -0.31711704 | 9.43E-07 |
| PDZD11 | 2166.910511 | -0.317188547 | 1.23E-02 |
| MAGEF1 | 1862.743773 | -0.317356062 | 1.15E-02 |
| CMTM3 | 5438.986239 | -0.317475139 | 1.64E-02 |
| GLT8D2 | 304.745755 | -0.317878155 | 4.52E-02 |
| TBC1D12 | 415.325258 | -0.317989473 | 4.33E-02 |
| RETREG2 | 4730.599494 | -0.318012606 | 1.31E-02 |
| KAT8 | 1737.846709 | -0.318233284 | 2.51E-02 |
| GMEB1 | 639.3399019 | -0.318329708 | 3.76E-02 |
| BRPF1 | 1247.041248 | -0.318510644 | 7.94E-03 |
| PMVK | 1036.769575 | -0.318655172 | 8.44E-04 |
| KMT2E | 3440.83109 | -0.318667864 | 6.95E-03 |
| FAM172A | 1449.935888 | -0.31871042 | 2.89E-03 |
| GSDMD | 3104.419199 | -0.318981817 | 4.16E-02 |
| GUCD1 | 4312.301395 | -0.320157847 | 1.64E-02 |

| | | | |
|---|---|---|---|
| AP000525.1 | 2001.229695 | -0.320504366 | 3.80E-04 |
| PCSK6 | 1048.029103 | -0.320763916 | 6.89E-04 |
| HCFC2 | 577.4168189 | -0.320995582 | 3.64E-02 |
| ARID1A | 5257.33245 | -0.321028263 | 3.95E-05 |
| UBR5 | 9785.128236 | -0.321120352 | 3.60E-06 |
| ARNTL2 | 885.1172776 | -0.321561256 | 3.68E-02 |
| PABPC1 | 53608.13181 | -0.321991458 | 1.21E-03 |
| SF3B4 | 3690.285838 | -0.322100202 | 9.49E-03 |
| DHX33 | 2973.454175 | -0.322822692 | 9.78E-04 |
| PDZD4 | 1438.460956 | -0.323133311 | 7.24E-03 |
| MFSD3 | 1352.000871 | -0.323202967 | 1.92E-03 |
| CDK19 | 1510.941392 | -0.323207504 | 1.12E-03 |
| EDC4 | 4898.051578 | -0.323484089 | 2.95E-03 |
| RIC8B | 960.5026178 | -0.323642922 | 1.89E-02 |
| PATZ1 | 2235.275444 | -0.324050119 | 2.37E-03 |
| PCDHGC3 | 8838.789475 | -0.324073268 | 4.03E-02 |
| XPO5 | 5961.325534 | -0.324184034 | 1.85E-04 |
| KRBA2 | 422.1868317 | -0.324316425 | 4.76E-03 |
| MICOS13 | 1664.422011 | -0.324428123 | 6.95E-03 |
| EHHADH | 803.6704901 | -0.32457811 | 8.31E-03 |
| RETREG3 | 2208.190505 | -0.324624842 | 2.05E-02 |
| LYSMD1 | 394.5452007 | -0.325053588 | 4.48E-02 |
| CHGB | 280.9084651 | -0.325261165 | 4.67E-02 |
| MEA1 | 2088.826949 | -0.325541097 | 2.64E-03 |
| TP53BP1 | 4052.421138 | -0.325830285 | 1.53E-05 |
| HOOK2 | 670.3343555 | -0.325946211 | 3.83E-02 |
| RAB11B | 2421.741717 | -0.326154029 | 2.08E-02 |
| RPARP-AS1 | 304.2994051 | -0.326187573 | 1.47E-02 |
| SLC35E2A | 570.8039971 | -0.326198352 | 3.89E-03 |
| UNC119 | 1445.579656 | -0.326208514 | 2.84E-02 |
| AC008894.2 | 658.6547525 | -0.326594964 | 1.16E-02 |
| PC | 2346.183981 | -0.326700121 | 4.19E-02 |
| UTP25 | 1455.765387 | -0.32672904 | 5.17E-06 |
| ZNF517 | 698.1262746 | -0.326902543 | 4.71E-03 |
| LPIN1 | 2699.104519 | -0.327183302 | 2.75E-02 |
| TRAPPC12 | 1553.179296 | -0.327494807 | 4.80E-03 |
| GARNL3 | 363.9609093 | -0.327506531 | 2.78E-02 |
| NUP153 | 5230.246428 | -0.32793765 | 3.91E-04 |
| ZNF470 | 625.7405749 | -0.328093878 | 3.21E-02 |
| POTEC | 621.7288682 | -0.328125111 | 1.46E-03 |
| TTC5 | 1049.690651 | -0.328352942 | 2.32E-03 |
| OFD1 | 1073.083644 | -0.328658917 | 2.68E-02 |
| RABL2A | 460.6065084 | -0.328934147 | 1.22E-02 |
| SPHK2 | 601.2242349 | -0.329006026 | 3.68E-02 |
| GNL3L | 1411.500738 | -0.329092238 | 1.65E-03 |
| ADCK5 | 414.4494723 | -0.329180061 | 1.81E-02 |
| ZNF710 | 1212.046584 | -0.329234287 | 1.68E-03 |

| | | | |
|---|---|---|---|
| ZNF266 | 2147.525112 | -0.32937097 | 1.61E-04 |
| BUB1 | 3783.438651 | -0.329421805 | 4.66E-02 |
| RPL17 | 27377.12479 | -0.330002415 | 1.87E-04 |
| NA | 887.2209421 | -0.330161845 | 2.10E-02 |
| GMDS | 1022.418347 | -0.330200924 | 3.66E-03 |
| DCBLD2 | 4378.874141 | -0.330206137 | 1.30E-03 |
| OAF | 6886.650965 | -0.330256478 | 8.58E-03 |
| IDH3G | 3348.08267 | -0.330578784 | 2.02E-02 |
| LRFN4 | 1416.02701 | -0.3307935 | 1.61E-02 |
| SLC25A6 | 13376.01443 | -0.330820459 | 1.51E-02 |
| MTG1 | 2244.95116 | -0.330839509 | 6.07E-04 |
| AC004475.1 | 1488.138642 | -0.331152302 | 4.89E-03 |
| CCM2 | 2359.251049 | -0.3311591 | 3.94E-03 |
| YLPM1 | 5289.212819 | -0.331278903 | 1.23E-03 |
| KIAA1958 | 673.3497986 | -0.3312901 | 1.58E-02 |
| NA | 455.9595989 | -0.331329686 | 1.51E-03 |
| ZNF552 | 298.9944392 | -0.331374097 | 3.98E-02 |
| PGD | 5301.957375 | -0.331461318 | 8.07E-05 |
| SVIL | 3315.265826 | -0.331901231 | 5.66E-03 |
| EME1 | 841.7569482 | -0.33191199 | 2.08E-02 |
| ALKBH4 | 894.7968205 | -0.332112486 | 1.97E-02 |
| CSPG4P12 | 639.7044511 | -0.332156911 | 1.70E-02 |
| ST6GALNAC4 | 1223.903519 | -0.332298809 | 2.85E-03 |
| EHMT1 | 4111.833905 | -0.332468337 | 2.06E-05 |
| NPIPA1 | 2359.416333 | -0.3324696 | 1.46E-03 |
| TIGD1 | 735.47436 | -0.332670773 | 7.18E-04 |
| CZIB | 1778.126786 | -0.332759256 | 2.04E-04 |
| VPS45 | 2279.513238 | -0.333179779 | 1.32E-04 |
| MSH3 | 1602.672514 | -0.333187163 | 2.81E-05 |
| CUL4B | 4541.326137 | -0.333238306 | 1.08E-02 |
| ANP32B | 12976.31036 | -0.333540336 | 8.42E-04 |
| NUDC | 5364.378525 | -0.33383526 | 8.29E-03 |
| FAM122C | 495.1261315 | -0.333960262 | 1.66E-02 |
| PHTF1 | 1158.451581 | -0.333970037 | 1.16E-03 |
| TSPAN14 | 2543.409951 | -0.334087421 | 1.87E-02 |
| PPP2R1B | 1715.618634 | -0.334144781 | 3.36E-02 |
| GNL1 | 4663.726163 | -0.334439276 | 2.46E-03 |
| GSPT2 | 941.9840154 | -0.334515706 | 2.84E-04 |
| SSBP2 | 2717.946164 | -0.33468087 | 7.48E-03 |
| AMDHD2 | 1252.631935 | -0.33503856 | 3.27E-03 |
| PAK1IP1 | 1899.848474 | -0.33516391 | 1.72E-02 |
| HNRNPDL | 6883.710354 | -0.335462331 | 7.29E-03 |
| BLOC1S1 | 1479.646415 | -0.335590302 | 2.70E-02 |
| ACCS | 1275.123585 | -0.335890104 | 2.91E-03 |
| ZNF282 | 2297.74343 | -0.336223982 | 5.79E-03 |
| ANKRD37 | 182.4951309 | -0.336259525 | 4.58E-02 |
| TSPO | 3048.132338 | -0.336263797 | 2.02E-02 |

| | | | |
|---|---|---|---|
| ATP5MC2 | 5676.214694 | -0.336721146 | 1.08E-03 |
| NCDN | 2694.254182 | -0.336955641 | 1.27E-03 |
| RIDA | 596.8398994 | -0.336973857 | 4.18E-02 |
| SLC35E2B | 7148.872086 | -0.336997191 | 1.61E-05 |
| RAB3B | 2500.739363 | -0.337057447 | 4.36E-03 |
| ZNF814 | 407.689954 | -0.337143019 | 1.21E-02 |
| PLCB3 | 4561.635039 | -0.337146691 | 2.04E-03 |
| THADA | 2754.68274 | -0.33731983 | 4.22E-05 |
| HARS2 | 2149.232641 | -0.33788057 | 9.18E-08 |
| CAPN15 | 2350.329042 | -0.338006714 | 4.94E-04 |
| NT5C | 924.5825339 | -0.338102009 | 2.18E-03 |
| CLDN15 | 890.7426039 | -0.338432789 | 1.08E-03 |
| REX1BD | 399.8892279 | -0.338505196 | 2.37E-02 |
| ZNF440 | 480.4141961 | -0.338701826 | 1.29E-02 |
| PHF23 | 4034.620571 | -0.338803149 | 1.02E-02 |
| FCHSD1 | 1323.690342 | -0.338995509 | 1.79E-02 |
| RPS27 | 26325.22995 | -0.339509911 | 3.80E-03 |
| PAQR7 | 1043.956206 | -0.339559536 | 6.59E-03 |
| ATP5IF1 | 2344.233637 | -0.339641331 | 9.93E-05 |
| MRPL9 | 1883.802852 | -0.339823975 | 3.05E-02 |
| TAB1 | 1987.787015 | -0.340461396 | 1.27E-03 |
| BANF1 | 7623.196999 | -0.340468991 | 1.16E-03 |
| TIRAP | 621.8397527 | -0.340929254 | 6.49E-04 |
| ABHD11 | 864.5830385 | -0.340942761 | 3.95E-03 |
| TRMT10C | 913.7961037 | -0.34136651 | 1.54E-02 |
| ZNF286A | 1332.71699 | -0.341402673 | 7.38E-03 |
| MTA2 | 8545.013148 | -0.341508122 | 1.49E-06 |
| JARID2 | 1429.283919 | -0.341525748 | 1.71E-05 |
| HDAC10 | 1350.08221 | -0.34155842 | 3.05E-03 |
| TTC31 | 2117.998023 | -0.341571219 | 1.50E-04 |
| PLPP7 | 799.2589398 | -0.341610153 | 1.95E-03 |
| IFT57 | 1167.083796 | -0.341687069 | 2.45E-02 |
| ITGB1BP1 | 3142.327611 | -0.341984416 | 3.42E-05 |
| FBXO41 | 1662.033975 | -0.342051559 | 6.08E-03 |
| PRPF6 | 7435.468691 | -0.342260052 | 2.30E-02 |
| MRPS2 | 2996.568875 | -0.342530335 | 8.02E-04 |
| MSL2 | 981.2483469 | -0.342555036 | 1.67E-03 |
| KIAA0753 | 1607.123434 | -0.342665788 | 2.73E-03 |
| RNF40 | 7519.534809 | -0.342673457 | 2.63E-04 |
| HOXC10 | 347.6687726 | -0.342731891 | 1.49E-02 |
| BOK | 3337.704686 | -0.342751671 | 5.00E-02 |
| PAN2 | 2947.00056 | -0.342943786 | 1.42E-02 |
| PIEZO2 | 3798.179918 | -0.342946703 | 1.19E-04 |
| ZNF212 | 1061.294324 | -0.343459694 | 2.12E-03 |
| TMEM198B | 935.5913628 | -0.343573081 | 2.47E-04 |
| CCT7 | 18359.8704 | -0.343674708 | 8.20E-06 |
| SLC25A53 | 297.7436183 | -0.343818253 | 2.19E-02 |

| | | | |
|---|---|---|---|
| ARFGAP2 | 4523.910026 | -0.3439705 | 2.44E-03 |
| ELL2 | 3965.592371 | -0.344001019 | 1.76E-02 |
| LLGL1 | 5186.655941 | -0.344012165 | 1.58E-03 |
| VRK3 | 1353.668644 | -0.344045829 | 3.86E-02 |
| DCTN1 | 12802.93387 | -0.344070719 | 3.87E-04 |
| ILF3-DT | 606.3224029 | -0.344557225 | 4.40E-03 |
| ADCY3 | 3045.328296 | -0.344894484 | 2.45E-03 |
| BPTF | 4182.268897 | -0.345040849 | 3.75E-02 |
| TMEM185B | 1772.296556 | -0.345088568 | 1.46E-02 |
| RACK1 | 52295.32079 | -0.345237219 | 2.54E-03 |
| WDR20 | 798.6054704 | -0.345451451 | 4.82E-04 |
| TYSND1 | 784.5755278 | -0.345471051 | 6.45E-04 |
| FAM225A | 1270.608866 | -0.345484547 | 6.36E-03 |
| ELOA | 3907.96247 | -0.345604433 | 1.76E-09 |
| CHD8 | 7116.522774 | -0.345998498 | 1.67E-04 |
| FIBP | 3013.116514 | -0.346015578 | 1.57E-03 |
| ANKS1A | 3891.174726 | -0.346232994 | 6.01E-04 |
| FBXL6 | 944.013837 | -0.346314853 | 5.57E-04 |
| UMPS | 1674.316527 | -0.346387371 | 2.78E-03 |
| MAPK7 | 1119.071792 | -0.346485328 | 1.92E-02 |
| GIT1 | 3699.953202 | -0.346608485 | 6.77E-05 |
| SNRPD3 | 3280.283592 | -0.346711889 | 8.35E-04 |
| CEP19 | 554.178459 | -0.34696656 | 5.18E-03 |
| ZNF786 | 452.9511319 | -0.347057206 | 8.12E-03 |
| RBM10 | 3973.238951 | -0.347324523 | 1.69E-06 |
| FKBP3 | 3306.432212 | -0.347338013 | 1.26E-06 |
| TRMT2B | 1049.45023 | -0.347377647 | 1.21E-04 |
| ZBTB45P2 | 717.7323167 | -0.347764107 | 1.54E-02 |
| SETDB1 | 2138.132053 | -0.347798429 | 1.02E-03 |
| DANCR | 709.3361258 | -0.347921078 | 2.00E-03 |
| VARS | 6546.556051 | -0.348150481 | 3.30E-04 |
| LRRC37A4P | 1314.268066 | -0.348468463 | 1.18E-03 |
| UPF1 | 5584.674034 | -0.348713732 | 1.55E-05 |
| ZNF3 | 2904.964969 | -0.348841135 | 5.71E-04 |
| CCNL2 | 6146.069744 | -0.348977325 | 1.36E-10 |
| SMAGP | 683.0418754 | -0.349057222 | 2.40E-02 |
| ARGLU1 | 2029.422784 | -0.349495471 | 1.10E-02 |
| COX10-AS1 | 646.976046 | -0.349602251 | 1.72E-02 |
| GPR137 | 1729.845174 | -0.349664458 | 1.18E-02 |
| PRKCZ | 1369.324973 | -0.349930584 | 1.29E-02 |
| SNX1 | 8206.680958 | -0.350129729 | 4.04E-04 |
| IGF2BP1 | 4568.368856 | -0.350381084 | 1.05E-06 |
| DPY19L3 | 1813.446008 | -0.350500069 | 3.59E-02 |
| CSTF1 | 2225.944012 | -0.350538663 | 5.40E-04 |
| SCAMP4 | 2956.628619 | -0.350631984 | 7.85E-04 |
| KCTD7 | 1741.806217 | -0.350962331 | 1.89E-03 |
| GYS1 | 3047.439824 | -0.351130365 | 9.73E-03 |

| | | | |
|---|---|---|---|
| RPL26 | 31543.39492 | -0.351342214 | 1.72E-05 |
| ZGPAT | 1521.912996 | -0.351357832 | 8.80E-04 |
| CEP78 | 2121.120832 | -0.351489651 | 9.63E-03 |
| AC018647.2 | 763.6559716 | -0.351675678 | 2.55E-04 |
| EEF1AKMT3 | 554.8192586 | -0.351772987 | 1.36E-03 |
| CLCN2 | 515.9349584 | -0.352007785 | 4.10E-02 |
| MLLT6 | 7194.777465 | -0.352163181 | 5.71E-05 |
| EEF2 | 104137.7217 | -0.352253911 | 5.03E-03 |
| MCCC2 | 8099.333828 | -0.352483194 | 4.90E-10 |
| FBXO45 | 1117.356984 | -0.353326937 | 1.02E-03 |
| PSKH1 | 2749.504495 | -0.353855342 | 2.48E-03 |
| ITFG2 | 735.0446452 | -0.353857349 | 1.14E-02 |
| ITPRID2 | 7029.408836 | -0.353987836 | 1.40E-02 |
| PRKD3 | 3907.703797 | -0.354072406 | 2.49E-03 |
| PHF6 | 2873.346761 | -0.354138474 | 3.34E-02 |
| NA | 4877.191813 | -0.354597224 | 4.34E-12 |
| FAM241A | 959.3567144 | -0.35468413 | 1.30E-02 |
| AC009163.6 | 182.0607473 | -0.354926177 | 2.10E-02 |
| YBEY | 319.8980182 | -0.354966158 | 3.70E-03 |
| NOP16 | 1938.308954 | -0.354980791 | 3.71E-03 |
| RNF31 | 2126.637285 | -0.355171654 | 2.28E-02 |
| TTF2 | 2656.857746 | -0.355190489 | 7.88E-03 |
| MTF1 | 1054.800846 | -0.355210286 | 4.60E-02 |
| HSD17B10 | 2671.357743 | -0.355575286 | 2.79E-04 |
| TOPORS | 1768.495239 | -0.355703681 | 6.07E-03 |
| TARDBP | 7443.761822 | -0.355743508 | 1.73E-05 |
| YWHAQ | 14732.7944 | -0.355778528 | 8.88E-06 |
| AC090114.2 | 835.2073606 | -0.355881674 | 6.45E-03 |
| SLC25A23 | 5173.267767 | -0.355991312 | 1.59E-03 |
| MRNIP | 1068.786205 | -0.356029693 | 1.57E-04 |
| PCYOX1 | 5760.612629 | -0.356067653 | 2.05E-03 |
| KCTD1 | 748.5229894 | -0.356124743 | 5.76E-03 |
| ERG28 | 2251.811792 | -0.356223639 | 2.48E-02 |
| NA | 1044.03691 | -0.356542939 | 4.51E-04 |
| ARID2 | 1422.741654 | -0.356842504 | 2.27E-02 |
| PEX16 | 1105.978028 | -0.357066672 | 1.46E-02 |
| HDAC8 | 1003.791273 | -0.357137739 | 4.66E-04 |
| ZNF512 | 2797.925537 | -0.357189137 | 2.94E-04 |
| CHMP1A | 5055.845608 | -0.357222657 | 4.19E-03 |
| SNUPN | 1526.134709 | -0.357264064 | 7.14E-07 |
| RPUSD2 | 434.9378424 | -0.357347508 | 3.10E-03 |
| AC078846.1 | 319.4735211 | -0.357378438 | 4.31E-03 |
| MSH5 | 896.9720349 | -0.357521853 | 7.81E-03 |
| TEPSIN | 793.317385 | -0.357570528 | 2.17E-03 |
| MSTO2P | 212.3275863 | -0.357722694 | 4.85E-02 |
| C2CD2 | 984.5173801 | -0.357738926 | 2.89E-02 |
| IP6K1 | 2965.032039 | -0.357800133 | 8.49E-04 |

| | | | |
|---|---|---|---|
| HNRNPA3 | 17215.90036 | -0.358081845 | 3.66E-06 |
| DDX54 | 3050.744023 | -0.358135696 | 5.77E-06 |
| TAF1C | 2965.008844 | -0.35817038 | 1.20E-06 |
| AC097374.1 | 1567.411536 | -0.358189631 | 2.05E-02 |
| PUS3 | 1045.525188 | -0.358391969 | 6.61E-03 |
| SRCAP | 10671.7796 | -0.358619827 | 6.20E-10 |
| TMEM168 | 1168.630788 | -0.359404898 | 4.34E-02 |
| CSPG4P11 | 285.6691582 | -0.359640636 | 1.12E-02 |
| VDAC3 | 5335.465565 | -0.359815482 | 2.03E-04 |
| E2F4 | 7133.361551 | -0.360252145 | 1.16E-05 |
| CNOT3 | 2190.48878 | -0.360360408 | 4.91E-05 |
| MRTO4 | 2839.374245 | -0.360384271 | 3.41E-04 |
| EPHB4 | 7703.29599 | -0.360399992 | 1.03E-02 |
| UBL5 | 4372.394973 | -0.360493323 | 1.34E-02 |
| RCC1L | 4660.194879 | -0.360564611 | 1.31E-04 |
| SLC39A4 | 422.1923132 | -0.36059991 | 1.33E-02 |
| TXN2 | 3663.452541 | -0.360859114 | 4.23E-03 |
| RPL35A | 14501.75282 | -0.360879277 | 3.50E-03 |
| APPL2 | 2709.579325 | -0.361164872 | 2.47E-06 |
| MYO1B | 8268.616834 | -0.361187786 | 2.60E-03 |
| RAB14 | 4903.793093 | -0.361509502 | 8.40E-05 |
| SMTN | 2139.458024 | -0.361853263 | 5.80E-03 |
| MAP4K2 | 2297.576008 | -0.362055975 | 8.45E-03 |
| NDOR1 | 1466.776151 | -0.362829231 | 7.44E-04 |
| RCOR2 | 339.1359079 | -0.363101935 | 2.34E-02 |
| TNK2 | 2521.478982 | -0.363122083 | 2.40E-06 |
| AP1G2 | 2614.075705 | -0.363156204 | 8.31E-03 |
| TADA3 | 4028.48948 | -0.363235688 | 1.26E-02 |
| POM121 | 7411.262456 | -0.363307153 | 4.62E-04 |
| CCDC80 | 5060.118978 | -0.36341076 | 4.28E-03 |
| DDX6 | 6778.175847 | -0.363455645 | 3.78E-08 |
| WIZ | 2939.743558 | -0.363472547 | 4.99E-04 |
| AC019205.1 | 379.8578792 | -0.363526917 | 1.58E-03 |
| NR6A1 | 312.4305955 | -0.364196893 | 2.45E-02 |
| ARSJ | 838.5835605 | -0.364392707 | 2.89E-02 |
| POLR1D | 2138.160546 | -0.364557481 | 8.44E-03 |
| PDCL | 1337.598657 | -0.364573987 | 2.85E-06 |
| SMAP2 | 1577.202322 | -0.364605994 | 1.15E-03 |
| PSME2 | 4940.514941 | -0.364827546 | 1.03E-07 |
| MATR3 | 12010.77523 | -0.364882224 | 1.61E-02 |
| BIN1 | 2819.19804 | -0.364885739 | 6.83E-03 |
| DMAC2 | 1815.547142 | -0.365083767 | 1.26E-03 |
| KEAP1 | 3402.522724 | -0.365191156 | 1.01E-03 |
| AC074117.1 | 550.5267816 | -0.365367461 | 2.93E-03 |
| RGS10 | 602.2226751 | -0.365422538 | 6.47E-03 |
| IGFBP4 | 22445.28338 | -0.36631166 | 1.04E-02 |
| TMEM80 | 695.471718 | -0.366795162 | 1.87E-03 |

| | | | |
|---|---|---|---|
| DUSP7 | 822.9598436 | -0.366807595 | 1.49E-03 |
| EXOG | 670.6987006 | -0.36703769 | 1.83E-04 |
| TAOK2 | 4230.053492 | -0.367228937 | 1.06E-03 |
| GABARAP | 9957.378077 | -0.367251154 | 4.35E-02 |
| PTPN1 | 5738.112123 | -0.367701773 | 6.02E-07 |
| HOXB4 | 680.793008 | -0.367757671 | 4.91E-04 |
| ARMCX4 | 791.492689 | -0.368149169 | 1.11E-02 |
| STK11 | 2145.068827 | -0.368251311 | 3.01E-04 |
| TENM3 | 2755.606116 | -0.368548512 | 7.74E-04 |
| OGFR | 4528.943543 | -0.368760471 | 1.60E-04 |
| DBR1 | 685.1075807 | -0.3688698 | 3.48E-04 |
| PLEKHM1P1 | 493.0380594 | -0.368998366 | 3.94E-04 |
| ZNF135 | 520.20737 | -0.369768085 | 2.60E-03 |
| PPP1R10 | 3168.508356 | -0.369823642 | 4.38E-03 |
| FP565260.1 | 1630.800222 | -0.369949378 | 7.17E-05 |
| STOML1 | 725.8469046 | -0.36996449 | 1.13E-02 |
| BCAT2 | 1673.235047 | -0.370108951 | 2.38E-03 |
| TBC1D10B | 3214.040616 | -0.370313944 | 2.13E-03 |
| IREB2 | 3969.02169 | -0.37053139 | 3.77E-03 |
| THAP12 | 4116.916326 | -0.370551992 | 4.79E-03 |
| RPL7 | 41042.59646 | -0.370736275 | 3.48E-06 |
| RBM38 | 948.7983218 | -0.370923253 | 1.64E-03 |
| FAAP100 | 1938.390015 | -0.371219356 | 1.73E-03 |
| MMS19 | 2806.241532 | -0.371371703 | 3.75E-04 |
| POLM | 811.6532932 | -0.37184191 | 6.09E-03 |
| LSS | 8142.1505 | -0.372277018 | 4.86E-02 |
| POLR3H | 2865.148515 | -0.372376433 | 2.26E-04 |
| RPS20 | 35647.71181 | -0.37268244 | 2.31E-03 |
| GAB1 | 870.6201106 | -0.372769276 | 7.46E-03 |
| PHKA2 | 1834.123583 | -0.372917922 | 1.55E-03 |
| CYBC1 | 1890.425223 | -0.373033499 | 3.19E-03 |
| MARS2 | 475.8319689 | -0.373121195 | 2.34E-03 |
| TTF1 | 1467.770898 | -0.37317202 | 5.13E-03 |
| HIBCH | 1071.09971 | -0.373188085 | 1.88E-02 |
| CSK | 3907.937769 | -0.37350667 | 1.57E-04 |
| LARS2 | 2509.944575 | -0.373540649 | 6.38E-05 |
| RPS17 | 42235.26551 | -0.373606408 | 5.88E-05 |
| GXYLT2 | 1174.465531 | -0.373756223 | 4.98E-03 |
| PTPA | 8374.95881 | -0.374286671 | 4.07E-03 |
| TMEM107 | 548.8358257 | -0.374351634 | 3.19E-03 |
| SSBP4 | 1404.437664 | -0.374453271 | 1.12E-02 |
| CABLES2 | 893.2707725 | -0.374586901 | 1.72E-02 |
| RAB35 | 2634.294251 | -0.374622581 | 3.25E-03 |
| FOXO3 | 3030.424993 | -0.37473617 | 2.63E-14 |
| LSM11 | 1292.011328 | -0.374778262 | 8.69E-04 |
| HCG11 | 485.9195478 | -0.375147515 | 1.26E-02 |
| AC008966.1 | 495.8421598 | -0.375202975 | 2.05E-03 |

| | | | |
|---|---|---|---|
| SESTD1 | 441.5668905 | -0.375268272 | 4.20E-02 |
| JCAD | 5110.161346 | -0.375626675 | 1.48E-04 |
| L3MBTL1 | 722.4876846 | -0.375754172 | 3.24E-03 |
| FANCC | 1644.315235 | -0.375784675 | 5.39E-03 |
| PCBP1-AS1 | 586.6476801 | -0.375846749 | 1.62E-02 |
| PLEKHG4 | 1389.84851 | -0.375850319 | 6.50E-03 |
| CAT | 4283.077306 | -0.375963645 | 8.75E-05 |
| TTN-AS1 | 253.0060172 | -0.376001239 | 2.08E-02 |
| ZMIZ1 | 3413.461136 | -0.376145131 | 1.30E-05 |
| CCDC134 | 408.6200617 | -0.376172689 | 3.39E-04 |
| AC009113.1 | 261.0991221 | -0.376384516 | 3.94E-03 |
| ZNF384 | 2497.210853 | -0.376461399 | 1.32E-04 |
| IPO7 | 18247.77458 | -0.376763142 | 6.45E-04 |
| SNHG6 | 2615.171978 | -0.377079364 | 3.30E-05 |
| ZNF229 | 1229.958577 | -0.377337174 | 2.94E-02 |
| TTLL11 | 281.7170078 | -0.377409667 | 1.62E-02 |
| SPATC1L | 320.1765012 | -0.377439386 | 1.76E-02 |
| IL17RC | 1364.10407 | -0.377487885 | 1.73E-02 |
| ALX4 | 946.9188903 | -0.377625005 | 4.32E-03 |
| ZNHIT3 | 1265.022696 | -0.377671025 | 7.03E-03 |
| TP53INP1 | 249.9843985 | -0.377770676 | 4.90E-02 |
| PHLDB2 | 5271.381741 | -0.377776172 | 4.38E-02 |
| RPP40 | 549.7937429 | -0.37788659 | 4.09E-02 |
| CRYZL2P | 328.6455636 | -0.377968649 | 9.89E-03 |
| TBC1D17 | 1298.653076 | -0.378035334 | 2.41E-02 |
| AC004943.2 | 695.5030925 | -0.378575598 | 3.84E-04 |
| CHERP | 3014.334714 | -0.379033718 | 3.47E-05 |
| PLCXD1 | 1585.892566 | -0.379054685 | 6.10E-05 |
| TRMT12 | 1056.448805 | -0.379144609 | 6.21E-05 |
| AARS2 | 1735.949022 | -0.379265107 | 2.92E-04 |
| ZNF599 | 320.7031162 | -0.379546116 | 1.45E-03 |
| LGR4 | 3291.633279 | -0.379860697 | 4.10E-03 |
| KDM6A | 1263.201627 | -0.380304532 | 4.47E-03 |
| HNRNPA2B1 | 37196.52692 | -0.380455172 | 3.49E-03 |
| AL031282.2 | 397.8015291 | -0.380462279 | 5.57E-03 |
| RUSC1-AS1 | 232.9101864 | -0.38059634 | 1.53E-02 |
| FGD5-AS1 | 5228.2829 | -0.380701853 | 2.84E-05 |
| IQCA1 | 679.4135308 | -0.381027997 | 7.83E-03 |
| SPAST | 681.3991413 | -0.381067199 | 1.39E-02 |
| OSGEP | 996.6215002 | -0.381080935 | 3.41E-04 |
| TAF5L | 1206.157522 | -0.381207186 | 2.03E-04 |
| INTS1 | 14293.109 | -0.381629796 | 1.45E-05 |
| FAXC | 492.2908407 | -0.381833086 | 4.89E-04 |
| HOXC13 | 1296.224136 | -0.38188205 | 7.37E-03 |
| ATIC | 7866.447087 | -0.382023271 | 1.01E-04 |
| DEPDC1B | 1923.476532 | -0.382056304 | 3.88E-02 |
| TCEANC2 | 825.1789663 | -0.382210283 | 5.50E-04 |

| | | | |
|---|---|---|---|
| NA | 2018.632065 | -0.382418887 | 1.97E-03 |
| AACS | 2808.450745 | -0.382619518 | 1.23E-04 |
| HERC2P2 | 2782.210698 | -0.382732539 | 4.90E-03 |
| DHPS | 2252.553779 | -0.382765168 | 4.66E-09 |
| COG8 | 3046.293015 | -0.382965649 | 1.20E-03 |
| RTN2 | 629.9105108 | -0.382966758 | 3.90E-02 |
| EXTL3 | 8190.652019 | -0.383307039 | 3.04E-02 |
| PDK2 | 1694.855698 | -0.383311805 | 4.46E-02 |
| MEIOB | 1375.792081 | -0.383341027 | 1.22E-02 |
| PHC2 | 12014.71889 | -0.38341802 | 3.80E-03 |
| NDN | 2108.156895 | -0.383453341 | 3.04E-02 |
| CHTF8 | 7617.256486 | -0.383514867 | 3.93E-04 |
| CPSF3 | 3527.478027 | -0.384514762 | 1.25E-09 |
| DHX34 | 1155.079236 | -0.384569886 | 1.29E-03 |
| SETD6 | 1072.763408 | -0.384640094 | 1.48E-03 |
| ZNF142 | 1567.042809 | -0.385119155 | 2.06E-05 |
| HDAC4 | 1112.550625 | -0.38522981 | 4.03E-02 |
| URB1-AS1 | 331.4038143 | -0.385360812 | 3.06E-02 |
| STARD8 | 1624.649461 | -0.385442416 | 1.24E-03 |
| INTS7 | 1984.401887 | -0.385520956 | 5.70E-04 |
| NUBP2 | 2876.219677 | -0.385689464 | 3.76E-04 |
| ZNF285 | 323.5126856 | -0.385696038 | 3.24E-02 |
| PRPF31 | 3047.900361 | -0.385801866 | 2.37E-04 |
| CDK2AP1 | 4145.313976 | -0.385970189 | 1.13E-07 |
| XAB2 | 2830.758882 | -0.386212629 | 8.08E-03 |
| RPL38 | 12269.32601 | -0.386418114 | 5.91E-03 |
| MOCS3 | 1071.09352 | -0.38657908 | 3.84E-04 |
| CAPN10-DT | 307.6229708 | -0.386624915 | 1.53E-02 |
| PLAGL1 | 554.3723733 | -0.387006398 | 2.82E-02 |
| TNFSF12 | 602.4423719 | -0.387059461 | 4.27E-02 |
| CCNO | 490.4138232 | -0.387104172 | 6.52E-04 |
| ACTR5 | 1029.980742 | -0.387253394 | 1.51E-04 |
| CREB3L4 | 569.1732261 | -0.387288602 | 1.07E-03 |
| RILPL2 | 1239.886752 | -0.38747605 | 1.36E-03 |
| BTBD11 | 910.1491859 | -0.387483732 | 3.08E-04 |
| ZSCAN12 | 304.742191 | -0.387659802 | 3.80E-03 |
| ZNF22 | 1306.917588 | -0.388085221 | 3.30E-03 |
| ZNF783 | 1538.74869 | -0.38819341 | 3.62E-05 |
| SFT2D2 | 1443.35455 | -0.388359155 | 1.14E-03 |
| ZNF251 | 1133.94733 | -0.388499327 | 3.62E-07 |
| NRDE2 | 1127.583602 | -0.388752733 | 1.19E-03 |
| PNP | 1512.768907 | -0.388788214 | 5.70E-04 |
| COLGALT1 | 8186.22967 | -0.38907044 | 5.30E-04 |
| TCTA | 1330.44691 | -0.389077176 | 1.20E-03 |
| ELOF1 | 2490.3667 | -0.389270323 | 1.98E-03 |
| NME2 | 13404.07001 | -0.389455811 | 4.62E-05 |
| LPCAT3 | 1832.190275 | -0.389533492 | 6.44E-05 |

| | | | |
|---|---|---|---|
| POLR2E | 5539.893555 | -0.389706753 | 3.49E-04 |
| ZNF821 | 353.8660785 | -0.389863134 | 1.39E-02 |
| BCLAF1 | 5637.954325 | -0.390418276 | 4.69E-02 |
| ALG12 | 1025.930514 | -0.390461913 | 3.46E-02 |
| SFMBT2 | 670.6519321 | -0.390511455 | 5.32E-04 |
| LAMA4 | 12155.3392 | -0.390516225 | 1.75E-08 |
| USP37 | 848.2020086 | -0.390528399 | 2.62E-02 |
| PDLIM7 | 2642.119675 | -0.39061502 | 2.23E-03 |
| C6orf47 | 1110.798322 | -0.390868072 | 3.26E-04 |
| WDR81 | 3131.426754 | -0.390958224 | 5.20E-03 |
| POLR2I | 806.7091448 | -0.391285123 | 2.91E-02 |
| LAP3 | 2191.36426 | -0.391342734 | 3.46E-05 |
| ACAA1 | 1616.455569 | -0.391859057 | 2.40E-04 |
| DAAM1 | 871.5055857 | -0.391900327 | 1.30E-02 |
| FOXP4 | 1453.076162 | -0.391935123 | 1.34E-03 |
| LSM3 | 1531.34614 | -0.391953395 | 4.46E-02 |
| PAXIP1 | 1709.812529 | -0.392090734 | 1.21E-03 |
| PKM | 127003.2279 | -0.392397793 | 4.38E-03 |
| DHX57 | 2397.342818 | -0.3924618 | 6.87E-07 |
| NOP56 | 9573.943269 | -0.39259102 | 1.70E-12 |
| HIRA | 2952.925862 | -0.392602754 | 2.43E-04 |
| SH3BGRL | 2014.089491 | -0.392634451 | 8.00E-04 |
| RBP5 | 155.6224082 | -0.392679369 | 8.89E-03 |
| C17orf49 | 1824.22033 | -0.392849591 | 1.58E-03 |
| ZBTB26 | 1136.425005 | -0.392899741 | 5.08E-03 |
| AGBL5 | 3091.418337 | -0.392958184 | 2.19E-05 |
| RNASEH1-AS1 | 526.78785 | -0.393010362 | 9.69E-03 |
| KANSL1L | 442.1730578 | -0.393059771 | 2.40E-02 |
| NOSIP | 2178.863594 | -0.39384155 | 1.43E-03 |
| ZSCAN9 | 524.7559014 | -0.393975214 | 2.30E-04 |
| AC092718.4 | 197.3954875 | -0.394121713 | 1.70E-02 |
| DOLPP1 | 1556.722068 | -0.394147248 | 1.57E-04 |
| CASC3 | 4319.53297 | -0.394174539 | 5.33E-09 |
| SLX1A-SULT1A3 | 269.3639749 | -0.394301984 | 1.53E-02 |
| BMS1P1 | 299.021387 | -0.394490129 | 1.01E-02 |
| RBM15B | 4182.730323 | -0.394490589 | 5.80E-06 |
| BEAN1 | 243.2446155 | -0.394696325 | 4.55E-02 |
| SCRIB | 5063.164234 | -0.394727063 | 1.52E-05 |
| AC011043.1 | 615.4699368 | -0.394809847 | 4.34E-03 |
| MRPS27 | 6656.267008 | -0.395234932 | 4.62E-04 |
| RPL14 | 17200.18652 | -0.395550512 | 2.55E-05 |
| AC125257.1 | 163.7034956 | -0.395995245 | 3.45E-02 |
| ZNF607 | 500.3212696 | -0.396136624 | 1.69E-02 |
| COTL1 | 13773.83675 | -0.396187834 | 3.22E-04 |
| TOR1AIP1 | 2235.13473 | -0.396286774 | 7.02E-04 |
| AGAP4 | 370.1450236 | -0.396647601 | 3.71E-02 |
| HMBS | 3683.837046 | -0.396764593 | 7.28E-04 |

| | | | |
|---|---|---|---|
| ECSIT | 1263.236298 | -0.396938519 | 5.70E-04 |
| CIZ1 | 7246.560402 | -0.396943568 | 3.61E-05 |
| SOD1 | 8136.187217 | -0.397154845 | 3.62E-05 |
| CTR9 | 4964.492339 | -0.397196144 | 7.92E-04 |
| CC2D2A | 1425.087652 | -0.39732855 | 3.84E-04 |
| PPIL2 | 3565.270233 | -0.397563437 | 1.54E-06 |
| LINC02482 | 133.6517581 | -0.397606491 | 3.11E-02 |
| ATXN7L3 | 3881.118133 | -0.397629888 | 8.50E-05 |
| CPSF6 | 3792.174803 | -0.397678978 | 7.96E-04 |
| HNRNPUL2 | 6953.721425 | -0.39773639 | 9.33E-09 |
| AC102953.2 | 378.0608071 | -0.397781375 | 2.18E-03 |
| TRIM62 | 875.1026027 | -0.398022264 | 3.04E-03 |
| IL11RA | 1015.928474 | -0.398184073 | 3.72E-02 |
| RSBN1 | 590.7070609 | -0.398529434 | 1.66E-03 |
| ERGIC1 | 7955.280628 | -0.398781957 | 1.17E-04 |
| MED4 | 1518.399422 | -0.399051631 | 1.06E-03 |
| ZNF689 | 997.135433 | -0.399192677 | 7.59E-03 |
| SYNGR1 | 3473.006517 | -0.399680833 | 4.23E-03 |
| TRABD | 1154.065959 | -0.399893507 | 1.80E-06 |
| CPSF7 | 6356.269338 | -0.399989851 | 5.74E-17 |
| FAM185A | 370.2486297 | -0.400498036 | 4.38E-03 |
| SURF6 | 3350.758565 | -0.400549541 | 5.41E-06 |
| NEU3 | 1544.659068 | -0.40069914 | 6.72E-08 |
| GPAA1 | 6075.963465 | -0.400945276 | 2.70E-04 |
| EMC10 | 4806.966245 | -0.400966913 | 5.96E-04 |
| RGS19 | 893.2063205 | -0.401261494 | 2.85E-03 |
| KDM5A | 4501.713184 | -0.401289793 | 9.52E-08 |
| GAS2L1 | 793.1223109 | -0.401358745 | 1.34E-02 |
| IVNS1ABP | 2139.15868 | -0.401393423 | 7.34E-04 |
| SMG6 | 4055.32213 | -0.401445474 | 1.09E-04 |
| HERC2P5 | 595.3123382 | -0.401612659 | 9.46E-03 |
| ZBTB34 | 895.6646402 | -0.401614765 | 4.62E-05 |
| NAPRT | 1467.532231 | -0.401858386 | 7.03E-03 |
| AC073508.3 | 191.8299393 | -0.401893134 | 9.24E-03 |
| THG1L | 1892.442873 | -0.401905902 | 6.38E-05 |
| BMP2K | 1364.424567 | -0.401971524 | 5.09E-03 |
| DCTN5 | 8916.310083 | -0.402206722 | 2.77E-05 |
| PA2G4 | 8850.309135 | -0.402359711 | 5.95E-09 |
| PABPN1 | 6487.789511 | -0.402393535 | 2.50E-07 |
| CDK2 | 2915.972948 | -0.402489766 | 4.49E-03 |
| ALG6 | 407.5561029 | -0.402643009 | 2.89E-02 |
| ABHD3 | 445.9021462 | -0.402680067 | 2.73E-02 |
| ST3GAL2 | 3203.355799 | -0.402735807 | 1.40E-03 |
| AL691432.2 | 135.8271995 | -0.402800311 | 4.20E-02 |
| SNRNP70 | 6284.377571 | -0.402890741 | 6.16E-06 |
| TADA1 | 504.945217 | -0.40301372 | 1.97E-04 |
| GOLGA2P7 | 5165.611373 | -0.403063104 | 9.49E-07 |

| | | | |
|---|---|---|---|
| CDKN3 | 1240.158803 | -0.403566268 | 2.95E-02 |
| IKBKE | 935.0775559 | -0.403986978 | 2.72E-02 |
| POLR1C | 1344.849713 | -0.404283516 | 1.29E-07 |
| EIF3L | 19700.69514 | -0.404430406 | 7.02E-04 |
| HMCES | 2773.444736 | -0.404730024 | 1.04E-02 |
| POTEI | 907.5491784 | -0.404843236 | 8.99E-03 |
| ZNF503 | 297.2788085 | -0.405030746 | 1.55E-03 |
| LRRC75B | 360.1727331 | -0.405371254 | 2.29E-03 |
| FGD1 | 1766.826493 | -0.405602703 | 3.69E-04 |
| TIMM21 | 780.7950079 | -0.405605649 | 1.98E-02 |
| SPRYD4 | 795.5056325 | -0.40581283 | 3.15E-03 |
| PIGCP1 | 268.8142606 | -0.406072198 | 1.23E-02 |
| PYGO2 | 3109.595671 | -0.40629863 | 7.67E-04 |
| NA | 309.2585295 | -0.406827178 | 9.62E-03 |
| ZBTB22 | 878.4057966 | -0.406836483 | 7.99E-04 |
| SRRM2 | 25535.81099 | -0.406886655 | 2.47E-05 |
| AC074143.1 | 1226.0207 | -0.40688976 | 3.63E-04 |
| TFAM | 1552.341068 | -0.407128204 | 6.66E-03 |
| INO80B | 868.9819165 | -0.407256052 | 3.58E-04 |
| TBCC | 863.9789017 | -0.407281372 | 2.74E-03 |
| RPS28 | 7175.876842 | -0.407497901 | 5.12E-03 |
| ZNF462 | 2159.207217 | -0.407666806 | 9.31E-06 |
| PRDM15 | 765.236804 | -0.407780945 | 5.41E-03 |
| RABL2B | 1249.900016 | -0.407886071 | 2.31E-06 |
| PEX6 | 2151.85644 | -0.408019627 | 3.91E-03 |
| CROCCP2 | 636.4083808 | -0.40809955 | 1.28E-04 |
| AGAP9 | 243.6519753 | -0.408276811 | 2.96E-02 |
| PRKD2 | 2127.037915 | -0.408317443 | 3.50E-04 |
| RPL34 | 13518.72636 | -0.408681307 | 1.82E-04 |
| TNFRSF10D | 1337.770238 | -0.408950627 | 3.70E-02 |
| ZNF200 | 1367.270175 | -0.409017861 | 1.54E-07 |
| RPL7L1 | 10467.69566 | -0.409234619 | 1.21E-09 |
| PIN1 | 1762.228934 | -0.409334771 | 3.74E-04 |
| RTEL1 | 1233.022643 | -0.409514847 | 8.56E-06 |
| EMILIN1 | 5275.584214 | -0.40961543 | 4.41E-02 |
| SPA17 | 608.872348 | -0.409970611 | 2.67E-03 |
| AC060766.1 | 170.7416219 | -0.409971621 | 4.45E-02 |
| RBX1 | 2129.996844 | -0.410179246 | 1.68E-03 |
| SPG7 | 3626.18426 | -0.410421203 | 1.74E-04 |
| MPND | 470.5937356 | -0.410721544 | 5.41E-03 |
| CARMIL1 | 2603.175438 | -0.411028653 | 2.47E-10 |
| CNP | 7610.34218 | -0.411075784 | 6.16E-03 |
| AC060234.2 | 135.2052896 | -0.41108579 | 1.82E-02 |
| AC105285.1 | 164.9294561 | -0.411179439 | 3.46E-02 |
| BMT2 | 620.0680209 | -0.411590961 | 4.14E-03 |
| SPATA2L | 395.8479867 | -0.41166165 | 2.93E-02 |
| PHPT1 | 3241.950508 | -0.411877213 | 8.91E-03 |

| | | | |
|---|---|---|---|
| POMGNT2 | 1580.571202 | -0.412419066 | 2.79E-02 |
| MORN1 | 444.3401887 | -0.412655273 | 6.42E-05 |
| RTL10 | 1453.480058 | -0.412732257 | 2.82E-02 |
| SNX32 | 323.3175144 | -0.41308492 | 2.12E-02 |
| B3GALT6 | 999.9083851 | -0.41316284 | 3.28E-03 |
| ITGB5 | 9170.149928 | -0.413263391 | 2.56E-02 |
| NUP50 | 3007.034293 | -0.413807683 | 5.10E-04 |
| SRD5A3 | 467.3307845 | -0.414044492 | 4.40E-03 |
| FARSA | 5326.766618 | -0.414090408 | 1.08E-04 |
| TBC1D13 | 4105.408839 | -0.41428447 | 4.67E-03 |
| AC132008.2 | 533.2068694 | -0.414866287 | 2.07E-03 |
| NPRL2 | 1001.425579 | -0.414931996 | 2.16E-04 |
| PUF60 | 8432.570762 | -0.415139889 | 5.21E-04 |
| INTS6L | 853.156252 | -0.415177783 | 9.70E-03 |
| CBX8 | 874.9361288 | -0.415179342 | 3.57E-03 |
| PLEKHO2 | 2012.556267 | -0.415259346 | 4.10E-03 |
| SIVA1 | 2030.777572 | -0.4154133 | 1.50E-04 |
| RGPD6 | 2488.190399 | -0.415611199 | 4.18E-02 |
| MALL | 777.0586729 | -0.415658195 | 2.50E-02 |
| MKS1 | 1373.870469 | -0.415693189 | 1.33E-05 |
| IQCE | 1179.997058 | -0.416104249 | 2.67E-03 |
| TAF6L | 1020.213158 | -0.416128944 | 7.53E-03 |
| SNRPB | 16462.47914 | -0.416185512 | 2.54E-04 |
| SENP8 | 218.9915801 | -0.416407594 | 7.42E-03 |
| ELOVL1 | 2900.615077 | -0.416594062 | 1.36E-03 |
| RPL37A | 41742.41886 | -0.417095616 | 1.49E-03 |
| SMN2 | 2795.058623 | -0.417138603 | 2.93E-05 |
| BAZ1A | 2148.548363 | -0.417222136 | 3.89E-02 |
| NUP160 | 7594.563443 | -0.417256273 | 7.41E-08 |
| LETM1 | 2668.77504 | -0.417341144 | 7.55E-08 |
| EVL | 1559.341381 | -0.417568075 | 6.54E-06 |
| EVC | 3307.119757 | -0.417590319 | 3.18E-06 |
| ISG20L2 | 2175.728689 | -0.417683611 | 2.94E-06 |
| COL6A2 | 15718.66709 | -0.4180731 | 1.74E-04 |
| SMG9 | 1877.07634 | -0.418112971 | 5.79E-04 |
| SRSF2 | 7548.406969 | -0.418169397 | 1.95E-02 |
| FAM214A | 916.7426868 | -0.418229676 | 2.81E-02 |
| RPL37 | 65992.22428 | -0.418471485 | 6.88E-04 |
| GTPBP6 | 1657.891782 | -0.418863327 | 1.24E-04 |
| JAZF1 | 445.0955495 | -0.418868176 | 1.58E-02 |
| MUC20-OT1 | 762.1769723 | -0.419180704 | 5.00E-03 |
| NIP7 | 3651.687745 | -0.419404692 | 2.01E-05 |
| DENND6A | 791.9044852 | -0.419715117 | 2.82E-02 |
| LMF2 | 2487.476587 | -0.41975433 | 6.06E-05 |
| PEBP1 | 5568.332282 | -0.419865949 | 2.09E-03 |
| TRMT2A | 1696.322086 | -0.420255091 | 7.80E-03 |
| MAU2 | 2566.508798 | -0.420386795 | 1.10E-05 |

| | | | |
|---|---|---|---|
| NDST2 | 1159.932404 | -0.420573636 | 7.56E-04 |
| AMOTL1 | 6561.397498 | -0.420653735 | 5.10E-05 |
| WDYHV1 | 642.0397183 | -0.420748033 | 2.50E-03 |
| RNF227 | 387.3186546 | -0.420834916 | 1.82E-03 |
| BUB3 | 4309.026554 | -0.420895702 | 1.59E-09 |
| BRF1 | 2155.441023 | -0.421065357 | 3.16E-04 |
| RNASEL | 490.163533 | -0.421067422 | 3.78E-04 |
| FADS2 | 16355.60439 | -0.421174417 | 2.80E-02 |
| MYO9A | 2981.552426 | -0.421523664 | 1.27E-02 |
| LSM2 | 1298.558421 | -0.42153921 | 1.28E-04 |
| ANAPC5 | 6108.373281 | -0.421586899 | 3.27E-05 |
| RPL23AP42 | 8204.968219 | -0.421623697 | 6.17E-04 |
| CCDC136 | 1824.450033 | -0.421719683 | 1.68E-04 |
| RPL31 | 25986.45476 | -0.421849737 | 3.28E-06 |
| WDR70 | 3681.263314 | -0.421868952 | 9.44E-04 |
| ARHGAP45 | 1747.783185 | -0.422053004 | 1.69E-03 |
| BBS1 | 1549.593752 | -0.422239065 | 1.61E-04 |
| SRSF8 | 1740.258175 | -0.422566028 | 6.56E-05 |
| C1orf74 | 515.7573069 | -0.422595386 | 1.27E-04 |
| IPO8 | 3229.342953 | -0.422690535 | 3.48E-06 |
| ZNF345 | 182.6037155 | -0.422717546 | 1.75E-02 |
| BRCC3 | 1658.547271 | -0.423031051 | 3.95E-03 |
| UBAC2-AS1 | 265.6527688 | -0.423037297 | 2.05E-02 |
| AMMECR1 | 735.4118195 | -0.423222685 | 2.88E-04 |
| SH3GLB2 | 3523.835095 | -0.423343444 | 6.95E-07 |
| TMEM147-AS1 | 1163.878717 | -0.424122445 | 5.57E-03 |
| ABCA2 | 4348.010355 | -0.424164051 | 9.65E-03 |
| SNRNP40 | 2625.67186 | -0.424331829 | 1.75E-06 |
| IPO5P1 | 378.5648789 | -0.424480683 | 3.36E-02 |
| PRKACA | 4284.940824 | -0.424759276 | 1.31E-05 |
| SDHAF1 | 427.5079267 | -0.424924512 | 3.03E-02 |
| FAM13B | 5560.921553 | -0.425321357 | 4.10E-04 |
| TSPAN15 | 535.6987614 | -0.425500951 | 1.74E-02 |
| TAF8 | 1218.431965 | -0.426071889 | 6.00E-05 |
| KDM3B | 13087.24983 | -0.426510735 | 8.82E-11 |
| SDHA | 10652.72361 | -0.426727716 | 4.39E-03 |
| DBNDD1 | 1129.439015 | -0.426869957 | 6.66E-03 |
| VPS13B | 2695.859008 | -0.426903565 | 1.29E-03 |
| PHF5A | 2143.181465 | -0.42702378 | 1.10E-07 |
| NA | 238.0695398 | -0.427149662 | 2.09E-02 |
| ZNF132 | 208.122955 | -0.427264802 | 9.54E-03 |
| TM2D3 | 1237.924129 | -0.427327776 | 3.73E-06 |
| DCAF12 | 4233.100473 | -0.427476622 | 4.28E-09 |
| N4BP2L1 | 189.5204037 | -0.427646904 | 1.56E-02 |
| STX1B | 399.8371736 | -0.42768066 | 8.83E-03 |
| ENC1 | 10344.19834 | -0.427812347 | 5.74E-06 |
| SMDT1 | 1180.252028 | -0.42795521 | 2.56E-03 |

| | | | |
|---|---|---|---|
| PIP4K2B | 6414.059097 | -0.428004437 | 7.16E-05 |
| BRD9 | 4818.224883 | -0.42805112 | 1.68E-11 |
| RAB11FIP4 | 856.5889553 | -0.428088253 | 1.49E-02 |
| MFSD10 | 2310.200077 | -0.428437727 | 2.25E-04 |
| ASB16-AS1 | 304.3514996 | -0.428488675 | 4.31E-03 |
| CACTIN | 1550.647358 | -0.428538751 | 1.31E-06 |
| PPA1 | 2590.012982 | -0.428585194 | 5.12E-05 |
| RBMX | 11615.11458 | -0.428613939 | 7.52E-10 |
| TMEM177 | 593.8295865 | -0.428652192 | 7.19E-05 |
| TLE2 | 901.3412673 | -0.428819956 | 4.98E-02 |
| PRIMPOL | 515.2385594 | -0.429086269 | 2.01E-02 |
| ATP11C | 2323.681458 | -0.429107003 | 2.86E-02 |
| MUTYH | 512.5623734 | -0.429185619 | 1.56E-02 |
| NSMCE4A | 1084.538538 | -0.429295219 | 7.93E-06 |
| TCF3 | 4011.452498 | -0.429440089 | 4.99E-06 |
| QTRT1 | 1398.971228 | -0.429452625 | 7.63E-05 |
| PLXNA3 | 2210.81147 | -0.429467947 | 1.53E-02 |
| ZNF318 | 3190.47659 | -0.429528512 | 8.76E-07 |
| EGFR | 7757.382542 | -0.429947064 | 3.75E-10 |
| CACYBP | 3900.728954 | -0.430451384 | 4.50E-03 |
| TIMM13 | 2178.951561 | -0.430517828 | 1.28E-04 |
| HAUS4 | 2423.93088 | -0.430997028 | 4.10E-04 |
| AL845472.2 | 297.2034584 | -0.431008751 | 6.21E-03 |
| HOXC9 | 348.2374019 | -0.431027031 | 4.17E-03 |
| AL049840.4 | 212.2357653 | -0.431373664 | 1.83E-02 |
| DUSP18 | 224.9387926 | -0.431447262 | 2.09E-02 |
| GFOD2 | 1499.238531 | -0.431493976 | 7.63E-05 |
| APRT | 3780.591045 | -0.431674476 | 3.89E-05 |
| ABRAXAS1 | 475.8088465 | -0.432052136 | 2.54E-02 |
| BTBD6 | 2124.514255 | -0.432282307 | 4.08E-04 |
| TCOF1 | 9862.217299 | -0.432421199 | 3.99E-09 |
| PLEK2 | 599.7952445 | -0.432556285 | 1.18E-02 |
| PKD1P1 | 1205.512043 | -0.432964218 | 7.89E-03 |
| PRPF8 | 28223.35904 | -0.433175035 | 3.76E-05 |
| PIK3R4 | 2327.394576 | -0.433515261 | 1.01E-02 |
| CCDC102A | 865.6513979 | -0.43387286 | 1.00E-02 |
| NXF1 | 3794.280569 | -0.434663116 | 1.04E-04 |
| TOB2 | 2449.947897 | -0.434704613 | 8.42E-04 |
| NISCH | 3398.117465 | -0.435429323 | 8.91E-09 |
| RALB | 5036.240045 | -0.435502374 | 1.43E-07 |
| LUC7L2 | 3487.963961 | -0.435579366 | 1.44E-04 |
| XYLB | 384.9369497 | -0.435886604 | 2.37E-02 |
| PSMB9 | 1826.833266 | -0.435963989 | 2.12E-05 |
| HOXB8 | 188.1046317 | -0.436029486 | 2.25E-02 |
| CCDC8 | 1342.593494 | -0.436218083 | 3.76E-03 |
| RPL9 | 21258.71788 | -0.436344197 | 1.37E-04 |
| DHRS13 | 519.4154562 | -0.436428563 | 9.71E-04 |

| | | | |
|---|---|---|---|
| TRIM34 | 459.8810898 | -0.436674737 | 1.04E-02 |
| GLRX5 | 2045.707253 | -0.436763682 | 1.75E-06 |
| ZDHHC12 | 1347.149644 | -0.436919895 | 1.69E-03 |
| AC099850.3 | 182.1025745 | -0.437138953 | 3.54E-02 |
| HDHD5 | 2611.229972 | -0.437168816 | 5.53E-05 |
| ZBTB12 | 788.0118232 | -0.43719773 | 2.02E-04 |
| SHLD3 | 185.7210417 | -0.437275919 | 7.74E-03 |
| CA5B | 553.48089 | -0.437654931 | 9.53E-03 |
| AL162231.1 | 198.401438 | -0.437659339 | 2.17E-02 |
| ZNF581 | 889.5295823 | -0.437771735 | 4.42E-03 |
| ABTB1 | 827.9921184 | -0.437803636 | 2.60E-03 |
| NUTM2D | 183.9396658 | -0.437964054 | 2.71E-02 |
| PITPNB | 3674.542116 | -0.438070338 | 3.34E-08 |
| TPPP3 | 274.850805 | -0.438112402 | 3.54E-02 |
| MRPL24 | 1793.201605 | -0.438185821 | 8.58E-06 |
| TNS2 | 4398.495576 | -0.438320659 | 1.48E-02 |
| F8A3 | 1246.56378 | -0.438443862 | 7.87E-04 |
| RPS15A | 28078.05717 | -0.438545945 | 1.58E-05 |
| EXOSC10 | 4054.640073 | -0.438807423 | 3.66E-10 |
| SLC4A3 | 1292.885625 | -0.438999827 | 1.79E-04 |
| JMJD7-PLA2G4 | 577.9573265 | -0.439160448 | 4.54E-03 |
| INA | 628.7562037 | -0.439215883 | 2.98E-04 |
| SNRPE | 2384.328884 | -0.439465336 | 5.79E-06 |
| CTDSP1 | 4035.144085 | -0.439701768 | 2.09E-04 |
| MAST3 | 693.4806052 | -0.439739325 | 5.18E-06 |
| ZNF614 | 555.4187751 | -0.439745936 | 8.46E-05 |
| ALAD | 2563.398008 | -0.439873114 | 1.85E-04 |
| KLF16 | 371.8280178 | -0.440671516 | 2.70E-02 |
| C2CD3 | 3048.904252 | -0.440923319 | 2.79E-06 |
| GTF3C2 | 4232.170189 | -0.441166381 | 3.82E-07 |
| ZNF71 | 490.1173141 | -0.441178149 | 4.93E-03 |
| CSRP1 | 17874.18646 | -0.441457129 | 1.31E-03 |
| SF3B3 | 19040.42579 | -0.441541105 | 2.17E-06 |
| TRAF2 | 1387.830748 | -0.4415697 | 1.53E-03 |
| LAMA5 | 8544.457214 | -0.441609367 | 3.04E-03 |
| FKBPL | 291.1264024 | -0.441759693 | 1.22E-02 |
| BICRAL | 968.5111093 | -0.441916977 | 7.49E-05 |
| HIGD2A | 893.9299553 | -0.441941329 | 4.47E-03 |
| OBSCN | 2244.375511 | -0.442303222 | 2.27E-03 |
| DTX3 | 1694.078695 | -0.442355217 | 5.11E-06 |
| NOA1 | 1315.983886 | -0.442944879 | 2.15E-04 |
| RAPGEF4 | 1768.711438 | -0.443490731 | 7.23E-04 |
| TCF7L2 | 706.3416085 | -0.443581541 | 7.28E-04 |
| AGPAT4 | 1218.159414 | -0.443643764 | 2.87E-06 |
| MAP3K4 | 1470.576972 | -0.443644118 | 7.46E-03 |
| DUSP16 | 1770.746084 | -0.443678144 | 2.66E-05 |
| ICMT | 5718.142823 | -0.443850631 | 3.45E-03 |

| | | | |
|---|---|---|---|
| RBM5 | 5755.397797 | -0.443886639 | 1.62E-12 |
| PTPRA | 8909.240941 | -0.4440887 | 5.16E-07 |
| EEF1B2 | 14457.85453 | -0.444128442 | 2.56E-05 |
| CSNK1G2 | 3597.310619 | -0.444166716 | 1.01E-04 |
| PPP1R3E | 786.1886405 | -0.444249119 | 2.03E-05 |
| NFATC4 | 3878.575819 | -0.444281118 | 2.96E-03 |
| HNRNPUL1 | 13088.87928 | -0.444323529 | 1.14E-05 |
| CAMK2G | 1503.621246 | -0.444336781 | 1.83E-04 |
| FAM32A | 4402.655163 | -0.444659726 | 5.94E-05 |
| RPL23A | 15036.58917 | -0.444804859 | 1.09E-03 |
| SHOX2 | 480.5175504 | -0.444957043 | 3.50E-03 |
| ELAC2 | 5596.559 | -0.445102037 | 7.74E-05 |
| GTF2IRD2B | 908.8548731 | -0.445172432 | 1.29E-02 |
| RBFA | 1166.399006 | -0.445174112 | 3.35E-11 |
| BAD | 1488.272155 | -0.445224186 | 1.97E-03 |
| NEURL4 | 2079.603323 | -0.445392277 | 1.23E-03 |
| HGH1 | 1789.133151 | -0.446304965 | 3.23E-06 |
| LRRC37B | 470.6949516 | -0.446381332 | 1.13E-03 |
| RAMP2 | 1068.842961 | -0.446574603 | 3.86E-03 |
| SH3BP1 | 1544.175936 | -0.446984981 | 1.34E-03 |
| GABPB1-AS1 | 2069.678936 | -0.447091634 | 2.30E-03 |
| TIAM2 | 690.2406023 | -0.447486351 | 9.93E-03 |
| ZNF276 | 1389.475469 | -0.447538382 | 1.36E-06 |
| ALDH7A1 | 9362.120636 | -0.447676626 | 1.38E-06 |
| TFAP2A-AS1 | 96.55106709 | -0.448086332 | 2.95E-02 |
| EMC9 | 672.5210451 | -0.448090535 | 1.35E-05 |
| KATNAL1 | 513.5647486 | -0.448217975 | 7.12E-04 |
| PKMYT1 | 1390.690032 | -0.448236553 | 4.78E-02 |
| HHIPL1 | 616.3811624 | -0.448252771 | 3.51E-04 |
| C3orf14 | 394.3980219 | -0.448446935 | 2.66E-03 |
| AC021092.1 | 246.9309242 | -0.448629426 | 8.60E-04 |
| LURAP1 | 118.0104379 | -0.44875469 | 3.78E-02 |
| GEMIN6 | 1396.151903 | -0.4490576 | 3.61E-08 |
| HSF4 | 604.4274697 | -0.449488632 | 1.06E-03 |
| KDM8 | 224.6049505 | -0.449610676 | 1.06E-02 |
| PPP4R1L | 870.1846927 | -0.4499044 | 3.31E-06 |
| GPHN | 1149.675776 | -0.450141145 | 2.71E-03 |
| SYT17 | 533.479335 | -0.45023862 | 7.13E-04 |
| CTBP1 | 3972.434996 | -0.450441727 | 1.96E-06 |
| TXNDC15 | 1878.519765 | -0.450570911 | 2.33E-02 |
| MPHOSPH6 | 905.9787589 | -0.450588091 | 3.63E-06 |
| SSRP1 | 19588.32243 | -0.450740151 | 2.42E-10 |
| TTC38 | 996.7800644 | -0.451112727 | 1.76E-03 |
| AMER1 | 999.99627 | -0.451231877 | 2.82E-05 |
| PRDM10 | 641.3958178 | -0.451610272 | 2.30E-02 |
| TRIM46 | 215.9672309 | -0.451788372 | 1.67E-02 |
| RPL27A | 44327.24657 | -0.45178893 | 1.02E-05 |

| | | | |
|---|---|---|---|
| MPST | 1990.715552 | -0.452023463 | 1.14E-04 |
| HDGF | 21317.82589 | -0.452048357 | 4.20E-06 |
| LMBR1 | 4188.33573 | -0.452112174 | 1.07E-05 |
| ZNF428 | 892.0978553 | -0.452365741 | 4.35E-04 |
| SDF2L1 | 443.8847136 | -0.452532784 | 3.25E-03 |
| AC232271.1 | 168.7313114 | -0.452664947 | 8.32E-03 |
| SNX22 | 156.8170892 | -0.452755707 | 4.80E-02 |
| MRPS21 | 2857.515873 | -0.452769923 | 1.44E-05 |
| EXOC3-AS1 | 613.1024133 | -0.45283667 | 1.02E-05 |
| SHPRH | 739.154152 | -0.452863364 | 1.85E-02 |
| LCMT2 | 524.7377588 | -0.452925564 | 5.35E-03 |
| RPS10 | 23450.91986 | -0.453107384 | 1.75E-04 |
| TBX6 | 135.3723593 | -0.453141351 | 2.68E-02 |
| POLR2L | 4348.331672 | -0.453147971 | 2.41E-03 |
| DOCK5 | 2093.893287 | -0.453204168 | 8.04E-03 |
| PMS2P3 | 353.3370009 | -0.45346666 | 1.44E-04 |
| CCDC77 | 870.8727388 | -0.453576238 | 2.48E-03 |
| PLCD1 | 580.6541518 | -0.453663998 | 1.21E-02 |
| SETD1A | 2873.426182 | -0.453784017 | 5.41E-05 |
| TSC22D1 | 5776.662557 | -0.453903833 | 7.98E-07 |
| BACE1 | 6341.742065 | -0.45391989 | 4.99E-07 |
| PM20D2 | 1660.566496 | -0.454018853 | 2.40E-03 |
| MAPK8IP3 | 3358.91059 | -0.454042974 | 3.19E-06 |
| GLMN | 563.3816818 | -0.454078547 | 1.06E-02 |
| USP35 | 816.1064056 | -0.454205256 | 1.54E-04 |
| FUBP1 | 4556.3948 | -0.454239922 | 1.13E-05 |
| SLC22A3 | 402.2075182 | -0.454313513 | 6.63E-04 |
| GPRASP2 | 355.0135181 | -0.454534063 | 3.34E-04 |
| UBALD2 | 914.1334782 | -0.454579764 | 1.37E-02 |
| AC135050.6 | 225.2054249 | -0.45464475 | 3.09E-02 |
| MAP7D2 | 1188.134455 | -0.454721827 | 3.76E-04 |
| PYM1 | 593.1831054 | -0.454729288 | 2.09E-04 |
| PPIH | 1924.711024 | -0.455025711 | 5.30E-06 |
| C1QBP | 10284.03275 | -0.455249397 | 2.29E-09 |
| WDR59 | 2811.589974 | -0.45535603 | 1.23E-08 |
| ACD | 1028.935391 | -0.455461451 | 6.22E-04 |
| TSPAN5 | 2843.317647 | -0.455528172 | 8.17E-05 |
| BUD13 | 1709.776183 | -0.455567413 | 1.40E-09 |
| PBX1 | 1185.042455 | -0.455675287 | 3.96E-06 |
| TCTN1 | 1339.541659 | -0.455806404 | 6.56E-08 |
| WDR89 | 465.2162936 | -0.456280682 | 2.80E-02 |
| PPP1R13L | 292.5531074 | -0.456555653 | 2.64E-02 |
| ATP13A1 | 3127.033339 | -0.456782643 | 2.22E-06 |
| CCDC120 | 662.5532422 | -0.456782861 | 7.99E-06 |
| HACD3 | 7474.997162 | -0.456953849 | 4.14E-04 |
| SNRK | 1621.869494 | -0.457235746 | 2.40E-05 |
| P3H4 | 2876.333403 | -0.457237151 | 1.08E-02 |

| | | | |
|---|---|---|---|
| HNRNPD | 10291.59296 | -0.457363019 | 2.01E-09 |
| GSTM2 | 1579.400594 | -0.457540784 | 1.09E-03 |
| AL139300.1 | 771.5096936 | -0.457542705 | 2.46E-02 |
| IPO5 | 38071.98562 | -0.458341495 | 4.94E-15 |
| SNHG20 | 472.8352944 | -0.458367667 | 6.22E-06 |
| AL590327.1 | 533.3376147 | -0.458531753 | 3.62E-03 |
| STEAP3 | 2478.209655 | -0.458620751 | 2.21E-04 |
| ACTR1B | 4339.501904 | -0.458868723 | 1.33E-04 |
| CEP89 | 891.0493953 | -0.458869329 | 1.70E-05 |
| IFT46 | 2480.303894 | -0.4590171 | 2.77E-08 |
| RIOX1 | 610.8090639 | -0.45937854 | 2.59E-04 |
| DIXDC1 | 1830.006981 | -0.459433526 | 4.09E-03 |
| TOMM22 | 3069.363182 | -0.459494556 | 2.42E-04 |
| CHD1L | 2797.267694 | -0.459497596 | 8.54E-16 |
| EIF3E | 11249.5084 | -0.459801391 | 2.21E-04 |
| LAMC3 | 1143.755153 | -0.460266899 | 3.59E-03 |
| TOMM40L | 1091.925039 | -0.460511186 | 9.55E-04 |
| KHSRP | 10535.99363 | -0.46075673 | 7.98E-10 |
| HOXA10 | 596.1595277 | -0.460812089 | 4.68E-05 |
| DNAAF5 | 2856.312045 | -0.460931957 | 6.10E-09 |
| PLEKHG5 | 1159.440594 | -0.461273433 | 1.51E-04 |
| SCO2 | 668.0786893 | -0.461439515 | 4.18E-03 |
| NEIL2 | 844.7899118 | -0.461447307 | 8.31E-04 |
| UBE3D | 288.116491 | -0.461592194 | 8.43E-03 |
| DOK4 | 2219.835095 | -0.461765279 | 9.94E-05 |
| JUN | 2696.774838 | -0.461828049 | 4.04E-02 |
| EVA1A | 932.7295082 | -0.462214291 | 2.31E-06 |
| RPL22 | 11894.40039 | -0.462290188 | 9.99E-08 |
| SCAF11 | 4087.643525 | -0.462606967 | 1.03E-02 |
| RPL8 | 63923.6203 | -0.462859016 | 4.97E-05 |
| EPC1 | 776.583326 | -0.463063073 | 2.54E-03 |
| TIMP1 | 7816.765979 | -0.463224062 | 1.42E-02 |
| ZNF784 | 232.483614 | -0.463225982 | 2.87E-02 |
| USP20 | 2227.906697 | -0.46327358 | 1.44E-05 |
| ZSWIM1 | 834.9451821 | -0.463334073 | 1.67E-04 |
| CNOT11 | 2756.127194 | -0.463414518 | 8.87E-07 |
| GPSM3 | 1935.741378 | -0.4635224 | 9.94E-04 |
| RPA3 | 2242.846147 | -0.464052383 | 8.40E-04 |
| PRPF4B | 3417.369769 | -0.464177458 | 6.26E-03 |
| PPM1D | 648.1218558 | -0.464181422 | 1.51E-05 |
| IQCK | 379.5866682 | -0.464344688 | 4.12E-04 |
| BRD3 | 2610.24856 | -0.464580776 | 4.63E-08 |
| GRK2 | 4126.550521 | -0.464634106 | 6.80E-06 |
| PLEC | 19438.93034 | -0.464702427 | 1.16E-05 |
| SEMA4D | 1038.790482 | -0.464734413 | 4.87E-03 |
| MAP4K4 | 13426.54364 | -0.464817149 | 7.74E-14 |
| ZNF446 | 728.7385639 | -0.464821699 | 6.18E-04 |

| | | | |
|---|---|---|---|
| RPP25L | 1196.735363 | -0.464885422 | 6.63E-04 |
| AC009053.1 | 341.7337856 | -0.464921747 | 9.46E-05 |
| SGK1 | 5483.663424 | -0.46501884 | 2.33E-03 |
| U2AF2 | 10147.28852 | -0.465295505 | 8.55E-08 |
| GATD1 | 2190.233517 | -0.465377568 | 3.60E-06 |
| UCKL1 | 1756.907951 | -0.465387357 | 1.82E-11 |
| MT-CO2 | 272442.4574 | -0.465500585 | 3.89E-03 |
| COPS9 | 1451.955329 | -0.4656357 | 1.41E-03 |
| TSC2 | 3389.517899 | -0.465685933 | 4.68E-09 |
| CARD6 | 1819.649875 | -0.466205719 | 5.70E-04 |
| DEPDC7 | 451.1246896 | -0.466271675 | 3.01E-03 |
| ZNF300 | 709.6145832 | -0.466654783 | 1.97E-02 |
| VARS2 | 1563.733782 | -0.467057082 | 3.37E-07 |
| PANK4 | 952.9214816 | -0.4671546 | 1.39E-02 |
| HSD11B1L | 516.8047837 | -0.467193691 | 5.75E-04 |
| AP006621.5 | 135.4954891 | -0.467456837 | 1.35E-02 |
| DCAF11 | 2586.756227 | -0.467568489 | 2.45E-03 |
| C2orf16 | 308.8560384 | -0.467669096 | 5.24E-04 |
| ATP2A1-AS1 | 206.3249085 | -0.467890389 | 4.11E-04 |
| RPL32 | 31619.84787 | -0.46802533 | 2.42E-05 |
| FARP2 | 2032.293464 | -0.468078095 | 2.42E-05 |
| PCDHGA7 | 577.8186976 | -0.468088688 | 2.74E-03 |
| BIN3 | 1269.359925 | -0.468364224 | 8.68E-07 |
| AL359258.1 | 351.9067717 | -0.468869249 | 2.28E-03 |
| CCDC86 | 3009.142193 | -0.469100658 | 6.24E-05 |
| GOLGA2P10 | 667.6520873 | -0.469279385 | 9.81E-05 |
| RPL36 | 12121.41414 | -0.46958096 | 1.71E-04 |
| SMARCD1 | 4173.785875 | -0.469593631 | 1.72E-08 |
| FAM111A | 2701.692 | -0.469690393 | 4.27E-03 |
| LINC01667 | 1112.755145 | -0.469700729 | 3.13E-07 |
| ZNF764 | 306.824933 | -0.469730317 | 1.76E-02 |
| CDK10 | 2722.429257 | -0.469889933 | 8.31E-13 |
| CYP4F12 | 335.7560523 | -0.469986924 | 1.45E-02 |
| PHLDB3 | 358.9219198 | -0.470012473 | 4.88E-03 |
| HARS | 4201.237933 | -0.470114173 | 3.10E-09 |
| AP1M1 | 4261.769271 | -0.470464593 | 3.87E-05 |
| RPS8 | 38197.62831 | -0.470653233 | 1.24E-07 |
| PHRF1 | 3416.257358 | -0.470776549 | 1.47E-05 |
| FMC1-LUC7L2 | 1583.580303 | -0.470789215 | 5.05E-06 |
| CRACR2A | 437.6015031 | -0.470910833 | 3.96E-05 |
| NSDHL | 2495.579896 | -0.471143575 | 2.97E-03 |
| SNHG3 | 1840.165392 | -0.471298068 | 2.24E-02 |
| LRFN1 | 256.0931305 | -0.471473113 | 1.89E-02 |
| UBTD2 | 1371.08928 | -0.471595725 | 6.54E-14 |
| SMARCB1 | 4122.10918 | -0.472026319 | 1.77E-07 |
| DPY19L1 | 3679.417338 | -0.472107743 | 1.76E-03 |
| SF3A2 | 2191.908115 | -0.472120219 | 1.27E-04 |

| | | | |
|---|---|---|---|
| MZT2B | 1646.035276 | -0.47229497 | 3.51E-03 |
| UBXN11 | 840.3304153 | -0.472325653 | 5.34E-05 |
| NTN4 | 1601.295178 | -0.472338067 | 9.15E-07 |
| FLT3LG | 325.6567815 | -0.472604341 | 6.29E-03 |
| SMARCD2 | 3933.648525 | -0.472693284 | 7.91E-08 |
| RPS12 | 23474.3027 | -0.47282677 | 2.29E-08 |
| N6AMT1 | 944.9122957 | -0.472960669 | 3.48E-06 |
| PARP10 | 2590.466427 | -0.473135837 | 8.26E-05 |
| SHQ1 | 1577.551243 | -0.473140076 | 2.13E-03 |
| AC008840.1 | 101.8367287 | -0.473321538 | 3.64E-02 |
| AL121845.3 | 393.5459396 | -0.473725586 | 2.34E-02 |
| DHX30 | 5148.154897 | -0.473747752 | 5.42E-09 |
| ZADH2 | 2400.126845 | -0.47377355 | 4.27E-13 |
| 9-Sep | 13119.5137 | -0.473983107 | 1.47E-04 |
| GSTM1 | 801.4680163 | -0.474083885 | 1.56E-04 |
| ZBTB46 | 1939.837957 | -0.474118801 | 2.15E-03 |
| UFC1 | 2361.765324 | -0.474224873 | 2.21E-05 |
| AC055713.1 | 109.6029545 | -0.474226602 | 1.66E-02 |
| MYO18A | 7255.058026 | -0.474255161 | 3.05E-04 |
| EIF3B | 14629.46064 | -0.474357972 | 2.91E-04 |
| SNRPD1 | 2850.34921 | -0.474438774 | 3.51E-04 |
| DELE1 | 3152.562817 | -0.474543605 | 4.05E-05 |
| LATS1 | 1070.227587 | -0.474615181 | 1.39E-06 |
| ZNF793 | 305.6492898 | -0.474646235 | 5.24E-03 |
| PCF11 | 2025.55404 | -0.474734712 | 2.80E-04 |
| ATP5F1D | 1814.642888 | -0.474793982 | 3.04E-04 |
| WBP1 | 3743.929191 | -0.474924504 | 3.66E-03 |
| AC245140.2 | 138.1623687 | -0.475078367 | 1.07E-02 |
| RPS4X | 33744.19624 | -0.475376025 | 2.46E-06 |
| GPANK1 | 1068.369362 | -0.475501028 | 8.87E-05 |
| RPL9P9 | 16918.13946 | -0.475586928 | 6.50E-06 |
| MHENCR | 187.7693423 | -0.475628346 | 9.11E-03 |
| HSD17B11 | 1994.692616 | -0.475648755 | 4.73E-07 |
| WDR4 | 1435.970238 | -0.47564884 | 6.46E-05 |
| CEP164 | 3691.315585 | -0.475665922 | 8.36E-07 |
| NSUN5P1 | 2362.073103 | -0.47585484 | 6.23E-07 |
| EBP | 1987.716815 | -0.475969459 | 6.67E-03 |
| ZBTB48 | 733.9219745 | -0.476029237 | 1.02E-03 |
| GDPD5 | 1564.667833 | -0.476066612 | 1.28E-03 |
| NUP62 | 3968.806784 | -0.476113105 | 1.12E-07 |
| GTF2IP4 | 7182.880039 | -0.476124513 | 6.50E-06 |
| TMEM256 | 1274.485443 | -0.476133666 | 8.70E-05 |
| TYK2 | 3847.834705 | -0.476331426 | 5.09E-06 |
| TERF1 | 1319.671707 | -0.476617448 | 7.43E-03 |
| AL513550.1 | 323.2667061 | -0.476769121 | 1.78E-04 |
| ARL10 | 1205.912163 | -0.476813568 | 3.27E-02 |
| HEXD | 641.0207043 | -0.476915718 | 1.87E-05 |

| | | | |
|---|---|---|---|
| E4F1 | 1257.798378 | -0.477037518 | 7.95E-09 |
| CIT | 2750.995169 | -0.477246687 | 1.38E-03 |
| ZNF233 | 106.6235961 | -0.477367945 | 6.47E-03 |
| EIF2B1 | 3995.522508 | -0.477631522 | 7.33E-17 |
| RAVER1 | 3064.395334 | -0.478123741 | 6.83E-06 |
| RPS6 | 79859.13227 | -0.478607326 | 2.77E-08 |
| MAP3K3 | 2379.952131 | -0.478806953 | 3.27E-05 |
| S100A10 | 12016.01031 | -0.478866175 | 5.63E-05 |
| PRR19 | 437.3132953 | -0.479605234 | 3.26E-03 |
| FBXL3 | 1749.761037 | -0.47962874 | 1.00E-05 |
| TFCP2 | 1699.603032 | -0.47970063 | 1.75E-12 |
| NAT14 | 1520.084379 | -0.47973006 | 3.62E-03 |
| STK25 | 5404.879135 | -0.480280079 | 3.26E-08 |
| SELENOW | 2630.523569 | -0.480308205 | 1.21E-02 |
| CDAN1 | 831.7323165 | -0.48047929 | 4.30E-04 |
| LDOC1 | 2258.350712 | -0.480499064 | 3.99E-03 |
| L3MBTL2 | 1878.053994 | -0.480509183 | 6.83E-07 |
| IBA57 | 513.5595263 | -0.480561269 | 1.32E-02 |
| ACIN1 | 10252.36296 | -0.480738669 | 2.56E-31 |
| AK1 | 2241.002632 | -0.48089067 | 3.39E-03 |
| SHPK | 1541.581613 | -0.4810591 | 2.87E-03 |
| ZNF136 | 332.4460914 | -0.481221053 | 1.39E-02 |
| PRKDC | 32717.12166 | -0.481437278 | 1.38E-08 |
| NRG2 | 272.570081 | -0.481727942 | 4.45E-03 |
| UBAC1 | 2818.103575 | -0.481979034 | 9.26E-08 |
| DNMT3A | 1317.672083 | -0.482155768 | 7.72E-05 |
| RPLP1 | 50415.89131 | -0.482682786 | 4.52E-04 |
| TADA2A | 885.8477906 | -0.483039833 | 7.92E-06 |
| PRRT3 | 215.8516639 | -0.483123249 | 2.47E-03 |
| ID3 | 8443.247581 | -0.483224517 | 1.28E-02 |
| OBSL1 | 7207.953164 | -0.483287084 | 6.88E-04 |
| SUV39H2 | 651.0796669 | -0.483630953 | 6.35E-03 |
| LPAL2 | 159.1006028 | -0.4838671 | 1.31E-02 |
| RAB9B | 263.064607 | -0.48398408 | 6.29E-04 |
| DACT3 | 383.0030205 | -0.484453609 | 1.12E-02 |
| DHRS12 | 99.15939236 | -0.48451269 | 4.85E-02 |
| MED11 | 514.0417921 | -0.484525659 | 9.79E-03 |
| PRCC | 2243.716349 | -0.48491283 | 1.27E-04 |
| AKT2 | 5598.789116 | -0.484932866 | 1.57E-07 |
| CXXC1 | 1916.272061 | -0.48541909 | 1.37E-09 |
| ATR | 1999.15314 | -0.485426901 | 1.58E-02 |
| ZNF365 | 213.2583861 | -0.485652287 | 1.31E-02 |
| HGSNAT | 3374.663359 | -0.485980711 | 2.55E-04 |
| RANBP6 | 1516.497612 | -0.485999812 | 4.57E-03 |
| RPL10A | 24437.36707 | -0.486028011 | 4.35E-07 |
| MORC2 | 2508.372783 | -0.486052171 | 1.23E-06 |
| AC240565.2 | 150.4166937 | -0.486333745 | 8.63E-03 |

| | | | |
|---|---|---|---|
| AC018665.1 | 157.0923195 | -0.486357063 | 3.00E-03 |
| STK36 | 1396.326232 | -0.486458695 | 4.10E-06 |
| UBB | 33616.13027 | -0.487005391 | 1.36E-04 |
| UBTF | 5711.417987 | -0.487150365 | 9.45E-12 |
| MEN1 | 3843.040286 | -0.487659264 | 1.28E-07 |
| P4HA3 | 312.285872 | -0.48780163 | 3.04E-03 |
| RPS16 | 19557.72628 | -0.487821068 | 7.15E-05 |
| PTPN18 | 1447.976884 | -0.487910078 | 1.30E-08 |
| ANKRD13B | 446.4179403 | -0.487958974 | 9.33E-03 |
| RAD51-AS1 | 223.6582899 | -0.488489633 | 1.43E-03 |
| AC004890.2 | 340.3912192 | -0.488661474 | 1.11E-03 |
| PPIA | 74455.26732 | -0.488845314 | 9.94E-04 |
| DGCR8 | 1736.836189 | -0.48916672 | 3.64E-06 |
| ZNF776 | 951.6721861 | -0.489922461 | 2.90E-05 |
| EFHC1 | 1062.860149 | -0.490026578 | 2.61E-04 |
| RUVBL1 | 5432.212495 | -0.490659344 | 8.36E-08 |
| ZIK1 | 840.9093282 | -0.491144769 | 2.80E-05 |
| KIFC2 | 1596.31756 | -0.491365677 | 4.04E-04 |
| HNRNPA0 | 10274.12928 | -0.49139195 | 8.36E-07 |
| SLC25A42 | 745.7633681 | -0.491552272 | 1.61E-04 |
| UBQLN4 | 3371.894706 | -0.491625804 | 1.97E-07 |
| MT-ATP8 | 15345.88142 | -0.491697494 | 1.94E-02 |
| JPT2 | 6701.70618 | -0.491797115 | 2.54E-07 |
| MIGA2 | 1872.139308 | -0.49263347 | 7.13E-07 |
| CCND3 | 1929.467615 | -0.492946236 | 2.36E-04 |
| U62317.2 | 110.991617 | -0.493029956 | 4.74E-02 |
| PRMT1 | 7046.38091 | -0.493148738 | 2.59E-06 |
| PCIF1 | 2716.372684 | -0.493620478 | 2.43E-06 |
| CERK | 2461.786519 | -0.493819082 | 2.98E-03 |
| DDX39B | 14330.28417 | -0.494168669 | 1.50E-08 |
| GTF2IP1 | 5488.638803 | -0.494352008 | 1.02E-06 |
| KIFAP3 | 1267.969376 | -0.494497541 | 4.13E-08 |
| HERC2P4 | 239.1856069 | -0.494533388 | 2.91E-02 |
| ZNF718 | 234.8439511 | -0.494645104 | 1.36E-03 |
| NR2F2 | 5643.974102 | -0.494763106 | 3.82E-05 |
| SFPQ | 12484.80377 | -0.494771649 | 1.18E-06 |
| AASDH | 769.6253677 | -0.49489752 | 3.25E-03 |
| ZDHHC17 | 996.3512409 | -0.495005334 | 6.43E-03 |
| DCLRE1B | 841.3736811 | -0.495324983 | 9.76E-06 |
| C12orf57 | 2937.498738 | -0.495354817 | 1.06E-03 |
| METRN | 2250.581151 | -0.495514275 | 1.63E-03 |
| KLHL25 | 490.3954555 | -0.495559484 | 3.47E-03 |
| POLR2G | 4145.462309 | -0.495594616 | 1.06E-07 |
| PLXNB1 | 1364.593559 | -0.495979001 | 2.38E-04 |
| NKRF | 953.9954924 | -0.496134005 | 1.75E-06 |
| NSUN5P2 | 311.8154007 | -0.496188085 | 6.74E-03 |
| RPS23 | 74273.86759 | -0.496310441 | 2.01E-06 |

| | | | |
|---|---|---|---|
| ZNF799 | 165.882558 | -0.49645888 | 1.11E-02 |
| PDE7A | 1118.518144 | -0.496786829 | 1.15E-04 |
| VIPR1 | 332.1102024 | -0.49686091 | 6.93E-03 |
| KIF9 | 221.1677954 | -0.49692556 | 1.16E-02 |
| ZNF436 | 1465.456013 | -0.497002877 | 7.91E-08 |
| GCSH | 1242.2651 | -0.497218649 | 1.31E-06 |
| ZMYND19 | 1227.019722 | -0.497370805 | 1.55E-07 |
| CNPY3 | 3245.507656 | -0.497949576 | 2.61E-04 |
| PALLD | 4841.86156 | -0.498200325 | 3.25E-06 |
| LRRC14 | 2220.494375 | -0.498244041 | 2.61E-06 |
| THAP11 | 2904.615872 | -0.498376451 | 4.63E-10 |
| AC020915.3 | 223.8893518 | -0.498399334 | 6.11E-04 |
| SFXN5 | 1044.332545 | -0.498656729 | 1.94E-04 |
| ORAI2 | 6379.575777 | -0.498985553 | 2.17E-05 |
| C16orf91 | 450.8027921 | -0.499139135 | 7.79E-05 |
| SETD1B | 1514.715378 | -0.499316228 | 1.21E-06 |
| UBA52 | 14827.05506 | -0.499518085 | 7.89E-05 |
| ZKSCAN2-DT | 185.4873067 | -0.499623207 | 2.80E-02 |
| TBRG4 | 6211.250692 | -0.499678834 | 3.94E-07 |
| DDIAS | 859.845589 | -0.499717633 | 1.65E-02 |
| ZMYM3 | 5158.185274 | -0.499820748 | 7.92E-04 |
| ZDHHC8 | 2492.861947 | -0.499930023 | 2.31E-05 |
| SKI | 2113.55652 | -0.500083807 | 5.94E-05 |
| SAFB | 4567.180569 | -0.500085239 | 3.15E-08 |
| TKFC | 1546.53106 | -0.50009454 | 9.05E-11 |
| LRIG3 | 486.7772918 | -0.500241243 | 3.41E-04 |
| AGMAT | 377.0349647 | -0.500902606 | 8.81E-05 |
| PCED1A | 3718.966878 | -0.500998195 | 3.18E-12 |
| NELFB | 5735.829947 | -0.501298931 | 5.71E-07 |
| CLK2 | 1573.991102 | -0.501481079 | 2.79E-07 |
| TET1 | 199.1206885 | -0.501715005 | 1.43E-02 |
| ATXN7L3B | 4368.840664 | -0.501785877 | 9.07E-06 |
| CHST10 | 1531.826562 | -0.50204381 | 1.20E-03 |
| MANSC1 | 461.6010775 | -0.502148609 | 2.29E-02 |
| FAM72B | 387.5188204 | -0.502216698 | 2.27E-02 |
| SMYD2 | 2015.433512 | -0.502529023 | 1.33E-11 |
| AC037459.3 | 258.5800368 | -0.502609459 | 1.47E-04 |
| AUNIP | 407.0831788 | -0.50283255 | 8.72E-04 |
| NDUFA2 | 1438.143049 | -0.502868536 | 1.25E-02 |
| PRRT1 | 226.6096598 | -0.502985425 | 1.09E-03 |
| USPL1 | 651.1617121 | -0.503261712 | 1.89E-03 |
| RBM26 | 3241.279658 | -0.503424707 | 5.60E-05 |
| COQ10A | 755.6505801 | -0.503547634 | 5.76E-07 |
| WDR18 | 1470.094668 | -0.503552024 | 1.28E-03 |
| ZNF324 | 851.7318929 | -0.50363822 | 3.06E-05 |
| MMS22L | 1204.062958 | -0.503780908 | 3.20E-03 |
| SHROOM2 | 932.6937715 | -0.503857 | 1.24E-02 |

| | | | |
|---|---|---|---|
| CIAO1 | 6123.071829 | -0.504023808 | 2.42E-08 |
| BRMS1L | 598.3185116 | -0.504141192 | 6.23E-04 |
| NONO | 22380.64719 | -0.504357511 | 3.47E-13 |
| SH2D2A | 145.948703 | -0.504380463 | 1.60E-02 |
| TIFA | 397.8935473 | -0.504825089 | 1.03E-03 |
| BAX | 1828.257783 | -0.504843088 | 3.44E-03 |
| RPS25 | 42066.75165 | -0.505140649 | 1.40E-06 |
| KDM4B | 1924.045144 | -0.505207703 | 4.80E-04 |
| NOC2L | 7230.373689 | -0.505279289 | 4.17E-12 |
| COL26A1 | 362.579038 | -0.505367234 | 3.10E-03 |
| SLC5A6 | 2109.913971 | -0.505398878 | 5.71E-07 |
| CBFB | 1922.228719 | -0.505444656 | 1.03E-04 |
| AC069281.2 | 301.6621336 | -0.505478928 | 7.04E-03 |
| MSH6 | 5759.369797 | -0.505544005 | 1.15E-03 |
| ADGRA2 | 6548.879872 | -0.505574183 | 2.95E-08 |
| PRR11 | 1991.797462 | -0.505612461 | 5.86E-05 |
| DBF4B | 1125.920736 | -0.505687123 | 1.32E-05 |
| THRA | 4147.420593 | -0.505692436 | 2.71E-06 |
| CYBA | 3231.264754 | -0.505725254 | 1.55E-03 |
| RPL4 | 113049.4838 | -0.505909192 | 1.03E-05 |
| KMT2A | 10311.72669 | -0.506036872 | 8.63E-09 |
| RABIF | 589.0444905 | -0.506060741 | 5.58E-04 |
| SUMO3 | 6194.823647 | -0.506456009 | 8.10E-06 |
| PSTPIP2 | 476.2849638 | -0.506697863 | 1.80E-04 |
| PPAN | 1600.837878 | -0.506806949 | 1.47E-04 |
| CCNG2 | 1339.046455 | -0.507343698 | 1.91E-04 |
| GID8 | 3607.78933 | -0.507366315 | 3.85E-08 |
| POLR2D | 1969.389092 | -0.507534145 | 1.43E-10 |
| AKT1S1 | 2743.622308 | -0.507646305 | 2.14E-05 |
| CLBA1 | 602.1476952 | -0.507711015 | 7.79E-05 |
| SCAND1 | 1159.837333 | -0.507891679 | 1.05E-03 |
| CPPED1 | 2575.427025 | -0.508107614 | 3.20E-08 |
| CSPG4P10 | 464.9625043 | -0.508203505 | 1.92E-05 |
| PITPNC1 | 453.1297312 | -0.508370612 | 6.06E-05 |
| MPZL2 | 259.3641946 | -0.508567858 | 1.84E-02 |
| NR2F6 | 1473.239063 | -0.508622784 | 2.82E-06 |
| RPS3A | 65602.08282 | -0.508750894 | 1.76E-05 |
| SCNN1D | 251.7293197 | -0.508910372 | 2.39E-02 |
| ZNHIT2 | 588.7238783 | -0.508941521 | 1.59E-03 |
| KRBA1 | 1022.141037 | -0.509762646 | 5.55E-07 |
| AC060766.7 | 170.9646921 | -0.510032502 | 1.21E-02 |
| ADAMTS13 | 316.9800591 | -0.510043415 | 1.05E-03 |
| STRA6LP | 197.9736375 | -0.510343266 | 3.43E-02 |
| AC008124.1 | 360.4932966 | -0.510518028 | 1.11E-03 |
| COL3A1 | 378.3612853 | -0.510559671 | 3.48E-02 |
| ACTL6A | 2808.889881 | -0.510640228 | 7.96E-13 |
| CIC | 4151.020787 | -0.510648083 | 7.00E-06 |

| | | | |
|---|---|---|---|
| AC006504.5 | 389.6895637 | -0.511296936 | 2.04E-06 |
| FTSJ3 | 5699.792994 | -0.511409082 | 4.65E-11 |
| AL157935.2 | 243.4140541 | -0.511569357 | 8.01E-03 |
| BX004987.1 | 445.7885911 | -0.512360264 | 1.16E-03 |
| MROH6 | 143.2816375 | -0.512549099 | 3.12E-02 |
| GTF3C5 | 4038.886948 | -0.51301612 | 2.08E-11 |
| 2-Mar | 358.2724243 | -0.513126574 | 2.65E-02 |
| NRBP2 | 1261.707177 | -0.513429596 | 5.52E-08 |
| PTTG1 | 6678.880036 | -0.513658453 | 1.34E-06 |
| MAGI3 | 464.1381935 | -0.514191076 | 9.42E-04 |
| MED25 | 1497.258246 | -0.514244755 | 1.78E-04 |
| RPL18 | 22761.13749 | -0.514386782 | 5.26E-05 |
| DCXR | 1850.066695 | -0.51459341 | 3.15E-06 |
| DNASE1 | 1176.911667 | -0.514648464 | 1.59E-07 |
| TWIST1 | 111.7034611 | -0.514786469 | 3.38E-02 |
| PEG3 | 8882.770975 | -0.514795691 | 5.19E-04 |
| ACBD7 | 133.8693676 | -0.5150469 | 2.14E-02 |
| CYB5B | 13391.33376 | -0.515257364 | 4.30E-04 |
| AL669831.1 | 130.1423115 | -0.515716376 | 1.17E-02 |
| SNAPC4 | 1363.319501 | -0.515948241 | 4.24E-08 |
| POLH | 851.2292712 | -0.516047877 | 6.62E-04 |
| ZBED4 | 1607.762094 | -0.516182044 | 7.24E-12 |
| NUMA1 | 17142.57348 | -0.51647631 | 1.18E-11 |
| PLK2 | 8444.080224 | -0.517061198 | 6.63E-06 |
| CD37 | 311.4740181 | -0.517763687 | 2.59E-03 |
| ADAM12 | 626.9604478 | -0.517849466 | 2.88E-03 |
| CD163L1 | 1044.256401 | -0.517880395 | 5.12E-03 |
| HELB | 231.1127368 | -0.517900555 | 3.55E-02 |
| HOXB7 | 1350.143893 | -0.518274404 | 1.45E-06 |
| ACTR3B | 853.544742 | -0.518320602 | 6.58E-07 |
| RPL13 | 48811.62292 | -0.518406965 | 2.12E-06 |
| VPS26B | 4900.815599 | -0.518558823 | 1.82E-07 |
| IGFBP2 | 14621.13362 | -0.518829309 | 1.95E-03 |
| UNC13A | 437.1254544 | -0.519074825 | 1.49E-02 |
| TRIM26 | 1999.723817 | -0.519120575 | 2.11E-06 |
| SYNJ1 | 938.8449281 | -0.519404197 | 1.54E-03 |
| AC010894.3 | 225.0721163 | -0.519414882 | 1.86E-03 |
| RPL36A | 17414.29202 | -0.519449417 | 3.42E-10 |
| PQBP1 | 2394.096602 | -0.519514677 | 1.31E-06 |
| GPX7 | 858.5942305 | -0.519645582 | 7.01E-04 |
| ZNF667 | 678.8257523 | -0.519731273 | 1.31E-03 |
| LANCL2 | 1253.792335 | -0.519742017 | 6.76E-04 |
| TUBG1 | 3738.175157 | -0.52002983 | 5.51E-06 |
| LHX4 | 142.5888116 | -0.520405398 | 9.49E-03 |
| DGCR2 | 4016.860652 | -0.520620183 | 2.59E-04 |
| SNX15 | 1870.19779 | -0.520693356 | 4.22E-10 |
| PPM1F | 3912.01717 | -0.520741988 | 1.74E-07 |

| | | | |
|---|---|---|---|
| NOVA2 | 439.1966609 | -0.520791766 | 1.53E-03 |
| SNHG10 | 266.7820462 | -0.521109204 | 7.65E-03 |
| RPS18 | 68722.47473 | -0.521581037 | 2.36E-06 |
| SREBF1 | 4307.962205 | -0.521796224 | 2.54E-06 |
| TCEA3 | 938.9865274 | -0.521957758 | 1.61E-04 |
| BOLA3-AS1 | 290.6439025 | -0.52203968 | 9.17E-05 |
| CBX6 | 5530.008558 | -0.522292763 | 1.31E-07 |
| DGCR6 | 250.3990308 | -0.522309084 | 1.18E-02 |
| IFT22 | 1788.73016 | -0.522418343 | 9.77E-08 |
| ZSCAN2 | 647.4750931 | -0.522534546 | 1.89E-04 |
| ZNF30 | 247.4851225 | -0.522602728 | 1.51E-02 |
| DNLZ | 303.6119897 | -0.522604151 | 6.60E-04 |
| CST3 | 10175.55565 | -0.522824719 | 4.42E-04 |
| IVD | 2392.483779 | -0.522850095 | 7.65E-08 |
| C11orf80 | 588.5319896 | -0.522995876 | 3.70E-04 |
| STX2 | 1209.839998 | -0.523029921 | 3.37E-05 |
| MTMR1 | 2691.207728 | -0.523239587 | 1.68E-06 |
| TGFBI | 26371.45713 | -0.523590726 | 2.63E-05 |
| AC122688.3 | 95.38874051 | -0.523902902 | 3.24E-02 |
| AC233723.2 | 80.15059256 | -0.52438837 | 2.13E-02 |
| TOE1 | 794.4759432 | -0.5244071 | 8.10E-06 |
| EIF4E2 | 3353.242311 | -0.524451473 | 3.85E-03 |
| MADD | 5953.472304 | -0.524558917 | 2.90E-07 |
| MAP6D1 | 244.6323378 | -0.524900662 | 1.33E-03 |
| CNTRL | 1364.911649 | -0.524961895 | 4.78E-02 |
| DHX9 | 14124.10549 | -0.525192905 | 1.24E-11 |
| ATP5MC1 | 1764.60637 | -0.525402524 | 6.41E-04 |
| AL031714.1 | 169.347552 | -0.525429574 | 9.54E-04 |
| SDHAP1 | 489.7331854 | -0.525560595 | 2.63E-03 |
| TARBP2 | 1194.823027 | -0.525640504 | 1.56E-06 |
| FAM122B | 3045.59739 | -0.525677886 | 1.90E-06 |
| SERPINF1 | 435.8532755 | -0.525735331 | 3.15E-03 |
| RPL5 | 35286.27166 | -0.525744681 | 1.78E-09 |
| RPL24 | 16195.05698 | -0.526099832 | 8.78E-07 |
| CACNB3 | 1731.645184 | -0.5261711 | 3.92E-05 |
| TEAD2 | 2552.04389 | -0.526571286 | 1.51E-04 |
| SLC35D1 | 878.4067731 | -0.526752009 | 7.71E-10 |
| SUMO2 | 6044.73372 | -0.526927758 | 5.70E-05 |
| ZNF326 | 1281.750808 | -0.526937739 | 2.76E-03 |
| ARF5 | 6943.705236 | -0.527147173 | 4.15E-04 |
| LEAP2 | 96.45515125 | -0.527183231 | 2.60E-02 |
| RBM4 | 5211.12219 | -0.527747754 | 8.34E-08 |
| SIPA1L2 | 2095.321232 | -0.527805463 | 1.58E-05 |
| SMC1A | 8629.188695 | -0.528480458 | 4.97E-06 |
| PI4KAP2 | 678.9268025 | -0.528802824 | 1.67E-08 |
| PKNOX2 | 537.5619923 | -0.529018734 | 1.46E-03 |
| PRR14 | 1269.197719 | -0.529094956 | 8.91E-06 |

| | | | |
|---|---|---|---|
| BHLHB9 | 316.3356222 | -0.529165248 | 4.43E-03 |
| BBS12 | 273.4155551 | -0.529200569 | 2.52E-04 |
| MIER3 | 2154.762199 | -0.529275839 | 1.64E-02 |
| SPIDR | 2126.955127 | -0.52931844 | 4.49E-06 |
| AC024075.2 | 79.91306537 | -0.529573466 | 4.48E-02 |
| CCDC85B | 1306.311525 | -0.529740511 | 8.78E-03 |
| MRPS26 | 2527.031429 | -0.529985649 | 3.99E-10 |
| ZNF107 | 246.4897489 | -0.530358981 | 2.40E-02 |
| WDR73 | 1944.452553 | -0.530472914 | 6.72E-10 |
| INTS5 | 1999.085661 | -0.530478255 | 7.93E-05 |
| ZNF691 | 543.1359512 | -0.531003508 | 1.75E-07 |
| NDE1 | 2540.9428 | -0.531373083 | 4.54E-08 |
| CPT1B | 640.0772637 | -0.531716955 | 6.97E-05 |
| KRI1 | 1878.252019 | -0.531758169 | 5.87E-14 |
| RPL3 | 72185.34694 | -0.532006015 | 1.44E-05 |
| AC002467.1 | 95.26917571 | -0.532010577 | 6.38E-03 |
| CCDC15 | 552.3203374 | -0.532389737 | 3.64E-02 |
| POLQ | 804.9903651 | -0.532491945 | 3.25E-02 |
| ARHGAP9 | 264.0659115 | -0.532695592 | 7.50E-03 |
| EVA1C | 367.5986656 | -0.532847506 | 1.00E-02 |
| DOT1L | 1539.184202 | -0.533047265 | 2.90E-06 |
| AFG3L1P | 627.678639 | -0.533133341 | 4.05E-05 |
| LY6G5B | 240.2465744 | -0.533289474 | 6.28E-03 |
| RMDN2 | 97.31966742 | -0.533371108 | 3.25E-02 |
| HECTD3 | 2870.664007 | -0.533680553 | 5.77E-03 |
| C12orf75 | 2759.517718 | -0.533709273 | 9.36E-08 |
| ACTB | 127290.7296 | -0.533792455 | 3.20E-03 |
| SLX4 | 472.5174403 | -0.533802785 | 1.58E-05 |
| RPA1 | 8159.632852 | -0.53381073 | 1.10E-06 |
| RPS13 | 31513.02511 | -0.533933047 | 4.56E-06 |
| TFPI2 | 2830.248721 | -0.533942618 | 1.49E-04 |
| DISC1 | 789.4212616 | -0.533995204 | 1.76E-07 |
| KMT2B | 2506.320285 | -0.534165155 | 1.66E-05 |
| DNAJC17 | 400.8681713 | -0.534306889 | 9.24E-05 |
| AC004816.1 | 101.0139062 | -0.53437147 | 3.41E-02 |
| PLCG1 | 10197.95981 | -0.534382414 | 2.13E-09 |
| PIDD1 | 1290.180358 | -0.534465762 | 1.16E-07 |
| ZNF574 | 810.6241985 | -0.534581427 | 8.77E-05 |
| PMF1-BGLAP | 147.0489674 | -0.534603929 | 2.37E-03 |
| NAB1 | 2310.147904 | -0.535505867 | 9.78E-05 |
| ABHD17B | 548.5468552 | -0.535544323 | 6.99E-05 |
| DOCK2 | 1547.341543 | -0.535808617 | 1.35E-03 |
| NMT2 | 1496.883168 | -0.535964131 | 3.40E-09 |
| AL137003.1 | 206.3088742 | -0.536013298 | 6.38E-03 |
| LINC00920 | 110.360093 | -0.536140702 | 1.81E-02 |
| PI4KA | 4563.049434 | -0.536247447 | 1.20E-12 |
| KMT2C | 4465.661044 | -0.536466971 | 6.45E-07 |

| | | | |
|---|---|---|---|
| MEX3B | 635.5673506 | -0.536469475 | 3.39E-03 |
| FAM156A | 1547.074356 | -0.53652816 | 1.05E-04 |
| SNX21 | 1691.955302 | -0.536777153 | 2.92E-03 |
| STK11IP | 891.7554413 | -0.537109762 | 1.16E-04 |
| DROSHA | 5892.566378 | -0.537412585 | 2.61E-12 |
| ILK | 12306.18143 | -0.537674239 | 1.30E-04 |
| NUDT15 | 1373.191692 | -0.537703122 | 1.92E-06 |
| NA | 75.43671028 | -0.53816587 | 2.62E-02 |
| BTBD2 | 6344.109791 | -0.538237598 | 1.28E-08 |
| APEX2 | 2147.285681 | -0.538984612 | 1.25E-05 |
| H2AFY | 11665.51269 | -0.539001008 | 5.55E-08 |
| PML | 4297.998273 | -0.539047012 | 6.09E-05 |
| NUTM2A | 89.88279148 | -0.539257773 | 1.82E-02 |
| DUBR | 562.7047495 | -0.54001485 | 1.52E-03 |
| PCMTD2 | 3440.324389 | -0.540060886 | 1.75E-05 |
| CKB | 5181.807168 | -0.540535662 | 1.50E-07 |
| GNAS | 41829.31542 | -0.540913554 | 1.03E-05 |
| ANXA11 | 5252.448292 | -0.54096758 | 2.04E-04 |
| AURKA | 3414.550899 | -0.541050144 | 4.04E-06 |
| TENM3-AS1 | 240.739707 | -0.541081103 | 3.20E-03 |
| ZNF41 | 598.0445322 | -0.541951925 | 8.35E-04 |
| TTC12 | 290.7928053 | -0.542816809 | 2.66E-02 |
| IL27RA | 1682.285814 | -0.542838396 | 6.38E-05 |
| LINC01089 | 367.5833516 | -0.542844581 | 1.04E-05 |
| PID1 | 439.8178603 | -0.543342092 | 2.70E-04 |
| FASN | 14089.9427 | -0.543393923 | 7.33E-04 |
| AC007406.5 | 157.0453592 | -0.543407433 | 9.54E-03 |
| THNSL1 | 293.0053922 | -0.543417834 | 1.35E-02 |
| ANLN | 11414.65258 | -0.543769168 | 1.93E-02 |
| QARS | 10369.56212 | -0.544125421 | 1.43E-04 |
| ING4 | 1112.173601 | -0.544175124 | 9.45E-05 |
| ODC1 | 5781.739414 | -0.5442782 | 5.97E-04 |
| SF3A1 | 5632.206611 | -0.544320224 | 7.30E-05 |
| CHKB-CPT1B | 289.6555874 | -0.544486108 | 2.40E-02 |
| FAM189B | 2509.143088 | -0.54460247 | 4.65E-07 |
| AL645608.2 | 67.27581695 | -0.544889766 | 4.50E-02 |
| DHX8 | 4969.121338 | -0.544918417 | 2.00E-05 |
| ILF3 | 17419.90848 | -0.544950534 | 2.46E-16 |
| RBM23 | 4441.381134 | -0.545094924 | 4.84E-09 |
| CSTF3 | 1829.155359 | -0.545260506 | 2.33E-06 |
| MSRB1 | 1234.914881 | -0.545391817 | 1.44E-05 |
| DUSP10 | 564.3339546 | -0.545519094 | 8.87E-03 |
| AL662795.2 | 323.8813937 | -0.54556355 | 2.22E-02 |
| TEN1-CDK3 | 280.7781373 | -0.545652507 | 4.01E-03 |
| CDH2 | 355.8677844 | -0.545944622 | 2.44E-02 |
| DMPK | 1237.876481 | -0.546072514 | 9.01E-08 |
| VIT | 826.8832205 | -0.546206138 | 9.13E-03 |

| | | | |
|---|---|---|---|
| TMCC1-AS1 | 380.6175541 | -0.546275207 | 4.42E-04 |
| CD3EAP | 786.1555899 | -0.546691217 | 2.45E-03 |
| INTS3 | 5212.447762 | -0.547257776 | 7.61E-10 |
| SZRD1 | 8977.152041 | -0.547322698 | 1.80E-07 |
| CBX5 | 9249.567894 | -0.547341043 | 1.85E-18 |
| PPM1G | 8237.17981 | -0.547631985 | 1.17E-11 |
| TAF5 | 278.1052842 | -0.547851162 | 7.14E-04 |
| HMGN3 | 1577.110129 | -0.548018885 | 2.71E-09 |
| SLC19A1 | 1452.445233 | -0.548120741 | 5.49E-04 |
| FAM193B | 1060.685105 | -0.548690819 | 6.58E-11 |
| CENPB | 6732.691902 | -0.548711681 | 3.17E-05 |
| OTX1 | 137.7147062 | -0.548972086 | 6.92E-03 |
| U47924.2 | 96.97950533 | -0.548985787 | 1.87E-02 |
| SMPD4 | 4560.855375 | -0.548995434 | 2.34E-06 |
| AC022400.8 | 288.4064756 | -0.549025869 | 4.94E-04 |
| AC093484.3 | 9569.147076 | -0.549582182 | 3.17E-05 |
| RDM1 | 181.5823242 | -0.549716773 | 1.87E-03 |
| CCAR1 | 2806.344201 | -0.549968791 | 1.83E-05 |
| MZT2A | 1203.994707 | -0.550295702 | 6.59E-05 |
| TTC30A | 485.3279646 | -0.550564562 | 3.63E-05 |
| CYHR1 | 3198.186788 | -0.550567771 | 2.06E-05 |
| ACTG1 | 90431.48433 | -0.550868705 | 9.25E-03 |
| SSBP3 | 3859.235303 | -0.5509484 | 3.29E-06 |
| CDR2L | 743.6601986 | -0.551253206 | 3.13E-07 |
| LRIG1 | 1253.713048 | -0.551575096 | 1.62E-08 |
| BZW2 | 4791.617903 | -0.551613716 | 2.06E-10 |
| ZNF224 | 508.5609643 | -0.551697585 | 1.87E-03 |
| RPS14 | 16221.32272 | -0.551819866 | 3.53E-04 |
| AC243964.2 | 208.6143951 | -0.551855516 | 4.63E-03 |
| PROSER3 | 399.7278117 | -0.551858164 | 2.05E-06 |
| KLF5 | 608.4056549 | -0.552045073 | 4.40E-02 |
| DDX25 | 1351.778353 | -0.552207079 | 2.37E-11 |
| NA | 94.5777297 | -0.552213618 | 4.13E-03 |
| C8orf44 | 173.4833532 | -0.55224385 | 6.06E-03 |
| FKBP5 | 2560.809448 | -0.552252559 | 4.78E-14 |
| HADH | 2162.047075 | -0.552263541 | 5.68E-10 |
| FCHSD2 | 1863.422345 | -0.55227234 | 3.94E-12 |
| TSPYL4 | 3260.739482 | -0.552286323 | 1.79E-08 |
| TDP1 | 1043.60661 | -0.552394525 | 2.24E-08 |
| HECW1 | 564.3576593 | -0.552565183 | 1.24E-03 |
| PAK4 | 2072.569239 | -0.55258059 | 1.80E-06 |
| MELTF-AS1 | 84.58329469 | -0.55299455 | 4.18E-02 |
| PIK3R1 | 4595.722417 | -0.553034134 | 2.41E-06 |
| PSPC1 | 1501.329787 | -0.553095361 | 9.93E-07 |
| HPS3 | 6068.142851 | -0.553155138 | 6.33E-05 |
| KCNJ14 | 169.2243641 | -0.553322302 | 6.90E-03 |
| RBBP4 | 6433.30477 | -0.553567898 | 5.16E-07 |

| | | | |
|---|---|---|---|
| SVBP | 540.4729975 | -0.553603773 | 4.61E-05 |
| ZBTB18 | 811.3954707 | -0.553767809 | 7.51E-07 |
| BOLA1 | 358.734382 | -0.553808745 | 5.68E-06 |
| NCAPD3 | 6605.075933 | -0.554260449 | 4.28E-06 |
| ERI1 | 2311.186083 | -0.554290997 | 1.08E-05 |
| RPL23 | 29548.49268 | -0.554297471 | 1.65E-08 |
| PNKP | 1324.117395 | -0.554529958 | 9.31E-08 |
| NAAA | 534.8626594 | -0.554689639 | 5.41E-08 |
| PYGL | 5642.226534 | -0.554916258 | 1.59E-20 |
| ZNF445 | 2350.177662 | -0.555002399 | 1.94E-13 |
| CNIH3 | 994.9039675 | -0.555161165 | 5.05E-04 |
| KBTBD7 | 448.7049977 | -0.555334945 | 2.94E-04 |
| RAPGEFL1 | 411.4915503 | -0.556023599 | 1.80E-04 |
| HSD3B7 | 136.8593015 | -0.556035644 | 4.04E-02 |
| WNK4 | 1114.048769 | -0.556259433 | 1.38E-04 |
| RHOU | 327.4778719 | -0.556771077 | 2.14E-02 |
| TRAF3IP2 | 955.3632333 | -0.556929962 | 1.86E-05 |
| RPL18A | 25731.2536 | -0.556974195 | 1.87E-05 |
| MT-ND3 | 19371.55593 | -0.557184818 | 2.91E-04 |
| PARPBP | 708.9645397 | -0.557190882 | 2.24E-03 |
| NHEJ1 | 722.382986 | -0.557789055 | 8.52E-05 |
| FOXA3 | 158.3876836 | -0.557889229 | 1.54E-03 |
| RRP8 | 1549.173165 | -0.558042083 | 2.30E-06 |
| COCH | 812.7057441 | -0.558279727 | 1.09E-05 |
| ZBED5 | 1813.585829 | -0.558295556 | 9.85E-04 |
| DFFB | 269.6359483 | -0.558388493 | 5.57E-04 |
| RHPN1 | 347.3131214 | -0.558564824 | 2.42E-03 |
| PPP2R1A | 11814.77005 | -0.55898172 | 5.59E-04 |
| SUOX | 1358.723531 | -0.559288433 | 1.27E-03 |
| MSANTD2 | 917.7133361 | -0.560164027 | 2.00E-05 |
| ALDH1A3 | 299.1188345 | -0.560193967 | 6.73E-03 |
| AC040162.1 | 268.8290852 | -0.560375829 | 3.87E-02 |
| UBL7 | 4320.443653 | -0.560380318 | 3.35E-05 |
| RTL8C | 5952.977431 | -0.560768532 | 9.81E-04 |
| HDAC3 | 3441.016786 | -0.560870263 | 2.18E-06 |
| SNRPD2 | 8421.120681 | -0.560906618 | 4.44E-05 |
| VWA1 | 1340.086787 | -0.561081144 | 5.42E-03 |
| DNM3 | 456.6554895 | -0.561656907 | 6.05E-05 |
| AL157392.3 | 363.3674805 | -0.561936227 | 1.82E-04 |
| SCN4B | 2747.623872 | -0.562178646 | 1.38E-02 |
| AC138035.1 | 323.6251374 | -0.562312387 | 8.82E-04 |
| DNMT1 | 10550.99223 | -0.562413866 | 2.85E-05 |
| ANKRD20A8P | 247.7582674 | -0.562662731 | 2.12E-02 |
| POLD2 | 6626.569676 | -0.562694084 | 8.43E-13 |
| MAGEA1 | 1398.25761 | -0.562815041 | 7.49E-08 |
| ZNF511 | 737.2524609 | -0.562989107 | 6.19E-05 |
| SCARNA12 | 64.51896559 | -0.563065771 | 2.40E-02 |

| | | | |
|---|---|---|---|
| FAAP20 | 1311.256067 | -0.563538876 | 5.64E-05 |
| HENMT1 | 1022.022023 | -0.563602509 | 9.53E-07 |
| EFCAB2 | 314.3175812 | -0.563760283 | 1.34E-03 |
| WHRN | 950.7847616 | -0.563900328 | 3.56E-07 |
| HOXC11 | 245.5107089 | -0.564202146 | 6.07E-04 |
| DSCR4 | 114.9837457 | -0.564688292 | 1.92E-02 |
| HPS6 | 665.5373466 | -0.564846698 | 1.61E-03 |
| NOD1 | 846.3666035 | -0.565428933 | 1.01E-08 |
| NENF | 1359.970011 | -0.565511077 | 7.05E-08 |
| RSRP1 | 1513.468845 | -0.565609529 | 2.44E-03 |
| RPL12 | 40443.63764 | -0.565718205 | 1.85E-04 |
| ANKRD54 | 1353.749219 | -0.56576093 | 1.04E-10 |
| CYB5D1 | 624.0858821 | -0.566001188 | 1.05E-04 |
| PPOX | 774.9046767 | -0.56647464 | 8.89E-06 |
| UBE2E3 | 2974.808693 | -0.566611407 | 1.18E-11 |
| AP003390.1 | 87.82873925 | -0.566641017 | 2.76E-02 |
| NSD2 | 9039.317095 | -0.566840903 | 7.38E-13 |
| MOB3A | 1958.109916 | -0.566905324 | 4.81E-06 |
| BAIAP2-DT | 1190.829781 | -0.566945073 | 1.27E-05 |
| FZD2 | 1127.496047 | -0.567055002 | 5.22E-05 |
| CPOX | 969.5245559 | -0.567616783 | 4.53E-05 |
| CDK3 | 153.5450677 | -0.567652783 | 2.42E-02 |
| CDK5 | 1685.224274 | -0.567675483 | 1.01E-05 |
| MT-CYB | 107218.2105 | -0.56767792 | 3.55E-03 |
| ELF2 | 1626.106279 | -0.56789494 | 2.10E-16 |
| ZNF668 | 589.6378144 | -0.568033941 | 1.03E-03 |
| HDX | 246.3347493 | -0.568118402 | 8.34E-03 |
| SLBP | 2440.223261 | -0.568198348 | 1.59E-03 |
| AC003665.1 | 447.9924771 | -0.568471721 | 1.05E-02 |
| AC126755.1 | 915.0766034 | -0.568561733 | 5.28E-04 |
| AL137003.2 | 405.2240052 | -0.568727438 | 1.92E-03 |
| PRKCZ-AS1 | 101.6789457 | -0.568922227 | 3.72E-03 |
| AC136475.2 | 116.3715542 | -0.568992016 | 4.77E-03 |
| AC139887.2 | 179.702569 | -0.569004438 | 3.42E-04 |
| MCRS1 | 3164.332928 | -0.569267889 | 3.38E-06 |
| GYG1 | 2317.222666 | -0.569295596 | 1.94E-05 |
| LINC01029 | 2160.242943 | -0.569307126 | 3.25E-08 |
| HNRNPL | 11795.62872 | -0.569644036 | 5.02E-11 |
| SF3A3 | 6221.122994 | -0.569826619 | 2.28E-03 |
| C17orf97 | 179.3364957 | -0.569993277 | 3.80E-03 |
| SETMAR | 386.4169727 | -0.570016876 | 1.36E-07 |
| TIE1 | 3876.156317 | -0.570112562 | 1.95E-05 |
| KDM2B | 1959.038517 | -0.570343264 | 1.26E-05 |
| ASGR1 | 173.8182471 | -0.570402579 | 4.27E-03 |
| AL031282.1 | 233.3734107 | -0.570607548 | 1.31E-04 |
| FAM160A2 | 1211.307237 | -0.57075077 | 5.54E-08 |
| AMPD2 | 4404.75949 | -0.570831991 | 6.39E-06 |

| | | | |
|---|---|---|---|
| URB2 | 1780.506995 | -0.570861755 | 4.60E-13 |
| AAAS | 2391.296236 | -0.570929519 | 2.55E-06 |
| RARA-AS1 | 119.2043537 | -0.571037423 | 7.14E-03 |
| NAA80 | 540.0798404 | -0.571110113 | 3.13E-04 |
| RPL39 | 26083.00633 | -0.571239501 | 1.66E-06 |
| PTBP2 | 289.7813242 | -0.57124069 | 3.13E-02 |
| CTDSPL2 | 1007.377581 | -0.571407962 | 8.02E-03 |
| TTYH2 | 540.5647135 | -0.571474973 | 1.22E-02 |
| GBP2 | 578.7100366 | -0.571601783 | 3.03E-03 |
| LIX1L | 2287.312237 | -0.571843207 | 1.21E-11 |
| NA | 104.0017355 | -0.572069293 | 1.94E-02 |
| AC016394.2 | 100.9367242 | -0.572179992 | 8.76E-03 |
| HMGA2 | 5164.377752 | -0.572270927 | 1.57E-04 |
| MPZL3 | 518.1093127 | -0.5725261 | 1.53E-04 |
| BID | 1337.216014 | -0.572600905 | 7.93E-04 |
| AP000892.3 | 436.2274301 | -0.572786734 | 1.03E-02 |
| NA | 115.5666921 | -0.572936081 | 1.30E-02 |
| VPS51 | 3467.254109 | -0.572967969 | 1.16E-03 |
| ATRNL1 | 133.6598312 | -0.573030163 | 1.95E-02 |
| EIF5A | 21651.00118 | -0.573206814 | 8.09E-12 |
| NUP188 | 7233.696582 | -0.573403027 | 8.92E-21 |
| ZYG11A | 209.58949 | -0.573890704 | 8.17E-03 |
| ZNF443 | 150.5584489 | -0.57422671 | 5.36E-03 |
| MEIS1 | 435.9633765 | -0.574398994 | 8.00E-03 |
| CBR3 | 396.1745623 | -0.574502044 | 1.07E-02 |
| RABEP2 | 542.7569312 | -0.574632242 | 4.85E-05 |
| DNM1P34 | 71.78782101 | -0.574667657 | 4.40E-02 |
| STAT6 | 5677.333921 | -0.574742853 | 1.06E-06 |
| PPIAP22 | 335.0730773 | -0.574796846 | 1.17E-02 |
| AC118282.4 | 232.0794676 | -0.575093402 | 1.46E-02 |
| F8A1 | 1206.55352 | -0.575547882 | 2.57E-04 |
| RPUSD4 | 2469.189106 | -0.575694094 | 1.05E-08 |
| BOP1 | 3358.256854 | -0.576117179 | 1.36E-08 |
| AP001453.4 | 83.2107295 | -0.576119491 | 7.01E-03 |
| MAGEE1 | 540.1230386 | -0.576204751 | 8.89E-03 |
| BAG1 | 1688.379934 | -0.576211566 | 9.74E-07 |
| RPL10 | 91368.98784 | -0.576851292 | 5.50E-07 |
| SUGT1P4-STRA | 208.5434078 | -0.576884437 | 1.47E-03 |
| ADORA2A-AS1 | 66.80349087 | -0.576926123 | 4.45E-02 |
| DPCD | 538.2956527 | -0.576941284 | 9.76E-05 |
| AP4M1 | 1444.153207 | -0.576980211 | 5.25E-07 |
| HDAC1 | 6926.946715 | -0.577164669 | 4.04E-09 |
| MCMBP | 2516.238275 | -0.577253731 | 4.16E-07 |
| CCDC88C | 617.6493224 | -0.577320122 | 2.77E-05 |
| AC133065.3 | 74.7330206 | -0.577623038 | 1.40E-02 |
| AC104581.4 | 154.1621748 | -0.577899713 | 1.27E-02 |
| NPDC1 | 1418.160007 | -0.578070494 | 4.18E-04 |

| | | | |
|---|---|---|---|
| PPME1 | 11264.96145 | -0.578079882 | 3.41E-07 |
| RPS21 | 23282.49507 | -0.578246571 | 2.28E-07 |
| NA | 83.64413233 | -0.57845034 | 1.75E-02 |
| ABCC6 | 487.0257148 | -0.578740403 | 1.51E-03 |
| LRFN3 | 657.5490135 | -0.579385371 | 7.99E-05 |
| ZNF10 | 240.692249 | -0.579451333 | 4.77E-02 |
| DNAAF3 | 180.1413094 | -0.580525963 | 2.89E-03 |
| YWHAH | 4738.809844 | -0.580767633 | 8.22E-05 |
| DOCK4 | 1431.176833 | -0.580798253 | 7.94E-06 |
| EEPD1 | 580.6320043 | -0.58146396 | 2.55E-03 |
| TMSB10 | 36922.00469 | -0.582093639 | 3.66E-05 |
| AC010422.6 | 130.2044948 | -0.582157841 | 2.80E-03 |
| ZNF75D | 1009.175721 | -0.582356911 | 2.56E-06 |
| MDC1 | 3115.594994 | -0.582457347 | 3.13E-10 |
| TPI1 | 33065.67208 | -0.582723046 | 9.76E-04 |
| RPS19 | 28332.18881 | -0.582886642 | 3.13E-08 |
| ZBTB9 | 785.6997796 | -0.582965912 | 8.12E-06 |
| CTCF | 3014.64559 | -0.58334523 | 6.65E-20 |
| AC010761.1 | 252.8642833 | -0.583493038 | 2.10E-03 |
| ARMC9 | 2187.710154 | -0.583695979 | 7.18E-10 |
| PODXL2 | 4334.400803 | -0.583728183 | 7.81E-08 |
| SHC3 | 593.2962704 | -0.583841941 | 3.18E-07 |
| CLK1 | 1374.728739 | -0.583996359 | 8.22E-03 |
| EPHB2 | 2544.09388 | -0.584076819 | 9.45E-09 |
| MCM6 | 6117.317718 | -0.584233404 | 4.98E-04 |
| KLHL23 | 726.4414864 | -0.584378134 | 7.07E-05 |
| AC073896.4 | 110.457009 | -0.584580409 | 3.36E-03 |
| PRMT2 | 3752.41591 | -0.584586957 | 2.96E-07 |
| ICK | 878.5792377 | -0.584694784 | 3.04E-06 |
| PLPP3 | 2030.88115 | -0.585054091 | 9.45E-07 |
| RPS24 | 15668.6721 | -0.585241933 | 1.38E-09 |
| C8orf48 | 96.08804137 | -0.585281717 | 1.65E-02 |
| INAFM2 | 266.0135533 | -0.585308537 | 2.10E-04 |
| APEX1 | 6343.691188 | -0.585561451 | 6.75E-07 |
| FGFR1OP | 667.7562339 | -0.585718577 | 3.38E-04 |
| SRGAP3 | 587.6223722 | -0.585796316 | 1.61E-05 |
| CCDC57 | 916.7750703 | -0.585952053 | 7.43E-06 |
| BVES | 1349.384071 | -0.586373339 | 1.40E-12 |
| RUVBL2 | 6983.216087 | -0.58651288 | 3.38E-12 |
| RAB40C | 1074.092055 | -0.586553886 | 1.14E-05 |
| ZNF214 | 149.0621941 | -0.586715369 | 4.80E-02 |
| CYTH2 | 1564.392588 | -0.587283427 | 1.05E-07 |
| TWNK | 1717.423571 | -0.587329579 | 2.92E-07 |
| ZNF662 | 318.1287865 | -0.587400533 | 1.31E-02 |
| PRKCD | 1783.577947 | -0.587654667 | 2.55E-09 |
| SERPINA5 | 1276.950546 | -0.587685117 | 1.80E-03 |
| NUDT16L1 | 643.9989348 | -0.587723299 | 3.77E-05 |

| | | | |
|---|---|---|---|
| AL353194.1 | 66.22251447 | -0.587949929 | 2.74E-02 |
| HOXB2 | 845.4751948 | -0.588055235 | 9.38E-06 |
| N4BP3 | 111.414007 | -0.588106225 | 3.32E-02 |
| TFDP1 | 3616.317136 | -0.588289725 | 6.37E-08 |
| PCSK1 | 1402.5926 | -0.588595603 | 7.59E-03 |
| ZNF75A | 1038.680233 | -0.58883728 | 7.24E-08 |
| MT-ATP6 | 93267.30604 | -0.58895542 | 2.59E-03 |
| DDT | 2299.728872 | -0.588978982 | 1.42E-07 |
| TUBB2A | 6585.09119 | -0.588996303 | 2.90E-03 |
| SELENOO | 1094.121601 | -0.589112347 | 2.50E-05 |
| AC046185.3 | 1008.471374 | -0.589339447 | 7.21E-12 |
| CCDC78 | 384.8868074 | -0.589396217 | 1.86E-04 |
| USP5 | 5161.921228 | -0.590241771 | 1.95E-06 |
| CHN1 | 893.6230543 | -0.590396283 | 3.18E-09 |
| LINC01238 | 72.75569433 | -0.590397144 | 1.76E-02 |
| BAHD1 | 1473.206705 | -0.590467205 | 2.25E-06 |
| PRORSD1P | 101.8748698 | -0.59049416 | 3.67E-02 |
| THOC1 | 1197.632741 | -0.590506472 | 5.95E-03 |
| METTL3 | 3919.669917 | -0.590510053 | 4.19E-18 |
| PUS7 | 1767.96461 | -0.590567837 | 3.13E-07 |
| NRSN2 | 5123.912099 | -0.590654826 | 3.78E-08 |
| RPS3 | 78001.32135 | -0.590877707 | 2.07E-09 |
| LZTR1 | 4101.887411 | -0.590907639 | 9.79E-06 |
| NA | 595.4944113 | -0.591034971 | 1.65E-10 |
| CFL1 | 47571.41666 | -0.591443903 | 7.39E-07 |
| ATAT1 | 537.2356922 | -0.591453055 | 5.96E-11 |
| PTPRN | 673.3985235 | -0.591758711 | 9.01E-03 |
| GNAI2 | 15461.96579 | -0.591807422 | 2.43E-05 |
| IRF2BP1 | 1114.399218 | -0.59183874 | 1.93E-06 |
| ARHGAP1 | 3525.834389 | -0.592015705 | 3.20E-08 |
| ZSWIM9 | 585.2098536 | -0.59209155 | 2.89E-07 |
| DHODH | 999.3050469 | -0.592142266 | 4.37E-08 |
| AF254983.1 | 196.670656 | -0.592311862 | 2.02E-02 |
| RPS11 | 38271.47838 | -0.59250666 | 5.94E-05 |
| PLLP | 455.6051462 | -0.592579281 | 4.83E-07 |
| NOTCH4 | 160.9558428 | -0.592679251 | 3.02E-03 |
| AC100810.1 | 387.3119823 | -0.592869855 | 6.32E-05 |
| KIF26A | 548.4895354 | -0.592897301 | 1.17E-04 |
| GPD1L | 1340.603692 | -0.592943934 | 1.46E-07 |
| ANKRD13D | 1126.817695 | -0.593037553 | 1.68E-07 |
| TMEM39B | 1129.7264 | -0.593267512 | 1.52E-09 |
| AP5S1 | 887.952493 | -0.594403935 | 1.07E-05 |
| AC139530.2 | 288.0852949 | -0.594413235 | 8.29E-05 |
| VASH1 | 4125.770126 | -0.594481677 | 5.45E-05 |
| DIPK2A | 160.7504337 | -0.594539838 | 1.01E-02 |
| NEIL1 | 519.4268325 | -0.594555731 | 1.10E-03 |
| MTA1 | 4030.972488 | -0.594592982 | 1.18E-12 |

| | | | |
|---|---|---|---|
| HDGFL2 | 2478.722792 | -0.594787662 | 1.33E-08 |
| ZNF362 | 968.369252 | -0.594791709 | 4.16E-03 |
| DCTN3 | 2523.316974 | -0.5949811 | 3.20E-07 |
| KIAA1586 | 323.4141024 | -0.595117597 | 6.84E-03 |
| AC008764.6 | 91.36670583 | -0.595127349 | 8.74E-03 |
| HCFC1 | 8795.74916 | -0.595371472 | 1.01E-07 |
| FRMD6-AS1 | 136.6120981 | -0.595464057 | 1.03E-02 |
| ACVR2B | 614.6329055 | -0.596428633 | 3.73E-06 |
| C8orf82 | 970.646144 | -0.596583471 | 4.90E-06 |
| PNRC2 | 3356.074831 | -0.596761783 | 3.15E-07 |
| AL512353.1 | 63.48155036 | -0.596771209 | 2.79E-02 |
| FAIM | 348.3492391 | -0.596823833 | 2.70E-06 |
| AC026362.1 | 273.9551125 | -0.596970718 | 4.90E-03 |
| SIPA1 | 2309.882455 | -0.597240275 | 9.23E-05 |
| FAM161A | 386.9162848 | -0.59785082 | 3.24E-02 |
| ALDH16A1 | 1223.646242 | -0.598001208 | 1.30E-05 |
| TET2 | 1173.805508 | -0.598262573 | 2.38E-05 |
| GINS4 | 1896.938065 | -0.598592108 | 1.55E-04 |
| EP400P1 | 318.6927657 | -0.598625207 | 3.28E-03 |
| PRRX2 | 217.4448498 | -0.599119695 | 1.14E-02 |
| UBTD1 | 536.0145794 | -0.599522395 | 4.77E-05 |
| DDX46 | 3171.764144 | -0.599708078 | 6.43E-04 |
| RMI1 | 821.7678593 | -0.599847399 | 3.43E-03 |
| CEP85L | 295.6479056 | -0.599960411 | 1.26E-02 |
| KCTD21 | 734.3641873 | -0.600223866 | 2.01E-04 |
| LAS1L | 2829.020455 | -0.600275233 | 1.51E-12 |
| ZNF618 | 3843.333358 | -0.600424698 | 4.68E-11 |
| ECT2 | 1973.1338 | -0.600723994 | 9.54E-03 |
| ASH1L-AS1 | 95.00739717 | -0.600761013 | 1.79E-02 |
| HERC2P9 | 445.6592513 | -0.600832652 | 3.27E-05 |
| RGS17 | 116.2456555 | -0.60095332 | 2.42E-02 |
| TOP1MT | 1570.636353 | -0.601448855 | 5.46E-17 |
| AL133372.2 | 107.2268163 | -0.601846441 | 4.98E-03 |
| MAPKBP1 | 1283.789127 | -0.602222089 | 4.10E-07 |
| AMACR | 1358.410533 | -0.602986526 | 1.45E-25 |
| FAM227A | 248.8095749 | -0.603002797 | 1.43E-02 |
| TRIM9 | 249.1688506 | -0.603241269 | 2.45E-03 |
| SPATA13 | 828.620463 | -0.603359385 | 1.48E-03 |
| EED | 1595.744461 | -0.603484129 | 1.44E-06 |
| ATP1B1 | 7211.224436 | -0.603697593 | 2.51E-05 |
| SH3D21 | 340.8390984 | -0.603739627 | 2.29E-02 |
| PTOV1-AS1 | 556.2368947 | -0.603945547 | 1.73E-06 |
| WWTR1 | 6889.014365 | -0.603963701 | 1.72E-16 |
| ABCB10 | 750.3101236 | -0.604084832 | 1.03E-05 |
| EXOSC4 | 1304.84018 | -0.604407878 | 3.89E-07 |
| RRN3P2 | 93.46092709 | -0.604853183 | 1.53E-02 |
| SALL2 | 1175.392959 | -0.60489122 | 2.14E-03 |

| | | | |
|---|---|---|---|
| KCNIP3 | 568.7111685 | -0.605150435 | 1.13E-04 |
| DPM3 | 388.6405479 | -0.60520892 | 7.79E-04 |
| AC132812.1 | 80.9606679 | -0.605374535 | 2.39E-02 |
| MCM3AP-AS1 | 539.3267377 | -0.605811554 | 8.92E-06 |
| RBM43 | 426.6895759 | -0.60595022 | 5.63E-06 |
| RBM12B | 1824.034248 | -0.606389615 | 8.92E-04 |
| USP49 | 781.7022822 | -0.606402757 | 2.03E-07 |
| POLR1E | 2520.906887 | -0.606695976 | 2.73E-13 |
| BATF2 | 455.2453915 | -0.606746653 | 1.81E-03 |
| FIGNL1 | 1385.407933 | -0.606970707 | 8.21E-04 |
| TUSC3 | 4463.581761 | -0.607136565 | 2.12E-24 |
| CALM1 | 7582.069497 | -0.607168043 | 1.35E-15 |
| IRAK1BP1 | 289.2275019 | -0.607578604 | 9.70E-04 |
| BTBD19 | 501.2512357 | -0.607626072 | 2.45E-06 |
| PCBD2 | 158.4686784 | -0.607908342 | 5.21E-03 |
| ARHGAP24 | 86.39537728 | -0.607933991 | 1.39E-02 |
| TAF4 | 859.8748059 | -0.608374104 | 2.42E-06 |
| ASPHD1 | 246.0080628 | -0.60851612 | 1.12E-05 |
| CCDC191 | 86.06487863 | -0.608517722 | 4.41E-02 |
| RCBTB2 | 954.637502 | -0.608676593 | 9.36E-07 |
| RPS26 | 3828.699601 | -0.608787448 | 4.19E-07 |
| RTEL1-TNFRSF6 | 821.9011062 | -0.608957414 | 1.61E-04 |
| ZNF341 | 271.7597439 | -0.609188888 | 2.30E-03 |
| GTF3C6 | 1875.255331 | -0.609704958 | 1.40E-08 |
| FAM229A | 264.9096072 | -0.609718919 | 1.73E-03 |
| TCEAL1 | 563.5790865 | -0.609763399 | 5.35E-05 |
| WNT6 | 159.8552245 | -0.610186934 | 3.36E-02 |
| FZR1 | 2317.725477 | -0.61056167 | 1.17E-05 |
| ARID3A | 498.0095121 | -0.611078853 | 8.26E-04 |
| AL671710.1 | 120.4166884 | -0.611237336 | 1.12E-02 |
| ANP32E | 1408.613357 | -0.612430796 | 5.68E-04 |
| NEK2 | 1071.988946 | -0.612568909 | 3.82E-03 |
| HMG20B | 2798.243651 | -0.612832346 | 3.20E-08 |
| NA | 101.4348357 | -0.613271785 | 3.11E-03 |
| THRB | 889.0566991 | -0.613353342 | 3.09E-07 |
| CNTROB | 2651.404963 | -0.613568212 | 2.22E-11 |
| DUT | 1804.676887 | -0.61361571 | 2.02E-07 |
| KPTN | 754.5468126 | -0.614152834 | 8.36E-05 |
| RAN | 15303.38853 | -0.614240885 | 3.97E-11 |
| TATDN2 | 2765.254714 | -0.614597135 | 2.56E-07 |
| ZNF418 | 240.3467327 | -0.614706612 | 3.71E-03 |
| FAM53B | 1491.038315 | -0.614907022 | 8.79E-10 |
| DHFRP1 | 909.4746823 | -0.614911947 | 3.51E-04 |
| CEP192 | 2010.196164 | -0.6150013 | 4.09E-05 |
| AL121832.2 | 88.90306509 | -0.615286693 | 1.78E-02 |
| GSTT2B | 1091.957125 | -0.615349935 | 1.56E-07 |
| GALNT11 | 2503.820462 | -0.61591769 | 1.65E-05 |

| | | | |
|---|---|---|---|
| AQP11 | 115.1563059 | -0.616321276 | 6.17E-03 |
| CCDC32 | 725.8717716 | -0.616440601 | 3.66E-04 |
| SMIM24 | 186.2624921 | -0.61656978 | 3.34E-02 |
| AC012435.1 | 67.2756116 | -0.61664933 | 1.45E-02 |
| VPS37D | 253.6964543 | -0.616650289 | 3.09E-03 |
| TUBGCP6 | 2476.919981 | -0.616998071 | 4.42E-06 |
| NMRAL1 | 1468.372654 | -0.617235906 | 9.78E-08 |
| PTOV1 | 4290.183077 | -0.617449805 | 8.64E-04 |
| NFYA | 1791.337203 | -0.617478841 | 2.12E-12 |
| GIPC3 | 892.0876321 | -0.617769197 | 2.03E-04 |
| GEMIN4 | 3368.788003 | -0.618010115 | 8.17E-07 |
| ASPRV1 | 67.17027102 | -0.618070597 | 2.97E-02 |
| RPL7P9 | 127.9709629 | -0.618125758 | 4.76E-03 |
| RCC1 | 5720.614234 | -0.61858768 | 1.91E-10 |
| AL450998.2 | 172.8671453 | -0.619035414 | 2.39E-04 |
| PAICS | 10941.13924 | -0.619172938 | 2.10E-08 |
| WDR77 | 2537.187295 | -0.619344438 | 4.84E-11 |
| CD207 | 40.47290139 | -0.619843478 | 4.72E-02 |
| GGT5 | 898.8201285 | -0.619877248 | 5.25E-03 |
| CD274 | 323.6243598 | -0.619900332 | 7.42E-03 |
| TUBB | 73676.804 | -0.621294675 | 2.38E-04 |
| AL928654.2 | 163.3762937 | -0.621410557 | 1.82E-04 |
| ZNF660 | 284.1719379 | -0.621565098 | 1.97E-02 |
| RFX5 | 4160.592771 | -0.621859086 | 1.02E-21 |
| CLASRP | 1076.251792 | -0.621968355 | 6.13E-11 |
| ZNRF2 | 314.0326182 | -0.622331604 | 1.26E-03 |
| SCD5 | 399.6806143 | -0.622912266 | 3.54E-05 |
| ZNF180 | 312.6482421 | -0.623559545 | 8.69E-03 |
| RBM4B | 1235.024212 | -0.624092042 | 1.20E-04 |
| ITPKA | 85.58454827 | -0.624207647 | 3.12E-02 |
| GTF2I | 19621.1412 | -0.624224071 | 3.87E-12 |
| LTBP3 | 6500.71372 | -0.62435633 | 1.01E-06 |
| MAPK12 | 3334.757022 | -0.624777248 | 4.37E-10 |
| CD6 | 246.0228803 | -0.625383404 | 3.74E-02 |
| AL627309.6 | 103.0867455 | -0.62559671 | 2.63E-02 |
| LDB2 | 2892.946269 | -0.62582967 | 1.14E-07 |
| ST8SIA6 | 68.51548135 | -0.625941378 | 3.65E-02 |
| MLPH | 3090.876894 | -0.626156886 | 4.88E-05 |
| WDR6 | 10006.27012 | -0.626731058 | 1.02E-04 |
| GLIPR1 | 1112.924389 | -0.626764679 | 9.67E-06 |
| TSEN2 | 547.339728 | -0.626860687 | 1.23E-05 |
| TRMU | 1734.138226 | -0.626884895 | 1.88E-20 |
| AC241952.1 | 387.9923078 | -0.627077941 | 1.95E-06 |
| MASTL | 798.0087179 | -0.627182563 | 3.35E-03 |
| TCTEX1D2 | 229.6803877 | -0.628005539 | 1.74E-05 |
| AL928654.1 | 326.5701445 | -0.628231055 | 2.34E-05 |
| HCG27 | 82.53773153 | -0.628837431 | 2.92E-02 |

| | | | |
|---|---|---|---|
| STON2 | 213.0074989 | -0.629132212 | 6.29E-03 |
| MAP3K7CL | 150.299139 | -0.629373161 | 3.59E-03 |
| POTEH | 157.7783814 | -0.629551536 | 1.88E-02 |
| MDK | 24505.83644 | -0.630363336 | 2.16E-04 |
| TCIRG1 | 2216.590153 | -0.630404431 | 1.45E-08 |
| AL161756.1 | 48.76356671 | -0.630526532 | 2.60E-02 |
| CTNNBIP1 | 1107.567502 | -0.630925878 | 8.20E-06 |
| XRCC1 | 1748.371832 | -0.631268276 | 2.22E-11 |
| AUTS2 | 1479.629524 | -0.631543365 | 6.78E-06 |
| AC002310.6 | 87.54832053 | -0.631671691 | 2.29E-02 |
| FUT8 | 3274.135694 | -0.631743486 | 7.72E-12 |
| LTA4H | 4008.592376 | -0.631862682 | 1.24E-07 |
| AC008969.1 | 361.7441285 | -0.631871031 | 2.35E-07 |
| CPSF1 | 6411.044963 | -0.632161566 | 3.17E-13 |
| PLEKHJ1 | 2128.603738 | -0.632208947 | 1.04E-09 |
| POLA1 | 2878.667761 | -0.632404017 | 2.12E-07 |
| AC002310.1 | 144.0995327 | -0.632920069 | 2.15E-04 |
| USF1 | 2130.590853 | -0.632990391 | 1.41E-09 |
| ALYREF | 2230.458412 | -0.633269406 | 1.29E-06 |
| NA | 205.8975033 | -0.633714712 | 6.25E-05 |
| TRPM2 | 1069.765725 | -0.633737201 | 5.86E-05 |
| PDCD4 | 2431.171143 | -0.633818465 | 4.15E-06 |
| MAGED4 | 331.0548708 | -0.634034567 | 3.01E-02 |
| TSPYL1 | 2490.20289 | -0.634155453 | 9.48E-09 |
| ZCCHC3 | 3514.471382 | -0.634158849 | 1.68E-09 |
| TXNDC16 | 735.2004753 | -0.63439481 | 2.60E-08 |
| HERC5 | 869.4560916 | -0.634728693 | 4.83E-06 |
| ACSL4 | 4513.913578 | -0.634831645 | 1.99E-07 |
| IQCC | 379.3166359 | -0.635138936 | 1.44E-05 |
| AP000787.1 | 90.97820889 | -0.635751626 | 4.03E-02 |
| ARHGDIB | 757.048678 | -0.635813515 | 4.74E-03 |
| C19orf48 | 2555.273303 | -0.636664224 | 5.73E-14 |
| AC078883.1 | 75.30345687 | -0.636972349 | 1.63E-02 |
| AC012510.1 | 46.483895 | -0.636991514 | 3.67E-02 |
| NA | 707.1082302 | -0.63707965 | 1.97E-10 |
| AC004812.2 | 138.0782283 | -0.637210397 | 1.64E-04 |
| BCORL1 | 1237.78771 | -0.637308905 | 8.40E-07 |
| H2AFV | 7403.940441 | -0.637597725 | 4.97E-16 |
| COQ4 | 1733.716167 | -0.637768672 | 6.28E-13 |
| RPUSD1 | 1588.871539 | -0.637939181 | 5.54E-08 |
| CSAD | 412.1667877 | -0.638014335 | 4.41E-06 |
| SRRT | 5581.018869 | -0.63846399 | 9.71E-08 |
| RASSF2 | 380.6043208 | -0.638668321 | 1.26E-05 |
| PHF14 | 2956.198094 | -0.638670002 | 4.92E-06 |
| TRPM2-AS | 87.9797349 | -0.639101836 | 1.10E-02 |
| AC132872.1 | 143.6391076 | -0.63967304 | 1.78E-04 |
| NA | 1367.280099 | -0.63982526 | 5.52E-10 |

| | | | |
|---|---|---|---|
| RNASEH2C | 2719.859161 | -0.639929167 | 3.72E-06 |
| EMC3-AS1 | 195.9109598 | -0.640501535 | 2.40E-03 |
| AC138932.1 | 1791.102047 | -0.640758911 | 5.51E-03 |
| SMARCA4 | 6882.643051 | -0.641312566 | 3.45E-10 |
| SLC2A4RG | 3962.621191 | -0.641446913 | 1.37E-07 |
| AC109439.2 | 140.6189782 | -0.641632904 | 8.59E-05 |
| ZNF724 | 94.80845695 | -0.642075713 | 3.02E-02 |
| GAMT | 1313.677418 | -0.642095893 | 1.58E-06 |
| AC009133.1 | 273.7952977 | -0.642134142 | 2.31E-06 |
| RCCD1 | 1438.649141 | -0.64293889 | 9.74E-15 |
| RAD9A | 799.115388 | -0.643013363 | 9.08E-08 |
| CCT3 | 14487.18287 | -0.643334019 | 5.18E-07 |
| REV3L | 1912.9575 | -0.643563774 | 2.54E-03 |
| AC104667.2 | 71.44130672 | -0.643603174 | 2.23E-02 |
| SOX12 | 4342.362057 | -0.643711447 | 1.76E-11 |
| GNG11 | 5111.312028 | -0.644750792 | 2.15E-07 |
| C3orf62 | 371.8425366 | -0.646127226 | 7.14E-03 |
| CMKLR1 | 1380.27909 | -0.646180959 | 3.28E-02 |
| ARL17A | 1881.979378 | -0.646416772 | 5.77E-13 |
| RPL29 | 36284.96416 | -0.646507345 | 1.26E-06 |
| PANO1 | 51.604583 | -0.646515827 | 2.88E-02 |
| MAGED4B | 1693.218086 | -0.64689198 | 1.30E-03 |
| SAMD10 | 302.1944671 | -0.64746298 | 1.90E-03 |
| AC026401.3 | 298.6534259 | -0.647621606 | 2.00E-05 |
| C11orf95 | 1680.071571 | -0.648059257 | 4.51E-06 |
| NCAPG2 | 4695.891488 | -0.648588858 | 6.29E-07 |
| RBM14 | 2760.301633 | -0.649206538 | 2.43E-07 |
| GTPBP3 | 853.3995395 | -0.649304852 | 5.80E-07 |
| NBPF4 | 231.6908026 | -0.649356944 | 9.79E-05 |
| SLIT2 | 273.2187921 | -0.649380317 | 2.71E-04 |
| SLC2A6 | 1907.900978 | -0.649689013 | 3.85E-05 |
| RNA5-8SN2 | 684.3031988 | -0.650336681 | 4.57E-02 |
| SH2B1 | 2927.777628 | -0.650479701 | 3.37E-11 |
| FAM102B | 808.5164647 | -0.650564415 | 2.55E-05 |
| TTC28 | 2641.399182 | -0.650674602 | 5.94E-05 |
| MTATP6P1 | 11442.5503 | -0.651239492 | 2.89E-04 |
| EFCAB7 | 208.1890505 | -0.651834984 | 3.73E-02 |
| TECR | 3400.127322 | -0.652303011 | 1.20E-05 |
| MYH10 | 12958.38845 | -0.652550994 | 8.77E-22 |
| EZH2 | 2381.538175 | -0.652618191 | 4.52E-08 |
| LIMCH1 | 5167.393543 | -0.652888451 | 4.62E-05 |
| LRRIQ3 | 42.99257129 | -0.6530736 | 4.67E-02 |
| LLGL2 | 1015.83685 | -0.653511697 | 3.50E-09 |
| U52111.1 | 82.8321453 | -0.653752314 | 2.70E-02 |
| PAXX | 1435.946098 | -0.653935474 | 4.05E-13 |
| PRR3 | 2046.71534 | -0.653944469 | 1.85E-12 |
| ZNF771 | 204.8286513 | -0.654354302 | 3.84E-04 |

| | | | |
|---|---|---|---|
| ZSWIM3 | 567.2068289 | -0.654421445 | 3.39E-04 |
| AP001062.1 | 128.0366995 | -0.654548544 | 1.75E-03 |
| ABCB9 | 492.6059606 | -0.654740435 | 8.47E-05 |
| FRMD5 | 262.6352491 | -0.654752312 | 6.53E-04 |
| SEMA6A-AS1 | 39.33274057 | -0.655076442 | 3.89E-02 |
| HAS3 | 161.7908666 | -0.655117705 | 4.77E-03 |
| EEF1G | 92857.80488 | -0.655272307 | 3.31E-08 |
| SAYSD1 | 553.4734968 | -0.655360798 | 9.48E-09 |
| CRMP1 | 630.7320603 | -0.655576102 | 1.41E-14 |
| IMPA2 | 418.8634436 | -0.655618093 | 8.64E-06 |
| PRXL2B | 1786.82515 | -0.65568168 | 4.92E-05 |
| GUSBP2 | 71.99712807 | -0.655723221 | 2.73E-02 |
| MT-ND2 | 53315.87307 | -0.65592122 | 4.10E-03 |
| AAR2 | 4059.122882 | -0.656361075 | 2.72E-04 |
| PRPH | 172.1192953 | -0.656814323 | 1.15E-02 |
| DSN1 | 2244.06107 | -0.656829257 | 7.46E-05 |
| RPL35 | 27266.11572 | -0.65717722 | 4.14E-09 |
| CYP27C1 | 309.7320071 | -0.657549847 | 8.63E-05 |
| BCS1L | 1798.308695 | -0.657683507 | 5.10E-18 |
| ZNF772 | 795.6098817 | -0.658779915 | 4.32E-09 |
| SIM2 | 439.4469742 | -0.658889143 | 1.87E-05 |
| AL359878.1 | 47.04444433 | -0.659078883 | 3.02E-02 |
| ZNF512B | 4994.652742 | -0.659236296 | 5.15E-16 |
| ZHX1 | 2058.307391 | -0.659489439 | 4.44E-04 |
| RPL21P16 | 5461.515123 | -0.659815738 | 1.74E-07 |
| WDR5 | 4924.709522 | -0.659913083 | 4.16E-19 |
| ZNF653 | 221.8018928 | -0.659950569 | 5.51E-03 |
| ZNF551 | 748.8878328 | -0.660284869 | 2.66E-05 |
| FOXD2 | 276.8095472 | -0.660578987 | 4.05E-07 |
| EEF1A1P19 | 59.86761148 | -0.660682188 | 3.54E-02 |
| MAST1 | 367.3052568 | -0.660720163 | 3.87E-04 |
| PLSCR3 | 1851.084236 | -0.662096476 | 6.04E-07 |
| DDX23 | 4531.807855 | -0.662491533 | 4.03E-08 |
| PECAM1 | 169.5651196 | -0.662529115 | 1.82E-03 |
| LINC00511 | 271.0163228 | -0.662798114 | 5.63E-05 |
| AP001816.1 | 121.3397646 | -0.663003963 | 5.58E-03 |
| CDKN2D | 386.7352033 | -0.663059937 | 1.71E-05 |
| HAUS7 | 1980.774412 | -0.663429891 | 1.27E-09 |
| AC140125.1 | 49.69899974 | -0.663440326 | 2.80E-02 |
| CTU1 | 196.5163835 | -0.663597856 | 1.59E-04 |
| AC134978.1 | 392.7299982 | -0.664139779 | 9.21E-04 |
| AC144652.1 | 365.0300178 | -0.664190841 | 7.25E-06 |
| NA | 1377.415725 | -0.664756104 | 3.16E-07 |
| LINC01011 | 116.1222739 | -0.665433507 | 1.49E-02 |
| RFC5 | 1626.892685 | -0.665470576 | 1.65E-09 |
| RAD54B | 595.254013 | -0.66547805 | 5.59E-03 |
| MIS18BP1 | 607.5874957 | -0.665516005 | 6.12E-03 |

| | | | |
|---|---|---|---|
| AL122010.1 | 52.24306951 | -0.665603962 | 3.67E-02 |
| NGEF | 148.4072051 | -0.666423519 | 3.36E-02 |
| TRIM41 | 1896.095923 | -0.666632848 | 2.50E-14 |
| AC067930.5 | 85.73263488 | -0.666659299 | 4.63E-02 |
| GRIN2A | 673.7685502 | -0.666889097 | 1.71E-03 |
| AC093627.4 | 67.41982005 | -0.667481289 | 3.99E-02 |
| PMF1 | 1385.681795 | -0.667516711 | 1.04E-06 |
| ERMAP | 476.2680804 | -0.667561308 | 4.56E-10 |
| TCTN2 | 617.1698295 | -0.667627023 | 5.47E-03 |
| TRUB2 | 3441.465998 | -0.668053209 | 9.39E-08 |
| CASP8AP2 | 923.2155655 | -0.668870001 | 1.21E-02 |
| AC127496.7 | 44.57894777 | -0.669052181 | 3.11E-02 |
| CNOT6 | 1530.549631 | -0.669838035 | 6.94E-06 |
| PDGFC | 565.39758 | -0.669870443 | 6.65E-05 |
| EMP2 | 4797.302819 | -0.670027076 | 3.13E-09 |
| CCHCR1 | 1287.575472 | -0.670235708 | 2.22E-08 |
| TBX1 | 62.73316178 | -0.670759058 | 3.08E-02 |
| TNXB | 1293.362643 | -0.670764912 | 7.94E-09 |
| PEG10 | 103399.184 | -0.671165908 | 2.85E-03 |
| CHST7 | 232.4396841 | -0.671455886 | 4.21E-03 |
| ZDHHC23 | 299.2083981 | -0.671514344 | 1.21E-06 |
| MMP2 | 12012.51999 | -0.671783867 | 4.04E-06 |
| LINC00665 | 1946.725265 | -0.671916743 | 3.33E-11 |
| GUSBP5 | 128.5868497 | -0.672073484 | 2.02E-03 |
| AL137802.2 | 105.9809884 | -0.672748862 | 3.92E-03 |
| AC009090.4 | 71.1676882 | -0.672800111 | 3.62E-02 |
| CAPS | 230.0045912 | -0.672861183 | 2.69E-04 |
| CPNE2 | 3888.561043 | -0.672909881 | 7.74E-11 |
| SLC27A5 | 255.6919464 | -0.672973348 | 7.48E-06 |
| ANAPC2 | 2613.355409 | -0.673030894 | 1.68E-06 |
| TSSC2 | 633.1595099 | -0.673063936 | 4.53E-07 |
| MTMR4 | 2791.918713 | -0.673518922 | 3.50E-13 |
| SPEG | 2203.384874 | -0.673522351 | 4.66E-06 |
| FHOD3 | 292.6071871 | -0.673554656 | 5.43E-05 |
| AGER | 190.6236298 | -0.673659113 | 3.28E-03 |
| AL049840.3 | 74.68180201 | -0.67419161 | 6.29E-03 |
| ENSA | 5268.579363 | -0.67423733 | 1.40E-15 |
| PPM1N | 138.511032 | -0.6747069 | 3.56E-02 |
| ARRB2 | 1284.031028 | -0.675185925 | 3.09E-11 |
| NADK2 | 3676.638597 | -0.675282101 | 5.50E-08 |
| AC107375.1 | 147.2864795 | -0.675609517 | 6.35E-05 |
| ARL3 | 697.8318546 | -0.675739676 | 9.88E-10 |
| RIMBP3 | 128.4407853 | -0.675836111 | 1.28E-02 |
| PRPF4 | 4611.579451 | -0.676049652 | 1.05E-14 |
| MON1A | 421.933622 | -0.676123405 | 4.36E-09 |
| SAFB2 | 1849.688607 | -0.676528649 | 1.27E-08 |
| DBNL | 8491.156225 | -0.676542423 | 1.02E-05 |

| | | | |
|---|---|---|---|
| MDFI | 1896.304313 | -0.676603884 | 8.30E-07 |
| CAPG | 110.5601278 | -0.676769272 | 1.68E-03 |
| RNFT2 | 124.865365 | -0.676846069 | 3.80E-03 |
| RPS5 | 32305.54264 | -0.676911374 | 1.55E-06 |
| MZF1 | 921.5328365 | -0.676980914 | 8.94E-12 |
| RALGDS | 3171.392639 | -0.677290124 | 4.14E-06 |
| NCOA5 | 2667.247924 | -0.677339317 | 2.16E-13 |
| RHOBTB2 | 2339.17076 | -0.677609321 | 1.29E-09 |
| SCARA3 | 3354.9439 | -0.677739121 | 1.80E-05 |
| ACAP3 | 1532.688697 | -0.677885637 | 1.82E-06 |
| TRIM36 | 230.900317 | -0.678037279 | 1.57E-04 |
| HDAC5 | 2924.778223 | -0.678208276 | 3.83E-06 |
| ZFP36L2 | 1694.899588 | -0.678392896 | 1.02E-12 |
| FAM168A | 5995.039326 | -0.67873482 | 8.86E-08 |
| LYG1 | 60.06585242 | -0.678860234 | 3.38E-02 |
| LIMD2 | 2211.717988 | -0.678891272 | 6.38E-10 |
| C2CD5 | 1692.653386 | -0.679599534 | 1.55E-04 |
| CRELD2 | 1562.4412 | -0.679681551 | 3.49E-07 |
| CDC6 | 2762.907527 | -0.680030106 | 2.75E-05 |
| Z92544.2 | 182.6149303 | -0.680038096 | 6.88E-03 |
| KLHDC3 | 5370.43767 | -0.680041836 | 6.29E-06 |
| RPLP2 | 26725.70048 | -0.680114544 | 2.28E-08 |
| CENPJ | 546.3815123 | -0.680174374 | 2.15E-03 |
| RAD51AP1 | 744.7072606 | -0.680194928 | 3.99E-03 |
| CHD3 | 11157.51814 | -0.680263288 | 1.78E-13 |
| MARVELD1 | 3273.994686 | -0.680761991 | 7.24E-06 |
| TNFRSF10A | 429.7567669 | -0.680813871 | 4.70E-06 |
| LRRC37A | 215.1036013 | -0.681106039 | 7.58E-06 |
| C1orf35 | 645.7215147 | -0.681150164 | 1.27E-05 |
| TSGA10 | 108.0443109 | -0.681411799 | 1.92E-04 |
| C1orf112 | 1218.550267 | -0.681620268 | 1.12E-07 |
| HOXA-AS2 | 648.9155533 | -0.68195397 | 9.43E-07 |
| AC114271.1 | 214.1344847 | -0.6819946 | 1.62E-03 |
| AKR7A2 | 2176.262776 | -0.682019975 | 1.43E-05 |
| EPS8 | 4496.746593 | -0.682126035 | 1.38E-06 |
| ARMC6 | 2014.71344 | -0.682511127 | 5.15E-08 |
| C1GALT1C1L | 229.7544466 | -0.682630468 | 3.02E-04 |
| ZNF92 | 182.2272354 | -0.682733567 | 4.12E-02 |
| SPATA6 | 471.5236482 | -0.682766281 | 2.05E-07 |
| CORO2B | 318.2158163 | -0.682867191 | 5.51E-05 |
| AL596244.1 | 271.7370762 | -0.682987216 | 3.12E-04 |
| MEPCE | 5159.706351 | -0.683284853 | 7.74E-17 |
| ZWILCH | 3027.212332 | -0.68345258 | 4.96E-05 |
| EFHD2 | 3883.799029 | -0.683869277 | 5.58E-07 |
| WDR27 | 1383.416826 | -0.68389925 | 1.54E-08 |
| AL662907.1 | 56.14748766 | -0.684473304 | 2.64E-02 |
| BX664615.1 | 109.2441282 | -0.684711819 | 5.95E-04 |

| | | | |
|---|---|---|---|
| BICD1 | 1218.161353 | -0.684893652 | 5.93E-14 |
| AL109918.1 | 846.5422957 | -0.685439483 | 1.33E-10 |
| CLEC4F | 53.65404115 | -0.686048452 | 1.79E-02 |
| HBE1 | 415.2750207 | -0.686132409 | 1.94E-02 |
| CEP83-DT | 49.55047776 | -0.686228936 | 1.78E-02 |
| PLEKHH3 | 964.0051629 | -0.686268956 | 1.43E-10 |
| GINS3 | 861.3140357 | -0.686357099 | 4.25E-04 |
| HEXIM2 | 229.1603445 | -0.686502455 | 5.42E-06 |
| SPDYE5 | 63.22792987 | -0.686573817 | 1.02E-02 |
| PXMP2 | 902.2577882 | -0.686640562 | 8.34E-08 |
| TGFB1I1 | 2716.369981 | -0.686706523 | 3.25E-06 |
| TMEM200A | 540.674388 | -0.686978703 | 2.31E-05 |
| MYL6 | 26337.83559 | -0.687102609 | 3.64E-04 |
| DOCK11 | 2817.853034 | -0.687203893 | 7.89E-03 |
| CROCC | 613.0046235 | -0.687643096 | 1.10E-07 |
| TRO | 3618.311323 | -0.687743701 | 4.85E-07 |
| AC245452.1 | 293.78634 | -0.688441532 | 3.63E-06 |
| GLYCTK | 459.0239151 | -0.688799512 | 2.02E-05 |
| ZFP14 | 308.8122709 | -0.689343722 | 1.56E-06 |
| ASB16 | 66.44886693 | -0.689380944 | 7.51E-03 |
| AL512625.1 | 409.7626639 | -0.68999651 | 4.90E-02 |
| VAT1L | 15308.2066 | -0.690008035 | 3.28E-09 |
| FBL | 5586.954134 | -0.69014019 | 9.18E-12 |
| FOXL1 | 378.206564 | -0.690643277 | 1.43E-05 |
| ZNF688 | 448.3177419 | -0.690711551 | 1.56E-05 |
| STOX1 | 103.1043832 | -0.690876494 | 2.85E-03 |
| PCOLCE | 18291.75263 | -0.69166729 | 1.45E-06 |
| METAP1 | 2773.955335 | -0.691715964 | 1.66E-17 |
| AC138969.1 | 3387.999623 | -0.691802156 | 3.06E-07 |
| SSSCA1-AS1 | 84.81215062 | -0.692269922 | 5.51E-03 |
| DDTL | 284.9097104 | -0.692304728 | 4.70E-09 |
| RFWD3 | 5369.116357 | -0.692853752 | 1.40E-08 |
| NCALD | 398.8456302 | -0.692950588 | 4.18E-02 |
| AL109811.1 | 48.68424945 | -0.69297415 | 3.99E-02 |
| TMC3-AS1 | 109.7734588 | -0.693170218 | 5.57E-04 |
| APOBEC3F | 829.4756475 | -0.693372425 | 8.09E-07 |
| CACTIN-AS1 | 46.52365665 | -0.69341278 | 2.65E-02 |
| RALY | 10373.36135 | -0.693499001 | 9.43E-12 |
| DNM1 | 2280.331942 | -0.693713665 | 1.38E-12 |
| CNBD2 | 95.00086923 | -0.694393711 | 5.50E-04 |
| APAF1 | 1252.400371 | -0.694759162 | 5.42E-05 |
| AC011448.1 | 195.9966694 | -0.694762027 | 2.65E-04 |
| CARHSP1 | 4813.82067 | -0.694821596 | 6.99E-07 |
| AC006058.1 | 319.7637519 | -0.69486456 | 6.75E-05 |
| PI4KAP1 | 543.6085889 | -0.695003706 | 1.93E-10 |
| SYCP2L | 81.04151592 | -0.695049016 | 3.56E-02 |
| JADE2 | 2540.950106 | -0.696100157 | 6.60E-11 |

| | | | |
|---|---|---|---|
| TSSK4 | 100.2316405 | -0.696136731 | 1.31E-03 |
| VILL | 408.5323477 | -0.696208969 | 8.28E-09 |
| CCNB2 | 4442.079969 | -0.696430237 | 6.63E-14 |
| RGS12 | 1044.633314 | -0.696505248 | 3.94E-07 |
| NA | 153.5758231 | -0.697160159 | 9.44E-05 |
| ZNF681 | 158.2520651 | -0.697290902 | 9.44E-05 |
| KBTBD4 | 1247.543525 | -0.697389824 | 1.68E-05 |
| AP003352.1 | 107.9554894 | -0.697942992 | 5.38E-03 |
| CEP68 | 1429.317071 | -0.698076565 | 1.94E-12 |
| LSP1 | 262.4820471 | -0.698223656 | 1.40E-02 |
| ELP6 | 1863.952023 | -0.698373024 | 5.79E-10 |
| SAMD9 | 759.2126716 | -0.698654016 | 2.07E-02 |
| ELK3 | 2668.256893 | -0.698957891 | 2.12E-11 |
| DAAM2 | 3400.637135 | -0.699331603 | 4.29E-05 |
| RASA4CP | 218.979591 | -0.699863957 | 2.30E-04 |
| MPP3 | 253.4525953 | -0.699937419 | 8.59E-03 |
| VAX2 | 144.6886263 | -0.700707282 | 1.30E-03 |
| AC009061.2 | 39.94211966 | -0.70094355 | 4.50E-02 |
| C2orf92 | 128.3120634 | -0.701309863 | 3.87E-05 |
| PGAP3 | 538.2201203 | -0.702147223 | 6.46E-05 |
| PRC1 | 7878.190155 | -0.70223769 | 3.49E-07 |
| AIP | 2244.172621 | -0.702439732 | 4.38E-06 |
| NAV1 | 6051.846112 | -0.702473464 | 1.87E-16 |
| GPM6B | 424.3810447 | -0.7027084 | 2.03E-03 |
| ZNF606 | 571.9029524 | -0.702766358 | 1.47E-06 |
| ZNF324B | 402.6473517 | -0.702803131 | 7.09E-09 |
| ZFP30 | 444.4963992 | -0.703033866 | 1.07E-05 |
| NRIP2 | 58.64968632 | -0.703167031 | 2.38E-02 |
| POTEG | 103.8770361 | -0.703206927 | 2.07E-03 |
| TTI1 | 3953.054055 | -0.703335309 | 1.54E-14 |
| ZNF726 | 144.2205326 | -0.703528105 | 3.04E-05 |
| FUZ | 1165.079848 | -0.703592402 | 1.13E-10 |
| SENCR | 247.900948 | -0.703956284 | 6.75E-07 |
| HMGN2 | 20275.41271 | -0.704111068 | 9.66E-18 |
| IGHV3-19 | 30.14242899 | -0.704331867 | 4.63E-02 |
| SOGA1 | 7215.020647 | -0.70454052 | 1.52E-12 |
| AL354836.1 | 91.21734912 | -0.705902708 | 1.85E-02 |
| TRIM28 | 15980.75068 | -0.705931618 | 2.71E-12 |
| AC011450.1 | 89.0077873 | -0.70621928 | 3.59E-03 |
| RPA2 | 2878.925117 | -0.706411446 | 3.25E-08 |
| NLE1 | 1547.763898 | -0.706492105 | 1.11E-09 |
| UBXN10 | 251.9791355 | -0.706555922 | 2.45E-05 |
| NR1H3 | 1127.557574 | -0.706588348 | 3.59E-05 |
| ACYP1 | 203.5393431 | -0.706631353 | 1.80E-06 |
| AC244197.3 | 405.6009799 | -0.70674724 | 8.89E-05 |
| ANKLE1 | 635.5348284 | -0.706769191 | 8.09E-10 |
| SPIN3 | 350.5108314 | -0.706777609 | 4.64E-05 |

| | | | |
|---|---|---|---|
| CHEK2 | 957.2903215 | -0.706908266 | 9.59E-08 |
| CENPBD1P1 | 1256.44121 | -0.707069296 | 6.03E-09 |
| ATAD3B | 666.0326199 | -0.707182051 | 1.83E-07 |
| GDF11 | 3916.115371 | -0.707238857 | 1.82E-06 |
| HRCT1 | 93.63600929 | -0.707565001 | 8.79E-03 |
| CRABP2 | 3933.139023 | -0.709030687 | 2.25E-05 |
| AC022150.4 | 176.7869855 | -0.709280532 | 6.87E-03 |
| MISP3 | 41.92976061 | -0.709548043 | 4.83E-02 |
| SMOC1 | 69.53858863 | -0.709789085 | 3.39E-02 |
| SYT13 | 311.0869747 | -0.709941678 | 3.55E-02 |
| DPH2 | 2221.522895 | -0.710693143 | 7.00E-12 |
| LINC00461 | 138.3576983 | -0.710700726 | 2.35E-02 |
| USP13 | 1845.289676 | -0.71118181 | 3.38E-20 |
| C1RL-AS1 | 573.792794 | -0.711590394 | 1.42E-04 |
| EXOSC5 | 1246.924041 | -0.711756779 | 3.19E-07 |
| TOR2A | 716.6961917 | -0.711820055 | 2.60E-09 |
| CYP3A5 | 68.36392406 | -0.71218531 | 5.34E-03 |
| STRA6 | 466.7762795 | -0.71231915 | 7.70E-04 |
| PLA2G6 | 356.9025369 | -0.712370572 | 1.46E-04 |
| IGFLR1 | 122.5102195 | -0.71272227 | 2.08E-02 |
| MAMSTR | 128.869854 | -0.713025402 | 1.67E-03 |
| ABHD14A | 756.7593186 | -0.713071601 | 2.37E-06 |
| ZNF692 | 657.41101 | -0.713091457 | 4.15E-10 |
| PAFAH1B3 | 1766.108722 | -0.713413489 | 7.47E-09 |
| MINDY1 | 1011.666293 | -0.713570084 | 4.19E-05 |
| ZNF284 | 227.9168793 | -0.713571364 | 8.08E-08 |
| ATAD2 | 3593.292633 | -0.713894358 | 8.42E-04 |
| SASS6 | 296.0806781 | -0.713998563 | 1.24E-02 |
| PRKRA-AS1 | 232.7012941 | -0.714658575 | 1.60E-04 |
| GPC6 | 245.0733971 | -0.714869069 | 1.14E-02 |
| MLKL | 890.2288898 | -0.715226319 | 1.90E-11 |
| PRMT6 | 1118.533194 | -0.715362297 | 2.59E-14 |
| AL162431.2 | 48.18395432 | -0.715531447 | 8.09E-03 |
| STIM2 | 1398.883632 | -0.716213836 | 1.24E-09 |
| PSD3 | 2830.348069 | -0.716824859 | 2.80E-10 |
| AL732372.3 | 267.4658968 | -0.716914376 | 3.51E-07 |
| LYPD3 | 55.2915181 | -0.716925128 | 2.90E-02 |
| ANKRD18EP | 104.7435619 | -0.717038692 | 2.68E-03 |
| SPOUT1 | 3020.906239 | -0.717057674 | 3.48E-09 |
| AC010970.1 | 17518.2263 | -0.717089852 | 3.15E-03 |
| NUP85 | 2328.574609 | -0.717175653 | 3.27E-08 |
| AC008755.1 | 1945.393163 | -0.717864115 | 8.97E-19 |
| GPR68 | 67.37087682 | -0.718263893 | 4.42E-02 |
| AC084125.2 | 187.0679811 | -0.718564281 | 5.70E-04 |
| AP001318.2 | 42.60818205 | -0.718789292 | 2.10E-02 |
| AXIN2 | 442.7244193 | -0.718895463 | 6.90E-04 |
| AP002495.2 | 126.5569363 | -0.719192906 | 1.14E-03 |

| | | | |
|---|---|---|---|
| MC1R | 850.7944033 | -0.719635186 | 8.24E-13 |
| ZNF578 | 160.044922 | -0.720044537 | 3.52E-05 |
| SLC25A10 | 1342.75035 | -0.720167187 | 2.58E-09 |
| DYSF | 2529.150232 | -0.720924002 | 9.53E-07 |
| TK1 | 5146.104526 | -0.721829613 | 1.32E-04 |
| RPL13A | 70398.25543 | -0.72197622 | 1.25E-08 |
| DTYMK | 2052.910667 | -0.722321535 | 3.48E-14 |
| MAP2K6 | 286.8384036 | -0.722753578 | 6.76E-06 |
| NHP2 | 3255.783206 | -0.72345532 | 2.62E-11 |
| AC021028.1 | 60.34363932 | -0.723479176 | 3.38E-02 |
| CENPS | 482.2845624 | -0.724526851 | 2.43E-10 |
| MB21D2 | 492.152103 | -0.724674556 | 1.54E-09 |
| MCAT | 555.0467995 | -0.725216441 | 1.79E-06 |
| TMPO | 5018.503462 | -0.725378094 | 4.24E-10 |
| REEP1 | 645.6708003 | -0.72602129 | 7.27E-04 |
| SYN1 | 187.3211338 | -0.72604805 | 7.55E-05 |
| TROAP | 1436.648234 | -0.726682963 | 3.04E-06 |
| ZNF521 | 1676.551674 | -0.726748722 | 2.44E-07 |
| SACS | 1170.373401 | -0.726783505 | 2.50E-05 |
| TULP3 | 1839.543258 | -0.727456992 | 1.94E-08 |
| EXOSC2 | 2807.209963 | -0.727714121 | 3.69E-35 |
| TLE6 | 48.19042002 | -0.728246426 | 3.69E-02 |
| C12orf76 | 572.4469284 | -0.728398475 | 7.77E-12 |
| RPS2P5 | 8753.092583 | -0.728935519 | 3.10E-10 |
| NA | 98.75516987 | -0.729462438 | 1.56E-03 |
| PGM5P2 | 53.12602511 | -0.729649058 | 2.74E-02 |
| BCL9 | 2022.851338 | -0.729733351 | 4.84E-14 |
| AC012313.2 | 95.14215469 | -0.730438394 | 9.00E-04 |
| DIP2A | 2335.099636 | -0.730697863 | 8.20E-19 |
| SMC2 | 3469.216138 | -0.731664645 | 6.06E-03 |
| NA | 96.45349882 | -0.73178671 | 1.73E-04 |
| PIK3R3 | 2884.989418 | -0.732989758 | 2.48E-06 |
| NECTIN1 | 3123.160443 | -0.732990844 | 4.03E-05 |
| MBOAT1 | 364.5205903 | -0.733110472 | 2.08E-07 |
| ROGDI | 517.5564016 | -0.733362157 | 4.37E-07 |
| SESN1 | 517.2971486 | -0.73337904 | 5.62E-06 |
| NA | 99.97772378 | -0.73346404 | 2.19E-02 |
| KIF3C | 1868.784399 | -0.734274838 | 1.60E-13 |
| EML6 | 415.0599144 | -0.734290707 | 1.12E-02 |
| C11orf1 | 496.8051139 | -0.734822048 | 4.17E-10 |
| LINC01776 | 61.85722524 | -0.734914947 | 1.83E-02 |
| NA | 36.61783023 | -0.735129099 | 3.43E-02 |
| NA | 400.6173156 | -0.735528646 | 7.85E-07 |
| BRCA1 | 2010.069614 | -0.73570057 | 9.92E-05 |
| SPTBN5 | 182.573012 | -0.735898535 | 9.54E-03 |
| AP001469.3 | 144.6518621 | -0.736036724 | 1.94E-04 |
| ARHGEF40 | 6406.672053 | -0.736557785 | 2.18E-07 |

| | | | |
|---|---|---|---|
| AC104447.1 | 94.69098291 | -0.736647196 | 1.31E-03 |
| KMT2E-AS1 | 139.9163431 | -0.736852904 | 3.58E-05 |
| AL358334.2 | 74.69140229 | -0.737019181 | 2.37E-03 |
| HOXA3 | 372.817861 | -0.737960696 | 1.15E-03 |
| S100A4 | 620.5299465 | -0.738093894 | 2.76E-08 |
| TAGLN2 | 18351.75881 | -0.738337204 | 5.82E-06 |
| LINC01515 | 119.0846713 | -0.738383189 | 4.13E-03 |
| IFFO2 | 2826.398817 | -0.738389391 | 1.77E-10 |
| AFMID | 694.8669109 | -0.739231394 | 4.96E-09 |
| ZNF891 | 383.1294511 | -0.739363151 | 7.62E-07 |
| RPLP0 | 75599.32004 | -0.739709917 | 4.39E-11 |
| MBD3 | 5402.244443 | -0.739968868 | 3.20E-12 |
| GLYATL2 | 43.13377158 | -0.740497575 | 4.18E-02 |
| TIGD5 | 583.978972 | -0.740813814 | 3.44E-09 |
| CIP2A | 1223.598363 | -0.740888941 | 4.73E-03 |
| MED16 | 2198.289011 | -0.741039958 | 1.38E-06 |
| LCTL | 194.6728429 | -0.741476186 | 6.82E-04 |
| DCTPP1 | 2007.818684 | -0.742820339 | 4.79E-10 |
| ADAMTS15 | 608.3231517 | -0.744073121 | 4.31E-08 |
| SPIN4 | 606.1825621 | -0.744127184 | 2.88E-05 |
| KBTBD6 | 801.5697179 | -0.744178847 | 3.88E-09 |
| LRRC37A2 | 983.0797025 | -0.744342644 | 5.50E-22 |
| SLC46A1 | 1659.182079 | -0.74513005 | 2.01E-05 |
| ZNF530 | 486.0653137 | -0.746975938 | 9.97E-08 |
| RPS15 | 13793.06285 | -0.747131838 | 7.88E-08 |
| FRMD4A | 2369.142518 | -0.747287179 | 1.48E-15 |
| PSIP1 | 5527.069374 | -0.747341015 | 1.80E-08 |
| ENKD1 | 782.8162869 | -0.747453038 | 1.34E-11 |
| AL606534.4 | 36.57530579 | -0.747630682 | 3.73E-02 |
| LINC01719 | 71.01155469 | -0.747636487 | 7.78E-03 |
| TET3 | 1452.642177 | -0.748124299 | 1.83E-12 |
| OLMALINC | 520.9366639 | -0.748316899 | 9.11E-10 |
| GEN1 | 694.1512265 | -0.748432674 | 1.56E-02 |
| CENPT | 2443.523131 | -0.748843094 | 8.42E-24 |
| TANC2 | 3184.769522 | -0.749162199 | 3.28E-07 |
| TBXAS1 | 153.5800672 | -0.750177208 | 1.12E-04 |
| LIMK2 | 1150.8064 | -0.7504173 | 1.28E-11 |
| SETD9 | 410.3844377 | -0.7509229 | 3.34E-07 |
| PKP2 | 1275.453997 | -0.751435091 | 7.51E-11 |
| AC004477.1 | 149.757773 | -0.751444443 | 1.65E-03 |
| SMIM10 | 1046.732466 | -0.751484188 | 6.54E-12 |
| ODF2 | 3894.579451 | -0.751551267 | 1.28E-30 |
| HNRNPA1P16 | 83.2465907 | -0.751668854 | 5.30E-03 |
| POMT2 | 1487.647994 | -0.7516902 | 1.16E-07 |
| POMK | 182.4422372 | -0.752388542 | 1.16E-07 |
| MAPK11 | 1201.739867 | -0.75258255 | 1.85E-17 |
| PTMA | 27507.72041 | -0.752909449 | 8.32E-11 |

| | | | |
|---|---|---|---|
| EEF1D | 14301.58075 | -0.752996731 | 1.19E-14 |
| YJEFN3 | 352.2178593 | -0.753309483 | 7.01E-06 |
| FLI1 | 3174.945059 | -0.753430252 | 5.94E-17 |
| TOPBP1 | 3698.866461 | -0.753593163 | 2.46E-06 |
| TPTE | 162.9454147 | -0.754137785 | 1.82E-03 |
| CLMP | 3028.617727 | -0.754417833 | 1.78E-06 |
| IGF2 | 2046.485533 | -0.7545325 | 1.40E-04 |
| SLC26A6 | 836.8867718 | -0.755564362 | 2.58E-05 |
| ANKRD36C | 322.2887014 | -0.755599986 | 9.51E-05 |
| ASIC1 | 1465.170741 | -0.755687433 | 1.63E-07 |
| ADORA2A | 130.0004362 | -0.755863134 | 3.11E-03 |
| GPR153 | 403.3307557 | -0.755914192 | 1.57E-04 |
| PPAN-P2RY11 | 179.025675 | -0.755921273 | 2.56E-03 |
| BMP4 | 3231.109292 | -0.756364181 | 8.56E-06 |
| HNRNPM | 10158.43398 | -0.756451249 | 1.75E-34 |
| SARM1 | 1278.675473 | -0.757345387 | 5.38E-07 |
| NFAM1 | 176.4578415 | -0.757357713 | 1.34E-02 |
| TBXA2R | 175.3616382 | -0.757694922 | 1.03E-03 |
| RCAN3 | 1155.641347 | -0.75774004 | 9.10E-12 |
| MIR762HG | 48.53944852 | -0.757887345 | 2.38E-02 |
| DCAF15 | 1202.53612 | -0.758034522 | 3.03E-11 |
| AP003392.4 | 253.0696273 | -0.75870314 | 1.37E-06 |
| LDHA | 37610.75825 | -0.758844919 | 4.52E-05 |
| KDM1A | 8012.116972 | -0.759503598 | 3.02E-12 |
| SLFN13 | 2137.894164 | -0.759800853 | 5.85E-06 |
| TMCO6 | 764.0004854 | -0.759823317 | 2.67E-22 |
| CEP170P1 | 71.64086969 | -0.760190052 | 2.94E-03 |
| C1orf53 | 34.30616792 | -0.760217285 | 3.55E-02 |
| ZSWIM5 | 539.9072653 | -0.760505045 | 8.33E-08 |
| EHBP1L1 | 5588.830947 | -0.76058486 | 1.48E-08 |
| PPP1R14B | 3533.962955 | -0.760639894 | 6.96E-12 |
| ARVCF | 806.5385148 | -0.760716498 | 5.65E-10 |
| CAPN5 | 3165.900756 | -0.761052619 | 7.82E-13 |
| AC016876.1 | 301.3414025 | -0.761747228 | 1.42E-06 |
| ANAPC15 | 955.6555701 | -0.761853117 | 3.59E-08 |
| RPL41 | 46211.12679 | -0.761917123 | 4.91E-11 |
| SLC25A12 | 2043.058397 | -0.762249148 | 1.62E-17 |
| NHLRC4 | 70.47882681 | -0.762387445 | 2.01E-02 |
| RNF32 | 194.6641487 | -0.762769196 | 2.81E-04 |
| AHDC1 | 1790.089992 | -0.762992316 | 1.46E-07 |
| AL591845.1 | 91.06227821 | -0.7632469 | 6.17E-03 |
| MTMR11 | 725.5845595 | -0.763346966 | 3.09E-07 |
| LPAR2 | 318.5182653 | -0.763844953 | 8.38E-06 |
| MYO19 | 2959.134123 | -0.764453846 | 3.84E-14 |
| METTL25 | 105.2190927 | -0.764573277 | 7.63E-04 |
| IRGQ | 1154.186896 | -0.764939231 | 4.72E-04 |
| LINC01948 | 68.24883831 | -0.765261733 | 1.23E-03 |

| | | | |
|---|---|---|---|
| FANCD2 | 2188.44749 | -0.765489338 | 9.94E-06 |
| FAM71F2 | 50.614311 | -0.766381664 | 2.56E-02 |
| NAA16 | 439.3530062 | -0.766592377 | 1.60E-05 |
| SH2D3C | 192.6025404 | -0.766689213 | 9.77E-03 |
| AC127024.5 | 44.36894571 | -0.766728061 | 1.04E-02 |
| 6-Sep | 2821.75174 | -0.767293537 | 1.10E-40 |
| IGFBP6 | 2175.730591 | -0.767348523 | 9.21E-08 |
| MPHOSPH9 | 1122.599799 | -0.767355772 | 5.59E-04 |
| ST3GAL5 | 3369.022929 | -0.767640594 | 2.80E-12 |
| AC022107.1 | 577.3918954 | -0.768357159 | 1.77E-17 |
| BCAS1 | 91.26170515 | -0.768390224 | 2.54E-02 |
| AL121603.2 | 41.03947167 | -0.768421561 | 3.45E-02 |
| BRICD5 | 122.1955978 | -0.768614345 | 1.65E-02 |
| TTC30B | 553.0113901 | -0.768655972 | 3.85E-14 |
| NUTF2 | 4592.139175 | -0.768661357 | 3.98E-08 |
| BCRP3 | 45.78330543 | -0.768688195 | 1.10E-02 |
| DEPTOR | 574.1833815 | -0.768856935 | 3.13E-04 |
| P3H2 | 6277.970193 | -0.768990817 | 1.26E-08 |
| FBXW9 | 659.5921597 | -0.769242718 | 4.20E-08 |
| HMGN2P15 | 98.57012837 | -0.769302295 | 1.81E-03 |
| CEP131 | 811.4527466 | -0.769374107 | 6.56E-12 |
| AP001107.1 | 45.2837746 | -0.769512762 | 6.78E-03 |
| SREBF2-AS1 | 139.3932799 | -0.770303923 | 1.26E-05 |
| ENO3 | 574.3900042 | -0.770375456 | 4.12E-11 |
| SLX1B-SULT1A4 | 282.9680088 | -0.770473731 | 6.50E-07 |
| CHTF18 | 1995.240987 | -0.771492696 | 7.55E-08 |
| STPG3 | 31.27329893 | -0.771907752 | 2.50E-02 |
| HPRT1 | 3591.552647 | -0.77192078 | 3.42E-22 |
| CYP4F29P | 60.47096418 | -0.772064626 | 4.24E-02 |
| AK5 | 674.3521065 | -0.772184069 | 3.41E-05 |
| SRM | 3029.639383 | -0.772590969 | 9.80E-14 |
| CBR3-AS1 | 84.35414603 | -0.773066267 | 9.47E-03 |
| AC138907.8 | 93.45109188 | -0.773914942 | 1.50E-02 |
| HAUS8 | 546.7861723 | -0.774152693 | 6.26E-05 |
| TRAIP | 640.6318974 | -0.774783334 | 6.76E-11 |
| ATOH8 | 972.7768433 | -0.774960046 | 1.50E-09 |
| ING5 | 1464.070134 | -0.775022574 | 4.21E-17 |
| RUNDC3A | 72.75626673 | -0.775553441 | 1.53E-02 |
| FUS | 18454.28601 | -0.776293952 | 5.29E-09 |
| HAUS1 | 1379.441748 | -0.776392038 | 1.10E-18 |
| MDFIC | 378.7965505 | -0.776881035 | 4.42E-04 |
| MATN2 | 4934.640224 | -0.777139559 | 2.31E-09 |
| ZBTB3 | 245.4977007 | -0.778297291 | 6.09E-05 |
| TMEM52 | 33.06314033 | -0.7795013 | 2.12E-02 |
| AC010643.1 | 279.3700195 | -0.780135551 | 2.88E-10 |
| SGO2 | 767.3696881 | -0.78029225 | 1.87E-02 |
| RAMP2-AS1 | 693.2912316 | -0.780644004 | 7.18E-11 |

| | | | |
|---|---|---|---|
| NAPA-AS1 | 80.1956042 | -0.781131474 | 1.36E-02 |
| AC022400.7 | 90.26412942 | -0.781156275 | 3.56E-02 |
| STARD9 | 896.3127905 | -0.781337609 | 4.03E-09 |
| ARHGAP27P1-E | 216.4464008 | -0.78166956 | 6.97E-05 |
| AC114811.2 | 115.9227166 | -0.781924824 | 1.38E-03 |
| CABIN1 | 5087.334398 | -0.782018493 | 3.89E-16 |
| JAG1 | 2378.045133 | -0.782287067 | 6.69E-07 |
| AC067968.1 | 221.9617431 | -0.782401608 | 3.50E-03 |
| LINC02166 | 38.8899247 | -0.78275025 | 1.95E-02 |
| SLC25A25-AS1 | 105.4103291 | -0.784170882 | 4.94E-04 |
| TGM1 | 90.89196127 | -0.785111593 | 1.92E-02 |
| CBX1 | 4941.429777 | -0.78514329 | 1.15E-07 |
| AC005224.1 | 53.6959204 | -0.78531379 | 1.80E-02 |
| HOXA6 | 98.81130786 | -0.785363727 | 1.84E-04 |
| AP001972.5 | 1533.814559 | -0.785431516 | 7.89E-05 |
| BTBD3 | 2563.138022 | -0.785710396 | 1.19E-23 |
| CETN2 | 2499.92649 | -0.786185352 | 1.29E-11 |
| USP21 | 1556.040878 | -0.7863156 | 2.54E-18 |
| PRSS23 | 21726.22584 | -0.786589931 | 6.71E-10 |
| TTC41P | 38.05365793 | -0.786885212 | 4.32E-02 |
| FBXL19 | 2478.860355 | -0.786933039 | 1.26E-10 |
| IL22RA1 | 124.5505552 | -0.787097121 | 8.07E-04 |
| ZIM2 | 133.2839755 | -0.787103371 | 3.15E-04 |
| LNX1 | 162.9220059 | -0.78752783 | 9.85E-06 |
| MOSPD3 | 923.006193 | -0.788201087 | 1.44E-06 |
| PRAC2 | 68.33346669 | -0.788853539 | 5.76E-03 |
| TMEM158 | 357.808961 | -0.788926028 | 3.79E-04 |
| VIM | 74772.47501 | -0.789068688 | 6.27E-13 |
| H1FX | 4099.276197 | -0.789531145 | 7.29E-17 |
| TRIM24 | 2116.562968 | -0.789595418 | 5.03E-26 |
| MEX3D | 606.3152883 | -0.789598692 | 5.42E-09 |
| AC135048.4 | 49.86613015 | -0.790358842 | 4.11E-03 |
| AC022497.1 | 36.62962661 | -0.791172048 | 3.05E-02 |
| DDX11L2 | 54.53949296 | -0.791312035 | 7.36E-03 |
| RANGAP1 | 7733.942925 | -0.791653745 | 3.87E-23 |
| ACOT11 | 447.5869448 | -0.792967489 | 2.46E-08 |
| ANKRD23 | 127.0011083 | -0.793041159 | 9.03E-05 |
| AC004918.1 | 38.88983206 | -0.793910343 | 3.14E-02 |
| CRTC1 | 1021.148261 | -0.794783166 | 7.71E-14 |
| HSPBP1 | 1969.006347 | -0.795096706 | 5.90E-08 |
| AC026356.1 | 50.63709256 | -0.795141349 | 5.27E-03 |
| CASP7 | 1127.428638 | -0.79532295 | 5.81E-10 |
| MAPK15 | 44.18083844 | -0.795904975 | 3.45E-02 |
| TNC | 30916.41729 | -0.795970757 | 4.74E-05 |
| DRC3 | 166.3603861 | -0.79599463 | 1.10E-02 |
| PFKFB4 | 995.5007413 | -0.796189184 | 6.87E-03 |
| CP | 65.43405638 | -0.796314694 | 2.14E-02 |

| | | | |
|---|---|---|---|
| FOXL2 | 28.92919324 | -0.796998565 | 4.68E-02 |
| HVCN1 | 174.3969288 | -0.797204359 | 3.93E-05 |
| LINC00893 | 98.4657195 | -0.797362758 | 4.41E-03 |
| H3F3AP4 | 420.9060945 | -0.797653141 | 2.93E-02 |
| CIB2 | 420.3516067 | -0.797904835 | 1.15E-06 |
| H2BFM | 64.76987712 | -0.798601619 | 1.26E-02 |
| CENPX | 954.3917312 | -0.799413902 | 7.99E-09 |
| RBBP7 | 8591.231336 | -0.800241022 | 4.15E-11 |
| PARD6A | 71.02224597 | -0.800394904 | 8.78E-03 |
| AC016526.3 | 48.75006189 | -0.800569471 | 3.71E-03 |
| DPYD | 377.6234819 | -0.800813085 | 3.41E-09 |
| MAMLD1 | 1047.30731 | -0.801209857 | 5.22E-06 |
| RBL1 | 1227.749753 | -0.801230018 | 1.53E-04 |
| RASA2 | 422.6520801 | -0.801630864 | 2.97E-03 |
| POLR3K | 902.123523 | -0.802066365 | 6.53E-07 |
| CKM | 89.28298883 | -0.80210832 | 6.94E-05 |
| DGAT2 | 873.0974125 | -0.802893603 | 5.32E-07 |
| ATP8B1 | 1504.624557 | -0.803163633 | 8.05E-10 |
| TELO2 | 1185.44392 | -0.803170288 | 2.32E-11 |
| NUDT8 | 148.3854709 | -0.803309139 | 1.09E-04 |
| CLEC18A | 39.39460018 | -0.803322197 | 4.21E-02 |
| PDS5B | 1981.632488 | -0.803702601 | 3.89E-08 |
| MYD88 | 2389.693906 | -0.803880206 | 1.13E-09 |
| CCP110 | 1115.965857 | -0.804410044 | 4.44E-05 |
| COMTD1 | 194.8452166 | -0.804476987 | 1.66E-05 |
| MCM3 | 8184.573665 | -0.804595893 | 6.37E-05 |
| SPIRE2 | 1116.819214 | -0.80502378 | 2.42E-18 |
| MGAT5B | 1657.991925 | -0.805079969 | 2.89E-09 |
| AC080038.2 | 35.49383439 | -0.805201091 | 8.46E-03 |
| IFT81 | 713.7989242 | -0.805432324 | 1.05E-05 |
| DAXX | 4618.188562 | -0.805574609 | 1.75E-18 |
| AC027031.2 | 487.0588179 | -0.805791486 | 4.86E-12 |
| AC007066.2 | 118.9970978 | -0.807344522 | 5.25E-05 |
| SHROOM4 | 459.884557 | -0.807448606 | 2.54E-14 |
| SPAG8 | 62.51808455 | -0.80795817 | 6.26E-03 |
| RDH5 | 60.9220358 | -0.808165259 | 1.09E-02 |
| FIRRE | 111.7449268 | -0.808793581 | 8.28E-05 |
| HMGB1P6 | 2288.973929 | -0.808879319 | 6.00E-07 |
| CUZD1 | 53.56285614 | -0.809289999 | 8.33E-03 |
| NADK2-AS1 | 55.41232106 | -0.809411915 | 6.10E-03 |
| ABHD17A | 2190.111595 | -0.809520905 | 5.43E-05 |
| MAEA | 2666.630086 | -0.809891231 | 2.15E-12 |
| CICP27 | 149.4400659 | -0.810031231 | 6.50E-07 |
| AL096870.2 | 67.05680501 | -0.810189766 | 1.63E-03 |
| EPS8L1 | 328.8165661 | -0.810622058 | 3.20E-06 |
| NA | 104.9217022 | -0.810673395 | 9.24E-05 |
| SCFD2 | 1398.46596 | -0.810822173 | 1.65E-15 |

| | | | |
|---|---|---|---|
| GRAPL | 65.18748568 | -0.810941443 | 1.29E-03 |
| AC106820.3 | 46.58828501 | -0.811577585 | 1.98E-02 |
| ANKDD1A | 276.2525082 | -0.81237496 | 2.02E-08 |
| TCAF1 | 6221.734152 | -0.812531991 | 7.28E-07 |
| RPSA | 54318.93969 | -0.813225232 | 9.12E-12 |
| AC002094.1 | 106.135426 | -0.813424647 | 4.91E-02 |
| PRDX2 | 4035.777858 | -0.813625067 | 5.47E-07 |
| LRRC46 | 47.82199529 | -0.813749346 | 2.89E-02 |
| AC131392.1 | 146.7752183 | -0.814475063 | 1.25E-03 |
| KIAA1841 | 487.9104014 | -0.815041359 | 4.38E-07 |
| SNHG19 | 151.0834963 | -0.815285415 | 1.78E-05 |
| NA | 30.75614378 | -0.816111582 | 2.92E-02 |
| PTMS | 13732.99283 | -0.81648824 | 9.50E-11 |
| FANCI | 5748.936805 | -0.817357075 | 8.95E-10 |
| ARL6IP6 | 1040.346011 | -0.817593941 | 7.46E-07 |
| H3F3A | 10452.75397 | -0.817797137 | 9.10E-14 |
| PLIN3 | 3764.213717 | -0.818192057 | 3.52E-05 |
| POLA2 | 2901.776547 | -0.81822668 | 1.51E-05 |
| DSCC1 | 574.0425517 | -0.818435228 | 1.10E-08 |
| HIRIP3 | 1467.028443 | -0.818474147 | 8.52E-26 |
| ANKRD36 | 491.341241 | -0.818481999 | 4.60E-04 |
| TGIF2 | 2230.264133 | -0.818797148 | 4.22E-12 |
| AC018653.3 | 136.7528489 | -0.819386739 | 4.96E-06 |
| VSTM4 | 321.1915059 | -0.8197838 | 6.42E-04 |
| PDE3B | 420.4228213 | -0.819998339 | 7.43E-06 |
| NCAPH2 | 1949.059842 | -0.820002018 | 4.43E-12 |
| ZBTB45 | 473.1003646 | -0.820261712 | 1.62E-07 |
| NOSTRIN | 322.9129418 | -0.820294157 | 3.76E-05 |
| SERINC5 | 1988.56082 | -0.821089463 | 2.90E-16 |
| AC118344.1 | 45.85678471 | -0.821717894 | 1.20E-02 |
| LINC00926 | 65.66827473 | -0.822243225 | 1.87E-02 |
| COL6A1 | 17941.64959 | -0.824761662 | 8.24E-18 |
| APOBEC3G | 2433.261157 | -0.824828884 | 6.21E-12 |
| TCTEX1D1 | 67.45108166 | -0.825221359 | 1.70E-02 |
| AP002490.1 | 78.86775456 | -0.825428888 | 8.51E-04 |
| NKX2-6 | 293.1226131 | -0.826054771 | 3.98E-07 |
| HAUS5 | 1590.911352 | -0.826126123 | 1.75E-09 |
| LINC01116 | 674.8071079 | -0.826909848 | 2.08E-11 |
| TSEN54 | 1095.294017 | -0.827033419 | 1.42E-11 |
| TTLL12 | 2960.526421 | -0.827530368 | 4.53E-17 |
| EME2 | 888.6364107 | -0.827566555 | 7.08E-07 |
| BRD8 | 7516.993316 | -0.827878941 | 4.56E-40 |
| MALINC1 | 67.02279075 | -0.828074701 | 2.10E-03 |
| IL17RD | 1697.446375 | -0.82830706 | 1.63E-05 |
| RAC3 | 1022.137781 | -0.828400143 | 5.42E-09 |
| HSBP1 | 10345.67201 | -0.828457229 | 2.93E-13 |
| C20orf27 | 2903.6291 | -0.828644451 | 1.57E-13 |

| | | | |
|---|---|---|---|
| PLEKHO1 | 792.133608 | -0.828652424 | 1.61E-07 |
| C20orf204 | 77.0840451 | -0.828689304 | 1.61E-02 |
| CYB561 | 2861.485413 | -0.829692494 | 3.21E-19 |
| UBE2T | 1400.850171 | -0.829841904 | 7.36E-05 |
| C17orf53 | 443.4942892 | -0.82988688 | 1.78E-04 |
| NPIPB11 | 99.95104275 | -0.830042574 | 2.74E-02 |
| STC1 | 1128.86879 | -0.830111013 | 1.90E-03 |
| TMEM14A | 661.6508551 | -0.830349721 | 4.47E-12 |
| FES | 826.5363836 | -0.83049032 | 7.78E-09 |
| ATP2A1 | 140.4236131 | -0.830849283 | 5.60E-03 |
| B3GNT3 | 176.5313853 | -0.831169411 | 2.40E-03 |
| LAT | 302.3278347 | -0.831501537 | 2.05E-07 |
| FAM86C2P | 198.0837881 | -0.831626811 | 1.66E-05 |
| RPS2 | 74595.74233 | -0.832014031 | 5.30E-13 |
| PAK1 | 4174.330043 | -0.83206819 | 1.09E-20 |
| BX088651.4 | 150.1509367 | -0.832791211 | 7.61E-06 |
| TBCD | 2586.174519 | -0.832834708 | 8.62E-13 |
| RPS9 | 22523.13304 | -0.832964439 | 7.22E-10 |
| RTKN | 1095.937803 | -0.832985578 | 1.00E-11 |
| SLC6A6 | 5652.703338 | -0.833208777 | 1.13E-14 |
| DCT | 247.3288116 | -0.833209039 | 6.13E-04 |
| LINC01297 | 1123.725903 | -0.83410954 | 3.96E-03 |
| CDC42BPG | 137.7976709 | -0.83419269 | 3.21E-03 |
| NYNRIN | 5607.528608 | -0.834939137 | 3.72E-04 |
| HNRNPA1P48 | 182.5805565 | -0.835025467 | 5.11E-04 |
| ROM1 | 228.2174532 | -0.835611151 | 2.27E-04 |
| SLC22A16 | 25.62185034 | -0.835737537 | 4.73E-02 |
| AC024361.1 | 32.63315248 | -0.836280544 | 1.41E-02 |
| CFAP45 | 61.16425464 | -0.836341902 | 3.17E-03 |
| CCDC142 | 726.9646701 | -0.836380426 | 5.35E-14 |
| NA | 432.2714357 | -0.83689734 | 1.30E-15 |
| DPF3 | 481.2364414 | -0.837571294 | 8.77E-06 |
| RF00017 | 34.86901515 | -0.83761836 | 1.67E-02 |
| TSHZ1 | 2579.087137 | -0.837986499 | 8.87E-09 |
| AC004597.1 | 144.9279273 | -0.838426814 | 1.16E-04 |
| DENND6B | 132.705302 | -0.838714448 | 6.84E-04 |
| RSBN1L | 1605.52993 | -0.839813375 | 3.09E-20 |
| AC092718.3 | 26.6206568 | -0.840121907 | 4.46E-02 |
| AL365205.1 | 676.4159332 | -0.840360052 | 5.59E-07 |
| AC007191.1 | 91.39415374 | -0.841218107 | 1.83E-04 |
| AL136295.5 | 119.402497 | -0.841864398 | 1.17E-03 |
| AC022075.1 | 57.59754974 | -0.842133872 | 8.25E-03 |
| CASP6 | 603.1914311 | -0.842649054 | 6.44E-10 |
| EFCAB12 | 77.01796851 | -0.843087263 | 8.06E-03 |
| TRIM59 | 116.3614613 | -0.843289946 | 4.39E-04 |
| AFF3 | 1285.889864 | -0.844636952 | 2.09E-06 |
| PPFIA3 | 924.0988512 | -0.845819553 | 6.04E-10 |

| | | | |
|---|---|---|---|
| SOX9 | 2034.137963 | -0.846583975 | 2.44E-11 |
| NDUFA6-DT | 233.1582508 | -0.84738034 | 3.82E-08 |
| PPP2R5D | 4036.10371 | -0.847524875 | 2.39E-11 |
| C5orf30 | 391.2718681 | -0.847577994 | 1.09E-07 |
| SNRNP25 | 1685.632207 | -0.84785517 | 1.51E-07 |
| FEN1 | 5803.451592 | -0.847936885 | 2.50E-04 |
| AC023794.4 | 102.4700055 | -0.848255111 | 6.51E-04 |
| CCDC3 | 1071.774426 | -0.84842761 | 1.50E-09 |
| AC092614.1 | 44.38551858 | -0.849845347 | 2.38E-02 |
| AC009630.2 | 148.3111538 | -0.85013353 | 5.82E-03 |
| TMEM59L | 339.1337644 | -0.850182963 | 9.55E-07 |
| RNASEH2A | 2104.874415 | -0.850561251 | 1.01E-09 |
| PLP2 | 4682.094735 | -0.850566569 | 2.99E-15 |
| MT-CO3 | 214402.6256 | -0.850595096 | 6.11E-06 |
| FNBP1L | 2268.026796 | -0.851002547 | 1.13E-22 |
| RFC2 | 3109.107604 | -0.851028097 | 3.35E-10 |
| MYCBP | 1295.260831 | -0.852018042 | 1.67E-11 |
| EGFL8 | 261.8969967 | -0.852512941 | 2.19E-07 |
| SGSM2 | 4223.108752 | -0.853047272 | 5.08E-14 |
| AC010761.2 | 141.550258 | -0.85332642 | 5.33E-06 |
| CPED1 | 752.439028 | -0.853676624 | 1.03E-09 |
| AC239804.1 | 24.72617584 | -0.853811852 | 4.91E-02 |
| ELMO3 | 151.9450077 | -0.854924482 | 2.15E-08 |
| RUSC1 | 898.0028328 | -0.855095609 | 4.80E-19 |
| SKAP1 | 149.3932027 | -0.855263379 | 1.06E-04 |
| NA | 33.09930813 | -0.856210558 | 2.37E-02 |
| LMCD1 | 1289.053248 | -0.856942149 | 5.71E-04 |
| GAB3 | 903.944943 | -0.856944293 | 4.42E-06 |
| AC007906.2 | 35.53760287 | -0.857633651 | 7.88E-03 |
| CDHR3 | 288.3718067 | -0.857991197 | 3.76E-05 |
| MPP2 | 1292.305072 | -0.858207465 | 1.55E-07 |
| DOK1 | 726.6865422 | -0.859063549 | 1.11E-11 |
| OIP5 | 389.6625071 | -0.859076672 | 4.08E-10 |
| ESAM | 145.5043028 | -0.859230211 | 3.27E-03 |
| TBCB | 2280.125148 | -0.859552173 | 7.56E-14 |
| NELFCD | 4856.196635 | -0.859667711 | 3.06E-13 |
| BX537318.1 | 32.49625477 | -0.859799951 | 2.43E-02 |
| POLD1 | 3115.56909 | -0.859869935 | 1.82E-05 |
| ASB9 | 574.9228447 | -0.861059044 | 4.71E-07 |
| AP000529.1 | 73.28206619 | -0.86144917 | 4.42E-03 |
| CDK1 | 1514.458408 | -0.861637041 | 2.07E-05 |
| ZNF436-AS1 | 269.1957982 | -0.861960059 | 5.04E-05 |
| AC021321.1 | 29.50129298 | -0.8623036 | 3.63E-02 |
| MTCL1 | 1412.753168 | -0.86248444 | 2.27E-17 |
| AL355312.4 | 60.37212687 | -0.862636912 | 2.30E-02 |
| CCDC157 | 226.1355338 | -0.863373296 | 3.59E-06 |
| PTGER4 | 362.933959 | -0.863395759 | 7.13E-03 |

| | | | |
|---|---|---|---|
| SLC6A16 | 260.3475046 | -0.863648666 | 4.27E-06 |
| NA | 175.0159474 | -0.864038133 | 2.85E-07 |
| ITGA9 | 2422.723682 | -0.86418104 | 4.86E-04 |
| FOXR2 | 367.8161971 | -0.864664865 | 1.41E-07 |
| AL596202.1 | 165.9496982 | -0.864897852 | 2.40E-06 |
| GMFG | 271.7051033 | -0.865061977 | 5.38E-05 |
| GPT | 53.48594233 | -0.865168306 | 5.54E-03 |
| CXCL14 | 171.4861655 | -0.865232968 | 2.82E-02 |
| RAD51 | 406.6156072 | -0.865635778 | 1.28E-03 |
| PASK | 805.208939 | -0.866129212 | 2.26E-06 |
| FIZ1 | 677.6044093 | -0.86631961 | 1.71E-10 |
| LINC01424 | 34.96071384 | -0.866802033 | 1.10E-02 |
| MND1 | 324.1099673 | -0.866921258 | 1.40E-06 |
| CCDC150 | 150.6043612 | -0.866965428 | 4.43E-03 |
| AADAT | 215.3693497 | -0.867273963 | 1.92E-05 |
| AKR7A3 | 21.31235145 | -0.867347602 | 4.80E-02 |
| AL390879.1 | 139.316736 | -0.867912242 | 3.58E-07 |
| AC020898.1 | 58.20265113 | -0.868449741 | 5.50E-03 |
| P2RY11 | 313.5632137 | -0.868942539 | 7.05E-08 |
| AC091132.3 | 70.90349035 | -0.869729133 | 2.34E-03 |
| ZNF253 | 186.1191461 | -0.870345801 | 2.99E-03 |
| AP000845.1 | 32.2697025 | -0.870507858 | 2.06E-02 |
| AC012513.3 | 165.7443447 | -0.870828668 | 4.16E-08 |
| PGM2L1 | 488.073069 | -0.870882308 | 2.99E-07 |
| SLF1 | 1156.522204 | -0.870967667 | 1.80E-04 |
| KIF24 | 939.5837656 | -0.871692637 | 2.07E-17 |
| AGAP2-AS1 | 607.7561161 | -0.872322557 | 2.00E-09 |
| ITGA4 | 921.9627008 | -0.872679399 | 6.45E-08 |
| CRYBB2 | 39.30274241 | -0.873722731 | 6.04E-03 |
| ARHGAP4 | 258.78952 | -0.87433528 | 1.25E-04 |
| ATP6V1B1 | 96.52137705 | -0.874966896 | 5.99E-04 |
| AC012511.1 | 21.46889958 | -0.875561987 | 2.81E-02 |
| NMB | 413.3741812 | -0.875654585 | 1.71E-08 |
| PPP1R14A | 175.0821301 | -0.875803827 | 1.65E-04 |
| AC096921.2 | 33.69750771 | -0.87664052 | 2.31E-02 |
| AL391988.1 | 50.15214144 | -0.876767442 | 9.51E-03 |
| HOXA2 | 98.19002439 | -0.877275651 | 8.81E-04 |
| FAM117B | 322.1446807 | -0.877646339 | 7.56E-09 |
| KIF20B | 638.9915225 | -0.877779234 | 7.98E-03 |
| CALHM2 | 962.5218492 | -0.8778027 | 4.34E-16 |
| PRMT7 | 1436.551168 | -0.877857515 | 2.05E-08 |
| WDR62 | 1308.836061 | -0.878428735 | 7.78E-09 |
| SFXN2 | 188.3754226 | -0.878851767 | 1.21E-04 |
| POLD3 | 1334.384664 | -0.878873745 | 6.69E-10 |
| ALDH1B1 | 2870.46866 | -0.879354659 | 2.90E-21 |
| U95743.1 | 74.02309161 | -0.879811812 | 4.83E-04 |
| THOP1 | 2596.814043 | -0.880238332 | 2.29E-14 |

| | | | |
|---|---|---|---|
| RPL28 | 25434.00515 | -0.880371007 | 1.43E-06 |
| AL929601.2 | 89.03516644 | -0.880415469 | 6.31E-05 |
| RTL5 | 2173.925386 | -0.880640808 | 2.64E-21 |
| PRR12 | 1625.764277 | -0.881539797 | 1.76E-12 |
| AC005332.3 | 49.48262873 | -0.88177165 | 4.14E-03 |
| PRICKLE1 | 152.583996 | -0.882085187 | 1.04E-03 |
| KCNN4 | 521.3339695 | -0.88220438 | 1.09E-06 |
| NPAS2 | 2199.95089 | -0.882377057 | 7.40E-11 |
| MAP3K12 | 1271.312178 | -0.882686634 | 4.12E-07 |
| FOXC1 | 860.7224109 | -0.882876777 | 7.22E-11 |
| AC009533.1 | 214.4304185 | -0.883053453 | 9.48E-07 |
| PIK3R5 | 80.61375241 | -0.883125319 | 1.83E-03 |
| TRMT9B | 213.1628729 | -0.883164895 | 9.45E-05 |
| F12 | 188.3703076 | -0.883709033 | 2.02E-06 |
| BRD3OS | 2561.136172 | -0.88385221 | 1.58E-16 |
| DEAF1 | 3949.248938 | -0.884072192 | 1.11E-11 |
| PDSS1 | 287.872663 | -0.88412029 | 8.72E-08 |
| Z97832.2 | 62.15081118 | -0.884143372 | 2.94E-04 |
| NA | 224.2178336 | -0.884534075 | 2.32E-04 |
| MIR31HG | 28.12104951 | -0.884798158 | 4.47E-02 |
| LINC01060 | 80.61008245 | -0.885039281 | 1.56E-04 |
| CDCA4 | 1712.782741 | -0.885086998 | 4.71E-10 |
| RANBP1 | 4498.47852 | -0.8853282 | 2.56E-15 |
| FAM84A | 44.9306386 | -0.885457595 | 1.25E-02 |
| PCDHB15 | 54.5526868 | -0.886336056 | 3.38E-03 |
| SETBP1 | 900.8322632 | -0.886523094 | 3.81E-16 |
| AC093627.5 | 69.5628685 | -0.886741613 | 9.65E-04 |
| KEL | 94.35231685 | -0.886755043 | 2.81E-04 |
| GMNN | 1365.264934 | -0.887169931 | 9.19E-11 |
| RPL7A | 72628.68302 | -0.887553677 | 4.12E-09 |
| AC004057.1 | 1236.708287 | -0.887574842 | 1.62E-05 |
| AC018845.3 | 27.32562862 | -0.887645199 | 2.04E-02 |
| KMT5C | 360.876878 | -0.888033697 | 3.78E-10 |
| ADAMTS9 | 372.2011459 | -0.889014799 | 1.25E-04 |
| ANO1 | 210.5349667 | -0.889028051 | 1.48E-02 |
| NPW | 37.18962742 | -0.8892185 | 2.40E-02 |
| AC092375.2 | 32.97172105 | -0.889657893 | 1.35E-02 |
| CDRT15P1 | 33.8116877 | -0.890827823 | 4.92E-02 |
| MIR647 | 20.76725866 | -0.891727873 | 4.48E-02 |
| MDH1B | 51.25257486 | -0.891761052 | 3.11E-03 |
| AL035587.1 | 56.86355169 | -0.891850194 | 1.01E-03 |
| RFX1 | 522.128581 | -0.892217108 | 6.90E-12 |
| PDXP | 978.5654494 | -0.892465562 | 8.58E-06 |
| TSTD3 | 24.12197618 | -0.893068614 | 3.14E-02 |
| CCDC188 | 246.0132551 | -0.893301289 | 8.06E-07 |
| RPL12P38 | 32.15082083 | -0.89364737 | 4.91E-02 |
| HIC2 | 492.267418 | -0.894003086 | 2.49E-11 |

| | | | |
|---|---|---|---|
| SAP30L-AS1 | 31.89079869 | -0.894120962 | 2.06E-02 |
| LRRC73 | 66.17494783 | -0.896505848 | 2.63E-03 |
| SFRP1 | 14944.91324 | -0.896567429 | 1.56E-07 |
| GLB1L2 | 252.2022969 | -0.896618081 | 8.67E-06 |
| FN3KRP | 1835.113705 | -0.897043044 | 1.46E-08 |
| ZNF296 | 43.61675919 | -0.897154568 | 4.25E-03 |
| SLC26A10 | 106.1172386 | -0.897722731 | 3.43E-04 |
| DCLK1 | 439.8908667 | -0.897742069 | 3.75E-03 |
| MANEAL | 1265.66944 | -0.899579787 | 3.09E-10 |
| CCDC96 | 51.5401067 | -0.899602672 | 3.64E-03 |
| PRDM13 | 87.34034449 | -0.900280026 | 5.18E-04 |
| PTRHD1 | 611.6881417 | -0.901654535 | 3.26E-05 |
| EMID1 | 737.0210262 | -0.90181162 | 8.36E-08 |
| CHEK1 | 2325.87416 | -0.901887732 | 3.55E-10 |
| AC005005.3 | 21.64510273 | -0.903072182 | 4.54E-02 |
| AL121761.1 | 44.35731965 | -0.903626216 | 2.75E-03 |
| PPP1R16A | 1407.164711 | -0.903715503 | 1.75E-12 |
| HMGB2 | 6919.321379 | -0.903917895 | 5.29E-08 |
| WDR54 | 1659.603232 | -0.903947066 | 7.37E-08 |
| AC034102.4 | 74.17309698 | -0.904131668 | 4.31E-04 |
| SKA3 | 713.8002196 | -0.904736444 | 6.08E-09 |
| HMGB1 | 9585.141429 | -0.904915836 | 8.15E-12 |
| CISH | 106.4716427 | -0.905142557 | 2.91E-05 |
| BRCA2 | 456.7661104 | -0.905259143 | 1.70E-02 |
| NBPF6 | 117.6512584 | -0.905344734 | 1.02E-03 |
| MAP3K1 | 6834.516553 | -0.90696653 | 3.68E-17 |
| TCAF1P1 | 529.6204086 | -0.907063151 | 4.95E-09 |
| KIF2C | 4458.272751 | -0.907338622 | 1.33E-18 |
| NA | 84.8373856 | -0.907594313 | 1.39E-02 |
| GCNT1 | 951.6921527 | -0.908077723 | 6.30E-06 |
| AC004408.2 | 62.99055256 | -0.908412709 | 3.37E-03 |
| LTBP2 | 2319.685725 | -0.908432164 | 4.07E-14 |
| ZGRF1 | 425.2006383 | -0.909834707 | 1.90E-03 |
| ATP6V1G2 | 37.63788298 | -0.910080327 | 2.34E-02 |
| TREX1 | 689.4221273 | -0.910495375 | 6.82E-05 |
| NA | 58.15628841 | -0.910867813 | 5.27E-04 |
| NA | 523.1918312 | -0.911336209 | 7.74E-17 |
| AC005944.1 | 200.5770361 | -0.912411677 | 7.51E-04 |
| ZDHHC8P1 | 115.4879533 | -0.912547681 | 3.21E-05 |
| PHETA1 | 683.9070219 | -0.912832144 | 1.74E-09 |
| NEFH | 4344.736666 | -0.913652255 | 1.42E-24 |
| AL031846.2 | 58.3896185 | -0.913795063 | 1.46E-02 |
| MCUB | 438.5499419 | -0.914220601 | 1.73E-06 |
| CALM3 | 16638.08905 | -0.915400473 | 2.97E-15 |
| GPR176 | 2600.86289 | -0.915426658 | 4.29E-10 |
| FUT2 | 72.47299879 | -0.915520959 | 3.54E-03 |
| ARAP3 | 3262.492922 | -0.915988076 | 8.31E-18 |

| | | | |
|---|---|---|---|
| MAP3K5 | 1322.675559 | -0.915997985 | 6.45E-29 |
| ZNF682 | 145.7408049 | -0.916628779 | 3.08E-06 |
| TWIST2 | 580.5860996 | -0.91714996 | 7.53E-12 |
| LRWD1 | 2037.680347 | -0.917424571 | 1.92E-13 |
| PCNT | 3658.598164 | -0.917489001 | 1.03E-20 |
| CEMIP | 1148.669696 | -0.917635748 | 1.99E-05 |
| NID2 | 2791.988157 | -0.917659663 | 1.37E-11 |
| AL355388.2 | 43.64300837 | -0.918340175 | 1.00E-03 |
| SIX1 | 639.7445453 | -0.918536117 | 5.24E-17 |
| ZNF785 | 729.5996148 | -0.918563386 | 1.78E-16 |
| CEL | 34.46526462 | -0.9186093 | 6.41E-03 |
| AC007686.3 | 43.47349693 | -0.918668025 | 1.49E-03 |
| TMTC1 | 1452.335955 | -0.920137409 | 5.58E-07 |
| FTCD | 30.49378675 | -0.920351765 | 1.13E-02 |
| KDM1B | 941.9864652 | -0.920662549 | 1.65E-21 |
| SPDYE2 | 89.92486882 | -0.920985411 | 4.71E-04 |
| ARHGAP11A | 1604.160296 | -0.921075322 | 2.86E-04 |
| AC113191.1 | 39.4017043 | -0.921611711 | 3.08E-03 |
| AC138907.9 | 36.3952997 | -0.922161522 | 3.28E-02 |
| HAS2 | 104.2657699 | -0.922215599 | 9.51E-03 |
| KCTD14 | 1366.012134 | -0.922405739 | 8.71E-12 |
| PBK | 2261.414972 | -0.922572057 | 6.53E-06 |
| PROCA1 | 284.7855565 | -0.922708432 | 3.78E-08 |
| TSHZ2 | 2202.117157 | -0.922950665 | 1.67E-11 |
| LEF1 | 105.2991748 | -0.923258185 | 1.59E-03 |
| NA | 105.2038393 | -0.923320135 | 6.28E-07 |
| NUSAP1 | 2530.933799 | -0.924495332 | 1.07E-06 |
| PCDH12 | 238.254452 | -0.924687063 | 5.89E-04 |
| GPRC5A | 3645.517891 | -0.925178432 | 3.91E-10 |
| GSN-AS1 | 75.19028754 | -0.92666624 | 1.36E-05 |
| FAIM2 | 317.161598 | -0.926996142 | 6.53E-04 |
| LINC02076 | 54.9376968 | -0.927275789 | 3.18E-04 |
| BUB1B | 1938.845997 | -0.927301021 | 4.53E-08 |
| AC090004.1 | 43.08723672 | -0.927354514 | 1.27E-02 |
| CDCA8 | 3450.166152 | -0.928084101 | 1.41E-18 |
| ZNF483 | 79.61362581 | -0.928144134 | 1.14E-03 |
| RPL13AP7 | 153.1708563 | -0.928648544 | 1.89E-03 |
| CDH12P3 | 37.29249809 | -0.928656768 | 4.88E-02 |
| MIR663AHG | 144.4925481 | -0.929214788 | 1.96E-02 |
| AC009159.3 | 20.44347584 | -0.929529379 | 2.45E-02 |
| DDX39A | 2714.807594 | -0.929718984 | 5.80E-13 |
| GDPGP1 | 127.8574598 | -0.929904684 | 7.04E-06 |
| AL929601.1 | 134.5228738 | -0.929920441 | 3.12E-07 |
| PXMP4 | 1242.073786 | -0.930002753 | 4.45E-05 |
| AL353593.1 | 40.48596538 | -0.930974446 | 5.97E-03 |
| NA | 791.4238412 | -0.931471193 | 8.60E-09 |
| FBXO46 | 574.6112734 | -0.931770608 | 1.97E-13 |

| | | | |
|---|---|---|---|
| SUV39H1 | 1125.56085 | -0.931805495 | 1.10E-09 |
| CROCCP3 | 208.8809749 | -0.932516836 | 4.59E-02 |
| PREP | 3723.278541 | -0.932611789 | 5.77E-20 |
| GAL3ST4 | 1047.330578 | -0.932636552 | 7.49E-10 |
| SAMD1 | 2077.502904 | -0.933162068 | 8.79E-14 |
| PIAS3 | 2689.601268 | -0.933766934 | 7.57E-15 |
| FMC1 | 495.7004804 | -0.93381561 | 2.05E-15 |
| AC097468.3 | 30.99750723 | -0.935133112 | 9.82E-03 |
| HBQ1 | 45.14721855 | -0.935748543 | 1.07E-03 |
| AL031666.1 | 32.03510505 | -0.935989556 | 3.81E-03 |
| FAM131B | 590.29546 | -0.93604052 | 1.11E-09 |
| LMNB2 | 11542.53977 | -0.936247655 | 5.15E-16 |
| PYGM | 80.99656343 | -0.936573549 | 1.45E-02 |
| RHOB | 1671.093319 | -0.936616411 | 2.63E-07 |
| RPL4P4 | 54.83496717 | -0.937041372 | 4.55E-03 |
| GALNT17 | 25.55343124 | -0.937163192 | 2.51E-02 |
| HES6 | 712.9778645 | -0.938437381 | 7.80E-15 |
| BAHCC1 | 1925.241566 | -0.938468572 | 1.56E-10 |
| APOBEC3C | 3158.950832 | -0.939156702 | 5.06E-08 |
| AL359513.1 | 23.59183929 | -0.939728775 | 2.59E-02 |
| NME1 | 4684.921094 | -0.939766362 | 1.21E-12 |
| HCAR1 | 765.884658 | -0.939880515 | 2.82E-14 |
| AC087292.1 | 79.73504335 | -0.940625242 | 2.49E-02 |
| IQCD | 101.3790606 | -0.940805407 | 5.90E-05 |
| RIPOR3 | 69.68518001 | -0.94122787 | 2.52E-02 |
| AC073857.1 | 101.7834547 | -0.941235351 | 6.41E-04 |
| ITIH4 | 62.83899631 | -0.941984173 | 4.31E-02 |
| EML4 | 8450.371969 | -0.942056948 | 6.53E-21 |
| CERKL | 136.7873575 | -0.942279602 | 1.53E-03 |
| MANEA | 94.93787047 | -0.942433159 | 3.29E-03 |
| SP4 | 489.826266 | -0.942709294 | 4.51E-06 |
| POLR2J4 | 326.6944127 | -0.943212856 | 1.23E-14 |
| ATF7 | 63.46670973 | -0.943293437 | 6.25E-05 |
| ENDOD1 | 5289.674656 | -0.943349381 | 2.99E-06 |
| ARL17B | 293.4542539 | -0.943670798 | 1.98E-06 |
| TYMSOS | 40.62691575 | -0.944606875 | 8.66E-03 |
| PPP1R1B | 37.78421622 | -0.944861297 | 1.42E-02 |
| DNA2 | 356.2619562 | -0.945261561 | 3.84E-08 |
| TOX | 45.34735463 | -0.945280152 | 2.88E-02 |
| ZCCHC24 | 4931.736738 | -0.945543081 | 7.07E-13 |
| HOXC5 | 53.48940178 | -0.946078045 | 1.42E-02 |
| DIRAS1 | 1073.655793 | -0.946091762 | 4.04E-17 |
| C17orf113 | 66.56775071 | -0.9461423 | 6.59E-05 |
| POM121L9P | 30.25348809 | -0.946233173 | 3.89E-02 |
| AC068134.3 | 53.58614978 | -0.946894286 | 2.98E-03 |
| TRIM14 | 2948.397171 | -0.946945295 | 8.29E-30 |
| CCDC85C | 2884.103042 | -0.947528274 | 2.19E-13 |

| | | | |
|---|---|---|---|
| TNNT1 | 3015.335765 | -0.947705566 | 2.56E-14 |
| HOTTIP | 31.11813955 | -0.948274771 | 7.40E-03 |
| FSD1 | 1426.531621 | -0.948290593 | 2.31E-19 |
| DNALI1 | 156.4162185 | -0.94835751 | 3.54E-04 |
| GLIDR | 39.68952859 | -0.949772309 | 9.09E-03 |
| AC112907.3 | 48.88332524 | -0.949996274 | 1.58E-02 |
| AC012640.2 | 113.968136 | -0.95143743 | 7.09E-06 |
| PTGIR | 176.9157293 | -0.951698325 | 1.77E-06 |
| CBX2 | 1976.013708 | -0.951947511 | 1.98E-15 |
| NA | 27.96683761 | -0.952274761 | 4.72E-02 |
| RADIL | 174.7008569 | -0.953534535 | 3.00E-06 |
| LINC00852 | 19.58380796 | -0.953657781 | 3.84E-02 |
| PLK1 | 5435.480583 | -0.95461683 | 7.61E-13 |
| NA | 21.30695445 | -0.954674747 | 4.29E-02 |
| MYLK2 | 26.74488626 | -0.954918208 | 2.51E-02 |
| GDA | 761.696661 | -0.955068466 | 1.53E-05 |
| AC015712.2 | 238.278475 | -0.955921758 | 3.95E-09 |
| AL359258.2 | 40.70092234 | -0.956631729 | 1.09E-02 |
| MAZ | 11428.16553 | -0.956666465 | 1.31E-18 |
| KANSL1-AS1 | 39.99049905 | -0.95749776 | 3.13E-03 |
| TUBA1B | 81573.23168 | -0.958844896 | 1.47E-07 |
| AC021188.1 | 55.39549357 | -0.958900846 | 4.47E-04 |
| XRCC2 | 856.2539064 | -0.959391024 | 9.98E-05 |
| KIAA0895L | 893.3242262 | -0.959599018 | 1.06E-17 |
| KAT6B | 1431.533217 | -0.959957805 | 5.48E-16 |
| KAT2A | 3508.866278 | -0.960316111 | 5.97E-34 |
| ZNF519 | 152.1448506 | -0.960586149 | 1.57E-03 |
| SLC29A1 | 2721.933491 | -0.960591419 | 5.30E-14 |
| FZD1 | 1665.798152 | -0.961331978 | 1.63E-17 |
| CHST2 | 604.9230275 | -0.962040176 | 1.06E-27 |
| B3GNT4 | 62.54714383 | -0.963223458 | 1.09E-02 |
| POLE | 4059.996044 | -0.963513347 | 3.60E-06 |
| S1PR5 | 103.3816303 | -0.964703466 | 1.39E-03 |
| POP1 | 1541.445619 | -0.965477692 | 3.54E-28 |
| PAK3 | 94.81753912 | -0.965586685 | 2.64E-02 |
| ARMC5 | 983.2956384 | -0.965983796 | 7.40E-16 |
| ESCO2 | 666.4800424 | -0.966348798 | 1.98E-04 |
| AHRR | 2060.829402 | -0.966647499 | 1.27E-22 |
| TSEN34 | 2579.236649 | -0.967322772 | 8.22E-12 |
| GDAP1 | 1189.14057 | -0.967461846 | 8.84E-12 |
| TPX2 | 13710.93008 | -0.967538205 | 1.48E-24 |
| CCNB1 | 16691.04164 | -0.967646405 | 5.64E-30 |
| TMSB15B | 331.5306934 | -0.967692952 | 1.22E-10 |
| AC012254.3 | 17.6987559 | -0.968561747 | 3.39E-02 |
| IP6K2 | 3247.021157 | -0.969602975 | 1.42E-15 |
| LINC02202 | 74.77782157 | -0.96960874 | 3.99E-03 |
| CYTL1 | 1870.470186 | -0.96967203 | 6.51E-14 |

| | | | |
|---|---|---|---|
| DTL | 1434.809302 | -0.969736549 | 6.90E-07 |
| C3orf70 | 235.4032375 | -0.969977177 | 3.64E-12 |
| COL1A1 | 951.6101759 | -0.970058926 | 1.53E-04 |
| LINC01670 | 119.7635046 | -0.972357672 | 7.11E-04 |
| FHOD1 | 2903.184332 | -0.972632009 | 3.69E-17 |
| TFEB | 289.5264894 | -0.973608681 | 1.24E-05 |
| SYCE1L | 52.13788359 | -0.974354816 | 1.79E-03 |
| AC129492.1 | 1124.648021 | -0.975377356 | 2.74E-09 |
| AC008467.1 | 33.37212616 | -0.975791765 | 2.76E-03 |
| NT5DC2 | 2832.960852 | -0.976123061 | 8.10E-08 |
| NAIP | 402.0713656 | -0.976333628 | 3.25E-12 |
| DOC2A | 97.79700132 | -0.97659083 | 2.50E-04 |
| AL773545.3 | 36.11612349 | -0.978280701 | 1.13E-02 |
| AC137767.1 | 53.94270911 | -0.979213662 | 1.23E-04 |
| EBPL | 1026.420101 | -0.979410515 | 1.56E-19 |
| MVK | 1449.963206 | -0.979569548 | 6.99E-05 |
| RPSAP52 | 35.61562351 | -0.979620194 | 4.21E-02 |
| ITPKB | 985.5116325 | -0.980608552 | 4.25E-09 |
| MCPH1-AS1 | 46.31687827 | -0.981094114 | 4.45E-03 |
| WAS | 120.5812993 | -0.981590951 | 4.11E-06 |
| NCAPD2 | 11336.63226 | -0.982757163 | 7.64E-34 |
| PLCD4 | 1142.359525 | -0.983092144 | 3.09E-07 |
| SOBP | 699.2504882 | -0.984253753 | 4.73E-19 |
| CU634019.7 | 82.61437957 | -0.984462734 | 1.20E-04 |
| MED15P9 | 38.96548172 | -0.985139054 | 1.29E-03 |
| LBR | 3086.654563 | -0.985557915 | 2.30E-28 |
| NA | 130.3090386 | -0.985568854 | 1.34E-06 |
| CDC25C | 1731.072231 | -0.986257773 | 7.26E-23 |
| FUT8-AS1 | 44.23740848 | -0.986495494 | 5.99E-03 |
| NPHP4 | 695.872442 | -0.986786757 | 7.73E-15 |
| HASPIN | 607.7763723 | -0.987762159 | 4.35E-07 |
| SHROOM3 | 3719.952372 | -0.988432921 | 3.60E-21 |
| LINC01963 | 343.5003131 | -0.988743666 | 2.25E-15 |
| AF186192.2 | 101.1648723 | -0.988868123 | 3.69E-06 |
| CLDN4 | 208.9669803 | -0.989052062 | 3.15E-07 |
| RNF144A | 711.0777741 | -0.989074785 | 4.88E-16 |
| DCLK2 | 707.8872453 | -0.989277721 | 6.50E-07 |
| AC148477.3 | 81.05410233 | -0.989646763 | 8.18E-05 |
| MARCKSL1 | 6894.580256 | -0.990824526 | 9.18E-12 |
| ITGB3BP | 726.7298882 | -0.991214085 | 6.23E-11 |
| SHOX | 186.8715178 | -0.991510586 | 2.49E-05 |
| RACGAP1 | 5057.43822 | -0.991564897 | 2.41E-28 |
| AQP3 | 243.1798854 | -0.991786334 | 1.20E-06 |
| AL590705.1 | 49.9269315 | -0.994016394 | 2.58E-04 |
| AC104964.4 | 21.11909575 | -0.994322023 | 3.39E-02 |
| HOXA4 | 390.6718035 | -0.995203444 | 2.99E-13 |
| ELN | 32.22984501 | -0.995236429 | 9.00E-03 |

| | | | |
|---|---|---|---|
| AL445623.1 | 26.5487547 | -0.995473886 | 4.25E-02 |
| AC109631.1 | 24.14286993 | -0.995547214 | 1.22E-02 |
| TPGS1 | 169.8498911 | -0.995578922 | 5.22E-07 |
| SOCS2-AS1 | 244.3080781 | -0.996293577 | 3.95E-05 |
| HJURP | 3243.06767 | -0.996779107 | 5.76E-16 |
| NA | 15.47188334 | -0.99818011 | 4.59E-02 |
| AL353763.1 | 33.38763978 | -0.99849593 | 1.43E-02 |
| NASP | 6492.894079 | -0.999075593 | 1.18E-13 |
| NDP | 64.67606061 | -0.999454485 | 3.47E-02 |
| LINC00894 | 215.4552753 | -1.000420572 | 6.96E-04 |
| MED15P7 | 23.90752639 | -1.000604981 | 3.09E-02 |
| MEIS2 | 693.3792644 | -1.000960644 | 6.50E-15 |
| AL035413.1 | 33.50113637 | -1.00099972 | 1.00E-02 |
| SNED1 | 2396.810234 | -1.002322223 | 1.23E-10 |
| NA | 209.1836675 | -1.002340197 | 3.84E-08 |
| AC091057.1 | 426.5278587 | -1.002843548 | 7.29E-06 |
| ZNF837 | 70.46561219 | -1.003308471 | 5.77E-05 |
| RAPGEF5 | 299.1615067 | -1.003590221 | 2.45E-06 |
| DRD2 | 181.3447405 | -1.003692111 | 3.52E-06 |
| ITGB8 | 92.24891433 | -1.003796873 | 1.46E-04 |
| FOXM1 | 5920.981055 | -1.004919409 | 1.45E-20 |
| KIRREL3 | 364.1978972 | -1.004993144 | 2.60E-05 |
| SFI1 | 851.4528004 | -1.005564828 | 2.09E-21 |
| AC090774.2 | 76.6145223 | -1.005965006 | 2.03E-02 |
| PIK3C2B | 986.2967161 | -1.006656362 | 1.33E-11 |
| SLC22A20P | 38.07364341 | -1.007461369 | 4.05E-03 |
| PYCR3 | 1124.038153 | -1.007571962 | 1.49E-16 |
| AL121832.3 | 132.6188102 | -1.007901575 | 2.93E-07 |
| TMSB15B-AS1 | 41.70051277 | -1.009229432 | 2.30E-02 |
| PRICKLE4 | 26.23740993 | -1.010402872 | 3.01E-02 |
| KIF14 | 937.9262382 | -1.011130268 | 1.67E-02 |
| CASP2 | 2799.212485 | -1.012190409 | 7.23E-13 |
| ZNF887P | 56.08603498 | -1.012341503 | 2.45E-03 |
| TSPY26P | 1120.670006 | -1.012786173 | 2.80E-10 |
| ZWINT | 1578.129043 | -1.012894109 | 5.31E-08 |
| TNFRSF1B | 1074.403305 | -1.013190528 | 9.03E-19 |
| AC254562.2 | 30.44230704 | -1.013211774 | 2.83E-02 |
| BNIPL | 32.83408996 | -1.013928893 | 2.08E-03 |
| PIK3R2 | 4034.204486 | -1.01435249 | 1.85E-21 |
| LZTS1 | 3910.639426 | -1.014718559 | 3.64E-19 |
| PLD2 | 1440.140448 | -1.014936439 | 1.76E-10 |
| MAD2L1 | 2085.432235 | -1.015985393 | 1.24E-08 |
| AC253536.6 | 33.45740573 | -1.016178474 | 8.22E-03 |
| ABCA11P | 119.0527732 | -1.016784459 | 2.35E-04 |
| SMC4 | 5575.897019 | -1.017256655 | 3.17E-05 |
| TTK | 1450.743421 | -1.018779873 | 6.36E-04 |
| RRM1 | 7162.102486 | -1.018819931 | 1.15E-07 |

| | | | |
|---|---|---|---|
| RIN1 | 1365.463973 | -1.018997375 | 3.18E-12 |
| NRP2 | 3214.243923 | -1.019516428 | 1.79E-35 |
| NA | 71.81406865 | -1.019583881 | 1.29E-05 |
| AL513165.1 | 119.8942597 | -1.020023949 | 1.26E-05 |
| TRHDE-AS1 | 181.9325313 | -1.020762136 | 2.40E-05 |
| NTHL1 | 700.6367838 | -1.020773605 | 5.63E-09 |
| EXOSC10-AS1 | 25.29981927 | -1.021313847 | 3.28E-02 |
| AC091153.3 | 18.33641116 | -1.024140474 | 3.63E-02 |
| RPL10P9 | 30.23878723 | -1.024338787 | 3.57E-02 |
| HNRNPA1 | 38484.91909 | -1.024365606 | 3.34E-44 |
| AC092535.1 | 18.53798512 | -1.024532941 | 4.56E-02 |
| AL133325.3 | 114.519684 | -1.024692812 | 1.06E-04 |
| PLEKHG4B | 728.7304898 | -1.024839424 | 1.94E-07 |
| SLC27A3 | 368.6739809 | -1.025338059 | 9.03E-07 |
| AC016773.1 | 28.24278393 | -1.025443971 | 2.39E-02 |
| AP001992.1 | 25.90380369 | -1.026234041 | 2.08E-02 |
| LINC01087 | 87.97856487 | -1.026669361 | 3.68E-05 |
| AC007996.1 | 132.0789388 | -1.026901735 | 1.93E-06 |
| FAM89B | 1379.792366 | -1.027089815 | 1.67E-15 |
| ACY1 | 1833.499762 | -1.027158848 | 2.32E-11 |
| RAMP1 | 108.3421471 | -1.028039228 | 2.63E-04 |
| KCTD12 | 316.8102465 | -1.028211885 | 1.09E-10 |
| LINC00342 | 173.0711505 | -1.028475594 | 1.58E-05 |
| ADGRF5 | 151.418413 | -1.029143192 | 5.39E-05 |
| C3AR1 | 62.64796635 | -1.029494292 | 1.27E-03 |
| ZNF395 | 5434.055165 | -1.031141441 | 1.23E-12 |
| AL390066.1 | 43.97765178 | -1.031211441 | 1.33E-03 |
| B4GALNT4 | 1110.705317 | -1.032053752 | 1.91E-08 |
| CDCA2 | 1941.804237 | -1.033063104 | 4.12E-18 |
| ZBTB7C | 327.535564 | -1.033182636 | 6.58E-05 |
| PANK1 | 348.7559744 | -1.033770879 | 2.57E-11 |
| EHMT2 | 4158.059463 | -1.033870642 | 1.09E-15 |
| HYAL3 | 410.5153546 | -1.033977248 | 1.29E-08 |
| CDC7 | 1042.852377 | -1.034339494 | 2.89E-10 |
| CRIP1 | 19.7420287 | -1.03456977 | 3.79E-02 |
| CORO2A | 1356.442854 | -1.034634599 | 7.52E-39 |
| BCL7C | 1987.222391 | -1.034736763 | 2.71E-18 |
| FOXA2 | 107.3111875 | -1.035150621 | 5.20E-04 |
| LINC01770 | 24.1553231 | -1.036074365 | 1.40E-02 |
| STAC2 | 256.3708383 | -1.037385226 | 2.01E-11 |
| LINC01126 | 42.41739339 | -1.037806405 | 8.70E-04 |
| AOAH | 33.00216439 | -1.037963694 | 4.32E-02 |
| SEMA3C | 1509.717254 | -1.038178383 | 7.30E-09 |
| AL450384.2 | 50.7999228 | -1.038435862 | 6.16E-03 |
| AC004846.1 | 97.33826787 | -1.038451727 | 3.60E-04 |
| DPYSL5 | 1435.744589 | -1.040957035 | 8.25E-21 |
| SPAG5 | 4370.625311 | -1.041587856 | 3.59E-13 |

| | | | |
|---|---|---|---|
| AC011379.1 | 25.8671817 | -1.041875928 | 5.26E-03 |
| CKAP2L | 1761.07225 | -1.042477533 | 1.59E-06 |
| MMP24 | 326.4706903 | -1.043048519 | 5.04E-07 |
| UBR7 | 3302.335545 | -1.043293426 | 1.17E-15 |
| AF186192.1 | 33.11822328 | -1.043889458 | 8.32E-03 |
| TUBB4B | 23483.62734 | -1.044293917 | 3.23E-11 |
| SYCP2 | 588.1176954 | -1.044492213 | 4.66E-04 |
| AL031283.2 | 36.90499216 | -1.04480669 | 1.36E-02 |
| BRI3BP | 787.7099777 | -1.047150456 | 8.13E-10 |
| AC138866.1 | 312.6038432 | -1.047409761 | 2.00E-09 |
| AP001486.2 | 32.71344285 | -1.048154597 | 1.32E-02 |
| FBXO5 | 1028.923715 | -1.048360753 | 8.56E-11 |
| AC021945.1 | 17.7759937 | -1.048366738 | 3.58E-02 |
| COL13A1 | 1295.802804 | -1.049024015 | 1.18E-11 |
| AC010973.2 | 51.84488928 | -1.049347273 | 7.12E-04 |
| EWSAT1 | 144.2678212 | -1.050096724 | 3.23E-03 |
| SYT11 | 517.0050096 | -1.051886786 | 3.90E-20 |
| TRIB2 | 5941.218021 | -1.052141373 | 7.69E-16 |
| SLC5A11 | 47.33680111 | -1.052538067 | 3.23E-02 |
| WRAP53 | 1064.570908 | -1.052623361 | 6.51E-08 |
| AC008608.2 | 35.30038961 | -1.053609858 | 9.52E-03 |
| BMPER | 162.7326693 | -1.053631654 | 1.24E-02 |
| ADAMTS3 | 506.1523499 | -1.054384243 | 3.51E-09 |
| TMTC4 | 1572.807064 | -1.054743199 | 2.63E-19 |
| IZUMO4 | 100.2450413 | -1.056288899 | 2.58E-06 |
| DLGAP5 | 2951.892619 | -1.05629848 | 2.21E-07 |
| AP003396.5 | 77.13102264 | -1.056396062 | 1.82E-02 |
| FAM57B | 70.39876574 | -1.057047055 | 4.09E-06 |
| BICC1 | 1248.921737 | -1.057338879 | 1.62E-19 |
| TFAP2A | 2148.867207 | -1.057437849 | 4.14E-19 |
| AC060780.3 | 14.00509407 | -1.058235775 | 3.12E-02 |
| CCDC180 | 210.7067393 | -1.059059869 | 1.26E-05 |
| KIF22 | 5070.961106 | -1.060232087 | 1.52E-25 |
| TUBB2B | 4106.075648 | -1.06063617 | 4.86E-12 |
| FAM171A1 | 2363.984497 | -1.061311993 | 2.34E-20 |
| HOXA5 | 769.5073558 | -1.062245521 | 5.10E-23 |
| TONSL | 1771.134022 | -1.062556156 | 1.33E-09 |
| AC004491.1 | 16.41975344 | -1.062759204 | 2.12E-02 |
| TNFRSF13C | 32.69246434 | -1.062903655 | 4.02E-02 |
| AC091078.2 | 27.36795842 | -1.062910199 | 2.37E-02 |
| SH3PXD2A | 3563.651804 | -1.063144191 | 1.23E-32 |
| FAM83D | 3176.229694 | -1.063165814 | 5.87E-23 |
| MAFB | 2006.013682 | -1.06357204 | 2.43E-08 |
| SELENBP1 | 2651.618252 | -1.0636722 | 7.24E-08 |
| PSRC1 | 2696.465216 | -1.063779567 | 1.03E-31 |
| POLN | 112.3528714 | -1.063967762 | 1.32E-04 |
| ELFN1-AS1 | 139.7402583 | -1.06474661 | 7.68E-08 |

| | | | |
|---|---|---|---|
| AP002807.1 | 119.5726333 | -1.064903521 | 5.60E-06 |
| BCAR3 | 3028.458632 | -1.064931901 | 4.72E-19 |
| TAL1 | 117.5851983 | -1.06606158 | 2.66E-05 |
| CHAF1A | 2043.033801 | -1.066470789 | 2.90E-05 |
| TMEM204 | 546.6914666 | -1.066704732 | 4.21E-16 |
| PDGFRA | 1613.900805 | -1.066707561 | 1.45E-12 |
| AL132780.5 | 20.48620529 | -1.066953087 | 3.12E-02 |
| AC120114.1 | 121.2192769 | -1.066973993 | 2.24E-08 |
| CDH5 | 1043.252914 | -1.067919552 | 1.49E-05 |
| NUF2 | 1217.065549 | -1.068390093 | 5.16E-05 |
| TSPOAP1 | 131.5695784 | -1.068775899 | 3.28E-05 |
| BOC | 1274.836958 | -1.069057177 | 3.87E-16 |
| COQ8A | 1679.746005 | -1.069303462 | 6.38E-15 |
| AC005837.1 | 17.76815366 | -1.069662372 | 4.02E-02 |
| IGHV3-66 | 19.3309565 | -1.070275333 | 3.38E-02 |
| AC073254.1 | 18.88198772 | -1.070615421 | 2.38E-02 |
| AC103563.3 | 94.0265393 | -1.07128824 | 1.67E-05 |
| TRIP13 | 4766.872522 | -1.071462933 | 1.25E-11 |
| AL357060.1 | 57.41758522 | -1.071745468 | 2.75E-04 |
| CYP2D6 | 86.52775803 | -1.07220321 | 9.24E-06 |
| ADAMTSL2 | 89.19405861 | -1.072390487 | 2.12E-06 |
| KIF15 | 1673.069654 | -1.072549742 | 1.22E-07 |
| C1orf220 | 69.7777801 | -1.073290211 | 1.09E-03 |
| MED15P6 | 29.92975537 | -1.073989588 | 2.11E-02 |
| AC234783.1 | 28.84823402 | -1.074024841 | 7.84E-03 |
| BX664727.3 | 36.41130282 | -1.074471194 | 1.35E-02 |
| GRWD1 | 2844.295269 | -1.074609663 | 2.48E-16 |
| WDR76 | 686.2599006 | -1.075376956 | 9.33E-04 |
| NA | 440.3647721 | -1.075727317 | 1.33E-09 |
| LAT2 | 82.5730175 | -1.076988103 | 2.60E-04 |
| AC090739.1 | 23.09489251 | -1.077151857 | 8.21E-03 |
| SP6 | 120.9574354 | -1.078108885 | 3.61E-05 |
| VPS9D1-AS1 | 411.3095622 | -1.079218283 | 5.23E-06 |
| CAMKK1 | 327.9390094 | -1.080475634 | 2.34E-12 |
| AC008013.1 | 35.41368548 | -1.081077241 | 1.93E-02 |
| AC009902.2 | 45.96325468 | -1.082308905 | 2.48E-04 |
| CDKN2C | 1684.89081 | -1.082362075 | 5.17E-10 |
| LECT2 | 122.0354449 | -1.082373481 | 7.47E-04 |
| ANKRD44 | 619.7998958 | -1.083770828 | 1.31E-04 |
| UBE2C | 4006.039724 | -1.083971216 | 2.93E-21 |
| AC092198.1 | 26.27954271 | -1.084462687 | 3.53E-02 |
| CYP4F32P | 226.164921 | -1.084617516 | 1.69E-11 |
| PRR34 | 41.40607072 | -1.084787162 | 7.20E-03 |
| CENPU | 1296.012335 | -1.085003043 | 3.89E-13 |
| HSD11B2 | 26.99250525 | -1.085479412 | 1.12E-02 |
| AC010542.5 | 24.71451806 | -1.085773318 | 3.11E-02 |
| AC006111.2 | 16.47899019 | -1.087400458 | 4.25E-02 |

| | | | |
|---|---|---|---|
| PARD3-AS1 | 19.68112082 | -1.087813413 | 3.68E-02 |
| ZP2 | 35.43078166 | -1.088113509 | 4.01E-02 |
| SSX2 | 555.8518566 | -1.088822419 | 4.08E-05 |
| HSPA1L | 100.3602291 | -1.089080696 | 5.33E-07 |
| ARHGEF19 | 358.8169912 | -1.090139692 | 6.63E-11 |
| AL136116.3 | 28.4971429 | -1.090496172 | 1.56E-02 |
| AL162171.4 | 23.20073892 | -1.090735527 | 1.45E-02 |
| TYRP1 | 289.0563429 | -1.090992783 | 3.32E-13 |
| ECSCR | 314.089158 | -1.092368531 | 4.92E-11 |
| WDHD1 | 1399.6444 | -1.093047416 | 1.53E-06 |
| AC005703.6 | 31.16690174 | -1.094053371 | 2.56E-03 |
| BMF | 718.4286987 | -1.094353524 | 2.83E-11 |
| CENPO | 2183.214192 | -1.09481837 | 4.38E-10 |
| AC025449.1 | 22.665462 | -1.094885359 | 2.09E-02 |
| AC073842.2 | 16.81019481 | -1.094924247 | 4.27E-02 |
| ATRIP | 655.0171284 | -1.095030431 | 3.91E-13 |
| AC007952.6 | 69.57289609 | -1.095193106 | 7.22E-05 |
| JAK3 | 626.4733155 | -1.095457854 | 6.70E-07 |
| BX119917.1 | 35.28777911 | -1.09630897 | 8.08E-04 |
| TNFRSF25 | 230.3667618 | -1.096624218 | 1.00E-11 |
| PTPRN2 | 749.2135336 | -1.097006669 | 2.42E-05 |
| CCDC39 | 40.01878636 | -1.09848388 | 4.40E-02 |
| LINC01977 | 90.72125105 | -1.098740112 | 1.43E-07 |
| ZNF367 | 499.3917033 | -1.099741221 | 1.36E-02 |
| SORBS1 | 666.9068481 | -1.100713718 | 1.14E-21 |
| PDE5A | 1577.849555 | -1.100884578 | 7.28E-09 |
| SRD5A1 | 2652.925209 | -1.100885501 | 1.80E-62 |
| KIAA0825 | 59.54724542 | -1.101344799 | 3.50E-03 |
| SIGLEC17P | 28.70313536 | -1.102925695 | 7.98E-03 |
| MN1 | 809.3964248 | -1.103648679 | 9.60E-11 |
| LARGE2 | 219.8105223 | -1.10478723 | 7.28E-07 |
| TMEM198 | 334.0886835 | -1.104961084 | 2.29E-12 |
| CRIP2 | 4276.416024 | -1.105069357 | 9.67E-20 |
| EFNB2 | 435.0140339 | -1.1051028 | 1.69E-23 |
| RUNX1 | 2054.795126 | -1.1051764 | 7.63E-21 |
| PLK4 | 998.2687593 | -1.105385661 | 5.52E-05 |
| MXD4 | 4041.346101 | -1.105648643 | 9.80E-07 |
| AC126335.2 | 44.10181047 | -1.10735458 | 1.87E-02 |
| TUBB4A | 872.2014509 | -1.107605302 | 2.23E-13 |
| FPR1 | 28.60611966 | -1.10821059 | 1.40E-02 |
| AL603839.2 | 21.50821683 | -1.108270151 | 3.76E-02 |
| TMEM249 | 28.67557228 | -1.109271408 | 4.66E-02 |
| CAMK4 | 96.91924048 | -1.109693511 | 4.83E-04 |
| ARHGAP19 | 772.317132 | -1.109835776 | 7.52E-18 |
| STARD4-AS1 | 775.5088185 | -1.110896727 | 2.51E-05 |
| PROCR | 2832.564645 | -1.112353662 | 1.67E-20 |
| LHX2 | 305.5215854 | -1.112749153 | 5.98E-06 |

| | | | |
|---|---|---|---|
| INKA1 | 307.9654041 | -1.113124607 | 3.15E-09 |
| CPXM1 | 3098.915536 | -1.113820866 | 6.05E-14 |
| FDXACB1 | 160.5580444 | -1.114214272 | 9.43E-07 |
| LETM2 | 181.2562158 | -1.115437792 | 3.94E-06 |
| AKR7L | 26.4332983 | -1.117891253 | 1.14E-02 |
| NRP1 | 5641.832865 | -1.117906832 | 7.00E-18 |
| AP001011.1 | 16.03711086 | -1.118359082 | 4.72E-02 |
| MMD | 3317.81114 | -1.118690566 | 3.89E-26 |
| STX1A | 710.3622965 | -1.119178916 | 2.31E-21 |
| DNMT3B | 895.4449263 | -1.120036051 | 3.67E-20 |
| EBF1 | 927.3327661 | -1.120886948 | 1.43E-07 |
| KCNMA1 | 1312.170868 | -1.122441448 | 1.25E-07 |
| PTX3 | 4981.988845 | -1.122728837 | 8.63E-09 |
| DPF1 | 212.3406063 | -1.123410356 | 5.87E-08 |
| NF1P6 | 29.64198724 | -1.123746233 | 3.54E-03 |
| FAM20A | 268.4477144 | -1.127063243 | 2.00E-04 |
| WDR90 | 1545.088941 | -1.127408525 | 1.72E-13 |
| AC007292.1 | 22.58854334 | -1.127939602 | 1.43E-02 |
| PBX4 | 93.55397893 | -1.127971518 | 2.01E-06 |
| FAM187A | 233.2630473 | -1.12809712 | 2.05E-13 |
| DPY19L2P1 | 178.5518984 | -1.128306291 | 6.45E-05 |
| SOX15 | 45.37839204 | -1.128402553 | 4.18E-04 |
| STK32B | 1361.46195 | -1.129364313 | 5.14E-18 |
| RAB36 | 529.2506332 | -1.129623703 | 6.05E-21 |
| AC097534.2 | 72.72643838 | -1.130395124 | 3.56E-05 |
| NRXN2 | 691.5951493 | -1.130501278 | 1.85E-14 |
| TERT | 319.3671589 | -1.13105592 | 2.85E-07 |
| AC112484.3 | 34.69759026 | -1.131193516 | 1.85E-03 |
| CYP4F35P | 52.7238723 | -1.131956297 | 3.11E-04 |
| PROM2 | 46.85997022 | -1.132685173 | 4.64E-02 |
| RPL13AP25 | 30.00146136 | -1.13360572 | 1.64E-02 |
| MSH5-SAPCD1 | 219.4053818 | -1.135043603 | 2.34E-05 |
| EVX1 | 13.13102745 | -1.13558089 | 3.87E-02 |
| RHOV | 18.9921507 | -1.136157176 | 2.36E-02 |
| PIF1 | 875.719037 | -1.136386761 | 7.86E-19 |
| SELE | 16.48521074 | -1.136558484 | 2.49E-02 |
| BEX1 | 1622.382863 | -1.13814903 | 8.62E-21 |
| GLYAT | 20.60510753 | -1.138546148 | 3.48E-02 |
| AC010809.1 | 20.89926275 | -1.139966057 | 3.24E-02 |
| TEDC2 | 938.6373295 | -1.140682571 | 3.11E-19 |
| AP002784.1 | 15.36806646 | -1.141382326 | 3.07E-02 |
| AC134043.2 | 93.25974137 | -1.141741811 | 7.87E-04 |
| SPATA33 | 423.1964117 | -1.143237074 | 4.43E-14 |
| TIMELESS | 5461.253155 | -1.146237272 | 6.88E-43 |
| KIF4A | 4156.483902 | -1.146265955 | 8.56E-30 |
| GVQW3 | 442.8556818 | -1.146616799 | 1.45E-11 |
| LINC01237 | 22.92773012 | -1.146772694 | 2.14E-02 |

| | | | |
|---|---|---|---|
| LGALS1 | 23005.81072 | -1.147783756 | 7.09E-19 |
| PLEKHA7 | 2443.226225 | -1.147909397 | 2.39E-12 |
| AC093249.2 | 23.67802045 | -1.1481207 | 4.59E-03 |
| GLDC | 119.3079729 | -1.148369253 | 2.02E-07 |
| HOXA-AS3 | 169.3705619 | -1.148572362 | 5.37E-08 |
| AL512310.11 | 40.74662864 | -1.149886549 | 2.45E-04 |
| NA | 39.83765738 | -1.150150627 | 1.47E-04 |
| PLXDC1 | 87.55184205 | -1.150593709 | 8.82E-03 |
| MTUS2 | 74.85023664 | -1.150710192 | 6.63E-05 |
| SLC12A9-AS1 | 29.85792199 | -1.150851974 | 2.23E-03 |
| LOXL1-AS1 | 19.08140033 | -1.151224418 | 4.90E-02 |
| DDX12P | 194.3383618 | -1.1512958 | 1.45E-09 |
| NA | 1596.217383 | -1.152947492 | 2.64E-33 |
| SAPCD1 | 48.82728607 | -1.153625642 | 2.52E-03 |
| HAPLN4 | 233.8119336 | -1.154149614 | 1.67E-06 |
| AC004076.2 | 16.73431622 | -1.155838759 | 1.83E-02 |
| LOXL1 | 102.2955771 | -1.156351554 | 3.81E-05 |
| AURKB | 2643.519517 | -1.156408832 | 7.65E-20 |
| TMSB15B | 161.5483094 | -1.156819876 | 4.58E-07 |
| KRTCAP3 | 21.62333393 | -1.15759845 | 6.27E-03 |
| STMN1 | 17626.57597 | -1.157631146 | 4.86E-32 |
| CKAP2 | 2416.929446 | -1.15768036 | 1.52E-12 |
| PRKAR1B | 1954.365488 | -1.158899336 | 1.88E-16 |
| LRRC45 | 550.2770296 | -1.1596361 | 1.81E-08 |
| LINC01711 | 47.1064077 | -1.160814389 | 5.92E-04 |
| CENPK | 922.9012945 | -1.161809924 | 4.81E-07 |
| PBX3 | 961.2103878 | -1.16211615 | 2.42E-36 |
| NDC80 | 1820.079162 | -1.162665998 | 5.70E-10 |
| RARRES1 | 136.06074 | -1.163220763 | 1.10E-04 |
| CCDC18 | 362.2515424 | -1.165325452 | 6.21E-04 |
| DNER | 848.2215533 | -1.166490973 | 2.64E-17 |
| TMSB15A | 178.0663652 | -1.167455939 | 1.00E-10 |
| AC007663.4 | 28.0616015 | -1.167487438 | 3.45E-03 |
| AL158013.1 | 39.60705562 | -1.168183993 | 4.55E-03 |
| PCDH19 | 17.98510115 | -1.169756098 | 1.41E-02 |
| MKI67 | 9082.160526 | -1.171332525 | 1.35E-13 |
| PAPPA | 2557.838757 | -1.171390157 | 1.59E-05 |
| AC008894.3 | 44.06928147 | -1.17364718 | 5.13E-04 |
| PSMC3IP | 718.5035865 | -1.174237562 | 3.19E-11 |
| SENP3-EIF4A1 | 170.1265395 | -1.176495223 | 9.44E-05 |
| AC022150.3 | 13.31789858 | -1.177136994 | 4.10E-02 |
| AC025588.1 | 23.81622268 | -1.177385402 | 4.36E-03 |
| AC106795.2 | 19.42540368 | -1.177692357 | 2.19E-02 |
| CFAP44 | 349.6453516 | -1.178155908 | 3.58E-07 |
| CU638689.5 | 79.83533743 | -1.179496834 | 8.03E-07 |
| FAT3 | 83.64781789 | -1.179803049 | 2.85E-02 |
| ATAD5 | 479.5242427 | -1.180234189 | 1.95E-04 |

| | | | |
|---|---|---|---|
| OXTR | 109.0439804 | -1.180770896 | 2.94E-03 |
| MAPK8IP1 | 1166.539598 | -1.181475484 | 8.63E-20 |
| CCNE2 | 275.1172858 | -1.181864488 | 8.23E-07 |
| GPRIN1 | 770.8989094 | -1.182739206 | 5.05E-14 |
| RPS2P46 | 29.36839468 | -1.183202931 | 3.63E-03 |
| SSX3 | 30.18188152 | -1.183351703 | 1.30E-03 |
| RN7SL268P | 15.40962059 | -1.184096912 | 1.87E-02 |
| ANKRD36B | 215.0634424 | -1.184468367 | 1.51E-03 |
| AL359921.2 | 32.19723881 | -1.18446879 | 1.82E-03 |
| BRIP1 | 600.5206492 | -1.185596938 | 4.69E-07 |
| ARHGEF39 | 1059.646998 | -1.186048968 | 4.48E-20 |
| MPP4 | 50.02754041 | -1.189105713 | 3.98E-02 |
| LRRC17 | 2568.192514 | -1.189645869 | 1.46E-11 |
| AC096537.1 | 58.48260446 | -1.189953134 | 1.79E-04 |
| RECQL4 | 2969.427958 | -1.190631969 | 1.85E-11 |
| PRPS2 | 2404.952892 | -1.191254372 | 2.06E-21 |
| RIPOR2 | 1126.46556 | -1.191708591 | 2.54E-08 |
| CHAF1B | 1422.02079 | -1.1918416 | 1.25E-07 |
| AP000640.1 | 28.79122399 | -1.191933463 | 1.37E-03 |
| TUBB3 | 3262.462153 | -1.192106827 | 5.17E-17 |
| BTN1A1 | 14.31306556 | -1.19217409 | 4.32E-02 |
| ERCC6L | 856.5915759 | -1.192192818 | 2.77E-09 |
| MCM7 | 17692.15162 | -1.19226555 | 2.31E-10 |
| MYO15B | 38.86608803 | -1.192704841 | 8.91E-03 |
| GJD4 | 14.77319841 | -1.192905791 | 4.21E-02 |
| DHFR | 7109.258431 | -1.193967877 | 6.57E-14 |
| FANCL | 998.7763732 | -1.194294028 | 2.45E-29 |
| LINC00242 | 19.69388058 | -1.194368794 | 3.15E-02 |
| NFATC2 | 610.572034 | -1.194790337 | 1.05E-08 |
| PRIM1 | 1111.63247 | -1.195185681 | 1.54E-10 |
| AC138409.2 | 185.7386091 | -1.195863774 | 5.23E-11 |
| AC244021.1 | 26.25435153 | -1.197259116 | 8.25E-03 |
| EPHB3 | 918.3399535 | -1.197543349 | 8.05E-19 |
| NA | 27.53951476 | -1.198077776 | 2.18E-03 |
| TACC3 | 5029.8107 | -1.19833088 | 6.56E-39 |
| BFSP2-AS1 | 17.70852641 | -1.198561718 | 3.00E-02 |
| CU634019.6 | 561.8181909 | -1.19935267 | 6.78E-13 |
| HOPX | 22.94674443 | -1.200026013 | 3.40E-03 |
| SLCO2A1 | 1257.747719 | -1.200191398 | 3.01E-31 |
| BLM | 908.9205934 | -1.200361365 | 1.59E-11 |
| COLEC10 | 58.18578616 | -1.200586529 | 1.18E-04 |
| NES | 7130.448034 | -1.200592608 | 1.12E-13 |
| YPEL1 | 157.0091846 | -1.200951675 | 8.72E-05 |
| CMBL | 2580.311176 | -1.201182212 | 7.13E-17 |
| AC118754.1 | 83.43697291 | -1.20141065 | 3.31E-06 |
| ANPEP | 1081.191916 | -1.202093162 | 1.53E-25 |
| AC015818.2 | 21.48308687 | -1.202450628 | 5.51E-03 |

| | | | |
|---|---|---|---|
| AC009902.3 | 19.06579071 | -1.202943112 | 1.48E-02 |
| SOX4 | 406.2559629 | -1.202951211 | 1.09E-09 |
| TBC1D4 | 1063.760572 | -1.203209739 | 7.11E-20 |
| CSGALNACT1 | 578.8533399 | -1.204592238 | 5.74E-17 |
| AP000527.1 | 63.32417353 | -1.204914194 | 3.81E-05 |
| AC008610.1 | 16.74355191 | -1.205902305 | 1.45E-02 |
| BRAT1 | 2764.474164 | -1.207167361 | 3.44E-19 |
| NHSL2 | 362.3466133 | -1.207714797 | 2.49E-12 |
| THY1 | 1129.874395 | -1.20799212 | 1.82E-06 |
| SLC23A3 | 29.17969002 | -1.208297715 | 2.45E-03 |
| E2F1 | 1910.727677 | -1.208330996 | 6.84E-03 |
| REM2 | 34.64299948 | -1.208988242 | 3.77E-04 |
| CCDC103 | 32.8956705 | -1.209146833 | 7.97E-03 |
| SERTAD4-AS1 | 147.6901043 | -1.209615835 | 1.68E-08 |
| KHK | 169.8357938 | -1.210261841 | 7.12E-10 |
| KIF18A | 852.0039837 | -1.211327368 | 3.60E-06 |
| NBL1 | 2346.529476 | -1.211629414 | 2.99E-19 |
| BX005266.2 | 176.8294464 | -1.213612828 | 3.57E-05 |
| WASIR2 | 30.59058839 | -1.214412395 | 1.54E-03 |
| IMPDH1P8 | 64.51239715 | -1.21463415 | 2.32E-05 |
| WEE1 | 1947.39113 | -1.214786507 | 1.12E-18 |
| COLQ | 55.80094762 | -1.215140172 | 4.22E-03 |
| NTNG2 | 150.2818736 | -1.215525323 | 1.38E-03 |
| NF1P7 | 19.5947773 | -1.215663848 | 6.44E-03 |
| PARP1 | 12306.40327 | -1.215840307 | 1.23E-26 |
| FBXO43 | 180.2093421 | -1.216425615 | 1.54E-08 |
| TEDC1 | 659.3959808 | -1.218469878 | 7.14E-14 |
| ACSL6 | 38.72385866 | -1.219240777 | 5.61E-03 |
| AL021877.2 | 56.88130604 | -1.221487451 | 6.00E-06 |
| RMI2 | 1653.217522 | -1.222547043 | 7.43E-12 |
| FXYD2 | 41.98422852 | -1.22298548 | 3.64E-03 |
| KIF11 | 2007.052058 | -1.223279688 | 2.79E-10 |
| AF274858.1 | 22.2363708 | -1.223591518 | 1.22E-02 |
| MTND2P28 | 235.219453 | -1.224273808 | 9.89E-04 |
| RIBC1 | 84.71474259 | -1.224693353 | 5.12E-08 |
| AL929601.4 | 26.25866688 | -1.22546022 | 1.04E-02 |
| HAND1 | 50.73723808 | -1.225988815 | 2.17E-02 |
| CNIH2 | 324.591191 | -1.226098657 | 2.32E-06 |
| NEIL3 | 593.7359702 | -1.228200754 | 3.67E-12 |
| SEC14L6 | 14.29596388 | -1.228255934 | 1.73E-02 |
| LINC00565 | 61.82331782 | -1.2282648 | 3.01E-05 |
| KDR | 808.088877 | -1.228971153 | 3.95E-06 |
| FAM96AP2 | 18.82771174 | -1.229688047 | 2.89E-02 |
| AC009506.1 | 50.51966715 | -1.229827703 | 6.52E-05 |
| DACT1 | 395.7859807 | -1.23021833 | 2.58E-10 |
| NA | 24.42026338 | -1.230680478 | 4.23E-03 |
| EXOC3L1 | 83.24562736 | -1.232639844 | 6.45E-05 |

| | | | |
|---|---|---|---|
| SAMD15 | 178.1043318 | -1.23603179 | 1.11E-08 |
| FERMT1 | 589.4759035 | -1.23618174 | 5.28E-16 |
| MIA | 176.8750646 | -1.237100047 | 2.67E-07 |
| RASSF10 | 29.88727853 | -1.237316955 | 1.52E-03 |
| MT-TT | 23.81572381 | -1.239031017 | 2.64E-02 |
| SUFU | 1234.590452 | -1.241650378 | 7.73E-16 |
| ZNF835 | 58.20874136 | -1.242366376 | 2.58E-02 |
| NTSR1 | 30.50159874 | -1.242366797 | 3.74E-03 |
| FAM95B1 | 154.0703667 | -1.243925992 | 1.85E-05 |
| LRRC18 | 14.9253248 | -1.244782145 | 3.13E-02 |
| SLC14A2 | 38.02249123 | -1.245069578 | 1.72E-03 |
| NPIPA5 | 155.7742575 | -1.245312139 | 7.82E-04 |
| MAF | 556.0220594 | -1.246593023 | 1.22E-19 |
| KCNAB3 | 147.848877 | -1.247419273 | 1.35E-10 |
| OPN3 | 842.1645973 | -1.247469058 | 4.04E-23 |
| ATP2A3 | 1211.316063 | -1.247548268 | 5.97E-25 |
| PTOV1-AS2 | 99.07661411 | -1.247944716 | 1.92E-08 |
| PAXBP1-AS1 | 94.88010745 | -1.248044719 | 3.94E-10 |
| ZNF488 | 398.2968029 | -1.248469859 | 9.96E-08 |
| AC092821.1 | 44.78592829 | -1.249776638 | 1.89E-03 |
| NME9 | 23.61128077 | -1.250197527 | 1.49E-02 |
| NAV2-AS4 | 43.2774696 | -1.252735046 | 9.24E-04 |
| GRAP | 220.9100729 | -1.255057863 | 6.98E-08 |
| ZNF29P | 10.7398875 | -1.256062015 | 3.94E-02 |
| FOXP3 | 47.74772429 | -1.256868342 | 2.52E-05 |
| AC110285.2 | 75.3928026 | -1.259727468 | 4.11E-08 |
| AL669942.1 | 573.7894056 | -1.261225021 | 6.31E-20 |
| CCDC34 | 1220.437521 | -1.261354631 | 4.77E-22 |
| SLC12A2 | 5479.546749 | -1.261640902 | 5.79E-21 |
| PTPN22 | 24.45633375 | -1.263760464 | 3.20E-02 |
| NA | 16.72335002 | -1.264791078 | 4.77E-02 |
| KIAA1549L | 2687.335253 | -1.264931896 | 2.29E-30 |
| NALCN | 1270.957975 | -1.265349143 | 5.03E-52 |
| AC011468.5 | 30.75586694 | -1.26601875 | 1.47E-04 |
| MMP9 | 160.911846 | -1.266456595 | 7.15E-05 |
| AL035448.1 | 10.52901536 | -1.268465151 | 4.81E-02 |
| SPC24 | 981.3362757 | -1.269024384 | 1.23E-12 |
| BIRC5 | 4253.014258 | -1.269130306 | 7.40E-29 |
| NKX2-4 | 192.9784856 | -1.26929047 | 7.91E-12 |
| AC021106.1 | 31.7070498 | -1.269602678 | 1.05E-03 |
| TNFRSF21 | 5207.604204 | -1.271446261 | 2.53E-57 |
| TRIM71 | 31.17178457 | -1.273335935 | 2.35E-03 |
| AC005329.1 | 103.148734 | -1.273427944 | 3.90E-03 |
| FAM222A | 185.6045506 | -1.273503175 | 3.18E-11 |
| AC005482.1 | 12.43280898 | -1.274659606 | 4.25E-02 |
| FAM110A | 439.3631602 | -1.27549312 | 9.95E-27 |
| HIC1 | 996.4257033 | -1.277081246 | 1.29E-08 |

| | | | |
|---|---|---|---|
| ADAMTS6 | 290.3621183 | -1.27708446 | 1.59E-11 |
| CDK5R1 | 277.1423831 | -1.278187965 | 2.45E-10 |
| JPH4 | 834.6588439 | -1.279779086 | 2.51E-07 |
| AC138649.1 | 32.3201417 | -1.279782898 | 3.65E-03 |
| NOTUM | 82.32665438 | -1.279803181 | 6.17E-05 |
| IQCH-AS1 | 120.9287616 | -1.28022277 | 3.20E-04 |
| NCAPH | 2363.792997 | -1.28026876 | 5.93E-16 |
| CLCN1 | 55.22163434 | -1.280438235 | 5.76E-05 |
| AC011445.1 | 28.48409379 | -1.282466627 | 8.86E-03 |
| NA | 36.98258257 | -1.283727756 | 4.72E-03 |
| AC140134.1 | 277.8328783 | -1.284976831 | 1.10E-08 |
| AC093677.2 | 17.96716077 | -1.285136984 | 1.37E-02 |
| SLC7A10 | 92.31430533 | -1.285181019 | 7.17E-06 |
| TIGD3 | 41.09771545 | -1.285268122 | 7.19E-04 |
| ADAMTS18 | 25.18789751 | -1.286176579 | 4.13E-02 |
| ARHGAP28 | 1096.61935 | -1.287148998 | 8.86E-11 |
| BAMBI | 4010.746087 | -1.287953756 | 6.98E-14 |
| FANCA | 2228.666704 | -1.288620023 | 2.36E-10 |
| PDCD4-AS1 | 327.1842865 | -1.288828619 | 4.31E-13 |
| TCF19 | 1894.285729 | -1.289116419 | 1.06E-09 |
| AGBL2 | 104.4122325 | -1.289954622 | 1.62E-06 |
| AC009533.2 | 17.73738798 | -1.290833987 | 3.22E-02 |
| AC027601.3 | 16.33985414 | -1.291061652 | 2.99E-02 |
| TMOD1 | 1497.162289 | -1.291799912 | 4.86E-10 |
| HEY1 | 1560.216718 | -1.292333862 | 1.77E-11 |
| TRERF1 | 1637.405227 | -1.293502391 | 8.57E-23 |
| CDC45 | 1432.239354 | -1.29386524 | 1.20E-10 |
| CDC20 | 4779.118958 | -1.294770646 | 2.70E-23 |
| APCDD1L | 1106.51774 | -1.296265104 | 4.23E-12 |
| FGD3 | 69.12946047 | -1.297371486 | 1.70E-05 |
| AC024075.1 | 11.31564554 | -1.298130613 | 2.38E-02 |
| FANCE | 547.4079603 | -1.298269483 | 1.88E-30 |
| AC120049.1 | 38.88637755 | -1.298965115 | 3.77E-04 |
| APCDD1 | 31.18290478 | -1.301479412 | 1.01E-02 |
| CD160 | 21.21898451 | -1.302267225 | 4.48E-03 |
| GPR162 | 140.5579805 | -1.302283286 | 1.02E-07 |
| ZFP2 | 31.7747584 | -1.303741281 | 8.14E-04 |
| RASL11B | 84.07592989 | -1.304105914 | 6.27E-08 |
| LTB | 51.92894334 | -1.304973095 | 2.48E-03 |
| C19orf57 | 543.2283454 | -1.308333184 | 1.71E-24 |
| NA | 45.78185892 | -1.309412325 | 2.27E-05 |
| SYCE2 | 183.0282083 | -1.309453656 | 5.09E-08 |
| ATXN7L2 | 355.4580206 | -1.310389849 | 5.36E-21 |
| SERPINE3 | 16.91027922 | -1.310392003 | 2.22E-02 |
| MYC | 932.6837416 | -1.310767782 | 1.29E-05 |
| NTRK1 | 102.2523374 | -1.310868243 | 1.92E-08 |
| AC239868.2 | 154.2788812 | -1.312631027 | 3.80E-15 |

| | | | |
|---|---|---|---|
| RAB40B | 390.349998 | -1.31455983 | 7.63E-17 |
| CD3E | 18.90742972 | -1.314659225 | 3.57E-02 |
| SH3TC2 | 503.4662709 | -1.315041591 | 1.27E-09 |
| FAM78A | 383.5158475 | -1.315704626 | 4.69E-25 |
| AC087742.1 | 12.64680862 | -1.315982618 | 2.39E-02 |
| SLA | 35.9612121 | -1.317075605 | 7.50E-03 |
| TICRR | 1882.00171 | -1.317964609 | 6.41E-22 |
| DUSP9 | 394.385755 | -1.31805684 | 1.37E-11 |
| MIR25 | 38.65376894 | -1.318346971 | 7.28E-04 |
| AC008264.2 | 36.20563716 | -1.318851814 | 1.02E-02 |
| U82695.1 | 9.740562148 | -1.319478966 | 4.12E-02 |
| UCP3 | 55.17028068 | -1.320218183 | 3.71E-04 |
| SUSD2 | 605.236627 | -1.320566853 | 3.06E-12 |
| AC004801.2 | 21.85766275 | -1.320615786 | 3.69E-02 |
| GLI3 | 3529.66754 | -1.321069577 | 3.97E-34 |
| CLDND2 | 28.29589719 | -1.32219374 | 3.40E-03 |
| ME3 | 73.97688032 | -1.32323664 | 8.71E-04 |
| AC139491.3 | 70.41335232 | -1.32375658 | 4.22E-05 |
| AL591848.4 | 12.13650127 | -1.324722336 | 2.61E-02 |
| NOX5 | 244.0562303 | -1.325136199 | 4.04E-10 |
| SH2D1B | 48.08555972 | -1.325244941 | 1.46E-03 |
| AC027307.3 | 30.32121398 | -1.329615389 | 7.92E-04 |
| AC103563.4 | 91.89867608 | -1.329647399 | 4.43E-06 |
| ZMAT4 | 504.9050422 | -1.332804513 | 1.93E-14 |
| NACAD | 1177.507431 | -1.334939485 | 6.97E-15 |
| DDX11 | 1754.937538 | -1.335214157 | 1.04E-19 |
| LINC01929 | 20.14398567 | -1.336796698 | 4.81E-02 |
| TCIM | 39.7471214 | -1.338025869 | 2.54E-03 |
| QRICH2 | 106.3556761 | -1.33844501 | 2.67E-08 |
| IPO4 | 3946.715919 | -1.339046981 | 8.04E-48 |
| COL9A3 | 4082.534637 | -1.339215519 | 8.64E-18 |
| PAQR8 | 949.7430723 | -1.339397701 | 3.14E-29 |
| AC099066.2 | 26.07679972 | -1.339617548 | 2.01E-02 |
| NEURL1B | 558.7832072 | -1.33975153 | 2.29E-12 |
| RCAN2 | 2097.152447 | -1.339861012 | 1.67E-12 |
| RPL13AP5 | 31.68415491 | -1.342736023 | 6.60E-03 |
| HSPA12B | 366.7055406 | -1.342803148 | 1.81E-13 |
| AC006027.1 | 33.73319651 | -1.342873789 | 3.18E-02 |
| PTGER2 | 910.6436438 | -1.344965018 | 1.58E-13 |
| ZCCHC18 | 174.0764069 | -1.345208116 | 8.72E-06 |
| CHRNA5 | 352.9120654 | -1.345428077 | 1.19E-12 |
| SLC38A4 | 1027.095872 | -1.346131305 | 2.74E-23 |
| IFNWP19 | 27.0852413 | -1.346312365 | 3.46E-03 |
| LPAR6 | 141.6397686 | -1.348237197 | 1.66E-05 |
| AC073655.2 | 12.2572459 | -1.348485152 | 3.89E-02 |
| LINC00327 | 20.45128611 | -1.34889619 | 1.59E-02 |
| AC009812.4 | 17.00709006 | -1.348997244 | 1.44E-02 |

| | | | |
|---|---|---|---|
| AC005041.1 | 156.4021284 | -1.35024876 | 6.63E-10 |
| MCM2 | 7582.703007 | -1.351642487 | 1.71E-10 |
| AC234782.3 | 13.05435962 | -1.351985703 | 2.89E-02 |
| CACNB2 | 46.1342047 | -1.35226728 | 7.51E-04 |
| AC092849.1 | 12.41354471 | -1.352387734 | 4.23E-02 |
| AL512310.10 | 21.73922699 | -1.352492345 | 8.96E-03 |
| FHDC1 | 144.9143232 | -1.352531051 | 3.14E-08 |
| CENPM | 685.6041869 | -1.353930351 | 2.04E-27 |
| CHI3L1 | 17.46316668 | -1.354346212 | 4.80E-02 |
| SHISAL2A | 14.73665807 | -1.355004594 | 3.24E-02 |
| SLC4A8 | 216.6168844 | -1.358985057 | 1.24E-06 |
| RPS2P7 | 39.60609226 | -1.359619701 | 1.77E-03 |
| DPY19L2P2 | 694.3173376 | -1.361777255 | 1.06E-16 |
| FANCG | 2394.608469 | -1.361886114 | 2.42E-21 |
| AC083843.4 | 12.60092653 | -1.362603539 | 3.90E-02 |
| ATP6V1G2-DD› | 701.8318688 | -1.362770228 | 3.52E-03 |
| KIF20A | 9603.437791 | -1.362854439 | 6.39E-46 |
| CFP | 142.794171 | -1.363871281 | 1.63E-07 |
| TMPO-AS1 | 252.5453917 | -1.36481495 | 7.35E-08 |
| H2AFX | 5031.626145 | -1.365901975 | 1.96E-40 |
| RSPH4A | 9.050537237 | -1.366010249 | 3.94E-02 |
| CCNA2 | 2526.037041 | -1.366469911 | 2.42E-17 |
| SLC43A2 | 1323.412679 | -1.366658554 | 1.96E-19 |
| ENHO | 164.5618616 | -1.368298685 | 7.07E-10 |
| AC012073.1 | 79.18141641 | -1.37058567 | 2.53E-07 |
| TCF4 | 3210.623798 | -1.372733444 | 2.64E-04 |
| BCL2L11 | 920.1726165 | -1.373678128 | 2.96E-27 |
| UBXN10-AS1 | 15.03613687 | -1.37419874 | 4.36E-03 |
| CASS4 | 18.80043433 | -1.375333186 | 2.94E-02 |
| TNFRSF11B | 3103.669719 | -1.37611782 | 2.34E-27 |
| GNB3 | 68.90723971 | -1.376580512 | 1.18E-05 |
| NWD1 | 21.24870833 | -1.37722693 | 1.94E-03 |
| ADGRF3 | 53.77481944 | -1.377936568 | 6.57E-04 |
| AL133215.2 | 51.61394003 | -1.38132197 | 1.53E-06 |
| LINC01993 | 11.62413837 | -1.381822154 | 3.26E-02 |
| ADM5 | 59.87160877 | -1.384526358 | 2.23E-05 |
| AC140125.2 | 68.76072382 | -1.385461337 | 6.27E-06 |
| HPDL | 404.0767023 | -1.38577034 | 1.22E-23 |
| SAC3D1 | 598.4358237 | -1.387582664 | 5.72E-22 |
| FGF17 | 24.00998586 | -1.38828244 | 3.11E-03 |
| RAD54L | 885.7258961 | -1.388644785 | 7.35E-10 |
| CYP26B1 | 155.6896713 | -1.39094621 | 3.56E-07 |
| TOP2A | 19561.07748 | -1.392397542 | 5.52E-14 |
| NA | 14.66958513 | -1.393677131 | 2.33E-02 |
| KIFC1 | 3153.021046 | -1.394919848 | 5.19E-26 |
| LINC02552 | 13.25560472 | -1.397100643 | 2.84E-02 |
| AL391069.2 | 56.01725847 | -1.397260369 | 6.41E-07 |

| | | | |
|---|---|---|---|
| ST3GAL5-AS1 | 58.4679868 | -1.397858652 | 1.87E-08 |
| CACNG4 | 946.3481827 | -1.398175243 | 4.37E-13 |
| KNTC1 | 2763.277137 | -1.398438402 | 4.23E-09 |
| KCNK17 | 7.93099649 | -1.39881302 | 4.70E-02 |
| TCAM1P | 268.7105326 | -1.399074071 | 1.48E-10 |
| CCNF | 2336.796344 | -1.399589939 | 1.74E-30 |
| AC018553.1 | 92.95374125 | -1.402131277 | 8.68E-13 |
| NREP | 12921.79755 | -1.404090773 | 4.36E-10 |
| PINLYP | 28.72624574 | -1.404496629 | 4.44E-03 |
| IQGAP3 | 3343.91172 | -1.406929766 | 8.71E-43 |
| AC138904.1 | 27.73708265 | -1.411263319 | 1.12E-03 |
| TAS1R3 | 9.427335722 | -1.412832449 | 4.45E-02 |
| AL133406.2 | 27.78123807 | -1.414794762 | 8.91E-05 |
| CCDC87 | 25.69209508 | -1.415449082 | 1.50E-03 |
| AC023300.2 | 9.952432568 | -1.416370669 | 3.15E-02 |
| LRRC37A6P | 65.4873206 | -1.416838187 | 2.33E-08 |
| AL353622.1 | 109.5816489 | -1.41686574 | 3.42E-11 |
| NCAPG | 2313.274187 | -1.416997633 | 1.15E-09 |
| MIR9-3HG | 38.19321297 | -1.418315489 | 4.09E-03 |
| LINC01639 | 38.42955259 | -1.418326642 | 6.51E-05 |
| ADGRF1 | 58.70965185 | -1.419856419 | 5.60E-07 |
| CCDC189 | 255.0331238 | -1.421058366 | 7.94E-16 |
| PLCB2 | 233.3908541 | -1.42299722 | 4.04E-08 |
| AC005393.1 | 10.54654791 | -1.424124058 | 2.05E-02 |
| RESF1 | 1525.676642 | -1.424554252 | 1.82E-07 |
| DYRK1B | 543.7450149 | -1.425743735 | 1.41E-17 |
| CEND1 | 59.0622389 | -1.426161129 | 7.04E-06 |
| FAM153B | 429.3182156 | -1.427388864 | 4.77E-12 |
| MEX3A | 1174.644861 | -1.427953264 | 1.39E-22 |
| NA | 139.7692151 | -1.429819505 | 1.61E-10 |
| IPO9-AS1 | 9.846607716 | -1.431460014 | 3.77E-02 |
| NF1P10 | 67.28745981 | -1.432263782 | 5.61E-06 |
| GNA14 | 109.087556 | -1.432360564 | 4.57E-08 |
| SLC49A3 | 8.681313307 | -1.433467503 | 4.29E-02 |
| FSIP1 | 21.52831629 | -1.43499243 | 2.00E-03 |
| ELL3 | 67.9189708 | -1.435108011 | 4.61E-08 |
| SBSPON | 767.0994634 | -1.435621348 | 2.25E-18 |
| CD93 | 23.58250152 | -1.436459909 | 3.86E-03 |
| UNG | 3547.676506 | -1.437347256 | 8.23E-17 |
| AL590999.1 | 37.94265019 | -1.440223671 | 7.15E-03 |
| FAM153A | 178.9188772 | -1.443095901 | 1.09E-09 |
| WFIKKN1 | 35.11312639 | -1.443516622 | 5.73E-04 |
| ARHGAP33 | 172.4402215 | -1.445020255 | 2.46E-08 |
| AC100793.4 | 11.90692052 | -1.445450396 | 1.33E-02 |
| CHRNA3 | 26.82226292 | -1.445561859 | 1.95E-02 |
| TNFRSF19 | 517.7426435 | -1.445936137 | 9.85E-16 |
| MCM4 | 9151.55886 | -1.448519534 | 7.16E-09 |

| | | | |
|---|---|---|---|
| PRSS2 | 39.46445423 | -1.450201158 | 2.22E-02 |
| APOBEC3B | 1479.583054 | -1.45116515 | 1.96E-10 |
| RPS2P55 | 23.60481798 | -1.452230593 | 8.90E-03 |
| NRGN | 1001.585436 | -1.452490476 | 1.91E-14 |
| SNX18P14 | 21.59630153 | -1.452514073 | 3.11E-03 |
| TGM2 | 27895.58294 | -1.453938788 | 1.20E-20 |
| SOCS2 | 977.7645512 | -1.454228917 | 2.90E-41 |
| PRC1-AS1 | 19.02665523 | -1.454689335 | 2.90E-02 |
| LINC01341 | 19.68380255 | -1.455005896 | 3.23E-03 |
| FAM110D | 49.12210533 | -1.455742214 | 9.60E-05 |
| MTND1P23 | 155.9045096 | -1.456208886 | 5.09E-08 |
| LINC01976 | 14.95955971 | -1.456258157 | 1.45E-02 |
| C4orf47 | 22.60644234 | -1.457129007 | 5.96E-03 |
| GNG2 | 1893.590023 | -1.458923672 | 1.81E-20 |
| AP001107.9 | 109.1125027 | -1.464321947 | 1.74E-10 |
| SPOCK2 | 70.07807607 | -1.46444285 | 4.59E-05 |
| ACTBL2 | 62.37567461 | -1.465368441 | 1.55E-03 |
| IGDCC3 | 90.23794552 | -1.467216245 | 4.61E-11 |
| AC012569.1 | 18.76329793 | -1.467665428 | 1.61E-02 |
| SCG3 | 17.83135647 | -1.468566966 | 1.58E-02 |
| TUBA3FP | 243.9884443 | -1.469372818 | 9.90E-20 |
| PRRT2 | 400.9286235 | -1.471007727 | 4.80E-19 |
| AC007952.5 | 15.85280466 | -1.472172408 | 3.40E-02 |
| EID3 | 84.86515522 | -1.472236388 | 1.23E-10 |
| DHRS2 | 1627.919122 | -1.473623662 | 2.93E-08 |
| B9D2 | 114.4858032 | -1.473815713 | 6.97E-10 |
| TRIM60P18 | 16.65121857 | -1.475961659 | 3.54E-03 |
| AL391832.2 | 14.02876155 | -1.476928417 | 2.24E-02 |
| SPC25 | 707.3497935 | -1.477281465 | 2.30E-16 |
| RAB27B | 854.5006837 | -1.477761528 | 7.50E-11 |
| ITPRIPL1 | 839.3701575 | -1.478358976 | 2.20E-33 |
| AC097534.1 | 25.37859066 | -1.481648374 | 3.70E-03 |
| AL031719.2 | 10.26048018 | -1.481684183 | 2.21E-02 |
| Z95152.1 | 9.983162855 | -1.481718874 | 4.89E-02 |
| SLC25A48 | 75.47198056 | -1.482897131 | 6.76E-07 |
| MCM5 | 5691.840419 | -1.483607272 | 6.64E-10 |
| RNF43 | 197.160326 | -1.484221235 | 8.02E-09 |
| TSPAN11 | 2754.87485 | -1.484654911 | 7.11E-11 |
| AL133415.1 | 10.82372751 | -1.484740105 | 4.85E-02 |
| BSN | 168.5919291 | -1.485081535 | 2.54E-13 |
| AC024405.2 | 18.19649933 | -1.48664596 | 1.35E-02 |
| TMEM119 | 206.1753641 | -1.486678147 | 1.18E-09 |
| POLE2 | 503.188362 | -1.487555907 | 7.92E-11 |
| A3GALT2 | 14.22687366 | -1.490229177 | 2.12E-02 |
| NA | 16.21782912 | -1.490898417 | 1.51E-02 |
| GINS1 | 2083.359063 | -1.493711798 | 3.65E-05 |
| SIRPG | 23.17823626 | -1.493933691 | 3.47E-03 |

| | | | |
|---|---|---|---|
| CCDC74A | 423.2740133 | -1.49555875 | 7.99E-29 |
| C1QL1 | 789.3243264 | -1.495859329 | 1.43E-19 |
| NR2E3 | 27.75191453 | -1.496218809 | 1.35E-03 |
| TFR2 | 211.3813313 | -1.496576765 | 2.05E-13 |
| DOK3 | 101.2399448 | -1.497115333 | 3.41E-10 |
| UCP2 | 2547.826664 | -1.498770578 | 1.33E-44 |
| AC126915.2 | 21.24752388 | -1.498936112 | 3.27E-03 |
| PAQR4 | 2220.425977 | -1.499919376 | 5.33E-13 |
| AC084876.2 | 15.49307114 | -1.502177163 | 6.09E-03 |
| HCRTR1 | 28.97834538 | -1.502241421 | 1.39E-03 |
| FBLN1 | 4794.018155 | -1.504248573 | 7.68E-24 |
| PCNA | 8363.090087 | -1.505977988 | 7.54E-07 |
| AC092683.1 | 166.0784964 | -1.506242465 | 6.01E-09 |
| SOWAHA | 80.37691478 | -1.508156982 | 1.64E-07 |
| AF001548.2 | 23.73397881 | -1.508447469 | 2.00E-03 |
| GRAMD2A | 20.24742378 | -1.511128191 | 9.78E-03 |
| INCENP | 3709.047014 | -1.511491105 | 2.10E-33 |
| AL050303.4 | 19.2713527 | -1.514478662 | 5.24E-03 |
| RIMBP3B | 55.67544132 | -1.517902634 | 9.28E-04 |
| AC008438.1 | 15.79731862 | -1.520010042 | 3.48E-03 |
| ELOVL3 | 25.0446582 | -1.521548042 | 1.64E-03 |
| AACSP1 | 270.037476 | -1.523023577 | 2.47E-13 |
| CARMIL3 | 83.73224846 | -1.525670922 | 7.46E-13 |
| PGM5P3-AS1 | 9.034595375 | -1.526219839 | 2.44E-02 |
| AJ011931.1 | 14.10779318 | -1.526671878 | 1.42E-02 |
| PLGLB2 | 28.97754901 | -1.531198365 | 3.44E-04 |
| USP24P1 | 23.05176201 | -1.532120773 | 2.99E-03 |
| AL121899.1 | 25.3059189 | -1.532413061 | 2.38E-04 |
| TLL2 | 230.6792461 | -1.534324912 | 1.47E-15 |
| VIM-AS1 | 75.42933042 | -1.535811675 | 2.63E-08 |
| MRVI1 | 377.0708102 | -1.536351014 | 2.20E-12 |
| CYP26A1 | 160.8035835 | -1.543091798 | 1.08E-13 |
| KANK4 | 25.95378457 | -1.543672953 | 1.82E-03 |
| FOXD4L5 | 21.56223852 | -1.545743905 | 4.78E-03 |
| ESPN | 28.24027471 | -1.545801176 | 3.89E-03 |
| AC104561.3 | 19.82206338 | -1.549228125 | 2.74E-03 |
| LAMP5 | 323.5776342 | -1.549976262 | 7.06E-17 |
| HELLS | 1113.450738 | -1.55155651 | 7.62E-14 |
| RPL5P30 | 9.877346493 | -1.551624978 | 2.63E-02 |
| GNRH1 | 35.48485024 | -1.55219827 | 4.35E-03 |
| HTR1D | 270.4186298 | -1.553798859 | 7.61E-15 |
| NPM2 | 23.96537148 | -1.554594895 | 1.45E-03 |
| CDCA5 | 3916.362872 | -1.554776308 | 1.47E-16 |
| WDR34 | 3834.872329 | -1.555725886 | 1.93E-52 |
| LHX1 | 640.0009217 | -1.556345692 | 1.50E-22 |
| AL162231.4 | 42.81852526 | -1.556562209 | 7.83E-05 |
| ENOX1 | 275.262444 | -1.557067158 | 2.50E-14 |

| | | | |
|---|---|---|---|
| OLFML2B | 1524.891652 | -1.559701186 | 2.19E-10 |
| FNDC5 | 839.7235065 | -1.560109518 | 1.58E-03 |
| EMILIN3 | 78.86613811 | -1.560441777 | 1.30E-05 |
| ESPL1 | 2910.925144 | -1.561000216 | 1.32E-30 |
| AC007787.2 | 9.107539871 | -1.561116383 | 3.05E-02 |
| PRDM1 | 260.727632 | -1.562399761 | 2.93E-08 |
| NRM | 1135.086586 | -1.563787271 | 1.06E-31 |
| MT-TL2 | 78.84596201 | -1.564095868 | 4.35E-04 |
| AC141557.1 | 12.84495437 | -1.564282452 | 9.46E-03 |
| A2M | 8519.613877 | -1.565125614 | 3.72E-11 |
| ENO1-AS1 | 7.014030703 | -1.566554277 | 4.08E-02 |
| TEKT2 | 47.2298313 | -1.569309296 | 1.16E-06 |
| NA | 221.0300842 | -1.570699751 | 1.55E-12 |
| THSD1 | 104.9408088 | -1.572736679 | 1.19E-16 |
| GTSE1 | 2245.155013 | -1.572770018 | 3.63E-37 |
| GPRACR | 10.49508947 | -1.573530485 | 2.40E-02 |
| AP002360.3 | 35.90771319 | -1.574004934 | 6.98E-05 |
| LGR6 | 65.23759692 | -1.574638394 | 2.18E-05 |
| GBX1 | 52.67623207 | -1.576837456 | 2.49E-07 |
| ROPN1L | 83.08733187 | -1.578526758 | 7.32E-13 |
| AC145098.2 | 70.74117309 | -1.578651933 | 6.87E-07 |
| TH | 17.682393 | -1.579506405 | 1.47E-02 |
| KNL1 | 1041.133435 | -1.579780772 | 5.56E-09 |
| OR7E104P | 10.84326186 | -1.580948455 | 3.88E-02 |
| PYY2 | 20.43247318 | -1.584842445 | 4.91E-04 |
| FXYD6 | 2385.941432 | -1.585214264 | 6.99E-26 |
| AC006213.5 | 7.711689508 | -1.586089816 | 4.57E-02 |
| PHF19 | 5086.402977 | -1.587087697 | 2.68E-38 |
| MTCO3P12 | 35.49839846 | -1.588277196 | 9.16E-04 |
| AP000757.1 | 13.68707276 | -1.588549663 | 3.83E-02 |
| MIR1-1HG-AS1 | 31.01433589 | -1.590361313 | 1.63E-03 |
| CDCA7 | 2462.20736 | -1.592086352 | 1.35E-23 |
| KCNJ6 | 165.3882011 | -1.592523812 | 5.26E-13 |
| AL031847.1 | 35.35801227 | -1.592751357 | 5.90E-06 |
| SFN | 28.73189058 | -1.596494261 | 6.11E-03 |
| LRRC74B | 22.31592487 | -1.596938794 | 7.23E-04 |
| AC100861.1 | 179.1552444 | -1.597391967 | 6.57E-14 |
| ZC3H12D | 48.81604572 | -1.59884254 | 1.57E-04 |
| AL513548.1 | 37.02258029 | -1.599494409 | 1.12E-04 |
| EXO1 | 1051.953812 | -1.600992553 | 9.53E-17 |
| NGFR | 2528.28695 | -1.603229943 | 3.65E-16 |
| FOXC2 | 196.881966 | -1.604387534 | 4.07E-10 |
| AC009121.1 | 31.56069955 | -1.604724173 | 1.71E-05 |
| LINC01845 | 13.01278341 | -1.604735683 | 1.23E-02 |
| NA | 19.75334832 | -1.60503758 | 2.34E-03 |
| EEF1A1P11 | 14.09671937 | -1.605203903 | 3.58E-02 |
| TAGLN | 4862.193472 | -1.606090213 | 2.99E-11 |

| | | | |
|---|---|---|---|
| PALM3 | 604.2618859 | -1.606091446 | 1.01E-19 |
| ITGA10 | 182.6699131 | -1.60623535 | 8.88E-13 |
| METTL7A | 666.3257155 | -1.606970961 | 1.26E-09 |
| CDCA3 | 1454.557961 | -1.607259209 | 1.33E-30 |
| LHX1-DT | 215.5440843 | -1.607652474 | 2.69E-21 |
| AF233439.1 | 16.16706051 | -1.607851971 | 9.19E-03 |
| LPAR1 | 468.0407392 | -1.609373756 | 3.19E-19 |
| MIMT1 | 17.01502868 | -1.611109223 | 3.88E-03 |
| ACOX2 | 57.51644016 | -1.612637247 | 2.15E-05 |
| RRS1-AS1 | 20.2277811 | -1.615150326 | 4.36E-04 |
| FAM107A | 21.23397353 | -1.621636729 | 1.02E-02 |
| DUSP27 | 48.58308619 | -1.621811386 | 7.16E-04 |
| ASF1B | 2796.164702 | -1.622335181 | 6.92E-13 |
| TGFB3 | 607.8050464 | -1.623244292 | 6.30E-18 |
| C1QTNF3-AMA | 63.15032994 | -1.623407425 | 1.55E-03 |
| CITED1 | 34.09631409 | -1.624917492 | 7.62E-05 |
| CSPG5 | 116.8237618 | -1.62594207 | 3.06E-10 |
| STEAP4 | 35.06140897 | -1.628632045 | 6.08E-04 |
| MYB | 40.76901378 | -1.630864777 | 5.03E-05 |
| 4-Mar | 1178.181843 | -1.631974672 | 1.29E-08 |
| AL049539.1 | 10.7466179 | -1.635657203 | 2.08E-02 |
| AC027228.2 | 92.37529988 | -1.635932236 | 6.35E-08 |
| AL390838.1 | 7.085227837 | -1.638933542 | 4.98E-02 |
| AL513365.2 | 8.494539678 | -1.6429587 | 2.18E-02 |
| GPR39 | 120.3391383 | -1.64415241 | 2.04E-06 |
| MT-TH | 24.1473077 | -1.646438076 | 1.70E-02 |
| KIAA1755 | 395.0065777 | -1.651095361 | 6.87E-04 |
| AL031651.2 | 13.37010399 | -1.652028428 | 3.22E-02 |
| AC009063.2 | 45.99358757 | -1.652883939 | 1.65E-08 |
| FAM78B | 308.7966026 | -1.65516592 | 4.45E-18 |
| H1FX-AS1 | 94.09648599 | -1.655871817 | 5.63E-18 |
| NA | 45.0838267 | -1.65640274 | 1.51E-08 |
| CD247 | 25.01308638 | -1.658171449 | 4.52E-04 |
| PCDHB12 | 24.55439884 | -1.658661004 | 3.51E-05 |
| ACTL8 | 26.23675356 | -1.660027418 | 1.14E-03 |
| AL136368.1 | 23.15736852 | -1.661704325 | 5.63E-05 |
| EPHB6 | 1812.119747 | -1.661739327 | 2.01E-18 |
| AC090627.1 | 17.38586849 | -1.661755956 | 7.07E-04 |
| AL359258.3 | 11.44985761 | -1.661803422 | 2.44E-02 |
| CDT1 | 1350.485546 | -1.662336802 | 2.19E-09 |
| ORC1 | 1063.331149 | -1.663495522 | 5.91E-09 |
| LMNB1 | 8403.611797 | -1.66750178 | 7.46E-22 |
| MYBL2 | 6940.146619 | -1.672228086 | 1.48E-24 |
| DQX1 | 18.43108469 | -1.673170559 | 4.20E-02 |
| RYR2 | 420.6593994 | -1.674795104 | 1.17E-02 |
| LINC02458 | 11.69928983 | -1.676987738 | 6.21E-03 |
| TBX3 | 4851.962125 | -1.681302388 | 2.78E-51 |

| | | | |
|---|---|---|---|
| CCDC74B | 422.4592504 | -1.681540923 | 7.38E-30 |
| RFLNA | 58.80224814 | -1.684084727 | 1.33E-05 |
| COLEC12 | 3956.507078 | -1.685390012 | 5.00E-55 |
| C11orf94 | 12.57013048 | -1.686868977 | 1.35E-03 |
| Z97192.1 | 5.753673836 | -1.686971768 | 4.45E-02 |
| AC078909.1 | 15.87716308 | -1.687869968 | 4.70E-02 |
| ALPK3 | 123.8681661 | -1.689838519 | 4.18E-05 |
| AC096711.2 | 11.37920918 | -1.692144689 | 2.14E-02 |
| NARF-IT1 | 9.897559897 | -1.693770163 | 3.58E-03 |
| TMEM121B | 53.36772833 | -1.693780432 | 9.10E-08 |
| NPIPA7 | 80.15957517 | -1.694022154 | 2.55E-07 |
| PPP1R16B | 527.7404052 | -1.694469623 | 3.80E-17 |
| CLDN5 | 29.88470087 | -1.699501768 | 6.95E-03 |
| CFAP74 | 16.55830783 | -1.700230284 | 1.04E-02 |
| CKMT1A | 221.78047 | -1.705325355 | 1.35E-10 |
| KCNH3 | 436.3653327 | -1.705458985 | 4.77E-22 |
| CNGA1 | 27.19220841 | -1.707636125 | 2.61E-04 |
| PARD6G-AS1 | 83.02067989 | -1.70787725 | 2.54E-10 |
| BARD1 | 329.1435678 | -1.708627021 | 3.66E-18 |
| ADCYAP1 | 83.8132783 | -1.709625794 | 5.74E-04 |
| AL359555.1 | 15.27189961 | -1.711940242 | 5.27E-03 |
| XKR5 | 178.029926 | -1.7188907 | 4.31E-23 |
| DISP2 | 460.1966062 | -1.719983947 | 1.20E-11 |
| LINC01778 | 11.93831883 | -1.723026848 | 1.42E-02 |
| FAM20C | 2024.588472 | -1.72665253 | 7.30E-24 |
| AC092073.1 | 35.30548196 | -1.729893553 | 1.50E-03 |
| CORO1A | 260.7712798 | -1.730407832 | 4.23E-23 |
| LINC01835 | 24.62093076 | -1.731339146 | 2.45E-03 |
| PIMREG | 1661.753605 | -1.735383318 | 4.57E-46 |
| PTPN6 | 55.81649397 | -1.736716821 | 4.97E-05 |
| AC137932.2 | 28.57861597 | -1.738286552 | 3.31E-04 |
| MAP1A | 1216.104392 | -1.73840131 | 8.50E-31 |
| HIST2H2AC | 8.790403176 | -1.739349598 | 2.40E-02 |
| AP003086.1 | 12.31159373 | -1.740088579 | 5.87E-03 |
| UNC5C | 16.04686619 | -1.741082381 | 6.45E-03 |
| LIG1 | 1872.991016 | -1.741254735 | 4.02E-12 |
| TRAF3IP3 | 99.96308918 | -1.742138122 | 1.17E-12 |
| AL157823.2 | 6.642025107 | -1.743467517 | 4.13E-02 |
| AP000866.2 | 16.58937699 | -1.745164853 | 2.83E-03 |
| ADRA1D | 1899.628549 | -1.748772302 | 7.52E-29 |
| FAM71F1 | 16.90444822 | -1.748965463 | 1.12E-02 |
| KRT71 | 12.44156458 | -1.750264887 | 1.49E-02 |
| SEMA6C | 475.9537505 | -1.754106176 | 1.50E-19 |
| CAMKV | 31.31748181 | -1.763680214 | 8.40E-06 |
| C21orf58 | 704.1135324 | -1.765357402 | 2.83E-33 |
| AC243773.1 | 15.24207664 | -1.767487979 | 1.99E-02 |
| ISM1 | 7.500361104 | -1.767550613 | 3.45E-02 |

| | | | |
|---|---|---|---|
| PHF2P2 | 183.8502381 | -1.768170042 | 5.83E-13 |
| PCLAF | 1284.567792 | -1.775801835 | 6.26E-22 |
| AL049796.1 | 51.65809039 | -1.775855291 | 1.29E-08 |
| GZMA | 20.76270662 | -1.77691412 | 4.32E-03 |
| FAM151A | 17.24186513 | -1.780157181 | 5.73E-03 |
| SAPCD2 | 1918.011644 | -1.780211167 | 3.38E-60 |
| LIPG | 311.095507 | -1.790450341 | 4.43E-26 |
| TRBC2 | 122.21731 | -1.7962205 | 2.22E-09 |
| AL023803.2 | 11.63335537 | -1.796378688 | 8.83E-03 |
| CRLF1 | 1052.296416 | -1.798648704 | 4.49E-40 |
| AC137800.2 | 32.96507486 | -1.799725341 | 3.92E-04 |
| ZNF285B | 8.246660278 | -1.800687968 | 2.33E-02 |
| ASPM | 1737.093246 | -1.807959049 | 2.73E-06 |
| BPI | 155.9343946 | -1.814031942 | 5.19E-12 |
| COL6A3 | 4751.934988 | -1.814048963 | 3.75E-15 |
| TXK | 110.571621 | -1.821149992 | 9.98E-12 |
| AL591135.1 | 25.44653968 | -1.821306368 | 4.05E-04 |
| KIF18B | 1701.1111 | -1.822319688 | 5.61E-37 |
| AMH | 31.73905909 | -1.82629329 | 1.51E-04 |
| C1QTNF3 | 44.26961409 | -1.827276975 | 6.24E-08 |
| TYMS | 5723.921539 | -1.830141396 | 2.52E-15 |
| BGLAP | 11.92623914 | -1.830141673 | 2.53E-02 |
| TF | 518.8802447 | -1.833792946 | 1.31E-14 |
| SLC26A4 | 67.63469275 | -1.83730419 | 5.82E-06 |
| SEMA3A | 14.92527264 | -1.838412758 | 1.05E-02 |
| AC159540.2 | 46.64017187 | -1.842956509 | 5.61E-04 |
| 1-Mar | 48.32321599 | -1.845326705 | 1.00E-02 |
| AL513534.2 | 18.4908913 | -1.848325813 | 1.24E-04 |
| CRNN | 18.66305069 | -1.849342685 | 3.48E-03 |
| BCO1 | 10.69519175 | -1.849349781 | 2.43E-03 |
| AC120053.1 | 12.97532409 | -1.851960765 | 4.37E-03 |
| AC124283.5 | 8.482519695 | -1.856097775 | 1.83E-02 |
| NOS2 | 73.8585863 | -1.858006965 | 1.95E-08 |
| BNC2-AS1 | 11.35366052 | -1.858254396 | 3.21E-03 |
| NA | 8.638736085 | -1.862820108 | 2.12E-02 |
| NF1P1 | 9.916285653 | -1.864672301 | 9.85E-03 |
| LRRN4 | 75.66148512 | -1.865511496 | 3.41E-07 |
| ADCY10P1 | 106.8816125 | -1.866877235 | 1.03E-07 |
| AC139495.1 | 32.95620183 | -1.870005384 | 1.09E-03 |
| AC254562.1 | 10.04046911 | -1.871887133 | 1.15E-02 |
| AC103808.2 | 11.94070214 | -1.872950485 | 2.09E-04 |
| DRAXIN | 93.60471832 | -1.873683961 | 8.67E-13 |
| GINS2 | 1477.928982 | -1.880209088 | 1.79E-08 |
| NA | 18.71341225 | -1.88349106 | 2.55E-04 |
| AC020658.5 | 8.914706664 | -1.886415713 | 2.45E-02 |
| CYP21A2 | 7.052953024 | -1.886706958 | 4.27E-02 |
| MT-TM | 13.86033428 | -1.889854257 | 2.23E-02 |

| | | | |
|---|---|---|---|
| ESPNP | 31.51150625 | -1.892382514 | 1.02E-04 |
| TMPRSS13 | 17.10204436 | -1.898043202 | 9.94E-04 |
| CTXN1 | 271.998645 | -1.899859489 | 6.86E-35 |
| AL021407.3 | 8.471258648 | -1.899925495 | 4.57E-02 |
| KRT15 | 92.32944513 | -1.900431343 | 1.30E-05 |
| AC002310.2 | 17.19992926 | -1.912861923 | 5.14E-03 |
| ELOVL2 | 1985.730308 | -1.913404133 | 4.72E-66 |
| AANAT | 8.096579261 | -1.913663514 | 4.59E-02 |
| RTN4RL1 | 146.9324584 | -1.915201542 | 2.70E-18 |
| STMN3 | 8539.398862 | -1.916781863 | 9.28E-62 |
| CKMT1B | 286.8831445 | -1.917376372 | 5.18E-23 |
| AC008013.2 | 40.82931398 | -1.920449948 | 2.02E-07 |
| ISLR | 87.15946199 | -1.921196646 | 1.56E-07 |
| TMEM145 | 253.2058034 | -1.921372512 | 8.55E-22 |
| LINC02657 | 19.24585261 | -1.92139535 | 2.60E-04 |
| BMX | 147.3418536 | -1.92263767 | 7.76E-08 |
| MT-TQ | 46.69971454 | -1.922651632 | 1.08E-04 |
| TACR1 | 14.86252692 | -1.929495208 | 3.85E-02 |
| DPYSL3 | 1641.152646 | -1.931216573 | 1.87E-50 |
| SASH3 | 23.27087915 | -1.932507479 | 5.82E-03 |
| LINC01132 | 37.93370493 | -1.933410814 | 4.44E-09 |
| NA | 50.17517098 | -1.935105098 | 6.51E-09 |
| ZMYND10 | 55.92339239 | -1.93890083 | 3.86E-05 |
| LINC00589 | 13.02961938 | -1.946015186 | 1.56E-03 |
| MT-TF | 12.06340084 | -1.951400128 | 3.07E-02 |
| IRX4 | 106.6038776 | -1.955144127 | 6.58E-10 |
| IGSF9 | 81.60414804 | -1.95822816 | 1.61E-11 |
| MCM10 | 945.6615549 | -1.960738445 | 3.52E-06 |
| AL445931.1 | 15.68498476 | -1.961285613 | 5.81E-04 |
| AC100812.1 | 7.009746242 | -1.965631508 | 4.99E-02 |
| RNF207 | 629.8990604 | -1.965997008 | 3.56E-35 |
| SHISA2 | 384.9045469 | -1.967489721 | 8.41E-41 |
| SKIDA1 | 152.151256 | -1.970042101 | 2.16E-23 |
| ANK1 | 275.01387 | -1.978652233 | 1.02E-10 |
| VSIG2 | 41.62470534 | -1.981805452 | 8.31E-08 |
| PLXNA4 | 515.4222672 | -1.98291078 | 2.26E-18 |
| POU3F2 | 6.773130487 | -1.983690186 | 2.24E-02 |
| PLAC9 | 94.03662567 | -1.987685057 | 4.39E-12 |
| RHBDL3 | 329.7370229 | -1.99042371 | 3.95E-23 |
| SMAD5-AS1 | 9.513127475 | -1.993688401 | 3.24E-02 |
| NKD1 | 470.6736306 | -1.995481873 | 2.47E-15 |
| AC009486.1 | 13.83506336 | -1.996881972 | 2.79E-03 |
| SOX17 | 62.40755451 | -2.001557463 | 1.09E-11 |
| SLC5A9 | 31.88326152 | -2.00366706 | 3.71E-03 |
| WNT7A | 74.42957687 | -2.008165649 | 1.15E-03 |
| CICP16 | 8.040177207 | -2.008531763 | 6.82E-03 |
| SERTAD4 | 765.4049029 | -2.012029454 | 2.75E-25 |

| | | | |
|---|---|---|---|
| ANKRD63 | 7.890383257 | -2.012056251 | 2.88E-02 |
| E2F2 | 349.0168751 | -2.014519704 | 1.79E-08 |
| SYNGAP1-AS1 | 10.96983291 | -2.016869223 | 1.10E-02 |
| PRKXP1 | 23.97776507 | -2.018148679 | 1.08E-06 |
| CYSLTR2 | 151.0712334 | -2.01848732 | 1.74E-11 |
| C22orf24 | 18.15540398 | -2.019704408 | 9.55E-05 |
| MDGA1 | 555.6681225 | -2.025824075 | 2.44E-10 |
| MXD3 | 586.6509012 | -2.026319209 | 9.39E-37 |
| GRM4 | 49.89927047 | -2.032187729 | 4.75E-03 |
| CLSPN | 1191.810265 | -2.033955233 | 8.67E-06 |
| HBG1 | 1243.158158 | -2.035631223 | 3.63E-03 |
| AC110285.7 | 11.64946197 | -2.045520828 | 7.20E-03 |
| AL022324.3 | 5.838635839 | -2.055887281 | 2.96E-02 |
| AC016957.2 | 7.752886535 | -2.058615183 | 1.71E-02 |
| AC067930.4 | 32.3071449 | -2.070107455 | 2.30E-06 |
| ATP5MC1P4 | 11.52322567 | -2.070958495 | 2.09E-02 |
| KCNQ2 | 12.26458218 | -2.07546811 | 3.49E-04 |
| NA | 10.22504646 | -2.084173528 | 1.25E-02 |
| AC141557.2 | 11.85635291 | -2.084864496 | 4.68E-03 |
| DPT | 7.418061856 | -2.085056856 | 2.78E-02 |
| ZAP70 | 104.1860247 | -2.088201654 | 1.37E-11 |
| GLYATL1 | 31.67131304 | -2.090801601 | 1.11E-05 |
| LFNG | 282.2624472 | -2.091155587 | 1.72E-29 |
| SYP | 149.7220486 | -2.092519672 | 1.55E-20 |
| AP000915.1 | 54.83911648 | -2.092622903 | 1.15E-10 |
| AC008728.1 | 6.476754203 | -2.095350223 | 3.08E-02 |
| AC104938.1 | 9.622931093 | -2.099517516 | 1.13E-02 |
| SNX18P9 | 5.651227874 | -2.111485135 | 3.36E-02 |
| RPS10P5 | 9.212638172 | -2.114830536 | 1.07E-02 |
| PAK6 | 23.09223846 | -2.117501989 | 1.29E-02 |
| COL8A1 | 2277.352965 | -2.121817013 | 3.54E-26 |
| HMGN1P17 | 7.596987093 | -2.127156584 | 2.86E-02 |
| SAMD7 | 13.89098362 | -2.132137776 | 2.60E-02 |
| CRIP3 | 31.00547865 | -2.140337628 | 7.11E-07 |
| LINC00452 | 9.072206744 | -2.140547927 | 1.42E-03 |
| DKK1 | 937.75661 | -2.140751387 | 1.99E-28 |
| RRM2 | 5728.927211 | -2.144886322 | 6.36E-17 |
| AL589993.1 | 9.839506058 | -2.145777364 | 3.29E-03 |
| BUD13P1 | 14.41791251 | -2.155577164 | 1.24E-03 |
| LINC02367 | 8.694116988 | -2.156163736 | 6.38E-03 |
| RN7SKP275 | 16.16681324 | -2.156343684 | 1.01E-03 |
| C21orf62 | 7.981432635 | -2.160905987 | 2.55E-02 |
| NA | 9.494831861 | -2.165762951 | 4.82E-03 |
| WNT7B | 200.4097672 | -2.165836941 | 2.00E-04 |
| HGFAC | 24.78828744 | -2.169461049 | 9.73E-07 |
| AC092364.2 | 6.569950109 | -2.173981795 | 1.18E-02 |
| TUBA1A | 19721.63494 | -2.17978925 | 7.24E-26 |

| | | | |
|---|---|---|---|
| AC010422.5 | 8.90689788 | -2.189272778 | 1.55E-02 |
| RGS14 | 562.7905907 | -2.189306453 | 3.95E-51 |
| AL021920.2 | 10.43786673 | -2.191437609 | 8.63E-03 |
| AL592424.1 | 19.3597784 | -2.199430761 | 1.27E-06 |
| MT-TS2 | 38.2703071 | -2.199975319 | 8.12E-04 |
| CCR1 | 28.49835598 | -2.200382718 | 6.80E-07 |
| DERL3 | 144.1694928 | -2.201375888 | 7.83E-15 |
| CLEC18B | 62.85794889 | -2.216547588 | 2.44E-12 |
| NPY | 6.722255392 | -2.219864428 | 1.45E-02 |
| AL807752.3 | 120.8915602 | -2.22043039 | 9.52E-03 |
| LAG3 | 53.74991194 | -2.22329682 | 8.05E-12 |
| CCNA1 | 17.31100278 | -2.225458044 | 6.22E-04 |
| GLP1R | 28.92280827 | -2.227625946 | 5.53E-08 |
| AC012508.2 | 9.930179044 | -2.228471442 | 4.64E-04 |
| KCNH4 | 17.34632279 | -2.240192116 | 3.47E-04 |
| CASC8 | 11.89497762 | -2.242099542 | 2.57E-04 |
| AC010343.3 | 9.445590034 | -2.242377583 | 4.45E-03 |
| CHRNB2 | 21.60480173 | -2.243998533 | 1.64E-06 |
| GABRR1 | 17.11856491 | -2.247451706 | 6.95E-05 |
| ELOVL2-AS1 | 21.42926749 | -2.251912024 | 1.08E-05 |
| RTN1 | 9.232349879 | -2.259007433 | 5.27E-04 |
| TREM2 | 9.532154477 | -2.266962597 | 3.20E-02 |
| AC093620.1 | 7.115466543 | -2.268152634 | 6.19E-03 |
| ANKRD35 | 7.132370749 | -2.278783858 | 2.26E-03 |
| UHRF1 | 3016.429635 | -2.279760589 | 1.06E-20 |
| AC008013.3 | 8.40054863 | -2.28483581 | 1.59E-02 |
| H19 | 25168.32442 | -2.291916797 | 1.11E-07 |
| DNAJC9-AS1 | 7.566559336 | -2.293412838 | 3.57E-02 |
| GRP | 8.013379437 | -2.297149791 | 1.70E-03 |
| GAL | 58.68265862 | -2.299845283 | 2.36E-10 |
| RIMBP3C | 27.06751474 | -2.310529524 | 4.55E-04 |
| AC093635.1 | 6.445982233 | -2.310982471 | 2.96E-02 |
| EDN1 | 1541.884744 | -2.315557281 | 1.49E-24 |
| AL135818.2 | 5.597184778 | -2.322412807 | 9.85E-03 |
| NPBWR1 | 390.8449915 | -2.327549899 | 1.65E-28 |
| CCR5 | 30.83657564 | -2.329168653 | 2.54E-07 |
| PLAC9P1 | 16.69223394 | -2.339879773 | 1.43E-03 |
| CD180 | 81.62185089 | -2.340887226 | 4.64E-11 |
| VWA5B1 | 9.228630555 | -2.342710542 | 4.67E-03 |
| NPM1P9 | 12.32308265 | -2.34694091 | 1.11E-03 |
| MIR4324 | 5.16723264 | -2.350635107 | 2.35E-02 |
| AL109613.1 | 11.43374656 | -2.360806845 | 2.27E-03 |
| ANGPT2 | 85.43122701 | -2.371703976 | 1.17E-14 |
| BX005266.1 | 12.39520324 | -2.37297806 | 3.99E-02 |
| CREG2 | 57.31823733 | -2.375884544 | 1.16E-10 |
| KMO | 12.61321509 | -2.379288802 | 2.76E-02 |
| MT-TI | 17.57118234 | -2.38079964 | 3.08E-03 |

| | | | |
|---|---|---|---|
| CD3D | 25.35654547 | -2.38908217 | 4.90E-04 |
| KCNC1 | 90.05506287 | -2.390141886 | 3.30E-16 |
| GNRH2 | 11.35355812 | -2.419635507 | 1.24E-04 |
| OR6W1P | 7.088915303 | -2.422721241 | 6.19E-03 |
| MIR5587 | 6.078807982 | -2.427587831 | 1.17E-02 |
| AC087045.2 | 9.78579654 | -2.437284412 | 3.16E-04 |
| FAM124B | 12.99155367 | -2.461619187 | 1.15E-03 |
| VCAN | 908.1972434 | -2.474868087 | 2.33E-25 |
| AC016722.1 | 5.455952972 | -2.490242585 | 3.69E-02 |
| AC112777.1 | 32.46563624 | -2.491767269 | 9.37E-07 |
| ZC3H11B | 16.96159532 | -2.507523038 | 1.40E-06 |
| LINC02102 | 11.0472963 | -2.535089092 | 1.10E-02 |
| AL358613.2 | 7.002776717 | -2.54299796 | 1.29E-02 |
| AL031710.2 | 12.472377 | -2.57070393 | 8.40E-04 |
| NA | 12.81908503 | -2.623263239 | 4.02E-04 |
| HHIP | 9.193770709 | -2.652578251 | 1.01E-02 |
| FAM111B | 742.562208 | -2.677767721 | 1.93E-09 |
| AC026333.4 | 5.069693882 | -2.72533983 | 2.39E-02 |
| SCG2 | 992.0410165 | -2.745403927 | 3.94E-29 |
| SIX2 | 191.1792146 | -2.750199728 | 4.98E-42 |
| AC016526.2 | 5.337631847 | -2.765322289 | 1.07E-02 |
| HSD17B2 | 349.7292265 | -2.774582418 | 7.08E-21 |
| BFSP2 | 15.52825355 | -2.78861666 | 3.35E-03 |
| AC034245.1 | 13.7967374 | -2.801041355 | 9.24E-04 |
| SNHG18 | 5.836971013 | -2.832803061 | 2.40E-02 |
| AP002768.1 | 6.270874198 | -2.908863563 | 1.01E-02 |
| GNG4 | 685.0541399 | -2.90907535 | 2.48E-70 |
| GNA15 | 12.58832049 | -2.909827829 | 2.11E-03 |
| SCARNA8 | 5.577300044 | -2.91366775 | 1.26E-02 |
| SLC1A2 | 13.68876523 | -2.942485404 | 3.03E-03 |
| RIBC2 | 202.8855199 | -2.947632199 | 1.38E-14 |
| KRT5 | 26.98052728 | -2.954997165 | 3.01E-04 |
| AC027020.2 | 13.79088975 | -2.961920778 | 2.10E-04 |
| CYP24A1 | 230.7349006 | -2.993609216 | 1.05E-24 |
| DISP3 | 10.62150296 | -3.007514332 | 1.50E-03 |
| AP000787.2 | 6.966646499 | -3.038536219 | 9.25E-04 |
| NRAP | 18.38544263 | -3.077274824 | 2.70E-07 |
| RPS23P8 | 7.341802009 | -3.102277714 | 1.36E-02 |
| LINC00892 | 26.51675302 | -3.128698255 | 1.11E-05 |
| SLC22A14 | 12.55910754 | -3.133455696 | 8.96E-05 |
| PNMT | 17.22271722 | -3.158197175 | 8.65E-07 |
| KISS1 | 240.3909755 | -3.17922589 | 4.20E-02 |
| F13A1 | 43.43632321 | -3.232532472 | 1.65E-09 |
| LRRTM2 | 15.22254316 | -3.303084675 | 5.47E-05 |
| LINC00273 | 6.012401461 | -3.392270859 | 1.96E-02 |
| AC132217.1 | 38.61820786 | -3.486595028 | 5.60E-05 |
| AC087623.3 | 5.389991956 | -3.507335903 | 1.59E-02 |

| | | | |
|---|---|---|---|
| ADAMTS7P3 | 14.41766203 | -3.750403417 | 1.55E-02 |
| RDH10-AS1 | 9.84593621 | -3.855289202 | 1.85E-04 |
| FP565260.2 | 17.2358532 | -3.863361426 | 3.54E-02 |
| ESX1 | 1145.550705 | -3.907313787 | 5.51E-83 |
| TRIM15 | 6.446979355 | -3.979853948 | 1.43E-03 |
| PRND | 26.16392127 | -4.057660812 | 1.09E-10 |
| GDF10 | 12.90454409 | -4.20419802 | 8.56E-04 |
| SPINK4 | 6.515444977 | -4.261637089 | 6.31E-05 |
| MS4A6E | 21.63496949 | -4.321626408 | 6.82E-06 |
| EVPLL | 5.603573818 | -4.382082038 | 2.39E-02 |
| NPTX1 | 3190.575678 | -4.760959476 | 6.20E-140 |
| DND1 | 12.25643446 | -5.023356347 | 5.42E-05 |
| TBX10 | 5.510975525 | -5.183092779 | 4.69E-04 |
| AL358333.3 | 10.81048479 | -5.55336953 | 3.26E-02 |
| PCDHGA11 | 39.1975755 | -6.302082138 | 3.63E-06 |
| AC114546.3 | 13.1763236 | -6.916115163 | 6.90E-03 |
| AC139530.3 | 18.27891962 | -7.480266174 | 3.18E-06 |
| AC093668.2 | 22.02253282 | -29.98837941 | 3.13E-06 |

| Gene | baseMean | log2 Fold Change | padj |
|---|---|---|---|
| AP003419.1 | 46.01432342 | 21.16535979 | 1.17E-08 |
| CXCR5 | 38.38259048 | 19.42048694 | 4.61E-03 |
| NMRK2 | 40.23946307 | 3.582906529 | 1.68E-02 |
| TFE3 | 7339.395033 | 2.138972877 | 1.73E-127 |
| PTGDS | 58.43516836 | 1.644368166 | 3.72E-02 |
| SLC9A3 | 209.2046598 | 1.630642611 | 2.67E-10 |
| RENBP | 435.6778357 | 1.185312426 | 1.72E-04 |
| MAT1A | 731.6526786 | 1.020656529 | 1.12E-02 |
| RRAGD | 701.8675929 | 1.012658656 | 9.92E-05 |
| GPNMB | 7636.758085 | 0.899695098 | 7.36E-05 |
| UAP1L1 | 2532.817372 | 0.663151668 | 3.02E-02 |
| COL12A1 | 7146.720884 | 0.56840206 | 2.21E-02 |
| AC010442.1 | 720.7233888 | 0.442177267 | 2.21E-02 |
| DAB2 | 9628.493215 | 0.358876294 | 1.68E-02 |
| OSBPL1A | 2509.624306 | 0.276254765 | 3.34E-02 |
| CDK2AP1 | 4145.313976 | -0.319665187 | 6.44E-03 |
| RNF139 | 1481.985634 | -0.396447993 | 2.02E-03 |
| MMP17 | 411.9505481 | -0.81035251 | 4.65E-02 |
| PCDHGA11 | 39.1975755 | -5.540257086 | 9.56E-03 |

| Gene | baseMean | log2 Fold Change | padj |
|---|---|---|---|
| CXCR5 | 38.38259048 | 23.81928614 | 1.85E-08 |
| AP003419.1 | 46.01432342 | 21.12221325 | 2.73E-11 |
| SALL3 | 117.0169671 | 12.06296652 | 4.53E-02 |
| NPPB | 2240.650956 | 10.67463235 | 2.75E-73 |
| AC089983.1 | 88.31094755 | 10.49058734 | 2.63E-16 |
| LINC01750 | 27.45105218 | 10.04584659 | 5.42E-03 |
| SFTA1P | 288.2208761 | 9.637580437 | 8.17E-26 |
| RGPD4 | 132.856446 | 9.101790444 | 2.02E-02 |
| SLC18A3 | 36.75075062 | 9.068957309 | 4.08E-12 |
| KCND3 | 287.4144475 | 8.87765951 | 8.68E-87 |
| KISS1 | 240.3909755 | 8.792092792 | 5.99E-54 |
| IMPDH1P10 | 18.85870357 | 8.683734076 | 3.08E-05 |
| XIRP1 | 24.70751638 | 8.583428205 | 4.78E-09 |
| HBG1 | 1243.158158 | 8.505035434 | 1.47E-62 |
| TNNI1 | 87.9705331 | 8.495083343 | 1.03E-24 |
| APCS | 10.98996835 | 8.476739544 | 1.38E-09 |
| MC5R | 25.26232457 | 8.417420863 | 2.06E-10 |
| SLC27A2 | 45.4861244 | 8.282755596 | 4.05E-09 |
| AC115522.1 | 25.34233385 | 8.255919686 | 6.22E-10 |
| PSG3 | 19.01407792 | 8.09901113 | 7.80E-10 |
| CEACAMP10 | 596.3616348 | 8.046529487 | 7.98E-102 |
| CHRND | 24.58768675 | 8.03157421 | 2.45E-05 |
| ANKRD1 | 2371.13122 | 7.931678425 | 1.24E-76 |
| INHBB | 89.82945103 | 7.68795146 | 7.84E-17 |
| PSG2 | 45.18656473 | 7.631610864 | 4.28E-16 |
| KRTAP3-1 | 7.798810147 | 7.618582463 | 7.12E-08 |
| KRTAP2-3 | 13.1496624 | 7.562411465 | 1.11E-07 |
| KRT34 | 47.26339011 | 7.454235422 | 6.00E-11 |
| KRT81 | 3638.578358 | 7.437935388 | 1.26E-200 |
| THAP12P8 | 7.763317663 | 7.412105763 | 2.95E-07 |
| NA | 9.834076694 | 7.352282717 | 7.36E-05 |
| PSG7 | 92.83788603 | 7.310502063 | 1.17E-29 |
| MYO7B | 109.193404 | 7.28434739 | 1.70E-15 |
| MARCO | 27.67416851 | 7.280673382 | 2.51E-08 |
| CAVIN2 | 121.5099942 | 7.27375915 | 9.54E-36 |
| PSG6 | 42.7531939 | 7.263658023 | 2.33E-13 |
| KRT80 | 1730.679913 | 6.984458242 | 8.65E-226 |
| PSG4 | 1067.986903 | 6.933426144 | 4.47E-182 |
| VSTM1 | 56.60755979 | 6.876642047 | 4.81E-21 |
| LCN2 | 20.89964453 | 6.852395855 | 2.76E-06 |
| LNCOG | 57.59267757 | 6.819224466 | 3.16E-23 |
| NPIPA2 | 22.55804979 | 6.809240805 | 4.73E-04 |
| PDGFB | 265.4831051 | 6.747696631 | 3.28E-82 |
| RFPL4AL1 | 48.58179161 | 6.732047494 | 3.16E-07 |
| CXCR4 | 15.34564949 | 6.710598189 | 3.91E-06 |
| LURAP1L-AS1 | 37.65792059 | 6.661974371 | 2.46E-12 |

| | | | |
|---|---|---|---|
| KRTAP1-5 | 5.600439711 | 6.640272946 | 1.39E-05 |
| FGF19 | 32.84708541 | 6.619362124 | 3.32E-07 |
| PSG5 | 429.1534162 | 6.602100157 | 1.74E-97 |
| AP003498.1 | 6.759281315 | 6.57010102 | 1.53E-04 |
| LCE2A | 7.928852411 | 6.569879344 | 8.58E-06 |
| AL365184.1 | 7.788440408 | 6.480446383 | 8.46E-06 |
| TGM4 | 11.45184573 | 6.434178557 | 6.72E-06 |
| PRAMEF1 | 8.708668192 | 6.42768725 | 5.31E-06 |
| AC062015.1 | 5.368918224 | 6.403434832 | 2.35E-05 |
| ADCY8 | 7.411685296 | 6.357646673 | 1.05E-05 |
| LINC01186 | 8.428108961 | 6.356599008 | 6.91E-06 |
| PSG11 | 67.8097526 | 6.317738201 | 9.54E-28 |
| TNNT2 | 85.46035183 | 6.313245986 | 4.08E-10 |
| SPNS3 | 24.62107375 | 6.279789239 | 8.76E-09 |
| RPL23AP47 | 7.779613667 | 6.222111127 | 2.52E-03 |
| SNX18P24 | 5.717837345 | 6.210523957 | 2.99E-04 |
| PSG9 | 110.5829147 | 6.17803222 | 5.03E-34 |
| PAPPA2 | 59.55621791 | 6.167556545 | 1.13E-27 |
| AC026316.2 | 27.22988646 | 6.161403425 | 1.03E-10 |
| DIRC3-AS1 | 14.94687545 | 6.143555174 | 9.02E-08 |
| PSG1 | 87.14686265 | 6.137198431 | 1.55E-43 |
| OR10Y1P | 5.655546436 | 6.052138752 | 8.48E-05 |
| AC067773.1 | 17.64783138 | 6.042997493 | 5.52E-10 |
| NA | 6.666296487 | 5.964809228 | 1.05E-03 |
| CSTA | 29.96038685 | 5.847990294 | 9.06E-10 |
| AC135977.1 | 22.06214483 | 5.833184665 | 1.09E-02 |
| RFPL4A | 89.139459 | 5.816692105 | 1.08E-13 |
| LINC00592 | 5.244624385 | 5.804665869 | 1.23E-04 |
| HBG2 | 280.0908711 | 5.799590666 | 7.53E-81 |
| LY6K | 276.1851787 | 5.786371904 | 9.78E-83 |
| CEACAMP8 | 13.78743094 | 5.769516794 | 2.45E-05 |
| LINC02331 | 15.15784713 | 5.763109222 | 5.65E-05 |
| REXO1L1P | 16.21114651 | 5.741275481 | 5.67E-04 |
| FOXB1 | 5.497409793 | 5.713649515 | 2.68E-04 |
| DRGX | 5.390330915 | 5.693403479 | 1.26E-04 |
| MYL2 | 7.674597299 | 5.679945465 | 4.10E-05 |
| MYOD1 | 190.9762848 | 5.64315215 | 1.25E-17 |
| AL159166.1 | 10.36203043 | 5.618215706 | 6.89E-05 |
| AC245041.1 | 18.51833017 | 5.615281656 | 2.27E-06 |
| CHAT | 14.31282578 | 5.602853305 | 1.80E-05 |
| IL7R | 8419.815253 | 5.601856492 | 1.61E-77 |
| GGTLC2 | 5.282843147 | 5.593794138 | 2.58E-04 |
| AC100800.1 | 63.02116095 | 5.567099965 | 2.62E-25 |
| CLDN9 | 13.60129441 | 5.565718917 | 1.29E-07 |
| AC114488.1 | 8.478510308 | 5.528183123 | 1.21E-04 |
| LGALS9C | 9.780642832 | 5.480594951 | 1.28E-06 |
| SPX | 445.5857189 | 5.457895833 | 1.10E-44 |

| | | | |
|---|---|---|---|
| PSG8 | 13.98429761 | 5.451482355 | 7.34E-09 |
| C17orf102 | 57.63053537 | 5.432595704 | 1.98E-12 |
| DEFB115 | 7.658037426 | 5.420669195 | 6.15E-07 |
| AL136452.1 | 5.435183264 | 5.397218799 | 4.77E-04 |
| LINC01293 | 57.56332832 | 5.339978219 | 1.17E-11 |
| IL1RAPL1 | 13.93100269 | 5.338975234 | 3.50E-10 |
| TINAGL1 | 244.1329187 | 5.327601074 | 3.18E-53 |
| AL009178.2 | 8.641259053 | 5.325573194 | 1.89E-05 |
| FRG2C | 27.16775073 | 5.317692149 | 5.86E-18 |
| TNFSF4 | 327.5208002 | 5.302612396 | 6.41E-99 |
| KRT86 | 789.1722789 | 5.302328522 | 9.12E-171 |
| FRG2B | 5.367969376 | 5.278447231 | 3.97E-04 |
| HBBP1 | 30.01402287 | 5.250355032 | 3.02E-14 |
| TCEA1P3 | 6.557677423 | 5.231828614 | 4.56E-05 |
| AC107871.1 | 14.44185302 | 5.228567503 | 1.33E-04 |
| AC132068.1 | 5.382205542 | 5.21280924 | 7.30E-04 |
| TMEFF2 | 21.7217407 | 5.200029761 | 7.81E-12 |
| FPGT-TNNI3K | 29.63087152 | 5.187584761 | 4.72E-06 |
| SAA1 | 78.53236772 | 5.182325854 | 1.52E-50 |
| AC018781.1 | 5.21339508 | 5.181368897 | 6.77E-04 |
| SAA2 | 14.69633019 | 5.147314499 | 7.87E-09 |
| MYH2 | 23.18872912 | 5.137359842 | 2.81E-04 |
| CCND2 | 40.55711956 | 5.1319368 | 4.18E-12 |
| GSX2 | 19.00376094 | 5.116301325 | 1.54E-13 |
| ADGRG4 | 32.72646161 | 5.087394078 | 3.88E-06 |
| TNFSF18 | 594.8574692 | 5.055025438 | 1.29E-17 |
| REN | 21.64424951 | 5.044876789 | 3.90E-15 |
| KCNU1 | 6.149298403 | 5.030626405 | 1.89E-03 |
| NGF | 85.12586789 | 4.966973251 | 4.94E-22 |
| KRT87P | 36.57107821 | 4.964156957 | 2.12E-18 |
| CHRNA4 | 6.453752451 | 4.952147498 | 1.03E-04 |
| PCDHGB8P | 11.70899875 | 4.919553304 | 2.95E-02 |
| SERPINB7 | 182.2948166 | 4.916494497 | 3.84E-37 |
| PLCE1-AS1 | 11.35185947 | 4.91086312 | 8.12E-05 |
| NA | 5.318823847 | 4.902548215 | 2.10E-02 |
| CPA4 | 8124.801267 | 4.893229893 | 1.35E-205 |
| LYPD5 | 426.1463919 | 4.889410278 | 9.85E-54 |
| GPR65 | 19.00070169 | 4.887892294 | 9.30E-11 |
| CDH4 | 706.3312431 | 4.837604468 | 1.31E-109 |
| CCN2 | 8122.707037 | 4.828236594 | 3.61E-137 |
| 4-Mar | 1178.181843 | 4.827438381 | 7.09E-83 |
| RPL13AP20 | 27.56620949 | 4.826081948 | 5.44E-14 |
| LINC01291 | 43.47043514 | 4.809781799 | 8.61E-11 |
| MEI4 | 8.544885276 | 4.799715403 | 1.72E-05 |
| KLHDC7B | 28.70749508 | 4.754819947 | 1.60E-03 |
| STXBP6 | 354.2817869 | 4.740371691 | 7.00E-98 |
| KCNQ3 | 69.36936743 | 4.738234531 | 3.77E-25 |

| | | | |
|---|---|---|---|
| AC112656.1 | 16.72979437 | 4.73238902 | 1.48E-08 |
| MME | 72.18384025 | 4.709525342 | 1.79E-15 |
| HSD17B2 | 349.7292265 | 4.698851287 | 1.46E-93 |
| AC114296.1 | 5.972914867 | 4.692457258 | 1.91E-03 |
| TMEM178B | 177.5366143 | 4.671916082 | 1.19E-40 |
| SYNPO2L | 54.0993148 | 4.662578175 | 5.15E-24 |
| MT3 | 7.356063884 | 4.644209081 | 5.48E-04 |
| CAPSL | 15.12718097 | 4.643573298 | 3.23E-06 |
| C4BPB | 51.0915616 | 4.627400142 | 1.26E-16 |
| ASS1 | 1006.358665 | 4.623584239 | 1.42E-29 |
| ASB18 | 18.35133093 | 4.621492373 | 1.96E-05 |
| LINC02535 | 49.64853035 | 4.590335069 | 1.01E-18 |
| OLR1 | 33.65778704 | 4.572896165 | 5.70E-09 |
| WFDC21P | 140.545174 | 4.567643265 | 1.54E-26 |
| VLDLR-AS1 | 123.3112072 | 4.555694014 | 8.03E-22 |
| C2orf78 | 9.859463982 | 4.540947117 | 2.47E-07 |
| PADI1 | 49.5811494 | 4.519912364 | 6.77E-22 |
| ALOXE3 | 48.23452377 | 4.515474789 | 5.12E-11 |
| HS3ST5 | 7.638299293 | 4.494994025 | 3.05E-05 |
| NXPH2 | 6.697336569 | 4.492832716 | 2.84E-03 |
| CRLF2 | 21.35257814 | 4.473773115 | 7.97E-10 |
| OSCAR | 347.1562098 | 4.466898158 | 2.63E-68 |
| AOX1 | 235.0610148 | 4.46129515 | 1.44E-34 |
| IL24 | 21.39327091 | 4.455662144 | 4.07E-12 |
| MYOSLID | 13.69503477 | 4.432133785 | 2.48E-07 |
| CEACAMP7 | 24.51020739 | 4.428098079 | 1.28E-11 |
| LAD1 | 18.12413109 | 4.426488618 | 1.90E-06 |
| CCN1 | 10596.59591 | 4.421427289 | 0.00E+00 |
| GPRIN2 | 25.95356115 | 4.394375442 | 2.21E-10 |
| FAM90A12P | 21.08359169 | 4.390080335 | 7.86E-06 |
| NA | 12.04352763 | 4.371464908 | 3.29E-03 |
| GFRA1 | 5.639523961 | 4.346310291 | 6.23E-05 |
| AC114550.1 | 11.42977472 | 4.34321379 | 3.95E-06 |
| FPR3 | 132.3685572 | 4.338164442 | 5.70E-43 |
| NA | 16.51338953 | 4.333846374 | 4.99E-03 |
| AL078612.2 | 11.28350549 | 4.330500423 | 3.68E-07 |
| AL451069.1 | 7.150260511 | 4.328282686 | 8.10E-04 |
| MYCT1 | 39.3580197 | 4.321826517 | 3.49E-23 |
| PHF24 | 57.9882298 | 4.301443058 | 6.35E-09 |
| ZNF536 | 5.604134159 | 4.297754662 | 5.75E-03 |
| ALCAM | 37.52071971 | 4.287095956 | 4.37E-10 |
| LHX5 | 8.739437811 | 4.268021598 | 1.17E-05 |
| AP000577.1 | 9.975781841 | 4.260929086 | 1.03E-08 |
| AL590004.3 | 1289.954072 | 4.247938251 | 2.18E-95 |
| LINC02154 | 47.14458306 | 4.235538444 | 1.25E-25 |
| LAMP3 | 296.1481714 | 4.214300393 | 1.32E-52 |
| HBE1 | 415.2750207 | 4.211041935 | 4.72E-71 |

| | | | |
|---|---|---|---|
| KRT73-AS1 | 27.90722466 | 4.207070173 | 8.49E-14 |
| HCN4 | 60.95695921 | 4.196742354 | 9.62E-16 |
| RBFOX3 | 23.02501928 | 4.195283352 | 3.90E-11 |
| LTF | 52.07108289 | 4.194473745 | 3.44E-20 |
| AC092132.1 | 5.105028167 | 4.182834376 | 3.31E-03 |
| ITIH3 | 47.92121289 | 4.162708509 | 4.72E-11 |
| AL137129.1 | 25.35779872 | 4.161376251 | 4.29E-05 |
| LINC00702 | 14.07835045 | 4.144396505 | 4.28E-03 |
| LINC01198 | 6.335604086 | 4.144197462 | 2.65E-04 |
| AC079054.1 | 7.797618888 | 4.136665053 | 4.34E-05 |
| TAGLN | 4862.193472 | 4.130792771 | 2.57E-73 |
| KRT4 | 36.4381841 | 4.121526567 | 5.79E-06 |
| AC034206.1 | 5.479961284 | 4.120348639 | 4.01E-04 |
| CST6 | 11.6176009 | 4.119025563 | 6.92E-06 |
| MYHAS | 12.21043099 | 4.117519072 | 2.39E-03 |
| SUSD4 | 14.80441536 | 4.114994724 | 2.27E-04 |
| CTNND2 | 5.418341109 | 4.113315237 | 2.35E-03 |
| AL353743.4 | 7.124459301 | 4.107809411 | 1.12E-02 |
| STX11 | 126.2269358 | 4.102672517 | 6.19E-47 |
| SPOCD1 | 189.9108213 | 4.078512568 | 3.56E-39 |
| FGF21 | 23.9633406 | 4.070042212 | 3.96E-03 |
| AC022028.2 | 14.49200066 | 4.058071187 | 3.14E-07 |
| AC103740.2 | 6.922522524 | 4.056188355 | 1.80E-03 |
| CSF2RB | 37.96421996 | 4.049175255 | 2.99E-21 |
| ANKRD34C | 8.916883708 | 4.048307935 | 4.91E-06 |
| AMOTL2 | 5194.214834 | 4.038764035 | 0.00E+00 |
| AP002478.1 | 8.764480814 | 4.037922708 | 1.93E-04 |
| AL161431.1 | 253.7190024 | 4.033662578 | 1.92E-98 |
| GDF6 | 21.29645127 | 4.017597586 | 1.23E-06 |
| PRR20G | 7.206833657 | 4.011162951 | 1.57E-04 |
| AL021153.1 | 5.180709888 | 4.009216772 | 3.49E-03 |
| AC010457.1 | 11.60862398 | 4.007257528 | 9.52E-07 |
| CA9 | 143.2752701 | 3.983716217 | 2.78E-06 |
| KCNE1B | 47.6023731 | 3.9637998 | 1.13E-08 |
| RNF144B | 69.84375385 | 3.962750329 | 3.56E-32 |
| DUSP13 | 11.52726261 | 3.96138273 | 8.98E-05 |
| PADI3 | 60.49052887 | 3.956003911 | 4.50E-16 |
| GUCY1A2 | 7.369270985 | 3.950202509 | 1.62E-02 |
| AC095350.1 | 25.6081823 | 3.943889293 | 5.76E-10 |
| SNX18P23 | 5.420301708 | 3.941794469 | 4.76E-03 |
| AP1M2 | 110.3798198 | 3.931041026 | 3.20E-11 |
| WNT7A | 74.42957687 | 3.924149434 | 8.08E-22 |
| AL049839.2 | 69.57473853 | 3.920643314 | 5.18E-06 |
| LINC01693 | 25.8599621 | 3.917611769 | 4.60E-11 |
| KRT7 | 53.45128578 | 3.908653037 | 2.33E-16 |
| FKBP9P1 | 272.6623261 | 3.885283653 | 9.25E-39 |
| MYH15 | 58.51415002 | 3.883154588 | 3.35E-09 |

| | | | |
|---|---|---|---|
| HBD | 5.800011754 | 3.87440689 | 6.25E-04 |
| GPA33 | 14.52145809 | 3.866687296 | 3.52E-05 |
| AC011510.1 | 10.81268386 | 3.863551203 | 3.27E-06 |
| C7orf69 | 6.788843734 | 3.861705243 | 1.53E-04 |
| CHRNA1 | 65.20886781 | 3.860482566 | 1.00E-05 |
| TNNC1 | 56.70589737 | 3.847901634 | 4.45E-20 |
| TMEM155 | 10.51064918 | 3.81252668 | 1.38E-06 |
| AC018464.1 | 5.840792081 | 3.812246004 | 3.84E-03 |
| TPO | 119.1349183 | 3.806336708 | 3.99E-24 |
| MGAM | 11.91207921 | 3.804194342 | 9.18E-05 |
| ANXA8L1 | 163.5609482 | 3.771284746 | 2.30E-28 |
| PPME1 | 11264.96145 | 3.766659503 | 1.10E-281 |
| AC245041.2 | 17.47365683 | 3.764055615 | 1.28E-07 |
| TM6SF1 | 340.5809889 | 3.762538469 | 6.38E-28 |
| ROBO4 | 76.28270049 | 3.756184797 | 1.07E-13 |
| CES1P1 | 26.62878007 | 3.751712519 | 3.57E-07 |
| LINC01511 | 8.69732256 | 3.750751154 | 1.15E-04 |
| AP003064.2 | 24.94135603 | 3.741975525 | 5.44E-08 |
| HYDIN2 | 17.04818201 | 3.738237437 | 3.79E-04 |
| LINC00707 | 90.45365985 | 3.724023029 | 1.49E-33 |
| CPNE4 | 26.75781287 | 3.7114427 | 2.24E-08 |
| OLIG2 | 29.16821453 | 3.707083982 | 1.42E-10 |
| NA | 53.49267065 | 3.704195485 | 3.84E-24 |
| ANKRD2 | 7.839182386 | 3.701862198 | 1.29E-03 |
| NCF4 | 7.267399721 | 3.701183191 | 3.94E-06 |
| PLAC8 | 321.3230234 | 3.683047137 | 3.00E-44 |
| TAGAP | 49.23096721 | 3.681140849 | 1.68E-02 |
| TMEM200C | 23.71648983 | 3.678159464 | 7.35E-05 |
| WNT7B | 200.4097672 | 3.660179928 | 4.20E-25 |
| SERPINE1 | 2089.772652 | 3.649129014 | 5.25E-112 |
| LRRC38 | 9.873393408 | 3.648211744 | 5.78E-06 |
| AP001783.1 | 27.85078654 | 3.641180276 | 1.25E-18 |
| IGFN1 | 141.2038353 | 3.639495745 | 1.96E-11 |
| CD177 | 28.34911627 | 3.636979245 | 9.41E-08 |
| AC010247.2 | 13.42529976 | 3.588167779 | 5.45E-07 |
| KRT8 | 14283.57687 | 3.561845464 | 3.44E-225 |
| GDF15 | 313.7269975 | 3.547432629 | 5.79E-11 |
| IRX1 | 12.32971566 | 3.545092412 | 1.82E-06 |
| DUX4L45 | 69.43942931 | 3.535368467 | 2.10E-33 |
| ITGB2 | 124.990794 | 3.53430358 | 8.08E-27 |
| INHBC | 11.53838661 | 3.529550562 | 1.90E-05 |
| MPP4 | 50.02754041 | 3.526990422 | 6.09E-18 |
| LINC00520 | 22.88133806 | 3.519527934 | 3.27E-06 |
| TENM2 | 351.6327637 | 3.493048385 | 2.55E-13 |
| PRELP | 480.5999178 | 3.468454992 | 8.22E-16 |
| RPL29P19 | 46.34515324 | 3.455415305 | 1.62E-18 |
| AC245100.6 | 52.1791593 | 3.455269513 | 8.61E-11 |

| | | | |
|---|---|---|---|
| GDAP1L1 | 48.33841871 | 3.443241797 | 1.67E-11 |
| FAM201A | 45.64172295 | 3.436907561 | 1.52E-12 |
| AC121757.1 | 28.94935304 | 3.430977974 | 4.02E-11 |
| AC016717.2 | 5.261098293 | 3.426082669 | 2.12E-02 |
| MEGF6 | 249.4909584 | 3.424798509 | 3.37E-20 |
| HRH2 | 76.37524624 | 3.419949347 | 1.15E-21 |
| FOSL1 | 1816.844697 | 3.419534893 | 4.01E-43 |
| CLDN1 | 777.3721858 | 3.419090483 | 2.89E-32 |
| AL590644.1 | 26.03942664 | 3.418009054 | 3.88E-11 |
| BX640514.2 | 77.90288435 | 3.414148728 | 4.36E-22 |
| SLAMF9 | 7.095931523 | 3.40197791 | 1.32E-04 |
| LINC00525 | 14.59269898 | 3.401969457 | 1.60E-07 |
| KCNJ1 | 6.187173029 | 3.387519657 | 1.27E-02 |
| LINC02243 | 8.297230953 | 3.383218686 | 1.03E-04 |
| SBSN | 9.307325147 | 3.379504451 | 3.80E-05 |
| INHBE | 431.5658161 | 3.376227348 | 4.40E-08 |
| TNFRSF12A | 3033.22403 | 3.376071254 | 5.09E-112 |
| AC245100.3 | 41.78719059 | 3.362574549 | 5.80E-09 |
| PTPRN | 673.3985235 | 3.357883403 | 1.36E-70 |
| DUX4L31 | 29.25806601 | 3.351854225 | 4.13E-16 |
| NEGR1 | 205.2653914 | 3.351377814 | 3.02E-24 |
| HCLS1 | 91.68229821 | 3.348208917 | 1.06E-32 |
| CSDC2 | 8.659595836 | 3.347783095 | 1.11E-03 |
| HNRNPA1P33 | 8.089797574 | 3.341848024 | 2.52E-03 |
| KRT17 | 109.932617 | 3.340107978 | 2.12E-08 |
| NDUFC2-KCTD1 | 12.13295113 | 3.337008252 | 2.27E-02 |
| ITGBL1 | 61.58578191 | 3.330055993 | 2.03E-18 |
| ESM1 | 570.3932995 | 3.329763757 | 2.56E-25 |
| EDN1 | 1541.884744 | 3.329648225 | 2.54E-56 |
| PSG8-AS1 | 10.13715312 | 3.321212599 | 1.21E-05 |
| GRIP1 | 17.48063495 | 3.31962677 | 8.55E-09 |
| TMEM52B | 10.55974503 | 3.316141059 | 6.51E-03 |
| PIWIL2 | 12.98599122 | 3.313069402 | 4.04E-06 |
| IL32 | 397.8318467 | 3.300352175 | 1.48E-33 |
| FAM19A5 | 5.778785156 | 3.286070856 | 4.21E-04 |
| CALCB | 23.00979005 | 3.27404512 | 5.82E-06 |
| ZP1 | 57.88032852 | 3.272951676 | 1.07E-08 |
| WDFY4 | 28.36478348 | 3.271866503 | 1.80E-06 |
| TMEM130 | 81.1847839 | 3.268741592 | 2.21E-18 |
| TMEM92-AS1 | 16.11057805 | 3.263809568 | 1.90E-05 |
| RXFP3 | 26.79648996 | 3.261170841 | 3.70E-09 |
| HDAC2-AS2 | 45.6529736 | 3.260352264 | 7.99E-17 |
| HNF4A | 25.93293679 | 3.259882421 | 8.10E-04 |
| EVI2B | 11.71834069 | 3.241657008 | 2.53E-05 |
| SNAPC1 | 2436.405541 | 3.235345229 | 1.35E-58 |
| FGF14 | 12.63602815 | 3.23501348 | 2.80E-04 |
| NKD2 | 30.11768859 | 3.227367815 | 3.42E-10 |

| | | | |
|---|---|---|---|
| ARC | 110.2728391 | 3.221588743 | 1.87E-22 |
| KLK8 | 6.266111381 | 3.220169318 | 2.43E-03 |
| AC004233.3 | 52.94805197 | 3.218322821 | 6.02E-22 |
| NA | 43.80094162 | 3.215749261 | 2.19E-10 |
| AXL | 11912.61544 | 3.206422493 | 4.27E-81 |
| EFEMP1 | 741.1103555 | 3.205095766 | 1.69E-63 |
| C5orf67 | 8.957053987 | 3.203686828 | 1.86E-04 |
| SERPINB2 | 585.2962819 | 3.19404869 | 3.52E-25 |
| AC005832.4 | 89.28362442 | 3.193766981 | 8.56E-03 |
| AC092384.2 | 8.422921767 | 3.192155351 | 1.01E-03 |
| PLAUR | 813.9719978 | 3.18995501 | 2.49E-80 |
| PROSER2-AS1 | 38.08475189 | 3.183125185 | 1.43E-12 |
| VCX3A | 15.59637735 | 3.18111045 | 1.26E-06 |
| FLT4 | 134.8507834 | 3.177110697 | 7.59E-18 |
| GNAT1 | 22.34118719 | 3.169158161 | 5.95E-04 |
| COL4A3 | 9.192934283 | 3.168542715 | 6.69E-03 |
| RGCC | 105.6600784 | 3.164433746 | 4.81E-23 |
| SLAMF7 | 176.448223 | 3.162214374 | 3.45E-09 |
| SZT2-AS1 | 13.31741173 | 3.16101076 | 4.30E-06 |
| AC010207.1 | 29.98635144 | 3.157755165 | 3.49E-12 |
| UCA1 | 6683.968826 | 3.156681468 | 8.76E-52 |
| AP000879.1 | 26.45327596 | 3.142305393 | 3.62E-02 |
| FMN2 | 287.7331264 | 3.134121937 | 1.57E-36 |
| SCG5 | 39.15796545 | 3.132687479 | 1.98E-16 |
| LINC01615 | 22.14922554 | 3.132027759 | 6.85E-10 |
| TMEM156 | 66.58521868 | 3.128890506 | 1.83E-12 |
| AL683807.1 | 11.74887014 | 3.127958284 | 2.32E-05 |
| NYAP2 | 8.4881526 | 3.124330498 | 2.88E-03 |
| MYO5B | 212.8051185 | 3.112466452 | 8.32E-25 |
| GBX2 | 105.7843387 | 3.104747908 | 5.02E-48 |
| TGFB2-AS1 | 227.6156955 | 3.093180397 | 1.99E-56 |
| IL21R | 123.0008977 | 3.087475827 | 8.49E-14 |
| RPL12P14 | 37.54375201 | 3.085101621 | 3.18E-12 |
| INKA2-AS1 | 103.6961057 | 3.084153918 | 2.63E-20 |
| MIR503 | 28.37757972 | 3.080558696 | 6.37E-08 |
| AC098613.1 | 9.120879227 | 3.078263559 | 3.32E-03 |
| ILDR2 | 71.76148004 | 3.047045061 | 5.03E-09 |
| SHANK2 | 32.47462154 | 3.046109302 | 2.16E-03 |
| C7orf65 | 7.884882127 | 3.044339887 | 7.14E-03 |
| TNFRSF9 | 304.3101605 | 3.043202088 | 1.41E-32 |
| CES5A | 10.8197907 | 3.041764752 | 1.42E-02 |
| RASGRF1 | 134.7074287 | 3.029697451 | 1.17E-24 |
| AC105411.1 | 36.104077 | 3.029044815 | 2.99E-09 |
| EXPH5 | 144.9258139 | 3.021341898 | 1.19E-34 |
| GRIK5 | 70.41443968 | 3.016648297 | 1.57E-09 |
| AC018816.1 | 5.312448188 | 3.015108248 | 7.94E-03 |
| CSF2RA | 877.8505258 | 3.013200921 | 7.90E-62 |

| | | | |
|---|---|---|---|
| RCSD1 | 77.4899084 | 3.01058697 | 8.67E-19 |
| AL365181.3 | 171.3149321 | 3.010407048 | 9.29E-31 |
| CD24 | 529.1194415 | 3.007781937 | 8.59E-49 |
| VN2R19P | 6.404195999 | 3.003509628 | 1.31E-03 |
| MOV10L1 | 49.20024885 | 3.002994979 | 3.87E-12 |
| BRS3 | 15.9865702 | 3.002798145 | 5.48E-06 |
| CLDN3 | 137.1034934 | 2.98218957 | 3.40E-39 |
| AC009238.1 | 10.03320931 | 2.977953973 | 3.63E-04 |
| LINC01583 | 42.75130781 | 2.974057003 | 2.34E-14 |
| AC092850.1 | 7.775102116 | 2.973287071 | 6.04E-04 |
| CD274 | 323.6243598 | 2.972020063 | 5.08E-54 |
| BDNF | 222.3395398 | 2.968174036 | 4.74E-64 |
| CDC42EP3 | 1854.372981 | 2.959094769 | 1.20E-40 |
| MFAP5 | 48.10545696 | 2.9590531 | 9.97E-08 |
| IL11 | 145.4784115 | 2.958361295 | 5.98E-35 |
| LINC01638 | 242.896981 | 2.95788181 | 2.12E-38 |
| FAM3B | 10.079902 | 2.954222196 | 2.91E-05 |
| C6orf132 | 13.19535717 | 2.953986706 | 1.07E-03 |
| NPY4R2 | 11.66202251 | 2.947974298 | 1.21E-02 |
| PLAC1 | 82.08555886 | 2.947634662 | 3.74E-32 |
| AL355312.4 | 60.37212687 | 2.946967286 | 1.45E-28 |
| LINC01425 | 7.678605227 | 2.940748292 | 2.91E-04 |
| AC105105.1 | 257.2624826 | 2.937123341 | 2.31E-09 |
| MYOZ2 | 10.39892218 | 2.936533786 | 1.54E-05 |
| TMEM178A | 50.3489753 | 2.93532176 | 1.03E-15 |
| CASP14 | 7.650197149 | 2.934437247 | 6.08E-03 |
| FAM90A24P | 23.25397222 | 2.932846266 | 3.99E-07 |
| MSLN | 108.5106732 | 2.931674978 | 3.81E-15 |
| LINC00882 | 70.53774856 | 2.92807953 | 7.85E-27 |
| ALDH2 | 6.145122017 | 2.920942874 | 9.21E-03 |
| RPLP0P2 | 306.6935964 | 2.919093324 | 1.39E-43 |
| NA | 26.57063328 | 2.918204246 | 1.49E-05 |
| AC005150.1 | 7.249293129 | 2.916986567 | 1.10E-03 |
| FLNC-AS1 | 13.89514009 | 2.91205391 | 5.22E-05 |
| KNDC1 | 17.66343112 | 2.910848817 | 2.11E-02 |
| MAGEB2 | 14.51957996 | 2.899640218 | 3.07E-07 |
| GADD45B | 977.0968945 | 2.896611578 | 1.75E-145 |
| KIF21B | 1486.646113 | 2.896141331 | 4.23E-50 |
| TDRG1 | 14.32537586 | 2.89460629 | 5.71E-07 |
| KCNK13 | 67.77061597 | 2.888238034 | 8.23E-34 |
| AC090505.1 | 11.22101694 | 2.884869906 | 3.62E-04 |
| MAP3K15 | 28.1466474 | 2.881242171 | 6.51E-09 |
| AC008429.1 | 55.50190177 | 2.870545866 | 5.04E-12 |
| FST | 410.4366751 | 2.868266875 | 7.23E-26 |
| AP001189.5 | 7.632947862 | 2.861266545 | 3.79E-03 |
| AC126768.2 | 6.376656943 | 2.858350833 | 9.36E-03 |
| UPB1 | 37.15425623 | 2.858344741 | 1.81E-09 |

| | | | |
|---|---|---|---|
| AL023881.1 | 23.0706953 | 2.85509356 | 2.79E-06 |
| TSPAN2 | 207.194681 | 2.848969028 | 9.48E-30 |
| NDRG1 | 8255.208812 | 2.843871188 | 6.90E-48 |
| EEF1A2 | 5981.383771 | 2.841416946 | 1.57E-128 |
| CDH1 | 23.87917036 | 2.837397602 | 9.44E-05 |
| FAM166B | 10.42880559 | 2.827919619 | 7.68E-05 |
| TRPM1 | 8.303459732 | 2.825984647 | 4.71E-02 |
| SLC12A8 | 358.0297075 | 2.822885445 | 7.34E-45 |
| GOLT1A | 20.85450575 | 2.813964164 | 1.15E-09 |
| AWAT2 | 6.560235983 | 2.804896236 | 1.80E-02 |
| AP000892.3 | 436.2274301 | 2.803204513 | 9.63E-52 |
| LINC02057 | 19.34808816 | 2.796740365 | 1.77E-06 |
| CES1 | 60.91092756 | 2.796732491 | 8.07E-13 |
| DNAJC6 | 116.1338755 | 2.796225863 | 2.15E-15 |
| SMPD4P1 | 9.902617318 | 2.795448521 | 1.96E-03 |
| KCNQ1 | 14.72470286 | 2.788702073 | 7.00E-04 |
| INHBA | 62.28707009 | 2.769666524 | 3.29E-26 |
| ADAMTS16 | 718.0575453 | 2.76793907 | 2.53E-33 |
| TMEM154 | 315.1802205 | 2.762915782 | 3.79E-20 |
| NEK11 | 62.08406486 | 2.762721317 | 1.12E-10 |
| AL590483.1 | 14.41730836 | 2.761769463 | 2.88E-06 |
| LOX | 1600.914318 | 2.755867261 | 1.43E-49 |
| PTPRR | 11.79398755 | 2.752227606 | 2.14E-04 |
| MUC3A | 74.07583469 | 2.750981772 | 8.08E-09 |
| SERPINB8 | 817.2619824 | 2.749801116 | 2.55E-38 |
| AC110048.1 | 5.151843255 | 2.749155931 | 1.29E-02 |
| CLSTN2 | 328.2908032 | 2.743963865 | 2.68E-31 |
| APLN | 612.3809223 | 2.743580287 | 5.38E-27 |
| TMEM132E | 67.2505779 | 2.743191172 | 2.45E-07 |
| HNF1B | 9.019723547 | 2.742417651 | 1.82E-03 |
| AC022239.1 | 18.69611026 | 2.735725194 | 8.62E-08 |
| SLC7A11-AS1 | 23.40959561 | 2.732596162 | 1.26E-02 |
| ARFGEF3 | 55.42513659 | 2.728455086 | 2.31E-13 |
| EPS8L2 | 349.987695 | 2.720567172 | 2.65E-38 |
| HRASLS | 31.71422465 | 2.71780002 | 1.72E-09 |
| GRIK4 | 72.0864243 | 2.716637086 | 3.42E-07 |
| TSKS | 10.55565234 | 2.715906609 | 4.71E-05 |
| AL355512.1 | 12.3124604 | 2.715200265 | 3.15E-03 |
| SLC6A1 | 26.45725069 | 2.713844911 | 1.46E-04 |
| CRYBG2 | 93.98396619 | 2.713835276 | 2.77E-10 |
| DCSTAMP | 13.89413438 | 2.709135347 | 2.49E-05 |
| HMGA1 | 9467.267639 | 2.705470539 | 2.11E-97 |
| ELFN2 | 490.7239094 | 2.704666578 | 9.26E-36 |
| RPSAP71 | 7.432948907 | 2.70463697 | 1.04E-03 |
| SLC7A3 | 41.66827949 | 2.701984579 | 4.39E-08 |
| AL391422.1 | 7.102066668 | 2.701694985 | 3.66E-03 |
| FOXG1-AS1 | 5.642949624 | 2.694280211 | 2.88E-02 |

| | | | |
|---|---|---|---|
| INSYN2B | 326.6455031 | 2.692499195 | 5.17E-29 |
| POU3F3 | 5.207919673 | 2.692143385 | 6.88E-03 |
| OXTR | 109.0439804 | 2.691931728 | 1.36E-16 |
| NCF2 | 11.76390963 | 2.691675683 | 6.77E-04 |
| KRT18 | 26282.33084 | 2.687268687 | 2.17E-122 |
| DMBT1 | 9.376975909 | 2.686486846 | 2.82E-03 |
| ADAM18 | 22.12303189 | 2.686011349 | 1.91E-05 |
| GAS6-DT | 47.94899173 | 2.677809612 | 2.95E-15 |
| SULF1 | 13.73868172 | 2.674797458 | 6.69E-03 |
| AC011591.1 | 33.09399697 | 2.674524352 | 1.48E-07 |
| MATN4 | 29.85064027 | 2.67446757 | 5.76E-09 |
| GALNT18 | 150.014102 | 2.67401514 | 1.52E-29 |
| AJUBA | 9629.715395 | 2.67219923 | 6.48E-56 |
| TGFB2 | 3830.689573 | 2.670272043 | 1.57E-149 |
| OGFRL1 | 1421.080525 | 2.665778775 | 8.28E-57 |
| AL590666.2 | 45.63170054 | 2.664395773 | 4.19E-17 |
| AC015909.1 | 13.01442352 | 2.662579896 | 6.00E-03 |
| AC073610.3 | 16.21540822 | 2.661100796 | 8.75E-05 |
| SPATA8 | 7.77623881 | 2.656141041 | 1.39E-03 |
| PSCA | 27.29284018 | 2.651427786 | 5.21E-08 |
| BEX2 | 181.7672279 | 2.650752278 | 2.99E-13 |
| AL353608.3 | 5.077825569 | 2.648638511 | 3.24E-02 |
| AC009299.1 | 54.0118018 | 2.647884646 | 3.78E-12 |
| AC087667.1 | 30.95840747 | 2.646008029 | 1.53E-09 |
| AC007952.3 | 21.04985286 | 2.643979711 | 3.40E-02 |
| AC004877.1 | 19.22436289 | 2.642887453 | 9.23E-06 |
| RBM24 | 126.6531076 | 2.641914033 | 1.65E-17 |
| ALPK2 | 2595.841625 | 2.637778074 | 3.86E-32 |
| CHAC1 | 753.1216115 | 2.637052523 | 1.92E-09 |
| AL138828.1 | 7.572296246 | 2.635840587 | 1.28E-02 |
| NPY4R | 21.74673346 | 2.635152016 | 6.43E-05 |
| GPR1 | 155.1271788 | 2.631531489 | 4.64E-04 |
| DUSP1 | 581.9843181 | 2.630878983 | 1.41E-38 |
| SUN3 | 13.02687629 | 2.630856794 | 1.47E-03 |
| AC015987.1 | 29.65507999 | 2.621724266 | 1.27E-09 |
| NEUROD2 | 17.25648078 | 2.616968691 | 4.56E-03 |
| CEACAMP11 | 8.440503374 | 2.613621296 | 3.94E-03 |
| RBPJL | 20.92174682 | 2.6073973 | 2.72E-05 |
| UGT3A1 | 24.54671501 | 2.606578874 | 2.08E-03 |
| NOC2LP1 | 35.01371521 | 2.605314466 | 5.02E-10 |
| S100P | 8.506367251 | 2.601402822 | 2.29E-02 |
| ANKFN1 | 98.2235767 | 2.600205326 | 2.57E-21 |
| HEPACAM | 11.78854112 | 2.595117496 | 1.95E-02 |
| GRIN2B | 12.85074978 | 2.592983726 | 4.73E-05 |
| GREM1 | 25.98707971 | 2.591279074 | 1.15E-06 |
| SMPD3 | 199.6240386 | 2.589841832 | 3.57E-14 |
| DHRS3 | 27.89732788 | 2.586422386 | 3.77E-08 |

| | | | |
|---|---|---|---|
| CES1P2 | 28.23427915 | 2.586116548 | 1.10E-07 |
| FBP1 | 21.31209753 | 2.585725918 | 7.93E-05 |
| AP004607.3 | 9.874376473 | 2.584856861 | 1.04E-03 |
| AC009292.1 | 5.752936398 | 2.583431589 | 7.08E-03 |
| CST7 | 33.33388902 | 2.580885904 | 1.71E-09 |
| DUXA | 17.7868599 | 2.576737263 | 5.01E-04 |
| LINC00513 | 40.82278957 | 2.571850096 | 1.50E-14 |
| LINC01204 | 6.565611826 | 2.571358289 | 8.90E-03 |
| MAGEA4 | 9.336679457 | 2.570129661 | 1.70E-04 |
| SBF2-AS1 | 216.8026323 | 2.568409915 | 9.79E-36 |
| AC084783.1 | 9.848268834 | 2.568406463 | 2.43E-03 |
| AC011944.1 | 11.91468553 | 2.566362237 | 1.14E-03 |
| MYRIP | 5.796576788 | 2.563668867 | 2.48E-02 |
| TCEAL2 | 28.40002448 | 2.561576559 | 3.07E-09 |
| ZNF185 | 2431.652507 | 2.558018728 | 5.08E-127 |
| F3 | 631.683175 | 2.557663625 | 1.89E-39 |
| GDA | 761.696661 | 2.555351994 | 9.99E-40 |
| AC026250.1 | 102.7984567 | 2.554147404 | 7.11E-28 |
| AL645608.7 | 11.78460553 | 2.55208187 | 1.04E-02 |
| AC069368.1 | 13.95312624 | 2.550907176 | 1.53E-02 |
| MUC5AC | 33.8582948 | 2.549452155 | 4.67E-06 |
| AC003092.1 | 11.66842353 | 2.547935739 | 1.34E-02 |
| FO681492.1 | 191.8394703 | 2.546773796 | 1.98E-21 |
| ANXA3 | 816.3716511 | 2.545434975 | 2.71E-40 |
| KIRREL3 | 364.1978972 | 2.543735382 | 1.09E-37 |
| COL8A1 | 2277.352965 | 2.538334684 | 4.53E-39 |
| PPL | 285.5073873 | 2.537692621 | 1.56E-31 |
| POU2F3 | 18.51370929 | 2.536346572 | 5.23E-05 |
| IL6 | 31.65351887 | 2.534766229 | 6.94E-06 |
| C6orf223 | 35.05250985 | 2.534329357 | 2.02E-06 |
| IL20RB | 107.3800024 | 2.530979956 | 1.78E-10 |
| AL035458.2 | 6.031236525 | 2.528057452 | 5.64E-03 |
| RBP7 | 234.865592 | 2.524755542 | 1.84E-48 |
| NA | 21.84005856 | 2.523809533 | 3.88E-09 |
| UGCG | 4030.044103 | 2.517555747 | 1.28E-103 |
| BMP4 | 3231.109292 | 2.515658314 | 4.91E-58 |
| WWC1 | 602.1446754 | 2.51432839 | 1.12E-58 |
| CRIM1 | 9816.058979 | 2.508939451 | 4.90E-112 |
| AC090152.1 | 15.60249365 | 2.508546756 | 4.18E-05 |
| BASP1 | 2291.351755 | 2.506771514 | 1.92E-63 |
| AL844175.1 | 5.243046801 | 2.50066062 | 1.63E-02 |
| NA | 8.854667232 | 2.49817624 | 4.14E-04 |
| ASNS | 14006.90675 | 2.491761177 | 5.18E-25 |
| GJB3 | 26.34134941 | 2.491022387 | 7.36E-05 |
| LPO | 43.94056569 | 2.490399887 | 6.33E-07 |
| IRX4 | 106.6038776 | 2.490006024 | 1.20E-26 |
| SLC35F3 | 102.7993497 | 2.48651918 | 3.81E-25 |

| | | | |
|---|---|---|---|
| LAMC2 | 82.01010143 | 2.481475456 | 5.47E-18 |
| ADM2 | 805.2354835 | 2.479977459 | 2.83E-10 |
| CELF2-AS1 | 7.070457865 | 2.471357889 | 3.95E-03 |
| ENO2 | 1006.576172 | 2.469399423 | 1.28E-30 |
| SH2D5 | 415.6853727 | 2.469313361 | 2.24E-85 |
| ADGRE4P | 80.48512716 | 2.463298847 | 3.25E-12 |
| DIO2 | 72.37432467 | 2.461866145 | 3.80E-08 |
| FO393418.1 | 6.878802365 | 2.457383643 | 2.03E-03 |
| NA | 225.5584139 | 2.456883629 | 1.52E-44 |
| LINC01605 | 45.79967183 | 2.453953641 | 2.77E-07 |
| AC012464.1 | 11.57719136 | 2.451222756 | 2.87E-04 |
| SHH | 5.381375728 | 2.45007984 | 1.44E-02 |
| NA | 17.66581051 | 2.448995102 | 5.35E-07 |
| NTN1 | 74.09737759 | 2.448296801 | 5.62E-12 |
| AC016705.2 | 94.42185277 | 2.445050806 | 2.50E-12 |
| ARHGEF28 | 3310.07867 | 2.443574885 | 2.34E-56 |
| ATF3 | 836.543072 | 2.441541232 | 2.14E-19 |
| NOX1 | 21.84778441 | 2.439283877 | 7.71E-05 |
| MATK | 61.01281891 | 2.436677559 | 2.24E-14 |
| S100A16 | 1005.645382 | 2.435121715 | 1.40E-82 |
| CYTOR | 2035.603011 | 2.433455629 | 4.75E-122 |
| CGB2 | 5.418755906 | 2.43278545 | 6.53E-03 |
| ARHGAP40 | 27.14455982 | 2.431614413 | 1.27E-06 |
| SSTR5 | 7.06647648 | 2.430680551 | 3.28E-03 |
| CATSPER1 | 133.9849911 | 2.429347592 | 4.40E-29 |
| SHOC1 | 8.469872835 | 2.429341631 | 3.35E-02 |
| AL391422.4 | 577.9328061 | 2.423121133 | 3.27E-30 |
| CREB5 | 770.2728268 | 2.422794793 | 5.55E-20 |
| FAM228A | 5.204431146 | 2.420572917 | 2.42E-02 |
| TFCP2L1 | 94.35599976 | 2.418527548 | 2.12E-12 |
| SFN | 28.73189058 | 2.418184934 | 8.87E-10 |
| CEACAM22P | 11.65861573 | 2.418178831 | 1.85E-03 |
| COX6B2 | 38.77772173 | 2.411248344 | 1.39E-08 |
| PIM1 | 1122.782319 | 2.410787368 | 6.28E-34 |
| KHDRBS3 | 71.81950796 | 2.409957694 | 6.93E-23 |
| MAEL | 7.774685191 | 2.409952409 | 5.35E-03 |
| AC010255.1 | 15.32312268 | 2.409835875 | 3.00E-04 |
| LINC01121 | 12.19044645 | 2.407817508 | 1.84E-03 |
| MS4A4A | 31.22211546 | 2.403890094 | 8.44E-06 |
| PPP1R15A | 3154.007942 | 2.402805393 | 5.79E-32 |
| STC2 | 5840.24853 | 2.401676519 | 7.45E-16 |
| JDP2 | 1542.857247 | 2.397867713 | 4.29E-31 |
| AC017002.1 | 16.76283255 | 2.39604205 | 4.30E-05 |
| CLGN | 74.76072684 | 2.395058164 | 2.12E-05 |
| UNC5A | 9.614455664 | 2.389043249 | 3.02E-03 |
| PTPRB | 576.9107832 | 2.386372098 | 3.29E-43 |
| KLHDC7A | 24.91322927 | 2.385451928 | 7.79E-03 |

| | | | |
|---|---|---|---|
| CDA | 83.82678768 | 2.385003609 | 5.45E-17 |
| IL31RA | 161.1209566 | 2.383272722 | 6.77E-20 |
| HAVCR2 | 347.7987367 | 2.378232112 | 2.59E-28 |
| RPS6P12 | 74.88269617 | 2.378198767 | 1.72E-21 |
| AKR1C7P | 9.921288973 | 2.377019279 | 3.34E-03 |
| PLCB4 | 1544.325878 | 2.374801506 | 8.09E-35 |
| AC011603.2 | 323.6017866 | 2.37264747 | 7.29E-05 |
| IGF2-AS | 6.867922769 | 2.368448625 | 3.72E-03 |
| ABL2 | 2982.838415 | 2.367685981 | 7.90E-111 |
| LCP1 | 1000.928624 | 2.365760834 | 1.26E-19 |
| TMEM92 | 202.8482872 | 2.360842559 | 6.03E-40 |
| NANOS1 | 479.3315027 | 2.358684918 | 3.26E-47 |
| ASNSP1 | 400.9698674 | 2.356697174 | 7.74E-13 |
| TEX19 | 43.55508042 | 2.353397611 | 3.63E-12 |
| AC011313.1 | 22.4197688 | 2.349578321 | 8.76E-06 |
| EPHB1 | 77.54038977 | 2.349055134 | 5.30E-08 |
| AP005901.1 | 10.95831507 | 2.347478954 | 2.61E-04 |
| MAFF | 607.0319197 | 2.347119424 | 2.83E-38 |
| AJAP1 | 312.8253769 | 2.346045217 | 1.45E-28 |
| AL136131.3 | 28.31490858 | 2.345888304 | 1.07E-04 |
| SH2D1B | 48.08555972 | 2.345766582 | 8.49E-14 |
| RIMS2 | 41.07355111 | 2.341595644 | 3.31E-04 |
| LINC02273 | 6.807350032 | 2.337858642 | 1.42E-02 |
| CYP2B7P | 34.19759031 | 2.335576948 | 2.82E-05 |
| NOG | 208.1326203 | 2.328005262 | 8.42E-14 |
| TMEM45A | 771.370834 | 2.327989867 | 5.75E-35 |
| NANOGNB | 5.163597743 | 2.32660986 | 2.75E-02 |
| TUNAR | 7.534935806 | 2.325256106 | 1.35E-02 |
| AC020934.1 | 20.74450596 | 2.321247729 | 5.54E-06 |
| SH2D4B | 5.657365801 | 2.317493116 | 2.80E-02 |
| SEMA3B | 1213.282702 | 2.315078322 | 1.29E-30 |
| DPP6 | 11.99597905 | 2.314086101 | 5.98E-03 |
| LNCOC1 | 35.33999473 | 2.31200141 | 1.20E-07 |
| PRKAG2 | 1255.564186 | 2.309249859 | 1.55E-67 |
| INSC | 8.952767796 | 2.308891553 | 4.89E-03 |
| SLC7A5 | 18550.72411 | 2.305827759 | 1.34E-12 |
| AL713998.1 | 83.75568185 | 2.305548483 | 4.46E-09 |
| FOXE1 | 349.40179 | 2.298111999 | 3.58E-37 |
| NLRP5 | 272.5607272 | 2.296725285 | 1.28E-26 |
| AC009139.1 | 7.51698613 | 2.294642136 | 1.25E-02 |
| HIF1A-AS1 | 59.94623766 | 2.294502453 | 1.40E-13 |
| GOLGA8IP | 12.87111558 | 2.289116089 | 2.53E-02 |
| LINC00958 | 7.133378578 | 2.285030662 | 4.92E-03 |
| UNC13D | 102.386188 | 2.283165003 | 3.30E-11 |
| ITGB2-AS1 | 20.00398093 | 2.282996901 | 3.16E-04 |
| PRKCE | 1501.609574 | 2.281798436 | 6.33E-29 |
| AC130304.1 | 149.9867397 | 2.281467934 | 3.13E-02 |

| | | | |
|---|---|---|---|
| KLC3 | 240.3054713 | 2.281408467 | 1.43E-37 |
| CNN1 | 297.6933427 | 2.280342001 | 1.96E-12 |
| USP41 | 16.73443121 | 2.277845267 | 2.60E-02 |
| SIK1B | 50.93325372 | 2.277506462 | 2.07E-07 |
| AC107956.1 | 22.9915758 | 2.27731425 | 4.36E-07 |
| MYCL | 127.9613386 | 2.273750965 | 3.68E-17 |
| OLFML1 | 58.53919204 | 2.272627589 | 2.68E-05 |
| CABP7 | 22.87835713 | 2.27244794 | 7.31E-06 |
| CELF2 | 578.2401139 | 2.272117432 | 2.21E-39 |
| HS1BP3-IT1 | 7.829249802 | 2.270151286 | 1.19E-02 |
| EPPK1 | 92.84336005 | 2.266764958 | 1.28E-23 |
| AC016571.1 | 5.653719177 | 2.262901178 | 1.31E-02 |
| ARL11 | 11.01279057 | 2.26277598 | 6.63E-04 |
| AL121890.4 | 11.68394512 | 2.261763175 | 3.97E-04 |
| AC093616.1 | 454.260492 | 2.26165619 | 1.31E-44 |
| KSR2 | 287.877213 | 2.261005241 | 6.19E-22 |
| SIGLEC6 | 13.02459145 | 2.258211796 | 1.17E-03 |
| SERINC2 | 3346.317931 | 2.249050216 | 1.13E-34 |
| TCL1B | 142.1677943 | 2.246221218 | 9.10E-23 |
| AC004917.1 | 9.77783445 | 2.244082331 | 5.82E-04 |
| TMEM184A | 186.8034924 | 2.241438151 | 2.56E-14 |
| MIR503HG | 2154.711243 | 2.240284436 | 2.51E-23 |
| AC017002.3 | 10.79083619 | 2.239236405 | 2.07E-03 |
| LIMS2 | 478.4740095 | 2.237549482 | 4.31E-34 |
| BMPER | 162.7326693 | 2.236491034 | 1.14E-09 |
| SLC5A4 | 15.92355371 | 2.235674399 | 5.39E-05 |
| KCNE1 | 13.69174082 | 2.234063494 | 3.16E-04 |
| ACTR3C | 65.52093073 | 2.233298684 | 3.22E-07 |
| NA | 13.87474242 | 2.225626611 | 3.62E-03 |
| C21orf59-TCP1( | 27.03478783 | 2.222473758 | 3.31E-02 |
| LINC01933 | 13.42810455 | 2.219760257 | 1.25E-02 |
| AC090241.2 | 9.365140553 | 2.217163236 | 2.49E-02 |
| AC243972.1 | 13.61856719 | 2.216849986 | 6.33E-04 |
| ADARB1 | 2219.524268 | 2.216144428 | 1.73E-22 |
| OLAH | 10.38694469 | 2.215920328 | 2.07E-03 |
| CRIM1-DT | 61.0346116 | 2.215266338 | 2.52E-17 |
| LINC01764 | 8.559247651 | 2.214113916 | 1.06E-02 |
| EMX1 | 5.979855963 | 2.213681407 | 4.61E-02 |
| LINC02487 | 24.11198403 | 2.211311276 | 4.92E-05 |
| SMIM34B | 6.97659786 | 2.208787207 | 4.04E-02 |
| AC124947.1 | 10.90404445 | 2.206013359 | 1.62E-02 |
| CD3D | 25.35654547 | 2.203296318 | 3.05E-09 |
| SLPI | 49.88731299 | 2.202943818 | 5.87E-10 |
| ADAP2 | 273.0761693 | 2.201190569 | 2.21E-32 |
| LINC00973 | 7.727607361 | 2.197943513 | 3.49E-02 |
| CAV1 | 23107.43986 | 2.196673475 | 4.64E-72 |
| AP003469.4 | 15.34615549 | 2.19625485 | 3.89E-06 |

| | | | |
|---|---|---|---|
| LINC00601 | 16.03766609 | 2.195586139 | 1.10E-04 |
| APBB1IP | 649.9227515 | 2.192806234 | 1.04E-50 |
| TRIM29 | 531.2890621 | 2.192686283 | 7.12E-19 |
| HYI | 1853.666591 | 2.188904225 | 2.93E-67 |
| RGS7 | 47.66134455 | 2.188878897 | 1.10E-13 |
| TBC1D2 | 462.5151675 | 2.188108678 | 4.24E-48 |
| GPRC5A | 3645.517891 | 2.185412993 | 4.57E-55 |
| AC008676.1 | 5.891633912 | 2.180396353 | 1.99E-02 |
| TUFT1 | 886.6362793 | 2.179837352 | 5.78E-76 |
| FAM180A | 81.88165982 | 2.175887889 | 7.34E-08 |
| HSPB3 | 33.71635809 | 2.175477586 | 1.29E-04 |
| TXNRD1 | 33083.72171 | 2.172909829 | 2.71E-33 |
| LINC00689 | 9.95143427 | 2.172020308 | 7.20E-03 |
| VXN | 179.481724 | 2.168912985 | 6.91E-25 |
| WNT5B | 481.1590035 | 2.167823205 | 1.76E-30 |
| ACKR3 | 110.6345116 | 2.16381406 | 1.88E-20 |
| MFNG | 16.54506896 | 2.161104482 | 2.46E-03 |
| PLEKHA7 | 2443.226225 | 2.160236453 | 2.25E-44 |
| MIR4435-2HG | 2097.286563 | 2.159708016 | 7.69E-106 |
| AC136628.1 | 42.07864024 | 2.15515025 | 3.57E-06 |
| SLC16A12 | 9.185038545 | 2.153364364 | 9.80E-03 |
| DUSP8 | 315.6478431 | 2.151044647 | 1.76E-18 |
| AC005481.1 | 58.82653227 | 2.14969616 | 3.24E-16 |
| TOX | 45.34735463 | 2.145757446 | 1.62E-11 |
| PPP2R2C | 56.81038469 | 2.143740999 | 2.79E-06 |
| IGFBPL1 | 580.1775613 | 2.141215968 | 6.93E-65 |
| NA | 13.14292939 | 2.141183123 | 3.40E-03 |
| AC124276.1 | 10.9518038 | 2.139498788 | 4.34E-03 |
| UNC5CL | 15.07775381 | 2.139409757 | 1.03E-03 |
| OR51B5 | 232.9156724 | 2.139285941 | 1.49E-53 |
| NA | 5.308877443 | 2.139150566 | 2.26E-02 |
| PICART1 | 40.09760807 | 2.13699361 | 2.67E-09 |
| NMNAT3 | 18.81273681 | 2.136134499 | 4.52E-05 |
| HSD11B1-AS1 | 25.91185308 | 2.132144786 | 5.57E-06 |
| HMX2 | 8.63397014 | 2.131892737 | 2.32E-03 |
| CAPN6 | 166.404887 | 2.128352791 | 1.95E-19 |
| LINC00239 | 13.37656744 | 2.127412554 | 1.07E-02 |
| SEMA7A | 526.884678 | 2.123932915 | 3.26E-33 |
| AC092162.3 | 14.46723367 | 2.123502691 | 1.68E-04 |
| AC106760.1 | 323.397699 | 2.12223638 | 7.01E-42 |
| BCL11A | 26.1444669 | 2.119703959 | 3.45E-05 |
| MYPN | 979.7692297 | 2.119507732 | 1.91E-30 |
| AC010247.1 | 21.51955622 | 2.119424862 | 1.36E-04 |
| LINC01436 | 88.90708551 | 2.116243335 | 2.52E-21 |
| CHST4 | 31.95928319 | 2.113992978 | 4.86E-04 |
| VSTM2L | 409.4949088 | 2.111362458 | 2.61E-30 |
| NA | 18.72396924 | 2.106377108 | 1.20E-04 |

| | | | |
|---|---|---|---|
| PEAR1 | 1120.147306 | 2.102784911 | 1.65E-59 |
| ATP6V0A4 | 28.12777374 | 2.102091736 | 1.73E-04 |
| AC122710.2 | 8.911190889 | 2.102009247 | 1.34E-02 |
| LURAP1L | 175.7105056 | 2.099682949 | 3.22E-14 |
| AC073283.1 | 6.951841042 | 2.098209558 | 1.75E-02 |
| DLGAP1-AS2 | 423.8719857 | 2.096867094 | 4.50E-19 |
| TEX37 | 13.34227782 | 2.095289544 | 4.90E-03 |
| AL591926.5 | 6.905487616 | 2.094957539 | 1.58E-02 |
| PNCK | 57.10837434 | 2.094511649 | 1.32E-05 |
| FCMR | 73.10846455 | 2.091688605 | 1.34E-11 |
| AC073610.1 | 10.58281692 | 2.091041232 | 9.00E-03 |
| AC148476.1 | 31.43670773 | 2.086468386 | 1.91E-07 |
| AC006273.1 | 24.67120854 | 2.086327706 | 1.99E-06 |
| MYL7 | 7.43415832 | 2.084253631 | 2.02E-02 |
| INHBA-AS1 | 14.99610622 | 2.078864428 | 6.44E-04 |
| AC139491.2 | 236.3875248 | 2.077964285 | 1.62E-15 |
| NMRK2 | 40.23946307 | 2.07643399 | 4.47E-02 |
| AL390195.2 | 16.80462659 | 2.074455887 | 6.56E-05 |
| AC090409.1 | 76.02384453 | 2.074381363 | 8.76E-16 |
| TRIB3 | 8474.977623 | 2.07348522 | 2.22E-09 |
| PTPRF | 14909.99649 | 2.073386543 | 1.33E-68 |
| NA | 1365.072079 | 2.073197126 | 3.52E-62 |
| SPECC1L-ADOR/ | 222.3063778 | 2.069944105 | 7.41E-14 |
| FAM90A2P | 9.664367904 | 2.069522802 | 1.52E-02 |
| OCLN | 724.0860483 | 2.069521979 | 6.07E-21 |
| PSAT1 | 16232.43473 | 2.06651447 | 1.59E-22 |
| AL157400.2 | 6.202634719 | 2.06647019 | 2.35E-02 |
| AC097059.2 | 16.24885375 | 2.061734863 | 3.20E-04 |
| EHD4 | 2189.688796 | 2.058895798 | 4.70E-90 |
| LINC01602 | 20.38071161 | 2.056103796 | 1.75E-05 |
| NA | 39.12265666 | 2.055876232 | 3.14E-07 |
| DUBR | 562.7047495 | 2.05532285 | 1.09E-44 |
| AC129492.2 | 18.58770482 | 2.055160872 | 1.45E-05 |
| CYFIP2 | 654.8815999 | 2.049032929 | 2.81E-17 |
| AREG | 9.792709533 | 2.046489083 | 1.04E-02 |
| ESR1 | 33.66109608 | 2.041839421 | 2.28E-02 |
| AP003717.1 | 9.464883929 | 2.041727934 | 4.66E-02 |
| LINC01203 | 6.584875129 | 2.041111527 | 1.64E-02 |
| FAM163B | 39.18434864 | 2.038818405 | 1.17E-05 |
| FGF23 | 33.33527346 | 2.037839168 | 1.94E-02 |
| AC104109.4 | 6.371128039 | 2.037781984 | 4.09E-02 |
| AC078905.1 | 7.967882544 | 2.035965028 | 6.85E-03 |
| AL355916.1 | 143.2120041 | 2.035940183 | 1.88E-22 |
| MYH9 | 39701.66481 | 2.035789822 | 1.82E-90 |
| LINC01018 | 72.32292642 | 2.035561688 | 4.18E-17 |
| OTOF | 23.81127999 | 2.035364243 | 9.99E-04 |
| CFAP52 | 12.90059407 | 2.032634561 | 8.08E-03 |

| | | | |
|---|---|---|---|
| GLI1 | 189.0118592 | 2.032475721 | 1.23E-09 |
| VENTX | 11.24059506 | 2.03164153 | 1.62E-03 |
| PAEP | 37.82752161 | 2.030920865 | 5.63E-11 |
| NA | 7.980453732 | 2.028130954 | 4.33E-03 |
| RPL24P8 | 15.75472878 | 2.026779961 | 2.93E-05 |
| INKA2 | 917.345134 | 2.01607587 | 1.65E-27 |
| PDE1C | 3542.718138 | 2.014273807 | 1.28E-26 |
| S100A3 | 298.3816144 | 2.013026669 | 1.13E-34 |
| APBA1 | 534.1102272 | 2.011431587 | 4.67E-30 |
| PSTPIP1 | 14.56411153 | 2.011116123 | 1.52E-04 |
| MET | 4328.732849 | 2.00928414 | 2.83E-42 |
| NA | 26.04020648 | 2.008798332 | 1.23E-06 |
| NLRP3 | 205.3152859 | 2.004044813 | 2.04E-15 |
| S100A2 | 104.4095725 | 2.000786902 | 3.25E-09 |
| CD55 | 1726.450139 | 1.9999417 | 5.17E-96 |
| PPP1R32 | 33.1700174 | 1.996026876 | 7.37E-08 |
| NA | 43.02971145 | 1.994065871 | 3.24E-05 |
| GAS6 | 1171.454117 | 1.992168872 | 1.87E-22 |
| DNAJB13 | 31.78302137 | 1.989982725 | 2.02E-06 |
| AL136295.4 | 47.25182977 | 1.987061429 | 2.60E-03 |
| AC005670.2 | 15.0788706 | 1.985558135 | 6.62E-03 |
| SHOX | 186.8715178 | 1.984090578 | 1.33E-24 |
| Z99943.1 | 6.154884971 | 1.983319557 | 4.27E-02 |
| FLNB | 29152.77127 | 1.98251782 | 3.71E-52 |
| GPR20 | 11.90770545 | 1.982290903 | 2.77E-02 |
| FUNDC2P2 | 18.74886774 | 1.979580796 | 4.00E-04 |
| MYL3 | 32.80788903 | 1.976433581 | 1.26E-05 |
| PRTN3 | 8.680751077 | 1.975379599 | 8.01E-03 |
| ARHGDIB | 757.048678 | 1.974080214 | 7.74E-23 |
| AC138969.2 | 22.23915958 | 1.973076448 | 1.33E-05 |
| AC009227.1 | 9.181000227 | 1.972514421 | 3.77E-03 |
| AL137918.1 | 5.869461034 | 1.971718228 | 2.14E-02 |
| CALB2 | 75.59255705 | 1.970132329 | 3.10E-10 |
| LINC00869 | 134.1130625 | 1.968485677 | 9.71E-07 |
| AC073569.2 | 42.59422281 | 1.96679908 | 8.28E-06 |
| XPNPEP2 | 24.90087282 | 1.965830596 | 2.15E-04 |
| CLIC6 | 238.7103255 | 1.964171494 | 2.06E-29 |
| AC091806.1 | 113.7234827 | 1.957942682 | 1.55E-10 |
| POU2F2 | 515.4171588 | 1.95759516 | 7.02E-18 |
| TCP10L | 15.48542202 | 1.956835291 | 1.61E-04 |
| PLEK2 | 599.7952445 | 1.955246241 | 1.84E-41 |
| NEUROG2 | 6.13522316 | 1.954750615 | 2.76E-02 |
| ZNF534 | 9.308285298 | 1.953845385 | 1.76E-02 |
| ADM | 794.9914859 | 1.952833353 | 9.54E-30 |
| MGLL | 4291.086206 | 1.952679343 | 1.31E-53 |
| CAV2 | 2124.493216 | 1.948515269 | 1.67E-58 |
| AC091182.2 | 17.00484763 | 1.948309626 | 2.51E-04 |

| | | | |
|---|---|---|---|
| ERBB3 | 99.31327608 | 1.945480316 | 5.24E-10 |
| DEPP1 | 122.3980038 | 1.938502549 | 1.24E-08 |
| C7orf57 | 122.2458911 | 1.936946746 | 4.74E-10 |
| DAW1 | 15.73967927 | 1.934207623 | 1.05E-02 |
| ZIC5 | 122.3265531 | 1.933003229 | 9.39E-18 |
| GNA15 | 12.58832049 | 1.927300337 | 2.03E-03 |
| GRIN1 | 70.83324153 | 1.927215425 | 1.10E-11 |
| LATS2 | 922.0682349 | 1.926800923 | 8.75E-69 |
| HRK | 25.95120166 | 1.926240547 | 8.75E-03 |
| EMP1 | 5374.264184 | 1.925836283 | 5.31E-27 |
| AC096564.1 | 10.91551866 | 1.924901086 | 2.70E-03 |
| ENPP7P4 | 5.157622277 | 1.921678179 | 4.94E-02 |
| WTIP | 955.4419514 | 1.920763905 | 2.32E-24 |
| VSIG1 | 26.36195593 | 1.91839969 | 7.96E-04 |
| AL356515.1 | 15.89611218 | 1.917936508 | 1.40E-03 |
| GRIN3B | 49.69900749 | 1.917775084 | 2.01E-11 |
| MYEOV | 22.27153524 | 1.917461805 | 8.65E-03 |
| THBS1 | 3896.636739 | 1.917327995 | 1.51E-65 |
| DBNDD2 | 498.5138506 | 1.913370243 | 1.01E-40 |
| PVALB | 24.08534313 | 1.912730508 | 4.47E-04 |
| DMKN | 409.4101885 | 1.905651787 | 8.68E-36 |
| SLC38A5 | 19.73587966 | 1.904182917 | 2.05E-05 |
| HES4 | 110.1398204 | 1.903237949 | 1.94E-11 |
| NA | 32.99010685 | 1.903047376 | 1.31E-06 |
| PCK2 | 6290.645737 | 1.903009822 | 4.71E-14 |
| PAWR | 1395.931334 | 1.90128469 | 5.58E-22 |
| SLC7A11 | 1423.958723 | 1.900425507 | 4.90E-04 |
| SNCB | 14.57954609 | 1.900163168 | 3.18E-03 |
| CACNA1H | 127.7563133 | 1.896317702 | 3.63E-10 |
| MAMDC2 | 4343.615439 | 1.892390248 | 1.07E-49 |
| IL1B | 73.1764465 | 1.891608421 | 4.04E-06 |
| AC142381.3 | 18.77409383 | 1.891075894 | 4.75E-03 |
| PRKG1-AS1 | 7.922083969 | 1.890497608 | 1.35E-02 |
| AC108463.2 | 15.81970056 | 1.88896558 | 7.13E-04 |
| TSHZ3 | 74.67057501 | 1.886257053 | 1.19E-09 |
| PXDC1 | 1898.914831 | 1.884426795 | 2.80E-99 |
| IGSF23 | 35.73858368 | 1.884307388 | 1.81E-06 |
| CRNN | 18.66305069 | 1.884002598 | 6.47E-05 |
| ARHGEF16 | 104.7080639 | 1.881638282 | 3.18E-16 |
| PLAU | 3831.106497 | 1.8790044 | 1.96E-34 |
| AC105345.1 | 9.725683969 | 1.877283964 | 2.59E-02 |
| AC147067.1 | 13.41097365 | 1.875887015 | 8.88E-04 |
| ADGRG3 | 68.61698108 | 1.875680254 | 3.95E-15 |
| RIMKLA | 73.27344799 | 1.875582321 | 1.51E-06 |
| ADTRP | 17.75775975 | 1.875520757 | 1.41E-04 |
| FADS3 | 3833.388893 | 1.874991526 | 1.29E-81 |
| ALS2CL | 852.7311576 | 1.874616722 | 6.60E-27 |

| | | | |
|---|---|---|---|
| KCNK9 | 51.84500957 | 1.874088211 | 1.06E-09 |
| ARLNC1 | 37.47117289 | 1.871999661 | 2.00E-06 |
| C1orf198 | 2749.169919 | 1.871800817 | 1.07E-127 |
| AL161908.1 | 16.68269058 | 1.870874875 | 1.88E-03 |
| OTOA | 11.52031922 | 1.870647276 | 2.29E-02 |
| HBEGF | 861.6443607 | 1.866966613 | 1.19E-47 |
| RAB33A | 27.18663417 | 1.865405891 | 2.91E-05 |
| LINC01169 | 5.898552156 | 1.863137127 | 3.23E-02 |
| PKD1L2 | 885.8875173 | 1.861773462 | 8.61E-11 |
| FAM129A | 2074.598291 | 1.860430869 | 4.27E-13 |
| ANXA1 | 9614.08568 | 1.860260597 | 3.73E-19 |
| TM4SF19 | 104.74363 | 1.857738234 | 1.64E-09 |
| MMP24 | 326.4706903 | 1.857604081 | 1.28E-24 |
| SDC4 | 4176.418327 | 1.856894944 | 8.79E-40 |
| GSG1 | 11.75434517 | 1.856364597 | 1.33E-02 |
| AC024475.4 | 21.46977055 | 1.856250644 | 2.83E-05 |
| CFAP57 | 85.99377228 | 1.855561444 | 3.43E-06 |
| MAFK | 1886.636662 | 1.853398636 | 2.55E-64 |
| LUCAT1 | 124.7362473 | 1.853284486 | 4.12E-05 |
| SDR42E1 | 246.9387746 | 1.852971993 | 4.40E-18 |
| AC000078.1 | 21.02708856 | 1.852781351 | 6.54E-06 |
| AC011731.1 | 14.19372554 | 1.849668723 | 9.40E-04 |
| MACC1 | 8.872948584 | 1.849349853 | 2.87E-02 |
| FLJ45513 | 22.04428444 | 1.848782685 | 4.05E-04 |
| GAL3ST1 | 22.23042295 | 1.84771658 | 3.17E-04 |
| SLC3A2 | 10651.30897 | 1.847066429 | 1.11E-14 |
| AP000781.2 | 21.60906742 | 1.845179796 | 8.66E-03 |
| FAM131B | 590.29546 | 1.844743843 | 9.64E-41 |
| VAV1 | 107.6228808 | 1.844552366 | 1.50E-11 |
| AL121772.1 | 44.2696685 | 1.842156561 | 1.86E-07 |
| PPP1R14A | 175.0821301 | 1.842081271 | 5.50E-21 |
| PVR | 6076.548739 | 1.841541968 | 1.73E-29 |
| NA | 17.7033213 | 1.840297457 | 9.19E-03 |
| CTH | 928.9606088 | 1.839830209 | 2.32E-09 |
| ERFE | 143.5214673 | 1.839399374 | 7.79E-13 |
| C19orf33 | 9.271186735 | 1.837692545 | 8.11E-03 |
| KCNH1 | 22.72830459 | 1.836435751 | 1.12E-03 |
| BEST1 | 300.9674433 | 1.835940939 | 1.62E-13 |
| GAS6-AS1 | 6.044418485 | 1.835432139 | 3.93E-02 |
| SOHLH1 | 91.92949427 | 1.833407076 | 1.47E-16 |
| ARNT2 | 978.2147149 | 1.83116949 | 7.65E-34 |
| HES2 | 24.47606085 | 1.827912521 | 3.31E-04 |
| CYTH3 | 8882.569858 | 1.826156102 | 7.91E-130 |
| ANOS1 | 67.41388281 | 1.823306206 | 1.05E-11 |
| ABI3 | 137.2170805 | 1.823001012 | 2.54E-13 |
| DDIT3 | 1960.790249 | 1.822772002 | 1.72E-05 |
| JUN | 2696.774838 | 1.821273799 | 7.10E-21 |

| | | | |
|---|---|---|---|
| SPANXB1 | 1009.08562 | 1.821131996 | 3.16E-45 |
| CFAP58 | 18.54124951 | 1.82046291 | 3.94E-04 |
| ANKRD33B | 166.908834 | 1.817442684 | 1.77E-20 |
| LAYN | 609.0277932 | 1.814902562 | 2.52E-29 |
| KCNG1 | 1345.743577 | 1.810826873 | 4.50E-16 |
| FMNL1 | 1214.864249 | 1.810510098 | 5.37E-60 |
| PARP8 | 150.9692421 | 1.809638966 | 3.33E-14 |
| COL13A1 | 1295.802804 | 1.807796912 | 2.22E-35 |
| KIAA1324 | 83.13749574 | 1.803824429 | 1.09E-08 |
| NUAK1 | 3145.049458 | 1.803511144 | 1.64E-58 |
| PRSS8 | 17.4747297 | 1.803257825 | 3.44E-03 |
| MCOLN2 | 40.13451401 | 1.801729001 | 5.05E-05 |
| VEGFA | 2322.712089 | 1.801027261 | 3.55E-11 |
| PROSER2 | 428.4427461 | 1.799736052 | 1.07E-21 |
| SLC7A1 | 3757.962617 | 1.798876233 | 3.43E-21 |
| IGF2 | 2046.485533 | 1.79814897 | 1.54E-22 |
| ATP2B3 | 7.381561982 | 1.796245917 | 3.59E-02 |
| CES3 | 106.6239909 | 1.795910276 | 5.40E-12 |
| NKX2-2 | 22.54412958 | 1.795498149 | 2.33E-05 |
| AC060766.6 | 9.190089575 | 1.793839517 | 1.27E-02 |
| NIPAL4 | 209.0449799 | 1.79325962 | 3.72E-10 |
| MAP3K20 | 5636.626502 | 1.792735325 | 1.80E-35 |
| PROC | 45.91681908 | 1.791249782 | 8.97E-07 |
| FAM83G | 1496.74242 | 1.789774577 | 1.18E-44 |
| RHCE | 20.12885806 | 1.788597983 | 4.28E-04 |
| CSF3R | 14.09332699 | 1.788276678 | 3.15E-03 |
| BNC1 | 1562.452238 | 1.787048218 | 3.01E-18 |
| EIF1AY | 6.426460361 | 1.783208771 | 4.76E-02 |
| RPS6KA2 | 1707.929685 | 1.780340794 | 1.63E-28 |
| NA | 310.8966264 | 1.780228411 | 9.66E-28 |
| AL590399.4 | 12.31094571 | 1.780049964 | 1.15E-02 |
| MTHFD1L | 2628.193976 | 1.779005718 | 4.20E-31 |
| AC004817.3 | 15.82350215 | 1.778179823 | 9.86E-03 |
| SHMT2 | 14299.48947 | 1.773410382 | 1.39E-18 |
| AC093799.1 | 7.917034643 | 1.773240517 | 3.72E-02 |
| SP8 | 10.28686341 | 1.772955855 | 4.94E-02 |
| FLNC | 24286.4346 | 1.772103803 | 1.43E-44 |
| SIK1 | 154.214165 | 1.771193845 | 2.82E-21 |
| FBP2 | 5.93318243 | 1.770849194 | 4.47E-02 |
| TBX1 | 62.73316178 | 1.764762012 | 6.73E-13 |
| SLC1A4 | 2242.247256 | 1.764551194 | 1.27E-14 |
| VAT1 | 16525.20215 | 1.764441723 | 1.23E-43 |
| NEDD9 | 1801.09962 | 1.762854807 | 2.41E-55 |
| CES4A | 56.32910085 | 1.76274649 | 2.37E-10 |
| CFD | 50.63983391 | 1.762249836 | 1.13E-07 |
| LINC00900 | 27.39226582 | 1.761190022 | 3.21E-03 |
| GATA3 | 89.10281909 | 1.758360062 | 1.68E-05 |

| | | | |
|---|---|---|---|
| AC079328.2 | 12.08875658 | 1.757571657 | 4.27E-02 |
| HPSE | 457.3034666 | 1.756955749 | 1.05E-44 |
| CAMK2B | 19.42018871 | 1.755799026 | 3.98E-03 |
| AC009108.3 | 8.624638713 | 1.754645157 | 1.62E-02 |
| ME3 | 73.97688032 | 1.754027493 | 1.76E-07 |
| AP001057.1 | 8.751761603 | 1.753875178 | 1.11E-02 |
| AL365181.2 | 32.29086124 | 1.751377668 | 1.51E-06 |
| TMEM45B | 21.77121324 | 1.74997882 | 4.56E-04 |
| ADRB1 | 31.51575746 | 1.749181523 | 5.13E-06 |
| SRMS | 9.882972601 | 1.747163152 | 1.49E-02 |
| IRF8 | 69.17208856 | 1.745937362 | 2.07E-06 |
| RAB11FIP1 | 3684.571645 | 1.745773047 | 2.28E-76 |
| BMP6 | 286.6928456 | 1.744226777 | 3.25E-12 |
| RAB3B | 2500.739363 | 1.743169978 | 3.86E-63 |
| PDLIM1 | 4528.312637 | 1.743139626 | 6.27E-54 |
| BICDL1 | 37.542591 | 1.742076645 | 1.39E-03 |
| VIP | 7.76050365 | 1.735021686 | 2.42E-02 |
| LRRC15 | 26.29205623 | 1.734996678 | 3.08E-02 |
| AL139352.1 | 15.41388576 | 1.734117111 | 2.17E-03 |
| TMEM71 | 20.65123729 | 1.733823752 | 1.26E-02 |
| ESRG | 287.4514818 | 1.733645612 | 1.70E-17 |
| NMNAT2 | 86.63767565 | 1.732943431 | 5.42E-08 |
| MINAR2 | 11.96470369 | 1.732790271 | 2.60E-02 |
| NA | 24.85123195 | 1.732227379 | 2.46E-04 |
| CITED4 | 281.9765894 | 1.731345645 | 1.26E-17 |
| TREML2 | 26.08468359 | 1.730872258 | 3.20E-05 |
| TTBK1 | 13.02622851 | 1.730596896 | 6.71E-03 |
| ATP6V1B1-AS1 | 56.91216974 | 1.730250002 | 6.37E-08 |
| RUSC2 | 2608.465768 | 1.730118326 | 4.62E-100 |
| HOXC13-AS | 178.5640571 | 1.726270105 | 1.16E-13 |
| C2CD3 | 3048.904252 | 1.726154266 | 1.20E-88 |
| QRFPR | 70.06962025 | 1.725631353 | 2.09E-07 |
| AC099518.6 | 18.71020804 | 1.723862777 | 7.15E-04 |
| JPH2 | 828.9286868 | 1.723733819 | 8.39E-23 |
| PHGDH | 8589.264013 | 1.723168749 | 7.28E-31 |
| NA | 39.88031917 | 1.722706451 | 1.26E-07 |
| UPP1 | 3542.442731 | 1.722678676 | 1.48E-17 |
| SHD | 7.266987622 | 1.721851599 | 4.30E-02 |
| LINC00211 | 14.16910301 | 1.720934519 | 9.21E-04 |
| TPM1-AS | 112.9196416 | 1.717328139 | 1.26E-05 |
| CDK15 | 14.04083495 | 1.716724486 | 2.30E-02 |
| PTH1R | 160.1294675 | 1.713761389 | 5.74E-16 |
| AC069528.2 | 12.19333953 | 1.712539171 | 8.43E-03 |
| DENND1C | 82.52985998 | 1.710308577 | 2.15E-11 |
| LMO7 | 2495.326444 | 1.707942037 | 8.96E-22 |
| CACNA2D4 | 7.946059862 | 1.706313515 | 4.09E-02 |
| NFE2L3 | 4016.428492 | 1.704795317 | 1.89E-54 |

| | | | |
|---|---|---|---|
| NMT2 | 1496.883168 | 1.70072784 | 4.86E-95 |
| ELF3 | 19.19275641 | 1.699270647 | 4.03E-03 |
| LINC01836 | 94.12756194 | 1.699263012 | 7.25E-15 |
| ZFP42 | 83.95063084 | 1.698752316 | 6.55E-11 |
| AC008514.1 | 9.822789484 | 1.69874671 | 3.90E-02 |
| AL136962.1 | 29.14557855 | 1.697427929 | 9.82E-03 |
| ACTN4 | 24964.65553 | 1.696706171 | 1.97E-61 |
| PTPN6 | 55.81649397 | 1.695329781 | 2.71E-07 |
| UCP2 | 2547.826664 | 1.695129316 | 8.54E-61 |
| VLDLR | 1352.477514 | 1.694572389 | 6.07E-71 |
| MEGF11 | 58.02669947 | 1.694249385 | 2.59E-05 |
| OR7E122P | 9.011555853 | 1.693769594 | 1.77E-02 |
| LINC01168 | 12.82812301 | 1.691836061 | 3.88E-03 |
| GAD1 | 53.22551783 | 1.691194665 | 6.19E-09 |
| LINC02561 | 10.77705008 | 1.690228286 | 1.77E-02 |
| LINC00632 | 86.52816748 | 1.689889739 | 1.83E-10 |
| NECAB2 | 101.487437 | 1.687327603 | 3.61E-09 |
| MYOF | 15695.67332 | 1.686211376 | 1.45E-41 |
| LINC00051 | 15.47321426 | 1.685312661 | 6.21E-04 |
| SH3BP4 | 4262.928303 | 1.685013308 | 8.64E-64 |
| SLC5A5 | 18.76023996 | 1.683804413 | 4.64E-03 |
| AL162727.1 | 20.90424433 | 1.683048916 | 1.43E-03 |
| AC104024.2 | 12.74438091 | 1.682994397 | 1.64E-02 |
| AC092490.1 | 224.8316177 | 1.682462848 | 7.87E-28 |
| CALHM6 | 28.56829687 | 1.679875059 | 6.17E-05 |
| IL2RG | 101.3761071 | 1.678775142 | 1.02E-09 |
| NAT16 | 35.39645748 | 1.677523499 | 5.06E-03 |
| C1QTNF1 | 954.1115381 | 1.675733684 | 2.40E-31 |
| TCAF2P1 | 28.17114646 | 1.675577133 | 2.16E-03 |
| WNT9A | 123.5863717 | 1.675147132 | 1.11E-13 |
| COL12A1 | 7146.720884 | 1.66992455 | 1.11E-35 |
| GABRR2 | 34.6200028 | 1.669644409 | 1.43E-03 |
| LOXL4 | 806.0990284 | 1.669022795 | 8.01E-39 |
| KCNK15 | 31.2411331 | 1.66784589 | 6.10E-06 |
| ANXA8 | 20.84443961 | 1.667433816 | 6.55E-04 |
| SSX4 | 18.60498698 | 1.667220502 | 1.88E-03 |
| SAMD7 | 13.89098362 | 1.667201382 | 5.77E-03 |
| GALNT8 | 42.41409185 | 1.667133062 | 5.74E-05 |
| AC084759.3 | 23.81259529 | 1.66490353 | 2.35E-04 |
| LINP1 | 60.87986528 | 1.663896844 | 3.47E-09 |
| LINC02133 | 8.747140843 | 1.663838325 | 2.87E-02 |
| PLCXD2 | 49.07100378 | 1.663665387 | 7.85E-07 |
| ITPK1 | 2503.186321 | 1.662501952 | 5.71E-50 |
| AC080023.1 | 19.56690321 | 1.662011086 | 1.07E-04 |
| TMC5 | 49.88234155 | 1.661230957 | 3.58E-03 |
| FOXO6 | 98.30035387 | 1.6605353 | 1.63E-13 |
| AL049650.1 | 34.41115523 | 1.659524501 | 2.61E-04 |

| | | | |
|---|---|---|---|
| PKP2 | 1275.453997 | 1.659408249 | 1.10E-54 |
| BIK | 12.27743953 | 1.653331628 | 7.55E-03 |
| MTHFD2 | 8169.227928 | 1.652331471 | 9.55E-17 |
| LINC01356 | 69.31803959 | 1.651459746 | 4.53E-08 |
| NNMT | 7823.226213 | 1.650169046 | 7.46E-18 |
| AL358473.1 | 37.43308023 | 1.649915879 | 3.14E-06 |
| LINC01219 | 18.91677116 | 1.64948722 | 3.89E-04 |
| SSR3 | 5937.31788 | 1.649286732 | 2.99E-109 |
| CARMIL2 | 174.2976759 | 1.648723758 | 7.38E-08 |
| AP002957.1 | 37.61139018 | 1.646696008 | 1.20E-05 |
| NOXA1 | 57.91642516 | 1.640930318 | 4.33E-07 |
| TRBV12-4 | 14.22207727 | 1.640483536 | 3.38E-02 |
| AC117402.1 | 30.025143 | 1.640019785 | 3.40E-07 |
| RAB32 | 1770.456458 | 1.635470657 | 3.08E-39 |
| LINC01133 | 24.06459905 | 1.632729423 | 4.23E-02 |
| SGK2 | 13.77953981 | 1.632593051 | 1.21E-02 |
| NPTX2 | 1549.746687 | 1.629435971 | 1.18E-21 |
| BACE2 | 2559.985201 | 1.629162144 | 6.42E-39 |
| P3H2-AS1 | 15.1095985 | 1.628460312 | 3.26E-03 |
| AC092667.1 | 12.34521846 | 1.62768447 | 9.16E-03 |
| NR2F2-AS1 | 143.3345313 | 1.624100778 | 1.76E-14 |
| CLIC3 | 15.9697621 | 1.622870076 | 2.09E-02 |
| SPTB | 236.3609432 | 1.62272722 | 7.18E-09 |
| UQCRHL | 16.37024274 | 1.62106881 | 3.26E-03 |
| FOXA2 | 107.3111875 | 1.619656406 | 1.60E-10 |
| PTK6 | 8.946125232 | 1.618132117 | 4.46E-02 |
| LPAR1 | 468.0407392 | 1.617784888 | 6.59E-24 |
| KLK1 | 68.0157263 | 1.616796148 | 2.81E-12 |
| CD99P1 | 133.5769418 | 1.615825343 | 7.29E-15 |
| CPZ | 462.7221784 | 1.615583285 | 4.97E-13 |
| CHMP4C | 55.12357223 | 1.615462409 | 9.69E-06 |
| NF2 | 3616.898035 | 1.614859228 | 6.41E-55 |
| MAP7D3 | 5669.030078 | 1.613653886 | 4.29E-39 |
| LINC01600 | 10.46319634 | 1.609923928 | 2.97E-02 |
| KCNS3 | 13.08495442 | 1.609602218 | 4.53E-02 |
| LINC02547 | 19.27199454 | 1.607554509 | 2.74E-04 |
| UAP1 | 2438.450767 | 1.60748835 | 2.88E-41 |
| LINC01405 | 15.01222138 | 1.607053022 | 4.78E-03 |
| AL442125.2 | 11.40234346 | 1.604300232 | 4.71E-03 |
| AC100803.2 | 22.40861951 | 1.602818615 | 2.09E-03 |
| TMEM225B | 15.79911079 | 1.602214159 | 2.91E-03 |
| AL158069.1 | 22.12769692 | 1.600086528 | 1.40E-03 |
| AC092437.1 | 11.49377109 | 1.599553665 | 8.73E-03 |
| PEA15 | 12565.60558 | 1.599245675 | 4.27E-34 |
| FAM155B | 469.1044739 | 1.598607096 | 5.77E-43 |
| CTPS1 | 4664.975413 | 1.598386733 | 5.64E-49 |
| PDCD1LG2 | 137.9408949 | 1.597674856 | 7.36E-10 |

| | | | |
|---|---|---|---|
| MAP6 | 112.0278279 | 1.596234331 | 5.37E-08 |
| SERTAD4-AS1 | 147.6901043 | 1.595515864 | 3.18E-19 |
| ADRB2 | 262.2256592 | 1.593172049 | 1.45E-27 |
| TRIM58 | 392.4915009 | 1.591478446 | 1.04E-28 |
| KLK6 | 11.46290878 | 1.591107569 | 6.12E-03 |
| NANOS3 | 11.09929873 | 1.590864506 | 1.23E-02 |
| ERICH2 | 16.08752266 | 1.590509745 | 2.81E-02 |
| LIPH | 201.1100145 | 1.586820176 | 1.31E-14 |
| AC005381.1 | 14.56912461 | 1.586560165 | 2.55E-02 |
| PAGE1 | 358.4447149 | 1.584649776 | 2.70E-24 |
| RBMS2 | 4001.840397 | 1.584250422 | 2.07E-79 |
| ELMOD1 | 26.26171552 | 1.580683211 | 1.93E-02 |
| AL451042.2 | 26.7035443 | 1.579436154 | 3.69E-05 |
| FAM131C | 149.5905496 | 1.578307267 | 1.43E-12 |
| AC106744.1 | 14.94496473 | 1.578118033 | 1.25E-02 |
| MICAL2 | 7480.93452 | 1.576973571 | 5.89E-33 |
| TNFRSF8 | 76.01422714 | 1.572654015 | 2.18E-14 |
| NXPH4 | 627.4375109 | 1.572356376 | 1.53E-23 |
| C19orf81 | 35.88473878 | 1.568729494 | 7.54E-05 |
| PHLDA2 | 934.0650038 | 1.568307972 | 5.05E-22 |
| MOCOS | 902.9342014 | 1.567509452 | 9.90E-11 |
| ADRA2C | 237.9240199 | 1.566164828 | 2.32E-20 |
| C1orf116 | 44.68469099 | 1.565010973 | 7.92E-07 |
| GOLGA7B | 57.16990062 | 1.564123793 | 9.89E-06 |
| NABP1 | 2078.385631 | 1.563518108 | 2.45E-18 |
| AL357033.1 | 14.66257818 | 1.563283292 | 4.28E-03 |
| LYPD1 | 36.76512797 | 1.561812725 | 1.12E-04 |
| AC007040.2 | 54.2009864 | 1.561433754 | 1.56E-06 |
| KLF4 | 330.1654842 | 1.560211204 | 7.42E-08 |
| AL096865.1 | 40.03954789 | 1.560089006 | 2.73E-06 |
| AC073365.1 | 60.71790693 | 1.557960575 | 1.36E-11 |
| AC117500.3 | 30.09938499 | 1.556087453 | 1.20E-02 |
| NA | 21.75015542 | 1.555866344 | 5.09E-04 |
| SLC45A2 | 191.8192019 | 1.555344528 | 6.51E-12 |
| ITGA3 | 22276.61878 | 1.554807782 | 2.30E-52 |
| SPINT1 | 91.07398261 | 1.554352078 | 4.65E-07 |
| AL450405.1 | 9.956692724 | 1.553188559 | 4.47E-02 |
| LINC01119 | 98.2556091 | 1.551499889 | 4.24E-09 |
| TRIM7 | 166.7515713 | 1.549620469 | 1.09E-21 |
| RUBCNL | 165.9995 | 1.548256184 | 1.09E-06 |
| AC092645.1 | 38.54954026 | 1.548118142 | 3.46E-03 |
| NUAK2 | 210.063746 | 1.547609825 | 1.90E-22 |
| AC016687.3 | 39.0356115 | 1.546294282 | 1.30E-04 |
| PPP1R13L | 292.5531074 | 1.545201619 | 7.80E-19 |
| DKK1 | 937.75661 | 1.544364148 | 5.01E-17 |
| FAM71F1 | 16.90444822 | 1.54395517 | 5.27E-03 |
| KCNK6 | 269.045259 | 1.543831786 | 1.82E-31 |

| | | | |
|---|---|---|---|
| GIT1 | 3699.953202 | 1.54336204 | 9.01E-85 |
| CRADD | 389.3041389 | 1.542717774 | 4.95E-46 |
| AL031728.1 | 20.12150877 | 1.542330618 | 9.00E-04 |
| IL12A | 40.27153141 | 1.541347261 | 2.13E-06 |
| GAS5 | 7842.947098 | 1.540425435 | 1.31E-23 |
| GCSAM | 14.9224264 | 1.537454105 | 1.49E-02 |
| VWF | 121.5698336 | 1.536353185 | 2.51E-06 |
| PCMTD1P2 | 10.24315486 | 1.535568241 | 3.38E-02 |
| CEBPG | 1838.694519 | 1.535024354 | 6.42E-09 |
| AC090358.1 | 11.99193219 | 1.534972144 | 2.66E-02 |
| METRNL | 538.6256251 | 1.533571619 | 1.74E-23 |
| TUBA3C | 15.7033151 | 1.532920917 | 3.49E-02 |
| AC002401.4 | 17.60670996 | 1.532776342 | 1.43E-02 |
| LINC02381 | 568.2933948 | 1.532742959 | 6.64E-39 |
| LINC00173 | 18.45505566 | 1.53255102 | 1.41E-02 |
| EPHA2 | 5020.238168 | 1.530699246 | 5.23E-38 |
| AC073130.2 | 9.878200911 | 1.52982243 | 4.59E-02 |
| FAM107A | 21.23397353 | 1.529578878 | 2.52E-03 |
| GLS2 | 26.64856311 | 1.529091615 | 3.61E-03 |
| KLF2 | 119.5906069 | 1.528805539 | 1.53E-11 |
| DLL4 | 11.35024759 | 1.528723973 | 1.27E-02 |
| CPNE7 | 438.1070185 | 1.528419897 | 3.86E-21 |
| KCNF1 | 83.96198227 | 1.526708312 | 1.85E-06 |
| CRYBA2 | 26.68788722 | 1.526583389 | 7.05E-05 |
| TIE1 | 3876.156317 | 1.52644077 | 2.33E-34 |
| USP43 | 231.4230637 | 1.524972215 | 2.82E-18 |
| PIM3 | 1569.131357 | 1.523745446 | 1.02E-70 |
| EZR | 9411.26178 | 1.522421305 | 4.90E-73 |
| AC116021.1 | 33.88852242 | 1.521869398 | 1.01E-04 |
| SHANK1 | 153.5495197 | 1.521822669 | 1.19E-09 |
| DUOXA1 | 48.41294367 | 1.521790277 | 8.88E-05 |
| WAS | 120.5812993 | 1.521224957 | 6.52E-18 |
| XDH | 47.86677611 | 1.521163133 | 6.19E-04 |
| FAM167A | 682.8611599 | 1.520731105 | 9.82E-12 |
| ANGPT4 | 46.20427387 | 1.51787632 | 8.92E-07 |
| AC009264.1 | 24.17787094 | 1.515100973 | 2.71E-03 |
| ADAMTS1 | 240.3419719 | 1.514853763 | 8.10E-08 |
| AC093673.1 | 597.9308383 | 1.514292294 | 3.13E-34 |
| CDC42EP1 | 3906.296252 | 1.513609279 | 9.87E-17 |
| SLC6A15 | 394.4907478 | 1.513455362 | 2.01E-09 |
| TPM1 | 23789.68 | 1.512667246 | 1.55E-45 |
| CHST3 | 1381.804227 | 1.512167809 | 9.92E-48 |
| CEACAM5 | 17.62300151 | 1.511928132 | 2.40E-03 |
| CBLN1 | 8.856461468 | 1.510899308 | 4.30E-02 |
| BST1 | 122.7096193 | 1.51061729 | 1.07E-08 |
| LINC01357 | 79.30204463 | 1.509463941 | 1.55E-10 |
| WWC2 | 2643.317265 | 1.509447136 | 1.28E-17 |

| | | | |
|---|---|---|---|
| AC129492.4 | 81.89087006 | 1.508402083 | 8.90E-04 |
| CCK | 19.10415853 | 1.508369595 | 6.99E-03 |
| AC245595.1 | 457.8250349 | 1.50826973 | 7.20E-27 |
| GABARAPL1 | 3444.554127 | 1.50774502 | 1.79E-63 |
| B3GALT5 | 20.11888974 | 1.507568089 | 2.88E-02 |
| LRRC20 | 1074.126572 | 1.506852108 | 6.18E-22 |
| GFPT2 | 1047.076629 | 1.50681855 | 9.61E-23 |
| OR51B4 | 93.62497888 | 1.506409316 | 2.90E-10 |
| RTBDN | 28.10301601 | 1.505485558 | 1.20E-04 |
| SYTL5 | 146.9965863 | 1.503599147 | 4.77E-05 |
| BCL2L1 | 5620.751235 | 1.502331064 | 7.26E-74 |
| PPP1R1C | 18.81541995 | 1.502299215 | 2.13E-03 |
| LMX1B | 158.1278197 | 1.500983293 | 3.70E-11 |
| PODXL | 5988.65413 | 1.5004302 | 2.41E-25 |
| TMEM255B | 84.91660562 | 1.500005087 | 1.26E-04 |
| C10orf55 | 41.45330799 | 1.499905316 | 1.20E-04 |
| RCAN1 | 3868.497247 | 1.498473915 | 3.89E-30 |
| PIK3CD | 607.3206334 | 1.49571269 | 1.04E-19 |
| SH2D3C | 192.6025404 | 1.495607962 | 1.32E-08 |
| CACNG6 | 238.4323998 | 1.495079717 | 3.73E-13 |
| C22orf34 | 161.5042601 | 1.494230769 | 6.08E-16 |
| ACOXL | 37.18565397 | 1.492264778 | 4.55E-04 |
| CD9 | 3271.154132 | 1.491840062 | 2.80E-22 |
| NA | 440.3647721 | 1.490837737 | 4.15E-20 |
| CHRD | 56.17107808 | 1.490832184 | 1.07E-04 |
| DUX4 | 10.46079859 | 1.487856525 | 2.30E-02 |
| CASS4 | 18.80043433 | 1.487769153 | 3.47E-03 |
| KLF15 | 216.4359999 | 1.487527171 | 5.51E-08 |
| SLC22A15 | 397.3289944 | 1.487036258 | 1.17E-05 |
| RASIP1 | 167.1458461 | 1.486677395 | 9.18E-09 |
| CEBPA | 131.3984501 | 1.484553953 | 3.03E-10 |
| WWC3 | 2266.235865 | 1.484163593 | 9.21E-33 |
| AC005336.2 | 28.78799943 | 1.483339745 | 5.24E-04 |
| CCNB3 | 41.40245284 | 1.483327156 | 1.20E-03 |
| PGM5P2 | 53.12602511 | 1.483265241 | 1.39E-08 |
| TTC9 | 226.5344015 | 1.4828177 | 8.52E-14 |
| CACNA1I | 43.84665629 | 1.480549688 | 1.20E-04 |
| ZNF469 | 430.1866375 | 1.478234164 | 2.83E-14 |
| RPS6KL1 | 347.9932618 | 1.477179356 | 1.06E-16 |
| ITPR3 | 5904.785391 | 1.475558712 | 8.87E-55 |
| FOXC2 | 196.881966 | 1.475373268 | 9.32E-11 |
| AC243964.3 | 235.8240355 | 1.471054315 | 1.86E-18 |
| FIBCD1 | 129.1126746 | 1.469153028 | 2.55E-08 |
| PSD2 | 18.55882236 | 1.468764342 | 3.36E-03 |
| CLEC4M | 17.52321515 | 1.468239324 | 1.87E-02 |
| AC015819.1 | 10.6041902 | 1.466737318 | 4.36E-02 |
| UBE2QL1 | 493.8624657 | 1.4657712 | 1.34E-16 |

| | | | |
|---|---|---|---|
| ALMS1-IT1 | 18.70546363 | 1.465345126 | 4.24E-03 |
| PFKP | 5727.138889 | 1.464185225 | 9.38E-46 |
| AC099568.2 | 26.25689458 | 1.459145072 | 7.87E-03 |
| NA | 42.13470809 | 1.458487366 | 5.57E-07 |
| AC016251.1 | 9.220325434 | 1.45800104 | 4.45E-02 |
| ANG | 121.0230573 | 1.457198123 | 1.73E-09 |
| NA | 23.5931674 | 1.452890428 | 3.11E-03 |
| AC226118.1 | 19.52819123 | 1.45276449 | 5.06E-03 |
| TUBB6 | 6630.385277 | 1.451846077 | 2.32E-19 |
| KCTD19 | 164.1371016 | 1.45089055 | 2.69E-11 |
| GPC4 | 272.3054783 | 1.450485063 | 1.17E-10 |
| FERMT3 | 650.0854354 | 1.450472648 | 1.22E-32 |
| DYSF | 2529.150232 | 1.450251707 | 3.00E-25 |
| SH3TC1 | 1122.413012 | 1.449261746 | 1.14E-10 |
| FLNA | 161615.4789 | 1.448408398 | 4.77E-61 |
| AC145124.1 | 226.6834811 | 1.447047109 | 4.30E-26 |
| DGAT2L7P | 9.133983121 | 1.446125031 | 4.15E-02 |
| HABP4 | 1578.110636 | 1.444762835 | 9.87E-73 |
| PAX7 | 19.03668264 | 1.444646893 | 8.79E-03 |
| PTPRH | 506.6079635 | 1.443656424 | 4.31E-07 |
| AC239859.6 | 53.28698724 | 1.443331434 | 1.69E-08 |
| SARS | 9946.604605 | 1.44261462 | 1.71E-23 |
| AC069209.1 | 17.56227689 | 1.440689509 | 8.85E-03 |
| ETS1 | 13708.86452 | 1.440477993 | 3.46E-84 |
| KIF1A | 33.08770637 | 1.439694162 | 6.42E-03 |
| HKDC1 | 536.4580666 | 1.439226775 | 3.74E-09 |
| CYP27B1 | 66.5197949 | 1.439012119 | 1.82E-05 |
| EIF4EBP1 | 6024.259061 | 1.435878023 | 1.91E-19 |
| AC124067.2 | 159.4351296 | 1.435528027 | 7.61E-12 |
| TMEM40 | 73.22422857 | 1.434797347 | 7.94E-05 |
| ADAMTS6 | 290.3621183 | 1.434549091 | 5.98E-17 |
| FENDRR | 1374.189237 | 1.43406324 | 2.53E-16 |
| CHURC1-FNTB | 112.0451196 | 1.433883339 | 1.19E-03 |
| SLC8B1 | 1452.09912 | 1.433440202 | 7.98E-31 |
| PTCH1 | 1620.679928 | 1.432984443 | 6.52E-18 |
| LINC00316 | 15.37048889 | 1.432287918 | 2.75E-02 |
| HIST1H2BJ | 99.37829003 | 1.431816362 | 3.76E-11 |
| VGF | 785.8211023 | 1.42822752 | 1.92E-12 |
| CDCP1 | 1149.145664 | 1.427523355 | 3.09E-23 |
| CXCL14 | 171.4861655 | 1.427472555 | 3.56E-05 |
| EBF2 | 15.16971605 | 1.426811899 | 1.45E-02 |
| MYLK | 2007.992478 | 1.426470309 | 4.22E-20 |
| ZMAT1 | 38.3723145 | 1.425728862 | 4.64E-04 |
| MESP1 | 46.744598 | 1.425430699 | 1.53E-05 |
| LINC01647 | 23.74194246 | 1.423425537 | 3.98E-03 |
| WFDC3 | 25.00327681 | 1.423415176 | 4.72E-03 |
| FRMD5 | 262.6352491 | 1.423347272 | 2.44E-17 |

| | | | |
|---|---|---|---|
| BAIAP2L1 | 1835.617301 | 1.422851603 | 2.18E-51 |
| ANO4 | 74.03198246 | 1.419734502 | 2.66E-06 |
| ARL4C | 1081.703425 | 1.418610678 | 1.37E-24 |
| NTRK1 | 102.2523374 | 1.417777176 | 7.53E-13 |
| MYT1 | 13.7927867 | 1.417501911 | 2.55E-02 |
| BOK | 3337.704686 | 1.412970148 | 4.32E-21 |
| PLAC9P1 | 16.69223394 | 1.41208427 | 7.97E-03 |
| CARS | 10244.17979 | 1.41103112 | 2.72E-57 |
| ARL4A | 624.3512308 | 1.408047184 | 7.44E-23 |
| VEPH1 | 23.61860105 | 1.407561792 | 2.25E-02 |
| SPANXA2-OT1 | 50.20779197 | 1.406476991 | 1.76E-04 |
| CCDC85C | 2884.103042 | 1.406442207 | 2.52E-29 |
| NA | 29.32595176 | 1.402255495 | 3.84E-03 |
| SPATA21 | 20.50351776 | 1.399773559 | 1.50E-02 |
| PIK3R5 | 80.61375241 | 1.39629209 | 3.45E-09 |
| GNB1L | 130.7836641 | 1.396289038 | 1.40E-11 |
| OXCT2 | 18.46815646 | 1.395370608 | 2.35E-02 |
| DENND2A | 3519.216737 | 1.394826611 | 7.79E-28 |
| KCNJ4 | 60.10916576 | 1.39413609 | 2.02E-06 |
| CCL26 | 75.15112479 | 1.391464579 | 2.26E-05 |
| CGN | 277.5497533 | 1.390853189 | 1.76E-09 |
| ADORA2B | 805.8301713 | 1.390534745 | 3.35E-31 |
| DNAH5 | 297.4360865 | 1.389376391 | 2.40E-20 |
| CCRL2 | 23.25096366 | 1.387703885 | 6.71E-03 |
| DPY19L2P5 | 27.43561117 | 1.387157268 | 6.91E-04 |
| CTIF | 3415.140153 | 1.384909661 | 4.59E-29 |
| SIGLEC14 | 107.3074879 | 1.384135881 | 2.20E-06 |
| SUGT1P1 | 47.60383308 | 1.38361813 | 3.33E-05 |
| YBX3 | 3730.363678 | 1.38291614 | 7.92E-92 |
| HAR1A | 46.38065491 | 1.382545337 | 1.60E-04 |
| MAP3K21 | 160.6828438 | 1.378918492 | 1.12E-09 |
| KCNS1 | 313.8676778 | 1.378395197 | 8.32E-13 |
| AC138150.2 | 38.54319126 | 1.377880609 | 6.43E-04 |
| DNAH10OS | 45.64178324 | 1.377723456 | 1.54E-04 |
| LINC02525 | 15.86356257 | 1.377517851 | 1.37E-02 |
| XBP1 | 4964.531383 | 1.376665336 | 3.75E-45 |
| OVAAL | 29.55126187 | 1.375426233 | 9.14E-05 |
| PRDM12 | 12.20491701 | 1.374876207 | 3.11E-02 |
| BRSK2 | 130.1529311 | 1.374681387 | 1.88E-05 |
| LINC00629 | 15.33200815 | 1.373879299 | 1.67E-02 |
| MYC | 932.6837416 | 1.373126449 | 3.67E-06 |
| AC010327.3 | 10.02375181 | 1.372963358 | 4.82E-02 |
| MTMR10 | 1017.928693 | 1.37274916 | 4.90E-37 |
| KRT8P33 | 61.62685347 | 1.372611015 | 1.40E-07 |
| AC138811.2 | 44.94366895 | 1.372278974 | 4.21E-03 |
| ARID3B | 403.876423 | 1.372255816 | 3.06E-22 |
| AC092687.3 | 14.56324383 | 1.372074292 | 1.05E-02 |

| | | | |
|---|---|---|---|
| SUSD3 | 356.0271866 | 1.372061999 | 5.14E-15 |
| SLC16A3 | 3928.098209 | 1.371703872 | 2.81E-32 |
| WARS | 18975.32743 | 1.370249839 | 3.60E-12 |
| PKP1 | 27.47331584 | 1.367513597 | 8.70E-04 |
| DOCK2 | 1547.341543 | 1.367340145 | 2.99E-19 |
| CD1D | 791.2080238 | 1.36648569 | 7.63E-21 |
| ETHE1 | 1360.286633 | 1.366038563 | 2.35E-39 |
| AC099548.2 | 253.6681035 | 1.36556899 | 6.08E-07 |
| MYO1E | 3124.761668 | 1.364170747 | 1.05E-33 |
| NLRP12 | 80.46937671 | 1.364093449 | 2.83E-04 |
| RRAD | 54.82459311 | 1.363337228 | 1.20E-04 |
| UPK1A-AS1 | 54.35396382 | 1.362782031 | 3.41E-03 |
| AC099489.1 | 53.33698487 | 1.361047285 | 1.62E-03 |
| SCGB3A2 | 67.32651415 | 1.360290332 | 1.37E-04 |
| SCARF2 | 1756.210107 | 1.359778203 | 4.53E-68 |
| CACNA2D3 | 94.70357472 | 1.358243893 | 1.81E-06 |
| PRDX6 | 12969.97135 | 1.354583613 | 3.15E-29 |
| CFLAR-AS1 | 19.01424679 | 1.353713908 | 8.85E-03 |
| IL18 | 12.86748055 | 1.353443409 | 4.92E-02 |
| DENND3 | 1469.517193 | 1.352424612 | 1.20E-43 |
| LUARIS | 40.9259168 | 1.351864719 | 2.84E-03 |
| NUPR1 | 3841.855504 | 1.349834299 | 1.23E-04 |
| AC009055.2 | 28.24339746 | 1.346127451 | 2.11E-02 |
| SLC13A5 | 229.8750143 | 1.345621946 | 6.29E-08 |
| N4BP3 | 111.414007 | 1.345069051 | 6.56E-09 |
| ATP1A3 | 357.0233239 | 1.344904442 | 5.81E-08 |
| SYNE3 | 2045.696794 | 1.343816318 | 8.45E-32 |
| AP000446.1 | 357.0586464 | 1.343190988 | 2.51E-15 |
| RELB | 233.1853472 | 1.342935799 | 4.14E-06 |
| NEDD4L | 4687.21832 | 1.341497269 | 1.22E-43 |
| DLGAP1-AS5 | 26.94921455 | 1.341380512 | 1.12E-02 |
| LIF | 154.155482 | 1.341300811 | 5.86E-06 |
| PCID2 | 2143.240142 | 1.341155154 | 1.65E-67 |
| HLA-DQA1 | 47.19583332 | 1.340356722 | 2.13E-03 |
| GUCA1B | 26.55959895 | 1.338243974 | 1.43E-04 |
| MMP23B | 44.95772942 | 1.337571083 | 5.79E-03 |
| ARL4D | 533.0645091 | 1.337020005 | 1.02E-14 |
| ZFAS1 | 5020.048184 | 1.335939348 | 4.28E-15 |
| AC117500.5 | 24.72259737 | 1.333726715 | 4.67E-02 |
| BATF3 | 241.6407464 | 1.333107586 | 8.39E-16 |
| SERP2 | 73.98911883 | 1.331886629 | 1.10E-07 |
| TGIF2-RAB5IF | 109.1225945 | 1.331832286 | 3.58E-05 |
| AL390719.1 | 186.2336545 | 1.329881877 | 6.32E-13 |
| WDR66 | 107.1767467 | 1.327118238 | 1.53E-04 |
| MAMLD1 | 1047.30731 | 1.326554463 | 1.25E-15 |
| SLFNL1 | 51.8325916 | 1.323011355 | 2.99E-05 |
| AZGP1 | 71.03585013 | 1.322884446 | 7.05E-05 |

| | | | |
|---|---|---|---|
| AC005736.1 | 74.59682991 | 1.322515884 | 1.96E-06 |
| AC073585.1 | 61.66117725 | 1.32239965 | 6.33E-06 |
| PRSS3 | 32.73514005 | 1.322127917 | 7.81E-04 |
| HLA-V | 43.93514239 | 1.318722734 | 5.13E-03 |
| MPP5 | 2731.596394 | 1.316704198 | 1.05E-19 |
| RASAL3 | 65.57954874 | 1.31663951 | 9.48E-07 |
| SEMA3F | 497.6585612 | 1.316482748 | 1.10E-23 |
| NOCT | 193.3207984 | 1.315970441 | 4.44E-13 |
| HES7 | 61.44827338 | 1.31463992 | 2.39E-06 |
| AL139289.2 | 61.65032174 | 1.314370726 | 6.41E-06 |
| TMEM183B | 23.23056025 | 1.314276285 | 4.72E-02 |
| LINC01611 | 16.09218297 | 1.312470219 | 1.30E-02 |
| HHLA1 | 19.15672691 | 1.310311827 | 2.30E-02 |
| NOTCH1 | 1196.391895 | 1.309417805 | 6.06E-26 |
| PPM1J | 142.1954387 | 1.309121714 | 1.61E-09 |
| DOP1B | 559.0709286 | 1.307357587 | 2.63E-09 |
| NT5M | 308.3312684 | 1.306649708 | 3.39E-16 |
| CACNA1B | 179.5369723 | 1.305660547 | 6.93E-08 |
| DCT | 247.3288116 | 1.304717432 | 6.48E-09 |
| LIN28A | 640.0340004 | 1.304152566 | 6.37E-21 |
| AC097652.1 | 33.6880894 | 1.303676304 | 1.41E-04 |
| AHNAK2 | 1587.711431 | 1.302930231 | 2.54E-14 |
| GARS | 17460.14145 | 1.302814821 | 3.86E-14 |
| LINC02256 | 15.471134 | 1.301447464 | 2.83E-02 |
| TEAD4 | 1502.038523 | 1.3010617 | 1.26E-38 |
| IL6R | 802.3266812 | 1.300698739 | 2.12E-16 |
| FBXL16 | 218.7308951 | 1.30058339 | 9.27E-05 |
| CD44 | 26250.0004 | 1.299837867 | 1.79E-30 |
| FAM189A2 | 29.59519028 | 1.298977367 | 9.38E-03 |
| TSC22D2 | 1341.340198 | 1.298595763 | 6.07E-22 |
| TMPRSS3 | 63.85416896 | 1.297645925 | 8.80E-04 |
| AC090617.9 | 15.7887937 | 1.297275736 | 1.33E-02 |
| LCN10 | 13.65945486 | 1.296967789 | 2.13E-02 |
| LINC01730 | 59.12212474 | 1.296418487 | 5.13E-04 |
| NBEAP3 | 75.26489319 | 1.292289623 | 4.89E-08 |
| FSTL3 | 2216.418323 | 1.29128766 | 1.20E-12 |
| PIK3R6 | 45.75679108 | 1.291159713 | 5.83E-05 |
| HS3ST3A1 | 154.9435874 | 1.290184724 | 4.39E-11 |
| ACTBL2 | 62.37567461 | 1.28889754 | 7.17E-04 |
| HEG1 | 10780.54503 | 1.288822295 | 6.37E-68 |
| ACTN1 | 17677.66597 | 1.288325148 | 2.87E-55 |
| AARS | 23738.39566 | 1.287689357 | 1.77E-15 |
| PAQR5 | 477.0127961 | 1.287347455 | 2.27E-10 |
| LINC00623 | 78.41347884 | 1.286012985 | 7.95E-04 |
| F11R | 1874.633537 | 1.285001489 | 1.55E-15 |
| CCND1 | 9113.412653 | 1.284825408 | 1.48E-18 |
| FBLN5 | 552.2943098 | 1.284592734 | 2.16E-07 |

| | | | |
|---|---|---|---|
| DUX4L26 | 66.00978395 | 1.283953723 | 4.30E-07 |
| LINC02495 | 16.81312737 | 1.283115934 | 1.62E-02 |
| LINC01503 | 111.7516947 | 1.282930773 | 2.66E-11 |
| BEX5 | 103.9003807 | 1.282811133 | 8.38E-08 |
| ATP8B3 | 1210.261948 | 1.281960472 | 1.84E-32 |
| LY96 | 154.5175083 | 1.281409302 | 3.16E-07 |
| CHRM4 | 157.7202575 | 1.281015243 | 6.23E-09 |
| SOX30 | 17.53791901 | 1.280583288 | 2.08E-02 |
| SMCO4 | 160.0160317 | 1.28047988 | 1.96E-05 |
| ERCC2 | 3374.31351 | 1.280412193 | 9.26E-30 |
| AC092111.1 | 85.97713132 | 1.280168595 | 4.87E-09 |
| AL353807.3 | 42.662876 | 1.280070853 | 1.10E-04 |
| EBI3 | 130.7138912 | 1.279994196 | 5.17E-07 |
| C4orf19 | 21.16468249 | 1.27827286 | 1.25E-02 |
| SHB | 1465.471817 | 1.277633999 | 4.32E-18 |
| QSOX1 | 8378.691381 | 1.275118426 | 1.07E-35 |
| DSP | 1365.602877 | 1.27347448 | 6.50E-27 |
| POLRMTP1 | 27.84699559 | 1.273054687 | 2.88E-03 |
| LINC01721 | 15.2378456 | 1.272996034 | 3.86E-02 |
| AC104129.1 | 41.03407315 | 1.272565962 | 8.20E-03 |
| SORBS2 | 136.3448756 | 1.271932231 | 4.47E-05 |
| LINC02577 | 30.04782846 | 1.271862058 | 2.98E-03 |
| AC022784.1 | 65.50596088 | 1.270386096 | 1.55E-04 |
| OPN3 | 842.1645973 | 1.269222872 | 3.92E-27 |
| CHRFAM7A | 55.5674447 | 1.268928794 | 2.14E-04 |
| EIF3CL | 2513.103972 | 1.268165857 | 2.46E-04 |
| UBASH3B | 966.8217515 | 1.267852682 | 5.09E-10 |
| HERC3 | 716.0548112 | 1.26676786 | 8.65E-06 |
| EYA2 | 133.8226664 | 1.265813297 | 2.27E-10 |
| AC108463.1 | 17.95471562 | 1.265097355 | 1.39E-02 |
| LINC01094 | 30.00020827 | 1.264770598 | 1.08E-03 |
| BHLHE41 | 332.2020079 | 1.26235703 | 8.42E-05 |
| LACTB | 732.0504195 | 1.2623442 | 2.55E-22 |
| FAM85B | 19.23518435 | 1.26190681 | 3.06E-03 |
| CLIP4 | 1736.450377 | 1.261208822 | 2.45E-10 |
| ANXA2P2 | 38.68847783 | 1.260386391 | 1.39E-03 |
| C3 | 235.9605219 | 1.257897646 | 1.44E-05 |
| MIR4527HG | 177.5777164 | 1.257742982 | 9.90E-16 |
| TMED10P2 | 29.80088923 | 1.257615392 | 1.55E-03 |
| KCNN4 | 521.3339695 | 1.257116413 | 1.12E-13 |
| ULBP1 | 1094.423407 | 1.256455197 | 1.65E-10 |
| STX3 | 2995.088942 | 1.256124066 | 3.40E-44 |
| LARP1B | 467.2784037 | 1.255430032 | 2.17E-12 |
| TXNRD2 | 1698.664478 | 1.253327426 | 8.68E-36 |
| STARD13-AS | 15.35750094 | 1.252795068 | 4.76E-02 |
| LINC02298 | 64.28724493 | 1.252490079 | 4.44E-06 |
| ACSM5 | 100.294989 | 1.251997685 | 2.64E-06 |

| | | | |
|---|---|---|---|
| CAMSAP3 | 101.4902695 | 1.251983121 | 5.72E-05 |
| CCM2L | 68.80571366 | 1.251976212 | 5.68E-04 |
| DDAH1 | 3716.229868 | 1.251462497 | 2.21E-19 |
| AL391056.1 | 11.43566403 | 1.250809316 | 4.33E-02 |
| LINC00941 | 153.5825482 | 1.250186842 | 3.33E-11 |
| IARS | 24186.3385 | 1.249698003 | 2.14E-21 |
| ADGRG1 | 794.4778913 | 1.249340522 | 3.61E-06 |
| NA | 40.08367415 | 1.248775079 | 7.42E-04 |
| AP002956.1 | 14354.37341 | 1.246025652 | 1.73E-21 |
| KLHL5 | 3640.730673 | 1.243159385 | 2.65E-17 |
| MTSS1L | 4140.209056 | 1.242834867 | 5.39E-32 |
| IZUMO1 | 20.46040962 | 1.241487701 | 4.80E-02 |
| KIFC3 | 5432.509691 | 1.239223874 | 6.35E-36 |
| PTTG1IP | 16185.65895 | 1.238772685 | 2.40E-33 |
| BCAN | 87.58170779 | 1.238618638 | 2.51E-06 |
| SCARF1 | 287.3364243 | 1.237969261 | 2.71E-10 |
| LCMT1-AS2 | 33.15623234 | 1.237855843 | 1.79E-02 |
| COLGALT2 | 105.74784 | 1.237449916 | 4.91E-08 |
| TUBA1C | 18892.83266 | 1.236960649 | 2.48E-14 |
| EFR3B | 66.27859465 | 1.236508234 | 1.28E-04 |
| SERPINE2 | 8188.667421 | 1.236232412 | 8.31E-30 |
| RBAKDN | 14.897235 | 1.234893794 | 3.77E-02 |
| AC012368.1 | 55.07852825 | 1.234704755 | 4.22E-05 |
| CAPN2 | 11550.35271 | 1.233138557 | 6.59E-26 |
| AC021106.1 | 31.7070498 | 1.232541264 | 8.62E-05 |
| FGFR4 | 381.567178 | 1.23154207 | 3.62E-07 |
| WNK4 | 1114.048769 | 1.230104646 | 6.99E-20 |
| AC006252.1 | 66.20634047 | 1.229504552 | 1.41E-04 |
| GOT1 | 2517.089131 | 1.226959899 | 5.29E-15 |
| KRT18P41 | 21.57738418 | 1.226678865 | 3.47E-03 |
| PARD3B | 69.05787518 | 1.226100481 | 5.56E-03 |
| MELTF | 431.330045 | 1.22438386 | 7.38E-11 |
| ASPDH | 17.12319354 | 1.223649273 | 4.61E-02 |
| PVT1 | 340.4968136 | 1.223464188 | 1.01E-08 |
| ACSBG1 | 82.41581756 | 1.223382789 | 8.33E-04 |
| TFPI2 | 2830.248721 | 1.222274398 | 1.16E-20 |
| PLS3-AS1 | 36.45111194 | 1.221334832 | 1.13E-03 |
| GALNT15 | 66.94304396 | 1.221054513 | 2.55E-03 |
| PYGB | 11837.43242 | 1.221025273 | 2.37E-51 |
| TRNP1 | 1601.234225 | 1.219030506 | 2.16E-17 |
| CALM2 | 26395.33632 | 1.218316546 | 1.76E-40 |
| SLC45A3 | 519.9116178 | 1.216165687 | 4.25E-35 |
| TDRKH-AS1 | 27.67236504 | 1.215874127 | 3.24E-02 |
| C6orf48 | 7131.05733 | 1.215790613 | 7.96E-42 |
| BLK | 325.3412728 | 1.215496158 | 3.41E-09 |
| CBX7 | 937.6131425 | 1.215292325 | 3.80E-16 |
| HCFC1R1 | 2331.026297 | 1.215109403 | 2.48E-12 |

| | | | |
|---|---|---|---|
| AL512785.2 | 18.65984878 | 1.213288632 | 3.27E-02 |
| SPANXD | 336.2169453 | 1.213252812 | 1.91E-07 |
| PLEKHN1 | 170.6434411 | 1.211983038 | 1.33E-10 |
| ARID3A | 498.0095121 | 1.211396298 | 3.82E-13 |
| AC015912.3 | 21.02346451 | 1.210843788 | 6.55E-03 |
| SESN2 | 1594.171601 | 1.210773084 | 1.65E-05 |
| CITED2 | 3299.680565 | 1.209913744 | 9.62E-15 |
| PM20D2 | 1660.566496 | 1.20902804 | 4.64E-19 |
| GSDME | 2169.264017 | 1.208243594 | 1.32E-23 |
| SAMD4A | 2581.581522 | 1.20754738 | 1.80E-21 |
| SOCS2-AS1 | 244.3080781 | 1.206094274 | 1.07E-07 |
| SH3GL3 | 111.9951372 | 1.204893816 | 1.20E-04 |
| GDPD5 | 1564.667833 | 1.204457969 | 4.98E-19 |
| CEBPB | 1453.556869 | 1.20423055 | 7.76E-07 |
| EXT1 | 6550.50112 | 1.203749863 | 1.29E-21 |
| AC125611.3 | 1591.780644 | 1.203345476 | 1.27E-12 |
| SV2B | 97.51491438 | 1.203133757 | 4.82E-02 |
| RASD2 | 211.4641263 | 1.202655757 | 7.84E-17 |
| DPYSL4 | 506.4894992 | 1.201813067 | 9.06E-10 |
| AC017104.1 | 30.59044047 | 1.201547904 | 8.00E-04 |
| SPANXC | 1478.430067 | 1.200251012 | 2.63E-09 |
| SLC30A2 | 14.29975304 | 1.199840774 | 4.77E-02 |
| WNT3 | 343.0286232 | 1.19973955 | 1.97E-15 |
| LONP1 | 5588.880683 | 1.19951565 | 1.41E-33 |
| NME1-NME2 | 345.9268082 | 1.196953357 | 4.65E-07 |
| MYADM | 3260.603661 | 1.196270027 | 1.60E-25 |
| NOS2 | 73.8585863 | 1.195742574 | 2.48E-05 |
| TNFSF15 | 89.03391984 | 1.195094316 | 2.30E-04 |
| AC092692.1 | 32.94360854 | 1.194974036 | 1.64E-04 |
| PRKCQ | 27.55521402 | 1.194909124 | 1.52E-03 |
| ARHGEF18 | 3169.607921 | 1.193680928 | 2.96E-30 |
| MMP19 | 140.8610266 | 1.193280636 | 4.20E-05 |
| NR4A1 | 297.1782047 | 1.193106659 | 1.49E-14 |
| F7 | 60.99527613 | 1.193092529 | 1.10E-04 |
| AIF1L | 1935.226753 | 1.192437355 | 2.46E-19 |
| AK5 | 674.3521065 | 1.191061621 | 6.88E-12 |
| ATF4 | 18601.19462 | 1.1887828 | 2.84E-33 |
| ADAMTS18 | 25.18789751 | 1.188012727 | 4.58E-02 |
| SLC2A3 | 3270.065993 | 1.187887146 | 2.28E-07 |
| SLC2A1 | 3250.842943 | 1.18788261 | 3.50E-19 |
| CXCL1 | 30.04715564 | 1.186565832 | 3.70E-02 |
| PANX2 | 146.5880724 | 1.184845852 | 1.00E-07 |
| ZNF385C | 35.7694324 | 1.183963215 | 2.96E-03 |
| NTSR1 | 30.50159874 | 1.183324844 | 8.28E-04 |
| AL161729.1 | 72.37469199 | 1.183132814 | 1.60E-06 |
| FBLN7 | 185.287182 | 1.183077939 | 1.26E-08 |
| EBNA1BP2 | 5155.502085 | 1.182165418 | 4.71E-35 |

| | | | |
|---|---|---|---|
| FGFRL1 | 1762.380023 | 1.181788048 | 7.37E-30 |
| ARHGAP30 | 309.6936613 | 1.181030022 | 2.20E-12 |
| PGRMC2 | 2270.365819 | 1.18049135 | 3.15E-80 |
| HOXB9 | 2074.166601 | 1.180379805 | 1.35E-16 |
| ZC3H12A | 238.2279361 | 1.180345308 | 3.15E-08 |
| GOLGA8S | 17.94810252 | 1.179495733 | 2.23E-02 |
| C12orf75 | 2759.517718 | 1.17720482 | 6.01E-36 |
| FOXF1 | 786.624986 | 1.17557083 | 7.62E-12 |
| TRHDE | 434.2884023 | 1.174991697 | 5.21E-04 |
| BEX1 | 1622.382863 | 1.173711477 | 2.55E-23 |
| CCDC9B | 1980.078368 | 1.173452175 | 7.34E-15 |
| AC019080.5 | 19.93720477 | 1.172524156 | 1.68E-02 |
| ITGB1-DT | 24.64010817 | 1.1723292 | 1.23E-02 |
| CARD10 | 1474.574126 | 1.171790719 | 7.01E-14 |
| HHAT | 280.182391 | 1.170771795 | 1.45E-18 |
| GPT2 | 3740.348619 | 1.170514273 | 1.49E-07 |
| NFIX | 4322.548575 | 1.16909868 | 1.92E-34 |
| POU4F1 | 102.5808769 | 1.168530557 | 2.07E-07 |
| PDE6A | 35.90315679 | 1.167461736 | 1.26E-03 |
| SERTAD2 | 1650.89911 | 1.165894049 | 4.88E-28 |
| PPP1R26 | 2889.950209 | 1.16526523 | 5.52E-33 |
| FZD2 | 1127.496047 | 1.165126102 | 3.73E-19 |
| LMCD1 | 1289.053248 | 1.164221417 | 1.18E-06 |
| CHST13 | 38.55414762 | 1.164149358 | 1.40E-03 |
| YARS | 12806.03362 | 1.163708017 | 1.55E-18 |
| GFI1 | 39.2406586 | 1.163073364 | 1.78E-02 |
| HR | 283.1205138 | 1.161259223 | 1.25E-03 |
| CNN2 | 8063.875845 | 1.160764746 | 1.11E-38 |
| TNNI3 | 204.9804876 | 1.160704024 | 2.84E-10 |
| CAVIN1 | 22717.39463 | 1.160284305 | 2.05E-18 |
| SLC22A18AS | 118.3495015 | 1.160212993 | 1.86E-05 |
| PSORS1C1 | 40.71724625 | 1.159680969 | 1.34E-02 |
| CEACAM19 | 174.7498355 | 1.159498642 | 1.67E-08 |
| SDR16C5 | 42.38694442 | 1.158611581 | 9.87E-03 |
| TIMP3 | 33487.39915 | 1.157498928 | 1.57E-15 |
| SOWAHD | 24.47470161 | 1.157218761 | 4.44E-03 |
| AL163195.2 | 16.13949065 | 1.156943588 | 3.98E-02 |
| MID1 | 1581.448112 | 1.156928429 | 2.40E-23 |
| AC104435.2 | 31.56962772 | 1.15515178 | 3.91E-03 |
| CDH2 | 355.8677844 | 1.155041046 | 5.75E-08 |
| TMEM151A | 32.37522276 | 1.154972382 | 1.90E-02 |
| STEAP3 | 2478.209655 | 1.154795708 | 7.76E-24 |
| DHRS2 | 1627.919122 | 1.153726804 | 1.94E-05 |
| C16orf95 | 250.6123435 | 1.153369532 | 2.13E-10 |
| ASMTL | 1137.639438 | 1.153195899 | 6.51E-27 |
| P3H2 | 6277.970193 | 1.152773498 | 1.69E-18 |
| SLC8A1 | 552.6757728 | 1.15116552 | 2.13E-08 |

| | | | |
|---|---|---|---|
| FOXF2 | 394.4896013 | 1.151121326 | 1.57E-23 |
| NRARP | 134.6099929 | 1.150719831 | 7.93E-09 |
| SMS | 6603.49377 | 1.150323421 | 2.25E-26 |
| AHNAK | 122265.4934 | 1.149081335 | 3.93E-42 |
| SPN | 423.4724567 | 1.148956313 | 1.51E-09 |
| VSIR | 1076.142038 | 1.147675268 | 6.42E-13 |
| AC125807.2 | 800.7685966 | 1.147148704 | 1.55E-23 |
| NBEAL2 | 1699.565215 | 1.146715788 | 1.30E-17 |
| KLF16 | 371.8280178 | 1.146241898 | 3.43E-11 |
| LSP1P4 | 1758.041799 | 1.145336433 | 2.60E-37 |
| MYH14 | 1638.03928 | 1.143828338 | 1.40E-29 |
| NOTUM | 82.32665438 | 1.143764851 | 5.18E-05 |
| PKP3 | 138.0497832 | 1.143387539 | 4.92E-04 |
| SEMA4A | 163.4203738 | 1.142663384 | 6.97E-06 |
| SNCAIP | 91.61249628 | 1.141512331 | 9.18E-06 |
| PCED1B | 73.68965833 | 1.140607045 | 1.78E-04 |
| ACO1 | 6073.137214 | 1.139635649 | 3.04E-37 |
| SLC1A5 | 12852.03691 | 1.138990209 | 3.15E-12 |
| RGS9 | 137.3520688 | 1.138478555 | 4.85E-05 |
| ZMYND12 | 26.09480414 | 1.137816126 | 4.81E-02 |
| LINC01777 | 21.41202174 | 1.137562194 | 1.95E-02 |
| CD200 | 73.55103506 | 1.134556434 | 3.22E-03 |
| SPATA6L | 55.35177118 | 1.134028838 | 4.25E-03 |
| TMEM158 | 357.808961 | 1.133618321 | 2.20E-08 |
| PTGFR | 15.23427547 | 1.132976352 | 4.59E-02 |
| ERVV-2 | 60.76003593 | 1.131862484 | 1.62E-02 |
| AC087203.3 | 32.47593761 | 1.13183781 | 1.43E-02 |
| PROM2 | 46.85997022 | 1.131612573 | 3.21E-02 |
| C16orf74 | 376.5939505 | 1.131188928 | 3.02E-18 |
| ADAP1 | 466.5208821 | 1.130732293 | 4.46E-17 |
| CCDC169-SOHLH | 38.91818841 | 1.130315818 | 2.99E-02 |
| AC079922.2 | 64.77901412 | 1.129637001 | 8.62E-05 |
| WDR1 | 12088.49163 | 1.128975414 | 1.10E-28 |
| MSX1 | 641.8880923 | 1.128055225 | 3.48E-16 |
| SYT15 | 456.2697294 | 1.127251822 | 7.27E-14 |
| AC092903.2 | 35.98719012 | 1.126625124 | 2.85E-03 |
| IGF2BP2 | 5316.699877 | 1.126361019 | 1.84E-42 |
| AC148477.4 | 20.92675249 | 1.12631584 | 4.71E-02 |
| S100A10 | 12016.01031 | 1.124928985 | 8.31E-24 |
| UBE2S | 5368.618216 | 1.124882137 | 6.86E-23 |
| NUP210 | 2474.763351 | 1.124549373 | 1.50E-23 |
| MAP3K7CL | 150.299139 | 1.123976784 | 2.90E-09 |
| FHL2 | 2309.54896 | 1.123703582 | 1.27E-23 |
| SRRM3 | 406.1814457 | 1.122823923 | 2.63E-15 |
| AC244153.1 | 322.5237666 | 1.122729093 | 3.89E-21 |
| BPIFA2 | 22.6494482 | 1.122562487 | 4.09E-02 |
| ESX1 | 1145.550705 | 1.120182263 | 3.08E-10 |

| | | | |
|---|---|---|---|
| MEI1 | 21.74231947 | 1.119986552 | 3.85E-02 |
| AC027117.1 | 17.37176545 | 1.119458266 | 2.87E-02 |
| DDIT4 | 5087.59468 | 1.118924712 | 4.28E-04 |
| C5AR1 | 91.91066328 | 1.117995893 | 2.79E-05 |
| CHRM3 | 17.48536067 | 1.117978364 | 4.70E-02 |
| SLC12A7 | 1143.958255 | 1.117752624 | 6.71E-11 |
| NPAS1 | 1188.241383 | 1.117640394 | 1.30E-25 |
| CYP24A1 | 230.7349006 | 1.116352939 | 3.95E-07 |
| DRAP1 | 3957.910986 | 1.114271285 | 1.27E-30 |
| MARS | 14346.5054 | 1.113140967 | 4.85E-54 |
| FOSL2 | 5985.00852 | 1.113051183 | 7.60E-25 |
| FOXL1 | 378.206564 | 1.111600767 | 3.13E-14 |
| RAB38 | 37.37841539 | 1.110642019 | 9.59E-03 |
| LINC02600 | 73.12744437 | 1.110249203 | 2.74E-06 |
| TLN1 | 34235.64152 | 1.109390658 | 2.70E-25 |
| TPBG | 2572.499946 | 1.109283425 | 1.01E-51 |
| OR2A1-AS1 | 214.8177946 | 1.109172913 | 5.68E-10 |
| ZP3 | 653.8483888 | 1.109159095 | 1.05E-13 |
| OTUD1 | 108.5394167 | 1.108340224 | 7.03E-07 |
| SLC37A2 | 644.4016214 | 1.108156805 | 3.83E-12 |
| ZYX | 10900.51147 | 1.108132009 | 8.77E-20 |
| PLEKHS1 | 100.4445389 | 1.107558303 | 2.28E-04 |
| DUSP23 | 365.8499249 | 1.107044383 | 4.93E-10 |
| AC139769.1 | 26.58478457 | 1.106759794 | 1.03E-02 |
| ARSI | 573.0458329 | 1.106555649 | 1.79E-07 |
| KIF17 | 286.1148535 | 1.103684043 | 1.39E-08 |
| TSEN15 | 2116.407831 | 1.102399237 | 2.54E-16 |
| ZSWIM4 | 566.8401579 | 1.102184271 | 3.50E-19 |
| GNGT2 | 60.85698961 | 1.101791007 | 5.68E-04 |
| NT5C2 | 2575.304732 | 1.101732047 | 1.25E-23 |
| FALEC | 20.55045138 | 1.101719152 | 3.50E-02 |
| WNT2B | 151.2588281 | 1.101532898 | 6.08E-07 |
| AC092198.1 | 26.27954271 | 1.101509073 | 1.21E-02 |
| RAC2 | 3122.912202 | 1.101253497 | 2.14E-19 |
| LEF1 | 105.2991748 | 1.100557426 | 4.27E-05 |
| RTN4R | 369.6459485 | 1.100495144 | 6.91E-10 |
| LINC01666 | 39.06634075 | 1.100046166 | 5.33E-04 |
| RAP1GAP2 | 2036.056692 | 1.099010683 | 2.16E-20 |
| NT5E | 5643.186994 | 1.098477548 | 2.45E-11 |
| EIF3D | 11749.83261 | 1.098466144 | 2.23E-18 |
| PLCG2 | 1243.876311 | 1.098032541 | 9.66E-13 |
| CLMN | 1499.986064 | 1.097289764 | 1.48E-24 |
| C1orf162 | 17.40202554 | 1.097104261 | 4.95E-02 |
| SPANXA1 | 416.0491261 | 1.097048503 | 2.91E-07 |
| SUPT3H | 446.3907523 | 1.097010644 | 1.44E-21 |
| PLPPR3 | 186.1600222 | 1.094968194 | 9.01E-11 |
| HIST3H2A | 214.9155915 | 1.094402006 | 2.21E-07 |

| | | | |
|---|---|---|---|
| PRAG1 | 3099.318571 | 1.093136851 | 1.14E-12 |
| FCRLB | 100.9744833 | 1.09281265 | 6.82E-06 |
| PIK3AP1 | 191.3237924 | 1.091944134 | 5.79E-08 |
| DNAH10 | 134.5372647 | 1.091925594 | 1.19E-02 |
| CEBPD | 169.0188464 | 1.091557297 | 6.18E-07 |
| FAM27C | 79.29343533 | 1.090259932 | 1.35E-06 |
| RIMS4 | 97.05276827 | 1.089922548 | 1.04E-03 |
| EIF1 | 14837.75257 | 1.089616623 | 1.04E-32 |
| TJP2 | 4658.735463 | 1.089168589 | 7.31E-31 |
| C3orf52 | 168.1696811 | 1.089051515 | 7.66E-08 |
| AZU1 | 16.26746334 | 1.08815208 | 4.14E-02 |
| NEK6 | 8959.611567 | 1.087212252 | 1.05E-25 |
| CXCL5 | 15.43725055 | 1.087108311 | 3.16E-02 |
| LINC02575 | 49.33029379 | 1.086869298 | 3.71E-03 |
| SPHK1 | 935.0801792 | 1.086233271 | 4.02E-22 |
| FAM24B | 116.4415182 | 1.084893582 | 1.92E-09 |
| AC098487.1 | 25.41401803 | 1.084268397 | 3.79E-02 |
| HIST1H2BN | 83.96504118 | 1.083994292 | 1.19E-04 |
| NFIL3 | 1500.966468 | 1.08380541 | 3.30E-08 |
| PCLO | 431.3526201 | 1.083571506 | 4.13E-05 |
| AC092802.1 | 33.0940247 | 1.082382932 | 1.20E-02 |
| TCAF2 | 445.3765845 | 1.082318441 | 1.62E-04 |
| ITGAX | 188.9758 | 1.081206134 | 3.57E-08 |
| ASAP1 | 7407.398524 | 1.080371131 | 4.67E-27 |
| MAPK13 | 160.0474767 | 1.079097515 | 1.67E-08 |
| CDT1 | 1350.485546 | 1.079052656 | 1.67E-04 |
| EVA1C | 367.5986656 | 1.078219684 | 5.61E-09 |
| MKNK2 | 4825.496095 | 1.077304034 | 3.39E-18 |
| ATG9B | 118.2688634 | 1.076563944 | 1.27E-02 |
| AP001453.2 | 57.55253509 | 1.075740504 | 7.89E-04 |
| PIPOX | 53.95477712 | 1.075636785 | 3.85E-04 |
| NFKBIE | 433.0629905 | 1.075619182 | 1.27E-08 |
| TYRP1 | 289.0563429 | 1.074546629 | 1.12E-15 |
| SYNPO | 2426.228123 | 1.073150399 | 2.22E-20 |
| BNIP3 | 2020.176455 | 1.07276197 | 5.36E-11 |
| CYP11A1 | 221.6021817 | 1.071087104 | 6.43E-11 |
| UPK1A | 68.1821967 | 1.071072481 | 8.08E-03 |
| RASAL2-AS1 | 121.4954753 | 1.069962639 | 1.32E-07 |
| NFKB2 | 732.7160331 | 1.069661751 | 3.46E-11 |
| HK2 | 2102.950957 | 1.067012598 | 4.13E-14 |
| AP000523.1 | 53.76492508 | 1.066600722 | 1.84E-04 |
| FGGY | 156.5456048 | 1.066205439 | 6.94E-05 |
| MYRF | 3061.445988 | 1.065886458 | 4.51E-25 |
| TRIO | 14689.77273 | 1.065849763 | 1.29E-32 |
| C19orf38 | 23.21258587 | 1.065549063 | 3.25E-02 |
| PDGFA | 1396.475027 | 1.065265045 | 1.54E-13 |
| ERN1 | 835.8781716 | 1.062729721 | 2.93E-04 |

| | | | |
|---|---|---|---|
| CACNG4 | 946.3481827 | 1.061721316 | 2.31E-08 |
| SMOC2 | 58.39393359 | 1.061209196 | 3.44E-02 |
| MSANTD3 | 2120.168871 | 1.061162819 | 3.00E-30 |
| STXBP2 | 1158.876155 | 1.060096677 | 2.88E-21 |
| SCN5A | 570.9436693 | 1.059368425 | 1.22E-07 |
| NXN | 1791.096427 | 1.059246125 | 4.32E-18 |
| AP000759.1 | 69.88374719 | 1.059083589 | 1.80E-04 |
| CD22 | 253.9919888 | 1.058553365 | 1.35E-05 |
| GALNT9 | 229.1882238 | 1.057942509 | 4.48E-05 |
| LAMB4 | 116.1014266 | 1.05779999 | 5.67E-05 |
| HIST1H1C | 590.5061429 | 1.057532286 | 2.37E-07 |
| NOS1AP | 114.7690526 | 1.057377662 | 4.77E-06 |
| CARD11 | 487.3641494 | 1.055375506 | 1.34E-06 |
| MAMDC2-AS1 | 35.10599911 | 1.054716252 | 3.40E-03 |
| TTC7A | 1548.824814 | 1.053526595 | 3.63E-27 |
| LINC02454 | 19.94867742 | 1.052997214 | 3.21E-02 |
| BRI3 | 3521.077973 | 1.051320439 | 1.13E-14 |
| TSPAN13 | 149.3123275 | 1.050946246 | 5.76E-04 |
| ZNF560 | 384.619609 | 1.048979998 | 6.31E-11 |
| MAD1L1 | 2125.818007 | 1.048831192 | 1.42E-28 |
| SLC1A6 | 42.11919478 | 1.047462054 | 7.61E-03 |
| RRAS2 | 2136.436366 | 1.047375541 | 3.79E-27 |
| RNASE4 | 101.2496272 | 1.047179697 | 5.34E-03 |
| OR2W3 | 22.54171169 | 1.047010776 | 1.85E-02 |
| GPR160 | 34.99439473 | 1.044657391 | 1.19E-02 |
| FTSJ1 | 3401.260365 | 1.044416175 | 1.04E-42 |
| KCMF1 | 2322.440073 | 1.043979105 | 6.66E-15 |
| PDE4C | 138.9413135 | 1.041767458 | 2.44E-02 |
| TBC1D30 | 168.1408535 | 1.041074257 | 2.90E-07 |
| PLEKHA6 | 883.8866704 | 1.040738194 | 3.18E-06 |
| ROBO3 | 1232.003209 | 1.040464134 | 6.20E-17 |
| ZBED1 | 2312.89623 | 1.039221769 | 3.14E-21 |
| ZNF438 | 178.9399402 | 1.038403521 | 7.85E-11 |
| ZIC2 | 467.9089163 | 1.037649717 | 2.37E-08 |
| DOCK9 | 3479.797232 | 1.037403126 | 3.87E-19 |
| ITGB4 | 449.0655626 | 1.036965505 | 7.36E-07 |
| KYAT1 | 621.6280245 | 1.036680961 | 8.57E-24 |
| AP003393.1 | 17.29893083 | 1.036573122 | 3.12E-02 |
| NTMT1 | 2667.508817 | 1.035891219 | 1.15E-29 |
| HOXC5 | 53.48940178 | 1.034482109 | 2.41E-03 |
| RUNX2 | 681.1290161 | 1.034458074 | 7.02E-12 |
| AC004951.3 | 24.48200905 | 1.03418652 | 4.07E-03 |
| RSPO4 | 543.3235221 | 1.03352516 | 1.72E-17 |
| AL359182.1 | 21.13015621 | 1.032799738 | 2.93E-02 |
| TCHH | 60.14085417 | 1.03212476 | 1.42E-02 |
| THEMIS2 | 327.7141705 | 1.030770998 | 4.44E-11 |
| XAGE3 | 258.2371969 | 1.030232493 | 1.59E-11 |

| | | | |
|---|---|---|---|
| RAET1E | 123.7618426 | 1.030217417 | 8.97E-07 |
| ASPHD2 | 456.4862353 | 1.029847201 | 1.05E-20 |
| ARHGEF2 | 9221.872532 | 1.029038244 | 2.09E-07 |
| COL9A2 | 63.23009237 | 1.02902886 | 8.29E-03 |
| CLU | 7930.293639 | 1.029027807 | 1.12E-16 |
| PPM1E | 322.4396099 | 1.028927693 | 3.89E-04 |
| TRHDE-AS1 | 181.9325313 | 1.028287606 | 3.69E-06 |
| LYG2 | 18.18221626 | 1.028105536 | 2.21E-02 |
| TMEM139 | 162.5650754 | 1.025578275 | 6.20E-08 |
| LINC01835 | 24.62093076 | 1.025308179 | 4.40E-02 |
| CRISPLD2 | 701.9003059 | 1.025306157 | 3.67E-06 |
| LRRC2 | 129.9918758 | 1.024864782 | 2.67E-07 |
| STAC | 1317.904973 | 1.022600008 | 5.09E-09 |
| AKAP2 | 11074.45609 | 1.022319256 | 2.12E-31 |
| PGM2 | 2123.780405 | 1.020857051 | 5.67E-28 |
| ACSL1 | 1243.219884 | 1.020817582 | 2.25E-32 |
| CDKN1A | 368.8877978 | 1.019905688 | 3.34E-13 |
| ARHGAP23 | 5404.484757 | 1.019736403 | 1.15E-30 |
| MB21D2 | 492.152103 | 1.019280921 | 5.98E-21 |
| RADIL | 174.7008569 | 1.018946706 | 7.59E-08 |
| MYL12A | 9438.49279 | 1.018748187 | 1.09E-33 |
| COMT | 3313.736764 | 1.018428303 | 1.57E-17 |
| GGT5 | 898.8201285 | 1.018126066 | 9.03E-07 |
| MESP2 | 29.91063221 | 1.017632326 | 1.01E-02 |
| ASGR1 | 173.8182471 | 1.017176063 | 9.53E-09 |
| PHLDB2 | 5271.381741 | 1.017082024 | 8.51E-10 |
| KIAA1211 | 73.81802387 | 1.016983383 | 1.21E-03 |
| AC015468.3 | 39.3845665 | 1.015473557 | 1.07E-03 |
| CCNB1IP1 | 4135.158345 | 1.015127684 | 6.85E-21 |
| PLXNA2 | 1668.333265 | 1.014504287 | 2.31E-11 |
| ALG1L6P | 33.26554236 | 1.014421414 | 2.22E-03 |
| LARGE2 | 219.8105223 | 1.014210273 | 1.97E-06 |
| DNAJB6 | 11954.55005 | 1.01403428 | 5.58E-52 |
| ADRA1B | 405.8511243 | 1.013794958 | 3.56E-03 |
| BLVRB | 2573.362732 | 1.013655635 | 1.12E-19 |
| CMIP | 2119.779756 | 1.012576464 | 1.38E-21 |
| CYS1 | 92.6595099 | 1.012319544 | 2.57E-05 |
| AL031587.5 | 395.7116765 | 1.011779401 | 1.47E-11 |
| AC020904.2 | 128.7665194 | 1.011335188 | 9.35E-08 |
| ACTG1 | 90431.48433 | 1.010042065 | 3.22E-07 |
| AC020916.1 | 458.5870598 | 1.009719648 | 2.07E-14 |
| DES | 57.98814614 | 1.008740085 | 4.28E-03 |
| AXIN2 | 442.7244193 | 1.00785911 | 3.68E-07 |
| AL844908.1 | 27.56971702 | 1.007758669 | 1.02E-02 |
| ANKRD20A19P | 42.14462645 | 1.0070286 | 3.89E-02 |
| CT55 | 51.54356207 | 1.006707617 | 1.28E-04 |
| XAGE2 | 41.30684287 | 1.006676569 | 3.84E-02 |

| | | | |
|---|---|---|---|
| EGFL7 | 1775.9885 | 1.006599626 | 5.24E-11 |
| HSBP1L1 | 284.4549299 | 1.004228522 | 7.63E-09 |
| MSRB3 | 1780.19995 | 1.003444394 | 1.43E-17 |
| SSUH2 | 29.24276905 | 1.003336043 | 5.28E-03 |
| TAX1BP3 | 4806.333429 | 1.001409951 | 3.72E-16 |
| LIMA1 | 10004.88249 | 1.000741833 | 1.40E-19 |
| SYDE1 | 3531.235541 | 0.999565018 | 3.87E-16 |
| FOXD3-AS1 | 24.57546857 | 0.997854165 | 3.34E-02 |
| AC010618.4 | 100.5779133 | 0.997518234 | 7.31E-04 |
| RANGAP1 | 7733.942925 | 0.997413858 | 5.31E-37 |
| ANKRD18A | 120.6515545 | 0.997301399 | 2.64E-03 |
| PORCN | 684.9381108 | 0.996140606 | 3.44E-13 |
| RAB31 | 4038.601522 | 0.995993784 | 7.18E-35 |
| TMEM56 | 329.4539423 | 0.99578093 | 1.22E-05 |
| SOCS3 | 897.8181479 | 0.995124761 | 3.90E-04 |
| GPR4 | 237.5154985 | 0.994979958 | 8.93E-05 |
| ESPNP | 31.51150625 | 0.994317215 | 2.65E-02 |
| CTNNAL1 | 7787.2537 | 0.992378794 | 1.82E-12 |
| L1CAM | 10255.39231 | 0.992326868 | 1.51E-09 |
| LINC00937 | 58.8563272 | 0.99153923 | 2.39E-02 |
| CKS1B | 3700.240863 | 0.991289344 | 1.56E-25 |
| SAMD11 | 775.4203297 | 0.990927035 | 3.81E-02 |
| ATXN2L | 9348.969127 | 0.989997511 | 1.34E-16 |
| AGT | 33.3491476 | 0.987561767 | 2.39E-02 |
| KLRG2 | 100.5493213 | 0.987462391 | 3.69E-04 |
| CD68 | 753.459441 | 0.987033848 | 4.13E-07 |
| TXNIP | 25810.30319 | 0.986548772 | 2.81E-07 |
| LHFPL4 | 27.81473221 | 0.985497033 | 2.59E-02 |
| RAB3IP | 248.084127 | 0.984704689 | 7.87E-05 |
| RPH3A | 45.77989665 | 0.984427942 | 5.01E-03 |
| MGAT5B | 1657.991925 | 0.983133725 | 8.44E-14 |
| TPK1 | 102.279142 | 0.982911647 | 1.07E-04 |
| SH3D21 | 340.8390984 | 0.982847893 | 5.35E-05 |
| RIN1 | 1365.463973 | 0.981293863 | 9.15E-12 |
| LDLR | 5914.413995 | 0.980577922 | 1.42E-12 |
| SH3RF2 | 238.2640979 | 0.980176005 | 1.66E-06 |
| AC105052.2 | 44.08002568 | 0.980110365 | 3.51E-03 |
| AJ011932.1 | 30.9980947 | 0.978857067 | 1.75E-02 |
| AL136162.1 | 32.3372411 | 0.977280258 | 6.63E-03 |
| FUT1 | 131.4538281 | 0.977241073 | 7.03E-03 |
| SYTL2 | 1943.873549 | 0.976594438 | 1.37E-06 |
| AL138478.1 | 119.427306 | 0.976290569 | 2.11E-04 |
| C1QTNF1-AS1 | 25.06605869 | 0.975104981 | 4.55E-02 |
| IER3 | 2184.936652 | 0.97502234 | 6.62E-14 |
| AC011558.1 | 173.9603832 | 0.974293861 | 3.23E-04 |
| MAP3K14 | 719.8879108 | 0.974135763 | 1.91E-09 |
| DPY19L2P3 | 201.8146384 | 0.973923962 | 9.77E-05 |

| | | | |
|---|---|---|---|
| UCHL1 | 173.5698622 | 0.97307644 | 1.65E-04 |
| CABLES1 | 2714.737389 | 0.972787794 | 1.57E-10 |
| ISL2 | 61.72259261 | 0.972757826 | 3.42E-04 |
| NRN1 | 401.8029362 | 0.971513187 | 4.52E-02 |
| UBALD2 | 914.1334782 | 0.971247113 | 6.08E-09 |
| LMNA | 16821.17365 | 0.970422999 | 7.43E-21 |
| DDX20 | 1407.651257 | 0.970269523 | 6.22E-42 |
| SEC14L5 | 39.34326117 | 0.970128755 | 1.52E-03 |
| IRS2 | 208.4093286 | 0.969818983 | 6.96E-09 |
| MEIOB | 1375.792081 | 0.969242924 | 1.77E-12 |
| HAX1 | 4016.217241 | 0.967746186 | 1.65E-17 |
| AC013394.1 | 193.2051003 | 0.967165809 | 9.39E-04 |
| PLCD3 | 2293.886567 | 0.966895943 | 2.93E-26 |
| LASP1 | 19014.37754 | 0.96651398 | 2.81E-21 |
| CDC20 | 4779.118958 | 0.966463558 | 1.64E-13 |
| ASB4 | 79.74807 | 0.965136026 | 1.53E-03 |
| JAG2 | 524.7348244 | 0.964721388 | 8.77E-06 |
| NAV3 | 102.7180682 | 0.964451749 | 3.94E-02 |
| AC005330.1 | 96.12826441 | 0.963699949 | 7.45E-06 |
| CASZ1 | 165.5766828 | 0.963643963 | 4.84E-06 |
| GEM | 363.9867785 | 0.963490697 | 6.32E-06 |
| CSRNP1 | 686.1282168 | 0.963236218 | 3.96E-19 |
| CYP2S1 | 1720.583561 | 0.962382723 | 3.64E-07 |
| FAM129B | 17789.76161 | 0.96232032 | 2.38E-23 |
| EEF1B2P6 | 25.72444092 | 0.961134036 | 4.36E-02 |
| LINC00346 | 364.435918 | 0.961019728 | 1.23E-18 |
| MYL12B | 10882.15315 | 0.960689433 | 4.82E-55 |
| SAMD12 | 473.3971852 | 0.960669417 | 1.72E-04 |
| NFASC | 750.1182267 | 0.96037014 | 1.91E-19 |
| AL161729.4 | 34.7473894 | 0.958845081 | 1.03E-02 |
| TLN2 | 3537.806415 | 0.958548283 | 2.69E-08 |
| TLE4 | 2216.44133 | 0.958435716 | 7.49E-07 |
| KTN1-AS1 | 455.5717954 | 0.958114672 | 7.11E-09 |
| ECSCR | 314.089158 | 0.957349315 | 1.37E-09 |
| MICB | 1091.298993 | 0.956793041 | 1.15E-06 |
| HEBP1 | 1059.86254 | 0.956422165 | 4.82E-18 |
| IFITM10 | 41.85794267 | 0.955940368 | 3.91E-02 |
| NA | 111.6290596 | 0.955875129 | 1.36E-02 |
| GRID2IP | 33.40003305 | 0.95582252 | 5.28E-03 |
| P2RX7 | 1431.080718 | 0.955742233 | 5.42E-10 |
| RPL12P38 | 32.15082083 | 0.955649669 | 1.85E-02 |
| ST8SIA2 | 194.4998905 | 0.954717785 | 2.18E-04 |
| AL353813.1 | 25.38625687 | 0.954453725 | 4.87E-02 |
| MYO18B | 38.53813517 | 0.952964297 | 4.17E-02 |
| HHEX | 480.0358831 | 0.952851828 | 8.88E-09 |
| LONRF3 | 116.1789548 | 0.952318758 | 1.62E-05 |
| ASTN1 | 2110.214754 | 0.950878306 | 1.11E-13 |

| | | | |
|---|---|---|---|
| SPRYD3 | 1098.046721 | 0.950867215 | 3.08E-15 |
| HOXC12 | 131.3367996 | 0.950805784 | 1.29E-06 |
| RNASET2 | 534.0947676 | 0.949976952 | 6.16E-12 |
| AL118516.1 | 162.6771387 | 0.949885517 | 2.22E-08 |
| ATP8B1 | 1504.624557 | 0.949410547 | 7.78E-14 |
| AC044860.1 | 85.8121818 | 0.94919549 | 3.79E-04 |
| RBCK1 | 7551.367995 | 0.949050644 | 7.00E-13 |
| SLC35G2 | 48.39117825 | 0.948832535 | 8.27E-03 |
| PYCR1 | 8366.040574 | 0.948014112 | 2.19E-15 |
| AC135506.1 | 22.3710912 | 0.947996682 | 3.35E-02 |
| AC025857.2 | 32.64328677 | 0.947753554 | 3.28E-02 |
| P4HA2 | 4210.481934 | 0.947205016 | 7.17E-21 |
| CD82 | 1265.274499 | 0.947036 | 1.46E-06 |
| CRY1 | 1344.096812 | 0.945921207 | 8.75E-14 |
| FPR1 | 28.60611966 | 0.945895434 | 1.79E-02 |
| APOPT1 | 868.812937 | 0.945876116 | 1.19E-16 |
| TAF7L | 37.70872258 | 0.944944058 | 8.17E-03 |
| SNX8 | 3274.668815 | 0.944702527 | 1.15E-11 |
| PGK1 | 28199.85894 | 0.944088131 | 8.54E-09 |
| LINC01881 | 83.25249054 | 0.944001335 | 1.50E-02 |
| INPP4B | 1588.237089 | 0.943519078 | 2.79E-12 |
| MEAK7 | 2844.519966 | 0.943034826 | 1.23E-15 |
| PATJ | 706.5965863 | 0.942639937 | 1.53E-06 |
| AC018630.2 | 44.03435817 | 0.942501518 | 3.18E-03 |
| CT45A1 | 1229.578091 | 0.942294757 | 1.53E-20 |
| MRPL23-AS1 | 189.6877581 | 0.941955576 | 8.61E-09 |
| SLC22A31 | 78.82971781 | 0.941440526 | 7.74E-05 |
| SOX7 | 73.75907218 | 0.94110678 | 5.46E-04 |
| IL17D | 149.0420822 | 0.941009651 | 8.57E-05 |
| PRICKLE1 | 152.583996 | 0.940580871 | 1.55E-04 |
| AC116351.1 | 130.5665582 | 0.939857841 | 1.17E-03 |
| AC105206.2 | 53.39233751 | 0.939841056 | 1.98E-03 |
| KIF6 | 184.9785171 | 0.938286597 | 4.75E-03 |
| NAMPTP1 | 291.5978854 | 0.937911155 | 4.52E-04 |
| CTSB | 51746.63918 | 0.937659077 | 5.88E-13 |
| ADAMTSL4 | 843.0356382 | 0.937435911 | 1.26E-10 |
| ELF4 | 1789.071958 | 0.936444218 | 3.77E-16 |
| EPAS1 | 9233.833509 | 0.93555635 | 7.00E-22 |
| LINC01747 | 30.54129894 | 0.935270169 | 3.72E-02 |
| TRABD2A | 897.350399 | 0.935223525 | 1.60E-18 |
| PHLDB3 | 358.9219198 | 0.93479143 | 6.14E-10 |
| LY6E-DT | 189.8540731 | 0.933822198 | 2.93E-07 |
| PDE2A | 399.4921856 | 0.932554817 | 6.89E-08 |
| FOXN3-AS1 | 91.62970335 | 0.932502038 | 1.57E-04 |
| IP6K3 | 192.4547338 | 0.931681158 | 1.05E-04 |
| ULBP2 | 214.0881506 | 0.931039613 | 7.97E-06 |
| CCDC71L | 1297.79424 | 0.930395053 | 3.89E-13 |

| | | | |
|---|---|---|---|
| FAM72D | 245.0786245 | 0.930246022 | 7.88E-05 |
| PTTG1 | 6678.880036 | 0.930229867 | 6.47E-20 |
| SLC20A1 | 3792.236403 | 0.927459258 | 2.70E-10 |
| CLCN1 | 55.22163434 | 0.927396226 | 1.19E-03 |
| PDXK | 5385.678385 | 0.926331339 | 2.36E-40 |
| SLC46A3 | 306.615023 | 0.926218437 | 1.46E-08 |
| NAMPT | 5630.167846 | 0.925857566 | 3.61E-05 |
| DTD1 | 3987.586041 | 0.925832414 | 1.00E-23 |
| FHAD1 | 349.9471983 | 0.925441513 | 2.76E-10 |
| MOGAT3 | 27.3877624 | 0.924784554 | 2.12E-02 |
| LINC00668 | 106.385289 | 0.92469303 | 6.48E-05 |
| CMTM7 | 1057.832695 | 0.924111447 | 1.45E-15 |
| LZTS3 | 1797.583311 | 0.923960757 | 3.51E-30 |
| ENPP7 | 27.35216795 | 0.923870479 | 2.68E-02 |
| PTPRU | 1753.900827 | 0.923280897 | 2.23E-11 |
| SERTAD4 | 765.4049029 | 0.92292931 | 2.42E-06 |
| NKX2-4 | 192.9784856 | 0.922317224 | 3.57E-08 |
| FBXO27 | 124.0869824 | 0.921772858 | 6.39E-03 |
| ANKRD13A | 3316.621453 | 0.921722159 | 7.02E-18 |
| PLEKHF1 | 202.8274138 | 0.921687272 | 6.36E-03 |
| AL122035.1 | 44.99986555 | 0.921682123 | 3.35E-03 |
| AP003774.1 | 35.71106587 | 0.921199436 | 8.46E-03 |
| SCN4A | 135.5757544 | 0.919673095 | 1.66E-02 |
| DICER1-AS1 | 124.6884539 | 0.919655363 | 5.01E-07 |
| FHL1 | 18557.55669 | 0.919286857 | 3.61E-37 |
| ROCK1P1 | 82.49286083 | 0.917304141 | 2.68E-03 |
| PYGL | 5642.226534 | 0.915717027 | 8.30E-57 |
| GADD45A | 1177.960422 | 0.914078593 | 6.89E-05 |
| NRTN | 83.69516378 | 0.914050764 | 1.93E-04 |
| NKX6-1 | 64.55961741 | 0.913265709 | 4.79E-04 |
| LINC01679 | 144.8361926 | 0.913187272 | 3.30E-05 |
| RAPGEF1 | 4833.32604 | 0.913070445 | 3.71E-23 |
| C4orf36 | 228.6572123 | 0.912162 | 1.91E-07 |
| AC244669.1 | 91.29498098 | 0.912136674 | 6.16E-05 |
| GSC | 33.42228245 | 0.911569494 | 2.55E-02 |
| MIR210HG | 249.23102 | 0.910366875 | 4.36E-03 |
| LINC00662 | 641.6290959 | 0.91035231 | 3.65E-03 |
| DLGAP4 | 3901.025781 | 0.909575984 | 1.48E-46 |
| INF2 | 4687.789987 | 0.909514552 | 2.52E-21 |
| DOCK6 | 1355.620577 | 0.909013291 | 6.99E-11 |
| MT-TC | 333.2264633 | 0.908901904 | 2.69E-03 |
| ST3GAL4 | 2762.325065 | 0.908762457 | 1.55E-14 |
| AC092338.1 | 1800.678811 | 0.90870594 | 1.29E-17 |
| SLC31A1 | 4679.004963 | 0.907871139 | 1.42E-29 |
| SERPINB1 | 1965.599407 | 0.907774676 | 9.82E-26 |
| MBP | 3639.442215 | 0.907771309 | 2.94E-07 |
| PPP1R18 | 5146.357623 | 0.907761265 | 9.22E-14 |

| | | | |
|---|---|---|---|
| WSCD1 | 732.1810093 | 0.907598192 | 4.98E-09 |
| EXOC3L1 | 83.24562736 | 0.906458424 | 1.49E-03 |
| KLHL29 | 1265.210865 | 0.90621111 | 3.60E-09 |
| AC004923.4 | 31.09287184 | 0.90567193 | 3.23E-02 |
| RNF41 | 3274.28356 | 0.90562831 | 3.08E-07 |
| CCNI | 9677.528827 | 0.905065761 | 1.05E-33 |
| MYO1D | 1277.296482 | 0.904826073 | 1.56E-18 |
| TGM2 | 27895.58294 | 0.904403718 | 1.88E-08 |
| GTPBP2 | 4056.078583 | 0.904391235 | 7.66E-07 |
| NXNL2 | 101.7112097 | 0.902929212 | 1.51E-05 |
| LARP6 | 2466.487265 | 0.902817529 | 9.85E-12 |
| GPR176 | 2600.86289 | 0.902543478 | 6.10E-10 |
| AC009404.1 | 51.36748406 | 0.902527154 | 3.64E-03 |
| C7orf61 | 25.21213737 | 0.902021699 | 4.99E-02 |
| SORCS2 | 333.4430111 | 0.901865997 | 3.64E-06 |
| RAP2B | 924.2648138 | 0.9012783 | 3.99E-09 |
| FAM219A | 2447.604118 | 0.900815542 | 2.90E-11 |
| SCN8A | 851.2309317 | 0.900193479 | 7.18E-07 |
| PFKFB4 | 995.5007413 | 0.900050498 | 1.81E-03 |
| NSD2 | 9039.317095 | 0.899804931 | 1.01E-31 |
| C11orf68 | 2384.928605 | 0.899347052 | 6.68E-12 |
| GSDMB | 99.2754146 | 0.899221875 | 4.60E-03 |
| ABRACL | 559.2061279 | 0.898996176 | 4.93E-11 |
| ANO2 | 401.3700442 | 0.898223581 | 6.15E-08 |
| PRELID2 | 136.0433833 | 0.898118519 | 2.37E-06 |
| CAP2 | 1928.403831 | 0.898013505 | 1.64E-19 |
| HSPA9 | 36794.30816 | 0.89785461 | 1.16E-07 |
| ALDH3B1 | 1300.36037 | 0.896978974 | 1.30E-18 |
| FTCDNL1 | 96.17166783 | 0.89672847 | 3.78E-04 |
| ANXA2 | 106890.7943 | 0.896447392 | 6.80E-28 |
| GAL | 58.68265862 | 0.896206319 | 5.24E-03 |
| MIR22HG | 876.9771294 | 0.895727341 | 3.63E-24 |
| CAVIN3 | 2094.464026 | 0.895491932 | 2.95E-11 |
| LYRM1 | 775.5876976 | 0.893848968 | 5.09E-08 |
| AC074183.2 | 32.94080807 | 0.893826906 | 2.40E-02 |
| ZC3H7B | 8780.439133 | 0.893369772 | 1.53E-31 |
| AL138724.1 | 134.917473 | 0.89278803 | 2.21E-04 |
| SNHG5 | 2995.960911 | 0.892221241 | 1.23E-14 |
| ADAM15 | 5546.526107 | 0.891795288 | 8.85E-15 |
| GCK | 86.15722206 | 0.891699952 | 1.16E-03 |
| SYDE2 | 235.870281 | 0.89039375 | 4.68E-04 |
| ZNRF2P2 | 85.72672436 | 0.88955167 | 1.75E-04 |
| ZNF710-AS1 | 199.9959931 | 0.889137587 | 2.31E-04 |
| HERPUD1 | 5168.49769 | 0.888890869 | 3.31E-11 |
| AL645608.8 | 27.87022459 | 0.888792817 | 3.13E-02 |
| SOHLH2 | 153.5629023 | 0.888746554 | 4.98E-05 |
| HIP1R | 1309.841327 | 0.888409907 | 7.13E-11 |

| | | | |
|---|---|---|---|
| LRRC59 | 6683.308593 | 0.88815241 | 5.41E-32 |
| SLC7A10 | 92.31430533 | 0.887579546 | 2.69E-04 |
| PCDHGA3 | 35.30244861 | 0.887478673 | 1.08E-02 |
| NXF3 | 122.3733355 | 0.886432014 | 1.22E-02 |
| FAM20C | 2024.588472 | 0.885841669 | 4.34E-07 |
| FAM171A1 | 2363.984497 | 0.885431593 | 8.21E-15 |
| RNF126 | 894.0071533 | 0.885120517 | 1.98E-18 |
| FAM43A | 796.7891411 | 0.884598746 | 2.21E-14 |
| SLC6A17 | 261.5610214 | 0.882671665 | 9.34E-05 |
| LENG1 | 325.7448315 | 0.88258274 | 1.24E-07 |
| MYO1C | 10838.86687 | 0.882507972 | 4.66E-14 |
| H2BFM | 64.76987712 | 0.8824773 | 2.82E-03 |
| PKN1 | 5938.711978 | 0.88239043 | 1.87E-19 |
| SBF1 | 7935.53834 | 0.8816403 | 2.02E-26 |
| SLC39A14 | 6805.43364 | 0.881304414 | 8.80E-15 |
| DLX4 | 530.3310774 | 0.880444292 | 1.04E-12 |
| INAFM2 | 266.0135533 | 0.880154189 | 7.30E-10 |
| PARVG | 194.1912916 | 0.879992264 | 3.81E-05 |
| WHAMM | 724.8447879 | 0.879233545 | 1.12E-10 |
| XAGE1A | 480.1985385 | 0.879171111 | 8.67E-19 |
| AL162231.2 | 108.8299862 | 0.879152438 | 1.24E-04 |
| AC006538.1 | 49.8630077 | 0.878684086 | 5.07E-04 |
| MMP17 | 411.9505481 | 0.878523948 | 2.76E-05 |
| TRIML2 | 231.9553741 | 0.877682944 | 1.35E-03 |
| RPSAP52 | 35.61562351 | 0.877329509 | 3.07E-02 |
| ZBTB25 | 608.6663508 | 0.877198553 | 1.63E-15 |
| PHACTR1 | 383.5787134 | 0.876401117 | 8.36E-11 |
| IGDCC4 | 762.010444 | 0.876280305 | 4.05E-09 |
| SLC19A3 | 219.3959215 | 0.875589123 | 1.98E-09 |
| ADCK2 | 1160.502158 | 0.875488705 | 1.24E-12 |
| HSPA2 | 2639.01931 | 0.875211573 | 3.33E-04 |
| RGS10 | 602.2226751 | 0.872922269 | 3.89E-13 |
| RAB20 | 447.2426815 | 0.872187353 | 2.72E-09 |
| ALAS1 | 2028.759906 | 0.872156365 | 4.63E-16 |
| TP53I11 | 1725.487319 | 0.871892642 | 4.33E-15 |
| PLA2G4C | 169.8016121 | 0.871824472 | 4.28E-03 |
| EEF2 | 104137.7217 | 0.871536044 | 4.20E-14 |
| AC156455.1 | 100.1937583 | 0.87133837 | 1.34E-04 |
| PNMA2 | 3733.387097 | 0.868538183 | 4.36E-13 |
| EHD1 | 3523.992995 | 0.868497378 | 1.67E-08 |
| LSP1 | 262.4820471 | 0.867379047 | 9.71E-04 |
| TPPP | 427.9957226 | 0.866630207 | 1.09E-05 |
| TCF15 | 164.4727977 | 0.86553988 | 1.46E-04 |
| CADM4 | 889.7866067 | 0.865028365 | 2.39E-06 |
| STXBP5-AS1 | 41.1795655 | 0.864984811 | 2.16E-02 |
| ANKZF1 | 2063.874003 | 0.864831605 | 6.87E-20 |
| HIVEP1 | 1633.6484 | 0.864382244 | 5.72E-11 |

| | | | |
|---|---|---|---|
| TMCO3 | 2286.119453 | 0.863695891 | 8.56E-21 |
| AC142086.6 | 63.49481042 | 0.862905622 | 4.80E-04 |
| P2RX5 | 672.8200669 | 0.859303738 | 5.07E-11 |
| CCDC85B | 1306.311525 | 0.859270223 | 5.70E-06 |
| SCARA3 | 3354.9439 | 0.858708097 | 2.66E-08 |
| EVC2 | 505.3089137 | 0.858346266 | 3.51E-11 |
| SNX18P3 | 247.9116142 | 0.857570516 | 5.50E-07 |
| LINC01224 | 480.4551308 | 0.856271404 | 1.06E-04 |
| KLHL36 | 2749.072336 | 0.856118582 | 4.35E-21 |
| AC006329.1 | 118.9663596 | 0.855801857 | 3.08E-06 |
| IGFL2-AS1 | 125.576691 | 0.855012881 | 4.43E-03 |
| FLNB-AS1 | 98.05334563 | 0.854186433 | 5.54E-04 |
| LINC00189 | 232.8947623 | 0.853219823 | 1.32E-05 |
| CDH13 | 11715.24575 | 0.853073668 | 4.42E-13 |
| LRP4-AS1 | 36.37525356 | 0.852953553 | 2.95E-02 |
| LYN | 763.4264982 | 0.852672769 | 4.60E-10 |
| LIMD1 | 1660.134219 | 0.852538862 | 5.88E-09 |
| TTC39B | 448.9058529 | 0.852102093 | 2.41E-07 |
| LAMA1 | 1777.372355 | 0.851952945 | 9.75E-06 |
| YBX2 | 1436.654908 | 0.851516786 | 6.67E-08 |
| IFITM2 | 1317.633411 | 0.851514739 | 9.59E-05 |
| AL442125.1 | 36.23637267 | 0.85148135 | 2.29E-02 |
| SLC12A4 | 2661.801366 | 0.850942898 | 7.14E-09 |
| IQGAP3 | 3343.91172 | 0.850672829 | 1.04E-16 |
| BX322562.1 | 38.96148071 | 0.850363566 | 1.92E-02 |
| ITGA7 | 128.2115376 | 0.849886194 | 1.41E-02 |
| RFLNB | 1500.315546 | 0.849830747 | 4.94E-15 |
| IGSF8 | 967.3449524 | 0.848658366 | 3.71E-15 |
| PROB1 | 216.9344506 | 0.848507845 | 2.07E-05 |
| DAPK3 | 2248.152994 | 0.847869395 | 5.14E-11 |
| PTGR1 | 3609.649913 | 0.847301228 | 5.90E-15 |
| SLC16A1-AS1 | 314.3438713 | 0.846721868 | 3.35E-03 |
| ACOT9 | 3014.294838 | 0.846632395 | 2.22E-22 |
| MYBL1 | 683.1035354 | 0.846423709 | 3.13E-04 |
| HCG4 | 34.57144931 | 0.84596241 | 2.45E-02 |
| TMED7-TICAM2 | 518.8254975 | 0.844081281 | 5.76E-04 |
| CSNK2A2 | 5047.769308 | 0.843982506 | 2.01E-42 |
| KCNAB2 | 1627.146031 | 0.843045894 | 1.75E-07 |
| DVL1 | 1980.054946 | 0.842462335 | 3.29E-19 |
| PTAFR | 29.50072821 | 0.841686254 | 4.92E-02 |
| TNFRSF10D | 1337.770238 | 0.84156862 | 2.02E-06 |
| PFDN2 | 1584.135587 | 0.841146101 | 1.23E-12 |
| S100A11 | 9916.271027 | 0.84094085 | 1.91E-17 |
| CLEC11A | 1129.023465 | 0.840727389 | 5.89E-07 |
| AL662907.1 | 56.14748766 | 0.840654693 | 1.99E-03 |
| MRPL58 | 996.859681 | 0.840348137 | 7.04E-10 |
| CA5B | 553.48089 | 0.840037363 | 4.17E-08 |

| | | | |
|---|---|---|---|
| ZMAT5 | 512.1863495 | 0.839568022 | 1.16E-11 |
| AL139020.1 | 290.5916199 | 0.839540046 | 7.05E-07 |
| RN7SL1 | 545.8515391 | 0.838910181 | 1.72E-04 |
| TPST2 | 341.6889748 | 0.8387371 | 1.91E-09 |
| BICD2 | 3147.686758 | 0.835571332 | 6.42E-18 |
| KLC2 | 1774.580244 | 0.835497033 | 1.30E-13 |
| ECHDC3 | 285.3637131 | 0.835410448 | 3.18E-11 |
| GYPC | 5168.208005 | 0.83508075 | 3.18E-11 |
| DGKD | 3157.094476 | 0.834228074 | 5.55E-11 |
| SBNO2 | 2829.160965 | 0.833648792 | 1.76E-21 |
| POC1A | 1050.629485 | 0.833480997 | 2.72E-14 |
| AURKA | 3414.550899 | 0.832807032 | 1.72E-13 |
| SIM2 | 439.4469742 | 0.832419773 | 1.26E-08 |
| PHLDB1 | 8195.744789 | 0.832353456 | 3.68E-14 |
| DAZAP1 | 4240.34988 | 0.831540556 | 3.80E-26 |
| DRP2 | 72.51218544 | 0.831535403 | 2.59E-03 |
| SNHG7 | 855.8523069 | 0.831453922 | 1.35E-17 |
| RP9P | 225.4968509 | 0.830902108 | 3.66E-09 |
| GRAMD1A | 1599.035645 | 0.830685508 | 7.01E-16 |
| LINC01271 | 81.39153448 | 0.830494565 | 4.56E-04 |
| NALT1 | 51.36180174 | 0.830291825 | 2.42E-03 |
| HOXB-AS1 | 138.8490023 | 0.830233561 | 3.57E-07 |
| GLIS1 | 41.15276081 | 0.830004122 | 4.71E-02 |
| JPH1 | 95.12718083 | 0.829666449 | 2.07E-03 |
| RALGPS1 | 346.6724631 | 0.829404117 | 3.57E-04 |
| MT2A | 3108.118129 | 0.829187058 | 1.12E-03 |
| AC006058.3 | 72.14677154 | 0.828591981 | 1.09E-03 |
| TMSB10 | 36922.00469 | 0.828450597 | 1.51E-09 |
| RHOBTB1 | 1185.631432 | 0.828248823 | 2.14E-17 |
| SH2D2A | 145.948703 | 0.827935274 | 7.66E-06 |
| LINC01257 | 134.8471246 | 0.827277503 | 5.61E-03 |
| GNG11 | 5111.312028 | 0.827226868 | 1.15E-11 |
| LINC01117 | 41.95177487 | 0.827216691 | 5.75E-03 |
| TRIM25 | 7151.293211 | 0.826951705 | 1.81E-12 |
| APOBR | 38.09673852 | 0.825158141 | 1.47E-02 |
| MRPS24 | 3240.035124 | 0.825041218 | 9.27E-23 |
| FER1L6 | 34.70666307 | 0.824619019 | 3.36E-02 |
| SLC6A11 | 59.07492838 | 0.823928114 | 3.24E-02 |
| SKAP1 | 149.3932027 | 0.823561294 | 8.04E-05 |
| FXYD5 | 4701.536881 | 0.82273275 | 4.65E-15 |
| IL4R | 1223.73997 | 0.821742063 | 7.51E-09 |
| LINC01029 | 2160.242943 | 0.821258899 | 1.30E-16 |
| CEMIP2 | 4440.133686 | 0.821038207 | 1.93E-14 |
| AURKB | 2643.519517 | 0.820591984 | 1.29E-10 |
| CIT | 2750.995169 | 0.819865273 | 5.91E-09 |
| IFRD1 | 3961.506108 | 0.819286204 | 1.95E-03 |
| CARD19 | 976.3250092 | 0.818630896 | 1.90E-12 |

| | | | |
|---|---|---|---|
| NUP50-DT | 172.5240029 | 0.818603122 | 4.53E-04 |
| STX8 | 1628.006817 | 0.81755664 | 2.19E-18 |
| AC012254.2 | 129.1850703 | 0.817440529 | 6.78E-04 |
| SYNGR3 | 539.8025067 | 0.817264633 | 5.41E-06 |
| KBTBD3 | 101.4109539 | 0.817025116 | 1.67E-02 |
| MTCH1 | 8829.71424 | 0.817003722 | 1.57E-08 |
| DGKA | 1692.88712 | 0.816982859 | 6.78E-10 |
| C10orf25 | 177.3571878 | 0.815173001 | 1.64E-08 |
| PCAT6 | 92.64316638 | 0.814203246 | 1.52E-03 |
| IRS1 | 1437.527567 | 0.814145444 | 1.56E-06 |
| ZNF793-AS1 | 53.5064077 | 0.813037703 | 3.22E-03 |
| HDAC9 | 214.4988075 | 0.812272145 | 2.16E-04 |
| DIAPH3 | 4667.695024 | 0.812271442 | 1.66E-06 |
| CMSS1 | 1886.108128 | 0.812257397 | 1.03E-10 |
| AC009549.1 | 45.61235665 | 0.812044985 | 1.00E-02 |
| C17orf51 | 1818.558345 | 0.811980574 | 1.05E-07 |
| MYDGF | 2881.007494 | 0.809595501 | 1.63E-15 |
| AC245297.3 | 91.66191408 | 0.809525336 | 2.28E-04 |
| GPC1 | 4941.093134 | 0.809206777 | 3.59E-12 |
| AC016831.1 | 169.9544083 | 0.808907943 | 9.30E-03 |
| PMAIP1 | 884.7431058 | 0.808806692 | 4.86E-03 |
| SLC41A1 | 2600.516694 | 0.808652235 | 1.55E-17 |
| CCND3 | 1929.467615 | 0.808624044 | 2.20E-10 |
| AC083899.1 | 108.9198831 | 0.808521836 | 1.78E-04 |
| AC091390.4 | 55.41285315 | 0.808416205 | 4.79E-03 |
| PNPLA4 | 1147.358362 | 0.808371113 | 2.59E-15 |
| C19orf73 | 29.56774817 | 0.808115786 | 4.41E-02 |
| GBE1 | 1827.158317 | 0.80809273 | 2.34E-07 |
| NDUFAF2 | 2164.104687 | 0.807772666 | 3.14E-17 |
| TIMM9 | 1153.293094 | 0.807294261 | 4.92E-12 |
| RPL23AP2 | 80.89177985 | 0.807108796 | 1.78E-02 |
| ZNF850 | 295.9530147 | 0.806967747 | 7.12E-03 |
| UBE2C | 4006.039724 | 0.806928475 | 2.56E-12 |
| DPP4 | 73.62921846 | 0.806363161 | 7.90E-03 |
| PLPP2 | 1427.512506 | 0.806339685 | 2.55E-08 |
| RASGRF2 | 415.7962532 | 0.805780325 | 9.03E-03 |
| TPM4 | 21265.75795 | 0.80575811 | 8.12E-19 |
| SOX17 | 62.40755451 | 0.805590956 | 1.53E-03 |
| RILPL1 | 800.4589162 | 0.804452083 | 8.69E-08 |
| FABP3 | 256.5391819 | 0.804177226 | 8.26E-03 |
| OAZ1 | 19674.06586 | 0.804143493 | 2.66E-11 |
| ST14 | 50.55600932 | 0.803469859 | 1.80E-02 |
| GTF2IRD1 | 2131.857364 | 0.80306606 | 3.01E-09 |
| RHEB | 6449.634271 | 0.802833209 | 5.95E-44 |
| FAM72C | 235.6365345 | 0.802385752 | 2.97E-02 |
| SDC2 | 2063.499858 | 0.801828282 | 1.03E-10 |
| KIAA1549L | 2687.335253 | 0.801270409 | 4.57E-13 |

| | | | |
|---|---|---|---|
| TTC7B | 1528.002513 | 0.800814267 | 3.95E-12 |
| TANGO6 | 1779.528087 | 0.800637521 | 4.00E-16 |
| THEM6 | 1224.94723 | 0.800560642 | 3.79E-05 |
| TSPO | 3048.132338 | 0.80042568 | 9.10E-10 |
| TTYH1 | 43.2858316 | 0.799952273 | 4.19E-02 |
| AVPI1 | 366.8041458 | 0.799711615 | 5.40E-07 |
| B3GNT5 | 183.3353908 | 0.799191405 | 9.90E-04 |
| TRIM8 | 1510.617956 | 0.797808163 | 2.00E-07 |
| VCL | 13667.92026 | 0.796080327 | 6.99E-07 |
| FBXO25 | 1326.467273 | 0.795691493 | 4.03E-21 |
| 5-Sep | 1245.849617 | 0.79506364 | 2.38E-14 |
| CNNM1 | 541.0648735 | 0.794572201 | 1.80E-11 |
| SLC6A9 | 461.5566515 | 0.794073154 | 9.28E-03 |
| BID | 1337.216014 | 0.792649641 | 1.35E-06 |
| MOSPD1 | 1494.579896 | 0.792480317 | 1.11E-06 |
| ACOT4 | 121.1949792 | 0.792341785 | 4.87E-05 |
| CERNA1 | 67.95049957 | 0.79174473 | 4.94E-03 |
| CBS | 6183.303293 | 0.791527743 | 9.28E-11 |
| CRYAB | 8154.737916 | 0.790879562 | 2.75E-03 |
| LRRC3 | 598.7888325 | 0.790798036 | 6.22E-05 |
| AL591845.1 | 91.06227821 | 0.790493027 | 2.26E-03 |
| TRIP6 | 1891.929213 | 0.790129086 | 6.42E-09 |
| NCOR1P1 | 48.21470094 | 0.789980463 | 2.09E-02 |
| RTN4 | 17887.31892 | 0.788518685 | 1.13E-14 |
| UCP3 | 55.17028068 | 0.788073548 | 2.82E-02 |
| C22orf23 | 284.8102428 | 0.787844543 | 1.09E-06 |
| CPPED1 | 2575.427025 | 0.787582544 | 4.50E-19 |
| TEX41 | 51.499136 | 0.787098522 | 2.60E-02 |
| SND1 | 22442.27026 | 0.786525047 | 2.24E-14 |
| SRF | 2490.360946 | 0.786273585 | 1.97E-21 |
| HSPA5 | 56281.23694 | 0.78582038 | 1.52E-10 |
| C2orf27A | 102.3823596 | 0.785696741 | 4.05E-04 |
| RBPMS | 1720.796055 | 0.785350661 | 1.22E-14 |
| LIMD1-AS1 | 173.1250136 | 0.784833132 | 3.14E-04 |
| TBL3 | 2640.040612 | 0.784708356 | 4.34E-10 |
| TIMM17B | 1538.405681 | 0.784609745 | 6.07E-14 |
| MAP4K3-DT | 286.7965894 | 0.784320526 | 2.06E-10 |
| G0S2 | 2016.718559 | 0.78413417 | 7.96E-03 |
| CT45A3 | 529.6423585 | 0.783409232 | 2.52E-10 |
| IER5 | 1734.988743 | 0.782936912 | 3.34E-11 |
| LNP1 | 176.8767432 | 0.782795523 | 2.43E-06 |
| PARVA | 12811.79845 | 0.782347494 | 2.05E-53 |
| FHL3 | 1027.877847 | 0.782100806 | 9.13E-08 |
| TUBB4B | 23483.62734 | 0.781774901 | 1.25E-06 |
| RPL39L | 913.2693129 | 0.781740023 | 1.34E-18 |
| AP003119.3 | 53.08071754 | 0.781468329 | 2.03E-03 |
| AC005077.4 | 26.51815576 | 0.781224623 | 3.75E-02 |

| | | | |
|---|---|---|---|
| RHPN2 | 791.741258 | 0.780154351 | 5.73E-11 |
| CDYL2 | 2093.83975 | 0.779765222 | 1.14E-10 |
| PLEKHB2 | 5020.256771 | 0.77965068 | 1.32E-23 |
| WDSUB1 | 397.7929314 | 0.778913807 | 4.74E-10 |
| FPR2 | 56.2775836 | 0.778806921 | 8.84E-03 |
| CYP4F2 | 99.75776246 | 0.778678729 | 3.75E-02 |
| LITAF | 5168.421173 | 0.77853017 | 2.88E-16 |
| CCDC155 | 74.62847622 | 0.777734409 | 4.65E-02 |
| SDHB | 2974.624177 | 0.776961534 | 9.97E-25 |
| NFKBIZ | 242.4207178 | 0.776831839 | 1.08E-05 |
| RELT | 599.7605953 | 0.776385787 | 8.93E-12 |
| PLEKHG4 | 1389.84851 | 0.775713022 | 1.09E-09 |
| IRAK1 | 19076.01415 | 0.775098394 | 7.99E-14 |
| PTPRM | 1734.337607 | 0.775062576 | 1.05E-06 |
| LINC01234 | 437.4067572 | 0.775017236 | 5.90E-11 |
| ITPKA | 85.58454827 | 0.774641171 | 3.22E-03 |
| AC018413.1 | 63.11440807 | 0.773648586 | 2.27E-02 |
| TOMM34 | 5597.477897 | 0.773589503 | 1.78E-18 |
| DAAM2 | 3400.637135 | 0.773496812 | 4.74E-06 |
| PLCB2 | 233.3908541 | 0.773109104 | 3.67E-03 |
| PHLPP1 | 1810.927214 | 0.772610865 | 1.43E-09 |
| PRICKLE3 | 449.079314 | 0.772261741 | 3.83E-11 |
| RNPEPL1 | 2127.531071 | 0.770562461 | 1.92E-09 |
| XPOT | 9329.971461 | 0.770399724 | 1.49E-05 |
| TLNRD1 | 501.8768901 | 0.768427028 | 3.16E-05 |
| CEBPB-AS1 | 71.98809642 | 0.768217571 | 4.30E-03 |
| RMC1 | 875.8919123 | 0.768215367 | 2.39E-12 |
| PABPC1 | 53608.13181 | 0.767981221 | 9.85E-17 |
| SLC35E4 | 401.9891485 | 0.767836454 | 8.15E-10 |
| PRXL2C | 968.9850378 | 0.767523267 | 9.63E-10 |
| TFAP2C | 755.5948606 | 0.767074068 | 4.89E-06 |
| CAMKK1 | 327.9390094 | 0.767064898 | 1.57E-07 |
| NSMF | 5938.632953 | 0.76640298 | 1.03E-15 |
| FOLR1 | 159.0086188 | 0.765963979 | 2.75E-02 |
| ATCAY | 1432.949226 | 0.76546505 | 1.71E-04 |
| GASK1A | 355.4300528 | 0.765131405 | 8.76E-11 |
| DNAH11 | 135.1776425 | 0.764887442 | 7.66E-04 |
| SMPDL3B | 107.507826 | 0.764659092 | 1.02E-02 |
| UBR4 | 15901.60597 | 0.764531287 | 1.04E-19 |
| SMYD3 | 591.4497563 | 0.76438485 | 4.91E-08 |
| CENPH | 2549.82782 | 0.764168404 | 2.11E-08 |
| ETS2 | 1570.480883 | 0.763972949 | 3.90E-14 |
| TIMM44 | 2236.04911 | 0.763699886 | 9.05E-09 |
| APCDD1L-DT | 254.384421 | 0.763584201 | 3.18E-03 |
| PDLIM4 | 2965.529172 | 0.76339305 | 3.61E-12 |
| PACS1 | 5469.333388 | 0.762779826 | 4.47E-17 |
| CD79B | 132.1895311 | 0.762197287 | 3.87E-04 |

| | | | |
|---|---|---|---|
| ARF1 | 11486.82838 | 0.761489037 | 7.26E-20 |
| INPP5B | 1336.420141 | 0.760489372 | 1.11E-07 |
| ALDH1L2 | 216.1567516 | 0.760472971 | 8.08E-03 |
| MAP1LC3B | 6359.449737 | 0.760343026 | 2.48E-13 |
| YDJC | 1016.804538 | 0.760171853 | 7.33E-13 |
| LAPTM5 | 330.5302971 | 0.759442716 | 1.77E-05 |
| TNNT1 | 3015.335765 | 0.759385877 | 1.39E-09 |
| GABRE | 2505.179407 | 0.759194748 | 1.19E-04 |
| TRIM6 | 699.1372443 | 0.757771647 | 3.01E-09 |
| EPB41L4A-AS1 | 1731.796056 | 0.757731873 | 3.00E-06 |
| UHRF1BP1 | 4704.980666 | 0.757505942 | 1.33E-05 |
| CHCHD6 | 358.4714213 | 0.75719372 | 5.88E-12 |
| CDHR3 | 288.3718067 | 0.757041863 | 1.92E-04 |
| PGF | 1251.66419 | 0.755405948 | 9.87E-06 |
| CDKN1C | 326.2645282 | 0.755343013 | 3.62E-04 |
| EML2 | 1514.807018 | 0.755170318 | 4.35E-10 |
| KDM7A | 570.1778443 | 0.755008516 | 3.26E-02 |
| DLC1 | 8727.154963 | 0.75450855 | 2.41E-33 |
| ANKRD11 | 12651.38724 | 0.754505636 | 2.76E-10 |
| MIF4GD | 956.5208067 | 0.754384784 | 6.15E-10 |
| KCTD20 | 6930.129421 | 0.754101611 | 1.05E-55 |
| AC145098.2 | 70.74117309 | 0.754001915 | 1.25E-02 |
| DCTN2 | 6303.198494 | 0.753841089 | 3.58E-14 |
| ID1 | 4464.244433 | 0.753527944 | 1.12E-03 |
| HS6ST1 | 1299.442214 | 0.75308794 | 1.30E-05 |
| MAD2L2 | 2128.707886 | 0.75291223 | 1.37E-17 |
| LGALS8 | 2152.324426 | 0.752254492 | 5.68E-14 |
| TIMP2 | 22643.24137 | 0.751889267 | 3.23E-08 |
| AL589843.1 | 84.43969269 | 0.751835767 | 1.25E-03 |
| PRKCH | 883.6995121 | 0.751349077 | 2.49E-11 |
| RFK | 851.8053873 | 0.750740058 | 4.83E-05 |
| MACROD1 | 1467.887466 | 0.750681968 | 5.88E-09 |
| SDCBP2 | 87.31079611 | 0.749918353 | 1.83E-03 |
| DGKQ | 1201.710352 | 0.749805165 | 1.41E-06 |
| CNN3 | 12063.72067 | 0.748419177 | 9.31E-20 |
| ADAMTS17 | 64.94372239 | 0.748014861 | 4.90E-02 |
| CYSTM1 | 1163.563966 | 0.746619861 | 9.83E-09 |
| BCAT1 | 1936.78461 | 0.746514136 | 1.73E-04 |
| PDCD2L | 340.1145273 | 0.746425117 | 2.03E-09 |
| RNF19B | 646.416233 | 0.746392344 | 8.81E-10 |
| FLII | 12054.38531 | 0.74604059 | 6.16E-13 |
| EIF3C | 53785.43486 | 0.745954571 | 2.15E-10 |
| FAM84A | 44.9306386 | 0.745949447 | 2.49E-02 |
| GPKOW | 1631.232436 | 0.74585772 | 1.38E-13 |
| GRPEL1 | 1981.801444 | 0.745652332 | 2.42E-18 |
| LINC00575 | 45.20874019 | 0.744980857 | 2.36E-02 |
| MROH1 | 1333.300415 | 0.744851124 | 9.51E-14 |

| | | | |
|---|---|---|---|
| TKTL1 | 81.13948348 | 0.744753372 | 6.44E-03 |
| ZPR1 | 5866.201251 | 0.744186699 | 3.56E-08 |
| CAPN15 | 2350.329042 | 0.744086735 | 1.34E-16 |
| ICAM5 | 825.1103563 | 0.744027649 | 2.18E-05 |
| OCIAD2 | 557.958916 | 0.743888949 | 7.40E-08 |
| PSPH | 3489.931624 | 0.743807028 | 9.21E-04 |
| NAT8L | 892.8349717 | 0.743568609 | 6.24E-11 |
| PRKCA | 7132.858038 | 0.743554904 | 1.58E-19 |
| P2RX4 | 1389.149782 | 0.743418445 | 4.20E-12 |
| TRAP1 | 2737.631956 | 0.743397031 | 1.10E-18 |
| SCRN1 | 19372.32534 | 0.743304208 | 2.41E-25 |
| FAM207A | 755.1657164 | 0.743149063 | 1.24E-11 |
| AC011476.3 | 154.9701394 | 0.742234506 | 1.46E-04 |
| ADCY3 | 3045.328296 | 0.74201743 | 2.13E-12 |
| SINHCAF | 2560.657428 | 0.741831261 | 1.64E-22 |
| HRH1 | 582.8781532 | 0.741593061 | 1.27E-05 |
| ARSJ | 838.5835605 | 0.7415737 | 8.02E-07 |
| BCAS4 | 1194.623206 | 0.741303564 | 9.25E-08 |
| NACC2 | 1511.713598 | 0.740652739 | 5.08E-15 |
| DHRSX | 467.6353937 | 0.740501207 | 2.16E-06 |
| EXD3 | 412.4310246 | 0.740341214 | 1.15E-10 |
| AP001505.1 | 63.01778494 | 0.739975456 | 3.90E-03 |
| CORO1C | 8847.320057 | 0.739651996 | 5.33E-15 |
| RAB43P1 | 75.2544523 | 0.739403274 | 4.17E-02 |
| KATNB1 | 1890.166344 | 0.738964198 | 5.53E-11 |
| LOXL2 | 22646.42599 | 0.738815089 | 5.92E-07 |
| GOLGA8N | 204.8886715 | 0.73875364 | 6.92E-06 |
| MYO1F | 331.6675034 | 0.738484091 | 1.67E-05 |
| CHIC2 | 379.1742009 | 0.737494702 | 6.57E-06 |
| TCEA1 | 7064.069542 | 0.737445551 | 1.26E-03 |
| AC092042.3 | 124.0100688 | 0.737419353 | 1.74E-03 |
| KIF12 | 74.36197482 | 0.737285888 | 1.00E-02 |
| ARID5A | 430.5825868 | 0.73705932 | 1.24E-06 |
| GRB10 | 4252.746946 | 0.736951034 | 1.15E-05 |
| SAMD4B | 3994.490781 | 0.73671316 | 1.48E-22 |
| GPRC5B | 2084.199969 | 0.736607196 | 2.89E-04 |
| AC023024.1 | 138.7335527 | 0.736443597 | 4.68E-03 |
| TAGLN2 | 18351.75881 | 0.736329932 | 6.53E-06 |
| ENO1 | 85505.67072 | 0.735296277 | 6.72E-08 |
| RALGPS2 | 1148.025275 | 0.735065146 | 4.19E-07 |
| SQSTM1 | 11897.91029 | 0.734901366 | 3.33E-08 |
| MS4A15 | 168.0965125 | 0.734234952 | 3.19E-05 |
| DUX4L46 | 65.62997736 | 0.733954419 | 4.64E-03 |
| RTKN2 | 192.5215251 | 0.733640199 | 1.23E-02 |
| KAZN | 2090.457273 | 0.733092795 | 1.44E-14 |
| FEZ2 | 2245.374732 | 0.732110831 | 1.58E-17 |
| CREB3L2 | 4451.672148 | 0.731370455 | 1.46E-19 |

| | | | |
|---|---|---|---|
| ZNF280A | 323.1856629 | 0.731229934 | 1.30E-05 |
| PRH1 | 47.6120715 | 0.730951147 | 3.78E-02 |
| PLP2 | 4682.094735 | 0.730318417 | 1.54E-11 |
| CDC20B | 40.89525028 | 0.730216323 | 2.32E-02 |
| PPFIA4 | 830.6144763 | 0.730194059 | 1.93E-02 |
| CCDC113 | 1000.208797 | 0.729902538 | 4.44E-08 |
| AC005520.2 | 60.31597595 | 0.729793733 | 4.83E-03 |
| DLGAP4-AS1 | 149.9801256 | 0.729312871 | 1.92E-04 |
| TLR6 | 73.63909273 | 0.728986096 | 3.77E-02 |
| THAP4 | 1823.13702 | 0.728890868 | 1.77E-10 |
| CNKSR1 | 91.16836387 | 0.728600877 | 1.40E-02 |
| C5AR2 | 63.75946987 | 0.728516539 | 4.66E-02 |
| ZDHHC9 | 2390.142154 | 0.727900383 | 1.07E-17 |
| AC040977.1 | 44.63630535 | 0.727622421 | 2.10E-02 |
| MAFG | 2434.517742 | 0.726895207 | 1.55E-15 |
| CNRIP1 | 1462.563083 | 0.726463938 | 3.71E-11 |
| SNX33 | 3457.403843 | 0.72636045 | 5.49E-08 |
| BHLHE40 | 1231.608194 | 0.726077204 | 5.33E-04 |
| LRRC23 | 334.2435134 | 0.724246029 | 4.09E-06 |
| DUSP14 | 1404.725443 | 0.724022392 | 3.88E-17 |
| AC104971.1 | 52.78632629 | 0.724013972 | 1.88E-02 |
| SLC43A1 | 1313.017204 | 0.72390582 | 3.94E-05 |
| WBP2 | 3411.363076 | 0.72348797 | 2.93E-11 |
| LINC02515 | 71.41042735 | 0.723029664 | 1.76E-02 |
| SNORD17 | 72.81252209 | 0.722544252 | 2.27E-02 |
| PFDN4 | 1460.258671 | 0.722092397 | 4.20E-12 |
| CRYL1 | 352.6870594 | 0.722084451 | 5.26E-06 |
| PAM16 | 284.7754864 | 0.721984513 | 4.78E-06 |
| HPCAL4 | 258.7617992 | 0.721378142 | 2.76E-06 |
| HGS | 3477.552588 | 0.720962593 | 2.00E-26 |
| LPP | 1835.54307 | 0.720331019 | 8.97E-04 |
| KCNC4 | 377.5097556 | 0.720087283 | 8.66E-03 |
| AC118282.4 | 232.0794676 | 0.719539868 | 1.23E-03 |
| C12orf66 | 315.7310445 | 0.719494653 | 1.52E-06 |
| ST3GAL3 | 868.2489115 | 0.718552331 | 1.36E-17 |
| TNIP1 | 3697.169382 | 0.718510289 | 2.28E-07 |
| FEZ1 | 659.6364878 | 0.718072757 | 4.14E-06 |
| FDFT1 | 15026.82734 | 0.717829907 | 2.86E-05 |
| LINC00899 | 351.447866 | 0.717358143 | 1.49E-05 |
| SOWAHA | 80.37691478 | 0.716562906 | 6.53E-03 |
| APLP2 | 27182.95191 | 0.716356439 | 8.25E-12 |
| LRG1 | 182.2253611 | 0.715733035 | 3.15E-02 |
| BEX4 | 216.329814 | 0.715197925 | 3.26E-05 |
| HCCS | 1607.400186 | 0.714149762 | 2.80E-05 |
| GSTO1 | 3403.432352 | 0.713584402 | 2.88E-10 |
| NATD1 | 1387.904722 | 0.712959299 | 6.33E-07 |
| HSPG2 | 6936.109191 | 0.71280857 | 2.34E-06 |

| | | | |
|---|---|---|---|
| RN7SL2 | 1281.448062 | 0.71246864 | 2.17E-02 |
| NOP53 | 6803.000535 | 0.711934767 | 7.66E-07 |
| PRR11 | 1991.797462 | 0.710890513 | 4.14E-09 |
| AFAP1 | 8267.209501 | 0.710789916 | 1.40E-14 |
| AC020928.2 | 89.02862678 | 0.710501836 | 6.82E-03 |
| WASF2 | 8001.080965 | 0.710308325 | 3.75E-17 |
| MED15 | 3949.807379 | 0.710244839 | 1.02E-14 |
| PRSS16 | 79.4703776 | 0.709982755 | 3.16E-02 |
| TMEM54 | 1494.946011 | 0.709116586 | 2.26E-10 |
| MAP4 | 17613.87065 | 0.707798015 | 8.99E-23 |
| TMCO4 | 1573.630919 | 0.707745423 | 1.83E-08 |
| SLC25A34-AS1 | 55.99924143 | 0.707268247 | 2.11E-02 |
| FILIP1 | 45.8162629 | 0.70690391 | 3.41E-02 |
| FSTL5 | 120.0386041 | 0.706865325 | 9.51E-03 |
| MIR621 | 113.5093563 | 0.706412442 | 2.80E-04 |
| RASA3 | 1805.923465 | 0.706315777 | 1.69E-07 |
| DMGDH | 358.5659548 | 0.705189305 | 8.42E-03 |
| TOR4A | 755.6938358 | 0.705081766 | 3.09E-05 |
| TENM3-AS1 | 240.739707 | 0.704828185 | 3.06E-05 |
| SSR4P1 | 122.1326671 | 0.704152333 | 2.03E-03 |
| ZNF788P | 519.4212818 | 0.703488529 | 5.42E-08 |
| TBC1D22B | 1030.373319 | 0.703394047 | 2.12E-14 |
| ANXA5 | 24506.2297 | 0.703380273 | 2.88E-11 |
| AGRN | 19104.86146 | 0.702667066 | 3.16E-08 |
| SYNE1 | 3339.578059 | 0.702113965 | 6.89E-04 |
| PRDM13 | 87.34034449 | 0.701731551 | 4.23E-03 |
| PKD1 | 2789.544658 | 0.70105394 | 3.59E-05 |
| FBXO41 | 1662.033975 | 0.700803191 | 1.05E-09 |
| NLK | 1029.221641 | 0.700315533 | 3.01E-17 |
| SDK1 | 1217.636645 | 0.699746638 | 6.19E-07 |
| SYT5 | 170.0209336 | 0.699717487 | 1.86E-05 |
| SAV1 | 1549.090313 | 0.699540918 | 3.10E-10 |
| SHROOM3 | 3719.952372 | 0.699532939 | 3.41E-11 |
| FAM86B3P | 213.3333834 | 0.699438274 | 5.43E-04 |
| OR2A20P | 429.4941753 | 0.69902879 | 6.22E-03 |
| PTP4A2 | 6807.446394 | 0.698914172 | 3.84E-08 |
| ZNF687-AS1 | 61.88834523 | 0.698895598 | 3.13E-02 |
| SMIM1 | 37.94733867 | 0.698855932 | 4.09E-02 |
| TWF2 | 2812.418961 | 0.698733562 | 1.66E-12 |
| HOXC8 | 845.3701195 | 0.698550675 | 4.75E-11 |
| WWTR1 | 6889.014365 | 0.698500753 | 8.93E-22 |
| NA | 264.1330256 | 0.698386255 | 1.77E-04 |
| SPECC1 | 1835.882639 | 0.698200806 | 2.17E-11 |
| HRCT1 | 93.63600929 | 0.69778522 | 5.55E-03 |
| AL138756.1 | 98.1393658 | 0.697613144 | 1.21E-03 |
| SRXN1 | 4583.879997 | 0.696695478 | 7.55E-06 |
| SRGN | 1547.782683 | 0.696342644 | 1.85E-24 |

| | | | |
|---|---|---|---|
| RRP9 | 1400.270047 | 0.696246366 | 4.37E-13 |
| MAP1B | 20810.43855 | 0.696189604 | 5.97E-04 |
| LRRC8A | 4409.169916 | 0.695853319 | 5.82E-12 |
| AC006547.1 | 93.48753224 | 0.695343442 | 1.22E-03 |
| PLEC | 19438.93034 | 0.695170264 | 1.36E-11 |
| KCTD13 | 506.4129761 | 0.695132541 | 2.04E-05 |
| SORT1 | 7230.535163 | 0.694916989 | 8.35E-13 |
| REXO2 | 5050.771619 | 0.694201072 | 9.10E-20 |
| VDAC2 | 3904.428237 | 0.694123367 | 1.27E-16 |
| GPR19 | 145.3282073 | 0.693933602 | 6.69E-03 |
| SIRT5 | 725.0325281 | 0.693582101 | 5.54E-08 |
| PPDPF | 3817.555591 | 0.693446341 | 2.03E-09 |
| TMEM206 | 691.1648412 | 0.693147068 | 1.65E-09 |
| HLA-L | 166.7642701 | 0.692907098 | 1.98E-02 |
| KIF1C | 10083.17981 | 0.692316258 | 9.79E-15 |
| GIHCG | 338.5023988 | 0.692195165 | 1.55E-08 |
| NCOR2 | 5055.339838 | 0.69189443 | 2.17E-09 |
| SMURF2 | 2056.949958 | 0.691813713 | 1.45E-10 |
| SNRPB | 16462.47914 | 0.691602934 | 2.13E-10 |
| ESRRA | 1871.490756 | 0.690692815 | 9.31E-13 |
| AC125232.1 | 781.9054721 | 0.690678601 | 3.87E-07 |
| COL7A1 | 231.364016 | 0.689640778 | 2.54E-02 |
| SDC1 | 3446.919541 | 0.689564283 | 7.18E-07 |
| PTPN14 | 5617.28597 | 0.689411603 | 1.50E-21 |
| CYP2R1 | 801.0685079 | 0.689244854 | 1.22E-04 |
| AK9 | 76.96140995 | 0.688783425 | 3.34E-02 |
| CT45A5 | 774.9288964 | 0.688320397 | 1.95E-14 |
| NA | 123.1612894 | 0.688168939 | 6.69E-03 |
| PIEZO1 | 9000.487395 | 0.688119053 | 6.80E-11 |
| RPL9 | 21258.71788 | 0.687858942 | 4.20E-10 |
| PLD6 | 169.4549217 | 0.687742456 | 2.40E-06 |
| TSSC2 | 633.1595099 | 0.687335963 | 1.12E-07 |
| TTC28-AS1 | 350.7466498 | 0.686972912 | 1.56E-06 |
| AXIN1 | 1792.469903 | 0.68661675 | 5.73E-08 |
| NEURL1 | 376.2572735 | 0.686176481 | 2.87E-06 |
| TMTC2 | 236.964812 | 0.686154686 | 7.72E-05 |
| TESMIN | 196.0399767 | 0.685579516 | 3.68E-03 |
| GOT2 | 7515.082647 | 0.684986472 | 2.06E-14 |
| CCNB1 | 16691.04164 | 0.684491813 | 2.03E-15 |
| GRK6 | 1476.105821 | 0.684099044 | 1.84E-10 |
| HACL1 | 1377.386205 | 0.683612447 | 1.65E-09 |
| MT-TN | 372.0712389 | 0.683440356 | 2.12E-02 |
| F12 | 188.3703076 | 0.683322304 | 1.17E-04 |
| PCBD1 | 805.7687183 | 0.683208216 | 4.04E-11 |
| H1F0 | 16936.24213 | 0.683038362 | 1.33E-04 |
| FGD1 | 1766.826493 | 0.682981182 | 2.32E-10 |
| MRPL2 | 1431.242601 | 0.682777083 | 1.76E-08 |

| | | | |
|---|---|---|---|
| GLIPR1 | 1112.924389 | 0.682127191 | 9.53E-07 |
| PLXNB2 | 10147.39702 | 0.681954848 | 9.71E-12 |
| CMC1 | 560.2089086 | 0.681326068 | 1.12E-10 |
| ARHGEF10L | 688.723664 | 0.680264627 | 9.82E-08 |
| SPTAN1 | 27548.29185 | 0.680007852 | 3.96E-21 |
| KIF9-AS1 | 229.0448139 | 0.67991349 | 6.68E-03 |
| MSH5 | 896.9720349 | 0.679757556 | 3.87E-08 |
| KLF10 | 1561.85262 | 0.679513083 | 9.18E-09 |
| YIF1A | 3045.656821 | 0.67875206 | 1.26E-10 |
| KLF13 | 1510.656057 | 0.678110413 | 3.51E-07 |
| AL450124.1 | 315.5916599 | 0.67757116 | 1.05E-06 |
| ABHD11 | 864.5830385 | 0.677536497 | 3.87E-10 |
| AC005253.1 | 114.5949675 | 0.677148408 | 8.10E-04 |
| SIMC1 | 762.4064736 | 0.677055171 | 9.38E-21 |
| COX7A2L | 3464.715754 | 0.677039458 | 2.21E-11 |
| C1GALT1 | 513.9821536 | 0.676974461 | 1.98E-03 |
| EIF4E3 | 382.3841659 | 0.676930182 | 2.30E-06 |
| ARHGAP45 | 1747.783185 | 0.676733399 | 9.67E-08 |
| CADM3 | 1193.863173 | 0.676306963 | 5.96E-05 |
| AC010601.1 | 55.14621582 | 0.674971057 | 2.83E-02 |
| SMKR1 | 71.51547513 | 0.674942765 | 3.89E-02 |
| AP5B1 | 930.0444665 | 0.67489882 | 3.15E-08 |
| PIP5K1A | 3765.167022 | 0.674854418 | 5.80E-12 |
| JUNB | 461.7586993 | 0.674263965 | 5.40E-06 |
| NFKBIB | 708.9637072 | 0.673375968 | 1.12E-04 |
| PKD1P6 | 914.9616331 | 0.673322827 | 4.19E-05 |
| LIN54 | 911.5084985 | 0.673289394 | 1.36E-04 |
| RAB11FIP5 | 2099.786999 | 0.673249656 | 1.49E-06 |
| CENPA | 1024.489308 | 0.672721729 | 9.26E-04 |
| FKBP9 | 12757.91606 | 0.672649027 | 8.85E-15 |
| AC020910.4 | 64.22503316 | 0.672563154 | 1.62E-02 |
| MAGEA10-MAG | 194.1530519 | 0.671723042 | 1.24E-04 |
| DGAT2 | 873.0974125 | 0.671105974 | 2.88E-05 |
| CLDN7 | 246.5956882 | 0.670680968 | 1.20E-04 |
| SNAPC2 | 1028.266326 | 0.670649587 | 9.75E-07 |
| LINC02210 | 631.473022 | 0.670130841 | 6.08E-07 |
| SQOR | 301.2361767 | 0.670002231 | 1.04E-04 |
| LDLRAD4 | 335.4509987 | 0.66976592 | 6.23E-05 |
| PTP4A1 | 2535.03776 | 0.669532878 | 2.60E-02 |
| TVP23C-CDRT4 | 503.1404065 | 0.669493165 | 2.17E-03 |
| APBA2 | 1434.218644 | 0.668874232 | 7.88E-08 |
| TUBA1B | 81573.23168 | 0.667430007 | 4.37E-04 |
| CACNG7 | 582.6298124 | 0.667383608 | 2.92E-05 |
| PLK3 | 284.7440062 | 0.667304196 | 7.16E-06 |
| NSUN7 | 276.9157398 | 0.667126109 | 9.46E-04 |
| MCRIP2 | 724.7230538 | 0.666283941 | 8.41E-08 |
| IMP4 | 3203.583644 | 0.665754675 | 5.62E-13 |

| | | | |
|---|---|---|---|
| CYTL1 | 1870.470186 | 0.665678436 | 4.01E-07 |
| AL671277.2 | 54.94515208 | 0.665455379 | 3.89E-02 |
| EVA1A | 932.7295082 | 0.665155494 | 6.43E-13 |
| ELL | 737.5986994 | 0.664818689 | 2.03E-13 |
| FCHO1 | 269.9933789 | 0.664661083 | 3.21E-04 |
| H1FX | 4099.276197 | 0.664606909 | 2.56E-12 |
| EPSTI1 | 1053.012588 | 0.663979532 | 1.05E-07 |
| ANKRD18B | 178.8942262 | 0.663895059 | 1.92E-02 |
| RRAS | 2730.533569 | 0.663832355 | 5.55E-06 |
| IGFBP4 | 22445.28338 | 0.663249854 | 6.66E-07 |
| FOXN3 | 4410.147751 | 0.662325697 | 5.12E-15 |
| CLTCL1 | 1082.315327 | 0.662081757 | 8.06E-07 |
| RASGRP2 | 175.0039685 | 0.661784432 | 7.18E-04 |
| GLRB | 86.56225061 | 0.661420506 | 8.98E-03 |
| AC105052.3 | 838.6690067 | 0.661371708 | 3.77E-08 |
| AC099850.3 | 182.1025745 | 0.66035103 | 3.99E-04 |
| PDZD7 | 149.1959024 | 0.659762416 | 1.50E-03 |
| TNS3 | 16971.90437 | 0.659291436 | 3.16E-08 |
| DUSP7 | 822.9598436 | 0.659154105 | 7.09E-10 |
| ACTB | 127290.7296 | 0.658963732 | 1.91E-04 |
| NBR2 | 171.9762602 | 0.65880701 | 2.46E-03 |
| TNXB | 1293.362643 | 0.658805234 | 1.03E-08 |
| EIF4EBP3 | 143.5782433 | 0.658553786 | 3.71E-03 |
| SNHG28 | 86.68670457 | 0.658320432 | 1.06E-02 |
| TGFB1I1 | 2716.369981 | 0.657706683 | 8.46E-06 |
| YRDC | 837.8103711 | 0.657685842 | 1.94E-12 |
| NES | 7130.448034 | 0.657630678 | 1.07E-04 |
| PRKCQ-AS1 | 135.4452557 | 0.656985363 | 4.79E-03 |
| TNFRSF10B | 5148.799909 | 0.656177425 | 6.07E-08 |
| NAV2 | 2387.667239 | 0.655951435 | 1.86E-07 |
| NUDT18 | 423.799999 | 0.655237633 | 3.89E-04 |
| SPINDOC | 2425.02415 | 0.655228969 | 1.88E-13 |
| AFAP1L1 | 1973.109642 | 0.654568072 | 4.56E-10 |
| ERICH5 | 54.53299792 | 0.654374762 | 4.33E-02 |
| MBOAT7 | 4718.094911 | 0.654337908 | 1.66E-07 |
| LHX6 | 463.3941 | 0.654264844 | 5.83E-05 |
| PTPN1 | 5738.112123 | 0.653825375 | 2.09E-20 |
| PPP1R35 | 1131.012291 | 0.653745528 | 3.27E-10 |
| BACH1 | 2018.700707 | 0.653490654 | 1.31E-03 |
| RPL12 | 40443.63764 | 0.653361005 | 1.24E-05 |
| TACC1 | 7595.804216 | 0.653349931 | 6.30E-09 |
| CGAS | 435.9810156 | 0.653348899 | 1.77E-05 |
| BAALC-AS1 | 90.98072801 | 0.65292246 | 4.79E-03 |
| AGPAT2 | 2074.660649 | 0.652628214 | 5.42E-07 |
| RIPOR1 | 6265.287925 | 0.652553796 | 1.90E-11 |
| CPM | 499.6059264 | 0.652552773 | 3.61E-03 |
| CCBE1 | 1908.550007 | 0.65252569 | 7.35E-07 |

| | | | |
|---|---|---|---|
| SMIM10L2B | 102.9373407 | 0.651900259 | 4.90E-02 |
| MED8 | 2065.090107 | 0.651636779 | 1.50E-09 |
| EPB41L4B | 494.6600528 | 0.651255495 | 2.68E-04 |
| NELFE | 4523.328843 | 0.65106974 | 2.02E-08 |
| NUDCD1 | 1948.346597 | 0.650717579 | 5.07E-05 |
| TCEA3 | 938.9865274 | 0.650160078 | 1.15E-06 |
| TCL6 | 482.6066254 | 0.649769993 | 7.54E-04 |
| MLPH | 3090.876894 | 0.649717834 | 2.23E-05 |
| BRSK1 | 458.4730848 | 0.649675416 | 1.82E-05 |
| NOD2 | 182.4262089 | 0.649605009 | 3.05E-04 |
| RPS6KA1 | 1985.693064 | 0.649321356 | 7.61E-08 |
| FADD | 1921.919656 | 0.649189404 | 6.13E-10 |
| DNAH3 | 118.1200945 | 0.647862647 | 9.57E-03 |
| GLIS2 | 746.7602133 | 0.645402699 | 4.10E-07 |
| EIF3F | 10611.45856 | 0.64530675 | 4.34E-07 |
| JUND | 1908.596123 | 0.644894187 | 9.09E-05 |
| SPSB1 | 702.3432713 | 0.644652867 | 2.34E-08 |
| AC090204.1 | 619.9733506 | 0.643794827 | 8.75E-11 |
| HOXC13 | 1296.224136 | 0.643620205 | 1.30E-06 |
| SYTL3 | 93.87811856 | 0.643580943 | 3.23E-02 |
| LLGL2 | 1015.83685 | 0.643186959 | 2.64E-09 |
| RAB34 | 4222.806935 | 0.643098139 | 2.38E-09 |
| CCT4 | 15099.11166 | 0.643043682 | 3.74E-28 |
| NPC1 | 3213.854759 | 0.64277825 | 1.17E-11 |
| CDC14B | 2116.738376 | 0.642757282 | 1.63E-12 |
| MRPL4 | 2091.730207 | 0.642593574 | 6.42E-11 |
| AKT1 | 9635.270702 | 0.642551258 | 1.50E-13 |
| GAS7 | 2811.772974 | 0.642512545 | 5.97E-07 |
| MICALL1 | 3271.141042 | 0.642107214 | 3.45E-13 |
| HOXA11-AS | 322.0818879 | 0.642022815 | 7.31E-06 |
| TRABD | 1154.065959 | 0.641960578 | 4.25E-16 |
| MINAR1 | 148.9115714 | 0.641727129 | 4.31E-04 |
| MIER2 | 921.5115839 | 0.641593443 | 2.01E-07 |
| RPL23AP7 | 323.4192596 | 0.641021799 | 2.12E-09 |
| ELOVL1 | 2900.615077 | 0.640829611 | 2.38E-07 |
| KIF2C | 4458.272751 | 0.640762924 | 8.58E-10 |
| AC008957.1 | 76.43432309 | 0.640083058 | 8.96E-03 |
| PRDM8 | 163.9781801 | 0.639825335 | 7.58E-03 |
| TEPP | 64.07188254 | 0.639681257 | 3.36E-02 |
| COPG1 | 7879.341187 | 0.639301363 | 2.84E-10 |
| AC015982.1 | 44.17162428 | 0.638851179 | 4.87E-02 |
| MAP2K1 | 3084.114411 | 0.638780927 | 2.42E-15 |
| IL3RA | 225.9906258 | 0.638776148 | 5.16E-04 |
| RPL4 | 113049.4838 | 0.638597299 | 1.50E-08 |
| PINK1 | 5036.849718 | 0.638333123 | 2.82E-06 |
| ARHGEF1 | 4120.570488 | 0.638326963 | 1.43E-09 |
| RPL9P9 | 16918.13946 | 0.637680854 | 6.33E-10 |

| | | | |
|---|---|---|---|
| SKA3 | 713.8002196 | 0.637556143 | 3.56E-05 |
| PRC1 | 7878.190155 | 0.637336291 | 4.59E-06 |
| CGNL1 | 895.5418779 | 0.637244658 | 3.26E-05 |
| HTT | 5698.107226 | 0.636910984 | 3.45E-05 |
| ZFAND2A | 542.0624813 | 0.636901294 | 1.35E-03 |
| CDK7 | 2722.817861 | 0.636517035 | 9.50E-22 |
| S100A6 | 9517.564517 | 0.636212629 | 2.29E-06 |
| TMEM18 | 1213.800047 | 0.636041673 | 4.39E-07 |
| CCDC169 | 311.1901616 | 0.635888789 | 2.24E-03 |
| PNO1 | 1549.618819 | 0.635293563 | 3.60E-14 |
| PSMD2 | 12961.65287 | 0.634945428 | 8.02E-10 |
| BORCS8-MEF2B | 221.7826825 | 0.634760977 | 6.30E-03 |
| HLA-E | 8048.516065 | 0.634364665 | 4.14E-06 |
| RFTN1 | 2508.131971 | 0.634363847 | 4.59E-08 |
| ALDOA | 42268.47046 | 0.633952749 | 1.50E-06 |
| CHPT1 | 1064.870081 | 0.633343114 | 1.47E-09 |
| AC023824.6 | 76.23987725 | 0.633171312 | 1.29E-02 |
| ACTR1B | 4339.501904 | 0.63299729 | 5.50E-08 |
| AC134312.5 | 157.3323084 | 0.632850881 | 2.66E-02 |
| SH2D4A | 2100.933704 | 0.632838028 | 3.95E-18 |
| SLC1A3 | 1565.100729 | 0.632466699 | 2.72E-06 |
| ADD2 | 5103.394758 | 0.631299458 | 5.19E-15 |
| DPP9 | 3238.309079 | 0.630547329 | 2.97E-12 |
| COX19 | 685.3613752 | 0.630384443 | 8.11E-04 |
| STK24 | 5540.39702 | 0.630303436 | 2.69E-13 |
| ZNF706 | 2431.618789 | 0.629598173 | 5.89E-24 |
| MAP4K2 | 2297.576008 | 0.629357375 | 9.77E-07 |
| IL1RAP | 613.1327742 | 0.629254594 | 2.89E-04 |
| DENND1A | 2021.521882 | 0.629016813 | 4.10E-06 |
| C2orf88 | 65.43652657 | 0.628969686 | 2.70E-02 |
| RAG1 | 104.3953946 | 0.628669845 | 2.64E-02 |
| NBPF3 | 1660.670122 | 0.628379529 | 1.67E-07 |
| HOXC4 | 316.5324271 | 0.62800188 | 1.20E-06 |
| UBE2D2 | 2275.060115 | 0.627654828 | 1.59E-12 |
| NRIP3 | 1681.226296 | 0.626907737 | 3.66E-14 |
| NA | 437.0559334 | 0.62680383 | 3.65E-04 |
| E2F7 | 1312.39899 | 0.626434964 | 6.70E-05 |
| AC064799.1 | 110.997097 | 0.626196231 | 2.53E-02 |
| WIPI1 | 1713.174141 | 0.625876948 | 3.14E-08 |
| NA | 53.61074811 | 0.625689874 | 2.29E-02 |
| PINX1 | 749.064652 | 0.625686888 | 4.23E-13 |
| PPP6R1 | 4169.616317 | 0.625650771 | 1.59E-13 |
| ZNF319 | 661.4451797 | 0.625573404 | 2.15E-05 |
| AP002387.2 | 105.8432616 | 0.625382556 | 3.37E-03 |
| HLA-DQB2 | 65.97511308 | 0.625183707 | 1.11E-02 |
| SPG21 | 4901.16491 | 0.624969441 | 9.70E-14 |
| RPL15 | 46122.89874 | 0.624528365 | 5.05E-07 |

| | | | |
|---|---|---|---|
| MXRA7 | 5655.55128 | 0.624258261 | 5.47E-08 |
| TMEM205 | 1463.331865 | 0.623340966 | 4.09E-06 |
| RPS16 | 19557.72628 | 0.623125162 | 2.24E-07 |
| THBS4 | 204.431369 | 0.623049631 | 9.06E-05 |
| SLC25A6 | 13376.01443 | 0.622964648 | 7.93E-07 |
| NDUFAF3 | 1905.99529 | 0.622857754 | 1.52E-08 |
| GAPDHP14 | 68.65157365 | 0.622706656 | 2.87E-02 |
| PKN3 | 1552.250126 | 0.622589527 | 6.27E-11 |
| CCDC50 | 3220.397041 | 0.622373845 | 8.00E-08 |
| UCK2 | 2195.040756 | 0.622277941 | 4.11E-14 |
| GALC | 2588.669822 | 0.622108514 | 2.13E-12 |
| RAD23A | 3349.438841 | 0.621692143 | 6.05E-08 |
| PPARD | 1678.792718 | 0.621660128 | 8.75E-10 |
| SHC2 | 1490.065423 | 0.621422574 | 3.60E-06 |
| DDX41 | 2155.136552 | 0.621177705 | 1.05E-07 |
| RHEBL1 | 167.6777451 | 0.62079582 | 4.97E-03 |
| RABEPK | 1331.448188 | 0.620625255 | 4.43E-07 |
| DNAJA3 | 2240.392464 | 0.620370026 | 7.01E-07 |
| KLF6 | 4088.683155 | 0.619715694 | 1.02E-10 |
| PDLIM5 | 3144.819155 | 0.619535959 | 2.62E-06 |
| SLC45A4 | 507.6338217 | 0.619303153 | 2.76E-06 |
| C11orf45 | 327.0867099 | 0.619131424 | 2.18E-04 |
| TCF3 | 4011.452498 | 0.618842775 | 1.08E-11 |
| RPL36 | 12121.41414 | 0.618834809 | 3.83E-07 |
| RNF187 | 6542.358528 | 0.618719488 | 8.10E-09 |
| STARD5 | 260.8407944 | 0.618493548 | 5.97E-05 |
| EFTUD2 | 8567.38897 | 0.618408346 | 6.92E-07 |
| C16orf58 | 6656.991018 | 0.618353545 | 2.70E-11 |
| LPXN | 808.5363228 | 0.617796969 | 7.24E-04 |
| XAB2 | 2830.758882 | 0.617638038 | 7.01E-06 |
| HDAC11 | 769.1429549 | 0.617582239 | 5.41E-05 |
| CRYBG1 | 722.0676876 | 0.61726314 | 6.90E-04 |
| ECSIT | 1263.236298 | 0.617183713 | 1.58E-08 |
| HAUS4 | 2423.93088 | 0.616302055 | 1.48E-07 |
| ADRM1 | 5929.584761 | 0.616198046 | 1.94E-11 |
| FLOT1 | 6068.185566 | 0.616080745 | 7.23E-07 |
| HAGH | 1298.627079 | 0.615717905 | 7.72E-07 |
| SZRD1 | 8977.152041 | 0.615705245 | 3.14E-09 |
| HTATIP2 | 3669.802027 | 0.615664245 | 2.46E-14 |
| SNHG1 | 3764.755585 | 0.615662898 | 1.03E-04 |
| AC105942.1 | 246.655278 | 0.615026109 | 1.10E-04 |
| EPG5 | 3554.316738 | 0.61497254 | 3.78E-14 |
| PITPNM3 | 292.5927778 | 0.614943333 | 4.58E-05 |
| ATP9A | 5110.679593 | 0.614790833 | 2.67E-13 |
| AC023024.2 | 59.16330905 | 0.614363996 | 3.78E-02 |
| CD3EAP | 786.1555899 | 0.614340145 | 4.79E-04 |
| SUSD2 | 605.236627 | 0.614128317 | 1.56E-03 |

| | | | |
|---|---|---|---|
| ATG4D | 562.7709515 | 0.61398423 | 7.98E-07 |
| RPS3 | 78001.32135 | 0.613975891 | 4.77E-10 |
| DNAAF3 | 180.1413094 | 0.6139155 | 1.02E-03 |
| IL15RA | 273.3091975 | 0.613808228 | 2.02E-03 |
| ZCCHC4 | 500.6188419 | 0.61376993 | 7.39E-06 |
| IGFBP6 | 2175.730591 | 0.613170476 | 2.52E-05 |
| OSBP2 | 391.2814485 | 0.61288387 | 4.55E-04 |
| GATA2-AS1 | 401.5088441 | 0.612876535 | 2.81E-05 |
| CHCHD4 | 918.971077 | 0.612871315 | 2.11E-06 |
| JOSD1 | 3246.967769 | 0.612369421 | 1.49E-13 |
| ACOT7 | 3467.030846 | 0.612233502 | 1.43E-09 |
| SPATC1L | 320.1765012 | 0.612066013 | 2.40E-05 |
| IDH2 | 4025.990048 | 0.611919059 | 1.25E-04 |
| ATXN10 | 5946.981823 | 0.611800246 | 9.52E-11 |
| HSPB8 | 1436.3874 | 0.611412761 | 4.30E-06 |
| MIR3936HG | 98.17743369 | 0.611360842 | 1.26E-02 |
| ACP6 | 1453.343694 | 0.610960483 | 4.84E-09 |
| TMEM268 | 1773.573704 | 0.610531552 | 2.22E-03 |
| EIF1AD | 1509.337044 | 0.610321147 | 4.35E-11 |
| FARP1 | 10323.71837 | 0.610288846 | 7.01E-15 |
| MYLK3 | 310.1611264 | 0.610153768 | 1.07E-03 |
| DIS3L | 2900.587099 | 0.609888805 | 6.91E-10 |
| RPS17 | 42235.26551 | 0.609602132 | 9.69E-12 |
| SCML1 | 645.1475496 | 0.608821533 | 2.68E-04 |
| OOEP | 184.8376679 | 0.608814421 | 1.31E-03 |
| RUVBL2 | 6983.216087 | 0.608708482 | 3.86E-13 |
| COL16A1 | 5086.360967 | 0.608518986 | 1.48E-05 |
| ARHGEF10 | 3349.754618 | 0.608478745 | 2.37E-08 |
| LINC01137 | 148.1694956 | 0.608421161 | 9.27E-05 |
| DIRAS1 | 1073.655793 | 0.608275537 | 7.63E-08 |
| AC008105.3 | 53.59516555 | 0.60824762 | 3.39E-02 |
| AL645568.1 | 62.75099522 | 0.608043155 | 4.83E-02 |
| SKA1 | 744.58815 | 0.607210908 | 2.74E-03 |
| CFAP20 | 1402.706065 | 0.606948811 | 2.00E-12 |
| LINC01694 | 305.0858403 | 0.606658611 | 1.70E-02 |
| ZG16B | 226.8742984 | 0.606208576 | 1.55E-03 |
| RNH1 | 8558.943629 | 0.606015987 | 5.88E-11 |
| ODC1-DT | 53.69138881 | 0.605324232 | 3.51E-02 |
| PREX1 | 4692.440271 | 0.604948447 | 8.69E-11 |
| CIAO2B | 2234.297862 | 0.604938503 | 3.96E-07 |
| CYB5R3 | 9520.782062 | 0.60490895 | 2.86E-06 |
| CCDC59 | 724.4599412 | 0.604868302 | 4.14E-06 |
| CYTH2 | 1564.392588 | 0.604816678 | 2.93E-08 |
| LRP5 | 4117.859614 | 0.604631881 | 3.32E-12 |
| TGFBR3L | 82.45148481 | 0.604544015 | 2.32E-02 |
| DTNB | 563.0925909 | 0.604536574 | 1.35E-08 |
| PDPK2P | 424.9377736 | 0.603854655 | 1.79E-05 |

| | | | |
|---|---|---|---|
| SH3BGRL3 | 7583.347441 | 0.60368892 | 5.14E-06 |
| INA | 628.7562037 | 0.603537586 | 1.42E-07 |
| FGF7P6 | 157.2628648 | 0.603037681 | 3.16E-02 |
| RPSA | 54318.93969 | 0.602874767 | 8.18E-07 |
| RTN4IP1 | 449.1760696 | 0.602807909 | 6.04E-06 |
| ESYT2 | 7763.823023 | 0.602799356 | 1.59E-15 |
| ULK2 | 882.4735602 | 0.602483447 | 4.29E-06 |
| ABCA2 | 4348.010355 | 0.602414678 | 1.20E-04 |
| RNASEH1 | 1425.965528 | 0.602230283 | 5.00E-18 |
| NLE1 | 1547.763898 | 0.601918571 | 2.10E-07 |
| FAM174B | 267.0840553 | 0.60189603 | 2.65E-05 |
| KDSR | 2931.962615 | 0.601870284 | 1.41E-12 |
| TTF2 | 2656.857746 | 0.601797127 | 1.52E-06 |
| RNF44 | 1765.055586 | 0.601428094 | 2.94E-06 |
| KLF11 | 805.9599835 | 0.601073753 | 3.80E-07 |
| FAHD2A | 1741.478709 | 0.600929935 | 1.74E-12 |
| ZXDC | 1312.034482 | 0.600789406 | 1.52E-11 |
| AC020898.1 | 58.20265113 | 0.600739395 | 4.55E-02 |
| MPST | 1990.715552 | 0.600545277 | 1.24E-07 |
| LEMD2 | 2746.266522 | 0.600520853 | 9.99E-17 |
| MRPS18B | 4057.860062 | 0.600383381 | 1.12E-10 |
| PCDHGA4 | 133.1061788 | 0.600376328 | 1.15E-02 |
| MAPKAPK5-AS1 | 536.4434689 | 0.600266564 | 1.02E-07 |
| RPS19 | 28332.18881 | 0.599298439 | 1.29E-08 |
| PPFIBP1 | 7522.570197 | 0.599174811 | 3.08E-17 |
| PDLIM7 | 2642.119675 | 0.59880831 | 8.05E-07 |
| RNF113A | 757.8717493 | 0.598371541 | 1.15E-07 |
| VAV2 | 2652.404696 | 0.598173881 | 1.39E-10 |
| GTPBP1 | 3068.954214 | 0.597831121 | 2.87E-10 |
| TXN | 6436.015601 | 0.597742013 | 2.59E-15 |
| SMIM10 | 1046.732466 | 0.59735381 | 3.27E-08 |
| CBARP | 284.9060585 | 0.596824751 | 4.62E-04 |
| BTBD11 | 910.1491859 | 0.596801889 | 2.65E-09 |
| TICAM1 | 498.5809167 | 0.596778743 | 1.94E-05 |
| SIX1 | 639.7445453 | 0.596531633 | 1.77E-08 |
| EPB41 | 2483.745836 | 0.595631096 | 7.46E-06 |
| CTNNA1 | 21483.2315 | 0.595613321 | 2.63E-19 |
| SDHAF4 | 156.402571 | 0.595577889 | 4.79E-04 |
| NOA1 | 1315.983886 | 0.595467919 | 2.36E-07 |
| SHKBP1 | 2696.583798 | 0.594987506 | 1.91E-08 |
| BBC3 | 539.5516664 | 0.594311556 | 6.28E-05 |
| PXN | 10227.78138 | 0.594015072 | 1.27E-07 |
| SLC2A8 | 513.2020919 | 0.593903796 | 1.63E-06 |
| ALDH1B1 | 2870.46866 | 0.593758071 | 2.33E-10 |
| IER5L | 924.1752821 | 0.593682972 | 1.48E-03 |
| AGPAT3 | 4177.175525 | 0.593244085 | 1.12E-19 |
| METTL21A | 721.6296818 | 0.593200912 | 2.67E-06 |

| | | | |
|---|---|---|---|
| SBDS | 6786.193804 | 0.593138252 | 5.33E-08 |
| LCMT1 | 1635.262159 | 0.593000211 | 4.58E-11 |
| SNHG17 | 1649.128822 | 0.592914515 | 6.56E-03 |
| NEMP2 | 430.5920123 | 0.592857036 | 5.37E-03 |
| AL357033.4 | 129.7848764 | 0.592442026 | 7.17E-03 |
| REPIN1 | 8966.942661 | 0.592292728 | 1.00E-15 |
| PPP2R5B | 1355.840148 | 0.592157373 | 1.08E-05 |
| RPLP0 | 75599.32004 | 0.591492703 | 2.37E-07 |
| PNP | 1512.768907 | 0.591397646 | 3.72E-08 |
| AL590617.2 | 125.1792277 | 0.591155813 | 8.34E-03 |
| YIF1B | 1626.377802 | 0.591001161 | 1.38E-09 |
| FSCN1 | 13820.42552 | 0.590950668 | 2.02E-05 |
| RFNG | 1336.381185 | 0.59077781 | 2.52E-05 |
| HILPDA | 944.6319639 | 0.590202546 | 1.04E-02 |
| ZNF593 | 490.6340416 | 0.590089931 | 7.89E-05 |
| TCP11L2 | 360.9121766 | 0.589900721 | 4.78E-03 |
| PRDM11 | 1207.530965 | 0.589886766 | 5.85E-05 |
| NA | 96.88619428 | 0.589803461 | 1.51E-02 |
| TROAP | 1436.648234 | 0.589723079 | 1.74E-04 |
| CREB3L1 | 742.9290274 | 0.589692211 | 1.51E-04 |
| C2orf49 | 1087.85998 | 0.58933199 | 8.45E-10 |
| AL445363.3 | 241.367527 | 0.589277838 | 1.21E-03 |
| HMG20B | 2798.243651 | 0.589271123 | 9.97E-08 |
| NARS | 10148.14967 | 0.588621565 | 1.03E-04 |
| NA | 141.9738255 | 0.587769056 | 3.98E-03 |
| EDEM1 | 2254.97478 | 0.587466838 | 3.58E-08 |
| C1orf226 | 396.125664 | 0.587132859 | 1.72E-04 |
| DNAJC15 | 2292.244546 | 0.586355506 | 3.82E-09 |
| EPB41L3 | 354.3759169 | 0.58587839 | 5.36E-03 |
| BCKDHA | 1382.500501 | 0.58563576 | 6.05E-04 |
| NUDC | 5364.378525 | 0.585579017 | 7.53E-07 |
| CDCA7 | 2462.20736 | 0.585397305 | 5.53E-04 |
| DHX35 | 1575.892245 | 0.585303782 | 3.43E-07 |
| POLRMT | 1932.426877 | 0.584696452 | 5.10E-07 |
| AC139256.2 | 279.1925693 | 0.584164732 | 3.81E-04 |
| DUSP9 | 394.385755 | 0.583836365 | 3.87E-03 |
| FAM210A | 907.4822041 | 0.583650304 | 8.49E-05 |
| TBC1D1 | 3064.045464 | 0.5834275 | 1.02E-08 |
| SLC2A13 | 99.18506297 | 0.583102612 | 2.19E-02 |
| ASPHD1 | 246.0080628 | 0.582983026 | 9.85E-06 |
| NSUN5 | 2016.221066 | 0.582972311 | 8.33E-11 |
| GMPS | 4155.036423 | 0.582695905 | 2.01E-13 |
| SUN2 | 5461.842658 | 0.582610212 | 2.62E-10 |
| PATL1 | 4544.71706 | 0.582331234 | 2.77E-23 |
| ALDH18A1 | 2818.954037 | 0.581914752 | 6.31E-06 |
| SPAG1 | 418.6973724 | 0.581316524 | 2.35E-04 |
| C8orf33 | 4642.341906 | 0.581276379 | 7.97E-11 |

| | | | |
|---|---|---|---|
| CHSY3 | 217.2220635 | 0.581260948 | 5.12E-03 |
| ATP11A | 1843.254605 | 0.58073024 | 1.38E-05 |
| ELP4 | 1116.525473 | 0.580488506 | 1.07E-09 |
| METTL22 | 1657.307153 | 0.580474813 | 1.03E-10 |
| PELI1 | 458.0342912 | 0.579941231 | 2.18E-04 |
| WDYHV1 | 642.0397183 | 0.579893923 | 1.06E-05 |
| ASRGL1 | 682.0832095 | 0.579430565 | 4.16E-04 |
| OLMALINC | 520.9366639 | 0.57898757 | 1.25E-06 |
| ZNF703 | 2016.919739 | 0.578372248 | 4.24E-07 |
| DNAH14 | 338.8482397 | 0.578178735 | 6.44E-03 |
| SRP14-AS1 | 113.3393621 | 0.578147698 | 4.16E-02 |
| PHB2 | 8408.780695 | 0.578128656 | 5.41E-11 |
| UQCRC1 | 6211.217267 | 0.57809125 | 1.89E-09 |
| PCBP2 | 21473.6083 | 0.577526352 | 1.79E-18 |
| GAGE2E | 1699.270948 | 0.576293921 | 3.53E-02 |
| TMEM42 | 1109.272036 | 0.574780677 | 5.92E-05 |
| AC141586.1 | 467.7885949 | 0.574276154 | 1.96E-04 |
| STK40 | 4424.176598 | 0.574188988 | 6.12E-15 |
| SETD3 | 2342.695916 | 0.574110748 | 1.02E-12 |
| NOC2L | 7230.373689 | 0.573700263 | 1.60E-15 |
| NDUFA4L2 | 114.6081155 | 0.572688312 | 2.99E-02 |
| STPG1 | 457.4197334 | 0.572363462 | 6.16E-05 |
| TMCC2 | 214.3735261 | 0.571906161 | 4.40E-03 |
| LRCH4 | 2232.681622 | 0.571640686 | 3.91E-06 |
| DDX19B | 2093.58396 | 0.571629381 | 2.57E-03 |
| MTFP1 | 587.7366352 | 0.57152596 | 1.42E-03 |
| AC122710.1 | 176.9889668 | 0.57121176 | 1.08E-03 |
| TSPAN14 | 2543.409951 | 0.571140199 | 1.50E-05 |
| TECPR1 | 2202.322973 | 0.57107264 | 5.50E-08 |
| NA | 158.6725789 | 0.570978809 | 1.34E-03 |
| ERO1A | 3104.479503 | 0.570780034 | 3.43E-04 |
| PSMB6 | 4576.834031 | 0.570730364 | 4.73E-13 |
| SOCS7 | 1710.926291 | 0.570694675 | 1.80E-12 |
| TFAP4 | 422.6089636 | 0.570611118 | 6.33E-04 |
| BNIP1 | 476.0777862 | 0.570344008 | 1.03E-07 |
| RNF181 | 2072.155686 | 0.569869557 | 1.32E-06 |
| PTBP3 | 2504.459584 | 0.569757336 | 1.28E-05 |
| FAM234B | 967.1611581 | 0.569690291 | 1.87E-08 |
| DSCR8 | 287.3163435 | 0.569430586 | 6.52E-04 |
| CLIC1 | 17383.67349 | 0.569384751 | 1.29E-07 |
| PLA2G16 | 1773.072902 | 0.569312802 | 1.03E-04 |
| DPH5 | 1379.944129 | 0.569025667 | 1.75E-06 |
| RPL18A | 25731.2536 | 0.568990031 | 1.24E-05 |
| TMBIM1 | 4490.409115 | 0.568734386 | 6.77E-07 |
| GID4 | 1136.933439 | 0.568638103 | 2.91E-08 |
| CETP | 1153.142862 | 0.568630261 | 2.80E-03 |
| NDUFC2 | 3541.592762 | 0.568314733 | 1.11E-14 |

| | | | |
|---|---|---|---|
| RPL8 | 63923.6203 | 0.568198095 | 4.10E-07 |
| SLC13A3 | 117.8644963 | 0.567786154 | 7.00E-03 |
| ZFAND3 | 4293.104697 | 0.567776518 | 1.14E-04 |
| OSBPL3 | 2070.819801 | 0.567726027 | 1.56E-05 |
| PQBP1 | 2394.096602 | 0.567692562 | 8.65E-08 |
| ANP32B | 12976.31036 | 0.56753427 | 2.32E-09 |
| FPGS | 3260.463423 | 0.567504099 | 5.61E-08 |
| HINT1 | 21385.80112 | 0.567458403 | 1.70E-09 |
| WDR45B | 3377.208925 | 0.567219385 | 3.38E-18 |
| AIFM2 | 829.2813547 | 0.567096822 | 3.20E-05 |
| SLC25A26 | 1028.574356 | 0.566830095 | 8.04E-07 |
| RNF220 | 2750.734764 | 0.566761394 | 2.07E-07 |
| LMNB2 | 11542.53977 | 0.566590098 | 2.21E-06 |
| CD33 | 476.0162093 | 0.566384302 | 3.88E-05 |
| RPS20 | 35647.71181 | 0.566131132 | 1.30E-06 |
| MORC4 | 2027.296062 | 0.566127911 | 3.34E-11 |
| MSN | 33120.58291 | 0.565748407 | 1.53E-10 |
| MIRLET7BHG | 367.6008969 | 0.565058731 | 1.28E-03 |
| EBF4 | 818.8381144 | 0.564565098 | 1.21E-03 |
| DLG3 | 615.1176337 | 0.563996521 | 3.32E-02 |
| AC022211.2 | 275.9392876 | 0.563792661 | 4.66E-05 |
| MAP3K10 | 677.0189669 | 0.563713389 | 1.05E-03 |
| HAUS2 | 847.5349321 | 0.563561509 | 1.45E-04 |
| PWWP2B | 214.222878 | 0.563083517 | 1.96E-04 |
| CASKIN2 | 1291.448053 | 0.562734924 | 8.89E-05 |
| FYCO1 | 2501.029106 | 0.562517051 | 1.17E-11 |
| ABCC1 | 5112.111329 | 0.562492559 | 1.13E-07 |
| IFT27 | 1262.90455 | 0.562179485 | 1.12E-05 |
| DNASE2 | 2894.136866 | 0.56194703 | 3.18E-04 |
| DCBLD1 | 748.5491359 | 0.561760056 | 6.98E-05 |
| STIP1 | 14572.5874 | 0.561492475 | 1.25E-12 |
| CPT1C | 384.7895272 | 0.561327845 | 1.86E-03 |
| PTBP1 | 14317.09673 | 0.561296758 | 1.08E-08 |
| KMT2B | 2506.320285 | 0.561132899 | 5.11E-06 |
| PLPBP | 2627.182497 | 0.560820203 | 7.65E-07 |
| BUD23 | 4063.891293 | 0.560380682 | 2.51E-20 |
| UBC | 59339.49899 | 0.560369969 | 2.18E-07 |
| RELL2 | 675.3055991 | 0.560038079 | 3.44E-03 |
| RPL38 | 12269.32601 | 0.560018327 | 3.05E-05 |
| TUBE1 | 792.6081694 | 0.560008792 | 3.02E-02 |
| SF3A2 | 2191.908115 | 0.559921546 | 3.45E-06 |
| OGDH | 7751.108466 | 0.558496576 | 1.63E-06 |
| AL691420.1 | 230.2834602 | 0.558465697 | 6.42E-05 |
| MAPKAPK2 | 3962.376668 | 0.558211802 | 9.61E-11 |
| POTEF | 330.2910566 | 0.558211344 | 4.05E-03 |
| DGKG | 127.0190231 | 0.558013565 | 1.47E-02 |
| TRADD | 922.3822232 | 0.557673334 | 9.32E-06 |

| | | | |
|---|---|---|---|
| PLIN3 | 3764.213717 | 0.557324663 | 7.26E-03 |
| BCAR3 | 3028.458632 | 0.557082249 | 6.90E-06 |
| AC027307.2 | 280.7966597 | 0.556717352 | 1.21E-04 |
| OSER1 | 2909.229889 | 0.556422255 | 2.94E-12 |
| VDAC1 | 14777.17519 | 0.55621746 | 4.80E-07 |
| SLC25A5 | 11977.73219 | 0.555952378 | 7.92E-04 |
| NR2F2 | 5643.974102 | 0.555867781 | 2.88E-06 |
| TRMT44 | 441.8763599 | 0.555513195 | 1.39E-04 |
| CEP85 | 1897.607299 | 0.555306071 | 2.49E-07 |
| MTLN | 469.6545991 | 0.555235325 | 5.78E-05 |
| BTG3 | 724.4575775 | 0.555225973 | 3.20E-05 |
| GCLM | 1084.260776 | 0.554896806 | 1.70E-03 |
| PNMA6A | 795.9455409 | 0.554806836 | 2.68E-04 |
| FAHD2B | 1473.812251 | 0.554723625 | 1.28E-12 |
| ZNF33B | 606.8960617 | 0.5547088 | 1.13E-02 |
| MEF2D | 1707.54484 | 0.554538262 | 6.43E-07 |
| SLC35F2 | 1705.247956 | 0.554529705 | 1.52E-06 |
| TRIP10 | 3342.101072 | 0.553792199 | 1.47E-19 |
| SLC35D2 | 1250.038697 | 0.55339666 | 1.26E-07 |
| FAM107B | 1775.665814 | 0.553330671 | 1.75E-03 |
| RPL3 | 72185.34694 | 0.553252918 | 6.33E-06 |
| TMEM39A | 1461.701079 | 0.553214521 | 7.92E-07 |
| FKBP1B | 606.1349468 | 0.553148677 | 7.92E-04 |
| PXK | 1077.365537 | 0.553098853 | 5.03E-04 |
| GMFG | 271.7051033 | 0.552847131 | 1.27E-02 |
| CASP4 | 3032.86953 | 0.552394729 | 1.23E-11 |
| IQGAP1 | 16665.12339 | 0.552256467 | 1.34E-07 |
| RPS6KA4 | 2634.070272 | 0.552208781 | 1.49E-08 |
| PARVB | 2787.345432 | 0.552077753 | 1.67E-06 |
| JADE1 | 1188.702036 | 0.55190921 | 3.20E-03 |
| MOK | 1296.653583 | 0.551883577 | 9.32E-08 |
| SKP2 | 5169.048372 | 0.551584415 | 1.72E-02 |
| NDUFA7 | 788.1011822 | 0.551460224 | 4.14E-03 |
| MVD | 2075.489569 | 0.551450802 | 2.96E-03 |
| CCDC86 | 3009.142193 | 0.551225884 | 1.65E-06 |
| FMOD | 1627.644591 | 0.551134991 | 5.06E-03 |
| MYZAP | 230.2645852 | 0.551088822 | 4.44E-03 |
| GAPDH | 113084.8488 | 0.551068039 | 1.94E-04 |
| PSRC1 | 2696.465216 | 0.550712804 | 2.49E-09 |
| FAH | 2365.929671 | 0.550364239 | 1.00E-10 |
| SFXN3 | 2024.371708 | 0.55035285 | 3.16E-05 |
| TRAPPC3 | 2005.937891 | 0.550343388 | 1.49E-10 |
| LYPLAL1 | 519.1981249 | 0.550239315 | 8.15E-05 |
| ILVBL | 2135.803995 | 0.550184976 | 2.34E-08 |
| SLC43A3 | 6224.28028 | 0.550032768 | 6.94E-13 |
| ZNF419 | 808.3470474 | 0.55002306 | 7.52E-05 |
| SPIRE1 | 1623.898975 | 0.549914011 | 4.40E-04 |

| | | | |
|---|---|---|---|
| STK32B | 1361.46195 | 0.549886092 | 4.29E-05 |
| MATN3 | 531.0146682 | 0.549466756 | 6.54E-04 |
| FSTL1 | 34023.92307 | 0.549081021 | 7.47E-07 |
| ACVR2B | 614.6329055 | 0.549064666 | 1.79E-05 |
| PMP22 | 6670.421425 | 0.548926083 | 7.93E-06 |
| FOXD1 | 1076.126703 | 0.548590793 | 1.25E-06 |
| PHYHIP | 101.0500315 | 0.548497832 | 3.91E-02 |
| RPL14 | 17200.18652 | 0.547911676 | 2.12E-09 |
| KIAA0930 | 2835.093006 | 0.547904596 | 1.24E-06 |
| PKP4 | 3485.234126 | 0.547866326 | 3.92E-06 |
| UBE2I | 4881.546543 | 0.54768286 | 1.45E-11 |
| SH2D3A | 372.6600886 | 0.547271833 | 6.94E-05 |
| RPL27 | 24671.76414 | 0.546051215 | 1.86E-06 |
| SLC38A1 | 10325.87367 | 0.545618829 | 3.04E-03 |
| ATP6V1E1 | 4139.268406 | 0.545399959 | 9.48E-16 |
| AC019080.1 | 129.0291347 | 0.545275843 | 3.71E-03 |
| MZT2A | 1203.994707 | 0.545176703 | 6.90E-05 |
| WDR31 | 314.5976903 | 0.544933153 | 4.52E-04 |
| PKMYT1 | 1390.690032 | 0.544413641 | 1.26E-02 |
| GMPPA | 1577.592379 | 0.544165243 | 3.23E-07 |
| SNHG29 | 16413.21366 | 0.543771289 | 5.13E-08 |
| STARD13 | 1275.002019 | 0.543765483 | 4.41E-04 |
| KDELR3 | 2158.686813 | 0.543585365 | 1.08E-03 |
| LCAT | 227.905104 | 0.543438014 | 4.65E-03 |
| ALG9 | 1298.005813 | 0.543381085 | 9.31E-08 |
| TMEM81 | 102.0745934 | 0.543184173 | 1.31E-02 |
| SIGIRR | 424.1308521 | 0.542689495 | 1.57E-04 |
| HLA-DQB1 | 303.1547478 | 0.54261456 | 3.22E-04 |
| ZFPL1 | 1177.567137 | 0.542315935 | 5.20E-06 |
| CSPG4 | 4172.53723 | 0.542240075 | 7.79E-05 |
| DTNBP1 | 753.5644288 | 0.542237214 | 7.85E-06 |
| CCDC61 | 301.5063114 | 0.541948614 | 1.27E-05 |
| NXT1 | 1469.60535 | 0.541893891 | 1.01E-08 |
| SP6 | 120.9574354 | 0.541321872 | 4.84E-02 |
| BEND7 | 460.5163483 | 0.541248317 | 9.36E-05 |
| MRPS30 | 2531.712345 | 0.540478806 | 3.66E-17 |
| UBE2T | 1400.850171 | 0.540245369 | 1.44E-02 |
| BMP8B | 199.0507767 | 0.540089495 | 2.58E-02 |
| SLC22A3 | 402.2075182 | 0.539772004 | 2.19E-05 |
| MMS19 | 2806.241532 | 0.53956826 | 7.63E-08 |
| PITPNM1 | 3508.712923 | 0.539338903 | 4.46E-06 |
| CEP170B | 2094.005391 | 0.539022706 | 6.98E-12 |
| CEMIP | 1148.669696 | 0.538783808 | 1.76E-02 |
| TBC1D16 | 6642.917329 | 0.538666561 | 1.36E-08 |
| AC008403.1 | 202.196078 | 0.538029632 | 1.53E-03 |
| GATAD2A | 4101.766037 | 0.537691313 | 1.94E-08 |
| METTL26 | 1644.085805 | 0.537552295 | 2.31E-06 |

| | | | |
|---|---|---|---|
| DAP3 | 4392.491215 | 0.537018308 | 2.43E-20 |
| CDCA8 | 3450.166152 | 0.536849698 | 7.49E-07 |
| PHLDA3 | 421.1813707 | 0.536738548 | 6.49E-03 |
| PSMD3 | 5874.943555 | 0.536584281 | 1.77E-10 |
| MRPS5 | 2726.626492 | 0.536577032 | 1.98E-23 |
| COPA | 14477.07068 | 0.536429742 | 3.50E-14 |
| CLPP | 1685.642619 | 0.536314426 | 5.11E-07 |
| HELZ2 | 4389.311923 | 0.536235674 | 2.06E-03 |
| DDX47 | 4767.009017 | 0.53574626 | 1.64E-13 |
| INPP5K | 1873.425128 | 0.53565852 | 8.89E-06 |
| RGS3 | 2410.104891 | 0.535435814 | 3.55E-05 |
| MED10 | 2839.473397 | 0.535305829 | 6.57E-05 |
| CD151 | 16569.59205 | 0.53526927 | 6.82E-05 |
| RDM1 | 181.5823242 | 0.535054368 | 1.63E-03 |
| RPL18 | 22761.13749 | 0.535017921 | 2.52E-05 |
| TRIB1 | 1261.800018 | 0.534730508 | 2.83E-03 |
| EEF1B2 | 14457.85453 | 0.534682742 | 2.68E-07 |
| GOLGA2 | 4842.364652 | 0.534305443 | 1.21E-17 |
| RPS15A | 28078.05717 | 0.533550632 | 9.74E-08 |
| PLK1 | 5435.480583 | 0.533048662 | 1.32E-04 |
| ICAM1 | 10019.94945 | 0.532852876 | 1.15E-03 |
| ATG7 | 1515.301373 | 0.532698704 | 1.61E-06 |
| TMEM147 | 1948.56684 | 0.532678592 | 1.64E-07 |
| SGF29 | 499.749353 | 0.53245468 | 7.80E-05 |
| TUBG1 | 3738.175157 | 0.532391159 | 2.97E-06 |
| CAMKK2 | 2052.006897 | 0.532340128 | 1.48E-11 |
| HLA-B | 7949.216215 | 0.532338777 | 1.50E-06 |
| CHCHD2 | 15154.40285 | 0.531606403 | 5.78E-07 |
| AZIN2 | 391.0001786 | 0.531373249 | 1.59E-02 |
| PRR14 | 1269.197719 | 0.531323346 | 6.70E-06 |
| INSIG2 | 868.8112722 | 0.531119856 | 1.11E-07 |
| PFKFB3 | 2049.457462 | 0.531106459 | 7.62E-06 |
| GARNL3 | 363.9609093 | 0.530998883 | 1.08E-04 |
| B4GALT1 | 11172.21506 | 0.530865128 | 3.83E-13 |
| EEF1D | 14301.58075 | 0.530658589 | 1.12E-07 |
| RBM42 | 2701.957043 | 0.530497161 | 4.96E-08 |
| SLC25A24 | 1865.58441 | 0.530436943 | 4.92E-03 |
| SLC35F6 | 2445.04949 | 0.530255979 | 1.18E-02 |
| RNPEP | 3309.936809 | 0.530136283 | 1.21E-04 |
| GYG2 | 1137.477831 | 0.530002332 | 1.39E-04 |
| MED30 | 472.5295775 | 0.529970295 | 2.87E-07 |
| RPL37A | 41742.41886 | 0.528986454 | 3.58E-05 |
| RPS2P5 | 8753.092583 | 0.528754342 | 8.81E-06 |
| CRTAP | 9964.199093 | 0.528732511 | 2.35E-06 |
| TRAF2 | 1387.830748 | 0.528705776 | 9.63E-05 |
| PUDP | 1113.873636 | 0.528612508 | 4.62E-06 |
| TXNDC11 | 2341.326044 | 0.528556286 | 2.62E-08 |

| | | | |
|---|---|---|---|
| BRD4 | 3830.430528 | 0.528531666 | 2.24E-04 |
| XPC | 3166.78906 | 0.528414589 | 5.24E-14 |
| FBL | 5586.954134 | 0.528403465 | 2.94E-07 |
| EEF1A1P6 | 301.9815671 | 0.528367702 | 1.46E-02 |
| SEC61G | 3651.15804 | 0.528331541 | 3.01E-06 |
| PROSER1 | 1189.283984 | 0.528232748 | 4.20E-03 |
| RTN3 | 7686.045019 | 0.528182508 | 5.64E-09 |
| SEMA4D | 1038.790482 | 0.528169301 | 1.06E-03 |
| CTSL | 7027.922032 | 0.528145419 | 3.17E-22 |
| FOXO1 | 731.2693503 | 0.528136517 | 1.46E-06 |
| ETNK2 | 2028.091551 | 0.528068025 | 2.47E-06 |
| LAMA5 | 8544.457214 | 0.528062636 | 2.93E-04 |
| PRR5L | 1135.682346 | 0.527454801 | 8.09E-06 |
| GALNT14 | 1213.5074 | 0.527414987 | 8.05E-06 |
| WDR54 | 1659.603232 | 0.526786211 | 2.93E-03 |
| HSF2BP | 157.1567785 | 0.526444684 | 4.09E-03 |
| MPRIP | 18080.17353 | 0.52623534 | 3.15E-08 |
| NMT1 | 6026.448678 | 0.526125845 | 1.69E-04 |
| RPS2 | 74595.74233 | 0.525795665 | 1.13E-05 |
| TGIF1 | 1706.310532 | 0.525753063 | 8.20E-03 |
| TNFAIP1 | 4428.834655 | 0.525380196 | 1.31E-06 |
| FAM221A | 241.7934268 | 0.525127587 | 4.39E-03 |
| FAM50B | 89.10699054 | 0.524823229 | 2.52E-02 |
| GRAMD2B | 728.6384239 | 0.524766056 | 6.83E-03 |
| NOTCH4 | 160.9558428 | 0.524412835 | 7.74E-03 |
| NECTIN2 | 5365.872754 | 0.523613706 | 8.46E-06 |
| PSMD9 | 1835.168441 | 0.523347647 | 1.66E-12 |
| ZDHHC24 | 651.9338299 | 0.523172674 | 7.54E-06 |
| CTSH | 1717.103531 | 0.523022774 | 7.16E-04 |
| MRPL45P2 | 92.66883192 | 0.522106079 | 3.00E-02 |
| OXNAD1 | 604.3305565 | 0.521571721 | 1.46E-04 |
| INPP5A | 820.5608163 | 0.521457559 | 4.75E-04 |
| ITPRIP | 1779.823694 | 0.521416137 | 7.87E-05 |
| HIST1H2BK | 512.5589084 | 0.521253572 | 4.80E-03 |
| DIAPH2 | 574.5007948 | 0.521102271 | 7.53E-05 |
| PSMD8 | 6091.348911 | 0.521038376 | 5.36E-08 |
| FAM72B | 387.5188204 | 0.520954061 | 1.54E-02 |
| KPNA2 | 10845.5826 | 0.520912064 | 2.86E-03 |
| CDKN2D | 386.7352033 | 0.520853088 | 6.76E-04 |
| FXR2 | 2852.945115 | 0.520710898 | 5.28E-11 |
| BOD1 | 2630.333907 | 0.520622563 | 3.74E-06 |
| MZT2B | 1646.035276 | 0.52055253 | 1.07E-03 |
| BUB1 | 3783.438651 | 0.519843853 | 7.37E-04 |
| EXOSC5 | 1246.924041 | 0.519764945 | 2.60E-04 |
| RPL22L1 | 2206.738069 | 0.519468776 | 1.91E-05 |
| NBPF9 | 4785.546614 | 0.519458153 | 3.66E-13 |
| KMT2E-AS1 | 139.9163431 | 0.519457561 | 2.57E-03 |

| | | | |
|---|---|---|---|
| NFKB1 | 2198.220305 | 0.519375217 | 3.88E-10 |
| CALCOCO2 | 4846.363505 | 0.519244778 | 1.80E-12 |
| ZNF385A | 632.1157056 | 0.519190488 | 6.94E-05 |
| RPS10 | 23450.91986 | 0.519120908 | 1.37E-05 |
| AC011462.1 | 3328.314682 | 0.519109434 | 1.92E-07 |
| SLC25A43 | 1114.800884 | 0.518653183 | 7.76E-08 |
| MRPS15 | 2177.285808 | 0.518633237 | 5.74E-10 |
| ZBED4 | 1607.762094 | 0.518587125 | 2.11E-12 |
| FLYWCH1 | 2934.327451 | 0.518450964 | 2.78E-08 |
| DRAM2 | 825.8159279 | 0.518387099 | 1.29E-04 |
| YARS2 | 1071.129074 | 0.518318391 | 8.05E-06 |
| BHMT2 | 687.4043628 | 0.518121918 | 5.01E-03 |
| HIST1H4I | 441.8190474 | 0.517773898 | 1.77E-02 |
| CISH | 106.4716427 | 0.517690526 | 1.70E-02 |
| ARF4 | 9639.726706 | 0.516890481 | 7.47E-06 |
| ANXA7 | 2952.787176 | 0.516562412 | 7.04E-12 |
| TRIM14 | 2948.397171 | 0.516305807 | 1.12E-09 |
| PPIF | 3038.649213 | 0.516074663 | 4.90E-07 |
| TTBK2 | 1066.932728 | 0.516006319 | 2.42E-02 |
| OLA1 | 3800.835144 | 0.515787771 | 3.37E-17 |
| PA2G4 | 8850.309135 | 0.515727419 | 3.02E-14 |
| UBE2R2 | 4405.95647 | 0.515623987 | 8.78E-11 |
| MRAS | 1665.819629 | 0.515009454 | 8.23E-06 |
| AK4 | 1671.133532 | 0.514954198 | 1.09E-02 |
| ZNF628 | 188.4749801 | 0.514649001 | 1.04E-02 |
| CARM1 | 4180.03102 | 0.514630322 | 4.42E-09 |
| CCDC80 | 5060.118978 | 0.514473719 | 2.44E-05 |
| UBE2J1 | 3574.235135 | 0.514437 | 5.32E-11 |
| OCEL1 | 585.363893 | 0.514083475 | 8.03E-03 |
| AC005586.1 | 92.41806109 | 0.513998537 | 2.69E-02 |
| TSPAN17 | 2270.832624 | 0.513790087 | 3.25E-06 |
| MSL3P1 | 461.1279657 | 0.513693475 | 5.72E-03 |
| ECM1 | 273.6904325 | 0.513063634 | 3.04E-02 |
| POLG | 4710.124758 | 0.513052948 | 6.38E-06 |
| PACSIN3 | 1919.961239 | 0.512944637 | 2.08E-08 |
| ABHD4 | 4610.109295 | 0.512722759 | 9.27E-05 |
| EDEM2 | 2837.570935 | 0.512686722 | 1.15E-04 |
| PCSK6 | 1048.029103 | 0.512388027 | 8.35E-09 |
| GPR3 | 101.6903154 | 0.511962232 | 4.73E-02 |
| NA | 504.5175673 | 0.511726246 | 6.41E-05 |
| DIPK1B | 1341.473757 | 0.511723013 | 7.29E-04 |
| DOCK5 | 2093.893287 | 0.511548921 | 2.23E-03 |
| LRFN4 | 1416.02701 | 0.511447719 | 7.24E-05 |
| LINC00957 | 360.1825963 | 0.511257374 | 9.61E-03 |
| LIG3 | 2382.176734 | 0.511050184 | 2.27E-07 |
| FAM222B | 821.712738 | 0.510904265 | 6.95E-09 |
| LRRC17 | 2568.192514 | 0.509930383 | 7.78E-03 |

| | | | |
|---|---|---|---|
| BOLA2-SMG1P6 | 948.6690809 | 0.509848278 | 4.63E-04 |
| IQSEC1 | 2478.420337 | 0.509755998 | 4.50E-10 |
| AP2B1 | 20918.25531 | 0.509575537 | 8.62E-10 |
| EEF1A1P5 | 932.495451 | 0.509472125 | 1.47E-02 |
| SPCS3 | 2551.177372 | 0.509228441 | 1.00E-05 |
| C2CD2L | 1120.873168 | 0.509199337 | 2.45E-06 |
| AHCY | 11764.42684 | 0.509050884 | 6.63E-07 |
| MAST4 | 393.8249765 | 0.508805795 | 3.43E-02 |
| IKBKG | 2243.828522 | 0.508390712 | 2.42E-10 |
| PTRH1 | 373.7101057 | 0.507773967 | 2.71E-03 |
| PFKFB2 | 1199.134629 | 0.507670701 | 9.84E-05 |
| RPL35 | 27266.11572 | 0.507376987 | 9.43E-06 |
| GSPT1 | 9602.68302 | 0.507249889 | 7.23E-19 |
| STK26 | 523.7241541 | 0.507168229 | 6.30E-03 |
| LZTS1 | 3910.639426 | 0.506952461 | 1.86E-05 |
| RHOB | 1671.093319 | 0.506938 | 9.11E-03 |
| MRPL9 | 1883.802852 | 0.506921128 | 5.87E-04 |
| ARNTL2 | 885.1172776 | 0.506828104 | 3.71E-04 |
| AL356056.2 | 88.00872493 | 0.506784593 | 2.54E-02 |
| TYRO3 | 2229.388693 | 0.506537197 | 6.24E-05 |
| MAP2K3 | 1746.737502 | 0.506043571 | 1.11E-02 |
| CWF19L1 | 851.5140741 | 0.506040408 | 2.13E-05 |
| CSNK1E | 5056.524669 | 0.506011721 | 1.60E-14 |
| GSTM3 | 4602.829408 | 0.504977565 | 1.47E-06 |
| AC004556.1 | 758.4552135 | 0.504961292 | 3.31E-03 |
| FOXP4 | 1453.076162 | 0.504914591 | 1.88E-05 |
| TESK1 | 1077.548336 | 0.504900195 | 2.39E-05 |
| RP9 | 827.2961611 | 0.504613773 | 6.08E-05 |
| SART1 | 4923.107164 | 0.504395428 | 4.54E-09 |
| AP3D1 | 8143.720888 | 0.50429421 | 1.43E-13 |
| ADK | 1521.24063 | 0.504280838 | 3.62E-08 |
| IMMP2L | 342.1216284 | 0.504159096 | 5.71E-03 |
| CEP250 | 4914.904169 | 0.504124569 | 5.60E-10 |
| SREBF2 | 12774.93631 | 0.50398113 | 1.06E-03 |
| ADAMTSL5 | 346.9843244 | 0.503925643 | 1.17E-02 |
| REEP4 | 1433.911046 | 0.503910408 | 3.51E-08 |
| SLC22A23 | 249.0103972 | 0.503856721 | 2.23E-02 |
| RHBDF1 | 1735.657285 | 0.503638358 | 2.00E-05 |
| GTF3A | 2319.291999 | 0.503620841 | 2.70E-13 |
| RABAC1 | 2206.457957 | 0.50358929 | 9.06E-04 |
| MRPL33 | 1615.604336 | 0.503416966 | 5.15E-04 |
| AL513165.1 | 119.8942597 | 0.502990452 | 3.38E-02 |
| NOC4L | 875.5322397 | 0.502700069 | 1.45E-08 |
| NCAPD2 | 11336.63226 | 0.502692099 | 1.88E-09 |
| ARHGAP10 | 734.6739707 | 0.502410561 | 2.70E-08 |
| LMO4 | 1106.20488 | 0.502257717 | 3.53E-03 |
| WDR4 | 1435.970238 | 0.502161116 | 1.89E-05 |

| | | | |
|---|---|---|---|
| KLF7 | 1229.558498 | 0.502075526 | 3.77E-04 |
| ATL3 | 3470.507866 | 0.501909218 | 9.72E-13 |
| GORASP2 | 3964.523769 | 0.501803443 | 2.45E-12 |
| SNRPA1 | 1845.883866 | 0.501628979 | 2.95E-09 |
| LGALS1 | 23005.81072 | 0.50157379 | 2.81E-04 |
| KCTD10 | 3551.12457 | 0.501532068 | 3.18E-05 |
| PIGBOS1 | 339.9707877 | 0.501064477 | 4.02E-03 |
| MDH2 | 11598.52765 | 0.500732163 | 6.75E-08 |
| AC093525.7 | 393.9993354 | 0.500681234 | 7.56E-05 |
| TPM2 | 11840.72667 | 0.500561925 | 1.30E-04 |
| PNPLA2 | 3504.906473 | 0.500400456 | 9.86E-04 |
| MYO18A | 7255.058026 | 0.500341605 | 1.27E-04 |
| STK39 | 1510.725526 | 0.499992764 | 1.78E-06 |
| GNB1 | 13180.59273 | 0.49960905 | 7.59E-15 |
| AC008443.1 | 461.6091636 | 0.499300741 | 3.01E-04 |
| SIRPA | 3488.218336 | 0.49904222 | 3.24E-04 |
| NPBWR1 | 390.8449915 | 0.498998354 | 2.12E-02 |
| TPD52L2 | 13159.04135 | 0.498981143 | 7.26E-06 |
| SIAH2 | 1088.269525 | 0.498964078 | 5.99E-06 |
| AL928654.4 | 120.2726713 | 0.498780792 | 2.43E-02 |
| FAU | 10785.67298 | 0.498648116 | 5.85E-05 |
| HAUS7 | 1980.774412 | 0.498574149 | 6.95E-06 |
| RPL5 | 35286.27166 | 0.498550629 | 1.46E-08 |
| GAB2 | 601.0589633 | 0.49823794 | 1.94E-02 |
| RPS7 | 42243.97316 | 0.497782989 | 1.37E-10 |
| CKAP4 | 11320.18536 | 0.497707544 | 1.33E-04 |
| BOLA2 | 1180.562019 | 0.497683521 | 5.17E-05 |
| MORN4 | 623.2349339 | 0.497613739 | 1.72E-06 |
| LCORL | 400.3564959 | 0.497521107 | 2.27E-02 |
| HSP90AB1 | 76450.15073 | 0.497431532 | 2.62E-08 |
| AIG1 | 902.4128692 | 0.497287562 | 3.63E-06 |
| ZDHHC18 | 1131.262007 | 0.49704486 | 1.05E-04 |
| ARHGDIA | 10479.86553 | 0.497040187 | 2.05E-04 |
| TIPARP | 910.2975068 | 0.496931602 | 6.80E-04 |
| RIPK2 | 724.4734466 | 0.496890625 | 4.56E-04 |
| CDC37 | 9701.163091 | 0.496664211 | 3.64E-10 |
| HMGCL | 1754.137431 | 0.496500583 | 1.45E-04 |
| RACK1 | 52295.32079 | 0.496484334 | 6.24E-06 |
| DDX59 | 684.8136277 | 0.496226087 | 1.37E-04 |
| MARS2 | 475.8319689 | 0.495950439 | 1.83E-05 |
| POLR1D | 2138.160546 | 0.495919161 | 1.86E-04 |
| EXOC2 | 2251.914411 | 0.495788779 | 6.11E-10 |
| EEF1A1 | 626427.2418 | 0.49526086 | 9.40E-09 |
| TRIM35 | 1131.368175 | 0.495250539 | 4.87E-06 |
| ADAMTS12 | 10377.03832 | 0.495158983 | 3.31E-06 |
| OSER1-DT | 334.8931331 | 0.494907041 | 2.14E-03 |
| DYRK4 | 644.2354557 | 0.494868985 | 1.10E-07 |

| | | | |
|---|---|---|---|
| ATF6 | 3656.141178 | 0.49474536 | 2.28E-08 |
| FAM200B | 455.2856727 | 0.494407501 | 3.59E-05 |
| AKAP8L | 2144.844662 | 0.494234802 | 1.42E-09 |
| QPCT | 409.1069256 | 0.49416696 | 2.23E-03 |
| GLRX3 | 1925.851693 | 0.493640378 | 1.58E-09 |
| MAP2K2 | 4092.995995 | 0.493580555 | 8.25E-07 |
| PPP2R3B | 555.037031 | 0.49351922 | 7.98E-05 |
| PKIG | 2587.098172 | 0.49320366 | 1.48E-03 |
| EIF3L | 19700.69514 | 0.492567984 | 2.56E-05 |
| MCFD2 | 14679.18902 | 0.492557911 | 1.69E-16 |
| ZNF496 | 1806.436781 | 0.49251515 | 5.87E-10 |
| PTPDC1 | 927.0040926 | 0.492454479 | 1.50E-02 |
| LRRK1 | 2208.045532 | 0.492449916 | 6.96E-07 |
| BTF3L4P2 | 254.3305076 | 0.492294154 | 1.78E-02 |
| REXO1 | 986.0877504 | 0.492259252 | 6.55E-10 |
| MRPL23 | 883.1208097 | 0.492235721 | 4.53E-03 |
| PLRG1 | 3630.724933 | 0.492031838 | 9.06E-11 |
| SNAP23 | 1810.39237 | 0.491855974 | 1.12E-07 |
| FOXK2 | 3237.994456 | 0.491593503 | 1.19E-04 |
| NME6 | 613.6357016 | 0.491526591 | 6.81E-06 |
| SLC41A3 | 2189.823322 | 0.491418996 | 5.31E-05 |
| IDS | 4286.442473 | 0.49122328 | 1.56E-07 |
| GHITM | 5118.84873 | 0.49118464 | 1.91E-15 |
| ALKBH3 | 1188.819224 | 0.490916875 | 9.76E-07 |
| EPS15L1 | 1774.911684 | 0.490236308 | 6.19E-10 |
| TTC9C | 1503.409275 | 0.490184545 | 2.94E-06 |
| CHKA | 2021.673855 | 0.490040432 | 3.35E-07 |
| AMZ2 | 2889.472315 | 0.489989517 | 8.77E-19 |
| PEMT | 720.8887767 | 0.489922943 | 1.61E-04 |
| TMEM79 | 312.6854783 | 0.489828031 | 9.98E-03 |
| KXD1 | 2873.498203 | 0.489551658 | 1.02E-06 |
| CORO7 | 454.3518415 | 0.48923778 | 5.16E-04 |
| PPIE | 2964.366664 | 0.489052233 | 1.85E-06 |
| KATNAL2 | 328.0324392 | 0.489006757 | 2.47E-02 |
| ATP6V1B2 | 7378.659165 | 0.488947075 | 4.84E-08 |
| MT-TP | 942.5677673 | 0.488764119 | 2.10E-02 |
| DGKZ | 4303.453503 | 0.488702787 | 4.14E-09 |
| VAC14 | 3528.466868 | 0.488544256 | 7.74E-06 |
| EIF4A1 | 39066.0312 | 0.488474059 | 2.09E-15 |
| STMN1 | 17626.57597 | 0.488385848 | 2.34E-06 |
| SAP30BP | 2342.287681 | 0.487963879 | 2.64E-08 |
| CTU2 | 1166.626706 | 0.487781509 | 1.86E-06 |
| AC068473.5 | 236.2554259 | 0.487747197 | 7.61E-05 |
| AL133372.2 | 107.2268163 | 0.487516925 | 2.07E-02 |
| CCT7 | 18359.8704 | 0.487455245 | 8.29E-11 |
| CHCHD7 | 1011.119218 | 0.487399523 | 8.94E-05 |
| PTGS1 | 2837.117892 | 0.487222191 | 2.28E-02 |

| | | | |
|---|---|---|---|
| STX18-AS1 | 165.4843433 | 0.487067457 | 1.22E-02 |
| ACAP1 | 278.1116252 | 0.486737424 | 3.85E-02 |
| CLK3 | 3618.729538 | 0.486503194 | 1.17E-06 |
| ME1 | 734.9613871 | 0.486478079 | 1.40E-03 |
| RNASEH2A | 2104.874415 | 0.486137836 | 8.76E-04 |
| DNPEP | 2813.70088 | 0.486103854 | 1.55E-06 |
| WDR74 | 2352.357433 | 0.485412457 | 3.86E-05 |
| UBTD1 | 536.0145794 | 0.485155195 | 1.05E-03 |
| TENT5C | 137.8763368 | 0.485088233 | 3.74E-02 |
| SH3BP2 | 2645.32467 | 0.484758086 | 1.36E-05 |
| RPS13 | 31513.02511 | 0.484746447 | 3.94E-05 |
| DKK3 | 22309.34202 | 0.484604396 | 1.83E-04 |
| CLTB | 1764.884731 | 0.484444141 | 2.10E-06 |
| MICALL2 | 1339.550355 | 0.484398191 | 2.54E-02 |
| LFNG | 282.2624472 | 0.484334013 | 8.06E-03 |
| PAFAH1B1 | 7948.894797 | 0.484314084 | 8.53E-13 |
| ZNF665 | 165.7284233 | 0.484294588 | 5.28E-03 |
| TTLL11 | 281.7170078 | 0.484221429 | 9.40E-04 |
| SERPINB6 | 5673.984218 | 0.4842059 | 1.85E-06 |
| TMEM43 | 7214.222135 | 0.484074264 | 1.44E-11 |
| CIDECP1 | 295.2909922 | 0.483894864 | 1.33E-02 |
| TNS1 | 3945.928745 | 0.483789186 | 1.61E-03 |
| FAF1 | 2836.781069 | 0.483633453 | 4.34E-09 |
| KIRREL1 | 10728.67019 | 0.483405545 | 7.94E-07 |
| SF3B2 | 19110.27107 | 0.483378674 | 3.86E-09 |
| ZNF524 | 171.9973563 | 0.483377651 | 3.21E-03 |
| TFE3 | 7339.395033 | 0.48326365 | 3.75E-07 |
| PSMG4 | 858.7223993 | 0.483258329 | 1.95E-06 |
| TAZ | 1646.042878 | 0.483226205 | 7.78E-06 |
| ARRDC1 | 1211.294091 | 0.483127729 | 3.88E-04 |
| SAPCD2 | 1918.011644 | 0.483033957 | 1.75E-05 |
| HDAC6 | 2892.832686 | 0.483029057 | 6.28E-05 |
| SMIM29 | 831.6872953 | 0.482977858 | 1.99E-03 |
| PRRG1 | 823.3572282 | 0.482652684 | 2.69E-04 |
| RHOA | 15149.00168 | 0.482486757 | 8.31E-06 |
| SYNGR2 | 1829.502754 | 0.48234664 | 1.23E-04 |
| YIPF1 | 1067.199585 | 0.482271958 | 9.92E-06 |
| EXOSC7 | 1415.908928 | 0.482100827 | 1.73E-05 |
| DLL3 | 453.7860412 | 0.481580788 | 8.30E-03 |
| KCTD5 | 3210.37524 | 0.480905367 | 7.48E-07 |
| CERS2 | 7121.847108 | 0.48089309 | 1.30E-07 |
| SH3GLB2 | 3523.835095 | 0.480743609 | 1.13E-08 |
| TP53INP2 | 2226.784429 | 0.480626913 | 1.86E-03 |
| DMTN | 300.3915311 | 0.480552679 | 4.09E-02 |
| ZNRF3 | 595.9671143 | 0.480443055 | 1.25E-05 |
| ACBD6 | 1499.904214 | 0.480313205 | 5.80E-06 |
| PRMT1 | 7046.38091 | 0.4797801 | 5.13E-06 |

| | | | |
|---|---|---|---|
| NA | 208.923837 | 0.479649746 | 4.47E-02 |
| TIMM50 | 2568.673392 | 0.479433704 | 9.65E-10 |
| VAMP2 | 1712.31311 | 0.479127722 | 4.63E-06 |
| RPL13 | 48811.62292 | 0.478870376 | 1.47E-05 |
| NSMCE3 | 513.061359 | 0.478715977 | 8.99E-06 |
| CLCF1 | 243.9135885 | 0.478476646 | 2.84E-03 |
| PLOD3 | 10152.46719 | 0.47847107 | 1.76E-07 |
| EIF2D | 2568.611508 | 0.477768122 | 7.49E-08 |
| TRAF7 | 3862.880105 | 0.477566654 | 1.49E-05 |
| UBXN1 | 5831.619773 | 0.477555953 | 9.04E-06 |
| ATP6V1D | 2512.368254 | 0.477470281 | 4.60E-04 |
| IFRD2 | 2613.653594 | 0.477176788 | 1.10E-06 |
| ERCC6 | 565.1973192 | 0.47670764 | 3.04E-02 |
| C2CD4C | 130.8640201 | 0.47667665 | 2.86E-02 |
| ABT1 | 1107.478359 | 0.476424186 | 2.68E-05 |
| FKBP4 | 6633.202591 | 0.476346563 | 8.74E-06 |
| CBL | 5340.886292 | 0.476135713 | 2.11E-11 |
| COPS4 | 2229.661381 | 0.475628637 | 7.39E-14 |
| ATF6B | 3233.231276 | 0.475313568 | 3.53E-10 |
| ZSWIM6 | 1861.149076 | 0.475298351 | 7.47E-08 |
| HIF3A | 567.7463239 | 0.475273861 | 4.70E-02 |
| PIGU | 1597.231878 | 0.475175451 | 6.57E-07 |
| AC011476.2 | 111.0196704 | 0.475134432 | 4.61E-02 |
| RAB11FIP3 | 2920.906787 | 0.475128775 | 3.80E-07 |
| PPTC7 | 1103.285373 | 0.474703512 | 8.84E-05 |
| SELENON | 6949.973863 | 0.474637526 | 4.28E-05 |
| SUCLG1 | 3552.834348 | 0.474576567 | 1.97E-09 |
| MFSD12 | 3145.648508 | 0.47441215 | 4.14E-08 |
| PRPS1 | 3124.775206 | 0.474355681 | 8.20E-09 |
| VOPP1 | 7094.186809 | 0.474307966 | 4.90E-04 |
| UBN1 | 2592.183221 | 0.474191213 | 3.21E-07 |
| SNRPA | 3244.309368 | 0.473983814 | 5.23E-04 |
| DGKH | 590.9709515 | 0.47385022 | 1.68E-02 |
| MRPL10 | 2784.242699 | 0.473843671 | 1.99E-06 |
| DNAJC25 | 601.7296739 | 0.473768558 | 6.49E-04 |
| ZNRF1 | 621.6109898 | 0.473721125 | 3.12E-03 |
| THOC5 | 3377.448188 | 0.473673333 | 6.06E-07 |
| COMMD7 | 2781.927981 | 0.473502264 | 1.32E-05 |
| ECI1 | 2757.268471 | 0.473401999 | 5.41E-08 |
| FAM241B | 259.3376186 | 0.473041744 | 3.32E-02 |
| PIP4K2C | 2211.016611 | 0.472973705 | 1.44E-06 |
| NA | 1744.479115 | 0.47294548 | 1.10E-05 |
| MRPL48 | 1187.5922 | 0.472711382 | 6.86E-05 |
| SPAG5 | 4370.625311 | 0.472603244 | 2.10E-03 |
| C9orf3 | 1301.867845 | 0.47248865 | 8.80E-06 |
| EFNA3 | 288.3960931 | 0.472395255 | 3.53E-02 |
| MARK4 | 1860.573636 | 0.472328566 | 6.75E-04 |

| | | | |
|---|---|---|---|
| PCDH1 | 2577.072522 | 0.472205683 | 3.59E-02 |
| HEIH | 828.3928894 | 0.47215542 | 3.10E-03 |
| NQO2 | 3009.345582 | 0.472136532 | 1.26E-06 |
| DIAPH1 | 8016.050664 | 0.472016513 | 1.07E-05 |
| CTF1 | 182.4779826 | 0.47189196 | 7.91E-03 |
| C20orf194 | 2675.497658 | 0.471573843 | 1.87E-08 |
| AATF | 3385.796557 | 0.471501888 | 4.31E-08 |
| SLC16A5 | 221.2180602 | 0.471479145 | 1.06E-02 |
| DAGLB | 1643.438416 | 0.471457796 | 2.14E-07 |
| TUBGCP3 | 960.1919668 | 0.471453807 | 1.55E-04 |
| KDM7A-DT | 360.7802649 | 0.471429787 | 3.21E-04 |
| COASY | 5076.517411 | 0.471427645 | 1.53E-05 |
| IPO5P1 | 378.5648789 | 0.471106424 | 1.66E-02 |
| STXBP1 | 3551.05391 | 0.471104873 | 1.80E-05 |
| ARRDC4 | 850.4171308 | 0.471052829 | 1.89E-02 |
| ATG2A | 2615.448837 | 0.470484825 | 5.56E-05 |
| HOXC9 | 348.2374019 | 0.470353128 | 1.17E-03 |
| SEC11C | 605.6753875 | 0.469975163 | 1.46E-05 |
| BCAM | 3742.931072 | 0.469482921 | 6.27E-04 |
| SLC6A8 | 4800.871325 | 0.469273072 | 1.32E-03 |
| C8orf58 | 1572.615118 | 0.468942676 | 5.31E-03 |
| LACTB2 | 789.8368156 | 0.468918794 | 2.58E-04 |
| RPS5 | 32305.54264 | 0.468828189 | 1.43E-03 |
| IAH1 | 2107.417709 | 0.468679745 | 2.60E-12 |
| CSRP2 | 540.994005 | 0.468616267 | 1.21E-03 |
| RIN3 | 1523.104933 | 0.468560125 | 1.92E-05 |
| MGAT4B | 6599.001212 | 0.46846804 | 4.01E-04 |
| ALPK1 | 381.0090307 | 0.468328578 | 3.68E-02 |
| CT45A10 | 1249.05887 | 0.468119825 | 1.16E-03 |
| STAG3 | 3593.012809 | 0.467892748 | 3.57E-08 |
| FRG1JP | 122.945195 | 0.467889555 | 1.88E-02 |
| SF3A1 | 5632.206611 | 0.467843464 | 8.13E-04 |
| FAM50A | 5466.553151 | 0.46765796 | 1.46E-07 |
| SEC31A | 8459.893365 | 0.46749758 | 1.99E-07 |
| NT5DC3 | 443.4467148 | 0.4670324 | 2.47E-02 |
| GALE | 2706.092626 | 0.466966265 | 1.89E-05 |
| SLC25A3 | 15226.79426 | 0.46677265 | 6.76E-05 |
| NDEL1 | 2052.460807 | 0.46628695 | 1.92E-14 |
| NA | 175.0159474 | 0.466247767 | 4.39E-03 |
| ITGA6 | 12300.09608 | 0.465762862 | 6.95E-05 |
| TPX2 | 13710.93008 | 0.465737801 | 2.56E-06 |
| SNHG15 | 1441.968187 | 0.465574097 | 3.02E-02 |
| CUL4B | 4541.326137 | 0.465397881 | 1.96E-04 |
| RPS3A | 65602.08282 | 0.465139695 | 1.06E-04 |
| FOPNL | 1315.094864 | 0.464907143 | 3.39E-03 |
| DDX54 | 3050.744023 | 0.464894327 | 1.45E-09 |
| APPL2 | 2709.579325 | 0.464445584 | 4.81E-10 |

| | | | |
|---|---|---|---|
| C5orf30 | 391.2718681 | 0.464266939 | 4.58E-03 |
| TDP1 | 1043.60661 | 0.46385567 | 2.17E-06 |
| NECAP2 | 4663.673728 | 0.463584546 | 3.50E-04 |
| ERGIC1 | 7955.280628 | 0.463428033 | 5.59E-06 |
| ZSCAN18 | 2383.204961 | 0.463385802 | 1.10E-04 |
| NPIPP1 | 408.711574 | 0.463164802 | 1.01E-03 |
| RPL10 | 91368.98784 | 0.463035505 | 8.79E-05 |
| ZNF674-AS1 | 251.204287 | 0.463035487 | 2.37E-04 |
| MRPL37 | 3539.378493 | 0.46274837 | 1.55E-06 |
| GSTM4 | 1524.811665 | 0.462476921 | 1.36E-06 |
| NELFB | 5735.829947 | 0.462036956 | 4.71E-06 |
| GTSF1 | 1663.890902 | 0.461682394 | 2.43E-09 |
| STIM1 | 3076.092754 | 0.461016193 | 3.76E-06 |
| NA | 225.0341295 | 0.460960475 | 1.93E-02 |
| CNIH4 | 1503.44764 | 0.460868722 | 1.54E-08 |
| RASAL2 | 1538.28984 | 0.460486725 | 2.28E-05 |
| NCS1 | 3791.736683 | 0.460236377 | 2.07E-07 |
| NPTXR | 2829.986031 | 0.459862451 | 1.41E-04 |
| MAGOH | 973.9237622 | 0.459641261 | 2.35E-06 |
| PRPF6 | 7435.468691 | 0.459474265 | 1.43E-03 |
| SUGCT | 319.5476151 | 0.459411286 | 2.52E-02 |
| SPTBN2 | 1745.729904 | 0.459385343 | 4.41E-03 |
| RNF20 | 5561.7265 | 0.459149383 | 5.65E-05 |
| FMNL3 | 2124.37167 | 0.458913844 | 5.86E-04 |
| POLR2J4 | 256.1167163 | 0.458848229 | 3.22E-02 |
| PDCD11 | 5588.612431 | 0.458733787 | 4.07E-04 |
| LRRC42 | 2265.315389 | 0.458704118 | 1.24E-07 |
| ZNF598 | 2854.685967 | 0.458577969 | 1.87E-08 |
| AC021054.1 | 291.9708505 | 0.45841201 | 4.95E-03 |
| NLN | 3580.468692 | 0.45824036 | 1.15E-03 |
| NDUFA9 | 2952.013751 | 0.458209263 | 5.22E-04 |
| ERO1B | 267.8671326 | 0.458129736 | 4.65E-02 |
| PLGRKT | 793.4817684 | 0.458126265 | 5.80E-06 |
| LRRC4B | 823.3866249 | 0.458093731 | 1.84E-02 |
| NA | 295.3263647 | 0.458070709 | 2.77E-02 |
| MAPK9 | 1850.100491 | 0.45753021 | 2.28E-05 |
| FGF2 | 458.0900956 | 0.457403135 | 2.77E-02 |
| SLC6A6 | 5652.703338 | 0.45724309 | 4.99E-05 |
| AKT3 | 2703.948896 | 0.45687713 | 1.85E-04 |
| NPLOC4 | 6856.1453 | 0.45685709 | 5.52E-05 |
| COX5A | 3745.18832 | 0.456834815 | 1.11E-05 |
| PRRC2B | 23100.42542 | 0.456281736 | 6.99E-11 |
| MCUR1 | 1934.263528 | 0.456125484 | 3.15E-08 |
| RPL23A | 15036.58917 | 0.45589371 | 8.00E-04 |
| DAP | 15213.24624 | 0.455820057 | 5.11E-05 |
| AGAP1 | 2415.072612 | 0.45576768 | 1.26E-04 |
| SAP30L | 1605.846888 | 0.455696625 | 2.10E-04 |

| | | | |
|---|---|---|---|
| GLUD1 | 5125.474283 | 0.45539362 | 1.35E-11 |
| SMAGP | 683.0418754 | 0.455092667 | 2.02E-03 |
| MRPL52 | 1805.217802 | 0.455082334 | 3.59E-08 |
| PDPR | 5125.12976 | 0.455076014 | 9.00E-10 |
| SNHG3 | 1840.165392 | 0.455068974 | 2.80E-02 |
| RPL26L1 | 1232.856647 | 0.454977807 | 2.58E-07 |
| WDR25 | 582.4176668 | 0.454881589 | 1.26E-02 |
| CHCHD3 | 3891.440959 | 0.454790598 | 1.84E-05 |
| PUS1 | 1053.427984 | 0.454617991 | 6.03E-04 |
| ACOT13 | 1368.242753 | 0.454430077 | 6.76E-04 |
| IFFO2 | 2826.398817 | 0.453962633 | 1.57E-04 |
| DNHD1 | 2181.427692 | 0.453888719 | 5.72E-05 |
| GNAO1 | 1149.018389 | 0.453779128 | 4.24E-03 |
| LONRF1 | 546.4227996 | 0.453770996 | 5.35E-03 |
| KRTCAP2 | 1559.895768 | 0.45358551 | 2.98E-05 |
| KARS | 13794.60913 | 0.453510515 | 7.65E-09 |
| ARIH2 | 4845.810584 | 0.453492898 | 1.58E-05 |
| RINL | 283.9051294 | 0.453405436 | 9.75E-04 |
| ECE1 | 7636.537465 | 0.453384547 | 1.92E-07 |
| AL590327.1 | 533.3376147 | 0.453361388 | 3.66E-03 |
| PACSIN2 | 4740.17459 | 0.45319019 | 6.43E-07 |
| BANF1 | 7623.196999 | 0.453160908 | 8.14E-06 |
| TNFRSF1B | 1074.403305 | 0.452726648 | 1.37E-04 |
| MECR | 1202.982923 | 0.451998083 | 3.56E-07 |
| DCAF4 | 983.4014373 | 0.451707092 | 6.09E-06 |
| ERCC1 | 2607.469316 | 0.451548801 | 7.41E-05 |
| RPL23AP42 | 8204.968219 | 0.451343614 | 2.24E-04 |
| TRAM2 | 9655.068286 | 0.451288656 | 4.28E-11 |
| CLCN4 | 1217.164316 | 0.451283012 | 1.47E-02 |
| LINC00963 | 2907.811582 | 0.451016915 | 1.66E-04 |
| MMGT1 | 2362.006228 | 0.450598355 | 1.00E-10 |
| ABHD14B | 2069.042673 | 0.450243642 | 2.47E-04 |
| FAM27E3 | 808.5898466 | 0.450193977 | 4.54E-03 |
| DUSP22 | 1490.329574 | 0.450028598 | 7.22E-09 |
| PANK4 | 952.9214816 | 0.449479735 | 1.83E-02 |
| SNHG16 | 2900.523027 | 0.449428556 | 2.74E-09 |
| KNSTRN | 2096.989694 | 0.449356278 | 7.58E-05 |
| CEP131 | 811.4527466 | 0.449254472 | 8.41E-05 |
| PSMB7 | 8583.989746 | 0.448975475 | 7.46E-08 |
| RPL32 | 31619.84787 | 0.448938862 | 5.80E-05 |
| OXA1L | 5881.365913 | 0.448719808 | 1.17E-04 |
| AC009303.4 | 97.14289571 | 0.448656145 | 4.30E-02 |
| MAP3K9 | 1540.316129 | 0.448500785 | 3.89E-04 |
| POLR3C | 1690.913297 | 0.448405135 | 4.03E-03 |
| SSH3 | 1977.929526 | 0.447960198 | 2.05E-03 |
| PRKX | 863.1302112 | 0.447860113 | 2.44E-03 |
| BYSL | 1631.402346 | 0.447722022 | 3.97E-04 |

| | | | |
|---|---|---|---|
| EIF2AK3-DT | 109.8054073 | 0.447625526 | 3.48E-02 |
| GOLGA3 | 4858.345823 | 0.447545975 | 4.60E-06 |
| RNF26 | 5061.4502 | 0.44752556 | 2.63E-04 |
| RPLP1 | 50415.89131 | 0.447430603 | 1.32E-03 |
| DHX32 | 1575.566691 | 0.447211177 | 1.49E-07 |
| GAS2L1 | 793.1223109 | 0.447173478 | 4.70E-03 |
| MICAL3 | 4913.970363 | 0.446938578 | 1.55E-08 |
| HYOU1 | 23967.33716 | 0.446605079 | 1.82E-05 |
| GNB2 | 11143.86627 | 0.446488786 | 1.76E-04 |
| FAXC | 492.2908407 | 0.44639505 | 2.04E-05 |
| CTDSPL | 1864.785257 | 0.446325188 | 8.18E-05 |
| MFSD14C | 181.1776867 | 0.445922989 | 9.87E-03 |
| ARPC1B | 9222.847879 | 0.445880121 | 1.09E-04 |
| BABAM2 | 1517.574927 | 0.44575199 | 8.92E-04 |
| ARL8A | 771.8332599 | 0.445229972 | 7.72E-03 |
| TP53 | 6389.292284 | 0.445006506 | 1.13E-03 |
| RHOBTB3 | 58008.22119 | 0.444969126 | 2.56E-03 |
| CDC42BPB | 7684.040986 | 0.444950224 | 1.80E-20 |
| BIRC5 | 4253.014258 | 0.444722935 | 2.53E-04 |
| PTGIR | 176.9157293 | 0.444608132 | 3.39E-02 |
| NOB1 | 4105.266417 | 0.444584659 | 6.11E-05 |
| DESI2 | 2337.936691 | 0.444044006 | 5.93E-03 |
| ELFN1 | 408.2898101 | 0.443815545 | 2.19E-03 |
| CALCOCO1 | 3982.64577 | 0.443628374 | 1.58E-03 |
| NBPF8 | 1942.365623 | 0.443335498 | 4.38E-07 |
| FBRSL1 | 1449.075711 | 0.443165484 | 2.86E-03 |
| ECPAS | 7118.119316 | 0.442780185 | 6.14E-10 |
| FAM86EP | 169.2220581 | 0.442686353 | 3.98E-02 |
| ZDHHC7 | 4188.484347 | 0.442644798 | 2.42E-03 |
| EP400 | 4945.291168 | 0.442356795 | 6.69E-14 |
| FAM83H | 345.1934365 | 0.442055172 | 2.03E-03 |
| 2-Mar | 481.9723773 | 0.442035298 | 2.20E-02 |
| MIB2 | 1313.54863 | 0.441834723 | 3.49E-06 |
| TPT1-AS1 | 345.5680486 | 0.441778778 | 2.67E-04 |
| TUBB | 73676.804 | 0.441647778 | 1.29E-02 |
| RPS6 | 79859.13227 | 0.44152668 | 3.84E-07 |
| MIIP | 828.9580369 | 0.441389142 | 1.59E-04 |
| GBGT1 | 427.374157 | 0.441350272 | 3.99E-03 |
| KCTD17 | 1231.905598 | 0.441241518 | 4.45E-05 |
| CLYBL | 500.0273141 | 0.441069753 | 2.10E-03 |
| NOVA2 | 439.1966609 | 0.440861007 | 7.96E-03 |
| WASH8P | 904.7545586 | 0.440860244 | 2.85E-03 |
| CARD8-AS1 | 133.2138116 | 0.44063211 | 2.55E-02 |
| GRHPR | 4130.066864 | 0.440487085 | 1.21E-06 |
| MFN2 | 7446.363905 | 0.440431531 | 1.05E-07 |
| MRPL14 | 1572.149765 | 0.440355003 | 9.63E-05 |
| PSMD7 | 7737.490292 | 0.439983176 | 1.91E-16 |

| | | | |
|---|---|---|---|
| YWHAZ | 33323.3696 | 0.439959136 | 1.16E-06 |
| FTL | 97215.3597 | 0.439816929 | 2.88E-03 |
| RAB11FIP4 | 856.5889553 | 0.439532481 | 1.14E-02 |
| MRPS9 | 1646.296068 | 0.439289207 | 6.32E-07 |
| CDC34 | 2227.878526 | 0.439102147 | 3.22E-06 |
| DOCK3 | 240.1180089 | 0.438902758 | 1.35E-02 |
| UXT | 1749.111555 | 0.438742709 | 5.97E-04 |
| KIAA2013 | 2549.440012 | 0.438530215 | 3.70E-04 |
| EEF1G | 92857.80488 | 0.438250161 | 3.97E-04 |
| TXNL4A | 1733.43983 | 0.438221675 | 5.93E-07 |
| AC069281.2 | 301.6621336 | 0.438051852 | 1.99E-02 |
| AC083798.2 | 648.5242041 | 0.437831529 | 3.21E-02 |
| DENND5A | 8765.225991 | 0.437743552 | 3.31E-06 |
| AHCYL1 | 7022.281892 | 0.436457207 | 2.06E-14 |
| GPI | 9672.754168 | 0.436287264 | 1.65E-04 |
| GOLGA2P7 | 5165.611373 | 0.436016684 | 9.33E-08 |
| RTEL1 | 1233.022643 | 0.435748005 | 1.39E-06 |
| SNX9 | 2199.738733 | 0.435268335 | 2.68E-10 |
| RPL21 | 8227.922657 | 0.435253455 | 1.97E-05 |
| AC002351.1 | 190.5787274 | 0.435214452 | 1.16E-02 |
| FASN | 14089.9427 | 0.43514192 | 8.88E-03 |
| MAP1S | 1324.327333 | 0.43499173 | 3.28E-03 |
| LARS | 10091.49174 | 0.434970265 | 3.70E-03 |
| RPS12 | 23474.3027 | 0.434770111 | 3.53E-07 |
| PITRM1 | 3121.362636 | 0.434723312 | 8.13E-08 |
| LSM4 | 3280.823502 | 0.434667935 | 6.20E-07 |
| STK32C | 480.8698462 | 0.434303143 | 4.30E-04 |
| APBB2 | 2409.951598 | 0.434166018 | 3.57E-04 |
| SLC35B1 | 1668.984745 | 0.43414772 | 2.42E-08 |
| VPS51 | 3467.254109 | 0.434007537 | 1.82E-02 |
| TAF7 | 3538.289254 | 0.433625131 | 1.71E-05 |
| FOXK1 | 3559.332545 | 0.433624878 | 2.55E-07 |
| RPUSD3 | 1681.748031 | 0.433621313 | 4.76E-06 |
| RPL30 | 35678.43918 | 0.433213127 | 3.00E-05 |
| SYNGR1 | 3473.006517 | 0.433176258 | 1.71E-03 |
| SCYL1 | 4860.578339 | 0.433136361 | 9.98E-04 |
| BAIAP2 | 994.5039872 | 0.432929114 | 2.78E-07 |
| MAGEA2B | 1301.131276 | 0.432889063 | 3.21E-02 |
| PLEKHG3 | 2080.800905 | 0.432781166 | 7.45E-05 |
| CCNY | 2484.643454 | 0.432768226 | 3.61E-09 |
| PGM1 | 2597.812756 | 0.432693662 | 1.49E-02 |
| MOAP1 | 1235.398058 | 0.432650678 | 1.21E-06 |
| ZNF721 | 490.7716122 | 0.432522431 | 2.70E-04 |
| EIF6 | 9045.030297 | 0.432511482 | 1.16E-04 |
| RRNAD1 | 1100.690946 | 0.432406866 | 1.14E-04 |
| SERINC3 | 7719.314089 | 0.432226575 | 1.03E-12 |
| LOXL3 | 3055.781083 | 0.432138388 | 3.21E-03 |

| | | | |
|---|---|---|---|
| BCL9L | 9610.645557 | 0.431978212 | 3.16E-04 |
| IGBP1 | 3196.48629 | 0.431721412 | 1.56E-02 |
| DUS2 | 833.3492684 | 0.431521214 | 7.94E-05 |
| LINC01006 | 968.1904904 | 0.431503628 | 3.21E-04 |
| IMPA2 | 418.8634436 | 0.4314831 | 4.03E-03 |
| PAQR7 | 1043.956206 | 0.431398698 | 3.34E-04 |
| EIF4H | 25471.60151 | 0.431315598 | 1.21E-10 |
| TMEM241 | 503.1300017 | 0.431268285 | 5.09E-03 |
| SPACA9 | 475.0875021 | 0.431107175 | 5.09E-03 |
| SGPL1 | 1970.346108 | 0.43105471 | 8.01E-03 |
| RPAP1 | 1524.937256 | 0.430788571 | 2.89E-04 |
| TTC27 | 1234.216433 | 0.430532527 | 4.78E-06 |
| CALML4 | 214.1852994 | 0.430399333 | 2.98E-02 |
| SNHG19 | 151.0834963 | 0.429752425 | 2.65E-02 |
| CRAT | 2562.999731 | 0.42973519 | 6.72E-05 |
| DHDDS | 1753.60419 | 0.429720091 | 1.18E-04 |
| VARS | 6546.556051 | 0.429661274 | 5.88E-06 |
| DNTTIP1 | 2150.378831 | 0.429621083 | 7.25E-04 |
| SLC37A1 | 253.5659223 | 0.429547916 | 2.02E-02 |
| ALG3 | 1585.689512 | 0.429445034 | 2.89E-04 |
| ADIPOR1 | 4348.452011 | 0.429117267 | 1.38E-04 |
| BCAR1 | 5994.364749 | 0.429076037 | 1.49E-04 |
| FARSB | 2957.701085 | 0.429024796 | 9.00E-10 |
| AL596244.1 | 271.7370762 | 0.429023609 | 2.98E-02 |
| RUNX1 | 2054.795126 | 0.428998132 | 6.38E-04 |
| CCDC28A | 816.7254079 | 0.428723288 | 4.46E-05 |
| RSL24D1 | 4258.03113 | 0.428618298 | 2.29E-03 |
| POLR1E | 2520.906887 | 0.428401142 | 3.66E-07 |
| NDST1 | 8021.325827 | 0.428400082 | 2.84E-05 |
| MCRIP1 | 1829.068626 | 0.428161443 | 4.72E-03 |
| SEL1L3 | 3269.858349 | 0.427773915 | 3.13E-03 |
| WDR62 | 1308.836061 | 0.427771597 | 8.61E-03 |
| SURF4 | 17447.89564 | 0.427263563 | 2.03E-04 |
| ASPH | 8470.783867 | 0.42719188 | 2.12E-04 |
| POC1B | 724.6914982 | 0.427189577 | 1.43E-03 |
| RPS8 | 38197.62831 | 0.427129277 | 2.08E-06 |
| DTX2 | 890.7965669 | 0.426831293 | 4.68E-04 |
| ZNF106 | 3656.174963 | 0.426698038 | 1.03E-06 |
| PWP2 | 996.9400637 | 0.426487197 | 3.99E-03 |
| NLRP1 | 4175.84274 | 0.42640248 | 3.58E-05 |
| MAST1 | 367.3052568 | 0.426235622 | 2.80E-02 |
| RPS21 | 23282.49507 | 0.426178464 | 2.22E-04 |
| LYPD6 | 673.2989076 | 0.425758463 | 1.92E-02 |
| BAZ2A | 7258.618219 | 0.425698945 | 6.29E-11 |
| CAB39 | 2508.723948 | 0.425543823 | 9.41E-04 |
| FBXO22 | 2360.772059 | 0.42548202 | 7.77E-07 |
| AL162171.1 | 1232.53051 | 0.425458221 | 5.20E-05 |

| | | | |
|---|---|---|---|
| TBCA | 6087.859397 | 0.425405594 | 9.12E-08 |
| SH3RF1 | 1138.990913 | 0.425328674 | 2.37E-03 |
| ADGRB2 | 2041.469019 | 0.425285436 | 1.98E-03 |
| PPM1F | 3912.01717 | 0.424547583 | 2.76E-05 |
| AC093525.6 | 218.5605161 | 0.424189951 | 5.88E-03 |
| DUSP3 | 4615.69139 | 0.424161962 | 5.24E-06 |
| CDC42EP2 | 1460.609892 | 0.424140986 | 4.50E-04 |
| GPR157 | 512.3955584 | 0.423889832 | 5.00E-02 |
| BSDC1 | 3231.619132 | 0.42382019 | 1.25E-04 |
| CYB5R1 | 1271.066427 | 0.423642253 | 4.23E-04 |
| AKAP13 | 5307.00127 | 0.423585581 | 1.38E-05 |
| MED27 | 1626.357315 | 0.423432243 | 3.09E-04 |
| PPP1R14B | 3533.962955 | 0.423135684 | 2.72E-04 |
| PEX10 | 1005.175471 | 0.422738658 | 1.01E-04 |
| XPO6 | 8367.253338 | 0.422195913 | 7.86E-08 |
| LINC00476 | 276.6276502 | 0.422184105 | 5.17E-03 |
| PSMA1 | 6765.463374 | 0.422139081 | 3.08E-09 |
| EEF1AKMT2 | 304.8747012 | 0.422130776 | 3.99E-03 |
| CIZ1 | 7246.560402 | 0.421959263 | 1.01E-05 |
| C14orf93 | 695.6090895 | 0.421510314 | 3.68E-04 |
| JTB | 2513.992003 | 0.421219564 | 3.71E-05 |
| TBL2 | 3247.426371 | 0.421093991 | 1.82E-06 |
| PLS1 | 372.3599433 | 0.420846813 | 4.11E-02 |
| SLC9A5 | 766.9386846 | 0.420712186 | 3.48E-04 |
| AC115618.2 | 149.3828333 | 0.420583063 | 4.44E-02 |
| RTL8A | 3282.957021 | 0.420495314 | 2.25E-04 |
| CCNK | 2190.032846 | 0.420407011 | 2.77E-07 |
| HEATR6 | 1746.01362 | 0.420008243 | 7.86E-07 |
| TMEM44 | 939.3230402 | 0.419989934 | 1.03E-04 |
| RAB3GAP1 | 4031.730799 | 0.41989205 | 2.21E-05 |
| KLHL23 | 726.4414864 | 0.419766036 | 5.48E-03 |
| AC147651.4 | 110.8532044 | 0.419688901 | 3.01E-02 |
| DARS2 | 3464.006142 | 0.419553673 | 5.29E-08 |
| ACO2 | 4130.560406 | 0.419481681 | 1.91E-03 |
| TMEM229B | 588.0733194 | 0.419377822 | 5.26E-03 |
| USP39 | 3397.710093 | 0.419368267 | 1.46E-05 |
| RFWD3 | 5369.116357 | 0.419324773 | 1.08E-03 |
| MORF4L2 | 11614.8889 | 0.419169784 | 9.28E-15 |
| TBRG4 | 6211.250692 | 0.418954398 | 2.84E-05 |
| CAD | 8898.770146 | 0.418864077 | 2.36E-07 |
| FOXM1 | 5920.981055 | 0.41872781 | 2.75E-04 |
| ATG101 | 1922.096598 | 0.418582764 | 4.57E-04 |
| SEH1L | 1842.288795 | 0.418300806 | 2.93E-05 |
| ABCF2 | 6738.907087 | 0.418293764 | 8.78E-09 |
| MAP7D1 | 4939.854588 | 0.418214994 | 8.48E-06 |
| ARSG | 201.1469179 | 0.418020214 | 3.78E-02 |
| UNC93B1 | 929.0976881 | 0.417755256 | 1.01E-02 |

| | | | |
|---|---|---|---|
| NSFL1C | 5125.345501 | 0.41753725 | 1.48E-07 |
| HSD11B1L | 516.8047837 | 0.417523936 | 2.15E-03 |
| PPT2 | 1542.529366 | 0.417184 | 8.41E-03 |
| AL160006.1 | 851.9614508 | 0.417071292 | 1.10E-04 |
| FDXR | 399.2808891 | 0.41700243 | 4.59E-04 |
| UBA52 | 14827.05506 | 0.416984811 | 1.30E-03 |
| RGMB | 1091.065118 | 0.416680639 | 4.78E-07 |
| RNF114 | 4494.094987 | 0.416678434 | 3.14E-07 |
| FITM2 | 2198.051981 | 0.416648168 | 2.88E-09 |
| SLC17A5 | 973.1245904 | 0.416411737 | 1.51E-02 |
| PPP1R3C | 1863.591223 | 0.416224953 | 9.30E-05 |
| GSK3B | 3261.575123 | 0.415337387 | 4.25E-06 |
| RPP21 | 594.9113622 | 0.415201282 | 1.37E-03 |
| MRPS10 | 1948.921082 | 0.414989081 | 3.25E-05 |
| COL18A1 | 7768.279912 | 0.41490533 | 1.09E-03 |
| RFT1 | 1944.490929 | 0.414846047 | 5.02E-09 |
| FRS3 | 243.2988226 | 0.414832233 | 1.70E-03 |
| SLC7A6OS | 1998.111305 | 0.414751443 | 3.06E-07 |
| EIF3M | 9035.068701 | 0.414715372 | 1.05E-11 |
| KANK2 | 5584.704178 | 0.414619378 | 5.96E-03 |
| C9orf40 | 891.5125514 | 0.414578579 | 3.63E-03 |
| TUSC1 | 473.2965741 | 0.414514001 | 2.31E-03 |
| NBAS | 3584.572694 | 0.414374819 | 1.37E-04 |
| COX7A2 | 5358.559036 | 0.414336382 | 6.12E-03 |
| URGCP | 3502.054904 | 0.414304487 | 1.51E-04 |
| APOL6 | 2761.451997 | 0.413992484 | 1.09E-02 |
| ANO10 | 1558.862414 | 0.413445727 | 2.73E-03 |
| LAS1L | 2829.020455 | 0.413335019 | 1.82E-06 |
| GALK1 | 1388.050362 | 0.413202997 | 1.44E-02 |
| DLG1 | 3174.228063 | 0.413129317 | 1.21E-02 |
| GLTP | 1149.725905 | 0.41300109 | 4.34E-05 |
| JMJD7 | 296.4011285 | 0.412925229 | 3.34E-03 |
| APEH | 7239.845918 | 0.412756736 | 2.76E-03 |
| GOLGA2P10 | 667.6520873 | 0.412681739 | 6.12E-04 |
| SAE1 | 11296.4598 | 0.412469953 | 3.67E-07 |
| PSMC3 | 7429.183721 | 0.412257813 | 2.07E-07 |
| EXOSC1 | 656.5367159 | 0.412194476 | 3.67E-05 |
| SLC27A1 | 950.222448 | 0.412083793 | 1.33E-03 |
| ANKRD37 | 182.4951309 | 0.412037417 | 8.87E-03 |
| DYNLRB1 | 5076.376126 | 0.411209484 | 2.94E-03 |
| FAHD2CP | 272.9451567 | 0.411133589 | 1.70E-02 |
| RRP12 | 2494.959313 | 0.410945399 | 5.85E-04 |
| MED13L | 4915.063541 | 0.410915864 | 5.05E-04 |
| GTF2E2 | 2020.494839 | 0.410905985 | 9.97E-08 |
| DAZAP2 | 5104.342531 | 0.410585366 | 3.22E-03 |
| TMEM106C | 4149.946741 | 0.410285274 | 2.47E-07 |
| NFKBIL1 | 706.4137555 | 0.410225722 | 8.21E-04 |

| | | | |
|---|---|---|---|
| COG5 | 3073.024123 | 0.410061068 | 2.76E-04 |
| AC010618.1 | 1835.153133 | 0.409919628 | 1.46E-03 |
| PLBD2 | 5126.821147 | 0.409568601 | 2.34E-04 |
| PFKL | 6261.863339 | 0.409332797 | 1.91E-03 |
| TUFM | 15387.79824 | 0.409252601 | 2.60E-05 |
| CEP41 | 1707.755942 | 0.40902867 | 1.10E-04 |
| NUDT21 | 7108.295312 | 0.408965645 | 1.90E-05 |
| STOML2 | 9108.069344 | 0.408652442 | 4.52E-07 |
| MFHAS1 | 1322.185779 | 0.40828078 | 6.08E-05 |
| RPS10-NUDT3 | 510.8586973 | 0.407825011 | 3.18E-02 |
| CHMP1B | 1645.111841 | 0.407660156 | 1.30E-07 |
| MFF | 2603.156878 | 0.407559623 | 3.76E-12 |
| ERICH1 | 745.9834042 | 0.407476554 | 1.11E-03 |
| KIAA1191 | 5326.813769 | 0.407329596 | 2.46E-05 |
| SNTA1 | 2042.451858 | 0.407208203 | 3.27E-03 |
| EMC6 | 677.2712484 | 0.407138573 | 1.49E-03 |
| VPS72 | 2156.237537 | 0.40713812 | 1.57E-03 |
| POLD2 | 6626.569676 | 0.407122802 | 4.06E-07 |
| ARMC4 | 198.3905927 | 0.407093397 | 4.61E-02 |
| UQCC1 | 3926.027674 | 0.407088009 | 4.66E-07 |
| NDUFS7 | 1677.92942 | 0.406622797 | 6.86E-06 |
| PFN1 | 38626.73183 | 0.406468117 | 1.74E-03 |
| CCDC130 | 833.3427174 | 0.406454077 | 2.38E-04 |
| SLX1A | 797.5724445 | 0.406359926 | 1.15E-02 |
| OGFOD2 | 630.4163552 | 0.406151609 | 2.19E-03 |
| PRMT5 | 6007.645453 | 0.406037724 | 6.66E-06 |
| RPL7A | 72628.68302 | 0.405860966 | 1.42E-02 |
| SLC25A25 | 1315.684058 | 0.405784417 | 5.59E-08 |
| CIB1 | 2207.068549 | 0.405321706 | 8.82E-04 |
| BAG6 | 12356.6706 | 0.405117605 | 1.98E-04 |
| THAP3 | 667.2347235 | 0.404653487 | 8.20E-03 |
| AMZ2P1 | 165.7323703 | 0.404513002 | 3.31E-02 |
| SLC25A19 | 457.3255449 | 0.404372883 | 3.43E-03 |
| CEP104 | 1693.378755 | 0.40430642 | 7.21E-04 |
| MVB12B | 642.8929973 | 0.40422073 | 3.34E-03 |
| PRDM4 | 1802.069444 | 0.404218641 | 2.64E-08 |
| EIF3J | 2363.847504 | 0.404190878 | 7.07E-03 |
| LDHA | 37610.75825 | 0.404025837 | 4.76E-02 |
| ATIC | 7866.447087 | 0.403836987 | 3.65E-05 |
| SPA17 | 608.872348 | 0.403777412 | 2.84E-03 |
| PET117 | 794.4960644 | 0.403733353 | 2.25E-06 |
| PURA | 1075.154351 | 0.403600964 | 3.47E-02 |
| MCTS1 | 2241.850458 | 0.403577699 | 7.78E-08 |
| ARHGEF39 | 1059.646998 | 0.403575374 | 3.55E-03 |
| RPLP2 | 26725.70048 | 0.403518123 | 1.72E-03 |
| SMIM13 | 1014.459578 | 0.403288931 | 1.26E-02 |
| WDR45 | 3949.598989 | 0.403256307 | 2.77E-03 |

| | | | |
|---|---|---|---|
| PDXDC1 | 6002.09432 | 0.403025869 | 4.80E-06 |
| UQCRFS1 | 3568.156548 | 0.402991702 | 5.84E-05 |
| STMN3 | 8539.398862 | 0.402794012 | 1.54E-03 |
| DHPS | 2252.553779 | 0.402691295 | 4.14E-10 |
| BTF3 | 37705.71318 | 0.402265476 | 1.17E-06 |
| TARS | 20704.77187 | 0.401929166 | 3.71E-03 |
| EIF2S2 | 12053.63184 | 0.401596109 | 9.65E-03 |
| GPRIN1 | 770.8989094 | 0.401480782 | 1.93E-02 |
| MRPS18C | 1122.775971 | 0.401418393 | 1.92E-05 |
| POLR3D | 3632.780335 | 0.401220178 | 4.70E-05 |
| PDP2 | 1653.370186 | 0.401075886 | 3.38E-05 |
| HTRA2 | 2044.632305 | 0.401072375 | 7.32E-04 |
| IMPDH1 | 3474.006773 | 0.40104016 | 6.20E-06 |
| BTBD2 | 6344.109791 | 0.401030769 | 3.64E-05 |
| STK3 | 890.5682343 | 0.400996694 | 4.66E-04 |
| RPS27L | 1580.599612 | 0.400960824 | 1.51E-04 |
| NA | 220.007574 | 0.4008426 | 3.65E-02 |
| LIPC | 496.2256483 | 0.400742367 | 1.65E-02 |
| VKORC1L1 | 3787.780139 | 0.400692384 | 5.43E-04 |
| CAP1 | 16233.00955 | 0.400520314 | 2.84E-05 |
| DHCR7 | 5667.85858 | 0.400314543 | 1.68E-02 |
| KLHL35 | 461.3989432 | 0.400039847 | 1.23E-02 |
| FAM114A1 | 1772.247523 | 0.400013339 | 9.09E-03 |
| SLC39A3 | 931.2260034 | 0.399809332 | 2.00E-04 |
| PERP | 2168.913583 | 0.399525661 | 9.82E-07 |
| ARHGAP22 | 180.5159699 | 0.3991416 | 4.72E-02 |
| MAPKBP1 | 1283.789127 | 0.399118131 | 1.20E-03 |
| ZNF787 | 811.1459574 | 0.398869831 | 5.02E-05 |
| SELENOM | 467.2787855 | 0.398820335 | 2.37E-02 |
| ZNF358 | 1642.217967 | 0.398785016 | 1.34E-03 |
| WDR13 | 3160.87114 | 0.398687497 | 6.40E-03 |
| ZBTB7A | 1167.70057 | 0.398269341 | 6.27E-03 |
| RUVBL1 | 5432.212495 | 0.398254217 | 1.94E-05 |
| MTX1 | 2583.38152 | 0.397991581 | 3.48E-03 |
| MRPL55 | 606.6771076 | 0.39773741 | 3.54E-04 |
| RPL39 | 26083.00633 | 0.397485651 | 1.40E-03 |
| PSMD13 | 8806.558774 | 0.397270099 | 2.10E-05 |
| DRG1 | 3037.512976 | 0.39711544 | 3.48E-05 |
| GART | 10888.60204 | 0.39658076 | 1.17E-04 |
| DDX51 | 1618.888558 | 0.396566457 | 1.12E-03 |
| MRPL38 | 1864.760936 | 0.396554138 | 6.23E-05 |
| HMGB2 | 6919.321379 | 0.39571141 | 3.16E-02 |
| ADAMTS2 | 1245.830925 | 0.395587143 | 2.76E-02 |
| NCAPH | 2363.792997 | 0.395362042 | 2.62E-02 |
| CDK2AP2 | 1734.372532 | 0.395354913 | 4.23E-04 |
| MAF1 | 4739.284522 | 0.395344094 | 2.95E-04 |
| CAMSAP1 | 3654.8622 | 0.395327786 | 5.24E-05 |

| | | | |
|---|---|---|---|
| FARSA | 5326.766618 | 0.395206538 | 2.38E-04 |
| MCMBP | 2516.238275 | 0.395205949 | 8.18E-04 |
| ELL2 | 3965.592371 | 0.394367142 | 5.36E-03 |
| FH | 3105.884307 | 0.394317289 | 6.30E-10 |
| THSD1 | 104.9408088 | 0.394054392 | 3.16E-02 |
| JRK | 2881.469205 | 0.393863796 | 4.05E-03 |
| TBL1X | 3544.334191 | 0.393814168 | 5.39E-04 |
| OGT | 16811.32788 | 0.393601043 | 1.94E-02 |
| NAA80 | 540.0798404 | 0.39345245 | 1.71E-02 |
| TRRAP | 9853.594491 | 0.393395408 | 1.67E-09 |
| SEC14L1 | 8187.091665 | 0.39321663 | 8.46E-04 |
| VIM | 74772.47501 | 0.393186021 | 7.73E-04 |
| PLXNA1 | 4699.089689 | 0.39305166 | 3.43E-04 |
| DHX37 | 2003.689058 | 0.392921738 | 4.79E-05 |
| BAG3 | 1430.928166 | 0.392739955 | 1.28E-03 |
| AL158212.3 | 189.4677657 | 0.392674957 | 2.90E-02 |
| JMJD6 | 1235.975837 | 0.392621236 | 2.67E-02 |
| SWAP70 | 3816.151683 | 0.392504547 | 9.88E-06 |
| AC011481.1 | 2788.474976 | 0.392148186 | 2.36E-04 |
| RPL13A | 70398.25543 | 0.392142442 | 3.86E-03 |
| CDK16 | 7022.735633 | 0.391682244 | 2.93E-04 |
| ZNF529-AS1 | 151.9124009 | 0.391616593 | 1.75E-02 |
| PLXND1 | 13090.28614 | 0.391336991 | 8.08E-03 |
| HDGF | 21317.82589 | 0.391325991 | 9.06E-05 |
| PARS2 | 429.7542574 | 0.391245529 | 3.03E-03 |
| PES1 | 4082.165987 | 0.391161455 | 9.43E-06 |
| MN1 | 809.3964248 | 0.391117914 | 3.79E-02 |
| MAPKAPK5 | 1373.001255 | 0.390850525 | 9.70E-07 |
| ARPC2 | 12473.43948 | 0.39069318 | 1.38E-06 |
| ITGA5 | 9206.177774 | 0.390685958 | 1.86E-02 |
| RPS23 | 74273.86759 | 0.390648275 | 2.74E-04 |
| ERF | 3013.278853 | 0.390540588 | 3.22E-05 |
| PHKA1 | 1733.661679 | 0.390423775 | 7.03E-10 |
| MRPS2 | 2996.568875 | 0.39040714 | 1.02E-04 |
| PLEKHM2 | 4411.871107 | 0.390245425 | 6.12E-06 |
| METTL23 | 915.3090469 | 0.389795828 | 2.46E-05 |
| RPL35A | 14501.75282 | 0.389669954 | 1.47E-03 |
| DOT1L | 1539.184202 | 0.389538536 | 8.76E-04 |
| GFPT1 | 3629.248179 | 0.389196114 | 4.49E-02 |
| AKAP1 | 1854.077959 | 0.388868946 | 1.31E-03 |
| RPS18 | 68722.47473 | 0.388376376 | 6.86E-04 |
| GAGE13 | 2562.719045 | 0.388143597 | 1.78E-05 |
| POLE4 | 2168.142337 | 0.387867689 | 1.21E-03 |
| PICALM | 8631.628285 | 0.387811388 | 9.98E-04 |
| SLC25A39 | 4759.181151 | 0.387746962 | 8.31E-05 |
| MARCKS | 3627.182142 | 0.387590238 | 1.15E-04 |
| SSSCA1 | 1070.342479 | 0.387494269 | 1.59E-03 |

| | | | |
|---|---|---|---|
| ORAI1 | 1063.716768 | 0.387404175 | 4.73E-04 |
| SOX13 | 1139.603012 | 0.38732783 | 4.47E-04 |
| TRIP4 | 1456.396546 | 0.387111778 | 1.99E-05 |
| TMEM109 | 5624.877036 | 0.3868355 | 2.30E-03 |
| SAP18 | 2287.019192 | 0.386814276 | 9.89E-05 |
| SPRYD7 | 559.7320342 | 0.38659979 | 3.61E-03 |
| H2AFZ | 12211.53725 | 0.386532135 | 4.83E-05 |
| EIF3B | 14629.46064 | 0.386379331 | 4.15E-03 |
| RPL17 | 27377.12479 | 0.386204892 | 9.28E-06 |
| GPN3 | 1009.88344 | 0.386184553 | 4.37E-03 |
| CALU | 26908.66603 | 0.386175702 | 4.93E-03 |
| SOCS2 | 977.7645512 | 0.385925099 | 5.51E-04 |
| CXorf40A | 833.72996 | 0.38578419 | 6.06E-04 |
| UBALD1 | 463.8856587 | 0.385735822 | 4.09E-03 |
| VASP | 4094.575671 | 0.385594205 | 5.88E-03 |
| ARL2 | 3980.279198 | 0.385006787 | 4.34E-04 |
| ZDHHC1 | 267.080948 | 0.384722275 | 2.91E-02 |
| PFAS | 3969.518652 | 0.384577453 | 1.65E-05 |
| PNPT1 | 1683.829152 | 0.384398062 | 3.08E-02 |
| CKAP5 | 15304.98664 | 0.38426024 | 4.05E-03 |
| ATP5MC2 | 5676.214694 | 0.384078372 | 1.53E-04 |
| TFIP11 | 2496.731842 | 0.383876214 | 4.81E-04 |
| ZNF710 | 1212.046584 | 0.383752747 | 1.70E-04 |
| RIOK3 | 2038.172139 | 0.383751324 | 9.86E-03 |
| TKT | 23860.21316 | 0.383547196 | 6.39E-04 |
| USP32 | 3325.072913 | 0.383384014 | 2.45E-06 |
| CARS2 | 1179.357325 | 0.383350305 | 9.08E-04 |
| LUZP1 | 6874.52116 | 0.383273091 | 6.04E-09 |
| TNFAIP8L1 | 574.1983135 | 0.383241928 | 3.12E-03 |
| ZNF853 | 558.4164668 | 0.383197864 | 3.51E-02 |
| UBE2H | 13961.67353 | 0.38305606 | 4.11E-11 |
| DYNC1H1 | 30557.26086 | 0.382998595 | 4.41E-05 |
| MYO9B | 6638.196893 | 0.382712251 | 2.22E-03 |
| LINC01547 | 217.7179449 | 0.38258138 | 4.97E-02 |
| ERP29 | 5731.028148 | 0.382516733 | 7.87E-05 |
| STX5 | 2654.151479 | 0.38187474 | 7.10E-03 |
| TMEM14C | 4616.94698 | 0.381526557 | 5.10E-05 |
| SMYD5 | 1789.083633 | 0.381396229 | 4.78E-04 |
| H3F3B | 13249.7928 | 0.38103713 | 4.90E-08 |
| CDCA3 | 1454.557961 | 0.380902559 | 1.36E-02 |
| MRPS23 | 2563.61453 | 0.380840404 | 3.75E-06 |
| LARP1 | 12477.89571 | 0.380748457 | 6.76E-10 |
| NYAP1 | 365.2685364 | 0.380445759 | 1.31E-02 |
| NAP1L1 | 25109.18802 | 0.380434226 | 8.40E-10 |
| UBXN6 | 3914.433524 | 0.380425563 | 3.22E-03 |
| OSBPL2 | 1833.379553 | 0.380370819 | 7.75E-04 |
| SHC1 | 17857.68695 | 0.380020481 | 1.10E-03 |

| | | | |
|---|---|---|---|
| KHSRP | 10535.99363 | 0.379824743 | 6.17E-07 |
| FAM216A | 1300.115299 | 0.379757342 | 3.66E-03 |
| RNMT | 3107.790568 | 0.379651584 | 1.73E-02 |
| DDA1 | 1367.310378 | 0.379537129 | 3.72E-03 |
| EIF2B5 | 3033.847097 | 0.3793796 | 2.88E-05 |
| RUNDC1 | 835.3750685 | 0.379350973 | 3.43E-03 |
| GPS1 | 5156.171516 | 0.379043484 | 2.48E-03 |
| CCT5 | 38883.96642 | 0.378799996 | 3.00E-07 |
| BUD31 | 4153.64177 | 0.378746626 | 4.39E-04 |
| ING1 | 361.767476 | 0.37865527 | 1.14E-02 |
| PDCD5 | 1993.549848 | 0.378227953 | 1.33E-06 |
| PRSS23 | 21726.22584 | 0.378164747 | 6.11E-03 |
| CMAS | 1197.058059 | 0.378017604 | 1.40E-03 |
| MRPL11 | 2079.489287 | 0.37790896 | 2.99E-05 |
| BOLA3 | 668.8711214 | 0.37783861 | 4.16E-04 |
| C12orf45 | 368.0857885 | 0.37766536 | 1.55E-02 |
| GAGE1 | 7974.436024 | 0.377537313 | 1.77E-05 |
| ULK1 | 1698.020532 | 0.377407932 | 1.93E-04 |
| AARSD1 | 1040.493228 | 0.37713761 | 9.58E-08 |
| PRRC2A | 20762.84586 | 0.37712251 | 2.27E-08 |
| NUTF2 | 4592.139175 | 0.376949691 | 1.31E-02 |
| ACLY | 17193.67914 | 0.376859963 | 1.50E-03 |
| SLC37A4 | 2303.912945 | 0.376519634 | 2.58E-03 |
| AC026464.4 | 1259.810063 | 0.376313823 | 3.73E-04 |
| SORBS3 | 1260.08882 | 0.376280467 | 2.65E-02 |
| DNM2 | 7737.278026 | 0.376064388 | 1.12E-07 |
| TBP | 646.4204299 | 0.37599379 | 1.13E-03 |
| ZNF622 | 2500.069831 | 0.375745274 | 2.08E-07 |
| SEC24C | 6118.579717 | 0.375679444 | 8.37E-04 |
| VAPA | 4644.133299 | 0.375294073 | 5.03E-06 |
| ILF2 | 9055.317368 | 0.37527355 | 2.26E-07 |
| CSNK2B | 5612.548763 | 0.375071322 | 1.64E-04 |
| UBE2M | 2402.768163 | 0.374861433 | 4.18E-05 |
| AC097374.1 | 1567.411536 | 0.374834308 | 1.44E-02 |
| SCRIB | 5063.164234 | 0.374724503 | 4.41E-05 |
| INTS1 | 14293.109 | 0.374232072 | 2.28E-05 |
| LAMB2 | 8298.833374 | 0.374174996 | 6.24E-03 |
| R3HDM4 | 1971.816613 | 0.374075669 | 7.54E-04 |
| TPM3 | 16136.67089 | 0.374004671 | 3.45E-04 |
| EPN1 | 6033.942424 | 0.374000477 | 4.81E-04 |
| RAC1 | 16484.60893 | 0.373934654 | 4.21E-07 |
| ATP5F1B | 32311.30569 | 0.373646971 | 4.76E-05 |
| GPN1 | 3418.273067 | 0.373491928 | 1.60E-06 |
| RNF5 | 1615.342707 | 0.373470188 | 2.52E-02 |
| TAF6 | 4881.790056 | 0.373227745 | 8.47E-06 |
| CMTM4 | 3789.945746 | 0.37306328 | 5.19E-08 |
| COTL1 | 13773.83675 | 0.372794213 | 8.01E-04 |

| | | | |
|---|---|---|---|
| ANKS6 | 4072.304987 | 0.372741897 | 4.61E-04 |
| AC233702.1 | 357.8868138 | 0.372594838 | 2.60E-02 |
| SLC25A28 | 399.421377 | 0.372225131 | 6.30E-03 |
| HMGXB3 | 4166.29827 | 0.372221695 | 3.03E-05 |
| CDKN2AIPNL | 855.1231238 | 0.372191504 | 1.34E-03 |
| EIF3G | 8552.070086 | 0.371570654 | 4.39E-04 |
| RNF157 | 2624.034765 | 0.371443243 | 9.48E-04 |
| CC2D1A | 2123.997049 | 0.371390068 | 1.38E-03 |
| ACP1 | 2949.308581 | 0.371379534 | 8.19E-06 |
| C1orf43 | 7614.203548 | 0.371376313 | 1.21E-04 |
| ATAD3A | 1829.111853 | 0.371212124 | 1.07E-04 |
| HOXA1 | 545.3612809 | 0.371125836 | 2.95E-03 |
| NUP37 | 1711.26298 | 0.370939771 | 7.99E-03 |
| AP5Z1 | 1954.951571 | 0.370932633 | 1.15E-02 |
| COQ5 | 1997.429849 | 0.370818755 | 4.53E-07 |
| MRPL21 | 1743.40577 | 0.370210959 | 1.32E-04 |
| MAN2C1 | 3192.371798 | 0.370061597 | 2.12E-04 |
| SIGMAR1 | 5799.589283 | 0.370056722 | 1.34E-04 |
| URB1 | 5309.208866 | 0.370026031 | 8.29E-06 |
| PSMB2 | 4850.469396 | 0.369957083 | 8.58E-04 |
| CSNK2A1 | 11719.97268 | 0.369914278 | 1.39E-09 |
| SFXN4 | 585.4507196 | 0.369560959 | 4.61E-04 |
| OSBPL6 | 864.281503 | 0.369496359 | 1.64E-02 |
| CTBS | 682.3989996 | 0.369388993 | 8.41E-04 |
| DLST | 6606.867678 | 0.369360969 | 3.50E-05 |
| FAM104A | 1453.554848 | 0.369258036 | 1.68E-05 |
| CDC25C | 1731.072231 | 0.369221201 | 5.02E-04 |
| AUP1 | 4484.134215 | 0.369213574 | 2.11E-03 |
| PTGES3 | 10958.19736 | 0.368970966 | 7.18E-06 |
| ERI3 | 2151.816879 | 0.36888176 | 8.80E-04 |
| RBMS3 | 446.5159869 | 0.368783664 | 4.54E-02 |
| CDK5RAP3 | 3684.848713 | 0.368435397 | 2.99E-08 |
| DFFA | 3664.020143 | 0.368336759 | 2.47E-07 |
| MRPS27 | 6656.267008 | 0.368299649 | 1.24E-03 |
| TXNRD3 | 920.3708362 | 0.368282757 | 5.88E-04 |
| ASXL1 | 11653.72846 | 0.368117896 | 2.34E-08 |
| ZNF330 | 1429.442326 | 0.368081999 | 1.75E-03 |
| UQCC2 | 1854.703094 | 0.367786056 | 8.20E-04 |
| TUBG2 | 1807.888365 | 0.367459623 | 4.56E-05 |
| TBRG1 | 2566.845992 | 0.367342714 | 3.13E-04 |
| DDR1 | 3398.200686 | 0.367123469 | 4.84E-03 |
| TBC1D9 | 1044.573317 | 0.366853631 | 1.43E-03 |
| HS1BP3 | 1553.181874 | 0.366804258 | 7.37E-03 |
| AHSA1 | 8364.353404 | 0.366703322 | 8.58E-08 |
| KIF13B | 1664.961045 | 0.366503112 | 1.44E-04 |
| STK35 | 2874.044657 | 0.365498695 | 5.05E-11 |
| CAMLG | 1170.821212 | 0.365486377 | 1.08E-03 |

| | | | |
|---|---|---|---|
| MSRA | 717.8651147 | 0.36540773 | 1.94E-02 |
| LENG8-AS1 | 110.7386974 | 0.36510303 | 4.72E-02 |
| JPT1 | 2871.360134 | 0.36502206 | 5.40E-05 |
| POLR3E | 2515.286688 | 0.364894833 | 7.83E-03 |
| RALA | 4010.82315 | 0.364880607 | 4.68E-02 |
| METTL2B | 1796.131861 | 0.364870689 | 4.39E-06 |
| PPFIA1 | 3610.881528 | 0.36486687 | 3.83E-04 |
| KSR1 | 1136.231107 | 0.364596042 | 4.17E-02 |
| SNX30 | 2405.921239 | 0.364387619 | 2.65E-03 |
| CCNB2 | 4442.079969 | 0.364373778 | 1.89E-04 |
| PRKCZ | 1369.324973 | 0.364359262 | 8.98E-03 |
| DNMT1 | 10550.99223 | 0.364237269 | 1.05E-02 |
| SORD | 1813.484548 | 0.364211724 | 4.32E-04 |
| SNRPN | 5362.88291 | 0.364175752 | 2.25E-04 |
| SLC25A37 | 6361.631733 | 0.364104871 | 5.36E-10 |
| ZMIZ2 | 2157.698902 | 0.363928299 | 8.41E-08 |
| MTCL1 | 1412.753168 | 0.363820652 | 7.25E-04 |
| DBN1 | 7236.824837 | 0.363549245 | 1.79E-03 |
| TFB1M | 1043.033676 | 0.363482187 | 5.94E-04 |
| PRELID3B | 2493.846453 | 0.36331064 | 1.83E-02 |
| MFGE8 | 7060.251313 | 0.363275259 | 4.47E-03 |
| GRK3 | 836.2779387 | 0.36262186 | 2.54E-02 |
| HOXC11 | 245.5107089 | 0.362446836 | 3.31E-02 |
| TMEM120B | 1090.727739 | 0.362325076 | 1.45E-03 |
| KLHL17 | 698.8288926 | 0.362229817 | 6.89E-03 |
| KIF20A | 9603.437791 | 0.361464689 | 5.13E-04 |
| DNAJC19 | 835.3569696 | 0.361150283 | 2.09E-03 |
| TRAF3 | 2982.287153 | 0.36114565 | 8.66E-05 |
| ANAPC16 | 1880.704335 | 0.361126298 | 2.91E-05 |
| TCOF1 | 9862.217299 | 0.361079955 | 1.29E-06 |
| TTLL12 | 2960.526421 | 0.360893053 | 5.64E-04 |
| RSPH3 | 647.1392878 | 0.360847248 | 1.07E-04 |
| RPL27A | 44327.24657 | 0.360385482 | 6.15E-04 |
| FBXW4 | 791.3351442 | 0.360171403 | 6.40E-03 |
| RSU1 | 1873.166407 | 0.359953294 | 6.79E-05 |
| SERPINB9 | 1503.778653 | 0.359720621 | 2.39E-05 |
| RPL37 | 65992.22428 | 0.35960285 | 4.36E-03 |
| APTX | 2086.750278 | 0.359496097 | 1.55E-05 |
| FAM86C1 | 447.2350556 | 0.359465345 | 4.20E-02 |
| SRP68 | 5045.523619 | 0.359157479 | 7.30E-09 |
| AFG1L | 293.8883154 | 0.358989704 | 3.19E-02 |
| METTL5 | 1304.863506 | 0.358502344 | 1.97E-03 |
| EIF3H | 12961.00924 | 0.358480318 | 2.78E-06 |
| LSM10 | 1191.931926 | 0.358419238 | 1.40E-03 |
| IL17RA | 1078.041984 | 0.358395213 | 5.00E-02 |
| LRP11 | 2422.615292 | 0.358281357 | 7.10E-03 |
| TRMT1 | 1512.289143 | 0.358075401 | 2.79E-03 |

| | | | |
|---|---|---|---|
| ATXN7L1 | 780.669098 | 0.357756772 | 1.57E-02 |
| GLE1 | 3667.136514 | 0.357694373 | 1.11E-04 |
| RANGRF | 1588.216267 | 0.357243163 | 6.96E-04 |
| RASSF1 | 1191.274444 | 0.357154394 | 1.46E-02 |
| IQCB1 | 903.2856941 | 0.357132572 | 1.26E-02 |
| TAMM41 | 651.7368094 | 0.356816703 | 2.76E-04 |
| PHF2 | 2669.663996 | 0.356584436 | 9.40E-09 |
| NUP93 | 5090.165098 | 0.356398096 | 1.47E-05 |
| CNDP2 | 7468.433963 | 0.356284131 | 3.26E-04 |
| SERP1 | 3386.382468 | 0.356251332 | 8.75E-04 |
| SELENOS | 1716.629402 | 0.35604028 | 5.13E-06 |
| CENPN | 2368.501306 | 0.355888623 | 2.80E-02 |
| SLC25A15 | 687.4465156 | 0.355869928 | 2.93E-04 |
| RNF123 | 2422.551839 | 0.355800454 | 5.43E-04 |
| SOGA1 | 7215.020647 | 0.355576074 | 7.75E-04 |
| MMAB | 1653.207457 | 0.355571069 | 1.53E-02 |
| MAP2K5 | 1236.431012 | 0.355488063 | 1.34E-03 |
| ITPKC | 871.852918 | 0.355454377 | 9.44E-04 |
| C9orf64 | 1570.97536 | 0.3550483 | 2.02E-05 |
| TRMT61A | 1134.142873 | 0.354964877 | 2.72E-02 |
| BOP1 | 3358.256854 | 0.354896813 | 7.78E-04 |
| PUSL1 | 454.6324044 | 0.354883229 | 1.53E-03 |
| RARA | 1421.915282 | 0.354877326 | 1.65E-02 |
| SLC22A5 | 1353.716616 | 0.354605015 | 2.66E-03 |
| WFS1 | 1309.328652 | 0.354506773 | 3.20E-02 |
| TCHP | 1059.989031 | 0.354186768 | 1.59E-02 |
| AC131212.3 | 157.8229129 | 0.353860881 | 3.98E-02 |
| NRBF2 | 506.3490214 | 0.353742116 | 1.08E-03 |
| STK11 | 2145.068827 | 0.35335654 | 5.33E-04 |
| ELMOD3 | 825.5341406 | 0.353115795 | 1.39E-02 |
| ZBTB39 | 920.5177444 | 0.353056547 | 5.27E-03 |
| ANAPC1 | 5455.149671 | 0.352948399 | 3.53E-05 |
| STK10 | 3765.459772 | 0.352578224 | 5.57E-04 |
| TMEM147-AS1 | 1163.878717 | 0.352534396 | 2.50E-02 |
| NIPSNAP1 | 5969.896051 | 0.352448281 | 7.98E-03 |
| RNF166 | 1068.399366 | 0.352327129 | 1.44E-04 |
| CCDC124 | 1599.856452 | 0.352321559 | 1.78E-05 |
| LIMK1 | 4481.843518 | 0.35217627 | 7.09E-03 |
| SELENOF | 4529.787408 | 0.352108474 | 5.33E-07 |
| EIF2S3 | 9547.502372 | 0.35208043 | 3.52E-11 |
| APBA3 | 709.0816651 | 0.351867062 | 3.48E-02 |
| EXOC4 | 8628.260536 | 0.351844867 | 2.31E-06 |
| PPID | 1757.95365 | 0.35177403 | 3.19E-03 |
| SLC4A3 | 1292.885625 | 0.351670626 | 3.31E-03 |
| STK16 | 1384.410778 | 0.351629616 | 5.29E-04 |
| RAN | 15303.38853 | 0.351490948 | 3.30E-04 |
| POLB | 1462.502926 | 0.351457594 | 5.12E-04 |

| | | | |
|---|---|---|---|
| RPL21P16 | 5461.515123 | 0.351453363 | 9.76E-03 |
| PAXX | 1435.946098 | 0.351311 | 1.66E-04 |
| TEAD3 | 1178.905986 | 0.351159975 | 2.71E-03 |
| DENND4B | 2722.961269 | 0.35110273 | 1.09E-04 |
| SETD7 | 2957.166962 | 0.351096107 | 5.77E-05 |
| TBC1D14 | 3382.105884 | 0.351025992 | 2.38E-05 |
| TMEM41A | 1259.205691 | 0.350585088 | 4.59E-03 |
| VANGL2 | 2032.007805 | 0.350375202 | 7.21E-04 |
| TMEM245 | 5277.538846 | 0.350144829 | 4.08E-06 |
| LAMC3 | 1143.755153 | 0.350120025 | 3.33E-02 |
| STAU2 | 1901.995673 | 0.3496272 | 1.85E-03 |
| PEX19 | 4514.041774 | 0.349585078 | 2.35E-04 |
| GTF2H4 | 781.8735298 | 0.349187206 | 8.32E-03 |
| CLTA | 9649.743942 | 0.348135134 | 2.57E-04 |
| RPS29 | 12227.15154 | 0.348045642 | 1.06E-02 |
| NSMCE2 | 1140.479055 | 0.347991871 | 4.10E-05 |
| EHD2 | 10312.92724 | 0.34782773 | 3.93E-03 |
| GPR108 | 2585.88666 | 0.347771067 | 8.20E-03 |
| GATA2 | 692.1570249 | 0.347501548 | 4.82E-02 |
| TRPT1 | 1109.105546 | 0.347414683 | 3.76E-02 |
| ZBTB17 | 1035.308393 | 0.347363212 | 2.78E-05 |
| PSMC1 | 6088.680726 | 0.347291916 | 7.15E-06 |
| FAM98C | 277.9088541 | 0.347121635 | 2.42E-02 |
| SLC39A13 | 2800.97386 | 0.346991006 | 3.45E-02 |
| FANCE | 547.4079603 | 0.346874054 | 2.96E-03 |
| BCL7A | 490.2871555 | 0.34687021 | 2.25E-02 |
| CNNM4 | 988.0553982 | 0.346761439 | 4.96E-03 |
| C7orf50 | 3262.29235 | 0.346733151 | 2.18E-03 |
| PRKCD | 1783.577947 | 0.346678205 | 7.61E-04 |
| DUSP12 | 1147.714159 | 0.346505272 | 2.76E-02 |
| RCN1 | 15497.19177 | 0.346479735 | 4.51E-07 |
| EXOSC9 | 1360.485009 | 0.346414671 | 4.38E-02 |
| RAB5C | 6053.456357 | 0.346303591 | 6.29E-05 |
| SRRD | 1362.097613 | 0.346290134 | 1.60E-03 |
| POGK | 3075.627708 | 0.34622703 | 1.00E-05 |
| TTL | 3635.258305 | 0.346125795 | 6.02E-04 |
| TCEAL3 | 1081.559372 | 0.346053329 | 2.05E-02 |
| RPTOR | 2416.020726 | 0.345846011 | 9.28E-04 |
| FLCN | 2181.401598 | 0.345601065 | 1.48E-03 |
| CDYL | 1827.428318 | 0.345487951 | 2.91E-03 |
| RASSF2 | 380.6043208 | 0.344811847 | 2.65E-02 |
| PAK4 | 2072.569239 | 0.344701172 | 4.56E-03 |
| PWP1 | 3685.791824 | 0.34468405 | 9.48E-08 |
| DERL1 | 3417.012654 | 0.344531995 | 2.12E-04 |
| INO80E | 2856.807769 | 0.344500229 | 4.55E-03 |
| CCNQ | 1123.60927 | 0.344460248 | 4.20E-03 |
| CUTA | 2762.537115 | 0.344434126 | 3.74E-02 |

| | | | |
|---|---|---|---|
| SNX24 | 1531.40493 | 0.344260248 | 6.92E-03 |
| IPO5 | 38071.98562 | 0.344133585 | 8.55E-09 |
| NA | 279.0168569 | 0.343883473 | 4.38E-02 |
| ARMC10 | 2257.733891 | 0.343538481 | 1.63E-04 |
| EMC8 | 1576.06358 | 0.343438319 | 1.03E-03 |
| MAN2A2 | 3877.249022 | 0.343288765 | 2.71E-03 |
| DESI1 | 2612.386155 | 0.342989047 | 7.40E-06 |
| C19orf54 | 722.8808513 | 0.342963601 | 2.00E-03 |
| BLVRA | 2172.864212 | 0.342409397 | 3.67E-04 |
| B4GALT3 | 1911.562604 | 0.342193699 | 5.21E-03 |
| ACD | 1028.935391 | 0.341986874 | 1.31E-02 |
| PTPN23 | 3372.476872 | 0.341976188 | 2.46E-03 |
| FAM174A | 424.3305524 | 0.341971829 | 1.20E-02 |
| DNAJA4 | 883.5233635 | 0.341879299 | 3.46E-03 |
| HDAC7 | 5153.464871 | 0.341720873 | 3.75E-05 |
| FAM89A | 912.360877 | 0.341341173 | 1.39E-03 |
| TRIM62 | 875.1026027 | 0.341278084 | 1.22E-02 |
| CYCS | 4213.819034 | 0.341270344 | 3.87E-03 |
| NOP14 | 2656.143157 | 0.341259797 | 2.34E-11 |
| PSMF1 | 6766.592909 | 0.34122548 | 8.62E-04 |
| ATPAF2 | 838.2054896 | 0.341025215 | 1.17E-02 |
| ZNF581 | 889.5295823 | 0.340689125 | 3.23E-02 |
| IDH3A | 2872.949552 | 0.340551891 | 2.54E-02 |
| MAVS | 12155.02289 | 0.340550966 | 6.99E-05 |
| GEMIN7 | 848.4478189 | 0.340480778 | 1.45E-02 |
| PEX6 | 2151.85644 | 0.340394157 | 1.92E-02 |
| LRRC75B | 360.1727331 | 0.340283956 | 1.08E-02 |
| COQ2 | 766.2534857 | 0.340231122 | 1.16E-03 |
| NDUFS6 | 3119.638103 | 0.340227621 | 3.44E-03 |
| VPS33A | 1967.463228 | 0.339714386 | 2.02E-04 |
| GOLT1B | 1313.786465 | 0.339621789 | 1.49E-02 |
| FAM83D | 3176.229694 | 0.339574957 | 3.84E-03 |
| ODC1 | 5781.739414 | 0.339409043 | 4.66E-02 |
| DIP2C | 1132.351831 | 0.339233396 | 1.80E-02 |
| EGFR | 7757.382542 | 0.339088305 | 1.23E-06 |
| CRYZL1 | 678.2921722 | 0.338758846 | 1.28E-02 |
| COPZ1 | 6355.496849 | 0.338592721 | 1.53E-03 |
| PEX14 | 1078.492463 | 0.338434755 | 5.35E-03 |
| RNF216 | 6139.992269 | 0.338261506 | 4.10E-07 |
| TRIM65 | 1666.174936 | 0.338159707 | 1.52E-03 |
| TRAPPC5 | 686.9271114 | 0.33812358 | 4.32E-02 |
| PSMD11 | 5804.341134 | 0.338084676 | 2.17E-07 |
| TSG101 | 4160.63018 | 0.338078578 | 2.75E-04 |
| AGFG1 | 2601.944699 | 0.337694507 | 9.92E-03 |
| XYLT2 | 1979.267738 | 0.337531959 | 2.58E-03 |
| UBE2V1 | 9433.955559 | 0.337411314 | 2.02E-03 |
| RRP1 | 1635.850816 | 0.337145257 | 1.63E-03 |

| | | | |
|---|---|---|---|
| 8-Sep | 9809.378699 | 0.337110132 | 1.95E-05 |
| RFX2 | 533.5043503 | 0.337089466 | 1.37E-02 |
| PYCR2 | 3197.901368 | 0.337041209 | 9.22E-03 |
| ZNF263 | 2605.294114 | 0.33691794 | 2.23E-04 |
| IFNGR2 | 3672.123779 | 0.336868784 | 4.22E-03 |
| WEE1 | 1947.39113 | 0.336821149 | 3.01E-02 |
| NME2 | 13404.07001 | 0.336767981 | 5.42E-04 |
| PHF7 | 290.484141 | 0.336727872 | 4.91E-02 |
| SEC16A | 7783.876163 | 0.336325909 | 4.86E-05 |
| HTR7P1 | 368.6181894 | 0.336303448 | 3.04E-02 |
| CUEDC2 | 1415.340345 | 0.336162428 | 8.08E-03 |
| PCGF1 | 527.1915392 | 0.336154065 | 1.75E-02 |
| SNRPC | 5495.320625 | 0.336141121 | 3.42E-04 |
| LLGL1 | 5186.655941 | 0.336036448 | 2.11E-03 |
| RCC1 | 5720.614234 | 0.335980359 | 1.09E-03 |
| RPL36A | 17414.29202 | 0.335970982 | 9.63E-05 |
| AC097532.2 | 288.9884563 | 0.335874631 | 3.79E-02 |
| RABL6 | 5448.593957 | 0.335747748 | 1.29E-04 |
| PITPNM2 | 1205.666963 | 0.335728499 | 4.14E-03 |
| MPP6 | 815.7351252 | 0.335681255 | 2.27E-02 |
| CYB5A | 1340.256396 | 0.335377725 | 4.65E-03 |
| COX10 | 905.009525 | 0.335064926 | 3.00E-03 |
| SUCLG2 | 2395.918286 | 0.334715064 | 1.09E-04 |
| NEK9 | 9405.283839 | 0.334625911 | 1.22E-04 |
| TSR3 | 1839.045511 | 0.334500236 | 8.52E-04 |
| NDUFV1 | 7369.650581 | 0.334360123 | 5.78E-05 |
| ADCY9 | 1485.946218 | 0.334325068 | 5.36E-03 |
| TGFBI | 26371.45713 | 0.334310071 | 1.15E-02 |
| PTDSS1 | 5001.392315 | 0.334298708 | 1.22E-03 |
| GAGE2A | 3777.159662 | 0.334056556 | 9.24E-04 |
| PHB | 7327.068528 | 0.334042318 | 2.31E-03 |
| NARS2 | 1356.554678 | 0.333985008 | 4.40E-03 |
| LYAR | 1412.981459 | 0.333938463 | 2.57E-02 |
| MIEN1 | 938.6749156 | 0.333882655 | 4.69E-02 |
| TARS2 | 1686.668846 | 0.333718816 | 4.35E-04 |
| BUD13 | 1709.776183 | 0.333462396 | 1.27E-05 |
| SCARB1 | 3539.739284 | 0.333273135 | 9.58E-04 |
| TRAPPC9 | 1714.828313 | 0.333074612 | 4.14E-02 |
| CSAG1 | 2662.965097 | 0.333001738 | 6.88E-05 |
| SURF1 | 3085.300649 | 0.332501497 | 4.17E-03 |
| YIPF2 | 2105.78297 | 0.332384873 | 6.01E-03 |
| COX17 | 554.2075333 | 0.332295139 | 3.90E-02 |
| VRK3 | 1353.668644 | 0.332269061 | 4.65E-02 |
| R3HDM2 | 1816.887832 | 0.332263483 | 2.40E-02 |
| HJURP | 3243.06767 | 0.332210221 | 1.49E-02 |
| PPP2CB | 3807.900051 | 0.331999437 | 4.95E-10 |
| TFAP2A | 2148.867207 | 0.331885015 | 1.10E-02 |

| | | | |
|---|---|---|---|
| TDRKH | 1007.401832 | 0.331857854 | 1.31E-03 |
| NDUFS8 | 2353.294169 | 0.3318397 | 2.04E-03 |
| COPS6 | 7614.260616 | 0.331576034 | 3.39E-05 |
| RRP7A | 1939.453049 | 0.331318822 | 4.49E-04 |
| MRPL45 | 2482.560395 | 0.331180424 | 6.96E-05 |
| POLR3A | 1936.558238 | 0.330832232 | 8.18E-04 |
| TP53I13 | 980.1633656 | 0.330776112 | 3.16E-03 |
| CBX4 | 1634.077251 | 0.330604283 | 2.17E-02 |
| SLC39A11 | 639.9957675 | 0.330443812 | 1.02E-02 |
| AGAP2-AS1 | 607.7561161 | 0.330377799 | 3.89E-02 |
| UROD | 4803.217769 | 0.329998845 | 5.70E-03 |
| HADHA | 12749.68248 | 0.329836334 | 1.17E-04 |
| USF2 | 2621.385513 | 0.329685664 | 3.70E-04 |
| UBXN11 | 840.3304153 | 0.329614445 | 6.39E-03 |
| DENND5B | 1132.684712 | 0.329586566 | 2.30E-03 |
| HEMK1 | 1344.047739 | 0.329431824 | 2.18E-03 |
| PSMB3 | 4278.148867 | 0.329431019 | 1.68E-02 |
| PRKRA | 1493.271741 | 0.329199632 | 6.28E-05 |
| RPS14 | 16221.32272 | 0.328887479 | 4.93E-02 |
| GPR107 | 9096.320411 | 0.328731826 | 1.97E-05 |
| MIEF2 | 859.0616354 | 0.328546244 | 3.46E-02 |
| AP1S1 | 3904.931659 | 0.328308747 | 1.99E-02 |
| SMC1A | 8629.188695 | 0.328200386 | 7.53E-03 |
| AK2 | 8587.731938 | 0.328036049 | 7.39E-04 |
| INTS4 | 2113.040451 | 0.328033447 | 1.66E-06 |
| PPP1R7 | 3054.404302 | 0.327911797 | 2.25E-05 |
| CYC1 | 5704.306008 | 0.327422226 | 6.30E-04 |
| MRTO4 | 2839.374245 | 0.327415689 | 1.27E-03 |
| SUPV3L1 | 1062.588736 | 0.327389534 | 1.73E-02 |
| GLUL | 5237.162868 | 0.327201632 | 7.65E-03 |
| SSH1 | 4488.68356 | 0.327182358 | 1.49E-05 |
| ZBTB18 | 811.3954707 | 0.32716257 | 5.11E-03 |
| PI4K2A | 1765.032642 | 0.327032553 | 2.20E-02 |
| DBNDD1 | 1129.439015 | 0.326935718 | 4.61E-02 |
| ZNRD1 | 1174.003622 | 0.326736291 | 1.81E-02 |
| CTDP1 | 1130.124063 | 0.326679841 | 2.02E-03 |
| TOMM7 | 3903.636684 | 0.326417678 | 2.57E-03 |
| CHST11 | 1000.865333 | 0.326343527 | 4.93E-02 |
| DNAAF5 | 2856.312045 | 0.326216963 | 5.88E-05 |
| RPL24 | 16195.05698 | 0.326181028 | 3.97E-03 |
| PDIA5 | 1145.144501 | 0.326043 | 2.75E-02 |
| NDUFA10 | 3977.783713 | 0.325838513 | 5.33E-04 |
| RECQL5 | 1712.668547 | 0.325758619 | 7.02E-03 |
| PELP1 | 3679.224611 | 0.325708768 | 4.06E-04 |
| INPPL1 | 9644.907944 | 0.325638596 | 1.10E-03 |
| EEF2K | 2141.191494 | 0.325546168 | 2.02E-02 |
| RAB5IF | 3288.76951 | 0.325534645 | 4.29E-03 |

| | | | |
|---|---|---|---|
| ST6GALNAC4 | 1223.903519 | 0.32526802 | 3.40E-03 |
| HDDC3 | 769.9143343 | 0.32511475 | 9.61E-03 |
| AGO2 | 1010.035818 | 0.324992669 | 4.74E-02 |
| SPR | 1452.076523 | 0.324878285 | 2.24E-02 |
| PFN2 | 10427.95587 | 0.32484007 | 8.35E-05 |
| ANO6 | 6166.89251 | 0.324814785 | 2.09E-03 |
| DDX10 | 2148.269443 | 0.324682901 | 6.63E-03 |
| SCLY | 1049.079079 | 0.324611657 | 1.93E-02 |
| ELP2 | 3950.602411 | 0.324415617 | 1.10E-03 |
| DBF4B | 1125.920736 | 0.324298387 | 7.56E-03 |
| LRIF1 | 1383.86094 | 0.324053572 | 2.16E-02 |
| AC245060.4 | 370.64184 | 0.32403285 | 4.19E-03 |
| SET | 25045.76652 | 0.323324417 | 1.71E-02 |
| PDE6D | 1308.982486 | 0.323213366 | 4.26E-04 |
| ADPRH | 512.9493166 | 0.323203121 | 2.24E-02 |
| PPCS | 1325.8896 | 0.323136779 | 8.64E-04 |
| RSL1D1 | 10055.13546 | 0.323131343 | 1.22E-02 |
| MEF2A | 1827.254252 | 0.322796789 | 1.32E-02 |
| CDK5RAP1 | 2109.566458 | 0.322738935 | 1.95E-04 |
| PTDSS2 | 1997.212239 | 0.32264454 | 1.25E-02 |
| SMCR8 | 1335.19602 | 0.322596985 | 3.88E-05 |
| NOP9 | 3695.923527 | 0.322590378 | 2.96E-04 |
| RPL19 | 41793.06361 | 0.322428816 | 2.00E-03 |
| TRMT2B | 1049.45023 | 0.322404298 | 3.49E-04 |
| ZNF692 | 657.41101 | 0.322290411 | 7.96E-03 |
| CTNNBL1 | 3608.026851 | 0.322245552 | 4.31E-07 |
| ESS2 | 772.4988779 | 0.321984049 | 1.00E-02 |
| WWP2 | 2287.978635 | 0.321782681 | 7.65E-04 |
| NR2C2AP | 655.2007936 | 0.321750276 | 4.07E-03 |
| CLIP2 | 3854.007439 | 0.321649968 | 2.22E-03 |
| RAB4B | 443.5158268 | 0.321397743 | 1.94E-02 |
| CHCHD10 | 1240.010778 | 0.32127809 | 1.39E-02 |
| KMT2D | 6520.858359 | 0.321240976 | 3.59E-05 |
| ABTB2 | 1119.031511 | 0.321009239 | 2.12E-04 |
| CUL4A | 3078.197308 | 0.320835586 | 3.82E-03 |
| MRPL46 | 1238.821764 | 0.320463202 | 1.16E-04 |
| IKBKB | 2090.625545 | 0.320432624 | 5.05E-04 |
| CFL1 | 47571.41666 | 0.320354091 | 1.31E-02 |
| EIF5A2 | 297.4165173 | 0.320351871 | 2.64E-02 |
| ATP6V0C | 2785.293244 | 0.319984433 | 2.10E-02 |
| EXOSC2 | 2807.209963 | 0.31990559 | 1.06E-07 |
| YWHAE | 30929.50022 | 0.319865462 | 5.56E-05 |
| KDELR2 | 11593.04683 | 0.319819935 | 7.85E-03 |
| USP31 | 1052.765356 | 0.319695386 | 3.26E-02 |
| SNAPC3 | 1889.170263 | 0.319533637 | 3.53E-03 |
| SHANK3 | 593.1335192 | 0.31936634 | 1.39E-02 |
| MAGOHB | 603.1906548 | 0.319290149 | 5.04E-03 |

| | | | |
|---|---|---|---|
| BRPF1 | 1247.041248 | 0.31910448 | 7.55E-03 |
| TMBIM6 | 24594.40474 | 0.319034294 | 2.13E-04 |
| RPS27A | 29292.20617 | 0.318744583 | 1.73E-04 |
| POM121 | 7411.262456 | 0.318716818 | 2.57E-03 |
| PELO | 2799.47239 | 0.318522391 | 1.30E-04 |
| AP2A1 | 5271.47238 | 0.318393779 | 9.23E-03 |
| NOP10 | 2112.578788 | 0.318295278 | 2.32E-03 |
| PTPN2 | 1323.929657 | 0.318167842 | 1.55E-03 |
| ISCA2 | 1204.879052 | 0.317736683 | 2.84E-04 |
| FAM192A | 4054.220044 | 0.317640114 | 4.11E-08 |
| XRCC6 | 20895.91892 | 0.317594372 | 2.47E-04 |
| MRPS35 | 1793.927643 | 0.317558877 | 1.64E-03 |
| STAG3L5P-PVRI( | 1727.688963 | 0.317548422 | 2.61E-02 |
| INTS11 | 4008.1811 | 0.317539604 | 5.32E-04 |
| TBC1D2B | 6479.875257 | 0.31744716 | 4.52E-04 |
| GDI1 | 9466.865383 | 0.317295124 | 6.26E-05 |
| RPL10A | 24437.36707 | 0.317211066 | 1.67E-03 |
| LIN52 | 827.3337377 | 0.316988311 | 7.10E-03 |
| DMWD | 1492.062478 | 0.316713788 | 1.55E-03 |
| LTB4R2 | 430.8196029 | 0.316698875 | 1.14E-02 |
| AC073869.1 | 886.8040614 | 0.316454914 | 2.00E-04 |
| DCAF7 | 7641.624779 | 0.316101831 | 3.47E-03 |
| LMTK2 | 2508.032902 | 0.315991142 | 4.46E-05 |
| RCCD1 | 1438.649141 | 0.315942886 | 2.57E-04 |
| FAM229B | 637.9454571 | 0.315766287 | 9.02E-03 |
| NADSYN1 | 1687.908599 | 0.315585302 | 2.93E-03 |
| NMNAT1 | 617.3995545 | 0.315581095 | 2.25E-02 |
| PPP6C | 3935.07532 | 0.315567353 | 6.41E-06 |
| 9-Mar | 716.9582159 | 0.315430724 | 3.66E-02 |
| CHTF18 | 1995.240987 | 0.315419306 | 4.91E-02 |
| NSUN4 | 1220.122466 | 0.315360548 | 2.10E-03 |
| PTGIS | 1269.187612 | 0.315202335 | 2.69E-02 |
| SLC44A2 | 6112.518187 | 0.315037188 | 1.46E-03 |
| MRPS12 | 1199.187636 | 0.314303229 | 7.21E-03 |
| TMEM134 | 704.4764652 | 0.314201997 | 3.94E-02 |
| USP36 | 1986.048346 | 0.314198961 | 7.24E-04 |
| EIF2B2 | 2814.549157 | 0.313914799 | 1.19E-02 |
| DGUOK | 3302.395692 | 0.313829397 | 3.17E-04 |
| NFE2L1 | 19179.84364 | 0.313610783 | 2.89E-03 |
| CANX | 22952.90753 | 0.313590287 | 4.77E-03 |
| GCN1 | 15271.39176 | 0.313419724 | 2.23E-03 |
| JAK1 | 6969.182078 | 0.313291085 | 5.23E-06 |
| NANS | 3649.193973 | 0.313261631 | 1.42E-05 |
| PRELID1 | 4349.040238 | 0.312634992 | 1.69E-03 |
| CS | 9835.018622 | 0.312625096 | 1.63E-04 |
| CLPB | 1753.166047 | 0.312582861 | 1.74E-03 |
| GGA3 | 1961.376462 | 0.312558098 | 3.36E-03 |

| | | | |
|---|---|---|---|
| PSMA7 | 11075.96575 | 0.312456507 | 1.09E-04 |
| CKS2 | 3853.770361 | 0.312160584 | 1.23E-03 |
| SIN3B | 3202.374621 | 0.31175183 | 5.33E-08 |
| EME1 | 841.7569482 | 0.311722712 | 3.09E-02 |
| TMEM8A | 4587.609126 | 0.311634246 | 1.88E-02 |
| PPM1M | 1735.27223 | 0.311433901 | 1.91E-02 |
| THOC6 | 2050.440282 | 0.311400177 | 3.15E-03 |
| EFHD2 | 3883.799029 | 0.311314172 | 3.96E-02 |
| EIF5A | 21651.00118 | 0.310944566 | 4.55E-04 |
| ATG3 | 1629.288593 | 0.310551812 | 5.82E-04 |
| GPX4 | 7821.112851 | 0.310268879 | 1.33E-02 |
| HLA-C | 3125.12319 | 0.31025686 | 4.77E-02 |
| SIPA1L3 | 2104.661638 | 0.309579636 | 2.62E-02 |
| FAM220A | 1262.311827 | 0.309522337 | 3.62E-02 |
| SLC22A18 | 622.2696081 | 0.308751293 | 2.14E-02 |
| QSOX2 | 2784.490846 | 0.308667435 | 9.14E-06 |
| TTLL4 | 2183.833793 | 0.308324916 | 1.94E-02 |
| NAGA | 2652.792476 | 0.308113759 | 2.47E-02 |
| LDLRAP1 | 1727.553505 | 0.307890883 | 4.03E-03 |
| OTUD5 | 3164.940614 | 0.307803862 | 1.38E-02 |
| LINC01578 | 1662.10461 | 0.307742592 | 2.22E-02 |
| TMEM167B | 1344.383655 | 0.307686105 | 1.20E-02 |
| TFG | 4603.610223 | 0.307424056 | 4.65E-06 |
| DEAF1 | 3949.248938 | 0.307085138 | 3.63E-02 |
| STX18 | 1205.829051 | 0.306834845 | 6.20E-03 |
| UBAP2 | 3251.690654 | 0.306685414 | 1.21E-04 |
| DNAJB1 | 5494.200964 | 0.30658342 | 3.61E-02 |
| YIPF6 | 1932.021176 | 0.306562747 | 5.37E-03 |
| MALSU1 | 1736.348579 | 0.306294583 | 2.04E-02 |
| UPF1 | 5584.674034 | 0.306140824 | 1.82E-04 |
| CLUH | 3865.37148 | 0.305743678 | 6.20E-04 |
| ELOVL5 | 4586.484995 | 0.305606661 | 1.07E-05 |
| SEC22B | 4612.525216 | 0.305556463 | 1.56E-04 |
| FAM57A | 872.282127 | 0.305554459 | 1.29E-02 |
| CAMK2N1 | 1039.323332 | 0.305453924 | 2.38E-02 |
| NR1H2 | 1443.06312 | 0.305430457 | 1.89E-02 |
| MCL1 | 7487.483715 | 0.305235231 | 8.62E-05 |
| RBM33 | 2975.818229 | 0.305230845 | 1.53E-02 |
| TMED3 | 5421.603628 | 0.305144652 | 3.50E-02 |
| RPIA | 1240.066785 | 0.305143727 | 1.15E-04 |
| CSTF2 | 2441.068491 | 0.30510762 | 9.72E-06 |
| CCDC144NL | 1842.281029 | 0.304899467 | 9.34E-03 |
| TNIP2 | 1383.159776 | 0.304859633 | 3.97E-04 |
| SNHG6 | 2615.171978 | 0.30446147 | 1.05E-03 |
| KAT8 | 1737.846709 | 0.304436247 | 3.31E-02 |
| ZNF277 | 1041.934133 | 0.304109965 | 5.67E-03 |
| TSPAN18 | 4006.491139 | 0.30405128 | 4.03E-03 |

| | | | |
|---|---|---|---|
| ELMSAN1 | 2538.049382 | 0.304049757 | 1.43E-03 |
| B4GALT5 | 5897.880734 | 0.303896396 | 2.30E-02 |
| QRSL1 | 944.9892823 | 0.303875364 | 3.31E-02 |
| AEBP1 | 10182.06945 | 0.303712992 | 2.00E-02 |
| PSMC4 | 5691.909785 | 0.303255386 | 5.91E-04 |
| DCTN1 | 12802.93387 | 0.302402483 | 2.20E-03 |
| PDPK1 | 4241.916576 | 0.3020265 | 2.62E-03 |
| PTPMT1 | 1669.947516 | 0.302020379 | 1.20E-03 |
| TRAF4 | 2091.976798 | 0.301607235 | 5.18E-03 |
| ELOC | 2405.537577 | 0.301369715 | 1.40E-04 |
| ATP6V1G1 | 2695.790776 | 0.30121877 | 5.20E-05 |
| NT5C3B | 3653.141291 | 0.301114583 | 9.10E-04 |
| TMEM184B | 4398.019052 | 0.301088554 | 1.52E-02 |
| TSEN54 | 1095.294017 | 0.300995061 | 2.57E-02 |
| RPL23 | 29548.49268 | 0.300871592 | 4.20E-03 |
| TAF3 | 497.5951653 | 0.30081478 | 1.36E-02 |
| MCU | 1122.807796 | 0.300600519 | 9.30E-04 |
| TCP1 | 10078.89212 | 0.300404028 | 3.83E-06 |
| PMM1 | 1793.70348 | 0.300293431 | 3.17E-02 |
| LSM6 | 664.6222914 | 0.300083281 | 3.72E-03 |
| DIABLO | 2319.996166 | 0.299829128 | 2.09E-03 |
| STX10 | 2145.396318 | 0.299718288 | 3.82E-03 |
| NDUFA8 | 3018.998062 | 0.299658971 | 8.24E-03 |
| RPL29 | 36284.96416 | 0.299606003 | 4.31E-02 |
| DTYMK | 2052.910667 | 0.299393125 | 3.43E-03 |
| MACF1 | 17521.76745 | 0.29922097 | 1.67E-05 |
| ZNF646 | 2349.21993 | 0.299112578 | 1.27E-02 |
| FCHSD1 | 1323.690342 | 0.298940068 | 4.06E-02 |
| CUL9 | 2107.237193 | 0.298439697 | 2.63E-02 |
| ABR | 5944.267889 | 0.298222099 | 6.59E-03 |
| BRD1 | 1179.651156 | 0.298216607 | 1.71E-03 |
| NOL7 | 2262.371265 | 0.29804629 | 5.53E-04 |
| NUBP2 | 2876.219677 | 0.297891145 | 7.96E-03 |
| DDHD2 | 3026.27877 | 0.29753555 | 1.25E-03 |
| RPN1 | 10565.1925 | 0.297429567 | 8.76E-03 |
| COX6A1 | 5248.247842 | 0.297423596 | 3.35E-03 |
| TMEM243 | 999.7284056 | 0.297346716 | 4.65E-03 |
| CPNE1 | 12635.24221 | 0.297315549 | 1.32E-02 |
| RAB7A | 10166.49167 | 0.297005386 | 5.85E-03 |
| CHCHD1 | 767.6253033 | 0.296800744 | 2.71E-02 |
| ZFYVE19 | 888.1861836 | 0.296267361 | 2.31E-02 |
| RPL26 | 31543.39492 | 0.296149139 | 3.91E-04 |
| DIS3L2 | 1432.177132 | 0.295902634 | 4.07E-03 |
| HOXB4 | 680.793008 | 0.295710731 | 5.88E-03 |
| USB1 | 2856.737564 | 0.295598618 | 3.18E-03 |
| ATP5PD | 3820.956853 | 0.29556107 | 4.40E-03 |
| HMOX2 | 1518.610831 | 0.295412973 | 4.77E-02 |

| | | | |
|---|---|---|---|
| PPAN | 1600.837878 | 0.295284434 | 4.05E-02 |
| MEX3D | 606.3152883 | 0.29518567 | 4.76E-02 |
| MPC2 | 1051.78884 | 0.295148417 | 2.36E-03 |
| NDE1 | 2540.9428 | 0.295092515 | 4.27E-03 |
| MYBBP1A | 5944.230185 | 0.295049391 | 3.03E-02 |
| ZNF480 | 608.6021465 | 0.294962849 | 2.56E-02 |
| NUP188 | 7233.696582 | 0.294896403 | 3.85E-06 |
| LETMD1 | 2616.411317 | 0.294790254 | 3.53E-03 |
| NCLN | 3912.468455 | 0.294460957 | 7.83E-03 |
| STRN4 | 4766.955253 | 0.294337042 | 2.25E-03 |
| RPUSD1 | 1588.871539 | 0.294283052 | 2.16E-02 |
| TMEM11 | 884.3596645 | 0.293983773 | 1.06E-03 |
| AFDN | 4425.730501 | 0.293630282 | 3.92E-02 |
| ATP6V1E2 | 503.4465659 | 0.292307393 | 4.08E-02 |
| PSMA5 | 4318.210261 | 0.292298141 | 2.45E-03 |
| CBLL1 | 1559.789929 | 0.292241258 | 1.91E-03 |
| NCK2 | 1845.894516 | 0.292067827 | 1.14E-02 |
| RPL31 | 25986.45476 | 0.291853614 | 2.07E-03 |
| SNAI1 | 1108.966077 | 0.291827927 | 4.50E-02 |
| IFT20 | 1159.347537 | 0.291758297 | 1.03E-04 |
| PRPF3 | 2440.32239 | 0.291636078 | 9.51E-07 |
| CHP1 | 2502.400603 | 0.291546414 | 3.64E-03 |
| C1orf109 | 1140.631876 | 0.291427021 | 1.78E-03 |
| CPNE2 | 3888.561043 | 0.291397227 | 9.81E-03 |
| ENTR1 | 4996.79375 | 0.291375917 | 2.00E-04 |
| PLCE1 | 1326.090032 | 0.291305082 | 2.66E-02 |
| STARD3 | 1832.710767 | 0.291278362 | 3.85E-02 |
| CANT1 | 3836.902629 | 0.290971485 | 2.13E-02 |
| MRPS18A | 1552.70699 | 0.290901184 | 6.98E-03 |
| BBS2 | 3444.019886 | 0.290591008 | 8.07E-04 |
| ZFYVE27 | 817.0816198 | 0.290337504 | 3.68E-03 |
| MRPL18 | 1831.355952 | 0.290270617 | 4.47E-05 |
| CERS5 | 1746.370704 | 0.289950927 | 1.43E-02 |
| COX8A | 6710.786952 | 0.289774923 | 1.47E-03 |
| ADIRF-AS1 | 624.3818623 | 0.289368213 | 4.07E-02 |
| RNF7 | 2168.713989 | 0.289356004 | 7.01E-04 |
| SCNM1 | 804.5286563 | 0.289271752 | 4.70E-02 |
| STYXL1 | 1931.959515 | 0.28924572 | 3.48E-02 |
| RUBCN | 1148.934787 | 0.28923639 | 3.84E-03 |
| PREB | 2909.820778 | 0.28917421 | 4.86E-03 |
| KRT10 | 834.2809818 | 0.289168085 | 9.21E-03 |
| EIF2AK4 | 3997.221069 | 0.289015148 | 1.32E-06 |
| NAPA | 3117.244375 | 0.288853077 | 1.68E-02 |
| NUDCD3 | 9536.33003 | 0.288720551 | 3.15E-02 |
| PPA1 | 2590.012982 | 0.288163619 | 9.55E-03 |
| CHID1 | 5044.44065 | 0.288147511 | 9.88E-03 |
| PEX5 | 2983.181293 | 0.288096034 | 8.84E-03 |

| | | | |
|---|---|---|---|
| RTCB | 5427.681484 | 0.287793589 | 1.08E-03 |
| BPNT1 | 1216.423968 | 0.287684513 | 2.54E-02 |
| CUX1 | 5903.96191 | 0.287286836 | 1.72E-03 |
| CCT3 | 14487.18287 | 0.287259749 | 4.42E-02 |
| C20orf27 | 2903.6291 | 0.287178073 | 2.16E-02 |
| TRIT1 | 1099.34828 | 0.287165339 | 1.63E-02 |
| DYM | 2560.880205 | 0.286731162 | 4.01E-03 |
| CAPN10 | 1051.764605 | 0.286648903 | 2.03E-03 |
| LAMP2 | 7955.899773 | 0.286501736 | 7.88E-04 |
| RELA | 4313.421176 | 0.286464611 | 2.42E-02 |
| RIC1 | 2976.271407 | 0.286320513 | 2.61E-06 |
| NIT2 | 1435.897603 | 0.286290414 | 6.36E-03 |
| HSF1 | 3249.795479 | 0.286012773 | 1.55E-02 |
| GNG5 | 2669.344981 | 0.285564138 | 6.53E-03 |
| PABPC4 | 6179.496463 | 0.285445041 | 1.38E-03 |
| NDUFS2 | 5203.207594 | 0.285297764 | 2.15E-04 |
| AAGAB | 3925.30352 | 0.285231711 | 1.18E-04 |
| FRG1HP | 786.3343413 | 0.28515111 | 7.43E-03 |
| FBXL5 | 1766.997965 | 0.285109044 | 1.47E-04 |
| NOD1 | 846.3666035 | 0.284950309 | 6.50E-03 |
| SNU13 | 7127.161613 | 0.284943804 | 4.44E-03 |
| TAP2 | 3870.967011 | 0.284937261 | 6.96E-03 |
| ATP8B2 | 4883.448368 | 0.284749792 | 3.02E-02 |
| TRIR | 3142.154741 | 0.284318785 | 1.70E-02 |
| LTBR | 4379.978864 | 0.284142699 | 5.71E-03 |
| DUS1L | 2081.113592 | 0.284119367 | 6.84E-05 |
| DIP2B | 4273.333063 | 0.28339177 | 8.48E-05 |
| STAG3L5P | 572.7572891 | 0.283176388 | 2.54E-02 |
| ZNF274 | 1260.893165 | 0.282809129 | 1.19E-02 |
| SNX15 | 1870.19779 | 0.282616645 | 1.28E-03 |
| NUP98 | 10252.36378 | 0.282361972 | 1.94E-04 |
| CDK9 | 5047.740811 | 0.282047387 | 3.35E-02 |
| TMEM70 | 998.0146355 | 0.281917958 | 1.37E-03 |
| KLHDC10 | 3849.172421 | 0.281620865 | 1.08E-03 |
| G6PD | 13437.0078 | 0.281314736 | 1.74E-02 |
| FAR2 | 1062.330359 | 0.281227206 | 1.07E-02 |
| COX4I1 | 10109.85953 | 0.28115771 | 3.94E-02 |
| TMED4 | 5466.26898 | 0.281153945 | 7.47E-03 |
| F8 | 532.5279691 | 0.281144478 | 4.54E-02 |
| ANKRD27 | 1949.923053 | 0.281134947 | 8.99E-04 |
| KLHDC4 | 1557.359036 | 0.281001118 | 3.20E-02 |
| ZNF275 | 1849.137419 | 0.280017059 | 1.72E-04 |
| TXNDC5 | 23972.78375 | 0.279828908 | 1.28E-02 |
| OCRL | 3249.787062 | 0.279440462 | 6.22E-05 |
| RAB2A | 3018.845323 | 0.279394872 | 4.27E-06 |
| C7orf31 | 557.6287 | 0.279273488 | 4.99E-02 |
| C19orf48 | 2555.273303 | 0.27918122 | 2.06E-03 |

| | | | |
|---|---|---|---|
| MPV17L2 | 689.7563318 | 0.278887534 | 3.04E-02 |
| PLLP | 455.6051462 | 0.278859467 | 2.85E-02 |
| ZNF281 | 1503.230845 | 0.278852453 | 4.37E-03 |
| DHX16 | 3120.149289 | 0.278790885 | 2.02E-03 |
| RHOG | 1700.297983 | 0.278701593 | 2.51E-02 |
| LRRC47 | 2958.002262 | 0.278604524 | 4.34E-02 |
| PCBP1 | 10328.69218 | 0.278602335 | 2.00E-03 |
| GUK1 | 4974.434092 | 0.278410965 | 2.62E-02 |
| CCDC144NL-AS1 | 1631.969598 | 0.278091566 | 1.91E-02 |
| CCDC22 | 951.556721 | 0.277844477 | 1.70E-02 |
| KIAA1958 | 673.3497986 | 0.277789667 | 4.81E-02 |
| ARFGAP1 | 6666.931935 | 0.277754914 | 1.27E-02 |
| CCT2 | 12555.94788 | 0.277592572 | 1.84E-05 |
| SSRP1 | 19588.32243 | 0.277372996 | 1.95E-04 |
| RBSN | 2154.092521 | 0.277369523 | 2.75E-03 |
| CDV3 | 9627.780627 | 0.276761657 | 2.62E-02 |
| ATP6V0B | 2681.18054 | 0.276653184 | 4.10E-02 |
| ITCH | 4592.164107 | 0.276601181 | 3.67E-02 |
| TOM1L2 | 4289.413806 | 0.27655237 | 4.74E-02 |
| RBPMS2 | 349.7867648 | 0.276518612 | 3.50E-02 |
| NACA | 14467.33557 | 0.27638214 | 1.92E-02 |
| RAB10 | 4799.435422 | 0.276122261 | 8.20E-03 |
| EMG1 | 2344.946931 | 0.275810451 | 6.49E-03 |
| TIMM10 | 1509.425945 | 0.275783807 | 1.75E-02 |
| PIGL | 581.9689132 | 0.275479081 | 2.45E-02 |
| SEC61B | 3153.007258 | 0.275426538 | 1.37E-03 |
| EMC4 | 1681.225534 | 0.275224306 | 4.52E-03 |
| ACTR3 | 8588.083485 | 0.275202988 | 3.89E-03 |
| ZZEF1 | 4289.269115 | 0.275069252 | 6.38E-04 |
| OPTN | 3436.814398 | 0.27464212 | 1.12E-02 |
| UBE2L3 | 4746.243312 | 0.274628175 | 1.92E-02 |
| GCDH | 1021.008364 | 0.27461066 | 2.80E-02 |
| MINK1 | 4742.302078 | 0.274302053 | 2.88E-02 |
| TOMM20 | 6224.484129 | 0.274213811 | 1.86E-03 |
| DHX34 | 1155.079236 | 0.274045885 | 2.91E-02 |
| SNX5 | 7442.90652 | 0.274002007 | 3.87E-03 |
| PASD1 | 4060.815916 | 0.273476315 | 8.65E-04 |
| POLR2J3 | 1905.043087 | 0.273446987 | 3.97E-02 |
| CLPTM1L | 15633.00909 | 0.273431623 | 2.71E-03 |
| YWHAG | 14945.9692 | 0.272858072 | 1.02E-05 |
| ARPC3 | 7377.140275 | 0.272755865 | 6.64E-03 |
| SSNA1 | 2077.769422 | 0.272652548 | 1.64E-02 |
| RPL11 | 36795.27254 | 0.272516375 | 1.38E-03 |
| TRIM27 | 4409.412563 | 0.27205627 | 4.30E-03 |
| GMPR2 | 4514.04944 | 0.27171441 | 2.03E-03 |
| GAGE12C | 12322.95692 | 0.270985636 | 1.34E-03 |
| FBXW5 | 3797.76075 | 0.270622803 | 3.26E-02 |

| | | | |
|---|---|---|---|
| RPL23AP82 | 694.2770221 | 0.270591544 | 1.37E-02 |
| NUP88 | 3423.063769 | 0.270478727 | 7.78E-04 |
| EXOSC8 | 1008.870093 | 0.270424639 | 1.65E-02 |
| SEC24D | 3034.081143 | 0.270408465 | 4.03E-02 |
| GADD45GIP1 | 1768.229665 | 0.270349697 | 3.39E-02 |
| NARF | 2110.426255 | 0.270306497 | 2.60E-02 |
| RPL7L1 | 10467.69566 | 0.270143819 | 1.13E-04 |
| YBX1 | 25603.98182 | 0.269944693 | 1.15E-03 |
| BRMS1 | 1809.448814 | 0.269854461 | 2.77E-02 |
| NDOR1 | 1466.776151 | 0.269725274 | 1.59E-02 |
| ZNF717 | 470.4797863 | 0.269650967 | 1.62E-02 |
| DNAJC18 | 922.7044842 | 0.269494726 | 1.48E-02 |
| RAP1A | 1573.438963 | 0.269283049 | 9.10E-04 |
| PFDN1 | 2381.99521 | 0.268982396 | 6.93E-03 |
| LRSAM1 | 1674.395831 | 0.26859434 | 3.16E-02 |
| GSTP1 | 12693.12152 | 0.268590475 | 5.00E-02 |
| UBE2F | 1396.145029 | 0.268377586 | 2.75E-02 |
| BAP1 | 5048.804008 | 0.268073087 | 6.00E-03 |
| TIAL1 | 2251.871338 | 0.268014557 | 1.14E-03 |
| DAGLA | 1136.539145 | 0.267978099 | 3.35E-02 |
| RAB29 | 1618.049604 | 0.267859014 | 9.93E-03 |
| SIRT6 | 725.8968626 | 0.267692729 | 1.85E-02 |
| PCGF2 | 2220.258047 | 0.267292881 | 3.14E-02 |
| CNOT3 | 2190.48878 | 0.266819049 | 3.65E-03 |
| ANKRD40 | 2527.947377 | 0.266376496 | 1.12E-02 |
| HEBP2 | 1235.697435 | 0.266341622 | 2.23E-02 |
| EIF3I | 10526.45264 | 0.266250151 | 5.88E-03 |
| FLAD1 | 987.6272258 | 0.266132954 | 2.67E-02 |
| FADS1 | 13379.58282 | 0.265583154 | 1.30E-02 |
| MYO19 | 2959.134123 | 0.265573334 | 1.78E-02 |
| NEFH | 4344.736666 | 0.26504974 | 7.09E-03 |
| SNAP29 | 1759.719219 | 0.264938875 | 8.57E-04 |
| QPCTL | 1074.333906 | 0.264862083 | 2.65E-02 |
| STX4 | 2129.364156 | 0.264600982 | 2.57E-02 |
| HUWE1 | 25381.83121 | 0.264506799 | 3.39E-04 |
| VPS11 | 3275.966273 | 0.264196279 | 2.93E-02 |
| TOP1MT | 1570.636353 | 0.264004974 | 4.74E-04 |
| NFS1 | 2756.988872 | 0.263838012 | 1.73E-02 |
| ANO8 | 402.3322437 | 0.26373522 | 3.84E-02 |
| MRM3 | 916.3975761 | 0.26357333 | 1.42E-03 |
| ARCN1 | 17474.68119 | 0.263469906 | 5.91E-04 |
| ITPRIPL2 | 3464.727928 | 0.263055876 | 1.95E-03 |
| REPS1 | 1371.046726 | 0.263041537 | 5.17E-04 |
| FKBP1A | 26088.7043 | 0.262757012 | 2.71E-02 |
| ESD | 3673.991792 | 0.262338061 | 9.63E-04 |
| GRIPAP1 | 2385.540867 | 0.261950738 | 2.36E-03 |
| PDE12 | 1424.591386 | 0.261847618 | 7.68E-03 |

| | | | |
|---|---|---|---|
| MANBAL | 3030.257961 | 0.261799535 | 8.20E-03 |
| KIF22 | 5070.961106 | 0.261690936 | 2.30E-02 |
| SENP2 | 2850.4508 | 0.261597058 | 4.67E-02 |
| CAMTA1 | 1421.581369 | 0.261512463 | 3.07E-02 |
| SECISBP2 | 2314.173011 | 0.261443306 | 5.71E-03 |
| CSNK1A1 | 6157.901358 | 0.260715624 | 1.06E-02 |
| HUS1 | 841.6123964 | 0.260590941 | 4.27E-02 |
| NETO2 | 4066.529492 | 0.260496879 | 4.14E-02 |
| ASPSCR1 | 867.8872405 | 0.259981094 | 2.55E-02 |
| TRIM11 | 1271.475392 | 0.259725954 | 1.17E-03 |
| RPS6KB2 | 2424.493208 | 0.259643334 | 1.94E-02 |
| DCAF8 | 5161.578887 | 0.259568306 | 2.45E-05 |
| PRKAR2A | 3963.018413 | 0.259449863 | 1.95E-04 |
| NIPAL3 | 1794.482202 | 0.259428217 | 4.98E-03 |
| CMTR1 | 5231.338082 | 0.259214846 | 4.02E-03 |
| NSMAF | 2623.86319 | 0.259041705 | 2.62E-02 |
| RRP1B | 3504.70681 | 0.258499616 | 1.11E-05 |
| ZNF544 | 1742.095957 | 0.258216862 | 3.72E-02 |
| CELSR1 | 3462.62622 | 0.258178279 | 1.77E-02 |
| MRPL22 | 1050.652787 | 0.257175564 | 1.01E-02 |
| SMNDC1 | 892.9026407 | 0.256980327 | 1.28E-02 |
| DCPS | 1817.087287 | 0.25670134 | 1.05E-02 |
| VPS16 | 3276.175891 | 0.256388285 | 1.47E-02 |
| KPNA1 | 3329.771721 | 0.256350771 | 1.08E-03 |
| LINC00847 | 682.5496375 | 0.256307594 | 4.69E-02 |
| DDX56 | 6339.961598 | 0.256212464 | 1.84E-02 |
| KMT2A | 10311.72669 | 0.255893603 | 6.96E-03 |
| RITA1 | 1304.280185 | 0.255855883 | 1.00E-02 |
| DDB2 | 1400.451522 | 0.255795229 | 1.38E-02 |
| ACAD9 | 2006.68078 | 0.255400574 | 8.37E-03 |
| SRI | 3330.530597 | 0.255388808 | 1.37E-03 |
| CENPV | 1153.791311 | 0.255108855 | 2.81E-02 |
| MRPS34 | 2965.552352 | 0.254810607 | 8.69E-03 |
| PARP12 | 3009.496546 | 0.254347571 | 1.56E-03 |
| SRR | 938.275198 | 0.253926308 | 4.32E-02 |
| OXSR1 | 3175.687546 | 0.253860138 | 4.72E-03 |
| EIF1B | 1124.244198 | 0.253810371 | 3.85E-02 |
| CACTIN | 1550.647358 | 0.253785348 | 6.73E-03 |
| HDLBP | 32128.14887 | 0.253509065 | 4.02E-03 |
| WDR12 | 2415.211772 | 0.252947751 | 1.36E-02 |
| MRGBP | 2498.724127 | 0.252917854 | 2.08E-03 |
| SHISA5 | 6497.044586 | 0.252781745 | 4.73E-02 |
| TRPC4AP | 7921.881414 | 0.252539774 | 4.86E-02 |
| ARHGAP35 | 10670.21885 | 0.252118132 | 2.28E-02 |
| SEPHS1 | 2733.449846 | 0.252077835 | 2.67E-04 |
| MKRN2 | 1621.236604 | 0.251938051 | 2.18E-04 |
| RBM38 | 948.7983218 | 0.251832149 | 4.33E-02 |

| | | | |
|---|---|---|---|
| PSMG3 | 1824.579345 | 0.251079938 | 3.44E-03 |
| RAD23B | 8479.786235 | 0.251077334 | 4.29E-02 |
| TMEM80 | 695.471718 | 0.250979383 | 4.20E-02 |
| SMIM26 | 1687.576725 | 0.250972389 | 4.27E-02 |
| PTK2 | 6063.525587 | 0.250972105 | 1.30E-02 |
| AC011529.1 | 1413.500826 | 0.250929524 | 1.57E-02 |
| EWSR1 | 12651.43444 | 0.250893525 | 2.72E-04 |
| UFD1 | 2667.109898 | 0.250638356 | 2.54E-02 |
| RANBP3 | 2474.200959 | 0.250562792 | 3.28E-02 |
| GRSF1 | 4306.976919 | 0.250499882 | 1.31E-02 |
| PPP2CA | 10438.57862 | 0.250357063 | 2.10E-02 |
| GPSM1 | 4074.013977 | 0.250221053 | 3.85E-02 |
| GTPBP6 | 1657.891782 | 0.250071846 | 3.26E-02 |
| SEC23B | 5037.762538 | 0.250051885 | 6.80E-04 |
| THOC7 | 1848.437883 | 0.250024666 | 2.82E-04 |
| HIRIP3 | 1467.028443 | 0.24984531 | 2.74E-03 |
| TCIRG1 | 2216.590153 | 0.249826683 | 4.51E-02 |
| METTL2A | 1073.794768 | 0.249533666 | 1.29E-02 |
| NPRL2 | 1001.425579 | 0.24936371 | 3.78E-02 |
| HM13 | 10209.03501 | 0.249240982 | 1.78E-03 |
| MAPKAP1 | 7259.855184 | 0.249075802 | 4.20E-03 |
| TBC1D20 | 4180.645879 | 0.249044574 | 7.60E-04 |
| NOM1 | 2171.555554 | 0.249002726 | 1.57E-02 |
| NELFA | 1190.639151 | 0.248717002 | 5.40E-03 |
| ILKAP | 1385.386995 | 0.248563636 | 2.09E-04 |
| MLLT6 | 7194.777465 | 0.248386568 | 6.68E-03 |
| ARL6IP4 | 4010.236254 | 0.248197598 | 2.00E-02 |
| RAB6A | 4718.383046 | 0.247907588 | 2.43E-02 |
| CPSF4 | 2678.248412 | 0.247894503 | 1.68E-03 |
| RHBDF2 | 1009.728671 | 0.247797286 | 4.85E-02 |
| PEX26 | 2862.067889 | 0.247626922 | 3.51E-02 |
| SLC33A1 | 1177.423913 | 0.247516704 | 4.46E-02 |
| LRRC28 | 807.0602025 | 0.2473671 | 6.88E-03 |
| TMEM14B | 2820.957355 | 0.246808208 | 6.10E-03 |
| POLDIP2 | 7076.671149 | 0.246779787 | 2.60E-02 |
| HPS5 | 2635.985793 | 0.246637735 | 2.37E-03 |
| PRPF8 | 28223.35904 | 0.246311324 | 3.06E-02 |
| HNRNPC | 25372.56747 | 0.246274678 | 1.87E-04 |
| TMX2 | 4622.100163 | 0.245887915 | 1.52E-03 |
| USP7 | 5820.658002 | 0.245785815 | 2.98E-05 |
| DDB1 | 22375.96724 | 0.24506117 | 4.27E-02 |
| HERC2 | 4605.110151 | 0.244775025 | 8.69E-03 |
| HDHD5 | 2611.229972 | 0.244691352 | 3.74E-02 |
| CYTH1 | 1067.619713 | 0.244671604 | 4.59E-02 |
| RNF212 | 2579.106999 | 0.244646478 | 3.11E-04 |
| PODXL2 | 4334.400803 | 0.244452739 | 4.41E-02 |
| SYMPK | 5323.531753 | 0.244133417 | 1.72E-02 |

| | | | |
|---|---|---|---|
| WRAP73 | 1404.388509 | 0.244039768 | 4.44E-03 |
| PPM1G | 8237.17981 | 0.243990893 | 5.31E-03 |
| METAP1 | 2773.955335 | 0.243850468 | 5.92E-03 |
| GALK2 | 1013.100879 | 0.243815582 | 1.81E-03 |
| POLR2E | 5539.893555 | 0.243681367 | 3.72E-02 |
| ATP5PO | 6020.894828 | 0.243243568 | 9.74E-03 |
| TCF20 | 3764.351559 | 0.243132307 | 1.44E-03 |
| SMIM19 | 1014.322732 | 0.242748609 | 9.30E-03 |
| BRPF3 | 3812.337617 | 0.242678217 | 1.53E-02 |
| GAK | 2571.379573 | 0.242405543 | 3.44E-03 |
| GATB | 1624.777046 | 0.242240015 | 1.13E-02 |
| UQCRC2 | 8408.55013 | 0.242132484 | 1.04E-04 |
| VPS37B | 1594.694277 | 0.241821408 | 4.90E-02 |
| NA | 1427.962526 | 0.241813161 | 2.41E-03 |
| TNKS | 4426.097578 | 0.241678825 | 2.78E-02 |
| USP10 | 5912.579759 | 0.241274568 | 2.89E-06 |
| BCAP31 | 10945.19398 | 0.241016273 | 4.00E-02 |
| VMP1 | 6417.676705 | 0.239271072 | 3.94E-02 |
| DGAT1 | 1372.034896 | 0.238918639 | 3.38E-02 |
| KIF3C | 1868.784399 | 0.238619289 | 3.21E-02 |
| PRPF31 | 3047.900361 | 0.238599098 | 3.40E-02 |
| IPO9 | 8938.980323 | 0.238400788 | 2.31E-02 |
| RAB8A | 1921.112499 | 0.238146413 | 4.84E-02 |
| TCP11L1 | 1101.58559 | 0.238029538 | 2.17E-03 |
| MORC2 | 2508.372783 | 0.237865258 | 3.02E-02 |
| YPEL5 | 2746.761684 | 0.237863758 | 3.27E-02 |
| TUBGCP4 | 1347.698251 | 0.237723141 | 3.79E-02 |
| MON1B | 3623.886187 | 0.237656442 | 3.49E-02 |
| KANSL3 | 5951.018254 | 0.237549756 | 1.98E-02 |
| ABHD12 | 4210.205228 | 0.237543717 | 1.60E-02 |
| UTP4 | 5907.120824 | 0.237429856 | 9.47E-03 |
| GUSB | 2894.849897 | 0.23741676 | 6.25E-03 |
| ALKBH1 | 826.8390791 | 0.237326658 | 3.01E-02 |
| DEDD2 | 826.3518582 | 0.23711003 | 4.83E-02 |
| PPP5C | 4020.73127 | 0.23649924 | 4.46E-02 |
| MRPS16 | 2989.27372 | 0.23628294 | 7.96E-03 |
| KAT7 | 3026.224646 | 0.236237164 | 1.50E-03 |
| EIF4G1 | 29771.85812 | 0.236124588 | 2.79E-02 |
| PDIA6 | 20872.47759 | 0.236087758 | 4.56E-03 |
| LTB4R | 770.8990611 | 0.236083719 | 4.63E-02 |
| PCCB | 2314.132373 | 0.235770449 | 4.04E-03 |
| PGS1 | 1688.179157 | 0.235366522 | 2.70E-02 |
| KHNYN | 5658.646905 | 0.235062982 | 4.82E-04 |
| RNF4 | 3198.021208 | 0.234874937 | 1.84E-03 |
| TMEM231 | 1533.842966 | 0.234116211 | 4.02E-02 |
| PDAP1 | 9323.361523 | 0.233809279 | 2.72E-03 |
| IGF2BP1 | 4568.368856 | 0.23334495 | 1.83E-03 |

| | | | |
|---|---|---|---|
| MYO10 | 14327.81064 | 0.233185113 | 3.42E-02 |
| ANKLE2 | 3008.70015 | 0.232666977 | 4.02E-02 |
| RPL22 | 11894.40039 | 0.232456629 | 1.37E-02 |
| RPS25 | 42066.75165 | 0.232239693 | 4.61E-02 |
| GLOD4 | 2925.096266 | 0.232085872 | 3.54E-02 |
| PACS2 | 2431.694362 | 0.231931852 | 1.83E-02 |
| FCHSD2 | 1863.422345 | 0.231112019 | 7.22E-03 |
| AAMP | 6026.877721 | 0.230981695 | 1.93E-02 |
| SMAD3 | 7634.379311 | 0.230052175 | 1.64E-02 |
| ATP2A2 | 12810.82753 | 0.229899281 | 6.99E-03 |
| WDR46 | 5956.684672 | 0.229802765 | 8.34E-03 |
| NOL6 | 8157.384097 | 0.2293015 | 2.31E-02 |
| AGAP3 | 4495.009324 | 0.229300339 | 7.32E-03 |
| LINC00221 | 1466.442818 | 0.229199667 | 3.78E-02 |
| CPSF1 | 6411.044963 | 0.229124729 | 1.75E-02 |
| CTDSP2 | 12475.24743 | 0.229042763 | 3.15E-03 |
| GIPC1 | 4074.727252 | 0.228956368 | 2.71E-02 |
| KANK1 | 4383.01843 | 0.228497188 | 5.00E-03 |
| HAGLR | 1176.304192 | 0.227952382 | 1.43E-02 |
| STT3A | 12540.12889 | 0.227338107 | 1.07E-02 |
| SSBP1 | 6663.456012 | 0.22729815 | 2.58E-04 |
| PHF5A | 2143.181465 | 0.22711466 | 8.11E-03 |
| HERPUD2 | 1607.403671 | 0.226916665 | 4.39E-03 |
| SRRM1 | 3120.305736 | 0.22672172 | 2.76E-02 |
| SMARCD1 | 4173.785875 | 0.22665897 | 1.22E-02 |
| CTDNEP1 | 6021.439003 | 0.226540244 | 4.66E-02 |
| EIF4A3 | 3461.150558 | 0.226434277 | 4.78E-02 |
| RAI1 | 4292.083718 | 0.226104532 | 1.92E-02 |
| FBXW8 | 1751.709567 | 0.225786575 | 2.35E-02 |
| RACGAP1 | 5057.43822 | 0.225535425 | 2.77E-02 |
| RPS4X | 33744.19624 | 0.225519451 | 4.37E-02 |
| DLAT | 2983.280395 | 0.225514784 | 3.00E-02 |
| CTPS2 | 1411.08626 | 0.225069545 | 4.06E-02 |
| ATXN2 | 1989.701599 | 0.225054786 | 2.83E-02 |
| ARID1A | 5257.33245 | 0.224311731 | 6.06E-03 |
| SEPHS2 | 6477.342138 | 0.224056778 | 3.18E-03 |
| ST13 | 13724.51909 | 0.223608337 | 1.13E-02 |
| AC007950.1 | 1503.756081 | 0.223305093 | 4.72E-02 |
| HNRNPUL2 | 6953.721425 | 0.223215866 | 2.41E-03 |
| TPT1 | 67889.21744 | 0.2229311 | 1.32E-02 |
| NRDC | 8374.393388 | 0.222851973 | 1.77E-02 |
| ELP3 | 3058.743108 | 0.22277291 | 1.39E-03 |
| AP1B1 | 6559.260492 | 0.222697863 | 1.78E-02 |
| TMEM161A | 1198.284385 | 0.222153932 | 2.56E-02 |
| CHD4 | 25148.89638 | 0.222075698 | 1.47E-02 |
| NFIC | 3080.234408 | 0.221832394 | 1.44E-02 |
| MTA1 | 4030.972488 | 0.220488034 | 1.74E-02 |

| | | | |
|---|---|---|---|
| SCAF1 | 3012.004697 | 0.22018419 | 2.38E-02 |
| SMAD2 | 2978.677024 | 0.219692159 | 1.42E-02 |
| UBE2D3 | 7342.501166 | 0.219608743 | 8.30E-03 |
| PDSS2 | 889.527782 | 0.219596736 | 1.93E-02 |
| PDCD6IP | 6347.384567 | 0.219455291 | 1.13E-02 |
| AK3 | 2865.631536 | 0.219380215 | 2.25E-03 |
| CHMP4B | 5505.366858 | 0.218628033 | 2.78E-02 |
| OSBP | 5786.050547 | 0.218107907 | 1.65E-02 |
| AFG3L2 | 4562.506854 | 0.218079107 | 1.94E-04 |
| XRCC3 | 1695.724094 | 0.217725787 | 6.50E-03 |
| TBC1D5 | 6004.939625 | 0.217673728 | 1.84E-03 |
| RER1 | 5729.27554 | 0.217650945 | 2.55E-02 |
| PITPNA | 4835.854441 | 0.217030425 | 4.54E-02 |
| PHF8 | 2614.998685 | 0.217017498 | 2.06E-02 |
| PTPN9 | 3529.233561 | 0.216594858 | 7.63E-03 |
| EIF2B4 | 2102.427611 | 0.216589031 | 4.32E-02 |
| RDH13 | 730.3832492 | 0.216420618 | 2.11E-02 |
| ATP5F1C | 4379.356069 | 0.216114749 | 4.46E-02 |
| TNKS1BP1 | 7024.243343 | 0.215806674 | 2.12E-02 |
| NA | 1855.274218 | 0.215504009 | 2.30E-02 |
| DCP1A | 1566.772076 | 0.215497476 | 4.53E-02 |
| PDCD7 | 2488.493922 | 0.215338966 | 6.48E-03 |
| KIAA0355 | 2133.566248 | 0.215273369 | 3.74E-02 |
| ARFGAP3 | 2468.560438 | 0.214708499 | 2.57E-03 |
| ODF2 | 3894.579451 | 0.214612997 | 2.62E-03 |
| TACC3 | 5029.8107 | 0.214555679 | 4.42E-02 |
| GNA11 | 4105.184978 | 0.21359343 | 3.31E-02 |
| BZW2 | 4791.617903 | 0.213550445 | 2.72E-02 |
| PLCXD1 | 1585.892566 | 0.213543236 | 3.65E-02 |
| INPP5E | 1157.460523 | 0.213511281 | 2.16E-02 |
| UBQLN1 | 7209.186043 | 0.213369953 | 2.32E-03 |
| STK25 | 5404.879135 | 0.213205742 | 2.62E-02 |
| RBFA | 1166.399006 | 0.212574962 | 2.77E-03 |
| NSUN2 | 8633.428321 | 0.212234527 | 1.36E-03 |
| KIAA1671 | 2181.6701 | 0.21191124 | 1.43E-02 |
| PSMB1 | 6788.226364 | 0.211792968 | 2.83E-02 |
| NBPF15 | 3036.575356 | 0.211399728 | 7.46E-03 |
| DNAJC7 | 4148.495269 | 0.210489838 | 3.17E-02 |
| DEF8 | 3273.679404 | 0.210425388 | 4.71E-02 |
| CDC42SE1 | 2555.88822 | 0.209861628 | 3.22E-02 |
| TBKBP1 | 1994.792743 | 0.209221954 | 4.97E-02 |
| EEFSEC | 1107.058862 | 0.208586696 | 4.90E-02 |
| LONP2 | 5795.25613 | 0.208557079 | 4.36E-03 |
| CHRAC1 | 1489.31076 | 0.208516442 | 3.29E-02 |
| SRPRA | 13337.9025 | 0.207997965 | 2.45E-02 |
| SNRNP70 | 6284.377571 | 0.207498243 | 3.34E-02 |
| LARP4B | 1886.896081 | 0.206661156 | 2.19E-02 |

| | | | |
|---|---|---|---|
| SMARCB1 | 4122.10918 | 0.206221763 | 4.01E-02 |
| UBE2Z | 8651.042852 | 0.205561944 | 4.33E-02 |
| KIF7 | 2072.428412 | 0.204256201 | 2.09E-02 |
| ATG16L1 | 1820.901969 | 0.203482603 | 4.79E-02 |
| TXLNA | 5726.03994 | 0.202382021 | 3.79E-02 |
| YTHDF1 | 3356.918214 | 0.202150114 | 2.12E-02 |
| LAPTM4A | 7258.494821 | 0.199784445 | 6.46E-03 |
| NGRN | 7742.66646 | 0.199385868 | 2.24E-02 |
| CLEC16A | 2148.484018 | 0.198726714 | 2.48E-02 |
| ATP6V1H | 2279.078079 | 0.198360644 | 3.98E-02 |
| COPS8 | 3379.611916 | 0.198241484 | 8.66E-03 |
| MTMR3 | 2790.3142 | 0.197686014 | 3.04E-02 |
| NRF1 | 837.4999078 | 0.197443469 | 4.01E-02 |
| MCM3AP | 8164.114642 | 0.197419346 | 1.39E-02 |
| ATXN1L | 3477.835901 | 0.195630297 | 4.65E-02 |
| GNB5 | 2160.045094 | 0.195573871 | 1.63E-02 |
| MOCS1 | 1229.149912 | 0.195491654 | 3.54E-02 |
| NKIRAS2 | 2781.961389 | 0.19532272 | 3.22E-02 |
| NUP205 | 10693.41834 | 0.195123657 | 3.11E-02 |
| TOLLIP | 3012.347413 | 0.195046364 | 1.10E-02 |
| MRPL15 | 2929.771809 | 0.194829689 | 1.45E-02 |
| SCAF4 | 1560.497146 | 0.194809623 | 4.96E-02 |
| PSME3 | 7880.901979 | 0.194809128 | 2.38E-02 |
| CALM1 | 7582.069497 | 0.194747876 | 2.27E-02 |
| YEATS2 | 5124.563984 | 0.194252074 | 3.33E-03 |
| TEX261 | 6210.156097 | 0.192952465 | 4.49E-02 |
| KIAA0556 | 2031.806147 | 0.192769796 | 4.72E-02 |
| MRPL44 | 1275.662237 | 0.192391315 | 4.83E-02 |
| NUP155 | 7161.690823 | 0.192015526 | 3.34E-02 |
| DCUN1D5 | 2156.25617 | 0.191853857 | 2.65E-02 |
| EDC3 | 4250.423147 | 0.190727759 | 3.53E-02 |
| EXOC6B | 3172.372247 | 0.189378329 | 1.47E-02 |
| EPN2 | 2916.468412 | 0.18885851 | 1.49E-02 |
| RHOQ | 3625.302826 | 0.188563892 | 1.83E-02 |
| TMED2 | 8167.232608 | 0.188379185 | 1.08E-02 |
| ZNF746 | 1278.679077 | 0.188000567 | 3.21E-02 |
| WDR59 | 2811.589974 | 0.187532209 | 3.47E-02 |
| PJA1 | 2541.974321 | 0.187318694 | 2.90E-02 |
| PPP6R2 | 3037.313945 | 0.186847382 | 3.51E-02 |
| BUB3 | 4309.026554 | 0.18649874 | 1.47E-02 |
| TMEM69 | 1083.687159 | 0.186158223 | 3.28E-02 |
| RALBP1 | 3901.009253 | 0.185874355 | 1.46E-02 |
| HNRNPAB | 5558.229272 | 0.185862091 | 2.37E-02 |
| FAM118A | 895.3172424 | 0.185765282 | 2.86E-02 |
| IL13RA1 | 5499.46665 | 0.184524407 | 2.45E-02 |
| MAGEA10 | 7039.294074 | 0.184504483 | 2.92E-02 |
| PPP2R2A | 3979.054954 | 0.183956259 | 1.68E-02 |

| | | | |
|---|---|---|---|
| E2F4 | 7133.361551 | 0.183891048 | 4.15E-02 |
| UCKL1 | 1756.907951 | 0.182130918 | 1.65E-02 |
| CFL2 | 3651.721601 | 0.180599494 | 2.02E-02 |
| GEMIN5 | 2211.92909 | 0.180307022 | 1.25E-02 |
| NONO | 22380.64719 | 0.179968882 | 2.02E-02 |
| MRPL35 | 2106.352528 | 0.178149843 | 3.03E-02 |
| ZC3H18 | 3644.307394 | 0.176851204 | 3.39E-02 |
| NBR1 | 8980.711034 | 0.176450051 | 3.98E-03 |
| YKT6 | 13205.20478 | 0.17544518 | 4.97E-02 |
| GFM2 | 5520.51197 | 0.175124822 | 4.06E-02 |
| MRFAP1 | 10526.41433 | 0.175048112 | 1.14E-02 |
| SFT2D1 | 1397.633585 | 0.173801059 | 1.80E-02 |
| ELAVL1 | 3947.132859 | 0.173625524 | 3.24E-02 |
| UBE2O | 3242.178472 | 0.171081329 | 2.55E-02 |
| NDUFV3 | 1383.346286 | 0.17030488 | 1.96E-02 |
| DIDO1 | 6933.614041 | 0.166656754 | 1.33E-03 |
| TNPO3 | 5287.867569 | 0.16663914 | 3.13E-03 |
| PSMD4 | 7480.110364 | 0.165856737 | 3.83E-02 |
| GDI2 | 6479.769526 | 0.164607394 | 1.68E-02 |
| ARHGEF7 | 1898.615677 | 0.164091986 | 1.63E-02 |
| CASC3 | 4319.53297 | 0.1632583 | 2.97E-02 |
| TTC1 | 6219.880764 | 0.16277632 | 2.12E-02 |
| VPS26C | 2551.853355 | 0.160455477 | 2.77E-02 |
| IMMT | 7144.203813 | 0.158124721 | 2.11E-02 |
| ISCU | 1888.172076 | 0.156264149 | 3.51E-02 |
| DNAJA2 | 4555.427379 | 0.154361795 | 1.03E-02 |
| CFLAR | 2074.937118 | 0.143625392 | 3.67E-02 |
| FBXW11 | 3631.29144 | 0.142186076 | 4.29E-02 |
| NFRKB | 3989.407039 | 0.134369406 | 1.97E-02 |
| NAA20 | 4011.398173 | 0.134152369 | 4.32E-02 |
| YWHAB | 18910.22592 | 0.133144132 | 3.70E-02 |
| ZNF207 | 9064.445444 | 0.132326146 | 3.19E-03 |
| RPN2 | 26174.83075 | -0.115445101 | 2.85E-02 |
| FOXO3 | 3030.424993 | -0.117915111 | 3.59E-02 |
| RNF145 | 10583.21953 | -0.131824876 | 1.39E-02 |
| ATP6AP2 | 6279.90048 | -0.144135168 | 1.75E-02 |
| AP3B1 | 6576.449798 | -0.144872668 | 1.42E-02 |
| UBQLN2 | 1858.80665 | -0.154181018 | 3.34E-02 |
| METTL9 | 4952.663648 | -0.156575135 | 4.10E-02 |
| RPRD2 | 2318.718644 | -0.157728242 | 1.75E-02 |
| INTS14 | 3408.072776 | -0.158640854 | 1.46E-02 |
| SRCAP | 10671.7796 | -0.159130536 | 1.25E-02 |
| CDC23 | 4764.631516 | -0.166777992 | 1.67E-02 |
| PWWP3A | 2910.551855 | -0.169019592 | 2.75E-02 |
| THRAP3 | 8007.967293 | -0.170994642 | 4.40E-02 |
| HNRNPA1 | 38484.91909 | -0.171568934 | 4.68E-02 |
| PCED1A | 3718.966878 | -0.171751066 | 3.46E-02 |

| | | | |
|---|---|---|---|
| VPS39 | 4054.900901 | -0.172640162 | 2.81E-02 |
| NCR3LG1 | 1883.189571 | -0.172751487 | 2.69E-02 |
| EIF2B1 | 3995.522508 | -0.173188825 | 6.15E-03 |
| CUL1 | 4998.254058 | -0.175233868 | 6.06E-03 |
| PTRH2 | 1318.331553 | -0.177109685 | 4.52E-02 |
| ABCF1 | 5978.512133 | -0.177947107 | 1.32E-02 |
| TRAK1 | 3100.499793 | -0.178255381 | 2.63E-02 |
| SUPT16H | 16245.5832 | -0.179341525 | 3.53E-02 |
| TMED10 | 13060.58854 | -0.18043311 | 1.26E-02 |
| YY1 | 5353.830341 | -0.183864369 | 3.23E-02 |
| RGP1 | 6625.186062 | -0.186735016 | 3.01E-02 |
| DOCK1 | 3315.886736 | -0.187217191 | 2.54E-02 |
| RARS | 5457.586302 | -0.187659645 | 8.62E-03 |
| ZCCHC14 | 2165.42474 | -0.187720258 | 3.67E-02 |
| RPL7 | 41042.59646 | -0.188899473 | 3.11E-02 |
| CAST | 16712.96545 | -0.189738119 | 1.72E-03 |
| LAMC1 | 24069.24177 | -0.190515512 | 4.47E-02 |
| SARAF | 7191.114511 | -0.191071696 | 8.23E-03 |
| TXNL1 | 2788.791101 | -0.194156917 | 8.10E-03 |
| NPTN | 4805.499864 | -0.194713741 | 3.54E-02 |
| EPS15 | 2833.132169 | -0.195021206 | 2.82E-02 |
| TRAM1 | 4228.399092 | -0.195105482 | 4.99E-02 |
| EZH1 | 1512.234473 | -0.197407253 | 2.61E-02 |
| DPYSL2 | 5563.378319 | -0.197910244 | 2.47E-02 |
| DERA | 1351.703334 | -0.199195103 | 2.01E-02 |
| TIMM8A | 916.3763145 | -0.200158066 | 4.24E-02 |
| CAPRIN1 | 19543.43926 | -0.200352995 | 2.44E-05 |
| HIPK1 | 2541.742476 | -0.20082882 | 2.09E-02 |
| DNAJA1 | 17897.86696 | -0.201542008 | 4.77E-02 |
| PEAK1 | 4818.635054 | -0.201943343 | 3.18E-02 |
| EVL | 1559.341381 | -0.203024929 | 4.85E-02 |
| THAP11 | 2904.615872 | -0.204568776 | 2.14E-02 |
| APOL3 | 1109.386461 | -0.205174181 | 4.93E-02 |
| KIAA1549 | 2899.370619 | -0.206152848 | 8.53E-03 |
| NCBP1 | 5885.144796 | -0.206725518 | 3.36E-02 |
| CFDP1 | 2477.404977 | -0.20697662 | 1.48E-02 |
| WASH4P | 1272.645148 | -0.20809605 | 4.59E-02 |
| NKAPD1 | 3040.424315 | -0.210303287 | 4.11E-02 |
| ELF1 | 1523.077728 | -0.212066125 | 1.14E-02 |
| MSL3 | 1482.073196 | -0.215204591 | 1.27E-02 |
| PANK2 | 3143.512483 | -0.215328203 | 4.06E-04 |
| GPATCH2L | 2485.997893 | -0.216272748 | 4.36E-02 |
| C6orf62 | 6945.794868 | -0.216933546 | 1.93E-02 |
| PUM1 | 5310.868167 | -0.217739014 | 9.96E-04 |
| ZNF3 | 2904.964969 | -0.218297269 | 4.59E-02 |
| UBE2N | 4070.400118 | -0.218329635 | 1.61E-02 |
| LSG1 | 3989.680754 | -0.218462098 | 2.82E-02 |

| | | | |
|---|---|---|---|
| ZNF618 | 3843.333358 | -0.218534347 | 3.35E-02 |
| INTS10 | 3396.001548 | -0.21894313 | 8.73E-03 |
| SIK3 | 2445.245228 | -0.218958155 | 1.82E-02 |
| C11orf49 | 1568.56911 | -0.21909624 | 3.83E-02 |
| WASHC5 | 4004.657705 | -0.219536834 | 2.28E-02 |
| TMEM59 | 6590.074609 | -0.220267746 | 1.40E-02 |
| DDX39B | 14330.28417 | -0.220319953 | 2.24E-02 |
| DNAJC3 | 5598.113646 | -0.220753776 | 3.66E-02 |
| UBR2 | 2838.907482 | -0.22078466 | 2.97E-02 |
| PPIL2 | 3565.270233 | -0.220785341 | 1.35E-02 |
| STRADA | 1484.97176 | -0.221315445 | 1.01E-02 |
| TMED8 | 2389.896747 | -0.221526071 | 2.00E-02 |
| ECD | 1457.84952 | -0.221591999 | 4.77E-03 |
| SLC9A6 | 2218.455759 | -0.221641422 | 2.34E-02 |
| G3BP1 | 7461.652877 | -0.221762889 | 1.04E-03 |
| SLC35B4 | 4181.027119 | -0.221772375 | 4.63E-02 |
| POP1 | 1541.445619 | -0.221879611 | 2.57E-02 |
| ARID1B | 3578.147371 | -0.222287453 | 9.27E-05 |
| ABI1 | 1233.020187 | -0.222729448 | 5.01E-03 |
| ZSCAN25 | 1601.665477 | -0.222767822 | 2.22E-02 |
| ABHD16A | 1665.899777 | -0.223103151 | 1.65E-02 |
| MED20 | 1921.206544 | -0.223441331 | 1.65E-03 |
| WLS | 5603.222117 | -0.223616686 | 4.73E-02 |
| PIDD1 | 1290.180358 | -0.223698047 | 4.74E-02 |
| ATP5MD | 1693.401647 | -0.224093401 | 3.26E-02 |
| CFAP298 | 1784.980603 | -0.224477744 | 1.47E-02 |
| VPS26B | 4900.815599 | -0.224707709 | 4.30E-02 |
| GID8 | 3607.78933 | -0.226017889 | 2.69E-02 |
| POTEC | 621.7288682 | -0.22623208 | 4.19E-02 |
| TIGD1 | 735.47436 | -0.226421647 | 3.27E-02 |
| ZNF789 | 838.5068154 | -0.226903093 | 1.01E-02 |
| KAT14 | 1554.577656 | -0.22734161 | 7.57E-03 |
| DDX17 | 25267.52818 | -0.227458524 | 2.07E-04 |
| UNK | 1262.706656 | -0.227596229 | 3.85E-02 |
| THUMPD3 | 1703.170376 | -0.227684375 | 4.34E-02 |
| STOM | 8123.48438 | -0.227969449 | 5.71E-03 |
| UTP3 | 1873.104973 | -0.229153851 | 1.79E-02 |
| MARF1 | 3255.495896 | -0.229953036 | 1.74E-02 |
| STX6 | 1556.556609 | -0.230849227 | 6.79E-04 |
| PPIL1 | 2099.664457 | -0.231007167 | 2.06E-03 |
| ELOA | 3907.96247 | -0.231563734 | 1.18E-04 |
| SETD5 | 6418.971384 | -0.231877939 | 5.71E-03 |
| RRP36 | 2595.60035 | -0.232290466 | 3.06E-02 |
| BPGM | 883.7041193 | -0.232832175 | 1.59E-02 |
| POLR3B | 989.7002471 | -0.232909119 | 1.79E-02 |
| SERBP1 | 15173.87957 | -0.233034896 | 2.72E-02 |
| ABCC5 | 1812.770361 | -0.233580938 | 4.07E-03 |

| | | | |
|---|---|---|---|
| WDR82 | 7613.00644 | -0.233719026 | 3.71E-03 |
| TRA2B | 6439.616221 | -0.233819852 | 8.98E-03 |
| SCAF8 | 2710.998518 | -0.233847507 | 2.75E-04 |
| SYPL1 | 4385.788923 | -0.235025381 | 1.21E-03 |
| SRRM2 | 25535.81099 | -0.235285719 | 2.39E-02 |
| COQ10A | 755.6505801 | -0.23534017 | 3.55E-02 |
| FBXW2 | 5336.381978 | -0.235544774 | 1.03E-05 |
| CHTF8 | 7617.256486 | -0.235689226 | 4.36E-02 |
| NA | 8125.450766 | -0.235747096 | 6.76E-03 |
| MLLT1 | 5500.18598 | -0.236224063 | 6.48E-03 |
| PSMA4 | 7281.516475 | -0.236417144 | 4.20E-03 |
| AGPAT4 | 1218.159414 | -0.236634172 | 2.33E-02 |
| PREP | 3723.278541 | -0.236678875 | 4.38E-02 |
| DNAH2 | 2042.058399 | -0.237077356 | 8.47E-03 |
| RECK | 1748.465281 | -0.238175884 | 3.02E-02 |
| NUP160 | 7594.563443 | -0.238349578 | 4.00E-03 |
| HDAC1 | 6926.946715 | -0.238777811 | 2.90E-02 |
| DDX42 | 5679.915558 | -0.239247274 | 4.88E-03 |
| UBA3 | 1913.314045 | -0.239568769 | 4.04E-02 |
| ATP13A1 | 3127.033339 | -0.239989401 | 2.27E-02 |
| ZNF595 | 2014.600462 | -0.240324367 | 2.76E-02 |
| TOR1B | 3507.737833 | -0.240529314 | 3.16E-03 |
| DUXAP9 | 5141.360363 | -0.240548371 | 7.37E-04 |
| FAM53B | 1491.038315 | -0.241168961 | 3.16E-02 |
| ACAD8 | 2448.858029 | -0.242699851 | 4.90E-03 |
| DUXAP8 | 8045.737965 | -0.243494749 | 1.23E-05 |
| SLC30A8 | 872.0711494 | -0.244603392 | 3.85E-02 |
| MTG2 | 2081.396989 | -0.244687319 | 9.70E-03 |
| SPG7 | 3626.18426 | -0.245102987 | 3.85E-02 |
| GSS | 4871.01293 | -0.245362979 | 3.19E-02 |
| HNRNPA3 | 17215.90036 | -0.245517314 | 2.46E-03 |
| MRI1 | 1491.264152 | -0.245649254 | 2.79E-02 |
| MCOLN1 | 614.275788 | -0.245663003 | 4.25E-02 |
| KDM2A | 7320.875344 | -0.245873878 | 1.44E-02 |
| HNRNPM | 10158.43398 | -0.246608313 | 2.17E-04 |
| MRM2 | 2951.072928 | -0.246771365 | 5.52E-05 |
| POLR2A | 10759.83302 | -0.247327659 | 5.67E-03 |
| SUMO1 | 4259.39701 | -0.247365232 | 7.68E-03 |
| CNOT4 | 1183.783448 | -0.247704758 | 2.18E-03 |
| UBE4A | 5703.241683 | -0.247760877 | 3.84E-03 |
| TMEM9B | 1478.565878 | -0.247816589 | 4.83E-02 |
| ARHGAP39 | 505.0589739 | -0.248470984 | 3.86E-02 |
| FRMD3 | 3496.851126 | -0.248764413 | 5.04E-03 |
| ISG20L2 | 2175.728689 | -0.249008743 | 9.36E-03 |
| CCDC120 | 662.5532422 | -0.24981369 | 2.54E-02 |
| RNF167 | 3862.807726 | -0.249946364 | 3.56E-02 |
| CENPT | 2443.523131 | -0.250031329 | 2.05E-03 |

| | | | |
|---|---|---|---|
| EAPP | 1334.720575 | -0.250638477 | 4.43E-02 |
| DTX3 | 1694.078695 | -0.250717449 | 1.69E-02 |
| FBXO11 | 2388.76673 | -0.251400186 | 4.31E-02 |
| PANX1 | 1303.080535 | -0.251507776 | 4.40E-03 |
| GOLM1 | 4855.950469 | -0.251903374 | 8.13E-04 |
| SRPRB | 2953.540073 | -0.252268361 | 4.13E-02 |
| RAP1B | 7934.008772 | -0.252425289 | 1.42E-02 |
| NSMCE4A | 1084.538538 | -0.252602756 | 1.51E-02 |
| MRPS21 | 2857.515873 | -0.252722991 | 2.59E-02 |
| DDX1 | 6598.918061 | -0.252905981 | 1.11E-02 |
| CDC42EP4 | 3656.079881 | -0.253044554 | 1.25E-02 |
| BAZ1B | 12077.77564 | -0.253134272 | 2.14E-03 |
| RBL2 | 4576.824447 | -0.253355527 | 8.08E-03 |
| PPM1A | 1396.887455 | -0.254454937 | 2.84E-02 |
| PI4KA | 4563.049434 | -0.254767534 | 1.71E-03 |
| SSR1 | 4631.397543 | -0.255565429 | 4.74E-06 |
| AC026412.1 | 1148.852394 | -0.255742755 | 1.35E-02 |
| NSD1 | 6067.037785 | -0.255894242 | 1.90E-03 |
| LINC00339 | 748.7352198 | -0.256675748 | 3.45E-02 |
| TIMELESS | 5461.253155 | -0.257583 | 5.71E-03 |
| NAA60 | 1498.058643 | -0.257750248 | 3.13E-02 |
| NDFIP1 | 3015.796015 | -0.257798369 | 1.22E-03 |
| MEA1 | 2088.826949 | -0.258666617 | 2.20E-02 |
| RHNO1 | 1266.859032 | -0.2592879 | 2.83E-02 |
| NUP50 | 3007.034293 | -0.259506341 | 4.32E-02 |
| DERL2 | 1918.396919 | -0.259681994 | 9.78E-04 |
| HRNR | 620.4357706 | -0.259692957 | 4.65E-02 |
| MORN2 | 607.2431179 | -0.259724182 | 4.95E-02 |
| SLC35D1 | 878.4067731 | -0.260068792 | 5.22E-03 |
| MTPN | 15377.19465 | -0.260550702 | 8.93E-04 |
| HOXB7 | 1350.143893 | -0.261892761 | 2.66E-02 |
| SP140L | 1768.433108 | -0.26221575 | 1.79E-03 |
| SV2A | 2278.471485 | -0.262563129 | 8.23E-03 |
| ITFG1 | 3584.265264 | -0.262639868 | 2.95E-02 |
| SUPT20H | 1593.510877 | -0.263066854 | 4.62E-02 |
| ASL | 1647.09413 | -0.263430809 | 1.12E-02 |
| MAGT1 | 4499.864434 | -0.263431133 | 2.43E-02 |
| NKRF | 953.9954924 | -0.263593679 | 2.02E-02 |
| PI4KAP2 | 678.9268025 | -0.26362194 | 9.64E-03 |
| ATP5PF | 3726.084089 | -0.263798577 | 1.76E-02 |
| FGFR1 | 4324.242565 | -0.263963473 | 8.73E-03 |
| CCDC97 | 2026.245674 | -0.264374656 | 1.63E-03 |
| IP6K1 | 2965.032039 | -0.266051429 | 1.82E-02 |
| RAB30-AS1 | 321.317745 | -0.266082408 | 3.63E-02 |
| ZMYND8 | 5516.740769 | -0.266517351 | 1.47E-02 |
| EFR3A | 3105.551569 | -0.26660296 | 3.61E-02 |
| ADGRE5 | 2806.008902 | -0.267351389 | 3.93E-02 |

| | | | |
|---|---|---|---|
| HSPA14 | 1041.680726 | -0.267668054 | 1.84E-02 |
| RNF14 | 1986.791575 | -0.268137122 | 1.36E-02 |
| MFAP1 | 1548.759448 | -0.268201819 | 1.45E-02 |
| USP13 | 1845.289676 | -0.268329772 | 1.38E-03 |
| PHYH | 939.2769998 | -0.268884625 | 4.73E-03 |
| TUB | 3581.772259 | -0.268920502 | 9.02E-03 |
| ACTR3B | 853.544742 | -0.268956731 | 1.78E-02 |
| HDDC2 | 1891.097272 | -0.269068035 | 1.28E-04 |
| BCL7C | 1987.222391 | -0.269262922 | 4.84E-02 |
| SIPA1L1 | 3181.582534 | -0.269296109 | 6.18E-04 |
| JADE2 | 2540.950106 | -0.269435267 | 2.33E-02 |
| SCAND2P | 815.8365651 | -0.270284008 | 2.27E-02 |
| SYNRG | 2863.187215 | -0.270623749 | 3.76E-03 |
| THSD4 | 3166.704389 | -0.270836108 | 4.99E-02 |
| TMEM223 | 1446.597385 | -0.271320412 | 5.00E-02 |
| STK19 | 688.7074119 | -0.271452404 | 2.02E-02 |
| DSE | 960.8994602 | -0.27148535 | 1.34E-02 |
| GTF2H5 | 972.3372066 | -0.271852678 | 2.44E-02 |
| BEX3 | 5302.998397 | -0.272324728 | 1.56E-02 |
| ST3GAL2 | 3203.355799 | -0.272422222 | 4.26E-02 |
| ZNF623 | 2360.212587 | -0.27266781 | 1.54E-02 |
| SMN2 | 2795.058623 | -0.273531778 | 1.00E-02 |
| MSH3 | 1602.672514 | -0.273706708 | 9.09E-04 |
| TRIM5 | 2409.462025 | -0.273751581 | 2.04E-03 |
| AC046185.3 | 1008.471374 | -0.274522561 | 3.16E-03 |
| TRAF5 | 901.765231 | -0.274820001 | 4.86E-02 |
| AC019205.1 | 379.8578792 | -0.274970622 | 2.55E-02 |
| TBC1D24 | 1209.420253 | -0.275180926 | 3.27E-02 |
| WNK1 | 11229.30839 | -0.27524928 | 5.86E-03 |
| PNMA8A | 2550.232377 | -0.276247635 | 2.37E-06 |
| HSD17B12 | 2891.077653 | -0.27647874 | 3.95E-02 |
| POLR2B | 7372.476681 | -0.276880122 | 4.89E-03 |
| ZFP91 | 6060.027481 | -0.277792289 | 3.58E-05 |
| MYH10 | 12958.38845 | -0.277925749 | 1.28E-04 |
| CDC40 | 1575.43891 | -0.277930085 | 1.94E-02 |
| ZFP1 | 767.8083205 | -0.278301677 | 4.71E-02 |
| MTMR4 | 2791.918713 | -0.278660998 | 5.86E-03 |
| HSDL2 | 2211.078784 | -0.278714947 | 1.29E-02 |
| MRPS25 | 2424.720312 | -0.2787691 | 4.74E-04 |
| ZNF783 | 1538.74869 | -0.278917225 | 4.71E-03 |
| ORAI2 | 6379.575777 | -0.279163776 | 2.87E-02 |
| NBPF19 | 1631.618351 | -0.279268552 | 1.20E-02 |
| GPC2 | 493.4075104 | -0.279894225 | 1.99E-02 |
| B3GALNT1 | 1058.702994 | -0.280078106 | 3.68E-02 |
| ENOX2 | 1065.289968 | -0.280174616 | 4.67E-03 |
| LSM1 | 1371.961436 | -0.280759913 | 8.16E-04 |
| MTFR1L | 2547.233496 | -0.280811606 | 2.87E-02 |

| | | | |
|---|---|---|---|
| GARS-DT | 1058.115265 | -0.280926669 | 1.90E-02 |
| DISC1 | 789.4212616 | -0.280940074 | 1.15E-02 |
| EYA3 | 2181.355695 | -0.281051093 | 1.32E-03 |
| TM9SF2 | 21150.58403 | -0.281123496 | 2.06E-02 |
| TCEANC2 | 825.1789663 | -0.281262946 | 1.62E-02 |
| DCK | 775.2998254 | -0.281693693 | 2.87E-02 |
| STAG3L1 | 415.1234277 | -0.282053292 | 1.92E-02 |
| AC010643.1 | 279.3700195 | -0.282650505 | 4.12E-02 |
| FUZ | 1165.079848 | -0.282846839 | 1.90E-02 |
| PPP4R3A | 2306.290159 | -0.282925747 | 3.78E-02 |
| NA | 432.2714357 | -0.283713049 | 1.40E-02 |
| POPDC3 | 714.293481 | -0.283746373 | 2.59E-02 |
| MKLN1 | 5424.076468 | -0.283830358 | 2.07E-03 |
| ATG14 | 977.1180919 | -0.284039894 | 4.37E-04 |
| VIRMA | 5473.931037 | -0.284712763 | 4.63E-04 |
| ZBTB44 | 2561.825986 | -0.285046213 | 4.66E-02 |
| FBXL4 | 893.34174 | -0.285065185 | 2.11E-02 |
| SOD1 | 8136.187217 | -0.286481048 | 4.37E-03 |
| RAD9A | 799.115388 | -0.286668128 | 3.15E-02 |
| CWC15 | 2318.485294 | -0.28700839 | 4.54E-02 |
| KDM5A | 4501.713184 | -0.287536078 | 2.40E-04 |
| NKX3-1 | 1379.272934 | -0.287831407 | 1.30E-02 |
| COA7 | 1675.441611 | -0.287831987 | 3.94E-02 |
| PINK1-AS | 414.9095693 | -0.288010768 | 4.64E-02 |
| HIP1 | 8387.051766 | -0.288242713 | 1.39E-03 |
| IFT46 | 2480.303894 | -0.288289099 | 9.52E-04 |
| STAG3L3 | 1698.480032 | -0.288786479 | 3.75E-03 |
| ARSA | 1347.401683 | -0.288898749 | 3.47E-02 |
| ZNF512 | 2797.925537 | -0.289253273 | 4.61E-03 |
| RAMP2-AS1 | 693.2912316 | -0.289256138 | 3.11E-02 |
| MAGEA11 | 992.8345221 | -0.289326585 | 1.98E-03 |
| SLC2A4RG | 3962.621191 | -0.289373404 | 3.20E-02 |
| KIAA0232 | 1363.339717 | -0.289542364 | 2.13E-02 |
| WRB | 849.4585615 | -0.28959542 | 6.20E-04 |
| AC026470.1 | 2189.407246 | -0.289831556 | 3.24E-02 |
| ZNF7 | 1564.127802 | -0.29007959 | 5.31E-05 |
| LRRC37A4P | 1314.268066 | -0.290147476 | 9.20E-03 |
| NDUFB1 | 1417.659011 | -0.290308412 | 4.77E-02 |
| MIGA2 | 1872.139308 | -0.290388902 | 6.23E-03 |
| CSTF2T | 1225.829139 | -0.290428426 | 6.63E-03 |
| RBM27 | 1405.851249 | -0.290480548 | 4.70E-02 |
| RXRA | 6547.240196 | -0.290667639 | 1.76E-02 |
| ZKSCAN5 | 2534.993214 | -0.290853696 | 2.66E-04 |
| ATP5F1E | 10326.63516 | -0.291171045 | 3.29E-02 |
| JHY | 621.7022963 | -0.291301002 | 1.46E-02 |
| PAXIP1 | 1709.812529 | -0.29136763 | 2.23E-02 |
| MCCC2 | 8099.333828 | -0.291576653 | 4.63E-07 |

| | | | |
|---|---|---|---|
| PRKD3 | 3907.703797 | -0.291673377 | 1.61E-02 |
| SPIDR | 2126.955127 | -0.292502555 | 1.92E-02 |
| DMPK | 1237.876481 | -0.292539142 | 7.95E-03 |
| TDP2 | 1202.826721 | -0.293200951 | 3.38E-02 |
| THADA | 2754.68274 | -0.293625492 | 5.10E-04 |
| LDB1 | 2533.864535 | -0.293946847 | 1.50E-02 |
| TAF2 | 2850.725467 | -0.29425006 | 2.87E-02 |
| BTBD9 | 821.1860443 | -0.29438199 | 4.49E-02 |
| NAA30 | 898.0928515 | -0.294429303 | 3.60E-02 |
| CHMP2B | 883.1991909 | -0.294737227 | 2.91E-02 |
| FGD5-AS1 | 5228.2829 | -0.294771164 | 1.76E-03 |
| TEDC2 | 938.6373295 | -0.295159044 | 4.12E-02 |
| MSANTD2 | 917.7133361 | -0.295206613 | 4.09E-02 |
| AQR | 2617.52052 | -0.295419413 | 8.83E-04 |
| PDP1 | 2884.757114 | -0.295463592 | 7.94E-03 |
| CDK13 | 3103.381094 | -0.295791309 | 3.78E-03 |
| TGS1 | 2101.273683 | -0.295808621 | 6.09E-03 |
| KIZ | 1237.572865 | -0.296565971 | 3.11E-02 |
| PNMA1 | 2633.572079 | -0.296634194 | 5.02E-05 |
| NA | 707.1082302 | -0.296638004 | 6.42E-03 |
| KLHL22 | 1002.038083 | -0.297318905 | 4.92E-02 |
| NCBP3 | 2535.90906 | -0.297411228 | 1.04E-02 |
| UBR7 | 3302.335545 | -0.297449733 | 4.67E-02 |
| AKT1S1 | 2743.622308 | -0.297485753 | 2.09E-02 |
| SSH2 | 1653.105872 | -0.297742879 | 1.04E-02 |
| ATMIN | 3490.625369 | -0.297971649 | 8.92E-04 |
| MKNK1 | 1050.384741 | -0.298065975 | 6.87E-03 |
| ZNF316 | 2117.898613 | -0.29808351 | 2.76E-03 |
| MRPS28 | 938.6174987 | -0.298262177 | 2.75E-02 |
| NAA40 | 3343.755419 | -0.298262507 | 4.22E-07 |
| ZNF22 | 1306.917588 | -0.299080127 | 3.11E-02 |
| AMD1 | 3417.285691 | -0.299173623 | 3.88E-04 |
| PARD3 | 3322.902457 | -0.300942189 | 1.95E-02 |
| NDUFB3 | 1402.880037 | -0.30106731 | 2.57E-02 |
| SREBF1 | 4307.962205 | -0.301142286 | 1.15E-02 |
| PABPN1 | 6487.789511 | -0.301245725 | 1.88E-04 |
| CTNNB1 | 13879.2314 | -0.301521787 | 2.24E-04 |
| RCN2 | 3584.929533 | -0.301615618 | 8.55E-03 |
| WDFY3 | 2746.737282 | -0.301902602 | 1.80E-02 |
| RNF149 | 1382.487106 | -0.301917362 | 4.93E-05 |
| PCF11 | 2025.55404 | -0.301925468 | 3.19E-02 |
| CCDC125 | 1183.595849 | -0.302102303 | 8.49E-03 |
| CSTB | 2993.214322 | -0.302233313 | 3.68E-02 |
| SLC35B3 | 642.8102402 | -0.302400814 | 3.35E-02 |
| PLSCR1 | 1248.540368 | -0.302800833 | 7.51E-03 |
| TCTN1 | 1339.541659 | -0.303023959 | 6.69E-04 |
| UFC1 | 2361.765324 | -0.303095367 | 1.09E-02 |

| | | | |
|---|---|---|---|
| NCOA5 | 2667.247924 | -0.3032502 | 2.35E-03 |
| LINC00888 | 561.0637431 | -0.303484311 | 5.65E-03 |
| TXNDC17 | 2660.628943 | -0.303817182 | 4.00E-03 |
| CABP4 | 464.2960878 | -0.304112855 | 3.94E-02 |
| VPS4B | 1727.894412 | -0.304125492 | 3.10E-04 |
| DYNC1LI2 | 8557.972053 | -0.304867482 | 1.76E-03 |
| PAIP2 | 3370.926984 | -0.30524122 | 1.66E-04 |
| LRRC27 | 266.4274175 | -0.305751057 | 1.45E-02 |
| HTATSF1 | 6072.239422 | -0.305811049 | 1.15E-02 |
| ITFG2 | 735.0446452 | -0.305881453 | 3.61E-02 |
| LCLAT1 | 1108.344898 | -0.30604716 | 4.02E-02 |
| RBM6 | 3881.850496 | -0.306096274 | 2.33E-03 |
| ZNF510 | 1009.980934 | -0.30673428 | 2.04E-02 |
| EPHX2 | 966.502786 | -0.307045919 | 1.45E-02 |
| COPS5 | 3660.217827 | -0.307392813 | 5.12E-06 |
| VCPIP1 | 1187.72626 | -0.308467268 | 2.62E-02 |
| MMUT | 1660.940296 | -0.308915849 | 1.66E-03 |
| RGS12 | 1044.633314 | -0.309184077 | 4.19E-02 |
| CFAP36 | 1119.165321 | -0.309253952 | 1.69E-03 |
| GSTT2B | 1091.957125 | -0.309316793 | 1.57E-02 |
| CMPK1 | 2535.089272 | -0.309623845 | 3.03E-02 |
| CUL3 | 3416.48265 | -0.309643386 | 5.88E-04 |
| RIOX1 | 610.8090639 | -0.309721873 | 2.14E-02 |
| MTERF4 | 1692.32588 | -0.309787205 | 8.54E-05 |
| PDF | 1295.291604 | -0.30981696 | 2.49E-05 |
| SFXN5 | 1044.332545 | -0.309898233 | 3.15E-02 |
| VPS41 | 3381.596704 | -0.310311017 | 1.33E-04 |
| DUT | 1804.676887 | -0.310402883 | 1.58E-02 |
| ROMO1 | 4188.143055 | -0.310437496 | 2.24E-02 |
| FAM160A2 | 1211.307237 | -0.310514228 | 5.90E-03 |
| NAGK | 1852.588991 | -0.310901519 | 2.18E-02 |
| NDFIP2 | 1600.419492 | -0.310925817 | 1.93E-02 |
| TEX10 | 3490.266542 | -0.311020349 | 1.20E-08 |
| BACE1 | 6341.742065 | -0.311057851 | 9.89E-04 |
| ANAPC13 | 1919.232172 | -0.311068772 | 1.82E-03 |
| RERE | 5375.12534 | -0.31122742 | 6.22E-05 |
| CD47 | 3412.049834 | -0.31183802 | 9.96E-05 |
| PRPS2 | 2404.952892 | -0.311875754 | 2.87E-02 |
| PPP4R1L | 870.1846927 | -0.312346604 | 2.35E-03 |
| FKBP10 | 18208.3748 | -0.312402808 | 1.21E-02 |
| KDM6A | 1263.201627 | -0.31264795 | 2.49E-02 |
| RNF103 | 872.356071 | -0.312933603 | 1.93E-03 |
| UFSP2 | 1095.762186 | -0.312958082 | 1.44E-02 |
| ZNF226 | 1041.694412 | -0.313117768 | 3.49E-02 |
| ZKSCAN7 | 396.0174361 | -0.313473183 | 4.23E-02 |
| TRIAP1 | 1088.600773 | -0.313565607 | 4.88E-03 |
| SLC39A6 | 2723.388463 | -0.31409485 | 7.83E-03 |

| | | | |
|---|---|---|---|
| SCAMP1 | 2340.239497 | -0.314114074 | 1.15E-02 |
| RIPK1 | 2531.004431 | -0.314384788 | 4.79E-03 |
| NELFCD | 4856.196635 | -0.314748722 | 1.65E-02 |
| SYNJ2BP | 3240.666765 | -0.314759857 | 8.46E-06 |
| ICK | 878.5792377 | -0.314801519 | 2.11E-02 |
| COL6A2 | 15718.66709 | -0.314889603 | 6.61E-03 |
| RPARP-AS1 | 304.2994051 | -0.315142171 | 2.29E-02 |
| SRP14 | 6129.603975 | -0.315259207 | 4.31E-03 |
| ZNF786 | 452.9511319 | -0.31544087 | 2.02E-02 |
| MAP3K7 | 3642.665461 | -0.315659262 | 8.98E-03 |
| SS18 | 2671.703371 | -0.315694351 | 2.13E-02 |
| KAT2A | 3508.866278 | -0.315792124 | 2.07E-04 |
| WSB1 | 2199.915048 | -0.315834125 | 2.22E-02 |
| TM9SF3 | 4388.18588 | -0.316365166 | 3.57E-02 |
| IFT22 | 1788.73016 | -0.316663377 | 2.35E-03 |
| LMBR1 | 4188.33573 | -0.316712621 | 3.28E-03 |
| PARG | 979.4293806 | -0.316855559 | 1.30E-02 |
| OSGEP | 996.6215002 | -0.316969155 | 4.17E-03 |
| DCAF16 | 2142.847711 | -0.317003865 | 2.96E-03 |
| MORN1 | 444.3401887 | -0.317057061 | 3.90E-03 |
| RMND5B | 1733.652011 | -0.317186663 | 3.73E-03 |
| OSGEPL1 | 614.0173471 | -0.317432741 | 1.44E-02 |
| SUZ12P1 | 346.4032018 | -0.318168702 | 2.41E-02 |
| YTHDF3 | 3985.073013 | -0.318215106 | 1.15E-04 |
| CYB561 | 2861.485413 | -0.318525855 | 1.49E-03 |
| PSME4 | 4808.399483 | -0.318721866 | 1.81E-03 |
| ZNF691 | 543.1359512 | -0.318751314 | 3.35E-03 |
| IFNGR1 | 1233.623812 | -0.319186972 | 6.54E-03 |
| PIGX | 1133.959941 | -0.319257229 | 5.14E-03 |
| WIPF1 | 1839.234446 | -0.319294896 | 2.01E-04 |
| CHD8 | 7116.522774 | -0.31946812 | 6.09E-04 |
| LRRC14 | 2220.494375 | -0.319536797 | 4.35E-03 |
| SPATA20 | 5233.898565 | -0.319628723 | 1.48E-02 |
| GNG10 | 1567.78708 | -0.319875734 | 1.47E-02 |
| ZNF649 | 459.1343401 | -0.319962126 | 1.17E-02 |
| TCF7L2 | 706.3416085 | -0.320044651 | 2.22E-02 |
| RBM7 | 1307.600618 | -0.320169574 | 4.11E-02 |
| MIS12 | 1596.761517 | -0.320439831 | 5.66E-04 |
| ZNF227 | 1191.452921 | -0.320590499 | 2.50E-03 |
| GSTCD | 1060.988874 | -0.320733937 | 3.38E-02 |
| GNA13 | 2404.762405 | -0.321574765 | 1.96E-02 |
| DBR1 | 685.1075807 | -0.321589355 | 2.75E-03 |
| C8orf82 | 970.646144 | -0.321628043 | 2.36E-02 |
| CTCF | 3014.64559 | -0.321976636 | 1.45E-06 |
| TOMM6 | 3943.972343 | -0.322242952 | 8.04E-03 |
| PARP10 | 2590.466427 | -0.32253298 | 1.12E-02 |
| HSD17B10 | 2671.357743 | -0.323174519 | 1.20E-03 |

| | | | |
|---|---|---|---|
| DGLUCY | 2153.555328 | -0.323278342 | 2.84E-03 |
| GNB4 | 1581.570179 | -0.323773683 | 1.98E-02 |
| HOTAIRM1 | 1189.090175 | -0.324498737 | 3.81E-03 |
| TTC5 | 1049.690651 | -0.324592437 | 2.89E-03 |
| ARMCX2 | 878.4276883 | -0.324639993 | 2.27E-03 |
| SUSD1 | 1885.99721 | -0.324683384 | 5.31E-03 |
| DHX15 | 7490.4444 | -0.324920643 | 6.91E-04 |
| SLC22A4 | 550.2859545 | -0.324931264 | 1.26E-02 |
| GOLPH3L | 1038.121567 | -0.325062816 | 2.28E-03 |
| CENPS | 482.2845624 | -0.325245258 | 9.22E-03 |
| KLHL7 | 2045.471299 | -0.325294479 | 3.74E-04 |
| CSPG4P12 | 639.7044511 | -0.325369587 | 2.16E-02 |
| UBL3 | 632.1579712 | -0.325437272 | 3.87E-02 |
| C2orf68 | 2322.099413 | -0.325453701 | 2.16E-09 |
| LAMTOR4 | 2886.284026 | -0.325814922 | 3.98E-03 |
| RAD51-AS1 | 223.6582899 | -0.326121605 | 4.94E-02 |
| WDR48 | 2718.702698 | -0.326315125 | 1.95E-04 |
| EIF4EBP2 | 3010.009684 | -0.326809619 | 1.57E-02 |
| BTG1 | 2988.2491 | -0.326849 | 1.58E-02 |
| FIP1L1 | 2275.696884 | -0.327096299 | 7.74E-03 |
| NEPRO | 1618.389282 | -0.327586808 | 3.13E-02 |
| USP8 | 1583.500754 | -0.327659554 | 4.98E-02 |
| CRBN | 1204.374755 | -0.327708906 | 3.61E-03 |
| TMEM30A | 6283.824916 | -0.327768169 | 8.22E-03 |
| NA | 2018.632065 | -0.327914061 | 9.97E-03 |
| AC006001.3 | 380.0272958 | -0.328224322 | 1.68E-02 |
| ZSCAN2 | 647.4750931 | -0.328477122 | 2.90E-02 |
| COA3 | 1646.581393 | -0.328534845 | 3.74E-02 |
| CSNK1G1 | 2578.916105 | -0.328643074 | 1.39E-05 |
| EIF2AK3 | 1083.717756 | -0.328749497 | 4.73E-02 |
| INTS8 | 2842.185022 | -0.328833752 | 3.46E-02 |
| LEPROT | 2664.218206 | -0.328964048 | 1.18E-03 |
| MFSD4B | 323.1202287 | -0.329219147 | 1.95E-02 |
| TLCD1 | 494.9847647 | -0.329647281 | 4.46E-02 |
| MYD88 | 2389.693906 | -0.329683837 | 2.47E-02 |
| TOB2 | 2449.947897 | -0.329960859 | 1.54E-02 |
| PAPOLG | 812.3893065 | -0.330233064 | 2.10E-02 |
| SETD2 | 3613.711957 | -0.33053483 | 6.24E-03 |
| YAP1 | 7292.138549 | -0.33090229 | 2.76E-04 |
| LTK | 604.491803 | -0.330932987 | 2.83E-02 |
| ZKSCAN8 | 2404.075644 | -0.330972702 | 1.80E-02 |
| RAVER2 | 998.6587245 | -0.33137311 | 1.49E-03 |
| TSC22D1 | 5776.662557 | -0.331440078 | 5.19E-04 |
| PSTPIP2 | 476.2849638 | -0.331447941 | 2.27E-02 |
| TRIM26 | 1999.723817 | -0.331917155 | 4.21E-03 |
| MIS18A | 904.3056736 | -0.332000128 | 3.00E-02 |
| KLHL8 | 892.952596 | -0.332379196 | 4.03E-02 |

| | | | |
|---|---|---|---|
| GTF2H2C | 2923.736029 | -0.332475798 | 1.34E-02 |
| ZBTB8OS | 542.9329176 | -0.332532689 | 1.17E-02 |
| ZNF251 | 1133.94733 | -0.332571209 | 2.61E-05 |
| NEK4 | 1172.213935 | -0.332614012 | 5.53E-03 |
| BIVM | 3286.082284 | -0.33332542 | 2.87E-02 |
| CCNL2 | 6146.069744 | -0.333425336 | 1.38E-09 |
| MRNIP | 1068.786205 | -0.333479205 | 5.86E-04 |
| ELK3 | 2668.256893 | -0.333648033 | 3.01E-03 |
| ENY2 | 2390.585504 | -0.333837311 | 1.27E-04 |
| FYTTD1 | 2885.666641 | -0.334213434 | 5.28E-03 |
| GNPDA2 | 591.1548042 | -0.334224895 | 4.12E-02 |
| INTS5 | 1999.085661 | -0.334281481 | 2.02E-02 |
| ZCCHC10 | 662.269274 | -0.334365538 | 4.56E-03 |
| PPP6R3 | 4508.605406 | -0.334509697 | 1.16E-02 |
| PRPF38A | 2292.914314 | -0.334716388 | 5.91E-04 |
| UBA5 | 1245.060766 | -0.335187687 | 3.91E-02 |
| OSTC | 3548.714486 | -0.335451125 | 5.58E-05 |
| LIPA | 3892.554295 | -0.335536113 | 1.70E-02 |
| ADAMTS7 | 1523.351905 | -0.335735754 | 1.01E-02 |
| NAGLU | 2962.678407 | -0.335812596 | 1.24E-02 |
| AEBP2 | 673.02458 | -0.335849017 | 1.20E-02 |
| KIAA0753 | 1607.123434 | -0.335965085 | 3.68E-03 |
| CCDC57 | 916.7750703 | -0.336170938 | 1.71E-02 |
| TNPO1 | 14951.12919 | -0.336244533 | 2.13E-02 |
| ALDH7A1 | 9362.120636 | -0.336303512 | 4.56E-04 |
| SCPEP1 | 3442.188977 | -0.336514092 | 4.41E-03 |
| CCNC | 1930.595924 | -0.336710231 | 4.82E-02 |
| VDAC3 | 5335.465565 | -0.337132842 | 5.94E-04 |
| STK36 | 1396.326232 | -0.337401563 | 2.45E-03 |
| IQCC | 379.3166359 | -0.337632854 | 3.54E-02 |
| SLC26A2 | 2274.410435 | -0.33780055 | 6.05E-03 |
| CDK12 | 4237.691112 | -0.337881754 | 2.37E-06 |
| DTWD1 | 275.7066094 | -0.337983546 | 4.55E-02 |
| GALM | 998.4623932 | -0.33800582 | 3.68E-03 |
| SLC16A4 | 951.4176711 | -0.338277095 | 1.44E-02 |
| C11orf54 | 1261.266506 | -0.339011843 | 4.18E-04 |
| TM9SF1 | 3915.379062 | -0.339421923 | 1.73E-03 |
| LAGE3 | 1873.462234 | -0.339633573 | 3.70E-02 |
| DNAJC30 | 848.6358511 | -0.340032966 | 4.22E-02 |
| APPL1 | 1651.578524 | -0.340350424 | 4.71E-02 |
| RAD21 | 11043.70758 | -0.340450078 | 1.04E-02 |
| CHD6 | 4186.747292 | -0.340803561 | 6.49E-03 |
| ATXN7L2 | 355.4580206 | -0.340815109 | 2.59E-02 |
| DNAJC4 | 1151.179245 | -0.341195811 | 4.33E-02 |
| ACBD5 | 839.2854957 | -0.341284703 | 3.06E-03 |
| VAMP7 | 2563.18571 | -0.341680273 | 7.54E-03 |
| LYRM2 | 2767.812549 | -0.341811934 | 5.60E-04 |

| | | | |
|---|---|---|---|
| FAM161B | 403.8397917 | -0.341826502 | 8.61E-03 |
| CHN1 | 893.6230543 | -0.341846829 | 1.28E-03 |
| BTBD1 | 3723.853847 | -0.341897147 | 2.71E-03 |
| SRSF1 | 10334.91913 | -0.342018798 | 1.01E-04 |
| THAP7 | 1127.050214 | -0.342039442 | 2.23E-02 |
| ANXA4 | 2729.466506 | -0.34206293 | 4.17E-05 |
| RBBP4 | 6433.30477 | -0.342150912 | 3.40E-03 |
| RYK | 3166.723865 | -0.342251499 | 1.11E-05 |
| EHBP1 | 2470.970071 | -0.342267469 | 3.34E-02 |
| OBSCN | 2244.375511 | -0.342833794 | 2.37E-02 |
| PHKB | 4756.907039 | -0.342855302 | 2.28E-07 |
| CDKN2A | 1458.259282 | -0.343000604 | 3.72E-02 |
| ACOT11 | 447.5869448 | -0.34340501 | 2.97E-02 |
| MREG | 275.7797383 | -0.343475014 | 9.36E-03 |
| SREK1IP1 | 1707.725758 | -0.343551479 | 5.10E-03 |
| NBDY | 1029.916909 | -0.343610666 | 8.61E-03 |
| WDR26 | 5431.769766 | -0.343672343 | 1.88E-09 |
| G6PC3 | 3386.25251 | -0.343808796 | 2.71E-03 |
| SASH1 | 1551.444222 | -0.344134986 | 3.81E-05 |
| NUP107 | 3600.901159 | -0.344373326 | 2.65E-02 |
| PGBD2 | 274.8945426 | -0.344634894 | 4.94E-02 |
| RPS6KA3 | 4793.968676 | -0.344638923 | 6.61E-03 |
| ENTPD5 | 1104.144112 | -0.345113085 | 1.30E-02 |
| PLPP5 | 1498.63886 | -0.345274957 | 6.64E-05 |
| C1GALT1C1 | 890.1808052 | -0.345366874 | 7.39E-04 |
| CCDC136 | 1824.450033 | -0.345692582 | 2.83E-03 |
| C9orf16 | 2033.48365 | -0.345961776 | 4.24E-02 |
| ADAMTS10 | 829.7042696 | -0.346149572 | 4.35E-02 |
| KDM5B | 4458.455445 | -0.346714328 | 1.27E-04 |
| UGDH | 1488.604689 | -0.346824907 | 4.92E-03 |
| PNMA3 | 556.7056313 | -0.347013677 | 4.26E-03 |
| RNGTT | 1700.91481 | -0.347624423 | 1.81E-02 |
| ZNF280D | 1224.036056 | -0.347742989 | 1.43E-02 |
| AFF4 | 7606.913083 | -0.347963557 | 4.03E-03 |
| NCBP2 | 4485.947445 | -0.348140351 | 1.08E-04 |
| KIFC2 | 1596.31756 | -0.348422013 | 1.75E-02 |
| ANGPTL2 | 1290.993455 | -0.348915866 | 3.24E-02 |
| TAPT1 | 725.1491562 | -0.349082029 | 1.33E-02 |
| TMEM260 | 833.9640819 | -0.349123691 | 4.22E-02 |
| NA | 887.2209421 | -0.349694385 | 1.46E-02 |
| OIP5 | 389.6625071 | -0.350154786 | 2.04E-02 |
| NLGN2 | 4393.574422 | -0.350222308 | 8.47E-03 |
| FBXL17 | 1252.937548 | -0.350252518 | 8.02E-03 |
| SS18L2 | 592.0948868 | -0.350335963 | 1.59E-03 |
| CHSY1 | 3084.922925 | -0.350361806 | 9.06E-06 |
| FLI1 | 3174.945059 | -0.350486257 | 2.58E-04 |
| ZNF891 | 383.1294511 | -0.350554218 | 3.36E-02 |

| | | | |
|---|---|---|---|
| TEX2 | 2687.825769 | -0.35056083 | 1.34E-06 |
| NFIB | 3177.725101 | -0.350656467 | 1.08E-04 |
| CDK5 | 1685.224274 | -0.35085353 | 1.06E-02 |
| OIP5-AS1 | 1986.522582 | -0.350874135 | 1.35E-05 |
| ZNF584 | 705.5673573 | -0.351138415 | 7.68E-03 |
| BRD3OS | 2561.136172 | -0.351292425 | 2.53E-03 |
| PIK3CG | 245.0376781 | -0.351618947 | 1.70E-02 |
| TRIM3 | 1257.094969 | -0.351768338 | 2.27E-03 |
| DROSHA | 5892.566378 | -0.351981826 | 1.06E-05 |
| RBPJ | 2693.98748 | -0.352833823 | 8.08E-04 |
| LATS1 | 1070.227587 | -0.352905227 | 6.09E-04 |
| PC | 2346.183981 | -0.35305214 | 2.65E-02 |
| SUGT1 | 954.8725404 | -0.353349506 | 1.18E-02 |
| FAM114A2 | 967.3843151 | -0.353616625 | 2.49E-02 |
| STIL | 1639.705519 | -0.353699794 | 2.97E-02 |
| ZFP62 | 704.7395684 | -0.353860537 | 1.25E-02 |
| ARMC7 | 495.6640675 | -0.353888087 | 2.85E-02 |
| RC3H2 | 3309.143174 | -0.35408069 | 4.05E-05 |
| RABGAP1L | 809.043815 | -0.354374525 | 3.46E-02 |
| SAMHD1 | 2674.03882 | -0.354869595 | 1.07E-02 |
| CBX1 | 4941.429777 | -0.354870781 | 3.04E-02 |
| CPT2 | 1235.342638 | -0.355029145 | 1.49E-03 |
| RNF34 | 1430.120546 | -0.355624494 | 2.56E-03 |
| CHUK | 797.0347497 | -0.356808301 | 3.62E-02 |
| ST6GALNAC2 | 968.4348124 | -0.356887122 | 4.32E-03 |
| WDR73 | 1944.452553 | -0.356973674 | 6.42E-05 |
| CHST2 | 604.9230275 | -0.356991709 | 1.23E-04 |
| PLEKHA1 | 499.4409068 | -0.357045121 | 1.50E-02 |
| AP1G1 | 6782.94064 | -0.35725348 | 2.38E-06 |
| IQCE | 1179.997058 | -0.357315839 | 1.22E-02 |
| GATD3B | 2946.051139 | -0.35737744 | 5.86E-03 |
| GGT7 | 2198.532399 | -0.357415417 | 1.10E-03 |
| ERAP1 | 6025.539569 | -0.35744374 | 5.00E-04 |
| ZFAT | 705.2663559 | -0.3575017 | 2.13E-02 |
| SLC25A44 | 2071.496953 | -0.358135434 | 1.15E-02 |
| MRFAP1L1 | 2334.33261 | -0.35922679 | 5.33E-04 |
| TFCP2 | 1699.603032 | -0.359276033 | 3.95E-07 |
| PWWP2A | 1799.661611 | -0.359723043 | 2.69E-04 |
| RAP2A | 2898.846822 | -0.360093935 | 1.11E-02 |
| GPAA1 | 6075.963465 | -0.360307779 | 1.30E-03 |
| ZSCAN26 | 550.8185622 | -0.360428294 | 4.65E-02 |
| ASB8 | 972.3934283 | -0.360706595 | 1.01E-04 |
| NRDE2 | 1127.583602 | -0.36089049 | 3.16E-03 |
| B4GALNT3 | 353.3939525 | -0.361112222 | 3.63E-02 |
| DOLPP1 | 1556.722068 | -0.361362367 | 6.87E-04 |
| ELK4 | 1399.151224 | -0.361454467 | 4.05E-02 |
| FBXL20 | 1050.705018 | -0.361589559 | 2.77E-03 |

| | | | |
|---|---|---|---|
| CD109 | 3127.492824 | -0.361778492 | 1.10E-02 |
| ZC3HAV1 | 3837.855423 | -0.361781243 | 2.21E-02 |
| ASB16-AS1 | 304.3514996 | -0.362082317 | 2.16E-02 |
| AC009163.6 | 182.0607473 | -0.36209932 | 2.29E-02 |
| INIP | 1067.234987 | -0.362778304 | 3.47E-04 |
| FZD3 | 812.4350586 | -0.363271551 | 3.70E-02 |
| SEC23IP | 2552.802326 | -0.363366809 | 6.86E-03 |
| SLC36A4 | 873.4201689 | -0.36342669 | 4.75E-02 |
| NAE1 | 2939.312569 | -0.363449484 | 4.27E-02 |
| TFAM | 1552.341068 | -0.363970952 | 1.81E-02 |
| WDR75 | 2797.494185 | -0.364028942 | 1.19E-03 |
| ERC1 | 4832.654921 | -0.364158086 | 7.79E-08 |
| FKBP14 | 2076.395674 | -0.364494169 | 4.39E-03 |
| 3-Mar | 427.8141635 | -0.364557122 | 1.29E-02 |
| MZF1 | 921.5328365 | -0.364812951 | 5.15E-04 |
| L3MBTL2 | 1878.053994 | -0.36486187 | 2.72E-04 |
| HARS2 | 2149.232641 | -0.364874032 | 1.16E-08 |
| SPART | 2174.26844 | -0.364957058 | 4.46E-04 |
| KIF13A | 2692.151826 | -0.365195287 | 8.08E-05 |
| KANSL1 | 2209.649256 | -0.365413797 | 1.23E-03 |
| TPRKB | 868.0736886 | -0.365759193 | 2.09E-02 |
| DCAF6 | 2454.745029 | -0.365887977 | 2.03E-03 |
| CECR7 | 1422.204956 | -0.365929999 | 1.41E-03 |
| CLTC | 25848.11783 | -0.365966142 | 7.28E-08 |
| CYB561A3 | 2743.50724 | -0.366160346 | 4.30E-03 |
| ADAM23 | 710.607505 | -0.366483602 | 1.26E-02 |
| SMIM3 | 518.8367427 | -0.366512787 | 2.09E-02 |
| CBWD2 | 1164.078601 | -0.366982038 | 8.50E-03 |
| ANKRD34A | 275.0037824 | -0.367026158 | 1.55E-02 |
| FBXO38 | 1593.343293 | -0.367269091 | 1.14E-02 |
| ACER3 | 1828.681258 | -0.367357253 | 1.45E-03 |
| G3BP2 | 7950.497687 | -0.3675206 | 1.21E-05 |
| SH2B1 | 2927.777628 | -0.367534408 | 3.91E-04 |
| KIAA1841 | 487.9104014 | -0.367545636 | 4.15E-02 |
| CDC26 | 1403.047321 | -0.36754798 | 4.12E-05 |
| ZNF671 | 336.4759413 | -0.367637867 | 1.38E-02 |
| NF1 | 6160.278563 | -0.367745522 | 1.69E-03 |
| PSEN1 | 3588.470478 | -0.368001692 | 4.10E-06 |
| FRYL | 2695.093653 | -0.368413527 | 1.66E-02 |
| CYP20A1 | 839.5415502 | -0.368489621 | 5.86E-04 |
| L3MBTL1 | 722.4876846 | -0.368737707 | 4.21E-03 |
| TMEM181 | 2526.062735 | -0.368843683 | 2.09E-03 |
| ARMC5 | 983.2956384 | -0.369307539 | 4.49E-03 |
| COLEC12 | 3956.507078 | -0.369324369 | 1.77E-03 |
| USP33 | 3250.439962 | -0.369672644 | 3.04E-03 |
| C19orf57 | 543.2283454 | -0.369684203 | 7.75E-03 |
| ZRSR2 | 304.3966028 | -0.369973854 | 3.08E-02 |

| | | | |
|---|---|---|---|
| SLC30A6 | 1921.760547 | -0.370350365 | 7.72E-04 |
| MAGI1 | 1754.890765 | -0.370522379 | 1.04E-03 |
| ZNF12 | 1848.754705 | -0.370527245 | 2.10E-02 |
| WWP1 | 1340.319215 | -0.370581375 | 3.21E-02 |
| CDC42SE2 | 2790.469273 | -0.370707783 | 8.67E-05 |
| MLYCD | 694.324879 | -0.370852677 | 6.96E-03 |
| D2HGDH | 937.3314087 | -0.37103912 | 1.47E-02 |
| MPHOSPH8 | 1367.059281 | -0.371222819 | 9.97E-03 |
| GTF3C4 | 5054.525674 | -0.3712628 | 2.00E-08 |
| REV1 | 1476.613051 | -0.371802003 | 1.70E-02 |
| DCAF12 | 4233.100473 | -0.371861989 | 5.37E-07 |
| ZNF346 | 892.9662731 | -0.372006926 | 1.50E-03 |
| ZNF528 | 1333.785262 | -0.372577207 | 6.68E-03 |
| SC5D | 2421.196059 | -0.37293032 | 7.19E-03 |
| CRELD2 | 1562.4412 | -0.373167001 | 9.39E-03 |
| FASTKD1 | 914.5231774 | -0.373714862 | 4.22E-02 |
| RNF227 | 387.3186546 | -0.37387736 | 7.70E-03 |
| JMJD7-PLA2G4E | 577.9573265 | -0.374253429 | 1.92E-02 |
| PALLD | 4841.86156 | -0.374352583 | 7.36E-04 |
| ING2 | 350.4735794 | -0.374380583 | 1.94E-02 |
| HCG18 | 1539.718788 | -0.374456298 | 2.13E-03 |
| IPP | 567.557776 | -0.374738697 | 2.71E-03 |
| DHX9 | 14124.10549 | -0.374780113 | 2.72E-06 |
| RNF139 | 1481.985634 | -0.37479263 | 2.01E-05 |
| TMA7 | 3015.831578 | -0.374819556 | 2.55E-03 |
| KPTN | 754.5468126 | -0.375436619 | 2.54E-02 |
| SACM1L | 1668.475358 | -0.375502886 | 1.26E-02 |
| PARP1 | 12306.40327 | -0.375560988 | 2.71E-03 |
| MFAP4 | 931.0022545 | -0.375638035 | 3.60E-02 |
| PTOV1-AS1 | 556.2368947 | -0.375809898 | 5.36E-03 |
| RBM5 | 5755.397797 | -0.37582106 | 4.55E-09 |
| SNX29 | 2506.115096 | -0.376043641 | 1.40E-04 |
| CLUAP1 | 987.7006588 | -0.376440641 | 2.20E-02 |
| AL513550.1 | 323.2667061 | -0.37681216 | 4.94E-03 |
| CC2D2A | 1425.087652 | -0.376849437 | 9.07E-04 |
| PYGO1 | 602.9014045 | -0.376969824 | 2.16E-02 |
| ZNF131 | 2746.314322 | -0.377055924 | 5.92E-03 |
| HDAC8 | 1003.791273 | -0.377149552 | 2.58E-04 |
| MGST2 | 872.3258944 | -0.377735646 | 2.22E-02 |
| LINC00355 | 589.7798369 | -0.378115527 | 5.25E-03 |
| PHAX | 2311.510306 | -0.378120652 | 1.26E-02 |
| ZNF17 | 345.8333988 | -0.378595482 | 1.03E-02 |
| ANKMY2 | 1123.608775 | -0.379175924 | 1.06E-04 |
| C16orf45 | 734.6864757 | -0.379372742 | 9.86E-04 |
| ATP2A3 | 1211.316063 | -0.379409355 | 4.10E-03 |
| SLC16A2 | 289.1988524 | -0.379411069 | 4.77E-02 |
| GSTZ1 | 1069.563483 | -0.379524139 | 3.38E-03 |

| | | | |
|---|---|---|---|
| DNM1 | 2280.331942 | -0.379939655 | 2.37E-04 |
| ZNF266 | 2147.525112 | -0.379951898 | 1.30E-05 |
| ENC1 | 10344.19834 | -0.379994642 | 7.41E-05 |
| PRAF2 | 2410.163933 | -0.380145612 | 2.78E-02 |
| MRPL32 | 2091.698119 | -0.380533756 | 6.13E-06 |
| USF1 | 2130.590853 | -0.380586428 | 5.52E-04 |
| FERMT2 | 5028.125652 | -0.380750479 | 2.44E-04 |
| AZI2 | 1519.970742 | -0.380770368 | 7.55E-03 |
| IFNAR1 | 2453.459114 | -0.380813578 | 2.20E-02 |
| HBS1L | 1899.075632 | -0.380966658 | 5.37E-03 |
| AC018647.2 | 763.6559716 | -0.381243847 | 9.53E-05 |
| PPP2R5E | 2442.538167 | -0.381491496 | 1.37E-04 |
| UBR1 | 1334.677991 | -0.381780194 | 1.62E-02 |
| ITPRID2 | 7029.408836 | -0.381820715 | 7.54E-03 |
| MTM1 | 703.2840345 | -0.382658608 | 8.25E-03 |
| PALM2-AKAP2 | 1579.919824 | -0.38302856 | 2.50E-02 |
| PSMC6 | 3382.500155 | -0.383029751 | 2.75E-03 |
| TRPC1 | 289.6312812 | -0.383383128 | 2.55E-02 |
| IPO4 | 3946.715919 | -0.383459712 | 1.08E-04 |
| MEPCE | 5159.706351 | -0.383484236 | 7.71E-06 |
| SLC38A7 | 1481.333423 | -0.383583571 | 1.82E-03 |
| TXN2 | 3663.452541 | -0.383636271 | 2.26E-03 |
| FBXW9 | 659.5921597 | -0.38373279 | 1.17E-02 |
| ZNF384 | 2497.210853 | -0.3841116 | 1.08E-04 |
| WDR7 | 1333.184344 | -0.384397974 | 2.04E-03 |
| LTBP3 | 6500.71372 | -0.384824844 | 4.48E-03 |
| ZNF586 | 548.3422355 | -0.384986643 | 4.06E-03 |
| MED11 | 514.0417921 | -0.385124711 | 4.99E-02 |
| KYAT3 | 811.592789 | -0.38546245 | 1.50E-02 |
| ACAA1 | 1616.455569 | -0.385602639 | 3.61E-04 |
| DEPDC7 | 451.1246896 | -0.385733902 | 1.89E-02 |
| RNF13 | 1302.496039 | -0.385767985 | 2.94E-02 |
| MDN1 | 7460.91208 | -0.385870194 | 3.38E-02 |
| RBM4 | 5211.12219 | -0.38620852 | 1.50E-04 |
| BLOC1S2 | 510.6569008 | -0.386754795 | 6.26E-03 |
| EPM2A | 231.7732836 | -0.387112136 | 2.15E-02 |
| QTRT1 | 1398.971228 | -0.387210618 | 4.77E-04 |
| MPPE1 | 700.4250634 | -0.387262274 | 3.27E-02 |
| POGLUT1 | 1622.036049 | -0.387644225 | 1.68E-05 |
| PYCR3 | 1124.038153 | -0.387773909 | 3.46E-03 |
| ZBTB26 | 1136.425005 | -0.387811114 | 6.30E-03 |
| MSL1 | 3890.623421 | -0.387899394 | 6.17E-06 |
| THAP8 | 365.5154966 | -0.387953626 | 3.91E-02 |
| ATP6V1A | 3354.193021 | -0.388218737 | 4.09E-05 |
| ADAT1 | 2374.903336 | -0.388273977 | 5.84E-07 |
| TTC30A | 485.3279646 | -0.388398109 | 5.86E-03 |
| LRCH1 | 1532.93364 | -0.388856847 | 9.07E-04 |

| | | | |
|---|---|---|---|
| FNIP1 | 3476.825664 | -0.388921417 | 1.05E-02 |
| SLC30A9 | 2686.38585 | -0.388928808 | 4.77E-02 |
| NA | 1377.415725 | -0.388945936 | 5.01E-03 |
| ZNF284 | 227.9168793 | -0.389362178 | 6.66E-03 |
| SLC36A1 | 875.5995958 | -0.389405066 | 2.80E-03 |
| RBM23 | 4441.381134 | -0.389435916 | 5.37E-05 |
| RSPRY1 | 1915.819271 | -0.389479673 | 3.90E-03 |
| RAPH1 | 2044.357541 | -0.389706011 | 2.20E-03 |
| DHX58 | 287.5296303 | -0.389924732 | 2.12E-02 |
| MIA3 | 2860.699507 | -0.390156487 | 1.88E-02 |
| SIN3A | 5272.090568 | -0.390264476 | 1.27E-10 |
| CLASP1 | 3823.442414 | -0.39027522 | 1.91E-05 |
| PRR3 | 2046.71534 | -0.39027753 | 6.09E-05 |
| SOCS5 | 1947.412593 | -0.390315007 | 3.43E-03 |
| PHKG2 | 1614.940655 | -0.390495003 | 1.53E-03 |
| HNRNPLL | 2217.846786 | -0.390713042 | 3.38E-06 |
| FAM230C | 171.8176099 | -0.390870635 | 1.57E-02 |
| SLC19A2 | 517.6445248 | -0.391415301 | 1.13E-02 |
| SLC30A7 | 941.1097818 | -0.391471933 | 7.36E-04 |
| TADA1 | 504.945217 | -0.39148871 | 4.59E-04 |
| TOMM40L | 1091.925039 | -0.391624367 | 6.40E-03 |
| TMED7 | 4176.355682 | -0.391715439 | 1.38E-03 |
| MAML3 | 1424.798106 | -0.391760097 | 3.84E-03 |
| TIMP1 | 7816.765979 | -0.391852396 | 4.42E-02 |
| WDPCP | 375.1359725 | -0.39185507 | 5.22E-03 |
| SDHAF2 | 1739.613726 | -0.392034849 | 1.23E-02 |
| ACADM | 1314.411771 | -0.392057767 | 1.89E-02 |
| PAPSS1 | 4577.996602 | -0.39229902 | 6.88E-04 |
| RFC3 | 1087.343074 | -0.392369403 | 2.33E-02 |
| OTUD6B-AS1 | 432.9170639 | -0.392475276 | 4.23E-02 |
| HEATR5B | 2084.146624 | -0.392697931 | 1.55E-02 |
| PRMT9 | 690.1474832 | -0.392870276 | 5.87E-03 |
| AF131215.5 | 146.1309368 | -0.393120094 | 3.17E-02 |
| ARSD | 1843.274745 | -0.39403485 | 3.49E-04 |
| SEZ6L2 | 3889.516264 | -0.394329367 | 9.18E-03 |
| SLC35F5 | 1967.09848 | -0.394521641 | 1.18E-02 |
| C6orf47 | 1110.798322 | -0.394609848 | 3.39E-04 |
| RELCH | 1303.408196 | -0.394849642 | 7.68E-03 |
| ZNF558 | 1415.933579 | -0.394994552 | 5.61E-04 |
| DICER1 | 1977.619445 | -0.395092062 | 4.96E-02 |
| ZNF736 | 672.2343097 | -0.395391036 | 2.24E-02 |
| TBC1D4 | 1063.760572 | -0.395946107 | 5.75E-03 |
| MFAP3 | 1637.248945 | -0.396106031 | 2.51E-04 |
| RBM34 | 1316.827311 | -0.396359994 | 1.50E-02 |
| BLM | 908.9205934 | -0.396438487 | 4.77E-02 |
| ZNF726 | 144.2205326 | -0.396498383 | 3.32E-02 |
| FERMT1 | 589.4759035 | -0.396611304 | 1.89E-02 |

| | | | |
|---|---|---|---|
| ORMDL3 | 2445.102458 | -0.396615809 | 1.02E-02 |
| SON | 15233.18633 | -0.396813638 | 1.51E-04 |
| C3orf14 | 394.3980219 | -0.39696323 | 1.06E-02 |
| RIT1 | 1170.596501 | -0.397161747 | 3.79E-04 |
| GPRASP2 | 355.0135181 | -0.39735325 | 2.75E-03 |
| AC009053.1 | 341.7337856 | -0.397474563 | 1.40E-03 |
| COL27A1 | 4166.309454 | -0.397808223 | 8.01E-04 |
| TNK1 | 762.1460284 | -0.397827612 | 1.26E-02 |
| AC008755.1 | 1945.393163 | -0.398283386 | 2.48E-06 |
| VPS26A | 1738.082387 | -0.39881133 | 2.88E-02 |
| EMSY | 1435.066954 | -0.398854714 | 3.67E-02 |
| PGAP2 | 928.1459418 | -0.399186931 | 1.20E-04 |
| CLCC1 | 1194.714376 | -0.399305893 | 6.89E-05 |
| CLCN5 | 680.0817195 | -0.399573875 | 4.10E-02 |
| LRRC29 | 210.1297555 | -0.399706161 | 2.02E-02 |
| TMEM106A | 336.5545532 | -0.400064328 | 1.09E-02 |
| UBE2Q2 | 1264.376967 | -0.400387035 | 1.17E-03 |
| ICE1 | 6660.02912 | -0.400614618 | 7.71E-03 |
| TVP23B | 715.0767378 | -0.400866097 | 4.32E-02 |
| ZKSCAN4 | 386.1799195 | -0.40093475 | 1.05E-02 |
| LAP3 | 2191.36426 | -0.400938361 | 2.54E-05 |
| CBX5 | 9249.567894 | -0.401025339 | 3.01E-10 |
| MESD | 4307.097829 | -0.40108958 | 2.40E-06 |
| ZNF76 | 1282.837094 | -0.401571158 | 3.69E-04 |
| HMBOX1 | 1002.827149 | -0.401712902 | 6.33E-03 |
| PYGO2 | 3109.595671 | -0.401730377 | 9.55E-04 |
| PATZ1 | 2235.275444 | -0.402118072 | 1.22E-04 |
| KIF2A | 4545.860791 | -0.402144932 | 6.76E-03 |
| RABL2A | 460.6065084 | -0.402358501 | 1.99E-03 |
| DHX57 | 2397.342818 | -0.402371017 | 4.52E-07 |
| SLC25A22 | 2360.771682 | -0.402441886 | 1.40E-05 |
| SSB | 4068.960498 | -0.402491738 | 1.51E-02 |
| BBS12 | 273.4155551 | -0.40252975 | 8.61E-03 |
| SYN1 | 187.3211338 | -0.402667976 | 4.45E-02 |
| Z97192.2 | 257.6198424 | -0.402960572 | 1.38E-02 |
| DENND6A | 791.9044852 | -0.40297359 | 3.71E-02 |
| ABCD3 | 2227.255344 | -0.403084216 | 1.22E-03 |
| CRTC1 | 1021.148261 | -0.40310122 | 3.33E-04 |
| CXXC5 | 1901.801502 | -0.403457095 | 2.61E-05 |
| SRPK2 | 3839.713398 | -0.403553375 | 8.46E-08 |
| TM2D3 | 1237.924129 | -0.403599042 | 1.90E-05 |
| IFFO1 | 1580.185706 | -0.403628274 | 7.60E-04 |
| LNPK | 1105.442779 | -0.403775147 | 1.79E-02 |
| FMC1-LUC7L2 | 1583.580303 | -0.403897684 | 1.36E-04 |
| NLRC5 | 1288.003349 | -0.404061 | 1.21E-02 |
| MIATNB | 181.1681693 | -0.404138885 | 3.12E-02 |
| NHLRC3 | 495.9969832 | -0.404168899 | 9.62E-04 |

| | | | |
|---|---|---|---|
| CCDC8 | 1342.593494 | -0.404265562 | 8.34E-03 |
| SLC24A1 | 908.7448531 | -0.404312338 | 1.90E-02 |
| PRXL2A | 2340.360732 | -0.404915541 | 1.57E-07 |
| PRRT3-AS1 | 120.3373431 | -0.405024561 | 3.31E-02 |
| ZNF814 | 407.689954 | -0.405054935 | 2.75E-03 |
| DCAF5 | 3256.108903 | -0.405255799 | 3.22E-06 |
| EP300 | 4842.321 | -0.405285193 | 8.11E-21 |
| SPAST | 681.3991413 | -0.405476389 | 9.28E-03 |
| SRP9 | 4972.131608 | -0.40564195 | 1.74E-03 |
| SS18L1 | 741.1511123 | -0.405766747 | 1.70E-02 |
| LMLN | 677.6147435 | -0.406127639 | 1.53E-02 |
| WDR83 | 712.4492011 | -0.406530206 | 1.95E-03 |
| ZNF324 | 851.7318929 | -0.406713762 | 1.17E-03 |
| MCM8 | 2658.923529 | -0.406750739 | 2.65E-02 |
| PRUNE1 | 1101.991682 | -0.406792159 | 1.62E-04 |
| NA | 191.9492736 | -0.406939763 | 4.07E-02 |
| BST2 | 6874.146056 | -0.406977231 | 5.05E-03 |
| FAM199X | 2323.398304 | -0.407140152 | 1.79E-02 |
| MAPRE2 | 4609.080376 | -0.407245883 | 1.38E-09 |
| UBA6-AS1 | 658.5300295 | -0.407281543 | 8.91E-03 |
| TSNAX | 1321.492617 | -0.407320254 | 2.07E-02 |
| ZMIZ1 | 3413.461136 | -0.407625869 | 2.37E-06 |
| COX20 | 1221.240216 | -0.40768153 | 4.11E-07 |
| STAG3L2 | 795.2782283 | -0.407742821 | 1.90E-05 |
| ZNF324B | 402.6473517 | -0.407907045 | 1.55E-03 |
| C1orf216 | 1881.654585 | -0.408157331 | 1.12E-02 |
| NOP14-AS1 | 749.5561467 | -0.408288818 | 6.83E-03 |
| ABHD14A | 756.7593186 | -0.408497145 | 1.20E-02 |
| ZNF445 | 2350.177662 | -0.408984322 | 1.46E-07 |
| IKZF4 | 437.2775442 | -0.409018035 | 1.28E-03 |
| MT-ND5 | 72234.1821 | -0.409175393 | 4.32E-02 |
| STEAP1B | 969.6948838 | -0.409195848 | 6.55E-05 |
| ZBTB11-AS1 | 138.5096279 | -0.409229877 | 3.33E-02 |
| CAAP1 | 1071.16947 | -0.409239713 | 2.58E-03 |
| MED13 | 2899.319374 | -0.409343368 | 4.93E-03 |
| NASP | 6492.894079 | -0.40998614 | 5.28E-03 |
| ERMP1 | 4149.788729 | -0.410212679 | 5.69E-06 |
| TECPR2 | 3050.458997 | -0.410239002 | 2.68E-03 |
| TIRAP | 621.8397527 | -0.410827352 | 4.30E-05 |
| PARPBP | 708.9645397 | -0.410926809 | 3.36E-02 |
| AL161457.2 | 446.413554 | -0.411064478 | 7.22E-03 |
| ORC5 | 1547.191739 | -0.411433906 | 4.30E-03 |
| RBM41 | 337.71327 | -0.411811662 | 4.08E-02 |
| CIPC | 1125.986789 | -0.411976375 | 1.27E-04 |
| APP | 42894.66391 | -0.412076121 | 2.10E-06 |
| SLC50A1 | 2499.022849 | -0.412392786 | 2.40E-04 |
| AP000866.1 | 162.9045803 | -0.412878407 | 2.36E-02 |

| | | | |
|---|---|---|---|
| ZNF200 | 1367.270175 | -0.413383999 | 1.97E-07 |
| ZNF530 | 486.0653137 | -0.413574956 | 6.09E-03 |
| FKBP3 | 3306.432212 | -0.413648453 | 7.95E-09 |
| MDP1 | 536.9280245 | -0.414615723 | 3.35E-03 |
| ARMCX4 | 791.492689 | -0.414919979 | 4.43E-03 |
| C11orf1 | 496.8051139 | -0.414986833 | 9.09E-04 |
| NOL8 | 2499.308708 | -0.415224194 | 4.85E-02 |
| DHX36 | 2895.374078 | -0.415283907 | 1.21E-02 |
| LPAR2 | 318.5182653 | -0.415797928 | 2.61E-02 |
| CTNND1 | 17473.49328 | -0.415810326 | 9.17E-08 |
| AAMDC | 728.8554866 | -0.415918549 | 8.74E-04 |
| RNF207 | 629.8990604 | -0.416485603 | 1.87E-02 |
| VSIG10 | 1369.992613 | -0.416834226 | 2.71E-04 |
| TMEM106B | 1864.784951 | -0.41702924 | 1.41E-03 |
| FAHD1 | 990.6586558 | -0.417129641 | 1.26E-03 |
| HMGB1 | 9585.141429 | -0.417158792 | 3.58E-03 |
| SLC12A9 | 3367.234538 | -0.41721679 | 8.80E-04 |
| EIF1AX | 3017.493396 | -0.417383989 | 5.56E-03 |
| GTF2IRD2B | 908.8548731 | -0.41749863 | 2.18E-02 |
| SLF2 | 1287.386998 | -0.417548334 | 1.01E-02 |
| C1RL-AS1 | 573.792794 | -0.417762861 | 4.01E-02 |
| AIDA | 4089.78514 | -0.418277759 | 1.15E-04 |
| GABPB1-AS1 | 2069.678936 | -0.418344189 | 4.83E-03 |
| PTPN12 | 6077.371603 | -0.418420689 | 3.77E-03 |
| NSF | 4379.616314 | -0.418508674 | 9.06E-11 |
| GGPS1 | 1335.938822 | -0.41853512 | 1.42E-04 |
| CCDC77 | 870.8727388 | -0.419653513 | 6.12E-03 |
| PCDHGB2 | 256.1679818 | -0.419733815 | 2.18E-02 |
| GALNS | 1902.302002 | -0.419873793 | 1.01E-03 |
| TEAD2 | 2552.04389 | -0.420122791 | 3.44E-03 |
| TICRR | 1882.00171 | -0.420332141 | 5.14E-03 |
| ANGEL2 | 967.7784368 | -0.421027257 | 8.77E-05 |
| ZNF572 | 210.6921965 | -0.421331754 | 4.66E-02 |
| CLINT1 | 9716.206222 | -0.421752142 | 1.52E-03 |
| SLX4IP | 516.7187441 | -0.421951044 | 1.79E-03 |
| AC139256.1 | 185.1133231 | -0.421969342 | 4.77E-02 |
| SARNP | 2010.285177 | -0.422057549 | 1.10E-06 |
| NPHP3 | 735.6109425 | -0.423060989 | 1.13E-02 |
| EOGT | 588.9739614 | -0.423155284 | 4.47E-03 |
| NBPF4 | 231.6908026 | -0.423825173 | 1.76E-02 |
| SERF1B | 1933.088998 | -0.423918254 | 7.11E-09 |
| USP6NL | 606.8158013 | -0.424015471 | 3.23E-03 |
| AC011043.1 | 615.4699368 | -0.424097739 | 2.38E-03 |
| AASDH | 769.6253677 | -0.424355791 | 1.49E-02 |
| ALDH3A2 | 1982.806079 | -0.424493214 | 2.96E-06 |
| LRRC37A17P | 728.9897898 | -0.42449452 | 8.09E-04 |
| AL031282.2 | 397.8015291 | -0.424960179 | 2.12E-03 |

| | | | |
|---|---|---|---|
| MICOS10 | 1821.045151 | -0.425076547 | 1.07E-06 |
| OST4 | 4570.163147 | -0.425277163 | 1.30E-02 |
| TMCC1-AS1 | 380.6175541 | -0.425343298 | 8.27E-03 |
| RLIM | 3157.917378 | -0.425781327 | 1.51E-02 |
| TAF11 | 1504.934528 | -0.426201787 | 1.89E-07 |
| ZBTB34 | 895.6646402 | -0.426224178 | 2.00E-05 |
| WDR6 | 10006.27012 | -0.426409495 | 1.23E-02 |
| UPF3A | 938.2065818 | -0.426468553 | 1.05E-02 |
| SLC25A53 | 297.7436183 | -0.427025091 | 3.93E-03 |
| TMEM184C | 1244.756484 | -0.427083016 | 2.44E-04 |
| UGDH-AS1 | 416.4032267 | -0.427190405 | 4.39E-02 |
| HCAR1 | 765.884658 | -0.427460771 | 1.23E-03 |
| PIGZ | 330.2327841 | -0.427488518 | 4.10E-02 |
| CASK | 2190.921557 | -0.42755619 | 2.95E-05 |
| CAPS2 | 142.4286161 | -0.427595046 | 4.43E-02 |
| B4GALT6 | 552.4881887 | -0.428102522 | 3.13E-02 |
| BOLA3-AS1 | 290.6439025 | -0.428363437 | 2.26E-03 |
| UTP14C | 661.9742037 | -0.428552748 | 1.17E-02 |
| RALB | 5036.240045 | -0.428757047 | 2.78E-07 |
| DYNLL2 | 4327.120161 | -0.429143246 | 2.85E-07 |
| LCMT2 | 524.7377588 | -0.429382868 | 9.65E-03 |
| TRAK2 | 3071.599689 | -0.429386923 | 1.10E-10 |
| MAT2A | 17360.67703 | -0.429460255 | 5.75E-08 |
| LEO1 | 1631.362175 | -0.42989749 | 1.70E-05 |
| ZNF225 | 359.5567769 | -0.429966819 | 3.86E-02 |
| FAM92A | 1091.539897 | -0.430127838 | 2.64E-04 |
| ATP2B4 | 5211.832201 | -0.430155979 | 2.47E-05 |
| BAIAP2-DT | 1190.829781 | -0.430563219 | 1.41E-03 |
| ZNF652 | 812.4173567 | -0.43061166 | 1.86E-05 |
| CRIPT | 678.109701 | -0.430700249 | 2.51E-05 |
| NA | 96.45349882 | -0.431221595 | 4.27E-02 |
| MTATP6P1 | 11442.5503 | -0.431338967 | 2.41E-02 |
| HSD17B4 | 7591.053364 | -0.431678593 | 8.35E-09 |
| CCDC40 | 197.5519158 | -0.431965373 | 1.63E-02 |
| TRIM34 | 459.8810898 | -0.432008282 | 1.26E-02 |
| C1orf112 | 1218.550267 | -0.432084055 | 1.47E-03 |
| ARHGEF40 | 6406.672053 | -0.432288891 | 4.26E-03 |
| NMB | 413.3741812 | -0.432369036 | 1.00E-02 |
| IRF2BPL | 3296.790079 | -0.432392921 | 6.50E-05 |
| IRAK1BP1 | 289.2275019 | -0.432497079 | 2.77E-02 |
| LINC01001 | 877.8552253 | -0.432523893 | 3.54E-02 |
| AASS | 1629.736665 | -0.432755699 | 9.96E-04 |
| COX11 | 1314.57339 | -0.433304654 | 6.03E-04 |
| DTNA | 635.6860956 | -0.433318666 | 6.63E-03 |
| RCOR1 | 3100.077146 | -0.433369153 | 6.83E-04 |
| CRK | 4662.264636 | -0.433718511 | 1.10E-06 |
| SGSH | 1798.745288 | -0.433880474 | 4.26E-03 |

| | | | |
|---|---|---|---|
| ING4 | 1112.173601 | -0.433902505 | 2.67E-03 |
| ZNHIT2 | 588.7238783 | -0.43399946 | 9.02E-03 |
| EMP3 | 3670.20766 | -0.434169863 | 2.45E-02 |
| HOXA7 | 998.7093727 | -0.434329753 | 3.83E-06 |
| CMTM6 | 3907.919114 | -0.434554233 | 3.64E-07 |
| DDX3X | 15192.34195 | -0.434681219 | 2.73E-06 |
| SYNJ2BP-COX16 | 287.8347004 | -0.434784209 | 2.80E-02 |
| DLD | 4716.307916 | -0.434848346 | 9.87E-05 |
| RBM43 | 426.6895759 | -0.434967437 | 2.01E-03 |
| EMP2 | 4797.302819 | -0.435349811 | 2.27E-04 |
| ADSS | 1748.129653 | -0.435350506 | 6.61E-04 |
| PGGT1B | 904.8288717 | -0.435587067 | 4.74E-03 |
| ERVK9-11 | 199.3391714 | -0.435719225 | 2.42E-02 |
| GOLGA8B | 542.4888917 | -0.43574529 | 1.01E-02 |
| APOC1 | 1307.212753 | -0.435830719 | 1.16E-02 |
| RPS6KC1 | 2016.543957 | -0.435900827 | 2.56E-08 |
| BAZ1A | 2148.548363 | -0.436019874 | 3.01E-02 |
| WDR90 | 1545.088941 | -0.436070307 | 9.34E-03 |
| RUFY2 | 371.2303911 | -0.436614334 | 3.94E-02 |
| CLDN12 | 1756.686322 | -0.436883969 | 4.18E-03 |
| NR2C1 | 844.9690927 | -0.436911632 | 1.70E-02 |
| SUB1 | 7877.607387 | -0.437174146 | 3.84E-06 |
| DALRD3 | 2136.868904 | -0.437407329 | 8.21E-03 |
| NECTIN3 | 569.8866024 | -0.43757194 | 9.14E-03 |
| ZNF268 | 934.5315959 | -0.437615571 | 2.23E-02 |
| ENTPD1-AS1 | 196.7310503 | -0.437783522 | 4.28E-02 |
| AHSA2P | 2124.375938 | -0.437924314 | 1.95E-03 |
| TRIM46 | 215.9672309 | -0.438775797 | 2.31E-02 |
| ZNF512B | 4994.652742 | -0.438914296 | 1.70E-07 |
| ACIN1 | 10252.36296 | -0.438947342 | 4.19E-26 |
| AL136295.7 | 229.180298 | -0.439131046 | 1.05E-03 |
| DDHD1 | 1115.601227 | -0.439257458 | 4.91E-02 |
| FUBP1 | 4556.3948 | -0.439515949 | 2.53E-05 |
| THTPA | 987.8526175 | -0.43976869 | 4.58E-03 |
| FDX1 | 969.3216053 | -0.439966212 | 1.19E-05 |
| DDT | 2299.728872 | -0.440089538 | 1.44E-04 |
| CDC73 | 1412.75039 | -0.440187636 | 3.15E-03 |
| CU633967.1 | 1209.838726 | -0.440416875 | 3.49E-03 |
| TMEM131 | 2558.958861 | -0.440543873 | 5.15E-08 |
| ATP1B3 | 8061.625662 | -0.440694851 | 2.03E-06 |
| PJA2 | 7418.704723 | -0.440726029 | 1.74E-04 |
| ISCA1 | 1370.129141 | -0.440880837 | 1.97E-05 |
| NKX2-6 | 293.1226131 | -0.440926561 | 1.19E-02 |
| SMAD5 | 6225.784563 | -0.441515313 | 2.40E-05 |
| PHLPP2 | 1636.290208 | -0.441687062 | 1.33E-07 |
| LPAL2 | 159.1006028 | -0.441689181 | 3.17E-02 |
| MINDY3 | 690.3990073 | -0.441781037 | 4.02E-04 |

| | | | |
|---|---|---|---|
| NMD3 | 1663.788087 | -0.442070446 | 1.54E-02 |
| KLHDC3 | 5370.43767 | -0.442151439 | 5.42E-03 |
| PARP14 | 3812.609189 | -0.442275479 | 7.96E-03 |
| EHHADH | 803.6704901 | -0.442383895 | 2.22E-04 |
| DPF3 | 481.2364414 | -0.44255261 | 3.11E-02 |
| MARVELD1 | 3273.994686 | -0.442643447 | 5.77E-03 |
| FMC1 | 495.7004804 | -0.443673355 | 3.40E-04 |
| TRAPPC13 | 1327.936223 | -0.444396477 | 8.01E-03 |
| TMEM53 | 424.5627033 | -0.444490335 | 1.09E-02 |
| GLT8D1 | 3116.387313 | -0.444571315 | 1.09E-04 |
| LINC01667 | 1112.755145 | -0.444581889 | 2.08E-06 |
| WDR86 | 1345.190106 | -0.444587896 | 1.04E-03 |
| OFD1 | 1073.083644 | -0.444850765 | 1.97E-03 |
| WAC-AS1 | 606.8259948 | -0.445192404 | 4.36E-06 |
| DDX6 | 6778.175847 | -0.445619186 | 1.18E-11 |
| SMIM30 | 1342.204583 | -0.446798328 | 1.23E-03 |
| ROGDI | 517.5564016 | -0.446918292 | 3.84E-03 |
| C5orf15 | 4632.545109 | -0.44737526 | 2.46E-05 |
| SORBS1 | 666.9068481 | -0.447407109 | 2.26E-04 |
| CCDC74B | 422.4592504 | -0.44767384 | 4.35E-03 |
| SHMT1 | 2837.59829 | -0.4481587 | 7.82E-04 |
| IGFBP7 | 8049.810997 | -0.448323667 | 2.62E-03 |
| MAZ | 11428.16553 | -0.448770072 | 9.94E-05 |
| SETD1B | 1514.715378 | -0.449482087 | 1.82E-05 |
| PAN3 | 841.5834156 | -0.449793885 | 1.44E-03 |
| TMCO1 | 3626.680009 | -0.449817227 | 2.08E-07 |
| GBP1 | 1438.088516 | -0.449936853 | 1.51E-04 |
| CAPN7 | 1738.504672 | -0.449967356 | 2.18E-04 |
| ENOX1 | 275.262444 | -0.45028081 | 4.61E-02 |
| ITPR1 | 1284.633357 | -0.450430186 | 2.73E-02 |
| TYK2 | 3847.834705 | -0.450514064 | 2.00E-05 |
| FGFR1OP | 667.7562339 | -0.450531717 | 8.53E-03 |
| ERMARD | 674.76412 | -0.450973575 | 2.40E-04 |
| CCT6P3 | 317.357428 | -0.450994588 | 8.40E-03 |
| AL590428.1 | 120.8243549 | -0.451246724 | 2.52E-02 |
| EIF3E | 11249.5084 | -0.45134923 | 3.14E-04 |
| OSBPL11 | 1303.92238 | -0.451610449 | 1.10E-06 |
| SLX1B-SULT1A4 | 282.9680088 | -0.452302502 | 6.30E-03 |
| SDR39U1 | 2016.387592 | -0.453443407 | 3.56E-05 |
| RAPGEFL1 | 411.4915503 | -0.453563471 | 3.21E-03 |
| APOBEC3D | 638.3823992 | -0.453604494 | 1.04E-02 |
| RABEP1 | 5406.597359 | -0.453874629 | 6.02E-04 |
| CRABP2 | 3933.139023 | -0.453907098 | 1.06E-02 |
| TUT7 | 1144.43367 | -0.453926018 | 1.11E-02 |
| IRF9 | 1259.63647 | -0.45394887 | 1.13E-03 |
| NRAS | 3533.096792 | -0.454278019 | 4.50E-07 |
| HELQ | 538.5349663 | -0.454315852 | 4.42E-04 |

| | | | |
|---|---|---|---|
| TSPYL1 | 2490.20289 | -0.454343453 | 7.27E-05 |
| CNTNAP1 | 3759.390506 | -0.454669894 | 3.13E-05 |
| IQCK | 379.5866682 | -0.454714081 | 6.63E-04 |
| ARL17A | 1881.979378 | -0.454716092 | 9.55E-07 |
| AC012467.2 | 176.2267427 | -0.454724777 | 1.13E-02 |
| NA | 284.9394827 | -0.455179763 | 9.69E-03 |
| UNG | 3547.676506 | -0.455233768 | 1.88E-02 |
| CBX2 | 1976.013708 | -0.455277216 | 3.54E-04 |
| THAP5 | 611.0550646 | -0.455280154 | 2.41E-02 |
| TARBP1 | 1916.655118 | -0.455436217 | 7.83E-03 |
| SCYL3 | 584.6494419 | -0.455561057 | 7.42E-04 |
| PIKFYVE | 2742.920494 | -0.455855657 | 1.06E-02 |
| ZNF714 | 190.8902153 | -0.455873239 | 3.62E-02 |
| SNAI3-AS1 | 316.1365721 | -0.455886949 | 1.28E-03 |
| VAMP1 | 983.9967304 | -0.45594558 | 6.20E-04 |
| ZNF664 | 5439.782785 | -0.455974796 | 2.71E-03 |
| IQSEC2 | 1053.000885 | -0.456044796 | 2.24E-03 |
| LMNB1 | 8403.611797 | -0.456150417 | 2.03E-02 |
| ZNF529 | 751.8925068 | -0.456185028 | 4.57E-03 |
| POLG2 | 631.5605938 | -0.456283217 | 1.01E-03 |
| SIX5 | 1314.827357 | -0.456305313 | 4.76E-04 |
| NA | 1596.217383 | -0.456945411 | 5.92E-06 |
| DENND4A | 1554.563487 | -0.456959814 | 2.41E-02 |
| PEX13 | 727.2580527 | -0.456966354 | 1.06E-04 |
| NSL1 | 1006.639425 | -0.457060276 | 5.13E-05 |
| HOXB5 | 349.4882309 | -0.457092894 | 1.59E-03 |
| AC060766.7 | 170.9646921 | -0.457285048 | 3.13E-02 |
| MTDH | 5816.793673 | -0.457582323 | 7.68E-06 |
| SLC12A6 | 634.7577138 | -0.457611386 | 2.13E-02 |
| SVBP | 540.4729975 | -0.457685861 | 1.17E-03 |
| SPATA5 | 574.9454006 | -0.458301306 | 8.62E-04 |
| CCDC134 | 408.6200617 | -0.458344107 | 1.94E-05 |
| CNBD2 | 95.00086923 | -0.458569411 | 3.61E-02 |
| MT-ND6 | 16357.53968 | -0.458592421 | 4.30E-02 |
| ZNF532 | 4172.318559 | -0.458796136 | 4.20E-05 |
| TRAPPC6B | 1174.086258 | -0.459179162 | 4.59E-02 |
| NIFK-AS1 | 296.8162705 | -0.459316592 | 1.23E-03 |
| KRBOX1 | 1196.380718 | -0.459422837 | 5.21E-05 |
| HERC1 | 6565.165456 | -0.459432401 | 1.97E-04 |
| SPTY2D1 | 1895.298232 | -0.459485777 | 8.43E-06 |
| MSX2 | 281.5175503 | -0.459746052 | 1.59E-02 |
| HOMER2 | 738.0435219 | -0.459871569 | 6.32E-04 |
| ARSB | 2222.90203 | -0.459976482 | 1.14E-03 |
| RAPGEF3 | 1404.252954 | -0.459985468 | 9.59E-03 |
| ZNF417 | 630.0890961 | -0.460069087 | 8.85E-03 |
| ZNF585B | 695.3659731 | -0.460662694 | 9.14E-04 |
| DOK4 | 2219.835095 | -0.461140225 | 1.14E-04 |

| | | | |
|---|---|---|---|
| SALL1 | 1727.630831 | -0.461496518 | 3.31E-03 |
| PIGW | 1124.188094 | -0.461566715 | 1.05E-04 |
| STMP1 | 2351.565466 | -0.461594225 | 4.69E-06 |
| SLC41A2 | 385.2660685 | -0.462081251 | 4.54E-03 |
| NUDT16L1 | 643.9989348 | -0.462326414 | 1.79E-03 |
| FUT8 | 3274.135694 | -0.462464818 | 1.15E-06 |
| NUMB | 4674.727443 | -0.462500899 | 1.21E-14 |
| TLE1 | 2878.688334 | -0.462609053 | 1.12E-03 |
| NCOA3 | 5143.412747 | -0.462716598 | 7.41E-04 |
| PIAS2 | 969.8538545 | -0.463067455 | 1.35E-04 |
| SMIM15 | 2643.17882 | -0.463651707 | 5.66E-03 |
| GPAM | 1106.907471 | -0.463882197 | 1.09E-04 |
| ACSL3 | 7289.522609 | -0.464106463 | 1.83E-06 |
| GTPBP8 | 382.465942 | -0.464163021 | 1.02E-03 |
| CASP1 | 701.6894892 | -0.464252016 | 2.05E-02 |
| DLK2 | 675.4898631 | -0.464291513 | 3.28E-03 |
| ADNP | 9338.900182 | -0.464642692 | 1.64E-16 |
| KIF3B | 5246.530593 | -0.464914828 | 1.54E-12 |
| SPRY2 | 1269.954807 | -0.465065873 | 1.19E-04 |
| TRUB1 | 555.5771618 | -0.465278877 | 1.88E-03 |
| KRAS | 1361.215751 | -0.465771337 | 4.64E-04 |
| ZNF81 | 350.8730169 | -0.465987571 | 2.82E-02 |
| EPHB4 | 7703.29599 | -0.466548794 | 5.97E-04 |
| DCAF13 | 3623.187632 | -0.467347127 | 4.45E-03 |
| SBNO1 | 2551.156243 | -0.468316874 | 1.31E-03 |
| N4BP2L2 | 1220.939384 | -0.468685157 | 2.98E-02 |
| MAP4K5 | 1453.252143 | -0.468793767 | 1.40E-02 |
| UBXN2B | 1528.527194 | -0.469055126 | 8.15E-03 |
| TENT2 | 2303.57107 | -0.469175383 | 4.44E-03 |
| FZD6 | 1565.705517 | -0.469352257 | 1.83E-04 |
| PDCL | 1337.598657 | -0.469588156 | 1.93E-09 |
| MTHFR | 2157.163497 | -0.469824337 | 2.19E-03 |
| ZIK1 | 840.9093282 | -0.469907954 | 8.61E-05 |
| LUC7L | 2156.329626 | -0.470289009 | 5.55E-10 |
| UHRF1BP1L | 1014.008429 | -0.470467151 | 1.82E-02 |
| SUV39H2 | 651.0796669 | -0.470739625 | 9.21E-03 |
| SUMF1 | 1209.276204 | -0.471015398 | 8.28E-04 |
| AC027290.2 | 87.57948417 | -0.471027917 | 4.54E-02 |
| BBS9 | 1146.378736 | -0.471233173 | 6.52E-08 |
| RSRC1 | 823.9064721 | -0.471514161 | 1.49E-03 |
| AC240274.1 | 1087.087042 | -0.471552263 | 1.91E-03 |
| SMAD1 | 441.7382654 | -0.471688385 | 2.74E-03 |
| ZNF138 | 290.8676414 | -0.471934706 | 9.87E-03 |
| SPPL2A | 1140.966053 | -0.472097857 | 1.43E-05 |
| ZNF551 | 748.8878328 | -0.472233461 | 4.10E-03 |
| SCP2 | 2504.721614 | -0.472361479 | 4.32E-04 |
| AHDC1 | 1790.089992 | -0.472529686 | 2.02E-03 |

| | | | |
|---|---|---|---|
| USP49 | 781.7022822 | -0.472670739 | 9.32E-05 |
| TNFRSF14 | 898.3680434 | -0.472677491 | 1.95E-03 |
| BCO2 | 146.8565289 | -0.473061099 | 3.52E-02 |
| TSPAN10 | 87.03663476 | -0.473128596 | 3.48E-02 |
| MPZL2 | 259.3641946 | -0.473241877 | 3.20E-02 |
| FAIM | 348.3492391 | -0.473455184 | 3.89E-04 |
| DNASE1 | 1176.911667 | -0.473958855 | 2.30E-06 |
| TNRC6C | 889.66465 | -0.474046676 | 4.94E-03 |
| MAFG-DT | 238.8518 | -0.474269206 | 5.66E-03 |
| ITGB1 | 24729.84288 | -0.474445647 | 7.88E-08 |
| KIF15 | 1673.069654 | -0.474552905 | 3.49E-02 |
| THOC1 | 1197.632741 | -0.474755175 | 3.38E-02 |
| BAG2 | 930.8421993 | -0.474813807 | 1.38E-02 |
| LYNX1 | 851.7560327 | -0.475493659 | 1.85E-02 |
| PNPLA8 | 1719.226756 | -0.476159796 | 1.59E-02 |
| WRN | 1840.206464 | -0.47685544 | 8.30E-03 |
| ARMC8 | 1212.528709 | -0.477346759 | 2.57E-03 |
| TFRC | 9364.404424 | -0.477433502 | 1.28E-05 |
| MEGF9 | 1097.538604 | -0.477789278 | 1.82E-05 |
| BCL9 | 2022.851338 | -0.478053982 | 1.87E-06 |
| PLBD1 | 1735.301632 | -0.47820621 | 6.19E-05 |
| NAA15 | 3518.266988 | -0.478315388 | 4.60E-03 |
| IPO8 | 3229.342953 | -0.478391349 | 1.43E-07 |
| PRKRA-AS1 | 232.7012941 | -0.479031557 | 1.82E-02 |
| SLC2A11 | 910.2474593 | -0.479269611 | 7.91E-05 |
| ZNF485 | 223.0510035 | -0.479677097 | 3.21E-02 |
| ZNF513 | 1599.06592 | -0.480135043 | 2.02E-05 |
| FOXRED2 | 5609.229125 | -0.480528027 | 3.04E-04 |
| DSN1 | 2244.06107 | -0.480560948 | 5.56E-03 |
| CACNA1C | 802.2770963 | -0.480708359 | 3.38E-02 |
| NBPF1 | 2185.792378 | -0.480908238 | 1.39E-03 |
| HIPK2 | 6086.845547 | -0.481248268 | 4.93E-13 |
| ABHD13 | 288.6171128 | -0.481617958 | 4.84E-03 |
| CACNB3 | 1731.645184 | -0.481689664 | 2.09E-04 |
| AC016582.3 | 142.7343043 | -0.481856345 | 2.73E-02 |
| ZNF35 | 293.7677343 | -0.482354004 | 5.34E-03 |
| BSN | 168.5919291 | -0.482365862 | 2.77E-02 |
| GBAP1 | 243.0011944 | -0.482706417 | 7.85E-03 |
| IFI16 | 7219.505499 | -0.482728235 | 8.28E-04 |
| CLIP3 | 2375.243907 | -0.482733268 | 4.81E-04 |
| MROH6 | 143.2816375 | -0.482755765 | 4.51E-02 |
| ANKRD23 | 127.0011083 | -0.482762194 | 2.70E-02 |
| FUS | 18454.28601 | -0.482794595 | 5.44E-04 |
| NA | 4877.191813 | -0.483364953 | 2.99E-21 |
| SIAE | 1123.263764 | -0.483703384 | 1.27E-05 |
| RCAN3 | 1155.641347 | -0.484255759 | 2.96E-05 |
| SUFU | 1234.590452 | -0.48436284 | 3.80E-03 |

| | | | |
|---|---|---|---|
| JAZF1 | 445.0955495 | -0.484634099 | 5.07E-03 |
| MSH2 | 2826.08355 | -0.484641634 | 9.89E-04 |
| UTP20 | 2908.602101 | -0.484654438 | 3.86E-05 |
| GOLGA8A | 1520.217999 | -0.484671516 | 5.07E-03 |
| PPP1R10 | 3168.508356 | -0.484814138 | 1.22E-04 |
| CNOT6 | 1530.549631 | -0.485226173 | 1.82E-03 |
| NOL11 | 2478.399255 | -0.485280415 | 4.54E-03 |
| NA | 115.5666921 | -0.485296156 | 4.47E-02 |
| AP001372.2 | 231.832704 | -0.485303382 | 9.87E-03 |
| H2AFY2 | 115.8074385 | -0.485531019 | 4.46E-02 |
| AC106782.2 | 196.6371558 | -0.485555405 | 2.01E-02 |
| OSBPL1A | 2509.624306 | -0.485722786 | 9.09E-12 |
| C7orf43 | 1455.089757 | -0.485753505 | 2.38E-06 |
| ETV5 | 3045.552529 | -0.48587477 | 1.78E-03 |
| AC148477.2 | 153.8592474 | -0.485959561 | 8.83E-03 |
| PCNX2 | 1178.527075 | -0.48600831 | 8.03E-07 |
| AC003665.1 | 447.9924771 | -0.486311497 | 3.39E-02 |
| PPP1CB | 4116.18422 | -0.486397025 | 2.20E-02 |
| MDK | 24505.83644 | -0.486450824 | 5.92E-03 |
| AC008124.1 | 360.4932966 | -0.48691172 | 2.44E-03 |
| PPIL4 | 1070.567896 | -0.487654128 | 6.63E-03 |
| GYG1 | 2317.222666 | -0.488329041 | 3.42E-04 |
| CD276 | 8333.957018 | -0.488693989 | 5.28E-03 |
| SNPH | 1189.696174 | -0.488909744 | 1.18E-04 |
| AC125257.1 | 163.7034956 | -0.488952765 | 9.65E-03 |
| NA | 389.3797951 | -0.489128959 | 9.21E-03 |
| DNAJC24 | 615.7328119 | -0.489130355 | 8.38E-03 |
| JAK2 | 767.597303 | -0.489148451 | 3.23E-02 |
| AC138035.1 | 323.6251374 | -0.489189965 | 5.14E-03 |
| ZNF681 | 158.2520651 | -0.489297715 | 1.11E-02 |
| ZSCAN30 | 924.1394521 | -0.489381274 | 1.09E-04 |
| TAF13 | 605.4475085 | -0.489665698 | 1.09E-03 |
| SDSL | 476.1971922 | -0.489702088 | 3.07E-02 |
| PRMT2 | 3752.41591 | -0.489739879 | 2.62E-05 |
| GABBR1 | 1499.290205 | -0.489877958 | 2.49E-08 |
| PYROXD2 | 332.6115709 | -0.490269318 | 7.78E-03 |
| ZNF808 | 291.7277947 | -0.490447876 | 4.05E-02 |
| CEP192 | 2010.196164 | -0.490661798 | 1.53E-03 |
| INTS3 | 5212.447762 | -0.490687513 | 5.41E-08 |
| AKAP12 | 6631.627566 | -0.491119917 | 3.41E-04 |
| DYRK1A | 2154.694965 | -0.491522399 | 8.45E-06 |
| SMARCA5 | 4168.205921 | -0.491541409 | 5.55E-03 |
| FANCL | 998.7763732 | -0.491628673 | 9.65E-06 |
| FAM98B | 1262.849122 | -0.491630052 | 8.82E-04 |
| TMEM263 | 1624.078972 | -0.491723982 | 5.89E-05 |
| MPP1 | 2061.247563 | -0.492209992 | 1.35E-05 |
| TMX3 | 2503.415976 | -0.49254587 | 4.85E-02 |

| | | | |
|---|---|---|---|
| RWDD2B | 1630.863067 | -0.492622945 | 6.94E-04 |
| PLXNB1 | 1364.593559 | -0.492901524 | 2.99E-04 |
| BFSP1 | 448.8421094 | -0.493051306 | 1.46E-04 |
| HSBP1 | 10345.67201 | -0.493137738 | 3.22E-05 |
| RBMS1 | 3419.859945 | -0.49318064 | 1.88E-05 |
| PRCP | 7324.04715 | -0.493748556 | 8.66E-05 |
| PRTG | 674.9816943 | -0.493836908 | 7.14E-04 |
| MLH3 | 531.7367748 | -0.493915749 | 2.24E-02 |
| LINC01560 | 138.5155022 | -0.493923311 | 1.18E-02 |
| CDC14A | 337.6127109 | -0.494541622 | 1.95E-03 |
| CCDC157 | 226.1355338 | -0.494857739 | 1.34E-02 |
| AGO3 | 1257.21731 | -0.495118224 | 1.96E-06 |
| SLC25A42 | 745.7633681 | -0.495488266 | 1.67E-04 |
| PGBD1 | 568.7505738 | -0.49565689 | 2.97E-05 |
| GTF2IP4 | 7182.880039 | -0.495738869 | 2.75E-06 |
| SCO2 | 668.0786893 | -0.49580228 | 2.11E-03 |
| GUSBP1 | 518.3979596 | -0.496109919 | 2.35E-04 |
| HOXD3 | 206.5031631 | -0.49642756 | 1.82E-02 |
| VEZT | 1666.310259 | -0.496654016 | 2.04E-05 |
| FAM102B | 808.5164647 | -0.497380838 | 1.98E-03 |
| HEXIM2 | 229.1603445 | -0.497795558 | 1.63E-03 |
| HMGN3-AS1 | 78.76644849 | -0.498127218 | 4.75E-02 |
| SAYSD1 | 553.4734968 | -0.498408475 | 2.40E-05 |
| FLG-AS1 | 202.2547421 | -0.498421396 | 8.66E-03 |
| CYB5D2 | 726.5046726 | -0.498480689 | 1.06E-05 |
| BAG4 | 1113.912056 | -0.498501528 | 2.96E-07 |
| FAM8A1 | 1246.636344 | -0.498575469 | 2.84E-05 |
| NUTM2D | 183.9396658 | -0.498794847 | 1.31E-02 |
| TMX4 | 4179.591097 | -0.498934556 | 2.46E-05 |
| DNAJC2 | 1522.385745 | -0.499285135 | 2.26E-02 |
| MYCBP2 | 9079.698849 | -0.499561558 | 4.53E-04 |
| FAM89B | 1379.792366 | -0.499673597 | 2.57E-04 |
| INTS13 | 1642.98614 | -0.499871307 | 3.04E-03 |
| ALG10B | 377.4116927 | -0.499881339 | 3.44E-02 |
| MORC3 | 970.0257404 | -0.50016199 | 2.87E-02 |
| LRBA | 3125.68714 | -0.500567395 | 3.85E-05 |
| CORO1A | 260.7712798 | -0.500768498 | 6.63E-03 |
| ZNF436 | 1465.456013 | -0.500934101 | 9.57E-08 |
| MYORG | 1497.758198 | -0.501063384 | 3.67E-03 |
| TOR3A | 1769.121677 | -0.501205429 | 1.53E-06 |
| GTF2IP20 | 291.374783 | -0.501775363 | 1.17E-02 |
| DCTN6 | 1034.325814 | -0.501853608 | 8.26E-06 |
| SLC37A3 | 2567.358471 | -0.501881291 | 4.02E-10 |
| TRIM61 | 105.2563199 | -0.501902988 | 4.52E-02 |
| MTBP | 545.7777279 | -0.502114197 | 4.32E-03 |
| PLEKHA3 | 389.0818929 | -0.502130808 | 2.24E-02 |
| BMF | 718.4286987 | -0.50215027 | 4.66E-03 |

| | | | |
|---|---|---|---|
| CPVL | 3426.438074 | -0.502304261 | 3.44E-04 |
| NSUN5P1 | 2362.073103 | -0.502468917 | 1.59E-07 |
| MTMR11 | 725.5845595 | -0.502900247 | 1.35E-03 |
| BBS4 | 1662.859635 | -0.503346199 | 3.94E-06 |
| ING5 | 1464.070134 | -0.503415366 | 1.18E-07 |
| NA | 213.9369198 | -0.503490098 | 2.86E-03 |
| ZMYM4 | 4083.321672 | -0.50351823 | 7.35E-04 |
| VBP1 | 3292.384531 | -0.50354981 | 4.82E-04 |
| ATG2B | 2630.035045 | -0.503723131 | 7.91E-03 |
| PTPRK | 2144.776039 | -0.503732002 | 7.66E-05 |
| BSCL2 | 1044.211796 | -0.503962679 | 8.84E-03 |
| ZMPSTE24 | 1890.659318 | -0.504315893 | 1.12E-06 |
| GON7 | 437.6675684 | -0.504321494 | 1.56E-05 |
| AC239868.1 | 580.739753 | -0.504482503 | 2.24E-03 |
| KLHL15 | 574.0645139 | -0.504696536 | 7.37E-04 |
| WTAP | 2928.312467 | -0.50489504 | 9.97E-13 |
| CCDC106 | 633.8088478 | -0.505394519 | 7.53E-03 |
| SLC12A5-AS1 | 153.3221957 | -0.50547525 | 8.74E-03 |
| TNKS2 | 1792.852115 | -0.505525493 | 3.51E-06 |
| ZNF610 | 237.0462589 | -0.506188121 | 4.62E-02 |
| AL139300.1 | 771.5096936 | -0.506370554 | 1.21E-02 |
| ACCS | 1275.123585 | -0.507008362 | 3.73E-06 |
| THBS3 | 1463.203303 | -0.507059502 | 1.14E-05 |
| GEMIN2 | 718.8377862 | -0.507197426 | 1.63E-05 |
| GPR158 | 359.1130548 | -0.507285103 | 3.03E-02 |
| ANPEP | 1081.191916 | -0.507708014 | 2.87E-05 |
| SETMAR | 386.4169727 | -0.508619267 | 8.15E-06 |
| TMEM87A | 1484.561536 | -0.508744497 | 1.28E-07 |
| SOCS4 | 860.6030037 | -0.508792958 | 3.18E-02 |
| ALDH16A1 | 1223.646242 | -0.508911771 | 2.89E-04 |
| TRIM52 | 1020.521846 | -0.508931148 | 6.62E-04 |
| TBCEL | 1876.676505 | -0.50899107 | 1.92E-04 |
| LHX1 | 640.0009217 | -0.509101293 | 2.88E-03 |
| IQCG | 313.8111129 | -0.509343138 | 2.89E-03 |
| SH3YL1 | 402.1516766 | -0.509498845 | 2.74E-03 |
| DOCK4 | 1431.176833 | -0.509962231 | 1.23E-04 |
| UBP1 | 4324.363057 | -0.510774949 | 1.55E-11 |
| GOLIM4 | 1524.415909 | -0.510911631 | 4.69E-04 |
| RWDD4 | 906.2997913 | -0.511382682 | 2.43E-03 |
| C16orf86 | 157.9991481 | -0.511598831 | 1.31E-02 |
| NA | 202.7798493 | -0.5129812 | 2.66E-03 |
| NIPSNAP3A | 968.4216957 | -0.513216979 | 1.09E-08 |
| TCTA | 1330.44691 | -0.513415076 | 1.32E-05 |
| HOXB13 | 1427.82097 | -0.513545962 | 9.21E-04 |
| HSD17B11 | 1994.692616 | -0.513695793 | 6.78E-08 |
| CHAC2 | 209.3736024 | -0.51371036 | 1.34E-03 |
| FAM168A | 5995.039326 | -0.513750744 | 8.59E-05 |

| | | | |
|---|---|---|---|
| DHFR | 7109.258431 | -0.514255202 | 2.86E-03 |
| PBX4 | 93.55397893 | -0.514386012 | 4.66E-02 |
| ITGB3BP | 726.7298882 | -0.514652437 | 1.53E-03 |
| EFNB3 | 2025.53239 | -0.514810469 | 4.77E-03 |
| METAP2 | 4028.775778 | -0.515137752 | 1.08E-02 |
| JARID2 | 1429.283919 | -0.515206197 | 5.69E-11 |
| ELF2 | 1626.106279 | -0.51521335 | 3.21E-13 |
| TTN-AS1 | 253.0060172 | -0.515850757 | 1.64E-03 |
| ZNF776 | 951.6721861 | -0.516088098 | 1.55E-05 |
| ADGRA3 | 1740.112729 | -0.516338519 | 1.30E-05 |
| SLC2A6 | 1907.900978 | -0.51668706 | 1.52E-03 |
| AC118549.1 | 1932.132923 | -0.51681282 | 1.30E-02 |
| ZCCHC2 | 526.4422095 | -0.516820789 | 4.84E-03 |
| ANKDD1A | 276.2525082 | -0.516988654 | 6.72E-04 |
| FICD | 254.4358081 | -0.517313475 | 4.42E-03 |
| PER3 | 895.4161079 | -0.517415717 | 1.32E-03 |
| KIF11 | 2007.052058 | -0.517561334 | 1.52E-02 |
| TRMT11 | 543.8145435 | -0.517807528 | 2.82E-02 |
| RAB36 | 529.2506332 | -0.518075219 | 3.68E-05 |
| NID1 | 12845.3558 | -0.518085197 | 1.96E-11 |
| NA | 126.8616337 | -0.518108066 | 3.85E-02 |
| RAB18 | 819.3678411 | -0.518202895 | 3.23E-06 |
| COMMD8 | 378.8613276 | -0.518309647 | 2.49E-02 |
| MAML2 | 2592.591056 | -0.518502092 | 5.92E-06 |
| ZNF549 | 563.6641424 | -0.518613967 | 3.15E-07 |
| ACP5 | 1625.175419 | -0.519014531 | 1.87E-02 |
| FLVCR1 | 455.1969064 | -0.519017075 | 2.64E-02 |
| CHEK1 | 2325.87416 | -0.519541053 | 6.25E-04 |
| NFYA | 1791.337203 | -0.519796875 | 7.41E-09 |
| TMSB15B | 161.5483094 | -0.519874762 | 3.95E-02 |
| HPF1 | 946.603226 | -0.519968323 | 1.43E-06 |
| TTYH2 | 540.5647135 | -0.519970465 | 2.54E-02 |
| OSCP1 | 227.0419356 | -0.52020369 | 9.39E-04 |
| FIZ1 | 677.6044093 | -0.520288689 | 2.65E-04 |
| LRRC37A | 215.1036013 | -0.520668785 | 1.15E-03 |
| EIF5B | 6141.87376 | -0.520731176 | 1.40E-03 |
| PABPC1L | 2421.736173 | -0.520903654 | 5.03E-03 |
| TRAFD1 | 2732.918766 | -0.52104427 | 1.85E-05 |
| CASP3 | 1463.757118 | -0.521391896 | 1.28E-07 |
| ERCC6L2 | 1307.127116 | -0.521455106 | 9.87E-03 |
| GPR89B | 811.0943084 | -0.521569581 | 2.86E-03 |
| FAM111A | 2701.692 | -0.521618817 | 1.38E-03 |
| GLI3 | 3529.66754 | -0.521850038 | 4.93E-06 |
| EIF3J-DT | 370.0506668 | -0.522174264 | 1.81E-03 |
| NA | 153.5758231 | -0.522249933 | 5.80E-03 |
| DZANK1 | 343.8105538 | -0.522300423 | 1.08E-03 |
| RAD51AP1 | 744.7072606 | -0.522380026 | 3.53E-02 |

| | | | |
|---|---|---|---|
| SERINC1 | 4518.27721 | -0.522731963 | 2.76E-05 |
| PRKAR1B | 1954.365488 | -0.523067434 | 4.92E-04 |
| CTSO | 316.7800897 | -0.523109711 | 3.44E-02 |
| GORAB | 305.0897635 | -0.523125204 | 9.22E-03 |
| MT-CYB | 107218.2105 | -0.523594166 | 8.03E-03 |
| ZNF117 | 141.3690325 | -0.523758965 | 3.46E-02 |
| AIFM3 | 126.0632533 | -0.524024236 | 3.80E-02 |
| GNAI3 | 4325.565096 | -0.524174411 | 3.52E-05 |
| NIPSNAP3B | 109.7404881 | -0.524196692 | 3.47E-02 |
| TBCCD1 | 605.9990156 | -0.524371576 | 1.74E-06 |
| ULK3 | 1952.350994 | -0.52444162 | 5.18E-08 |
| RBM26 | 3241.279658 | -0.524675142 | 2.80E-05 |
| TNFRSF21 | 5207.604204 | -0.524688098 | 2.18E-10 |
| OPRL1 | 280.5460774 | -0.52474006 | 2.34E-02 |
| LINC00680 | 465.4090567 | -0.524921942 | 8.86E-06 |
| AC008840.1 | 101.8367287 | -0.524961469 | 2.38E-02 |
| CNTLN | 1036.894075 | -0.52517482 | 4.61E-03 |
| CYHR1 | 3198.186788 | -0.525583985 | 5.71E-05 |
| ZNF557 | 384.8904144 | -0.525717532 | 3.91E-07 |
| SYNC | 139.3471639 | -0.525940743 | 1.19E-02 |
| IDUA | 285.6584141 | -0.525953609 | 4.42E-02 |
| RBBP9 | 1634.768975 | -0.526319174 | 2.65E-07 |
| ISYNA1 | 5628.667813 | -0.526402677 | 9.19E-03 |
| DENND4C | 2174.261154 | -0.526639794 | 5.08E-06 |
| TBCK | 1057.417717 | -0.526692537 | 1.75E-07 |
| JRKL | 493.6218218 | -0.526844548 | 2.22E-02 |
| AC090114.2 | 835.2073606 | -0.527003529 | 3.20E-05 |
| ZNF561-AS1 | 275.8020633 | -0.527266836 | 4.47E-03 |
| PHACTR2 | 1004.303295 | -0.527306593 | 1.11E-02 |
| ARHGAP1 | 3525.834389 | -0.527608144 | 1.21E-06 |
| SPON2 | 97.66510103 | -0.527631667 | 3.37E-02 |
| AC018521.5 | 171.255319 | -0.527643269 | 1.93E-02 |
| SOWAHC | 557.8818764 | -0.527707276 | 2.22E-04 |
| EDRF1 | 901.3236475 | -0.527838478 | 2.29E-03 |
| HECW1 | 564.3576593 | -0.528676367 | 2.30E-03 |
| BRD8 | 7516.993316 | -0.52911011 | 9.84E-17 |
| BATF2 | 455.2453915 | -0.529193346 | 8.23E-03 |
| MYO9A | 2981.552426 | -0.529244337 | 1.28E-03 |
| BTN3A1 | 1442.820939 | -0.529249419 | 2.12E-05 |
| GPD1L | 1340.603692 | -0.529327296 | 4.43E-06 |
| AC104581.4 | 154.1621748 | -0.529467086 | 2.76E-02 |
| ARID2 | 1422.741654 | -0.529599967 | 3.98E-04 |
| SLC35G1 | 581.9719221 | -0.529794249 | 2.36E-06 |
| KCNQ1OT1 | 858.4641324 | -0.530625894 | 2.30E-02 |
| RNF146 | 744.1983505 | -0.530678977 | 1.05E-02 |
| SEPSECS | 589.7060353 | -0.530696413 | 1.77E-02 |
| ARMCX5 | 434.7492399 | -0.530973625 | 9.24E-04 |

| | | | |
|---|---|---|---|
| RMI1 | 821.7678593 | -0.531087348 | 1.17E-02 |
| SNX21 | 1691.955302 | -0.531093111 | 3.43E-03 |
| RPS6KB1 | 1346.706903 | -0.53120153 | 1.49E-02 |
| AL050341.2 | 228.804696 | -0.531426156 | 4.66E-03 |
| PSMA3 | 2944.602149 | -0.531453303 | 2.82E-06 |
| AL031282.1 | 233.3734107 | -0.531515513 | 5.44E-04 |
| ZNF420 | 350.8025213 | -0.531666229 | 3.31E-03 |
| RUNDC3A-AS1 | 128.8383308 | -0.531736167 | 1.84E-02 |
| TTLL3 | 1264.197568 | -0.531840999 | 6.24E-06 |
| BMS1P1 | 299.021387 | -0.531889068 | 4.76E-04 |
| ANTXR1 | 11230.02048 | -0.532007147 | 4.54E-08 |
| SGCE | 1697.348195 | -0.532243943 | 1.31E-07 |
| SPTSSA | 1145.077999 | -0.532293829 | 3.26E-05 |
| CHKB-CPT1B | 289.6555874 | -0.532447345 | 2.95E-02 |
| SSBP3 | 3859.235303 | -0.532466686 | 8.50E-06 |
| FBXO16 | 183.9879856 | -0.532475504 | 3.28E-03 |
| APTR | 399.1142399 | -0.532513689 | 4.29E-05 |
| RFX5 | 4160.592771 | -0.533033476 | 5.35E-16 |
| HERC2P3 | 1250.815945 | -0.53318451 | 5.02E-06 |
| KCTD1 | 748.5229894 | -0.533550568 | 2.25E-05 |
| PPM1D | 648.1218558 | -0.533763618 | 9.43E-07 |
| MFSD6 | 998.7542204 | -0.533850816 | 1.48E-07 |
| CD2AP | 647.6098537 | -0.534006672 | 3.61E-02 |
| TRIM38 | 1546.239524 | -0.534066539 | 5.05E-10 |
| WASF1 | 1399.641949 | -0.534460744 | 6.13E-06 |
| HERC2P4 | 239.1856069 | -0.534592165 | 1.94E-02 |
| PCSK9 | 829.7416089 | -0.534602868 | 1.32E-02 |
| MED4 | 1518.399422 | -0.534700217 | 7.57E-06 |
| SLC26A11 | 414.2417322 | -0.534744039 | 6.57E-03 |
| BTN3A2 | 884.7388591 | -0.535087088 | 9.66E-04 |
| ZNF503 | 297.2788085 | -0.535151122 | 3.20E-05 |
| PCNX1 | 5764.565374 | -0.535727606 | 2.25E-07 |
| DNM1P41 | 127.8426165 | -0.535922425 | 1.02E-02 |
| ZNF215 | 464.4303147 | -0.535967125 | 5.19E-04 |
| U62317.2 | 110.991617 | -0.5359934 | 3.33E-02 |
| ATR | 1999.15314 | -0.536110735 | 7.13E-03 |
| DNM1L | 3372.045611 | -0.536198345 | 7.66E-04 |
| BBS10 | 716.8808489 | -0.536411766 | 9.38E-03 |
| XYLT1 | 2293.659344 | -0.536585161 | 1.14E-10 |
| PTPRG | 3213.930849 | -0.537249884 | 2.58E-05 |
| PBRM1 | 2780.16346 | -0.537379245 | 9.97E-06 |
| CDH5 | 1043.252914 | -0.537397638 | 4.80E-02 |
| SF3B1 | 12797.31495 | -0.537892417 | 6.95E-05 |
| ZNF322 | 1703.612534 | -0.538156756 | 9.63E-04 |
| H3F3A | 10452.75397 | -0.538771056 | 2.06E-06 |
| NA | 130.178278 | -0.53899161 | 2.53E-02 |
| ZC3H13 | 3640.434787 | -0.539045859 | 1.08E-03 |

| | | | |
|---|---|---|---|
| NA | 365.8632067 | -0.53927134 | 4.88E-03 |
| HERC2P5 | 595.3123382 | -0.539371536 | 3.72E-04 |
| CASTOR3 | 877.9701155 | -0.539697989 | 7.61E-05 |
| AC016394.2 | 100.9367242 | -0.539723989 | 1.83E-02 |
| BTBD7 | 2004.166963 | -0.539773353 | 1.73E-12 |
| PCSK7 | 5369.388781 | -0.539996037 | 6.71E-11 |
| NEDD1 | 792.5713997 | -0.54014554 | 4.10E-02 |
| PNPLA7 | 124.432657 | -0.540220908 | 2.24E-02 |
| SOCS6 | 907.8110569 | -0.540259367 | 1.09E-07 |
| GTF3C6 | 1875.255331 | -0.540374272 | 8.04E-07 |
| MTMR9 | 1043.343317 | -0.540595613 | 5.67E-07 |
| HAUS6 | 2447.994906 | -0.541239472 | 1.45E-02 |
| MRE11 | 2199.237406 | -0.541493243 | 3.25E-03 |
| FLVCR1-DT | 243.0547659 | -0.541510862 | 9.31E-04 |
| NUDT11 | 434.1893591 | -0.541593313 | 3.91E-07 |
| ARHGEF5 | 1397.221325 | -0.541751955 | 5.19E-07 |
| NA | 79.65493325 | -0.542051527 | 4.62E-02 |
| RPS6KA6 | 346.3436661 | -0.542130685 | 2.57E-02 |
| ALX4 | 946.9188903 | -0.54236869 | 2.54E-05 |
| ACAP2 | 1696.026809 | -0.542385975 | 3.06E-03 |
| ZSWIM5 | 539.9072653 | -0.542474928 | 2.74E-04 |
| SGSM2 | 4223.108752 | -0.542595952 | 3.85E-06 |
| C1orf131 | 519.7394715 | -0.542751924 | 4.81E-07 |
| LINC01842 | 95.88066794 | -0.542971935 | 1.86E-02 |
| ANKRD13C | 893.5313821 | -0.543118204 | 2.12E-09 |
| SCD5 | 399.6806143 | -0.543801381 | 4.52E-04 |
| PUS10 | 251.3955229 | -0.54401151 | 3.57E-02 |
| PDIK1L | 258.2112201 | -0.544307822 | 3.53E-03 |
| SPTY2D1OS | 81.55678222 | -0.544326439 | 4.68E-02 |
| CHPF | 6703.093104 | -0.544848249 | 2.76E-04 |
| BX088651.4 | 150.1509367 | -0.54534467 | 5.68E-03 |
| BORCS8 | 545.5502618 | -0.54538278 | 1.95E-04 |
| TNRC6A | 4408.836177 | -0.545428876 | 4.02E-07 |
| MAGI3 | 464.1381935 | -0.545638239 | 5.34E-04 |
| AC145350.2 | 120.1467579 | -0.546556796 | 4.66E-02 |
| HBP1 | 3602.158838 | -0.546742296 | 1.42E-12 |
| KLHDC2 | 2270.566987 | -0.546886735 | 2.28E-11 |
| POLR2L | 4348.331672 | -0.547026568 | 1.88E-04 |
| TMEM234 | 454.68688 | -0.547058099 | 5.32E-06 |
| SLC38A9 | 1524.406775 | -0.547327123 | 7.72E-07 |
| DUSP5P1 | 154.5629042 | -0.547375705 | 4.39E-03 |
| SENP1 | 1760.284492 | -0.547828247 | 2.24E-06 |
| CBX3 | 8096.996929 | -0.547948717 | 1.63E-04 |
| MGA | 1832.066296 | -0.549044568 | 1.44E-04 |
| UPF3B | 556.6795719 | -0.549295461 | 2.90E-02 |
| VAT1L | 15308.2066 | -0.549480499 | 4.07E-06 |
| SMCHD1 | 2551.389501 | -0.549760378 | 4.76E-02 |

| | | | |
|---|---|---|---|
| ACTR6 | 773.9997215 | -0.54998445 | 3.31E-04 |
| WBP1L | 2089.230149 | -0.5502076 | 5.15E-04 |
| DCLRE1C | 292.8190817 | -0.550730458 | 1.57E-02 |
| MBD5 | 732.2921705 | -0.550757613 | 2.40E-04 |
| MOB1B | 708.4653324 | -0.550773203 | 2.64E-02 |
| USP37 | 848.2020086 | -0.550891869 | 1.09E-03 |
| MIR4453HG | 190.5201642 | -0.551018003 | 8.01E-03 |
| HDHD2 | 1699.767881 | -0.551370887 | 1.41E-05 |
| ZMYM5 | 236.1654716 | -0.551400391 | 2.50E-02 |
| THNSL1 | 293.0053922 | -0.551702226 | 1.34E-02 |
| NA | 221.0300842 | -0.551792837 | 2.21E-02 |
| EXOSC4 | 1304.84018 | -0.552079035 | 5.18E-06 |
| DNAAF2 | 775.7030037 | -0.552142222 | 9.01E-08 |
| TENT5A | 439.8439513 | -0.552229482 | 1.63E-02 |
| PHF14 | 2956.198094 | -0.552247825 | 1.13E-04 |
| FAM135A | 586.9742666 | -0.552324728 | 3.36E-02 |
| CMTM1 | 94.02007502 | -0.553161855 | 3.90E-02 |
| 7-Mar | 2762.733203 | -0.553305852 | 1.20E-03 |
| PGM2L1 | 488.073069 | -0.553548504 | 2.03E-03 |
| AC021087.3 | 146.4656657 | -0.553915489 | 4.38E-03 |
| PIGF | 619.1242501 | -0.554094532 | 2.61E-06 |
| ATAT1 | 537.2356922 | -0.554097637 | 2.42E-09 |
| SFRP1 | 14944.91324 | -0.554148076 | 2.14E-03 |
| SORL1 | 4958.874962 | -0.554304934 | 4.06E-05 |
| KDELC1 | 2977.176079 | -0.554311252 | 8.64E-08 |
| C4orf3 | 2769.442182 | -0.554461637 | 4.57E-04 |
| DLX2 | 377.9123392 | -0.554656298 | 1.84E-02 |
| ZADH2 | 2400.126845 | -0.554766191 | 3.10E-17 |
| FEM1B | 2626.446844 | -0.554794639 | 6.85E-07 |
| ZNF414 | 209.8551727 | -0.555170454 | 1.04E-02 |
| PXYLP1 | 564.4135188 | -0.555282267 | 1.58E-04 |
| FDXACB1 | 160.5580444 | -0.555372856 | 2.37E-02 |
| GLMP | 3840.216257 | -0.555550393 | 2.70E-04 |
| PRPF40A | 3930.555096 | -0.555756894 | 4.87E-03 |
| SHLD1 | 405.9876561 | -0.555888648 | 2.11E-04 |
| ZNF792 | 217.2366505 | -0.556076044 | 4.84E-03 |
| ASF1A | 781.5517459 | -0.556254694 | 2.96E-03 |
| VWA1 | 1340.086787 | -0.556316328 | 6.14E-03 |
| AF186192.2 | 101.1648723 | -0.556417465 | 1.53E-02 |
| FIGNL1 | 1385.407933 | -0.556430017 | 2.62E-03 |
| SCRN3 | 689.0235599 | -0.557353731 | 1.02E-03 |
| CCDC102A | 865.6513979 | -0.557519914 | 6.87E-04 |
| MYCBP | 1295.260831 | -0.557590419 | 2.27E-05 |
| ERGIC2 | 921.606192 | -0.557672923 | 1.03E-02 |
| CDKN2C | 1684.89081 | -0.557755977 | 2.76E-03 |
| ASAH1 | 3432.713782 | -0.557867676 | 1.86E-07 |
| MED21 | 921.6848111 | -0.55789631 | 2.90E-06 |

| | | | |
|---|---|---|---|
| ZNF33A | 507.7272835 | -0.558010109 | 1.02E-02 |
| TAOK1 | 4239.749078 | -0.558118999 | 4.88E-03 |
| LGALSL | 1263.152514 | -0.558266414 | 6.54E-06 |
| CERS6 | 1849.566839 | -0.558713303 | 1.76E-06 |
| ADAM1A | 87.26285511 | -0.559080951 | 1.65E-02 |
| CDC27 | 3449.786474 | -0.559100212 | 5.07E-06 |
| CKAP2 | 2416.929446 | -0.559145748 | 1.43E-03 |
| AC009283.1 | 58.64454459 | -0.559481277 | 4.06E-02 |
| ICE2 | 2369.142308 | -0.559487177 | 2.21E-05 |
| XPO1 | 12353.51509 | -0.559661399 | 1.51E-03 |
| LINC00265 | 516.0461094 | -0.559760908 | 6.04E-05 |
| ZNF45 | 1552.966555 | -0.559997667 | 3.33E-10 |
| OMA1 | 317.5654528 | -0.560011086 | 3.17E-02 |
| ZFHX4 | 573.6192349 | -0.560250858 | 2.01E-02 |
| LSM11 | 1292.011328 | -0.560434841 | 3.25E-07 |
| ZNF799 | 165.882558 | -0.561286025 | 4.99E-03 |
| TBC1D8B | 395.7373092 | -0.561538698 | 2.15E-02 |
| TMEM200A | 540.674388 | -0.561623963 | 8.13E-04 |
| CSRP1 | 17874.18646 | -0.56184357 | 2.77E-05 |
| CDK6 | 3470.812973 | -0.561848271 | 4.93E-03 |
| ZKSCAN3 | 370.8912552 | -0.561970419 | 8.96E-05 |
| LEAP2 | 96.45515125 | -0.562085401 | 2.09E-02 |
| MANBA | 1597.878041 | -0.562290558 | 2.08E-06 |
| POC1B-AS1 | 131.0831049 | -0.562396962 | 1.68E-02 |
| DHFR2 | 351.7586684 | -0.562568545 | 1.82E-04 |
| GTF2IP1 | 5488.638803 | -0.562575659 | 2.29E-08 |
| ABCA7 | 1472.770357 | -0.563488805 | 1.34E-04 |
| TGFBR2 | 6562.551738 | -0.563646065 | 5.38E-08 |
| CSKMT | 343.0414308 | -0.563810287 | 1.07E-05 |
| ZNF502 | 192.5176735 | -0.563937757 | 4.95E-02 |
| PCMTD1 | 851.989544 | -0.564009154 | 1.74E-03 |
| DEPDC1B | 1923.476532 | -0.564092386 | 1.28E-03 |
| CEP135 | 329.7092278 | -0.564397285 | 4.75E-02 |
| ZNF383 | 287.0882816 | -0.564479366 | 2.29E-03 |
| ZNF785 | 729.5996148 | -0.564598235 | 1.15E-06 |
| KIAA0040 | 1749.222652 | -0.564599246 | 2.55E-05 |
| CLEC2B | 380.8367918 | -0.564827007 | 2.42E-02 |
| UBLCP1 | 2106.228726 | -0.565062442 | 1.62E-02 |
| ZSCAN12 | 304.742191 | -0.565115276 | 2.46E-05 |
| ZNF462 | 2159.207217 | -0.565820505 | 4.34E-10 |
| ZC2HC1A | 281.9343025 | -0.565995993 | 1.07E-02 |
| RNFT2 | 124.865365 | -0.566101008 | 2.04E-02 |
| TUT4 | 1942.183595 | -0.566219988 | 4.19E-03 |
| AC009690.2 | 70.56743767 | -0.566322011 | 4.80E-02 |
| ANP32E | 1408.613357 | -0.566514383 | 1.70E-03 |
| NA | 111.8505229 | -0.566525365 | 2.33E-02 |
| BIRC6 | 6959.61733 | -0.566527902 | 1.21E-05 |

| | | | |
|---|---|---|---|
| SAFB2 | 1849.688607 | -0.56670304 | 2.94E-06 |
| AL732372.2 | 291.0174159 | -0.566899335 | 6.34E-03 |
| KCNH3 | 436.3653327 | -0.567085006 | 2.65E-03 |
| NBL1 | 2346.529476 | -0.567494352 | 7.10E-05 |
| ACADSB | 511.8617066 | -0.567534109 | 4.05E-03 |
| TUG1 | 9645.076116 | -0.567696894 | 6.08E-09 |
| HLTF | 3216.029954 | -0.568123771 | 4.34E-04 |
| CARD8 | 1192.652439 | -0.568532494 | 1.58E-12 |
| AC233723.2 | 80.15059256 | -0.568980958 | 1.55E-02 |
| TYW3 | 1219.576547 | -0.569661213 | 6.23E-07 |
| AC024075.2 | 79.91306537 | -0.569703289 | 3.49E-02 |
| NLRP11 | 220.6543597 | -0.570541021 | 3.07E-03 |
| FUT10 | 580.7966807 | -0.570588085 | 4.28E-04 |
| LNPEP | 4087.045638 | -0.571071864 | 7.83E-07 |
| FAM241A | 959.3567144 | -0.571403478 | 2.81E-05 |
| IRAK3 | 959.1155894 | -0.571407374 | 6.20E-05 |
| MTMR2 | 2934.886144 | -0.572067146 | 1.46E-19 |
| TOPORS | 1768.495239 | -0.57228946 | 3.95E-06 |
| NFYB | 983.766186 | -0.572348206 | 8.11E-04 |
| KIDINS220 | 3754.609944 | -0.572534903 | 1.14E-04 |
| LINC01089 | 367.5833516 | -0.572601115 | 4.84E-06 |
| HELB | 231.1127368 | -0.572946451 | 1.98E-02 |
| RABEP2 | 542.7569312 | -0.57371674 | 6.57E-05 |
| MATR3 | 12010.77523 | -0.573978383 | 6.08E-05 |
| SMARCA2 | 4976.288903 | -0.574211775 | 1.37E-09 |
| EFNB1 | 5451.422566 | -0.57423031 | 1.48E-05 |
| R3HCC1L | 545.7380229 | -0.574278962 | 5.44E-04 |
| LBHD1 | 835.1787337 | -0.574332111 | 2.46E-05 |
| CADPS2 | 265.7565386 | -0.574385895 | 1.75E-02 |
| RASL12 | 186.42904 | -0.574835985 | 3.17E-02 |
| SREBF2-AS1 | 139.3932799 | -0.574925918 | 1.96E-03 |
| TCF12 | 5252.519598 | -0.574935271 | 1.07E-02 |
| HECTD1 | 6357.213805 | -0.575750294 | 1.93E-04 |
| MED7 | 1176.800016 | -0.57595245 | 1.13E-04 |
| TMEM50B | 1177.523805 | -0.576379111 | 4.59E-05 |
| CRACR2B | 125.8372559 | -0.576742574 | 2.69E-03 |
| MBIP | 613.297196 | -0.576756147 | 1.20E-04 |
| KLHL9 | 2682.692212 | -0.576812455 | 6.37E-11 |
| RAPGEF4 | 1768.711438 | -0.577172766 | 7.73E-06 |
| FAM126B | 379.8792403 | -0.577319826 | 2.40E-02 |
| ZNF639 | 844.3296518 | -0.578212114 | 1.39E-04 |
| TBXA2R | 175.3616382 | -0.578423939 | 1.62E-02 |
| DDX46 | 3171.764144 | -0.578620783 | 1.10E-03 |
| SNX16 | 332.872172 | -0.578874942 | 1.51E-03 |
| CSE1L | 9704.638476 | -0.578897803 | 1.32E-03 |
| TRBV20-1 | 248.8926942 | -0.578938135 | 1.85E-03 |
| IFT57 | 1167.083796 | -0.578946787 | 5.21E-05 |

| | | | |
|---|---|---|---|
| RASA1 | 3077.667782 | -0.578965096 | 4.07E-03 |
| OXCT1 | 3184.337989 | -0.578978241 | 1.81E-07 |
| RNF138 | 715.7112521 | -0.579422841 | 1.50E-03 |
| CSNK1G3 | 2107.355913 | -0.580304644 | 5.25E-03 |
| TSBP1-AS1 | 166.0090922 | -0.580647138 | 6.48E-03 |
| TCTEX1D2 | 229.6803877 | -0.580865148 | 1.36E-04 |
| ACOX1 | 1603.338045 | -0.580866424 | 7.34E-07 |
| SPTBN1 | 22759.67806 | -0.58114179 | 3.09E-25 |
| EXO1 | 1051.953812 | -0.581569472 | 5.40E-03 |
| STON1 | 979.0446458 | -0.581639826 | 6.40E-07 |
| KCNN3 | 360.2014472 | -0.582066666 | 5.20E-03 |
| NDUFA5 | 1338.229747 | -0.582148088 | 1.35E-04 |
| UTRN | 4972.912532 | -0.582205124 | 1.69E-04 |
| CUL7 | 4517.472227 | -0.582499777 | 6.49E-04 |
| STT3B | 6050.763375 | -0.582875405 | 4.72E-11 |
| ASH1L | 3747.874399 | -0.583496413 | 9.48E-08 |
| UGT3A2 | 1561.296932 | -0.58353628 | 1.42E-02 |
| RRM2B | 1392.151338 | -0.583718077 | 1.65E-03 |
| CD58 | 338.3302354 | -0.583778941 | 1.18E-04 |
| AGGF1 | 3409.272125 | -0.583958276 | 3.26E-07 |
| SRSF10 | 4121.068345 | -0.584140457 | 2.55E-06 |
| CRISPLD1 | 185.2859406 | -0.584377038 | 3.69E-02 |
| ZNF160 | 881.0002724 | -0.584561985 | 1.37E-04 |
| CACNG8 | 671.2693113 | -0.584763891 | 1.56E-03 |
| AC004918.3 | 104.4916378 | -0.584787639 | 2.37E-02 |
| PRKD1 | 819.8096354 | -0.585342065 | 8.51E-07 |
| FAM171A2 | 602.8195332 | -0.585360041 | 1.12E-04 |
| SELENOI | 2088.637171 | -0.585949859 | 2.46E-06 |
| AC010168.2 | 137.9560396 | -0.586221857 | 3.35E-03 |
| PHF3 | 3186.201128 | -0.586375801 | 7.83E-03 |
| CCDC18-AS1 | 304.8541376 | -0.586798415 | 3.50E-04 |
| RBL1 | 1227.749753 | -0.586802326 | 8.07E-03 |
| HAPLN4 | 233.8119336 | -0.587157367 | 2.47E-02 |
| HAS3 | 161.7908666 | -0.587365209 | 1.43E-02 |
| DDTL | 284.9097104 | -0.58741894 | 1.78E-06 |
| PRPF38B | 1919.634009 | -0.587560769 | 7.34E-04 |
| AC073073.2 | 98.76486797 | -0.587581763 | 7.97E-03 |
| DBT | 1105.356408 | -0.587686942 | 6.00E-04 |
| KLHL28 | 332.9547999 | -0.587783759 | 3.25E-04 |
| C3orf62 | 371.8425366 | -0.587954643 | 1.68E-02 |
| ENTPD1 | 1129.684048 | -0.588032529 | 1.91E-07 |
| SKAP2 | 3278.228177 | -0.588229859 | 3.13E-09 |
| PRR22 | 99.43339654 | -0.588500216 | 2.42E-02 |
| CHML | 800.3653271 | -0.588737773 | 3.18E-02 |
| ZDHHC20 | 1478.023608 | -0.588967006 | 5.81E-04 |
| ARHGAP42 | 723.8663377 | -0.588985742 | 7.33E-04 |
| REC8 | 133.8922578 | -0.589037731 | 3.06E-03 |

| | | | |
|---|---|---|---|
| KYNU | 102.2786772 | -0.589290627 | 1.40E-02 |
| AL671710.1 | 120.4166884 | -0.589356484 | 1.75E-02 |
| ATF2 | 1408.171597 | -0.589374566 | 2.65E-03 |
| AC034236.2 | 124.9206715 | -0.58992967 | 9.64E-03 |
| PPIC | 3752.098519 | -0.590111704 | 9.54E-13 |
| GOPC | 1413.898633 | -0.590423123 | 3.09E-03 |
| NME3 | 701.7911262 | -0.590525542 | 6.96E-04 |
| MYNN | 726.2999099 | -0.59069112 | 1.53E-04 |
| HERC2P8 | 188.6041447 | -0.590704466 | 4.18E-03 |
| ADAM12 | 626.9604478 | -0.591280612 | 6.24E-04 |
| AC008687.4 | 62.22366419 | -0.591365575 | 3.83E-02 |
| AP1AR | 523.019085 | -0.591578577 | 2.16E-02 |
| PLEKHF2 | 603.1114698 | -0.591643457 | 1.40E-03 |
| SLC2A12 | 396.0274564 | -0.59181898 | 5.91E-04 |
| TAPT1-AS1 | 124.3582695 | -0.59208026 | 4.87E-02 |
| TRAPPC8 | 1683.150941 | -0.592146218 | 1.20E-06 |
| CYP46A1 | 69.16182907 | -0.592267086 | 4.58E-02 |
| ELMOD2 | 619.7299235 | -0.592275736 | 4.69E-03 |
| HNMT | 837.2644328 | -0.592935409 | 3.74E-05 |
| GUSBP9 | 127.4596327 | -0.593054351 | 3.92E-02 |
| UBR5 | 9785.128236 | -0.593069528 | 7.53E-19 |
| GSPT2 | 941.9840154 | -0.593294988 | 5.72E-11 |
| F2RL2 | 2246.079531 | -0.593340877 | 1.97E-06 |
| Z97832.2 | 62.15081118 | -0.593692646 | 2.35E-02 |
| A1BG-AS1 | 468.9123604 | -0.593717034 | 4.29E-06 |
| FCGR2A | 305.3492327 | -0.594133541 | 5.43E-04 |
| SNX1 | 8206.680958 | -0.594254938 | 3.55E-10 |
| SETD6 | 1072.763408 | -0.59527118 | 3.82E-07 |
| WDR19 | 1924.718523 | -0.595435004 | 1.03E-04 |
| ZNF790 | 195.3086188 | -0.596522831 | 4.68E-03 |
| MLLT3 | 633.9740889 | -0.596594137 | 1.27E-04 |
| GTF2A1 | 961.5536787 | -0.596667963 | 3.84E-03 |
| ZNF75D | 1009.175721 | -0.596798207 | 1.93E-06 |
| NA | 100.5426996 | -0.596821775 | 1.18E-02 |
| OBSCN-AS1 | 87.8409625 | -0.596939406 | 1.33E-02 |
| ZNF141 | 264.4869761 | -0.597001433 | 1.12E-02 |
| ATP1B1 | 7211.224436 | -0.597031395 | 3.39E-05 |
| PHF23 | 4034.620571 | -0.597133739 | 1.44E-06 |
| LIMS1 | 1656.651267 | -0.597247762 | 2.66E-05 |
| RANBP6 | 1516.497612 | -0.597737547 | 3.82E-04 |
| ERMAP | 476.2680804 | -0.597802198 | 6.49E-08 |
| MYO5A | 3183.631342 | -0.597920673 | 1.45E-05 |
| XRCC4 | 617.0391537 | -0.598318251 | 2.80E-03 |
| IP6K2 | 3247.021157 | -0.598423736 | 2.06E-06 |
| PAPSS2 | 187.5959033 | -0.598755615 | 6.26E-03 |
| FAM161A | 386.9162848 | -0.599137437 | 3.46E-02 |
| FAM215B | 156.6873607 | -0.599498655 | 6.66E-03 |

| | | | |
|---|---|---|---|
| BRCC3 | 1658.547271 | -0.599695206 | 2.51E-05 |
| LRRC56 | 715.9976465 | -0.599944931 | 1.29E-04 |
| MAN1A2 | 2382.046761 | -0.600008424 | 6.88E-05 |
| FRG1-DT | 78.2155875 | -0.600057251 | 3.55E-02 |
| CDK1 | 1514.458408 | -0.600146146 | 4.72E-03 |
| LMNTD2 | 174.1655401 | -0.600289505 | 1.01E-02 |
| SENP6 | 2495.39091 | -0.600648455 | 1.55E-02 |
| TTC8 | 1064.914823 | -0.60103963 | 2.11E-06 |
| MAPK8IP2 | 998.1611941 | -0.601228744 | 2.89E-05 |
| NFATC3 | 2721.717658 | -0.601233439 | 3.11E-19 |
| RMDN1 | 2329.178846 | -0.601234904 | 5.14E-12 |
| AC015813.6 | 283.6121634 | -0.601258042 | 4.58E-03 |
| AC073343.1 | 62.96201287 | -0.601390509 | 2.34E-02 |
| GPATCH11 | 944.2607256 | -0.601412933 | 9.77E-05 |
| OLFML3 | 780.6792935 | -0.601637647 | 4.03E-02 |
| CDK2AP1 | 4145.313976 | -0.601685772 | 4.15E-17 |
| RSRP1 | 1513.468845 | -0.601793229 | 1.30E-03 |
| ZNF701 | 285.7463812 | -0.602460663 | 4.70E-04 |
| PHF20L1 | 2256.175305 | -0.602540627 | 2.75E-03 |
| CAMSAP2 | 2288.391164 | -0.602829495 | 1.11E-03 |
| CCDC82 | 785.0585224 | -0.603175689 | 2.62E-02 |
| SLX4 | 472.5174403 | -0.603320738 | 1.58E-06 |
| ZNF829 | 399.8671455 | -0.603500636 | 2.02E-03 |
| SLC25A5-AS1 | 91.85803581 | -0.603518692 | 8.20E-03 |
| SPHK2 | 601.2242349 | -0.603810207 | 3.66E-05 |
| PGAM2 | 238.1250556 | -0.603854479 | 4.05E-02 |
| MFSD11 | 952.6038169 | -0.604008005 | 8.69E-09 |
| C5orf51 | 3030.487835 | -0.604209105 | 7.97E-03 |
| 4-Sep | 536.272959 | -0.604239688 | 3.58E-04 |
| DONSON | 1377.01341 | -0.604381539 | 4.05E-08 |
| BMS1P2 | 130.4097474 | -0.604579639 | 1.45E-02 |
| STC1 | 1128.86879 | -0.604602 | 3.18E-02 |
| BPHL | 824.4779233 | -0.604821623 | 1.97E-04 |
| PIR | 1579.361561 | -0.604945515 | 5.00E-06 |
| PMEPA1 | 367.6116401 | -0.60509383 | 6.26E-03 |
| CSPP1 | 914.3606557 | -0.605138007 | 5.37E-04 |
| ZBED9 | 232.3704514 | -0.605293106 | 4.20E-02 |
| POTEG | 103.8770361 | -0.605464818 | 1.17E-02 |
| SNX7 | 1214.597973 | -0.606280028 | 1.21E-08 |
| MRPL50 | 1319.50401 | -0.606575875 | 3.23E-08 |
| PIGN | 1278.798581 | -0.606791845 | 5.20E-07 |
| AP000873.2 | 168.6041013 | -0.607049506 | 6.11E-04 |
| LYSMD3 | 1057.37091 | -0.607113103 | 1.10E-02 |
| LRTOMT | 373.8084847 | -0.607230274 | 6.88E-04 |
| C7orf25 | 138.5021058 | -0.607342009 | 2.03E-02 |
| ZNF467 | 538.7262357 | -0.607373267 | 6.30E-03 |
| MIA | 176.8750646 | -0.607837953 | 1.98E-02 |

| | | | |
|---|---|---|---|
| SH2B3 | 6559.800718 | -0.608002963 | 1.21E-07 |
| TTC12 | 290.7928053 | -0.608065296 | 1.23E-02 |
| NUF2 | 1217.065549 | -0.608104511 | 3.36E-02 |
| VAX2 | 144.6886263 | -0.608900665 | 7.05E-03 |
| ZNF26 | 696.8708955 | -0.608960238 | 2.87E-03 |
| AC073508.3 | 191.8299393 | -0.609063281 | 7.97E-05 |
| C5orf63 | 487.5369067 | -0.609279639 | 2.52E-05 |
| DCUN1D4 | 1235.219247 | -0.609315105 | 2.79E-04 |
| DAB2 | 9628.493215 | -0.609802781 | 1.23E-12 |
| FP671120.2 | 584.4415846 | -0.609858873 | 3.08E-09 |
| PER2 | 398.8658407 | -0.610018748 | 7.18E-04 |
| LRIG1 | 1253.713048 | -0.610122816 | 6.43E-10 |
| FBLN1 | 4794.018155 | -0.610902982 | 1.27E-04 |
| PRRT1 | 226.6096598 | -0.611548129 | 9.60E-05 |
| CMTM8 | 58.12048249 | -0.61173132 | 2.37E-02 |
| SFXN2 | 188.3754226 | -0.611858524 | 1.20E-02 |
| CILP2 | 791.2075924 | -0.612218599 | 4.50E-04 |
| FRRS1L | 848.3939876 | -0.612227595 | 1.30E-07 |
| SLC27A3 | 368.6739809 | -0.612276978 | 5.70E-03 |
| ZNF597 | 311.6737611 | -0.612405542 | 8.49E-06 |
| ZNF229 | 1229.958577 | -0.612950478 | 1.72E-04 |
| DPYSL3 | 1641.152646 | -0.613145311 | 5.42E-06 |
| ARRDC3 | 2277.994295 | -0.613224031 | 2.07E-04 |
| ZNF354C | 587.1740906 | -0.613357433 | 6.84E-04 |
| APLP1 | 955.4105536 | -0.613452361 | 5.04E-05 |
| AC007637.1 | 50.37287676 | -0.613653115 | 4.38E-02 |
| AC099778.1 | 87.65766901 | -0.613888936 | 1.62E-02 |
| RCHY1 | 708.5633167 | -0.614371955 | 2.67E-03 |
| STK38L | 684.3303227 | -0.614481519 | 2.66E-03 |
| ADGRG6 | 2889.841449 | -0.61501493 | 6.05E-04 |
| AC026412.3 | 60.51370205 | -0.615548536 | 2.93E-02 |
| XXYLT1 | 1532.304719 | -0.615820362 | 3.04E-09 |
| EFNA4 | 501.216771 | -0.616002408 | 3.65E-04 |
| IRX3 | 770.2032991 | -0.616023589 | 3.48E-03 |
| RTKN | 1095.937803 | -0.616100109 | 9.67E-07 |
| TBC1D8 | 2589.114755 | -0.6162048 | 1.45E-16 |
| CTSA | 8271.579524 | -0.616426587 | 3.89E-06 |
| B3GNT2 | 692.6789905 | -0.616471931 | 4.84E-08 |
| AC005519.1 | 52.47188024 | -0.616955914 | 2.47E-02 |
| F2R | 9742.24398 | -0.617117906 | 1.80E-19 |
| PMS2P7 | 61.88986737 | -0.617318549 | 3.29E-02 |
| RNF141 | 2039.07365 | -0.617344904 | 1.35E-11 |
| RSBN1L | 1605.52993 | -0.617598384 | 3.48E-11 |
| SRD5A3 | 467.3307845 | -0.617737192 | 1.29E-05 |
| HGSNAT | 3374.663359 | -0.617784104 | 2.22E-06 |
| U91328.1 | 298.4707967 | -0.61821154 | 1.27E-05 |
| ZNF407 | 755.8998402 | -0.61823764 | 1.70E-10 |

| | | | |
|---|---|---|---|
| MBNL1 | 2385.858077 | -0.618253685 | 1.43E-04 |
| PIAS3 | 2689.601268 | -0.618357709 | 6.29E-07 |
| FAM225B | 439.9626523 | -0.618771244 | 4.59E-03 |
| TRIM66 | 654.3240807 | -0.61895965 | 1.29E-02 |
| TASOR | 4044.453702 | -0.61976985 | 5.74E-04 |
| FKBP5 | 2560.809448 | -0.619956278 | 4.28E-17 |
| WBP1 | 3743.929191 | -0.620016203 | 9.48E-05 |
| ZNF470 | 625.7405749 | -0.6200393 | 1.94E-05 |
| CLBA1 | 602.1476952 | -0.620150009 | 1.39E-06 |
| ZNF563 | 145.358265 | -0.62022791 | 1.90E-02 |
| ZBTB1 | 1574.403671 | -0.620339922 | 4.60E-07 |
| FER1L4 | 915.777995 | -0.620452214 | 3.65E-02 |
| APOBEC3G | 2433.261157 | -0.620692098 | 4.56E-07 |
| USP24 | 5365.564556 | -0.620833494 | 4.20E-14 |
| GCC2 | 1356.216985 | -0.620912883 | 1.79E-02 |
| EIF4E | 1732.254152 | -0.62221585 | 3.99E-07 |
| DCBLD2 | 4378.874141 | -0.622455871 | 1.74E-10 |
| ZNF451 | 1376.809275 | -0.622479011 | 4.63E-04 |
| B3GALT6 | 999.9083851 | -0.622767591 | 4.59E-06 |
| EPC1 | 776.583326 | -0.622820276 | 3.81E-05 |
| RUSC1 | 898.0028328 | -0.622994727 | 2.73E-10 |
| ARG2 | 2408.765317 | -0.623164524 | 5.00E-07 |
| TASOR2 | 3918.209476 | -0.623235088 | 3.18E-03 |
| ITGB8 | 92.24891433 | -0.623644548 | 3.17E-02 |
| AP5M1 | 1165.533304 | -0.623694094 | 6.56E-08 |
| MIER1 | 835.1737643 | -0.623741098 | 2.76E-04 |
| ITGAV | 3581.278314 | -0.624079486 | 5.84E-04 |
| HOXB6 | 896.7173835 | -0.62416314 | 2.69E-07 |
| TDRD1 | 1309.35831 | -0.624248971 | 1.73E-08 |
| IKZF5 | 323.6334647 | -0.624551284 | 1.51E-02 |
| NA | 75.43671028 | -0.625017358 | 1.17E-02 |
| RSF1 | 2839.203396 | -0.625255558 | 3.31E-03 |
| SLC25A12 | 2043.058397 | -0.625558686 | 5.93E-12 |
| AC004148.2 | 154.6999536 | -0.625759301 | 1.14E-03 |
| C5orf24 | 1607.435982 | -0.625889565 | 5.99E-08 |
| AC009902.2 | 45.96325468 | -0.626063631 | 4.74E-02 |
| ACACB | 1090.006831 | -0.626174814 | 1.88E-08 |
| DLX1 | 957.5198547 | -0.62622807 | 1.11E-03 |
| FBXO3 | 1527.174634 | -0.626525595 | 1.13E-07 |
| RAD50 | 5501.443131 | -0.626602309 | 2.94E-03 |
| ABCC6P1 | 93.64657159 | -0.626619759 | 4.24E-02 |
| ZNF134 | 1247.108248 | -0.626658522 | 5.42E-07 |
| TRIM41 | 1896.095923 | -0.626928773 | 1.46E-12 |
| FAM187A | 233.2630473 | -0.627029306 | 7.50E-05 |
| WDR89 | 465.2162936 | -0.627172646 | 1.83E-03 |
| ZNF286A | 1332.71699 | -0.627567345 | 2.09E-07 |
| HAP1 | 265.2190354 | -0.627586201 | 2.51E-02 |

| | | | |
|---|---|---|---|
| AC022150.4 | 176.7869855 | -0.627685106 | 2.12E-02 |
| RUSC1-AS1 | 232.9101864 | -0.627776148 | 4.45E-05 |
| CPT1B | 640.0772637 | -0.628009664 | 2.72E-06 |
| ZNF561 | 1131.693688 | -0.628421499 | 6.29E-12 |
| GTF2H2B | 334.5807758 | -0.628523628 | 4.52E-05 |
| ZNF337-AS1 | 269.830778 | -0.628926257 | 1.53E-05 |
| TNFSF13 | 541.1227978 | -0.628981318 | 7.07E-04 |
| SEMA5A | 1442.927626 | -0.6295027 | 1.15E-04 |
| CCDC200 | 85.92076755 | -0.62954429 | 4.02E-02 |
| PCLAF | 1284.567792 | -0.629755313 | 1.55E-03 |
| COLEC11 | 893.9691598 | -0.629755789 | 4.55E-04 |
| CFB | 187.4850491 | -0.630019411 | 9.20E-03 |
| LINC00641 | 237.5139497 | -0.630083388 | 5.41E-03 |
| MGAT2 | 2529.203773 | -0.630351623 | 1.01E-15 |
| SETDB2 | 563.5613098 | -0.630480759 | 2.99E-02 |
| SOS2 | 1401.00858 | -0.630506998 | 2.80E-06 |
| GSTM2 | 1579.400594 | -0.630733211 | 3.90E-06 |
| CD27-AS1 | 940.6269923 | -0.631810532 | 5.12E-11 |
| ZFP30 | 444.4963992 | -0.63199087 | 1.21E-04 |
| AMACR | 1358.410533 | -0.632247529 | 1.38E-26 |
| STAG1 | 1643.158292 | -0.632430394 | 1.11E-03 |
| RAB42 | 462.7888802 | -0.632589021 | 9.52E-06 |
| EEPD1 | 580.6320043 | -0.632874374 | 1.13E-03 |
| AC135048.1 | 219.3732508 | -0.632942288 | 1.39E-02 |
| ALX1 | 332.6739069 | -0.633163044 | 4.75E-06 |
| EPC2 | 1225.634859 | -0.633380834 | 4.99E-05 |
| ZNF527 | 289.327272 | -0.633490891 | 2.62E-04 |
| BARD1 | 329.1435678 | -0.63390522 | 2.16E-03 |
| SMIM8 | 188.533859 | -0.634037903 | 1.80E-03 |
| RRN3P2 | 93.46092709 | -0.634113315 | 1.34E-02 |
| NAA16 | 439.3530062 | -0.634227291 | 6.30E-04 |
| TRIM33 | 1174.386367 | -0.634242354 | 1.31E-03 |
| SRPX | 4619.232277 | -0.634316157 | 4.66E-08 |
| CD163L1 | 1044.256401 | -0.634618775 | 4.48E-04 |
| FAM172A | 1449.935888 | -0.634781739 | 4.11E-10 |
| MAN1A1 | 698.1988176 | -0.635033793 | 3.36E-06 |
| PCAT19 | 298.9928344 | -0.635189647 | 1.71E-02 |
| NA | 96.70339755 | -0.635207681 | 4.44E-03 |
| PHF6 | 2873.346761 | -0.63529587 | 3.86E-05 |
| SCAMP5 | 945.5999557 | -0.63541946 | 4.44E-04 |
| NAPEPLD | 824.1140459 | -0.635545408 | 2.20E-04 |
| KCTD3 | 2461.517131 | -0.636429047 | 4.24E-06 |
| POLE2 | 503.188362 | -0.638420594 | 9.92E-03 |
| STS | 1112.178348 | -0.638498901 | 5.98E-12 |
| SLIT2 | 273.2187921 | -0.638993882 | 4.79E-04 |
| AL731571.1 | 87.35052019 | -0.639071791 | 1.31E-02 |
| PCYOX1L | 784.1777684 | -0.639572743 | 1.12E-05 |

| | | | |
|---|---|---|---|
| FAM230E | 99.91948562 | -0.639575916 | 1.02E-02 |
| MATN2 | 4934.640224 | -0.639659973 | 1.45E-06 |
| NDUFB2-AS1 | 48.92324914 | -0.640020484 | 3.24E-02 |
| ARHGEF34P | 257.9797001 | -0.640523716 | 1.38E-04 |
| EPHX1 | 3232.5919 | -0.640792485 | 3.44E-08 |
| OTUD6B | 390.2730135 | -0.640891274 | 1.93E-02 |
| CASP6 | 603.1914311 | -0.640896325 | 5.40E-06 |
| AFG3L1P | 627.678639 | -0.640929419 | 9.79E-07 |
| SLC26A1 | 198.7225818 | -0.641145926 | 5.52E-03 |
| NA | 443.9465271 | -0.642022957 | 3.16E-04 |
| TP53BP1 | 4052.421138 | -0.64254642 | 9.63E-19 |
| EBF3 | 94.37900685 | -0.643209613 | 3.46E-02 |
| KBTBD6 | 801.5697179 | -0.643557236 | 6.61E-07 |
| AL360270.2 | 99.39925113 | -0.643747097 | 3.03E-02 |
| ABHD3 | 445.9021462 | -0.644215271 | 2.32E-04 |
| CCDC183 | 146.8971672 | -0.644226159 | 5.50E-04 |
| APH1B | 999.3437867 | -0.644307009 | 2.66E-06 |
| ASAH2B | 169.8476485 | -0.644381854 | 3.02E-02 |
| HAPLN3 | 305.6184811 | -0.644683236 | 5.43E-04 |
| HENMT1 | 1022.022023 | -0.644745443 | 2.62E-08 |
| ST3GAL1 | 3868.982245 | -0.644932898 | 1.56E-07 |
| LRRC37A6P | 65.4873206 | -0.645050821 | 1.56E-02 |
| FAM78A | 383.5158475 | -0.645772971 | 8.18E-07 |
| LINC00565 | 61.82331782 | -0.647172931 | 4.05E-02 |
| OSTM1 | 1362.323596 | -0.64723567 | 2.49E-05 |
| MRTFB | 2701.886717 | -0.647396122 | 3.33E-04 |
| RLF | 1290.363741 | -0.647403642 | 1.65E-05 |
| SSBP3-AS1 | 46.60351791 | -0.648119891 | 4.26E-02 |
| ZNF837 | 70.46561219 | -0.648170562 | 1.36E-02 |
| SRSF11 | 4244.378978 | -0.648288235 | 1.04E-05 |
| SULT1C4 | 67.64718645 | -0.648304289 | 3.78E-02 |
| CTDSPL2 | 1007.377581 | -0.648358834 | 2.39E-03 |
| TTPAL | 3969.055166 | -0.648389107 | 6.85E-24 |
| CCDC144B | 1018.828952 | -0.64848868 | 1.46E-03 |
| CEP170P1 | 71.64086969 | -0.648856662 | 1.65E-02 |
| TOPBP1 | 3698.866461 | -0.649323433 | 6.95E-05 |
| PDCD10 | 979.8580687 | -0.649403987 | 2.35E-03 |
| BACH2 | 445.9839007 | -0.649485433 | 1.14E-04 |
| SLC7A8 | 1004.591585 | -0.649898803 | 4.36E-05 |
| ZNF182 | 399.0536131 | -0.649970707 | 2.85E-03 |
| ZNRF2 | 314.0326182 | -0.649999317 | 8.57E-04 |
| RPAP3 | 1271.227255 | -0.650201287 | 4.37E-03 |
| AL354822.1 | 208.7467612 | -0.650361836 | 1.10E-02 |
| AL353194.1 | 66.22251447 | -0.650800018 | 1.72E-02 |
| MIB1 | 2496.134306 | -0.651026947 | 3.89E-04 |
| BLOC1S6 | 1357.049245 | -0.651245235 | 2.59E-05 |
| HNRNPH2 | 2907.243069 | -0.651578209 | 2.43E-09 |

| | | | |
|---|---|---|---|
| FAR1 | 2147.702616 | -0.652024281 | 3.43E-03 |
| GTF2I | 19621.1412 | -0.652328888 | 4.15E-13 |
| COL5A2 | 264.4417449 | -0.652430904 | 8.85E-03 |
| ZNF135 | 520.20737 | -0.652439295 | 5.14E-08 |
| TLK1 | 1635.937326 | -0.652680363 | 2.65E-02 |
| REL | 362.8332223 | -0.652773549 | 7.46E-04 |
| LMBRD1 | 897.240277 | -0.653036701 | 5.73E-10 |
| APOBEC3F | 829.4756475 | -0.65320802 | 4.72E-06 |
| TRPV4 | 204.8151447 | -0.653257062 | 2.10E-03 |
| PDSS1 | 287.872663 | -0.653415058 | 1.34E-04 |
| AC245452.1 | 293.78634 | -0.653789345 | 1.86E-05 |
| LAMP5 | 323.5776342 | -0.654076009 | 8.55E-04 |
| DPYD | 377.6234819 | -0.654083431 | 2.80E-06 |
| CACHD1 | 561.4740815 | -0.654659802 | 2.89E-07 |
| ZNF546 | 127.0354486 | -0.655060635 | 2.37E-02 |
| NEAT1 | 4927.823974 | -0.655127021 | 8.61E-03 |
| ZNF883 | 505.0140234 | -0.655286425 | 5.53E-05 |
| SLC25A13 | 3952.548398 | -0.655791765 | 7.98E-18 |
| AP5S1 | 887.952493 | -0.656510486 | 1.31E-06 |
| GPR137 | 1729.845174 | -0.65688732 | 4.34E-07 |
| LINC01948 | 68.24883831 | -0.65700131 | 8.93E-03 |
| TAPBPL | 874.5015255 | -0.657094147 | 1.75E-05 |
| GXYLT1 | 1016.912635 | -0.65716295 | 5.35E-05 |
| ARFIP1 | 1104.574144 | -0.657268018 | 1.34E-05 |
| AL121761.1 | 44.35731965 | -0.657359654 | 4.12E-02 |
| AL023806.3 | 69.85019706 | -0.657497213 | 1.75E-02 |
| PRKAR2B | 977.5424306 | -0.65764898 | 7.20E-09 |
| TOX2 | 3322.582474 | -0.657912316 | 8.82E-04 |
| DHX29 | 3842.598765 | -0.658348183 | 2.17E-04 |
| AC084018.2 | 65.15693012 | -0.658402388 | 2.68E-02 |
| METTL14 | 856.0442512 | -0.658743336 | 8.20E-03 |
| NAP1L5 | 362.4858112 | -0.659110858 | 8.17E-05 |
| ZNF594 | 820.0920276 | -0.659425618 | 8.66E-03 |
| PALD1 | 72.23173115 | -0.659597342 | 8.09E-03 |
| ZNF320 | 1074.743161 | -0.659729062 | 9.03E-08 |
| APPBP2 | 2193.313553 | -0.659812385 | 1.06E-05 |
| FAM217B | 1604.216906 | -0.660137513 | 8.37E-04 |
| PCYOX1 | 5760.612629 | -0.660184276 | 1.50E-09 |
| FAM186B | 70.38773791 | -0.660492523 | 2.65E-02 |
| MYO1B | 8268.616834 | -0.6606158 | 5.53E-09 |
| PPARG | 1847.312041 | -0.660760266 | 6.57E-06 |
| MTX3 | 1901.727013 | -0.660766026 | 8.29E-03 |
| ZNF192P1 | 85.4674867 | -0.661029492 | 1.82E-02 |
| NPR3 | 210.5822672 | -0.661066637 | 3.04E-02 |
| AL157392.3 | 363.3674805 | -0.661095081 | 1.24E-05 |
| ORAI3 | 3085.151218 | -0.661273973 | 1.40E-03 |
| ZFP69 | 265.9010029 | -0.661450662 | 2.88E-03 |

| | | | |
|---|---|---|---|
| NUP35 | 774.1365033 | -0.661636531 | 1.65E-05 |
| ZSCAN16-AS1 | 82.51571009 | -0.662109184 | 3.86E-02 |
| DNTTIP2 | 1683.644075 | -0.662127498 | 2.46E-03 |
| POT1 | 1255.972173 | -0.662323666 | 2.25E-03 |
| LIN7C | 545.565844 | -0.662662892 | 8.74E-04 |
| ZRANB2 | 2901.416846 | -0.662716089 | 1.38E-04 |
| RNF219 | 1318.110589 | -0.66305755 | 7.03E-04 |
| HIPK3 | 1459.481618 | -0.663100464 | 1.87E-07 |
| POLR2J4 | 326.6944127 | -0.663470758 | 1.45E-07 |
| CATSPER2 | 113.6358509 | -0.663638744 | 2.67E-02 |
| PAXIP1-AS2 | 428.289815 | -0.664005983 | 2.18E-05 |
| GLRX | 2880.011914 | -0.66450482 | 2.94E-12 |
| AL132639.2 | 45.55583347 | -0.665240547 | 3.66E-02 |
| NSUN5P2 | 311.8154007 | -0.665387124 | 2.20E-04 |
| ANAPC4 | 975.3825386 | -0.665784844 | 2.00E-05 |
| CROT | 779.6871208 | -0.665977191 | 5.31E-06 |
| CLCN3 | 2983.363341 | -0.666002004 | 3.63E-08 |
| SPIN4 | 606.1825621 | -0.666034981 | 2.52E-04 |
| LINC01524 | 71.63097704 | -0.666076737 | 2.36E-02 |
| STX17 | 1044.005854 | -0.666123708 | 8.50E-05 |
| AC113191.1 | 39.4017043 | -0.666149088 | 4.68E-02 |
| GNPTAB | 2239.063169 | -0.666274426 | 3.32E-09 |
| ZNF816 | 264.0347429 | -0.666460609 | 3.86E-02 |
| STRN3 | 913.2813553 | -0.666803837 | 3.70E-06 |
| BPTF | 4182.268897 | -0.66687572 | 1.18E-05 |
| LRIG3 | 486.7772918 | -0.667197865 | 1.60E-06 |
| KPNA5 | 218.1163122 | -0.667450951 | 1.00E-02 |
| CKMT1B | 286.8831445 | -0.668579255 | 8.64E-04 |
| POP5 | 1281.270367 | -0.668703906 | 6.60E-17 |
| ZNF112 | 266.0353975 | -0.668704322 | 7.59E-03 |
| AP4S1 | 353.5657067 | -0.669075909 | 2.85E-03 |
| TM7SF3 | 2482.855936 | -0.669230099 | 8.38E-09 |
| NCOA1 | 2266.228525 | -0.669266415 | 2.61E-08 |
| SYT16 | 160.8716979 | -0.669993616 | 5.83E-03 |
| AC022031.2 | 97.99760851 | -0.669998455 | 6.07E-03 |
| MED23 | 1471.504237 | -0.670497599 | 7.00E-08 |
| U47924.2 | 96.97950533 | -0.670803635 | 4.41E-03 |
| ZNF439 | 282.7368766 | -0.670888726 | 4.65E-04 |
| ERAP2 | 8236.42801 | -0.670930865 | 4.98E-22 |
| U2SURP | 3767.063815 | -0.670972696 | 1.20E-03 |
| JAK3 | 626.4733155 | -0.670983179 | 4.07E-03 |
| MTF1 | 1054.800846 | -0.671453718 | 4.37E-05 |
| ZNF680 | 151.8603691 | -0.67180808 | 9.29E-03 |
| BVES | 1349.384071 | -0.672005203 | 8.30E-16 |
| CENPJ | 546.3815123 | -0.67205029 | 2.77E-03 |
| ZNF578 | 160.044922 | -0.67219599 | 1.18E-04 |
| ZBED5 | 1813.585829 | -0.672303229 | 5.71E-05 |

| | | | |
|---|---|---|---|
| LIG4 | 286.7947973 | -0.672637613 | 1.47E-02 |
| PIP4K2A | 1913.589145 | -0.672659264 | 1.90E-10 |
| ZNF793 | 305.6492898 | -0.67280527 | 8.12E-05 |
| GOLGA6L4 | 124.8781029 | -0.672821338 | 1.40E-02 |
| AC015818.9 | 37.31164793 | -0.672924461 | 4.77E-02 |
| MTF2 | 777.0723014 | -0.673351914 | 1.05E-04 |
| FAM230B | 183.8312062 | -0.673745805 | 6.97E-04 |
| ADAT2 | 484.6253686 | -0.673759207 | 3.49E-03 |
| AC072061.1 | 35.64870075 | -0.67448315 | 4.61E-02 |
| DNAJC13 | 3775.927102 | -0.674582236 | 5.46E-05 |
| LINC01297 | 1123.725903 | -0.674879135 | 2.48E-02 |
| MTERF1 | 599.7245431 | -0.675066578 | 4.55E-04 |
| AC132812.1 | 80.9606679 | -0.675328272 | 1.31E-02 |
| NEK1 | 848.2975352 | -0.675444024 | 1.52E-02 |
| Z99129.4 | 56.19699926 | -0.675454377 | 4.71E-02 |
| TEFM | 462.4051509 | -0.675497836 | 8.11E-07 |
| TXNDC9 | 936.8841297 | -0.675870406 | 1.03E-05 |
| EID1 | 3347.970584 | -0.676087307 | 6.08E-08 |
| ZNF564 | 315.0327144 | -0.676275571 | 1.90E-06 |
| SPOPL | 436.6793134 | -0.676409288 | 4.25E-03 |
| FZD7 | 2542.472438 | -0.676538443 | 8.85E-15 |
| DHRS11 | 725.2445982 | -0.676745119 | 1.65E-10 |
| MAP4K4 | 13426.54364 | -0.676999752 | 2.69E-28 |
| RWDD3 | 366.8926154 | -0.677086033 | 1.93E-03 |
| FNDC3A | 2717.186824 | -0.677489628 | 2.16E-06 |
| ANKRD36 | 491.341241 | -0.677522198 | 5.12E-03 |
| VAMP8 | 385.8872751 | -0.677762967 | 7.39E-03 |
| TNRC6B | 2313.664522 | -0.677991071 | 2.59E-07 |
| ZNF449 | 394.9404058 | -0.678056728 | 4.97E-04 |
| GLDC | 119.3079729 | -0.678192273 | 3.60E-03 |
| FOXD2 | 276.8095472 | -0.67868768 | 4.73E-07 |
| AL355388.2 | 43.64300837 | -0.678701564 | 2.25E-02 |
| SNRNP25 | 1685.632207 | -0.678920549 | 4.17E-05 |
| RGS14 | 562.7905907 | -0.679237192 | 3.95E-06 |
| TICAM2 | 385.835864 | -0.680091112 | 3.41E-02 |
| AC012640.2 | 113.968136 | -0.680156087 | 2.23E-03 |
| SOX18 | 172.617791 | -0.680453102 | 4.86E-03 |
| AL390728.4 | 395.6292296 | -0.680726386 | 1.10E-03 |
| DPY19L2P2 | 694.3173376 | -0.68089088 | 7.51E-05 |
| PPIL3 | 802.2648949 | -0.681507441 | 1.76E-07 |
| AC010132.3 | 1353.721702 | -0.681571476 | 5.72E-07 |
| ALMS1 | 1742.721093 | -0.681583501 | 7.80E-04 |
| AC008906.1 | 67.2150028 | -0.681746872 | 2.63E-02 |
| DCP2 | 2102.574905 | -0.681955035 | 2.04E-07 |
| AC091132.3 | 70.90349035 | -0.681999512 | 2.41E-02 |
| SRSF6 | 9770.517034 | -0.682063587 | 4.04E-11 |
| TEX9 | 137.6996453 | -0.682249041 | 3.26E-03 |

| | | | |
|---|---|---|---|
| CYP4F11 | 312.2638214 | -0.682267585 | 1.13E-03 |
| GLI4 | 718.4885791 | -0.682361951 | 1.05E-06 |
| CCP110 | 1115.965857 | -0.682892267 | 7.38E-04 |
| ASB16 | 66.44886693 | -0.682902199 | 1.14E-02 |
| COL4A3BP | 2975.02318 | -0.683236235 | 3.38E-14 |
| TERF1 | 1319.671707 | -0.683241808 | 7.11E-05 |
| BCAP29 | 2042.475783 | -0.683556858 | 1.35E-07 |
| SPRY3 | 329.3369305 | -0.683834795 | 8.47E-09 |
| NEO1 | 7967.414653 | -0.683921784 | 6.20E-08 |
| TUSC3 | 4463.581761 | -0.685265083 | 2.68E-30 |
| KBTBD4 | 1247.543525 | -0.685267129 | 2.82E-05 |
| NDRG4 | 400.4921828 | -0.685596751 | 1.83E-05 |
| MYLIP | 1308.827892 | -0.686372802 | 2.85E-07 |
| KLF12 | 261.469215 | -0.686381513 | 2.02E-03 |
| STOX1 | 103.1043832 | -0.686990016 | 4.81E-03 |
| ZNF195 | 778.8841383 | -0.687075313 | 3.26E-04 |
| ZBED6 | 1064.580852 | -0.687311166 | 6.96E-07 |
| CCDC96 | 51.5401067 | -0.687861171 | 3.58E-02 |
| IFIT5 | 791.850336 | -0.688308368 | 1.84E-07 |
| CREBRF | 445.0915497 | -0.688628934 | 3.06E-03 |
| PCMTD2 | 3440.324389 | -0.688976762 | 2.82E-08 |
| DIPK1A | 488.9177531 | -0.689141537 | 1.09E-04 |
| SMARCA1 | 5733.867665 | -0.689148103 | 8.69E-18 |
| FTX | 118.6607101 | -0.689191907 | 2.56E-03 |
| SUCO | 1308.492429 | -0.689282544 | 7.55E-04 |
| SNTB2 | 3318.817684 | -0.689285717 | 6.37E-12 |
| STRA6LP | 197.9736375 | -0.689664961 | 3.38E-03 |
| TUBD1 | 294.8775476 | -0.690303458 | 2.68E-05 |
| EXOC3-AS1 | 613.1024133 | -0.690473399 | 1.84E-11 |
| LUC7L2 | 3487.963961 | -0.690640264 | 4.66E-10 |
| SERPINA5 | 1276.950546 | -0.690815816 | 2.00E-04 |
| RBM4B | 1235.024212 | -0.690913056 | 1.98E-05 |
| PLSCR4 | 241.6913682 | -0.690945684 | 5.10E-04 |
| ZNF75A | 1038.680233 | -0.691168791 | 3.27E-10 |
| STAC2 | 256.3708383 | -0.691767629 | 1.58E-05 |
| PCDHGB1 | 109.8113584 | -0.692269279 | 1.34E-03 |
| RAPGEF6 | 3147.194901 | -0.692279829 | 5.44E-06 |
| CNFN | 49.70271958 | -0.69237906 | 2.65E-02 |
| SLC25A46 | 2798.417499 | -0.692441644 | 4.52E-06 |
| LINC01106 | 1890.349103 | -0.692502707 | 1.01E-14 |
| HOTAIR | 192.0254968 | -0.693051076 | 2.34E-03 |
| BCKDHB | 817.3871201 | -0.693133206 | 2.89E-08 |
| CU633906.2 | 132.698479 | -0.693303761 | 1.69E-02 |
| CETN2 | 2499.92649 | -0.693373779 | 4.15E-09 |
| TMX1 | 2060.293302 | -0.693588491 | 2.11E-05 |
| ZNF254 | 243.3316025 | -0.694139386 | 1.32E-03 |
| HIBCH | 1071.09971 | -0.694253793 | 3.33E-06 |

| | | | |
|---|---|---|---|
| GUSBP2 | 71.99712807 | -0.694673661 | 2.36E-02 |
| METTL18 | 221.5883232 | -0.694835802 | 8.26E-03 |
| AC092072.1 | 204.0970607 | -0.694898722 | 3.49E-03 |
| DNM3 | 456.6554895 | -0.695754265 | 9.30E-07 |
| EXOG | 670.6987006 | -0.695756794 | 8.34E-13 |
| ENAH | 5040.728136 | -0.696037488 | 3.78E-04 |
| CTR9 | 4964.492339 | -0.696044561 | 7.28E-10 |
| PBDC1 | 1042.742786 | -0.696202254 | 2.27E-16 |
| PCDHB13 | 90.25920142 | -0.696901288 | 8.70E-03 |
| LECT2 | 122.0354449 | -0.697351531 | 4.24E-02 |
| AC010894.3 | 225.0721163 | -0.697615193 | 3.04E-05 |
| SYNJ1 | 938.8449281 | -0.697731082 | 1.56E-05 |
| MPHOSPH10 | 1233.545703 | -0.69805931 | 4.39E-06 |
| RGS2 | 189.9077921 | -0.698208803 | 1.59E-03 |
| AC011468.5 | 30.75586694 | -0.698411433 | 4.88E-02 |
| OXR1 | 901.5233758 | -0.698450119 | 9.17E-07 |
| NCAPG | 2313.274187 | -0.69883071 | 5.34E-03 |
| AC243964.2 | 208.6143951 | -0.69895863 | 3.21E-04 |
| LINC01977 | 90.72125105 | -0.698988701 | 1.40E-03 |
| MR1 | 1383.354792 | -0.699138012 | 1.32E-06 |
| FAM234A | 7638.339532 | -0.699146066 | 9.10E-10 |
| HFE | 563.7342057 | -0.69919264 | 4.27E-09 |
| FAM156A | 1547.074356 | -0.6995581 | 2.82E-07 |
| FBXL21P | 1580.257631 | -0.699698566 | 4.58E-03 |
| VPS45 | 2279.513238 | -0.700037288 | 7.31E-17 |
| UBTD2 | 1371.08928 | -0.700574631 | 1.47E-27 |
| DNMT3B | 895.4449263 | -0.700606811 | 2.21E-08 |
| AC005041.1 | 156.4021284 | -0.700642293 | 2.21E-03 |
| PHF2P2 | 183.8502381 | -0.70098912 | 7.31E-03 |
| AL845472.2 | 297.2034584 | -0.701157877 | 5.91E-06 |
| CEP350 | 2166.233297 | -0.702199083 | 3.25E-03 |
| CXADR | 430.9583909 | -0.702220695 | 9.23E-04 |
| RBM12B | 1824.034248 | -0.702231164 | 1.01E-04 |
| ZEB1 | 1261.88444 | -0.70396139 | 6.64E-03 |
| SRGAP3 | 587.6223722 | -0.704132644 | 1.83E-07 |
| QSER1 | 3135.860873 | -0.704623191 | 1.81E-04 |
| LHX4 | 142.5888116 | -0.704857094 | 4.91E-04 |
| PIH1D2 | 92.08842891 | -0.704861578 | 8.33E-03 |
| ZNF845 | 255.5422777 | -0.704888322 | 5.76E-03 |
| AC245060.6 | 212.7784863 | -0.705107843 | 1.91E-02 |
| C22orf46 | 2615.757336 | -0.705126485 | 1.82E-22 |
| CAND1 | 6229.939359 | -0.70523916 | 4.20E-12 |
| AC055713.1 | 109.6029545 | -0.705330653 | 3.94E-04 |
| AL732372.3 | 267.4658968 | -0.7058274 | 1.04E-06 |
| LCTL | 194.6728429 | -0.706241286 | 1.44E-03 |
| TTC34 | 117.7523898 | -0.706389722 | 1.62E-03 |
| ZNF606 | 571.9029524 | -0.706408299 | 2.07E-06 |

| | | | |
|---|---|---|---|
| RAB30 | 961.6113688 | -0.706844919 | 1.62E-08 |
| TMEM221 | 72.69405852 | -0.70886264 | 1.87E-02 |
| THRA | 4147.420593 | -0.708915565 | 1.98E-11 |
| AL162258.2 | 44.64770892 | -0.708981333 | 2.54E-02 |
| CGB3 | 47.16534887 | -0.709122592 | 3.94E-02 |
| C11orf71 | 513.7332104 | -0.709268655 | 1.34E-08 |
| DPF1 | 212.3406063 | -0.709345742 | 1.10E-03 |
| KRIT1 | 1969.030056 | -0.709997093 | 5.39E-05 |
| ZNF614 | 555.4187751 | -0.710204691 | 2.33E-10 |
| SYT13 | 311.0869747 | -0.710411814 | 3.61E-02 |
| PSMG3-AS1 | 1134.425137 | -0.710473251 | 1.32E-05 |
| CLOCK | 1168.238529 | -0.710602758 | 2.88E-04 |
| AC004908.1 | 54.79869408 | -0.710720034 | 3.81E-02 |
| FLG | 765.867577 | -0.710767017 | 9.07E-08 |
| MFN1 | 2306.740901 | -0.71086058 | 2.05E-07 |
| LARGE1 | 1127.72581 | -0.711012275 | 1.04E-07 |
| TMF1 | 1309.052105 | -0.711075842 | 2.82E-03 |
| RANBP17 | 424.6546508 | -0.711583807 | 4.59E-04 |
| AC138866.1 | 312.6038432 | -0.711645907 | 8.91E-05 |
| AC092919.2 | 104.0774614 | -0.711659962 | 3.80E-03 |
| CUL5 | 1325.527547 | -0.711711772 | 3.30E-03 |
| DNA2 | 356.2619562 | -0.711982024 | 5.81E-05 |
| GIMAP2 | 643.0757087 | -0.712375584 | 1.00E-10 |
| HIST1H3H | 87.21353459 | -0.712909932 | 3.17E-02 |
| KNL1 | 1041.133435 | -0.712960261 | 1.63E-02 |
| MANEAL | 1265.66944 | -0.713021028 | 1.12E-06 |
| NA | 243.5273505 | -0.713418759 | 7.14E-07 |
| SNX4 | 778.4362423 | -0.713476552 | 1.25E-04 |
| AL121832.2 | 88.90306509 | -0.714174904 | 6.49E-03 |
| ALKBH8 | 742.8315573 | -0.714448151 | 7.05E-03 |
| FIRRE | 111.7449268 | -0.714554517 | 8.74E-04 |
| SCOC | 836.9758959 | -0.715017014 | 7.03E-03 |
| ZNF487 | 57.49883629 | -0.715897982 | 3.11E-02 |
| DPY19L4 | 1724.934286 | -0.715917698 | 7.13E-06 |
| DST | 4851.010378 | -0.716248115 | 3.38E-03 |
| TMEM198B | 935.5913628 | -0.716308273 | 3.39E-15 |
| FAM200A | 562.1375119 | -0.71640348 | 3.59E-08 |
| THUMPD3-AS1 | 493.6368638 | -0.716547105 | 6.40E-04 |
| TIAM1 | 2164.633176 | -0.716664241 | 7.76E-14 |
| RRAGB | 446.9069824 | -0.716760574 | 2.08E-05 |
| ASIC1 | 1465.170741 | -0.716803884 | 9.30E-07 |
| PEX12 | 557.9871048 | -0.717376484 | 1.66E-10 |
| AP003392.4 | 253.0696273 | -0.717517161 | 8.84E-06 |
| FBXL3 | 1749.761037 | -0.717531445 | 2.12E-11 |
| HLA-DMA | 431.2072074 | -0.717776489 | 7.22E-03 |
| CA11 | 828.0280318 | -0.718060655 | 3.72E-04 |
| CCDC142 | 726.9646701 | -0.718117235 | 2.55E-10 |

| | | | |
|---|---|---|---|
| PLA2G6 | 356.9025369 | -0.718338701 | 1.57E-04 |
| LAPTM4B | 16786.53935 | -0.718619674 | 4.10E-09 |
| AC073578.2 | 49.81234939 | -0.718764355 | 3.61E-02 |
| GALNT7 | 2610.119088 | -0.719231372 | 1.77E-08 |
| MAP3K12 | 1271.312178 | -0.71933592 | 5.65E-05 |
| KCNH2 | 218.6381721 | -0.719436937 | 3.95E-03 |
| TCAF1P1 | 529.6204086 | -0.719721469 | 6.40E-06 |
| TKFC | 1546.53106 | -0.720434944 | 8.54E-21 |
| ZNF559-ZNF177 | 287.9463382 | -0.720475995 | 4.87E-06 |
| PPIP5K2 | 3322.167985 | -0.720545867 | 1.16E-04 |
| YAE1 | 670.6135456 | -0.721553573 | 7.13E-08 |
| ZBTB33 | 1269.78301 | -0.721614372 | 1.31E-04 |
| MON2 | 1932.049245 | -0.721635407 | 8.89E-05 |
| ARMT1 | 767.7113099 | -0.72210701 | 5.77E-03 |
| ZFX | 1304.403397 | -0.722118235 | 6.29E-08 |
| INO80D | 402.4125441 | -0.722370084 | 5.80E-03 |
| AC146944.4 | 132.6466037 | -0.722375059 | 3.04E-03 |
| RBBP8 | 1615.696352 | -0.722752312 | 6.59E-04 |
| FAM133B | 963.4062515 | -0.724120579 | 6.87E-07 |
| CD164 | 6736.973925 | -0.724672392 | 3.94E-12 |
| TPD52L1 | 2117.343253 | -0.724701693 | 1.69E-05 |
| AC060234.2 | 135.2052896 | -0.724795906 | 3.24E-05 |
| SCFD2 | 1398.46596 | -0.725276021 | 2.28E-12 |
| PANK3 | 2532.270152 | -0.725549973 | 2.96E-04 |
| ANKRD17 | 5883.493827 | -0.725819052 | 9.16E-24 |
| TIMM8B | 2524.292083 | -0.725820337 | 7.18E-07 |
| MSRB1 | 1234.914881 | -0.7258218 | 5.54E-09 |
| KIF26A | 548.4895354 | -0.725971811 | 2.16E-06 |
| AL365295.1 | 41.34757873 | -0.726282103 | 2.75E-02 |
| USP32P2 | 439.998467 | -0.726482101 | 4.55E-03 |
| AC007066.2 | 118.9970978 | -0.726681018 | 4.93E-04 |
| SECISBP2L | 1068.391519 | -0.726696558 | 2.95E-04 |
| AL627309.6 | 103.0867455 | -0.727148652 | 9.87E-03 |
| DSCR4 | 114.9837457 | -0.727310784 | 2.65E-03 |
| HNRNPDL | 6883.710354 | -0.727327501 | 3.56E-10 |
| SUCLA2 | 1298.223595 | -0.727379683 | 1.47E-04 |
| USP15 | 1224.130539 | -0.727504721 | 1.42E-03 |
| VASH1 | 4125.770126 | -0.727900463 | 5.39E-07 |
| SEC62 | 2051.27731 | -0.728291497 | 1.26E-04 |
| ICAM4 | 123.6028687 | -0.728464324 | 8.06E-03 |
| HERC6 | 1046.33378 | -0.728854477 | 7.85E-11 |
| ZNF566 | 241.2730474 | -0.729217348 | 1.82E-02 |
| MAPK6 | 2936.015372 | -0.729537584 | 1.49E-03 |
| ASNSD1 | 1743.763404 | -0.730170617 | 6.60E-04 |
| NA | 105.2038393 | -0.730248378 | 1.78E-04 |
| BTN2A3P | 353.6383477 | -0.730487194 | 2.91E-10 |
| HERC2P2 | 2782.210698 | -0.730577559 | 1.20E-08 |

| | | | |
|---|---|---|---|
| ELAC1 | 322.3283585 | -0.730623564 | 2.14E-07 |
| CPEB4 | 593.1823739 | -0.730650099 | 6.01E-04 |
| PAG1 | 564.5162007 | -0.731536676 | 1.07E-03 |
| B3GNT3 | 176.5313853 | -0.73308103 | 9.19E-03 |
| CRYZ | 1531.82456 | -0.733222115 | 4.49E-07 |
| ZBTB45P2 | 717.7323167 | -0.733244759 | 5.46E-08 |
| GPBP1 | 4156.015503 | -0.733290365 | 1.40E-04 |
| AC005224.1 | 53.6959204 | -0.733443023 | 3.47E-02 |
| C1R | 3793.027515 | -0.733763635 | 5.39E-06 |
| MRPL1 | 701.2691386 | -0.734057542 | 3.24E-05 |
| AC106820.4 | 312.6824286 | -0.734105358 | 1.60E-04 |
| ZDHHC21 | 343.4902923 | -0.734167171 | 3.19E-02 |
| CHKB-DT | 40.59241617 | -0.734842339 | 4.62E-02 |
| CLN5 | 1590.052287 | -0.735244764 | 2.52E-08 |
| C2orf69 | 301.3506584 | -0.735995658 | 5.73E-04 |
| FAM106A | 885.1586983 | -0.736121497 | 4.72E-04 |
| ARHGEF3 | 994.1096122 | -0.736525021 | 7.53E-10 |
| LINC01366 | 70.30869446 | -0.736608718 | 4.29E-03 |
| AHI1 | 978.9390272 | -0.736701156 | 1.08E-03 |
| UBN2 | 1275.398508 | -0.736714147 | 7.39E-04 |
| IFT88 | 335.7827756 | -0.737496117 | 5.74E-04 |
| RIOK2 | 1734.119616 | -0.737683131 | 2.04E-03 |
| HELZ | 2055.418843 | -0.737754106 | 6.95E-13 |
| ACER2 | 79.7112544 | -0.737834756 | 1.09E-02 |
| UBAC2-AS1 | 265.6527688 | -0.737884313 | 2.66E-05 |
| RASA2 | 422.6520801 | -0.737890675 | 6.98E-03 |
| KANSL1L | 442.1730578 | -0.738008291 | 8.09E-06 |
| MTHFD2L | 161.4402153 | -0.738057014 | 1.07E-03 |
| SPAG8 | 62.51808455 | -0.738922504 | 1.64E-02 |
| POMK | 182.4422372 | -0.739176367 | 6.15E-07 |
| CPNE8 | 535.1743269 | -0.739196087 | 1.22E-03 |
| SH3BP5 | 1459.805366 | -0.739814851 | 3.10E-08 |
| AC093297.2 | 88.64698431 | -0.73987203 | 3.23E-02 |
| CCDC186 | 315.7125108 | -0.739991931 | 2.62E-02 |
| PDE8B | 904.5066421 | -0.740065852 | 1.11E-04 |
| RASSF5 | 662.6694676 | -0.740300222 | 4.61E-08 |
| PIK3C2B | 986.2967161 | -0.740310631 | 1.36E-06 |
| PCBD2 | 158.4686784 | -0.740375802 | 6.19E-04 |
| RHBDL1 | 79.67093928 | -0.740565733 | 1.26E-02 |
| C1GALT1C1L | 229.7544466 | -0.740896799 | 1.17E-04 |
| TSPAN6 | 4046.676907 | -0.741774045 | 9.59E-13 |
| IGFLR1 | 122.5102195 | -0.74186558 | 1.70E-02 |
| ZNF615 | 310.8107361 | -0.742102145 | 5.97E-04 |
| TIGD7 | 862.8036905 | -0.742137252 | 5.52E-05 |
| AL354714.1 | 37.61021756 | -0.742417523 | 3.24E-02 |
| SLC4A11 | 468.4280727 | -0.742825202 | 2.60E-05 |
| AP002490.1 | 78.86775456 | -0.74302386 | 4.07E-03 |

| | | | |
|---|---|---|---|
| ABCB10 | 750.3101236 | -0.743043318 | 6.56E-08 |
| KMT2C | 4465.661044 | -0.743093971 | 2.29E-12 |
| SGK1 | 5483.663424 | -0.744744472 | 3.26E-07 |
| ACVR2A | 430.0205324 | -0.745122545 | 4.19E-08 |
| MEIS3P1 | 287.1832009 | -0.745544518 | 4.45E-06 |
| CTSK | 134.8465847 | -0.745735879 | 2.09E-02 |
| EFCAB2 | 314.3175812 | -0.74620631 | 2.04E-05 |
| SEMA6A-AS1 | 39.33274057 | -0.746419759 | 2.68E-02 |
| AC107068.1 | 89.06929586 | -0.746619769 | 5.59E-03 |
| LIN9 | 409.9594675 | -0.746664869 | 1.12E-04 |
| UNC13A | 437.1254544 | -0.747566862 | 2.83E-04 |
| PKN2 | 2294.834694 | -0.747628762 | 1.33E-04 |
| HACD4 | 437.4754155 | -0.747770168 | 4.31E-10 |
| RASL11B | 84.07592989 | -0.749090406 | 3.02E-03 |
| LINC01087 | 87.97856487 | -0.749094748 | 4.12E-03 |
| LINC00920 | 110.360093 | -0.749562871 | 9.93E-04 |
| IGFBP2 | 14621.13362 | -0.749608222 | 3.31E-06 |
| GCNT1 | 951.6921527 | -0.74995667 | 2.99E-04 |
| HIST4H4 | 75.16508419 | -0.751109578 | 1.32E-02 |
| FGD4 | 160.9786872 | -0.75125379 | 9.66E-03 |
| DIS3 | 4813.73347 | -0.751256457 | 1.04E-05 |
| INTU | 674.7119295 | -0.752254544 | 5.01E-04 |
| USP46 | 1234.24571 | -0.752874199 | 1.36E-06 |
| LYPD6B | 296.0477993 | -0.753536791 | 1.05E-02 |
| ZWILCH | 3027.212332 | -0.753933396 | 7.07E-06 |
| MXD4 | 4041.346101 | -0.754016816 | 1.41E-03 |
| FMR1 | 2502.389221 | -0.754043465 | 2.54E-07 |
| WDR5B | 519.3509633 | -0.754264352 | 1.71E-09 |
| NA | 138.6279895 | -0.754535551 | 4.35E-04 |
| PLK4 | 998.2687593 | -0.755092543 | 8.98E-03 |
| RAD54B | 595.254013 | -0.755137531 | 1.52E-03 |
| SAMD14 | 319.8574801 | -0.755236124 | 2.41E-03 |
| GCFC2 | 532.1667663 | -0.755299888 | 1.30E-04 |
| HDX | 246.3347493 | -0.755624889 | 4.79E-04 |
| NA | 151.0137063 | -0.755747014 | 3.80E-02 |
| UBXN8 | 546.4351699 | -0.755931604 | 4.24E-09 |
| TSSK4 | 100.2316405 | -0.75601606 | 8.75E-04 |
| C14orf132 | 1060.585725 | -0.756263579 | 9.95E-05 |
| NPAT | 1282.054313 | -0.757259555 | 7.45E-05 |
| AC010654.1 | 46.46695817 | -0.757317953 | 2.30E-02 |
| AC138907.8 | 93.45109188 | -0.757444139 | 2.07E-02 |
| G2E3 | 721.7049318 | -0.757595911 | 4.53E-04 |
| SLK | 1599.19667 | -0.758163625 | 1.82E-03 |
| RECQL | 2422.766935 | -0.758555416 | 5.46E-04 |
| FBXO2 | 682.1087411 | -0.758984943 | 1.07E-04 |
| SLC9A7P1 | 124.8698123 | -0.759049436 | 6.20E-04 |
| CENPC | 439.1903264 | -0.759115605 | 7.56E-03 |

| | | | |
|---|---|---|---|
| AL049796.1 | 51.65809039 | -0.759207611 | 1.61E-02 |
| PAXBP1 | 1660.356288 | -0.759719413 | 4.06E-04 |
| C6orf226 | 100.1026753 | -0.76031868 | 1.94E-03 |
| AC025171.2 | 287.6890618 | -0.76032672 | 8.29E-06 |
| FRK | 159.893411 | -0.760616874 | 1.37E-03 |
| SUZ12 | 1749.24164 | -0.761036857 | 1.12E-05 |
| USP1 | 2030.465933 | -0.761343495 | 1.86E-04 |
| HOXA-AS2 | 648.9155533 | -0.761360099 | 5.05E-08 |
| LINC01347 | 72.27855982 | -0.761913264 | 4.74E-03 |
| ZNF397 | 629.7096529 | -0.761935999 | 2.28E-06 |
| CNPY4 | 2550.890741 | -0.762461246 | 8.68E-10 |
| BTN3A3 | 848.1382446 | -0.762756891 | 3.68E-05 |
| ANXA9 | 166.5640661 | -0.762782929 | 8.07E-03 |
| AP001453.1 | 87.62101159 | -0.763179573 | 2.29E-02 |
| NUDT15 | 1373.191692 | -0.763274685 | 9.46E-12 |
| AMDHD2 | 1252.631935 | -0.76343537 | 1.37E-12 |
| DBP | 393.6402923 | -0.763861039 | 9.86E-04 |
| AC021092.1 | 246.9309242 | -0.764494839 | 1.87E-08 |
| HSPB6 | 899.0889698 | -0.764695085 | 1.45E-05 |
| PAK1 | 4174.330043 | -0.764796572 | 1.78E-17 |
| RAB22A | 2724.769956 | -0.764845233 | 2.92E-19 |
| ZNF613 | 252.1276893 | -0.76517453 | 8.15E-07 |
| ZNF189 | 953.5826802 | -0.765403177 | 3.37E-07 |
| AC016205.1 | 101.1451885 | -0.765441977 | 1.02E-02 |
| CCDC18 | 362.2515424 | -0.765564207 | 3.70E-02 |
| CARMIL1 | 2603.175438 | -0.765587487 | 2.10E-32 |
| SLC6A16 | 260.3475046 | -0.766158558 | 1.07E-04 |
| ITGB5 | 9170.149928 | -0.766682993 | 7.44E-06 |
| ESCO2 | 666.4800424 | -0.766874185 | 4.38E-03 |
| C18orf54 | 375.9919352 | -0.767129898 | 8.01E-03 |
| KRCC1 | 655.4614552 | -0.767343439 | 1.75E-03 |
| GNAQ | 3447.817147 | -0.767407256 | 3.33E-10 |
| AC247036.1 | 655.3655526 | -0.76843002 | 1.34E-04 |
| JKAMP | 1240.07522 | -0.768594986 | 1.07E-08 |
| LRRC49 | 476.3752078 | -0.768791115 | 3.36E-02 |
| ZDHHC17 | 996.3512409 | -0.768906662 | 1.06E-05 |
| SOCS1 | 103.8450996 | -0.769001126 | 6.52E-04 |
| GVINP1 | 117.3805028 | -0.769325922 | 1.85E-03 |
| KIAA2026 | 1257.987222 | -0.769832797 | 8.76E-08 |
| ELOVL6 | 1116.973458 | -0.769885772 | 1.61E-05 |
| C4orf33 | 225.6424876 | -0.770542635 | 2.21E-04 |
| ROR1 | 803.2210795 | -0.770597183 | 1.80E-07 |
| NA | 43.48618695 | -0.77061247 | 3.85E-02 |
| UFL1 | 1471.591244 | -0.770694083 | 2.77E-03 |
| VILL | 408.5323477 | -0.770763566 | 3.22E-10 |
| DDX58 | 932.5016077 | -0.770891549 | 6.55E-15 |
| PIK3CA | 1376.042323 | -0.771424718 | 2.18E-04 |

| | | | |
|---|---|---|---|
| APOL4 | 164.0383577 | -0.771750709 | 7.22E-05 |
| BMP2K | 1364.424567 | -0.771781573 | 1.55E-08 |
| TTC37 | 7042.255014 | -0.771787724 | 1.87E-03 |
| KLRG1 | 35.37736487 | -0.771892708 | 3.49E-02 |
| HECTD2 | 302.8748965 | -0.772207542 | 1.44E-03 |
| ZNF418 | 240.3467327 | -0.772260594 | 2.30E-04 |
| SMC2 | 3469.216138 | -0.772747796 | 3.62E-03 |
| TGFBR1 | 4359.922317 | -0.772749796 | 3.51E-07 |
| AC018553.1 | 92.95374125 | -0.772969268 | 1.06E-04 |
| ARL6 | 147.678687 | -0.772992688 | 5.91E-04 |
| ANKRD6 | 555.8492612 | -0.773529807 | 2.52E-05 |
| USP27X-AS1 | 60.68377957 | -0.773530715 | 1.96E-02 |
| BTBD19 | 501.2512357 | -0.774209298 | 2.17E-09 |
| AC084018.1 | 34.87481708 | -0.774447158 | 2.54E-02 |
| LINC00652 | 176.6260923 | -0.77451418 | 1.13E-06 |
| CCSAP | 674.7629138 | -0.774589065 | 3.90E-08 |
| PIK3C2A | 3270.990908 | -0.774599441 | 2.43E-02 |
| BHLHB9 | 316.3356222 | -0.775009942 | 1.90E-05 |
| NA | 104.9217022 | -0.775389845 | 3.39E-04 |
| ZNF684 | 75.25162679 | -0.775396704 | 3.97E-02 |
| CMTR2 | 1344.960866 | -0.775458064 | 4.02E-06 |
| PDCD4 | 2431.171143 | -0.775680988 | 1.33E-08 |
| ZNF300 | 709.6145832 | -0.776054087 | 4.28E-05 |
| CRYGS | 33.5931757 | -0.776335716 | 4.49E-02 |
| THY1 | 1129.874395 | -0.776942754 | 3.61E-03 |
| CDC37L1 | 623.5580681 | -0.77770769 | 1.50E-05 |
| SETBP1 | 900.8322632 | -0.777719161 | 2.65E-12 |
| AP000347.1 | 429.715956 | -0.778311781 | 6.03E-05 |
| MZF1-AS1 | 123.7415297 | -0.778376853 | 7.57E-05 |
| GPATCH2 | 545.3141153 | -0.778478048 | 7.28E-07 |
| PROS1 | 404.9411068 | -0.779092693 | 2.88E-05 |
| HLA-DMB | 409.4052964 | -0.779378814 | 3.32E-02 |
| VSIG2 | 41.62470534 | -0.780232236 | 4.05E-02 |
| TMEM150A | 656.5324793 | -0.780722598 | 1.22E-08 |
| IFIT3 | 1284.613263 | -0.780751977 | 1.22E-09 |
| AC009126.1 | 120.3707254 | -0.780869579 | 9.58E-04 |
| IGSF1 | 60.08272131 | -0.781282998 | 2.23E-02 |
| AL928654.1 | 326.5701445 | -0.781442929 | 1.65E-07 |
| HCFC2 | 577.4168189 | -0.781686242 | 3.96E-08 |
| S1PR2 | 135.2910706 | -0.78255249 | 1.54E-04 |
| SMPDL3A | 94.57710573 | -0.782628791 | 3.58E-03 |
| AL355482.1 | 38.9604449 | -0.78292415 | 4.57E-02 |
| CYB5D1 | 624.0858821 | -0.783115485 | 6.08E-08 |
| DSCC1 | 574.0425517 | -0.784613788 | 9.32E-08 |
| AL512625.1 | 409.7626639 | -0.784843665 | 2.32E-02 |
| AC018653.3 | 136.7528489 | -0.785188637 | 2.33E-05 |
| PARP9 | 1481.054975 | -0.78551921 | 1.42E-07 |

| | | | |
|---|---|---|---|
| ACSL4 | 4513.913578 | -0.785534477 | 8.61E-11 |
| TAF9B | 1307.227021 | -0.786118687 | 1.74E-05 |
| CRLF1 | 1052.296416 | -0.786504229 | 1.73E-08 |
| NPIPA5 | 155.7742575 | -0.787274878 | 4.81E-02 |
| ZNF771 | 204.8286513 | -0.787682017 | 2.12E-05 |
| GPR75 | 82.90617554 | -0.787764877 | 1.39E-03 |
| MSI2 | 3793.636912 | -0.787957099 | 1.07E-23 |
| AL929601.2 | 89.03516644 | -0.788131145 | 6.30E-04 |
| AC073857.1 | 101.7834547 | -0.78824616 | 6.19E-03 |
| EP400P1 | 318.6927657 | -0.78847227 | 8.97E-05 |
| LRRC37A2 | 983.0797025 | -0.788518379 | 3.04E-23 |
| SLC9A3R1 | 2447.527257 | -0.788736096 | 1.15E-10 |
| RBAK | 798.7510512 | -0.788943633 | 3.62E-04 |
| AC124312.3 | 178.0884694 | -0.789093542 | 1.76E-04 |
| RAB9B | 263.064607 | -0.789121025 | 3.78E-08 |
| AC018892.3 | 49.79431166 | -0.789445286 | 2.75E-02 |
| RNF125 | 78.72295167 | -0.789577275 | 4.75E-02 |
| PTX3 | 4981.988845 | -0.790575265 | 9.17E-05 |
| NFATC2 | 610.572034 | -0.790600345 | 2.76E-04 |
| PLTP | 6996.727233 | -0.790962239 | 4.96E-08 |
| P2RX6 | 798.6419577 | -0.79149792 | 8.32E-11 |
| AL136295.5 | 119.402497 | -0.791644872 | 3.02E-03 |
| DDIAS | 859.845589 | -0.792068245 | 6.59E-05 |
| ZNF84 | 831.139286 | -0.792193861 | 9.95E-06 |
| USP38 | 1006.367779 | -0.792339708 | 3.07E-09 |
| ZNF211 | 653.8157566 | -0.792912568 | 7.02E-11 |
| TRMT10A | 167.6673084 | -0.792957538 | 1.58E-02 |
| TBXAS1 | 153.5800672 | -0.793026473 | 8.78E-05 |
| C17orf97 | 179.3364957 | -0.793201228 | 5.90E-05 |
| CCNE1 | 424.1412281 | -0.793221963 | 3.76E-08 |
| PDE3B | 420.4228213 | -0.793346073 | 2.07E-05 |
| SEMA6C | 475.9537505 | -0.793818296 | 9.95E-05 |
| TOR1AIP1 | 2235.13473 | -0.7954106 | 9.82E-13 |
| TTLL7 | 129.1930967 | -0.795645239 | 3.82E-02 |
| AP000662.1 | 132.6091028 | -0.7960012 | 2.44E-03 |
| AC022893.1 | 38.04951431 | -0.796377865 | 2.56E-02 |
| ZNF440 | 480.4141961 | -0.796741047 | 1.09E-09 |
| SH3BGRL | 2014.089491 | -0.796777788 | 8.90E-13 |
| SLC44A1 | 3595.730604 | -0.796780737 | 8.80E-27 |
| MAJIN | 69.31755048 | -0.797014646 | 2.58E-02 |
| KALRN | 216.1013953 | -0.797127794 | 2.57E-05 |
| 6-Mar | 13289.27483 | -0.797144683 | 6.54E-25 |
| PRICKLE2 | 161.0608885 | -0.797243323 | 1.90E-02 |
| AC009275.1 | 248.9884021 | -0.797489427 | 9.65E-06 |
| RASSF8-AS1 | 287.0778321 | -0.798038387 | 8.28E-08 |
| NPAS2 | 2199.95089 | -0.798505236 | 6.33E-09 |
| SP3 | 2625.874883 | -0.798679976 | 1.50E-06 |

| | | | |
|---|---|---|---|
| LINC01355 | 100.5912123 | -0.798930393 | 2.55E-02 |
| CU634019.6 | 561.8181909 | -0.799343041 | 3.55E-06 |
| AC244197.3 | 405.6009799 | -0.800256122 | 1.10E-05 |
| GLCE | 3567.17982 | -0.800574496 | 1.64E-13 |
| XRN1 | 1398.276899 | -0.800911351 | 4.02E-03 |
| PRAP1 | 135.3358466 | -0.801063095 | 4.70E-02 |
| ANKIB1 | 5079.40403 | -0.801082529 | 2.39E-12 |
| RIC3 | 102.3729829 | -0.801151876 | 9.76E-03 |
| H1FX-AS1 | 94.09648599 | -0.801173248 | 3.41E-05 |
| METTL25 | 105.2190927 | -0.801617963 | 8.14E-04 |
| AC020917.4 | 47.82870504 | -0.801876687 | 8.77E-03 |
| FLVCR2 | 215.8963486 | -0.802040568 | 1.55E-03 |
| BOC | 1274.836958 | -0.802053394 | 2.54E-09 |
| PLEKHG2 | 2307.085248 | -0.802291254 | 4.86E-10 |
| LHB | 40.4321548 | -0.802336082 | 2.87E-02 |
| CCDC34 | 1220.437521 | -0.803282617 | 2.44E-09 |
| PCM1 | 4712.755582 | -0.803330164 | 5.01E-04 |
| LMO2 | 1198.776963 | -0.803583203 | 2.20E-11 |
| ORC3 | 1863.49752 | -0.804116037 | 1.61E-10 |
| RNFT1 | 531.588277 | -0.804324308 | 2.00E-07 |
| LTN1 | 2205.287303 | -0.805752104 | 4.06E-04 |
| TMEM67 | 702.5993906 | -0.805853932 | 6.61E-04 |
| CD37 | 311.4740181 | -0.805868247 | 1.61E-06 |
| CLUHP3 | 309.9195558 | -0.806371098 | 7.78E-06 |
| STXBP5 | 998.2124252 | -0.806651326 | 9.87E-08 |
| MBTD1 | 632.4450253 | -0.807107905 | 4.60E-06 |
| ERI2 | 682.0807066 | -0.807402425 | 6.25E-06 |
| HELLS | 1113.450738 | -0.807697826 | 2.18E-04 |
| IGHV3-19 | 30.14242899 | -0.807926699 | 2.63E-02 |
| LINC02503 | 75.87403053 | -0.808075094 | 3.88E-02 |
| PHC2 | 12014.71889 | -0.808694855 | 5.72E-11 |
| TMEM135 | 1235.723969 | -0.809233866 | 9.46E-06 |
| AL358334.2 | 74.69140229 | -0.809286428 | 1.30E-03 |
| SETX | 6083.649708 | -0.809842353 | 2.73E-09 |
| SMC6 | 1775.052863 | -0.8099945 | 3.93E-03 |
| KLHL13 | 144.3735064 | -0.810207306 | 1.12E-03 |
| HAUS1 | 1379.441748 | -0.810811152 | 1.31E-19 |
| CIP2A | 1223.598363 | -0.810882693 | 1.89E-03 |
| PIGM | 786.4326629 | -0.810885599 | 1.45E-23 |
| RC3H1 | 637.8260933 | -0.81092172 | 4.67E-07 |
| TUFMP1 | 318.1004999 | -0.811142321 | 1.93E-03 |
| PRRG4 | 102.5023206 | -0.811461729 | 5.92E-03 |
| AC007390.2 | 108.6961985 | -0.811945185 | 2.61E-03 |
| ABCA11P | 119.0527732 | -0.812093429 | 4.91E-03 |
| GCNA | 103.4698222 | -0.8122492 | 4.15E-04 |
| TDG | 846.6086933 | -0.812613193 | 5.97E-09 |
| AC069061.2 | 47.29811414 | -0.813292542 | 6.09E-03 |

| | | | |
|---|---|---|---|
| KIAA0586 | 1386.15036 | -0.813430096 | 1.16E-04 |
| KRR1 | 1171.71655 | -0.813565096 | 3.86E-03 |
| PYM1 | 593.1831054 | -0.813778536 | 1.78E-11 |
| CHST15 | 1086.666004 | -0.813859144 | 2.30E-11 |
| PAQR8 | 949.7430723 | -0.813970887 | 2.63E-11 |
| NA | 140.364548 | -0.814103769 | 1.97E-02 |
| NA | 50.17517098 | -0.814313923 | 1.34E-02 |
| ETFRF1 | 176.6651362 | -0.815138857 | 5.35E-04 |
| AC004951.1 | 210.411599 | -0.815154259 | 7.41E-08 |
| MICE | 40.47422612 | -0.81524214 | 2.65E-02 |
| ANKAR | 93.50273591 | -0.816846351 | 1.02E-02 |
| SCAF11 | 4087.643525 | -0.817106652 | 1.40E-06 |
| ELL3 | 67.9189708 | -0.817322527 | 3.00E-03 |
| CEP126 | 263.8688328 | -0.817598861 | 4.89E-03 |
| USP46-AS1 | 131.0591345 | -0.817933935 | 4.09E-05 |
| TTC41P | 38.05365793 | -0.81801374 | 4.52E-02 |
| YES1 | 1836.174064 | -0.818240686 | 3.16E-05 |
| MINDY4 | 770.5683414 | -0.818249973 | 1.70E-15 |
| YTHDC2 | 2799.831602 | -0.818611082 | 4.67E-05 |
| ZNF124 | 122.9671591 | -0.818984004 | 3.39E-04 |
| BRCA1 | 2010.069614 | -0.819211435 | 1.35E-05 |
| LINC01123 | 472.9229212 | -0.819352248 | 1.49E-07 |
| AC244102.1 | 41.47769037 | -0.819900161 | 1.66E-02 |
| GCNT2 | 168.7839907 | -0.819926035 | 4.56E-05 |
| AC093010.3 | 72.5179379 | -0.820458521 | 2.75E-02 |
| AL031123.2 | 49.34209947 | -0.820580792 | 1.74E-02 |
| CCDC171 | 280.7352548 | -0.820625632 | 4.72E-04 |
| ZNF583 | 197.0848784 | -0.820952034 | 3.86E-03 |
| KIAA0895 | 489.2677565 | -0.821164064 | 2.03E-06 |
| FBXW7 | 632.8639789 | -0.821261902 | 4.64E-10 |
| NA | 209.1836675 | -0.821985091 | 1.28E-05 |
| AL136304.1 | 55.96859215 | -0.822180091 | 7.99E-03 |
| VTA1 | 2431.669244 | -0.822976083 | 8.52E-15 |
| C12orf4 | 972.3065059 | -0.823077156 | 3.27E-06 |
| LINC02557 | 375.7815063 | -0.823078647 | 1.74E-05 |
| TTC30B | 553.0113901 | -0.823319682 | 3.18E-15 |
| NA | 213.3305447 | -0.823351269 | 2.67E-06 |
| OAS2 | 1649.202108 | -0.823561074 | 7.38E-05 |
| ZNF429 | 118.5701516 | -0.82407288 | 1.25E-04 |
| BMPR1A | 1056.786776 | -0.824286734 | 5.92E-05 |
| DDAH2 | 1845.955686 | -0.824355205 | 2.89E-11 |
| CCDC183-AS1 | 300.2302564 | -0.82450952 | 1.57E-06 |
| SBSPON | 767.0994634 | -0.824871261 | 1.13E-06 |
| NEB | 43.88871341 | -0.825270585 | 4.36E-02 |
| ZEB2P1 | 76.62255836 | -0.825645435 | 1.65E-03 |
| CYP2T1P | 71.51101725 | -0.826309025 | 2.12E-03 |
| TRMT10C | 913.7961037 | -0.826456011 | 3.53E-10 |

| | | | |
|---|---|---|---|
| LPIN2 | 5759.958478 | -0.826497244 | 3.60E-15 |
| C6orf120 | 1436.962007 | -0.826527919 | 4.20E-12 |
| ENTPD7 | 448.2118662 | -0.826582506 | 7.53E-13 |
| AC008622.2 | 52.92455998 | -0.826787171 | 4.39E-03 |
| NDUFA6-DT | 233.1582508 | -0.827002592 | 1.70E-07 |
| ADAM22 | 711.1995431 | -0.827247934 | 8.42E-06 |
| CCAR1 | 2806.344201 | -0.827337645 | 4.02E-11 |
| PPP1R16B | 527.7404052 | -0.827574369 | 9.09E-05 |
| FNDC3B | 3823.40014 | -0.827714093 | 5.86E-05 |
| SLC25A20 | 850.5040378 | -0.828333991 | 1.75E-06 |
| IFT74 | 706.580519 | -0.829490465 | 1.99E-03 |
| NTF3 | 90.45861571 | -0.829894713 | 1.20E-03 |
| AC027228.2 | 92.37529988 | -0.82991334 | 8.58E-03 |
| CCT6B | 86.79091986 | -0.830346513 | 1.18E-03 |
| HOXA-AS3 | 169.3705619 | -0.830668822 | 1.42E-04 |
| AF254983.1 | 196.670656 | -0.831675653 | 8.62E-04 |
| CENPK | 922.9012945 | -0.832957878 | 5.14E-04 |
| CICP27 | 149.4400659 | -0.833007145 | 8.51E-07 |
| NA | 83.64413233 | -0.833031229 | 6.24E-04 |
| HAUS3 | 535.9848163 | -0.833067689 | 3.07E-05 |
| ATAD2B | 555.2963023 | -0.833087626 | 1.84E-04 |
| ZNF552 | 298.9944392 | -0.833575329 | 3.12E-08 |
| AC027796.3 | 301.6582265 | -0.833689136 | 1.70E-06 |
| ITPKB | 985.5116325 | -0.833696135 | 9.57E-07 |
| TANC2 | 3184.769522 | -0.833709811 | 1.23E-08 |
| ZNF708 | 103.7787774 | -0.833974287 | 2.17E-03 |
| SESN1 | 517.2971486 | -0.834624913 | 2.98E-07 |
| ZNF569 | 229.4674729 | -0.834636034 | 5.88E-04 |
| LRRC58 | 1475.387555 | -0.834812838 | 2.02E-04 |
| NA | 921.6526783 | -0.835107593 | 5.61E-06 |
| LINC01776 | 61.85722524 | -0.83513025 | 9.03E-03 |
| AP000527.1 | 63.32417353 | -0.83604119 | 6.49E-03 |
| AL606760.1 | 46.42269404 | -0.836071697 | 1.97E-02 |
| RALGAPA1 | 1361.744049 | -0.836473339 | 6.89E-06 |
| CYP3A5 | 68.36392406 | -0.836935668 | 1.67E-03 |
| PLAG1 | 287.0523866 | -0.837096256 | 2.93E-06 |
| GPC3 | 69.20991286 | -0.837212153 | 1.55E-02 |
| LUC7L3 | 3823.723063 | -0.837366629 | 5.83E-04 |
| AC114811.2 | 115.9227166 | -0.837371325 | 8.80E-04 |
| AP006284.1 | 314.2504097 | -0.837575571 | 2.14E-03 |
| ZNF146 | 2240.559327 | -0.837986911 | 3.32E-03 |
| CNTF | 43.98519678 | -0.838147926 | 1.22E-02 |
| NA | 202.4228237 | -0.838292032 | 1.59E-07 |
| AL929601.1 | 134.5228738 | -0.838504853 | 8.77E-06 |
| MMP2 | 12012.51999 | -0.839753835 | 5.03E-09 |
| PDE7A | 1118.518144 | -0.840261558 | 2.14E-11 |
| AC068338.2 | 87.31231627 | -0.840389839 | 4.83E-05 |

| | | | |
|---|---|---|---|
| RAB40B | 390.349998 | -0.840903769 | 2.02E-07 |
| PPM1L | 96.48854693 | -0.841524487 | 8.08E-03 |
| GLP1R | 28.92280827 | -0.841560716 | 3.79E-02 |
| DZIP3 | 1549.200322 | -0.841809766 | 1.69E-04 |
| TET3 | 1452.642177 | -0.842076366 | 2.87E-15 |
| HAT1 | 2459.082348 | -0.842678155 | 8.51E-07 |
| AL450384.2 | 50.7999228 | -0.842969763 | 3.41E-02 |
| BX664615.1 | 109.2441282 | -0.843419521 | 4.13E-05 |
| ZNF654 | 394.779115 | -0.844289277 | 1.39E-06 |
| BDH2 | 851.1838392 | -0.84460115 | 8.25E-16 |
| AL049697.1 | 38.98059825 | -0.845870235 | 4.13E-02 |
| SMARCAD1 | 2144.28931 | -0.846401347 | 2.88E-07 |
| SLC35E3 | 1051.924473 | -0.846669687 | 4.24E-10 |
| CRYBG3 | 1054.976464 | -0.846740227 | 2.58E-03 |
| ZNF772 | 795.6098817 | -0.847063544 | 5.47E-14 |
| FAM227A | 248.8095749 | -0.847250975 | 3.83E-04 |
| CPEB3 | 39.77725826 | -0.848500397 | 1.39E-02 |
| SHC3 | 593.2962704 | -0.848832514 | 1.82E-13 |
| RNF168 | 803.0081132 | -0.849847822 | 2.13E-21 |
| AC104109.2 | 39.74171862 | -0.849946399 | 1.64E-02 |
| TMSB15B | 331.5306934 | -0.85003675 | 2.43E-08 |
| PSPC1 | 1501.329787 | -0.850598309 | 2.32E-14 |
| PRPF4B | 3417.369769 | -0.851098816 | 1.02E-07 |
| AC093627.4 | 67.41982005 | -0.851194068 | 5.89E-03 |
| WNK3 | 384.5832355 | -0.851519575 | 4.85E-09 |
| ZNF30 | 247.4851225 | -0.851861616 | 5.40E-05 |
| AC007485.2 | 149.7281123 | -0.851867592 | 3.67E-06 |
| NID2 | 2791.988157 | -0.8521485 | 5.19E-10 |
| CCDC24 | 384.6898969 | -0.852298999 | 5.00E-08 |
| ZNF137P | 75.91940324 | -0.852787026 | 2.86E-02 |
| P3H4 | 2876.333403 | -0.853880804 | 3.50E-07 |
| AC007952.6 | 69.57289609 | -0.854016296 | 3.04E-03 |
| THUMPD1 | 2204.022143 | -0.854212954 | 1.97E-09 |
| PRKAR1A | 9864.643983 | -0.854708249 | 8.95E-15 |
| NA | 99.97772378 | -0.855378986 | 7.90E-03 |
| KTN1 | 7762.162086 | -0.855564366 | 1.38E-04 |
| RGS11 | 225.5788872 | -0.855893509 | 1.50E-04 |
| SCUBE3 | 863.7147381 | -0.856164605 | 1.49E-08 |
| AC109439.2 | 140.6189782 | -0.857121348 | 6.07E-07 |
| PLGLB1 | 77.66513972 | -0.857170017 | 1.85E-03 |
| AC083799.1 | 89.31240053 | -0.857369885 | 1.17E-02 |
| ALG6 | 407.5561029 | -0.857493955 | 7.18E-07 |
| ANGPT1 | 607.9227096 | -0.857608656 | 2.08E-06 |
| RBM12B-AS1 | 53.27711546 | -0.857674143 | 1.10E-02 |
| ASB9 | 574.9228447 | -0.857781314 | 7.10E-07 |
| OSBPL8 | 2211.075378 | -0.85807387 | 1.69E-03 |
| MDFI | 1896.304313 | -0.858242758 | 2.71E-10 |

| | | | |
|---|---|---|---|
| IL2RB | 1447.939605 | -0.858845951 | 1.05E-10 |
| SWT1 | 219.6789585 | -0.859995387 | 3.28E-04 |
| AC008494.3 | 51.81223161 | -0.860660251 | 1.18E-02 |
| AC012313.3 | 61.80867221 | -0.860881187 | 1.12E-02 |
| AC022400.5 | 62.08055453 | -0.861071957 | 4.84E-03 |
| BRMS1L | 598.3185116 | -0.861260497 | 2.52E-09 |
| C3orf38 | 678.6615537 | -0.861261555 | 2.56E-16 |
| AL353150.1 | 115.1320375 | -0.861567551 | 1.83E-05 |
| ROBO1 | 184.8300666 | -0.861569276 | 4.52E-03 |
| KIAA1324L | 213.9158876 | -0.862637295 | 3.90E-03 |
| AL590705.1 | 49.9269315 | -0.86266684 | 2.50E-03 |
| ZNF724 | 94.80845695 | -0.863323228 | 3.75E-03 |
| CPLANE1 | 2794.991298 | -0.863481826 | 4.46E-05 |
| NA | 53.08899995 | -0.863573807 | 1.88E-02 |
| NRG1 | 723.2771418 | -0.863659632 | 6.31E-09 |
| FZD9 | 356.4608133 | -0.864386134 | 3.07E-06 |
| ADAMTS13 | 316.9800591 | -0.864457968 | 1.90E-08 |
| SPIN3 | 350.5108314 | -0.864641667 | 7.00E-07 |
| PRPF39 | 861.6590797 | -0.865200333 | 3.95E-04 |
| POLR2M | 3888.908804 | -0.865213498 | 6.28E-19 |
| DCUN1D1 | 609.5664724 | -0.865747228 | 1.46E-06 |
| IBTK | 2116.937579 | -0.866127159 | 2.55E-06 |
| AC109466.1 | 33.83691283 | -0.866406555 | 2.53E-02 |
| GPR162 | 140.5579805 | -0.867023663 | 6.71E-04 |
| IGSF9 | 81.60414804 | -0.868011248 | 3.61E-03 |
| PAX9 | 66.37338436 | -0.868473464 | 5.72E-03 |
| AC139491.3 | 70.41335232 | -0.868647574 | 1.08E-02 |
| AC006213.1 | 86.97093276 | -0.868783686 | 1.15E-02 |
| PPP1R1B | 37.78421622 | -0.86911994 | 2.97E-02 |
| AC006333.2 | 155.503166 | -0.869236803 | 2.57E-06 |
| GCSH | 1242.2651 | -0.869829449 | 1.53E-17 |
| SLC29A2 | 765.8787347 | -0.870052198 | 4.34E-07 |
| C2CD6 | 329.5712679 | -0.87041568 | 9.43E-08 |
| TSPAN9 | 2172.099223 | -0.87053125 | 1.59E-10 |
| NDRG2 | 3068.619395 | -0.8705771 | 1.86E-06 |
| TSPY26P | 1120.670006 | -0.870894847 | 9.48E-08 |
| C2CD5 | 1692.653386 | -0.872219475 | 8.03E-07 |
| AL109976.1 | 46.47939616 | -0.872326487 | 1.30E-02 |
| ZNF709 | 132.4124894 | -0.872456452 | 5.36E-03 |
| AC004846.1 | 97.33826787 | -0.872939087 | 3.83E-03 |
| SHLD2 | 793.2501601 | -0.873083955 | 4.18E-07 |
| KIF18A | 852.0039837 | -0.873122321 | 1.32E-03 |
| ZNF43 | 401.036937 | -0.873255797 | 7.09E-11 |
| TMEM173 | 1611.363547 | -0.873291481 | 7.10E-13 |
| MAF | 556.0220594 | -0.87343848 | 6.71E-10 |
| CYP4F12 | 335.7560523 | -0.873916147 | 1.47E-06 |
| SULT1B1 | 66.07862671 | -0.875287751 | 3.94E-03 |

| | | | |
|---|---|---|---|
| ZNF354B | 411.7844575 | -0.875705877 | 1.23E-04 |
| ZNF682 | 145.7408049 | -0.875799113 | 4.02E-05 |
| ZFP14 | 308.8122709 | -0.875931421 | 2.91E-09 |
| PBX1 | 1185.042455 | -0.876170608 | 2.83E-19 |
| SNRK | 1621.869494 | -0.876964483 | 6.01E-17 |
| NCAM1 | 1326.577166 | -0.877251011 | 7.10E-18 |
| ETAA1 | 484.0096387 | -0.877345762 | 3.47E-04 |
| SHOC2 | 769.3246051 | -0.877404203 | 1.14E-03 |
| PJVK | 37.82220968 | -0.877476529 | 2.04E-02 |
| AL512310.11 | 40.74662864 | -0.877480688 | 7.83E-03 |
| NR4A3 | 62.52509383 | -0.877576653 | 3.61E-03 |
| AC253536.6 | 33.45740573 | -0.877906127 | 3.02E-02 |
| CCR1 | 28.49835598 | -0.878337348 | 4.32E-02 |
| PPP1R12A-AS1 | 32.66543125 | -0.878583562 | 4.41E-02 |
| GSAP | 508.953102 | -0.878692529 | 4.24E-10 |
| TMSB15A | 178.0663652 | -0.879625892 | 2.53E-06 |
| DPY19L2P1 | 178.5518984 | -0.880130683 | 2.77E-03 |
| AKAP11 | 1962.018282 | -0.880198283 | 9.94E-05 |
| REST | 1531.355795 | -0.880244916 | 3.92E-11 |
| CBLN3 | 46.02617143 | -0.880699356 | 1.16E-02 |
| CXCR6 | 36.33111727 | -0.881077422 | 5.00E-02 |
| LRRC40 | 617.8751783 | -0.881455952 | 4.48E-03 |
| FAM110D | 49.12210533 | -0.881500522 | 2.49E-02 |
| AL136368.1 | 23.15736852 | -0.88185362 | 3.86E-02 |
| HIST1H4H | 35.41351939 | -0.882008274 | 1.10E-02 |
| CAMK1 | 633.1255623 | -0.882325327 | 2.04E-06 |
| UBXN10 | 251.9791355 | -0.882401804 | 1.97E-07 |
| DMTF1 | 2324.734966 | -0.88358786 | 5.36E-08 |
| AMMECR1 | 735.4118195 | -0.883598267 | 1.16E-14 |
| LRRC8C | 1363.062089 | -0.883681102 | 3.20E-15 |
| CAPZA2 | 3179.046091 | -0.883887556 | 5.28E-08 |
| STRA6 | 466.7762795 | -0.884189012 | 2.45E-05 |
| LRRC39 | 48.17217601 | -0.88430702 | 1.24E-02 |
| AC024896.1 | 158.5161316 | -0.884404582 | 1.90E-06 |
| EEF1A1P19 | 59.86761148 | -0.884643889 | 5.25E-03 |
| SRSF8 | 1740.258175 | -0.884673016 | 4.68E-18 |
| PCDHGC3 | 8838.789475 | -0.884696687 | 3.07E-10 |
| WASIR2 | 30.59058839 | -0.884741827 | 2.87E-02 |
| AC112702.1 | 125.4890082 | -0.884806868 | 1.58E-03 |
| ZNF700 | 334.5364929 | -0.884986636 | 4.53E-05 |
| AL132655.1 | 28.77482108 | -0.885516463 | 2.68E-02 |
| TSPYL2 | 2040.348022 | -0.885807901 | 1.15E-07 |
| ESCO1 | 556.9806551 | -0.886201987 | 2.35E-04 |
| AC010761.1 | 252.8642833 | -0.886443762 | 2.66E-06 |
| SGO2 | 767.3696881 | -0.886603898 | 6.80E-03 |
| AC080038.1 | 8743.391557 | -0.886931299 | 2.88E-10 |
| VPS50 | 1142.578612 | -0.887566041 | 3.56E-05 |

| | | | |
|---|---|---|---|
| AC093110.1 | 87.1898104 | -0.888146188 | 4.17E-04 |
| SLCO2A1 | 1257.747719 | -0.888172807 | 2.28E-17 |
| HSPA7 | 29.38649302 | -0.888336103 | 1.47E-02 |
| ZNF334 | 302.7358372 | -0.888862233 | 3.92E-06 |
| TRMT13 | 719.993185 | -0.888995313 | 4.50E-04 |
| SETD9 | 410.3844377 | -0.889570678 | 3.35E-09 |
| HERC2P9 | 445.6592513 | -0.891050983 | 7.30E-10 |
| ZNF14 | 181.160133 | -0.89142552 | 1.65E-04 |
| NRG2 | 272.570081 | -0.892178454 | 6.19E-08 |
| USP34 | 5304.481039 | -0.892413174 | 1.95E-05 |
| AKAP9 | 1494.681228 | -0.892718042 | 1.29E-03 |
| AC135776.4 | 95.35057604 | -0.894574216 | 4.01E-03 |
| AC102945.2 | 89.66476115 | -0.894920008 | 3.04E-04 |
| GPX7 | 858.5942305 | -0.895154462 | 1.50E-09 |
| NR3C1 | 4920.832451 | -0.895467843 | 3.79E-17 |
| NA | 149.0104143 | -0.89550673 | 2.44E-04 |
| IZUMO4 | 100.2450413 | -0.895912768 | 9.07E-05 |
| MYO5C | 156.771633 | -0.896502494 | 1.20E-02 |
| TNRC6C-AS1 | 1271.337069 | -0.896638794 | 9.62E-11 |
| AC026691.1 | 44.42698353 | -0.896858234 | 5.89E-03 |
| NA | 34.38690557 | -0.896952145 | 3.31E-02 |
| ANKRD36C | 322.2887014 | -0.896983222 | 4.58E-06 |
| BORCS7 | 376.7341782 | -0.89698474 | 7.67E-09 |
| PPWD1 | 2051.396356 | -0.897313302 | 6.85E-07 |
| THOC2 | 2817.232692 | -0.899115014 | 9.73E-05 |
| SUOX | 1358.723531 | -0.899154323 | 7.15E-08 |
| THNSL2 | 53.591121 | -0.899376762 | 3.04E-02 |
| GSKIP | 690.2447286 | -0.899568933 | 3.30E-05 |
| ADAMTS14 | 70.35092211 | -0.900056165 | 1.84E-02 |
| ZSCAN31 | 211.2724114 | -0.900308975 | 2.44E-05 |
| ARL15 | 968.3590657 | -0.900310024 | 2.35E-12 |
| GPRASP1 | 51.27077291 | -0.900357578 | 1.14E-02 |
| RFX7 | 2054.624167 | -0.900705751 | 5.31E-16 |
| CYP27C1 | 309.7320071 | -0.900895441 | 1.06E-07 |
| ZNF503-AS2 | 106.5020803 | -0.901177671 | 6.74E-05 |
| ST8SIA6 | 68.51548135 | -0.901660033 | 2.75E-03 |
| PUS7L | 453.9353949 | -0.902262261 | 1.13E-03 |
| ACP2 | 3153.085252 | -0.902682867 | 9.27E-17 |
| TGFA | 403.9619259 | -0.902991799 | 3.29E-04 |
| CCDC151 | 67.50158075 | -0.903106273 | 4.07E-02 |
| ZBTB6 | 524.4678034 | -0.903544626 | 7.46E-06 |
| DBI | 4740.869507 | -0.903571981 | 2.86E-16 |
| TOP2B | 4796.333484 | -0.904218743 | 7.39E-10 |
| AL133355.1 | 44.43468705 | -0.904406163 | 7.53E-03 |
| TTC33 | 553.6499434 | -0.904522928 | 3.00E-07 |
| ARHGAP24 | 86.39537728 | -0.905373955 | 3.17E-04 |
| JUP | 7928.88656 | -0.905419778 | 2.49E-09 |

| | | | |
|---|---|---|---|
| HLA-DRA | 272.0727889 | -0.905722514 | 7.75E-04 |
| CASP8AP2 | 923.2155655 | -0.906325578 | 4.55E-04 |
| CCNT2 | 1129.848901 | -0.906789901 | 3.64E-05 |
| CLK1 | 1374.728739 | -0.907160843 | 1.80E-05 |
| ZNF41 | 598.0445322 | -0.907668534 | 1.20E-08 |
| SMPD1 | 2976.700773 | -0.908423194 | 9.80E-11 |
| AC010226.1 | 53.78791204 | -0.909827651 | 6.36E-03 |
| ZNF260 | 617.7063357 | -0.910002118 | 2.95E-04 |
| NA | 138.1124267 | -0.910269732 | 4.38E-03 |
| EMID1 | 737.0210262 | -0.910313855 | 7.90E-08 |
| CYP26A1 | 160.8035835 | -0.910952296 | 1.64E-05 |
| MINPP1 | 1359.435503 | -0.911072045 | 5.83E-15 |
| RBP5 | 155.6224082 | -0.912582045 | 2.16E-09 |
| ERVW-1 | 167.7450715 | -0.912589208 | 2.92E-06 |
| AC005703.6 | 31.16690174 | -0.913158612 | 1.64E-02 |
| NAB1 | 2310.147904 | -0.913454952 | 6.13E-12 |
| C8orf48 | 96.08804137 | -0.913743892 | 2.14E-04 |
| AC015922.3 | 166.9699555 | -0.913827347 | 1.76E-07 |
| GPT | 53.48594233 | -0.914409397 | 4.22E-03 |
| FAM126A | 3391.101266 | -0.914589473 | 8.48E-09 |
| MTERF2 | 526.1795391 | -0.914740963 | 2.48E-10 |
| ZNF432 | 386.5222351 | -0.914968386 | 9.25E-13 |
| GLMN | 563.3816818 | -0.915018929 | 5.50E-08 |
| CD207 | 40.47290139 | -0.916042527 | 4.03E-03 |
| CARF | 206.4300318 | -0.916124937 | 1.98E-04 |
| CENPQ | 301.7951478 | -0.916533077 | 5.70E-07 |
| IL10RB-DT | 121.8730826 | -0.916705475 | 1.89E-03 |
| SHLD3 | 185.7210417 | -0.916790001 | 2.66E-08 |
| DENND1B | 124.7418185 | -0.916836682 | 6.22E-03 |
| SNX22 | 156.8170892 | -0.916892304 | 2.68E-05 |
| AC011450.1 | 89.0077873 | -0.917140865 | 2.02E-04 |
| ZNF235 | 385.1857377 | -0.917421812 | 2.59E-08 |
| ZC3H12C | 987.4220162 | -0.917522502 | 1.19E-15 |
| NQO1 | 36091.71617 | -0.91815824 | 1.29E-08 |
| DKKL1 | 62.86708431 | -0.918294741 | 2.14E-03 |
| SPDYE2 | 89.92486882 | -0.918730377 | 7.02E-04 |
| TRANK1 | 2119.181798 | -0.918788435 | 2.08E-07 |
| RASSF8 | 897.5966724 | -0.919162912 | 3.92E-05 |
| FAM222A | 185.6045506 | -0.919565712 | 3.04E-06 |
| PRMT6 | 1118.533194 | -0.919774532 | 3.34E-22 |
| NUDT12 | 724.1957269 | -0.92026827 | 2.38E-11 |
| AC112907.3 | 48.88332524 | -0.920438214 | 2.41E-02 |
| AC090617.5 | 55.59406078 | -0.920553887 | 1.74E-03 |
| FPGT | 345.5712041 | -0.920627892 | 1.49E-05 |
| PANK1 | 348.7559744 | -0.920694278 | 8.59E-09 |
| ZNF638 | 3723.556478 | -0.920804482 | 1.55E-05 |
| SYNE2 | 2558.962612 | -0.920810477 | 6.97E-11 |

| | | | |
|---|---|---|---|
| CEP85L | 295.6479056 | -0.921045738 | 8.32E-05 |
| NA | 92.09315874 | -0.921654497 | 1.77E-04 |
| SLC35A1 | 1520.866646 | -0.921892071 | 1.91E-10 |
| A2M | 8519.613877 | -0.922374136 | 2.05E-04 |
| RNF152 | 191.2408013 | -0.923047765 | 1.69E-02 |
| WNT6 | 159.8552245 | -0.923212164 | 8.18E-04 |
| AC021504.1 | 106.4338101 | -0.923336508 | 2.59E-04 |
| LAMTOR3 | 1013.807565 | -0.923707854 | 1.31E-10 |
| CLK4 | 476.8723821 | -0.923734631 | 1.20E-05 |
| MEIS2 | 693.3792644 | -0.924726495 | 1.70E-12 |
| AC020915.3 | 223.8893518 | -0.925468302 | 3.55E-10 |
| HOMER1 | 828.6769183 | -0.925740585 | 1.28E-09 |
| TRA2A | 2138.299833 | -0.925903905 | 2.53E-17 |
| KDM8 | 224.6049505 | -0.926272201 | 4.84E-08 |
| AC137767.1 | 53.94270911 | -0.926559743 | 6.84E-04 |
| ATP11C | 2323.681458 | -0.926664222 | 2.47E-07 |
| USP32P3 | 29.91988469 | -0.927149651 | 2.28E-02 |
| TTK | 1450.743421 | -0.927898646 | 2.23E-03 |
| E2F8 | 533.6095413 | -0.92790071 | 3.35E-08 |
| DACT1 | 395.7859807 | -0.928368855 | 3.21E-06 |
| GUSBP3 | 41.01851898 | -0.928417456 | 1.10E-02 |
| HMGN5 | 66.92264094 | -0.928463924 | 3.00E-02 |
| CCNG2 | 1339.046455 | -0.929008652 | 1.46E-12 |
| UBE2D1 | 283.642357 | -0.929577243 | 2.03E-07 |
| ZBTB46 | 1939.837957 | -0.929985553 | 1.84E-10 |
| AC003102.1 | 62.22034967 | -0.930219676 | 1.30E-03 |
| AC010595.1 | 65.32033256 | -0.93035236 | 1.73E-03 |
| AC023355.1 | 61.08893835 | -0.930889231 | 3.57E-02 |
| ZNF37BP | 389.2242676 | -0.930927468 | 2.54E-05 |
| DM1-AS | 32.87255979 | -0.931171723 | 2.78E-02 |
| MPZL3 | 518.1093127 | -0.931229847 | 3.64E-10 |
| TCAF1 | 6221.734152 | -0.931443247 | 1.08E-08 |
| DMRTA1 | 131.8968401 | -0.931919914 | 1.29E-03 |
| AP000808.1 | 86.69107461 | -0.932279429 | 6.35E-04 |
| NA | 25.92113813 | -0.932279661 | 4.25E-02 |
| ZNF641 | 822.7066855 | -0.932918949 | 7.14E-08 |
| EML6 | 415.0599144 | -0.933066991 | 9.20E-04 |
| DNAJC12 | 164.2504545 | -0.933961673 | 7.10E-07 |
| FMO5 | 105.7304208 | -0.934095901 | 1.09E-03 |
| TOGARAM1 | 1182.064204 | -0.934659959 | 7.28E-07 |
| CLDN2 | 59.31407275 | -0.935158541 | 1.52E-02 |
| ODR4 | 586.5015614 | -0.935562787 | 1.25E-04 |
| AC240565.1 | 50.67336097 | -0.935946383 | 2.53E-02 |
| ODF2L | 553.6489667 | -0.936553149 | 3.47E-03 |
| NCOA2 | 1947.675613 | -0.937364484 | 3.16E-14 |
| MDM1 | 410.0232745 | -0.937432918 | 1.68E-05 |
| STYX | 677.5133932 | -0.937882154 | 6.68E-10 |

| | | | |
|---|---|---|---|
| ZSCAN1 | 53.59253485 | -0.938141706 | 1.09E-02 |
| AL157838.1 | 39.49941962 | -0.938390112 | 8.36E-03 |
| ZNF181 | 409.5785145 | -0.938604341 | 1.88E-05 |
| CTXN1 | 271.998645 | -0.939064464 | 4.82E-10 |
| NA | 25.22626024 | -0.939684356 | 4.11E-02 |
| FNDC10 | 630.1756976 | -0.939686836 | 1.78E-09 |
| FRS2 | 817.1134476 | -0.939934157 | 1.54E-08 |
| HOXB-AS3 | 267.2395337 | -0.940551505 | 1.19E-08 |
| NEMF | 1174.676923 | -0.940748006 | 2.71E-03 |
| ROCK1 | 2028.443087 | -0.940863405 | 1.46E-04 |
| C2orf76 | 103.0839047 | -0.941283222 | 2.19E-05 |
| BCLAF1 | 5637.954325 | -0.941461204 | 8.97E-08 |
| MFSD1 | 1429.551435 | -0.941594765 | 9.05E-16 |
| TBL1XR1 | 2738.467211 | -0.941999614 | 3.80E-08 |
| KRBA2 | 422.1868317 | -0.94202702 | 4.37E-17 |
| PPP4R2 | 1886.522598 | -0.94452032 | 2.97E-09 |
| ZNF184 | 394.1955242 | -0.944946065 | 8.88E-10 |
| RBM45 | 475.9013446 | -0.946330186 | 2.99E-11 |
| EIF3EP1 | 83.17454681 | -0.946336486 | 1.20E-04 |
| ZNF224 | 508.5609643 | -0.946598249 | 4.41E-08 |
| FNIP2 | 1389.398079 | -0.947070313 | 1.87E-18 |
| CASP7 | 1127.428638 | -0.947095395 | 1.93E-13 |
| TESC | 28.26307814 | -0.947607475 | 2.45E-02 |
| AC020951.1 | 35.13358648 | -0.947802068 | 8.56E-03 |
| AL133406.2 | 27.78123807 | -0.948459301 | 1.19E-02 |
| TMEM14A | 661.6508551 | -0.948705902 | 7.59E-15 |
| CEP83 | 365.6925099 | -0.949206461 | 4.05E-03 |
| LMBRD2 | 905.4643708 | -0.949623246 | 2.03E-04 |
| KIAA1109 | 3318.943946 | -0.949839585 | 1.38E-04 |
| AC015799.1 | 53.59283895 | -0.949877898 | 1.51E-03 |
| STXBP3 | 829.6540019 | -0.949985738 | 9.63E-05 |
| CSAD | 412.1667877 | -0.950022297 | 9.77E-12 |
| MUC13 | 39.74750968 | -0.950165682 | 1.39E-02 |
| CDKL5 | 533.1492249 | -0.950622845 | 9.36E-07 |
| SRBD1 | 1161.090299 | -0.951265869 | 1.29E-07 |
| NIPBL | 5744.833362 | -0.951839199 | 3.85E-04 |
| AC138409.2 | 185.7386091 | -0.952199751 | 5.86E-07 |
| EBLN2 | 45.02589026 | -0.952279545 | 1.11E-02 |
| ZNF252P | 1467.129611 | -0.95336298 | 3.09E-07 |
| ZNF107 | 246.4897489 | -0.954579584 | 2.68E-05 |
| AL359878.1 | 47.04444433 | -0.954628079 | 1.97E-03 |
| RARRES3 | 1750.500845 | -0.956268581 | 2.72E-07 |
| SPRED2 | 2021.337216 | -0.956699874 | 1.32E-18 |
| AL359644.1 | 20.06275388 | -0.956780231 | 4.05E-02 |
| SNORA26 | 19.2884351 | -0.957302059 | 4.19E-02 |
| NEK7 | 933.9272831 | -0.957571888 | 4.39E-07 |
| AKR7L | 26.4332983 | -0.958504229 | 3.90E-02 |

| | | | |
|---|---|---|---|
| AC007996.1 | 132.0789388 | -0.959320179 | 1.78E-05 |
| PHOSPHO2 | 116.2748943 | -0.959780187 | 1.81E-04 |
| EPM2AIP1 | 1443.908971 | -0.960939947 | 2.18E-05 |
| GSTA4 | 526.6998491 | -0.960972369 | 5.18E-14 |
| C6orf163 | 32.03509104 | -0.961240027 | 1.68E-02 |
| ZNF443 | 150.5584489 | -0.961641258 | 3.22E-06 |
| MINDY1 | 1011.666293 | -0.961656661 | 2.04E-08 |
| AL355488.1 | 83.60491341 | -0.961895766 | 1.47E-03 |
| GALNT1 | 2272.25254 | -0.962046666 | 1.48E-09 |
| KIAA0895L | 893.3242262 | -0.962428518 | 2.23E-17 |
| MYL5 | 176.9455721 | -0.962627088 | 1.55E-06 |
| CCDC189 | 255.0331238 | -0.962948658 | 9.48E-08 |
| TMSB15B-AS1 | 41.70051277 | -0.963078552 | 3.16E-02 |
| CAMKV | 31.31748181 | -0.963086674 | 1.35E-02 |
| POLI | 891.7794694 | -0.963753097 | 1.06E-07 |
| ARHGAP18 | 3596.763254 | -0.964427598 | 7.33E-31 |
| SLC38A4 | 1027.095872 | -0.965219942 | 2.81E-12 |
| ZNF292 | 1527.571376 | -0.965269762 | 1.33E-04 |
| AC087501.4 | 28.25252368 | -0.965388278 | 3.64E-02 |
| LINC01238 | 72.75569433 | -0.967096258 | 1.23E-04 |
| MIER3 | 2154.762199 | -0.967144315 | 2.63E-06 |
| CX3CL1 | 347.8028956 | -0.96745404 | 2.04E-05 |
| PAPLN | 1234.774959 | -0.967550549 | 5.19E-15 |
| DACT3 | 383.0030205 | -0.967935961 | 9.19E-08 |
| ZNF271P | 785.0492098 | -0.967967286 | 4.91E-08 |
| PHTF1 | 1158.451581 | -0.967981266 | 7.41E-23 |
| PKD2 | 2184.663901 | -0.968041023 | 5.64E-23 |
| PLEKHH2 | 523.7251953 | -0.968159716 | 2.00E-04 |
| TTC28 | 2641.399182 | -0.968577042 | 8.17E-10 |
| MMP9 | 160.911846 | -0.968922244 | 3.42E-03 |
| WASHC4 | 2069.96772 | -0.96916739 | 5.34E-05 |
| NKTR | 2044.428997 | -0.97026753 | 1.84E-04 |
| KBTBD7 | 448.7049977 | -0.970642658 | 1.41E-10 |
| FRMD6-AS1 | 136.6120981 | -0.970953682 | 2.33E-05 |
| NFAM1 | 176.4578415 | -0.971271784 | 1.11E-03 |
| DNM1P47 | 297.1681704 | -0.971400042 | 2.12E-06 |
| CCDC69 | 1021.222201 | -0.971501259 | 1.29E-08 |
| ADAM10 | 7600.927323 | -0.971990573 | 2.66E-18 |
| MIR3142HG | 184.9959843 | -0.972066044 | 2.97E-07 |
| COL4A5 | 2752.108464 | -0.972312658 | 2.09E-15 |
| ASPM | 1737.093246 | -0.972354577 | 1.99E-02 |
| CWC22 | 1209.979763 | -0.972626663 | 1.09E-04 |
| NAIP | 402.0713656 | -0.973345474 | 1.05E-11 |
| CLSTN3 | 6610.167495 | -0.97349107 | 3.81E-08 |
| AC126768.3 | 60.52388494 | -0.974355227 | 1.39E-02 |
| AL133215.2 | 51.61394003 | -0.974861318 | 1.07E-03 |
| AC126335.2 | 44.10181047 | -0.974956816 | 4.54E-02 |

| | | | |
|---|---|---|---|
| PFN1P6 | 39.48164717 | -0.97562741 | 5.75E-03 |
| PNISR | 4426.738011 | -0.97571381 | 2.11E-06 |
| DNASE1L2 | 80.57937059 | -0.976050633 | 1.62E-04 |
| NA | 130.3090386 | -0.976385858 | 4.37E-06 |
| SCFD1 | 2000.414519 | -0.977473073 | 1.37E-07 |
| ATXN1 | 3041.292894 | -0.977893044 | 6.62E-24 |
| IFT81 | 713.7989242 | -0.977910725 | 1.02E-07 |
| CEACAM1 | 714.1367716 | -0.978307672 | 3.67E-06 |
| AC022007.1 | 130.4691102 | -0.978761676 | 9.48E-07 |
| NA | 27.8812932 | -0.97888999 | 4.46E-02 |
| AL353803.2 | 202.4477704 | -0.979783182 | 1.42E-05 |
| XKR5 | 178.029926 | -0.980308883 | 1.73E-08 |
| KCP | 42.91701193 | -0.980773121 | 1.53E-02 |
| AC007406.5 | 157.0453592 | -0.98094455 | 2.13E-06 |
| AC109347.1 | 25.84637001 | -0.980986479 | 4.57E-02 |
| KCND1 | 722.1343561 | -0.98100457 | 8.54E-09 |
| TACR2 | 92.22274491 | -0.982179324 | 5.59E-04 |
| COPS2 | 1367.152129 | -0.982257324 | 1.17E-04 |
| BMT2 | 620.0680209 | -0.982931901 | 1.27E-12 |
| AC005046.1 | 173.6432524 | -0.983093235 | 4.86E-04 |
| HS2ST1 | 2383.667486 | -0.983286821 | 8.07E-12 |
| SLC18B1 | 776.1425478 | -0.983305438 | 2.65E-22 |
| ZNF148 | 1250.193518 | -0.983613213 | 2.82E-08 |
| FAM153B | 429.3182156 | -0.983720434 | 3.23E-06 |
| C1orf21 | 2338.356182 | -0.984472751 | 5.47E-14 |
| ZNF136 | 332.4460914 | -0.984611125 | 1.57E-07 |
| WDHD1 | 1399.6444 | -0.985057137 | 1.94E-05 |
| ANKRD12 | 1010.776964 | -0.985377383 | 4.08E-04 |
| DYRK1B | 543.7450149 | -0.985473828 | 7.71E-09 |
| LIMCH1 | 5167.393543 | -0.985620012 | 2.35E-10 |
| ZNF484 | 334.2493911 | -0.985732105 | 1.08E-05 |
| NXT2 | 357.2870437 | -0.986011538 | 1.14E-10 |
| FOXC1 | 860.7224109 | -0.987818474 | 4.56E-13 |
| UPF2 | 1004.08589 | -0.988065079 | 2.35E-06 |
| AP005131.6 | 30.74446118 | -0.989262966 | 8.19E-03 |
| CHM | 1261.466858 | -0.989357679 | 1.10E-05 |
| PANO1 | 51.604583 | -0.989399436 | 9.25E-04 |
| NF1P10 | 67.28745981 | -0.989739489 | 2.60E-03 |
| SLC7A14 | 53.52073558 | -0.990120168 | 3.00E-02 |
| SSX3 | 30.18188152 | -0.990385219 | 1.15E-02 |
| PSD3 | 2830.348069 | -0.990506577 | 1.56E-18 |
| AC004542.2 | 51.77550836 | -0.990610674 | 4.03E-02 |
| SPATA6 | 471.5236482 | -0.990687075 | 1.61E-13 |
| NFATC4 | 3878.575819 | -0.991016453 | 1.10E-12 |
| MAN1C1 | 266.1644314 | -0.99102178 | 5.38E-05 |
| ZNF441 | 190.1687265 | -0.991420558 | 5.10E-03 |
| MST1 | 313.337359 | -0.99150229 | 2.34E-08 |

| | | | |
|---|---|---|---|
| PGBD5 | 220.9593528 | -0.991701934 | 8.76E-05 |
| BCL2L11 | 920.1726165 | -0.992537489 | 1.38E-14 |
| ZNF362 | 968.369252 | -0.992570267 | 5.34E-07 |
| ANKRD50 | 1173.25523 | -0.993158093 | 6.90E-13 |
| LOXL1 | 102.2955771 | -0.993274353 | 7.30E-04 |
| PCDHGA7 | 577.8186976 | -0.993335364 | 2.88E-11 |
| ZNF345 | 182.6037155 | -0.994087705 | 1.42E-08 |
| AC015922.2 | 73.21311534 | -0.994621269 | 1.33E-04 |
| SPRN | 304.4446147 | -0.994629851 | 9.40E-09 |
| SPSB4 | 329.8186146 | -0.994759548 | 2.32E-04 |
| SYNE4 | 22.45815938 | -0.995474296 | 2.64E-02 |
| AC087292.1 | 79.73504335 | -0.99556275 | 1.85E-02 |
| AC007731.1 | 33.03692063 | -0.995610981 | 1.12E-02 |
| AC092279.1 | 230.0617854 | -0.995730785 | 1.09E-13 |
| AL391987.4 | 23.88118373 | -0.995981117 | 1.86E-02 |
| UGT8 | 101.0890748 | -0.996019655 | 1.43E-05 |
| POLK | 1122.115659 | -0.996055292 | 2.01E-04 |
| NA | 39.83765738 | -0.996254191 | 1.71E-03 |
| IFT80 | 1020.407039 | -0.996891618 | 2.67E-04 |
| AC007382.1 | 86.90946211 | -0.997659036 | 4.35E-04 |
| AL365205.1 | 676.4159332 | -0.997695497 | 2.95E-09 |
| AP002495.2 | 126.5569363 | -0.997722068 | 8.81E-06 |
| AL137003.1 | 206.3088742 | -0.997771962 | 2.11E-07 |
| NA | 27.53951476 | -0.998286651 | 1.54E-02 |
| RGPD6 | 2488.190399 | -0.998399925 | 5.69E-08 |
| LINC02482 | 133.6517581 | -0.998838227 | 3.72E-08 |
| KRTCAP3 | 21.62333393 | -0.999548715 | 2.62E-02 |
| APOE | 6958.007481 | -0.999792441 | 1.28E-06 |
| XAF1 | 1197.67208 | -1.000484249 | 1.06E-09 |
| NA | 94.5777297 | -1.000649501 | 5.82E-07 |
| AC012313.10 | 41.82089725 | -1.000791336 | 3.71E-03 |
| CEP19 | 554.178459 | -1.00098733 | 4.08E-17 |
| ROM1 | 228.2174532 | -1.001206218 | 1.09E-05 |
| AC138907.9 | 36.3952997 | -1.00153401 | 2.56E-02 |
| NACAD | 1177.507431 | -1.002184415 | 1.10E-08 |
| AL928970.1 | 299.1134828 | -1.003145932 | 1.87E-12 |
| NPIPA7 | 80.15957517 | -1.004044771 | 3.13E-03 |
| NEIL1 | 519.4268325 | -1.004698744 | 1.05E-08 |
| GAL3ST4 | 1047.330578 | -1.004751222 | 4.09E-11 |
| RBM25 | 4151.031687 | -1.004971442 | 1.44E-14 |
| C11orf96 | 147.5207521 | -1.005160124 | 3.92E-06 |
| DNAAF4 | 158.9747061 | -1.005211664 | 2.91E-06 |
| SUGT1P4-STRA6 | 208.5434078 | -1.006795285 | 3.96E-08 |
| PCDHB5 | 79.84968752 | -1.007816577 | 4.32E-04 |
| LIFR-AS1 | 357.2237635 | -1.008420064 | 1.46E-12 |
| LGR6 | 65.23759692 | -1.008902983 | 9.22E-03 |
| FBXO6 | 243.709265 | -1.009047682 | 1.69E-07 |

| | | | |
|---|---|---|---|
| AC109322.1 | 83.15132005 | -1.009524051 | 1.87E-05 |
| RIDA | 596.8398994 | -1.009592836 | 2.52E-11 |
| HOTTIP | 31.11813955 | -1.011210742 | 7.19E-03 |
| LRRC46 | 47.82199529 | -1.011662538 | 6.96E-03 |
| CYP2D6 | 86.52775803 | -1.012940421 | 4.71E-05 |
| AC106820.3 | 46.58828501 | -1.013070411 | 4.19E-03 |
| ZNF625 | 116.0250302 | -1.013087853 | 1.67E-03 |
| MLLT11 | 927.8788135 | -1.013116764 | 1.95E-17 |
| 7-Sep | 8237.371082 | -1.013361412 | 1.91E-07 |
| DNAJC28 | 72.00157702 | -1.013430704 | 1.08E-03 |
| RICTOR | 2048.459973 | -1.013959136 | 1.11E-04 |
| AL096870.2 | 67.05680501 | -1.015692405 | 1.33E-04 |
| AC092375.2 | 32.97172105 | -1.016632589 | 7.19E-03 |
| LGI4 | 122.8848522 | -1.0166502 | 2.93E-05 |
| AC010319.4 | 27.61844253 | -1.016968187 | 1.32E-02 |
| TTC3 | 9113.083029 | -1.017621866 | 9.41E-16 |
| RGPD5 | 665.842043 | -1.017687557 | 3.46E-03 |
| ATM | 4856.471657 | -1.017742035 | 1.70E-05 |
| KIF20B | 638.9915225 | -1.018012108 | 1.88E-03 |
| AC127024.5 | 44.36894571 | -1.018190152 | 1.21E-03 |
| SHBG | 42.47836588 | -1.018961635 | 4.33E-02 |
| INTS6L | 853.156252 | -1.018994396 | 1.17E-11 |
| TRIM23 | 971.8216866 | -1.020196956 | 4.07E-05 |
| AP001107.9 | 109.1125027 | -1.02028474 | 1.39E-05 |
| TRIM22 | 535.1244149 | -1.021606312 | 1.55E-08 |
| SPOCK2 | 70.07807607 | -1.021745726 | 6.48E-03 |
| SLC9A3-AS1 | 632.0318409 | -1.021990786 | 3.64E-09 |
| CFP | 142.794171 | -1.022237422 | 1.50E-04 |
| CHD1 | 2713.459699 | -1.022266027 | 5.89E-06 |
| ARRB1 | 3818.529655 | -1.022362035 | 3.64E-14 |
| AP001486.2 | 32.71344285 | -1.0226045 | 2.10E-02 |
| ZFP3 | 574.7828544 | -1.022832531 | 1.90E-22 |
| TBC1D19 | 280.9976935 | -1.025045793 | 4.40E-08 |
| MAP9 | 380.48986 | -1.026079845 | 2.56E-04 |
| PRDM1 | 260.727632 | -1.026156601 | 4.32E-04 |
| VPS13B | 2695.859008 | -1.026334406 | 1.68E-16 |
| LINC01126 | 42.41739339 | -1.027460066 | 1.62E-03 |
| AL592435.1 | 78.35341732 | -1.027941404 | 1.20E-03 |
| AL121594.1 | 307.8997471 | -1.028004904 | 1.18E-05 |
| SLC4A7 | 1002.091398 | -1.028261847 | 4.19E-06 |
| ICA1L | 195.1054069 | -1.028544769 | 9.93E-09 |
| IGDCC3 | 90.23794552 | -1.028877452 | 6.99E-06 |
| ESAM | 145.5043028 | -1.029392054 | 4.08E-04 |
| BMI1 | 922.8108579 | -1.030502821 | 1.16E-04 |
| GBP4 | 532.7411693 | -1.030629263 | 9.78E-06 |
| BLZF1 | 801.6579964 | -1.031426302 | 2.56E-06 |
| RNF32 | 194.6641487 | -1.032245593 | 8.03E-07 |

| | | | |
|---|---|---|---|
| SERINC5 | 1988.56082 | -1.032434926 | 1.13E-24 |
| NA | 25.35615402 | -1.032584385 | 2.65E-02 |
| ARHGEF35 | 220.0362306 | -1.032790835 | 5.43E-08 |
| AC010542.5 | 24.71451806 | -1.033692237 | 4.90E-02 |
| ARID4B | 1086.545711 | -1.034454496 | 6.59E-06 |
| CITED1 | 34.09631409 | -1.034727987 | 1.45E-02 |
| FAXDC2 | 500.5299023 | -1.037291224 | 1.34E-09 |
| ZNF326 | 1281.750808 | -1.03734896 | 5.24E-10 |
| TRO | 3618.311323 | -1.038158719 | 7.81E-15 |
| USP53 | 1004.853788 | -1.038396118 | 4.03E-05 |
| ZNF518A | 469.4591773 | -1.039054846 | 2.23E-04 |
| AL773545.3 | 36.11612349 | -1.039322453 | 9.48E-03 |
| CCN4 | 138.2887904 | -1.039506029 | 2.40E-02 |
| APOD | 125.6884614 | -1.039588246 | 5.40E-04 |
| PPIG | 1943.519505 | -1.03966403 | 5.42E-07 |
| LRP6 | 2588.5102 | -1.04007746 | 2.46E-17 |
| ATPSCKMT | 793.0329605 | -1.040175056 | 9.17E-21 |
| AC135178.2 | 138.3094543 | -1.040369955 | 4.98E-07 |
| SPOCK1 | 3684.499858 | -1.040415468 | 3.21E-25 |
| PSMA3-AS1 | 956.8211118 | -1.040454139 | 3.10E-10 |
| KCNC1 | 90.05506287 | -1.04135757 | 2.58E-04 |
| CSGALNACT1 | 578.8533399 | -1.04165399 | 1.13E-12 |
| DRAXIN | 93.60471832 | -1.042351994 | 6.70E-05 |
| EFNB2 | 435.0140339 | -1.042367053 | 7.29E-20 |
| AL021068.1 | 38.23277759 | -1.04274668 | 6.03E-03 |
| CCDC180 | 210.7067393 | -1.043044466 | 2.29E-05 |
| ACY3 | 92.98786841 | -1.043366329 | 2.49E-04 |
| NR1H3 | 1127.557574 | -1.043571842 | 4.45E-10 |
| ZNF302 | 1153.441241 | -1.044268572 | 3.40E-08 |
| ZMYM2 | 1599.798682 | -1.044288182 | 7.07E-15 |
| PINLYP | 28.72624574 | -1.044300751 | 4.64E-02 |
| AP003396.5 | 77.13102264 | -1.04696666 | 2.17E-02 |
| TEX14 | 48.118117 | -1.047351762 | 1.67E-02 |
| CHN2 | 294.5890476 | -1.047747655 | 8.85E-14 |
| DNAJB14 | 680.7839592 | -1.047828157 | 6.55E-05 |
| ZNF879 | 144.5956642 | -1.047898845 | 5.70E-04 |
| ZNF658 | 263.7837688 | -1.048438151 | 2.05E-07 |
| SREK1 | 3158.431512 | -1.048827124 | 3.15E-05 |
| AC112484.3 | 34.69759026 | -1.049504652 | 5.72E-03 |
| TOB1 | 570.1577494 | -1.049680553 | 5.65E-10 |
| TEN1-CDK3 | 280.7781373 | -1.049773724 | 1.10E-08 |
| AC126335.1 | 71.26480399 | -1.049812503 | 7.52E-04 |
| AUTS2 | 1479.629524 | -1.050081239 | 1.77E-14 |
| AC083805.2 | 26.76387639 | -1.050920298 | 3.71E-02 |
| ING3 | 694.3881223 | -1.051070954 | 1.12E-07 |
| CD4 | 153.7742743 | -1.052239634 | 8.27E-05 |
| NPW | 37.18962742 | -1.052372375 | 8.29E-03 |

| | | | |
|---|---|---|---|
| ANK3 | 152.7292105 | -1.052891599 | 6.64E-03 |
| ZNF519 | 152.1448506 | -1.053099381 | 6.05E-04 |
| ASPG | 144.3937042 | -1.053852025 | 2.90E-03 |
| RTP4 | 188.0812098 | -1.054018451 | 2.49E-08 |
| STPG3 | 31.27329893 | -1.054723829 | 2.50E-03 |
| EHD3 | 6392.771573 | -1.054822664 | 4.22E-13 |
| TSC22D3 | 3742.290724 | -1.055876242 | 1.17E-11 |
| ZNF567 | 254.7474616 | -1.056163005 | 2.61E-05 |
| CCDC78 | 384.8868074 | -1.05647462 | 8.72E-12 |
| ZNF239 | 420.1033766 | -1.057046981 | 1.91E-07 |
| EFHC2 | 108.0319244 | -1.057143852 | 1.89E-07 |
| AL592424.1 | 19.3597784 | -1.057842193 | 1.70E-02 |
| TET1 | 199.1206885 | -1.057990453 | 1.15E-07 |
| MICU3 | 114.4954291 | -1.058024733 | 4.30E-03 |
| MINDY2 | 1015.699395 | -1.058403645 | 3.95E-09 |
| AC105450.1 | 36.06228179 | -1.058431433 | 6.94E-03 |
| RIBC1 | 84.71474259 | -1.058543592 | 6.21E-06 |
| NCR1 | 49.57956449 | -1.059032534 | 4.02E-03 |
| DYNC2LI1 | 454.8326072 | -1.059049711 | 4.01E-08 |
| EDEM3 | 1391.633374 | -1.059536816 | 1.80E-09 |
| BRIP1 | 600.5206492 | -1.059815086 | 8.80E-06 |
| ZDHHC8P1 | 115.4879533 | -1.060662452 | 2.95E-06 |
| SPRED1 | 1716.526315 | -1.061716642 | 4.35E-23 |
| ZNF253 | 186.1191461 | -1.062148516 | 2.95E-04 |
| TNFSF13B | 34.70978961 | -1.06237253 | 1.21E-02 |
| AC092535.1 | 18.53798512 | -1.063681385 | 4.55E-02 |
| GMPR | 1897.304968 | -1.063712911 | 5.82E-12 |
| HTR2B | 48.42776478 | -1.064105675 | 3.66E-03 |
| KEL | 94.35231685 | -1.064347463 | 2.42E-05 |
| AL137802.2 | 105.9809884 | -1.064433985 | 7.47E-06 |
| CPXM1 | 3098.915536 | -1.064476957 | 1.03E-12 |
| ARHGEF6 | 2573.200329 | -1.064810083 | 6.52E-48 |
| ARHGAP12 | 906.8280344 | -1.066142122 | 1.99E-07 |
| FP236241.1 | 248.492216 | -1.066682206 | 1.08E-06 |
| SNX13 | 1939.795473 | -1.067395648 | 4.93E-09 |
| COLEC10 | 58.18578616 | -1.06791845 | 1.18E-03 |
| CASD1 | 796.3896898 | -1.068154952 | 5.88E-11 |
| 6-Sep | 2821.75174 | -1.068195283 | 3.70E-74 |
| MTUS1 | 3968.754988 | -1.069253079 | 7.04E-29 |
| CEBPZ | 2916.110111 | -1.069732464 | 1.91E-07 |
| TUBA1A | 19721.63494 | -1.070268651 | 8.10E-07 |
| SHPRH | 739.154152 | -1.070694145 | 1.76E-09 |
| SLCO4C1 | 24.23165258 | -1.071038488 | 2.79E-02 |
| GABPA | 1270.757625 | -1.071513297 | 3.52E-10 |
| UBQLNL | 35.35447442 | -1.071669616 | 7.04E-03 |
| LINC02166 | 38.8899247 | -1.072562505 | 2.04E-03 |
| EFCAB12 | 77.01796851 | -1.072876768 | 6.76E-04 |

| | | | |
|---|---|---|---|
| CCDC138 | 483.0823704 | -1.073137429 | 1.27E-09 |
| CCDC188 | 246.0132551 | -1.073398575 | 4.77E-09 |
| PDS5B | 1981.632488 | -1.073682762 | 1.24E-13 |
| CMAHP | 193.4739888 | -1.074593118 | 1.05E-06 |
| AC138915.3 | 16.81197511 | -1.074747171 | 3.84E-02 |
| AC107982.3 | 18.97466292 | -1.075029395 | 3.79E-02 |
| ZBTB41 | 298.4376423 | -1.075119228 | 4.41E-03 |
| AL138899.1 | 26.53149751 | -1.075333322 | 2.90E-02 |
| NA | 137.0107692 | -1.076039543 | 2.37E-05 |
| FBN1 | 3258.584951 | -1.076041658 | 3.26E-19 |
| EPHX4 | 58.81641382 | -1.076072442 | 2.13E-04 |
| FAM117B | 322.1446807 | -1.076081103 | 4.39E-12 |
| NMI | 638.8842373 | -1.076259906 | 4.52E-14 |
| KAZALD1 | 613.8092983 | -1.07756842 | 4.79E-10 |
| SPDYE1 | 47.23973887 | -1.078624849 | 1.52E-04 |
| KCNMB2-AS1 | 30.03868167 | -1.079083885 | 2.16E-02 |
| NCCRP1 | 25.68808836 | -1.079324153 | 5.67E-03 |
| AP000254.1 | 23.68196743 | -1.079487199 | 2.60E-02 |
| WFDC1 | 322.5959827 | -1.080101759 | 1.96E-12 |
| FHOD3 | 292.6071871 | -1.080223382 | 1.53E-10 |
| MED15P9 | 38.96548172 | -1.081671747 | 6.06E-04 |
| NLGN3 | 48.82620059 | -1.082381076 | 6.41E-04 |
| MEIS3P2 | 55.25686464 | -1.083060155 | 2.83E-03 |
| AC008467.1 | 33.37212616 | -1.084277222 | 1.65E-03 |
| ZGRF1 | 425.2006383 | -1.084784242 | 1.85E-04 |
| TMCO1-AS1 | 41.50881023 | -1.084828229 | 7.52E-04 |
| AC080038.2 | 35.49383439 | -1.085384023 | 7.12E-04 |
| EXOC5 | 2309.050245 | -1.085494144 | 1.06E-06 |
| DNALI1 | 156.4162185 | -1.085881564 | 4.52E-05 |
| MANEA | 94.93787047 | -1.086007715 | 8.03E-04 |
| SASS6 | 296.0806781 | -1.086047272 | 9.29E-05 |
| CCDC181 | 83.36213419 | -1.086053933 | 3.75E-03 |
| ZNF396 | 54.79317222 | -1.086193054 | 1.12E-03 |
| REV3L | 1912.9575 | -1.086313331 | 1.10E-07 |
| AC022107.1 | 577.3918954 | -1.086422022 | 5.99E-31 |
| C22orf42 | 36.40816898 | -1.087487225 | 1.82E-02 |
| TPR | 4477.96525 | -1.088195643 | 2.61E-06 |
| ZNF442 | 63.06922939 | -1.088203367 | 1.57E-02 |
| THRB | 889.0566991 | -1.088319875 | 5.30E-20 |
| BNIPL | 32.83408996 | -1.089237454 | 1.39E-03 |
| RIF1 | 2064.822231 | -1.090883458 | 6.13E-03 |
| CEP295 | 1241.620742 | -1.091397395 | 2.12E-05 |
| TMC7 | 119.7353915 | -1.092214993 | 8.92E-09 |
| NINJ2 | 86.32428491 | -1.092378411 | 1.44E-04 |
| AL031123.3 | 37.00347853 | -1.093059238 | 5.21E-03 |
| TRIM24 | 2116.562968 | -1.09363979 | 4.41E-47 |
| PTPN13 | 4334.427018 | -1.094402843 | 6.35E-07 |

| | | | |
|---|---|---|---|
| AL137003.2 | 405.2240052 | -1.094675144 | 8.23E-10 |
| AC093726.1 | 112.1297998 | -1.094744669 | 2.97E-06 |
| ETV4 | 2074.53517 | -1.095779347 | 8.92E-07 |
| DSCR9 | 20.72678461 | -1.096711668 | 4.88E-02 |
| BDP1 | 2234.630743 | -1.096851199 | 6.19E-03 |
| AC009902.3 | 19.06579071 | -1.096862711 | 3.59E-02 |
| AC015818.2 | 21.48308687 | -1.09716772 | 1.71E-02 |
| TBK1 | 788.5180789 | -1.097359205 | 4.56E-06 |
| KMT2E | 3440.83109 | -1.098102669 | 1.75E-24 |
| EVI5 | 724.956478 | -1.098356473 | 6.83E-11 |
| NECAB1 | 84.7509274 | -1.099006382 | 4.67E-04 |
| SLC5A3 | 5415.192645 | -1.099014809 | 3.61E-14 |
| AL031666.1 | 32.03510505 | -1.100226357 | 1.39E-03 |
| PALMD | 192.6518655 | -1.102328177 | 1.30E-05 |
| AC040162.1 | 268.8290852 | -1.102899086 | 1.11E-05 |
| HMGN3 | 1577.110129 | -1.10371471 | 3.04E-33 |
| AL604028.1 | 19.48662782 | -1.103779016 | 2.22E-02 |
| LINC00926 | 65.66827473 | -1.104455741 | 1.44E-03 |
| RARB | 1624.994019 | -1.105056729 | 7.28E-16 |
| NBN | 2080.013296 | -1.10537215 | 2.08E-07 |
| ZSCAN23 | 119.0486742 | -1.10560107 | 1.32E-06 |
| QRICH2 | 106.3556761 | -1.106006497 | 8.09E-06 |
| GDPGP1 | 127.8574598 | -1.106381139 | 2.28E-07 |
| C1QTNF3 | 44.26961409 | -1.106701911 | 9.37E-04 |
| FAM153A | 178.9188772 | -1.107084715 | 3.93E-06 |
| TRIP11 | 1200.792687 | -1.107189577 | 1.60E-08 |
| STXBP4 | 406.927206 | -1.107817103 | 1.37E-06 |
| C1orf53 | 34.30616792 | -1.10837682 | 2.88E-03 |
| COL6A3 | 4751.934988 | -1.109753083 | 3.63E-06 |
| TBC1D10C | 46.80751802 | -1.109960526 | 7.00E-03 |
| ZNF608 | 2639.516423 | -1.112463541 | 4.05E-31 |
| FRAS1 | 577.0625188 | -1.112808838 | 2.55E-09 |
| GLTPD2 | 18.81306677 | -1.113490279 | 3.20E-02 |
| TP53INP1 | 249.9843985 | -1.113812702 | 4.37E-10 |
| ZCCHC24 | 4931.736738 | -1.115009717 | 1.72E-17 |
| FAP | 92.15692023 | -1.115015052 | 5.60E-03 |
| ZNF10 | 240.692249 | -1.115036772 | 4.36E-05 |
| INSYN1 | 368.9945437 | -1.115038291 | 7.95E-09 |
| TRIM2 | 1676.098241 | -1.115313627 | 1.70E-05 |
| SPIN2B | 409.744545 | -1.115363575 | 1.35E-26 |
| AL589666.2 | 15.697453 | -1.116235107 | 3.97E-02 |
| KIAA1217 | 1220.659916 | -1.11631571 | 6.41E-21 |
| PNRC2 | 3356.074831 | -1.116405285 | 7.33E-23 |
| COL11A2 | 524.1032316 | -1.116550669 | 2.38E-06 |
| AC016526.3 | 48.75006189 | -1.11719083 | 1.20E-04 |
| FCGR2C | 56.33897174 | -1.117645971 | 9.62E-05 |
| AFAP1L2 | 2158.646464 | -1.118491926 | 4.38E-20 |

| | | | |
|---|---|---|---|
| AL161891.1 | 53.9731438 | -1.120111119 | 1.36E-02 |
| MPHOSPH9 | 1122.599799 | -1.120248946 | 2.25E-07 |
| TMEM249 | 28.67557228 | -1.12052289 | 4.98E-02 |
| RAB37 | 569.4378075 | -1.120791755 | 7.69E-11 |
| CACNA1G | 138.963321 | -1.121505211 | 1.49E-08 |
| ACSL6 | 38.72385866 | -1.122364251 | 1.50E-02 |
| KCNIP4 | 24.49992728 | -1.123156426 | 1.59E-02 |
| AC016727.1 | 32.90193357 | -1.123489551 | 4.91E-03 |
| ITGA11 | 5052.132686 | -1.123754752 | 1.45E-14 |
| AC004943.1 | 20.64228712 | -1.124140677 | 2.67E-02 |
| ZBTB10 | 1109.922168 | -1.124579517 | 4.91E-18 |
| FXYD6 | 2385.941432 | -1.124993255 | 2.27E-13 |
| SERPINF1 | 435.8532755 | -1.125659344 | 4.72E-11 |
| AL365203.2 | 100.0752658 | -1.126530088 | 1.92E-07 |
| DCLK1 | 439.8908667 | -1.126774196 | 1.91E-04 |
| ZNF180 | 312.6482421 | -1.127611541 | 9.66E-07 |
| AC137894.1 | 39.95266791 | -1.128408632 | 1.06E-03 |
| SLF1 | 1156.522204 | -1.128610369 | 8.91E-07 |
| CCL5 | 161.9253664 | -1.129222631 | 3.28E-04 |
| RAB26 | 425.0303471 | -1.130846606 | 4.42E-08 |
| LINC02289 | 34.71716412 | -1.132411138 | 1.98E-02 |
| LRGUK | 25.47846217 | -1.133279159 | 3.27E-02 |
| HOXA5 | 769.5073558 | -1.133511933 | 8.26E-25 |
| AC048341.2 | 122.453961 | -1.133879834 | 3.08E-04 |
| NCALD | 398.8456302 | -1.135520861 | 3.79E-04 |
| LINC02076 | 54.9376968 | -1.136835133 | 2.83E-05 |
| KCNC3 | 168.2532864 | -1.13781967 | 7.08E-06 |
| CAMK1G | 592.4143727 | -1.138386468 | 3.12E-07 |
| AL132780.5 | 20.48620529 | -1.139200509 | 2.73E-02 |
| NRROS | 203.4804337 | -1.139317937 | 2.36E-08 |
| SKIDA1 | 152.151256 | -1.13974611 | 5.40E-09 |
| USP16 | 1293.924721 | -1.141640525 | 6.07E-08 |
| PTCH2 | 132.6977029 | -1.14195404 | 1.45E-04 |
| C4A | 1312.82826 | -1.142758396 | 4.81E-02 |
| ACTL8 | 26.23675356 | -1.145016921 | 3.02E-02 |
| CCDC28B | 704.535287 | -1.145726869 | 1.55E-11 |
| ACOX2 | 57.51644016 | -1.146168963 | 4.73E-03 |
| IREB2 | 3969.02169 | -1.146227252 | 1.50E-22 |
| ZNF252P-AS1 | 45.88853292 | -1.146424353 | 3.86E-04 |
| MCC | 3773.783562 | -1.146431225 | 2.27E-39 |
| NT5C1B | 24.42320539 | -1.14722091 | 3.91E-02 |
| AC018692.2 | 18.65657282 | -1.147598831 | 2.34E-02 |
| GSN-AS1 | 75.19028754 | -1.148204406 | 3.28E-07 |
| CCR5 | 30.83657564 | -1.14825732 | 8.08E-03 |
| TSHZ1 | 2579.087137 | -1.149129528 | 1.38E-15 |
| SH3BP5-AS1 | 201.3828996 | -1.150681225 | 8.19E-07 |
| AL121899.1 | 25.3059189 | -1.15160587 | 7.05E-03 |

| | | | |
|---|---|---|---|
| AC012513.3 | 165.7443447 | -1.152362442 | 8.07E-12 |
| FMO4 | 51.19801347 | -1.152493067 | 1.32E-02 |
| GNG4 | 685.0541399 | -1.152537121 | 9.23E-13 |
| LONRF2 | 240.7121002 | -1.152663989 | 1.49E-07 |
| AC009159.3 | 20.44347584 | -1.153095634 | 8.24E-03 |
| MMP28 | 50.44223442 | -1.153414703 | 2.19E-02 |
| LINC01139 | 24.08748545 | -1.154026861 | 1.92E-02 |
| AC097376.2 | 104.004178 | -1.154554278 | 1.76E-06 |
| CARNS1 | 82.0911703 | -1.155277858 | 2.23E-04 |
| CASC15 | 47.77630025 | -1.155950586 | 1.36E-03 |
| ZNF624 | 132.3153627 | -1.156073054 | 1.55E-04 |
| SPDYE5 | 63.22792987 | -1.156343566 | 2.27E-05 |
| FBXL22 | 54.18573168 | -1.156602559 | 1.28E-04 |
| MPZ | 270.8214232 | -1.157955735 | 4.14E-05 |
| C8orf31 | 287.7631837 | -1.1582894 | 1.51E-11 |
| BRWD3 | 947.0454821 | -1.159111803 | 1.47E-09 |
| RAET1G | 88.40783674 | -1.159740005 | 1.29E-03 |
| CLEC18B | 62.85794889 | -1.159805006 | 3.94E-04 |
| AC021945.1 | 17.7759937 | -1.16017814 | 2.92E-02 |
| RHOJ | 2033.908093 | -1.1604512 | 1.51E-15 |
| ZFPM2-AS1 | 556.3674849 | -1.160828185 | 1.91E-07 |
| PCOLCE | 18291.75263 | -1.160906314 | 6.33E-17 |
| AL390728.6 | 38.95737465 | -1.161112369 | 2.12E-02 |
| EXOSC10-AS1 | 25.29981927 | -1.161150316 | 1.97E-02 |
| CAT | 4283.077306 | -1.162906507 | 6.12E-38 |
| EXTL2 | 2261.873093 | -1.162914621 | 3.04E-25 |
| SEMA3C | 1509.717254 | -1.162996514 | 9.18E-11 |
| GOLGB1 | 3111.727492 | -1.163647757 | 5.06E-04 |
| TMEM198 | 334.0886835 | -1.164172948 | 3.37E-13 |
| SCN4B | 2747.623872 | -1.164842727 | 3.47E-08 |
| FSIP1 | 21.52831629 | -1.165098368 | 1.71E-02 |
| AL450326.1 | 41.15195871 | -1.165326196 | 2.26E-04 |
| AVIL | 167.3458292 | -1.167374295 | 2.00E-06 |
| STAG2 | 4891.459207 | -1.167451096 | 2.87E-04 |
| HAND1 | 50.73723808 | -1.167862401 | 3.37E-02 |
| LIX1L-AS1 | 18.03875301 | -1.17022318 | 3.14E-02 |
| SEMA3F-AS1 | 37.36207118 | -1.170453603 | 1.62E-02 |
| AL031283.2 | 36.90499216 | -1.170852612 | 7.15E-03 |
| ABCC6P2 | 34.63096341 | -1.172080711 | 8.61E-03 |
| SPRY4 | 1960.932788 | -1.172321342 | 1.48E-19 |
| ZIM2 | 133.2839755 | -1.172683285 | 3.65E-07 |
| CENPE | 900.3298078 | -1.172708353 | 3.46E-02 |
| FMNL2 | 4188.966288 | -1.174598724 | 2.51E-27 |
| AC002128.1 | 38.25855138 | -1.174667507 | 2.78E-03 |
| ERVK13-1 | 1300.392176 | -1.175056968 | 4.61E-43 |
| ARHGAP5-AS1 | 297.6510154 | -1.1757463 | 5.56E-14 |
| MTSS1 | 1117.759609 | -1.175994341 | 6.83E-24 |

| | | | |
|---|---|---|---|
| TMEM117 | 26.69301115 | -1.176489505 | 4.12E-02 |
| MIS18BP1 | 607.5874957 | -1.176695761 | 4.31E-07 |
| AC018665.1 | 157.0923195 | -1.176875368 | 1.52E-12 |
| PGAP3 | 538.2201203 | -1.177460203 | 8.07E-12 |
| AC027031.2 | 487.0588179 | -1.177605932 | 7.44E-23 |
| AC010422.6 | 130.2044948 | -1.178149922 | 2.38E-09 |
| AC140134.1 | 277.8328783 | -1.178570717 | 2.90E-07 |
| KAT6B | 1431.533217 | -1.179619408 | 4.54E-23 |
| LIPG | 311.095507 | -1.181546641 | 1.00E-11 |
| LRRCC1 | 377.3329368 | -1.183031512 | 3.69E-04 |
| MAP7 | 29.05664401 | -1.183254394 | 1.01E-02 |
| AL513327.1 | 97.34746054 | -1.183520271 | 4.05E-07 |
| LINC01854 | 22.5958603 | -1.185347082 | 1.05E-02 |
| ZNF711 | 258.9397943 | -1.185484718 | 2.17E-04 |
| C2orf16 | 308.8560384 | -1.185492271 | 3.62E-18 |
| CCNE2 | 275.1172858 | -1.186352785 | 1.47E-06 |
| EPCAM | 29.64192716 | -1.186939723 | 6.56E-03 |
| GIN1 | 179.1267096 | -1.187651493 | 1.68E-03 |
| PRRT3 | 215.8516639 | -1.188161141 | 6.94E-14 |
| CYB5B | 13391.33376 | -1.188960644 | 4.68E-18 |
| AC106820.5 | 64.56184719 | -1.189436737 | 1.91E-05 |
| CNIH3 | 994.9039675 | -1.190769806 | 8.46E-15 |
| DOK3 | 101.2399448 | -1.191224312 | 1.11E-06 |
| AC026356.1 | 50.63709256 | -1.191431145 | 7.35E-05 |
| NA | 26.93695681 | -1.193310553 | 3.01E-02 |
| ITGA1 | 11661.05432 | -1.193445108 | 2.96E-21 |
| PELI2 | 304.1972169 | -1.193730749 | 1.47E-08 |
| GRIA1 | 156.935266 | -1.193777607 | 1.79E-04 |
| ATP2B1-AS1 | 26.62582568 | -1.194213128 | 7.40E-03 |
| ITIH4 | 62.83899631 | -1.195095901 | 9.98E-03 |
| SCUBE2 | 118.6778719 | -1.196274079 | 1.55E-03 |
| AC244034.3 | 16.18034274 | -1.19825971 | 4.56E-02 |
| AL121672.1 | 18.75379888 | -1.198644724 | 1.42E-02 |
| C21orf91 | 509.9850167 | -1.201118705 | 5.97E-10 |
| ZNF704 | 2053.150087 | -1.202075132 | 9.78E-09 |
| OTULINL | 778.6536385 | -1.203657376 | 8.36E-12 |
| CDH12P3 | 37.29249809 | -1.204535302 | 1.16E-02 |
| VWA5A | 131.4514081 | -1.204795716 | 1.16E-04 |
| AGL | 980.3410774 | -1.205353617 | 5.15E-10 |
| INKA1 | 307.9654041 | -1.20554673 | 2.50E-10 |
| EFHB | 27.47713761 | -1.205714693 | 4.22E-02 |
| AP003774.3 | 29.72754542 | -1.206600621 | 1.04E-02 |
| PDCD4-AS1 | 327.1842865 | -1.20674221 | 2.69E-11 |
| KAT2B | 1031.112119 | -1.207269669 | 6.35E-27 |
| HLA-DPA1 | 429.1936339 | -1.207928236 | 3.10E-06 |
| AC022150.2 | 15.74442329 | -1.208698376 | 2.54E-02 |
| AC008103.1 | 69.65190709 | -1.208931094 | 2.47E-05 |

| | | | |
|---|---|---|---|
| NA | 51.51323304 | -1.209111352 | 2.51E-04 |
| AC092171.4 | 13.85223688 | -1.2093506 | 3.89E-02 |
| PKDCC | 957.8413157 | -1.209513689 | 1.02E-19 |
| AC016355.1 | 46.93499633 | -1.210134884 | 3.90E-03 |
| SHE | 294.9360923 | -1.210416771 | 4.01E-10 |
| NFAT5 | 5313.455934 | -1.21176142 | 9.29E-12 |
| EPHB3 | 918.3399535 | -1.211858913 | 7.23E-19 |
| AC090948.2 | 15.23363767 | -1.211890985 | 3.31E-02 |
| ADNP-AS1 | 43.48973831 | -1.212101363 | 2.23E-04 |
| RASSF3 | 2924.247126 | -1.212868068 | 7.12E-46 |
| COL6A1 | 17941.64959 | -1.213153066 | 1.25E-37 |
| WDR35 | 1138.627068 | -1.21349221 | 4.18E-08 |
| TMEM63C | 137.7208928 | -1.214052192 | 1.72E-03 |
| TTC14 | 1432.076423 | -1.214230378 | 5.22E-10 |
| AC024075.1 | 11.31564554 | -1.214438781 | 4.61E-02 |
| LARP7 | 955.0836205 | -1.215449164 | 1.87E-06 |
| AL139384.2 | 17.88755738 | -1.215953559 | 4.22E-02 |
| SIPA1L2 | 2095.321232 | -1.216704749 | 6.09E-25 |
| AC140125.2 | 68.76072382 | -1.217836121 | 1.25E-04 |
| COL9A3 | 4082.534637 | -1.218373659 | 8.69E-15 |
| C3orf18 | 20.94029741 | -1.220607832 | 3.66E-02 |
| RDX | 9248.951892 | -1.221383796 | 1.08E-21 |
| RN7SKP181 | 58.13072017 | -1.221861302 | 1.23E-04 |
| ZNF658B | 53.47303321 | -1.22373026 | 3.49E-03 |
| AC104667.2 | 71.44130672 | -1.224063957 | 1.25E-05 |
| PHIP | 1703.51194 | -1.225139262 | 1.32E-05 |
| DLEU2 | 124.6313412 | -1.225811545 | 6.00E-06 |
| ZNF644 | 1095.23322 | -1.225883308 | 1.29E-10 |
| EPS8 | 4496.746593 | -1.226294637 | 3.06E-19 |
| KBTBD8 | 99.19555138 | -1.226314051 | 5.85E-04 |
| AC105339.2 | 180.435808 | -1.227008915 | 6.52E-05 |
| AC090627.1 | 17.38586849 | -1.228619849 | 1.57E-02 |
| ABCA5 | 556.3366742 | -1.232138589 | 1.28E-06 |
| ELOVL3 | 25.0446582 | -1.232177928 | 1.43E-02 |
| PTOV1-AS2 | 99.07661411 | -1.232227543 | 7.74E-08 |
| AC006330.1 | 20.83963745 | -1.232609443 | 1.28E-02 |
| SALL2 | 1175.392959 | -1.23305614 | 3.35E-11 |
| SLC44A5 | 50.24975973 | -1.234965309 | 1.48E-04 |
| GDAP1 | 1189.14057 | -1.23507453 | 3.48E-18 |
| TBC1D32 | 228.4429051 | -1.235101181 | 5.29E-06 |
| NRXN2 | 691.5951493 | -1.235670722 | 7.73E-17 |
| NA | 14.84169264 | -1.236008502 | 2.90E-02 |
| CCDC150 | 150.6043612 | -1.23689496 | 3.71E-05 |
| LPP-AS2 | 110.5716903 | -1.236955609 | 1.51E-09 |
| LINC00893 | 98.4657195 | -1.237242531 | 1.25E-05 |
| OLFM5P | 19.59671498 | -1.238444245 | 1.31E-02 |
| AC027419.2 | 33.09885901 | -1.238957064 | 9.44E-04 |

| | | | |
|---|---|---|---|
| ST6GAL1 | 1260.199539 | -1.24081436 | 2.57E-13 |
| AC148477.3 | 81.05410233 | -1.241464691 | 2.88E-06 |
| ZNF660 | 284.1719379 | -1.241939394 | 8.27E-07 |
| AC099786.2 | 59.76050552 | -1.242684261 | 7.21E-04 |
| C20orf204 | 77.0840451 | -1.243048684 | 2.04E-04 |
| AARD | 73.18662623 | -1.247053264 | 8.15E-06 |
| AOAH | 33.00216439 | -1.248361927 | 1.71E-02 |
| AF131216.4 | 16.971124 | -1.248771521 | 3.14E-02 |
| CHIC1 | 312.476867 | -1.253081254 | 9.80E-11 |
| NF1P6 | 29.64198724 | -1.255407275 | 2.17E-03 |
| CD160 | 21.21898451 | -1.25625847 | 8.77E-03 |
| SENP7 | 434.7236845 | -1.258218026 | 4.08E-06 |
| ST3GAL5 | 3369.022929 | -1.261410437 | 2.40E-31 |
| PID1 | 439.8178603 | -1.262073984 | 1.03E-17 |
| AC012501.2 | 74.6821662 | -1.262665189 | 1.41E-05 |
| AL845552.2 | 20.43738086 | -1.262682465 | 1.31E-02 |
| MXRA5 | 2340.291441 | -1.262963878 | 3.30E-47 |
| FOXR2 | 367.8161971 | -1.263733117 | 3.30E-14 |
| AC012569.1 | 18.76329793 | -1.264298814 | 4.81E-02 |
| KIAA1586 | 323.4141024 | -1.264605591 | 2.49E-09 |
| HIST1H2AI | 17.08190837 | -1.26512467 | 2.19E-02 |
| CEP162 | 363.5001121 | -1.265180524 | 5.94E-05 |
| ATP7A | 1068.071532 | -1.265603699 | 9.31E-20 |
| N4BP2 | 343.3328604 | -1.265718624 | 2.99E-05 |
| HIC1 | 996.4257033 | -1.266207329 | 2.24E-08 |
| AC008555.4 | 16.62721988 | -1.266299524 | 3.81E-02 |
| AL033529.1 | 69.57657961 | -1.268302827 | 3.24E-04 |
| AL353622.1 | 109.5816489 | -1.268895014 | 8.95E-09 |
| EGFLAM | 33.98079973 | -1.26946438 | 3.15E-02 |
| AC008937.3 | 41.52574998 | -1.269586179 | 6.92E-04 |
| VIPR1 | 332.1102024 | -1.270947026 | 4.90E-12 |
| PTBP2 | 289.7813242 | -1.271837456 | 3.11E-07 |
| PARD6G-AS1 | 83.02067989 | -1.273240194 | 4.93E-06 |
| JCAD | 5110.161346 | -1.273834574 | 6.41E-43 |
| AC139100.2 | 16.33863358 | -1.273887388 | 2.22E-02 |
| STKLD1 | 37.49942356 | -1.276428842 | 7.07E-03 |
| LRRC24 | 60.46001221 | -1.277125974 | 9.86E-04 |
| TEX35 | 14.82468883 | -1.277397219 | 3.18E-02 |
| SELP | 136.3642521 | -1.278363633 | 8.07E-04 |
| EME2 | 888.6364107 | -1.279073469 | 5.97E-15 |
| AC011290.2 | 43.40247706 | -1.280529599 | 5.77E-03 |
| TCN2 | 1055.694199 | -1.280855535 | 1.53E-11 |
| ENDOD1 | 5289.674656 | -1.281886304 | 9.76E-11 |
| LAG3 | 53.74991194 | -1.283857601 | 7.63E-05 |
| ANKRD18EP | 104.7435619 | -1.283886109 | 1.39E-07 |
| NA | 120.1022931 | -1.284415392 | 1.87E-04 |
| STIM2 | 1398.883632 | -1.284517783 | 1.51E-27 |

| | | | |
|---|---|---|---|
| BRCA2 | 456.7661104 | -1.28515401 | 4.26E-04 |
| AC034238.3 | 15.59327677 | -1.286463904 | 4.14E-02 |
| LINC01963 | 343.5003131 | -1.28841411 | 8.20E-23 |
| LINC01389 | 27.21805161 | -1.288712516 | 8.79E-03 |
| AC048341.1 | 119.2109249 | -1.289812842 | 4.79E-08 |
| AC135050.3 | 16.30602706 | -1.289937564 | 2.95E-02 |
| AL132656.3 | 17.11838999 | -1.290347149 | 4.09E-02 |
| IFI30 | 2930.300089 | -1.290355396 | 2.98E-17 |
| TRPV6 | 109.5943365 | -1.291397441 | 1.98E-02 |
| PON2 | 4494.206146 | -1.291657145 | 9.97E-64 |
| SELENBP1 | 2651.618252 | -1.293455507 | 3.98E-11 |
| AL596202.1 | 165.9496982 | -1.294043596 | 1.26E-11 |
| SCN10A | 31.43251154 | -1.294631753 | 1.58E-03 |
| AC024405.2 | 18.19649933 | -1.2955955 | 4.48E-02 |
| USP51 | 104.9721947 | -1.295839904 | 3.42E-06 |
| ATRX | 2948.433124 | -1.297224658 | 1.34E-04 |
| AC131392.1 | 146.7752183 | -1.297627795 | 2.93E-07 |
| PBLD | 150.0356914 | -1.298197811 | 1.87E-08 |
| EEA1 | 1040.930789 | -1.300573625 | 3.57E-05 |
| HESX1 | 24.15406492 | -1.300705333 | 4.67E-03 |
| EREG | 68.09509223 | -1.30157719 | 4.41E-03 |
| NA | 168.0298735 | -1.301718201 | 6.47E-12 |
| AP002414.2 | 19.53611069 | -1.303112397 | 9.50E-03 |
| AC011008.2 | 45.63903778 | -1.303380767 | 9.07E-05 |
| TET2 | 1173.805508 | -1.304278107 | 2.45E-21 |
| IGSF11 | 16.86279637 | -1.30462557 | 3.21E-02 |
| AL358852.1 | 15.31578304 | -1.306523615 | 1.94E-02 |
| TM7SF2 | 2521.278961 | -1.306540167 | 5.95E-10 |
| SYNPO2 | 528.338954 | -1.307508767 | 1.73E-18 |
| TEK | 960.4093854 | -1.307944022 | 1.48E-08 |
| BBOX1-AS1 | 25.44159368 | -1.3079759 | 1.10E-02 |
| NARF-IT1 | 9.897559897 | -1.30817554 | 3.09E-02 |
| C3orf70 | 235.4032375 | -1.308256084 | 5.31E-19 |
| ZNF521 | 1676.551674 | -1.310105601 | 1.35E-21 |
| RN7SL832P | 28.57002741 | -1.312782683 | 2.78E-03 |
| JAG1 | 2378.045133 | -1.313617054 | 1.13E-17 |
| AC145207.8 | 18.98836007 | -1.315554828 | 2.75E-02 |
| SLC25A45 | 87.16993623 | -1.315712953 | 2.46E-04 |
| MAOB | 49.54773042 | -1.317480666 | 2.34E-04 |
| AC024060.1 | 46.15207851 | -1.317591275 | 1.70E-03 |
| AC104964.4 | 21.11909575 | -1.317771515 | 7.05E-03 |
| SCN9A | 28.85194318 | -1.317959743 | 3.19E-02 |
| CD101 | 54.93496383 | -1.318034364 | 7.08E-03 |
| TPTE2P6 | 16.97085291 | -1.318098671 | 2.10E-02 |
| AL121672.2 | 17.12607123 | -1.318806819 | 1.27E-02 |
| CLEC4F | 53.65404115 | -1.319712271 | 9.27E-06 |
| TMTC3 | 861.6792463 | -1.320021199 | 1.32E-06 |

| | | | |
|---|---|---|---|
| PDE5A | 1577.849555 | -1.320313316 | 3.94E-12 |
| FNBP1L | 2268.026796 | -1.320776674 | 1.93E-51 |
| MARCKSL1 | 6894.580256 | -1.322896735 | 2.97E-20 |
| AC010136.1 | 15.27942093 | -1.325395959 | 2.88E-02 |
| FAS | 239.2567132 | -1.325634444 | 2.52E-08 |
| GPCPD1 | 2138.032665 | -1.325641964 | 5.55E-10 |
| AL353763.1 | 33.38763978 | -1.326361195 | 1.82E-03 |
| SSBP2 | 2717.946164 | -1.326376344 | 1.56E-31 |
| CHGB | 280.9084651 | -1.326665258 | 4.60E-18 |
| ZFYVE16 | 2545.697203 | -1.326890944 | 4.89E-05 |
| AC016738.1 | 27.13772455 | -1.327817871 | 2.18E-02 |
| AF111169.3 | 28.42239266 | -1.32989715 | 2.65E-03 |
| TNFAIP8L3 | 111.5799797 | -1.330204663 | 3.45E-06 |
| AKR1C3 | 83.72430564 | -1.331348698 | 1.63E-03 |
| AL122023.1 | 13.6568424 | -1.331527755 | 2.76E-02 |
| EPN3 | 37.87810074 | -1.332576709 | 3.97E-04 |
| BICD1 | 1218.161353 | -1.332822457 | 8.89E-47 |
| FRMD4A | 2369.142518 | -1.334049305 | 2.00E-46 |
| SVEP1 | 143.2123067 | -1.334103582 | 1.25E-04 |
| RNASEL | 490.163533 | -1.334562009 | 2.02E-29 |
| C1QTNF6 | 3661.584941 | -1.334619154 | 1.74E-10 |
| RN7SL268P | 15.40962059 | -1.335484049 | 1.31E-02 |
| ANKRD26 | 244.6223901 | -1.335709334 | 3.08E-06 |
| SLC25A27 | 441.4577795 | -1.336888673 | 1.13E-12 |
| UST | 680.7892115 | -1.338570908 | 2.44E-14 |
| AC138932.5 | 14.49975663 | -1.339772905 | 3.64E-02 |
| C4A-AS1 | 29.21201907 | -1.340170304 | 1.11E-02 |
| CDC7 | 1042.852377 | -1.34069781 | 3.40E-16 |
| AC012321.1 | 316.3933628 | -1.341271075 | 9.96E-08 |
| CFAP300 | 138.3243478 | -1.343461055 | 3.14E-09 |
| PDE7B | 23.34403892 | -1.34349913 | 1.80E-02 |
| IMPG2 | 21.46058389 | -1.343631255 | 4.35E-03 |
| AC099786.1 | 15.87473704 | -1.343669027 | 4.77E-02 |
| LHFPL2 | 4882.279716 | -1.343914854 | 8.65E-96 |
| SATB1 | 25.96066018 | -1.345389759 | 5.07E-03 |
| AL158071.5 | 12.03021384 | -1.345798008 | 3.97E-02 |
| FAM86HP | 34.88382065 | -1.347028739 | 4.88E-04 |
| ITGA2 | 6584.396695 | -1.347438973 | 8.41E-24 |
| AC108515.1 | 38.85659606 | -1.347580091 | 5.48E-03 |
| CCDC103 | 32.8956705 | -1.347740472 | 4.05E-03 |
| AC010809.1 | 20.89926275 | -1.347937045 | 1.44E-02 |
| DIXDC1 | 1830.006981 | -1.348254147 | 8.24E-20 |
| CA12 | 123.9598698 | -1.34898374 | 8.99E-05 |
| SULF2 | 7414.451633 | -1.349013983 | 7.08E-60 |
| STON2 | 213.0074989 | -1.350642891 | 1.55E-09 |
| AC006449.5 | 65.52035688 | -1.350646017 | 9.36E-07 |
| KCNRG | 21.37247774 | -1.350877427 | 1.61E-02 |

| | | | |
|---|---|---|---|
| CASKIN1 | 97.09318984 | -1.351884366 | 4.49E-03 |
| GIMAP4 | 15.60675452 | -1.352707156 | 4.59E-02 |
| FBXO40 | 30.39114751 | -1.353966007 | 8.59E-04 |
| AL390198.1 | 33.74893479 | -1.354373735 | 7.18E-04 |
| C2orf50 | 15.27994356 | -1.354751642 | 3.59E-02 |
| SPIN2A | 48.63188211 | -1.355392012 | 3.52E-05 |
| TPTE2P1 | 16.81761646 | -1.355596701 | 2.07E-02 |
| TDRD10 | 80.80343204 | -1.356688056 | 6.07E-08 |
| ZNF233 | 106.6235961 | -1.357943878 | 3.38E-13 |
| ITGA10 | 182.6699131 | -1.359650921 | 4.32E-09 |
| TPTE | 162.9454147 | -1.360142771 | 1.86E-08 |
| ADD3 | 3597.694114 | -1.361560511 | 1.30E-50 |
| C15orf65 | 39.22759693 | -1.361610553 | 4.48E-03 |
| CMBL | 2580.311176 | -1.36170677 | 2.96E-21 |
| CMPK2 | 34.3634101 | -1.362988368 | 4.40E-03 |
| JMJD1C-AS1 | 18.31117869 | -1.364415235 | 7.57E-03 |
| RANBP2 | 6080.340109 | -1.365925759 | 6.43E-09 |
| PLCL2 | 39.70894612 | -1.36614644 | 2.14E-03 |
| PLA2R1 | 868.557516 | -1.366602713 | 1.43E-20 |
| RFLNA | 58.80224814 | -1.367008349 | 5.59E-04 |
| AC103760.1 | 21.44567501 | -1.367345802 | 9.93E-03 |
| EFCAB7 | 208.1890505 | -1.368659516 | 2.96E-06 |
| KIAA0825 | 59.54724542 | -1.368760844 | 4.39E-04 |
| ITGA8 | 483.5758338 | -1.369401666 | 9.70E-11 |
| AC004801.2 | 21.85766275 | -1.369798828 | 3.68E-02 |
| AC114956.3 | 27.47485298 | -1.370343405 | 1.72E-03 |
| AC010809.2 | 18.6237117 | -1.370826832 | 8.52E-03 |
| COLQ | 55.80094762 | -1.37086306 | 1.09E-03 |
| NYNRIN | 5607.528608 | -1.374199332 | 9.99E-10 |
| SPARC | 56594.83605 | -1.37442457 | 1.22E-13 |
| CCDC194 | 12.17004459 | -1.376398769 | 4.25E-02 |
| MEIS1 | 435.9633765 | -1.379152734 | 1.41E-11 |
| OR7E126P | 11.34187201 | -1.380824755 | 4.23E-02 |
| LINC02552 | 13.25560472 | -1.381582149 | 3.99E-02 |
| SNX25P1 | 19.83565817 | -1.382148765 | 5.71E-03 |
| ATF7 | 63.46670973 | -1.384764549 | 6.56E-08 |
| VCAM1 | 2482.479407 | -1.386462607 | 9.55E-05 |
| FAM13B | 5560.921553 | -1.386510814 | 3.98E-35 |
| TIA1 | 2320.581695 | -1.387328559 | 2.69E-13 |
| AC093503.3 | 11.79264787 | -1.391362853 | 2.98E-02 |
| SLC43A2 | 1323.412679 | -1.392337752 | 8.03E-20 |
| ABAT | 360.9278879 | -1.392345886 | 4.16E-18 |
| NAAA | 534.8626594 | -1.395444555 | 7.81E-40 |
| ADAMTS3 | 506.1523499 | -1.397228089 | 1.17E-14 |
| STARD4 | 2398.647591 | -1.399159879 | 1.43E-23 |
| AJ011931.1 | 14.10779318 | -1.399211277 | 3.19E-02 |
| LRRN2 | 226.4845914 | -1.399441072 | 4.41E-10 |

| | | | |
|---|---|---|---|
| AC104447.1 | 94.69098291 | -1.400784595 | 6.61E-09 |
| TCIM | 39.7471214 | -1.401858198 | 2.34E-03 |
| AC067930.4 | 32.3071449 | -1.402609138 | 1.47E-03 |
| MBL1P | 30.16663922 | -1.402689962 | 9.95E-03 |
| RHBDL3 | 329.7370229 | -1.405883723 | 2.02E-12 |
| AP000924.1 | 15.43826193 | -1.407537315 | 2.24E-02 |
| GALNT12 | 1294.735295 | -1.407751743 | 4.08E-12 |
| THRB-AS1 | 13.0541707 | -1.407864143 | 3.10E-02 |
| CD226 | 25.91145121 | -1.409837446 | 4.19E-02 |
| AL356585.1 | 10.47909926 | -1.410033068 | 4.74E-02 |
| PGAP1 | 1273.05056 | -1.410334878 | 6.92E-13 |
| AL391069.2 | 56.01725847 | -1.412022525 | 1.69E-06 |
| AC015802.1 | 42.72525233 | -1.413110094 | 1.06E-04 |
| AC006511.3 | 14.06890334 | -1.413118787 | 2.77E-02 |
| LOXL1-AS1 | 19.08140033 | -1.415217817 | 1.21E-02 |
| ZNF750 | 11.54075097 | -1.417476196 | 3.31E-02 |
| JMJD1C | 1264.147845 | -1.417896925 | 1.81E-08 |
| PCDHB12 | 24.55439884 | -1.41922538 | 7.96E-04 |
| DDX60 | 693.0435674 | -1.419739934 | 1.06E-07 |
| GRIN2A | 673.7685502 | -1.419796474 | 3.59E-12 |
| FZD4 | 3295.196726 | -1.419802807 | 1.29E-14 |
| DMXL2 | 3319.662109 | -1.419872901 | 1.61E-09 |
| KCNAB3 | 147.848877 | -1.420015866 | 3.37E-12 |
| ST3GAL6 | 110.4910764 | -1.42047607 | 1.18E-07 |
| METTL7B | 30.52122426 | -1.421706849 | 3.16E-03 |
| ZFP37 | 238.528989 | -1.421867681 | 5.19E-10 |
| ZFHX2 | 54.06332065 | -1.422080855 | 1.69E-03 |
| ARID4A | 694.162518 | -1.422572933 | 9.56E-10 |
| SCN2B | 23.74475667 | -1.423269137 | 3.67E-03 |
| LINC01354 | 15.60851471 | -1.42769758 | 3.63E-02 |
| AL049597.2 | 16.72602794 | -1.428118334 | 8.06E-03 |
| LINC01063 | 10.14136766 | -1.4287934 | 2.70E-02 |
| DSTNP1 | 29.83426616 | -1.429805406 | 1.43E-03 |
| PLPP3 | 2030.88115 | -1.430298283 | 4.90E-35 |
| ZNF471 | 354.7592985 | -1.430714286 | 1.20E-14 |
| MEGF10 | 176.8006285 | -1.431642982 | 2.07E-07 |
| LAMA4 | 12155.3392 | -1.434097546 | 3.38E-103 |
| AC044802.1 | 26.95094169 | -1.43471285 | 1.82E-03 |
| CCDC88A | 3210.87298 | -1.435967083 | 4.15E-08 |
| TLE2 | 901.3412673 | -1.436258961 | 1.26E-13 |
| CRHBP | 46.71371443 | -1.436458244 | 2.55E-03 |
| AC112484.1 | 14.85166309 | -1.436794865 | 2.21E-02 |
| AC007496.2 | 13.47156693 | -1.438020825 | 3.23E-02 |
| PYCARD-AS1 | 20.17777663 | -1.438435681 | 1.02E-02 |
| MYB | 40.76901378 | -1.440825872 | 5.22E-04 |
| SLC25A48 | 75.47198056 | -1.441843432 | 3.22E-06 |
| TNMD | 57.18971116 | -1.444345613 | 2.51E-02 |

| | | | |
|---|---|---|---|
| DOP1A | 788.5080826 | -1.444412667 | 1.02E-10 |
| AC004540.1 | 39.99496744 | -1.444999346 | 1.05E-04 |
| KANK4 | 25.95378457 | -1.445089108 | 5.55E-03 |
| BX571846.1 | 39.76887745 | -1.445358133 | 8.34E-05 |
| GALNT4 | 422.8417552 | -1.445392931 | 4.62E-16 |
| A2M-AS1 | 30.00575528 | -1.44585593 | 5.78E-04 |
| SEMA6A | 182.063587 | -1.446428256 | 1.54E-09 |
| AC090181.3 | 12.66189901 | -1.446721904 | 3.27E-02 |
| AC010976.1 | 25.86267738 | -1.447594232 | 3.42E-03 |
| ZFP2 | 31.7747584 | -1.449708827 | 7.01E-04 |
| AC025766.1 | 17.13948499 | -1.451684715 | 1.28E-02 |
| GMFB | 1903.598644 | -1.452419128 | 3.82E-10 |
| SLC26A10 | 106.1172386 | -1.452764522 | 1.37E-08 |
| TXNDC16 | 735.2004753 | -1.454385269 | 4.11E-36 |
| RMDN2 | 97.31966742 | -1.454943062 | 1.47E-08 |
| CHD9 | 2148.656633 | -1.455586749 | 1.04E-10 |
| SLIT3 | 447.7973117 | -1.456778628 | 2.01E-14 |
| SSPO | 225.5519511 | -1.457038239 | 1.18E-04 |
| NR2E3 | 27.75191453 | -1.457231703 | 2.04E-03 |
| PAXBP1-AS1 | 94.88010745 | -1.459382614 | 8.31E-12 |
| PCDHB14 | 50.50684348 | -1.459902753 | 1.48E-05 |
| EML4 | 8450.371969 | -1.46358505 | 3.78E-49 |
| GJA3 | 323.6162197 | -1.466012694 | 9.09E-24 |
| MSC-AS1 | 127.9931706 | -1.466903017 | 1.32E-03 |
| AC109631.1 | 24.14286993 | -1.468514631 | 5.79E-04 |
| AC073130.3 | 40.02148935 | -1.469247122 | 2.48E-05 |
| BAZ2B | 879.1717393 | -1.469397507 | 5.79E-09 |
| LINGO1 | 53.96417116 | -1.470408911 | 6.15E-04 |
| 2-Mar | 358.2724243 | -1.471303782 | 4.07E-12 |
| AC113382.1 | 25.08330705 | -1.471924836 | 2.06E-03 |
| LINC00271 | 18.92309629 | -1.473548393 | 8.08E-03 |
| CPA1 | 26.12893113 | -1.474483083 | 1.21E-02 |
| MFSD4A | 25.6331144 | -1.474891135 | 1.02E-02 |
| FANCM | 393.9957169 | -1.476426584 | 1.05E-06 |
| AC012501.1 | 23.96649049 | -1.476557463 | 8.85E-03 |
| AC021087.1 | 21.45280627 | -1.476904104 | 2.85E-03 |
| AL022069.2 | 10.01820396 | -1.477197156 | 3.06E-02 |
| HLF | 27.2149322 | -1.478000481 | 6.35E-03 |
| VPS13A | 2553.723416 | -1.478087897 | 4.71E-05 |
| PRRT2 | 400.9286235 | -1.478140481 | 9.11E-19 |
| DCLK2 | 707.8872453 | -1.478217781 | 5.92E-14 |
| IQGAP2 | 1477.33195 | -1.478304548 | 2.54E-14 |
| SLC22A17 | 525.7143164 | -1.478560474 | 1.20E-09 |
| Z83843.1 | 106.7766771 | -1.480007063 | 3.54E-07 |
| RTL5 | 2173.925386 | -1.481213931 | 6.61E-57 |
| MDGA1 | 555.6681225 | -1.481242004 | 7.45E-06 |
| SVIL | 3315.265826 | -1.481703739 | 3.21E-42 |

| | | | |
|---|---|---|---|
| CCDC121 | 148.176966 | -1.481903229 | 8.23E-10 |
| SNCA | 34.72054321 | -1.482140218 | 2.51E-03 |
| LINC01237 | 22.92773012 | -1.482193281 | 3.14E-03 |
| IPO9-AS1 | 9.846607716 | -1.484218191 | 4.66E-02 |
| PECAM1 | 169.5651196 | -1.484290788 | 1.28E-11 |
| AC009506.1 | 50.51966715 | -1.485508291 | 4.54E-06 |
| RCBTB2 | 954.637502 | -1.485853553 | 8.34E-34 |
| FAM78B | 308.7966026 | -1.487008912 | 2.44E-14 |
| C1orf54 | 51.82230516 | -1.487225622 | 1.14E-04 |
| AL662795.2 | 323.8813937 | -1.48871551 | 2.28E-11 |
| MED15P6 | 29.92975537 | -1.488931303 | 1.70E-03 |
| AC134312.1 | 26.72549384 | -1.490195487 | 1.46E-02 |
| LTBP1 | 2228.493436 | -1.490613685 | 7.40E-18 |
| AC021242.3 | 17.8700325 | -1.491623025 | 2.37E-02 |
| CFAP44 | 349.6453516 | -1.497076691 | 2.15E-10 |
| AL359715.2 | 10.4764427 | -1.499917004 | 3.86E-02 |
| IL1R1 | 1260.323681 | -1.500857384 | 5.11E-32 |
| MANSC1 | 461.6010775 | -1.501286475 | 1.45E-13 |
| AP001992.1 | 25.90380369 | -1.50147492 | 1.66E-03 |
| NTRK3 | 38.86279822 | -1.502598621 | 4.20E-02 |
| AC018638.6 | 17.14440965 | -1.504920337 | 1.05E-02 |
| LINC00342 | 173.0711505 | -1.505172931 | 9.20E-10 |
| NMUR1 | 23.97944588 | -1.506868399 | 8.43E-04 |
| TRGV5 | 22.90767887 | -1.506875653 | 2.03E-03 |
| AC010245.2 | 16.07305884 | -1.508001742 | 7.66E-03 |
| AC017083.1 | 17.17452587 | -1.509806146 | 1.23E-02 |
| GATA6-AS1 | 51.19632388 | -1.510186505 | 1.96E-06 |
| SNED1 | 2396.810234 | -1.510984315 | 7.62E-23 |
| CEND1 | 59.0622389 | -1.511667339 | 7.21E-06 |
| SAMD5 | 74.58486099 | -1.514210051 | 1.16E-05 |
| TENM4 | 2010.113719 | -1.514485576 | 1.86E-08 |
| MIS18A-AS1 | 12.05924686 | -1.51464892 | 2.62E-02 |
| RGL1 | 1815.256907 | -1.514666657 | 3.12E-32 |
| ZNF492 | 38.86841734 | -1.515175601 | 1.87E-04 |
| AL162171.4 | 23.20073892 | -1.515253106 | 1.48E-03 |
| FRRS1 | 170.6658154 | -1.515795312 | 2.06E-06 |
| ADGRF1 | 58.70965185 | -1.516939408 | 3.61E-07 |
| AC025171.4 | 21.72926794 | -1.520655242 | 1.66E-03 |
| KCNJ6 | 165.3882011 | -1.520884992 | 1.71E-11 |
| CDH12P2 | 21.78894892 | -1.521548114 | 4.75E-03 |
| AC091060.1 | 13.56319733 | -1.523644058 | 2.61E-02 |
| GDF11 | 3916.115371 | -1.524347359 | 9.62E-27 |
| PCDHGC5 | 14.16082268 | -1.524513409 | 1.76E-02 |
| TCEANC | 77.99028319 | -1.528798263 | 6.85E-08 |
| Z92544.2 | 182.6149303 | -1.52902172 | 3.61E-10 |
| EPHX3 | 10.35639321 | -1.52943021 | 2.22E-02 |
| MDH1B | 51.25257486 | -1.529652391 | 2.82E-06 |

| | | | |
|---|---|---|---|
| NHLRC4 | 70.47882681 | -1.530021607 | 2.07E-06 |
| AC005839.1 | 24.1898049 | -1.531446782 | 1.05E-03 |
| CFAP221 | 20.09340792 | -1.531802364 | 4.66E-03 |
| ARHGAP20 | 387.949941 | -1.533093468 | 1.82E-12 |
| ITM2B | 18797.51183 | -1.534058602 | 7.16E-24 |
| PLEKHB1 | 266.0658083 | -1.535925629 | 2.28E-08 |
| AC091563.1 | 20.66601571 | -1.536914977 | 4.50E-03 |
| SNAP25 | 73.26660247 | -1.538472187 | 3.11E-04 |
| RFTN2 | 36.09268729 | -1.538693716 | 2.49E-02 |
| GVQW2 | 19.57657147 | -1.544030176 | 2.91E-03 |
| MSC | 323.6871895 | -1.54432538 | 2.79E-04 |
| CEACAM21 | 17.26810419 | -1.545600533 | 6.36E-03 |
| ABCA8 | 289.3288945 | -1.548659076 | 3.07E-06 |
| AC008972.2 | 38.5054591 | -1.550679046 | 8.01E-06 |
| TNC | 30916.41729 | -1.553319415 | 7.79E-17 |
| NRIP1 | 1058.701831 | -1.553621143 | 3.92E-10 |
| LNX1 | 162.9220059 | -1.554027715 | 9.54E-17 |
| TGFBR3 | 1252.634429 | -1.554459451 | 1.19E-40 |
| SYPL2 | 83.161266 | -1.554948951 | 4.62E-07 |
| AL445931.1 | 15.68498476 | -1.556163337 | 7.73E-03 |
| AC008972.1 | 18.70068471 | -1.557323111 | 7.05E-03 |
| BTBD8 | 29.23506076 | -1.55857688 | 6.19E-03 |
| MEF2C | 193.607049 | -1.558863571 | 3.15E-13 |
| AP001972.5 | 1533.814559 | -1.559758293 | 2.95E-16 |
| AL441883.1 | 18.77579353 | -1.560188684 | 2.29E-02 |
| NA | 35.14152274 | -1.560331263 | 6.32E-04 |
| RB1 | 3726.742893 | -1.561949813 | 1.02E-26 |
| TRPS1 | 933.1259396 | -1.562607356 | 1.47E-15 |
| AL049830.3 | 11.08898876 | -1.563348146 | 4.81E-02 |
| PLCD4 | 1142.359525 | -1.565879183 | 1.06E-16 |
| AC092683.1 | 166.0784964 | -1.569327814 | 3.54E-09 |
| MT-TH | 24.1473077 | -1.569485636 | 3.23E-02 |
| KCNE4 | 2965.933508 | -1.569596654 | 8.62E-33 |
| NHSL2 | 362.3466133 | -1.572856787 | 3.90E-19 |
| AC010834.3 | 26.81611228 | -1.573235087 | 1.70E-03 |
| TRPM2-AS | 87.9797349 | -1.573550529 | 9.19E-10 |
| ABCC6 | 487.0257148 | -1.573908394 | 2.76E-19 |
| IQCN | 27.27984108 | -1.574210647 | 9.50E-03 |
| CD247 | 25.01308638 | -1.574449038 | 1.36E-03 |
| FAM110B | 546.9806472 | -1.576007251 | 4.80E-08 |
| SCG2 | 992.0410165 | -1.578204733 | 2.34E-10 |
| GPER1 | 115.2248845 | -1.578280164 | 7.85E-07 |
| ZNF214 | 149.0621941 | -1.579291213 | 1.35E-08 |
| RN7SKP74 | 10.63960034 | -1.579411978 | 1.61E-02 |
| AC112694.1 | 23.75344823 | -1.579929346 | 4.48E-04 |
| ARHGAP5 | 3812.275453 | -1.581683894 | 2.60E-11 |
| IGLV1-51 | 9.959366418 | -1.584842911 | 4.26E-02 |

| | | | |
|---|---|---|---|
| AC012511.1 | 21.46889958 | -1.585265031 | 2.49E-04 |
| RTCA-AS1 | 15.71405731 | -1.586203186 | 8.76E-03 |
| NADK2 | 3676.638597 | -1.589400141 | 8.70E-40 |
| MAP2K6 | 286.8384036 | -1.591504986 | 6.63E-22 |
| TSNAXIP1 | 79.41687888 | -1.592153673 | 2.67E-08 |
| ZBP1 | 10.18674428 | -1.59222335 | 4.13E-02 |
| TNIK | 2274.191439 | -1.592990105 | 1.80E-35 |
| CCDC87 | 25.69209508 | -1.594026126 | 7.10E-04 |
| RASEF | 196.4761225 | -1.598132424 | 2.12E-21 |
| AC011379.1 | 25.8671817 | -1.59876955 | 9.16E-05 |
| RN7SL521P | 9.532866364 | -1.60151429 | 3.14E-02 |
| AGBL2 | 104.4122325 | -1.60258396 | 1.00E-08 |
| PCDHA1 | 13.97894304 | -1.603429271 | 2.15E-02 |
| AQP6 | 26.44147426 | -1.607738268 | 5.78E-03 |
| AC253536.5 | 9.237527182 | -1.608417454 | 3.29E-02 |
| F2RL3 | 45.08989024 | -1.610051018 | 2.42E-04 |
| MAP3K5 | 1322.675559 | -1.613872015 | 1.12E-79 |
| AL929601.4 | 26.25866688 | -1.614635877 | 1.67E-03 |
| AC139100.1 | 11.20839737 | -1.617042634 | 1.17E-02 |
| GPR137C | 168.6258987 | -1.617905502 | 3.49E-16 |
| NOX3 | 28.38747355 | -1.620005192 | 1.43E-04 |
| NEK2P2 | 32.61469177 | -1.620469696 | 1.45E-03 |
| AC007364.1 | 9.772733044 | -1.622454856 | 3.34E-02 |
| PEAK3 | 21.50540919 | -1.622932147 | 3.02E-03 |
| AC106820.2 | 28.77241803 | -1.623118936 | 2.20E-04 |
| ZNF663P | 25.43069399 | -1.623917149 | 1.18E-03 |
| SELE | 16.48521074 | -1.624467515 | 4.20E-03 |
| AC140479.2 | 142.8946411 | -1.625369379 | 8.29E-13 |
| EDNRA | 37.02980821 | -1.626319408 | 1.16E-02 |
| SLC44A4 | 43.19448386 | -1.627023781 | 4.55E-04 |
| TMEM150C | 53.30532729 | -1.627189984 | 4.54E-04 |
| NOSTRIN | 322.9129418 | -1.629277191 | 3.08E-16 |
| ZHX1 | 2058.307391 | -1.630699886 | 3.55E-20 |
| AL358472.3 | 18.74929553 | -1.63080533 | 4.19E-03 |
| ATP2B1 | 2269.200867 | -1.631839445 | 2.47E-16 |
| AL139274.2 | 45.84750634 | -1.632098808 | 2.53E-05 |
| TNFRSF19 | 517.7426435 | -1.632431124 | 3.01E-19 |
| AL022068.1 | 12.71267123 | -1.636698546 | 1.23E-02 |
| GAB1 | 870.6201106 | -1.638064367 | 1.39E-36 |
| MAGEB3 | 36.6029307 | -1.641784526 | 5.26E-04 |
| AC092070.3 | 6.920013472 | -1.642243518 | 4.95E-02 |
| LINC02199 | 16.66390552 | -1.650249177 | 2.11E-02 |
| AL031673.1 | 19.97831105 | -1.650475443 | 1.99E-03 |
| TBX3 | 4851.962125 | -1.650826241 | 3.10E-49 |
| LINC00327 | 20.45128611 | -1.650911752 | 5.47E-03 |
| TRAC | 26.18792344 | -1.651795545 | 5.05E-03 |
| NFIA | 845.235121 | -1.654756724 | 8.52E-20 |

| | | | |
|---|---|---|---|
| PLXDC1 | 87.55184205 | -1.655649435 | 1.45E-04 |
| FAM111B | 742.562208 | -1.656490606 | 3.50E-04 |
| DOCK11 | 2817.853034 | -1.656813163 | 3.86E-12 |
| LINC01359 | 12.98320562 | -1.65836421 | 1.77E-02 |
| LRP3 | 178.7844076 | -1.658674455 | 5.21E-08 |
| SIRPB1 | 258.9285303 | -1.660168709 | 6.44E-09 |
| LRRC37A7P | 23.79231287 | -1.661673613 | 3.02E-03 |
| AC073655.2 | 12.2572459 | -1.668182478 | 2.16E-02 |
| AC026740.1 | 48.90043332 | -1.669231549 | 2.07E-07 |
| C3orf80 | 32.31383395 | -1.669380885 | 3.90E-04 |
| ZNF92 | 182.2272354 | -1.669453857 | 1.47E-07 |
| KITLG | 103.264813 | -1.669468007 | 2.02E-06 |
| RAB11FIP1P1 | 56.13576336 | -1.673958903 | 2.00E-04 |
| IFIT1 | 1401.914147 | -1.674562527 | 1.48E-16 |
| SIRPG-AS1 | 14.55610769 | -1.676372369 | 4.11E-02 |
| EDRF1-DT | 13.63250119 | -1.677660121 | 6.99E-03 |
| CARD6 | 1819.649875 | -1.678732095 | 8.39E-40 |
| FAM171B | 279.1691248 | -1.67905723 | 2.13E-07 |
| AKAP5 | 222.2346906 | -1.680582122 | 6.33E-17 |
| AC008434.1 | 11.24641032 | -1.682809871 | 8.04E-03 |
| GPRC5C | 1051.716387 | -1.683632102 | 3.89E-15 |
| BTG2 | 1018.081876 | -1.687062645 | 1.02E-26 |
| ELN | 32.22984501 | -1.687551103 | 2.42E-05 |
| AC006058.1 | 319.7637519 | -1.688866586 | 3.85E-21 |
| AC005837.1 | 17.76815366 | -1.691966166 | 3.01E-03 |
| SCN11A | 27.86262684 | -1.694365385 | 1.89E-04 |
| ACPP | 27.55663385 | -1.697074135 | 6.82E-05 |
| MAPK10 | 51.86224909 | -1.698043374 | 4.69E-06 |
| DUSP6 | 3860.577203 | -1.698905118 | 1.47E-35 |
| AF131216.3 | 48.22332352 | -1.699579129 | 4.92E-06 |
| PKNOX2 | 537.5619923 | -1.699686787 | 1.12E-25 |
| ZMAT4 | 504.9050422 | -1.701267498 | 6.49E-22 |
| CASC8 | 11.89497762 | -1.701661606 | 5.17E-03 |
| CNIH2 | 324.591191 | -1.702381508 | 4.72E-11 |
| AC138123.1 | 17.09661336 | -1.702646439 | 9.48E-03 |
| AC025470.2 | 12.21345488 | -1.703773256 | 2.62E-02 |
| AC021321.1 | 29.50129298 | -1.703813786 | 9.65E-05 |
| RF02271 | 10.29439901 | -1.703824208 | 2.37E-02 |
| AC079465.1 | 12.33074335 | -1.704255773 | 2.47E-02 |
| TMEM233 | 90.45126575 | -1.706495031 | 1.94E-04 |
| PDZRN3 | 211.3588518 | -1.707250335 | 3.15E-18 |
| VASH2 | 115.4401106 | -1.710722297 | 7.27E-07 |
| KIAA1614-AS1 | 20.95491454 | -1.712406363 | 1.20E-03 |
| CHRM1 | 20.60036107 | -1.713850917 | 1.95E-02 |
| LINC00933 | 28.50852274 | -1.714524642 | 7.50E-03 |
| TRAF3IP3 | 99.96308918 | -1.717329863 | 1.35E-11 |
| SIGLEC15 | 122.8361965 | -1.722256469 | 4.50E-11 |

| | | | |
|---|---|---|---|
| ADCY10P1 | 106.8816125 | -1.723109291 | 1.26E-06 |
| AP002784.1 | 15.36806646 | -1.723659501 | 2.85E-03 |
| LGR4 | 3291.633279 | -1.723776142 | 2.84E-46 |
| HAS2-AS1 | 12.5539903 | -1.724406452 | 9.32E-03 |
| SOBP | 699.2504882 | -1.726968916 | 5.13E-51 |
| OASL | 87.89182137 | -1.730985592 | 7.19E-07 |
| NF1P7 | 19.5947773 | -1.732171751 | 4.14E-04 |
| AP000787.1 | 90.97820889 | -1.732837836 | 1.03E-08 |
| C2CD4A | 10.978227 | -1.737223376 | 7.23E-03 |
| AC083843.3 | 226.7278591 | -1.739821221 | 2.23E-09 |
| LIX1-AS1 | 19.51395383 | -1.740156929 | 1.36E-03 |
| NR2F1-AS1 | 377.042445 | -1.741285588 | 2.50E-26 |
| AC023300.2 | 9.952432568 | -1.743850268 | 1.39E-02 |
| GALNT5 | 69.69342666 | -1.745129159 | 1.06E-06 |
| ETV1 | 1912.447401 | -1.745447199 | 3.15E-50 |
| NT5C1A | 27.98056727 | -1.746358619 | 1.48E-04 |
| RN7SL81P | 11.13600973 | -1.748923355 | 1.94E-02 |
| EMILIN1 | 5275.584214 | -1.749823413 | 3.17E-23 |
| FAM214A | 916.7426868 | -1.750598646 | 2.00E-24 |
| PROX1 | 36.00156087 | -1.750668868 | 1.79E-03 |
| GFRA3 | 106.5806063 | -1.750988983 | 2.15E-10 |
| LRRC73 | 66.17494783 | -1.752630681 | 1.01E-08 |
| NAP1L2 | 60.86264806 | -1.757928915 | 8.37E-04 |
| PLPPR1 | 23.86116289 | -1.75805722 | 3.42E-04 |
| AC097717.1 | 10.81695375 | -1.760910905 | 2.49E-02 |
| LINC00922 | 15.62938541 | -1.761873998 | 1.46E-02 |
| BCRP3 | 45.78330543 | -1.76407183 | 4.87E-08 |
| MMP1 | 74.36681813 | -1.764342424 | 9.65E-05 |
| NA | 11.84966354 | -1.76883986 | 7.12E-03 |
| NTNG2 | 150.2818736 | -1.770862287 | 3.22E-06 |
| AC120049.1 | 38.88637755 | -1.773587991 | 7.89E-06 |
| CD38 | 109.1153045 | -1.774772931 | 3.94E-06 |
| ADGRF3 | 53.77481944 | -1.775424281 | 1.67E-05 |
| EMILIN3 | 78.86613811 | -1.779624305 | 1.17E-06 |
| LINC00578 | 8.053509764 | -1.780382297 | 2.83E-02 |
| GIMAP5 | 56.13836726 | -1.78109877 | 1.88E-04 |
| PEG3 | 8882.770975 | -1.781532614 | 1.33E-38 |
| AC120053.1 | 12.97532409 | -1.783157 | 9.49E-03 |
| VPS13C | 3551.851662 | -1.784961282 | 1.17E-08 |
| RYR3 | 63.81883754 | -1.78624712 | 1.69E-03 |
| BCAS1 | 91.26170515 | -1.786434792 | 8.54E-08 |
| LINC01132 | 37.93370493 | -1.788515634 | 1.89E-07 |
| NA | 224.2178336 | -1.78958763 | 8.97E-14 |
| APC | 2832.53114 | -1.793232579 | 6.69E-14 |
| AC091181.1 | 14.26359752 | -1.797348322 | 4.92E-03 |
| AC079594.2 | 20.72220822 | -1.799014825 | 2.90E-02 |
| CCDC39 | 40.01878636 | -1.802210276 | 9.66E-04 |

| | | | |
|---|---|---|---|
| PRKCB | 1065.831152 | -1.804948329 | 6.72E-24 |
| CPE | 2379.926947 | -1.806774467 | 4.78E-44 |
| NGFR | 2528.28695 | -1.80884021 | 3.42E-20 |
| TNN | 27.80903769 | -1.810137355 | 2.83E-04 |
| SCTR | 23.25023017 | -1.812815605 | 3.38E-03 |
| COL5A3 | 625.5339231 | -1.814643662 | 7.85E-19 |
| SNTB1 | 288.127424 | -1.815762356 | 2.02E-07 |
| AMBP | 14.23918813 | -1.816604208 | 9.61E-03 |
| LPL | 5043.224616 | -1.822855458 | 1.35E-39 |
| LINC01262 | 74.40238564 | -1.8232092 | 4.22E-12 |
| CACNA1E | 31.3844744 | -1.824959625 | 7.94E-04 |
| LINC02108 | 7.599326394 | -1.825940659 | 4.31E-02 |
| IL6ST | 22565.49008 | -1.826434537 | 1.23E-09 |
| UBXN10-AS1 | 15.03613687 | -1.830346027 | 4.68E-04 |
| FZD1 | 1665.798152 | -1.83100439 | 1.64E-58 |
| FXYD2 | 41.98422852 | -1.837066802 | 1.92E-05 |
| FLG2 | 11.11265756 | -1.841883127 | 1.42E-02 |
| BAMBI | 4010.746087 | -1.845232245 | 1.72E-27 |
| NA | 17.82293091 | -1.848754956 | 5.89E-04 |
| LINC02260 | 53.3414811 | -1.849080697 | 6.78E-06 |
| AC006111.2 | 16.47899019 | -1.849631128 | 1.52E-03 |
| TMEM191C | 53.90717508 | -1.850075131 | 6.55E-05 |
| GAS1 | 64.47266057 | -1.851430253 | 2.54E-10 |
| PLAT | 9788.326261 | -1.853103805 | 9.86E-22 |
| DYNC2H1 | 1286.378422 | -1.853540705 | 2.98E-10 |
| AC006213.5 | 7.711689508 | -1.854478403 | 3.17E-02 |
| ENHO | 164.5618616 | -1.855724907 | 1.39E-15 |
| NA | 791.4238412 | -1.856054429 | 9.29E-31 |
| BX119917.1 | 35.28777911 | -1.861058352 | 5.54E-07 |
| EXTL3-AS1 | 56.09264869 | -1.861298322 | 1.04E-06 |
| AL445670.1 | 7.507883969 | -1.866276738 | 4.59E-02 |
| EYA1 | 61.04274887 | -1.866537825 | 1.77E-09 |
| CHRNB2 | 21.60480173 | -1.867394567 | 8.80E-05 |
| NA | 33.09930813 | -1.867935108 | 6.46E-06 |
| NWD1 | 21.24870833 | -1.872791666 | 1.46E-04 |
| NRP1 | 5641.832865 | -1.873269319 | 2.60E-48 |
| VMO1 | 25.53474 | -1.873789076 | 1.62E-04 |
| LINC01686 | 57.01245041 | -1.875011013 | 1.03E-08 |
| BPI | 155.9343946 | -1.876830801 | 6.42E-12 |
| SELL | 65.28807749 | -1.877872595 | 3.85E-06 |
| PDGFRL | 578.6213394 | -1.878180826 | 2.79E-21 |
| TNFSF10 | 463.9076168 | -1.881096485 | 2.47E-08 |
| AC008555.7 | 8.255115474 | -1.8837816 | 4.49E-02 |
| AC103691.1 | 22.88767344 | -1.885811059 | 1.71E-03 |
| NRP2 | 3214.243923 | -1.888015491 | 8.18E-114 |
| AC009318.2 | 8.898131022 | -1.890295608 | 3.20E-02 |
| RAB17 | 2197.833679 | -1.891338868 | 5.28E-34 |

| | | | |
|---|---|---|---|
| CLMP | 3028.617727 | -1.897634737 | 8.73E-36 |
| PDE4D | 2698.714731 | -1.898669985 | 5.45E-23 |
| TAL2 | 5.354113806 | -1.899137311 | 4.52E-02 |
| IL17RD | 1697.446375 | -1.900386844 | 6.64E-25 |
| AC006058.4 | 114.8916737 | -1.902821346 | 2.29E-13 |
| PLAC9 | 94.03662567 | -1.905499331 | 1.22E-10 |
| HPS3 | 6068.142851 | -1.905850449 | 5.67E-49 |
| FN1 | 116782.2439 | -1.906988369 | 6.78E-16 |
| MBNL3 | 313.5751306 | -1.907434407 | 4.34E-08 |
| AC008264.2 | 36.20563716 | -1.911929862 | 3.06E-04 |
| AC020612.3 | 8.979197569 | -1.912829884 | 9.55E-03 |
| PTHLH | 279.5986255 | -1.918039233 | 1.14E-14 |
| HTRA1 | 853.162337 | -1.918282373 | 6.64E-30 |
| LIFR | 13533.60101 | -1.923858768 | 2.87E-21 |
| PARP15 | 35.78849082 | -1.926600347 | 7.40E-06 |
| NA | 10.45776167 | -1.928074701 | 1.00E-02 |
| LGALS9 | 267.2477864 | -1.931070933 | 1.11E-07 |
| PCDH12 | 238.254452 | -1.931454147 | 1.59E-13 |
| ALMS1P1 | 12.52298362 | -1.93495525 | 1.59E-02 |
| ARHGAP28 | 1096.61935 | -1.939459991 | 1.09E-22 |
| SLC35E1P1 | 14.23395906 | -1.94568589 | 5.62E-03 |
| ABCA9 | 44.27440312 | -1.946623069 | 4.67E-06 |
| MMD | 3317.81114 | -1.949093103 | 1.68E-75 |
| COL3A1 | 378.3612853 | -1.950330771 | 3.80E-18 |
| AP001351.1 | 15.36296917 | -1.950556056 | 5.37E-03 |
| RAPGEF5 | 299.1615067 | -1.953419173 | 5.62E-20 |
| LHFPL3-AS2 | 9.327085567 | -1.955304731 | 2.94E-02 |
| AC012254.3 | 17.6987559 | -1.956436289 | 1.84E-04 |
| ST3GAL5-AS1 | 58.4679868 | -1.957964088 | 1.90E-12 |
| AC073210.1 | 9.605607766 | -1.958934792 | 4.24E-02 |
| CR383656.4 | 19.28138109 | -1.96214103 | 1.26E-03 |
| CAMK4 | 96.91924048 | -1.964718594 | 2.57E-09 |
| RGS4 | 778.1227395 | -1.966891048 | 7.34E-12 |
| GTF2IRD1P1 | 11.75515321 | -1.967581097 | 1.47E-02 |
| NA | 14.66958513 | -1.967623001 | 3.93E-03 |
| NA | 6.194262225 | -1.96878696 | 4.37E-02 |
| PTGER2 | 910.6436438 | -1.969609908 | 5.82E-27 |
| MARVELD3 | 105.4161867 | -1.973600343 | 1.13E-13 |
| AL499616.1 | 13.79995614 | -1.975275857 | 4.47E-03 |
| SLCO6A1 | 11.45394403 | -1.975407183 | 1.49E-02 |
| DDN | 37.56274831 | -1.975663801 | 9.18E-05 |
| ZAP70 | 104.1860247 | -1.975841728 | 3.71E-10 |
| NBPF22P | 6.433062377 | -1.976546412 | 2.65E-02 |
| C3AR1 | 62.64796635 | -1.977120089 | 5.61E-09 |
| AL356488.3 | 34.82499263 | -1.978138764 | 1.93E-08 |
| TPI1P2 | 29.76508584 | -1.978687878 | 3.31E-06 |
| S1PR3 | 4057.598699 | -1.979663934 | 5.63E-53 |

| | | | |
|---|---|---|---|
| IL1R2 | 12.0580723 | -1.980544093 | 5.99E-03 |
| SLC30A10 | 9.108997572 | -1.981887444 | 1.17E-02 |
| NPTX1 | 3190.575678 | -1.984936771 | 1.36E-27 |
| LINC00461 | 138.3576983 | -1.991332893 | 1.93E-10 |
| NA | 6.372103817 | -1.995106543 | 4.71E-02 |
| TCF4 | 3210.623798 | -1.995282679 | 4.73E-08 |
| DUSP27 | 48.58308619 | -1.996962141 | 5.61E-05 |
| AC093677.2 | 17.96716077 | -1.997003996 | 4.39E-04 |
| HEY2 | 65.51168791 | -1.997920517 | 9.78E-10 |
| RNF150 | 107.7773362 | -1.999686726 | 2.48E-13 |
| CTNNA2 | 10.07638971 | -2.001882555 | 2.70E-02 |
| NA | 9.251941923 | -2.003168198 | 1.19E-02 |
| SGPP2 | 21.98751951 | -2.005520647 | 2.86E-03 |
| SLC14A2 | 38.02249123 | -2.008664832 | 1.32E-06 |
| OR7E128P | 14.04561412 | -2.010295042 | 1.65E-03 |
| SPAG17 | 14.86134536 | -2.011859175 | 2.26E-03 |
| RTN1 | 9.232349879 | -2.012663618 | 9.73E-03 |
| DOCK10 | 51.60308168 | -2.014827811 | 7.91E-04 |
| SLC14A1 | 545.9822054 | -2.016596351 | 4.36E-03 |
| AC068279.1 | 52.80865783 | -2.018644812 | 3.37E-09 |
| CFI | 1692.65687 | -2.019975386 | 3.94E-09 |
| TSPAN7 | 39.81758527 | -2.019987588 | 4.23E-08 |
| TNFRSF11B | 3103.669719 | -2.020588432 | 1.40E-56 |
| AC024909.1 | 67.99772793 | -2.021114743 | 7.78E-09 |
| IFI44 | 187.1328943 | -2.022281343 | 4.74E-13 |
| PLVAP | 42.61214939 | -2.024169026 | 4.71E-05 |
| AL008726.1 | 10.79443075 | -2.024609519 | 9.89E-03 |
| COL1A2 | 3558.442137 | -2.024866967 | 2.98E-17 |
| MCTP1 | 299.6263461 | -2.026069498 | 1.53E-07 |
| GATM | 9.70851274 | -2.027032332 | 1.21E-02 |
| IDI2-AS1 | 13.37064553 | -2.032419225 | 1.21E-02 |
| LGALS12 | 389.1815947 | -2.033313404 | 5.26E-13 |
| KCNK2 | 141.6576935 | -2.037557717 | 1.61E-11 |
| STK32A | 19.69460038 | -2.040398777 | 4.88E-03 |
| PCDH19 | 17.98510115 | -2.043270628 | 9.71E-05 |
| NAP1L3 | 138.9603458 | -2.046545869 | 3.85E-11 |
| CCT8P1 | 14.76212583 | -2.047696575 | 8.62E-04 |
| ELMO1 | 314.1325128 | -2.050130074 | 6.60E-18 |
| METTL7A | 666.3257155 | -2.05048237 | 9.40E-15 |
| CYP39A1 | 219.4277118 | -2.051187971 | 1.73E-23 |
| SYNGR4 | 26.26163114 | -2.052914033 | 5.45E-04 |
| PCDH18 | 57.24297472 | -2.057970717 | 1.16E-05 |
| AC017116.1 | 20.85556617 | -2.060147563 | 1.90E-02 |
| LDB2 | 2892.946269 | -2.0604715 | 4.43E-72 |
| EPHB6 | 1812.119747 | -2.063246561 | 1.15E-27 |
| SLC7A4 | 76.41360638 | -2.065065502 | 1.21E-07 |
| WSCD2 | 53.14963086 | -2.065183984 | 5.26E-04 |

| | | | |
|---|---|---|---|
| SLC1A2 | 13.68876523 | -2.068370483 | 4.67E-02 |
| RF00017 | 8.264416105 | -2.070813647 | 4.39E-02 |
| EVX1 | 13.13102745 | -2.075439666 | 5.10E-04 |
| ZACN | 24.35531445 | -2.080947895 | 4.39E-03 |
| GNG13 | 36.65081425 | -2.086910421 | 1.67E-08 |
| AL158211.5 | 9.05234245 | -2.088025121 | 2.45E-02 |
| SOX4 | 406.2559629 | -2.090173782 | 1.07E-25 |
| AKR1B10 | 66.94033728 | -2.102689388 | 2.99E-03 |
| AC004846.2 | 7.837252615 | -2.1033091 | 1.75E-02 |
| TEX26-AS1 | 9.224930339 | -2.103909486 | 1.41E-02 |
| BUD13P1 | 14.41791251 | -2.105335774 | 2.75E-03 |
| AC010680.3 | 9.42834952 | -2.110618358 | 1.62E-02 |
| AC066613.1 | 23.66354476 | -2.110655594 | 1.87E-04 |
| MYO7A | 246.39633 | -2.111835217 | 6.69E-18 |
| AC073621.1 | 9.543323116 | -2.113591888 | 8.29E-03 |
| NPB | 18.04693315 | -2.114072126 | 4.95E-02 |
| AC023421.2 | 9.598318077 | -2.114779993 | 2.77E-03 |
| NA | 45.0838267 | -2.116208323 | 3.68E-10 |
| ABCC11 | 26.48989666 | -2.116932911 | 4.14E-05 |
| NPM1P9 | 12.32308265 | -2.116940963 | 5.30E-03 |
| AL035448.1 | 10.52901536 | -2.117781786 | 4.21E-03 |
| DEPDC1-AS1 | 8.111181073 | -2.118070816 | 3.68E-02 |
| RRS1-AS1 | 20.2277811 | -2.119094198 | 4.02E-05 |
| CP | 65.43405638 | -2.119776435 | 4.43E-09 |
| GALR1 | 16.93168639 | -2.125543726 | 8.96E-03 |
| GJA4 | 10.57516355 | -2.133765242 | 1.76E-02 |
| DRC7 | 9.706805279 | -2.136989822 | 1.92E-02 |
| IFIT2 | 1178.033649 | -2.137268174 | 4.59E-53 |
| CLDN10 | 8.452333829 | -2.139706987 | 3.06E-02 |
| ATRNL1 | 133.6598312 | -2.148363584 | 2.59E-18 |
| CDH11 | 43.29998427 | -2.151224261 | 5.51E-06 |
| FAT4 | 26.78071554 | -2.151379514 | 2.78E-02 |
| PIEZO2 | 3798.179918 | -2.153969332 | 1.03E-140 |
| RSPH4A | 9.050537237 | -2.155789582 | 6.32E-03 |
| AF230666.1 | 12.29874585 | -2.158429572 | 2.05E-02 |
| TMOD1 | 1497.162289 | -2.15929985 | 4.56E-26 |
| TMEM37 | 476.6411504 | -2.15947589 | 6.96E-23 |
| HGFAC | 24.78828744 | -2.16034324 | 5.92E-06 |
| RYR2 | 420.6593994 | -2.160490351 | 7.88E-04 |
| CFAP206 | 10.33592942 | -2.165304854 | 6.76E-03 |
| SAMD15 | 178.1043318 | -2.168386646 | 1.82E-21 |
| TRIB2 | 5941.218021 | -2.171414657 | 2.72E-64 |
| NR2F1 | 1908.419877 | -2.173853472 | 7.79E-30 |
| MYADML2 | 7.200107882 | -2.17495406 | 9.35E-03 |
| PAMR1 | 17.14486306 | -2.176459005 | 5.93E-03 |
| AL513534.2 | 18.4908913 | -2.182519049 | 5.85E-05 |
| SLC5A9 | 31.88326152 | -2.184668416 | 2.31E-03 |

| | | | |
|---|---|---|---|
| FKBP6 | 8.466083031 | -2.185105675 | 1.55E-02 |
| ANGPTL1 | 9.294027893 | -2.187512427 | 3.70E-02 |
| LINC01060 | 80.61008245 | -2.189884307 | 9.07E-17 |
| AL137847.1 | 17.62640037 | -2.190832711 | 2.76E-04 |
| Z95152.1 | 9.983162855 | -2.194357047 | 1.07E-02 |
| LRRC10B | 64.95964827 | -2.195063796 | 2.80E-07 |
| PDGFRA | 1613.900805 | -2.199902952 | 9.76E-48 |
| AL139393.2 | 31.78021253 | -2.201277631 | 3.79E-06 |
| SGSM1 | 82.61987736 | -2.20216291 | 2.21E-10 |
| DRD2 | 181.3447405 | -2.202616435 | 1.79E-20 |
| LANCL1-AS1 | 14.20954401 | -2.20456154 | 6.98E-04 |
| AC005614.1 | 9.40582381 | -2.204687348 | 3.42E-02 |
| CPXM2 | 397.4177067 | -2.207727289 | 3.95E-24 |
| AC104117.3 | 6.324456609 | -2.207786678 | 3.31E-02 |
| ZC3H12D | 48.81604572 | -2.211111467 | 5.68E-07 |
| NOX5 | 244.0562303 | -2.21117259 | 3.20E-24 |
| CABP5 | 17.04454047 | -2.213970613 | 1.43E-03 |
| AL355922.2 | 5.830788662 | -2.21515749 | 3.53E-02 |
| DEPTOR | 574.1833815 | -2.216022243 | 5.32E-27 |
| AC084083.1 | 13.67494269 | -2.218045047 | 1.34E-03 |
| TNKS2-AS1 | 15.76441824 | -2.22902879 | 1.63E-03 |
| NA | 24.42026338 | -2.229380523 | 3.98E-06 |
| NA | 13.3543947 | -2.233734367 | 2.99E-03 |
| MIR3153 | 9.557373238 | -2.236765444 | 1.11E-02 |
| AL136084.3 | 71.27286851 | -2.237425627 | 3.61E-06 |
| RESF1 | 1525.676642 | -2.242580709 | 8.98E-17 |
| AL121574.1 | 8.127876777 | -2.246338405 | 1.31E-02 |
| CLEC3A | 27.21407875 | -2.246533793 | 4.57E-02 |
| NA | 6.062339289 | -2.24675438 | 4.05E-02 |
| DCLK3 | 37.85752614 | -2.246931312 | 6.28E-04 |
| AC137932.2 | 28.57861597 | -2.255027694 | 3.20E-05 |
| CCN3 | 283.7792535 | -2.259331895 | 9.63E-30 |
| EID3 | 84.86515522 | -2.262652335 | 2.08E-17 |
| BMP7 | 617.2385659 | -2.268383657 | 6.98E-21 |
| AC106786.2 | 8.560531846 | -2.268474581 | 2.93E-02 |
| TLR3 | 366.6552858 | -2.268786046 | 4.72E-19 |
| Z99289.1 | 8.782062838 | -2.270345575 | 1.43E-02 |
| NPAS3 | 33.85604416 | -2.270907655 | 1.41E-06 |
| PAK3 | 94.81753912 | -2.271409973 | 2.13E-08 |
| SAMD9L | 932.6396089 | -2.271416343 | 2.91E-14 |
| AC010680.4 | 7.049811061 | -2.279412833 | 1.81E-02 |
| AC005790.1 | 5.466737288 | -2.284824861 | 2.91E-02 |
| CADPS | 45.90525098 | -2.287877602 | 2.23E-05 |
| AC092881.1 | 9.303526403 | -2.293551774 | 3.23E-02 |
| EWSAT1 | 144.2678212 | -2.297652232 | 1.06E-10 |
| AC026412.2 | 5.929610817 | -2.299267718 | 3.94E-02 |
| AC004067.1 | 10.2570936 | -2.301432184 | 1.24E-02 |

| | | | |
|---|---|---|---|
| COL2A1 | 456.0622899 | -2.301562419 | 7.27E-19 |
| AC106875.2 | 5.368267226 | -2.30170699 | 4.10E-02 |
| TMEM119 | 206.1753641 | -2.307189208 | 2.76E-19 |
| TTC23L | 18.95128468 | -2.310859759 | 2.07E-04 |
| BX088651.2 | 39.56877275 | -2.311214913 | 1.46E-04 |
| ATP2B2 | 75.43341306 | -2.311981531 | 3.74E-08 |
| JPH4 | 834.6588439 | -2.313656288 | 1.82E-21 |
| PYGM | 80.99656343 | -2.321637379 | 2.07E-09 |
| AC090360.1 | 8.70452416 | -2.322819984 | 6.37E-03 |
| AL355516.1 | 10.65810529 | -2.326353552 | 6.37E-03 |
| PLXNA4 | 515.4222672 | -2.329661968 | 1.34E-23 |
| PRKXP1 | 23.97776507 | -2.330812499 | 4.17E-07 |
| HAS2 | 104.2657699 | -2.331566402 | 8.72E-11 |
| HEPH | 29.34641488 | -2.337736399 | 1.93E-04 |
| BFSP2 | 15.52825355 | -2.342138779 | 2.19E-02 |
| AC073626.2 | 8.616285844 | -2.344215404 | 1.53E-02 |
| SPRY4-AS1 | 9.432133691 | -2.34869554 | 1.08E-02 |
| ADAMTS9 | 372.2011459 | -2.350847882 | 3.48E-24 |
| AC016957.2 | 7.752886535 | -2.355274551 | 1.63E-02 |
| TSPAN11 | 2754.87485 | -2.357362487 | 5.42E-26 |
| SULT2A1 | 70.46590007 | -2.35894833 | 1.44E-07 |
| ECT2L | 5.984631906 | -2.361372622 | 3.02E-02 |
| SLC18A1 | 15.55119098 | -2.363037535 | 8.98E-05 |
| ADCY5 | 17.41633914 | -2.368059965 | 1.85E-03 |
| AL109613.1 | 11.43374656 | -2.370719971 | 1.49E-03 |
| SCN2A | 55.63473643 | -2.377115579 | 1.89E-06 |
| SLC25A15P4 | 6.66330282 | -2.378826563 | 2.15E-02 |
| FAIM2 | 317.161598 | -2.380235064 | 2.11E-19 |
| AC012360.1 | 6.488712197 | -2.382738147 | 1.68E-02 |
| PTENP1 | 23.71083276 | -2.385125003 | 5.49E-06 |
| SAMD9 | 759.2126716 | -2.392046679 | 6.13E-18 |
| C16orf89 | 40.33127915 | -2.392919321 | 1.49E-05 |
| AC104446.1 | 5.961573802 | -2.39300344 | 2.28E-02 |
| AC025175.1 | 9.155338254 | -2.393076266 | 6.16E-03 |
| LRRTM2 | 15.22254316 | -2.394401591 | 1.22E-03 |
| PIK3R1 | 4595.722417 | -2.394735788 | 4.62E-100 |
| LMOD2 | 18.51326571 | -2.398108004 | 3.98E-06 |
| VNN2 | 32.72060292 | -2.403699655 | 6.90E-05 |
| SERPINE3 | 16.91027922 | -2.40708467 | 3.08E-05 |
| C7 | 11.38273905 | -2.408867476 | 1.14E-02 |
| STARD4-AS1 | 775.5088185 | -2.422089648 | 2.80E-21 |
| MAP1A | 1216.104392 | -2.425436404 | 4.80E-55 |
| DNER | 848.2215533 | -2.428317071 | 8.72E-63 |
| TGFB3 | 607.8050464 | -2.431853681 | 1.81E-36 |
| AP001931.1 | 8.404150489 | -2.434350502 | 5.79E-03 |
| NADK2-AS1 | 55.41232106 | -2.440411209 | 2.56E-12 |
| PRSS35 | 16.35980289 | -2.444402943 | 1.45E-03 |

| | | | |
|---|---|---|---|
| NREP | 12921.79755 | -2.445731665 | 5.99E-29 |
| CLEC3B | 312.9420554 | -2.44620358 | 4.06E-13 |
| TTC3P1 | 64.45909731 | -2.446844363 | 2.04E-11 |
| AL513548.1 | 37.02258029 | -2.461897789 | 1.38E-07 |
| GIMAP8 | 25.83136293 | -2.471815831 | 7.73E-03 |
| AC008555.1 | 13.9843821 | -2.474176683 | 9.48E-05 |
| TEX15 | 71.56116553 | -2.484999408 | 6.12E-08 |
| AC026355.2 | 13.91176656 | -2.491363864 | 1.45E-04 |
| C22orf15 | 18.48934264 | -2.492348612 | 1.14E-03 |
| RXFP4 | 11.38414106 | -2.492593002 | 6.61E-04 |
| ECM2 | 30.17136374 | -2.49534957 | 2.52E-05 |
| PRODH | 16.67697053 | -2.496337939 | 2.33E-02 |
| AC025164.1 | 16.69760917 | -2.496995028 | 4.32E-03 |
| TLL2 | 230.6792461 | -2.498513464 | 1.90E-34 |
| AC002451.1 | 7.972422639 | -2.50151507 | 6.45E-03 |
| C3orf49 | 7.959344906 | -2.504240677 | 8.03E-03 |
| CACNA1D | 40.85940222 | -2.507621505 | 3.09E-05 |
| COL15A1 | 15563.86287 | -2.508095071 | 6.36E-26 |
| RTN4RL1 | 146.9324584 | -2.508104186 | 4.24E-26 |
| GALNT17 | 25.55343124 | -2.529068216 | 6.34E-08 |
| BFSP2-AS1 | 17.70852641 | -2.531395095 | 6.39E-05 |
| C1orf115 | 406.0582373 | -2.532760817 | 7.37E-16 |
| KCNK17 | 7.93099649 | -2.53765231 | 1.65E-03 |
| GABRA2 | 53.37695158 | -2.539564296 | 4.88E-07 |
| AC087045.2 | 9.78579654 | -2.541106245 | 6.50E-04 |
| RGS5 | 1408.439858 | -2.550282559 | 1.71E-12 |
| TMEM100 | 132.9671535 | -2.552378994 | 4.63E-27 |
| GNG2 | 1893.590023 | -2.564734169 | 6.22E-59 |
| LINC00589 | 13.02961938 | -2.568320212 | 5.09E-04 |
| AL135818.2 | 5.597184778 | -2.57552378 | 8.97E-03 |
| CNTFR | 700.6352166 | -2.581350776 | 5.92E-32 |
| AL357060.1 | 57.41758522 | -2.591992294 | 1.15E-15 |
| AC093835.1 | 6.606173093 | -2.592840821 | 1.36E-02 |
| MGP | 1888.148666 | -2.595957629 | 2.64E-09 |
| LPAR6 | 141.6397686 | -2.604473994 | 8.99E-15 |
| SMAD5-AS1 | 9.513127475 | -2.608158807 | 9.90E-03 |
| AC096921.2 | 33.69750771 | -2.61254028 | 5.11E-09 |
| TDH | 10.90157754 | -2.624132997 | 3.16E-03 |
| OLFML2B | 1524.891652 | -2.626843399 | 1.90E-27 |
| AC010420.1 | 5.023018429 | -2.627987117 | 1.66E-02 |
| VSTM4 | 321.1915059 | -2.635101687 | 3.45E-28 |
| GPRIN3 | 52.6600565 | -2.636205797 | 1.55E-07 |
| INTS6L-AS1 | 6.545704255 | -2.638359943 | 1.20E-02 |
| SUSD5 | 41.31474875 | -2.639367296 | 1.25E-08 |
| ATP8A1 | 225.1808029 | -2.643340519 | 7.87E-07 |
| SLC26A4 | 67.63469275 | -2.645148394 | 1.96E-10 |
| HLX | 777.8677108 | -2.65265016 | 3.72E-34 |

| | | | |
|---|---|---|---|
| AL589993.1 | 9.839506058 | -2.653328669 | 1.53E-03 |
| RGS16 | 706.038458 | -2.661413876 | 6.14E-18 |
| PLEKHG1 | 194.9176734 | -2.666632374 | 7.44E-16 |
| SPINK4 | 6.515444977 | -2.670579156 | 1.40E-03 |
| AC092364.2 | 6.569950109 | -2.672873647 | 5.16E-03 |
| ENPP2 | 849.7703046 | -2.679435849 | 1.09E-30 |
| SPP1 | 3358.989557 | -2.692775455 | 4.33E-29 |
| SOD3 | 336.2709669 | -2.698459633 | 1.15E-24 |
| CLEC7A | 23.75435793 | -2.700255384 | 5.09E-05 |
| KCNMB2 | 23.37913114 | -2.708180294 | 6.48E-05 |
| SLC12A2 | 5479.546749 | -2.727142984 | 2.29E-93 |
| KDR | 808.088877 | -2.728267291 | 5.78E-26 |
| SEMA5B | 29.336469 | -2.731411633 | 1.25E-04 |
| MTUS2 | 74.85023664 | -2.732914218 | 1.08E-15 |
| SLC9A9 | 84.18489021 | -2.736673736 | 1.71E-09 |
| ITGA9 | 2422.723682 | -2.737313062 | 2.25E-32 |
| LRFN2 | 11.69458686 | -2.738142303 | 4.62E-04 |
| NA | 12.81908503 | -2.74669525 | 6.43E-04 |
| MEP1A | 8.233482334 | -2.75614497 | 2.19E-02 |
| TMTC1 | 1452.335955 | -2.75861769 | 6.16E-53 |
| PCSK1 | 1402.5926 | -2.766095149 | 9.47E-43 |
| AC117453.1 | 8.645326187 | -2.772505023 | 2.05E-02 |
| C1QTNF4 | 9.09059817 | -2.772942761 | 1.89E-03 |
| TSHZ2 | 2202.117157 | -2.780733787 | 1.03E-91 |
| RCAN2 | 2097.152447 | -2.781273734 | 6.18E-50 |
| AC159540.2 | 46.64017187 | -2.797670425 | 1.42E-06 |
| ANKRD44 | 619.7998958 | -2.798435346 | 1.83E-24 |
| RAB27B | 854.5006837 | -2.807664216 | 1.14E-34 |
| GJA5 | 604.0411684 | -2.807950829 | 3.07E-12 |
| FAM124B | 12.99155367 | -2.809199809 | 7.85E-04 |
| RGPD1 | 11.87981957 | -2.815931465 | 1.52E-02 |
| GIMAP6 | 27.79027384 | -2.821467479 | 1.13E-07 |
| GPM6B | 424.3810447 | -2.838799347 | 1.09E-36 |
| F13A1 | 43.43632321 | -2.839167552 | 2.80E-06 |
| AL355987.2 | 12.14815253 | -2.852247492 | 1.90E-02 |
| STEAP4 | 35.06140897 | -2.881460884 | 6.85E-08 |
| SESN3 | 604.4270253 | -2.898947654 | 4.80E-77 |
| DPT | 7.418061856 | -2.908212969 | 7.04E-03 |
| AP002768.1 | 6.270874198 | -2.910496274 | 1.82E-02 |
| AC020663.2 | 9.161786122 | -2.924208436 | 4.50E-04 |
| LYVE1 | 7.222638394 | -2.934998126 | 8.14E-03 |
| TF | 518.8802447 | -2.938117131 | 2.43E-33 |
| DLK1 | 278.9725961 | -2.944584069 | 9.02E-14 |
| PIK3R3 | 2884.989418 | -2.946807451 | 5.11E-86 |
| PLA2G7 | 21.73822793 | -2.961663449 | 5.20E-07 |
| MRVI1 | 377.0708102 | -2.969613939 | 1.16E-37 |
| AC092040.1 | 5.510916573 | -2.972623826 | 1.02E-02 |

| | | | |
|---|---|---|---|
| OTX2-AS1 | 6.633732328 | -2.975632836 | 1.19E-02 |
| CLCA2 | 6.610713691 | -2.984676893 | 2.36E-02 |
| 1-Mar | 48.32321599 | -2.984859169 | 5.81E-05 |
| AC013268.3 | 14.76815553 | -2.986938972 | 2.20E-03 |
| AP003068.4 | 67.36195346 | -2.992744613 | 1.07E-11 |
| MIR3125 | 7.117429883 | -3.00182304 | 8.03E-03 |
| PHBP6 | 8.166162406 | -3.009212483 | 5.22E-03 |
| RSAD2 | 15.70848725 | -3.017786239 | 1.79E-03 |
| HEY1 | 1560.216718 | -3.025902551 | 4.52E-57 |
| SPRY1 | 121.128746 | -3.032310974 | 1.05E-11 |
| AC003965.1 | 5.997039492 | -3.037984739 | 2.82E-03 |
| CRIP3 | 31.00547865 | -3.039790553 | 9.44E-10 |
| LINC01585 | 5.893009501 | -3.041455875 | 1.30E-02 |
| PPP1R9A | 8.422797227 | -3.049160206 | 3.71E-02 |
| AC007906.2 | 35.53760287 | -3.053390063 | 4.95E-13 |
| AC009486.1 | 13.83506336 | -3.072445732 | 1.50E-04 |
| IFI44L | 45.59889335 | -3.08018235 | 1.99E-05 |
| CCL15-CCL14 | 48.89464838 | -3.085333171 | 6.73E-05 |
| GAB3 | 903.944943 | -3.097034016 | 8.31E-63 |
| ERG | 136.5897879 | -3.106784885 | 3.01E-18 |
| UNC5C | 16.04686619 | -3.108794772 | 9.41E-06 |
| AMIGO3 | 93.09257147 | -3.117874373 | 2.37E-02 |
| WDR11-AS1 | 9.780944371 | -3.122884805 | 3.41E-02 |
| LINC02202 | 74.77782157 | -3.127473467 | 8.34E-17 |
| VCAN | 908.1972434 | -3.136869035 | 3.79E-38 |
| PEG10 | 103399.184 | -3.140743786 | 2.04E-54 |
| LINC01352 | 18.42901211 | -3.141664342 | 2.39E-06 |
| ZBTB7C | 327.535564 | -3.144869086 | 5.74E-32 |
| PDK4 | 122.2626061 | -3.148126459 | 1.51E-23 |
| MAFB | 2006.013682 | -3.14980632 | 4.84E-64 |
| CYP4F8 | 24.3394758 | -3.157677669 | 3.60E-06 |
| NA | 8.171653104 | -3.161586931 | 3.51E-02 |
| STUM | 6.712616375 | -3.17375407 | 1.14E-02 |
| OFCC1 | 6.101181922 | -3.174105935 | 2.32E-02 |
| EVX1-AS | 5.588263269 | -3.184061449 | 1.25E-02 |
| AC025188.1 | 5.405294077 | -3.18473433 | 2.49E-02 |
| AC104809.2 | 6.881124712 | -3.185110592 | 1.02E-03 |
| AL133481.1 | 5.83759897 | -3.215296596 | 2.27E-02 |
| APLNR | 28.83062591 | -3.217689776 | 6.17E-08 |
| ABCA6 | 33.80518392 | -3.218669288 | 1.91E-08 |
| LINC02620 | 7.019322023 | -3.229117399 | 2.27E-03 |
| PCYT1B | 54.97020519 | -3.232531774 | 2.07E-17 |
| NA | 106.8658323 | -3.233772795 | 1.94E-14 |
| PDE11A | 1409.744978 | -3.240567607 | 1.39E-38 |
| FAM20A | 268.4477144 | -3.241655511 | 7.01E-25 |
| RGS8 | 9.840947122 | -3.264059694 | 5.29E-03 |
| LINC02198 | 13.44948038 | -3.27160061 | 6.38E-04 |

| | | | |
|---|---|---|---|
| CXCL12 | 179.5598405 | -3.271665628 | 6.36E-21 |
| KLKB1 | 7.057022666 | -3.27406772 | 1.18E-02 |
| GNA14 | 109.087556 | -3.275791244 | 1.17E-25 |
| BAALC | 163.1251946 | -3.289267622 | 1.68E-23 |
| ISLR | 87.15946199 | -3.294361299 | 2.04E-16 |
| SHISA2 | 384.9045469 | -3.298265976 | 9.17E-80 |
| SCARA5 | 29.23635387 | -3.314130927 | 2.07E-07 |
| PI15 | 14.13753266 | -3.350802698 | 2.09E-03 |
| CNTFR-AS1 | 13.20071498 | -3.35511571 | 3.51E-06 |
| AC005336.1 | 13.39810125 | -3.371039431 | 3.23E-02 |
| AC243960.1 | 7.082069548 | -3.386017134 | 1.13E-02 |
| AC009063.2 | 45.99358757 | -3.387473658 | 3.06E-16 |
| EBF1 | 927.3327661 | -3.426735739 | 3.30E-57 |
| SFRP5 | 9.335463047 | -3.458162143 | 9.66E-03 |
| LINC01091 | 34.16317966 | -3.482201704 | 2.04E-13 |
| LINC02068 | 5.24218061 | -3.502826061 | 1.83E-02 |
| REEP1 | 645.6708003 | -3.515056445 | 6.60E-60 |
| AL445423.1 | 7.941162631 | -3.52468749 | 3.46E-03 |
| AC010273.2 | 19.41594626 | -3.53776569 | 1.16E-05 |
| ZC3H11B | 16.96159532 | -3.575996903 | 3.25E-07 |
| RPRM | 7.716992075 | -3.593000889 | 7.95E-03 |
| SPARCL1 | 344.9499686 | -3.597764701 | 8.72E-25 |
| AC083843.4 | 12.60092653 | -3.59920619 | 1.62E-04 |
| LINC01197 | 9.765364234 | -3.631365205 | 2.08E-03 |
| AP000787.2 | 6.966646499 | -3.632282964 | 1.36E-03 |
| AC134043.2 | 93.25974137 | -3.640587495 | 3.99E-20 |
| GNRH2 | 11.35355812 | -3.641832282 | 5.40E-07 |
| A3GALT2 | 14.22687366 | -3.651286401 | 1.13E-04 |
| FNDC5 | 839.7235065 | -3.668267771 | 3.06E-15 |
| MRAP2 | 60.19755106 | -3.696607708 | 3.39E-13 |
| GPR88 | 5.493524169 | -3.700279685 | 1.24E-02 |
| TRPA1 | 331.5429719 | -3.731197981 | 1.57E-33 |
| TMEM72 | 5.552220368 | -3.763611321 | 1.97E-02 |
| GRM6 | 10.24229563 | -3.780315667 | 1.59E-03 |
| LINC01929 | 20.14398567 | -3.783578888 | 5.69E-06 |
| CREG2 | 57.31823733 | -3.799309242 | 3.76E-21 |
| TDO2 | 14.91077846 | -3.844802145 | 2.60E-04 |
| H19 | 25168.32442 | -3.865864692 | 2.68E-20 |
| MMP16 | 8.240602331 | -3.886168501 | 2.49E-02 |
| RIPPLY1 | 5.620511406 | -3.896671897 | 1.25E-02 |
| NRAP | 18.38544263 | -3.910393147 | 1.48E-07 |
| TM4SF1 | 371.1106316 | -3.952797867 | 5.86E-15 |
| MAST4-AS1 | 5.212654004 | -4.004166298 | 1.04E-02 |
| AC064875.1 | 87.04053269 | -4.007456606 | 3.03E-23 |
| AC112206.2 | 10.72136123 | -4.060441641 | 1.80E-03 |
| EDN3 | 6.521416936 | -4.082876147 | 7.40E-03 |
| NAV2-AS4 | 43.2774696 | -4.099484602 | 2.38E-13 |

| | | | |
|---|---|---|---|
| CACNA2D1 | 5.59929796 | -4.124816059 | 2.84E-03 |
| NA | 5.349206308 | -4.135897278 | 4.34E-02 |
| AC013275.1 | 5.193056198 | -4.157016883 | 5.29E-03 |
| ZP2 | 35.43078166 | -4.161516527 | 1.60E-10 |
| RANBP3L | 670.2879169 | -4.161944702 | 3.11E-40 |
| CYP17A1 | 10.76100124 | -4.186194951 | 5.80E-03 |
| LINC02387 | 53.29999358 | -4.223349394 | 1.14E-12 |
| CMKLR1 | 1380.27909 | -4.236961089 | 1.23E-53 |
| AC103808.2 | 11.94070214 | -4.254919813 | 1.23E-06 |
| ADGRF5 | 151.418413 | -4.370871469 | 7.92E-33 |
| CYP4X1 | 528.1658392 | -4.379601941 | 9.06E-69 |
| AC110741.1 | 5.364867705 | -4.414035238 | 6.40E-03 |
| LINC02458 | 11.69928983 | -4.43561754 | 2.08E-05 |
| UGT1A9 | 6.632546904 | -4.44575998 | 1.64E-02 |
| AC016747.4 | 5.08681378 | -4.461150914 | 6.20E-03 |
| COL14A1 | 56.31998566 | -4.470580499 | 2.90E-12 |
| SELENOP | 510.7451053 | -4.583657674 | 1.36E-20 |
| SMTNL2 | 14.55934715 | -4.618420914 | 2.63E-05 |
| CYSLTR2 | 151.0712334 | -4.671143771 | 5.32E-34 |
| PRND | 26.16392127 | -4.679114546 | 4.72E-09 |
| ALX3 | 5.060448825 | -4.706067005 | 1.20E-02 |
| ITIH6 | 6.419265348 | -4.732272904 | 4.21E-03 |
| HLX-AS1 | 5.864391477 | -4.800004616 | 2.16E-03 |
| BMX | 147.3418536 | -4.807845872 | 3.11E-25 |
| AC012499.1 | 8.245841383 | -4.807942284 | 1.03E-03 |
| PCDHGA11 | 39.1975755 | -4.827862527 | 2.89E-04 |
| KCNA3 | 8.28462565 | -4.90301095 | 1.04E-03 |
| GABRR1 | 17.11856491 | -4.956365201 | 1.31E-06 |
| CLEC10A | 5.718442853 | -5.011209137 | 8.18E-04 |
| OVCH1 | 6.504097974 | -5.364660649 | 1.92E-03 |
| ANKRD35 | 7.132370749 | -5.374180304 | 1.61E-04 |
| POSTN | 840.5191803 | -5.550452058 | 3.94E-39 |
| KRT127P | 5.310953075 | -5.832309266 | 1.18E-04 |
| CPHL1P | 12.06318751 | -5.990381904 | 8.80E-05 |
| NA | 13.76245846 | -6.022456168 | 1.15E-02 |
| CD180 | 81.62185089 | -6.059454466 | 4.23E-16 |
| OR6W1P | 7.088915303 | -6.110397211 | 2.61E-05 |
| IL1RAPL2 | 7.265012822 | -6.131363936 | 9.62E-05 |
| AP001180.4 | 9.219709272 | -6.141570358 | 1.35E-05 |
| AC136475.4 | 7.580164663 | -6.436261184 | 4.53E-02 |
| LINC02126 | 16.96698924 | -7.221565112 | 3.14E-07 |
| LINC00892 | 26.51675302 | -7.321152348 | 3.51E-09 |
| PRSS2 | 39.46445423 | -7.43261712 | 6.42E-08 |
| CCL14 | 36.66059148 | -7.854527892 | 6.83E-06 |
| NA | 19.37100855 | -22.17679345 | 8.80E-04 |

| Gene | baseMean | log2 Fold Change | padj |
|---|---|---|---|
| CXCR5 | 38.38259048 | 30.65672614 | 1.52E-12 |
| AP003419.1 | 46.01432342 | 21.49639377 | 1.33E-10 |
| AC010323.1 | 29.99683061 | 8.805027769 | 3.89E-02 |
| BRD7P2 | 96.67767377 | 8.130172938 | 3.04E-02 |
| AC135977.1 | 22.06214483 | 5.802842552 | 4.36E-02 |
| NPIPA2 | 22.55804979 | 5.795465655 | 1.68E-02 |
| AC005832.4 | 89.28362442 | 4.637043961 | 3.67E-04 |
| MRO | 19.70905316 | 4.544863668 | 2.69E-03 |
| FBLN5 | 552.2943098 | 4.508474311 | 6.72E-99 |
| GPR1 | 155.1271788 | 4.315170567 | 6.92E-09 |
| AC245041.1 | 18.51833017 | 4.067230964 | 5.75E-03 |
| WFDC21P | 140.545174 | 3.962585841 | 5.66E-19 |
| CRYAB | 8154.737916 | 3.775009 | 8.42E-59 |
| AL157896.1 | 13.64846009 | 3.652029825 | 1.87E-04 |
| PI3 | 17.94074282 | 3.558883406 | 1.42E-07 |
| AP1M2 | 110.3798198 | 3.353705062 | 2.11E-07 |
| MYO7B | 109.193404 | 3.298030762 | 5.37E-03 |
| NA | 179.9563504 | 3.243432438 | 4.08E-02 |
| UQCRFS1P1 | 23.22662339 | 3.239924288 | 1.93E-02 |
| INMT | 744.6966048 | 3.073439578 | 2.53E-140 |
| TRIML2 | 231.9553741 | 3.052083975 | 4.90E-43 |
| KLHDC7A | 24.91322927 | 3.050873691 | 1.59E-03 |
| LIMS2 | 478.4740095 | 3.031266294 | 2.37E-63 |
| NR1H4 | 131.3960236 | 2.959386092 | 4.29E-05 |
| LRRC15 | 26.29205623 | 2.916779535 | 1.67E-04 |
| PKD1P5 | 347.9474761 | 2.892371127 | 5.00E-09 |
| KIAA1210 | 25.70861216 | 2.873269919 | 3.58E-11 |
| AC005336.1 | 13.39810125 | 2.772929624 | 3.00E-04 |
| TNMD | 57.18971116 | 2.643686414 | 4.13E-07 |
| ZNF185 | 2431.652507 | 2.636625261 | 5.29E-135 |
| MYL4 | 57.40459191 | 2.612892431 | 8.79E-13 |
| AL355312.3 | 14.17484134 | 2.610982674 | 2.03E-02 |
| HSPB3 | 33.71635809 | 2.5973643 | 1.52E-05 |
| MFAP5 | 48.10545696 | 2.576574259 | 3.61E-05 |
| SLCO1B3 | 13.29089053 | 2.554189173 | 4.78E-03 |
| TNNI1 | 87.9705331 | 2.554167827 | 4.91E-02 |
| LINC02535 | 49.64853035 | 2.552523581 | 3.81E-05 |
| CCN2 | 8122.707037 | 2.542250351 | 4.01E-37 |
| ITIH3 | 47.92121289 | 2.492169322 | 1.30E-03 |
| AC002401.4 | 17.60670996 | 2.490089433 | 4.49E-05 |
| MT-TE | 43.64527229 | 2.489954658 | 1.36E-09 |
| C1S | 1174.241073 | 2.478127578 | 2.58E-20 |
| CHST4 | 31.95928319 | 2.477585239 | 1.57E-04 |
| CSF2RA | 877.8505258 | 2.472038024 | 1.18E-40 |
| WTIP | 955.4419514 | 2.455370831 | 7.66E-39 |
| MT-TS2 | 38.2703071 | 2.443526992 | 5.43E-06 |

| | | | |
|---|---|---|---|
| SYNPO2L | 54.0993148 | 2.435579092 | 7.16E-06 |
| COL12A1 | 7146.720884 | 2.424618059 | 3.61E-74 |
| GPR17 | 17.22731363 | 2.360222298 | 5.21E-05 |
| AMOTL2 | 5194.214834 | 2.324098712 | 3.44E-118 |
| MT-TA | 109.576612 | 2.269963798 | 1.50E-08 |
| LINC01220 | 13.87085948 | 2.261687262 | 3.91E-02 |
| PIWIL2 | 12.98599122 | 2.260090733 | 1.47E-02 |
| STXBP6 | 354.2817869 | 2.258014088 | 2.82E-20 |
| CD177 | 28.34911627 | 2.245565811 | 1.08E-02 |
| AL590399.4 | 12.31094571 | 2.220511145 | 3.89E-03 |
| AKR1B10 | 66.94033728 | 2.210995282 | 1.31E-03 |
| FRMD7 | 28.169423 | 2.185749297 | 6.98E-03 |
| MT-TL2 | 78.84596201 | 2.157392747 | 5.59E-08 |
| CXADR | 430.9583909 | 2.152101105 | 6.16E-33 |
| GPC4 | 272.3054783 | 2.142224094 | 8.00E-23 |
| SELP | 136.3642521 | 2.14136542 | 6.74E-10 |
| TINAGL1 | 244.1329187 | 2.123649587 | 2.89E-07 |
| KCNJ5 | 31.40138865 | 2.113789311 | 2.35E-07 |
| SLC5A1 | 28.6409774 | 2.111241644 | 7.04E-06 |
| NPPB | 2240.650956 | 2.092351514 | 1.21E-02 |
| VTN | 154.8509553 | 2.021817643 | 5.28E-10 |
| PKD1L2 | 885.8875173 | 1.997103682 | 2.67E-11 |
| LOX | 1600.914318 | 1.978880774 | 1.28E-24 |
| TMEM255A | 638.2751286 | 1.977874356 | 1.25E-25 |
| WWC1 | 602.1446754 | 1.93228055 | 1.46E-33 |
| CPA4 | 8124.801267 | 1.929860219 | 8.19E-31 |
| CXCL14 | 171.4861655 | 1.921781447 | 7.34E-08 |
| STX11 | 126.2269358 | 1.914438561 | 7.27E-09 |
| LINC01693 | 25.8599621 | 1.905346002 | 2.04E-02 |
| MT-TH | 24.1473077 | 1.888763195 | 4.60E-03 |
| TCAF2P1 | 28.17114646 | 1.880806378 | 2.34E-03 |
| ANXA8L1 | 163.5609482 | 1.87313477 | 2.39E-06 |
| PTMAP5 | 14.19092898 | 1.868162748 | 4.27E-02 |
| AC005307.1 | 131.4519514 | 1.861142728 | 1.07E-07 |
| MMP28 | 50.44223442 | 1.833835157 | 9.01E-06 |
| MEGF6 | 249.4909584 | 1.825406097 | 2.74E-05 |
| VSTM2L | 409.4949088 | 1.774107765 | 1.76E-20 |
| NA | 111.6290596 | 1.753136896 | 6.64E-06 |
| ITGB2 | 124.990794 | 1.726472885 | 8.09E-06 |
| CALB2 | 75.59255705 | 1.705661943 | 5.90E-07 |
| MATN3 | 531.0146682 | 1.681473147 | 6.61E-31 |
| CARD11 | 487.3641494 | 1.678464366 | 1.06E-14 |
| ARSI | 573.0458329 | 1.676202747 | 2.55E-15 |
| IGFL2-AS1 | 125.576691 | 1.675403735 | 3.70E-09 |
| BICDL1 | 37.542591 | 1.670842929 | 1.00E-02 |
| MT-TI | 17.57118234 | 1.662235181 | 1.80E-02 |
| EDNRB | 780.2042551 | 1.657933588 | 1.23E-33 |

| | | | |
|---|---|---|---|
| OCLN | 724.0860483 | 1.652703285 | 1.59E-12 |
| AC002480.1 | 40.60269188 | 1.647735343 | 4.36E-06 |
| ALPP | 37.81297781 | 1.636765281 | 3.02E-03 |
| CCN1 | 10596.59591 | 1.636580835 | 1.78E-45 |
| XDH | 47.86677611 | 1.621294717 | 1.25E-03 |
| C17orf47 | 23.56653353 | 1.616985359 | 1.20E-03 |
| SYDE2 | 235.870281 | 1.614464483 | 8.52E-11 |
| LINC02582 | 131.1487649 | 1.608313135 | 1.90E-10 |
| MT-TL1 | 29.60530902 | 1.605664924 | 3.78E-03 |
| CYP4F11 | 312.2638214 | 1.600900542 | 5.98E-20 |
| PPP4R4 | 23.02234291 | 1.590469961 | 3.89E-02 |
| SLIT3 | 447.7973117 | 1.578487458 | 2.97E-20 |
| AC025423.2 | 28.53963443 | 1.577694488 | 2.06E-02 |
| DMKN | 409.4101885 | 1.572975798 | 1.41E-23 |
| UGT3A2 | 1561.296932 | 1.567863682 | 1.00E-14 |
| FLJ45513 | 22.04428444 | 1.566641579 | 1.48E-02 |
| HES4 | 110.1398204 | 1.550874704 | 5.39E-07 |
| IL6 | 31.65351887 | 1.548168252 | 4.20E-02 |
| AP000350.4 | 21.66484245 | 1.547643133 | 3.05E-02 |
| AJUBA | 9629.715395 | 1.539807686 | 7.72E-18 |
| LATS2 | 922.0682349 | 1.524222948 | 2.64E-42 |
| AC140479.2 | 142.8946411 | 1.49188987 | 8.20E-16 |
| MAGEB3 | 36.6029307 | 1.481465182 | 2.27E-04 |
| TMEM179 | 21.2981889 | 1.479083901 | 1.61E-02 |
| KRT8 | 14283.57687 | 1.477447241 | 1.15E-37 |
| CEACAMP10 | 596.3616348 | 1.475456254 | 5.28E-03 |
| SEPT4-AS1 | 108.7083786 | 1.474169391 | 1.07E-07 |
| AL390195.1 | 23.24237934 | 1.472454809 | 2.17E-02 |
| AC020934.1 | 20.74450596 | 1.466733258 | 3.20E-02 |
| TMPRSS3 | 63.85416896 | 1.466679632 | 6.39E-04 |
| SEMA3B | 1213.282702 | 1.464998026 | 1.64E-11 |
| AC027237.3 | 72.9435641 | 1.46452766 | 4.91E-04 |
| CPA1 | 26.12893113 | 1.460397091 | 6.23E-03 |
| C7orf57 | 122.2458911 | 1.459224484 | 3.64E-05 |
| AC108515.1 | 38.85659606 | 1.451128462 | 2.78E-04 |
| XAGE2 | 41.30684287 | 1.44569597 | 4.36E-03 |
| POSTN | 840.5191803 | 1.441406884 | 1.14E-03 |
| MMP19 | 140.8610266 | 1.441008126 | 2.33E-06 |
| AC083805.2 | 26.76387639 | 1.440990162 | 2.95E-04 |
| MSC | 323.6871895 | 1.428840718 | 2.35E-03 |
| TCAF2 | 445.3765845 | 1.41636532 | 3.13E-06 |
| AL603750.1 | 23.85849569 | 1.41272066 | 3.69E-03 |
| BPIFA2 | 22.6494482 | 1.405799021 | 2.49E-02 |
| CYP4F8 | 24.3394758 | 1.395403593 | 2.71E-02 |
| RIMS4 | 97.05276827 | 1.384526422 | 9.62E-05 |
| NA | 26.04020648 | 1.381308999 | 7.59E-03 |
| CD70 | 205.6557704 | 1.377096503 | 1.40E-14 |

| | | | |
|---|---|---|---|
| TEX14 | 48.118117 | 1.373256378 | 1.96E-03 |
| HHLA1 | 19.15672691 | 1.372639048 | 4.92E-02 |
| CLCNKB | 120.2619914 | 1.371945031 | 2.27E-08 |
| STON1-GTF2A1 | 83.30213425 | 1.365065876 | 4.41E-04 |
| NR0B1 | 84.72388409 | 1.361655478 | 1.57E-04 |
| AC005736.1 | 74.59682991 | 1.349592703 | 5.21E-06 |
| ANO4 | 74.03198246 | 1.347948188 | 4.15E-05 |
| MT-TS1 | 975.9977628 | 1.339160393 | 9.84E-03 |
| ELFN2 | 490.7239094 | 1.334397029 | 4.47E-08 |
| SLC45A2 | 191.8192019 | 1.334167919 | 3.76E-08 |
| AC116351.1 | 130.5665582 | 1.320528406 | 8.62E-06 |
| LINC01638 | 242.896981 | 1.314542796 | 9.34E-07 |
| FAM189A2 | 29.59519028 | 1.313089279 | 3.23E-02 |
| FMOD | 1627.644591 | 1.311709567 | 3.34E-12 |
| AC105460.1 | 37.50005812 | 1.306259354 | 5.59E-04 |
| MSC-AS1 | 127.9931706 | 1.304680368 | 7.59E-03 |
| MT-TT | 23.81572381 | 1.291060072 | 2.23E-02 |
| MT-TY | 275.5489081 | 1.277712092 | 9.12E-04 |
| AL139002.1 | 119.7836048 | 1.270756934 | 3.91E-10 |
| BMP2 | 67.08671781 | 1.269755184 | 2.80E-03 |
| IGFBP5 | 6426.850593 | 1.267052795 | 2.11E-02 |
| CLDN2 | 59.31407275 | 1.266233311 | 4.05E-04 |
| MYPN | 979.7692297 | 1.264222282 | 3.83E-10 |
| LINC01389 | 27.21805161 | 1.234361283 | 2.98E-03 |
| C4orf48 | 140.9285521 | 1.232757661 | 2.11E-02 |
| COL14A1 | 56.31998566 | 1.22395072 | 2.29E-04 |
| DNAJC6 | 116.1338755 | 1.222131397 | 9.07E-03 |
| SPP1 | 3358.989557 | 1.221346105 | 6.76E-06 |
| SEMA3F | 497.6585612 | 1.220861792 | 1.22E-19 |
| GFRA3 | 106.5806063 | 1.217646775 | 3.13E-06 |
| ASS1 | 1006.358665 | 1.215132134 | 3.52E-02 |
| IP6K3 | 192.4547338 | 1.212070378 | 4.82E-07 |
| CREB3L1 | 742.9290274 | 1.210055102 | 8.88E-16 |
| MAFK | 1886.636662 | 1.206600486 | 2.82E-26 |
| PEAR1 | 1120.147306 | 1.204280732 | 1.25E-18 |
| GFI1 | 39.2406586 | 1.202224191 | 4.57E-02 |
| H19 | 25168.32442 | 1.196363723 | 4.13E-02 |
| SYNPO | 2426.228123 | 1.19331499 | 4.53E-24 |
| MTND4P12 | 341.7639862 | 1.19149666 | 1.17E-04 |
| AC104435.2 | 31.56962772 | 1.187598358 | 1.09E-02 |
| POLRMTP1 | 27.84699559 | 1.18735175 | 2.27E-02 |
| CAVIN1 | 22717.39463 | 1.186499356 | 3.85E-18 |
| LINC01123 | 472.9229212 | 1.181174594 | 1.90E-15 |
| ATP1A2 | 143.772855 | 1.178499551 | 4.24E-02 |
| FEZ1 | 659.6364878 | 1.177769676 | 1.74E-14 |
| NEXN | 357.8514529 | 1.175558011 | 8.74E-10 |
| TMEM56 | 329.4539423 | 1.168581995 | 9.51E-07 |

| | | | |
|---|---|---|---|
| LINC01106 | 1890.349103 | 1.162064975 | 1.53E-42 |
| ACKR1 | 26.76636376 | 1.160262502 | 1.62E-02 |
| WWC2 | 2643.317265 | 1.156036946 | 1.36E-09 |
| KCNIP1 | 288.8772349 | 1.154563781 | 1.30E-06 |
| C1QTNF2 | 299.2471779 | 1.153695141 | 2.63E-04 |
| AC096637.2 | 78.23099629 | 1.153274777 | 7.17E-07 |
| INSYN1 | 368.9945437 | 1.151619082 | 1.12E-10 |
| DNAJB4 | 526.3585122 | 1.15087875 | 4.16E-07 |
| CD24 | 529.1194415 | 1.13927053 | 2.19E-06 |
| AC125807.2 | 800.7685966 | 1.132348765 | 3.62E-22 |
| NA | 40.08367415 | 1.132171782 | 9.84E-03 |
| LINC01730 | 59.12212474 | 1.130802706 | 1.22E-02 |
| LAMP3 | 296.1481714 | 1.126537521 | 2.06E-03 |
| ESRG | 287.4514818 | 1.120451704 | 9.19E-07 |
| CHRD | 56.17107808 | 1.11924485 | 2.18E-02 |
| C19orf81 | 35.88473878 | 1.118376046 | 3.14E-02 |
| MSLN | 108.5106732 | 1.11147753 | 4.13E-02 |
| AL391121.1 | 39.07113611 | 1.110699712 | 3.86E-02 |
| PLCE1 | 1326.090032 | 1.109904896 | 1.25E-21 |
| NPR3 | 210.5822672 | 1.103224413 | 1.70E-04 |
| MAMDC2-AS1 | 35.10599911 | 1.102587904 | 1.01E-02 |
| LMOD1 | 281.9537545 | 1.100357597 | 2.29E-04 |
| OLFM2 | 487.2499097 | 1.097628585 | 1.03E-07 |
| TEK | 960.4093854 | 1.096278245 | 1.55E-05 |
| GPNMB | 7636.758085 | 1.093491426 | 7.51E-10 |
| GABRE | 2505.179407 | 1.086310424 | 1.09E-07 |
| JPH2 | 828.9286868 | 1.083546886 | 2.09E-08 |
| MT-TC | 333.2264633 | 1.083037597 | 1.57E-03 |
| MYO5C | 156.771633 | 1.079870275 | 6.03E-03 |
| DTX4 | 442.4496529 | 1.078847647 | 3.38E-06 |
| NPAS1 | 1188.241383 | 1.076973119 | 3.96E-23 |
| RBM47 | 230.5145896 | 1.076151617 | 2.13E-10 |
| LINC01252 | 31.49057911 | 1.066489295 | 3.38E-02 |
| GALNT9 | 229.1882238 | 1.062944268 | 2.76E-04 |
| LINC00623 | 78.41347884 | 1.061940699 | 2.88E-02 |
| TMED10P2 | 29.80088923 | 1.061139599 | 3.20E-02 |
| GHR | 39.67772673 | 1.052536144 | 3.73E-02 |
| TRIM29 | 531.2890621 | 1.050170147 | 4.83E-04 |
| PLS3-AS1 | 36.45111194 | 1.04905019 | 2.46E-02 |
| PALMD | 192.6518655 | 1.048506729 | 1.44E-05 |
| PTCH1 | 1620.679928 | 1.047777632 | 6.47E-09 |
| LAMA1 | 1777.372355 | 1.045386708 | 1.99E-07 |
| NXNL2 | 101.7112097 | 1.044096847 | 1.34E-06 |
| IL6R | 802.3266812 | 1.041678923 | 8.67E-10 |
| COL1A1 | 951.6101759 | 1.036450981 | 2.66E-04 |
| MAN1C1 | 266.1644314 | 1.036420521 | 4.58E-05 |
| GAD1 | 53.22551783 | 1.03146616 | 4.32E-03 |

| | | | |
|---|---|---|---|
| PPFIBP2 | 404.1679279 | 1.030413291 | 2.65E-03 |
| CACNA1I | 43.84665629 | 1.029101608 | 4.33E-02 |
| MYCL | 127.9613386 | 1.02752704 | 2.86E-03 |
| SPTAN1 | 27548.29185 | 1.023460204 | 4.39E-46 |
| C2orf72 | 79.40764904 | 1.02151909 | 4.79E-02 |
| PANX2 | 146.5880724 | 1.014373592 | 4.58E-05 |
| CDH3 | 1193.523897 | 1.009906298 | 1.75E-14 |
| TEAD4 | 1502.038523 | 1.0095767 | 1.99E-22 |
| LINC01836 | 94.12756194 | 1.009496561 | 9.58E-05 |
| SLC47A1 | 75.64051756 | 1.00912195 | 6.69E-03 |
| RCAN1 | 3868.497247 | 1.005103508 | 8.31E-13 |
| PTHLH | 279.5986255 | 1.005074176 | 2.07E-04 |
| CLU | 7930.293639 | 1.004966719 | 7.44E-15 |
| HOXA1 | 545.3612809 | 1.001835386 | 9.01E-19 |
| MORC4 | 2027.296062 | 1.001768409 | 8.28E-34 |
| CALHM5 | 354.1155895 | 1.001459262 | 6.22E-07 |
| KRT15 | 92.32944513 | 0.997651699 | 4.14E-02 |
| SPTB | 236.3609432 | 0.994838677 | 4.31E-03 |
| KRT80 | 1730.679913 | 0.987493874 | 8.17E-04 |
| ADAMTSL1 | 298.7360119 | 0.984631112 | 2.18E-04 |
| ALS2CL | 852.7311576 | 0.982908332 | 6.50E-07 |
| NA | 98.28641465 | 0.971789126 | 4.91E-03 |
| TP63 | 52.16142317 | 0.952598822 | 3.26E-02 |
| SSPN | 336.9878138 | 0.937349642 | 4.04E-04 |
| PTH1R | 160.1294675 | 0.936237932 | 2.18E-04 |
| AC245060.6 | 212.7784863 | 0.935273807 | 3.99E-03 |
| PLA2R1 | 868.557516 | 0.934671386 | 3.34E-10 |
| RAB3D | 372.3493686 | 0.929028573 | 3.13E-06 |
| HUNK | 113.2340518 | 0.926572998 | 2.42E-04 |
| FAM167B | 176.2258737 | 0.921792967 | 8.81E-04 |
| MYLK3 | 310.1611264 | 0.921675844 | 1.32E-06 |
| SMPDL3A | 94.57710573 | 0.921658917 | 1.65E-04 |
| SNX10 | 285.7625478 | 0.919976056 | 3.05E-03 |
| RGS9 | 137.3520688 | 0.917508357 | 7.02E-03 |
| TNFRSF12A | 3033.22403 | 0.915043687 | 1.06E-07 |
| HSBP1L1 | 284.4549299 | 0.913162524 | 9.32E-07 |
| C1R | 3793.027515 | 0.911878056 | 6.62E-08 |
| MT-TN | 372.0712389 | 0.905954067 | 7.15E-03 |
| AC126335.1 | 71.26480399 | 0.904438373 | 4.77E-03 |
| DPP4 | 73.62921846 | 0.902798775 | 7.90E-03 |
| RASGRF1 | 134.7074287 | 0.901625976 | 3.81E-02 |
| GATA4 | 1468.75688 | 0.893446717 | 9.50E-07 |
| MALL | 777.0586729 | 0.889828126 | 7.39E-07 |
| DAG1 | 8402.798769 | 0.88971037 | 7.44E-15 |
| PIM1 | 1122.782319 | 0.888365247 | 2.27E-04 |
| BHMT2 | 687.4043628 | 0.887698782 | 2.26E-06 |
| CAP2 | 1928.403831 | 0.88728159 | 3.48E-18 |

| | | | |
|---|---|---|---|
| CAV1 | 23107.43986 | 0.885928362 | 3.88E-11 |
| HDAC9 | 214.4988075 | 0.883595999 | 2.07E-04 |
| ANXA1 | 9614.08568 | 0.883008961 | 4.17E-04 |
| FLNC | 24286.4346 | 0.882200608 | 1.89E-10 |
| SHISA3 | 161.7304846 | 0.881219917 | 2.92E-02 |
| SPTBN5 | 182.573012 | 0.878704594 | 4.36E-03 |
| GAREM1 | 159.1268217 | 0.87614807 | 9.17E-05 |
| PCSK5 | 671.7166073 | 0.874528236 | 1.46E-03 |
| PDCD1LG2 | 137.9408949 | 0.872331774 | 8.42E-03 |
| GRHL1 | 65.06406828 | 0.871529692 | 4.88E-02 |
| ABCG2 | 617.2687721 | 0.867135775 | 1.32E-03 |
| COL3A1 | 378.3612853 | 0.866946839 | 5.67E-04 |
| PLS3 | 10420.30682 | 0.866436307 | 1.29E-20 |
| ZBTB45P2 | 717.7323167 | 0.86568346 | 1.29E-10 |
| ADAMTS15 | 608.3231517 | 0.85762205 | 2.00E-10 |
| LRRN2 | 226.4845914 | 0.849245875 | 4.23E-04 |
| NSUN7 | 276.9157398 | 0.849150248 | 8.17E-05 |
| CAV2 | 2124.493216 | 0.845750155 | 2.20E-10 |
| AC090409.1 | 76.02384453 | 0.84422103 | 1.72E-02 |
| CYP39A1 | 219.4277118 | 0.843563484 | 1.59E-05 |
| TNFSF9 | 133.0035316 | 0.842243453 | 6.81E-03 |
| LINC01666 | 39.06634075 | 0.841497591 | 3.72E-02 |
| TP53I11 | 1725.487319 | 0.841343478 | 4.43E-13 |
| GPC3 | 69.20991286 | 0.837035146 | 2.68E-02 |
| SYDE1 | 3531.235541 | 0.836981807 | 1.80E-10 |
| C9orf3 | 1301.867845 | 0.835327157 | 9.38E-16 |
| AC099548.2 | 253.6681035 | 0.830666655 | 1.90E-02 |
| PDE11A | 1409.744978 | 0.826967472 | 9.62E-03 |
| TMEM139 | 162.5650754 | 0.82644982 | 1.01E-04 |
| CEBPD | 169.0188464 | 0.825307521 | 1.37E-03 |
| AC068279.1 | 52.80865783 | 0.823684338 | 1.00E-02 |
| RTN4R | 369.6459485 | 0.8175232 | 5.56E-05 |
| GLT8D2 | 304.745755 | 0.817505994 | 9.10E-09 |
| SERPINA1 | 129.2485732 | 0.815322689 | 1.61E-02 |
| AC104563.1 | 50.11300233 | 0.814244132 | 3.07E-02 |
| PHACTR2 | 1004.303295 | 0.810111486 | 1.99E-04 |
| OTUD1 | 108.5394167 | 0.807704586 | 2.29E-03 |
| FIGN | 438.4513192 | 0.807660867 | 9.82E-07 |
| AL161431.1 | 253.7190024 | 0.802473317 | 2.51E-03 |
| TEAD1 | 10939.96256 | 0.802215226 | 5.71E-07 |
| SORL1 | 4958.874962 | 0.793940204 | 1.01E-08 |
| FAS | 239.2567132 | 0.789846645 | 2.57E-03 |
| PARP8 | 150.9692421 | 0.783028594 | 1.18E-02 |
| CX3CL1 | 347.8028956 | 0.779612947 | 1.49E-03 |
| MTND1P23 | 155.9045096 | 0.777501763 | 1.32E-02 |
| RIPOR2 | 1126.46556 | 0.772211996 | 2.60E-03 |
| WNT5B | 481.1590035 | 0.771410967 | 1.18E-03 |

| | | | |
|---|---|---|---|
| B3GNT5 | 183.3353908 | 0.765181814 | 7.15E-03 |
| MYORG | 1497.758198 | 0.765034493 | 2.03E-05 |
| CYS1 | 92.6595099 | 0.760408012 | 9.89E-03 |
| ETS2 | 1570.480883 | 0.75967052 | 5.00E-13 |
| IGF2 | 2046.485533 | 0.758069547 | 7.77E-04 |
| AC008957.3 | 81.64226081 | 0.75795751 | 2.47E-02 |
| GPX8 | 5676.176415 | 0.757084506 | 8.61E-09 |
| ITGA3 | 22276.61878 | 0.756760512 | 8.92E-12 |
| SYTL4 | 710.6228113 | 0.75648724 | 2.27E-04 |
| HSPG2 | 6936.109191 | 0.755478228 | 4.13E-06 |
| FBLN7 | 185.287182 | 0.755245805 | 2.89E-03 |
| PLIN4 | 68.6546174 | 0.753886907 | 3.78E-02 |
| THBS1 | 3896.636739 | 0.753663846 | 1.53E-09 |
| PLEKHA6 | 883.8866704 | 0.753249003 | 5.86E-03 |
| PPP1R12B | 1810.477078 | 0.752705016 | 1.24E-18 |
| STON1 | 979.0446458 | 0.752072266 | 6.81E-11 |
| AC020898.1 | 58.20265113 | 0.749136421 | 3.02E-02 |
| SRPX2 | 1641.567739 | 0.747562791 | 3.23E-04 |
| TUFT1 | 886.6362793 | 0.746585741 | 3.37E-08 |
| FGD4 | 160.9786872 | 0.746557915 | 2.09E-02 |
| EFEMP1 | 741.1103555 | 0.746253972 | 2.89E-03 |
| SLC9A3 | 209.2046598 | 0.745640709 | 1.19E-02 |
| PARVA | 12811.79845 | 0.745270121 | 1.38E-47 |
| PDPN | 1362.464471 | 0.744927062 | 5.74E-04 |
| IDH1 | 9760.191328 | 0.744572742 | 2.64E-08 |
| ITGB3 | 1822.991296 | 0.744312622 | 5.52E-07 |
| ITGA1 | 11661.05432 | 0.74243683 | 8.84E-08 |
| DLGAP1-AS2 | 423.8719857 | 0.74213227 | 2.05E-02 |
| AL162171.1 | 1232.53051 | 0.738052637 | 1.05E-12 |
| AC093616.1 | 454.260492 | 0.736590477 | 2.24E-04 |
| KLF2 | 119.5906069 | 0.733948017 | 1.42E-02 |
| ASPH | 8470.783867 | 0.733410211 | 2.70E-10 |
| IFI6 | 1717.788303 | 0.728928122 | 8.85E-04 |
| NXN | 1791.096427 | 0.728752886 | 5.77E-08 |
| ARHGAP23 | 5404.484757 | 0.727259915 | 8.19E-15 |
| APOLD1 | 451.9599406 | 0.724031017 | 7.65E-07 |
| MSRB3 | 1780.19995 | 0.718677032 | 1.86E-08 |
| MMP24 | 326.4706903 | 0.715378476 | 1.91E-03 |
| MYOF | 15695.67332 | 0.711984947 | 5.10E-07 |
| NECTIN2 | 5365.872754 | 0.711641622 | 5.30E-09 |
| ZNF219 | 899.6886662 | 0.708355956 | 7.74E-08 |
| SPANXB1 | 1009.08562 | 0.708125423 | 2.19E-06 |
| LHX8 | 280.7864281 | 0.707924216 | 3.15E-02 |
| PRRG4 | 102.5023206 | 0.704191438 | 3.42E-02 |
| HNRNPA1P48 | 182.5805565 | 0.703506809 | 1.09E-02 |
| SGMS1 | 501.840403 | 0.701851238 | 9.11E-04 |
| MAP3K21 | 160.6828438 | 0.701784344 | 1.83E-02 |

| | | | |
|---|---|---|---|
| L3HYPDH | 792.4822493 | 0.700027354 | 1.67E-11 |
| TENT5C | 137.8763368 | 0.699826032 | 5.23E-03 |
| TSPAN33 | 331.3822405 | 0.699668773 | 1.45E-04 |
| ROCK2 | 3281.835884 | 0.699030354 | 2.26E-02 |
| MIR22HG | 876.9771294 | 0.698978589 | 1.94E-14 |
| MID1IP1 | 1888.372417 | 0.698671995 | 5.17E-04 |
| CARD16 | 266.3336459 | 0.697486759 | 9.62E-05 |
| EHD2 | 10312.92724 | 0.696997666 | 5.36E-09 |
| ACKR3 | 110.6345116 | 0.6939674 | 4.24E-02 |
| RALGPS1 | 346.6724631 | 0.693920315 | 1.61E-02 |
| TENT5B | 961.9812951 | 0.693644736 | 3.06E-04 |
| PLEKHA7 | 2443.226225 | 0.691691723 | 2.39E-04 |
| PKNOX2 | 537.5619923 | 0.68996343 | 1.02E-04 |
| SORT1 | 7230.535163 | 0.688219068 | 1.56E-11 |
| ADORA1 | 215.6833986 | 0.687320012 | 6.33E-05 |
| AP000662.1 | 132.6091028 | 0.686705048 | 1.92E-02 |
| SLC13A5 | 229.8750143 | 0.684344232 | 4.42E-02 |
| DSE | 960.8994602 | 0.68414389 | 4.66E-12 |
| MIR503HG | 2154.711243 | 0.683780501 | 2.74E-02 |
| EMP1 | 5374.264184 | 0.683002985 | 2.58E-03 |
| LMBRD2 | 905.4643708 | 0.682950608 | 3.81E-02 |
| PROSER2 | 428.4427461 | 0.682499738 | 5.00E-03 |
| NOS1AP | 114.7690526 | 0.680182159 | 2.15E-02 |
| ADAM23 | 710.607505 | 0.678920237 | 2.16E-06 |
| RFK | 851.8053873 | 0.677732193 | 1.66E-03 |
| GCLM | 1084.260776 | 0.677585816 | 5.21E-04 |
| AL592183.1 | 230.8207847 | 0.677529858 | 1.43E-02 |
| TMEM92 | 202.8482872 | 0.676403884 | 4.51E-03 |
| CGNL1 | 895.5418779 | 0.676046783 | 6.57E-05 |
| USP43 | 231.4230637 | 0.675463923 | 2.26E-03 |
| SGMS2 | 606.7809837 | 0.674704379 | 4.71E-03 |
| COL1A2 | 3558.442137 | 0.664780499 | 4.72E-02 |
| BASP1 | 2291.351755 | 0.664324012 | 2.93E-04 |
| RNF122 | 196.3554192 | 0.664198765 | 3.19E-03 |
| ME1 | 734.9613871 | 0.663507933 | 3.81E-05 |
| WWC3 | 2266.235865 | 0.662406663 | 3.75E-06 |
| PTPRK | 2144.776039 | 0.662403431 | 5.46E-07 |
| SKA2 | 1597.49907 | 0.65904316 | 2.84E-07 |
| AP002761.3 | 81.18544798 | 0.653017968 | 1.92E-02 |
| KRT18 | 26282.33084 | 0.65241719 | 7.14E-07 |
| LIPH | 201.1100145 | 0.651676302 | 2.01E-02 |
| PTPN14 | 5617.28597 | 0.6509988 | 2.76E-18 |
| RHBDF1 | 1735.657285 | 0.65059796 | 1.39E-07 |
| FZD5 | 364.7941564 | 0.649237661 | 4.83E-06 |
| ADAM9 | 9482.944296 | 0.649174371 | 6.71E-04 |
| CADM3 | 1193.863173 | 0.649144841 | 7.92E-04 |
| CHRM4 | 157.7202575 | 0.648240927 | 2.93E-02 |

| | | | |
|---|---|---|---|
| AC243964.3 | 235.8240355 | 0.647791146 | 2.29E-03 |
| LINC01503 | 111.7516947 | 0.645800451 | 9.47E-03 |
| B3GALNT1 | 1058.702994 | 0.644485694 | 3.49E-07 |
| ICA1L | 195.1054069 | 0.643929035 | 6.54E-04 |
| MBNL2 | 5478.880166 | 0.642446825 | 1.85E-02 |
| KCTD11 | 1277.421524 | 0.641682389 | 4.07E-05 |
| NANOS1 | 479.3315027 | 0.641233674 | 2.82E-03 |
| ELFN1 | 408.2898101 | 0.640455968 | 1.94E-05 |
| GLI2 | 1186.161085 | 0.639347398 | 3.37E-03 |
| RFTN1 | 2508.131971 | 0.636016352 | 3.49E-07 |
| CHST3 | 1381.804227 | 0.635956518 | 6.73E-08 |
| RRAS2 | 2136.436366 | 0.635847069 | 1.86E-09 |
| BIRC3 | 138.5705745 | 0.635552555 | 3.77E-02 |
| TBC1D8B | 395.7373092 | 0.63553395 | 2.53E-02 |
| CASC10 | 644.5185033 | 0.634116357 | 3.63E-08 |
| QKI | 4899.422462 | 0.632590523 | 4.55E-05 |
| SLC25A30 | 977.3527418 | 0.632483545 | 1.34E-03 |
| DDR2 | 12220.85219 | 0.632095975 | 2.49E-03 |
| CALD1 | 17077.04623 | 0.632073133 | 2.86E-05 |
| AC087473.1 | 176.7463438 | 0.631386917 | 3.11E-02 |
| USP48 | 3298.933719 | 0.630220477 | 1.15E-07 |
| PARD6B | 133.5606062 | 0.630188308 | 3.51E-02 |
| BEND7 | 460.5163483 | 0.6301256 | 2.07E-05 |
| GADD45B | 977.0968945 | 0.629185014 | 3.59E-06 |
| COL4A5 | 2752.108464 | 0.629061171 | 3.58E-06 |
| SBDS | 6786.193804 | 0.627615032 | 6.73E-08 |
| HOTAIRM1 | 1189.090175 | 0.625805586 | 6.53E-09 |
| ARHGAP29 | 4982.779967 | 0.62425706 | 9.98E-03 |
| SBDSP1 | 833.1214146 | 0.622927964 | 1.93E-02 |
| KLHL5 | 3640.730673 | 0.622847211 | 4.09E-04 |
| TGFB2 | 3830.689573 | 0.620925466 | 1.42E-07 |
| SBF2-AS1 | 216.8026323 | 0.620636869 | 3.76E-02 |
| STAC | 1317.904973 | 0.620512895 | 3.99E-03 |
| LPCAT2 | 1115.439456 | 0.619671578 | 2.32E-06 |
| ANO6 | 6166.89251 | 0.617676595 | 4.16E-09 |
| SLC12A7 | 1143.958255 | 0.617591094 | 3.73E-03 |
| UGT8 | 101.0890748 | 0.617244811 | 1.18E-02 |
| RAB37 | 569.4378075 | 0.616369045 | 2.35E-03 |
| FLNB | 29152.77127 | 0.615989659 | 8.17E-05 |
| RALGPS2 | 1148.025275 | 0.614118479 | 2.18E-04 |
| WHAMMP2 | 146.3450519 | 0.613364674 | 7.86E-03 |
| LGR4 | 3291.633279 | 0.612666617 | 8.24E-06 |
| PPIAP22 | 335.0730773 | 0.612447534 | 2.29E-02 |
| CEACAM19 | 174.7498355 | 0.612100827 | 2.45E-02 |
| NFE2L3 | 4016.428492 | 0.609402544 | 1.34E-06 |
| SLC2A13 | 99.18506297 | 0.608470708 | 4.44E-02 |
| NUDCD1 | 1948.346597 | 0.604688791 | 1.17E-03 |

| | | | |
|---|---|---|---|
| CLIC6 | 238.7103255 | 0.603495591 | 1.19E-02 |
| WNT5A | 1223.951769 | 0.603186515 | 5.06E-04 |
| TLCD2 | 1283.167735 | 0.60087806 | 8.05E-05 |
| VCL | 13667.92026 | 0.598748926 | 1.83E-03 |
| RARRES2 | 5285.601134 | 0.59792186 | 1.25E-03 |
| CEP112 | 429.4679967 | 0.597527126 | 1.36E-02 |
| WHAMM | 724.8447879 | 0.597112787 | 1.54E-04 |
| PLEKHF2 | 603.1114698 | 0.596614726 | 4.26E-03 |
| SQOR | 301.2361767 | 0.594530376 | 3.34E-03 |
| C1orf226 | 396.125664 | 0.594321146 | 8.26E-04 |
| JDP2 | 1542.857247 | 0.594268308 | 4.38E-02 |
| ADAM33 | 189.1758325 | 0.594100709 | 4.24E-03 |
| SVIP | 315.9252992 | 0.592628241 | 1.05E-02 |
| NFE2 | 135.7777788 | 0.590329007 | 1.16E-02 |
| NMRK1 | 236.4018946 | 0.58918883 | 2.33E-03 |
| SMIM3 | 518.8367427 | 0.588826656 | 2.50E-04 |
| NA | 243.5273505 | 0.588092308 | 3.61E-05 |
| SLC46A3 | 306.615023 | 0.586320281 | 3.19E-03 |
| KIF23 | 4117.414188 | 0.585718769 | 2.97E-03 |
| CTNNA1 | 21483.2315 | 0.58450458 | 1.42E-17 |
| MILR1 | 125.3341487 | 0.584139359 | 3.70E-03 |
| ERBB2 | 4218.931161 | 0.583344126 | 2.26E-06 |
| SLC16A4 | 951.4176711 | 0.582763824 | 2.17E-05 |
| MCC | 3773.783562 | 0.581919992 | 3.97E-10 |
| KIAA1522 | 1610.958184 | 0.576804587 | 2.31E-05 |
| CNN2 | 8063.875845 | 0.576527736 | 5.21E-09 |
| FHOD3 | 292.6071871 | 0.575301575 | 2.73E-03 |
| PEG10 | 103399.184 | 0.574252278 | 4.91E-02 |
| CSF1 | 3189.285983 | 0.574134878 | 7.97E-07 |
| CRIM1 | 9816.058979 | 0.573925416 | 1.43E-05 |
| LAYN | 609.0277932 | 0.571351486 | 7.24E-03 |
| MICAL3 | 4913.970363 | 0.570448548 | 2.01E-12 |
| USP18 | 291.189923 | 0.569341356 | 5.81E-03 |
| ZSWIM4 | 566.8401579 | 0.568672312 | 9.00E-05 |
| AHNAK | 122265.4934 | 0.565724144 | 1.15E-09 |
| SMO | 2973.87738 | 0.564732509 | 1.42E-04 |
| RAB6B | 890.8632009 | 0.562406741 | 1.53E-03 |
| CREB3L2 | 4451.672148 | 0.561035353 | 8.94E-11 |
| CXCL16 | 487.4807416 | 0.560560676 | 8.76E-03 |
| ATP13A3 | 4319.212019 | 0.559975843 | 1.29E-02 |
| LINC01138 | 170.844276 | 0.559327769 | 4.07E-02 |
| AGFG2 | 3050.773205 | 0.558988245 | 4.09E-03 |
| EID2 | 658.2350262 | 0.556617388 | 1.13E-08 |
| MYO1C | 10838.86687 | 0.554021412 | 4.06E-05 |
| ASPA | 207.1723575 | 0.552813005 | 4.22E-03 |
| PKP4 | 3485.234126 | 0.552511901 | 2.57E-05 |
| ADGRB2 | 2041.469019 | 0.552458719 | 2.31E-04 |

| | | | |
|---|---|---|---|
| SAMD4A | 2581.581522 | 0.5514013 | 2.92E-04 |
| SDC2 | 2063.499858 | 0.550361527 | 1.22E-04 |
| ARHGEF10L | 688.723664 | 0.548431074 | 1.67E-04 |
| TPM1 | 23789.68 | 0.547885298 | 1.12E-05 |
| MAFF | 607.0319197 | 0.546795482 | 3.66E-02 |
| SLC39A8 | 375.9403401 | 0.544462436 | 2.14E-02 |
| CYTOR | 2035.603011 | 0.542089518 | 1.22E-05 |
| KCNK6 | 269.045259 | 0.542032349 | 1.54E-03 |
| NQO1 | 36091.71617 | 0.541719704 | 7.49E-03 |
| NUAK1 | 3145.049458 | 0.540339088 | 5.55E-05 |
| PCSK9 | 829.7416089 | 0.538387005 | 4.23E-02 |
| B4GALNT3 | 353.3939525 | 0.537239582 | 2.57E-03 |
| TXNRD1 | 33083.72171 | 0.535282031 | 3.49E-02 |
| C21orf91 | 509.9850167 | 0.533822275 | 2.97E-02 |
| 4-Sep | 536.272959 | 0.53359811 | 6.81E-03 |
| EVI5 | 724.956478 | 0.532970098 | 1.00E-02 |
| RGPD6 | 2488.190399 | 0.532523846 | 2.78E-02 |
| LHFPL6 | 422.6305817 | 0.530427699 | 4.97E-03 |
| ARHGEF10 | 3349.754618 | 0.529286319 | 1.27E-05 |
| COL26A1 | 362.579038 | 0.528535417 | 6.75E-03 |
| FBXL7 | 181.7020041 | 0.527571473 | 1.00E-02 |
| NBPF10 | 2314.64115 | 0.526586951 | 2.59E-06 |
| PLAUR | 813.9719978 | 0.525772156 | 2.93E-02 |
| PIP5K1A | 3765.167022 | 0.524071969 | 1.32E-06 |
| TRNP1 | 1601.234225 | 0.52386682 | 3.90E-03 |
| CYBRD1 | 6049.057482 | 0.521122077 | 5.77E-10 |
| SERPINB9 | 1503.778653 | 0.519138544 | 1.74E-09 |
| TTC33 | 553.6499434 | 0.519025955 | 1.64E-02 |
| C11orf96 | 147.5207521 | 0.518959831 | 4.25E-02 |
| OSMR | 15770.68321 | 0.517522014 | 2.97E-04 |
| PDGFRB | 11707.28027 | 0.517158491 | 4.98E-05 |
| FRG1JP | 122.945195 | 0.516283491 | 2.57E-02 |
| DCUN1D1 | 609.5664724 | 0.516039234 | 2.07E-02 |
| TMEM64 | 564.8664716 | 0.515382066 | 3.97E-03 |
| OTULINL | 778.6536385 | 0.512981605 | 2.46E-02 |
| C16orf87 | 427.9873414 | 0.512279889 | 3.64E-02 |
| NACC2 | 1511.713598 | 0.512043638 | 1.08E-06 |
| ITPR2 | 1727.732014 | 0.511395634 | 1.86E-02 |
| RBMS3 | 446.5159869 | 0.511214076 | 1.31E-02 |
| MPP5 | 2731.596394 | 0.510699333 | 6.10E-03 |
| SCARF2 | 1756.210107 | 0.509988962 | 5.36E-09 |
| SEL1L3 | 3269.858349 | 0.509688507 | 1.92E-03 |
| CASP4 | 3032.86953 | 0.507849445 | 4.69E-09 |
| PHACTR4 | 2601.10134 | 0.507388623 | 9.71E-12 |
| LAMB2 | 8298.833374 | 0.506447896 | 7.84E-04 |
| PRKAG2 | 1255.564186 | 0.505865994 | 3.81E-03 |
| BAIAP2L1 | 1835.617301 | 0.505085315 | 4.08E-06 |

| | | | |
|---|---|---|---|
| RAPH1 | 2044.357541 | 0.501727266 | 2.66E-04 |
| EPHA2 | 5020.238168 | 0.50133251 | 6.13E-04 |
| NR4A1 | 297.1782047 | 0.50055455 | 1.54E-02 |
| TUBB6 | 6630.385277 | 0.499508955 | 2.21E-02 |
| EIF4E3 | 382.3841659 | 0.499003126 | 3.58E-03 |
| FAM43A | 796.7891411 | 0.497516767 | 2.80E-04 |
| SLC16A5 | 221.2180602 | 0.497449921 | 2.49E-02 |
| FSCN1 | 13820.42552 | 0.497156683 | 2.62E-03 |
| GABARAPL1 | 3444.554127 | 0.496189755 | 1.67E-06 |
| PRDM5 | 571.6075791 | 0.494123453 | 3.81E-03 |
| TBC1D9 | 1044.573317 | 0.493857617 | 5.39E-05 |
| RAPGEF2 | 1880.163938 | 0.493689197 | 7.52E-03 |
| REEP3 | 450.3834737 | 0.49040323 | 7.51E-03 |
| MBP | 3639.442215 | 0.490004043 | 3.97E-02 |
| TMOD3 | 4530.510323 | 0.48923353 | 5.23E-03 |
| LYN | 763.4264982 | 0.488772849 | 3.81E-03 |
| NCOA3 | 5143.412747 | 0.4885963 | 1.98E-03 |
| RAPGEF1 | 4833.32604 | 0.486859146 | 3.41E-06 |
| MFAP4 | 931.0022545 | 0.482777561 | 2.01E-02 |
| TVP23B | 715.0767378 | 0.482712712 | 3.89E-02 |
| IGFBP7 | 8049.810997 | 0.482706004 | 5.51E-03 |
| NA | 175.0159474 | 0.482332237 | 1.04E-02 |
| PAPLN | 1234.774959 | 0.482306704 | 8.87E-04 |
| ABCC4 | 5268.937555 | 0.478623849 | 4.31E-07 |
| OSTM1 | 1362.323596 | 0.477994746 | 1.01E-02 |
| PLXNB2 | 10147.39702 | 0.47798829 | 2.85E-05 |
| STXBP5 | 998.2124252 | 0.477667935 | 1.06E-02 |
| F11R | 1874.633537 | 0.477285706 | 3.05E-02 |
| PHLPP2 | 1636.290208 | 0.477109458 | 2.03E-08 |
| MFSD6 | 998.7542204 | 0.476167477 | 8.38E-06 |
| RNF41 | 3274.28356 | 0.474767811 | 4.94E-02 |
| TGFBR3 | 1252.634429 | 0.47461049 | 4.25E-04 |
| CAMTA1 | 1421.581369 | 0.474388666 | 7.61E-05 |
| SIRPA | 3488.218336 | 0.473370583 | 3.87E-03 |
| TPD52L1 | 2117.343253 | 0.472919061 | 3.02E-02 |
| PLOD2 | 2680.245608 | 0.472908755 | 7.09E-03 |
| FAM114A1 | 1772.247523 | 0.472682428 | 7.79E-03 |
| HEG1 | 10780.54503 | 0.472636644 | 1.13E-08 |
| ACSS3 | 176.7728467 | 0.472563597 | 4.77E-02 |
| PTPRJ | 3731.301994 | 0.472317818 | 3.81E-09 |
| NBPF14 | 4716.123144 | 0.471092337 | 1.93E-05 |
| INSR | 813.980436 | 0.470812909 | 4.77E-02 |
| SPART | 2174.26844 | 0.470444057 | 1.59E-05 |
| BCAM | 3742.931072 | 0.470122106 | 3.49E-03 |
| MID2 | 540.4827019 | 0.467821056 | 2.99E-03 |
| USP12 | 609.8723212 | 0.46495009 | 4.28E-02 |
| LIN28A | 640.0340004 | 0.46493065 | 1.16E-02 |

| | | | |
|---|---|---|---|
| LOXL2 | 22646.42599 | 0.462989512 | 1.41E-02 |
| SHTN1 | 750.5620927 | 0.462278616 | 2.22E-03 |
| MYH14 | 1638.03928 | 0.461443972 | 1.50E-04 |
| AASS | 1629.736665 | 0.461368272 | 2.01E-03 |
| WWP1 | 1340.319215 | 0.461237853 | 2.21E-02 |
| E2F8 | 533.6095413 | 0.459507125 | 3.09E-02 |
| AC244153.1 | 322.5237666 | 0.457986316 | 2.44E-03 |
| PPIC | 3752.098519 | 0.454106232 | 3.37E-07 |
| TJP1 | 3258.84199 | 0.453761723 | 3.58E-04 |
| HOXC6 | 874.2215878 | 0.453201984 | 9.48E-08 |
| FAM83H | 345.1934365 | 0.453192519 | 6.27E-03 |
| NA | 148.9071066 | 0.451897025 | 4.44E-02 |
| MYADM | 3260.603661 | 0.450077541 | 1.69E-03 |
| CTSV | 629.8242174 | 0.449881145 | 1.16E-02 |
| RBPMS | 1720.796055 | 0.449438093 | 1.67E-04 |
| CASP10 | 374.7130969 | 0.448851986 | 2.94E-02 |
| EPDR1 | 1988.545424 | 0.447304566 | 2.46E-03 |
| P4HA1 | 2000.926193 | 0.447119724 | 5.32E-04 |
| SMG1P3 | 308.3634136 | 0.446345275 | 3.97E-02 |
| SLC17A5 | 973.1245904 | 0.444971521 | 3.35E-02 |
| FCHO2 | 1078.451211 | 0.444933666 | 3.58E-02 |
| SH3BGRL2 | 622.823889 | 0.444244207 | 3.66E-02 |
| NKX3-1 | 1379.272934 | 0.44388655 | 2.31E-04 |
| AC145124.1 | 226.6834811 | 0.442592032 | 2.01E-02 |
| PGM3 | 2990.489355 | 0.442462458 | 1.03E-02 |
| PPM1K | 858.4480811 | 0.441445894 | 1.57E-02 |
| CLTB | 1764.884731 | 0.440797771 | 1.26E-04 |
| PRRG1 | 823.3572282 | 0.440327645 | 4.78E-03 |
| NBPF19 | 1631.618351 | 0.44013207 | 1.34E-04 |
| COL5A1 | 25458.72823 | 0.439994496 | 1.54E-04 |
| METRNL | 538.6256251 | 0.437518222 | 4.75E-02 |
| C8orf58 | 1572.615118 | 0.436951022 | 3.99E-02 |
| SLFN12 | 950.4862695 | 0.436484744 | 1.34E-02 |
| CTBS | 682.3989996 | 0.435662946 | 2.78E-04 |
| EHD4 | 2189.688796 | 0.435211361 | 7.84E-04 |
| ATG5 | 1481.378423 | 0.434956799 | 4.27E-02 |
| PPARG | 1847.312041 | 0.433636391 | 1.94E-02 |
| SLC44A1 | 3595.730604 | 0.433560286 | 7.07E-08 |
| PAM | 16151.86896 | 0.43353328 | 2.63E-04 |
| SEC24D | 3034.081143 | 0.432326945 | 2.55E-03 |
| NEDD4L | 4687.21832 | 0.432062252 | 2.43E-04 |
| FTH1 | 76509.70998 | 0.431451632 | 6.43E-03 |
| B4GALNT1 | 1140.587409 | 0.430943685 | 1.68E-02 |
| ARHGAP31 | 2564.11328 | 0.430837434 | 8.34E-03 |
| SCD5 | 399.6806143 | 0.430650938 | 2.06E-02 |
| INF2 | 4687.789987 | 0.430400586 | 1.63E-04 |
| GBP1 | 1438.088516 | 0.429590325 | 1.48E-03 |

| | | | |
|---|---|---|---|
| GNE | 4850.27859 | 0.429185933 | 4.80E-04 |
| NOTCH2 | 14889.49296 | 0.429060824 | 1.43E-07 |
| FZD6 | 1565.705517 | 0.429046797 | 3.17E-03 |
| MLLT3 | 633.9740889 | 0.42765089 | 2.64E-02 |
| FAM174B | 267.0840553 | 0.427290722 | 1.91E-02 |
| LRP6 | 2588.5102 | 0.426104522 | 5.61E-03 |
| FZD7 | 2542.472438 | 0.42568701 | 1.12E-05 |
| PEA15 | 12565.60558 | 0.424797386 | 1.68E-02 |
| ADAMTS12 | 10377.03832 | 0.424654142 | 5.67E-04 |
| TENM3 | 2755.606116 | 0.424273952 | 5.01E-04 |
| COL16A1 | 5086.360967 | 0.42348382 | 1.78E-02 |
| LLGL2 | 1015.83685 | 0.422492835 | 1.07E-03 |
| S100A10 | 12016.01031 | 0.421688877 | 2.84E-03 |
| TUB | 3581.772259 | 0.420893695 | 9.90E-05 |
| HOXC4 | 316.5324271 | 0.420748515 | 8.92E-03 |
| CDKN1B | 2344.932369 | 0.420069318 | 6.05E-05 |
| RAB18 | 819.3678411 | 0.419879283 | 5.25E-04 |
| DNAJC3 | 5598.113646 | 0.417711576 | 1.05E-04 |
| RAB11FIP1 | 3684.571645 | 0.417542658 | 3.39E-04 |
| MCUR1 | 1934.263528 | 0.417071437 | 2.99E-06 |
| TAX1BP3 | 4806.333429 | 0.416893117 | 8.72E-03 |
| PPP2R5A | 1252.576734 | 0.415114742 | 1.28E-03 |
| LAMA5 | 8544.457214 | 0.414166967 | 2.59E-02 |
| HOOK3 | 1766.319764 | 0.413433035 | 3.72E-02 |
| TNIK | 2274.191439 | 0.4123974 | 1.31E-02 |
| PRTG | 674.9816943 | 0.412195172 | 2.04E-02 |
| MIR4435-2HG | 2097.286563 | 0.412179471 | 1.10E-03 |
| ALDH3B1 | 1300.36037 | 0.41210987 | 9.29E-04 |
| FLNA | 161615.4789 | 0.411596496 | 1.00E-04 |
| NEO1 | 7967.414653 | 0.41140708 | 9.72E-03 |
| PCDHGB5 | 192.1740464 | 0.410642735 | 3.77E-02 |
| SYNE2 | 2558.962612 | 0.408983293 | 2.84E-02 |
| ARNTL2 | 885.1172776 | 0.40865459 | 2.18E-02 |
| SH3BP5 | 1459.805366 | 0.407683506 | 1.60E-02 |
| CDYL2 | 2093.83975 | 0.40754212 | 7.85E-03 |
| SHC1 | 17857.68695 | 0.407000289 | 2.57E-03 |
| FOXD2 | 276.8095472 | 0.406688238 | 7.43E-03 |
| PNPLA6 | 3135.262182 | 0.4064402 | 1.90E-02 |
| NT5DC1 | 1286.054086 | 0.40610638 | 6.15E-04 |
| MYH9 | 39701.66481 | 0.404890982 | 1.63E-03 |
| FAM160B1 | 711.6560439 | 0.404690162 | 1.57E-02 |
| LOXL3 | 3055.781083 | 0.402309559 | 2.80E-02 |
| NUCB2 | 1933.069367 | 0.402126269 | 1.97E-02 |
| SGCD | 2815.580324 | 0.401799975 | 6.74E-10 |
| TRAM1 | 4228.399092 | 0.399801224 | 5.71E-05 |
| MYL12B | 10882.15315 | 0.39869597 | 5.36E-09 |
| EGLN1 | 1845.305623 | 0.398506352 | 1.23E-02 |

| | | | |
|---|---|---|---|
| SYNJ2 | 2575.00764 | 0.396260877 | 7.83E-03 |
| MAPRE3 | 973.794316 | 0.395359246 | 2.88E-03 |
| FAM122B | 3045.59739 | 0.39445119 | 2.86E-03 |
| MGLL | 4291.086206 | 0.392534049 | 2.67E-02 |
| BCOR | 2863.827776 | 0.391356426 | 2.07E-04 |
| UBR3 | 2018.764591 | 0.39073088 | 3.07E-02 |
| PICALM | 8631.628285 | 0.388763734 | 5.20E-03 |
| ATP11A | 1843.254605 | 0.387479762 | 2.44E-02 |
| BCL2L1 | 5620.751235 | 0.38719234 | 1.12E-04 |
| SGCE | 1697.348195 | 0.387188642 | 8.71E-04 |
| MEGF9 | 1097.538604 | 0.386842308 | 2.65E-03 |
| RBPMS2 | 349.7867648 | 0.385735883 | 7.09E-03 |
| INTS6 | 1176.900312 | 0.385164012 | 3.08E-02 |
| SEC23A | 5558.829986 | 0.385083444 | 8.87E-04 |
| NFIB | 3177.725101 | 0.383729263 | 1.06E-04 |
| DCAF6 | 2454.745029 | 0.38266641 | 5.37E-03 |
| HLA-C | 3125.12319 | 0.382044423 | 4.44E-02 |
| RRBP1 | 17700.10449 | 0.379413144 | 1.86E-05 |
| SERTAD2 | 1650.89911 | 0.37876893 | 6.01E-03 |
| PDLIM5 | 3144.819155 | 0.378311227 | 2.87E-02 |
| RAP1A | 1573.438963 | 0.377248203 | 7.74E-06 |
| SOWAHC | 557.8818764 | 0.376689656 | 3.32E-02 |
| SPECC1 | 1835.882639 | 0.376329602 | 3.54E-03 |
| LRRC8C | 1363.062089 | 0.375978493 | 6.32E-03 |
| PLLP | 455.6051462 | 0.375079009 | 8.15E-03 |
| 10-Sep | 3710.919446 | 0.374336572 | 3.89E-02 |
| PLEKHA8 | 1554.564933 | 0.373204128 | 2.42E-02 |
| FBN1 | 3258.584951 | 0.372420598 | 1.91E-02 |
| MYO1E | 3124.761668 | 0.372219003 | 1.43E-02 |
| LINC01578 | 1662.10461 | 0.371786081 | 1.89E-02 |
| BCL6 | 700.8646573 | 0.368661744 | 7.91E-03 |
| SLC1A3 | 1565.100729 | 0.368509182 | 4.19E-02 |
| GSTA4 | 526.6998491 | 0.367928931 | 2.07E-02 |
| SERPINH1 | 45395.88893 | 0.366315275 | 3.23E-04 |
| PDLIM1 | 4528.312637 | 0.3652555 | 1.82E-02 |
| AMIGO1 | 1025.261971 | 0.365237782 | 2.92E-02 |
| PKD2 | 2184.663901 | 0.365067638 | 2.27E-03 |
| ULK2 | 882.4735602 | 0.364889304 | 3.52E-02 |
| ADCY6 | 3447.816817 | 0.36431315 | 1.42E-02 |
| NEDD9 | 1801.09962 | 0.362619291 | 2.01E-02 |
| XYLT1 | 2293.659344 | 0.361790524 | 1.13E-04 |
| SLC9A3R1 | 2447.527257 | 0.361532682 | 2.49E-02 |
| GJC1 | 1901.807077 | 0.361058749 | 1.15E-02 |
| LINC00265 | 516.0461094 | 0.360683079 | 3.98E-02 |
| GINM1 | 1586.931872 | 0.360669163 | 3.21E-05 |
| GPC1 | 4941.093134 | 0.360525631 | 1.94E-02 |
| JOSD1 | 3246.967769 | 0.360186578 | 2.25E-04 |

| | | | |
|---|---|---|---|
| STS | 1112.178348 | 0.358847122 | 6.16E-04 |
| SH3BP4 | 4262.928303 | 0.357852577 | 6.80E-03 |
| UBE2J1 | 3574.235135 | 0.356731651 | 7.13E-05 |
| NDST1 | 8021.325827 | 0.356177179 | 3.69E-03 |
| KIF13A | 2692.151826 | 0.355827314 | 6.42E-04 |
| ARHGAP18 | 3596.763254 | 0.355814596 | 3.05E-04 |
| VASN | 2614.474492 | 0.355527233 | 4.11E-02 |
| HACD1 | 437.3528397 | 0.355050904 | 2.40E-03 |
| LDLRAP1 | 1727.553505 | 0.353187522 | 3.91E-03 |
| CREG1 | 2653.32401 | 0.3516012 | 1.04E-03 |
| BLVRB | 2573.362732 | 0.350754678 | 2.01E-02 |
| MYO10 | 14327.81064 | 0.35039615 | 3.97E-03 |
| SLC44A2 | 6112.518187 | 0.350080797 | 2.07E-03 |
| NOTCH2NLA | 1200.993235 | 0.349806981 | 3.31E-03 |
| PIR | 1579.361561 | 0.349531749 | 4.63E-02 |
| NTN4 | 1601.295178 | 0.348702075 | 2.27E-03 |
| ARHGEF12 | 14026.13928 | 0.348109143 | 3.92E-03 |
| AC092338.1 | 1800.678811 | 0.347251693 | 1.33E-02 |
| DAB2 | 9628.493215 | 0.346702397 | 7.04E-04 |
| CD47 | 3412.049834 | 0.345183282 | 6.89E-05 |
| G6PD | 13437.0078 | 0.344762866 | 1.26E-02 |
| SMURF2 | 2056.949958 | 0.343894484 | 1.38E-02 |
| OPTN | 3436.814398 | 0.343604809 | 5.28E-03 |
| DPYSL5 | 1435.744589 | 0.3432241 | 1.82E-02 |
| TSPAN18 | 4006.491139 | 0.34208141 | 5.31E-03 |
| AFAP1 | 8267.209501 | 0.341407265 | 3.04E-03 |
| TCF7L1 | 1427.682103 | 0.341369894 | 2.39E-02 |
| P2RX6 | 798.6419577 | 0.340548039 | 3.20E-02 |
| PEAK1 | 4818.635054 | 0.3396588 | 6.07E-04 |
| STK3 | 890.5682343 | 0.339249515 | 1.62E-02 |
| IFNGR2 | 3672.123779 | 0.3392467 | 1.77E-02 |
| MRAS | 1665.819629 | 0.338673896 | 2.27E-02 |
| TRAF5 | 901.765231 | 0.338306849 | 3.84E-02 |
| GSTM3 | 4602.829408 | 0.338108689 | 1.00E-02 |
| KANK1 | 4383.01843 | 0.334647919 | 1.14E-04 |
| SOCS6 | 907.8110569 | 0.334086727 | 4.91E-03 |
| PGRMC1 | 3576.645105 | 0.333916127 | 1.73E-04 |
| SWAP70 | 3816.151683 | 0.331996191 | 1.44E-03 |
| CRYZL1 | 678.2921722 | 0.331613016 | 4.94E-02 |
| FOSL2 | 5985.00852 | 0.329457511 | 2.68E-02 |
| PQLC3 | 634.4680855 | 0.329359457 | 3.48E-02 |
| FGD5 | 1364.669439 | 0.32678218 | 3.80E-02 |
| MEF2A | 1827.254252 | 0.325924557 | 4.58E-02 |
| FMNL2 | 4188.966288 | 0.325770822 | 2.70E-02 |
| YAP1 | 7292.138549 | 0.325414487 | 2.02E-03 |
| NUDCD2 | 2588.159888 | 0.32500894 | 4.49E-03 |
| NRBP2 | 1261.707177 | 0.324971666 | 4.57E-03 |

| | | | |
|---|---|---|---|
| UBE2W | 670.5578627 | 0.324082871 | 2.70E-02 |
| ITGB1 | 24729.84288 | 0.322494778 | 2.60E-03 |
| INPPL1 | 9644.907944 | 0.321077488 | 6.81E-03 |
| STOM | 8123.48438 | 0.321014301 | 2.99E-04 |
| TRIM37 | 2440.200354 | 0.320850348 | 2.01E-02 |
| TRAM2 | 9655.068286 | 0.320771606 | 4.04E-05 |
| DZIP1 | 7215.684019 | 0.320364558 | 3.70E-03 |
| EPHX1 | 3232.5919 | 0.319861524 | 3.94E-02 |
| DENND3 | 1469.517193 | 0.31952924 | 1.71E-02 |
| ADAM10 | 7600.927323 | 0.319506009 | 3.77E-02 |
| PON2 | 4494.206146 | 0.319352252 | 5.08E-04 |
| ITSN1 | 4401.632807 | 0.319083966 | 5.31E-06 |
| SAT1 | 2227.264793 | 0.317888579 | 4.58E-02 |
| ENY2 | 2390.585504 | 0.317312687 | 1.28E-03 |
| BPGM | 883.7041193 | 0.316144583 | 2.04E-03 |
| TGFBR2 | 6562.551738 | 0.316056405 | 1.82E-02 |
| KIRREL1 | 10728.67019 | 0.31373626 | 1.10E-02 |
| OSBPL1A | 2509.624306 | 0.313476165 | 8.42E-05 |
| SHROOM4 | 459.884557 | 0.313182531 | 1.75E-02 |
| LGMN | 4006.816281 | 0.310908663 | 8.76E-03 |
| SPPL2A | 1140.966053 | 0.310612679 | 2.06E-02 |
| CTSZ | 14220.45977 | 0.308660711 | 2.78E-02 |
| TFE3 | 7339.395033 | 0.308206544 | 1.00E-02 |
| TTC39C | 430.9709289 | 0.30731013 | 2.67E-02 |
| ZHX3 | 3668.787153 | 0.306729343 | 4.01E-02 |
| NFASC | 750.1182267 | 0.305486103 | 4.49E-02 |
| SETD7 | 2957.166962 | 0.305386131 | 3.06E-03 |
| CACUL1 | 1176.624673 | 0.303259366 | 3.62E-02 |
| MROH1 | 1333.300415 | 0.302734529 | 2.46E-02 |
| KLHL42 | 1254.378234 | 0.302442369 | 1.92E-03 |
| CPD | 5957.815368 | 0.301769797 | 4.42E-02 |
| MARCKS | 3627.182142 | 0.30141564 | 1.63E-02 |
| FERMT2 | 5028.125652 | 0.30021096 | 2.10E-02 |
| RNF11 | 2230.717256 | 0.299429835 | 2.85E-04 |
| PFN2 | 10427.95587 | 0.299413638 | 2.00E-03 |
| PWWP2A | 1799.661611 | 0.299410918 | 1.17E-02 |
| SPTBN1 | 22759.67806 | 0.298872879 | 2.33E-06 |
| ACSL3 | 7289.522609 | 0.297871658 | 1.57E-02 |
| SLC35G1 | 581.9719221 | 0.294200915 | 3.75E-02 |
| FBXO9 | 2299.674237 | 0.293036411 | 8.81E-03 |
| COG5 | 3073.024123 | 0.292994319 | 4.91E-02 |
| TRAK1 | 3100.499793 | 0.291896049 | 5.08E-04 |
| EIF2AK2 | 3388.221179 | 0.291352917 | 2.30E-02 |
| USP32 | 3325.072913 | 0.290303865 | 2.90E-03 |
| PDP2 | 1653.370186 | 0.289004458 | 1.75E-02 |
| COPB2 | 9595.875253 | 0.287976955 | 3.81E-03 |
| CASP3 | 1463.757118 | 0.287459904 | 2.11E-02 |

| | | | |
|---|---|---|---|
| TIAL1 | 2251.871338 | 0.287450489 | 2.31E-03 |
| FADS3 | 3833.388893 | 0.287321594 | 4.28E-02 |
| JCAD | 5110.161346 | 0.285348825 | 2.36E-02 |
| GTF2A2 | 1816.018377 | 0.282644738 | 1.37E-02 |
| NXPE3 | 1902.101291 | 0.281225515 | 3.58E-02 |
| PPFIBP1 | 7522.570197 | 0.279075052 | 1.40E-03 |
| GALNT10 | 5382.628084 | 0.27677623 | 4.36E-02 |
| SCRN1 | 19372.32534 | 0.276714205 | 2.04E-03 |
| CUL1 | 4998.254058 | 0.2756548 | 2.99E-05 |
| TRIM5 | 2409.462025 | 0.274963234 | 7.85E-03 |
| PBX1 | 1185.042455 | 0.274459818 | 3.23E-02 |
| ZNF281 | 1503.230845 | 0.274400904 | 2.05E-02 |
| CCNYL1 | 1291.923322 | 0.274348297 | 3.49E-02 |
| AHCYL1 | 7022.281892 | 0.271438608 | 3.48E-05 |
| NFKB1 | 2198.220305 | 0.27005536 | 1.09E-02 |
| ARHGAP10 | 734.6739707 | 0.268943206 | 2.18E-02 |
| FYCO1 | 2501.029106 | 0.267774992 | 1.23E-02 |
| TXNDC11 | 2341.326044 | 0.267535233 | 3.64E-02 |
| SNRPE | 2384.328884 | 0.266082459 | 3.81E-02 |
| EIF2AK1 | 12276.79952 | 0.26573513 | 2.62E-02 |
| ADGRA2 | 6548.879872 | 0.262225244 | 3.12E-02 |
| LUZP1 | 6874.52116 | 0.2599448 | 8.87E-04 |
| ZNF618 | 3843.333358 | 0.259930276 | 3.57E-02 |
| SNAP23 | 1810.39237 | 0.259806471 | 3.62E-02 |
| KLHDC10 | 3849.172421 | 0.259338083 | 1.32E-02 |
| KLHL7 | 2045.471299 | 0.25698527 | 2.27E-02 |
| MAP2K4 | 1534.37162 | 0.255645197 | 4.28E-02 |
| TRIM24 | 2116.562968 | 0.254894645 | 7.15E-03 |
| MGME1 | 2856.925219 | 0.254718371 | 6.63E-03 |
| CEBPZOS | 1243.790674 | 0.253048662 | 2.76E-02 |
| PHF10 | 1866.438536 | 0.252923032 | 2.60E-02 |
| DAPK1 | 5388.257129 | 0.252180932 | 7.53E-03 |
| PIEZO2 | 3798.179918 | 0.249503773 | 3.02E-02 |
| ANKRD13C | 893.5313821 | 0.247389997 | 3.03E-02 |
| RAB2A | 3018.845323 | 0.246695216 | 2.94E-04 |
| KPNA4 | 3635.1137 | 0.246681773 | 1.17E-02 |
| TRIP10 | 3342.101072 | 0.244375767 | 1.18E-03 |
| STT3B | 6050.763375 | 0.242942928 | 4.45E-02 |
| TMEM106C | 4149.946741 | 0.242333429 | 1.80E-02 |
| HDAC7 | 5153.464871 | 0.241597769 | 2.27E-02 |
| PDCD6IP | 6347.384567 | 0.238035485 | 2.30E-02 |
| FRMD3 | 3496.851126 | 0.237969896 | 2.96E-02 |
| PDXDC1 | 6002.09432 | 0.235784432 | 4.67E-02 |
| PRPS1 | 3124.775206 | 0.235608254 | 3.24E-02 |
| C1orf198 | 2749.169919 | 0.233781624 | 3.94E-02 |
| PHKB | 4756.907039 | 0.233590221 | 3.29E-03 |
| CYP2U1 | 1376.719402 | 0.233263826 | 1.14E-02 |

| | | | |
|---|---|---|---|
| TRAK2 | 3071.599689 | 0.231813018 | 3.96E-03 |
| RCN1 | 15497.19177 | 0.231275277 | 6.46E-03 |
| CFL2 | 3651.721601 | 0.228913549 | 9.81E-03 |
| DOCK1 | 3315.886736 | 0.224104086 | 2.27E-02 |
| FARP1 | 10323.71837 | 0.22375544 | 3.89E-02 |
| KCTD20 | 6930.129421 | 0.222533078 | 1.20E-04 |
| PAFAH1B1 | 7948.894797 | 0.221933799 | 1.06E-02 |
| PGRMC2 | 2270.365819 | 0.217993313 | 9.25E-03 |
| NNT | 4515.752462 | 0.217319322 | 4.31E-02 |
| CSDE1 | 17721.67776 | 0.2169896 | 3.99E-02 |
| REPIN1 | 8966.942661 | 0.216620257 | 3.22E-02 |
| PPP2CB | 3807.900051 | 0.215699531 | 5.58E-04 |
| SEC22B | 4612.525216 | 0.215045689 | 4.23E-02 |
| RIC1 | 2976.271407 | 0.215025853 | 3.15E-03 |
| SGCB | 2046.737694 | 0.211122733 | 3.34E-03 |
| SIPA1L1 | 3181.582534 | 0.207890288 | 3.86E-02 |
| COX20 | 1221.240216 | 0.207847301 | 4.70E-02 |
| PYGL | 5642.226534 | 0.206096631 | 6.69E-03 |
| MTMR12 | 3615.825772 | 0.204511642 | 4.13E-02 |
| EXOC6B | 3172.372247 | 0.201030661 | 3.43E-02 |
| MACF1 | 17521.76745 | 0.198553047 | 2.80E-02 |
| EIF2AK4 | 3997.221069 | 0.19743322 | 7.21E-03 |
| UBE2G1 | 2388.742941 | 0.196806312 | 4.91E-02 |
| ELOVL5 | 4586.484995 | 0.195530937 | 3.05E-02 |
| HSPA4 | 19032.1917 | 0.195150519 | 3.83E-02 |
| WDR26 | 5431.769766 | 0.19460005 | 5.64E-03 |
| PNMA8A | 2550.232377 | 0.193720114 | 5.23E-03 |
| EGFR | 7757.382542 | 0.188500467 | 4.57E-02 |
| UBR5 | 9785.128236 | 0.187632789 | 4.40E-02 |
| UBTD2 | 1371.08928 | 0.180107559 | 2.92E-02 |
| 2-Sep | 17670.70577 | 0.17167224 | 3.15E-02 |
| FNBP1 | 5457.462751 | 0.164580397 | 3.28E-02 |
| CFLAR | 2074.937118 | 0.163760252 | 4.58E-02 |
| DCAF8 | 5161.578887 | 0.162091576 | 4.87E-02 |
| GOLPH3 | 5308.918063 | 0.157075555 | 2.73E-02 |
| RNF145 | 10583.21953 | 0.153704102 | 1.42E-02 |
| UBE2K | 2831.245226 | 0.14387574 | 4.58E-02 |
| C2orf68 | 2322.099413 | -0.146193691 | 4.37E-02 |
| STK35 | 2874.044657 | -0.16369264 | 2.95E-02 |
| CSNK2A1 | 11719.97268 | -0.165707266 | 4.98E-02 |
| SSBP1 | 6663.456012 | -0.183445395 | 1.86E-02 |
| TSR1 | 7944.401475 | -0.186133778 | 1.10E-02 |
| HNRNPF | 6684.601055 | -0.18640393 | 7.68E-03 |
| PA2G4 | 8850.309135 | -0.187327314 | 4.96E-02 |
| MRM2 | 2951.072928 | -0.188868738 | 1.16E-02 |
| PDF | 1295.291604 | -0.191813333 | 4.32E-02 |
| QSOX2 | 2784.490846 | -0.195805247 | 3.29E-02 |

| | | | |
|---|---|---|---|
| NFX1 | 3764.769076 | -0.198935765 | 6.27E-03 |
| ELOA | 3907.96247 | -0.201629367 | 4.58E-03 |
| BUB3 | 4309.026554 | -0.201716173 | 3.12E-02 |
| DUXAP8 | 8045.737965 | -0.207063387 | 1.49E-03 |
| NOP56 | 9573.943269 | -0.207685537 | 2.59E-03 |
| MAP4K4 | 13426.54364 | -0.208572109 | 9.33E-03 |
| PRXL2A | 2340.360732 | -0.212282417 | 3.81E-02 |
| CCT7 | 18359.8704 | -0.213969743 | 3.66E-02 |
| TMEM230 | 10289.00671 | -0.215146281 | 2.59E-02 |
| ZNF7 | 1564.127802 | -0.216088763 | 1.36E-02 |
| BFAR | 4729.283267 | -0.21810905 | 3.82E-03 |
| ADAT1 | 2374.903336 | -0.220494534 | 2.95E-02 |
| NEU3 | 1544.659068 | -0.220559022 | 2.13E-02 |
| MRPS33 | 2202.644019 | -0.220954026 | 3.76E-02 |
| DHPS | 2252.553779 | -0.222261702 | 6.46E-03 |
| RNF149 | 1382.487106 | -0.225928776 | 1.21E-02 |
| RPIA | 1240.066785 | -0.227027617 | 2.54E-02 |
| UTP4 | 5907.120824 | -0.231766727 | 4.51E-02 |
| POLD2 | 6626.569676 | -0.231953044 | 2.94E-02 |
| MLH1 | 2396.676964 | -0.232697682 | 4.68E-02 |
| YBX3 | 3730.363678 | -0.234442203 | 1.17E-02 |
| FBXO17 | 1499.143492 | -0.234818168 | 3.03E-02 |
| WIPF1 | 1839.234446 | -0.235004124 | 3.23E-02 |
| TCOF1 | 9862.217299 | -0.238038069 | 1.15E-02 |
| ACTL6A | 2808.889881 | -0.238233185 | 8.76E-03 |
| TMEM43 | 7214.222135 | -0.238447092 | 9.67E-03 |
| GUSB | 2894.849897 | -0.238535312 | 2.63E-02 |
| ISG20L2 | 2175.728689 | -0.238560349 | 4.76E-02 |
| ODF2 | 3894.579451 | -0.238880683 | 3.84E-03 |
| ENTR1 | 4996.79375 | -0.239345748 | 1.39E-02 |
| RPL6 | 45697.77222 | -0.239895914 | 5.47E-03 |
| RPL17 | 27377.12479 | -0.24005779 | 3.79E-02 |
| METTL2B | 1796.131861 | -0.240345297 | 1.86E-02 |
| SPAG7 | 2103.845243 | -0.240569677 | 5.47E-03 |
| WDR46 | 5956.684672 | -0.240808236 | 2.39E-02 |
| CSNK1E | 5056.524669 | -0.241894281 | 3.50E-03 |
| SCAMP1-AS1 | 798.8270424 | -0.243207273 | 4.76E-02 |
| EXOSC2 | 2807.209963 | -0.243708045 | 5.77E-04 |
| TRIM14 | 2948.397171 | -0.244285868 | 3.37E-02 |
| SSRP1 | 19588.32243 | -0.245069998 | 6.32E-03 |
| VPS37C | 1181.127109 | -0.24527444 | 4.24E-02 |
| RPL31 | 25986.45476 | -0.245838833 | 4.44E-02 |
| U2AF2 | 10147.28852 | -0.246225022 | 3.51E-02 |
| PSMC4 | 5691.909785 | -0.246374364 | 2.88E-02 |
| TOMM5 | 5140.633679 | -0.246559011 | 3.28E-02 |
| KAT2A | 3508.866278 | -0.246570636 | 2.12E-02 |
| EIF2B1 | 3995.522508 | -0.247161214 | 2.66E-04 |

| | | | |
|---|---|---|---|
| SLC25A22 | 2360.771682 | -0.247380597 | 4.44E-02 |
| C9orf78 | 3511.313378 | -0.247971506 | 5.53E-05 |
| POP5 | 1281.270367 | -0.248578409 | 1.52E-02 |
| AC022107.1 | 577.3918954 | -0.249351539 | 4.13E-02 |
| PHB2 | 8408.780695 | -0.249499599 | 3.88E-02 |
| COQ9 | 2727.947584 | -0.250851943 | 3.68E-02 |
| LARS2 | 2509.944575 | -0.251137615 | 4.20E-02 |
| PRPF19 | 12095.58731 | -0.252570938 | 1.67E-02 |
| CCDC137 | 2193.678733 | -0.252614187 | 1.42E-02 |
| DNAAF5 | 2856.312045 | -0.254377769 | 1.18E-02 |
| FKBP5 | 2560.809448 | -0.256269248 | 5.47E-03 |
| MRPS23 | 2563.61453 | -0.257188724 | 1.38E-02 |
| ZBED4 | 1607.762094 | -0.257542468 | 6.34E-03 |
| TIMM8A | 916.3763145 | -0.260607392 | 2.11E-02 |
| MRPS25 | 2424.720312 | -0.260752619 | 5.47E-03 |
| RPS8 | 38197.62831 | -0.261103188 | 2.68E-02 |
| HARS2 | 2149.232641 | -0.261269433 | 3.63E-04 |
| MRPS26 | 2527.031429 | -0.261310585 | 1.83E-02 |
| MTG1 | 2244.95116 | -0.261454711 | 3.60E-02 |
| CLP1 | 1304.941902 | -0.262487696 | 1.98E-03 |
| RPP25 | 1596.165225 | -0.263342698 | 1.94E-02 |
| VARS | 6546.556051 | -0.263819173 | 3.64E-02 |
| RPAIN | 1766.99041 | -0.263954536 | 1.65E-03 |
| RPL36A | 17414.29202 | -0.264449208 | 1.37E-02 |
| STOML2 | 9108.069344 | -0.264543298 | 9.37E-03 |
| HSD17B10 | 2671.357743 | -0.265113196 | 3.76E-02 |
| COQ4 | 1733.716167 | -0.265321352 | 2.49E-02 |
| DAXX | 4618.188562 | -0.265622722 | 3.64E-02 |
| MRPS16 | 2989.27372 | -0.266788501 | 1.10E-02 |
| SNX1 | 8206.680958 | -0.267033231 | 3.81E-02 |
| AATF | 3385.796557 | -0.268491464 | 1.60E-02 |
| SPATA2 | 1058.007254 | -0.268510366 | 2.49E-02 |
| EEF1A1 | 626427.2418 | -0.268655257 | 1.62E-02 |
| RRS1 | 2053.440274 | -0.269098198 | 1.55E-02 |
| CHN1 | 893.6230543 | -0.269857605 | 4.62E-02 |
| TOP1MT | 1570.636353 | -0.270059359 | 2.24E-03 |
| RCC1 | 5720.614234 | -0.271190223 | 4.19E-02 |
| MRPL15 | 2929.771809 | -0.274408521 | 1.92E-03 |
| AHRR | 2060.829402 | -0.275267895 | 4.54E-02 |
| UTP14A | 3064.894564 | -0.276174706 | 3.35E-02 |
| NDUFS2 | 5203.207594 | -0.277264899 | 2.18E-03 |
| GTF3C2 | 4232.170189 | -0.278670985 | 1.12E-02 |
| THG1L | 1892.442873 | -0.278809877 | 3.34E-02 |
| SNHG20 | 472.8352944 | -0.279190732 | 3.58E-02 |
| MOK | 1296.653583 | -0.279383279 | 4.92E-02 |
| SFXN4 | 585.4507196 | -0.279384973 | 4.51E-02 |
| WDR12 | 2415.211772 | -0.279650484 | 2.46E-02 |

| | | | |
|---|---|---|---|
| PRR3 | 2046.71534 | -0.280016386 | 2.34E-02 |
| RUVBL2 | 6983.216087 | -0.281050765 | 9.37E-03 |
| PATZ1 | 2235.275444 | -0.281552265 | 3.88E-02 |
| ZNF3 | 2904.964969 | -0.281891892 | 2.94E-02 |
| RUVBL1 | 5432.212495 | -0.282923927 | 1.68E-02 |
| RPS27A | 29292.20617 | -0.283469696 | 5.41E-03 |
| GEMIN6 | 1396.151903 | -0.283742329 | 4.56E-03 |
| RPL11 | 36795.27254 | -0.284297849 | 4.41E-03 |
| PFDN6 | 2605.546708 | -0.284460152 | 2.97E-02 |
| EXOC3-AS1 | 613.1024133 | -0.28463193 | 3.42E-02 |
| NOC2L | 7230.373689 | -0.284640032 | 1.29E-03 |
| RCE1 | 673.4495861 | -0.28483757 | 1.51E-02 |
| FBL | 5586.954134 | -0.28516283 | 4.04E-02 |
| TSC22D1 | 5776.662557 | -0.285200961 | 1.55E-02 |
| UTP25 | 1455.765387 | -0.286546908 | 5.10E-04 |
| EEF1D | 14301.58075 | -0.28931603 | 2.96E-02 |
| EBNA1BP2 | 5155.502085 | -0.289742721 | 3.07E-02 |
| WRAP73 | 1404.388509 | -0.291273296 | 3.08E-03 |
| GFOD2 | 1499.238531 | -0.293039649 | 4.14E-02 |
| IRAK1 | 19076.01415 | -0.293389455 | 4.13E-02 |
| FKBP1A | 26088.7043 | -0.293638196 | 4.60E-02 |
| NPRL2 | 1001.425579 | -0.293763098 | 4.69E-02 |
| PHB | 7327.068528 | -0.293852509 | 3.52E-02 |
| RPS24 | 15668.6721 | -0.293864197 | 2.12E-02 |
| RPL27A | 44327.24657 | -0.294453208 | 2.80E-02 |
| CORO2A | 1356.442854 | -0.294857165 | 2.64E-03 |
| NA | 707.1082302 | -0.295833706 | 2.45E-02 |
| RPL34 | 13518.72636 | -0.295907456 | 3.86E-02 |
| TIMM13 | 2178.951561 | -0.296412011 | 4.65E-02 |
| MRPS18B | 4057.860062 | -0.296770181 | 1.45E-02 |
| MTMR14 | 1785.623778 | -0.297816864 | 2.89E-02 |
| RALB | 5036.240045 | -0.297839555 | 3.15E-03 |
| PNP | 1512.768907 | -0.298121163 | 4.26E-02 |
| ZNF691 | 543.1359512 | -0.298727017 | 2.36E-02 |
| QTRT1 | 1398.971228 | -0.299355301 | 3.49E-02 |
| RPL26 | 31543.39492 | -0.299679596 | 2.05E-03 |
| COA4 | 2229.22841 | -0.301019777 | 3.32E-02 |
| CRMP1 | 630.7320603 | -0.302590872 | 3.55E-03 |
| PHF5A | 2143.181465 | -0.303823542 | 1.53E-03 |
| ATAT1 | 537.2356922 | -0.304091183 | 6.04E-03 |
| VARS2 | 1563.733782 | -0.304437354 | 7.40E-03 |
| NFYA | 1791.337203 | -0.304689806 | 5.75E-03 |
| BCS1L | 1798.308695 | -0.305228503 | 8.69E-04 |
| PPM1F | 3912.01717 | -0.305385152 | 1.78E-02 |
| MAX | 2278.562027 | -0.308311246 | 1.61E-02 |
| C19orf48 | 2555.273303 | -0.308656077 | 3.37E-03 |
| POLR3D | 3632.780335 | -0.309056377 | 1.18E-02 |

| | | | |
|---|---|---|---|
| FZD1 | 1665.798152 | -0.310663979 | 4.92E-02 |
| RPS15A | 28078.05717 | -0.311253958 | 1.57E-02 |
| MRTO4 | 2839.374245 | -0.311307071 | 1.17E-02 |
| RPS17 | 42235.26551 | -0.311794645 | 5.70E-03 |
| GAGE12C | 12322.95692 | -0.312015921 | 1.14E-03 |
| ALDH1B1 | 2870.46866 | -0.312234223 | 9.37E-03 |
| ZADH2 | 2400.126845 | -0.312285422 | 2.55E-05 |
| CLIC1 | 17383.67349 | -0.312503047 | 2.91E-02 |
| TRMT1 | 1512.289143 | -0.31265047 | 4.27E-02 |
| PDLIM3 | 622.703077 | -0.31277845 | 2.69E-02 |
| TOE1 | 794.4759432 | -0.313026417 | 4.58E-02 |
| RALY | 10373.36135 | -0.313036688 | 2.05E-02 |
| RPS3 | 78001.32135 | -0.31381299 | 1.40E-02 |
| ZNF772 | 795.6098817 | -0.314838122 | 3.62E-02 |
| PES1 | 4082.165987 | -0.315205332 | 2.99E-03 |
| RPS4X | 33744.19624 | -0.315993645 | 1.37E-02 |
| TMEM177 | 593.8295865 | -0.316443221 | 2.06E-02 |
| POLR1E | 2520.906887 | -0.317225785 | 1.76E-03 |
| DCAF4 | 983.4014373 | -0.317759775 | 1.15E-02 |
| WDR77 | 2537.187295 | -0.318467002 | 7.43E-03 |
| SORD | 1813.484548 | -0.31905401 | 1.21E-02 |
| TXN2 | 3663.452541 | -0.319303828 | 5.00E-02 |
| ATIC | 7866.447087 | -0.319653591 | 7.63E-03 |
| RRP12 | 2494.959313 | -0.319780948 | 3.98E-02 |
| AHCY | 11764.42684 | -0.320183091 | 1.42E-02 |
| RPS12 | 23474.3027 | -0.320605781 | 1.72E-03 |
| BZW2 | 4791.617903 | -0.320934699 | 2.62E-03 |
| PPIH | 1924.711024 | -0.322127069 | 9.66E-03 |
| ASTN2 | 485.4395629 | -0.322597061 | 3.85E-02 |
| 6-Sep | 2821.75174 | -0.322662645 | 5.46E-07 |
| RPL23AP42 | 8204.968219 | -0.322858642 | 4.55E-02 |
| EEF1AKNMT | 2434.565317 | -0.323338587 | 2.76E-03 |
| CIAO1 | 6123.071829 | -0.323495664 | 3.53E-03 |
| SPRED2 | 2021.337216 | -0.323498105 | 2.69E-02 |
| C6orf48 | 7131.05733 | -0.324232557 | 5.70E-03 |
| RPLP0 | 75599.32004 | -0.324308642 | 3.39E-02 |
| FARSA | 5326.766618 | -0.324394013 | 1.57E-02 |
| CRADD | 389.3041389 | -0.324776103 | 4.91E-02 |
| LDLRAD3 | 1672.485159 | -0.324928777 | 4.44E-02 |
| TKFC | 1546.53106 | -0.325392487 | 2.97E-04 |
| SP110 | 1100.571643 | -0.327608006 | 8.85E-04 |
| TMEM208 | 1264.667512 | -0.327855129 | 3.05E-02 |
| C1QBP | 10284.03275 | -0.327955433 | 2.18E-04 |
| NOP10 | 2112.578788 | -0.328002722 | 8.50E-03 |
| FAM86DP | 1360.276101 | -0.328568096 | 2.30E-02 |
| SLC43A3 | 6224.28028 | -0.329069326 | 2.89E-04 |
| TBX3 | 4851.962125 | -0.329271192 | 3.55E-02 |

| | | | |
|---|---|---|---|
| SNHG6 | 2615.171978 | -0.329751814 | 2.17E-03 |
| NMRAL1 | 1468.372654 | -0.330072711 | 3.22E-02 |
| BCAR3 | 3028.458632 | -0.330701278 | 4.86E-02 |
| PPIF | 3038.649213 | -0.33187974 | 1.06E-02 |
| AC090204.1 | 619.9733506 | -0.331926024 | 1.06E-02 |
| RPL30 | 35678.43918 | -0.331964346 | 9.81E-03 |
| RPL9P9 | 16918.13946 | -0.332017665 | 1.25E-02 |
| VILL | 408.5323477 | -0.332354716 | 3.88E-02 |
| FAM89A | 912.360877 | -0.332931226 | 1.00E-02 |
| IMPDH2 | 17822.23971 | -0.333263578 | 4.29E-03 |
| CLDN15 | 890.7426039 | -0.333544733 | 6.81E-03 |
| WDR4 | 1435.970238 | -0.334024125 | 3.04E-02 |
| ZIK1 | 840.9093282 | -0.334183642 | 2.67E-02 |
| RPS27 | 26325.22995 | -0.334674106 | 2.05E-02 |
| SMN2 | 2795.058623 | -0.33516889 | 5.70E-03 |
| RANGAP1 | 7733.942925 | -0.335705807 | 5.08E-04 |
| DCTPP1 | 2007.818684 | -0.335771191 | 3.83E-02 |
| CYB561 | 2861.485413 | -0.336084441 | 3.91E-03 |
| TMEM101 | 1425.19861 | -0.336157593 | 4.87E-02 |
| PMVK | 1036.769575 | -0.33642029 | 2.41E-03 |
| EIF3K | 7247.367878 | -0.336771943 | 1.99E-02 |
| RPL8 | 63923.6203 | -0.337312661 | 2.06E-02 |
| RPLP2 | 26725.70048 | -0.337577352 | 4.19E-02 |
| RPL24 | 16195.05698 | -0.338230128 | 1.29E-02 |
| RPL27 | 24671.76414 | -0.338481823 | 2.29E-02 |
| RPS7 | 42243.97316 | -0.338803274 | 1.75E-04 |
| MPV17 | 3093.539863 | -0.33897147 | 1.16E-02 |
| CALHM2 | 962.5218492 | -0.340390049 | 1.57E-02 |
| FBXL19-AS1 | 285.6288051 | -0.341352818 | 2.10E-02 |
| GPANK1 | 1068.369362 | -0.343124313 | 2.83E-02 |
| ZNF619 | 453.2585032 | -0.343985004 | 3.05E-02 |
| PIM3 | 1569.131357 | -0.344053065 | 2.30E-03 |
| SAPCD2 | 1918.011644 | -0.344182887 | 1.60E-02 |
| CSKMT | 343.0414308 | -0.344959501 | 3.26E-02 |
| FANCE | 547.4079603 | -0.345267736 | 1.61E-02 |
| PLP2 | 4682.094735 | -0.345408514 | 1.48E-02 |
| EXOSC4 | 1304.84018 | -0.346135032 | 2.70E-02 |
| RAB36 | 529.2506332 | -0.346303121 | 2.98E-02 |
| AMOTL1 | 6561.397498 | -0.346688068 | 5.97E-03 |
| MON1A | 421.933622 | -0.347203854 | 1.77E-02 |
| PUS3 | 1045.525188 | -0.34727954 | 3.55E-02 |
| ABHD5 | 550.9249173 | -0.347732539 | 5.67E-03 |
| DPM2 | 4556.786876 | -0.347860964 | 2.01E-03 |
| RAMP2-AS1 | 693.2912316 | -0.348109365 | 2.90E-02 |
| LIMD2 | 2211.717988 | -0.349323044 | 1.38E-02 |
| DDTL | 284.9097104 | -0.34941304 | 2.06E-02 |
| FAM171A1 | 2363.984497 | -0.349660581 | 2.36E-02 |

| | | | |
|---|---|---|---|
| DDX25 | 1351.778353 | -0.350005569 | 2.87E-04 |
| CROCC | 613.0046235 | -0.350049693 | 4.54E-02 |
| GPX1 | 6093.658858 | -0.351374016 | 4.37E-02 |
| EIF3I | 10526.45264 | -0.352490427 | 1.04E-03 |
| RPL18 | 22761.13749 | -0.352939238 | 3.46E-02 |
| UBIAD1 | 1239.930969 | -0.355740213 | 1.19E-02 |
| RPS2 | 74595.74233 | -0.356828666 | 2.02E-02 |
| SERPINE2 | 8188.667421 | -0.357318214 | 1.47E-02 |
| ITPA | 3951.47197 | -0.358843137 | 7.57E-03 |
| WDR3 | 4591.267674 | -0.359671858 | 2.50E-02 |
| C11orf1 | 496.8051139 | -0.360776941 | 1.69E-02 |
| APRT | 3780.591045 | -0.363626161 | 3.91E-03 |
| CENPBD1P1 | 1256.44121 | -0.363906862 | 2.21E-02 |
| RPL35 | 27266.11572 | -0.364036853 | 1.10E-02 |
| NHP2 | 3255.783206 | -0.364812518 | 8.23E-03 |
| RPS6 | 79859.13227 | -0.366261784 | 2.54E-04 |
| POLR1C | 1344.849713 | -0.366465357 | 1.68E-05 |
| ZNF135 | 520.20737 | -0.366963861 | 1.34E-02 |
| IPO4 | 3946.715919 | -0.367178768 | 1.36E-03 |
| SYT11 | 517.0050096 | -0.367258677 | 1.11E-02 |
| MAP3K1 | 6834.516553 | -0.367286133 | 8.10E-03 |
| PLEKHJ1 | 2128.603738 | -0.367336415 | 4.14E-03 |
| RMND1 | 749.3812191 | -0.368194473 | 2.73E-02 |
| NAV2 | 2387.667239 | -0.368433665 | 2.75E-02 |
| RPL10A | 24437.36707 | -0.368787653 | 1.29E-03 |
| RPS19 | 28332.18881 | -0.370376048 | 4.45E-03 |
| SNHG5 | 2995.960911 | -0.370578499 | 1.55E-02 |
| CCNB1IP1 | 4135.158345 | -0.371953758 | 8.53E-03 |
| RPL22L1 | 2206.738069 | -0.371961647 | 1.57E-02 |
| RPL29 | 36284.96416 | -0.372000854 | 3.76E-02 |
| CCDC51 | 1006.545882 | -0.37267821 | 3.65E-03 |
| SRM | 3029.639383 | -0.373695642 | 3.95E-03 |
| KLF13 | 1510.656057 | -0.373908302 | 3.75E-02 |
| RUNX1 | 2054.795126 | -0.375076057 | 1.61E-02 |
| RPS28 | 7175.876842 | -0.375083381 | 4.36E-02 |
| TSSC2 | 633.1595099 | -0.375590364 | 3.23E-02 |
| TNFRSF21 | 5207.604204 | -0.376423574 | 6.60E-05 |
| UBL7 | 4320.443653 | -0.376995978 | 3.34E-02 |
| TSEN2 | 547.339728 | -0.377565679 | 4.91E-02 |
| GLI3 | 3529.66754 | -0.377901972 | 7.15E-03 |
| PROCR | 2832.564645 | -0.37910169 | 1.77E-02 |
| MGAT5B | 1657.991925 | -0.379816491 | 3.80E-02 |
| RPL37A | 41742.41886 | -0.382255787 | 1.89E-02 |
| LGALS3 | 3865.647832 | -0.382444275 | 1.31E-03 |
| PPFIA3 | 924.0988512 | -0.383169086 | 3.89E-02 |
| ILF3-DT | 606.3224029 | -0.383528464 | 6.81E-03 |
| UCP2 | 2547.826664 | -0.38413298 | 4.67E-03 |

| | | | |
|---|---|---|---|
| TRAIP | 640.6318974 | -0.385654774 | 1.01E-02 |
| RPL10 | 91368.98784 | -0.385749445 | 7.18E-03 |
| EEF1B2 | 14457.85453 | -0.385834021 | 2.07E-03 |
| RAB3IL1 | 1180.86828 | -0.388611065 | 4.83E-02 |
| TTLL12 | 2960.526421 | -0.389375247 | 1.20E-03 |
| RPL19 | 41793.06361 | -0.391658665 | 8.13E-04 |
| TGFBI | 26371.45713 | -0.392035023 | 1.18E-02 |
| MANEAL | 1265.66944 | -0.394521312 | 4.38E-02 |
| CDK5 | 1685.224274 | -0.394711119 | 1.55E-02 |
| ALKBH2 | 710.5069644 | -0.396261445 | 5.57E-03 |
| RPL41 | 46211.12679 | -0.3964868 | 7.02E-03 |
| THTPA | 987.8526175 | -0.396845405 | 4.34E-02 |
| MDFI | 1896.304313 | -0.399775356 | 2.67E-02 |
| PLK2 | 8444.080224 | -0.400474434 | 3.99E-03 |
| NA | 646.9206097 | -0.401850323 | 5.32E-03 |
| GAGE13 | 2562.719045 | -0.40364513 | 6.55E-05 |
| CDK4 | 12333.38824 | -0.403996131 | 7.91E-03 |
| ANKH | 6322.166104 | -0.405436363 | 4.83E-08 |
| DPH2 | 2221.522895 | -0.405535672 | 1.22E-03 |
| C20orf27 | 2903.6291 | -0.406226408 | 3.70E-03 |
| RPS21 | 23282.49507 | -0.406334572 | 2.79E-03 |
| ATP5MC2 | 5676.214694 | -0.40905947 | 3.53E-04 |
| SRD5A1 | 2652.925209 | -0.410209694 | 1.01E-08 |
| SNHG29 | 16413.21366 | -0.415347147 | 3.44E-04 |
| ITPRIPL1 | 839.3701575 | -0.415449707 | 7.53E-03 |
| ARG2 | 2408.765317 | -0.415728151 | 6.44E-03 |
| AGMAT | 377.0349647 | -0.416854381 | 6.79E-03 |
| CHGB | 280.9084651 | -0.418719761 | 3.19E-02 |
| RPL13A | 70398.25543 | -0.418912973 | 9.34E-03 |
| SETD9 | 410.3844377 | -0.418931292 | 2.97E-02 |
| TRIB2 | 5941.218021 | -0.419351064 | 1.50E-02 |
| ABHD11 | 864.5830385 | -0.421072911 | 1.61E-03 |
| RPS18 | 68722.47473 | -0.42179982 | 1.28E-03 |
| IL2RB | 1447.939605 | -0.424599451 | 1.19E-02 |
| MICOS13 | 1664.422011 | -0.425621593 | 1.61E-03 |
| SPOUT1 | 3020.906239 | -0.425882802 | 4.73E-03 |
| RPS5 | 32305.54264 | -0.426362898 | 1.93E-02 |
| TIMP3 | 33487.39915 | -0.426779198 | 3.20E-02 |
| NIP7 | 3651.687745 | -0.428553607 | 1.02E-04 |
| AC055811.4 | 242.529967 | -0.430655235 | 2.84E-02 |
| PSENEN | 1598.522587 | -0.43274193 | 6.10E-04 |
| PRAG1 | 3099.318571 | -0.434931068 | 4.13E-02 |
| PRKAR1B | 1954.365488 | -0.435226758 | 2.04E-02 |
| PDSS1 | 287.872663 | -0.436649992 | 4.77E-02 |
| CIB2 | 420.3516067 | -0.437163488 | 4.65E-02 |
| KDM8 | 224.6049505 | -0.437324331 | 4.62E-02 |
| LINC01029 | 2160.242943 | -0.437439316 | 2.24E-04 |

| | | | |
|---|---|---|---|
| CALM3 | 16638.08905 | -0.437575638 | 2.46E-03 |
| RAMP2 | 1068.842961 | -0.438833666 | 2.07E-02 |
| LINC00665 | 1946.725265 | -0.439859823 | 2.01E-04 |
| SORBS1 | 666.9068481 | -0.443867115 | 1.26E-03 |
| GAS5 | 7842.947098 | -0.445377942 | 3.90E-02 |
| FRMD4A | 2369.142518 | -0.448008317 | 3.30E-05 |
| ZNF551 | 748.8878328 | -0.448569995 | 2.68E-02 |
| NRXN2 | 691.5951493 | -0.449855864 | 2.06E-02 |
| RPS11 | 38271.47838 | -0.451288381 | 1.51E-02 |
| PAK1 | 4174.330043 | -0.451542717 | 9.58E-06 |
| SOCS2 | 977.7645512 | -0.451694414 | 3.14E-04 |
| SRD5A3 | 467.3307845 | -0.452511204 | 7.85E-03 |
| TMEM204 | 546.6914666 | -0.452655246 | 5.54E-03 |
| RPL18A | 25731.2536 | -0.456362624 | 3.58E-03 |
| NOP16 | 1938.308954 | -0.45750422 | 7.91E-04 |
| AC046185.3 | 1008.471374 | -0.460503439 | 1.08E-06 |
| LAT | 302.3278347 | -0.460561263 | 2.68E-02 |
| MRPL24 | 1793.201605 | -0.46135147 | 2.21E-05 |
| BCL2L11 | 920.1726165 | -0.464063247 | 3.29E-03 |
| TNFRSF1B | 1074.403305 | -0.46408554 | 7.04E-04 |
| FAM78A | 383.5158475 | -0.471732101 | 2.05E-03 |
| RPS29 | 12227.15154 | -0.473078174 | 1.87E-03 |
| HPCAL4 | 258.7617992 | -0.473931901 | 2.17E-02 |
| CDCA7L | 2435.713917 | -0.47506564 | 4.67E-05 |
| ANKLE1 | 635.5348284 | -0.475131462 | 3.61E-04 |
| MLPH | 3090.876894 | -0.481143282 | 1.21E-02 |
| SPRY4 | 1960.932788 | -0.481353171 | 2.64E-03 |
| GALNT6 | 1395.585243 | -0.482163883 | 3.28E-02 |
| NA | 131.4282674 | -0.482641668 | 2.28E-02 |
| RPS23 | 74273.86759 | -0.483683926 | 3.47E-05 |
| FMC1 | 495.7004804 | -0.488266305 | 4.23E-04 |
| PAFAH2 | 548.3130673 | -0.488776657 | 4.57E-02 |
| NA | 437.0559334 | -0.489611563 | 3.53E-02 |
| RTL5 | 2173.925386 | -0.489733465 | 2.99E-06 |
| SPRY2 | 1269.954807 | -0.490081167 | 2.73E-04 |
| ZNF561-AS1 | 275.8020633 | -0.490583447 | 3.23E-02 |
| P2RX5 | 672.8200669 | -0.490975217 | 2.86E-03 |
| HHAT | 280.182391 | -0.491848217 | 5.93E-03 |
| PLIN2 | 5843.809196 | -0.491910029 | 3.80E-03 |
| PRRT3-AS1 | 120.3373431 | -0.49494675 | 2.18E-02 |
| MAP1A | 1216.104392 | -0.499513505 | 1.01E-02 |
| RAP1GAP | 474.2442089 | -0.502482312 | 2.89E-02 |
| LINC01410 | 734.486561 | -0.502501529 | 1.24E-03 |
| DPYSL3 | 1641.152646 | -0.502975692 | 1.49E-03 |
| TNFRSF10A | 429.7567669 | -0.503551676 | 5.50E-03 |
| SENCR | 247.900948 | -0.506408427 | 2.60E-03 |
| P2RY11 | 313.5632137 | -0.506983055 | 1.21E-02 |

| | | | |
|---|---|---|---|
| MEX3A | 1174.644861 | -0.510327215 | 5.87E-03 |
| HOXB9 | 2074.166601 | -0.513437297 | 4.89E-03 |
| STX1A | 710.3622965 | -0.51472337 | 1.63E-04 |
| AC027031.2 | 487.0588179 | -0.515105798 | 1.25E-04 |
| LINC00957 | 360.1825963 | -0.521280864 | 3.64E-02 |
| TRIM9 | 249.1688506 | -0.527002956 | 3.77E-02 |
| ACY1 | 1833.499762 | -0.528092935 | 6.44E-03 |
| CDKN1A | 368.8877978 | -0.531218001 | 3.44E-03 |
| CCDC78 | 384.8868074 | -0.534793845 | 4.20E-03 |
| STMN3 | 8539.398862 | -0.535219529 | 1.11E-04 |
| TLL2 | 230.6792461 | -0.536611988 | 3.88E-02 |
| FAM20C | 2024.588472 | -0.538804303 | 1.78E-02 |
| MARCKSL1 | 6894.580256 | -0.538956426 | 2.73E-03 |
| ADM | 794.9914859 | -0.539508648 | 2.60E-02 |
| NME1 | 4684.921094 | -0.539721311 | 6.58E-04 |
| TWNK | 1717.423571 | -0.541307348 | 2.21E-05 |
| CYTL1 | 1870.470186 | -0.54156467 | 4.08E-04 |
| IL17RD | 1697.446375 | -0.550188143 | 2.80E-02 |
| SERPINA5 | 1276.950546 | -0.553117741 | 1.68E-02 |
| BOLA2 | 1180.562019 | -0.554407293 | 5.31E-05 |
| ZMAT4 | 504.9050422 | -0.560012467 | 1.18E-02 |
| NPAS2 | 2199.95089 | -0.560365266 | 4.67E-04 |
| TUBB3 | 3262.462153 | -0.564961739 | 1.14E-03 |
| BTBD11 | 910.1491859 | -0.567577586 | 3.87E-07 |
| REEP1 | 645.6708003 | -0.567681109 | 4.24E-02 |
| THSD1 | 104.9408088 | -0.569177856 | 8.65E-03 |
| SNHG19 | 151.0834963 | -0.58307177 | 1.12E-02 |
| ATP2A3 | 1211.316063 | -0.58387507 | 2.74E-05 |
| AC018553.1 | 92.95374125 | -0.585627553 | 1.17E-02 |
| SNED1 | 2396.810234 | -0.587180038 | 2.07E-03 |
| FBLN1 | 4794.018155 | -0.588508557 | 1.50E-03 |
| CPNE7 | 438.1070185 | -0.589122823 | 6.65E-03 |
| NPBWR1 | 390.8449915 | -0.589899282 | 2.56E-02 |
| TTYH2 | 540.5647135 | -0.593324488 | 3.64E-02 |
| NALCN | 1270.957975 | -0.595332917 | 5.42E-12 |
| SLC37A2 | 644.4016214 | -0.599355607 | 2.99E-03 |
| DHRS12 | 99.15939236 | -0.602341408 | 4.44E-02 |
| SBSPON | 767.0994634 | -0.603918206 | 2.77E-03 |
| DNER | 848.2215533 | -0.607726052 | 1.57E-04 |
| TUBA1A | 19721.63494 | -0.610621832 | 3.47E-02 |
| CNIH3 | 994.9039675 | -0.613015728 | 6.78E-04 |
| CDKN2B | 1954.875598 | -0.622042924 | 3.97E-02 |
| TUBB2B | 4106.075648 | -0.624528363 | 6.71E-04 |
| ENOX1 | 275.262444 | -0.633412782 | 1.31E-02 |
| PALD1 | 72.23173115 | -0.63673947 | 3.29E-02 |
| HECW1 | 564.3576593 | -0.639365493 | 9.35E-04 |
| EXOSC5 | 1246.924041 | -0.641629607 | 4.13E-05 |

| | | | |
|---|---|---|---|
| CDK5R1 | 277.1423831 | -0.6451981 | 1.04E-02 |
| NTRK1 | 102.2523374 | -0.646559165 | 2.23E-02 |
| LRRC17 | 2568.192514 | -0.648522933 | 2.87E-03 |
| BBC3 | 539.5516664 | -0.649620528 | 1.11E-04 |
| PMEPA1 | 367.6116401 | -0.655120792 | 1.17E-02 |
| ID3 | 8443.247581 | -0.656389005 | 2.60E-03 |
| UBXN10 | 251.9791355 | -0.66105977 | 5.67E-04 |
| GMFG | 271.7051033 | -0.661181672 | 1.27E-02 |
| SHISA2 | 384.9045469 | -0.66140389 | 5.18E-05 |
| ARL9 | 148.4194161 | -0.66737686 | 6.81E-03 |
| CHST2 | 604.9230275 | -0.667888556 | 2.45E-13 |
| UNC13A | 437.1254544 | -0.668344814 | 5.89E-03 |
| AL645608.2 | 67.27581695 | -0.681410107 | 3.86E-02 |
| LCTL | 194.6728429 | -0.687905665 | 7.72E-03 |
| USP2-AS1 | 184.375442 | -0.693635796 | 1.03E-02 |
| DUSP5 | 787.2594332 | -0.695421139 | 9.34E-03 |
| RAB7B | 81.30781683 | -0.698526144 | 3.77E-02 |
| AC011476.2 | 111.0196704 | -0.699642591 | 1.36E-02 |
| SPANXC | 1478.430067 | -0.70053865 | 5.72E-03 |
| HLA-DRA | 272.0727889 | -0.701275039 | 4.19E-02 |
| TMEM145 | 253.2058034 | -0.70468042 | 3.29E-03 |
| FXYD6 | 2385.941432 | -0.705265758 | 6.61E-05 |
| SIGLEC15 | 122.8361965 | -0.705965594 | 3.32E-02 |
| PPP1R16B | 527.7404052 | -0.709130323 | 4.97E-03 |
| NRROS | 203.4804337 | -0.7119011 | 2.86E-03 |
| FAM78B | 308.7966026 | -0.713459386 | 1.60E-03 |
| VCAN | 908.1972434 | -0.721264152 | 2.51E-02 |
| BSN | 168.5919291 | -0.721703789 | 2.60E-03 |
| COL6A3 | 4751.934988 | -0.722641989 | 1.86E-02 |
| LINC01366 | 70.30869446 | -0.723247619 | 1.57E-02 |
| PLXNA4 | 515.4222672 | -0.726102815 | 1.21E-02 |
| PRR7-AS1 | 73.11687539 | -0.727335538 | 2.50E-02 |
| PRDM13 | 87.34034449 | -0.730881351 | 2.16E-02 |
| BEX1 | 1622.382863 | -0.737128577 | 2.52E-08 |
| C2orf70 | 148.9513194 | -0.74744398 | 2.14E-02 |
| AC008622.2 | 52.92455998 | -0.75133838 | 3.12E-02 |
| NKD1 | 470.6736306 | -0.753340125 | 2.53E-02 |
| FAM155B | 469.1044739 | -0.753399289 | 1.24E-07 |
| LFNG | 282.2624472 | -0.759871787 | 1.63E-04 |
| NOX5 | 244.0562303 | -0.764870005 | 2.91E-03 |
| CEMIP | 1148.669696 | -0.765869481 | 2.65E-03 |
| TMEM59L | 339.1337644 | -0.779309681 | 6.33E-05 |
| RGS14 | 562.7905907 | -0.789017061 | 4.40E-07 |
| CRYBB2 | 39.30274241 | -0.795640467 | 4.68E-02 |
| LAMP5 | 323.5776342 | -0.805617486 | 1.67E-04 |
| COL8A1 | 2277.352965 | -0.811096789 | 8.41E-04 |
| MYC | 932.6837416 | -0.812317059 | 4.24E-02 |

| | | | |
|---|---|---|---|
| ARHGAP9 | 264.0659115 | -0.81251587 | 1.49E-04 |
| TMEM200A | 540.674388 | -0.814662664 | 3.13E-06 |
| PLP1 | 431.1818229 | -0.819297984 | 5.62E-03 |
| AC010970.1 | 17518.2263 | -0.827386866 | 3.11E-03 |
| AC096537.1 | 58.48260446 | -0.832188382 | 3.68E-02 |
| ANGPT2 | 85.43122701 | -0.838797087 | 2.31E-02 |
| PTX3 | 4981.988845 | -0.84369839 | 1.93E-04 |
| LPAR1 | 468.0407392 | -0.849132651 | 1.37E-05 |
| NPIPA7 | 80.15957517 | -0.863881339 | 4.40E-02 |
| GAL | 58.68265862 | -0.866899667 | 4.44E-02 |
| ESX1 | 1145.550705 | -0.876877928 | 1.71E-05 |
| PQLC2L | 60.00408119 | -0.882353018 | 7.62E-03 |
| BPI | 155.9343946 | -0.888862106 | 6.17E-03 |
| AC247036.1 | 655.3655526 | -0.897272201 | 4.13E-05 |
| PCDH12 | 238.254452 | -0.898720682 | 4.72E-03 |
| IGHV3-19 | 30.14242899 | -0.902735714 | 3.15E-02 |
| ADCY1 | 191.0931613 | -0.915555707 | 2.87E-03 |
| PLPP4 | 110.7522772 | -0.916486916 | 2.84E-03 |
| QRICH2 | 106.3556761 | -0.923919469 | 9.68E-04 |
| ITGA10 | 182.6699131 | -0.924342477 | 2.90E-04 |
| KCNK13 | 67.77061597 | -0.925498794 | 2.80E-02 |
| SMOC1 | 69.53858863 | -0.925724046 | 1.52E-02 |
| PLCB4 | 1544.325878 | -0.932508761 | 5.84E-05 |
| DRD2 | 181.3447405 | -0.933802011 | 1.26E-04 |
| THBS2 | 4772.078864 | -0.934731478 | 2.04E-07 |
| SLC22A18AS | 118.3495015 | -0.935539324 | 7.49E-03 |
| EIF4EBP3 | 143.5782433 | -0.942447391 | 2.78E-04 |
| TIE1 | 3876.156317 | -0.944181516 | 2.96E-12 |
| PIK3R5 | 80.61375241 | -0.950916747 | 2.73E-03 |
| SYP | 149.7220486 | -0.957441304 | 7.49E-05 |
| GPM6B | 424.3810447 | -0.959226263 | 1.03E-04 |
| UBE2QL1 | 493.8624657 | -0.966033767 | 6.53E-06 |
| MIR4527HG | 177.5777164 | -0.967579175 | 4.78E-07 |
| CUX2 | 98.45321618 | -0.978325948 | 4.47E-02 |
| EDN1 | 1541.884744 | -0.98516332 | 1.54E-04 |
| SPOCK2 | 70.07807607 | -0.988753482 | 3.05E-02 |
| ADAMTS14 | 70.35092211 | -1.001471883 | 2.53E-02 |
| SYT13 | 311.0869747 | -1.018111625 | 7.23E-03 |
| CLEC3B | 312.9420554 | -1.025308735 | 1.85E-02 |
| FST | 410.4366751 | -1.033380058 | 5.77E-03 |
| RGCC | 105.6600784 | -1.051195268 | 3.67E-02 |
| IGSF9B | 83.02754075 | -1.0529525 | 2.06E-02 |
| CFP | 142.794171 | -1.079614605 | 3.14E-04 |
| ATP2B2 | 75.43341306 | -1.083585066 | 1.57E-02 |
| SIGLEC17P | 28.70313536 | -1.119389876 | 2.64E-02 |
| RUBCNL | 165.9995 | -1.13609578 | 5.81E-03 |
| CXCL8 | 99.7196793 | -1.136251872 | 3.86E-02 |

| | | | |
|---|---|---|---|
| SLC4A5 | 33.49355513 | -1.15362045 | 1.52E-02 |
| FLT4 | 134.8507834 | -1.157731034 | 3.10E-02 |
| TMEM163 | 27.6127971 | -1.188630452 | 3.57E-02 |
| ME3 | 73.97688032 | -1.195588483 | 1.10E-02 |
| KCNC1 | 90.05506287 | -1.202149636 | 9.15E-05 |
| CHD5 | 93.8602919 | -1.211000548 | 1.06E-02 |
| CRLF1 | 1052.296416 | -1.21134106 | 4.89E-18 |
| GNG4 | 685.0541399 | -1.217976116 | 2.60E-13 |
| LCP2 | 54.88249637 | -1.248252786 | 2.60E-03 |
| VGF | 785.8211023 | -1.249620893 | 5.79E-08 |
| ATP6V1G2-DDX | 701.8318688 | -1.265441435 | 3.12E-02 |
| TMEM233 | 90.45126575 | -1.287043347 | 1.11E-02 |
| RGS7 | 47.66134455 | -1.31654479 | 3.44E-03 |
| NPTX1 | 3190.575678 | -1.349426196 | 3.28E-12 |
| GALNT17 | 25.55343124 | -1.349558807 | 4.33E-03 |
| SNORA71C | 20.68944665 | -1.37676549 | 1.36E-02 |
| SNORA26 | 19.2884351 | -1.389640613 | 5.50E-03 |
| CREG2 | 57.31823733 | -1.421807933 | 5.10E-04 |
| SCG2 | 992.0410165 | -1.426123802 | 9.85E-08 |
| KRT75 | 23.29573459 | -1.440458306 | 2.12E-02 |
| PRICKLE1 | 152.583996 | -1.451909084 | 1.86E-07 |
| LRRC18 | 14.9253248 | -1.459553454 | 4.30E-02 |
| F13A1 | 43.43632321 | -1.485234863 | 2.01E-02 |
| DHRS2 | 1627.919122 | -1.497639562 | 1.72E-07 |
| GLP1R | 28.92280827 | -1.498047229 | 6.10E-04 |
| DCT | 247.3288116 | -1.502137754 | 2.88E-09 |
| TF | 518.8802447 | -1.502371483 | 3.40E-09 |
| LAIR1 | 55.86959733 | -1.505658186 | 3.19E-02 |
| KCNQ2 | 12.26458218 | -1.506615731 | 3.65E-02 |
| LYZ | 14.04044372 | -1.50850841 | 4.78E-02 |
| TRBC2 | 122.21731 | -1.527674713 | 2.58E-06 |
| CASC8 | 11.89497762 | -1.527913951 | 3.12E-02 |
| GABRA2 | 53.37695158 | -1.530042351 | 2.06E-03 |
| ACTL8 | 26.23675356 | -1.536462664 | 1.12E-02 |
| CAPN6 | 166.404887 | -1.567632057 | 6.25E-07 |
| ACTBL2 | 62.37567461 | -1.594293207 | 2.99E-03 |
| DRAXIN | 93.60471832 | -1.627299244 | 1.98E-09 |
| IRX4 | 106.6038776 | -1.629917011 | 1.46E-07 |
| MMP9 | 160.911846 | -1.634569282 | 1.82E-06 |
| DKK1 | 937.75661 | -1.656533485 | 6.65E-17 |
| ANK2 | 166.3824866 | -1.658104412 | 1.01E-03 |
| UBXN10-AS1 | 15.03613687 | -1.666489875 | 2.99E-03 |
| C16orf89 | 40.33127915 | -1.707097121 | 5.37E-03 |
| HSD17B2 | 349.7292265 | -1.714320742 | 1.49E-08 |
| AC099066.2 | 26.07679972 | -1.727998295 | 1.43E-02 |
| CHI3L1 | 17.46316668 | -1.736781109 | 3.20E-02 |
| PTPN22 | 24.45633375 | -1.740335826 | 1.37E-02 |

| | | | |
|---|---|---|---|
| TRIM67 | 26.95075226 | -1.74175667 | 3.18E-03 |
| UNC5C | 16.04686619 | -1.756358412 | 2.04E-02 |
| TYRP1 | 289.0563429 | -1.82644991 | 2.85E-31 |
| STK32A | 19.69460038 | -1.842022577 | 3.64E-02 |
| CYP24A1 | 230.7349006 | -1.950777274 | 5.04E-15 |
| GABRR1 | 17.11856491 | -1.953454356 | 2.56E-03 |
| KIAA1755 | 395.0065777 | -1.961837578 | 2.78E-04 |
| LINC00589 | 13.02961938 | -1.972573115 | 8.10E-03 |
| MMP1 | 74.36681813 | -2.004251259 | 3.55E-05 |
| EPHB1 | 77.54038977 | -2.099437221 | 3.53E-04 |
| TNFSF18 | 594.8574692 | -2.0999391 | 1.04E-02 |
| LRRTM2 | 15.22254316 | -2.138390453 | 6.65E-03 |
| AL441883.1 | 18.77579353 | -2.194248123 | 5.08E-03 |
| RAB3C | 13.43604499 | -2.7588066 | 3.41E-03 |
| AC018521.2 | 17.20774676 | -3.052749127 | 2.18E-02 |
| PCDHGA11 | 39.1975755 | -6.088929535 | 5.85E-05 |
| AC022149.1 | 85.0235935 | -9.813450773 | 1.01E-02 |
| AC087164.2 | 292.3801368 | -15.28098829 | 4.15E-04 |
| AC093668.2 | 22.02253282 | -30.1133505 | 2.57E-05 |

| Gene | baseMean | log2 Fold Change | padj |
|---|---|---|---|
| AP003419.1 | 46.01432342 | 19.16194236 | 5.10E-08 |
| NA | 14.62702115 | 6.813647723 | 2.50E-04 |
| NA | 14.44229704 | 6.357907239 | 1.96E-03 |
| NA | 14.13000731 | 6.332410087 | 4.53E-02 |
| MAGEL2 | 23.89919631 | 6.279538582 | 2.15E-18 |
| NEFL | 99.35758727 | 5.058186958 | 4.48E-41 |
| AC233968.1 | 53.18545577 | 5.04359733 | 7.78E-03 |
| KRTAP2-3 | 13.1496624 | 4.958708413 | 1.50E-02 |
| ATP1A3 | 357.0233239 | 4.621605854 | 5.49E-97 |
| SV2C | 14.29657055 | 4.225484757 | 3.91E-02 |
| CDS1 | 52.31633794 | 4.026255113 | 1.20E-23 |
| NAP1L2 | 60.86264806 | 3.667746739 | 2.00E-23 |
| S100B | 57.72904892 | 3.420190979 | 3.98E-34 |
| LINC00632 | 86.52816748 | 3.366794358 | 6.16E-37 |
| CDR1 | 15.96781407 | 3.335398982 | 1.83E-06 |
| SNORD116-24 | 8.382697762 | 3.274394858 | 5.14E-05 |
| SNORD116-6 | 8.657214572 | 3.041070733 | 1.20E-03 |
| VGF | 785.8211023 | 2.740050719 | 1.05E-45 |
| AC124312.2 | 11.88291938 | 2.692173834 | 9.58E-05 |
| BEX5 | 103.9003807 | 2.683385634 | 1.04E-33 |
| MAGEE1 | 540.1230386 | 2.674409046 | 7.95E-45 |
| H3F3AP4 | 420.9060945 | 2.674257779 | 4.80E-14 |
| PNMA3 | 556.7056313 | 2.658796731 | 8.60E-164 |
| MAP1A | 1216.104392 | 2.643201796 | 2.78E-74 |
| SNORD116-1 | 8.986201518 | 2.632002525 | 1.57E-03 |
| MAGEE2 | 17.34802969 | 2.626955003 | 1.36E-04 |
| MUC5AC | 33.8582948 | 2.53571313 | 1.70E-06 |
| NA | 106.0004149 | 2.532855727 | 1.34E-15 |
| DNAH8 | 13.86457734 | 2.521823661 | 1.85E-04 |
| AC124312.3 | 178.0884694 | 2.414566349 | 6.44E-49 |
| SNHG14 | 4449.319971 | 2.398138559 | 3.12E-79 |
| MCF2L | 207.9155546 | 2.387394159 | 4.03E-12 |
| PWAR5 | 61.86062971 | 2.356018921 | 3.07E-07 |
| ELF3 | 19.19275641 | 2.314123568 | 4.10E-04 |
| SNCAIP | 91.61249628 | 2.306125637 | 3.31E-10 |
| ALDOC | 1821.65617 | 2.285357054 | 9.44E-06 |
| MIAT | 450.5079603 | 2.269948101 | 6.92E-19 |
| NA | 82.95313313 | 2.253870783 | 1.70E-25 |
| SNRPN | 5362.88291 | 2.166072102 | 3.32E-129 |
| MT-TF | 12.06340084 | 1.987757125 | 2.45E-02 |
| STARD13-AS | 15.35750094 | 1.970603537 | 3.71E-03 |
| SNORD116-4 | 18.00845193 | 1.922104872 | 1.95E-04 |
| NPPB | 2240.650956 | 1.914817686 | 3.61E-04 |
| MAP6 | 112.0278279 | 1.841098644 | 9.90E-09 |
| HAND1 | 50.73723808 | 1.782918312 | 1.57E-03 |
| LGI3 | 21.16732851 | 1.761934739 | 1.38E-02 |

| | | | |
|---|---|---|---|
| KRT15 | 92.32944513 | 1.75359024 | 3.53E-07 |
| ABCA3 | 301.3116936 | 1.676351755 | 1.03E-10 |
| NPIPA7 | 80.15957517 | 1.659875693 | 4.28E-06 |
| SNURF | 237.1015219 | 1.653400973 | 5.24E-09 |
| PHOSPHO1 | 77.15191877 | 1.61158585 | 9.47E-06 |
| EPB41L3 | 354.3759169 | 1.599992093 | 6.96E-17 |
| AC006449.7 | 76.39049316 | 1.579176483 | 2.29E-03 |
| SCARNA9 | 19.28743193 | 1.568793168 | 2.22E-02 |
| NEFH | 4344.736666 | 1.567556641 | 2.78E-74 |
| MT-TQ | 46.69971454 | 1.537326175 | 3.77E-03 |
| AHNAK2 | 1587.711431 | 1.516378839 | 2.01E-18 |
| TMEM151A | 32.37522276 | 1.47204666 | 6.55E-03 |
| CUX2 | 98.45321618 | 1.464652909 | 2.58E-03 |
| LRRC15 | 26.29205623 | 1.461233334 | 3.56E-02 |
| GDF5 | 28.69338378 | 1.454666229 | 8.45E-03 |
| LBP | 19.22947916 | 1.453807144 | 4.34E-02 |
| PRND | 26.16392127 | 1.439646422 | 2.30E-02 |
| CORO2B | 318.2158163 | 1.422163948 | 7.57E-20 |
| SDR16C5 | 42.38694442 | 1.384186429 | 3.61E-03 |
| AIFM3 | 126.0632533 | 1.37908591 | 6.76E-08 |
| CXCL14 | 171.4861655 | 1.374492644 | 2.26E-03 |
| WNT7B | 200.4097672 | 1.342954954 | 9.65E-04 |
| KIF5A | 324.9960343 | 1.337820972 | 4.87E-06 |
| NFASC | 750.1182267 | 1.322091559 | 6.86E-34 |
| PAK3 | 94.81753912 | 1.317770111 | 1.35E-02 |
| AC022966.1 | 180.2258423 | 1.30869393 | 1.83E-02 |
| SNAP25 | 73.26660247 | 1.304352303 | 1.74E-02 |
| AL645608.2 | 67.27581695 | 1.299769782 | 1.69E-04 |
| EEF1A2 | 5981.383771 | 1.296312557 | 3.46E-26 |
| NEFM | 79.5989356 | 1.288333501 | 3.26E-02 |
| RASGRP2 | 175.0039685 | 1.263467661 | 3.53E-08 |
| ACKR1 | 26.76636376 | 1.243035966 | 1.71E-02 |
| TRO | 3618.311323 | 1.242593231 | 8.29E-20 |
| SLIT2 | 273.2187921 | 1.223578695 | 2.32E-11 |
| SCN1B | 334.4603313 | 1.216469956 | 1.03E-08 |
| ALPL | 124.5448661 | 1.213966146 | 5.04E-08 |
| AL139289.1 | 27.90948198 | 1.210968238 | 2.21E-02 |
| LYNX1 | 851.7560327 | 1.210031371 | 1.44E-10 |
| AC100793.3 | 34.38183358 | 1.206983809 | 2.84E-02 |
| COL7A1 | 231.364016 | 1.193936181 | 6.37E-04 |
| AP001107.4 | 33.31533398 | 1.182652275 | 6.93E-03 |
| SYT2 | 35.32637237 | 1.170781078 | 7.59E-03 |
| REEP2 | 551.7378508 | 1.161754952 | 5.85E-17 |
| ASPHD1 | 246.0080628 | 1.159822642 | 4.67E-21 |
| LYPD1 | 36.76512797 | 1.149504902 | 7.78E-03 |
| CNTNAP3 | 149.2453626 | 1.144737947 | 1.65E-06 |
| MTND1P23 | 155.9045096 | 1.14221542 | 4.28E-05 |

| | | | |
|---|---|---|---|
| PTPRN | 673.3985235 | 1.140531821 | 6.65E-08 |
| MAP1B | 20810.43855 | 1.138659784 | 7.95E-08 |
| NEGR1 | 205.2653914 | 1.132072769 | 1.66E-02 |
| CNTNAP1 | 3759.390506 | 1.13179363 | 1.75E-27 |
| VCAN | 908.1972434 | 1.124174996 | 1.27E-04 |
| TUBB3 | 3262.462153 | 1.119768209 | 4.30E-14 |
| NTSR1 | 30.50159874 | 1.116542906 | 1.40E-02 |
| HLA-DRA | 272.0727889 | 1.109765077 | 3.16E-04 |
| ADAMTS6 | 290.3621183 | 1.085236637 | 4.25E-09 |
| AC016026.1 | 401.4028746 | 1.084565471 | 2.41E-09 |
| HLA-DMB | 409.4052964 | 1.084028415 | 2.69E-02 |
| TRIM36 | 230.900317 | 1.083632051 | 5.88E-11 |
| CADM3 | 1193.863173 | 1.080454244 | 4.86E-10 |
| ARL4C | 1081.703425 | 1.058288691 | 1.88E-13 |
| ATP13A2 | 2347.326166 | 1.057505846 | 9.90E-29 |
| VWA7 | 89.11714845 | 1.055728465 | 6.48E-03 |
| CALB2 | 75.59255705 | 1.017316808 | 2.73E-04 |
| CDK5R2 | 57.6406424 | 0.998953519 | 4.80E-02 |
| LINC02600 | 73.12744437 | 0.998101082 | 1.07E-03 |
| BEND6 | 355.9990232 | 0.994667713 | 3.46E-03 |
| CD74 | 214.5084595 | 0.99284946 | 4.63E-02 |
| BMF | 718.4286987 | 0.992820475 | 7.81E-08 |
| RALGPS1 | 346.6724631 | 0.992759623 | 1.93E-04 |
| AL662797.1 | 102.1259687 | 0.991012467 | 3.24E-02 |
| AC008735.2 | 41.98639427 | 0.985584179 | 3.56E-02 |
| CDH2 | 355.8677844 | 0.976479592 | 8.05E-05 |
| RARRES1 | 136.06074 | 0.97414216 | 9.69E-03 |
| SLC25A48 | 75.47198056 | 0.966246708 | 1.53E-02 |
| GPRASP2 | 355.0135181 | 0.961450992 | 3.49E-16 |
| CCNDBP1 | 1210.781179 | 0.958572573 | 6.44E-29 |
| FHOD3 | 292.6071871 | 0.957896746 | 6.20E-08 |
| MAGED2 | 15306.61468 | 0.953784175 | 1.37E-10 |
| SCN4B | 2747.623872 | 0.949727645 | 1.69E-04 |
| CD24 | 529.1194415 | 0.926166391 | 1.36E-04 |
| GPT | 53.48594233 | 0.918659191 | 4.53E-02 |
| LINC02593 | 656.2390751 | 0.917404468 | 4.04E-02 |
| FO681492.1 | 191.8394703 | 0.916430009 | 1.94E-02 |
| RN7SL2 | 1281.448062 | 0.902843404 | 2.64E-02 |
| CCN1 | 10596.59591 | 0.901349922 | 1.14E-13 |
| B4GAT1 | 2676.554409 | 0.900267072 | 3.37E-06 |
| WNT9A | 123.5863717 | 0.889646042 | 1.12E-03 |
| LINC00707 | 90.45365985 | 0.8865359 | 3.86E-02 |
| AP006284.1 | 314.2504097 | 0.881578346 | 4.15E-02 |
| CDH4 | 706.3312431 | 0.880276858 | 1.34E-03 |
| RF00017 | 34.86901515 | 0.87610418 | 3.36E-02 |
| THBS1 | 3896.636739 | 0.87184867 | 3.62E-13 |
| SCN8A | 851.2309317 | 0.87168522 | 2.98E-05 |

| | | | |
|---|---|---|---|
| IDS | 4286.442473 | 0.870580293 | 4.12E-21 |
| COL12A1 | 7146.720884 | 0.862309274 | 9.90E-09 |
| MT-ND6 | 16357.53968 | 0.855453563 | 6.28E-04 |
| AHI1 | 978.9390272 | 0.854902068 | 1.10E-03 |
| PRICKLE1 | 152.583996 | 0.8533124 | 1.40E-02 |
| ADAM22 | 711.1995431 | 0.852908438 | 4.40E-05 |
| C2orf27A | 102.3823596 | 0.847681401 | 1.27E-03 |
| LINC00893 | 98.4657195 | 0.847349833 | 2.98E-02 |
| KBTBD11 | 254.4879915 | 0.843919982 | 1.72E-11 |
| ACKR3 | 110.6345116 | 0.843122197 | 3.13E-02 |
| PTCH2 | 132.6977029 | 0.841455734 | 3.30E-02 |
| CCN2 | 8122.707037 | 0.839558614 | 9.45E-04 |
| PRDM1 | 260.727632 | 0.833436101 | 4.61E-02 |
| DYNC1H1 | 30557.26086 | 0.82625451 | 1.96E-19 |
| CFB | 187.4850491 | 0.82095444 | 1.62E-02 |
| PDZD7 | 149.1959024 | 0.820774185 | 1.07E-03 |
| DPYSL3 | 1641.152646 | 0.81759028 | 5.24E-09 |
| ZNF497 | 162.7110401 | 0.815698438 | 3.83E-02 |
| KRT81 | 3638.578358 | 0.815031651 | 1.57E-02 |
| ANXA6 | 16028.14149 | 0.807150444 | 2.12E-06 |
| TNFRSF11A | 99.87689972 | 0.804211763 | 2.79E-02 |
| SLC26A10 | 106.1172386 | 0.796685156 | 2.88E-02 |
| ESYT1 | 9950.813896 | 0.79227514 | 6.12E-13 |
| AC009336.2 | 199.8657579 | 0.791500033 | 3.30E-02 |
| MT-ND5 | 72234.1821 | 0.785502025 | 3.72E-04 |
| SYP | 149.7220486 | 0.776975641 | 2.89E-03 |
| SH3RF2 | 238.2640979 | 0.772802683 | 2.93E-03 |
| VIPR1 | 332.1102024 | 0.771865383 | 1.07E-05 |
| IL2RB | 1447.939605 | 0.770893377 | 6.86E-08 |
| FBLN5 | 552.2943098 | 0.767556657 | 2.83E-02 |
| TMOD2 | 1004.63622 | 0.76342473 | 2.84E-03 |
| SPTAN1 | 27548.29185 | 0.762042849 | 3.93E-25 |
| AC012313.2 | 95.14215469 | 0.753383826 | 1.51E-02 |
| EDN1 | 1541.884744 | 0.752815244 | 1.35E-02 |
| PREPL | 3920.905607 | 0.751536908 | 5.26E-05 |
| LRIG1 | 1253.713048 | 0.747993145 | 4.77E-14 |
| AC139768.1 | 342.3986867 | 0.738801516 | 1.57E-03 |
| LY6G5B | 240.2465744 | 0.730346878 | 1.12E-03 |
| STRA6 | 466.7762795 | 0.729949547 | 6.93E-03 |
| KRT80 | 1730.679913 | 0.729907703 | 3.58E-03 |
| AC015813.1 | 221.2712838 | 0.727373485 | 2.61E-02 |
| CPNE7 | 438.1070185 | 0.723567674 | 3.80E-04 |
| PPL | 285.5073873 | 0.721314796 | 2.32E-02 |
| NA | 45.17878706 | 0.711776856 | 3.75E-02 |
| LINC01638 | 242.896981 | 0.706090966 | 8.45E-03 |
| KLC2 | 1774.580244 | 0.702457431 | 1.35E-08 |
| PDZRN3 | 211.3588518 | 0.698825324 | 2.17E-03 |

| | | | |
|---|---|---|---|
| MMP25 | 133.6636292 | 0.693962252 | 4.30E-02 |
| SYNC | 139.3471639 | 0.68988744 | 3.00E-03 |
| CD99P1 | 133.5769418 | 0.683443301 | 2.21E-02 |
| NA | 221.0300842 | 0.680128493 | 2.55E-02 |
| AC139256.1 | 185.1133231 | 0.678979467 | 6.17E-03 |
| MT-ND1 | 34633.62567 | 0.663098677 | 1.86E-02 |
| FHDC1 | 144.9143232 | 0.656917968 | 4.28E-02 |
| MLLT11 | 927.8788135 | 0.653501394 | 3.33E-07 |
| LPCAT4 | 1464.730274 | 0.652874689 | 1.72E-03 |
| HOXA-AS3 | 169.3705619 | 0.652867464 | 2.57E-02 |
| AC244197.3 | 405.6009799 | 0.652589558 | 4.44E-03 |
| LZTS3 | 1797.583311 | 0.643712233 | 1.31E-13 |
| SH3BP4 | 4262.928303 | 0.642109373 | 1.16E-08 |
| KIAA1841 | 487.9104014 | 0.639271121 | 5.18E-04 |
| SEMA5A | 1442.927626 | 0.632504696 | 1.57E-03 |
| AMOTL2 | 5194.214834 | 0.631484347 | 1.74E-08 |
| MAST1 | 367.3052568 | 0.628330504 | 9.83E-03 |
| AGRN | 19104.86146 | 0.625131926 | 2.35E-05 |
| PDGFC | 565.39758 | 0.624198935 | 2.29E-03 |
| MFAP4 | 931.0022545 | 0.621725143 | 4.66E-03 |
| NACC2 | 1511.713598 | 0.62172395 | 6.76E-10 |
| CLU | 7930.293639 | 0.608336708 | 4.68E-05 |
| NPIPP1 | 408.711574 | 0.602956417 | 1.40E-04 |
| TMEM158 | 357.808961 | 0.600880015 | 2.63E-02 |
| YJEFN3 | 352.2178593 | 0.599980379 | 1.02E-02 |
| UBE2QL1 | 493.8624657 | 0.597220444 | 1.18E-02 |
| CKMT1B | 286.8831445 | 0.590465999 | 2.87E-02 |
| MT-RNR1 | 32125.31546 | 0.590022154 | 6.19E-05 |
| NA | 437.0559334 | 0.584873067 | 5.04E-03 |
| AC011448.1 | 195.9966694 | 0.583067749 | 3.23E-02 |
| TMSB15A | 178.0663652 | 0.581503432 | 7.83E-03 |
| DENND2A | 3519.216737 | 0.580409479 | 3.52E-04 |
| CALM3 | 16638.08905 | 0.579585128 | 2.82E-05 |
| PEG3 | 8882.770975 | 0.577439928 | 1.27E-03 |
| MDN1 | 7460.91208 | 0.570824103 | 1.03E-02 |
| ADRA2C | 237.9240199 | 0.569354016 | 2.18E-02 |
| AC010761.2 | 141.550258 | 0.569265885 | 1.73E-02 |
| NA | 365.8632067 | 0.565066484 | 2.52E-02 |
| DNM1 | 2280.331942 | 0.563410432 | 2.91E-07 |
| MOAP1 | 1235.398058 | 0.562395209 | 2.62E-09 |
| AKAP12 | 6631.627566 | 0.562343788 | 4.65E-04 |
| PLA2R1 | 868.557516 | 0.554377434 | 2.61E-03 |
| CYP26A1 | 160.8035835 | 0.551328599 | 4.09E-02 |
| ARMC5 | 983.2956384 | 0.547552042 | 1.30E-04 |
| FAM20C | 2024.588472 | 0.546331375 | 2.77E-02 |
| PYGL | 5642.226534 | 0.544213258 | 1.05E-19 |
| KIAA0408 | 379.6278526 | 0.544137627 | 4.21E-02 |

| | | | |
|---|---|---|---|
| FHL2 | 2309.54896 | 0.538808728 | 9.25E-05 |
| SLC6A6 | 5652.703338 | 0.535450033 | 2.65E-05 |
| SULF2 | 7414.451633 | 0.532454419 | 1.07E-08 |
| TRNP1 | 1601.234225 | 0.519353986 | 8.88E-03 |
| UBR4 | 15901.60597 | 0.519128887 | 3.92E-08 |
| LPXN | 808.5363228 | 0.518267832 | 4.15E-02 |
| PALM3 | 604.2618859 | 0.510055583 | 4.76E-02 |
| NBL1 | 2346.529476 | 0.508894808 | 5.23E-03 |
| DOCK4 | 1431.176833 | 0.508010413 | 1.59E-03 |
| PIK3CB | 1228.520412 | 0.50537344 | 1.37E-05 |
| TRIO | 14689.77273 | 0.504550344 | 1.47E-06 |
| PPARGC1B | 475.6106913 | 0.503947394 | 3.09E-02 |
| NA | 105.2038393 | 0.501283792 | 3.62E-02 |
| AC138969.1 | 3387.999623 | 0.50035398 | 4.72E-03 |
| ATP9A | 5110.679593 | 0.50020468 | 8.74E-08 |
| HSF4 | 604.4274697 | 0.494682984 | 4.51E-03 |
| MMP2 | 12012.51999 | 0.492733388 | 1.34E-02 |
| PKP2 | 1275.453997 | 0.490399816 | 4.65E-04 |
| GFPT2 | 1047.076629 | 0.488853911 | 3.13E-02 |
| NAIP | 402.0713656 | 0.48211772 | 1.27E-02 |
| PLK2 | 8444.080224 | 0.480980487 | 5.66E-04 |
| NBPF20 | 948.1141545 | 0.48052856 | 9.81E-05 |
| PLEC | 19438.93034 | 0.480375208 | 1.09E-04 |
| LAMA5 | 8544.457214 | 0.477393644 | 1.43E-02 |
| PYGB | 11837.43242 | 0.476871577 | 3.94E-07 |
| GZF1 | 2171.95145 | 0.475476546 | 3.80E-04 |
| AHNAK | 122265.4934 | 0.475197849 | 1.78E-06 |
| DSP | 1365.602877 | 0.475137802 | 2.50E-03 |
| STMN3 | 8539.398862 | 0.471322607 | 2.35E-03 |
| CDCP1 | 1149.145664 | 0.467445773 | 2.58E-02 |
| C11orf45 | 327.0867099 | 0.467121998 | 4.25E-02 |
| NA | 213.9369198 | 0.465416033 | 3.05E-02 |
| PNMA6A | 795.9455409 | 0.465023172 | 2.61E-02 |
| TYRP1 | 289.0563429 | 0.464692105 | 2.23E-02 |
| MYH9 | 39701.66481 | 0.463504363 | 3.72E-04 |
| CARD19 | 976.3250092 | 0.461495421 | 1.67E-03 |
| DBNDD1 | 1129.439015 | 0.460621849 | 3.07E-02 |
| GPRC5A | 3645.517891 | 0.457434097 | 3.07E-02 |
| PRCP | 7324.04715 | 0.455604213 | 4.47E-03 |
| CHST11 | 1000.865333 | 0.454357931 | 3.26E-02 |
| ADAMTS15 | 608.3231517 | 0.453842498 | 6.55E-03 |
| CAMKMT | 213.8627465 | 0.451920408 | 4.46E-02 |
| SLC9A7 | 825.0912289 | 0.451043299 | 2.14E-02 |
| STARD13 | 1275.002019 | 0.44755826 | 4.03E-02 |
| ELOVL4 | 381.9483083 | 0.444249575 | 9.27E-03 |
| SDK1 | 1217.636645 | 0.442015011 | 2.84E-02 |
| NA | 400.6173156 | 0.44016618 | 2.16E-02 |

| | | | |
|---|---|---|---|
| PXDN | 21810.67112 | 0.436604177 | 3.56E-06 |
| CNNM1 | 541.0648735 | 0.435709791 | 4.56E-03 |
| TACC2 | 880.4470114 | 0.434621556 | 3.16E-02 |
| SRRM2 | 25535.81099 | 0.433821735 | 1.24E-04 |
| NBEAL2 | 1699.565215 | 0.432111153 | 3.13E-02 |
| DENND5B | 1132.684712 | 0.430470898 | 3.09E-04 |
| MT-RNR2 | 147786.8412 | 0.426784729 | 2.20E-02 |
| CALM1 | 7582.069497 | 0.422419726 | 7.77E-07 |
| SIPA1L2 | 2095.321232 | 0.421722747 | 8.27E-03 |
| MC1R | 850.7944033 | 0.420522067 | 1.06E-03 |
| MYH10 | 12958.38845 | 0.419228646 | 3.60E-08 |
| UGCG | 4030.044103 | 0.418715899 | 1.35E-02 |
| VLDLR | 1352.477514 | 0.418609186 | 1.64E-03 |
| ANPEP | 1081.191916 | 0.418545675 | 6.99E-03 |
| FLNB | 29152.77127 | 0.418535202 | 3.73E-02 |
| EPDR1 | 1988.545424 | 0.415064065 | 1.13E-02 |
| KLF6 | 4088.683155 | 0.414716368 | 4.02E-04 |
| HIVEP1 | 1633.6484 | 0.41450076 | 3.18E-02 |
| MACF1 | 17521.76745 | 0.413542251 | 1.90E-08 |
| TBC1D4 | 1063.760572 | 0.412379517 | 2.84E-02 |
| TSC22D1 | 5776.662557 | 0.412318175 | 1.67E-04 |
| TMEM131L | 1289.718128 | 0.411867254 | 1.13E-02 |
| DDX24 | 7689.582107 | 0.411186391 | 2.30E-04 |
| PLXNB1 | 1364.593559 | 0.41035785 | 3.36E-02 |
| VAMP1 | 983.9967304 | 0.405146668 | 2.46E-02 |
| TTC19 | 2471.205667 | 0.404286979 | 3.19E-08 |
| RWDD2A | 631.5284039 | 0.403826428 | 2.04E-03 |
| NBPF19 | 1631.618351 | 0.401671005 | 1.68E-03 |
| PPP3CA | 1295.712804 | 0.398243014 | 2.92E-02 |
| FAT1 | 14154.47872 | 0.397143003 | 8.68E-03 |
| PIF1 | 875.719037 | 0.39680806 | 3.33E-02 |
| CAMTA1 | 1421.581369 | 0.396603765 | 3.22E-03 |
| NPIPA1 | 2359.416333 | 0.39531236 | 1.67E-03 |
| GUSBP11 | 471.3827146 | 0.394558042 | 3.58E-03 |
| DOCK9 | 3479.797232 | 0.394409463 | 1.81E-02 |
| SH3PXD2B | 1746.191447 | 0.392788324 | 1.55E-03 |
| COLEC12 | 3956.507078 | 0.39250772 | 8.32E-03 |
| MIR4435-2HG | 2097.286563 | 0.390896279 | 3.61E-03 |
| CEP250 | 4914.904169 | 0.390090194 | 4.24E-05 |
| ELMSAN1 | 2538.049382 | 0.389296392 | 3.57E-04 |
| HOTAIRM1 | 1189.090175 | 0.38919109 | 3.81E-03 |
| SYT17 | 533.479335 | 0.38792152 | 3.57E-02 |
| NEURL4 | 2079.603323 | 0.387498983 | 4.81E-02 |
| XYLT1 | 2293.659344 | 0.385915938 | 6.93E-05 |
| MAP2K4 | 1534.37162 | 0.385224837 | 6.37E-04 |
| PDLIM5 | 3144.819155 | 0.384875093 | 4.63E-02 |
| SLC25A28 | 399.421377 | 0.383218322 | 3.57E-02 |

| | | | |
|---|---|---|---|
| PFKP | 5727.138889 | 0.382195107 | 8.65E-03 |
| MB21D2 | 492.152103 | 0.377954631 | 7.86E-03 |
| FOSL2 | 5985.00852 | 0.377751174 | 1.48E-02 |
| PRKDC | 32717.12166 | 0.373470962 | 3.09E-04 |
| NAV2 | 2387.667239 | 0.372280263 | 4.20E-02 |
| NBPF10 | 2314.64115 | 0.371272718 | 6.59E-03 |
| EZR | 9411.26178 | 0.365516258 | 9.80E-04 |
| CCDC92 | 1103.160319 | 0.360501287 | 8.65E-03 |
| CTSZ | 14220.45977 | 0.359495404 | 1.38E-02 |
| RAB30 | 961.6113688 | 0.357794801 | 4.25E-02 |
| ABL2 | 2982.838415 | 0.356465277 | 2.52E-02 |
| MLST8 | 2699.894163 | 0.352288607 | 2.56E-02 |
| SERPINB9 | 1503.778653 | 0.348187966 | 4.76E-04 |
| HOXA1 | 545.3612809 | 0.346421109 | 3.82E-02 |
| NAT8L | 892.8349717 | 0.344183193 | 3.71E-02 |
| ATP2A2 | 12810.82753 | 0.344078093 | 3.61E-04 |
| LUZP1 | 6874.52116 | 0.342264562 | 4.03E-06 |
| ANXA4 | 2729.466506 | 0.342125872 | 4.47E-04 |
| SON | 15233.18633 | 0.341940207 | 1.49E-02 |
| MICAL3 | 4913.970363 | 0.339813507 | 3.60E-04 |
| PTPN14 | 5617.28597 | 0.333005817 | 1.95E-04 |
| ARID5B | 1968.657859 | 0.328189461 | 2.58E-03 |
| AKAP13 | 5307.00127 | 0.327011704 | 1.28E-02 |
| CCNO | 490.4138232 | 0.323679575 | 3.71E-02 |
| STK36 | 1396.326232 | 0.321263051 | 3.58E-02 |
| TSPAN18 | 4006.491139 | 0.321259195 | 2.11E-02 |
| ZMIZ2 | 2157.698902 | 0.318314649 | 6.87E-05 |
| NOTCH2 | 14889.49296 | 0.317295328 | 7.73E-04 |
| NBPF15 | 3036.575356 | 0.315476535 | 4.12E-04 |
| CRMP1 | 630.7320603 | 0.315466415 | 3.07E-03 |
| ZMIZ1 | 3413.461136 | 0.315416738 | 4.11E-03 |
| PDP1 | 2884.757114 | 0.315195922 | 3.25E-02 |
| MAN2C1 | 3192.371798 | 0.307175152 | 2.83E-02 |
| EPHB2 | 2544.09388 | 0.30659324 | 3.73E-02 |
| IFI27L1 | 506.8812782 | 0.305384069 | 3.57E-02 |
| USP22 | 19092.274 | 0.303702525 | 1.26E-03 |
| INPP5E | 1157.460523 | 0.303027909 | 7.74E-03 |
| PRRC2C | 9875.15191 | 0.302951842 | 1.48E-02 |
| STK39 | 1510.725526 | 0.300306275 | 4.48E-02 |
| HEG1 | 10780.54503 | 0.299554158 | 2.89E-03 |
| HUWE1 | 25381.83121 | 0.298947521 | 6.54E-04 |
| ACOT9 | 3014.294838 | 0.297684561 | 1.74E-02 |
| BTBD11 | 910.1491859 | 0.293064664 | 4.02E-02 |
| GON4L | 2736.602273 | 0.289771892 | 3.71E-03 |
| KMT2A | 10311.72669 | 0.289034832 | 1.98E-02 |
| SRRM1 | 3120.305736 | 0.288431294 | 3.07E-02 |
| NRBP2 | 1261.707177 | 0.287506424 | 4.45E-02 |

| | | | |
|---|---|---|---|
| BCOR | 2863.827776 | 0.286850052 | 3.01E-02 |
| FAM193B | 1060.685105 | 0.285694125 | 1.72E-02 |
| EP400 | 4945.291168 | 0.285517364 | 5.14E-05 |
| IQSEC1 | 2478.420337 | 0.285394526 | 1.12E-02 |
| FOXK1 | 3559.332545 | 0.283829653 | 1.40E-02 |
| EGFR | 7757.382542 | 0.282225005 | 9.92E-04 |
| MSTO1 | 1555.257013 | 0.279132596 | 2.96E-03 |
| ABCC4 | 5268.937555 | 0.279110063 | 2.52E-02 |
| CAST | 16712.96545 | 0.277754924 | 2.91E-05 |
| IFT172 | 1439.982738 | 0.276485827 | 3.54E-02 |
| ENC1 | 10344.19834 | 0.276441311 | 4.55E-02 |
| ANKMY2 | 1123.608775 | 0.273504972 | 3.71E-02 |
| PTPRJ | 3731.301994 | 0.264915425 | 1.05E-02 |
| DPYSL2 | 5563.378319 | 0.263600337 | 1.73E-02 |
| PARN | 3180.356521 | 0.262763794 | 5.12E-03 |
| SZT2 | 2700.467225 | 0.260634543 | 5.75E-03 |
| IMPDH1 | 3474.006773 | 0.258633059 | 4.53E-02 |
| NID1 | 12845.3558 | 0.254054683 | 2.17E-02 |
| AKAP8L | 2144.844662 | 0.246416967 | 4.67E-02 |
| TRRAP | 9853.594491 | 0.242874222 | 5.03E-03 |
| PITRM1 | 3121.362636 | 0.241890901 | 3.97E-02 |
| PNMA1 | 2633.572079 | 0.241486459 | 1.12E-02 |
| SPTBN1 | 22759.67806 | 0.23677937 | 1.07E-03 |
| SLC23A2 | 3962.862943 | 0.236396169 | 6.93E-03 |
| ESYT2 | 7763.823023 | 0.234676686 | 3.75E-02 |
| EP300 | 4842.321 | 0.231901959 | 2.09E-06 |
| TRAM2 | 9655.068286 | 0.212162088 | 3.54E-02 |
| HIPK2 | 6086.845547 | 0.205072102 | 3.60E-02 |
| NDUFS1 | 6300.475034 | 0.200995099 | 5.71E-03 |
| RNF145 | 10583.21953 | 0.200695629 | 8.86E-04 |
| NOP56 | 9573.943269 | 0.197883715 | 8.96E-03 |
| RIC1 | 2976.271407 | 0.187302634 | 2.79E-02 |
| SPEN | 5159.913258 | 0.157797677 | 3.19E-02 |
| TUSC3 | 4463.581761 | -0.181482283 | 4.75E-02 |
| PRKAG1 | 3564.972411 | -0.182889067 | 2.83E-02 |
| DYNLL1 | 9168.515016 | -0.18761319 | 3.91E-02 |
| TMED10 | 13060.58854 | -0.203928684 | 3.57E-02 |
| FUCA2 | 6047.718625 | -0.206051712 | 4.20E-02 |
| TXNDC12 | 4250.368422 | -0.224333534 | 4.74E-02 |
| CNIH1 | 3853.018552 | -0.228704863 | 1.94E-02 |
| ZCRB1 | 2337.235148 | -0.230063474 | 4.34E-02 |
| SPCS1 | 3376.39699 | -0.24731742 | 2.68E-02 |
| MAGEA10 | 7039.294074 | -0.250297442 | 1.96E-02 |
| PHYH | 939.2769998 | -0.25211204 | 4.33E-02 |
| HIP1 | 8387.051766 | -0.263547604 | 3.47E-02 |
| BIVM-ERCC5 | 1830.656259 | -0.265490613 | 4.73E-02 |
| SDCBP | 7542.179046 | -0.268590462 | 1.56E-02 |

| | | | |
|---|---|---|---|
| NEK9 | 9405.283839 | -0.271913436 | 2.46E-02 |
| AC046185.3 | 1008.471374 | -0.272799566 | 2.84E-02 |
| PLPP5 | 1498.63886 | -0.274239289 | 1.81E-02 |
| TPT1 | 67889.21744 | -0.274582437 | 1.81E-02 |
| DDX25 | 1351.778353 | -0.275164222 | 1.90E-02 |
| PPIC | 3752.098519 | -0.275498043 | 1.65E-02 |
| GALK2 | 1013.100879 | -0.276709329 | 3.55E-03 |
| CYBRD1 | 6049.057482 | -0.280861324 | 1.01E-02 |
| SMARCA1 | 5733.867665 | -0.281977529 | 1.15E-02 |
| MAGEC2 | 3875.244618 | -0.284194121 | 1.38E-02 |
| TMC6 | 4347.104689 | -0.288622108 | 1.85E-02 |
| FAR2 | 1062.330359 | -0.290997545 | 4.72E-02 |
| MRPL52 | 1805.217802 | -0.30517712 | 4.93E-03 |
| TOMM7 | 3903.636684 | -0.305906781 | 4.28E-02 |
| RAD51D | 1062.319911 | -0.312408541 | 2.77E-02 |
| SRGN | 1547.782683 | -0.312506763 | 1.44E-04 |
| FP671120.2 | 584.4415846 | -0.314345929 | 4.72E-02 |
| PLCXD1 | 1585.892566 | -0.317521047 | 1.17E-02 |
| CMC4 | 890.8548752 | -0.318437466 | 3.75E-02 |
| DHRS4 | 1323.074625 | -0.323086694 | 1.51E-02 |
| PMVK | 1036.769575 | -0.323155898 | 7.73E-03 |
| MED28 | 3238.863894 | -0.328832567 | 3.59E-02 |
| KRBA2 | 422.1868317 | -0.332295758 | 4.69E-02 |
| HEXB | 16325.57173 | -0.332670067 | 8.89E-03 |
| PDLIM3 | 622.703077 | -0.332740533 | 2.56E-02 |
| KHDC4 | 2471.678525 | -0.333617293 | 2.27E-02 |
| ELOVL2 | 1985.730308 | -0.336525705 | 4.88E-02 |
| ZNF721 | 490.7716122 | -0.33660825 | 2.11E-02 |
| CKB | 5181.807168 | -0.337513591 | 1.98E-02 |
| C19orf25 | 1676.958754 | -0.339528236 | 3.03E-02 |
| KAT2B | 1031.112119 | -0.341807021 | 3.24E-02 |
| MTSS1 | 1117.759609 | -0.343490396 | 4.72E-02 |
| GALNT14 | 1213.5074 | -0.348559794 | 4.21E-02 |
| CT45A5 | 774.9288964 | -0.349640899 | 2.76E-03 |
| PRICKLE3 | 449.079314 | -0.35235028 | 4.63E-02 |
| LPIN2 | 5759.958478 | -0.3525039 | 1.91E-02 |
| GSR | 4194.454514 | -0.355531372 | 1.03E-03 |
| TTYH3 | 13891.95316 | -0.356033249 | 2.18E-02 |
| P2RX4 | 1389.149782 | -0.358168186 | 2.46E-02 |
| AC008969.1 | 361.7441285 | -0.366152658 | 4.30E-02 |
| AC074117.1 | 550.5267816 | -0.370764002 | 2.74E-02 |
| BLVRB | 2573.362732 | -0.37209612 | 2.48E-02 |
| ARMC9 | 2187.710154 | -0.372218073 | 1.57E-03 |
| AGMAT | 377.0349647 | -0.373783424 | 3.23E-02 |
| DAPK1 | 5388.257129 | -0.374889939 | 2.19E-05 |
| HACD4 | 437.4754155 | -0.376799775 | 1.51E-02 |
| CSKMT | 343.0414308 | -0.3772899 | 3.19E-02 |

| | | | |
|---|---|---|---|
| HSPB8 | 1436.3874 | -0.37946893 | 4.96E-02 |
| TXNDC16 | 735.2004753 | -0.383514555 | 1.34E-02 |
| ACVR2B | 614.6329055 | -0.385246117 | 3.49E-02 |
| FKBP14 | 2076.395674 | -0.388231035 | 1.92E-02 |
| CRACR2A | 437.6015031 | -0.388234028 | 7.31E-03 |
| CHST2 | 604.9230275 | -0.391673743 | 2.37E-04 |
| SPCS3 | 2551.177372 | -0.395331541 | 8.89E-03 |
| DPP7 | 4715.09999 | -0.395740113 | 2.88E-02 |
| DDT | 2299.728872 | -0.396975759 | 8.15E-03 |
| PROCR | 2832.564645 | -0.399694165 | 2.46E-02 |
| TMEM35B | 784.729291 | -0.400253451 | 1.65E-04 |
| FRG1BP | 884.4544904 | -0.403681468 | 1.50E-03 |
| ITGB3 | 1822.991296 | -0.404325396 | 4.48E-02 |
| CPE | 2379.926947 | -0.404619204 | 4.53E-02 |
| HES6 | 712.9778645 | -0.40684402 | 1.92E-02 |
| PFN1 | 38626.73183 | -0.406979346 | 1.86E-02 |
| LDB2 | 2892.946269 | -0.408118912 | 1.15E-02 |
| TUB | 3581.772259 | -0.409796293 | 4.65E-04 |
| C11orf71 | 513.7332104 | -0.412276666 | 2.01E-02 |
| TMSB4X | 28768.79103 | -0.415159299 | 1.36E-02 |
| LINC01234 | 437.4067572 | -0.417055012 | 1.44E-02 |
| CORO7 | 454.3518415 | -0.417966089 | 4.72E-02 |
| TFPT | 1099.521882 | -0.418306019 | 3.07E-02 |
| NUDT14 | 692.5963255 | -0.423662794 | 3.32E-02 |
| ZKSCAN7 | 396.0174361 | -0.424812666 | 3.86E-02 |
| ZNF219 | 899.6886662 | -0.425623721 | 1.52E-02 |
| AC010442.1 | 720.7233888 | -0.427681187 | 2.58E-03 |
| CAPN10-DT | 307.6229708 | -0.429795762 | 4.81E-02 |
| BRI3 | 3521.077973 | -0.430627427 | 3.30E-02 |
| HMGA2 | 5164.377752 | -0.431388569 | 4.77E-02 |
| SYNGR2 | 1829.502754 | -0.431596902 | 1.08E-02 |
| NMT1 | 6026.448678 | -0.433058721 | 2.54E-02 |
| CMBL | 2580.311176 | -0.433411483 | 4.98E-02 |
| IL10RB | 1854.424156 | -0.433764326 | 9.79E-03 |
| TLCD1 | 494.9847647 | -0.436522767 | 4.30E-02 |
| HPSE | 457.3034666 | -0.440533008 | 1.63E-02 |
| PPP1R3B | 1558.68049 | -0.440573976 | 1.03E-07 |
| ELFN1 | 408.2898101 | -0.441130761 | 3.30E-02 |
| ACYP2 | 296.8828932 | -0.445000139 | 4.20E-02 |
| RAMP2 | 1068.842961 | -0.448730424 | 3.61E-02 |
| AS3MT | 520.8533791 | -0.450796774 | 3.88E-02 |
| MAMDC2 | 4343.615439 | -0.452529541 | 1.52E-02 |
| MAGEA1 | 1398.25761 | -0.4557078 | 2.50E-04 |
| AC027031.2 | 487.0588179 | -0.456832214 | 2.52E-03 |
| MR1 | 1383.354792 | -0.458230057 | 2.75E-02 |
| HMOX1 | 1720.384188 | -0.461972328 | 3.19E-02 |
| MEIS3P1 | 287.1832009 | -0.462486724 | 4.29E-02 |

| | | | |
|---|---|---|---|
| FAM49A | 399.0533098 | -0.466479559 | 2.35E-02 |
| CLIP3 | 2375.243907 | -0.467940535 | 9.79E-03 |
| FEZ1 | 659.6364878 | -0.46839502 | 4.45E-02 |
| AC016065.1 | 209.0021807 | -0.470550238 | 3.60E-02 |
| EHD3 | 6392.771573 | -0.47193002 | 2.52E-02 |
| GNG7 | 471.4326956 | -0.476773543 | 1.90E-02 |
| HAGHL | 852.8669522 | -0.478495214 | 4.02E-02 |
| SCAPER | 932.7521639 | -0.482141561 | 2.88E-02 |
| XAGE3 | 258.2371969 | -0.482490165 | 3.84E-02 |
| AL391244.2 | 314.9832061 | -0.48519255 | 2.95E-02 |
| AC245297.1 | 206.1312181 | -0.485710704 | 3.03E-02 |
| ZNF524 | 171.9973563 | -0.487868754 | 4.52E-02 |
| FTH1 | 76509.70998 | -0.492267159 | 2.58E-03 |
| SLC27A5 | 255.6919464 | -0.492646094 | 2.21E-02 |
| GPX7 | 858.5942305 | -0.494490942 | 1.74E-02 |
| MAGEA11 | 992.8345221 | -0.495315937 | 3.55E-07 |
| CPVL | 3426.438074 | -0.495935705 | 5.75E-03 |
| NA | 202.7798493 | -0.496464016 | 2.28E-02 |
| AL050341.2 | 228.804696 | -0.496886233 | 4.53E-02 |
| ANKRD34A | 275.0037824 | -0.499662134 | 5.12E-03 |
| SYT7 | 257.2568568 | -0.509164269 | 2.46E-02 |
| SEL1L3 | 3269.858349 | -0.509220976 | 5.11E-03 |
| IFITM3 | 20144.00674 | -0.512698784 | 2.52E-02 |
| PYCARD | 961.5747621 | -0.517207258 | 9.27E-03 |
| CASP10 | 374.7130969 | -0.520430511 | 2.83E-02 |
| TSHZ2 | 2202.117157 | -0.522717192 | 4.46E-03 |
| ZNF521 | 1676.551674 | -0.528332791 | 4.66E-03 |
| NA | 791.4238412 | -0.528412361 | 2.19E-02 |
| PRKCB | 1065.831152 | -0.529396015 | 4.41E-02 |
| DUSP15 | 247.1476797 | -0.529675447 | 2.84E-02 |
| COL9A3 | 4082.534637 | -0.530677828 | 1.88E-02 |
| PLTP | 6996.727233 | -0.531078835 | 6.00E-03 |
| FTL | 97215.3597 | -0.532884389 | 3.07E-03 |
| ECSCR | 314.089158 | -0.545140413 | 2.27E-02 |
| AP006621.5 | 135.4954891 | -0.547879724 | 3.77E-02 |
| SLC7A8 | 1004.591585 | -0.548893436 | 7.21E-03 |
| DYSF | 2529.150232 | -0.54889548 | 4.24E-03 |
| USP2 | 233.5515925 | -0.550652418 | 3.19E-02 |
| IFI30 | 2930.300089 | -0.554946807 | 8.27E-03 |
| PRR7 | 203.9707106 | -0.566697366 | 1.34E-02 |
| COCH | 812.7057441 | -0.572112946 | 1.72E-04 |
| FAM133A | 452.9836405 | -0.577165461 | 1.88E-03 |
| CD33 | 476.0162093 | -0.59404095 | 7.78E-04 |
| OLFML2A | 2480.21215 | -0.594791233 | 3.85E-02 |
| FBXO2 | 682.1087411 | -0.596821526 | 3.27E-02 |
| FXYD6 | 2385.941432 | -0.598701658 | 2.96E-03 |
| SLC2A4 | 550.3986875 | -0.598860012 | 8.26E-03 |

| | | | |
|---|---|---|---|
| ITGA1 | 11661.05432 | -0.605869963 | 8.05E-05 |
| ABHD8 | 415.0651293 | -0.605929976 | 1.83E-02 |
| COL8A2 | 659.7014155 | -0.606844213 | 2.51E-02 |
| OOEP | 184.8376679 | -0.611635137 | 1.72E-02 |
| MIR4527HG | 177.5777164 | -0.614512124 | 1.09E-02 |
| AC132872.1 | 143.6391076 | -0.615191313 | 1.31E-02 |
| AC021078.1 | 620.0647068 | -0.628114319 | 1.00E-02 |
| CLIC6 | 238.7103255 | -0.630700978 | 3.73E-02 |
| PDE6B | 188.8286447 | -0.631160739 | 3.32E-02 |
| ID3 | 8443.247581 | -0.633545579 | 8.92E-03 |
| IFIT2 | 1178.033649 | -0.645845061 | 5.26E-05 |
| IFITM1 | 3198.919102 | -0.646880618 | 1.07E-02 |
| LRRN2 | 226.4845914 | -0.648227194 | 4.98E-02 |
| AC005253.1 | 114.5949675 | -0.648448479 | 1.93E-02 |
| ARHGAP9 | 264.0659115 | -0.651789617 | 1.30E-02 |
| TBXA2R | 175.3616382 | -0.652109888 | 3.48E-02 |
| PPP1R16B | 527.7404052 | -0.657020165 | 3.07E-02 |
| CAPG | 110.5601278 | -0.657597942 | 4.04E-02 |
| NGFR | 2528.28695 | -0.661733709 | 1.92E-02 |
| ANO7 | 227.8760111 | -0.665373924 | 1.24E-03 |
| MS4A15 | 168.0965125 | -0.668690605 | 1.05E-02 |
| FAM83H | 345.1934365 | -0.671561185 | 1.13E-04 |
| COLEC11 | 893.9691598 | -0.672327913 | 2.56E-03 |
| CAMK1G | 592.4143727 | -0.673210937 | 3.77E-02 |
| ZNF467 | 538.7262357 | -0.676830905 | 3.06E-02 |
| ID1 | 4464.244433 | -0.678920822 | 3.54E-02 |
| GPNMB | 7636.758085 | -0.686854918 | 1.21E-03 |
| SLC45A2 | 191.8192019 | -0.691482943 | 3.25E-02 |
| PTGER2 | 910.6436438 | -0.691946313 | 3.88E-03 |
| TPPP | 427.9957226 | -0.708053752 | 5.11E-03 |
| CKM | 89.28298883 | -0.7099069 | 1.47E-02 |
| RARRES2 | 5285.601134 | -0.711876487 | 1.26E-04 |
| GZMM | 799.9992698 | -0.711906934 | 2.60E-02 |
| REEP1 | 645.6708003 | -0.713376534 | 1.49E-02 |
| AC020916.1 | 458.5870598 | -0.723661884 | 4.04E-06 |
| IFITM2 | 1317.633411 | -0.73417128 | 1.34E-02 |
| HOXC12 | 131.3367996 | -0.734637807 | 7.32E-03 |
| LINC02557 | 375.7815063 | -0.740555766 | 3.30E-03 |
| DMTN | 300.3915311 | -0.759426466 | 9.83E-03 |
| VAMP8 | 385.8872751 | -0.773951699 | 2.12E-02 |
| SHE | 294.9360923 | -0.778443837 | 1.02E-03 |
| PDGFRL | 578.6213394 | -0.778819138 | 2.96E-03 |
| TM7SF2 | 2521.278961 | -0.779488746 | 6.05E-03 |
| WFDC1 | 322.5959827 | -0.78276321 | 6.00E-07 |
| ST6GAL1 | 1260.199539 | -0.787648907 | 1.65E-04 |
| GREB1 | 310.3314579 | -0.790380427 | 1.51E-02 |
| NA | 224.2178336 | -0.805959089 | 1.05E-02 |

| | | | |
|---|---|---|---|
| RPL4P5 | 84.9281609 | -0.807145417 | 3.41E-02 |
| GAB3 | 903.944943 | -0.808916533 | 4.12E-04 |
| ELMO1 | 314.1325128 | -0.80993627 | 6.99E-03 |
| TRBV20-1 | 248.8926942 | -0.810056449 | 8.24E-05 |
| LINC01311 | 79.3079992 | -0.814911298 | 1.74E-02 |
| TMEM119 | 206.1753641 | -0.815905454 | 1.40E-02 |
| SYPL2 | 83.161266 | -0.817744341 | 4.96E-02 |
| SDR42E1 | 246.9387746 | -0.82216737 | 7.59E-03 |
| KCNIP1 | 288.8772349 | -0.831400554 | 7.59E-03 |
| TDRD10 | 80.80343204 | -0.839101347 | 1.93E-02 |
| AC008622.2 | 52.92455998 | -0.839816113 | 4.76E-02 |
| ENPP2 | 849.7703046 | -0.845069169 | 6.24E-03 |
| SOX17 | 62.40755451 | -0.85748193 | 1.96E-02 |
| AC132192.2 | 139.1062199 | -0.870749297 | 2.38E-03 |
| FPR2 | 56.2775836 | -0.87282445 | 4.67E-02 |
| MAFB | 2006.013682 | -0.88108257 | 1.15E-04 |
| RGS4 | 778.1227395 | -0.887522091 | 3.57E-02 |
| AARD | 73.18662623 | -0.890727879 | 5.21E-03 |
| AC023509.1 | 209.0395314 | -0.892320814 | 1.08E-04 |
| CD79B | 132.1895311 | -0.901985716 | 2.61E-03 |
| MRVI1 | 377.0708102 | -0.903494127 | 1.65E-03 |
| PCSK9 | 829.7416089 | -0.907833036 | 8.36E-05 |
| SLC7A10 | 92.31430533 | -0.924545256 | 9.23E-03 |
| CNTFR | 700.6352166 | -0.92522386 | 6.25E-04 |
| RTN4RL1 | 146.9324584 | -0.929058931 | 6.59E-04 |
| SPACA6P-AS | 74.7839744 | -0.931242089 | 6.03E-03 |
| BAIAP2L2 | 100.9679441 | -0.957158689 | 1.74E-02 |
| IL4I1 | 44.42488749 | -0.958395962 | 3.25E-02 |
| CFP | 142.794171 | -0.960071447 | 5.40E-03 |
| CHAD | 59.05109073 | -0.964930938 | 4.04E-02 |
| TRAF3IP3 | 99.96308918 | -0.966518184 | 1.13E-03 |
| AC018892.3 | 49.79431166 | -0.976541202 | 4.67E-02 |
| JPH4 | 834.6588439 | -0.983125044 | 2.58E-03 |
| MAT1A | 731.6526786 | -0.992804551 | 4.75E-03 |
| RENBP | 435.6778357 | -0.994014946 | 8.69E-04 |
| RRAGD | 701.8675929 | -0.99633554 | 1.11E-05 |
| AL139089.1 | 33.40727554 | -1.004135632 | 2.88E-02 |
| FCRLA | 42.33965956 | -1.006739333 | 1.71E-02 |
| APOC1 | 1307.212753 | -1.022424168 | 2.33E-09 |
| SLC6A12 | 262.088674 | -1.04334047 | 2.30E-02 |
| TMEM255A | 638.2751286 | -1.063444865 | 2.00E-07 |
| AC009086.2 | 455.0561247 | -1.065339961 | 4.17E-02 |
| AC018755.2 | 97.94631105 | -1.114966086 | 3.92E-02 |
| AC006277.1 | 45.89973383 | -1.12974499 | 6.03E-03 |
| CAPN6 | 166.404887 | -1.156940767 | 2.20E-04 |
| RPLP0P2 | 306.6935964 | -1.168994936 | 4.25E-05 |
| RASAL3 | 65.57954874 | -1.173632055 | 1.79E-03 |

| | | | |
|---|---|---|---|
| KDR | 808.088877 | -1.181117213 | 3.57E-04 |
| CYSLTR2 | 151.0712334 | -1.182607466 | 1.54E-03 |
| AC048341.2 | 122.453961 | -1.196178175 | 5.48E-04 |
| SMPDL3B | 107.507826 | -1.196705941 | 2.83E-02 |
| LYPD9P | 50.79022482 | -1.218283009 | 1.17E-02 |
| SSX6P | 45.48695315 | -1.253017582 | 1.44E-02 |
| CRYZL2P-SEC16B | 94.80652887 | -1.279886266 | 4.15E-02 |
| NA | 39.88031917 | -1.28315263 | 8.88E-03 |
| ZNF252P-AS1 | 45.88853292 | -1.289416548 | 9.04E-04 |
| APOE | 6958.007481 | -1.292046753 | 4.27E-09 |
| AL121944.1 | 25.22065618 | -1.304685523 | 1.43E-02 |
| HTR2B | 48.42776478 | -1.309464687 | 1.70E-05 |
| LPL | 5043.224616 | -1.398618308 | 1.07E-22 |
| C1orf115 | 406.0582373 | -1.430482376 | 5.13E-04 |
| RNA5-8SN2 | 684.3031988 | -1.442507797 | 1.53E-05 |
| PGAM2 | 238.1250556 | -1.530905276 | 4.54E-06 |
| FABP6 | 25.52013632 | -1.606444698 | 1.01E-02 |
| NOS1 | 28.20981896 | -1.687894024 | 6.88E-03 |
| SLC9A3 | 209.2046598 | -1.730870937 | 1.14E-12 |
| RNA5S1 | 26.79550676 | -1.81163295 | 1.17E-02 |
| AC079298.3 | 24.94196526 | -1.898307303 | 3.13E-02 |
| CYP4F8 | 24.3394758 | -2.073295356 | 5.14E-03 |
| UNC5CL | 15.07775381 | -2.098175619 | 4.26E-02 |
| MS4A6E | 21.63496949 | -2.112362086 | 4.09E-02 |
| PSG6 | 42.7531939 | -2.65822914 | 3.89E-02 |
| AMIGO3 | 93.09257147 | -3.849986197 | 4.34E-02 |
| FPGT-TNNI3K | 29.63087152 | -5.73211694 | 2.71E-06 |

| rank | motif_ID | consensus | p-value |
|---|---|---|---|
| 1 | TEAD4_HUMAN.H11MO.0.A | RCATTCCWGRVAK | 6.81e-421 |
| 2 | TEAD1_HUMAN.H11MO.0.A | RCATTCCWGSCW | 1.42e-389 |
| 3 | TEAD2_HUMAN.H11MO.0.D | HYACATTCCWGG | 1.66e-365 |
| 4 | JUN_HUMAN.H11MO.0.A | DRTGAGTCAYH | 1.33E-207 |
| 5 | FOS_HUMAN.H11MO.0.A | VTGAGTCAB | 1.11E-206 |
| 6 | FOSL2_HUMAN.H11MO.0.A | NDRTGAGTCAYH | 8.43E-204 |
| 7 | FOSL1_HUMAN.H11MO.0.A | KRVTGAGTCAYH | 8.63E-204 |
| 8 | JUND_HUMAN.H11MO.0.A | RRTGAGTCAYY | 7.22E-188 |
| 9 | FOSB_HUMAN.H11MO.0.A | RTGAGTCAB | 6.19E-186 |
| 10 | JUNB_HUMAN.H11MO.0.A | DVTGAGTCABH | 5.68E-183 |
| 11 | ETV6_HUMAN.H11MO.0.D | RCAGGAARK | 1.71E-126 |
| 12 | ETV4_HUMAN.H11MO.0.B | VRCAGGAARBR | 6.94E-112 |
| 13 | BATF_HUMAN.H11MO.1.A | DATGAYKCAAH | 3.53E-100 |
| 14 | ETV5_HUMAN.H11MO.0.C | RRRSAGGAARDGRV | 3.40E-99 |
| 15 | ZN394_HUMAN.H11MO.1.D | GAAWGGAAW | 1.03E-94 |
| 16 | ERG_HUMAN.H11MO.0.A | RRRSAGGAAGYGG | 9.17E-88 |
| 17 | ETV7_HUMAN.H11MO.0.D | VAGGAAR | 3.59E-81 |
| 18 | ETV2_HUMAN.H11MO.0.B | RRARRCAGGAARYRGS | 4.20E-79 |
| 19 | FLI1_HUMAN.H11MO.1.A | RRRCAGGAAGBRG | 8.39E-79 |
| 20 | ETS2_HUMAN.H11MO.0.B | RDVRGGAARVRRR | 2.86E-76 |
| 21 | VEZF1_HUMAN.H11MO.0.C | GGRRRRGRRGGAGGGGGRGRRR | 1.98E-73 |
| 22 | FLI1_HUMAN.H11MO.0.A | SRRGGMAGGAAGGRRRGR | 7.41E-73 |
| 23 | ETS1_HUMAN.H11MO.0.A | RRRCMGGAAGTGG | 1.06E-72 |
| 24 | BACH2_HUMAN.H11MO.0.A | TGCTGAGTCAB | 2.82E-72 |
| 25 | IRF3_HUMAN.H11MO.0.B | RAAARGGAAAVDGAAASDGA | 5.89E-71 |
| 26 | MAZ_HUMAN.H11MO.0.A | GGGGGAGGGGGDGRGRRRGRRG | 2.38E-68 |
| 27 | NFAT5_HUMAN.H11MO.0.D | RRRGGAARRDN | 3.55E-68 |
| 28 | EGR2_HUMAN.H11MO.0.A | GRGRRKGWGKGGGHGGRG | 3.92E-67 |
| 29 | E2F7_HUMAN.H11MO.0.B | GDGGCGGGAARDR | 1.34E-66 |
| 30 | BC11A_HUMAN.H11MO.0.A | RAAAGAGGAAGTGARAV | 1.21E-66 |
| 31 | ZBT17_HUMAN.H11MO.0.A | SRRGGWGGGGGAGGGGMRR | 8.43E-63 |
| 32 | NFAC1_HUMAN.H11MO.0.B | AWGGAAARWVWGAMW | 1.71E-62 |
| 33 | PATZ1_HUMAN.H11MO.0.C | GGGGMGGGGGMKGGRRVGGVRG | 1.14E-61 |
| 34 | WT1_HUMAN.H11MO.0.C | RGGVGGGGGAGGRGGVGGRG | 6.98E-61 |
| 35 | FOXG1_HUMAN.H11MO.0.D | DTTGTTTAYDTWTDTTT | 1.19E-60 |
| 36 | ZN467_HUMAN.H11MO.0.C | GGGGAGGGGRRGGRGRRGGRRR | 1.60E-60 |
| 37 | ZN770_HUMAN.H11MO.0.C | GGGAGGCYGAGGBRGGAGGATC | 3.53E-60 |
| 38 | KLF15_HUMAN.H11MO.0.A | RGGGMGGRGVDGGGGGRGG | 2.45E-59 |
| 39 | EGR2_HUMAN.H11MO.1.A | GRDGHGKGGGHGGRGG | 3.66E-59 |
| 40 | FOXP1_HUMAN.H11MO.0.A | TBTGTTTMY | 2.67E-59 |
| 41 | ELF5_HUMAN.H11MO.0.A | VGRASSAGGAAGTRV | 8.43E-59 |
| 42 | ZN341_HUMAN.H11MO.0.C | GRGGRRRRGGRGRRGGAASAGC | 3.09E-58 |
| 43 | STAT1_HUMAN.H11MO.1.A | RRRAAAADGAAASTRAAAV | 6.08E-58 |
| 44 | EGR1_HUMAN.H11MO.0.A | VDRDGCGKGGGYGGRRR | 8.53E-57 |
| 45 | NFE2_HUMAN.H11MO.0.A | VVRTGACTCAGCA | 6.86E-56 |
| 46 | KLF6_HUMAN.H11MO.0.A | GSRRGGGHGGGGMHGGGRV | 1.30E-55 |

| | | | |
|---|---|---|---|
| 47 | RUNX1_HUMAN.H11MO.0.A | BYTGTGGTTWB | 2.83E-55 |
| 48 | RUNX2_HUMAN.H11MO.0.A | KGKYTGTGGTTTKK | 1.65E-54 |
| 49 | CPEB1_HUMAN.H11MO.0.D | TTTTTATTYTTTTTTTT | 1.37E-51 |
| 50 | FOXK1_HUMAN.H11MO.0.A | TGTTTMCHTT | 3.73E-52 |
| 51 | RUNX3_HUMAN.H11MO.0.A | BYTGTGGTTW | 3.89E-51 |
| 52 | NF2L2_HUMAN.H11MO.0.A | WWYTGCTGAGTCAT | 1.24E-50 |
| 53 | SP4_HUMAN.H11MO.0.A | SRGVARGRGGCGGRGCHDRR | 5.14E-51 |
| 54 | TAF1_HUMAN.H11MO.0.A | RARRWGGCGGMGGMGR | 4.76E-50 |
| 55 | ELF2_HUMAN.H11MO.0.C | RAVCCGGAAGTGR | 7.66E-50 |
| 56 | PRDM6_HUMAN.H11MO.0.C | RRRARGAAAAAAA | 5.37E-49 |
| 57 | ELF3_HUMAN.H11MO.0.A | RAABVAGGAAGTRR | 2.37E-48 |
| 58 | ETV1_HUMAN.H11MO.0.A | SRCCGGAAGYG | 7.89E-48 |
| 59 | BACH1_HUMAN.H11MO.0.A | TGCTGAGTCABSS | 8.60E-48 |
| 60 | ZN263_HUMAN.H11MO.0.A | GGGAGGAGGRRGRGGRGGRR | 1.47E-47 |
| 61 | ZN394_HUMAN.H11MO.0.C | NRARWRGAAWNGAMWGRAAK | 1.04E-47 |
| 62 | FOXA1_HUMAN.H11MO.0.A | TGTTTACWYWGB | 1.76E-46 |
| 63 | HMGA1_HUMAN.H11MO.0.D | NWATTTT | 5.10E-46 |
| 64 | SP4_HUMAN.H11MO.1.A | GRRRGGGGMGGGGCBK | 2.10E-45 |
| 65 | FOXJ2_HUMAN.H11MO.0.C | TGTTTRTTTW | 1.03E-44 |
| 66 | FEV_HUMAN.H11MO.0.B | GCVGGAAGYG | 1.03E-44 |
| 67 | TBX15_HUMAN.H11MO.0.D | GGGGGGGGGGGGTGGGDR | 6.68E-44 |
| 68 | PURA_HUMAN.H11MO.0.D | GGWRKKGGDGGGCVKGG | 1.85E-43 |
| 69 | E2F3_HUMAN.H11MO.0.A | DDGGCGGGAAA | 2.02E-43 |
| 70 | PEBB_HUMAN.H11MO.0.C | TYTGTGGTYWB | 6.57E-43 |
| 71 | ELF1_HUMAN.H11MO.0.A | VRACCCGGAAGTGS | 1.41E-42 |
| 72 | FOXF1_HUMAN.H11MO.0.D | WTGTTTATTTW | 4.56E-42 |
| 73 | ZN281_HUMAN.H11MO.0.A | RGGWGGGGGAGGGGV | 8.37E-42 |
| 74 | KLF3_HUMAN.H11MO.0.B | GGRGVDGGGCGGGGCYGGG | 6.50E-41 |
| 75 | FOXD3_HUMAN.H11MO.0.D | KBTGTTTNCWTWRBH | 4.59E-41 |
| 76 | KLF16_HUMAN.H11MO.0.D | GGGGGYGKGKBGGGGGGGG | 1.57E-40 |
| 77 | MAFK_HUMAN.H11MO.1.A | TGCTGAGTCAGCA | 4.51E-40 |
| 78 | RXRA_HUMAN.H11MO.0.A | VRGDVRGNVARAGGTCARRG | 5.05E-40 |
| 79 | EHF_HUMAN.H11MO.0.B | VWACSAGGAAGTDVS | 2.13E-39 |
| 80 | FOXA2_HUMAN.H11MO.0.A | TGTTTACWYWGB | 2.38E-39 |
| 81 | SOX10_HUMAN.H11MO.1.A | BBYCTTTGTTYYY | 8.15E-40 |
| 82 | KLF5_HUMAN.H11MO.0.A | WGGGTGKGGCDGGG | 4.92E-39 |
| 83 | FOXP2_HUMAN.H11MO.0.C | BTGTTTACH | 6.29E-39 |
| 84 | SP2_HUMAN.H11MO.0.A | GGSSVGGGGGCGGGGCCDGSGS | 1.25E-38 |
| 85 | ZSC22_HUMAN.H11MO.0.C | RGKCWGAGGGMGGAGG | 1.35E-38 |
| 86 | E2F4_HUMAN.H11MO.0.A | SRGGGCGGGAARD | 4.34E-38 |
| 87 | GABPA_HUMAN.H11MO.0.A | GVVRCCGGAAGTGV | 1.20E-37 |
| 88 | TFDP1_HUMAN.H11MO.0.C | RRRRGGCGGGAARN | 6.25E-37 |
| 89 | FOXM1_HUMAN.H11MO.0.A | TGTTTRCTYWKB | 8.48E-37 |
| 90 | FOXO1_HUMAN.H11MO.0.A | KBBYTGTTTHCN | 1.26E-36 |
| 91 | ZN770_HUMAN.H11MO.1.C | DGGAGGCYGRG | 2.93E-36 |
| 92 | SP3_HUMAN.H11MO.0.B | SGVVGGGGGCGGGGCBRGSS | 2.38E-36 |
| 93 | FOXF2_HUMAN.H11MO.0.D | TGTTTACHTWD | 5.71E-36 |

| | | | |
|---|---|---|---|
| 94 | ELK1_HUMAN.H11MO.0.B | RCCGGAAGTGV | 1.27E-35 |
| 95 | NFKB1_HUMAN.H11MO.0.A | AGGGAAAK | 3.40E-36 |
| 96 | E2F6_HUMAN.H11MO.0.A | DGGGCGGGARRVR | 8.13E-36 |
| 97 | NF2L1_HUMAN.H11MO.0.C | HGTCATN | 1.77E-35 |
| 98 | ZFX_HUMAN.H11MO.1.A | DGVRGVVSAGGCCKVGG | 3.74E-35 |
| 99 | FUBP1_HUMAN.H11MO.0.D | TYGTNTTTTTTT | 6.65E-35 |
| 100 | PO3F3_HUMAN.H11MO.0.D | WWWAWTTATGCWWATTW | 1.06E-34 |
| 101 | HSF1_HUMAN.H11MO.1.A | TTCTGGAAR | 1.64E-34 |
| 102 | VEZF1_HUMAN.H11MO.1.C | RRRRGGAGGGGR | 1.06E-34 |
| 103 | ZFP82_HUMAN.H11MO.0.C | TTCCADTTCAYTAATTYTCTCTTC | 6.47E-34 |
| 104 | NFAC1_HUMAN.H11MO.1.B | AATGGAAAAW | 6.24E-34 |
| 105 | STAT2_HUMAN.H11MO.0.A | RRGRAAAHGAAACTGAAAV | 2.72E-33 |
| 106 | SP2_HUMAN.H11MO.1.B | RGGRGGCGGGGC | 5.79E-33 |
| 107 | MAZ_HUMAN.H11MO.1.A | RRGGGAGGGGS | 7.73E-33 |
| 108 | ZN713_HUMAN.H11MO.0.D | RWAGAARAADGVSANGMAA | 9.59E-33 |
| 109 | KLF1_HUMAN.H11MO.0.A | DGGGYGKGGCYGGG | 2.12E-32 |
| 110 | SALL4_HUMAN.H11MO.0.B | GSDGGGWGGG | 1.17E-32 |
| 111 | ELK4_HUMAN.H11MO.0.A | SVCCGGAAGTGV | 1.33E-31 |
| 112 | HXD11_HUMAN.H11MO.0.D | DWTTTATKRHN | 8.71E-32 |
| 113 | FOXO3_HUMAN.H11MO.0.B | BYTGTTTACH | 1.23E-31 |
| 114 | TBX1_HUMAN.H11MO.0.D | GGGYGGGYAGGGGTGGGMDD | 1.67E-31 |
| 115 | TBX21_HUMAN.H11MO.0.A | GDGDRGGTGTGRV | 7.52E-32 |
| 116 | FOXJ3_HUMAN.H11MO.0.A | TTGTTTATKKTTW | 8.63E-31 |
| 117 | ANDR_HUMAN.H11MO.0.A | TGTTCTTKKYTGTTTRYW | 7.09E-31 |
| 118 | BATF_HUMAN.H11MO.0.A | RBTYTCRDWATGASTCAB | 8.99E-31 |
| 119 | HXC10_HUMAN.H11MO.0.D | WWWWWWWNNWTTTWATKRC | 1.29E-30 |
| 120 | FOXJ3_HUMAN.H11MO.1.B | TKKTTWWTGTTTA | 5.07E-30 |
| 121 | NFAC2_HUMAN.H11MO.0.B | TGGAAAAWY | 5.33E-30 |
| 122 | HXD12_HUMAN.H11MO.0.D | DDTTTTAHKRN | 8.06E-30 |
| 123 | ZN148_HUMAN.H11MO.0.D | KGRGKGGGGGAGGGG | 2.41E-29 |
| 124 | ZN121_HUMAN.H11MO.0.C | CTGGGCAACABAGYRAGACC | 1.60E-28 |
| 125 | ELK3_HUMAN.H11MO.0.D | VMCHGGAARTSC | 8.69E-29 |
| 126 | RARA_HUMAN.H11MO.1.A | VAGGTCAGRG | 3.88E-29 |
| 127 | NFAC4_HUMAN.H11MO.0.C | WGGAAAAYTW | 1.88E-28 |
| 128 | SPIC_HUMAN.H11MO.0.D | KVDRRWGGGKDRG | 3.50E-28 |
| 129 | FOXA3_HUMAN.H11MO.0.B | CHTGTTTACWTWG | 1.33E-27 |
| 130 | HAND1_HUMAN.H11MO.1.D | KSYCTGG | 6.75E-28 |
| 131 | TEAD3_HUMAN.H11MO.0.D | GAHATTYYWGCYBYW | 2.26E-27 |
| 132 | IRF1_HUMAN.H11MO.0.A | RAAANWGAAASTGAAASYRR | 8.50E-27 |
| 133 | MEF2B_HUMAN.H11MO.0.A | DKKKCTATTTTTRG | 4.90E-27 |
| 134 | ZN554_HUMAN.H11MO.1.D | GCWGAGYCA | 2.71E-27 |
| 135 | SP1_HUMAN.H11MO.0.A | VGGGGGCGGGGCCKGGGGGGG | 7.16E-26 |
| 136 | COT1_HUMAN.H11MO.1.C | BSARAGGTCAGRG | 4.24E-26 |
| 137 | WT1_HUMAN.H11MO.1.B | GWGGGAGGAGS | 2.95E-25 |
| 138 | SUH_HUMAN.H11MO.0.A | VSYSTGGGAAV | 3.63E-25 |
| 139 | MEF2A_HUMAN.H11MO.0.A | DKCTATTTTTRGM | 8.85E-25 |
| 140 | RORG_HUMAN.H11MO.0.C | RRAASTRGGTCA | 8.76E-25 |

| | | | |
|---|---|---|---|
| 141 | KLF4_HUMAN.H11MO.0.A | WGGGYGKGGC | 1.25E-24 |
| 142 | MAFF_HUMAN.H11MO.1.B | TGCTGASTC | 7.46E-25 |
| 143 | GATA3_HUMAN.H11MO.0.A | RVAGATAAVAD | 8.99E-25 |
| 144 | FOXL1_HUMAN.H11MO.0.D | TKTTTGTTTRTTTAYWTTT | 2.49E-23 |
| 145 | SMAD3_HUMAN.H11MO.0.B | SYCTSHCWSCWS | 8.16E-25 |
| 146 | MEF2C_HUMAN.H11MO.0.A | DKCTATTTTTRGM | 5.74E-24 |
| 147 | STAT3_HUMAN.H11MO.0.A | SAYTTCCMRGAA | 6.70E-24 |
| 148 | MEF2D_HUMAN.H11MO.0.A | DKCTATTTTTAG | 2.31E-23 |
| 149 | COT2_HUMAN.H11MO.1.A | RGRGYCARAGGTCARVR | 8.72E-24 |
| 150 | STA5B_HUMAN.H11MO.0.A | YTTCYKRGAAAY | 3.49E-23 |
| 151 | PATZ1_HUMAN.H11MO.1.C | AGGGAGGGG | 1.57E-23 |
| 152 | ARNT2_HUMAN.H11MO.0.D | GNCGTGGSASRC | 4.84E-23 |
| 153 | NFIC_HUMAN.H11MO.0.A | CYTGGCHBYMWGCCARG | 1.02E-22 |
| 154 | COT1_HUMAN.H11MO.0.C | GRGGKSARAGGTCAGGG | 2.49E-22 |
| 155 | E2F1_HUMAN.H11MO.0.A | RRNKGGCGGGAARR | 4.95E-22 |
| 156 | ARI3A_HUMAN.H11MO.0.D | RATTAAWBRRWATCAAAWTWWA | 1.33E-21 |
| 157 | MLX_HUMAN.H11MO.0.D | YGATCACGTGN | 3.78E-21 |
| 158 | ZFP28_HUMAN.H11MO.0.C | TTCTATTTCTTCTTGWGTCA | 2.13E-21 |
| 159 | SP1_HUMAN.H11MO.1.A | VGGGGGCGGGGCCK | 9.82E-22 |
| 160 | SPI1_HUMAN.H11MO.0.A | RRAAAGAGGAAGTGRVD | 1.47E-21 |
| 161 | PROX1_HUMAN.H11MO.0.D | KGGKARGGCGKSKKGGG | 8.93E-22 |
| 162 | ATF2_HUMAN.H11MO.0.B | RRTGABGTCAY | 2.94E-21 |
| 163 | MAFG_HUMAN.H11MO.1.A | TGCTGAGT | 1.64E-21 |
| 164 | NR1D1_HUMAN.H11MO.0.B | RRRNMASTGGGTCAGDRDR | 2.28E-21 |
| 165 | SMCA5_HUMAN.H11MO.0.C | GAATBGAATGGAAWC | 5.81E-21 |
| 166 | STA5A_HUMAN.H11MO.0.A | TTCYRRGAAR | 7.04E-21 |
| 167 | FOXB1_HUMAN.H11MO.0.D | RTMAATRTTTACATAAB | 2.17E-20 |
| 168 | BCL6_HUMAN.H11MO.0.A | YKCTTTCYAGGAA | 5.13E-21 |
| 169 | MAFF_HUMAN.H11MO.0.B | TGCTGASTCAGCABTTTT | 1.87E-20 |
| 170 | Z354A_HUMAN.H11MO.0.C | RTTTARWYCATTTACATTTAATGT | 5.74E-20 |
| 171 | KLF12_HUMAN.H11MO.0.C | VGGGGCGGGGC | 5.50E-20 |
| 172 | FOXD2_HUMAN.H11MO.0.D | WAWGTAAATRTTWWYTTAA | 2.78E-19 |
| 173 | SRY_HUMAN.H11MO.0.B | YWTTGTTTH | 3.49E-20 |
| 174 | RARA_HUMAN.H11MO.0.A | GGRRGRTSMRRAGKTCAAGG | 1.26E-19 |
| 175 | SRBP2_HUMAN.H11MO.0.B | VSGTGGGGWGABG | 1.05E-19 |
| 176 | PAX5_HUMAN.H11MO.0.A | SCSKGGGCARCVRAGCGWGAC | 6.78E-19 |
| 177 | FOXC1_HUMAN.H11MO.0.C | CDWTGTTTACTTWRG | 2.00E-18 |
| 178 | ZF64A_HUMAN.H11MO.0.D | GGGGCCCGGGAMYCDG | 9.57E-19 |
| 179 | HXA11_HUMAN.H11MO.0.D | TWAADTTTTAYGR | 1.36E-18 |
| 180 | IRF8_HUMAN.H11MO.0.B | RAAARRGGAASTGAAASTDR | 2.96E-18 |
| 181 | STAT4_HUMAN.H11MO.0.A | YTTCYYRGMAV | 1.74E-18 |
| 182 | XBP1_HUMAN.H11MO.0.D | GSKGACGTGGC | 4.52E-18 |
| 183 | KLF9_HUMAN.H11MO.0.C | GRRGKGGGCGTGGCC | 8.66E-18 |
| 184 | MAFB_HUMAN.H11MO.0.B | TGCTGASTYAD | 2.66E-18 |
| 185 | RREB1_HUMAN.H11MO.0.D | DKGKKKGKGGKTGKTTTGGGKT | 1.43E-17 |
| 186 | SOX4_HUMAN.H11MO.0.B | BYCTTTGTYYYB | 5.35E-18 |
| 187 | RARG_HUMAN.H11MO.1.B | VAGGTCAS | 7.19E-18 |

| | | | |
|---|---|---|---|
| 188 | PRDM1_HUMAN.H11MO.0.A | RAAAGTGAAAGTGR | 1.33E-17 |
| 189 | PO2F1_HUMAN.H11MO.0.C | AWKBRWATGSAAWYAW | 1.68E-17 |
| 190 | IRF4_HUMAN.H11MO.0.A | RAAWGRGGAASTGAAASY | 4.82E-17 |
| 191 | ZSC31_HUMAN.H11MO.0.C | RGCAGCAGGGCARTTAKR | 5.31E-17 |
| 192 | STAT1_HUMAN.H11MO.0.A | YCCVTTTCCHGGAAAWCHC | 8.17E-17 |
| 193 | RXRB_HUMAN.H11MO.0.C | TSAGGTCASR | 5.85E-17 |
| 194 | ATF3_HUMAN.H11MO.0.A | GGTSACGTGAB | 2.09E-16 |
| 195 | MAFK_HUMAN.H11MO.0.A | RAWWHTGCTGASTCAGCA | 4.05E-16 |
| 196 | SPIB_HUMAN.H11MO.0.A | RRAAAGAGGAAGTGARA | 2.42E-16 |
| 197 | Z324A_HUMAN.H11MO.0.C | KVTCAAACCATCCTTKSCTKSCW | 9.64E-17 |
| 198 | UBIP1_HUMAN.H11MO.0.D | SCAGARA | 8.57E-17 |
| 199 | RXRA_HUMAN.H11MO.1.A | SARAGGTCAS | 1.06E-16 |
| 200 | ZNF76_HUMAN.H11MO.0.C | RRKGCMTKCTGGGARWTGTAGT | 4.10E-16 |
| 201 | USF1_HUMAN.H11MO.0.A | RGTCACGTGRBS | 1.02E-15 |
| 202 | BMAL1_HUMAN.H11MO.0.A | KSCACGTGHSY | 4.00E-16 |
| 203 | HIF1A_HUMAN.H11MO.0.C | GDACGTGM | 7.09E-16 |
| 204 | IKZF1_HUMAN.H11MO.0.C | BTGGGARR | 2.80E-16 |
| 205 | USF2_HUMAN.H11MO.0.A | GGGTCACGTGRBSVSSSB | 4.41E-15 |
| 206 | SMCA1_HUMAN.H11MO.0.C | CMARGAAGMATD | 6.57E-16 |
| 207 | ATF1_HUMAN.H11MO.0.B | HRRTGACGYMA | 2.61E-15 |
| 208 | HNF1B_HUMAN.H11MO.0.A | DGTTAATRATTAACH | 1.04E-14 |
| 209 | HNF1A_HUMAN.H11MO.0.C | DGTTAATKATTAACH | 7.43E-15 |
| 210 | FOXP3_HUMAN.H11MO.0.D | AATKTGWTT | 4.03E-15 |
| 211 | TFEB_HUMAN.H11MO.0.C | RGTCACGTG | 8.62E-14 |
| 212 | NFAC3_HUMAN.H11MO.0.B | TGGAAAAHY | 4.36E-15 |
| 213 | SMAD1_HUMAN.H11MO.0.D | RSCCTGTCTGCC | 4.68E-15 |
| 214 | ZN335_HUMAN.H11MO.1.A | YGCCTGW | 6.20E-15 |
| 215 | IRF7_HUMAN.H11MO.0.C | GAAASYGAAA | 7.43E-15 |
| 216 | RARB_HUMAN.H11MO.0.D | SAGGTCAVVVV | 1.40E-14 |
| 217 | SOX2_HUMAN.H11MO.0.A | BBCCTTTGTYYYB | 9.12E-15 |
| 218 | TFE3_HUMAN.H11MO.0.B | RGTCACGTGA | 4.47E-14 |
| 219 | CR3L2_HUMAN.H11MO.0.D | TGATGACGTGGCA | 8.65E-14 |
| 220 | MNX1_HUMAN.H11MO.0.D | MBATTTDHWTHWCYHYTHH | 1.08E-14 |
| 221 | MYCN_HUMAN.H11MO.0.A | RGCCACGTGSDS | 3.74E-14 |
| 222 | ZN563_HUMAN.H11MO.1.C | SHRGCAGCWSWG | 3.15E-14 |
| 223 | LMX1B_HUMAN.H11MO.0.D | TAATTAAHWRMYTWWWWW | 1.27E-13 |
| 224 | LMX1A_HUMAN.H11MO.0.D | TAATTAATTWKKTWATTWW | 2.03E-13 |
| 225 | NFIA_HUMAN.H11MO.0.C | TTGGCHNRNWGCCAR | 7.70E-14 |
| 226 | MITF_HUMAN.H11MO.0.A | TCACGTGACC | 2.73E-13 |
| 227 | LYL1_HUMAN.H11MO.0.A | CCAKCTGYTYYCTB | 6.50E-14 |
| 228 | AP2C_HUMAN.H11MO.0.A | VTGSCCTGRGGSHR | 2.10E-13 |
| 229 | FOXO4_HUMAN.H11MO.0.C | TTGTTTAYK | 3.40E-13 |
| 230 | ESR2_HUMAN.H11MO.1.A | AGGTCAS | 2.38E-13 |
| 231 | BHE40_HUMAN.H11MO.0.A | DGCACGTGAS | 8.71E-13 |
| 232 | HXA10_HUMAN.H11MO.0.C | KATRRTTTATGW | 7.35E-13 |
| 233 | ZN322_HUMAN.H11MO.0.B | GWGCCTGSTACWSWGCCTGR | 9.04E-13 |
| 234 | COT2_HUMAN.H11MO.0.A | CARAGGTCARRGD | 3.85E-13 |

| | | | |
|---|---|---|---|
| 235 | HXC12_HUMAN.H11MO.0.D | DTTTTAYKACYT | 9.32E-13 |
| 236 | IRX2_HUMAN.H11MO.0.D | RCDKGKBDKGDCRKGTDVK | 5.87E-13 |
| 237 | STAT6_HUMAN.H11MO.0.B | TTCTSWGGAAR | 7.86E-13 |
| 238 | AP2A_HUMAN.H11MO.0.A | ATKSCCTGRGGSHAD | 8.98E-13 |
| 239 | TF65_HUMAN.H11MO.0.A | HWGGGRATTTCCMV | 2.22E-12 |
| 240 | ZN263_HUMAN.H11MO.1.A | SRGGGAGGASG | 4.72E-12 |
| 241 | HXA9_HUMAN.H11MO.0.B | RTGATTTATKR | 5.36E-12 |
| 242 | MAF_HUMAN.H11MO.0.A | VAAWBTGCTGASTHWGCMD | 4.07E-12 |
| 243 | AP2D_HUMAN.H11MO.0.D | CGCCYGVGGCSCGT | 1.47E-11 |
| 244 | HEY1_HUMAN.H11MO.0.D | KGRCACGTGBCK | 2.33E-11 |
| 245 | CREB1_HUMAN.H11MO.0.A | NRRTGACGTMA | 1.29E-11 |
| 246 | ZFHX3_HUMAN.H11MO.0.D | ATTAWTAATTA | 1.81E-11 |
| 247 | SPDEF_HUMAN.H11MO.0.D | RWCCMGGAWGWAGT | 1.34E-11 |
| 248 | IRF2_HUMAN.H11MO.0.A | RAAAVHGAAAGTGAAASTRV | 4.50E-11 |
| 249 | KLF14_HUMAN.H11MO.0.D | DRGKGGGCGTGGY | 4.37E-11 |
| 250 | ZN335_HUMAN.H11MO.0.A | RGCTGYCYNBVSCKMTGCCTGW | 2.61E-11 |
| 251 | EGR4_HUMAN.H11MO.0.D | GGSGGYRGGGM | 2.51E-11 |
| 252 | ESR1_HUMAN.H11MO.1.A | CHRGGTCASV | 1.55E-11 |
| 253 | PO4F1_HUMAN.H11MO.0.D | CATTAATTATTCAT | 8.72E-11 |
| 254 | OSR2_HUMAN.H11MO.0.C | GBYSCWGCTNCTGYDG | 2.02E-11 |
| 255 | BHE23_HUMAN.H11MO.0.D | RRSRTRTGKTDB | 2.62E-11 |
| 256 | MAFG_HUMAN.H11MO.0.A | RWWWNTGCTGASTCAGCA | 3.64E-10 |
| 257 | ZN282_HUMAN.H11MO.0.D | MGKRTTRTGGGARA | 5.29E-11 |
| 258 | MAX_HUMAN.H11MO.0.A | VRGCACGTGG | 7.74E-11 |
| 259 | PTF1A_HUMAN.H11MO.0.B | SCAGCTGBYBBYBYYCHC | 5.26E-11 |
| 260 | ZN768_HUMAN.H11MO.0.C | GAGGCHCAGAGAGGTTAAGT | 2.73E-10 |
| 261 | NR4A1_HUMAN.H11MO.0.A | AAAGGKCAV | 3.70E-11 |
| 262 | REL_HUMAN.H11MO.0.B | DRWRGGGRAATKCCA | 5.98E-11 |
| 263 | FOXO6_HUMAN.H11MO.0.D | WWSWTGTTTACRT | 2.57E-10 |
| 264 | AP2B_HUMAN.H11MO.0.B | GCCYGVGGGS | 1.13E-10 |
| 265 | EGR3_HUMAN.H11MO.0.D | WGAGTGGGYGT | 3.25E-10 |
| 266 | SMAD2_HUMAN.H11MO.0.A | TGTCTGNCWCCT | 1.33E-10 |
| 267 | NKX61_HUMAN.H11MO.0.B | GAYWAATKRSHWWTTAATK | 2.77E-10 |
| 268 | NRL_HUMAN.H11MO.0.D | TGCTGACKYWR | 2.04E-10 |
| 269 | BARX1_HUMAN.H11MO.0.D | BWAATKGTTTYY | 1.82E-10 |
| 270 | EPAS1_HUMAN.H11MO.0.B | VTACGTGMC | 9.74E-10 |
| 271 | MXI1_HUMAN.H11MO.0.A | VCACGTGGSNGSBGS | 3.85E-10 |
| 272 | ARNT_HUMAN.H11MO.0.B | STACGTGMC | 9.33E-10 |
| 273 | GSX2_HUMAN.H11MO.0.D | YTAATTRSHWWTTAAT | 4.96E-10 |
| 274 | CREB3_HUMAN.H11MO.0.D | KGRTGACGTGGCA | 5.87E-09 |
| 275 | NKX61_HUMAN.H11MO.1.B | WAATGRV | 2.58E-10 |
| 276 | JDP2_HUMAN.H11MO.0.D | BGATGACGTCAY | 7.90E-09 |
| 277 | ZN554_HUMAN.H11MO.0.C | GCWGAGYCANGTRGDKKRCT | 5.85E-10 |
| 278 | RARG_HUMAN.H11MO.0.B | RGGTCASVRTGVCMTGRV | 5.05E-10 |
| 279 | HAND1_HUMAN.H11MO.0.D | RGSYCTGGYYWNTSTCA | 5.15E-10 |
| 280 | PO4F3_HUMAN.H11MO.0.D | TCATTAATTATKCAT | 4.30E-09 |
| 281 | BPTF_HUMAN.H11MO.0.D | KKTBTTGTKKKS | 5.00E-10 |

| | | | |
|---|---|---|---|
| 282 | ZEP2_HUMAN.H11MO.0.D | GGYDGGGWAACCSS | 9.97E-10 |
| 283 | E2F4_HUMAN.H11MO.1.A | GGCGGGAADTTBNR | 2.23E-09 |
| 284 | THA_HUMAN.H11MO.1.D | BTSAGGTCAS | 1.11E-09 |
| 285 | HXB13_HUMAN.H11MO.0.A | TTTTATDGSH | 1.46E-09 |
| 286 | ATF6A_HUMAN.H11MO.0.B | GKGSTGACGTGG | 1.32E-08 |
| 287 | HSF1_HUMAN.H11MO.0.A | BRGAANSTTCYRGAA | 2.93E-09 |
| 288 | THAP1_HUMAN.H11MO.0.C | SGCCGCCATSTYGGSYBCGGGC | 1.80E-09 |
| 289 | BATF3_HUMAN.H11MO.0.B | BRSTTTCADTWTGASWM | 1.68E-09 |
| 290 | EVX1_HUMAN.H11MO.0.D | TGATTTATKGCYW | 2.33E-09 |
| 291 | ZFX_HUMAN.H11MO.0.A | SCAGGCCBCR | 1.71E-09 |
| 292 | FOXD1_HUMAN.H11MO.0.D | CWWTGTTTACHTWVR | 3.29E-09 |
| 293 | SRF_HUMAN.H11MO.0.A | TTKCCWTWTWTGGBCWTD | 6.21E-09 |
| 294 | LEF1_HUMAN.H11MO.0.A | BCCTTTSWWNTBYH | 1.16E-09 |
| 295 | NR1H3_HUMAN.H11MO.1.B | MVAGGTCAVR | 1.55E-09 |
| 296 | FOXQ1_HUMAN.H11MO.0.C | HATTGTTTATTT | 1.90E-08 |
| 297 | ATF2_HUMAN.H11MO.1.B | WDDWGATGATGTCATCWWTW | 6.05E-08 |
| 298 | PITX2_HUMAN.H11MO.0.D | TKGGATTAVH | 5.90E-09 |
| 299 | HSFY1_HUMAN.H11MO.0.D | WWTCGAACGWWW | 3.61E-08 |
| 300 | CREM_HUMAN.H11MO.0.C | CRVTGACGTCA | 1.56E-08 |
| 301 | MXI1_HUMAN.H11MO.1.A | SCCACGTGSYB | 8.42E-09 |
| 302 | NFIB_HUMAN.H11MO.0.D | YCYTGGCAS | 4.77E-09 |
| 303 | GATA2_HUMAN.H11MO.0.A | STGBNNNNNDVAGATAAGR | 1.78E-08 |
| 304 | RARG_HUMAN.H11MO.2.D | GGRGKTCRVCNRVAGGTCAV | 1.38E-08 |
| 305 | THB_HUMAN.H11MO.0.C | GRSSYSASVTSAGGTCA | 1.65E-08 |
| 306 | PTF1A_HUMAN.H11MO.1.B | RGCAGCTGG | 8.99E-09 |
| 307 | NR1H4_HUMAN.H11MO.1.B | RGGTCAKT | 9.03E-09 |
| 308 | CREB5_HUMAN.H11MO.0.D | RRTGACGTMAT | 4.26E-07 |
| 309 | GATA1_HUMAN.H11MO.0.A | STGBNNNDNVSAGATAAGV | 4.74E-08 |
| 310 | PPARG_HUMAN.H11MO.0.A | MWSTRGGKSARAGGKCA | 1.98E-08 |
| 311 | PO6F1_HUMAN.H11MO.0.D | CATAAWTTATGCR | 4.84E-07 |
| 312 | THA_HUMAN.H11MO.0.C | RGGTCABYTSAGGTCAK | 2.33E-08 |
| 313 | CLOCK_HUMAN.H11MO.0.C | CWGHCACGTGMVVM | 2.40E-08 |
| 314 | ITF2_HUMAN.H11MO.0.C | DGCAGSTGKS | 1.87E-08 |
| 315 | RORA_HUMAN.H11MO.0.C | AWSTRGGTCARRG | 2.14E-08 |
| 316 | GATA2_HUMAN.H11MO.1.A | DSAGATAAGRR | 4.68E-08 |
| 317 | RFX4_HUMAN.H11MO.0.D | SGTTGCCATGGYWAC | 3.07E-07 |
| 318 | THB_HUMAN.H11MO.1.D | SGTSAGGTCA | 6.87E-08 |
| 319 | ZNF41_HUMAN.H11MO.1.C | CACAAGGRARWV | 5.79E-08 |
| 320 | TEF_HUMAN.H11MO.0.D | TGTTTATGTAAMTK | 1.64E-07 |
| 321 | NKX22_HUMAN.H11MO.0.D | DSAGWGSH | 2.92E-08 |
| 322 | ZEB1_HUMAN.H11MO.0.A | BVCAGGTGWG | 6.68E-08 |
| 323 | ZEP1_HUMAN.H11MO.0.D | TGGGAAATCCCB | 1.98E-07 |
| 324 | TBX3_HUMAN.H11MO.0.C | DRAGGTGBSAR | 5.86E-08 |
| 325 | GATA4_HUMAN.H11MO.0.A | DSAGATAAGA | 1.85E-07 |
| 326 | HXC13_HUMAN.H11MO.0.D | DDTTTTACKAG | 1.97E-07 |
| 327 | BHA15_HUMAN.H11MO.0.B | SSAGCAGCTGG | 1.44E-07 |
| 328 | NR4A2_HUMAN.H11MO.0.C | AAAGGTCAS | 1.61E-07 |

| | | | |
|---|---|---|---|
| 329 | MYC_HUMAN.H11MO.0.A | RCCACGTGCTB | 3.46E-07 |
| 330 | PO5F1_HUMAN.H11MO.1.A | WHATGCAVAT | 1.29E-07 |
| 331 | HSF4_HUMAN.H11MO.0.D | TRGAABVTTCTAGAA | 1.57E-06 |
| 332 | ZKSC1_HUMAN.H11MO.0.B | WGHRSMCCTACTRTGTGCH | 9.09E-07 |
| 333 | MAF_HUMAN.H11MO.1.B | TGCTGAST | 1.81E-07 |
| 334 | MEIS1_HUMAN.H11MO.0.A | RTGATTKATKRB | 3.97E-07 |
| 335 | ATF2_HUMAN.H11MO.2.C | RRTGACGTCAS | 9.70E-06 |
| 336 | DLX1_HUMAN.H11MO.0.D | WWWWWTATGSTAATTA | 3.75E-07 |
| 337 | ZNF85_HUMAN.H11MO.0.C | AWAGAARWCTGMWGKAATCT | 8.01E-07 |
| 338 | PO3F2_HUMAN.H11MO.0.A | YYCATGCATWTYCATTM | 4.05E-07 |
| 339 | TAL1_HUMAN.H11MO.0.A | CTGSNVVSBRSWGATAAGV | 1.01E-06 |
| 340 | DMBX1_HUMAN.H11MO.0.D | YMMTWWWAAYCYAATTAABD | 1.01E-06 |
| 341 | SRBP1_HUMAN.H11MO.0.A | DGTSRGGTGAB | 2.87E-07 |
| 342 | FOXH1_HUMAN.H11MO.0.A | TGTGGATTS | 4.40E-07 |
| 343 | E4F1_HUMAN.H11MO.0.D | YGTKACGTC | 4.75E-06 |
| 344 | RFX3_HUMAN.H11MO.0.B | BGTTRCCATGGYRA | 5.59E-06 |
| 345 | ZN418_HUMAN.H11MO.0.C | TGCTTYTRGCYTCTKVY | 6.26E-07 |
| 346 | HSF2_HUMAN.H11MO.0.A | VGAABRTTCTAGAA | 7.71E-07 |
| 347 | HEY2_HUMAN.H11MO.0.D | GKBGGCWCGTGGCHTBV | 2.47E-06 |
| 348 | NFIC_HUMAN.H11MO.1.A | TYYTGGCWS | 3.58E-07 |
| 349 | HNF1B_HUMAN.H11MO.1.A | DGTTAATVATT | 2.31E-06 |
| 350 | HIC2_HUMAN.H11MO.0.D | HCRTGCCAGVN | 1.07E-06 |
| 351 | TBP_HUMAN.H11MO.0.A | MTATAAAARS | 1.31E-06 |
| 352 | GATA1_HUMAN.H11MO.1.A | DSAGATAAGVV | 1.74E-06 |
| 353 | HXC11_HUMAN.H11MO.0.D | DTTTWAYGACY | 2.22E-06 |
| 354 | ZN436_HUMAN.H11MO.0.C | TCMRGGARGGCTTCCTGGAGGAGG | 5.92E-05 |
| 355 | HES1_HUMAN.H11MO.0.D | VMGVCACGMGMCM | 1.59E-06 |
| 356 | PO3F4_HUMAN.H11MO.0.D | TMATTTGCATAATTWAW | 4.88E-06 |
| 357 | ZIM3_HUMAN.H11MO.0.C | KTRGGTTTCTGTTKYYTKY | 5.45E-06 |
| 358 | PAX4_HUMAN.H11MO.0.D | TWAYTAATTMRH | 2.50E-06 |
| 359 | RFX2_HUMAN.H11MO.0.A | CKGTTGCCATGGCRACVS | 1.53E-05 |
| 360 | ATF7_HUMAN.H11MO.0.D | KRTGACGTCAT | 5.19E-05 |
| 361 | MAFA_HUMAN.H11MO.0.D | STGCTGACBMYGCAGYHTYCV | 2.21E-06 |
| 362 | HEN1_HUMAN.H11MO.0.C | BGGGKMKCAGCTGCGKCCYY | 1.07E-05 |
| 363 | ZNF8_HUMAN.H11MO.0.C | TGTGGTATATCCATACAATGGA | 1.98E-04 |
| 364 | ZN528_HUMAN.H11MO.0.C | CCMAGGGAAGYCATTTCTSA | 1.01E-05 |
| 365 | NR1D1_HUMAN.H11MO.1.D | AWAARTAGGTCARW | 1.10E-05 |
| 366 | HXA2_HUMAN.H11MO.0.D | NGTAATTARBD | 5.38E-06 |
| 367 | HXB2_HUMAN.H11MO.0.D | WAGAAYVTTCTAGAA | 1.20E-05 |
| 368 | CTCFL_HUMAN.H11MO.0.A | GCCDSYAGGGGGCGCYV | 7.48E-06 |
| 369 | SIX2_HUMAN.H11MO.0.A | KGAAAYHTGAKMC | 3.35E-06 |
| 370 | MECP2_HUMAN.H11MO.0.C | SCCGGRR | 5.73E-06 |

### AME (Analysis of Motif Enrichment): Version 5.0.5 compiled on Jul 15 2019 at 00:31:14
### The format of this file is described at http://meme-suite.org//doc/ame-output-format.html.
### ame --verbose 1 --oc . --scoring avg --method fisher --hit-lo-fraction 0.25 --evalue-report-threshold 1

| adj_p-value | E-value | tests | FASTA_max | pos | neg | PWM_min | TP |
|---|---|---|---|---|---|---|---|
| 2.57e-417 | 3.35e-414 | 3773 | 3469 | 3469 | 3469 | 1.8 | 2397 |
| 4.96e-386 | 6.46e-383 | 3489 | 3469 | 3469 | 3469 | 1.24 | 2484 |
| 4.76e-362 | 6.20e-359 | 2867 | 3469 | 3469 | 3469 | 1.26 | 2132 |
| 2.68E-204 | 3.50E-201 | 2024 | 3469 | 3469 | 3469 | 1.62 | 1371 |
| 2.85E-203 | 3.71E-200 | 2572 | 3469 | 3469 | 3469 | 2.38 | 1373 |
| 1.49E-200 | 1.94E-197 | 1771 | 3469 | 3469 | 3469 | 1.07 | 1422 |
| 1.57E-200 | 2.04E-197 | 1819 | 3469 | 3469 | 3469 | 1.21 | 1414 |
| 1.56E-184 | 2.03E-181 | 2157 | 3469 | 3469 | 3469 | 2.3 | 1251 |
| 1.78E-182 | 2.32E-179 | 2877 | 3469 | 3469 | 3469 | 3.36 | 1347 |
| 1.31E-179 | 1.71E-176 | 2311 | 3469 | 3469 | 3469 | 1.08 | 1680 |
| 7.83E-123 | 1.02E-119 | 4573 | 3469 | 3469 | 3469 | 1.62 | 2315 |
| 2.99E-108 | 3.90E-105 | 4317 | 3469 | 3469 | 3469 | 1.66 | 2170 |
| 8.27E-97 | 1.08E-93 | 2341 | 3469 | 3469 | 3469 | 2.06 | 1166 |
| 1.48E-95 | 1.93E-92 | 4357 | 3469 | 3469 | 3469 | 2.28 | 1848 |
| 4.38E-91 | 5.70E-88 | 4245 | 3469 | 3469 | 3469 | 1.02 | 2535 |
| 2.55E-84 | 3.32E-81 | 2786 | 3469 | 3469 | 3469 | 1.03 | 1796 |
| 1.63E-77 | 2.12E-74 | 4543 | 3469 | 3469 | 3469 | 1.3 | 2450 |
| 6.99E-76 | 9.10E-73 | 1664 | 3469 | 3469 | 3469 | 1.17 | 1127 |
| 2.27E-75 | 2.96E-72 | 2706 | 3469 | 3469 | 3469 | 1.05 | 1721 |
| 1.29E-72 | 1.67E-69 | 4502 | 3469 | 3469 | 3469 | 2.34 | 1762 |
| 6.10E-70 | 7.94E-67 | 3083 | 3469 | 3469 | 3469 | 4.54 | 1348 |
| 1.93E-69 | 2.51E-66 | 2605 | 3469 | 3469 | 3469 | 1.43 | 1514 |
| 2.24E-69 | 2.91E-66 | 2109 | 3469 | 3469 | 3469 | 1.11 | 1354 |
| 5.29E-69 | 6.89E-66 | 1877 | 3469 | 3469 | 3469 | 1.23 | 1179 |
| 1.39E-67 | 1.80E-64 | 2352 | 3469 | 3469 | 3469 | 1.22 | 1478 |
| 6.64E-65 | 8.64E-62 | 2791 | 3469 | 3469 | 3469 | 3.01 | 1426 |
| 1.66E-64 | 2.15E-61 | 4656 | 3469 | 3469 | 3469 | 2.49 | 1775 |
| 1.06E-63 | 1.39E-60 | 2717 | 3469 | 3469 | 3469 | 2.21 | 1357 |
| 3.87E-63 | 5.04E-60 | 2896 | 3469 | 3469 | 3469 | 1.71 | 1414 |
| 4.67E-63 | 6.08E-60 | 3855 | 3469 | 3469 | 3469 | 1.28 | 2098 |
| 2.92E-59 | 3.80E-56 | 3458 | 3469 | 3469 | 3469 | 3.14 | 1499 |
| 6.63E-59 | 8.63E-56 | 3870 | 3469 | 3469 | 3469 | 1.52 | 1856 |
| 3.07E-58 | 3.99E-55 | 2696 | 3469 | 3469 | 3469 | 1.53 | 1579 |
| 1.32E-57 | 1.72E-54 | 1894 | 3469 | 3469 | 3469 | 1.28 | 1228 |
| 2.15E-57 | 2.79E-54 | 1804 | 3469 | 3469 | 3469 | 1.32 | 1090 |
| 3.04E-57 | 3.95E-54 | 1893 | 3469 | 3469 | 3469 | 5.43 | 893 |
| 5.73E-57 | 7.46E-54 | 1622 | 3469 | 3469 | 3469 | 4.13 | 779 |
| 4.87E-56 | 6.34E-53 | 1992 | 3469 | 3469 | 3469 | 1.45 | 1218 |
| 8.34E-56 | 1.09E-52 | 2280 | 3469 | 3469 | 3469 | 1.22 | 1400 |
| 8.40E-56 | 1.09E-52 | 3149 | 3469 | 3469 | 3469 | 1.09 | 1832 |
| 1.83E-55 | 2.38E-52 | 2166 | 3469 | 3469 | 3469 | 1.04 | 1392 |
| 9.01E-55 | 1.17E-51 | 2917 | 3469 | 3469 | 3469 | 4.62 | 1143 |
| 1.31E-54 | 1.71E-51 | 2154 | 3469 | 3469 | 3469 | 1.1 | 1374 |
| 2.20E-53 | 2.87E-50 | 2584 | 3469 | 3469 | 3469 | 2.67 | 1112 |
| 9.89E-53 | 1.29E-49 | 1442 | 3469 | 3469 | 3469 | 1.83 | 744 |
| 3.79E-52 | 4.94E-49 | 2923 | 3469 | 3469 | 3469 | 1.45 | 1645 |

| | | | | | | | |
|---|---|---|---|---|---|---|---|
| 5.89E-52 | 7.67E-49 | 2085 | 3469 | 3469 | 3469 | 1.51 | 1126 |
| 4.39E-51 | 5.72E-48 | 2666 | 3469 | 3469 | 3469 | 1.63 | 1318 |
| 1.20E-48 | 1.56E-45 | 872 | 3469 | 3469 | 3469 | 2.21 | 517 |
| 1.23E-48 | 1.60E-45 | 3286 | 3469 | 3469 | 3469 | 1.64 | 1403 |
| 8.03E-48 | 1.04E-44 | 2064 | 3469 | 3469 | 3469 | 6.73 | 530 |
| 9.88E-48 | 1.29E-44 | 798 | 3469 | 3469 | 3469 | 1.07 | 594 |
| 1.33E-47 | 1.73E-44 | 2582 | 3469 | 3469 | 3469 | 1.35 | 1486 |
| 1.19E-46 | 1.55E-43 | 2507 | 3469 | 3469 | 3469 | 1.35 | 1341 |
| 1.23E-46 | 1.61E-43 | 1611 | 3469 | 3469 | 3469 | 1.05 | 1051 |
| 1.86E-45 | 2.43E-42 | 3469 | 3469 | 3469 | 3469 | 2.26 | 1447 |
| 4.30E-45 | 5.59E-42 | 1814 | 3469 | 3469 | 3469 | 1.03 | 1172 |
| 1.23E-44 | 1.60E-41 | 1562 | 3469 | 3469 | 3469 | 1.05 | 1016 |
| 1.51E-44 | 1.96E-41 | 1755 | 3469 | 3469 | 3469 | 1.04 | 1138 |
| 3.23E-44 | 4.20E-41 | 2195 | 3469 | 3469 | 3469 | 1.84 | 1190 |
| 3.77E-44 | 4.91E-41 | 3635 | 3469 | 3469 | 3469 | 1.92 | 1477 |
| 2.62E-43 | 3.41E-40 | 1486 | 3469 | 3469 | 3469 | 1.63 | 808 |
| 1.67E-42 | 2.17E-39 | 3262 | 3469 | 3469 | 3469 | 2.09 | 858 |
| 4.27E-42 | 5.56E-39 | 2038 | 3469 | 3469 | 3469 | 3.53 | 841 |
| 1.54E-41 | 2.00E-38 | 1497 | 3469 | 3469 | 3469 | 2.2 | 769 |
| 4.16E-41 | 5.42E-38 | 4027 | 3469 | 3469 | 3469 | 1.54 | 1606 |
| 1.39E-40 | 1.81E-37 | 2080 | 3469 | 3469 | 3469 | 5.04 | 885 |
| 5.39E-40 | 7.02E-37 | 2915 | 3469 | 3469 | 3469 | 2.73 | 1185 |
| 6.62E-40 | 8.62E-37 | 3285 | 3469 | 3469 | 3469 | 2.04 | 1231 |
| 1.60E-39 | 2.08E-36 | 2434 | 3469 | 3469 | 3469 | 3.08 | 802 |
| 2.01E-39 | 2.62E-36 | 1423 | 3469 | 3469 | 3469 | 1.17 | 886 |
| 5.86E-39 | 7.63E-36 | 1285 | 3469 | 3469 | 3469 | 1.14 | 829 |
| 1.45E-38 | 1.89E-35 | 1734 | 3469 | 3469 | 3469 | 2.03 | 948 |
| 1.20E-37 | 1.56E-34 | 1839 | 3469 | 3469 | 3469 | 2.15 | 960 |
| 1.67E-37 | 2.17E-34 | 3631 | 3469 | 3469 | 3469 | 1.59 | 1454 |
| 3.16E-37 | 4.11E-34 | 2015 | 3469 | 3469 | 3469 | 6.72 | 728 |
| 7.75E-37 | 1.01E-33 | 1719 | 3469 | 3469 | 3469 | 2.01 | 762 |
| 1.89E-36 | 2.46E-33 | 3739 | 3469 | 3469 | 3469 | 2.62 | 1368 |
| 2.98E-36 | 3.87E-33 | 1399 | 3469 | 3469 | 3469 | 1.04 | 916 |
| 3.29E-36 | 4.29E-33 | 1381 | 3469 | 3469 | 3469 | 1.06 | 894 |
| 3.35E-36 | 4.37E-33 | 4113 | 3469 | 3469 | 3469 | 1.82 | 1699 |
| 1.27E-35 | 1.65E-32 | 2583 | 3469 | 3469 | 3469 | 8.54 | 658 |
| 1.32E-35 | 1.72E-32 | 2097 | 3469 | 3469 | 3469 | 1.04 | 1278 |
| 2.19E-35 | 2.85E-32 | 1759 | 3469 | 3469 | 3469 | 1.44 | 1061 |
| 2.44E-35 | 3.18E-32 | 1808 | 3469 | 3469 | 3469 | 2.06 | 884 |
| 5.99E-35 | 7.80E-32 | 1381 | 3469 | 3469 | 3469 | 1.05 | 896 |
| 1.31E-34 | 1.70E-31 | 1091 | 3469 | 3469 | 3469 | 1.81 | 543 |
| 1.28E-33 | 1.66E-30 | 2045 | 3469 | 3469 | 3469 | 1.27 | 1133 |
| 2.26E-33 | 2.94E-30 | 2662 | 3469 | 3469 | 3469 | 1.1 | 1499 |
| 4.49E-33 | 5.85E-30 | 3568 | 3469 | 3469 | 3469 | 1.54 | 1513 |
| 4.91E-33 | 6.39E-30 | 1677 | 3469 | 3469 | 3469 | 24.2 | 339 |
| 5.65E-33 | 7.36E-30 | 2380 | 3469 | 3469 | 3469 | 2.63 | 1117 |
| 7.39E-33 | 9.62E-30 | 1294 | 3469 | 3469 | 3469 | 1.2 | 771 |

| | | | | | | | |
|---|---|---|---|---|---|---|---|
| 1.24E-32 | 1.62E-29 | 981 | 3469 | 3469 | 3469 | 1.04 | 664 |
| 1.52E-32 | 1.97E-29 | 4467 | 3469 | 3469 | 3469 | 1.88 | 1811 |
| 2.10E-32 | 2.73E-29 | 2577 | 3469 | 3469 | 3469 | 1.06 | 1521 |
| 5.22E-32 | 6.80E-29 | 2953 | 3469 | 3469 | 3469 | 1.26 | 1374 |
| 1.15E-31 | 1.50E-28 | 3085 | 3469 | 3469 | 3469 | 2.02 | 1348 |
| 1.38E-31 | 1.79E-28 | 2072 | 3469 | 3469 | 3469 | 2.17 | 785 |
| 2.10E-31 | 2.73E-28 | 1990 | 3469 | 3469 | 3469 | 1.33 | 1073 |
| 4.51E-31 | 5.87E-28 | 2741 | 3469 | 3469 | 3469 | 1.02 | 1610 |
| 4.63E-31 | 6.02E-28 | 4378 | 3469 | 3469 | 3469 | 4.39 | 1201 |
| 8.60E-31 | 1.12E-27 | 1329 | 3469 | 3469 | 3469 | 1.48 | 835 |
| 1.75E-30 | 2.28E-27 | 2814 | 3469 | 3469 | 3469 | 1.06 | 1631 |
| 3.19E-30 | 4.15E-27 | 1171 | 3469 | 3469 | 3469 | 1.17 | 760 |
| 1.11E-29 | 1.44E-26 | 1912 | 3469 | 3469 | 3469 | 2.92 | 785 |
| 1.66E-29 | 2.16E-26 | 2148 | 3469 | 3469 | 3469 | 1.65 | 1130 |
| 2.28E-29 | 2.97E-26 | 2381 | 3469 | 3469 | 3469 | 1.12 | 1384 |
| 3.85E-29 | 5.01E-26 | 1813 | 3469 | 3469 | 3469 | 40.4 | 252 |
| 4.54E-29 | 5.92E-26 | 3877 | 3469 | 3469 | 3469 | 1.25 | 2050 |
| 1.14E-28 | 1.48E-25 | 859 | 3469 | 3469 | 3469 | 1.03 | 588 |
| 2.22E-28 | 2.89E-25 | 2544 | 3469 | 3469 | 3469 | 1.03 | 1486 |
| 2.22E-28 | 2.90E-25 | 1815 | 3469 | 3469 | 3469 | 1.02 | 1115 |
| 2.28E-28 | 2.97E-25 | 1363 | 3469 | 3469 | 3469 | 1.33 | 856 |
| 3.16E-28 | 4.12E-25 | 4204 | 3469 | 3469 | 3469 | 1.39 | 1873 |
| 6.64E-28 | 8.65E-25 | 770 | 3469 | 3469 | 3469 | 1.22 | 508 |
| 1.22E-27 | 1.59E-24 | 1717 | 3469 | 3469 | 3469 | 1.18 | 1010 |
| 1.35E-27 | 1.76E-24 | 1502 | 3469 | 3469 | 3469 | 5.54 | 323 |
| 2.55E-27 | 3.32E-24 | 1981 | 3469 | 3469 | 3469 | 2.48 | 712 |
| 3.73E-27 | 4.86E-24 | 736 | 3469 | 3469 | 3469 | 1.05 | 512 |
| 1.50E-26 | 1.95E-23 | 2813 | 3469 | 3469 | 3469 | 1.02 | 1637 |
| 2.10E-26 | 2.74E-23 | 2612 | 3469 | 3469 | 3469 | 1.13 | 1389 |
| 5.01E-26 | 6.52E-23 | 2080 | 3469 | 3469 | 3469 | 3.66 | 886 |
| 5.08E-26 | 6.62E-23 | 317 | 3469 | 3469 | 3469 | 197 | 97 |
| 8.83E-26 | 1.15E-22 | 1016 | 3469 | 3469 | 3469 | 1.2 | 627 |
| 1.22E-25 | 1.58E-22 | 3135 | 3469 | 3469 | 3469 | 5.2 | 648 |
| 5.73E-25 | 7.46E-22 | 3046 | 3469 | 3469 | 3469 | 2.04 | 1234 |
| 1.68E-24 | 2.19E-21 | 4804 | 3469 | 3469 | 3469 | 3.14 | 1014 |
| 1.94E-24 | 2.53E-21 | 1460 | 3469 | 3469 | 3469 | 1.16 | 852 |
| 3.20E-24 | 4.17E-21 | 4742 | 3469 | 3469 | 3469 | 1.34 | 2329 |
| 4.19E-24 | 5.46E-21 | 1854 | 3469 | 3469 | 3469 | 1.2 | 1031 |
| 8.85E-24 | 1.15E-20 | 1041 | 3469 | 3469 | 3469 | 1.26 | 649 |
| 9.00E-24 | 1.17E-20 | 1837 | 3469 | 3469 | 3469 | 1.53 | 855 |
| 9.01E-24 | 1.17E-20 | 3322 | 3469 | 3469 | 3469 | 1.02 | 1883 |
| 4.14E-23 | 5.38E-20 | 578 | 3469 | 3469 | 3469 | 1.66 | 404 |
| 1.18E-22 | 1.54E-19 | 2786 | 3469 | 3469 | 3469 | 3.28 | 872 |
| 1.02E-21 | 1.32E-18 | 3447 | 3469 | 3469 | 3469 | 5.86 | 781 |
| 1.21E-21 | 1.57E-18 | 3323 | 3469 | 3469 | 3469 | 2.07 | 1154 |
| 1.27E-21 | 1.66E-18 | 1439 | 3469 | 3469 | 3469 | 1.2 | 808 |
| 1.87E-21 | 2.43E-18 | 2132 | 3469 | 3469 | 3469 | 1.47 | 992 |

| | | | | | | | |
|---|---|---|---|---|---|---|---|
| 2.02E-21 | 2.63E-18 | 1609 | 3469 | 3469 | 3469 | 7.84 | 417 |
| 2.05E-21 | 2.67E-18 | 2743 | 3469 | 3469 | 3469 | 1.09 | 1495 |
| 2.38E-21 | 3.10E-18 | 2644 | 3469 | 3469 | 3469 | 1.46 | 1190 |
| 3.01E-21 | 3.92E-18 | 121 | 3469 | 3469 | 3469 | 1.24 | 111 |
| 4.11E-21 | 5.35E-18 | 5035 | 3469 | 3469 | 3469 | 1.28 | 2449 |
| 8.21E-21 | 1.07E-17 | 1431 | 3469 | 3469 | 3469 | 1.46 | 705 |
| 1.21E-20 | 1.58E-17 | 1807 | 3469 | 3469 | 3469 | 1.03 | 1086 |
| 2.25E-20 | 2.92E-17 | 971 | 3469 | 3469 | 3469 | 1.03 | 628 |
| 2.55E-20 | 3.32E-17 | 2927 | 3469 | 3469 | 3469 | 1.06 | 1662 |
| 6.07E-20 | 7.90E-17 | 1741 | 3469 | 3469 | 3469 | 1.03 | 1043 |
| 6.27E-20 | 8.17E-17 | 4005 | 3469 | 3469 | 3469 | 1.37 | 1951 |
| 1.05E-19 | 1.37E-16 | 2172 | 3469 | 3469 | 3469 | 1.86 | 877 |
| 2.92E-19 | 3.80E-16 | 2852 | 3469 | 3469 | 3469 | 8.9 | 509 |
| 7.07E-19 | 9.21E-16 | 2836 | 3469 | 3469 | 3469 | 2.2 | 1170 |
| 7.27E-19 | 9.47E-16 | 1470 | 3469 | 3469 | 3469 | 1.11 | 866 |
| 8.51E-19 | 1.11E-15 | 639 | 3469 | 3469 | 3469 | 1.18 | 433 |
| 1.09E-18 | 1.43E-15 | 290 | 3469 | 3469 | 3469 | 1.29 | 216 |
| 1.23E-18 | 1.60E-15 | 577 | 3469 | 3469 | 3469 | 8.33 | 179 |
| 1.27E-18 | 1.66E-15 | 1298 | 3469 | 3469 | 3469 | 1.12 | 788 |
| 1.97E-18 | 2.57E-15 | 1342 | 3469 | 3469 | 3469 | 1.11 | 819 |
| 2.60E-18 | 3.39E-15 | 2915 | 3469 | 3469 | 3469 | 1.47 | 1396 |
| 4.22E-18 | 5.49E-15 | 1435 | 3469 | 3469 | 3469 | 1.04 | 864 |
| 5.82E-18 | 7.58E-15 | 3554 | 3469 | 3469 | 3469 | 1.16 | 1791 |
| 8.07E-18 | 1.05E-14 | 3536 | 3469 | 3469 | 3469 | 1.98 | 1266 |
| 9.30E-18 | 1.21E-14 | 1600 | 3469 | 3469 | 3469 | 1.28 | 846 |
| 1.07E-17 | 1.39E-14 | 1522 | 3469 | 3469 | 3469 | 1.03 | 917 |
| 1.07E-17 | 1.40E-14 | 494 | 3469 | 3469 | 3469 | 1.09 | 344 |
| 1.18E-17 | 1.54E-14 | 2303 | 3469 | 3469 | 3469 | 1.33 | 1124 |
| 3.56E-17 | 4.64E-14 | 1905 | 3469 | 3469 | 3469 | 1.11 | 1113 |
| 5.68E-17 | 7.40E-14 | 990 | 3469 | 3469 | 3469 | 1.35 | 627 |
| 7.48E-17 | 9.73E-14 | 1360 | 3469 | 3469 | 3469 | 1.07 | 816 |
| 8.97E-17 | 1.17E-13 | 322 | 3469 | 3469 | 3469 | 1.12 | 238 |
| 1.02E-16 | 1.33E-13 | 2922 | 3469 | 3469 | 3469 | 2.34 | 832 |
| 2.42E-16 | 3.15E-13 | 1920 | 3469 | 3469 | 3469 | 7.24 | 396 |
| 3.89E-16 | 5.07E-13 | 3692 | 3469 | 3469 | 3469 | 1.02 | 2033 |
| 6.83E-16 | 8.89E-13 | 1007 | 3469 | 3469 | 3469 | 36.9 | 121 |
| 1.02E-15 | 1.33E-12 | 512 | 3469 | 3469 | 3469 | 1.07 | 350 |
| 1.68E-15 | 2.19E-12 | 1760 | 3469 | 3469 | 3469 | 1.08 | 1031 |
| 2.55E-15 | 3.32E-12 | 1876 | 3469 | 3469 | 3469 | 1.14 | 1034 |
| 3.09E-15 | 4.03E-12 | 1046 | 3469 | 3469 | 3469 | 1.13 | 645 |
| 4.46E-15 | 5.80E-12 | 2557 | 3469 | 3469 | 3469 | 1.16 | 1357 |
| 6.19E-15 | 8.06E-12 | 1369 | 3469 | 3469 | 3469 | 1.24 | 691 |
| 6.56E-15 | 8.54E-12 | 758 | 3469 | 3469 | 3469 | 1.22 | 469 |
| 9.49E-15 | 1.24E-11 | 3568 | 3469 | 3469 | 3469 | 1.14 | 1801 |
| 1.20E-14 | 1.56E-11 | 837 | 3469 | 3469 | 3469 | 3.01 | 417 |
| 1.61E-14 | 2.09E-11 | 3001 | 3469 | 3469 | 3469 | 1.4 | 1386 |
| 2.34E-14 | 3.04E-11 | 3252 | 3469 | 3469 | 3469 | 3.32 | 861 |

| | | | | | | | |
|---|---|---|---|---|---|---|---|
| 2.34E-14 | 3.05E-11 | 1762 | 3469 | 3469 | 3469 | 1.26 | 939 |
| 2.97E-14 | 3.87E-11 | 1774 | 3469 | 3469 | 3469 | 1.15 | 971 |
| 5.69E-14 | 7.40E-11 | 1179 | 3469 | 3469 | 3469 | 1.3 | 660 |
| 7.93E-14 | 1.03E-10 | 1493 | 3469 | 3469 | 3469 | 1.12 | 886 |
| 8.05E-14 | 1.05E-10 | 986 | 3469 | 3469 | 3469 | 1.36 | 547 |
| 1.47E-13 | 1.92E-10 | 2519 | 3469 | 3469 | 3469 | 4.78 | 376 |
| 1.66E-13 | 2.16E-10 | 795 | 3469 | 3469 | 3469 | 2.63 | 217 |
| 2.09E-13 | 2.72E-10 | 515 | 3469 | 3469 | 3469 | 1.24 | 337 |
| 2.14E-13 | 2.79E-10 | 886 | 3469 | 3469 | 3469 | 1.11 | 553 |
| 2.15E-13 | 2.79E-10 | 2226 | 3469 | 3469 | 3469 | 1.24 | 1228 |
| 2.73E-13 | 3.55E-10 | 3183 | 3469 | 3469 | 3469 | 2.34 | 1087 |
| 3.48E-13 | 4.53E-10 | 3278 | 3469 | 3469 | 3469 | 9.96 | 225 |
| 3.52E-13 | 4.58E-10 | 858 | 3469 | 3469 | 3469 | 1.31 | 526 |
| 4.36E-13 | 5.68E-10 | 428 | 3469 | 3469 | 3469 | 14.8 | 101 |
| 4.77E-13 | 6.21E-10 | 1193 | 3469 | 3469 | 3469 | 2.99 | 309 |
| 6.21E-13 | 8.08E-10 | 876 | 3469 | 3469 | 3469 | 1.03 | 534 |
| 1.16E-12 | 1.51E-09 | 4144 | 3469 | 3469 | 3469 | 2.45 | 1353 |
| 1.80E-12 | 2.34E-09 | 407 | 3469 | 3469 | 3469 | 20.6 | 87 |
| 2.10E-12 | 2.74E-09 | 3199 | 3469 | 3469 | 3469 | 1.34 | 1462 |
| 2.28E-12 | 2.97E-09 | 872 | 3469 | 3469 | 3469 | 1.02 | 544 |
| 6.11E-12 | 7.96E-09 | 588 | 3469 | 3469 | 3469 | 1.31 | 331 |
| 7.18E-12 | 9.35E-09 | 966 | 3469 | 3469 | 3469 | 1.06 | 589 |
| 9.96E-12 | 1.30E-08 | 2470 | 3469 | 3469 | 3469 | 1.02 | 1382 |
| 1.01E-11 | 1.31E-08 | 117 | 3469 | 3469 | 3469 | 6.68 | 84 |
| 1.08E-11 | 1.41E-08 | 2483 | 3469 | 3469 | 3469 | 1.13 | 1337 |
| 1.10E-11 | 1.44E-08 | 2355 | 3469 | 3469 | 3469 | 1.11 | 1280 |
| 1.60E-11 | 2.08E-08 | 2573 | 3469 | 3469 | 3469 | 1.01 | 1442 |
| 1.91E-11 | 2.48E-08 | 2569 | 3469 | 3469 | 3469 | 2.06 | 898 |
| 2.43E-11 | 3.16E-08 | 1739 | 3469 | 3469 | 3469 | 8.6 | 402 |
| 2.75E-11 | 3.58E-08 | 3017 | 3469 | 3469 | 3469 | 1.44 | 1401 |
| 2.76E-11 | 3.59E-08 | 617 | 3469 | 3469 | 3469 | 33.8 | 68 |
| 2.87E-11 | 3.74E-08 | 332 | 3469 | 3469 | 3469 | 1.08 | 230 |
| 3.03E-11 | 3.94E-08 | 2815 | 3469 | 3469 | 3469 | 1.18 | 1417 |
| 3.28E-11 | 4.26E-08 | 876 | 3469 | 3469 | 3469 | 2.36 | 305 |
| 1.03E-10 | 1.34E-07 | 3266 | 3469 | 3469 | 3469 | 1.61 | 1350 |
| 1.06E-10 | 1.38E-07 | 833 | 3469 | 3469 | 3469 | 2.14 | 351 |
| 1.21E-10 | 1.58E-07 | 597 | 3469 | 3469 | 3469 | 1.28 | 356 |
| 1.95E-10 | 2.54E-07 | 2538 | 3469 | 3469 | 3469 | 6.31 | 358 |
| 2.03E-10 | 2.65E-07 | 744 | 3469 | 3469 | 3469 | 54.7 | 65 |
| 2.98E-10 | 3.88E-07 | 4590 | 3469 | 3469 | 3469 | 1.69 | 1637 |
| 4.33E-10 | 5.63E-07 | 2061 | 3469 | 3469 | 3469 | 4.55 | 635 |
| 6.36E-10 | 8.28E-07 | 1868 | 3469 | 3469 | 3469 | 1.91 | 775 |
| 6.46E-10 | 8.42E-07 | 2719 | 3469 | 3469 | 3469 | 1.42 | 1225 |
| 7.37E-10 | 9.60E-07 | 846 | 3469 | 3469 | 3469 | 6.68 | 119 |
| 8.11E-10 | 1.06E-06 | 1104 | 3469 | 3469 | 3469 | 1.08 | 633 |
| 9.82E-10 | 1.28E-06 | 1086 | 3469 | 3469 | 3469 | 1.47 | 575 |
| 9.96E-10 | 1.30E-06 | 2588 | 3469 | 3469 | 3469 | 3.34 | 663 |

| | | | | | | | |
|---|---|---|---|---|---|---|---|
| 1.14E-09 | 1.48E-06 | 1223 | 3469 | 3469 | 3469 | 1.02 | 723 |
| 1.57E-09 | 2.04E-06 | 2674 | 3469 | 3469 | 3469 | 1.57 | 1017 |
| 1.86E-09 | 2.43E-06 | 2372 | 3469 | 3469 | 3469 | 1.12 | 1242 |
| 2.10E-09 | 2.73E-06 | 2333 | 3469 | 3469 | 3469 | 4.2 | 700 |
| 2.77E-09 | 3.61E-06 | 1251 | 3469 | 3469 | 3469 | 2.99 | 413 |
| 6.84E-09 | 8.90E-06 | 1450 | 3469 | 3469 | 3469 | 1.04 | 837 |
| 7.27E-09 | 9.46E-06 | 1355 | 3469 | 3469 | 3469 | 4.13 | 252 |
| 9.01E-09 | 1.17E-05 | 2214 | 3469 | 3469 | 3469 | 2.38 | 658 |
| 9.12E-09 | 1.19E-05 | 620 | 3469 | 3469 | 3469 | 1.07 | 388 |
| 9.19E-09 | 1.20E-05 | 395 | 3469 | 3469 | 3469 | 1.66 | 208 |
| 9.60E-09 | 1.25E-05 | 746 | 3469 | 3469 | 3469 | 1.02 | 457 |
| 1.23E-08 | 1.60E-05 | 678 | 3469 | 3469 | 3469 | 1.43 | 372 |
| 1.86E-08 | 2.42E-05 | 1386 | 3469 | 3469 | 3469 | 1.09 | 774 |
| 2.81E-08 | 3.65E-05 | 623 | 3469 | 3469 | 3469 | 1.25 | 371 |
| 2.81E-08 | 3.66E-05 | 643 | 3469 | 3469 | 3469 | 1.05 | 400 |
| 2.92E-08 | 3.80E-05 | 1120 | 3469 | 3469 | 3469 | 2.62 | 422 |
| 2.94E-08 | 3.83E-05 | 1173 | 3469 | 3469 | 3469 | 1.05 | 684 |
| 3.95E-08 | 5.14E-05 | 2543 | 3469 | 3469 | 3469 | 1.11 | 1333 |
| 4.28E-08 | 5.57E-05 | 490 | 3469 | 3469 | 3469 | 1.3 | 282 |
| 5.04E-08 | 6.56E-05 | 2492 | 3469 | 3469 | 3469 | 1.06 | 1364 |
| 8.46E-08 | 1.10E-04 | 3225 | 3469 | 3469 | 3469 | 1.15 | 1501 |
| 8.89E-08 | 1.16E-04 | 244 | 3469 | 3469 | 3469 | 1.16 | 168 |
| 1.00E-07 | 1.30E-04 | 1893 | 3469 | 3469 | 3469 | 1.26 | 900 |
| 1.00E-07 | 1.31E-04 | 1296 | 3469 | 3469 | 3469 | 1.64 | 487 |
| 1.24E-07 | 1.61E-04 | 2352 | 3469 | 3469 | 3469 | 1.86 | 976 |
| 1.29E-07 | 1.68E-04 | 471 | 3469 | 3469 | 3469 | 51.3 | 69 |
| 1.54E-07 | 2.01E-04 | 4169 | 3469 | 3469 | 3469 | 2.32 | 872 |
| 1.58E-07 | 2.06E-04 | 2641 | 3469 | 3469 | 3469 | 1.08 | 1418 |
| 2.10E-07 | 2.73E-04 | 817 | 3469 | 3469 | 3469 | 1.04 | 490 |
| 3.01E-07 | 3.93E-04 | 2658 | 3469 | 3469 | 3469 | 1.12 | 1390 |
| 3.20E-07 | 4.17E-04 | 987 | 3469 | 3469 | 3469 | 9.82 | 180 |
| 3.95E-07 | 5.14E-04 | 2974 | 3469 | 3469 | 3469 | 1.59 | 1211 |
| 4.31E-07 | 5.62E-04 | 1558 | 3469 | 3469 | 3469 | 1.19 | 813 |
| 4.32E-07 | 5.62E-04 | 2116 | 3469 | 3469 | 3469 | 1.04 | 1154 |
| 4.70E-07 | 6.11E-04 | 2586 | 3469 | 3469 | 3469 | 1.03 | 1411 |
| 5.21E-07 | 6.79E-04 | 535 | 3469 | 3469 | 3469 | 1.04 | 326 |
| 5.60E-07 | 7.30E-04 | 1456 | 3469 | 3469 | 3469 | 5.02 | 234 |
| 5.65E-07 | 7.36E-04 | 606 | 3469 | 3469 | 3469 | 1.03 | 370 |
| 6.63E-07 | 8.63E-04 | 1337 | 3469 | 3469 | 3469 | 1.15 | 712 |
| 6.69E-07 | 8.71E-04 | 114 | 3469 | 3469 | 3469 | 1.06 | 87 |
| 9.42E-07 | 1.23E-03 | 3657 | 3469 | 3469 | 3469 | 2.43 | 703 |
| 1.13E-06 | 1.47E-03 | 143 | 3469 | 3469 | 3469 | 4.26 | 52 |
| 1.14E-06 | 1.48E-03 | 1946 | 3469 | 3469 | 3469 | 1.61 | 851 |
| 1.17E-06 | 1.52E-03 | 2320 | 3469 | 3469 | 3469 | 1.31 | 1097 |
| 1.18E-06 | 1.54E-03 | 2297 | 3469 | 3469 | 3469 | 1.05 | 1266 |
| 1.23E-06 | 1.60E-03 | 286 | 3469 | 3469 | 3469 | 1.14 | 189 |
| 1.65E-06 | 2.14E-03 | 3293 | 3469 | 3469 | 3469 | 5.95 | 385 |

| | | | | | | | |
|---|---|---|---|---|---|---|---|
| 1.88E-06 | 2.45E-03 | 1883 | 3469 | 3469 | 3469 | 2.29 | 467 |
| 2.13E-06 | 2.77E-03 | 955 | 3469 | 3469 | 3469 | 1.1 | 527 |
| 2.46E-06 | 3.20E-03 | 2221 | 3469 | 3469 | 3469 | 3.4 | 501 |
| 2.47E-06 | 3.21E-03 | 1696 | 3469 | 3469 | 3469 | 1.02 | 951 |
| 2.50E-06 | 3.26E-03 | 190 | 3469 | 3469 | 3469 | 1.43 | 119 |
| 2.54E-06 | 3.31E-03 | 866 | 3469 | 3469 | 3469 | 4.92 | 219 |
| 2.60E-06 | 3.39E-03 | 1443 | 3469 | 3469 | 3469 | 1.19 | 819 |
| 2.83E-06 | 3.68E-03 | 1685 | 3469 | 3469 | 3469 | 1.13 | 890 |
| 3.36E-06 | 4.37E-03 | 1439 | 3469 | 3469 | 3469 | 1.83 | 508 |
| 3.41E-06 | 4.44E-03 | 1989 | 3469 | 3469 | 3469 | 2.93 | 686 |
| 3.69E-06 | 4.81E-03 | 1123 | 3469 | 3469 | 3469 | 1.09 | 629 |
| 4.59E-06 | 5.97E-03 | 738 | 3469 | 3469 | 3469 | 16.5 | 92 |
| 5.34E-06 | 6.96E-03 | 4615 | 3469 | 3469 | 3469 | 1.92 | 1488 |
| 5.49E-06 | 7.15E-03 | 3538 | 3469 | 3469 | 3469 | 2.98 | 811 |
| 6.13E-06 | 7.98E-03 | 322 | 3469 | 3469 | 3469 | 1.42 | 180 |
| 7.33E-06 | 9.54E-03 | 121 | 3469 | 3469 | 3469 | 2.71 | 73 |
| 1.08E-05 | 1.41E-02 | 1832 | 3469 | 3469 | 3469 | 15.6 | 61 |
| 1.12E-05 | 1.46E-02 | 310 | 3469 | 3469 | 3469 | 1.23 | 171 |
| 1.23E-05 | 1.60E-02 | 788 | 3469 | 3469 | 3469 | 1.41 | 370 |
| 1.37E-05 | 1.78E-02 | 1622 | 3469 | 3469 | 3469 | 3.92 | 230 |
| 1.76E-05 | 2.30E-02 | 3694 | 3469 | 3469 | 3469 | 7 | 536 |
| 1.86E-05 | 2.42E-02 | 1041 | 3469 | 3469 | 3469 | 4.73 | 202 |
| 1.98E-05 | 2.58E-02 | 1438 | 3469 | 3469 | 3469 | 3.12 | 414 |
| 2.12E-05 | 2.76E-02 | 1289 | 3469 | 3469 | 3469 | 2.03 | 511 |
| 2.54E-05 | 3.31E-02 | 2824 | 3469 | 3469 | 3469 | 2.22 | 984 |
| 3.14E-05 | 4.08E-02 | 3473 | 3469 | 3469 | 3469 | 1.93 | 1153 |
| 4.43E-05 | 5.77E-02 | 104 | 3469 | 3469 | 3469 | 2.44 | 58 |
| 5.02E-05 | 6.53E-02 | 1059 | 3469 | 3469 | 3469 | 2.93 | 295 |
| 5.07E-05 | 6.60E-02 | 2559 | 3469 | 3469 | 3469 | 12.5 | 215 |
| 5.90E-05 | 7.69E-02 | 122 | 3469 | 3469 | 3469 | 1.3 | 83 |
| 6.21E-05 | 8.09E-02 | 2661 | 3469 | 3469 | 3469 | 30.8 | 50 |
| 6.30E-05 | 8.20E-02 | 2628 | 3469 | 3469 | 3469 | 6.6 | 97 |
| 6.48E-05 | 8.44E-02 | 3463 | 3469 | 3469 | 3469 | 1.46 | 1484 |
| 7.50E-05 | 9.77E-02 | 3500 | 3469 | 3469 | 3469 | 2.53 | 934 |
| 9.38E-05 | 1.22E-01 | 2004 | 3469 | 3469 | 3469 | 2.11 | 555 |
| 1.15E-04 | 1.50E-01 | 375 | 3469 | 3469 | 3469 | 9.45 | 71 |
| 1.30E-04 | 1.70E-01 | 1897 | 3469 | 3469 | 3469 | 2.19 | 639 |
| 1.50E-04 | 1.95E-01 | 2589 | 3469 | 3469 | 3469 | 1.74 | 963 |
| 1.61E-04 | 2.09E-01 | 980 | 3469 | 3469 | 3469 | 1.34 | 457 |
| 1.70E-04 | 2.22E-01 | 5841 | 3469 | 3469 | 3469 | 1.6 | 2066 |
| 1.88E-04 | 2.45E-01 | 2813 | 3469 | 3469 | 3469 | 1.04 | 1491 |
| 1.94E-04 | 2.53E-01 | 983 | 3469 | 3469 | 3469 | 1.16 | 540 |
| 2.36E-04 | 3.07E-01 | 4021 | 3469 | 3469 | 3469 | 6.16 | 336 |
| 2.53E-04 | 3.30E-01 | 1372 | 3469 | 3469 | 3469 | 4.59 | 263 |
| 2.54E-04 | 3.31E-01 | 1294 | 3469 | 3469 | 3469 | 1.05 | 715 |
| 2.58E-04 | 3.36E-01 | 1797 | 3469 | 3469 | 3469 | 1.11 | 961 |
| 2.88E-04 | 3.75E-01 | 1792 | 3469 | 3469 | 3469 | 2.32 | 556 |

| | | | | | | | |
|---|---|---|---|---|---|---|---|
| 3.21E-04 | 4.18E-01 | 927 | 3469 | 3469 | 3469 | 2.14 | 335 |
| 3.35E-04 | 4.36E-01 | 2596 | 3469 | 3469 | 3469 | 1.09 | 1302 |
| 4.48E-04 | 5.84E-01 | 285 | 3469 | 3469 | 3469 | 1.28 | 166 |
| 4.80E-04 | 6.25E-01 | 528 | 3469 | 3469 | 3469 | 33.6 | 41 |
| 5.03E-04 | 6.54E-01 | 2771 | 3469 | 3469 | 3469 | 1.33 | 1241 |
| 5.38E-04 | 7.01E-01 | 1355 | 3469 | 3469 | 3469 | 1.65 | 520 |
| 5.43E-04 | 7.07E-01 | 56 | 3469 | 3469 | 3469 | 1.11 | 44 |
| 5.61E-04 | 7.31E-01 | 1497 | 3469 | 3469 | 3469 | 1.13 | 792 |
| 6.47E-04 | 8.42E-01 | 808 | 3469 | 3469 | 3469 | 1.18 | 461 |
| 6.82E-04 | 8.88E-01 | 1683 | 3469 | 3469 | 3469 | 1.06 | 930 |
| 7.13E-04 | 9.28E-01 | 708 | 3469 | 3469 | 3469 | 2.51 | 232 |
| 7.51E-04 | 9.78E-01 | 744 | 3469 | 3469 | 3469 | 1.07 | 432 |
| 7.99E-04 | 1.04E+00 | 2789 | 3469 | 3469 | 3469 | 2.06 | 820 |
| 8.13E-04 | 1.06E+00 | 1850 | 3469 | 3469 | 3469 | 1.02 | 1016 |
| 8.31E-04 | 1.08E+00 | 175 | 3469 | 3469 | 3469 | 1.91 | 92 |
| 1.07E-03 | 1.40E+00 | 192 | 3469 | 3469 | 3469 | 13.6 | 47 |
| 1.13E-03 | 1.47E+00 | 1805 | 3469 | 3469 | 3469 | 1.08 | 975 |
| 1.19E-03 | 1.55E+00 | 1549 | 3469 | 3469 | 3469 | 1.76 | 475 |
| 1.42E-03 | 1.85E+00 | 575 | 3469 | 3469 | 3469 | 1.68 | 270 |
| 1.50E-03 | 1.95E+00 | 4185 | 3469 | 3469 | 3469 | 4.8 | 689 |
| 1.73E-03 | 2.25E+00 | 749 | 3469 | 3469 | 3469 | 1.34 | 354 |
| 1.78E-03 | 2.32E+00 | 1664 | 3469 | 3469 | 3469 | 1.04 | 910 |
| 1.98E-03 | 2.58E+00 | 1509 | 3469 | 3469 | 3469 | 1.22 | 676 |
| 2.07E-03 | 2.70E+00 | 1192 | 3469 | 3469 | 3469 | 1.79 | 501 |
| 2.14E-03 | 2.78E+00 | 962 | 3469 | 3469 | 3469 | 1.04 | 530 |
| 2.54E-03 | 3.31E+00 | 43 | 3469 | 3469 | 3469 | 28.4 | 20 |
| 2.63E-03 | 3.42E+00 | 1652 | 3469 | 3469 | 3469 | 1.46 | 678 |
| 2.96E-03 | 3.86E+00 | 608 | 3469 | 3469 | 3469 | 1.12 | 350 |
| 3.06E-03 | 3.98E+00 | 562 | 3469 | 3469 | 3469 | 1.23 | 320 |
| 3.37E-03 | 4.39E+00 | 1353 | 3469 | 3469 | 3469 | 1.08 | 710 |
| 3.50E-03 | 4.56E+00 | 229 | 3469 | 3469 | 3469 | 12.9 | 50 |
| 3.68E-03 | 4.79E+00 | 71 | 3469 | 3469 | 3469 | 1.08 | 52 |
| 3.73E-03 | 4.86E+00 | 1693 | 3469 | 3469 | 3469 | 1.98 | 615 |
| 3.96E-03 | 5.16E+00 | 370 | 3469 | 3469 | 3469 | 6.03 | 101 |
| 4.34E-03 | 5.65E+00 | 22 | 3469 | 3469 | 3469 | 2.38 | 18 |
| 4.75E-03 | 6.18E+00 | 472 | 3469 | 3469 | 3469 | 1.27 | 280 |
| 5.33E-03 | 6.94E+00 | 487 | 3469 | 3469 | 3469 | 2.73 | 174 |
| 5.38E-03 | 7.00E+00 | 1002 | 3469 | 3469 | 3469 | 1.15 | 521 |
| 5.66E-03 | 7.37E+00 | 472 | 3469 | 3469 | 3469 | 37.4 | 36 |
| 5.68E-03 | 7.39E+00 | 761 | 3469 | 3469 | 3469 | 347 | 17 |
| 5.88E-03 | 7.65E+00 | 1761 | 3469 | 3469 | 3469 | 1.03 | 960 |
| 6.56E-03 | 8.54E+00 | 1149 | 3469 | 3469 | 3469 | 4.94 | 296 |

0.0 --control --shuffle-- --kmer 2 YAP_chip_seq.fasta db/HUMAN/HOCOMOCOv11_full_HUMAN_mono_m

| %TP | FP | %FP |
|---|---|---|
| 69.1 | 630 | 18.16 |
| 71.61 | 768 | 22.14 |
| 61.46 | 520 | 14.99 |
| 39.52 | 307 | 8.85 |
| 39.58 | 310 | 8.94 |
| 40.99 | 346 | 9.97 |
| 40.76 | 341 | 9.83 |
| 36.06 | 274 | 7.9 |
| 38.83 | 336 | 9.69 |
| 48.43 | 573 | 16.52 |
| 66.73 | 1327 | 38.25 |
| 62.55 | 1240 | 35.75 |
| 33.61 | 432 | 12.45 |
| 53.27 | 988 | 28.48 |
| 73.08 | 1702 | 49.06 |
| 51.77 | 990 | 28.54 |
| 70.63 | 1674 | 48.26 |
| 32.49 | 474 | 13.66 |
| 49.61 | 963 | 27.76 |
| 50.79 | 1012 | 29.17 |
| 38.86 | 668 | 19.26 |
| 43.64 | 809 | 23.32 |
| 39.03 | 676 | 19.49 |
| 33.99 | 539 | 15.54 |
| 42.61 | 787 | 22.69 |
| 41.11 | 755 | 21.76 |
| 51.17 | 1064 | 30.67 |
| 39.12 | 703 | 20.27 |
| 40.76 | 753 | 21.71 |
| 60.48 | 1383 | 39.87 |
| 43.21 | 844 | 24.33 |
| 53.5 | 1170 | 33.73 |
| 45.52 | 920 | 26.52 |
| 35.4 | 626 | 18.05 |
| 31.42 | 518 | 14.93 |
| 25.74 | 372 | 10.72 |
| 22.46 | 294 | 8.48 |
| 35.11 | 625 | 18.02 |
| 40.36 | 777 | 22.4 |
| 52.81 | 1165 | 33.58 |
| 40.13 | 772 | 22.25 |
| 32.95 | 570 | 16.43 |
| 39.61 | 761 | 21.94 |
| 32.06 | 552 | 15.91 |
| 21.45 | 285 | 8.22 |
| 47.42 | 1012 | 29.17 |

| | | |
|---|---|---|
| 32.46 | 570 | 16.43 |
| 37.99 | 731 | 21.07 |
| 14.9 | 154 | 4.44 |
| 40.44 | 816 | 23.52 |
| 15.28 | 163 | 4.7 |
| 17.12 | 204 | 5.88 |
| 42.84 | 895 | 25.8 |
| 38.66 | 774 | 22.31 |
| 30.3 | 536 | 15.45 |
| 41.71 | 872 | 25.14 |
| 33.78 | 641 | 18.48 |
| 29.29 | 518 | 14.93 |
| 32.8 | 616 | 17.76 |
| 34.3 | 660 | 19.03 |
| 42.58 | 906 | 26.12 |
| 23.29 | 366 | 10.55 |
| 24.73 | 405 | 11.67 |
| 24.24 | 395 | 11.39 |
| 22.17 | 345 | 9.95 |
| 46.3 | 1041 | 30.01 |
| 25.51 | 435 | 12.54 |
| 34.16 | 678 | 19.54 |
| 35.49 | 717 | 20.67 |
| 23.12 | 377 | 10.87 |
| 25.54 | 442 | 12.74 |
| 23.9 | 401 | 11.56 |
| 27.33 | 494 | 14.24 |
| 27.67 | 508 | 14.64 |
| 41.91 | 926 | 26.69 |
| 20.99 | 333 | 9.6 |
| 21.97 | 360 | 10.38 |
| 39.43 | 856 | 24.68 |
| 26.41 | 481 | 13.87 |
| 25.77 | 464 | 13.38 |
| 48.98 | 1160 | 33.44 |
| 18.97 | 289 | 8.33 |
| 36.84 | 784 | 22.6 |
| 30.59 | 602 | 17.35 |
| 25.48 | 460 | 13.26 |
| 25.83 | 472 | 13.61 |
| 15.65 | 215 | 6.2 |
| 32.66 | 672 | 19.37 |
| 43.21 | 995 | 28.68 |
| 43.61 | 1009 | 29.09 |
| 9.77 | 93 | 2.68 |
| 32.2 | 662 | 19.08 |
| 22.23 | 386 | 11.13 |

| | | |
|---|---|---|
| 19.14 | 308 | 8.88 |
| 52.21 | 1293 | 37.27 |
| 43.85 | 1022 | 29.46 |
| 39.61 | 891 | 25.68 |
| 38.86 | 870 | 25.08 |
| 22.63 | 402 | 11.59 |
| 30.93 | 635 | 18.3 |
| 46.41 | 1114 | 32.11 |
| 34.62 | 744 | 21.45 |
| 24.07 | 446 | 12.86 |
| 47.02 | 1138 | 32.8 |
| 21.91 | 391 | 11.27 |
| 22.63 | 412 | 11.88 |
| 32.57 | 695 | 20.03 |
| 39.9 | 919 | 26.49 |
| 7.26 | 56 | 1.61 |
| 59.09 | 1557 | 44.88 |
| 16.95 | 271 | 7.81 |
| 42.84 | 1019 | 29.37 |
| 32.14 | 690 | 19.89 |
| 24.68 | 476 | 13.72 |
| 53.99 | 1387 | 39.98 |
| 14.64 | 218 | 6.28 |
| 29.12 | 606 | 17.47 |
| 9.31 | 99 | 2.85 |
| 20.52 | 368 | 10.61 |
| 14.76 | 224 | 6.46 |
| 47.19 | 1174 | 33.84 |
| 40.04 | 945 | 27.24 |
| 25.54 | 513 | 14.79 |
| 2.8 | 1 | 0.03 |
| 18.07 | 313 | 9.02 |
| 18.68 | 327 | 9.43 |
| 35.57 | 816 | 23.52 |
| 29.23 | 627 | 18.07 |
| 24.56 | 496 | 14.3 |
| 67.14 | 1886 | 54.37 |
| 29.72 | 647 | 18.65 |
| 18.71 | 340 | 9.8 |
| 24.65 | 502 | 14.47 |
| 54.28 | 1435 | 41.37 |
| 11.65 | 167 | 4.81 |
| 25.14 | 522 | 15.05 |
| 22.51 | 453 | 13.06 |
| 33.27 | 770 | 22.2 |
| 23.29 | 478 | 13.78 |
| 28.6 | 632 | 18.22 |

| | | |
|---|---|---|
| 12.02 | 181 | 5.22 |
| 43.1 | 1083 | 31.22 |
| 34.3 | 805 | 23.21 |
| 3.2 | 10 | 0.29 |
| 70.6 | 2042 | 58.86 |
| 20.32 | 400 | 11.53 |
| 31.31 | 720 | 20.76 |
| 18.1 | 343 | 9.89 |
| 47.91 | 1251 | 36.06 |
| 30.07 | 688 | 19.83 |
| 56.24 | 1537 | 44.31 |
| 25.28 | 547 | 15.77 |
| 14.67 | 256 | 7.38 |
| 33.73 | 807 | 23.26 |
| 24.96 | 545 | 15.71 |
| 12.48 | 206 | 5.94 |
| 6.23 | 65 | 1.87 |
| 5.16 | 44 | 1.27 |
| 22.72 | 482 | 13.89 |
| 23.61 | 509 | 14.67 |
| 40.24 | 1018 | 29.35 |
| 24.91 | 549 | 15.83 |
| 51.63 | 1398 | 40.3 |
| 36.49 | 902 | 26 |
| 24.39 | 536 | 15.45 |
| 26.43 | 597 | 17.21 |
| 9.92 | 149 | 4.3 |
| 32.4 | 777 | 22.4 |
| 32.08 | 772 | 22.25 |
| 18.07 | 363 | 10.46 |
| 23.52 | 518 | 14.93 |
| 6.86 | 84 | 2.42 |
| 23.98 | 530 | 15.28 |
| 11.42 | 189 | 5.45 |
| 58.6 | 1658 | 47.79 |
| 3.49 | 21 | 0.61 |
| 10.09 | 162 | 4.67 |
| 29.72 | 714 | 20.58 |
| 29.81 | 718 | 20.7 |
| 18.59 | 389 | 11.21 |
| 39.12 | 1013 | 29.2 |
| 19.92 | 428 | 12.34 |
| 13.52 | 253 | 7.29 |
| 51.92 | 1441 | 41.54 |
| 12.02 | 215 | 6.2 |
| 39.95 | 1045 | 30.12 |
| 24.82 | 573 | 16.52 |

| | | |
|---|---|---|
| 27.07 | 643 | 18.54 |
| 27.99 | 672 | 19.37 |
| 19.03 | 410 | 11.82 |
| 25.54 | 602 | 17.35 |
| 15.77 | 320 | 9.22 |
| 10.84 | 188 | 5.42 |
| 6.26 | 81 | 2.33 |
| 9.71 | 164 | 4.73 |
| 15.94 | 328 | 9.46 |
| 35.4 | 911 | 26.26 |
| 31.33 | 782 | 22.54 |
| 6.49 | 85 | 2.45 |
| 15.16 | 308 | 8.88 |
| 2.91 | 18 | 0.52 |
| 8.91 | 144 | 4.15 |
| 15.39 | 316 | 9.11 |
| 39 | 1032 | 29.75 |
| 2.51 | 13 | 0.37 |
| 42.14 | 1139 | 32.83 |
| 15.68 | 328 | 9.46 |
| 9.54 | 167 | 4.81 |
| 16.98 | 368 | 10.61 |
| 39.84 | 1072 | 30.9 |
| 2.42 | 14 | 0.4 |
| 38.54 | 1030 | 29.69 |
| 36.9 | 977 | 28.16 |
| 41.57 | 1131 | 32.6 |
| 25.89 | 632 | 18.22 |
| 11.59 | 221 | 6.37 |
| 40.39 | 1094 | 31.54 |
| 1.96 | 7 | 0.2 |
| 6.63 | 100 | 2.88 |
| 40.85 | 1110 | 32 |
| 8.79 | 151 | 4.35 |
| 38.92 | 1052 | 30.33 |
| 10.12 | 188 | 5.42 |
| 10.26 | 193 | 5.56 |
| 10.32 | 192 | 5.53 |
| 1.87 | 7 | 0.2 |
| 47.19 | 1331 | 38.37 |
| 18.3 | 418 | 12.05 |
| 22.34 | 540 | 15.57 |
| 35.31 | 945 | 27.24 |
| 3.43 | 34 | 0.98 |
| 18.25 | 421 | 12.14 |
| 16.58 | 373 | 10.75 |
| 19.11 | 444 | 12.8 |

| | | |
|---|---|---|
| 20.84 | 499 | 14.38 |
| 29.32 | 758 | 21.85 |
| 35.8 | 967 | 27.88 |
| 20.18 | 479 | 13.81 |
| 11.91 | 244 | 7.03 |
| 24.13 | 606 | 17.47 |
| 7.26 | 124 | 3.57 |
| 18.97 | 449 | 12.94 |
| 11.18 | 230 | 6.63 |
| 6 | 96 | 2.77 |
| 13.17 | 285 | 8.22 |
| 10.72 | 218 | 6.28 |
| 22.31 | 555 | 16 |
| 10.69 | 220 | 6.34 |
| 11.53 | 243 | 7 |
| 12.16 | 259 | 7.47 |
| 19.72 | 479 | 13.81 |
| 38.43 | 1069 | 30.82 |
| 8.13 | 153 | 4.41 |
| 39.32 | 1100 | 31.71 |
| 43.27 | 1233 | 35.54 |
| 4.84 | 74 | 2.13 |
| 25.94 | 674 | 19.43 |
| 14.04 | 316 | 9.11 |
| 28.13 | 743 | 21.42 |
| 1.99 | 14 | 0.4 |
| 25.14 | 647 | 18.65 |
| 40.88 | 1158 | 33.38 |
| 14.13 | 323 | 9.31 |
| 40.07 | 1135 | 32.72 |
| 5.19 | 82 | 2.36 |
| 34.91 | 966 | 27.85 |
| 23.44 | 604 | 17.41 |
| 33.27 | 915 | 26.38 |
| 40.67 | 1158 | 33.38 |
| 9.4 | 194 | 5.59 |
| 6.75 | 121 | 3.49 |
| 10.67 | 229 | 6.6 |
| 20.52 | 517 | 14.9 |
| 2.51 | 27 | 0.78 |
| 20.27 | 506 | 14.59 |
| 1.5 | 9 | 0.26 |
| 24.53 | 642 | 18.51 |
| 31.62 | 867 | 24.99 |
| 36.49 | 1026 | 29.58 |
| 5.45 | 94 | 2.71 |
| 11.1 | 239 | 6.89 |

| | | |
|---|---|---|
| 13.46 | 309 | 8.91 |
| 15.19 | 363 | 10.46 |
| 14.44 | 338 | 9.74 |
| 27.41 | 738 | 21.27 |
| 3.43 | 48 | 1.38 |
| 6.31 | 115 | 3.32 |
| 23.61 | 619 | 17.84 |
| 25.66 | 683 | 19.69 |
| 14.64 | 347 | 10 |
| 19.78 | 500 | 14.41 |
| 18.13 | 453 | 13.06 |
| 2.65 | 30 | 0.86 |
| 42.89 | 1244 | 35.86 |
| 23.38 | 611 | 17.61 |
| 5.19 | 91 | 2.62 |
| 2.1 | 22 | 0.63 |
| 1.76 | 13 | 0.37 |
| 4.93 | 86 | 2.48 |
| 10.67 | 239 | 6.89 |
| 6.63 | 126 | 3.63 |
| 15.45 | 374 | 10.78 |
| 5.82 | 107 | 3.08 |
| 11.93 | 275 | 7.93 |
| 14.73 | 358 | 10.32 |
| 28.37 | 779 | 22.46 |
| 33.24 | 937 | 27.01 |
| 1.67 | 16 | 0.46 |
| 8.5 | 182 | 5.25 |
| 6.2 | 117 | 3.37 |
| 2.39 | 31 | 0.89 |
| 1.44 | 9 | 0.26 |
| 2.8 | 35 | 1.01 |
| 42.78 | 1259 | 36.29 |
| 26.92 | 738 | 21.27 |
| 16 | 401 | 11.56 |
| 2.05 | 23 | 0.66 |
| 18.42 | 477 | 13.75 |
| 27.76 | 771 | 22.23 |
| 13.17 | 322 | 9.28 |
| 59.56 | 1841 | 53.07 |
| 42.98 | 1275 | 36.75 |
| 15.57 | 395 | 11.39 |
| 9.69 | 216 | 6.23 |
| 7.58 | 161 | 4.64 |
| 20.61 | 551 | 15.88 |
| 27.7 | 775 | 22.34 |
| 16.03 | 408 | 11.76 |

| | | |
|---|---|---|
| 9.66 | 222 | 6.4 |
| 37.53 | 1097 | 31.62 |
| 4.79 | 92 | 2.65 |
| 1.18 | 8 | 0.23 |
| 35.77 | 1041 | 30.01 |
| 14.99 | 381 | 10.98 |
| 1.27 | 12 | 0.35 |
| 22.83 | 625 | 18.02 |
| 13.29 | 333 | 9.6 |
| 26.81 | 753 | 21.71 |
| 6.69 | 142 | 4.09 |
| 12.45 | 309 | 8.91 |
| 23.64 | 649 | 18.71 |
| 29.29 | 834 | 24.04 |
| 2.65 | 41 | 1.18 |
| 1.35 | 13 | 0.37 |
| 28.11 | 798 | 23 |
| 13.69 | 345 | 9.95 |
| 7.78 | 176 | 5.07 |
| 19.86 | 531 | 15.31 |
| 10.2 | 246 | 7.09 |
| 26.23 | 741 | 21.36 |
| 19.49 | 527 | 15.19 |
| 14.44 | 372 | 10.72 |
| 15.28 | 399 | 11.5 |
| 0.58 | 2 | 0.06 |
| 19.54 | 530 | 15.28 |
| 10.09 | 246 | 7.09 |
| 9.22 | 221 | 6.37 |
| 20.47 | 562 | 16.2 |
| 1.44 | 16 | 0.46 |
| 1.5 | 19 | 0.55 |
| 17.73 | 475 | 13.69 |
| 2.91 | 49 | 1.41 |
| 0.52 | 2 | 0.06 |
| 8.07 | 190 | 5.48 |
| 5.02 | 104 | 3 |
| 15.02 | 396 | 11.42 |
| 1.04 | 8 | 0.23 |
| 0.49 | 0 | 0 |
| 27.67 | 796 | 22.95 |
| 8.53 | 201 | 5.79 |

ıeme_format.meme db/MOUSE/HOCOMOCOv11_full_MOUSE_mono_meme_format.meme

| rank | motif_ID | consensus | p-value |
|---|---|---|---|
| 1 | TEAD4_HUMAN.H11MO.0.A | RCATTCCWGRVAK | 1.36E-235 |
| 2 | TEAD1_HUMAN.H11MO.0.A | RCATTCCWGSCW | 1.41E-216 |
| 3 | TEAD2_HUMAN.H11MO.0.D | HYACATTCCWGG | 2.42E-205 |
| 4 | MITF_HUMAN.H11MO.0.A | TCACGTGACC | 2.05E-172 |
| 5 | TFE3_HUMAN.H11MO.0.B | RGTCACGTGA | 8.03E-155 |
| 6 | FOSL1_HUMAN.H11MO.0.A | KRVTGAGTCAYH | 2.52E-153 |
| 7 | FOSL2_HUMAN.H11MO.0.A | NDRTGAGTCAYH | 5.02E-151 |
| 8 | FOS_HUMAN.H11MO.0.A | VTGAGTCAB | 1.20E-148 |
| 9 | USF1_HUMAN.H11MO.0.A | RGTCACGTGRBS | 2.56E-145 |
| 10 | JUND_HUMAN.H11MO.0.A | RRTGAGTCAYY | 2.24E-143 |
| 11 | JUN_HUMAN.H11MO.0.A | DRTGAGTCAYH | 2.16E-142 |
| 12 | MLX_HUMAN.H11MO.0.D | YGATCACGTGN | 1.98E-140 |
| 13 | FOSB_HUMAN.H11MO.0.A | RTGAGTCAB | 4.13E-139 |
| 14 | USF2_HUMAN.H11MO.0.A | GGGTCACGTGRBSVSSSB | 7.91E-131 |
| 15 | JUNB_HUMAN.H11MO.0.A | DVTGAGTCABH | 1.36E-125 |
| 16 | BHE40_HUMAN.H11MO.0.A | DGCACGTGAS | 1.02E-120 |
| 17 | BMAL1_HUMAN.H11MO.0.A | KSCACGTGHSY | 7.18E-117 |
| 18 | TFEB_HUMAN.H11MO.0.C | RGTCACGTG | 6.07E-105 |
| 19 | MYCN_HUMAN.H11MO.0.A | RGCCACGTGSDS | 8.88E-90 |
| 20 | ETV6_HUMAN.H11MO.0.D | RCAGGAARK | 4.43E-89 |
| 21 | ETV5_HUMAN.H11MO.0.C | RRRSAGGAARDGRV | 1.20E-87 |
| 22 | ETV4_HUMAN.H11MO.0.B | VRCAGGAARBR | 5.34E-85 |
| 23 | MAX_HUMAN.H11MO.0.A | VRGCACGTGG | 1.53E-83 |
| 24 | ZBT17_HUMAN.H11MO.0.A | SRRGGWGGGGGAGGGGMRR | 8.73E-84 |
| 25 | ATF3_HUMAN.H11MO.0.A | GGTSACGTGAB | 4.03E-82 |
| 26 | FLI1_HUMAN.H11MO.0.A | SRRGGMAGGAAGGRRRGR | 9.12E-80 |
| 27 | ERG_HUMAN.H11MO.0.A | RRRSAGGAAGYGG | 1.23E-78 |
| 28 | VEZF1_HUMAN.H11MO.0.C | GGRRRRGRRGGAGGGGGRGRRR | 4.56E-77 |
| 29 | MAZ_HUMAN.H11MO.0.A | GGGGGAGGGGGDGRGRRRGRRG | 7.80E-77 |
| 30 | FLI1_HUMAN.H11MO.1.A | RRRCAGGAAGBRG | 5.06E-76 |
| 31 | HEY1_HUMAN.H11MO.0.D | KGRCACGTGBCK | 2.07E-74 |
| 32 | CR3L2_HUMAN.H11MO.0.D | TGATGACGTGGCA | 6.85E-74 |
| 33 | CLOCK_HUMAN.H11MO.0.C | CWGHCACGTGMVVM | 3.61E-74 |
| 34 | ZN770_HUMAN.H11MO.0.C | GGGAGGCYGAGGBRGGAGGATC | 2.55E-71 |
| 35 | EGR2_HUMAN.H11MO.0.A | GRGRRKGWGKGGGHGGRG | 8.35E-71 |
| 36 | ETS2_HUMAN.H11MO.0.B | RDVRGGAARVRRR | 6.02E-71 |
| 37 | E2F7_HUMAN.H11MO.0.B | GDGGCGGGAARDR | 1.36E-70 |
| 38 | IRF3_HUMAN.H11MO.0.B | RAAARGGAAAVDGAAASDGA | 4.28E-70 |
| 39 | ZN394_HUMAN.H11MO.1.D | GAAWGGAAW | 4.70E-70 |
| 40 | BC11A_HUMAN.H11MO.0.A | RAAAGAGGAAGTGARAV | 5.99E-70 |
| 41 | ZN467_HUMAN.H11MO.0.C | GGGGAGGGGRRGGRGRRGGRRR | 1.72E-68 |
| 42 | KLF15_HUMAN.H11MO.0.A | RGGGMGGRGVDGGGGGRGG | 1.67E-65 |
| 43 | WT1_HUMAN.H11MO.0.C | RGGVGGGGGAGGRGGVGGRG | 1.88E-63 |
| 44 | ETV7_HUMAN.H11MO.0.D | VAGGAAR | 1.21E-63 |
| 45 | MYC_HUMAN.H11MO.0.A | RCCACGTGCTB | 4.60E-62 |
| 46 | NFAC1_HUMAN.H11MO.0.B | AWGGAAARWVWGAMW | 1.67E-61 |

| | | | |
|---|---|---|---|
| 47 | ZN263_HUMAN.H11MO.0.A | GGGAGGAGGRRGRGGRGGRR | 3.25E-61 |
| 48 | STAT1_HUMAN.H11MO.1.A | RRRAAAADGAAASTRAAAV | 4.22E-61 |
| 49 | PATZ1_HUMAN.H11MO.0.C | GGGGMGGGGGMKGGRRVGGVRG | 3.17E-61 |
| 50 | ZN341_HUMAN.H11MO.0.C | GRGGRRRRGGRGRRGGAASAGC | 8.05E-60 |
| 51 | NFAT5_HUMAN.H11MO.0.D | RRRGGAARRDN | 2.71E-59 |
| 52 | TAF1_HUMAN.H11MO.0.A | RARRWGGCGGMGGMGR | 5.05E-59 |
| 53 | MXI1_HUMAN.H11MO.1.A | SCCACGTGSYB | 8.45E-59 |
| 54 | EGR1_HUMAN.H11MO.0.A | VDRDGCGKGGGYGGRRR | 1.30E-58 |
| 55 | BATF_HUMAN.H11MO.1.A | DATGAYKCAAH | 3.47E-58 |
| 56 | PRDM6_HUMAN.H11MO.0.C | RRRARGAAAAAAA | 8.31E-58 |
| 57 | MXI1_HUMAN.H11MO.0.A | VCACGTGGSNGSBGS | 3.86E-57 |
| 58 | SP4_HUMAN.H11MO.0.A | SRGVARGRGGCGGRGCHDRR | 7.34E-56 |
| 59 | ETS1_HUMAN.H11MO.0.A | RRRCMGGAAGTGG | 1.94E-55 |
| 60 | E2F3_HUMAN.H11MO.0.A | DDGGCGGGAAA | 1.52E-55 |
| 61 | NFE2_HUMAN.H11MO.0.A | VVRTGACTCAGCA | 7.34E-54 |
| 62 | ELF5_HUMAN.H11MO.0.A | VGRASSAGGAAGTRV | 8.68E-53 |
| 63 | SP4_HUMAN.H11MO.1.A | GRRRGGGGMGGGGCBK | 3.47E-52 |
| 64 | FOXG1_HUMAN.H11MO.0.D | DTTGTTTAYDTWTDTTT | 1.41E-51 |
| 65 | EGR2_HUMAN.H11MO.1.A | GRDGHGKGGGHGGRGG | 1.26E-51 |
| 66 | ETV2_HUMAN.H11MO.0.B | RRARRCAGGAARYRGS | 3.88E-51 |
| 67 | KLF6_HUMAN.H11MO.0.A | GSRRGGGHGGGGMHGGGRV | 8.45E-51 |
| 68 | BACH2_HUMAN.H11MO.0.A | TGCTGAGTCAB | 3.49E-50 |
| 69 | TBX15_HUMAN.H11MO.0.D | GGGGGGGGGGGGGTGGGDR | 5.42E-50 |
| 70 | FOXP1_HUMAN.H11MO.0.A | TBTGTTTMY | 2.80E-48 |
| 71 | STAT2_HUMAN.H11MO.0.A | RRGRAAAHGAAACTGAAAV | 9.40E-47 |
| 72 | PURA_HUMAN.H11MO.0.D | GGWRKKGGDGGGCVKGG | 2.07E-46 |
| 73 | ZN281_HUMAN.H11MO.0.A | RGGWGGGGGAGGGGV | 1.94E-45 |
| 74 | SP2_HUMAN.H11MO.0.A | GGSSVGGGGGCGGGGCCDGSGS | 2.18E-45 |
| 75 | FOXK1_HUMAN.H11MO.0.A | TGTTTMCHTT | 4.24E-44 |
| 76 | SP3_HUMAN.H11MO.0.B | SGVVGGGGGCGGGGCBRGSS | 5.21E-43 |
| 77 | MEF2C_HUMAN.H11MO.0.A | DKCTATTTTTRGM | 1.28E-42 |
| 78 | MEF2A_HUMAN.H11MO.0.A | DKCTATTTTTRGM | 1.25E-42 |
| 79 | ZSC22_HUMAN.H11MO.0.C | RGKCWGAGGGMGGAGG | 1.42E-42 |
| 80 | BACH1_HUMAN.H11MO.0.A | TGCTGAGTCABSS | 1.63E-42 |
| 81 | TFDP1_HUMAN.H11MO.0.C | RRRRGGCGGGAARN | 1.28E-42 |
| 82 | HMGA1_HUMAN.H11MO.0.D | NWATTTT | 1.05E-42 |
| 83 | NF2L2_HUMAN.H11MO.0.A | WWYTGCTGAGTCAT | 1.28E-41 |
| 84 | KLF16_HUMAN.H11MO.0.D | GGGGGYGKGKBGGGGGGGG | 6.84E-42 |
| 85 | ZN394_HUMAN.H11MO.0.C | NRARWRGAAWNGAMWGRAAK | 9.32E-42 |
| 86 | KLF1_HUMAN.H11MO.0.A | DGGGYGKGGCYGGG | 1.98E-41 |
| 87 | ZN121_HUMAN.H11MO.0.C | CTGGGCAACABAGYRAGACC | 1.79E-40 |
| 88 | ZN770_HUMAN.H11MO.1.C | DGGAGGCYGRG | 5.71E-41 |
| 89 | SMAD3_HUMAN.H11MO.0.B | SYCTSHCWSCWS | 3.59E-41 |
| 90 | KLF3_HUMAN.H11MO.0.B | GGRGVDGGGCGGGGCYGGG | 1.18E-40 |
| 91 | SP2_HUMAN.H11MO.1.B | RGGRGGCGGGGC | 4.69E-40 |
| 92 | ELF3_HUMAN.H11MO.0.A | RAABVAGGAAGTRR | 1.13E-39 |
| 93 | CPEB1_HUMAN.H11MO.0.D | TTTTTATTYTTTTTTTT | 3.42E-39 |

| | | | |
|---|---|---|---|
| 94 | ELF2_HUMAN.H11MO.0.C | RAVCCGGAAGTGR | 4.60E-39 |
| 95 | E2F6_HUMAN.H11MO.0.A | DGGGCGGGARRVR | 2.74E-39 |
| 96 | SRBP1_HUMAN.H11MO.0.A | DGTSRGGTGAB | 8.05E-39 |
| 97 | VEZF1_HUMAN.H11MO.1.C | RRRRGGAGGGGR | 1.73E-38 |
| 98 | IRF1_HUMAN.H11MO.0.A | RAAANWGAAASTGAAASYRR | 2.47E-37 |
| 99 | MEF2D_HUMAN.H11MO.0.A | DKCTATTTTTAG | 2.97E-37 |
| 100 | HXC10_HUMAN.H11MO.0.D | WWWWWWWNNWTTTWATKRC | 1.97E-37 |
| 101 | SP1_HUMAN.H11MO.0.A | VGGGGGCGGGGCCKGGGGGGGG | 1.42E-36 |
| 102 | KLF5_HUMAN.H11MO.0.A | WGGGTGKGGCDGGG | 5.58E-37 |
| 103 | PO3F3_HUMAN.H11MO.0.D | WWWAWTTATGCWWATTW | 1.14E-36 |
| 104 | FEV_HUMAN.H11MO.0.B | GCVGGAAGYG | 8.78E-37 |
| 105 | HXD12_HUMAN.H11MO.0.D | DDTTTTAHKRN | 3.10E-36 |
| 106 | XBP1_HUMAN.H11MO.0.D | GSKGACGTGGC | 1.15E-35 |
| 107 | MAZ_HUMAN.H11MO.1.A | RRGGGAGGGGS | 7.79E-36 |
| 108 | MEF2B_HUMAN.H11MO.0.A | DKKKCTATTTTTRG | 1.18E-35 |
| 109 | ZFP28_HUMAN.H11MO.0.C | TTCTATTTCTTCTTGWGTCA | 9.19E-35 |
| 110 | ELF1_HUMAN.H11MO.0.A | VRACCCGGAAGTGS | 2.30E-34 |
| 111 | TBX1_HUMAN.H11MO.0.D | GGGYGGGYAGGGGTGGGMDD | 2.56E-34 |
| 112 | ARNT_HUMAN.H11MO.0.B | STACGTGMC | 9.38E-34 |
| 113 | ZN713_HUMAN.H11MO.0.D | RWAGAARAADGVSANGMAA | 5.22E-34 |
| 114 | HSF1_HUMAN.H11MO.1.A | TTCTGGAAR | 5.47E-34 |
| 115 | FOXD3_HUMAN.H11MO.0.D | KBTGTTTNCWTWRBH | 8.42E-34 |
| 116 | E2F4_HUMAN.H11MO.0.A | SRGGGCGGGAARD | 2.85E-33 |
| 117 | SALL4_HUMAN.H11MO.0.B | GSDGGGWGGG | 1.89E-33 |
| 118 | EPAS1_HUMAN.H11MO.0.B | VTACGTGMC | 1.29E-32 |
| 119 | ZN148_HUMAN.H11MO.0.D | KGRGKGGGGGAGGGG | 5.71E-33 |
| 120 | ZFX_HUMAN.H11MO.1.A | DGVRGVVSAGGCCKVGG | 4.82E-33 |
| 121 | ARI3A_HUMAN.H11MO.0.D | RATTAAWBRRWATCAAAWTWWA | 1.10E-31 |
| 122 | SOX10_HUMAN.H11MO.1.A | BBYCTTTGTTYYY | 2.58E-32 |
| 123 | MAFK_HUMAN.H11MO.1.A | TGCTGAGTCAGCA | 8.37E-32 |
| 124 | SPIC_HUMAN.H11MO.0.D | KVDRRWGGGKDRG | 5.11E-32 |
| 125 | HXA11_HUMAN.H11MO.0.D | TWAADTTTTAYGR | 2.29E-31 |
| 126 | FOXF1_HUMAN.H11MO.0.D | WTGTTTATTTW | 7.34E-31 |
| 127 | HXD11_HUMAN.H11MO.0.D | DWTTTATKRHN | 4.67E-31 |
| 128 | ZFP82_HUMAN.H11MO.0.C | TTCCADTTCAYTAATTYTCTCTTC | 9.06E-31 |
| 129 | EHF_HUMAN.H11MO.0.B | VWACSAGGAAGTDVS | 1.54E-30 |
| 130 | FOXO1_HUMAN.H11MO.0.A | KBBYTGTTTHCN | 6.24E-31 |
| 131 | FOXJ2_HUMAN.H11MO.0.C | TGTTTRTTTW | 1.85E-30 |
| 132 | PAX5_HUMAN.H11MO.0.A | SCSKGGGCARCVRAGCGWGAC | 3.22E-30 |
| 133 | SP1_HUMAN.H11MO.1.A | VGGGGGCGGGGCCK | 2.39E-30 |
| 134 | FOXJ3_HUMAN.H11MO.0.A | TTGTTTATKKTTW | 7.79E-30 |
| 135 | ATF6A_HUMAN.H11MO.0.B | GKGSTGACGTGG | 1.48E-29 |
| 136 | NF2L1_HUMAN.H11MO.0.C | HGTCATN | 2.68E-30 |
| 137 | E2F1_HUMAN.H11MO.0.A | RRNKGGCGGGAARR | 5.84E-30 |
| 138 | NFKB1_HUMAN.H11MO.0.A | AGGGAAAK | 2.61E-30 |
| 139 | PROX1_HUMAN.H11MO.0.D | KGGKARGGCGKSKKGGG | 1.33E-29 |
| 140 | KLF12_HUMAN.H11MO.0.C | VGGGGCGGGGC | 7.20E-29 |

| | | | |
|---|---|---|---|
| 141 | FUBP1_HUMAN.H11MO.0.D | TYGTNTTTTTTT | 1.05E-28 |
| 142 | ZN554_HUMAN.H11MO.1.D | GCWGAGYCA | 6.96E-29 |
| 143 | GABPA_HUMAN.H11MO.0.A | GVVRCCGGAAGTGV | 3.33E-28 |
| 144 | ANDR_HUMAN.H11MO.0.A | TGTTCTTKKYTGTTTRYW | 3.94E-28 |
| 145 | SUH_HUMAN.H11MO.0.A | VSYSTGGGAAV | 2.86E-28 |
| 146 | NFAC2_HUMAN.H11MO.0.B | TGGAAAAWY | 3.20E-27 |
| 147 | ETV1_HUMAN.H11MO.0.A | SRCCGGAAGYG | 1.74E-26 |
| 148 | WT1_HUMAN.H11MO.1.B | GWGGGAGGAGS | 2.04E-26 |
| 149 | LYL1_HUMAN.H11MO.0.A | CCAKCTGYTYYCTB | 6.44E-26 |
| 150 | NFAC1_HUMAN.H11MO.1.B | AATGGAAAAW | 1.41E-25 |
| 151 | STAT3_HUMAN.H11MO.0.A | SAYTTCCMRGAA | 2.81E-25 |
| 152 | FOXM1_HUMAN.H11MO.0.A | TGTTTRCTYWKB | 2.11E-25 |
| 153 | PATZ1_HUMAN.H11MO.1.C | AGGGAGGGG | 1.78E-25 |
| 154 | BATF_HUMAN.H11MO.0.A | RBTYTCRDWATGASTCAB | 1.26E-24 |
| 155 | TBX21_HUMAN.H11MO.0.A | GDGDRGGTGTGRV | 4.93E-25 |
| 156 | GATA3_HUMAN.H11MO.0.A | RVAGATAAVAD | 8.66E-25 |
| 157 | NFAC4_HUMAN.H11MO.0.C | WGGAAAAYTW | 3.25E-24 |
| 158 | Z354A_HUMAN.H11MO.0.C | RTTTARWYCATTTACATTTAATGT | 9.22E-24 |
| 159 | FOXP2_HUMAN.H11MO.0.C | BTGTTTACH | 5.25E-24 |
| 160 | SPI1_HUMAN.H11MO.0.A | RRAAAGAGGAAGTGRVD | 1.72E-23 |
| 161 | KLF4_HUMAN.H11MO.0.A | WGGGYGKGGC | 1.77E-23 |
| 162 | IRF7_HUMAN.H11MO.0.C | GAAASYGAAA | 1.72E-23 |
| 163 | FOXO3_HUMAN.H11MO.0.B | BYTGTTTACH | 4.58E-23 |
| 164 | RXRA_HUMAN.H11MO.0.A | VRGDVRGNVARAGGTCARRG | 2.83E-23 |
| 165 | RARA_HUMAN.H11MO.1.A | VAGGTCAGRG | 8.96E-23 |
| 166 | FOXL1_HUMAN.H11MO.0.D | TKTTTGTTTRTTTAYWTTT | 3.95E-21 |
| 167 | IRF8_HUMAN.H11MO.0.B | RAAARRGGAASTGAAASTDR | 5.98E-22 |
| 168 | IRF2_HUMAN.H11MO.0.A | RAAAVHGAAAGTGAAASTRV | 1.11E-21 |
| 169 | STAT1_HUMAN.H11MO.0.A | YCCVTTTCCHGGAAAWCHC | 1.38E-21 |
| 170 | COT1_HUMAN.H11MO.0.C | GRGGKSARAGGTCAGGG | 8.80E-22 |
| 171 | STA5A_HUMAN.H11MO.0.A | TTCYRRGAAR | 1.84E-21 |
| 172 | FOXA1_HUMAN.H11MO.0.A | TGTTTACWYWGB | 2.81E-21 |
| 173 | PRDM1_HUMAN.H11MO.0.A | RAAAGTGAAAGTGR | 3.31E-21 |
| 174 | FOXJ3_HUMAN.H11MO.1.B | TKKTTWWTGTTTA | 1.74E-20 |
| 175 | ITF2_HUMAN.H11MO.0.C | DGCAGSTGKS | 1.25E-20 |
| 176 | HXC12_HUMAN.H11MO.0.D | DTTTTAYKACYT | 4.05E-20 |
| 177 | ELK1_HUMAN.H11MO.0.B | RCCGGAAGTGV | 6.19E-20 |
| 178 | FOXF2_HUMAN.H11MO.0.D | TGTTTACHTWD | 1.71E-19 |
| 179 | KLF9_HUMAN.H11MO.0.C | GRRGKGGGCGTGGCC | 2.18E-19 |
| 180 | NFIC_HUMAN.H11MO.0.A | CYTGGCHBYMWGCCARG | 1.11E-19 |
| 181 | STA5B_HUMAN.H11MO.0.A | YTTCYKRGAAAY | 2.03E-19 |
| 182 | HIF1A_HUMAN.H11MO.0.C | GDACGTGM | 3.40E-19 |
| 183 | MNX1_HUMAN.H11MO.0.D | MBATTTDHWTHWCYHYTHH | 1.97E-19 |
| 184 | RORG_HUMAN.H11MO.0.C | RRAASTRGGTCA | 2.87E-19 |
| 185 | FOXD2_HUMAN.H11MO.0.D | WAWGTAAATRTTWWYTTAA | 5.22E-18 |
| 186 | ARNT2_HUMAN.H11MO.0.D | GNCGTGGSASRC | 9.26E-19 |
| 187 | IRX2_HUMAN.H11MO.0.D | RCDKGKBDKGDCRKGTDVK | 1.39E-18 |

| | | | |
|---|---|---|---|
| 188 | SRY_HUMAN.H11MO.0.B | YWTTGTTTH | 1.88E-18 |
| 189 | RUNX2_HUMAN.H11MO.0.A | KGKYTGTGGTTTKK | 2.91E-18 |
| 190 | RUNX1_HUMAN.H11MO.0.A | BYTGTGGTTWB | 7.04E-18 |
| 191 | SMCA5_HUMAN.H11MO.0.C | GAATBGAATGGAAWC | 1.02E-17 |
| 192 | HES1_HUMAN.H11MO.0.D | VMGVCACGMGMCM | 8.68E-18 |
| 193 | SMAD2_HUMAN.H11MO.0.A | TGTCTGNCWCCT | 7.76E-18 |
| 194 | Z324A_HUMAN.H11MO.0.C | KVTCAAACCATCCTTKSCTKSCW | 1.06E-17 |
| 195 | HEY2_HUMAN.H11MO.0.D | GKBGGCWCGTGGCHTBV | 3.22E-17 |
| 196 | HES5_HUMAN.H11MO.0.D | YGGCACGTGYCR | 1.64E-16 |
| 197 | KLF14_HUMAN.H11MO.0.D | DRGKGGGCGTGGY | 3.97E-17 |
| 198 | ZN554_HUMAN.H11MO.0.C | GCWGAGYCANGTRGDKKRCT | 2.88E-17 |
| 199 | SMCA1_HUMAN.H11MO.0.C | CMARGAAGMATD | 2.74E-17 |
| 200 | ESR2_HUMAN.H11MO.1.A | AGGTCAS | 3.37E-17 |
| 201 | ZF64A_HUMAN.H11MO.0.D | GGGGCCCGGGAMYCDG | 4.44E-17 |
| 202 | ELK4_HUMAN.H11MO.0.A | SVCCGGAAGTGV | 1.31E-16 |
| 203 | NR1D1_HUMAN.H11MO.0.B | RRRNMASTGGGTCAGDRDR | 7.10E-17 |
| 204 | BHE41_HUMAN.H11MO.0.D | WSVRKSYCACGTGMWGRRRV | 2.52E-16 |
| 205 | FOXA2_HUMAN.H11MO.0.A | TGTTTACWYWGB | 2.85E-16 |
| 206 | IRF4_HUMAN.H11MO.0.A | RAAWGRGGAASTGAAASY | 3.32E-16 |
| 207 | PITX2_HUMAN.H11MO.0.D | TKGGATTAVH | 2.99E-16 |
| 208 | DLX1_HUMAN.H11MO.0.D | WWWWWTATGSTAATTA | 3.80E-16 |
| 209 | SRBP2_HUMAN.H11MO.0.B | VSGTGGGGWGABG | 3.11E-16 |
| 210 | EGR4_HUMAN.H11MO.0.D | GGSGGYRGGGM | 8.92E-16 |
| 211 | OSR2_HUMAN.H11MO.0.C | GBYSCWGCTNCTGYDG | 5.94E-16 |
| 212 | COT1_HUMAN.H11MO.1.C | BSARAGGTCAGRG | 8.26E-16 |
| 213 | CREB3_HUMAN.H11MO.0.D | KGRTGACGTGGCA | 1.40E-14 |
| 214 | TEAD3_HUMAN.H11MO.0.D | GAHATTYYWGCYBYW | 1.78E-15 |
| 215 | PTF1A_HUMAN.H11MO.0.B | SCAGCTGBYBBYBYYCHC | 1.24E-15 |
| 216 | AP2D_HUMAN.H11MO.0.D | CGCCYGVGGCSCGT | 5.72E-15 |
| 217 | BATF3_HUMAN.H11MO.0.B | BRSTTTCADTWTGASWM | 3.19E-15 |
| 218 | ZN563_HUMAN.H11MO.1.C | SHRGCAGCWSWG | 1.87E-15 |
| 219 | HAND1_HUMAN.H11MO.1.D | KSYCTGG | 1.26E-15 |
| 220 | RXRB_HUMAN.H11MO.0.C | TSAGGTCASR | 2.57E-15 |
| 221 | FOXB1_HUMAN.H11MO.0.D | RTMAATRTTTACATAAB | 1.50E-14 |
| 222 | LMX1B_HUMAN.H11MO.0.D | TAATTAAHWRMYTWWWWW | 1.03E-14 |
| 223 | SPIB_HUMAN.H11MO.0.A | RRAAAGAGGAAGTGARA | 1.19E-14 |
| 224 | HNF1A_HUMAN.H11MO.0.C | DGTTAATKATTAACH | 9.87E-15 |
| 225 | E2F4_HUMAN.H11MO.1.A | GGCGGGAADTTBNR | 8.44E-15 |
| 226 | ESR1_HUMAN.H11MO.1.A | CHRGGTCASV | 4.55E-15 |
| 227 | IKZF1_HUMAN.H11MO.0.C | BTGGGARR | 3.34E-15 |
| 228 | PTF1A_HUMAN.H11MO.1.B | RGCAGCTGG | 5.13E-15 |
| 229 | RUNX3_HUMAN.H11MO.0.A | BYTGTGGTTW | 1.09E-14 |
| 230 | RARG_HUMAN.H11MO.1.B | VAGGTCAS | 5.97E-15 |
| 231 | ELK3_HUMAN.H11MO.0.D | VMCHGGAARTSC | 1.80E-14 |
| 232 | RREB1_HUMAN.H11MO.0.D | DKGKKKGKGGKTGKTTTGGGKT | 2.58E-14 |
| 233 | CR3L1_HUMAN.H11MO.0.D | GCCACGTGGCA | 2.80E-13 |
| 234 | NFAC3_HUMAN.H11MO.0.B | TGGAAAAHY | 1.44E-14 |

| | | | |
|---|---|---|---|
| 235 | BCL6_HUMAN.H11MO.0.A | YKCTTTCYAGGAA | 1.95E-14 |
| 236 | MAFF_HUMAN.H11MO.1.B | TGCTGASTC | 2.62E-14 |
| 237 | ZN350_HUMAN.H11MO.0.C | CAGYYYTTTTATDWCCHR | 3.77E-14 |
| 238 | PO4F1_HUMAN.H11MO.0.D | CATTAATTATTCAT | 2.04E-13 |
| 239 | FOXC1_HUMAN.H11MO.0.C | CDWTGTTTACTTWRG | 3.79E-13 |
| 240 | LMX1A_HUMAN.H11MO.0.D | TAATTAATTWKKTWATTWW | 3.22E-13 |
| 241 | RARB_HUMAN.H11MO.0.D | SAGGTCAVVVV | 1.36E-13 |
| 242 | LEF1_HUMAN.H11MO.0.A | BCCTTTSWWNTBYH | 7.23E-14 |
| 243 | COT2_HUMAN.H11MO.1.A | RGRGYCARAGGTCARVR | 1.15E-13 |
| 244 | GSX2_HUMAN.H11MO.0.D | YTAATTRSHWWTTAAT | 2.47E-13 |
| 245 | MAFF_HUMAN.H11MO.0.B | TGCTGASTCAGCABTTTT | 2.41E-13 |
| 246 | TFAP4_HUMAN.H11MO.0.A | YCAGCTGWKB | 4.09E-13 |
| 247 | SOX4_HUMAN.H11MO.0.B | BYCTTTGTYYYB | 2.21E-13 |
| 248 | NKX61_HUMAN.H11MO.0.B | GAYWAATKRSHWWTTAATK | 5.07E-13 |
| 249 | ZFHX3_HUMAN.H11MO.0.D | ATTAWTAATTA | 1.77E-12 |
| 250 | ZN322_HUMAN.H11MO.0.B | GWGCCTGSTACWSWGCCTGR | 1.10E-12 |
| 251 | BHA15_HUMAN.H11MO.0.B | SSAGCAGCTGG | 9.39E-13 |
| 252 | HAND1_HUMAN.H11MO.0.D | RGSYCTGGYYWNTSTCA | 1.07E-12 |
| 253 | MAFB_HUMAN.H11MO.0.B | TGCTGASTYAD | 6.71E-13 |
| 254 | HNF1B_HUMAN.H11MO.0.A | DGTTAATRATTAACH | 3.63E-12 |
| 255 | HXB13_HUMAN.H11MO.0.A | TTTTATDGSH | 1.62E-12 |
| 256 | SOX2_HUMAN.H11MO.0.A | BBCCTTTGTYYYB | 1.09E-12 |
| 257 | HXA10_HUMAN.H11MO.0.C | KATRRTTTATGW | 3.00E-12 |
| 258 | FOXP3_HUMAN.H11MO.0.D | AATKTGWTT | 1.42E-12 |
| 259 | STAT4_HUMAN.H11MO.0.A | YTTCYYRGMAV | 1.52E-12 |
| 260 | ZN282_HUMAN.H11MO.0.D | MGKRTTRTGGGARA | 2.13E-12 |
| 261 | PAX4_HUMAN.H11MO.0.D | TWAYTAATTMRH | 3.25E-12 |
| 262 | NRF1_HUMAN.H11MO.0.A | CWSTGCGCATGCGCRDS | 2.71E-11 |
| 263 | MAFG_HUMAN.H11MO.1.A | TGCTGAGT | 2.28E-12 |
| 264 | TAL1_HUMAN.H11MO.1.A | VARSAARCAGVTGKB | 2.84E-12 |
| 265 | MAF_HUMAN.H11MO.0.A | VAAWBTGCTGASTHWGCMD | 4.42E-12 |
| 266 | HNF1B_HUMAN.H11MO.1.A | DGTTAATVATT | 1.41E-11 |
| 267 | ZN263_HUMAN.H11MO.1.A | SRGGGAGGASG | 7.67E-12 |
| 268 | GSX1_HUMAN.H11MO.0.D | TAATTAGNDDTTAAT | 6.88E-12 |
| 269 | HESX1_HUMAN.H11MO.0.D | ABSYGGCWMRTRRMMT | 1.08E-11 |
| 270 | PEBB_HUMAN.H11MO.0.C | TYTGTGGTYWB | 7.46E-12 |
| 271 | PO3F4_HUMAN.H11MO.0.D | TMATTTGCATAATTWAW | 3.05E-11 |
| 272 | PO2F1_HUMAN.H11MO.0.C | AWKBRWATGSAAWYAW | 2.39E-11 |
| 273 | MYOG_HUMAN.H11MO.0.B | CHSCAGCTGYYYB | 3.68E-11 |
| 274 | HXC13_HUMAN.H11MO.0.D | DDTTTTACKAG | 3.77E-11 |
| 275 | BARX1_HUMAN.H11MO.0.D | BWAATKGTTTYY | 3.85E-11 |
| 276 | PIT1_HUMAN.H11MO.0.C | YTMATGAATATWY | 7.62E-11 |
| 277 | VENTX_HUMAN.H11MO.0.D | YVYTAATYGSTTHHHWWTT | 1.56E-10 |
| 278 | HSF1_HUMAN.H11MO.0.A | BRGAANSTTCYRGAA | 2.25E-10 |
| 279 | FOXA3_HUMAN.H11MO.0.B | CHTGTTTACWTWG | 1.35E-10 |
| 280 | FOXD1_HUMAN.H11MO.0.D | CWWTGTTTACHTWVR | 1.94E-10 |
| 281 | BPTF_HUMAN.H11MO.0.D | KKTBTTGTKKKS | 6.58E-11 |

| | | | |
|---|---|---|---|
| 282 | TYY1_HUMAN.H11MO.0.A | CAARATGGCGGC | 4.92E-10 |
| 283 | ZN335_HUMAN.H11MO.1.A | YGCCTGW | 8.41E-11 |
| 284 | REL_HUMAN.H11MO.0.B | DRWRGGGRAATKCCA | 8.81E-11 |
| 285 | RARA_HUMAN.H11MO.0.A | GGRRGRTSMRRAGKTCAAGG | 2.25E-10 |
| 286 | STAT6_HUMAN.H11MO.0.B | TTCTSWGGAAR | 2.03E-10 |
| 287 | HTF4_HUMAN.H11MO.0.A | DSCAGSTGKG | 2.01E-10 |
| 288 | MAFK_HUMAN.H11MO.0.A | RAWWHTGCTGASTCAGCA | 1.77E-09 |
| 289 | NRL_HUMAN.H11MO.0.D | TGCTGACKYWR | 4.24E-10 |
| 290 | MSX2_HUMAN.H11MO.0.D | TAATTDK | 4.81E-10 |
| 291 | PO4F3_HUMAN.H11MO.0.D | TCATTAATTATKCAT | 3.56E-09 |
| 292 | FOXO4_HUMAN.H11MO.0.C | TTGTTTAYK | 9.50E-10 |
| 293 | ATF2_HUMAN.H11MO.0.B | RRTGABGTCAY | 1.38E-09 |
| 294 | MYOD1_HUMAN.H11MO.1.A | CHGCAGCTGTY | 1.47E-09 |
| 295 | KLF13_HUMAN.H11MO.0.D | DRGKGGGCGKD | 2.83E-09 |
| 296 | CTCF_HUMAN.H11MO.0.A | YGGCCACCAGRKGGCRSYV | 3.46E-09 |
| 297 | ZNF76_HUMAN.H11MO.0.C | RRKGCMTKCTGGGARWTGTAGT | 4.03E-09 |
| 298 | HES7_HUMAN.H11MO.0.D | TGGCACGTGCCG | 3.72E-08 |
| 299 | THB_HUMAN.H11MO.0.C | GRSSYSASVTSAGGTCA | 3.09E-09 |
| 300 | OLIG2_HUMAN.H11MO.0.B | CCAKCTGYTYYYYWBCHB | 2.20E-09 |
| 301 | FOXO6_HUMAN.H11MO.0.D | WWSWTGTTTACRT | 8.46E-09 |
| 302 | TBX3_HUMAN.H11MO.0.C | DRAGGTGBSAR | 1.90E-09 |
| 303 | TBP_HUMAN.H11MO.0.A | MTATAAAARS | 5.02E-09 |
| 304 | NR2E3_HUMAN.H11MO.0.C | AAAGTCAAAAGTCA | 5.66E-09 |
| 305 | THAP1_HUMAN.H11MO.0.C | SGCCGCCATSTYGGSYBCGGGC | 4.75E-09 |
| 306 | ZSC31_HUMAN.H11MO.0.C | RGCAGCAGGGCARTTAKR | 6.13E-09 |
| 307 | SMAD4_HUMAN.H11MO.0.B | CWGTCTGDCACCT | 4.73E-09 |
| 308 | MYOD1_HUMAN.H11MO.0.A | VCAGCTGYYBCYGBC | 8.62E-09 |
| 309 | ASCL2_HUMAN.H11MO.0.D | VCASCTGCT | 7.67E-09 |
| 310 | SIX2_HUMAN.H11MO.0.A | KGAAAYHTGAKMC | 7.62E-09 |
| 311 | ATOH1_HUMAN.H11MO.0.B | RRCAGMTGG | 8.67E-09 |
| 312 | JDP2_HUMAN.H11MO.0.D | BGATGACGTCAY | 1.68E-07 |
| 313 | IRF9_HUMAN.H11MO.0.C | GAAAGCGAAAYT | 6.82E-08 |
| 314 | UBIP1_HUMAN.H11MO.0.D | SCAGARA | 9.75E-09 |
| 315 | ASCL1_HUMAN.H11MO.0.A | CHVCASCTGCYBYY | 1.71E-08 |
| 316 | TFE2_HUMAN.H11MO.0.A | SMCAGCTGCWB | 1.67E-08 |
| 317 | FEZF1_HUMAN.H11MO.0.C | DYTGYYCTTTTH | 1.09E-08 |
| 318 | ZN418_HUMAN.H11MO.0.C | TGCTTYTRGCYTCTKVY | 2.24E-08 |
| 319 | HXD10_HUMAN.H11MO.0.D | TGCTTTAATT | 2.06E-08 |
| 320 | HXB3_HUMAN.H11MO.0.D | YTAATTASYTTTTW | 3.87E-08 |
| 321 | FOXQ1_HUMAN.H11MO.0.C | HATTGTTTATTT | 1.92E-07 |
| 322 | ATF1_HUMAN.H11MO.0.B | HRRTGACGYMA | 6.03E-08 |
| 323 | HXC11_HUMAN.H11MO.0.D | DTTTWAYGACY | 7.15E-08 |
| 324 | RARG_HUMAN.H11MO.0.B | RGGTCASVRTGVCMTGRV | 3.91E-08 |
| 325 | HSF4_HUMAN.H11MO.0.D | TRGAABVTTCTAGAA | 3.85E-07 |
| 326 | ZN335_HUMAN.H11MO.0.A | RGCTGYCYNBVSCKMTGCCTGW | 1.23E-07 |
| 327 | NR4A1_HUMAN.H11MO.0.A | AAAGGKCAV | 3.46E-08 |
| 328 | THA_HUMAN.H11MO.0.C | RGGTCABYTSAGGTCAK | 6.17E-08 |

| | | | |
|---|---|---|---|
| 329 | THB_HUMAN.H11MO.1.D | SGTSAGGTCA | 8.63E-08 |
| 330 | ZN768_HUMAN.H11MO.0.C | GAGGCHCAGAGAGGTTAAGT | 3.81E-07 |
| 331 | HXA1_HUMAN.H11MO.0.C | TKGATGGATG | 2.30E-07 |
| 332 | ZN219_HUMAN.H11MO.0.D | GDGGGGGGYGGA | 8.41E-08 |
| 333 | TYY2_HUMAN.H11MO.0.D | AWAATGGCGGH | 4.61E-07 |
| 334 | RFX1_HUMAN.H11MO.0.B | SSSSCBGTTGCCAKGGVRACSS | 3.37E-07 |
| 335 | CDX1_HUMAN.H11MO.0.C | HTTTATGD | 9.34E-08 |
| 336 | TF65_HUMAN.H11MO.0.A | HWGGGRATTTCCMV | 2.15E-07 |
| 337 | NR1H3_HUMAN.H11MO.1.B | MVAGGTCAVR | 8.03E-08 |
| 338 | PO6F1_HUMAN.H11MO.0.D | CATAAWTTATGCR | 2.11E-06 |
| 339 | E4F1_HUMAN.H11MO.0.D | YGTKACGTC | 9.37E-07 |
| 340 | THA_HUMAN.H11MO.1.D | BTSAGGTCAS | 1.38E-07 |
| 341 | ZEP2_HUMAN.H11MO.0.D | GGYDGGGWAACCSS | 2.63E-07 |
| 342 | MAFA_HUMAN.H11MO.0.D | STGCTGACBMYGCAGYHTYCV | 3.02E-07 |
| 343 | CDC5L_HUMAN.H11MO.0.D | GBGWKTTAAYRTAWY | 7.29E-07 |
| 344 | BSH_HUMAN.H11MO.0.D | TAATKVGTTAATBR | 4.87E-07 |
| 345 | EVX2_HUMAN.H11MO.0.A | KTTTTATKRCYBY | 7.05E-07 |
| 346 | ZBT7B_HUMAN.H11MO.0.D | SSCSGTTGCCATGGVRACSGS | 2.28E-06 |
| 347 | MYF6_HUMAN.H11MO.0.C | CASCTGC | 4.31E-07 |
| 348 | ZBT14_HUMAN.H11MO.0.C | GGAGCGCGC | 1.59E-06 |
| 349 | HIC2_HUMAN.H11MO.0.D | HCRTGCCAGVN | 6.19E-07 |
| 350 | VSX1_HUMAN.H11MO.0.D | TAATTAGCHWWWWW | 1.22E-06 |
| 351 | NANOG_HUMAN.H11MO.0.A | BYYWTTGWNATGCAAAT | 5.87E-07 |
| 352 | HXD8_HUMAN.H11MO.0.D | WAATWTGCAYN | 4.81E-07 |
| 353 | ATF2_HUMAN.H11MO.2.C | RRTGACGTCAS | 1.91E-05 |
| 354 | AHR_HUMAN.H11MO.0.B | DTYGCGTGM | 1.40E-06 |
| 355 | HSF2_HUMAN.H11MO.0.A | VGAABRTTCTAGAA | 9.53E-07 |
| 356 | ZEB1_HUMAN.H11MO.0.A | BVCAGGTGWG | 5.43E-07 |
| 357 | HXA9_HUMAN.H11MO.0.B | RTGATTTATKR | 1.15E-06 |
| 358 | MSX1_HUMAN.H11MO.0.D | TAATTGSWWKTTAATTGS | 3.93E-06 |
| 359 | IRF5_HUMAN.H11MO.0.D | KAAAGGRAARCMAAAWSWGA | 1.41E-05 |
| 360 | ATF2_HUMAN.H11MO.1.B | WDDWGATGATGTCATCWWTW | 2.03E-05 |
| 361 | ATF7_HUMAN.H11MO.0.D | KRTGACGTCAT | 2.67E-05 |
| 362 | LHX9_HUMAN.H11MO.0.D | YTAATTAGYNHWWWWWWD | 3.84E-06 |
| 363 | ZNF85_HUMAN.H11MO.1.C | TGMWGKAATCTYY | 1.88E-06 |
| 364 | ZNF41_HUMAN.H11MO.1.C | CACAAGGRARWV | 1.05E-06 |
| 365 | COT2_HUMAN.H11MO.0.A | CARAGGTCARRGD | 1.12E-06 |
| 366 | TWST1_HUMAN.H11MO.1.A | VMCATCTGGW | 1.36E-06 |
| 367 | GBX2_HUMAN.H11MO.0.D | BTAATTRSYBHHWWWWW | 3.09E-06 |
| 368 | ZN563_HUMAN.H11MO.0.C | RGGNTCMTNDCHGGCAGCTGY | 1.31E-06 |
| 369 | RORA_HUMAN.H11MO.0.C | AWSTRGGTCARRG | 9.83E-07 |
| 370 | EVX1_HUMAN.H11MO.0.D | TGATTTATKGCYW | 2.23E-06 |
| 371 | ZN143_HUMAN.H11MO.0.A | DGCMTKCTGGGARWTGTAGTYY | 1.07E-05 |
| 372 | CREB5_HUMAN.H11MO.0.D | RRTGACGTMAT | 3.33E-05 |
| 373 | SOX2_HUMAN.H11MO.1.A | BCCWTTGTYATGYWRA | 2.77E-06 |
| 374 | ZNF85_HUMAN.H11MO.0.C | AWAGAARWCTGMWGKAATCT | 7.79E-06 |
| 375 | IRX3_HUMAN.H11MO.0.D | WNWCATGTWMT | 3.22E-06 |

| | | | |
|---|---|---|---|
| 376 | NR0B1_HUMAN.H11MO.0.D | GMGKGGGASR | 2.72E-06 |
| 377 | MEOX1_HUMAN.H11MO.0.D | GGTAATTAGBGKHAWW | 1.63E-05 |
| 378 | HLTF_HUMAN.H11MO.0.D | KANKGCTGSMAM | 2.93E-06 |
| 379 | PAX2_HUMAN.H11MO.0.D | RHTCAGTSAYGMGTGAYW | 7.33E-06 |
| 380 | SMAD1_HUMAN.H11MO.0.D | RSCCTGTCTGCC | 3.51E-06 |
| 381 | BHE23_HUMAN.H11MO.0.D | RRSRTRTGKTDB | 2.71E-06 |
| 382 | NDF2_HUMAN.H11MO.0.B | RRCAGATGG | 3.89E-06 |

### AME (Analysis of Motif Enrichment): Version 5.0.5 compiled on Jul 15 2019 at 00:31:14
### The format of this file is described at http://meme-suite.org//doc/ame-output-format.html.
### ame --verbose 1 --oc . --scoring avg --method fisher --hit-lo-fraction 0.25 --evalue-report-thresho

| adj_p-value | E-value | tests | FASTA_max | pos | neg | PWM_min | TP |
|---|---|---|---|---|---|---|---|
| 3.95E-232 | 5.14E-229 | 2893 | 3131 | 3131 | 3131 | 1.82 | 1720 |
| 3.80E-213 | 4.95E-210 | 2695 | 3131 | 3131 | 3131 | 1.65 | 1677 |
| 5.20E-202 | 6.77E-199 | 2145 | 3131 | 3131 | 3131 | 1.36 | 1496 |
| 2.75E-169 | 3.58E-166 | 1343 | 3131 | 3131 | 3131 | 1.08 | 1092 |
| 9.57E-152 | 1.25E-148 | 1191 | 3131 | 3131 | 3131 | 1.03 | 992 |
| 3.50E-150 | 4.55E-147 | 1385 | 3131 | 3131 | 3131 | 1.32 | 1068 |
| 6.82E-148 | 8.88E-145 | 1358 | 3131 | 3131 | 3131 | 1.86 | 990 |
| 2.41E-145 | 3.13E-142 | 2010 | 3131 | 3131 | 3131 | 2.5 | 1062 |
| 2.38E-142 | 3.10E-139 | 929 | 3131 | 3131 | 3131 | 1.6 | 722 |
| 3.78E-140 | 4.92E-137 | 1691 | 3131 | 3131 | 3131 | 1.63 | 1106 |
| 3.39E-139 | 4.41E-136 | 1567 | 3131 | 3131 | 3131 | 1.77 | 1018 |
| 1.52E-137 | 1.98E-134 | 768 | 3131 | 3131 | 3131 | 1.56 | 654 |
| 9.31E-136 | 1.21E-132 | 2253 | 3131 | 3131 | 3131 | 5.12 | 905 |
| 7.16E-128 | 9.32E-125 | 905 | 3131 | 3131 | 3131 | 1.07 | 775 |
| 2.56E-122 | 3.33E-119 | 1886 | 3131 | 3131 | 3131 | 5.98 | 783 |
| 1.38E-117 | 1.80E-114 | 1356 | 3131 | 3131 | 3131 | 1.14 | 989 |
| 1.12E-113 | 1.46E-110 | 1560 | 3131 | 3131 | 3131 | 1.16 | 1097 |
| 2.56E-102 | 3.33E-99 | 422 | 3131 | 3131 | 3131 | 1.42 | 399 |
| 1.24E-86 | 1.61E-83 | 1392 | 3131 | 3131 | 3131 | 1.03 | 1019 |
| 1.66E-85 | 2.16E-82 | 3741 | 3131 | 3131 | 3131 | 1.72 | 1820 |
| 4.59E-84 | 5.98E-81 | 3832 | 3131 | 3131 | 3131 | 2.06 | 1652 |
| 1.93E-81 | 2.51E-78 | 3613 | 3131 | 3131 | 3131 | 1.46 | 1885 |
| 2.59E-80 | 3.37E-77 | 1695 | 3131 | 3131 | 3131 | 1.03 | 1178 |
| 2.90E-80 | 3.77E-77 | 3319 | 3131 | 3131 | 3131 | 3.87 | 1336 |
| 4.93E-79 | 6.42E-76 | 1224 | 3131 | 3131 | 3131 | 1.02 | 908 |
| 2.13E-76 | 2.78E-73 | 2339 | 3131 | 3131 | 3131 | 1.06 | 1528 |
| 2.99E-75 | 3.89E-72 | 2425 | 3131 | 3131 | 3131 | 1.49 | 1349 |
| 1.39E-73 | 1.81E-70 | 3041 | 3131 | 3131 | 3131 | 5.8 | 1195 |
| 2.16E-73 | 2.82E-70 | 2777 | 3131 | 3131 | 3131 | 7.06 | 1099 |
| 1.14E-72 | 1.49E-69 | 2256 | 3131 | 3131 | 3131 | 1.03 | 1474 |
| 1.41E-71 | 1.84E-68 | 682 | 3131 | 3131 | 3131 | 1.31 | 506 |
| 4.26E-71 | 5.55E-68 | 622 | 3131 | 3131 | 3131 | 1.05 | 518 |
| 9.90E-71 | 1.29E-67 | 2746 | 3131 | 3131 | 3131 | 2.08 | 832 |
| 4.10E-68 | 5.34E-65 | 1608 | 3131 | 3131 | 3131 | 65.5 | 325 |
| 2.25E-67 | 2.93E-64 | 2698 | 3131 | 3131 | 3131 | 3.53 | 1136 |
| 2.38E-67 | 3.10E-64 | 3953 | 3131 | 3131 | 3131 | 2.69 | 1396 |
| 3.98E-67 | 5.18E-64 | 2923 | 3131 | 3131 | 3131 | 2.92 | 1018 |
| 8.99E-67 | 1.17E-63 | 2099 | 3131 | 3131 | 3131 | 1.14 | 1374 |
| 1.60E-66 | 2.08E-63 | 3398 | 3131 | 3131 | 3131 | 1.01 | 2046 |
| 1.97E-66 | 2.57E-63 | 3295 | 3131 | 3131 | 3131 | 2.54 | 1297 |
| 3.27E-65 | 4.26E-62 | 1902 | 3131 | 3131 | 3131 | 4.08 | 938 |
| 3.45E-62 | 4.49E-59 | 2061 | 3131 | 3131 | 3131 | 1.17 | 1331 |
| 3.72E-60 | 4.84E-57 | 1977 | 3131 | 3131 | 3131 | 7.12 | 804 |
| 4.70E-60 | 6.12E-57 | 3883 | 3131 | 3131 | 3131 | 1.73 | 1856 |
| 5.94E-59 | 7.74E-56 | 1293 | 3131 | 3131 | 3131 | 1.12 | 870 |
| 5.29E-58 | 6.88E-55 | 3174 | 3131 | 3131 | 3131 | 2 | 1355 |

| | | | | | | | |
|---|---|---|---|---|---|---|---|
| 6.95E-58 | 9.05E-55 | 2142 | 3131 | 3131 | 3131 | 1.18 | 1365 |
| 7.81E-58 | 1.02E-54 | 1850 | 3131 | 3131 | 3131 | 2.26 | 942 |
| 8.86E-58 | 1.15E-54 | 2795 | 3131 | 3131 | 3131 | 14 | 749 |
| 2.31E-56 | 3.01E-53 | 2872 | 3131 | 3131 | 3131 | 9.4 | 792 |
| 1.08E-55 | 1.41E-52 | 4001 | 3131 | 3131 | 3131 | 2.66 | 1449 |
| 1.27E-55 | 1.65E-52 | 2512 | 3131 | 3131 | 3131 | 1.06 | 1555 |
| 1.61E-55 | 2.10E-52 | 1910 | 3131 | 3131 | 3131 | 1.18 | 1120 |
| 3.35E-55 | 4.36E-52 | 2579 | 3131 | 3131 | 3131 | 5.78 | 754 |
| 6.54E-55 | 8.52E-52 | 1883 | 3131 | 3131 | 3131 | 1.89 | 912 |
| 2.45E-54 | 3.20E-51 | 2953 | 3131 | 3131 | 3131 | 3.55 | 1063 |
| 6.94E-54 | 9.04E-51 | 1800 | 3131 | 3131 | 3131 | 2.26 | 732 |
| 1.93E-52 | 2.51E-49 | 2630 | 3131 | 3131 | 3131 | 3.84 | 996 |
| 3.54E-52 | 4.61E-49 | 1821 | 3131 | 3131 | 3131 | 1.47 | 1013 |
| 4.72E-52 | 6.14E-49 | 3101 | 3131 | 3131 | 3131 | 2.16 | 1082 |
| 8.95E-51 | 1.17E-47 | 1219 | 3131 | 3131 | 3131 | 1.61 | 689 |
| 1.66E-49 | 2.15E-46 | 1906 | 3131 | 3131 | 3131 | 1.15 | 1173 |
| 7.19E-49 | 9.36E-46 | 2074 | 3131 | 3131 | 3131 | 9.42 | 617 |
| 2.30E-48 | 2.99E-45 | 1628 | 3131 | 3131 | 3131 | 1.38 | 985 |
| 2.93E-48 | 3.82E-45 | 2327 | 3131 | 3131 | 3131 | 5.34 | 784 |
| 5.27E-48 | 6.87E-45 | 1360 | 3131 | 3131 | 3131 | 1.11 | 902 |
| 2.53E-47 | 3.29E-44 | 2993 | 3131 | 3131 | 3131 | 7.16 | 840 |
| 5.53E-47 | 7.20E-44 | 1583 | 3131 | 3131 | 3131 | 3.53 | 576 |
| 1.22E-46 | 1.59E-43 | 2259 | 3131 | 3131 | 3131 | 2.8 | 1102 |
| 7.50E-45 | 9.76E-42 | 2682 | 3131 | 3131 | 3131 | 1.38 | 1360 |
| 9.80E-44 | 1.28E-40 | 1042 | 3131 | 3131 | 3131 | 1.39 | 683 |
| 5.89E-43 | 7.66E-40 | 2838 | 3131 | 3131 | 3131 | 2.02 | 1304 |
| 3.54E-42 | 4.61E-39 | 1828 | 3131 | 3131 | 3131 | 4.21 | 779 |
| 4.38E-42 | 5.70E-39 | 2011 | 3131 | 3131 | 3131 | 26.1 | 462 |
| 1.16E-40 | 1.51E-37 | 2733 | 3131 | 3131 | 3131 | 2 | 1054 |
| 1.31E-39 | 1.70E-36 | 2510 | 3131 | 3131 | 3131 | 44.5 | 385 |
| 1.70E-39 | 2.22E-36 | 1334 | 3131 | 3131 | 3131 | 1.3 | 795 |
| 1.71E-39 | 2.23E-36 | 1371 | 3131 | 3131 | 3131 | 2.29 | 616 |
| 2.56E-39 | 3.33E-36 | 1805 | 3131 | 3131 | 3131 | 1.12 | 1126 |
| 2.61E-39 | 3.39E-36 | 1602 | 3131 | 3131 | 3131 | 1.38 | 887 |
| 2.71E-39 | 3.52E-36 | 2111 | 3131 | 3131 | 3131 | 2.48 | 791 |
| 2.94E-39 | 3.83E-36 | 2811 | 3131 | 3131 | 3131 | 2.34 | 839 |
| 8.56E-39 | 1.11E-35 | 669 | 3131 | 3131 | 3131 | 1.14 | 483 |
| 1.51E-38 | 1.97E-35 | 2212 | 3131 | 3131 | 3131 | 4.49 | 915 |
| 2.81E-38 | 3.66E-35 | 3018 | 3131 | 3131 | 3131 | 1.93 | 1328 |
| 3.72E-38 | 4.84E-35 | 1882 | 3131 | 3131 | 3131 | 24.5 | 343 |
| 5.71E-38 | 7.44E-35 | 319 | 3131 | 3131 | 3131 | 44.5 | 150 |
| 9.08E-38 | 1.18E-34 | 1592 | 3131 | 3131 | 3131 | 45.4 | 222 |
| 1.68E-37 | 2.19E-34 | 4685 | 3131 | 3131 | 3131 | 2.97 | 1144 |
| 2.40E-37 | 3.13E-34 | 2035 | 3131 | 3131 | 3131 | 21.7 | 426 |
| 9.47E-37 | 1.23E-33 | 2018 | 3131 | 3131 | 3131 | 8.02 | 540 |
| 1.65E-36 | 2.15E-33 | 1467 | 3131 | 3131 | 3131 | 1.42 | 831 |
| 2.75E-36 | 3.58E-33 | 803 | 3131 | 3131 | 3131 | 1.15 | 571 |

| | | | | | | | |
|---|---|---|---|---|---|---|---|
| 6.73E-36 | 8.76E-33 | 1462 | 3131 | 3131 | 3131 | 1.21 | 872 |
| 6.96E-36 | 9.06E-33 | 2536 | 3131 | 3131 | 3131 | 2.68 | 910 |
| 2.23E-35 | 2.90E-32 | 2765 | 3131 | 3131 | 3131 | 2.42 | 873 |
| 7.11E-35 | 9.26E-32 | 4101 | 3131 | 3131 | 3131 | 2.65 | 1494 |
| 2.36E-34 | 3.07E-31 | 954 | 3131 | 3131 | 3131 | 1.58 | 601 |
| 2.83E-34 | 3.69E-31 | 954 | 3131 | 3131 | 3131 | 1.37 | 569 |
| 3.44E-34 | 4.48E-31 | 1749 | 3131 | 3131 | 3131 | 2.84 | 724 |
| 1.11E-33 | 1.45E-30 | 785 | 3131 | 3131 | 3131 | 11.5 | 334 |
| 1.43E-33 | 1.87E-30 | 2568 | 3131 | 3131 | 3131 | 2.23 | 1146 |
| 2.16E-33 | 2.81E-30 | 1894 | 3131 | 3131 | 3131 | 1.05 | 1175 |
| 3.33E-33 | 4.34E-30 | 3797 | 3131 | 3131 | 3131 | 1.55 | 1416 |
| 7.04E-33 | 9.17E-30 | 2272 | 3131 | 3131 | 3131 | 1.38 | 1135 |
| 1.72E-32 | 2.24E-29 | 1500 | 3131 | 3131 | 3131 | 1.02 | 959 |
| 1.72E-32 | 2.24E-29 | 2208 | 3131 | 3131 | 3131 | 3.19 | 965 |
| 1.98E-32 | 2.58E-29 | 1686 | 3131 | 3131 | 3131 | 2.24 | 685 |
| 5.42E-32 | 7.06E-29 | 590 | 3131 | 3131 | 3131 | 3.01 | 322 |
| 2.94E-31 | 3.83E-28 | 1281 | 3131 | 3131 | 3131 | 1.27 | 747 |
| 3.66E-31 | 4.76E-28 | 1430 | 3131 | 3131 | 3131 | 1.51 | 850 |
| 8.36E-31 | 1.09E-27 | 891 | 3131 | 3131 | 3131 | 1.02 | 608 |
| 1.07E-30 | 1.39E-27 | 2054 | 3131 | 3131 | 3131 | 1.24 | 1156 |
| 1.24E-30 | 1.62E-27 | 2274 | 3131 | 3131 | 3131 | 1.02 | 1365 |
| 2.58E-30 | 3.36E-27 | 3071 | 3131 | 3131 | 3131 | 1.44 | 1419 |
| 4.66E-30 | 6.07E-27 | 1637 | 3131 | 3131 | 3131 | 1.03 | 1025 |
| 7.03E-30 | 9.15E-27 | 3727 | 3131 | 3131 | 3131 | 2.01 | 1636 |
| 9.53E-30 | 1.24E-26 | 741 | 3131 | 3131 | 3131 | 1.02 | 518 |
| 1.21E-29 | 1.58E-26 | 2126 | 3131 | 3131 | 3131 | 16.9 | 533 |
| 1.44E-29 | 1.88E-26 | 2996 | 3131 | 3131 | 3131 | 1.99 | 1314 |
| 6.54E-29 | 8.51E-26 | 593 | 3131 | 3131 | 3131 | 1.31 | 425 |
| 8.99E-29 | 1.17E-25 | 3477 | 3131 | 3131 | 3131 | 3.06 | 1051 |
| 1.21E-28 | 1.57E-25 | 1441 | 3131 | 3131 | 3131 | 1.52 | 729 |
| 2.23E-28 | 2.91E-25 | 4365 | 3131 | 3131 | 3131 | 2.38 | 1180 |
| 3.83E-28 | 4.99E-25 | 1676 | 3131 | 3131 | 3131 | 1.28 | 947 |
| 7.81E-28 | 1.02E-24 | 1065 | 3131 | 3131 | 3131 | 1.09 | 689 |
| 1.05E-27 | 1.36E-24 | 2243 | 3131 | 3131 | 3131 | 1.5 | 1023 |
| 1.15E-27 | 1.49E-24 | 1266 | 3131 | 3131 | 3131 | 1.51 | 792 |
| 1.86E-27 | 2.42E-24 | 1208 | 3131 | 3131 | 3131 | 1.08 | 769 |
| 1.90E-27 | 2.47E-24 | 3040 | 3131 | 3131 | 3131 | 1.55 | 1311 |
| 2.30E-27 | 2.99E-24 | 1242 | 3131 | 3131 | 3131 | 1.4 | 729 |
| 3.01E-27 | 3.92E-24 | 934 | 3131 | 3131 | 3131 | 15.3 | 192 |
| 3.51E-27 | 4.57E-24 | 1470 | 3131 | 3131 | 3131 | 48.5 | 188 |
| 5.26E-27 | 6.85E-24 | 676 | 3131 | 3131 | 3131 | 4.65 | 302 |
| 5.26E-27 | 6.85E-24 | 355 | 3131 | 3131 | 3131 | 1.04 | 276 |
| 7.01E-27 | 9.13E-24 | 2615 | 3131 | 3131 | 3131 | 1.63 | 874 |
| 9.53E-27 | 1.24E-23 | 1632 | 3131 | 3131 | 3131 | 1.84 | 732 |
| 9.99E-27 | 1.30E-23 | 3829 | 3131 | 3131 | 3131 | 2.53 | 1176 |
| 3.76E-26 | 4.90E-23 | 2824 | 3131 | 3131 | 3131 | 2 | 1132 |
| 1.02E-25 | 1.33E-22 | 1415 | 3131 | 3131 | 3131 | 1.8 | 713 |

| | | | | | | | |
|---|---|---|---|---|---|---|---|
| 1.97E-25 | 2.57E-22 | 1883 | 3131 | 3131 | 3131 | 2.04 | 780 |
| 2.04E-25 | 2.66E-22 | 2932 | 3131 | 3131 | 3131 | 1.02 | 1674 |
| 3.47E-25 | 4.52E-22 | 1041 | 3131 | 3131 | 3131 | 1.2 | 632 |
| 6.35E-25 | 8.27E-22 | 1612 | 3131 | 3131 | 3131 | 1.43 | 863 |
| 8.69E-25 | 1.13E-21 | 3043 | 3131 | 3131 | 3131 | 1.07 | 1671 |
| 7.38E-24 | 9.61E-21 | 2309 | 3131 | 3131 | 3131 | 1.02 | 1355 |
| 2.49E-23 | 3.24E-20 | 1428 | 3131 | 3131 | 3131 | 1.41 | 734 |
| 6.61E-23 | 8.61E-20 | 3246 | 3131 | 3131 | 3131 | 5.32 | 731 |
| 2.63E-22 | 3.42E-19 | 4082 | 3131 | 3131 | 3131 | 1.45 | 1708 |
| 3.21E-22 | 4.18E-19 | 2283 | 3131 | 3131 | 3131 | 1.04 | 1327 |
| 4.45E-22 | 5.80E-19 | 1583 | 3131 | 3131 | 3131 | 1.04 | 963 |
| 4.86E-22 | 6.33E-19 | 2300 | 3131 | 3131 | 3131 | 2.21 | 773 |
| 6.53E-22 | 8.50E-19 | 3669 | 3131 | 3131 | 3131 | 1.7 | 1635 |
| 1.63E-21 | 2.13E-18 | 1293 | 3131 | 3131 | 3131 | 1.35 | 709 |
| 1.86E-21 | 2.43E-18 | 3784 | 3131 | 3131 | 3131 | 1.56 | 1444 |
| 2.00E-21 | 2.61E-18 | 2312 | 3131 | 3131 | 3131 | 1.09 | 1292 |
| 8.16E-21 | 1.06E-17 | 2511 | 3131 | 3131 | 3131 | 1.03 | 1442 |
| 9.38E-21 | 1.22E-17 | 1018 | 3131 | 3131 | 3131 | 1.77 | 596 |
| 9.42E-21 | 1.23E-17 | 1793 | 3131 | 3131 | 3131 | 1.79 | 718 |
| 1.97E-20 | 2.56E-17 | 1144 | 3131 | 3131 | 3131 | 1.11 | 719 |
| 2.97E-20 | 3.86E-17 | 1678 | 3131 | 3131 | 3131 | 8.09 | 397 |
| 3.84E-20 | 5.00E-17 | 2237 | 3131 | 3131 | 3131 | 1.05 | 1287 |
| 7.13E-20 | 9.28E-17 | 1555 | 3131 | 3131 | 3131 | 1.64 | 712 |
| 9.72E-20 | 1.27E-16 | 3430 | 3131 | 3131 | 3131 | 1.95 | 1356 |
| 2.50E-19 | 3.25E-16 | 2788 | 3131 | 3131 | 3131 | 3.85 | 680 |
| 4.63E-19 | 6.02E-16 | 117 | 3131 | 3131 | 3131 | 2.13 | 97 |
| 5.60E-19 | 7.29E-16 | 935 | 3131 | 3131 | 3131 | 1.59 | 519 |
| 7.24E-19 | 9.42E-16 | 651 | 3131 | 3131 | 3131 | 1.19 | 439 |
| 1.20E-18 | 1.56E-15 | 868 | 3131 | 3131 | 3131 | 1.26 | 523 |
| 2.24E-18 | 2.91E-15 | 2544 | 3131 | 3131 | 3131 | 1.07 | 1448 |
| 2.25E-18 | 2.94E-15 | 1226 | 3131 | 3131 | 3131 | 1.07 | 742 |
| 3.41E-18 | 4.44E-15 | 1215 | 3131 | 3131 | 3131 | 2.53 | 501 |
| 5.46E-18 | 7.10E-15 | 1646 | 3131 | 3131 | 3131 | 1.09 | 971 |
| 1.08E-17 | 1.41E-14 | 620 | 3131 | 3131 | 3131 | 1.73 | 371 |
| 4.02E-17 | 5.23E-14 | 3211 | 3131 | 3131 | 3131 | 1.01 | 1788 |
| 4.70E-17 | 6.12E-14 | 1161 | 3131 | 3131 | 3131 | 1.12 | 675 |
| 5.51E-17 | 7.17E-14 | 889 | 3131 | 3131 | 3131 | 1.04 | 565 |
| 1.91E-16 | 2.48E-13 | 1114 | 3131 | 3131 | 3131 | 2.29 | 447 |
| 2.04E-16 | 2.66E-13 | 937 | 3131 | 3131 | 3131 | 15.3 | 210 |
| 2.73E-16 | 3.55E-13 | 2450 | 3131 | 3131 | 3131 | 14.5 | 276 |
| 3.07E-16 | 4.00E-13 | 1511 | 3131 | 3131 | 3131 | 2.36 | 547 |
| 3.43E-16 | 4.46E-13 | 1008 | 3131 | 3131 | 3131 | 1.01 | 633 |
| 5.07E-16 | 6.60E-13 | 2575 | 3131 | 3131 | 3131 | 3.01 | 630 |
| 5.12E-16 | 6.67E-13 | 1781 | 3131 | 3131 | 3131 | 1.78 | 685 |
| 1.39E-15 | 1.81E-12 | 266 | 3131 | 3131 | 3131 | 2.13 | 159 |
| 1.98E-15 | 2.58E-12 | 2141 | 3131 | 3131 | 3131 | 1.05 | 1214 |
| 3.18E-15 | 4.14E-12 | 2293 | 3131 | 3131 | 3131 | 1.5 | 907 |

| | | | | | | | |
|---|---|---|---|---|---|---|---|
| 4.80E-15 | 6.25E-12 | 2558 | 3131 | 3131 | 3131 | 2.29 | 839 |
| 5.95E-15 | 7.75E-12 | 2046 | 3131 | 3131 | 3131 | 1.03 | 1183 |
| 1.04E-14 | 1.36E-11 | 1479 | 3131 | 3131 | 3131 | 3.64 | 418 |
| 1.37E-14 | 1.79E-11 | 1348 | 3131 | 3131 | 3131 | 1.83 | 550 |
| 1.55E-14 | 2.02E-11 | 1784 | 3131 | 3131 | 3131 | 1.25 | 880 |
| 1.98E-14 | 2.57E-11 | 2548 | 3131 | 3131 | 3131 | 1.21 | 1284 |
| 2.01E-14 | 2.62E-11 | 1893 | 3131 | 3131 | 3131 | 1.46 | 953 |
| 2.22E-14 | 2.88E-11 | 689 | 3131 | 3131 | 3131 | 1.11 | 437 |
| 2.27E-14 | 2.95E-11 | 138 | 3131 | 3131 | 3131 | 1.04 | 115 |
| 3.11E-14 | 4.05E-11 | 784 | 3131 | 3131 | 3131 | 11.9 | 162 |
| 4.85E-14 | 6.32E-11 | 1682 | 3131 | 3131 | 3131 | 1.07 | 988 |
| 7.46E-14 | 9.71E-11 | 2723 | 3131 | 3131 | 3131 | 1.03 | 1517 |
| 7.90E-14 | 1.03E-10 | 2345 | 3131 | 3131 | 3131 | 3.49 | 588 |
| 7.91E-14 | 1.03E-10 | 1781 | 3131 | 3131 | 3131 | 1.45 | 888 |
| 1.10E-13 | 1.43E-10 | 836 | 3131 | 3131 | 3131 | 1.09 | 512 |
| 2.09E-13 | 2.72E-10 | 2942 | 3131 | 3131 | 3131 | 1.11 | 1572 |
| 2.19E-13 | 2.85E-10 | 867 | 3131 | 3131 | 3131 | 2.76 | 365 |
| 3.13E-13 | 4.07E-10 | 1098 | 3131 | 3131 | 3131 | 1.24 | 613 |
| 3.48E-13 | 4.53E-10 | 1048 | 3131 | 3131 | 3131 | 1.11 | 639 |
| 4.74E-13 | 6.17E-10 | 1582 | 3131 | 3131 | 3131 | 9.11 | 145 |
| 5.42E-13 | 7.06E-10 | 1426 | 3131 | 3131 | 3131 | 1.07 | 840 |
| 1.08E-12 | 1.41E-09 | 3492 | 3131 | 3131 | 3131 | 1.88 | 1252 |
| 1.14E-12 | 1.48E-09 | 1274 | 3131 | 3131 | 3131 | 1.46 | 665 |
| 1.35E-12 | 1.76E-09 | 2273 | 3131 | 3131 | 3131 | 3.9 | 565 |
| 2.01E-12 | 2.62E-09 | 2434 | 3131 | 3131 | 3131 | 1.42 | 1141 |
| 2.42E-12 | 3.16E-09 | 173 | 3131 | 3131 | 3131 | 1.05 | 135 |
| 2.61E-12 | 3.39E-09 | 1463 | 3131 | 3131 | 3131 | 1.13 | 823 |
| 2.69E-12 | 3.50E-09 | 2167 | 3131 | 3131 | 3131 | 1.5 | 1031 |
| 4.37E-12 | 5.69E-09 | 764 | 3131 | 3131 | 3131 | 1.06 | 482 |
| 5.02E-12 | 6.53E-09 | 1574 | 3131 | 3131 | 3131 | 1.22 | 833 |
| 5.39E-12 | 7.02E-09 | 2877 | 3131 | 3131 | 3131 | 1.26 | 1392 |
| 5.42E-12 | 7.06E-09 | 4293 | 3131 | 3131 | 3131 | 1.9 | 1610 |
| 5.75E-12 | 7.48E-09 | 2241 | 3131 | 3131 | 3131 | 3.16 | 500 |
| 6.38E-12 | 8.31E-09 | 425 | 3131 | 3131 | 3131 | 1.23 | 275 |
| 7.84E-12 | 1.02E-08 | 762 | 3131 | 3131 | 3131 | 1.13 | 466 |
| 9.10E-12 | 1.18E-08 | 762 | 3131 | 3131 | 3131 | 1.31 | 449 |
| 9.49E-12 | 1.24E-08 | 962 | 3131 | 3131 | 3131 | 1.07 | 587 |
| 9.87E-12 | 1.28E-08 | 1169 | 3131 | 3131 | 3131 | 1.08 | 670 |
| 1.00E-11 | 1.31E-08 | 2204 | 3131 | 3131 | 3131 | 1.37 | 1060 |
| 1.22E-11 | 1.59E-08 | 3663 | 3131 | 3131 | 3131 | 3.85 | 750 |
| 1.32E-11 | 1.72E-08 | 2576 | 3131 | 3131 | 3131 | 2.6 | 792 |
| 1.59E-11 | 2.06E-08 | 1459 | 3131 | 3131 | 3131 | 4.39 | 365 |
| 1.73E-11 | 2.25E-08 | 2888 | 3131 | 3131 | 3131 | 5.13 | 422 |
| 1.73E-11 | 2.25E-08 | 959 | 3131 | 3131 | 3131 | 1.03 | 587 |
| 2.17E-11 | 2.82E-08 | 840 | 3131 | 3131 | 3131 | 2.35 | 422 |
| 2.78E-11 | 3.61E-08 | 99 | 3131 | 3131 | 3131 | 1.08 | 84 |
| 2.91E-11 | 3.79E-08 | 2021 | 3131 | 3131 | 3131 | 1.04 | 1138 |

| | | | | | | | |
|---|---|---|---|---|---|---|---|
| 3.62E-11 | 4.71E-08 | 1858 | 3131 | 3131 | 3131 | 1.37 | 869 |
| 5.98E-11 | 7.79E-08 | 2283 | 3131 | 3131 | 3131 | 1.52 | 937 |
| 9.17E-11 | 1.19E-07 | 2429 | 3131 | 3131 | 3131 | 1.06 | 1357 |
| 9.66E-11 | 1.26E-07 | 473 | 3131 | 3131 | 3131 | 1.47 | 270 |
| 1.45E-10 | 1.89E-07 | 384 | 3131 | 3131 | 3131 | 1.37 | 237 |
| 1.73E-10 | 2.26E-07 | 539 | 3131 | 3131 | 3131 | 1.17 | 347 |
| 2.14E-10 | 2.78E-07 | 1569 | 3131 | 3131 | 3131 | 15.3 | 183 |
| 2.77E-10 | 3.61E-07 | 3833 | 3131 | 3131 | 3131 | 1.67 | 1517 |
| 2.83E-10 | 3.69E-07 | 2469 | 3131 | 3131 | 3131 | 4.07 | 514 |
| 3.20E-10 | 4.17E-07 | 1297 | 3131 | 3131 | 3131 | 1.25 | 694 |
| 3.95E-10 | 5.15E-07 | 1640 | 3131 | 3131 | 3131 | 1.74 | 672 |
| 5.35E-10 | 6.96E-07 | 1306 | 3131 | 3131 | 3131 | 1.05 | 763 |
| 5.69E-10 | 7.41E-07 | 2574 | 3131 | 3131 | 3131 | 2.34 | 890 |
| 7.59E-10 | 9.88E-07 | 1497 | 3131 | 3131 | 3131 | 1.34 | 762 |
| 9.69E-10 | 1.26E-06 | 547 | 3131 | 3131 | 3131 | 1.09 | 345 |
| 9.71E-10 | 1.26E-06 | 886 | 3131 | 3131 | 3131 | 1.16 | 540 |
| 1.55E-09 | 2.02E-06 | 1648 | 3131 | 3131 | 3131 | 1.13 | 904 |
| 1.98E-09 | 2.58E-06 | 1859 | 3131 | 3131 | 3131 | 1.2 | 975 |
| 2.00E-09 | 2.60E-06 | 2974 | 3131 | 3131 | 3131 | 1.54 | 1098 |
| 2.12E-09 | 2.76E-06 | 584 | 3131 | 3131 | 3131 | 1.59 | 295 |
| 2.27E-09 | 2.96E-06 | 1405 | 3131 | 3131 | 3131 | 1.08 | 786 |
| 2.90E-09 | 3.77E-06 | 2649 | 3131 | 3131 | 3131 | 1.36 | 1271 |
| 2.91E-09 | 3.79E-06 | 968 | 3131 | 3131 | 3131 | 1.1 | 562 |
| 3.05E-09 | 3.97E-06 | 2146 | 3131 | 3131 | 3131 | 1.1 | 1169 |
| 3.36E-09 | 4.37E-06 | 2217 | 3131 | 3131 | 3131 | 2.05 | 828 |
| 3.65E-09 | 4.75E-06 | 1711 | 3131 | 3131 | 3131 | 1.03 | 978 |
| 4.22E-09 | 5.49E-06 | 1296 | 3131 | 3131 | 3131 | 1.03 | 752 |
| 6.10E-09 | 7.94E-06 | 225 | 3131 | 3131 | 3131 | 9.31 | 69 |
| 6.74E-09 | 8.77E-06 | 2956 | 3131 | 3131 | 3131 | 1.18 | 1416 |
| 7.31E-09 | 9.51E-06 | 2576 | 3131 | 3131 | 3131 | 1.03 | 1422 |
| 8.43E-09 | 1.10E-05 | 1906 | 3131 | 3131 | 3131 | 1.9 | 698 |
| 9.98E-09 | 1.30E-05 | 707 | 3131 | 3131 | 3131 | 1.03 | 437 |
| 1.14E-08 | 1.49E-05 | 1488 | 3131 | 3131 | 3131 | 13.1 | 315 |
| 1.19E-08 | 1.54E-05 | 1725 | 3131 | 3131 | 3131 | 1.11 | 939 |
| 1.21E-08 | 1.57E-05 | 1121 | 3131 | 3131 | 3131 | 1.05 | 650 |
| 1.44E-08 | 1.87E-05 | 1928 | 3131 | 3131 | 3131 | 21.2 | 80 |
| 1.69E-08 | 2.19E-05 | 552 | 3131 | 3131 | 3131 | 1.12 | 345 |
| 3.61E-08 | 4.70E-05 | 1511 | 3131 | 3131 | 3131 | 1.05 | 863 |
| 4.45E-08 | 5.80E-05 | 1211 | 3131 | 3131 | 3131 | 3.32 | 466 |
| 4.60E-08 | 5.99E-05 | 1221 | 3131 | 3131 | 3131 | 1.32 | 600 |
| 8.28E-08 | 1.08E-04 | 2151 | 3131 | 3131 | 3131 | 1.12 | 1123 |
| 1.22E-07 | 1.59E-04 | 1600 | 3131 | 3131 | 3131 | 1.19 | 838 |
| 1.43E-07 | 1.86E-04 | 914 | 3131 | 3131 | 3131 | 1.31 | 473 |
| 1.58E-07 | 2.06E-04 | 704 | 3131 | 3131 | 3131 | 4.43 | 198 |
| 1.63E-07 | 2.12E-04 | 1210 | 3131 | 3131 | 3131 | 1.19 | 647 |
| 1.73E-07 | 2.26E-04 | 893 | 3131 | 3131 | 3131 | 1.2 | 484 |
| 1.83E-07 | 2.38E-04 | 2773 | 3131 | 3131 | 3131 | 8.84 | 293 |

| | | | | | | | |
|---|---|---|---|---|---|---|---|
| 1.90E-07 | 2.47E-04 | 386 | 3131 | 3131 | 3131 | 111 | 35 |
| 1.91E-07 | 2.49E-04 | 2269 | 3131 | 3131 | 3131 | 1.82 | 864 |
| 1.92E-07 | 2.49E-04 | 2175 | 3131 | 3131 | 3131 | 1.1 | 1166 |
| 3.64E-07 | 4.74E-04 | 1618 | 3131 | 3131 | 3131 | 13.5 | 198 |
| 4.17E-07 | 5.43E-04 | 2055 | 3131 | 3131 | 3131 | 1.02 | 1142 |
| 5.70E-07 | 7.42E-04 | 2832 | 3131 | 3131 | 3131 | 1.03 | 1530 |
| 7.56E-07 | 9.84E-04 | 427 | 3131 | 3131 | 3131 | 1.34 | 255 |
| 7.85E-07 | 1.02E-03 | 1849 | 3131 | 3131 | 3131 | 1.18 | 900 |
| 8.40E-07 | 1.09E-03 | 1744 | 3131 | 3131 | 3131 | 1.04 | 958 |
| 9.58E-07 | 1.25E-03 | 269 | 3131 | 3131 | 3131 | 1.11 | 181 |
| 1.47E-06 | 1.92E-03 | 1553 | 3131 | 3131 | 3131 | 1.21 | 823 |
| 1.74E-06 | 2.27E-03 | 1260 | 3131 | 3131 | 3131 | 4.34 | 232 |
| 1.91E-06 | 2.48E-03 | 1299 | 3131 | 3131 | 3131 | 2.79 | 496 |
| 1.97E-06 | 2.57E-03 | 698 | 3131 | 3131 | 3131 | 2.61 | 367 |
| 2.00E-06 | 2.61E-03 | 578 | 3131 | 3131 | 3131 | 51.8 | 62 |
| 2.88E-06 | 3.75E-03 | 715 | 3131 | 3131 | 3131 | 1.27 | 426 |
| 3.05E-06 | 3.97E-03 | 82 | 3131 | 3131 | 3131 | 1.12 | 65 |
| 3.53E-06 | 4.59E-03 | 1141 | 3131 | 3131 | 3131 | 15.1 | 135 |
| 5.47E-06 | 7.13E-03 | 2484 | 3131 | 3131 | 3131 | 10.7 | 221 |
| 5.80E-06 | 7.55E-03 | 686 | 3131 | 3131 | 3131 | 1.03 | 413 |
| 6.75E-06 | 8.79E-03 | 3546 | 3131 | 3131 | 3131 | 1.23 | 1657 |
| 7.25E-06 | 9.44E-03 | 1446 | 3131 | 3131 | 3131 | 1.2 | 697 |
| 7.52E-06 | 9.79E-03 | 1329 | 3131 | 3131 | 3131 | 1.08 | 752 |
| 7.57E-06 | 9.85E-03 | 1594 | 3131 | 3131 | 3131 | 1.45 | 817 |
| 8.14E-06 | 1.06E-02 | 1329 | 3131 | 3131 | 3131 | 1.14 | 746 |
| 8.59E-06 | 1.12E-02 | 1817 | 3131 | 3131 | 3131 | 1.05 | 997 |
| 1.03E-05 | 1.34E-02 | 1191 | 3131 | 3131 | 3131 | 3.34 | 432 |
| 1.13E-05 | 1.48E-02 | 1479 | 3131 | 3131 | 3131 | 7.14 | 370 |
| 1.15E-05 | 1.50E-02 | 1515 | 3131 | 3131 | 3131 | 1.06 | 835 |
| 1.42E-05 | 1.85E-02 | 1644 | 3131 | 3131 | 3131 | 1.06 | 906 |
| 2.37E-05 | 3.08E-02 | 141 | 3131 | 3131 | 3131 | 1.54 | 84 |
| 2.62E-05 | 3.41E-02 | 384 | 3131 | 3131 | 3131 | 1.49 | 185 |
| 2.69E-05 | 3.51E-02 | 2763 | 3131 | 3131 | 3131 | 1.48 | 1254 |
| 2.75E-05 | 3.57E-02 | 1601 | 3131 | 3131 | 3131 | 1.04 | 892 |
| 2.83E-05 | 3.69E-02 | 1691 | 3131 | 3131 | 3131 | 1.11 | 917 |
| 3.00E-05 | 3.91E-02 | 2747 | 3131 | 3131 | 3131 | 1.11 | 1404 |
| 3.66E-05 | 4.76E-02 | 1635 | 3131 | 3131 | 3131 | 1.06 | 913 |
| 3.70E-05 | 4.82E-02 | 1793 | 3131 | 3131 | 3131 | 1.94 | 661 |
| 4.93E-05 | 6.42E-02 | 1273 | 3131 | 3131 | 3131 | 1.88 | 460 |
| 5.20E-05 | 6.76E-02 | 271 | 3131 | 3131 | 3131 | 1.09 | 172 |
| 5.41E-05 | 7.05E-02 | 898 | 3131 | 3131 | 3131 | 1.04 | 511 |
| 6.79E-05 | 8.84E-02 | 949 | 3131 | 3131 | 3131 | 1.05 | 533 |
| 7.61E-05 | 9.91E-02 | 1945 | 3131 | 3131 | 3131 | 1.16 | 993 |
| 8.04E-05 | 1.05E-01 | 209 | 3131 | 3131 | 3131 | 83.7 | 25 |
| 1.17E-04 | 1.53E-01 | 958 | 3131 | 3131 | 3131 | 3.25 | 304 |
| 1.21E-04 | 1.58E-01 | 3505 | 3131 | 3131 | 3131 | 1.08 | 1756 |
| 1.40E-04 | 1.82E-01 | 2267 | 3131 | 3131 | 3131 | 30.8 | 46 |

| | | | | | | | |
|---|---|---|---|---|---|---|---|
| 1.44E-04 | 1.87E-01 | 1664 | 3131 | 3131 | 3131 | 3.37 | 396 |
| 1.51E-04 | 1.97E-01 | 397 | 3131 | 3131 | 3131 | 5.82 | 134 |
| 1.55E-04 | 2.01E-01 | 673 | 3131 | 3131 | 3131 | 816 | 22 |
| 1.62E-04 | 2.11E-01 | 1923 | 3131 | 3131 | 3131 | 1.59 | 885 |
| 1.79E-04 | 2.33E-01 | 389 | 3131 | 3131 | 3131 | 45.7 | 21 |
| 1.94E-04 | 2.53E-01 | 577 | 3131 | 3131 | 3131 | 36.3 | 49 |
| 2.20E-04 | 2.87E-01 | 2362 | 3131 | 3131 | 3131 | 2.23 | 481 |
| 2.23E-04 | 2.90E-01 | 1037 | 3131 | 3131 | 3131 | 1.98 | 415 |
| 2.37E-04 | 3.09E-01 | 2956 | 3131 | 3131 | 3131 | 6.05 | 233 |
| 2.43E-04 | 3.16E-01 | 115 | 3131 | 3131 | 3131 | 1.34 | 76 |
| 2.55E-04 | 3.32E-01 | 272 | 3131 | 3131 | 3131 | 3 | 104 |
| 2.68E-04 | 3.49E-01 | 1935 | 3131 | 3131 | 3131 | 2.45 | 543 |
| 4.23E-04 | 5.51E-01 | 1609 | 3131 | 3131 | 3131 | 1.12 | 818 |
| 4.51E-04 | 5.87E-01 | 1490 | 3131 | 3131 | 3131 | 2.18 | 493 |
| 5.28E-04 | 6.88E-01 | 725 | 3131 | 3131 | 3131 | 1.49 | 338 |
| 5.51E-04 | 7.17E-01 | 1132 | 3131 | 3131 | 3131 | 3.7 | 208 |
| 6.76E-04 | 8.80E-01 | 958 | 3131 | 3131 | 3131 | 1.2 | 503 |
| 7.25E-04 | 9.44E-01 | 318 | 3131 | 3131 | 3131 | 52.4 | 35 |
| 7.87E-04 | 1.02E+00 | 1825 | 3131 | 3131 | 3131 | 3.29 | 709 |
| 8.84E-04 | 1.15E+00 | 555 | 3131 | 3131 | 3131 | 2.02 | 231 |
| 9.24E-04 | 1.20E+00 | 1493 | 3131 | 3131 | 3131 | 2.51 | 558 |
| 9.42E-04 | 1.23E+00 | 770 | 3131 | 3131 | 3131 | 1.06 | 437 |
| 9.50E-04 | 1.24E+00 | 1619 | 3131 | 3131 | 3131 | 1.07 | 890 |
| 9.63E-04 | 1.25E+00 | 2002 | 3131 | 3131 | 3131 | 1.15 | 982 |
| 1.09E-03 | 1.41E+00 | 57 | 3131 | 3131 | 3131 | 1.22 | 37 |
| 1.15E-03 | 1.50E+00 | 822 | 3131 | 3131 | 3131 | 1.04 | 466 |
| 1.27E-03 | 1.65E+00 | 1332 | 3131 | 3131 | 3131 | 10.2 | 58 |
| 1.32E-03 | 1.72E+00 | 2435 | 3131 | 3131 | 3131 | 1.01 | 1312 |
| 1.45E-03 | 1.89E+00 | 1258 | 3131 | 3131 | 3131 | 11.2 | 96 |
| 1.68E-03 | 2.19E+00 | 428 | 3131 | 3131 | 3131 | 1.85 | 192 |
| 1.69E-03 | 2.20E+00 | 120 | 3131 | 3131 | 3131 | 1.59 | 77 |
| 1.70E-03 | 2.22E+00 | 84 | 3131 | 3131 | 3131 | 2.61 | 51 |
| 1.92E-03 | 2.50E+00 | 72 | 3131 | 3131 | 3131 | 1.83 | 40 |
| 2.00E-03 | 2.61E+00 | 522 | 3131 | 3131 | 3131 | 4.73 | 152 |
| 2.00E-03 | 2.61E+00 | 1069 | 3131 | 3131 | 3131 | 1.38 | 504 |
| 2.25E-03 | 2.93E+00 | 2143 | 3131 | 3131 | 3131 | 1.07 | 1136 |
| 2.36E-03 | 3.07E+00 | 2101 | 3131 | 3131 | 3131 | 1.62 | 850 |
| 2.67E-03 | 3.48E+00 | 1973 | 3131 | 3131 | 3131 | 1.03 | 1065 |
| 2.74E-03 | 3.56E+00 | 888 | 3131 | 3131 | 3131 | 2.04 | 326 |
| 2.76E-03 | 3.60E+00 | 2115 | 3131 | 3131 | 3131 | 5.77 | 248 |
| 2.91E-03 | 3.79E+00 | 2966 | 3131 | 3131 | 3131 | 2.41 | 757 |
| 2.98E-03 | 3.88E+00 | 1338 | 3131 | 3131 | 3131 | 9.91 | 100 |
| 3.00E-03 | 3.91E+00 | 282 | 3131 | 3131 | 3131 | 1.93 | 139 |
| 3.02E-03 | 3.94E+00 | 91 | 3131 | 3131 | 3131 | 9.33 | 30 |
| 5.73E-03 | 7.47E+00 | 2073 | 3131 | 3131 | 3131 | 2.75 | 561 |
| 5.74E-03 | 7.48E+00 | 740 | 3131 | 3131 | 3131 | 1.18 | 424 |
| 5.77E-03 | 7.52E+00 | 1796 | 3131 | 3131 | 3131 | 1.52 | 678 |

| | | | | | | | |
|---|---|---|---|---|---|---|---|
| 5.98E-03 | 7.78E+00 | 2204 | 3131 | 3131 | 3131 | 8.61 | 129 |
| 6.26E-03 | 8.15E+00 | 386 | 3131 | 3131 | 3131 | 1.74 | 176 |
| 6.77E-03 | 8.81E+00 | 2321 | 3131 | 3131 | 3131 | 1.15 | 1155 |
| 6.96E-03 | 9.06E+00 | 953 | 3131 | 3131 | 3131 | 5.42 | 95 |
| 7.05E-03 | 9.17E+00 | 2016 | 3131 | 3131 | 3131 | 1.14 | 1024 |
| 7.35E-03 | 9.57E+00 | 2723 | 3131 | 3131 | 3131 | 1.28 | 1068 |
| 7.36E-03 | 9.58E+00 | 1899 | 3131 | 3131 | 3131 | 1.47 | 858 |

old 10.0 --control --shuffle-- --kmer 2 YT_chip_seq.fasta db/HUMAN/HOCOMOCOv11_full_HUMAN_monc

| %TP | FP | %FP |
|---|---|---|
| 54.93 | 504 | 16.1 |
| 53.56 | 514 | 16.42 |
| 47.78 | 408 | 13.03 |
| 34.88 | 221 | 7.06 |
| 31.68 | 198 | 6.32 |
| 34.11 | 244 | 7.79 |
| 31.62 | 204 | 6.52 |
| 33.92 | 250 | 7.98 |
| 23.06 | 82 | 2.62 |
| 35.32 | 288 | 9.2 |
| 32.51 | 237 | 7.57 |
| 20.89 | 61 | 1.95 |
| 28.9 | 180 | 5.75 |
| 24.75 | 128 | 4.09 |
| 25.01 | 141 | 4.5 |
| 31.59 | 267 | 8.53 |
| 35.04 | 346 | 11.05 |
| 12.74 | 14 | 0.45 |
| 32.55 | 369 | 11.79 |
| 58.13 | 1036 | 33.09 |
| 52.76 | 886 | 28.3 |
| 60.2 | 1117 | 35.68 |
| 37.62 | 506 | 16.16 |
| 42.67 | 630 | 20.12 |
| 29 | 316 | 10.09 |
| 48.8 | 809 | 25.84 |
| 43.09 | 661 | 21.11 |
| 38.17 | 543 | 17.34 |
| 35.1 | 470 | 15.01 |
| 47.08 | 778 | 24.85 |
| 16.16 | 97 | 3.1 |
| 16.54 | 104 | 3.32 |
| 26.57 | 289 | 9.23 |
| 10.38 | 25 | 0.8 |
| 36.28 | 521 | 16.64 |
| 44.59 | 733 | 23.41 |
| 32.51 | 432 | 13.8 |
| 43.88 | 718 | 22.93 |
| 65.35 | 1352 | 43.18 |
| 41.42 | 654 | 20.89 |
| 29.96 | 381 | 12.17 |
| 42.51 | 702 | 22.42 |
| 25.68 | 304 | 9.71 |
| 59.28 | 1193 | 38.1 |
| 27.79 | 355 | 11.34 |
| 43.28 | 741 | 23.67 |

| | | |
|---|---|---|
| 43.6 | 751 | 23.99 |
| 30.09 | 411 | 13.13 |
| 23.92 | 274 | 8.75 |
| 25.3 | 308 | 9.84 |
| 46.28 | 834 | 26.64 |
| 49.66 | 931 | 29.73 |
| 35.77 | 558 | 17.82 |
| 24.08 | 286 | 9.13 |
| 29.13 | 400 | 12.78 |
| 33.95 | 517 | 16.51 |
| 23.38 | 276 | 8.82 |
| 31.81 | 473 | 15.11 |
| 32.35 | 488 | 15.59 |
| 34.56 | 542 | 17.31 |
| 22.01 | 258 | 8.24 |
| 37.46 | 629 | 20.09 |
| 19.71 | 216 | 6.9 |
| 31.46 | 483 | 15.43 |
| 25.04 | 332 | 10.6 |
| 28.81 | 421 | 13.45 |
| 26.83 | 376 | 12.01 |
| 18.4 | 196 | 6.26 |
| 35.2 | 584 | 18.65 |
| 43.44 | 813 | 25.97 |
| 21.81 | 279 | 8.91 |
| 41.65 | 774 | 24.72 |
| 24.88 | 353 | 11.27 |
| 14.76 | 140 | 4.47 |
| 33.66 | 574 | 18.33 |
| 12.3 | 102 | 3.26 |
| 25.39 | 377 | 12.04 |
| 19.67 | 248 | 7.92 |
| 35.96 | 642 | 20.5 |
| 28.33 | 448 | 14.31 |
| 25.26 | 374 | 11.95 |
| 26.8 | 410 | 13.09 |
| 15.43 | 164 | 5.24 |
| 29.22 | 473 | 15.11 |
| 42.41 | 822 | 26.25 |
| 10.95 | 83 | 2.65 |
| 4.79 | 4 | 0.13 |
| 7.09 | 27 | 0.86 |
| 36.54 | 665 | 21.24 |
| 13.61 | 132 | 4.22 |
| 17.25 | 206 | 6.58 |
| 26.54 | 418 | 13.35 |
| 18.24 | 230 | 7.35 |

| | | |
|---|---|---|
| 27.85 | 453 | 14.47 |
| 29.06 | 482 | 15.39 |
| 27.88 | 455 | 14.53 |
| 47.72 | 994 | 31.75 |
| 19.2 | 258 | 8.24 |
| 18.17 | 236 | 7.54 |
| 23.12 | 347 | 11.08 |
| 10.67 | 90 | 2.87 |
| 36.6 | 691 | 22.07 |
| 37.53 | 718 | 22.93 |
| 45.23 | 933 | 29.8 |
| 36.25 | 686 | 21.91 |
| 30.63 | 541 | 17.28 |
| 30.82 | 545 | 17.41 |
| 21.88 | 326 | 10.41 |
| 10.28 | 88 | 2.81 |
| 23.86 | 379 | 12.1 |
| 27.15 | 460 | 14.69 |
| 19.42 | 278 | 8.88 |
| 36.92 | 718 | 22.93 |
| 43.6 | 905 | 28.9 |
| 45.32 | 956 | 30.53 |
| 32.74 | 610 | 19.48 |
| 52.25 | 1164 | 37.18 |
| 16.54 | 219 | 6.99 |
| 17.02 | 228 | 7.28 |
| 41.97 | 865 | 27.63 |
| 13.57 | 160 | 5.11 |
| 33.57 | 638 | 20.38 |
| 23.28 | 378 | 12.07 |
| 37.69 | 752 | 24.02 |
| 30.25 | 556 | 17.76 |
| 22.01 | 352 | 11.24 |
| 32.67 | 622 | 19.87 |
| 25.3 | 433 | 13.83 |
| 24.56 | 416 | 13.29 |
| 41.87 | 877 | 28.01 |
| 23.28 | 385 | 12.3 |
| 6.13 | 32 | 1.02 |
| 6 | 30 | 0.96 |
| 9.65 | 90 | 2.87 |
| 8.82 | 76 | 2.43 |
| 27.91 | 502 | 16.03 |
| 23.38 | 390 | 12.46 |
| 37.56 | 760 | 24.27 |
| 36.15 | 726 | 23.19 |
| 22.77 | 381 | 12.17 |

| | | |
|---|---|---|
| 24.91 | 435 | 13.89 |
| 53.47 | 1236 | 39.48 |
| 20.19 | 322 | 10.28 |
| 27.56 | 506 | 16.16 |
| 53.37 | 1238 | 39.54 |
| 43.28 | 945 | 30.18 |
| 23.44 | 411 | 13.13 |
| 23.35 | 409 | 13.06 |
| 54.55 | 1294 | 41.33 |
| 42.38 | 932 | 29.77 |
| 30.76 | 609 | 19.45 |
| 24.69 | 449 | 14.34 |
| 52.22 | 1226 | 39.16 |
| 22.64 | 402 | 12.84 |
| 46.12 | 1046 | 33.41 |
| 41.26 | 906 | 28.94 |
| 46.06 | 1051 | 33.57 |
| 19.04 | 318 | 10.16 |
| 22.93 | 413 | 13.19 |
| 22.96 | 417 | 13.32 |
| 12.68 | 173 | 5.53 |
| 41.11 | 912 | 29.13 |
| 22.74 | 414 | 13.22 |
| 43.31 | 978 | 31.24 |
| 21.72 | 390 | 12.46 |
| 3.1 | 8 | 0.26 |
| 16.58 | 269 | 8.59 |
| 14.02 | 211 | 6.74 |
| 16.7 | 274 | 8.75 |
| 46.25 | 1078 | 34.43 |
| 23.7 | 449 | 14.34 |
| 16 | 259 | 8.27 |
| 31.01 | 646 | 20.63 |
| 11.85 | 168 | 5.37 |
| 57.11 | 1422 | 45.42 |
| 21.56 | 403 | 12.87 |
| 18.05 | 316 | 10.09 |
| 14.28 | 228 | 7.28 |
| 6.71 | 67 | 2.14 |
| 8.82 | 107 | 3.42 |
| 17.47 | 305 | 9.74 |
| 20.22 | 375 | 11.98 |
| 20.12 | 371 | 11.85 |
| 21.88 | 417 | 13.32 |
| 5.08 | 42 | 1.34 |
| 38.77 | 886 | 28.3 |
| 28.97 | 611 | 19.51 |

| | | |
|---|---|---|
| 26.8 | 553 | 17.66 |
| 37.78 | 862 | 27.53 |
| 13.35 | 215 | 6.87 |
| 17.57 | 318 | 10.16 |
| 28.11 | 594 | 18.97 |
| 41.01 | 960 | 30.66 |
| 30.44 | 659 | 21.05 |
| 13.96 | 233 | 7.44 |
| 3.67 | 23 | 0.73 |
| 5.17 | 46 | 1.47 |
| 31.56 | 694 | 22.17 |
| 48.45 | 1188 | 37.94 |
| 18.78 | 352 | 11.24 |
| 28.36 | 607 | 19.39 |
| 16.35 | 295 | 9.42 |
| 50.21 | 1246 | 39.8 |
| 11.66 | 184 | 5.88 |
| 19.58 | 379 | 12.1 |
| 20.41 | 401 | 12.81 |
| 4.63 | 39 | 1.25 |
| 26.83 | 573 | 18.3 |
| 39.99 | 946 | 30.21 |
| 21.24 | 426 | 13.61 |
| 18.05 | 342 | 10.92 |
| 36.44 | 847 | 27.05 |
| 4.31 | 38 | 1.21 |
| 26.29 | 564 | 18.01 |
| 32.93 | 748 | 23.89 |
| 15.39 | 282 | 9.01 |
| 26.6 | 575 | 18.36 |
| 44.46 | 1087 | 34.72 |
| 51.42 | 1297 | 41.42 |
| 15.97 | 294 | 9.39 |
| 8.78 | 128 | 4.09 |
| 14.88 | 271 | 8.66 |
| 14.34 | 258 | 8.24 |
| 18.75 | 369 | 11.79 |
| 21.4 | 438 | 13.99 |
| 33.85 | 780 | 24.91 |
| 23.95 | 503 | 16.07 |
| 25.3 | 541 | 17.28 |
| 11.66 | 193 | 6.16 |
| 13.48 | 235 | 7.51 |
| 18.75 | 371 | 11.85 |
| 13.48 | 239 | 7.63 |
| 2.68 | 15 | 0.48 |
| 36.35 | 857 | 27.37 |

| | | |
|---|---|---|
| 27.75 | 614 | 19.61 |
| 29.93 | 676 | 21.59 |
| 43.34 | 1068 | 34.11 |
| 8.62 | 130 | 4.15 |
| 7.57 | 108 | 3.45 |
| 11.08 | 188 | 6 |
| 5.84 | 70 | 2.24 |
| 48.45 | 1226 | 39.16 |
| 16.42 | 317 | 10.12 |
| 22.17 | 471 | 15.04 |
| 21.46 | 452 | 14.44 |
| 24.37 | 533 | 17.02 |
| 28.43 | 643 | 20.54 |
| 24.34 | 533 | 17.02 |
| 11.02 | 191 | 6.1 |
| 17.25 | 346 | 11.05 |
| 28.87 | 662 | 21.14 |
| 31.14 | 727 | 23.22 |
| 35.07 | 838 | 26.76 |
| 9.42 | 155 | 4.95 |
| 25.1 | 559 | 17.85 |
| 40.59 | 1003 | 32.03 |
| 17.95 | 368 | 11.75 |
| 37.34 | 908 | 29 |
| 26.45 | 596 | 19.04 |
| 31.24 | 733 | 23.41 |
| 24.02 | 532 | 16.99 |
| 2.2 | 12 | 0.38 |
| 45.23 | 1146 | 36.6 |
| 45.42 | 1153 | 36.83 |
| 22.29 | 486 | 15.52 |
| 13.96 | 270 | 8.62 |
| 10.06 | 172 | 5.49 |
| 29.99 | 703 | 22.45 |
| 20.76 | 448 | 14.31 |
| 2.56 | 16 | 0.51 |
| 11.02 | 199 | 6.36 |
| 27.56 | 640 | 20.44 |
| 14.88 | 297 | 9.49 |
| 19.16 | 410 | 13.09 |
| 35.87 | 882 | 28.17 |
| 26.76 | 623 | 19.9 |
| 15.11 | 308 | 9.84 |
| 6.32 | 94 | 3 |
| 20.66 | 456 | 14.56 |
| 15.46 | 318 | 10.16 |
| 9.36 | 161 | 5.14 |

| | | |
|---|---|---|
| 1.12 | 1 | 0.03 |
| 27.6 | 647 | 20.66 |
| 37.24 | 927 | 29.61 |
| 6.32 | 94 | 3 |
| 36.47 | 909 | 29.03 |
| 48.87 | 1283 | 40.98 |
| 8.14 | 141 | 4.5 |
| 28.74 | 688 | 21.97 |
| 30.6 | 742 | 23.7 |
| 5.78 | 88 | 2.81 |
| 26.29 | 622 | 19.87 |
| 7.41 | 123 | 3.93 |
| 15.84 | 336 | 10.73 |
| 11.72 | 231 | 7.38 |
| 1.98 | 13 | 0.42 |
| 13.61 | 281 | 8.97 |
| 2.08 | 17 | 0.54 |
| 4.31 | 56 | 1.79 |
| 7.06 | 116 | 3.7 |
| 13.19 | 273 | 8.72 |
| 52.92 | 1423 | 45.45 |
| 22.26 | 517 | 16.51 |
| 24.02 | 567 | 18.11 |
| 26.09 | 625 | 19.96 |
| 23.83 | 562 | 17.95 |
| 31.84 | 791 | 25.26 |
| 13.8 | 289 | 9.23 |
| 11.82 | 237 | 7.57 |
| 26.67 | 644 | 20.57 |
| 28.94 | 710 | 22.68 |
| 2.68 | 30 | 0.96 |
| 5.91 | 98 | 3.13 |
| 40.05 | 1039 | 33.18 |
| 28.49 | 701 | 22.39 |
| 29.29 | 724 | 23.12 |
| 44.84 | 1185 | 37.85 |
| 29.16 | 722 | 23.06 |
| 21.11 | 492 | 15.71 |
| 14.69 | 319 | 10.19 |
| 5.49 | 91 | 2.91 |
| 16.32 | 365 | 11.66 |
| 17.02 | 385 | 12.3 |
| 31.72 | 800 | 25.55 |
| 0.8 | 1 | 0.03 |
| 9.71 | 193 | 6.16 |
| 56.08 | 1542 | 49.25 |
| 1.47 | 8 | 0.26 |

| | | |
|---|---|---|
| 12.65 | 268 | 8.56 |
| 4.28 | 65 | 2.08 |
| 0.7 | 0 | 0 |
| 28.27 | 704 | 22.48 |
| 0.67 | 0 | 0 |
| 1.56 | 11 | 0.35 |
| 15.36 | 341 | 10.89 |
| 13.25 | 288 | 9.2 |
| 7.44 | 135 | 4.31 |
| 2.43 | 29 | 0.93 |
| 3.32 | 46 | 1.47 |
| 17.34 | 397 | 12.68 |
| 26.13 | 649 | 20.73 |
| 15.75 | 357 | 11.4 |
| 10.8 | 228 | 7.28 |
| 6.64 | 121 | 3.86 |
| 16.07 | 370 | 11.82 |
| 1.12 | 6 | 0.19 |
| 22.64 | 552 | 17.63 |
| 7.38 | 143 | 4.57 |
| 17.82 | 418 | 13.35 |
| 13.96 | 315 | 10.06 |
| 28.43 | 721 | 23.03 |
| 31.36 | 806 | 25.74 |
| 1.18 | 9 | 0.29 |
| 14.88 | 341 | 10.89 |
| 1.85 | 17 | 0.54 |
| 41.9 | 1123 | 35.87 |
| 3.07 | 41 | 1.31 |
| 6.13 | 115 | 3.67 |
| 2.46 | 33 | 1.05 |
| 1.63 | 17 | 0.54 |
| 1.28 | 11 | 0.35 |
| 4.85 | 84 | 2.68 |
| 16.1 | 376 | 12.01 |
| 36.28 | 958 | 30.6 |
| 27.15 | 688 | 21.97 |
| 34.01 | 892 | 28.49 |
| 10.41 | 224 | 7.15 |
| 7.92 | 156 | 4.98 |
| 24.18 | 601 | 19.2 |
| 3.19 | 45 | 1.44 |
| 4.44 | 77 | 2.46 |
| 0.96 | 6 | 0.19 |
| 17.92 | 429 | 13.7 |
| 13.54 | 313 | 10 |
| 21.65 | 536 | 17.12 |

| | | |
|---|---|---|
| 4.12 | 66 | 2.11 |
| 5.62 | 107 | 3.42 |
| 36.89 | 984 | 31.43 |
| 3.03 | 44 | 1.41 |
| 32.71 | 860 | 27.47 |
| 34.11 | 900 | 28.74 |
| 27.4 | 704 | 22.48 |

›_meme_format.meme db/MOUSE/HOCOMOCOv11_full_MOUSE_mono_meme_format.meme

| rank | motif_ID | consensus | p-value | adj_p-value |
|---|---|---|---|---|
| 1 | MITF_HUMAN.H11MO.0.A | TCACGTGACC | 8.05E-137 | 1.96E-134 |
| 2 | USF1_HUMAN.H11MO.0.A | RGTCACGTGRBS | 8.63E-136 | 1.88E-133 |
| 3 | USF2_HUMAN.H11MO.0.A | GGGTCACGTGRBSVSSSSB | 1.51E-134 | 3.44E-132 |
| 4 | TFE3_HUMAN.H11MO.0.B | RGTCACGTGA | 1.48E-132 | 3.64E-130 |
| 5 | TFEB_HUMAN.H11MO.0.C | RGTCACGTG | 3.33E-124 | 6.13E-122 |
| 6 | MLX_HUMAN.H11MO.0.D | YGATCACGTGN | 5.42E-119 | 1.19E-116 |
| 7 | BHE40_HUMAN.H11MO.0.A | DGCACGTGAS | 1.48E-116 | 4.24E-114 |
| 8 | ATF3_HUMAN.H11MO.0.A | GGTSACGTGAB | 3.29E-110 | 1.16E-107 |
| 9 | CR3L2_HUMAN.H11MO.0.D | TGATGACGTGGCA | 3.12E-82 | 6.62E-80 |
| 10 | CLOCK_HUMAN.H11MO.0.C | CWGHCACGTGMVVM | 1.88E-62 | 9.10E-60 |
| 11 | BMAL1_HUMAN.H11MO.0.A | KSCACGTGHSY | 3.53E-58 | 9.40E-56 |
| 12 | SRBP1_HUMAN.H11MO.0.A | DGTSRGGTGAB | 7.19E-58 | 4.24E-55 |
| 13 | HEY1_HUMAN.H11MO.0.D | KGRCACGTGBCK | 5.74E-56 | 1.14E-53 |
| 14 | MYCN_HUMAN.H11MO.0.A | RGCCACGTGSDS | 2.12E-48 | 6.52E-46 |
| 15 | ATF6A_HUMAN.H11MO.0.B | GKGSTGACGTGG | 2.72E-39 | 3.62E-37 |
| 16 | BHE41_HUMAN.H11MO.0.D | WSVRKSYCACGTGMWGRRRV | 5.54E-33 | 1.06E-30 |
| 17 | SP1_HUMAN.H11MO.1.A | VGGGGGCGGGGCCK | 8.80E-33 | 4.44E-30 |
| 18 | SP2_HUMAN.H11MO.0.A | GGSSVGGGGGCGGGGCCDGSGS | 1.72E-31 | 1.17E-28 |
| 19 | MAX_HUMAN.H11MO.0.A | VRGCACGTGG | 7.16E-31 | 2.24E-28 |
| 20 | SP4_HUMAN.H11MO.0.A | SRGVARGRGGCGGRGCHDRR | 1.08E-30 | 6.95E-28 |
| 21 | SP2_HUMAN.H11MO.1.B | RGGRGGCGGGGC | 3.83E-30 | 1.99E-27 |
| 22 | KLF12_HUMAN.H11MO.0.C | VGGGGCGGGGC | 8.34E-30 | 3.18E-27 |
| 23 | SP1_HUMAN.H11MO.0.A | VGGGGGCGGGGCCKGGGGGGG | 9.94E-29 | 2.91E-26 |
| 24 | SP3_HUMAN.H11MO.0.B | SGVVGGGGGCGGGGCBRGSS | 1.63E-28 | 1.25E-25 |
| 25 | KLF3_HUMAN.H11MO.0.B | GGRGVDGGGCGGGGCYGGG | 9.55E-28 | 5.61E-25 |
| 26 | XBP1_HUMAN.H11MO.0.D | GSKGACGTGGC | 3.18E-27 | 1.29E-24 |
| 27 | SP4_HUMAN.H11MO.1.A | GRRRGGGGMGGGGCBK | 9.23E-27 | 4.83E-24 |
| 28 | MXI1_HUMAN.H11MO.0.A | VCACGTGGSNGSBGS | 1.75E-26 | 6.21E-24 |
| 29 | ARNT_HUMAN.H11MO.0.B | STACGTGMC | 1.77E-25 | 5.15E-23 |
| 30 | MXI1_HUMAN.H11MO.1.A | SCCACGTGSYB | 2.23E-25 | 7.77E-23 |
| 31 | TAF1_HUMAN.H11MO.0.A | RARRWGGCGGMGGMGR | 2.73E-23 | 1.39E-20 |
| 32 | ELF2_HUMAN.H11MO.0.C | RAVCCGGAAGTGR | 7.18E-23 | 1.95E-20 |
| 33 | E2F7_HUMAN.H11MO.0.B | GDGGCGGGAARDR | 1.86E-22 | 1.13E-19 |
| 34 | MAZ_HUMAN.H11MO.1.A | RRGGGAGGGGS | 5.84E-22 | 3.29E-19 |
| 35 | CREB3_HUMAN.H11MO.0.D | KGRTGACGTGGCA | 7.25E-20 | 4.35E-18 |
| 36 | GABPA_HUMAN.H11MO.0.A | GVVRCCGGAAGTGV | 4.63E-20 | 1.01E-17 |
| 37 | KLF9_HUMAN.H11MO.0.C | GRRGKGGGCGTGGCC | 1.09E-19 | 3.41E-17 |
| 38 | E2F4_HUMAN.H11MO.0.A | SRGGGCGGGAARD | 1.18E-19 | 6.04E-17 |
| 39 | WT1_HUMAN.H11MO.0.C | RGGVGGGGGAGGRGGVGGRG | 4.12E-19 | 2.13E-16 |
| 40 | ELK4_HUMAN.H11MO.0.A | SVCCGGAAGTGV | 1.10E-18 | 2.13E-16 |
| 41 | KLF1_HUMAN.H11MO.0.A | DGGGYGKGGCYGGG | 5.72E-19 | 3.04E-16 |
| 42 | TYY1_HUMAN.H11MO.0.A | CAARATGGCGGC | 5.66E-18 | 4.47E-16 |
| 43 | ETV1_HUMAN.H11MO.0.A | SRCCGGAAGYG | 2.95E-18 | 8.19E-16 |
| 44 | ELF1_HUMAN.H11MO.0.A | VRACCCGGAAGTGS | 1.23E-17 | 2.91E-15 |
| 45 | ELK1_HUMAN.H11MO.0.B | RCCGGAAGTGV | 6.54E-17 | 1.12E-14 |
| 46 | KLF6_HUMAN.H11MO.0.A | GSRRGGGHGGGGMHGGGRV | 2.54E-16 | 1.92E-13 |

| | | | | |
|---|---|---|---|---|
| 47 | FLI1_HUMAN.H11MO.1.A | RRRCAGGAAGBRG | 4.66E-15 | 1.58E-12 |
| 48 | TYY2_HUMAN.H11MO.0.D | AWAATGGCGGH | 2.04E-14 | 2.30E-12 |
| 49 | TFDP1_HUMAN.H11MO.0.C | RRRRGGCGGGAARN | 1.26E-14 | 6.38E-12 |
| 50 | NRF1_HUMAN.H11MO.0.A | CWSTGCGCATGCGCRDS | 7.12E-14 | 6.41E-12 |
| 51 | ESR2_HUMAN.H11MO.1.A | AGGTCAS | 2.84E-14 | 1.07E-11 |
| 52 | ETS1_HUMAN.H11MO.0.A | RRRCMGGAAGTGG | 5.55E-14 | 1.58E-11 |
| 53 | KLF4_HUMAN.H11MO.0.A | WGGGYGKGGC | 5.70E-14 | 2.51E-11 |
| 54 | EPAS1_HUMAN.H11MO.0.B | VTACGTGMC | 1.38E-13 | 3.04E-11 |
| 55 | FLI1_HUMAN.H11MO.0.A | SRRGGMAGGAAGGRRRGR | 9.83E-14 | 3.55E-11 |
| 56 | HESX1_HUMAN.H11MO.0.D | ABSYGGCWMRTRRMMT | 2.06E-13 | 4.14E-11 |
| 57 | KLF15_HUMAN.H11MO.0.A | RGGGMGGRGVDGGGGGRGG | 1.09E-13 | 5.57E-11 |
| 58 | MYC_HUMAN.H11MO.0.A | RCCACGTGCTB | 3.84E-13 | 7.37E-11 |
| 59 | PATZ1_HUMAN.H11MO.0.C | GGGGMGGGGGMKGGRRVGGVRG | 1.08E-13 | 7.80E-11 |
| 60 | ERG_HUMAN.H11MO.0.A | RRRSAGGAAGYGG | 2.30E-13 | 7.93E-11 |
| 61 | EGR2_HUMAN.H11MO.0.A | GRGRRKGWGKGGGHGGRG | 1.77E-13 | 1.16E-10 |
| 62 | HES1_HUMAN.H11MO.0.D | VMGVCACGMGMCM | 3.78E-13 | 1.60E-10 |
| 63 | ELF5_HUMAN.H11MO.0.A | VGRASSAGGAAGTRV | 6.19E-13 | 1.72E-10 |
| 64 | ZN467_HUMAN.H11MO.0.C | GGGGAGGGGRRGGRGRRGGRRR | 6.68E-13 | 2.78E-10 |
| 65 | FEV_HUMAN.H11MO.0.B | GCVGGAAGYG | 9.64E-13 | 7.10E-10 |
| 66 | ESR1_HUMAN.H11MO.1.A | CHRGGTCASV | 3.05E-12 | 9.90E-10 |
| 67 | KLF5_HUMAN.H11MO.0.A | WGGGTGKGGCDGGG | 2.30E-12 | 1.38E-09 |
| 68 | RXRB_HUMAN.H11MO.0.C | TSAGGTCASR | 4.77E-12 | 1.46E-09 |
| 69 | MAZ_HUMAN.H11MO.0.A | GGGGGAGGGGGDGRGRRRGRRG | 3.50E-12 | 2.26E-09 |
| 70 | ZBT17_HUMAN.H11MO.0.A | SRRGGWGGGGGAGGGGMRR | 1.92E-11 | 1.37E-08 |
| 71 | ETV2_HUMAN.H11MO.0.B | RRARRCAGGAARYRGS | 8.12E-11 | 1.54E-08 |
| 72 | ZFP42_HUMAN.H11MO.0.A | GGCMGCCATTTT | 8.30E-11 | 1.60E-08 |
| 73 | EHF_HUMAN.H11MO.0.B | VWACSAGGAAGTDVS | 1.10E-10 | 1.70E-08 |
| 74 | KLF14_HUMAN.H11MO.0.D | DRGKGGGCGTGGY | 7.55E-11 | 2.05E-08 |
| 75 | VEZF1_HUMAN.H11MO.1.C | RRRRGGAGGGGR | 2.55E-11 | 2.17E-08 |
| 76 | ETS2_HUMAN.H11MO.0.B | RDVRGGAARVRRR | 4.78E-11 | 2.97E-08 |
| 77 | ZN341_HUMAN.H11MO.0.C | GRGGRRRRGGRGRRGGAASAGC | 1.14E-10 | 6.60E-08 |
| 78 | VEZF1_HUMAN.H11MO.0.C | GGRRRRGRRGGAGGGGGRGRRR | 1.02E-10 | 7.06E-08 |
| 79 | IRF3_HUMAN.H11MO.0.B | RAAARGGAAAVDGAAASDGA | 3.34E-10 | 9.47E-08 |
| 80 | EGR1_HUMAN.H11MO.0.A | VDRDGCGKGGGYGGRRR | 1.61E-10 | 1.09E-07 |
| 81 | E2F6_HUMAN.H11MO.0.A | DGGGCGGGARRVR | 2.80E-10 | 1.54E-07 |
| 82 | NFYC_HUMAN.H11MO.0.A | BBRRCCAATCAGMR | 1.57E-09 | 3.78E-07 |
| 83 | ETV5_HUMAN.H11MO.0.C | RRRSAGGAARDGRV | 8.56E-10 | 4.90E-07 |
| 84 | NFYA_HUMAN.H11MO.0.A | BYRRCCAATCAGAR | 4.31E-09 | 5.77E-07 |
| 85 | ZSC22_HUMAN.H11MO.0.C | RGKCWGAGGGMGGAGG | 2.48E-09 | 9.07E-07 |
| 86 | ZFX_HUMAN.H11MO.1.A | DGVRGVVSAGGCCKVGG | 1.63E-09 | 1.16E-06 |
| 87 | E2F1_HUMAN.H11MO.0.A | RRNKGGCGGGAARR | 2.77E-09 | 1.27E-06 |
| 88 | ZN770_HUMAN.H11MO.0.C | GGGAGGCYGAGGBRGGAGGATC | 7.69E-09 | 2.27E-06 |
| 89 | SPI1_HUMAN.H11MO.0.A | RRAAAGAGGAAGTGRVD | 2.39E-08 | 3.32E-06 |
| 90 | NFYB_HUMAN.H11MO.0.A | YRRCCAATSAGMR | 2.80E-08 | 3.36E-06 |
| 91 | E2F3_HUMAN.H11MO.0.A | DDGGCGGGAAA | 8.33E-09 | 4.70E-06 |
| 92 | PATZ1_HUMAN.H11MO.1.C | AGGGAGGGG | 8.17E-09 | 6.38E-06 |
| 93 | TBX1_HUMAN.H11MO.0.D | GGGYGGGYAGGGGTGGGMDD | 1.63E-08 | 6.63E-06 |

| | | | | |
|---|---|---|---|---|
| 94 | TBX15_HUMAN.H11MO.0.D | GGGGGGGGGGGGGTGGGDR | 1.45E-08 | 9.27E-06 |
| 95 | ZN148_HUMAN.H11MO.0.D | KGRGKGGGGGAGGGG | 1.81E-08 | 1.10E-05 |
| 96 | CEBPZ_HUMAN.H11MO.0.D | DCTSATTGGCT | 1.65E-07 | 1.12E-05 |
| 97 | THA_HUMAN.H11MO.1.D | BTSAGGTCAS | 4.09E-08 | 1.20E-05 |
| 98 | KLF16_HUMAN.H11MO.0.D | GGGGGYGKGKBGGGGGGGG | 2.23E-08 | 1.47E-05 |
| 99 | ETV7_HUMAN.H11MO.0.D | VAGGAAR | 1.02E-07 | 6.03E-05 |
| 100 | ZN121_HUMAN.H11MO.0.C | CTGGGCAACABAGYRAGACC | 2.36E-06 | 7.79E-05 |
| 101 | ZN263_HUMAN.H11MO.0.A | GGGAGGAGGRRGRGGRGGRR | 3.38E-07 | 1.43E-04 |
| 102 | ELF3_HUMAN.H11MO.0.A | RAABVAGGAAGTRR | 8.51E-07 | 1.56E-04 |
| 103 | RFX2_HUMAN.H11MO.0.A | CKGTTGCCATGGCRACVS | 3.80E-06 | 1.83E-04 |
| 104 | ZBT7B_HUMAN.H11MO.0.D | SSCSGTTGCCATGGVRACSGS | 2.81E-06 | 2.50E-04 |
| 105 | ZN281_HUMAN.H11MO.0.A | RGGWGGGGGAGGGGV | 6.10E-07 | 2.76E-04 |
| 106 | RARA_HUMAN.H11MO.1.A | VAGGTCAGRG | 9.28E-07 | 3.74E-04 |
| 107 | ETV4_HUMAN.H11MO.0.B | VRCAGGAARBR | 1.59E-06 | 7.48E-04 |
| 108 | NFAT5_HUMAN.H11MO.0.D | RRRGGAARRDN | 1.48E-06 | 9.34E-04 |
| 109 | HXB13_HUMAN.H11MO.0.A | TTTTATDGSH | 3.80E-06 | 9.43E-04 |
| 110 | PURA_HUMAN.H11MO.0.D | GGWRKKGGDGGGCVKGG | 1.34E-06 | 9.45E-04 |
| 111 | GSC2_HUMAN.H11MO.0.D | RKMGGRTTRGB | 2.36E-06 | 1.09E-03 |
| 112 | PRDM6_HUMAN.H11MO.0.C | RRRARGAAAAAAA | 3.03E-06 | 1.12E-03 |
| 113 | HES7_HUMAN.H11MO.0.D | TGGCACGTGCCG | 6.71E-05 | 1.14E-03 |
| 114 | THAP1_HUMAN.H11MO.0.C | SGCCGCCATSTYGGSYBCGGGC | 2.38E-06 | 1.14E-03 |
| 115 | FOXI1_HUMAN.H11MO.0.B | YTBTGATTGGYY | 3.43E-05 | 1.65E-03 |
| 116 | HXA1_HUMAN.H11MO.0.C | TKGATGGATG | 1.20E-05 | 1.91E-03 |
| 117 | EGR2_HUMAN.H11MO.1.A | GRDGHGKGGGHGGRGG | 3.05E-06 | 1.95E-03 |
| 118 | RARG_HUMAN.H11MO.1.B | VAGGTCAS | 4.37E-06 | 1.96E-03 |
| 119 | THB_HUMAN.H11MO.1.D | SGTSAGGTCA | 9.68E-06 | 2.25E-03 |
| 120 | BC11A_HUMAN.H11MO.0.A | RAAAGAGGAAGTGARAV | 4.99E-06 | 2.30E-03 |
| 121 | WT1_HUMAN.H11MO.1.B | GWGGGAGGAGS | 3.50E-06 | 2.34E-03 |
| 122 | FOS_HUMAN.H11MO.0.A | VTGAGTCAB | 1.34E-05 | 2.50E-03 |
| 123 | SPIC_HUMAN.H11MO.0.D | KVDRRWGGGKDRG | 3.31E-06 | 2.83E-03 |
| 124 | EGR4_HUMAN.H11MO.0.D | GGSGGYRGGGM | 6.54E-06 | 2.84E-03 |
| 125 | FOSL1_HUMAN.H11MO.0.A | KRVTGAGTCAYH | 4.30E-05 | 3.52E-03 |
| 126 | SPIB_HUMAN.H11MO.0.A | RRAAAGAGGAAGTGARA | 3.69E-05 | 3.79E-03 |
| 127 | ATF7_HUMAN.H11MO.0.D | KRTGACGTCAT | 2.20E-04 | 4.83E-03 |
| 128 | OSR2_HUMAN.H11MO.0.C | GBYSCWGCTNCTGYDG | 1.77E-05 | 5.59E-03 |
| 129 | SPDEF_HUMAN.H11MO.0.D | RWCCMGGAWGWAGT | 3.10E-05 | 5.90E-03 |
| 130 | HXA7_HUMAN.H11MO.0.D | VMTCRATMGATTGGM | 2.56E-05 | 6.48E-03 |
| 131 | FOSB_HUMAN.H11MO.0.A | RTGAGTCAB | 3.06E-05 | 7.01E-03 |
| 132 | IRF8_HUMAN.H11MO.0.B | RAAARRGGAASTGAAASTDR | 5.47E-05 | 7.03E-03 |

### AME (Analysis of Motif Enrichment): Version 5.0.5 compiled on Jul 15 2019 at 00:31:14
### The format of this file is described at http://meme-suite.org//doc/ame-output-format.html.
### ame --verbose 1 --oc . --scoring avg --method fisher --hit-lo-fraction 0.25 --evalue-report-threshold 10.0 --

| E-value | tests | FASTA_max | pos | neg | PWM_min | TP | %TP | FP |
|---|---|---|---|---|---|---|---|---|
| 2.55E-131 | 243 | 279 | 279 | 1116 | 27.8 | 167 | 59.86 | 2 |
| 2.45E-130 | 218 | 279 | 279 | 1116 | 30.4 | 166 | 59.5 | 2 |
| 4.48E-129 | 228 | 279 | 279 | 1116 | 14.3 | 168 | 60.22 | 4 |
| 4.73E-127 | 246 | 279 | 279 | 1116 | 29.7 | 161 | 57.71 | 1 |
| 7.99E-119 | 184 | 279 | 279 | 1116 | 2.77 | 168 | 60.22 | 12 |
| 1.55E-113 | 219 | 279 | 279 | 1116 | 8.19 | 166 | 59.5 | 15 |
| 5.52E-111 | 286 | 279 | 279 | 1116 | 10.6 | 158 | 56.63 | 10 |
| 1.51E-104 | 354 | 279 | 279 | 1116 | 11.1 | 157 | 56.27 | 15 |
| 8.61E-77 | 212 | 279 | 279 | 1116 | 1.59 | 149 | 53.41 | 42 |
| 1.18E-56 | 484 | 279 | 279 | 1116 | 1.76 | 164 | 58.78 | 114 |
| 1.22E-52 | 266 | 279 | 279 | 1116 | 1.04 | 156 | 55.91 | 109 |
| 5.52E-52 | 590 | 279 | 279 | 1116 | 2.71 | 176 | 63.08 | 158 |
| 1.48E-50 | 198 | 279 | 279 | 1116 | 1.07 | 130 | 46.59 | 64 |
| 8.49E-43 | 308 | 279 | 279 | 1116 | 1.03 | 159 | 56.99 | 149 |
| 4.72E-34 | 133 | 279 | 279 | 1116 | 1.04 | 92 | 32.97 | 41 |
| 1.38E-27 | 192 | 279 | 279 | 1116 | 2.41 | 89 | 31.9 | 52 |
| 5.78E-27 | 505 | 279 | 279 | 1116 | 5.73 | 107 | 38.35 | 87 |
| 1.52E-25 | 681 | 279 | 279 | 1116 | 27.7 | 86 | 30.82 | 51 |
| 2.92E-25 | 313 | 279 | 279 | 1116 | 1.03 | 138 | 49.46 | 170 |
| 9.05E-25 | 646 | 279 | 279 | 1116 | 5.77 | 121 | 43.37 | 127 |
| 2.59E-24 | 518 | 279 | 279 | 1116 | 10.1 | 92 | 32.97 | 66 |
| 4.14E-24 | 381 | 279 | 279 | 1116 | 6.17 | 94 | 33.69 | 71 |
| 3.79E-23 | 293 | 279 | 279 | 1116 | 9.44 | 88 | 31.54 | 63 |
| 1.63E-22 | 766 | 279 | 279 | 1116 | 16.4 | 100 | 35.84 | 88 |
| 7.30E-22 | 587 | 279 | 279 | 1116 | 9.3 | 100 | 35.84 | 91 |
| 1.68E-21 | 407 | 279 | 279 | 1116 | 1.03 | 157 | 56.27 | 247 |
| 6.28E-21 | 523 | 279 | 279 | 1116 | 10.7 | 94 | 33.69 | 82 |
| 8.08E-21 | 354 | 279 | 279 | 1116 | 1.65 | 109 | 39.07 | 117 |
| 6.71E-20 | 291 | 279 | 279 | 1116 | 1.02 | 125 | 44.8 | 163 |
| 1.01E-19 | 349 | 279 | 279 | 1116 | 1.12 | 132 | 47.31 | 183 |
| 1.81E-17 | 510 | 279 | 279 | 1116 | 1.97 | 128 | 45.88 | 184 |
| 2.54E-17 | 272 | 279 | 279 | 1116 | 2.57 | 81 | 29.03 | 70 |
| 1.47E-16 | 605 | 279 | 279 | 1116 | 2.77 | 118 | 42.29 | 161 |
| 4.28E-16 | 563 | 279 | 279 | 1116 | 13.3 | 84 | 30.11 | 80 |
| 5.66E-15 | 60 | 279 | 279 | 1116 | 1.06 | 44 | 15.77 | 16 |
| 1.32E-14 | 219 | 279 | 279 | 1116 | 1.33 | 89 | 31.9 | 100 |
| 4.44E-14 | 312 | 279 | 279 | 1116 | 2.78 | 94 | 33.69 | 114 |
| 7.86E-14 | 512 | 279 | 279 | 1116 | 2.45 | 114 | 40.86 | 167 |
| 2.77E-13 | 516 | 279 | 279 | 1116 | 5.9 | 104 | 37.28 | 143 |
| 2.77E-13 | 194 | 279 | 279 | 1116 | 4.11 | 51 | 18.28 | 29 |
| 3.96E-13 | 532 | 279 | 279 | 1116 | 15.7 | 64 | 22.94 | 51 |
| 5.82E-13 | 79 | 279 | 279 | 1116 | 64.1 | 26 | 9.32 | 1 |
| 1.07E-12 | 278 | 279 | 279 | 1116 | 4.01 | 60 | 21.51 | 46 |
| 3.79E-12 | 237 | 279 | 279 | 1116 | 4.39 | 55 | 19.71 | 39 |
| 1.46E-11 | 172 | 279 | 279 | 1116 | 1.82 | 61 | 21.86 | 53 |
| 2.51E-10 | 759 | 279 | 279 | 1116 | 5.52 | 114 | 40.86 | 190 |

| | | | | | | | | |
|---|---|---|---|---|---|---|---|---|
| 2.06E-09 | 339 | 279 | 279 | 1116 | 3.92 | 73 | 26.16 | 88 |
| 3.00E-09 | 113 | 279 | 279 | 1116 | 19.7 | 21 | 7.53 | 1 |
| 8.31E-09 | 506 | 279 | 279 | 1116 | 1.59 | 126 | 45.16 | 241 |
| 8.34E-09 | 90 | 279 | 279 | 1116 | 17.5 | 26 | 9.32 | 6 |
| 1.39E-08 | 376 | 279 | 279 | 1116 | 1.43 | 106 | 37.99 | 182 |
| 2.06E-08 | 285 | 279 | 279 | 1116 | 5.11 | 50 | 17.92 | 43 |
| 3.27E-08 | 441 | 279 | 279 | 1116 | 1.32 | 130 | 46.59 | 260 |
| 3.96E-08 | 220 | 279 | 279 | 1116 | 1.01 | 87 | 31.18 | 133 |
| 4.62E-08 | 361 | 279 | 279 | 1116 | 9.23 | 52 | 18.64 | 48 |
| 5.39E-08 | 201 | 279 | 279 | 1116 | 1.31 | 72 | 25.81 | 95 |
| 7.25E-08 | 512 | 279 | 279 | 1116 | 8.09 | 88 | 31.54 | 135 |
| 9.59E-08 | 192 | 279 | 279 | 1116 | 1.03 | 78 | 27.96 | 112 |
| 1.02E-07 | 725 | 279 | 279 | 1116 | 15.6 | 79 | 28.32 | 111 |
| 1.03E-07 | 344 | 279 | 279 | 1116 | 4.78 | 64 | 22.94 | 76 |
| 1.51E-07 | 653 | 279 | 279 | 1116 | 3.55 | 131 | 46.95 | 268 |
| 2.09E-07 | 424 | 279 | 279 | 1116 | 1.27 | 120 | 43.01 | 235 |
| 2.24E-07 | 278 | 279 | 279 | 1116 | 1.05 | 100 | 35.84 | 175 |
| 3.62E-07 | 416 | 279 | 279 | 1116 | 6.04 | 92 | 32.97 | 152 |
| 9.25E-07 | 737 | 279 | 279 | 1116 | 2.3 | 80 | 28.67 | 120 |
| 1.29E-06 | 325 | 279 | 279 | 1116 | 1.81 | 82 | 29.39 | 129 |
| 1.80E-06 | 600 | 279 | 279 | 1116 | 1.68 | 147 | 52.69 | 335 |
| 1.91E-06 | 307 | 279 | 279 | 1116 | 1.36 | 90 | 32.26 | 153 |
| 2.94E-06 | 645 | 279 | 279 | 1116 | 2.31 | 147 | 52.69 | 337 |
| 1.79E-05 | 714 | 279 | 279 | 1116 | 4.63 | 114 | 40.86 | 232 |
| 2.01E-05 | 190 | 279 | 279 | 1116 | 1.18 | 70 | 25.09 | 107 |
| 2.08E-05 | 193 | 279 | 279 | 1116 | 17.7 | 22 | 7.89 | 7 |
| 2.22E-05 | 155 | 279 | 279 | 1116 | 3.2 | 42 | 15.05 | 41 |
| 2.67E-05 | 272 | 279 | 279 | 1116 | 4.35 | 52 | 18.64 | 62 |
| 2.83E-05 | 851 | 279 | 279 | 1116 | 8.34 | 54 | 19.35 | 64 |
| 3.87E-05 | 622 | 279 | 279 | 1116 | 2.7 | 104 | 37.28 | 204 |
| 8.60E-05 | 579 | 279 | 279 | 1116 | 13.9 | 52 | 18.64 | 63 |
| 9.19E-05 | 695 | 279 | 279 | 1116 | 4.26 | 122 | 43.73 | 266 |
| 1.23E-04 | 283 | 279 | 279 | 1116 | 3.94 | 63 | 22.58 | 93 |
| 1.42E-04 | 676 | 279 | 279 | 1116 | 2.65 | 124 | 44.44 | 275 |
| 2.00E-04 | 549 | 279 | 279 | 1116 | 2.06 | 106 | 37.99 | 218 |
| 4.92E-04 | 241 | 279 | 279 | 1116 | 4.13 | 35 | 12.54 | 32 |
| 6.38E-04 | 572 | 279 | 279 | 1116 | 5.03 | 58 | 20.79 | 83 |
| 7.52E-04 | 134 | 279 | 279 | 1116 | 4.09 | 32 | 11.47 | 28 |
| 1.18E-03 | 366 | 279 | 279 | 1116 | 1.1 | 112 | 40.14 | 248 |
| 1.52E-03 | 714 | 279 | 279 | 1116 | 1.85 | 136 | 48.75 | 329 |
| 1.65E-03 | 457 | 279 | 279 | 1116 | 1.17 | 123 | 44.09 | 286 |
| 2.95E-03 | 295 | 279 | 279 | 1116 | 2.11 | 76 | 27.24 | 140 |
| 4.33E-03 | 139 | 279 | 279 | 1116 | 1.15 | 54 | 19.35 | 83 |
| 4.37E-03 | 120 | 279 | 279 | 1116 | 7.53 | 26 | 9.32 | 20 |
| 6.12E-03 | 565 | 279 | 279 | 1116 | 1.34 | 127 | 45.52 | 306 |
| 8.30E-03 | 780 | 279 | 279 | 1116 | 7.03 | 61 | 21.86 | 98 |
| 8.63E-03 | 406 | 279 | 279 | 1116 | 1.58 | 110 | 39.43 | 251 |

| | | | | | | | | |
|---|---|---|---|---|---|---|---|---|
| 1.21E-02 | 639 | 279 | 279 | 1116 | 1.1 | 169 | 60.57 | 467 |
| 1.43E-02 | 607 | 279 | 279 | 1116 | 15.7 | 70 | 25.09 | 126 |
| 1.46E-02 | 68 | 279 | 279 | 1116 | 1.96 | 27 | 9.68 | 25 |
| 1.56E-02 | 294 | 279 | 279 | 1116 | 1.41 | 78 | 27.96 | 153 |
| 1.91E-02 | 657 | 279 | 279 | 1116 | 4.33 | 120 | 43.01 | 287 |
| 7.85E-02 | 592 | 279 | 279 | 1116 | 1.03 | 155 | 55.56 | 425 |
| 1.01E-01 | 33 | 279 | 279 | 1116 | 729 | 8 | 2.87 | 0 |
| 1.86E-01 | 422 | 279 | 279 | 1116 | 10.3 | 55 | 19.71 | 95 |
| 2.03E-01 | 183 | 279 | 279 | 1116 | 1.49 | 53 | 19 | 93 |
| 2.38E-01 | 48 | 279 | 279 | 1116 | 65.2 | 9 | 3.23 | 1 |
| 3.25E-01 | 89 | 279 | 279 | 1116 | 13.4 | 12 | 4.3 | 4 |
| 3.60E-01 | 453 | 279 | 279 | 1116 | 31.1 | 39 | 13.98 | 55 |
| 4.87E-01 | 403 | 279 | 279 | 1116 | 2.34 | 68 | 24.37 | 137 |
| 9.74E-01 | 471 | 279 | 279 | 1116 | 13.4 | 16 | 5.73 | 9 |
| 1.22E+00 | 631 | 279 | 279 | 1116 | 1.37 | 142 | 50.9 | 394 |
| 1.23E+00 | 248 | 279 | 279 | 1116 | 38.6 | 9 | 3.23 | 1 |
| 1.23E+00 | 705 | 279 | 279 | 1116 | 1.05 | 176 | 63.08 | 527 |
| 1.42E+00 | 461 | 279 | 279 | 1116 | 40.3 | 8 | 2.87 | 0 |
| 1.46E+00 | 371 | 279 | 279 | 1116 | 2.6 | 77 | 27.6 | 171 |
| 1.48E+00 | 17 | 279 | 279 | 1116 | 1.08 | 11 | 3.94 | 6 |
| 1.49E+00 | 480 | 279 | 279 | 1116 | 2.05 | 90 | 32.26 | 212 |
| 2.14E+00 | 48 | 279 | 279 | 1116 | 7.02 | 14 | 5.02 | 10 |
| 2.49E+00 | 159 | 279 | 279 | 1116 | 1.11E+03 | 7 | 2.51 | 0 |
| 2.54E+00 | 639 | 279 | 279 | 1116 | 1.03 | 162 | 58.06 | 477 |
| 2.55E+00 | 448 | 279 | 279 | 1116 | 2.9 | 51 | 18.28 | 94 |
| 2.93E+00 | 233 | 279 | 279 | 1116 | 2.06 | 47 | 16.85 | 86 |
| 2.99E+00 | 461 | 279 | 279 | 1116 | 3.24 | 73 | 26.16 | 161 |
| 3.05E+00 | 669 | 279 | 279 | 1116 | 17.7 | 22 | 7.89 | 21 |
| 3.26E+00 | 187 | 279 | 279 | 1116 | 4.36 | 27 | 9.68 | 35 |
| 3.69E+00 | 856 | 279 | 279 | 1116 | 2.6 | 116 | 41.58 | 304 |
| 3.70E+00 | 435 | 279 | 279 | 1116 | 1.02 | 118 | 42.29 | 316 |
| 4.59E+00 | 82 | 279 | 279 | 1116 | 10.3 | 18 | 6.45 | 18 |
| 4.94E+00 | 103 | 279 | 279 | 1116 | 1.27 | 36 | 12.9 | 61 |
| 6.29E+00 | 22 | 279 | 279 | 1116 | 2.85 | 8 | 2.87 | 3 |
| 7.28E+00 | 316 | 279 | 279 | 1116 | 5.89 | 32 | 11.47 | 48 |
| 7.68E+00 | 191 | 279 | 279 | 1116 | 1.6 | 48 | 17.2 | 94 |
| 8.43E+00 | 254 | 279 | 279 | 1116 | 7.96 | 17 | 6.09 | 15 |
| 9.13E+00 | 230 | 279 | 279 | 1116 | 3.15 | 34 | 12.19 | 55 |
| 9.15E+00 | 129 | 279 | 279 | 1116 | 1.22 | 44 | 15.77 | 85 |

-control --shuffle-- --kmer 2 TFE3_chip_seq.fasta db/HUMAN/HOCOMOCOv11_full_HUMAN_mono_meme_form

| %FP |
|---|
| 0.18 |
| 0.18 |
| 0.36 |
| 0.09 |
| 1.08 |
| 1.34 |
| 0.9 |
| 1.34 |
| 3.76 |
| 10.22 |
| 9.77 |
| 14.16 |
| 5.73 |
| 13.35 |
| 3.67 |
| 4.66 |
| 7.8 |
| 4.57 |
| 15.23 |
| 11.38 |
| 5.91 |
| 6.36 |
| 5.65 |
| 7.89 |
| 8.15 |
| 22.13 |
| 7.35 |
| 10.48 |
| 14.61 |
| 16.4 |
| 16.49 |
| 6.27 |
| 14.43 |
| 7.17 |
| 1.43 |
| 8.96 |
| 10.22 |
| 14.96 |
| 12.81 |
| 2.6 |
| 4.57 |
| 0.09 |
| 4.12 |
| 3.49 |
| 4.75 |
| 17.03 |

| |
|---|
| 7.89 |
| 0.09 |
| 21.59 |
| 0.54 |
| 16.31 |
| 3.85 |
| 23.3 |
| 11.92 |
| 4.3 |
| 8.51 |
| 12.1 |
| 10.04 |
| 9.95 |
| 6.81 |
| 24.01 |
| 21.06 |
| 15.68 |
| 13.62 |
| 10.75 |
| 11.56 |
| 30.02 |
| 13.71 |
| 30.2 |
| 20.79 |
| 9.59 |
| 0.63 |
| 3.67 |
| 5.56 |
| 5.73 |
| 18.28 |
| 5.65 |
| 23.84 |
| 8.33 |
| 24.64 |
| 19.53 |
| 2.87 |
| 7.44 |
| 2.51 |
| 22.22 |
| 29.48 |
| 25.63 |
| 12.54 |
| 7.44 |
| 1.79 |
| 27.42 |
| 8.78 |
| 22.49 |

| |
|---|
| 41.85 |
| 11.29 |
| 2.24 |
| 13.71 |
| 25.72 |
| 38.08 |
| 0 |
| 8.51 |
| 8.33 |
| 0.09 |
| 0.36 |
| 4.93 |
| 12.28 |
| 0.81 |
| 35.3 |
| 0.09 |
| 47.22 |
| 0 |
| 15.32 |
| 0.54 |
| 19 |
| 0.9 |
| 0 |
| 42.74 |
| 8.42 |
| 7.71 |
| 14.43 |
| 1.88 |
| 3.14 |
| 27.24 |
| 28.32 |
| 1.61 |
| 5.47 |
| 0.27 |
| 4.3 |
| 8.42 |
| 1.34 |
| 4.93 |
| 7.62 |

ıat.meme db/MOUSE/HOCOMOCOv11_full_MOUSE_mono_meme_format.meme

| rank | motif_ID | consensus | p-value |
|---|---|---|---|
| 1 | TEAD4_HUMAN.H11MO.0.A | RCATTCCWGRVAK | 2.37E-155 |
| 2 | TEAD1_HUMAN.H11MO.0.A | RCATTCCWGSCW | 5.07E-142 |
| 3 | FOSL2_HUMAN.H11MO.0.A | NDRTGAGTCAYH | 1.47E-58 |
| 4 | FOSL1_HUMAN.H11MO.0.A | KRVTGAGTCAYH | 2.37E-57 |
| 5 | FOS_HUMAN.H11MO.0.A | VTGAGTCAB | 5.73E-57 |
| 6 | FOSB_HUMAN.H11MO.0.A | RTGAGTCAB | 8.17E-56 |
| 7 | JUN_HUMAN.H11MO.0.A | DRTGAGTCAYH | 8.14E-55 |
| 8 | JUND_HUMAN.H11MO.0.A | RRTGAGTCAYY | 1.48E-54 |
| 9 | JUNB_HUMAN.H11MO.0.A | DVTGAGTCABH | 2.19E-52 |
| 10 | ETV4_HUMAN.H11MO.0.B | VRCAGGAARBR | 5.02E-44 |
| 11 | ERG_HUMAN.H11MO.0.A | RRRSAGGAAGYGG | 1.58E-38 |
| 12 | MAZ_HUMAN.H11MO.0.A | GGGGGAGGGGGDGRGRRRGRRG | 6.64E-38 |
| 13 | ETV5_HUMAN.H11MO.0.C | RRRSAGGAARDGRV | 1.11E-37 |
| 14 | ZBT17_HUMAN.H11MO.0.A | SRRGGWGGGGGAGGGGMRR | 2.27E-36 |
| 15 | KLF15_HUMAN.H11MO.0.A | RGGGMGGRGVDGGGGGRGG | 1.57E-35 |
| 16 | ZN467_HUMAN.H11MO.0.C | GGGGAGGGGRRGGRGRRGGRRR | 2.89E-35 |
| 17 | ETS1_HUMAN.H11MO.0.A | RRRCMGGAAGTGG | 3.47E-35 |
| 18 | VEZF1_HUMAN.H11MO.0.C | GGRRRRGRRGGAGGGGGRGRRR | 2.97E-35 |
| 19 | EGR2_HUMAN.H11MO.0.A | GRGRRKGWGKGGGHGGRG | 1.11E-34 |
| 20 | FLI1_HUMAN.H11MO.0.A | SRRGGMAGGAAGGRRRGR | 4.52E-34 |
| 21 | ETV2_HUMAN.H11MO.0.B | RRARRCAGGAARYRGS | 9.52E-32 |
| 22 | E2F7_HUMAN.H11MO.0.B | GDGGCGGGAARDR | 5.22E-31 |
| 23 | PATZ1_HUMAN.H11MO.0.C | GGGGMGGGGGMKGGRRVGGVRG | 5.64E-31 |
| 24 | SP4_HUMAN.H11MO.0.A | SRGVARGRGGCGGRGCHDRR | 2.06E-28 |
| 25 | WT1_HUMAN.H11MO.0.C | RGGVGGGGGAGGRGGVGGRG | 1.58E-27 |
| 26 | E2F3_HUMAN.H11MO.0.A | DDGGCGGGAAA | 1.65E-27 |
| 27 | ETS2_HUMAN.H11MO.0.B | RDVRGGAARVRRR | 3.03E-27 |
| 28 | BC11A_HUMAN.H11MO.0.A | RAAAGAGGAAGTGARAV | 1.78E-26 |
| 29 | ZN341_HUMAN.H11MO.0.C | GRGGRRRRGGRGRRGGAASAGC | 2.14E-26 |
| 30 | IRF3_HUMAN.H11MO.0.B | RAAARGGAAAVDGAAASDGA | 5.32E-26 |
| 31 | ELF5_HUMAN.H11MO.0.A | VGRASSAGGAAGTRV | 6.12E-25 |
| 32 | TAF1_HUMAN.H11MO.0.A | RARRWGGCGGMGGMGR | 7.89E-25 |
| 33 | ZN263_HUMAN.H11MO.0.A | GGGAGGAGGRRGRGGRGGRR | 1.14E-24 |
| 34 | KLF6_HUMAN.H11MO.0.A | GSRRGGGHGGGGMHGGGRV | 1.25E-24 |
| 35 | EGR1_HUMAN.H11MO.0.A | VDRDGCGKGGGYGGRRR | 6.67E-24 |
| 36 | ELF3_HUMAN.H11MO.0.A | RAABVAGGAAGTRR | 5.09E-23 |
| 37 | ZN770_HUMAN.H11MO.0.C | GGGAGGCYGAGGBRGGAGGATC | 5.95E-23 |
| 38 | ELF1_HUMAN.H11MO.0.A | VRACCCGGAAGTGS | 8.81E-23 |
| 39 | ETV1_HUMAN.H11MO.0.A | SRCCGGAAGYG | 1.17E-22 |
| 40 | BACH2_HUMAN.H11MO.0.A | TGCTGAGTCAB | 4.39E-22 |
| 41 | KLF5_HUMAN.H11MO.0.A | WGGGTGKGGCDGGG | 8.66E-22 |
| 42 | RUNX1_HUMAN.H11MO.0.A | BYTGTGGTTWB | 2.01E-21 |
| 43 | FEV_HUMAN.H11MO.0.B | GCVGGAAGYG | 1.12E-21 |
| 44 | ELF2_HUMAN.H11MO.0.C | RAVCCGGAAGTGR | 3.22E-21 |
| 45 | NFAC1_HUMAN.H11MO.0.B | AWGGAAARWVWGAMW | 2.13E-21 |

| | | | |
|---|---|---|---|
| 46 | EHF_HUMAN.H11MO.0.B | VWACSAGGAAGTDVS | 5.75E-21 |
| 47 | RUNX3_HUMAN.H11MO.0.A | BYTGTGGTTW | 1.09E-20 |
| 48 | PRDM6_HUMAN.H11MO.0.C | RRRARGAAAAAAA | 1.01E-20 |
| 49 | RXRA_HUMAN.H11MO.0.A | VRGDVRGNVARAGGTCARRG | 1.54E-20 |
| 50 | KLF1_HUMAN.H11MO.0.A | DGGGYGKGGCYGGG | 3.39E-20 |
| 51 | RUNX2_HUMAN.H11MO.0.A | KGKYTGTGGTTTKK | 6.39E-20 |
| 52 | KLF3_HUMAN.H11MO.0.B | GGRGVDGGGCGGGGCYGGG | 1.04E-19 |
| 53 | SP2_HUMAN.H11MO.0.A | GGSSVGGGGGCGGGGCCDGSGS | 3.00E-19 |
| 54 | ZN394_HUMAN.H11MO.0.C | NRARWRGAAWNGAMWGRAAK | 2.42E-19 |
| 55 | FOXP1_HUMAN.H11MO.0.A | TBTGTTTMY | 7.35E-19 |
| 56 | ELK1_HUMAN.H11MO.0.B | RCCGGAAGTGV | 5.22E-18 |
| 57 | ZSC22_HUMAN.H11MO.0.C | RGKCWGAGGGMGGAGG | 2.90E-18 |
| 58 | NFE2_HUMAN.H11MO.0.A | VVRTGACTCAGCA | 5.85E-18 |
| 59 | Z354A_HUMAN.H11MO.0.C | RTTTARWYCATTTACATTTAATGT | 9.99E-18 |
| 60 | TBX21_HUMAN.H11MO.0.A | GDGDRGGTGTGRV | 3.51E-18 |
| 61 | FOXJ3_HUMAN.H11MO.0.A | TTGTTTATKKTTW | 2.32E-17 |
| 62 | BACH1_HUMAN.H11MO.0.A | TGCTGAGTCABSS | 2.37E-17 |
| 63 | FOXM1_HUMAN.H11MO.0.A | TGTTTRCTYWKB | 1.49E-17 |
| 64 | E2F6_HUMAN.H11MO.0.A | DGGGCGGGARRVR | 3.69E-17 |
| 65 | ZN281_HUMAN.H11MO.0.A | RGGWGGGGGAGGGGV | 6.23E-17 |
| 66 | FOXA2_HUMAN.H11MO.0.A | TGTTTACWYWGB | 1.03E-16 |
| 67 | SP3_HUMAN.H11MO.0.B | SGVVGGGGGCGGGGCBRGSS | 5.51E-17 |
| 68 | NF2L2_HUMAN.H11MO.0.A | WWYTGCTGAGTCAT | 3.31E-16 |
| 69 | STAT2_HUMAN.H11MO.0.A | RRGRAAAHGAAACTGAAAV | 2.48E-16 |
| 70 | PEBB_HUMAN.H11MO.0.C | TYTGTGGTYWB | 1.61E-16 |
| 71 | FOXJ2_HUMAN.H11MO.0.C | TGTTTRTTTW | 4.17E-16 |
| 72 | FOXO1_HUMAN.H11MO.0.A | KBBYTGTTTHCN | 1.79E-16 |
| 73 | MEF2B_HUMAN.H11MO.0.A | DKKKCTATTTTTRG | 3.55E-16 |
| 74 | SP1_HUMAN.H11MO.0.A | VGGGGGCGGGGCCKGGGGGGGG | 1.04E-15 |
| 75 | ZFP82_HUMAN.H11MO.0.C | TTCCADTTCAYTAATTYTCTCTTC | 8.19E-16 |
| 76 | FOXA1_HUMAN.H11MO.0.A | TGTTTACWYWGB | 8.84E-16 |
| 77 | NFKB1_HUMAN.H11MO.0.A | AGGGAAAK | 3.03E-16 |
| 78 | GABPA_HUMAN.H11MO.0.A | GVVRCCGGAAGTGV | 1.73E-15 |
| 79 | SMAD3_HUMAN.H11MO.0.B | SYCTSHCWSCWS | 4.74E-16 |
| 80 | ANDR_HUMAN.H11MO.0.A | TGTTCTTKKYTGTTTRYW | 1.49E-15 |
| 81 | SRBP2_HUMAN.H11MO.0.B | VSGTGGGGWGABG | 7.93E-16 |
| 82 | MEF2C_HUMAN.H11MO.0.A | DKCTATTTTTRGM | 4.21E-15 |
| 83 | MEF2A_HUMAN.H11MO.0.A | DKCTATTTTTRGM | 7.43E-15 |
| 84 | ZN121_HUMAN.H11MO.0.C | CTGGGCAACABAGYRAGACC | 4.78E-14 |
| 85 | KLF4_HUMAN.H11MO.0.A | WGGGYGKGGC | 1.13E-14 |
| 86 | FOXK1_HUMAN.H11MO.0.A | TGTTTMCHTT | 1.68E-14 |
| 87 | TFDP1_HUMAN.H11MO.0.C | RRRRGGCGGGAARN | 2.61E-14 |
| 88 | STA5A_HUMAN.H11MO.0.A | TTCYRRGAAR | 8.53E-14 |
| 89 | SALL4_HUMAN.H11MO.0.B | GSDGGGWGGG | 5.41E-14 |
| 90 | FOXO3_HUMAN.H11MO.0.B | BYTGTTTACH | 3.62E-13 |
| 91 | MEF2D_HUMAN.H11MO.0.A | DKCTATTTTTAG | 9.09E-13 |
| 92 | E2F4_HUMAN.H11MO.0.A | SRGGGCGGGAARD | 7.95E-13 |

| | | | |
|---|---|---|---|
| 93 | BATF_HUMAN.H11MO.0.A | RBTYTCRDWATGASTCAB | 3.07E-12 |
| 94 | IRF1_HUMAN.H11MO.0.A | RAAANWGAAASTGAAASYRR | 5.09E-12 |
| 95 | SUH_HUMAN.H11MO.0.A | VSYSTGGGAAV | 1.77E-12 |
| 96 | COT1_HUMAN.H11MO.0.C | GRGGKSARAGGTCAGGG | 2.23E-12 |
| 97 | ELK4_HUMAN.H11MO.0.A | SVCCGGAAGTGV | 8.56E-12 |
| 98 | ZFP28_HUMAN.H11MO.0.C | TTCTATTTCTTCTTGWGTCA | 1.29E-11 |
| 99 | FOXP2_HUMAN.H11MO.0.C | BTGTTTACH | 5.73E-12 |
| 100 | SPI1_HUMAN.H11MO.0.A | RRAAAGAGGAAGTGRVD | 9.60E-12 |
| 101 | IRF8_HUMAN.H11MO.0.B | RAAARRGGAASTGAAASTDR | 1.86E-11 |
| 102 | IKZF1_HUMAN.H11MO.0.C | BTGGGARR | 5.25E-12 |
| 103 | FOXA3_HUMAN.H11MO.0.B | CHTGTTTACWTWG | 6.29E-11 |
| 104 | NFAC2_HUMAN.H11MO.0.B | TGGAAAAWY | 3.74E-11 |
| 105 | RORG_HUMAN.H11MO.0.C | RRAASTRGGTCA | 8.21E-11 |
| 106 | PAX5_HUMAN.H11MO.0.A | SCSKGGGCARCVRAGCGWGAC | 2.12E-10 |
| 107 | STAT3_HUMAN.H11MO.0.A | SAYTTCCMRGAA | 1.16E-10 |
| 108 | BCL6_HUMAN.H11MO.0.A | YKCTTTCYAGGAA | 1.20E-10 |
| 109 | STA5B_HUMAN.H11MO.0.A | YTTCYKRGAAAY | 1.77E-10 |
| 110 | HXB13_HUMAN.H11MO.0.A | TTTTATDGSH | 2.13E-10 |
| 111 | NFIC_HUMAN.H11MO.0.A | CYTGGCHBYMWGCCARG | 1.46E-10 |
| 112 | STAT1_HUMAN.H11MO.0.A | YCCVTTTCCHGGAAAWCHC | 4.62E-10 |
| 113 | NFAC4_HUMAN.H11MO.0.C | WGGAAAAYTW | 1.63E-10 |
| 114 | SMCA5_HUMAN.H11MO.0.C | GAATBGAATGGAAWC | 3.72E-10 |
| 115 | Z324A_HUMAN.H11MO.0.C | KVTCAAACCATCCTTKSCTKSCW | 2.86E-10 |
| 116 | E2F1_HUMAN.H11MO.0.A | RRNKGGCGGGAARR | 7.78E-10 |
| 117 | MAFF_HUMAN.H11MO.0.B | TGCTGASTCAGCABTTTT | 9.67E-10 |
| 118 | GATA3_HUMAN.H11MO.0.A | RVAGATAAVAD | 1.10E-09 |
| 119 | HIF1A_HUMAN.H11MO.0.C | GDACGTGM | 4.74E-09 |
| 120 | SOX4_HUMAN.H11MO.0.B | BYCTTTGTYYYB | 1.40E-09 |
| 121 | KLF12_HUMAN.H11MO.0.C | VGGGGCGGGGC | 2.68E-09 |
| 122 | PO2F1_HUMAN.H11MO.0.C | AWKBRWATGSAAWYAW | 3.50E-09 |
| 123 | KLF9_HUMAN.H11MO.0.C | GRRGKGGGCGTGGCC | 6.08E-09 |
| 124 | SOX2_HUMAN.H11MO.0.A | BBCCTTTGTYYYB | 2.71E-09 |
| 125 | PRDM1_HUMAN.H11MO.0.A | RAAAGTGAAAGTGR | 6.32E-09 |
| 126 | NR1D1_HUMAN.H11MO.0.B | RRRNMASTGGGTCAGDRDR | 5.40E-09 |
| 127 | SRY_HUMAN.H11MO.0.B | YWTTGTTTH | 6.86E-09 |
| 128 | SPIB_HUMAN.H11MO.0.A | RRAAAGAGGAAGTGARA | 2.94E-08 |
| 129 | ZN322_HUMAN.H11MO.0.B | GWGCCTGSTACWSWGCCTGR | 3.73E-08 |
| 130 | TBP_HUMAN.H11MO.0.A | MTATAAAARS | 2.72E-08 |
| 131 | RARA_HUMAN.H11MO.0.A | GGRRGRTSMRRAGKTCAAGG | 2.89E-08 |
| 132 | ARNT_HUMAN.H11MO.0.B | STACGTGMC | 1.21E-07 |
| 133 | EVX2_HUMAN.H11MO.0.A | KTTTTATKRCYBY | 6.40E-08 |
| 134 | MAF_HUMAN.H11MO.0.A | VAAWBTGCTGASTHWGCMD | 3.78E-08 |
| 135 | MYCN_HUMAN.H11MO.0.A | RGCCACGTGSDS | 8.99E-08 |
| 136 | MXI1_HUMAN.H11MO.0.A | VCACGTGGSNGSBGS | 8.19E-08 |
| 137 | PTF1A_HUMAN.H11MO.0.B | SCAGCTGBYBBYBYYCHC | 5.41E-08 |
| 138 | PPARG_HUMAN.H11MO.0.A | MWSTRGGKSARAGGKCA | 6.90E-08 |
| 139 | LYL1_HUMAN.H11MO.0.A | CCAKCTGYTYYCTB | 4.84E-08 |

| | | | |
|---|---|---|---|
| 140 | IRF7_HUMAN.H11MO.0.C | GAAASYGAAA | 1.03E-07 |
| 141 | HNF1A_HUMAN.H11MO.0.C | DGTTAATKATTAACH | 3.08E-07 |
| 142 | MAFB_HUMAN.H11MO.0.B | TGCTGASTYAD | 1.12E-07 |
| 143 | MAX_HUMAN.H11MO.0.A | VRGCACGTGG | 3.73E-07 |
| 144 | COT2_HUMAN.H11MO.0.A | CARAGGTCARRGD | 2.07E-07 |
| 145 | IRF2_HUMAN.H11MO.0.A | RAAAVHGAAAGTGAAASTRV | 8.70E-07 |
| 146 | ATF2_HUMAN.H11MO.0.B | RRTGABGTCAY | 4.75E-07 |
| 147 | ATF6A_HUMAN.H11MO.0.B | GKGSTGACGTGG | 3.79E-06 |
| 148 | IRF4_HUMAN.H11MO.0.A | RAAWGRGGAASTGAAASY | 6.80E-07 |
| 149 | FOXC1_HUMAN.H11MO.0.C | CDWTGTTTACTTWRG | 1.91E-06 |
| 150 | MAFK_HUMAN.H11MO.0.A | RAWWHTGCTGASTCAGCA | 1.95E-06 |
| 151 | NKX61_HUMAN.H11MO.0.B | GAYWAATKRSHWWTTAATK | 5.35E-07 |
| 152 | SMCA1_HUMAN.H11MO.0.C | CMARGAAGMATD | 3.59E-07 |
| 153 | OLIG2_HUMAN.H11MO.0.B | CCAKCTGYTYYYYWBCHB | 5.62E-07 |
| 154 | TBX3_HUMAN.H11MO.0.C | DRAGGTGBSAR | 3.81E-07 |
| 155 | ZN768_HUMAN.H11MO.0.C | GAGGCHCAGAGAGGTTAAGT | 3.57E-06 |
| 156 | RXRB_HUMAN.H11MO.0.C | TSAGGTCASR | 8.10E-07 |
| 157 | ATF3_HUMAN.H11MO.0.A | GGTSACGTGAB | 3.01E-06 |
| 158 | EPAS1_HUMAN.H11MO.0.B | VTACGTGMC | 5.44E-06 |
| 159 | BMAL1_HUMAN.H11MO.0.A | KSCACGTGHSY | 3.00E-06 |
| 160 | STAT4_HUMAN.H11MO.0.A | YTTCYYRGMAV | 1.20E-06 |
| 161 | REL_HUMAN.H11MO.0.B | DRWRGGGRAATKCCA | 1.45E-06 |
| 162 | HNF1B_HUMAN.H11MO.0.A | DGTTAATRATTAACH | 5.66E-06 |
| 163 | MYC_HUMAN.H11MO.0.A | RCCACGTGCTB | 4.95E-06 |
| 164 | ZNF76_HUMAN.H11MO.0.C | RRKGCMTKCTGGGARWTGTAGT | 5.53E-06 |
| 165 | LEF1_HUMAN.H11MO.0.A | BCCTTTSWWNTBYH | 1.50E-06 |
| 166 | ITF2_HUMAN.H11MO.0.C | DGCAGSTGKS | 2.20E-06 |
| 167 | NFAC3_HUMAN.H11MO.0.B | TGGAAAAHY | 3.56E-06 |
| 168 | FOXO4_HUMAN.H11MO.0.C | TTGTTTAYK | 5.47E-06 |
| 169 | BHE40_HUMAN.H11MO.0.A | DGCACGTGAS | 1.29E-05 |
| 170 | TFAP4_HUMAN.H11MO.0.A | YCAGCTGWKB | 7.26E-06 |
| 171 | ZSC31_HUMAN.H11MO.0.C | RGCAGCAGGGCARTTAKR | 6.85E-06 |
| 172 | TF65_HUMAN.H11MO.0.A | HWGGGRATTTCCMV | 9.16E-06 |
| 173 | SRBP1_HUMAN.H11MO.0.A | DGTSRGGTGAB | 4.61E-06 |
| 174 | ATF1_HUMAN.H11MO.0.B | HRRTGACGYMA | 1.90E-05 |
| 175 | BHA15_HUMAN.H11MO.0.B | SSAGCAGCTGG | 9.53E-06 |
| 176 | TWST1_HUMAN.H11MO.0.A | MCATCTGKWWYYMATTA | 2.01E-05 |
| 177 | HXC9_HUMAN.H11MO.0.C | WTTTATKGCH | 1.22E-05 |
| 178 | OSR2_HUMAN.H11MO.0.C | GBYSCWGCTNCTGYDG | 9.14E-06 |
| 179 | PO3F2_HUMAN.H11MO.0.A | YYCATGCATWTYCATTM | 1.31E-05 |
| 180 | HTF4_HUMAN.H11MO.0.A | DSCAGSTGKG | 1.16E-05 |
| 181 | HXA9_HUMAN.H11MO.0.B | RTGATTTATKR | 2.45E-05 |
| 182 | ZN418_HUMAN.H11MO.0.C | TGCTTYTRGCYTCTKVY | 1.97E-05 |
| 183 | NF2L1_HUMAN.H11MO.0.C | HGTCATN | 1.38E-05 |
| 184 | CREB1_HUMAN.H11MO.0.A | NRRTGACGTMA | 7.07E-05 |
| 185 | ZN436_HUMAN.H11MO.0.C | TCMRGGARGGCTTCCTGGAGGAGG | 7.31E-04 |
| 186 | RORA_HUMAN.H11MO.0.C | AWSTRGGTCARRG | 1.67E-05 |

| | | | |
|---|---|---|---|
| 187 | MYOG_HUMAN.H11MO.0.B | CHSCAGCTGYYYB | 4.84E-05 |

### AME (Analysis of Motif Enrichment): Version 5.0.5 compiled on Jul 15 2019 at 00:31:14
### The format of this file is described at http://meme-suite.org//doc/ame-output-format.html.
### ame --verbose 1 --oc . --scoring avg --method fisher --hit-lo-fraction 0.25 --evalue-report-threshold 10.0 --con

| adj_p-value | E-value | tests | FASTA_max | pos | neg | PWM_min |
|---|---|---|---|---|---|---|
| 3.01E-152 | 1.21E-149 | 1274 | 1150 | 1150 | 1150 | 2.94 |
| 5.78E-139 | 2.32E-136 | 1140 | 1150 | 1150 | 1150 | 1.38 |
| 7.46E-56 | 3.00E-53 | 507 | 1150 | 1150 | 1150 | 1.37 |
| 1.24E-54 | 4.97E-52 | 522 | 1150 | 1150 | 1150 | 1.34 |
| 4.22E-54 | 1.70E-51 | 737 | 1150 | 1150 | 1150 | 2.35 |
| 6.99E-53 | 2.81E-50 | 856 | 1150 | 1150 | 1150 | 4.44 |
| 4.65E-52 | 1.87E-49 | 572 | 1150 | 1150 | 1150 | 1.99 |
| 9.40E-52 | 3.78E-49 | 635 | 1150 | 1150 | 1150 | 3.4 |
| 1.54E-49 | 6.19E-47 | 702 | 1150 | 1150 | 1150 | 3.23 |
| 7.31E-41 | 2.94E-38 | 1458 | 1150 | 1150 | 1150 | 2.35 |
| 1.57E-35 | 6.33E-33 | 994 | 1150 | 1150 | 1150 | 1.99 |
| 7.31E-35 | 2.94E-32 | 1101 | 1150 | 1150 | 1150 | 16.9 |
| 1.69E-34 | 6.78E-32 | 1518 | 1150 | 1150 | 1150 | 2.18 |
| 2.94E-33 | 1.18E-30 | 1294 | 1150 | 1150 | 1150 | 5.15 |
| 1.27E-32 | 5.12E-30 | 809 | 1150 | 1150 | 1150 | 1.82 |
| 2.23E-32 | 8.96E-30 | 772 | 1150 | 1150 | 1150 | 6.31 |
| 2.68E-32 | 1.08E-29 | 771 | 1150 | 1150 | 1150 | 1.33 |
| 3.58E-32 | 1.44E-29 | 1205 | 1150 | 1150 | 1150 | 20.1 |
| 1.19E-31 | 4.77E-29 | 1065 | 1150 | 1150 | 1150 | 4.83 |
| 4.11E-31 | 1.65E-28 | 909 | 1150 | 1150 | 1150 | 4.78 |
| 5.60E-29 | 2.25E-26 | 588 | 1150 | 1150 | 1150 | 1.07 |
| 5.63E-28 | 2.26E-25 | 1080 | 1150 | 1150 | 1150 | 2.2 |
| 6.14E-28 | 2.47E-25 | 1088 | 1150 | 1150 | 1150 | 9.24 |
| 2.13E-25 | 8.56E-23 | 1032 | 1150 | 1150 | 1150 | 5.17 |
| 1.24E-24 | 4.97E-22 | 783 | 1150 | 1150 | 1150 | 3.83 |
| 1.99E-24 | 8.01E-22 | 1205 | 1150 | 1150 | 1150 | 1.25 |
| 4.65E-24 | 1.87E-21 | 1534 | 1150 | 1150 | 1150 | 1.52 |
| 2.31E-23 | 9.29E-21 | 1295 | 1150 | 1150 | 1150 | 1.15 |
| 2.43E-23 | 9.75E-21 | 1131 | 1150 | 1150 | 1150 | 5.95 |
| 4.37E-23 | 1.76E-20 | 821 | 1150 | 1150 | 1150 | 2.12 |
| 4.60E-22 | 1.85E-19 | 751 | 1150 | 1150 | 1150 | 1.04 |
| 7.59E-22 | 3.05E-19 | 962 | 1150 | 1150 | 1150 | 3.35 |
| 9.79E-22 | 3.94E-19 | 856 | 1150 | 1150 | 1150 | 1.12 |
| 1.43E-21 | 5.73E-19 | 1138 | 1150 | 1150 | 1150 | 13.1 |
| 6.64E-21 | 2.67E-18 | 995 | 1150 | 1150 | 1150 | 2.7 |
| 3.19E-20 | 1.28E-17 | 627 | 1150 | 1150 | 1150 | 1.03 |
| 4.09E-20 | 1.64E-17 | 687 | 1150 | 1150 | 1150 | 1.33 |
| 4.63E-20 | 1.86E-17 | 526 | 1150 | 1150 | 1150 | 1.08 |
| 6.65E-20 | 2.67E-17 | 569 | 1150 | 1150 | 1150 | 1.03 |
| 2.46E-19 | 9.89E-17 | 560 | 1150 | 1150 | 1150 | 1.97 |
| 8.63E-19 | 3.47E-16 | 996 | 1150 | 1150 | 1150 | 3.95 |
| 1.27E-18 | 5.10E-16 | 631 | 1150 | 1150 | 1150 | 1.11 |
| 1.60E-18 | 6.42E-16 | 1433 | 1150 | 1150 | 1150 | 1.89 |
| 1.88E-18 | 7.56E-16 | 584 | 1150 | 1150 | 1150 | 1.59 |
| 2.68E-18 | 1.08E-15 | 1257 | 1150 | 1150 | 1150 | 1.54 |

| | | | | | | |
|---|---|---|---|---|---|---|
| 2.89E-18 | 1.16E-15 | 503 | 1150 | 1150 | 1150 | 1.23 |
| 6.73E-18 | 2.70E-15 | 615 | 1150 | 1150 | 1150 | 2.75 |
| 1.15E-17 | 4.60E-15 | 1136 | 1150 | 1150 | 1150 | 4.04 |
| 2.08E-17 | 8.34E-15 | 1345 | 1150 | 1150 | 1150 | 4.08 |
| 2.41E-17 | 9.69E-15 | 712 | 1150 | 1150 | 1150 | 17.9 |
| 5.37E-17 | 2.16E-14 | 840 | 1150 | 1150 | 1150 | 1.34 |
| 7.95E-17 | 3.20E-14 | 768 | 1150 | 1150 | 1150 | 30 |
| 2.30E-16 | 9.23E-14 | 765 | 1150 | 1150 | 1150 | 2.14 |
| 2.91E-16 | 1.17E-13 | 1202 | 1150 | 1150 | 1150 | 1.81 |
| 7.52E-16 | 3.02E-13 | 1024 | 1150 | 1150 | 1150 | 1.29 |
| 1.95E-15 | 7.83E-13 | 373 | 1150 | 1150 | 1150 | 1.16 |
| 2.04E-15 | 8.20E-13 | 703 | 1150 | 1150 | 1150 | 2.14 |
| 2.56E-15 | 1.03E-12 | 437 | 1150 | 1150 | 1150 | 1.08 |
| 4.08E-15 | 1.64E-12 | 408 | 1150 | 1150 | 1150 | 1.47 |
| 5.10E-15 | 2.05E-12 | 1452 | 1150 | 1150 | 1150 | 1.3 |
| 6.74E-15 | 2.71E-12 | 291 | 1150 | 1150 | 1150 | 1.37 |
| 1.35E-14 | 5.41E-12 | 568 | 1150 | 1150 | 1150 | 1.05 |
| 1.38E-14 | 5.53E-12 | 924 | 1150 | 1150 | 1150 | 2.35 |
| 3.58E-14 | 1.44E-11 | 970 | 1150 | 1150 | 1150 | 1.03 |
| 4.71E-14 | 1.89E-11 | 756 | 1150 | 1150 | 1150 | 1.56 |
| 4.73E-14 | 1.90E-11 | 458 | 1150 | 1150 | 1150 | 1.04 |
| 5.23E-14 | 2.10E-11 | 950 | 1150 | 1150 | 1150 | 2.98 |
| 8.00E-14 | 3.22E-11 | 242 | 1150 | 1150 | 1150 | 1.27 |
| 9.55E-14 | 3.84E-11 | 386 | 1150 | 1150 | 1150 | 1.81 |
| 1.19E-13 | 4.78E-11 | 741 | 1150 | 1150 | 1150 | 4.2 |
| 2.01E-13 | 8.08E-11 | 482 | 1150 | 1150 | 1150 | 1.11 |
| 2.06E-13 | 8.29E-11 | 1152 | 1150 | 1150 | 1150 | 1.92 |
| 2.35E-13 | 9.44E-11 | 662 | 1150 | 1150 | 1150 | 2.46 |
| 3.21E-13 | 1.29E-10 | 308 | 1150 | 1150 | 1150 | 8.17 |
| 4.21E-13 | 1.69E-10 | 514 | 1150 | 1150 | 1150 | 1.49 |
| 4.35E-13 | 1.75E-10 | 492 | 1150 | 1150 | 1150 | 1.11 |
| 4.56E-13 | 1.83E-10 | 1504 | 1150 | 1150 | 1150 | 2.17 |
| 7.20E-13 | 2.90E-10 | 416 | 1150 | 1150 | 1150 | 1.37 |
| 8.40E-13 | 3.38E-10 | 1775 | 1150 | 1150 | 1150 | 2.08 |
| 9.47E-13 | 3.81E-10 | 635 | 1150 | 1150 | 1150 | 1.91 |
| 1.05E-12 | 4.21E-10 | 1320 | 1150 | 1150 | 1150 | 3.95 |
| 2.26E-12 | 9.07E-10 | 536 | 1150 | 1150 | 1150 | 1.66 |
| 4.00E-12 | 1.61E-09 | 538 | 1150 | 1150 | 1150 | 2.4 |
| 6.40E-12 | 2.57E-09 | 134 | 1150 | 1150 | 1150 | 44.4 |
| 7.32E-12 | 2.94E-09 | 647 | 1150 | 1150 | 1150 | 5.86 |
| 1.78E-11 | 7.15E-09 | 1060 | 1150 | 1150 | 1150 | 1.42 |
| 2.05E-11 | 8.24E-09 | 787 | 1150 | 1150 | 1150 | 2.42 |
| 4.32E-11 | 1.74E-08 | 507 | 1150 | 1150 | 1150 | 1.11 |
| 7.68E-11 | 3.09E-08 | 1420 | 1150 | 1150 | 1150 | 2.25 |
| 2.22E-10 | 8.91E-08 | 612 | 1150 | 1150 | 1150 | 1.12 |
| 3.48E-10 | 1.40E-07 | 383 | 1150 | 1150 | 1150 | 2 |
| 4.40E-10 | 1.77E-07 | 554 | 1150 | 1150 | 1150 | 1.04 |

| | | | | | | |
|---|---|---|---|---|---|---|
| 1.57E-09 | 6.33E-07 | 512 | 1150 | 1150 | 1150 | 1.5 |
| 1.86E-09 | 7.46E-07 | 365 | 1150 | 1150 | 1150 | 14.9 |
| 2.16E-09 | 8.70E-07 | 1222 | 1150 | 1150 | 1150 | 2.32 |
| 2.38E-09 | 9.58E-07 | 1069 | 1150 | 1150 | 1150 | 1.21 |
| 2.87E-09 | 1.15E-06 | 335 | 1150 | 1150 | 1150 | 1.08 |
| 3.10E-09 | 1.25E-06 | 240 | 1150 | 1150 | 1150 | 1.41 |
| 3.94E-09 | 1.58E-06 | 687 | 1150 | 1150 | 1150 | 1.41 |
| 4.40E-09 | 1.77E-06 | 458 | 1150 | 1150 | 1150 | 1.26 |
| 6.66E-09 | 2.68E-06 | 358 | 1150 | 1150 | 1150 | 6.53 |
| 7.73E-09 | 3.11E-06 | 1472 | 1150 | 1150 | 1150 | 3.12 |
| 3.22E-08 | 1.29E-05 | 512 | 1150 | 1150 | 1150 | 1.26 |
| 3.48E-08 | 1.40E-05 | 931 | 1150 | 1150 | 1150 | 1.16 |
| 5.71E-08 | 2.30E-05 | 696 | 1150 | 1150 | 1150 | 1.46 |
| 7.44E-08 | 2.99E-05 | 350 | 1150 | 1150 | 1150 | 71.1 |
| 7.84E-08 | 3.15E-05 | 674 | 1150 | 1150 | 1150 | 1.06 |
| 9.44E-08 | 3.80E-05 | 790 | 1150 | 1150 | 1150 | 2.58 |
| 1.05E-07 | 4.21E-05 | 593 | 1150 | 1150 | 1150 | 1.11 |
| 1.17E-07 | 4.71E-05 | 549 | 1150 | 1150 | 1150 | 1.17 |
| 1.45E-07 | 5.83E-05 | 996 | 1150 | 1150 | 1150 | 16.2 |
| 1.58E-07 | 6.33E-05 | 341 | 1150 | 1150 | 1150 | 1.12 |
| 1.65E-07 | 6.61E-05 | 1009 | 1150 | 1150 | 1150 | 1.08 |
| 1.95E-07 | 7.83E-05 | 523 | 1150 | 1150 | 1150 | 3.47 |
| 2.14E-07 | 8.59E-05 | 748 | 1150 | 1150 | 1150 | 1.25 |
| 4.31E-07 | 1.73E-04 | 554 | 1150 | 1150 | 1150 | 4.16 |
| 5.91E-07 | 2.37E-04 | 611 | 1150 | 1150 | 1150 | 1.12 |
| 9.71E-07 | 3.90E-04 | 884 | 1150 | 1150 | 1150 | 1.31 |
| 1.27E-06 | 5.09E-04 | 267 | 1150 | 1150 | 1150 | 1.04 |
| 1.39E-06 | 5.60E-04 | 992 | 1150 | 1150 | 1150 | 3.24 |
| 1.49E-06 | 6.01E-04 | 557 | 1150 | 1150 | 1150 | 1.17 |
| 1.97E-06 | 7.91E-04 | 563 | 1150 | 1150 | 1150 | 1.05 |
| 1.99E-06 | 8.02E-04 | 328 | 1150 | 1150 | 1150 | 8.12 |
| 2.77E-06 | 1.12E-03 | 1025 | 1150 | 1150 | 1150 | 4.17 |
| 3.65E-06 | 1.47E-03 | 577 | 1150 | 1150 | 1150 | 1.05 |
| 6.41E-06 | 2.58E-03 | 1189 | 1150 | 1150 | 1150 | 1.69 |
| 6.82E-06 | 2.74E-03 | 995 | 1150 | 1150 | 1150 | 2.89 |
| 9.19E-06 | 3.69E-03 | 313 | 1150 | 1150 | 1150 | 1.4 |
| 1.35E-05 | 5.42E-03 | 362 | 1150 | 1150 | 1150 | 1.6 |
| 1.53E-05 | 6.16E-03 | 563 | 1150 | 1150 | 1150 | 3.88 |
| 1.95E-05 | 7.85E-03 | 676 | 1150 | 1150 | 1150 | 2.7 |
| 2.19E-05 | 8.80E-03 | 181 | 1150 | 1150 | 1150 | 1.01 |
| 2.43E-05 | 9.77E-03 | 380 | 1150 | 1150 | 1150 | 1.69 |
| 2.57E-05 | 1.03E-02 | 679 | 1150 | 1150 | 1150 | 1.15 |
| 2.57E-05 | 1.03E-02 | 286 | 1150 | 1150 | 1150 | 2.31 |
| 4.14E-05 | 1.67E-02 | 506 | 1150 | 1150 | 1150 | 2.38 |
| 4.47E-05 | 1.80E-02 | 827 | 1150 | 1150 | 1150 | 2.54 |
| 6.09E-05 | 2.45E-02 | 882 | 1150 | 1150 | 1150 | 1.69 |
| 7.44E-05 | 2.99E-02 | 1537 | 1150 | 1150 | 1150 | 5.15 |

| | | | | | | |
|---|---|---|---|---|---|---|
| 8.98E-05 | 3.61E-02 | 871 | 1150 | 1150 | 1150 | 1.48 |
| 1.20E-04 | 4.81E-02 | 388 | 1150 | 1150 | 1150 | 1.12 |
| 1.26E-04 | 5.07E-02 | 1123 | 1150 | 1150 | 1150 | 1.98 |
| 1.49E-04 | 5.97E-02 | 398 | 1150 | 1150 | 1150 | 1.88 |
| 1.92E-04 | 7.71E-02 | 927 | 1150 | 1150 | 1150 | 5.19 |
| 2.03E-04 | 8.14E-02 | 233 | 1150 | 1150 | 1150 | 1.45 |
| 2.36E-04 | 9.47E-02 | 496 | 1150 | 1150 | 1150 | 1.48 |
| 2.50E-04 | 1.00E-01 | 66 | 1150 | 1150 | 1150 | 1.04 |
| 2.80E-04 | 1.13E-01 | 412 | 1150 | 1150 | 1150 | 1.29 |
| 2.86E-04 | 1.15E-01 | 150 | 1150 | 1150 | 1150 | 1.08 |
| 3.30E-04 | 1.33E-01 | 169 | 1150 | 1150 | 1150 | 2.45 |
| 3.32E-04 | 1.33E-01 | 621 | 1150 | 1150 | 1150 | 2.51 |
| 3.89E-04 | 1.57E-01 | 1084 | 1150 | 1150 | 1150 | 3.31 |
| 4.96E-04 | 1.99E-01 | 883 | 1150 | 1150 | 1150 | 4.88 |
| 5.36E-04 | 2.15E-01 | 1406 | 1150 | 1150 | 1150 | 2.96 |
| 6.60E-04 | 2.65E-01 | 185 | 1150 | 1150 | 1150 | 152 |
| 7.04E-04 | 2.83E-01 | 870 | 1150 | 1150 | 1150 | 10.4 |
| 7.19E-04 | 2.89E-01 | 239 | 1150 | 1150 | 1150 | 1.67 |
| 9.02E-04 | 3.63E-01 | 166 | 1150 | 1150 | 1150 | 1.01 |
| 1.05E-03 | 4.23E-01 | 351 | 1150 | 1150 | 1150 | 5.54 |
| 1.07E-03 | 4.28E-01 | 889 | 1150 | 1150 | 1150 | 1.03 |
| 1.28E-03 | 5.17E-01 | 887 | 1150 | 1150 | 1150 | 4.34 |
| 1.37E-03 | 5.50E-01 | 242 | 1150 | 1150 | 1150 | 1.73 |
| 1.50E-03 | 6.04E-01 | 304 | 1150 | 1150 | 1150 | 2.78 |
| 1.62E-03 | 6.53E-01 | 294 | 1150 | 1150 | 1150 | 1.25 |
| 2.24E-03 | 8.99E-01 | 1494 | 1150 | 1150 | 1150 | 2.59 |
| 2.66E-03 | 1.07E+00 | 1210 | 1150 | 1150 | 1150 | 3.69 |
| 2.80E-03 | 1.12E+00 | 786 | 1150 | 1150 | 1150 | 1.19 |
| 3.18E-03 | 1.28E+00 | 583 | 1150 | 1150 | 1150 | 1.34 |
| 3.26E-03 | 1.31E+00 | 254 | 1150 | 1150 | 1150 | 4.22 |
| 3.48E-03 | 1.40E+00 | 480 | 1150 | 1150 | 1150 | 1.24 |
| 3.75E-03 | 1.51E+00 | 548 | 1150 | 1150 | 1150 | 1.63 |
| 4.15E-03 | 1.67E+00 | 454 | 1150 | 1150 | 1150 | 1.15 |
| 4.30E-03 | 1.73E+00 | 936 | 1150 | 1150 | 1150 | 1.51 |
| 5.26E-03 | 2.11E+00 | 278 | 1150 | 1150 | 1150 | 1.42 |
| 5.80E-03 | 2.33E+00 | 610 | 1150 | 1150 | 1150 | 4.1 |
| 6.21E-03 | 2.50E+00 | 310 | 1150 | 1150 | 1150 | 4.51 |
| 6.95E-03 | 2.79E+00 | 574 | 1150 | 1150 | 1150 | 1.5 |
| 7.41E-03 | 2.98E+00 | 813 | 1150 | 1150 | 1150 | 4.54 |
| 8.21E-03 | 3.30E+00 | 632 | 1150 | 1150 | 1150 | 1.16 |
| 1.21E-02 | 4.87E+00 | 1048 | 1150 | 1150 | 1150 | 3.24 |
| 1.22E-02 | 4.90E+00 | 500 | 1150 | 1150 | 1150 | 5.14 |
| 1.23E-02 | 4.95E+00 | 628 | 1150 | 1150 | 1150 | 1.2 |
| 1.29E-02 | 5.18E+00 | 938 | 1150 | 1150 | 1150 | 2.36 |
| 1.72E-02 | 6.90E+00 | 245 | 1150 | 1150 | 1150 | 1.14 |
| 1.74E-02 | 7.00E+00 | 24 | 1150 | 1150 | 1150 | 3.28 |
| 2.06E-02 | 8.26E+00 | 1246 | 1150 | 1150 | 1150 | 2.77 |

| | | | | | | |
|---|---|---|---|---|---|---|
| 2.24E-02 | 9.02E+00 | 469 | 1150 | 1150 | 1150 | 22.1 |

trol --shuffle-- --kmer 2 taz4sa_chip_seq.fasta db/HUMAN/HOCOMOCOv11_core_HUMAN_mono_meme_form

| TP | %TP | FP | %FP |
|---|---|---|---|
| 731 | 63.57 | 135 | 11.74 |
| 817 | 71.04 | 226 | 19.65 |
| 395 | 34.35 | 89 | 7.74 |
| 401 | 34.87 | 95 | 8.26 |
| 398 | 34.61 | 94 | 8.17 |
| 352 | 30.61 | 70 | 6.09 |
| 363 | 31.57 | 78 | 6.78 |
| 327 | 28.43 | 59 | 5.13 |
| 342 | 29.74 | 71 | 6.17 |
| 621 | 54 | 297 | 25.83 |
| 515 | 44.78 | 227 | 19.74 |
| 357 | 31.04 | 113 | 9.83 |
| 654 | 56.87 | 351 | 30.52 |
| 483 | 42 | 209 | 18.17 |
| 478 | 41.57 | 208 | 18.09 |
| 361 | 31.39 | 123 | 10.7 |
| 479 | 41.65 | 210 | 18.26 |
| 326 | 28.35 | 100 | 8.7 |
| 411 | 35.74 | 160 | 13.91 |
| 344 | 29.91 | 115 | 10 |
| 414 | 36 | 172 | 14.96 |
| 461 | 40.09 | 211 | 18.35 |
| 361 | 31.39 | 136 | 11.83 |
| 352 | 30.61 | 138 | 12 |
| 386 | 33.57 | 166 | 14.43 |
| 655 | 56.96 | 397 | 34.52 |
| 767 | 66.7 | 511 | 44.43 |
| 745 | 64.78 | 492 | 42.78 |
| 404 | 35.13 | 184 | 16 |
| 430 | 37.39 | 206 | 17.91 |
| 490 | 42.61 | 260 | 22.61 |
| 289 | 25.13 | 106 | 9.22 |
| 546 | 47.48 | 310 | 26.96 |
| 267 | 23.22 | 92 | 8 |
| 424 | 36.87 | 210 | 18.26 |
| 418 | 36.35 | 209 | 18.17 |
| 447 | 38.87 | 233 | 20.26 |
| 354 | 30.78 | 160 | 13.91 |
| 382 | 33.22 | 182 | 15.83 |
| 280 | 24.35 | 109 | 9.48 |
| 367 | 31.91 | 174 | 15.13 |
| 398 | 34.61 | 200 | 17.39 |
| 429 | 37.3 | 224 | 19.48 |
| 308 | 26.78 | 132 | 11.48 |
| 606 | 52.7 | 382 | 33.22 |

| | | | |
|---|---|---|---|
| 323 | 28.09 | 144 | 12.52 |
| 239 | 20.78 | 86 | 7.48 |
| 379 | 32.96 | 188 | 16.35 |
| 383 | 33.3 | 192 | 16.7 |
| 140 | 12.17 | 29 | 2.52 |
| 457 | 39.74 | 256 | 22.26 |
| 138 | 12 | 29 | 2.52 |
| 404 | 35.13 | 215 | 18.7 |
| 568 | 49.39 | 358 | 31.13 |
| 540 | 46.96 | 335 | 29.13 |
| 246 | 21.39 | 100 | 8.7 |
| 342 | 29.74 | 170 | 14.78 |
| 293 | 25.48 | 134 | 11.65 |
| 274 | 23.83 | 121 | 10.52 |
| 686 | 59.65 | 479 | 41.65 |
| 201 | 17.48 | 72 | 6.26 |
| 368 | 32 | 195 | 16.96 |
| 308 | 26.78 | 147 | 12.78 |
| 584 | 50.78 | 386 | 33.57 |
| 415 | 36.09 | 236 | 20.52 |
| 307 | 26.7 | 150 | 13.04 |
| 440 | 38.26 | 257 | 22.35 |
| 170 | 14.78 | 56 | 4.87 |
| 224 | 19.48 | 91 | 7.91 |
| 189 | 16.43 | 67 | 5.83 |
| 315 | 27.39 | 159 | 13.83 |
| 439 | 38.17 | 259 | 22.52 |
| 272 | 23.65 | 126 | 10.96 |
| 132 | 11.48 | 35 | 3.04 |
| 331 | 28.78 | 173 | 15.04 |
| 313 | 27.22 | 159 | 13.83 |
| 548 | 47.65 | 358 | 31.13 |
| 242 | 21.04 | 107 | 9.3 |
| 646 | 56.17 | 453 | 39.39 |
| 300 | 26.09 | 150 | 13.04 |
| 255 | 22.17 | 115 | 10 |
| 271 | 23.57 | 130 | 11.3 |
| 230 | 20 | 101 | 8.78 |
| 52 | 4.52 | 2 | 0.17 |
| 184 | 16 | 70 | 6.09 |
| 512 | 44.52 | 336 | 29.22 |
| 278 | 24.17 | 139 | 12.09 |
| 313 | 27.22 | 169 | 14.7 |
| 609 | 52.96 | 431 | 37.48 |
| 360 | 31.3 | 211 | 18.35 |
| 178 | 15.48 | 73 | 6.35 |
| 348 | 30.26 | 203 | 17.65 |

| | | | |
|---|---|---|---|
| 271 | 23.57 | 144 | 12.52 |
| 97 | 8.43 | 25 | 2.17 |
| 381 | 33.13 | 233 | 20.26 |
| 576 | 50.09 | 411 | 35.74 |
| 217 | 18.87 | 105 | 9.13 |
| 167 | 14.52 | 70 | 6.09 |
| 343 | 29.83 | 204 | 17.74 |
| 279 | 24.26 | 153 | 13.3 |
| 98 | 8.52 | 27 | 2.35 |
| 420 | 36.52 | 270 | 23.48 |
| 293 | 25.48 | 169 | 14.7 |
| 509 | 44.26 | 357 | 31.04 |
| 320 | 27.83 | 192 | 16.7 |
| 36 | 3.13 | 1 | 0.09 |
| 398 | 34.61 | 260 | 22.61 |
| 246 | 21.39 | 133 | 11.57 |
| 347 | 30.17 | 217 | 18.87 |
| 318 | 27.65 | 193 | 16.78 |
| 104 | 9.04 | 33 | 2.87 |
| 220 | 19.13 | 116 | 10.09 |
| 569 | 49.48 | 419 | 36.43 |
| 154 | 13.39 | 67 | 5.83 |
| 424 | 36.87 | 286 | 24.87 |
| 139 | 12.09 | 58 | 5.04 |
| 363 | 31.57 | 236 | 20.52 |
| 440 | 38.26 | 305 | 26.52 |
| 172 | 14.96 | 85 | 7.39 |
| 290 | 25.22 | 175 | 15.22 |
| 321 | 27.91 | 203 | 17.65 |
| 341 | 29.65 | 221 | 19.22 |
| 84 | 7.3 | 26 | 2.26 |
| 261 | 22.7 | 153 | 13.3 |
| 348 | 30.26 | 229 | 19.91 |
| 489 | 42.52 | 356 | 30.96 |
| 270 | 23.48 | 163 | 14.17 |
| 185 | 16.09 | 99 | 8.61 |
| 192 | 16.7 | 105 | 9.13 |
| 79 | 6.87 | 25 | 2.17 |
| 222 | 19.3 | 128 | 11.13 |
| 124 | 10.78 | 57 | 4.96 |
| 176 | 15.3 | 94 | 8.17 |
| 375 | 32.61 | 259 | 22.52 |
| 100 | 8.7 | 40 | 3.48 |
| 150 | 13.04 | 75 | 6.52 |
| 284 | 24.7 | 181 | 15.74 |
| 377 | 32.78 | 263 | 22.87 |
| 115 | 10 | 49 | 4.26 |

| | | | |
|---|---|---|---|
| 403 | 35.04 | 288 | 25.04 |
| 232 | 20.17 | 143 | 12.43 |
| 323 | 28.09 | 217 | 18.87 |
| 144 | 12.52 | 74 | 6.43 |
| 167 | 14.52 | 90 | 7.83 |
| 140 | 12.17 | 73 | 6.35 |
| 217 | 18.87 | 132 | 11.48 |
| 51 | 4.43 | 15 | 1.3 |
| 229 | 19.91 | 143 | 12.43 |
| 102 | 8.87 | 47 | 4.09 |
| 83 | 7.22 | 34 | 2.96 |
| 219 | 19.04 | 134 | 11.65 |
| 229 | 19.91 | 141 | 12.26 |
| 165 | 14.35 | 91 | 7.91 |
| 295 | 25.65 | 197 | 17.13 |
| 18 | 1.57 | 0 | 0 |
| 76 | 6.61 | 28 | 2.43 |
| 95 | 8.26 | 43 | 3.74 |
| 110 | 9.57 | 55 | 4.78 |
| 60 | 5.22 | 20 | 1.74 |
| 499 | 43.39 | 388 | 33.74 |
| 177 | 15.39 | 103 | 8.96 |
| 122 | 10.61 | 64 | 5.57 |
| 95 | 8.26 | 44 | 3.83 |
| 182 | 15.83 | 111 | 9.65 |
| 379 | 32.96 | 277 | 24.09 |
| 254 | 22.09 | 168 | 14.61 |
| 414 | 36 | 313 | 27.22 |
| 304 | 26.43 | 215 | 18.7 |
| 50 | 4.35 | 16 | 1.39 |
| 272 | 23.65 | 188 | 16.35 |
| 248 | 21.57 | 167 | 14.52 |
| 256 | 22.26 | 175 | 15.22 |
| 366 | 31.83 | 270 | 23.48 |
| 118 | 10.26 | 64 | 5.57 |
| 186 | 16.17 | 116 | 10.09 |
| 81 | 7.04 | 37 | 3.22 |
| 242 | 21.04 | 164 | 14.26 |
| 164 | 14.26 | 98 | 8.52 |
| 340 | 29.57 | 251 | 21.83 |
| 271 | 23.57 | 189 | 16.43 |
| 79 | 6.87 | 36 | 3.13 |
| 328 | 28.52 | 242 | 21.04 |
| 123 | 10.7 | 67 | 5.83 |
| 140 | 12.17 | 85 | 7.39 |
| 20 | 1.74 | 4 | 0.35 |
| 292 | 25.39 | 209 | 18.17 |

| | | | | |
|---|---|---|---|---|
| | 64 | 5.57 | 27 | 2.35 |

ıat.meme

| rank | motif_ID | consensus | p-value | adj_p-value |
|---|---|---|---|---|
| 1 | TEAD4_HUMAN.H11MO.0.A | RCATTCCWGRVAK | 7.68E-166 | 9.93E-163 |
| 2 | TEAD1_HUMAN.H11MO.0.A | RCATTCCWGSCW | 1.24E-154 | 1.46E-151 |
| 3 | ZBT17_HUMAN.H11MO.0.A | SRRGGWGGGGGAGGGGMRR | 1.18E-53 | 1.67E-50 |
| 4 | MAZ_HUMAN.H11MO.0.A | GGGGGAGGGGGDGRGRRRGRRG | 7.39E-52 | 9.28E-49 |
| 5 | VEZF1_HUMAN.H11MO.0.C | GGRRRRGRRGGAGGGGGRGRRR | 6.84E-48 | 9.24E-45 |
| 6 | EGR2_HUMAN.H11MO.0.A | GRGRRKGWGKGGGHGGRG | 1.25E-47 | 1.39E-44 |
| 7 | ZN467_HUMAN.H11MO.0.C | GGGGAGGGGRRGGRGRRGGRRR | 1.51E-46 | 1.38E-43 |
| 8 | KLF15_HUMAN.H11MO.0.A | RGGGMGGRGVDGGGGGRGG | 9.52E-45 | 8.79E-42 |
| 9 | ETV4_HUMAN.H11MO.0.B | VRCAGGAARBR | 9.34E-45 | 1.43E-41 |
| 10 | ERG_HUMAN.H11MO.0.A | RRRSAGGAAGYGG | 3.10E-40 | 3.19E-37 |
| 11 | ETV5_HUMAN.H11MO.0.C | RRRSAGGAARDGRV | 2.53E-40 | 4.06E-37 |
| 12 | KLF6_HUMAN.H11MO.0.A | GSRRGGGHGGGGMHGGGRV | 3.58E-40 | 4.47E-37 |
| 13 | E2F7_HUMAN.H11MO.0.B | GDGGCGGGAARDR | 2.19E-39 | 2.72E-36 |
| 14 | SP4_HUMAN.H11MO.0.A | SRGVARGRGGCGGRGCHDRR | 4.14E-38 | 4.66E-35 |
| 15 | EGR1_HUMAN.H11MO.0.A | VDRDGCGKGGGYGGRRR | 6.76E-38 | 7.18E-35 |
| 16 | ZN341_HUMAN.H11MO.0.C | GRGGRRRRGGRGRRGGAASAGC | 1.92E-36 | 2.46E-33 |
| 17 | ZN263_HUMAN.H11MO.0.A | GGGAGGAGGRRGRGGRGGRR | 1.05E-35 | 1.02E-32 |
| 18 | ZN770_HUMAN.H11MO.0.C | GGGAGGCYGAGGBRGGAGGATC | 1.87E-35 | 1.49E-32 |
| 19 | PATZ1_HUMAN.H11MO.0.C | GGGGMGGGGGMKGGRRVGGVRG | 1.41E-35 | 1.72E-32 |
| 20 | FLI1_HUMAN.H11MO.0.A | SRRGGMAGGAAGGRRRGR | 6.82E-35 | 7.12E-32 |
| 21 | IRF3_HUMAN.H11MO.0.B | RAAARGGAAAVDGAAASDGA | 1.54E-34 | 1.41E-31 |
| 22 | WT1_HUMAN.H11MO.0.C | RGGVGGGGGAGGRGGVGGRG | 2.61E-34 | 2.40E-31 |
| 23 | ZN281_HUMAN.H11MO.0.A | RGGWGGGGGAGGGGV | 3.57E-34 | 3.01E-31 |
| 24 | BC11A_HUMAN.H11MO.0.A | RAAAGAGGAAGTGARAV | 2.56E-34 | 3.46E-31 |
| 25 | ETS1_HUMAN.H11MO.0.A | RRRCMGGAAGTGG | 1.03E-33 | 7.65E-31 |
| 26 | IRF1_HUMAN.H11MO.0.A | RAAANWGAAASTGAAASYRR | 4.36E-30 | 1.84E-27 |
| 27 | ETS2_HUMAN.H11MO.0.B | RDVRGGAARVRRR | 3.46E-30 | 5.83E-27 |
| 28 | KLF3_HUMAN.H11MO.0.B | GGRGVDGGGCGGGGCYGGG | 9.80E-30 | 8.76E-27 |
| 29 | ETV2_HUMAN.H11MO.0.B | RRARRCAGGAARYRGS | 9.48E-29 | 5.33E-26 |
| 30 | NFAC1_HUMAN.H11MO.0.B | AWGGAAARWVWGAMW | 1.31E-28 | 1.73E-25 |
| 31 | KLF5_HUMAN.H11MO.0.A | WGGGTGKGGCDGGG | 1.78E-28 | 1.96E-25 |
| 32 | TAF1_HUMAN.H11MO.0.A | RARRWGGCGGMGGMGR | 3.04E-28 | 3.18E-25 |
| 33 | FOXP1_HUMAN.H11MO.0.A | TBTGTTTMY | 2.64E-27 | 2.73E-24 |
| 34 | TBX21_HUMAN.H11MO.0.A | GDGDRGGTGTGRV | 2.01E-27 | 2.97E-24 |
| 35 | SALL4_HUMAN.H11MO.0.B | GSDGGGWGGG | 1.95E-27 | 3.00E-24 |
| 36 | ZSC22_HUMAN.H11MO.0.C | RGKCWGAGGGMGGAGG | 6.73E-27 | 5.31E-24 |
| 37 | SP2_HUMAN.H11MO.0.A | GGSSVGGGGGCGGGGCCDGSGS | 1.87E-26 | 1.66E-23 |
| 38 | SP3_HUMAN.H11MO.0.B | SGVVGGGGGCGGGGCBRGSS | 3.81E-26 | 4.23E-23 |
| 39 | STAT2_HUMAN.H11MO.0.A | RRGRAAAHGAAACTGAAAV | 3.77E-25 | 1.68E-22 |
| 40 | E2F3_HUMAN.H11MO.0.A | DDGGCGGGAAA | 4.46E-25 | 5.84E-22 |
| 41 | FOSL1_HUMAN.H11MO.0.A | KRVTGAGTCAYH | 1.18E-23 | 4.69E-21 |
| 42 | FEV_HUMAN.H11MO.0.B | GCVGGAAGYG | 4.10E-24 | 6.59E-21 |
| 43 | KLF1_HUMAN.H11MO.0.A | DGGGYGKGGCYGGG | 1.28E-23 | 1.05E-20 |
| 44 | ELF5_HUMAN.H11MO.0.A | VGRASSAGGAAGTRV | 1.27E-23 | 1.06E-20 |
| 45 | JUND_HUMAN.H11MO.0.A | RRTGAGTCAYY | 2.21E-23 | 1.11E-20 |

| | | | | |
|---|---|---|---|---|
| 46 | ZFP82_HUMAN.H11MO.0.C | TTCCADTTCAYTAATTYTCTCTTC | 2.56E-23 | 1.50E-20 |
| 47 | TFDP1_HUMAN.H11MO.0.C | RRRRGGCGGGAARN | 2.86E-23 | 2.73E-20 |
| 48 | JUN_HUMAN.H11MO.0.A | DRTGAGTCAYH | 6.97E-23 | 3.23E-20 |
| 49 | ELF3_HUMAN.H11MO.0.A | RAABVAGGAAGTRR | 5.59E-23 | 3.66E-20 |
| 50 | FOSL2_HUMAN.H11MO.0.A | NDRTGAGTCAYH | 1.08E-22 | 4.13E-20 |
| 51 | FOS_HUMAN.H11MO.0.A | VTGAGTCAB | 3.92E-22 | 2.30E-19 |
| 52 | PRDM6_HUMAN.H11MO.0.C | RRRARGAAAAAAA | 2.31E-22 | 2.84E-19 |
| 53 | ZN121_HUMAN.H11MO.0.C | CTGGGCAACABAGYRAGACC | 2.38E-21 | 3.03E-19 |
| 54 | FOXK1_HUMAN.H11MO.0.A | TGTTTMCHTT | 3.40E-22 | 3.54E-19 |
| 55 | KLF4_HUMAN.H11MO.0.A | WGGGYGKGGC | 6.54E-22 | 4.85E-19 |
| 56 | FOSB_HUMAN.H11MO.0.A | RTGAGTCAB | 2.20E-21 | 1.61E-18 |
| 57 | E2F6_HUMAN.H11MO.0.A | DGGGCGGGARRVR | 2.76E-21 | 3.16E-18 |
| 58 | SP1_HUMAN.H11MO.0.A | VGGGGGCGGGGCCKGGGGGGGG | 8.47E-21 | 3.45E-18 |
| 59 | PRDM1_HUMAN.H11MO.0.A | RAAAGTGAAAGTGR | 5.60E-21 | 3.97E-18 |
| 60 | SUH_HUMAN.H11MO.0.A | VSYSTGGGAAV | 9.56E-21 | 1.22E-17 |
| 61 | FOXO1_HUMAN.H11MO.0.A | KBBYTGTTTHCN | 1.24E-20 | 1.48E-17 |
| 62 | JUNB_HUMAN.H11MO.0.A | DVTGAGTCABH | 3.31E-20 | 1.79E-17 |
| 63 | FOXJ3_HUMAN.H11MO.0.A | TTGTTTATKKTTW | 1.08E-19 | 2.93E-17 |
| 64 | ZN394_HUMAN.H11MO.0.C | NRARWRGAAWNGAMWGRAAK | 1.48E-19 | 1.89E-16 |
| 65 | IRF2_HUMAN.H11MO.0.A | RAAAVHGAAAGTGAAASTRV | 7.81E-19 | 2.19E-16 |
| 66 | NFKB1_HUMAN.H11MO.0.A | AGGGAAAK | 5.70E-19 | 9.20E-16 |
| 67 | E2F4_HUMAN.H11MO.0.A | SRGGGCGGGAARD | 3.23E-18 | 2.36E-15 |
| 68 | EHF_HUMAN.H11MO.0.B | VWACSAGGAAGTDVS | 4.75E-18 | 2.42E-15 |
| 69 | ANDR_HUMAN.H11MO.0.A | TGTTCTTKKYTGTTTRYW | 1.55E-17 | 1.01E-14 |
| 70 | ELF1_HUMAN.H11MO.0.A | VRACCCGGAAGTGS | 3.04E-17 | 1.61E-14 |
| 71 | ELF2_HUMAN.H11MO.0.C | RAVCCGGAAGTGR | 5.34E-17 | 3.15E-14 |
| 72 | KLF12_HUMAN.H11MO.0.C | VGGGGCGGGGC | 1.13E-16 | 7.13E-14 |
| 73 | LYL1_HUMAN.H11MO.0.A | CCAKCTGYTYYCTB | 8.92E-17 | 1.45E-13 |
| 74 | SPI1_HUMAN.H11MO.0.A | RRAAAGAGGAAGTGRVD | 3.64E-16 | 1.83E-13 |
| 75 | MEF2B_HUMAN.H11MO.0.A | DKKKCTATTTTTRG | 2.59E-16 | 1.87E-13 |
| 76 | ETV1_HUMAN.H11MO.0.A | SRCCGGAAGYG | 8.22E-16 | 4.74E-13 |
| 77 | RXRA_HUMAN.H11MO.0.A | VRGDVRGNVARAGGTCARRG | 4.91E-16 | 6.92E-13 |
| 78 | MEF2C_HUMAN.H11MO.0.A | DKCTATTTTTRGM | 1.95E-15 | 1.14E-12 |
| 79 | MEF2A_HUMAN.H11MO.0.A | DKCTATTTTTRGM | 2.07E-15 | 1.21E-12 |
| 80 | FOXP2_HUMAN.H11MO.0.C | BTGTTTACH | 2.74E-15 | 1.97E-12 |
| 81 | E2F1_HUMAN.H11MO.0.A | RRNKGGCGGGAARR | 3.74E-15 | 2.74E-12 |
| 82 | FOXO3_HUMAN.H11MO.0.B | BYTGTTTACH | 7.39E-15 | 4.69E-12 |
| 83 | SRY_HUMAN.H11MO.0.B | YWTTGTTTH | 6.53E-15 | 6.74E-12 |
| 84 | Z354A_HUMAN.H11MO.0.C | RTTTARWYCATTTACATTTAATGT | 1.48E-14 | 6.85E-12 |
| 85 | SMCA5_HUMAN.H11MO.0.C | GAATBGAATGGAAWC | 1.71E-14 | 1.04E-11 |
| 86 | MEF2D_HUMAN.H11MO.0.A | DKCTATTTTTAG | 2.97E-14 | 1.22E-11 |
| 87 | SOX4_HUMAN.H11MO.0.B | BYCTTTGTYYYB | 3.96E-14 | 4.37E-11 |
| 88 | SMAD3_HUMAN.H11MO.0.B | SYCTSHCWSCWS | 8.92E-14 | 1.61E-10 |
| 89 | RUNX2_HUMAN.H11MO.0.A | KGKYTGTGGTTTKK | 2.60E-13 | 2.08E-10 |
| 90 | IRF8_HUMAN.H11MO.0.B | RAAARRGGAASTGAAASTDR | 6.38E-13 | 2.67E-10 |
| 91 | NFIC_HUMAN.H11MO.0.A | CYTGGCHBYMWGCCARG | 2.86E-13 | 3.07E-10 |
| 92 | SOX2_HUMAN.H11MO.0.A | BBCCTTTGTYYYB | 3.74E-13 | 4.21E-10 |

| | | | | |
|---|---|---|---|---|
| 93 | IRF7_HUMAN.H11MO.0.C | GAAASYGAAA | 4.63E-13 | 4.35E-10 |
| 94 | FOXM1_HUMAN.H11MO.0.A | TGTTTRCTYWKB | 4.90E-13 | 4.39E-10 |
| 95 | ZFP28_HUMAN.H11MO.0.C | TTCTATTTCTTCTTGWGTCA | 1.69E-12 | 4.65E-10 |
| 96 | GATA3_HUMAN.H11MO.0.A | RVAGATAAVAD | 4.97E-13 | 4.95E-10 |
| 97 | KLF9_HUMAN.H11MO.0.C | GRRGKGGGCGTGGCC | 2.36E-12 | 9.73E-10 |
| 98 | PAX5_HUMAN.H11MO.0.A | SCSKGGGCARCVRAGCGWGAC | 3.60E-12 | 1.33E-09 |
| 99 | IRF4_HUMAN.H11MO.0.A | RAAWGRGGAASTGAAASY | 3.28E-12 | 1.52E-09 |
| 100 | SMCA1_HUMAN.H11MO.0.C | CMARGAAGMATD | 1.36E-12 | 1.61E-09 |
| 101 | SRBP2_HUMAN.H11MO.0.B | VSGTGGGGWGABG | 1.79E-12 | 2.45E-09 |
| 102 | NFE2_HUMAN.H11MO.0.A | VVRTGACTCAGCA | 1.05E-11 | 3.98E-09 |
| 103 | NFAC2_HUMAN.H11MO.0.B | TGGAAAAWY | 4.06E-12 | 4.18E-09 |
| 104 | STAT3_HUMAN.H11MO.0.A | SAYTTCCMRGAA | 5.97E-12 | 4.25E-09 |
| 105 | NFIA_HUMAN.H11MO.0.C | TTGGCHNRNWGCCAR | 7.10E-12 | 6.25E-09 |
| 106 | NF2L2_HUMAN.H11MO.0.A | WWYTGCTGAGTCAT | 3.80E-11 | 6.80E-09 |
| 107 | FOXJ2_HUMAN.H11MO.0.C | TGTTTRTTTW | 2.64E-11 | 1.25E-08 |
| 108 | TBX3_HUMAN.H11MO.0.C | DRAGGTGBSAR | 8.97E-12 | 1.28E-08 |
| 109 | IKZF1_HUMAN.H11MO.0.C | BTGGGARR | 8.80E-12 | 1.34E-08 |
| 110 | NFAC4_HUMAN.H11MO.0.C | DGGAAAAYTW | 1.52E-11 | 1.71E-08 |
| 111 | FOXA1_HUMAN.H11MO.0.A | TGTTTACWYWGB | 5.31E-11 | 2.69E-08 |
| 112 | OSR2_HUMAN.H11MO.0.C | GBYSCWGCTNCTGYDG | 3.35E-11 | 2.97E-08 |
| 113 | STA5A_HUMAN.H11MO.0.A | TTCYRRGAAR | 5.46E-11 | 3.07E-08 |
| 114 | PTF1A_HUMAN.H11MO.0.B | SCAGCTGBYBBYBYYCHC | 1.14E-10 | 9.74E-08 |
| 115 | FOXA2_HUMAN.H11MO.0.A | TGTTTACWYWGB | 2.96E-10 | 1.36E-07 |
| 116 | MAF_HUMAN.H11MO.0.A | VAAWBTGCTGASTHWGCMD | 1.96E-10 | 1.50E-07 |
| 117 | STAT1_HUMAN.H11MO.0.A | YCCVTTTCCHGGAAAWCHC | 5.60E-10 | 2.25E-07 |
| 118 | Z324A_HUMAN.H11MO.0.C | KVTCAAACCATCCTTKSCTKSCW | 3.48E-10 | 2.70E-07 |
| 119 | BACH2_HUMAN.H11MO.0.A | TGCTGAGTCAB | 6.34E-10 | 3.26E-07 |
| 120 | COT1_HUMAN.H11MO.0.C | GRGGKSARAGGTCAGGG | 3.42E-10 | 3.74E-07 |
| 121 | RUNX1_HUMAN.H11MO.0.A | BYTGTGGTTWB | 6.09E-10 | 3.75E-07 |
| 122 | BHA15_HUMAN.H11MO.0.B | SSAGCAGCTGG | 6.89E-10 | 4.37E-07 |
| 123 | GABPA_HUMAN.H11MO.0.A | GVVRCCGGAAGTGV | 1.89E-09 | 7.48E-07 |
| 124 | MXI1_HUMAN.H11MO.0.A | VCACGTGGSNGSBGS | 1.83E-09 | 9.88E-07 |
| 125 | MAFF_HUMAN.H11MO.0.B | TGCTGASTCAGCABTTTT | 1.48E-09 | 1.01E-06 |
| 126 | LEF1_HUMAN.H11MO.0.A | BCCTTTSWWNTBYH | 1.73E-09 | 2.66E-06 |
| 127 | ATF2_HUMAN.H11MO.0.B | RRTGABGTCAY | 7.18E-09 | 3.30E-06 |
| 128 | RUNX3_HUMAN.H11MO.0.A | BYTGTGGTTW | 5.86E-09 | 3.43E-06 |
| 129 | BACH1_HUMAN.H11MO.0.A | TGCTGAGTCABSS | 6.79E-09 | 3.73E-06 |
| 130 | MYOG_HUMAN.H11MO.0.B | CHSCAGCTGYYYB | 1.05E-08 | 5.17E-06 |
| 131 | ZN768_HUMAN.H11MO.0.C | GAGGCHCAGAGAGGTTAAGT | 3.23E-08 | 6.19E-06 |
| 132 | ELK1_HUMAN.H11MO.0.B | RCCGGAAGTGV | 2.05E-08 | 6.98E-06 |
| 133 | SPIB_HUMAN.H11MO.0.A | RRAAAGAGGAAGTGARA | 2.73E-08 | 9.00E-06 |
| 134 | STA5B_HUMAN.H11MO.0.A | YTTCYKRGAAAY | 1.52E-08 | 1.06E-05 |
| 135 | BMAL1_HUMAN.H11MO.0.A | KSCACGTGHSY | 3.23E-08 | 1.15E-05 |
| 136 | ITF2_HUMAN.H11MO.0.C | DGCAGSTGKS | 1.05E-08 | 1.29E-05 |
| 137 | REL_HUMAN.H11MO.0.B | DRWRGGGRAATKCCA | 1.67E-08 | 1.60E-05 |
| 138 | MAX_HUMAN.H11MO.0.A | VRGCACGTGG | 4.01E-08 | 1.72E-05 |
| 139 | ZN322_HUMAN.H11MO.0.B | GWGCCTGSTACWSWGCCTGR | 7.14E-08 | 2.54E-05 |

| | | | | |
|---|---|---|---|---|
| 140 | PEBB_HUMAN.H11MO.0.C | TYTGTGGTYWB | 3.34E-08 | 2.55E-05 |
| 141 | MYCN_HUMAN.H11MO.0.A | RGCCACGTGSDS | 8.35E-08 | 2.72E-05 |
| 142 | MZF1_HUMAN.H11MO.0.B | RRDRGGGAWTKRR | 5.65E-08 | 7.13E-05 |
| 143 | NFAC3_HUMAN.H11MO.0.B | TGGAAAAHY | 8.19E-08 | 7.26E-05 |
| 144 | PO2F1_HUMAN.H11MO.0.C | AWKBRWATGSAAWYAW | 1.41E-07 | 8.75E-05 |
| 145 | BCL6_HUMAN.H11MO.0.A | YKCTTTCYAGGAA | 1.14E-07 | 9.68E-05 |
| 146 | HTF4_HUMAN.H11MO.0.A | DSCAGSTGKG | 1.05E-07 | 1.12E-04 |
| 147 | NDF2_HUMAN.H11MO.0.B | RRCAGATGG | 1.69E-07 | 1.34E-04 |
| 148 | FOXA3_HUMAN.H11MO.0.B | CHTGTTTACWTWG | 3.21E-07 | 1.66E-04 |
| 149 | HIF1A_HUMAN.H11MO.0.C | GDACGTGM | 5.72E-07 | 1.66E-04 |
| 150 | HNF1B_HUMAN.H11MO.0.A | DGTTAATRATTAACH | 6.93E-07 | 1.67E-04 |
| 151 | STAT4_HUMAN.H11MO.0.A | YTTCYYRGMAV | 2.03E-07 | 2.00E-04 |
| 152 | FOXO4_HUMAN.H11MO.0.C | TTGTTTAYK | 3.78E-07 | 2.13E-04 |
| 153 | MYC_HUMAN.H11MO.0.A | RCCACGTGCTB | 6.70E-07 | 2.14E-04 |
| 154 | NANOG_HUMAN.H11MO.0.A | BYYWTTGWNATGCAAAT | 3.50E-07 | 2.41E-04 |
| 155 | SRF_HUMAN.H11MO.0.A | TTKCCWTWTWTGGBCWTD | 1.17E-06 | 2.85E-04 |
| 156 | RARA_HUMAN.H11MO.0.A | GGRRGRTSMRRAGKTCAAGG | 4.65E-07 | 3.27E-04 |
| 157 | CLOCK_HUMAN.H11MO.0.C | CWGHCACGTGMVVM | 5.74E-07 | 4.55E-04 |
| 158 | PRGR_HUMAN.H11MO.0.A | RGAACATTYTGTTCTB | 1.03E-06 | 5.07E-04 |
| 159 | BHE40_HUMAN.H11MO.0.A | DGCACGTGAS | 2.48E-06 | 5.69E-04 |
| 160 | HXB13_HUMAN.H11MO.0.A | TTTTATDGSH | 9.22E-07 | 5.84E-04 |
| 161 | OLIG2_HUMAN.H11MO.0.B | CCAKCTGYTYYYYWBCHB | 6.25E-07 | 6.18E-04 |
| 162 | RXRB_HUMAN.H11MO.0.C | TSAGGTCASR | 8.40E-07 | 6.59E-04 |
| 163 | HSF1_HUMAN.H11MO.0.A | BRGAANSTTCYRGAA | 2.47E-06 | 7.36E-04 |
| 164 | ZN418_HUMAN.H11MO.0.C | TGCTTYTRGCYTCTKVY | 1.20E-06 | 7.94E-04 |
| 165 | TF65_HUMAN.H11MO.0.A | HWGGGRATTTCCMV | 2.01E-06 | 1.03E-03 |
| 166 | ARNT_HUMAN.H11MO.0.B | STACGTGMC | 4.91E-06 | 1.05E-03 |
| 167 | TFE2_HUMAN.H11MO.0.A | SMCAGCTGCWB | 1.78E-06 | 1.20E-03 |
| 168 | TBP_HUMAN.H11MO.0.A | MTATAAAARS | 1.96E-06 | 1.33E-03 |
| 169 | HSF2_HUMAN.H11MO.0.A | VGAABRTTCTAGAA | 2.39E-06 | 1.35E-03 |
| 170 | ZN554_HUMAN.H11MO.0.C | GCWGAGYCANGTRGDKKRCT | 2.25E-06 | 1.47E-03 |
| 171 | ELK4_HUMAN.H11MO.0.A | SVCCGGAAGTGV | 5.51E-06 | 1.69E-03 |
| 172 | FOXH1_HUMAN.H11MO.0.A | TGTGGATTS | 3.30E-06 | 2.00E-03 |
| 173 | CREM_HUMAN.H11MO.0.C | CRVTGACGTCA | 8.81E-06 | 2.26E-03 |
| 174 | ZNF76_HUMAN.H11MO.0.C | RRKGCMTKCTGGGARWTGTAGT | 8.68E-06 | 2.65E-03 |
| 175 | ZN335_HUMAN.H11MO.0.A | RGCTGYCYNBVSCKMTGCCTGW | 7.72E-06 | 3.03E-03 |
| 176 | HXA10_HUMAN.H11MO.0.C | KATRRTTTATGW | 7.58E-06 | 3.06E-03 |
| 177 | SOX3_HUMAN.H11MO.0.B | BCCWTTGTBYB | 3.63E-06 | 3.22E-03 |
| 178 | FOXC1_HUMAN.H11MO.0.C | CDWTGTTTACTTWRG | 2.10E-05 | 3.43E-03 |
| 179 | PIT1_HUMAN.H11MO.0.C | YTMATGAATATWY | 5.62E-06 | 3.69E-03 |
| 180 | ZNF85_HUMAN.H11MO.0.C | AWAGAARWCTGMWGKAATCT | 1.18E-05 | 3.78E-03 |
| 181 | NKX61_HUMAN.H11MO.0.B | GAYWAATKRSHWWTTAATK | 6.59E-06 | 4.17E-03 |
| 182 | ZBT18_HUMAN.H11MO.0.C | KCCAGMTGTKB | 6.99E-06 | 5.21E-03 |
| 183 | ZEB1_HUMAN.H11MO.0.A | BVCAGGTGWG | 5.72E-06 | 5.80E-03 |
| 184 | ZN350_HUMAN.H11MO.0.C | CAGYYYTTTTATDWCCHR | 6.77E-06 | 7.19E-03 |
| 185 | SIX1_HUMAN.H11MO.0.A | TGWAAYHTGAKMCY | 1.96E-05 | 7.71E-03 |
| 186 | SMAD2_HUMAN.H11MO.0.A | TGTCTGNCWCCT | 7.79E-06 | 7.93E-03 |

| | | | | |
|---|---|---|---|---|
| 187 | SIX2_HUMAN.H11MO.0.A | KGAAAYHTGAKMC | 1.28E-05 | 8.30E-03 |
| 188 | EPAS1_HUMAN.H11MO.0.B | VTACGTGMC | 4.81E-05 | 9.23E-03 |
| 189 | HNF1A_HUMAN.H11MO.0.C | DGTTAATKATTAACH | 2.40E-05 | 1.01E-02 |
| 190 | MAFK_HUMAN.H11MO.0.A | RAWWHTGCTGASTCAGCA | 8.74E-05 | 1.41E-02 |
| 191 | IRF9_HUMAN.H11MO.0.C | GAAAGCGAAAYT | 9.77E-05 | 1.67E-02 |
| 192 | USF1_HUMAN.H11MO.0.A | RGTCACGTGRBS | 1.77E-04 | 1.91E-02 |
| 193 | ZSC31_HUMAN.H11MO.0.C | RGCAGCAGGGCARTTAKR | 3.77E-05 | 2.22E-02 |

| E-value | tests | FASTA_max | pos | neg | PWM_min | TP | %TP |
|---|---|---|---|---|---|---|---|
| 3.99E-160 | 1294 | 1166 | 1166 | 1166 | 2.69 | 762 | 65.35 |
| 5.86E-149 | 1179 | 1166 | 1166 | 1166 | 2.34 | 745 | 63.89 |
| 6.70E-48 | 1413 | 1166 | 1166 | 1166 | 4.93 | 579 | 49.66 |
| 3.73E-46 | 1256 | 1166 | 1166 | 1166 | 9.09 | 506 | 43.4 |
| 3.71E-42 | 1350 | 1166 | 1166 | 1166 | 22.1 | 376 | 32.25 |
| 5.61E-42 | 1113 | 1166 | 1166 | 1166 | 7.43 | 391 | 33.53 |
| 5.56E-41 | 917 | 1166 | 1166 | 1166 | 6.35 | 451 | 38.68 |
| 3.53E-39 | 923 | 1166 | 1166 | 1166 | 2.01 | 552 | 47.34 |
| 5.75E-39 | 1531 | 1166 | 1166 | 1166 | 1.6 | 777 | 66.64 |
| 1.28E-34 | 1029 | 1166 | 1166 | 1166 | 1.02 | 672 | 57.63 |
| 1.63E-34 | 1605 | 1166 | 1166 | 1166 | 1.7 | 780 | 66.9 |
| 1.80E-34 | 1247 | 1166 | 1166 | 1166 | 2.31 | 619 | 53.09 |
| 1.09E-33 | 1242 | 1166 | 1166 | 1166 | 2.02 | 573 | 49.14 |
| 1.87E-32 | 1127 | 1166 | 1166 | 1166 | 4.09 | 459 | 39.37 |
| 2.89E-32 | 1063 | 1166 | 1166 | 1166 | 2.75 | 466 | 39.97 |
| 9.90E-31 | 1279 | 1166 | 1166 | 1166 | 10.1 | 365 | 31.3 |
| 4.11E-30 | 979 | 1166 | 1166 | 1166 | 2.39 | 511 | 43.83 |
| 5.97E-30 | 793 | 1166 | 1166 | 1166 | 1.43 | 516 | 44.25 |
| 6.91E-30 | 1220 | 1166 | 1166 | 1166 | 5.92 | 493 | 42.28 |
| 2.86E-29 | 1043 | 1166 | 1166 | 1166 | 2.78 | 476 | 40.82 |
| 5.68E-29 | 919 | 1166 | 1166 | 1166 | 1.99 | 502 | 43.05 |
| 9.67E-29 | 920 | 1166 | 1166 | 1166 | 6.8 | 410 | 35.16 |
| 1.21E-28 | 845 | 1166 | 1166 | 1166 | 12.3 | 288 | 24.7 |
| 1.39E-28 | 1352 | 1166 | 1166 | 1166 | 1.49 | 720 | 61.75 |
| 3.07E-28 | 740 | 1166 | 1166 | 1166 | 1.08 | 492 | 42.2 |
| 7.39E-25 | 422 | 1166 | 1166 | 1166 | 1.15 | 315 | 27.02 |
| 2.35E-24 | 1687 | 1166 | 1166 | 1166 | 1.21 | 915 | 78.47 |
| 3.52E-24 | 894 | 1166 | 1166 | 1166 | 4.95 | 374 | 32.08 |
| 2.14E-23 | 562 | 1166 | 1166 | 1166 | 1.89 | 312 | 26.76 |
| 6.97E-23 | 1324 | 1166 | 1166 | 1166 | 2.74 | 517 | 44.34 |
| 7.87E-23 | 1100 | 1166 | 1166 | 1166 | 4.02 | 411 | 35.25 |
| 1.28E-22 | 1047 | 1166 | 1166 | 1166 | 2.17 | 442 | 37.91 |
| 1.10E-21 | 1036 | 1166 | 1166 | 1166 | 2.34 | 400 | 34.31 |
| 1.19E-21 | 1479 | 1166 | 1166 | 1166 | 1.62 | 574 | 49.23 |
| 1.21E-21 | 1539 | 1166 | 1166 | 1166 | 2.59 | 634 | 54.37 |
| 2.13E-21 | 789 | 1166 | 1166 | 1166 | 2.38 | 370 | 31.73 |
| 6.68E-21 | 888 | 1166 | 1166 | 1166 | 3.28 | 443 | 37.99 |
| 1.70E-20 | 1111 | 1166 | 1166 | 1166 | 22 | 257 | 22.04 |
| 6.76E-20 | 446 | 1166 | 1166 | 1166 | 1.17 | 317 | 27.19 |
| 2.35E-19 | 1310 | 1166 | 1166 | 1166 | 1.3 | 683 | 58.58 |
| 1.88E-18 | 396 | 1166 | 1166 | 1166 | 2.65 | 228 | 19.55 |
| 2.65E-18 | 1605 | 1166 | 1166 | 1166 | 1.21 | 794 | 68.1 |
| 4.22E-18 | 823 | 1166 | 1166 | 1166 | 1.5 | 467 | 40.05 |
| 4.24E-18 | 832 | 1166 | 1166 | 1166 | 1.15 | 507 | 43.48 |
| 4.46E-18 | 501 | 1166 | 1166 | 1166 | 8.11 | 157 | 13.46 |

| | | | | | | | |
|---|---|---|---|---|---|---|---|
| 6.04E-18 | 586 | 1166 | 1166 | 1166 | 1.4 | 395 | 33.88 |
| 1.10E-17 | 955 | 1166 | 1166 | 1166 | 2.45 | 353 | 30.27 |
| 1.30E-17 | 463 | 1166 | 1166 | 1166 | 7.47 | 157 | 13.46 |
| 1.47E-17 | 655 | 1166 | 1166 | 1166 | 1.3 | 397 | 34.05 |
| 1.66E-17 | 383 | 1166 | 1166 | 1166 | 2.69 | 222 | 19.04 |
| 9.25E-17 | 587 | 1166 | 1166 | 1166 | 8.88 | 173 | 14.84 |
| 1.14E-16 | 1231 | 1166 | 1166 | 1166 | 10 | 264 | 22.64 |
| 1.22E-16 | 127 | 1166 | 1166 | 1166 | 3.8 | 87 | 7.46 |
| 1.43E-16 | 1042 | 1166 | 1166 | 1166 | 1.67 | 465 | 39.88 |
| 1.95E-16 | 742 | 1166 | 1166 | 1166 | 2.21 | 359 | 30.79 |
| 6.46E-16 | 729 | 1166 | 1166 | 1166 | 3.84 | 238 | 20.41 |
| 1.27E-15 | 1147 | 1166 | 1166 | 1166 | 2.91 | 399 | 34.22 |
| 1.39E-15 | 407 | 1166 | 1166 | 1166 | 1.28 | 288 | 24.7 |
| 1.60E-15 | 708 | 1166 | 1166 | 1166 | 1.35 | 401 | 34.39 |
| 4.92E-15 | 1279 | 1166 | 1166 | 1166 | 1.01 | 751 | 64.41 |
| 5.94E-15 | 1189 | 1166 | 1166 | 1166 | 1.32 | 609 | 52.23 |
| 7.19E-15 | 540 | 1166 | 1166 | 1166 | 3.3 | 223 | 19.13 |
| 1.18E-14 | 272 | 1166 | 1166 | 1166 | 5.64 | 138 | 11.84 |
| 7.58E-14 | 1276 | 1166 | 1166 | 1166 | 2.11 | 566 | 48.54 |
| 8.80E-14 | 280 | 1166 | 1166 | 1166 | 1.31 | 199 | 17.07 |
| 3.70E-13 | 1613 | 1166 | 1166 | 1166 | 1.28 | 816 | 69.98 |
| 9.47E-13 | 729 | 1166 | 1166 | 1166 | 5.17 | 196 | 16.81 |
| 9.72E-13 | 509 | 1166 | 1166 | 1166 | 1.14 | 323 | 27.7 |
| 4.06E-12 | 651 | 1166 | 1166 | 1166 | 1.94 | 320 | 27.44 |
| 6.46E-12 | 529 | 1166 | 1166 | 1166 | 1.04 | 349 | 29.93 |
| 1.27E-11 | 590 | 1166 | 1166 | 1166 | 1.03 | 381 | 32.68 |
| 2.87E-11 | 634 | 1166 | 1166 | 1166 | 1.56 | 343 | 29.42 |
| 5.82E-11 | 1624 | 1166 | 1166 | 1166 | 1.64 | 614 | 52.66 |
| 7.36E-11 | 503 | 1166 | 1166 | 1166 | 1.09 | 331 | 28.39 |
| 7.53E-11 | 723 | 1166 | 1166 | 1166 | 5.93 | 169 | 14.49 |
| 1.90E-10 | 576 | 1166 | 1166 | 1166 | 1.02 | 371 | 31.82 |
| 2.78E-10 | 1408 | 1166 | 1166 | 1166 | 2.45 | 518 | 44.43 |
| 4.56E-10 | 581 | 1166 | 1166 | 1166 | 1.43 | 315 | 27.02 |
| 4.88E-10 | 585 | 1166 | 1166 | 1166 | 1.4 | 325 | 27.87 |
| 7.90E-10 | 718 | 1166 | 1166 | 1166 | 1.11 | 427 | 36.62 |
| 1.10E-09 | 733 | 1166 | 1166 | 1166 | 1.43 | 380 | 32.59 |
| 1.88E-09 | 634 | 1166 | 1166 | 1166 | 1.17 | 372 | 31.9 |
| 2.71E-09 | 1032 | 1166 | 1166 | 1166 | 4.35 | 212 | 18.18 |
| 2.75E-09 | 464 | 1166 | 1166 | 1166 | 1.65 | 285 | 24.44 |
| 4.18E-09 | 609 | 1166 | 1166 | 1166 | 5.01 | 130 | 11.15 |
| 4.91E-09 | 411 | 1166 | 1166 | 1166 | 1.07 | 270 | 23.16 |
| 1.76E-08 | 1105 | 1166 | 1166 | 1166 | 3.34 | 323 | 27.7 |
| 6.47E-08 | 1803 | 1166 | 1166 | 1166 | 2.41 | 541 | 46.4 |
| 8.36E-08 | 801 | 1166 | 1166 | 1166 | 1.33 | 420 | 36.02 |
| 1.08E-07 | 419 | 1166 | 1166 | 1166 | 1.65 | 231 | 19.81 |
| 1.24E-07 | 1076 | 1166 | 1166 | 1166 | 9.78 | 172 | 14.75 |
| 1.69E-07 | 1124 | 1166 | 1166 | 1166 | 1.91 | 483 | 41.42 |

| | | | | | | | |
|---|---|---|---|---|---|---|---|
| 1.75E-07 | 938 | 1166 | 1166 | 1166 | 2.42 | 350 | 30.02 |
| 1.76E-07 | 896 | 1166 | 1166 | 1166 | 3.5 | 229 | 19.64 |
| 1.87E-07 | 276 | 1166 | 1166 | 1166 | 1.38 | 191 | 16.38 |
| 1.99E-07 | 997 | 1166 | 1166 | 1166 | 3 | 288 | 24.7 |
| 3.91E-07 | 413 | 1166 | 1166 | 1166 | 1.13 | 263 | 22.56 |
| 5.36E-07 | 370 | 1166 | 1166 | 1166 | 27.7 | 60 | 5.15 |
| 6.09E-07 | 462 | 1166 | 1166 | 1166 | 1.16 | 290 | 24.87 |
| 6.49E-07 | 1186 | 1166 | 1166 | 1166 | 1.63 | 518 | 44.43 |
| 9.86E-07 | 1373 | 1166 | 1166 | 1166 | 2.26 | 432 | 37.05 |
| 1.60E-06 | 378 | 1166 | 1166 | 1166 | 1.77 | 182 | 15.61 |
| 1.68E-06 | 1029 | 1166 | 1166 | 1166 | 2.17 | 427 | 36.62 |
| 1.71E-06 | 712 | 1166 | 1166 | 1166 | 1.18 | 396 | 33.96 |
| 2.51E-06 | 880 | 1166 | 1166 | 1166 | 7.29 | 100 | 8.58 |
| 2.73E-06 | 179 | 1166 | 1166 | 1166 | 1.56 | 112 | 9.61 |
| 5.02E-06 | 472 | 1166 | 1166 | 1166 | 2.85 | 222 | 19.04 |
| 5.14E-06 | 1426 | 1166 | 1166 | 1166 | 1.27 | 678 | 58.15 |
| 5.40E-06 | 1527 | 1166 | 1166 | 1166 | 1.17 | 776 | 66.55 |
| 6.86E-06 | 1123 | 1166 | 1166 | 1166 | 1.7 | 501 | 42.97 |
| 1.08E-05 | 507 | 1166 | 1166 | 1166 | 1.1 | 306 | 26.24 |
| 1.20E-05 | 889 | 1166 | 1166 | 1166 | 3.48 | 241 | 20.67 |
| 1.23E-05 | 562 | 1166 | 1166 | 1166 | 1.11 | 330 | 28.3 |
| 3.92E-05 | 853 | 1166 | 1166 | 1166 | 1.91 | 357 | 30.62 |
| 5.47E-05 | 459 | 1166 | 1166 | 1166 | 1.23 | 266 | 22.81 |
| 6.02E-05 | 764 | 1166 | 1166 | 1166 | 1.62 | 334 | 28.64 |
| 9.06E-05 | 403 | 1166 | 1166 | 1166 | 1.29 | 235 | 20.15 |
| 1.09E-04 | 776 | 1166 | 1166 | 1166 | 1.15 | 458 | 39.28 |
| 1.31E-04 | 515 | 1166 | 1166 | 1166 | 16.6 | 56 | 4.8 |
| 1.50E-04 | 1095 | 1166 | 1166 | 1166 | 1.42 | 545 | 46.74 |
| 1.51E-04 | 616 | 1166 | 1166 | 1166 | 1.56 | 287 | 24.61 |
| 1.76E-04 | 634 | 1166 | 1166 | 1166 | 1.48 | 319 | 27.36 |
| 3.01E-04 | 396 | 1166 | 1166 | 1166 | 1.03 | 250 | 21.44 |
| 3.97E-04 | 539 | 1166 | 1166 | 1166 | 2.04 | 191 | 16.38 |
| 4.07E-04 | 685 | 1166 | 1166 | 1166 | 1.09 | 406 | 34.82 |
| 1.07E-03 | 1542 | 1166 | 1166 | 1166 | 2.38 | 508 | 43.57 |
| 1.33E-03 | 459 | 1166 | 1166 | 1166 | 1.26 | 248 | 21.27 |
| 1.38E-03 | 585 | 1166 | 1166 | 1166 | 5.36 | 125 | 10.72 |
| 1.50E-03 | 550 | 1166 | 1166 | 1166 | 1.7 | 242 | 20.75 |
| 2.08E-03 | 491 | 1166 | 1166 | 1166 | 6.74 | 145 | 12.44 |
| 2.49E-03 | 192 | 1166 | 1166 | 1166 | 1.27 | 132 | 11.32 |
| 2.81E-03 | 340 | 1166 | 1166 | 1166 | 1.02 | 217 | 18.61 |
| 3.62E-03 | 330 | 1166 | 1166 | 1166 | 1.1 | 211 | 18.1 |
| 4.25E-03 | 694 | 1166 | 1166 | 1166 | 1.02 | 408 | 34.99 |
| 4.62E-03 | 356 | 1166 | 1166 | 1166 | 2.08 | 132 | 11.32 |
| 5.18E-03 | 1232 | 1166 | 1166 | 1166 | 1.56 | 506 | 43.4 |
| 6.44E-03 | 957 | 1166 | 1166 | 1166 | 1.04 | 544 | 46.66 |
| 6.91E-03 | 428 | 1166 | 1166 | 1166 | 1.78 | 167 | 14.32 |
| 1.02E-02 | 356 | 1166 | 1166 | 1166 | 1.25 | 209 | 17.92 |

| | | | | | | | |
|---|---|---|---|---|---|---|---|
| 1.03E-02 | 764 | 1166 | 1166 | 1166 | 7.31 | 101 | 8.66 |
| 1.09E-02 | 326 | 1166 | 1166 | 1166 | 5.12 | 59 | 5.06 |
| 2.86E-02 | 1262 | 1166 | 1166 | 1166 | 8.79 | 119 | 10.21 |
| 2.92E-02 | 887 | 1166 | 1166 | 1166 | 1.07 | 494 | 42.37 |
| 3.52E-02 | 621 | 1166 | 1166 | 1166 | 4.04 | 143 | 12.26 |
| 3.89E-02 | 846 | 1166 | 1166 | 1166 | 1.57 | 364 | 31.22 |
| 4.51E-02 | 1067 | 1166 | 1166 | 1166 | 3.1 | 284 | 24.36 |
| 5.38E-02 | 792 | 1166 | 1166 | 1166 | 1.21 | 410 | 35.16 |
| 6.67E-02 | 517 | 1166 | 1166 | 1166 | 2.24 | 200 | 17.15 |
| 6.69E-02 | 291 | 1166 | 1166 | 1166 | 1.05 | 174 | 14.92 |
| 6.72E-02 | 241 | 1166 | 1166 | 1166 | 1.46 | 130 | 11.15 |
| 8.02E-02 | 983 | 1166 | 1166 | 1166 | 4.35 | 219 | 18.78 |
| 8.57E-02 | 564 | 1166 | 1166 | 1166 | 4.89 | 152 | 13.04 |
| 8.59E-02 | 319 | 1166 | 1166 | 1166 | 1.3 | 176 | 15.09 |
| 9.70E-02 | 689 | 1166 | 1166 | 1166 | 1.07 | 398 | 34.13 |
| 1.14E-01 | 244 | 1166 | 1166 | 1166 | 15.9 | 36 | 3.09 |
| 1.31E-01 | 703 | 1166 | 1166 | 1166 | 8.92 | 106 | 9.09 |
| 1.83E-01 | 793 | 1166 | 1166 | 1166 | 2.1 | 136 | 11.66 |
| 2.04E-01 | 493 | 1166 | 1166 | 1166 | 1.15 | 276 | 23.67 |
| 2.29E-01 | 229 | 1166 | 1166 | 1166 | 1.6 | 97 | 8.32 |
| 2.35E-01 | 633 | 1166 | 1166 | 1166 | 1.44 | 303 | 25.99 |
| 2.48E-01 | 988 | 1166 | 1166 | 1166 | 5.28 | 166 | 14.24 |
| 2.65E-01 | 785 | 1166 | 1166 | 1166 | 3.24 | 165 | 14.15 |
| 2.96E-01 | 298 | 1166 | 1166 | 1166 | 2.54 | 122 | 10.46 |
| 3.19E-01 | 659 | 1166 | 1166 | 1166 | 1.07 | 377 | 32.33 |
| 4.14E-01 | 512 | 1166 | 1166 | 1166 | 1.73 | 238 | 20.41 |
| 4.20E-01 | 213 | 1166 | 1166 | 1166 | 1.03 | 137 | 11.75 |
| 4.82E-01 | 673 | 1166 | 1166 | 1166 | 5.62 | 156 | 13.38 |
| 5.33E-01 | 678 | 1166 | 1166 | 1166 | 1.02 | 389 | 33.36 |
| 5.41E-01 | 565 | 1166 | 1166 | 1166 | 1.14 | 300 | 25.73 |
| 5.92E-01 | 654 | 1166 | 1166 | 1166 | 11.3 | 59 | 5.06 |
| 6.78E-01 | 306 | 1166 | 1166 | 1166 | 1.04 | 187 | 16.04 |
| 8.05E-01 | 608 | 1166 | 1166 | 1166 | 1.29 | 294 | 25.21 |
| 9.10E-01 | 257 | 1166 | 1166 | 1166 | 1.37 | 135 | 11.58 |
| 1.07E+00 | 306 | 1166 | 1166 | 1166 | 1.54 | 158 | 13.55 |
| 1.22E+00 | 393 | 1166 | 1166 | 1166 | 3.34 | 113 | 9.69 |
| 1.23E+00 | 404 | 1166 | 1166 | 1166 | 1.06 | 236 | 20.24 |
| 1.29E+00 | 889 | 1166 | 1166 | 1166 | 1.72 | 374 | 32.08 |
| 1.38E+00 | 164 | 1166 | 1166 | 1166 | 1.38 | 99 | 8.49 |
| 1.48E+00 | 658 | 1166 | 1166 | 1166 | 1.28 | 331 | 28.39 |
| 1.52E+00 | 321 | 1166 | 1166 | 1166 | 1.39 | 173 | 14.84 |
| 1.68E+00 | 634 | 1166 | 1166 | 1166 | 1.57 | 292 | 25.04 |
| 2.09E+00 | 747 | 1166 | 1166 | 1166 | 1.73 | 294 | 25.21 |
| 2.33E+00 | 1018 | 1166 | 1166 | 1166 | 1.01 | 561 | 48.11 |
| 2.89E+00 | 1066 | 1166 | 1166 | 1166 | 2.73 | 329 | 28.22 |
| 3.10E+00 | 395 | 1166 | 1166 | 1166 | 1.12 | 226 | 19.38 |
| 3.19E+00 | 1021 | 1166 | 1166 | 1166 | 1.22 | 490 | 42.02 |

| | | | | | | | |
|---|---|---|---|---|---|---|---|
| 3.34E+00 | 653 | 1166 | 1166 | 1166 | 2.16 | 203 | 17.41 |
| 3.71E+00 | 193 | 1166 | 1166 | 1166 | 1.02 | 120 | 10.29 |
| 4.05E+00 | 422 | 1166 | 1166 | 1166 | 1.07 | 246 | 21.1 |
| 5.65E+00 | 162 | 1166 | 1166 | 1166 | 1.22 | 101 | 8.66 |
| 6.70E+00 | 172 | 1166 | 1166 | 1166 | 1.43 | 90 | 7.72 |
| 7.67E+00 | 109 | 1166 | 1166 | 1166 | 3.9 | 39 | 3.34 |
| 8.94E+00 | 596 | 1166 | 1166 | 1166 | 1.1 | 337 | 28.9 |

| FP | %FP |
|---|---|
| 139 | 11.92 |
| 144 | 12.35 |
| 229 | 19.64 |
| 178 | 15.27 |
| 100 | 8.58 |
| 110 | 9.43 |
| 153 | 13.12 |
| 235 | 20.15 |
| 441 | 37.82 |
| 356 | 30.53 |
| 462 | 39.62 |
| 308 | 26.42 |
| 271 | 23.24 |
| 184 | 15.78 |
| 190 | 16.3 |
| 122 | 10.46 |
| 233 | 19.98 |
| 238 | 20.41 |
| 219 | 18.78 |
| 208 | 17.84 |
| 230 | 19.73 |
| 160 | 13.72 |
| 79 | 6.78 |
| 427 | 36.62 |
| 225 | 19.3 |
| 107 | 9.18 |
| 659 | 56.52 |
| 149 | 12.78 |
| 109 | 9.35 |
| 266 | 22.81 |
| 181 | 15.52 |
| 206 | 17.67 |
| 177 | 15.18 |
| 321 | 27.53 |
| 376 | 32.25 |
| 156 | 13.38 |
| 214 | 18.35 |
| 81 | 6.95 |
| 124 | 10.63 |
| 435 | 37.31 |
| 70 | 6 |
| 554 | 47.51 |
| 246 | 21.1 |
| 280 | 24.01 |
| 31 | 2.66 |

| | |
|---|---|
| 189 | 16.21 |
| 157 | 13.46 |
| 32 | 2.74 |
| 192 | 16.47 |
| 69 | 5.92 |
| 42 | 3.6 |
| 97 | 8.32 |
| 5 | 0.43 |
| 251 | 21.53 |
| 167 | 14.32 |
| 83 | 7.12 |
| 201 | 17.24 |
| 119 | 10.21 |
| 204 | 17.5 |
| 528 | 45.28 |
| 388 | 33.28 |
| 77 | 6.6 |
| 29 | 2.49 |
| 354 | 30.36 |
| 66 | 5.66 |
| 608 | 52.14 |
| 66 | 5.66 |
| 156 | 13.38 |
| 156 | 13.38 |
| 180 | 15.44 |
| 207 | 17.75 |
| 178 | 15.27 |
| 416 | 35.68 |
| 171 | 14.67 |
| 55 | 4.72 |
| 205 | 17.58 |
| 331 | 28.39 |
| 162 | 13.89 |
| 170 | 14.58 |
| 255 | 21.87 |
| 216 | 18.52 |
| 211 | 18.1 |
| 88 | 7.55 |
| 143 | 12.26 |
| 37 | 3.17 |
| 133 | 11.41 |
| 175 | 15.01 |
| 367 | 31.48 |
| 261 | 22.38 |
| 110 | 9.43 |
| 67 | 5.75 |
| 318 | 27.27 |

| | |
|---|---|
| 203 | 17.41 |
| 108 | 9.26 |
| 83 | 7.12 |
| 153 | 13.12 |
| 137 | 11.75 |
| 7 | 0.6 |
| 159 | 13.64 |
| 354 | 30.36 |
| 277 | 23.76 |
| 80 | 6.86 |
| 275 | 23.58 |
| 249 | 21.36 |
| 27 | 2.32 |
| 36 | 3.09 |
| 111 | 9.52 |
| 515 | 44.17 |
| 616 | 52.83 |
| 346 | 29.67 |
| 179 | 15.35 |
| 126 | 10.81 |
| 199 | 17.07 |
| 224 | 19.21 |
| 151 | 12.95 |
| 206 | 17.67 |
| 128 | 10.98 |
| 317 | 27.19 |
| 9 | 0.77 |
| 398 | 34.13 |
| 170 | 14.58 |
| 197 | 16.9 |
| 143 | 12.26 |
| 97 | 8.32 |
| 275 | 23.58 |
| 369 | 31.65 |
| 145 | 12.44 |
| 52 | 4.46 |
| 140 | 12.01 |
| 67 | 5.75 |
| 60 | 5.15 |
| 123 | 10.55 |
| 119 | 10.21 |
| 285 | 24.44 |
| 60 | 5.15 |
| 374 | 32.08 |
| 412 | 35.33 |
| 86 | 7.38 |
| 120 | 10.29 |

| | |
|---|---|
| 39 | 3.34 |
| 15 | 1.29 |
| 52 | 4.46 |
| 371 | 31.82 |
| 71 | 6.09 |
| 253 | 21.7 |
| 183 | 15.69 |
| 296 | 25.39 |
| 117 | 10.03 |
| 98 | 8.4 |
| 65 | 5.57 |
| 131 | 11.23 |
| 80 | 6.86 |
| 100 | 8.58 |
| 288 | 24.7 |
| 6 | 0.51 |
| 47 | 4.03 |
| 69 | 5.92 |
| 184 | 15.78 |
| 44 | 3.77 |
| 207 | 17.75 |
| 92 | 7.89 |
| 92 | 7.89 |
| 62 | 5.32 |
| 274 | 23.5 |
| 154 | 13.21 |
| 75 | 6.43 |
| 87 | 7.46 |
| 287 | 24.61 |
| 208 | 17.84 |
| 19 | 1.63 |
| 115 | 9.86 |
| 204 | 17.5 |
| 75 | 6.43 |
| 93 | 7.98 |
| 58 | 4.97 |
| 157 | 13.46 |
| 276 | 23.67 |
| 50 | 4.29 |
| 239 | 20.5 |
| 106 | 9.09 |
| 205 | 17.58 |
| 207 | 17.75 |
| 455 | 39.02 |
| 238 | 20.41 |
| 152 | 13.04 |
| 388 | 33.28 |

| | |
|---|---|
| 131 | 11.23 |
| 68 | 5.83 |
| 170 | 14.58 |
| 55 | 4.72 |
| 47 | 4.03 |
| 13 | 1.11 |
| 253 | 21.7 |

| rank | motif_ID | consensus | p-value |
|---|---|---|---|
| 1 | SP3_HUMAN.H11MO.0.B | SGVVGGGGGCGGGGCBRGSS | 5.20E-23 |
| 2 | SP2_HUMAN.H11MO.0.A | GGSSVGGGGGCGGGGCCDGSGS | 1.49E-20 |
| 3 | EGR2_HUMAN.H11MO.0.A | GRGRRKGWGKGGGHGGRG | 4.29E-20 |
| 4 | KLF1_HUMAN.H11MO.0.A | DGGGYGKGGCYGGG | 4.02E-18 |
| 5 | KLF6_HUMAN.H11MO.0.A | GSRRGGGHGGGGMHGGGRV | 1.24E-17 |
| 6 | KLF3_HUMAN.H11MO.0.B | GGRGVDGGGCGGGGCYGGG | 1.76E-17 |
| 7 | SP4_HUMAN.H11MO.0.A | SRGVARGRGGCGGRGCHDRR | 2.45E-17 |
| 8 | SP1_HUMAN.H11MO.0.A | VGGGGGCGGGGCCKGGGGGGG | 2.36E-16 |
| 9 | KLF5_HUMAN.H11MO.0.A | WGGGTGKGGCDGGG | 2.74E-16 |
| 10 | EGR1_HUMAN.H11MO.0.A | VDRDGCGKGGGYGGRRR | 2.71E-16 |
| 11 | KLF15_HUMAN.H11MO.0.A | RGGGMGGRGVDGGGGGRGG | 5.13E-16 |
| 12 | PATZ1_HUMAN.H11MO.0.C | GGGGMGGGGGMKGGRRVGGVRG | 7.61E-16 |
| 13 | ZN281_HUMAN.H11MO.0.A | RGGWGGGGGAGGGGV | 1.78E-15 |
| 14 | E2F7_HUMAN.H11MO.0.B | GDGGCGGGAARDR | 2.99E-15 |
| 15 | KLF12_HUMAN.H11MO.0.C | VGGGGCGGGGC | 7.32E-15 |
| 16 | ZBT17_HUMAN.H11MO.0.A | SRRGGWGGGGGAGGGGMRR | 5.87E-15 |
| 17 | ZN467_HUMAN.H11MO.0.C | GGGGAGGGGRRGGRGRRGGRRR | 2.27E-14 |
| 18 | WT1_HUMAN.H11MO.0.C | RGGVGGGGGAGGRGGVGGRG | 2.42E-14 |
| 19 | SALL4_HUMAN.H11MO.0.B | GSDGGGWGGG | 2.02E-14 |
| 20 | ZN263_HUMAN.H11MO.0.A | GGGAGGAGGRRGRGGRGGRR | 1.50E-12 |
| 21 | MAZ_HUMAN.H11MO.0.A | GGGGGAGGGGGDGRGRRRGRRG | 1.21E-12 |
| 22 | KLF4_HUMAN.H11MO.0.A | WGGGYGKGGC | 1.83E-12 |
| 23 | TAF1_HUMAN.H11MO.0.A | RARRWGGCGGMGGMGR | 3.28E-12 |
| 24 | VEZF1_HUMAN.H11MO.0.C | GGRRRRGRRGGAGGGGGRGRRR | 4.74E-12 |
| 25 | KLF9_HUMAN.H11MO.0.C | GRRGKGGGCGTGGCC | 1.48E-10 |
| 26 | ZN341_HUMAN.H11MO.0.C | GRGGRRRRGGRGRRGGAASAGC | 4.88E-10 |
| 27 | E2F4_HUMAN.H11MO.0.A | SRGGGCGGGAARD | 1.18E-09 |
| 28 | BC11A_HUMAN.H11MO.0.A | RAAAGAGGAAGTGARAV | 2.23E-09 |
| 29 | ETV5_HUMAN.H11MO.0.C | RRRSAGGAARDGRV | 2.07E-09 |
| 30 | ELF5_HUMAN.H11MO.0.A | VGRASSAGGAAGTRV | 1.68E-08 |
| 31 | ETS2_HUMAN.H11MO.0.B | RDVRGGAARVRRR | 1.32E-08 |
| 32 | FLI1_HUMAN.H11MO.0.A | SRRGGMAGGAAGGRRRGR | 2.92E-08 |
| 33 | ERG_HUMAN.H11MO.0.A | RRRSAGGAAGYGG | 4.33E-08 |
| 34 | FEV_HUMAN.H11MO.0.B | GCVGGAAGYG | 2.72E-08 |
| 35 | TYY1_HUMAN.H11MO.0.A | CAARATGGCGGC | 5.04E-07 |
| 36 | RXRA_HUMAN.H11MO.0.A | VRGDVRGNVARAGGTCARRG | 1.03E-07 |
| 37 | MZF1_HUMAN.H11MO.0.B | RRDRGGGAWTKRR | 1.19E-07 |
| 38 | SPI1_HUMAN.H11MO.0.A | RRAAAGAGGAAGTGRVD | 4.99E-07 |
| 39 | ELF3_HUMAN.H11MO.0.A | RAABVAGGAAGTRR | 5.21E-07 |
| 40 | ETS1_HUMAN.H11MO.0.A | RRRCMGGAAGTGG | 4.14E-07 |
| 41 | GATA3_HUMAN.H11MO.0.A | RVAGATAAVAD | 7.41E-07 |
| 42 | ELK4_HUMAN.H11MO.0.A | SVCCGGAAGTGV | 1.10E-06 |
| 43 | GABPA_HUMAN.H11MO.0.A | GVVRCCGGAAGTGV | 1.45E-06 |
| 44 | NRF1_HUMAN.H11MO.0.A | CWSTGCGCATGCGCRDS | 3.29E-06 |
| 45 | ELF2_HUMAN.H11MO.0.C | RAVCCGGAAGTGR | 1.52E-06 |

| | | | |
|---|---|---|---|
| 46 | CREB1_HUMAN.H11MO.0.A | NRRTGACGTMA | 2.83E-06 |
| 47 | ETV1_HUMAN.H11MO.0.A | SRCCGGAAGYG | 1.61E-06 |
| 48 | GATA2_HUMAN.H11MO.0.A | STGBNNNNNDVAGATAAGR | 4.44E-06 |
| 49 | ELK1_HUMAN.H11MO.0.B | RCCGGAAGTGV | 3.44E-06 |
| 50 | SOX3_HUMAN.H11MO.0.B | BCCWTTGTBYB | 1.95E-06 |
| 51 | NFKB1_HUMAN.H11MO.0.A | AGGGAAAK | 1.24E-06 |
| 52 | SOX4_HUMAN.H11MO.0.B | BYCTTTGTYYYB | 2.19E-06 |
| 53 | ETV2_HUMAN.H11MO.0.B | RRARRCAGGAARYRGS | 5.48E-06 |
| 54 | TFDP1_HUMAN.H11MO.0.C | RRRRGGCGGGAARN | 2.71E-06 |
| 55 | ETV4_HUMAN.H11MO.0.B | VRCAGGAARBR | 2.10E-06 |
| 56 | FOXJ2_HUMAN.H11MO.0.C | TGTTTRTTTW | 1.22E-05 |
| 57 | MYCN_HUMAN.H11MO.0.A | RGCCACGTGSDS | 1.03E-05 |
| 58 | ATF1_HUMAN.H11MO.0.B | HRRTGACGYMA | 9.53E-06 |
| 59 | PRDM6_HUMAN.H11MO.0.C | RRRARGAAAAAAA | 6.85E-06 |
| 60 | ELF1_HUMAN.H11MO.0.A | VRACCCGGAAGTGS | 1.18E-05 |
| 61 | GATA1_HUMAN.H11MO.0.A | STGBNNNDNVSAGATAAGV | 2.07E-05 |
| 62 | E2F6_HUMAN.H11MO.0.A | DGGGCGGGARRVR | 6.13E-06 |
| 63 | SOX2_HUMAN.H11MO.0.A | BBCCTTTGTYYYB | 8.24E-06 |
| 64 | MEF2C_HUMAN.H11MO.0.A | DKCTATTTTTRGM | 2.07E-05 |
| 65 | MEF2A_HUMAN.H11MO.0.A | DKCTATTTTTRGM | 2.20E-05 |
| 66 | SPIB_HUMAN.H11MO.0.A | RRAAAGAGGAAGTGARA | 4.95E-05 |
| 67 | IRF3_HUMAN.H11MO.0.B | RAAARGGAAAVDGAAASDGA | 1.47E-05 |
| 68 | SUH_HUMAN.H11MO.0.A | VSYSTGGGAAV | 8.23E-06 |
| 69 | AIRE_HUMAN.H11MO.0.C | HTTGGWWWWAWTGGDTWH | 2.30E-05 |
| 70 | ZN394_HUMAN.H11MO.0.C | NRARWRGAAWNGAMWGRAAK | 1.60E-05 |
| 71 | NKX61_HUMAN.H11MO.0.B | GAYWAATKRSHWWTTAATK | 3.33E-05 |
| 72 | ATF2_HUMAN.H11MO.0.B | RRTGABGTCAY | 3.73E-05 |
| 73 | IRF4_HUMAN.H11MO.0.A | RAAWGRGGAASTGAAASY | 4.63E-05 |
| 74 | FOXQ1_HUMAN.H11MO.0.C | HATTGTTTATTT | 2.36E-04 |
| 75 | ZFP42_HUMAN.H11MO.0.A | GGCMGCCATTTT | 4.24E-05 |
| 76 | TBX21_HUMAN.H11MO.0.A | GDGDRGGTGTGRV | 1.70E-05 |
| 77 | RXRG_HUMAN.H11MO.0.B | GGTCAAAGGTCAH | 2.12E-04 |
| 78 | FOXP1_HUMAN.H11MO.0.A | TBTGTTTMY | 2.64E-05 |
| 79 | STAT2_HUMAN.H11MO.0.A | RRGRAAAHGAAACTGAAAV | 7.30E-05 |
| 80 | FOSL2_HUMAN.H11MO.0.A | NDRTGAGTCAYH | 1.55E-04 |
| 81 | FOSL1_HUMAN.H11MO.0.A | KRVTGAGTCAYH | 1.55E-04 |
| 82 | SOX17_HUMAN.H11MO.0.C | SCCATTGTKBY | 3.01E-05 |
| 83 | PPARG_HUMAN.H11MO.0.A | MWSTRGGKSARAGGKCA | 4.13E-05 |
| 84 | PPARA_HUMAN.H11MO.0.B | AWSTRGGKSAAAGGTCA | 4.52E-05 |
| 85 | NFAC2_HUMAN.H11MO.0.B | TGGAAAAWY | 4.56E-05 |
| 86 | ZN770_HUMAN.H11MO.0.C | GGGAGGCYGAGGBRGGAGGATC | 7.59E-05 |
| 87 | NFAC4_HUMAN.H11MO.0.C | DGGAAAAYTW | 4.70E-05 |
| 88 | IRF8_HUMAN.H11MO.0.B | RAAARRGGAASTGAAASTDR | 1.40E-04 |
| 89 | RORG_HUMAN.H11MO.0.C | RRAASTRGGTCA | 8.30E-05 |
| 90 | NFAC1_HUMAN.H11MO.0.B | AWGGAAARWVWGAMW | 4.70E-05 |
| 91 | PO2F1_HUMAN.H11MO.0.C | AWKBRWATGSAAWYAW | 9.34E-05 |
| 92 | SRY_HUMAN.H11MO.0.B | YWTTGTTTH | 5.98E-05 |

| | | | |
|---|---|---|---|
| 93 | JUND_HUMAN.H11MO.0.A | RRTGAGTCAYY | 1.55E-04 |
| 94 | HXB13_HUMAN.H11MO.0.A | TTTTATDGSY | 9.68E-05 |
| 95 | MEF2B_HUMAN.H11MO.0.A | DKKKCTATTTTTRG | 1.20E-04 |
| 96 | FOS_HUMAN.H11MO.0.A | VTGAGTCAB | 1.55E-04 |
| 97 | EHF_HUMAN.H11MO.0.B | VWACSAGGAAGTDVS | 1.71E-04 |

### AME (Analysis of Motif Enrichment): Version 5.0.5 compiled on Jul 15 2019 at 00:31:14
### The format of this file is described at http://meme-suite.org//doc/ame-output-format.html.
### ame --verbose 1 --oc . --scoring avg --method fisher --hit-lo-fraction 0.25 --evalue-report-threshold 10.0 --co

| adj_p-value | E-value | tests | FASTA_max | pos | neg | PWM_min |
|---|---|---|---|---|---|---|
| 2.91E-20 | 1.17E-17 | 560 | 156 | 156 | 1092 | 3.66 |
| 7.63E-18 | 3.07E-15 | 513 | 156 | 156 | 1092 | 4.51 |
| 1.53E-17 | 6.15E-15 | 357 | 156 | 156 | 1092 | 1.32 |
| 1.36E-15 | 5.45E-13 | 337 | 156 | 156 | 1092 | 1.22 |
| 5.97E-15 | 2.40E-12 | 483 | 156 | 156 | 1092 | 1.75 |
| 6.83E-15 | 2.75E-12 | 388 | 156 | 156 | 1092 | 1.28 |
| 1.08E-14 | 4.34E-12 | 440 | 156 | 156 | 1092 | 1.09 |
| 4.13E-14 | 1.66E-11 | 175 | 156 | 156 | 1092 | 2.2 |
| 9.63E-14 | 3.87E-11 | 352 | 156 | 156 | 1092 | 1.13 |
| 1.00E-13 | 4.03E-11 | 370 | 156 | 156 | 1092 | 1.04 |
| 1.52E-13 | 6.12E-11 | 297 | 156 | 156 | 1092 | 1.43 |
| 3.62E-13 | 1.46E-10 | 476 | 156 | 156 | 1092 | 3.3 |
| 4.53E-13 | 1.82E-10 | 255 | 156 | 156 | 1092 | 1.09 |
| 1.33E-12 | 5.34E-10 | 444 | 156 | 156 | 1092 | 1.11 |
| 2.43E-12 | 9.76E-10 | 332 | 156 | 156 | 1092 | 2.56 |
| 2.64E-12 | 1.06E-09 | 450 | 156 | 156 | 1092 | 2.97 |
| 5.54E-12 | 2.23E-09 | 244 | 156 | 156 | 1092 | 4.49 |
| 7.42E-12 | 2.98E-09 | 306 | 156 | 156 | 1092 | 1.96 |
| 1.25E-11 | 5.01E-09 | 618 | 156 | 156 | 1092 | 2.49 |
| 3.97E-10 | 1.60E-07 | 264 | 156 | 156 | 1092 | 2.37 |
| 5.00E-10 | 2.01E-07 | 415 | 156 | 156 | 1092 | 1.78 |
| 5.05E-10 | 2.03E-07 | 276 | 156 | 156 | 1092 | 1.09 |
| 1.42E-09 | 5.73E-07 | 435 | 156 | 156 | 1092 | 1.07 |
| 2.01E-09 | 8.09E-07 | 425 | 156 | 156 | 1092 | 3.66 |
| 2.67E-08 | 1.07E-05 | 180 | 156 | 156 | 1092 | 1.06 |
| 2.11E-07 | 8.47E-05 | 432 | 156 | 156 | 1092 | 2.7 |
| 4.29E-07 | 1.72E-04 | 364 | 156 | 156 | 1092 | 1.02 |
| 8.24E-07 | 3.31E-04 | 370 | 156 | 156 | 1092 | 1.5 |
| 1.02E-06 | 4.09E-04 | 493 | 156 | 156 | 1092 | 2.49 |
| 3.78E-06 | 1.52E-03 | 225 | 156 | 156 | 1092 | 1.3 |
| 6.74E-06 | 2.71E-03 | 512 | 156 | 156 | 1092 | 2.99 |
| 7.86E-06 | 3.16E-03 | 269 | 156 | 156 | 1092 | 1.11 |
| 1.27E-05 | 5.10E-03 | 293 | 156 | 156 | 1092 | 1.26 |
| 1.90E-05 | 7.65E-03 | 701 | 156 | 156 | 1092 | 2.2 |
| 3.28E-05 | 1.32E-02 | 65 | 156 | 156 | 1092 | 11.8 |
| 4.46E-05 | 1.79E-02 | 433 | 156 | 156 | 1092 | 1.34 |
| 4.52E-05 | 1.82E-02 | 381 | 156 | 156 | 1092 | 1.24 |
| 4.89E-05 | 1.97E-02 | 98 | 156 | 156 | 1092 | 4.11 |
| 8.59E-05 | 3.45E-02 | 165 | 156 | 156 | 1092 | 1.06 |
| 1.07E-04 | 4.29E-02 | 258 | 156 | 156 | 1092 | 1.43 |
| 1.93E-04 | 7.74E-02 | 260 | 156 | 156 | 1092 | 1.06 |
| 2.11E-04 | 8.50E-02 | 192 | 156 | 156 | 1092 | 4.54 |
| 2.88E-04 | 1.16E-01 | 199 | 156 | 156 | 1092 | 1.45 |
| 3.12E-04 | 1.25E-01 | 95 | 156 | 156 | 1092 | 1.09 |
| 3.76E-04 | 1.51E-01 | 248 | 156 | 156 | 1092 | 3.19 |

| | | | | | | |
|---|---|---|---|---|---|---|
| 3.96E-04 | 1.59E-01 | 140 | 156 | 156 | 1092 | 1.66 |
| 4.41E-04 | 1.77E-01 | 274 | 156 | 156 | 1092 | 1.28 |
| 5.46E-04 | 2.20E-01 | 123 | 156 | 156 | 1092 | 1.22 |
| 5.94E-04 | 2.39E-01 | 173 | 156 | 156 | 1092 | 4.05 |
| 6.07E-04 | 2.44E-01 | 311 | 156 | 156 | 1092 | 2.85 |
| 6.10E-04 | 2.45E-01 | 492 | 156 | 156 | 1092 | 1.13 |
| 7.85E-04 | 3.15E-01 | 358 | 156 | 156 | 1092 | 5.1 |
| 8.71E-04 | 3.50E-01 | 159 | 156 | 156 | 1092 | 2.08 |
| 8.88E-04 | 3.57E-01 | 328 | 156 | 156 | 1092 | 1.4 |
| 9.10E-04 | 3.66E-01 | 434 | 156 | 156 | 1092 | 4.61 |
| 1.23E-03 | 4.95E-01 | 101 | 156 | 156 | 1092 | 113 |
| 1.62E-03 | 6.52E-01 | 157 | 156 | 156 | 1092 | 2.04 |
| 1.94E-03 | 7.81E-01 | 204 | 156 | 156 | 1092 | 1.16 |
| 2.22E-03 | 8.92E-01 | 324 | 156 | 156 | 1092 | 1.1 |
| 2.32E-03 | 9.32E-01 | 196 | 156 | 156 | 1092 | 7.06 |
| 2.57E-03 | 1.03E+00 | 124 | 156 | 156 | 1092 | 1.4 |
| 2.59E-03 | 1.04E+00 | 424 | 156 | 156 | 1092 | 1.45 |
| 2.94E-03 | 1.18E+00 | 357 | 156 | 156 | 1092 | 3.12 |
| 3.02E-03 | 1.21E+00 | 146 | 156 | 156 | 1092 | 2.25 |
| 3.08E-03 | 1.24E+00 | 140 | 156 | 156 | 1092 | 1.2 |
| 3.16E-03 | 1.27E+00 | 64 | 156 | 156 | 1092 | 1.58 |
| 3.20E-03 | 1.29E+00 | 218 | 156 | 156 | 1092 | 1.46 |
| 3.86E-03 | 1.55E+00 | 470 | 156 | 156 | 1092 | 1.21 |
| 4.77E-03 | 1.92E+00 | 208 | 156 | 156 | 1092 | 10.4 |
| 4.84E-03 | 1.94E+00 | 304 | 156 | 156 | 1092 | 1.52 |
| 5.71E-03 | 2.29E+00 | 172 | 156 | 156 | 1092 | 10.3 |
| 5.74E-03 | 2.31E+00 | 154 | 156 | 156 | 1092 | 15.1 |
| 5.86E-03 | 2.36E+00 | 127 | 156 | 156 | 1092 | 13.8 |
| 5.88E-03 | 2.37E+00 | 25 | 156 | 156 | 1092 | 82.8 |
| 8.11E-03 | 3.26E+00 | 192 | 156 | 156 | 1092 | 1.67 |
| 8.25E-03 | 3.32E+00 | 489 | 156 | 156 | 1092 | 1.18 |
| 8.45E-03 | 3.40E+00 | 40 | 156 | 156 | 1092 | 110 |
| 8.64E-03 | 3.47E+00 | 329 | 156 | 156 | 1092 | 3.53 |
| 8.65E-03 | 3.48E+00 | 119 | 156 | 156 | 1092 | 86.5 |
| 1.05E-02 | 4.23E+00 | 68 | 156 | 156 | 1092 | 153 |
| 1.08E-02 | 4.35E+00 | 70 | 156 | 156 | 1092 | 187 |
| 1.09E-02 | 4.39E+00 | 365 | 156 | 156 | 1092 | 1.34 |
| 1.11E-02 | 4.44E+00 | 269 | 156 | 156 | 1092 | 1.87 |
| 1.13E-02 | 4.53E+00 | 251 | 156 | 156 | 1092 | 1.64 |
| 1.22E-02 | 4.91E+00 | 270 | 156 | 156 | 1092 | 1.15 |
| 1.34E-02 | 5.39E+00 | 178 | 156 | 156 | 1092 | 1.37 |
| 1.43E-02 | 5.75E+00 | 306 | 156 | 156 | 1092 | 1.67 |
| 1.55E-02 | 6.25E+00 | 112 | 156 | 156 | 1092 | 1.5 |
| 1.66E-02 | 6.65E+00 | 201 | 156 | 156 | 1092 | 5.6 |
| 1.74E-02 | 6.98E+00 | 373 | 156 | 156 | 1092 | 1.59 |
| 1.75E-02 | 7.03E+00 | 189 | 156 | 156 | 1092 | 2.03 |
| 1.90E-02 | 7.65E+00 | 321 | 156 | 156 | 1092 | 72 |

| | | | | | | |
|---|---|---|---|---|---|---|
| 1.91E-02 | 7.68E+00 | 124 | 156 | 156 | 1092 | 178 |
| 1.95E-02 | 7.82E+00 | 203 | 156 | 156 | 1092 | 12.7 |
| 2.39E-02 | 9.60E+00 | 202 | 156 | 156 | 1092 | 44.7 |
| 2.40E-02 | 9.63E+00 | 156 | 156 | 156 | 1092 | 84.5 |
| 2.46E-02 | 9.87E+00 | 145 | 156 | 156 | 1092 | 1.05 |

ntrol --shuffle-- --kmer 2 camta1_chip_seq.fasta db/HUMAN/HOCOMOCOv11_core_HUMAN_mono_meme_fo

| TP | %TP | FP | %FP |
|---|---|---|---|
| 74 | 47.44 | 131 | 12 |
| 63 | 40.38 | 102 | 9.34 |
| 89 | 57.05 | 224 | 20.51 |
| 86 | 55.13 | 226 | 20.7 |
| 89 | 57.05 | 247 | 22.62 |
| 88 | 56.41 | 243 | 22.25 |
| 102 | 65.38 | 327 | 29.95 |
| 51 | 32.69 | 82 | 7.51 |
| 86 | 55.13 | 244 | 22.34 |
| 92 | 58.97 | 278 | 25.46 |
| 73 | 46.79 | 179 | 16.39 |
| 67 | 42.95 | 152 | 13.92 |
| 72 | 46.15 | 179 | 16.39 |
| 97 | 62.18 | 320 | 29.3 |
| 61 | 39.1 | 133 | 12.18 |
| 70 | 44.87 | 174 | 15.93 |
| 49 | 31.41 | 87 | 7.97 |
| 68 | 43.59 | 170 | 15.57 |
| 72 | 46.15 | 189 | 17.31 |
| 53 | 33.97 | 117 | 10.71 |
| 80 | 51.28 | 251 | 22.99 |
| 70 | 44.87 | 199 | 18.22 |
| 93 | 59.62 | 334 | 30.59 |
| 63 | 40.38 | 168 | 15.38 |
| 51 | 32.69 | 126 | 11.54 |
| 60 | 38.46 | 175 | 16.03 |
| 79 | 50.64 | 285 | 26.1 |
| 73 | 46.79 | 254 | 23.26 |
| 65 | 41.67 | 209 | 19.14 |
| 51 | 32.69 | 148 | 13.55 |
| 65 | 41.67 | 220 | 20.15 |
| 61 | 39.1 | 203 | 18.59 |
| 60 | 38.46 | 200 | 18.32 |
| 56 | 35.9 | 176 | 16.12 |
| 12 | 7.69 | 8 | 0.73 |
| 77 | 49.36 | 304 | 27.84 |
| 71 | 45.51 | 269 | 24.63 |
| 22 | 14.1 | 37 | 3.39 |
| 42 | 26.92 | 122 | 11.17 |
| 51 | 32.69 | 166 | 15.2 |
| 57 | 36.54 | 202 | 18.5 |
| 23 | 14.74 | 43 | 3.94 |
| 39 | 25 | 113 | 10.35 |
| 28 | 17.95 | 67 | 6.14 |
| 29 | 18.59 | 68 | 6.23 |

| | | | |
|---|---|---|---|
| 25 | 16.03 | 54 | 4.95 |
| 52 | 33.33 | 180 | 16.48 |
| 32 | 20.51 | 86 | 7.88 |
| 22 | 14.1 | 43 | 3.94 |
| 48 | 30.77 | 160 | 14.65 |
| 85 | 54.49 | 375 | 34.34 |
| 42 | 26.92 | 130 | 11.9 |
| 30 | 19.23 | 78 | 7.14 |
| 53 | 33.97 | 189 | 17.31 |
| 39 | 25 | 115 | 10.53 |
| 7 | 4.49 | 2 | 0.18 |
| 24 | 15.38 | 55 | 5.04 |
| 42 | 26.92 | 139 | 12.73 |
| 64 | 41.03 | 259 | 23.72 |
| 15 | 9.62 | 22 | 2.01 |
| 29 | 18.59 | 80 | 7.33 |
| 66 | 42.31 | 270 | 24.73 |
| 51 | 32.69 | 186 | 17.03 |
| 29 | 18.59 | 80 | 7.33 |
| 34 | 21.79 | 104 | 9.52 |
| 18 | 11.54 | 37 | 3.39 |
| 46 | 29.49 | 163 | 14.93 |
| 75 | 48.08 | 328 | 30.04 |
| 22 | 14.1 | 50 | 4.58 |
| 57 | 36.54 | 225 | 20.6 |
| 24 | 15.38 | 60 | 5.49 |
| 8 | 5.13 | 5 | 0.46 |
| 13 | 8.33 | 19 | 1.74 |
| 4 | 2.56 | 0 | 0 |
| 33 | 21.15 | 103 | 9.43 |
| 78 | 50 | 354 | 32.42 |
| 6 | 3.85 | 3 | 0.27 |
| 37 | 23.72 | 120 | 10.99 |
| 10 | 6.41 | 11 | 1.01 |
| 5 | 3.21 | 1 | 0.09 |
| 5 | 3.21 | 1 | 0.09 |
| 60 | 38.46 | 248 | 22.71 |
| 42 | 26.92 | 149 | 13.64 |
| 43 | 27.56 | 155 | 14.19 |
| 52 | 33.33 | 205 | 18.77 |
| 37 | 23.72 | 127 | 11.63 |
| 53 | 33.97 | 211 | 19.32 |
| 27 | 17.31 | 81 | 7.42 |
| 17 | 10.9 | 35 | 3.21 |
| 61 | 39.1 | 258 | 23.63 |
| 34 | 21.79 | 113 | 10.35 |
| 9 | 5.77 | 8 | 0.73 |

| | | | |
|---|---|---|---|
| 5 | 3.21 | 1 | 0.09 |
| 15 | 9.62 | 28 | 2.56 |
| 10 | 6.41 | 12 | 1.1 |
| 5 | 3.21 | 1 | 0.09 |
| 33 | 21.15 | 112 | 10.26 |

rmat.meme

| | seqnames | start | end | width | strand | V5 | geneId |
|---|---|---|---|---|---|---|---|
| 1 | chr1 | 8964070 | 8964581 | 512 | * | 335 | 2023 |
| 2 | chr1 | 15298307 | 15298697 | 391 | * | 86 | 23254 |
| 3 | chr1 | 15664780 | 15665109 | 330 | * | 86 | 114827 |
| 4 | chr1 | 16470691 | 16473814 | 3124 | * | 143 | 1969 |
| 5 | chr1 | 16500171 | 16500446 | 276 | * | 130 | 1969 |
| 6 | chr1 | 17516511 | 17516973 | 463 | * | 179 | 29943 |
| 7 | chr1 | 19379010 | 19379343 | 334 | * | 133 | 23352 |
| 8 | chr1 | 20091968 | 20092502 | 535 | * | 30 | 255104 |
| 9 | chr1 | 26218626 | 26219157 | 532 | * | 95 | 3925 |
| 10 | chr1 | 26745568 | 26745877 | 310 | * | 20 | 79727 |
| 11 | chr1 | 27319850 | 27320229 | 380 | * | 60 | 388610 |
| 12 | chr1 | 27325825 | 27326224 | 400 | * | 98 | 388610 |
| 13 | chr1 | 28764542 | 28764887 | 346 | * | 69 | 65979 |
| 14 | chr1 | 29450908 | 29451362 | 455 | * | 342 | 399474 |
| 15 | chr1 | 29598235 | 29598672 | 438 | * | 153 | 10076 |
| 16 | chr1 | 33564956 | 33565296 | 341 | * | 104 | 113451 |
| 17 | chr1 | 43672425 | 43672985 | 561 | * | 229 | 10969 |
| 18 | chr1 | 43938555 | 43938888 | 334 | * | 114 | 81888 |
| 19 | chr1 | 44287576 | 44288112 | 537 | * | 52 | 6487 |
| 20 | chr1 | 48035652 | 48035948 | 297 | * | 123 | 2306 |
| 21 | chr1 | 53611742 | 53612084 | 343 | * | 76 | 6512 |
| 22 | chr1 | 54789786 | 54790127 | 342 | * | 59 | 23648 |
| 23 | chr1 | 59228823 | 59229084 | 262 | * | 153 | 3725 |
| 24 | chr1 | 64197014 | 64197317 | 304 | * | 45 | 4919 |
| 25 | chr1 | 64386261 | 64386568 | 308 | * | 114 | 4919 |
| 26 | chr1 | 85666871 | 85668161 | 1291 | * | 392 | 84144 |
| 27 | chr1 | 86042753 | 86043047 | 295 | * | 77 | 23576 |
| 28 | chr1 | 86043944 | 86044422 | 479 | * | 69 | 23576 |
| 29 | chr1 | 86045969 | 86047024 | 1056 | * | 462 | 3491 |
| 30 | chr1 | 86072754 | 86073248 | 495 | * | 69 | 3491 |
| 31 | chr1 | 86078137 | 86078727 | 591 | * | 45 | 3491 |
| 32 | chr1 | 92022749 | 92023151 | 403 | * | 37 | 8317 |
| 33 | chr1 | 92202283 | 92202618 | 336 | * | 45 | 7049 |
| 34 | chr1 | 94219904 | 94220360 | 457 | * | 60 | 8412 |
| 35 | chr1 | 94511228 | 94511545 | 318 | * | 76 | 24 |
| 36 | chr1 | 94701686 | 94701982 | 297 | * | 37 | 9411 |
| 37 | chr1 | 94791431 | 94791703 | 273 | * | 30 | 9411 |
| 38 | chr1 | 145092201 | 145092948 | 748 | * | 211 | 9554 |
| 39 | chr1 | 150533666 | 150534069 | 404 | * | 50 | 54507 |
| 40 | chr1 | 151167717 | 151168396 | 680 | * | 218 | 8394 |
| 41 | chr1 | 151960675 | 151961361 | 687 | * | 69 | 6281 |
| 42 | chr1 | 152080915 | 152081601 | 687 | * | 60 | 7062 |
| 43 | chr1 | 153936471 | 153936966 | 496 | * | 185 | 27173 |
| 44 | chr1 | 154944978 | 154945721 | 744 | * | 114 | 6464 |
| 45 | chr1 | 156866462 | 156866916 | 455 | * | 69 | 375033 |
| 46 | chr1 | 157989200 | 157989745 | 546 | * | 398 | 55243 |

| | | | | | | | |
|---|---|---|---|---|---|---|---|
| 47 | chr1 | 202558413 | 202558703 | 291 | * | 111 | 4660 |
| 48 | chr1 | 205270625 | 205270897 | 273 | * | 46 | 81788 |
| 49 | chr1 | 214578011 | 214578393 | 383 | * | 30 | 5784 |
| 50 | chr1 | 214725070 | 214725779 | 710 | * | 505 | 5784 |
| 51 | chr10 | 14116505 | 14116793 | 289 | * | 60 | 55691 |
| 52 | chr10 | 65027392 | 65027699 | 308 | * | 23 | 221037 |
| 53 | chr10 | 73526579 | 73526884 | 306 | * | 185 | 64115 |
| 54 | chr10 | 88855004 | 88855272 | 269 | * | 30 | 54537 |
| 55 | chr10 | 92690523 | 92691144 | 622 | * | 115 | 27063 |
| 56 | chr10 | 95048585 | 95049037 | 453 | * | 164 | 26509 |
| 57 | chr10 | 95218439 | 95218982 | 544 | * | 30 | 26509 |
| 58 | chr10 | 99334390 | 99334663 | 274 | * | 153 | 26287 |
| 59 | chr10 | 99447240 | 99447501 | 262 | * | 52 | 60370 |
| 60 | chr10 | 112170369 | 112170880 | 512 | * | 18 | 1847 |
| 61 | chr11 | 355489 | 356436 | 948 | * | 263 | 338707 |
| 62 | chr11 | 843827 | 844806 | 980 | * | 493 | 7106 |
| 63 | chr11 | 6340835 | 6341474 | 640 | * | 52 | 112464 |
| 64 | chr11 | 8939027 | 8939344 | 318 | * | 67 | 56672 |
| 65 | chr11 | 9685672 | 9685988 | 317 | * | 90 | 23075 |
| 66 | chr11 | 9787186 | 9787560 | 375 | * | 339 | 440028 |
| 67 | chr11 | 9927781 | 9928133 | 353 | * | 24 | 81846 |
| 68 | chr11 | 10315832 | 10316481 | 650 | * | 74 | 81846 |
| 69 | chr11 | 12065501 | 12065883 | 383 | * | 159 | 27122 |
| 70 | chr11 | 12204273 | 12204587 | 315 | * | 120 | 9645 |
| 71 | chr11 | 12221949 | 12222713 | 765 | * | 58 | 9645 |
| 72 | chr11 | 12307969 | 12308403 | 435 | * | 36 | 84953 |
| 73 | chr11 | 12398728 | 12399341 | 614 | * | 444 | 55742 |
| 74 | chr11 | 12439588 | 12440294 | 707 | * | 39 | 55742 |
| 75 | chr11 | 14443839 | 14444215 | 377 | * | 191 | 22800 |
| 76 | chr11 | 19391844 | 19392220 | 377 | * | 77 | 89797 |
| 77 | chr11 | 35103927 | 35104185 | 259 | * | 52 | 960 |
| 78 | chr11 | 35104949 | 35105468 | 520 | * | 69 | 960 |
| 79 | chr11 | 35121076 | 35121919 | 844 | * | 111 | 960 |
| 80 | chr11 | 36469026 | 36469974 | 949 | * | 105 | 79899 |
| 81 | chr11 | 44009067 | 44009525 | 459 | * | 37 | 387763 |
| 82 | chr11 | 46312294 | 46312889 | 596 | * | 153 | 90993 |
| 83 | chr11 | 46316827 | 46317184 | 358 | * | 38 | 90993 |
| 84 | chr11 | 57545545 | 57546305 | 761 | * | 174 | 1500 |
| 85 | chr11 | 62276155 | 62276573 | 419 | * | 51 | 79026 |
| 86 | chr11 | 62312607 | 62313996 | 1390 | * | 146 | 79026 |
| 87 | chr11 | 62320800 | 62321240 | 441 | * | 38 | 79026 |
| 88 | chr11 | 64622417 | 64622721 | 305 | * | 133 | 10938 |
| 89 | chr11 | 64649077 | 64649334 | 258 | * | 23 | 10938 |
| 90 | chr11 | 65325933 | 65326245 | 313 | * | 52 | 4054 |
| 91 | chr11 | 65836942 | 65837240 | 299 | * | 26 | 55690 |
| 92 | chr11 | 67780335 | 67781497 | 1163 | * | 373 | 221 |
| 93 | chr11 | 68121673 | 68122150 | 478 | * | 81 | 4041 |

| | | | | | | | |
|---|---|---|---|---|---|---|---|
| 94 | chr11 | 69311795 | 69312508 | 714 | * | 340 | 595 |
| 95 | chr11 | 69451331 | 69451765 | 435 | * | 287 | 595 |
| 96 | chr11 | 76841869 | 76842265 | 397 | * | 20 | 4647 |
| 97 | chr11 | 78076388 | 78076753 | 366 | * | 29 | 9846 |
| 98 | chr11 | 85780367 | 85780966 | 600 | * | 54 | 8301 |
| 99 | chr11 | 86451743 | 86452412 | 670 | * | 164 | 11098 |
| 100 | chr11 | 101881810 | 101882194 | 385 | * | 138 | 85016 |
| 101 | chr11 | 101980790 | 101981077 | 288 | * | 11 | 10413 |
| 102 | chr11 | 101982302 | 101982654 | 353 | * | 45 | 10413 |
| 103 | chr11 | 113423910 | 113424358 | 449 | * | 146 | 1813 |
| 104 | chr11 | 114033451 | 114033746 | 296 | * | 60 | 4837 |
| 105 | chr11 | 114163101 | 114163752 | 652 | * | 40 | 4837 |
| 106 | chr11 | 114165660 | 114166745 | 1086 | * | 215 | 4837 |
| 107 | chr11 | 114178170 | 114178516 | 347 | * | 58 | 4837 |
| 108 | chr11 | 116064234 | 116064588 | 355 | * | 38 | 283143 |
| 109 | chr11 | 116165568 | 116165954 | 387 | * | 105 | 84811 |
| 110 | chr11 | 117069784 | 117070058 | 275 | * | 49 | 6876 |
| 111 | chr11 | 118790380 | 118790702 | 323 | * | 19 | 283149 |
| 112 | chr11 | 119454860 | 119455274 | 415 | * | 58 | 5818 |
| 113 | chr11 | 121337186 | 121337565 | 380 | * | 52 | 6653 |
| 114 | chr11 | 125260776 | 125261290 | 515 | * | 101 | 63876 |
| 115 | chr11 | 128078002 | 128078509 | 508 | * | 60 | 2113 |
| 116 | chr11 | 128080300 | 128081133 | 834 | * | 56 | 2113 |
| 117 | chr11 | 128351195 | 128351501 | 307 | * | 78 | 2113 |
| 118 | chr11 | 128372581 | 128373071 | 491 | * | 56 | 2113 |
| 119 | chr11 | 128391683 | 128392123 | 441 | * | 39 | 2113 |
| 120 | chr11 | 128392416 | 128392706 | 291 | * | 152 | 2113 |
| 121 | chr11 | 130348299 | 130349056 | 758 | * | 38 | 170689 |
| 122 | chr11 | 130350333 | 130350607 | 275 | * | 12 | 170689 |
| 123 | chr12 | 1714807 | 1715268 | 462 | * | 52 | 81029 |
| 124 | chr12 | 8122893 | 8123555 | 663 | * | 347 | 6515 |
| 125 | chr12 | 12419925 | 12420296 | 372 | * | 378 | 4040 |
| 126 | chr12 | 14413562 | 14413821 | 260 | * | 174 | 55729 |
| 127 | chr12 | 48162361 | 48162681 | 321 | * | 17 | 10411 |
| 128 | chr12 | 48214295 | 48214642 | 348 | * | 89 | 51564 |
| 129 | chr12 | 52585646 | 52586238 | 593 | * | 174 | 144501 |
| 130 | chr12 | 52674686 | 52675158 | 473 | * | 196 | 3892 |
| 131 | chr12 | 53318538 | 53319368 | 831 | * | 69 | 3856 |
| 132 | chr12 | 53339108 | 53339865 | 758 | * | 69 | 3875 |
| 133 | chr12 | 54238921 | 54239402 | 482 | * | 74 | 3229 |
| 134 | chr12 | 56614848 | 56615309 | 462 | * | 30 | 10193 |
| 135 | chr12 | 57230122 | 57230526 | 405 | * | 85 | 8630 |
| 136 | chr12 | 58258971 | 58259397 | 427 | * | 95 | 10106 |
| 137 | chr12 | 65672464 | 65672880 | 417 | * | 23 | 253827 |
| 138 | chr12 | 66285880 | 66286286 | 407 | * | 37 | 8091 |
| 139 | chr12 | 66290051 | 66290336 | 286 | * | 37 | 8091 |
| 140 | chr12 | 76371820 | 76372207 | 388 | * | 30 | 22822 |

| | | | | | | | |
|---|---|---|---|---|---|---|---|
| 141 | chr12 | 94179180 | 94179599 | 420 | * | 30 | 8738 |
| 142 | chr12 | 104612982 | 104613669 | 688 | * | 77 | 7296 |
| 143 | chr12 | 105879193 | 105879720 | 528 | * | 23 | 387882 |
| 144 | chr12 | 106424495 | 106424846 | 352 | * | 114 | 9891 |
| 145 | chr12 | 106468366 | 106469154 | 789 | * | 77 | 9891 |
| 146 | chr12 | 106477725 | 106478039 | 315 | * | 86 | 9891 |
| 147 | chr12 | 106517472 | 106517733 | 262 | * | 60 | 9891 |
| 148 | chr12 | 107487861 | 107488260 | 400 | * | 86 | 1407 |
| 149 | chr12 | 110547203 | 110547467 | 265 | * | 45 | 28981 |
| 150 | chr12 | 120524622 | 120524935 | 314 | * | 95 | 92558 |
| 151 | chr12 | 122362573 | 122363015 | 443 | * | 143 | 144406 |
| 152 | chr12 | 124985044 | 124985492 | 449 | * | 164 | 9612 |
| 153 | chr12 | 125399323 | 125399731 | 409 | * | 52 | 7316 |
| 154 | chr13 | 21635103 | 21635912 | 810 | * | 361 | 26524 |
| 155 | chr13 | 30716638 | 30716998 | 361 | * | 95 | 84056 |
| 156 | chr13 | 47149937 | 47150222 | 286 | * | 22 | 23143 |
| 157 | chr13 | 49888203 | 49888509 | 307 | * | 37 | 81602 |
| 158 | chr13 | 60737079 | 60737981 | 903 | * | 48 | 81624 |
| 159 | chr13 | 78549376 | 78549892 | 517 | * | 33 | 1910 |
| 160 | chr13 | 80510585 | 80511317 | 733 | * | 52 | 10253 |
| 161 | chr13 | 81030328 | 81030675 | 348 | * | 20 | 10253 |
| 162 | chr13 | 97774389 | 97775392 | 1004 | * | 60 | 10150 |
| 163 | chr13 | 97845914 | 97847006 | 1093 | * | 94 | 10150 |
| 164 | chr13 | 97874681 | 97876342 | 1662 | * | 25 | 10150 |
| 165 | chr13 | 97908526 | 97908826 | 301 | * | 22 | 10150 |
| 166 | chr13 | 97910015 | 97911140 | 1126 | * | 44 | 10150 |
| 167 | chr13 | 97927088 | 97927361 | 274 | * | 67 | 10150 |
| 168 | chr13 | 97959831 | 97961450 | 1620 | * | 345 | 10150 |
| 169 | chr13 | 97980281 | 97980794 | 514 | * | 65 | 10150 |
| 170 | chr13 | 102108183 | 102108524 | 342 | * | 53 | 9358 |
| 171 | chr13 | 102164992 | 102165294 | 303 | * | 22 | 9358 |
| 172 | chr13 | 114884318 | 114885047 | 730 | * | 123 | 22821 |
| 173 | chr14 | 23082912 | 23083540 | 629 | * | 130 | 63874 |
| 174 | chr14 | 23449613 | 23450968 | 1356 | * | 1429 | 84962 |
| 175 | chr14 | 23451833 | 23452477 | 645 | * | 885 | 84962 |
| 176 | chr14 | 55031972 | 55032386 | 415 | * | 12 | 23034 |
| 177 | chr14 | 55118581 | 55119512 | 932 | * | 311 | 23034 |
| 178 | chr14 | 55240189 | 55240475 | 287 | * | 52 | 23034 |
| 179 | chr14 | 61645457 | 61645774 | 318 | * | 23 | 5583 |
| 180 | chr14 | 62028132 | 62028842 | 711 | * | 86 | 400221 |
| 181 | chr14 | 73084087 | 73084440 | 354 | * | 23 | 8110 |
| 182 | chr14 | 89021184 | 89021544 | 361 | * | 104 | 11099 |
| 183 | chr14 | 100040747 | 100041237 | 491 | * | 104 | 317762 |
| 184 | chr15 | 57719653 | 57720169 | 517 | * | 102 | 84952 |
| 185 | chr15 | 57843980 | 57844277 | 298 | * | 196 | 1.01E+08 |
| 186 | chr15 | 58762023 | 58762459 | 437 | * | 164 | 3990 |
| 187 | chr15 | 58839441 | 58839708 | 268 | * | 17 | 102 |

| | | | | | | | |
|---|---|---|---|---|---|---|---|
| 188 | chr15 | 59063573 | 59064048 | 476 | * | 23 | 54629 |
| 189 | chr15 | 60689716 | 60690450 | 735 | * | 302 | 302 |
| 190 | chr15 | 63268526 | 63269190 | 665 | * | 60 | 7168 |
| 191 | chr15 | 63337020 | 63337462 | 443 | * | 96 | 7168 |
| 192 | chr15 | 63414111 | 63414391 | 281 | * | 22 | 114294 |
| 193 | chr15 | 66088669 | 66089001 | 333 | * | 30 | 10260 |
| 194 | chr15 | 67055636 | 67055984 | 349 | * | 45 | 4091 |
| 195 | chr15 | 67058182 | 67058687 | 506 | * | 57 | 4091 |
| 196 | chr15 | 67095516 | 67095881 | 366 | * | 114 | 4091 |
| 197 | chr15 | 67175511 | 67176123 | 613 | * | 1228 | 4091 |
| 198 | chr15 | 67223905 | 67224284 | 380 | * | 58 | 4088 |
| 199 | chr15 | 67336643 | 67337123 | 481 | * | 229 | 4088 |
| 200 | chr15 | 67397813 | 67398320 | 508 | * | 23 | 4088 |
| 201 | chr15 | 67401605 | 67402067 | 463 | * | 152 | 4088 |
| 202 | chr15 | 67442228 | 67442870 | 643 | * | 30 | 4088 |
| 203 | chr15 | 67470904 | 67471304 | 401 | * | 192 | 4088 |
| 204 | chr15 | 74488207 | 74488593 | 387 | * | 19 | 64220 |
| 205 | chr15 | 74501838 | 74502186 | 349 | * | 72 | 64220 |
| 206 | chr15 | 79296668 | 79296972 | 305 | * | 60 | 5923 |
| 207 | chr15 | 80773028 | 80773502 | 475 | * | 69 | 9915 |
| 208 | chr15 | 83478872 | 83479421 | 550 | * | 23 | 123720 |
| 209 | chr15 | 83989428 | 83989742 | 315 | * | 12 | 646 |
| 210 | chr15 | 90955823 | 90956124 | 302 | * | 60 | 8826 |
| 211 | chr15 | 96703904 | 96704280 | 377 | * | 17 | 7026 |
| 212 | chr15 | 96808034 | 96808347 | 314 | * | 33 | 7026 |
| 213 | chr15 | 101663625 | 101663995 | 371 | * | 23 | 22856 |
| 214 | chr16 | 122632 | 123022 | 391 | * | 105 | 64285 |
| 215 | chr16 | 2885818 | 2886111 | 294 | * | 84 | 124220 |
| 216 | chr16 | 3070424 | 3071127 | 704 | * | 69 | 51330 |
| 217 | chr16 | 3071665 | 3072737 | 1073 | * | 174 | 51330 |
| 218 | chr16 | 23916392 | 23916936 | 545 | * | 139 | 5579 |
| 219 | chr16 | 24508293 | 24508567 | 275 | * | 45 | 5930 |
| 220 | chr16 | 24551207 | 24551508 | 302 | * | 24 | 5930 |
| 221 | chr16 | 24647313 | 24647791 | 479 | * | 135 | 5930 |
| 222 | chr16 | 24657768 | 24658027 | 260 | * | 45 | 27327 |
| 223 | chr16 | 24712329 | 24712742 | 414 | * | 74 | 27327 |
| 224 | chr16 | 30064485 | 30065116 | 632 | * | 76 | 226 |
| 225 | chr16 | 31150057 | 31150570 | 514 | * | 30 | 5652 |
| 226 | chr16 | 46782372 | 46782661 | 290 | * | 40 | 91807 |
| 227 | chr16 | 53089042 | 53089337 | 296 | * | 37 | 80205 |
| 228 | chr16 | 56848498 | 56849047 | 550 | * | 185 | 9688 |
| 229 | chr16 | 57570068 | 57570480 | 413 | * | 65 | 92922 |
| 230 | chr16 | 66908556 | 66909073 | 518 | * | 52 | 57546 |
| 231 | chr16 | 68002219 | 68002601 | 383 | * | 43 | 6560 |
| 232 | chr16 | 68679683 | 68680140 | 458 | * | 52 | 1001 |
| 233 | chr16 | 68741230 | 68741915 | 686 | * | 484 | 999 |
| 234 | chr16 | 68824045 | 68824395 | 351 | * | 36 | 999 |

| | | | | | | | |
|---|---|---|---|---|---|---|---|
| 235 | chr16 | 70728005 | 70728910 | 906 | * | 165 | 92154 |
| 236 | chr16 | 71663055 | 71663326 | 272 | * | 30 | 91862 |
| 237 | chr16 | 75278976 | 75279622 | 647 | * | 210 | 9564 |
| 238 | chr16 | 75287317 | 75287687 | 371 | * | 126 | 9564 |
| 239 | chr16 | 77868278 | 77868626 | 349 | * | 17 | 57687 |
| 240 | chr16 | 77930456 | 77930776 | 321 | * | 60 | 57687 |
| 241 | chr16 | 80966773 | 80967083 | 311 | * | 34 | 55839 |
| 242 | chr16 | 82038351 | 82038610 | 260 | * | 33 | 93517 |
| 243 | chr16 | 82667758 | 82668084 | 327 | * | 30 | 1012 |
| 244 | chr16 | 82687616 | 82687913 | 298 | * | 133 | 1012 |
| 245 | chr16 | 84565888 | 84566173 | 286 | * | 263 | 57707 |
| 246 | chr16 | 88365647 | 88366127 | 481 | * | 146 | 84627 |
| 247 | chr17 | 783017 | 783542 | 526 | * | 93 | 64359 |
| 248 | chr17 | 836637 | 836916 | 280 | * | 174 | 64359 |
| 249 | chr17 | 854803 | 855462 | 660 | * | 261 | 64359 |
| 250 | chr17 | 1391299 | 1391915 | 617 | * | 995 | 4641 |
| 251 | chr17 | 1682984 | 1683275 | 292 | * | 23 | 5176 |
| 252 | chr17 | 3571288 | 3571708 | 421 | * | 120 | 30851 |
| 253 | chr17 | 7254529 | 7255474 | 946 | * | 573 | 147040 |
| 254 | chr17 | 8760323 | 8760592 | 270 | * | 11 | 146850 |
| 255 | chr17 | 9551502 | 9552052 | 551 | * | 153 | 124739 |
| 256 | chr17 | 17843351 | 17844209 | 859 | * | 37 | 146691 |
| 257 | chr17 | 36607100 | 36607720 | 621 | * | 744 | 57636 |
| 258 | chr17 | 36620868 | 36621397 | 530 | * | 75 | 57636 |
| 259 | chr17 | 38269796 | 38270102 | 307 | * | 30 | 339287 |
| 260 | chr17 | 40575233 | 40575965 | 733 | * | 398 | 284119 |
| 261 | chr17 | 41607973 | 41608311 | 339 | * | 36 | 2118 |
| 262 | chr17 | 42620499 | 42620814 | 316 | * | 95 | 2535 |
| 263 | chr17 | 43448803 | 43449797 | 995 | * | 844 | 201176 |
| 264 | chr17 | 47099626 | 47100213 | 588 | * | 69 | 10642 |
| 265 | chr17 | 47913737 | 47914249 | 513 | * | 52 | 729220 |
| 266 | chr17 | 48105588 | 48106198 | 611 | * | 423 | 284080 |
| 267 | chr17 | 48287766 | 48288048 | 283 | * | 139 | 1277 |
| 268 | chr17 | 48296272 | 48296659 | 388 | * | 30 | 1277 |
| 269 | chr17 | 53511290 | 53511717 | 428 | * | 37 | 23531 |
| 270 | chr17 | 54618156 | 54618426 | 271 | * | 69 | 9241 |
| 271 | chr17 | 54674927 | 54675249 | 323 | * | 11 | 9241 |
| 272 | chr17 | 54776778 | 54777253 | 476 | * | 335 | 9241 |
| 273 | chr17 | 55550823 | 55551166 | 344 | * | 69 | 124540 |
| 274 | chr17 | 55952432 | 55953028 | 597 | * | 275 | 51649 |
| 275 | chr17 | 57833365 | 57833963 | 599 | * | 37 | 81671 |
| 276 | chr17 | 64948358 | 64948702 | 345 | * | 20 | 27092 |
| 277 | chr17 | 70335322 | 70335750 | 429 | * | 185 | 6662 |
| 278 | chr17 | 73549106 | 73549585 | 480 | * | 174 | 3993 |
| 279 | chr17 | 76349080 | 76349568 | 489 | * | 287 | 9021 |
| 280 | chr17 | 76879382 | 76879698 | 317 | * | 33 | 7077 |
| 281 | chr17 | 79393021 | 79393284 | 264 | * | 45 | 57597 |

| | | | | | | | |
|---|---|---|---|---|---|---|---|
| 282 | chr17 | 81066619 | 81067385 | 767 | * | 152 | 284207 |
| 283 | chr18 | 3247456 | 3247778 | 323 | * | 95 | 10627 |
| 284 | chr18 | 3261453 | 3262365 | 913 | * | 1036 | 103910 |
| 285 | chr18 | 10925846 | 10926263 | 418 | * | 45 | 63895 |
| 286 | chr18 | 12308451 | 12308825 | 375 | * | 67 | 84617 |
| 287 | chr18 | 21032635 | 21033036 | 402 | * | 60 | 8780 |
| 288 | chr18 | 43169234 | 43169991 | 758 | * | 95 | 8170 |
| 289 | chr18 | 46456244 | 46456505 | 262 | * | 60 | 4092 |
| 290 | chr18 | 56251878 | 56252268 | 391 | * | 60 | 115701 |
| 291 | chr18 | 74919591 | 74919935 | 345 | * | 30 | 2587 |
| 292 | chr18 | 77725352 | 77725768 | 417 | * | 45 | 440498 |
| 293 | chr19 | 782252 | 782992 | 741 | * | 299 | 5725 |
| 294 | chr19 | 1027936 | 1028849 | 914 | * | 164 | 1265 |
| 295 | chr19 | 1361554 | 1361831 | 278 | * | 37 | 84939 |
| 296 | chr19 | 2462456 | 2463941 | 1486 | * | 4037 | 84823 |
| 297 | chr19 | 2540320 | 2541524 | 1205 | * | 889 | 4616 |
| 298 | chr19 | 2578785 | 2579220 | 436 | * | 46 | 4616 |
| 299 | chr19 | 6744832 | 6745182 | 351 | * | 69 | 9322 |
| 300 | chr19 | 10119513 | 10120072 | 560 | * | 95 | 50509 |
| 301 | chr19 | 13147874 | 13148322 | 449 | * | 10 | 4784 |
| 302 | chr19 | 13164686 | 13165028 | 343 | * | 299 | 4784 |
| 303 | chr19 | 15217867 | 15218127 | 261 | * | 69 | 85360 |
| 304 | chr19 | 17403331 | 17403630 | 300 | * | 60 | 126549 |
| 305 | chr19 | 34971987 | 34972691 | 705 | * | 1010 | 126374 |
| 306 | chr19 | 39420787 | 39421056 | 270 | * | 45 | 54938 |
| 307 | chr19 | 39919116 | 39919508 | 393 | * | 335 | 6217 |
| 308 | chr19 | 41731721 | 41732368 | 648 | * | 169 | 558 |
| 309 | chr19 | 45129235 | 45129594 | 360 | * | 164 | 147710 |
| 310 | chr19 | 45879617 | 45879985 | 369 | * | 252 | 2068 |
| 311 | chr19 | 46714594 | 46714921 | 328 | * | 69 | 93429 |
| 312 | chr19 | 47660872 | 47661161 | 290 | * | 69 | 10055 |
| 313 | chr19 | 48222464 | 48223086 | 623 | * | 323 | 30846 |
| 314 | chr19 | 48246324 | 48246698 | 375 | * | 30 | 29997 |
| 315 | chr19 | 50143944 | 50144269 | 326 | * | 30 | 6237 |
| 316 | chr19 | 54362249 | 54362538 | 290 | * | 52 | 91663 |
| 317 | chr19 | 57439175 | 57439468 | 294 | * | 12 | 1E+08 |
| 318 | chr2 | 1759470 | 1759855 | 386 | * | 10 | 7837 |
| 319 | chr2 | 1795916 | 1796463 | 548 | * | 69 | 7837 |
| 320 | chr2 | 12959365 | 12959825 | 461 | * | 52 | 1E+08 |
| 321 | chr2 | 20212343 | 20212760 | 418 | * | 82 | 4148 |
| 322 | chr2 | 23787410 | 23787677 | 268 | * | 15 | 114818 |
| 323 | chr2 | 28619533 | 28620004 | 472 | * | 174 | 2355 |
| 324 | chr2 | 31510648 | 31510982 | 335 | * | 45 | 30845 |
| 325 | chr2 | 36598512 | 36599788 | 1277 | * | 104 | 51232 |
| 326 | chr2 | 36608141 | 36608427 | 287 | * | 69 | 51232 |
| 327 | chr2 | 37817298 | 37817610 | 313 | * | 52 | 10602 |
| 328 | chr2 | 37995788 | 37996157 | 370 | * | 119 | 10602 |

| | | | | | | | |
|---|---|---|---|---|---|---|---|
| 329 | chr2 | 42365721 | 42365987 | 267 | * | 171 | 27436 |
| 330 | chr2 | 43269100 | 43269362 | 263 | * | 196 | 678 |
| 331 | chr2 | 43313576 | 43314014 | 439 | * | 69 | 678 |
| 332 | chr2 | 43452061 | 43452327 | 267 | * | 23 | 678 |
| 333 | chr2 | 46166084 | 46166380 | 297 | * | 52 | 5581 |
| 334 | chr2 | 46294732 | 46295067 | 336 | * | 38 | 2034 |
| 335 | chr2 | 46324833 | 46325342 | 510 | * | 60 | 2034 |
| 336 | chr2 | 47082393 | 47082726 | 334 | * | 30 | 1E+08 |
| 337 | chr2 | 47261357 | 47262006 | 650 | * | 385 | 57217 |
| 338 | chr2 | 47297012 | 47297694 | 683 | * | 157 | 57217 |
| 339 | chr2 | 54892808 | 54893067 | 260 | * | 17 | 6711 |
| 340 | chr2 | 64881252 | 64882185 | 934 | * | 524 | 9792 |
| 341 | chr2 | 64995020 | 64995286 | 267 | * | 38 | 9792 |
| 342 | chr2 | 71970378 | 71970822 | 445 | * | 60 | 8291 |
| 343 | chr2 | 85034081 | 85034438 | 358 | * | 23 | 129293 |
| 344 | chr2 | 100323892 | 100324162 | 271 | * | 11 | 51455 |
| 345 | chr2 | 101680065 | 101680357 | 293 | * | 23 | 11138 |
| 346 | chr2 | 102422721 | 102423005 | 285 | * | 30 | 9448 |
| 347 | chr2 | 102649874 | 102650159 | 286 | * | 114 | 7850 |
| 348 | chr2 | 121672588 | 121673530 | 943 | * | 123 | 2736 |
| 349 | chr2 | 121692915 | 121693341 | 427 | * | 60 | 2736 |
| 350 | chr2 | 121694657 | 121695161 | 505 | * | 23 | 2736 |
| 351 | chr2 | 128432859 | 128433243 | 385 | * | 34 | 55679 |
| 352 | chr2 | 129120439 | 129120787 | 349 | * | 142 | 9394 |
| 353 | chr2 | 159314442 | 159315041 | 600 | * | 211 | 8502 |
| 354 | chr2 | 172960330 | 172960646 | 317 | * | 17 | 1746 |
| 355 | chr2 | 173647752 | 173648188 | 437 | * | 37 | 11069 |
| 356 | chr2 | 192731708 | 192732269 | 562 | * | 52 | 8436 |
| 357 | chr2 | 220233574 | 220233832 | 259 | * | 17 | 23549 |
| 358 | chr2 | 233250324 | 233250763 | 440 | * | 120 | 347694 |
| 359 | chr2 | 235357742 | 235358019 | 278 | * | 17 | 10123 |
| 360 | chr2 | 235890038 | 235890450 | 413 | * | 287 | 23677 |
| 361 | chr2 | 235905190 | 235905827 | 638 | * | 45 | 23677 |
| 362 | chr2 | 238223793 | 238224104 | 312 | * | 133 | 1293 |
| 363 | chr2 | 238341290 | 238341615 | 326 | * | 45 | 1293 |
| 364 | chr20 | 304994 | 305383 | 390 | * | 20 | 6666 |
| 365 | chr20 | 2820877 | 2821281 | 405 | * | 20 | 64773 |
| 366 | chr20 | 5637706 | 5638516 | 811 | * | 94 | 56261 |
| 367 | chr20 | 5730964 | 5731922 | 959 | * | 236 | 149840 |
| 368 | chr20 | 15458413 | 15458985 | 573 | * | 30 | 140733 |
| 369 | chr20 | 17830666 | 17831141 | 476 | * | 94 | 27131 |
| 370 | chr20 | 22744025 | 22744375 | 351 | * | 56 | 3170 |
| 371 | chr20 | 22756320 | 22757008 | 689 | * | 24 | 3170 |
| 372 | chr20 | 30300377 | 30301599 | 1223 | * | 152 | 598 |
| 373 | chr20 | 30307845 | 30308569 | 725 | * | 180 | 598 |
| 374 | chr20 | 32978385 | 32978861 | 477 | * | 113 | 83737 |
| 375 | chr20 | 36536186 | 36536704 | 519 | * | 54 | 128434 |

| | | | | | | | |
|---|---|---|---|---|---|---|---|
| 376 | chr20 | 42838826 | 42839323 | 498 | * | 26 | 51526 |
| 377 | chr20 | 42844950 | 42845318 | 369 | * | 17 | 1.01E+08 |
| 378 | chr20 | 43201625 | 43201969 | 345 | * | 28 | 11142 |
| 379 | chr20 | 43966362 | 43966660 | 299 | * | 33 | 6385 |
| 380 | chr20 | 44600466 | 44600869 | 404 | * | 49 | 63925 |
| 381 | chr20 | 45902156 | 45902451 | 296 | * | 29 | 23613 |
| 382 | chr20 | 45987762 | 45988519 | 758 | * | 179 | 23613 |
| 383 | chr20 | 47155146 | 47155546 | 401 | * | 37 | 57580 |
| 384 | chr20 | 49157486 | 49157795 | 310 | * | 147 | 5770 |
| 385 | chr20 | 49407284 | 49407715 | 432 | * | 96 | 55653 |
| 386 | chr20 | 52228074 | 52228382 | 309 | * | 61 | 7764 |
| 387 | chr20 | 52237521 | 52238282 | 762 | * | 56 | 7764 |
| 388 | chr20 | 52790525 | 52790851 | 327 | * | 37 | 1591 |
| 389 | chr20 | 55606089 | 55606573 | 485 | * | 81 | 655 |
| 390 | chr20 | 61447250 | 61447555 | 306 | * | 22 | 1299 |
| 391 | chr21 | 18899958 | 18900225 | 268 | * | 45 | 1525 |
| 392 | chr21 | 30631119 | 30631588 | 470 | * | 82 | 571 |
| 393 | chr21 | 35899149 | 35899617 | 469 | * | 86 | 1827 |
| 394 | chr21 | 44144420 | 44144744 | 325 | * | 196 | 5152 |
| 395 | chr21 | 46289913 | 46290181 | 269 | * | 86 | 754 |
| 396 | chr21 | 46348705 | 46349043 | 339 | * | 37 | 3689 |
| 397 | chr22 | 19868621 | 19869072 | 452 | * | 207 | 10587 |
| 398 | chr22 | 20923381 | 20923702 | 322 | * | 19 | 51586 |
| 399 | chr22 | 24603833 | 24604126 | 294 | * | 45 | 56241 |
| 400 | chr22 | 26812912 | 26813292 | 381 | * | 123 | 57168 |
| 401 | chr22 | 30602839 | 30603333 | 495 | * | 133 | 3976 |
| 402 | chr22 | 36025786 | 36026155 | 370 | * | 229 | 4151 |
| 403 | chr22 | 36725465 | 36726120 | 656 | * | 95 | 4627 |
| 404 | chr22 | 36764587 | 36765371 | 785 | * | 143 | 4627 |
| 405 | chr22 | 37893173 | 37893471 | 299 | * | 30 | 29775 |
| 406 | chr22 | 37942123 | 37942481 | 359 | * | 69 | 11135 |
| 407 | chr22 | 37956847 | 37957348 | 502 | * | 69 | 11135 |
| 408 | chr22 | 38614686 | 38615080 | 395 | * | 153 | 23764 |
| 409 | chr22 | 39570745 | 39572785 | 2041 | * | 770 | 23492 |
| 410 | chr22 | 39867482 | 39867846 | 365 | * | 23 | 4248 |
| 411 | chr22 | 41686719 | 41687035 | 317 | * | 114 | 5905 |
| 412 | chr22 | 42063285 | 42063611 | 327 | * | 30 | 4809 |
| 413 | chr22 | 46084606 | 46084919 | 314 | * | 54 | 25814 |
| 414 | chr22 | 50731767 | 50732557 | 791 | * | 584 | 23654 |
| 415 | chr3 | 11641936 | 11642260 | 325 | * | 113 | 9686 |
| 416 | chr3 | 11760965 | 11761840 | 876 | * | 347 | 9686 |
| 417 | chr3 | 13034848 | 13035176 | 329 | * | 23 | 9922 |
| 418 | chr3 | 16525302 | 16526428 | 1127 | * | 133 | 23180 |
| 419 | chr3 | 49506637 | 49507329 | 693 | * | 107 | 1605 |
| 420 | chr3 | 50301808 | 50302548 | 741 | * | 102 | 7869 |
| 421 | chr3 | 58033690 | 58034187 | 498 | * | 68 | 2317 |
| 422 | chr3 | 111454057 | 111454368 | 312 | * | 52 | 90102 |

| | | | | | | | |
|---|---|---|---|---|---|---|---|
| 423 | chr3 | 111578322 | 111578600 | 279 | * | 37 | 90102 |
| 424 | chr3 | 124493126 | 124493603 | 478 | * | 263 | 3693 |
| 425 | chr3 | 124597900 | 124598363 | 464 | * | 45 | 3693 |
| 426 | chr3 | 124782026 | 124782362 | 337 | * | 114 | 57493 |
| 427 | chr3 | 129330739 | 129330997 | 259 | * | 52 | 23129 |
| 428 | chr3 | 134032225 | 134032601 | 377 | * | 123 | 51421 |
| 429 | chr3 | 134045636 | 134046173 | 538 | * | 261 | 51421 |
| 430 | chr3 | 134076332 | 134076949 | 618 | * | 192 | 51421 |
| 431 | chr3 | 134077317 | 134077625 | 309 | * | 10 | 51421 |
| 432 | chr3 | 134082696 | 134083239 | 544 | * | 645 | 51421 |
| 433 | chr3 | 134092213 | 134093797 | 1585 | * | 961 | 51421 |
| 434 | chr3 | 136505277 | 136505744 | 468 | * | 143 | 80723 |
| 435 | chr3 | 170469452 | 170469769 | 318 | * | 114 | 200916 |
| 436 | chr3 | 171757589 | 171758116 | 528 | * | 67 | 64778 |
| 437 | chr3 | 177196207 | 177196989 | 783 | * | 45 | 1.01E+08 |
| 438 | chr3 | 177233844 | 177234429 | 586 | * | 23 | 1.01E+08 |
| 439 | chr3 | 184286822 | 184287144 | 323 | * | 37 | 2049 |
| 440 | chr3 | 192596895 | 192597905 | 1011 | * | 196 | 151963 |
| 441 | chr3 | 192607022 | 192607376 | 355 | * | 164 | 151963 |
| 442 | chr3 | 194823637 | 194824260 | 624 | * | 123 | 152002 |
| 443 | chr4 | 1779774 | 1780320 | 547 | * | 62 | 2261 |
| 444 | chr4 | 7912518 | 7912776 | 259 | * | 23 | 60312 |
| 445 | chr4 | 41145281 | 41145556 | 276 | * | 45 | 323 |
| 446 | chr4 | 77725068 | 77725345 | 278 | * | 17 | 57619 |
| 447 | chr4 | 79490187 | 79490652 | 466 | * | 114 | 306 |
| 448 | chr4 | 84176435 | 84176722 | 288 | * | 23 | 27235 |
| 449 | chr4 | 124753629 | 124753926 | 298 | * | 30 | 285419 |
| 450 | chr4 | 129731925 | 129732353 | 429 | * | 60 | 79960 |
| 451 | chr4 | 129733455 | 129733772 | 318 | * | 45 | 79960 |
| 452 | chr4 | 143329330 | 143329597 | 268 | * | 52 | 8821 |
| 453 | chr4 | 159731308 | 159731853 | 546 | * | 77 | 57600 |
| 454 | chr4 | 187354862 | 187355272 | 411 | * | 27 | 285441 |
| 455 | chr5 | 956680 | 957627 | 948 | * | 233 | 1.01E+08 |
| 456 | chr5 | 1327633 | 1328504 | 872 | * | 384 | 81037 |
| 457 | chr5 | 10513926 | 10514734 | 809 | * | 141 | 651746 |
| 458 | chr5 | 10551236 | 10552097 | 862 | * | 294 | 651746 |
| 459 | chr5 | 10552688 | 10552969 | 282 | * | 53 | 651746 |
| 460 | chr5 | 14035822 | 14036352 | 531 | * | 77 | 1767 |
| 461 | chr5 | 14036749 | 14037478 | 730 | * | 60 | 1767 |
| 462 | chr5 | 14270035 | 14270392 | 358 | * | 29 | 7204 |
| 463 | chr5 | 17169474 | 17169762 | 289 | * | 28 | 10409 |
| 464 | chr5 | 21111593 | 21111994 | 402 | * | 26 | 728411 |
| 465 | chr5 | 31362426 | 31363063 | 638 | * | 39 | 29102 |
| 466 | chr5 | 31853831 | 31854099 | 269 | * | 11 | 23037 |
| 467 | chr5 | 33736408 | 33736681 | 274 | * | 39 | 81792 |
| 468 | chr5 | 33892176 | 33893068 | 893 | * | 71 | 81792 |
| 469 | chr5 | 34609234 | 34609521 | 288 | * | 81 | 26064 |

| | | | | | | | |
|---|---|---|---|---|---|---|---|
| 470 | chr5 | 34610042 | 34610300 | 259 | * | 41 | 26064 |
| 471 | chr5 | 38696955 | 38697581 | 627 | * | 65 | 3977 |
| 472 | chr5 | 40405575 | 40405875 | 301 | * | 11 | 5734 |
| 473 | chr5 | 52095156 | 52095617 | 462 | * | 195 | 53918 |
| 474 | chr5 | 54455975 | 54456254 | 280 | * | 52 | 493869 |
| 475 | chr5 | 54930125 | 54930574 | 450 | * | 28 | 153129 |
| 476 | chr5 | 55290455 | 55291121 | 667 | * | 53 | 3572 |
| 477 | chr5 | 55575888 | 55576220 | 333 | * | 108 | 79722 |
| 478 | chr5 | 55642918 | 55643204 | 287 | * | 58 | 79722 |
| 479 | chr5 | 55684433 | 55684739 | 307 | * | 15 | 79722 |
| 480 | chr5 | 55864863 | 55865176 | 314 | * | 88 | 4214 |
| 481 | chr5 | 56988528 | 56989496 | 969 | * | 76 | 345651 |
| 482 | chr5 | 57137449 | 57138090 | 642 | * | 113 | 345651 |
| 483 | chr5 | 57312239 | 57312566 | 328 | * | 81 | 10769 |
| 484 | chr5 | 58422841 | 58423809 | 969 | * | 82 | 5144 |
| 485 | chr5 | 59086266 | 59086604 | 339 | * | 15 | 5144 |
| 486 | chr5 | 60597535 | 60597796 | 262 | * | 124 | 57688 |
| 487 | chr5 | 60746113 | 60746445 | 333 | * | 76 | 57688 |
| 488 | chr5 | 66539951 | 66540268 | 318 | * | 20 | 4064 |
| 489 | chr5 | 66563843 | 66564454 | 612 | * | 52 | 4064 |
| 490 | chr5 | 67729695 | 67730150 | 456 | * | 61 | 5295 |
| 491 | chr5 | 68797782 | 68798216 | 435 | * | 126 | 1.01E+08 |
| 492 | chr5 | 71475069 | 71475701 | 633 | * | 113 | 4131 |
| 493 | chr5 | 73234342 | 73234911 | 570 | * | 178 | 64283 |
| 494 | chr5 | 92414022 | 92414405 | 384 | * | 36 | 441094 |
| 495 | chr5 | 92695060 | 92695597 | 538 | * | 118 | 441094 |
| 496 | chr5 | 95048073 | 95048748 | 676 | * | 57 | 22836 |
| 497 | chr5 | 95590853 | 95591421 | 569 | * | 111 | 5122 |
| 498 | chr5 | 109496359 | 109496704 | 346 | * | 37 | 642987 |
| 499 | chr5 | 111016339 | 111016604 | 266 | * | 154 | 9315 |
| 500 | chr5 | 122188300 | 122188599 | 300 | * | 15 | 28966 |
| 501 | chr5 | 123996222 | 123997206 | 985 | * | 164 | 57507 |
| 502 | chr5 | 124379437 | 124380387 | 951 | * | 390 | 57507 |
| 503 | chr5 | 125690129 | 125690443 | 315 | * | 51 | 65983 |
| 504 | chr5 | 130971229 | 130971873 | 645 | * | 18 | 51735 |
| 505 | chr5 | 131724146 | 131724537 | 392 | * | 104 | 6584 |
| 506 | chr5 | 135329471 | 135329793 | 323 | * | 37 | 7045 |
| 507 | chr5 | 135347543 | 135348276 | 734 | * | 47 | 7045 |
| 508 | chr5 | 135393248 | 135394350 | 1103 | * | 292 | 7045 |
| 509 | chr5 | 149318184 | 149318575 | 392 | * | 23 | 5145 |
| 510 | chr5 | 150016653 | 150017378 | 726 | * | 89 | 11346 |
| 511 | chr5 | 150024418 | 150024971 | 554 | * | 93 | 11346 |
| 512 | chr5 | 153747565 | 153748260 | 696 | * | 111 | 55568 |
| 513 | chr5 | 156717240 | 156717707 | 468 | * | 63 | 26999 |
| 514 | chr5 | 159510283 | 159510830 | 548 | * | 87 | 114825 |
| 515 | chr5 | 161616955 | 161617227 | 273 | * | 23 | 2566 |
| 516 | chr5 | 162886932 | 162887286 | 355 | * | 23 | 134492 |

| | | | | | | | |
|---|---|---|---|---|---|---|---|
| 517 | chr5 | 174715190 | 174715507 | 318 | * | 77 | 1812 |
| 518 | chr6 | 4582105 | 4582582 | 478 | * | 120 | 9425 |
| 519 | chr6 | 16769689 | 16770169 | 481 | * | 45 | 6310 |
| 520 | chr6 | 30712449 | 30712855 | 407 | * | 52 | 8870 |
| 521 | chr6 | 30811884 | 30812556 | 673 | * | 77 | 780 |
| 522 | chr6 | 34203148 | 34203491 | 344 | * | 95 | 3159 |
| 523 | chr6 | 37093895 | 37094450 | 556 | * | 104 | 5292 |
| 524 | chr6 | 37105305 | 37105858 | 554 | * | 77 | 5292 |
| 525 | chr6 | 45413507 | 45414146 | 640 | * | 252 | 860 |
| 526 | chr6 | 74289898 | 74290319 | 422 | * | 104 | 26503 |
| 527 | chr6 | 74295491 | 74296051 | 561 | * | 77 | 26503 |
| 528 | chr6 | 75915126 | 75915419 | 294 | * | 19 | 1303 |
| 529 | chr6 | 75915687 | 75915967 | 281 | * | 18 | 1303 |
| 530 | chr6 | 76027250 | 76027550 | 301 | * | 37 | 27145 |
| 531 | chr6 | 86113030 | 86114258 | 1229 | * | 45 | 4907 |
| 532 | chr6 | 86388763 | 86389022 | 260 | * | 52 | 387066 |
| 533 | chr6 | 90927742 | 90928211 | 470 | * | 133 | 60468 |
| 534 | chr6 | 91005482 | 91005836 | 355 | * | 30 | 60468 |
| 535 | chr6 | 99419064 | 99419410 | 347 | * | 17 | 26235 |
| 536 | chr6 | 119027409 | 119027695 | 287 | * | 23 | 387119 |
| 537 | chr6 | 132272297 | 132273024 | 728 | * | 385 | 1490 |
| 538 | chr6 | 157469879 | 157470346 | 468 | * | 69 | 57492 |
| 539 | chr6 | 158402258 | 158402799 | 542 | * | 174 | 8871 |
| 540 | chr7 | 500185 | 500446 | 262 | * | 37 | 5154 |
| 541 | chr7 | 500861 | 501374 | 514 | * | 297 | 5154 |
| 542 | chr7 | 1576493 | 1577559 | 1067 | * | 394 | 7975 |
| 543 | chr7 | 2215151 | 2215523 | 373 | * | 130 | 8379 |
| 544 | chr7 | 2348700 | 2350144 | 1445 | * | 412 | 29886 |
| 545 | chr7 | 3988324 | 3989052 | 729 | * | 138 | 221935 |
| 546 | chr7 | 5631069 | 5631877 | 809 | * | 425 | 6624 |
| 547 | chr7 | 5813226 | 5813825 | 600 | * | 30 | 54476 |
| 548 | chr7 | 6322698 | 6323466 | 769 | * | 149 | 9265 |
| 549 | chr7 | 23373716 | 23374140 | 425 | * | 23 | 10643 |
| 550 | chr7 | 27135175 | 27135707 | 533 | * | 160 | 3198 |
| 551 | chr7 | 29762982 | 29763513 | 532 | * | 52 | 442524 |
| 552 | chr7 | 30027444 | 30028753 | 1310 | * | 98 | 9805 |
| 553 | chr7 | 30805396 | 30805765 | 370 | * | 11 | 84182 |
| 554 | chr7 | 32996897 | 32997393 | 497 | * | 74 | 11328 |
| 555 | chr7 | 33756017 | 33756425 | 409 | * | 17 | 168667 |
| 556 | chr7 | 35840996 | 35841294 | 299 | * | 23 | 989 |
| 557 | chr7 | 36087542 | 36088137 | 596 | * | 48 | 80820 |
| 558 | chr7 | 36428631 | 36429056 | 426 | * | 20 | 54443 |
| 559 | chr7 | 39773417 | 39773784 | 368 | * | 100 | 349114 |
| 560 | chr7 | 41067410 | 41067749 | 340 | * | 33 | 79783 |
| 561 | chr7 | 42533048 | 42533614 | 567 | * | 29 | 2737 |
| 562 | chr7 | 43921821 | 43922187 | 367 | * | 36 | 55665 |
| 563 | chr7 | 47521973 | 47522485 | 513 | * | 28 | 64759 |

| | | | | | | | |
|---|---|---|---|---|---|---|---|
| 564 | chr7 | 47618572 | 47619039 | 468 | * | 35 | 64759 |
| 565 | chr7 | 47619710 | 47619991 | 282 | * | 47 | 64759 |
| 566 | chr7 | 47644438 | 47644824 | 387 | * | 193 | 64759 |
| 567 | chr7 | 47823274 | 47824126 | 853 | * | 80 | 80099 |
| 568 | chr7 | 47826043 | 47826981 | 939 | * | 251 | 80099 |
| 569 | chr7 | 48123033 | 48123377 | 345 | * | 26 | 7378 |
| 570 | chr7 | 66056857 | 66057377 | 521 | * | 76 | 493754 |
| 571 | chr7 | 66119560 | 66119997 | 438 | * | 52 | 27342 |
| 572 | chr7 | 68164229 | 68164599 | 371 | * | 37 | 26053 |
| 573 | chr7 | 68428485 | 68429035 | 551 | * | 61 | 26053 |
| 574 | chr7 | 73895574 | 73895864 | 291 | * | 114 | 9569 |
| 575 | chr7 | 73896804 | 73898105 | 1302 | * | 77 | 9569 |
| 576 | chr7 | 74066702 | 74066976 | 275 | * | 16 | 2969 |
| 577 | chr7 | 75849398 | 75849763 | 366 | * | 45 | 222183 |
| 578 | chr7 | 75987692 | 75987968 | 277 | * | 36 | 7532 |
| 579 | chr7 | 76052654 | 76052980 | 327 | * | 52 | 7784 |
| 580 | chr7 | 90254986 | 90255560 | 575 | * | 174 | 5218 |
| 581 | chr7 | 92324853 | 92325376 | 524 | * | 102 | 257415 |
| 582 | chr7 | 92327440 | 92328167 | 728 | * | 114 | 257415 |
| 583 | chr7 | 93543595 | 93544227 | 633 | * | 38 | 2791 |
| 584 | chr7 | 93577012 | 93577341 | 330 | * | 11 | 2791 |
| 585 | chr7 | 100463203 | 100464075 | 873 | * | 53 | 7205 |
| 586 | chr7 | 100760671 | 100760984 | 314 | * | 120 | 5054 |
| 587 | chr7 | 100765346 | 100765637 | 292 | * | 46 | 5054 |
| 588 | chr7 | 101000819 | 101001099 | 281 | * | 60 | 136227 |
| 589 | chr7 | 101045057 | 101046005 | 949 | * | 230 | 136227 |
| 590 | chr7 | 106358737 | 106359081 | 345 | * | 53 | 168455 |
| 591 | chr7 | 107731824 | 107732430 | 607 | * | 23 | 22798 |
| 592 | chr7 | 115994461 | 115995253 | 793 | * | 218 | 858 |
| 593 | chr7 | 116151772 | 116153031 | 1260 | * | 67 | 858 |
| 594 | chr7 | 116165247 | 116165638 | 392 | * | 38 | 857 |
| 595 | chr7 | 129609830 | 129610365 | 536 | * | 61 | 7328 |
| 596 | chr7 | 129934234 | 129934492 | 259 | * | 19 | 51200 |
| 597 | chr7 | 130070754 | 130071032 | 279 | * | 39 | 95681 |
| 598 | chr7 | 131546885 | 131547278 | 394 | * | 70 | 91584 |
| 599 | chr7 | 138575623 | 138575983 | 361 | * | 45 | 57670 |
| 600 | chr7 | 140325921 | 140326646 | 726 | * | 222 | 27147 |
| 601 | chr7 | 141062494 | 141062872 | 379 | * | 97 | 55750 |
| 602 | chr7 | 150008900 | 150009212 | 313 | * | 51 | 65999 |
| 603 | chr7 | 150009673 | 150010046 | 374 | * | 26 | 65999 |
| 604 | chr7 | 150065922 | 150066506 | 585 | * | 50 | 29803 |
| 605 | chr7 | 151217693 | 151217966 | 274 | * | 45 | 6009 |
| 606 | chr7 | 151384460 | 151385166 | 707 | * | 265 | 51422 |
| 607 | chr7 | 151451083 | 151451720 | 638 | * | 649 | 51422 |
| 608 | chr7 | 157102105 | 157102943 | 839 | * | 97 | 10049 |
| 609 | chr8 | 1771745 | 1772089 | 345 | * | 45 | 9639 |
| 610 | chr8 | 6451629 | 6452008 | 380 | * | 60 | 285 |

| | | | | | | | |
|---|---|---|---|---|---|---|---|
| 611 | chr8 | 11368114 | 11368472 | 359 | * | 23 | 640 |
| 612 | chr8 | 11748677 | 11748955 | 279 | * | 23 | 1508 |
| 613 | chr8 | 13051438 | 13051855 | 418 | * | 77 | 10395 |
| 614 | chr8 | 13224167 | 13224555 | 389 | * | 174 | 10395 |
| 615 | chr8 | 17533644 | 17533914 | 271 | * | 45 | 57509 |
| 616 | chr8 | 18523702 | 18524358 | 657 | * | 86 | 23362 |
| 617 | chr8 | 18577622 | 18577889 | 268 | * | 76 | 23362 |
| 618 | chr8 | 22454683 | 22455558 | 876 | * | 683 | 541565 |
| 619 | chr8 | 23571892 | 23572451 | 560 | * | 60 | 137814 |
| 620 | chr8 | 27471827 | 27472407 | 581 | * | 54 | 1191 |
| 621 | chr8 | 27862149 | 27862489 | 341 | * | 123 | 286133 |
| 622 | chr8 | 29877616 | 29877918 | 303 | * | 45 | 51669 |
| 623 | chr8 | 37000279 | 37000782 | 504 | * | 17 | 157855 |
| 624 | chr8 | 37002190 | 37002479 | 290 | * | 36 | 157855 |
| 625 | chr8 | 37378906 | 37379175 | 270 | * | 52 | 80139 |
| 626 | chr8 | 37659128 | 37659509 | 382 | * | 57 | 25960 |
| 627 | chr8 | 38857155 | 38857665 | 511 | * | 23 | 8754 |
| 628 | chr8 | 67434445 | 67434904 | 460 | * | 25 | 254778 |
| 629 | chr8 | 87520578 | 87520848 | 271 | * | 45 | 51115 |
| 630 | chr8 | 101800788 | 101801622 | 835 | * | 49 | 26986 |
| 631 | chr8 | 102063535 | 102063910 | 376 | * | 114 | 441374 |
| 632 | chr8 | 107072179 | 107072635 | 457 | * | 30 | 55074 |
| 633 | chr8 | 118662650 | 118662928 | 279 | * | 56 | 90390 |
| 634 | chr8 | 119214273 | 119214617 | 345 | * | 37 | 2131 |
| 635 | chr8 | 119221383 | 119221775 | 393 | * | 60 | 2131 |
| 636 | chr8 | 128866269 | 128866857 | 589 | * | 32 | 5820 |
| 637 | chr8 | 128919590 | 128920372 | 783 | * | 102 | 5820 |
| 638 | chr8 | 131154432 | 131154696 | 265 | * | 23 | 50807 |
| 639 | chr8 | 142164055 | 142164415 | 361 | * | 101 | 22898 |
| 640 | chr8 | 143757658 | 143758007 | 350 | * | 96 | 8000 |
| 641 | chr8 | 144097296 | 144097653 | 358 | * | 60 | 1E+08 |
| 642 | chr8 | 144120539 | 144120934 | 396 | * | 21 | 286122 |
| 643 | chr8 | 145014188 | 145014897 | 710 | * | 107 | 5339 |
| 644 | chr9 | 707011 | 707400 | 390 | * | 58 | 23189 |
| 645 | chr9 | 5509461 | 5509868 | 408 | * | 22 | 80380 |
| 646 | chr9 | 35518166 | 35518583 | 418 | * | 196 | 9853 |
| 647 | chr9 | 35907883 | 35908233 | 351 | * | 42 | 158376 |
| 648 | chr9 | 73034611 | 73034989 | 379 | * | 77 | 687 |
| 649 | chr9 | 73169044 | 73169380 | 337 | * | 67 | 687 |
| 650 | chr9 | 75169179 | 75169690 | 512 | * | 53 | 117531 |
| 651 | chr9 | 78987063 | 78987379 | 317 | * | 37 | 55312 |
| 652 | chr9 | 80129858 | 80130639 | 782 | * | 252 | 9630 |
| 653 | chr9 | 84174793 | 84175055 | 263 | * | 24 | 7088 |
| 654 | chr9 | 89990731 | 89991127 | 397 | * | 41 | 1612 |
| 655 | chr9 | 91149434 | 91150423 | 990 | * | 185 | 158046 |
| 656 | chr9 | 97675333 | 97675718 | 386 | * | 27 | 84909 |
| 657 | chr9 | 97684284 | 97685128 | 845 | * | 93 | 84909 |

| | | | | | | | |
|---|---|---|---|---|---|---|---|
| 658 | chr9 | 98269647 | 98270217 | 571 | * | 37 | 5727 |
| 659 | chr9 | 100639252 | 100639765 | 514 | * | 60 | 2304 |
| 660 | chr9 | 100893014 | 100893273 | 260 | * | 10 | 7464 |
| 661 | chr9 | 100985598 | 100985883 | 286 | * | 67 | 55357 |
| 662 | chr9 | 103110905 | 103111182 | 278 | * | 30 | 54881 |
| 663 | chr9 | 109679615 | 109679892 | 278 | * | 37 | 58499 |
| 664 | chr9 | 116372433 | 116372744 | 312 | * | 36 | 5998 |
| 665 | chr9 | 116382853 | 116383559 | 707 | * | 85 | 5998 |
| 666 | chr9 | 116939571 | 116939897 | 327 | * | 37 | 85301 |
| 667 | chr9 | 124113550 | 124113911 | 362 | * | 50 | 2040 |
| 668 | chr9 | 127042288 | 127042783 | 496 | * | 28 | 10783 |
| 669 | chr9 | 127048011 | 127048705 | 695 | * | 28 | 10783 |
| 670 | chr9 | 127059621 | 127060063 | 443 | * | 192 | 10783 |
| 671 | chr9 | 127071709 | 127072485 | 777 | * | 91 | 10783 |
| 672 | chr9 | 130860793 | 130861640 | 848 | * | 98 | 114789 |
| 673 | chr9 | 131314982 | 131315253 | 272 | * | 30 | 6709 |
| 674 | chr9 | 131316389 | 131317843 | 1455 | * | 255 | 6709 |
| 675 | chr9 | 131320799 | 131321311 | 513 | * | 93 | 6709 |
| 676 | chr9 | 131988204 | 131988566 | 363 | * | 46 | 389792 |
| 677 | chr9 | 132319341 | 132319928 | 588 | * | 60 | 1.01E+08 |
| 678 | chr9 | 132945366 | 132945762 | 397 | * | 93 | 23413 |
| 679 | chr9 | 134483402 | 134483665 | 264 | * | 32 | 2889 |
| 680 | chr9 | 134483975 | 134484312 | 338 | * | 168 | 2889 |
| 681 | chr9 | 134609440 | 134609876 | 437 | * | 96 | 2889 |
| 682 | chr9 | 136579908 | 136580245 | 338 | * | 62 | 1757 |
| 683 | chr9 | 137409818 | 137410210 | 393 | * | 21 | 6256 |
| 684 | chr9 | 137494456 | 137494750 | 295 | * | 183 | 1289 |
| 685 | chr9 | 140306252 | 140306592 | 341 | * | 81 | 54932 |
| 686 | chrX | 68209948 | 68210316 | 369 | * | 30 | 1947 |
| 687 | chrX | 128800119 | 128800525 | 407 | * | 45 | 8862 |
| 688 | chrX | 133738181 | 133738449 | 269 | * | 60 | 10761 |
| 689 | chrX | 133803948 | 133804345 | 398 | * | 45 | 10761 |
| 690 | chrX | 135296457 | 135297045 | 589 | * | 196 | 2273 |
| 691 | chrX | 139805527 | 139805829 | 303 | * | 86 | 286411 |
| 692 | chrX | 152066053 | 152066588 | 536 | * | 557 | 7739 |
| 693 | chrX | 153597502 | 153597835 | 334 | * | 52 | 2316 |

| ENSEMBL | baseMean | log2 Fold Change | padj | Gene |
|---|---|---|---|---|
| ENSG00000074800 | 85505.67072 | 0.789447698 | 6.16E-09 | ENO1 |
| ENSG00000189337 | 2090.457273 | 1.387914469 | 2.25E-51 | KAZN |
| ENSG00000142621 | 349.9471983 | 0.606337834 | 9.70E-05 | FHAD1 |
| ENSG00000142627 | 5020.238168 | 1.629833058 | 5.31E-43 | EPHA2 |
| ENSG00000142627 | 5020.238168 | 1.629833058 | 5.31E-43 | EPHA2 |
| ENSG00000142623 | 49.5811494 | 3.962982254 | 1.13E-16 | PADI1 |
| ENSG00000127481 | 15901.60597 | 0.771680083 | 4.76E-20 | UBR4 |
| ENSG00000162542 | 1573.630919 | 0.874906788 | 1.83E-12 | TMCO4 |
| ENSG00000117632 | 17626.57597 | 0.685300807 | 1.12E-11 | STMN1 |
| ENSG00000131914 | 640.0340004 | 2.374330869 | 7.25E-70 | LIN28A |
| ENSG00000253368 | 1601.234225 | 1.388374179 | 2.40E-22 | TRNP1 |
| ENSG00000253368 | 1601.234225 | 1.388374179 | 2.40E-22 | TRNP1 |
| ENSG00000204138 | 2601.10134 | 0.200137755 | 1.33E-02 | PHACTR4 |
| ENSG00000253304 | 55.71415746 | 0.750407845 | 1.10E-02 | TMEM200B |
| ENSG00000060656 | 1753.900827 | 0.823606809 | 3.97E-09 | PTPRU |
| ENSG00000142920 | 391.0001786 | 0.50744707 | 2.49E-02 | AZIN2 |
| ENSG00000117395 | 5155.502085 | 0.946285368 | 1.41E-22 | EBNA1BP2 |
| ENSG00000178922 | 1853.666591 | 1.624499058 | 6.43E-37 | HYI |
| ENSG00000126091 | 868.2489115 | 0.507866547 | 6.67E-09 | ST3GAL3 |
| ENSG00000186564 | 276.8095472 | -0.59811741 | 1.98E-05 | FOXD2 |
| ENSG00000162383 | 15.21491097 | 1.621882739 | 2.08E-03 | SLC1A7 |
| ENSG00000157216 | 3859.235303 | -0.542267744 | 6.46E-06 | SSBP3 |
| ENSG00000177606 | 2696.774838 | 1.677887543 | 8.72E-18 | JUN |
| ENSG00000185483 | 803.2210795 | -0.89125324 | 1.72E-09 | ROR1 |
| ENSG00000185483 | 803.2210795 | -0.89125324 | 1.72E-09 | ROR1 |
| ENSG00000097096 | 235.870281 | 1.291897068 | 1.36E-07 | SYDE2 |
| ENSG00000153904 | 3716.229868 | 1.204092801 | 5.88E-18 | DDAH1 |
| ENSG00000153904 | 3716.229868 | 1.204092801 | 5.88E-18 | DDAH1 |
| ENSG00000142871 | 10596.59591 | 4.163417313 | 2.09E-303 | CCN1 |
| ENSG00000142871 | 10596.59591 | 4.163417313 | 2.09E-303 | CCN1 |
| ENSG00000142871 | 10596.59591 | 4.163417313 | 2.09E-303 | CCN1 |
| ENSG00000097046 | 1042.852377 | -1.013545796 | 1.46E-09 | CDC7 |
| ENSG00000069702 | 1252.634429 | -0.764091099 | 1.27E-10 | TGFBR3 |
| ENSG00000137936 | 3028.458632 | 0.353489729 | 7.72E-03 | BCAR3 |
| ENSG00000198691 | 54.74305486 | 1.026348836 | 2.48E-02 | ABCA4 |
| ENSG00000137962 | 4982.779967 | 0.443426328 | 4.01E-02 | ARHGAP29 |
| ENSG00000137962 | 4982.779967 | 0.443426328 | 4.01E-02 | ARHGAP29 |
| ENSG00000265808 | 4612.525216 | 0.398501777 | 5.03E-07 | SEC22B |
| ENSG00000143382 | 843.0356382 | 1.054531604 | 3.38E-13 | ADAMTSL4 |
| ENSG00000143398 | 3765.167022 | 0.923842394 | 7.89E-22 | PIP5K1A |
| ENSG00000197747 | 12016.01031 | 1.448456552 | 4.10E-39 | S100A10 |
| ENSG00000159450 | 60.14085417 | 1.364748755 | 7.62E-04 | TCHH |
| ENSG00000143570 | 6731.336194 | 0.30402251 | 3.39E-02 | SLC39A1 |
| ENSG00000160691 | 17857.68695 | 0.471747848 | 3.67E-05 | SHC1 |
| ENSG00000187800 | 1120.147306 | 1.902616643 | 1.68E-48 | PEAR1 |
| ENSG00000183853 | 10728.67019 | 0.646225205 | 1.64E-11 | KIRREL1 |

| | | | | |
|---|---|---|---|---|
| ENSG00000077157 | 1810.477078 | 0.449403283 | 3.32E-07 | PPP1R12B |
| ENSG00000163545 | 210.063746 | 1.357489241 | 6.41E-17 | NUAK2 |
| ENSG00000152104 | 5617.28597 | 0.751215291 | 2.26E-25 | PTPN14 |
| ENSG00000152104 | 5617.28597 | 0.751215291 | 2.26E-25 | PTPN14 |
| ENSG00000151474 | 2369.142518 | -1.620589534 | 2.97E-66 | FRMD4A |
| ENSG00000171988 | 1264.147845 | -1.147539346 | 8.55E-06 | JMJD1C |
| ENSG00000107738 | 1076.142038 | 1.239144648 | 6.50E-15 | VSIR |
| ENSG00000122376 | 793.2501601 | -0.719260164 | 4.96E-05 | SHLD2 |
| ENSG00000148677 | 2371.13122 | 7.508110522 | 9.13E-69 | ANKRD1 |
| ENSG00000138119 | 15695.67332 | 1.567003655 | 6.04E-36 | MYOF |
| ENSG00000138119 | 15695.67332 | 1.567003655 | 6.04E-36 | MYOF |
| ENSG00000165887 | 7.839182386 | 4.452964021 | 7.83E-05 | ANKRD2 |
| ENSG00000119986 | 366.8041458 | 0.991259613 | 3.20E-10 | AVPI1 |
| ENSG00000138166 | 787.2594332 | -0.593307411 | 1.10E-02 | DUSP5 |
| ENSG00000182272 | 1110.705317 | -0.603243977 | 2.09E-03 | B4GALNT4 |
| ENSG00000214063 | 4504.292425 | 0.286714485 | 2.20E-02 | TSPAN4 |
| ENSG00000170955 | 2094.464026 | 0.830231468 | 1.09E-09 | CAVIN3 |
| ENSG00000166452 | 922.1010123 | 0.283238444 | 3.36E-02 | AKIP1 |
| ENSG00000133789 | 3816.151683 | 0.670327923 | 3.69E-15 | SWAP70 |
| ENSG00000245522 | 102.7984567 | 2.265063254 | 1.46E-21 | AC026250.1 |
| ENSG00000133812 | 3829.910888 | 0.306651837 | 5.44E-03 | SBF2 |
| ENSG00000133812 | 3829.910888 | 0.306651837 | 5.44E-03 | SBF2 |
| ENSG00000050165 | 22309.34202 | 0.307103942 | 2.83E-02 | DKK3 |
| ENSG00000133816 | 7480.93452 | 1.532830488 | 3.87E-31 | MICAL2 |
| ENSG00000133816 | 7480.93452 | 1.532830488 | 3.87E-31 | MICAL2 |
| ENSG00000133816 | 7480.93452 | 1.532830488 | 3.87E-31 | MICAL2 |
| ENSG00000197702 | 12811.79845 | 0.983629944 | 3.10E-84 | PARVA |
| ENSG00000197702 | 12811.79845 | 0.983629944 | 3.10E-84 | PARVA |
| ENSG00000133818 | 2136.436366 | 1.200576442 | 1.87E-35 | RRAS2 |
| ENSG00000166833 | 2387.667239 | 0.836171488 | 1.46E-11 | NAV2 |
| ENSG00000026508 | 26250.0004 | 1.043074392 | 7.92E-20 | CD44 |
| ENSG00000026508 | 26250.0004 | 1.043074392 | 7.92E-20 | CD44 |
| ENSG00000026508 | 26250.0004 | 1.043074392 | 7.92E-20 | CD44 |
| ENSG00000135362 | 1135.682346 | 1.408426387 | 1.53E-37 | PRR5L |
| ENSG00000187479 | 147.5207521 | -0.629921572 | 6.50E-03 | C11orf96 |
| ENSG00000157613 | 742.9290274 | 1.331661607 | 4.26E-20 | CREB3L1 |
| ENSG00000157613 | 742.9290274 | 1.331661607 | 4.26E-20 | CREB3L1 |
| ENSG00000198561 | 17473.49328 | -0.493167374 | 1.58E-10 | CTNND1 |
| ENSG00000124942 | 122265.4934 | 1.312324699 | 7.24E-55 | AHNAK |
| ENSG00000124942 | 122265.4934 | 1.312324699 | 7.24E-55 | AHNAK |
| ENSG00000124942 | 122265.4934 | 1.312324699 | 7.24E-55 | AHNAK |
| ENSG00000110047 | 3523.992995 | 1.149829284 | 2.59E-14 | EHD1 |
| ENSG00000110047 | 3523.992995 | 1.149829284 | 2.59E-14 | EHD1 |
| ENSG00000168056 | 6500.71372 | -0.624296133 | 1.33E-06 | LTBP3 |
| ENSG00000175115 | 5469.333388 | 0.739178707 | 5.12E-16 | PACS1 |
| ENSG00000006534 | 1300.36037 | 0.874665701 | 1.49E-17 | ALDH3B1 |
| ENSG00000162337 | 4117.859614 | 0.760848971 | 6.74E-19 | LRP5 |

| | | | | |
|---|---|---|---|---|
| ENSG00000110092 | 9113.412653 | 1.172440106 | 1.55E-15 | CCND1 |
| ENSG00000110092 | 9113.412653 | 1.172440106 | 1.55E-15 | CCND1 |
| ENSG00000137474 | 246.39633 | -0.882477911 | 5.23E-04 | MYO7A |
| ENSG00000033327 | 601.0589633 | 0.609388694 | 3.56E-03 | GAB2 |
| ENSG00000073921 | 8631.628285 | 0.508904921 | 9.70E-06 | PICALM |
| ENSG00000150687 | 21726.22584 | 0.524920452 | 8.21E-05 | PRSS23 |
| ENSG00000137691 | 138.3243478 | -1.030916047 | 7.84E-06 | CFAP300 |
| ENSG00000137693 | 7292.138549 | 1.496556748 | 1.76E-73 | YAP1 |
| ENSG00000137693 | 7292.138549 | 1.496556748 | 1.76E-73 | YAP1 |
| ENSG00000149295 | 181.3447405 | -1.699514989 | 1.07E-12 | DRD2 |
| ENSG00000166741 | 7823.226213 | 1.644060024 | 1.03E-17 | NNMT |
| ENSG00000166741 | 7823.226213 | 1.644060024 | 1.03E-17 | NNMT |
| ENSG00000166741 | 7823.226213 | 1.644060024 | 1.03E-17 | NNMT |
| ENSG00000166741 | 7823.226213 | 1.644060024 | 1.03E-17 | NNMT |
| ENSG00000246100 | 27.39226582 | 1.883359434 | 1.53E-03 | LINC00900 |
| ENSG00000137656 | 1709.776183 | 0.296364772 | 1.63E-04 | BUD13 |
| ENSG00000149591 | 4862.193472 | 4.154828998 | 3.49E-74 | TAGLN |
| ENSG00000186174 | 9610.645557 | 0.323326848 | 1.05E-02 | BCL9L |
| ENSG00000110400 | 3123.160443 | -0.393502595 | 4.76E-02 | NECTIN1 |
| ENSG00000137642 | 4958.874962 | 0.501110345 | 2.54E-04 | SORL1 |
| ENSG00000165495 | 537.5619923 | -1.525324899 | 9.88E-21 | PKNOX2 |
| ENSG00000134954 | 13708.86452 | 0.899227179 | 6.01E-33 | ETS1 |
| ENSG00000134954 | 13708.86452 | 0.899227179 | 6.01E-33 | ETS1 |
| ENSG00000134954 | 13708.86452 | 0.899227179 | 6.01E-33 | ETS1 |
| ENSG00000134954 | 13708.86452 | 0.899227179 | 6.01E-33 | ETS1 |
| ENSG00000134954 | 13708.86452 | 0.899227179 | 6.01E-33 | ETS1 |
| ENSG00000134954 | 13708.86452 | 0.899227179 | 6.01E-33 | ETS1 |
| ENSG00000166106 | 608.3231517 | 0.845501582 | 7.59E-11 | ADAMTS15 |
| ENSG00000166106 | 608.3231517 | 0.845501582 | 7.59E-11 | ADAMTS15 |
| ENSG00000111186 | 481.1590035 | 2.338572567 | 2.38E-35 | WNT5B |
| ENSG00000059804 | 3270.065993 | 0.878659474 | 2.30E-04 | SLC2A3 |
| ENSG00000070018 | 2588.5102 | -0.812213603 | 7.70E-11 | LRP6 |
| ENSG00000171681 | 4087.275155 | -0.217907536 | 3.94E-02 | ATF7IP |
| ENSG00000079337 | 1404.252954 | -0.529698791 | 2.64E-03 | RAPGEF3 |
| ENSG00000061273 | 5153.464871 | 0.267199119 | 1.96E-03 | HDAC7 |
| ENSG00000167767 | 1730.679913 | 7.592655617 | 1.93E-267 | KRT80 |
| ENSG00000170442 | 789.1722789 | 5.14302061 | 2.71E-160 | KRT86 |
| ENSG00000170421 | 14283.57687 | 4.642117003 | 0.00E+00 | KRT8 |
| ENSG00000111057 | 26282.33084 | 3.175387246 | 3.95E-171 | KRT18 |
| ENSG00000123364 | 1296.224136 | 0.955539546 | 1.35E-13 | HOXC13 |
| ENSG00000181852 | 3274.28356 | 1.127984306 | 9.81E-11 | RNF41 |
| ENSG00000025423 | 125.5224641 | 0.741781196 | 1.09E-03 | HSD17B6 |
| ENSG00000175215 | 12475.24743 | 0.387778399 | 1.37E-07 | CTDSP2 |
| ENSG00000174099 | 1780.19995 | 1.505181526 | 4.87E-39 | MSRB3 |
| ENSG00000149948 | 5164.377752 | 1.294155138 | 7.88E-20 | HMGA2 |
| ENSG00000149948 | 5164.377752 | 1.294155138 | 7.88E-20 | HMGA2 |
| ENSG00000139289 | 5771.939055 | -0.571282357 | 7.23E-04 | PHLDA1 |

| | | | | |
|---|---|---|---|---|
| ENSG00000169372 | 389.3041389 | 0.800483045 | 5.77E-12 | CRADD |
| ENSG00000198431 | 33083.72171 | 2.593612209 | 2.98E-47 | TXNRD1 |
| ENSG00000235162 | 2759.517718 | 1.056364505 | 7.55E-29 | C12orf75 |
| ENSG00000074590 | 3145.049458 | 2.221828755 | 8.47E-89 | NUAK1 |
| ENSG00000074590 | 3145.049458 | 2.221828755 | 8.47E-89 | NUAK1 |
| ENSG00000074590 | 3145.049458 | 2.221828755 | 8.47E-89 | NUAK1 |
| ENSG00000074590 | 3145.049458 | 2.221828755 | 8.47E-89 | NUAK1 |
| ENSG00000008405 | 1344.096812 | 1.135342776 | 1.51E-19 | CRY1 |
| ENSG00000122970 | 713.7989242 | -0.704327772 | 2.41E-04 | IFT81 |
| ENSG00000135127 | 37.542591 | 3.718293734 | 1.48E-14 | BICDL1 |
| ENSG00000158023 | 107.1767467 | 2.385939353 | 5.73E-13 | WDR66 |
| ENSG00000196498 | 5055.339838 | 0.359340854 | 4.11E-03 | NCOR2 |
| ENSG00000150991 | 59339.49899 | 0.474249903 | 1.75E-05 | UBC |
| ENSG00000150457 | 922.0682349 | 1.529782224 | 1.93E-42 | LATS2 |
| ENSG00000102781 | 513.5647486 | 0.338351508 | 1.53E-02 | KATNAL1 |
| ENSG00000136141 | 1532.93364 | -0.309852978 | 1.18E-02 | LRCH1 |
| ENSG00000102543 | 324.1905473 | 0.393955673 | 4.37E-03 | CDADC1 |
| ENSG00000139734 | 4667.695024 | 1.187855396 | 5.75E-13 | DIAPH3 |
| ENSG00000136160 | 780.2042551 | 0.738363229 | 4.28E-07 | EDNRB |
| ENSG00000136158 | 1269.954807 | -0.588032876 | 9.34E-07 | SPRY2 |
| ENSG00000136158 | 1269.954807 | -0.588032876 | 9.34E-07 | SPRY2 |
| ENSG00000139793 | 5478.880166 | 0.497481869 | 3.49E-02 | MBNL2 |
| ENSG00000139793 | 5478.880166 | 0.497481869 | 3.49E-02 | MBNL2 |
| ENSG00000139793 | 5478.880166 | 0.497481869 | 3.49E-02 | MBNL2 |
| ENSG00000139793 | 5478.880166 | 0.497481869 | 3.49E-02 | MBNL2 |
| ENSG00000139793 | 5478.880166 | 0.497481869 | 3.49E-02 | MBNL2 |
| ENSG00000139793 | 5478.880166 | 0.497481869 | 3.49E-02 | MBNL2 |
| ENSG00000139793 | 5478.880166 | 0.497481869 | 3.49E-02 | MBNL2 |
| ENSG00000139793 | 5478.880166 | 0.497481869 | 3.49E-02 | MBNL2 |
| ENSG00000198542 | 61.58578191 | 3.195801831 | 8.07E-17 | ITGBL1 |
| ENSG00000198542 | 61.58578191 | 3.195801831 | 8.07E-17 | ITGBL1 |
| ENSG00000185989 | 1805.923465 | 0.462363101 | 1.21E-03 | RASA3 |
| ENSG00000100439 | 4610.109295 | 0.519024066 | 8.36E-05 | ABHD4 |
| ENSG00000129474 | 9629.715395 | 2.98002213 | 1.52E-69 | AJUBA |
| ENSG00000129474 | 9629.715395 | 2.98002213 | 1.52E-69 | AJUBA |
| ENSG00000020577 | 2581.581522 | 1.425541248 | 1.11E-29 | SAMD4A |
| ENSG00000020577 | 2581.581522 | 1.425541248 | 1.11E-29 | SAMD4A |
| ENSG00000020577 | 2581.581522 | 1.425541248 | 1.11E-29 | SAMD4A |
| ENSG00000027075 | 883.6995121 | 0.703860761 | 6.26E-10 | PRKCH |
| ENSG00000232774 | 143.2120041 | 2.445557406 | 2.38E-32 | AL355916.1 |
| ENSG00000205683 | 481.2364414 | -0.726174595 | 2.18E-04 | DPF3 |
| ENSG00000070778 | 1232.53051 | 0.957847939 | 2.03E-22 | AL162171.1 |
| ENSG00000205476 | 2884.103042 | 1.49178026 | 7.03E-33 | CCDC85C |
| ENSG00000128849 | 895.5418779 | 1.068789398 | 3.38E-13 | CGNL1 |
| ENSG00000263155 | 230.2645852 | 0.545945604 | 5.89E-03 | MYZAP |
| ENSG00000166035 | 496.2256483 | 0.52290644 | 1.25E-03 | LIPC |
| ENSG00000137845 | 7600.927323 | -0.6431847 | 2.12E-08 | ADAM10 |

| | | | | |
|---|---|---|---|---|
| ENSG00000128923 | 1015.699395 | -0.817667972 | 9.90E-06 | MINDY2 |
| ENSG00000182718 | 106890.7943 | 1.133497065 | 2.60E-44 | ANXA2 |
| ENSG00000140416 | 23789.68 | 1.806728813 | 7.03E-65 | TPM1 |
| ENSG00000140416 | 23789.68 | 1.806728813 | 7.03E-65 | TPM1 |
| ENSG00000103642 | 732.0504195 | 1.298021657 | 1.48E-23 | LACTB |
| ENSG00000174485 | 1554.563487 | -0.447264797 | 3.00E-02 | DENND4A |
| ENSG00000137834 | 1299.992546 | 1.17263253 | 1.02E-14 | SMAD6 |
| ENSG00000137834 | 1299.992546 | 1.17263253 | 1.02E-14 | SMAD6 |
| ENSG00000137834 | 1299.992546 | 1.17263253 | 1.02E-14 | SMAD6 |
| ENSG00000137834 | 1299.992546 | 1.17263253 | 1.02E-14 | SMAD6 |
| ENSG00000166949 | 7634.379311 | 0.367479333 | 5.13E-05 | SMAD3 |
| ENSG00000166949 | 7634.379311 | 0.367479333 | 5.13E-05 | SMAD3 |
| ENSG00000166949 | 7634.379311 | 0.367479333 | 5.13E-05 | SMAD3 |
| ENSG00000166949 | 7634.379311 | 0.367479333 | 5.13E-05 | SMAD3 |
| ENSG00000166949 | 7634.379311 | 0.367479333 | 5.13E-05 | SMAD3 |
| ENSG00000166949 | 7634.379311 | 0.367479333 | 5.13E-05 | SMAD3 |
| ENSG00000137868 | 466.7762795 | -1.596373508 | 1.13E-14 | STRA6 |
| ENSG00000137868 | 466.7762795 | -1.596373508 | 1.13E-14 | STRA6 |
| ENSG00000058335 | 134.7074287 | 2.893702412 | 2.93E-22 | RASGRF1 |
| ENSG00000172379 | 978.2147149 | 2.345531993 | 1.77E-55 | ARNT2 |
| ENSG00000156232 | 724.8447879 | 0.950625558 | 3.26E-12 | WHAMM |
| ENSG00000169594 | 1562.452238 | 1.845566869 | 2.10E-19 | BNC1 |
| ENSG00000140575 | 16665.12339 | 0.553503957 | 1.43E-07 | IQGAP1 |
| ENSG00000185551 | 5643.974102 | 0.577229389 | 1.27E-06 | NR2F2 |
| ENSG00000185551 | 5643.974102 | 0.577229389 | 1.27E-06 | NR2F2 |
| ENSG00000131873 | 3084.922925 | -0.45602812 | 5.55E-09 | CHSY1 |
| ENSG00000007384 | 1735.657285 | 0.559783851 | 2.08E-06 | RHBDF1 |
| ENSG00000162078 | 226.8742984 | 0.518549863 | 9.79E-03 | ZG16B |
| ENSG00000006327 | 3033.22403 | 3.33949236 | 1.75E-109 | TNFRSF12A |
| ENSG00000006327 | 3033.22403 | 3.33949236 | 1.75E-109 | TNFRSF12A |
| ENSG00000166501 | 1065.831152 | -1.655506666 | 3.34E-20 | PRKCB |
| ENSG00000122257 | 2157.766372 | -0.350759871 | 4.18E-02 | RBBP6 |
| ENSG00000122257 | 2157.766372 | -0.350759871 | 4.18E-02 | RBBP6 |
| ENSG00000122257 | 2157.766372 | -0.350759871 | 4.18E-02 | RBBP6 |
| ENSG00000090905 | 4408.836177 | -0.471882334 | 1.73E-05 | TNRC6A |
| ENSG00000090905 | 4408.836177 | -0.471882334 | 1.73E-05 | TNRC6A |
| ENSG00000149925 | 42268.47046 | 0.537814681 | 6.63E-05 | ALDOA |
| ENSG00000052344 | 17.4747297 | 2.062348289 | 7.72E-04 | PRSS8 |
| ENSG00000140795 | 310.1611264 | 1.711528806 | 2.27E-24 | MYLK3 |
| ENSG00000177200 | 2148.656633 | -1.257208278 | 3.85E-08 | CHD9 |
| ENSG00000102900 | 5090.165098 | 0.412150592 | 4.54E-07 | NUP93 |
| ENSG00000135736 | 865.6513979 | -0.371371302 | 3.63E-02 | CCDC102A |
| ENSG00000172840 | 1653.370186 | 0.601947145 | 1.28E-10 | PDP2 |
| ENSG00000124067 | 2661.801366 | 1.224363279 | 1.41E-17 | SLC12A4 |
| ENSG00000062038 | 1193.523897 | 0.906880617 | 1.79E-12 | CDH3 |
| ENSG00000039068 | 23.87917036 | 3.136967277 | 1.17E-05 | CDH1 |
| ENSG00000039068 | 23.87917036 | 3.136967277 | 1.17E-05 | CDH1 |

| | | | | |
|---|---|---|---|---|
| ENSG00000132613 | 4140.209056 | 1.124432269 | 3.26E-26 | MTSS1L |
| ENSG00000140832 | 105.4161867 | -1.47190961 | 1.15E-08 | MARVELD3 |
| ENSG00000050820 | 5994.364749 | 0.407616191 | 3.93E-04 | BCAR1 |
| ENSG00000050820 | 5994.364749 | 0.407616191 | 3.93E-04 | BCAR1 |
| ENSG00000171724 | 15308.2066 | -0.34407068 | 6.94E-03 | VAT1L |
| ENSG00000171724 | 15308.2066 | -0.34407068 | 6.94E-03 | VAT1L |
| ENSG00000166451 | 2368.501306 | 0.553452105 | 3.08E-04 | CENPN |
| ENSG00000184860 | 246.9387746 | 2.085926158 | 9.95E-23 | SDR42E1 |
| ENSG00000140945 | 11715.24575 | 0.779044434 | 5.43E-11 | CDH13 |
| ENSG00000140945 | 11715.24575 | 0.779044434 | 5.43E-11 | CDH13 |
| ENSG00000140950 | 2844.519966 | 0.881368943 | 1.07E-13 | MEAK7 |
| ENSG00000225614 | 430.1866375 | 1.345121251 | 9.16E-12 | ZNF469 |
| ENSG00000167693 | 1791.096427 | 1.934863089 | 1.05E-59 | NXN |
| ENSG00000167693 | 1791.096427 | 1.934863089 | 1.05E-59 | NXN |
| ENSG00000167693 | 1791.096427 | 1.934863089 | 1.05E-59 | NXN |
| ENSG00000197879 | 10838.86687 | 1.230741327 | 1.01E-26 | MYO1C |
| ENSG00000132386 | 435.8532755 | -1.455416043 | 4.74E-17 | SERPINF1 |
| ENSG00000213977 | 4806.333429 | 1.056596244 | 7.03E-18 | TAX1BP3 |
| ENSG00000213859 | 1277.421524 | 0.691127389 | 9.27E-07 | KCTD11 |
| ENSG00000276231 | 45.75679108 | 1.067170092 | 1.56E-03 | PIK3R6 |
| ENSG00000154914 | 231.4230637 | 1.889042835 | 8.08E-28 | USP43 |
| ENSG00000175662 | 4289.413806 | 0.454992127 | 4.91E-04 | TOM1L2 |
| ENSG00000275832 | 5404.484757 | 1.122243191 | 5.86E-37 | ARHGAP23 |
| ENSG00000275832 | 5404.484757 | 1.122243191 | 5.86E-37 | ARHGAP23 |
| ENSG00000188895 | 3890.623421 | -0.379501794 | 1.24E-05 | MSL1 |
| ENSG00000177469 | 22717.39463 | 1.689564489 | 2.24E-38 | CAVIN1 |
| ENSG00000175832 | 2074.53517 | -1.30621195 | 3.43E-09 | ETV4 |
| ENSG00000180340 | 1127.496047 | 1.089366535 | 1.05E-16 | FZD2 |
| ENSG00000159314 | 474.1617742 | -0.487731324 | 2.06E-02 | ARHGAP27 |
| ENSG00000159217 | 4568.368856 | 0.444338346 | 2.89E-10 | IGF2BP1 |
| ENSG00000204584 | 22.04428444 | 1.974876834 | 1.69E-04 | FLJ45513 |
| ENSG00000246640 | 40.09760807 | 2.306534101 | 1.26E-10 | PICART1 |
| ENSG00000108821 | 951.6101759 | 1.445112198 | 3.52E-09 | COL1A1 |
| ENSG00000108821 | 951.6101759 | 1.445112198 | 3.52E-09 | COL1A1 |
| ENSG00000108960 | 3317.81114 | -1.8052227 | 1.26E-64 | MMD |
| ENSG00000183691 | 208.1326203 | 2.661632609 | 7.78E-18 | NOG |
| ENSG00000183691 | 208.1326203 | 2.661632609 | 7.78E-18 | NOG |
| ENSG00000183691 | 208.1326203 | 2.661632609 | 7.78E-18 | NOG |
| ENSG00000153944 | 3793.636912 | -0.60881878 | 2.18E-14 | MSI2 |
| ENSG00000181610 | 2563.61453 | 0.432014944 | 1.40E-07 | MRPS23 |
| ENSG00000062716 | 6417.676705 | 0.333750315 | 2.72E-03 | VMP1 |
| ENSG00000075461 | 946.3481827 | 1.503253182 | 4.89E-16 | CACNG4 |
| ENSG00000125398 | 2034.137963 | -0.714625467 | 4.09E-08 | SOX9 |
| ENSG00000073350 | 1015.83685 | 1.443722946 | 7.88E-45 | LLGL2 |
| ENSG00000184557 | 897.8181479 | 1.175959172 | 2.23E-05 | SOCS3 |
| ENSG00000035862 | 22643.24137 | 0.561238916 | 6.59E-05 | TIMP2 |
| ENSG00000266074 | 1925.241566 | -0.481660967 | 2.39E-03 | BAHCC1 |

| | | | | |
|---|---|---|---|---|
| ENSG00000176845 | 538.6256251 | 1.649327975 | 6.20E-27 | METRNL |
| ENSG00000101608 | 9438.49279 | 1.075116123 | 2.05E-37 | MYL12A |
| ENSG00000118680 | 10882.15315 | 1.265432765 | 2.55E-95 | MYL12B |
| ENSG00000154864 | 3798.179918 | -1.241257333 | 7.72E-49 | PIEZO2 |
| ENSG00000176014 | 6630.385277 | 1.622740739 | 5.13E-24 | TUBB6 |
| ENSG00000101782 | 2038.172139 | 0.42904334 | 3.69E-03 | RIOK3 |
| ENSG00000132874 | 38.02249123 | -1.709996199 | 5.25E-05 | SLC14A2 |
| ENSG00000101665 | 443.250496 | 0.732494727 | 1.20E-03 | SMAD7 |
| ENSG00000198796 | 2595.841625 | 3.05642035 | 5.31E-43 | ALPK2 |
| ENSG00000166573 | 16.93168639 | -1.920086837 | 2.20E-02 | GALR1 |
| ENSG00000226742 | 284.4549299 | 1.298451394 | 2.24E-14 | HSBP1L1 |
| ENSG00000011304 | 14317.09673 | 0.343686763 | 9.48E-04 | PTBP1 |
| ENSG00000064666 | 8063.875845 | 1.385288766 | 5.99E-55 | CNN2 |
| ENSG00000160953 | 2910.551855 | -0.197067128 | 9.69E-03 | PWWP3A |
| ENSG00000176619 | 11542.53977 | 0.515534353 | 2.23E-05 | LMNB2 |
| ENSG00000099860 | 977.0968945 | 2.948432547 | 2.81E-150 | GADD45B |
| ENSG00000099860 | 977.0968945 | 2.948432547 | 2.81E-150 | GADD45B |
| ENSG00000125733 | 3342.101072 | 0.588537466 | 8.29E-22 | TRIP10 |
| ENSG00000080573 | 625.5339231 | -0.583078072 | 9.22E-03 | COL5A3 |
| ENSG00000008441 | 4322.548575 | 0.823560494 | 2.65E-17 | NFIX |
| ENSG00000008441 | 4322.548575 | 0.823560494 | 2.65E-17 | NFIX |
| ENSG00000105137 | 3531.235541 | 1.114608601 | 7.12E-20 | SYDE1 |
| ENSG00000160117 | 635.5348284 | -0.377979304 | 2.49E-03 | ANKLE1 |
| ENSG00000142279 | 955.4419514 | 2.308478611 | 5.49E-35 | WTIP |
| ENSG00000104835 | 871.2070724 | -0.493160923 | 9.16E-04 | SARS2 |
| ENSG00000105193 | 19557.72628 | 0.342074846 | 8.72E-03 | RPS16 |
| ENSG00000167601 | 11912.61544 | 3.198676255 | 1.04E-80 | AXL |
| ENSG00000216588 | 35.73858368 | 3.030178483 | 4.17E-16 | IGSF23 |
| ENSG00000104884 | 3374.31351 | 1.213072723 | 1.05E-26 | ERCC2 |
| ENSG00000268460 | 30.28041748 | 0.99870528 | 1.27E-02 | AC006262.1 |
| ENSG00000142230 | 11296.4598 | 0.535864296 | 1.77E-11 | SAE1 |
| ENSG00000024422 | 10312.92724 | 0.797585316 | 7.83E-13 | EHD2 |
| ENSG00000105373 | 6803.000535 | 0.344297525 | 3.16E-02 | NOP53 |
| ENSG00000126458 | 2730.533569 | 0.700994899 | 1.66E-06 | RRAS |
| ENSG00000179820 | 3260.603661 | 1.269940585 | 1.20E-28 | MYADM |
| ENSG00000268654 | 17.01502868 | -1.403360214 | 2.26E-02 | MIMT1 |
| ENSG00000130508 | 21810.67112 | 0.271883229 | 2.14E-03 | PXDN |
| ENSG00000130508 | 21810.67112 | 0.271883229 | 2.14E-03 | PXDN |
| ENSG00000264370 | 7.117429883 | -3.876098581 | 6.03E-03 | MIR3125 |
| ENSG00000132031 | 531.0146682 | 1.772885735 | 8.01E-35 | MATN3 |
| ENSG00000119771 | 1265.210865 | 0.984978247 | 1.28E-10 | KLHL29 |
| ENSG00000075426 | 5985.00852 | 1.169947569 | 2.27E-27 | FOSL2 |
| ENSG00000013016 | 6392.771573 | -1.189668697 | 2.04E-16 | EHD3 |
| ENSG00000150938 | 9816.058979 | 2.438982844 | 7.79E-106 | CRIM1 |
| ENSG00000150938 | 9816.058979 | 2.438982844 | 7.79E-106 | CRIM1 |
| ENSG00000163171 | 1854.372981 | 3.105214755 | 1.04E-44 | CDC42EP3 |
| ENSG00000163171 | 1854.372981 | 3.105214755 | 1.04E-44 | CDC42EP3 |

| | | | | |
|---|---|---|---|---|
| ENSG00000143924 | 8450.371969 | -1.266649032 | 6.79E-37 | EML4 |
| ENSG00000152518 | 1694.899588 | -0.250930154 | 2.02E-02 | ZFP36L2 |
| ENSG00000152518 | 1694.899588 | -0.250930154 | 2.02E-02 | ZFP36L2 |
| ENSG00000152518 | 1694.899588 | -0.250930154 | 2.02E-02 | ZFP36L2 |
| ENSG00000171132 | 1501.609574 | 1.827231423 | 1.06E-18 | PRKCE |
| ENSG00000116016 | 9233.833509 | 0.864745175 | 9.52E-19 | EPAS1 |
| ENSG00000116016 | 9233.833509 | 0.864745175 | 9.52E-19 | EPAS1 |
| ENSG00000239332 | 98.2556091 | 1.473124067 | 3.98E-08 | LINC01119 |
| ENSG00000068724 | 1548.824814 | 1.099712786 | 2.23E-29 | TTC7A |
| ENSG00000068724 | 1548.824814 | 1.099712786 | 2.23E-29 | TTC7A |
| ENSG00000115306 | 22759.67806 | -0.445719781 | 4.34E-15 | SPTBN1 |
| ENSG00000179833 | 1650.89911 | 1.079206742 | 5.94E-24 | SERTAD2 |
| ENSG00000179833 | 1650.89911 | 1.079206742 | 5.94E-24 | SERTAD2 |
| ENSG00000135636 | 2529.150232 | 0.837814203 | 8.09E-09 | DYSF |
| ENSG00000186854 | 897.350399 | 0.987950091 | 2.10E-20 | TRABD2A |
| ENSG00000135945 | 1476.613051 | -0.388212075 | 1.33E-02 | REV1 |
| ENSG00000204634 | 2589.114755 | -0.454573307 | 2.87E-09 | TBC1D8 |
| ENSG00000071054 | 13426.54364 | -0.626051304 | 3.36E-24 | MAP4K4 |
| ENSG00000115590 | 12.0580723 | -2.205566462 | 2.99E-03 | IL1R2 |
| ENSG00000074047 | 1186.161085 | 0.899172303 | 6.97E-07 | GLI2 |
| ENSG00000074047 | 1186.161085 | 0.899172303 | 6.97E-07 | GLI2 |
| ENSG00000074047 | 1186.161085 | 0.899172303 | 6.97E-07 | GLI2 |
| ENSG00000072163 | 478.4740095 | 3.741328975 | 6.07E-98 | LIMS2 |
| ENSG00000136720 | 1299.442214 | 1.365340025 | 1.19E-16 | HS6ST1 |
| ENSG00000144283 | 3485.234126 | 0.500713642 | 3.38E-05 | PKP4 |
| ENSG00000115844 | 377.9123392 | -0.792234399 | 5.51E-04 | DLX2 |
| ENSG00000091428 | 1768.711438 | -0.861659202 | 8.30E-12 | RAPGEF4 |
| ENSG00000168497 | 121.5099942 | 7.175347267 | 9.06E-35 | CAVIN2 |
| ENSG00000123992 | 2813.70088 | 0.428262244 | 3.45E-05 | DNPEP |
| ENSG00000244280 | 5.851314691 | 2.230888476 | 2.90E-02 | ECEL1P2 |
| ENSG00000188042 | 1081.703425 | 1.426287135 | 9.91E-25 | ARL4C |
| ENSG00000130147 | 4262.928303 | 1.704774805 | 2.86E-65 | SH3BP4 |
| ENSG00000130147 | 4262.928303 | 1.704774805 | 2.86E-65 | SH3BP4 |
| ENSG00000163359 | 4751.934988 | -1.023217403 | 2.55E-05 | COL6A3 |
| ENSG00000163359 | 4751.934988 | -1.023217403 | 2.55E-05 | COL6A3 |
| ENSG00000177732 | 4342.362057 | -0.268085631 | 1.20E-02 | SOX12 |
| ENSG00000132635 | 3718.966878 | -0.214872424 | 7.03E-03 | PCED1A |
| ENSG00000125772 | 2138.032665 | -1.195336303 | 3.40E-08 | GPCPD1 |
| ENSG00000171984 | 405.9876561 | -0.756336997 | 4.31E-07 | SHLD1 |
| ENSG00000172264 | 76.02264208 | -0.668063508 | 4.19E-02 | MACROD2 |
| ENSG00000089006 | 7442.90652 | 0.314058688 | 8.37E-04 | SNX5 |
| ENSG00000125798 | 107.3111875 | 1.094613206 | 5.20E-05 | FOXA2 |
| ENSG00000125798 | 107.3111875 | 1.094613206 | 5.20E-05 | FOXA2 |
| ENSG00000171552 | 5620.751235 | 1.818596542 | 3.80E-108 | BCL2L1 |
| ENSG00000171552 | 5620.751235 | 1.818596542 | 3.80E-108 | BCL2L1 |
| ENSG00000078747 | 4592.164107 | 0.326966382 | 1.22E-02 | ITCH |
| ENSG00000132821 | 409.4949088 | 3.096817337 | 8.25E-66 | VSTM2L |

| | | | | |
|---|---|---|---|---|
| ENSG00000132823 | 2909.229889 | 0.595781932 | 7.42E-14 | OSER1 |
| ENSG00000223891 | 334.8931331 | 0.425466188 | 1.17E-02 | OSER1-DT |
| ENSG00000168734 | 2587.098172 | 0.727044228 | 1.17E-06 | PKIG |
| ENSG00000124145 | 4176.418327 | 1.721108012 | 3.32E-34 | SDC4 |
| ENSG00000198026 | 1385.685245 | -0.200729708 | 3.33E-02 | ZNF335 |
| ENSG00000101040 | 5516.740769 | -0.473603966 | 3.95E-06 | ZMYND8 |
| ENSG00000101040 | 5516.740769 | -0.473603966 | 3.95E-06 | ZMYND8 |
| ENSG00000124126 | 4692.440271 | 0.448252634 | 3.20E-06 | PREX1 |
| ENSG00000196396 | 5738.112123 | 0.453820411 | 4.29E-10 | PTPN1 |
| ENSG00000124243 | 1194.623206 | 0.751132225 | 7.13E-08 | BCAS4 |
| ENSG00000171940 | 4777.839968 | -0.284547502 | 3.10E-03 | ZNF217 |
| ENSG00000171940 | 4777.839968 | -0.284547502 | 3.10E-03 | ZNF217 |
| ENSG00000019186 | 230.7349006 | 1.084260869 | 1.24E-06 | CYP24A1 |
| ENSG00000101144 | 617.2385659 | -2.465790858 | 3.97E-24 | BMP7 |
| ENSG00000092758 | 4082.534637 | -0.769084839 | 2.54E-06 | COL9A3 |
| ENSG00000154639 | 430.9583909 | 1.088675926 | 1.43E-08 | CXADR |
| ENSG00000156273 | 2018.700707 | 0.637138201 | 1.99E-03 | BACH1 |
| ENSG00000159200 | 3868.497247 | 1.691601081 | 3.12E-38 | RCAN1 |
| ENSG00000160191 | 550.0523679 | 0.41613068 | 2.37E-03 | PDE9A |
| ENSG00000183255 | 16185.65895 | 1.262106492 | 1.36E-34 | PTTG1IP |
| ENSG00000160255 | 124.990794 | 4.512183942 | 1.05E-43 | ITGB2 |
| ENSG00000184470 | 1698.664478 | 0.969891323 | 2.00E-21 | TXNRD2 |
| ENSG00000099917 | 3949.807379 | 0.580186947 | 5.55E-10 | MED15 |
| ENSG00000099994 | 605.236627 | 2.137551043 | 1.31E-34 | SUSD2 |
| ENSG00000128203 | 456.4862353 | 0.96477011 | 7.38E-18 | ASPHD2 |
| ENSG00000128342 | 154.155482 | 1.243210766 | 3.53E-05 | LIF |
| ENSG00000198125 | 5.190958042 | 4.128701746 | 3.91E-03 | MB |
| ENSG00000100345 | 39701.66481 | 1.695973263 | 6.55E-63 | MYH9 |
| ENSG00000100345 | 39701.66481 | 1.695973263 | 6.55E-63 | MYH9 |
| ENSG00000100065 | 1474.574126 | 1.095791657 | 3.79E-12 | CARD10 |
| ENSG00000128283 | 3906.296252 | 1.66022256 | 5.55E-20 | CDC42EP1 |
| ENSG00000128283 | 3906.296252 | 1.66022256 | 5.55E-20 | CDC42EP1 |
| ENSG00000185022 | 607.0319197 | 2.042904178 | 1.07E-28 | MAFF |
| ENSG00000100307 | 937.6131425 | 0.978174434 | 1.42E-10 | CBX7 |
| ENSG00000128268 | 1251.709054 | 0.968942726 | 4.77E-08 | MGAT3 |
| ENSG00000100401 | 7733.942925 | 0.651366095 | 4.83E-16 | RANGAP1 |
| ENSG00000100138 | 7127.161613 | 0.229772096 | 2.89E-02 | SNU13 |
| ENSG00000130638 | 5946.981823 | 0.632411445 | 2.22E-11 | ATXN10 |
| ENSG00000196576 | 10147.39702 | 0.693129203 | 4.58E-12 | PLXNB2 |
| ENSG00000144560 | 1747.13902 | -0.334142349 | 1.69E-02 | VGLL4 |
| ENSG00000144560 | 1747.13902 | -0.334142349 | 1.69E-02 | VGLL4 |
| ENSG00000144711 | 2478.420337 | 0.382229125 | 6.27E-06 | IQSEC1 |
| ENSG00000131378 | 2508.131971 | 0.976858737 | 4.01E-18 | RFTN1 |
| ENSG00000173402 | 8402.798769 | 0.953859743 | 3.36E-18 | DAG1 |
| ENSG00000012171 | 1213.282702 | 3.544449289 | 1.25E-71 | SEMA3B |
| ENSG00000136068 | 29152.77127 | 1.894065626 | 1.20E-47 | FLNB |
| ENSG00000144824 | 5271.381741 | 0.979027565 | 4.40E-09 | PHLDB2 |

| | | | | |
|---|---|---|---|---|
| ENSG00000144824 | 5271.381741 | 0.979027565 | 4.40E-09 | PHLDB2 |
| ENSG00000082781 | 9170.149928 | -0.532078806 | 3.15E-03 | ITGB5 |
| ENSG00000082781 | 9170.149928 | -0.532078806 | 3.15E-03 | ITGB5 |
| ENSG00000173706 | 10780.54503 | 1.561382908 | 1.49E-99 | HEG1 |
| ENSG00000004399 | 13090.28614 | 0.343019483 | 2.48E-02 | PLXND1 |
| ENSG00000114019 | 5194.214834 | 3.908633168 | 0.00E+00 | AMOTL2 |
| ENSG00000114019 | 5194.214834 | 3.908633168 | 0.00E+00 | AMOTL2 |
| ENSG00000114019 | 5194.214834 | 3.908633168 | 0.00E+00 | AMOTL2 |
| ENSG00000114019 | 5194.214834 | 3.908633168 | 0.00E+00 | AMOTL2 |
| ENSG00000114019 | 5194.214834 | 3.908633168 | 0.00E+00 | AMOTL2 |
| ENSG00000114019 | 5194.214834 | 3.908633168 | 0.00E+00 | AMOTL2 |
| ENSG00000168917 | 48.39117825 | 0.768444574 | 4.63E-02 | SLC35G2 |
| ENSG00000163584 | 2206.738069 | 0.305701355 | 2.11E-02 | RPL22L1 |
| ENSG00000075420 | 3823.40014 | -0.510424873 | 2.18E-02 | FNDC3B |
| ENSG00000228221 | 8.053509764 | -2.63823571 | 9.54E-03 | LINC00578 |
| ENSG00000228221 | 8.053509764 | -2.63823571 | 9.54E-03 | LINC00578 |
| ENSG00000182580 | 918.3399535 | -0.711419928 | 5.03E-07 | EPHB3 |
| ENSG00000180611 | 492.152103 | 1.204761574 | 6.54E-29 | MB21D2 |
| ENSG00000180611 | 492.152103 | 1.204761574 | 6.54E-29 | MB21D2 |
| ENSG00000173950 | 1532.304719 | -0.264792978 | 2.18E-02 | XXYLT1 |
| ENSG00000068078 | 194.2592122 | 1.963993087 | 3.99E-10 | FGFR3 |
| ENSG00000196526 | 8267.209501 | 0.579882137 | 6.87E-10 | AFAP1 |
| ENSG00000163697 | 2409.951598 | 0.768676197 | 2.93E-11 | APBB2 |
| ENSG00000138771 | 3719.952372 | 1.151313416 | 3.16E-29 | SHROOM3 |
| ENSG00000138772 | 816.3716511 | 3.398650038 | 7.32E-72 | ANXA3 |
| ENSG00000173085 | 766.2534857 | 0.405794626 | 9.92E-05 | COQ2 |
| ENSG00000249464 | 34.16317966 | -3.874173877 | 5.74E-15 | LINC01091 |
| ENSG00000077684 | 1188.702036 | 0.751534371 | 3.48E-05 | JADE1 |
| ENSG00000077684 | 1188.702036 | 0.751534371 | 3.48E-05 | JADE1 |
| ENSG00000109452 | 1588.237089 | 1.049844951 | 5.30E-15 | INPP4B |
| ENSG00000052795 | 1389.398079 | -0.625273139 | 1.71E-08 | FNIP2 |
| ENSG00000251165 | 217.3634243 | 0.765455208 | 1.05E-02 | F11-AS1 |
| ENSG00000215246 | 130.5665582 | 2.11330432 | 5.12E-16 | AC116351.1 |
| ENSG00000049656 | 15633.00909 | 0.453293676 | 1.74E-07 | CLPTM1L |
| ENSG00000164236 | 166.908834 | 1.699949145 | 1.49E-17 | ANKRD33B |
| ENSG00000164236 | 166.908834 | 1.699949145 | 1.49E-17 | ANKRD33B |
| ENSG00000164236 | 166.908834 | 1.699949145 | 1.49E-17 | ANKRD33B |
| ENSG00000039139 | 297.4360865 | 0.404079956 | 2.36E-02 | DNAH5 |
| ENSG00000039139 | 297.4360865 | 0.404079956 | 2.36E-02 | DNAH5 |
| ENSG00000038382 | 14689.77273 | 0.745652083 | 3.10E-16 | TRIO |
| ENSG00000176788 | 2291.351755 | 2.817583098 | 3.99E-80 | BASP1 |
| ENSG00000183666 | 518.3979596 | -0.580532905 | 1.66E-05 | GUSBP1 |
| ENSG00000113360 | 5892.566378 | -0.294455051 | 3.45E-04 | DROSHA |
| ENSG00000133401 | 5704.21073 | 0.328400238 | 3.11E-02 | PDZD2 |
| ENSG00000151388 | 10377.03832 | 0.99066368 | 1.61E-22 | ADAMTS12 |
| ENSG00000151388 | 10377.03832 | 0.99066368 | 1.61E-22 | ADAMTS12 |
| ENSG00000039560 | 6837.00494 | 0.414674154 | 3.15E-02 | RAI14 |

| | | | | |
|---|---|---|---|---|
| ENSG00000039560 | 6837.00494 | 0.414674154 | 3.15E-02 | RAI14 |
| ENSG00000113594 | 13533.60101 | -1.751473269 | 9.88E-18 | LIFR |
| ENSG00000171522 | 362.933959 | -0.685882544 | 4.46E-02 | PTGER4 |
| ENSG00000152684 | 2799.47239 | 0.614980176 | 7.47E-15 | PELO |
| ENSG00000164294 | 5676.176415 | 0.640834267 | 3.51E-07 | GPX8 |
| ENSG00000177058 | 1524.406775 | -0.327726088 | 5.87E-03 | SLC38A9 |
| ENSG00000134352 | 22565.49008 | -1.208308474 | 1.20E-04 | IL6ST |
| ENSG00000164512 | 43.24059336 | 1.138823993 | 2.10E-03 | ANKRD55 |
| ENSG00000164512 | 43.24059336 | 1.138823993 | 2.10E-03 | ANKRD55 |
| ENSG00000164512 | 43.24059336 | 1.138823993 | 2.10E-03 | ANKRD55 |
| ENSG00000095015 | 6834.516553 | 0.372893754 | 1.45E-03 | MAP3K1 |
| ENSG00000169067 | 62.37567461 | 2.235655734 | 2.75E-10 | ACTBL2 |
| ENSG00000169067 | 62.37567461 | 2.235655734 | 2.75E-10 | ACTBL2 |
| ENSG00000145632 | 8444.080224 | -0.457675495 | 1.00E-04 | PLK2 |
| ENSG00000113448 | 2698.714731 | -1.949685301 | 4.98E-24 | PDE4D |
| ENSG00000113448 | 2698.714731 | -1.949685301 | 4.98E-24 | PDE4D |
| ENSG00000130449 | 1861.149076 | 0.426135224 | 2.30E-06 | ZSWIM6 |
| ENSG00000130449 | 1861.149076 | 0.426135224 | 2.30E-06 | ZSWIM6 |
| ENSG00000134061 | 81.62185089 | -5.177747191 | 7.87E-17 | CD180 |
| ENSG00000134061 | 81.62185089 | -5.177747191 | 7.87E-17 | CD180 |
| ENSG00000145675 | 4595.722417 | -1.977630202 | 5.60E-69 | PIK3R1 |
| ENSG00000197822 | 724.0860483 | 2.636221107 | 9.49E-34 | OCLN |
| ENSG00000131711 | 20810.43855 | 0.746524055 | 2.33E-04 | MAP1B |
| ENSG00000214944 | 3310.07867 | 1.910476554 | 1.52E-34 | ARHGEF28 |
| ENSG00000237187 | 377.042445 | -1.6087087 | 3.32E-22 | NR2F1-AS1 |
| ENSG00000237187 | 377.042445 | -1.6087087 | 3.32E-22 | NR2F1-AS1 |
| ENSG00000164292 | 58008.22119 | 1.079446649 | 2.11E-15 | RHOBTB3 |
| ENSG00000175426 | 1402.5926 | -3.123620584 | 4.26E-53 | PCSK1 |
| ENSG00000186952 | 61.95177943 | -0.788433286 | 1.17E-02 | TMEM232 |
| ENSG00000134986 | 12921.79755 | -2.31527136 | 5.50E-26 | NREP |
| ENSG00000064652 | 1531.40493 | 0.785184449 | 1.99E-11 | SNX24 |
| ENSG00000168916 | 2639.516423 | -0.59642762 | 1.62E-09 | ZNF608 |
| ENSG00000168916 | 2639.516423 | -0.59642762 | 1.62E-09 | ZNF608 |
| ENSG00000155324 | 728.6384239 | 0.8068512 | 1.29E-05 | GRAMD2B |
| ENSG00000158987 | 3147.194901 | -0.428639178 | 8.62E-03 | RAPGEF6 |
| ENSG00000197375 | 1353.716616 | 0.848112162 | 5.87E-15 | SLC22A5 |
| ENSG00000120708 | 26371.45713 | 0.389657595 | 2.85E-03 | TGFBI |
| ENSG00000120708 | 26371.45713 | 0.389657595 | 2.85E-03 | TGFBI |
| ENSG00000120708 | 26371.45713 | 0.389657595 | 2.85E-03 | TGFBI |
| ENSG00000132915 | 35.90315679 | 1.714067358 | 5.84E-07 | PDE6A |
| ENSG00000171992 | 2426.228123 | 1.330966963 | 5.06E-31 | SYNPO |
| ENSG00000171992 | 2426.228123 | 1.330966963 | 5.06E-31 | SYNPO |
| ENSG00000164574 | 5382.628084 | 0.386925517 | 2.41E-04 | GALNT10 |
| ENSG00000055163 | 654.8815999 | 2.330770875 | 3.38E-22 | CYFIP2 |
| ENSG00000170234 | 1799.661611 | -0.226820279 | 3.43E-02 | PWWP2A |
| ENSG00000113327 | 26.96306728 | -1.134571235 | 2.12E-02 | GABRG2 |
| ENSG00000170584 | 2588.159888 | 0.228541499 | 2.87E-02 | NUDCD2 |

| | | | | |
|---|---|---|---|---|
| ENSG00000184845 | 26.45660106 | 1.361070962 | 7.17E-03 | DRD1 |
| ENSG00000153046 | 1827.428318 | 0.398023778 | 5.52E-04 | CDYL |
| ENSG00000124788 | 3041.292894 | -0.947733958 | 2.04E-22 | ATXN1 |
| ENSG00000137331 | 2184.936652 | 0.643325579 | 2.09E-06 | IER3 |
| ENSG00000204580 | 3398.200686 | 0.286001906 | 3.78E-02 | DDR1 |
| ENSG00000137309 | 9467.267639 | 2.413551812 | 1.59E-77 | HMGA1 |
| ENSG00000137193 | 1122.782319 | 2.747930724 | 5.91E-44 | PIM1 |
| ENSG00000137193 | 1122.782319 | 2.747930724 | 5.91E-44 | PIM1 |
| ENSG00000124813 | 681.1290161 | 0.9044351 | 4.10E-09 | RUNX2 |
| ENSG00000119899 | 973.1245904 | 0.542953893 | 1.08E-03 | SLC17A5 |
| ENSG00000119899 | 973.1245904 | 0.542953893 | 1.08E-03 | SLC17A5 |
| ENSG00000111799 | 7146.720884 | 3.914485629 | 2.64E-196 | COL12A1 |
| ENSG00000111799 | 7146.720884 | 3.914485629 | 2.64E-196 | COL12A1 |
| ENSG00000118407 | 45.8162629 | 1.797486879 | 2.56E-10 | FILIP1 |
| ENSG00000135318 | 5643.186994 | 0.848971581 | 4.86E-07 | NT5E |
| ENSG00000203875 | 2995.960911 | 0.630920536 | 1.27E-07 | SNHG5 |
| ENSG00000112182 | 445.9839007 | -0.422281714 | 1.93E-02 | BACH2 |
| ENSG00000112182 | 445.9839007 | -0.422281714 | 1.93E-02 | BACH2 |
| ENSG00000112234 | 893.34174 | -0.419912959 | 4.63E-04 | FBXL4 |
| ENSG00000111860 | 295.6479056 | -0.975434364 | 4.92E-05 | CEP85L |
| ENSG00000118523 | 8122.707037 | 4.572167702 | 6.21E-123 | CCN2 |
| ENSG00000049618 | 3578.147371 | -0.302278691 | 7.11E-08 | ARID1B |
| ENSG00000078269 | 2575.00764 | 0.338964492 | 9.15E-03 | SYNJ2 |
| ENSG00000197461 | 1396.475027 | 1.258517676 | 1.32E-18 | PDGFA |
| ENSG00000197461 | 1396.475027 | 1.258517676 | 1.32E-18 | PDGFA |
| ENSG00000198517 | 1886.636662 | 1.611584072 | 2.45E-48 | MAFK |
| ENSG00000002822 | 2125.818007 | 0.620671877 | 2.65E-10 | MAD1L1 |
| ENSG00000106266 | 3274.668815 | 0.837603861 | 2.73E-09 | SNX8 |
| ENSG00000146555 | 1217.636645 | 1.050973017 | 1.21E-14 | SDK1 |
| ENSG00000075618 | 13820.42552 | 0.698810655 | 3.53E-07 | FSCN1 |
| ENSG00000011275 | 6139.992269 | 0.312206186 | 4.19E-06 | RNF216 |
| ENSG00000008256 | 8882.569858 | 1.714966959 | 2.69E-114 | CYTH3 |
| ENSG00000136231 | 3445.067623 | 0.331742981 | 1.83E-03 | IGF2BP3 |
| ENSG00000105991 | 545.3612809 | 0.839147808 | 2.14E-13 | HOXA1 |
| ENSG00000227855 | 201.8146384 | 0.792722292 | 2.30E-03 | DPY19L2P3 |
| ENSG00000136193 | 19372.32534 | 0.816428257 | 1.92E-30 | SCRN1 |
| ENSG00000106125 | 770.5683414 | -0.655953281 | 4.08E-10 | MINDY4 |
| ENSG00000122642 | 12757.91606 | 1.122308259 | 4.37E-40 | FKBP9 |
| ENSG00000164619 | 162.7326693 | 0.95837238 | 2.13E-02 | BMPER |
| ENSG00000122545 | 8237.371082 | -0.587655631 | 4.86E-03 | 7-Sep |
| ENSG00000122547 | 580.6320043 | -0.72640529 | 1.67E-04 | EEPD1 |
| ENSG00000011426 | 11414.65258 | 0.855314117 | 1.02E-04 | ANLN |
| ENSG00000188185 | 516.0461094 | -0.393566491 | 7.45E-03 | LINC00265 |
| ENSG00000175600 | 319.5476151 | 0.871727421 | 4.40E-06 | SUGCT |
| ENSG00000106571 | 3529.66754 | -0.551298388 | 1.49E-06 | GLI3 |
| ENSG00000106608 | 3502.054904 | 0.632415521 | 2.08E-09 | URGCP |
| ENSG00000136205 | 16971.90437 | 0.918192476 | 3.17E-15 | TNS3 |

| | | | | |
|---|---|---|---|---|
| ENSG00000136205 | 16971.90437 | 0.918192476 | 3.17E-15 | TNS3 |
| ENSG00000136205 | 16971.90437 | 0.918192476 | 3.17E-15 | TNS3 |
| ENSG00000136205 | 16971.90437 | 0.918192476 | 3.17E-15 | TNS3 |
| ENSG00000136275 | 6.788843734 | 3.806163813 | 2.63E-04 | C7orf69 |
| ENSG00000136275 | 6.788843734 | 3.806163813 | 2.63E-04 | C7orf69 |
| ENSG00000183696 | 3542.442731 | 2.125364727 | 2.08E-26 | UPP1 |
| ENSG00000229180 | 380.0272958 | -0.284169068 | 4.92E-02 | AC006001.3 |
| ENSG00000154710 | 2589.467863 | -0.188625002 | 2.98E-02 | RABGEF1 |
| ENSG00000158321 | 1479.629524 | -1.089612099 | 2.40E-15 | AUTS2 |
| ENSG00000158321 | 1479.629524 | -1.089612099 | 2.40E-15 | AUTS2 |
| ENSG00000006704 | 2131.857364 | 0.742953693 | 6.04E-08 | GTF2IRD1 |
| ENSG00000006704 | 2131.857364 | 0.742953693 | 6.04E-08 | GTF2IRD1 |
| ENSG00000263001 | 19621.1412 | -0.630247002 | 3.04E-12 | GTF2I |
| ENSG00000177679 | 406.1814457 | 0.808692874 | 4.88E-08 | SRRM3 |
| ENSG00000170027 | 14945.9692 | 0.34856001 | 9.94E-09 | YWHAG |
| ENSG00000188372 | 653.8483888 | 0.885842955 | 7.03E-09 | ZP3 |
| ENSG00000058091 | 3306.003124 | 0.540644783 | 2.14E-07 | CDK14 |
| ENSG00000234545 | 963.4062515 | -0.364513108 | 2.31E-02 | FAM133B |
| ENSG00000234545 | 963.4062515 | -0.364513108 | 2.31E-02 | FAM133B |
| ENSG00000127920 | 5111.312028 | 0.936478497 | 9.99E-15 | GNG11 |
| ENSG00000127920 | 5111.312028 | 0.936478497 | 9.99E-15 | GNG11 |
| ENSG00000087077 | 1891.929213 | 1.306659749 | 3.65E-23 | TRIP6 |
| ENSG00000106366 | 2089.772652 | 2.879923768 | 1.30E-69 | SERPINE1 |
| ENSG00000106366 | 2089.772652 | 2.879923768 | 1.30E-69 | SERPINE1 |
| ENSG00000160963 | 362.579038 | 0.926516967 | 3.46E-09 | COL26A1 |
| ENSG00000160963 | 362.579038 | 0.926516967 | 3.46E-09 | COL26A1 |
| ENSG00000253276 | 1297.79424 | 0.818889506 | 3.08E-10 | CCDC71L |
| ENSG00000091128 | 116.1014266 | 1.341597788 | 2.27E-07 | LAMB4 |
| ENSG00000105971 | 2124.493216 | 1.994195648 | 3.31E-61 | CAV2 |
| ENSG00000105971 | 2124.493216 | 1.994195648 | 3.31E-61 | CAV2 |
| ENSG00000105974 | 23107.43986 | 2.211744154 | 4.56E-73 | CAV1 |
| ENSG00000186591 | 13961.67353 | 0.427879274 | 1.22E-13 | UBE2H |
| ENSG00000128510 | 8124.801267 | 6.169402228 | 0.00E+00 | CPA4 |
| ENSG00000106477 | 1707.755942 | 0.239685783 | 3.95E-02 | CEP41 |
| ENSG00000221866 | 515.4222672 | -2.898214396 | 4.74E-33 | PLXNA4 |
| ENSG00000122778 | 2899.370619 | -0.319773893 | 2.23E-05 | KIAA1549 |
| ENSG00000146966 | 3519.216737 | 1.488625991 | 1.28E-31 | DENND2A |
| ENSG00000006530 | 2711.497476 | 0.191961864 | 3.64E-02 | AGK |
| ENSG00000127399 | 24.33443373 | 1.416282635 | 4.24E-03 | LRRC61 |
| ENSG00000127399 | 24.33443373 | 1.416282635 | 4.24E-03 | LRRC61 |
| ENSG00000214022 | 8966.942661 | 0.7976291 | 5.92E-28 | REPIN1 |
| ENSG00000106615 | 6449.634271 | 0.810435783 | 1.48E-44 | RHEB |
| ENSG00000106617 | 1255.564186 | 2.129482436 | 3.13E-57 | PRKAG2 |
| ENSG00000106617 | 1255.564186 | 2.129482436 | 3.13E-57 | PRKAG2 |
| ENSG00000105993 | 11954.55005 | 0.84205073 | 7.17E-36 | DNAJB6 |
| ENSG00000104728 | 3349.754618 | 0.981261274 | 1.48E-20 | ARHGEF10 |
| ENSG00000091879 | 85.43122701 | -2.635816037 | 1.09E-14 | ANGPT2 |

| | | | | |
|---|---|---|---|---|
| ENSG00000136573 | 325.3412728 | 1.223563511 | 3.33E-09 | BLK |
| ENSG00000164733 | 51746.63918 | 1.060123044 | 2.30E-16 | CTSB |
| ENSG00000164741 | 8727.154963 | 0.962947777 | 4.65E-54 | DLC1 |
| ENSG00000164741 | 8727.154963 | 0.962947777 | 4.65E-54 | DLC1 |
| ENSG00000129422 | 3968.754988 | -0.89726615 | 1.54E-20 | MTUS1 |
| ENSG00000156011 | 2830.348069 | -1.040706938 | 3.32E-20 | PSD3 |
| ENSG00000156011 | 2830.348069 | -1.040706938 | 3.32E-20 | PSD3 |
| ENSG00000241852 | 1572.615118 | 0.701178051 | 1.38E-05 | C8orf58 |
| ENSG00000180053 | 293.1226131 | -0.409231347 | 2.56E-02 | NKX2-6 |
| ENSG00000120885 | 7930.293639 | 1.642905719 | 1.24E-41 | CLU |
| ENSG00000168079 | 29.23635387 | -2.953056641 | 1.72E-06 | SCARA5 |
| ENSG00000133872 | 7191.114511 | -0.23368088 | 1.03E-03 | SARAF |
| ENSG00000215262 | 6.149298403 | 3.688246302 | 3.77E-02 | KCNU1 |
| ENSG00000215262 | 6.149298403 | 3.688246302 | 3.77E-02 | KCNU1 |
| ENSG00000183779 | 2016.919739 | 0.457197591 | 1.14E-04 | ZNF703 |
| ENSG00000020181 | 6548.879872 | 0.350126644 | 2.40E-04 | ADGRA2 |
| ENSG00000168615 | 9482.944296 | 0.426628452 | 1.72E-02 | ADAM9 |
| ENSG00000169085 | 179.481724 | 1.786199157 | 1.14E-16 | VXN |
| ENSG00000176623 | 2329.178846 | -0.337581627 | 2.55E-04 | RMDN1 |
| ENSG00000070756 | 53608.13181 | 0.495211903 | 2.35E-07 | PABPC1 |
| ENSG00000248599 | 5.21339508 | 4.774231062 | 2.34E-03 | AC018781.1 |
| ENSG00000164830 | 901.5233758 | -0.598421295 | 4.39E-05 | OXR1 |
| ENSG00000164758 | 472.5295775 | 0.485628576 | 5.20E-06 | MED30 |
| ENSG00000182197 | 6550.50112 | 1.207486276 | 9.86E-22 | EXT1 |
| ENSG00000182197 | 6550.50112 | 1.207486276 | 9.86E-22 | EXT1 |
| ENSG00000249859 | 340.4968136 | 0.872603845 | 9.44E-05 | PVT1 |
| ENSG00000249859 | 340.4968136 | 0.872603845 | 9.44E-05 | PVT1 |
| ENSG00000153317 | 7407.398524 | 1.004462889 | 1.99E-23 | ASAP1 |
| ENSG00000105339 | 1469.517193 | 1.133550485 | 1.75E-30 | DENND3 |
| ENSG00000167653 | 27.29284018 | 2.599935796 | 1.31E-07 | PSCA |
| ENSG00000247317 | 189.8540731 | 0.96110214 | 1.77E-07 | LY6E-DT |
| ENSG00000177335 | 287.7631837 | -1.049506788 | 1.59E-09 | C8orf31 |
| ENSG00000178209 | 19438.93034 | 0.527868266 | 5.68E-07 | PLEC |
| ENSG00000107104 | 4383.01843 | 0.188996789 | 2.66E-02 | KANK1 |
| ENSG00000197646 | 137.9408949 | 1.414650157 | 1.12E-07 | PDCD1LG2 |
| ENSG00000198853 | 2608.465768 | 1.40352794 | 2.30E-65 | RUSC2 |
| ENSG00000235387 | 28.07979697 | 1.069205389 | 8.15E-03 | SPAAR |
| ENSG00000119138 | 1741.781689 | 0.461335251 | 8.07E-05 | KLF9 |
| ENSG00000119138 | 1741.781689 | 0.461335251 | 8.07E-05 | KLF9 |
| ENSG00000165091 | 14.90855238 | -2.324723512 | 1.10E-03 | TMC1 |
| ENSG00000135002 | 851.8053873 | 0.842902488 | 4.49E-06 | RFK |
| ENSG00000156049 | 109.087556 | -2.952194286 | 7.38E-22 | GNA14 |
| ENSG00000196781 | 2878.688334 | -0.434228303 | 2.69E-03 | TLE1 |
| ENSG00000196730 | 5388.257129 | 0.946048752 | 1.83E-39 | DAPK1 |
| ENSG00000130045 | 101.7112097 | 1.106723649 | 8.29E-08 | NXNL2 |
| ENSG00000148120 | 1301.867845 | 1.195648926 | 1.72E-33 | C9orf3 |
| ENSG00000148120 | 1301.867845 | 1.195648926 | 1.72E-33 | C9orf3 |

| | | | | |
|---|---|---|---|---|
| ENSG00000185920 | 1620.679928 | 2.003098833 | 1.34E-34 | PTCH1 |
| ENSG00000178919 | 349.40179 | 2.32085975 | 1.18E-37 | FOXE1 |
| ENSG00000106789 | 1356.442854 | -0.399021698 | 2.18E-06 | CORO2A |
| ENSG00000095383 | 462.5151675 | 1.639217343 | 1.29E-26 | TBC1D2 |
| ENSG00000136891 | 3490.266542 | -0.36393643 | 2.95E-11 | TEX10 |
| ENSG00000148143 | 2159.207217 | -0.739220654 | 2.15E-16 | ZNF462 |
| ENSG00000138835 | 2410.104891 | 0.321411029 | 2.26E-02 | RGS3 |
| ENSG00000138835 | 2410.104891 | 0.321411029 | 2.26E-02 | RGS3 |
| ENSG00000196739 | 4166.309454 | -0.740578715 | 5.36E-11 | COL27A1 |
| ENSG00000148175 | 8123.48438 | -0.238974626 | 4.00E-03 | STOM |
| ENSG00000119408 | 8959.611567 | 0.911296503 | 3.24E-18 | NEK6 |
| ENSG00000119408 | 8959.611567 | 0.911296503 | 3.24E-18 | NEK6 |
| ENSG00000119408 | 8959.611567 | 0.911296503 | 3.24E-18 | NEK6 |
| ENSG00000119408 | 8959.611567 | 0.911296503 | 3.24E-18 | NEK6 |
| ENSG00000148339 | 1315.684058 | 0.374604317 | 9.33E-07 | SLC25A25 |
| ENSG00000197694 | 27548.29185 | 1.017738029 | 1.13E-46 | SPTAN1 |
| ENSG00000197694 | 27548.29185 | 1.017738029 | 1.13E-46 | SPTAN1 |
| ENSG00000197694 | 27548.29185 | 1.017738029 | 1.13E-46 | SPTAN1 |
| ENSG00000188483 | 924.1752821 | 0.46661864 | 1.78E-02 | IER5L |
| ENSG00000204054 | 2907.811582 | 0.470267326 | 9.23E-05 | LINC00963 |
| ENSG00000107130 | 3791.736683 | 0.342412313 | 2.07E-04 | NCS1 |
| ENSG00000107263 | 4833.32604 | 0.917301875 | 2.78E-23 | RAPGEF1 |
| ENSG00000107263 | 4833.32604 | 0.917301875 | 2.78E-23 | RAPGEF1 |
| ENSG00000107263 | 4833.32604 | 0.917301875 | 2.78E-23 | RAPGEF1 |
| ENSG00000123453 | 781.3490579 | -0.313157891 | 4.29E-02 | SARDH |
| ENSG00000186350 | 6547.240196 | -0.412632166 | 4.57E-04 | RXRA |
| ENSG00000130635 | 25458.72823 | 0.872894659 | 5.90E-19 | COL5A1 |
| ENSG00000187609 | 412.4310246 | 0.601449401 | 5.44E-07 | EXD3 |
| ENSG00000090776 | 5451.422566 | -0.452485923 | 9.87E-04 | EFNB1 |
| ENSG00000171388 | 612.3809223 | 1.987565964 | 2.28E-14 | APLN |
| ENSG00000170965 | 82.08555886 | 2.601999582 | 1.62E-24 | PLAC1 |
| ENSG00000170965 | 82.08555886 | 2.601999582 | 1.62E-24 | PLAC1 |
| ENSG00000022267 | 18557.55669 | 0.968986433 | 3.33E-41 | FHL1 |
| ENSG00000203930 | 86.52816748 | 1.348295974 | 1.23E-06 | LINC00632 |
| ENSG00000147394 | 2431.652507 | 2.905150958 | 3.22E-164 | ZNF185 |
| ENSG00000196924 | 161615.4789 | 1.566609371 | 2.04E-71 | FLNA |

| | seqnames | start | end | width | strand |
|---|---|---|---|---|---|
| 1 | chr1 | 15298307 | 15298697 | 391 | * |
| 2 | chr1 | 16470691 | 16473814 | 3124 | * |
| 3 | chr1 | 16500171 | 16500446 | 276 | * |
| 4 | chr1 | 85666871 | 85668161 | 1291 | * |
| 5 | chr1 | 86045969 | 86047024 | 1056 | * |
| 6 | chr1 | 86072754 | 86073248 | 495 | * |
| 7 | chr1 | 86078137 | 86078727 | 591 | * |
| 8 | chr1 | 152080915 | 152081601 | 687 | * |
| 9 | chr10 | 92690523 | 92691144 | 622 | * |
| 10 | chr10 | 95048585 | 95049037 | 453 | * |
| 11 | chr10 | 95218439 | 95218982 | 544 | * |
| 12 | chr10 | 99447240 | 99447501 | 262 | * |
| 13 | chr10 | 112170369 | 112170880 | 512 | * |
| 14 | chr11 | 11371436 | 11371752 | 317 | * |
| 15 | chr11 | 12065501 | 12065883 | 383 | * |
| 16 | chr11 | 12398728 | 12399341 | 614 | * |
| 17 | chr11 | 12439588 | 12440294 | 707 | * |
| 18 | chr11 | 12578393 | 12578738 | 346 | * |
| 19 | chr11 | 12695648 | 12696339 | 692 | * |
| 20 | chr11 | 12824088 | 12824355 | 268 | * |
| 21 | chr11 | 12915216 | 12915827 | 612 | * |
| 22 | chr11 | 14443839 | 14444215 | 377 | * |
| 23 | chr11 | 44009067 | 44009525 | 459 | * |
| 24 | chr11 | 69311795 | 69312508 | 714 | * |
| 25 | chr11 | 69451331 | 69451765 | 435 | * |
| 26 | chr11 | 114033451 | 114033746 | 296 | * |
| 27 | chr11 | 114163101 | 114163752 | 652 | * |
| 28 | chr11 | 114165660 | 114166745 | 1086 | * |
| 29 | chr11 | 114178170 | 114178516 | 347 | * |
| 30 | chr11 | 116064234 | 116064588 | 355 | * |
| 31 | chr11 | 121337186 | 121337565 | 380 | * |
| 32 | chr12 | 1714807 | 1715268 | 462 | * |
| 33 | chr12 | 8122893 | 8123555 | 663 | * |
| 34 | chr12 | 52585646 | 52586238 | 593 | * |
| 35 | chr12 | 52674686 | 52675158 | 473 | * |
| 36 | chr12 | 56614848 | 56615309 | 462 | * |
| 37 | chr12 | 65672464 | 65672880 | 417 | * |
| 38 | chr12 | 98884556 | 98884886 | 331 | * |
| 39 | chr12 | 104612982 | 104613669 | 688 | * |
| 40 | chr12 | 120524622 | 120524935 | 314 | * |
| 41 | chr13 | 21635103 | 21635912 | 810 | * |
| 42 | chr13 | 78549376 | 78549892 | 517 | * |
| 43 | chr13 | 97774389 | 97775392 | 1004 | * |
| 44 | chr13 | 97845914 | 97847006 | 1093 | * |
| 45 | chr13 | 97874681 | 97876342 | 1662 | * |
| 46 | chr13 | 97908526 | 97908826 | 301 | * |

| | | | | | |
|---|---|---|---|---|---|
| 47 | chr13 | 97910015 | 97911140 | 1126 | * |
| 48 | chr13 | 97927088 | 97927361 | 274 | * |
| 49 | chr13 | 97959831 | 97961450 | 1620 | * |
| 50 | chr13 | 97980281 | 97980794 | 514 | * |
| 51 | chr13 | 102108183 | 102108524 | 342 | * |
| 52 | chr13 | 102164992 | 102165294 | 303 | * |
| 53 | chr14 | 23082912 | 23083540 | 629 | * |
| 54 | chr14 | 23449613 | 23450968 | 1356 | * |
| 55 | chr14 | 23451833 | 23452477 | 645 | * |
| 56 | chr14 | 55031972 | 55032386 | 415 | * |
| 57 | chr14 | 55118581 | 55119512 | 932 | * |
| 58 | chr14 | 55240189 | 55240475 | 287 | * |
| 59 | chr14 | 75761947 | 75762206 | 260 | * |
| 60 | chr14 | 89021184 | 89021544 | 361 | * |
| 61 | chr14 | 103576347 | 103576719 | 373 | * |
| 62 | chr15 | 63268526 | 63269190 | 665 | * |
| 63 | chr15 | 63337020 | 63337462 | 443 | * |
| 64 | chr15 | 79296668 | 79296972 | 305 | * |
| 65 | chr15 | 83478872 | 83479421 | 550 | * |
| 66 | chr15 | 83989428 | 83989742 | 315 | * |
| 67 | chr16 | 122632 | 123022 | 391 | * |
| 68 | chr16 | 3070424 | 3071127 | 704 | * |
| 69 | chr16 | 3071665 | 3072737 | 1073 | * |
| 70 | chr16 | 23916392 | 23916936 | 545 | * |
| 71 | chr16 | 68741230 | 68741915 | 686 | * |
| 72 | chr16 | 68824045 | 68824395 | 351 | * |
| 73 | chr16 | 71663055 | 71663326 | 272 | * |
| 74 | chr16 | 77953028 | 77953311 | 284 | * |
| 75 | chr16 | 88365647 | 88366127 | 481 | * |
| 76 | chr17 | 783017 | 783542 | 526 | * |
| 77 | chr17 | 836637 | 836916 | 280 | * |
| 78 | chr17 | 854803 | 855462 | 660 | * |
| 79 | chr17 | 3571288 | 3571708 | 421 | * |
| 80 | chr17 | 47913737 | 47914249 | 513 | * |
| 81 | chr17 | 76349080 | 76349568 | 489 | * |
| 82 | chr17 | 81066619 | 81067385 | 767 | * |
| 83 | chr18 | 3623983 | 3624832 | 850 | * |
| 84 | chr18 | 56251878 | 56252268 | 391 | * |
| 85 | chr18 | 60905271 | 60905622 | 352 | * |
| 86 | chr18 | 77725352 | 77725768 | 417 | * |
| 87 | chr19 | 2540320 | 2541524 | 1205 | * |
| 88 | chr19 | 2578785 | 2579220 | 436 | * |
| 89 | chr19 | 34971987 | 34972691 | 705 | * |
| 90 | chr19 | 45129235 | 45129594 | 360 | * |
| 91 | chr19 | 54362249 | 54362538 | 290 | * |
| 92 | chr2 | 20212343 | 20212760 | 418 | * |
| 93 | chr2 | 23787410 | 23787677 | 268 | * |

| | | | | | |
|---|---|---|---|---|---|
| 94 | chr2 | 26981036 | 26981590 | 555 | * |
| 95 | chr2 | 46166084 | 46166380 | 297 | * |
| 96 | chr2 | 46294732 | 46295067 | 336 | * |
| 97 | chr2 | 46324833 | 46325342 | 510 | * |
| 98 | chr2 | 128432859 | 128433243 | 385 | * |
| 99 | chr2 | 192731708 | 192732269 | 562 | * |
| 100 | chr20 | 4066133 | 4066421 | 289 | * |
| 101 | chr20 | 4115377 | 4115882 | 506 | * |
| 102 | chr20 | 42844950 | 42845318 | 369 | * |
| 103 | chr20 | 55606089 | 55606573 | 485 | * |
| 104 | chr21 | 18899958 | 18900225 | 268 | * |
| 105 | chr21 | 30631119 | 30631588 | 470 | * |
| 106 | chr21 | 35899149 | 35899617 | 469 | * |
| 107 | chr21 | 46348705 | 46349043 | 339 | * |
| 108 | chr22 | 30602839 | 30603333 | 495 | * |
| 109 | chr22 | 36025786 | 36026155 | 370 | * |
| 110 | chr22 | 37942123 | 37942481 | 359 | * |
| 111 | chr22 | 37956847 | 37957348 | 502 | * |
| 112 | chr22 | 38614686 | 38615080 | 395 | * |
| 113 | chr3 | 16525302 | 16526428 | 1127 | * |
| 114 | chr3 | 49506637 | 49507329 | 693 | * |
| 115 | chr3 | 50301808 | 50302548 | 741 | * |
| 116 | chr3 | 58033690 | 58034187 | 498 | * |
| 117 | chr3 | 86914672 | 86914984 | 313 | * |
| 118 | chr3 | 134032225 | 134032601 | 377 | * |
| 119 | chr3 | 134045636 | 134046173 | 538 | * |
| 120 | chr3 | 134076332 | 134076949 | 618 | * |
| 121 | chr3 | 134077317 | 134077625 | 309 | * |
| 122 | chr3 | 134082696 | 134083239 | 544 | * |
| 123 | chr3 | 134092213 | 134093797 | 1585 | * |
| 124 | chr4 | 184019710 | 184020366 | 657 | * |
| 125 | chr4 | 187354862 | 187355272 | 411 | * |
| 126 | chr5 | 956680 | 957627 | 948 | * |
| 127 | chr5 | 17169474 | 17169762 | 289 | * |
| 128 | chr5 | 34609234 | 34609521 | 288 | * |
| 129 | chr5 | 34610042 | 34610300 | 259 | * |
| 130 | chr5 | 38765596 | 38766327 | 732 | * |
| 131 | chr5 | 38783291 | 38783609 | 319 | * |
| 132 | chr5 | 38845886 | 38846339 | 454 | * |
| 133 | chr5 | 52160527 | 52160818 | 292 | * |
| 134 | chr5 | 68797782 | 68798216 | 435 | * |
| 135 | chr5 | 73680967 | 73681251 | 285 | * |
| 136 | chr5 | 73838716 | 73839702 | 987 | * |
| 137 | chr5 | 73890729 | 73891358 | 630 | * |
| 138 | chr5 | 150016653 | 150017378 | 726 | * |
| 139 | chr5 | 150024418 | 150024971 | 554 | * |
| 140 | chr5 | 156717240 | 156717707 | 468 | * |

| | | | | | |
|---|---|---|---|---|---|
| 141 | chr5 | 174715190 | 174715507 | 318 | * |
| 142 | chr6 | 30712449 | 30712855 | 407 | * |
| 143 | chr6 | 37093895 | 37094450 | 556 | * |
| 144 | chr6 | 37105305 | 37105858 | 554 | * |
| 145 | chr6 | 74289898 | 74290319 | 422 | * |
| 146 | chr6 | 74295491 | 74296051 | 561 | * |
| 147 | chr6 | 75915126 | 75915419 | 294 | * |
| 148 | chr6 | 75915687 | 75915967 | 281 | * |
| 149 | chr6 | 76027250 | 76027550 | 301 | * |
| 150 | chr6 | 132272297 | 132273024 | 728 | * |
| 151 | chr7 | 500185 | 500446 | 262 | * |
| 152 | chr7 | 500861 | 501374 | 514 | * |
| 153 | chr7 | 1576493 | 1577559 | 1067 | * |
| 154 | chr7 | 22600871 | 22601348 | 478 | * |
| 155 | chr7 | 22616853 | 22617235 | 383 | * |
| 156 | chr7 | 22626202 | 22626955 | 754 | * |
| 157 | chr7 | 27135175 | 27135707 | 533 | * |
| 158 | chr7 | 48123033 | 48123377 | 345 | * |
| 159 | chr7 | 100760671 | 100760984 | 314 | * |
| 160 | chr7 | 100765346 | 100765637 | 292 | * |
| 161 | chr7 | 105470608 | 105471680 | 1073 | * |
| 162 | chr7 | 129934234 | 129934492 | 259 | * |
| 163 | chr7 | 134427070 | 134427416 | 347 | * |
| 164 | chr7 | 134464064 | 134464569 | 506 | * |
| 165 | chr7 | 134855248 | 134855779 | 532 | * |
| 166 | chr7 | 135442942 | 135443311 | 370 | * |
| 167 | chr7 | 151384460 | 151385166 | 707 | * |
| 168 | chr7 | 151451083 | 151451720 | 638 | * |
| 169 | chr8 | 1771745 | 1772089 | 345 | * |
| 170 | chr8 | 11748677 | 11748955 | 279 | * |
| 171 | chr8 | 17533644 | 17533914 | 271 | * |
| 172 | chr8 | 27206199 | 27206514 | 316 | * |
| 173 | chr8 | 38857155 | 38857665 | 511 | * |
| 174 | chr8 | 67434445 | 67434904 | 460 | * |
| 175 | chr8 | 102063535 | 102063910 | 376 | * |
| 176 | chr8 | 143757658 | 143758007 | 350 | * |
| 177 | chr8 | 144097296 | 144097653 | 358 | * |
| 178 | chr9 | 707011 | 707400 | 390 | * |
| 179 | chr9 | 5509461 | 5509868 | 408 | * |
| 180 | chr9 | 78987063 | 78987379 | 317 | * |
| 181 | chr9 | 98269647 | 98270217 | 571 | * |
| 182 | chr9 | 100639252 | 100639765 | 514 | * |
| 183 | chr9 | 124113550 | 124113911 | 362 | * |
| 184 | chr9 | 127421345 | 127422015 | 671 | * |
| 185 | chr9 | 131314982 | 131315253 | 272 | * |
| 186 | chr9 | 131316389 | 131317843 | 1455 | * |
| 187 | chr9 | 131320799 | 131321311 | 513 | * |

| | | | | | |
|---|---|---|---|---|---|
| 188 | chrX | 133738181 | 133738449 | 269 | * |
| 189 | chrX | 133803948 | 133804345 | 398 | * |
| 190 | chrX | 139805527 | 139805829 | 303 | * |
| 191 | chrX | 152066053 | 152066588 | 536 | * |

| V5 | geneId | ENSEMBL | baseMean | log2 Fold Change |
|---|---|---|---|---|
| 86 | 23254 | ENSG00000189337 | 2090.457273 | 1.819562339 |
| 143 | 1969 | ENSG00000142627 | 5020.238168 | 1.064095 |
| 130 | 1969 | ENSG00000142627 | 5020.238168 | 1.064095 |
| 392 | 84144 | ENSG00000097096 | 235.870281 | 1.598244045 |
| 462 | 3491 | ENSG00000142871 | 10596.59591 | 1.740805097 |
| 69 | 3491 | ENSG00000142871 | 10596.59591 | 1.740805097 |
| 45 | 3491 | ENSG00000142871 | 10596.59591 | 1.740805097 |
| 60 | 7062 | ENSG00000159450 | 60.14085417 | 4.470135267 |
| 115 | 27063 | ENSG00000148677 | 2371.13122 | 2.922114295 |
| 164 | 26509 | ENSG00000138119 | 15695.67332 | 2.025635732 |
| 30 | 26509 | ENSG00000138119 | 15695.67332 | 2.025635732 |
| 52 | 60370 | ENSG00000119986 | 366.8041458 | 1.359429399 |
| 18 | 1847 | ENSG00000138166 | 787.2594332 | 1.302938139 |
| 87 | 283106 | ENSG00000254598 | 12.69435714 | 2.069371384 |
| 159 | 27122 | ENSG00000050165 | 22309.34202 | 1.093975211 |
| 444 | 55742 | ENSG00000197702 | 12811.79845 | 1.125315857 |
| 39 | 55742 | ENSG00000197702 | 12811.79845 | 1.125315857 |
| 142 | 7003 | ENSG00000187079 | 10939.96256 | 1.187366447 |
| 554 | 7003 | ENSG00000187079 | 10939.96256 | 1.187366447 |
| 45 | 7003 | ENSG00000187079 | 10939.96256 | 1.187366447 |
| 37 | 7003 | ENSG00000187079 | 10939.96256 | 1.187366447 |
| 191 | 22800 | ENSG00000133818 | 2136.436366 | 1.285937127 |
| 37 | 387763 | ENSG00000187479 | 147.5207521 | 1.078930128 |
| 340 | 595 | ENSG00000110092 | 9113.412653 | 1.078201034 |
| 287 | 595 | ENSG00000110092 | 9113.412653 | 1.078201034 |
| 60 | 4837 | ENSG00000166741 | 7823.226213 | 1.112634451 |
| 40 | 4837 | ENSG00000166741 | 7823.226213 | 1.112634451 |
| 215 | 4837 | ENSG00000166741 | 7823.226213 | 1.112634451 |
| 58 | 4837 | ENSG00000166741 | 7823.226213 | 1.112634451 |
| 38 | 283143 | ENSG00000246100 | 27.39226582 | 2.43778509 |
| 52 | 6653 | ENSG00000137642 | 4958.874962 | 1.536327103 |
| 52 | 81029 | ENSG00000111186 | 481.1590035 | 1.668509358 |
| 347 | 6515 | ENSG00000059804 | 3270.065993 | 2.168291612 |
| 174 | 144501 | ENSG00000167767 | 1730.679913 | 3.075015596 |
| 196 | 3892 | ENSG00000170442 | 789.1722789 | 1.011367361 |
| 30 | 10193 | ENSG00000181852 | 3274.28356 | 1.444728516 |
| 23 | 253827 | ENSG00000174099 | 1780.19995 | 1.5565635 |
| 23 | 643770 | ENSG00000245017 | 7.064652985 | 1.93138767 |
| 77 | 7296 | ENSG00000198431 | 33083.72171 | 2.000025042 |
| 95 | 92558 | ENSG00000135127 | 37.542591 | 2.742854188 |
| 361 | 26524 | ENSG00000150457 | 922.0682349 | 1.856232447 |
| 33 | 1910 | ENSG00000136160 | 780.2042551 | 2.082298048 |
| 60 | 10150 | ENSG00000139793 | 5478.880166 | 1.231430013 |
| 94 | 10150 | ENSG00000139793 | 5478.880166 | 1.231430013 |
| 25 | 10150 | ENSG00000139793 | 5478.880166 | 1.231430013 |
| 22 | 10150 | ENSG00000139793 | 5478.880166 | 1.231430013 |

| | | | | |
|---|---|---|---|---|
| 44 | 10150 | ENSG00000139793 | 5478.880166 | 1.231430013 |
| 67 | 10150 | ENSG00000139793 | 5478.880166 | 1.231430013 |
| 345 | 10150 | ENSG00000139793 | 5478.880166 | 1.231430013 |
| 65 | 10150 | ENSG00000139793 | 5478.880166 | 1.231430013 |
| 53 | 9358 | ENSG00000198542 | 61.58578191 | 1.360256698 |
| 22 | 9358 | ENSG00000198542 | 61.58578191 | 1.360256698 |
| 130 | 63874 | ENSG00000100439 | 4610.109295 | 1.143863046 |
| 1429 | 84962 | ENSG00000129474 | 9629.715395 | 2.41294946 |
| 885 | 84962 | ENSG00000129474 | 9629.715395 | 2.41294946 |
| 12 | 23034 | ENSG00000020577 | 2581.581522 | 1.573953635 |
| 311 | 23034 | ENSG00000020577 | 2581.581522 | 1.573953635 |
| 52 | 23034 | ENSG00000020577 | 2581.581522 | 1.573953635 |
| 15 | 731223 | ENSG00000259687 | 13.87085948 | 2.196446389 |
| 104 | 11099 | ENSG00000070778 | 1232.53051 | 1.375748917 |
| 30 | 91828 | ENSG00000205436 | 86.63631761 | 1.261960233 |
| 60 | 7168 | ENSG00000140416 | 23789.68 | 1.33657541 |
| 96 | 7168 | ENSG00000140416 | 23789.68 | 1.33657541 |
| 60 | 5923 | ENSG00000058335 | 134.7074287 | 2.459396605 |
| 23 | 123720 | ENSG00000156232 | 724.8447879 | 1.514378994 |
| 12 | 646 | ENSG00000169594 | 1562.452238 | 1.226491598 |
| 105 | 64285 | ENSG00000007384 | 1735.657285 | 1.044026739 |
| 69 | 51330 | ENSG00000006327 | 3033.22403 | 2.936404904 |
| 174 | 51330 | ENSG00000006327 | 3033.22403 | 2.936404904 |
| 139 | 5579 | ENSG00000166501 | 1065.831152 | 2.338768189 |
| 484 | 999 | ENSG00000039068 | 23.87917036 | 1.703672002 |
| 36 | 999 | ENSG00000039068 | 23.87917036 | 1.703672002 |
| 30 | 91862 | ENSG00000140832 | 105.4161867 | 1.33581724 |
| 17 | 10143 | ENSG00000166509 | 27.21407875 | 4.850407839 |
| 146 | 84627 | ENSG00000225614 | 430.1866375 | 1.286291157 |
| 93 | 64359 | ENSG00000167693 | 1791.096427 | 1.527285754 |
| 174 | 64359 | ENSG00000167693 | 1791.096427 | 1.527285754 |
| 261 | 64359 | ENSG00000167693 | 1791.096427 | 1.527285754 |
| 120 | 30851 | ENSG00000213977 | 4806.333429 | 1.032618893 |
| 52 | 729220 | ENSG00000204584 | 22.04428444 | 1.290335976 |
| 287 | 9021 | ENSG00000184557 | 897.8181479 | 1.11022357 |
| 152 | 284207 | ENSG00000176845 | 538.6256251 | 1.789221388 |
| 143 | 649446 | ENSG00000177337 | 142.860099 | 1.182975185 |
| 60 | 115701 | ENSG00000198796 | 2595.841625 | 1.585857684 |
| 76 | 596 | ENSG00000171791 | 308.996463 | 1.164816339 |
| 45 | 440498 | ENSG00000226742 | 284.4549299 | 2.970641489 |
| 889 | 4616 | ENSG00000099860 | 977.0968945 | 1.064075522 |
| 46 | 4616 | ENSG00000099860 | 977.0968945 | 1.064075522 |
| 1010 | 126374 | ENSG00000142279 | 955.4419514 | 2.230720722 |
| 164 | 147710 | ENSG00000216588 | 35.73858368 | 1.644353837 |
| 52 | 91663 | ENSG00000179820 | 3260.603661 | 1.622775922 |
| 82 | 4148 | ENSG00000132031 | 531.0146682 | 2.011154986 |
| 15 | 114818 | ENSG00000119771 | 1265.210865 | 1.307548046 |

| | | | | |
|---|---|---|---|---|
| 45 | 54978 | ENSG00000213699 | 2445.04949 | 1.466724367 |
| 52 | 5581 | ENSG00000171132 | 1501.609574 | 1.487310913 |
| 38 | 2034 | ENSG00000116016 | 9233.833509 | 1.044855878 |
| 60 | 2034 | ENSG00000116016 | 9233.833509 | 1.044855878 |
| 34 | 55679 | ENSG00000072163 | 478.4740095 | 3.512004826 |
| 52 | 8436 | ENSG00000168497 | 121.5099942 | 2.41419518 |
| 90 | 54498 | ENSG00000088826 | 2279.030744 | 1.036663145 |
| 37 | 54498 | ENSG00000088826 | 2279.030744 | 1.036663145 |
| 17 | 100505783 | ENSG00000223891 | 334.8931331 | 1.018728451 |
| 81 | 655 | ENSG00000101144 | 617.2385659 | 1.520959265 |
| 45 | 1525 | ENSG00000154639 | 430.9583909 | 2.673380869 |
| 82 | 571 | ENSG00000156273 | 2018.700707 | 1.171362318 |
| 86 | 1827 | ENSG00000159200 | 3868.497247 | 1.916557243 |
| 37 | 3689 | ENSG00000160255 | 124.990794 | 1.34529355 |
| 133 | 3976 | ENSG00000128342 | 154.155482 | 1.002146009 |
| 229 | 4151 | ENSG00000198125 | 5.190958042 | 3.025789133 |
| 69 | 11135 | ENSG00000128283 | 3906.296252 | 1.29127043 |
| 69 | 11135 | ENSG00000128283 | 3906.296252 | 1.29127043 |
| 153 | 23764 | ENSG00000185022 | 607.0319197 | 3.699491821 |
| 133 | 23180 | ENSG00000131378 | 2508.131971 | 1.553417001 |
| 107 | 1605 | ENSG00000173402 | 8402.798769 | 1.172653743 |
| 102 | 7869 | ENSG00000012171 | 1213.282702 | 1.7631064 |
| 68 | 2317 | ENSG00000136068 | 29152.77127 | 1.634231923 |
| 23 | 389136 | ENSG00000206538 | 29.00567059 | 1.824417818 |
| 123 | 51421 | ENSG00000114019 | 5194.214834 | 2.617020907 |
| 261 | 51421 | ENSG00000114019 | 5194.214834 | 2.617020907 |
| 192 | 51421 | ENSG00000114019 | 5194.214834 | 2.617020907 |
| 10 | 51421 | ENSG00000114019 | 5194.214834 | 2.617020907 |
| 645 | 51421 | ENSG00000114019 | 5194.214834 | 2.617020907 |
| 961 | 51421 | ENSG00000114019 | 5194.214834 | 2.617020907 |
| 655 | 152641 | ENSG00000251359 | 13.98774191 | 1.617468807 |
| 27 | 285441 | ENSG00000251165 | 217.3634243 | 1.691148918 |
| 233 | 100506688 | ENSG00000215246 | 130.5665582 | 3.170881367 |
| 28 | 10409 | ENSG00000176788 | 2291.351755 | 2.289317224 |
| 81 | 26064 | ENSG00000039560 | 6837.00494 | 1.284856627 |
| 41 | 26064 | ENSG00000039560 | 6837.00494 | 1.284856627 |
| 108 | 9180 | ENSG00000145623 | 15770.68321 | 1.237806056 |
| 56 | 9180 | ENSG00000145623 | 15770.68321 | 1.237806056 |
| 122 | 9180 | ENSG00000145623 | 15770.68321 | 1.237806056 |
| 30 | 3672 | ENSG00000213949 | 11661.05432 | 1.677980974 |
| 126 | 100506658 | ENSG00000197822 | 724.0860483 | 2.52656351 |
| 82 | 3074 | ENSG00000049860 | 16325.57173 | 1.383755226 |
| 89 | 3074 | ENSG00000049860 | 16325.57173 | 1.383755226 |
| 127 | 3074 | ENSG00000049860 | 16325.57173 | 1.383755226 |
| 89 | 11346 | ENSG00000171992 | 2426.228123 | 1.408445403 |
| 93 | 11346 | ENSG00000171992 | 2426.228123 | 1.408445403 |
| 63 | 26999 | ENSG00000055163 | 654.8815999 | 1.87876886 |

| | | | | |
|---|---|---|---|---|
| 77 | 1812 | ENSG00000184845 | 26.45660106 | 1.683372035 |
| 52 | 8870 | ENSG00000137331 | 2184.936652 | 1.618783256 |
| 104 | 5292 | ENSG00000137193 | 1122.782319 | 1.496184939 |
| 77 | 5292 | ENSG00000137193 | 1122.782319 | 1.496184939 |
| 104 | 26503 | ENSG00000119899 | 973.1245904 | 2.192337972 |
| 77 | 26503 | ENSG00000119899 | 973.1245904 | 2.192337972 |
| 19 | 1303 | ENSG00000111799 | 7146.720884 | 2.615659848 |
| 18 | 1303 | ENSG00000111799 | 7146.720884 | 2.615659848 |
| 37 | 27145 | ENSG00000118407 | 45.8162629 | 1.020757796 |
| 385 | 1490 | ENSG00000118523 | 8122.707037 | 3.123332247 |
| 37 | 5154 | ENSG00000197461 | 1396.475027 | 2.048728128 |
| 297 | 5154 | ENSG00000197461 | 1396.475027 | 2.048728128 |
| 394 | 7975 | ENSG00000198517 | 1886.636662 | 1.979224901 |
| 409 | 100506178 | ENSG00000232759 | 40.60269188 | 1.929100963 |
| 60 | 100506178 | ENSG00000232759 | 40.60269188 | 1.929100963 |
| 138 | 100506178 | ENSG00000232759 | 40.60269188 | 1.929100963 |
| 160 | 3198 | ENSG00000105991 | 545.3612809 | 1.242660095 |
| 26 | 7378 | ENSG00000183696 | 3542.442731 | 4.034437348 |
| 120 | 5054 | ENSG00000106366 | 2089.772652 | 2.557298491 |
| 46 | 5054 | ENSG00000106366 | 2089.772652 | 2.557298491 |
| 72 | 222255 | ENSG00000146776 | 780.669098 | 1.121204671 |
| 19 | 51200 | ENSG00000128510 | 8124.801267 | 3.172005188 |
| 90 | 800 | ENSG00000122786 | 17077.04623 | 1.130978105 |
| 335 | 800 | ENSG00000122786 | 17077.04623 | 1.130978105 |
| 85 | 78996 | ENSG00000122783 | 3033.595272 | 1.333068503 |
| 53 | 389558 | ENSG00000189320 | 81.88165982 | 2.531951624 |
| 265 | 51422 | ENSG00000106617 | 1255.564186 | 1.178963824 |
| 649 | 51422 | ENSG00000106617 | 1255.564186 | 1.178963824 |
| 45 | 9639 | ENSG00000104728 | 3349.754618 | 1.589110071 |
| 23 | 1508 | ENSG00000164733 | 51746.63918 | 1.20430749 |
| 45 | 57509 | ENSG00000129422 | 3968.754988 | 1.019009773 |
| 23 | 2185 | ENSG00000120899 | 244.6017206 | 1.122408158 |
| 23 | 8754 | ENSG00000168615 | 9482.944296 | 1.069424991 |
| 25 | 254778 | ENSG00000169085 | 179.481724 | 1.56034171 |
| 114 | 441374 | ENSG00000248599 | 5.21339508 | 3.794824329 |
| 96 | 8000 | ENSG00000167653 | 27.29284018 | 2.767445721 |
| 60 | 100133669 | ENSG00000247317 | 189.8540731 | 1.381054605 |
| 58 | 23189 | ENSG00000107104 | 4383.01843 | 1.019399234 |
| 22 | 80380 | ENSG00000197646 | 137.9408949 | 2.244504139 |
| 37 | 55312 | ENSG00000135002 | 851.8053873 | 1.014500019 |
| 37 | 5727 | ENSG00000185920 | 1620.679928 | 1.455093934 |
| 60 | 2304 | ENSG00000178919 | 349.40179 | 1.499902651 |
| 50 | 2040 | ENSG00000148175 | 8123.48438 | 1.450155304 |
| 476 | 100379345 | ENSG00000224020 | 24.1636788 | 1.530872184 |
| 30 | 6709 | ENSG00000197694 | 27548.29185 | 1.129892226 |
| 255 | 6709 | ENSG00000197694 | 27548.29185 | 1.129892226 |
| 93 | 6709 | ENSG00000197694 | 27548.29185 | 1.129892226 |

| | | | | |
|---|---|---|---|---|
| 60 | 10761 | ENSG00000170965 | 82.08555886 | 1.842281053 |
| 45 | 10761 | ENSG00000170965 | 82.08555886 | 1.842281053 |
| 86 | 286411 | ENSG00000203930 | 86.52816748 | 2.76982743 |
| 557 | 7739 | ENSG00000147394 | 2431.652507 | 2.623454688 |

| padj | Gene |
|---|---|
| 6.02E-90 | KAZN |
| 1.06E-18 | EPHA2 |
| 1.06E-18 | EPHA2 |
| 1.15E-11 | SYDE2 |
| 1.11E-52 | CCN1 |
| 1.11E-52 | CCN1 |
| 1.11E-52 | CCN1 |
| 8.26E-41 | TCHH |
| 6.95E-11 | ANKRD1 |
| 7.56E-60 | MYOF |
| 7.56E-60 | MYOF |
| 1.72E-19 | AVPI1 |
| 2.80E-10 | DUSP5 |
| 1.34E-02 | CSNK2A3 |
| 2.03E-19 | DKK3 |
| 3.16E-111 | PARVA |
| 3.16E-111 | PARVA |
| 1.64E-16 | TEAD1 |
| 1.64E-16 | TEAD1 |
| 1.64E-16 | TEAD1 |
| 1.64E-16 | TEAD1 |
| 3.00E-41 | RRAS2 |
| 5.42E-09 | C11orf96 |
| 2.55E-13 | CCND1 |
| 2.55E-13 | CCND1 |
| 1.43E-08 | NNMT |
| 1.43E-08 | NNMT |
| 1.43E-08 | NNMT |
| 1.43E-08 | NNMT |
| 1.30E-05 | LINC00900 |
| 1.22E-34 | SORL1 |
| 2.86E-18 | WNT5B |
| 7.52E-23 | SLC2A3 |
| 1.85E-42 | KRT80 |
| 2.63E-06 | KRT86 |
| 2.65E-17 | RNF41 |
| 3.11E-42 | MSRB3 |
| 1.45E-02 | LINC02453 |
| 2.58E-28 | TXNRD1 |
| 3.52E-08 | BICDL1 |
| 2.44E-64 | LATS2 |
| 9.89E-55 | EDNRB |
| 3.16E-09 | MBNL2 |
| 3.16E-09 | MBNL2 |
| 3.16E-09 | MBNL2 |
| 3.16E-09 | MBNL2 |

| | |
|---|---|
| 3.16E-09 | MBNL2 |
| 3.16E-09 | MBNL2 |
| 3.16E-09 | MBNL2 |
| 3.16E-09 | MBNL2 |
| 1.45E-03 | ITGBL1 |
| 1.45E-03 | ITGBL1 |
| 1.36E-20 | ABHD4 |
| 1.04E-45 | AJUBA |
| 1.04E-45 | AJUBA |
| 2.48E-36 | SAMD4A |
| 2.48E-36 | SAMD4A |
| 2.48E-36 | SAMD4A |
| 1.18E-02 | LINC01220 |
| 3.73E-47 | AL162171.1 |
| 1.02E-10 | EXOC3L4 |
| 1.10E-35 | TPM1 |
| 1.10E-35 | TPM1 |
| 1.99E-16 | RASGRF1 |
| 2.75E-31 | WHAMM |
| 4.97E-09 | BNC1 |
| 5.26E-21 | RHBDF1 |
| 7.07E-85 | TNFRSF12A |
| 7.07E-85 | TNFRSF12A |
| 3.79E-45 | PRKCB |
| 2.89E-02 | CDH1 |
| 2.89E-02 | CDH1 |
| 2.94E-10 | MARVELD3 |
| 3.00E-15 | CLEC3A |
| 4.49E-11 | ZNF469 |
| 2.00E-37 | NXN |
| 2.00E-37 | NXN |
| 2.00E-37 | NXN |
| 3.28E-17 | TAX1BP3 |
| 2.00E-02 | FLJ45513 |
| 5.69E-05 | SOCS3 |
| 3.04E-32 | METRNL |
| 1.65E-09 | DLGAP1-AS1 |
| 4.62E-12 | ALPK2 |
| 7.95E-08 | BCL2 |
| 1.98E-80 | HSBP1L1 |
| 3.12E-19 | GADD45B |
| 3.12E-19 | GADD45B |
| 5.28E-33 | WTIP |
| 4.25E-05 | IGSF23 |
| 7.29E-47 | MYADM |
| 9.00E-46 | MATN3 |
| 1.73E-18 | KLHL29 |

| | |
|---|---|
| 8.71E-15 | SLC35F6 |
| 1.08E-12 | PRKCE |
| 3.23E-27 | EPAS1 |
| 3.23E-27 | EPAS1 |
| 6.61E-87 | LIMS2 |
| 2.85E-04 | CAVIN2 |
| 1.06E-04 | SMOX |
| 1.06E-04 | SMOX |
| 3.39E-12 | OSER1-DT |
| 7.39E-11 | BMP7 |
| 1.93E-52 | CXADR |
| 8.09E-10 | BACH1 |
| 3.42E-49 | RCAN1 |
| 2.51E-04 | ITGB2 |
| 9.57E-04 | LIF |
| 4.48E-02 | MB |
| 2.02E-12 | CDC42EP1 |
| 2.02E-12 | CDC42EP1 |
| 1.47E-96 | MAFF |
| 1.40E-45 | RFTN1 |
| 2.71E-27 | DAG1 |
| 5.08E-18 | SEMA3B |
| 1.33E-35 | FLNB |
| 5.46E-05 | VGLL3 |
| 2.42E-151 | AMOTL2 |
| 2.42E-151 | AMOTL2 |
| 2.42E-151 | AMOTL2 |
| 2.42E-151 | AMOTL2 |
| 2.42E-151 | AMOTL2 |
| 2.42E-151 | AMOTL2 |
| 3.53E-03 | WWC2-AS2 |
| 1.43E-10 | F11-AS1 |
| 4.07E-37 | AC116351.1 |
| 5.22E-53 | BASP1 |
| 1.90E-14 | RAI14 |
| 1.90E-14 | RAI14 |
| 1.92E-25 | OSMR |
| 1.92E-25 | OSMR |
| 1.92E-25 | OSMR |
| 4.70E-42 | ITGA1 |
| 3.30E-31 | OCLN |
| 4.72E-54 | HEXB |
| 4.72E-54 | HEXB |
| 4.72E-54 | HEXB |
| 5.63E-35 | SYNPO |
| 5.63E-35 | SYNPO |
| 1.01E-14 | CYFIP2 |

| | |
|---|---|
| 2.73E-04 | DRD1 |
| 1.32E-37 | IER3 |
| 1.92E-13 | PIM1 |
| 1.92E-13 | PIM1 |
| 2.18E-50 | SLC17A5 |
| 2.18E-50 | SLC17A5 |
| 1.55E-87 | COL12A1 |
| 1.55E-87 | COL12A1 |
| 6.44E-04 | FILIP1 |
| 1.80E-57 | CCN2 |
| 2.93E-49 | PDGFA |
| 2.93E-49 | PDGFA |
| 9.59E-74 | MAFK |
| 1.93E-09 | AC002480.1 |
| 1.93E-09 | AC002480.1 |
| 1.93E-09 | AC002480.1 |
| 1.85E-30 | HOXA1 |
| 1.27E-94 | UPP1 |
| 5.61E-55 | SERPINE1 |
| 5.61E-55 | SERPINE1 |
| 3.10E-18 | ATXN7L1 |
| 3.78E-86 | CPA4 |
| 4.02E-18 | CALD1 |
| 4.02E-18 | CALD1 |
| 4.40E-18 | CYREN |
| 1.46E-10 | FAM180A |
| 7.48E-18 | PRKAG2 |
| 7.48E-18 | PRKAG2 |
| 1.15E-53 | ARHGEF10 |
| 5.76E-21 | CTSB |
| 3.29E-27 | MTUS1 |
| 8.46E-06 | PTK2B |
| 1.96E-11 | ADAM9 |
| 3.32E-13 | VXN |
| 1.91E-02 | AC018781.1 |
| 6.70E-09 | PSCA |
| 8.98E-16 | LY6E-DT |
| 1.04E-45 | KANK1 |
| 1.67E-19 | PDCD1LG2 |
| 1.19E-08 | RFK |
| 1.33E-18 | PTCH1 |
| 4.77E-16 | FOXE1 |
| 5.04E-90 | STOM |
| 1.88E-04 | MIR181A2HG |
| 1.62E-57 | SPTAN1 |
| 1.62E-57 | SPTAN1 |
| 1.62E-57 | SPTAN1 |

| | |
|---|---|
| 1.21E-12 | PLAC1 |
| 1.21E-12 | PLAC1 |
| 1.60E-29 | LINC00632 |
| 1.95E-134 | ZNF185 |

| | seqnames | start | end | width | strand | V5 | geneId |
|---|---|---|---|---|---|---|---|
| 1 | chr5 | 39423971 | 39425270 | 1300 | * | 246 | 1601 |
| 2 | chr6 | 75915126 | 75915419 | 294 | * | 19 | 1303 |
| 3 | chr6 | 75915687 | 75915967 | 281 | * | 18 | 1303 |

| ENSEMBL | baseMean | log2 Fold Change | padj | Gene |
|---|---|---|---|---|
| ENSG00000153071 | 9628.493215 | 0.358876294 | 1.68E-02 | DAB2 |
| ENSG00000111799 | 7146.720884 | 0.56840206 | 2.21E-02 | COL12A1 |
| ENSG00000111799 | 7146.720884 | 0.56840206 | 2.21E-02 | COL12A1 |

| | seqnames | start | end | width | strand | V5 | geneId |
|---|---|---|---|---|---|---|---|
| 1 | chr1 | 11918805 | 11919783 | 979 | * | 32 | 4879 |
| 2 | chr1 | 16126741 | 16127095 | 355 | * | 35 | 440567 |
| 3 | chr1 | 16471673 | 16472390 | 718 | * | 133 | 1969 |
| 4 | chr1 | 16472812 | 16473079 | 268 | * | 143 | 1969 |
| 5 | chr1 | 16500060 | 16500469 | 410 | * | 428 | 1969 |
| 6 | chr1 | 26745520 | 26745805 | 286 | * | 68 | 79727 |
| 7 | chr1 | 31869995 | 31870851 | 857 | * | 173 | 347735 |
| 8 | chr1 | 32041442 | 32041720 | 279 | * | 104 | 64129 |
| 9 | chr1 | 41331964 | 41332315 | 352 | * | 154 | 163732 |
| 10 | chr1 | 43672519 | 43672978 | 460 | * | 326 | 10969 |
| 11 | chr1 | 43938233 | 43938717 | 485 | * | 124 | 81888 |
| 12 | chr1 | 86072824 | 86073265 | 442 | * | 428 | 3491 |
| 13 | chr1 | 95111719 | 95112144 | 426 | * | 68 | 2152 |
| 14 | chr1 | 162508731 | 162509087 | 357 | * | 28 | 6675 |
| 15 | chr1 | 204273366 | 204273663 | 298 | * | 35 | 22874 |
| 16 | chr1 | 210612302 | 210612835 | 534 | * | 254 | 55733 |
| 17 | chr1 | 212686475 | 212686772 | 298 | * | 104 | 467 |
| 18 | chr1 | 218653748 | 218654207 | 460 | * | 85 | 728463 |
| 19 | chr1 | 218879344 | 218880109 | 766 | * | 87 | 728463 |
| 20 | chr10 | 10827768 | 10828129 | 362 | * | 77 | 207107 |
| 21 | chr10 | 47057731 | 47058040 | 310 | * | 72 | 5540 |
| 22 | chr10 | 71598801 | 71599065 | 265 | * | 68 | 1305 |
| 23 | chr10 | 73526553 | 73526989 | 437 | * | 219 | 64115 |
| 24 | chr10 | 92690792 | 92691202 | 411 | * | 186 | 27063 |
| 25 | chr10 | 100028822 | 100029417 | 596 | * | 133 | 84171 |
| 26 | chr10 | 101155708 | 101156050 | 343 | * | 77 | 2805 |
| 27 | chr11 | 707713 | 708231 | 519 | * | 92 | 64787 |
| 28 | chr11 | 9787192 | 9787471 | 280 | * | 174 | 440028 |
| 29 | chr11 | 12108006 | 12108314 | 309 | * | 59 | 9645 |
| 30 | chr11 | 12204300 | 12204763 | 464 | * | 84 | 9645 |
| 31 | chr11 | 12221919 | 12222589 | 671 | * | 71 | 9645 |
| 32 | chr11 | 14443880 | 14444276 | 397 | * | 301 | 22800 |
| 33 | chr11 | 35188198 | 35188474 | 277 | * | 175 | 960 |
| 34 | chr11 | 62320766 | 62321499 | 734 | * | 66 | 79026 |
| 35 | chr11 | 65670612 | 65671088 | 477 | * | 265 | 8061 |
| 36 | chr11 | 65782763 | 65783279 | 517 | * | 347 | 1474 |
| 37 | chr11 | 69064692 | 69065140 | 449 | * | 142 | 26579 |
| 38 | chr11 | 69068965 | 69069447 | 483 | * | 102 | 26579 |
| 39 | chr11 | 69235335 | 69235901 | 567 | * | 327 | 26579 |
| 40 | chr11 | 69251856 | 69252129 | 274 | * | 339 | 26579 |
| 41 | chr11 | 69259075 | 69259410 | 336 | * | 15 | 595 |
| 42 | chr11 | 69264482 | 69264887 | 406 | * | 105 | 595 |
| 43 | chr11 | 69312001 | 69312382 | 382 | * | 519 | 595 |
| 44 | chr11 | 69382580 | 69383128 | 549 | * | 364 | 595 |
| 45 | chr11 | 69402593 | 69403055 | 463 | * | 47 | 595 |
| 46 | chr11 | 69451291 | 69451559 | 269 | * | 100 | 595 |

| | | | | | | | |
|---|---|---|---|---|---|---|---|
| 47 | chr11 | 71010432 | 71010721 | 290 | * | 256 | 22941 |
| 48 | chr11 | 73850844 | 73851313 | 470 | * | 166 | 26005 |
| 49 | chr11 | 75201801 | 75202124 | 324 | * | 143 | 81544 |
| 50 | chr11 | 114162084 | 114162403 | 320 | * | 22 | 4837 |
| 51 | chr11 | 114165658 | 114166091 | 434 | * | 84 | 4837 |
| 52 | chr11 | 114166532 | 114166994 | 463 | * | 32 | 4837 |
| 53 | chr11 | 115547383 | 115547745 | 363 | * | 138 | 283143 |
| 54 | chr11 | 120436804 | 120437119 | 316 | * | 49 | 2900 |
| 55 | chr11 | 120581233 | 120581564 | 332 | * | 140 | 2900 |
| 56 | chr11 | 122577411 | 122577702 | 292 | * | 68 | 84959 |
| 57 | chr11 | 124768577 | 124768904 | 328 | * | 131 | 54538 |
| 58 | chr11 | 128050405 | 128050708 | 304 | * | 31 | 2113 |
| 59 | chr11 | 128077756 | 128078447 | 692 | * | 170 | 2113 |
| 60 | chr11 | 128080241 | 128080896 | 656 | * | 92 | 2113 |
| 61 | chr11 | 128392416 | 128392689 | 274 | * | 122 | 2113 |
| 62 | chr11 | 129731767 | 129732255 | 489 | * | 266 | 120224 |
| 63 | chr12 | 4225529 | 4226142 | 614 | * | 174 | 894 |
| 64 | chr12 | 6225691 | 6226037 | 347 | * | 29 | 7450 |
| 65 | chr12 | 33045015 | 33045321 | 307 | * | 114 | 5318 |
| 66 | chr12 | 50595583 | 50595897 | 315 | * | 51 | 51474 |
| 67 | chr12 | 52541124 | 52541411 | 288 | * | 146 | 144501 |
| 68 | chr12 | 52585667 | 52586263 | 597 | * | 133 | 144501 |
| 69 | chr12 | 53370603 | 53370879 | 277 | * | 144 | 3856 |
| 70 | chr12 | 104613389 | 104614010 | 622 | * | 114 | 7296 |
| 71 | chr12 | 105879187 | 105879861 | 675 | * | 85 | 387882 |
| 72 | chr12 | 106410492 | 106410751 | 260 | * | 186 | 9891 |
| 73 | chr12 | 106424406 | 106424821 | 416 | * | 114 | 9891 |
| 74 | chr12 | 113767506 | 113767933 | 428 | * | 85 | 80024 |
| 75 | chr12 | 118125055 | 118125341 | 287 | * | 21 | 283455 |
| 76 | chr13 | 21635502 | 21635893 | 392 | * | 197 | 26524 |
| 77 | chr13 | 99762286 | 99762554 | 269 | * | 18 | 23348 |
| 78 | chr13 | 102102592 | 102103143 | 552 | * | 304 | 9358 |
| 79 | chr13 | 102107201 | 102108511 | 1311 | * | 151 | 9358 |
| 80 | chr13 | 102237633 | 102237947 | 315 | * | 72 | 9358 |
| 81 | chr14 | 23446951 | 23447252 | 302 | * | 174 | 84962 |
| 82 | chr14 | 23449667 | 23450184 | 518 | * | 606 | 84962 |
| 83 | chr14 | 23451856 | 23452210 | 355 | * | 242 | 84962 |
| 84 | chr14 | 67688136 | 67688636 | 501 | * | 289 | 64398 |
| 85 | chr14 | 75919704 | 75920208 | 505 | * | 424 | 122953 |
| 86 | chr14 | 80892031 | 80892336 | 306 | * | 123 | 1734 |
| 87 | chr14 | 100040819 | 100041679 | 861 | * | 104 | 317762 |
| 88 | chr14 | 100046219 | 100046595 | 377 | * | 610 | 317762 |
| 89 | chr14 | 105434344 | 105434661 | 318 | * | 133 | 113146 |
| 90 | chr14 | 105439591 | 105439991 | 401 | * | 186 | 113146 |
| 91 | chr15 | 63400188 | 63400604 | 417 | * | 508 | 114294 |
| 92 | chr15 | 74714954 | 74715596 | 643 | * | 212 | 8482 |
| 93 | chr16 | 3070872 | 3071210 | 339 | * | 338 | 51330 |

| | | | | | | | |
|---|---|---|---|---|---|---|---|
| 94 | chr16 | 4994834 | 4995111 | 278 | * | 68 | 5493 |
| 95 | chr16 | 57682614 | 57683027 | 414 | * | 35 | 9289 |
| 96 | chr16 | 67005020 | 67005415 | 396 | * | 157 | 23491 |
| 97 | chr16 | 70727806 | 70728577 | 772 | * | 606 | 92154 |
| 98 | chr16 | 77332420 | 77332843 | 424 | * | 123 | 170692 |
| 99 | chr16 | 81544985 | 81545304 | 320 | * | 82 | 80790 |
| 100 | chr16 | 81551292 | 81551837 | 546 | * | 95 | 80790 |
| 101 | chr16 | 84839999 | 84840262 | 264 | * | 42 | 83716 |
| 102 | chr16 | 84871900 | 84872211 | 312 | * | 51 | 83716 |
| 103 | chr16 | 84884903 | 84885246 | 344 | * | 96 | 83716 |
| 104 | chr16 | 86806149 | 86806730 | 582 | * | 208 | 2300 |
| 105 | chr16 | 87212597 | 87213073 | 477 | * | 42 | 1.01E+08 |
| 106 | chr16 | 88365689 | 88366172 | 484 | * | 114 | 84627 |
| 107 | chr16 | 89647827 | 89648138 | 312 | * | 35 | 27132 |
| 108 | chr17 | 831197 | 831588 | 392 | * | 59 | 64359 |
| 109 | chr17 | 836585 | 836917 | 333 | * | 144 | 64359 |
| 110 | chr17 | 854946 | 855343 | 398 | * | 29 | 64359 |
| 111 | chr17 | 3571343 | 3571701 | 359 | * | 14 | 30851 |
| 112 | chr17 | 9551279 | 9552360 | 1082 | * | 186 | 124739 |
| 113 | chr17 | 15824088 | 15824437 | 350 | * | 94 | 136 |
| 114 | chr17 | 15857520 | 15857792 | 273 | * | 28 | 136 |
| 115 | chr17 | 36607203 | 36607464 | 262 | * | 363 | 57636 |
| 116 | chr17 | 36621050 | 36621428 | 379 | * | 43 | 57636 |
| 117 | chr17 | 42620333 | 42621215 | 883 | * | 400 | 2535 |
| 118 | chr17 | 48105528 | 48106199 | 672 | * | 428 | 284080 |
| 119 | chr17 | 54776711 | 54777165 | 455 | * | 95 | 9241 |
| 120 | chr17 | 81066625 | 81066952 | 328 | * | 93 | 284207 |
| 121 | chr18 | 3247445 | 3247787 | 343 | * | 124 | 10627 |
| 122 | chr18 | 55880870 | 55881205 | 336 | * | 28 | 23327 |
| 123 | chr18 | 56252034 | 56252358 | 325 | * | 254 | 115701 |
| 124 | chr18 | 76252022 | 76252617 | 596 | * | 522 | 27164 |
| 125 | chr18 | 76610946 | 76611303 | 358 | * | 85 | 27164 |
| 126 | chr19 | 2540426 | 2541577 | 1152 | * | 949 | 4616 |
| 127 | chr19 | 6801190 | 6801484 | 295 | * | 77 | 7409 |
| 128 | chr19 | 13148148 | 13148413 | 266 | * | 54 | 4784 |
| 129 | chr19 | 16449517 | 16449963 | 447 | * | 314 | 10365 |
| 130 | chr19 | 34972216 | 34972661 | 446 | * | 197 | 126374 |
| 131 | chr19 | 39150272 | 39150540 | 269 | * | 77 | 81 |
| 132 | chr19 | 39156768 | 39157026 | 259 | * | 144 | 81 |
| 133 | chr19 | 41729361 | 41729778 | 418 | * | 606 | 558 |
| 134 | chr19 | 41731738 | 41732350 | 613 | * | 389 | 558 |
| 135 | chr19 | 54545376 | 54545939 | 564 | * | 133 | 284415 |
| 136 | chr2 | 36599212 | 36599778 | 567 | * | 208 | 51232 |
| 137 | chr2 | 37817283 | 37817662 | 380 | * | 123 | 10602 |
| 138 | chr2 | 37995707 | 37996071 | 365 | * | 315 | 10602 |
| 139 | chr2 | 46191293 | 46191638 | 346 | * | 59 | 5581 |
| 140 | chr2 | 47082424 | 47082853 | 430 | * | 153 | 1E+08 |

| | | | | | | | |
|---|---|---|---|---|---|---|---|
| 141 | chr2 | 47261340 | 47262054 | 715 | * | 116 | 57217 |
| 142 | chr2 | 47270488 | 47271211 | 724 | * | 133 | 57217 |
| 143 | chr2 | 47296935 | 47297372 | 438 | * | 387 | 57217 |
| 144 | chr2 | 47497063 | 47497428 | 366 | * | 28 | 805 |
| 145 | chr2 | 64881278 | 64881930 | 653 | * | 856 | 9792 |
| 146 | chr2 | 71944204 | 71944554 | 351 | * | 55 | 8291 |
| 147 | chr2 | 71970201 | 71970583 | 383 | * | 301 | 8291 |
| 148 | chr2 | 72060855 | 72061164 | 310 | * | 31 | 8291 |
| 149 | chr2 | 85185189 | 85185534 | 346 | * | 133 | 56888 |
| 150 | chr2 | 173918142 | 173918437 | 296 | * | 123 | 51776 |
| 151 | chr2 | 192500571 | 192501167 | 597 | * | 123 | 64859 |
| 152 | chr2 | 202016018 | 202016307 | 290 | * | 59 | 65072 |
| 153 | chr2 | 217237552 | 217238307 | 756 | * | 258 | 57574 |
| 154 | chr2 | 235336300 | 235336569 | 270 | * | 114 | 10123 |
| 155 | chr2 | 235889990 | 235890375 | 386 | * | 231 | 23677 |
| 156 | chr2 | 236007477 | 236007859 | 383 | * | 123 | 23677 |
| 157 | chr2 | 237149120 | 237149456 | 337 | * | 59 | 401036 |
| 158 | chr2 | 237569826 | 237570096 | 271 | * | 51 | 57007 |
| 159 | chr20 | 8995838 | 8996913 | 1076 | * | 404 | 5332 |
| 160 | chr20 | 9181402 | 9181921 | 520 | * | 256 | 5332 |
| 161 | chr20 | 22743670 | 22744411 | 742 | * | 72 | 3170 |
| 162 | chr20 | 22756082 | 22756691 | 610 | * | 169 | 3170 |
| 163 | chr20 | 30300584 | 30301410 | 827 | * | 221 | 598 |
| 164 | chr20 | 30307711 | 30308503 | 793 | * | 258 | 598 |
| 165 | chr20 | 43966332 | 43966805 | 474 | * | 181 | 6385 |
| 166 | chr20 | 45503618 | 45503902 | 285 | * | 30 | 2139 |
| 167 | chr20 | 52757538 | 52757830 | 293 | * | 103 | 1591 |
| 168 | chr20 | 59760991 | 59761669 | 679 | * | 841 | 1002 |
| 169 | chr20 | 60163773 | 60164312 | 540 | * | 17 | 1002 |
| 170 | chr21 | 46348349 | 46349309 | 961 | * | 144 | 3689 |
| 171 | chr22 | 19868703 | 19869020 | 318 | * | 707 | 10587 |
| 172 | chr22 | 24870319 | 24870743 | 425 | * | 95 | 51733 |
| 173 | chr22 | 26812797 | 26813143 | 347 | * | 77 | 57168 |
| 174 | chr22 | 30018125 | 30018465 | 341 | * | 154 | 4771 |
| 175 | chr22 | 33157409 | 33157784 | 376 | * | 28 | 7078 |
| 176 | chr22 | 33184852 | 33185296 | 445 | * | 154 | 7078 |
| 177 | chr22 | 36725492 | 36725792 | 301 | * | 35 | 4627 |
| 178 | chr22 | 37828296 | 37828586 | 291 | * | 59 | 114794 |
| 179 | chr22 | 37851225 | 37851615 | 391 | * | 35 | 114794 |
| 180 | chr22 | 37893077 | 37893368 | 292 | * | 254 | 29775 |
| 181 | chr22 | 37956958 | 37957853 | 896 | * | 54 | 11135 |
| 182 | chr22 | 39570767 | 39572655 | 1889 | * | 825 | 23492 |
| 183 | chr22 | 39610309 | 39611133 | 825 | * | 51 | 5155 |
| 184 | chr22 | 46357860 | 46358120 | 261 | * | 85 | 7477 |
| 185 | chr22 | 50345813 | 50346430 | 618 | * | 314 | 415116 |
| 186 | chr3 | 13812040 | 13812473 | 434 | * | 85 | 7476 |
| 187 | chr3 | 13891832 | 13892170 | 339 | * | 240 | 7476 |

| | | | | | | | |
|---|---|---|---|---|---|---|---|
| 188 | chr3 | 58033638 | 58034235 | 598 | * | 59 | 2317 |
| 189 | chr3 | 58546541 | 58546843 | 303 | * | 363 | 11170 |
| 190 | chr3 | 99282083 | 99282401 | 319 | * | 28 | 1295 |
| 191 | chr3 | 100215632 | 100215982 | 351 | * | 87 | 55076 |
| 192 | chr3 | 111578200 | 111578601 | 402 | * | 35 | 90102 |
| 193 | chr3 | 123469624 | 123469902 | 279 | * | 137 | 4638 |
| 194 | chr3 | 124781958 | 124782238 | 281 | * | 145 | 57493 |
| 195 | chr3 | 127487152 | 127487561 | 410 | * | 77 | 11343 |
| 196 | chr3 | 134044958 | 134045329 | 372 | * | 35 | 51421 |
| 197 | chr3 | 134045737 | 134046377 | 641 | * | 751 | 51421 |
| 198 | chr3 | 134076653 | 134077052 | 400 | * | 289 | 51421 |
| 199 | chr3 | 134082797 | 134083226 | 430 | * | 536 | 51421 |
| 200 | chr3 | 134092703 | 134093716 | 1014 | * | 463 | 51421 |
| 201 | chr3 | 159646507 | 159646862 | 356 | * | 105 | 3592 |
| 202 | chr3 | 192597396 | 192597697 | 302 | * | 92 | 151963 |
| 203 | chr4 | 184086259 | 184086638 | 380 | * | 35 | 80014 |
| 204 | chr5 | 5145916 | 5146231 | 316 | * | 16 | 170690 |
| 205 | chr5 | 5165658 | 5165914 | 257 | * | 19 | 170690 |
| 206 | chr5 | 5222025 | 5222447 | 423 | * | 153 | 170690 |
| 207 | chr5 | 6427875 | 6428154 | 280 | * | 31 | 134111 |
| 208 | chr5 | 10513757 | 10514503 | 747 | * | 181 | 651746 |
| 209 | chr5 | 10551317 | 10551977 | 661 | * | 1211 | 651746 |
| 210 | chr5 | 14035982 | 14036418 | 437 | * | 65 | 1767 |
| 211 | chr5 | 14036749 | 14037034 | 286 | * | 16 | 1767 |
| 212 | chr5 | 14038434 | 14038782 | 349 | * | 39 | 1767 |
| 213 | chr5 | 14265590 | 14266278 | 689 | * | 481 | 7204 |
| 214 | chr5 | 17128206 | 17128493 | 288 | * | 21 | 10409 |
| 215 | chr5 | 17249446 | 17249702 | 257 | * | 147 | 10409 |
| 216 | chr5 | 35919307 | 35919575 | 269 | * | 107 | 133690 |
| 217 | chr5 | 52658546 | 52658816 | 271 | * | 99 | 10468 |
| 218 | chr5 | 52766264 | 52766773 | 510 | * | 130 | 10468 |
| 219 | chr5 | 56988697 | 56989141 | 445 | * | 66 | 345651 |
| 220 | chr5 | 64494601 | 64494913 | 313 | * | 131 | 11174 |
| 221 | chr5 | 68797814 | 68798074 | 261 | * | 35 | 1.01E+08 |
| 222 | chr5 | 72934427 | 72934767 | 341 | * | 314 | 64283 |
| 223 | chr5 | 73060351 | 73060727 | 377 | * | 155 | 64283 |
| 224 | chr5 | 73234304 | 73234652 | 349 | * | 62 | 64283 |
| 225 | chr5 | 150017173 | 150017606 | 434 | * | 314 | 11346 |
| 226 | chr5 | 159253189 | 159253468 | 280 | * | 28 | 147 |
| 227 | chr5 | 159332516 | 159333064 | 549 | * | 210 | 147 |
| 228 | chr5 | 165499698 | 165499975 | 278 | * | 35 | 57451 |
| 229 | chr5 | 165767142 | 165767458 | 317 | * | 134 | 57451 |
| 230 | chr5 | 165784591 | 165786011 | 1421 | * | 3304 | 57451 |
| 231 | chr5 | 169213301 | 169213653 | 353 | * | 197 | 1794 |
| 232 | chr5 | 169397953 | 169398353 | 401 | * | 40 | 1E+08 |
| 233 | chr6 | 3797982 | 3798238 | 257 | * | 19 | 221749 |
| 234 | chr6 | 11277458 | 11277725 | 268 | * | 142 | 4739 |

| | | | | | | | |
|---|---|---|---|---|---|---|---|
| 235 | chr6 | 11653213 | 11653675 | 463 | * | 133 | 84830 |
| 236 | chr6 | 17278821 | 17279118 | 298 | * | 35 | 221662 |
| 237 | chr6 | 26055282 | 26055574 | 293 | * | 95 | 3006 |
| 238 | chr6 | 71959848 | 71960195 | 348 | * | 62 | 79627 |
| 239 | chr6 | 75915759 | 75916078 | 320 | * | 85 | 1303 |
| 240 | chr6 | 86113468 | 86113781 | 314 | * | 155 | 4907 |
| 241 | chr6 | 114338324 | 114338625 | 302 | * | 242 | 222537 |
| 242 | chr6 | 132272505 | 132272761 | 257 | * | 277 | 1490 |
| 243 | chr6 | 168962029 | 168962401 | 373 | * | 12 | 64094 |
| 244 | chr7 | 501135 | 501578 | 444 | * | 338 | 5154 |
| 245 | chr7 | 1576704 | 1577115 | 412 | * | 213 | 7975 |
| 246 | chr7 | 6322756 | 6323770 | 1015 | * | 164 | 9265 |
| 247 | chr7 | 6324193 | 6324739 | 547 | * | 47 | 9265 |
| 248 | chr7 | 6327785 | 6328055 | 271 | * | 42 | 9265 |
| 249 | chr7 | 6346258 | 6346537 | 280 | * | 34 | 9265 |
| 250 | chr7 | 12700114 | 12700403 | 290 | * | 160 | 10124 |
| 251 | chr7 | 24739878 | 24740240 | 363 | * | 102 | 1687 |
| 252 | chr7 | 32155441 | 32155729 | 289 | * | 335 | 5137 |
| 253 | chr7 | 34019283 | 34019725 | 443 | * | 57 | 168667 |
| 254 | chr7 | 34026776 | 34027130 | 355 | * | 57 | 168667 |
| 255 | chr7 | 41296451 | 41296799 | 349 | * | 21 | 285954 |
| 256 | chr7 | 41956330 | 41956671 | 342 | * | 59 | 3624 |
| 257 | chr7 | 47803070 | 47803413 | 344 | * | 21 | 84847 |
| 258 | chr7 | 47826082 | 47826630 | 549 | * | 515 | 80099 |
| 259 | chr7 | 48122982 | 48123264 | 283 | * | 34 | 7378 |
| 260 | chr7 | 76050343 | 76050697 | 355 | * | 133 | 7784 |
| 261 | chr7 | 98450303 | 98450698 | 396 | * | 24 | 222865 |
| 262 | chr7 | 98462978 | 98463509 | 532 | * | 565 | 222865 |
| 263 | chr7 | 100765284 | 100765588 | 305 | * | 153 | 5054 |
| 264 | chr7 | 115994831 | 115995441 | 611 | * | 288 | 858 |
| 265 | chr7 | 116152279 | 116152885 | 607 | * | 41 | 857 |
| 266 | chr7 | 128470372 | 128470710 | 339 | * | 46 | 2318 |
| 267 | chr7 | 131222771 | 131223571 | 801 | * | 116 | 5420 |
| 268 | chr7 | 131338876 | 131339512 | 637 | * | 101 | 5420 |
| 269 | chr7 | 135443008 | 135443310 | 303 | * | 35 | 389558 |
| 270 | chr7 | 140325982 | 140326484 | 503 | * | 320 | 27147 |
| 271 | chr7 | 143067173 | 143067524 | 352 | * | 174 | 9715 |
| 272 | chr7 | 150723100 | 150723397 | 298 | * | 19 | 285973 |
| 273 | chr7 | 151297124 | 151297379 | 256 | * | 147 | 51422 |
| 274 | chr7 | 151384709 | 151385121 | 413 | * | 262 | 51422 |
| 275 | chr7 | 151439876 | 151440200 | 325 | * | 81 | 51422 |
| 276 | chr7 | 151451244 | 151451638 | 395 | * | 877 | 51422 |
| 277 | chr7 | 157101776 | 157102060 | 285 | * | 153 | 10049 |
| 278 | chr7 | 157102547 | 157102999 | 453 | * | 155 | 10049 |
| 279 | chr8 | 8395155 | 8395490 | 336 | * | 51 | 157285 |
| 280 | chr8 | 26014830 | 26015136 | 307 | * | 31 | 64641 |
| 281 | chr8 | 37000437 | 37001059 | 623 | * | 95 | 157855 |

| | | | | | | | |
|---|---|---|---|---|---|---|---|
| 282 | chr8 | 86777505 | 86777898 | 394 | * | 23 | 254958 |
| 283 | chr8 | 86796533 | 86796824 | 292 | * | 12 | 254958 |
| 284 | chr8 | 86831537 | 86831838 | 302 | * | 14 | 254958 |
| 285 | chr8 | 102039342 | 102039846 | 505 | * | 69 | 441374 |
| 286 | chr8 | 118859969 | 118860266 | 298 | * | 46 | 2131 |
| 287 | chr8 | 118953013 | 118953333 | 321 | * | 143 | 2131 |
| 288 | chr8 | 119037362 | 119037626 | 265 | * | 11 | 2131 |
| 289 | chr8 | 119345170 | 119345503 | 334 | * | 353 | 2131 |
| 290 | chr8 | 128866268 | 128866659 | 392 | * | 133 | 5820 |
| 291 | chr8 | 128919904 | 128920393 | 490 | * | 208 | 5820 |
| 292 | chr8 | 128930920 | 128931185 | 266 | * | 66 | 5820 |
| 293 | chr8 | 131495578 | 131495937 | 360 | * | 394 | 50807 |
| 294 | chr8 | 131536926 | 131537264 | 339 | * | 95 | 50807 |
| 295 | chr8 | 142164137 | 142164445 | 309 | * | 77 | 22898 |
| 296 | chr9 | 5446945 | 5447411 | 467 | * | 51 | 29126 |
| 297 | chr9 | 5509580 | 5509899 | 320 | * | 108 | 80380 |
| 298 | chr9 | 12814390 | 12814655 | 266 | * | 26 | 286343 |
| 299 | chr9 | 35518187 | 35518497 | 311 | * | 642 | 9853 |
| 300 | chr9 | 38017295 | 38017614 | 320 | * | 150 | 6461 |
| 301 | chr9 | 38046868 | 38047547 | 680 | * | 86 | 6461 |
| 302 | chr9 | 38443304 | 38443719 | 416 | * | 54 | 347252 |
| 303 | chr9 | 38681842 | 38682957 | 1116 | * | 301 | 158228 |
| 304 | chr9 | 74755450 | 74755735 | 286 | * | 33 | 9615 |
| 305 | chr9 | 100639312 | 100639728 | 417 | * | 62 | 2304 |
| 306 | chr9 | 100985523 | 100985914 | 392 | * | 1108 | 55357 |
| 307 | chr9 | 110517251 | 110517857 | 607 | * | 127 | 9314 |
| 308 | chr9 | 112813243 | 112813522 | 280 | * | 33 | 11217 |
| 309 | chr9 | 112878372 | 112878672 | 301 | * | 38 | 11217 |
| 310 | chr9 | 114490989 | 114491312 | 324 | * | 38 | 158401 |
| 311 | chr9 | 117566629 | 117567069 | 441 | * | 300 | 9966 |
| 312 | chr9 | 127048280 | 127048972 | 693 | * | 125 | 10783 |
| 313 | chr9 | 127071997 | 127072265 | 269 | * | 395 | 10783 |
| 314 | chr9 | 133837918 | 133838229 | 312 | * | 21 | 84929 |
| 315 | chrX | 10415805 | 10416114 | 310 | * | 43 | 4281 |
| 316 | chrX | 48345644 | 48346099 | 456 | * | 197 | 24140 |
| 317 | chrX | 128756857 | 128757159 | 303 | * | 51 | 8862 |
| 318 | chrX | 133803925 | 133804379 | 455 | * | 67 | 10761 |
| 319 | chrX | 139805465 | 139805789 | 325 | * | 70 | 286411 |
| 320 | chrX | 139806485 | 139806748 | 264 | * | 55 | 286411 |
| 321 | chrX | 153597303 | 153598110 | 808 | * | 35 | 2316 |

| ENSEMBL | baseMean | log2 Fold Change | padj | Gene |
|---|---|---|---|---|
| ENSG00000120937 | 2240.650956 | 10.67463235 | 2.75E-73 | NPPB |
| ENSG00000233954 | 16.37024274 | 1.62106881 | 3.26E-03 | UQCRHL |
| ENSG00000142627 | 5020.238168 | 1.530699246 | 5.23E-38 | EPHA2 |
| ENSG00000142627 | 5020.238168 | 1.530699246 | 5.23E-38 | EPHA2 |
| ENSG00000142627 | 5020.238168 | 1.530699246 | 5.23E-38 | EPHA2 |
| ENSG00000131914 | 640.0340004 | 1.304152566 | 6.37E-21 | LIN28A |
| ENSG00000168528 | 3346.317931 | 2.249050216 | 1.13E-34 | SERINC2 |
| ENSG00000142910 | 244.1329187 | 5.327601074 | 3.18E-53 | TINAGL1 |
| ENSG00000179862 | 281.9765894 | 1.731345645 | 1.26E-17 | CITED4 |
| ENSG00000117395 | 5155.502085 | 1.182165418 | 4.71E-35 | EBNA1BP2 |
| ENSG00000178922 | 1853.666591 | 2.188904225 | 2.93E-67 | HYI |
| ENSG00000142871 | 10596.59591 | 4.421427289 | 0.00E+00 | CCN1 |
| ENSG00000117525 | 631.683175 | 2.557663625 | 1.89E-39 | F3 |
| ENSG00000117143 | 2438.450767 | 1.60748835 | 2.88E-41 | UAP1 |
| ENSG00000143850 | 883.8866704 | 1.040738194 | 3.18E-06 | PLEKHA6 |
| ENSG00000054392 | 280.182391 | 1.170771795 | 1.45E-18 | HHAT |
| ENSG00000162772 | 836.543072 | 2.441541232 | 2.14E-19 | ATF3 |
| ENSG00000232480 | 227.6156955 | 3.093180397 | 1.99E-56 | TGFB2-AS1 |
| ENSG00000232480 | 227.6156955 | 3.093180397 | 1.99E-56 | TGFB2-AS1 |
| ENSG00000225383 | 288.2208761 | 9.637580437 | 8.17E-26 | SFTA1P |
| ENSG00000204174 | 21.74673346 | 2.635152016 | 6.43E-05 | NPY4R |
| ENSG00000197467 | 1295.802804 | 1.807796912 | 2.22E-35 | COL13A1 |
| ENSG00000107738 | 1076.142038 | 1.147675268 | 6.42E-13 | VSIR |
| ENSG00000148677 | 2371.13122 | 7.931678425 | 1.24E-76 | ANKRD1 |
| ENSG00000138131 | 806.0990284 | 1.669022795 | 8.01E-39 | LOXL4 |
| ENSG00000120053 | 2517.089131 | 1.226959899 | 5.29E-15 | GOT1 |
| ENSG00000177106 | 349.987695 | 2.720567172 | 2.65E-38 | EPS8L2 |
| ENSG00000245522 | 102.7984567 | 2.554147404 | 7.11E-28 | AC026250.1 |
| ENSG00000133816 | 7480.93452 | 1.576973571 | 5.89E-33 | MICAL2 |
| ENSG00000133816 | 7480.93452 | 1.576973571 | 5.89E-33 | MICAL2 |
| ENSG00000133816 | 7480.93452 | 1.576973571 | 5.89E-33 | MICAL2 |
| ENSG00000133818 | 2136.436366 | 1.047375541 | 3.79E-27 | RRAS2 |
| ENSG00000026508 | 26250.0004 | 1.299837867 | 1.79E-30 | CD44 |
| ENSG00000124942 | 122265.4934 | 1.149081335 | 3.93E-42 | AHNAK |
| ENSG00000175592 | 1816.844697 | 3.419534893 | 4.01E-43 | FOSL1 |
| ENSG00000175315 | 11.6176009 | 4.119025563 | 6.92E-06 | CST6 |
| ENSG00000172927 | 22.27153524 | 1.917461805 | 8.65E-03 | MYEOV |
| ENSG00000172927 | 22.27153524 | 1.917461805 | 8.65E-03 | MYEOV |
| ENSG00000172927 | 22.27153524 | 1.917461805 | 8.65E-03 | MYEOV |
| ENSG00000172927 | 22.27153524 | 1.917461805 | 8.65E-03 | MYEOV |
| ENSG00000110092 | 9113.412653 | 1.284825408 | 1.48E-18 | CCND1 |
| ENSG00000110092 | 9113.412653 | 1.284825408 | 1.48E-18 | CCND1 |
| ENSG00000110092 | 9113.412653 | 1.284825408 | 1.48E-18 | CCND1 |
| ENSG00000110092 | 9113.412653 | 1.284825408 | 1.48E-18 | CCND1 |
| ENSG00000110092 | 9113.412653 | 1.284825408 | 1.48E-18 | CCND1 |
| ENSG00000110092 | 9113.412653 | 1.284825408 | 1.48E-18 | CCND1 |

| | | | | |
|---|---|---|---|---|
| ENSG00000162105 | 32.47462154 | 3.046109302 | 2.16E-03 | SHANK2 |
| ENSG00000168014 | 3048.904252 | 1.726154266 | 1.20E-88 | C2CD3 |
| ENSG00000158555 | 1564.667833 | 1.204457969 | 4.98E-19 | GDPD5 |
| ENSG00000166741 | 7823.226213 | 1.650169046 | 7.46E-18 | NNMT |
| ENSG00000166741 | 7823.226213 | 1.650169046 | 7.46E-18 | NNMT |
| ENSG00000166741 | 7823.226213 | 1.650169046 | 7.46E-18 | NNMT |
| ENSG00000246100 | 27.39226582 | 1.761190022 | 3.21E-03 | LINC00900 |
| ENSG00000149403 | 72.0864243 | 2.716637086 | 3.42E-07 | GRIK4 |
| ENSG00000149403 | 72.0864243 | 2.716637086 | 3.42E-07 | GRIK4 |
| ENSG00000154127 | 966.8217515 | 1.267852682 | 5.09E-10 | UBASH3B |
| ENSG00000154133 | 76.28270049 | 3.756184797 | 1.07E-13 | ROBO4 |
| ENSG00000134954 | 13708.86452 | 1.440477993 | 3.46E-84 | ETS1 |
| ENSG00000134954 | 13708.86452 | 1.440477993 | 3.46E-84 | ETS1 |
| ENSG00000134954 | 13708.86452 | 1.440477993 | 3.46E-84 | ETS1 |
| ENSG00000134954 | 13708.86452 | 1.440477993 | 3.46E-84 | ETS1 |
| ENSG00000151715 | 21.77121324 | 1.74997882 | 4.56E-04 | TMEM45B |
| ENSG00000118971 | 40.55711956 | 5.1319368 | 4.18E-12 | CCND2 |
| ENSG00000110799 | 121.5698336 | 1.536353185 | 2.51E-06 | VWF |
| ENSG00000057294 | 1275.453997 | 1.659408249 | 1.10E-54 | PKP2 |
| ENSG00000050405 | 10004.88249 | 1.000741833 | 1.40E-19 | LIMA1 |
| ENSG00000167767 | 1730.679913 | 6.984458242 | 8.65E-226 | KRT80 |
| ENSG00000167767 | 1730.679913 | 6.984458242 | 8.65E-226 | KRT80 |
| ENSG00000170421 | 14283.57687 | 3.561845464 | 3.44E-225 | KRT8 |
| ENSG00000198431 | 33083.72171 | 2.172909829 | 2.71E-33 | TXNRD1 |
| ENSG00000235162 | 2759.517718 | 1.17720482 | 6.01E-36 | C12orf75 |
| ENSG00000074590 | 3145.049458 | 1.803511144 | 1.64E-58 | NUAK1 |
| ENSG00000074590 | 3145.049458 | 1.803511144 | 1.64E-58 | NUAK1 |
| ENSG00000089060 | 1452.09912 | 1.433440202 | 7.98E-31 | SLC8B1 |
| ENSG00000171435 | 287.877213 | 2.261005241 | 6.19E-22 | KSR2 |
| ENSG00000150457 | 922.0682349 | 1.926800923 | 8.75E-69 | LATS2 |
| ENSG00000088387 | 3479.797232 | 1.037403126 | 3.87E-19 | DOCK9 |
| ENSG00000198542 | 61.58578191 | 3.330055993 | 2.03E-18 | ITGBL1 |
| ENSG00000198542 | 61.58578191 | 3.330055993 | 2.03E-18 | ITGBL1 |
| ENSG00000198542 | 61.58578191 | 3.330055993 | 2.03E-18 | ITGBL1 |
| ENSG00000129474 | 9629.715395 | 2.67219923 | 6.48E-56 | AJUBA |
| ENSG00000129474 | 9629.715395 | 2.67219923 | 6.48E-56 | AJUBA |
| ENSG00000129474 | 9629.715395 | 2.67219923 | 6.48E-56 | AJUBA |
| ENSG00000072415 | 2731.596394 | 1.316704198 | 1.05E-19 | MPP5 |
| ENSG00000140044 | 1542.857247 | 2.397867713 | 4.29E-31 | JDP2 |
| ENSG00000211448 | 72.37432467 | 2.461866145 | 3.80E-08 | DIO2 |
| ENSG00000205476 | 2884.103042 | 1.406442207 | 2.52E-29 | CCDC85C |
| ENSG00000205476 | 2884.103042 | 1.406442207 | 2.52E-29 | CCDC85C |
| ENSG00000185567 | 1587.711431 | 1.302930231 | 2.54E-14 | AHNAK2 |
| ENSG00000185567 | 1587.711431 | 1.302930231 | 2.54E-14 | AHNAK2 |
| ENSG00000103642 | 732.0504195 | 1.2623442 | 2.55E-22 | LACTB |
| ENSG00000138623 | 526.884678 | 2.123932915 | 3.26E-33 | SEMA7A |
| ENSG00000006327 | 3033.22403 | 3.376071254 | 5.09E-112 | TNFRSF12A |

| | | | | |
|---|---|---|---|---|
| ENSG00000118898 | 285.5073873 | 2.537692621 | 1.56E-31 | PPL |
| ENSG00000205336 | 794.4778913 | 1.249340522 | 3.61E-06 | ADGRG1 |
| ENSG00000172828 | 106.6239909 | 1.795910276 | 5.40E-12 | CES3 |
| ENSG00000132613 | 4140.209056 | 1.242834867 | 5.39E-32 | MTSS1L |
| ENSG00000140873 | 25.18789751 | 1.188012727 | 4.58E-02 | ADAMTS18 |
| ENSG00000153815 | 2119.779756 | 1.012576464 | 1.38E-21 | CMIP |
| ENSG00000153815 | 2119.779756 | 1.012576464 | 1.38E-21 | CMIP |
| ENSG00000103196 | 701.9003059 | 1.025306157 | 3.67E-06 | CRISPLD2 |
| ENSG00000103196 | 701.9003059 | 1.025306157 | 3.67E-06 | CRISPLD2 |
| ENSG00000103196 | 701.9003059 | 1.025306157 | 3.67E-06 | CRISPLD2 |
| ENSG00000176678 | 378.206564 | 1.111600767 | 3.13E-14 | FOXL1 |
| ENSG00000260456 | 250.6123435 | 1.153369532 | 2.13E-10 | C16orf95 |
| ENSG00000225614 | 430.1866375 | 1.478234164 | 2.83E-14 | ZNF469 |
| ENSG00000178773 | 438.1070185 | 1.528419897 | 3.86E-21 | CPNE7 |
| ENSG00000167693 | 1791.096427 | 1.059246125 | 4.32E-18 | NXN |
| ENSG00000167693 | 1791.096427 | 1.059246125 | 4.32E-18 | NXN |
| ENSG00000167693 | 1791.096427 | 1.059246125 | 4.32E-18 | NXN |
| ENSG00000213977 | 4806.333429 | 1.001409951 | 3.72E-16 | TAX1BP3 |
| ENSG00000154914 | 231.4230637 | 1.524972215 | 2.82E-18 | USP43 |
| ENSG00000170425 | 805.8301713 | 1.390534745 | 3.35E-31 | ADORA2B |
| ENSG00000170425 | 805.8301713 | 1.390534745 | 3.35E-31 | ADORA2B |
| ENSG00000275832 | 5404.484757 | 1.019736403 | 1.15E-30 | ARHGAP23 |
| ENSG00000275832 | 5404.484757 | 1.019736403 | 1.15E-30 | ARHGAP23 |
| ENSG00000180340 | 1127.496047 | 1.165126102 | 3.73E-19 | FZD2 |
| ENSG00000246640 | 40.09760807 | 2.13699361 | 2.67E-09 | PICART1 |
| ENSG00000183691 | 208.1326203 | 2.328005262 | 8.42E-14 | NOG |
| ENSG00000176845 | 538.6256251 | 1.533571619 | 1.74E-23 | METRNL |
| ENSG00000101608 | 9438.49279 | 1.018748187 | 1.09E-33 | MYL12A |
| ENSG00000049759 | 4687.21832 | 1.341497269 | 1.22E-43 | NEDD4L |
| ENSG00000198796 | 2595.841625 | 2.637778074 | 3.86E-32 | ALPK2 |
| ENSG00000256463 | 117.0169671 | 12.06296652 | 4.53E-02 | SALL3 |
| ENSG00000256463 | 117.0169671 | 12.06296652 | 4.53E-02 | SALL3 |
| ENSG00000099860 | 977.0968945 | 2.896611578 | 1.75E-145 | GADD45B |
| ENSG00000141968 | 107.6228808 | 1.844552366 | 1.50E-11 | VAV1 |
| ENSG00000008441 | 4322.548575 | 1.16909868 | 1.92E-34 | NFIX |
| ENSG00000127528 | 119.5906069 | 1.528805539 | 1.53E-11 | KLF2 |
| ENSG00000142279 | 955.4419514 | 1.920763905 | 2.32E-24 | WTIP |
| ENSG00000130402 | 24964.65553 | 1.696706171 | 1.97E-61 | ACTN4 |
| ENSG00000130402 | 24964.65553 | 1.696706171 | 1.97E-61 | ACTN4 |
| ENSG00000167601 | 11912.61544 | 3.206422493 | 4.27E-81 | AXL |
| ENSG00000167601 | 11912.61544 | 3.206422493 | 4.27E-81 | AXL |
| ENSG00000189068 | 56.60755979 | 6.876642047 | 4.81E-21 | VSTM1 |
| ENSG00000150938 | 9816.058979 | 2.508939451 | 4.90E-112 | CRIM1 |
| ENSG00000163171 | 1854.372981 | 2.959094769 | 1.20E-40 | CDC42EP3 |
| ENSG00000163171 | 1854.372981 | 2.959094769 | 1.20E-40 | CDC42EP3 |
| ENSG00000171132 | 1501.609574 | 2.281798436 | 6.33E-29 | PRKCE |
| ENSG00000239332 | 98.2556091 | 1.551499889 | 4.24E-09 | LINC01119 |

| | | | | |
|---|---|---|---|---|
| ENSG00000068724 | 1548.824814 | 1.053526595 | 3.63E-27 | TTC7A |
| ENSG00000068724 | 1548.824814 | 1.053526595 | 3.63E-27 | TTC7A |
| ENSG00000068724 | 1548.824814 | 1.053526595 | 3.63E-27 | TTC7A |
| ENSG00000143933 | 26395.33632 | 1.218316546 | 1.76E-40 | CALM2 |
| ENSG00000179833 | 1650.89911 | 1.165894049 | 4.88E-28 | SERTAD2 |
| ENSG00000135636 | 2529.150232 | 1.450251707 | 3.00E-25 | DYSF |
| ENSG00000135636 | 2529.150232 | 1.450251707 | 3.00E-25 | DYSF |
| ENSG00000135636 | 2529.150232 | 1.450251707 | 3.00E-25 | DYSF |
| ENSG00000176407 | 2322.440073 | 1.043979105 | 6.66E-15 | KCMF1 |
| ENSG00000091436 | 5636.626502 | 1.792735325 | 1.80E-35 | MAP3K20 |
| ENSG00000173559 | 2078.385631 | 1.563518108 | 2.45E-18 | NABP1 |
| ENSG00000226312 | 19.01424679 | 1.353713908 | 8.85E-03 | CFLAR-AS1 |
| ENSG00000144583 | 1178.181843 | 4.827438381 | 7.09E-83 | 4-Mar |
| ENSG00000188042 | 1081.703425 | 1.418610678 | 1.37E-24 | ARL4C |
| ENSG00000130147 | 4262.928303 | 1.685013308 | 8.64E-64 | SH3BP4 |
| ENSG00000130147 | 4262.928303 | 1.685013308 | 8.64E-64 | SH3BP4 |
| ENSG00000182177 | 18.35133093 | 4.621492373 | 1.96E-05 | ASB18 |
| ENSG00000144476 | 110.6345116 | 2.16381406 | 1.88E-20 | ACKR3 |
| ENSG00000101333 | 1544.325878 | 2.374801506 | 8.09E-35 | PLCB4 |
| ENSG00000101333 | 1544.325878 | 2.374801506 | 8.09E-35 | PLCB4 |
| ENSG00000125798 | 107.3111875 | 1.619656406 | 1.60E-10 | FOXA2 |
| ENSG00000125798 | 107.3111875 | 1.619656406 | 1.60E-10 | FOXA2 |
| ENSG00000171552 | 5620.751235 | 1.502331064 | 7.26E-74 | BCL2L1 |
| ENSG00000171552 | 5620.751235 | 1.502331064 | 7.26E-74 | BCL2L1 |
| ENSG00000124145 | 4176.418327 | 1.856894944 | 8.79E-40 | SDC4 |
| ENSG00000064655 | 133.8226664 | 1.265813297 | 2.27E-10 | EYA2 |
| ENSG00000019186 | 230.7349006 | 1.116352939 | 3.95E-07 | CYP24A1 |
| ENSG00000179242 | 706.3312431 | 4.837604468 | 1.31E-109 | CDH4 |
| ENSG00000179242 | 706.3312431 | 4.837604468 | 1.31E-109 | CDH4 |
| ENSG00000160255 | 124.990794 | 3.53430358 | 8.08E-27 | ITGB2 |
| ENSG00000184470 | 1698.664478 | 1.253327426 | 8.68E-36 | TXNRD2 |
| ENSG00000100024 | 37.15425623 | 2.858344741 | 1.81E-09 | UPB1 |
| ENSG00000128203 | 456.4862353 | 1.029847201 | 1.05E-20 | ASPHD2 |
| ENSG00000186575 | 3616.898035 | 1.614859228 | 6.41E-55 | NF2 |
| ENSG00000100234 | 33487.39915 | 1.157498928 | 1.57E-15 | TIMP3 |
| ENSG00000100234 | 33487.39915 | 1.157498928 | 1.57E-15 | TIMP3 |
| ENSG00000100345 | 39701.66481 | 2.035789822 | 1.82E-90 | MYH9 |
| ENSG00000166897 | 490.7239094 | 2.704666578 | 9.26E-36 | ELFN2 |
| ENSG00000166897 | 490.7239094 | 2.704666578 | 9.26E-36 | ELFN2 |
| ENSG00000100065 | 1474.574126 | 1.171790719 | 7.01E-14 | CARD10 |
| ENSG00000128283 | 3906.296252 | 1.513609279 | 9.87E-17 | CDC42EP1 |
| ENSG00000100307 | 937.6131425 | 1.215292325 | 3.80E-16 | CBX7 |
| ENSG00000100311 | 265.4831051 | 6.747696631 | 3.28E-82 | PDGFB |
| ENSG00000188064 | 200.4097672 | 3.660179928 | 4.20E-25 | WNT7B |
| ENSG00000198355 | 1569.131357 | 1.523745446 | 1.02E-70 | PIM3 |
| ENSG00000154764 | 74.42957687 | 3.924149434 | 8.08E-22 | WNT7A |
| ENSG00000154764 | 74.42957687 | 3.924149434 | 8.08E-22 | WNT7A |

| | | | | |
|---|---|---|---|---|
| ENSG00000136068 | 29152.77127 | 1.98251782 | 3.71E-52 | FLNB |
| ENSG00000168309 | 21.23397353 | 1.529578878 | 2.52E-03 | FAM107A |
| ENSG00000144810 | 2277.352965 | 2.538334684 | 4.53E-39 | COL8A1 |
| ENSG00000181458 | 771.370834 | 2.327989867 | 5.75E-35 | TMEM45A |
| ENSG00000144824 | 5271.381741 | 1.017082024 | 8.51E-10 | PHLDB2 |
| ENSG00000065534 | 2007.992478 | 1.426470309 | 4.22E-20 | MYLK |
| ENSG00000173706 | 10780.54503 | 1.288822295 | 6.37E-68 | HEG1 |
| ENSG00000074416 | 4291.086206 | 1.952679343 | 1.31E-53 | MGLL |
| ENSG00000114019 | 5194.214834 | 4.038764035 | 0.00E+00 | AMOTL2 |
| ENSG00000114019 | 5194.214834 | 4.038764035 | 0.00E+00 | AMOTL2 |
| ENSG00000114019 | 5194.214834 | 4.038764035 | 0.00E+00 | AMOTL2 |
| ENSG00000114019 | 5194.214834 | 4.038764035 | 0.00E+00 | AMOTL2 |
| ENSG00000114019 | 5194.214834 | 4.038764035 | 0.00E+00 | AMOTL2 |
| ENSG00000168811 | 40.27153141 | 1.541347261 | 2.13E-06 | IL12A |
| ENSG00000180611 | 492.152103 | 1.019280921 | 5.98E-21 | MB21D2 |
| ENSG00000151718 | 2643.317265 | 1.509447136 | 1.28E-17 | WWC2 |
| ENSG00000145536 | 718.0575453 | 2.76793907 | 2.53E-33 | ADAMTS16 |
| ENSG00000145536 | 718.0575453 | 2.76793907 | 2.53E-33 | ADAMTS16 |
| ENSG00000145536 | 718.0575453 | 2.76793907 | 2.53E-33 | ADAMTS16 |
| ENSG00000215218 | 493.8624657 | 1.4657712 | 1.34E-16 | UBE2QL1 |
| ENSG00000164236 | 166.908834 | 1.817442684 | 1.77E-20 | ANKRD33B |
| ENSG00000164236 | 166.908834 | 1.817442684 | 1.77E-20 | ANKRD33B |
| ENSG00000039139 | 297.4360865 | 1.389376391 | 2.40E-20 | DNAH5 |
| ENSG00000039139 | 297.4360865 | 1.389376391 | 2.40E-20 | DNAH5 |
| ENSG00000039139 | 297.4360865 | 1.389376391 | 2.40E-20 | DNAH5 |
| ENSG00000038382 | 14689.77273 | 1.065849763 | 1.29E-32 | TRIO |
| ENSG00000176788 | 2291.351755 | 2.506771514 | 1.92E-63 | BASP1 |
| ENSG00000176788 | 2291.351755 | 2.506771514 | 1.92E-63 | BASP1 |
| ENSG00000152611 | 15.12718097 | 4.643573298 | 3.23E-06 | CAPSL |
| ENSG00000134363 | 410.4366751 | 2.868266875 | 7.23E-26 | FST |
| ENSG00000134363 | 410.4366751 | 2.868266875 | 7.23E-26 | FST |
| ENSG00000169067 | 62.37567461 | 1.28889754 | 7.17E-04 | ACTBL2 |
| ENSG00000049192 | 290.3621183 | 1.434549091 | 5.98E-17 | ADAMTS6 |
| ENSG00000197822 | 724.0860483 | 2.069521979 | 6.07E-21 | OCLN |
| ENSG00000214944 | 3310.07867 | 2.443574885 | 2.34E-56 | ARHGEF28 |
| ENSG00000214944 | 3310.07867 | 2.443574885 | 2.34E-56 | ARHGEF28 |
| ENSG00000214944 | 3310.07867 | 2.443574885 | 2.34E-56 | ARHGEF28 |
| ENSG00000171992 | 2426.228123 | 1.073150399 | 2.22E-20 | SYNPO |
| ENSG00000170214 | 405.8511243 | 1.013794958 | 3.56E-03 | ADRA1B |
| ENSG00000170214 | 405.8511243 | 1.013794958 | 3.56E-03 | ADRA1B |
| ENSG00000145934 | 351.6327637 | 3.493048385 | 2.55E-13 | TENM2 |
| ENSG00000145934 | 351.6327637 | 3.493048385 | 2.55E-13 | TENM2 |
| ENSG00000145934 | 351.6327637 | 3.493048385 | 2.55E-13 | TENM2 |
| ENSG00000134516 | 1547.341543 | 1.367340145 | 2.99E-19 | DOCK2 |
| ENSG00000204767 | 326.6455031 | 2.692499195 | 5.17E-29 | INSYN2B |
| ENSG00000168994 | 1898.914831 | 1.884426795 | 2.80E-99 | PXDC1 |
| ENSG00000111859 | 1801.09962 | 1.762854807 | 2.41E-55 | NEDD9 |

| | | | | |
|---|---|---|---|---|
| ENSG00000111863 | 17.75775975 | 1.875520757 | 1.41E-04 | ADTRP |
| ENSG00000112183 | 126.6531076 | 2.641914033 | 1.65E-17 | RBM24 |
| ENSG00000187837 | 590.5061429 | 1.057532286 | 2.37E-07 | HIST1H1C |
| ENSG00000119900 | 1421.080525 | 2.665778775 | 8.28E-57 | OGFRL1 |
| ENSG00000111799 | 7146.720884 | 1.66992455 | 1.11E-35 | COL12A1 |
| ENSG00000135318 | 5643.186994 | 1.098477548 | 2.45E-11 | NT5E |
| ENSG00000249853 | 7.638299293 | 4.494994025 | 3.05E-05 | HS3ST5 |
| ENSG00000118523 | 8122.707037 | 4.828236594 | 3.61E-137 | CCN2 |
| ENSG00000112562 | 58.39393359 | 1.061209196 | 3.44E-02 | SMOC2 |
| ENSG00000197461 | 1396.475027 | 1.065265045 | 1.54E-13 | PDGFA |
| ENSG00000198517 | 1886.636662 | 1.853398636 | 2.55E-64 | MAFK |
| ENSG00000008256 | 8882.569858 | 1.826156102 | 7.91E-130 | CYTH3 |
| ENSG00000008256 | 8882.569858 | 1.826156102 | 7.91E-130 | CYTH3 |
| ENSG00000008256 | 8882.569858 | 1.826156102 | 7.91E-130 | CYTH3 |
| ENSG00000008256 | 8882.569858 | 1.826156102 | 7.91E-130 | CYTH3 |
| ENSG00000122644 | 624.3512308 | 1.408047184 | 7.44E-23 | ARL4A |
| ENSG00000105928 | 2169.264017 | 1.208243594 | 1.32E-23 | GSDME |
| ENSG00000154678 | 3542.718138 | 2.014273807 | 1.28E-26 | PDE1C |
| ENSG00000164619 | 162.7326693 | 2.236491034 | 1.14E-09 | BMPER |
| ENSG00000164619 | 162.7326693 | 2.236491034 | 1.14E-09 | BMPER |
| ENSG00000224116 | 14.99610622 | 2.078864428 | 6.44E-04 | INHBA-AS1 |
| ENSG00000122641 | 62.28707009 | 2.769666524 | 3.29E-26 | INHBA |
| ENSG00000146666 | 14.59269898 | 3.401969457 | 1.60E-07 | LINC00525 |
| ENSG00000136275 | 6.788843734 | 3.861705243 | 1.53E-04 | C7orf69 |
| ENSG00000183696 | 3542.442731 | 1.722678676 | 1.48E-17 | UPP1 |
| ENSG00000188372 | 653.8483888 | 1.109159095 | 1.05E-13 | ZP3 |
| ENSG00000166448 | 81.1847839 | 3.268741592 | 2.21E-18 | TMEM130 |
| ENSG00000166448 | 81.1847839 | 3.268741592 | 2.21E-18 | TMEM130 |
| ENSG00000106366 | 2089.772652 | 3.649129014 | 5.25E-112 | SERPINE1 |
| ENSG00000105971 | 2124.493216 | 1.948515269 | 1.67E-58 | CAV2 |
| ENSG00000105974 | 23107.43986 | 2.196673475 | 4.64E-72 | CAV1 |
| ENSG00000128591 | 24286.4346 | 1.772103803 | 1.43E-44 | FLNC |
| ENSG00000128567 | 5988.65413 | 1.5004302 | 2.41E-25 | PODXL |
| ENSG00000128567 | 5988.65413 | 1.5004302 | 2.41E-25 | PODXL |
| ENSG00000189320 | 81.88165982 | 2.175887889 | 7.34E-08 | FAM180A |
| ENSG00000146966 | 3519.216737 | 1.394826611 | 7.79E-28 | DENND2A |
| ENSG00000159784 | 590.29546 | 1.844743843 | 9.64E-41 | FAM131B |
| ENSG00000181652 | 118.2688634 | 1.076563944 | 1.27E-02 | ATG9B |
| ENSG00000106617 | 1255.564186 | 2.309249859 | 1.55E-67 | PRKAG2 |
| ENSG00000106617 | 1255.564186 | 2.309249859 | 1.55E-67 | PRKAG2 |
| ENSG00000106617 | 1255.564186 | 2.309249859 | 1.55E-67 | PRKAG2 |
| ENSG00000106617 | 1255.564186 | 2.309249859 | 1.55E-67 | PRKAG2 |
| ENSG00000105993 | 11954.55005 | 1.01403428 | 5.58E-52 | DNAJB6 |
| ENSG00000105993 | 11954.55005 | 1.01403428 | 5.58E-52 | DNAJB6 |
| ENSG00000275342 | 3099.318571 | 1.093136851 | 1.14E-12 | PRAG1 |
| ENSG00000221818 | 15.16971605 | 1.426811899 | 1.45E-02 | EBF2 |
| ENSG00000215262 | 6.149298403 | 5.030626405 | 1.89E-03 | KCNU1 |

| | | | | |
|---|---|---|---|---|
| ENSG00000205176 | 16.21114651 | 5.741275481 | 5.67E-04 | REXO1L1P |
| ENSG00000205176 | 16.21114651 | 5.741275481 | 5.67E-04 | REXO1L1P |
| ENSG00000205176 | 16.21114651 | 5.741275481 | 5.67E-04 | REXO1L1P |
| ENSG00000248599 | 5.21339508 | 5.181368897 | 6.77E-04 | AC018781.1 |
| ENSG00000182197 | 6550.50112 | 1.203749863 | 1.29E-21 | EXT1 |
| ENSG00000182197 | 6550.50112 | 1.203749863 | 1.29E-21 | EXT1 |
| ENSG00000182197 | 6550.50112 | 1.203749863 | 1.29E-21 | EXT1 |
| ENSG00000182197 | 6550.50112 | 1.203749863 | 1.29E-21 | EXT1 |
| ENSG00000249859 | 340.4968136 | 1.223464188 | 1.01E-08 | PVT1 |
| ENSG00000249859 | 340.4968136 | 1.223464188 | 1.01E-08 | PVT1 |
| ENSG00000249859 | 340.4968136 | 1.223464188 | 1.01E-08 | PVT1 |
| ENSG00000153317 | 7407.398524 | 1.080371131 | 4.67E-27 | ASAP1 |
| ENSG00000153317 | 7407.398524 | 1.080371131 | 4.67E-27 | ASAP1 |
| ENSG00000105339 | 1469.517193 | 1.352424612 | 1.20E-43 | DENND3 |
| ENSG00000120217 | 323.6243598 | 2.972020063 | 5.08E-54 | CD274 |
| ENSG00000197646 | 137.9408949 | 1.597674856 | 7.36E-10 | PDCD1LG2 |
| ENSG00000153714 | 175.7105056 | 2.099682949 | 3.22E-14 | LURAP1L |
| ENSG00000198853 | 2608.465768 | 1.730118326 | 4.62E-100 | RUSC2 |
| ENSG00000107338 | 1465.471817 | 1.277633999 | 4.32E-18 | SHB |
| ENSG00000107338 | 1465.471817 | 1.277633999 | 4.32E-18 | SHB |
| ENSG00000137142 | 580.1775613 | 2.141215968 | 6.93E-65 | IGFBPL1 |
| ENSG00000204860 | 45.64172295 | 3.436907561 | 1.52E-12 | FAM201A |
| ENSG00000119125 | 761.696661 | 2.555351994 | 9.99E-40 | GDA |
| ENSG00000178919 | 349.40179 | 2.298111999 | 3.58E-37 | FOXE1 |
| ENSG00000095383 | 462.5151675 | 2.188108678 | 4.24E-48 | TBC1D2 |
| ENSG00000136826 | 330.1654842 | 1.560211204 | 7.42E-08 | KLF4 |
| ENSG00000241978 | 11074.45609 | 1.022319256 | 2.12E-31 | AKAP2 |
| ENSG00000241978 | 11074.45609 | 1.022319256 | 2.12E-31 | AKAP2 |
| ENSG00000165181 | 8.469872835 | 2.429341631 | 3.35E-02 | SHOC1 |
| ENSG00000181634 | 89.03391984 | 1.195094316 | 2.30E-04 | TNFSF15 |
| ENSG00000119408 | 8959.611567 | 1.087212252 | 1.05E-25 | NEK6 |
| ENSG00000119408 | 8959.611567 | 1.087212252 | 1.05E-25 | NEK6 |
| ENSG00000130720 | 129.1126746 | 1.469153028 | 2.55E-08 | FIBCD1 |
| ENSG00000101871 | 1581.448112 | 1.156928429 | 2.40E-23 | MID1 |
| ENSG00000068438 | 3401.260365 | 1.044416175 | 1.04E-42 | FTSJ1 |
| ENSG00000171388 | 612.3809223 | 2.743580287 | 5.38E-27 | APLN |
| ENSG00000170965 | 82.08555886 | 2.947634662 | 3.74E-32 | PLAC1 |
| ENSG00000203930 | 86.52816748 | 1.689889739 | 1.83E-10 | LINC00632 |
| ENSG00000203930 | 86.52816748 | 1.689889739 | 1.83E-10 | LINC00632 |
| ENSG00000196924 | 161615.4789 | 1.448408398 | 4.77E-61 | FLNA |

| | seqnames | start | end | width | strand | V5 | geneId |
|---|---|---|---|---|---|---|---|
| 1 | chr1 | 16470691 | 16473814 | 3124 | * | 143 | 1969 |
| 2 | chr1 | 16500171 | 16500446 | 276 | * | 130 | 1969 |
| 3 | chr1 | 26745568 | 26745877 | 310 | * | 20 | 79727 |
| 4 | chr1 | 27319850 | 27320229 | 380 | * | 60 | 388610 |
| 5 | chr1 | 27325825 | 27326224 | 400 | * | 98 | 388610 |
| 6 | chr1 | 28764542 | 28764887 | 346 | * | 69 | 65979 |
| 7 | chr1 | 43672425 | 43672985 | 561 | * | 229 | 10969 |
| 8 | chr1 | 48035652 | 48035948 | 297 | * | 123 | 2306 |
| 9 | chr1 | 85666871 | 85668161 | 1291 | * | 392 | 84144 |
| 10 | chr1 | 86045969 | 86047024 | 1056 | * | 462 | 3491 |
| 11 | chr1 | 86072754 | 86073248 | 495 | * | 69 | 3491 |
| 12 | chr1 | 86078137 | 86078727 | 591 | * | 45 | 3491 |
| 13 | chr1 | 92202283 | 92202618 | 336 | * | 45 | 7049 |
| 14 | chr1 | 94219904 | 94220360 | 457 | * | 60 | 8412 |
| 15 | chr1 | 94701686 | 94701982 | 297 | * | 37 | 9411 |
| 16 | chr1 | 94791431 | 94791703 | 273 | * | 30 | 9411 |
| 17 | chr1 | 110419465 | 110420459 | 995 | * | 180 | 1435 |
| 18 | chr1 | 145092201 | 145092948 | 748 | * | 211 | 9554 |
| 19 | chr1 | 151167717 | 151168396 | 680 | * | 218 | 8394 |
| 20 | chr1 | 151960675 | 151961361 | 687 | * | 69 | 6281 |
| 21 | chr1 | 154944978 | 154945721 | 744 | * | 114 | 6464 |
| 22 | chr1 | 156866462 | 156866916 | 455 | * | 69 | 375033 |
| 23 | chr1 | 157989200 | 157989745 | 546 | * | 398 | 55243 |
| 24 | chr1 | 202558413 | 202558703 | 291 | * | 111 | 4660 |
| 25 | chr1 | 214578011 | 214578393 | 383 | * | 30 | 5784 |
| 26 | chr1 | 214725070 | 214725779 | 710 | * | 505 | 5784 |
| 27 | chr1 | 231275423 | 231276229 | 807 | * | 30 | 440730 |
| 28 | chr10 | 14116505 | 14116793 | 289 | * | 60 | 55691 |
| 29 | chr10 | 95048585 | 95049037 | 453 | * | 164 | 26509 |
| 30 | chr10 | 95218439 | 95218982 | 544 | * | 30 | 26509 |
| 31 | chr10 | 112170369 | 112170880 | 512 | * | 18 | 1847 |
| 32 | chr11 | 9685672 | 9685988 | 317 | * | 90 | 23075 |
| 33 | chr11 | 12398728 | 12399341 | 614 | * | 444 | 55742 |
| 34 | chr11 | 12439588 | 12440294 | 707 | * | 39 | 55742 |
| 35 | chr11 | 12578393 | 12578738 | 346 | * | 142 | 7003 |
| 36 | chr11 | 12695648 | 12696339 | 692 | * | 554 | 7003 |
| 37 | chr11 | 12824088 | 12824355 | 268 | * | 45 | 7003 |
| 38 | chr11 | 12915216 | 12915827 | 612 | * | 37 | 7003 |
| 39 | chr11 | 14443839 | 14444215 | 377 | * | 191 | 22800 |
| 40 | chr11 | 19391844 | 19392220 | 377 | * | 77 | 89797 |
| 41 | chr11 | 36171351 | 36171636 | 286 | * | 76 | 143458 |
| 42 | chr11 | 44009067 | 44009525 | 459 | * | 37 | 387763 |
| 43 | chr11 | 46312294 | 46312889 | 596 | * | 153 | 90993 |
| 44 | chr11 | 46316827 | 46317184 | 358 | * | 38 | 90993 |
| 45 | chr11 | 62276155 | 62276573 | 419 | * | 51 | 79026 |
| 46 | chr11 | 62312607 | 62313996 | 1390 | * | 146 | 79026 |

| | | | | | | | |
|---|---|---|---|---|---|---|---|
| 47 | chr11 | 62320800 | 62321240 | 441 | * | 38 | 79026 |
| 48 | chr11 | 67780335 | 67781497 | 1163 | * | 373 | 221 |
| 49 | chr11 | 85780367 | 85780966 | 600 | * | 54 | 8301 |
| 50 | chr11 | 94470403 | 94470880 | 478 | * | 69 | 154810 |
| 51 | chr11 | 101980790 | 101981077 | 288 | * | 11 | 10413 |
| 52 | chr11 | 101982302 | 101982654 | 353 | * | 45 | 10413 |
| 53 | chr11 | 102139149 | 102139523 | 375 | * | 90 | 330 |
| 54 | chr11 | 113423910 | 113424358 | 449 | * | 146 | 1813 |
| 55 | chr11 | 121337186 | 121337565 | 380 | * | 52 | 6653 |
| 56 | chr11 | 125260776 | 125261290 | 515 | * | 101 | 63876 |
| 57 | chr11 | 128584864 | 128585765 | 902 | * | 77 | 100507392 |
| 58 | chr11 | 130348299 | 130349056 | 758 | * | 38 | 170689 |
| 59 | chr11 | 130350333 | 130350607 | 275 | * | 12 | 170689 |
| 60 | chr12 | 1714807 | 1715268 | 462 | * | 52 | 81029 |
| 61 | chr12 | 12419925 | 12420296 | 372 | * | 378 | 4040 |
| 62 | chr12 | 48214295 | 48214642 | 348 | * | 89 | 51564 |
| 63 | chr12 | 52585646 | 52586238 | 593 | * | 174 | 144501 |
| 64 | chr12 | 53318538 | 53319368 | 831 | * | 69 | 3856 |
| 65 | chr12 | 53339108 | 53339865 | 758 | * | 69 | 3875 |
| 66 | chr12 | 56614848 | 56615309 | 462 | * | 30 | 10193 |
| 67 | chr12 | 65672464 | 65672880 | 417 | * | 23 | 253827 |
| 68 | chr12 | 94179180 | 94179599 | 420 | * | 30 | 8738 |
| 69 | chr12 | 104612982 | 104613669 | 688 | * | 77 | 7296 |
| 70 | chr12 | 106424495 | 106424846 | 352 | * | 114 | 9891 |
| 71 | chr12 | 106468366 | 106469154 | 789 | * | 77 | 9891 |
| 72 | chr12 | 106477725 | 106478039 | 315 | * | 86 | 9891 |
| 73 | chr12 | 106517472 | 106517733 | 262 | * | 60 | 9891 |
| 74 | chr12 | 120524622 | 120524935 | 314 | * | 95 | 92558 |
| 75 | chr13 | 21635103 | 21635912 | 810 | * | 361 | 26524 |
| 76 | chr13 | 78549376 | 78549892 | 517 | * | 33 | 1910 |
| 77 | chr13 | 80510585 | 80511317 | 733 | * | 52 | 10253 |
| 78 | chr13 | 81030328 | 81030675 | 348 | * | 20 | 10253 |
| 79 | chr13 | 97774389 | 97775392 | 1004 | * | 60 | 10150 |
| 80 | chr13 | 97845914 | 97847006 | 1093 | * | 94 | 10150 |
| 81 | chr13 | 97874681 | 97876342 | 1662 | * | 25 | 10150 |
| 82 | chr13 | 97908526 | 97908826 | 301 | * | 22 | 10150 |
| 83 | chr13 | 97910015 | 97911140 | 1126 | * | 44 | 10150 |
| 84 | chr13 | 97927088 | 97927361 | 274 | * | 67 | 10150 |
| 85 | chr13 | 97959831 | 97961450 | 1620 | * | 345 | 10150 |
| 86 | chr13 | 97980281 | 97980794 | 514 | * | 65 | 10150 |
| 87 | chr14 | 23449613 | 23450968 | 1356 | * | 1429 | 84962 |
| 88 | chr14 | 23451833 | 23452477 | 645 | * | 885 | 84962 |
| 89 | chr14 | 55031972 | 55032386 | 415 | * | 12 | 23034 |
| 90 | chr14 | 55118581 | 55119512 | 932 | * | 311 | 23034 |
| 91 | chr14 | 55240189 | 55240475 | 287 | * | 52 | 23034 |
| 92 | chr14 | 75761947 | 75762206 | 260 | * | 15 | 731223 |
| 93 | chr14 | 89021184 | 89021544 | 361 | * | 104 | 11099 |

| | | | | | | | |
|---|---|---|---|---|---|---|---|
| 94 | chr14 | 102746509 | 102747063 | 555 | * | 30 | 5891 |
| 95 | chr15 | 57719653 | 57720169 | 517 | * | 102 | 84952 |
| 96 | chr15 | 58839441 | 58839708 | 268 | * | 17 | 102 |
| 97 | chr15 | 63268526 | 63269190 | 665 | * | 60 | 7168 |
| 98 | chr15 | 63337020 | 63337462 | 443 | * | 96 | 7168 |
| 99 | chr15 | 69706738 | 69707872 | 1135 | * | 69 | 9493 |
| 100 | chr15 | 77436810 | 77437535 | 726 | * | 69 | 79834 |
| 101 | chr15 | 79296668 | 79296972 | 305 | * | 60 | 5923 |
| 102 | chr15 | 83478872 | 83479421 | 550 | * | 23 | 123720 |
| 103 | chr16 | 122632 | 123022 | 391 | * | 105 | 64285 |
| 104 | chr16 | 3070424 | 3071127 | 704 | * | 69 | 51330 |
| 105 | chr16 | 3071665 | 3072737 | 1073 | * | 174 | 51330 |
| 106 | chr16 | 46782372 | 46782661 | 290 | * | 40 | 91807 |
| 107 | chr16 | 66908556 | 66909073 | 518 | * | 52 | 57546 |
| 108 | chr16 | 68679683 | 68680140 | 458 | * | 52 | 1001 |
| 109 | chr17 | 783017 | 783542 | 526 | * | 93 | 64359 |
| 110 | chr17 | 836637 | 836916 | 280 | * | 174 | 64359 |
| 111 | chr17 | 854803 | 855462 | 660 | * | 261 | 64359 |
| 112 | chr17 | 1391299 | 1391915 | 617 | * | 995 | 4641 |
| 113 | chr17 | 3571288 | 3571708 | 421 | * | 120 | 30851 |
| 114 | chr17 | 7254529 | 7255474 | 946 | * | 573 | 147040 |
| 115 | chr17 | 8287787 | 8288752 | 966 | * | 30 | 6154 |
| 116 | chr17 | 9551502 | 9552052 | 551 | * | 153 | 124739 |
| 117 | chr17 | 33760201 | 33760832 | 632 | * | 133 | 55106 |
| 118 | chr17 | 36607100 | 36607720 | 621 | * | 744 | 57636 |
| 119 | chr17 | 36620868 | 36621397 | 530 | * | 75 | 57636 |
| 120 | chr17 | 40575233 | 40575965 | 733 | * | 398 | 284119 |
| 121 | chr17 | 47913737 | 47914249 | 513 | * | 52 | 729220 |
| 122 | chr17 | 48287766 | 48288048 | 283 | * | 139 | 1277 |
| 123 | chr17 | 48296272 | 48296659 | 388 | * | 30 | 1277 |
| 124 | chr17 | 55952432 | 55953028 | 597 | * | 275 | 51649 |
| 125 | chr17 | 57232047 | 57232686 | 640 | * | 185 | 348235 |
| 126 | chr17 | 73549106 | 73549585 | 480 | * | 174 | 3993 |
| 127 | chr17 | 81066619 | 81067385 | 767 | * | 152 | 284207 |
| 128 | chr18 | 3261453 | 3262365 | 913 | * | 1036 | 103910 |
| 129 | chr18 | 10925846 | 10926263 | 418 | * | 45 | 63895 |
| 130 | chr18 | 12308451 | 12308825 | 375 | * | 67 | 84617 |
| 131 | chr18 | 77725352 | 77725768 | 417 | * | 45 | 440498 |
| 132 | chr19 | 1027936 | 1028849 | 914 | * | 164 | 1265 |
| 133 | chr19 | 2540320 | 2541524 | 1205 | * | 889 | 4616 |
| 134 | chr19 | 2578785 | 2579220 | 436 | * | 46 | 4616 |
| 135 | chr19 | 6744832 | 6745182 | 351 | * | 69 | 9322 |
| 136 | chr19 | 15217867 | 15218127 | 261 | * | 69 | 85360 |
| 137 | chr19 | 17403331 | 17403630 | 300 | * | 60 | 126549 |
| 138 | chr19 | 34971987 | 34972691 | 705 | * | 1010 | 126374 |
| 139 | chr19 | 40029827 | 40030125 | 299 | * | 30 | 163126 |
| 140 | chr19 | 48222464 | 48223086 | 623 | * | 323 | 30846 |

| | | | | | | | |
|---|---|---|---|---|---|---|---|
| 141 | chr19 | 54362249 | 54362538 | 290 | * | 52 | 91663 |
| 142 | chr2 | 11294959 | 11295219 | 261 | * | 23 | 130814 |
| 143 | chr2 | 11484979 | 11485525 | 547 | * | 114 | 9475 |
| 144 | chr2 | 20212343 | 20212760 | 418 | * | 82 | 4148 |
| 145 | chr2 | 28619533 | 28620004 | 472 | * | 174 | 2355 |
| 146 | chr2 | 36598512 | 36599788 | 1277 | * | 104 | 51232 |
| 147 | chr2 | 36608141 | 36608427 | 287 | * | 69 | 51232 |
| 148 | chr2 | 54892808 | 54893067 | 260 | * | 17 | 6711 |
| 149 | chr2 | 64881252 | 64882185 | 934 | * | 524 | 9792 |
| 150 | chr2 | 64995020 | 64995286 | 267 | * | 38 | 9792 |
| 151 | chr2 | 102422721 | 102423005 | 285 | * | 30 | 9448 |
| 152 | chr2 | 121672588 | 121673530 | 943 | * | 123 | 2736 |
| 153 | chr2 | 121692915 | 121693341 | 427 | * | 60 | 2736 |
| 154 | chr2 | 121694657 | 121695161 | 505 | * | 23 | 2736 |
| 155 | chr2 | 128432859 | 128433243 | 385 | * | 34 | 55679 |
| 156 | chr2 | 159314442 | 159315041 | 600 | * | 211 | 8502 |
| 157 | chr2 | 201983011 | 201983757 | 747 | * | 86 | 8837 |
| 158 | chr2 | 202897750 | 202898370 | 621 | * | 143 | 8324 |
| 159 | chr2 | 235890038 | 235890450 | 413 | * | 287 | 23677 |
| 160 | chr2 | 235905190 | 235905827 | 638 | * | 45 | 23677 |
| 161 | chr2 | 238223793 | 238224104 | 312 | * | 133 | 1293 |
| 162 | chr2 | 238341290 | 238341615 | 326 | * | 45 | 1293 |
| 163 | chr20 | 17951957 | 17952911 | 955 | * | 94 | 92667 |
| 164 | chr20 | 30300377 | 30301599 | 1223 | * | 152 | 598 |
| 165 | chr20 | 30307845 | 30308569 | 725 | * | 180 | 598 |
| 166 | chr20 | 32497279 | 32497654 | 376 | * | 123 | 22913 |
| 167 | chr20 | 36536186 | 36536704 | 519 | * | 54 | 128434 |
| 168 | chr20 | 52790525 | 52790851 | 327 | * | 37 | 1591 |
| 169 | chr20 | 62066261 | 62066597 | 337 | * | 85 | 3785 |
| 170 | chr21 | 18899958 | 18900225 | 268 | * | 45 | 1525 |
| 171 | chr21 | 35899149 | 35899617 | 469 | * | 86 | 1827 |
| 172 | chr21 | 46348705 | 46349043 | 339 | * | 37 | 3689 |
| 173 | chr22 | 36725465 | 36726120 | 656 | * | 95 | 4627 |
| 174 | chr22 | 36764587 | 36765371 | 785 | * | 143 | 4627 |
| 175 | chr22 | 38614686 | 38615080 | 395 | * | 153 | 23764 |
| 176 | chr22 | 41686719 | 41687035 | 317 | * | 114 | 5905 |
| 177 | chr22 | 50731767 | 50732557 | 791 | * | 584 | 23654 |
| 178 | chr3 | 16525302 | 16526428 | 1127 | * | 133 | 23180 |
| 179 | chr3 | 49506637 | 49507329 | 693 | * | 107 | 1605 |
| 180 | chr3 | 50301808 | 50302548 | 741 | * | 102 | 7869 |
| 181 | chr3 | 58033690 | 58034187 | 498 | * | 68 | 2317 |
| 182 | chr3 | 124782026 | 124782362 | 337 | * | 114 | 57493 |
| 183 | chr3 | 134032225 | 134032601 | 377 | * | 123 | 51421 |
| 184 | chr3 | 134045636 | 134046173 | 538 | * | 261 | 51421 |
| 185 | chr3 | 134076332 | 134076949 | 618 | * | 192 | 51421 |
| 186 | chr3 | 134077317 | 134077625 | 309 | * | 10 | 51421 |
| 187 | chr3 | 134082696 | 134083239 | 544 | * | 645 | 51421 |

| | | | | | | | |
|---|---|---|---|---|---|---|---|
| 188 | chr3 | 134092213 | 134093797 | 1585 | * | 961 | 51421 |
| 189 | chr3 | 170469452 | 170469769 | 318 | * | 114 | 200916 |
| 190 | chr4 | 7912518 | 7912776 | 259 | * | 23 | 60312 |
| 191 | chr5 | 956680 | 957627 | 948 | * | 233 | 100506688 |
| 192 | chr5 | 16890183 | 16890440 | 258 | * | 21 | 4651 |
| 193 | chr5 | 17169474 | 17169762 | 289 | * | 28 | 10409 |
| 194 | chr5 | 32224101 | 32224370 | 270 | * | 94 | 64083 |
| 195 | chr5 | 33736408 | 33736681 | 274 | * | 39 | 81792 |
| 196 | chr5 | 33892176 | 33893068 | 893 | * | 71 | 81792 |
| 197 | chr5 | 38765596 | 38766327 | 732 | * | 108 | 9180 |
| 198 | chr5 | 38783291 | 38783609 | 319 | * | 56 | 9180 |
| 199 | chr5 | 38845886 | 38846339 | 454 | * | 122 | 9180 |
| 200 | chr5 | 39423971 | 39425270 | 1300 | * | 246 | 1601 |
| 201 | chr5 | 52160527 | 52160818 | 292 | * | 30 | 3672 |
| 202 | chr5 | 54455975 | 54456254 | 280 | * | 52 | 493869 |
| 203 | chr5 | 55864863 | 55865176 | 314 | * | 88 | 4214 |
| 204 | chr5 | 56988528 | 56989496 | 969 | * | 76 | 345651 |
| 205 | chr5 | 57137449 | 57138090 | 642 | * | 113 | 345651 |
| 206 | chr5 | 57312239 | 57312566 | 328 | * | 81 | 10769 |
| 207 | chr5 | 68797782 | 68798216 | 435 | * | 126 | 100506658 |
| 208 | chr5 | 135329471 | 135329793 | 323 | * | 37 | 7045 |
| 209 | chr5 | 135347543 | 135348276 | 734 | * | 47 | 7045 |
| 210 | chr5 | 135393248 | 135394350 | 1103 | * | 292 | 7045 |
| 211 | chr5 | 150016653 | 150017378 | 726 | * | 89 | 11346 |
| 212 | chr5 | 150024418 | 150024971 | 554 | * | 93 | 11346 |
| 213 | chr5 | 153747565 | 153748260 | 696 | * | 111 | 55568 |
| 214 | chr5 | 158635479 | 158636405 | 927 | * | 129 | 153830 |
| 215 | chr5 | 159510283 | 159510830 | 548 | * | 87 | 114825 |
| 216 | chr5 | 162886932 | 162887286 | 355 | * | 23 | 134492 |
| 217 | chr6 | 37093895 | 37094450 | 556 | * | 104 | 5292 |
| 218 | chr6 | 37105305 | 37105858 | 554 | * | 77 | 5292 |
| 219 | chr6 | 74289898 | 74290319 | 422 | * | 104 | 26503 |
| 220 | chr6 | 74295491 | 74296051 | 561 | * | 77 | 26503 |
| 221 | chr6 | 75915126 | 75915419 | 294 | * | 19 | 1303 |
| 222 | chr6 | 75915687 | 75915967 | 281 | * | 18 | 1303 |
| 223 | chr6 | 86388763 | 86389022 | 260 | * | 52 | 387066 |
| 224 | chr6 | 132272297 | 132273024 | 728 | * | 385 | 1490 |
| 225 | chr6 | 143998853 | 143999499 | 647 | * | 143 | 9749 |
| 226 | chr6 | 158402258 | 158402799 | 542 | * | 174 | 8871 |
| 227 | chr6 | 163837611 | 163838004 | 394 | * | 37 | 9444 |
| 228 | chr7 | 1576493 | 1577559 | 1067 | * | 394 | 7975 |
| 229 | chr7 | 5631069 | 5631877 | 809 | * | 425 | 6624 |
| 230 | chr7 | 22600871 | 22601348 | 478 | * | 409 | 100506178 |
| 231 | chr7 | 22616853 | 22617235 | 383 | * | 60 | 100506178 |
| 232 | chr7 | 22626202 | 22626955 | 754 | * | 138 | 100506178 |
| 233 | chr7 | 27135175 | 27135707 | 533 | * | 160 | 3198 |
| 234 | chr7 | 30027444 | 30028753 | 1310 | * | 98 | 9805 |

| | | | | | | | |
|---|---|---|---|---|---|---|---|
| 235 | chr7 | 39773417 | 39773784 | 368 | * | 100 | 349114 |
| 236 | chr7 | 42533048 | 42533614 | 567 | * | 29 | 2737 |
| 237 | chr7 | 55132663 | 55133075 | 413 | * | 61 | 1956 |
| 238 | chr7 | 55133348 | 55133893 | 546 | * | 45 | 1956 |
| 239 | chr7 | 101000819 | 101001099 | 281 | * | 60 | 136227 |
| 240 | chr7 | 101045057 | 101046005 | 949 | * | 230 | 136227 |
| 241 | chr7 | 115994461 | 115995253 | 793 | * | 218 | 858 |
| 242 | chr7 | 116151772 | 116153031 | 1260 | * | 67 | 858 |
| 243 | chr7 | 116165247 | 116165638 | 392 | * | 38 | 857 |
| 244 | chr7 | 129934234 | 129934492 | 259 | * | 19 | 51200 |
| 245 | chr7 | 130013296 | 130014253 | 958 | * | 262 | 1357 |
| 246 | chr7 | 131546885 | 131547278 | 394 | * | 70 | 91584 |
| 247 | chr7 | 134427070 | 134427416 | 347 | * | 90 | 800 |
| 248 | chr7 | 134464064 | 134464569 | 506 | * | 335 | 800 |
| 249 | chr7 | 150065922 | 150066506 | 585 | * | 50 | 29803 |
| 250 | chr7 | 151384460 | 151385166 | 707 | * | 265 | 51422 |
| 251 | chr7 | 151451083 | 151451720 | 638 | * | 649 | 51422 |
| 252 | chr8 | 1771745 | 1772089 | 345 | * | 45 | 9639 |
| 253 | chr8 | 6451629 | 6452008 | 380 | * | 60 | 285 |
| 254 | chr8 | 22454683 | 22455558 | 876 | * | 683 | 541565 |
| 255 | chr8 | 27471827 | 27472407 | 581 | * | 54 | 1191 |
| 256 | chr8 | 37659128 | 37659509 | 382 | * | 57 | 25960 |
| 257 | chr8 | 38857155 | 38857665 | 511 | * | 23 | 8754 |
| 258 | chr8 | 74048447 | 74048874 | 428 | * | 68 | 157869 |
| 259 | chr8 | 142164055 | 142164415 | 361 | * | 101 | 22898 |
| 260 | chr9 | 707011 | 707400 | 390 | * | 58 | 23189 |
| 261 | chr9 | 5509461 | 5509868 | 408 | * | 22 | 80380 |
| 262 | chr9 | 78987063 | 78987379 | 317 | * | 37 | 55312 |
| 263 | chr9 | 86046814 | 86047642 | 829 | * | 436 | 257019 |
| 264 | chr9 | 86124470 | 86124739 | 270 | * | 66 | 257019 |
| 265 | chr9 | 86153128 | 86153406 | 279 | * | 29 | 257019 |
| 266 | chr9 | 89990731 | 89991127 | 397 | * | 41 | 1612 |
| 267 | chr9 | 91149434 | 91150423 | 990 | * | 185 | 158046 |
| 268 | chr9 | 97675333 | 97675718 | 386 | * | 27 | 84909 |
| 269 | chr9 | 97684284 | 97685128 | 845 | * | 93 | 84909 |
| 270 | chr9 | 98269647 | 98270217 | 571 | * | 37 | 5727 |
| 271 | chr9 | 100893014 | 100893273 | 260 | * | 10 | 7464 |
| 272 | chr9 | 124113550 | 124113911 | 362 | * | 50 | 2040 |
| 273 | chr9 | 131314982 | 131315253 | 272 | * | 30 | 6709 |
| 274 | chr9 | 131316389 | 131317843 | 1455 | * | 255 | 6709 |
| 275 | chr9 | 131320799 | 131321311 | 513 | * | 93 | 6709 |
| 276 | chr9 | 134483402 | 134483665 | 264 | * | 32 | 2889 |
| 277 | chr9 | 134483975 | 134484312 | 338 | * | 168 | 2889 |
| 278 | chr9 | 134609440 | 134609876 | 437 | * | 96 | 2889 |
| 279 | chr9 | 137494456 | 137494750 | 295 | * | 183 | 1289 |
| 280 | chrX | 152066053 | 152066588 | 536 | * | 557 | 7739 |
| 281 | chrX | 153597502 | 153597835 | 334 | * | 52 | 2316 |

| ENSEMBL | baseMean | log2FoldChange | padj | Gene |
|---|---|---|---|---|
| ENSG00000142627 | 5020.238168 | 0.50133251 | 6.13E-04 | EPHA2 |
| ENSG00000142627 | 5020.238168 | 0.50133251 | 6.13E-04 | EPHA2 |
| ENSG00000131914 | 640.0340004 | 0.46493065 | 1.16E-02 | LIN28A |
| ENSG00000253368 | 1601.234225 | 0.52386682 | 3.90E-03 | TRNP1 |
| ENSG00000253368 | 1601.234225 | 0.52386682 | 3.90E-03 | TRNP1 |
| ENSG00000204138 | 2601.10134 | 0.507388623 | 9.71E-12 | PHACTR4 |
| ENSG00000117395 | 5155.502085 | -0.289742721 | 3.07E-02 | EBNA1BP2 |
| ENSG00000186564 | 276.8095472 | 0.406688238 | 7.43E-03 | FOXD2 |
| ENSG00000097096 | 235.870281 | 1.614464483 | 8.52E-11 | SYDE2 |
| ENSG00000142871 | 10596.59591 | 1.636580835 | 1.78E-45 | CCN1 |
| ENSG00000142871 | 10596.59591 | 1.636580835 | 1.78E-45 | CCN1 |
| ENSG00000142871 | 10596.59591 | 1.636580835 | 1.78E-45 | CCN1 |
| ENSG00000069702 | 1252.634429 | 0.47461049 | 4.25E-04 | TGFBR3 |
| ENSG00000137936 | 3028.458632 | -0.330701278 | 4.86E-02 | BCAR3 |
| ENSG00000137962 | 4982.779967 | 0.62425706 | 9.98E-03 | ARHGAP29 |
| ENSG00000137962 | 4982.779967 | 0.62425706 | 9.98E-03 | ARHGAP29 |
| ENSG00000184371 | 3189.285983 | 0.574134878 | 7.97E-07 | CSF1 |
| ENSG00000265808 | 4612.525216 | 0.215045689 | 4.23E-02 | SEC22B |
| ENSG00000143398 | 3765.167022 | 0.524071969 | 1.32E-06 | PIP5K1A |
| ENSG00000197747 | 12016.01031 | 0.421688877 | 2.84E-03 | S100A10 |
| ENSG00000160691 | 17857.68695 | 0.407000289 | 2.57E-03 | SHC1 |
| ENSG00000187800 | 1120.147306 | 1.204280732 | 1.25E-18 | PEAR1 |
| ENSG00000183853 | 10728.67019 | 0.31373626 | 1.10E-02 | KIRREL1 |
| ENSG00000077157 | 1810.477078 | 0.752705016 | 1.24E-18 | PPP1R12B |
| ENSG00000152104 | 5617.28597 | 0.6509988 | 2.76E-18 | PTPN14 |
| ENSG00000152104 | 5617.28597 | 0.6509988 | 2.76E-18 | PTPN14 |
| ENSG00000119283 | 26.95075226 | -1.74175667 | 3.18E-03 | TRIM67 |
| ENSG00000151474 | 2369.142518 | -0.448008317 | 3.30E-05 | FRMD4A |
| ENSG00000138119 | 15695.67332 | 0.711984947 | 5.10E-07 | MYOF |
| ENSG00000138119 | 15695.67332 | 0.711984947 | 5.10E-07 | MYOF |
| ENSG00000138166 | 787.2594332 | -0.695421139 | 9.34E-03 | DUSP5 |
| ENSG00000133789 | 3816.151683 | 0.331996191 | 1.44E-03 | SWAP70 |
| ENSG00000197702 | 12811.79845 | 0.745270121 | 1.38E-47 | PARVA |
| ENSG00000197702 | 12811.79845 | 0.745270121 | 1.38E-47 | PARVA |
| ENSG00000187079 | 10939.96256 | 0.802215226 | 5.71E-07 | TEAD1 |
| ENSG00000187079 | 10939.96256 | 0.802215226 | 5.71E-07 | TEAD1 |
| ENSG00000187079 | 10939.96256 | 0.802215226 | 5.71E-07 | TEAD1 |
| ENSG00000187079 | 10939.96256 | 0.802215226 | 5.71E-07 | TEAD1 |
| ENSG00000133818 | 2136.436366 | 0.635847069 | 1.86E-09 | RRAS2 |
| ENSG00000166833 | 2387.667239 | -0.368433665 | 2.75E-02 | NAV2 |
| ENSG00000179241 | 1672.485159 | -0.324928777 | 4.44E-02 | LDLRAD3 |
| ENSG00000187479 | 147.5207521 | 0.518959831 | 4.25E-02 | C11orf96 |
| ENSG00000157613 | 742.9290274 | 1.210055102 | 8.88E-16 | CREB3L1 |
| ENSG00000157613 | 742.9290274 | 1.210055102 | 8.88E-16 | CREB3L1 |
| ENSG00000124942 | 122265.4934 | 0.565724144 | 1.15E-09 | AHNAK |
| ENSG00000124942 | 122265.4934 | 0.565724144 | 1.15E-09 | AHNAK |

| | | | | |
|---|---|---|---|---|
| ENSG00000124942 | 122265.4934 | 0.565724144 | 1.15E-09 | AHNAK |
| ENSG00000006534 | 1300.36037 | 0.41210987 | 9.29E-04 | ALDH3B1 |
| ENSG00000073921 | 8631.628285 | 0.388763734 | 5.20E-03 | PICALM |
| ENSG00000166025 | 6561.397498 | -0.346688068 | 5.97E-03 | AMOTL1 |
| ENSG00000137693 | 7292.138549 | 0.325414487 | 2.02E-03 | YAP1 |
| ENSG00000137693 | 7292.138549 | 0.325414487 | 2.02E-03 | YAP1 |
| ENSG00000023445 | 138.5705745 | 0.635552555 | 3.77E-02 | BIRC3 |
| ENSG00000149295 | 181.3447405 | -0.933802011 | 1.26E-04 | DRD2 |
| ENSG00000137642 | 4958.874962 | 0.793940204 | 1.01E-08 | SORL1 |
| ENSG00000165495 | 537.5619923 | 0.68996343 | 1.02E-04 | PKNOX2 |
| ENSG00000254703 | 247.900948 | -0.506408427 | 2.60E-03 | SENCR |
| ENSG00000166106 | 608.3231517 | 0.85762205 | 2.00E-10 | ADAMTS15 |
| ENSG00000166106 | 608.3231517 | 0.85762205 | 2.00E-10 | ADAMTS15 |
| ENSG00000111186 | 481.1590035 | 0.771410967 | 1.18E-03 | WNT5B |
| ENSG00000070018 | 2588.5102 | 0.426104522 | 5.61E-03 | LRP6 |
| ENSG00000061273 | 5153.464871 | 0.241597769 | 2.27E-02 | HDAC7 |
| ENSG00000167767 | 1730.679913 | 0.987493874 | 8.17E-04 | KRT80 |
| ENSG00000170421 | 14283.57687 | 1.477447241 | 1.15E-37 | KRT8 |
| ENSG00000111057 | 26282.33084 | 0.65241719 | 7.14E-07 | KRT18 |
| ENSG00000181852 | 3274.28356 | 0.474767811 | 4.94E-02 | RNF41 |
| ENSG00000174099 | 1780.19995 | 0.718677032 | 1.86E-08 | MSRB3 |
| ENSG00000169372 | 389.3041389 | -0.324776103 | 4.91E-02 | CRADD |
| ENSG00000198431 | 33083.72171 | 0.535282031 | 3.49E-02 | TXNRD1 |
| ENSG00000074590 | 3145.049458 | 0.540339088 | 5.55E-05 | NUAK1 |
| ENSG00000074590 | 3145.049458 | 0.540339088 | 5.55E-05 | NUAK1 |
| ENSG00000074590 | 3145.049458 | 0.540339088 | 5.55E-05 | NUAK1 |
| ENSG00000074590 | 3145.049458 | 0.540339088 | 5.55E-05 | NUAK1 |
| ENSG00000135127 | 37.542591 | 1.670842929 | 1.00E-02 | BICDL1 |
| ENSG00000150457 | 922.0682349 | 1.524222948 | 2.64E-42 | LATS2 |
| ENSG00000136160 | 780.2042551 | 1.657933588 | 1.23E-33 | EDNRB |
| ENSG00000136158 | 1269.954807 | -0.490081167 | 2.73E-04 | SPRY2 |
| ENSG00000136158 | 1269.954807 | -0.490081167 | 2.73E-04 | SPRY2 |
| ENSG00000139793 | 5478.880166 | 0.642446825 | 1.85E-02 | MBNL2 |
| ENSG00000139793 | 5478.880166 | 0.642446825 | 1.85E-02 | MBNL2 |
| ENSG00000139793 | 5478.880166 | 0.642446825 | 1.85E-02 | MBNL2 |
| ENSG00000139793 | 5478.880166 | 0.642446825 | 1.85E-02 | MBNL2 |
| ENSG00000139793 | 5478.880166 | 0.642446825 | 1.85E-02 | MBNL2 |
| ENSG00000139793 | 5478.880166 | 0.642446825 | 1.85E-02 | MBNL2 |
| ENSG00000139793 | 5478.880166 | 0.642446825 | 1.85E-02 | MBNL2 |
| ENSG00000139793 | 5478.880166 | 0.642446825 | 1.85E-02 | MBNL2 |
| ENSG00000129474 | 9629.715395 | 1.539807686 | 7.72E-18 | AJUBA |
| ENSG00000129474 | 9629.715395 | 1.539807686 | 7.72E-18 | AJUBA |
| ENSG00000020577 | 2581.581522 | 0.5514013 | 2.92E-04 | SAMD4A |
| ENSG00000020577 | 2581.581522 | 0.5514013 | 2.92E-04 | SAMD4A |
| ENSG00000020577 | 2581.581522 | 0.5514013 | 2.92E-04 | SAMD4A |
| ENSG00000259687 | 13.87085948 | 2.261687262 | 3.91E-02 | LINC01220 |
| ENSG00000070778 | 1232.53051 | 0.738052637 | 1.05E-12 | AL162171.1 |

| | | | | |
|---|---|---|---|---|
| ENSG00000080823 | 1296.653583 | -0.279383279 | 4.92E-02 | MOK |
| ENSG00000128849 | 895.5418779 | 0.676046783 | 6.57E-05 | CGNL1 |
| ENSG00000137845 | 7600.927323 | 0.319506009 | 3.77E-02 | ADAM10 |
| ENSG00000140416 | 23789.68 | 0.547885298 | 1.12E-05 | TPM1 |
| ENSG00000140416 | 23789.68 | 0.547885298 | 1.12E-05 | TPM1 |
| ENSG00000137807 | 4117.414188 | 0.585718769 | 2.97E-03 | KIF23 |
| ENSG00000173517 | 4818.635054 | 0.3396588 | 6.07E-04 | PEAK1 |
| ENSG00000058335 | 134.7074287 | 0.901625976 | 3.81E-02 | RASGRF1 |
| ENSG00000156232 | 724.8447879 | 0.597112787 | 1.54E-04 | WHAMM |
| ENSG00000007384 | 1735.657285 | 0.65059796 | 1.39E-07 | RHBDF1 |
| ENSG00000006327 | 3033.22403 | 0.915043687 | 1.06E-07 | TNFRSF12A |
| ENSG00000006327 | 3033.22403 | 0.915043687 | 1.06E-07 | TNFRSF12A |
| ENSG00000140795 | 310.1611264 | 0.921675844 | 1.32E-06 | MYLK3 |
| ENSG00000172840 | 1653.370186 | 0.289004458 | 1.75E-02 | PDP2 |
| ENSG00000062038 | 1193.523897 | 1.009906298 | 1.75E-14 | CDH3 |
| ENSG00000167693 | 1791.096427 | 0.728752886 | 5.77E-08 | NXN |
| ENSG00000167693 | 1791.096427 | 0.728752886 | 5.77E-08 | NXN |
| ENSG00000167693 | 1791.096427 | 0.728752886 | 5.77E-08 | NXN |
| ENSG00000197879 | 10838.86687 | 0.554021412 | 4.06E-05 | MYO1C |
| ENSG00000213977 | 4806.333429 | 0.416893117 | 8.72E-03 | TAX1BP3 |
| ENSG00000213859 | 1277.421524 | 0.641682389 | 4.07E-05 | KCTD11 |
| ENSG00000161970 | 31543.39492 | -0.299679596 | 2.05E-03 | RPL26 |
| ENSG00000154914 | 231.4230637 | 0.675463923 | 2.26E-03 | USP43 |
| ENSG00000172123 | 950.4862695 | 0.436484744 | 1.34E-02 | SLFN12 |
| ENSG00000275832 | 5404.484757 | 0.727259915 | 8.19E-15 | ARHGAP23 |
| ENSG00000275832 | 5404.484757 | 0.727259915 | 8.19E-15 | ARHGAP23 |
| ENSG00000177469 | 22717.39463 | 1.186499356 | 3.85E-18 | CAVIN1 |
| ENSG00000204584 | 22.04428444 | 1.566641579 | 1.48E-02 | FLJ45513 |
| ENSG00000108821 | 951.6101759 | 1.036450981 | 2.66E-04 | COL1A1 |
| ENSG00000108821 | 951.6101759 | 1.036450981 | 2.66E-04 | COL1A1 |
| ENSG00000181610 | 2563.61453 | -0.257188724 | 1.38E-02 | MRPS23 |
| ENSG00000182628 | 1597.49907 | 0.65904316 | 2.84E-07 | SKA2 |
| ENSG00000073350 | 1015.83685 | 0.422492835 | 1.07E-03 | LLGL2 |
| ENSG00000176845 | 538.6256251 | 0.437518222 | 4.75E-02 | METRNL |
| ENSG00000118680 | 10882.15315 | 0.39869597 | 5.36E-09 | MYL12B |
| ENSG00000154864 | 3798.179918 | 0.249503773 | 3.02E-02 | PIEZO2 |
| ENSG00000176014 | 6630.385277 | 0.499508955 | 2.21E-02 | TUBB6 |
| ENSG00000226742 | 284.4549299 | 0.913162524 | 9.32E-07 | HSBP1L1 |
| ENSG00000064666 | 8063.875845 | 0.576527736 | 5.21E-09 | CNN2 |
| ENSG00000099860 | 977.0968945 | 0.629185014 | 3.59E-06 | GADD45B |
| ENSG00000099860 | 977.0968945 | 0.629185014 | 3.59E-06 | GADD45B |
| ENSG00000125733 | 3342.101072 | 0.244375767 | 1.18E-03 | TRIP10 |
| ENSG00000105137 | 3531.235541 | 0.836981807 | 1.80E-10 | SYDE1 |
| ENSG00000160117 | 635.5348284 | -0.475131462 | 3.61E-04 | ANKLE1 |
| ENSG00000142279 | 955.4419514 | 2.455370831 | 7.66E-39 | WTIP |
| ENSG00000176396 | 658.2350262 | 0.556617388 | 1.13E-08 | EID2 |
| ENSG00000024422 | 10312.92724 | 0.696997666 | 5.36E-09 | EHD2 |

| | | | | |
|---|---|---|---|---|
| ENSG00000179820 | 3260.603661 | 0.450077541 | 1.69E-03 | MYADM |
| ENSG00000162976 | 634.4680855 | 0.329359457 | 3.48E-02 | PQLC3 |
| ENSG00000134318 | 3281.835884 | 0.699030354 | 2.26E-02 | ROCK2 |
| ENSG00000132031 | 531.0146682 | 1.681473147 | 6.61E-31 | MATN3 |
| ENSG00000075426 | 5985.00852 | 0.329457511 | 2.68E-02 | FOSL2 |
| ENSG00000150938 | 9816.058979 | 0.573925416 | 1.43E-05 | CRIM1 |
| ENSG00000150938 | 9816.058979 | 0.573925416 | 1.43E-05 | CRIM1 |
| ENSG00000115306 | 22759.67806 | 0.298872879 | 2.33E-06 | SPTBN1 |
| ENSG00000179833 | 1650.89911 | 0.37876893 | 6.01E-03 | SERTAD2 |
| ENSG00000179833 | 1650.89911 | 0.37876893 | 6.01E-03 | SERTAD2 |
| ENSG00000071054 | 13426.54364 | -0.208572109 | 9.33E-03 | MAP4K4 |
| ENSG00000074047 | 1186.161085 | 0.639347398 | 3.37E-03 | GLI2 |
| ENSG00000074047 | 1186.161085 | 0.639347398 | 3.37E-03 | GLI2 |
| ENSG00000074047 | 1186.161085 | 0.639347398 | 3.37E-03 | GLI2 |
| ENSG00000072163 | 478.4740095 | 3.031266294 | 2.37E-63 | LIMS2 |
| ENSG00000144283 | 3485.234126 | 0.552511901 | 2.57E-05 | PKP4 |
| ENSG00000003402 | 2074.937118 | 0.163760252 | 4.58E-02 | CFLAR |
| ENSG00000155760 | 2542.472438 | 0.42568701 | 1.12E-05 | FZD7 |
| ENSG00000130147 | 4262.928303 | 0.357852577 | 6.80E-03 | SH3BP4 |
| ENSG00000130147 | 4262.928303 | 0.357852577 | 6.80E-03 | SH3BP4 |
| ENSG00000163359 | 4751.934988 | -0.722641989 | 1.86E-02 | COL6A3 |
| ENSG00000163359 | 4751.934988 | -0.722641989 | 1.86E-02 | COL6A3 |
| ENSG00000125871 | 2856.925219 | 0.254718371 | 6.63E-03 | MGME1 |
| ENSG00000171552 | 5620.751235 | 0.38719234 | 1.12E-04 | BCL2L1 |
| ENSG00000171552 | 5620.751235 | 0.38719234 | 1.12E-04 | BCL2L1 |
| ENSG00000125970 | 10373.36135 | -0.313036688 | 2.05E-02 | RALY |
| ENSG00000132821 | 409.4949088 | 1.774107765 | 1.76E-20 | VSTM2L |
| ENSG00000019186 | 230.7349006 | -1.950777274 | 5.04E-15 | CYP24A1 |
| ENSG00000075043 | 12.26458218 | -1.506615731 | 3.65E-02 | KCNQ2 |
| ENSG00000154639 | 430.9583909 | 2.152101105 | 6.16E-33 | CXADR |
| ENSG00000159200 | 3868.497247 | 1.005103508 | 8.31E-13 | RCAN1 |
| ENSG00000160255 | 124.990794 | 1.726472885 | 8.09E-06 | ITGB2 |
| ENSG00000100345 | 39701.66481 | 0.404890982 | 1.63E-03 | MYH9 |
| ENSG00000100345 | 39701.66481 | 0.404890982 | 1.63E-03 | MYH9 |
| ENSG00000185022 | 607.0319197 | 0.546795482 | 3.66E-02 | MAFF |
| ENSG00000100401 | 7733.942925 | -0.335705807 | 5.08E-04 | RANGAP1 |
| ENSG00000196576 | 10147.39702 | 0.47798829 | 2.85E-05 | PLXNB2 |
| ENSG00000131378 | 2508.131971 | 0.636016352 | 3.49E-07 | RFTN1 |
| ENSG00000173402 | 8402.798769 | 0.88971037 | 7.44E-15 | DAG1 |
| ENSG00000012171 | 1213.282702 | 1.464998026 | 1.64E-11 | SEMA3B |
| ENSG00000136068 | 29152.77127 | 0.615989659 | 8.17E-05 | FLNB |
| ENSG00000173706 | 10780.54503 | 0.472636644 | 1.13E-08 | HEG1 |
| ENSG00000114019 | 5194.214834 | 2.324098712 | 3.44E-118 | AMOTL2 |
| ENSG00000114019 | 5194.214834 | 2.324098712 | 3.44E-118 | AMOTL2 |
| ENSG00000114019 | 5194.214834 | 2.324098712 | 3.44E-118 | AMOTL2 |
| ENSG00000114019 | 5194.214834 | 2.324098712 | 3.44E-118 | AMOTL2 |
| ENSG00000114019 | 5194.214834 | 2.324098712 | 3.44E-118 | AMOTL2 |

| | | | | |
|---|---|---|---|---|
| ENSG00000114019 | 5194.214834 | 2.324098712 | 3.44E-118 | AMOTL2 |
| ENSG00000163584 | 2206.738069 | -0.371961647 | 1.57E-02 | RPL22L1 |
| ENSG00000196526 | 8267.209501 | 0.341407265 | 3.04E-03 | AFAP1 |
| ENSG00000215246 | 130.5665582 | 1.320528406 | 8.62E-06 | AC116351.1 |
| ENSG00000145555 | 14327.81064 | 0.35039615 | 3.97E-03 | MYO10 |
| ENSG00000176788 | 2291.351755 | 0.664324012 | 2.93E-04 | BASP1 |
| ENSG00000113384 | 5308.918063 | 0.157075555 | 2.73E-02 | GOLPH3 |
| ENSG00000151388 | 10377.03832 | 0.424654142 | 5.67E-04 | ADAMTS12 |
| ENSG00000151388 | 10377.03832 | 0.424654142 | 5.67E-04 | ADAMTS12 |
| ENSG00000145623 | 15770.68321 | 0.517522014 | 2.97E-04 | OSMR |
| ENSG00000145623 | 15770.68321 | 0.517522014 | 2.97E-04 | OSMR |
| ENSG00000145623 | 15770.68321 | 0.517522014 | 2.97E-04 | OSMR |
| ENSG00000153071 | 9628.493215 | 0.346702397 | 7.04E-04 | DAB2 |
| ENSG00000213949 | 11661.05432 | 0.74243683 | 8.84E-08 | ITGA1 |
| ENSG00000164294 | 5676.176415 | 0.757084506 | 8.61E-09 | GPX8 |
| ENSG00000095015 | 6834.516553 | -0.367286133 | 8.10E-03 | MAP3K1 |
| ENSG00000169067 | 62.37567461 | -1.594293207 | 2.99E-03 | ACTBL2 |
| ENSG00000169067 | 62.37567461 | -1.594293207 | 2.99E-03 | ACTBL2 |
| ENSG00000145632 | 8444.080224 | -0.400474434 | 3.99E-03 | PLK2 |
| ENSG00000197822 | 724.0860483 | 1.652703285 | 1.59E-12 | OCLN |
| ENSG00000120708 | 26371.45713 | -0.392035023 | 1.18E-02 | TGFBI |
| ENSG00000120708 | 26371.45713 | -0.392035023 | 1.18E-02 | TGFBI |
| ENSG00000120708 | 26371.45713 | -0.392035023 | 1.18E-02 | TGFBI |
| ENSG00000171992 | 2426.228123 | 1.19331499 | 4.53E-24 | SYNPO |
| ENSG00000171992 | 2426.228123 | 1.19331499 | 4.53E-24 | SYNPO |
| ENSG00000164574 | 5382.628084 | 0.27677623 | 4.36E-02 | GALNT10 |
| ENSG00000145860 | 10583.21953 | 0.153704102 | 1.42E-02 | RNF145 |
| ENSG00000170234 | 1799.661611 | 0.299410918 | 1.17E-02 | PWWP2A |
| ENSG00000170584 | 2588.159888 | 0.32500894 | 4.49E-03 | NUDCD2 |
| ENSG00000137193 | 1122.782319 | 0.888365247 | 2.27E-04 | PIM1 |
| ENSG00000137193 | 1122.782319 | 0.888365247 | 2.27E-04 | PIM1 |
| ENSG00000119899 | 973.1245904 | 0.444971521 | 3.35E-02 | SLC17A5 |
| ENSG00000119899 | 973.1245904 | 0.444971521 | 3.35E-02 | SLC17A5 |
| ENSG00000111799 | 7146.720884 | 2.424618059 | 3.61E-74 | COL12A1 |
| ENSG00000111799 | 7146.720884 | 2.424618059 | 3.61E-74 | COL12A1 |
| ENSG00000203875 | 2995.960911 | -0.370578499 | 1.55E-02 | SNHG5 |
| ENSG00000118523 | 8122.707037 | 2.542250351 | 4.01E-37 | CCN2 |
| ENSG00000112419 | 1004.303295 | 0.810111486 | 1.99E-04 | PHACTR2 |
| ENSG00000078269 | 2575.00764 | 0.396260877 | 7.83E-03 | SYNJ2 |
| ENSG00000112531 | 4899.422462 | 0.632590523 | 4.55E-05 | QKI |
| ENSG00000198517 | 1886.636662 | 1.206600486 | 2.82E-26 | MAFK |
| ENSG00000075618 | 13820.42552 | 0.497156683 | 2.62E-03 | FSCN1 |
| ENSG00000232759 | 40.60269188 | 1.647735343 | 4.36E-06 | AC002480.1 |
| ENSG00000232759 | 40.60269188 | 1.647735343 | 4.36E-06 | AC002480.1 |
| ENSG00000232759 | 40.60269188 | 1.647735343 | 4.36E-06 | AC002480.1 |
| ENSG00000105991 | 545.3612809 | 1.001835386 | 9.01E-19 | HOXA1 |
| ENSG00000136193 | 19372.32534 | 0.276714205 | 2.04E-03 | SCRN1 |

| | | | | |
|---|---|---|---|---|
| ENSG00000188185 | 516.0461094 | 0.360683079 | 3.98E-02 | LINC00265 |
| ENSG00000106571 | 3529.66754 | -0.377901972 | 7.15E-03 | GLI3 |
| ENSG00000146648 | 7757.382542 | 0.188500467 | 4.57E-02 | EGFR |
| ENSG00000146648 | 7757.382542 | 0.188500467 | 4.57E-02 | EGFR |
| ENSG00000160963 | 362.579038 | 0.528535417 | 6.75E-03 | COL26A1 |
| ENSG00000160963 | 362.579038 | 0.528535417 | 6.75E-03 | COL26A1 |
| ENSG00000105971 | 2124.493216 | 0.845750155 | 2.20E-10 | CAV2 |
| ENSG00000105971 | 2124.493216 | 0.845750155 | 2.20E-10 | CAV2 |
| ENSG00000105974 | 23107.43986 | 0.885928362 | 3.88E-11 | CAV1 |
| ENSG00000128510 | 8124.801267 | 1.929860219 | 8.19E-31 | CPA4 |
| ENSG00000091704 | 26.12893113 | 1.460397091 | 6.23E-03 | CPA1 |
| ENSG00000221866 | 515.4222672 | -0.726102815 | 1.21E-02 | PLXNA4 |
| ENSG00000122786 | 17077.04623 | 0.632073133 | 2.86E-05 | CALD1 |
| ENSG00000122786 | 17077.04623 | 0.632073133 | 2.86E-05 | CALD1 |
| ENSG00000214022 | 8966.942661 | 0.216620257 | 3.22E-02 | REPIN1 |
| ENSG00000106617 | 1255.564186 | 0.505865994 | 3.81E-03 | PRKAG2 |
| ENSG00000106617 | 1255.564186 | 0.505865994 | 3.81E-03 | PRKAG2 |
| ENSG00000104728 | 3349.754618 | 0.529286319 | 1.27E-05 | ARHGEF10 |
| ENSG00000091879 | 85.43122701 | -0.838797087 | 2.31E-02 | ANGPT2 |
| ENSG00000241852 | 1572.615118 | 0.436951022 | 3.99E-02 | C8orf58 |
| ENSG00000120885 | 7930.293639 | 1.004966719 | 7.44E-15 | CLU |
| ENSG00000020181 | 6548.879872 | 0.262225244 | 3.12E-02 | ADGRA2 |
| ENSG00000168615 | 9482.944296 | 0.649174371 | 6.71E-04 | ADAM9 |
| ENSG00000164764 | 767.0994634 | -0.603918206 | 2.77E-03 | SBSPON |
| ENSG00000105339 | 1469.517193 | 0.31952924 | 1.71E-02 | DENND3 |
| ENSG00000107104 | 4383.01843 | 0.334647919 | 1.14E-04 | KANK1 |
| ENSG00000197646 | 137.9408949 | 0.872331774 | 8.42E-03 | PDCD1LG2 |
| ENSG00000135002 | 851.8053873 | 0.677732193 | 1.66E-03 | RFK |
| ENSG00000172159 | 3496.851126 | 0.237969896 | 2.96E-02 | FRMD3 |
| ENSG00000172159 | 3496.851126 | 0.237969896 | 2.96E-02 | FRMD3 |
| ENSG00000172159 | 3496.851126 | 0.237969896 | 2.96E-02 | FRMD3 |
| ENSG00000196730 | 5388.257129 | 0.252180932 | 7.53E-03 | DAPK1 |
| ENSG00000130045 | 101.7112097 | 1.044096847 | 1.34E-06 | NXNL2 |
| ENSG00000148120 | 1301.867845 | 0.835327157 | 9.38E-16 | C9orf3 |
| ENSG00000148120 | 1301.867845 | 0.835327157 | 9.38E-16 | C9orf3 |
| ENSG00000185920 | 1620.679928 | 1.047777632 | 6.47E-09 | PTCH1 |
| ENSG00000106789 | 1356.442854 | -0.294857165 | 2.64E-03 | CORO2A |
| ENSG00000148175 | 8123.48438 | 0.321014301 | 2.99E-04 | STOM |
| ENSG00000197694 | 27548.29185 | 1.023460204 | 4.39E-46 | SPTAN1 |
| ENSG00000197694 | 27548.29185 | 1.023460204 | 4.39E-46 | SPTAN1 |
| ENSG00000197694 | 27548.29185 | 1.023460204 | 4.39E-46 | SPTAN1 |
| ENSG00000107263 | 4833.32604 | 0.486859146 | 3.41E-06 | RAPGEF1 |
| ENSG00000107263 | 4833.32604 | 0.486859146 | 3.41E-06 | RAPGEF1 |
| ENSG00000107263 | 4833.32604 | 0.486859146 | 3.41E-06 | RAPGEF1 |
| ENSG00000130635 | 25458.72823 | 0.439994496 | 1.54E-04 | COL5A1 |
| ENSG00000147394 | 2431.652507 | 2.636625261 | 5.29E-135 | ZNF185 |
| ENSG00000196924 | 161615.4789 | 0.411596496 | 1.00E-04 | FLNA |

| | seqnames | start | end | width | strand | V5 | geneId |
|---|---|---|---|---|---|---|---|
| 1 | chr1 | 7740313 | 7740827 | 515 | * | 69 | 23261 |
| 2 | chr1 | 17338253 | 17338505 | 253 | * | 72 | 23400 |
| 3 | chr11 | 66025420 | 66025979 | 560 | * | 155 | 64837 |
| 4 | chr11 | 118023704 | 118024144 | 441 | * | 21 | 6330 |
| 5 | chr12 | 6579427 | 6579824 | 398 | * | 152 | 6843 |
| 6 | chr14 | 74180690 | 74181113 | 424 | * | 63 | 9240 |
| 7 | chr14 | 90848967 | 90850029 | 1063 | * | 2104 | 801 |
| 8 | chr14 | 93650620 | 93651214 | 595 | * | 390 | 64112 |
| 9 | chr14 | 94547159 | 94547611 | 453 | * | 72 | 57062 |
| 10 | chr14 | 102438663 | 102438987 | 325 | * | 734 | 1778 |
| 11 | chr15 | 25199719 | 25200233 | 515 | * | 998 | 6638 |
| 12 | chr15 | 43477510 | 43478110 | 601 | * | 354 | 23582 |
| 13 | chr15 | 43810677 | 43810950 | 274 | * | 163 | 4130 |
| 14 | chr15 | 68871784 | 68872166 | 383 | * | 370 | 10391 |
| 15 | chr16 | 2255245 | 2256208 | 964 | * | 201 | 64223 |
| 16 | chr16 | 14723865 | 14724136 | 272 | * | 80 | 5073 |
| 17 | chr16 | 29911504 | 29911843 | 340 | * | 108 | 253982 |
| 18 | chr16 | 89984285 | 89984736 | 452 | * | 229 | 4157 |
| 19 | chr16 | 89988993 | 89989671 | 679 | * | 145 | 10381 |
| 20 | chr17 | 8534378 | 8534812 | 435 | * | 81 | 4628 |
| 21 | chr17 | 11924100 | 11924347 | 248 | * | 20 | 6416 |
| 22 | chr17 | 20946725 | 20947072 | 348 | * | 208 | 23326 |
| 23 | chr17 | 40834596 | 40835299 | 704 | * | 354 | 8506 |
| 24 | chr2 | 44588278 | 44589158 | 881 | * | 630 | 79823 |
| 25 | chr2 | 219827819 | 219828174 | 356 | * | 101 | 8941 |
| 26 | chr20 | 23342130 | 23342711 | 582 | * | 302 | 64412 |
| 27 | chr20 | 57581374 | 57581627 | 254 | * | 50 | 1522 |
| 28 | chr3 | 158450219 | 158450618 | 400 | * | 63 | 5918 |
| 29 | chr4 | 85503927 | 85504225 | 299 | * | 81 | 1040 |
| 30 | chr5 | 71404225 | 71405041 | 817 | * | 338 | 4131 |
| 31 | chr5 | 95998014 | 95998608 | 595 | * | 60 | 831 |
| 32 | chr5 | 149109565 | 149109830 | 266 | * | 54 | 133522 |
| 33 | chr5 | 150535279 | 150535547 | 269 | * | 109 | 309 |
| 34 | chr7 | 4681569 | 4681855 | 287 | * | 86 | 221937 |
| 35 | chr7 | 100812238 | 100812524 | 287 | * | 34 | 7425 |
| 36 | chr9 | 129677046 | 129677295 | 250 | * | 94 | 9649 |
| 37 | chr9 | 131314330 | 131314659 | 330 | * | 234 | 6709 |
| 38 | chrX | 75647964 | 75648287 | 324 | * | 111 | 57692 |
| 39 | chrX | 148586639 | 148586884 | 246 | * | 30 | 3423 |

| ENSEMBL | baseMean | log2 Fold Change | padj | Gene |
|---|---|---|---|---|
| ENSG00000171735 | 1421.581369 | 0.396603765 | 3.22E-03 | CAMTA1 |
| ENSG00000159363 | 2347.326166 | 1.057505846 | 9.90E-29 | ATP13A2 |
| ENSG00000174996 | 1774.580244 | 0.702457431 | 1.35E-08 | KLC2 |
| ENSG00000177098 | 2747.623872 | 0.949727645 | 1.69E-04 | SCN4B |
| ENSG00000139190 | 983.9967304 | 0.405146668 | 2.46E-02 | VAMP1 |
| ENSG00000176903 | 2633.572079 | 0.241486459 | 1.12E-02 | PNMA1 |
| ENSG00000198668 | 7582.069497 | 0.422419726 | 7.77E-07 | CALM1 |
| ENSG00000165943 | 1235.398058 | 0.562395209 | 2.62E-09 | MOAP1 |
| ENSG00000089737 | 7689.582107 | 0.411186391 | 2.30E-04 | DDX24 |
| ENSG00000197102 | 30557.26086 | 0.82625451 | 1.96E-19 | DYNC1H1 |
| ENSG00000128739 | 5362.88291 | 2.166072102 | 3.32E-129 | SNRPN |
| ENSG00000166946 | 1210.781179 | 0.958572573 | 6.44E-29 | CCNDBP1 |
| ENSG00000166963 | 1216.104392 | 2.643201796 | 2.78E-74 | MAP1A |
| ENSG00000103647 | 318.2158163 | 1.422163948 | 7.57E-20 | CORO2B |
| ENSG00000167965 | 2699.894163 | 0.352288607 | 2.56E-02 | MLST8 |
| ENSG00000140694 | 3180.356521 | 0.262763794 | 5.12E-03 | PARN |
| ENSG00000174939 | 246.0080628 | 1.159822642 | 4.67E-21 | ASPHD1 |
| ENSG00000258839 | 850.7944033 | 0.420522067 | 1.06E-03 | MC1R |
| ENSG00000258947 | 3262.462153 | 1.119768209 | 4.30E-14 | TUBB3 |
| ENSG00000133026 | 12958.38845 | 0.419228646 | 3.60E-08 | MYH10 |
| ENSG00000065559 | 1534.37162 | 0.385224837 | 6.37E-04 | MAP2K4 |
| ENSG00000124422 | 19092.274 | 0.303702525 | 1.26E-03 | USP22 |
| ENSG00000108797 | 3759.390506 | 1.13179363 | 1.75E-27 | CNTNAP1 |
| ENSG00000143919 | 213.8627465 | 0.451920408 | 4.46E-02 | CAMKMT |
| ENSG00000171450 | 57.6406424 | 0.998953519 | 4.80E-02 | CDK5R2 |
| ENSG00000125812 | 2171.95145 | 0.475476546 | 3.80E-04 | GZF1 |
| ENSG00000101160 | 14220.45977 | 0.359495404 | 1.38E-02 | CTSZ |
| ENSG00000118849 | 136.06074 | 0.97414216 | 9.69E-03 | RARRES1 |
| ENSG00000163624 | 52.31633794 | 4.026255113 | 1.20E-23 | CDS1 |
| ENSG00000131711 | 20810.43855 | 1.138659784 | 7.95E-08 | MAP1B |
| ENSG00000153113 | 16712.96545 | 0.277754924 | 2.91E-05 | CAST |
| ENSG00000155846 | 475.6106913 | 0.503947394 | 3.09E-02 | PPARGC1B |
| ENSG00000197043 | 16028.14149 | 0.807150444 | 2.12E-06 | ANXA6 |
| ENSG00000164916 | 3559.332545 | 0.283829653 | 1.40E-02 | FOXK1 |
| ENSG00000128564 | 785.8211023 | 2.740050719 | 1.05E-45 | VGF |
| ENSG00000136828 | 346.6724631 | 0.992759623 | 1.93E-04 | RALGPS1 |
| ENSG00000197694 | 27548.29185 | 0.762042849 | 3.93E-25 | SPTAN1 |
| ENSG00000198934 | 540.1230386 | 2.674409046 | 7.95E-45 | MAGEE1 |
| ENSG00000010404 | 4286.442473 | 0.870580293 | 4.12E-21 | IDS |

| short hairpin number | Gene name | Gene description |
|---|---|---|
| sh#1 | YEATS2 | YEATS domain containing 2 |
| sh#2 | YEATS2 | YEATS domain containing 2 |
| sh#3 | YEATS2 | YEATS domain containing 2 |
| sh#4 | YEATS2 | YEATS domain containing 2 |
| sh#5 | YEATS2 | YEATS domain containing 2 |
| sh#1 | ZZZ3 | zinc finger, ZZ-type containing 3 |
| sh#2 | ZZZ3 | zinc finger, ZZ-type containing 3 |
| sh#3 | ZZZ3 | zinc finger, ZZ-type containing 3 |
| sh#4 | ZZZ3 | zinc finger, ZZ-type containing 3 |
| sh#5 | ZZZ3 | zinc finger, ZZ-type containing 3 |
| sh#1 | EP400 | E1A binding protein p400 |
| sh#2 | EP400 | E1A binding protein p400 |
| sh#3 | EP400 | E1A binding protein p400 |
| sh#4 | EP400 | E1A binding protein p400 |
| sh#5 | EP400 | E1A binding protein p400 |
| sh#1 | NCOA2 | nuclear receptor coactivator 2 |
| sh#2 | NCOA2 | nuclear receptor coactivator 2 |
| sh#3 | NCOA2 | nuclear receptor coactivator 2 |
| sh#4 | NCOA2 | nuclear receptor coactivator 2 |
| sh#5 | NCOA2 | nuclear receptor coactivator 2 |
| sh#1 | TCF20 | transcription factor 20 |
| sh#2 | TCF20 | transcription factor 20 |
| sh#3 | TCF20 | transcription factor 20 |
| sh#4 | TCF20 | transcription factor 20 |
| sh#5 | TCF20 | transcription factor 20 |
| sh#1 | NCOR2 | nuclear receptor co-repressor 2 |
| sh#2 | NCOR2 | nuclear receptor co-repressor 2 |
| sh#3 | NCOR2 | nuclear receptor co-repressor 2 |
| sh#4 | NCOR2 | nuclear receptor co-repressor 2 |
| sh#5 | NCOR2 | nuclear receptor co-repressor 2 |
| sh#1 | EP300 | E1A binding protein p300 |
| sh#2 | EP300 | E1A binding protein p300 |
| sh#3 | EP300 | E1A binding protein p300 |
| sh#4 | EP300 | E1A binding protein p300 |
| sh#5 | EP300 | E1A binding protein p300 |
| sh#1 | CSRP2BP (KAT14) | CSRP2 binding protein (KAT14) |
| sh#2 | CSRP2BP (KAT14) | CSRP2 binding protein (KAT14) |
| sh#3 | CSRP2BP (KAT14) | CSRP2 binding protein (KAT14) |
| sh#4 | CSRP2BP (KAT14) | CSRP2 binding protein (KAT14) |
| sh#5 | CSRP2BP (KAT14) | CSRP2 binding protein (KAT14) |
| sh#1 | NFRKB | nuclear factor related to kappaB binding protein |
| sh#2 | NFRKB | nuclear factor related to kappaB binding protein |
| sh#3 | NFRKB | nuclear factor related to kappaB binding protein |
| sh#4 | NFRKB | nuclear factor related to kappaB binding protein |
| sh#5 | NFRKB | nuclear factor related to kappaB binding protein |
| sh#1 | TRIM24 | tripartite motif-containing 24 |

| | | |
|---|---|---|
| sh#2 | TRIM24 | tripartite motif-containing 24 |
| sh#3 | TRIM24 | tripartite motif-containing 24 |
| sh#4 | TRIM24 | tripartite motif-containing 24 |
| sh#5 | TRIM24 | tripartite motif-containing 24 |
| sh#1 | CREBBP | CREB binding protein |
| sh#2 | CREBBP | CREB binding protein |
| sh#3 | CREBBP | CREB binding protein |
| sh#4 | CREBBP | CREB binding protein |
| sh#5 | CREBBP | CREB binding protein |
| sh#1 | NCOA6 | nuclear receptor coactivator 6 |
| sh#2 | NCOA6 | nuclear receptor coactivator 6 |
| sh#3 | NCOA6 | nuclear receptor coactivator 6 |
| sh#4 | NCOA6 | nuclear receptor coactivator 6 |
| sh#5 | NCOA6 | nuclear receptor coactivator 6 |
| sh#1 | EPC2 | enhancer of polycomb homolog 2 |
| sh#2 | EPC2 | enhancer of polycomb homolog 2 |
| sh#3 | EPC2 | enhancer of polycomb homolog 2 |
| sh#4 | EPC2 | enhancer of polycomb homolog 2 |
| sh#5 | EPC2 | enhancer of polycomb homolog 2 |
| sh#1 | IRF2BP2 | interferon regulatory factor 2 binding protein 2 |
| sh#2 | IRF2BP2 | interferon regulatory factor 2 binding protein 2 |
| sh#3 | IRF2BP2 | interferon regulatory factor 2 binding protein 2 |
| sh#4 | IRF2BP2 | interferon regulatory factor 2 binding protein 2 |
| sh#5 | IRF2BP2 | interferon regulatory factor 2 binding protein 2 |
| sh#1 | NCOR1 | nuclear receptor co-repressor 1 |
| sh#2 | NCOR1 | nuclear receptor co-repressor 1 |
| sh#3 | NCOR1 | nuclear receptor co-repressor 1 |
| sh#4 | NCOR1 | nuclear receptor co-repressor 1 |
| sh#5 | NCOR1 | nuclear receptor co-repressor 1 |
| sh#1 | MLL2 (KMT2D) | myeloid/lymphoid or mixed-lineage leukemia 2 |
| sh#2 | MLL2 (KMT2D) | myeloid/lymphoid or mixed-lineage leukemia 2 |
| sh#3 | MLL2 (KMT2D) | myeloid/lymphoid or mixed-lineage leukemia 2 |
| sh#4 | MLL2 (KMT2D) | myeloid/lymphoid or mixed-lineage leukemia 2 |
| sh#5 | MLL2 (KMT2D) | myeloid/lymphoid or mixed-lineage leukemia 2 |
| sh#1 | EPC1 | enhancer of polycomb homolog 1 |
| sh#2 | EPC1 | enhancer of polycomb homolog 1 |
| sh#3 | EPC1 | enhancer of polycomb homolog 1 |
| sh#4 | EPC1 | enhancer of polycomb homolog 1 |
| sh#5 | EPC1 | enhancer of polycomb homolog 1 |
| sh#1 | C11orf30 (EMSY) | chromosome 11 open reading frame 30 |
| sh#2 | C11orf30 (EMSY) | chromosome 11 open reading frame 30 |
| sh#3 | C11orf30 (EMSY) | chromosome 11 open reading frame 30 |
| sh#4 | C11orf30 (EMSY) | chromosome 11 open reading frame 30 |
| sh#5 | C11orf30 (EMSY) | chromosome 11 open reading frame 30 |

| Clone ID |
|---|
| TRCN0000144252 |
| TRCN0000144134 |
| TRCN0000144016 |
| TRCN0000140296 |
| TRCN0000431997 |
| TRCN0000376579 |
| TRCN0000370301 |
| TRCN0000135859 |
| TRCN0000133685 |
| TRCN0000370272 |
| TRCN0000370638 |
| TRCN0000312678 |
| TRCN0000312679 |
| TRCN0000312676 |
| TRCN0000050260 |
| TRCN0000276447 |
| TRCN0000019642 |
| TRCN0000019639 |
| TRCN0000319399 |
| TRCN0000019640 |
| TRCN0000015490 |
| TRCN0000297821 |
| TRCN0000280429 |
| TRCN0000280378 |
| TRCN0000015492 |
| TRCN0000060707 |
| TRCN0000060704 |
| TRCN0000060703 |
| TRCN0000060705 |
| TRCN0000060706 |
| TRCN0000039885 |
| TRCN0000039886 |
| TRCN0000039883 |
| TRCN0000231133 |
| TRCN0000231134 |
| TRCN0000107700 |
| TRCN0000107701 |
| TRCN0000107702 |
| TRCN0000107703 |
| TRCN0000107704 |
| TRCN0000014868 |
| TRCN0000014870 |
| TRCN0000232106 |
| TRCN0000014872 |
| TRCN0000232107 |
| TRCN0000021260 |

TRCN0000021259
TRCN0000021262
TRCN0000195174
TRCN0000194983
TRCN0000006485
TRCN0000006486
TRCN0000006488
TRCN0000011027
TRCN0000367481
TRCN0000063263
TRCN0000063264
TRCN0000303657
TRCN0000303658
TRCN0000299538
TRCN0000062734
TRCN0000062735
TRCN0000062737
TRCN0000062736
TRCN0000363358
TRCN0000364917
TRCN0000364916
TRCN0000364854
TRCN0000034174
TRCN0000364851
TRCN0000060656
TRCN0000299678
TRCN0000299677
TRCN0000303750
TRCN0000310464
TRCN0000013138
TRCN0000013140
TRCN0000013139
TRCN0000013141
TRCN0000235741
TRCN0000073264
TRCN0000073265
TRCN0000073266
TRCN0000073263
TRCN0000229403
TRCN0000230140
TRCN0000230141
TRCN0000218503
TRCN0000218618
TRCN0000230142

**Oligo Sequence**

CCGGCGTCAGAGTTCAAGTTCATTTCTCGAGAAATGAACTTGAACTCTGACGTTTTTG
CCGGCCTAACTACAAACAGCAAGAACTCGAGTTCTTGCTGTTTGTAGTTAGGTTTTTG
CCGGCCAGTCAGAAATCTGTTCTATCTCGAGATAGAACAGATTTCTGACTGGTTTTTG
CCGGGCGAGCTACGAACAATGCTAACTCGAGTTAGCATTGTTCGTAGCTCGCTTTTTG
CCGGATAACAGCAATATGGATATAGCTCGAGCTATATCCATATTGCTGTTATTTTTTG
CCGGTCAAAGAACTTGGTCATAAATCTCGAGATTTATGACCAAGTTCTTTGATTTTTG
CCGGTTCCCGATCTACTCGTGTTACCTCGAGGTAACACGAGTAGATCGGGAATTTTTG
CCGGCGCATCCTGAAGAAATCTCTTCTCGAGAAGAGATTTCTTCAGGATGCGTTTTTG
CCGGGCATCAGATGACGAAAGTATTCTCGAGAATACTTTCGTCATCTGATGCTTTTTG
CCGGGACGACAGCACCCTCTTAATACTCGAGTATTAAGAGGGTGCTGTCGTCTTTTTG
CCGGGCCTTATACGAGGACGTTATCCTCGAGGATAACGTCCTCGTATAAGGCTTTTTG
CCGGTCACAACCTCGGTCAAGTTTACTCGAGTAAACTTGACCGAGGTTGTGATTTTTG
CCGGGGAGGTCTTCAAGGAGTATTTCTCGAGAAATACTCCTTGAAGACCTCCTTTTTG
CCGGCAGCCGTACCACAGGTATAAACTCGAGTTTATACCTGTGGTACGGCTGTTTTTG
CCGGGCGGAAACTCATGGAGGAAATCTCGAGATTTCCTCCATGAGTTTCCGCTTTTTG
CCGGATCCGTTCTCAGACTACTAATCTCGAGATTAGTAGTCTGAGAACGGATTTTTTG
CCGGGCACAGGAAATAGCCATAGTTCTCGAGAACTATGGCTATTTCCTGTGCTTTTT
CCGGGCACTCTTGTTGCTGCACAAACTCGAGTTTGTGCAGCAACAAGAGTGCTTTTT
CCGGGCACTCTTGTTGCTGCACAAACTCGAGTTTGTGCAGCAACAAGAGTGCTTTTTG
CCGGCCAGGAATGATGGGTAATCAACTCGAGTTGATTACCCATCATTCCTGGTTTTT
CCGGCGCTTGATGATATACTGTCTTCTCGAGAAGACAGTATATCATCAAGCGTTTTT
CCGGGCACAGGCTTATGGAACACAACTCGAGTTGTGTTCCATAAGCCTGTGCTTTTTG
CCGGGCTAGTTTCAACAACAAGAAACTCGAGTTTCTTGTTGTTGAAACTAGCTTTTTG
CCGGCGCTTGATGATATACTGTCTTCTCGAGAAGACAGTATATCATCAAGCGTTTTTG
CCGGGCACAGGCTTATGGAACACAACTCGAGTTGTGTTCCATAAGCCTGTGCTTTTT
CCGGGCAGTGTAAGAACTTCTACTTCTCGAGAAGTAGAAGTTCTTACACTGCTTTTTG
CCGGCCTCTATTACTACCTGACTAACTCGAGTTAGTCAGGTAGTAATAGAGGTTTTTG
CCGGGCCTGTCTAAAGCCTTAACTACTCGAGTAGTTAAGGCTTTAGACAGGCTTTTTG
CCGGGCTTCACAACACAGGCATGAACTCGAGTTCATGCCTGTGTTGTGAAGCTTTTTG
CCGGGCAGCGCATCAAGTTCATCAACTCGAGTTGATGAACTTGATGCGCTGCTTTTTG
CCGGGCCTTCACAATTCCGAGACATCTCGAGATGTCTCGGAATTGTGAAGGCTTTTTG
CCGGCCCGGTGAACTCTCCTATAATCTCGAGATTATAGGAGAGTTCACCGGGTTTTTG
CCGGCCTCACTTTATGGAAGAGTTACTCGAGTAACTCTTCCATAAAGTGAGGTTTTTG
CCGGTAACCAATGGTGGTGATATTACTCGAGTAATATCACCACCATTGGTTATTTTTG
CCGGATACTCAGCCGGAGGATATTTCTCGAGAAATATCCTCCGGCTGAGTATTTTTTG
CCGGCGGCCATTGATGTGTTGTCATCTCGAGATGACAACACATCAATGGCCGTTTTTG
CCGGGCTGTCCTACTTCTGTGACAACTCGAGTTGTCACAGAAGTAGGACAGCTTTTTG
CCGGCCCTTATACCTCTCGGATCTTCTCGAGAAGATCCGAGAGGTATAAGGGTTTTTG
CCGGCAAGGTTATTTCAGGTGGAAACTCGAGTTTCCACCTGAAATAACCTTGTTTTTG
CCGGGCACCTCTCGATTACTGTTATCTCGAGATAACAGTAATCGAGAGGTGCTTTTTG
CCGGCAGGGAGGTTGCATCATTGTTCTCGAGAACAATGATGCAACCTCCCTGTTTTT
CCGGCCCTGTGTGAAATACGACATTCTCGAGAATGTCGTATTTCACACAGGGTTTTT
CCGGTCACCAGCACTCAGGTAAATACTCGAGTATTTACCTGAGTGCTGGTGATTTTTG
CCGGGCCACGTTGGATTTACAAGAACTCGAGTTCTTGTAAATCCAACGTGGCTTTTT
CCGGAGAGGATGGCTTCCGTGAAAGCTCGAGCTTTCACGGAAGCCATCCTCTTTTTG
CCGGCCATGAAATGAGCCTGGCTTTCTCGAGAAAGCCAGGCTCATTTCATGGTTTTT

CCGGCGCATGAAACTTATGCAACAACTCGAGTTGTTGCATAAGTTTCATGCGTTTTT
CCGGGCAGCAGTACAGCATTACTTTCTCGAGAAAGTAATGCTGTACTGCTGCTTTTT
CCGGCCTTGTTAAGTTAACACCTATCTCGAGATAGGTGTTAACTTAACAAGGTTTTTG
CCGGCCATTTCTGTTAACCTCTTATCTCGAGATAAGAGGTTAACAGAAATGGTTTTTG
CCGGCCCGATAACTTTGTGATGTTTCTCGAGAAACATCACAAAGTTATCGGGTTTTT
CCGGGCTATCAGAATAGGTATCATTCTCGAGAATGATACCTATTCTGATAGCTTTTT
CCGGCCGTTTACCATGAGATCCTTACTCGAGTAAGGATCTCATGGTAAACGGTTTTT
CCGGCGTGCCAAATATGTCTCAGATCTCGAGATCTGAGACATATTTGGCACGTTTTT
CCGGTAACTCTGGCCATAGCTTAATCTCGAGATTAAGCTATGGCCAGAGTTATTTTTG
CCGGGCAGATTATGACCAATCAAATCTCGAGATTTGATTGGTCATAATCTGCTTTTTG
CCGGCCCTAAACTTACCTCACAGTACTCGAGTACTGTGAGGTAAGTTTAGGGTTTTTG
CCGGACAAATGAACCCAGCTAATTTCTCGAGAAATTAGCTGGGTTCATTTGTTTTTTG
CCGGGCCCATTGTTGGTCAACTTATCTCGAGATAAGTTGACCAACAATGGGCTTTTTG
CCGGGCAGATTATGACCAATCAAATCTCGAGATTTGATTGGTCATAATCTGCTTTTTG
CCGGGCCAGTATTATGCTCCTCGTTCTCGAGAACGAGGAGCATAATACTGGCTTTTTG
CCGGCGAGAATTTAGTAGAGCCATACTCGAGTATGGCTCTACTAAATTCTCGTTTTTG
CCGGCCACACCACTACAAACCACTTCTCGAGAAGTGGTTTGTAGTGGTGTGGTTTTTG
CCGGGCAGAGAGCAACGTCAACTATCTCGAGATAGTTGACGTTGCTCTCTGCTTTTTG
CCGGGACATTGCCTGTGATCAATAACTCGAGTTATTGATCACAGGCAATGTCTTTTTG
CCGGGGCCCTTCGAGAGCAAGTTTACTCGAGTAAACTTGCTCTCGAAGGGCCTTTTTG
CCGGAGCTAGTGGAGAGGTCTATTGCTCGAGCAATAGACCTCTCCACTAGCTTTTTTG
CCGGCCCAATGATTACCCTTAATTACTCGAGTAATTAAGGGTAATCATTGGGTTTTTG
CCGGCCTGATCTTAGTAGCAGACAACTCGAGTTGTCTGCTACTAAGATCAGGTTTTTG
CCGGTGATCTTAGTAGCAGACAATGCTCGAGCATTGTCTGCTACTAAGATCATTTTTG
CCGGGCCATCAAACACAATGTCAAACTCGAGTTTGACATTGTGTTTGATGGCTTTTTG
CCGGGCCCACAGATGATGAAGAAATCTCGAGATTTCTTCATCATCTGTGGGCTTTTTG
CCGGGCTCTCAAAGTTCAGACTCTTCTCGAGAAGAGTCTGAACTTTGAGAGCTTTTTG
CCGGCTATCAGCCAGAGGTTGTTAACTCGAGTTAACAACCTCTGGCTGATAGTTTTTG
CCGGGCCATCAAACACAATGTCAAACTCGAGTTTGACATTGTGTTTGATGGCTTTTTG
CCGGCCCACCTGAATCATCACCTTTCTCGAGAAAGGTGATGATTCAGGTGGGTTTTT
CCGGCCTCGCCTCAAGAAATGGAAACTCGAGTTTCCATTTCTTGAGGCGAGGTTTTT
CCGGCCTGAATTGAACAACAGTCTTCTCGAGAAGACTGTTGTTCAATTCAGGTTTTT
CCGGCGCTTTGAGTTGCCATTTGATCTCGAGATCAAATGGCAACTCAAAGCGTTTTT
CCGGCTGTTGTATAGTTGGACTATGCTCGAGCATAGTCCAACTATACAACAGTTTTTG
CCGGCCAGTCTTCAATGCTAAAGATCTCGAGATCTTTAGCATTGAAGACTGGTTTTTG
CCGGGCTTTGTTACAGCCTTCAAATCTCGAGATTTGAAGGCTGTAACAAAGCTTTTTG
CCGGGCTCATTCACATACAGCCTTTCTCGAGAAAGGCTGTATGTGAATGAGCTTTTTG
CCGGGCCGAACAATACCAGCAACATCTCGAGATGTTGCTGGTATTGTTCGGCTTTTTG
CCGGATCCGACCGAAACGGAAATATCTCGAGATATTTCCGTTTCGGTCGGATTTTTTG
CCGGCTACGAGTGGTAGCATCATTACTCGAGTAATGATGCTACCACTCGTAGTTTTTG
CCGGAGACTAGCCCTACCATAATTTCTCGAGAAATTATGGTAGGGCTAGTCTTTTTTG
GTACCGGCGTACTCCTTCTGACAAATTTCTCGAGAAATTTGTCAGAAGGAGTACGTTTTTTG
GTACCGGTAATATGTCTGGACCTAATAGCTCGAGCTATTAGGTCCAGACATATTATTTTTTG
CCGGCATTATGCAGCAGGCATTAAGCTCGAGCTTAATGCCTGCTGCATAATGTTTTTG

| RefSeq ID |
|---|
| NM_018023 |
| NM_018023 |
| NM_018023 |
| NM_018023 |
| NM_018023 |
| NM_015534 |
| NM_015534 |
| NM_015534 |
| NM_015534 |
| NM_015534 |
| NM_015409 |
| NM_015409 |
| NM_015409 |
| NM_015409 |
| NM_015409 |
| NM_006540 |
| NM_006540 |
| NM_006540 |
| NM_006540 |
| NM_006540 |
| NM_005650 |
| NM_005650 |
| NM_005650 |
| NM_005650 |
| NM_005650 |
| NM_006312 |
| NM_006312 |
| NM_006312 |
| NM_006312 |
| NM_006312 |
| NM_001429 |
| NM_001429 |
| NM_001429 |
| NM_001429 |
| NM_001429 |
| NM_020536 |
| NM_020536 |
| NM_020536 |
| NM_020536 |
| NM_020536 |
| NM_006165 |
| NM_006165 |
| NM_006165 |
| NM_006165 |
| NM_006165 |
| NM_003852 |

NM_003852
NM_003852
NM_003852
NM_003852
NM_004380
NM_004380
NM_004380
NM_004380
NM_004380
NM_014071
NM_014071
NM_014071
NM_014071
NM_014071
NM_015630
NM_015630
NM_015630
NM_015630
NM_015630
NM_182972
NM_182972
NM_182972
NM_182972
NM_182972
NM_006311
NM_006311
NM_006311
NM_006311
NM_006311
NM_003482
NM_003482
NM_003482
NM_003482
NM_003482
NM_025209
NM_025209
NM_025209
NM_025209
NM_025209
NM_020193
NM_020193
NM_020193
NM_020193
NM_020193
